# Proton-secreting cells modulate mucosal immune surveillance in the male reproductive tract

**DOI:** 10.1101/2025.03.26.645301

**Authors:** AAS Da Silva, F Barrachina, MC Avenatti, ML Elizagaray, I Bastepe, E Sasso-Cerri, MA Battistone

## Abstract

Proton-secreting cells in various organs, such as the kidney and epididymis, regulate pH balance, maintaining cellular homeostasis, and supporting key physiological processes. More recently, these specialized cells have emerged as key contributors to mucosal immunity, orchestrating immune activation. Epididymitis is an inflammatory condition that significantly impacts male fertility, often due to a lack of diagnosis and treatment. This study explores the involvement of region-specific epididymal proton-secreting clear cells (CCs) in the immune response by interacting with the immune system during LPS-induced mouse epididymitis. We found that in response to LPS, CCs rapidly shifted to a proinflammatory phenotype, marked by the upregulation of cytokines and chemokines, alongside the downregulation of genes involved in sperm maturation. Morphological changes in CCs, including increased apical blebs and altered shape across different epididymal segments, suggest their active role in immune responses. Moreover, mononuclear phagocytes (MPs) reduced their luminal-reaching projections in the proximal epididymis after the LPS challenge. This bacteria antigen triggered the migration of dendritic cells and neutrophil infiltration in the distal epididymis. These immune landscape alterations contributed to epithelial damage and impaired sperm maturation, as evidenced by decreased sperm motility following LPS injection. Our findings indicate that proton-secreting cells are immune gatekeepers in the epididymis, initiating immune responses and disrupting sperm maturation. This research enhances the understanding of epithelial immunoregulation and will help to develop novel diagnostic and therapeutic strategies for epididymitis and male infertility. Furthermore, insights into CC-mediated immune responses could inform the development of new approaches for male contraception.

## Introduction

Epithelia provides a barrier to the environment and constitutes the first line of defense against stressors. Some epithelia have an even more complex function, as they also need to provide a balance between tolerance and immune activation. This relates in particular – but not exclusively – to the epididymal epithelium, where control of autoimmune responses against antigenic spermatozoa, while protecting against pathogens, is a key determinant of male fertility (Breton et al., 2019). This balance is maintained through intricate interactions between different epithelial cell types and immune cells (Voisin et al., 2018; Battistone et al., 2024). The epididymis exhibits differential immune responses along its length, shaped by a region-specific immune cell distribution that protects developing sperm and combats infections; however, severe inflammation can compromise the epididymis, potentially leading to fertility issues (Jhonson et al 2005; Michel et al., 2016; Klein et al., 2020; Pleuger et al., 2020, da Silva 2024).

Epididymitis, the most common intrascrotal inflammation, affects approximately 400 per 100,000 men annually worldwide (Pilatz et al., 2015, Redshaw et al., 2014). It is frequently caused by bacteria ascending the urogenital tract, from sexually transmitted diseases or urinary tract infections, caused by *Escherichia Coli* among others (Michel et al., 2016; Pleuger et al., 2020). Persistent oligozoospermia and azoospermia occur in up to 40% of males with epididymitis (Michel et al., 2016), underscoring the need for studies on the molecular mechanisms of this inflammatory process. Lipopolysaccharide (LPS)-induced epididymitis is a widely used mouse model that mimics this inflammatory male disease (Battistone et al., 2019; Barrachina et al., 2022, Zang et al., 2009; Liu et al., 2023). The model is characterized by high levels of proinflammatory cytokines in the distal epididymis (Liu et al., 2023; Andrade et al., 2021; Battistone et al., 2019) and immune infiltration (Battistone et al., 2019). Additionally, LPS alters the transcriptional profile of *Wfdc* genes, which are involved in innate immunity and fertility (Andrade et al., 2021). RNA sequencing (RNA-seq) of whole rat epididymis following proximal LPS injection revealed upregulated genes enriched in inflammatory-related processes (Song et al., 2019).

In the epididymis, epithelial proton-secreting clear cells (CCs), strategically located within the epididymal epithelium, work with resident mononuclear phagocytes (MPs) to balance inflammation and immune tolerance (Battistone et al., 2019). These proton-secreting cells not only contribute to sperm maturation by establishing an acidic environment (Breton and Brown, 2013) but also function as immune sensors and mediators, expressing chemokines in response to bacterial antigens and viral infection (Battistone et al., 2019; da Silva et al., 2024). A previous study by our group demonstrated that a 6-h intravasal-epididymal injection of LPS upregulates the expression of chemokines and cytokines in isolated CCs, including *Cxcl10*, *Cxcl1*, *Cxcl2*, *Il6*, and *Ccl5*. This discovery highlighted the involvement of these cells in the epididymal immune responses (Battistone et al., 2019). However, the specific mechanisms by which these cells contribute to the epididymal mucosal immunity remain poorly understood. Similar proton-secreting cells are found in the kidney (Battistone et al., 2020; Brown et al., 2009) and the respiratory tract (Paranjapye et al., 2022), making studying their role crucial for advancing our understanding of mucosal functions across other systems. Importantly, renal proton-secreting intercalated cells play a significant role in inflammation-induced acute kidney injury (Battistone et al., 2020).

Given the impact of inflammation on male fertility and the role of CCs in modulating epididymal immune responses, we aimed to characterize the molecular mechanisms by which epithelial CCs respond to LPS-induced epididymitis. RNA-seq analysis revealed an early activation of CCs following LPS-triggered inflammation, leading to the upregulation of several pro-inflammatory genes, activation of MPs, and neutrophil infiltration. LPS challenge also altered the distal epididymis/cauda luminal pH and sperm motility. Our findings demonstrate that CCs, in collaboration with MPs, are likely modulators of the early immune response in epididymitis.

## Materials and Methods

### Animals

Adult male transgenic mice (12 -15 weeks) that express EGFP under the control of the ATP6V1B1 promoter (Miller et al., 2005) were used in this study; they are referred to as B1-EGFP transgenic mice. Adult C57BL/6 wild-type male mice (12-15 weeks) were purchased from the Jackson Laboratories (Bar Harbor, ME, USA). All procedures performed were approved by the Massachusetts General Hospital Subcommittee on Research Animal Care and were executed following the National Institutes of Health (NIH) Guide for the Care and Use of Laboratory Animals.

### Animal model of epididymitis

Intravasal-epididymal injection was performed as previously reported (Silva et al., 2018; Battistone et al., 2019). Males were anesthetized with 2% isoflurane (Baxter, Deerfield, IL) (mixed with oxygen). A single incision in the scrotum was made to expose the proximal portion of the vas deferens, adjacent to the cauda epididymis. LPS (25 μg, 1 mg/ml endotoxin units; purity ≥99.9%, S-form; Innaxon, Oakield Close, UK) or sterile saline (Hospira, Lake Forest, IL) were injected into the vas deferens using a 31-G needle (25 μl). The injection was made bilaterally in a retrograde direction into the cauda epididymis. Mice were euthanized at 1 h, 4 h, 24 h, or 48 h post-injections.

### Isolation of EGFP^+^ CCs from B1-EGFP mice, RNA extraction, and RNA-seq

Double fluorescence-activated Cell Sorting (FACS) isolation was performed at the HSCI-CRM Flow Cytometry Core (Boston, MA). EGFP+ CCs, 1 h after intravasal-epididymal injections of saline or LPS from the epididymis of B1-EGFP mice, were isolated, as we reported (Battistone et al., 2020). The CC RNA, from the three anatomical regions (IS/caput, corpus, and cauda), was isolated using a PicoRNA kit (Thermo Fisher Scientific). Each sample was obtained from 6 epididymides. DNA contamination was removed by digestion with an RNase-free DNase set (Qiagen, Hilden, Germany). RNA was analyzed by a Bioanalyzer (Agilent RNA 6000 Pico Kit, Agilent Technologies, Santa Clara, CA). RNA-seq libraries were set using the Clontech SMARTER Kit v4, and sequencing on an Illumina HiSeq2500 instrument. Transcriptome mapping was made with STAR (Ensembl annotation of mm9 reference genome) (Dobin et al., 2013). HTSeq was used for gene reading counts (Anders et al., 2015). Differential expression analysis was made using the EdgeR package (Robinson et al., 2010) and including only those genes with counts per million (CPM) value of >1. Differentially expressed genes were defined based on the criteria of >2-fold change in expression value and P<0.05. Multiplot studio software was used to obtain volcano plots, and Morpheus was used to plot heatmaps.

### Sperm collection

Sperm were collected from the distal cauda 4 and 24 h after saline or LPS intravasal-epididymal injections. Sperm recovered by incising the cauda region three times and placing it in modified Human Tubal Fluid (HTF) medium (90126, FUJIFILM Irvine Scientific, Inc., Santa Ana, CA) supplemented with 0.3% Bovine Serum Albumin (BSA; A0281, Sigma-Aldrich). After 15 min of incubation at 37 °C, tissues were removed from the tubes, and sperm was collected and analyzed.

### Sperm Capacitation and Computer-Assisted Sperm Analysis (CASA)

The CASA analysis was performed as previously described (Barrachina et al., 2023). Sperm was obtained from the distal cauda, as indicated above. After 15 min of incubation at 37 °C, sperm was diluted (1:5 ratio) in HTF medium with 0.3 % BSA and incubated for 45 min at 37 °C to achieve sperm capacitation. Sperm analysis was performed using Hamilton Thorne’s CASA version 14 (Hamilton Thorne Inc., Beverly, MA) and 100µm thickness slides (Leja, IMV technologies). Sperm were considered hyperactivated when presenting curvilinear velocity (VCL) ≥ 238.5 µm/s, linearity (LIN) < 33%, and amplitude of lateral head displacement (ALH) ≥ 4.22 µm, as previously reported (Barrachina et al., 2023).

### Distal cauda luminal pH measurement

For the pH assessment in the distal portion of the cauda, the luminal content from the cauda distal portion was recovered by an incision in the respective region 4 h and 24 h post injections of saline or LPS using a pH strip. Two independent researchers blinded to the experimental conditions conducted the pH strip test with at least four mice per group.

### Confocal microscopy

Epididymides from B1-EGFP mice after 1 h, 24 h, and 48h post injections were removed and fixed for 4 h by immersion in 4% paraformaldehyde (PFA) at room temperature. After several washes in phosphate-buffered saline (PBS), the epididymides were soaked in 30% sucrose in PBS (with 0.02% NaAzide) for 48 h at 4 °C. Then, the tissues were embedded in Tissue-Tek OCT compound (Sakura Finetek, Torrance, CA, USA), and frozen on a cutting block in a Reichert Frigocut microtome. Epididymides were cut at 25-µm thickness and sections were placed onto Fisherbrand Superfrost Plus microscope slides (Fisher Scientific, Pittsburgh, PA, USA).

For immunofluorescence (IF) antigen retrieval, the slides were exposed to 1% SDS in PBS for 4 min (Brown et al., 1996). For the blocking: 1% bovine serum albumin in PBS for 30 min at room temperature. Slides were incubated with the primary antibodies for 18 h at 4 °C. The antibodies used were rat antibody against Ly6G (0.2 mg/ml; Clone 1A8, 551461, BD Biosciences), rabbit antibody against AQP9 (0.5 μg/mL; (Pastor-Soler et al., 2001), rabbit monoclonal antibody against cleaved caspase-3 (ASP175) (5A1E) (Cell Signaling Technology, Danvers, MA), rat antibody against F4/80 (20 μg/mL; clone BM8, 14–4801, eBioscience, San Diego, CA) and rabbit antibody against neutrophil elastase (1mg/ml; Ab68672, Abcam, Waltham, MA). The secondary antibodies (Jackson ImmunoResearch Laboratories, West Grove, PA) were donkey Cy3-conjugated anti-rat IgG (3 μg/ml; 712–166–153), Cy3 donkey anti-rat IgG (3 μg/mL;712-166-153, Jackson ImmunoResearch, West Grove, PA) and donkey Alexa Fluor 647-conjugated anti-rabbit IgG (1.5 μg/ml; 711–606–152). All antibodies were diluted in DAKO (Dako, Carpinteria, CA). Slides were mounted with DAPI in the SlowFade Diamond Mounting medium (Thermo Fisher Scientific, Waltham, MA). For negative controls, incubations were achieved with secondary antibodies alone. To evaluate the EGFP^+^ apical cellular protrusions in the CCs (CC blebs), the epididymal sections were only mounted with SlowFade Diamond Antifade Mounting medium containing DAPI.

Images were taken using a Nikon CSA-W1 SoRa spinning disk confocal microscope (Nikon, Yokogawa Electric Corporation, Tokyo, Japan) at the Molecular Imaging Core (MGH, Charlestown, MA) and the stitching images were taken using a Nikon E800. The number of F4/80^+^ luminal projections and CC blebs per tissue area (110,000 µm2) was evaluated in epididymal sections using Fiji software.

### Flow cytometry analysis

Epididymides from mice injected with LPS or saline were removed, and single-cell suspensions were made as previously described (Battistone et al., 2019^b^). Briefly, the epididymides were divided into proximal and distal regions. They were incubated for 30 min at 37 °C with gentle shaking (10 s/min at 1400 rpm) in a medium containing RPMI 1640 with collagenase type I (0.5 mg/mL, Gibco) and collagenase type II (0.5 mg/mL, Sigma). After digestion, the cell suspensions were passed through a 70 μm nylon mesh strainer, washed with 2% fetal bovine serum (FBS) with 2 mM EDTA in PBS, and centrifuged for 5 min at 400 g. Cells were incubated with different antibodies (0.8 μg/mL for each one) against CD64 Alexa Fluor 647 (Clone X54-5/7.1; 558,539, BD Biosciences), CD45 Brilliant Violet 711 (Clone 30-F11; 563,709, BD Biosciences), F4/80 PE/Cyanine 7 (Clone BM8; 123,113, Biolegend, San Diego, CA), Ly6C R718 (Clone AL-21; 566,987, BD Biosciences, San Jose, CA), Ly6G PE (0.2 mg/ml; Clone 1A8, 551461, BD Biosciences), CD11c Brilliant Violet 421 (clone N418, 117329, Biolegend), CD11b APC/Cyanine 7 (Clone M1-70; 101,225, Biolegend), which were diluted in 2% FBS in PBS with BD Horizon Brilliant Stain buffer (BD Biosciences). After 30 min, the cell suspension was washed in 2% FBS in PBS and passed through a 40 μm strainer. Flow cytometry analyses were performed at the HSCI-CRM Flow Cytometry Core (Boston, MA). DAPI was used as the viability dye for flow cytometry analyses. Data were acquired on the Flow and Mass Cytometry Core (CNY) cytometer (BD Biosciences) and analyzed using FlowJo software version 10.10 (BD Biosciences).

### Statistical Analysis

Data analysis was performed using GraphPad Prism version 10.2.3 (GraphPad Software; https://www.graphpad.com). To examine whether the samples were normally distributed, we performed a test of normality (Shapiro–Wilk test). Student’s *t-test* (two-tailed) or one-way ANOVA were used as parametric tests. Mann–Whitney *U* test (two-tailed) was used as a nonparametric test. *P*-values < 0.05 were determined statistically significant. Data were expressed as the means ± SEM.

## Results

### CCs change their transcriptomics profile after saline and LPS intravasal-epididymal injection

We characterized the transcriptomic profiles of CCs that were isolated by fluorescence-activated double-cell sorting (FACS) from the proximal, middle, and distal epididymal regions of B1 EGFP mice after LPS and saline intravasal-epididymal injections. The complete transcriptome dataset is listed in Suppl. Table 1. Initially, we analyzed the transcriptomic profiles of saline-treated CCs (saline CCs) from the 3 epididymal regions and compared them to non-injected controls (Battistone et al., 2020). We observed region-specific variations in the gene expression profile of the CCs (**Fig. 1**). CCs from the proximal epididymis of saline-injected mice showed increased expression of acid-base transport-related genes (*Adcy7, Adcy8, Atp4a*) and sperm-related genes (*Wfdc6b, Wfdc10, Lcn12*). EnrichR analysis identified pathways related to epithelial secretion and gap junctions. Notably, there was downregulation of V-ATPase subunits (*Atp6v0b, Atp6v1e1, Atp6v1c2, Atp6v0d2*) and pro-inflammatory genes (*Tnfrsf21, Nfkbid, Il18, Nfkbia*) in the CCs of this region (**Fig. 1A**). In contrast, CCs from the corpus of saline-injected mice displayed upregulation of pro-inflammatory genes (*Cacnb3, C1qa, C1qb, C1qc, Fcer1g, Vegfa, Cmtm3*) and downregulation of sperm maturation genes (*Mfge8, Hsp1b, Abcc3*) (**Fig. 1B**). These upregulated genes were associated with the complement cascade, VEGF signaling, high-affinity IgE receptor signaling, and natural killer cell cytotoxicity. CCs from the distal regions of saline-injected mice showed elevated expression of sperm maturation-related genes (*Defb37, Defb22, Abcc3, Defb26, Nqo1, Hsp1a, Hsp1b*) and acid-base transport genes (*Slc26a3, P2rx4, Atp6v0d2*). The pathway-enriched analysis identified FoxO signaling, MAPK signaling, apoptosis, infection, Toll-like receptor signaling, and TNF signaling pathways. Interestingly, under the same condition, pro-inflammatory (*Bcl2, Cd1d1, Anxa1*) and other sperm maturation-related genes (*Cdc14a, Cdc14b, Gpx5*) were downregulated in cauda CCs after saline injections compared to CCs from non-injected epididymis (**Fig. 1C**). The disruption in genes related to acidification following intravasal-epididymal saline injections may reflect a compensatory response to pH disturbances caused by the saline solution. These findings confirm that the saline-injected epididymis serves as an appropriate control for assessing LPS-induced effects on CC functionality, consistent with our previous report using flow cytometry (Battistone et al. 2020).

**Figure 1:**
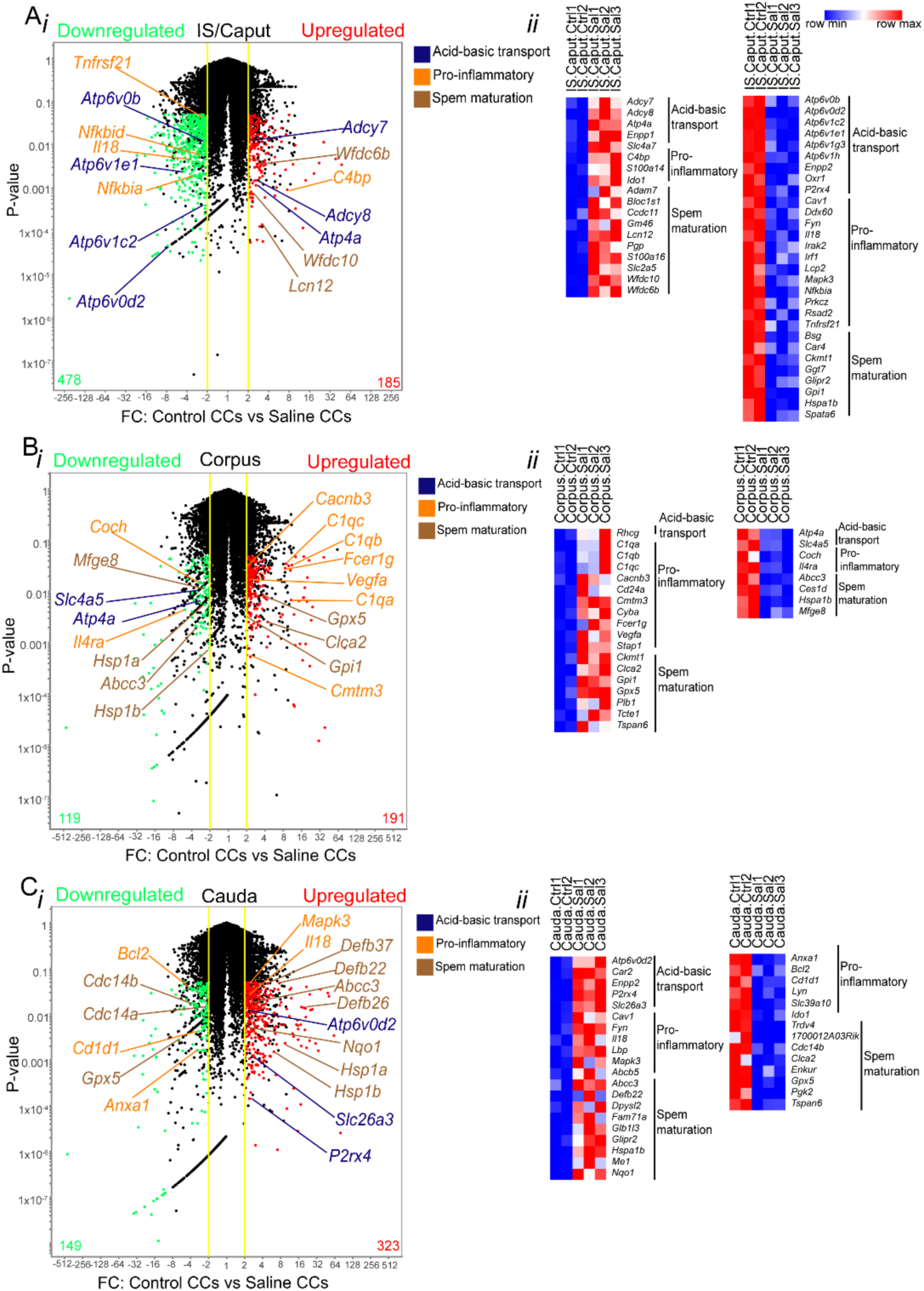
Saline injection changed the transcriptomics profile of CCs. Volcano plots show the differential gene expression profiles of EGFP^+^ CCs from control (non-injected) mice compared to those subjected to saline (25 µl) intravasal-epididymal injection in the IS/caput (**Ai**), corpus (**Bi**), and cauda (**Ci**) regions of the mouse epididymis. Upregulated genes are represented in red, while downregulated genes are shown in green. The most significant genes associated with acid-base transport, pro-inflammatory responses, and sperm maturation are highlighted. FC: Fold Change. The yellow lines show ±2FC. The black dots represent transcripts that are not significantly differentially expressed. Data was analyzed using the student’s *t*-test, two-tailed; the value of *P*<0.05 was considered significant. Heat-map of the genes related to acid-base transport, pro-inflammation, and sperm maturation genes upregulated (right) and downregulated (left) from IS/caput (**Aii**), corpus (**Bii**), and cauda (**Cii**). The complete list is provided in Suppl. Table 1. Heatmaps represent row-normalized gene expression of the indicated genes using a color gradient scale ranging from higher (red) to lower (blue) relative levels.

Following the LPS challenge, CCs from all three evaluated epididymal regions changed their transcriptomic profile compared to saline injection (**Fig. 2)**. CCs isolated from the distal region after LPS injection showed the highest number of distinct genes compared to saline CCs, as illustrated in the volcano plots comparing the gene expression profiles of proximal, middle and distal CCs. Middle CCs showed fewer genes differentially expressed between LPS and saline. Specifically, IS/Caput CCs showed an upregulation of immune-related genes, including *Cxcl1*, *Ptgr1*, and *Mir130a* (**Fig. 2A**), distal CCs exhibited an increased expression of genes such as *Cxcl1*, *Cxcl16*, *Mir150*, *Dusp4*, *Dusp10,* and *Atf3* (**Fig. 2C**). EnrichR analysis of upregulated genes in proximal LPS-treated CCs revealed significant enrichment in pathways related to epithelial cell signaling during bacterial infections, even in regions far from the injection site. In distal LPS-treated CCs, upregulated genes were associated with epithelial cell signaling during bacterial infections, chemokine signaling pathways, and NOD-like receptor signaling pathways, as expected. Following LPS exposure, CCs downregulated genes associated with sperm maturation in all epididymal regions. In proximal CCs, genes such as *Spag11a* and *Rnase9* were downregulated (**Fig. 2A**). In corpus CCs, downregulated genes included *Spag11a*, *Defb43*, *Defb40*, *Wfdc9*, and *Adam7* (**Fig. 2B**). Similarly, in distal CCs, genes such as *Defb8*, *Defb29*, *Defb47*, *Wfdc13*, *Adam7*, and *Gpx5* showed decreased expression (**Fig. 2C**). Interestingly, enriched pathways downregulated in CCs were primarily linked to pathways involved in tyrosine, arginine, proline, and glutathione metabolism. We also found downregulation of several beta-defensins in CCs across all epididymal regions comparing LPS and saline injections (**Fig. 2D**). Our findings demonstrated that CCs quickly shift to pro-inflammatory phenotype after the LPS challenge.

**Figure 2:**
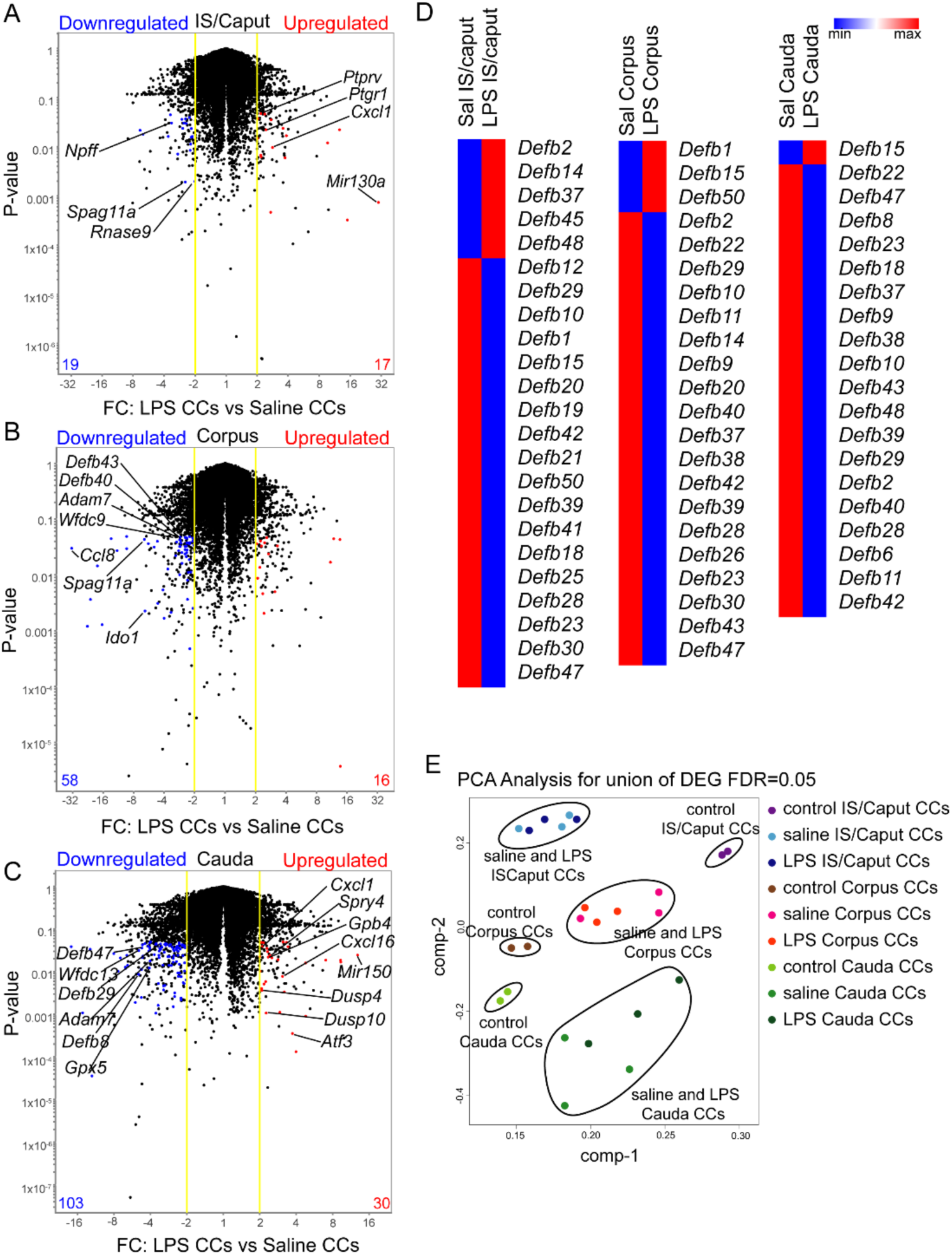
LPS injections changed the CC transcriptomics profile. Volcano plots show the differential gene expression profiles of EGFP^+^ CCs from saline versus LPS (25 µg) intravasal-epididymal injected epididymis in the IS/caput (**A**), corpus (**B**), and cauda (**C**) regions of the mouse epididymis. Upregulated genes are in red, and downregulated genes are in blue, highlighted are some significant genes related to pro-inflammation and sperm maturation genes. FC: Fold Change. The yellow lines show ±2FC. The black dots represent transcripts that are not significantly differentially expressed. Data was analyzed using the student’s *t*-test, two-tailed; the value of *P*<0.05 was considered significant. Each sample of RNA was obtained from double cell sorting live EGFP^+^ CCs from a pool of 4-12 epididymides. Heat-map of the up and downregulated beta-defensins (**D**) from IS/caput, corpus, and cauda. Heatmaps represent row-normalized gene expression of the indicated genes using a color gradient scale ranging from higher (red) to lower (blue) relative levels. Three-dimensional PCA plot of the transcriptomes (population-level RNA-seq) from cauda, corpus, and caput CCs, showing differential gene expression patterns in cells from the three regions (**E**).

Principal Component Analysis (PCA) of all the RNA-seq data revealed a significant separation of CCs from different epididymal regions based on global transcriptome expression profiles, consistent with our previous findings (Battistone et al., 2020). Interestingly, while LPS- and saline-treated CCs clustered closely together, they were distinctly separated from control (non-injected) CCs, indicating transcriptomic differences induced by intravasal injections of saline and LPS (**Fig. 2E**). Additionally, this analysis highlights that CCs from regions distant from the injection site also exhibit altered responses. These findings suggest that region-specific effects may be induced following cauda injections, offering new insights into the molecular mechanisms governing distant tissue responses.

Using previous transcriptomics data from whole epididymis (Johnston et al., 2005), we made Venn diagrams comparing segment 7 with other segments grouped into proximal (1, 2, and 3), middle (4, 5, and 6), and distal (8, 9, and 10) regions and analyzed the distinct immune response- and sperm maturation-related gene expression. Notably, we revealed that segment 7 shows significant enrichment in immune response genes compared to the other segments (**Suppl. Fig. 1A, Suppl. Table 2**). However, concerning sperm maturation, segment 7 displays similar expression profiles to the other regions (**Suppl. Fig. 1B, Suppl. Table 3**), suggesting that immune response may play a more prominent role in this segment than sperm maturation.

### CC morphological alterations after the LPS challenge

In previous work, we demonstrated that, under physiological conditions, CCs in the proximal epididymis exhibit apical membrane protrusions (Battistone et al., 2020). To examine the effects of the LPS challenge on these CC membrane blebs, we performed intravasal-epididymal injections on B1-EGFP mice. Following 1, 24, and 48 h of LPS-induced epididymitis, we observed a marked increase in the number of EGFP^+^ apical blebs in CCs within the proximal epididymis (segments 1-2) compared to CCs in saline-injected controls only 48 h (**Fig. 3A**, arrows) while no difference was observed at 1 and 24 h (**Fig. 3B**). The number of CCs did not change in the epididymal regions post-intravasal epididymal injections (**Fig. 3C**). These cells were negative for the cleaved caspase-3 (**Fig. 3D**), indicating that their morphology changes might not be related to apoptosis. In the distal portion, CCs also showed morphological changes at 1 and 24 h post-injection with either LPS or saline. Specifically, CCs at segments 7-8 displayed irregular shapes with vacuoles (**Fig. 4A, B**). Importantly, we did not observe changes in CC morphology in the epididymal segments 8-9 (injection site) and no differences in the luminal pH of the distal portion (segments 9-10) 4 h post-LPS injections (**Fig. 4C**). However, the luminal pH increased 24h after LPS injections (**Fig. 4D**). Interestingly, CCs from all epididymal portions showed upregulated genes related to vesicle formation, endocytosis, tight junctions and regulation of actin cytoskeleton after the LPS challenge: CCs from IS/Caput showed upregulation of *Rnd1*, *Pdcd6ip;* CCs from corpus *Sytl5, Inf2, Rab22a, Rhoq, Rhog, Rhoc,* and CCs, from cauda, upregulated *Sytl5, Sytl4, Rnd3, cd81, Anxa6* (**Fig. 4E**).

**Figure 3:**
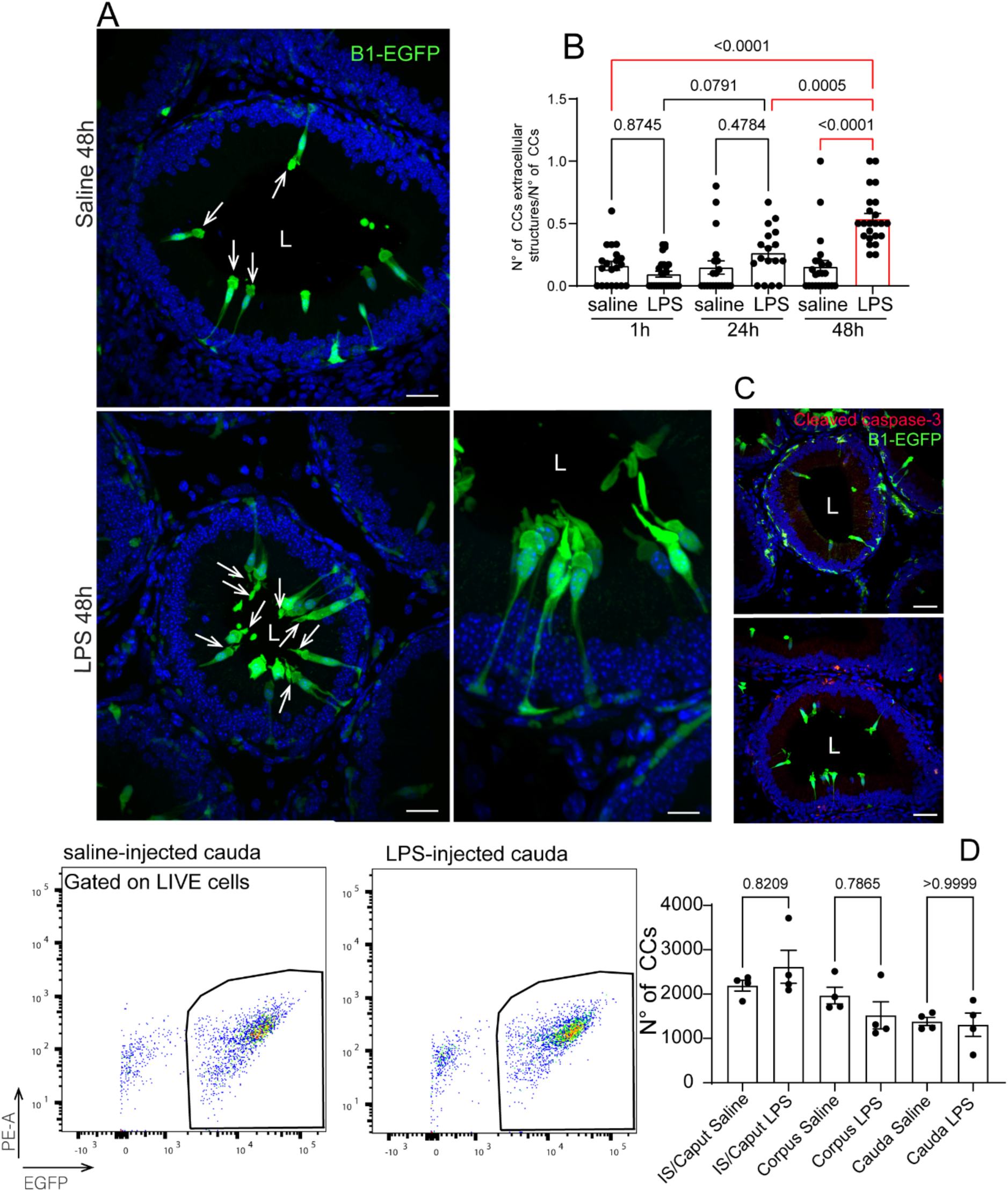
Altered morphology of proximal CCs after LPS injections. Confocal microscopy images of EGFP^+^ CCs from IS. In panel **A**, note the apical blebs in the CCs from both saline and LPS (arrows). The number of EGFP^+^ blebs in the CCs showed no difference between 1- and 24-h post-intravasal-epididymal injection of LPS while increasing after 48-h post-injections (**B**). The cell sorting analysis showed no difference in the number of EGFP^+^ CCs between saline and LPS in the epididymal regions (**C**). Data was analyzed using one-way ANOVA. Data are shown as means ± SEM. **D**) Immunolabeling for cleaved caspase-3 (red) in IS shows that the EGFP^+^ CCs are negative for this marker. Lumen (L).

**Figure 4:**
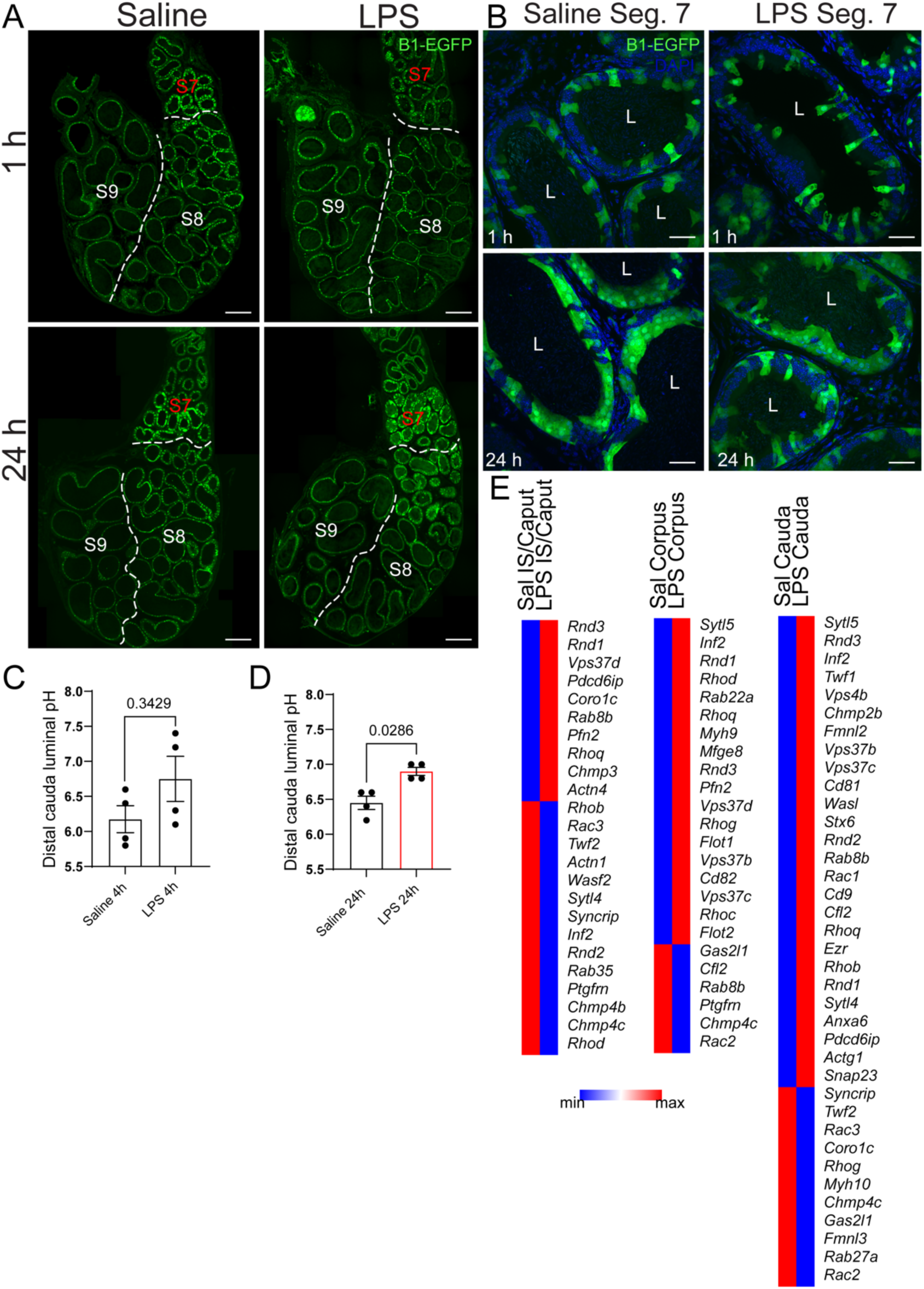
Altered morphology of distal CCs post-LPS injections. Confocal microscopy images of EGFP^+^ CCs from distal epididymis 1 h and 24 h post-intravasal-epididymal injections of saline or LPS. **A**) segmental division of distal epididymis: segments 7, 8 and 9. **B**) CCs from segment 7 showed irregular shapes 1 h and 24 h post-intravasal-epididymal injections. In segment 9 (near the injection site), the luminal pH was similar at 4 h but showed a marked increase at 24 h following LPS injection compared to saline (**C, D**). (**E**) Heatmap of genes related to GTPases and vesicle formation from IS/Caput, Corpus, and Cauda CCs. Data was analyzed using the student’s *t*-test, two-tailed; the value of *P*<0.05 was considered significant. Data are shown as means ± SEM. Lumen (L).

Analysis using aquaporin 9 immunolabeling revealed epithelial damage 24 h after both saline and LPS injections, with localized damage observed in segments 7 and 8. No damage was detected in the proximal regions (segments 1, 2, and 3) (**Suppl. Fig. 2, arrows**). These observations are consistent with our previous study using CX3CR1 heterozygous mice in which epithelial damage occurred in the distal epididymis, 48 h after intravasal epididymal LPS injection (Barrachina et al., 2022).

### Immune response induced by LPS-mediated epididymitis

The epididymis contains a complex network of MPs, that include DCs and macrophages and are strategically positioned in the interstitium and the epithelium, with some extending luminal projections between epithelial cells in the IS (Da Silva & Smith, 2015; Battistone et al., 2020). LPS injections in the distal epididymal segments caused morphological changes in F4/80^+^ MPs in the proximal region at 1-, 24-, and 48-h post-injection (**Fig. 5A**). 1 h and 24 h post-LPS challenges, the luminal reaching projections in the Segments 1-2 significantly decreased in the LPS-treated epididymis compared to saline group. 48 h after injections, the MP projections in both saline- and LPS-treated groups had been reduced compared to earlier points (**Fig. 5A-B**).

**Figure 5:**
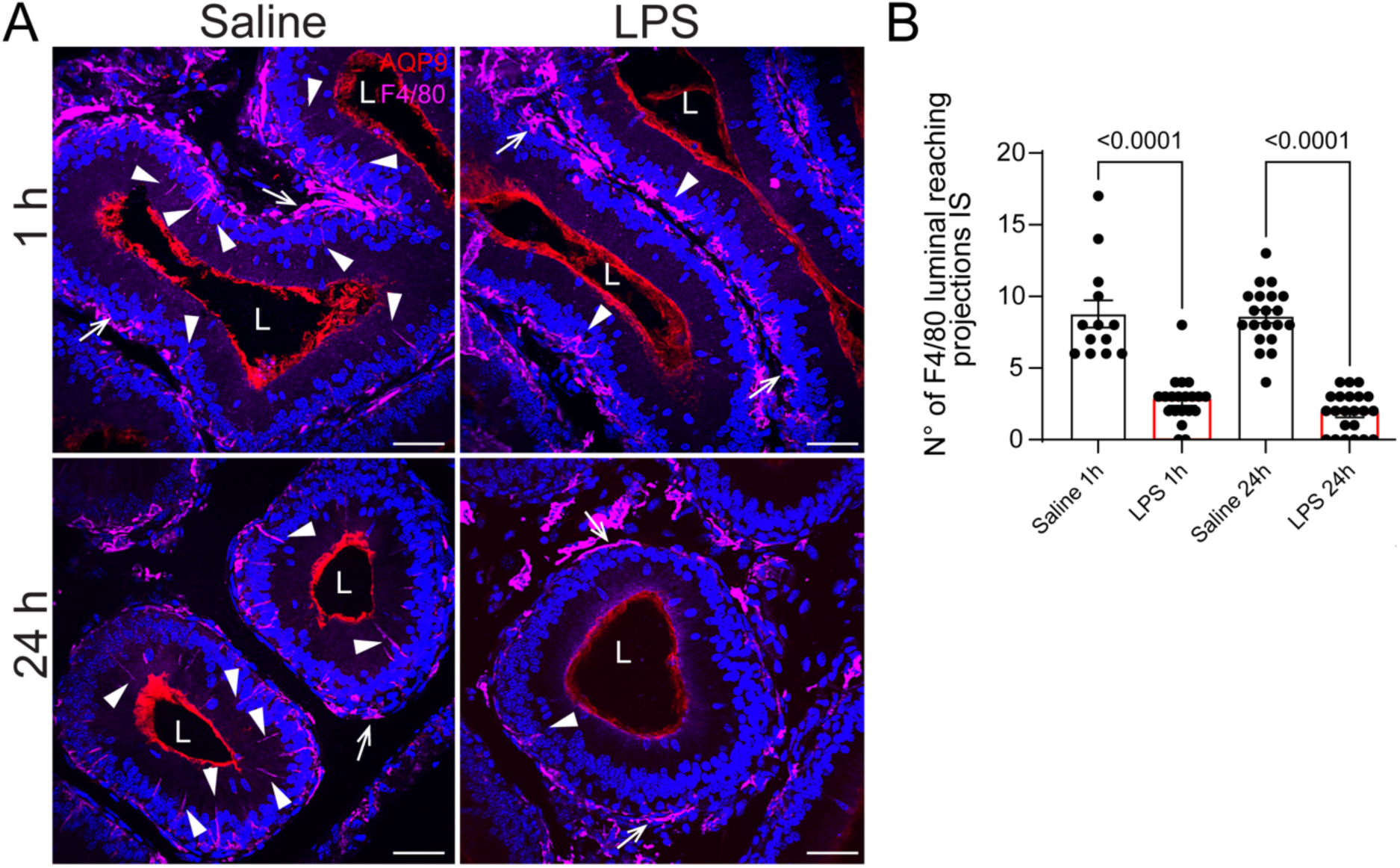
Response of proximal F4/80^+^ MPs post-LPS intravasal-epididymal injection. **A**) Confocal microscopy images of B1-EGFP epididymis immunolabeled for aquaporin 9 (red) and F4/80 (magenta) 1 and 24 h post-intravasal-epididymal injection with saline or LPS. Notably, F4/80^+^ MPs are observed in the basal portion of the epithelium in both saline- and LPS-treated groups (arrows). However, luminal-reaching projections were more numerous in the saline-treated group (arrowheads). (**B**) Quantification of the number of F4/80+ luminal projections in the IS of saline and LPS injected epididymis. Luminal reaching projections decreased in the IS/S1 after the LPS challenge compared to saline at 1- and 24-h post-intravasal-epididymal injections. The quantifications were performed in a standardized area of tissue (110,000 µm^2^). Each image quantification is represented as a dot. Data was analyzed using one-way ANOVA. Data are shown as means ± SEM. Nuclei are labeled with DAPI (blue). Lumen (L).

Flow cytometry analysis revealed no significant differences in the number of CD45^+^ cells in the distal epididymis (**Fig. 6A-B**) after saline or LPS challenges. Interestingly, 48 h after LPS challenge, we found a decrease in CD64^−^ CD11c^+^ cells (DCs) in the distal portion, indicating DC migration in response to LPS (**Fig. 6C**). In contrast, we observed neutrophil infiltration in the distal epididymis, as evidenced by an increased number of Ly6G^+^ cells following LPS injection compared to saline (**Fig. 7A**). Ly6G^+^ neutrophils predominantly localized to the epididymal interstitial compartment by IF (**Fig. 7B**). Furthermore, N-elastase-positive structures were detected in the LPS-injected cauda (**Fig. 7C**), suggesting the formation of neutrophil extracellular traps (NETs), which are formed during neutrophil activation and play a critical role in trapping and neutralizing pathogens.

**Figure 6:**
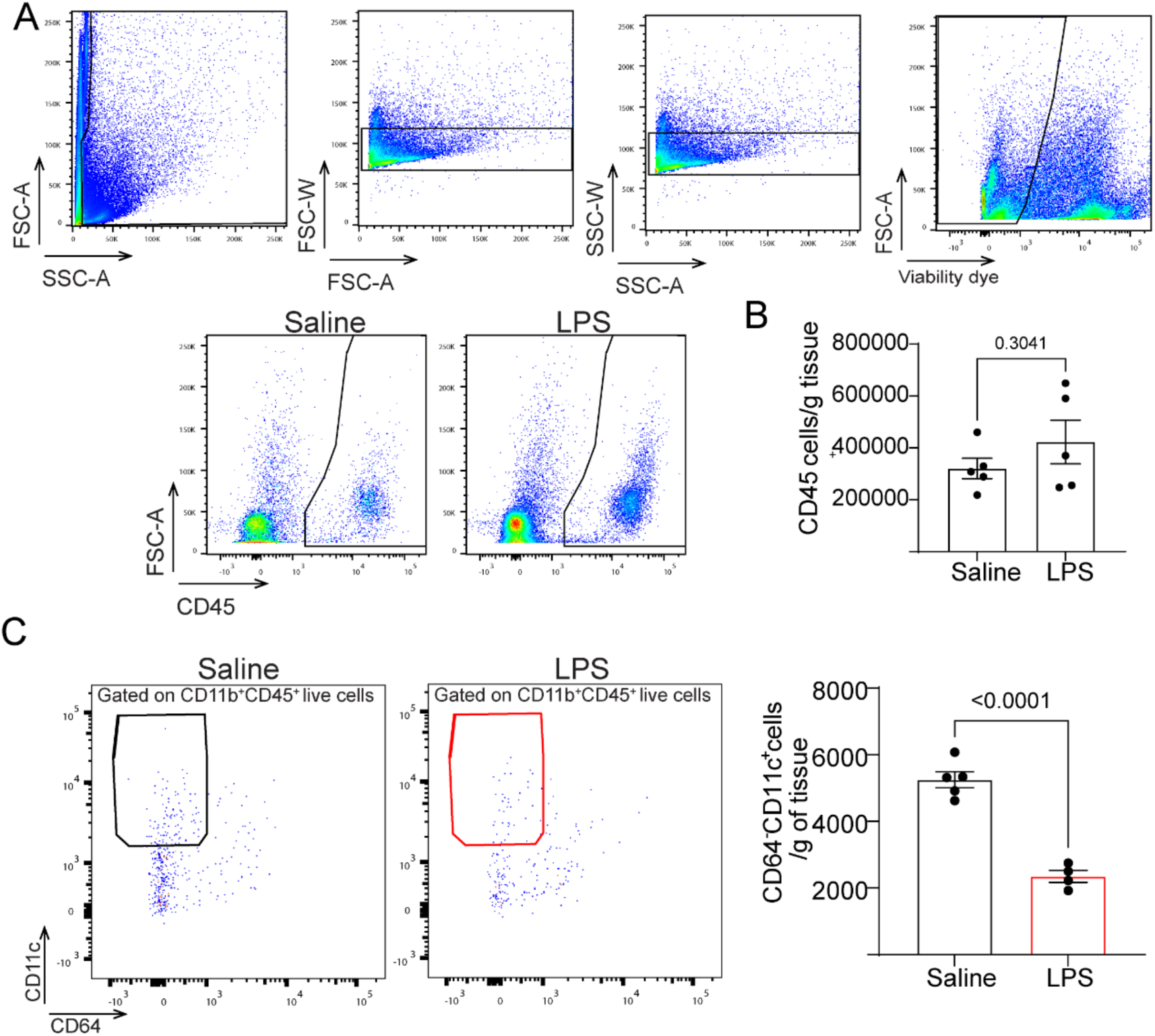
Reduced number of dendritic cells (DCs) 48 h post-LPS injections. **A**) Flow cytometry analysis conducted 48 h after intravasal-epididymal injection of saline or LPS, illustrating the gating strategy used to analyze myeloid cells in the distal epididymis. Key parameters include forward scatter area (FSC-A), side scatter area (SSC-A), and forward scatter width (FSC-W). (**B**) The absolute number of CD45^+^ live cells per gram of tissue 48 h after intravasal-epididymal injection of saline or LPS. **C**) The number of CD64^−^ CD11c^+^ CD45^+^ cells (DCs) per gram of tissue 48 h after intravasal-epididymal injection of saline or LPS, each dot represents a distal epididymis. Data was analyzed using the student’s *t*-test, two-tailed; the value of *P*<0.05 was considered significant. Data are shown as means ± SEM.

**Figure 7:**
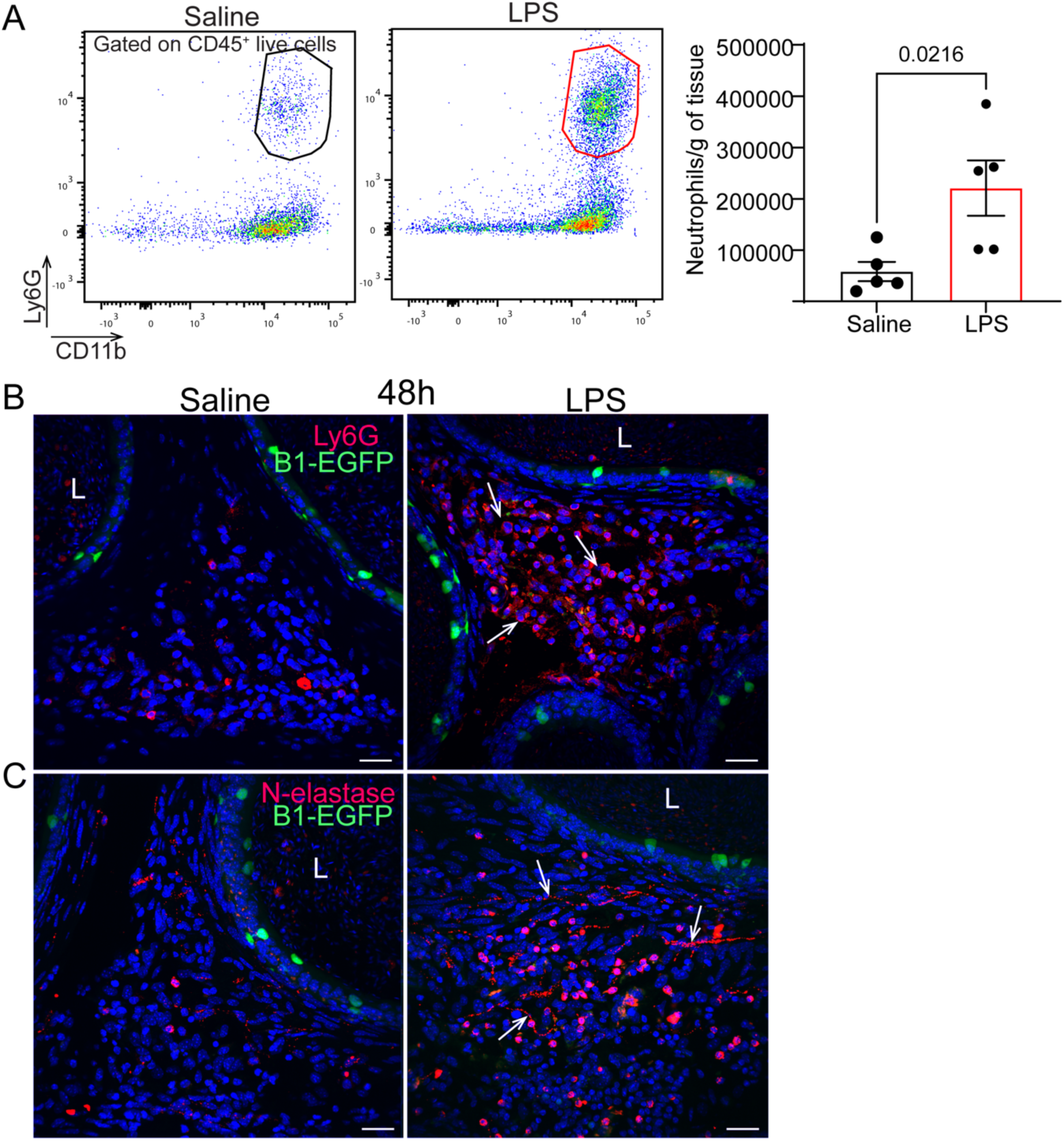
LPS caused infiltration of neutrophils in the distal epididymis. **A**) Flow cytometry analysis of Ly6G^+^CD11b^+^CD45^+^ (neutrophils) cells showed an increased number of neutrophils in the distal epididymis, each dot represents a distal epididymis. Data was analyzed using the student’s *t*-test, two-tailed; the value of *P*<0.05 was considered significant. Data are shown as means ± SEM. **B**) Confocal microscopy images of B1-EGFP^+^ epididymis subjected to immunolabeling for Ly6G (red) and **C**) N-elastase (red). Note the abundance of Ly6G-positive cells in the interstitial compartment and the presence of N-elastase-positive structures (arrows). Lumen (L).

### LPS exposure impaired sperm motility

We then evaluated the impact of the LPS injection on sperm functionality. Sperm count and motility under capacitated conditions (60 min) were analyzed after 4 h and 24 h post-intravasal-epididymal injections. The total and progressive motility decreased 24 h after the LPS treatment and no significant changes were observed between groups at 4 h (**Fig. 8**, **Suppl. Table 4)**. No change was observed in the sperm concentration under the different conditions. The motility decline may be linked to the downregulation of sperm maturation-related genes in CCs and the inflammatory environment induced by LPS.

**Figure 8:**
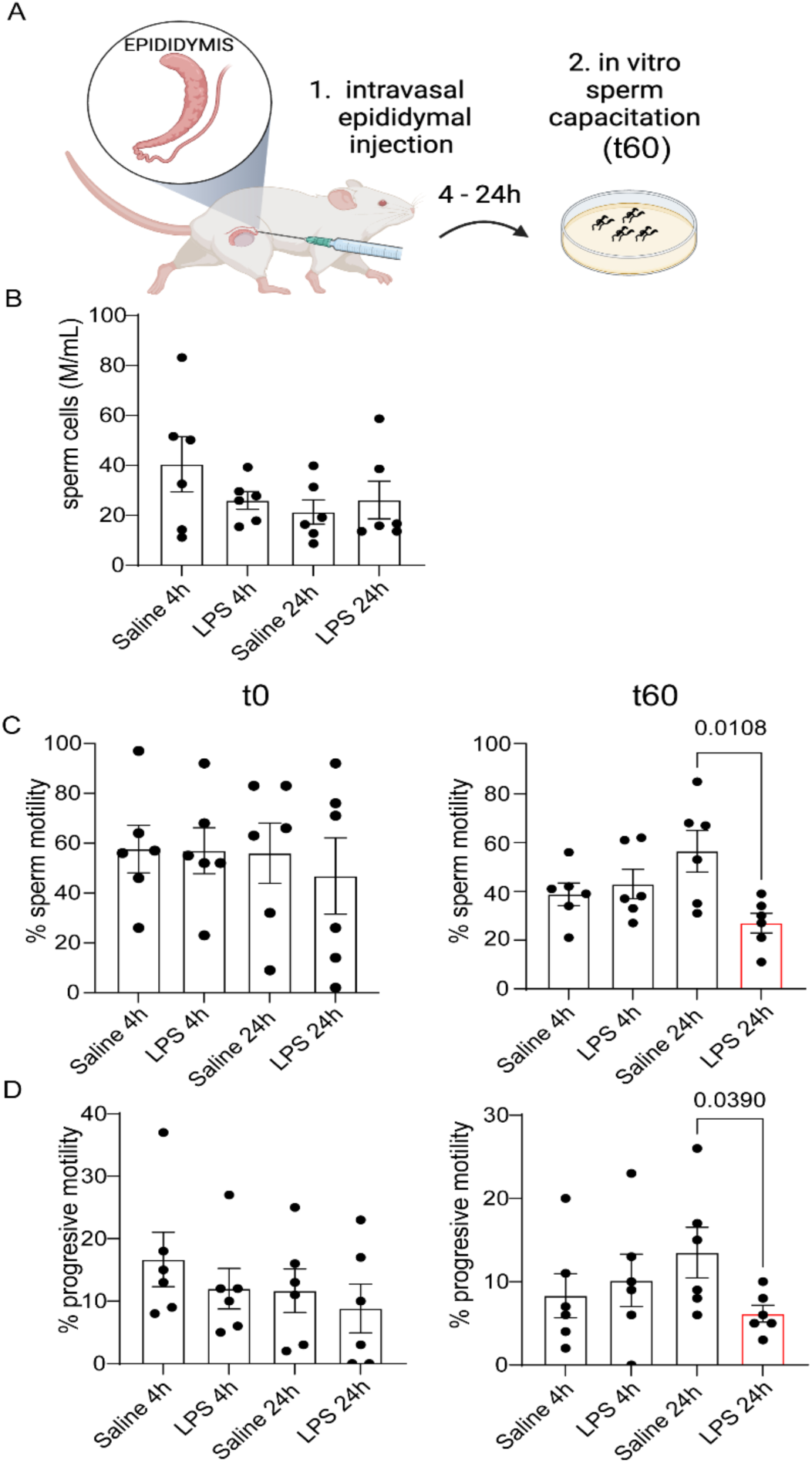
LPS-induced epididymitis disrupted sperm motility. **A.** Computer-Assisted Sperm Analysis (CASA) was performed on sperm isolated from the distal epididymis at 4 and 24 h following intravasal-epididymal injections of saline or LPS, assessed at time 0 (t0) and after 60 min of *in vitro* incubation in capacitating medium (t60). No difference was observed in the sperm concentration at t0 and t1h. However, after 1h in capacitating condition (t60) sperm motility (**B**) and progressive motility (**C**) decrease 24 h post-LPS injections. Data was analyzed using the student’s *t*-test, two-tailed; the value of *P*<0.05 was considered significant. Data are shown as means ± SEM.

We provide insights into the underlying mechanisms of epididymal cell response to bacterial infections and their impact on sperm function following LPS intravasal-epididymal injections. Our results demonstrate that all regions rapidly respond to inflammatory stimuli, highlighting the cellular dynamics of the epididymal mucosa.

## Discussion

Our findings demonstrated that CCs exhibit significant functional plasticity within the epididymis, particularly in response to inflammatory stimuli. The shift after the LPS challenge towards a more pro-inflammatory profile, along with the downregulation of genes associated with sperm maturation, suggests that these cells play an active role in initiating early immune responses. Consequently, the CC immune response triggers immune cell infiltration, which disrupts the epididymal environment and impairs sperm maturation. These insights underscore the importance of CCs in modulating immune responses that could impact male fertility under inflammatory conditions. Our findings demonstrate that CCs are key immune regulators beyond acid secretion. This cellular heterogeneity extends to other tissues containing proton-secreting cells, including the kidney, inner ear, olfactory mucosa, and lung, highlighting broad implications for epithelial physiology, reproduction, and immunology.

pH homeostasis is essential for sperm maturation, and successful fertilization, and the epididymal acidic luminal environment provides optimal sperm storage and maturation (Breton and Brown, 2013; Battistone et al., 2024). CCs are pivotal in maintaining this acidity through their V-ATPase-mediated proton secretion (Breton and Brown, 2013). Our results demonstrate that CCs dynamically respond to external stimuli and stressors, exhibiting region-specific profiles within the epididymis. Extracellular acidosis also impacts immune cell function (Lardner, 2001). For example, acidosis exacerbates neutrophil recruitment and IL-1β production during Pseudomonas aeruginosa infections (Torres et al., 2018). In our study, luminal pH increased, consistent with altered expression of V ATPase subunits in the different epididymal regions. This pH imbalance likely affects sperm maturation, as evidenced by reduced motility after LPS exposure. Recent studies have demonstrated that sperm incubated *in vitro* with LPS exhibit a reduction in motility in both mouse and human sperm (Bhagwat et al.,2025). Kidney proton-secreting intercalated cells are critical for urinary acidification (Brown & Breton, 1996) and recently, it was reported their role in defense against bacterial infections and injury (Battistone et al., 2020; Saxena et al., 2021). Further research is needed to clarify how luminal pH changes link to immune responses in the epididymis.

LPS intravasal-epididymal injection led to the upregulation of multiple pro-inflammatory mediators and the downregulation of sperm maturation-related genes in both distal and proximal CCs. Although certain cytokines/chemokines are essential for normal sperm maturation (Li et al., 2009), elevated levels of some of these mediators induce detrimental effects (Azenabor et al., 2015). We observed an upregulation of *Cxcl1, Cxcl16, Mir130a,* and *Mir150* in CCs following LPS treatment. CXCL1 plays a key role in recruiting several immune cells, particularly neutrophils and macrophages, while CXCL16 facilitates the migration of lymphocytes (Zhou et al., 2023). MicroRNAs, such as *Mir130a* and *Mir150*, are also involved in immune responses. *Mir130a* contributes to the production of pro-inflammatory cytokines (Lin et al., 2015). *Mir150* is crucial for the maturation and differentiation of both myeloid and lymphoid cells (Tsitsiou & Lindsay, 2009) (Huang et al., 2015). This pro-inflammatory response aligns with the immune cell infiltration observed at 24 h (Battistone et al., 2019, Barrachina et al., 2022) and 48 h (current study) following the LPS challenge. Additionally, in response to the LPS challenge, CCs exhibited upregulation of several dual-specificity phosphatases (DUSPs) in the proximal and distal epididymal regions. DUSPs regulate mitogen-activated protein kinase (MAPK) signaling pathways, which control the production of pro-inflammatory soluble mediators (Chen et al., 2002). In the epididymis, DUSP6 modulates cell proliferation in the caput and corpus regions (Xu et al., 2010); however, its role, in the epididymal immune response remains largely unexplored.

LPS exposure downregulated the expression of important mediators involved in sperm maturation, like Spag*11a* (Sperm Associated Antigen 11A) and β-defensins genes, contributing to decreased sperm motility. It was reported that decreased SPAG11A levels result in lower sperm counts and subfertility (Pujianto et al., 2013, Sangeeta & Yenugu, 2020; Sangeeta, Aisha, and Yenugu, 2024). In addition, deletion of multiple β-defensin genes in experimental models leads to male sterility or subfertility, marked by reduced sperm motility and premature capacitation (Zhou et al., 2013; Dorin, 2015). LPS exposure also led to the downregulation of *Adam7* in CCs. ADAM7 is also crucial for sperm maturation and male fertility (Cornwall & Hsia, 1997; Choi et al., 2015). This protein is secreted into the epididymal lumen and is transferred to sperm membranes during epididymal transit (Oh et al., 2009). Studies on *Adam7*-null mice revealed reduced fertility and impaired sperm motility (Choi et al., 2015), consistent with the motility defects observed following LPS injections.

We observed that CCs changed their cellular morphology after LPS injections, increasing apical blebs in the proximal epididymis and showing irregular shapes in the distal regions. Interestingly, LPS induces the upregulation of genes related to GTPases and vesicle formation in CCs. These results provide novel evidence that proximal CCs actively respond during the LPS-induced inflammatory process, further supporting their previously reported role in immune activation (Battistone et al., 2019, Barrachina et al., 2023). During the sperm transit through the epididymis, CCs (also principal cells) transfer proteins and RNAs via extracellular vesicles to sperm, contributing to their maturation (Sullivan, 2015; Nixon et al., 2019; Barrachina et al., 2022). Studies from our research group using Regulatory T cell-depleted mice revealed an increase in bleb formation in CCs, accompanied by the loss of extracellular vesicle-associated proteins crucial for sperm maturation, sperm-egg interaction, and embryogenesis (Barrachina et al., 2023, Elizagaray et al., 2024). Extracellular vesicles are crucial in the pathogenesis of inflammatory diseases, as they facilitate antigen presentation and enhance immune responses (Lu et al., 2021). Therefore, the increased formation of CC blebs, coupled with their altered morphology, may indicate CC involvement in responding to stressors and influencing the initiation of immune responses by producing extracellular vesicles. The cellular morphological changes may also be associated with disruptions in the sperm maturation process.

Several studies have highlighted the complexity of the epididymal segments, each expressing distinct and overlapping genes, proteins, and signal transduction pathways (Jhonson et al., 2005; Breton, Nair, Battistone, 2019; Battistone et al., 2019; Battistone et al., 2020^a^; Mendelson et al., 2020; Pleuger et al., 2022; Barrachina et al., 2022; Barrachina et al., 2023; Battistone et al., 2024; Elizagaray et al., 2024; da Silva et al., 2024). Through microdissection and transcriptomics analysis, ten anatomically distinct segments could be categorized (Johnston et al., 2005). In our study, intravasal-epididymal injection of LPS resulted in epithelial damage and altered the morphology of CCs, particularly in segments 7 and 8 of the epididymis. These changes were not limited to LPS, as similar alterations were observed in the saline group, suggesting that this region is susceptible to external stimuli. These results are consistent with a previous study using CX3CR1-EGFP transgenic mice (Barrachina et al., 2022). Epididymal segment 7 presents an enrichment of immune-related genes compared to the other segments. In contrast, sperm maturation-related genes displayed similar expression patterns across regions (Johnston et al., 2005), suggesting that the noted alterations may be associated with an immune response in this specific segment.

Our results revealed important inter-regional communication within the epididymis. Specifically, we observed signaling pathways facilitating coordinated immune responses by CCs and MPs across different epididymal regions, underscoring the epididymis’ capacity to maintain an integrated defense system. This regional communication likely supports a rapid and efficient immune response to localized inflammatory challenges, enhancing the overall protection of the reproductive tract. High-resolution three-dimensional imaging by Damon-Soubeyrand et al. (2023) revealed notable connections between epididymal segments, particularly through lymphatic and vascular structures. The epididymal segments are drained by initial and collecting lymphatic vessels that connect to the proximal lymph nodes, supporting immune surveillance and fluid reabsorption across the epididymis. These lymphatic connections suggest a mechanism by which immune responses in the proximal epididymis may be activated following LPS challenge in distal regions. Moreover, the reaction of CCs and MPs in the proximal region following intravasal-epididymal injections may be associated with the release of soluble mediators (Dellière et al., 2020; Canesi et al., 2022), which could play a pivotal role in facilitating dynamic signaling across the different epididymal regions.

The luminal-reaching projections of epididymal MPs play a crucial role in sensing and capturing luminal antigens, contributing to the immune surveillance within the epididymis (Battistone et al., 2019). These highly dynamic projections that extend into the lumen, are in close interaction with CCs, particularly under physiological conditions where they help maintain tissue homeostasis and coordinate local immune responses (Battistone et al., 2019, Battistone et al., 2024). Furthermore, during autoimmune-associated epididymitis, the activity of epididymal MPs becomes more pronounced (Barrachina et al., 2023; Elizagaray et al., 2024). However, using the LPS-induced epididymitis model, the number of luminal reaching projections of F4/80^+^MPs decreased, similar to what we previously reported for CX3CR1^+^ MPs (Barrachina et al., 2022), suggesting that these cells perceive the inflammatory challenge initiated on distal epididymal portions.

DCs efficiently capture and present antigens, initiating antigen-specific immune responses and activating T and B lymphocytes (Granucci et al., 2004). Their migration is influenced by environmental signals, including bacterial antigens and soluble mediators (Lambrecht et al., 1996). The decreased number of DCs after LPS injections in the distal epididymis suggests their activation and migration to nearby lymph nodes. We observed that 48 h after the LPS challenge, neutrophils accumulated in the interstitial compartment of the cauda, with N-elastase-positive structures indicative of NET formation. Neutrophils transmigrate across the epithelium, disrupting barrier function (Chin & Parkos, 2007; Gough, 2017). Late neutrophil infiltration was also observed after intravasal epididymal injection of LPS (Barrachina et al., 2022) and uropathogenic *E. coli* intravasal injection, another mouse model of epididymitis (Pleuger et al., 2020). Neutrophil infiltration releases mediators like reactive oxygen species and elastases, exacerbating inflammation and epithelial damage (Pettersen & Adler, 2002). The persistent presence of neutrophils may contribute to epididymal damage observed during epididymitis (Chin & Parkos, 2007; Gough, 2017). Notably, our group showed that renal intercalated cells sense DAMPs, such as UDP-glucose, triggering chemokine production and neutrophil recruitment, highlighting the role of proton-secreting cells in immune activation (Battistone et al., 2020^b^).

In conclusion, our findings indicate that CCs, together with MPs, serve as key gatekeepers of the epididymal mucosal barrier, monitoring potential threats and initiating immune responses during epididymitis. The distinct transcriptomic and morphological profiles exhibited by CCs in response to various stressors, such as saline and LPS, demonstrate their remarkable adaptability and underscore their dual role in the epididymal immune response and sperm maturation. CCs not only contribute to maintaining the delicate balance required for sperm storage and function but also actively participate in the immune defense, highlighting their functional plasticity in adapting to both physiological and pathological stimuli. The shift of CCs to a pro-inflammatory profile disrupts sperm maturation, impairs motility, and highlights their role in early immune defense. Sustained neutrophil infiltration likely worsens epididymal damage, contributing to the pathogenesis of epididymitis. Additionally, our findings reveal inter-regional communication within the epididymis, coordinating immune responses across regions to ensure rapid and effective defense against localized inflammation, thereby protecting the reproductive tract. By shedding light on the immunoregulatory mechanisms in the epididymis, our study may help identify new diagnostic and therapeutic targets for epididymitis and male infertility and support the development of novel male contraceptive methods.

## Author Contributions

A.A.S.S., F.B., and M.A.B. were involved in the study design and conceptualization. A.A.S.S., F.B., M.C.A., I.B., and M.L.E. performed the experiments and data analysis. A.A.S.S., F.B., M.C.A., M.L.E., S-C.E., and M.A.B. were involved in data interpretation. A.A.S.S. and M.A.B. wrote the original manuscript. All authors contributed to the writing of the manuscript, made critical comments, and approved the final version.

## Classification

Biological Sciences. Immunology and Inflammation

---

**Supplementary Figure 1:**
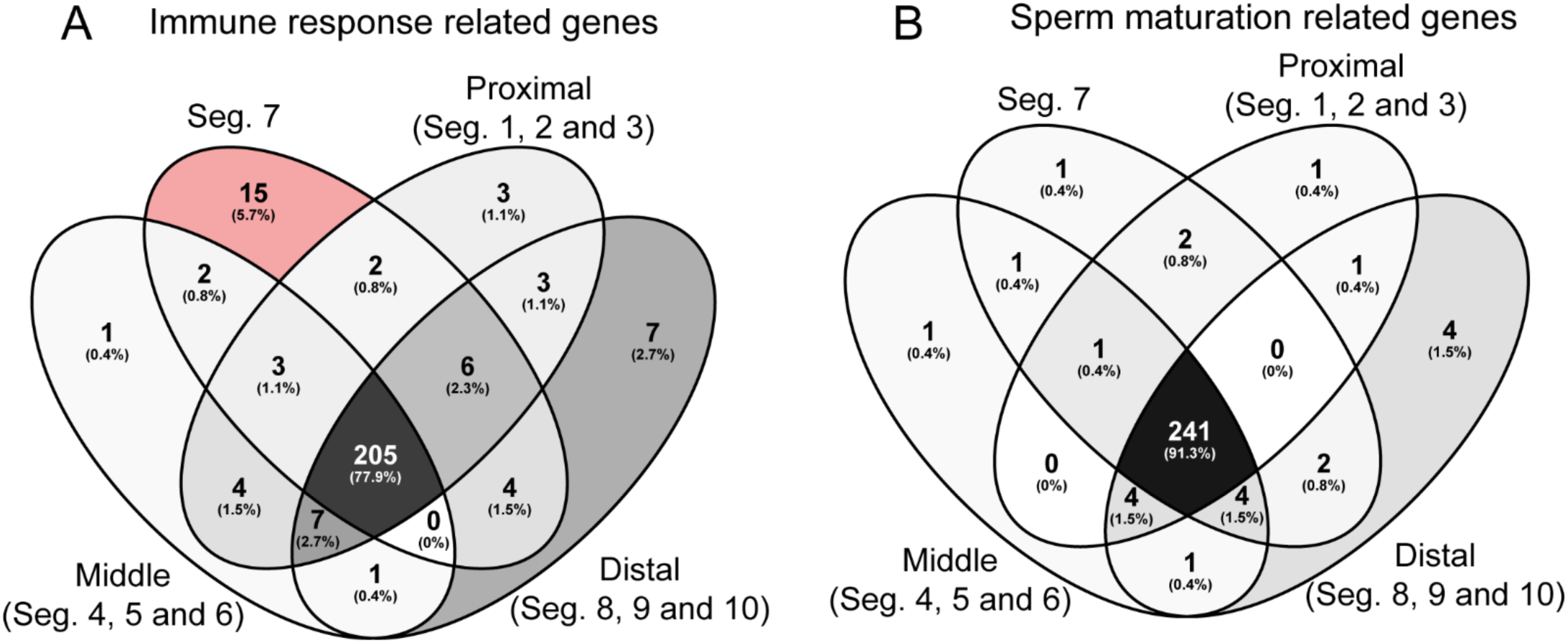
Enrichment of immune response-related genes on segment 7 of epididymis. Venn diagrams comparing segment 7 with other segments grouped into proximal (1, 2, and 3), middle (4, 5, and 6), and distal (8, 9, and 10) regions highlighting the distinct patterns of immune response (**A**) and sperm maturation-related gene expression (**B**).

**Supplementary Figure 2:**
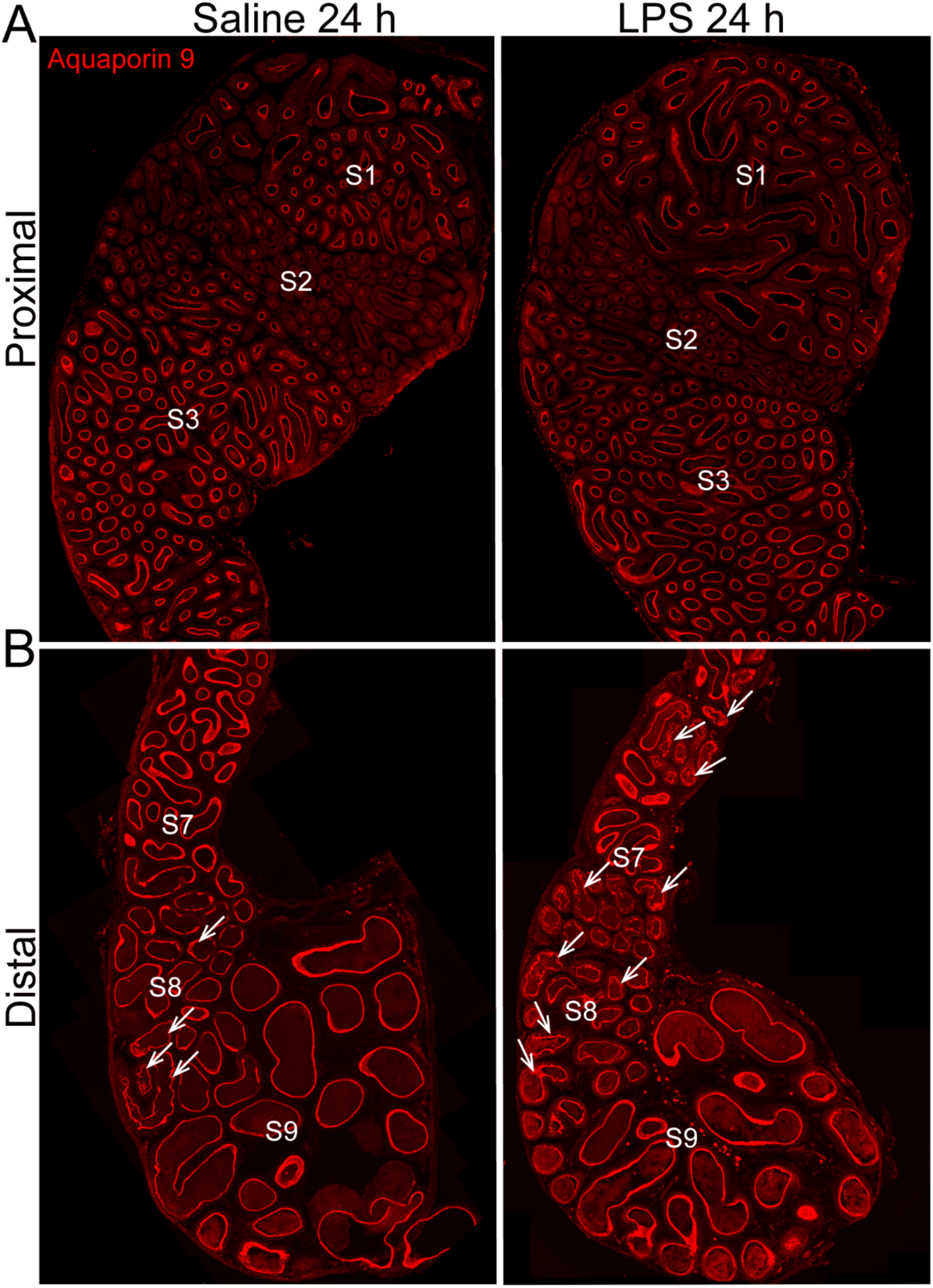
LPS or saline injections caused epididymal epithelial damage in segments 7/8. Epididymal segments S1, S2, S3, S7, S8, and S9 were analyzed using aquaporin 9 (red) immunolabeling 24 h after saline and LPS injections. **A.** No epithelial damage was observed in the proximal segments (1, 2, and 3). **B.** In segments 7 and 8, saline injections caused mild epithelial damage in a few regions (indicated by arrows), whereas LPS injections resulted in more extensive epithelial damage across several regions (arrows).

**Supplementary Table 4:** Computer-assisted sperm analysis (CASA) 4 and 24 h post-LPS injections. Sperm was isolated from distal epididymides 4 and 24 h post-intravasal-epididymal injections of saline or LPS. Distal sperm were capacitated in HTF medium (dilution ratio 1:5) with 0.3 % BSA and incubated for 60 min at 37 ^C^. Data are shown as mean ±SEM. Sperm analyses were performed using Hamilton Thorne’s CASA version 14.

| **Sperm parameters** | **Units** | **Saline 4h** | **LPS 4h** | **Saline 24h** | **LPS 24h** |
| --- | --- | --- | --- | --- | --- |
| Path Velocity (VAP) t0 | µm/s | 56.4±14.2 | 49.0±10.8 | 45.7±14.3 | 34.5±16.7 |
| Path Velocity (VAP) t60 | µm/s | 45.9±9.2 | 51.1±16.0 | 48.5±10.6 | 56.9±17.1 |
| Progressive Velocity (VSL) t0 | µm/s | 37.4±9.9 | 34.7±7.18 | 31.2±9.0 | 22.7±13.3 |
| Progressive Velocity (VSL) t60 | µm/s | 28.4±9.2 | 33.9±11.1 | 34.5±6.8 | 34.5±13.7 |
| Curvilinear Velocity (VCL) t0 | µm/s | 83.0±45.7 | 67.8±32.2 | 80.9±28.3 | 68.9±24.2 |
| Curvilinear Velocity (VCL) t60 | µm/s | 78.7±7.9 | 82.9±25.3 | 58.3±22.8 | 93.6±27.2 |
| Lateral Amplitude (ALH) t0 | µm | 8.7±0.7 | 7.8±1.9 | 7.9±1.6 | 5.0±4.0 |
| Lateral Amplitude (ALH) t60 | µm | 7.5±0.7 | 7.1±3.7 | 6.9±3.1 | 6.7±1.7 |
| Beat Frequency (BCF) t0 | Hz | 30.9±2.6 | 33.0±2.7 | 32.7±5.9 | 41.3±11.7 |
| Beat Frequency (BCF) t60 | Hz | 30.9±2.67 | 33.0±2.7 | 32.7±5.9 | 41.3±11.7 |
| Straightness (STR) t0 | % | 60.3±4.0 | 66.5±12.9 | 64.8±16.6 | 60.0±4.1 |
| Straightness (STR) t60 | % | 59.5±6.0 | 64.0±11.9 | 74.5±14.5 | 57.5±6.9 |
| Linearity (LIN) t0 | % | 33.6±3.0 | 41.6±15.4 | 39.5±17.4 | 29.0±6.8 |
| Linearity (LIN) t60 | % | 36.6±10.3 | 38.1±8.6 | 49.5±23.3 | 32.8±4.0 |
| Elongation t0 | % | 78.3±5.2 | 80.0±5.8 | 68.2±20.7 | 77.3±4.9 |
| Elongation t60 | % | 83.3±3.0 | 84.3±1.0 | 80.0±1.6 | 86.1±7.0 |
| Area t0 | µm sq | 2.2±0.3 | 2.3±0.4 | 2.6±0.7 | 2.1±0.7 |
| Area t60 | µm sq | 2.7±0.7 | 2.4±0.5 | 3.2±0.3 | 2.8±0.7 |

## Acknowledgments

The authors thank the Microscopy Core of the Program in Membrane Biology (PMB) (MGH, Boston, MA), the MGB Molecular Imaging Core (MGH, Charlestown, MA), and the MGH Pathology Flow Cytometry Facility (MGH, Boston, MA). This work was supported by the National Institutes of Health (grant HD104672 to M.A.B.), the MGH Physician/Scientist Development and Claflin Distinguished Scholar Awards (to M.A.B), and the Lalor Foundation (to F.B. and M.L.E).

