## Supplementary material for "Proton-secreting cells modulate mucosal immune surveillance in the male reproductive tract": Suppl Tables 1-3

| ID | Name | Length | IS.Caput.Sal | IS.Caput.Sa | IS.Caput.Sal | IS.Caput.LPS | IS.Caput.LPS | IS.Caput.LPS | Corpus.Sal1 | Corpus.Sal2 | Corpus.Sal3 | Corpus.LPS1 | Corpus.LPS2 | Corpus.LPS3 | Cauda.Sal1 | Cauda.Sal2 | Cauda.Sal3 | Cauda.LPS1 | Cauda.LPS2 | Cauda.LPS3 |
|---|---|---|---|---|---|---|---|---|---|---|---|---|---|---|---|---|---|---|---|---|
| ENSMUSG00 | Gnai3 | 3253 | 71.4212395 | 76.6821248 | 81.6535675 | 81.4580491 | 76.9696486 | 86.2184052 | 67.8928943 | 62.5190725 | 63.3814011 | 68.6605845 | 73.990237 | 67.7406197 | 50.681607 | 46.9295159 | 66.520311 | 58.4305488 | 74.8499736 | 59.169033 |
| ENSMUSG00 | Pbsn | 1107 | 0 | 0 | 0 | 0 | 0 | 0 | 0 | 0 | 0 | 0 | 0 | 0 | 0 | 0 | 0 | 0 | 0 | 0 |
| ENSMUSG00 | Cdc45 | 2338 | 1.4544115 | 0.76941672 | 1.38623889 | 1.52000062 | 1.60576991 | 1.20808632 | 2.12482091 | 1.04547209 | 1.97300674 | 1.18985949 | 0.98083747 | 0.88331031 | 1.73020411 | 3.35716457 | 3.04870367 | 1.58861801 | 1.95073966 | 3.31748676 |
| ENSMUSG00 | H19 | 2893 | 0 | 0 | 0 | 0 | 0 | 0 | 0 | 0 | 0 | 0 | 0 | 0 | 0 | 0 | 0 | 0 | 0 | 0 |
| ENSMUSG00 | Scml2 | 6101 | 2.6056279 | 1.7745764 | 2.50549674 | 2.88257058 | 1.96440731 | 1.73000037 | 2.13259903 | 1.20192448 | 1.69444607 | 1.76979365 | 2.49015519 | 2.43352995 | 1.56057368 | 2.51213716 | 2.16779207 | 1.95187328 | 2.18612544 | 2.49754508 |
| ENSMUSG00 | Apoh | 1584 | 0.02683408 | 0.10515434 | 0.15269422 | 0.11505314 | 0.06077263 | 0.25026633 | 0 | 0.14466819 | 0 | 0.26790172 | 0.30160959 | 0 | 0.2491511 | 0.38116991 | 0.13005542 | 0.02417336 | 0.14517537 | 0.03472797 |
| ENSMUSG00 | Narf | 4795 | 12.6318823 | 13.6447392 | 15.2838162 | 17.3977627 | 13.9125852 | 16.6588126 | 18.7573311 | 16.9257258 | 16.3199958 | 17.6704513 | 20.8635585 | 15.8891302 | 13.3849477 | 13.4053424 | 17.1594384 | 15.0686916 | 16.7053098 | 15.8316094 |
| ENSMUSG00 | Cav2 | 3515 | 241.402796 | 293.845316 | 272.708358 | 231.279008 | 268.908881 | 261.071714 | 308.010251 | 345.611362 | 349.458505 | 323.683206 | 331.366568 | 297.561793 | 342.502449 | 280.606852 | 315.054173 | 253.850932 | 371.279868 | 302.808309 |
| ENSMUSG00 | Klf6 | 4213 | 94.1308551 | 63.9847947 | 72.2904693 | 76.2414817 | 89.2034322 | 78.1928727 | 66.269195 | 83.8365387 | 60.5919831 | 55.0856305 | 91.5242558 | 60.7573294 | 78.113748 | 90.0880803 | 70.0612263 | 137.075467 | 141.00584 | 166.202449 |
| ENSMUSG00 | Scmh1 | 5784 | 10.1486252 | 13.4253728 | 9.60110872 | 11.7526069 | 12.5489179 | 12.1911376 | 10.4948422 | 10.2678399 | 12.2762021 | 12.4724371 | 9.84573833 | 9.4570106 | 12.7679599 | 12.1168962 | 12.6226067 | 9.69843358 | 12.338099 | 13.2957696 |
| ENSMUSG00 | Cox5a | 685 | 266.076197 | 231.099085 | 236.924371 | 228.935934 | 241.854133 | 252.538092 | 292.716983 | 260.210134 | 254.634882 | 266.040099 | 268.655679 | 234.425783 | 279.283467 | 242.729683 | 227.179966 | 241.09108 | 228.782815 | 243.164343 |
| ENSMUSG00 | Tbx2 | 3619 | 60.7804051 | 58.3412886 | 60.2563898 | 62.3804849 | 51.6164675 | 64.9018948 | 51.7053454 | 47.06773 | 56.9031718 | 57.8602579 | 47.1413033 | 44.6648684 | 40.7714188 | 45.9949183 | 47.2468795 | 39.2851686 | 44.2462748 | 48.062668 |
| ENSMUSG00 | Tbx4 | 3391 | 0 | 0 | 0 | 0.01343587 | 0 | 0 | 0 | 0 | 0 | 0 | 0 | 0 | 0 | 0 | 0 | 0 | 0 | 0 |
| ENSMUSG00 | Zfy2 | 2807 | 0 | 0 | 0 | 0 | 0 | 0 | 0 | 0 | 0 | 0 | 0 | 0 | 0 | 0 | 0 | 0 | 0 | 0 |
| ENSMUSG00 | Ngfr | 3440 | 0 | 0.00968398 | 0 | 0.02648898 | 0 | 0 | 0 | 0 | 0.01197282 | 0.0137066 | 0 | 0 | 0 | 0.28352495 | 0 | 0 | 0.02228273 | 0 |
| ENSMUSG00 | Wnt3 | 3071 | 0 | 0 | 0 | 0 | 0 | 0.1452213 | 0.01540628 | 0 | 0 | 0 | 0 | 0 | 0 | 0 | 0.10733066 | 0.12468449 | 0 | 0 |
| ENSMUSG00 | Wnt9a | 3462 | 2.08719813 | 1.34714188 | 1.73261634 | 1.47395637 | 1.45980704 | 1.30251264 | 2.81525473 | 1.09215709 | 0.95173924 | 1.81139328 | 2.62196479 | 1.72509414 | 1.6956956 | 1.98547803 | 2.23740396 | 1.88024358 | 2.3580304 | 1.50949311 |
| ENSMUSG00 | Fer | 4416 | 7.94084471 | 9.14294108 | 10.7241137 | 10.2140928 | 11.291819 | 10.2337367 | 8.72113239 | 10.8108023 | 8.37533767 | 7.8477733 | 7.73530238 | 8.6580002 | 6.63567909 | 6.75206162 | 6.54970386 | 8.0986022 | 8.03671896 | 9.42977667 |
| ENSMUSG00 | Xpo6 | 9659 | 7.00131847 | 8.46704165 | 7.94790275 | 9.19334045 | 8.49123076 | 8.22886268 | 8.08219487 | 7.94373501 | 7.40240101 | 8.23514198 | 5.49023749 | 7.40818712 | 7.27286988 | 6.81827476 | 8.88100625 | 7.70648523 | 6.45186445 | 8.98119488 |
| ENSMUSG00 | Tfe3 | 8492 | 6.46186791 | 7.77510084 | 5.86155944 | 6.78157796 | 7.06788846 | 6.47710013 | 5.40429799 | 6.98005282 | 5.36899116 | 5.15259687 | 6.78480922 | 6.80935309 | 5.32124779 | 5.66057824 | 5.91435429 | 7.61573668 | 6.19214312 | 8.05185111 |
| ENSMUSG00 | Axin2 | 5492 | 5.00743789 | 4.87683302 | 5.08221541 | 4.83650562 | 4.88155162 | 5.57603593 | 5.26366475 | 4.81925252 | 5.50453434 | 5.46027809 | 5.92402526 | 6.1486664 | 6.01827833 | 4.99789797 | 5.91165806 | 5.82167829 | 6.67151126 | 6.70085441 |
| ENSMUSG00 | Baat1 | 6223 | 2.7731161 | 3.29221114 | 3.11711163 | 3.67534055 | 3.82858785 | 3.10550636 | 3.56574746 | 3.87877203 | 3.19008526 | 4.47791509 | 3.23976096 | 2.80737625 | 3.35327029 | 3.36594623 | 3.44284253 | 3.31035622 | 4.18799323 | 1.2552295 |
| ENSMUSG00 | Gna12 | 4017 | 27.1622571 | 32.9728821 | 31.1652219 | 36.589477 | 32.7110161 | 35.045884 | 27.2427748 | 25.4330785 | 21.1315432 | 27.2668866 | 26.8191514 | 28.6373116 | 27.0177292 | 27.0894938 | 27.7651511 | 29.0539635 | 27.4209055 | 33.6326528 |
| ENSMUSG00 | Slc22a18 | 1807 | 9.78536427 | 11.4668623 | 12.2071554 | 11.497419 | 10.5746396 | 8.96720746 | 10.9183105 | 10.7792613 | 15.0660138 | 12.0812261 | 13.3251683 | 14.239635 | 9.39135559 | 14.7274278 | 10.237686 | 10.3831752 | 14.7620933 | 12.9379471 |
| ENSMUSG00 | Itgb2l | 3359 | 0.03796235 | 0.35703013 | 0.25922106 | 0.29840517 | 0.22926787 | 0.44256684 | 0.08451208 | 0.21603324 | 0.19618465 | 0.19651975 | 0.18489862 | 0.21601775 | 0.26435681 | 0.16592112 | 0.20852231 | 0.1709911 | 0.15974045 | 0.01637663 |
| ENSMUSG00 | Igsf5 | 2802 | 25.9399897 | 27.2257415 | 25.0154338 | 26.0325596 | 26.2990632 | 29.6750489 | 20.8533756 | 25.6251665 | 26.7374276 | 26.385547 | 23.3077688 | 23.3262459 | 25.2293496 | 20.8849215 | 23.9533671 | 27.5495591 | 24.7438512 | 24.1474665 |
| ENSMUSG00 | Pih1d2 | 2882 | 0.97340098 | 0.64730121 | 0.43640243 | 0.553309 | 0.6179324 | 0.67056093 | 0.67308114 | 0.42406554 | 0.41443753 | 0.40901026 | 0.48073345 | 0.65847763 | 0.35946227 | 0.56403298 | 0.7576969 | 0.31886696 | 0.69152236 | 0.85892082 |
| ENSMUSG00 | Dlat | 4667 | 36.2482538 | 35.2972492 | 39.2004911 | 38.1025833 | 35.4878829 | 35.5161547 | 38.7159033 | 36.4656835 | 35.8208844 | 39.1693329 | 40.332834 | 34.5633737 | 29.6604585 | 29.1482177 | 31.419828 | 31.1362449 | 36.5524636 | 29.8324511 |
| ENSMUSG00 | Sdhd | 1283 | 362.370706 | 348.759778 | 333.901781 | 347.655981 | 333.096921 | 334.432673 | 437.762129 | 387.669257 | 405.701626 | 422.959816 | 419.660002 | 362.823559 | 410.45847 | 385.561944 | 371.873747 | 389.919552 | 360.67943 | 376.445797 |
| ENSMUSG00 | Fgf23 | 1811 | 0 | 0 | 0 | 0 | 0 | 0 | 0 | 0 | 0 | 0 | 0 | 0 | 0 | 0 | 0 | 0 | 0 | 0.21262493 |
| ENSMUSG00 | Fgf6 | 1229 | 0 | 0 | 0 | 0 | 0 | 0 | 0 | 0 | 0 | 0 | 0 | 0 | 0 | 0 | 0 | 0 | 0 | 0 |
| ENSMUSG00 | Ccnd2 | 5696 | 7.90255994 | 7.60886155 | 6.42036323 | 5.90309879 | 9.36274077 | 7.38593237 | 12.7335563 | 8.60938285 | 8.87216553 | 8.17852788 | 9.13394132 | 9.45612963 | 18.759293 | 25.0158813 | 11.5590039 | 5.45856271 | 8.92216813 | 13.7329617 |
| ENSMUSG00 | Gpr107 | 10361 | 11.1708903 | 11.2436243 | 12.5357522 | 10.8219023 | 11.1724039 | 11.8991604 | 10.890912 | 11.5377106 | 11.1224661 | 10.3302863 | 12.2238601 | 9.81528204 | 8.92745342 | 10.7402793 | 10.2834884 | 9.05433724 | 11.844494 | 10.4167423 |
| ENSMUSG00 | Nalcn | 7257 | 0 | 0 | 0 | 0.00627822 | 0 | 0 | 0 | 0.12104523 | 0 | 0 | 0.07241622 | 0 | 0.04078703 | 0.40319381 | 0.1362598 | 0.03693458 | 0 | 0.01516029 |
| ENSMUSG00 | Btbd17 | 3162 | 0.43015991 | 0.71640633 | 0.88730701 | 0.77808235 | 0.66976744 | 0.70520968 | 0 | 0.19325694 | 0.27353473 | 0.32805676 | 0.07554548 | 0.26478018 | 0.23402238 | 0 | 0.18242308 | 0.09687693 | 0.18181355 | 0.5567019 |
| ENSMUSG00 | Slfn4 | 3572 | 0 | 0 | 0 | 0 | 0 | 0 | 0 | 0 | 0 | 0 | 0 | 0 | 0 | 0 | 0 | 0 | 0 | 0 |
| ENSMUSG00 | Th | 2574 | 0 | 0 | 0 | 0.01770048 | 0 | 0 | 0 | 0.16321539 | 0.30401857 | 0 | 0 | 0.02168441 | 1.62906487 | 2.09305134 | 0.64027282 | 0 | 0.11911825 | 0.5770186 |
| ENSMUSG00 | Ins2 | 822 | 0 | 0 | 0 | 0 | 0 | 0 | 0 | 0 | 0 | 0 | 0 | 0 | 0 | 0 | 0 | 0 | 0 | 0 |
| ENSMUSG00 | Scnn1g | 2978 | 0 | 0 | 0.09746178 | 0 | 0.0161625 | 0 | 0.30186061 | 0.37192064 | 0 | 0.3166602 | 0.88234477 | 0 | 0 | 0.10917012 | 0 | 0.3857348 | 0.01286981 | 0 |
| ENSMUSG00 | Drp2 | 8620 | 0 | 0 | 0 | 0 | 0 | 0 | 0 | 0 | 0 | 0 | 0 | 0 | 0 | 0 | 0 | 0 | 0 | 0 |
| ENSMUSG00 | Tspan32 | 2962 | 0.14350161 | 0.02249351 | 0 | 0.18458222 | 0.03249961 | 0.05018845 | 0.25557153 | 0.21920015 | 0.13904968 | 0.11142975 | 0.59678375 | 0.1884391 | 0.61623259 | 0.54879914 | 0.25038083 | 0.40074572 | 0 | 0 |
| ENSMUSG00 | Lhx2 | 4702 | 0.48814962 | 0.12752703 | 0.38065092 | 0.86238472 | 0.30709437 | 0.36885276 | 0.02012449 | 0.14620656 | 0 | 0.09025018 | 0.15240842 | 0.08309435 | 0.11540857 | 0.03951007 | 0.12267584 | 0.2361607 | 0.16302125 | 0.1754863 |
| ENSMUSG00 | Clec2g | 3079 | 0 | 0 | 0 | 0.04439205 | 0 | 0 | 0 | 0.02480832 | 0 | 0 | 0.03103278 | 0 | 0.04806617 | 0 | 0 | 0.01243605 | 0 | 0 |
| ENSMUSG00 | Gmpr | 1861 | 21.6294474 | 18.6523567 | 16.8436581 | 20.7607553 | 23.173672 | 18.4790957 | 24.6605581 | 22.1027498 | 25.7830687 | 30.3527902 | 30.31823 | 25.853359 | 29.8483282 | 22.8851306 | 26.9437364 | 37.035515 | 34.2279986 | 27.3715383 |
| ENSMUSG00 | Glra1 | 3270 | 0 | 0.01018743 | 0 | 0.02786608 | 0.11775394 | 0.01515374 | 0 | 0 | 0 | 0.01441918 | 0 | 0 | 0 | 0 | 0.02519973 | 0.035129 | 0.01172058 | 0 |
| ENSMUSG00 | Mid2 | 8103 | 5.08824148 | 6.05987981 | 6.28026088 | 5.97697027 | 6.35581369 | 7.19165922 | 5.93817037 | 4.72279245 | 6.577237 | 4.67259221 | 4.51630485 | 5.6690473 | 4.17644024 | 4.59110379 | 4.56608629 | 6.11477798 | 4.83394736 | 4.85394438 |
| ENSMUSG00 | Trim25 | 6232 | 5.47001784 | 4.30843914 | 3.54728873 | 5.21261627 | 4.27873647 | 4.40504082 | 3.90223315 | 3.46256537 | 4.23628957 | 4.05532364 | 4.86795549 | 4.06616027 | 3.44341849 | 3.19712989 | 3.94032852 | 4.13504152 | 3.29020681 | 3.83969214 |
| ENSMUSG00 | Dgke | 11704 | 0.46122329 | 1.20397764 | 0.99193842 | 0.86030338 | 1.21316793 | 0.92720853 | 1.09954278 | 0.78969253 | 0.95365224 | 0.86211983 | 0.7510759 | 1.06824235 | 0.83034755 | 0.63491575 | 0.88359454 | 1.07962239 | 0.97584044 | 1.08570605 |
| ENSMUSG00 | Scpep1 | 2786 | 23.6021204 | 21.0447573 | 26.8606789 | 19.7551118 | 21.6991013 | 30.1300541 | 19.2918892 | 18.2599713 | 20.8741422 | 19.1581465 | 24.0589615 | 17.8105255 | 17.2289862 | 16.1203993 | 17.569077 | 17.4135677 | 19.3970131 | 20.0212614 |
| ENSMUSG00 | Mnt | 2671 | 7.78174132 | 10.962948 | 9.41753492 | 10.3028344 | 9.94714409 | 7.94031073 | 7.61679219 | 9.06551221 | 8.80475496 | 8.96763656 | 8.33512941 | 8.06620957 | 8.07115232 | 7.96383078 | 7.06488678 | 9.96329919 | 9.72865193 | 8.79404289 |
| ENSMUSG00 | Itgb2 | 3148 | 0 | 0 | 0.06146573 | 0 | 0 | 0 | 0.79656035 | 1.64998943 | 0.28783461 | 0.01497799 | 0.4249361 | 0.10638309 | 0.50146803 | 0.6048944 | 0.61514394 | 0.75413522 | 0 | 0 |
| ENSMUSG00 | Hddc2 | 2549 | 15.4079179 | 12.9905912 | 14.0612709 | 13.2804456 | 15.7103807 | 13.8996487 | 14.7562102 | 16.4216842 | 18.1938081 | 14.169258 | 14.8441613 | 17.9775045 | 18.405121 | 19.7692565 | 16.0345012 | 10.5603362 | 16.5093268 | 20.0700154 |
| ENSMUSG00 | Tpd52l1 | 1274 | 26.0235772 | 28.9722816 | 25.6675875 | 25.7487849 | 28.1839993 | 23.6872957 | 50.2465107 | 33.9654568 | 38.6648923 | 41.081068 | 25.9874774 | 32.7708914 | 79.9610747 | 70.212961 | 73.0244217 | 67.8951979 | 70.0643214 | 60.8381722 |
| ENSMUSG00 | Pemt | 2147 | 11.3043575 | 10.4888293 | 9.82340905 | 9.35837008 | 9.774345 | 8.12415567 | 8.10948572 | 10.5665054 | 6.77170008 | 6.56639976 | 7.38765078 | 8.8389964 | 13.395672 | 8.93405128 | 11.4757943 | 11.1287094 | 12.0494878 | 9.1468333 |
| ENSMUSG00 | Cdh1 | 3837 | 129.287962 | 170.158826 | 150.781189 | 171.33185 | 156.300174 | 169.553776 | 83.2934965 | 105.07141 | 92.6240408 | 101.096908 | 104.116289 | 114.817051 | 56.5317801 | 72.1535588 | 73.6087218 | 76.9005527 | 85.9519898 | 73.0730823 |
| ENSMUSG00 | Cdh4 | 7257 | 0.12299968 | 0.14689439 | 0.1066524 | 0.16323373 | 0.17244454 | 0.12290881 | 0.09127429 | 0.1736736 | 0.18161341 | 0.01949182 | 0.05924964 | 0.14613443 | 0.10196758 | 0.10239843 | 0.07380739 | 0.1846729 | 0.08978187 | 0.14402274 |
| ENSMUSG00 | Ckmt1 | 2658 | 75.1116673 | 87.6938754 | 91.6876772 | 79.3632939 | 87.8253686 | 93.9227509 | 141.012431 | 129.994983 | 139.860606 | 148.707429 | 143.900044 | 130.509535 | 122.123407 | 94.5656136 | 114.645112 | 113.877822 | 127.898515 | 100.022202 |
| ENSMUSG00 | Bcl6b | 3710 | 0.14893997 | 0.26039729 | 0.11734819 | 0.28245391 | 0.29839198 | 0.22706104 | 0.16578567 | 0.13382783 | 0.64388623 | 0.17792718 | 0.29617899 | 0.37611632 | 0.11967311 | 0.13770484 | 0.26653298 | 0.06192551 | 0.59917122 | 0.2817178 |
| ENSMUSG00 | Clec10a | 1990 | 0.10679692 | 0 | 0.07292492 | 0 | 0 | 0 | 0.04755043 | 1.26668269 | 0.72438596 | 0 | 0 | 0 | 0 | 0 | 0.31056449 | 0 | 0.01925945 | 0 |
| ENSMUSG00 | Alox12 | 3971 | 105.540427 | 133.763309 | 102.606458 | 106.347348 | 110.117987 | 106.842232 | 56.6298572 | 72.7492651 | 72.0841815 | 71.812504 | 80.7638621 | 67.6928308 | 143.90857 | 126.010727 | 133.637885 | 170.654034 | 175.088731 | 148.91662 |
| ENSMUSG00 | Arvcf | 5943 | 10.3062357 | 12.3150645 | 10.4590617 | 11.6068351 | 10.6014956 | 11.9400159 | 10.4290105 | 14.0096647 | 13.2853022 | 14.5426955 | 11.6563503 | 12.2845174 | 13.8873124 | 17.4897977 | 16.8813375 | 11.8099753 | 13.1559222 | 14.7912759 |
| ENSMUSG00 | Comt | 3219 | 52.6726148 | 62.2378831 | 56.2930241 | 55.0865438 | 65.1627597 | 55.3869945 | 63.9213547 | 69.8829066 | 62.9504994 | 51.1055619 | 63.7444693 | 61.5550178 | 56.9638821 | 73.8574972 | 62.4871065 | 46.4863913 | 60.0909663 | 62.4598587 |
| ENSMUSG00 | Rtcd1 | 1696 | 19.523309 | 23.000825 | 22.3613489 | 20.5508248 | 19.9509094 | 23.3739309 | 18.4954635 | 20.559943 | 19.8647228 | 18.5155474 | 19.0705465 | 20.108119 | 23.8806038 | 30.232472 | 23.9289697 | 23.7735906 | 25.2194265 | 23.1907496 |
| ENSMUSG00 | Dbt | 3285 | 7.78938081 | 10.9319026 | 12.1780547 | 11.6364433 | 11.6923212 | 10.1066458 | 10.4131102 | 13.2656107 | 9.57884263 | 11.6549075 | 11.8964737 | 12.4884359 | 8.54485873 | 8.75156472 | 8.87997004 | 9.89611132 | 10.6170259 | 12.1405182 |
| ENSMUSG00 | Dazap2 | 1834 | 610.809114 | 648.420773 | 631.150604 | 675.739349 | 680.04057 | 634.053435 | 644.783586 | 675.176702 | 653.819095 | 703.404166 | 706.699481 | 652.10589 | 669.53149 | 682.657178 | 676.073949 | 681.151072 | 751.45873 | 689.413478 |
| ENSMUSG00 | Mcts1 | 1781 | 28.0424379 | 22.0714291 | 22.4348883 | 26.2468453 | 23.5119447 | 22.4531477 | 27.6544069 | 27.5131119 | 27.1262117 | 25.2829431 | 25.4835545 | 24.9149083 | 28.529969 | 28.0594262 | 28.8711633 | 30.6152863 | 29.9767162 | 26.3154177 |
| ENSMUSG00 | Rem1 | 2112 | 0.56351559 | 1.71927342 | 0.89326121 | 0.9060435 | 0.04557947 | 1.22004836 | 0 | 0 | 0 | 0 | 0 | 0 | 0 | 0 | 0 | 0 | 0 | 0.02604598 |
| ENSMUSG00 | Rnf17 | 5120 | 0.03320717 | 0.04554497 | 0 | 0.07118913 | 0.01880153 | 0.15485229 | 0.0646853 | 0.00745945 | 0.30568117 | 0.04604561 | 0.04665523 | 0.08721197 | 0.07708112 | 0.03628444 | 0.02414154 | 0.25427356 | 0.55393476 | 0.01074397 |
| ENSMUSG00 | Trappc10 | 5025 | 11.1317038 | 9.6657114 | 10.9935466 | 10.0642307 | 10.6321264 | 9.77248935 | 8.85053098 | 9.73621243 | 11.1715863 | 9.18617681 | 7.81512754 | 10.4300313 | 10.249258 | 10.0744389 | 9.32262071 | 12.4206348 | 9.70170142 | 8.30883821 |
| ENSMUSG00 | Ccm2 | 3312 | 22.2279483 | 18.9396678 | 19.7028661 | 19.4239718 | 18.7761007 | 19.7642394 | 17.4708352 | 16.0980054 | 15.0096994 | 17.7099864 | 17.3963164 | 16.4480938 | 22.1487201 | 21.1606866 | 16.9807137 | 16.9833642 | 16.5016001 | 17.2235639 |
| ENSMUSG00 | Wap | 2761 | 0 | 0 | 0 | 0 | 0.08716393 | 0 | 0.0514082 | 0.01383282 | 0.02983449 | 0.0170774 | 0 | 0.22237316 | 0.10720446 | 0 | 0 | 0 | 0 | 0 |
| ENSMUSG00 | Tbrg4 | 3249 | 1.47832714 | 2.28648058 | 2.53108654 | 2.91680432 | 2.53325914 | 2.22674642 | 2.51926545 | 1.89257519 | 2.77619174 | 2.78637582 | 2.92619783 | 1.83818895 | 2.97600978 | 1.92980973 | 1.86415165 | 1.75601739 | 1.89921016 | 2.86135397 |
| ENSMUSG00 | Tmprss2 | 3145 | 12.1906737 | 15.9308721 | 10.3668552 | 14.848989 | 12.7637597 | 13.07751 | 6.51395548 | 7.517042 | 5.14667749 | 8.65054238 | 6.82065385 | 5.57269254 | 12.2192571 | 13.0250191 | 9.1966589 | 12.3698751 | 13.2831999 | 11.7189512 |
| ENSMUSG00 | Mx1 | 4963 | 0 | 0.08054699 | 0 | 0 | 0.0193963 | 0.01996886 | 0 | 0.02308628 | 0 | 0 | 0.00962623 | 0 | 0 | 0.03743227 | 0 | 0 | 0 | 0 |
| ENSMUSG00 | Fap | 5810 | 0.04389519 | 0 | 0 | 0 | 0.02485297 | 0 | 0 | 0 | 0 | 0 | 0 | 0 | 0.05094518 | 0.00799382 | 0 | 0.05931419 | 0 | 0 |
| ENSMUSG00 | Gcg | 1487 | 0 | 0 | 0 | 0 | 0 | 0 | 0 | 0 | 0 | 0 | 0 | 0 | 0 | 0 | 0 | 0 | 0 | 0 |
| ENSMUSG00 | Ndufa9 | 1311 | 141.52181 | 143.593566 | 139.032389 | 135.119253 | 146.635319 | 155.272791 | 153.883496 | 152.420022 | 157.803103 | 150.69523 | 163.768622 | 135.558406 | 157.177283 | 132.034412 | 128.350232 | 132.600578 | 139.915833 | 133.893257 |
| ENSMUSG00 | Egfl6 | 2692 | 1.23157642 | 1.69534417 | 1.72505903 | 1.26934559 | 1.18005459 | 0.93877764 | 0.22847876 | 0.73774329 | 0.90267624 | 1.48878521 | 0.37268727 | 0.82935606 | 1.44770485 | 2.01855803 | 3.41305608 | 0.48361094 | 0.69761833 | 1.34866309 |
| ENSMUSG00 | Lck | 3548 | 0.88652284 | 0.76052549 | 0.7362375 | 0.9374172 | 1.15310415 | 0.87988225 | 0.50673106 | 0.82886555 | 1.00992866 | 0.47841751 | 0.75405797 | 0.69218973 | 0.45883689 | 0.92940536 | 0.59224333 | 0.33455717 | 0.46449571 | 0.89924695 |
| ENSMUSG00 | Tssk3 | 1278 | 0.79821927 | 1.40758708 | 0.26495674 | 0.42780323 | 0.71557633 | 0.11632097 | 0.296167 | 1.0160733 | 1.67582067 | 0.51651788 | 0.93456492 | 0.78613609 | 0.69481574 | 1.41730783 | 0.25791271 | 1.19845407 | 0.23991422 | 1.9799835 |
| ENSMUSG00 | Cttnbp2 | 9995 | 1.04615041 | 1.53982563 | 0.89535799 | 1.14415429 | 1.07387884 | 1.16011402 | 0.16567721 | 0.28276516 | 0.56865825 | 0.27361088 | 0.34893166 | 0.55843584 | 0.40472408 | 0.30668432 | 0.21847748 | 0.34095688 | 0.11503641 | 0.59989921 |
| ENSMUSG00 | Galnt1 | 5503 | 13.5787933 | 14.3409497 | 14.2052744 | 12.7335792 | 16.0498055 | 15.4880432 | 12.526726 | 13.949978 | 13.3371563 | 12.4409997 | 14.6372124 | 15.822721 | 10.6947079 | 13.8834315 | 13.9110469 | 10.9521018 | 15.9559415 | 12.7451592 |
| ENSMUSG00 | Myf5 | 2080 | 0 | 0 | 0 | 0 | 0 | 0 | 0 | 0 | 0 | 0 | 0 | 0 | 0 | 0 | 0 | 0 | 0 | 0 |
| ENSMUSG00 | Mkrn2 | 2314 | 24.889591 | 25.9852092 | 23.2878621 | 21.6385427 | 25.7923878 | 24.9476813 | 21.4276956 | 22.6117508 | 20.130488 | 23.8606195 | 20.7905718 | 21.732892 | 17.5667674 | 24.4062274 | 21.6869037 | 20.5352825 | 20.3887949 | 22.9164991 |
| ENSMUSG00 | Pparg | 1975 | 0 | 0.01686729 | 0 | 0.11534442 | 0.02437059 | 0 | 0 | 0 | 0.1547034 | 0.35451684 | 0 | 0 | 0 | 0 | 0 | 0 | 0 | 0 |
| ENSMUSG00 | Raf1 | 6697 | 19.1422444 | 21.4442119 | 19.538659 | 21.1511553 | 23.3364725 | 19.7189839 | 19.3715642 | 21.6938777 | 19.274084 | 18.5800666 | 20.9376591 | 20.7693939 | 19.9920597 | 20.3266567 | 20.7515078 | 17.981777 | 22.8115006 | 20.9785364 |
| ENSMUSG00 | 1-Sep | 2354 | 0.66809325 | 0.67927737 | 0.41099006 | 0.77418937 | 0.67474659 | 0.65256361 | 1.06523874 | 1.15193737 | 1.15476212 | 0.76114134 | 1.09594212 | 0.42679776 | 1.25739806 | 1.61784841 | 0.40256406 | 0.37412233 | 1.09085042 | 0.44399874 |
| ENSMUSG00 | Pdgfb | 2911 | 3.76720557 | 3.27292606 | 4.83569129 | 3.80327509 | 4.150159 | 3.54069687 | 1.33275152 | 2.11232456 | 1.99494975 | 1.63593989 | 2.03507209 | 1.82153485 | 0.50840176 | 1.81883406 | 1.75506456 | 2.88067435 | 1.15861014 | 1.30389157 |
| ENSMUSG00 | Acvrl1 | 4297 | 0.13848556 | 0.79076454 | 0.76551084 | 0.50894348 | 0.32483728 | 0.78417172 | 0.1101063 | 0.2577565 | 0.20128388 | 0.02194587 | 0 | 0.23381084 | 0.18368878 | 0.41072267 | 0.04794223 | 0.34752937 | 0 | 0.30724193 |
| ENSMUSG00 | Grasp | 2095 | 1.9291626 | 2.80791976 | 2.6056565 | 3.20404352 | 3.04875526 | 3.16346628 | 1.27425669 | 2.57155827 | 1.27359799 | 1.36396049 | 0.38124655 | 0.89082354 | 0.91036456 | 1.389848 | 1.78779819 | 0.80209751 | 1.20426768 | 0.38410349 |
| ENSMUSG00 | Acvr1b | 3268 | 2.17208213 | 2.81345129 | 2.32394255 | 2.07729364 | 2.22396587 | 1.94086594 | 1.72283011 | 1.70627012 | 2.64662433 | 2.52489996 | 1.92971069 | 1.77626344 | 2.30960323 | 1.44960126 | 1.19771965 | 3.12839418 | 2.42764488 | 2.01991824 |
| ENSMUSG00 | Tom1l2 | 12272 | 5.33046495 | 5.23364244 | 5.3371708 | 6.3782492 | 5.14578584 | 6.66652242 | 5.38975922 | 5.15062141 | 4.40325201 | 6.38947356 | 5.51249521 | 5.8308083 | 4.49422128 | 4.18572039 | 5.35835098 | 5.57572649 | 6.78018176 | 4.50490159 |
| ENSMUSG00 | Gpa33 | 2997 | 0 | 0 | 0 | 0.03040443 | 0.01606003 | 0.01653411 | 0 | 0 | 0 | 0.01573263 | 0 | 0 | 0.11522303 | 0 | 0 | 0.01277631 | 0 | 0 |
| ENSMUSG00 | Zfp385a | 3711 | 0.02290767 | 0.26930391 | 0.20856278 | 0.14732755 | 0.36316191 | 0.26705866 | 0.1784903 | 0.07204172 | 0.15537893 | 0.24140753 | 0.30897305 | 0.15040599 | 0.25257516 | 0.50060994 | 0.86599875 | 0.11349951 | 0.4544212 | 0.54846051 |
| ENSMUSG00 | Itga5 | 4349 | 0 | 0.11489846 | 0.15572072 | 0.13619075 | 0.2656165 | 0.29624536 | 0.21757958 | 0.03512753 | 0 | 0.18430949 | 0 | 0 | 0 | 0 | 0 | 0 | 0.04406334 | 0 |
| ENSMUSG00 | Gabra2 | 2382 | 0 | 0 | 0 | 0 | 0 | 0.02080299 | 0 | 0 | 0 | 0 | 0 | 0 | 0 | 0 | 0 | 0 | 0.01608997 | 0.23093664 |
| ENSMUSG00 | Wdr77 | 6811 | 9.23618566 | 8.64736411 | 8.09658268 | 8.86336569 | 8.60735326 | 9.01421772 | 10.6767857 | 7.95698415 | 8.68962313 | 8.75725145 | 9.64477579 | 6.7690115 | 8.11937737 | 8.27824432 | 9.10414647 | 8.16296608 | 9.22284549 | 7.91497938 |
| ENSMUSG00 | Adora3 | 2521 | 0 | 0 | 0 | 0 | 0 | 0 | 0 | 0.09089822 | 0 | 0 | 0 | 0 | 0 | 0 | 0 | 0 | 0 | 0 |
| ENSMUSG00 | Atp5f1 | 2743 | 144.096111 | 127.033494 | 129.495747 | 129.507642 | 140.500296 | 125.44447 | 157.375458 | 154.022733 | 165.677001 | 154.791135 | 153.078421 | 137.453445 | 154.470229 | 137.876122 | 137.288451 | 145.275374 | 140.57633 | 137.412468 |
| ENSMUSG00 | Sox9 | 1698 | 8.68627567 | 12.301051 | 9.51517028 | 14.5698746 | 13.549505 | 12.5486899 | 13.597519 | 36.3929959 | 16.1301725 | 27.2963141 | 23.8593432 | 20.9719827 | 34.5149103 | 25.273461 | 24.2162266 | 47.4460765 | 27.7177224 | 20.4421357 |
| ENSMUSG00 | Hnrnpd | 5344 | 26.8043494 | 30.0215005 | 28.4411737 | 30.4962305 | 28.1099801 | 28.4946763 | 30.2698445 | 31.2242901 | 33.0478629 | 30.1926846 | 29.2782539 | 30.1116308 | 29.7431582 | 29.8444978 | 29.9605625 | 29.7139126 | 28.7518945 | 29.738269 |
| ENSMUSG00 | Dynlt1c | 6502 | 0 | 0 | 0 | 0 | 0 | 0 | 0 | 0 | 0 | 0 | 0 | 0 | 0 | 0 | 0 | 0 | 0 | 0 |
| ENSMUSG00 | C1d | 4056 | 27.1735507 | 22.9560008 | 21.6225864 | 23.5331331 | 25.3475811 | 24.3121153 | 28.2056355 | 25.7064247 | 26.7367101 | 25.737588 | 23.2632009 | 22.8574496 | 26.6606417 | 28.7641882 | 25.1313238 | 25.6214765 | 27.5446636 | 24.2224518 |
| ENSMUSG00 | Gm2a | 4123 | 26.1752709 | 26.275414 | 25.1429725 | 27.537745 | 26.64008 | 27.5706939 | 27.0358165 | 25.6406912 | 26.4121144 | 26.3828941 | 24.8086796 | 25.2341487 | 26.5145595 | 28.3192907 | 28.7701381 | 29.3285807 | 32.1725233 | 28.4985331 |
| ENSMUSG00 | Krit1 | 4763 | 15.8937053 | 16.4431271 | 17.6919357 | 18.6816543 | 17.6237823 | 16.1673206 | 17.5026119 | 18.1540204 | 15.7205723 | 17.6208801 | 18.3757556 | 18.9372477 | 18.9953257 | 16.0989268 | 16.8508568 | 16.0863963 | 18.7889782 | 17.5086722 |
| ENSMUSG00 | Clcn4-2 | 4759 | 44.7380598 | 43.6728613 | 46.5134009 | 44.5749574 | 43.2974922 | 46.8559153 | 34.1697158 | 39.5968299 | 40.9528457 | 37.0745811 | 44.4018998 | 41.0730091 | 29.4809777 | 32.0882873 | 32.2236165 | 29.5849051 | 35.4914878 | 27.0248564 |
| ENSMUSG00 | Vmn2r88 | 3174 | 0 | 0 | 0 | 0 | 0 | 0 | 0 | 0 | 0 | 0 | 0 | 0 | 0 | 0 | 0 | 0 | 0 | 0 |
| ENSMUSG00 | Grm6 | 5410 | 0 | 0 | 0 | 0 | 0 | 0 | 0 | 0 | 0 | 0 | 0 | 0 | 0 | 0 | 0 | 0 | 0 | 0 |
| ENSMUSG00 | Sema4f | 5143 | 0.15702865 | 0.05829595 | 0.01881141 | 0.06201192 | 0.12166343 | 0.15415978 | 0.00919943 | 0.17080017 | 0.0960992 | 0 | 0 | 0 | 0.09592051 | 0 | 0 | 0 | 0.06706916 | 0.12835102 |
| ENSMUSG00 | Hk2 | 6685 | 9.90621754 | 8.62597156 | 7.84396513 | 6.04527245 | 6.27119005 | 7.62748883 | 14.4450528 | 12.2375656 | 14.0409977 | 10.8760485 | 10.0981326 | 12.1567098 | 14.8327829 | 9.71261543 | 11.2171775 | 8.87242347 | 8.94949132 | 10.9524489 |
| ENSMUSG00 | Myo18a | 10638 | 6.92437207 | 8.60536128 | 7.00274833 | 8.62567619 | 7.87267787 | 9.95899082 | 5.40817988 | 6.049464 | 5.06797792 | 6.55092493 | 5.0568343 | 5.46193878 | 4.15968507 | 4.23489047 | 5.49586462 | 5.99663019 | 5.22762331 | 5.0572388 |
| ENSMUSG00 | Sez6 | 5750 | 0 | 0 | 0 | 0 | 0 | 0 | 0 | 0 | 0 | 0.00820012 | 0 | 0 | 0 | 0 | 0 | 0 | 0 | 0 |
| ENSMUSG00 | Haao | 1212 | 0.14028111 | 0.08245766 | 0.23947292 | 0.11277486 | 0.07942562 | 0.36796584 | 0.15614746 | 0.25209507 | 0.06796455 | 0.11670966 | 1.18254848 | 0.32236769 | 0.52913773 | 0.22992122 | 0.54391493 | 0 | 0 | 0.0907741 |
| ENSMUSG00 | Cd52 | 1183 | 15.198385 | 14.9527868 | 15.0068349 | 9.32017982 | 6.91667542 | 8.67068119 | 650.299374 | 528.979301 | 430.909134 | 252.413697 | 140.01334 | 400.051566 | 2926.80469 | 4307.16892 | 2011.52752 | 639.061498 | 853.286566 | 1970.51641 |
| ENSMUSG00 | Abhd15 | 1892 | 0.02246574 | 0.03521448 | 0 | 0.02408089 | 0.17807794 | 0 | 0.0500134 | 0.1009313 | 0.10884386 | 0 | 0.17675725 | 0.02950088 | 0.13036976 | 0 | 0.02177672 | 0.16190532 | 0.06077108 | 0 |
| ENSMUSG00 | Hoxb6 | 2073 | 65.9824681 | 62.4958252 | 65.0580813 | 67.0338568 | 59.4161084 | 59.8314 | 56.1452919 | 55.6028319 | 56.6041406 | 59.9107344 | 58.5145295 | 62.3315286 | 66.5612024 | 64.2779 | 54.5577766 | 57.9069244 | 50.3621962 | 60.0244088 |
| ENSMUSG00 | Loxl3 | 4183 | 0.01016141 | 0.27077179 | 0.1503361 | 0 | 0.03451967 | 0.10661597 | 0 | 0.08217347 | 0.1476925 | 0.24798362 | 0.2398456 | 0.04003036 | 0.01179343 | 0.02220611 | 0.08864786 | 0.03661545 | 0.04581197 | 0 |
| ENSMUSG00 | Tcl1b5 | 1035 | 0 | 0 | 0 | 0 | 0 | 0 | 0 | 0 | 0 | 0 | 0 | 0 | 0 | 0 | 0 | 0 | 0 | 0 |
| ENSMUSG00 | Btn1a1 | 3925 | 0.27073361 | 0.10184834 | 0.35740952 | 0.51074801 | 0.30657276 | 0.1514988 | 0.27724627 | 0.20434152 | 0 | 0.30032295 | 0.1704075 | 0 | 0.01256864 | 0 | 0.41988847 | 0 | 0.23435188 | 0.19621083 |
| ENSMUSG00 | Kat2b | 4198 | 8.79871319 | 6.87207394 | 8.52701583 | 6.09940613 | 7.62451864 | 7.3184123 | 4.64335963 | 5.03106206 | 5.86696913 | 4.56007278 | 4.90495643 | 6.40856345 | 5.26457812 | 5.21085402 | 3.2682511 | 5.20817933 | 5.33171925 | 5.12352566 |
| ENSMUSG00 | Rab5b | 3160 | 22.3959234 | 25.1111753 | 24.737856 | 24.409762 | 25.8785248 | 25.8270102 | 26.6657852 | 26.8434575 | 22.3397747 | 27.9323153 | 21.7406298 | 23.8629621 | 26.0241475 | 19.7681321 | 24.4862411 | 23.3378829 | 23.3596354 | 25.728943 |
| ENSMUSG00 | Cryba1 | 1908 | 0 | 0 | 0 | 0 | 0 | 0 | 0 | 0 | 0 | 0 | 0 | 0 | 0.0258553 | 0 | 0 | 0 | 0 | 0 |
| ENSMUSG00 | Dnmt3l | 2970 | 0 | 0 | 0 | 0 | 0.05005327 | 0 | 0 | 0 | 0 | 0.15875658 | 0.03217169 | 0 | 0 | 0.02578492 | 0 | 0 | 0 | 0 |
| ENSMUSG00 | Aire | 4111 | 0.66172008 | 1.33705365 | 1.10608413 | 0.58738393 | 0.85468996 | 0.81965115 | 0.41431682 | 0.48309534 | 0.45083756 | 0.4817148 | 0.65079 | 0.48877739 | 0.28799952 | 0.61006585 | 1.20267251 | 0.70787793 | 0.84838069 | 0.54861916 |
| ENSMUSG00 | Icosl | 2834 | 4.09453531 | 5.07804174 | 6.77639081 | 4.01914647 | 4.99322003 | 5.2280407 | 2.05344372 | 4.08337965 | 3.2699245 | 3.44396456 | 5.02361949 | 3.30876193 | 3.65550551 | 4.96561697 | 4.01257213 | 5.0801935 | 4.28702756 | 4.50321551 |
| ENSMUSG00 | Spg7 | 3939 | 23.0384744 | 23.5279187 | 20.7297586 | 24.9261362 | 21.4449165 | 21.6124895 | 25.0436495 | 23.8617675 | 22.2819155 | 26.0711429 | 26.9984914 | 20.603192 | 28.8176546 | 30.2434842 | 23.4406415 | 20.4819268 | 22.5151183 | 30.5280293 |
| ENSMUSG00 | Sult5a1 | 2053 | 60.7246377 | 57.847281 | 61.2385803 | 68.6411642 | 69.068021 | 65.1933431 | 65.6109094 | 72.8874002 | 77.2372556 | 75.4227518 | 92.2454629 | 66.7449838 | 49.6923551 | 43.7293818 | 42.4458308 | 43.9045737 | 55.6132676 | 46.3008945 |
| ENSMUSG00 | Rpl13 | 2348 | 68.3920592 | 59.9433467 | 56.0786838 | 57.1259432 | 62.8297896 | 54.1112474 | 75.0192094 | 59.3706426 | 62.7445344 | 60.8059325 | 55.3033804 | 61.4732979 | 97.8864577 | 99.5343474 | 70.9621346 | 61.1703869 | 53.9147183 | 79.3742988 |
| ENSMUSG00 | Chmp1a | 3395 | 55.3506283 | 59.3450322 | 52.9472868 | 64.0136028 | 58.5238824 | 58.3248081 | 59.7296441 | 52.1306606 | 46.0391244 | 54.5948202 | 52.0247486 | 53.9578806 | 59.125647 | 64.2145909 | 57.8035378 | 61.1070724 | 64.3249602 | 68.2955478 |
| ENSMUSG00 | Rpa1 | 6612 | 9.03202808 | 12.2278273 | 11.5958932 | 12.0724364 | 14.5880785 | 11.391433 | 14.4972002 | 11.5928844 | 11.536211 | 13.4777419 | 11.9220631 | 11.708458 | 11.0720758 | 11.5688766 | 13.7837026 | 12.9141038 | 14.7694199 | 13.5193283 |
| ENSMUSG00 | Serpinf1 | 2712 | 0.015673 | 0 | 0 | 0.03359959 | 0 | 0 | 0.06978271 | 0.04224823 | 0.03037354 | 0 | 0 | 0 | 0.0545707 | 0 | 0 | 0.2541412 | 0.04239635 | 0 |
| ENSMUSG00 | Tubgcp3 | 9566 | 5.75415042 | 6.50169073 | 4.90005759 | 6.60125522 | 6.80770361 | 5.39765298 | 6.2368062 | 5.94884796 | 6.01479851 | 5.68312365 | 5.48368225 | 6.63416353 | 5.45611211 | 5.3212126 | 5.38816082 | 5.97214993 | 7.40002826 | 6.67630911 |
| ENSMUSG00 | Oprm1 | 8203 | 0 | 0 | 0 | 0 | 0 | 0 | 0 | 0 | 0 | 0 | 0 | 0 | 0 | 0 | 0 | 0 | 0 | 0 |
| ENSMUSG00 | Polr3d | 2040 | 20.3358097 | 15.3337293 | 14.6069092 | 14.7626717 | 16.4214798 | 12.1938589 | 13.451644 | 17.5984599 | 15.8689222 | 15.4164309 | 18.6650205 | 16.5805351 | 16.3714255 | 19.9891709 | 20.338219 | 13.4017126 | 20.5909909 | 23.2710094 |
| ENSMUSG00 | Tcf7 | 4136 | 10.965431 | 10.4465241 | 13.0407365 | 11.4563554 | 11.3114676 | 9.60862967 | 12.0111975 | 10.9978606 | 13.3637099 | 10.4652673 | 10.2688438 | 10.7690761 | 13.7881277 | 14.0028474 | 14.0360233 | 14.5071039 | 14.4001609 | 13.1936736 |
| ENSMUSG00 | Ddx3x | 3287 | 87.350917 | 91.2532095 | 80.8383944 | 82.140173 | 86.9654684 | 87.9949212 | 72.3004522 | 85.7266641 | 65.7831481 | 77.317399 | 73.5155887 | 79.2830327 | 64.7303192 | 71.3546216 | 75.3836803 | 73.7970788 | 85.0944067 | 78.1708359 |
| ENSMUSG00 | Il12rb1 | 2991 | 0.4547528 | 0.11137711 | 0.12908423 | 0.10662899 | 0.43449079 | 0.49701839 | 0.44291374 | 0.11492197 | 0.30294328 | 0.29951968 | 0.04791871 | 0.33590168 | 0.21441489 | 0.46583839 | 0.44080568 | 0.10241553 | 0.03844162 | 0.2942647 |
| ENSMUSG00 | Slc5a5 | 2913 | 0.02918309 | 0.06861564 | 0.26569739 | 0.09384355 | 0.36350921 | 0.01701089 | 0.06496763 | 0.1311102 | 0.67866555 | 0.22660825 | 0.24600905 | 0.45986128 | 0.52498779 | 0.01594373 | 0 | 0.18402626 | 0.19735477 | 0.03776801 |
| ENSMUSG00 | Kcnn3 | 3072 | 0.08301792 | 0 | 0 | 0 | 0.03133589 | 0 | 0 | 0 | 0 | 0 | 0 | 0 | 0 | 0 | 0 | 0.17466411 | 0 | 0 |
| ENSMUSG00 | Usp32 | 8579 | 4.00328503 | 4.34904319 | 3.67072715 | 3.84499313 | 3.36065069 | 3.68511995 | 3.16557614 | 3.57928564 | 3.71586003 | 3.51198268 | 2.68974297 | 3.66292319 | 3.68594929 | 2.21420116 | 3.35702151 | 4.78911548 | 2.92618306 | 2.71871561 |
| ENSMUSG00 | Car4 | 2211 | 0.80742532 | 0.4670736 | 0.78762872 | 0.26788493 | 0.21769301 | 0.94130023 | 43.5250968 | 30.7819363 | 28.7430094 | 14.6719511 | 6.35270818 | 20.3218491 | 3.68148636 | 14.4730783 | 5.10593682 | 1.52400427 | 1.43875289 | 7.93663729 |
| ENSMUSG00 | Txnrd3 | 2820 | 21.0415694 | 24.6657174 | 26.2795207 | 24.4123184 | 24.0830299 | 21.3849918 | 28.068834 | 23.5655249 | 23.8502058 | 25.1470417 | 28.6819166 | 23.0981838 | 28.0247275 | 27.2406101 | 33.1804019 | 32.2890299 | 32.1016714 | 33.2200386 |
| ENSMUSG00 | Fasl | 2382 | 0 | 0 | 0 | 0 | 0 | 0 | 0 | 0 | 0 | 0 | 0 | 0 | 0 | 0 | 0 | 0 | 0 | 0 |
| ENSMUSG00 | Znf512b | 9357 | 6.05075286 | 6.74659982 | 7.3669211 | 8.22893715 | 6.17788183 | 7.37174363 | 7.02333131 | 6.7511204 | 7.51806871 | 6.93881782 | 7.0868486 | 6.72269443 | 5.87849607 | 6.13993119 | 6.83830462 | 6.27332491 | 6.53312424 | 7.2839888 |
| ENSMUSG00 | Dnajc5 | 2094 | 76.9518254 | 101.434104 | 93.6053223 | 101.087207 | 94.5629102 | 98.3718785 | 94.0379034 | 88.0395064 | 95.1380978 | 111.481916 | 99.1090829 | 90.7604256 | 87.2143065 | 85.0144159 | 84.4100644 | 86.4190117 | 101.251707 | 86.953268 |
| ENSMUSG00 | Tpd52l2 | 5140 | 47.7066849 | 50.2933967 | 48.8817348 | 46.8551907 | 47.598165 | 49.0128594 | 45.4993331 | 48.3943807 | 48.2138649 | 50.352181 | 52.1899595 | 50.7336139 | 47.4699733 | 50.4921324 | 46.4519487 | 53.6217494 | 51.2110909 | 42.316344 |
| ENSMUSG00 | Fmr1 | 8279 | 8.36857554 | 10.0192181 | 10.5347838 | 9.43799868 | 9.64498869 | 7.691178 | 10.5952055 | 9.87217552 | 6.01952942 | 8.57699711 | 9.18682617 | 9.84308097 | 7.40068139 | 6.94501512 | 8.69915687 | 6.56753986 | 11.4113204 | 8.033097 |
| ENSMUSG00 | Bcl11a | 13496 | 0.06298929 | 0.05183542 | 0.00716857 | 0.01687946 | 0.0677613 | 0.04405993 | 0.09815908 | 0.0537682 | 0.07019042 | 0.05939256 | 0.05663895 | 0 | 0.20469675 | 0.20303802 | 0.08853328 | 0.07944109 | 0.12211253 | 0.04483552 |
| ENSMUSG00 | Il4 | 1220 | 0 | 0 | 0.07930087 | 0 | 0 | 0 | 0 | 0 | 0 | 0.2318887 | 0 | 0 | 0.20217999 | 0 | 0 | 0 | 0.09424499 | 0 |
| ENSMUSG00 | Pxmp4 | 3109 | 12.1951164 | 11.6579057 | 11.3426515 | 10.6392146 | 9.68044399 | 10.2962579 | 10.9417228 | 10.9331741 | 10.8762076 | 12.1781971 | 13.2921644 | 11.3462524 | 12.4400846 | 10.292691 | 11.5958062 | 10.1114791 | 12.5247725 | 11.8723418 |
| ENSMUSG00 | Dlg3 | 6526 | 7.80933283 | 7.85604416 | 7.59033023 | 10.3395505 | 8.18670476 | 8.42836872 | 8.85209638 | 7.15741774 | 7.74377181 | 7.88253406 | 8.023499 | 6.74817006 | 7.21156131 | 5.8144069 | 8.66204945 | 7.25209788 | 7.54662757 | 6.2629124 |
| ENSMUSG00 | Gnb1l | 4178 | 1.80072191 | 1.62657502 | 1.57462904 | 1.72298827 | 1.67044733 | 1.61301383 | 1.65333919 | 1.85568638 | 1.53784022 | 2.14423974 | 1.48653533 | 1.83024073 | 1.97185979 | 2.28996697 | 2.83026485 | 2.01625984 | 2.49515384 | 2.51477726 |
| ENSMUSG00 | Dbh | 2398 | 0 | 0 | 0 | 0 | 0 | 0 | 0.80893238 | 0.07963387 | 0 | 0.11797507 | 0.17930552 | 0.06982777 | 0 | 0.13745306 | 0 | 0 | 0 | 0 |
| ENSMUSG00 | Mmp11 | 2264 | 6.23309118 | 4.517252 | 4.70060807 | 6.25860636 | 3.72044463 | 3.78649422 | 3.17646061 | 1.92311567 | 2.0556874 | 3.49881544 | 3.88277048 | 3.7473413 | 1.78675672 | 2.89249638 | 2.42040942 | 4.6171978 | 3.87664399 | 1.60362237 |
| ENSMUSG00 | Smarcb1 | 7032 | 20.5937336 | 19.5035816 | 20.7059622 | 18.9772894 | 19.2883619 | 19.4067446 | 18.1930438 | 17.5591634 | 21.7588035 | 19.1097134 | 20.3070749 | 18.0813537 | 16.556218 | 18.7242588 | 19.7219052 | 18.8567084 | 18.1221469 | 19.6818704 |
| ENSMUSG00 | Vpreb3 | 856 | 0 | 0.15566773 | 0 | 0 | 0 | 0.05788871 | 0.33163093 | 0.17846917 | 0.09623018 | 0.0550826 | 0.27905934 | 0 | 0 | 0.43405689 | 0 | 0.13419605 | 0.62683196 | 0 |
| ENSMUSG00 | Hip1r | 4878 | 21.871257 | 29.8436547 | 25.1288491 | 26.4511844 | 25.7532433 | 30.2415924 | 29.272174 | 28.1784453 | 31.9748535 | 35.860826 | 25.3663679 | 24.7726342 | 32.7969266 | 40.0934916 | 36.7925333 | 36.42239 | 34.4213831 | 34.9699139 |
| ENSMUSG00 | Nsun5 | 3239 | 5.52475428 | 6.70577556 | 6.34725236 | 7.28639116 | 4.87411885 | 6.54787588 | 5.82826764 | 5.36509512 | 6.10358986 | 5.99755781 | 7.19795608 | 5.34203625 | 6.44254431 | 11.901387 | 5.17722545 | 6.77385552 | 6.79200198 | 5.82529287 |
| ENSMUSG00 | Top1mt | 1889 | 5.08532016 | 6.15468506 | 4.09728155 | 4.72735029 | 3.36337423 | 5.3776127 | 6.51206804 | 4.54912151 | 6.21395288 | 4.59276305 | 7.3091387 | 7.62331442 | 4.62241893 | 4.1551354 | 5.38739243 | 4.52027816 | 5.76213251 | 6.72689447 |
| ENSMUSG00 | Hoxa10 | 4974 | 2.93109678 | 5.11681769 | 2.71335243 | 2.03348449 | 5.36089379 | 2.99866763 | 0 | 0 | 0.00828036 | 0.00947943 | 0.00960494 | 0.17954375 | 0.00991796 | 0.02801212 | 0 | 0.03849076 | 0.05393729 | 0 |
| ENSMUSG00 | Hoxa4 | 1982 | 0.0643368 | 0.03361543 | 0.19525138 | 0.11493704 | 0.12142261 | 0.22501241 | 0.1193559 | 0 | 0.02078028 | 0.02378946 | 0.43387954 | 0 | 0.74669905 | 0.30492822 | 0.04157574 | 0.11591506 | 0 | 0.27754342 |
| ENSMUSG00 | Snrpn | 2185 | 15.698708 | 14.3314053 | 14.8773512 | 15.2009159 | 16.2128585 | 16.6914471 | 19.3581396 | 17.654154 | 16.436907 | 14.7817994 | 16.8578915 | 16.4764771 | 21.4486597 | 20.9795489 | 17.9702896 | 17.8397429 | 18.4527526 | 21.4497753 |
| ENSMUSG00 | Mmp14 | 2587 | 107.09267 | 112.558179 | 103.815168 | 110.811786 | 105.808369 | 116.612695 | 75.8429376 | 107.431818 | 88.0725957 | 97.7278975 | 78.966263 | 97.4994121 | 64.2630697 | 59.7830734 | 69.646078 | 76.4776067 | 80.6081876 | 62.1324352 |
| ENSMUSG00 | Slc7a7 | 2259 | 0.97842813 | 1.25347322 | 0.66382445 | 0.76640978 | 1.470165 | 0.789685 | 1.42419751 | 0.86224547 | 1.23978888 | 1.06449131 | 1.83993864 | 0.81536824 | 2.05276681 | 1.00741934 | 0.89370351 | 1.10176604 | 0.373253 | 0.46267066 |
| ENSMUSG00 | Oxa1l | 2524 | 21.084184 | 19.8108773 | 20.1236953 | 20.9934606 | 21.319925 | 20.7909449 | 20.3571986 | 24.0140026 | 24.6890643 | 21.3336383 | 25.7613782 | 23.3744673 | 22.965532 | 24.5653156 | 21.2863823 | 24.2881388 | 24.3563313 | 22.0777437 |
| ENSMUSG00 | Heatr6 | 6626 | 6.67150363 | 5.98787459 | 5.7601442 | 6.78671155 | 6.16722052 | 6.24457177 | 6.71917158 | 5.66603253 | 6.48317778 | 4.95986118 | 5.94843657 | 5.51754588 | 6.79747064 | 6.28740513 | 6.3549638 | 7.33335056 | 7.06254279 | 6.32613029 |
| ENSMUSG00 | Ccl3 | 763 | 0 | 0 | 0 | 0 | 0 | 0 | 0 | 0.70077803 | 0.37785794 | 0.1853894 | 0 | 0 | 0.3879312 | 1.40001836 | 0.21599767 | 0 | 0 | 0 |
| ENSMUSG00 | Expi | 823 | 1.18787248 | 1.74052789 | 3.29151604 | 10.1861873 | 2.10540614 | 3.79322262 | 0 | 0 | 0.05004437 | 0 | 0 | 0 | 0 | 0.05006264 | 0 | 0 | 0 | 0 |
| ENSMUSG00 | Icam4 | 904 | 0 | 0 | 0 | 0 | 0 | 0 | 0 | 0 | 0 | 0 | 0 | 0 | 0 | 0 | 0 | 0 | 0 | 0 |
| ENSMUSG00 | Ilf2 | 1959 | 76.5049774 | 80.586935 | 81.4127257 | 76.0281031 | 77.8366178 | 79.5524998 | 77.9125586 | 81.6291929 | 88.1967502 | 89.0544166 | 82.18561 | 86.6439152 | 77.9892536 | 78.2840105 | 92.8139184 | 91.7880002 | 84.9282567 | 82.49962 |
| ENSMUSG00 | 2500003M10 | 4779 | 25.6596428 | 26.3979766 | 26.5603983 | 25.2735569 | 27.9284002 | 25.3933134 | 27.0768313 | 27.6832982 | 25.7340314 | 25.2674097 | 27.9112124 | 25.9398591 | 21.8323928 | 24.9470543 | 24.9847476 | 26.9452422 | 25.598983 | 26.3937815 |
| ENSMUSG00 | Snapin | 3836 | 33.9952763 | 34.0510579 | 31.6647379 | 35.6791911 | 34.3172084 | 34.6972478 | 33.7454352 | 31.063685 | 34.7444121 | 29.6597098 | 30.2142986 | 30.4832672 | 27.9967633 | 30.8376149 | 30.5789441 | 32.5709729 | 31.4023856 | 28.1068429 |
| ENSMUSG00 | S100a4 | 619 | 0.20600247 | 0.16145183 | 0.39073934 | 0.22081282 | 0.07775755 | 0 | 1.98728538 | 5.36791439 | 1.53035519 | 0 | 2.70133045 | 0.0901707 | 1.19544226 | 1.27552421 | 0.86529924 | 0.43301172 | 0 | 0.2666031 |
| ENSMUSG00 | S100a3 | 709 | 0.11990177 | 0.04698575 | 0.40936697 | 0 | 0 | 0.13978204 | 0 | 0.16160396 | 0.75518294 | 0.33251554 | 0.40430148 | 0 | 0.13915915 | 0.32753235 | 0.34867325 | 0 | 0.05405684 | 0 |
| ENSMUSG00 | S100a5 | 429 | 0 | 0 | 0 | 0 | 0 | 0 | 0 | 0.08902658 | 0 | 0 | 0 | 0 | 0 | 0 | 0 | 0 | 0 | 0 |
| ENSMUSG00 | S100a6 | 716 | 332.205258 | 320.566817 | 344.35738 | 274.893452 | 362.804458 | 361.748797 | 41.9602669 | 42.8336993 | 36.8722858 | 27.7899396 | 39.6345326 | 46.7729016 | 76.547151 | 50.8549784 | 52.42265 | 93.7478139 | 52.8324791 | 54.6249649 |
| ENSMUSG00 | Scn4a | 7015 | 0 | 0 | 0 | 0 | 0 | 0 | 0.04046701 | 0 | 0.03522724 | 0.04704988 | 0 | 0 | 0 | 0 | 0 | 0 | 0 | 0 |
| ENSMUSG00 | Icam2 | 1897 | 0 | 0 | 0 | 0 | 0 | 0 | 0 | 0 | 0 | 0.07456621 | 0.5036896 | 0.08826937 | 0 | 0.22034622 | 0 | 0.14129375 | 0 | 0 |
| ENSMUSG00 | Mapk7 | 4299 | 10.1640663 | 9.57774767 | 8.10163802 | 7.29146278 | 9.27034957 | 7.37700616 | 4.74337395 | 8.27102271 | 9.47510214 | 9.52007686 | 7.30126721 | 8.11462878 | 4.99171529 | 5.58539037 | 5.97082329 | 7.74897149 | 6.4278344 | 6.84574834 |
| ENSMUSG00 | Epn2 | 5209 | 20.4243532 | 18.7253128 | 18.1365914 | 19.0588435 | 16.6599843 | 17.7986494 | 14.3691031 | 10.7633801 | 11.5360197 | 13.6500788 | 11.6754698 | 15.1191975 | 12.4726694 | 11.8762765 | 13.8577704 | 14.8928337 | 13.1408651 | 12.5140702 |
| ENSMUSG00 | B9d1 | 1654 | 11.3843972 | 8.92237735 | 8.68617806 | 12.1753214 | 9.9814086 | 8.98779932 | 13.2155125 | 11.2221932 | 13.4715266 | 12.2295355 | 12.6802943 | 13.0939961 | 14.2269193 | 12.8605756 | 11.9071002 | 12.2928129 | 10.3114886 | 11.740154 |
| ENSMUSG00 | Sec24b | 6548 | 8.29590943 | 9.40168426 | 9.10882146 | 7.36157375 | 8.20330281 | 9.44438172 | 10.0073397 | 8.22991445 | 8.07628073 | 7.96406197 | 7.12830398 | 8.41326496 | 8.8071184 | 6.82333721 | 7.58843561 | 7.23942761 | 10.1493279 | 8.27488853 |
| ENSMUSG00 | N4bp3 | 2721 | 9.27896897 | 11.434856 | 9.493372 | 12.3070076 | 10.7726356 | 14.4050762 | 9.58077401 | 10.5130869 | 11.1859005 | 12.7017514 | 12.8348014 | 15.4052049 | 11.6395042 | 11.231242 | 11.37169 | 15.1839638 | 11.1133584 | 11.7255722 |
| ENSMUSG00 | Rmnd5b | 2624 | 31.1661428 | 34.1634902 | 34.0310736 | 34.4138681 | 34.4664194 | 33.6709314 | 28.596764 | 30.609231 | 27.1071314 | 25.965231 | 29.0582446 | 28.8650359 | 28.7080174 | 28.5319622 | 27.0071174 | 29.1703217 | 32.3962372 | 34.7580408 |
| ENSMUSG00 | Nhp2 | 1349 | 44.8053079 | 41.5114716 | 37.9027588 | 39.4141874 | 41.3527795 | 39.5981073 | 42.7883382 | 41.1651246 | 43.751132 | 39.77576 | 39.5586572 | 37.4035278 | 51.489503 | 57.2890744 | 47.218388 | 44.5635683 | 38.297886 | 48.6477867 |
| ENSMUSG00 | 1300018I17R | 5563 | 4.50800896 | 5.94039027 | 5.66952037 | 6.18346609 | 5.74502918 | 5.60285267 | 6.43819844 | 4.63416711 | 6.12282016 | 6.66195448 | 5.41043031 | 5.78925711 | 4.90392778 | 6.64560373 | 5.03632177 | 5.00400352 | 5.17401563 | 5.93303326 |
| ENSMUSG00 | Zfp276 | 5861 | 3.44479758 | 3.45576517 | 3.39690053 | 3.38928772 | 3.00567891 | 3.06058515 | 3.55995417 | 2.90629781 | 4.24443875 | 2.94440495 | 3.70070485 | 2.84744636 | 3.72872195 | 2.64670281 | 2.66428761 | 2.99216826 | 3.32845677 | 3.04094021 |
| ENSMUSG00 | C1ql4 | 872 | 0 | 0 | 0 | 0 | 0 | 0 | 0 | 0 | 0 | 0 | 0 | 0 | 0 | 0 | 0 | 0 | 0 | 0 |
| ENSMUSG00 | Mfsd10 | 2961 | 20.9008903 | 17.607119 | 18.7547494 | 22.2496691 | 17.9783545 | 16.902486 | 18.5192148 | 20.0313411 | 18.5833114 | 18.2488031 | 21.8948391 | 17.5496055 | 13.7782829 | 17.8655237 | 18.130911 | 16.0869691 | 14.6522689 | 13.3760746 |
| ENSMUSG00 | Luzp1 | 4366 | 14.5935087 | 13.3984064 | 12.3094348 | 10.0910513 | 9.87773789 | 11.4745336 | 9.09192341 | 10.3660094 | 10.0183417 | 9.9463577 | 7.98802558 | 10.3423892 | 8.92629745 | 8.5101397 | 8.88013378 | 12.3221034 | 10.2355617 | 10.2307363 |
| ENSMUSG00 | Slc13a2 | 2400 | 0.15939441 | 0.5552149 | 0.1612451 | 0.37967537 | 0.32087949 | 0.53682128 | 0.53226764 | 0.47740503 | 0.22309363 | 0.07858451 | 0.03981247 | 0.11628263 | 0.88386352 | 1.12239878 | 0.48068482 | 0 | 0.14372361 | 0.06876138 |
| ENSMUSG00 | Kctd10 | 4695 | 5.64019695 | 7.81912874 | 5.80070236 | 8.00593431 | 8.05786837 | 8.51737082 | 6.83237408 | 5.27941835 | 6.63195014 | 5.85492226 | 5.43382914 | 5.61128706 | 5.68446587 | 6.33103641 | 6.39743008 | 5.47241695 | 5.52649691 | 5.88169789 |
| ENSMUSG00 | Poldip2 | 3524 | 25.3052949 | 27.1588947 | 25.4633652 | 27.758105 | 26.2786099 | 25.4372574 | 26.8114131 | 30.0423778 | 25.2214816 | 26.9069422 | 33.3773972 | 25.9279908 | 26.7237883 | 30.3389016 | 28.4342179 | 29.2286419 | 23.7527334 | 27.442114 |
| ENSMUSG00 | Sebox | 1432 | 0 | 0 | 0 | 0 | 0 | 0 | 0 | 0 | 0 | 0 | 0 | 0 | 0 | 0 | 0 | 0 | 0 | 0 |
| ENSMUSG00 | Ift20 | 1668 | 51.6534723 | 47.4129562 | 46.6624762 | 44.4684533 | 45.3040289 | 47.562306 | 45.9795279 | 46.8704123 | 53.9770524 | 49.5251991 | 42.2470406 | 43.3006399 | 56.7257898 | 67.6891945 | 55.380029 | 50.3195512 | 49.0337635 | 53.2941945 |
| ENSMUSG00 | Col6a1 | 4358 | 0 | 0 | 0 | 0 | 0 | 0 | 0 | 0.02629124 | 0 | 0.07573541 | 0 | 0 | 0.03395956 | 0 | 0 | 0 | 0 | 0 |
| ENSMUSG00 | Pcbp3 | 3578 | 5.88039747 | 7.69045574 | 6.3001859 | 6.32863025 | 6.7269033 | 5.62280879 | 6.99508305 | 7.55735632 | 5.01881522 | 6.45719525 | 6.75632448 | 7.8466401 | 5.34957627 | 8.35941651 | 6.13762697 | 6.88117959 | 6.41628056 | 4.68914968 |
| ENSMUSG00 | Lgals9 | 2688 | 6.98931838 | 6.41967232 | 5.25486267 | 7.50875842 | 7.10876998 | 8.12974535 | 2.88663666 | 1.66239252 | 2.72738088 | 2.57855408 | 2.04394354 | 3.17700758 | 4.88180431 | 2.66085917 | 4.19986098 | 5.84045723 | 3.27940765 | 4.93199208 |
| ENSMUSG00 | Araf | 4311 | 41.4699073 | 44.9658389 | 41.4950859 | 47.4316787 | 46.12224 | 42.2077564 | 48.9479351 | 47.0959896 | 41.8457293 | 41.1679998 | 46.67783 | 44.435016 | 44.5486323 | 49.6545578 | 47.3564624 | 48.3628757 | 48.4879664 | 48.9480244 |
| ENSMUSG00 | Cfp | 1835 | 0.41694451 | 0.19969582 | 0.34270076 | 0.3476047 | 0.23606938 | 0.32405057 | 0.74772081 | 2.35190271 | 1.43647793 | 0.59098974 | 1.53608858 | 0.12166902 | 0.91405187 | 1.34143886 | 0.35925062 | 0.37560269 | 0.16709012 | 0.56957658 |
| ENSMUSG00 | Timp1 | 985 | 2.24392807 | 0.77786454 | 1.71885641 | 1.06386195 | 1.46594691 | 0.65399547 | 0.28819906 | 0.42651414 | 0 | 0.33508114 | 0 | 0.67998777 | 1.20199595 | 0 | 0.12548697 | 0.46648456 | 0.38909946 | 0 |
| ENSMUSG00 | Ust | 1631 | 13.0043426 | 12.5408443 | 12.6719265 | 11.9279987 | 13.2502966 | 11.4235609 | 13.575925 | 10.3501176 | 12.6766591 | 12.8645389 | 13.9721982 | 11.6696143 | 14.8812405 | 15.7471369 | 12.7570728 | 13.311327 | 12.2662949 | 13.0524367 |
| ENSMUSG00 | Cnnm3 | 5605 | 4.71689911 | 5.85427311 | 6.47281851 | 6.13712551 | 5.56458279 | 6.21508862 | 6.76136589 | 6.22117099 | 6.27534067 | 6.8475776 | 5.01189573 | 5.00897026 | 4.55913169 | 4.085091 | 5.99829966 | 4.30385061 | 5.96262817 | 4.7893745 |
| ENSMUSG00 | Lman2l | 7302 | 6.28089358 | 6.14066771 | 7.86351418 | 6.76364997 | 6.96732995 | 6.67760748 | 6.6154814 | 6.51185168 | 7.50178937 | 5.80505098 | 6.71283311 | 6.61960611 | 6.1614227 | 5.52724075 | 6.1051881 | 5.34872905 | 6.4874422 | 6.76502025 |
| ENSMUSG00 | Mcm3ap | 6640 | 3.6487877 | 5.09727418 | 3.89756608 | 4.76195253 | 3.820109 | 4.44034284 | 3.5128239 | 2.60559613 | 5.31578976 | 3.78483808 | 3.28093088 | 4.04327315 | 4.45770337 | 3.42035523 | 4.3187173 | 3.02172866 | 3.15730187 | 4.63103775 |
| ENSMUSG00 | Pcnt | 10004 | 5.32801787 | 5.1947336 | 4.46309162 | 5.06436239 | 4.37344248 | 4.918619 | 4.92800994 | 4.75305288 | 4.4010781 | 4.96769704 | 4.65619597 | 5.38405782 | 4.48247829 | 2.88302462 | 3.90646427 | 5.81784519 | 4.07245638 | 3.1947512 |
| ENSMUSG00 | Ftcd | 3285 | 0.12939171 | 0.05070456 | 0.04417674 | 0.01386942 | 0 | 0.10559182 | 1.35384835 | 0.96498307 | 0.51404784 | 0.47366003 | 0.18906376 | 0.69663384 | 1.56180195 | 3.84559871 | 1.60541831 | 0.31471732 | 0.35001184 | 2.69603232 |
| ENSMUSG00 | Mxd1 | 4656 | 4.16287806 | 4.49323401 | 4.62332925 | 5.68534509 | 5.5512979 | 5.01274431 | 3.07898233 | 3.23191721 | 3.43220965 | 4.12163579 | 3.84785426 | 2.57739851 | 2.70181248 | 4.20949417 | 4.30067787 | 3.12509253 | 3.30086878 | 2.64648623 |
| ENSMUSG00 | Gmcl1 | 3808 | 16.0175755 | 16.7089359 | 19.2198402 | 18.9638275 | 17.0130171 | 18.3480447 | 19.021983 | 16.7492941 | 17.4783113 | 16.0470881 | 18.7310959 | 15.7421275 | 12.9807198 | 12.5623451 | 12.5508941 | 15.9477162 | 14.5233315 | 11.2820673 |
| ENSMUSG00 | Snrnp27 | 1936 | 51.6385196 | 44.497492 | 48.7483253 | 38.100894 | 46.1927243 | 46.6348557 | 49.1944332 | 44.6235611 | 43.0160821 | 43.9115279 | 45.356598 | 40.1607531 | 43.471204 | 40.8304942 | 41.9464187 | 42.2858051 | 42.7805411 | 39.6088299 |
| ENSMUSG00 | Oas1c | 3433 | 4.28394725 | 3.57097147 | 3.98768106 | 3.78237624 | 3.2947865 | 3.34874285 | 1.52977202 | 3.41539981 | 4.22302754 | 4.32638262 | 2.65803002 | 2.42252657 | 2.85961303 | 3.65275374 | 2.43632853 | 2.71034594 | 3.36038898 | 2.48366197 |
| ENSMUSG00 | Oas1h | 1981 | 0 | 0.0168162 | 0 | 0 | 0 | 0 | 0 | 0 | 0 | 0 | 0 | 0 | 0 | 0 | 0 | 0 | 0 | 0 |
| ENSMUSG00 | Ocrl | 9714 | 3.70618531 | 3.77573569 | 3.85932531 | 2.94547414 | 3.16618275 | 3.37697237 | 4.86569552 | 4.52143944 | 3.59968621 | 3.92679833 | 3.10827401 | 3.17748213 | 8.54192762 | 8.41482333 | 6.51488469 | 3.55155829 | 4.43470843 | 6.1555383 |
| ENSMUSG00 | Calm1 | 4878 | 257.270464 | 233.763913 | 229.471811 | 208.041554 | 233.604611 | 237.828741 | 184.85698 | 176.328638 | 190.126683 | 162.862279 | 175.20423 | 185.525862 | 177.698056 | 138.570572 | 145.801816 | 175.251353 | 148.103874 | 152.103901 |
| ENSMUSG00 | Agpat3 | 9772 | 11.4005389 | 10.8202135 | 9.77173034 | 11.8425146 | 11.3483373 | 10.2127717 | 14.1473238 | 13.5854268 | 13.5925617 | 16.9553389 | 11.1615053 | 11.9205166 | 13.5041832 | 15.5320587 | 14.7485717 | 12.0529992 | 12.8564881 | 14.5065972 |
| ENSMUSG00 | Slc26a3 | 5215 | 177.37251 | 175.577797 | 196.147013 | 167.977411 | 180.22475 | 187.882301 | 124.618784 | 127.371479 | 139.189099 | 121.036714 | 127.925124 | 125.116989 | 213.46578 | 209.002846 | 221.78525 | 258.437737 | 205.5436 | 234.213271 |
| ENSMUSG00 | Sema6b | 3959 | 1.09510683 | 1.00132216 | 0.70868016 | 0.93216585 | 0.69298299 | 0.86363693 | 0.7289051 | 0.71387668 | 0.56177617 | 0.48829978 | 0.55510182 | 1.07148026 | 0.36136034 | 1.16139547 | 0.62442368 | 0.40621508 | 0.54212494 | 1.19494397 |
| ENSMUSG00 | Uhrf1 | 4188 | 1.46149602 | 1.90904838 | 1.97513698 | 2.15403218 | 2.0572145 | 2.23626233 | 1.63809418 | 0.93930694 | 1.90787585 | 1.89143222 | 1.31187207 | 1.54599256 | 2.01426883 | 1.56366197 | 0.89525826 | 1.86515853 | 1.61065622 | 1.56305723 |
| ENSMUSG00 | Dpp9 | 3611 | 10.6880919 | 12.3897028 | 9.57825373 | 10.6994643 | 9.18385362 | 12.2818323 | 10.285365 | 11.8247316 | 12.5236207 | 10.3676706 | 11.6295177 | 11.2527838 | 7.18596407 | 7.49845 | 11.0677123 | 9.10873214 | 7.69497792 | 10.2980216 |
| ENSMUSG00 | Ramp2 | 3294 | 0.03871145 | 0.23260369 | 0.22028019 | 0.42877728 | 0.014612 | 0.25573663 | 0.48835188 | 0.85799568 | 0.75020976 | 0.58687882 | 0.84121056 | 0.5930626 | 0.13478666 | 0 | 0.02501612 | 0.29060874 | 0.442137 | 0 |
| ENSMUSG00 | Lsr | 2351 | 31.458531 | 33.5254393 | 31.2956783 | 37.7705127 | 35.7867301 | 34.3770772 | 20.2452382 | 25.7161093 | 25.7700403 | 27.1552753 | 26.092321 | 24.026138 | 21.6968109 | 26.2149315 | 27.3742366 | 28.5347274 | 22.5784437 | 26.9780944 |
| ENSMUSG00 | Gramd1a | 2663 | 36.45581 | 34.7890195 | 33.4781887 | 41.129835 | 36.6547282 | 37.2157218 | 32.2109977 | 31.795928 | 27.4989204 | 34.4556022 | 33.5303783 | 29.5322075 | 37.9576036 | 42.0316375 | 37.3490634 | 43.7113956 | 38.0672382 | 40.776559 |
| ENSMUSG00 | Hpn | 3272 | 13.6141273 | 12.9199458 | 13.7492003 | 15.5397694 | 12.4448678 | 16.5832039 | 29.9753006 | 28.7026643 | 31.3933895 | 32.8123934 | 28.4576388 | 27.0037267 | 45.7134392 | 40.1559868 | 39.2623629 | 61.2392263 | 48.7043834 | 41.6266957 |
| ENSMUSG00 | Gabrg1 | 5023 | 0 | 0.05305657 | 0 | 0 | 0.01916461 | 0 | 0 | 0 | 0 | 0 | 0 | 0 | 0.20624532 | 0 | 0 | 0 | 0 | 0 |
| ENSMUSG00 | Ckb | 1470 | 24.3175871 | 19.0132777 | 15.7954385 | 23.2144369 | 19.9731112 | 17.0906367 | 28.6772768 | 25.1238457 | 20.0609152 | 27.5206146 | 33.344971 | 17.8078543 | 41.2441676 | 42.6527333 | 32.4567359 | 54.3882913 | 45.1572423 | 40.3025904 |
| ENSMUSG00 | Sp1 | 8316 | 4.8556899 | 4.45453804 | 4.14164906 | 4.35010452 | 4.89653769 | 4.2187753 | 3.78342129 | 4.79012454 | 5.9035987 | 4.0993216 | 5.06704105 | 4.60432233 | 3.43472584 | 3.57992545 | 3.11142101 | 2.97907924 | 4.1663026 | 3.300811 |
| ENSMUSG00 | Itgb7 | 2727 | 0.20262827 | 0.0977276 | 0.10643241 | 0.18378126 | 0.01765014 | 0.12719807 | 0.26024576 | 0.42015844 | 2.00872996 | 0.62245153 | 2.01471222 | 0.388888 | 1.30249287 | 0.68124807 | 0.51369743 | 0.37911492 | 0.01405438 | 0.10086012 |
| ENSMUSG00 | Myg1 | 3050 | 15.3394162 | 11.3591508 | 12.4819568 | 12.5180837 | 13.0035099 | 11.0478226 | 12.4409077 | 10.0928119 | 12.0723754 | 12.1045904 | 12.9697264 | 12.7369512 | 10.9985914 | 9.88292469 | 13.1980066 | 11.0352275 | 9.67581922 | 10.6591417 |
| ENSMUSG00 | Rarg | 5378 | 0.88519519 | 0.70614912 | 0.20687824 | 0.27956758 | 0.08054803 | 0.41462867 | 0.18474642 | 0.41189277 | 0.27569999 | 0.6049458 | 0.32868603 | 0.70573913 | 0.26601443 | 0.31089385 | 0.62821264 | 0.45567104 | 0.22092138 | 0.39891319 |
| ENSMUSG00 | Plin5 | 1501 | 81.3856602 | 74.2604557 | 78.9252337 | 67.6586061 | 76.3825731 | 70.8462134 | 85.5788963 | 69.1330828 | 77.4888211 | 63.9879964 | 69.2593669 | 64.2568054 | 87.9166414 | 78.592926 | 70.874362 | 82.1168989 | 69.017975 | 91.9505988 |
| ENSMUSG00 | Efnb2 | 7055 | 5.25967664 | 7.6116776 | 6.03383511 | 5.62490002 | 7.3886457 | 7.00972758 | 3.92316331 | 5.67878528 | 5.41175055 | 5.42434664 | 4.49646668 | 5.02380521 | 5.41924475 | 4.28562734 | 5.52684515 | 5.68795982 | 5.99748147 | 3.50873112 |
| ENSMUSG00 | Rrp15 | 3689 | 5.73802593 | 4.66868157 | 4.30103496 | 4.79199924 | 5.51905766 | 3.97603934 | 4.00150605 | 4.51392991 | 4.7449903 | 4.56297128 | 4.9471494 | 5.29560366 | 4.2124028 | 6.01796524 | 4.83607311 | 5.52211367 | 4.47780807 | 4.65243733 |
| ENSMUSG00 | Rnd2 | 1983 | 3.57960887 | 3.5950375 | 2.70774678 | 1.95294441 | 2.79131176 | 4.29811359 | 5.65461669 | 2.65786764 | 3.24008899 | 2.99595996 | 4.023408 | 4.47538596 | 3.880877 | 3.27895725 | 3.44904603 | 4.69219243 | 4.36799955 | 3.43980294 |
| ENSMUSG00 | Srr | 6885 | 3.16705233 | 3.42080119 | 3.21085016 | 3.7322337 | 3.92885129 | 3.67777006 | 4.69347272 | 4.69846984 | 4.63610018 | 3.67070108 | 3.60133674 | 4.33716478 | 3.62555558 | 4.07439779 | 3.71621877 | 3.63719059 | 4.5980423 | 4.10670746 |
| ENSMUSG00 | Sync | 2356 | 6.29639492 | 5.76895621 | 8.29495172 | 8.95363477 | 7.23204626 | 10.2007962 | 8.95647487 | 9.38599363 | 9.89455345 | 8.40547342 | 10.7878939 | 9.73694284 | 5.13001686 | 4.82971197 | 5.47372113 | 5.75334254 | 5.84004269 | 6.42084225 |
| ENSMUSG00 | Fndc5 | 2757 | 0 | 0 | 0 | 0.01652559 | 0 | 0 | 0 | 0.02770577 | 0 | 0.06840871 | 0.0173286 | 0 | 0.08946666 | 0 | 0 | 0 | 0 | 0.01995252 |
| ENSMUSG00 | Acp5 | 1605 | 0.31779571 | 0.45662534 | 0.60278542 | 0.76644747 | 0.2399099 | 0.71010148 | 1.2675671 | 0.57110134 | 1.53968283 | 1.05758587 | 2.79803495 | 0.41731337 | 1.93639301 | 2.37284434 | 0.48774428 | 0.97814013 | 0.382069 | 0.23991511 |
| ENSMUSG00 | Cnn1 | 2361 | 0 | 0.08465793 | 0 | 0.0771894 | 0.16308996 | 0.29383239 | 0 | 0 | 0 | 0.0798826 | 0 | 0 | 0.083578 | 0.05901409 | 0.05235268 | 0.34057719 | 0 | 0 |
| ENSMUSG00 | Fbxo9 | 3675 | 35.5424234 | 33.829584 | 32.9993036 | 35.3454524 | 34.9955102 | 32.253099 | 35.2366809 | 31.613956 | 32.2655725 | 32.3062506 | 32.9159714 | 31.1960193 | 31.2965304 | 29.4967347 | 32.6249055 | 30.5282938 | 30.6923243 | 35.6548822 |
| ENSMUSG00 | Ccdc132 | 10045 | 5.37820595 | 6.50007692 | 5.30206639 | 6.84436196 | 7.01015474 | 7.24668647 | 6.89554624 | 6.63091586 | 7.18355158 | 6.43540201 | 6.31609207 | 4.77864309 | 6.36477499 | 6.71809447 | 5.94746203 | 5.8589013 | 6.94413742 | 6.66461751 |
| ENSMUSG00 | Apbb3 | 2048 | 8.11500187 | 12.4272711 | 13.1562775 | 11.3457679 | 12.9495556 | 11.7832916 | 14.6234989 | 12.2148553 | 12.9914498 | 11.7416302 | 12.9701547 | 14.0356769 | 11.0804111 | 13.3118551 | 16.1949475 | 14.3776744 | 12.8565264 | 13.5105374 |
| ENSMUSG00 | Hars | 1974 | 43.9047892 | 40.4682474 | 41.38951 | 38.1752621 | 41.1584028 | 39.6873717 | 40.3859493 | 36.7413233 | 44.5456997 | 38.2651604 | 37.9731053 | 36.0228744 | 36.1867357 | 33.7390179 | 40.7423694 | 37.4953111 | 31.6279319 | 37.7595439 |
| ENSMUSG00 | Zmat2 | 2251 | 41.9575749 | 40.1353039 | 47.0841426 | 39.2055721 | 38.488433 | 41.0334498 | 46.8923086 | 39.6176187 | 46.3462658 | 43.0242312 | 37.8209578 | 40.6157509 | 51.3043173 | 49.6421654 | 42.7024566 | 50.1639277 | 44.6260436 | 50.1460182 |
| ENSMUSG00 | Ube2c | 2770 | 0.16879312 | 0.14431579 | 0.03492674 | 0 | 0 | 0.21466888 | 0.03416078 | 0.15166658 | 0.5501448 | 0.05106574 | 0 | 0.12090035 | 0.21371228 | 0.20120182 | 1.97826965 | 1.35468575 | 1.25909495 | 0.63548427 |
| ENSMUSG00 | Smg5 | 4425 | 6.70477127 | 8.52207825 | 8.90947508 | 6.90880469 | 8.10356681 | 7.69327186 | 5.94482474 | 6.99978268 | 7.39030362 | 7.05395832 | 4.25386071 | 4.8688917 | 4.74924221 | 4.45023572 | 6.21049441 | 4.96696241 | 4.90230148 | 4.10237406 |
| ENSMUSG00 | Cct3 | 4575 | 47.6243786 | 50.5700657 | 49.0819515 | 49.6241932 | 51.067182 | 50.0293063 | 42.4521414 | 45.0294685 | 39.746113 | 39.6168969 | 45.6551567 | 45.0551348 | 35.2494071 | 46.6775768 | 44.4256339 | 41.7974412 | 44.6009191 | 45.8709819 |
| ENSMUSG00 | Rhbg | 2327 | 20.1657561 | 28.3023601 | 33.5725189 | 29.9665256 | 24.1176721 | 25.6813909 | 29.2815194 | 45.6766033 | 60.4786674 | 60.8886391 | 63.1935205 | 53.1532051 | 73.3300824 | 49.0986055 | 66.892771 | 69.2258639 | 72.4856178 | 69.6655941 |
| ENSMUSG00 | 0610031J06R | 2546 | 72.4057142 | 74.3325028 | 71.5343055 | 74.5332874 | 71.5929281 | 75.9445273 | 78.4394414 | 77.2698605 | 63.947485 | 78.356098 | 77.5920869 | 68.4651666 | 88.2781678 | 87.8533876 | 90.4298092 | 83.905459 | 83.3965769 | 88.3473046 |
| ENSMUSG00 | Mef2d | 3903 | 19.6135849 | 26.3481692 | 23.5980532 | 25.2611068 | 22.8019285 | 23.6527652 | 17.9528766 | 25.8334467 | 20.0286976 | 24.8619388 | 21.4332442 | 18.9627386 | 14.6238863 | 20.1103017 | 18.4525544 | 19.9056219 | 23.2923331 | 23.1565366 |
| ENSMUSG00 | Tmem79 | 3547 | 3.99047748 | 4.40477795 | 4.25501592 | 4.18745431 | 3.41957845 | 3.98154189 | 3.26800291 | 4.1562637 | 4.15697001 | 4.58612704 | 4.3235866 | 3.91826896 | 5.59104335 | 6.27198135 | 5.37815629 | 4.64194 | 4.6138508 | 4.90072673 |
| ENSMUSG00 | Snd1 | 7518 | 30.281687 | 38.8916298 | 36.3798804 | 37.1918235 | 34.2198899 | 41.6432789 | 31.3592816 | 36.5007198 | 32.618318 | 35.2720877 | 37.9314614 | 36.1116498 | 31.1162479 | 33.2484808 | 40.7905268 | 35.1582951 | 35.0024414 | 37.3678516 |
| ENSMUSG00 | Col18a1 | 7962 | 71.0181307 | 104.465755 | 80.373453 | 104.884248 | 89.6927093 | 103.026369 | 59.0487428 | 71.0315367 | 65.023173 | 79.573474 | 66.5861234 | 65.4127038 | 43.9972299 | 34.9176472 | 47.5924026 | 42.3735923 | 44.2807844 | 42.0479055 |
| ENSMUSG00 | Slc19a1 | 4637 | 1.32914611 | 1.602065 | 2.15943946 | 1.55243584 | 1.39091606 | 2.05178453 | 1.17337893 | 2.4791704 | 2.43370829 | 2.09468296 | 1.69999313 | 1.57684964 | 1.37239968 | 2.29365883 | 2.79889802 | 1.54417589 | 1.89275868 | 2.58615366 |
| ENSMUSG00 | Kpnb1 | 6740 | 8.3938263 | 7.01844359 | 7.88608149 | 7.36142096 | 7.52686165 | 6.93297148 | 7.5461617 | 7.1454925 | 9.34335053 | 5.62450526 | 6.38653377 | 8.15703674 | 7.23876353 | 5.19567387 | 6.52256085 | 8.59550282 | 6.25503362 | 6.35787747 |
| ENSMUSG00 | Npepps | 6811 | 13.0492326 | 12.5944359 | 12.9971459 | 13.1712959 | 10.5648548 | 11.909826 | 13.3581385 | 11.3943564 | 12.9527994 | 11.4363473 | 11.9314645 | 13.0053526 | 10.8137649 | 8.98059948 | 10.5075764 | 12.5142992 | 10.1738283 | 8.64994176 |
| ENSMUSG00 | Tbx21 | 2482 | 0 | 0 | 0 | 0 | 0 | 0 | 0 | 0 | 0 | 0 | 0 | 0 | 0 | 0 | 0 | 0 | 0 | 0 |
| ENSMUSG00 | Mrpl10 | 1065 | 71.5603574 | 65.2494811 | 65.224779 | 65.7961373 | 72.0397055 | 70.816207 | 77.210738 | 66.7380853 | 83.2238328 | 66.4979871 | 71.9091629 | 66.4022953 | 59.2446218 | 69.6879358 | 62.9435976 | 71.0803106 | 67.4758748 | 70.246371 |
| ENSMUSG00 | Cyp51 | 3896 | 10.5171945 | 10.5257117 | 10.8765946 | 9.87000036 | 12.2924189 | 10.7220109 | 10.7109299 | 9.92061377 | 9.51433886 | 10.1175534 | 9.51575148 | 9.03995973 | 13.9284161 | 20.0629874 | 16.1591318 | 13.7791149 | 16.7726736 | 15.0936178 |
| ENSMUSG00 | Tcf25 | 3984 | 124.346844 | 129.66442 | 122.815828 | 126.779553 | 123.060184 | 124.20522 | 108.080495 | 124.058203 | 122.453383 | 112.610176 | 126.500511 | 123.581817 | 103.232981 | 105.571701 | 114.793492 | 108.076777 | 115.103951 | 110.832856 |
| ENSMUSG00 | Tubb6 | 1753 | 0.48494211 | 0.93116476 | 0.52429953 | 0.90966147 | 0.38439642 | 0.2544965 | 2.80691307 | 3.72556585 | 1.76211562 | 0.3227658 | 1.062877 | 1.75120447 | 1.74476832 | 0.7948218 | 0.86962782 | 0.87371608 | 0.98384675 | 1.53761907 |
| ENSMUSG00 | Def8 | 4028 | 5.48726205 | 6.26064075 | 6.07671459 | 7.46531511 | 6.38094519 | 7.51656411 | 6.23709844 | 7.88879811 | 4.69329959 | 6.87126684 | 9.96672171 | 9.58898672 | 7.3665964 | 4.56600263 | 6.66916942 | 6.274032 | 6.850778 | 6.35035616 |
| ENSMUSG00 | Meox1 | 2225 | 0 | 0 | 0 | 0 | 0 | 0 | 0 | 0 | 0 | 0 | 0 | 0 | 0 | 0 | 0 | 0 | 0 | 0 |
| ENSMUSG00 | Sost | 2064 | 0 | 0 | 0 | 0 | 0 | 0 | 0 | 0 | 0 | 0 | 0 | 0 | 0 | 0 | 0 | 0 | 0 | 0 |
| ENSMUSG00 | Nkx2-1 | 2829 | 0 | 0 | 0 | 0 | 0 | 0 | 0 | 0 | 0 | 0 | 0 | 0 | 0 | 0 | 0 | 0 | 0 | 0 |
| ENSMUSG00 | Pax9 | 5052 | 0 | 0 | 0.076601 | 0 | 0 | 0 | 0 | 0.02267957 | 0 | 0 | 0 | 0.01104823 | 0.09764829 | 0 | 0.01631099 | 0 | 0 | 0 |
| ENSMUSG00 | Irx2 | 3777 | 0.07877581 | 0.03527974 | 0 | 0.04825104 | 0.02548685 | 0 | 0.02505305 | 0.05055918 | 0.03271367 | 0.11235275 | 0 | 0 | 0.14367252 | 0.03688966 | 0.01090854 | 0 | 0 | 0.16020656 |
| ENSMUSG00 | Col1a1 | 6702 | 0 | 0 | 0 | 0 | 0 | 0 | 0 | 0.00569866 | 0 | 0.03517659 | 0 | 0 | 0.02208233 | 0 | 0.0122953 | 0 | 0 | 0.06566291 |
| ENSMUSG00 | Itga3 | 6963 | 20.816121 | 21.7397374 | 22.5506936 | 23.9877622 | 19.6661386 | 23.4348918 | 12.325894 | 14.2172493 | 12.6582139 | 13.4077829 | 10.9711559 | 12.8577226 | 9.91881146 | 9.79174498 | 9.78115351 | 13.236463 | 10.287498 | 12.5060198 |
| ENSMUSG00 | Sgca | 2007 | 0 | 0 | 0 | 0.04540214 | 0.04796405 | 0 | 0 | 0 | 0 | 0 | 0 | 0 | 0 | 0 | 0 | 0 | 0 | 0 |
| ENSMUSG00 | Dlx3 | 2634 | 0 | 0 | 0 | 0 | 0 | 0 | 0 | 0 | 0 | 0 | 0 | 0 | 0 | 0 | 0 | 0 | 0 | 0 |
| ENSMUSG00 | Foxm1 | 5295 | 0.17149062 | 0.11520309 | 0.08364299 | 0.08753323 | 0.24042844 | 0.11424261 | 0.11816807 | 0 | 0.52224977 | 0.15399845 | 0.47729065 | 0.05361735 | 0.24642264 | 0 | 0.22164141 | 0.20598213 | 0.55961548 | 0.23250727 |
| ENSMUSG00 | Itfg2 | 3622 | 17.5207697 | 13.9156292 | 15.6526167 | 16.4281402 | 15.694037 | 14.392456 | 16.889921 | 15.5743176 | 14.7598281 | 16.8581338 | 17.2264207 | 16.935785 | 14.410041 | 13.5921404 | 18.4849609 | 16.9569669 | 14.9940261 | 15.9620572 |
| ENSMUSG00 | Nrip2 | 3639 | 0.30369183 | 0.26547786 | 0.25256858 | 0.38812651 | 0.62165441 | 0.19063981 | 0.26003121 | 0.23089663 | 0.28295216 | 0.44053968 | 0.07877157 | 0.03067637 | 0.43380636 | 0.40841187 | 0.3509888 | 0.74708247 | 0.33702707 | 0 |
| ENSMUSG00 | Tulp3 | 6894 | 0.57955999 | 0.64267696 | 0.7367596 | 0.70053245 | 0.53759184 | 0.4887708 | 0.40490978 | 0.64263398 | 0.64521957 | 0.8617622 | 0.54746471 | 0.68818267 | 1.62436107 | 0.39073935 | 0.70521955 | 1.0497425 | 0.80610865 | 0.72611383 |
| ENSMUSG00 | Gtf2h4 | 2952 | 5.71631263 | 5.98097356 | 6.11223808 | 5.18580992 | 4.54905376 | 5.47228697 | 6.13846749 | 6.70178335 | 7.46435838 | 5.79800314 | 7.33131985 | 5.59669245 | 4.92984946 | 5.22338652 | 6.21093915 | 6.18720181 | 6.3357835 | 6.07485393 |
| ENSMUSG00 | Tubb5 | 3753 | 43.3546 | 40.094416 | 35.0846655 | 35.2664092 | 34.2425356 | 36.6001006 | 24.8854858 | 25.3496601 | 29.0599237 | 25.6546059 | 25.3195823 | 26.6808683 | 27.5380673 | 21.7803341 | 22.4176864 | 30.2509062 | 25.571289 | 21.9420817 |
| ENSMUSG00 | Ell2 | 3635 | 13.2835983 | 13.9208491 | 12.1366327 | 12.972677 | 11.5860889 | 13.8638597 | 10.230472 | 10.7380014 | 10.7640137 | 10.6364723 | 8.59554961 | 8.75238726 | 6.50068456 | 12.1764004 | 9.91784358 | 10.3547915 | 11.6929473 | 11.1834195 |
| ENSMUSG00 | Jup | 6380 | 35.4498498 | 41.3800449 | 38.5926755 | 43.0044842 | 38.2792133 | 43.8906736 | 31.1611092 | 32.9544162 | 30.2701524 | 36.5159287 | 32.9782002 | 33.3056783 | 26.9701767 | 26.6580326 | 29.9776838 | 26.6173736 | 34.2293479 | 29.8756355 |
| ENSMUSG00 | Fkbp10 | 3185 | 0.97421597 | 1.07730867 | 0.41007388 | 0.7295489 | 0.83116352 | 0.93348949 | 0.65361315 | 0.79142812 | 1.43538564 | 1.05108318 | 1.43999875 | 0.94632521 | 1.13068445 | 1.31239179 | 1.52646267 | 0.42077591 | 1.35977129 | 1.07082092 |
| ENSMUSG00 | Klhl10 | 2453 | 0.12129484 | 0.35309223 | 0.0394403 | 0 | 0.01962166 | 0 | 0.964384 | 0.2646844 | 0.36938579 | 0.11532989 | 0.52585564 | 0.36406466 | 0.12066511 | 0.07573435 | 0.38631708 | 0.09365823 | 0.24998808 | 0 |
| ENSMUSG00 | Mnx1 | 3262 | 0 | 0 | 0 | 0 | 0 | 0 | 0 | 0 | 0 | 0 | 0 | 0 | 0 | 0 | 0 | 0 | 0 | 0 |
| ENSMUSG00 | Nom1 | 4884 | 7.84135209 | 8.16452383 | 8.50795712 | 8.98347373 | 8.96806932 | 8.75593961 | 4.47552368 | 8.35947547 | 6.99934645 | 7.22127888 | 7.19950234 | 6.6055391 | 4.7069356 | 5.78173733 | 11.886362 | 7.69888953 | 7.89440104 | 7.7940831 |
| ENSMUSG00 | Ergic1 | 6555 | 46.7329983 | 44.2596485 | 46.5360004 | 40.8277001 | 41.3839087 | 42.484571 | 36.3271873 | 38.303105 | 40.7215574 | 39.9288412 | 36.2885611 | 40.1566232 | 43.0026812 | 32.6773655 | 36.669899 | 41.7370541 | 35.841373 | 33.9453603 |
| ENSMUSG00 | Tnk1 | 4467 | 12.46514 | 19.0913385 | 14.1969327 | 11.8517872 | 13.8781996 | 16.7394392 | 12.2015236 | 13.8422878 | 15.5359926 | 11.895868 | 12.0105839 | 12.5450918 | 14.1800273 | 12.9132654 | 11.7507907 | 11.0320152 | 14.7573832 | 12.4623273 |
| ENSMUSG00 | Acap1 | 4708 | 0.16250917 | 0.7146564 | 0.05137376 | 0.28064365 | 0.11245777 | 0.30523137 | 1.46721562 | 0.8112235 | 0.87481979 | 0.16024028 | 0.44649494 | 0.13041043 | 1.25739806 | 2.30839347 | 1.1639352 | 0.07319785 | 0.27678294 | 0.5141038 |
| ENSMUSG00 | Tcea3 | 1388 | 22.5081444 | 21.5525785 | 20.4228305 | 19.0713017 | 20.7022755 | 23.7767439 | 11.4873003 | 12.7956051 | 15.0146808 | 13.5880989 | 13.2861196 | 13.7900636 | 22.4268298 | 15.7936866 | 18.9087832 | 28.0558701 | 17.8101308 | 19.0629541 |
| ENSMUSG00 | Csn3 | 4218 | 0 | 0 | 0 | 0.02160315 | 0 | 0 | 0 | 0 | 0 | 0 | 0 | 0 | 0.01169557 | 0 | 0 | 0 | 0 | 0 |
| ENSMUSG00 | Ifrd1 | 5303 | 17.9462736 | 18.230062 | 16.8937919 | 19.1162217 | 20.7031714 | 17.6700394 | 12.0177595 | 23.0105083 | 13.4440616 | 14.5906664 | 16.1802424 | 13.1776748 | 12.8190424 | 15.957218 | 11.5998346 | 23.2357486 | 23.6332248 | 20.7464102 |
| ENSMUSG00 | Stk38l | 4568 | 7.31371899 | 8.55429617 | 8.85294908 | 6.76234415 | 9.83079789 | 9.68708211 | 8.14092943 | 7.09836901 | 8.13271393 | 7.46277549 | 8.19312358 | 8.02777809 | 5.71291249 | 4.50410042 | 8.68587967 | 5.78382642 | 6.92189031 | 5.50331913 |
| ENSMUSG00 | Brpf1 | 6772 | 5.7242647 | 8.5348304 | 6.21455573 | 6.76822366 | 6.26880636 | 7.39778701 | 7.6432473 | 6.8184605 | 5.08445469 | 5.79984285 | 5.46039838 | 7.59099605 | 7.05157943 | 7.24231487 | 6.72293551 | 6.47412071 | 7.30078603 | 8.83784822 |
| ENSMUSG00 | Akr1b3 | 3257 | 18.7273342 | 19.6890763 | 17.3176348 | 21.234776 | 18.9749432 | 15.8532233 | 17.0831165 | 14.8454349 | 15.6298844 | 14.9110299 | 15.0644404 | 14.6522541 | 17.7213212 | 16.1420654 | 13.8645702 | 14.3192999 | 15.9565424 | 13.4440433 |
| ENSMUSG00 | Hoxc13 | 2459 | 0 | 0 | 0 | 0 | 0 | 0 | 0 | 0 | 0 | 0 | 0 | 0 | 0 | 0 | 0 | 0 | 0 | 0 |
| ENSMUSG00 | Hoxc11 | 2019 | 0 | 0 | 0 | 0 | 0 | 0 | 0 | 0.01891649 | 0 | 0 | 0 | 0 | 0 | 0 | 0 | 0 | 0 | 0 |
| ENSMUSG00 | Hoxc8 | 2297 | 0 | 0 | 0 | 0 | 0 | 0.02157281 | 0 | 0 | 0.07172227 | 0 | 0 | 0 | 0 | 0 | 0 | 0 | 0 | 0 |
| ENSMUSG00 | Hoxc6 | 2498 | 0 | 0 | 0 | 0 | 0 | 0 | 0 | 0 | 0 | 0 | 0 | 0 | 0 | 0 | 0 | 0 | 0 | 0 |
| ENSMUSG00 | Gstt1 | 1771 | 24.4806773 | 23.3622895 | 22.5615675 | 20.4523039 | 24.8677076 | 23.223472 | 19.315114 | 16.7132196 | 19.3025455 | 22.2574748 | 26.6255697 | 21.0214833 | 10.9193177 | 12.0633993 | 13.4701868 | 10.3780302 | 14.1099636 | 10.8403039 |
| ENSMUSG00 | Gstt3 | 2457 | 48.715741 | 43.2782899 | 47.0150551 | 50.3266888 | 45.6244405 | 46.2653524 | 47.7170608 | 47.8143385 | 44.3211834 | 49.5303073 | 49.3110683 | 50.0682622 | 33.8717723 | 31.2651691 | 35.2485431 | 32.8049363 | 30.9480558 | 31.4337753 |
| ENSMUSG00 | Ddt | 1043 | 19.9625047 | 10.7955496 | 10.8527384 | 9.52282614 | 13.7981258 | 8.64678573 | 17.3283254 | 11.1458029 | 14.097398 | 13.833284 | 15.9402546 | 15.4121868 | 17.7367871 | 16.7430267 | 16.3147101 | 16.2634123 | 13.1918893 | 15.0312516 |
| ENSMUSG00 | Tat | 2407 | 0 | 0.17992008 | 0.08038809 | 0.0946428 | 0 | 0.18528234 | 0.03931257 | 0 | 0 | 0.13712294 | 0 | 0 | 0 | 0.01929542 | 0 | 0 | 0 | 0 |
| ENSMUSG00 | Marveld3 | 3280 | 31.7752109 | 29.9003538 | 29.7025275 | 31.7816065 | 28.1014128 | 26.4533038 | 20.5117773 | 23.625727 | 21.0076344 | 24.4378645 | 23.9166103 | 23.0239808 | 27.237836 | 27.5832568 | 26.0650087 | 26.4298583 | 22.3297419 | 24.3851345 |
| ENSMUSG00 | Ddx18 | 2469 | 9.19310004 | 8.1899258 | 10.0704717 | 8.59920885 | 9.57180013 | 8.44945356 | 8.41242043 | 9.15751406 | 9.47506721 | 9.0138242 | 8.33981657 | 10.0599311 | 9.99026271 | 11.117236 | 10.2461544 | 9.73936875 | 9.57769347 | 13.6130272 |
| ENSMUSG00 | Ubl3 | 3657 | 63.7751985 | 63.155126 | 59.3524831 | 71.462442 | 64.7550081 | 70.3927949 | 60.6254344 | 57.6384656 | 55.8050822 | 57.865014 | 67.7105851 | 63.6301606 | 55.483231 | 53.8863174 | 51.5103947 | 62.236634 | 65.2186343 | 60.0181394 |
| ENSMUSG00 | Gramd3 | 3197 | 35.8176241 | 37.6372142 | 36.9799525 | 38.7632282 | 37.3372445 | 41.0123657 | 29.1246019 | 31.2993726 | 25.1215845 | 27.122347 | 30.619609 | 31.4257719 | 17.6218485 | 13.9463009 | 16.7409512 | 17.8697486 | 16.9513243 | 14.935222 |
| ENSMUSG00 | Eef1e1 | 1028 | 23.5680451 | 20.9663838 | 21.786911 | 17.5064324 | 17.7451353 | 20.7273107 | 31.8946367 | 20.9909605 | 23.1974636 | 19.2180396 | 20.0301626 | 18.9490917 | 24.8579116 | 26.3845768 | 20.2400639 | 18.2886072 | 16.8143579 | 18.7287815 |
| ENSMUSG00 | Akt1 | 4183 | 60.5315165 | 69.4768561 | 61.568414 | 71.2660562 | 65.4492904 | 70.7456192 | 65.4324283 | 69.1352768 | 62.0013124 | 70.4498913 | 76.08816 | 63.1412181 | 72.1404107 | 79.2647239 | 80.1672157 | 73.9357485 | 74.9942004 | 81.99421 |
| ENSMUSG00 | Cldn15 | 2117 | 0.02007802 | 0.04720769 | 0 | 0 | 0 | 0 | 0.44697854 | 0.03608163 | 0.23346159 | 0 | 0 | 0 | 0.02330275 | 0 | 0 | 0 | 0.10862437 | 0 |
| ENSMUSG00 | Il16 | 8831 | 0.20696666 | 0.1735243 | 0.30675096 | 0.23732398 | 0.2071128 | 0.28056128 | 0.49825378 | 0.32003598 | 0.22386511 | 0.1868729 | 0.254266 | 0.15169017 | 0.11172442 | 0.21036842 | 0.03265892 | 0.09972641 | 0.0781195 | 0.19933093 |
| ENSMUSG00 | Tcirg1 | 4391 | 5.08203539 | 6.6079551 | 5.44215987 | 6.83778825 | 5.22863302 | 5.46197291 | 4.3207434 | 4.67076295 | 5.58095579 | 6.47503395 | 6.49547944 | 4.13119798 | 8.54966728 | 6.20876322 | 9.56146332 | 10.2201302 | 5.74327109 | 9.54610324 |
| ENSMUSG00 | Naglu | 2651 | 27.9466321 | 27.5701584 | 31.002123 | 30.1105053 | 29.5945061 | 32.0756284 | 26.342329 | 31.8966348 | 29.9538296 | 32.3883179 | 28.5820763 | 28.7394868 | 27.0199713 | 23.7914261 | 32.4670125 | 30.4476048 | 33.3240718 | 27.4734279 |
| ENSMUSG00 | Coasy | 2930 | 32.0021906 | 25.7634875 | 24.6820573 | 31.0219397 | 27.4992629 | 28.074245 | 21.1857456 | 23.1891754 | 22.6314984 | 26.3915199 | 34.5186304 | 25.4694679 | 24.6996323 | 25.9484543 | 23.9334635 | 25.4704416 | 25.7557947 | 26.4719592 |
| ENSMUSG00 | Smo | 3767 | 4.37802185 | 5.7128032 | 4.956778 | 7.98255621 | 7.62802188 | 6.93238402 | 3.35346039 | 5.92098832 | 5.68542294 | 4.28073812 | 3.91889094 | 5.70454741 | 3.07751541 | 4.15493959 | 3.06249951 | 3.04942453 | 3.55080372 | 5.78274655 |
| ENSMUSG00 | Tspan33 | 5497 | 0.1469162 | 0.28482918 | 0.08799987 | 0.02486504 | 0.07004828 | 0.02704351 | 0.232389 | 0.06947863 | 0.17232852 | 0 | 0.15643974 | 0.15230761 | 0.09871768 | 0 | 0.05996224 | 0.03482864 | 0.0348611 | 0 |
| ENSMUSG00 | Crnkl1 | 3830 | 8.27907606 | 8.8631434 | 9.19475982 | 7.87504213 | 8.53304861 | 8.55205141 | 5.67010956 | 9.19409792 | 9.00081297 | 7.8666578 | 8.44481642 | 8.02987731 | 5.51280955 | 8.52485468 | 7.73470462 | 6.84832012 | 9.23633735 | 7.36806051 |
| ENSMUSG00 | Rin2 | 5384 | 12.3710273 | 18.5312255 | 13.7735182 | 18.3124257 | 16.0469545 | 16.419405 | 9.47308096 | 10.846245 | 10.2277808 | 11.3935856 | 12.8577108 | 8.27282666 | 7.23852427 | 9.38543222 | 8.63212389 | 10.5967691 | 10.3859709 | 7.98980711 |
| ENSMUSG00 | Folh1 | 2652 | 1.58673168 | 2.04751198 | 1.71459723 | 2.62852179 | 2.06901934 | 1.38269317 | 6.60093937 | 5.60212841 | 4.61251704 | 5.10265906 | 1.11691079 | 6.58759517 | 6.64083502 | 14.6757712 | 3.15381442 | 1.37364695 | 2.76030268 | 4.60483473 |
| ENSMUSG00 | Chordc1 | 2194 | 35.5723817 | 28.9054004 | 33.6204348 | 37.3791613 | 32.2927036 | 33.4943956 | 38.8809299 | 27.0863164 | 38.2017136 | 37.329431 | 29.3312986 | 28.4420742 | 32.9628946 | 31.244974 | 29.0514161 | 33.9275026 | 36.9637394 | 31.7167366 |
| ENSMUSG00 | D10Wsu52e | 2603 | 104.246271 | 104.25157 | 99.3302035 | 104.424583 | 104.214953 | 105.539898 | 86.391534 | 94.740047 | 94.319178 | 86.7116468 | 87.2723502 | 95.4848811 | 93.7552804 | 91.906836 | 96.9969783 | 94.5864793 | 92.0243395 | 89.2233763 |
| ENSMUSG00 | Pwp1 | 2845 | 35.2591267 | 33.9451952 | 35.3152276 | 33.8705127 | 34.918907 | 32.7274469 | 30.2335502 | 31.6412979 | 29.619547 | 33.0966448 | 33.8538897 | 30.2719042 | 30.032647 | 32.4047499 | 30.3110216 | 34.4951939 | 31.4019677 | 30.7238912 |
| ENSMUSG00 | Fbxo7 | 6326 | 8.92971529 | 8.12548147 | 8.80144379 | 9.35564284 | 8.65096383 | 9.22749291 | 7.65110189 | 9.63564247 | 7.99510613 | 8.50441866 | 10.2482799 | 8.48793351 | 11.0813554 | 10.0288687 | 7.7375037 | 10.5804587 | 10.1601644 | 9.26963449 |
| ENSMUSG00 | Capns1 | 2928 | 68.6063736 | 73.190522 | 70.825589 | 67.0500478 | 69.7978647 | 64.0733092 | 50.5928954 | 51.3275627 | 53.410246 | 52.8835073 | 52.7188835 | 52.2128073 | 51.1852338 | 40.6861626 | 43.847011 | 43.4431003 | 43.588309 | 46.7802854 |
| ENSMUSG00 | Lrp3 | 4942 | 0.08600805 | 0.08088926 | 0.039153 | 0 | 0.01947872 | 0.06016115 | 0.01914718 | 0.10046565 | 0 | 0.05724488 | 0.23201113 | 0.18070631 | 0 | 0.00939783 | 0.08337021 | 0 | 0.18612528 | 0.17809504 |
| ENSMUSG00 | Dsg4 | 3717 | 0 | 0 | 0 | 0 | 0 | 0 | 0 | 0 | 0 | 0 | 0 | 0 | 0 | 0 | 0 | 0 | 0 | 0 |
| ENSMUSG00 | Evx2 | 7406 | 0 | 0 | 0 | 0 | 0 | 0 | 0 | 0 | 0 | 0 | 0 | 0 | 0 | 0 | 0 | 0 | 0 | 0 |
| ENSMUSG00 | Hoxd13 | 2481 | 0 | 0 | 0 | 0 | 0 | 0 | 0 | 0 | 0 | 0 | 0 | 0 | 0 | 0 | 0 | 0 | 0 | 0 |
| ENSMUSG00 | Hoxd12 | 2617 | 0 | 0 | 0 | 0 | 0 | 0 | 0 | 0 | 0 | 0 | 0 | 0 | 0 | 0 | 0 | 0 | 0 | 0 |
| ENSMUSG00 | Folr1 | 2131 | 0.39892235 | 0.84415593 | 0.95339666 | 0.47036273 | 0.29362506 | 0.95338436 | 1.1989135 | 1.6130062 | 1.37223961 | 0.70803496 | 0.85192324 | 0.65480599 | 0.16204759 | 0.37050656 | 0.29001564 | 0.68279825 | 0.55753881 | 0.07744126 |
| ENSMUSG00 | Clpb | 10260 | 8.86974482 | 9.09773193 | 8.2838492 | 9.7960686 | 9.85625454 | 9.75599628 | 8.07912412 | 9.17957742 | 8.45006001 | 10.0183755 | 9.79246956 | 7.69233401 | 8.18832896 | 8.74561178 | 7.67810671 | 7.78501043 | 8.37127011 | 7.92431381 |
| ENSMUSG00 | 7-Sep | 3625 | 84.1895624 | 80.7688351 | 79.4260007 | 87.1378539 | 85.3362409 | 83.754372 | 72.1763074 | 73.3399485 | 77.2943081 | 72.7486575 | 71.8798961 | 75.1856222 | 63.1040277 | 60.6271508 | 69.8864728 | 77.3734343 | 86.5592808 | 64.1444122 |
| ENSMUSG00 | Zdhhc4 | 3175 | 28.2207594 | 28.4130134 | 27.2872418 | 27.3940263 | 29.6674566 | 29.1385681 | 35.1231446 | 33.0559754 | 30.7828667 | 32.2555362 | 35.0600483 | 32.1357579 | 34.5090357 | 33.0740414 | 30.4048017 | 34.1298957 | 30.1299014 | 31.8446419 |
| ENSMUSG00 | Rac1 | 4266 | 88.4378674 | 95.8077588 | 94.3091074 | 97.5194317 | 97.3244195 | 98.8963839 | 94.9580771 | 92.8488995 | 88.6292111 | 101.076695 | 92.8285586 | 92.9606851 | 88.8575838 | 80.2047801 | 92.0612355 | 104.594573 | 95.4743453 | 98.9416024 |
| ENSMUSG00 | Nup214 | 9661 | 3.14136173 | 3.84127566 | 3.07906251 | 4.23494647 | 3.59208332 | 3.81095964 | 3.03631725 | 4.17463792 | 3.41906486 | 4.0117874 | 3.6445652 | 3.7726558 | 3.23228482 | 2.60079198 | 4.65708509 | 3.9634207 | 3.67354838 | 3.34233989 |
| ENSMUSG00 | Aif1l | 4009 | 29.7504425 | 39.4287062 | 30.6120345 | 30.9460523 | 34.337067 | 33.2988437 | 3.35165897 | 5.00149944 | 5.28058596 | 5.2102174 | 5.00510914 | 7.42074037 | 2.31339497 | 1.00789115 | 1.74713507 | 3.23784383 | 3.87182595 | 0.75467719 |
| ENSMUSG00 | Cpa3 | 1441 | 0 | 0 | 0 | 0 | 0 | 0 | 0 | 0 | 0 | 0 | 0 | 0 | 0 | 0 | 0 | 0 | 0 | 0 |
| ENSMUSG00 | Ltbp1 | 11195 | 0.2050272 | 0.32435064 | 0.13395082 | 0.03255814 | 0.14618003 | 0.30984291 | 0.81566298 | 0.36844837 | 0.3347899 | 0.22322352 | 0.04694279 | 0.45370481 | 0.30846219 | 0.40656727 | 0.08832848 | 0.08892861 | 0.09585854 | 0.27516838 |
| ENSMUSG00 | Ugp2 | 3514 | 41.3681565 | 36.3465101 | 35.5161378 | 36.6796228 | 37.5850878 | 38.045895 | 34.8449667 | 36.5838436 | 43.9877614 | 36.8188759 | 37.5374674 | 33.3400899 | 40.2348534 | 39.3069766 | 41.2953557 | 41.2871689 | 43.299772 | 39.0730593 |
| ENSMUSG00 | Kcnh6 | 3759 | 0.01130758 | 0 | 0 | 0 | 0 | 0 | 0 | 0 | 0 | 0 | 0 | 0.01484854 | 0 | 0.01096078 | 0 | 0 | 0 | 0 |
| ENSMUSG00 | Trmt1 | 6253 | 7.51810726 | 6.8085525 | 6.04959231 | 6.60136033 | 6.11944336 | 6.41102854 | 6.87028826 | 6.59648082 | 7.29145577 | 6.49990504 | 7.02145959 | 7.26594424 | 7.60530437 | 6.89270959 | 6.22009736 | 8.75056421 | 6.49702138 | 7.00259858 |
| ENSMUSG00 | Nacc1 | 4294 | 6.65195116 | 7.03651492 | 7.5590682 | 6.98164117 | 6.99448541 | 8.19339563 | 6.84237858 | 7.1243862 | 7.13618252 | 6.44561314 | 7.58791842 | 5.68035504 | 6.36466748 | 5.3431251 | 8.44372827 | 5.33250657 | 6.75663874 | 7.23804036 |
| ENSMUSG00 | Nfix | 6243 | 32.6941943 | 30.85298 | 30.4746268 | 30.7535225 | 29.1042787 | 29.8284755 | 17.4684807 | 18.5303199 | 20.8736402 | 22.6652668 | 18.9630542 | 21.6807595 | 20.4502646 | 16.7088612 | 18.9277691 | 22.9694577 | 20.9772476 | 21.8432766 |
| ENSMUSG00 | Slc1a5 | 5242 | 3.64075239 | 3.30459843 | 4.60480573 | 3.33755075 | 3.60851698 | 3.66777196 | 4.69335999 | 8.94701834 | 6.14419216 | 4.33549007 | 4.70276203 | 6.63356892 | 3.54790782 | 2.62255816 | 4.29936114 | 4.33161583 | 4.10168878 | 3.44200439 |
| ENSMUSG00 | Uba1 | 4895 | 101.065933 | 121.355573 | 107.133288 | 109.597813 | 109.085916 | 116.061714 | 99.0908242 | 108.038855 | 100.143393 | 110.70542 | 120.467071 | 102.018944 | 94.733406 | 85.9334218 | 89.4061137 | 81.9160859 | 91.5837987 | 89.520438 |
| ENSMUSG00 | Vwf | 16008 | 3.17036358 | 3.47529569 | 4.01904019 | 3.79675369 | 3.55998236 | 3.77341218 | 4.60181416 | 4.03205648 | 4.11402041 | 4.33569685 | 3.27691807 | 3.65409885 | 3.03547842 | 2.67500303 | 3.53898921 | 2.0499157 | 2.78445048 | 2.46386366 |
| ENSMUSG00 | Siae | 3423 | 4.48272525 | 6.08254715 | 6.04845778 | 6.16265367 | 4.96364857 | 6.44200013 | 9.74450442 | 5.86888801 | 5.9439494 | 8.20970469 | 6.07131374 | 7.54970838 | 5.90887703 | 5.92932112 | 4.20080135 | 5.10093981 | 4.37790888 | 4.46758156 |
| ENSMUSG00 | Vsig2 | 1310 | 0.0324467 | 0 | 0 | 0 | 0 | 0 | 0.03611655 | 0.11661802 | 0 | 0.14397161 | 0 | 0 | 0 | 0 | 0 | 0 | 0 | 0 |
| ENSMUSG00 | Esam | 2857 | 0.56534711 | 0.75790624 | 0.64340012 | 1.35550884 | 0.60649256 | 0.53767404 | 0.4802477 | 0.5748244 | 0.85054408 | 0.90769642 | 0.90299186 | 0.6642396 | 0.86335172 | 1.43054942 | 0.59127189 | 0.16082859 | 0.38903137 | 0.63538695 |
| ENSMUSG00 | Spa17 | 572 | 11.3693915 | 7.9787876 | 11.5013988 | 7.64660884 | 7.90979165 | 8.66306531 | 23.408196 | 16.6924836 | 20.5212535 | 14.1781835 | 16.2869177 | 20.3941844 | 21.1299296 | 20.8673609 | 16.7111206 | 10.9784259 | 8.84453005 | 11.2518627 |
| ENSMUSG00 | Fam50a | 2532 | 34.7998538 | 33.2602672 | 32.3636495 | 32.4433503 | 32.3350719 | 32.8003086 | 36.045086 | 31.9024214 | 30.7434893 | 31.3593145 | 31.8688408 | 32.8677539 | 34.1348811 | 32.4668383 | 33.4233791 | 31.3493006 | 27.9273372 | 30.6113078 |
| ENSMUSG00 | Emd | 2791 | 19.2042375 | 12.2461589 | 14.9401588 | 9.12526829 | 15.1932013 | 12.7654659 | 11.8663114 | 13.0683426 | 17.619742 | 12.6703789 | 12.5984842 | 13.0989821 | 15.1301043 | 11.1326028 | 12.5331856 | 21.360973 | 11.9057325 | 9.85473069 |
| ENSMUSG00 | Taco1 | 1378 | 6.19995674 | 4.06136882 | 3.1242701 | 4.36433806 | 4.22638807 | 4.06346799 | 3.77677408 | 4.01879416 | 5.29028541 | 5.13251487 | 4.16037374 | 4.41502701 | 4.51075586 | 4.28040573 | 4.15603503 | 2.94543134 | 2.16941303 | 2.75444729 |
| ENSMUSG00 | Grik3 | 8962 | 0.01897129 | 0.02601989 | 0.05937389 | 0 | 0.01074134 | 0.0055292 | 0.02111702 | 0.02130797 | 0.0275741 | 0.01052236 | 0.01599251 | 0.03736822 | 0.02201826 | 0.02072934 | 0.03677889 | 0.02563531 | 0.01710614 | 0.01227608 |
| ENSMUSG00 | Gria3 | 6707 | 3.58698817 | 5.48840735 | 3.92354264 | 5.03365646 | 4.53546679 | 3.45768292 | 1.3050314 | 1.26415884 | 0.73075822 | 1.18808243 | 1.34627511 | 1.9556703 | 1.68436097 | 0.57475164 | 0.8600295 | 0.49097842 | 1.42859315 | 1.41070022 |
| ENSMUSG00 | Npas1 | 2079 | 0.30687515 | 0.3525174 | 0.09307077 | 0.30680838 | 0.23151478 | 0.19067911 | 0.20481679 | 0.09185282 | 0.43583615 | 0.49894924 | 0.06893933 | 0.18793152 | 0.11864338 | 0.08935851 | 0.15854375 | 0.0368356 | 0.0553049 | 0.05291881 |
| ENSMUSG00 | Sipa1l2 | 6344 | 2.31822051 | 3.08238791 | 2.85178127 | 3.59805851 | 3.98317464 | 4.33508309 | 3.49773746 | 2.39605551 | 1.61006556 | 3.55265387 | 3.10266124 | 3.7065653 | 2.59723524 | 2.17432124 | 2.253616 | 2.92732418 | 2.6461094 | 2.83543157 |
| ENSMUSG00 | Ap4e1 | 7188 | 1.55521164 | 1.48089258 | 2.03238817 | 2.31354774 | 1.52002597 | 1.51663904 | 1.45466083 | 1.83310779 | 1.26057783 | 1.63991038 | 1.10331707 | 1.46760715 | 1.48929133 | 1.33749667 | 1.27250214 | 1.06007664 | 1.8981861 | 1.9667975 |
| ENSMUSG00 | Blvra | 2047 | 20.9099718 | 19.6101795 | 18.5979328 | 16.8266485 | 18.8812578 | 18.5429379 | 14.3532846 | 15.46727 | 13.0179168 | 15.2024739 | 16.5240208 | 14.3970053 | 17.9542151 | 15.7460658 | 12.5597317 | 18.593485 | 15.5402181 | 14.9682816 |
| ENSMUSG00 | Pdzd4 | 3962 | 0.05364106 | 0.26905922 | 0.13430309 | 0.02299901 | 0.07289034 | 0.062535 | 0.01294162 | 0.0867571 | 0.10395385 | 0.05950366 | 0.10852464 | 0.18314074 | 0.01245127 | 0 | 0.10399181 | 0.11597357 | 0.05804083 | 0.19437847 |
| ENSMUSG00 | Srpk3 | 2192 | 0.42660304 | 1.04862669 | 0.33102325 | 0.64439959 | 0.21957994 | 1.40158279 | 0.97129131 | 1.55069517 | 1.1461576 | 0.81739358 | 2.61541743 | 1.73150778 | 2.16052192 | 0.29663195 | 0.24435229 | 1.29265737 | 0.57699261 | 0.5270033 |
| ENSMUSG00 | Idh3g | 2398 | 89.9202494 | 80.2121148 | 83.3727277 | 80.8813035 | 88.1950261 | 82.4294638 | 97.5256764 | 96.3091982 | 94.6362393 | 88.2650154 | 103.319823 | 97.1769772 | 93.1299372 | 86.9228792 | 81.9563893 | 75.112184 | 85.5868724 | 87.078636 |
| ENSMUSG00 | Pnck | 2254 | 0.22629198 | 0.25125075 | 0.10730597 | 0.36384153 | 0.40572606 | 0.35174966 | 0.16792433 | 0.35582984 | 0.31063477 | 0.23010547 | 0.23315197 | 0.56954758 | 0.24075026 | 0.06181556 | 0.38386543 | 0.25481771 | 0.18704049 | 0.12202553 |
| ENSMUSG00 | Ssr4 | 1639 | 100.726116 | 91.9712302 | 96.0977516 | 88.8704446 | 98.6131786 | 84.0491759 | 114.889849 | 109.474013 | 113.281765 | 102.241366 | 107.471795 | 102.572773 | 168.70372 | 169.227633 | 140.950045 | 135.874419 | 125.221062 | 155.898292 |
| ENSMUSG00 | Bcap31 | 1211 | 319.087164 | 275.789799 | 285.327933 | 294.05873 | 284.657999 | 288.682523 | 265.591735 | 266.9999 | 282.659895 | 277.88156 | 299.826329 | 272.395182 | 284.381597 | 252.240101 | 266.772419 | 297.598016 | 274.930257 | 268.09558 |
| ENSMUSG00 | Fam98a | 3471 | 10.7640593 | 9.88622162 | 11.6090898 | 10.1728059 | 8.44492697 | 9.67927207 | 9.29623942 | 8.38450321 | 8.32985712 | 8.02825285 | 9.46965469 | 6.88248447 | 7.60373833 | 9.20585771 | 7.3239297 | 5.93498899 | 8.43598123 | 7.08414596 |
| ENSMUSG00 | Ltbp2 | 8635 | 94.2743638 | 111.442915 | 88.3720256 | 104.318171 | 92.4233267 | 109.044707 | 50.1070583 | 60.2099796 | 48.2981654 | 54.0253686 | 44.1344347 | 55.964332 | 21.7266105 | 15.8184205 | 21.6481134 | 17.7196604 | 19.2008756 | 21.6660072 |
| ENSMUSG00 | Mll1 | 16633 | 4.58707335 | 4.11579376 | 4.55728504 | 4.0129219 | 4.3059161 | 3.86101982 | 3.63243498 | 4.2869726 | 4.58095678 | 3.98001487 | 4.12749446 | 3.97652619 | 5.33566519 | 5.21040504 | 3.44811915 | 6.18800894 | 3.97940929 | 3.92567845 |
| ENSMUSG00 | Ift46 | 5301 | 11.265756 | 10.4947243 | 8.7512197 | 8.82686145 | 11.2498497 | 9.60019947 | 7.80065674 | 7.87479642 | 9.05155459 | 6.74216809 | 8.82318135 | 8.11806751 | 6.05830563 | 6.51846841 | 6.05470893 | 6.55151498 | 7.51198311 | 5.71779246 |
| ENSMUSG00 | Tmem25 | 4167 | 1.85647757 | 3.29371548 | 2.41461347 | 1.82594058 | 2.26394457 | 2.62806674 | 2.21405625 | 2.49300059 | 2.6785567 | 2.74960905 | 2.84333806 | 1.28588999 | 2.14280706 | 1.5603965 | 1.48313736 | 1.85618015 | 1.92229327 | 1.70294571 |
| ENSMUSG00 | Cd3g | 1226 | 0 | 0 | 0 | 0 | 0 | 0 | 0 | 0 | 0 | 0 | 0 | 0.22763321 | 0 | 0 | 0 | 0 | 0 | 0 |
| ENSMUSG00 | Trappc6a | 1430 | 93.7789024 | 77.8776261 | 76.9578892 | 80.4168363 | 79.4008447 | 76.5121928 | 113.583515 | 97.9114316 | 101.785417 | 108.908233 | 108.378997 | 100.390129 | 149.651648 | 150.439869 | 107.901977 | 97.0653509 | 100.747238 | 115.711464 |
| ENSMUSG00 | Supt6h | 7798 | 8.23067656 | 6.56603211 | 6.78643783 | 6.9527626 | 6.2896166 | 6.74852566 | 5.81245786 | 7.36616738 | 5.77287275 | 6.92930315 | 5.55679499 | 7.20063562 | 7.62942963 | 5.67001426 | 6.50940201 | 5.95620759 | 7.30843851 | 6.1513139 |
| ENSMUSG00 | Spag5 | 5247 | 0.03240341 | 0 | 0 | 0 | 0 | 0 | 0 | 0.02911561 | 0.12559258 | 0 | 0.10926234 | 0.04255053 | 0.15983278 | 0.00885155 | 0 | 0.14595238 | 0.07304421 | 0.27258181 |
| ENSMUSG00 | Fosl1 | 3205 | 0 | 0.03118212 | 0 | 0 | 0 | 0 | 0 | 0 | 0 | 0 | 0 | 0 | 0 | 0 | 0 | 0 | 0 | 0 |
| ENSMUSG00 | Unc119 | 2732 | 11.0619254 | 10.766942 | 7.61369621 | 9.15556857 | 8.61512104 | 9.05081042 | 7.10036543 | 6.96186545 | 7.43226657 | 8.85370087 | 8.5687151 | 7.3344886 | 6.44637426 | 6.54501232 | 7.75168365 | 6.68966932 | 6.46721189 | 5.6982347 |
| ENSMUSG00 | Rab34 | 2266 | 17.4259967 | 18.3764862 | 16.3308697 | 18.2163752 | 17.2688676 | 20.2715728 | 23.6563395 | 20.4277104 | 22.7561861 | 22.3476855 | 21.9688909 | 21.4050357 | 19.3104195 | 18.8768775 | 19.9280253 | 15.1743007 | 16.6768441 | 21.4841392 |
| ENSMUSG00 | Sdf2 | 1511 | 60.789997 | 50.0905994 | 52.8555252 | 50.8679562 | 53.9931242 | 57.259479 | 56.3618942 | 56.2649158 | 53.3980042 | 59.0085902 | 70.7613217 | 54.1164432 | 72.1205859 | 58.6468026 | 58.7892485 | 67.3063225 | 64.9847602 | 65.7852123 |
| ENSMUSG00 | Ccne1 | 2651 | 0.30463913 | 0.25132323 | 0.12773094 | 0.2234227 | 0 | 0.22430509 | 0.12492974 | 0.28813582 | 0.21750706 | 0.03557201 | 0.5046016 | 0.2737094 | 0.26052314 | 0.42046703 | 0.06216757 | 0.80885478 | 0.27468866 | 0.33200517 |
| ENSMUSG00 | Hsf2bp | 3447 | 0.28361446 | 0.08697884 | 0.09823461 | 0.27756946 | 0.06981712 | 0.27313662 | 0.21961209 | 0.06863917 | 0.13143362 | 0.05471506 | 0 | 0.3400432 | 0.07155776 | 0.21558033 | 0 | 0.01110839 | 0.04447496 | 0 |
| ENSMUSG00 | Bbc3 | 2003 | 14.748426 | 14.4195104 | 12.4858289 | 12.2830574 | 11.9428687 | 14.00242 | 7.06265152 | 9.2477859 | 13.9207545 | 9.062916 | 9.51682897 | 10.4497621 | 5.44301234 | 5.63450486 | 7.73429107 | 10.246513 | 7.57724093 | 7.2503266 |
| ENSMUSG00 | Mybpc3 | 4131 | 0 | 0.00806412 | 0 | 0 | 0 | 0 | 0 | 0 | 0 | 0 | 0 | 0 | 0 | 0 | 0 | 0 | 0 | 0 |
| ENSMUSG00 | Psmc3 | 2869 | 94.4180853 | 91.6830299 | 96.9156683 | 86.215026 | 93.3615041 | 91.1777207 | 79.3711829 | 78.9540393 | 83.8947362 | 78.935109 | 78.3316154 | 79.4337227 | 74.401954 | 75.9228895 | 73.470603 | 64.6362536 | 67.9293202 | 73.9909874 |
| ENSMUSG00 | Acp2 | 4354 | 16.5861952 | 17.6740436 | 18.9761116 | 17.9983914 | 18.0632897 | 17.8908813 | 17.4733091 | 15.9383544 | 13.8959558 | 13.6990445 | 17.0954031 | 15.332001 | 12.7578638 | 13.8030689 | 15.0271178 | 15.1086963 | 15.1755939 | 14.8577652 |
| ENSMUSG00 | Rapsn | 2155 | 0.11834388 | 0.13912578 | 0.06734134 | 0.33827225 | 0.08934 | 0.298926 | 0.7245097 | 0.2481177 | 0.40135352 | 0.35007483 | 0.73158869 | 0.20720432 | 0.20602657 | 0.08620712 | 0.34414293 | 0.07107305 | 0.23120272 | 0 |
| ENSMUSG00 | Slc39a13 | 4475 | 5.79400165 | 5.1365133 | 4.58332444 | 5.3044255 | 4.09793585 | 5.12690852 | 4.79999022 | 5.13686823 | 4.58343796 | 5.85827732 | 6.84329273 | 5.78736703 | 4.98279924 | 5.23079778 | 4.22603668 | 4.50930729 | 4.66767191 | 5.59310471 |
| ENSMUSG00 | Celf2 | 18155 | 3.2402734 | 3.73405341 | 3.31993494 | 3.37786647 | 3.64005129 | 3.58919552 | 3.04385618 | 2.68850952 | 2.91061965 | 2.28027086 | 2.28414563 | 3.01905705 | 2.99442428 | 2.8958803 | 2.56445928 | 2.32422194 | 2.23983481 | 2.46942561 |
| ENSMUSG00 | Nr1h3 | 2896 | 0.13209482 | 0.03450921 | 0.26725707 | 0.61356379 | 0.21606181 | 0.35932576 | 0.8658743 | 0.30332364 | 0.31288099 | 0.52100225 | 0.676372 | 0.5396542 | 0.18737952 | 0.24055984 | 0.51217404 | 0.42310063 | 0.19851328 | 0.531856 |
| ENSMUSG00 | Ddb2 | 2775 | 6.46385019 | 5.29404914 | 5.66536842 | 5.86136678 | 4.89124413 | 3.94636185 | 8.6271047 | 6.04196925 | 4.25965046 | 6.09985675 | 6.55937268 | 6.91913079 | 5.97316188 | 5.25529493 | 6.47346968 | 7.39595156 | 7.844806 | 7.31472446 |
| ENSMUSG00 | Sfpi1 | 2803 | 0.04549252 | 0.01188473 | 0 | 0.01625439 | 0 | 0 | 0.35446531 | 0.20438317 | 0.51428043 | 0 | 0.22157491 | 0 | 0.21119622 | 0.29824958 | 0.11759274 | 0 | 0 | 0 |
| ENSMUSG00 | Sf3a1 | 6327 | 11.004185 | 10.2881927 | 10.6808633 | 10.3983164 | 11.3426107 | 11.0822061 | 9.205296 | 11.9521032 | 10.0834381 | 9.53894423 | 11.1754289 | 9.27173406 | 12.1789876 | 8.59586318 | 12.6072724 | 7.07471472 | 11.1398862 | 8.04226706 |
| ENSMUSG00 | Stat6 | 4430 | 19.1417215 | 23.695018 | 20.7471123 | 26.719547 | 21.7299881 | 26.6891247 | 16.2977761 | 16.7080984 | 18.501392 | 20.7016067 | 15.8530901 | 17.0848845 | 13.585765 | 12.0356235 | 14.9367266 | 17.070869 | 19.05933 | 16.2171317 |
| ENSMUSG00 | Clgn | 4109 | 5.26530411 | 4.88059438 | 3.62595458 | 4.97856143 | 6.44257922 | 5.00471756 | 5.03179379 | 5.40028859 | 2.37556686 | 5.23259204 | 3.51132574 | 5.56934088 | 5.52267752 | 4.80378125 | 3.35909497 | 3.50383752 | 4.04809269 | 2.38296933 |
| ENSMUSG00 | Napsa | 2149 | 41.6348979 | 42.2573056 | 33.7646792 | 41.2572323 | 43.1821074 | 47.0162977 | 189.867168 | 135.601689 | 151.617741 | 206.923352 | 158.753364 | 118.929697 | 343.831297 | 212.877739 | 179.932272 | 234.269384 | 205.274861 | 182.305658 |
| ENSMUSG00 | Vrk3 | 3169 | 30.2861115 | 30.6743532 | 27.644198 | 31.8163758 | 30.6804941 | 29.9912095 | 24.9627009 | 28.2616547 | 25.161595 | 30.9180061 | 34.20681 | 32.1613758 | 28.0206535 | 23.4198494 | 24.9757617 | 34.4845041 | 26.6675559 | 29.0754982 |
| ENSMUSG00 | 1500002O20l | 3855 | 8.1151257 | 9.55747367 | 6.33686974 | 6.72483378 | 7.1792104 | 7.68677942 | 7.90385609 | 8.97595761 | 8.8676545 | 8.25595971 | 8.0306545 | 9.19401965 | 7.5373539 | 8.55390455 | 7.27840715 | 6.35693612 | 6.57164261 | 7.16331299 |
| ENSMUSG00 | Paxip1 | 5031 | 8.59228066 | 11.3890237 | 10.9419753 | 10.5593675 | 11.4135132 | 10.9033743 | 10.7396302 | 11.485809 | 7.71991341 | 10.749723 | 8.49902362 | 9.13064804 | 6.68741156 | 7.96685498 | 8.46798028 | 8.95806895 | 9.78154745 | 10.5294712 |
| ENSMUSG00 | Rmnd5a | 9948 | 4.8538279 | 6.0042239 | 5.36835319 | 5.7432197 | 5.59797303 | 5.51914242 | 5.40281496 | 5.98531919 | 4.95579688 | 5.5976056 | 5.28271555 | 6.45796418 | 6.52601559 | 5.71914023 | 6.26638059 | 6.61656153 | 6.93094171 | 6.72407255 |
| ENSMUSG00 | Mov10 | 4201 | 8.41806866 | 8.79409657 | 9.23484202 | 9.88005508 | 8.28359458 | 8.88198248 | 10.654081 | 9.29127238 | 7.67651555 | 9.02384323 | 6.83474174 | 10.1640043 | 8.26700066 | 6.01418318 | 6.74760068 | 10.9831426 | 9.88948009 | 7.24114164 |
| ENSMUSG00 | Ppm1j | 1711 | 2.21096474 | 2.74524727 | 1.78114109 | 1.75746869 | 2.53177857 | 1.94040511 | 0.58069331 | 1.02679749 | 1.56465431 | 0.96450883 | 1.39611217 | 1.30486645 | 0.37481878 | 0.4885994 | 0.69833146 | 0 | 0.13439964 | 0.90020747 |
| ENSMUSG00 | Rhoc | 910 | 125.507042 | 102.09853 | 100.361786 | 92.9745707 | 109.486903 | 99.9222705 | 67.6415333 | 61.7793586 | 59.1999589 | 61.1404724 | 67.5149413 | 84.0301733 | 60.7162059 | 61.0408006 | 62.6173097 | 75.529275 | 55.2993712 | 52.167977 |
| ENSMUSG00 | 4930432K09l | 1525 | 0.13936123 | 0.13106712 | 0 | 0.05975219 | 0 | 0.0324936 | 3.89294019 | 3.28078998 | 2.61973248 | 1.20582126 | 1.06514662 | 2.78163302 | 3.81715819 | 5.63420087 | 2.53963689 | 1.05456099 | 0.57803595 | 2.12822118 |
| ENSMUSG00 | Tead3 | 5375 | 0.98849247 | 1.5308437 | 1.1159661 | 1.27994748 | 0.99398019 | 1.14317004 | 1.03867835 | 1.13689012 | 1.27199286 | 1.57022807 | 1.06660372 | 1.27727005 | 1.32163648 | 1.51213308 | 1.56374277 | 0.85486007 | 0.56330743 | 1.17693903 |
| ENSMUSG00 | Ppard | 4719 | 1.11689804 | 1.12243064 | 1.08658492 | 0.97513572 | 1.01996024 | 0.77705071 | 1.20311962 | 1.61866507 | 1.27425965 | 1.33888414 | 1.19482706 | 1.79783444 | 1.57853772 | 1.34834498 | 1.67635065 | 1.5822565 | 0.86090008 | 1.45711768 |
| ENSMUSG00 | Def6 | 2612 | 4.31235515 | 4.030197 | 4.62993208 | 5.00613313 | 3.96185436 | 3.25501807 | 6.75636648 | 5.67329714 | 5.51886697 | 8.33982577 | 6.36511698 | 5.94056439 | 6.98805873 | 7.55694378 | 8.89650747 | 5.43867356 | 7.11648315 | 10.3615928 |
| ENSMUSG00 | Peg3 | 13121 | 4.10765667 | 3.25994636 | 2.42586563 | 3.46195878 | 2.53113538 | 2.9646289 | 1.97241334 | 1.74064908 | 1.39370574 | 2.28189136 | 1.07412642 | 1.36125387 | 9.08359974 | 9.39076014 | 5.79666734 | 13.7800662 | 10.2497505 | 8.73706108 |
| ENSMUSG00 | Zim1 | 5888 | 0 | 0 | 0 | 0 | 0 | 0 | 0 | 0 | 0 | 0 | 0 | 0 | 0 | 0 | 0 | 0 | 0 | 0 |
| ENSMUSG00 | Metrn | 1771 | 0 | 0.01881022 | 0.08194274 | 0.02572617 | 0 | 0.05596017 | 0.02671523 | 0.04313089 | 0 | 0.05324755 | 0.02697626 | 0 | 0 | 0 | 0.0216209 | 0 | 0 | 0 |
| ENSMUSG00 | Lmf1 | 14147 | 1.71559027 | 1.94032833 | 1.87380325 | 1.54264086 | 1.67391719 | 1.95100534 | 1.95310698 | 2.20294058 | 2.33488271 | 1.99308126 | 2.07687845 | 2.09895614 | 2.10271762 | 1.68744334 | 1.45328171 | 1.58878819 | 1.6688343 | 1.2481744 |
| ENSMUSG00 | Narfl | 3347 | 14.9726927 | 12.2124055 | 15.1898955 | 14.1161646 | 13.9635787 | 13.8427863 | 12.8636205 | 14.9140932 | 14.237466 | 14.9608745 | 15.2445939 | 14.1582006 | 13.1620562 | 13.2796509 | 13.0116655 | 12.469902 | 15.7450428 | 14.2001399 |
| ENSMUSG00 | Angptl4 | 3850 | 0.07728214 | 0.173054 | 0.50258213 | 0.35502112 | 0.0375054 | 0.21880428 | 0 | 0.3075234 | 0.19256033 | 0 | 0.11168172 | 0 | 0 | 0.09650719 | 0 | 0.11945859 | 0 | 0 |
| ENSMUSG00 | Rab11b-ps2 | 1544 | 1.2112874 | 1.16508827 | 0.84591018 | 1.09181259 | 0.71699109 | 0.8986247 | 0.85800195 | 0.51945625 | 0.69355532 | 0.79398853 | 0.74261593 | 1.04835118 | 0.70291592 | 0.69184845 | 0.64043869 | 0.7687881 | 0.96808652 | 0.5344149 |
| ENSMUSG00 | Dbf4 | 7708 | 0.46321157 | 0.82547519 | 0.489509 | 0.41967231 | 0.49955292 | 0.75216267 | 0.6015364 | 0.38648254 | 0.9511157 | 0.45266833 | 0.45865944 | 0.57205985 | 0.275204 | 0.42178076 | 0.37951615 | 0.36263808 | 0.64639577 | 0.72793577 |
| ENSMUSG00 | Daxx | 3002 | 8.22635152 | 8.0896402 | 7.38010556 | 8.40800085 | 9.28327242 | 8.50088534 | 7.86443264 | 6.74282921 | 6.68149007 | 8.98407804 | 7.35244198 | 7.58587286 | 5.86658171 | 8.47813454 | 6.50550882 | 8.99229767 | 9.44752156 | 9.10710394 |
| ENSMUSG00 | Cd320 | 2774 | 27.3510235 | 23.8858495 | 26.5234822 | 21.9757308 | 25.4019958 | 22.7399531 | 27.8691122 | 26.5584514 | 26.3391777 | 25.7850097 | 28.6236413 | 26.036578 | 28.9162572 | 27.4244467 | 25.1605752 | 23.2587143 | 23.169863 | 21.9322538 |
| ENSMUSG00 | Ipo4 | 5645 | 7.49205494 | 5.62394831 | 7.40384946 | 7.42536064 | 5.37167616 | 6.30272147 | 5.51492343 | 6.08236843 | 7.56606144 | 7.58420524 | 8.62403591 | 6.23909355 | 6.9038467 | 6.60666115 | 8.79501769 | 7.00016064 | 7.25110453 | 7.18187983 |
| ENSMUSG00 | Tm9sf1 | 4075 | 24.5956331 | 26.9200885 | 29.1190847 | 33.1840932 | 26.9420657 | 30.0478048 | 23.4415457 | 25.4084915 | 23.4586265 | 26.5085303 | 26.1795049 | 23.9562194 | 24.7930716 | 21.4497601 | 23.5581902 | 23.105915 | 25.2060063 | 24.1500103 |
| ENSMUSG00 | Rec8 | 2812 | 4.79165747 | 3.4355403 | 2.97603867 | 5.86525535 | 5.03228505 | 6.09717134 | 2.74252015 | 3.36831998 | 3.83743497 | 3.43737346 | 4.99496365 | 4.30725419 | 2.66659012 | 4.87233491 | 2.27106727 | 2.27401545 | 3.16205579 | 3.54077109 |
| ENSMUSG00 | Irf9 | 3689 | 18.5506461 | 15.9565964 | 16.9418816 | 15.7592552 | 15.4872847 | 17.6503909 | 14.9928224 | 14.6081539 | 13.1966554 | 12.0017648 | 13.3521231 | 14.0560451 | 11.6877462 | 19.8668392 | 12.1069359 | 10.8882752 | 10.6594689 | 17.6852265 |
| ENSMUSG00 | Gmpr2 | 1533 | 17.8006021 | 16.6238513 | 19.2484367 | 16.4947029 | 14.5683057 | 16.8408181 | 19.6596063 | 15.8948159 | 18.8603614 | 15.3478153 | 18.2934773 | 17.7678039 | 13.3868738 | 15.844917 | 14.0294926 | 14.936584 | 16.0255423 | 14.3892053 |
| ENSMUSG00 | Mdp1 | 1727 | 38.3457234 | 38.9839949 | 41.1749216 | 35.2985972 | 36.7887315 | 28.8651129 | 34.6009922 | 36.8655094 | 43.3328889 | 28.5852845 | 35.824303 | 28.5703797 | 34.2209826 | 35.7407019 | 31.0144891 | 28.7567561 | 32.3343454 | 27.1701031 |
| ENSMUSG00 | Dhrs1 | 3150 | 24.3156594 | 28.4798807 | 27.5191639 | 27.0328863 | 28.7569144 | 26.1292351 | 22.9353844 | 22.0667214 | 25.0780118 | 24.653082 | 25.3586446 | 25.3031004 | 22.6456992 | 22.1899529 | 23.9753816 | 23.2904139 | 22.2414193 | 20.4319539 |
| ENSMUSG00 | Ncan | 7180 | 0 | 0.00463968 | 0 | 0 | 0 | 0 | 0 | 0 | 0 | 0 | 0 | 0 | 0 | 0 | 0 | 0 | 0 | 0 |
| ENSMUSG00 | Tmem161a | 3697 | 12.5664478 | 10.6777873 | 9.68255679 | 12.5949114 | 8.9311603 | 10.5753602 | 9.66217814 | 10.6922198 | 10.6949027 | 13.8888601 | 11.6820564 | 9.70772817 | 12.2095494 | 11.0802502 | 11.3340499 | 9.40434718 | 11.1029115 | 10.936352 |
| ENSMUSG00 | Armc6 | 3933 | 1.8264365 | 1.13499309 | 1.56202348 | 1.59863313 | 1.43184212 | 1.81428773 | 1.407471 | 1.90330813 | 1.68599772 | 1.3307216 | 2.53876591 | 2.05778568 | 1.12887683 | 0.82661737 | 2.99609585 | 1.62586612 | 1.12065196 | 1.55250721 |
| ENSMUSG00 | 2310045N01l | 2858 | 10.9163048 | 10.09411 | 11.627927 | 10.6649191 | 9.21209316 | 10.9924538 | 11.2570405 | 11.2118354 | 13.3445464 | 9.68420672 | 10.1133135 | 9.13986357 | 13.1873985 | 12.0579121 | 11.6194731 | 11.8971516 | 8.74343792 | 11.2019944 |
| ENSMUSG00 | Slc25a42 | 6877 | 0.9394775 | 1.11898772 | 1.29427506 | 0.78176577 | 1.23881787 | 1.505965 | 1.80939866 | 1.81048759 | 2.85077527 | 2.24200668 | 1.63950708 | 2.14269811 | 2.56810039 | 1.55331396 | 2.19877432 | 1.22494309 | 1.54932537 | 1.73578249 |
| ENSMUSG00 | Snx9 | 3288 | 24.794686 | 25.0251973 | 24.5251445 | 24.6096152 | 22.9973394 | 27.2329044 | 19.8287322 | 20.3739276 | 20.5807316 | 20.5925821 | 24.5268035 | 20.5404354 | 18.4544581 | 17.1622768 | 17.831452 | 19.7275818 | 22.3570065 | 19.9424742 |
| ENSMUSG00 | Ranbp3 | 2463 | 27.3703651 | 32.7583556 | 29.7547294 | 34.7396026 | 31.5993993 | 29.3332869 | 22.57101 | 28.5628929 | 26.9560144 | 24.6952526 | 32.3155018 | 29.2108766 | 24.9764112 | 30.9250115 | 25.0086586 | 23.6770585 | 22.8899646 | 25.9523272 |
| ENSMUSG00 | Ndufa11 | 2657 | 75.411893 | 64.3564494 | 67.4352866 | 61.8683658 | 68.9462181 | 65.0134849 | 77.513019 | 66.6821809 | 63.4616468 | 68.4989517 | 71.078062 | 64.932711 | 75.0468986 | 71.8773381 | 70.400852 | 66.1763151 | 63.7569561 | 68.9218353 |
| ENSMUSG00 | Bmp8b | 3870 | 0.01098325 | 0 | 0 | 0 | 0 | 0.01280432 | 0.0122255 | 0.06908186 | 0 | 0.1096528 | 0 | 0 | 0.08923086 | 0 | 0 | 0.01978843 | 0.05942062 | 0 |
| ENSMUSG00 | Nr2f6 | 3217 | 11.3629006 | 13.0890576 | 11.8039233 | 14.7007659 | 12.2686284 | 13.0004685 | 10.6038053 | 10.4355367 | 11.7145359 | 12.7366991 | 8.55404914 | 11.6246484 | 6.6092808 | 9.25416849 | 9.08047968 | 7.24867265 | 7.10055112 | 9.3876281 |
| ENSMUSG00 | Use1 | 2334 | 42.2502178 | 34.6973633 | 35.4200357 | 34.5709552 | 36.0680096 | 29.3834547 | 37.8866311 | 36.9815036 | 37.5866661 | 37.4741022 | 37.4380459 | 39.410545 | 38.4679046 | 41.5887501 | 32.0750759 | 33.3689355 | 35.3212788 | 36.0128171 |
| ENSMUSG00 | Ocel1 | 7775 | 2.80452158 | 2.94353162 | 3.01129115 | 3.46322536 | 2.42671564 | 3.37787123 | 2.68358732 | 3.11433867 | 3.501516 | 3.06252156 | 2.5561907 | 3.05819579 | 2.91867292 | 2.63432058 | 3.06825735 | 2.21617663 | 2.27246596 | 2.14376326 |

| | | | | | | | | | | | | | | | | | | | |
|---|---|---|---|---|---|---|---|---|---|---|---|---|---|---|---|---|---|---|---|
| ENSMUSG00 Dyrk1b | 2829 | 7.52742778 | 8.14864715 | 7.8143172 | 7.90755488 | 7.26487501 | 9.02073446 | 9.91743323 | 7.35767385 | 6.17288183 | 7.30010888 | 8.17358781 | 8.18787556 | 5.24881835 | 4.76097043 | 5.92754798 | 5.4681532 | 5.94741758 | 6.41675688 |
| ENSMUSG00 Braf | 10697 | 5.40801576 | 6.01668801 | 5.78384083 | 6.11626247 | 5.66046181 | 5.71637592 | 4.86528436 | 6.62306314 | 5.14013261 | 5.37316138 | 5.89538611 | 5.65618194 | 4.96685752 | 5.07554811 | 5.61576783 | 5.50537106 | 7.10489359 | 5.46131358 |
| ENSMUSG00 Ndufb2 | 4711 | 10.4751665 | 8.54920168 | 10.2271357 | 9.34238353 | 10.2680075 | 9.30888754 | 10.6556469 | 10.0284444 | 10.4649245 | 9.61830308 | 10.1715736 | 9.27693989 | 9.07891579 | 8.24182914 | 8.53591988 | 9.00572978 | 9.51038868 | 8.4889876 |
| ENSMUSG00 Hltf | 6079 | 7.55150355 | 7.61719411 | 7.77445948 | 7.26248593 | 7.86231291 | 7.52379883 | 7.86080041 | 7.07429595 | 6.57195596 | 5.35186466 | 4.5110752 | 5.16012541 | 4.54447667 | 6.07386897 | 4.43908462 | 6.90982009 | 7.87457556 | 5.24844249 |
| ENSMUSG00 Prpf6 | 3837 | 37.4537399 | 39.3121671 | 35.7916218 | 40.9895035 | 34.9480165 | 34.7269483 | 29.0633266 | 28.1491045 | 27.14639 | 30.0083444 | 31.71301 | 30.082562 | 31.5765413 | 32.1247349 | 33.3306306 | 34.2290288 | 33.0622994 | 32.070725 |
| ENSMUSG00 Rgs19 | 2429 | 31.568274 | 28.8419171 | 23.5395277 | 24.7969126 | 28.3758397 | 29.376672 | 12.8946041 | 13.3020883 | 13.1240688 | 14.9274972 | 13.9056796 | 18.865648 | 10.8859233 | 8.6999011 | 9.83816485 | 18.4753363 | 10.5085689 | 9.80606966 |
| ENSMUSG00 Rgs20 | 4922 | 0.11226479 | 0.04060897 | 0 | 0.02776984 | 0 | 0 | 0.11534989 | 0.00775953 | 0.07531057 | 0.00957958 | 0 | 0 | 0.06013643 | 0.12268825 | 0 | 0 | 0.0233602 | 0 |
| ENSMUSG00 Abhd3 | 2775 | 0.09190308 | 0 | 0.40093377 | 0.04925518 | 0.26017256 | 0.0892842 | 1.10822491 | 0.19268239 | 0.07420994 | 0.27185991 | 0.22381062 | 0.4223888 | 3.59100803 | 5.22182172 | 1.87077335 | 2.24914198 | 3.71523383 | 4.20249752 |
| ENSMUSG00 Snrpd1 | 833 | 65.2120228 | 61.0669742 | 61.4976814 | 48.0223252 | 60.9016177 | 51.8727295 | 72.3037698 | 56.7154883 | 55.6240459 | 53.2072762 | 58.1558318 | 55.6816514 | 67.5130681 | 62.3900766 | 67.168923 | 61.8717378 | 54.0617033 | 58.3770112 |
| ENSMUSG00 Tchp | 3978 | 3.21620362 | 3.34133856 | 2.61445677 | 4.20334421 | 2.97648397 | 3.21382737 | 2.04569652 | 3.33151424 | 3.19925424 | 3.41362558 | 2.49803704 | 3.59195818 | 3.53433796 | 3.56094687 | 3.92543733 | 2.87805219 | 3.39136664 | 2.93160649 |
| ENSMUSG00 Tiam1 | 10002 | 0.29322707 | 0.19317615 | 0.25632845 | 0.32341873 | 0.23579927 | 0.30221123 | 0.89876115 | 0.45058023 | 0.63002769 | 0.63640721 | 0.52064292 | 0.64175177 | 0.30579673 | 0.43184364 | 0.39545585 | 0.28712213 | 0.31038093 | 0.23099205 |
| ENSMUSG00 Tsc2 | 6299 | 11.073344 | 16.1936946 | 11.3119877 | 11.5728958 | 13.4637958 | 14.1208377 | 9.89217307 | 12.0537381 | 8.55900129 | 11.3329361 | 10.0874257 | 10.6066595 | 8.84199629 | 10.1603354 | 11.1196452 | 9.97537494 | 10.6113767 | 9.52769256 |
| ENSMUSG00 Rpl3l | 1584 | 2.28089644 | 2.20824109 | 1.52694225 | 2.07095656 | 1.64086103 | 2.84677952 | 9.43864049 | 7.52274593 | 6.96842565 | 4.55432929 | 3.86060271 | 4.72177954 | 7.2876696 | 9.17739858 | 4.05772899 | 2.73159005 | 3.62938418 | 5.17446774 |
| ENSMUSG00 Slc9a3r2 | 3350 | 4.80879454 | 5.0217945 | 3.60996495 | 5.22251971 | 2.68676708 | 6.36050013 | 2.01961585 | 2.9299843 | 1.93023372 | 2.4490216 | 1.7113418 | 2.76579104 | 2.25306965 | 2.60641446 | 2.15232008 | 2.01168564 | 1.56737393 | 1.75700729 |
| ENSMUSG00 Pdl60 | 1900 | 203.644535 | 196.581142 | 191.177284 | 202.770627 | 182.090667 | 202.122992 | 166.690038 | 170.277811 | 170.989072 | 165.350071 | 167.514091 | 168.827691 | 154.123295 | 158.081007 | 161.9438 | 155.480019 | 169.946869 | 176.058094 |
| ENSMUSG00 Golga2 | 5394 | 15.8941305 | 18.2992409 | 16.8060801 | 18.7177001 | 16.6061383 | 17.9966268 | 16.095433 | 16.4976451 | 16.7983564 | 19.3882573 | 17.1118316 | 15.8734198 | 17.6146223 | 18.2624971 | 18.8133912 | 19.3866609 | 19.2697288 | 22.6808034 |
| ENSMUSG00 Uck1 | 3952 | 12.90643 | 12.9475216 | 11.285525 | 12.2433778 | 11.2048 | 11.2220892 | 12.1514092 | 13.7036505 | 14.2047624 | 13.7801271 | 14.0471917 | 10.7902749 | 17.4134678 | 16.0650474 | 17.4105765 | 14.6205786 | 14.8766547 | 15.7148992 |
| ENSMUSG00 Scin | 3848 | 0.12150648 | 0.42420266 | 0.11313975 | 0.21312339 | 0.56287333 | 0.37344836 | 0.19672632 | 0.07940208 | 0.32110069 | 0.44111885 | 0.29797271 | 0.18856643 | 0.80766912 | 2.42600353 | 0.81375214 | 1.04483206 | 0.52788299 | 0.64329777 |
| ENSMUSG00 Ikzf4 | 8382 | 0.4716036 | 0.41333107 | 0.42129178 | 0.38592629 | 0.35602234 | 0.48476785 | 0.44592002 | 0.56500333 | 0.36361276 | 0.20813362 | 0.38188048 | 0.42617542 | 0.38255483 | 1.25779144 | 0.61443504 | 0.31977362 | 0.74530976 | 0.74159259 |
| ENSMUSG00 1810046J19R | 743 | 160.695881 | 118.410974 | 135.094314 | 123.560571 | 133.836547 | 118.246294 | 209.245576 | 169.578379 | 170.511067 | 183.081802 | 166.794404 | 151.445997 | 237.297809 | 231.032768 | 184.325669 | 156.87296 | 162.486992 | 180.130763 |
| ENSMUSG00 Pon1 | 2890 | 0.04412302 | 0.02305391 | 0 | 0 | 0 | 0 | 0 | 0 | 0 | 0.01631512 | 0 | 0 | 0.01706987 | 0.03214124 | 0 | 0 | 0 | 0 |
| ENSMUSG00 Axl | 6694 | 1.86682428 | 1.68704379 | 2.232958 | 2.14396907 | 2.05642817 | 2.49466257 | 1.3004979 | 1.55188728 | 1.73507581 | 1.80319391 | 0.65660236 | 1.7760287 | 0.56745707 | 0.86727083 | 0.53548482 | 0.68069649 | 0.90462428 | 0.62454319 |
| ENSMUSG00 Tgfb1 | 2551 | 5.79843248 | 6.62079082 | 5.63188495 | 6.10814471 | 5.07545569 | 6.33249359 | 4.91488041 | 4.9406087 | 4.23005376 | 5.37861805 | 3.65194704 | 6.3670552 | 3.28750527 | 2.05730375 | 4.02163364 | 3.52735897 | 2.55408486 | 2.11324675 |
| ENSMUSG00 Ccdc97 | 3902 | 13.1371713 | 13.9415572 | 14.4302382 | 17.98155 | 15.0612708 | 14.959795 | 13.8348966 | 14.0554305 | 12.1912931 | 15.720929 | 12.7946775 | 14.9766783 | 14.1851387 | 12.9500684 | 13.895758 | 13.3555912 | 10.6865738 | 14.7320647 |
| ENSMUSG00 Zfp40 | 7290 | 2.6645906 | 3.16221163 | 3.8221061 | 2.94990575 | 3.75679898 | 2.87528207 | 5.55550798 | 5.76815296 | 4.73447191 | 3.54438757 | 4.09593265 | 5.41312393 | 4.03317184 | 3.51038301 | 4.95096886 | 4.38582403 | 3.60130499 | 3.51635717 |
| ENSMUSG00 Akap8l | 2075 | 48.2203298 | 50.9727418 | 47.3244658 | 55.9028525 | 49.918024 | 48.8124277 | 51.7361294 | 51.389488 | 47.8160561 | 61.3981688 | 48.6728975 | 47.1272486 | 49.5934359 | 61.6419357 | 57.7020344 | 60.545293 | 58.9209095 | 62.8031681 |
| ENSMUSG00 Shh | 2689 | 0 | 0.0619429 | 0.39576708 | 0.1694349 | 0.1431965 | 0.05528382 | 0.87974487 | 0.78117595 | 0.45949999 | 0.22795059 | 0.17766812 | 0.47741177 | 0.66044962 | 2.19352886 | 0.55160134 | 0.45567104 | 0.08551796 | 0.06137126 |
| ENSMUSG00 Pdcd2l | 1466 | 26.9933962 | 21.7238246 | 19.2372225 | 20.542872 | 20.5529224 | 22.6130824 | 20.0417282 | 21.4408917 | 22.1384546 | 20.1982549 | 22.4535787 | 24.4812217 | 21.1662865 | 22.1132149 | 18.1837699 | 16.4289168 | 15.4769221 | 18.9867722 |
| ENSMUSG00 Gtf2f1 | 1718 | 50.9913666 | 56.4457713 | 50.2881986 | 46.1179605 | 48.4120862 | 46.9856828 | 47.6982304 | 42.6385494 | 52.0465225 | 47.2055935 | 48.4979788 | 47.2711229 | 48.0397541 | 52.5807625 | 45.5663307 | 57.9039569 | 50.796844 | 56.2898775 |
| ENSMUSG00 Clpp | 1201 | 94.813794 | 77.9704793 | 86.959577 | 80.8414534 | 90.1722134 | 84.3345441 | 97.5799381 | 85.2571448 | 92.0438038 | 98.1881006 | 101.039463 | 85.2339094 | 97.8423205 | 94.7057202 | 96.3999767 | 103.808674 | 90.6619563 | 96.8728232 |
| ENSMUSG00 Alkbh7 | 1250 | 27.4073375 | 22.5994674 | 26.6634899 | 23.5823965 | 22.2562015 | 20.4950106 | 27.4035036 | 22.0599316 | 24.6789602 | 25.4991003 | 21.4796213 | 22.996053 | 30.7041853 | 23.4449753 | 25.5779259 | 28.8864342 | 17.8447238 | 21.6955917 |
| ENSMUSG00 Pspn | 472 | 0.27016002 | 0.35289083 | 0.61491776 | 0.67569345 | 0.40789766 | 0.10498461 | 1.5035809 | 0.48549664 | 0.17451913 | 0 | 0.70852693 | 0 | 1.56775161 | 1.08238339 | 0.43645716 | 0 | 0.08119978 | 0.69926831 |
| ENSMUSG00 Dennd1c | 2496 | 5.27908837 | 5.12506065 | 5.01952899 | 5.36656532 | 3.99171002 | 3.95071874 | 4.15123265 | 3.04498721 | 3.67972477 | 3.06026199 | 3.92382471 | 2.52691101 | 2.62866386 | 2.93997028 | 2.24495657 | 2.13236956 | 1.96545112 | 2.97525217 |
| ENSMUSG00 Med6 | 1838 | 22.8251408 | 18.6320214 | 19.738927 | 19.5084559 | 22.0233666 | 24.1562836 | 21.6227695 | 23.5845358 | 23.9320995 | 19.2399496 | 21.236203 | 17.8865208 | 20.6399589 | 25.5467749 | 23.4476754 | 21.6244018 | 24.5638617 | 24.9606066 |
| ENSMUSG00 Prkd1 | 3759 | 1.41344693 | 2.37506136 | 1.80162124 | 1.90292204 | 2.34321416 | 1.70053276 | 0.85588246 | 0.47753203 | 0.20817872 | 0.57699717 | 0.26689921 | 0.08909124 | 0.81366823 | 0.19768699 | 0.35074482 | 0.29540506 | 0.42822678 | 0.45365318 |
| ENSMUSG00 Lcp2 | 4725 | 3.45438891 | 2.4535211 | 1.83256337 | 2.48777766 | 2.58740923 | 2.26526782 | 8.89177966 | 4.02535797 | 3.86150825 | 5.85766409 | 3.98377432 | 4.1581192 | 14.7317112 | 5.80919696 | 6.85384612 | 24.9435957 | 5.86453178 | 8.91787847 |
| ENSMUSG00 Cse1l | 5087 | 23.8052382 | 24.0662249 | 25.6844713 | 22.9193459 | 23.2569822 | 24.1383279 | 18.4618965 | 25.7218735 | 22.6942803 | 26.3421071 | 23.3662466 | 20.5399883 | 18.0958045 | 18.150353 | 20.102666 | 18.6974383 | 20.6210094 | 15.9068992 |
| ENSMUSG00 Naa20 | 3592 | 22.9447484 | 20.5608815 | 19.9985224 | 17.9479058 | 21.0108175 | 21.3689266 | 20.0999855 | 21.9989088 | 21.4073566 | 19.4798562 | 20.1633733 | 21.0707234 | 20.8342201 | 20.3774726 | 22.1836885 | 22.119435 | 19.6859738 | 21.6851044 |
| ENSMUSG00 Prkra | 2020 | 8.37478221 | 7.68505381 | 9.12391835 | 7.66869063 | 6.24285347 | 8.09524855 | 8.52565105 | 8.39476568 | 12.6617952 | 8.26304404 | 8.94006649 | 7.95787665 | 8.84067032 | 6.96661311 | 6.89412173 | 7.86663469 | 8.65187692 | 11.3286081 |
| ENSMUSG00 Fkbp7 | 1768 | 6.82775452 | 6.21790446 | 6.21085486 | 5.10242465 | 7.78604648 | 7.2871667 | 3.96056362 | 6.56701942 | 3.82046866 | 5.84050001 | 5.99889185 | 5.61945019 | 3.90637354 | 4.15054627 | 5.56966734 | 4.33151667 | 4.79078711 | 4.72928972 |
| ENSMUSG00 Plekha3 | 3118 | 45.5860515 | 53.377556 | 49.3509301 | 49.6524787 | 53.1951921 | 54.161551 | 50.3323143 | 51.9847838 | 53.2069549 | 50.8404953 | 53.4901796 | 51.2508794 | 44.0791281 | 56.0963544 | 51.0064162 | 46.5062637 | 54.195203 | 54.3739791 |
| ENSMUSG00 Ykt6 | 2977 | 40.8632226 | 35.6404327 | 36.1054701 | 35.4907833 | 34.0496578 | 38.4004556 | 41.8455773 | 44.183619 | 39.4156478 | 42.0349232 | 36.7660161 | 36.5417287 | 53.1266801 | 42.0758158 | 47.4156972 | 42.7537719 | 50.054633 | 44.790754 |
| ENSMUSG00 Baz1b | 10776 | 6.93430767 | 5.61397698 | 4.84812665 | 5.4033978 | 5.64576387 | 4.75938003 | 5.22476688 | 5.11075021 | 5.43114689 | 5.03623417 | 5.09847831 | 5.2003457 | 5.87807923 | 6.10720785 | 4.88255261 | 6.50968752 | 5.60525649 | 4.02767123 |
| ENSMUSG00 Pex6 | 4686 | 12.9801338 | 13.4929307 | 11.6753015 | 13.5633071 | 12.2332738 | 11.6003732 | 13.6001235 | 14.3934682 | 13.8343098 | 14.4088363 | 12.2241091 | 13.316669 | 12.232968 | 9.57426129 | 12.2215456 | 10.9249983 | 8.70234313 | 11.2225152 |
| ENSMUSG00 Mrpl2 | 1209 | 65.8496234 | 60.2332397 | 60.0567982 | 61.162593 | 62.8621236 | 64.9636747 | 71.4190563 | 65.6441788 | 69.7002574 | 66.572581 | 70.2991325 | 68.9270341 | 74.9159637 | 66.3816237 | 62.4668751 | 65.9079851 | 54.937529 | 68.7045088 |
| ENSMUSG00 Mea1 | 836 | 137.379169 | 129.625413 | 129.323972 | 115.810071 | 136.450549 | 112.027113 | 166.61307 | 130.840958 | 140.063713 | 150.137766 | 149.49676 | 132.862642 | 181.100065 | 160.998762 | 142.529784 | 159.94115 | 148.308099 | 152.39365 |
| ENSMUSG00 Gnmt | 1201 | 85.4704414 | 77.9982169 | 81.5220861 | 79.9309912 | 84.8420336 | 78.7645032 | 128.149995 | 111.460758 | 102.057511 | 105.529634 | 111.740099 | 115.814014 | 130.209973 | 143.160709 | 133.965092 | 116.976052 | 112.425932 | 134.110462 |
| ENSMUSG00 Grin2d | 6606 | 1.29330009 | 1.62378927 | 1.3913065 | 1.21385768 | 1.13663035 | 1.50023414 | 2.82185823 | 2.26633693 | 1.87664869 | 2.69085909 | 1.70676509 | 2.4587281 | 0.67209697 | 1.70140314 | 0.97920743 | 1.00856277 | 1.18935678 | 1.24907145 |
| ENSMUSG00 Kdelr1 | 2143 | 65.9693026 | 61.4026748 | 61.1497404 | 52.7258003 | 58.1491136 | 58.3857453 | 59.654157 | 56.8876101 | 65.4602299 | 53.7073573 | 57.0490521 | 55.7375259 | 68.2773993 | 49.001438 | 51.6413342 | 56.5693248 | 56.3537851 | 55.6765061 |
| ENSMUSG00 Tmem143 | 5232 | 1.70605638 | 2.00565017 | 2.16349505 | 1.85483609 | 1.79390771 | 1.96998635 | 3.62622023 | 2.56951943 | 2.84180661 | 3.11814666 | 2.21890576 | 2.63502841 | 1.85749 | 2.36126285 | 2.58297216 | 1.95405049 | 2.6810827 | 2.4918116 |
| ENSMUSG00 Pdxk-ps | 2639 | 0.25770474 | 0.03786991 | 0.12831175 | 0.10358707 | 0.07295479 | 0.03755417 | 0.23306738 | 0 | 0.04682059 | 0.303737 | 0 | 0.06345092 | 0.03738683 | 0 | 0.35908897 | 0.08705708 | 0.07261519 | 0.18760211 |
| [illegible] | | | | | | | | | | | | | | | | | | | |

| | | | | | | | | | | | | | | | | | | | | |
|---|---|---|---|---|---|---|---|---|---|---|---|---|---|---|---|---|---|---|---|---|
| ENSMUSG00 | Ndufs3 | 2411 | 57.3493727 | 55.931396 | 55.1550539 | 52.0049747 | 56.5165806 | 50.9297693 | 63.3452209 | 60.9716121 | 59.0379919 | 59.1583896 | 61.4277774 | 51.2550298 | 68.0743628 | 67.9035288 | 62.5286159 | 68.5768738 | 61.5191908 | 61.488819 |
| ENSMUSG00 | Por | 4002 | 74.9629017 | 88.3182928 | 72.789955 | 81.3313595 | 85.1148485 | 77.2636326 | 54.5597235 | 60.5047055 | 56.5105342 | 75.0264064 | 81.7379162 | 58.8700429 | 60.6602611 | 60.4167714 | 65.9410209 | 59.4164598 | 66.089399 | 71.214938 |
| ENSMUSG00 | Igf1r | 4454 | 4.66659881 | 7.7485762 | 7.15719724 | 7.45711749 | 6.7540306 | 6.65301632 | 3.51605225 | 5.04201451 | 5.33556794 | 4.13918387 | 4.08672186 | 4.82465876 | 3.53320198 | 3.86005269 | 3.89444433 | 2.579071 | 3.03753543 | 2.58125355 |
| ENSMUSG00 | Insr | 9647 | 13.4384562 | 11.7097568 | 10.7056585 | 10.0832207 | 10.6970918 | 9.01472452 | 9.84311668 | 10.2379551 | 12.4323763 | 10.1759888 | 8.84483009 | 10.9178144 | 11.5467471 | 9.46502289 | 9.23799786 | 11.066053 | 10.1347967 | 8.02869517 |
| ENSMUSG00 | Fcer2a | 2970 | 0.02862301 | 0.02243293 | 0.04886215 | 0.04602126 | 0 | 0.25026633 | 0 | 0 | 0.24961525 | 0 | 0.1447726 | 0 | 0.06644029 | 0 | 0.29132413 | 0 | 0 | 0 |
| ENSMUSG00 | Cyp2a5 | 2148 | 0 | 0 | 0 | 0 | 0 | 0 | 0 | 0 | 0 | 0 | 0 | 0 | 0 | 0 | 0 | 0 | 0 | 0 |
| ENSMUSG00 | Atp4a | 3771 | 5.03840193 | 4.10779523 | 4.00226504 | 3.91455274 | 3.62489163 | 5.05908312 | 2.45910503 | 2.4205739 | 1.84580248 | 1.71297967 | 1.29224231 | 2.48661663 | 0.26163838 | 1.17003137 | 0.83036814 | 0.40615866 | 0.35572007 | 0.5251466 |
| ENSMUSG00 | Trim28 | 3829 | 43.049118 | 41.5867418 | 43.2064439 | 38.0766106 | 39.7474909 | 39.9501929 | 33.7206322 | 37.7435494 | 36.2708099 | 37.3609908 | 35.6347039 | 38.5709698 | 37.7623007 | 35.7822037 | 37.9303955 | 33.9705389 | 33.3515856 | 35.097218 |
| ENSMUSG00 | Ube2m | 2676 | 86.8211107 | 83.9171225 | 82.8821512 | 87.410464 | 81.1191987 | 85.6062359 | 93.4407728 | 90.0718429 | 91.330637 | 93.0503976 | 82.0529556 | 80.3653767 | 105.798159 | 95.1270714 | 88.0229055 | 80.3015279 | 79.1589847 | 87.5088817 |
| ENSMUSG00 | Adcy9 | 6730 | 0.51789368 | 0.46034163 | 0.6253339 | 0.50096839 | 0.77239937 | 0.59639991 | 0.78034285 | 0.59586958 | 0.47122759 | 0.60952618 | 0.42592831 | 0.56396212 | 0.54243119 | 1.03515797 | 1.04075253 | 0.80791475 | 0.67199154 | 0.9890196 |
| ENSMUSG00 | Mef2c | 7382 | 0 | 0 | 0 | 0 | 0 | 0.07383908 | 0.45505286 | 0.09312696 | 0.82015955 | 0.08303429 | 0.4012527 | 0.06048839 | 0.38759837 | 0.27053587 | 0.24557958 | 0.18154582 | 0 | 0.02235537 |
| ENSMUSG00 | Ctr9 | 5287 | 23.1298263 | 24.5105274 | 26.2316837 | 25.3959519 | 23.4787651 | 25.2028183 | 25.0299911 | 22.9212205 | 20.3946091 | 25.2831934 | 29.9192146 | 29.0638394 | 22.2168247 | 26.8896069 | 25.2492985 | 27.1010881 | 28.3224592 | 27.4681373 |
| ENSMUSG00 | Eif4g2 | 12073 | 84.0456856 | 99.2626569 | 99.078991 | 96.1676545 | 97.5196707 | 102.048341 | 87.8221763 | 94.6980507 | 88.6603461 | 90.7357151 | 88.6327773 | 88.9407535 | 70.7064035 | 79.5123699 | 84.2869235 | 75.6709684 | 90.6174228 | 80.3471041 |
| ENSMUSG00 | Mrvi1 | 9724 | 0 | 0 | 0 | 0 | 0 | 0 | 0 | 0.12961223 | 0 | 0 | 0.0049131 | 0 | 0 | 0 | 0 | 0 | 0 | 0 |
| ENSMUSG00 | Pcyt1a | 5849 | 7.84845104 | 7.25035294 | 6.74864092 | 6.98722119 | 6.05660725 | 7.04023279 | 8.67142961 | 7.07822948 | 6.99938394 | 6.71508562 | 6.57528493 | 8.34038111 | 7.48961233 | 5.78069681 | 6.54406658 | 6.33703333 | 8.15801669 | 6.45174341 |
| ENSMUSG00 | Zfp592 | 5829 | 7.45973496 | 7.72671693 | 8.35686139 | 8.32432874 | 6.7875178 | 7.37042718 | 8.15735844 | 7.05665078 | 5.81514882 | 8.20223248 | 7.16337514 | 7.60295696 | 7.71842658 | 7.42595461 | 8.69410083 | 7.66600426 | 8.98159574 | 7.27603728 |
| ENSMUSG00 | Psmd4 | 2517 | 92.0692172 | 91.8520007 | 80.872394 | 82.5601602 | 89.8959766 | 84.5566113 | 82.5763204 | 77.5987072 | 77.0385841 | 81.3757072 | 80.9156329 | 84.3111439 | 81.1810891 | 87.4447875 | 82.7468673 | 77.7221745 | 72.0692742 | 90.3049777 |
| ENSMUSG00 | Tmod4 | 1770 | 0.48028447 | 0.24467097 | 0.08198903 | 0.25740703 | 0.05438635 | 0.33595074 | 0 | 0.28050917 | 0.74461497 | 0.10655526 | 0.78275356 | 0.12613709 | 0.89187647 | 0.76094832 | 0.58194288 | 0.7571589 | 0.28149258 | 1.02559352 |
| ENSMUSG00 | Insrr | 6185 | 10.2947055 | 16.7937921 | 12.5450514 | 13.7824922 | 14.7625318 | 14.7897084 | 10.6482214 | 9.65153193 | 8.1174394 | 13.0436303 | 8.78255906 | 5.80266306 | 4.16350215 | 4.50548948 | 4.44323974 | 2.14823618 | 3.29661923 | 2.63261044 |
| ENSMUSG00 | Cabp5 | 2595 | 0 | 0 | 0 | 0 | 0 | 0 | 0.03646449 | 0.11774151 | 0 | 0 | 0 | 0.06452678 | 0 | 0 | 0 | 0 | 0 | 0 |
| ENSMUSG00 | Snx6 | 1914 | 74.5284069 | 83.0907821 | 70.4373193 | 80.076987 | 66.4643021 | 65.9408633 | 59.3756817 | 70.2987637 | 73.2491639 | 76.9586191 | 68.4674562 | 63.3394039 | 68.0182496 | 70.0058475 | 68.0665202 | 73.2002782 | 73.0282157 | 55.6988761 |
| ENSMUSG00 | Mthfd2 | 2965 | 19.3674512 | 16.549711 | 18.5826142 | 16.5494923 | 18.8144686 | 15.8100796 | 20.1058939 | 18.175204 | 20.5863124 | 18.2400866 | 18.8521759 | 16.8482354 | 22.0787367 | 25.3758589 | 14.8547937 | 14.4122489 | 15.8217158 | 22.7457554 |
| ENSMUSG00 | Kit | 5973 | 68.1235644 | 81.6397364 | 83.951116 | 90.6186517 | 87.7623942 | 94.5508965 | 66.6955896 | 88.7383493 | 82.6971176 | 84.939154 | 84.4400195 | 86.6904239 | 56.798191 | 63.1851087 | 85.7210338 | 78.9018577 | 95.356956 | 77.5542755 |
| ENSMUSG00 | Tomm40l | 4632 | 2.12892937 | 1.43838058 | 1.83802706 | 1.39673322 | 1.43398218 | 1.27305166 | 1.53214634 | 1.2368006 | 1.62718748 | 1.02811335 | 1.217065 | 1.62675182 | 1.0863246 | 1.53409874 | 1.00513295 | 1.15731542 | 1.07565168 | 0.95007093 |
| ENSMUSG00 | Nr1i3 | 3574 | 0.05946443 | 0.0279627 | 0.14888328 | 0 | 0.04040173 | 0.01386478 | 0 | 0.09617561 | 0.09219142 | 0 | 0.08020418 | 0.01561714 | 0.013803 | 0.31187971 | 0 | 0.01071366 | 0.10723642 | 0.0461744 |
| ENSMUSG00 | Apoa2 | 572 | 0 | 0.17471798 | 0 | 0.15930435 | 0.25244016 | 0 | 0 | 0 | 0 | 0.4121565 | 0 | 0.09757983 | 0.51746766 | 0.32478383 | 0.50421484 | 0.53553297 | 0 | 0 |
| ENSMUSG00 | Pan2 | 4418 | 8.54336721 | 10.4658889 | 8.21189402 | 10.8797876 | 10.1318898 | 8.53545275 | 8.83498418 | 10.1316197 | 10.897926 | 8.64465134 | 11.7328723 | 9.55107309 | 10.0271756 | 11.973702 | 8.81291768 | 8.45028117 | 8.31068185 | 12.8869229 |
| ENSMUSG00 | Cs | 2817 | 95.7083179 | 105.520396 | 105.14417 | 112.972629 | 109.86449 | 104.11879 | 92.8114532 | 100.219467 | 94.0257305 | 110.63761 | 99.5862819 | 99.7033774 | 84.4264016 | 84.3808446 | 99.6179614 | 85.5931682 | 94.0266375 | 80.5511414 |
| ENSMUSG00 | Ampd3 | 5623 | 7.37774352 | 5.2490173 | 5.37674844 | 6.76568949 | 7.30153493 | 5.41969254 | 5.57015711 | 4.69339322 | 6.60683571 | 6.06259265 | 6.10037706 | 4.744778 | 3.85145148 | 2.39530239 | 3.63435385 | 3.615919 | 2.44693946 | 3.19900016 |
| ENSMUSG00 | Bcas2 | 4548 | 38.6079336 | 39.5608931 | 40.3112753 | 36.7553808 | 37.2525003 | 38.7988752 | 41.2581541 | 38.964984 | 40.3261984 | 39.893559 | 39.6969147 | 39.5545632 | 40.2530222 | 35.8338396 | 38.6015457 | 41.969806 | 39.4302426 | 42.7929243 |
| ENSMUSG00 | Sh2d1a | 2724 | 0 | 0 | 0 | 0 | 0 | 0 | 0 | 0 | 0 | 0 | 0 | 0 | 0 | 0 | 0 | 0 | 0 | 0 |
| ENSMUSG00 | Ctcf | 4628 | 21.1974819 | 19.5069022 | 16.410208 | 18.4291865 | 16.7858524 | 15.9643746 | 16.2138956 | 16.8102687 | 20.1126892 | 15.8629311 | 17.6420493 | 17.4514399 | 15.7546616 | 17.3814087 | 15.9713896 | 16.5059987 | 16.074188 | 17.2349189 |
| ENSMUSG00 | Pard6a | 1424 | 9.28308269 | 8.23464801 | 6.08066147 | 8.15875443 | 5.61088436 | 7.34243454 | 8.53887534 | 7.45610103 | 10.817245 | 8.94009121 | 13.4870318 | 12.1116852 | 10.2197441 | 10.3716431 | 6.91514878 | 9.00797295 | 5.3290778 | 8.92352785 |
| ENSMUSG00 | Agrp | 654 | 0.32496312 | 0.81499435 | 0.96155336 | 0.27866082 | 0.22078864 | 1.288068 | 0.28937418 | 0.52558352 | 0.37785794 | 0.64886289 | 1.89931028 | 0.93879555 | 1.65948346 | 0 | 0.06299932 | 0.93677327 | 0.5274261 | 0 |
| ENSMUSG00 | Pvalb | 1613 | 0 | 0 | 0 | 0 | 0 | 0 | 0 | 0 | 0 | 0 | 0 | 0 | 0 | 0 | 0 | 0 | 0 | 0 |
| ENSMUSG00 | Tfap4 | 2106 | 0.18164605 | 0.77508633 | 0.52829592 | 0.41104456 | 0.61707594 | 0.42352764 | 0.67396979 | 0.38083592 | 0.48891875 | 0.64927369 | 0.83935112 | 0.47705695 | 0.30451801 | 0.44106446 | 0.48909729 | 0.32727015 | 0.2547807 | 0.52240367 |
| ENSMUSG00 | Ranbp1 | 2049 | 97.6229177 | 88.102769 | 93.7960157 | 84.9183154 | 87.0086116 | 86.674962 | 94.7869922 | 86.5247106 | 87.6193373 | 84.2913743 | 83.8218525 | 82.3750921 | 86.2164938 | 91.5508389 | 85.8013855 | 89.1952509 | 79.2338671 | 85.8292407 |
| ENSMUSG00 | Cd247 | 5557 | 0.04589366 | 0.03596857 | 0.08704972 | 0.06559085 | 0.02598448 | 0.06242022 | 0.00851407 | 0 | 0.01482329 | 0.06787936 | 0.0601808 | 0 | 0.02663231 | 0.01671553 | 0.02224306 | 0 | 0.04827858 | 0 |
| ENSMUSG00 | Rfx5 | 5528 | 3.54466103 | 4.54978928 | 4.77784869 | 5.22534409 | 5.4679537 | 5.14530917 | 3.91134122 | 5.63766287 | 3.97113113 | 3.82971522 | 5.85951915 | 5.35135694 | 3.75700745 | 5.11657909 | 3.50302664 | 3.83044607 | 5.75449103 | 4.81628214 |
| ENSMUSG00 | Psmb4 | 1495 | 125.667477 | 114.199721 | 119.882227 | 112.51597 | 118.993706 | 109.977237 | 106.524734 | 113.887445 | 108.407317 | 108.557004 | 112.263163 | 112.527363 | 122.950317 | 108.825176 | 108.777619 | 123.042193 | 108.15963 | 112.04196 |
| ENSMUSG00 | Mmp8 | 2443 | 0 | 0 | 0 | 0 | 0 | 0 | 0.04056712 | 0 | 0.13487293 | 0.03860066 | 0 | 0.18277745 | 0.02019317 | 0.13307761 | 0.0337303 | 0 | 0.12550568 | 0 |
| [illegible] | | | | | | | | | | | | | | | | | | | | |

| | | | | | | | | | | | | | | | | | | | | |
|---|---|---|---|---|---|---|---|---|---|---|---|---|---|---|---|---|---|---|---|---|
| ENSMUSG00 | Zmynd10 | 1743 | 0 | 0.11467433 | 0 | 0 | 0 | 0 | 0.84147622 | 1.073682 | 1.08696485 | 0.16230879 | 0.1918673 | 0.67247786 | 1.04720651 | 1.09248857 | 0.73278728 | 0.48330084 | 0.08795478 | 0.50496025 |
| ENSMUSG00 | Tmem115 | 2175 | 35.7238905 | 33.0065688 | 31.6707619 | 34.7730269 | 31.9551714 | 32.1466239 | 32.0638569 | 31.4494679 | 38.2513848 | 30.263164 | 36.3748649 | 33.9513203 | 32.366274 | 37.0698555 | 34.1736123 | 33.2556126 | 30.1323989 | 28.6047355 |
| ENSMUSG00 | Hyal2 | 1889 | 35.372227 | 33.9653966 | 34.7756772 | 29.8353689 | 29.531445 | 40.5813017 | 26.0232257 | 27.2138558 | 28.7150033 | 29.2788644 | 27.9972891 | 28.9567757 | 23.8955555 | 26.922327 | 26.56617 | 30.9932974 | 22.8050597 | 22.6268269 |
| ENSMUSG00 | Ifrd2 | 1968 | 8.61766526 | 7.60035036 | 6.53829221 | 5.83403616 | 5.69854584 | 4.93512997 | 5.86600288 | 7.21929558 | 8.03639332 | 7.11573113 | 5.09793764 | 6.72170327 | 8.52279059 | 5.09752179 | 7.41125548 | 6.32339807 | 6.56298885 | 8.05011318 |
| ENSMUSG00 | Hyal1 | 3939 | 6.45293101 | 7.67913378 | 6.82804846 | 8.36268978 | 8.12585155 | 8.27765897 | 4.48023083 | 6.24420085 | 4.67386965 | 4.91976112 | 5.44578735 | 4.93116288 | 6.13674523 | 7.59329583 | 6.35962071 | 7.85448357 | 5.91581328 | 6.36815249 |
| ENSMUSG00 | Tusc2 | 1658 | 39.0955328 | 33.5941852 | 33.0853941 | 35.0638677 | 38.6971377 | 37.0898326 | 33.5868656 | 32.4566315 | 27.9213774 | 38.7329661 | 38.8423667 | 39.2527519 | 43.6191741 | 43.0826335 | 33.1252557 | 38.6601187 | 39.0891243 | 39.6476975 |
| ENSMUSG00 | Nprl2 | 1445 | 28.8858705 | 21.6476178 | 23.4670206 | 24.7826857 | 20.7849968 | 24.4848801 | 22.0028514 | 21.2238748 | 24.5978983 | 23.754818 | 24.8297535 | 19.9700329 | 22.941902 | 24.8451762 | 25.0630918 | 24.1138081 | 19.09684 | 23.3360432 |
| ENSMUSG00 | Slc38a3 | 4246 | 0.09009576 | 0.15691425 | 0.20506913 | 0.15022483 | 0.13602993 | 0.38512487 | 0.28971494 | 0.55768463 | 0.29100223 | 0.44418939 | 0.46132202 | 0.48638236 | 0.5693038 | 0.01093832 | 0.37844105 | 0.26152323 | 0.48742818 | 0.01295551 |
| ENSMUSG00 | Cacna2d2 | 6399 | 0.02656989 | 0.08329525 | 0.00755955 | 0.05696021 | 0.05265252 | 0 | 0.04436257 | 0.07162195 | 0 | 0.00736845 | 0.02986402 | 0.05233536 | 0.20815155 | 0.04354814 | 0.01931625 | 0.02991921 | 0.02395768 | 0 |
| ENSMUSG00 | Rassf1 | 5579 | 4.78459411 | 4.63956243 | 3.81508395 | 3.61777068 | 4.63288098 | 4.05907855 | 2.56110935 | 5.21645647 | 3.25564679 | 2.81433665 | 3.1513147 | 3.27150415 | 3.66076604 | 5.05315667 | 3.19036963 | 5.20927962 | 4.08749716 | 4.49617363 |
| ENSMUSG00 | Epn3 | 5052 | 3.17190249 | 5.15650896 | 4.00240216 | 4.13945355 | 3.62990161 | 4.05092616 | 2.65033415 | 2.86518617 | 1.7283336 | 3.36924067 | 3.40439134 | 1.92239218 | 2.71462253 | 1.90299408 | 3.73521627 | 3.16814606 | 2.81454001 | 3.26657403 |
| ENSMUSG00 | Rnf112 | 4258 | 0 | 0 | 0 | 0 | 0 | 0 | 0 | 0 | 0 | 0 | 0 | 0 | 0 | 0 | 0 | 0 | 0 | 0 |
| ENSMUSG00 | Slc3a2 | 2948 | 97.3524249 | 104.198846 | 91.151615 | 102.805993 | 98.8924904 | 107.762067 | 66.0098536 | 73.5087148 | 76.7008045 | 80.3225345 | 79.5060876 | 85.9008349 | 73.0607703 | 85.0897273 | 81.3269517 | 92.8300445 | 92.3315334 | 104.793468 |
| ENSMUSG00 | Nxf1 | 2262 | 54.2683238 | 58.3638372 | 58.6812411 | 60.788343 | 58.8349995 | 54.8322246 | 53.8385657 | 68.3816223 | 68.4074888 | 62.7216915 | 61.6089982 | 60.3312533 | 54.2607472 | 64.2250687 | 65.5545533 | 65.4935279 | 59.0990821 | 57.5625798 |
| ENSMUSG00 | Stx5a | 4420 | 39.649059 | 33.6671264 | 32.7342148 | 35.5520457 | 33.1478678 | 31.5254043 | 32.6907062 | 36.3519088 | 33.9928528 | 31.03199 | 33.9288676 | 32.9842784 | 31.9429745 | 34.8015424 | 37.0627571 | 33.5692542 | 37.6672927 | 36.3035213 |
| ENSMUSG00 | Slc47a1 | 4491 | 0.13250333 | 0 | 0 | 0.01014497 | 0.06430451 | 0 | 0 | 0 | 0 | 0 | 0 | 0.01242834 | 0 | 0.26888138 | 0 | 0 | 0 | 0 |
| ENSMUSG00 | 1700027A23l | 1079 | 0 | 0.9879635 | 0.22415908 | 0.46447775 | 0.0446079 | 0 | 0.08769727 | 0 | 0 | 0 | 0.39849363 | 0 | 0 | 0 | 0 | 0.1419485 | 0.0710404 | 0 |
| ENSMUSG00 | Tnfrsf13b | 2885 | 0.13259847 | 0.24248554 | 0.36887961 | 0.1579239 | 0.05005053 | 0.51527973 | 0.36078993 | 0.23828882 | 0.59959572 | 0.35955476 | 0.48023355 | 0.1934685 | 0.23939232 | 0.04829541 | 1.04253504 | 0.75652153 | 0.53138713 | 0.6482757 |
| ENSMUSG00 | Spire2 | 2315 | 25.6499918 | 29.4419791 | 22.4210791 | 29.521195 | 23.535783 | 27.2486522 | 19.0068213 | 17.8506175 | 19.2856082 | 17.8215401 | 23.0310383 | 18.5891472 | 13.8086749 | 13.3413901 | 14.5406787 | 14.3899907 | 14.4861813 | 19.4135811 |
| ENSMUSG00 | Prox1 | 9269 | 0.29807276 | 0.09703832 | 0.00521885 | 0.00491542 | 0 | 0.13899785 | 0.05614839 | 0.12361334 | 0 | 0 | 0.01546282 | 0 | 0 | 0.07516035 | 0.04000583 | 0 | 0.03721401 | 0.13649902 |
| ENSMUSG00 | Raver1 | 3560 | 11.6889234 | 14.7568635 | 17.0530283 | 15.3576554 | 13.2092386 | 16.6196528 | 10.5523222 | 14.7083662 | 13.3393406 | 13.3770254 | 16.1978581 | 12.527167 | 12.5962676 | 9.78456897 | 12.0132626 | 10.5729402 | 11.0134275 | 10.7391374 |
| ENSMUSG00 | 2610507B11l | 8888 | 15.4325161 | 16.6264625 | 16.632483 | 16.2908415 | 15.173903 | 18.186433 | 15.8258995 | 18.5805522 | 15.8851683 | 17.0077807 | 15.2279993 | 15.7876435 | 15.2524875 | 16.172868 | 17.1147777 | 13.2345574 | 13.7988468 | 14.2040842 |
| ENSMUSG00 | AI597479 | 3454 | 9.6971855 | 10.6284914 | 10.041639 | 12.1619232 | 12.3325861 | 10.1716526 | 12.4514259 | 9.26613585 | 11.3638534 | 10.7024179 | 10.4014965 | 10.6007744 | 5.35595512 | 10.5554744 | 7.94448067 | 4.97755724 | 9.86452749 | 7.50124183 |
| ENSMUSG00 | Tmem86a | 1788 | 5.99066239 | 9.09210983 | 7.65643685 | 9.96329325 | 7.18748525 | 6.87308609 | 6.61530746 | 8.5014408 | 4.63002778 | 8.06941566 | 9.1381632 | 9.08409274 | 5.10425319 | 5.06521087 | 8.06518152 | 7.96650221 | 4.93011649 | 6.52233256 |
| ENSMUSG00 | Optc | 2971 | 0.11445352 | 0.2691045 | 0.47217515 | 0.1380173 | 0.11340406 | 0.05003642 | 0.74846715 | 0.2699564 | 0.34657115 | 0.50785005 | 0 | 0.39452336 | 0.76380619 | 0.56276915 | 0.74886705 | 0.46397235 | 0.10320107 | 0.55546052 |
| ENSMUSG00 | Tex14 | 4982 | 0.0511905 | 0 | 0 | 0 | 0 | 0.1491953 | 0.02849017 | 0 | 0.08267065 | 0 | 0.04794757 | 0.02240693 | 0 | 0.03728951 | 0 | 0.0768579 | 0.03846477 | 0.02208314 |
| ENSMUSG00 | Ifi35 | 1901 | 31.2807687 | 24.1128577 | 28.6017086 | 27.3462134 | 27.5473608 | 27.8652688 | 22.4741447 | 20.9947714 | 24.6339129 | 19.6192563 | 21.9649204 | 24.5753338 | 26.7549746 | 20.0093149 | 22.4105254 | 21.5321722 | 17.9232393 | 17.2753481 |
| ENSMUSG00 | Rdm1 | 1567 | 16.2288649 | 15.0513906 | 15.1881155 | 14.7121177 | 16.6173393 | 15.7480927 | 12.2282291 | 16.2080074 | 16.7689834 | 19.6486338 | 15.4879891 | 19.3057365 | 15.6779163 | 12.566875 | 11.5953326 | 16.1030697 | 15.7267447 | 15.6918128 |
| ENSMUSG00 | Nedd8 | 733 | 350.711193 | 286.909006 | 289.911213 | 282.751965 | 332.129973 | 287.108403 | 378.565978 | 321.691531 | 314.939183 | 310.628576 | 313.307265 | 300.018705 | 434.228554 | 417.299808 | 361.764275 | 371.622621 | 331.080644 | 356.546066 |
| ENSMUSG00 | Gosr1 | 4280 | 18.6407045 | 20.6882413 | 17.9818427 | 19.5337655 | 20.6584915 | 18.4549199 | 15.0007723 | 17.1330403 | 16.1859158 | 17.262886 | 17.4133026 | 13.7583 | 14.9839936 | 14.9749628 | 21.3227678 | 19.395803 | 19.0556915 | 19.7544386 |
| ENSMUSG00 | Mrpl52 | 407 | 130.335282 | 103.785626 | 120.517837 | 118.884101 | 123.936747 | 125.768977 | 127.058374 | 99.6568338 | 112.225666 | 109.01428 | 113.040012 | 119.722539 | 149.08663 | 137.734677 | 108.622284 | 119.011347 | 96.8045042 | 134.887195 |
| ENSMUSG00 | Spaca7 | 754 | 0 | 0 | 0 | 0 | 0 | 0 | 0 | 0 | 0 | 0 | 0 | 0 | 0 | 0.3695816 | 0 | 0 | 0 | 0 |
| ENSMUSG00 | 4632411B12l | 4797 | 16.6671328 | 19.7849564 | 17.0219969 | 20.619768 | 18.3417045 | 20.2156972 | 16.6782864 | 18.112928 | 17.4293573 | 19.1669525 | 19.7095357 | 17.4183963 | 20.2593031 | 21.5325517 | 20.6737847 | 18.6783451 | 18.5439515 | 17.9579448 |
| ENSMUSG00 | Eya4 | 4172 | 0 | 0 | 0 | 0 | 0 | 0 | 0 | 0 | 0 | 0 | 0 | 0 | 0 | 0 | 0 | 0 | 0 | 0 |
| ENSMUSG00 | Ebf3 | 6951 | 0 | 0 | 0 | 0 | 0 | 0 | 0 | 0 | 0 | 0 | 0 | 0 | 0 | 0 | 0 | 0 | 0 | 0 |
| ENSMUSG00 | Gm16119 | 1673 | 2.15955766 | 1.41375702 | 1.35896947 | 0.84422736 | 1.43849144 | 2.01409796 | 5.06214556 | 3.83522033 | 3.88970083 | 4.14294882 | 6.16819548 | 5.23793128 | 2.00512276 | 2.41520359 | 2.8813999 | 3.02113579 | 2.35959867 | 2.46603885 |
| ENSMUSG00 | Myt1 | 6879 | 0 | 0 | 0 | 0 | 0 | 0 | 0 | 0 | 0 | 0.0068543 | 0 | 0 | 0.00717138 | 0 | 0.00598947 | 0 | 0 | 0 |
| ENSMUSG00 | Faf1 | 5148 | 18.9572425 | 22.5709742 | 20.757018 | 21.5149376 | 22.9159567 | 22.3699598 | 19.7779497 | 20.7728684 | 20.0652256 | 19.5728541 | 22.3469503 | 18.9738558 | 18.5617568 | 21.3906238 | 22.2574839 | 19.9857929 | 22.7739204 | 24.6515169 |
| ENSMUSG00 | Gm266 | 1212 | 0.63126499 | 0.87954836 | 0.75833092 | 1.16534024 | 0.71483055 | 1.06301244 | 0.858811 | 1.10291591 | 0.40778728 | 1.28380628 | 0 | 0.78289296 | 0.28492031 | 0.68976367 | 0.30595215 | 0.34752201 | 0.50595771 | 0.13616116 |
| ENSMUSG00 | 1700021C14l | 1139 | 2.87348424 | 2.3982944 | 2.33585967 | 2.40005501 | 2.07064465 | 2.47981195 | 3.07387027 | 2.51486407 | 3.10978082 | 3.10474341 | 3.3975168 | 2.15618012 | 3.24837032 | 2.81355753 | 2.09805991 | 2.82389027 | 2.79287326 | 2.02843963 |
| ENSMUSG00 | Mettl16 | 13643 | 2.87251868 | 2.05595961 | 2.36850533 | 2.12059394 | 2.08502286 | 2.56789435 | 2.46568311 | 2.33191169 | 3.7162355 | 2.82358166 | 2.33919756 | 1.82465627 | 2.29610551 | 3.14892478 | 1.82104656 | 2.09934577 | 2.53392361 | 2.14907673 |

| | | | | | | | | | | | | | | | | | | | |
|---|---|---|---|---|---|---|---|---|---|---|---|---|---|---|---|---|---|---|---|
| ENSMUSG00 Pls3 | 5453 | 212.51029 | 226.622983 | 252.601555 | 239.352143 | 235.840247 | 246.745897 | 174.570163 | 182.914502 | 168.024066 | 174.638319 | 172.753829 | 184.847799 | 157.268669 | 152.116523 | 186.876597 | 178.153494 | 209.111312 | 183.578897 |
| ENSMUSG00 Mpped2 | 3413 | 30.0637254 | 28.413078 | 32.7828818 | 27.3526457 | 28.9806924 | 30.1701085 | 21.4175473 | 20.9146206 | 24.8349984 | 22.7533572 | 21.4308413 | 27.0656084 | 21.1174718 | 18.1666735 | 20.4378067 | 23.7282844 | 21.2125329 | 21.194543 |
| ENSMUSG00 Nkap | 4317 | 7.21711699 | 7.5854934 | 7.18263446 | 7.16607577 | 9.4546841 | 6.61162255 | 6.08258902 | 6.90064255 | 7.67059501 | 7.84206739 | 6.94988972 | 7.4731186 | 6.33075797 | 5.34693339 | 5.9172955 | 5.87175865 | 5.80620759 | 7.77288745 |
| ENSMUSG00 Ndufa1 | 416 | 391.231536 | 293.890197 | 377.918206 | 311.589354 | 372.906707 | 333.170663 | 517.709891 | 401.020225 | 442.755044 | 425.603103 | 407.580115 | 414.592301 | 666.09947 | 761.303442 | 515.910828 | 456.541857 | 493.819595 | 624.935188 |
| ENSMUSG00 Wt1 | 4845 | 0 | 0 | 0 | 0 | 0 | 0 | 0 | 0 | 0 | 0 | 0 | 0 | 0 | 0 | 0 | 0 | 0 | 0 |
| ENSMUSG00 E2f3 | 4786 | 1.80287312 | 2.14383022 | 1.46555827 | 1.33275098 | 1.39789749 | 1.93613898 | 1.04787797 | 1.17306376 | 1.41132231 | 1.06399414 | 1.4074946 | 2.04089865 | 1.96874137 | 1.81467705 | 0.91252924 | 2.17614818 | 2.09008849 | 1.19535042 |
| ENSMUSG00 Cr1l | 1776 | 41.1888558 | 43.9857753 | 42.5175005 | 44.9453545 | 45.0965771 | 45.6465496 | 46.5933983 | 47.9340456 | 46.9841672 | 46.0356528 | 47.04865 | 52.767172 | 49.2208094 | 53.3479382 | 53.4738662 | 58.3198722 | 55.1156319 | 55.5356579 |
| ENSMUSG00 Ppfibp1 | 7148 | 12.2675123 | 15.6777527 | 15.0710481 | 15.0489124 | 13.8375914 | 16.8179815 | 8.28038071 | 8.00929088 | 8.12436867 | 9.21509967 | 11.1082794 | 8.20680769 | 8.75110116 | 9.34989392 | 10.6981096 | 11.8653743 | 11.6083426 | 7.85734443 |
| ENSMUSG00 Cd46 | 3358 | 0.4050523 | 0.33729553 | 0.27370372 | 0.35276568 | 0.25800316 | 0.23610594 | 0.05635816 | 0.52318359 | 0.56419885 | 0.56165221 | 0.41258898 | 0.74797642 | 0.80799745 | 0.44258809 | 0.14723605 | 0.49032046 | 0.59349834 | 0.62249734 |
| ENSMUSG00 Cd34 | 2613 | 0.03253362 | 0.50995628 | 0.03702528 | 0.12205408 | 0.34998337 | 0.43617025 | 0.23538646 | 0 | 0 | 0 | 0 | 0 | 0 | 0.46213022 | 0 | 0 | 0.05867018 | 0 |
| ENSMUSG00 5033414D02l | 3045 | 27.1781864 | 24.1450107 | 24.7030672 | 25.1969782 | 24.7377539 | 23.9382828 | 24.0059405 | 25.4365163 | 30.7850607 | 23.412763 | 22.0753256 | 25.277438 | 29.0969207 | 26.6614991 | 25.5327869 | 26.8725872 | 25.777424 | 26.953559 |
| ENSMUSG00 Cd274 | 3615 | 4.08002658 | 6.46905996 | 3.89396883 | 6.71757583 | 4.85979312 | 7.29240782 | 5.94189661 | 4.6802861 | 5.69661352 | 4.60420422 | 7.04399803 | 4.90992551 | 6.93239667 | 5.64009803 | 5.08323483 | 5.75153522 | 4.67549016 | 4.21508231 |
| ENSMUSG00 Pdcd1lg2 | 1639 | 1.24481296 | 1.42275936 | 1.0329918 | 0.83394224 | 1.6738985 | 1.33027473 | 0.54846913 | 1.00199713 | 0.92977491 | 0.94934302 | 0.40808384 | 1.12380529 | 0.48158064 | 0.28336844 | 0.37707342 | 0.16353524 | 0.6313687 | 1.0404407 |
| ENSMUSG00 Glt8a | 2818 | 7.82831312 | 7.89674001 | 7.51866795 | 7.19470005 | 7.85687895 | 6.52379849 | 8.91519961 | 8.00983315 | 8.6231527 | 7.49592444 | 8.4258887 | 6.87297194 | 6.79232928 | 7.96043088 | 7.29580425 | 7.4325629 | 9.22116015 | 8.1596191 |
| ENSMUSG00 Mtif3 | 6417 | 4.06703726 | 4.43341307 | 4.45515918 | 4.23163198 | 4.26039156 | 4.4479312 | 4.93255138 | 4.88043789 | 4.78808489 | 6.23091731 | 6.32085706 | 5.3232329 | 5.75808104 | 6.13753874 | 6.14459856 | 5.34648344 | 5.78746793 | 5.44347558 |
| ENSMUSG00 Lnx2 | 5236 | 7.32232024 | 8.74177171 | 6.85507248 | 8.89293111 | 8.59498637 | 9.07583489 | 6.15353979 | 7.60784486 | 7.66937603 | 7.28512585 | 8.28488585 | 7.61122672 | 6.62344115 | 6.12927127 | 6.86168005 | 5.85767313 | 6.48531304 | 6.47165961 |
| ENSMUSG00 Il19 | 1459 | 0 | 0 | 0 | 0 | 0 | 0 | 0 | 0.02617711 | 0.16937566 | 0 | 0.032745 | 0.03825611 | 0 | 0 | 0.11295834 | 0 | 0 | 0 |
| ENSMUSG00 Dyrk3 | 2069 | 0.84229687 | 0.43472602 | 0.1870412 | 0.30829126 | 0.2791605 | 0.40715151 | 1.32630999 | 1.29215475 | 0.21897133 | 0.22789127 | 0.16163592 | 0.45861105 | 0.66761415 | 0.98769446 | 1.03551518 | 0 | 0.62981832 | 0 |
| ENSMUSG00 Mapkapk2 | 2581 | 65.6269378 | 95.6277541 | 68.9899129 | 104.626234 | 86.4360582 | 102.86848 | 61.3726653 | 73.6620609 | 52.9951255 | 85.6604601 | 69.765137 | 56.702312 | 57.0537672 | 57.4207992 | 63.7259245 | 74.4597282 | 78.4345216 | 72.2727163 |
| ENSMUSG00 Il10 | 1305 | 0 | 0 | 0 | 0 | 0 | 0 | 0.76135345 | 0.02926621 | 0 | 0 | 0 | 0.17108249 | 0 | 0.21353603 | 0 | 0.08802438 | 0 | 0 |
| ENSMUSG00 Lamp2 | 9540 | 77.8905123 | 79.9823734 | 79.0608056 | 79.540796 | 79.8263413 | 83.3410493 | 71.3558516 | 79.103111 | 74.4681562 | 80.1561955 | 72.4036006 | 75.4506064 | 78.2071192 | 64.5494921 | 73.3854128 | 66.6152213 | 80.6057464 | 73.0629341 |
| ENSMUSG00 Atxn10 | 2666 | 139.218003 | 129.25303 | 142.671069 | 125.865376 | 135.657642 | 137.468874 | 110.615127 | 113.4598 | 111.478581 | 108.131808 | 108.23734 | 101.058591 | 124.62508 | 112.852512 | 124.810115 | 125.787374 | 123.029426 | 114.351264 |
| ENSMUSG00 Foxred2 | 8797 | 0.72476713 | 0.31052146 | 1.13826541 | 1.00993562 | 0.37752674 | 0.45626593 | 3.07637289 | 1.99276034 | 2.72485531 | 2.19754329 | 1.81389521 | 2.36028492 | 3.93107582 | 5.40624593 | 4.18713095 | 1.4973251 | 2.09559495 | 3.26415297 |
| ENSMUSG00 Eif3d | 1895 | 35.2826606 | 38.0241637 | 33.5679116 | 32.8424203 | 37.8451537 | 35.7459561 | 28.5124429 | 36.761447 | 34.8835661 | 33.4658026 | 38.749399 | 32.2817764 | 32.7751366 | 28.5281888 | 34.1570682 | 32.7743352 | 33.2498322 | 33.6731207 |
| ENSMUSG00 H3f3b | 4546 | 457.328018 | 412.879102 | 470.742765 | 524.072277 | 490.435292 | 504.814386 | 415.344125 | 483.277553 | 460.009712 | 484.544524 | 501.185167 | 536.215889 | 357.900564 | 410.865688 | 466.576352 | 598.271789 | 533.153693 | 610.146104 |
| ENSMUSG00 Nup50 | 5088 | 9.36483929 | 10.0370807 | 7.94816654 | 9.70679482 | 9.68692803 | 9.86574668 | 7.56928472 | 9.03766755 | 9.55999904 | 9.98986991 | 10.6761257 | 8.40306555 | 8.73586035 | 11.8666104 | 9.28006742 | 12.1539566 | 10.9299247 | 8.49786908 |
| ENSMUSG00 Phf21b | 6107 | 0.0278403 | 0.07091332 | 0.007921 | 0 | 0.02364431 | 0.01622818 | 0.00774729 | 0.0687926 | 0.02023245 | 0 | 0.06258387 | 0.22849051 | 0 | 0.02281517 | 0 | 0.13166903 | 0.05020638 | 0.0540453 |
| ENSMUSG00 Nlrp14 | 3634 | 0 | 0 | 0 | 0 | 0 | 0 | 0 | 0 | 0 | 0 | 0 | 0 | 0 | 0 | 0 | 0 | 0 | 0 |
| ENSMUSG00 Ift27 | 1005 | 39.8406725 | 40.3069446 | 48.0847332 | 39.1688978 | 40.5649148 | 40.4803923 | 38.6975344 | 33.6321156 | 45.8174374 | 44.6641488 | 44.542424 | 34.3224671 | 45.6995174 | 40.2053294 | 43.210388 | 47.2060324 | 38.8983312 | 44.1167562 |
| ENSMUSG00 Pacsin2 | 5187 | 12.3082272 | 10.6739985 | 10.5942415 | 11.4890777 | 9.55771764 | 10.9480302 | 10.0335351 | 9.73401889 | 11.2752752 | 9.3083324 | 8.79604499 | 11.7829479 | 7.6085471 | 5.39922589 | 7.2203999 | 6.50357145 | 6.58352237 | 3.23458214 |
| ENSMUSG00 Cmah | 9953 | 0.12811768 | 0.06359339 | 0.16524666 | 0.1419062 | 0.09671842 | 0.06970142 | 0.35176713 | 0.07674551 | 0.04138101 | 0.02368668 | 0.22560264 | 0.02243169 | 0.2726068 | 0.21465166 | 0.02069806 | 0.01923571 | 0 | 0.01105377 |
| ENSMUSG00 Ttll12 | 5401 | 4.35990882 | 5.89035623 | 5.37384155 | 5.66876908 | 4.5271278 | 5.48649041 | 5.1333512 | 5.26109006 | 4.4152921 | 5.29910699 | 4.89160378 | 5.90089675 | 4.66739673 | 4.7467388 | 4.20330629 | 3.93469489 | 4.63378483 | 4.48139362 |
| ENSMUSG00 Bik | 938 | 155.927837 | 144.580802 | 139.602502 | 164.078047 | 168.820926 | 159.329472 | 161.60962 | 157.818499 | 153.681029 | 179.403902 | 162.271874 | 183.1563 | 143.893524 | 124.329535 | 126.196237 | 174.430454 | 147.176249 | 143.797793 |
| ENSMUSG00 Scube1 | 10802 | 0.428908 | 0.15728176 | 0.20599726 | 0.29946622 | 0.31636425 | 0.26606726 | 0.06569989 | 0.14142715 | 0.22495875 | 0.07420496 | 0.21671682 | 0.13951332 | 0 | 0 | 0 | 0 | 0.11708645 | 0.19860722 |
| ENSMUSG00 Tox4 | 6057 | 14.9613728 | 18.2101689 | 16.1884012 | 18.9480387 | 17.8001501 | 18.1619727 | 16.0599088 | 17.8508653 | 14.5720164 | 16.8300843 | 17.8258884 | 16.025002 | 15.3118713 | 14.6302326 | 17.7812229 | 18.5795105 | 21.2860596 | 19.2718053 |
| ENSMUSG00 Mrps18c | 1194 | 68.2076359 | 52.5082638 | 59.919976 | 54.8335182 | 66.0705384 | 54.9894238 | 64.1534566 | 62.1826049 | 62.6076433 | 60.932609 | 63.5398945 | 52.8706151 | 72.0146833 | 68.965969 | 63.1481133 | 70.0711702 | 70.6500666 | 66.9876392 |
| ENSMUSG00 Sulf1 | 4586 | 0 | 0 | 0.07383661 | 0.06954368 | 0.05247702 | 0 | 0.19601851 | 0 | 0 | 0 | 0 | 0 | 0 | 0.59751443 | 0.03593681 | 0.10854293 | 0.24235992 | 0 |
| ENSMUSG00 Snf8 | 4003 | 60.4289175 | 55.3578247 | 57.1588305 | 60.8011738 | 49.8874223 | 55.8412144 | 56.4490019 | 57.2075056 | 56.4346837 | 56.0555076 | 48.6223285 | 59.6919439 | 45.9059185 | 58.6613305 | 56.9905109 | 49.6065676 | 53.1761811 | 50.9002576 |
| ENSMUSG00 Plcg1 | 7278 | 10.3664039 | 11.1546473 | 10.1293295 | 10.3667339 | 8.99414897 | 10.6826379 | 8.0804699 | 7.17353862 | 8.2961572 | 9.36146829 | 8.44825111 | 8.3899883 | 9.37428436 | 8.3277733 | 9.20496426 | 8.35469697 | 10.1476744 | 9.47807362 |
| ENSMUSG00 Kctd2 | 2620 | 19.7275931 | 20.4200413 | 17.5954101 | 18.3983149 | 18.8869792 | 19.9913127 | 19.9002184 | 26.0641281 | 23.1870652 | 21.0198555 | 22.9757434 | 24.307508 | 15.9481427 | 18.4535477 | 19.6100535 | 21.5128909 | 16.3545038 | 18.0354285 |
| ENSMUSG00 Tmprss6 | 3173 | 0 | 0 | 0 | 0.02871796 | 0 | 0.031234 | 0 | 0 | 0 | 0 | 0 | 0 | 0 | 0.04391184 | 0 | 0 | 0 | 0 |
| ENSMUSG00 Kctd5 | 2744 | 18.975525 | 16.8628317 | 17.5406934 | 16.0891589 | 18.2775016 | 17.8960492 | 16.5525407 | 13.7514918 | 16.7808034 | 16.9426361 | 12.1874894 | 16.4558568 | 13.2139064 | 13.1851108 | 12.2073123 | 10.0750068 | 14.7355114 | 17.3607458 |
| ENSMUSG00 Pom121l2 | 4900 | 0 | 0 | 0 | 0 | 0 | 0 | 0 | 0 | 0 | 0 | 0 | 0 | 0 | 0 | 0 | 0 | 0 | 0 |
| ENSMUSG00 Etaa1 | 4179 | 2.81740458 | 4.84667138 | 2.89384604 | 2.84552107 | 3.75472771 | 3.28454348 | 5.26451201 | 4.67009276 | 5.1741854 | 2.86582403 | 4.61858895 | 3.91337381 | 5.22949015 | 4.48992852 | 4.90001753 | 3.57342352 | 4.06282592 | 2.79060316 |
| ENSMUSG00 Matn4 | 5221 | 0 | 0 | 0 | 0 | 0 | 0 | 0 | 0.0146303 | 0 | 0 | 0 | 0.19243094 | 0 | 0.16901698 | 0.00789151 | 0 | 0 | 0.05268062 |
| ENSMUSG00 Svs4 | 728 | 0 | 0 | 0 | 0 | 0 | 0 | 0 | 0 | 0 | 0 | 0 | 0 | 2.03290868 | 0.89315553 | 1.69786631 | 0.10519398 | 0.4211681 | 0 |
| ENSMUSG00 Svs6 | 531 | 0 | 0.06273615 | 0 | 0 | 0 | 0 | 0 | 0 | 0 | 0 | 0 | 0.10511424 | 0.2787114 | 0 | 0 | 0 | 0 | 0 |
| ENSMUSG00 Slpi | 1325 | 0 | 0 | 0 | 0 | 0 | 0.44877947 | 0 | 0 | 0 | 0 | 0 | 0 | 0 | 0 | 0.09328654 | 0 | 0.11570203 | 0 |
| ENSMUSG00 Svs3a | 1288 | 0 | 0 | 0 | 0 | 0 | 0 | 0 | 0 | 0 | 0 | 0 | 0 | 0 | 0 | 0 | 0.05945746 | 0 | 0 |
| ENSMUSG00 Svs5 | 643 | 0 | 0.31085127 | 0 | 0 | 0 | 0 | 0 | 0.11879441 | 0.19216104 | 0 | 0 | 0.43402537 | 0.30688596 | 1.22791522 | 0.32038535 | 0 | 0.17881632 | 0 |
| ENSMUSG00 Rbpjl | 4128 | 0.03089039 | 0 | 0 | 0 | 0 | 0 | 0 | 0.00925204 | 0 | 0 | 0.02314678 | 0 | 0.13145617 | 0.3375297 | 0 | 0.27827476 | 0.11141365 | 0 |
| ENSMUSG00 Sdc4 | 2878 | 118.698436 | 190.015452 | 155.810503 | 142.303067 | 174.532577 | 164.378022 | 98.0118804 | 165.933912 | 153.898468 | 124.167887 | 158.530526 | 167.621533 | 79.5858554 | 98.487931 | 102.488512 | 83.4862965 | 149.762868 | 122.346523 |
| ENSMUSG00 Ccdc70 | 1112 | 0 | 0 | 0 | 0 | 0 | 0 | 0 | 0 | 0 | 0 | 0 | 0 | 0 | 0 | 0 | 0 | 0 | 0 |
| ENSMUSG00 Il13ra1 | 3815 | 50.2262997 | 50.1658132 | 53.4834996 | 55.2703835 | 51.2229646 | 58.359222 | 37.1804486 | 40.5950699 | 39.4375733 | 35.4835306 | 41.5009731 | 39.707394 | 29.1853571 | 27.8299302 | 36.6224052 | 40.3180524 | 41.9127944 | 44.9589501 |
| ENSMUSG00 Prl5a1 | 1056 | 0 | 0 | 0 | 0 | 0 | 0 | 0 | 0 | 0.31201906 | 0 | 0 | 0 | 0 | 0 | 0 | 0 | 0 | 0 |
| ENSMUSG00 Nbr1 | 7212 | 19.8263189 | 24.0700904 | 21.6446731 | 27.5817415 | 25.0336724 | 25.8069977 | 19.5758505 | 23.8411253 | 18.7486594 | 27.4130475 | 24.8347642 | 21.2288356 | 19.467365 | 17.8512216 | 24.4684226 | 21.1893808 | 23.7652801 | 22.8060495 |
| ENSMUSG00 Cyth1 | 3950 | 45.4428755 | 43.1633905 | 39.0417253 | 41.1548876 | 43.5136961 | 44.7103652 | 32.6517374 | 42.5917804 | 36.5465159 | 38.950062 | 35.8735511 | 39.0568333 | 35.868675 | 33.6513869 | 32.5127214 | 30.9233007 | 33.4651641 | 30.150561 |
| ENSMUSG00 Rnd3 | 5138 | 0.49636251 | 1.31617702 | 0.69669935 | 0.75373468 | 1.48948538 | 1.53345361 | 0.42358568 | 0.83996525 | 0.2404818 | 0.64238015 | 0.31614414 | 0.71697815 | 0.64329281 | 0.79546121 | 0.73774682 | 1.01353106 | 1.32031034 | 0.9207441 |
| ENSMUSG00 Brca1 | 8453 | 0.08045461 | 0.3783317 | 0.00572265 | 0.09701867 | 0.02847032 | 0.09379436 | 0.2406773 | 0.02710924 | 0.02436207 | 0.02788992 | 0.19216238 | 0.12545813 | 0.16924472 | 0.39010176 | 0.33144514 | 0.10871579 | 0.04080642 | 0.24729044 |
| ENSMUSG00 Gast | 457 | 0 | 0 | 0 | 0 | 0 | 0 | 0 | 0 | 0 | 0 | 0 | 0 | 0 | 0.3048846 | 0 | 0 | 0.50318989 | 0 |
| ENSMUSG00 Cntnap1 | 5850 | 0.0871901 | 0.00569451 | 0 | 0.02336464 | 0.05759375 | 0.12705829 | 0 | 0.00652862 | 0.09856602 | 0.06447959 | 0.08983326 | 0.19082278 | 0.16022332 | 0.06351328 | 0.14790302 | 0.09818104 | 0.05896353 | 0.00940327 |
| ENSMUSG00 Nt5c3l | 3406 | 18.7816471 | 20.4122167 | 22.141613 | 21.0816811 | 20.77332 | 20.2662824 | 21.2254179 | 20.6099928 | 19.1058999 | 23.1323621 | 22.232328 | 17.6001238 | 21.7402254 | 23.9311137 | 23.0322262 | 21.6185667 | 20.4797006 | 23.0792758 |
| ENSMUSG00 Ccdc56 | 944 | 244.945082 | 207.464518 | 220.396773 | 209.223652 | 234.286217 | 218.472963 | 309.587307 | 241.129998 | 260.993366 | 275.661792 | 280.627272 | 250.1062 | 271.377803 | 231.38405 | 232.849895 | 218.791882 | 211.647231 | 220.735695 |
| ENSMUSG00 Zpbp2 | 1747 | 0 | 0 | 0.02768949 | 0.07823877 | 0 | 0 | 0.1895757 | 0.17489366 | 0.37720907 | 0 | 0 | 0 | 0.08471423 | 0 | 0.02358418 | 0 | 0 | 0 |
| ENSMUSG00 Gsdma | 2714 | 0.32889046 | 0.89603584 | 0.85553775 | 0.889733 | 0.69165255 | 0.9311678 | 0.10459693 | 0.09850656 | 0 | 0.22585083 | 0.01760315 | 0 | 0 | 0.15401503 | 0 | 0.12697696 | 0.32479913 | 0.32429834 |
| ENSMUSG00 Med24 | 6749 | 8.77310864 | 10.0397728 | 9.85532734 | 10.4839683 | 11.0969437 | 10.7930832 | 8.18805882 | 7.79240452 | 7.49400524 | 10.1301704 | 9.51393454 | 9.83328238 | 10.8546299 | 9.70309131 | 9.42587084 | 8.12448508 | 8.29674319 | 10.3024909 |
| ENSMUSG00 Gsdma2 | 3163 | 0 | 0 | 0 | 0 | 0 | 0 | 0 | 0 | 0 | 0 | 0 | 0 | 0 | 0 | 0 | 0 | 0 | 0 |
| ENSMUSG00 Psmd3 | 3525 | 70.1667005 | 85.4133157 | 76.3821475 | 75.4956198 | 77.1203134 | 76.2057784 | 54.8558635 | 52.7651063 | 51.7840107 | 55.0693462 | 65.5567303 | 60.6926056 | 44.937528 | 50.9106252 | 52.4691587 | 48.6643748 | 52.0367821 | 52.4654232 |
| ENSMUSG00 Exosc10 | 3692 | 24.5567553 | 25.3004755 | 22.6800057 | 23.3481842 | 24.3658086 | 23.4207801 | 21.8878807 | 19.0341334 | 24.6204334 | 24.1628198 | 23.6545569 | 22.4199966 | 21.2186036 | 17.4354023 | 21.024846 | 23.3249123 | 23.0144638 | 19.6524951 |
| ENSMUSG00 Glod4 | 5898 | 29.4826509 | 32.1437235 | 33.5530879 | 32.6760287 | 32.9366639 | 32.4050345 | 31.6861787 | 30.9139588 | 33.8053447 | 33.99999 | 30.5782415 | 31.6932272 | 31.006005 | 26.9339525 | 29.4097237 | 31.2791025 | 31.8606025 | 29.0621188 |

[illegible]

| | | | | | | | | | | | | | | | | | | | | |
|---|---|---|---|---|---|---|---|---|---|---|---|---|---|---|---|---|---|---|---|---|
| ENSMUSG00 | Katna1 | 1889 | 31.8845074 | 29.2744332 | 30.6783956 | 28.3158634 | 28.35936 | 30.0621666 | 29.1790741 | 29.3772158 | 30.4592708 | 29.2289431 | 26.7580233 | 29.252253 | 28.9619356 | 35.1834246 | 34.4836738 | 34.2973572 | 35.9321714 | 40.0119178 |
| ENSMUSG00 | Pcmt1 | 6301 | 23.7519005 | 23.2413082 | 24.397886 | 22.8058298 | 25.5440649 | 24.9139914 | 26.9038769 | 24.227112 | 25.0805683 | 25.330127 | 25.7640547 | 23.367392 | 25.7659641 | 23.0046019 | 26.057483 | 26.4406337 | 25.5163967 | 23.9120685 |
| ENSMUSG00 | Lrp11 | 4207 | 0.40413764 | 0.25339021 | 0.20697018 | 0.44402254 | 0.1716137 | 0.25913005 | 0.32613922 | 0.21787917 | 0.63634978 | 0.56038392 | 0.39746222 | 0.50415859 | 1.33678121 | 0.88317732 | 0.33298144 | 0.73723299 | 0.96567328 | 0.61455384 |
| ENSMUSG00 | 1700021F05f | 1608 | 23.7109098 | 22.5400677 | 23.8859348 | 18.0204131 | 21.5516076 | 23.5745281 | 28.2757988 | 20.3075274 | 28.2260823 | 29.5278347 | 25.3433082 | 21.8333655 | 24.6659586 | 21.5757048 | 22.9580809 | 25.8604475 | 22.2616675 | 21.8257525 |
| ENSMUSG00 | Sec63 | 8030 | 14.6200867 | 13.6943791 | 13.8313357 | 12.828583 | 13.0669481 | 14.1746736 | 10.51131 | 12.4470133 | 13.3048346 | 12.0842027 | 10.899592 | 13.512534 | 14.1360824 | 16.7383797 | 11.6575261 | 14.8632408 | 16.1466827 | 12.8377417 |
| ENSMUSG00 | Nr2e1 | 3953 | 0 | 0 | 0 | 0 | 0 | 0 | 0 | 0 | 0 | 0 | 0 | 0 | 0 | 0 | 0 | 0 | 0 | 0 |
| ENSMUSG00 | Snx3 | 1804 | 147.849213 | 138.145101 | 138.845976 | 137.693356 | 144.689258 | 134.045089 | 152.349973 | 148.281367 | 153.033933 | 139.152075 | 136.54528 | 129.267094 | 153.683689 | 136.963717 | 128.195307 | 132.870955 | 138.051151 | 134.534467 |
| ENSMUSG00 | Afg1 | 9222 | 14.3020561 | 13.3620034 | 14.0945214 | 13.0329684 | 14.0815365 | 13.9759987 | 14.0162914 | 13.6212114 | 12.2773511 | 12.6287396 | 14.671295 | 13.5393231 | 12.4479908 | 11.9006049 | 13.8544811 | 12.0825024 | 12.6590653 | 14.280178 |
| ENSMUSG00 | Adat2 | 1460 | 2.50372955 | 2.14480278 | 2.21988119 | 1.74754691 | 3.62637781 | 1.96853325 | 3.14337661 | 1.41259571 | 0.95913804 | 1.96999514 | 2.25785763 | 2.33202426 | 2.23007297 | 5.47149523 | 2.17295877 | 3.27830543 | 2.83509594 | 1.09264664 |
| ENSMUSG00 | Pex3 | 4633 | 12.3029227 | 11.137853 | 13.2601734 | 12.4597115 | 11.3966588 | 11.9255985 | 12.2034646 | 10.4116133 | 10.4899824 | 11.3373372 | 10.6934237 | 12.4329298 | 9.98777 | 11.3278262 | 12.6103618 | 13.8186695 | 15.287502 | 10.7691042 |
| ENSMUSG00 | Fuca2 | 2704 | 103.716402 | 99.4828479 | 109.895931 | 109.538591 | 105.697989 | 124.010114 | 84.581912 | 83.7718708 | 73.7518895 | 81.7464855 | 80.9206027 | 79.677684 | 83.1746342 | 92.4244209 | 86.2733763 | 98.2896102 | 85.6955588 | 86.2161961 |
| ENSMUSG00 | Cep57l1 | 2966 | 1.84867421 | 1.49380139 | 1.30474795 | 1.84333288 | 1.80129586 | 1.15277769 | 2.79154377 | 2.24055227 | 1.80520804 | 1.70098626 | 1.72350659 | 1.9759422 | 1.23080306 | 3.00649521 | 2.08369298 | 1.89774757 | 2.0933446 | 3.44966077 |
| ENSMUSG00 | Ltv1 | 1755 | 116.011278 | 111.973084 | 111.245338 | 113.032927 | 127.775859 | 117.006569 | 144.31041 | 139.472995 | 146.347073 | 153.515167 | 164.639857 | 140.286546 | 110.329217 | 109.560411 | 127.478318 | 124.406293 | 145.290515 | 123.684294 |
| ENSMUSG00 | Fam164b | 1151 | 0 | 0 | 0 | 0 | 0 | 0.0430519 | 0 | 0 | 0 | 0 | 0 | 0 | 0 | 0 | 0 | 0 | 0 | 0 |
| ENSMUSG00 | Plagl1 | 5750 | 66.1897993 | 75.1423023 | 68.5221573 | 70.0451534 | 69.6866547 | 76.0871452 | 69.125881 | 69.8887753 | 68.0544498 | 70.734255 | 77.254185 | 72.8612806 | 69.8797333 | 60.5711672 | 71.7122037 | 76.2349345 | 89.3302663 | 82.2840569 |
| ENSMUSG00 | Cd164 | 2881 | 219.932197 | 231.802704 | 252.328155 | 248.174061 | 243.496339 | 245.906085 | 238.287737 | 245.367295 | 244.888933 | 255.916538 | 249.255606 | 259.549612 | 162.019646 | 154.921097 | 186.343726 | 166.400002 | 205.293792 | 179.118511 |
| ENSMUSG00 | Utrn | 13154 | 3.39645465 | 3.05169816 | 2.55952388 | 2.93372393 | 2.67480889 | 2.74999966 | 3.15801521 | 4.00680518 | 2.83990952 | 4.40178376 | 4.42374178 | 4.11594897 | 2.58773171 | 2.77873626 | 3.13224539 | 3.91813574 | 3.32740087 | 4.03556241 |
| ENSMUSG00 | Smpd2 | 4245 | 12.9468063 | 11.8498163 | 11.8284392 | 11.3231806 | 10.9870045 | 13.9494739 | 11.4129996 | 9.83375638 | 11.8950927 | 10.0632596 | 12.4473743 | 13.9243314 | 10.3893366 | 11.7614591 | 11.2782586 | 11.6630755 | 10.8162317 | 10.5612301 |
| ENSMUSG00 | Mical1 | 5962 | 0 | 0.01117507 | 0.02434092 | 0.02292572 | 0 | 0.11635999 | 0.15077841 | 0.16014929 | 0 | 0 | 0.24039731 | 0 | 0.41371953 | 0.12464029 | 0 | 0.01926733 | 0 | 0 |
| ENSMUSG00 | Zbtb24 | 2858 | 11.8978799 | 10.1523901 | 10.1892461 | 12.0358952 | 9.885738 | 12.8476472 | 10.2306633 | 8.940069 | 12.1484369 | 9.68420672 | 11.4840436 | 7.90949732 | 7.30139993 | 10.2378499 | 8.26049389 | 9.63294147 | 12.6726209 | 6.54412046 |
| ENSMUSG00 | Grm1 | 8849 | 0 | 0 | 0 | 0 | 0 | 0 | 0 | 0 | 0 | 0.00532837 | 0 | 0.00630757 | 0 | 0 | 0 | 0 | 0 | 0 |
| ENSMUSG00 | Wasf1 | 2844 | 1.2703727 | 2.09669763 | 2.16014007 | 1.69812613 | 1.32007386 | 2.10825625 | 1.81331997 | 1.36976971 | 1.88264664 | 2.83501064 | 1.24308964 | 1.47193203 | 2.13355338 | 2.40059101 | 2.04269317 | 1.91183764 | 1.75190527 | 2.92066636 |
| ENSMUSG00 | Rab32 | 2105 | 0.88846924 | 1.23439703 | 1.26391647 | 1.08221008 | 1.44052788 | 1.5772129 | 2.2476332 | 0.94347027 | 0.93916996 | 0.31359138 | 0.4312229 | 1.14017743 | 1.99202516 | 0.90461168 | 2.0551845 | 1.49160561 | 2.29411563 | 1.30662961 |
| ENSMUSG00 | Slc22a16 | 2407 | 0 | 0 | 0 | 0 | 0 | 0 | 0 | 0 | 0 | 0 | 0 | 0 | 0 | 0 | 0 | 0 | 0 | 0 |
| ENSMUSG00 | Amd-ps4 | 1679 | 0 | 0 | 0 | 0 | 0 | 0.05902648 | 0 | 0 | 0 | 0 | 0 | 0 | 0 | 0 | 0 | 0 | 0.02282686 | 0 |
| ENSMUSG00 | Glt8d6 | 1014 | 43.1758691 | 43.8258391 | 37.4966419 | 40.4837287 | 43.3377183 | 43.9817162 | 50.9521158 | 43.0888643 | 39.0337689 | 40.9197425 | 34.2528548 | 39.3021529 | 39.5044545 | 39.1614346 | 41.5266175 | 44.3702798 | 41.0854879 | 34.2315053 |
| ENSMUSG00 | Slc16a10 | 6139 | 9.04939975 | 7.81949512 | 8.58888224 | 7.80007454 | 7.98931914 | 9.60543296 | 5.98826381 | 5.14499378 | 6.5479784 | 5.56069542 | 4.80162068 | 6.57350123 | 9.90816973 | 9.72912468 | 9.81884285 | 13.6908101 | 10.1949573 | 11.8100672 |
| ENSMUSG00 | Rev3l | 13681 | 5.25683487 | 4.49739899 | 3.88940015 | 5.25845253 | 3.86293781 | 4.41160949 | 4.01505886 | 3.74358684 | 4.50670376 | 3.30168655 | 3.73301883 | 3.76157013 | 4.3594977 | 2.98401819 | 3.22240076 | 3.94633115 | 3.53819991 | 3.79165175 |
| ENSMUSG00 | Traf3ip2 | 3677 | 10.6118443 | 13.7437206 | 12.4979205 | 14.0511896 | 12.9067383 | 12.6678188 | 11.6576794 | 11.5709373 | 11.4363428 | 10.1944001 | 12.473201 | 12.7357468 | 9.52560818 | 11.1657801 | 11.5301607 | 8.1642198 | 10.2460565 | 11.0706388 |
| ENSMUSG00 | Fyn | 4343 | 12.9482726 | 11.9275962 | 12.697634 | 12.4734243 | 12.7228755 | 11.6722188 | 11.6021191 | 13.3141371 | 11.1050909 | 13.0280552 | 14.5205953 | 12.1193115 | 18.0380117 | 18.415086 | 17.7309942 | 21.7065336 | 19.8294242 | 20.8611556 |
| ENSMUSG00 | Tube1 | 2917 | 1.18029457 | 0.61669396 | 0.84574907 | 0.93714867 | 0.57751708 | 0.84937836 | 0.25951418 | 1.13909462 | 0.39534537 | 1.05066702 | 2.32569219 | 1.79865351 | 1.97868849 | 1.60810861 | 1.10172141 | 0.68258882 | 1.30075536 | 1.88581099 |
| ENSMUSG00 | Lama4 | 11964 | 0 | 0 | 0.01212977 | 0 | 0 | 0 | 0 | 0 | 0 | 0 | 0 | 0 | 0 | 0 | 0 | 0 | 0 | 0 |
| ENSMUSG00 | Popdc3 | 2077 | 0 | 0.08019474 | 0 | 0 | 0.23173772 | 0 | 0.34169003 | 0.58842411 | 0.7733626 | 0.02270135 | 0.09200762 | 0.29560534 | 1.16382472 | 2.8622259 | 0.55543744 | 0.38714625 | 0.46131797 | 0.84751633 |
| ENSMUSG00 | Prep | 2692 | 30.947305 | 31.6299248 | 28.1759642 | 32.4783225 | 34.7937309 | 37.5326983 | 27.8568336 | 27.0127542 | 28.5949472 | 27.9891619 | 28.6969197 | 26.0625141 | 25.5272514 | 27.1038861 | 26.7687672 | 32.1743514 | 29.9264027 | 26.2376274 |
| ENSMUSG00 | Tnfaip3 | 8636 | 7.10716468 | 11.6764857 | 12.9167857 | 15.6635874 | 16.1851675 | 14.3218646 | 16.7862492 | 21.8160168 | 21.0081753 | 29.7995065 | 26.4322315 | 28.2891565 | 20.7986928 | 24.5988022 | 26.0348414 | 50.2643246 | 48.2274049 | 42.9193332 |
| ENSMUSG00 | Perp | 1898 | 176.716725 | 192.891837 | 185.134523 | 184.644654 | 195.672246 | 193.328763 | 199.520907 | 200.399439 | 217.520355 | 225.021639 | 206.756327 | 214.822663 | 200.082771 | 201.975499 | 233.967529 | 210.658863 | 227.675973 | 234.353866 |
| ENSMUSG00 | D10Bwg1379 | 8182 | 0.19740854 | 0.43157746 | 0.28378507 | 0.21160104 | 0.51767408 | 0.2301398 | 0.56090563 | 0.47612137 | 0.61915686 | 0.48983253 | 0.20436611 | 0.402484 | 0.95263297 | 0.88551425 | 0.61938296 | 1.31504042 | 1.0445813 | 0.72610407 |
| ENSMUSG00 | Hebp2 | 1906 | 40.6096147 | 33.6100187 | 37.8409936 | 39.3938096 | 36.3388448 | 36.0076264 | 46.9651566 | 45.0253558 | 49.8300658 | 48.4370813 | 42.3357318 | 48.6117523 | 64.1366689 | 52.6577507 | 62.6021542 | 59.6055783 | 47.7570594 | 52.729086 |
| ENSMUSG00 | Reps1 | 5490 | 31.0698126 | 33.6709016 | 32.1168521 | 30.2246491 | 32.0178115 | 30.4537201 | 36.0920764 | 37.9976142 | 36.4226838 | 38.5536442 | 38.1242428 | 39.0912973 | 33.2204186 | 45.0397671 | 38.9125806 | 36.1494022 | 45.7262741 | 37.9552817 |
| ENSMUSG00 | Fam184a | 4682 | 0.07262738 | 0.47671164 | 0.26862764 | 0.1362355 | 0.21588432 | 0.16933869 | 0.04042091 | 0.08973602 | 0.01759356 | 0.28197772 | 0.12244757 | 0.10729296 | 0.04214602 | 0.52574469 | 0 | 0 | 0.32743526 | 0 |
| ENSMUSG00 | Aif1a | 2176 | 27.3861474 | 25.59704 | 26.387583 | 26.2562268 | 26.9636098 | 26.2338001 | 27.6787869 | 33.1901806 | 36.4546691 | 28.2124153 | 27.6638087 | 31.9349724 | 34.4824661 | 36.412506 | 33.2111806 | 36.3373641 | 35.8252242 | 36.5800448 |
| ENSMUSG00 | Gopc | 4198 | 9.96309986 | 10.3001755 | 10.0019591 | 11.6127483 | 9.08062972 | 10.1041305 | 11.1688577 | 9.99843979 | 10.9883034 | 9.44586504 | 7.48835906 | 8.93476066 | 9.49504268 | 11.0633843 | 8.61719058 | 11.5108972 | 10.489975 | 10.5877461 |
| ENSMUSG00 | Qrsl1 | 3240 | 13.7879444 | 14.5281233 | 12.7652372 | 12.8949005 | 13.7859337 | 15.1564071 | 12.3830715 | 12.9665564 | 15.0000273 | 12.0787295 | 12.9611693 | 10.0606009 | 13.0942743 | 11.1953186 | 12.2205849 | 11.2390024 | 10.7171682 | 10.3226966 |
| ENSMUSG00 | Rtn4ip1 | 3125 | 73.0408945 | 63.8434955 | 67.1501999 | 59.0179544 | 68.5552615 | 62.3492315 | 72.6419947 | 57.9669007 | 48.2771863 | 58.9044306 | 49.5483649 | 48.1354274 | 36.2925049 | 29.5458706 | 37.602188 | 26.6135038 | 29.7044132 | 41.4020541 |
| ENSMUSG00 | Nmbr | 1265 | 0 | 0 | 0 | 0 | 0 | 0 | 0 | 0.09057487 | 0 | 0 | 0 | 0 | 0 | 0 | 0 | 0 | 0 | 0 |
| ENSMUSG00 | Aim1 | 7523 | 10.9384582 | 11.9471206 | 10.9632951 | 11.2464255 | 11.3116099 | 12.8113873 | 8.33301869 | 8.39694717 | 6.76131157 | 8.11019673 | 9.8814084 | 6.52901543 | 5.31811577 | 6.17361257 | 7.84817621 | 6.16884437 | 6.83688561 | 6.89533267 |
| ENSMUSG00 | Gje1 | 1477 | 0 | 0 | 0 | 0 | 0 | 0 | 0 | 0 | 0 | 0 | 0 | 0 | 0 | 0 | 0 | 0 | 0 | 0 |
| ENSMUSG00 | Vta1 | 2768 | 27.2260394 | 24.3107104 | 27.000399 | 23.8668765 | 27.2480941 | 26.5128607 | 25.6220035 | 23.0147858 | 24.9827168 | 27.1695707 | 23.0762715 | 23.330463 | 29.1749814 | 25.9402309 | 24.7536804 | 30.9036188 | 30.8908845 | 27.1070887 |
| ENSMUSG00 | Smpdl3a | 2092 | 28.6686441 | 34.6982775 | 32.5573283 | 33.8441027 | 29.449743 | 35.6722834 | 32.2504207 | 37.6447102 | 33.0555258 | 35.9264919 | 32.6568789 | 38.47332 | 35.8434581 | 30.0154905 | 33.363019 | 50.572155 | 40.744591 | 39.2847062 |
| ENSMUSG00 | Reep3 | 3015 | 52.8530365 | 58.1401159 | 60.0698168 | 59.5089196 | 54.4058361 | 59.1510077 | 31.0082433 | 40.9919118 | 42.5798904 | 35.45295 | 34.876512 | 44.874682 | 34.4750745 | 34.2438495 | 39.3977067 | 34.0487955 | 43.5000292 | 36.946415 |
| ENSMUSG00 | Fabp7 | 1404 | 0 | 0 | 0 | 0 | 0 | 0 | 0.13479396 | 0 | 0 | 0 | 0.03402775 | 0 | 0 | 0 | 0 | 0 | 0 | 0 |

[illegible]

| | | | | | | | | | | | | | | | | | | | | |
|---|---|---|---|---|---|---|---|---|---|---|---|---|---|---|---|---|---|---|---|---|
| ENSMUSG00 | Acaca | 10128 | 0.78060454 | 0.64797 | 0.65911801 | 0.72426226 | 0.54176928 | 0.60668829 | 0.78947894 | 1.08981085 | 0.82551962 | 0.48882542 | 0.73587021 | 0.57314661 | 1.35409489 | 1.39864205 | 1.0495657 | 1.20981382 | 1.01416346 | 1.06455222 |
| ENSMUSG00 | Shmt1 | 4275 | 4.8719383 | 5.06511842 | 6.29138781 | 5.01970804 | 5.9897505 | 5.26244235 | 6.99452938 | 6.0839827 | 7.66888106 | 5.73529022 | 5.59888987 | 7.82072091 | 2.55025818 | 3.59602174 | 2.85278609 | 2.33774235 | 2.95852116 | 2.12315851 |
| ENSMUSG00 | Llgl1 | 4937 | 18.7515239 | 20.4317285 | 21.467775 | 22.0745429 | 21.4872918 | 22.262095 | 13.3207664 | 13.476031 | 17.0852713 | 16.9520961 | 12.0864742 | 16.1443723 | 11.3512372 | 10.9313408 | 13.3193606 | 12.6808068 | 13.717352 | 13.6046424 |
| ENSMUSG00 | Drg2 | 2822 | 28.9794326 | 27.8709225 | 24.0667741 | 23.4263201 | 27.6306578 | 22.9677447 | 21.6947578 | 24.8074676 | 28.7517491 | 27.7524161 | 28.1198462 | 27.1760171 | 32.6548623 | 32.504278 | 31.6384733 | 32.5917927 | 31.2233014 | 27.8359335 |
| ENSMUSG00 | Srebf1 | 5159 | 33.3845655 | 37.1355795 | 33.8023991 | 36.394081 | 33.6149101 | 40.8697192 | 24.6605531 | 28.0576091 | 24.948219 | 29.2555536 | 27.1981107 | 24.8298014 | 31.163543 | 41.7537657 | 43.030429 | 34.6463569 | 36.3278914 | 35.1230854 |
| ENSMUSG00 | Tom1l1 | 7978 | 4.57125107 | 4.28833572 | 4.1534054 | 3.90050054 | 4.223157 | 3.54036828 | 4.66721964 | 3.59519857 | 4.07838399 | 4.23162491 | 5.81467594 | 3.51209108 | 4.29134502 | 4.27299576 | 4.57049097 | 5.87941766 | 5.95695765 | 4.06810892 |
| ENSMUSG00 | Myocd | 9063 | 0 | 0 | 0 | 0 | 0 | 0 | 0 | 0 | 0 | 0 | 0 | 0 | 0 | 0 | 0 | 0 | 0 | 0 |
| ENSMUSG00 | Cox11 | 1419 | 7.27889907 | 5.79865037 | 4.84076202 | 6.03627649 | 6.20728825 | 5.55242046 | 6.43505781 | 7.72460852 | 5.31158026 | 4.78484938 | 6.26225671 | 5.74283773 | 6.22298322 | 6.74241156 | 6.01037496 | 5.36985967 | 5.48290223 | 6.47393997 |
| ENSMUSG00 | 1700108M19 | 1956 | 0 | 0.03406226 | 0.14838506 | 0 | 0 | 0 | 0 | 0 | 0 | 0 | 0.02442483 | 0 | 0.07566245 | 0.09497768 | 0 | 0.09787988 | 0 | 0 |
| ENSMUSG00 | Stxbp4 | 11814 | 2.10115311 | 1.59599612 | 1.73610774 | 1.6891601 | 1.80484528 | 1.38415457 | 1.70203898 | 1.31252035 | 1.69431578 | 1.7041942 | 1.42750638 | 2.33392054 | 1.69116527 | 1.38774021 | 1.00440563 | 1.03715883 | 1.21655336 | 1.63434878 |
| ENSMUSG00 | Bzw2 | 1848 | 33.1669177 | 28.6079887 | 24.7626405 | 30.3493753 | 29.2750445 | 27.2969063 | 21.3777527 | 23.0849099 | 20.5041096 | 21.2790511 | 26.7053745 | 25.2801459 | 22.396904 | 15.2803058 | 22.4512807 | 23.0199485 | 19.4742385 | 18.7531046 |
| ENSMUSG00 | 1700086D15 | 1089 | 0 | 0 | 0.13326041 | 0 | 0.04419828 | 0.22751485 | 0 | 0 | 0 | 0 | 0 | 0 | 0 | 0.21324191 | 0 | 0.10548377 | 0 | 0 |
| ENSMUSG00 | Elac2 | 3715 | 9.16464226 | 9.81004207 | 8.34654992 | 8.46221281 | 8.48624765 | 8.16319595 | 7.88332389 | 10.6506942 | 6.49671554 | 10.1662755 | 9.68359185 | 11.4185474 | 9.72031318 | 8.55121286 | 10.3807958 | 11.4614147 | 9.33655416 | 10.7649046 |
| ENSMUSG00 | Pctp | 2717 | 8.18189439 | 5.93428075 | 8.95542354 | 7.86460427 | 5.70426184 | 7.34992699 | 10.8312427 | 10.3598824 | 9.12561004 | 11.0371171 | 10.0051348 | 11.0932859 | 13.1999646 | 11.5041851 | 8.70434048 | 9.75233726 | 10.142292 | 8.98934243 |
| ENSMUSG00 | Twistnb | 2320 | 10.4614032 | 6.90668195 | 7.11007494 | 5.91115273 | 7.36501418 | 6.49311681 | 8.46325938 | 10.4864484 | 7.42067396 | 7.47907707 | 7.86639403 | 6.78449002 | 7.08083123 | 7.80741124 | 7.3701059 | 9.17654212 | 10.7544911 | 8.44105258 |
| ENSMUSG00 | Efcab10 | 671 | 0 | 0.19858655 | 0.1441834 | 0.47530151 | 0 | 0 | 0 | 0 | 0 | 0 | 0.21359892 | 0 | 0.29407998 | 0.5537298 | 0 | 0.39945492 | 0.79965448 | 0.16396157 |
| ENSMUSG00 | Atxn7l1 | 12068 | 2.8564549 | 2.95642275 | 3.39913438 | 3.42425151 | 2.88360796 | 2.65666379 | 3.36771555 | 3.06032923 | 3.3275483 | 3.32883652 | 2.99286283 | 2.70105626 | 3.64225541 | 2.68242699 | 2.53327432 | 3.75037272 | 2.77887884 | 3.42781306 |
| ENSMUSG00 | Atp6v1c2 | 2189 | 154.156506 | 148.04378 | 143.728388 | 141.345387 | 151.212078 | 145.602185 | 167.528814 | 165.244515 | 187.568924 | 181.386968 | 193.718836 | 174.178525 | 240.867762 | 246.372199 | 226.053306 | 288.832576 | 243.807074 | 261.550837 |
| ENSMUSG00 | Sypl | 6631 | 33.8003006 | 37.5529911 | 37.5914345 | 39.2741561 | 39.6901461 | 36.1613147 | 36.7955791 | 36.9022353 | 39.335576 | 37.9921893 | 38.1925886 | 35.7570404 | 30.9561315 | 31.5323905 | 30.6075802 | 30.3911125 | 35.5924199 | 30.3873259 |
| ENSMUSG00 | Pdia6 | 3433 | 135.352923 | 136.114176 | 122.124494 | 137.598866 | 134.427289 | 138.337722 | 161.232459 | 167.654968 | 154.212489 | 163.688343 | 150.686644 | 162.683227 | 180.155621 | 212.170878 | 170.879042 | 164.260348 | 169.258662 | 187.764845 |
| ENSMUSG00 | Nampt | 4536 | 25.0383224 | 27.2833779 | 29.2096781 | 25.3317799 | 28.1512331 | 27.3435829 | 30.3944326 | 31.827002 | 37.7452262 | 32.5876223 | 31.1653664 | 28.8184043 | 34.4322862 | 31.2698948 | 37.5864282 | 34.0360955 | 38.0305693 | 34.501964 |
| ENSMUSG00 | Pik3cg | 6801 | 0 | 0 | 0 | 0 | 0 | 0 | 0 | 0 | 0 | 0 | 0 | 0 | 0.13781891 | 0.30047638 | 0.01211632 | 0 | 0 | 0.11323739 |
| ENSMUSG00 | Nbas | 8166 | 1.37415705 | 2.46807507 | 2.26880506 | 1.88024393 | 2.48145235 | 2.83990685 | 1.73236481 | 1.70242891 | 1.88128464 | 1.53589114 | 1.56207616 | 2.07787919 | 1.51632516 | 1.89962346 | 1.4228311 | 1.13005589 | 1.26721775 | 1.50894439 |
| ENSMUSG00 | Tspan13 | 2466 | 73.2378317 | 102.653967 | 85.350053 | 72.73878 | 91.2867019 | 94.6043267 | 60.0329977 | 69.957454 | 72.8864375 | 48.0111176 | 77.7844519 | 65.9104661 | 59.3343336 | 78.7063427 | 69.0535403 | 67.963499 | 69.1769455 | 69.5978966 |
| ENSMUSG00 | Rock2 | 8161 | 6.51562005 | 6.91484411 | 5.96889414 | 7.86055023 | 5.6559876 | 7.24984241 | 5.21187334 | 6.91684484 | 5.45048855 | 5.85845033 | 5.75453795 | 5.34150625 | 6.2866308 | 6.88039476 | 6.76511272 | 6.49828345 | 7.84277856 | 5.55416051 |
| ENSMUSG00 | Agr2 | 1283 | 1.78899416 | 1.45403124 | 3.54412459 | 1.56249879 | 1.01290876 | 1.58352461 | 0.73753202 | 0.92280941 | 0 | 0 | 0 | 0 | 0 | 0.0361996 | 0 | 0.02984459 | 0.08961722 | 0 |
| ENSMUSG00 | Matn3 | 3006 | 0 | 0 | 0 | 0 | 0 | 0 | 0 | 0 | 0 | 0 | 0 | 0 | 0 | 0 | 0.02741288 | 0 | 0.01274993 | 0 |
| ENSMUSG00 | Laptm4a | 2780 | 559.12654 | 485.589357 | 513.298839 | 487.437619 | 506.804917 | 490.162094 | 432.723804 | 430.296488 | 461.866773 | 431.632463 | 424.217572 | 430.423084 | 508.278456 | 416.944277 | 448.133613 | 496.235253 | 471.027432 | 450.619564 |
| ENSMUSG00 | Fam49a | 4833 | 0.13192171 | 0.0068928 | 0.05004503 | 0.1036978 | 0.19918032 | 0.06151798 | 0.45031724 | 0 | 0.32383356 | 0 | 0.21747343 | 0 | 0.02041462 | 0 | 0.06820038 | 0 | 0.01586025 | 0 |
| ENSMUSG00 | Snx13 | 7618 | 12.9390264 | 15.6987779 | 14.5920678 | 14.365664 | 13.375595 | 16.3592971 | 10.5767252 | 11.9670864 | 12.9755366 | 13.5609334 | 12.5489225 | 13.6938138 | 11.2094446 | 10.8824752 | 12.8504721 | 13.1438617 | 13.0756164 | 11.8134548 |
| ENSMUSG00 | Ntsr2 | 2192 | 8.64840717 | 13.0698399 | 14.8077732 | 7.23323149 | 13.7237465 | 8.56773998 | 1.14396532 | 2.82261368 | 0.20688416 | 0.06453107 | 0.10897573 | 0 | 0.04501087 | 0.04237599 | 0 | 0.03493669 | 0 | 0.35133554 |
| ENSMUSG00 | Sdc1 | 3526 | 54.5840718 | 67.7878661 | 51.9542143 | 57.6684461 | 59.5438035 | 60.5988052 | 16.9874792 | 24.4362053 | 24.763305 | 25.4742172 | 26.3941064 | 24.3144804 | 21.4620423 | 19.2573045 | 23.0429575 | 18.0702129 | 22.1849041 | 19.3140312 |
| ENSMUSG00 | Lpin1 | 5703 | 7.17736008 | 8.16612767 | 6.63301784 | 9.75452133 | 7.2835087 | 7.05537781 | 6.13082057 | 7.80188476 | 6.73081408 | 8.66455147 | 6.55931834 | 5.53948186 | 9.11727877 | 10.0413045 | 11.8410223 | 9.379621 | 9.84534893 | 10.7838298 |
| ENSMUSG00 | Pum2 | 7257 | 6.14998413 | 5.9170898 | 6.13251315 | 5.21092281 | 5.25955842 | 5.13485678 | 5.33954582 | 5.09968829 | 5.91946203 | 5.14583946 | 5.32588417 | 5.85306865 | 5.03719866 | 4.01913833 | 4.25811863 | 5.77762369 | 4.40987432 | 5.54108544 |
| ENSMUSG00 | Nrcam | 8372 | 0 | 0.06764443 | 0 | 0 | 0 | 0.01775659 | 0 | 0 | 0 | 0 | 0.01711955 | 0.02000083 | 0.08838733 | 0.01664265 | 0 | 0 | 0 | 0 |
| ENSMUSG00 | Rgs9 | 6784 | 25.719892 | 30.3174836 | 30.4901556 | 30.262097 | 29.1884926 | 29.0567179 | 19.9809589 | 24.8779815 | 19.391175 | 21.7543781 | 22.3240888 | 23.3991369 | 14.4272588 | 14.8150066 | 18.1714409 | 13.6816675 | 20.6941665 | 21.2122381 |
| ENSMUSG00 | Slc7a15 | 3211 | 0.02647473 | 0.08299693 | 0.01506494 | 0 | 0.0299794 | 0.03086436 | 0.01473456 | 0.45198109 | 0.02565339 | 0.02936824 | 0 | 0 | 8.44987683 | 20.6257454 | 7.67316425 | 1.3355802 | 4.72663145 | 7.50357793 |
| ENSMUSG00 | Trib2 | 4102 | 17.5222459 | 15.3327022 | 15.4483971 | 18.5598501 | 16.004862 | 15.9216243 | 22.9066261 | 26.1164526 | 26.8385072 | 29.5410306 | 32.0867895 | 23.1862236 | 22.573381 | 27.2980728 | 23.2323742 | 20.7601928 | 28.9923206 | 22.2208898 |
| ENSMUSG00 | Ang | 3891 | 7.37368127 | 6.57525129 | 8.11819977 | 7.99747964 | 9.75998144 | 9.19482746 | 7.79425011 | 10.924735 | 12.8396925 | 8.39769657 | 9.61392765 | 6.97157852 | 8.50725173 | 8.06892909 | 8.90529645 | 7.62663088 | 9.7416365 | 9.38731608 |
| ENSMUSG00 | Hs1bp3 | 2951 | 8.05164127 | 7.75532305 | 8.21251735 | 7.7041549 | 7.22549718 | 8.32875495 | 7.87208585 | 5.6427948 | 6.15494864 | 6.37517133 | 6.58909119 | 4.97442198 | 6.33574946 | 6.35832304 | 4.56554008 | 6.72129266 | 5.74050262 | 4.80933566 |
| ENSMUSG00 | Fam84a | 3377 | 23.4741348 | 23.6554102 | 17.6333479 | 22.2610966 | 17.2744719 | 19.1490427 | 2.74601275 | 3.01965397 | 2.14652851 | 2.77849865 | 1.88157225 | 2.28088879 | 5.39042863 | 4.4559918 | 6.23452622 | 2.89135472 | 3.14373238 | 4.43070093 |

[illegible]

| | | | | | | | | | | | | | | | | | | | |
|---|---|---|---|---|---|---|---|---|---|---|---|---|---|---|---|---|---|---|---|
| ENSMUSG00 1700020O03 | 4018 | 4.79214653 | 5.15694878 | 4.18964374 | 6.05515124 | 5.17496043 | 5.24139168 | 4.46279376 | 5.32298291 | 3.31091204 | 5.32763048 | 5.51706838 | 4.47303219 | 5.04614684 | 4.98193172 | 5.73212288 | 5.74645004 | 5.97119509 | 6.47568628 |
| ENSMUSG00 Esrrb | 6037 | 0.02816311 | 0.06069933 | 0.00801284 | 0.12829845 | 0.03986411 | 0.19699612 | 0.14106812 | 0.0856943 | 0.34111741 | 0.23430861 | 0.3956846 | 0.03698238 | 0.44126611 | 0.11539859 | 0.10919743 | 0.02537062 | 0.0571371 | 0.10904393 |
| ENSMUSG00 Vash1 | 5074 | 0.1675411 | 0.05252329 | 0.06673526 | 0.01795863 | 0.16126186 | 0.07812808 | 0.01864907 | 0 | 0.05682018 | 0 | 0 | 0 | 0.05833494 | 0 | 0.04060067 | 0 | 0 | 0 |
| ENSMUSG00 Angel1 | 4695 | 1.43041913 | 1.9157575 | 1.90609225 | 1.85349509 | 1.81455814 | 1.72036114 | 1.6627459 | 2.13129061 | 1.29831828 | 1.84786569 | 3.04253729 | 1.78324801 | 1.37646031 | 0.99911668 | 1.59716361 | 2.33250558 | 1.19999268 | 0.87873972 |
| ENSMUSG00 Ccnk | 2626 | 1.6024419 | 2.0043554 | 1.98946736 | 2.0472975 | 2.30945254 | 1.75491402 | 1.67558231 | 2.38520716 | 2.24283007 | 2.76512121 | 1.23712764 | 1.31781077 | 2.57367581 | 2.49376098 | 2.41623747 | 2.34759626 | 2.30599959 | 2.95364968 |
| ENSMUSG00 Cyp46a1 | 2104 | 0 | 0.20583062 | 0.06897366 | 0.3897808 | 0.27451667 | 0.02355168 | 0.11243507 | 0.14521826 | 0.11745204 | 0.08964012 | 0.15894711 | 0.26528357 | 0.44548785 | 0.68429977 | 0.09791244 | 0 | 0.05464776 | 0.26145013 |
| ENSMUSG00 Hhipl1 | 3093 | 0 | 0.01077042 | 0.03127936 | 0.01473037 | 0.06224626 | 0 | 0.03059339 | 0 | 0 | 0 | 0 | 0.10827481 | 0 | 0.06006348 | 0 | 0 | 0 | 0 |
| ENSMUSG00 Evl | 2678 | 8.60261591 | 9.87693727 | 8.54394319 | 12.6747491 | 10.4962821 | 9.3443355 | 7.93255894 | 11.2523546 | 7.55137388 | 10.2118775 | 8.88421558 | 8.60786731 | 7.95794931 | 10.6658378 | 9.56959214 | 10.0087473 | 8.4008724 | 12.653327 |
| ENSMUSG00 Degs2 | 3570 | 3.27420824 | 2.78074019 | 3.14360197 | 2.32271991 | 2.7773603 | 3.03978954 | 3.31321281 | 2.93129363 | 3.17263077 | 3.61884949 | 2.89058572 | 4.62785326 | 3.7448038 | 2.6929765 | 3.03529669 | 2.87447697 | 3.14554762 | 4.71506629 |
| ENSMUSG00 Yy1 | 3289 | 10.5842928 | 9.01443474 | 9.38349418 | 9.44744063 | 9.90736161 | 9.08491893 | 9.55172357 | 8.65106105 | 10.806921 | 9.66238188 | 10.7054255 | 11.9301948 | 12.1792389 | 10.6190191 | 11.6126003 | 14.1334136 | 9.75345409 | 8.73054233 |
| ENSMUSG00 Slc25a29 | 2866 | 8.42391486 | 8.20617701 | 7.9328539 | 8.36186648 | 5.2397628 | 10.0108279 | 11.4897504 | 9.54143759 | 13.0055118 | 9.78878871 | 10.9685704 | 13.5352008 | 18.7791772 | 21.9904489 | 16.0723446 | 13.8679869 | 11.5941729 | 14.4144589 |
| ENSMUSG00 Wars | 3891 | 52.0090319 | 57.4050259 | 50.0767361 | 53.1721157 | 56.1229858 | 53.3605638 | 60.9921351 | 60.4835733 | 56.7465718 | 55.1969811 | 62.1037625 | 59.7174514 | 79.3798854 | 95.6215842 | 74.8320215 | 73.0680442 | 75.9630948 | 80.9090511 |
| ENSMUSG00 Meg3 | 16216 | 0.00262119 | 0 | 0 | 0 | 0 | 0 | 0 | 0 | 0 | 0 | 0 | 0 | 0 | 0 | 0 | 0 | 0 | 0 |
| ENSMUSG00 Hsp90aa1 | 4440 | 57.6117453 | 52.2427213 | 48.1665265 | 48.1469414 | 50.12658 | 49.7871046 | 43.3379877 | 46.7856929 | 51.993338 | 44.8037876 | 45.4184909 | 50.1460987 | 62.2982117 | 70.5238822 | 67.2309172 | 63.5417107 | 63.6699923 | 78.3260299 |
| ENSMUSG00 Zfp839 | 2821 | 4.56542656 | 5.00697169 | 4.47553755 | 4.50603735 | 5.23803636 | 5.23456739 | 6.50738016 | 4.06158126 | 3.54779274 | 6.11739007 | 6.72338125 | 4.76837103 | 4.80903128 | 5.30131022 | 3.73895722 | 3.5833819 | 5.21704999 | 5.55746027 |
| ENSMUSG00 Fdft1 | 2852 | 54.3982793 | 48.6027184 | 47.779886 | 54.9224651 | 49.2795291 | 54.7304035 | 42.4520145 | 40.4555567 | 42.1540812 | 44.7204251 | 44.1064045 | 36.9299282 | 52.929757 | 61.4097663 | 51.0541019 | 59.3155342 | 57.1131703 | 60.2361291 |
| ENSMUSG00 Tecpr2 | 7809 | 1.0396323 | 1.46749079 | 0.99113393 | 1.27773962 | 1.41147529 | 0.93914772 | 0.52711014 | 1.34008429 | 1.00210501 | 1.5879927 | 0.96663381 | 1.20794557 | 1.44034795 | 0.80291354 | 0.60148257 | 0.94635513 | 1.5607328 | 1.63428259 |
| ENSMUSG00 Cirip | 3139 | 15.4908956 | 13.4036908 | 14.5320927 | 13.7307257 | 13.0794938 | 11.5238915 | 14.8765894 | 13.298597 | 15.1808852 | 13.5639009 | 11.6735881 | 13.8338915 | 14.7885741 | 12.517266 | 12.9550607 | 13.6011555 | 13.5527842 | 12.3547054 |
| ENSMUSG00 Traf3 | 7546 | 8.22953382 | 8.63945587 | 8.72467199 | 9.3706256 | 9.32532102 | 10.2966719 | 5.49870387 | 5.87108227 | 6.78436776 | 5.77980394 | 4.90032042 | 5.98394792 | 5.00771914 | 4.41914322 | 5.68937467 | 6.2109334 | 5.00791529 | 5.45279775 |
| ENSMUSG00 Amn | 1673 | 0.07621968 | 0.29868106 | 0.23131395 | 0.51742967 | 0.28769829 | 0.38504814 | 0 | 0 | 0 | 0 | 0 | 0.1668131 | 0.0884613 | 0.1388048 | 0.1477641 | 0.06866218 | 0 | 0 |
| ENSMUSG00 Cdc42bpb | 6681 | 4.5361758 | 5.05602077 | 5.69824403 | 4.18716977 | 4.35139678 | 4.24250316 | 4.02947378 | 4.00731538 | 3.40909193 | 4.57321594 | 3.18928775 | 3.95997965 | 2.70251186 | 3.47583351 | 3.18832576 | 3.10634773 | 4.03283741 | 3.54047536 |
| ENSMUSG00 1200009I06R | 4578 | 17.8451176 | 16.60551 | 17.8785525 | 22.6710483 | 19.0719331 | 19.1261862 | 21.8787552 | 22.525008 | 20.3233594 | 26.4797855 | 22.3429852 | 18.6052209 | 18.9655252 | 21.6495593 | 24.9027316 | 24.53175 | 22.9639651 | 23.2268683 |
| ENSMUSG00 Tnfaip2 | 4997 | 0 | 0.10666526 | 0.25169337 | 0.11852983 | 0 | 0.38674337 | 0.03787287 | 0.14521826 | 0.09890698 | 0 | 0.01912146 | 0 | 0 | 0.03717758 | 0.00824526 | 0.00766272 | 0.00766986 | 0 |
| ENSMUSG00 Eif5 | 4117 | 77.9588497 | 71.0275894 | 73.0714076 | 74.1349128 | 74.6003898 | 69.9780406 | 83.1103459 | 86.6170661 | 80.3022011 | 81.966865 | 84.2008984 | 80.2867024 | 87.0647828 | 99.0363476 | 89.2583622 | 86.2631488 | 97.1516236 | 94.3051434 |
| ENSMUSG00 Ppp1r13b | 3826 | 12.7537747 | 13.0865865 | 11.4928226 | 11.9320874 | 14.5175745 | 12.8738675 | 11.9703798 | 17.888339 | 16.1581182 | 13.568459 | 12.1622607 | 16.7767935 | 11.1789786 | 19.3132627 | 14.1179403 | 10.2181678 | 19.0529578 | 16.6637623 |
| ENSMUSG00 Zfyve21 | 1300 | 75.0706801 | 67.7276765 | 74.2719743 | 68.6219422 | 76.2705867 | 72.4613435 | 133.931276 | 122.950157 | 136.549141 | 114.503669 | 120.833895 | 120.13248 | 125.986127 | 117.432089 | 122.558782 | 104.356632 | 93.8699456 | 118.608097 |
| ENSMUSG00 Xrcc3 | 4182 | 1.77867188 | 1.7604375 | 1.78133039 | 1.57971101 | 1.77243334 | 2.38165936 | 1.36892256 | 1.83564631 | 2.51137291 | 2.32258366 | 2.36475763 | 1.721717 | 1.38016124 | 1.81023105 | 1.96057141 | 1.77627399 | 1.83291712 | 1.85468294 |
| ENSMUSG00 Klc1 | 4138 | 18.9824952 | 17.1233751 | 16.0621631 | 17.5505811 | 15.7958319 | 19.423522 | 14.6008436 | 14.7490235 | 14.3724816 | 15.6105518 | 15.4477754 | 14.2304311 | 14.5802162 | 15.9714684 | 18.4102651 | 19.3396253 | 12.5778422 | 10.9539642 |
| ENSMUSG00 2010107E04f | 361 | 463.671422 | 352.876752 | 412.715993 | 380.26434 | 471.1862 | 379.812781 | 586.886915 | 472.062326 | 450.998052 | 502.070092 | 519.039854 | 402.769532 | 612.754341 | 534.171332 | 460.863941 | 458.532675 | 476.477619 | 472.072611 |
| ENSMUSG00 Kif26a | 6874 | 0 | 0.00484622 | 0 | 0 | 0 | 0.02883488 | 0 | 0 | 0 | 0 | 0.0069501 | 0 | 0 | 0 | 0 | 0 | 0.01115109 | 0 |
| ENSMUSG00 Gpr132 | 2436 | 0 | 0 | 0 | 0.05610966 | 0.01975859 | 0 | 0.27191195 | 0.26653154 | 0.5072231 | 0 | 0.45107719 | 0.11456417 | 0.16200968 | 0.07626287 | 0.30444499 | 0 | 0 | 0 |
| ENSMUSG00 Hecw1 | 9482 | 0.08965445 | 0.01756639 | 0 | 0 | 0 | 0.03658185 | 0 | 0 | 0 | 0.03978123 | 0 | 0 | 0 | 0 | 0 | 0 | 0.03637805 | 0 |
| ENSMUSG00 Ggps1 | 2386 | 32.796324 | 31.4559726 | 35.1549462 | 36.2426078 | 33.3857224 | 34.2673975 | 33.1743973 | 37.3760518 | 32.9008797 | 33.3374838 | 33.9390422 | 33.1946459 | 27.4985121 | 28.1272868 | 33.4309355 | 33.0268525 | 36.270234 | 29.3489492 |
| ENSMUSG00 Gng4 | 3224 | 80.1322766 | 93.4083634 | 81.517801 | 94.6267844 | 95.2934456 | 89.9449742 | 150.317547 | 140.212554 | 105.585159 | 139.726371 | 139.916612 | 103.563677 | 85.9330172 | 86.7801434 | 91.3105201 | 64.2531594 | 84.0230355 | 77.5826328 |
| ENSMUSG00 Gpr137b | 4203 | 0.75847923 | 0.91148771 | 0.73659432 | 0.79132911 | 0.81307794 | 0.81349955 | 0.63038544 | 0.71786814 | 0.53896226 | 1.02086938 | 1.39812512 | 0.38511878 | 0.91551024 | 1.23762498 | 0.99009211 | 1.05679526 | 1.50460123 | 1.11248491 |
| ENSMUSG00 Mtr | 4271 | 1.04496452 | 1.91874783 | 1.81216691 | 2.48553578 | 1.65661269 | 1.72871864 | 2.42600718 | 2.22662332 | 2.28546107 | 1.74427795 | 1.83448682 | 1.973347 | 1.61706121 | 1.46802899 | 1.25408622 | 2.30407073 | 2.36903357 | 2.47289827 |
| ENSMUSG00 Ryr2 | 18086 | 0 | 0 | 0 | 0.00503827 | 0.01064512 | 0 | 0.01307992 | 0 | 0 | 0.04171244 | 0.00792463 | 0 | 0 | 0 | 0 | 0 | 0.01059557 | 0 |
| ENSMUSG00 Amph | 3172 | 0 | 0.01050217 | 0 | 0 | 0 | 0.10935345 | 0.04474717 | 0 | 0 | 0 | 0.33135217 | 0.07038545 | 0 | 0 | 0 | 0.37421464 | 0 | 0 |
| ENSMUSG00 Gli3 | 9730 | 0.02184233 | 0.04793222 | 0.04971586 | 0.03746026 | 0.07914808 | 0.13750502 | 0.02431278 | 0.02747655 | 0.03809647 | 0.12599366 | 0.02455034 | 0.00573645 | 0.10140168 | 0.04295959 | 0.00423449 | 0 | 0.1339254 | 0 |
| ENSMUSG00 Sfrp4 | 4573 | 0 | 0 | 0 | 0 | 0 | 0 | 0 | 0 | 0 | 0 | 0 | 0 | 0 | 0 | 0 | 0 | 0 | 0 |
| ENSMUSG00 Aoah | 3541 | 0 | 0 | 0 | 0 | 0 | 0 | 0 | 0 | 0.08141926 | 0 | 0 | 0 | 0 | 0 | 0 | 0 | 0 | 0 |
| [illegible] | | | | | | | | | | | | | | | | | | | |

| | | | | | | | | | | | | | | | | | | | | |
|---|---|---|---|---|---|---|---|---|---|---|---|---|---|---|---|---|---|---|---|---|
| ENSMUSG00 | 1190002H23 | 913 | 2.18810873 | 1.2405678 | 1.21861029 | 1.99610271 | 0.10543685 | 1.4654149 | 3.62747812 | 2.63540126 | 1.30822449 | 6.4038195 | 1.30818616 | 4.09600153 | 4.59278529 | 2.8995761 | 2.34663845 | 2.34860243 | 0.75561155 | 3.19329973 |
| ENSMUSG00 | Tdrd3 | 6027 | 5.02840393 | 4.8253122 | 5.45777346 | 4.30134963 | 5.39856984 | 4.52198498 | 5.19678006 | 5.77923859 | 5.17992018 | 5.79702524 | 4.20914268 | 4.51007594 | 5.03386911 | 5.6639131 | 4.76480578 | 4.79029666 | 6.2509955 | 4.65482735 |
| ENSMUSG00 | Naa16 | 6807 | 2.32913628 | 1.52690216 | 1.99690936 | 1.94773967 | 1.81016196 | 2.03830842 | 2.2241894 | 1.91326711 | 1.45214688 | 1.63472396 | 2.58982807 | 2.21393109 | 1.75383047 | 1.42600474 | 1.31951509 | 1.52442406 | 1.73417063 | 2.44053918 |
| ENSMUSG00 | Diap3 | 5069 | 0.11739445 | 0.04600321 | 0 | 0 | 0 | 0.01955129 | 0.18667461 | 0.03767252 | 0 | 0 | 0.02827478 | 0 | 0 | 0.1191109 | 0.01625629 | 0.0679849 | 0.08317011 | 0 |
| ENSMUSG00 | Mtrf1 | 1851 | 7.73865171 | 6.80295733 | 7.73558346 | 7.16275738 | 5.82471686 | 5.1935334 | 6.13454508 | 6.60268437 | 8.01034342 | 7.25983275 | 7.40757055 | 4.79453828 | 5.41025349 | 8.98270302 | 9.43784965 | 4.92337362 | 8.13734991 | 5.94371764 |
| ENSMUSG00 | Wbp4 | 3442 | 21.5366145 | 23.5087217 | 20.8700443 | 19.8551907 | 20.653937 | 23.8693876 | 21.9931105 | 23.1018541 | 27.6531197 | 21.3561727 | 22.4161993 | 23.3997679 | 22.8170867 | 22.76327 | 25.1374978 | 24.9856782 | 23.7952634 | 25.3789836 |
| ENSMUSG00 | Sugt1 | 1938 | 51.6510266 | 43.5921051 | 40.4610385 | 44.1269764 | 46.4430842 | 46.1520558 | 48.5577494 | 42.3506051 | 45.2881657 | 44.8150647 | 38.8017465 | 37.6424515 | 46.124579 | 46.8035617 | 44.0291136 | 48.5449032 | 49.6185134 | 44.3933142 |
| ENSMUSG00 | Lect1 | 2183 | 0 | 0 | 0.0443184 | 0 | 0 | 0 | 0 | 0.01749537 | 0 | 0 | 0 | 0 | 0 | 0 | 0 | 0 | 0 | 0 |
| ENSMUSG00 | Olfm4 | 2563 | 0 | 0 | 0 | 0 | 0.11267715 | 0 | 0 | 0 | 0 | 0 | 0 | 0 | 0.34645904 | 0 | 0 | 0 | 0 | 0 |
| ENSMUSG00 | Elp3 | 2755 | 15.6752301 | 17.3275417 | 15.5743453 | 15.8099304 | 17.593048 | 16.5835283 | 14.7004186 | 15.7067121 | 13.9182019 | 18.5864478 | 15.8324999 | 18.6795067 | 16.7245048 | 15.0374323 | 15.7328627 | 14.5518207 | 19.4344235 | 13.7972024 |
| ENSMUSG00 | Scara5 | 6390 | 0 | 0 | 0 | 0.01426011 | 0.03766191 | 0.00775473 | 0.03702088 | 0.14942254 | 0.21914578 | 0.14019771 | 0.12710083 | 0.13102268 | 0.26248594 | 0.49424068 | 0 | 0.05393043 | 0.24591208 | 0.00860862 |
| ENSMUSG00 | Pbk | 1979 | 0.21478108 | 0.0168332 | 0.56219869 | 0.13813354 | 0 | 0 | 0.83675784 | 0 | 1.08221264 | 0.26208072 | 0 | 0.08461192 | 0.1994216 | 0.32855848 | 0.87441402 | 1.12221083 | 1.64615222 | 0.19457491 |
| ENSMUSG00 | Esco2 | 3999 | 0 | 0.11662429 | 0.1814461 | 0 | 0 | 0 | 0.08281789 | 0 | 0 | 0 | 0.20309435 | 0 | 0.12336063 | 0.03484178 | 0 | 0.23937614 | 0.08625573 | 0 |
| ENSMUSG00 | Ccdc25 | 2218 | 25.1428273 | 22.1685446 | 20.7626695 | 19.8636293 | 21.1363813 | 19.8836487 | 26.8560247 | 21.3002713 | 20.5932579 | 21.555822 | 15.745489 | 21.6417808 | 27.112537 | 30.3415161 | 23.2385692 | 20.6990254 | 22.5845221 | 26.338896 |
| ENSMUSG00 | Clu | 3470 | 109.460015 | 128.125039 | 144.590852 | 127.767874 | 134.866482 | 110.915297 | 97.5841046 | 111.374329 | 65.9102945 | 52.613119 | 50.0053744 | 76.2599586 | 96.943615 | 145.221426 | 88.6130293 | 100.636985 | 90.9227885 | 114.472841 |
| ENSMUSG00 | Adam2 | 2530 | 0 | 0 | 0 | 0 | 0 | 0 | 0 | 0 | 0 | 0 | 0 | 0 | 0 | 0 | 0 | 0 | 0 | 0 |
| ENSMUSG00 | Epha2 | 2147 | 1.36602569 | 2.06363062 | 2.6360983 | 3.18311907 | 2.30907692 | 2.79267851 | 10.269077 | 7.45347769 | 8.76676186 | 5.09499914 | 6.29730473 | 8.57902592 | 4.61840492 | 5.3431251 | 3.97239034 | 1.51592996 | 2.35634429 | 3.6638576 |
| ENSMUSG00 | Chrna2 | 3520 | 0.01207533 | 0.17981392 | 0.2061372 | 0.25886957 | 0.27347684 | 0.2956271 | 2.13713521 | 2.00727115 | 2.10612865 | 0.32148207 | 0.97721506 | 2.80663985 | 0.40642773 | 0.35624726 | 0.43308454 | 0 | 0 | 0 |
| ENSMUSG00 | Trim35 | 5092 | 38.1978958 | 37.3101798 | 35.6816536 | 37.6424418 | 40.0217134 | 40.4245984 | 33.8677759 | 37.9673883 | 36.8025622 | 39.3817637 | 35.1556892 | 33.0706705 | 37.4058391 | 33.1365612 | 38.6042072 | 46.9984871 | 40.1101407 | 44.357304 |
| ENSMUSG00 | Stmn4 | 2013 | 0 | 0 | 0 | 0 | 0 | 0.02461636 | 0 | 0 | 0 | 0 | 0 | 0.0554552 | 0 | 0 | 0 | 0 | 0.34270906 | 0.02732693 |
| ENSMUSG00 | Dpysl2 | 4506 | 9.00853154 | 8.23581095 | 7.30004453 | 8.04851116 | 8.84448131 | 7.86289492 | 7.93794613 | 11.2644702 | 9.47856566 | 8.29793779 | 9.73311404 | 10.6156287 | 10.5758172 | 9.18368439 | 11.0821762 | 14.2336368 | 13.0816344 | 9.70533507 |
| ENSMUSG00 | Bnip3l | 4351 | 36.5558191 | 36.1534099 | 39.1235241 | 39.7180053 | 40.0785933 | 37.3780905 | 40.81004 | 39.798748 | 41.9437948 | 43.7588002 | 44.480891 | 42.9361119 | 38.1979382 | 35.5562526 | 38.1713334 | 42.7348172 | 38.2470193 | 39.685954 |
| ENSMUSG00 | Ppp2r2a | 2960 | 40.2219588 | 45.1637313 | 44.6801459 | 40.6817028 | 40.39179 | 43.4423458 | 35.6283653 | 36.8376043 | 33.3388148 | 39.871017 | 37.3322564 | 37.0155897 | 34.3990126 | 32.6206952 | 36.9700447 | 40.0499056 | 37.1350742 | 32.8753749 |
| ENSMUSG00 | Ebf2 | 7281 | 0 | 0 | 0 | 0 | 0 | 0 | 0 | 0 | 0 | 0 | 0 | 0 | 0 | 0 | 0 | 0 | 0 | 0 |
| ENSMUSG00 | Nefm | 5777 | 0 | 0.02306588 | 0 | 0 | 0 | 0 | 0 | 0 | 0 | 0 | 0 | 0 | 0 | 0 | 0 | 0.06628113 | 0.00663429 | 0.07617671 |
| ENSMUSG00 | Nefl | 3376 | 0 | 0 | 0 | 0 | 0 | 0 | 0 | 0 | 0 | 0 | 0 | 0 | 0 | 0 | 0 | 0 | 0 | 0 |
| ENSMUSG00 | Adam7 | 4268 | 29.9468286 | 67.6014472 | 42.8878723 | 39.3801998 | 23.558479 | 55.102454 | 74.0618575 | 69.1543781 | 37.4229869 | 25.6080905 | 14.7421791 | 32.9689047 | 143.615059 | 218.628903 | 117.73528 | 26.053403 | 31.609317 | 96.0597171 |
| ENSMUSG00 | Adamdec1 | 2411 | 0 | 0.17962158 | 0 | 0 | 0 | 0.10276386 | 0 | 0 | 0 | 0 | 0 | 0 | 0 | 0 | 0 | 0 | 0 | 0 |
| ENSMUSG00 | Nkx3-1 | 3193 | 0 | 0 | 0.04544961 | 0 | 0 | 0 | 0 | 0 | 0 | 0 | 0 | 0 | 0.13905019 | 0.10181917 | 0 | 0 | 0 | 0.55129703 |
| ENSMUSG00 | Pibf1 | 3274 | 9.06188542 | 7.99753661 | 7.23977699 | 7.33374171 | 7.4829411 | 6.6443647 | 9.95676108 | 10.0672093 | 7.30890826 | 8.8281555 | 9.57250237 | 9.4446784 | 6.78050176 | 8.76678253 | 8.80912924 | 9.59019489 | 7.3515316 | 10.4171186 |
| ENSMUSG00 | Entpd4 | 4302 | 9.16894546 | 9.40070981 | 9.79389701 | 9.95522587 | 11.9826337 | 10.2284583 | 11.8556643 | 11.7098509 | 10.3875801 | 10.0285666 | 12.9487684 | 10.7816866 | 10.9282467 | 13.1602377 | 13.8679342 | 10.7252864 | 11.3054558 | 12.5183439 |
| ENSMUSG00 | 6720463M24 | 2661 | 2.06056659 | 2.59142018 | 1.9996574 | 2.20870903 | 1.80879082 | 2.42084005 | 3.76936788 | 3.35852017 | 6.20661136 | 3.22488839 | 2.62124913 | 3.25119418 | 4.85718238 | 4.53794165 | 2.32252288 | 2.93546934 | 4.07603984 | 3.76236656 |
| ENSMUSG00 | Tnfrsf10b | 3163 | 1.2228805 | 0.78990423 | 0.48939392 | 0.46094006 | 0.92824766 | 0.43865841 | 0.31412149 | 0.65203596 | 0.45574709 | 0.46211565 | 0.30208636 | 0.40586792 | 0.85781077 | 0.16151912 | 0.50801792 | 0.71424149 | 0.23022436 | 0.67826594 |
| ENSMUSG00 | Rhobtb2 | 5498 | 1.06688146 | 1.0664004 | 1.60130639 | 1.34246802 | 0.79665425 | 1.39699413 | 1.78132494 | 1.31985385 | 1.36339503 | 0.86617334 | 1.39032255 | 0.76139954 | 1.1485059 | 2.04428322 | 1.22900239 | 1.40682115 | 1.41510336 | 0.71037588 |
| ENSMUSG00 | Klhl1 | 4111 | 0 | 0 | 0 | 0 | 0 | 0 | 0 | 0.00929029 | 0 | 0 | 0 | 0 | 0 | 0 | 0 | 0 | 0 | 0 |
| ENSMUSG00 | Pebp4 | 1040 | 0 | 0 | 0 | 0.26285218 | 0 | 0.04764686 | 0 | 0 | 0 | 0 | 0 | 0 | 0 | 0 | 0 | 0.11045368 | 0 | 0 |
| ENSMUSG00 | Bin3 | 1703 | 36.4650982 | 32.7847391 | 34.7675873 | 32.8531782 | 31.7393719 | 32.4435102 | 36.4499323 | 34.828421 | 35.3096377 | 39.3153251 | 36.5816476 | 34.2497753 | 40.3809117 | 44.3714211 | 42.0241354 | 42.1129229 | 32.5424694 | 47.9997255 |
| ENSMUSG00 | Pdlim2 | 4901 | 2.80130011 | 2.54213884 | 1.81610479 | 2.83536391 | 1.58115481 | 2.50746336 | 1.18740247 | 1.30139384 | 1.49585795 | 1.11599298 | 1.97884443 | 1.85634626 | 2.18425343 | 2.88084117 | 2.05125069 | 2.34384457 | 1.37633713 | 2.05400256 |
| ENSMUSG00 | Sorbs3 | 2893 | 20.7750843 | 23.2718145 | 20.1653915 | 23.0561248 | 20.9797289 | 20.3657795 | 17.8587782 | 24.5154826 | 20.315644 | 26.1586169 | 23.037009 | 21.9944194 | 27.2834662 | 26.92248 | 25.2222452 | 31.9110454 | 29.5561591 | 30.7085058 |
| ENSMUSG00 | Ppp3cc | 3291 | 7.03899147 | 6.36700409 | 8.77514361 | 8.74949256 | 8.90681906 | 6.760613 | 10.3797491 | 12.4870936 | 11.4135832 | 10.9173005 | 13.2829192 | 7.7846822 | 10.4030236 | 12.1084859 | 8.88881941 | 11.3906122 | 11.3080627 | 9.3604071 |
| ENSMUSG00 | Slc39a14 | 10975 | 1.27031414 | 1.74835782 | 1.56470189 | 1.68129594 | 1.69722592 | 1.68863074 | 1.66402679 | 2.64127867 | 1.5611472 | 1.89462051 | 2.68584505 | 2.15634086 | 1.63615653 | 1.61231866 | 1.50165124 | 2.58178127 | 2.45148614 | 2.5462074 |
| ENSMUSG00 | Fam160b2 | 4016 | 12.5102386 | 15.4453708 | 13.8038012 | 13.0693036 | 16.0359845 | 13.8811816 | 9.77826781 | 15.9578813 | 14.829607 | 13.4313756 | 14.1445283 | 14.8573066 | 15.3056696 | 16.4566575 | 16.1995161 | 12.2613847 | 19.1345182 | 13.0400074 |
| ENSMUSG00 | Hr | 6577 | 0.85953906 | 0.62806734 | 1.27976194 | 0.8520615 | 0.90014089 | 1.08493137 | 0.5467179 | 0.5168198 | 0.29432359 | 0.50183202 | 0.58111551 | 0.90805472 | 0.38253425 | 0.46606504 | 0.53874621 | 0.34931374 | 0.80417044 | 0.30109896 |
| ENSMUSG00 | Sftpc | 798 | 0 | 0 | 0 | 0 | 0 | 0 | 0 | 0 | 0 | 0 | 0 | 0 | 0 | 0 | 0 | 0 | 0 | 0 |
| ENSMUSG00 | Bmp1 | 4045 | 3.22597999 | 3.37658508 | 3.252806 | 2.57935208 | 3.46264272 | 3.6138582 | 1.1111754 | 1.7467477 | 1.22184966 | 1.95829868 | 1.65352143 | 0.74512875 | 0.24391554 | 0.80372957 | 1.77232898 | 0.70049566 | 0.45479907 | 0.42157783 |
| ENSMUSG00 | Epb4.9 | 6685 | 0.78206981 | 0.70263547 | 0.6946685 | 0.63383353 | 1.02959837 | 0.63006468 | 0.77144084 | 0.57131492 | 0.75780725 | 0.89575756 | 0.29301022 | 0.31727677 | 1.48327829 | 0.97265104 | 1.23265687 | 1.16275143 | 1.35303008 | 0.73235759 |
| ENSMUSG00 | Xpo7 | 15093 | 4.00184557 | 3.81179145 | 3.95500805 | 3.57110683 | 3.64186423 | 3.95292462 | 2.7366308 | 2.94040758 | 3.34283985 | 3.39580033 | 3.36479036 | 3.49471948 | 2.50369367 | 2.31097261 | 2.8472287 | 2.77037986 | 3.73029419 | 2.6533247 |
| ENSMUSG00 | Fgf17 | 1682 | 0 | 0 | 0 | 0 | 0 | 0 | 0 | 0 | 0 | 0 | 0 | 0 | 0 | 0 | 0 | 0 | 0 | 0 |
| ENSMUSG00 | Dok2 | 2746 | 0.30957885 | 0.01213143 | 0.08807999 | 0.232285 | 0.21033616 | 0.09022712 | 0.01722967 | 0.30598429 | 0.56995178 | 0.22321892 | 0.03479604 | 0.22358787 | 0.07186004 | 0.13530688 | 0.01500421 | 0.25099451 | 0 | 0 |
| ENSMUSG00 | Gfra2 | 3869 | 0.05493044 | 0 | 0.01250285 | 0 | 0 | 0 | 0.04891463 | 0.4343411 | 0 | 0.15842831 | 0.11113327 | 0.15869018 | 0.34426512 | 0.25208732 | 0.07454404 | 0.26721282 | 0 | 0 |
| ENSMUSG00 | Rb1 | 5984 | 7.87027658 | 8.26698662 | 9.44189229 | 7.75847329 | 7.5045262 | 9.04271141 | 7.74839996 | 9.77148531 | 6.09813719 | 9.74689504 | 10.2831127 | 8.91707444 | 5.55643587 | 7.60615068 | 8.64103487 | 7.31386435 | 11.029058 | 7.29900246 |
| ENSMUSG00 | Rcbtb2 | 7949 | 5.94077879 | 5.47322177 | 6.17676856 | 5.9494734 | 6.37601153 | 6.62656382 | 4.57115831 | 5.97702209 | 6.00517949 | 6.35873115 | 5.80583846 | 6.95150408 | 5.18205446 | 4.31195578 | 5.83114233 | 5.4287979 | 5.33260591 | 5.07946714 |
| ENSMUSG00 | Itm2b | 1767 | 2046.50275 | 2038.96782 | 2160.02733 | 2199.46135 | 2150.57357 | 2156.98815 | 3108.63311 | 2997.85502 | 2935.05787 | 3060.60632 | 2785.16923 | 2923.00971 | 3745.06586 | 3974.63611 | 3396.66839 | 3215.28419 | 3229.60326 | 3504.55018 |
| ENSMUSG00 | Med4 | 1297 | 39.8178789 | 35.2135528 | 36.848919 | 33.5121328 | 34.2527105 | 33.7737984 | 42.6799031 | 35.659984 | 44.01273 | 37.9532261 | 42.9127422 | 39.548646 | 37.8071903 | 41.6457006 | 41.0427218 | 38.3201298 | 40.3061442 | 40.6312446 |
| ENSMUSG00 | Sucla2 | 3414 | 61.0311578 | 57.9901964 | 58.5044835 | 56.504236 | 60.2706426 | 60.9611837 | 60.6860636 | 59.2687018 | 58.6672601 | 61.6936706 | 62.0066904 | 56.6004171 | 54.1870796 | 43.9137417 | 52.8596411 | 53.4655318 | 58.3426377 | 48.2578426 |
| ENSMUSG00 | Uchl3 | 1662 | 18.38822 | 15.393684 | 18.016375 | 16.3109635 | 18.9689591 | 18.6046364 | 24.9089014 | 17.5795354 | 25.3512669 | 17.2772433 | 19.5469144 | 20.1500587 | 21.4899967 | 21.4615227 | 22.9062802 | 25.273644 | 22.5760798 | 19.8257852 |
| ENSMUSG00 | Gpc5 | 4983 | 0.02559011 | 0.00668531 | 0.06795399 | 0 | 0 | 0.02983307 | 0.07595855 | 0.12263264 | 0.03306162 | 0.151397 | 0.02876277 | 0 | 0.1584007 | 0.27029471 | 0.00826842 | 0 | 0.09229692 | 0 |
| ENSMUSG00 | Trim52 | 3082 | 0 | 0 | 0 | 0 | 0.01561711 | 0 | 0 | 0 | 0 | 0 | 0 | 0 | 0 | 0 | 0 | 0 | 0 | 0 |
| ENSMUSG00 | Spry2 | 2083 | 3.22410841 | 1.72721679 | 2.15973995 | 1.90293368 | 2.21827397 | 2.37891184 | 5.56486142 | 3.13533404 | 4.98271818 | 2.35413977 | 4.72474385 | 3.93898339 | 3.6945651 | 2.2742664 | 3.44170462 | 2.75736491 | 2.31834537 | 2.85212843 |
| ENSMUSG00 | 4930449E01f | 592 | 0 | 0.05627178 | 0 | 0.15392245 | 0 | 0 | 0 | 0.19354258 | 0 | 0 | 0 | 0.28284964 | 0 | 0 | 0 | 0.12936016 | 0 | 0 |
| ENSMUSG00 | Rbm26 | 8072 | 10.0154664 | 9.87170998 | 10.63115 | 10.1936628 | 9.34970965 | 9.74848128 | 9.77670323 | 10.8729114 | 8.8832661 | 8.75021723 | 11.9259838 | 10.6071886 | 12.6018847 | 10.742209 | 9.17234836 | 9.84778258 | 11.124692 | 10.0927263 |
| ENSMUSG00 | Rnf219 | 3450 | 3.52361749 | 2.32707464 | 2.90241182 | 2.16580037 | 2.38566926 | 3.20297379 | 1.53594784 | 3.23251638 | 1.99366613 | 1.7630263 | 2.93573658 | 2.81505081 | 1.37271422 | 2.74625908 | 1.50475348 | 1.67590774 | 2.32179596 | 1.13207147 |
| ENSMUSG00 | Ednrb | 6800 | 0 | 0 | 0 | 0 | 0 | 0 | 0 | 0.01736018 | 0.06864419 | 0 | 0 | 0 | 0 | 0 | 0 | 0 | 0 | 0 |
| ENSMUSG00 | Scel | 3129 | 0.04075281 | 0.28745546 | 0 | 0.29121792 | 0.13844273 | 0.36424189 | 3.11486477 | 2.53883659 | 1.30312082 | 1.7178588 | 0.39697952 | 1.80165609 | 1.50152041 | 2.34521119 | 0.7900586 | 0.35488259 | 0.02449747 | 1.7228803 |
| ENSMUSG00 | Fbxl3 | 7229 | 12.7944755 | 15.8384863 | 12.647112 | 13.6639015 | 14.1352987 | 14.374337 | 11.1589594 | 12.8593592 | 11.9645432 | 13.227504 | 12.1403512 | 13.2493674 | 9.77903409 | 9.61133694 | 11.3419693 | 12.6856493 | 12.8037083 | 13.3394611 |
| ENSMUSG00 | Cln5 | 2392 | 30.0130612 | 28.7449054 | 29.7683264 | 29.2565903 | 30.3037948 | 35.424404 | 22.9047166 | 25.3072566 | 26.8607711 | 21.5253211 | 27.462612 | 28.3978517 | 21.0361855 | 17.8436941 | 17.5175511 | 23.4193807 | 20.5891687 | 21.5253026 |
| ENSMUSG00 | Irg1 | 2618 | 0 | 0 | 0 | 0 | 0 | 0 | 0 | 0 | 0 | 0 | 0 | 0 | 0 | 0 | 0 | 0 | 0 | 0 |
| ENSMUSG00 | Dct | 2475 | 0 | 0 | 0 | 0 | 0 | 0 | 0 | 0 | 0 | 0 | 0 | 0 | 0 | 0 | 0 | 0 | 0 | 0 |
| ENSMUSG00 | Tgds | 2253 | 7.35775351 | 5.91440643 | 5.58238699 | 5.19715419 | 7.1354028 | 5.89442193 | 6.42595639 | 6.91633386 | 7.03808636 | 5.5459105 | 4.68631417 | 8.59655345 | 7.09433697 | 6.67904319 | 5.54108808 | 6.76416411 | 8.64170387 | 6.25047995 |
| ENSMUSG00 | Gpr180 | 2868 | 18.3477712 | 19.258291 | 17.0859087 | 15.4729628 | 16.480317 | 15.9474104 | 17.882477 | 16.3795868 | 17.7211159 | 19.7940888 | 16.1415393 | 20.3956614 | 20.692572 | 18.1695488 | 16.1186003 | 17.1159549 | 18.6553384 | 19.103581 |
| ENSMUSG00 | Cldn10 | 2622 | 58.3919314 | 37.5055926 | 43.3000289 | 48.5671697 | 45.6537354 | 50.9323482 | 101.734891 | 67.4557649 | 81.6033369 | 101.152824 | 70.3505395 | 98.8162875 | 103.113196 | 142.573783 | 154.670829 | 192.957207 | 138.73184 | 166.600807 |
| ENSMUSG00 | Dnajc3 | 5178 | 46.2155545 | 42.5578979 | 42.2731218 | 46.0889823 | 35.3413623 | 48.366071 | 41.8486036 | 38.7455948 | 35.9765567 | 46.2856363 | 34.2396429 | 45.8878477 | 48.2648692 | 41.8786103 | 45.9200809 | 52.1633728 | 48.8738004 | 51.0146254 |
| ENSMUSG00 | Mbnl2 | 4752 | 59.8042102 | 62.0480696 | 60.8842772 | 66.1651446 | 58.5848162 | 64.9649685 | 49.3437788 | 50.9875002 | 47.738916 | 52.9056292 | 59.517625 | 52.8674868 | 51.4497016 | 40.5408406 | 42.5194508 | 47.9760683 | 56.4328913 | 46.3618418 |
| ENSMUSG00 | Nipbl | 8768 | 9.97669392 | 11.0371758 | 10.7748067 | 8.67779927 | 8.6020443 | 9.43242613 | 9.08157375 | 10.7372573 | 11.2971682 | 8.08789435 | 9.46999063 | 11.4648732 | 7.89378193 | 8.1097055 | 8.97054861 | 10.0748667 | 9.66899735 | 9.50488191 |
| ENSMUSG00 | Nup155 | 6367 | 4.16573423 | 3.12357434 | 3.19856389 | 5.35254612 | 3.50765079 | 3.8913722 | 2.44477326 | 2.72931413 | 2.85918642 | 2.8659215 | 3.42911199 | 2.78771489 | 3.6260035 | 4.55176858 | 3.13201713 | 2.62226767 | 2.20840525 | 3.95699244 |
| [illegible] | | | | | | | | | | | | | | | | | | | | |

| | | | | | | | | | | | | | | | | | | | |
|---|---|---|---|---|---|---|---|---|---|---|---|---|---|---|---|---|---|---|---|
| ENSMUSG00 Nit2 | 1270 | 15.5629188 | 15.0039177 | 14.207344 | 11.4081985 | 13.5678966 | 11.5883164 | 15.3859342 | 18.5849643 | 18.6474382 | 13.8481987 | 14.6710502 | 11.7344739 | 14.9549513 | 10.166501 | 11.7116234 | 12.6328854 | 14.2742822 | 13.5140483 |
| ENSMUSG00 Tomm70a | 6354 | 17.2255002 | 18.5648345 | 19.8777601 | 17.0083093 | 19.2027741 | 20.0503743 | 16.9995036 | 19.7092993 | 16.7105505 | 17.9356704 | 17.0753747 | 16.4794119 | 14.1070339 | 16.5339197 | 16.1719673 | 16.2587439 | 16.7926362 | 15.9036401 |
| ENSMUSG00 Tmem30c | 3941 | 0.07549765 | 0.0084529 | 0 | 0.01156078 | 0.02442625 | 0.17603103 | 0 | 0 | 0 | 0 | 0.12122547 | 0.11330254 | 0.02503523 | 0.07070909 | 0.01045459 | 0 | 0.00972502 | 0 |
| ENSMUSG00 Tmem45a | 3334 | 0 | 0 | 0 | 0 | 0.04331007 | 0 | 0 | 0 | 0 | 0.01414238 | 0 | 0 | 0.02959323 | 0.01393044 | 0.16065394 | 0 | 0.1149559 | 0.06599773 |
| ENSMUSG00 Gpr128 | 2682 | 0 | 0 | 0 | 0 | 0 | 0 | 0 | 0.04272081 | 0 | 0 | 0 | 0 | 0.42305522 | 0.65804449 | 0.33796951 | 0.12849197 | 0.74309002 | 0.38969912 |
| ENSMUSG00 Slc7a4 | 4891 | 22.6387208 | 25.3098988 | 24.023575 | 21.9840656 | 26.0292247 | 27.8005931 | 24.6478646 | 23.9180799 | 21.8269166 | 20.4181536 | 21.8020256 | 23.6911297 | 31.8120767 | 46.4820707 | 28.7425289 | 24.3240477 | 21.948877 | 32.6275646 |
| ENSMUSG00 Tfg | 3010 | 56.3440705 | 54.6508544 | 50.6073911 | 57.7762479 | 49.9388696 | 54.0141259 | 52.2797243 | 46.2876691 | 52.5983278 | 57.3954076 | 43.0450789 | 46.822109 | 48.0863273 | 49.7615222 | 54.2874986 | 57.2832575 | 52.2561869 | 54.2049869 |

[illegible]

| | | | | | | | | | | | | | | | | | | | |
|---|---|---|---|---|---|---|---|---|---|---|---|---|---|---|---|---|---|---|---|
| ENSMUSG00 Qpct | 1917 | 10.3103322 | 9.12324959 | 6.43466366 | 8.05696089 | 8.23540478 | 11.7354935 | 8.49012079 | 10.7783462 | 9.53928691 | 16.5039759 | 12.7599263 | 8.96777469 | 72.9813862 | 70.6231167 | 69.7438961 | 90.742947 | 68.5754817 | 82.2410537 |
| ENSMUSG00 Man2a1 | 6929 | 8.95007242 | 9.25010946 | 8.55909196 | 7.90364775 | 8.55802135 | 7.92386041 | 5.40111546 | 7.24272143 | 8.51191955 | 6.19920489 | 6.32954422 | 7.97481685 | 7.26914238 | 5.21482176 | 4.81051504 | 6.15059114 | 6.32779386 | 3.63604718 |
| ENSMUSG00 Cyp1b1 | 6752 | 0 | 0 | 0 | 0 | 0 | 0 | 0 | 0 | 0 | 0 | 0 | 0 | 0 | 0.01375713 | 0 | 0 | 0 | 0 |
| ENSMUSG00 4930583I09R | 626 | 0 | 0 | 0 | 0 | 0 | 0 | 0 | 0 | 0 | 0 | 0 | 0 | 0 | 0 | 0 | 0 | 0.18367235 | 0 |
| ENSMUSG00 Vapa | 1634 | 191.533422 | 192.7418 | 181.889211 | 205.331414 | 202.277797 | 207.066135 | 216.729132 | 207.533868 | 205.655313 | 219.998141 | 192.006213 | 195.047389 | 174.986409 | 167.130498 | 199.930928 | 190.117318 | 198.316304 | 182.297621 |
| ENSMUSG00 Hnrpll | 2753 | 25.3054789 | 27.0326937 | 31.4348877 | 28.928698 | 26.3300679 | 27.3053022 | 19.7809275 | 24.7494536 | 22.9645121 | 25.1082205 | 22.2649007 | 29.1952612 | 27.8645592 | 21.7964987 | 19.3511122 | 21.8644514 | 28.3583969 | 18.3230479 |
| ENSMUSG00 Ralbp1 | 5189 | 47.8705432 | 44.6247693 | 44.3649318 | 39.2567796 | 43.113737 | 41.0440642 | 24.7368082 | 24.0827791 | 28.0979506 | 24.3250015 | 26.2951014 | 27.0311737 | 32.8372408 | 34.2982738 | 33.1502208 | 46.001859 | 32.5725514 | 41.8531419 |
| ENSMUSG00 Snf7 | 2656 | 95.9567152 | 98.1200194 | 96.4920798 | 91.4994768 | 95.4121153 | 86.3254888 | 91.4901803 | 110.824113 | 107.773451 | 90.9994371 | 96.9529486 | 97.7824842 | 87.68674 | 107.839114 | 97.5589552 | 93.1171059 | 109.62533 | 95.16824 |
| ENSMUSG00 Twsg1 | 4041 | 162.64725 | 187.767333 | 183.642249 | 201.275863 | 196.184339 | 192.472088 | 98.418802 | 126.911057 | 113.041239 | 117.240848 | 128.392984 | 116.880014 | 110.334538 | 104.737681 | 130.038269 | 145.942078 | 141.098817 | 112.604635 |
| ENSMUSG00 Ndufv2 | 2126 | 166.661876 | 142.230076 | 143.18656 | 139.726251 | 152.183815 | 138.47262 | 180.660549 | 164.626141 | 179.081915 | 159.882135 | 162.448342 | 149.331839 | 179.460512 | 167.906517 | 159.573665 | 174.865243 | 165.79819 | 169.529477 |
| ENSMUSG00 ORF19 | 2979 | 4.58011464 | 4.13755316 | 4.38430789 | 4.28234052 | 3.68381289 | 4.19177202 | 3.60522931 | 2.74359655 | 4.72836133 | 3.24467746 | 3.80082751 | 3.39128396 | 3.59349649 | 3.99116696 | 5.22799199 | 3.12340302 | 3.71812682 | 3.87778194 |
| ENSMUSG00 D6Wsu116e | 4268 | 44.2081716 | 49.5010251 | 49.5409328 | 48.3792533 | 47.827434 | 49.7036674 | 59.6507791 | 54.0044865 | 49.176777 | 52.0778855 | 62.9201128 | 51.4215523 | 51.0541421 | 56.6078122 | 55.7495279 | 48.6348132 | 62.3565618 | 60.8348134 |
| ENSMUSG00 9130404H23 | 1793 | 0 | 0 | 0 | 0 | 0.08053306 | 0 | 0 | 0 | 0 | 0 | 0 | 0.12451905 | 0.16508171 | 0 | 0 | 0.08542244 | 0 | 0 |
| ENSMUSG00 Lhcgr | 2542 | 0 | 0 | 0 | 0 | 0 | 0 | 0 | 0 | 0 | 0 | 0 | 0 | 0 | 0 | 0 | 0 | 0 | 0 |
| ENSMUSG00 Nrxn1 | 15676 | 0.00271148 | 0 | 0 | 0 | 0 | 0 | 0 | 0 | 0 | 0 | 0 | 0 | 0 | 0 | 0 | 0 | 0 | 0 |
| ENSMUSG00 Cacna1h | 11518 | 0.11440011 | 0.07519841 | 0.00839964 | 0.00791128 | 0 | 0 | 0 | 0.02321122 | 0 | 0 | 0 | 0 | 0 | 0 | 0 | 0 | 0 | 0 |
| ENSMUSG00 Prss41 | 2564 | 1.19359309 | 0.71459016 | 1.3772495 | 0.85293687 | 0.91983785 | 1.10160133 | 1.27323512 | 0.13406069 | 0.75497903 | 0.95625451 | 1.6024362 | 1.43675263 | 1.07745217 | 0.45284797 | 1.33374771 | 0.88110212 | 0.91181908 | 0.8796308 |
| ENSMUSG00 Prss21 | 1088 | 0 | 0 | 0 | 0 | 0 | 0 | 0 | 0 | 0 | 0 | 0 | 0 | 0 | 0 | 0 | 0 | 0 | 0 |
| ENSMUSG00 1600002H07 | 3309 | 1.4643669 | 0.7047152 | 2.09048499 | 0.95004898 | 1.44003035 | 1.01830942 | 1.10095989 | 0.56555688 | 1.80478839 | 0.69821229 | 1.42935053 | 1.33255888 | 1.01377165 | 1.40356867 | 2.24124511 | 1.27288208 | 1.49413487 | 1.67903287 |
| [illegible] | | | | | | | | | | | | | | | | | | | |

| | | | | | | | | | | | | | | | | | | | |
|---|---|---|---|---|---|---|---|---|---|---|---|---|---|---|---|---|---|---|---|
| ENSMUSG00 Pold4 | 906 | 56.1105856 | 44.0127311 | 39.6171739 | 48.3771795 | 45.369385 | 45.5053801 | 54.4148029 | 49.9957939 | 58.3248341 | 56.5704354 | 54.7355485 | 49.9014202 | 64.3057332 | 47.5718688 | 46.0675233 | 61.2818772 | 51.482454 | 45.6587729 |
| ENSMUSG00 Pacs1 | 4353 | 15.9455439 | 21.4203517 | 18.8915694 | 21.4146328 | 17.9126387 | 19.5228436 | 12.70584 | 16.9948733 | 12.196053 | 16.8650689 | 17.3298092 | 15.9509957 | 19.1978562 | 18.1487233 | 19.2898624 | 17.9445989 | 17.3538091 | 19.4231558 |
| ENSMUSG00 Cdk2ap2 | 1512 | 67.3841977 | 60.5007986 | 56.9796677 | 60.4768624 | 56.9179097 | 57.8442955 | 72.6587569 | 74.086188 | 70.9868122 | 70.5701332 | 71.6940349 | 71.9106552 | 88.2232171 | 90.1850797 | 82.6483591 | 73.770198 | 71.8111105 | 87.0249889 |
| ENSMUSG00 Cabp2 | 1338 | 0 | 0 | 0 | 0 | 0 | 0 | 0 | 0 | 0 | 0 | 0 | 0 | 0.18434947 | 0 | 0 | 0 | 0 | 0 |
| ENSMUSG00 Adrbk1 | 4504 | 16.5717338 | 21.0572402 | 17.7856497 | 16.7313427 | 16.0831583 | 16.9319842 | 14.6539048 | 19.5880217 | 16.6794416 | 19.5553982 | 17.7883227 | 15.6516834 | 19.375702 | 16.9834396 | 16.4019515 | 17.5385264 | 16.0572207 | 15.6697788 |
| ENSMUSG00 Klc2 | 4529 | 2.27119731 | 2.46408027 | 2.91586968 | 2.16286698 | 3.13511095 | 2.6805961 | 2.32959316 | 2.95993227 | 2.9191591 | 2.32161776 | 2.28906293 | 3.17960719 | 2.28741502 | 3.09695615 | 3.03848966 | 1.9699076 | 3.18186965 | 1.9433555 |
| ENSMUSG00 Mbl2 | 1095 | 0.27172259 | 0.36507281 | 0.08835348 | 0 | 0 | 0.04525364 | 0.04320793 | 0.17439453 | 0.41374582 | 0.08612001 | 0 | 0 | 0.63072771 | 0.12724408 | 0 | 0 | 0 | 0 |
| ENSMUSG00 Acy3 | 1499 | 5.27415793 | 5.91142753 | 6.00231931 | 7.56817881 | 6.19710558 | 7.96678372 | 11.2679295 | 11.2870142 | 11.6223457 | 11.0720798 | 9.24265374 | 9.12263998 | 11.8475585 | 13.8185877 | 14.0178743 | 12.0057275 | 9.48569456 | 10.0183363 |
| ENSMUSG00 Pip5k1b | 4175 | 46.8116885 | 45.8880131 | 48.8138164 | 47.143404 | 48.4662532 | 49.5764714 | 45.0235385 | 44.0927855 | 44.9450936 | 44.2369592 | 49.0565861 | 48.5161771 | 37.7522097 | 38.3122005 | 40.7179927 | 38.37315 | 39.8960686 | 40.8582611 |
| ENSMUSG00 Dkk1 | 2423 | 0 | 0.02749723 | 0.01996431 | 0.0564107 | 0 | 0 | 0.03905297 | 0.141862 | 0 | 0.05837891 | 0.01971728 | 0.04607153 | 0 | 0.01916801 | 0.01700436 | 0.01580297 | 0.0158177 | 0 |
| ENSMUSG00 Nudt8 | 4511 | 3.34500942 | 4.34227041 | 3.53874197 | 4.11069498 | 3.74513527 | 4.0863704 | 4.4680118 | 3.95385766 | 4.59251107 | 4.50497741 | 4.26807986 | 3.67485076 | 4.05722485 | 3.7476497 | 4.29278015 | 3.42077472 | 3.17757372 | 2.99983158 |
| ENSMUSG00 Rab1b | 1852 | 195.932337 | 204.176384 | 190.124691 | 204.434276 | 196.893873 | 201.635745 | 236.767769 | 195.375173 | 198.57149 | 221.419906 | 221.255842 | 185.409264 | 198.393153 | 206.365224 | 206.341485 | 194.120147 | 192.728295 | 191.165557 |
| ENSMUSG00 Doc2g | 2817 | 1.50888094 | 0.83962211 | 0.7727401 | 0.84102744 | 0.90557045 | 0.96748326 | 1.36042847 | 1.03039495 | 1.0965526 | 0.78668195 | 1.40763988 | 0.97087947 | 1.03322084 | 0.89030199 | 0.90681451 | 0.91071022 | 0.50339829 | 0.70299178 |
| ENSMUSG00 Cnih2 | 1231 | 1.17398537 | 2.00256228 | 0.74662638 | 0.25907986 | 1.01659628 | 1.65041598 | 2.61353547 | 1.79947956 | 2.07438178 | 1.64701888 | 2.21216298 | 1.08820138 | 2.48462946 | 1.05640491 | 1.80737938 | 1.86631718 | 1.27650542 | 1.16184953 |
| ENSMUSG00 Yif1a | 1447 | 52.7863174 | 49.0139266 | 46.7020193 | 48.6782132 | 52.8553053 | 43.970774 | 52.1518472 | 49.4099359 | 54.9627933 | 52.9182737 | 61.3447635 | 50.4925379 | 54.7185399 | 47.2786045 | 47.0386803 | 61.7095345 | 51.5696614 | 58.316496 |
| ENSMUSG00 Cbwd1 | 1360 | 18.1584612 | 21.138991 | 17.179717 | 21.0384822 | 19.6420718 | 20.2583234 | 19.7600012 | 20.1071766 | 19.5938792 | 23.4366731 | 24.1685084 | 19.2071545 | 26.8786402 | 23.1537436 | 19.6616248 | 24.9452045 | 25.8702504 | 18.9700522 |
| ENSMUSG00 Rin1 | 4166 | 0.51014374 | 0.6157208 | 0.34834515 | 0.43745602 | 0.17330265 | 0.51146605 | 0.38613324 | 0.66007032 | 0.06920442 | 0.35085737 | 0.1834852 | 0.62970143 | 0.09473244 | 0.27870912 | 0.77141655 | 0.62500271 | 0.47838873 | 0.52817193 |
| ENSMUSG00 Aldh3b1 | 1855 | 7.67613691 | 4.05860598 | 4.3549216 | 8.12976047 | 8.01766276 | 5.74330874 | 13.2628534 | 7.92672503 | 11.2791105 | 11.3619215 | 8.96263372 | 10.0498282 | 11.462025 | 14.471527 | 14.3261475 | 15.2130338 | 14.6073811 | 10.9721668 |
| ENSMUSG00 Asah2 | 4910 | 0.51941152 | 0.7666715 | 0.90638794 | 0.59387105 | 1.04890343 | 0.80737651 | 1.38758162 | 0.89452674 | 1.39245654 | 0.74903358 | 1.23572703 | 1.07993645 | 0.45212551 | 1.02158074 | 0.91465776 | 1.04499825 | 1.53773533 | 1.17636583 |
| ENSMUSG00 Rce1 | 1438 | 44.2786882 | 38.5484395 | 35.2541445 | 38.8441171 | 37.621893 | 38.766916 | 36.9815376 | 39.0157435 | 40.9860251 | 40.2322065 | 38.4060167 | 41.4930065 | 46.3816079 | 46.799362 | 39.0813089 | 48.808542 | 35.7410043 | 40.4343712 |
| ENSMUSG00 Slc29a2 | 2460 | 1.43411773 | 0.69063317 | 0.27529651 | 1.07420348 | 0.99785695 | 1.22874665 | 0.96163982 | 1.00914884 | 0.58598701 | 0.97751458 | 0.69914573 | 0.74874669 | 0.84225225 | 1.51037683 | 0.8709272 | 0.38913219 | 0.49855346 | 0.15652998 |
| ENSMUSG00 Pcx | 4304 | 3.99967386 | 3.97835214 | 3.06830246 | 4.26605503 | 3.13125897 | 5.41119535 | 10.2781958 | 9.06892083 | 6.50716327 | 9.93626577 | 7.6479894 | 7.67724727 | 10.9919397 | 11.9887038 | 13.9189271 | 12.7665013 | 13.5620237 | 13.5094391 |
| ENSMUSG00 2700046G09 | 1639 | 7.15767455 | 6.686969 | 6.58163348 | 6.50474947 | 6.28446105 | 7.34674451 | 10.622981 | 7.41011834 | 9.64955586 | 10.7016849 | 8.2491234 | 7.38987113 | 9.02963709 | 11.7597903 | 10.6083323 | 8.03658874 | 7.59978432 | 9.16259069 |
| ENSMUSG00 Minpp1 | 2614 | 21.3176304 | 21.6648509 | 23.4650484 | 25.9004254 | 21.2855791 | 24.8142801 | 21.5929709 | 19.9874547 | 24.0753619 | 23.9361068 | 16.8875523 | 25.4309312 | 21.6086631 | 20.9478114 | 23.4852021 | 23.7595123 | 25.6437235 | 22.7906896 |
| ENSMUSG00 Apba1 | 6883 | 0.0432277 | 0.10163748 | 0.12650349 | 0.17872268 | 0.04195722 | 0.10798939 | 0.57052918 | 0.29963529 | 0.44878522 | 0.21921001 | 0.15270218 | 0.21894855 | 0.45153433 | 0.14170069 | 0.26338348 | 0.21139664 | 0.35080004 | 0.11988037 |
| ENSMUSG00 Papss2 | 3622 | 0.18776444 | 0.17475014 | 0.3205314 | 0.31447435 | 0.35879678 | 0.08208625 | 0 | 0.02108912 | 0.06822725 | 0.26035728 | 0.0923315 | 0.6164071 | 0.29964168 | 0.07693664 | 0.06825216 | 0.2114335 | 0.49733185 | 0.24299992 |
| ENSMUSG00 Cpt1a | 4281 | 22.7965158 | 25.3523498 | 20.8703365 | 19.7739828 | 22.1490048 | 24.052927 | 18.2354404 | 16.0941588 | 16.9517965 | 20.0894377 | 20.1656973 | 24.0681414 | 13.2404515 | 15.6549447 | 17.1600967 | 10.6258448 | 16.4191611 | 13.0294871 |
| ENSMUSG00 Peli3 | 4755 | 3.11972796 | 3.06156357 | 3.16386287 | 3.43025318 | 2.53059536 | 3.94962903 | 0.69650631 | 2.32929478 | 0.70159995 | 1.39815965 | 1.75827923 | 1.29121407 | 2.38619158 | 2.38325075 | 3.41396699 | 2.6251815 | 2.71629065 | 2.88060306 |
| ENSMUSG00 Mrpl11 | 816 | 80.4264605 | 75.8929784 | 90.5225256 | 72.026651 | 84.5847776 | 81.130456 | 102.336861 | 83.6862935 | 84.0891363 | 87.8297414 | 94.3204143 | 88.9220115 | 95.0969435 | 88.7901671 | 97.0963137 | 78.3643558 | 85.85842 | 74.5589118 |
| ENSMUSG00 Lao1 | 2497 | 0 | 0 | 0 | 0 | 0 | 0 | 0 | 0 | 0 | 0 | 0 | 0 | 0 | 0 | 0 | 0 | 0 | 0 |
| ENSMUSG00 Mtl5 | 4349 | 0 | 0.20681723 | 0.38930181 | 0.05238106 | 0.08853883 | 0.07975837 | 0.08703183 | 0.07903693 | 0.2272882 | 0.21683469 | 0.01098527 | 0 | 0.12477606 | 0.16018885 | 0.0663166 | 0.11445801 | 0.1145647 | 0.08854076 |
| ENSMUSG00 Mus81 | 3532 | 8.13519224 | 8.53572176 | 9.65553196 | 9.04255155 | 8.36721436 | 10.059261 | 10.8503029 | 10.2185222 | 7.46301531 | 8.81072031 | 9.48194957 | 7.09547919 | 9.69319099 | 9.38875379 | 8.4806147 | 9.76779085 | 8.72433259 | 12.3349979 |
| ENSMUSG00 Gal | 706 | 0 | 0.56622483 | 0.1370355 | 0 | 0 | 0 | 0 | 0 | 0 | 0 | 0.20300974 | 0 | 0 | 0 | 0.05835914 | 0 | 0.2714327 | 0 |
| ENSMUSG00 Ppp6r3 | 4928 | 13.8952595 | 15.1287048 | 15.8431165 | 16.2718015 | 15.7249182 | 14.0473151 | 13.6715209 | 12.0823773 | 14.9184112 | 15.4808924 | 14.464334 | 13.3196468 | 12.5732322 | 10.5648989 | 14.3804132 | 13.5275867 | 17.1566175 | 16.6657054 |
| ENSMUSG00 Efemp2 | 2392 | 2.52330058 | 1.14199721 | 2.26498135 | 2.3428129 | 1.40854292 | 1.94730642 | 2.45266396 | 3.28914502 | 2.42780047 | 2.64138555 | 2.49660109 | 3.40680884 | 1.54677834 | 0.42716134 | 1.0679333 | 2.11302683 | 1.65033804 | 2.09273776 |
| ENSMUSG00 Ctsw | 1222 | 0.03478329 | 0.24534865 | 0.35626986 | 0 | 0.23632696 | 0.08110104 | 0 | 0 | 0 | 0 | 0 | 0 | 0 | 0.03800662 | 0 | 0.34467814 | 0.06272716 | 0 |
| ENSMUSG00 Fibp | 1199 | 91.3559955 | 81.3790385 | 79.8828942 | 83.8262417 | 81.4107933 | 84.268577 | 72.1725519 | 78.7419673 | 73.3731423 | 80.2230481 | 77.3006 | 82.2571108 | 83.3581855 | 76.1930942 | 79.1042382 | 86.6408823 | 72.5930111 | 81.7107749 |
| ENSMUSG00 Fosl1 | 1703 | 0.124795 | 0.1369291 | 0.08521467 | 0.0267534 | 0.2826302 | 0.14548659 | 0.11112784 | 1.12132714 | 0.09673873 | 0.16612109 | 0.11221364 | 0.06554981 | 0.40554718 | 1.39086815 | 0.1693546 | 0.56210522 | 0.99022728 | 0.7429298 |
| ENSMUSG00 Lrp5 | 10175 | 1.86730356 | 1.58788734 | 1.6069045 | 1.19107988 | 1.59887864 | 1.40744374 | 1.39961832 | 1.62528848 | 1.14553159 | 1.52457802 | 1.10809732 | 1.71149747 | 1.53207723 | 1.36935884 | 1.98415355 | 1.24938395 | 1.46148434 | 1.04341598 |
| ENSMUSG00 Drap1 | 1907 | 91.1398347 | 77.4040033 | 78.6862565 | 76.4288311 | 78.1922913 | 80.0327317 | 103.97873 | 81.872334 | 90.8392686 | 90.6420966 | 77.9881729 | 88.1284531 | 107.666198 | 106.063975 | 98.7584226 | 107.000864 | 85.0735264 | 102.633666 |
| ENSMUSG00 Smarca2 | 7924 | 10.5351042 | 12.2548065 | 8.81516631 | 12.2699733 | 10.666287 | 11.2062719 | 11.3325927 | 10.0108051 | 10.5980951 | 12.4422161 | 10.4485112 | 12.7141937 | 10.7703454 | 11.3238234 | 13.2693552 | 12.6072936 | 13.1027181 | 14.2451649 |
| ENSMUSG00 Ovol1 | 2896 | 0.36693004 | 0.47162592 | 0.26725707 | 0.40904253 | 0.39888335 | 0.29088276 | 1.50302709 | 0.81765502 | 1.59284868 | 1.10712977 | 1.02280643 | 1.67678268 | 3.03214132 | 3.48009909 | 1.74992796 | 3.01459202 | 2.19688028 | 2.90621317 |
| ENSMUSG00 Vldlr | 9715 | 0.10500507 | 0.13373164 | 0.1443986 | 0.08441573 | 0.16349495 | 0.18872374 | 0.56492751 | 0.31450254 | 0.54265301 | 0.34944422 | 0.17703536 | 0.37344499 | 0.03554539 | 0.1912263 | 0.02120512 | 0.12612449 | 0.00789013 | 0.27745201 |
| ENSMUSG00 Rnaseh2c | 1610 | 16.7644638 | 14.6700882 | 17.4865805 | 13.6966121 | 13.1540661 | 13.6346963 | 12.6069188 | 11.4102767 | 11.7164126 | 12.4173281 | 13.0268365 | 12.0645035 | 15.412394 | 15.1159638 | 12.7443322 | 9.94128813 | 9.14117889 | 12.5735101 |
| ENSMUSG00 Kat5 | 2476 | 14.3171716 | 16.8313533 | 12.2301414 | 15.6777099 | 13.393738 | 13.2887783 | 13.1466571 | 16.6281946 | 16.8671757 | 15.0059589 | 15.3204027 | 14.6076532 | 13.5284216 | 13.6931275 | 13.8947089 | 15.6038865 | 14.2717471 | 11.3084149 |
| ENSMUSG00 Rela | 2787 | 76.3476538 | 95.9822535 | 84.7361231 | 102.99052 | 91.566365 | 93.4514416 | 60.1127363 | 83.236689 | 65.8659134 | 75.4546597 | 73.3852161 | 63.8064948 | 51.5445078 | 61.5921589 | 62.7706517 | 82.351597 | 77.6152206 | 78.1812242 |
| ENSMUSG00 Slc1a1 | 6416 | 6.02200826 | 7.06133668 | 6.41612755 | 7.2573858 | 7.6518956 | 6.82740282 | 16.0240728 | 18.7866618 | 15.008428 | 18.1297982 | 17.9230557 | 15.9721877 | 22.3823584 | 27.7318109 | 28.1591058 | 13.9650906 | 19.8799745 | 17.5504429 |
| ENSMUSG00 Kcnk7 | 1223 | 0.93838083 | 0.87163746 | 1.54257374 | 0.40978862 | 0.23613372 | 0.81034724 | 0.96714389 | 0.62456913 | 0.47147279 | 0.69395966 | 2.34382462 | 1.50606449 | 2.17818768 | 1.44307058 | 1.24649624 | 1.34627647 | 2.38168321 | 1.25940719 |
| ENSMUSG00 Ehbp1l1 | 6420 | 0.45683133 | 0.26982407 | 0.15823117 | 0.25548249 | 0.35236767 | 0.32417676 | 0.39795711 | 0.29744862 | 0.21812173 | 0.34518428 | 0.47626127 | 0.48686559 | 0.23820708 | 0.80300525 | 0.69311029 | 0.79324779 | 0.39997849 | 0.7882925 |
| ENSMUSG00 Fam89b | 1949 | 0 | 0.0170923 | 0 | 0.02337663 | 0 | 0 | 0 | 0 | 0 | 0 | 0 | 0.0490251 | 0 | 0.0238297 | 0 | 0 | 0.0196646 | 0 |
| ENSMUSG00 Ltbp3 | 5165 | 18.6561925 | 27.9402629 | 23.6689722 | 25.8987853 | 25.5336826 | 22.6225258 | 11.1938226 | 12.4448816 | 13.1972282 | 15.3456597 | 11.0719507 | 11.7250714 | 11.614252 | 9.28881748 | 12.9946437 | 15.783292 | 14.2694035 | 12.0455566 |
| ENSMUSG00 Scyl1 | 2620 | 35.7075925 | 42.111567 | 38.5880452 | 47.2130669 | 39.993587 | 42.4980887 | 39.0961641 | 41.2390482 | 38.1682673 | 40.0421049 | 46.8085185 | 30.2086296 | 36.4528977 | 37.0311827 | 35.6975312 | 36.0545528 | 41.4275087 | 40.458988 |
| ENSMUSG00 Capn1 | 3628 | 9.30240071 | 11.2389698 | 11.9333819 | 13.8642208 | 10.2684975 | 13.3032973 | 10.4979897 | 12.3588425 | 13.9293977 | 14.5948842 | 12.0622552 | 13.4000667 | 12.2921867 | 11.0861576 | 19.8853154 | 16.2851178 | 14.5255398 | 18.4526138 |
| ENSMUSG00 Smc5 | 3437 | 9.89355276 | 7.91871824 | 7.54384995 | 7.35710783 | 7.95430103 | 8.83788935 | 11.4530547 | 6.82284989 | 6.57881791 | 9.63043168 | 5.72687893 | 8.73692944 | 9.77452303 | 8.37804311 | 6.01780071 | 8.0101678 | 8.32986435 | 6.83412528 |
| ENSMUSG00 Arl2 | 3305 | 47.0450632 | 46.8598019 | 47.1001575 | 45.8505396 | 47.665896 | 47.9334612 | 53.2106589 | 48.0380686 | 48.1527071 | 50.6031905 | 57.4311377 | 47.4897559 | 54.6755864 | 44.0410801 | 42.2986019 | 47.9993908 | 45.4233298 | 47.103108 |
| ENSMUSG00 Men1 | 3201 | 13.4380625 | 15.652169 | 13.3741251 | 15.2439483 | 15.0365272 | 16.6259406 | 14.1598083 | 13.398959 | 15.2213834 | 15.7169011 | 12.820584 | 14.734219 | 11.3582078 | 8.54594423 | 14.1457388 | 10.6940965 | 12.7874055 | 11.3592688 |
| ENSMUSG00 Map4k2 | 4502 | 5.57986651 | 6.00105669 | 4.35168365 | 4.34155665 | 4.84312781 | 4.08353269 | 4.21421241 | 6.11655312 | 6.84307281 | 5.47752506 | 7.52386618 | 5.6906684 | 5.9719891 | 5.64302883 | 4.77725726 | 5.28175637 | 6.82756393 | 3.76339513 |
| ENSMUSG00 Sf1 | 5763 | 28.9268264 | 29.9486561 | 30.1589614 | 32.5876496 | 29.5573274 | 28.6069659 | 27.1742091 | 31.6315004 | 30.016201 | 28.4311459 | 27.9288279 | 29.7141164 | 22.5730116 | 26.9493368 | 30.3417323 | 28.2246226 | 31.616036 | 30.0388094 |
| ENSMUSG00 Rps6ka4 | 3764 | 7.02396905 | 9.29291683 | 9.84434757 | 9.51407568 | 9.5266427 | 9.63671652 | 10.8351565 | 10.1771997 | 5.93068319 | 7.92943548 | 7.29824685 | 7.2364621 | 7.20843637 | 3.9114707 | 5.40743056 | 8.94193515 | 6.39450437 | 6.05041716 |
| ENSMUSG00 Prdx5 | 1645 | 73.2536012 | 67.2738204 | 64.3706127 | 69.2971021 | 73.5877131 | 69.0424713 | 114.815936 | 96.6070434 | 87.530735 | 98.6868519 | 92.9070469 | 95.5143405 | 116.807184 | 111.917546 | 85.1832776 | 101.417736 | 90.8181795 | 96.8093398 |
| ENSMUSG00 Esrra | 2531 | 13.7037628 | 11.7799448 | 13.072894 | 10.1346772 | 9.60356438 | 10.5527157 | 11.6832969 | 13.3092449 | 16.8260993 | 16.0396086 | 11.5520642 | 13.4963198 | 11.9285395 | 11.7807602 | 12.6160434 | 16.8381849 | 10.5544958 | 10.5845259 |
| ENSMUSG00 Kcnk4 | 1708 | 0 | 0.07801615 | 0.02832174 | 0.29342593 | 0 | 0.11604856 | 0.0831019 | 0.02236089 | 0 | 0.49690437 | 0.16782772 | 0.13071584 | 0 | 0.2447288 | 0 | 0.04483678 | 0.29171069 | 0 |
| ENSMUSG00 Gpr137 | 3048 | 14.7959291 | 13.5962075 | 13.7280524 | 16.3379992 | 15.5228612 | 12.2906386 | 19.7912289 | 16.4648349 | 20.4311062 | 16.938983 | 19.1695453 | 17.7628585 | 17.5121832 | 19.0621894 | 17.1808325 | 16.8463598 | 14.6364205 | 14.7990379 |
| ENSMUSG00 Bad | 2112 | 22.6412514 | 16.956137 | 17.2010044 | 17.3658337 | 18.2773687 | 18.2068756 | 22.3122293 | 17.2516818 | 24.376489 | 20.9186595 | 19.9514741 | 21.036585 | 25.6236332 | 21.4188168 | 17.10879 | 19.6529443 | 17.2395748 | 20.0293575 |
| ENSMUSG00 Plcb3 | 4200 | 16.7389431 | 18.5362461 | 15.9863 | 20.0685552 | 15.1959359 | 16.0574453 | 15.8046877 | 15.4133624 | 14.0524469 | 15.9414282 | 14.6396123 | 15.5486489 | 11.8279145 | 13.247623 | 16.3334739 | 13.2649604 | 12.2917909 | 14.4333418 |
| ENSMUSG00 Vegfb | 1598 | 55.4322821 | 46.8631954 | 46.284811 | 45.6465782 | 45.9331562 | 46.3277747 | 53.2637731 | 51.4091724 | 52.8620424 | 57.5958528 | 50.286283 | 49.2141856 | 53.1290548 | 40.8928885 | 41.1095128 | 50.7266683 | 40.9645276 | 51.6011583 |
| ENSMUSG00 Dnajc4 | 1080 | 43.6071621 | 37.6312324 | 42.0580972 | 35.3098094 | 45.9035939 | 35.0080886 | 42.537603 | 41.834826 | 39.8899032 | 40.5146784 | 46.4478763 | 41.7583846 | 42.800006 | 45.4980041 | 41.2397055 | 41.836034 | 40.952358 | 43.2942044 |
| ENSMUSG00 Majeed1 | 4425 | 59.8530513 | 59.8352279 | 63.5360358 | 67.9348634 | 65.6552071 | 62.7867959 | 65.3503036 | 65.5528353 | 55.3621233 | 54.9718595 | 56.2179003 | 60.5331898 | 57.8047437 | 65.5989005 | 64.4978933 | 53.3732128 | 54.1245264 | 66.80654 |
| ENSMUSG00 Fasn | 10251 | 17.8711646 | 19.5080777 | 16.492585 | 18.9026555 | 15.9829351 | 19.5001197 | 14.6585766 | 17.503454 | 15.942649 | 18.3340848 | 16.8803921 | 14.3908688 | 15.6114271 | 18.5531692 | 18.6454022 | 16.8873121 | 18.6939308 | 15.3581176 |
| ENSMUSG00 Arhgap19 | 8733 | 0.17035167 | 0.12588177 | 0.31019326 | 0.50084281 | 0.28108647 | 0.2723613 | 0.40632668 | 0.28426728 | 0.03301335 | 0.35634334 | 0.08752998 | 0.41543777 | 0.84733626 | 0.32973015 | 0.71240524 | 0.45599716 | 0.45642218 | 0.46612549 |
| ENSMUSG00 Dus1l | 4099 | 12.7235548 | 13.1983753 | 12.3795641 | 15.89468 | 11.1434973 | 13.3583241 | 15.3976857 | 12.4947577 | 12.7005992 | 14.8043316 | 16.5155199 | 14.1615733 | 11.8666188 | 14.5031549 | 12.9464757 | 12.0691546 | 13.3146247 | 12.4672994 |
| ENSMUSG00 Gps1 | 2935 | 48.8628497 | 51.6434987 | 48.769089 | 49.2090326 | 52.1333511 | 49.5020834 | 46.5550313 | 46.5726093 | 49.536082 | 46.0422199 | 50.9491039 | 48.6651723 | 44.1718155 | 47.9473883 | 46.5219612 | 44.0439832 | 44.9338288 | 48.0369133 |
| ENSMUSG00 Zdhhc16 | 1819 | 25.797534 | 25.7309601 | 24.2532489 | 25.8738641 | 28.0218291 | 27.3234699 | 22.6549441 | 28.4080926 | 25.6311907 | 28.5392657 | 30.0990118 | 26.7571511 | 26.089777 | 26.1200118 | 25.0742839 | 23.6606062 | 28.3391255 | 25.2213277 |
| ENSMUSG00 Rfng | 3348 | 13.7494342 | 14.1689251 | 11.8622068 | 14.9692798 | 12.9386892 | 12.6249945 | 17.3536349 | 17.7957431 | 19.1293405 | 16.7168115 | 15.9677953 | 17.3715413 | 13.6738409 | 12.5682029 | 16.7981254 | 12.9007761 | 12.8441174 | 16.4797294 |
| ENSMUSG00 Mms19 | 10357 | 4.57596517 | 5.73816774 | 4.86210728 | 5.08091206 | 5.8091054 | 5.56912058 | 6.34520721 | 6.23572004 | 4.64476686 | 6.03667399 | 6.08430725 | 5.59396136 | 6.48264357 | 6.03140847 | 6.11837337 | 5.30530285 | 6.20947438 | 6.90999786 |
| ENSMUSG00 Slc16a3 | 5309 | 5.15602437 | 5.52182085 | 8.06377366 | 5.85282205 | 6.93556632 | 5.94558133 | 11.1575577 | 10.7548769 | 12.653081 | 12.2916883 | 14.1102157 | 12.0273343 | 8.79964695 | 4.715269 | 10.1820383 | 7.69562594 | 6.98810612 | 6.24797351 |
| ENSMUSG00 Csnk1d | 5838 | 32.581477 | 39.8636312 | 33.6742082 | 43.7816819 | 37.9003859 | 42.8302661 | 38.5195551 | 36.9101635 | 31.9375397 | 40.5037302 | 38.814085 | 38.1378345 | 32.6175404 | 31.2093448 | 37.1929085 | 35.280092 | 42.9808609 | 38.8870475 |
| ENSMUSG00 Cd7 | 1129 | 0 | 0 | 0 | 0 | 0 | 0 | 0 | 0.10148557 | 0 | 0 | 0 | 0 | 0 | 0.08227473 | 0 | 0 | 0 | 0 |
| ENSMUSG00 Sectm1a | 2047 | 0.22841081 | 0.50449425 | 0.87436279 | 0.6232092 | 1.41080382 | 1.35561948 | 0.18490544 | 0.31718165 | 0.3621677 | 0.09213621 | 0 | 0.81801166 | 0 | 0 | 0.10063888 | 0 | 0.11233893 | 0 |
| ENSMUSG00 1110031I02R | 1299 | 37.7278429 | 39.5189915 | 35.0419492 | 34.3724584 | 38.2017039 | 36.4302237 | 39.955357 | 43.4846522 | 48.922859 | 41.8149424 | 39.9779676 | 44.2142547 | 46.9773531 | 42.2966554 | 43.1680659 | 38.2316534 | 37.9427389 | 44.9728032 |
| ENSMUSG00 Rab40b | 3189 | 3.10558357 | 2.94582508 | 2.18431746 | 2.8431006 | 2.82241133 | 2.62603072 | 3.3084752 | 2.63478474 | 2.72510343 | 2.6318047 | 3.25091439 | 2.15281483 | 4.47065666 | 3.07297035 | 2.81653784 | 2.6655738 | 2.60796689 | 5.34738886 |
| ENSMUSG00 Ubtd1 | 1524 | 11.3793385 | 11.2135923 | 9.80801895 | 13.9313954 | 9.38004024 | 11.9329746 | 7.29559025 | 7.41794692 | 7.70220275 | 9.86947135 | 12.8528432 | 10.6210906 | 9.80811741 | 10.2396413 | 6.38029343 | 6.45714311 | 8.95286201 | 14.0049432 |
| ENSMUSG00 Ankrd2 | 1091 | 0.38959831 | 0.3664113 | 0.62074291 | 0.12528243 | 0.13235176 | 0.22709777 | 0 | 0.28005428 | 0.11325348 | 0 | 0 | 0.40608075 | 0 | 0.0425702 | 0.18882475 | 0.21058079 | 0.49181316 | 0.60504975 |
| ENSMUSG00 Wdr45l | 4783 | 29.5127931 | 28.3469537 | 27.822618 | 30.472461 | 30.269878 | 29.9098352 | 29.3688781 | 24.7456105 | 30.1385752 | 27.8389266 | 26.4295505 | 24.2494139 | 30.8388946 | 27.0138932 | 30.93347 | 32.198374 | 29.8805689 | 24.1635236 |
| ENSMUSG00 Fn3k | 4031 | 0.4006938 | 0.32230287 | 0.03600114 | 0.18084265 | 0.28657062 | 0.23356535 | 0.59859753 | 0.43583491 | 0.47000241 | 0.30412262 | 0.21333397 | 0.6092506 | 0.47728722 | 0.2419563 | 0.26575055 | 0.32296717 | 0.36129975 | 0.32751639 |
| ENSMUSG00 Hoga1 | 2569 | 12.7565164 | 15.0290558 | 10.6764468 | 10.144382 | 17.2180762 | 15.9903527 | 6.85103798 | 12.5920455 | 8.00002778 | 6.2953255 | 12.5899754 | 14.3612896 | 10.2158739 | 10.594097 | 7.81049347 | 6.30475941 | 13.7550976 | 13.3400831 |
| ENSMUSG00 Pi4k2a | 3762 | 15.4790248 | 17.5596675 | 17.3074885 | 15.6351165 | 17.1826617 | 19.7051806 | 20.2481162 | 20.273851 | 19.4984542 | 19.6022594 | 19.8236869 | 21.112623 | 15.9718967 | 13.5801425 | 16.8332779 | 20.5091903 | 20.7320612 | 20.9245167 |
| ENSMUSG00 D19Ertd386e | 6615 | 4.25373039 | 5.43380391 | 4.84832209 | 4.93836263 | 5.26795355 | 5.91786236 | 4.62755907 | 4.88447051 | 5.06192552 | 5.01087595 | 6.0233281 | 4.7335732 | 5.67522132 | 4.27580487 | 6.91986827 | 4.24293503 | 5.85178532 | 5.62980578 |
| ENSMUSG00 Loxl4 | 4492 | 0.06623692 | 0.48945927 | 0.77535489 | 0.16228333 | 0.60004179 | 0.72806777 | 0.54769798 | 0.41661347 | 0.62348243 | 0.61929909 | 0.65940504 | 0.83251322 | 0.16473258 | 0.17576792 | 0.66957114 | 0.32391876 | 1.24568997 | 0.62454685 |
| ENSMUSG00 Hps1 | 6435 | 4.40574241 | 6.57457275 | 5.96118253 | 6.06772572 | 6.24555653 | 6.12189949 | 5.43342187 | 6.8669168 | 6.8612191 | 6.7190668 | 7.03075147 | 7.70230122 | 7.82717755 | 7.44837579 | 8.4259903 | 6.93217679 | 8.07621734 | 8.1039057 |
| ENSMUSG00 Cnnm1 | 6447 | 0.00659302 | 0 | 0 | 0 | 0 | 0 | 0.08072584 | 0.02962029 | 0 | 0.00731359 | 0 | 0 | 0.06886726 | 0 | 0.00639081 | 0 | 0.05350344 | 0 |
| ENSMUSG00 Got1 | 3066 | 46.761239 | 50.9254844 | 45.2338263 | 45.1449617 | 51.1460559 | 51.8639015 | 53.0222297 | 52.9910241 | 47.2852366 | 53.3636465 | 53.2910496 | 54.1044192 | 50.4743068 | 50.8370376 | 43.2844788 | 47.5447953 | 51.626747 | 51.8154926 |
| ENSMUSG00 Entpd7 | 9045 | 0.82237765 | 0.81394689 | 0.98404971 | 0.95705897 | 0.42571077 | 0.91490398 | 1.12462421 | 0.80649518 | 0.91070239 | 0.93832245 | 0.59685686 | 0.79604428 | 1.38532968 | 1.65339924 | 1.02491433 | 1.3673705 | 1.45339081 | 1.42312117 |
| ENSMUSG00 Cutc | 1535 | 8.4179632 | 8.57237343 | 10.6831445 | 9.82456396 | 8.68569567 | 10.168802 | 11.3427139 | 11.8682579 | 14.6500571 | 12.1639599 | 11.4224168 | 11.6358385 | 16.2297187 | 16.7319742 | 12.6154601 | 11.5245998 | 12.1096117 | 15.6964096 |
| ENSMUSG00 Abcc2 | 5818 | 0 | 0 | 0 | 0 | 0 | 0 | 0 | 0 | 0 | 0 | 0 | 0 | 0 | 0 | 0 | 0 | 0.01317508 | 0 |
| ENSMUSG00 Dnmbp | 4884 | 5.73523976 | 6.9094926 | 6.30916029 | 7.15506163 | 6.56344414 | 7.59930333 | 7.03296579 | 8.15615801 | 7.64025047 | 8.06118698 | 7.91358342 | 8.63977087 | 7.13110844 | 6.12407704 | 6.27640409 | 7.55776936 | 7.40786738 | 6.60019176 |
| ENSMUSG00 Cpn1 | 3679 | 0.28883648 | 0.13582316 | 0.22352542 | 0.12384084 | 0.01308288 | 0.22897431 | 2.39199736 | 2.63682256 | 2.94429292 | 1.84552902 | 0.70123614 | 2.13917054 | 0.46931696 | 1.91886417 | 1.53427919 | 0 | 0.19793412 | 0.88217921 |
| ENSMUSG00 Cyp2c44 | 1955 | 0 | 0 | 0 | 0 | 0 | 0 | 0 | 0 | 0 | 0 | 0 | 0 | 0 | 0 | 0 | 0 | 0 | 0 |
| ENSMUSG00 Erlin1 | 4725 | 4.77677216 | 3.79308721 | 4.97556312 | 4.23307904 | 2.36330292 | 2.34916663 | 5.90782658 | 3.51612594 | 4.65473004 | 4.55041708 | 4.3578851 | 3.89623675 | 3.63333486 | 6.29083935 | 4.81338812 | 3.06324859 | 4.7857175 | 3.05024042 |
| ENSMUSG00 Chuk | 6271 | 19.8258077 | 22.4334799 | 20.6962497 | 18.7228211 | 23.647657 | 23.2394498 | 23.0565375 | 23.0335937 | 25.6668839 | 24.6919007 | 30.8773572 | 24.7792704 | 24.5597585 | 27.5213249 | 30.9915068 | 30.7252968 | 29.7577324 | 32.1142305 |
| ENSMUSG00 Cwf19l1 | 3560 | 6.99663853 | 8.03813936 | 5.96516287 | 6.62938791 | 7.23330875 | 7.23803974 | 6.60516894 | 8.50746492 | 7.01096308 | 7.81430194 | 9.71603088 | 6.50660112 | 6.73463815 | 5.23148288 | 8.54122142 | 8.6799214 | 9.58157423 | 7.8805181 |
| ENSMUSG00 Gm13540 | 869 | 1.32064413 | 0.4983517 | 1.22464634 | 0.7864392 | 0.72004029 | 1.14045417 | 1.25223431 | 1.84589287 | 1.5166496 | 0.59684435 | 1.20949262 | 0.38537857 | 1.64628953 | 1.81714496 | 1.99132951 | 2.29126765 | 1.41132509 | 0.56971457 |
| ENSMUSG00 Scd3 | 1990 | 0.93981294 | 1.25551109 | 1.87173962 | 1.48817481 | 1.20934482 | 1.44425053 | 3.8278097 | 3.70408727 | 3.47705259 | 2.77217703 | 1.34442094 | 1.90726887 | 4.61092292 | 9.56888233 | 5.30030066 | 4.90658537 | 3.60151634 | 4.00820124 |
| ENSMUSG00 Scd2 | 5447 | 111.198597 | 120.720253 | 119.792648 | 126.612361 | 111.833601 | 135.394407 | 112.909402 | 102.671443 | 109.533297 | 111.994823 | 105.969717 | 116.211755 | 112.402869 | 102.403798 | 116.49443 | 101.276442 | 114.964818 | 109.129504 |
| ENSMUSG00 Ndufb8 | 3113 | 44.0890502 | 37.6147842 | 39.1277062 | 37.6284757 | 40.7567462 | 36.9775137 | 48.6653382 | 44.4985042 | 47.3651547 | 45.4694543 | 47.099694 | 40.0732431 | 48.8565053 | 49.1743373 | 42.8692064 | 41.7961722 | 42.2044799 | 47.1456141 |
| ENSMUSG00 Sema4g | 4921 | 0.29367527 | 0.3249378 | 0.18677039 | 0.14813589 | 0.10759016 | 0.14097506 | 0.00961444 | 0.08537217 | 0.17576038 | 0.01916306 | 0.08737546 | 0.12476576 | 0.0400991 | 0.0755035 | 0.05860819 | 0.02334319 | 0.04672969 | 0 |
| ENSMUSG00 Mrpl43 | 1715 | 35.1937901 | 27.5632635 | 27.3881621 | 25.6098232 | 26.2690849 | 27.766867 | 28.2496695 | 30.2644168 | 29.7311408 | 28.53786 | 29.0271176 | 28.2821054 | 31.583933 | 30.5204003 | 23.6878916 | 29.2035653 | 22.7946489 | 28.0337955 |
| ENSMUSG00 Peo1 | 3597 | 5.80206879 | 4.89922742 | 3.98069641 | 5.26922281 | 5.27216512 | 3.85731593 | 3.99862507 | 3.94983978 | 4.74039965 | 4.28642757 | 4.68850718 | 4.50001172 | 2.83895104 | 4.23510166 | 5.15448989 | 3.45967401 | 3.71862041 | 5.5972569 |
| ENSMUSG00 Sbn3 | 3016 | 63.137662 | 67.664055 | 63.9634082 | 70.5471078 | 68.1602938 | 67.7571198 | 71.5808203 | 64.7472657 | 62.1348298 | 70.0850805 | 76.0027357 | 65.3280136 | 78.0216331 | 62.9212669 | 72.1710278 | 72.6201175 | 74.2889694 | 77.1878447 |
| ENSMUSG00 Kazald1 | 1675 | 1.67483082 | 1.01430305 | 1.1840685 | 1.16962681 | 0.91953526 | 1.0945977 | 3.02236218 | 2.12053339 | 2.21300682 | 2.28012356 | 2.62405742 | 2.06601259 | 4.09381284 | 3.43824886 | 3.34532036 | 5.4178352 | 3.13474787 | 1.51069786 |
| ENSMUSG00 Tlx1 | 3581 | 0 | 0 | 0 | 0 | 0 | 0 | 0 | 0 | 0 | 0 | 0 | 0 | 0 | 0 | 0 | 0 | 0 | 0 |
| ENSMUSG00 Lbx1 | 2477 | 0 | 0 | 0 | 0 | 0 | 0 | 0 | 0 | 0 | 0 | 0.11572457 | 0 | 0 | 0 | 0 | 0 | 0 | 0 |
| ENSMUSG00 Btrc | 6306 | 12.7353809 | 14.2535671 | 12.8295682 | 13.3404009 | 13.5666709 | 14.7762669 | 11.3406421 | 12.4361437 | 12.11829 | 13.5143423 | 13.8826092 | 13.3522249 | 12.739647 | 11.6478355 | 12.2141581 | 11.4119122 | 15.4386684 | 12.6796514 |
| ENSMUSG00 Poll | 2315 | 5.23281865 | 4.71992627 | 5.72542001 | 6.08136618 | 5.3225799 | 6.72119151 | 6.13123269 | 5.8567183 | 7.24099867 | 7.98404997 | 5.40692835 | 4.65331442 | 5.22087245 | 5.27636932 | 5.05453213 | 5.92139847 | 5.39713727 | 2.39996535 |
| ENSMUSG00 Fgf8 | 1811 | 0 | 0.0367895 | 0.02671095 | 0 | 0 | 0 | 0 | 0.10544562 | 0 | 0 | 0 | 0 | 0 | 0 | 0.13650433 | 0.10571675 | 0 | 0 |
| ENSMUSG00 Mgea5 | 5059 | 30.7341241 | 38.3766649 | 38.5439219 | 39.6981782 | 38.3418961 | 42.8333868 | 24.2502029 | 27.8497277 | 24.4481334 | 31.4555492 | 33.9873642 | 28.3105337 | 25.275416 | 34.1881363 | 29.0911164 | 23.1605572 | 34.636851 | 31.3265935 |
| ENSMUSG00 Kcnip2 | 3422 | 0.01242115 | 0.20443331 | 0.38167309 | 0 | 0.1547198 | 0 | 0.30417269 | 0 | 0.30089506 | 0.0137787 | 0.18149458 | 0.08155415 | 0.30273824 | 0.16286647 | 0.18060296 | 0.12308494 | 0.16799955 | 0.0482254 |
| ENSMUSG00 Ldb1 | 4110 | 16.5056596 | 17.5156118 | 15.2417815 | 14.9098795 | 13.3621715 | 14.9261032 | 16.1277526 | 16.6522592 | 15.7931749 | 17.8507285 | 14.7974506 | 15.671843 | 18.9405755 | 21.3575 | 20.771198 | 23.048896 | 18.5570148 | 20.0495479 |
| ENSMUSG00 Gbf1 | 8684 | 7.17066823 | 8.03284207 | 7.79301808 | 7.81211366 | 8.74622012 | 8.51937255 | 7.95990602 | 7.8900472 | 6.42650033 | 7.61773799 | 7.60856375 | 8.36849294 | 8.24849653 | 7.73889849 | 7.90914252 | 8.70842349 | 8.2487159 | 8.35525239 |
| ENSMUSG00 Nfkb2 | 3266 | 67.8573141 | 80.8444579 | 80.6028023 | 90.2990325 | 80.907612 | 96.6778991 | 58.8583633 | 77.8348532 | 63.3939421 | 70.0042131 | 66.8937187 | 70.7009786 | 39.2268797 | 39.6324775 | 50.7261042 | 51.6911476 | 51.9270863 | 44.2969843 |
| ENSMUSG00 Fbxl15 | 1344 | 7.87484288 | 9.02224218 | 7.27042644 | 9.89867928 | 7.41317574 | 8.81183283 | 9.61038791 | 8.95134431 | 10.8175893 | 9.96339265 | 10.1308506 | 10.0916711 | 8.11187033 | 8.98471929 | 8.52234564 | 10.512823 | 9.06827561 | 8.67703172 |
| ENSMUSG00 Tmem180 | 2794 | 6.86107172 | 7.72611147 | 7.58325207 | 9.94711419 | 9.40588057 | 10.6944518 | 9.70299392 | 7.25847018 | 9.81754456 | 9.11287751 | 10.9434407 | 8.82982207 | 11.335394 | 9.20902806 | 10.23403 | 10.5251204 | 10.9327339 | 11.005759 |
| ENSMUSG00 Actr1a | 2731 | 62.1313302 | 64.6009109 | 64.3681469 | 58.8072799 | 62.8658263 | 62.9978363 | 47.8324081 | 57.7850629 | 60.8975286 | 56.0076461 | 60.282866 | 64.1951652 | 52.8001443 | 60.4742493 | 60.9499399 | 53.9237188 | 59.7418696 | 55.8953757 |
| ENSMUSG00 Pitx3 | 1576 | 0 | 0 | 0 | 0 | 0 | 0 | 0 | 0 | 0 | 0 | 0 | 0 | 0 | 0 | 0 | 0 | 0 | 0 |
| ENSMUSG00 Sufu | 7297 | 4.25809013 | 5.1359471 | 4.88574645 | 4.05847318 | 4.26769283 | 5.60244007 | 4.66837451 | 4.51693069 | 4.59446674 | 5.53117747 | 5.2508588 | 4.54358004 | 3.88733074 | 5.08549073 | 5.92868764 | 4.72270065 | 5.43616792 | 5.61625113 |
| ENSMUSG00 Hexa | 2144 | 192.522278 | 144.267828 | 170.209847 | 175.677777 | 158.359417 | 158.527145 | 231.730616 | 167.768678 | 181.478085 | 184.512314 | 206.54202 | 195.901521 | 175.123704 | 126.508148 | 141.95704 | 134.356455 | 129.029483 | 145.502166 |
| ENSMUSG00 Arih1 | 7687 | 17.0584712 | 18.2274012 | 17.236256 | 16.6490144 | 14.9836989 | 18.1398975 | 18.4031365 | 16.6442758 | 15.2433508 | 16.4693168 | 16.2585263 | 14.3405664 | 13.9453956 | 12.7423677 | 13.3675921 | 13.8079308 | 15.9447766 | 15.714843 |
| ENSMUSG00 Bbs4 | 2526 | 10.6683617 | 9.85462584 | 8.90486604 | 10.6778061 | 11.6804708 | 10.4362843 | 12.7178579 | 11.3851461 | 10.3210865 | 10.7890366 | 12.6340745 | 13.4125523 | 11.834973 | 11.0134633 | 11.5644905 | 15.5678756 | 11.7436872 | 8.79797273 |
| ENSMUSG00 Adpgk | 2803 | 10.1145032 | 11.7658813 | 13.150421 | 12.0282457 | 10.6807194 | 12.7108154 | 11.8323896 | 13.0668976 | 11.9166123 | 12.8179935 | 15.5954645 | 11.5295286 | 11.0878016 | 16.0557691 | 13.6995541 | 11.8027416 | 12.2786353 | 17.6625744 |

| | | | | | | | | | | | | | | | | | | | | |
|---|---|---|---|---|---|---|---|---|---|---|---|---|---|---|---|---|---|---|---|---|
| ENSMUSG00 | Tmem89 | 536 | 0 | 0 | 0 | 0 | 0 | 0 | 0 | 0 | 0 | 0 | 0 | 0 | 0 | 0 | 0 | 0 | 0 | 0 |
| ENSMUSG00 | Arhgef9 | 9209 | 0.1476996 | 0.17001911 | 0.15233276 | 0.2918994 | 0.12543883 | 0.31209236 | 0.89908989 | 0.62209364 | 0.43382474 | 0.66048873 | 1.01163176 | 0.54548916 | 1.25887725 | 0.87249724 | 0.73821878 | 0.84406486 | 0.99051565 | 1.13494736 |
| ENSMUSG00 | Cnksr2 | 9521 | 0.36607756 | 0.48984405 | 0.10669511 | 0.12920368 | 0.15671564 | 0.33309263 | 0.33294291 | 0.12034157 | 0.05191032 | 0.12380712 | 0.04516066 | 0.02931187 | 0.11399036 | 0.02439034 | 0.05625672 | 0.1045642 | 0.17711974 | 0.0288883 |
| ENSMUSG00 | Rps6ka6 | 6702 | 0.08879028 | 0.33800012 | 0.28149324 | 0.09517377 | 0.17954314 | 0.02218117 | 0 | 0.12537046 | 0.01229081 | 0.35176592 | 0.09267002 | 0.00832821 | 0.12513318 | 0 | 0 | 0.00571331 | 0.0686363 | 0.13953369 |
| ENSMUSG00 | Tmem47 | 1993 | 1.96210547 | 2.60753211 | 4.1990069 | 4.25205933 | 6.27912702 | 6.26557394 | 2.18402733 | 4.36922617 | 3.09983812 | 2.46044814 | 3.93130617 | 4.53694799 | 1.88119705 | 0.06991082 | 1.73654325 | 0.21133802 | 1.17305776 | 1.32485556 |
| ENSMUSG00 | Alox5 | 3878 | 0.07672414 | 0.19757518 | 0.3243197 | 0.25846905 | 0.23581912 | 0.19166864 | 0.12200278 | 0.36439373 | 0.23365223 | 0.13374361 | 0.14783381 | 0.30225088 | 0.2035355 | 0 | 0.07437104 | 0.01974761 | 0.15812809 | 0.01418492 |
| ENSMUSG00 | 8-Mar | 6538 | 23.3459907 | 29.2927239 | 28.2043282 | 29.3728666 | 28.2401437 | 30.8700342 | 21.7024662 | 22.4668063 | 22.7603061 | 25.0104985 | 25.772756 | 25.5003647 | 22.4023344 | 25.3531581 | 27.450899 | 25.13663 | 24.8552308 | 22.8685763 |
| ENSMUSG00 | Myo3a | 8046 | 0 | 0.00414031 | 0 | 0 | 0 | 0 | 0 | 0 | 0 | 0.01758043 | 0 | 0 | 0 | 0.15585264 | 0 | 0 | 0.05239737 | 0 |
| ENSMUSG00 | Wdr73 | 3712 | 8.50790565 | 11.7205388 | 8.88759367 | 8.26039408 | 9.2711006 | 8.86395881 | 9.50842093 | 11.0502802 | 11.2064604 | 11.1779684 | 9.7943813 | 9.42791499 | 11.2299219 | 10.4223935 | 11.0218602 | 12.460952 | 11.3161696 | 11.5293871 |
| ENSMUSG00 | Nmb | 1060 | 17.4832611 | 17.2849923 | 19.0299643 | 15.4305801 | 19.2981769 | 17.9979268 | 24.5490315 | 17.5468868 | 22.2251764 | 26.333223 | 26.6367929 | 15.0596976 | 37.9762684 | 33.1242736 | 32.0283793 | 22.7576252 | 26.9368785 | 29.9436364 |
| ENSMUSG00 | Sec11a | 3888 | 81.6759697 | 73.9087 | 74.488767 | 72.2203488 | 76.0603238 | 75.361449 | 74.753546 | 68.938858 | 74.3327626 | 77.3353999 | 74.7958262 | 71.1620986 | 78.8574758 | 72.4972136 | 77.6344132 | 79.3387688 | 79.9351702 | 78.8775129 |
| ENSMUSG00 | Slc28a1 | 2959 | 0 | 0 | 0.03269586 | 0 | 0 | 0 | 0.33577771 | 0 | 0 | 0.09560805 | 0 | 0.07545206 | 0 | 0 | 0.12531734 | 0 | 0.12952449 | 0 |
| ENSMUSG00 | A930017K11f | 2536 | 0.06704286 | 0.06568 | 0.2098221 | 0.05389713 | 0.22775358 | 0.15631777 | 0.7835696 | 0.36144229 | 0.01624074 | 0.24170313 | 0.37677412 | 1.71672779 | 0.31124238 | 0 | 0.19496004 | 0 | 0.27203208 | 0.34706061 |
| ENSMUSG00 | Pigq | 5356 | 20.6256446 | 20.3696282 | 18.0000833 | 20.2370658 | 20.2107349 | 20.0671918 | 17.605334 | 19.1175935 | 17.732656 | 19.7987176 | 19.1866945 | 15.892249 | 14.5527314 | 15.920714 | 18.0391427 | 17.1006596 | 15.9430526 | 18.7848499 |
| ENSMUSG00 | Rab40c | 4876 | 6.97377786 | 6.98231704 | 7.03380497 | 7.64334173 | 6.40640247 | 7.9572991 | 6.18092217 | 6.40717497 | 7.15442552 | 7.2234568 | 7.19171852 | 6.57058866 | 7.891488 | 9.98224028 | 8.2386212 | 7.6015808 | 7.91521349 | 9.17194499 |
| ENSMUSG00 | 0610011F06f | 1329 | 112.899377 | 100.013881 | 96.4194747 | 96.1958543 | 103.434743 | 98.6207527 | 102.635405 | 91.0985069 | 95.2031425 | 95.8624531 | 104.177449 | 92.3961307 | 93.0216589 | 82.6837553 | 85.937331 | 90.7853303 | 74.8646101 | 94.0411516 |
| ENSMUSG00 | Fam195a | 811 | 21.7504908 | 13.7605667 | 14.6731054 | 16.1233289 | 10.9201898 | 13.0755672 | 29.0526684 | 21.9924191 | 20.5170806 | 23.5462821 | 26.7445514 | 20.1652147 | 24.0272593 | 28.9201777 | 21.0326068 | 18.8384123 | 14.980809 | 23.061771 |
| ENSMUSG00 | Rhot2 | 4186 | 4.4576618 | 3.87562816 | 3.62859258 | 4.32100683 | 4.04740495 | 3.35007731 | 4.83751232 | 6.25895558 | 4.4866343 | 5.03496521 | 5.2728215 | 3.56014857 | 5.17360528 | 4.77089288 | 5.05914948 | 6.15613444 | 4.22083681 | 4.91481269 |
| ENSMUSG00 | Rhbdl1 | 1561 | 0 | 0.29877035 | 0.03098881 | 0 | 0.03083403 | 0 | 0 | 0.02446663 | 0.0527694 | 0.06041089 | 0 | 0.10726905 | 0.25282213 | 0.02975278 | 0.02639433 | 0.02452954 | 0.0982096 | 0.31715693 |
| ENSMUSG00 | Jmjd8 | 2537 | 21.8976212 | 20.9830528 | 18.6286713 | 19.6287826 | 18.7443203 | 18.6335466 | 18.537171 | 20.0521403 | 19.1240503 | 18.4179531 | 18.3604984 | 19.404578 | 16.7810188 | 19.3501775 | 18.4489426 | 14.9419398 | 16.2399563 | 16.9125358 |
| ENSMUSG00 | Wdr24 | 4461 | 7.49867149 | 6.55653914 | 6.74475137 | 8.39524287 | 6.8513274 | 6.65368469 | 8.93012232 | 9.29767967 | 8.17082861 | 6.51083461 | 7.44308365 | 8.74583012 | 8.37127581 | 7.6625977 | 8.40471102 | 7.0813161 | 6.70130275 | 10.7650639 |
| ENSMUSG00 | Fbxl16 | 3472 | 0 | 0 | 0 | 0 | 0 | 0 | 0.05450769 | 0 | 0 | 0.08148163 | 0 | 0.03215188 | 0 | 0 | 0 | 0 | 0 | 0 |
| ENSMUSG00 | Gng13 | 769 | 0.05527331 | 0 | 0 | 0 | 0 | 0 | 0 | 0 | 0 | 0 | 0 | 0 | 0 | 0 | 0 | 0 | 0 | 0 |
| ENSMUSG00 | Prps2 | 3865 | 15.3084618 | 24.3662489 | 21.4270334 | 20.3580656 | 21.6188925 | 22.0647489 | 13.4409629 | 13.3698112 | 15.1532278 | 12.9557689 | 10.2472033 | 10.8165411 | 18.5329738 | 27.4218389 | 19.5933919 | 18.0307647 | 21.9149072 | 23.0568572 |
| ENSMUSG00 | Sdc3 | 6080 | 7.2776132 | 12.7279364 | 8.64835979 | 10.0638952 | 9.8005463 | 11.279767 | 2.68468326 | 2.30536376 | 1.94416612 | 1.76815137 | 1.93299996 | 2.57045815 | 1.17650132 | 0.72568887 | 1.41630348 | 1.05802994 | 1.17878578 | 1.0042781 |
| ENSMUSG00 | Hadha | 5857 | 15.9367196 | 16.7787328 | 16.3943073 | 15.6122616 | 16.7808414 | 16.4724556 | 18.6520361 | 16.2498663 | 18.2058886 | 17.2035262 | 16.4850932 | 17.6204815 | 17.6961513 | 16.9774272 | 15.3142884 | 18.5994158 | 17.0004643 | 14.2383141 |
| ENSMUSG00 | Il6 | 1082 | 0 | 0 | 0 | 0 | 0 | 0 | 0 | 0 | 0 | 0 | 0 | 0 | 0 | 0 | 0 | 0 | 0 | 0.25420105 |
| ENSMUSG00 | Tyms | 8387 | 1.78393013 | 1.66822649 | 1.97254649 | 1.68402572 | 1.62410092 | 2.0265396 | 2.43135384 | 2.09928431 | 2.21475124 | 1.87208587 | 2.6430882 | 1.65710027 | 1.9057519 | 1.77757864 | 1.71939246 | 2.5886222 | 2.5087799 | 2.05292124 |
| ENSMUSG00 | Agbl1 | 4573 | 0.08365331 | 0 | 0 | 0 | 0 | 0 | 0.04138437 | 0 | 0.09006454 | 0 | 0 | 0 | 0 | 0.13202999 | 0 | 0 | 0 | 0 |
| ENSMUSG00 | Hspa4l | 11771 | 2.97547065 | 3.14422101 | 2.83148096 | 3.75062883 | 2.87867422 | 3.46881764 | 2.94624022 | 2.76441482 | 2.98113257 | 2.83200639 | 3.23478395 | 3.00629769 | 3.27314819 | 3.75624596 | 2.96822015 | 3.85150617 | 4.24255924 | 3.7012329 |
| ENSMUSG00 | Plk4 | 6100 | 1.76988766 | 1.63833906 | 1.60187756 | 1.18010574 | 1.48341011 | 1.43784161 | 1.90026333 | 1.45882455 | 0.91825674 | 1.8087319 | 1.66037561 | 1.86662216 | 2.07032309 | 1.11161258 | 1.19552056 | 1.56301004 | 1.63357987 | 1.82161304 |
| ENSMUSG00 | Mfsd8 | 3172 | 4.94464374 | 5.52414324 | 6.02381604 | 6.88011987 | 7.32904134 | 4.60846671 | 4.3404759 | 6.55002109 | 3.73950711 | 6.27288675 | 9.21761494 | 4.6278434 | 4.61903513 | 5.49071021 | 5.31255875 | 5.13036196 | 5.90843606 | 5.60149479 |
| ENSMUSG00 | Larp1b | 4635 | 0.28428489 | 0.07187248 | 0.09392919 | 0.12778718 | 0.1038445 | 0.22451077 | 0.0204154 | 0.28840002 | 0.21326351 | 0.3153553 | 0.39168251 | 0.2528865 | 0.46830731 | 0.0501015 | 0.23111984 | 0.52045486 | 0.33902442 | 0.3916506 |
| ENSMUSG00 | Phf17 | 5936 | 14.8653547 | 16.2299343 | 16.7139674 | 16.0492156 | 16.1277622 | 17.9227963 | 15.279381 | 19.1605415 | 14.2931642 | 17.2128862 | 18.704347 | 19.6614809 | 12.8399279 | 11.845746 | 15.8462183 | 14.0106468 | 17.1551501 | 18.5711337 |
| ENSMUSG00 | D3Ertd751e | 3783 | 5.7302775 | 7.59953151 | 6.70043085 | 7.12982772 | 5.85267906 | 8.25223953 | 10.8932972 | 9.46985817 | 12.0630874 | 11.2174552 | 10.6840113 | 10.1952479 | 6.67669618 | 7.63632629 | 9.52983384 | 9.62579108 | 11.5900829 | 9.45173655 |
| ENSMUSG00 | Crisp4 | 3281 | 1.96915171 | 3.91916403 | 1.59230152 | 2.34678954 | 0.86552377 | 2.56749915 | 4.09533704 | 4.26041429 | 3.41442618 | 1.75324163 | 0.53876058 | 1.54807232 | 1.8644187 | 5.39323295 | 2.13479564 | 0.59519079 | 0.91114025 | 1.45863831 |
| ENSMUSG00 | Crispld1 | 4575 | 0 | 0 | 0 | 0 | 0 | 0 | 0 | 0 | 0 | 0 | 0 | 0 | 0 | 0 | 0 | 0 | 0 | 0 |
| ENSMUSG00 | Gdap1 | 5402 | 0 | 0 | 0 | 0.06747285 | 0 | 0 | 0 | 0.09191063 | 0 | 0 | 0 | 0 | 0.02739647 | 0.14615873 | 0 | 0.01417646 | 0 | 0 |
| ENSMUSG00 | Ly96 | 598 | 18.8359058 | 13.2583091 | 18.0389586 | 13.6378377 | 14.7293345 | 18.9758796 | 18.116066 | 19.3516688 | 22.383976 | 16.5579393 | 19.1738964 | 15.3073055 | 31.2655462 | 19.8824187 | 15.8467523 | 18.9532104 | 21.1499632 | 24.1009799 |
| ENSMUSG00 | Itih5 | 7886 | 9.93368494 | 13.3572624 | 14.7095772 | 13.4383704 | 16.2901476 | 16.6076433 | 13.3730613 | 14.689013 | 15.3757421 | 15.0372836 | 16.1572171 | 20.7592364 | 14.8258488 | 14.6823624 | 21.0605468 | 15.3288673 | 22.8081805 | 19.2176121 |
| ENSMUSG00 | Atp5c1 | 22595 | 25.1814245 | 23.8166184 | 24.1021998 | 23.0940757 | 25.2578101 | 24.2884056 | 29.6523499 | 28.1976392 | 27.1490138 | 27.6435214 | 28.491594 | 25.1497792 | 27.730236 | 25.2539083 | 23.6742554 | 24.306359 | 24.3120519 | 25.392564 |
| [illegible] | [illegible] | [illegible] | [illegible] | [illegible] | [illegible] | [illegible] | [illegible] | [illegible] | [illegible] | [illegible] | [illegible] | [illegible] | [illegible] | [illegible] | [illegible] | [illegible] | [illegible] | [illegible] | [illegible] | [illegible] |

| | | | | | | | | | | | | | | | | | | | |
|---|---|---|---|---|---|---|---|---|---|---|---|---|---|---|---|---|---|---|---|
| ENSMUSG00 Adck3 | 5708 | 5.92749126 | 5.32259277 | 5.47464972 | 4.78917775 | 7.0578872 | 8.00413811 | 7.29417615 | 7.3132964 | 8.68755518 | 8.5826175 | 9.28213542 | 9.13311648 | 8.67716274 | 6.7941158 | 7.91837891 | 5.99045417 | 9.07798763 | 8.68311233 |
| ENSMUSG00 Cdc42bpa | 19461 | 2.2824063 | 2.31089909 | 2.44838022 | 2.08127889 | 2.16409399 | 2.78306037 | 2.16372665 | 1.89970945 | 1.39044966 | 1.72263245 | 1.93937707 | 2.10230105 | 1.56911036 | 1.56555785 | 1.54974251 | 1.97739378 | 1.6582263 | 1.47550248 |
| ENSMUSG00 Ahctf1 | 10810 | 2.70130027 | 2.74268972 | 3.01607396 | 2.79435452 | 2.31977171 | 2.910822 | 3.26943304 | 3.74504593 | 3.38712419 | 3.03578986 | 2.60309441 | 2.51970799 | 2.56927562 | 3.19652183 | 2.83189232 | 2.49367138 | 3.0490856 | 3.12447655 |
| ENSMUSG00 Tfb2m | 3215 | 2.81605054 | 2.71476774 | 2.69326965 | 3.52867808 | 2.2456574 | 2.97470531 | 2.84023236 | 3.00549854 | 2.79274042 | 2.47852841 | 3.15032386 | 2.49998613 | 2.68525211 | 2.99033471 | 3.20385349 | 3.02513663 | 3.13524606 | 2.82317344 |
| ENSMUSG00 Kif26b | 7804 | 1.06753133 | 1.44708754 | 0.99176895 | 1.58214288 | 0.96831266 | 0.75560934 | 0.33950667 | 0.93474486 | 0.55414969 | 0.5800189 | 0.67340408 | 0.49350086 | 1.31484352 | 1.67232042 | 1.42547669 | 1.27569938 | 1.17375512 | 1.22649725 |
| ENSMUSG00 Efcab2 | 2529 | 5.76483803 | 7.00769463 | 5.50872944 | 5.0803537 | 7.44151133 | 5.85854778 | 10.3268481 | 9.09127175 | 10.7811283 | 9.21014132 | 7.17852282 | 7.8790793 | 7.97815507 | 9.01702132 | 8.22727787 | 7.50974348 | 8.71396828 | 13.1813043 |
| ENSMUSG00 Parp1 | 3823 | 12.3190518 | 12.2603302 | 10.2238589 | 13.6456698 | 13.2321872 | 11.3415229 | 14.0589075 | 12.3478445 | 14.6194355 | 12.6910996 | 13.496457 | 14.4831643 | 11.0458073 | 9.70672922 | 14.3661193 | 10.6869155 | 11.8698236 | 11.5687475 |
| ENSMUSG00 Mixl1 | 2292 | 0 | 0.02906884 | 0 | 0 | 0 | 0 | 0 | 0 | 0 | 0 | 0 | 0 | 0.30132934 | 0 | 0 | 0 | 0 | 0 |
| ENSMUSG00 Acbd3 | 3462 | 24.5184392 | 26.625297 | 25.8914361 | 26.9523451 | 23.5237478 | 28.2974449 | 25.214296 | 25.7815264 | 27.9900305 | 24.923682 | 23.1974885 | 26.392328 | 21.5738078 | 25.7843836 | 23.0762036 | 24.9298178 | 26.04904 | 20.1159818 |
| ENSMUSG00 Fam36a | 550 | 2.85943911 | 2.11991145 | 1.14337435 | 2.07095656 | 1.83776436 | 3.06325989 | 1.634438 | 4.30532536 | 2.77072924 | 2.65758509 | 2.86649751 | 2.13114348 | 3.04960943 | 1.94220729 | 2.24735759 | 3.06324859 | 2.36926199 | 2.00033115 |
| ENSMUSG00 Pppde1 | 4869 | 19.9998682 | 20.3886881 | 16.8795704 | 19.19197 | 20.3045741 | 18.9601032 | 17.0438363 | 19.6492027 | 14.5662724 | 18.1572332 | 15.89555 | 15.4871555 | 12.2696553 | 14.7278026 | 12.4730116 | 14.4386024 | 17.4825848 | 15.4441259 |
| ENSMUSG00 Sdccag8 | 4282 | 4.98309163 | 5.6480993 | 4.58656079 | 4.16029153 | 5.02451422 | 4.50163787 | 6.11020818 | 5.16426926 | 6.28089556 | 4.78994767 | 4.79757874 | 5.14880587 | 5.54148814 | 4.39277333 | 5.38833986 | 4.96293487 | 4.86910451 | 5.06155723 |
| ENSMUSG00 Capn2 | 4178 | 30.1442883 | 39.3966037 | 38.543677 | 41.918778 | 37.0378494 | 39.1986081 | 21.3122216 | 24.5992712 | 19.6764685 | 24.1960526 | 23.5673024 | 22.4705468 | 14.8184673 | 13.4952422 | 16.9421429 | 17.0007364 | 16.1359397 | 17.9852656 |
| ENSMUSG00 Trp53bp2 | 4343 | 15.4146102 | 16.3151108 | 15.1369164 | 17.708276 | 14.2301151 | 16.0079404 | 14.3691972 | 12.9799645 | 10.5645357 | 14.6457054 | 14.7956065 | 16.8230952 | 13.2331761 | 11.3570391 | 12.5226926 | 13.3307387 | 15.7170469 | 15.4147094 |
| ENSMUSG00 Srp9 | 1602 | 30.0348685 | 34.518979 | 33.4870444 | 34.4125241 | 23.8256027 | 22.4874141 | 28.3226336 | 39.4082654 | 34.6049003 | 40.6755755 | 25.885558 | 23.3436292 | 28.3304394 | 43.3130253 | 31.1712146 | 32.8410078 | 21.7708678 | 21.7358081 |
| ENSMUSG00 Cnih3 | 3597 | 0 | 0 | 0 | 0 | 0 | 0.01377613 | 0.01315337 | 0 | 0 | 0 | 0 | 0 | 0 | 0 | 0 | 0.12774181 | 0 | 0 |
| ENSMUSG00 Nvl | 3037 | 10.4688415 | 8.99459113 | 11.1496448 | 10.2613613 | 11.6962001 | 8.74555983 | 14.0053007 | 10.3875286 | 12.9919928 | 9.08237122 | 8.82507465 | 8.58278381 | 11.3867876 | 12.4024218 | 13.2409347 | 12.2676197 | 13.0993402 | 11.8096601 |
| ENSMUSG00 Tmem63a | 4524 | 12.9187924 | 12.7095613 | 12.5317811 | 13.0620412 | 15.6184049 | 19.0806503 | 14.8192012 | 13.5496918 | 14.9942952 | 16.373509 | 20.3075255 | 13.0656027 | 12.74735 | 7.69961661 | 11.4388051 | 11.1384702 | 12.7246017 | 14.0562616 |
| ENSMUSG00 Pycr2 | 1591 | 230.853065 | 124.918119 | 159.988383 | 129.69577 | 172.016416 | 220.88497 | 244.949426 | 154.065895 | 195.733027 | 156.388599 | 230.106541 | 275.254361 | 170.661768 | 118.139046 | 125.495122 | 101.995974 | 145.644746 | 170.421048 |
| ENSMUSG00 Wdr64 | 3967 | 0 | 0 | 0 | 0 | 0 | 0 | 0 | 0 | 0 | 0 | 0 | 0 | 0 | 0 | 0 | 0 | 0 | 0 |
| ENSMUSG00 Opn3 | 2208 | 0.53901491 | 0.60349446 | 0.70106566 | 1.09363014 | 0.89375402 | 0.58350139 | 0.81425806 | 1.60864738 | 1.38034518 | 0.96095183 | 1.55787908 | 1.13754747 | 1.00540592 | 1.28310205 | 1.17558787 | 1.00582211 | 0.74639075 | 0.7224928 |
| ENSMUSG00 Fh1 | 3571 | 34.625471 | 33.3595379 | 35.2337027 | 35.7496629 | 37.3895145 | 35.0657401 | 39.4559389 | 34.4277075 | 34.7161614 | 37.7232034 | 36.363018 | 34.8554824 | 37.6171411 | 34.9858849 | 33.4712164 | 31.5031657 | 32.8526449 | 35.4455207 |
| ENSMUSG00 Rgs7 | 1613 | 0.15810977 | 0 | 0 | 0 | 0 | 0 | 0 | 0 | 0 | 0 | 0 | 0 | 0 | 0.20155525 | 0 | 0 | 0 | 0.2728288 |
| ENSMUSG00 1810030J14R | 797 | 0 | 0 | 0 | 0 | 0 | 0 | 0 | 0 | 0 | 0 | 0 | 0 | 0 | 0 | 0 | 0 | 0 | 0 |
| ENSMUSG00 Spna1 | 9640 | 0 | 0 | 0 | 0 | 0 | 0 | 0 | 0 | 0 | 0 | 0 | 0 | 0 | 0 | 0 | 0 | 0 | 0 |
| ENSMUSG00 Ifi202b | 1789 | 0 | 0.01862096 | 0 | 0.02546732 | 0.13452187 | 0.11079426 | 0.13223219 | 0.23483311 | 0.04604418 | 0 | 0.10681936 | 0.06239873 | 0.11030054 | 0.1298046 | 0.04606099 | 0 | 0.02142331 | 0.06149704 |
| ENSMUSG00 Mnda | 1644 | 0 | 0 | 0 | 0 | 0 | 0 | 0.028779 | 0.02323139 | 0.07515788 | 0 | 0 | 0 | 0.0600145 | 0 | 0 | 0.23291124 | 0 | 0 |
| ENSMUSG00 Apcs | 981 | 0 | 0 | 0 | 0 | 0 | 0 | 0 | 0 | 0 | 0 | 0 | 0 | 0 | 0 | 0 | 0 | 0 | 0 |
| ENSMUSG00 Dusp23 | 1415 | 6.24811068 | 6.56840812 | 6.63213066 | 5.92454569 | 4.8982311 | 4.09729317 | 7.95789229 | 7.4225517 | 10.0128349 | 9.36349655 | 6.65135464 | 6.03519178 | 6.72866434 | 10.8314833 | 6.52236644 | 7.60400048 | 3.87325827 | 6.8809978 |
| ENSMUSG00 Ccdc19 | 2610 | 1.51455225 | 2.27191386 | 0.51894975 | 1.50124514 | 1.93634176 | 1.27204335 | 1.95776602 | 1.56574217 | 1.56224715 | 1.55362471 | 2.43459308 | 1.36865993 | 1.54989165 | 0.74737612 | 1.05766446 | 0.71886581 | 1.73275978 | 1.79148432 |
| ENSMUSG00 Tagln2 | 2077 | 377.307622 | 431.608081 | 371.70994 | 404.762826 | 372.147599 | 373.995499 | 369.435256 | 425.283526 | 419.777252 | 422.245104 | 388.939226 | 388.909133 | 382.565812 | 358.292543 | 350.084284 | 441.97353 | 371.416324 | 338.000106 |
| ENSMUSG00 Slamf9 | 1309 | 0.4870723 | 1.14521027 | 1.03472792 | 0.38286592 | 0.47800994 | 1.40065022 | 0.83131521 | 2.50920291 | 0.25171285 | 0.360204 | 0.76644318 | 0.21319963 | 1.31903522 | 0.42576703 | 0.78688991 | 0.26326621 | 0.70269757 | 0 |
| ENSMUSG00 Copa | 7807 | 27.592703 | 34.2815156 | 32.8397222 | 35.3598524 | 29.8458617 | 30.2571898 | 28.4469982 | 33.442201 | 27.2114831 | 38.2182209 | 29.0798774 | 22.492132 | 25.6991043 | 35.9678433 | 34.9740888 | 35.671524 | 33.554533 | 28.3254271 |
| ENSMUSG00 Dcaf8 | 3901 | 66.966627 | 73.5855446 | 72.3184899 | 74.1520304 | 72.1053478 | 70.0292797 | 70.8659274 | 61.5327478 | 61.2993875 | 74.3944062 | 69.5498562 | 73.7865604 | 63.1539592 | 67.5171545 | 71.3132288 | 76.7578954 | 75.1592339 | 61.1995702 |
| ENSMUSG00 Vangl2 | 7411 | 1.14708342 | 1.95535137 | 1.18796148 | 2.3422963 | 1.30542662 | 0.90934716 | 0.88100792 | 0.75240733 | 1.02813459 | 1.13248215 | 0.70266772 | 0.95649563 | 0.53918301 | 0.72069492 | 0.32801131 | 0.64584077 | 0.4447526 | 0.17814311 |
| ENSMUSG00 Uck2 | 1324 | 4.91185191 | 5.63601836 | 5.58999256 | 5.02410308 | 5.12583175 | 4.49118431 | 7.54001152 | 7.15386389 | 7.15475727 | 6.48144108 | 5.66515746 | 5.26960561 | 8.30892564 | 6.55970116 | 7.65527398 | 7.31686078 | 6.77368087 | 5.19345796 |
| ENSMUSG00 Fmo9 | 4319 | 0 | 0 | 0 | 0 | 0 | 0 | 0 | 0 | 0 | 0 | 0 | 0 | 0 | 0 | 0 | 0 | 0 | 0 |
| ENSMUSG00 Tada1 | 4260 | 7.04428504 | 5.94314545 | 6.68826511 | 7.7539336 | 6.12382691 | 6.3976534 | 6.4527386 | 7.5398616 | 6.73873274 | 7.08367373 | 5.75318162 | 6.14496379 | 5.41956274 | 6.74856575 | 4.94225236 | 6.02223168 | 4.99321474 | 4.44204993 |
| ENSMUSG00 Dusp27 | 4014 | 0.11648155 | 0 | 0 | 0 | 0 | 0 | 0 | 0 | 0 | 0 | 0 | 0 | 0 | 0 | 0 | 0 | 0 | 0 |
| ENSMUSG00 Pou2f1 | 17172 | 2.04704056 | 1.86429603 | 1.90710925 | 2.2791135 | 1.83591952 | 1.73140229 | 1.85701989 | 1.61470325 | 1.74368722 | 1.43604809 | 2.55122507 | 2.3012747 | 1.64612093 | 1.80669988 | 1.39881826 | 1.91096265 | 1.99978814 | 1.87079655 |
| ENSMUSG00 Mpzl1 | 2350 | 112.141315 | 117.502375 | 120.007524 | 128.113779 | 117.64671 | 123.185987 | 98.712368 | 102.794445 | 105.490059 | 108.366361 | 109.028554 | 114.220222 | 78.0703831 | 66.2470557 | 93.2031792 | 87.286291 | 97.5934301 | 103.065999 |
| [illegible] | | | | | | | | | | | | | | | | | | | |

| | | | | | | | | | | | | | | | | | | | | |
|---|---|---|---|---|---|---|---|---|---|---|---|---|---|---|---|---|---|---|---|---|
| ENSMUSG00 | Hao1 | 2275 | 0 | 0.07321515 | 0.19136781 | 0.02002683 | 0 | 0 | 0 | 0 | 0.03620793 | 0 | 0 | 0 | 0 | 0 | 0.01811057 | 0.28612762 | 0 | 0 |
| ENSMUSG00 | Tubgcp4 | 8376 | 2.86209638 | 3.24538224 | 3.1994909 | 3.61181154 | 3.75815163 | 3.47271426 | 3.39480898 | 3.49732243 | 2.51269216 | 4.23883471 | 2.99449059 | 3.87164517 | 3.38656309 | 3.40456897 | 3.87617312 | 3.9726048 | 3.935126 | 2.68609416 |
| ENSMUSG00 | 6330527O06 | 2927 | 0 | 0 | 0 | 0 | 0 | 0 | 0 | 0 | 0 | 0 | 0 | 0 | 0 | 0 | 0 | 0 | 0 | 0 |
| ENSMUSG00 | Ubr1 | 10058 | 6.63483062 | 8.34312791 | 8.41655181 | 7.52857981 | 7.05373389 | 7.6018958 | 6.08225232 | 9.33735509 | 5.89256275 | 6.68491776 | 7.39091623 | 8.04105141 | 6.81759445 | 8.25631619 | 7.45544161 | 7.95658873 | 8.16977337 | 6.77632563 |
| ENSMUSG00 | Snap25 | 6928 | 0 | 0 | 0 | 0 | 0 | 0 | 0 | 0 | 0.10106391 | 0 | 0 | 0 | 0 | 0.04692676 | 0.00594711 | 0 | 0 | 0 |
| ENSMUSG00 | Mkks | 4136 | 14.1204332 | 9.76995665 | 10.9822884 | 9.38539888 | 9.89171545 | 9.57268717 | 13.1894388 | 11.5611431 | 12.8956813 | 12.1410781 | 11.1698223 | 10.6476204 | 10.5080801 | 9.84803304 | 8.33793575 | 10.7854346 | 8.35839454 | 8.17954561 |
| ENSMUSG00 | Jag1 | 6390 | 6.37910232 | 5.33319105 | 6.25297904 | 8.11400132 | 7.7131596 | 7.22740965 | 6.98213713 | 8.67846139 | 7.03200136 | 9.49655009 | 9.12135357 | 9.11917868 | 8.67747653 | 8.26399489 | 9.83937 | 10.1449137 | 10.1243802 | 8.22123582 |
| ENSMUSG00 | 2210009G21 | 5892 | 1.41395358 | 1.3569407 | 1.89651825 | 1.144439 | 1.40507313 | 1.32039761 | 1.61402723 | 1.93165918 | 2.55143894 | 1.66451906 | 1.79196637 | 1.13677521 | 1.18892265 | 1.26544566 | 1.97196856 | 2.04710477 | 1.44406618 | 1.2977369 |
| ENSMUSG00 | Mtch2 | 6694 | 42.8417124 | 42.2557193 | 41.8553164 | 39.2516496 | 43.1274691 | 39.8775883 | 42.2732496 | 42.5057362 | 40.7927397 | 39.6209599 | 40.2311684 | 40.9153654 | 44.7996303 | 39.8805818 | 36.9422973 | 42.8266773 | 40.4332699 | 34.4977935 |
| ENSMUSG00 | Cdan1 | 13193 | 0.63147233 | 0.63883592 | 0.59765675 | 0.5249207 | 0.59831998 | 0.63100578 | 0.54151554 | 0.41976035 | 0.42144923 | 0.40742668 | 0.67717102 | 0.57537559 | 0.44870993 | 0.70759202 | 0.37163535 | 0.5775662 | 0.50547833 | 0.75052219 |
| ENSMUSG00 | Haus2 | 3211 | 9.54413949 | 10.1360005 | 10.8919534 | 10.6985448 | 12.9660897 | 11.6358646 | 7.52936124 | 11.0735368 | 13.0190948 | 8.63426145 | 11.6350101 | 12.9153028 | 9.18732063 | 8.37468907 | 10.3292596 | 10.5057693 | 14.0963428 | 12.6601463 |
| ENSMUSG00 | Lrrc57 | 2932 | 10.2493723 | 11.6231551 | 9.61861125 | 11.452418 | 10.3749576 | 10.6305148 | 13.8936619 | 12.2705467 | 8.1474008 | 12.028897 | 8.9618783 | 9.53739664 | 10.7682221 | 11.0882883 | 9.40103714 | 10.852488 | 10.7188143 | 9.41833955 |
| ENSMUSG00 | Snap23 | 4395 | 20.4160243 | 22.0949003 | 20.2959704 | 19.5513378 | 20.7969336 | 19.787269 | 18.6451786 | 21.7075361 | 20.0637839 | 18.6028031 | 20.664436 | 18.6433203 | 16.4664215 | 13.9702124 | 21.5804281 | 18.2348672 | 20.7720681 | 17.572875 |
| ENSMUSG00 | Zfp106 | 11534 | 6.08059133 | 6.23280958 | 7.01235848 | 7.13792328 | 6.77702827 | 6.47871703 | 4.7993614 | 5.67554861 | 4.32076331 | 5.33889531 | 5.06992797 | 6.0925888 | 4.09317191 | 5.79846418 | 4.72599778 | 4.82367371 | 5.20697999 | 4.92668695 |
| ENSMUSG00 | Vps39 | 8115 | 6.51588897 | 8.66586811 | 9.1322549 | 10.3305387 | 8.95615585 | 9.09840703 | 7.97581575 | 7.75143398 | 6.8517309 | 7.16992826 | 9.41369792 | 7.77223644 | 9.93325358 | 7.54322947 | 9.63146661 | 9.47002453 | 11.3349492 | 9.32748378 |
| ENSMUSG00 | Ehd4 | 3998 | 32.7878847 | 33.3712209 | 31.9687001 | 33.8915823 | 32.4089891 | 34.8901813 | 32.7685863 | 36.1480867 | 31.7500854 | 37.8809552 | 32.0013354 | 31.7749995 | 31.4771688 | 25.045886 | 29.8345433 | 33.1379444 | 32.171849 | 29.4033169 |
| ENSMUSG00 | Itpka | 1811 | 0 | 0.073579 | 0 | 0.25157948 | 0 | 0 | 0.02612517 | 0 | 0 | 0 | 0 | 0.09246106 | 0.13620077 | 0 | 0 | 0 | 0.23279363 | 0 |
| ENSMUSG00 | Ltk | 4058 | 0.03142324 | 0 | 0 | 0.01122746 | 0.02372199 | 0 | 0 | 0 | 0 | 0.0116192 | 0.08241121 | 0.06877238 | 0 | 0 | 0 | 0 | 0 | 0 |
| ENSMUSG00 | Tyro3 | 5849 | 0.03633542 | 0.13669165 | 0.02481118 | 0.08568499 | 0.19749806 | 0.025416 | 0.03235608 | 0.19589196 | 0.09154124 | 0 | 0.17969723 | 0.07634216 | 0 | 0.03970259 | 0.11270728 | 0 | 0 | 0.0376195 |
| ENSMUSG00 | Ubox5 | 5578 | 4.12249915 | 4.75983806 | 4.59626588 | 3.64292323 | 4.53016492 | 4.45956835 | 5.31822332 | 5.27901439 | 4.46716422 | 5.00416212 | 5.46440006 | 4.04258295 | 4.63426401 | 4.22975912 | 4.90459539 | 4.83952639 | 4.30810114 | 6.63698975 |
| ENSMUSG00 | Oxt | 534 | 0 | 0 | 0 | 0 | 0 | 0 | 0.08860052 | 0.07152135 | 0.30851323 | 0 | 0 | 0 | 0 | 0 | 0 | 0 | 0 | 0 |
| ENSMUSG00 | Ptpra | 3604 | 55.4076906 | 58.0572388 | 62.6681502 | 63.2974886 | 64.4519044 | 63.7008365 | 49.623176 | 61.6546607 | 57.6428373 | 63.8313766 | 69.5149507 | 63.125594 | 49.8794413 | 40.4903781 | 52.1878753 | 47.3639089 | 56.8832858 | 48.6290272 |
| ENSMUSG00 | Rtf1 | 5884 | 7.72230339 | 6.07490406 | 6.58517978 | 5.35830093 | 6.11874217 | 6.65306926 | 6.47292769 | 5.99758325 | 7.25173867 | 6.40268727 | 6.39002249 | 6.5074005 | 7.63789542 | 7.53019364 | 6.84125087 | 7.21690748 | 6.52667395 | 7.40435291 |
| ENSMUSG00 | Ndufaf1 | 1831 | 23.8873982 | 16.1925046 | 19.6294555 | 20.4539478 | 22.5018715 | 20.3515323 | 19.1989735 | 14.955736 | 20.2221069 | 19.0559915 | 20.7172676 | 17.7719996 | 14.333468 | 16.3606971 | 16.6966436 | 19.2812343 | 22.3761941 | 17.214756 |
| ENSMUSG00 | Nusap1 | 4249 | 0.61021787 | 0.28224622 | 0.13661623 | 0.31096029 | 0.29452342 | 0.12828432 | 0.16702523 | 0.3595425 | 0.51374096 | 0.41058508 | 0.86577355 | 0.30213232 | 0.17415363 | 1.47562998 | 0.79513476 | 0.74796903 | 0.88396731 | 0.60647918 |
| ENSMUSG00 | 4930402H24 | 10803 | 6.29137985 | 6.17350868 | 5.08676576 | 5.95081307 | 5.9390775 | 6.62813106 | 4.99272924 | 5.92524914 | 4.10988281 | 5.09348058 | 4.81597054 | 5.64718312 | 4.34274306 | 3.44364658 | 3.76813267 | 4.6786635 | 4.69011983 | 2.68858728 |
| ENSMUSG00 | Atrn | 13645 | 1.94692085 | 2.87598844 | 2.11290759 | 2.14699535 | 2.24345207 | 2.41862371 | 1.94521164 | 2.08245858 | 1.56656663 | 2.36358234 | 1.88718964 | 2.34798024 | 1.95592285 | 1.49083989 | 2.09555733 | 1.31891429 | 1.4746285 | 1.45938414 |
| ENSMUSG00 | Chac1 | 1634 | 31.2155516 | 25.9734561 | 23.8019084 | 20.3546893 | 20.8257467 | 27.5683616 | 5.99371147 | 5.32917243 | 4.94036542 | 4.96323192 | 4.18103982 | 7.17337157 | 3.68328881 | 4.77515709 | 3.40404532 | 2.69487139 | 2.46282814 | 3.87150996 |
| ENSMUSG00 | Dll4 | 3841 | 5.53308722 | 5.70681708 | 6.37256089 | 5.2429007 | 4.1352603 | 4.48954733 | 5.34592623 | 6.01572597 | 7.43094387 | 6.59201449 | 4.45285997 | 7.46921389 | 5.02181194 | 5.0543162 | 6.99385953 | 7.35706021 | 3.48239459 | 4.94093773 |
| ENSMUSG00 | Spint1 | 2756 | 327.471844 | 333.612438 | 321.185498 | 341.327606 | 341.359427 | 335.793488 | 284.56276 | 299.427881 | 306.851498 | 304.460782 | 314.714939 | 313.588432 | 277.858981 | 262.773097 | 273.326577 | 313.118802 | 291.507923 | 284.167145 |
| ENSMUSG00 | Gfra4 | 2165 | 11.701194 | 14.3099268 | 10.9482817 | 13.7209242 | 11.3382361 | 11.146504 | 12.3908956 | 14.8006585 | 15.7136545 | 16.7477556 | 14.365588 | 20.0833262 | 11.8032024 | 8.28056246 | 9.15378678 | 13.0170379 | 8.74512271 | 11.2813133 |
| ENSMUSG00 | Ppp1r14d | 1130 | 4.1000568 | 2.59427849 | 3.08220724 | 3.9109923 | 3.06681284 | 4.25364173 | 6.40605299 | 4.900795 | 6.05040851 | 7.51073149 | 6.0881363 | 5.63096065 | 12.7040601 | 7.72698091 | 7.94861875 | 5.92996132 | 5.83373723 | 6.1337588 |
| ENSMUSG00 | Adam33 | 5926 | 0.49622282 | 0.48906881 | 0.82445605 | 0.46129981 | 0.91780415 | 0.56024859 | 1.38121726 | 2.79063622 | 2.17539309 | 1.51970711 | 1.24958614 | 1.299791 | 0.0582726 | 0 | 0.07647943 | 0.08399897 | 0.07760978 | 0.25991478 |
| ENSMUSG00 | Siglec1 | 7776 | 0.14758742 | 0.00856813 | 0.1804054 | 0.02343675 | 0.02475922 | 0.0573527 | 0.26163133 | 0.26522502 | 0.47139916 | 0 | 0.13516578 | 0.22969408 | 0 | 0 | 0 | 0.06401465 | 0 | 0 |
| ENSMUSG00 | Rad51 | 4882 | 0.35696686 | 0.73695055 | 0.75304963 | 0.23331137 | 0.67041598 | 0.60900533 | 0.7268437 | 0.58673293 | 0.55680255 | 0.63743269 | 0.40122353 | 1.02896551 | 1.09132468 | 0.84668656 | 1.12245124 | 0.86275437 | 1.56225585 | 1.0140966 |
| ENSMUSG00 | Rpusd2 | 2703 | 1.68259483 | 1.70076929 | 1.14535921 | 1.29789139 | 1.24647971 | 2.4932193 | 3.53576069 | 1.8368525 | 1.34088546 | 2.16303633 | 1.18421096 | 1.1357238 | 1.29580693 | 1.90724887 | 2.19497745 | 1.18994118 | 1.50299204 | 1.6484416 |
| ENSMUSG00 | Casc5 | 8594 | 0.05440504 | 0.06202075 | 0.04503005 | 0.04241196 | 0.00560064 | 0.02306387 | 0.26976045 | 0.08888155 | 0.17732151 | 0.04389174 | 0.05559106 | 0.01948417 | 0.03444165 | 0.10268067 | 0.37874365 | 0.04009954 | 0.24974082 | 0.10881485 |
| ENSMUSG00 | 1700037H04 | 2194 | 34.8333211 | 35.7422757 | 35.4974402 | 34.970282 | 32.2488276 | 35.6174389 | 36.7244723 | 35.5290307 | 33.433539 | 35.3093006 | 34.5138146 | 33.3520208 | 35.6835701 | 32.239902 | 30.7227646 | 46.6154113 | 37.8895798 | 34.1236984 |
| ENSMUSG00 | Spef1 | 4213 | 0.32284966 | 0.67210919 | 0.43631478 | 0.24873108 | 0.35416322 | 0.42342711 | 0.28075409 | 0.54392218 | 0.43014638 | 0.87295392 | 0.26081748 | 0.66242182 | 0.17564177 | 0.70553563 | 0.54765894 | 0.40899058 | 0.62770341 | 0.70507756 |
| ENSMUSG00 | Cdc25b | 4089 | 1.49688074 | 1.43386387 | 1.69171309 | 1.44850471 | 1.65972151 | 1.92684877 | 2.77697308 | 1.61586833 | 3.27356753 | 1.94875736 | 2.03297696 | 3.41254968 | 1.60458425 | 1.05632187 | 2.84148661 | 2.10696666 | 2.0714384 | 1.9103187 |
| ENSMUSG00 | D2Ertd750e | 1885 | 0.69902412 | 1.66122656 | 1.25745517 | 0.60425788 | 0.3319443 | 0.4731826 | 1.4808743 | 0.16208977 | 0.43699221 | 0.87547725 | 0.25344806 | 0 | 1.36088048 | 1.55224271 | 2.33876206 | 0.95472602 | 2.17555107 | 0.9338416 |
| ENSMUSG00 | Ivd | 5629 | 9.20479829 | 7.74677181 | 7.70848405 | 8.37727498 | 8.44809746 | 9.7098357 | 10.6661555 | 10.957671 | 11.7874359 | 12.757243 | 11.0249904 | 12.0674474 | 14.2763712 | 14.4637565 | 13.4693534 | 12.9177231 | 12.0446295 | 10.6812851 |
| ENSMUSG00 | Smox | 3724 | 6.00368491 | 7.4605139 | 6.80658698 | 7.80557097 | 6.1005016 | 5.10962666 | 10.468756 | 8.25587646 | 7.8966628 | 8.40710713 | 3.98979916 | 6.81958283 | 6.63675901 | 8.82986409 | 5.97444688 | 6.52914492 | 4.74447446 | 6.45515028 |
| ENSMUSG00 | Adra1d | 2486 | 0 | 0 | 0 | 0 | 0 | 0 | 0 | 0 | 0 | 0 | 0 | 0 | 0 | 0 | 0 | 0 | 0 | 0.04425511 |
| ENSMUSG00 | Prnd | 3453 | 0 | 0.22189301 | 0.16810958 | 0.5541743 | 0.12545245 | 0.37311644 | 0.20552858 | 0.32075866 | 0.46518219 | 0.46426988 | 0.49808819 | 1.00219261 | 0.62861406 | 0.17485466 | 1.20514253 | 0.64316688 | 0.83245649 | 1.16294955 |
| ENSMUSG00 | Rassf2 | 13121 | 0.10690274 | 0.11425046 | 0.28387789 | 0.30904146 | 0.08437118 | 0.32478737 | 0.47236956 | 0.34929413 | 0.48340244 | 0.66120947 | 0.53888376 | 0.47643885 | 0.4474124 | 0.74333194 | 0.38623515 | 0.14591345 | 0.57835583 | 0.91395361 |
| ENSMUSG00 | Slc23a2 | 8331 | 3.17347491 | 4.04665094 | 3.60580511 | 3.08990398 | 4.05576887 | 3.78292385 | 2.69757802 | 2.98442612 | 3.22333553 | 2.79587653 | 2.56910112 | 3.12208604 | 3.41077714 | 2.62018018 | 2.40354773 | 3.18513875 | 4.07139272 | 3.67123555 |
| ENSMUSG00 | 5730494N06 | 3225 | 11.6114915 | 10.8770473 | 13.9195771 | 11.6551509 | 12.9844259 | 11.4163352 | 16.2843639 | 13.0623938 | 14.3163052 | 16.3894382 | 15.0805915 | 16.9783457 | 16.413379 | 16.4750499 | 15.9695953 | 16.2067222 | 13.999497 | 12.7245872 |
| ENSMUSG00 | Pcna | 2382 | 25.3389379 | 21.6491857 | 22.2981261 | 26.2234222 | 23.7022445 | 23.881838 | 30.0918172 | 26.8244707 | 26.5412686 | 31.4733913 | 28.9618136 | 28.3999089 | 31.6453273 | 34.8818104 | 30.7022509 | 34.7380364 | 34.3681655 | 37.5041097 |
| ENSMUSG00 | Fsip1 | 1963 | 0 | 0.1187928 | 0 | 0 | 0.19615659 | 0 | 0 | 0 | 0 | 0 | 0 | 0 | 0 | 0 | 0 | 0 | 0 | 0 |
| ENSMUSG00 | 4921508D12 | 1071 | 0 | 0 | 0 | 0 | 0 | 0 | 0 | 0 | 0 | 0 | 0 | 0 | 0 | 0 | 0 | 0 | 0 | 0 |

[illegible]

| | | | | | | | | | | | | | | | | | | | | |
|---|---|---|---|---|---|---|---|---|---|---|---|---|---|---|---|---|---|---|---|---|
| ENSMUSG00 | Tdo2 | 4066 | 0.08363045 | 0 | 0.03569124 | 0.08964298 | 0.0591883 | 0.0121871 | 0.11636173 | 0.03757246 | 0.14181289 | 0 | 0.19974774 | 0.06863707 | 0 | 0.0228451 | 0 | 0.09417267 | 0 | 0 |
| ENSMUSG00 | Rrh | 2119 | 0.06017722 | 0.06288418 | 0.47939789 | 0.51602882 | 0.18171562 | 0.21046465 | 0 | 0 | 0 | 0 | 0 | 0 | 0 | 0 | 0.23332641 | 0 | 0 | 0 |
| ENSMUSG00 | Ppa2 | 2094 | 78.1900373 | 63.4917673 | 63.1812825 | 62.314628 | 66.405505 | 58.3794621 | 65.3430122 | 56.5225669 | 63.7663248 | 61.6066495 | 59.638845 | 54.9360461 | 65.3282744 | 57.3564518 | 62.9632182 | 64.2016822 | 61.2598451 | 66.016182 |
| ENSMUSG00 | Ctso | 4697 | 20.1621316 | 20.6459091 | 19.5574482 | 22.3294707 | 23.6509431 | 21.2685354 | 14.1424316 | 14.7744507 | 16.0203884 | 18.3202115 | 14.2907849 | 17.87242 | 14.2208678 | 16.5822301 | 14.7279992 | 16.1330875 | 18.1635804 | 16.8880417 |
| ENSMUSG00 | Ints12 | 4866 | 19.6889985 | 14.6984759 | 18.6097922 | 16.9753747 | 17.8343318 | 14.674371 | 15.01249 | 12.9898122 | 13.5595557 | 15.5424842 | 12.1646473 | 13.5237703 | 12.3177722 | 16.5980822 | 15.6474466 | 16.6036132 | 14.6578789 | 12.525707 |
| ENSMUSG00 | Egf | 5725 | 0.27470594 | 0.38404385 | 0.45627435 | 0.62870262 | 0.59691993 | 0.36353097 | 1.84292182 | 1.64110585 | 1.14387004 | 0.79064935 | 1.10988113 | 1.05294176 | 1.34424089 | 2.29583873 | 1.28822332 | 0.54175357 | 0.91715331 | 1.5854153 |
| ENSMUSG00 | Gstcd | 5163 | 0.98791809 | 0.76136384 | 1.39602092 | 1.22660956 | 0.75512024 | 1.13252422 | 0.33906045 | 0.55479957 | 0.84558797 | 1.08675841 | 1.31397329 | 1.00539543 | 1.261246 | 0.72864053 | 0.5506309 | 1.0234561 | 1.11348916 | 1.31050167 |
| ENSMUSG00 | Pdgfc | 5464 | 8.02806399 | 10.4316182 | 10.3758744 | 11.6821053 | 11.4515924 | 10.7739106 | 6.58948548 | 7.89151214 | 8.76645641 | 7.77503361 | 7.69435642 | 6.57856638 | 5.9227192 | 6.03501136 | 6.47733099 | 3.60200809 | 5.66757831 | 5.99019372 |
| ENSMUSG00 | Glrb | 3622 | 8.56675276 | 9.7032312 | 10.7378019 | 4.54100966 | 5.86034742 | 9.34415158 | 6.0087886 | 6.69579667 | 6.94780815 | 4.51719879 | 5.57946092 | 6.11784043 | 4.16774343 | 6.82171576 | 7.79212197 | 2.48434366 | 3.11096943 | 6.39393538 |
| ENSMUSG00 | Pitx2 | 4478 | 0 | 0 | 0 | 0 | 0 | 0 | 0 | 0 | 0 | 0 | 0 | 0 | 0 | 0 | 0 | 0 | 0 | 0 |
| ENSMUSG00 | Enpep | 5342 | 0 | 0.04365224 | 0 | 0.0938172 | 0 | 0 | 0 | 0 | 0 | 0 | 0 | 0 | 0.1985207 | 0 | 0 | 0 | 0 | 0.10761783 | 0 |
| ENSMUSG00 | Alpk1 | 4169 | 2.23282167 | 2.86863253 | 1.92612282 | 3.12555977 | 2.66694036 | 2.47228798 | 2.06546116 | 3.05062845 | 2.7365471 | 2.79352931 | 3.27743778 | 2.22245142 | 3.53807706 | 6.18289002 | 2.53988963 | 3.11358021 | 3.45662932 | 3.16675117 |
| ENSMUSG00 | Aimp1 | 1115 | 39.2648711 | 41.6784641 | 38.1782123 | 30.2786851 | 34.6204509 | 39.4644192 | 43.1542551 | 32.5406119 | 41.3712087 | 32.0117331 | 37.8771868 | 27.8828018 | 47.8276256 | 39.2796183 | 35.6587457 | 36.1957849 | 38.7731506 | 38.580378 |
| ENSMUSG00 | Tbck | 4081 | 2.54135333 | 2.48969192 | 2.32325703 | 2.15468796 | 2.14653519 | 2.0763336 | 4.06928456 | 2.45195036 | 2.01845213 | 2.95775056 | 1.95501545 | 2.7217145 | 2.76819629 | 2.23059082 | 1.48410407 | 3.50911227 | 2.44176359 | 3.35635079 |
| ENSMUSG00 | Dkk2 | 3890 | 0.05759509 | 0.10833462 | 0.20974973 | 0.04938867 | 0.45653586 | 0.1611471 | 0.01282186 | 0 | 0 | 0 | 0 | 0 | 0 | 0 | 0 | 0 | 0 | 0 |
| ENSMUSG00 | Papss1 | 2587 | 78.0767671 | 81.8335689 | 93.9422561 | 86.0323547 | 88.7473042 | 81.9813296 | 78.1107274 | 88.1067868 | 82.1501435 | 80.0668881 | 85.1158808 | 74.6078706 | 76.2194331 | 80.0339163 | 77.7526076 | 66.7680441 | 75.6599916 | 68.1287893 |
| ENSMUSG00 | Kcnq5 | 12414 | 0 | 0 | 0 | 0 | 0 | 0 | 0 | 0 | 0 | 0 | 0 | 0 | 0.04768673 | 0 | 0 | 0 | 0 | 0 |
| ENSMUSG00 | Fubp1 | 5751 | 25.6243167 | 24.8731643 | 27.74055 | 25.5335152 | 23.9948227 | 24.0482837 | 22.7884056 | 25.8667027 | 21.7713455 | 21.578969 | 21.6237553 | 21.0897391 | 23.1776805 | 21.3201862 | 21.8222812 | 23.775997 | 21.1990872 | 20.4024386 |
| ENSMUSG00 | Dnajb4 | 4309 | 17.1243875 | 16.5288856 | 16.9739563 | 17.3510498 | 17.235451 | 18.9057076 | 14.3398372 | 16.220027 | 14.423405 | 14.9254025 | 15.1451829 | 14.8315 | 16.4859505 | 18.4741624 | 16.1784712 | 18.8120714 | 19.2120681 | 19.5321265 |
| ENSMUSG00 | Ptgfr | 4940 | 0.2581286 | 0.026974 | 0.04896106 | 0.19368055 | 0 | 0.02006184 | 0.22985917 | 0.38656278 | 0.46889168 | 0.50586787 | 0.25144715 | 0.0677923 | 0.04993109 | 0.03760655 | 0 | 0.29454313 | 0.22499243 | 0 |
| ENSMUSG00 | Ifi44 | 3116 | 1.82788626 | 2.64065625 | 1.73871483 | 2.96819384 | 2.93487339 | 4.03927931 | 0.71363797 | 0.96829262 | 2.22058236 | 1.16514896 | 1.33389647 | 0.93145522 | 0.5224497 | 0.68563159 | 1.38837079 | 2.38394667 | 0.81178934 | 0.82972658 |
| ENSMUSG00 | Efna3 | 2428 | 0.36763126 | 0.10976242 | 0.49808001 | 0.2064133 | 0.53523968 | 0.36735964 | 0.6625334 | 0.53481947 | 0.47496806 | 0.56316738 | 0.37385676 | 0.52873156 | 0.65017354 | 0.30605659 | 0.30544811 | 0.22078604 | 0.33148774 | 0.1132807 |
| ENSMUSG00 | Efna4 | 1595 | 20.9994224 | 16.6460042 | 18.4698934 | 14.653803 | 19.0717089 | 15.4716372 | 14.890887 | 15.7319175 | 13.9182019 | 16.7318483 | 14.0179815 | 16.79719 | 14.9387561 | 23.9645667 | 18.8313067 | 16.6366087 | 14.4654738 | 21.9691542 |
| ENSMUSG00 | Adam15 | 5858 | 10.1220076 | 9.91197928 | 9.57065922 | 9.58975208 | 9.67075353 | 10.4637643 | 7.48700126 | 10.9139777 | 9.23849124 | 11.1477849 | 10.5940033 | 13.0153969 | 11.7982274 | 9.97382416 | 13.9190644 | 13.6612102 | 13.6870285 | 13.4001358 |
| ENSMUSG00 | Zbtb7b | 5835 | 4.90976869 | 4.96696109 | 4.81682744 | 4.97384494 | 4.66058901 | 4.95951952 | 4.08664785 | 4.44432583 | 4.55980963 | 4.69486865 | 4.66696252 | 5.05067178 | 5.57995979 | 5.16576053 | 4.88628563 | 4.33107127 | 4.48618008 | 4.44032378 |
| ENSMUSG00 | Cks1b | 1528 | 22.7547343 | 18.6621973 | 19.7229774 | 18.337724 | 20.4434676 | 18.1931175 | 24.5542895 | 17.7464697 | 17.2508967 | 20.2427103 | 16.6962224 | 18.8487447 | 23.6005442 | 22.1582066 | 19.5222032 | 23.5557402 | 18.0595116 | 19.2603875 |
| [illegible] | | | | | | | | | | | | | | | | | | | | |

| | | | | | | | | | | | | | | |
|---|---|---|---|---|---|---|---|---|---|---|---|---|---|---|
| 22.4968448 | 21.746665 | 23.9849181 | 24.0277709 | 22.0531658 | 25.8224067 | 23.511531 | 23.0338931 | 23.7677415 | 22.1820018 | 27.8488505 | 24.2257118 | 22.4933996 | 22.8048729 | 22.3074288 |
| 0.01081953 | 0.01143005 | 0.01176745 | 0.05617749 | 0 | 0.03912279 | 0.05598516 | 0 | 0 | 0.02343002 | 0 | 0.00978427 | 0.009093 | 0 | 0 |
| 0 | 0 | 0 | 0 | 0 | 0 | 0 | 0 | 0 | 0 | 0 | 0 | 0 | 0 | 0 |
| 5.14908222 | 5.27282722 | 5.5715819 | 6.07577144 | 6.0957839 | 6.23269183 | 6.39993181 | 6.91697512 | 5.36234417 | 6.37306825 | 8.86467476 | 6.08424978 | 5.52906345 | 6.5746497 | 6.56635467 |
| 0 | 0 | 0 | 0 | 0 | 0 | 0 | 0 | 0 | 0 | 0 | 0 | 0 | 0 | 0 |
| 12.1440893 | 12.3042699 | 12.7936149 | 13.3679824 | 12.8881998 | 9.06103347 | 12.2934743 | 12.4909801 | 12.0764797 | 11.2476771 | 14.5918878 | 14.5478948 | 11.6681924 | 13.1842461 | 15.9026327 |
| 64.0303591 | 65.406595 | 67.3650824 | 85.4595197 | 74.2891411 | 67.1759913 | 66.8452299 | 74.3909068 | 71.441548 | 81.0413467 | 87.6872691 | 82.1032124 | 73.6440294 | 63.9272332 | 83.1296635 |
| 6.11802374 | 6.78426848 | 6.14727108 | 7.97310152 | 6.58899606 | 5.03614577 | 5.13647056 | 8.02976181 | 5.85704596 | 8.37918737 | 7.70282107 | 6.79669953 | 8.85331963 | 7.15742317 | 8.53631758 |
| 165.372206 | 161.651683 | 156.100615 | 172.581687 | 152.194864 | 160.11577 | 160.081437 | 157.464233 | 161.204988 | 168.447609 | 160.615138 | 169.729986 | 179.17157 | 164.693379 | 163.436617 |
| 0 | 0 | 0 | 0 | 0 | 0 | 0.01689384 | 0 | 0 | 0 | 0 | 0 | 0 | 0 | 0 |
| 0.05556225 | 0 | 0.04028678 | 0 | 0.15525367 | 0.30136475 | 0 | 0 | 0.38571799 | 0.58155512 | 0 | 0 | 0 | 0.03115959 | 0.20125283 |
| 0.02527658 | 0 | 0 | 0 | 0.05297143 | 0 | 0.01307925 | 0.23854348 | 0 | 0 | 0 | 0 | 0 | 0 | 0 |
| 0.14571763 | 0.05131335 | 0.05282807 | 0 | 0 | 0 | 0 | 0 | 0 | 0.15777799 | 0 | 0 | 0 | 0 | 0 |
| 0.0761205 | 0 | 0 | 0.09728863 | 0 | 0 | 0 | 0 | 0 | 0 | 0 | 0 | 0 | 0 | 0 |
| 4.21356298 | 4.32322668 | 3.79145999 | 2.39239095 | 2.00745162 | 1.86339526 | 1.0979871 | 1.49396078 | 2.63666803 | 0.2954007 | 0.18540554 | 0.30154166 | 0.07642836 | 0.35699811 | 0.51239354 |
| 36.5533886 | 32.130431 | 37.0734338 | 34.1308293 | 40.6419298 | 36.254797 | 37.9397392 | 33.0025309 | 28.8684716 | 31.1898471 | 33.5179013 | 34.6073675 | 28.5799562 | 30.9406763 | 28.7426224 |
| 180.667147 | 203.482329 | 185.382204 | 185.908094 | 212.783655 | 199.820023 | 186.545978 | 213.956606 | 206.324754 | 188.298689 | 200.486947 | 186.90151 | 187.531382 | 204.62499 | 193.118742 |
| 1.09761605 | 1.8131167 | 0.36898663 | 0.63207915 | 1.76491391 | 1.19969483 | 1.29080988 | 1.46484761 | 0.61120962 | 0.85353076 | 1.88176876 | 1.20915648 | 1.40885283 | 1.41016598 | 0.59032988 |
| 33.2810029 | 37.7410907 | 33.3693938 | 29.0945144 | 32.9128644 | 29.1309803 | 28.7120047 | 30.851382 | 29.1948748 | 20.4524325 | 24.7628946 | 24.7112391 | 22.0959329 | 27.549844 | 22.340169 |
| 10.9531169 | 12.5465838 | 12.3437495 | 10.2669192 | 10.7227773 | 10.411118 | 11.1196511 | 12.7534121 | 9.739713 | 10.3025498 | 11.9775103 | 11.5343667 | 11.4233188 | 12.7742867 | 11.7496137 |
| 1.17317792 | 1.09572952 | 1.74713969 | 0.87676883 | 1.45792763 | 0.91474771 | 0.93921792 | 1.12407766 | 1.22028968 | 0.92445986 | 0.52220587 | 0.80641579 | 1.12947724 | 1.2927943 | 0.72922955 |
| 1.00994594 | 1.01898219 | 0.66646267 | 0.21211164 | 0.8846559 | 0.04103264 | 0.03523091 | 0.14278942 | 0.36144638 | 0.58977136 | 1.58476712 | 0.94409543 | 0.94415196 | 0.63002132 | 0.09590629 |
| 3.15769614 | 2.97278736 | 2.8502751 | 1.13764575 | 1.29648891 | 2.19428396 | 1.53389841 | 1.70061733 | 2.32894207 | 1.58159848 | 1.59849994 | 1.25294689 | 2.58159211 | 1.16550974 | 2.15264303 |
| 9.93376416 | 10.3620675 | 10.5070606 | 18.090837 | 16.444481 | 16.9107472 | 14.3665979 | 11.8124897 | 13.0199373 | 24.1361703 | 30.4368844 | 19.5923483 | 16.1903093 | 14.3962913 | 15.8680116 |
| 0 | 0 | 0 | 0 | 0 | 0 | 0 | 0 | 0 | 0.12046866 | 0 | 0 | 0 | 0 | 0 |
| 15.1835195 | 14.1681773 | 16.4111355 | 13.6003535 | 14.3540145 | 13.3654746 | 14.0963783 | 13.1174972 | 12.2935763 | 17.4052541 | 14.7830087 | 16.5185524 | 19.6696515 | 15.4628305 | 15.4582553 |
| 0 | 0 | 0 | 0 | 0 | 0 | 0 | 0 | 0 | 0 | 0 | 0 | 0 | 0 | 0 |
| 0.01362064 | 0 | 0 | 0 | 0.02283552 | 0 | 0 | 0 | 0.0166863 | 0 | 0 | 0.35720332 | 0 | 0 | 0 |
| 82.3631032 | 82.1949037 | 83.1156904 | 72.220611 | 73.6439147 | 71.5424234 | 73.3071175 | 77.6812316 | 84.8199252 | 83.9340321 | 72.2815001 | 76.273598 | 81.4552942 | 80.1860816 | 76.7649347 |
| 27.2765973 | 30.3429134 | 29.3998673 | 12.2448114 | 14.7470529 | 15.9142305 | 14.3201719 | 16.3018076 | 15.5185252 | 8.27725635 | 7.57417021 | 7.83815563 | 8.21101355 | 9.43472031 | 6.80403041 |
| 0.02070329 | 0.00729051 | 0 | 0 | 0 | 0.00623849 | 0.03570941 | 0 | 0.01690871 | 0 | 0 | 0 | 0.0347991 | 0.02322102 | 0 |
| 0.08946695 | 0 | 0 | 0.04645329 | 0.01874934 | 0 | 0.09258852 | 0.09381435 | 0.02740091 | 0.04843585 | 0 | 0.10113293 | 0.0375951 | 0.03763014 | 0 |
| 0 | 0 | 0 | 0 | 0 | 0 | 0 | 0.07844821 | 0 | 0 | 0 | 0 | 0.01047909 | 0 | 0 |
| 0.66415517 | 0.4560605 | 0.07223431 | 0.31036014 | 0 | 0.06003865 | 0 | 0 | 0 | 0.97081761 | 0.27081101 | 0.42042404 | 0.08372582 | 0.53075776 | 0.80188202 |
| 44.468065 | 52.1090617 | 50.5763685 | 43.3401795 | 50.7818127 | 50.3990483 | 44.6999559 | 52.7625708 | 47.5586143 | 37.4317702 | 41.2756094 | 39.0508749 | 50.385064 | 53.1988071 | 44.2321116 |
| 3.10024055 | 3.07122943 | 3.94001047 | 3.85624277 | 2.61787404 | 2.63831868 | 3.4199538 | 2.85792374 | 2.4624557 | 4.54955369 | 3.72756634 | 2.83440414 | 3.84623998 | 3.8880366 | 2.49542807 |
| 20.1807577 | 20.558607 | 19.6882963 | 15.9845907 | 18.0209258 | 18.3237209 | 14.0486663 | 18.0612648 | 18.0842075 | 13.2205379 | 15.0268509 | 15.8677477 | 14.8273776 | 16.9762068 | 18.1210367 |
| 31.1508407 | 29.0462946 | 34.1125576 | 30.7055068 | 35.3069135 | 31.2329467 | 28.8524247 | 33.6285489 | 31.4044953 | 26.0433632 | 29.5751415 | 27.0487924 | 25.8744602 | 25.8266365 | 22.9742398 |
| 10.1108497 | 12.5382922 | 10.1507819 | 5.66976794 | 9.36228915 | 5.18874166 | 8.06504006 | 6.80169262 | 9.29943439 | 6.97282818 | 5.7401023 | 5.87990794 | 7.307767 | 7.55011038 | 8.60162884 |
| 0 | 0.14780681 | 0.04150145 | 0.11887608 | 0.18125931 | 0.54041492 | 0.14479557 | 0.21340015 | 0.06232905 | 0.30298777 | 0.02593193 | 0.17253583 | 0.1496562 | 0.19259446 | 0.47606988 |
| 27.3239766 | 25.3021126 | 26.071938 | 27.1703075 | 27.9948012 | 27.1590861 | 29.0846309 | 27.3915657 | 27.9982315 | 29.5854534 | 26.843436 | 25.4340007 | 31.5574141 | 25.219803 | 29.4009802 |

| | | | | | | | | | | | | | | | | | | | | |
|---|---|---|---|---|---|---|---|---|---|---|---|---|---|---|---|---|---|---|---|---|
| ENSMUSG00 | Piwil1 | 3922 | 0 | 0 | 0.08633725 | 0 | 0 | 0 | 0 | 0 | 0 | 0 | 0 | 0 | 0 | 0 | 0 | 0 | 0 | 0 |
| ENSMUSG00 | Scarb2 | 2098 | 28.0801592 | 29.3750497 | 27.1150008 | 28.904552 | 29.0443353 | 30.7519824 | 20.3638461 | 26.0866123 | 21.6729806 | 25.1260659 | 22.6122659 | 22.6934033 | 28.9454672 | 21.0968614 | 23.4483593 | 25.2593902 | 27.7673838 | 23.1258494 |
| ENSMUSG00 | Zcchc8 | 4272 | 17.9492831 | 19.4013298 | 20.3028417 | 21.618033 | 19.2550228 | 20.1597966 | 19.6028656 | 16.9505606 | 18.4047422 | 17.9353682 | 18.5977893 | 18.0826023 | 15.3700332 | 16.7859717 | 18.7393781 | 14.6995777 | 16.8843845 | 17.0744559 |
| ENSMUSG00 | Stx2 | 5773 | 1.28111911 | 1.6965167 | 2.17023114 | 1.6731234 | 1.47572328 | 1.23602869 | 1.82759872 | 1.1378994 | 0.34958241 | 1.16794571 | 0.93514123 | 1.0635238 | 1.4868792 | 1.5205149 | 1.32747088 | 1.61174738 | 0.94272201 | 0.78135229 |
| ENSMUSG00 | Ran | 2716 | 48.2173959 | 41.8864263 | 42.5494713 | 42.5079847 | 45.0129183 | 44.5901623 | 47.3126789 | 40.400137 | 42.5816407 | 42.9842199 | 43.6588538 | 39.9094318 | 40.5226463 | 42.8701499 | 38.1827379 | 42.1535037 | 39.9914306 | 39.0491744 |
| ENSMUSG00 | B3gnt4 | 1422 | 0 | 0 | 0 | 0 | 0 | 0 | 0 | 0 | 0 | 0 | 0 | 0 | 0 | 0 | 0 | 0 | 0 | 0 |
| ENSMUSG00 | Gbas | 2785 | 49.5257796 | 44.0663204 | 44.2222651 | 46.2481409 | 45.0383458 | 43.681135 | 45.7837234 | 48.354187 | 52.0562066 | 46.0840985 | 51.4974597 | 44.612447 | 46.0903586 | 39.4232756 | 42.6513772 | 48.396028 | 48.812702 | 42.7629142 |
| ENSMUSG00 | Diablo | 5015 | 21.5195697 | 19.6489613 | 20.1307194 | 21.1588579 | 23.8692112 | 19.9791879 | 22.0761054 | 20.8135266 | 21.3529294 | 22.517634 | 25.2830987 | 20.2004841 | 24.2675653 | 24.4491307 | 22.7656055 | 26.3949411 | 24.256763 | 24.2303324 |
| ENSMUSG00 | Vps33a | 4094 | 12.1576847 | 14.1909349 | 14.6278531 | 16.9379359 | 13.2145291 | 13.7135435 | 15.4396041 | 12.6872681 | 14.6476715 | 15.8589444 | 17.0608181 | 13.4562919 | 12.037759 | 13.8855796 | 15.9009424 | 14.1508766 | 17.2440251 | 16.5269178 |
| ENSMUSG00 | Mmp17 | 3479 | 0 | 0.1723576 | 0.02780887 | 0.01309602 | 0 | 0 | 0 | 0.0658679 | 0.07103165 | 0.27105894 | 0.20598574 | 0.33691547 | 0.05671965 | 0.01334964 | 0.04737172 | 0 | 0.0813175 | 0.03162352 |
| ENSMUSG00 | Il31 | 1627 | 0 | 0 | 0 | 0 | 0 | 0 | 0 | 0 | 0 | 0 | 0 | 0 | 0 | 0 | 0 | 0.04706897 | 0 | 0.10143044 |
| ENSMUSG00 | Bcl7a | 8572 | 8.21641119 | 7.03410389 | 6.7887724 | 6.3887512 | 7.36128697 | 6.6825665 | 6.20385571 | 6.07727915 | 5.3765412 | 5.62156075 | 6.2477518 | 4.94214745 | 4.92053071 | 6.01410838 | 5.26795442 | 4.96723698 | 5.10599989 | 5.31352547 |
| ENSMUSG00 | Shwap | 3949 | 6.61957034 | 8.22488525 | 8.06021347 | 7.33776252 | 6.31358229 | 7.42851818 | 8.41061043 | 8.06595685 | 7.73876797 | 8.66837442 | 9.44852939 | 9.07410873 | 9.05688529 | 6.59790657 | 8.00242932 | 9.5120501 | 7.53132601 | 7.68929524 |
| ENSMUSG00 | Psmd9 | 4555 | 11.1791879 | 10.5972302 | 9.88710357 | 12.4730236 | 9.98565705 | 9.75824413 | 13.0148818 | 11.6799158 | 11.1036315 | 10.6412564 | 12.0617349 | 11.9473701 | 12.3573474 | 12.9696771 | 11.5237722 | 13.819925 | 11.3674703 | 13.0306973 |
| ENSMUSG00 | Wdr66 | 7560 | 0.03935664 | 0.07050348 | 0.04479031 | 0.04821275 | 0.00636666 | 0.08520973 | 0 | 0.03031143 | 0 | 0 | 0.06319439 | 0 | 0.00652539 | 0.04914718 | 0.00544994 | 0 | 0.00506962 | 0.16007941 |
| ENSMUSG00 | Hpd | 2328 | 0 | 0.01430966 | 0 | 0 | 0.02067522 | 0.02128554 | 0 | 0 | 0 | 0 | 0 | 0 | 0 | 0 | 0.01769826 | 0.08223928 | 0 | 0 |
| ENSMUSG00 | Psph | 1808 | 14.011662 | 10.2628773 | 12.1736484 | 9.6010829 | 9.79676322 | 11.6481813 | 27.1890893 | 22.328191 | 18.7480655 | 16.3254094 | 12.6571931 | 18.3067964 | 32.3331427 | 45.9817016 | 28.8502045 | 17.8534193 | 16.8737458 | 25.4052014 |
| ENSMUSG00 | Cct6a | 1584 | 131.433303 | 128.793033 | 129.698474 | 128.1692 | 141.509071 | 130.044642 | 110.217036 | 120.388046 | 117.943204 | 114.959606 | 125.017174 | 108.001898 | 106.574382 | 109.190519 | 110.313004 | 111.801805 | 118.87443 | 99.8776458 |
| ENSMUSG00 | Rhof | 4595 | 0.27750931 | 0.23199404 | 0.23160341 | 0.26771452 | 0.23044664 | 0.02156811 | 0.33978638 | 0.14961115 | 0.17926666 | 0.17444221 | 0.86296443 | 0.02429409 | 0.93403195 | 0.25268818 | 0.34969765 | 0.43332135 | 0.40036175 | 0.1795271 |
| ENSMUSG00 | Gm13103 | 2037 | 0 | 0 | 0 | 0 | 0 | 0 | 0 | 0 | 0 | 0 | 0 | 0 | 0 | 0 | 0 | 0 | 0 | 0 |
| ENSMUSG00 | Tmem116 | 3714 | 0.78967613 | 0.57405095 | 1.05499622 | 0.5765668 | 0.38878775 | 0.93395028 | 0.53503837 | 0.85351896 | 0.60992417 | 1.04102253 | 0.81039913 | 0.72136559 | 0.62428651 | 0.37515418 | 0.80983134 | 0.65982738 | 0.74299768 | 0.51839492 |
| ENSMUSG00 | Mapkapk5 | 5866 | 5.12293888 | 4.46367795 | 5.59932273 | 4.06989213 | 4.46364925 | 4.15614472 | 4.41993659 | 4.70080371 | 4.93592236 | 4.71828551 | 5.96983371 | 5.76615948 | 3.91055942 | 5.64518144 | 5.37319964 | 5.28078783 | 6.0958805 | 4.99827035 |
| ENSMUSG00 | Aldh2 | 4691 | 242.89837 | 254.03301 | 236.835402 | 265.800277 | 255.352629 | 241.520721 | 253.437006 | 228.397243 | 205.045809 | 268.812281 | 238.528523 | 219.71691 | 299.840466 | 281.100091 | 284.678071 | 317.621854 | 292.859969 | 301.582649 |
| ENSMUSG00 | Acad10 | 6397 | 3.58141158 | 4.18689494 | 3.05501114 | 4.34457356 | 3.76959415 | 3.39285561 | 3.96429512 | 4.07775689 | 2.30495117 | 4.58460801 | 5.36226672 | 3.80422524 | 3.47798885 | 3.46315924 | 4.90786081 | 3.41185653 | 4.09804394 | 3.44828072 |
| ENSMUSG00 | Brap | 5142 | 17.3261083 | 18.7231164 | 20.6213105 | 20.2552601 | 19.9473219 | 17.0283314 | 18.2644239 | 18.3831575 | 16.4922279 | 18.9904913 | 16.9284275 | 16.7816053 | 15.80118 | 19.681382 | 19.5591206 | 16.6879135 | 18.4699657 | 20.5936465 |
| ENSMUSG00 | Fam168a | 6858 | 3.81170651 | 4.20175758 | 4.38028031 | 3.66720567 | 3.72675 | 3.48993443 | 3.9599705 | 4.06539133 | 3.55532478 | 3.84328421 | 4.91821532 | 3.79266532 | 3.34490253 | 2.60054383 | 2.99789682 | 3.46725973 | 3.749191 | 2.3341572 |
| ENSMUSG00 | Vps29 | 3147 | 61.5224267 | 56.9611324 | 56.489585 | 53.5092532 | 56.0239073 | 56.4967296 | 68.5710609 | 57.416033 | 62.1265937 | 56.3950579 | 60.8003523 | 56.5960532 | 69.6791777 | 66.500495 | 56.9123494 | 68.6116095 | 58.8960826 | 59.7286678 |
| ENSMUSG00 | 1500011H22I | 1381 | 14.8352606 | 13.3637534 | 15.5874156 | 13.1965371 | 12.1636795 | 14.2809471 | 15.2455772 | 10.4538219 | 13.7188974 | 13.8959712 | 15.6366989 | 14.9946494 | 10.0735704 | 12.9814755 | 12.7990351 | 13.5306419 | 12.1001198 | 13.224492 |
| ENSMUSG00 | Gpn3 | 2614 | 15.8052912 | 13.1263508 | 14.1937635 | 16.2792714 | 14.7305046 | 14.7482887 | 18.37122 | 19.6367976 | 18.2298388 | 16.197908 | 15.6264688 | 18.9183921 | 17.9285851 | 21.6940439 | 20.2540166 | 17.8416067 | 19.9109071 | 16.2459948 |
| ENSMUSG00 | Arpc3 | 2647 | 85.2512201 | 67.5696747 | 68.274843 | 63.6684182 | 71.9342237 | 66.3261561 | 77.8594747 | 68.3625246 | 75.1222131 | 72.6409399 | 71.1300005 | 62.837429 | 75.8895229 | 77.2547476 | 66.5419915 | 73.0949903 | 68.385002 | 68.6211071 |
| ENSMUSG00 | Anapc7 | 3272 | 24.2404213 | 27.9779442 | 21.4812774 | 23.1286353 | 24.6837921 | 23.6405309 | 20.2582711 | 19.1662362 | 28.3220234 | 22.076674 | 24.7781493 | 21.1696935 | 30.0484447 | 22.3136019 | 25.1969173 | 26.8806617 | 24.2936246 | 24.0580782 |
| ENSMUSG00 | Atp2a2 | 5653 | 37.8132903 | 51.357133 | 48.2023875 | 46.8505839 | 46.0458948 | 53.3921281 | 35.1350833 | 43.0162792 | 33.3470162 | 42.4297944 | 40.5829384 | 37.0853757 | 33.7984284 | 39.353826 | 44.6635653 | 35.9537492 | 47.0925983 | 44.6164434 |
| ENSMUSG00 | P2rx7 | 5767 | 4.0758388 | 6.00752731 | 4.26948621 | 5.02459237 | 4.07289384 | 4.89770386 | 3.24879848 | 5.07289409 | 4.74212701 | 3.54018632 | 4.22493997 | 3.66813527 | 5.10684145 | 3.88980305 | 4.06514397 | 4.01696159 | 5.37644775 | 4.5491224 |
| ENSMUSG00 | Ift81 | 4427 | 5.84722209 | 5.2674556 | 4.81877725 | 5.63981303 | 5.47966784 | 5.63022927 | 5.38639941 | 4.69317075 | 4.53079584 | 6.11350885 | 5.70882156 | 6.68224558 | 4.78052688 | 3.53549976 | 5.48175207 | 5.15500384 | 5.80911345 | 4.77151502 |
| ENSMUSG00 | P2rx4 | 3493 | 32.6242133 | 35.7543774 | 30.6887327 | 34.7349159 | 31.6378176 | 34.3592101 | 38.847055 | 37.2958158 | 37.177522 | 40.4148885 | 46.7218031 | 39.2451381 | 47.51004 | 46.4174947 | 47.5003337 | 50.0639004 | 44.9206583 | 41.1977736 |
| ENSMUSG00 | Camkk2 | 3402 | 4.27300711 | 3.99519719 | 4.0382371 | 3.01329658 | 4.59814086 | 3.91819087 | 3.89404765 | 3.48020128 | 2.50605784 | 3.03527455 | 3.55292901 | 2.44460133 | 3.50920752 | 2.27988318 | 4.19039928 | 2.53244758 | 2.15177034 | 4.07474865 |
| ENSMUSG00 | Anapc5 | 2733 | 97.0324893 | 100.158087 | 94.3221893 | 98.0237617 | 100.173575 | 93.0314951 | 87.9430695 | 100.197411 | 103.290296 | 92.93847 | 89.0820302 | 90.9225503 | 79.2234852 | 98.3941698 | 86.9559365 | 88.0838085 | 93.7332486 | 99.9543447 |
| ENSMUSG00 | Rnf34 | 1948 | 36.6574413 | 33.2273888 | 31.5123255 | 35.1063283 | 36.4695684 | 34.1120204 | 33.8815129 | 30.5656855 | 38.3745001 | 36.0165536 | 40.1231171 | 33.4951281 | 30.9464081 | 34.1416495 | 40.6939392 | 42.3594758 | 36.3785026 | 35.2137352 |
| ENSMUSG00 | Kdm2b | 12990 | 1.49864285 | 1.29250952 | 1.05386521 | 1.12938078 | 1.57846031 | 1.26266011 | 1.32941708 | 1.34070327 | 1.2206935 | 1.06352241 | 1.22839385 | 1.12146943 | 0.67598778 | 1.0618856 | 1.50660039 | 1.12307324 | 1.02085441 | 1.52026708 |
| ENSMUSG00 | Morn3 | 1546 | 0.08248094 | 0.17238238 | 0.31289476 | 0.20629192 | 0.0311332 | 0.09615666 | 0.64266899 | 0.34585617 | 0.47953253 | 0.15249257 | 0.27812072 | 0.10830982 | 0 | 0.03004145 | 0.02665042 | 0.12383767 | 0.22311557 | 0.10674471 |
| ENSMUSG00 | Ncor2 | 11609 | 9.68439922 | 10.2816866 | 10.5547551 | 10.6043537 | 8.66532177 | 11.2303594 | 7.86982367 | 9.07353311 | 8.27349081 | 8.96387304 | 6.58042541 | 7.26483471 | 5.79625594 | 6.05705473 | 6.28546434 | 6.43179296 | 5.87654479 | 5.84255565 |
| ENSMUSG00 | Dhx37 | 5491 | 2.57771328 | 3.36708364 | 3.1802667 | 3.38534076 | 3.7516779 | 3.10437814 | 2.69693471 | 3.33166286 | 2.06270112 | 2.7993315 | 1.9837353 | 3.13080023 | 2.17416208 | 1.77622625 | 3.4215825 | 1.72241487 | 3.16186715 | 3.51633518 |
| ENSMUSG00 | Aacs | 3651 | 33.959901 | 26.9167455 | 29.8111321 | 32.1710141 | 27.3288079 | 31.8407869 | 27.0061964 | 26.5390646 | 23.2611875 | 25.0282287 | 19.9160468 | 29.3678137 | 26.5914032 | 24.7930776 | 28.0319542 | 23.9014775 | 24.9630058 | 26.2163368 |
| ENSMUSG00 | Tmem132b | 1805 | 1.01258868 | 3.11904661 | 1.79558257 | 2.42319128 | 1.59995314 | 0.85104418 | 0.18348407 | 0.16927381 | 0 | 0.54856774 | 0 | 0.12369122 | 0.13665351 | 0 | 0.82174848 | 0.61519535 | 0.82810281 | 1.06665858 |
| ENSMUSG00 | Anxa3 | 1584 | 8.56007017 | 15.562842 | 11.0656007 | 7.30587453 | 8.75125885 | 11.1994183 | 5.64526283 | 4.7740503 | 4.21225729 | 1.6371772 | 1.77949656 | 2.74849854 | 2.05549655 | 3.19596308 | 2.08088666 | 3.79521803 | 1.47594957 | 2.56986988 |
| ENSMUSG00 | Mrpl1 | 4285 | 7.30077236 | 5.86182549 | 6.27670546 | 6.90061092 | 6.8069885 | 6.46440562 | 6.73529385 | 7.96826319 | 6.86281733 | 8.98999406 | 8.16132313 | 7.72431457 | 8.8417532 | 8.54094303 | 7.58647085 | 9.09681175 | 8.02303111 | 6.71406367 |
| ENSMUSG00 | Mfsd7a | 2498 | 1.32722327 | 1.38692594 | 0.77459616 | 0.43773621 | 0.46243642 | 1.01168511 | 0.81442962 | 1.07024346 | 0.90682881 | 1.28352595 | 0.86063776 | 2.41316716 | 0.98742829 | 1.33866065 | 1.1380734 | 1.0729964 | 0.39891262 | 0.99095669 |
| ENSMUSG00 | Pde6b | 5275 | 0.43512408 | 0.32207727 | 0.45851688 | 0.22456628 | 0.14599256 | 0.27242261 | 0.22423071 | 0.02896106 | 0.00780787 | 0.2860327 | 0 | 0.16929869 | 0.22444853 | 0 | 0.01562144 | 0 | 0.15257862 | 0.33370453 |
| ENSMUSG00 | Pxmp2 | 1527 | 0.8907437 | 0.58902956 | 1.39386728 | 1.84989178 | 1.22930257 | 1.26559045 | 4.36875424 | 3.17644735 | 2.83207869 | 2.62462985 | 3.09739416 | 3.03385723 | 3.97369078 | 3.40650865 | 1.6728857 | 1.980981 | 2.40951179 | 3.20616274 |
| ENSMUSG00 | Pgam5 | 3649 | 4.81080781 | 4.17209993 | 4.98450189 | 4.8195569 | 4.48475036 | 4.19616187 | 5.30306541 | 4.8041416 | 4.71799495 | 4.43208858 | 5.10612053 | 4.26762671 | 3.58261389 | 3.75472891 | 3.8841697 | 3.49431961 | 3.87569295 | 3.72355422 |
| ENSMUSG00 | Ankla2 | 5205 | 9.74230067 | 10.5090821 | 10.3066753 | 11.8957655 | 10.3014338 | 9.78678388 | 8.54446075 | 9.9791868 | 9.25015119 | 10.522473 | 11.253045 | 9.42593034 | 7.60119071 | 11.124856 | 8.39862647 | 10.5345149 | 10.8683217 | 14.2205532 |

[illegible]

| | | | | | | | | | | | | | | | | | | | | |
|---|---|---|---|---|---|---|---|---|---|---|---|---|---|---|---|---|---|---|---|---|
| ENSMUSG00 | Plcz1 | 2643 | 0 | 0 | 0 | 0 | 0 | 0 | 0 | 0 | 0 | 0 | 0 | 0 | 0.1679861 | 0.01757249 | 0 | 0.1593631 | 0 | 0 |
| ENSMUSG00 | Plekha5 | 5139 | 16.2692511 | 15.7067801 | 14.496057 | 15.8962741 | 15.9128504 | 13.5476923 | 14.0492692 | 13.7341038 | 16.9186096 | 15.0379456 | 14.4933178 | 14.5757189 | 15.9927951 | 14.5053046 | 12.5953774 | 16.913734 | 15.5796136 | 13.6907273 |
| ENSMUSG00 | Aebp2 | 6883 | 6.89172983 | 7.11462364 | 5.37639848 | 6.70541013 | 6.97888418 | 7.84003005 | 5.45783337 | 6.84167255 | 7.43786708 | 6.45299469 | 4.62271137 | 6.4468185 | 4.37199906 | 5.56681274 | 4.44160315 | 5.43511891 | 5.71302929 | 6.42558794 |
| ENSMUSG00 | Slco1c1 | 2898 | 0 | 0.01149513 | 0 | 0 | 0 | 0 | 0 | 0 | 0 | 0 | 0 | 0 | 0 | 0 | 0 | 0 | 0 | 0 |
| ENSMUSG00 | Slco1b2 | 4914 | 0.06054868 | 0.12202525 | 0.03937609 | 0.15761859 | 0.09794856 | 0.06050395 | 0.47177885 | 0.06217729 | 0 | 0 | 0 | 0 | 0 | 0.00945138 | 0 | 0 | 0.05459586 | 0 |
| ENSMUSG00 | Slco1a4 | 3836 | 0.0110806 | 0.00868428 | 0.03783123 | 0.09501782 | 0.02509485 | 0 | 0 | 0.01991262 | 0.01073684 | 0 | 0 | 0.01455049 | 0.01286025 | 0 | 0.01074076 | 0.05989146 | 0.00999121 | 0.01434023 |
| ENSMUSG00 | Recql | 5420 | 6.64241404 | 6.06024238 | 5.43532841 | 8.15391384 | 6.05645239 | 5.53125532 | 4.47811887 | 6.52512262 | 4.55939289 | 4.57588006 | 5.87931685 | 5.26232538 | 5.2608576 | 4.94432443 | 6.69715327 | 4.64856451 | 5.16202892 | 5.02389443 |
| ENSMUSG00 | Gys2 | 2840 | 0 | 0 | 0 | 0 | 0 | 0 | 0 | 0 | 0 | 0 | 0 | 0 | 0 | 0.0327071 | 0 | 0 | 0 | 0 |
| ENSMUSG00 | Golt1b | 2929 | 22.3772555 | 22.5763383 | 21.8002936 | 24.2971496 | 22.8581447 | 22.3486381 | 18.8346137 | 18.6593813 | 24.9312709 | 21.2492073 | 19.5569052 | 20.3139284 | 23.0069645 | 26.4805824 | 22.4505576 | 25.2961165 | 25.7122477 | 21.5040721 |
| ENSMUSG00 | Ldhb | 1914 | 632.159008 | 599.475452 | 574.568359 | 630.499154 | 631.021087 | 632.043305 | 659.460631 | 608.085303 | 586.89709 | 648.999622 | 630.933974 | 582.68169 | 737.37556 | 737.257423 | 675.412757 | 868.320545 | 718.147499 | 746.387932 |
| ENSMUSG00 | Kcnj8 | 2185 | 12.8196389 | 14.3161591 | 10.9144853 | 7.86110467 | 11.4767653 | 8.27768776 | 6.58263359 | 7.70839792 | 5.82454616 | 9.94804311 | 7.95884892 | 4.31709244 | 4.06395657 | 3.35842829 | 6.82607013 | 2.92655901 | 3.68353426 | 7.37650722 |
| ENSMUSG00 | Abcc9 | 7630 | 0.63507078 | 0.82954782 | 0.23457675 | 0.7165564 | 0.30279585 | 0.47409562 | 0.3782534 | 0.54060019 | 0.83128747 | 0.29862303 | 0.69502233 | 0.33650334 | 0.04525864 | 0.09739258 | 0.03239965 | 0.22582927 | 0.14567007 | 0.00720958 |
| ENSMUSG00 | Rad18 | 6703 | 2.24479074 | 3.19064271 | 2.56914468 | 2.88877276 | 3.59750765 | 2.87572928 | 3.18335345 | 4.46708093 | 3.89560659 | 3.7774023 | 3.8915601 | 2.9893813 | 3.69455803 | 3.84551223 | 3.27006232 | 5.63819626 | 4.97447088 | 5.33431588 |
| ENSMUSG00 | Sspn | 4990 | 1.55880706 | 1.90931618 | 1.56074918 | 3.2047949 | 2.44035605 | 1.99601191 | 1.63081779 | 2.30379021 | 2.33582845 | 2.87250777 | 2.19247805 | 2.5391093 | 2.00688962 | 1.07035472 | 1.44494434 | 2.12555075 | 2.28114432 | 1.60948489 |
| ENSMUSG00 | Bhlhe41 | 6490 | 0.68113071 | 0.76994363 | 0.90933293 | 0.74414032 | 0.5265588 | 0.73298342 | 1.21015481 | 0.63555924 | 1.52942223 | 1.08250459 | 1.40601188 | 0.96322869 | 2.04472816 | 1.94650104 | 2.4822509 | 2.76117169 | 2.07871441 | 2.91573694 |
| ENSMUSG00 | Srgap3 | 15055 | 0.15810627 | 0.16153047 | 0.08996738 | 0.15131532 | 0.20141556 | 0.18761248 | 0.13199153 | 0.20802238 | 0.14225831 | 0.14719914 | 0.13962791 | 0.17054271 | 0.33095474 | 0.25913672 | 0.15873067 | 0.07121468 | 0.08146407 | 0.31788723 |
| ENSMUSG00 | Rassf8 | 5553 | 0.57408395 | 0.45593012 | 0.11324615 | 0.40203335 | 0.73675734 | 0.42833265 | 0.49417169 | 0.31637863 | 0.00741698 | 0.19529375 | 0.52481046 | 0.54277792 | 0.31981794 | 0.36800646 | 0.20033171 | 0.28271473 | 0.10352862 | 0.0099062 |
| ENSMUSG00 | Lrmp | 3610 | 0 | 0 | 0 | 0.02524158 | 0 | 0 | 0.02621201 | 0.11637574 | 0.14831709 | 0.15673364 | 0.07940436 | 0 | 0.35529913 | 0 | 0.05706587 | 0.14849543 | 0 | 0.12190384 |
| ENSMUSG00 | Thumpd3 | 3897 | 12.455969 | 11.9591324 | 12.6985172 | 12.088817 | 11.6964156 | 11.7619396 | 14.6539398 | 13.9362577 | 12.5768421 | 12.413811 | 14.8584166 | 13.1339396 | 11.2538041 | 10.7737888 | 13.4272456 | 14.0900003 | 14.5555349 | 14.0592944 |
| ENSMUSG00 | Kras | 6083 | 18.5798558 | 19.1564202 | 17.7812286 | 16.9421473 | 16.7350022 | 16.2433258 | 17.2279441 | 18.1638369 | 16.5206475 | 17.3239885 | 14.4982037 | 14.8370749 | 18.3200396 | 17.3774031 | 13.7361097 | 18.1350057 | 15.8774072 | 13.2209952 |
| ENSMUSG00 | Bcat1 | 10448 | 0.00406826 | 0 | 0.0138898 | 0.05232892 | 0 | 0 | 0.06339754 | 0.0731095 | 0 | 0 | 0 | 0.02136894 | 0 | 0.02222631 | 0 | 0.01832437 | 0 | 0.01579511 |
| ENSMUSG00 | Mtmr14 | 3129 | 14.6574257 | 16.5978914 | 14.1765827 | 16.6139826 | 12.752114 | 16.7392903 | 14.8334094 | 12.8772721 | 13.6630244 | 16.0936245 | 16.2456235 | 13.7532361 | 13.8110449 | 17.470339 | 14.5765811 | 12.8247225 | 14.1350421 | 16.1212371 |
| ENSMUSG00 | Cpne9 | 5270 | 0.07258948 | 0.04424863 | 0 | 0.02593608 | 0.02739958 | 0.0094028 | 0.03591095 | 0.19567265 | 0.03907639 | 0.03578801 | 0.05439274 | 0.21182415 | 0.0093609 | 0 | 0.06254506 | 0.10898655 | 0.13090576 | 0 |
| ENSMUSG00 | Ogg1 | 2403 | 3.9798854 | 5.00455881 | 5.73718525 | 4.87273758 | 3.94589636 | 4.16548156 | 4.62691617 | 5.1018565 | 4.64483802 | 4.59145425 | 4.85105696 | 4.92422824 | 4.57803477 | 4.40667995 | 3.8406777 | 4.49352946 | 3.30151621 | 4.44101819 |
| ENSMUSG00 | Camk1 | 6009 | 45.5963329 | 42.3715178 | 42.7062038 | 47.691624 | 43.3099203 | 43.5988188 | 42.2656782 | 39.9720451 | 45.4772062 | 45.094873 | 41.040828 | 48.6819583 | 40.6953426 | 35.7779465 | 41.4003985 | 41.196335 | 36.5856622 | 37.5332564 |
| ENSMUSG00 | Etnk1 | 6421 | 17.7474501 | 19.6058912 | 23.5426386 | 18.2641533 | 19.677044 | 22.7428603 | 18.4579132 | 20.8597968 | 16.2924389 | 18.1744262 | 18.0653479 | 22.401022 | 12.692155 | 12.8677825 | 12.2494581 | 11.9445704 | 12.4511222 | 14.0756833 |
| ENSMUSG00 | Tll3 | 4645 | 0.39348172 | 0.58091376 | 0.37490788 | 0.33349311 | 0.27977652 | 0.17068756 | 0.61114332 | 0.41933531 | 0.36354083 | 0.29437468 | 0.63768513 | 0.45661898 | 1.24259081 | 0.60992235 | 0.71847708 | 0.40392675 | 0.76735105 | 0.0710559 |
| ENSMUSG00 | Cidec | 2933 | 0.01449205 | 0.03407388 | 0.0329857 | 0.06213576 | 0 | 0.06757959 | 0.04839347 | 0.13021617 | 0.01404245 | 0.03215186 | 0.04886831 | 0.11418138 | 0 | 0.09501007 | 0.01404758 | 0 | 0.0392018 | 0 |
| ENSMUSG00 | 5730419I09R | 4447 | 12.0815252 | 11.1242743 | 11.0844676 | 11.6079746 | 10.7152247 | 10.1512346 | 9.97960262 | 11.5255687 | 9.7617692 | 12.4795093 | 13.6008764 | 11.5597538 | 9.08541494 | 8.12536541 | 10.5621259 | 8.43822695 | 7.23088894 | 10.4649661 |
| ENSMUSG00 | Il17rc | 2854 | 6.3296075 | 8.53248972 | 8.64418377 | 7.88617936 | 8.17939137 | 8.66391523 | 6.56475853 | 8.21658552 | 9.33695715 | 6.36055387 | 9.90987226 | 8.07703877 | 6.44737391 | 7.76237901 | 8.01221567 | 7.91571772 | 5.82817549 | 7.34354228 |
| ENSMUSG00 | Cmas | 3047 | 17.8279012 | 18.0285404 | 19.5113452 | 18.3619831 | 17.1707913 | 17.8402851 | 20.1703872 | 18.6762979 | 20.6676018 | 20.2096549 | 18.8936085 | 15.9002281 | 17.6474532 | 21.2176467 | 18.4440178 | 15.9093892 | 16.7921255 | 18.5229221 |
| ENSMUSG00 | St8sia1 | 11347 | 0 | 0.017615 | 0 | 0 | 0.00424182 | 0 | 0 | 0 | 0 | 0 | 0 | 0.00983796 | 0 | 0.00409307 | 0 | 0 | 0 | 0 |
| ENSMUSG00 | Creld1 | 4459 | 7.93099494 | 9.05477188 | 8.68966087 | 10.3097317 | 9.46662863 | 7.79019202 | 9.65564876 | 7.7429764 | 7.43555622 | 9.67546389 | 9.18213488 | 8.87492818 | 8.61842645 | 7.07233356 | 6.40338153 | 7.87452052 | 5.9135439 | 9.83230726 |
| ENSMUSG00 | Tmem111 | 2414 | 92.1943255 | 101.649867 | 91.2364058 | 98.8225469 | 104.478575 | 93.3167883 | 90.5879047 | 89.7537278 | 84.2839221 | 93.6959087 | 96.5196855 | 84.5789949 | 99.1951642 | 94.8121221 | 97.3373957 | 92.8396542 | 104.563791 | 96.1406053 |
| ENSMUSG00 | Itpr2 | 13711 | 49.3563495 | 48.228499 | 45.7945025 | 41.467163 | 46.7979487 | 48.2082936 | 38.7445671 | 41.0559273 | 45.0585469 | 38.4915631 | 47.0815578 | 44.9220215 | 34.0476941 | 31.048538 | 33.5207757 | 36.1150999 | 38.8518125 | 34.7081746 |
| ENSMUSG00 | Med21 | 2170 | 33.3577487 | 30.8873471 | 32.9475013 | 28.7223542 | 34.0250557 | 28.7040489 | 30.4807028 | 30.5363217 | 31.1271363 | 28.4859779 | 29.457555 | 28.9624129 | 35.6008214 | 31.7831658 | 28.9550105 | 31.4441762 | 32.2152837 | 34.7292517 |
| ENSMUSG00 | 1700023A16f | 1657 | 0 | 0 | 0 | 0 | 0 | 0 | 0 | 0 | 0 | 0 | 0 | 0 | 0 | 0 | 0 | 0.06932518 | 0 | 0 |
| ENSMUSG00 | Sec13 | 1655 | 100.522815 | 91.4846551 | 84.7925145 | 86.4421022 | 95.6531099 | 91.3207477 | 86.3063309 | 85.3386732 | 85.13539 | 84.2444891 | 91.8838023 | 84.2462388 | 87.7541779 | 93.7861873 | 80.6604478 | 87.9642836 | 81.3999599 | 83.3279943 |
| ENSMUSG00 | Ccdc91 | 2910 | 24.4514312 | 23.3762646 | 26.8132318 | 30.7811042 | 22.4615644 | 24.2484854 | 23.1848385 | 23.5454192 | 24.216539 | 21.8578346 | 23.4770415 | 22.6331553 | 23.3436598 | 23.8125699 | 22.4413973 | 24.6454666 | 22.5479795 | 23.1945615 |
| ENSMUSG00 | Atp2b2 | 11839 | 0 | 0 | 0.00408595 | 0 | 0 | 0 | 0 | 0.00322598 | 0 | 0 | 0 | 0 | 0.12084007 | 0 | 0.01044047 | 0 | 0.00647458 | 0.00929286 |
| ENSMUSG00 | Far2 | 5336 | 0.23897213 | 0.31839535 | 0.37168567 | 0.58915144 | 0.4600315 | 0.43646523 | 0.4433347 | 0.74437966 | 0.74098679 | 0.53901666 | 0.83265951 | 0.61715219 | 0.76734429 | 0.34815658 | 0.79530739 | 0.61712748 | 0.43095536 | 0.60823413 |
| ENSMUSG00 | Ergic2 | 7060 | 13.7449457 | 12.4333536 | 13.1896666 | 12.5841411 | 13.0624314 | 13.4690787 | 14.2273112 | 15.0172959 | 14.2986048 | 13.6977468 | 12.8775844 | 12.9182426 | 13.0457067 | 12.8740905 | 13.5801729 | 11.747515 | 11.9918966 | 14.8976504 |
| ENSMUSG00 | Tmtc1 | 8556 | 4.75922845 | 4.92918701 | 5.37672129 | 5.35699049 | 4.68605568 | 4.64484483 | 4.18603295 | 4.71825261 | 4.46235392 | 3.95126861 | 5.12592472 | 4.90572443 | 3.615137 | 4.97770315 | 2.99525102 | 5.77760334 | 5.15586346 | 4.41693038 |
| ENSMUSG00 | Slc6a11 | 4052 | 0 | 0 | 0 | 0 | 0 | 0 | 0 | 0 | 0 | 0 | 0 | 0 | 0 | 0 | 0 | 0 | 0 | 0 |
| ENSMUSG00 | Caprin2 | 7128 | 4.09070577 | 4.79971133 | 4.92014724 | 4.80666461 | 4.67274006 | 4.80372319 | 4.30115263 | 3.74529872 | 3.98113207 | 3.93584013 | 4.75202659 | 4.14232401 | 3.03133835 | 4.47630304 | 3.2080336 | 3.18551209 | 4.21546251 | 4.1750739 |
| ENSMUSG00 | Slc6a1 | 4305 | 0.00987344 | 0 | 0 | 0 | 0 | 0 | 0 | 0 | 0 | 0 | 0 | 0 | 0 | 0 | 0 | 0 | 0 | 0 |
| ENSMUSG00 | Dennd5b | 10599 | 0.85820433 | 0.74803933 | 0.82151481 | 0.76945249 | 0.89007048 | 0.73401068 | 0.48656307 | 0.98372732 | 0.42356168 | 0.66728988 | 1.05926175 | 0.7267253 | 1.00534901 | 1.03413573 | 0.67639124 | 1.41977627 | 1.34877901 | 1.07952582 |
| ENSMUSG00 | Atg7 | 3652 | 10.3935165 | 13.8013168 | 12.3582979 | 13.5111202 | 12.599704 | 13.4872446 | 11.517243 | 10.9808386 | 9.33801617 | 13.9825333 | 10.9233544 | 10.5456756 | 10.549898 | 11.7254788 | 11.2593518 | 12.5922835 | 10.0328422 | 9.89621663 |
| ENSMUSG00 | Vgll4 | 5157 | 15.5613287 | 13.1068184 | 14.670584 | 15.3195367 | 12.6839799 | 13.0680081 | 9.64235516 | 9.92395176 | 9.85537337 | 11.6939596 | 10.7185625 | 11.7973768 | 11.8140232 | 9.14111862 | 11.0414098 | 12.5407865 | 9.65403523 | 9.02418156 |
| ENSMUSG00 | 1500001M20 | 5032 | 9.97587697 | 8.35470439 | 9.81505852 | 8.19410674 | 8.48430371 | 9.53240185 | 9.96650232 | 10.6941763 | 10.2311496 | 10.4008904 | 12.1146357 | 9.49487424 | 13.7545071 | 12.0355902 | 13.1579691 | 13.4458472 | 14.2200311 | 11.6314963 |
| ENSMUSG00 | Timp4 | 1411 | 4.45837424 | 3.39975675 | 3.08548386 | 3.58417854 | 2.96773732 | 3.51188757 | 3.52078759 | 3.08570792 | 3.06490727 | 1.97157441 | 3.41975252 | 3.28327427 | 3.18157652 | 2.73200514 | 3.88363355 | 3.69066095 | 2.66192655 | 4.01554783 |
| [illegible] | | | | | | | | | | | | | | | | | | | | |

| | | | | | | | | | | | | | | | | | | | |
|---|---|---|---|---|---|---|---|---|---|---|---|---|---|---|---|---|---|---|---|
| ENSMUSG00 Ano1 | 6731 | 40.3644459 | 46.8488867 | 44.4496041 | 45.6287327 | 49.7122468 | 51.6435041 | 31.5042976 | 31.6501591 | 30.631362 | 30.6749264 | 33.4304048 | 32.3152038 | 35.3187505 | 36.6115477 | 36.5923193 | 44.9122486 | 37.4323393 | 41.8840695 |
| ENSMUSG00 Fadd | 3901 | 2.27725757 | 2.4593985 | 2.36845534 | 1.85701257 | 2.33194913 | 2.03241153 | 2.03756218 | 2.88817193 | 2.62892654 | 1.99432608 | 2.85351585 | 1.73127279 | 2.51654743 | 1.40487114 | 1.64763976 | 1.81588371 | 1.91582361 | 2.4677246 |
| ENSMUSG00 Cttn | 5537 | 41.2538949 | 45.7548421 | 41.6202815 | 43.2899863 | 41.0994645 | 42.0442013 | 33.1966331 | 34.5021486 | 30.6909092 | 39.6314562 | 35.7470928 | 34.6466376 | 35.949844 | 35.0196975 | 42.0200806 | 42.0110963 | 40.4028526 | 36.4111208 |
| ENSMUSG00 Zfp300 | 5320 | 0.33556718 | 0.60739676 | 0.2636903 | 0.29974371 | 0.40713093 | 0.28874713 | 0.75593566 | 0.48817356 | 0.65805523 | 0.19498411 | 1.06865039 | 0.61900829 | 0.35237084 | 0.13968147 | 0.03097861 | 0.47503385 | 0.63396882 | 0.42394236 |
| ENSMUSG00 Gm498 | 1779 | 0 | 0 | 0 | 0 | 0 | 0 | 0 | 0 | 0 | 0 | 0 | 0 | 0 | 0 | 0 | 0 | 0 | 0.40197773 |
| ENSMUSG00 Slc6a14 | 3420 | 0 | 0 | 0 | 0 | 0 | 0 | 0 | 0 | 0 | 0 | 0.01396929 | 0 | 0 | 0 | 0.01204724 | 0.01119608 | 0 | 0.01608453 |
| ENSMUSG00 Nadsyn1 | 3491 | 3.62834215 | 4.50406361 | 3.71357963 | 3.69343327 | 3.1435344 | 2.52660744 | 2.466602 | 3.29301436 | 3.50398029 | 3.37659004 | 5.28247894 | 2.59012814 | 4.70567988 | 5.77391404 | 2.92695098 | 5.63774626 | 4.84156314 | 5.24721642 |
| ENSMUSG00 Dock11 | 8119 | 2.97886995 | 3.22501969 | 4.43875848 | 4.23680114 | 3.03529313 | 3.92442514 | 3.31579188 | 2.86478299 | 2.6480307 | 2.49139693 | 2.74210256 | 2.68113172 | 3.83402387 | 3.70111153 | 5.41471365 | 5.27268125 | 4.88578856 | 4.97988588 |
| ENSMUSG00 Cul4b | 5958 | 9.5526067 | 9.24799044 | 9.93776454 | 9.32173767 | 8.63595609 | 10.2049688 | 8.43337781 | 8.50002108 | 8.22624266 | 10.3592263 | 9.26151008 | 8.93725111 | 8.26338594 | 9.05807814 | 11.1544326 | 9.58871874 | 11.1479477 | 9.37130637 |
| ENSMUSG00 Tnni2 | 1343 | 0.34814366 | 0.71934023 | 0.39620911 | 0.67849656 | 0.3942302 | 0.33207342 | 0.95118565 | 0.3412575 | 1.28803698 | 0.10532547 | 0.60474631 | 0.45716477 | 0.03673263 | 0 | 0.03067875 | 0.114045 | 0.05707565 | 0 |
| ENSMUSG00 Syt8 | 1811 | 0.11735278 | 0 | 0.0534219 | 0.02515795 | 0 | 0 | 0.18287617 | 0 | 0 | 0 | 0 | 0.24656284 | 0 | 0.15387329 | 0 | 0.23257685 | 0 | 0 |
| ENSMUSG00 Smarca1 | 6842 | 0.04969912 | 0.05842659 | 0.11312138 | 0.31297414 | 0.21104322 | 0.09415164 | 0.97502013 | 0.96011301 | 0.79459516 | 0.47550402 | 0.81696436 | 1.01972491 | 10.4258919 | 15.0084591 | 6.67824108 | 1.41029422 | 2.44790876 | 7.11532585 |
| ENSMUSG00 Sash3 | 2669 | 0.09555304 | 0 | 0 | 0.01707045 | 0.09016846 | 0.25992442 | 0.90406393 | 0.2861926 | 0.38578602 | 0.26499084 | 1.98689411 | 0.52281437 | 0.27724944 | 0.24361829 | 0.41680105 | 0.57385699 | 0 | 0.02061038 |
| ENSMUSG00 Elf4 | 10535 | 1.08935904 | 1.6790837 | 1.54281027 | 1.46608391 | 1.29752884 | 1.90967345 | 0.43113594 | 1.09846207 | 0.78580823 | 0.84589302 | 1.16092922 | 0.56689852 | 1.21281126 | 0.89052735 | 0.94644295 | 1.16307493 | 1.07320907 | 1.31583245 |
| ENSMUSG00 Rab33a | 1156 | 0.11030755 | 0.92215624 | 0.16738246 | 0.51236469 | 0.70782241 | 0.68585098 | 0.49113508 | 1.18938277 | 0 | 0.20393903 | 1.11585111 | 1.25536957 | 1.15221606 | 0.56247165 | 1.21181047 | 1.85490832 | 0.56362201 | 1.80825783 |
| ENSMUSG00 Slc25a14 | 2350 | 8.95321793 | 7.47059372 | 6.31943567 | 6.76629978 | 8.17218619 | 7.88626483 | 8.85854414 | 5.21691965 | 8.57030051 | 5.95904633 | 7.86762082 | 5.41530684 | 10.3282142 | 7.25318301 | 6.90783533 | 10.8028394 | 6.001735 | 8.26307007 |
| ENSMUSG00 Rbmx2 | 1866 | 6.26416045 | 5.17724508 | 5.54766104 | 3.95546044 | 3.92071404 | 4.83311764 | 4.79212021 | 5.05547877 | 5.78288699 | 4.27034772 | 5.35100017 | 4.96538049 | 6.60931368 | 6.09796431 | 6.16036125 | 6.85373144 | 5.46344854 | 7.51732266 |
| ENSMUSG00 Enox2 | 5498 | 4.80869762 | 5.4107702 | 5.28783044 | 5.04668534 | 5.45401754 | 6.40814725 | 4.83625419 | 6.43255086 | 5.55845666 | 6.96369062 | 4.7531652 | 5.46177273 | 5.0426587 | 3.68308878 | 4.42141105 | 4.57565098 | 6.14838011 | 4.17220762 |
| ENSMUSG00 Igsf1 | 6160 | 0.1518042 | 0.11897462 | 0.15705692 | 0.17751056 | 0.03906812 | 0 | 0 | 0.02480026 | 0 | 0.05358034 | 0.1240908 | 0.06342689 | 0 | 0 | 0.1872798 | 0.05594407 | 0 | 0 |
| ENSMUSG00 2610018G03 | 5363 | 0.63405073 | 0.68327771 | 0.80276789 | 0.91750751 | 0.49361473 | 0.41578837 | 1.27037587 | 0.89018279 | 0.7372563 | 0.3868415 | 1.04226555 | 0.82219604 | 1.73852925 | 1.22107334 | 1.55187662 | 1.82778212 | 1.49360359 | 1.9488589 |
| ENSMUSG00 1700080O16 | 2515 | 0 | 0 | 0 | 0 | 0 | 0 | 0 | 0 | 0 | 0 | 0 | 0 | 0 | 0 | 0 | 0 | 0 | 0 |
| ENSMUSG00 Gpc4 | 2642 | 64.0795292 | 68.7567041 | 67.6899855 | 78.3436126 | 76.1511888 | 81.1374434 | 65.3996607 | 78.654376 | 63.7284165 | 81.86233 | 76.9426375 | 83.7023676 | 72.7655116 | 65.9920909 | 72.1884944 | 67.5230658 | 78.5239384 | 67.1269342 |
| ENSMUSG00 3830403N18 | 1014 | 0.96412135 | 0 | 0.1908226 | 1.52768788 | 0.52214118 | 0 | 3.45279905 | 0 | 0 | 1.85998829 | 0 | 0 | 0.8270637 | 0 | 0.60949047 | 1.17062015 | 0 | 0 |
| ENSMUSG00 Slc9a9 | 5964 | 0.42049051 | 0.32396845 | 0.37310235 | 0.51947535 | 0.59721032 | 0.29080243 | 0.22212525 | 0.13448029 | 0.08977611 | 0.28461189 | 0.24832725 | 0.20589279 | 0.44666726 | 0.91891387 | 0.39377745 | 0.87315939 | 1.04105635 | 0.37816455 |
| ENSMUSG00 Brs3 | 2769 | 0 | 0 | 0 | 0 | 0 | 0 | 0 | 0 | 0 | 0 | 0 | 0 | 0 | 0 | 0 | 0 | 0 | 0 |
| ENSMUSG00 Vgll1 | 2445 | 0 | 0 | 0 | 0 | 0 | 0 | 0 | 0 | 0 | 0 | 0 | 0 | 0.04035331 | 0 | 0 | 0 | 0 | 0 |
| ENSMUSG00 Cd40lg | 1245 | 0 | 0 | 0 | 0 | 0 | 0 | 0 | 0 | 0 | 0 | 0 | 0 | 0 | 0 | 0 | 0 | 0 | 0 |
| ENSMUSG00 Arhgef6 | 10690 | 0.8349941 | 0.87255476 | 0.53396413 | 0.74585433 | 0.63935764 | 0.43109488 | 0.23457175 | 0.07145445 | 0.19264039 | 0.00882146 | 0.31730796 | 0.24540095 | 0.06922159 | 0.11730499 | 0.00385421 | 0.36177281 | 0.02151149 | 0 |
| ENSMUSG00 Rbmx | 3651 | 37.5456567 | 41.7802637 | 45.4454147 | 43.36473 | 40.9998035 | 40.8392702 | 33.9002925 | 37.5229108 | 43.9503329 | 36.1863244 | 42.1482173 | 41.5063062 | 34.7255621 | 31.3443526 | 34.6788226 | 31.0016531 | 37.0980918 | 35.0153832 |
| ENSMUSG00 Fgf13 | 4186 | 0.5990935 | 0.5093228 | 0.10400425 | 0.64216474 | 0.73589181 | 0.66291282 | 0.19214418 | 0 | 0 | 0.55193131 | 0 | 0.37334891 | 0 | 0 | 0.46260705 | 0.27441907 | 0.25636319 | 0 |
| ENSMUSG00 F9 | 2726 | 0 | 0 | 0 | 0 | 0 | 0 | 0.15620474 | 0 | 0 | 0 | 0 | 0 | 0 | 0 | 0 | 0 | 0 | 0.04035885 |
| ENSMUSG00 Mcf2 | 5206 | 102.466377 | 130.819979 | 129.519524 | 124.212044 | 127.938832 | 136.455626 | 87.3094327 | 95.9798695 | 91.3603312 | 92.6983188 | 94.3202119 | 95.9459017 | 46.574409 | 51.9752695 | 64.4219487 | 43.5568531 | 65.8747032 | 64.5506402 |
| ENSMUSG00 Cacna1f | 8854 | 0 | 0 | 0 | 0 | 0 | 0 | 0 | 0 | 0 | 0 | 0.00539586 | 0 | 0 | 0 | 0 | 0 | 0 | 0 |
| ENSMUSG00 Ccdc22 | 3165 | 0 | 0 | 0 | 0 | 0 | 0 | 0 | 0 | 0 | 0 | 0 | 0 | 0 | 0 | 0 | 0 | 0 | 0 |
| ENSMUSG00 Syp | 5346 | 0.00795084 | 0.05608231 | 0.09048547 | 0.04261227 | 0.00900335 | 0 | 0.08850108 | 0.00714411 | 0.03852087 | 0.06173867 | 0.11617554 | 0.25057537 | 0.00922782 | 0.0781887 | 0.04624193 | 0.10027469 | 0.05018408 | 0 |
| ENSMUSG00 Prickle3 | 4921 | 4.56060413 | 4.19711327 | 4.67908971 | 4.63850502 | 4.4894438 | 4.99454498 | 4.15343981 | 4.09010284 | 3.35618631 | 4.49373701 | 3.52414345 | 3.62954928 | 5.08256087 | 4.66234082 | 4.00210195 | 4.22511681 | 4.11223019 | 4.13602306 |
| ENSMUSG00 Plp2 | 1876 | 44.2271341 | 46.6664638 | 44.9698491 | 36.4051202 | 38.4850136 | 43.9265437 | 35.0557696 | 35.1386389 | 36.1809051 | 33.0507328 | 32.3932554 | 34.3938731 | 41.4956106 | 41.4926921 | 43.1342515 | 40.3112738 | 37.4682455 | 37.3569307 |
| ENSMUSG00 Magix | 3468 | 0.58830693 | 0.41304915 | 0.36266199 | 0.28902623 | 0.70782241 | 0.48581111 | 0.47749243 | 0.60570419 | 0.8194549 | 0.55743334 | 0.57858946 | 0.65987375 | 1.20911562 | 3.30786897 | 0.85539562 | 2.39592324 | 1.66876322 | 1.80825783 |
| ENSMUSG00 Gpkow | 3726 | 15.2179025 | 18.623392 | 18.8508766 | 18.2317545 | 18.0720776 | 18.6587454 | 18.2850933 | 20.2852293 | 16.8681221 | 18.1339124 | 16.8866399 | 22.0955723 | 16.1659342 | 18.9216652 | 17.75891 | 19.751623 | 23.3187641 | 20.0932352 |
| ENSMUSG00 Praf2 | 1336 | 18.8664442 | 16.6813819 | 15.8589867 | 17.6651355 | 17.4730412 | 19.5837151 | 28.3309455 | 21.4689328 | 23.4911115 | 26.5752092 | 26.0688209 | 21.3904336 | 21.2319255 | 22.2139011 | 20.8783333 | 25.2786794 | 18.6754635 | 25.7340507 |
| ENSMUSG00 Ccdc120 | 4128 | 8.64930908 | 10.4183494 | 8.53099082 | 7.99084208 | 7.11251779 | 9.57923483 | 4.03441448 | 6.80024607 | 5.98641217 | 6.1679699 | 6.49267241 | 7.58541345 | 3.11909651 | 6.55932726 | 7.04658393 | 7.20731636 | 5.96063048 | 5.3836432 |
| ENSMUSG00 Gripap1 | 5376 | 10.3100353 | 9.38784128 | 9.51993957 | 9.65290727 | 9.0336888 | 8.81183283 | 10.4464565 | 11.2460143 | 11.0244413 | 10.708893 | 9.16219907 | 10.693849 | 9.99301673 | 12.6649985 | 10.8138756 | 12.8988635 | 10.5083634 | 15.9113023 |
| ENSMUSG00 Otud5 | 5061 | 16.5115938 | 19.3189774 | 19.4794023 | 20.4984189 | 19.0777789 | 18.3876771 | 17.6873323 | 16.2096978 | 17.3909473 | 17.3100191 | 17.6241548 | 15.2525314 | 17.2920018 | 19.95967 | 15.8505096 | 18.9750728 | 18.0991602 | 18.3254997 |
| ENSMUSG00 Pim2 | 3043 | 2.65395447 | 1.64210783 | 1.66914909 | 2.94956482 | 1.77153317 | 3.51738102 | 2.13008117 | 2.00814472 | 2.93706011 | 2.6341175 | 1.7897947 | 3.54006667 | 4.73378897 | 5.02139492 | 4.18379256 | 2.95704658 | 3.02277728 | 4.10353836 |
| ENSMUSG00 Slc35a2 | 2462 | 13.3454513 | 12.6107301 | 13.7536439 | 11.3439968 | 13.3135012 | 10.2647823 | 11.8185611 | 14.2717344 | 15.0727257 | 12.371793 | 12.61321 | 10.51928 | 10.6398246 | 13.5634845 | 13.6557553 | 12.333246 | 13.185367 | 12.9590909 |
| ENSMUSG00 Pqbp1 | 1675 | 43.6724824 | 48.2291155 | 40.8648033 | 40.5561296 | 39.4825451 | 39.3167659 | 37.9066359 | 42.5018735 | 39.2193986 | 42.4778574 | 42.0134411 | 39.1875936 | 44.8846817 | 40.9262525 | 41.6935148 | 47.5032132 | 44.5729113 | 39.8364456 |
| ENSMUSG00 Timm17b | 2551 | 26.8760678 | 23.3751786 | 24.5186103 | 24.968381 | 26.1508609 | 21.6392572 | 27.3378631 | 27.4877502 | 26.8979754 | 26.8006741 | 29.3653998 | 27.984411 | 33.551892 | 31.4785681 | 29.7685494 | 31.5811203 | 28.410438 | 28.960106 |
| ENSMUSG00 Eras | 1075 | 0 | 0 | 0 | 0 | 0 | 0 | 0 | 0 | 0.11493911 | 0 | 0 | 0 | 0.13767047 | 0 | 0.11498109 | 0 | 0 | 0 |
| ENSMUSG00 Hdac6 | 6047 | 13.3272389 | 15.8934513 | 11.2874237 | 14.0970256 | 11.2230879 | 14.1356814 | 28.4721083 | 28.5290362 | 28.7290762 | 20.3823281 | 18.5585212 | 23.6757358 | 34.647371 | 50.2382629 | 28.4942793 | 15.1718696 | 16.0353119 | 28.5279574 |
| ENSMUSG00 Gata1 | 2428 | 0.14005 | 0 | 0 | 0 | 0 | 0.06122661 | 0 | 0 | 0 | 0.07767826 | 0.01967667 | 0 | 0 | 0 | 0 | 0 | 0 | 0 |
| ENSMUSG00 Glod5 | 669 | 0 | 0 | 0 | 0 | 0 | 0 | 0 | 0.28544396 | 0 | 0.07047938 | 0.28564998 | 0 | 0 | 0 | 0.06158678 | 0.28617793 | 0.11457787 | 0 |
| ENSMUSG00 Was | 2370 | 0.05380402 | 0.04216822 | 0.08164309 | 0.19224069 | 0 | 0.08363331 | 0.41922627 | 0.1289195 | 0.24329587 | 0.05968443 | 0.04031642 | 0.07065274 | 0.14570608 | 0.11757997 | 0 | 0.22619924 | 0 | 0 |
| ENSMUSG00 Wdr13 | 7694 | 7.68452039 | 7.1223955 | 7.63891856 | 8.34950254 | 6.98144357 | 7.50955125 | 6.21693767 | 8.3096025 | 9.12698328 | 8.00348563 | 7.22771662 | 6.96426255 | 6.18091607 | 4.84723189 | 7.28283286 | 6.34029553 | 7.24284322 | 7.42129618 |
| ENSMUSG00 Rbm3 | 6617 | 35.1436932 | 34.9742597 | 34.3885578 | 30.7160071 | 33.4966765 | 33.3996058 | 34.6854776 | 33.5922294 | 33.1073111 | 32.6000404 | 34.6344984 | 31.1089865 | 33.6533587 | 34.9120282 | 33.2688398 | 35.0269075 | 34.0053935 | 33.9598309 |
| ENSMUSG00 Ebp | 2332 | 33.5193048 | 25.4560795 | 26.4063054 | 26.0627929 | 27.0587272 | 27.8362268 | 32.0760486 | 26.9410386 | 28.6468819 | 29.0344811 | 26.8580491 | 25.7058412 | 34.3969543 | 30.5511278 | 32.4029389 | 35.0723574 | 30.3224776 | 33.0951015 |
| ENSMUSG00 Porcn | 3414 | 10.6823202 | 11.2799372 | 11.916268 | 11.1033359 | 11.2082248 | 10.8133528 | 14.0524477 | 12.5406219 | 14.3802949 | 15.1920837 | 16.0509307 | 12.9811471 | 11.3865117 | 12.2300043 | 13.528689 | 10.9914456 | 15.8289627 | 13.357513 |
| ENSMUSG00 Slc38a5 | 2254 | 25.0618363 | 28.8938368 | 33.3077725 | 53.2219298 | 32.4794392 | 36.6039492 | 1.32240407 | 1.08443379 | 1.13290327 | 1.33879548 | 1.25054239 | 0.39620701 | 0.43772775 | 1.85466867 | 0.65805502 | 0.5775868 | 1.47932024 | 1.85478798 |
| ENSMUSG00 Ftsj1 | 5067 | 14.3613304 | 13.3330471 | 14.6065722 | 13.9551492 | 14.9420793 | 14.2291745 | 14.0621461 | 13.6880605 | 14.5172917 | 13.7627571 | 16.2455602 | 13.0754276 | 17.0670714 | 13.409848 | 16.1569946 | 18.0986787 | 15.0975505 | 17.3701541 |
| ENSMUSG00 Otc | 4376 | 0 | 0 | 0 | 0 | 0 | 0 | 0 | 0 | 0 | 0 | 0 | 0 | 0 | 0 | 0 | 0 | 0 | 0 |
| ENSMUSG00 Rpgr | 7194 | 1.0753325 | 0.99095904 | 1.45914041 | 1.2286409 | 1.32473184 | 1.67379959 | 2.0190426 | 1.83688786 | 2.06676844 | 1.32394246 | 1.74656854 | 2.70000703 | 1.80348822 | 1.66563449 | 1.42607554 | 1.24548264 | 1.8433276 | 1.1852115 |

| | | | | | | | | | | | | | | | | | | | | |
|---|---|---|---|---|---|---|---|---|---|---|---|---|---|---|---|---|---|---|---|---|
| [illegible] | Mthfsd | 4866 | 7.10166628 | 7.84557681 | 8.65872276 | 8.22083784 | 6.10304088 | 7.74961575 | 9.35363689 | 9.19882771 | 6.46660461 | 9.04060956 | 7.5206778 | 6.8708553 | 10.006289 | 6.52851537 | 11.6170437 | 9.36412306 | 7.77395295 | 8.46728749 |
| [illegible] | Cox4i1 | 674 | 940.411254 | 837.418942 | 814.096372 | 776.092507 | 882.085343 | 802.474907 | 1210.68512 | 1118.17608 | 1051.35609 | 1059.07195 | 1062.95738 | 999.795986 | 1373.0229 | 1527.48677 | 1137.44414 | 1076.00015 | 1065.744 | 1237.94975 |
| [illegible] | Cox4nb | 4928 | 11.6354469 | 12.1475793 | 13.9486174 | 11.3810157 | 12.4822643 | 13.0015594 | 14.477987 | 11.5708719 | 10.5975045 | 13.3185424 | 14.0571611 | 12.6174205 | 13.7544753 | 14.7870887 | 12.6999114 | 15.7575795 | 10.9503705 | 15.4936364 |
| [illegible] | 5430437P03Rik | 1583 | 54.1853729 | 63.0903707 | 61.3607384 | 60.1820238 | 58.92588 | 62.0113488 | 60.5829439 | 58.2657308 | 55.3923513 | 58.6479689 | 60.4203834 | 56.5561104 | 60.8935985 | 54.6590871 | 50.8057088 | 64.0515025 | 52.514048 | 58.5535975 |
| [illegible] | Gins2 | 4359 | 1.37490934 | 1.73480775 | 1.68680354 | 1.65144414 | 1.71150451 | 1.65971533 | 2.37703067 | 2.90889598 | 2.50388316 | 2.42297718 | 1.50153001 | 2.15118865 | 2.62560331 | 2.65303459 | 2.5331537 | 1.74806856 | 2.49705628 | 2.59965038 |
| [illegible] | Gse1 | 7308 | 2.40792869 | 2.69402305 | 3.31624777 | 3.2418915 | 2.81889208 | 2.75971026 | 3.04282418 | 3.49626672 | 3.53365427 | 4.01955229 | 2.46458119 | 3.14669581 | 3.44945399 | 2.60564803 | 3.81683817 | 2.95510435 | 3.25154681 | 3.07864322 |
| [illegible] | Zdhhc7 | 3776 | 9.62445061 | 10.9572602 | 9.81306257 | 11.7039759 | 9.40713974 | 10.7215528 | 11.4648043 | 7.90954944 | 9.39131093 | 10.6388769 | 10.1218132 | 11.3079931 | 9.65473699 | 11.0698302 | 9.54750037 | 8.04143318 | 11.0634702 | 12.018674 |
| [illegible] | 6430548M08Rik | 6773 | 5.3719815 | 5.73003422 | 4.77093139 | 4.0562985 | 4.3704611 | 4.54337038 | 4.98763513 | 5.60508608 | 4.69452092 | 6.33502731 | 5.51602208 | 4.06276711 | 6.43143112 | 8.48240604 | 7.96291697 | 6.25833492 | 5.92464681 | 7.33400611 |
| [illegible] | Crispld2 | 4852 | 3.04859878 | 3.21319755 | 2.70181919 | 3.51191892 | 2.53952441 | 2.34895481 | 0.14626756 | 0.74779024 | 0.34803115 | 0.04858894 | 0.07877157 | 0.26458373 | 0.16267739 | 0.01914431 | 0.35664991 | 0.41826096 | 0.04739443 | 0.21541076 |
| [illegible] | Usp10 | 5259 | 30.6240943 | 27.8525948 | 31.7431172 | 33.0596967 | 28.3904216 | 35.5509534 | 18.2449349 | 22.6946677 | 18.2006965 | 20.8094281 | 21.4028907 | 22.4684841 | 15.2714131 | 14.6247212 | 15.2380731 | 15.180817 | 16.1642378 | 19.2777683 |
| [illegible] | Cotl1 | 2448 | 72.4914665 | 53.7388151 | 58.9848808 | 35.548037 | 52.5754754 | 36.4155914 | 27.8116605 | 28.3790768 | 51.2476009 | 27.3504896 | 24.7071476 | 48.7019017 | 38.8730672 | 18.6876741 | 24.2193786 | 37.3051874 | 20.8853268 | 19.0329712 |
| [illegible] | Klhl36 | 2136 | 26.7846287 | 28.5249455 | 25.6814529 | 30.9926018 | 28.3698932 | 26.6554733 | 15.4164909 | 14.0897065 | 20.5546937 | 18.3216684 | 16.2157514 | 19.8072436 | 15.9358722 | 11.2848695 | 13.3287805 | 12.9786516 | 13.7264125 | 10.4816041 |
| [illegible] | Lrrc50 | 2809 | 0 | 0 | 0.08610454 | 0.12975734 | 0 | 0.24696984 | 0.13474597 | 0 | 0 | 0.08392792 | 0 | 0.15896237 | 0.40392812 | 0 | 0 | 0.28625944 | 0 | 0 |
| [illegible] | Taf1c | 3643 | 1.73847687 | 2.09405786 | 1.89882373 | 3.26418682 | 3.05201053 | 2.23075715 | 4.58451157 | 3.29190622 | 2.17068653 | 3.59810472 | 3.22608833 | 2.74252089 | 3.45309879 | 3.86290378 | 3.9810452 | 3.05862387 | 2.79846143 | 2.61229082 |
| [illegible] | Mast3 | 7307 | 2.93760967 | 3.55149098 | 2.00590936 | 3.58527446 | 2.63483912 | 3.47893148 | 1.89716914 | 3.43402332 | 2.35609191 | 3.1231614 | 1.60840835 | 2.79574826 | 1.08021168 | 2.11022814 | 2.2103476 | 3.07603484 | 2.61732888 | 3.29738453 |
| [illegible] | Pik3r2 | 4184 | 15.1267225 | 17.4526444 | 16.0474331 | 20.7768816 | 16.4159309 | 19.4586858 | 16.7245344 | 15.627484 | 15.9371341 | 23.7331217 | 15.734678 | 16.4485449 | 14.1723147 | 16.5729021 | 16.5042561 | 17.8183109 | 15.3708237 | 17.8279992 |
| [illegible] | Mbtps1 | 6776 | 13.9007482 | 17.3201485 | 15.9198602 | 17.3610709 | 15.5278018 | 18.2229392 | 14.3488128 | 14.3390602 | 11.6095403 | 16.9160802 | 15.7369698 | 15.8567223 | 12.0053618 | 12.6802777 | 14.8729347 | 14.613269 | 13.6031204 | 13.1840008 |
| [illegible] | Necab2 | 2220 | 0 | 0 | 0.10894939 | 0.12313796 | 0 | 0 | 0 | 0 | 0.11131491 | 0.02123906 | 0 | 0.05028438 | 0 | 0 | 0 | 0 | 0.0172641 | 0.02477888 |
| [illegible] | Il10 | 984 | 86.9541863 | 74.6831755 | 83.4246758 | 79.3151107 | 77.4317431 | 80.0699658 | 125.061258 | 119.622951 | 115.816147 | 119.170527 | 122.010641 | 116.4528 | 120.772956 | 123.048512 | 92.4941433 | 99.5789272 | 101.580267 | 109.682792 |
| [illegible] | Hsbp1 | 1159 | 330.469492 | 293.751319 | 321.919793 | 291.999515 | 314.830124 | 303.473083 | 366.907988 | 316.248909 | 336.457231 | 344.619161 | 327.829374 | 308.454977 | 341.067 | 337.49103 | 317.561259 | 337.016813 | 330.221226 | 342.822069 |
| [illegible] | Rab3a | 2381 | 17.1198924 | 15.1663912 | 16.0500164 | 15.9970739 | 15.5453378 | 18.8138056 | 28.1173627 | 22.5689669 | 26.6216076 | 25.6051488 | 23.2353641 | 26.770889 | 29.0894632 | 31.3073332 | 27.4619358 | 29.6225531 | 24.3221481 | 26.0836966 |
| [illegible] | Cdh13 | 11275 | 0.04146846 | 0 | 0 | 0 | 0 | 0 | 0 | 0.02709882 | 0 | 0 | 0 | 0 | 0 | 0 | 0 | 0 | 0 | 0 |
| [illegible] | Pde4c | 4454 | 1.02111671 | 1.76511968 | 1.12951216 | 2.42433038 | 0.5511289 | 2.33634352 | 0.88166869 | 0.71171293 | 0.72127262 | 1.45030228 | 0.91173585 | 1.31581602 | 2.68036012 | 1.48070508 | 1.77608862 | 3.1034821 | 1.11863911 | 3.01352089 |
| [illegible] | Mphosph6 | 1049 | 44.2880433 | 47.0000795 | 43.3009962 | 37.4391041 | 42.0294014 | 42.6087375 | 32.7896259 | 39.6851464 | 42.7570216 | 39.9589849 | 32.9278312 | 34.7983254 | 36.6815019 | 47.5509532 | 40.6124011 | 38.4366154 | 37.8878645 | 40.9553025 |
| [illegible] | Hsd17b2 | 1275 | 0.933447 | 1.22800473 | 1.13820071 | 0.17867076 | 0.67950951 | 0.77729778 | 0.14843193 | 0 | 0.35533465 | 0 | 0 | 0 | 0 | 0.36426735 | 0 | 0 | 0 | 0.43144397 |
| [illegible] | Bcmo1 | 2760 | 0.30800852 | 0.20518812 | 0.05257992 | 0.01650762 | 0.29646475 | 0.08976944 | 0 | 0.05535131 | 0.11938121 | 0.01708359 | 0.01730977 | 0 | 0.28598213 | 0.21875838 | 0 | 0 | 0 | 0.07972334 |
| [illegible] | 1700030J22Rik | 2968 | 0.45827683 | 0.50508094 | 0.34226554 | 0.16885832 | 0.2919052 | 0.25043497 | 0.28693673 | 0.24449314 | 0.43018261 | 0.33361347 | 0.33803037 | 0.03761163 | 0.41553165 | 0.09388966 | 0.18046504 | 0.29672641 | 0.27117663 | 0 |
| [illegible] | Lsm4 | 1058 | 72.5962695 | 70.6560819 | 65.8392096 | 63.0878355 | 76.2467902 | 61.6364815 | 86.9336936 | 72.991529 | 69.721219 | 78.257689 | 84.7124973 | 72.2754799 | 77.5415675 | 92.8442768 | 60.3614477 | 61.7426996 | 61.5104463 | 71.3870544 |
| [illegible] | Comp | 2490 | 0.4267588 | 0 | 0.03885424 | 0 | 0.13531063 | 0.03980139 | 0.4750269 | 0.01533831 | 0 | 0 | 0.26861422 | 0.02241593 | 0.25755619 | 0.1678702 | 0.01654681 | 0 | 0.18470505 | 0.4197482 |
| [illegible] | Ntpcr | 2898 | 8.57035425 | 6.96385187 | 10.3811246 | 8.02131062 | 7.13594124 | 8.20981168 | 10.6620122 | 8.09712905 | 9.57151423 | 10.2410349 | 7.7381975 | 9.16469182 | 13.4026299 | 12.2909853 | 9.95678815 | 9.70747793 | 8.60850108 | 8.27787151 |
| [illegible] | BC021891 | 5682 | 1.24927216 | 1.17257635 | 1.00458933 | 0.8178857 | 1.20287454 | 1.2907083 | 2.13997861 | 2.43323648 | 3.2111275 | 1.56007255 | 1.78252221 | 2.32810842 | 2.1705349 | 2.56660391 | 1.44299712 | 1.46908701 | 1.23437387 | 1.76199532 |
| [illegible] | Mau2 | 5353 | 16.9528397 | 17.6179345 | 15.2539733 | 20.1718055 | 16.2028258 | 16.9773423 | 17.7124245 | 18.6788174 | 16.5192321 | 20.7787239 | 20.0274625 | 17.0168431 | 21.021129 | 20.241744 | 20.1505087 | 22.2175652 | 19.1810479 | 21.6008112 |
| [illegible] | Pbx4 | 3160 | 0.94157036 | 1.15962606 | 0.68886357 | 0.80741091 | 0.77681269 | 1.34858705 | 0.83845254 | 0.87020664 | 1.5640449 | 1.32797867 | 1.37579785 | 1.1304438 | 1.38941159 | 1.70490954 | 0.95180809 | 0.92091334 | 1.3098861 | 2.5067441 |
| [illegible] | Lpar2 | 5974 | 0.6403525 | 0.74164964 | 0.32389374 | 0.37370123 | 0.75734865 | 0.52256816 | 0.57022311 | 0.54341383 | 0.53775498 | 0.42620321 | 0.43184596 | 0.6633599 | 0.94964354 | 0.74633954 | 0.61381628 | 0.52558249 | 0.33360687 | 0.45119622 |
| [illegible] | Atp13a1 | 4291 | 19.1377302 | 22.2344743 | 22.1744894 | 23.9007017 | 23.6901941 | 23.8582959 | 18.5457763 | 20.711657 | 17.708954 | 18.2958838 | 26.4649444 | 20.981278 | 16.9804805 | 26.2580648 | 23.8702093 | 17.6238522 | 22.4545119 | 26.3059163 |
| [illegible] | Ints10 | 6901 | 4.86582945 | 5.55134449 | 4.65440141 | 6.41063238 | 5.15425179 | 5.76595349 | 5.59442776 | 4.80933165 | 4.39259174 | 5.62993472 | 5.7944704 | 5.84766324 | 4.1961796 | 5.59267304 | 5.02705551 | 5.13796587 | 5.19273839 | 5.57982535 |
| [illegible] | Dctn1 | 6268 | 22.527472 | 29.9167648 | 27.2429104 | 29.1262133 | 28.2817286 | 32.270941 | 26.200113 | 27.9679526 | 23.4976038 | 34.5731704 | 29.2762628 | 24.6130331 | 22.202511 | 24.4891048 | 27.2726955 | 21.8087856 | 24.183233 | 26.4601877 |
| [illegible] | Pgr | 6382 | 0 | 0.00521982 | 0 | 0 | 0 | 0 | 0 | 0 | 0.01795318 | 0.03872126 | 0 | 0 | 0 | 0 | 0 | 0 | 0 | 0 |
| [illegible] | Cdh5 | 3974 | 0 | 0 | 0 | 0 | 0 | 0 | 0 | 0 | 0 | 0 | 0 | 0 | 0 | 0 | 0 | 0 | 0 | 0 |
| [illegible] | Bean1 | 3379 | 0 | 0 | 0 | 0 | 0 | 0.05865964 | 0 | 0 | 0 | 0 | 0 | 0 | 0 | 0 | 0 | 0 | 0 | 0 |
| [illegible] | Cmtm3 | 1611 | 35.3022505 | 28.4741498 | 31.3482096 | 32.3819961 | 35.5238096 | 33.6810945 | 37.0336984 | 38.3820605 | 37.6073027 | 34.4484033 | 30.0409888 | 51.0344326 | 30.9893857 | 26.3500285 | 24.807889 | 38.6946672 | 31.4270877 | 28.9557557 |
| [illegible] | Cmtm1 | 2269 | 0 | 0 | 0 | 0 | 0 | 0 | 0 | 0 | 0 | 0 | 0 | 0 | 0 | 0 | 0 | 0 | 0 | 0 |
| [illegible] | Ces2g | 2847 | 0.01492981 | 0 | 0.20389265 | 0 | 0 | 0.12183672 | 0 | 0 | 0 | 0 | 0 | 0.01960508 | 0.31189832 | 0.01631334 | 0 | 0 | 0.04038598 | 0.23186136 |
| [illegible] | Nae1 | 4554 | 29.6248197 | 29.2091319 | 31.0593331 | 33.1253004 | 30.3229005 | 30.4345621 | 28.5912368 | 27.7846792 | 26.4447457 | 28.4933543 | 28.9125572 | 29.2560357 | 26.9299838 | 25.6288958 | 29.3947859 | 31.1437 | 32.3172991 | 22.2621396 |
| [illegible] | Fam96b | 1316 | 43.7648279 | 39.4388216 | 39.8824319 | 37.5982478 | 40.2318511 | 35.0559232 | 49.2181288 | 41.558906 | 45.6306535 | 45.395092 | 43.0918508 | 39.6562648 | 50.6814226 | 44.1854084 | 46.3360963 | 53.9443664 | 44.7625519 | 48.5718708 |
| [illegible] | Rrad | 1570 | 12.0205721 | 10.2272707 | 8.01090312 | 10.3310394 | 11.3431922 | 11.6464705 | 10.4570061 | 26.7833344 | 15.582669 | 19.1005397 | 19.3534227 | 16.2114281 | 23.2834081 | 23.0445504 | 16.0869769 | 61.3137497 | 45.16156 | 25.4022945 |
| [illegible] | Cdh16 | 3298 | 15.9426631 | 14.8281773 | 14.9902207 | 22.5180419 | 17.2358405 | 16.9633221 | 34.5592006 | 44.9090772 | 36.7156933 | 49.6812292 | 56.0754591 | 37.3345517 | 53.4304453 | 36.5581719 | 35.6422192 | 64.2627386 | 60.8479353 | 41.7155173 |
| [illegible] | Car7 | 2450 | 1.19708455 | 1.50927806 | 0.43437456 | 0.85543185 | 1.04122121 | 1.39556678 | 1.37110212 | 0.81061426 | 0.23535152 | 1.42414369 | 1.48199871 | 0.56954758 | 0.08054191 | 1.38384424 | 1.24445516 | 1.17216145 | 1.03246351 | 0.80829708 |
| [illegible] | Ces2d-ps | 1867 | 0 | 0 | 0 | 0 | 0 | 0 | 0 | 0 | 0 | 0 | 0 | 0 | 0 | 0 | 0 | 0 | 0 | 0 |
| [illegible] | Cbfb | 2905 | 14.5878349 | 17.2699548 | 16.801684 | 16.765837 | 13.6028604 | 15.8125246 | 15.3257249 | 14.5407219 | 22.5568835 | 14.0884029 | 19.6855853 | 15.4285635 | 15.8609331 | 12.0866541 | 12.2824604 | 15.4612332 | 12.9689328 | 12.8764863 |
| [illegible] | Ces2e | 2687 | 0 | 0 | 0.14402242 | 0.10173661 | 0.08956443 | 0 | 0 | 0 | 0 | 0 | 0 | 0.33235973 | 0.60586277 | 0 | 0.39867527 | 0.52726181 | 0.07131801 | 0.65511404 |
| [illegible] | Tradd | 2320 | 14.8768116 | 11.4010509 | 11.884876 | 13.7075901 | 11.7217831 | 11.512467 | 10.5229924 | 9.16946903 | 10.0658424 | 11.8689701 | 13.1792989 | 10.9706647 | 10.9083076 | 10.5900527 | 9.34138724 | 11.4376685 | 9.76329374 | 10.1482318 |
| [illegible] | D230025D16Rik | 4969 | 5.88520046 | 4.03589501 | 4.40998375 | 6.21662066 | 6.81925423 | 4.83660209 | 4.42752983 | 5.73385635 | 7.24431773 | 5.9211187 | 6.98020122 | 5.47036178 | 6.01632962 | 5.29027037 | 5.57203573 | 5.60218788 | 6.32472599 | 6.42066575 |
| [illegible] | Hsd11b2 | 1840 | 9.91017419 | 9.99386827 | 8.64939755 | 8.04746707 | 9.60022608 | 8.91410588 | 2.8799022 | 4.33815875 | 4.85732278 | 2.46003669 | 2.85611165 | 2.66944473 | 2.65427164 | 1.23682624 | 3.53795969 | 3.97473153 | 3.02027683 | 1.58450144 |
| [illegible] | Tsnaxip1 | 2293 | 0 | 0.0871685 | 0.02109618 | 0.07947849 | 0 | 0.25902525 | 0.63963936 | 0.29980953 | 0.82624497 | 0.12337733 | 0 | 0.36512645 | 0.08605655 | 0.58738706 | 0.23358928 | 0.08349456 | 0.26743164 | 0.04798003 |
| [illegible] | Ctrl | 881 | 0.28947906 | 0.30250074 | 0.43926021 | 0.62058176 | 0.16389985 | 0.22498403 | 0.10740676 | 0.26010717 | 0.04674973 | 0.10703905 | 1.68107118 | 0 | 0.44796293 | 0.26358733 | 0.28060083 | 0.78232796 | 0.65254762 | 0.31219697 |
| [illegible] | Psmb10 | 1262 | 35.5669302 | 33.8672292 | 32.312985 | 29.8565639 | 30.3971023 | 35.4957775 | 16.7956261 | 22.546228 | 27.1857113 | 15.9161645 | 21.6539431 | 23.8388606 | 21.6950983 | 15.7144416 | 17.9563041 | 22.3007896 | 20.0134942 | 19.527797 |
| [illegible] | Dpep3 | 1671 | 0 | 0 | 0 | 0 | 0 | 0 | 0 | 0 | 0 | 0 | 0 | 0 | 0 | 0 | 0 | 0 | 0 | 0 |
| [illegible] | [illegible] | [illegible] | [illegible] | [illegible] | [illegible] | [illegible] | [illegible] | [illegible] | [illegible] | [illegible] | [illegible] | [illegible] | [illegible] | [illegible] | [illegible] | [illegible] | [illegible] | [illegible] | [illegible] | [illegible] |

| | | | | | | | | | | | | | | | | | | | | |
|---|---|---|---|---|---|---|---|---|---|---|---|---|---|---|---|---|---|---|---|---|
| ENSMUSG00 | Scn5a | 11226 | 0 | 0.00593495 | 0 | 0.00811706 | 0 | 0 | 0 | 0 | 0 | 0 | 0 | 0 | 0.07031095 | 0.06619503 | 0 | 0 | 0 | 0 |
| ENSMUSG00 | Wdr48 | 3817 | 15.211441 | 13.8069161 | 14.3333673 | 18.4297229 | 17.1872706 | 15.4746813 | 12.370462 | 16.589731 | 12.9591316 | 14.9716144 | 15.4451922 | 14.2134724 | 11.8644747 | 13.116774 | 13.2876907 | 13.8536361 | 16.7382596 | 10.9384103 |
| ENSMUSG00 | Gorasp1 | 3474 | 13.1895739 | 11.8714344 | 12.1421182 | 13.8755282 | 11.4164379 | 11.4396351 | 13.8914601 | 10.1252742 | 13.0886337 | 12.8259685 | 13.9584291 | 12.5159474 | 17.5089965 | 18.4225539 | 12.6545942 | 12.4879845 | 14.3089255 | 13.6018488 |
| ENSMUSG00 | Ttc21a | 5473 | 0.07766339 | 0.12173541 | 0.04419288 | 0.09989632 | 0.05276659 | 0.42553965 | 0.14696063 | 0.07676163 | 0.037627 | 0.07753633 | 0 | 0.02039673 | 0.16224639 | 0.15274869 | 0.07528148 | 0.09095156 | 0.1330531 | 0.3316829 |
| ENSMUSG00 | Csrnp1 | 2866 | 5.23528511 | 5.53277657 | 6.43067518 | 10.349002 | 8.46423222 | 9.95895832 | 8.66683756 | 14.605329 | 6.28000956 | 11.4175115 | 9.88504898 | 9.8738803 | 11.5670092 | 14.3253919 | 14.7353785 | 22.819385 | 18.8020842 | 22.2454831 |
| ENSMUSG00 | Mobp | 6489 | 0 | 0 | 0 | 0 | 0 | 0 | 0 | 0 | 0 | 0 | 0 | 0 | 0 | 0 | 0 | 0 | 0 | 0 |
| ENSMUSG00 | Rpsa | 1854 | 36.1775446 | 28.1021395 | 30.9183568 | 26.95818 | 31.6206489 | 25.6583734 | 36.1352499 | 25.9354016 | 31.3452933 | 26.4237221 | 31.8242037 | 23.6328453 | 42.546783 | 39.8306898 | 32.4901158 | 27.9641221 | 28.2795977 | 32.2815038 |
| ENSMUSG00 | Slc25a38 | 2905 | 12.68571 | 10.951399 | 13.0716769 | 12.2646254 | 12.1448193 | 11.6674939 | 17.3452678 | 16.6968506 | 17.140963 | 15.2894879 | 18.3370323 | 15.7744093 | 18.8157536 | 20.4322009 | 20.3951247 | 16.7266027 | 18.5628571 | 15.9062477 |
| ENSMUSG00 | Hhatl | 2116 | 6.16686628 | 3.66819676 | 5.25799243 | 5.98580828 | 4.13989136 | 5.22223988 | 56.7483833 | 40.2861258 | 47.3177787 | 57.066612 | 47.0976197 | 33.420815 | 40.7524534 | 38.0376198 | 37.8719406 | 46.3070047 | 43.2891539 | 32.07998 |
| ENSMUSG00 | Nktr | 8841 | 9.16354095 | 7.63020142 | 7.18407933 | 6.44172666 | 7.21352775 | 7.10138145 | 9.4721685 | 8.39792221 | 8.68825381 | 8.18110834 | 7.97600257 | 9.24263432 | 12.4208628 | 10.3121529 | 9.43241139 | 11.4858829 | 10.3521318 | 12.636975 |
| ENSMUSG00 | Deb1 | 709 | 47.9607064 | 33.4068657 | 45.1668224 | 42.3479947 | 39.2387192 | 33.827254 | 50.9161834 | 52.6290228 | 53.5598978 | 54.0005233 | 63.8796342 | 45.4240301 | 55.8028176 | 55.4184738 | 54.334915 | 49.2539266 | 47.6781296 | 49.2676767 |
| ENSMUSG00 | Pccb | 3522 | 21.807738 | 14.7174513 | 20.2586477 | 23.0780531 | 18.7088596 | 24.6919499 | 23.8041076 | 19.8986537 | 25.6450963 | 25.5299237 | 19.031308 | 25.7525417 | 26.178692 | 19.0022515 | 27.7367658 | 23.1896268 | 18.608168 | 20.0231899 |
| ENSMUSG00 | Vipr1 | 5551 | 0.73509222 | 1.19424715 | 0.65357859 | 1.29681949 | 0.87575649 | 1.27653412 | 0.38354721 | 0.7224288 | 0.73454604 | 0.39072822 | 0.2065572 | 0.24132155 | 0.81760699 | 1.51439017 | 0.54925511 | 0.12416338 | 0.22784504 | 0.67386403 |
| ENSMUSG00 | Lyzl4 | 1524 | 0 | 0.04371771 | 0.03174116 | 0.0896871 | 0 | 0 | 0 | 0.10024253 | 0.21620218 | 0 | 0 | 0 | 0 | 0 | 0.02703514 | 0 | 0 | 0 |
| ENSMUSG00 | Amotl2 | 4625 | 12.5539612 | 12.2519423 | 9.60149208 | 12.3531999 | 11.3851512 | 11.3462367 | 6.87440438 | 8.18349639 | 5.13829613 | 6.82028551 | 7.7163014 | 8.19434268 | 9.0450737 | 7.6821031 | 10.8504854 | 9.97625555 | 9.42205396 | 9.63402734 |
| ENSMUSG00 | Cck | 844 | 0 | 0.11841076 | 0 | 0 | 0 | 0 | 0.11211535 | 0.13575498 | 0.63438946 | 0.67038915 | 0.62265941 | 0.85971992 | 0.05845014 | 0.44022832 | 0 | 0.18147207 | 0.13623091 | 0 |
| ENSMUSG00 | Cep63 | 7862 | 5.78485606 | 5.65242952 | 5.74058812 | 5.48795586 | 6.04250937 | 5.93094916 | 6.94464913 | 5.86342276 | 6.3649983 | 6.53704739 | 6.41698755 | 5.65824003 | 5.03233244 | 5.06856105 | 4.9838056 | 5.95642494 | 5.68410865 | 5.09369495 |
| ENSMUSG00 | Trak1 | 5496 | 29.1565709 | 34.9312608 | 33.2435997 | 38.3820297 | 33.5855035 | 38.2645199 | 33.0655021 | 38.5954518 | 30.3653133 | 42.0633002 | 31.4935725 | 32.8739628 | 18.8315798 | 24.3544141 | 25.1212543 | 18.2674623 | 25.6763874 | 22.9705058 |
| ENSMUSG00 | Ephb1 | 7494 | 0 | 0.04000748 | 0.07100465 | 0.01823901 | 0 | 0.00661232 | 0 | 0 | 0 | 0 | 0 | 0 | 0.00658286 | 0 | 0.06597527 | 0 | 0 | 0 |
| ENSMUSG00 | Abhd5 | 3717 | 8.94243951 | 8.02126455 | 8.13383279 | 7.3422291 | 7.29036132 | 9.41195316 | 9.57200284 | 6.86374087 | 8.67609412 | 8.99377148 | 7.51906126 | 7.9136008 | 9.17093216 | 10.4833439 | 8.94529341 | 8.87987721 | 8.93970929 | 8.68720625 |
| ENSMUSG00 | Ryk | 4662 | 11.1414254 | 10.897075 | 10.1063532 | 9.67512525 | 10.6133886 | 11.1924128 | 10.0673911 | 10.5434624 | 11.1933324 | 8.31357316 | 8.70033028 | 10.5836649 | 8.63467278 | 8.89630417 | 10.8615856 | 10.1845459 | 11.8711224 | 7.457262 |
| ENSMUSG00 | Slco2a1 | 4247 | 2.69222801 | 4.62003642 | 5.29637294 | 5.09571369 | 3.93260596 | 5.64716813 | 2.49541796 | 3.65107497 | 3.88881394 | 5.36232392 | 7.46940721 | 2.19477647 | 29.7362157 | 28.9359678 | 24.3115373 | 34.9907811 | 29.8795312 | 28.728561 |
| ENSMUSG00 | Rab6b | 4726 | 2.69817029 | 2.1710477 | 2.18018662 | 2.13055243 | 2.16929745 | 1.56228466 | 2.71302073 | 2.79614288 | 0.95863663 | 1.18724792 | 0.90980666 | 0.96844781 | 2.46346434 | 3.12509941 | 2.00515401 | 3.01398772 | 1.6624822 | 1.53643717 |
| ENSMUSG00 | 1110059G10 | 3931 | 6.24980711 | 6.89816736 | 7.48183833 | 6.47891727 | 7.35876014 | 7.65161773 | 6.55950395 | 8.5400971 | 8.19329823 | 6.77693903 | 10.4397582 | 7.6673767 | 5.91079446 | 7.63237864 | 7.87137331 | 7.51006317 | 7.79005628 | 6.60501103 |
| ENSMUSG00 | Srprb | 5556 | 11.582584 | 13.5446055 | 12.5374161 | 11.3492594 | 11.5825021 | 10.7828033 | 11.8281697 | 13.2326088 | 12.0312662 | 11.5500553 | 12.279093 | 11.1711693 | 11.1365516 | 11.3184475 | 10.4635341 | 11.4954523 | 11.851076 | 11.4453793 |
| ENSMUSG00 | Trf | 2828 | 0 | 0.01177967 | 0 | 0.03222139 | 0 | 0 | 0 | 0.02701019 | 0.17476598 | 0 | 0 | 0 | 0 | 0 | 0.07284575 | 0.01353982 | 0 | 0 |
| ENSMUSG00 | Topbp1 | 5200 | 4.3976509 | 5.27240614 | 5.09782589 | 5.50237228 | 5.95169748 | 5.06962582 | 4.59478901 | 6.63225757 | 5.49681576 | 5.28631923 | 5.50330772 | 7.11649698 | 4.81934884 | 4.77838207 | 7.28950604 | 3.3651553 | 4.79078711 | 5.51148935 |
| ENSMUSG00 | Bfsp2 | 1822 | 0.13997314 | 0 | 0 | 0 | 0 | 0 | 0.05193488 | 0.0628854 | 0.27126136 | 0 | 0.05244233 | 0 | 0.08122709 | 0 | 0 | 0.12609421 | 0 | 0 |
| ENSMUSG00 | Uba5 | 2565 | 37.6332377 | 35.9623408 | 38.2839246 | 34.4594253 | 35.0152708 | 35.3727311 | 32.0029232 | 31.9386757 | 31.8573284 | 31.856596 | 29.5031323 | 31.4874323 | 41.4080382 | 39.1108104 | 37.6355733 | 42.4704788 | 33.4253123 | 36.308155 |
| ENSMUSG00 | Nphp3 | 6804 | 2.21146859 | 3.02087827 | 2.76562365 | 2.10261139 | 2.95695827 | 2.86217288 | 2.22517008 | 4.58039394 | 2.3668324 | 2.95904619 | 4.31126169 | 3.34697095 | 4.65477526 | 4.12290253 | 4.08745594 | 4.77225677 | 4.79923647 | 3.72710144 |
| ENSMUSG00 | Dnajc13 | 7761 | 16.0797831 | 14.345019 | 14.8779303 | 13.4904368 | 14.7354014 | 15.3427675 | 12.9239633 | 12.1353517 | 12.8160592 | 13.0984301 | 12.6624262 | 13.7219797 | 12.1343422 | 11.962599 | 12.3748005 | 10.2818742 | 12.0988825 | 10.9578764 |
| ENSMUSG00 | Acpp | 7778 | 33.2696724 | 46.4787256 | 43.2799431 | 42.0522933 | 42.5315906 | 43.283784 | 6.52693552 | 6.49634204 | 7.13271236 | 7.73518865 | 5.65093363 | 7.57795573 | 11.7843536 | 15.3639286 | 16.2570809 | 22.2369084 | 24.0906744 | 19.1874141 |
| ENSMUSG00 | Gnai2 | 3118 | 182.903126 | 178.14599 | 158.338759 | 171.138855 | 160.604405 | 172.973686 | 121.832424 | 129.080031 | 129.649023 | 135.841811 | 113.308473 | 126.900972 | 148.043217 | 138.185311 | 130.938492 | 161.979831 | 135.776868 | 140.998326 |
| ENSMUSG00 | Mrpl3 | 3769 | 13.5330887 | 10.9245787 | 15.1063001 | 14.2400929 | 12.0297883 | 13.6207567 | 13.6954451 | 13.3962207 | 13.4082926 | 16.4758493 | 15.1855665 | 11.950979 | 18.3898498 | 15.329383 | 14.8015141 | 14.9037732 | 15.6193134 | 16.9011795 |
| ENSMUSG00 | Cpne4 | 5034 | 0.05066171 | 0.02647032 | 0.00960936 | 0 | 0 | 0.02953083 | 0.19737113 | 0.06069512 | 0 | 0 | 0.18031868 | 0.19957925 | 0.25479337 | 0.08303472 | 0.00818466 | 0.04563839 | 0 | 0.04371006 |
| ENSMUSG00 | Nudt16 | 1603 | 18.9854685 | 16.1472981 | 21.697173 | 18.2471556 | 19.8472873 | 21.4532733 | 27.9802992 | 24.3497413 | 27.7231756 | 23.1488481 | 25.9886237 | 26.6020999 | 34.6215889 | 27.4955951 | 26.0112131 | 25.4394865 | 31.2252924 | 29.5120598 |
| ENSMUSG00 | 1700080E11Rik | 868 | 0 | 0 | 0 | 0 | 0 | 0 | 0 | 0 | 0 | 0 | 0 | 0.12860752 | 0 | 0 | 0 | 0 | 0 | 0 |
| ENSMUSG00 | Aste1 | 2765 | 4.08910554 | 5.87945475 | 4.07632281 | 4.69616551 | 5.23967777 | 5.28681968 | 4.82537991 | 5.08311178 | 4.97515236 | 4.50191163 | 6.16841236 | 4.84475914 | 4.67448909 | 6.68526104 | 5.75182661 | 6.50871084 | 8.03951255 | 8.75370957 |
| ENSMUSG00 | Atp2c1 | 6892 | 18.4772936 | 21.0453194 | 20.6492928 | 20.2750614 | 20.3436294 | 22.5906397 | 12.4116836 | 16.1203858 | 15.3941475 | 13.9974374 | 14.1134381 | 17.4039261 | 12.4618207 | 11.6312384 | 15.0351006 | 16.0173855 | 17.322463 | 15.0612572 |
| ENSMUSG00 | Pik3r4 | 4864 | 6.57152393 | 8.94462168 | 11.2281488 | 10.9312785 | 8.97525797 | 9.17907339 | 8.07350401 | 8.97490048 | 6.13055868 | 9.39330416 | 9.08549272 | 9.46708915 | 8.01237966 | 6.5598454 | 6.57327454 | 5.96716291 | 7.4698457 | 8.86659944 |
| ENSMUSG00 | Col6a4 | 8712 | 4.94234887 | 1.54481282 | 4.79737491 | 5.65852272 | 1.11600649 | 4.69249371 | 1.8899004 | 0.73210872 | 2.26450196 | 3.48001632 | 1.52998317 | 2.5627026 | 5.39638627 | 3.37988423 | 4.94210582 | 4.14902817 | 2.70994019 | 5.97952526 |
| ENSMUSG00 | Manf | 3517 | 35.036254 | 28.5579687 | 29.2414346 | 26.4919919 | 29.3554098 | 27.2068037 | 44.0168 | 36.5417782 | 32.2980979 | 32.4705724 | 27.6570416 | 33.7560177 | 53.9747562 | 58.540415 | 44.6809025 | 38.1164107 | 38.6532201 | 45.7184017 |
| ENSMUSG00 | Mapkapk3 | 6697 | 7.29258582 | 7.9439588 | 6.2263675 | 6.47664838 | 5.84310198 | 5.99338722 | 6.13220924 | 6.12492761 | 5.68259528 | 6.0478504 | 4.95084682 | 7.07592915 | 5.70886007 | 6.2970332 | 4.54035363 | 7.50145989 | 6.81025733 | 6.94903752 |
| ENSMUSG00 | Cish | 1821 | 26.6795256 | 37.2460476 | 32.6740485 | 33.226286 | 30.4491906 | 32.4364956 | 14.0820824 | 27.9154792 | 17.3928779 | 15.4838663 | 22.40517 | 18.9730341 | 22.9998889 | 26.3208667 | 24.9109901 | 34.0010501 | 28.8131249 | 22.6863477 |
| ENSMUSG00 | Hemk1 | 3039 | 2.96515213 | 4.08874944 | 4.09081846 | 4.55760365 | 3.91200564 | 3.45678826 | 4.06337914 | 4.43630077 | 4.36395462 | 3.42885996 | 4.00875762 | 6.57519158 | 3.31152061 | 5.11970032 | 3.74189846 | 4.56110559 | 4.09873198 | 3.89172687 |
| ENSMUSG00 | Rbm5 | 3062 | 54.0823533 | 41.0699463 | 42.8758202 | 47.2424284 | 43.5576618 | 39.0498844 | 54.3585906 | 42.1338783 | 49.6336522 | 49.8146718 | 53.1578326 | 43.839539 | 54.0524437 | 58.9120952 | 55.8414294 | 56.135381 | 56.9887752 | 53.2125977 |
| ENSMUSG00 | Rbm6 | 4202 | 14.5763824 | 15.5227622 | 17.1989658 | 16.5568098 | 16.8381551 | 18.0781391 | 14.8513621 | 15.5787191 | 17.6625658 | 15.7991885 | 12.7339252 | 16.0061574 | 14.0059441 | 15.5513639 | 15.1882937 | 15.1084786 | 15.049593 | 11.4547759 |
| ENSMUSG00 | Mon1a | 2075 | 7.43584525 | 8.23591072 | 10.5838953 | 8.78285192 | 8.60575599 | 9.55233418 | 11.5830559 | 9.16617658 | 10.6787207 | 10.4754093 | 12.4099772 | 11.3245272 | 9.24824859 | 11.7061483 | 11.2385931 | 9.57726518 | 9.73395589 | 8.8544779 |
| ENSMUSG00 | Mst1r | 4687 | 0.00906874 | 0.08529011 | 0.01032079 | 0.03888291 | 0.14376905 | 0 | 0.01009445 | 0.07333724 | 0 | 0.13077857 | 0.09173771 | 0 | 0.22103057 | 0 | 0.35162412 | 0 | 0.13083438 | 0.14083834 |
| ENSMUSG00 | Traip | 2682 | 1.9176459 | 2.76986406 | 2.63330926 | 1.85166064 | 2.62015692 | 2.86378587 | 2.89309446 | 0.78321481 | 1.87351041 | 1.70530135 | 1.62099971 | 0.70758111 | 2.07848868 | 2.16462003 | 2.27361306 | 1.42768857 | 2.27214064 | 2.09206894 |
| ENSMUSG00 | Bsn | 9812 | 0.12562679 | 0.30556059 | 0.10353079 | 0.0278604 | 0.12263535 | 0.13130565 | 0.06750688 | 0.22985264 | 0.10074158 | 0.06247036 | 0.12172584 | 0.07963914 | 0.12066511 | 0.31240418 | 0.20995493 | 0.31609653 | 0.24217595 | 0.25789023 |
| ENSMUSG00 | Apeh | 2366 | 29.5883453 | 27.0332869 | 20.3430527 | 31.1378735 | 29.5586158 | 27.205833 | 23.7963178 | 22.6474812 | 21.1677106 | 25.2294126 | 27.9865141 | 24.3928128 | 30.0870484 | 26.5198487 | 29.1684726 | 29.3572113 | 28.8500147 | 24.063578 |
| ENSMUSG00 | Mst1 | 2344 | 0.03626722 | 0 | 0.08254869 | 0 | 0.08213639 | 0.08456098 | 0.18166131 | 0.29328637 | 0 | 0 | 0 | 0.64292785 | 0 | 0 | 0 | 0 | 0 | 0 |
| ENSMUSG00 | Amigo3 | 2542 | 0.53507696 | 0.65524969 | 1.0276045 | 0.77070217 | 0.68164801 | 1.13062885 | 0.98645633 | 1.72782308 | 1.63644299 | 1.00162784 | 0.35709056 | 0.90025262 | 1.10618383 | 0.80391025 | 1.42633238 | 1.9582136 | 0.69355219 | 1.14692473 |
| ENSMUSG00 | Ip6k1 | 4572 | 19.811577 | 22.4490435 | 24.4089533 | 25.6206135 | 20.8234788 | 27.1066025 | 23.9150483 | 24.592833 | 23.9353833 | 25.9060732 | 23.9919759 | 25.8934865 | 23.2740476 | 26.8689001 | 26.2601343 | 27.9055783 | 28.5603004 | 25.4832214 |
| ENSMUSG00 | Cdhr4 | 3105 | 0 | 0 | 0 | 0.10271411 | 0 | 0 | 0 | 0 | 0 | 0 | 0.01538646 | 0 | 0 | 0 | 0 | 0 | 0 | 0 |
| ENSMUSG00 | Uba7 | 5028 | 3.19549653 | 4.91610332 | 4.42558154 | 3.22588142 | 5.36075522 | 3.88301055 | 4.10269054 | 3.81316349 | 5.38177026 | 2.65386814 | 2.31843474 | 1.97597214 | 2.53135136 | 4.7848125 | 5.3017913 | 3.91435406 | 3.62072216 | 5.34993102 |
| ENSMUSG00 | Nckipsd | 2924 | 7.31193692 | 7.2572892 | 6.68362047 | 8.64787264 | 8.26341508 | 7.94809578 | 10.8411405 | 8.67296689 | 7.62035056 | 10.1751347 | 7.95704678 | 9.25806989 | 9.21177388 | 10.8009506 | 10.4272063 | 7.25478676 | 10.2893786 | 9.97087092 |
| ENSMUSG00 | Ip6k2 | 2835 | 55.5490925 | 38.1423826 | 53.4582965 | 52.8411689 | 42.6141546 | 49.238113 | 77.9699598 | 67.2105452 | 77.9856143 | 69.7364722 | 63.666241 | 76.9803318 | 78.3742347 | 58.7800302 | 69.1197883 | 61.0218569 | 56.2524591 | 69.8140268 |
| ENSMUSG00 | Prkar2a | 4178 | 30.0730733 | 29.9321697 | 31.6430968 | 29.9996249 | 30.1717348 | 34.7746804 | 24.528306 | 28.5391767 | 20.9777179 | 26.4418616 | 23.8417397 | 28.3353328 | 15.4678822 | 11.6499291 | 17.7113438 | 15.3602341 | 17.1174892 | 15.8786459 |
| ENSMUSG00 | Slc25a20 | 1774 | 49.3337979 | 50.8707049 | 45.0740956 | 48.5659159 | 52.8799996 | 50.3907167 | 34.457712 | 40.4744738 | 32.0855495 | 31.6552918 | 34.6328053 | 35.9908658 | 33.5924217 | 28.327228 | 23.7826343 | 33.7147287 | 29.4900763 | 25.5200084 |
| ENSMUSG00 | Qars | 3751 | 21.2015954 | 23.0108528 | 20.891794 | 21.5962508 | 21.4418674 | 19.4987563 | 20.2822681 | 21.443135 | 20.3132642 | 21.369287 | 20.3403382 | 19.9841202 | 22.5025088 | 23.673979 | 19.2112827 | 21.8861803 | 21.9065797 | 15.8383992 |
| ENSMUSG00 | Nicn1 | 2106 | 23.2487988 | 25.047883 | 22.3968528 | 24.6392745 | 30.1852007 | 23.8360872 | 31.6016375 | 32.2135159 | 21.3747857 | 31.1132012 | 31.1398449 | 24.6775131 | 28.5975346 | 24.5219494 | 26.347105 | 28.4454892 | 28.5447278 | 37.2381382 |
| ENSMUSG00 | Amt | 1617 | 1.57718649 | 1.4627183 | 1.43594895 | 1.40881399 | 1.42877695 | 1.5628877 | 3.1600305 | 1.72420864 | 1.65561133 | 2.47854655 | 1.09318087 | 2.51981655 | 2.44066381 | 1.86695466 | 1.50333444 | 1.82336226 | 2.41761428 | 2.007135 |
| ENSMUSG00 | Klhdc8b | 1891 | 34.6604873 | 43.9180567 | 38.8830069 | 46.2356658 | 51.8480849 | 45.0455574 | 44.260248 | 51.2194249 | 45.8474964 | 47.4499145 | 52.3478128 | 52.3622345 | 55.9060279 | 68.2293362 | 55.472851 | 82.7166215 | 65.6269431 | 69.1177354 |
| ENSMUSG00 | 1700102P08Rik | 1090 | 0 | 0.12224915 | 0 | 0.04179912 | 0.13247318 | 0.13638367 | 0.04340613 | 0 | 0.26450056 | 0.30280268 | 0.08766047 | 0.05120703 | 0.27155184 | 0.17043702 | 0.4913947 | 0.070258 | 0 | 0.25233535 |
| ENSMUSG00 | Usp4 | 3751 | 35.3322151 | 39.7694322 | 43.6535325 | 39.8886882 | 42.7425854 | 43.7136751 | 35.7083962 | 36.2985643 | 31.0298836 | 38.2510237 | 40.5278373 | 39.8194383 | 35.8909097 | 38.6435608 | 41.3333657 | 31.1959734 | 39.6545877 | 31.3688295 |
| ENSMUSG00 | Nt5m | 3354 | 9.99898148 | 7.45914835 | 7.31227784 | 8.44930518 | 7.60581978 | 7.84511077 | 8.92931597 | 8.87056693 | 9.17302542 | 8.47700479 | 8.19039986 | 7.73830742 | 11.6637479 | 9.09772374 | 8.46388551 | 9.97793407 | 9.00450863 | 8.49574159 |
| ENSMUSG00 | Srek1 | 4268 | 22.7068076 | 24.9144232 | 22.7700146 | 24.7660785 | 22.588623 | 22.9419404 | 26.1727355 | 27.1498943 | 26.7500051 | 23.9841089 | 29.5739082 | 21.4605207 | 28.5265164 | 29.5009187 | 30.7177486 | 28.6820693 | 31.9685138 | 29.2574209 |
| ENSMUSG00 | Oas1d | 2065 | 0.26758706 | 0.56462533 | 0.32795614 | 0.04412692 | 0.74587 | 0.28795777 | 0 | 0 | 0 | 0.13699962 | 0.02313557 | 0 | 0 | 0.4048395 | 0 | 0.166884 | 0 | 0 |
| ENSMUSG00 | Eml4 | 5457 | 10.0557417 | 10.341221 | 11.50611 | 10.0439685 | 10.0374068 | 10.6424416 | 8.14988422 | 9.58134486 | 7.23047134 | 9.02922575 | 7.04761619 | 10.2589536 | 6.83432829 | 6.20466027 | 6.22138209 | 7.11502216 | 6.39122781 | 5.82650975 |
| ENSMUSG00 | Thsd7a | 14070 | 0.28397204 | 0.24623603 | 0.30598751 | 0.42743837 | 0.29419655 | 0.25709663 | 0 | 0 | 0.00292726 | 0 | 0.00679104 | 0 | 0 | 0 | 0.00292833 | 0 | 0.02723973 | 0.01172902 |
| ENSMUSG00 | Flcn | 5557 | 20.8586675 | 16.521565 | 16.9659908 | 18.824574 | 18.960011 | 21.1515357 | 24.3246938 | 16.5773033 | 21.6049475 | 20.8304798 | 22.5849948 | 20.0984598 | 24.5017261 | 17.4760818 | 25.7500456 | 23.2899502 | 22.4150558 | 28.1133459 |
| ENSMUSG00 | Atxn2l | 4908 | 24.1971194 | 23.9869128 | 24.4627348 | 23.5510125 | 22.4576416 | 23.3830748 | 24.8227686 | 26.3020212 | 26.6940315 | 24.478401 | 22.1256073 | 25.8153126 | 22.3441019 | 21.272679 | 22.1622061 | 21.7432606 | 22.0758845 | 23.794689 |
| ENSMUSG00 | Chsy1 | 4208 | 0.43434472 | 0.87082185 | 0.43683321 | 0.70377088 | 0.54903335 | 0.78898126 | 0.69709746 | 0.52641619 | 0.38171913 | 0.39217553 | 0.7720288 | 0.35813282 | 0.3634243 | 0.26489023 | 0.1762424 | 0.30028281 | 0.10018736 | 0.01307251 |

[illegible]

[illegible]

| | | | | | | | | | | | | | | | | | | | |
|---|---|---|---|---|---|---|---|---|---|---|---|---|---|---|---|---|---|---|---|
| MUSG00 Dysf | 8358 | 0.21359385 | 0.16740148 | 0.23150768 | 0.20169402 | 0.12093448 | 0.24900871 | 0.48116508 | 0.29245199 | 0.35972908 | 0.0789794 | 0.17148226 | 0.2738026 | 0.0295118 | 0.07779579 | 0.01971838 | 0.30236861 | 0.03668466 | 0 |
| MUSG00 Tubgcp5 | 3658 | 5.55425756 | 7.12156459 | 6.59879487 | 8.14568699 | 8.19742713 | 7.12540674 | 5.22534781 | 7.16238055 | 4.15468133 | 7.70807013 | 7.66645725 | 7.2172795 | 8.23996759 | 6.06896495 | 6.90447068 | 7.74604961 | 7.58563121 | 7.95511686 |
| MUSG00 Atp7a | 8767 | 7.44216325 | 10.3962677 | 9.32488494 | 9.14131139 | 8.07048338 | 10.8239403 | 4.009732 | 4.00787159 | 4.5146848 | 7.20141344 | 5.55294658 | 4.60939201 | 3.24677954 | 2.93487218 | 4.82650826 | 3.17960103 | 4.90499659 | 3.65179652 |
| MUSG00 Atp6v1h | 1962 | 302.020723 | 307.303809 | 293.076532 | 309.243847 | 324.313981 | 317.344596 | 326.052361 | 347.138181 | 338.623693 | 348.030828 | 350.617547 | 331.850002 | 309.569096 | 340.045523 | 352.145204 | 363.448513 | 374.453 | 324.447188 |
| MUSG00 Lpcat2b | 3210 | 0.03972446 | 0 | 0 | 0 | 0.04498311 | 0 | 0 | 0 | 0 | 0 | 0 | 0.05216417 | 0 | 0 | 0 | 0 | 0 | 0 |
| MUSG00 BC016423 | 8321 | 2.55409062 | 2.55021195 | 2.27305016 | 2.76509162 | 2.3831694 | 2.775096 | 2.57572931 | 2.12970493 | 2.55899876 | 2.08259536 | 1.81431161 | 3.09900686 | 1.54736575 | 2.40564856 | 2.30245445 | 1.95571544 | 2.7267357 | 2.10225886 |
| MUSG00 Epha4 | 1637 | 0 | 0.0610499 | 0.08865033 | 0.11132815 | 0 | 0.09081136 | 0 | 0 | 0 | 0 | 0 | 0 | 0.3314912 | 0 | 0 | 0 | 0 | 0.16801804 |
| MUSG00 Tmem87a | 8736 | 22.2986746 | 21.8692134 | 22.7858376 | 22.1546836 | 22.7877331 | 21.656632 | 16.7349105 | 19.3934864 | 17.7975157 | 21.5027932 | 19.7859203 | 19.4294219 | 16.9126708 | 15.911991 | 19.6245048 | 17.9837869 | 21.400604 | 16.6047407 |
| MUSG00 Atg3 | 1671 | 20.2223907 | 23.464568 | 23.6222626 | 25.1117426 | 22.0928698 | 21.1436858 | 21.915029 | 19.9990138 | 19.1513601 | 18.7925604 | 21.5287515 | 19.1730642 | 20.1047491 | 20.1229918 | 21.8706045 | 20.5087335 | 18.1883623 | 19.6531638 |
| MUSG00 Tcea1 | 2833 | 9.57229167 | 8.29000721 | 9.08391039 | 9.26338212 | 8.32497091 | 7.71364473 | 10.9221645 | 8.42578592 | 9.78415994 | 8.87092299 | 9.05582517 | 7.78227558 | 8.34097719 | 8.37731333 | 10.4712744 | 7.86626668 | 10.173447 | 8.67951666 |
| MUSG00 Ppp1r16a | 3879 | 14.8368158 | 16.6435651 | 13.0068554 | 17.1015418 | 15.5228246 | 17.2329563 | 16.4661296 | 14.5326079 | 17.264572 | 17.6374505 | 19.1149099 | 13.3819454 | 15.8843941 | 12.9310684 | 14.4561272 | 14.0073142 | 15.8976571 | 13.7841845 |
| MUSG00 Tpsb2 | 1182 | 0 | 0 | 0 | 0 | 0 | 0 | 0 | 0 | 0 | 0 | 0 | 0 | 0 | 0 | 0 | 0 | 0 | 0 |
| MUSG00 Dnahc8 | 14717 | 0 | 0 | 0.00328692 | 0 | 0 | 0.00673408 | 0 | 0 | 0.00559713 | 0 | 0.03246243 | 0.0037926 | 0.01340814 | 0 | 0.00559918 | 0.00260179 | 0.00260422 | 0.00373779 |
| MUSG00 Fgb | 1636 | 0 | 0 | 0 | 0 | 0 | 0 | 0 | 0 | 0 | 0 | 0 | 0 | 0 | 0 | 0 | 0 | 0 | 0 |
| MUSG00 Tpbpa | 1367 | 0 | 0 | 0 | 0 | 0 | 0 | 0 | 0 | 0 | 0 | 0 | 0 | 0 | 0 | 0 | 0 | 0 | 0 |
| MUSG00 Foxh1 | 1303 | 0.13048404 | 0 | 0 | 0 | 0 | 0.07605945 | 0 | 0 | 0 | 0.03618626 | 0 | 0 | 0 | 0 | 0 | 0 | 0.08824167 | 0 |
| MUSG00 Mrpl15 | 5106 | 16.3161271 | 10.9346672 | 12.9602408 | 12.9116395 | 12.5278744 | 13.130601 | 15.27979 | 13.3366732 | 13.9466286 | 13.9993079 | 12.7343749 | 11.0625638 | 14.3184296 | 12.3978243 | 13.5643979 | 14.8482966 | 14.2616459 | 14.1993738 |
| MUSG00 Pla2g4c | 3728 | 0.3648513 | 0.24126828 | 0.37629624 | 0.13443441 | 0.16784201 | 0.21267268 | 0.05076468 | 0.08195795 | 0.58553791 | 0.15177265 | 0.2691186 | 0.4791044 | 0.83366706 | 0.34882898 | 0.8509978 | 0.47246994 | 0.56543625 | 1.09191896 |
| MUSG00 B3galt2 | 5078 | 0.09207502 | 0.07216263 | 0.0190522 | 0.07177794 | 0.08530668 | 0.06830822 | 0.93171877 | 0.51143824 | 0.41364953 | 0.71496734 | 0.03763289 | 0.06594998 | 2.04982957 | 2.80786428 | 1.64708859 | 0.12064784 | 0.48304116 | 0.74746522 |
| MUSG00 Olfr1015 | 1015 | 0 | 0 | 0 | 0 | 0 | 0 | 0 | 0 | 0 | 0 | 0 | 0 | 0 | 0 | 0 | 0 | 0 | 0 |
| MUSG00 Pla2g4b | 6613 | 3.77938054 | 3.06782891 | 3.14541328 | 3.37591286 | 3.79203572 | 3.39443344 | 5.1154643 | 3.82905834 | 4.50292619 | 4.16392117 | 4.42133292 | 4.15262453 | 2.32001002 | 3.06911631 | 3.7943063 | 4.56846957 | 3.77873062 | 3.41883304 |
| MUSG00 Kcnk10 | 3880 | 0.05477471 | 0.06010568 | 0 | 0 | 0 | 0.12771323 | 0 | 0.18702465 | 0 | 0 | 0 | 0 | 0 | 0.23940251 | 0 | 0.02960614 | 0 | 0.11342084 |
| MUSG00 Ston1 | 4558 | 0.47559543 | 0.44582855 | 0.54125714 | 0.49979206 | 0.54911365 | 0.50009341 | 0.08304112 | 0.33516808 | 0.75903188 | 0.43447335 | 0.59744902 | 0.31838684 | 0.46539545 | 0.27511855 | 0.49716665 | 0.6888613 | 0.47088035 | 0.69998425 |
| MUSG00 Engase | 5190 | 1.13019543 | 1.67527272 | 1.44468154 | 1.08854904 | 1.22416453 | 1.66130552 | 1.6955989 | 0.94929093 | 1.5395345 | 2.00776598 | 2.08037391 | 1.97882118 | 1.54934538 | 1.28862208 | 2.16724947 | 1.18782038 | 1.27754323 | 1.17649535 |
| MUSG00 Fgg | 2075 | 0.12290653 | 0 | 0.09325018 | 0 | 0 | 0.09552334 | 0.27361549 | 0.75464506 | 0.3374317 | 0 | 0 | 0.51108318 | 0 | 0 | 0.05956851 | 0 | 0 | 0 |
| MUSG00 Cdk20 | 1630 | 38.56758 | 28.223992 | 32.4963287 | 30.7187655 | 31.832033 | 34.0181036 | 26.0945389 | 27.7656422 | 26.3037809 | 26.8730082 | 30.3942527 | 27.6681321 | 30.2347149 | 27.9519323 | 26.2628323 | 28.6357365 | 27.2751561 | 27.4033096 |
| MUSG00 Kif9 | 3261 | 27.2418945 | 31.5660359 | 27.264811 | 32.8749271 | 31.5713538 | 33.0047646 | 30.5261811 | 32.359892 | 29.2258809 | 34.6003167 | 33.9156769 | 30.8432456 | 28.6218913 | 30.1366725 | 26.7854335 | 34.9324002 | 34.0599842 | 28.3226894 |
| MUSG00 Ppargc1b | 4249 | 5.3319037 | 5.28427647 | 5.30526363 | 6.15486925 | 6.93262821 | 6.18097171 | 7.5495402 | 6.6942275 | 7.4831702 | 7.37943461 | 5.07099305 | 6.29223403 | 3.78493882 | 3.70547085 | 5.44958212 | 5.02851469 | 4.98810124 | 4.59595973 |
| MUSG00 Best4-ps | 1371 | 0.15501523 | 0 | 0 | 0 | 0 | 0 | 0 | 0 | 0 | 0 | 0 | 0.04071164 | 0 | 0 | 0 | 0 | 0 | 0 |
| MUSG00 Lgals3bp | 3495 | 18.7776801 | 18.4817459 | 17.5916329 | 19.7626733 | 18.75699 | 22.7559764 | 17.2599902 | 13.9765616 | 15.3079496 | 14.1114837 | 10.9766214 | 14.6925349 | 14.2279178 | 9.36854982 | 10.4448007 | 18.1976249 | 10.6699533 | 15.8810268 |
| MUSG00 Rbm46 | 1784 | 2.00136479 | 2.44618225 | 2.98267284 | 1.17478029 | 1.86160467 | 2.83317199 | 5.01238583 | 6.336856 | 5.65621993 | 2.9072743 | 4.74000429 | 5.60033834 | 6.60892838 | 6.66462241 | 8.63754591 | 4.29266899 | 6.68132194 | 7.27698945 |
| MUSG00 D3Ertd254e | 2385 | 8.7327196 | 8.11521658 | 5.90217918 | 6.80072611 | 6.23595992 | 7.95752493 | 6.34803239 | 7.07800497 | 9.96420109 | 8.42188662 | 6.59034018 | 6.17833749 | 8.04617016 | 4.45941133 | 7.82570433 | 5.33018099 | 5.39942799 | 6.50422142 |
| MUSG00 Pxk | 3026 | 95.0537099 | 95.3369676 | 103.221377 | 107.488531 | 102.769451 | 111.763849 | 99.5974107 | 101.905306 | 96.3650137 | 108.356243 | 97.9023521 | 98.192558 | 99.2670338 | 87.7925247 | 97.8026358 | 112.163894 | 113.72499 | 108.454835 |
| MUSG00 Cfhr2 | 4227 | 0 | 0 | 0 | 0 | 0 | 0 | 0 | 0 | 0 | 0 | 0 | 0 | 0 | 0 | 0 | 0 | 0 | 0 |
| MUSG00 Mtap9 | 6204 | 1.47987073 | 1.10076456 | 1.23195 | 0.98407156 | 1.17924765 | 1.02236459 | 1.76926846 | 1.57595987 | 0.95597329 | 1.0108065 | 0.82397172 | 1.84432799 | 1.47105169 | 1.52717502 | 0.90983449 | 1.16032144 | 0.80309778 | 1.31227398 |
| MUSG00 Mapkbp1 | 7572 | 1.28548406 | 1.25385299 | 1.10520612 | 1.24552776 | 1.11239919 | 0.85729109 | 0.93100755 | 1.18027234 | 0.64183952 | 1.30143912 | 1.11676805 | 1.4668934 | 2.13041957 | 2.71721219 | 3.81435429 | 2.41718309 | 2.18660331 | 2.52088748 |
| MUSG00 6330503K22f | 5956 | 4.01786671 | 2.99234359 | 2.74517348 | 3.38877479 | 3.6042374 | 2.29626502 | 2.94711281 | 4.48228497 | 2.95276062 | 3.27743303 | 4.96519464 | 4.08590143 | 5.99669377 | 5.51309096 | 4.09525203 | 7.0782335 | 4.74896023 | 5.46766138 |
| MUSG00 Zdhhc15 | 5497 | 1.14439986 | 1.44838671 | 1.75119748 | 1.20181034 | 0.88435952 | 1.59556737 | 1.747221 | 1.38262472 | 1.56594175 | 1.03788159 | 1.46010424 | 1.05599944 | 1.15768916 | 2.07845106 | 1.54402774 | 2.37531328 | 1.88249958 | 2.14152244 |
| MUSG00 Usp36 | 9559 | 2.8547257 | 3.34557783 | 2.59098729 | 3.06472973 | 1.79254784 | 2.67488341 | 2.40052822 | 2.87671616 | 2.41285164 | 2.99901952 | 2.01915297 | 3.04799413 | 2.4100853 | 3.45937757 | 3.36198489 | 2.13103914 | 2.99906581 | 2.82555407 |
| MUSG00 Gucy1a3 | 4644 | 0.03661083 | 0 | 0 | 0.19621466 | 0.02072865 | 0 | 0 | 0 | 0 | 0 | 0 | 0.01201888 | 0.1487181 | 0 | 0 | 0 | 0 | 0 |
| MUSG00 Chmp2a | 921 | 259.000052 | 255.398856 | 247.855255 | 228.893542 | 270.134542 | 260.1384 | 284.646638 | 258.887262 | 285.219976 | 270.975757 | 276.845769 | 257.624736 | 293.90253 | 284.009881 | 266.221997 | 286.451992 | 268.367306 | 283.347668 |
| MUSG00 Gde1 | 1685 | 421.368819 | 402.799351 | 395.112699 | 402.262111 | 412.849076 | 406.714721 | 450.467245 | 416.761842 | 422.180119 | 458.495117 | 468.790008 | 399.819019 | 510.212061 | 453.002122 | 482.950701 | 521.72941 | 484.753733 | 503.112964 |
| MUSG00 Parl | 3963 | 22.5342859 | 23.5031169 | 24.0709054 | 22.3838893 | 24.387813 | 23.4071959 | 25.8470729 | 23.3799567 | 26.1274036 | 24.485528 | 24.4480484 | 20.9431971 | 21.6846328 | 24.9975368 | 23.8880917 | 28.9184426 | 24.8835634 | 25.5820802 |
| MUSG00 Rbm34 | 3935 | 10.5101744 | 10.658433 | 11.5801437 | 10.6984511 | 10.4703753 | 12.0513255 | 10.1719254 | 10.7249313 | 13.3973927 | 11.3353406 | 12.6752368 | 10.0567484 | 11.6842051 | 11.8618317 | 10.6694753 | 11.1320089 | 10.2170992 | 10.9459036 |
| MUSG00 Vhl | 2992 | 15.9678536 | 17.1797446 | 16.0382828 | 18.1208699 | 15.8294829 | 17.5057618 | 16.7144056 | 17.436772 | 20.5932579 | 18.6270492 | 18.9535012 | 20.8562536 | 19.8020163 | 23.5480216 | 22.5286582 | 23.650081 | 24.6456535 | 17.705137 |
| MUSG00 Ndufb7 | 4692 | 36.3087693 | 28.6198373 | 33.3006187 | 27.6259956 | 32.8060298 | 31.0285446 | 43.5716295 | 36.1004486 | 39.3080174 | 36.4483377 | 35.3119258 | 34.8431534 | 44.9580729 | 48.9385269 | 36.205033 | 35.2058142 | 30.3784097 | 40.3423564 |
| MUSG00 6720456B07f | 1089 | 189.068019 | 168.981109 | 162.400025 | 153.292623 | 172.417479 | 159.442404 | 189.294162 | 157.819845 | 173.596058 | 177.042448 | 161.399515 | 147.662924 | 191.030941 | 193.239816 | 173.65945 | 194.547228 | 187.478589 | 158.35955 |
| MUSG00 Mga | 15430 | 5.50391067 | 4.94188042 | 4.77778744 | 4.62401785 | 4.72584993 | 4.4093262 | 4.45530282 | 5.2919868 | 6.56635313 | 4.52255611 | 3.82695066 | 5.31026524 | 4.35770598 | 4.76781817 | 3.95727193 | 4.25837215 | 5.67318614 | 4.61320701 |
| MUSG00 Zswim5 | 5568 | 5.45818981 | 7.25728101 | 5.96850132 | 6.51339642 | 6.40548769 | 5.62452005 | 3.35641311 | 5.83037752 | 5.2074905 | 4.58974159 | 4.81353299 | 5.01218234 | 4.05783371 | 4.27906193 | 5.29078888 | 4.4837421 | 5.07988634 | 4.1197553 |
| MUSG00 Trim36 | 4838 | 0.44807027 | 0.53708263 | 0.06999064 | 0.16951195 | 0.26861553 | 0.14339361 | 0.68455716 | 0.30787592 | 0.20431508 | 0.06822136 | 0.01974988 | 0.4384033 | 0.04078703 | 0.08639867 | 0.28955207 | 0 | 0.01584386 | 0.31836606 |
| MUSG00 Aspm | 11894 | 0.03573665 | 0.04201223 | 0.04880464 | 0 | 0.07688806 | 0 | 0.01591144 | 0.00642213 | 0.0034628 | 0 | 0.21288656 | 0 | 0 | 0.07809667 | 0.05196003 | 0.00965796 | 0.11922591 | 0 |
| MUSG00 Ppp3r1 | 3222 | 52.9929522 | 47.384542 | 47.7279494 | 50.0860394 | 52.0308784 | 50.4447443 | 49.2656852 | 54.8467554 | 53.9822024 | 50.1799384 | 46.2477019 | 47.4138636 | 49.9290448 | 48.2603366 | 41.6363333 | 52.8842964 | 54.5870194 | 46.7628626 |
| MUSG00 Tnks1bp1 | 6247 | 4.66080448 | 5.74323468 | 5.14941535 | 5.84920083 | 5.07746722 | 6.45684731 | 4.55934572 | 4.01059964 | 6.96880857 | 5.27586707 | 4.2979673 | 4.35124503 | 4.94345769 | 5.24884356 | 5.24335472 | 6.33171081 | 5.5400426 | 4.25314528 |
| MUSG00 9430020K01f | 6726 | 2.26239266 | 1.77807449 | 2.05691788 | 1.55798992 | 1.76755652 | 1.41452942 | 0.97073293 | 0.93124502 | 1.52474612 | 1.2968897 | 1.33536904 | 0.4647156 | 2.1783496 | 1.9196337 | 2.45028581 | 1.40615225 | 1.33338589 | 2.83796611 |
| MUSG00 Zfp446 | 3211 | 4.81840052 | 6.00690307 | 5.4233793 | 5.6756206 | 6.325653 | 5.20064504 | 4.77399812 | 6.68456249 | 7.91407045 | 5.06602075 | 4.4784374 | 6.32728159 | 6.06854791 | 5.40955736 | 8.93063933 | 4.55528248 | 6.09926432 | 7.04102861 |
| MUSG00 4931417G12f | 1538 | 0 | 0 | 0 | 0 | 0 | 0 | 0 | 0 | 0 | 0 | 0 | 0 | 0 | 0 | 0 | 0 | 0 | 0 |
| MUSG00 Zbtb41 | 8349 | 2.86117007 | 2.53367922 | 3.27357104 | 2.93044446 | 3.44746561 | 2.8607519 | 1.83039829 | 2.58916035 | 2.21989844 | 2.16297992 | 2.12295001 | 2.00559334 | 1.89669966 | 3.18750294 | 3.11366726 | 1.92163906 | 3.25927306 | 1.26503156 |
| MUSG00 Slc16a2 | 4294 | 2.27670948 | 2.49807917 | 2.59103679 | 2.50405367 | 2.85832337 | 3.09271835 | 3.37160684 | 3.6377952 | 3.53931633 | 2.0533725 | 2.1806921 | 2.2097491 | 3.09042519 | 4.08846414 | 1.83267284 | 1.23057844 | 0.87470356 | 2.02408916 |
| MUSG00 Cdkl4 | 2921 | 0.29103167 | 0.9237742 | 0.29809091 | 0.59271472 | 0.29660206 | 0.32232179 | 0.1295794 | 0.16997646 | 0.49350498 | 0.61339498 | 0 | 0.19108409 | 0.05066613 | 0.06360026 | 0.19747408 | 0 | 0.31490282 | 0.28248429 |
| MUSG00 2310014L17f | 3237 | 0.24948976 | 0.23669969 | 0.11955151 | 0.15482592 | 0.2676474 | 0.04592468 | 0.482335 | 0.11798703 | 0.02544734 | 0.07283087 | 0.05903609 | 0.22415929 | 0.19812015 | 0.15782668 | 0.33093619 | 0.02365808 | 0.20128114 | 0.13595084 |
| MUSG00 Rfc3 | 3576 | 9.15240088 | 7.37802355 | 7.52116412 | 7.21128386 | 8.53345627 | 8.85464115 | 9.89649995 | 8.61892303 | 7.20994375 | 7.04096072 | 8.60376769 | 8.61584053 | 8.82548235 | 7.98744792 | 6.85540429 | 8.67320805 | 7.50235118 | 9.12203587 |
| MUSG00 Zfp944 | 5021 | 4.14808529 | 3.90784599 | 3.59357236 | 3.98352887 | 4.08368841 | 4.02659138 | 4.48532865 | 4.29769117 | 4.26548261 | 3.3524798 | 4.50060852 | 4.68002269 | 2.50540508 | 3.14498899 | 3.33977958 | 3.2029586 | 3.89293197 | 2.73895174 |
| MUSG00 Gria2 | 6939 | 0.11638541 | 0.08161395 | 0.09759755 | 0.10505501 | 0 | 0.19995339 | 0 | 0 | 0 | 0 | 0 | 0 | 0.09353118 | 0.03346598 | 0.13062894 | 0 | 0 | 0 |
| MUSG00 Coil | 3509 | 2.51954306 | 2.11706338 | 2.75711202 | 2.77858749 | 2.16723965 | 3.00790318 | 2.15731793 | 2.5577699 | 2.51180233 | 2.67397833 | 2.40985114 | 1.98830374 | 0.99816647 | 1.56181314 | 2.38356108 | 1.39675057 | 2.00970038 | 1.5990108 |
| MUSG00 Tesk2 | 4546 | 9.29391662 | 6.82965627 | 6.85273945 | 7.53671477 | 8.02551653 | 8.73113497 | 6.15085641 | 6.6890827 | 6.38726212 | 6.61727864 | 6.15840864 | 7.08439008 | 7.55279684 | 6.69178998 | 6.34427829 | 7.42901798 | 6.38209563 | 3.78747259 |
| MUSG00 Dnahc17 | 16723 | 0.3126315 | 0.33466281 | 0.42232467 | 0.37869911 | 0.29645328 | 0.4148408 | 0.46984249 | 0.60521358 | 0.81274593 | 0.53570733 | 0.59422301 | 1.01131052 | 0.21239598 | 0.09720403 | 0.43855032 | 0.20836245 | 0.47899277 | 0.13815598 |
| MUSG00 Ttc37 | 7235 | 3.09021736 | 2.44494082 | 2.38023176 | 3.21162856 | 2.6144915 | 2.47249991 | 2.92311921 | 2.60246778 | 3.55223025 | 2.48296797 | 2.19890272 | 2.88528788 | 1.94327525 | 2.63193861 | 2.05580673 | 1.89468382 | 2.57980741 | 2.3037677 |
| MUSG00 1700018A14f | 710 | 0 | 0 | 0 | 0 | 0 | 0 | 0 | 0 | 0 | 0.26563776 | 0 | 0.07861361 | 0 | 0 | 0 | 0 | 0 | 0 |
| MUSG00 Kcnk1 | 2298 | 1203.8879 | 1113.72051 | 1230.00963 | 1214.24743 | 1205.8534 | 1218.82905 | 1113.00092 | 1138.80836 | 1229.43004 | 1178.70603 | 1254.60201 | 1253.71499 | 904.761961 | 789.185694 | 951.68781 | 861.005464 | 976.503344 | 956.459473 |
| MUSG00 Neu4 | 4602 | 0 | 0 | 0 | 0 | 0 | 0 | 0 | 0 | 0 | 0 | 0 | 0 | 0 | 0 | 0 | 0 | 0 | 0 |
| MUSG00 Gal3st2 | 2842 | 0 | 0 | 0 | 0 | 0 | 0 | 0 | 0 | 0 | 0 | 0 | 0 | 0 | 0 | 0 | 0 | 0 | 0 |
| MUSG00 Pqlc1 | 5761 | 24.2370254 | 28.5249969 | 22.4864284 | 26.0981507 | 25.9583209 | 23.6194769 | 21.1556086 | 25.1853735 | 24.9578418 | 26.1982882 | 24.5301731 | 25.5196069 | 20.3116313 | 22.2022247 | 16.871111 | 27.1776243 | 22.9186066 | 21.4269112 |

[illegible]

| | | | | | | | | | | | | | | | | | | | | |
|---|---|---|---|---|---|---|---|---|---|---|---|---|---|---|---|---|---|---|---|---|
| ENSMUSG00 | Tmem106a | 3211 | 3.60056303 | 6.04840154 | 5.2124701 | 4.54049648 | 4.55686851 | 3.93520619 | 5.09815849 | 4.94800355 | 3.93779518 | 4.15560543 | 4.56770858 | 5.40600158 | 4.04057747 | 4.32475308 | 4.72194723 | 5.28269669 | 5.03696583 | 4.06015518 |
| ENSMUSG00 | Zfr2 | 7303 | 8.11922779 | 8.82200978 | 10.0019212 | 10.5246449 | 8.85131774 | 8.69191452 | 8.01393726 | 10.1822001 | 9.81866684 | 11.3244329 | 9.26976806 | 9.28605094 | 8.49781623 | 5.39293251 | 7.99433174 | 6.58537627 | 8.09769631 | 8.18771724 |
| ENSMUSG00 | Cog7 | 4392 | 14.0812912 | 16.6185226 | 15.0671652 | 16.6393249 | 14.8165667 | 15.9083229 | 12.5176077 | 15.3482674 | 11.0374611 | 15.0620302 | 15.3593446 | 16.2541513 | 11.7713682 | 9.21056469 | 13.339848 | 12.1619757 | 15.8471209 | 13.3514817 |
| ENSMUSG00 | 1700047I17R | 3832 | 0 | 0 | 0 | 0 | 0 | 0 | 0 | 0 | 0.01074805 | 0 | 0 | 0 | 0 | 0 | 0 | 0 | 0 | 0 |
| ENSMUSG00 | Cebpa | 2633 | 0 | 0.08856447 | 0 | 0.13843082 | 0.12796182 | 0.03763975 | 0 | 0.18856864 | 0.14078186 | 0.35815194 | 0 | 0.12719103 | 0.44966427 | 0.28222765 | 0.34425911 | 0.23268125 | 0.11644906 | 0.27159832 |
| ENSMUSG00 | Atcay | 3983 | 0.01067165 | 0 | 0 | 0 | 0 | 0.01244106 | 0 | 0 | 0 | 0 | 0 | 0 | 0 | 0 | 0.0206887 | 0 | 0 | 0 |
| ENSMUSG00 | 5031414D18 | 2637 | 0 | 0.08843013 | 0 | 0 | 0.14602025 | 0 | 0 | 0.18828261 | 0.23428052 | 0.01788043 | 0 | 0 | 0.09353795 | 0 | 0.15624405 | 0 | 0.05813621 | 0 |
| ENSMUSG00 | Lbx2 | 933 | 0.09111506 | 0 | 0 | 0.14649853 | 0 | 0 | 0 | 0 | 0 | 0 | 0 | 0 | 0.05287451 | 0.2488965 | 0 | 0 | 0 | 0 |
| ENSMUSG00 | Dopey1 | 9334 | 0.78325373 | 1.67742236 | 1.25934946 | 1.0152879 | 0.89725249 | 1.25288677 | 1.91602664 | 1.74717761 | 2.44895182 | 1.01030005 | 1.00320247 | 1.3873188 | 0.76635183 | 1.63206135 | 1.26244322 | 1.12812482 | 1.76562113 | 1.16689556 |
| ENSMUSG00 | Dapk3 | 1475 | 38.5284206 | 39.7270379 | 36.9606568 | 41.6999389 | 39.3539661 | 40.9859898 | 41.1219352 | 41.6102988 | 35.7135957 | 36.6976321 | 43.6290638 | 40.0737537 | 49.6998163 | 47.1683003 | 43.5200163 | 42.4441648 | 37.3129236 | 47.5875391 |
| ENSMUSG00 | Parm1 | 5064 | 94.033469 | 92.9195559 | 90.0890136 | 97.4380154 | 88.0137467 | 97.9899435 | 86.795969 | 102.163168 | 94.8574908 | 103.649611 | 102.295394 | 101.28162 | 101.352541 | 91.1089187 | 103.695464 | 115.045733 | 119.38369 | 101.436619 |
| ENSMUSG00 | Hrh2 | 2786 | 0 | 0 | 0 | 0 | 0 | 0 | 0 | 0 | 0 | 0 | 0 | 0 | 0 | 0 | 0 | 0 | 0 | 0 |
| ENSMUSG00 | Otoa | 4357 | 0 | 0 | 0.01110249 | 0 | 0 | 0 | 0 | 0 | 0 | 0.02264366 | 0 | 0 | 0 | 0.0213193 | 0 | 0.01757659 | 0 | 0 |
| ENSMUSG00 | Vat1 | 2760 | 6.88399047 | 6.07115427 | 6.43227741 | 5.16688655 | 4.76087506 | 6.57112337 | 6.49892222 | 5.02313119 | 5.67060724 | 4.81757185 | 6.57771167 | 8.65547231 | 9.1871759 | 4.91364983 | 6.79228548 | 5.7297177 | 5.62396748 | 5.7400807 |
| ENSMUSG00 | Eef2 | 3084 | 177.642417 | 154.401592 | 152.461321 | 165.742333 | 155.898115 | 153.366032 | 166.223371 | 154.862514 | 172.839133 | 162.749428 | 154.122264 | 156.334531 | 211.852112 | 207.447245 | 160.504375 | 173.276173 | 148.707026 | 159.194643 |
| ENSMUSG00 | Htr2a | 2968 | 0 | 0 | 0 | 0 | 0.1459526 | 0 | 0 | 0 | 0 | 0 | 0 | 0.01880582 | 0.03324253 | 0 | 0 | 0 | 0.16787125 | 0 |
| ENSMUSG00 | Foxn2 | 7498 | 1.14511144 | 1.49725865 | 1.00643781 | 1.20313241 | 1.15547429 | 1.0111454 | 0.43539275 | 1.27341966 | 0.75803403 | 0.42761374 | 0.7964617 | 0.75929549 | 0.65793435 | 0.42120538 | 0.23628526 | 0.53110472 | 0.19423837 | 0.33014268 |
| ENSMUSG00 | Dpp4 | 6291 | 7.82403336 | 8.72140149 | 12.5182177 | 8.73416301 | 8.04111457 | 8.52266058 | 19.4109083 | 20.1312997 | 16.4654407 | 21.9976656 | 14.3985569 | 18.4366471 | 15.5108142 | 17.8511848 | 21.278629 | 8.46033131 | 17.3019684 | 18.61618 |
| ENSMUSG00 | Igsf6 | 2127 | 0.2198199 | 0.14095724 | 0.27291131 | 0.12852199 | 0.15840313 | 0.18637605 | 0.31141397 | 0.12569197 | 0.13554566 | 0.17734162 | 0 | 0.0262415 | 0.23193191 | 0.74240666 | 0.40678546 | 0.45005415 | 0 | 0.05172459 |
| ENSMUSG00 | Rundc1 | 3218 | 17.4088943 | 21.6564869 | 16.9863547 | 21.4213363 | 19.9527614 | 22.0682111 | 15.6875787 | 16.5088974 | 16.382455 | 19.6485684 | 21.4675544 | 18.7844508 | 15.8512127 | 14.6057851 | 17.7968187 | 12.1487601 | 16.7096938 | 17.094191 |
| ENSMUSG00 | Zbtb7a | 6801 | 15.2183655 | 14.5624518 | 13.3861174 | 12.8825008 | 12.9795542 | 14.6669097 | 12.0838293 | 14.0898012 | 11.7969965 | 15.4118532 | 11.8085142 | 14.3704198 | 14.3549278 | 11.957594 | 12.8917675 | 13.5461258 | 12.1837162 | 13.192156 |
| ENSMUSG00 | Epgn | 1755 | 0.29063368 | 0.47454265 | 0.24806939 | 0 | 0 | 0.05647035 | 0 | 0 | 0 | 0 | 0 | 0.31803796 | 0 | 0 | 0 | 0.32727015 | 0 | 0 |
| ENSMUSG00 | Baz1a | 8079 | 4.27734969 | 4.38729044 | 4.53857359 | 3.51901122 | 4.38483671 | 4.13399461 | 4.38633451 | 4.62336546 | 4.94503284 | 3.44336117 | 5.01462617 | 5.2713641 | 4.40256373 | 5.57053109 | 4.12576541 | 5.327224 | 5.0238332 | 4.5279188 |
| ENSMUSG00 | Ncapd3 | 5467 | 2.65900315 | 3.91808871 | 4.87540063 | 3.81689378 | 3.70651909 | 4.18755495 | 4.84636916 | 5.00895418 | 4.75373906 | 4.47616883 | 4.35191683 | 4.05319518 | 3.53724375 | 6.83026271 | 4.39372729 | 3.45995245 | 6.16922284 | 3.12929069 |
| ENSMUSG00 | Map2k2 | 3160 | 63.9460781 | 60.1740507 | 62.1048774 | 59.4456284 | 59.4033235 | 60.7648236 | 54.1101334 | 54.6296389 | 53.3730323 | 50.5377949 | 55.4552366 | 53.3958063 | 57.6683868 | 55.35077 | 55.4265234 | 52.6495852 | 48.8660286 | 57.5506667 |
| ENSMUSG00 | C8a | 3855 | 0.88207888 | 0.54441306 | 1.00386055 | 0.70912131 | 0.28716842 | 1.14401901 | 1.23957991 | 0.32693885 | 1.28207053 | 0.99071517 | 0.12392985 | 1.41891957 | 1.11332732 | 0.92767697 | 1.13290919 | 1.04293483 | 0.68599598 | 0.84190332 |
| ENSMUSG00 | Nek11 | 3796 | 0.17915775 | 0.02632737 | 0.03822987 | 0.08401668 | 0.177515 | 0.09137754 | 0.08724677 | 0.37226525 | 0 | 0.08694808 | 0.28946893 | 0.07351905 | 0.05198305 | 0.48940029 | 0.01085394 | 0.26226443 | 0.38366683 | 0.52168805 |
| ENSMUSG00 | Tbr1 | 5478 | 0 | 0 | 0 | 0 | 0 | 0 | 0 | 0 | 0 | 0 | 0 | 0 | 0 | 0 | 0 | 0 | 0 | 0 |
| ENSMUSG00 | Creb3l3 | 3004 | 0 | 0 | 0 | 0 | 0.11215828 | 0 | 0 | 0 | 0 | 0 | 0 | 0 | 0 | 0 | 0 | 0 | 0.02551684 | 0 |
| ENSMUSG00 | Ccl5 | 957 | 0 | 0.13923885 | 0 | 0.14282459 | 0.10058918 | 0 | 0 | 0.3591762 | 0 | 0 | 0 | 0.17497073 | 0 | 1.06768017 | 0.55968676 | 0 | 0 | 0 |
| ENSMUSG00 | Zc3h12b | 7717 | 0.08812788 | 0 | 0.01253687 | 0.07084781 | 0.01247426 | 0.07063367 | 0 | 0.04454213 | 0.12275364 | 0.10386963 | 0.29097098 | 0.10125946 | 0.10867469 | 0.00601841 | 0.12813753 | 0.04961851 | 0.0844301 | 0.44908303 |
| ENSMUSG00 | Kri1 | 2580 | 4.48116584 | 4.557927 | 3.59989063 | 3.74377574 | 3.37669697 | 3.34192854 | 4.25447345 | 4.14491189 | 5.58731803 | 3.08855382 | 4.33307763 | 4.3700635 | 5.37297239 | 3.8883422 | 4.34372992 | 5.29835149 | 3.93661577 | 3.34745339 |
| ENSMUSG00 | Anapc13 | 402 | 217.495391 | 154.548626 | 203.722355 | 169.550553 | 200.789144 | 177.872126 | 260.219734 | 230.389492 | 259.003761 | 213.585648 | 216.294588 | 244.36708 | 375.634325 | 406.790129 | 279.391645 | 222.218873 | 248.548896 | 327.591546 |
| ENSMUSG00 | Rrp12 | 4285 | 1.26969954 | 2.60439196 | 2.17878445 | 1.53110627 | 1.3029879 | 1.97748365 | 1.85496618 | 1.64000048 | 1.3937094 | 1.32043976 | 1.23757769 | 0.96391109 | 0.69076197 | 1.96181559 | 1.34614185 | 1.23316309 | 0.5366576 | 1.56618694 |
| ENSMUSG00 | Dhx57 | 5053 | 10.3970706 | 9.08473545 | 10.9517749 | 8.86334981 | 9.64924573 | 9.75756381 | 5.88014296 | 7.57347857 | 7.14019153 | 7.95953886 | 7.75291232 | 9.17926293 | 8.92328763 | 7.0314124 | 7.57495457 | 8.10824261 | 9.39764137 | 8.57850309 |
| ENSMUSG00 | 4933434I20R | 2104 | 0 | 0 | 0 | 0 | 0 | 0 | 0 | 0 | 0 | 0 | 0 | 0 | 0 | 0.08829674 | 0 | 0 | 0 | 0 |
| ENSMUSG00 | Zc4h2 | 3707 | 2.60282572 | 2.77682339 | 2.62289711 | 2.08939238 | 2.90843024 | 2.47295811 | 4.2756265 | 2.62720869 | 3.42202505 | 3.00177124 | 3.04151341 | 3.16193395 | 3.79271552 | 1.85425544 | 3.36770203 | 2.21046398 | 2.51235235 | 2.65622609 |
| [illegible] | | | | | | | | | | | | | | | | | | | | |

| | | | | | | | | | | | | | | | | | | | |
|---|---|---|---|---|---|---|---|---|---|---|---|---|---|---|---|---|---|---|---|
| ENSMUSG00 Lrrk2 | 11780 | 0.11185573 | 0.00565584 | 0.09444747 | 0 | 0 | 0.13040193 | 0.20885054 | 0.12644344 | 0.04545201 | 0.02801824 | 0.34878153 | 0.16109784 | 0.02931438 | 0.22078683 | 0.13990342 | 0.12026761 | 0.03578856 | 0 |
| ENSMUSG00 9530068E07F | 3425 | 176.809122 | 194.119341 | 192.166498 | 187.85783 | 198.472163 | 208.020205 | 154.840239 | 161.389384 | 162.413163 | 172.427024 | 159.51954 | 170.901855 | 132.785677 | 144.390261 | 142.767902 | 159.5796 | 165.054855 | 148.982328 |
| ENSMUSG00 Macrod1 | 1263 | 7.60583514 | 8.83596165 | 8.73251379 | 9.84811172 | 7.73616825 | 8.55304507 | 17.0070912 | 14.0310964 | 14.870191 | 14.3356057 | 12.596248 | 14.8488224 | 12.3818035 | 12.6866272 | 10.7326302 | 7.54897961 | 9.68019692 | 13.3276221 |
| ENSMUSG00 Zbtb20 | 3078 | 0 | 0 | 0 | 0 | 0 | 0 | 0 | 0 | 0 | 0 | 0 | 0 | 0 | 0 | 0 | 0 | 0 | 0 |
| ENSMUSG00 Snapc4 | 6442 | 3.23308542 | 4.17316138 | 3.62688841 | 4.37787744 | 3.72831883 | 4.93065853 | 3.60610453 | 5.03935766 | 2.6340957 | 4.49402853 | 3.2408657 | 2.73793067 | 3.60685083 | 4.80878707 | 4.214813 | 3.67927445 | 4.01587246 | 2.80083623 |
| ENSMUSG00 Naa30 | 6941 | 5.22971046 | 6.95918407 | 6.35595155 | 4.89021438 | 5.56141806 | 6.63225609 | 5.0850394 | 6.96608291 | 5.05559882 | 5.30538816 | 6.1121111 | 5.58076211 | 5.6005692 | 5.39985283 | 5.63916987 | 5.45590127 | 6.08494153 | 6.95836273 |
| ENSMUSG00 Noa1 | 3625 | 4.9364632 | 3.27155596 | 4.08339318 | 5.5427367 | 3.06716534 | 3.28073271 | 4.3331888 | 3.91933067 | 5.37413019 | 4.60451006 | 5.12674727 | 4.09571483 | 2.5584553 | 3.90770942 | 3.8416899 | 3.68646124 | 2.87579386 | 5.81199666 |
| ENSMUSG00 Mudeng | 3157 | 9.41118722 | 10.9213955 | 10.4959988 | 10.7949513 | 11.1906341 | 10.1397105 | 11.2998923 | 10.4524028 | 11.0109025 | 11.2313363 | 10.0937274 | 9.03446423 | 8.26626045 | 9.32706728 | 9.93170226 | 10.1275442 | 10.646868 | 10.4198435 |
| ENSMUSG00 Gramd1c | 3606 | 6.10584614 | 8.03722074 | 9.13543377 | 7.64404654 | 9.39680373 | 8.87716747 | 6.86204412 | 8.64254031 | 7.25275583 | 8.48608054 | 7.63127457 | 7.97145486 | 6.37511743 | 8.11419164 | 10.5231927 | 6.80651119 | 8.34335637 | 5.53752239 |
| ENSMUSG00 Lrrn3 | 3178 | 0 | 0 | 0 | 0 | 0.01514535 | 0 | 0 | 0 | 0 | 0 | 0 | 0 | 0 | 0 | 0 | 0 | 0 | 0 |
| ENSMUSG00 Slc2a13 | 12952 | 0.02953572 | 0.00771608 | 0 | 0 | 0 | 0.0153035 | 0.0292234 | 0.00589753 | 0 | 0.01092126 | 0 | 0.06464136 | 0 | 0.00358586 | 0.05407863 | 0.00886904 | 0 | 0 |
| ENSMUSG00 BC031181 | 1044 | 240.455526 | 219.373705 | 218.0979 | 217.593263 | 241.950513 | 221.515908 | 273.407464 | 242.946115 | 230.74706 | 242.347389 | 225.420925 | 223.529967 | 241.887054 | 222.433389 | 220.215213 | 222.004834 | 215.897464 | 194.534121 |
| ENSMUSG00 Zdhhc23 | 1652 | 1.54377153 | 2.2585013 | 2.16685305 | 3.00614639 | 2.09775939 | 1.88972289 | 1.40334217 | 1.34089548 | 1.62053482 | 2.05499433 | 2.19787945 | 1.7906962 | 1.70213032 | 2.50213304 | 0.99761637 | 0.69535001 | 2.22719401 | 0.96565623 |
| ENSMUSG00 Gm15430 | 563 | 0.60398119 | 1.77510982 | 0.60144709 | 0.64740383 | 0.76942684 | 1.14420166 | 2.26899171 | 1.01755957 | 0.73155445 | 1.59123155 | 0.93343613 | 1.58623553 | 2.19058247 | 1.97985452 | 1.53682535 | 0.61210562 | 1.42957768 | 0.87936405 |
| ENSMUSG00 Lzts1 | 1910 | 0.40057234 | 0.20929567 | 0.05065291 | 0 | 0.02519996 | 0 | 0.12385518 | 0.01999602 | 0.06469086 | 0 | 0 | 0 | 0 | 0 | 0 | 0 | 0 | 0 |
| ENSMUSG00 Skp1a | 2310 | 324.750585 | 285.366039 | 313.527488 | 275.003308 | 311.523979 | 303.859078 | 349.70419 | 313.326501 | 338.317808 | 325.44191 | 322.491311 | 307.203608 | 362.344 | 333.894788 | 334.856282 | 354.212982 | 366.721273 | 357.559194 |
| ENSMUSG00 Znrd1 | 954 | 28.7377762 | 25.5608371 | 29.5616019 | 25.7892704 | 30.0193864 | 28.1006592 | 35.0133662 | 31.4266624 | 32.0771292 | 35.9314059 | 40.914194 | 29.7215478 | 36.0940024 | 39.9204944 | 33.7731831 | 38.9327979 | 34.6704342 | 36.0384609 |
| ENSMUSG00 H2-Ea-ps | 1038 | 0 | 0 | 0 | 0 | 0 | 0 | 0 | 0.11038267 | 0.51582342 | 0 | 0 | 0.10754463 | 0 | 0 | 0 | 0 | 0 | 0 |
| ENSMUSG00 Srp72 | 5591 | 25.2400616 | 25.2215134 | 23.3691478 | 25.6367457 | 27.126398 | 25.7468594 | 23.0005118 | 23.3690232 | 21.2820326 | 21.2604047 | 26.4466994 | 22.2124574 | 21.4851044 | 24.6051488 | 24.8491587 | 24.1824602 | 27.3994292 | 25.4236329 |
| ENSMUSG00 Qsox2 | 6966 | 3.69769404 | 4.17008954 | 4.40264402 | 4.44753232 | 5.41708702 | 4.51708094 | 5.75277621 | 5.89937195 | 5.81199325 | 4.83284555 | 4.99084665 | 5.937325 | 5.01392441 | 3.81366896 | 4.19349743 | 3.38052305 | 5.13878285 | 2.91391909 |
| ENSMUSG00 Slc18a1 | 5180 | 1.63292086 | 1.29907425 | 1.35408531 | 1.37210867 | 0.9756471 | 1.12880744 | 1.74454086 | 2.16030384 | 2.06727685 | 1.90241254 | 1.20820841 | 1.59473322 | 2.59040183 | 2.54635537 | 2.19529526 | 3.42989225 | 2.36764769 | 1.50797165 |
| ENSMUSG00 Kidins220 | 7429 | 12.7475477 | 11.5915778 | 11.5382818 | 12.5785034 | 12.7051692 | 13.6671895 | 9.23453822 | 12.3846409 | 10.0568501 | 12.1034313 | 11.1575653 | 11.8483376 | 8.18103676 | 11.0155448 | 11.0809946 | 11.4062612 | 12.2474935 | 13.7134023 |
| ENSMUSG00 Igsf10 | 10077 | 0.05061646 | 0.05289335 | 0 | 0.0497342 | 0 | 0.03933927 | 0.11268277 | 0 | 0 | 0.03743233 | 0.0474099 | 0.00553892 | 0 | 0 | 0 | 0 | 0 | 0 |
| ENSMUSG00 6720456H20 | 8171 | 3.55293541 | 3.18818789 | 2.90679273 | 4.57785062 | 3.72284613 | 3.7842254 | 3.3699644 | 4.28618688 | 4.66756989 | 4.77796871 | 3.97588689 | 3.81849901 | 3.63453851 | 4.0640708 | 2.86913295 | 3.70206582 | 4.16518806 | 3.93835852 |
| ENSMUSG00 Ubac1 | 2558 | 24.1106374 | 24.0535245 | 24.1111224 | 22.317431 | 20.9048347 | 24.0789866 | 30.814278 | 23.1573167 | 25.1498583 | 25.4001834 | 21.6462771 | 26.7295491 | 25.630226 | 24.2024896 | 22.2758998 | 22.9773582 | 20.0171746 | 20.364982 |
| ENSMUSG00 P2ry12 | 4270 | 0 | 0 | 0 | 0 | 0 | 0.06962913 | 0 | 0.07155485 | 0 | 0 | 0 | 0 | 0 | 0 | 0 | 0 | 0 | 0 |
| ENSMUSG00 Csgalnact1 | 5507 | 0.4862586 | 0.42344336 | 0.64123256 | 1.00934218 | 0.69047067 | 0.84582476 | 0.73885789 | 0.75594187 | 0.80024645 | 0.57365119 | 0.73740176 | 1.47976879 | 0.55539838 | 0.18554021 | 0.56112524 | 1.52272435 | 0.25054416 | 1.48835244 |
| ENSMUSG00 Gpr101 | 5408 | 0 | 0 | 0 | 0 | 0 | 0 | 0 | 0 | 0 | 0 | 0 | 0 | 0 | 0 | 0 | 0 | 0 | 0 |
| ENSMUSG00 P2ry13 | 2522 | 0.01685376 | 0 | 0 | 0 | 0 | 0 | 0 | 0 | 0 | 0 | 0 | 0 | 0 | 0 | 0 | 0.16700899 | 0 | 0 |
| ENSMUSG00 Fam82a1 | 1791 | 9.80158442 | 5.63584978 | 4.99670749 | 4.73163275 | 5.93922676 | 7.13824973 | 6.8683956 | 4.37154801 | 7.93375094 | 3.97529659 | 6.50870456 | 10.4712801 | 8.20821404 | 9.28363109 | 6.80941404 | 8.08143473 | 8.6025524 | 9.42925503 |
| ENSMUSG00 Serbp1 | 6677 | 36.2156577 | 34.1211448 | 34.4200453 | 33.7698979 | 33.9237431 | 32.1569559 | 34.8272902 | 32.7698477 | 33.4204796 | 34.5950719 | 34.5808558 | 35.8367149 | 35.2275845 | 33.5410198 | 35.2221778 | 34.7637655 | 34.9683691 | 34.5361952 |
| ENSMUSG00 1810006K21F | 1653 | 50.5793595 | 42.4220462 | 47.8760409 | 41.2612725 | 53.3440316 | 42.8377836 | 61.3948556 | 49.1672307 | 51.7011616 | 52.3706541 | 54.8848434 | 48.6909773 | 64.9104175 | 65.4937494 | 58.4000275 | 60.2735393 | 51.1249151 | 61.684776 |
| ENSMUSG00 Abt1 | 1275 | 26.0031665 | 28.217981 | 24.7748355 | 23.6202747 | 23.8960845 | 27.4386117 | 28.6844712 | 30.7636057 | 29.3958662 | 28.9560789 | 30.1637973 | 24.5151145 | 30.3729844 | 30.5620308 | 26.4659406 | 30.4823267 | 27.5949338 | 36.586449 |
| ENSMUSG00 C530008M17 | 9682 | 0 | 0.04128845 | 0.03497363 | 0 | 0.00497128 | 0 | 0 | 0.05128085 | 0.00850785 | 0.03895947 | 0.00493441 | 0.02305956 | 0.24457055 | 0.32139577 | 0.19149659 | 0.14632849 | 0.15834041 | 0.14772121 |
| ENSMUSG00 P2ry14 | 3230 | 7.52723241 | 7.51860678 | 5.60114562 | 5.48707314 | 5.8712006 | 5.89109898 | 2.57802833 | 2.82600129 | 2.05295017 | 2.26264985 | 2.66238159 | 1.76259987 | 2.56587062 | 3.07710053 | 3.558896 | 5.65468103 | 4.0580785 | 1.83931379 |
| ENSMUSG00 Gadd45a | 2075 | 17.3298212 | 16.5199847 | 10.1176444 | 13.6573347 | 17.211512 | 12.5135578 | 21.1139955 | 17.4120543 | 19.3328513 | 21.9733639 | 20.9519095 | 15.0635041 | 23.8932901 | 17.7942407 | 20.1738703 | 20.224822 | 23.6237753 | 23.1966113 |
| ENSMUSG00 Sec24a | 13655 | 1.72448682 | 1.82482906 | 1.94131781 | 1.90185245 | 1.87522398 | 2.06121953 | 2.02001405 | 2.46971009 | 2.35566963 | 1.80937155 | 1.819332 | 2.56290153 | 1.82081921 | 2.10877585 | 1.95220847 | 2.3582864 | 2.86570115 | 1.73225309 |
| ENSMUSG00 Glb1l2 | 6787 | 51.2479528 | 65.3054453 | 61.1956876 | 60.3161903 | 65.5777071 | 63.0525132 | 66.9989918 | 77.7803722 | 67.0805577 | 67.5268654 | 68.5757545 | 77.5596749 | 54.5289588 | 72.0714791 | 61.1254555 | 41.229963 | 61.9647055 | 64.8567662 |
| ENSMUSG00 Ppp1r11 | 2235 | 34.650752 | 31.5540926 | 32.5953184 | 30.4963411 | 33.810792 | 34.2545742 | 38.7392405 | 34.1766464 | 35.1974251 | 33.902094 | 40.2079181 | 32.4904595 | 37.0817096 | 33.3316851 | 34.177935 | 42.5394086 | 32.6330841 | 30.2734681 |
| ENSMUSG00 Gltl6d1 | 1397 | 0.18255623 | 0 | 0.17313361 | 0 | 0.03445377 | 0 | 0 | 0 | 0 | 0 | 0 | 0 | 0 | 0 | 0 | 0 | 0 | 0 |
| ENSMUSG00 Gng12 | 4780 | 54.3230377 | 66.1797581 | 60.4669129 | 60.0204804 | 63.2763616 | 66.4711564 | 46.8375725 | 48.3317661 | 47.0026032 | 51.0766405 | 47.4751156 | 45.2830836 | 40.7968764 | 42.1203177 | 45.6061574 | 39.4601532 | 56.7838566 | 43.9267281 |
| [illegible] | | | | | | | | | | | | | | | | | | | |

| | | | | | | | | | | | | | | | | | | | |
|---|---|---|---|---|---|---|---|---|---|---|---|---|---|---|---|---|---|---|---|
| ENSMUSG00 Azin1 | 8364 | 7.90238507 | 9.21640809 | 9.91877147 | 9.85412831 | 11.3309132 | 10.3145994 | 13.1914356 | 15.1874618 | 10.7053426 | 11.4099741 | 10.5443058 | 12.0853856 | 11.4246681 | 14.3208156 | 11.9210623 | 15.2539818 | 19.4426683 | 15.5148832 |
| ENSMUSG00 Ints7 | 4184 | 9.84405248 | 10.8680929 | 7.74623933 | 10.8675722 | 9.73221975 | 10.5998319 | 9.00116932 | 9.07343403 | 10.3753794 | 10.8748634 | 11.6696959 | 8.75121287 | 7.55778178 | 8.99132667 | 10.1231356 | 9.93871884 | 8.92203948 | 9.084917 |
| ENSMUSG00 Fbxo27 | 3053 | 0.15314672 | 0 | 0 | 0.13431032 | 0.06306181 | 0.21100083 | 0.18596533 | 0.02501959 | 0 | 0 | 0.1095397 | 0 | 0 | 0 | 0 | 0 | 0 | 0 |
| ENSMUSG00 Klf10 | 3809 | 22.7200153 | 36.9074333 | 41.5918383 | 44.3768639 | 39.0463755 | 49.2665272 | 39.7481157 | 48.3396093 | 53.0266929 | 62.8345943 | 59.51488 | 64.8129891 | 29.347893 | 35.8481536 | 39.4816695 | 57.1091994 | 62.5254946 | 48.4958205 |
| ENSMUSG00 4930427A07l | 3783 | 4.75275957 | 6.18177418 | 5.98435427 | 6.00977032 | 6.38705411 | 6.78517473 | 2.41378457 | 3.10950567 | 2.29721248 | 2.13131648 | 2.51314215 | 2.3754485 | 2.50376107 | 2.77461373 | 3.28915408 | 6.75121204 | 2.73542167 | 3.94064708 |
| ENSMUSG00 C330005M16 | 2217 | 0 | 0 | 0 | 0 | 0 | 0 | 0 | 0 | 0 | 0 | 0 | 0 | 0 | 0 | 0 | 0 | 0 | 0 |
| ENSMUSG00 Uggt1 | 9726 | 21.8906464 | 23.9588418 | 24.5946029 | 25.9050561 | 23.5562365 | 25.1890515 | 24.5465533 | 26.0035049 | 24.1630947 | 24.5497801 | 26.3238744 | 24.9121727 | 22.3276887 | 24.2152958 | 23.2484206 | 22.6609846 | 23.6987816 | 22.5273773 |
| ENSMUSG00 Dtl | 4373 | 0.11663895 | 0.28186076 | 0.11061864 | 0.09376844 | 0.14308598 | 0.03399456 | 0.9845538 | 0.17467369 | 0.03767347 | 0.463636 | 0.05462492 | 0.22974661 | 0.64301836 | 0.79654849 | 0.4993557 | 0.12258598 | 0.54338678 | 0.23900595 |
| ENSMUSG00 Thoc2 | 12160 | 10.1963486 | 9.51718705 | 9.6070786 | 8.79373117 | 9.72534017 | 10.0327985 | 9.68820481 | 10.603417 | 10.4558202 | 9.44177323 | 9.77107909 | 9.63003784 | 11.566225 | 11.1106785 | 9.93106582 | 10.438599 | 10.5996165 | 8.87564699 |
| ENSMUSG00 Tbx10 | 1769 | 0 | 0 | 0 | 0 | 0 | 0.02801172 | 0 | 0 | 0.06984712 | 0.1066155 | 0 | 0.0631042 | 0.08366069 | 0 | 0 | 0 | 0 | 0 |
| ENSMUSG00 Erv3 | 3376 | 0 | 0 | 0 | 0 | 0 | 0.01467794 | 0 | 0 | 0 | 0 | 0 | 0 | 0 | 0 | 0 | 0 | 0 | 0 |
| ENSMUSG00 Asxl2 | 14883 | 4.06687971 | 3.823048 | 4.09532005 | 3.51435053 | 3.77410165 | 3.74567125 | 3.58906232 | 3.8826248 | 4.3226055 | 4.63174952 | 4.12489563 | 4.57536171 | 4.501293 | 3.65735876 | 4.27158508 | 4.47405538 | 4.37006826 | 4.41314745 |
| ENSMUSG00 Ubr5 | 9731 | 6.66996237 | 8.29484562 | 7.362162 | 7.47255439 | 7.35012214 | 8.7739554 | 6.47139818 | 7.19811592 | 6.06094857 | 7.00646562 | 6.41679897 | 6.78552347 | 6.85404914 | 5.48871652 | 6.88880191 | 8.03115093 | 7.78262899 | 7.34886843 |
| ENSMUSG00 Slc2a12 | 4231 | 0.52239876 | 0.75585863 | 0.19436304 | 0.61379804 | 0.48916869 | 0.52703215 | 0.91695572 | 0.48744735 | 0.92477405 | 0.60178161 | 0.01129165 | 0.48810672 | 0.71123776 | 0.37322122 | 0.33109263 | 0.60635091 | 0.53444848 | 0.3900433 |
| ENSMUSG00 Zmat4 | 6026 | 0 | 0.00552819 | 0 | 0.00756074 | 0.11182325 | 0.00822316 | 0.01570285 | 0 | 0 | 0 | 0 | 0 | 0.00818651 | 0.02312185 | 0 | 0 | 0 | 0.09128627 |
| ENSMUSG00 Cib2 | 1427 | 10.0677992 | 7.82047622 | 8.94927261 | 8.5885921 | 5.80146522 | 9.063254 | 12.4332548 | 7.4671901 | 10.3904314 | 11.5316016 | 7.43240419 | 8.44862167 | 27.690866 | 15.2318381 | 16.5730155 | 40.1689133 | 12.9723906 | 17.8095356 |
| ENSMUSG00 Nenf | 738 | 93.4768302 | 80.9349855 | 85.7352001 | 81.1208838 | 95.2203368 | 88.2280378 | 93.9201553 | 88.4428397 | 94.4276215 | 100.434831 | 90.4357954 | 92.8748424 | 132.688151 | 114.03345 | 111.824819 | 108.905129 | 105.059754 | 105.247776 |
| ENSMUSG00 Fam168b | 9394 | 7.89562829 | 8.17041817 | 8.07427034 | 9.44297992 | 9.15603019 | 9.85357742 | 6.99063214 | 7.67181854 | 7.15963171 | 7.89023904 | 8.74229866 | 7.63499323 | 6.86886819 | 8.55314788 | 7.20171969 | 8.6983347 | 8.44125062 | 9.55076145 |
| ENSMUSG00 Arhgef4 | 8945 | 0.11879591 | 0.26441761 | 0.09734193 | 0.14261702 | 0.05918962 | 0.13295311 | 0.37553943 | 0.44831775 | 0.47425502 | 0.24247427 | 0.34182195 | 0.64894678 | 0.57907784 | 0.76844326 | 0.23030495 | 0.27824365 | 0.11568586 | 0.70721602 |
| ENSMUSG00 Samd4b | 4299 | 15.5723778 | 21.9916245 | 18.0936582 | 21.6836233 | 16.4246411 | 22.6381877 | 17.4327247 | 20.4065942 | 16.8137556 | 18.6891831 | 16.3472817 | 18.2806357 | 16.5357741 | 15.3085071 | 19.2829799 | 13.5473398 | 19.1943515 | 17.6453962 |
| ENSMUSG00 Pank2 | 5238 | 5.83452111 | 6.72236143 | 6.68622298 | 7.39344935 | 6.91930861 | 7.6249529 | 8.59902068 | 7.52473449 | 8.04387278 | 7.44437411 | 8.06282232 | 6.99027771 | 7.16716095 | 7.29734325 | 7.60632007 | 7.48560175 | 8.1218384 | 6.64772137 |
| ENSMUSG00 Ppfia1 | 6538 | 7.02785183 | 9.10525267 | 7.54680345 | 9.78398688 | 7.8183088 | 10.663918 | 9.21938711 | 8.79745458 | 8.58629871 | 10.5580651 | 7.02229047 | 7.31630817 | 5.65151515 | 7.91353829 | 8.03487057 | 8.43939512 | 8.51760621 | 7.26106747 |
| ENSMUSG00 Mavs | 3395 | 15.4120388 | 18.7808187 | 18.0528021 | 17.6205158 | 18.9125143 | 18.3761094 | 19.3152792 | 20.5080264 | 22.7102084 | 22.1240266 | 22.4310114 | 18.4956172 | 15.853055 | 11.6965227 | 14.0048882 | 15.5530921 | 18.909145 | 19.9620676 |
| ENSMUSG00 Bcdin3d | 1269 | 3.78493687 | 4.64647934 | 6.70901603 | 4.27246988 | 5.57556563 | 4.6077404 | 4.95869685 | 6.89208838 | 6.45872075 | 6.7995104 | 7.41660112 | 7.60922744 | 5.17032703 | 9.14974141 | 7.27277267 | 6.45720251 | 4.62090105 | 8.62632958 |
| ENSMUSG00 Atg14 | 3440 | 8.55045983 | 8.88021046 | 11.7137066 | 9.48305458 | 8.87082547 | 10.0257856 | 8.71983676 | 7.43863651 | 9.94941703 | 8.56662487 | 11.4715453 | 9.37832935 | 8.70478889 | 10.0178816 | 9.80932392 | 9.43907996 | 9.88239109 | 13.6083575 |
| ENSMUSG00 Prss40 | 2092 | 0 | 0 | 0 | 0 | 0 | 0 | 0 | 0 | 0 | 0 | 0 | 0 | 0 | 0 | 0 | 0 | 0 | 0 |
| ENSMUSG00 Mrpl47 | 3554 | 5.40583556 | 5.44591771 | 6.60133996 | 5.38425397 | 5.97247562 | 5.66077936 | 7.21538322 | 5.7815173 | 6.61719204 | 6.35486406 | 5.99539434 | 6.10925513 | 5.48286644 | 5.35792792 | 7.74413037 | 5.86102713 | 5.32729057 | 5.8042811 |
| ENSMUSG00 Rapgef6 | 10270 | 7.5946444 | 7.85626386 | 7.53628516 | 8.3136706 | 8.78278724 | 7.57524749 | 6.08108434 | 8.60165791 | 7.70793411 | 8.64506078 | 8.03847403 | 8.54354641 | 7.12838999 | 8.44314617 | 7.72278433 | 7.08022039 | 7.8331935 | 8.75753549 |
| ENSMUSG00 1700021A07l | 442 | 0 | 0 | 0 | 0 | 0 | 0 | 0 | 0 | 0.55909297 | 0 | 0 | 0 | 0 | 0 | 0 | 0 | 0 | 0 |
| ENSMUSG00 Fbxo34 | 3801 | 6.68721265 | 5.92462944 | 5.29423537 | 6.1371362 | 6.09088537 | 6.27068268 | 5.27771004 | 4.56178655 | 6.14384489 | 6.40088474 | 5.46753668 | 6.02063185 | 4.85402185 | 3.5434847 | 4.61769608 | 4.93617407 | 6.40284097 | 7.0335243 |
| ENSMUSG00 H2-M11 | 1001 | 0 | 0 | 0 | 0 | 0 | 0 | 0 | 0 | 0 | 0 | 0 | 0 | 0 | 0 | 0 | 0 | 0 | 0 |
| ENSMUSG00 Shank2 | 10654 | 0.08777115 | 0.0969307 | 0.18161641 | 0.05986997 | 0.00451773 | 0.01395328 | 0.04440837 | 0 | 0.00386583 | 0 | 0.00448423 | 0 | 0.01852146 | 0.01743724 | 0.0502741 | 0.03594012 | 0.00719472 | 0.02581618 |
| ENSMUSG00 Aldh8a1 | 4560 | 0 | 0 | 0 | 0 | 0.04222099 | 0 | 0.09338029 | 0 | 0 | 0.0310202 | 0 | 0.02448055 | 0.0108184 | 0 | 0.01807086 | 0.15114713 | 0.02521467 | 0 |
| ENSMUSG00 Dlgap5 | 3710 | 0.32079378 | 0.65548282 | 0.78232125 | 0.51578541 | 0.99896445 | 0.74796579 | 1.32628534 | 0.73090581 | 0.73269812 | 0.82609049 | 1.1975933 | 0.84250057 | 2.10092801 | 0.83874767 | 1.0106042 | 0.87727807 | 1.33263943 | 1.31962547 |
| ENSMUSG00 H2-DMb2 | 1702 | 1.09884121 | 1.42881391 | 1.33581429 | 0.53538242 | 0.5373129 | 2.96967028 | 4.16974256 | 2.26641166 | 2.58928154 | 1.27434333 | 9.79639278 | 1.37735478 | 4.52160934 | 3.95675245 | 2.10607248 | 1.6423116 | 0 | 1.26049069 |
| ENSMUSG00 Plekhg2 | 8117 | 2.86963613 | 3.06985892 | 2.46724671 | 2.60445048 | 1.83229819 | 2.17941677 | 1.39309231 | 1.98090445 | 2.37975556 | 1.40574968 | 2.07768392 | 2.59239926 | 1.37353866 | 2.22579155 | 1.12178685 | 2.44357948 | 1.44012819 | 1.35540487 |
| ENSMUSG00 Zdhhc18 | 5477 | 5.78169731 | 5.62006833 | 5.26394451 | 7.03754674 | 5.49250544 | 7.1655587 | 7.74003292 | 6.08763325 | 6.97095127 | 7.1109149 | 4.03867186 | 7.01135217 | 7.7371037 | 5.55429564 | 7.56778624 | 7.26381981 | 6.0739868 | 7.81396477 |
| ENSMUSG00 Rps16 | 891 | 51.3306054 | 41.7626295 | 44.627432 | 40.9077839 | 40.3530269 | 39.6533098 | 47.9498867 | 42.6503259 | 47.6117971 | 39.5833064 | 42.8419661 | 41.2822914 | 61.6787383 | 61.5084435 | 42.357604 | 43.8773402 | 43.7891922 | 50.3787105 |
| ENSMUSG00 Vash2 | 4862 | 0.39340455 | 0.22610562 | 0.40792158 | 0.42168799 | 0.42568341 | 0.55035944 | 1.62509613 | 2.01095331 | 1.29607917 | 0.96008219 | 0.72713841 | 1.71051701 | 1.2277174 | 3.26694321 | 0.77962631 | 0.74029558 | 1.34008031 | 1.75368399 |
| ENSMUSG00 Mcrs1 | 1900 | 30.4247576 | 31.0686571 | 31.2391167 | 35.2978196 | 31.6404067 | 30.2010871 | 32.8947626 | 29.2272385 | 35.8539458 | 28.5137674 | 36.3844026 | 30.9336277 | 23.9129978 | 24.3220352 | 25.2847443 | 23.9417061 | 28.6640357 | 24.3777199 |
| ENSMUSG00 Wdhd1 | 8035 | 0.38617024 | 0.31094799 | 0.16856986 | 0.26650518 | 0.26956273 | 0.43786485 | 0.20609132 | 0.09981835 | 0.67661731 | 0.37556254 | 0.29729283 | 0.09725193 | 0.11665295 | 0.56068158 | 0.40509308 | 0.60998105 | 0.50561138 | 0.23961652 |
| ENSMUSG00 Tob1 | 2286 | 45.2384485 | 53.8456449 | 53.9811356 | 53.0748299 | 48.5740804 | 55.1452993 | 48.161244 | 51.1403953 | 56.3747191 | 55.1741606 | 66.2705133 | 59.7466869 | 56.6475428 | 67.9392074 | 69.2279861 | 71.3384501 | 76.6861251 | 84.3184208 |
| ENSMUSG00 Ephx3 | 2955 | 0.01438415 | 0 | 0 | 0.03083658 | 0.0325766 | 0 | 0 | 0 | 0 | 0.03191249 | 0 | 0 | 0.08347194 | 0.06286848 | 0 | 0.09070533 | 0.02593996 | 0 |
| ENSMUSG00 Pkd2l1 | 4426 | 0.02881056 | 0.06021309 | 0.0218588 | 0.08235164 | 0.14137257 | 0.03358748 | 0.01068972 | 0.09492011 | 0.10236143 | 0.21306237 | 0.26985404 | 0.02522172 | 0.24521061 | 0 | 0.82849943 | 0.23358482 | 0.13854965 | 0.18642942 |
| ENSMUSG00 Kcnh3 | 3758 | 0.24883286 | 0.71802672 | 0.33467584 | 0.35158869 | 0.17930999 | 0.84389967 | 0.84352035 | 0.96548117 | 0.49318606 | 0.45168316 | 0.49580186 | 2.39125111 | 0.38068803 | 0.54378389 | 0.12060062 | 0.09170183 | 0.10198589 | 0.26348162 |
| ENSMUSG00 Gch1 | 2769 | 2.9165704 | 3.4167071 | 2.53310289 | 1.41504146 | 2.43353894 | 1.93271767 | 1.00810692 | 0.67584966 | 0.80320399 | 1.36224495 | 1.397534 | 1.53195751 | 1.83502618 | 1.59342301 | 0.50590571 | 1.02329539 | 0.30450651 | 1.09263303 |
| ENSMUSG00 Nr0b2 | 1130 | 4.5890544 | 5.66024397 | 6.84934943 | 4.91898001 | 5.19654397 | 4.38519766 | 9.42066616 | 5.880954 | 6.77937339 | 6.71793206 | 6.04585758 | 6.22369335 | 5.15147455 | 3.61688468 | 2.33353945 | 2.40587002 | 2.61161492 | 3.60236628 |
| ENSMUSG00 BC030499 | 4185 | 0 | 0.01592014 | 0.01155879 | 0.09798074 | 0 | 0 | 0 | 0.03650409 | 0.08857315 | 0.0112666 | 0 | 0 | 0.40078499 | 0 | 0 | 0.01829898 | 0.01831603 | 0.11829915 |
| ENSMUSG00 BC022687 | 1910 | 8.65681334 | 7.18581802 | 7.29401924 | 10.3049064 | 7.43398821 | 8.87279313 | 8.74417574 | 7.79844866 | 7.31006746 | 10.6644523 | 8.15425993 | 7.68561218 | 14.6962623 | 5.86022254 | 9.42674342 | 10.7855219 | 8.54816884 | 11.232226 |
| ENSMUSG00 1810019J16R | 2436 | 12.5980037 | 11.5692564 | 12.5501113 | 11.8391384 | 10.9264999 | 13.0187806 | 9.45865133 | 10.3006603 | 8.87640426 | 9.83274107 | 13.2773592 | 10.7919446 | 10.6116275 | 10.0285673 | 9.42088121 | 10.3428652 | 10.4311719 | 11.1553771 |
| ENSMUSG00 Nme1 | 3378 | 13.1869226 | 10.9859574 | 10.768767 | 9.94034626 | 11.769381 | 13.1436499 | 16.5972542 | 13.3978676 | 15.5455321 | 12.3669399 | 11.8235244 | 12.4916048 | 17.3786209 | 19.2898326 | 16.3562127 | 15.631364 | 14.0234763 | 13.6301428 |
| ENSMUSG00 Gm5519 | 511 | 0 | 0 | 0 | 0 | 0 | 0 | 0 | 0 | 0 | 0 | 0 | 0 | 0 | 0 | 0 | 0 | 0 | 0 |
| ENSMUSG00 Lphn3 | 13688 | 0 | 0.00243373 | 0 | 0.0166427 | 0.00703272 | 0.01086048 | 0 | 0 | 0 | 0 | 0 | 0.02854395 | 0 | 0 | 0 | 0 | 0 | 0 |
| ENSMUSG00 Osbpl5 | 4714 | 2.04681268 | 1.85150155 | 2.49358674 | 2.03932549 | 1.2252505 | 1.93417543 | 0.19069599 | 0.42129931 | 1.36298185 | 0.85019299 | 0.76010222 | 0.82882825 | 0.44999415 | 0.85715647 | 0.95268765 | 0.80415149 | 0.73985851 | 1.54034834 |
| ENSMUSG00 Bclaf1 | 5753 | 33.6391563 | 33.7008594 | 33.6167868 | 35.1864627 | 32.7544727 | 33.8850085 | 27.616196 | 35.6298868 | 32.8031662 | 32.3408092 | 37.5190792 | 32.3659048 | 29.8068651 | 35.3436841 | 30.8385016 | 32.2005838 | 38.4928461 | 31.4201155 |
| ENSMUSG00 Kcnmb2 | 3331 | 0 | 0 | 0 | 0 | 0 | 0.07438117 | 0 | 0 | 0 | 0 | 0 | 0 | 0 | 0 | 0 | 0 | 0 | 0 |
| ENSMUSG00 Tnfrsf23 | 4325 | 0 | 0.11553605 | 0 | 0 | 0 | 0.08020096 | 0 | 0 | 0 | 0 | 0 | 0 | 0 | 0 | 0 | 0 | 0.07089257 | 0 |
| ENSMUSG00 Spag1 | 3847 | 0.17678264 | 0.66677745 | 0.18861527 | 0.50926044 | 0.20018476 | 0.30914105 | 0.34436054 | 0.58574259 | 0.24624119 | 0.47800297 | 0.55884407 | 0.44977529 | 0.65399734 | 0.79680524 | 0.19278087 | 0.45785494 | 0.38854317 | 0.02859844 |
| ENSMUSG00 Atoh8 | 2352 | 0 | 0 | 0.02056698 | 0 | 0 | 0 | 0 | 0 | 0 | 0.02004707 | 0 | 0 | 0.02097445 | 0.53315917 | 0.03503534 | 0.06512008 | 0.01629519 | 0.04677645 |
| ENSMUSG00 Wdtc1 | 6684 | 1.02383802 | 1.22107407 | 1.17966569 | 1.09744586 | 1.9154835 | 1.68289505 | 1.53603401 | 1.34850493 | 0.82570214 | 1.24155053 | 1.30087409 | 1.3945565 | 1.74182114 | 1.2298928 | 1.48557375 | 1.4665463 | 1.09520088 | 1.32502486 |
| ENSMUSG00 Kcnk2 | 3926 | 0.67124832 | 0.63638998 | 0.09857062 | 0.23209905 | 0.47813169 | 0.25243369 | 0.13256227 | 0.45721929 | 0.38815616 | 0.16813801 | 0.52326113 | 0.17060315 | 1.35706752 | 2.21218654 | 1.2173664 | 0.88753064 | 0.99574179 | 2.52206806 |
| ENSMUSG00 Cldn11 | 1841 | 0.09235237 | 0.34380499 | 0.13137841 | 0.32172383 | 0.36602223 | 0.5114079 | 0.23129501 | 0.64310944 | 0.96198815 | 0.25611463 | 0.70066479 | 0.5457262 | 0.08038878 | 0.12613821 | 0.15665991 | 0.12479285 | 0.27063653 | 0.53784026 |
| ENSMUSG00 Rgs22 | 4765 | 0 | 0.06991164 | 0 | 0 | 0.06060683 | 0 | 0 | 0 | 0 | 0 | 0 | 0 | 0 | 0 | 0 | 0 | 0 | 0 |
| ENSMUSG00 Cdkn3 | 1144 | 0 | 0.49503427 | 0 | 0 | 0 | 0.17326131 | 0.16542895 | 0.2003098 | 1.04406377 | 0.12364695 | 0.04176133 | 0.3415294 | 0.04312231 | 0.77136159 | 2.30498215 | 0.43512054 | 1.17257027 | 0.8655279 |
| ENSMUSG00 Slc25a43 | 2082 | 0 | 0 | 0 | 0.10941653 | 0 | 0 | 0 | 0 | 0.31651501 | 0.04529366 | 0.25241333 | 0 | 0 | 0.24538183 | 0 | 0.07356505 | 0.14726723 | 0.15852769 |
| [illegible] | | | | | | | | | | | | | | | | | | | |

| | | | | | | | | | | | | | | | | | | | | |
|---|---|---|---|---|---|---|---|---|---|---|---|---|---|---|---|---|---|---|---|---|
| ENSMUSG00 | Rpf2 | 1820 | 19.5710646 | 15.9059918 | 17.1699454 | 15.7210636 | 17.6924489 | 17.4251371 | 19.6789549 | 14.4795372 | 15.9541174 | 18.2644208 | 17.7974845 | 14.321931 | 14.3387825 | 15.745056 | 20.1027372 | 18.4510235 | 18.152345 | 16.2911585 |
| ENSMUSG00 | Grtp1 | 3081 | 1.90383456 | 0.97311278 | 1.44445466 | 1.3604726 | 1.34350712 | 1.67266611 | 2.6719916 | 2.15692243 | 2.64684522 | 1.86705154 | 2.06233998 | 2.35509125 | 4.78748564 | 3.5424734 | 4.39964683 | 4.44921696 | 4.68971565 | 2.85668844 |
| ENSMUSG00 | Tbkbp1 | 5463 | 0.00778056 | 0.00609791 | 0 | 0 | 0 | 0 | 0 | 0.02097331 | 0 | 0 | 0 | 0 | 0 | 0 | 0 | 0 | 0 | 0 |
| ENSMUSG00 | Jarid2 | 7988 | 5.74681899 | 5.55492928 | 5.02629321 | 5.96035194 | 6.07975851 | 6.50736324 | 6.56265001 | 6.34468177 | 4.5888832 | 6.8117065 | 6.62677191 | 6.04413472 | 3.99571238 | 4.34323151 | 4.25529339 | 4.25184887 | 5.0666712 | 4.12499436 |
| ENSMUSG00 | Tbc1d17 | 3828 | 23.3289903 | 23.1223511 | 24.9702445 | 26.4344513 | 23.311543 | 24.7893116 | 22.0372274 | 26.1599998 | 27.0380653 | 27.9110485 | 28.2431377 | 26.4934846 | 24.4211032 | 25.3331386 | 26.6604634 | 21.9860907 | 22.1367405 | 26.6998224 |
| ENSMUSG00 | C1s | 4862 | 0.04371162 | 0 | 0 | 0.01874169 | 0 | 0 | 0.03892446 | 0 | 0 | 0.0678846 | 0 | 0 | 0.14204995 | 0 | 0.0169484 | 0.03150194 | 0 | 0 |
| ENSMUSG00 | AI317395 | 4733 | 0.08980599 | 0.02815372 | 0.06132288 | 0 | 0.02033886 | 0.0418785 | 0.18993047 | 0 | 0 | 0.04981059 | 0.06056407 | 0 | 0.01042297 | 0 | 0.01741033 | 0 | 0 | 0.10460215 |
| ENSMUSG00 | 1700003F12l | 973 | 0.21842331 | 0 | 0 | 0 | 0 | 0 | 0 | 0 | 0 | 0 | 0 | 0 | 0.15210252 | 0 | 0.1270346 | 0 | 0 | 0 |
| ENSMUSG00 | Fchsd1 | 7519 | 0.92144483 | 1.14306779 | 1.08726249 | 1.29672343 | 0.88338948 | 1.07422471 | 0.67958097 | 1.31049871 | 0.61349777 | 1.51128069 | 0.85142232 | 0.64582559 | 0.85948679 | 0.84623487 | 1.09045214 | 1.1101679 | 0.902215 | 0.76818143 |
| ENSMUSG00 | Armc10 | 3757 | 10.3179985 | 8.44127636 | 8.76826568 | 8.29485077 | 8.99350823 | 9.00838888 | 7.11516199 | 8.88479098 | 6.90644262 | 7.79360839 | 9.66435146 | 8.7504459 | 9.12581382 | 8.75230606 | 8.89392117 | 8.57130711 | 8.27325706 | 11.1130987 |
| ENSMUSG00 | Car14 | 2530 | 0.35280976 | 0.63202329 | 0.86039876 | 0.59427449 | 0.53268532 | 0.23503273 | 0.22440796 | 0.30191622 | 0.21163032 | 0.95046872 | 0.26436736 | 0.39710748 | 2.67133307 | 4.95648363 | 3.22446959 | 0.9383469 | 0.53020569 | 3.87020593 |
| ENSMUSG00 | C1rl | 3004 | 0.0565981 | 0 | 0.11272128 | 0.04550038 | 0.04806783 | 0 | 0 | 0.44498472 | 0 | 0.03139195 | 0.20674915 | 0 | 0.06568831 | 0 | 0.06857782 | 0.03823962 | 0 | 0 |
| ENSMUSG00 | BC021785 | 6014 | 0.02120311 | 0 | 0.01608697 | 0.01515166 | 0.01600663 | 0.00823956 | 0.03146836 | 0 | 0.06163596 | 0 | 0.04766374 | 0.09280955 | 0 | 0.02316799 | 0.00685094 | 0.05730221 | 0.02549138 | 0.07317473 |
| ENSMUSG00 | Rgs4 | 5338 | 0 | 0 | 0 | 0 | 0 | 0 | 0 | 0 | 0 | 0 | 0 | 0 | 0 | 0 | 0 | 0 | 0 | 0 |
| ENSMUSG00 | Cbfa2t2 | 8789 | 5.33914146 | 5.14721929 | 5.91666232 | 5.8577745 | 5.07660635 | 6.49502208 | 4.37113384 | 4.9016967 | 4.36279966 | 5.34328143 | 4.70193868 | 5.13130679 | 3.56420156 | 3.61449036 | 3.5158911 | 4.06039892 | 4.17320129 | 5.06341647 |
| ENSMUSG00 | Osbpl7 | 4980 | 1.22053016 | 2.00680086 | 2.11755615 | 2.93676611 | 1.97166916 | 1.98011927 | 3.24918397 | 4.37141955 | 1.55483232 | 3.07710413 | 2.05298014 | 2.57783181 | 3.76409486 | 5.24128054 | 3.24317468 | 2.15288556 | 1.50072849 | 2.58476526 |
| ENSMUSG00 | Zfp280d | 4452 | 7.36107159 | 8.23094871 | 9.13794677 | 8.91367242 | 7.75170469 | 7.46852746 | 7.18404559 | 7.68652124 | 7.42874487 | 5.78263342 | 7.66202164 | 7.77307071 | 7.87848005 | 7.58419846 | 8.91219639 | 8.31694011 | 10.8040213 | 8.30326139 |
| ENSMUSG00 | Mc3r | 2622 | 0 | 0 | 0 | 0 | 0 | 0 | 0 | 0 | 0 | 0 | 0 | 0 | 0 | 0 | 0 | 0 | 0 | 0 |
| ENSMUSG00 | Ubn2 | 19377 | 5.22951642 | 4.09856736 | 4.33881384 | 3.5692659 | 3.73589375 | 3.93312196 | 3.57463807 | 4.69693425 | 4.63154347 | 3.63296692 | 4.03117392 | 4.68947199 | 4.67936543 | 4.10103904 | 3.64662581 | 4.2999619 | 4.04881714 | 3.806947 |
| ENSMUSG00 | Atf5 | 1988 | 3.52784409 | 3.6697806 | 2.65227103 | 2.63557349 | 3.17167103 | 3.09081437 | 2.18952035 | 1.90193553 | 2.44467246 | 2.30061278 | 2.11478689 | 1.82495879 | 2.65518871 | 2.38294613 | 1.96888723 | 3.52473901 | 2.50624678 | 2.6287048 |
| ENSMUSG00 | Tmc3 | 9253 | 1.68128114 | 1.86851746 | 1.98136475 | 1.98434031 | 1.38887103 | 1.6333712 | 0.98685259 | 0.92044804 | 2.42587821 | 1.63572633 | 0.84676248 | 1.57439619 | 1.29554262 | 1.79190956 | 0.88610284 | 1.21248762 | 1.13906102 | 1.31384552 |
| ENSMUSG00 | Srd5a2 | 849 | 2.75357442 | 4.23768266 | 2.4500139 | 2.84421125 | 2.09762212 | 4.26071796 | 3.62228991 | 5.17329361 | 3.20177861 | 3.22113167 | 0.95662461 | 1.70931358 | 2.78908366 | 3.17285874 | 5.19265781 | 0.270605 | 1.17371463 | 1.36064928 |
| ENSMUSG00 | Pcid2 | 4078 | 10.8816586 | 12.4086038 | 10.7351753 | 10.8260549 | 11.4369382 | 10.1584319 | 9.00309928 | 10.7890259 | 10.4834731 | 12.556563 | 11.937882 | 11.2507294 | 12.8833967 | 10.8081017 | 12.8009738 | 13.0139239 | 11.99224 | 13.5431935 |
| ENSMUSG00 | BC028528 | 1675 | 0 | 0.03977659 | 0.23103776 | 0.05440125 | 0.02873548 | 0.02958372 | 0.05649275 | 0 | 0.12294482 | 0.05629935 | 0.19965654 | 0 | 0.58903782 | 0 | 0.19678355 | 0.09144026 | 0.04576274 | 0 |
| ENSMUSG00 | 1110054O05l | 7150 | 6.19446062 | 5.9497295 | 7.26617785 | 6.1109149 | 7.15583704 | 7.24925305 | 6.53113484 | 7.09897941 | 6.75113237 | 6.04056562 | 6.82213043 | 5.83917701 | 5.94742835 | 7.54148048 | 6.26378899 | 5.9497713 | 7.70278163 | 6.95499755 |
| ENSMUSG00 | Cul7 | 6558 | 8.76288472 | 9.47370353 | 8.37205809 | 7.36424321 | 7.58162201 | 8.28145715 | 10.4682368 | 12.614325 | 9.10022283 | 11.9063075 | 8.27574761 | 9.48985419 | 11.9606204 | 10.6726502 | 13.2500887 | 13.5984789 | 13.2137477 | 10.7199815 |
| ENSMUSG00 | Ranbp9 | 3029 | 24.2485785 | 25.823267 | 30.7584746 | 28.2782315 | 31.6695028 | 31.0502107 | 24.3045653 | 30.1857416 | 21.9326015 | 29.9342365 | 28.2801256 | 29.7966083 | 22.6057118 | 22.7083834 | 28.7145872 | 26.3066207 | 25.8629749 | 24.6805467 |
| ENSMUSG00 | Gm129 | 2818 | 5.05295741 | 4.8468015 | 4.46313623 | 4.15514137 | 6.2001023 | 5.55665856 | 3.223575 | 4.77066205 | 5.30543124 | 5.13671608 | 6.2897479 | 3.96136711 | 7.14244934 | 8.4713488 | 7.20807915 | 7.63638089 | 8.98995112 | 7.8668098 |
| ENSMUSG00 | Fndc4 | 2604 | 3.00343794 | 2.55859403 | 3.26948592 | 2.44951851 | 2.03322258 | 2.22644771 | 2.23481548 | 3.62270484 | 1.0122646 | 2.49876999 | 2.53185264 | 3.23662252 | 1.66713084 | 0.48156312 | 0.74365327 | 0.64699951 | 0.41211072 | 0.80274426 |
| ENSMUSG00 | Reep2 | 1838 | 0.83252793 | 0.50748696 | 0.81587565 | 1.06590038 | 0.62849084 | 1.40192717 | 0.28315531 | 0.95584903 | 0.42575832 | 0.66698492 | 0.44187938 | 0.66808737 | 0.29523998 | 1.28870972 | 0.33624774 | 0.60414995 | 0.91749568 | 0.9876499 |
| ENSMUSG00 | Sp6 | 3838 | 0 | 0 | 0 | 0 | 0 | 0 | 0 | 0 | 0 | 0 | 0 | 0 | 0 | 0 | 0 | 0 | 0 | 0 |
| ENSMUSG00 | Eftud1 | 4397 | 7.97515811 | 7.81871886 | 8.13009755 | 9.13914967 | 8.8776416 | 9.79334216 | 8.16700553 | 6.957497 | 6.88471437 | 7.39913242 | 5.64998372 | 8.74618866 | 6.63069436 | 5.50315432 | 4.91944891 | 8.14230564 | 7.1126355 | 6.99342521 |
| ENSMUSG00 | Ift172 | 5359 | 8.10604402 | 10.2568157 | 8.93633048 | 8.68882017 | 8.30789875 | 9.03396542 | 6.77156647 | 8.40959784 | 10.3677197 | 9.66945753 | 8.88815704 | 7.11365881 | 7.70494771 | 5.39926599 | 6.55042464 | 6.51633401 | 7.41637803 | 7.41119146 |
| ENSMUSG00 | Cyp24a1 | 3548 | 0 | 0 | 0 | 0.06420666 | 0 | 0 | 0 | 0 | 0 | 0 | 0 | 0 | 0 | 0 | 0 | 0 | 0 | 0 |
| ENSMUSG00 | Rad9b | 2907 | 0.90654314 | 0.73341081 | 1.38114999 | 0.89335381 | 1.00999221 | 0.95457622 | 1.36713623 | 1.44518895 | 1.48764505 | 1.42733467 | 1.7256177 | 1.74723952 | 1.42548368 | 1.98110314 | 2.1401565 | 1.97577954 | 0.87015328 | 1.04076397 |
| ENSMUSG00 | Fam154b | 4881 | 0.71407999 | 0.70980147 | 0.45588658 | 0.90543358 | 1.03541323 | 0.78171696 | 1.30858669 | 0.76682144 | 1.43448221 | 0.68586354 | 1.04731009 | 0.80047047 | 0.86919584 | 0.54237103 | 0.62464969 | 0.7452587 | 0.83232687 | 1.61161694 |
| ENSMUSG00 | Bpifb5 | 1622 | 0 | 0 | 0.02982338 | 0.11235769 | 0 | 0 | 0.29169346 | 0 | 0 | 0.08720845 | 0 | 0.61940933 | 0.48662804 | 0.68721214 | 0.22861529 | 1.58167121 | 0.49620976 | 0 |
| ENSMUSG00 | Susd4 | 3624 | 35.9370473 | 36.9622372 | 33.0232103 | 41.902583 | 38.1442839 | 34.4708723 | 12.5853815 | 13.089118 | 12.9219284 | 14.9752923 | 13.4070455 | 15.540288 | 6.92879302 | 8.50962308 | 8.56091931 | 6.48742631 | 9.2220008 | 9.21372179 |
| ENSMUSG00 | Susd1 | 5403 | 89.1011704 | 92.8297123 | 93.085248 | 86.3914305 | 90.3932314 | 90.1267283 | 54.9310448 | 60.3599712 | 60.9908037 | 55.5458498 | 56.2105147 | 59.834225 | 38.8866622 | 35.063008 | 48.926371 | 37.22757 | 41.8021225 | 50.2544791 |
| ENSMUSG00 | Sct | 697 | 0.42688125 | 0.14338405 | 0.34701241 | 0.39220411 | 0 | 0 | 0 | 0 | 0 | 0 | 0 | 0 | 0 | 0 | 0.05911271 | 0 | 0.05498751 | 0 |
| ENSMUSG00 | Pptc7 | 4712 | 8.85825188 | 8.63222492 | 9.80405804 | 8.51852293 | 9.10134635 | 10.0220193 | 10.7236717 | 10.9016908 | 9.29144021 | 10.9071024 | 10.5851139 | 9.71325198 | 8.18708814 | 9.61014117 | 8.54285231 | 10.3446399 | 7.87348374 | 10.4484615 |
| ENSMUSG00 | Pln | 2477 | 1.01243657 | 1.03556433 | 0.56634331 | 0.44144734 | 0.64124081 | 0.6401645 | 0.28651198 | 0.55507731 | 0.01662758 | 0.17131866 | 1.00294624 | 0.56333935 | 0.51781585 | 0.35625259 | 0.53227686 | 0.83475688 | 0.23209304 | 0.31091138 |
| ENSMUSG00 | Akap12 | 6240 | 0.16348145 | 0.06940186 | 0.06976951 | 0.08761739 | 0.13112864 | 0.07941143 | 0.24262912 | 0.34887291 | 0.15840968 | 0.3098043 | 0.2296873 | 0.35779271 | 0.38738204 | 0.35726221 | 0 | 0.34976997 | 0.62648755 | 0.27328242 |
| ENSMUSG00 | Colec10 | 4581 | 0 | 0 | 0 | 0 | 0 | 0 | 0 | 0 | 0 | 0 | 0 | 0 | 0 | 0 | 0 | 0 | 0 | 0 |
| ENSMUSG00 | Tctn1 | 5162 | 2.84904899 | 3.53651027 | 2.50207916 | 2.99209493 | 2.92782333 | 2.81266 | 3.27210895 | 3.60319643 | 3.75003146 | 3.90030033 | 3.97970402 | 2.77888905 | 2.83835324 | 3.29301356 | 3.38424248 | 3.48635906 | 3.15549713 | 3.51686122 |
| ENSMUSG00 | Gm9766 | 1338 | 4.06626497 | 3.28647387 | 3.36228574 | 2.38361218 | 4.17287231 | 3.33314351 | 1.80339808 | 1.99810775 | 0.70798943 | 2.32581944 | 1.78531235 | 2.62809174 | 1.99097424 | 2.2215408 | 2.06315713 | 2.48974801 | 2.49206863 | 1.39783978 |
| ENSMUSG00 | AI481877 | 5753 | 0 | 0 | 0 | 0 | 0 | 0 | 0 | 0 | 0.02863655 | 0 | 0 | 0 | 0.00857499 | 0 | 0 | 0 | 0 | 0 |
| ENSMUSG00 | Capn8 | 3683 | 2.60824594 | 4.18785502 | 3.5593882 | 2.33804979 | 3.28023699 | 3.13488648 | 0.61661922 | 1.53474764 | 1.77807657 | 1.60028181 | 1.94576263 | 1.12146593 | 1.55376117 | 0.45397425 | 1.0739477 | 0.44705298 | 0.5411261 | 1.13513226 |
| ENSMUSG00 | Atp6v0a4 | 5065 | 40.7428686 | 32.1750599 | 39.4724187 | 35.8101713 | 39.1327269 | 29.6533732 | 30.7228827 | 29.3625301 | 40.2513826 | 35.0488446 | 32.3718968 | 38.701798 | 23.5897144 | 18.1558328 | 20.3526738 | 22.3468974 | 23.2227848 | 17.8874627 |
| ENSMUSG00 | Slc35f1 | 4933 | 0.08616496 | 0.12830833 | 0 | 0 | 0 | 0.06027091 | 0 | 0.01548445 | 0 | 0.17204797 | 0 | 0 | 0.07000272 | 0.2165445 | 0 | 0.0465728 | 0 | 0 |
| [illegible] | [illegible] | [illegible] | [illegible] | [illegible] | [illegible] | [illegible] | [illegible] | [illegible] | [illegible] | [illegible] | [illegible] | [illegible] | [illegible] | [illegible] | [illegible] | [illegible] | [illegible] | [illegible] | [illegible] | [illegible] |

| | | | | | | | | | | | | | | | | | | | |
|---|---|---|---|---|---|---|---|---|---|---|---|---|---|---|---|---|---|---|---|
| ENSMUSG00 Fam100a | 3152 | 17.989183 | 16.9735115 | 19.3064407 | 17.3600299 | 17.9273091 | 17.5446861 | 17.8473272 | 20.320006 | 19.783117 | 22.2440025 | 20.71966 | 22.4006387 | 18.2960061 | 19.6414875 | 22.3392954 | 24.4418471 | 22.7866372 | 28.6389417 |
| ENSMUSG00 Nphp4 | 6019 | 0.36015351 | 0.30993887 | 0.3134354 | 0.46931131 | 0.34385657 | 0.18935253 | 0.36944607 | 0.44417149 | 0.02052825 | 0.44651771 | 0.4365547 | 0.4729355 | 0.37701748 | 0.23148739 | 0.31488147 | 0.19084868 | 0.09551328 | 0.37470898 |
| ENSMUSG00 Fam190a | 8020 | 1.57936938 | 2.6625393 | 2.79263648 | 3.18700073 | 3.63690097 | 3.3982548 | 3.01456096 | 3.02396204 | 3.07614999 | 3.56863801 | 3.18102591 | 3.51457726 | 2.74954701 | 2.04423477 | 3.16974563 | 2.84553641 | 3.58413001 | 3.66270112 |
| ENSMUSG00 Grin3a | 19722 | 0 | 0.00506737 | 0.00490554 | 0 | 0 | 0.02010049 | 0.00239898 | 0.03292115 | 0 | 0 | 0 | 0 | 0.00250136 | 0.00235494 | 0 | 0 | 0 | 0 |
| ENSMUSG00 Myo9a | 12989 | 2.51974637 | 2.65190027 | 2.45051836 | 2.22385881 | 2.36046157 | 2.35765566 | 1.93417757 | 2.43168195 | 2.78085875 | 2.52651393 | 2.41283964 | 3.05956977 | 2.08129114 | 1.77709688 | 1.77633931 | 2.32001756 | 2.46086162 | 1.68978625 |
| ENSMUSG00 Fam149b | 4492 | 8.73381066 | 8.4542964 | 9.19657055 | 9.69642885 | 10.039985 | 8.95743982 | 9.17394108 | 7.41401934 | 7.60098432 | 8.21883363 | 8.30637635 | 7.14470302 | 6.05117685 | 7.39259183 | 8.60353059 | 6.8449149 | 7.26936887 | 7.05370558 |
| ENSMUSG00 Rcan2 | 3816 | 27.2563327 | 24.8537238 | 24.3769127 | 34.6841808 | 29.527472 | 27.8409488 | 58.2357581 | 44.4377009 | 52.1091452 | 57.4927207 | 54.0848587 | 44.8163496 | 25.6226048 | 23.9279549 | 24.8224258 | 22.8780359 | 27.7805391 | 20.9599689 |
| ENSMUSG00 Rbms3 | 11796 | 6.58332965 | 6.76368105 | 7.60706161 | 6.77853788 | 7.3201654 | 6.83891576 | 6.26905029 | 6.62441975 | 8.05155183 | 6.06371794 | 7.82478974 | 7.3483998 | 3.47531564 | 3.37030678 | 4.85854224 | 4.27507036 | 5.54945025 | 4.62606255 |
| ENSMUSG00 Olfr303 | 1039 | 0 | 0 | 0 | 0 | 0 | 0 | 0 | 0 | 0 | 0 | 0 | 0 | 0 | 0 | 0 | 0 | 0 | 0 |
| ENSMUSG00 2810432L12F | 3796 | 2.95610286 | 1.57086619 | 1.65662775 | 2.23244316 | 1.44547927 | 1.56647208 | 1.27131013 | 0.82502028 | 0.61844874 | 1.25453663 | 0.93133481 | 1.07337813 | 0.51983053 | 0.07341004 | 0.79233761 | 0.87757715 | 0.38366683 | 0.52168805 |
| ENSMUSG00 Stub1 | 1665 | 57.3883698 | 53.9808941 | 47.8796264 | 56.3698206 | 57.58486 | 55.2371613 | 53.8483643 | 55.1209267 | 55.2616674 | 53.8339261 | 54.4605833 | 58.8327254 | 60.9760275 | 66.5559114 | 54.2920202 | 50.3872197 | 52.3907818 | 64.7224265 |
| ENSMUSG00 Mocos | 2869 | 0.22222992 | 1.30046851 | 0.5058229 | 1.30219785 | 0.52007307 | 0.74268649 | 0.758586 | 0.23961772 | 1.24894628 | 1.13398345 | 0.81595433 | 1.12837519 | 1.47875388 | 0.46945923 | 0.74676922 | 0.77408687 | 0.78816714 | 0.30677787 |
| ENSMUSG00 Gm7488 | 683 | 0.87126276 | 0.97548739 | 0.63742573 | 0.60036515 | 0.70471338 | 0.58041269 | 1.24689344 | 0.50326738 | 0.60302366 | 0.89745116 | 0.62953825 | 0.49032793 | 0.28891313 | 0.34000064 | 0.78421792 | 1.17731004 | 0.72949028 | 0.16108084 |
| ENSMUSG00 6430573F11F | 4210 | 0.25240603 | 0.43520408 | 0.56301734 | 0.17315361 | 0.40014663 | 0.55320154 | 0.53943197 | 0.26308306 | 0.479368 | 1.04157135 | 0.69222624 | 0.35796268 | 0.21092031 | 1.26866272 | 0.59698216 | 1.25513155 | 1.45658135 | 1.26743072 |
| ENSMUSG00 Prex1 | 9011 | 0 | 0.06654446 | 0.05368275 | 0.07078622 | 0.10682926 | 0.1154819 | 0.30453173 | 0.09324524 | 0.33823129 | 0.08895372 | 0.09013143 | 0.13007756 | 0.79929641 | 0.96898107 | 0.60812417 | 0.06798909 | 0.11909181 | 0.53110557 |
| ENSMUSG00 C330006K01F | 3906 | 9.23883626 | 7.47109456 | 6.26651469 | 8.53832167 | 7.45514947 | 6.19092012 | 9.75082093 | 10.2863306 | 9.86957981 | 10.3572205 | 6.56813945 | 9.17400291 | 10.6595336 | 11.2126919 | 12.2887385 | 9.71479568 | 10.243895 | 9.32310001 |
| ENSMUSG00 Hs3st6 | 1849 | 0.34482295 | 0.70265164 | 0.39242994 | 0.59138186 | 1.14537839 | 0.93799117 | 0.92117709 | 2.39606202 | 1.04692604 | 0.89252278 | 1.36942823 | 2.65645122 | 0.29348355 | 0.42701432 | 0.80219362 | 0 | 0.72548426 | 0.56526394 |
| ENSMUSG00 Fam40b | 8329 | 0.05613602 | 0.26397539 | 0.13358041 | 0.30085934 | 0.19648042 | 0.02379769 | 0.14769236 | 0.01375642 | 0.15329355 | 0.02264411 | 0.04588782 | 0.11392319 | 0.06515201 | 0.14498094 | 0.22755091 | 0.05516716 | 0.10583561 | 0.01320905 |
| ENSMUSG00 Hnrnpu | 5357 | 118.327364 | 124.974657 | 130.799064 | 133.332181 | 117.845494 | 123.821708 | 114.709179 | 111.668396 | 107.383249 | 124.535244 | 107.295226 | 107.588675 | 99.4005442 | 100.214151 | 110.414325 | 110.947696 | 120.237399 | 108.796245 |
| ENSMUSG00 Ccdc151 | 2187 | 0.01943538 | 0.10662563 | 0.33178004 | 0.02083267 | 0.06602459 | 0.24923643 | 0.0216336 | 0.0698535 | 0.65913491 | 0.38807163 | 0.26213969 | 0.22969408 | 0.22556889 | 0.06370931 | 0.0188393 | 0 | 0.0525738 | 0 |
| ENSMUSG00 Lonrf1 | 3918 | 5.17482618 | 4.21725256 | 5.25960284 | 5.05846204 | 3.31690132 | 5.95695189 | 4.62500154 | 5.10791701 | 3.86846294 | 4.40458331 | 3.18255849 | 3.91763839 | 3.19813859 | 2.63159454 | 3.69110416 | 2.58984455 | 2.48465529 | 5.36331771 |
| ENSMUSG00 Zfp189 | 3037 | 2.54723149 | 2.79709846 | 4.10944051 | 3.93052144 | 2.09200327 | 3.26326859 | 5.09425288 | 2.67862421 | 3.55313373 | 4.64210085 | 4.01139757 | 4.22706697 | 3.4598941 | 2.69152433 | 2.19777809 | 2.38291893 | 2.39775977 | 2.42713872 |
| ENSMUSG00 Coro7 | 6960 | 4.31769532 | 5.04479749 | 4.43424028 | 5.3220013 | 4.64722022 | 6.1015363 | 4.09907261 | 4.72466357 | 4.39086131 | 4.85733538 | 4.69512523 | 5.26078659 | 5.94676417 | 4.17062566 | 4.25039039 | 5.26495852 | 5.72691793 | 7.25551149 |
| ENSMUSG00 Kcne1 | 3752 | 0 | 0 | 0 | 0 | 0 | 0 | 0 | 0 | 0.03293165 | 0 | 0 | 0 | 0 | 0 | 0 | 0.0714377 | 0.11236388 | 0.02932255 |
| ENSMUSG00 Mrpl12 | 1527 | 36.2699701 | 29.6914532 | 28.7009944 | 26.8831047 | 27.6750681 | 31.9318205 | 35.9725083 | 35.0409665 | 31.3146986 | 36.3434039 | 35.7608235 | 32.6413796 | 32.5325741 | 29.1378099 | 34.1592467 | 30.4669862 | 26.7807196 | 35.5920088 |
| ENSMUSG00 Vasn | 3184 | 5.05950431 | 4.62446584 | 4.55780751 | 3.93507764 | 5.47228529 | 3.57950023 | 2.77872832 | 4.01835892 | 3.69954256 | 5.37553558 | 3.8111933 | 2.92751748 | 2.58744667 | 7.01620793 | 3.94675708 | 4.23313248 | 4.00837213 | 5.73587419 |
| ENSMUSG00 Ccbl1 | 3253 | 12.6613943 | 10.0460957 | 9.24941158 | 10.770502 | 11.7037663 | 12.0187848 | 8.27571912 | 8.53546774 | 8.53357259 | 10.1316759 | 10.3539335 | 10.2262942 | 10.6610322 | 10.3796039 | 8.24537746 | 8.00418476 | 8.88350073 | 9.55430227 |
| ENSMUSG00 Cpeb3 | 11181 | 0.99980872 | 0.52437791 | 0.58839103 | 0.95351797 | 0.90831195 | 0.4653463 | 1.19329 | 0.9256901 | 0.95037305 | 0.93196543 | 0.94857712 | 0.91852982 | 1.05890888 | 1.21292134 | 1.00599452 | 1.34244863 | 1.1208925 | 0.69370218 |
| ENSMUSG00 Baat | 2530 | 0 | 0 | 0 | 0 | 0 | 0 | 0 | 0 | 0.03255851 | 0 | 0 | 0 | 0 | 0 | 0 | 0 | 0.07574367 | 0 |
| ENSMUSG00 Rxrb | 3779 | 11.7088881 | 12.0504701 | 10.6373124 | 13.2379007 | 10.8389169 | 11.4998008 | 9.87819626 | 13.8559893 | 13.2856213 | 12.6018021 | 10.8975957 | 11.5205654 | 9.72539782 | 12.5235578 | 13.5739447 | 9.54478756 | 11.6530604 | 11.3977588 |
| ENSMUSG00 D2Wsu81e | 3629 | 5.32925189 | 6.31558865 | 7.17138587 | 6.21458169 | 5.38483357 | 5.74860866 | 9.75196579 | 6.90388977 | 8.89783091 | 7.56178267 | 7.02998838 | 6.01375698 | 7.95237575 | 9.88008693 | 8.58318442 | 6.9216419 | 6.80135994 | 10.8987456 |
| ENSMUSG00 Dusp26 | 2557 | 1.3132221 | 1.66759893 | 2.66744927 | 1.5679984 | 1.88235916 | 3.10067946 | 0.12952239 | 0 | 0.64429434 | 0.79291366 | 0.22420786 | 0.52388577 | 0.07717156 | 0 | 0.06445296 | 0.17969781 | 0.2398204 | 0.04302629 |
| ENSMUSG00 Icmt | 4187 | 11.0653551 | 16.1591804 | 10.8600713 | 17.3887148 | 11.6450057 | 14.4149103 | 14.0344751 | 13.7098593 | 10.672885 | 12.4774252 | 11.8781304 | 9.74474549 | 8.98979051 | 8.9959768 | 10.6964632 | 10.9467058 | 12.1834968 | 10.2608341 |
| ENSMUSG00 1810049H13F | 956 | 25.2541213 | 20.1410595 | 21.8591685 | 19.3491464 | 21.0451298 | 20.4223609 | 23.161437 | 22.7316705 | 26.9263309 | 24.1178806 | 24.7870077 | 23.879295 | 26.4720435 | 17.9266404 | 22.6694753 | 20.3068389 | 21.7690159 | 21.8080036 |
| ENSMUSG00 Zmynd8 | 12576 | 4.78588814 | 4.45284472 | 4.63887386 | 4.70971355 | 4.17939414 | 4.84652213 | 6.23010467 | 4.78316108 | 4.36886228 | 5.36519214 | 4.43711446 | 4.961984 | 5.22111815 | 4.07345964 | 5.2255472 | 4.92029434 | 4.74812107 | 3.52117771 |
| ENSMUSG00 Kcne2 | 1764 | 0 | 0.0944243 | 0.21938109 | 0.05165651 | 0 | 0 | 0.42913994 | 0.1948592 | 0.23348365 | 0.05345885 | 0.67708274 | 0.8859629 | 0.41948909 | 0.18430193 | 0.16349824 | 0.28218702 | 0.02172693 | 0 |
| ENSMUSG00 Capsl | 941 | 4.60736233 | 4.92082072 | 5.19205799 | 3.87341503 | 4.85922715 | 3.68617572 | 3.720657 | 4.82985748 | 3.72035477 | 4.55973857 | 2.79237271 | 4.86385158 | 4.92794923 | 5.67595117 | 5.9985262 | 5.5340304 | 3.99147407 | 7.19035083 |
| ENSMUSG00 Tbc1d13 | 4742 | 2.07954467 | 1.14508683 | 1.10171684 | 1.26825345 | 1.38041769 | 0.68968377 | 1.5564694 | 0.9825966 | 1.11174062 | 1.17329882 | 0.92688658 | 1.37714731 | 1.93499296 | 1.17530377 | 1.07739201 | 1.86527421 | 1.09919367 | 1.28764463 |
| ENSMUSG00 Mrps6 | 1741 | 183.765916 | 152.692071 | 145.120591 | 139.090724 | 150.33969 | 144.445791 | 178.190256 | 157.661572 | 151.853099 | 161.574437 | 162.86294 | 149.397466 | 232.690238 | 209.065085 | 194.672027 | 223.145608 | 194.845522 | 203.290404 |
| ENSMUSG00 Lap3 | 3116 | 47.8115026 | 60.1684752 | 49.196315 | 49.8598078 | 46.7417205 | 52.3357016 | 29.9120595 | 29.9557867 | 35.8994149 | 32.9116751 | 36.7511475 | 29.3766645 | 30.0962692 | 27.2762131 | 27.0930642 | 34.6901106 | 29.8517081 | 29.3758516 |
| ENSMUSG00 Sdk1 | 11214 | 0.23500276 | 0.04753044 | 0.12509652 | 0.33721836 | 0.27040407 | 0.21210373 | 0.37549745 | 0.415505 | 0.37829598 | 0.62648964 | 0.28543983 | 0.58732372 | 0.70386185 | 0.63366732 | 0.49600589 | 0.54632577 | 0.43405027 | 0.92711978 |
| [illegible] | | | | | | | | | | | | | | | | | | | |

| | | | | | | | | | | | | | | | | | | | | |
|---|---|---|---|---|---|---|---|---|---|---|---|---|---|---|---|---|---|---|---|---|
| ENSMUSG00 | Col16a1 | 6609 | 1.35702711 | 1.67849807 | 1.84447415 | 1.5993588 | 1.66775768 | 2.03939227 | 0.05727061 | 0.24271159 | 0.00623188 | 0.50653653 | 0.06505896 | 0.18579885 | 1.20922539 | 0.32326041 | 0.44885944 | 0.26651051 | 0.12178124 | 0.11652708 |
| ENSMUSG00 | Slco4c1 | 2381 | 0 | 1.53902493 | 0.46727896 | 0.36356986 | 0.18193503 | 0.9781514 | 0.01987093 | 0.22456683 | 0 | 0.75251017 | 0.80260325 | 0.35163164 | 0.78732165 | 0.35111028 | 0.01730431 | 0.25730774 | 0.28974101 | 0.34655044 |
| ENSMUSG00 | Apobec2 | 1215 | 0.17491842 | 0.10967208 | 0 | 0.0374988 | 0.11884426 | 0.04078414 | 0.19470238 | 0.31434076 | 0.03389837 | 0.31045731 | 0 | 0 | 0.52783121 | 0.03822559 | 0 | 0.12605961 | 0 | 0 |
| ENSMUSG00 | Dnajc16 | 6494 | 5.02024484 | 4.89382524 | 6.61467431 | 7.57713703 | 4.33587548 | 7.00483668 | 4.67006886 | 4.69906522 | 3.62141985 | 6.05538751 | 4.85547776 | 6.3860544 | 3.78307589 | 4.18382986 | 4.34602183 | 4.20406576 | 5.92540839 | 3.80337064 |
| ENSMUSG00 | Limd2 | 4220 | 48.2363242 | 41.3095676 | 42.1950155 | 42.2789217 | 42.1325417 | 41.40354 | 41.3481422 | 51.143428 | 45.9688363 | 51.9104662 | 57.5337296 | 48.6866005 | 44.6909762 | 39.1693145 | 44.0524692 | 49.5963175 | 45.2922376 | 38.402092 |
| ENSMUSG00 | Ap1g2 | 4349 | 6.42122342 | 7.26924273 | 7.75266743 | 8.42287414 | 6.83962493 | 8.2948701 | 8.11571821 | 8.000294 | 4.79199287 | 7.67594801 | 7.34914859 | 7.31545819 | 6.30686273 | 7.60363077 | 6.61271235 | 6.77943612 | 6.86506905 | 8.55050728 |
| ENSMUSG00 | Cyp2s1 | 5858 | 0.58772947 | 0.84163679 | 0.94137632 | 0.54443037 | 0.4190386 | 0.48216214 | 0.36346666 | 0.8084428 | 0.51324951 | 0.57147489 | 0.68506257 | 1.07667632 | 0.22737483 | 0.20613627 | 0.40090281 | 0.2549221 | 0.51031942 | 0.54464462 |
| ENSMUSG00 | A930016O22l | 1210 | 0.52692367 | 0.24778186 | 0.39978124 | 0.48949882 | 0.67623364 | 0.08190534 | 0.3128111 | 0.63127938 | 0.61269197 | 0.38967523 | 0.11845031 | 0.55354376 | 0.16308072 | 0.30706835 | 0.03405087 | 0.91770877 | 0.50679401 | 0.13638622 |
| ENSMUSG00 | Agmat | 1400 | 0.24288672 | 0.11897462 | 0 | 0.0325436 | 0.10313984 | 0 | 0 | 0 | 0 | 0 | 0 | 0 | 0 | 0.39809218 | 0 | 0.13675217 | 0.24638334 | 0 |
| ENSMUSG00 | St8sia4 | 5406 | 0 | 0.01232442 | 0.02684436 | 0.01685573 | 0.06232399 | 0.06416373 | 0.04375941 | 0.2048797 | 0.06094934 | 0 | 0.7335038 | 0.18584571 | 0.18250802 | 0.16323301 | 0.0152429 | 0.03541492 | 0 | 0 |
| ENSMUSG00 | Sh3pxd2b | 9824 | 2.50946683 | 3.83857861 | 2.87562518 | 2.10552872 | 2.89555852 | 2.85493152 | 1.83972347 | 2.78356679 | 2.39807482 | 2.63014916 | 3.38470797 | 2.66465246 | 3.77622168 | 2.87911739 | 3.34259365 | 3.6326187 | 3.49945932 | 3.05170635 |
| ENSMUSG00 | Camta2 | 5786 | 7.05236242 | 9.68987227 | 10.3920322 | 12.5674778 | 10.315172 | 10.8423368 | 13.6393966 | 11.8748932 | 9.52429279 | 12.5414677 | 8.65332812 | 10.8718029 | 9.71120874 | 11.9923032 | 12.9956515 | 9.19874599 | 10.551986 | 10.9618908 |
| ENSMUSG00 | Creg1 | 2356 | 17.9149575 | 22.2981469 | 21.332809 | 22.8385371 | 22.5133191 | 24.5450085 | 21.8690608 | 26.5207005 | 22.6910431 | 25.67672 | 23.7252553 | 24.3778934 | 25.8594728 | 25.9227398 | 22.6293775 | 26.4101176 | 27.8988112 | 28.7887218 |
| ENSMUSG00 | Klc3 | 2682 | 7.24267914 | 8.5704314 | 6.51112769 | 7.15182687 | 6.67601626 | 8.79459403 | 5.73326646 | 4.88441239 | 5.82016759 | 5.04558234 | 5.84272423 | 7.28392316 | 4.67200111 | 4.69289623 | 5.13099167 | 5.22534016 | 4.04412454 | 5.53782954 |
| ENSMUSG00 | Rac1a1 | 1748 | 0 | 0 | 0 | 0 | 0 | 0 | 0 | 0 | 0 | 0 | 0 | 0 | 0 | 0 | 0 | 0 | 0 | 0 |
| ENSMUSG00 | Il17rd | 9146 | 0.17195403 | 0.02549642 | 0.18511629 | 0.05977832 | 0.13682812 | 0.08126952 | 0.11380701 | 0.08769303 | 0.00450323 | 0.01546601 | 0.0574596 | 0.01830822 | 0.01618147 | 0.05078076 | 0.0630682 | 0.17583703 | 0.04190498 | 0.07217464 |
| ENSMUSG00 | 1110037F02l | 7175 | 5.24872278 | 5.78971137 | 4.94185334 | 4.97202756 | 4.83667135 | 4.89656977 | 6.15889089 | 4.70019391 | 4.77591374 | 5.72379966 | 5.04050781 | 6.1299989 | 3.84342045 | 5.27553048 | 4.45034227 | 6.14250719 | 4.98909565 | 5.24407373 |
| ENSMUSG00 | Zfhx2 | 10552 | 3.28295285 | 3.16017884 | 3.36029167 | 2.88426626 | 3.18842064 | 2.84111105 | 3.04447583 | 3.3190327 | 3.70022965 | 3.29322103 | 3.02894686 | 4.01479226 | 3.78685112 | 2.147907 | 2.74885286 | 3.34208201 | 2.66961981 | 2.70562229 |
| ENSMUSG00 | Scamp5 | 3183 | 8.07905482 | 6.48884525 | 4.8327938 | 7.58635045 | 5.18669488 | 5.44877686 | 9.31983968 | 5.15051399 | 5.0981738 | 6.05863576 | 4.65291773 | 7.83839182 | 6.40090538 | 9.79075798 | 5.86374641 | 7.06144723 | 6.68271906 | 5.04638894 |
| ENSMUSG00 | Rcsd1 | 2663 | 2.12286459 | 1.62623967 | 2.54310712 | 2.78875337 | 1.59054047 | 1.65609962 | 2.43403568 | 1.44852897 | 2.44366109 | 0.8675871 | 3.01396659 | 1.76061422 | 0.94477198 | 0.645299 | 0.26302157 | 0.67580118 | 0.44615667 | 0.47510682 |
| ENSMUSG00 | Kcna2 | 3453 | 0 | 0 | 0 | 0 | 0 | 0 | 0 | 0 | 0 | 0 | 0 | 0 | 0 | 0 | 0 | 0 | 0 | 0 |
| ENSMUSG00 | Hnrnpul1 | 3573 | 39.2337169 | 51.5962905 | 52.3268108 | 49.4757492 | 48.562996 | 51.757851 | 49.8815733 | 45.9954961 | 46.4774787 | 49.3676969 | 42.6671963 | 45.1461699 | 44.2509921 | 46.8470448 | 41.3399322 | 42.5129694 | 46.8110718 | 46.1565357 |
| ENSMUSG00 | Hesx1 | 1137 | 0.14953448 | 0 | 0 | 0 | 0 | 0 | 0.16644742 | 0 | 0 | 0.0414694 | 0.04201843 | 0.09818059 | 0.34710232 | 0 | 0.65226738 | 0.50515313 | 0.60674876 | 0 |
| ENSMUSG00 | Esrp1 | 7723 | 4.9698529 | 6.25883847 | 5.96291608 | 5.8286044 | 5.0668463 | 6.19810186 | 5.28078845 | 5.84037644 | 4.98632555 | 4.67050214 | 6.96550605 | 4.64708935 | 5.51255271 | 6.30840964 | 7.52223149 | 4.7448027 | 6.36207596 | 6.33213724 |
| ENSMUSG00 | AK129341 | 3636 | 0.78323619 | 1.55753356 | 1.46344564 | 0.72677133 | 1.17814665 | 0.87221533 | 1.75665887 | 1.83819319 | 2.39008659 | 1.42645142 | 1.06429363 | 1.1973657 | 0.7190846 | 1.23901993 | 1.60908167 | 1.84291977 | 1.96058615 | 1.55828878 |
| ENSMUSG00 | Eif4h | 4189 | 107.637839 | 117.712452 | 109.738281 | 115.572131 | 106.237232 | 117.736538 | 100.927691 | 106.152815 | 104.564 | 116.666756 | 116.192714 | 104.502803 | 99.8061587 | 90.1829092 | 103.471083 | 97.5683799 | 104.310841 | 92.4741298 |
| ENSMUSG00 | Erg | 5606 | 0.00758209 | 0 | 0 | 0 | 0 | 0 | 0 | 0 | 0 | 0 | 0 | 0 | 0 | 0 | 0 | 0 | 0 | 0 |
| ENSMUSG00 | Ppp1r13l | 3617 | 1.57470102 | 1.54729506 | 1.57812457 | 2.17917077 | 1.98276379 | 1.97279337 | 1.36038668 | 1.07703208 | 1.56000903 | 1.30358593 | 0.81892381 | 1.29624431 | 1.5548351 | 1.36109297 | 0.97963334 | 0.92100715 | 0.91126943 | 1.50564047 |
| ENSMUSG00 | Ints8 | 7190 | 4.28598785 | 5.11508139 | 5.04591763 | 4.81591011 | 3.85590932 | 5.87189555 | 5.32349892 | 4.87099207 | 4.54254618 | 4.78064849 | 3.98678374 | 4.02121185 | 3.63642784 | 5.05135971 | 4.11442519 | 7.16284659 | 4.87207724 | 4.621071 |
| ENSMUSG00 | Slc25a34 | 2665 | 0.07974705 | 0.03750044 | 0.01815142 | 0 | 0.10836456 | 0.11156338 | 0 | 0.01433111 | 0.06181841 | 0.08846286 | 0 | 0 | 0.01851104 | 0 | 0 | 0.05747183 | 0.0287627 | 0 |
| ENSMUSG00 | Rnf167 | 2524 | 50.6559309 | 47.4748338 | 49.082651 | 52.1858079 | 49.3905239 | 43.741478 | 53.0111949 | 56.4563603 | 56.8027977 | 57.3318179 | 63.7693483 | 56.2137141 | 52.3027776 | 51.2835466 | 50.0001449 | 59.2715543 | 55.6217217 | 55.2924341 |
| ENSMUSG00 | Cd53 | 2794 | 0.01521302 | 0.03576903 | 0.12119353 | 0.06522698 | 0.01722689 | 0.01773541 | 2.70938748 | 1.7770266 | 2.88924735 | 0.23625979 | 5.91629765 | 0.99884865 | 1.9951706 | 3.60714637 | 1.09123663 | 1.52120881 | 0 | 0.63002556 |
| ENSMUSG00 | Siah1b | 2456 | 2.95944019 | 4.80161423 | 2.63927242 | 4.21105744 | 4.25269847 | 3.57118642 | 4.85455826 | 4.69629704 | 3.27010203 | 2.53415832 | 6.5359878 | 4.81796436 | 1.72742056 | 5.57858542 | 2.51638167 | 2.75954296 | 3.54237353 | 2.88932197 |
| ENSMUSG00 | Lat2 | 1492 | 1.3104813 | 1.49595437 | 1.6535188 | 2.07650872 | 2.00012015 | 2.78983219 | 2.0612092 | 1.71507437 | 3.36779821 | 0.8532634 | 4.16269745 | 2.09495785 | 2.67820731 | 2.73932955 | 1.04936939 | 1.74514833 | 1.97796572 | 1.62225248 |
| ENSMUSG00 | Myh6 | 6258 | 0 | 0.0106465 | 0 | 0 | 0 | 0 | 0.0151207 | 0.04272081 | 0 | 0 | 0 | 0 | 0 | 0 | 0 | 0 | 0 | 0 |
| ENSMUSG00 | Cmtm5 | 873 | 0 | 0 | 0.05541069 | 0 | 0 | 0 | 0 | 0 | 0 | 0 | 0 | 0 | 0 | 0 | 0 | 0 | 0 | 0 |
| ENSMUSG00 | Appl1 | 8697 | 3.51399581 | 3.59673539 | 4.36624369 | 3.24276031 | 3.5917694 | 4.21058642 | 2.69829697 | 4.14991609 | 3.77436508 | 3.94142363 | 3.75190257 | 3.07412928 | 3.05169271 | 2.94781376 | 2.50137076 | 2.72089173 | 4.14683746 | 4.31369562 |
| ENSMUSG00 | Spen | 14377 | 3.37924575 | 3.09564072 | 3.47231574 | 3.67923576 | 3.01640559 | 3.21574045 | 2.95848218 | 3.61814371 | 2.98506986 | 3.97814586 | 2.99403475 | 3.63382209 | 4.03178708 | 3.2239827 | 2.67378811 | 3.56619067 | 3.19363592 | 2.89642441 |
| ENSMUSG00 | Snrnp25 | 1253 | 19.0306494 | 15.8987316 | 16.639259 | 16.4354286 | 15.2885121 | 15.3838894 | 22.0138003 | 17.343557 | 18.571733 | 19.266688 | 20.5512391 | 15.0564198 | 28.4652643 | 20.4976699 | 23.3793346 | 18.9161101 | 17.4043599 | 24.8045852 |
| ENSMUSG00 | Il25 | 982 | 0 | 0 | 0 | 0 | 0 | 0 | 0 | 0 | 0 | 0 | 0 | 0 | 0 | 0 | 0 | 0 | 0.07805763 | 0 |
| ENSMUSG00 | AI314976 | 1487 | 47.8790651 | 39.6528734 | 44.3722229 | 41.210225 | 37.4503266 | 38.0559662 | 50.335345 | 53.577237 | 52.8749553 | 61.1659089 | 46.6825922 | 44.1795796 | 58.5215212 | 60.1555564 | 57.4106414 | 61.0538199 | 57.0641704 | 58.0425612 |
| ENSMUSG00 | Cept1 | 3842 | 32.1056796 | 30.9891421 | 34.4481986 | 35.4811673 | 33.3866675 | 32.1925099 | 26.4271236 | 28.9673765 | 30.0590813 | 28.3984194 | 29.6821515 | 28.1983865 | 21.1092327 | 20.2845337 | 26.5525904 | 32.1214544 | 32.6401987 | 26.6168478 |
| ENSMUSG00 | Gm9772 | 404 | 0 | 0.24737298 | 0 | 0 | 0.23827685 | 0 | 0 | 0 | 0 | 0 | 0 | 0 | 0 | 0 | 0 | 0 | 0 | 0.13616116 |
| ENSMUSG00 | Rfwd2 | 5036 | 5.24984501 | 5.29857591 | 4.96606735 | 4.27021702 | 5.2662212 | 5.20520176 | 5.43964279 | 5.30112972 | 4.84162378 | 4.1008753 | 4.37336296 | 4.20058303 | 5.22118981 | 5.67178589 | 5.73516495 | 6.20435577 | 4.22380754 | 4.3692698 |
| ENSMUSG00 | Ttc3 | 15207 | 24.1748713 | 24.9753605 | 23.9593234 | 23.8036758 | 24.8746491 | 24.5173123 | 20.7115476 | 21.3578082 | 21.3258779 | 21.8808781 | 22.5946933 | 23.4868432 | 18.922409 | 17.1550233 | 17.8656579 | 20.4231681 | 20.7572421 | 18.632992 |
| ENSMUSG00 | C1qtnf4 | 1506 | 17.3294675 | 15.4619609 | 15.7711842 | 15.9130872 | 15.1171314 | 14.4446547 | 24.6930715 | 21.0235734 | 24.2579279 | 23.7945116 | 24.9343409 | 21.5701963 | 28.3019764 | 31.9804241 | 30.0120231 | 27.8662057 | 22.6242217 | 29.5500447 |
| ENSMUSG00 | Iqcc | 3583 | 6.6907394 | 7.77269873 | 7.80348885 | 5.36610681 | 5.89726891 | 6.37560987 | 10.3921514 | 10.1263696 | 9.58681388 | 8.79058604 | 9.48032249 | 8.31860559 | 9.2247794 | 7.66074676 | 7.43996837 | 6.42273376 | 6.95285877 | 7.63034497 |
| ENSMUSG00 | Iqsec3 | 17189 | 0 | 0 | 0 | 0 | 0 | 0 | 0 | 0 | 0 | 0 | 0 | 0 | 0 | 0 | 0 | 0 | 0 | 0 |
| ENSMUSG00 | S100g | 477 | 528.685974 | 514.708659 | 560.301374 | 429.916689 | 554.072102 | 570.220031 | 129.638724 | 222.108437 | 282.088778 | 200.860019 | 214.336292 | 361.57316 | 123.691767 | 100.093337 | 100.283032 | 247.403883 | 138.601388 | 117.052921 |
| ENSMUSG00 | Chi3l3 | 2005 | 0 | 0 | 0 | 0 | 0 | 0 | 0 | 0 | 0 | 0 | 0 | 0 | 0 | 0 | 0 | 0.0954878 | 0 | 0 |
| ENSMUSG00 | Emi2 | 3517 | 16.5089765 | 14.9751737 | 14.0843034 | 16.4522395 | 10.1409597 | 17.1891768 | 15.2417587 | 16.137023 | 11.0548965 | 18.6216454 | 13.8828569 | 11.8868158 | 11.4037102 | 12.8094298 | 12.5350198 | 9.68969024 | 14.5044928 | 14.1863121 |
| ENSMUSG00 | Agbl2 | 5788 | 0.19827916 | 0.14964327 | 0.25072666 | 0.14956113 | 0.26610601 | 0.18834833 | 0.13078833 | 0.07258404 | 0.02134754 | 0.20365715 | 0.04952484 | 0 | 0 | 0.04814522 | 0.07118444 | 0.11246377 | 0.0728385 | 0.00950399 |
| ENSMUSG00 | Tex264 | 2649 | 42.9063196 | 43.8135611 | 43.2421744 | 43.2907318 | 42.6627998 | 40.6298744 | 44.9015005 | 41.0471007 | 43.8141191 | 43.377223 | 47.9372743 | 44.0794134 | 45.4583654 | 45.0765378 | 49.2583343 | 55.188199 | 44.2148597 | 45.0621977 |
| ENSMUSG00 | Fam116a | 5690 | 15.8217861 | 13.547634 | 13.3898612 | 13.5241835 | 16.3344015 | 13.6465964 | 14.6843921 | 12.827009 | 15.4612298 | 15.4959253 | 15.5835365 | 15.7539462 | 13.2476572 | 17.5736591 | 13.2945618 | 15.2624073 | 15.101504 | 14.0664763 |
| ENSMUSG00 | Hlcs | 8367 | 3.81007315 | 4.67025516 | 4.83330601 | 5.32552903 | 4.52728863 | 5.0399637 | 4.61987076 | 5.97968772 | 4.69113775 | 6.60459235 | 4.64788051 | 6.22397672 | 3.50223005 | 4.58501448 | 4.30675599 | 5.03859911 | 5.44181197 | 4.06306059 |
| ENSMUSG00 | 1700123O20l | 1887 | 35.7250711 | 34.33682 | 34.6074541 | 32.9333675 | 30.8381005 | 31.0393911 | 35.6036058 | 28.072529 | 29.7276282 | 33.007991 | 34.0273154 | 32.3594779 | 40.077281 | 39.2079657 | 39.7386496 | 44.6825212 | 34.3047777 | 34.9236406 |
| ENSMUSG00 | Snrpd2 | 610 | 56.0928962 | 47.2933875 | 50.1181495 | 43.245647 | 48.0530189 | 40.2108248 | 50.8029585 | 44.077789 | 45.9803561 | 44.8318162 | 45.6603292 | 38.5219573 | 55.3973169 | 51.3930475 | 46.3348645 | 48.4595884 | 40.9651567 | 61.8627003 |
| ENSMUSG00 | Tmem146 | 2543 | 1.18673515 | 1.17898564 | 1.7880896 | 2.4366111 | 2.32804822 | 1.85116386 | 1.89771658 | 1.41175219 | 1.79775983 | 1.13102355 | 0.97691618 | 1.3169248 | 1.18334524 | 1.62545174 | 1.53918514 | 1.27986694 | 1.25091728 | 1.66563162 |
| ENSMUSG00 | Zmynd15 | 3995 | 0.36174618 | 0.52533475 | 0.36325555 | 0.43337164 | 0.38553731 | 0.28528482 | 0.01184297 | 0.28680152 | 0.42399967 | 0.10622186 | 0.23917376 | 0.19559932 | 0.07409049 | 0 | 0.16501249 | 0.11501559 | 0.47008474 | 0.24785079 |
| ENSMUSG00 | Gpr161 | 3039 | 0.1678388 | 0.05480897 | 0.01591758 | 0.01499212 | 0.0791904 | 0.09783363 | 0 | 0 | 0.27105308 | 0.06206081 | 0.01572062 | 0.09183228 | 0 | 0.13754419 | 0 | 0 | 0 | 0 |
| ENSMUSG00 | Gm11639 | 3751 | 0 | 0 | 0 | 0 | 0 | 0 | 0 | 0 | 0 | 0 | 0 | 0 | 0 | 0 | 0 | 0 | 0 | 0 |
| ENSMUSG00 | 4930579G18l | 962 | 0.39765757 | 0.8657197 | 0.55312769 | 0.14208226 | 0.90059732 | 0.61812142 | 0.59017895 | 0.59551563 | 0.68501482 | 0.39210564 | 1.0429045 | 0.46416351 | 0.46152521 | 0.38622942 | 0.25697436 | 0.87566878 | 0.99600564 | 0.45745619 |
| ENSMUSG00 | Six5 | 3004 | 1.71209264 | 1.57471071 | 1.72302522 | 1.88068226 | 1.53817066 | 1.96297447 | 1.95298675 | 1.9197912 | 2.09771535 | 2.05617248 | 1.59037811 | 2.21107319 | 2.18413615 | 1.88621127 | 0.80921831 | 1.81000874 | 0.84205579 | 1.02546937 |
| ENSMUSG00 | D4Ertd22e | 3322 | 15.3284771 | 14.7110222 | 15.5663167 | 15.9916248 | 12.5618235 | 13.4845488 | 16.3928036 | 13.4972548 | 13.3403645 | 14.7895945 | 11.4044377 | 12.0805122 | 12.8750067 | 13.2537612 | 10.3562008 | 10.8347896 | 10.9602595 | 9.72015221 |
| ENSMUSG00 | Tiprl | 1879 | 63.3186513 | 56.9634535 | 53.316435 | 47.9372989 | 56.1240259 | 56.5676499 | 54.1363809 | 54.920634 | 61.2865876 | 55.9083379 | 57.9706787 | 57.7761914 | 50.4871085 | 53.7110175 | 48.4156655 | 54.5320199 | 51.9720088 | 58.2001619 |
| ENSMUSG00 | Sft2d2 | 5518 | 16.5306466 | 20.3028748 | 23.4328464 | 22.1777755 | 20.6553815 | 19.6307132 | 20.9211556 | 22.1139409 | 17.7046784 | 20.8495317 | 20.2511105 | 19.9674009 | 15.3949912 | 16.5895789 | 17.4050067 | 14.801353 | 17.6767716 | 16.9872269 |
| ENSMUSG00 | Psme4 | 8332 | 8.70815356 | 10.1713878 | 9.5214342 | 9.4927956 | 8.94819368 | 8.65329181 | 8.09744121 | 8.31505256 | 9.76770945 | 8.46018979 | 9.14559035 | 8.80912102 | 9.1416803 | 9.0970656 | 8.56470173 | 9.63699036 | 8.86859103 | 8.78746052 |
| ENSMUSG00 | Plekhh2 | 6856 | 0.24178849 | 0.55877813 | 0.26811467 | 0.21265365 | 0.82840825 | 0.69385391 | 0.26913572 | 0.12255438 | 0.19223578 | 0.16505497 | 0.17420857 | 0.1791051 | 0.98577489 | 0.43355041 | 0.43869801 | 1.34597964 | 0.58696925 | 0.16843347 |
| ENSMUSG00 | Reps2 | 8278 | 18.6236136 | 20.8537591 | 23.6257771 | 21.0027718 | 23.7054666 | 24.6985478 | 27.1084847 | 27.5623837 | 29.7977824 | 23.5867434 | 25.7169866 | 24.6376457 | 17.1809516 | 15.24946 | 19.5555585 | 12.4382029 | 23.501363 | 19.8492633 |
| ENSMUSG00 | Dlk1 | 5889 | 0 | 0 | 0 | 0 | 0 | 0 | 0 | 0 | 0 | 0 | 0 | 0 | 0 | 0 | 0 | 0 | 0 | 0.02802298 |
| ENSMUSG00 | Erf | 3688 | 2.40878031 | 2.31239179 | 2.07240179 | 2.22369522 | 1.22678981 | 1.5989087 | 1.19308003 | 1.6569372 | 1.85383992 | 1.25291999 | 1.47677475 | 2.04314383 | 1.083483 | 1.64972219 | 1.5528786 | 2.12840958 | 1.28987081 | 1.64072715 |
| ENSMUSG00 | Bsdc1 | 5652 | 12.8598463 | 14.8705648 | 14.0190805 | 15.7915934 | 14.391888 | 14.7991887 | 14.0799586 | 14.1903831 | 17.2266319 | 15.5417127 | 15.1895967 | 12.9170005 | 14.4626658 | 15.3416686 | 14.3461893 | 14.4504362 | 16.4439614 | 13.0223257 |
| ENSMUSG00 | Crocc | 8380 | 0.99922669 | 1.35557243 | 1.65670683 | 1.73436434 | 1.02811627 | 1.04072567 | 1.71635494 | 1.46297866 | 1.09109863 | 1.99743433 | 1.56779398 | 1.1123169 | 1.36575236 | 1.69592968 | 1.07183045 | 1.22456835 | 1.36462187 | 2.45506037 |
| ENSMUSG00 | Ino80d | 14395 | 3.27462593 | 2.61273064 | 3.06136062 | 2.46241698 | 2.74179767 | 2.82617536 | 2.31058099 | 2.47010258 | 2.75530494 | 2.16837562 | 2.6915937 | 2.56298388 | 2.42632841 | 2.3423694 | 1.94630483 | 3.00047274 | 1.83976875 | 2.59473313 |
| ENSMUSG00 | Raph6a | 3164 | 0 | 0 | 0 | 0 | 0 | 0 | 0 | 0 | 0 | 0.01490225 | 0.12079636 | 0 | 0 | 0 | 0.06050981 | 0 | 0 | 0 |
| ENSMUSG00 | Begain | 2644 | 0.03215218 | 0.06299715 | 0.14636469 | 0.1378549 | 0.07281683 | 0.20615736 | 0 | 0.10111453 | 0.29569967 | 0.44582738 | 0.21683037 | 0.12666187 | 0 | 0 | 0.37399294 | 0.04344623 | 0 | 0.04161052 |
| ENSMUSG00 | Osbpl10 | 1862 | 12.6237177 | 13.1498267 | 10.4696739 | 15.0728267 | 11.5806132 | 13.998248 | 10.3925272 | 12.7581495 | 11.0181753 | 12.3574346 | 10.85328 | 10.6415468 | 9.21992869 | 10.9999154 | 15.0688827 | 12.4413628 | 12.4117922 | 13.471618 |
| ENSMUSG00 | Wdr25 | 2734 | 1.05718799 | 1.23065191 | 0.81389261 | 1.13319349 | 0.96827206 | 1.64934117 | 3.02842678 | 2.40274075 | 1.12984224 | 2.50067739 | 1.83481004 | 1.2249231 | 3.3298512 | 1.54587123 | 1.82347778 | 2.08679609 | 1.40183968 | 1.20722253 |
| ENSMUSG00 | Tmem205 | 781 | 106.235728 | 94.9908008 | 99.8439577 | 92.8721928 | 104.460449 | 101.453036 | 96.6244851 | 84.600328 | 99.723049 | 99.0708118 | 92.6748554 | 95.0510003 | 129.046232 | 145.397943 | 114.583584 | 105.066289 | 97.8032135 | 113.046884 |
| ENSMUSG00 | Gfer | 1772 | 18.5180564 | 20.5291651 | 15.3419436 | 15.8897999 | 15.7542414 | 16.0515062 | 19.5445152 | 16.488255 | 16.2700683 | 18.7325593 | 16.7428043 | 17.0407864 | 16.6759698 | 20.6534655 | 17.4153303 | 15.5366516 | 11.4849118 | 16.7014105 |
| ENSMUSG00 | Foxa3 | 3550 | 0.01197329 | 0.05630348 | 0 | 0 | 0.06779144 | 0 | 0 | 0.02151685 | 0 | 0 | 0 | 0.18867266 | 0 | 0.02616568 | 0 | 0 | 0 | 0 |
| ENSMUSG00 | Kcnd3 | 4994 | 0 | 0 | 0 | 0 | 0.02891385 | 0 | 0.10421295 | 0.04588595 | 0 | 0.13218059 | 0.01913294 | 0.02235309 | 0.57293777 | 0.78119811 | 0.58576501 | 0.04600393 | 0.08441916 | 0.04406016 |
| ENSMUSG00 | Ccr6 | 1880 | 0 | 0 | 0 | 0 | 0 | 0 | 0.57882533 | 0.14220575 | 0.26289265 | 0 | 0.86401521 | 0.14844591 | 0.62976916 | 0.22233872 | 0.24107293 | 0.24440813 | 0 | 0 |
| ENSMUSG00 | Kcnk18 | 3029 | 0 | 0 | 0 | 0 | 0 | 0 | 0 | 0 | 0 | 0 | 0 | 0 | 0 | 0 | 0 | 0 | 0 | 0 |
| ENSMUSG00 | Rnasek | 2176 | 296.149345 | 288.211648 | 299.577985 | 266.163601 | 291.423757 | 285.224259 | 416.682068 | 376.693631 | 379.536771 | 410.943514 | 399.127284 | 347.052351 | 484.930933 | 451.698631 | 438.504978 | 432.81056 | 431.118001 | 471.521075 |
| ENSMUSG00 | Atp1a3 | 6359 | 0 | 0 | 0 | 0 | 0 | 0 | 0 | 0.00600604 | 0 | 0 | 0.00751297 | 0 | 0 | 0 | 0.00647925 | 0 | 0.00602709 | 0 |
| ENSMUSG00 | Fbxw4 | 3344 | 2.22440365 | 3.2575707 | 2.92208527 | 2.84756527 | 2.24538878 | 3.11186425 | 2.90044832 | 3.14082257 | 2.68500611 | 4.34282793 | 3.04308198 | 2.38685399 | 2.90621642 | 4.5832981 | 3.06793882 | 2.31300918 | 2.91114815 | 2.43461358 |
| ENSMUSG00 | Slc19a2 | 4233 | 4.62907776 | 6.64998715 | 5.53101552 | 7.62041564 | 6.35618836 | 7.05889401 | 8.18163973 | 5.33231983 | 4.2033014 | 7.24924501 | 5.30456662 | 6.40831845 | 5.60563482 | 7.21951559 | 4.5260391 | 7.02853814 | 7.89523527 | 8.45994058 |
| ENSMUSG00 | 4930505A04l | 984 | 0 | 0.06770913 | 0.09832018 | 0 | 0 | 0.05035847 | 0 | 0.07762683 | 0 | 0 | 0 | 0.1701697 | 0 | 0 | 0 | 0 | 0.07789898 | 0 |
| ENSMUSG00 | S100pbp | 11032 | 1.72609852 | 1.57022344 | 1.65308384 | 1.61892036 | 1.63610147 | 1.71583976 | 1.85270824 | 1.87292329 | 1.50081394 | 1.65830972 | 1.95309158 | 1.49759211 | 1.17606003 | 1.84816483 | 1.63954705 | 1.33281302 | 1.47996759 | 1.8399529 |
| ENSMUSG00 | Rfx3 | 12855 | 1.30937765 | 0.64008439 | 0.98967238 | 0.66631477 | 0.87989125 | 0.92128381 | 0.52262936 | 0.96557999 | 0.78496277 | 0.48416125 | 1.00344137 | 0.92917556 | 0.64471117 | 0.44077625 | 0.84614631 | 0.85785258 | 0.62013767 | 0.62476309 |
| ENSMUSG00 | Padi6 | 2514 | 0 | 0 | 0 | 0.07249172 | 0 | 0 | 0 | 0 | 0.06553145 | 0.13128676 | 0 | 0 | 0 | 0 | 0.01638884 | 0.07615475 | 0 | 0 |
| ENSMUSG00 | Ulk4 | 7937 | 1.31740876 | 1.41864158 | 1.37130456 | 1.61303433 | 2.11642199 | 1.39648965 | 1.04914092 | 2.05951219 | 1.28172728 | 1.6396112 | 1.16171941 | 1.125174 | 1.00068523 | 1.26394392 | 1.23028458 | 0.97451212 | 1.376212 | 1.46931216 |
| ENSMUSG00 | Slc16a11 | 2443 | 17.1029832 | 22.1995054 | 19.0880406 | 21.7081685 | 20.4703515 | 19.6142011 | 19.9863629 | 17.8377123 | 21.8998297 | 26.7116551 | 21.8047805 | 21.2707253 | 17.9921156 | 16.7487683 | 16.2580024 | 19.6390221 | 16.958955 | 15.6043025 |
| ENSMUSG00 | Arhgef1 | 6479 | 11.6120021 | 15.9134755 | 13.1255852 | 12.5734137 | 12.807445 | 15.4570265 | 17.9859744 | 18.1441912 | 19.0453467 | 19.4455896 | 20.6171915 | 17.5226204 | 19.7434267 | 23.6915741 | 23.4020259 | 20.460269 | 21.2956735 | 22.6098551 |
| ENSMUSG00 | Tet2 | 10644 | 4.27288034 | 3.83392479 | 3.68118748 | 4.25048074 | 3.41861465 | 3.0819156 | 3.79603794 | 5.43606629 | 4.3957048 | 4.31018736 | 5.09986817 | 4.50447709 | 5.14916949 | 3.87034062 | 4.07602765 | 4.21973717 | 5.23547874 | 4.27400707 |
| ENSMUSG00 | Rcc2 | 4139 | 19.9432355 | 21.7471467 | 18.652852 | 22.488817 | 19.8621226 | 21.8491758 | 13.5342382 | 17.4214196 | 16.1701107 | 17.7484406 | 14.5090898 | 15.2249053 | 13.6351083 | 14.9577116 | 13.418627 | 14.6076031 | 11.0839762 | 14.8719958 |
| ENSMUSG00 | Mgl2 | 1998 | 0 | 0 | 0.16947683 | 0 | 0.0481801 | 0 | 0 | 0.15292253 | 0 | 0 | 0 | 0 | 0 | 0 | 0 | 0 | 0 | 0 |
| ENSMUSG00 | Rps19 | 2568 | 26.3836646 | 22.0010392 | 24.7330395 | 20.1902136 | 24.7969373 | 22.6537808 | 33.5868433 | 26.4580545 | 29.1898204 | 27.8350725 | 28.4268445 | 26.277701 | 46.4311698 | 49.5367427 | 36.532688 | 31.5360727 | 27.9387959 | 38.4721011 |
| ENSMUSG00 | Cables1 | 4646 | 0.34765318 | 0.3441711 | 0.19782546 | 0.27458227 | 0.24863671 | 0.30930462 | 0.79431532 | 0.17263032 | 0.70919528 | 0.33490599 | 0.52443454 | 0.37242478 | 1.35912299 | 0.18993492 | 1.00210414 | 0.78295474 | 1.0146652 | 0.36704311 |
| ENSMUSG00 | Asgr2 | 1544 | 0 | 0 | 0 | 0 | 0 | 0 | 0 | 0 | 0 | 0 | 0 | 0 | 0 | 0 | 0 | 0 | 0 | 0 |
| ENSMUSG00 | Arhgef10l | 5604 | 17.9076946 | 23.3618262 | 18.5328283 | 22.2520661 | 20.1494456 | 27.6943543 | 13.6264568 | 15.6613385 | 15.2207841 | 18.089581 | 15.9420365 | 16.6929069 | 15.9333994 | 13.923281 | 17.6966711 | 15.5512888 | 15.5589446 | 14.1645505 |
| ENSMUSG00 | Slc22a2 | 2137 | 0 | 0 | 0 | 0 | 0 | 0 | 0 | 0 | 0 | 0 | 0 | 0 | 0 | 0 | 0 | 0 | 0 | 0 |
| ENSMUSG00 | D630013G24 | 8385 | 8.1157766 | 5.20451895 | 5.87866755 | 5.25432676 | 5.34990493 | 4.01858782 | 4.62123841 | 4.57306762 | 5.70274834 | 4.35799583 | 4.80313536 | 5.55626813 | 4.67726586 | 3.11842829 | 3.87201265 | 4.14643968 | 3.69321979 | 2.9521882 |
| ENSMUSG00 | Igsf21 | 1969 | 0.17269752 | 0 | 0 | 0.04627836 | 0.12222429 | 0 | 0.1682015 | 0 | 0 | 0 | 0 | 0 | 0 | 0.47175305 | 0 | 0.01946485 | 0.1955631 | 0 |
| ENSMUSG00 | Gm11992 | 2393 | 0.08881148 | 0 | 0.24257516 | 0 | 0 | 0.22778106 | 0.17794154 | 0 | 0 | 0.33496112 | 0 | 0.27969467 | 0.22676602 | 1.10627371 | 0 | 0.11200763 | 0.56056013 | 0.02298751 |
| ENSMUSG00 | Sun3 | 1288 | 0 | 0 | 0 | 0 | 0 | 0 | 0 | 0 | 0 | 0 | 0 | 0 | 0 | 0 | 0 | 0 | 0 | 0 |
| ENSMUSG00 | Mill2 | 1115 | 1.60109183 | 1.16520437 | 1.60522029 | 0.531205 | 1.46769991 | 0.84439636 | 0.21216448 | 0.51379914 | 0.81264874 | 0.08457525 | 0.47132246 | 0.05005889 | 0.75214582 | 0.24992334 | 0 | 0.13736541 | 0.37810696 | 0 |
| ENSMUSG00 | Sh3kbp1 | 8958 | 0.03321458 | 0.06321938 | 0.07560052 | 0.04577466 | 0.10746132 | 0.04425339 | 0.12147707 | 0.06821595 | 0.22528905 | 0.00526353 | 0.37865841 | 0.01869245 | 0.00550702 | 0.3473715 | 0.18397658 | 0.11113594 | 0 | 0.22720886 |
| ENSMUSG00 | Abhd4 | 2384 | 294.790512 | 338.257228 | 304.282492 | 316.825081 | 324.002144 | 321.219775 | 274.528644 | 291.072886 | 262.425828 | 321.669059 | 326.128219 | 277.696969 | 294.853393 | 280.651646 | 293.18087 | 309.007129 | 337.107433 | 320.409587 |
| ENSMUSG00 | Npnt | 6841 | 68.4084181 | 83.2067568 | 60.7692033 | 73.3332348 | 75.7615194 | 75.8755861 | 2.88399168 | 4.24856282 | 2.80557175 | 2.30205158 | 2.13698835 | 4.22635773 | 1.69463537 | 3.53032092 | 2.74636888 | 3.36952867 | 2.14573479 | 2.30779325 |
| ENSMUSG00 | Trim62 | 5997 | 0.76547591 | 0.74991508 | 0.86309284 | 1.53465582 | 0.73036603 | 0.57014151 | 0.90727999 | 1.30555986 | 0.80353883 | 0.39311909 | 0.89224535 | 0.79111746 | 2.40292098 | 1.44823151 | 1.23666501 | 2.03680236 | 1.53381879 | 0.96314094 |
| ENSMUSG00 | Iqcf4 | 995 | 0 | 0 | 0 | 0.04578999 | 0 | 0 | 0 | 0 | 0 | 0 | 0 | 0 | 0 | 0 | 0 | 0 | 0 | 0 |
| ENSMUSG00 | Cmtm8 | 1060 | 91.3861285 | 80.2652187 | 84.0604178 | 78.0125429 | 79.2360442 | 82.3229847 | 97.6605108 | 82.3658791 | 79.4977465 | 89.0970364 | 85.093511 | 76.6148953 | 81.3045844 | 67.5191874 | 71.8306371 | 70.729254 | 69.7104721 | 74.3660848 |
| ENSMUSG00 | Nrg3 | 8915 | 0 | 0 | 0 | 0 | 0 | 0 | 0 | 0 | 0 | 0 | 0 | 0 | 0 | 0 | 0 | 0 | 0 | 0 |
| ENSMUSG00 | Mtap7d2 | 4607 | 0.36904863 | 0.20246604 | 0.09450006 | 0.04944763 | 0.15671345 | 0.29041107 | 3.30685535 | 1.64143601 | 1.40357781 | 1.05416159 | 0.518504 | 1.36904056 | 5.9964996 | 7.94398543 | 4.45373884 | 1.47942872 | 0.89846756 | 4.32239996 |
| ENSMUSG00 | Iffo2 | 6201 | 0.67174766 | 0.90789858 | 0.62407393 | 0.72738968 | 0.82276793 | 0.56736721 | 0.25941479 | 0.61590715 | 0.67083343 | 0.43341237 | 0.96304948 | 0.4590548 | 0.54892796 | 0.51679439 | 0.45181516 | 0.48164286 | 0.5995244 | 0.33709822 |
| ENSMUSG00 | Ghitm | 2843 | 165.042082 | 167.490155 | 165.657647 | 168.910801 | 182.335849 | 185.818393 | 179.415403 | 176.923651 | 171.424566 | 185.020487 | 185.403136 | 162.519189 | 154.103678 | 150.179562 | 165.835176 | 163.600425 | 192.682997 | 154.598228 |
| ENSMUSG00 | Dpcd | 1339 | 28.5378292 | 24.3564776 | 26.011159 | 18.5102376 | 26.7080054 | 25.9420958 | 25.7233983 | 23.3889245 | 24.54581 | 25.1071333 | 21.8715829 | 24.8023295 | 25.3843846 | 26.9854369 | 28.2164502 | 28.5392279 | 25.2741749 | 27.6894234 |
| ENSMUSG00 | Irgq | 6120 | 5.26452997 | 5.78621022 | 6.07830798 | 6.40236898 | 5.8513319 | 6.05644522 | 4.70034457 | 5.39810916 | 6.17797736 | 5.58566338 | 3.13815904 | 3.85517245 | 3.78856227 | 1.73785883 | 3.17090406 | 4.41718445 | 3.89525436 | 4.79981421 |
| ENSMUSG00 | Fam117b | 5681 | 10.1231303 | 13.1996699 | 13.623948 | 13.1606537 | 12.4968469 | 11.7492575 | 11.5595843 | 13.4322162 | 11.4620457 | 12.6736708 | 13.749702 | 15.0322063 | 6.61695493 | 12.0749722 | 10.8062521 | 11.2492561 | 12.1435195 | 12.1037464 |
| ENSMUSG00 | Lrit1 | 4321 | 0 | 0 | 0 | 0 | 0 | 0 | 0 | 0 | 0 | 0 | 0 | 0 | 0 | 0 | 0 | 0 | 0 | 0 |
| ENSMUSG00 | Ramp3 | 1438 | 6.62111226 | 7.11200176 | 7.40067919 | 5.51294973 | 6.22568693 | 5.85811176 | 31.1908342 | 28.5247846 | 28.1832625 | 14.1648844 | 19.1033388 | 25.6565737 | 9.12537552 | 7.46076766 | 6.5613048 | 2.39649142 | 5.70363528 | 7.4977642 |
| ENSMUSG00 | Slc7a13 | 2137 | 0 | 0 | 0 | 0 | 0 | 0 | 0 | 0 | 0 | 0.02206397 | 0 | 0 | 0 | 0.10866656 | 0 | 0 | 0.10760776 | 0 |
| ENSMUSG00 | Wdr43 | 3432 | 16.9921624 | 16.1614128 | 16.4909763 | 15.5720003 | 14.6555537 | 16.2143706 | 13.6341024 | 15.5128814 | 15.5049471 | 17.1182332 | 17.6232801 | 16.214515 | 10.7662022 | 16.9970203 | 18.9560771 | 17.5498617 | 15.9022864 | 17.3426146 |
| ENSMUSG00 | Wwp1 | 8568 | 12.2435544 | 11.9557831 | 12.494237 | 12.5867171 | 12.280157 | 13.1689513 | 10.7071994 | 10.1275838 | 10.6859973 | 11.5455387 | 11.8433721 | 12.9050919 | 12.9663256 | 11.4375612 | 12.9548309 | 15.8426941 | 16.2734673 | 14.4264079 |
| ENSMUSG00 | Msln | 2612 | 0 | 0 | 0 | 0 | 0 | 0 | 0 | 0 | 0 | 0 | 0 | 0 | 0 | 0 | 0 | 0 | 0 | 0 |
| ENSMUSG00 | Pif1 | 4050 | 0 | 0.06580325 | 0.10749673 | 0 | 0 | 0 | 0 | 0 | 0 | 0.2033992 | 0 | 0 | 0 | 0 | 0.03440303 | 0.12207868 | 0.02836341 | 0.12302268 | 0 |
| ENSMUSG00 | 4930596D02l | 1159 | 0 | 0 | 0 | 0 | 0 | 0 | 0 | 0 | 0 | 0 | 0 | 0 | 0 | 0 | 0 | 0 | 0 | 0 |
| ENSMUSG00 | Nacad | 4901 | 1.89933351 | 1.71288499 | 1.51013062 | 2.65873468 | 2.08201751 | 2.89167146 | 1.47701283 | 1.83130271 | 0.83196594 | 1.08713109 | 1.11127224 | 1.7082941 | 0.61400672 | 0.19900548 | 0.92474416 | 0.62502522 | 0.83675041 | 1.14485388 |
| ENSMUSG00 | Fzd7 | 8930 | 0 | 0 | 0 | 0 | 0.10240835 | 0.04439215 | 0 | 0.11547535 | 0 | 0.0316802 | 0 | 0.08750496 | 0.00552429 | 0.13522355 | 0.01845534 | 0.22296882 | 0.11588018 | 0 |
| ENSMUSG00 | Grid1 | 6077 | 0 | 0 | 0 | 0 | 0.02376103 | 0 | 0.03114213 | 0.05656272 | 0.01355488 | 0 | 0 | 0.0459237 | 0 | 0 | 0 | 0 | 0 | 0 |
| ENSMUSG00 | Rwdd2b | 2004 | 6.61757232 | 5.53552584 | 4.97252857 | 4.68342073 | 5.40403335 | 4.50029816 | 6.70499043 | 6.78467827 | 4.82975609 | 3.95275357 | 4.88715893 | 7.49222217 | 4.89872831 | 5.16817939 | 3.76241753 | 5.40739633 | 3.86322953 | 3.43120676 |
| ENSMUSG00 | Ostc | 1054 | 91.2207857 | 78.7941606 | 81.8770191 | 81.2231521 | 81.7877375 | 79.3595961 | 94.5355805 | 89.7920049 | 85.6554483 | 92.8251511 | 84.3087503 | 86.794944 | 98.0085179 | 92.7119164 | 83.4976503 | 84.973179 | 85.5250952 | 83.9750926 |
| ENSMUSG00 | Tspyl2 | 3511 | 14.4670138 | 18.2646885 | 17.1532456 | 19.2184297 | 18.4247523 | 16.3717377 | 16.5749345 | 19.4388559 | 14.3231945 | 21.7153766 | 20.6557639 | 19.9194034 | 23.5348794 | 25.8213781 | 24.8429774 | 25.7050013 | 22.0722792 | 28.7971342 |
| ENSMUSG00 | Elmo1 | 3811 | 8.1195928 | 9.52795977 | 10.3068241 | 8.69138789 | 8.49981229 | 8.34763972 | 8.70275201 | 10.8033088 | 9.16456714 | 12.2485427 | 10.743411 | 9.87136095 | 13.488286 | 12.2112242 | 12.6599375 | 11.8257793 | 13.184407 | 13.5249365 |
| ENSMUSG00 | Iqsec2 | 7869 | 0.85345251 | 0.98639018 | 0.78071399 | 0.63689349 | 0.80128123 | 0.58564039 | 0.94998135 | 0.62610496 | 0.68565683 | 0.71304279 | 0.75891089 | 0.53198306 | 1.04067839 | 0.63743315 | 0.72255874 | 1.06078948 | 0.84747435 | 0.41943657 |
| ENSMUSG00 | Ccdc8 | 4807 | 0.65433389 | 0.81774943 | 0.58366232 | 0.72981078 | 0.52066986 | 0.54634801 | 0.80708127 | 1.05670679 | 0.25704087 | 0.60814304 | 0.4869925 | 0.84762708 | 0.01026252 | 0.08695585 | 0.10285389 | 0.10355271 | 0.15946036 | 0.08010479 |
| ENSMUSG00 | Pde9a | 2770 | 0.81327593 | 0.56523683 | 0.69853473 | 1.44742668 | 1.09469718 | 0.78711923 | 1.622637 | 0.77212077 | 1.04081448 | 1.0553587 | 1.41427675 | 1.02765299 | 1.35351109 | 0.31856955 | 0.40160361 | 1.71409217 | 1.85405191 | 0.93336751 |
| ENSMUSG00 | Nbl1 | 2423 | 117.832137 | 91.8957429 | 114.495335 | 118.876154 | 87.8611219 | 117.850019 | 140.551656 | 110.27406 | 126.092272 | 147.387299 | 104.856471 | 173.228965 | 115.786469 | 95.6292002 | 134.011334 | 127.387778 | 92.3437562 | 130.201084 |
| ENSMUSG00 | 8430410K20l | 2738 | 21.0197255 | 19.0898214 | 18.6568474 | 21.3494594 | 20.9895971 | 21.5729943 | 24.9005005 | 21.8023101 | 22.6992156 | 18.5640826 | 18.3736418 | 17.1850269 | 14.2878782 | 17.8278801 | 23.7006759 | 20.6277012 | 22.956584 | 18.0015192 |
| ENSMUSG00 | H2afv | 2837 | 32.7216441 | 29.0269561 | 28.4921287 | 30.304438 | 30.0294342 | 29.5185477 | 28.534365 | 26.7629524 | 27.6560848 | 29.267673 | 29.1162859 | 29.2161646 | 35.1948539 | 30.4661426 | 29.1910918 | 30.5299097 | 25.9786637 | 30.3063919 |
| ENSMUSG00 | Zfp598 | 3274 | 11.1391084 | 12.9731033 | 13.1793491 | 12.4270045 | 11.1288535 | 14.2271135 | 9.24866051 | 10.347178 | 8.11402036 | 12.3709389 | 12.286657 | 11.3199756 | 11.3309718 | 10.6818564 | 12.307612 | 11.0404195 | 9.54060231 | 11.1563979 |
| ENSMUSG00 | N4bp2l1 | 1901 | 4.1141254 | 4.41601754 | 3.71516856 | 4.36197268 | 2.60788434 | 2.91946668 | 3.6585817 | 3.09396632 | 2.68654811 | 2.90196333 | 4.82524567 | 3.6407902 | 1.81653562 | 4.666397 | 0.45514607 | 4.59245581 | 2.64110726 | 1.76515282 |
| ENSMUSG00 | Smc1a | 5655 | 18.174627 | 18.5974902 | 16.8259477 | 17.9424307 | 16.3333619 | 19.0937942 | 14.9677069 | 16.1009173 | 14.20953 | 12.0565724 | 15.5279175 | 15.1704109 | 10.8521494 | 13.2145953 | 11.6282375 | 14.2734926 | 13.3786225 | 12.4706764 |
| ENSMUSG00 | Cyyr1 | 8887 | 0.00478285 | 0 | 0.02177272 | 0 | 0 | 0 | 0.01597142 | 0 | 0.05097915 | 0.01061116 | 0 | 0.01884179 | 0 | 0 | 0 | 0 | 0 | 0 |
| ENSMUSG00 | Ripk2 | 3954 | 3.60122255 | 2.74658663 | 2.37341044 | 2.54653283 | 3.85883151 | 3.70956225 | 2.28546325 | 2.55002383 | 2.20828056 | 1.45482696 | 3.62480717 | 2.51269295 | 2.08356859 | 2.98350991 | 3.1469069 | 3.77676704 | 3.00484371 | 3.14417251 |
| ENSMUSG00 | Txndc3 | 2163 | 0 | 0 | 0 | 0 | 0 | 0 | 0 | 0 | 0 | 0 | 0 | 0 | 0 | 0 | 0 | 0 | 0 | 0 |
| ENSMUSG00 | Pnmal1 | 1683 | 0.1010224 | 0 | 0 | 0 | 0 | 0 | 0 | 0 | 0 | 0 | 0 | 0 | 0 | 0 | 0 | 0 | 0.13663564 | 0 |
| ENSMUSG00 | Tmco4 | 3682 | 22.9495627 | 23.4149295 | 24.5152112 | 26.4555083 | 24.2228285 | 27.5891102 | 28.449286 | 28.0374969 | 27.8752844 | 30.2215263 | 37.0703575 | 25.7097674 | 25.0411063 | 21.7336129 | 31.7572014 | 22.1819298 | 30.1863827 | 24.4717319 |
| ENSMUSG00 | Dnahc7b | 12292 | 0.2731784 | 0.2222305 | 0.06296587 | 0.11490338 | 0.18795414 | 0.16931458 | 0.13856626 | 0.13981924 | 0.18093653 | 0.21864547 | 0.11660013 | 0.25428548 | 0.25685346 | 0.16624958 | 0.20446591 | 0.16821451 | 0.23696702 | 0.14768146 |
| ENSMUSG00 | Brca2 | 10697 | 0.6834524 | 0.63218823 | 0.66023515 | 0.57073758 | 0.42745936 | 0.67169733 | 0.84479028 | 0.89259611 | 0.83551219 | 1.08873738 | 0.65206543 | 0.89225748 | 0.53035155 | 0.55574864 | 0.80885545 | 0.91278909 | 0.82406734 | 0.7405171 |
| ENSMUSG00 | Osgin2 | 7238 | 7.86328139 | 7.60332982 | 8.90211971 | 7.9753859 | 7.30823212 | 8.6282012 | 5.47122303 | 6.89130665 | 5.66755591 | 6.70324311 | 5.88111191 | 7.10226931 | 8.59457691 | 6.14720029 | 7.001646 | 9.2843349 | 9.62128765 | 7.80524352 |
| ENSMUSG00 | Otud3 | 2240 | 3.88998263 | 4.65488209 | 4.42704184 | 4.08829014 | 3.86774387 | 5.4861955 | 4.81575482 | 3.6828388 | 3.56704645 | 3.80994522 | 2.94327869 | 4.43535176 | 3.72191697 | 2.71614975 | 4.43284597 | 3.57265042 | 4.63679753 | 4.10112537 |
| ENSMUSG00 | Zmiz2 | 4637 | 13.603123 | 17.8597919 | 16.7434799 | 18.1084763 | 15.9540148 | 16.9806542 | 15.2437227 | 15.4021549 | 17.1869545 | 18.0183408 | 16.7526596 | 16.9842355 | 16.5007125 | 19.4510282 | 16.0292445 | 15.9372164 | 15.1420694 | 16.252421 |
| ENSMUSG00 | Spem1 | 1038 | 0.12284733 | 0 | 0 | 0 | 0 | 0 | 0.13674185 | 0 | 0.67453831 | 0 | 0 | 0.10754463 | 0 | 0.26846293 | 0.03969321 | 0.18444416 | 0 | 0 |
| ENSMUSG00 | Lonp1 | 2926 | 29.7797713 | 36.4096499 | 30.7004941 | 31.6404792 | 32.7514217 | 34.3109495 | 36.834683 | 35.2555294 | 32.3749099 | 30.8107124 | 34.3535587 | 29.6818766 | 31.7133754 | 33.650535 | 35.4001065 | 29.051657 | 29.3930999 | 38.5214148 |
| ENSMUSG00 | Hectd2 | 9293 | 0.07318227 | 0.31904095 | 0.07808059 | 0.19610909 | 0.18645747 | 0.1493034 | 0.07636825 | 0.02876862 | 0.03545595 | 0.07610681 | 0.07197347 | 0.01801861 | 0.0053085 | 0.12994149 | 0.00886722 | 0 | 0.13609898 | 0.05327472 |
| ENSMUSG00 | Prkd2 | 3785 | 7.16467696 | 6.44254652 | 6.2240183 | 7.54736454 | 6.24379776 | 8.13005219 | 5.5750197 | 5.21676906 | 6.22422378 | 6.36565637 | 5.47802694 | 6.14930814 | 5.12217794 | 5.32542508 | 6.0849854 | 7.06125336 | 6.70330477 | 6.06684813 |
| ENSMUSG00 | Chrnb1 | 2302 | 24.8900857 | 26.0193674 | 24.8381898 | 22.6815625 | 25.9059311 | 26.8213319 | 15.6818306 | 20.6059791 | 21.774105 | 16.6113034 | 17.3500718 | 23.0827588 | 7.00761814 | 5.44739513 | 6.51492891 | 9.84710668 | 8.64089839 | 8.19640469 |
| ENSMUSG00 | Pla2g5 | 3145 | 0.33787898 | 0.72027879 | 1.30739271 | 0.86920911 | 0.65808354 | 0.92960613 | 0.34600687 | 0.06071924 | 0.43216376 | 0.49474506 | 0.12152613 | 0.07098971 | 0.17254407 | 0.11814076 | 0.02620131 | 0.06087537 | 0 | 0.26236458 |
| ENSMUSG00 | Rpusd1 | 1697 | 6.36200042 | 5.45726847 | 7.01230906 | 5.15481468 | 5.21878253 | 4.64283125 | 8.41981676 | 6.16659886 | 4.61133647 | 7.25181706 | 7.51674361 | 5.55854271 | 7.18030853 | 12.5347036 | 7.30799548 | 5.05426992 | 6.73025131 | 7.68247395 |
| ENSMUSG00 | Pla2g2d | 2238 | 0.01899248 | 0 | 0 | 0.10178964 | 0 | 0.0685661 | 0 | 0 | 0 | 0 | 0 | 0 | 0 | 0 | 0.18409989 | 0 | 0 | 0 |
| ENSMUSG00 | 2310036O22l | 888 | 153.028207 | 142.480149 | 137.22176 | 121.803963 | 133.067424 | 130.187531 | 136.343632 | 133.974475 | 138.308773 | 133.487464 | 135.14718 | 125.773807 | 154.162241 | 154.656719 | 137.941696 | 152.515629 | 133.667276 | 138.823658 |
| ENSMUSG00 | Map6d1 | 3270 | 0 | 0 | 0.08875877 | 0 | 0 | 0 | 0 | 0 | 0 | 0.02883835 | 0.08766047 | 0 | 0.12068971 | 0 | 0 | 0 | 0 | 0.03364471 |
| ENSMUSG00 | Yeats2 | 5430 | 5.46383294 | 6.37423519 | 5.14024439 | 5.81469682 | 5.15002719 | 4.27084334 | 5.17564112 | 4.70547278 | 6.46241463 | 5.95679235 | 4.61912582 | 4.63588652 | 4.64246956 | 6.33795005 | 3.83940834 | 3.54699365 | 4.06555194 | 4.14341154 |
| ENSMUSG00 | Clvs1 | 7710 | 0 | 0 | 0 | 0 | 0 | 0 | 0 | 0 | 0.00534196 | 0 | 0 | 0 | 0 | 0 | 0 | 0 | 0 | 0 |
| ENSMUSG00 | Arhgap11a | 7555 | 0.11252197 | 0.5158979 | 0.48021374 | 0.44626304 | 0.43321917 | 0.52471458 | 0.58866867 | 0.73806628 | 0.26167475 | 0.40566455 | 0.54383143 | 0.40633507 | 0.5811437 | 0.94056193 | 0.61079739 | 0.81091955 | 0.81674835 | 0.34221416 |
| ENSMUSG00 | Elovl6 | 5946 | 59.0826236 | 53.9191547 | 55.125974 | 54.8249701 | 54.5511325 | 55.9364191 | 51.5299207 | 55.2395998 | 52.5671827 | 45.0413714 | 51.5995263 | 49.6389546 | 38.9898871 | 40.5467974 | 39.0188297 | 41.2657605 | 42.7480661 | 40.1235277 |
| ENSMUSG00 | Arhgap12 | 6536 | 12.1090327 | 11.9469743 | 13.4921888 | 12.0733979 | 13.8887405 | 13.9878815 | 12.2407803 | 9.85195692 | 9.47113421 | 12.8264918 | 12.8428094 | 12.2203509 | 8.67988141 | 9.79902027 | 6.93416638 | 6.93050621 | 11.252779 | 7.05288119 |
| ENSMUSG00 | Phf8 | 8856 | 4.12284849 | 4.50641907 | 5.46769466 | 4.62954598 | 4.853411 | 4.20772986 | 4.45559781 | 5.89532679 | 5.45526005 | 4.8183589 | 5.4431948 | 5.36979952 | 4.64018599 | 6.19883822 | 4.56399349 | 4.43233837 | 5.59141549 | 4.42880455 |
| ENSMUSG00 | Ublcp1 | 3573 | 7.88719051 | 9.57524275 | 9.3281171 | 8.47973537 | 8.97169358 | 8.72338915 | 4.95240468 | 7.81378286 | 7.79235507 | 7.60112093 | 6.3111616 | 8.73242524 | 5.61939276 | 7.89016542 | 7.48385384 | 5.69054367 | 7.02595143 | 6.52780891 |
| ENSMUSG00 | Chd7 | 11292 | 3.05651815 | 3.84992268 | 3.56418502 | 3.16732375 | 2.87717397 | 3.24734528 | 5.06981416 | 5.18837631 | 3.9574362 | 4.51798272 | 3.16468907 | 4.14218254 | 4.1546806 | 4.24873736 | 2.84236734 | 2.96708125 | 2.73565314 | 3.13725334 |
| ENSMUSG00 | Vps41 | 3166 | 29.2139176 | 31.6609282 | 33.4764387 | 35.6026607 | 34.2822135 | 38.5184072 | 34.3861258 | 31.907424 | 29.1531844 | 33.1812277 | 30.6478019 | 33.6022277 | 30.7895984 | 27.6082667 | 34.6296084 | 28.506305 | 32.6366698 | 28.3038013 |
| ENSMUSG00 | Pldr | 3325 | 0.20453619 | 0.35066204 | 0.81471209 | 0.12332313 | 0.1302819 | 0.50670464 | 0.28458754 | 0.14932368 | 0.35922074 | 0.1134453 | 0.10057886 | 0.33573331 | 0 | 0.34920366 | 0.60718082 | 0.01151597 | 0.25358753 | 0.62867551 |
| ENSMUSG00 | Rbbp8 | 3900 | 15.639725 | 15.0933036 | 16.0872997 | 15.81786 | 15.365191 | 15.4248766 | 11.7068552 | 11.9473669 | 12.081378 | 12.1382836 | 11.8702397 | 11.0772623 | 11.6246236 | 11.3847558 | 14.4416736 | 13.3329859 | 13.9645302 | 12.5553603 |
| ENSMUSG00 | Mul1 | 4197 | 10.3098092 | 10.6042475 | 11.1453905 | 10.3128406 | 12.3512227 | 11.5233424 | 13.4148411 | 13.2585967 | 10.1686024 | 13.5936028 | 10.8481142 | 11.8892548 | 13.6112366 | 12.0619622 | 13.4000771 | 13.7579785 | 12.0357539 | 15.0465701 |
| ENSMUSG00 | Wnk3-ps | 10505 | 0.28727915 | 0.49152771 | 0.49731949 | 0.50743857 | 0.31614495 | 0.33962845 | 0.0540459 | 0.04726333 | 0.25484279 | 0.06283768 | 0.11824359 | 0.09563845 | 0.01408812 | 0.00442114 | 0 | 0.0984147 | 0.14593545 | 0 |
| ENSMUSG00 | Lamp3 | 3337 | 0 | 0 | 0 | 0 | 0 | 0 | 0 | 0.01144513 | 0 | 0 | 0 | 0 | 0 | 0 | 0.01234689 | 0 | 0 | 0 |
| ENSMUSG00 | Kcnj1 | 3263 | 0 | 0 | 0 | 0 | 0 | 0 | 0 | 0 | 0 | 0 | 0 | 0 | 0 | 0 | 0 | 0 | 0 | 0 |
| ENSMUSG00 | Tmco5b | 1225 | 0 | 0 | 0 | 0 | 0 | 0 | 0 | 0 | 0 | 0 | 0 | 0 | 0 | 0 | 0 | 0 | 0 | 0 |
| ENSMUSG00 | Zfp236 | 11802 | 1.95922858 | 1.82907604 | 1.63950281 | 1.58664542 | 1.56606157 | 1.82222389 | 1.5955301 | 1.94489356 | 1.88797704 | 1.51016487 | 1.62326371 | 1.52284726 | 1.68870484 | 1.49933887 | 1.31264066 | 1.70980767 | 1.11387221 | 1.50550258 |
| ENSMUSG00 | Car8 | 3839 | 0 | 0 | 0 | 0 | 0 | 0.09035403 | 0.17253907 | 0 | 0 | 0 | 0 | 0 | 0 | 0 | 0 | 0.04991703 | 0 | 0 |
| ENSMUSG00 | Rusc1 | 3828 | 2.59827879 | 3.4548639 | 2.41362181 | 2.91600205 | 2.89193899 | 3.19736813 | 2.66968094 | 2.60402744 | 1.80755869 | 3.36262852 | 1.53508879 | 1.77886908 | 9.96174819 | 9.68978517 | 9.01956736 | 10.5329179 | 7.03850228 | 8.4640449 |
| ENSMUSG00 | Uspl1 | 6884 | 6.32267581 | 7.19585615 | 5.58642601 | 6.26100348 | 6.46746505 | 6.19769082 | 4.7353916 | 5.18737598 | 6.16840467 | 5.96575574 | 5.73243982 | 4.88103267 | 4.47169035 | 5.35011059 | 5.21902335 | 5.25077475 | 5.85138554 | 5.90524838 |
| ENSMUSG00 | Dmxl2 | 13961 | 2.82535659 | 2.52453564 | 2.16903016 | 2.65318595 | 2.24783427 | 2.15446668 | 3.45669597 | 2.44840629 | 2.49874499 | 2.71536175 | 2.13876864 | 3.94599663 | 4.86216734 | 2.83435015 | 3.52962258 | 7.73711077 | 3.46625727 | 3.97565996 |
| ENSMUSG00 | Tox | 8013 | 0.05304527 | 0.00415736 | 0 | 0.02274356 | 0.09610767 | 0.08039255 | 0 | 0.04289675 | 0.00513996 | 0.01765283 | 0.04173527 | 0.04179383 | 0.11697322 | 0.14490231 | 0.05141839 | 0.02867136 | 0.02869809 | 0.05491986 |
| ENSMUSG00 | Ttc1 | 2622 | 33.7674606 | 29.2345422 | 27.7105426 | 28.3931146 | 30.1971832 | 30.1814323 | 33.5266809 | 32.0163617 | 30.5992878 | 33.9693663 | 34.346223 | 27.2479207 | 35.3902885 | 35.373319 | 31.5847168 | 36.8594558 | 38.3116797 | 36.2321614 |
| ENSMUSG00 | Sox15 | 859 | 1.48446482 | 1.35733562 | 0.84470658 | 1.00775302 | 1.5689102 | 1.32679031 | 0.33047273 | 1.15599621 | 2.01377609 | 0.60379247 | 1.3904237 | 2.01430214 | 1.72288419 | 0.81101433 | 1.67875955 | 1.20354645 | 0.26770405 | 1.0886552 |
| ENSMUSG00 | Gpr110 | 3837 | 0.11077711 | 0.2778245 | 0.28986383 | 0.47498528 | 0.96589995 | 0.37451897 | 0.29593544 | 0.21898172 | 0.06440425 | 0.66357518 | 0.56030053 | 0.10182685 | 45.0634272 | 49.4822594 | 44.8524626 | 82.4690069 | 58.3934146 | 53.0306713 |
| ENSMUSG00 | Cdk13 | 7067 | 8.49862937 | 10.5166361 | 9.16543896 | 8.95490174 | 9.20139081 | 10.0760263 | 8.71003188 | 11.3599023 | 8.89936459 | 12.0628939 | 8.89706229 | 9.4302959 | 9.29118182 | 8.97766568 | 10.0919617 | 8.84796404 | 8.57962384 | 10.1736101 |
| ENSMUSG00 | Katnal1 | 12096 | 2.21344527 | 2.04866865 | 2.32711148 | 2.43660073 | 2.16430538 | 1.83498303 | 2.32691717 | 2.18774466 | 2.09711471 | 1.96039046 | 1.84813032 | 2.72204008 | 2.09185597 | 1.80816376 | 1.95143755 | 1.43059634 | 1.69170462 | 1.7733139 |
| ENSMUSG00 | Cftr | 7843 | 0 | 0.2221997 | 0.2548481 | 0.14185786 | 0.18589231 | 0.13501241 | 0.28431378 | 0 | 0.21005491 | 0.04208274 | 0 | 0.00711662 | 0 | 0 | 0 | 0 | 0 | 0 |
| ENSMUSG00 | Gtf3c3 | 2987 | 6.48890535 | 7.31612273 | 7.660087 | 6.69611599 | 7.36400707 | 7.69751201 | 7.46041907 | 7.47993151 | 7.12870059 | 6.23519511 | 7.9651588 | 7.36236058 | 7.25032196 | 4.80456076 | 5.71056048 | 6.88384874 | 7.1340546 | 5.54326787 |
| ENSMUSG00 | Sntb2 | 4986 | 2.85584315 | 2.16473681 | 2.18292104 | 2.50375575 | 2.00791017 | 2.40508655 | 1.53723506 | 3.13291067 | 1.56122172 | 2.04262974 | 1.52350951 | 1.84708871 | 1.75125338 | 2.04927782 | 1.39652285 | 2.33460583 | 2.55201175 | 1.68800508 |
| ENSMUSG00 | Nlrp6-2 | 1902 | 0.02234762 | 0.03502933 | 0 | 0.02395428 | 0.02530595 | 0.02605296 | 0.07462568 | 0.02008013 | 0.02165432 | 0.07437019 | 0.02511827 | 0.02934577 | 0.05187373 | 0.04883711 | 0.06498668 | 0.16105408 | 0.04030105 | 0 |
| ENSMUSG00 | Slc7a1 | 7512 | 7.72924261 | 6.73175898 | 6.24631582 | 6.64735152 | 6.94555448 | 6.41837191 | 7.16744257 | 8.06860258 | 7.36335073 | 6.46501922 | 7.26291302 | 7.28159602 | 8.20228497 | 8.30330263 | 7.09180135 | 8.72140965 | 6.38261414 | 7.02991779 |
| ENSMUSG00 | Thoc6 | 2103 | 7.70065243 | 8.11041457 | 7.79773029 | 7.25770321 | 8.99469615 | 6.85679766 | 8.54912886 | 7.15540017 | 9.89024748 | 8.20597118 | 8.56450754 | 9.58129063 | 8.23371515 | 6.38247326 | 7.85631189 | 6.59115543 | 9.16696497 | 8.99616106 |
| ENSMUSG00 | Ak7 | 3075 | 0.02764564 | 0.43333846 | 0 | 0.222249 | 0 | 0 | 0.10770366 | 0.13662323 | 0 | 0.03066712 | 0 | 0 | 0.0320858 | 0 | 0 | 0.17433122 | 0 | 0.0894457 |
| ENSMUSG00 | Inhba | 1975 | 0 | 0 | 0 | 0 | 0 | 0.07526998 | 0 | 0 | 0 | 0 | 0 | 0 | 0 | 0.07054798 | 0 | 0 | 0 | 0 |
| ENSMUSG00 | Pcf11 | 6913 | 4.74055992 | 4.96826182 | 4.96820578 | 4.14550801 | 3.92686315 | 4.12879713 | 6.24174211 | 6.44735044 | 6.82767931 | 3.96275981 | 5.40431325 | 6.9355785 | 4.65987878 | 5.95919362 | 5.41168997 | 6.35868903 | 5.32232678 | 4.185562 |
| ENSMUSG00 | Atp1b2 | 5456 | 0 | 0.01831721 | 0 | 0.01670126 | 0.0264655 | 0.01816449 | 0.13874686 | 0.00700007 | 0 | 0 | 0.04378204 | 0 | 0 | 0 | 0.09061926 | 0.02105422 | 0.08429537 | 0 |
| ENSMUSG00 | Mup4 | 915 | 0.23226872 | 0.65533562 | 0.31720348 | 0.99586982 | 0.36822237 | 0.97480787 | 0 | 0 | 0 | 0 | 0 | 0 | 0 | 0 | 0 | 0 | 0 | 0 |
| ENSMUSG00 | Atg2b | 11438 | 1.54219689 | 2.45230433 | 1.92005445 | 1.68095477 | 3.00035509 | 2.38275953 | 2.99065106 | 2.59112644 | 2.94549483 | 1.81380569 | 2.41422679 | 2.24960836 | 2.65679848 | 1.90031762 | 2.88173148 | 2.57100774 | 3.05256657 | 2.65474969 |
| ENSMUSG00 | Ankrd42 | 4732 | 2.52408167 | 4.30139019 | 2.66810892 | 3.48543915 | 3.82451464 | 4.19920671 | 2.83956061 | 2.4051851 | 2.8548557 | 3.94583231 | 2.95817051 | 2.44164035 | 2.50204145 | 3.65114127 | 2.4553759 | 3.60087073 | 3.02107116 | 3.40610065 |
| ENSMUSG00 | Wrap53 | 2311 | 4.46938891 | 5.2470331 | 5.08644218 | 6.34818532 | 5.41510175 | 5.23187667 | 3.86936232 | 4.13158832 | 3.01190877 | 3.71329641 | 4.34129869 | 5.82067273 | 4.48277916 | 3.6174538 | 3.65483295 | 5.94821638 | 3.71488122 | 3.52286793 |
| ENSMUSG00 | Bdkrb1 | 1337 | 1.4306155 | 1.71921444 | 0.79597432 | 1.39715992 | 1.69199732 | 1.593693 | 0.21232317 | 0.25709171 | 0.61610345 | 0.56425673 | 0.50026135 | 1.12716746 | 0.11069241 | 0.03473754 | 0.73959412 | 0.14319599 | 0.74531318 | 0 |
| ENSMUSG00 | Rap1gap | 5566 | 0.35128245 | 0.3291788 | 0.41716304 | 0.22101117 | 0.30266211 | 0.38281846 | 0.79052805 | 0.87830174 | 0.69556818 | 0.77087927 | 0.96133585 | 0.1203356 | 0.66473119 | 0.93455584 | 0.54777491 | 1.13509706 | 1.52175918 | 0.89035837 |
| ENSMUSG00 | Tmem29 | 7734 | 0.23632306 | 0.26705449 | 0.33149691 | 0.26509529 | 0.21159625 | 0.17299248 | 0.10399735 | 0.23703586 | 0.29289609 | 0.22557228 | 0.19767244 | 0.27424298 | 0.36357891 | 0.33629026 | 0.36225831 | 0.17328307 | 0.34688916 | 0.56189804 |
| ENSMUSG00 | Rgl2 | 4709 | 11.4995953 | 12.7478945 | 13.3543405 | 14.2614099 | 12.1326382 | 14.4059657 | 12.4887789 | 12.3198682 | 10.8979398 | 13.3171449 | 9.56716624 | 10.9995615 | 10.9789444 | 10.720901 | 12.0568579 | 9.48117397 | 10.8166593 | 12.3241893 |
| ENSMUSG00 | Ssr2 | 1162 | 108.4577 | 96.2691039 | 90.5858882 | 97.1995085 | 105.500869 | 98.8922454 | 116.897333 | 106.163046 | 111.189415 | 118.891188 | 123.014351 | 106.731844 | 131.990474 | 122.065613 | 105.166794 | 132.79789 | 107.722621 | 116.077736 |
| ENSMUSG00 | BC125332 | 3724 | 0 | 0.10734552 | 0 | 0 | 0 | 0 | 0 | 0 | 0 | 0 | 0 | 0 | 0.02649405 | 0 | 0 | 0.04112847 | 0 | 0 |
| ENSMUSG00 | Tcl1 | 2785 | 0 | 0 | 0 | 0 | 0 | 0 | 0 | 0 | 0 | 0 | 0 | 0 | 0 | 0 | 0 | 0 | 0 | 0 |
| ENSMUSG00 | D19Bwg1357 | 3444 | 17.7351736 | 18.7844485 | 15.8435953 | 17.5814831 | 19.9291879 | 19.6541911 | 17.007287 | 15.6362623 | 14.7453467 | 18.5508719 | 18.477423 | 14.877694 | 14.9399506 | 17.301906 | 16.9041227 | 17.4442401 | 16.6481243 | 15.6050805 |
| ENSMUSG00 | Myzap | 5792 | 9.32002306 | 8.35698123 | 7.10736781 | 8.42470278 | 8.50951137 | 7.58006249 | 7.42528058 | 9.45578471 | 7.82913568 | 6.12991705 | 6.77196839 | 9.24157811 | 5.5873166 | 5.9979588 | 5.03637803 | 6.6373912 | 7.23250043 | 7.44598406 |
| ENSMUSG00 | 4930506M07 | 4116 | 12.1133556 | 11.6951244 | 12.9513194 | 10.8035907 | 13.436244 | 14.1458848 | 9.448742 | 9.3439624 | 7.64492178 | 8.74052152 | 9.72677726 | 9.35685306 | 6.84366491 | 10.629332 | 8.71879378 | 9.6470748 | 7.64477399 | 8.35293773 |
| ENSMUSG00 | B4galnt3 | 4601 | 0 | 0.02896144 | 0.0210274 | 0.00990242 | 0.01046119 | 0.33386975 | 0.03084939 | 0 | 0 | 0 | 0 | 0 | 0.15010799 | 0 | 0 | 0 | 0 | 0 |
| ENSMUSG00 | Ccdc9 | 5429 | 5.8093278 | 6.71289487 | 5.91637947 | 6.56267023 | 7.44719394 | 6.87294315 | 5.62976434 | 6.07110762 | 5.54565169 | 6.17501382 | 7.12796397 | 7.02193729 | 4.7796258 | 5.63762269 | 6.70681846 | 6.56632078 | 4.98404229 | 6.17066605 |
| ENSMUSG00 | Ninj2 | 932 | 0 | 0.46466483 | 0 | 0.34219668 | 0.10328739 | 0.10633634 | 0 | 0 | 0 | 0 | 0.30756411 | 0 | 0 | 0 | 0 | 0 | 0 | 0 |
| ENSMUSG00 | Cldn5 | 1413 | 0 | 0 | 0 | 0 | 0 | 0 | 0 | 0 | 0 | 0 | 0 | 0 | 0 | 0 | 0 | 0 | 0 | 0 |
| ENSMUSG00 | Htr2c | 8402 | 0 | 0 | 0 | 0 | 0 | 0 | 0 | 0 | 0 | 0 | 0 | 0 | 0 | 0 | 0 | 0 | 0 | 0 |
| ENSMUSG00 | Mdfic | 7676 | 2.50844773 | 2.62128558 | 1.84646227 | 1.79252676 | 1.98146012 | 1.98185112 | 1.59023856 | 1.29862129 | 1.21799623 | 1.08724264 | 1.84850999 | 1.15616081 | 1.87661801 | 0.95598825 | 1.19697068 | 2.12005057 | 1.66267025 | 1.90625617 |
| ENSMUSG00 | 2810422O20l | 1599 | 7.84179295 | 5.87506952 | 5.3244161 | 7.66474104 | 5.80954426 | 6.29093491 | 4.43833761 | 7.81045378 | 7.34093621 | 7.87319435 | 8.18657825 | 8.44739859 | 5.09055355 | 7.72615838 | 8.55466952 | 5.53166373 | 6.25588711 | 3.23380615 |
| ENSMUSG00 | 1700013G24l | 1046 | 0 | 0 | 0 | 0 | 0 | 0 | 0 | 0 | 0 | 0 | 0 | 0 | 0.13320484 | 0 | 0 | 0 | 0 | 0.05258997 |
| ENSMUSG00 | BC055324 | 4068 | 0.62692 | 0.74519994 | 0.85616868 | 0.94078853 | 0.75723770 | 0.36543314 | 0.76760984 | 0.92946112 | 1.27568854 | 0.81134445 | 1.90254259 | 0.53510591 | 0.77611669 | 1.11885953 | 1.43820574 | 1.40248292 | 0.81024128 | 1.27110522 |
| ENSMUSG00 | Wapal | 4906 | 21.0966375 | 20.4250277 | 21.6921661 | 21.8239817 | 20.0729547 | 23.0087907 | 19.7795158 | 23.0275431 | 22.5241382 | 24.1520011 | 20.6349647 | 20.512768 | 17.2651655 | 18.1478425 | 19.4838177 | 22.4077321 | 23.3816972 | 17.9962528 |
| ENSMUSG00 | Dicer1 | 10027 | 3.99320593 | 4.7343048 | 5.88085504 | 4.64380047 | 4.22420393 | 4.51198222 | 3.51058473 | 4.51742189 | 3.93093603 | 3.86535136 | 5.80332098 | 5.29934285 | 3.59645273 | 4.17334424 | 4.4706585 | 4.27700013 | 3.86435486 | 4.04325438 |
| ENSMUSG00 | Pik3r1 | 7378 | 3.38750928 | 3.48119293 | 3.1536552 | 3.30993762 | 2.70733916 | 3.11635516 | 3.95020469 | 3.38544743 | 4.35422639 | 3.77052249 | 3.14053332 | 4.85682507 | 4.51328872 | 3.12229159 | 4.63503543 | 4.6449029 | 3.68302311 | 3.01215494 |
| ENSMUSG00 | Meis3 | 2749 | 3.5098854 | 4.13230154 | 3.66012889 | 3.62963576 | 4.91999656 | 3.8214549 | 8.53659103 | 8.89164698 | 7.61104038 | 7.90704773 | 8.01173367 | 11.8169209 | 6.87308997 | 10.3565753 | 8.91776128 | 11.7977972 | 9.92663638 | 12.1864482 |
| ENSMUSG00 | Paqr6 | 2083 | 0.32649199 | 0.19191298 | 0.09289204 | 0.13123681 | 0.09242808 | 0.11894559 | 0 | 0.03667057 | 0 | 0.02263596 | 0.11467825 | 0.05359361 | 0.26051421 | 0.02229673 | 0.61317726 | 0.14705946 | 0.03679913 | 0 |
| ENSMUSG00 | Hibch | 2510 | 8.04380821 | 8.75956581 | 10.8117731 | 7.98679662 | 7.90053886 | 7.2058756 | 12.9308756 | 8.11017947 | 8.94288888 | 10.6699599 | 7.5754715 | 10.562725 | 7.44892297 | 9.17779575 | 7.04201892 | 7.16995437 | 9.39070621 | 8.65681162 |
| ENSMUSG00 | Nthl1 | 1074 | 9.65666941 | 6.85488792 | 5.90030957 | 7.80561653 | 6.05010214 | 6.32097253 | 7.70923309 | 6.93437474 | 12.7701208 | 6.89260746 | 7.91809988 | 7.84745349 | 6.01720779 | 5.31901559 | 5.71604453 | 6.77394358 | 6.06654066 | 4.55848277 |
| ENSMUSG00 | Ccnb1 | 4963 | 0.00856441 | 0.24835323 | 0.13645566 | 0.03672057 | 0 | 0.07987545 | 0.07626464 | 0 | 0.26555884 | 0 | 0.02887868 | 0 | 0.07951951 | 0.0842226 | 0.55621685 | 0.16973471 | 0.67184925 | 0.76478508 |
| ENSMUSG00 | Cirh1a | 2231 | 18.6138758 | 21.7556642 | 21.6390781 | 20.9119271 | 22.9333191 | 22.7218496 | 23.306425 | 25.4558953 | 21.0455527 | 24.7271729 | 19.5939431 | 20.7651277 | 24.6549026 | 19.2354714 | 22.7153356 | 27.7697009 | 23.1572605 | 23.4238688 |
| ENSMUSG00 | Mfsd6 | 9048 | 13.2100525 | 16.965718 | 15.9053109 | 17.6040463 | 15.9269429 | 14.6226568 | 10.1182619 | 12.452209 | 9.25877244 | 11.1988131 | 11.5741279 | 9.96884513 | 10.6754967 | 11.0566496 | 10.1432476 | 10.7152761 | 13.2074927 | 9.52688663 |
| ENSMUSG00 | Gk5 | 6284 | 3.74727363 | 3.21784322 | 4.65722245 | 3.25539607 | 2.66548527 | 3.75351706 | 3.53866313 | 4.72238967 | 3.30331062 | 3.83418354 | 3.99138339 | 4.21903879 | 4.28631872 | 3.42935337 | 4.14375928 | 2.3946863 | 3.92777454 | 4.13181069 |
| ENSMUSG00 | Arhgap32 | 14367 | 0.86389026 | 1.04573782 | 0.75757252 | 1.34777224 | 1.11560733 | 0.81742868 | 0.95501336 | 1.19891233 | 0.83135586 | 0.91564322 | 0.67504118 | 0.68764337 | 0.62149906 | 0.71119226 | 0.82018828 | 0.31182579 | 0.63757117 | 0.5743277 |
| ENSMUSG00 | Mmrn2 | 3928 | 0 | 0 | 0.03694156 | 0.08119331 | 0 | 0.03784577 | 0 | 0.06806182 | 0.01048537 | 0.06001872 | 0.01216267 | 0.02841938 | 0.06279521 | 0.01182365 | 0.02097839 | 0.01949624 | 0.05854322 | 0.04201307 |
| ENSMUSG00 | Serpina3h | 1429 | 1.39800089 | 0.88585723 | 1.25249869 | 1.40285931 | 0.74100932 | 1.35238391 | 0.13243577 | 1.63032648 | 1.35462998 | 0.69290747 | 0.06686488 | 0.82024417 | 0.55235177 | 1.46254999 | 0.9514705 | 1.31297394 | 0.90871266 | 1.73226718 |
| ENSMUSG00 | Rpl21 | 3419 | 31.7514535 | 24.4658313 | 27.2923487 | 23.3868479 | 27.353123 | 25.783654 | 28.8664078 | 24.9887114 | 27.3211517 | 24.0787154 | 28.1283976 | 25.0427688 | 33.0707091 | 30.496337 | 30.0304994 | 26.5984769 | 24.6391344 | 31.8566924 |
| ENSMUSG00 | Tardbp | 11765 | 10.3110729 | 10.1425233 | 10.25856 | 10.2081524 | 10.7596225 | 8.8491537 | 10.1823802 | 10.0374763 | 11.1254354 | 9.3660173 | 9.48595861 | 10.3234069 | 10.1640941 | 9.13485918 | 9.23809392 | 10.7727931 | 10.632982 | 10.4126885 |
| ENSMUSG00 | Cacna2d4 | 6039 | 0 | 0 | 0 | 0 | 0 | 0.01641091 | 0 | 0 | 0.03410044 | 0 | 0 | 0 | 0 | 0 | 0 | 0 | 0 | 0 |
| ENSMUSG00 | Gpr12 | 2466 | 0.15512838 | 0.09456215 | 0 | 0 | 0 | 0.06028313 | 0 | 0 | 0 | 0 | 0 | 0 | 0 | 0 | 0 | 0 | 0 | 0 |
| ENSMUSG00 | Fam35a | 3690 | 2.39595572 | 2.44655673 | 2.77918386 | 2.2101428 | 4.0044717 | 3.33037342 | 4.5132962 | 3.78818951 | 3.97355003 | 3.88450238 | 2.65416436 | 3.50927743 | 2.41980407 | 3.14661839 | 3.38321719 | 3.27908725 | 4.16499866 | 4.74062221 |
| ENSMUSG00 | Smpx | 2452 | 0 | 0 | 0.01972819 | 0 | 0 | 0.02020911 | 0 | 0 | 0.08398555 | 0 | 0 | 0.04552664 | 8.20857351 | 8.65617779 | 5.93154536 | 2.17063394 | 1.73499957 | 4.35227027 |
| ENSMUSG00 | Dcp1b | 4494 | 3.63195096 | 3.15783109 | 3.48754424 | 4.02486306 | 3.44870481 | 3.06534489 | 3.16891775 | 3.22944213 | 4.29828124 | 3.6511893 | 3.11483374 | 3.82537251 | 3.1504807 | 2.66634948 | 4.26317835 | 3.57856144 | 2.20883643 | 3.89250921 |
| ENSMUSG00 | Syt15 | 5291 | 0 | 0 | 0 | 0 | 0 | 0 | 0 | 0.03609186 | 0 | 0.02673448 | 0.00902948 | 0 | 0.03511177 | 0 | 0 | 0 | 0 | 0 |
| ENSMUSG00 | Serpina3g | 2365 | 6.18257106 | 5.25399981 | 7.28159696 | 5.64456067 | 5.59673531 | 6.2019489 | 4.62123841 | 9.20493419 | 5.64246558 | 6.3598623 | 10.1610165 | 5.87657505 | 4.31784646 | 3.96689457 | 3.46685396 | 6.86478543 | 3.9865831 | 3.55872868 |
| ENSMUSG00 | Fam38b | 14570 | 0 | 0 | 0 | 0 | 0 | 0 | 0 | 0 | 0 | 0 | 0 | 0 | 0.01015757 | 0 | 0.00565567 | 0 | 0 | 0.0037755 |
| ENSMUSG00 | Zfp281 | 4935 | 4.33234114 | 5.15050421 | 4.66581569 | 4.69920396 | 4.78880943 | 4.86992417 | 4.77441015 | 7.82421861 | 7.22746153 | 5.75176699 | 6.31190941 | 5.91521612 | 5.66792241 | 5.52435244 | 4.64195848 | 7.02964343 | 6.34500194 | 4.92685414 |
| ENSMUSG00 | Stx3 | 4964 | 16.200603 | 14.8377939 | 12.8145029 | 15.5480276 | 14.1758405 | 16.7305362 | 13.2769061 | 12.9026307 | 14.2377238 | 13.4499186 | 13.7531055 | 14.617317 | 12.7106209 | 14.2026843 | 12.7572102 | 14.64824 | 13.3802322 | 14.6277238 |
| ENSMUSG00 | Cep78 | 2533 | 3.62460246 | 3.47201108 | 3.43751894 | 3.12973617 | 4.84549567 | 3.40393827 | 3.43684679 | 3.95041825 | 2.78045566 | 2.32682112 | 2.99890185 | 4.14265478 | 2.86292611 | 3.48376494 | 2.22842999 | 2.91752358 | 2.54197311 | 4.47369759 |
| ENSMUSG00 | Kif14 | 8740 | 0 | 0 | 0 | 0 | 0 | 0 | 0 | 0 | 0 | 0 | 0 | 0 | 0 | 0 | 0 | 0 | 0 | 0 |
| ENSMUSG00 | Gm3402 | 956 | 0 | 0 | 0 | 0 | 0 | 0 | 0 | 0 | 0 | 0 | 0 | 0 | 0 | 0 | 0 | 0 | 0 | 0 |
| ENSMUSG00 | Rrp9 | 2774 | 7.61538302 | 8.55038958 | 6.13824178 | 8.0643377 | 7.56507524 | 7.85984238 | 8.27204372 | 7.84775392 | 8.71538744 | 7.98876371 | 7.93952987 | 6.59968902 | 9.70988711 | 11.2343124 | 5.71831255 | 8.14400085 | 7.86145023 | 8.5468367 |
| ENSMUSG00 | 4930569F06l | 432 | 0 | 0 | 0 | 0 | 0 | 0 | 0 | 0 | 0 | 0 | 0 | 0 | 0 | 0 | 0 | 0 | 0 | 0 |
| ENSMUSG00 | Irf8 | 3851 | 0.27593596 | 0.2941154 | 0.28890969 | 0.2011264 | 0.2874667 | 0.14154247 | 1.10572348 | 2.13226864 | 1.83954316 | 1.08969426 | 2.58041863 | 0.85513479 | 1.92152365 | 1.13366508 | 1.17688163 | 0.67612602 | 0.95542054 | 1.09969645 |
| ENSMUSG00 | Upk2 | 868 | 0.29381458 | 0 | 0 | 0 | 0.05545152 | 0 | 0.59958464 | 0.35200371 | 0 | 0.21728435 | 0 | 0.19291128 | 1.19351413 | 9.68476937 | 3.27523889 | 1.98511213 | 0.92724912 | 1.90123641 |
| ENSMUSG00 | Rnf123 | 6998 | 6.20752928 | 8.86376237 | 8.31571263 | 10.4690139 | 8.26730098 | 10.0337559 | 7.16653896 | 7.21496942 | 8.7104949 | 8.19982936 | 6.64944406 | 9.01281777 | 6.54187185 | 6.95533061 | 7.66567959 | 8.43728445 | 9.1133121 | 8.26158491 |
| ENSMUSG00 | Eif2c1 | 9497 | 4.368227 | 2.59922656 | 2.72505409 | 2.32674597 | 2.29078967 | 1.90968732 | 2.60551027 | 2.37672 | 4.03321677 | 2.0008143 | 2.78188397 | 3.33236608 | 2.8102103 | 2.0197334 | 1.63556771 | 2.36670386 | 1.63039107 | 1.74926295 |
| ENSMUSG00 | Rbp3 | 5272 | 0 | 0 | 0 | 0 | 0 | 0 | 0 | 0 | 0 | 0 | 0.00906202 | 0 | 0 | 0 | 0 | 0 | 0 | 0 |
| ENSMUSG00 | Serpina3a | 1842 | 0 | 0 | 0 | 0 | 0 | 0 | 0 | 0 | 0 | 0 | 0 | 0 | 0 | 0 | 0 | 0 | 0 | 0 |
| ENSMUSG00 | H2-Ob | 3256 | 0.18276181 | 0 | 0 | 0.09795065 | 0 | 0.0304378 | 2.23776184 | 0.46919413 | 0.240339 | 0.24617996 | 1.54065437 | 0.34284805 | 1.27269074 | 1.04128329 | 0.55677778 | 0.67032083 | 0 | 0 |
| ENSMUSG00 | Sox5 | 14713 | 0 | 0 | 0 | 0 | 0 | 0 | 0.054667 | 0.01557496 | 0.14556507 | 0.0032047 | 0.13313215 | 0 | 0.1408238 | 0.07891675 | 0.04480961 | 0.02342252 | 0 | 0 |
| ENSMUSG00 | Ptchd2 | 5429 | 0.00782928 | 0.06136101 | 0.04455105 | 0 | 0.06205995 | 0.03650966 | 0 | 0.00703489 | 0 | 0.01736994 | 0 | 0 | 0.07269393 | 0.16254147 | 0 | 0.11284762 | 0.09177415 | 0.12158948 |
| ENSMUSG00 | Hspb8 | 2496 | 22.2232591 | 20.1665798 | 20.5820069 | 22.0321236 | 22.3111521 | 20.547708 | 15.5055174 | 17.2294251 | 20.0157226 | 18.7582726 | 14.4511593 | 18.4710486 | 30.4173961 | 25.7898658 | 27.3686617 | 44.1814701 | 25.9193867 | 28.518343 |
| ENSMUSG00 | Serpina5 | 2157 | 0 | 0 | 0 | 0 | 0 | 0 | 0 | 0.19476886 | 0 | 0 | 0 | 0 | 0 | 0 | 0 | 0 | 0 | 0 |
| ENSMUSG00 | Ptchd1 | 4654 | 0 | 0 | 0 | 0 | 0 | 0 | 0 | 0 | 0 | 0 | 0 | 0 | 0 | 0 | 0 | 0 | 0 | 0 |
| ENSMUSG00 | Fbxo2 | 1282 | 0.79572872 | 0.49371684 | 0.60372581 | 1.06617108 | 0.90106565 | 1.23688571 | 1.07025561 | 0.59582531 | 1.44570453 | 2.46419432 | 2.79494687 | 0.91429713 | 5.50270216 | 3.26050535 | 3.18171141 | 4.3607088 | 3.82665055 | 2.96070855 |
| ENSMUSG00 | Fmod | 3389 | 0 | 0.04914856 | 0.05709475 | 0 | 0.09941678 | 0.05848655 | 0 | 0.1690428 | 0 | 0.0417386 | 0 | 0 | 0.17467778 | 0.31520036 | 0.57139957 | 0 | 0.04523611 | 0.3084016 |
| ENSMUSG00 | Gltscr2 | 1542 | 82.5295052 | 76.1746206 | 73.6896387 | 67.6028985 | 73.290374 | 75.6787857 | 68.483722 | 71.2330411 | 70.7542674 | 79.1654804 | 77.1463337 | 73.6969449 | 76.0773666 | 75.2080974 | 62.2565663 | 63.2217162 | 58.7321917 | 74.2016485 |
| ENSMUSG00 | L3mbtl4 | 3495 | 0 | 0 | 0 | 0 | 0 | 0 | 0.09476073 | 0.07649408 | 0 | 0 | 0.06834758 | 0 | 0.07057499 | 0 | 0.10609843 | 0 | 0 | 0 |
| ENSMUSG00 | Tssk1 | 1448 | 0.38160724 | 0.39110442 | 0.60132842 | 0.22025367 | 0.03324028 | 0.136886 | 0.81686255 | 0.05275194 | 0.48354335 | 0.22793848 | 0.03299376 | 0.19273364 | 0 | 0.03207465 | 0.11381645 | 0.10577516 | 0 | 0 |
| ENSMUSG00 | Serpina12 | 3685 | 0 | 0 | 0 | 0 | 0 | 0 | 0 | 0 | 0 | 0 | 0 | 0 | 0 | 0 | 0 | 0 | 0 | 0 |
| ENSMUSG00 | Camsap2 | 8404 | 5.143713 | 4.506992 | 4.9559269 | 6.30503267 | 5.22326444 | 4.55786091 | 5.01613481 | 4.04464995 | 3.73932788 | 4.93724641 | 4.23516707 | 5.05422647 | 4.61973095 | 4.64772459 | 5.05949615 | 6.11446872 | 5.00741004 | 4.07135464 |
| ENSMUSG00 | Sepw1 | 730 | 402.285289 | 337.829255 | 350.343637 | 344.828452 | 355.78063 | 370.830936 | 413.953535 | 300.72593 | 369.437404 | 376.172186 | 337.506631 | 318.913875 | 376.476865 | 337.896641 | 337.344794 | 361.40039 | 295.059984 | 365.019333 |
| ENSMUSG00 | Prelp | 3673 | 0.09257866 | 0.00906967 | 0 | 0 | 0 | 0 | 0 | 0.0207963 | 0 | 0 | 0 | 0.04558862 | 0 | 0 | 0 | 0 | 0.03130381 | 0 |
| ENSMUSG00 | Crx | 5633 | 0 | 0 | 0 | 0 | 0.00854463 | 0 | 0 | 0 | 0.02193495 | 0 | 0 | 0 | 0 | 0 | 0 | 0 | 0 | 0 |
| ENSMUSG00 | Obox6 | 2020 | 0 | 0 | 0 | 0 | 0.02382768 | 0 | 0 | 0 | 0 | 0 | 0 | 0 | 0 | 0 | 0 | 0 | 0 | 0 |
| ENSMUSG00 | Sdk2 | 16393 | 0.79342304 | 1.03232764 | 0.73476539 | 0.83378962 | 0.66062849 | 0.79499597 | 0.27129823 | 0.62671605 | 0.48992683 | 0.46020329 | 0.32932168 | 0.65373069 | 0.00902798 | 0.1246593 | 0.07037416 | 0.0700737 | 0.08182885 | 0.01677823 |
| ENSMUSG00 | Tmtc4 | 8481 | 8.70551713 | 10.3540607 | 8.2761458 | 9.07352952 | 9.71037866 | 10.0203912 | 3.8939099 | 5.5795763 | 3.94333814 | 5.14816074 | 5.01352589 | 5.3900516 | 3.18758291 | 2.89693694 | 3.51240713 | 2.97981381 | 4.45581049 | 3.39226068 |
| ENSMUSG00 | Vmn1r90 | 4515 | 0 | 0.02213481 | 0 | 0.03027312 | 0.01066045 | 0.06585081 | 0 | 0 | 0 | 0 | 0.01058139 | 0.02472455 | 0.01092623 | 0 | 0 | 0 | 0 | 0 |
| ENSMUSG00 | Cdc42ep4 | 6288 | 20.678011 | 23.3423365 | 19.2786366 | 24.6354247 | 19.2742023 | 25.1309904 | 18.4269642 | 19.6367743 | 17.9274004 | 19.3911766 | 19.8834393 | 20.1586148 | 15.1024079 | 14.8461539 | 18.5367687 | 20.2840352 | 19.0961018 | 15.5719163 |
| ENSMUSG00 | 5730559C18l | 3980 | 8.21268351 | 8.13571185 | 7.63280831 | 8.59707143 | 8.67100233 | 9.41252929 | 8.36887587 | 7.34100197 | 7.25420793 | 11.5625845 | 7.46633773 | 8.2601571 | 9.06071144 | 14.3299848 | 11.6461684 | 11.5064237 | 9.17712587 | 12.2181031 |
| ENSMUSG00 | Mbp | 4683 | 6.12662691 | 8.08102666 | 6.02215849 | 6.04172721 | 5.5192917 | 6.67686844 | 1.68721277 | 3.28668336 | 3.24531803 | 4.07773539 | 2.55044621 | 3.49220353 | 2.99172849 | 2.28104636 | 3.1497239 | 3.8347843 | 3.45370431 | 4.63988835 |
| ENSMUSG00 | Entpd3 | 3735 | 7.35163152 | 9.38290889 | 7.42116008 | 8.53888381 | 8.28616517 | 8.19908684 | 5.07962095 | 7.38284191 | 6.4975785 | 7.24618572 | 7.12467252 | 9.35491426 | 3.68503478 | 2.68592313 | 4.97507408 | 3.57789076 | 5.54115136 | 3.40216965 |
| ENSMUSG00 | Ccdc64 | 7078 | 1.36919753 | 1.05426509 | 1.41471048 | 1.69292945 | 1.04723315 | 1.23916839 | 2.12566147 | 2.2339156 | 1.89697713 | 1.65207324 | 1.20821102 | 1.58504495 | 1.91668229 | 1.96196413 | 0.954656 | 1.40654958 | 1.75441088 | 1.63208709 |
| ENSMUSG00 | 5330439B14l | 311 | 1.36672592 | 0.96403874 | 0.46662569 | 0.14649853 | 0.30953006 | 0.63733419 | 1.521308 | 1.59646698 | 0.79459516 | 0.15166998 | 2.15064122 | 0.89735792 | 0 | 3.28543383 | 0 | 1.60057201 | 0.86264977 | 0 |
| ENSMUSG00 | Nppa | 862 | 0 | 0 | 0 | 0 | 0 | 0 | 0 | 0 | 0 | 0 | 0 | 0 | 0 | 0 | 0 | 0.04446206 | 0 | 0 |
| ENSMUSG00 | Ccdc74a | 1270 | 0.26774914 | 1.33776189 | 0.26662576 | 0.50224773 | 0.26529407 | 0.58526851 | 4.61950566 | 3.57865818 | 3.47004503 | 2.30184536 | 1.20377848 | 2.5930111 | 4.19515517 | 3.29131328 | 2.72514228 | 1.7487049 | 1.59944389 | 3.63839761 |
| ENSMUSG00 | Mmp1b | 1439 | 0 | 0 | 0 | 0 | 0 | 0 | 0 | 0.05308187 | 0 | 0.0327663 | 0 | 0 | 0.44566708 | 0.16137626 | 0.08589622 | 0.05321836 | 0 | 0 |
| ENSMUSG00 | D11Wsu47e | 2789 | 17.3282127 | 13.2463247 | 14.4305404 | 16.4503305 | 14.5137855 | 15.8128121 | 18.1175838 | 14.6799051 | 14.6936476 | 15.8239721 | 14.0806798 | 16.9508305 | 14.3980569 | 13.854959 | 16.0876638 | 13.0015078 | 14.0854981 | 16.7453322 |
| ENSMUSG00 | Gucy1a2 | 5273 | 0.43526912 | 0.24007017 | 0.59629802 | 0.25057278 | 0.49291179 | 0.46987231 | 0.26020628 | 0.40560867 | 0.14059497 | 0.33979267 | 0.08154269 | 0.41282208 | 0.36486721 | 0.27304508 | 0.5860263 | 0.23237236 | 0.58147236 | 0.76155221 |
| ENSMUSG00 | A2ld1 | 1626 | 18.9260439 | 17.2301009 | 17.3145109 | 15.1589945 | 17.7016546 | 14.7804894 | 15.2471364 | 13.5998776 | 13.8808184 | 14.9339558 | 13.6331985 | 11.7054987 | 17.536192 | 15.0814749 | 14.0125833 | 18.1326985 | 16.1696431 | 14.5811347 |
| ENSMUSG00 | D11Wsu99e | 5113 | 14.5563394 | 13.3955233 | 12.0531738 | 12.1810967 | 12.1906593 | 12.7637297 | 14.4815847 | 12.2651916 | 13.0253464 | 13.9063682 | 11.8759969 | 13.2088699 | 10.0190252 | 14.4609793 | 11.8616644 | 17.1195636 | 12.2707115 | 13.685035 |
| ENSMUSG00 | Mrps27 | 1837 | 12.4715786 | 11.6604197 | 10.7174888 | 12.2769281 | 11.1355839 | 12.4623641 | 12.5686376 | 10.5200629 | 12.5106564 | 14.116977 | 13.7577316 | 10.6952368 | 10.1778969 | 12.9952427 | 11.4386464 | 13.7571045 | 11.1828498 | 11.0497335 |
| ENSMUSG00 | Kctd12b | 4903 | 2.6441115 | 5.04143739 | 6.5510961 | 3.25236906 | 4.84951464 | 4.81074876 | 4.17833775 | 2.25898362 | 3.03249688 | 1.89449083 | 3.67349773 | 2.44755608 | 3.06878131 | 0.87147788 | 2.9495709 | 1.06391907 | 4.18986236 | 2.77121137 |
| ENSMUSG00 | Gcn1l1 | 13930 | 3.99725632 | 6.90650671 | 5.45895116 | 6.34845564 | 6.09855314 | 7.87223255 | 4.72107851 | 5.43412359 | 5.52011665 | 5.0772473 | 4.81522195 | 5.15283145 | 4.58631301 | 4.94447822 | 6.20241656 | 5.12648836 | 5.45867718 | 5.84052181 |
| ENSMUSG00 | Kif21b | 10871 | 0.02736972 | 0.03064382 | 0.040048 | 0.00419106 | 0.06198574 | 0.0273495 | 0.03916973 | 0.10539712 | 0.00378866 | 0 | 0.08349961 | 0 | 0.22689687 | 0.09826272 | 0.04548051 | 0.13736857 | 0.01057666 | 0.01012034 |
| ENSMUSG00 | Slc5a12 | 4677 | 0 | 0 | 0 | 0 | 0 | 0 | 0 | 0 | 0 | 0 | 0 | 0 | 0 | 0.03972126 | 0 | 0 | 0 | 0 |
| ENSMUSG00 | Ddx24 | 3148 | 47.0014351 | 50.7418449 | 47.6205751 | 47.2978059 | 52.9787533 | 50.4971948 | 47.2525612 | 51.5015083 | 52.4120655 | 49.8617189 | 47.7446059 | 45.8333824 | 49.4102717 | 59.574722 | 56.7241243 | 55.2829767 | 64.7821556 | 58.538916 |
| ENSMUSG00 | Klf8 | 5195 | 0.24545819 | 0.09618738 | 0.15829644 | 0.16663327 | 0.35207182 | 0.28615631 | 0.39161601 | 0.0955729 | 0.04756864 | 0.05445702 | 0.22990837 | 0 | 0.06647227 | 0.19668333 | 0.13482704 | 0.38327461 | 0.19919346 | 0.70945335 |
| ENSMUSG00 | Pcca | 3783 | 15.7189377 | 13.9222008 | 17.2881346 | 17.2464751 | 14.8479923 | 16.6485658 | 15.4707332 | 15.0427385 | 13.870373 | 13.4110909 | 10.5956093 | 15.6543532 | 9.2717402 | 11.0739008 | 10.9565861 | 10.8707672 | 12.2891351 | 11.2984605 |
| ENSMUSG00 | Pnpla3 | 4632 | 0 | 0 | 0.01044334 | 0.0196723 | 0.0415647 | 0 | 0 | 0 | 0.15115949 | 0 | 0 | 0.07230008 | 0 | 0 | 0.00889498 | 0 | 0 | 0 |
| ENSMUSG00 | Slc39a11 | 3333 | 5.90455941 | 6.60660758 | 5.63124206 | 6.06933804 | 6.77283894 | 5.96179003 | 6.64336456 | 5.22524317 | 4.21380194 | 7.93625698 | 7.52530849 | 5.07415114 | 6.60127066 | 7.51076001 | 6.7494898 | 7.35253187 | 5.19756562 | 6.8658231 |
| ENSMUSG00 | Rragb | 2835 | 5.54741274 | 6.70958116 | 4.65819181 | 4.21057976 | 5.75545754 | 5.36602794 | 11.5486327 | 12.771216 | 9.79178541 | 13.0891018 | 11.5772122 | 9.70621575 | 14.3210469 | 14.1052414 | 11.4220769 | 14.1411873 | 12.5726477 | 14.0287775 |
| ENSMUSG00 | Bbox1 | 4395 | 0 | 0.19707287 | 0.03301947 | 0.18659813 | 0.03285456 | 0 | 0 | 0.14772943 | 0.16868198 | 0.1609239 | 0 | 0.46989295 | 0.2469402 | 0 | 0.8062193 | 0.28750627 | 0.00872043 | 0.43807025 |
| ENSMUSG00 | Prima1 | 1674 | 0 | 0.01990018 | 0.02889697 | 0 | 0 | 0.02960139 | 0 | 0 | 0 | 0 | 0 | 0 | 0.05893897 | 0.05548875 | 0.14767583 | 0.04574744 | 0.18316032 | 0.1314435 |
| ENSMUSG00 | Rims1 | 8739 | 0 | 0.01143594 | 0.03321217 | 0.01564059 | 0.02203086 | 0.0056703 | 0 | 0 | 0 | 0 | 0 | 0.00638696 | 0 | 0 | 0.00876316 | 0 | 0 | 0 |
| ENSMUSG00 | Pyroxd1 | 4526 | 8.09532959 | 7.66947322 | 6.28449754 | 7.92234686 | 7.10387207 | 6.48148879 | 6.25121122 | 8.13496105 | 8.69958221 | 7.88623119 | 8.08564255 | 8.62022716 | 7.98945987 | 6.64952908 | 9.06689122 | 9.28923706 | 9.61121233 | 7.79072855 |
| ENSMUSG00 | Lrrc18 | 3271 | 7.03005204 | 4.62367899 | 5.35347539 | 5.11186282 | 6.75406694 | 6.24143266 | 5.78571433 | 5.60451029 | 8.86435558 | 5.27580477 | 8.31059351 | 6.89377184 | 5.80641643 | 4.9553612 | 5.75637757 | 6.22763777 | 5.83506446 | 6.86142328 |
| ENSMUSG00 | BC006965 | 5866 | 0 | 0 | 0.00824643 | 0 | 0 | 0 | 0 | 0.01953243 | 0 | 0 | 0 | 0 | 0.00840981 | 0 | 0 | 0 | 0 | 0 |
| ENSMUSG00 | Lrrc29 | 1903 | 3.0600153 | 4.53391466 | 2.51654204 | 3.59125415 | 3.33863054 | 4.45271542 | 3.20721786 | 3.91356725 | 3.09494049 | 3.7165557 | 3.61513086 | 3.51964241 | 5.21057035 | 4.10016116 | 4.76318565 | 4.46690218 | 2.85987081 | 3.00627803 |
| ENSMUSG00 | Iapp | 1034 | 197.521635 | 142.530217 | 99.3669037 | 205.33314 | 83.4628038 | 98.1949237 | 26.1729713 | 36.4933207 | 21.70856 | 43.6394807 | 18.6202208 | 26.8282247 | 22.9961162 | 11.274145 | 16.5762546 | 22.9225203 | 8.00626705 | 10.3208575 |
| ENSMUSG00 | Bivm | 3709 | 6.57804558 | 7.76013497 | 9.71644112 | 9.20065306 | 9.64195721 | 8.92456889 | 10.2432141 | 7.69202604 | 8.73922564 | 9.90304606 | 9.57044868 | 8.23164396 | 6.0650726 | 8.35217209 | 9.19809875 | 7.32982779 | 8.90832484 | 4.77566254 |
| ENSMUSG00 | Fcho2 | 5221 | 16.29053 | 16.7553458 | 19.0584853 | 17.6275253 | 17.7881365 | 21.4023011 | 17.2449776 | 14.9082774 | 16.2653466 | 17.1949701 | 15.9127854 | 18.8261601 | 16.0912191 | 13.725958 | 15.562051 | 16.1347129 | 19.629289 | 16.2677764 |
| ENSMUSG00 | Amot | 8388 | 4.50490003 | 4.3924727 | 4.80967147 | 4.03031651 | 4.52743059 | 4.81467309 | 3.81863181 | 3.47865945 | 4.35532183 | 4.04364946 | 2.83072894 | 3.68644219 | 2.73478082 | 2.45287681 | 2.04828908 | 2.05878206 | 2.99281408 | 3.65284602 |
| ENSMUSG00 | Kcnj2 | 5442 | 0.1171587 | 0.18976474 | 0.00888893 | 0 | 0.03537811 | 0.10926733 | 0 | 0.01403616 | 0 | 0 | 0.03511574 | 0.01025646 | 0 | 0 | 0 | 0 | 0 | 0 |
| ENSMUSG00 | Rasl12 | 3871 | 0 | 0.02581728 | 0 | 0 | 0 | 0 | 0 | 0 | 0 | 0 | 0 | 0 | 0 | 0 | 0 | 0 | 0 | 0 |
| ENSMUSG00 | Cox6a1 | 969 | 495.937585 | 410.927786 | 432.216745 | 396.225924 | 450.125379 | 434.212859 | 599.73337 | 459.412428 | 493.600627 | 532.914863 | 512.804276 | 471.927475 | 616.21623 | 590.20897 | 490.124184 | 526.189606 | 474.668616 | 557.073647 |
| ENSMUSG00 | Slco1a1 | 6539 | 0.02600103 | 0.00509449 | 0.0073977 | 0.00696759 | 0.07360747 | 0 | 0 | 0.01168142 | 0 | 0.00721069 | 0 | 0 | 0 | 0 | 0 | 0.01758356 | 0 | 0 |
| ENSMUSG00 | Lhfpl1 | 1897 | 0 | 0.03512166 | 0 | 0 | 0 | 0 | 0 | 0 | 0 | 0 | 0 | 0 | 0 | 0 | 0 | 0 | 0 | 0 |
| ENSMUSG00 | Btbd7 | 4286 | 6.41643698 | 6.96415147 | 7.09914853 | 8.21714589 | 7.50166239 | 8.91394251 | 5.65191125 | 5.07033994 | 5.8426042 | 6.14961342 | 6.08612397 | 6.84998566 | 5.49027637 | 5.8840736 | 5.26795442 | 7.60273143 | 4.76619605 | 8.53501073 |
| ENSMUSG00 | Zic5 | 5468 | 0 | 0 | 0 | 0 | 0 | 0 | 0 | 0 | 0 | 0.02586908 | 0 | 0 | 0 | 0.05945659 | 0.04521019 | 0 | 0 | 0 |
| ENSMUSG00 | 1810011H11l | 1468 | 0 | 0 | 0 | 0 | 0 | 0 | 0 | 0 | 0 | 0 | 0 | 0 | 0 | 0 | 0 | 0 | 0 | 0 |
| ENSMUSG00 | Mpped1 | 4284 | 0 | 0 | 0 | 0 | 0 | 0.01156693 | 0 | 0.10698152 | 0 | 0 | 0 | 0 | 0 | 0.02168258 | 0 | 0 | 0 | 0 |
| ENSMUSG00 | Trpc5 | 5003 | 0.5182522 | 0.25968476 | 0.8218569 | 0.59193841 | 0.45216878 | 0.68341767 | 1.2577626 | 0.49820351 | 0.58449783 | 1.2063342 | 1.29869965 | 0.92598441 | 1.45934914 | 1.12327295 | 0.93883218 | 1.13272235 | 1.96112972 | 1.0885272 |
| ENSMUSG00 | Ubr7 | 3244 | 14.4260785 | 17.5087807 | 15.1353679 | 16.2497313 | 12.9973999 | 16.6957885 | 20.7831589 | 16.8121919 | 13.4072012 | 18.9823731 | 16.8037077 | 16.7756692 | 14.3707342 | 17.1803036 | 20.1054448 | 21.3171507 | 22.6129878 | 20.8064655 |
| ENSMUSG00 | AK010878 | 4786 | 0 | 0 | 0 | 0 | 0 | 0 | 0 | 0 | 0 | 0 | 0 | 0 | 0 | 0 | 0 | 0 | 0 | 0 |
| ENSMUSG00 | Golt1a | 1837 | 11.1295534 | 10.318474 | 9.45350974 | 9.32550499 | 9.74691105 | 10.1155553 | 7.28877961 | 7.65095486 | 9.77535159 | 8.54718791 | 7.67208098 | 7.65680292 | 9.18427638 | 12.4390261 | 10.833071 | 12.5064586 | 9.72240302 | 8.74396253 |
| ENSMUSG00 | Alg13 | 6050 | 1.05384734 | 1.16732786 | 0.87951873 | 0.92628239 | 0.89899295 | 0.99924521 | 0.82112914 | 1.23730758 | 1.22538394 | 0.70920893 | 1.16081304 | 0.86721856 | 0.54632041 | 0.49130935 | 1.23264159 | 0.60125747 | 1.01992294 | 1.15473662 |
| ENSMUSG00 | Pi4ka | 11465 | 3.25508457 | 4.60830791 | 4.30783903 | 4.50245645 | 4.29891756 | 3.95903218 | 3.70578157 | 4.19733337 | 3.71810238 | 4.48276969 | 3.95034109 | 4.06994278 | 5.45168244 | 4.14006605 | 4.25132496 | 5.46388432 | 5.21157408 | 4.99472133 |
| ENSMUSG00 | Khdc1c | 1943 | 0 | 0 | 0 | 0 | 0 | 0 | 0 | 0 | 0 | 0 | 0 | 0 | 0.33006429 | 0.40635589 | 0 | 0.13794866 | 0.05917596 | 0.02831143 |
| ENSMUSG00 | Coro2b | 7823 | 0.48356905 | 0.3747328 | 0.42047809 | 0.44844694 | 0.34454656 | 0.48773622 | 1.07047733 | 0.70789957 | 0.66862936 | 0.58463738 | 0.42748908 | 0.64213341 | 0.55492889 | 0.36808557 | 0.57933927 | 0.66566824 | 0.5291116 | 0.7031715 |
| ENSMUSG00 | Prrxl1 | 3248 | 0 | 0 | 0 | 0 | 0 | 0 | 0 | 0.02351749 | 0 | 0 | 0 | 0 | 0 | 0 | 0 | 0 | 0 | 0 |
| ENSMUSG00 | Pgm5 | 3622 | 0 | 0 | 0 | 0 | 0 | 0.13681042 | 0 | 0 | 0.09096966 | 0 | 0.07914129 | 0 | 0.19068107 | 0.14105051 | 0 | 0.10571675 | 0 | 0 |
| ENSMUSG00 | Coq5 | 2030 | 30.9052413 | 35.2821293 | 28.7381663 | 32.7007101 | 30.5151654 | 31.0497917 | 28.8071286 | 33.0373688 | 34.1259702 | 31.1241095 | 26.7822181 | 29.1451243 | 36.3549499 | 40.0380064 | 29.2673121 | 35.4801145 | 34.0971883 | 31.000206 |
| ENSMUSG00 | Kirrel | 12466 | 0 | 0 | 0 | 0.01827412 | 0 | 0.00397503 | 0 | 0.19607843 | 0.03964689 | 0.01512938 | 0 | 0 | 0.00395732 | 0 | 0 | 0.0030716 | 0.0092234 | 0 |
| ENSMUSG00 | Gm13178 | 1223 | 0 | 0 | 0 | 0 | 0 | 0 | 0 | 0 | 0 | 0 | 0 | 0 | 0 | 0 | 0 | 0 | 0 | 0 |
| ENSMUSG00 | Tspo | 850 | 228.427817 | 195.801435 | 208.746011 | 196.180497 | 218.632185 | 195.529257 | 169.546376 | 147.692267 | 153.892205 | 191.653741 | 186.659573 | 175.85217 | 185.429971 | 137.638419 | 157.72925 | 173.614119 | 151.546687 | 172.405012 |
| ENSMUSG00 | Tmem45b | 1566 | 0.02714251 | 0 | 0.30889866 | 0.49459627 | 0.39956258 | 0.37971443 | 0.24169951 | 0.48777015 | 0.52600914 | 0.75275215 | 0 | 0.46334841 | 1.03956148 | 1.09733795 | 1.15764269 | 0.39121949 | 0.95448632 | 0.03512714 |
| ENSMUSG00 | Rnf10 | 4379 | 41.1559594 | 51.1142581 | 41.2704977 | 49.9101004 | 44.8784299 | 46.2710956 | 40.397832 | 45.4401499 | 39.0138542 | 47.0645636 | 42.6908912 | 42.8527649 | 46.5943845 | 44.7789306 | 46.7622135 | 44.8574597 | 50.8858393 | 50.1224791 |
| ENSMUSG00 | Pde3a | 4740 | 0.12554271 | 0.1335327 | 0.02041077 | 0.09612035 | 0.1421618 | 0.16726661 | 0 | 0.07251722 | 0.13902621 | 0 | 0.05039553 | 0.10597911 | 0 | 0 | 0.17384623 | 0 | 0 | 0.23210594 |
| ENSMUSG00 | Utp15 | 4058 | 6.97596038 | 7.73305726 | 9.21457343 | 8.90337806 | 8.65852742 | 11.6127771 | 7.71833254 | 9.80692048 | 8.44435217 | 9.78336425 | 10.2543097 | 10.0820307 | 10.5884919 | 9.31628564 | 8.18345344 | 8.73758069 | 11.2674402 | 8.89255151 |
| ENSMUSG00 | Ccrl1 | 2695 | 1.56141463 | 3.33746992 | 2.13597407 | 1.4200845 | 1.08944243 | 1.41579239 | 1.5449038 | 0.65189258 | 1.42127865 | 1.04973736 | 1.50681687 | 0.51777052 | 0.58575931 | 0.82720452 | 0.88671252 | 0.7246089 | 0.91016067 | 0.46946547 |
| ENSMUSG00 | Cd1d2 | 1236 | 0.72217532 | 0.16171308 | 0.15654864 | 0 | 0.38941686 | 0.20045604 | 0.2296732 | 0.09270001 | 0.06664485 | 0 | 0 | 0 | 0.15965022 | 0.30060898 | 0 | 0 | 0 | 0 |
| ENSMUSG00 | Trem3 | 990 | 0 | 0 | 0 | 0 | 0 | 0 | 0 | 0 | 0 | 0 | 0 | 0 | 0 | 0 | 0 | 0 | 0 | 0 |
| ENSMUSG00 | Plekha6 | 7443 | 9.73684336 | 9.50198501 | 7.68855118 | 8.15973023 | 8.58136003 | 6.95056481 | 5.32688767 | 5.58800567 | 5.92094208 | 5.41000948 | 4.3583497 | 5.53432205 | 3.54595939 | 4.43037781 | 3.64796793 | 3.78636129 | 3.0277929 | 3.08192899 |
| ENSMUSG00 | Gpr155 | 7694 | 27.7990702 | 24.9911652 | 26.8839636 | 25.4156489 | 25.1419549 | 26.1803824 | 25.7655478 | 23.4445953 | 23.6338013 | 27.6996593 | 22.1488532 | 24.0412147 | 19.2736864 | 21.7733069 | 21.2166057 | 19.8320861 | 22.4209335 | 22.8716055 |
| ENSMUSG00 | Tpp2 | 4652 | 26.2779205 | 26.5815699 | 29.8123198 | 23.7305267 | 29.7772305 | 27.8760756 | 22.2019729 | 18.2013233 | 23.3909662 | 22.2273199 | 23.7128932 | 22.460645 | 19.8916858 | 22.4333328 | 24.2231847 | 29.7468304 | 32.1472937 | 23.2949141 |
| ENSMUSG00 | Ubac2 | 2144 | 59.2177989 | 59.2764419 | 57.9399375 | 63.432704 | 58.9301771 | 63.6743382 | 77.9202749 | 66.3735501 | 71.384838 | 70.9899581 | 74.4478248 | 71.1493497 | 56.6028528 | 59.7014482 | 66.2222769 | 59.4540261 | 58.6156378 | 59.9609526 |
| ENSMUSG00 | Ppp2r2d | 4313 | 23.9085456 | 22.3450505 | 21.4332985 | 22.8914405 | 22.0851094 | 20.5311233 | 20.5134963 | 21.3941211 | 22.7084476 | 18.8143659 | 21.8437788 | 20.3824875 | 24.4543564 | 22.2690407 | 21.3793374 | 20.4015339 | 21.3091723 | 20.7766832 |
| ENSMUSG00 | Slc24a4 | 4509 | 0 | 0 | 0 | 0 | 0 | 0 | 0 | 0 | 0 | 0 | 0 | 0 | 0 | 0 | 0 | 0 | 0 | 0 |
| ENSMUSG00 | Enc1 | 4739 | 2.79839943 | 2.69230607 | 2.47022459 | 2.02856728 | 1.89927616 | 1.84032098 | 0.72880894 | 1.48288712 | 2.42478116 | 1.49242572 | 1.35088509 | 1.80202496 | 1.85293971 | 0.68602788 | 1.00852089 | 1.65637783 | 1.01901528 | 0.71968023 |
| ENSMUSG00 | Ydjc | 5278 | 0.52346276 | 0.44812722 | 0.51324701 | 0.34529022 | 0.24622242 | 0.49759282 | 0.57370433 | 0.58612819 | 0.18728238 | 0.46453895 | 0.38017208 | 0.61335893 | 0.49602874 | 0.92395399 | 0.57766486 | 0.42077591 | 0.39938354 | 0.52111696 |
| ENSMUSG00 | Mapk1ip1 | 2973 | 12.5385265 | 10.3311431 | 12.5937143 | 11.0646061 | 10.6042415 | 10.1338924 | 11.8401053 | 11.2791555 | 12.1218302 | 10.8955712 | 12.6146459 | 13.1794804 | 14.7514546 | 12.6225438 | 10.7819746 | 15.5326177 | 14.3610813 | 13.9511828 |
| ENSMUSG00 | Cir1 | 3150 | 2.73922246 | 3.6591306 | 2.94848185 | 2.76259031 | 2.97959527 | 2.81585375 | 2.95891992 | 4.0132334 | 3.28184618 | 3.20325413 | 3.70066345 | 2.87051979 | 3.69597856 | 3.71552699 | 2.56365237 | 3.05109284 | 3.54061981 | 3.61488416 |
| ENSMUSG00 | Tram2 | 6484 | 0.83253506 | 1.05836771 | 1.34288024 | 0.8150958 | 1.12090075 | 0.97057328 | 0.97047907 | 1.0720261 | 1.09254792 | 1.70161391 | 0.97996136 | 0.87803788 | 0.84451617 | 0.95982537 | 1.11200991 | 1.14564741 | 1.19991337 | 1.00109108 |
| ENSMUSG00 | Cpsf2 | 5300 | 11.3400602 | 11.9360729 | 11.801316 | 10.7627222 | 12.0602254 | 12.0515988 | 11.1407968 | 10.6938727 | 11.2446959 | 11.0047962 | 11.2406365 | 8.36181813 | 10.6761715 | 11.0414245 | 10.5180576 | 12.9393354 | 9.82744104 | 9.48647614 |
| ENSMUSG00 | Lad1 | 2982 | 15.5652757 | 15.3829159 | 16.5624998 | 16.2259655 | 13.5744292 | 15.3211336 | 11.9947637 | 15.9199048 | 13.9360142 | 10.4199576 | 12.7047425 | 11.2117981 | 15.6002676 | 21.4776648 | 18.1137625 | 10.8759706 | 10.9760756 | 8.30117305 |
| ENSMUSG00 | 2700046A07l | 2463 | 1.05270635 | 0.51396264 | 0.45172196 | 0.33296744 | 0.97709955 | 0.8449918 | 0.96046851 | 1.27152943 | 0.58527326 | 1.18690361 | 0.95045593 | 0.67984973 | 2.34341629 | 1.28225657 | 2.45904536 | 1.10378933 | 1.26042633 | 3.17145479 |
| ENSMUSG00 | Capza3 | 1264 | 0 | 0 | 0 | 0 | 0 | 0.03920311 | 0 | 0 | 0 | 0 | 0 | 0 | 0 | 0 | 0 | 0 | 0 | 0 |
| ENSMUSG00 | Myrip | 5421 | 0.02352251 | 0.04916125 | 0 | 0.06723637 | 0.01775758 | 0 | 0 | 0 | 0 | 0.08697787 | 0 | 0.17503528 | 0.0182003 | 0.16278134 | 0 | 0.09888738 | 0 | 0 |
| ENSMUSG00 | Abca9 | 8009 | 0 | 0 | 0 | 0 | 0 | 0 | 0 | 0 | 0.05142529 | 0 | 0 | 0.00696912 | 0 | 0.01159797 | 0 | 0 | 0 | 0 |
| ENSMUSG00 | Gck | 3289 | 0.29723899 | 0.17218583 | 0.05883068 | 0.15237807 | 0.23414739 | 0.28625781 | 0.97818856 | 0.76640272 | 0.76387275 | 0.44441222 | 0.45029605 | 0.91640188 | 0.49496907 | 0 | 0.02505415 | 0.12806223 | 0.13983447 | 0.33450354 |
| ENSMUSG00 | Phlda3 | 1556 | 13.8223773 | 15.3504789 | 11.2850845 | 13.4984842 | 12.4969776 | 13.1524929 | 7.44961847 | 6.03813046 | 7.64968063 | 7.42411459 | 6.17144387 | 7.46121968 | 5.70677706 | 7.0740673 | 9.18826471 | 9.52343513 | 9.87714978 | 10.7472805 |
| ENSMUSG00 | Pramel1 | 2790 | 0 | 0 | 0 | 0 | 0 | 0 | 0 | 0 | 0 | 0 | 0 | 0 | 0 | 0 | 0.01476758 | 0 | 0 | 0 |
| ENSMUSG00 | Efhc1 | 2273 | 0.26180047 | 1.11384952 | 0.44691779 | 0.36080018 | 0.33880809 | 0.47961291 | 0.31222621 | 0.9913558 | 0.05435968 | 0.26966966 | 0.65157224 | 0.41745106 | 0.69451005 | 0.85818375 | 0.47128924 | 0.15161261 | 0.75876963 | 1.06484852 |
| ENSMUSG00 | Poldip3 | 5671 | 24.0970095 | 27.0333167 | 22.7494631 | 29.9750055 | 25.0632288 | 27.5768695 | 23.1933076 | 29.2622092 | 24.9617448 | 27.7117429 | 32.2571532 | 30.5701725 | 30.4116351 | 26.8378195 | 29.2864524 | 34.820078 | 32.9534515 | 29.9635215 |
| ENSMUSG00 | Fam169a | 4575 | 2.40630395 | 4.4125932 | 2.52705437 | 5.97521893 | 4.30294137 | 4.84154577 | 4.25038493 | 4.23246951 | 4.21317801 | 6.23522961 | 3.35207905 | 5.1972617 | 4.85231973 | 6.84225462 | 7.11458561 | 6.11812765 | 7.31341141 | 4.96584942 |
| ENSMUSG00 | Oasl1 | 2666 | 1.49868213 | 1.72437337 | 1.48785802 | 0.8544832 | 1.3179409 | 1.82151834 | 0.78085442 | 0.27218891 | 1.25135325 | 0.3006609 | 0.62720313 | 0.48153047 | 0.27756143 | 1.58530081 | 0.80363125 | 1.1777061 | 0.53191035 | 1.87765518 |
| ENSMUSG00 | Abca8a | 7016 | 0 | 0 | 0 | 0 | 0 | 0 | 0 | 0.01088723 | 0 | 0 | 0 | 0 | 0 | 0.1191553 | 0.01761754 | 0 | 0.04370159 | 0 |
| ENSMUSG00 | Sytl3 | 5020 | 0.47416133 | 0.65696744 | 0.28908484 | 0.39933983 | 0.26846491 | 0.37510037 | 0.35814378 | 0.57821167 | 0.81224404 | 0.5071988 | 0.39971081 | 0.48922095 | 0.65841404 | 0.76790025 | 0.6565985 | 0.35849772 | 0.27484994 | 0.19724381 |
| ENSMUSG00 | Ptprc | 6825 | 0.94040756 | 0.93227294 | 0.82217282 | 0.56742693 | 1.15657663 | 0.62440075 | 1.96876202 | 2.08169578 | 1.08020312 | 1.38170559 | 1.27399889 | 1.68469253 | 0.99025241 | 1.08879912 | 1.47299335 | 1.08840701 | 1.30281332 | 1.13645188 |
| ENSMUSG00 | Pdcd7 | 3285 | 7.90583335 | 9.01527031 | 5.77242737 | 7.36466196 | 7.60440438 | 7.30092026 | 7.34534741 | 8.8476159 | 9.80452217 | 8.61200059 | 8.55149942 | 8.47854357 | 5.69156671 | 7.36601812 | 8.50370012 | 10.1059229 | 6.79022977 | 7.53549407 |
| ENSMUSG00 | Haus1 | 995 | 22.8545419 | 17.0079902 | 17.4533642 | 18.5907377 | 19.4946384 | 18.8748603 | 25.1066276 | 22.7235198 | 25.9951074 | 22.3195791 | 19.4460886 | 21.8213997 | 21.2201614 | 20.7247988 | 18.5096437 | 12.6224314 | 18.4890679 | 18.1336553 |
| ENSMUSG00 | Rpl37 | 863 | 104.218948 | 81.6030804 | 93.664159 | 88.0600486 | 96.4869385 | 93.6506471 | 98.9015907 | 85.014606 | 89.9135524 | 86.3245784 | 86.8031691 | 90.4174928 | 142.679566 | 147.404815 | 102.025174 | 92.7315985 | 93.4397783 | 128.057121 |
| ENSMUSG00 | Fhdc1 | 5595 | 1.67133847 | 2.20299747 | 1.21906484 | 1.55534575 | 1.79795747 | 1.93959761 | 1.63205488 | 1.13314366 | 0.27236838 | 1.61804022 | 1.01612512 | 0.80805517 | 2.00149155 | 2.77253355 | 1.45070715 | 2.80592476 | 2.87704105 | 2.86106346 |
| ENSMUSG00 | Rhod | 2006 | 7.69161461 | 4.69967551 | 4.94345649 | 4.56518939 | 4.36690434 | 4.64402488 | 6.29735059 | 4.55034106 | 6.54960046 | 6.32280118 | 6.78756888 | 8.34730747 | 16.2800245 | 12.3634809 | 12.0975665 | 14.5641742 | 12.0175677 | 12.3126066 |
| ENSMUSG00 | Smek1 | 4579 | 17.2842625 | 18.3042677 | 19.4170231 | 20.0392975 | 19.0046993 | 19.3925526 | 15.2094919 | 19.342254 | 18.4030599 | 19.2865838 | 17.2987292 | 19.4300428 | 16.9144158 | 16.2082664 | 16.4752236 | 15.0018235 | 19.8118246 | 17.2872034 |
| ENSMUSG00 | Card6 | 6098 | 1.16404795 | 1.95572583 | 1.35648962 | 1.62878118 | 1.28656995 | 1.47081744 | 0.62069766 | 1.25888371 | 1.41160738 | 1.01291278 | 1.25634438 | 1.40957888 | 1.28628646 | 1.75936113 | 1.45266227 | 1.51956822 | 1.53983974 | 0.77579258 |
| ENSMUSG00 | Tcf20 | 2807 | 2.86194452 | 2.82453467 | 1.77501732 | 2.56453331 | 3.12077311 | 2.57737766 | 3.47218693 | 1.94567636 | 2.68512019 | 2.75479706 | 2.28067133 | 2.66451685 | 2.75921304 | 1.34021057 | 1.82009011 | 2.57817057 | 1.93884366 | 1.23461836 |
| ENSMUSG00 | Oosp1 | 1133 | 0 | 0 | 0 | 0 | 0 | 0 | 0 | 0 | 0 | 0 | 0.5481681 | 0 | 0.04354097 | 0 | 0 | 0 | 0 | 0.48551727 |
| ENSMUSG00 | Mcm3 | 2974 | 2.7726981 | 2.85635105 | 1.90306088 | 2.58904388 | 2.24966908 | 2.31601546 | 1.82950843 | 2.42715671 | 1.38488621 | 2.63181464 | 1.10843044 | 1.05100104 | 4.24645959 | 1.43673707 | 3.75441212 | 3.28315564 | 4.02078165 | 3.21842117 |
| ENSMUSG00 | Ankrd13a | 4462 | 39.9807763 | 41.7792371 | 43.3106799 | 43.1819042 | 41.0771773 | 46.554249 | 38.3421441 | 41.5477188 | 47.0201953 | 45.5127922 | 42.1965736 | 43.8319323 | 45.8713861 | 42.4575151 | 48.5701891 | 41.9720665 | 48.8655991 | 47.1806032 |
| ENSMUSG00 | Il17f | 1170 | 0 | 0.17083535 | 0 | 0 | 0 | 0 | 0.20219094 | 0 | 0 | 0 | 0 | 0 | 0 | 0 | 0 | 0 | 0 | 0 |
| ENSMUSG00 | 8430432A02l | 1247 | 2.93139145 | 2.80501515 | 2.94818629 | 2.33833748 | 2.54747952 | 2.98031677 | 1.7832365 | 2.72584107 | 1.94868038 | 1.39901846 | 1.76234811 | 2.19323774 | 1.14725389 | 2.42090271 | 1.94939198 | 0.76765452 | 1.26012684 | 1.23516839 |
| ENSMUSG00 | Ipo9 | 8973 | 8.70662137 | 9.90513783 | 9.97875902 | 11.2366646 | 10.5189683 | 10.2441013 | 8.28354158 | 10.7771273 | 7.89488545 | 7.88209683 | 9.22169041 | 9.50477348 | 8.04880496 | 9.52898305 | 8.24673959 | 8.90161674 | 10.2681843 | 7.23400713 |
| ENSMUSG00 | Ndufa7 | 1406 | 169.990516 | 140.041918 | 153.331723 | 138.883524 | 162.787088 | 153.514932 | 202.154174 | 162.399086 | 166.618177 | 187.933059 | 190.998041 | 168.239824 | 243.856637 | 222.621552 | 190.557196 | 190.310846 | 183.43815 | 201.869357 |
| ENSMUSG00 | Olfr680-ps1 | 898 | 0 | 0 | 0 | 0 | 0 | 0 | 0 | 0 | 0 | 0 | 0 | 0 | 0 | 0 | 0 | 0 | 0 | 0 |
| ENSMUSG00 | Macc1 | 3814 | 1.2036075 | 1.91282743 | 1.43319584 | 1.66045757 | 1.22412076 | 2.28664948 | 0.94278018 | 0.37050836 | 1.01508455 | 0.32142587 | 0.28810279 | 0.19024741 | 0.8536724 | 0.94982664 | 1.17749596 | 1.10434368 | 0.41200267 | 0.73557012 |
| ENSMUSG00 | Shisa4 | 3349 | 15.1160539 | 17.3975073 | 16.3796905 | 14.6927226 | 16.1685321 | 16.9121453 | 15.6249098 | 17.0491614 | 19.5540639 | 16.2190535 | 17.3325096 | 15.6163857 | 14.2442413 | 15.3657954 | 13.9635013 | 21.8035797 | 16.3765097 | 13.1732767 |
| ENSMUSG00 | Git2 | 9965 | 3.43795001 | 3.75083465 | 3.76211601 | 4.12404034 | 3.31827713 | 3.45103834 | 3.56091412 | 2.59470712 | 3.24036411 | 4.20168836 | 3.15942776 | 3.78078999 | 3.40595676 | 2.58203958 | 3.78731813 | 3.72339173 | 3.07302672 | 2.89812153 |
| ENSMUSG00 | Lman1 | 4893 | 32.5152001 | 38.9229136 | 41.1466653 | 40.7097662 | 39.1606761 | 39.8102995 | 36.4248644 | 40.4950304 | 34.747177 | 36.5025269 | 37.4740018 | 36.6172648 | 29.1777169 | 27.8683508 | 28.7139394 | 29.9094015 | 35.9137689 | 28.9492929 |
| ENSMUSG00 | Wipi1 | 4080 | 5.82362583 | 5.73177739 | 5.41830965 | 4.80177673 | 5.35585621 | 5.82973336 | 4.69647916 | 5.46675564 | 5.02717766 | 4.8884185 | 4.51988578 | 6.99063813 | 4.93319173 | 5.14527635 | 5.07950554 | 4.25138362 | 5.08199182 | 4.82677946 |
| ENSMUSG00 | Gpr45 | 1545 | 0 | 0 | 0 | 0 | 0 | 0 | 0 | 0 | 0 | 0 | 0 | 0 | 0 | 0 | 0 | 0 | 0 | 0 |
| ENSMUSG00 | Dlx1 | 5316 | 0 | 0 | 0 | 0 | 0 | 0 | 0 | 0 | 0 | 0.106435 | 0 | 0 | 0 | 0.05241996 | 0 | 0 | 0 | 0 |
| ENSMUSG00 | Tdrkh | 3837 | 4.71910477 | 4.39309994 | 4.65202833 | 4.3459323 | 4.3151894 | 4.17136642 | 3.68686239 | 3.34444806 | 3.34902083 | 5.67725434 | 3.82249472 | 2.80751183 | 3.90849696 | 4.56331012 | 4.71396483 | 4.55056476 | 5.57364443 | 3.91386139 |
| ENSMUSG00 | Ammecr1l | 3472 | 9.9652112 | 10.9475842 | 7.95543947 | 11.1803024 | 8.87224399 | 10.4471777 | 8.27154266 | 9.73510257 | 9.89330477 | 8.25680518 | 8.86148423 | 8.84176683 | 8.88031345 | 6.35395781 | 8.14062999 | 9.18665782 | 8.86406003 | 8.96749839 |
| ENSMUSG00 | Slc16a6 | 5542 | 5.02361788 | 4.68255947 | 6.53767128 | 4.27494461 | 6.56581276 | 5.32902007 | 7.3589912 | 5.9542107 | 5.81159425 | 5.60669676 | 5.79299388 | 6.06297887 | 4.41512648 | 5.74056295 | 6.43820776 | 5.32005913 | 6.22404677 | 5.30040833 |
| ENSMUSG00 | Metap1d | 4398 | 8.24395524 | 8.91939884 | 8.22723981 | 8.36010977 | 8.91939922 | 8.06724812 | 10.0585163 | 10.0040126 | 10.6759045 | 7.80484583 | 11.2865352 | 8.14771609 | 8.09860034 | 9.16631829 | 9.15277856 | 7.53971487 | 11.8604116 | 7.70477711 |
| ENSMUSG00 | Nol4 | 7655 | 0 | 0.04786961 | 0 | 0 | 0.0691641 | 0 | 0 | 0 | 0 | 0 | 0 | 0 | 0 | 0 | 0 | 0 | 0 | 0 |
| ENSMUSG00 | Rnpep | 3169 | 16.698941 | 18.8797596 | 17.2642672 | 20.5161282 | 15.4769423 | 18.6076847 | 13.1531935 | 14.0886458 | 11.6450357 | 10.8912322 | 13.1008012 | 10.9024598 | 12.033314 | 10.8452587 | 11.2722464 | 13.2065743 | 11.658741 | 12.6022756 |
| ENSMUSG00 | Gm6363 | 889 | 0 | 0.11241697 | 0 | 0 | 0 | 0 | 0 | 0.08592217 | 0.09265808 | 0.05303791 | 0 | 0 | 0.05549147 | 0 | 0.3244217 | 0 | 0.0431117 | 0.06187751 |
| ENSMUSG00 | BC057022 | 3545 | 0.11990177 | 0.11276579 | 0.05458226 | 0.0257044 | 0.05430965 | 0.16773845 | 0.01334631 | 0.10773597 | 0.0116182 | 0.02660124 | 0.09433701 | 0.04723469 | 0.08349549 | 0.03930388 | 0.08135709 | 0.0432052 | 0.08649094 | 0.03103476 |
| ENSMUSG00 | AW549877 | 5077 | 10.6995499 | 12.3553634 | 11.9671369 | 12.5725868 | 13.8698059 | 14.0157033 | 12.3849813 | 13.9820049 | 15.4540698 | 13.9492528 | 14.5050287 | 16.446766 | 8.85195522 | 12.3314221 | 10.3957442 | 8.90707255 | 13.8524236 | 10.6290986 |
| ENSMUSG00 | Agrn | 9141 | 7.9188617 | 8.96142729 | 7.00122313 | 6.4845114 | 8.30369146 | 9.27521356 | 7.0443667 | 9.3256145 | 8.76356779 | 8.86171187 | 7.62016076 | 8.31647877 | 10.49133 | 8.01758778 | 11.7055509 | 9.58417129 | 11.7523914 | 10.7177791 |
| ENSMUSG00 | Mvk | 2727 | 3.46026736 | 4.40995776 | 3.72513435 | 4.44416494 | 5.34799152 | 3.74325747 | 4.61502479 | 3.36126754 | 4.06276961 | 3.37161244 | 5.25577101 | 3.27484636 | 6.60291522 | 5.50107819 | 5.0312131 | 4.94253532 | 4.04766172 | 5.84968667 |
| ENSMUSG00 | Mfsd9 | 7624 | 2.3415758 | 1.82644147 | 1.78291737 | 2.04379291 | 1.79926525 | 1.75488432 | 2.27130646 | 1.98877017 | 1.44239241 | 2.48617297 | 2.77601084 | 2.05721422 | 2.16118315 | 2.61948552 | 1.28079338 | 1.30079582 | 2.47331297 | 1.30596122 |
| ENSMUSG00 | Tmco7 | 4391 | 2.1683351 | 2.67049391 | 2.44566699 | 2.84302577 | 2.75133519 | 1.96360183 | 2.8984538 | 2.13098123 | 3.21725687 | 3.23214796 | 2.20868061 | 2.72023498 | 1.61780826 | 1.48079532 | 1.95170203 | 1.00282848 | 1.6758481 | 1.1400202 |
| ENSMUSG00 | Tnfrsf18 | 1664 | 0.58751145 | 2.10207566 | 2.15122671 | 1.56068481 | 0.95453935 | 1.31028863 | 1.25105641 | 0.87218227 | 0.29701814 | 0.96341581 | 1.17714741 | 1.81132559 | 1.1858634 | 1.25599996 | 0.94090092 | 0.41420128 | 0.46065261 | 0.76034222 |
| ENSMUSG00 | Pkp2 | 5597 | 8.16384925 | 8.98739865 | 9.31689579 | 9.80901526 | 9.06397133 | 9.79191055 | 8.9012419 | 8.2840051 | 8.80097782 | 7.48916821 | 7.99805898 | 6.65160746 | 6.09928297 | 6.97864526 | 7.48650748 | 7.55276587 | 8.26511352 | 7.36141164 |
| ENSMUSG00 | Pigs | 2548 | 23.5213101 | 25.1023379 | 24.6234179 | 26.8037698 | 23.9904015 | 22.578777 | 21.6165004 | 23.6379194 | 17.7806779 | 24.0194713 | 24.112479 | 22.8914315 | 21.7230813 | 26.7764381 | 25.7913978 | 30.2056989 | 26.7441742 | 25.1081598 |
| ENSMUSG00 | S100a10 | 1256 | 251.376152 | 203.855816 | 216.294384 | 181.228485 | 219.812671 | 198.01367 | 221.872356 | 196.161774 | 194.291486 | 180.531634 | 182.883758 | 170.113341 | 270.579282 | 287.945946 | 243.371292 | 265.869735 | 223.549722 | 258.096072 |
| ENSMUSG00 | Znrf3 | 9042 | 6.20513519 | 7.90637812 | 6.82643494 | 8.56096155 | 6.70184604 | 7.53539136 | 6.05928801 | 6.15842985 | 5.46602731 | 5.60051485 | 6.52533441 | 6.95072286 | 5.91961184 | 5.91722945 | 6.44771086 | 6.50457564 | 5.65866029 | 6.27841165 |
| ENSMUSG00 | Dcaf17 | 9789 | 5.72293615 | 5.18971945 | 6.19679304 | 4.97080349 | 5.15786985 | 5.16838707 | 5.19090992 | 6.44928401 | 5.3644711 | 5.88601077 | 5.9346562 | 5.17160139 | 5.12015813 | 5.74086686 | 4.65090604 | 5.20633348 | 5.78672661 | 5.11934786 |
| ENSMUSG00 | 2310008H04 | 3327 | 3.59000735 | 3.63468007 | 3.27142901 | 5.1353747 | 4.57159239 | 4.06609446 | 4.06715545 | 4.15559052 | 3.13200736 | 3.13204251 | 4.02073591 | 3.87180099 | 3.42520977 | 4.59275766 | 4.91644656 | 7.22768303 | 6.15156073 | 4.81143675 |
| ENSMUSG00 | Mettl8 | 5845 | 2.18161725 | 2.21706003 | 2.03590906 | 2.37743687 | 2.00927106 | 2.09401629 | 1.50558739 | 1.88838397 | 3.44571535 | 2.35551845 | 2.49296191 | 2.16769126 | 2.0171648 | 2.20102861 | 2.24159021 | 1.8473826 | 2.22285965 | 2.10813343 |
| ENSMUSG00 | Arhgef11 | 9094 | 5.44986092 | 7.00032338 | 6.25015375 | 4.70440077 | 5.80610515 | 5.52523332 | 4.86965774 | 6.11901587 | 4.86412117 | 5.59441487 | 4.56525609 | 5.33360576 | 3.99797922 | 4.15719014 | 6.52863889 | 4.15158774 | 5.17957102 | 6.77481851 |
| ENSMUSG00 | Rptn | 4143 | 0 | 0 | 0 | 0 | 0 | 0 | 0 | 0 | 0 | 0 | 0 | 0 | 0 | 0 | 0 | 0 | 0 | 0 |
| ENSMUSG00 | Elmod1 | 2648 | 0 | 0.6667611 | 0.0730718 | 0.15485249 | 0 | 0.01871327 | 0 | 0 | 0 | 0 | 0 | 0 | 0 | 0 | 0 | 0 | 0 | 0 |
| ENSMUSG00 | Hrnr | 11187 | 0 | 0 | 0 | 0 | 0 | 0 | 0 | 0 | 0 | 0 | 0 | 0 | 0 | 0 | 0 | 0 | 0 | 0 |
| ENSMUSG00 | Rapgef5 | 3959 | 5.79762441 | 4.6111306 | 6.87908502 | 5.78863483 | 4.52262582 | 7.30962274 | 5.16268686 | 3.41503169 | 3.96364297 | 5.09737333 | 3.82537556 | 3.66559037 | 4.87213429 | 2.59261008 | 2.58095122 | 2.92087988 | 3.03009117 | 3.72377888 |
| ENSMUSG00 | Zbed3 | 1892 | 51.9183202 | 54.7585101 | 53.7908202 | 56.3733641 | 54.5681693 | 52.0670372 | 51.6888517 | 54.906625 | 51.9185201 | 52.5585799 | 56.7138249 | 58.1167311 | 38.615523 | 41.9027786 | 41.2015559 | 48.8346912 | 40.0886583 | 40.151996 |
| ENSMUSG00 | Tlk1 | 6491 | 8.70927193 | 7.66745709 | 8.75657035 | 8.97042284 | 7.82301332 | 8.9394933 | 9.11120762 | 8.6493347 | 8.18527274 | 8.43351818 | 7.78707535 | 8.40975474 | 8.1168522 | 6.1176544 | 7.72489501 | 7.12012989 | 7.36885203 | 5.94074624 |
| ENSMUSG00 | Foxn4 | 3530 | 0 | 0 | 0 | 0 | 0 | 0 | 0 | 0 | 0 | 0.01335714 | 0 | 0 | 0 | 0 | 0 | 0 | 0.05428654 | 0 |
| ENSMUSG00 | Acacb | 9577 | 0.02662953 | 0.1426155 | 0.01515303 | 0.08563212 | 0.07538674 | 0.07761209 | 0.29147416 | 0.56628601 | 0.58917747 | 0.49233271 | 0.34919569 | 0.25060807 | 0.44814418 | 0.27642403 | 0.17638757 | 0.31185835 | 0.52425028 | 0.35037648 |
| ENSMUSG00 | Wdr41 | 5066 | 13.3657216 | 16.2027184 | 14.8959154 | 14.9202078 | 17.1587553 | 15.7089795 | 14.6439559 | 14.1807953 | 15.9672951 | 14.2122234 | 14.0891804 | 15.2154422 | 12.9610703 | 13.458334 | 11.508133 | 12.781172 | 14.3893835 | 13.4971022 |
| ENSMUSG00 | Ncapg2 | 6575 | 0.59474923 | 0.59785879 | 0.66950437 | 0.61671984 | 0.77596713 | 1.1681633 | 0.69799694 | 0.56925558 | 0.77048539 | 0.51632709 | 0.39963844 | 0.97624353 | 0.69777465 | 0.61454534 | 0.43864774 | 0.78619498 | 0.96180061 | 0.66931233 |
| ENSMUSG00 | Lce3b | 561 | 0 | 0 | 0 | 0 | 0 | 0 | 0 | 0 | 0 | 0 | 0 | 0 | 0 | 0 | 0 | 0 | 0 | 0 |
| ENSMUSG00 | Mat2b | 4602 | 47.086351 | 46.2413664 | 46.5445442 | 47.7985054 | 50.0981996 | 46.7854471 | 51.4353178 | 47.9438307 | 52.4720865 | 54.3739205 | 53.1109708 | 53.7295492 | 52.4620607 | 49.259798 | 50.9961022 | 53.0094437 | 58.022449 | 52.6662591 |

| | | | | | | | | | | | | | | | | | | | |
|---|---|---|---|---|---|---|---|---|---|---|---|---|---|---|---|---|---|---|---|
| ENSMUSG00 Igsf3 | 7319 | 3.24639888 | 3.33629614 | 3.20551472 | 3.22456906 | 2.93960512 | 3.76435577 | 1.64194841 | 2.26994057 | 2.05397981 | 1.90045873 | 2.42171193 | 2.5471282 | 2.48041338 | 2.0940769 | 2.20672358 | 3.58369532 | 2.73347821 | 3.1191118 |
| ENSMUSG00 2010003K11f | 1363 | 0 | 0 | 0 | 0 | 0 | 0 | 0 | 0 | 0 | 0 | 0 | 0 | 0 | 0 | 0 | 0 | 0 | 0 |
| ENSMUSG00 Csgalnact2 | 5489 | 2.84193835 | 3.2833441 | 3.36649455 | 3.11266016 | 3.22691709 | 3.00620519 | 4.09428357 | 3.54857446 | 3.51912478 | 3.55627458 | 2.97668716 | 4.66740556 | 2.0041934 | 3.03760746 | 2.30440475 | 3.03450796 | 2.70916436 | 3.35726922 |
| ENSMUSG00 Tbca | 499 | 203.581905 | 165.029007 | 166.350657 | 156.861471 | 161.468618 | 156.801135 | 206.127784 | 172.286769 | 202.961207 | 178.01989 | 172.621744 | 181.764432 | 221.647612 | 208.486484 | 188.751016 | 211.864463 | 200.080167 | 208.79208 |
| ENSMUSG00 Sln | 513 | 0 | 0 | 0 | 0 | 0 | 0 | 0 | 0 | 0 | 0 | 0 | 0 | 0 | 0 | 0 | 0 | 0 | 0 |
| ENSMUSG00 Dstyk | 3328 | 7.03736538 | 8.91880672 | 11.2794049 | 8.43317407 | 6.551611 | 7.75750427 | 6.80972753 | 7.87259256 | 6.1507507 | 7.32479375 | 8.28309827 | 6.44026876 | 6.55189526 | 6.22417758 | 8.13384081 | 5.39612226 | 8.82149742 | 5.3554539 |
| ENSMUSG00 Wdr60 | 3703 | 6.2328681 | 6.03644397 | 7.7857478 | 5.96735255 | 6.83699483 | 6.90499879 | 6.98893745 | 7.07534109 | 5.78368571 | 7.10507492 | 6.94110928 | 6.58694155 | 7.23392684 | 5.88233243 | 7.79970043 | 9.36842838 | 7.96955134 | 6.68487659 |
| ENSMUSG00 Wdr11 | 9839 | 7.61897553 | 9.39009582 | 8.2905978 | 7.44934183 | 8.32409137 | 7.89635842 | 7.28415972 | 6.39115521 | 7.87496095 | 8.1935292 | 8.1186204 | 6.96030983 | 7.0525347 | 5.98921056 | 8.38649784 | 6.15335106 | 7.65014994 | 6.44312496 |
| ENSMUSG00 Zfp386 | 1212 | 42.6103869 | 55.1641738 | 45.938889 | 40.5237672 | 50.3955537 | 41.0486341 | 51.0992547 | 59.7465305 | 48.682616 | 44.81651 | 57.2747646 | 43.4735854 | 39.8481411 | 63.2283368 | 51.3659661 | 29.63415 | 52.7777141 | 40.2129279 |
| ENSMUSG00 Myo3b | 7055 | 6.21762462 | 6.59175058 | 5.73214336 | 6.39339956 | 7.33406349 | 6.53913465 | 4.84192122 | 6.04690482 | 6.33997961 | 5.56050817 | 4.57095634 | 7.32605295 | 5.52405594 | 6.05649332 | 4.74212096 | 4.09771915 | 5.78561392 | 5.70753595 |
| ENSMUSG00 Tmcc2 | 7185 | 0.05915821 | 0.05100095 | 0.22217488 | 0.24730421 | 0.14067786 | 0.14483054 | 0.21071757 | 0.11694264 | 0.17196875 | 0.11156047 | 0.30586612 | 0.02330508 | 0.00686596 | 0 | 0.04587508 | 0.01598773 | 0 | 0.06890494 |
| ENSMUSG00 Abhd14b | 1573 | 60.9610153 | 59.3829341 | 57.0764605 | 55.7854872 | 55.8428231 | 56.4831858 | 71.7059292 | 56.1838647 | 67.2651996 | 62.6777626 | 64.084655 | 73.1316469 | 75.4561939 | 59.5830695 | 58.7246588 | 65.6271313 | 61.2538528 | 60.9891177 |
| ENSMUSG00 Svop | 4877 | 0 | 0.00683061 | 0 | 0 | 0 | 0 | 0 | 0 | 0 | 0 | 0 | 0 | 0 | 0 | 0 | 0.10991768 | 0.00785858 | 0 |
| ENSMUSG00 Hnrnpf | 2807 | 30.5122657 | 32.7313724 | 30.0546623 | 33.5337077 | 30.7962005 | 31.7229292 | 24.4569637 | 28.722537 | 26.5430734 | 28.3374551 | 27.1808367 | 25.1936033 | 24.4287015 | 26.2085623 | 25.4372271 | 27.2276638 | 28.6457324 | 26.7892586 |
| ENSMUSG00 Arsb | 4659 | 4.61635953 | 5.80598199 | 5.32638357 | 6.19997899 | 5.02084458 | 7.73231108 | 6.13368922 | 7.00071081 | 4.9770355 | 7.62062236 | 7.6804988 | 5.90622916 | 4.82836537 | 6.77870346 | 7.28698907 | 6.22150457 | 7.31317406 | 7.71001217 |
| ENSMUSG00 4933440N22 | 650 | 0.32696289 | 1.07626274 | 0.37210408 | 0.21028174 | 0.22214734 | 0.22870492 | 0.50952116 | 0.70509051 | 0.38018322 | 0.5077768 | 0.51449955 | 0.343481 | 0.5312668 | 0.57161954 | 0.12677402 | 0.47126901 | 0.3537812 | 1.18481153 |
| ENSMUSG00 Lce1c | 454 | 0 | 0 | 0 | 0 | 0 | 0 | 0 | 0 | 0 | 0 | 0 | 0 | 0 | 0 | 0 | 0 | 0 | 0 |
| ENSMUSG00 Dao | 2471 | 0 | 0 | 0.0587295 | 0.0184383 | 0.07791489 | 0 | 0 | 0 | 0 | 0 | 0 | 0 | 0 | 0.07985741 | 0.15036532 | 0.08337021 | 0 | 0.03102088 | 0.06678564 |
| ENSMUSG00 Zfp239 | 3832 | 0.6322534 | 0.34773376 | 0.75741436 | 0.87983228 | 0.2763315 | 0.71122138 | 0.93835167 | 0.36876798 | 0.73086719 | 0.7382678 | 0.74804215 | 0.83024342 | 0.18023143 | 0.35148187 | 0.33331113 | 0.94927132 | 0.22003615 | 0.35887987 |
| ENSMUSG00 Kank3 | 2991 | 18.6022318 | 18.2769841 | 19.6340575 | 18.8124005 | 18.0072292 | 19.5659573 | 16.2770801 | 16.0890762 | 19.5673818 | 15.4489098 | 19.5029169 | 21.8336092 | 10.6052901 | 6.27329031 | 11.4747229 | 9.07657661 | 9.63603319 | 10.9613599 |
| ENSMUSG00 Dmgdh | 3395 | 0 | 0 | 0 | 0 | 0 | 0 | 0 | 0 | 0 | 0 | 0 | 0 | 0 | 0.05472057 | 0 | 0.05639265 | 0 | 0 |
| ENSMUSG00 Uggt2 | 13554 | 0.24460703 | 0.54562952 | 0.60315233 | 0.26891571 | 0.41193029 | 0.20473315 | 0.37699346 | 0.40576258 | 0.27652154 | 0.40353265 | 0.35600345 | 0.48592653 | 0.44403821 | 0.31867346 | 0.43469253 | 0.41528105 | 0.80023182 | 0.30438859 |
| ENSMUSG00 Inpp5f | 7960 | 4.17041991 | 4.12644644 | 3.97440815 | 3.80629328 | 3.83362307 | 3.97791417 | 3.44146245 | 3.24347538 | 2.70092478 | 3.60146075 | 2.44276484 | 3.52003299 | 3.27226768 | 3.38411692 | 2.60874173 | 3.71361167 | 3.52929342 | 3.30331068 |
| ENSMUSG00 6230427J02R | 1580 | 0.24211809 | 0.42168221 | 0.15308079 | 0.37486935 | 0.12185297 | 0 | 0.0898342 | 0.45927573 | 0.05213483 | 0.65652878 | 0 | 0 | 1.31135476 | 0.47031987 | 0.28684628 | 0.55739492 | 0 | 0.38297479 |
| ENSMUSG00 Csdc2 | 4623 | 1.35155978 | 0.92956162 | 0.97312099 | 1.13335931 | 1.05155187 | 1.17906137 | 1.22810328 | 1.93316508 | 1.39872009 | 1.77465333 | 2.44920293 | 2.19737196 | 1.70735599 | 2.52162361 | 2.9499708 | 1.82217902 | 2.66120296 | 3.12943869 |
| ENSMUSG00 Ccdc115 | 1699 | 24.9927433 | 22.4896349 | 23.3183763 | 21.1044979 | 18.9524761 | 21.1452218 | 27.5410474 | 30.1447979 | 28.8232885 | 28.5013374 | 23.9858973 | 26.807287 | 31.4167949 | 26.5160475 | 27.6697912 | 23.9344468 | 27.8593153 | 22.146103 |
| ENSMUSG00 Klhdc8a | 4264 | 0.73766018 | 0.96876146 | 1.23656539 | 1.30357585 | 0.92561392 | 0.45322622 | 0.95424259 | 1.51372327 | 0.88863965 | 1.54810001 | 1.32210251 | 0.9162984 | 0.28923497 | 0.22873495 | 1.71995238 | 0.24245929 | 1.24937975 | 0.56763619 |
| ENSMUSG00 Vwa1 | 3619 | 0.09396005 | 0.23932999 | 0.41436293 | 0.42803965 | 0.15959743 | 0.41077149 | 0.5360099 | 0.35881229 | 0.27313522 | 0.40388831 | 0.03960345 | 0.50895741 | 0.02726273 | 0.12833404 | 0.66031783 | 0.02116088 | 0.16944481 | 0.25840144 |
| ENSMUSG00 Bhmt2 | 2459 | 0 | 0 | 0 | 0 | 0 | 0 | 0 | 0 | 0 | 0 | 0 | 0 | 0 | 0 | 0.13404329 | 0 | 0 | 0 |
| ENSMUSG00 Ssh1 | 12576 | 1.59191618 | 2.23834253 | 1.65014667 | 1.41291407 | 1.83709632 | 1.90708676 | 1.40327841 | 1.50935305 | 1.54580434 | 1.66467178 | 1.28022909 | 1.22052379 | 1.01990287 | 1.65449675 | 1.36945375 | 1.08392623 | 1.18550648 | 0.44616165 |
| ENSMUSG00 Lce1f | 1141 | 0 | 0 | 0 | 0 | 0 | 0 | 0 | 0 | 0 | 0 | 0 | 0 | 0 | 0 | 0 | 0 | 0 | 0 |
| ENSMUSG00 Rassf4 | 9992 | 18.9001698 | 20.0570828 | 17.1427813 | 15.0107044 | 17.1197815 | 18.5426012 | 6.70010415 | 11.3445807 | 7.83170336 | 4.94534998 | 8.83109971 | 9.74763123 | 4.15707309 | 3.33270723 | 3.83068909 | 2.95456949 | 3.80117696 | 1.99842932 |
| ENSMUSG00 Ppig | 8059 | 15.2795006 | 14.2858124 | 14.813981 | 13.3760306 | 14.6922115 | 15.5009854 | 15.0585707 | 15.2930739 | 15.7356101 | 13.9889976 | 14.7848054 | 13.7132413 | 15.7440987 | 14.49973 | 14.0133347 | 13.5744218 | 15.8460381 | 15.9655417 |
| ENSMUSG00 BC024479 | 2285 | 6.62225062 | 4.98599992 | 4.97495826 | 5.50321586 | 5.350332 | 5.59501324 | 5.92184953 | 5.14803499 | 4.93877694 | 4.35395991 | 5.47791646 | 8.59829901 | 5.20305998 | 5.24401512 | 4.72420466 | 3.75365254 | 7.0782045 | 6.30738992 |
| ENSMUSG00 Cox10 | 3030 | 9.18841264 | 8.04786751 | 6.54559322 | 8.13482673 | 7.38658235 | 7.91535414 | 6.58942261 | 8.89895581 | 7.34017113 | 8.16967631 | 8.08863158 | 9.85524075 | 6.70783826 | 6.74435592 | 6.4997834 | 8.85865208 | 8.00678084 | 8.13335967 |
| ENSMUSG00 Klhl23 | 4919 | 49.927812 | 36.5568211 | 47.8818295 | 47.2004639 | 40.3920677 | 45.1706561 | 61.7113535 | 46.8573183 | 52.2722948 | 50.6110415 | 41.0540251 | 52.2981071 | 49.5926674 | 41.1189267 | 51.4370308 | 52.5123882 | 49.7174427 | 42.0703923 |
| ENSMUSG00 Dzip1 | 6035 | 0.73248356 | 0.85559215 | 0.61719335 | 1.07957404 | 0.76564452 | 0.87035456 | 0.72909348 | 0.53792116 | 1.10558667 | 0.79691329 | 0.91037618 | 1.22082311 | 1.07900797 | 1.36985046 | 0.97627548 | 1.65598153 | 1.079614 | 1.92326786 |
| ENSMUSG00 Sprr2i | 615 | 0 | 0 | 0 | 0 | 0 | 0 | 0 | 0 | 0 | 0 | 0 | 0 | 0 | 0 | 0 | 0 | 0 | 0 |
| ENSMUSG00 Gm9774 | 930 | 2.3766335 | 1.86265645 | 1.9245383 | 1.8126437 | 1.5526427 | 1.65175779 | 2.08582778 | 1.39628138 | 1.15145098 | 2.23078596 | 1.28427308 | 1.62045472 | 2.44007333 | 1.59807613 | 1.81641268 | 2.05863481 | 1.81328717 | 1.65618816 |
| ENSMUSG00 Papd4 | 1858 | 11.873082 | 13.0526302 | 14.6838919 | 13.1680736 | 15.4654243 | 11.8681197 | 14.7183684 | 13.3200628 | 16.1598331 | 13.7797803 | 11.5708995 | 14.0890989 | 19.9864165 | 17.6477533 | 14.1921398 | 18.1148783 | 18.9568713 | 16.3724629 |
| ENSMUSG00 Armc5 | 3828 | 4.53033224 | 7.10988364 | 4.76405981 | 6.45091067 | 5.74615703 | 6.9513631 | 4.21463519 | 5.67697935 | 4.48661888 | 6.08475638 | 4.53038399 | 4.89918042 | 3.82747634 | 4.67110074 | 5.31702419 | 3.85106684 | 5.17625274 | 6.13607329 |
| ENSMUSG00 Pnliprp1 | 1606 | 1.93205346 | 1.86684961 | 1.02409716 | 1.21987852 | 1.10889239 | 1.69701142 | 0 | 0 | 0.89758907 | 0 | 0.17848677 | 0 | 0 | 0.20243376 | 0 | 0.14305333 | 0.07159333 | 0.58228818 |
| ENSMUSG00 Bend6 | 4709 | 0 | 0 | 0 | 0.14512968 | 0 | 0 | 0 | 0.00811051 | 0 | 0 | 0 | 0 | 0 | 0 | 0 | 0 | 0 | 0 |
| ENSMUSG00 1700069L16f | 1514 | 0 | 0 | 0 | 0 | 0 | 0 | 0 | 0.12613079 | 0 | 0 | 0 | 0 | 0 | 0 | 0 | 0 | 0 | 0 |
| ENSMUSG00 Nfrkb | 10002 | 2.94926936 | 4.16660974 | 3.07594134 | 3.51205411 | 2.88252572 | 3.04192946 | 2.87603567 | 3.50536147 | 3.47544684 | 2.96047207 | 3.63017081 | 3.2701438 | 2.7028485 | 3.51511439 | 4.24291167 | 4.17666993 | 3.46017258 | 3.65737412 |
| ENSMUSG00 Tekt3 | 1740 | 0 | 0 | 0 | 0 | 0 | 0 | 0 | 0 | 0 | 0 | 0 | 0 | 0 | 0 | 0 | 0 | 0 | 0 |
| ENSMUSG00 Cmklr1 | 4303 | 0 | 0 | 0 | 0 | 0 | 0 | 0 | 0 | 0 | 0 | 0 | 0 | 0 | 0 | 0 | 0 | 0 | 0 |
| ENSMUSG00 Slc35f2 | 2560 | 0.96300789 | 0.67666816 | 1.56836055 | 1.70853916 | 0.9776797 | 1.31624449 | 16.1343628 | 13.9193404 | 12.2111584 | 6.29903926 | 4.9081305 | 10.1819978 | 28.8090689 | 41.1102742 | 20.42374 | 4.98077042 | 8.26410777 | 21.3804927 |
| ENSMUSG00 Zfp451 | 17179 | 2.12043401 | 2.07296606 | 2.71729768 | 2.28879337 | 2.39272733 | 2.46912742 | 1.64970573 | 2.26099734 | 2.28241342 | 1.92950372 | 1.71588273 | 2.06965348 | 2.48396929 | 1.91951231 | 2.32881488 | 2.97337856 | 2.78874621 | 2.71219008 |
| ENSMUSG00 Chchd7 | 3816 | 20.127582 | 18.2539994 | 19.4711066 | 15.843686 | 19.638733 | 17.4785061 | 24.3258585 | 21.7484514 | 22.6547423 | 20.251573 | 21.6464631 | 21.8869852 | 20.5678932 | 20.5687913 | 19.5642608 | 16.6467814 | 17.4959144 | 21.8537227 |
| ENSMUSG00 Cdrt4 | 721 | 0.05895309 | 0.18481495 | 0 | 0.37914877 | 0.53405741 | 0.20618336 | 0.19686274 | 0.58268575 | 0.39986909 | 0.13079252 | 0.06626208 | 0.54189964 | 0.13684304 | 0 | 0.11429003 | 0 | 0 | 0.22888671 |
| ENSMUSG00 Slc35e2 | 2455 | 15.2187575 | 19.2414191 | 17.5563403 | 16.2943368 | 17.3902307 | 18.1659716 | 21.5267871 | 20.2240827 | 17.0282336 | 19.6861388 | 19.7521722 | 16.7560665 | 19.0696495 | 15.2291203 | 18.9476809 | 18.3420588 | 21.2941218 | 19.695725 |
| ENSMUSG00 Tbc1d22b | 4126 | 11.5792094 | 15.6714318 | 14.8075058 | 15.7796249 | 13.3336861 | 17.4263249 | 10.6872072 | 15.0325888 | 14.5240864 | 13.9303702 | 17.9011357 | 17.0720713 | 10.1031193 | 11.0088021 | 13.2611891 | 14.0040055 | 14.2121266 | 15.4654784 |
| ENSMUSG00 Kdm5b | 6609 | 23.9184067 | 25.3790925 | 25.5445031 | 26.0309432 | 24.3973285 | 24.6151648 | 21.3189829 | 18.3825135 | 20.1227507 | 19.5480295 | 17.6960355 | 18.4025312 | 15.08546 | 17.6458017 | 17.7050112 | 19.7565398 | 19.2704319 | 15.2234311 |
| ENSMUSG00 0610010F05f | 5105 | 7.24378122 | 7.21072441 | 6.91727271 | 5.97960817 | 8.05184384 | 7.50328365 | 7.43286357 | 5.56614053 | 6.53498095 | 5.19996982 | 6.82232022 | 5.72916889 | 3.49816926 | 4.75813079 | 4.01119947 | 4.09533235 | 5.06762985 | 4.73045991 |
| ENSMUSG00 Abhd14a | 1823 | 3.07771982 | 3.92883832 | 5.51930571 | 3.92379811 | 3.74916794 | 4.24038751 | 3.14033689 | 3.98055757 | 3.59224136 | 3.64687282 | 3.87860332 | 4.22521198 | 3.08493613 | 4.3310449 | 2.66691366 | 2.43648407 | 2.8802538 | 3.98310592 |
| ENSMUSG00 Fbxo38 | 4366 | 18.9744819 | 21.0284763 | 20.8628854 | 19.0550772 | 19.468845 | 23.2555087 | 20.5462298 | 16.4981381 | 18.6593972 | 20.4434909 | 17.6721388 | 19.7387501 | 16.9938825 | 16.0948017 | 19.1946781 | 17.7859257 | 19.9005303 | 17.6643994 |
| ENSMUSG00 Sprr2d | 679 | 0 | 0 | 0 | 0 | 0 | 0 | 0 | 0 | 0 | 0 | 0 | 0 | 0 | 0 | 0 | 0 | 0 | 0 |

| | | | | | | | | | | | | | | | | | | | | |
|---|---|---|---|---|---|---|---|---|---|---|---|---|---|---|---|---|---|---|---|---|
| ENSMUSG00 | Hfm1 | 7141 | 0 | 0 | 0 | 0 | 0 | 0 | 0 | 0.01604498 | 0 | 0 | 0 | 0 | 0 | 0 | 0 | 0 | 0 | 0 |
| ENSMUSG00 | Usp48 | 7848 | 12.4731677 | 12.3352791 | 10.2935519 | 12.4990989 | 11.5975367 | 11.7883478 | 11.2916218 | 13.5240426 | 12.647745 | 14.3290555 | 11.8706892 | 13.4062849 | 11.710673 | 12.3448479 | 12.9831101 | 12.7049875 | 10.6901458 | 12.336395 |
| ENSMUSG00 | Otud1 | 2528 | 1.34510051 | 3.729252 | 2.37275228 | 2.5772268 | 3.52231245 | 3.68509253 | 3.7805226 | 3.79204628 | 3.89382012 | 3.20803835 | 3.00483323 | 3.70926872 | 3.14178745 | 2.73740863 | 3.78115545 | 2.46889597 | 4.16919765 | 3.63390855 |
| ENSMUSG00 | Lrit2 | 2778 | 0 | 0 | 0 | 0 | 0 | 0 | 0 | 0 | 0 | 0 | 0 | 0 | 0 | 0 | 0 | 0 | 0 | 0 |
| ENSMUSG00 | A030009H04 | 2250 | 1.73798942 | 1.61382465 | 0.47298563 | 1.11371442 | 1.04820634 | 1.25533592 | 2.12381359 | 1.6634913 | 0.8054252 | 1.63455771 | 0.55206619 | 1.09150629 | 1.64439724 | 1.79583805 | 1.04377275 | 0.714758 | 0.76652594 | 2.24926125 |
| ENSMUSG00 | Hilpda | 1100 | 11.7855261 | 13.3857266 | 11.0379601 | 10.1476871 | 11.0265861 | 14.2351489 | 10.5808355 | 8.54120999 | 10.6710518 | 12.6878256 | 11.9003078 | 7.71270974 | 19.3739893 | 13.2998978 | 14.1208969 | 12.287804 | 11.8114679 | 10.2516972 |
| ENSMUSG00 | Gm9781 | 2376 | 2.86230142 | 2.74803336 | 3.03352526 | 2.97220618 | 3.54507014 | 2.58608542 | 4.32106537 | 3.40773961 | 3.17219376 | 3.23466525 | 2.67427167 | 3.00690438 | 3.96565497 | 3.44030277 | 2.87855988 | 3.07807492 | 3.19385808 | 3.5191011 |
| ENSMUSG00 | 4922505E12 | 2637 | 0 | 0 | 0 | 0 | 0 | 0 | 0 | 0 | 0 | 0 | 0 | 0 | 0 | 0 | 0 | 0 | 0 | 0 |
| ENSMUSG00 | Psapl1 | 2547 | 0.50064989 | 0.83707312 | 0.37984712 | 0.64397236 | 1.00156732 | 0.44747266 | 0 | 0 | 0 | 0 | 0 | 0 | 0 | 0.03646964 | 0 | 0 | 0.37619059 | 0 |
| ENSMUSG00 | Lengl9 | 1689 | 9.2107131 | 8.54025056 | 9.45131144 | 8.52415039 | 8.77716548 | 7.07057951 | 9.3841015 | 9.29371071 | 11.0464723 | 9.85446906 | 8.88178624 | 9.81483232 | 11.7123143 | 13.8864797 | 9.70883318 | 11.9020538 | 12.253523 | 11.6597159 |
| ENSMUSG00 | E130012A19 | 2323 | 0.10978522 | 1.43404624 | 0.74965437 | 1.23562023 | 0.99454685 | 0.6826033 | 1.91450358 | 1.80850808 | 1.32974115 | 1.42081327 | 1.99490787 | 2.35468572 | 0.67956149 | 1.0796301 | 0.58529976 | 0.75822985 | 1.12190624 | 1.72865467 |
| ENSMUSG00 | Gpr149 | 4284 | 0 | 0 | 0 | 0 | 0 | 0 | 0 | 0 | 0 | 0 | 0 | 0 | 0 | 0 | 0 | 0 | 0 | 0 |
| [illegible] | [illegible] | [illegible] | [illegible] | [illegible] | [illegible] | [illegible] | [illegible] | [illegible] | [illegible] | [illegible] | [illegible] | [illegible] | [illegible] | [illegible] | [illegible] | [illegible] | [illegible] | [illegible] | [illegible] | [illegible] |

| | | | | | | | | | | | | | | | | | | | |
|---|---|---|---|---|---|---|---|---|---|---|---|---|---|---|---|---|---|---|---|
| ENSMUSG00 Gm3345 | 1720 | 0 | 0 | 0 | 0 | 0.11193471 | 0 | 0 | 0 | 0.31129343 | 0 | 0.08332842 | 0 | 0.17208808 | 0.13501188 | 0 | 0.22261981 | 0 | 0.73558689 |
| ENSMUSG00 Zfp444 | 3413 | 9.26570494 | 8.16961396 | 8.12131055 | 11.3869238 | 9.81535859 | 10.5551823 | 8.08183177 | 8.50460763 | 6.54060696 | 11.9914475 | 8.16079719 | 8.71659776 | 6.70671246 | 6.01473385 | 8.37793138 | 10.2878661 | 8.83761959 | 9.6543964 |
| ENSMUSG00 Chchd8 | 1422 | 3.67660805 | 4.333956 | 3.94608265 | 3.04381098 | 3.4863489 | 3.17109617 | 5.42332396 | 5.55965351 | 4.11285882 | 5.60370524 | 6.58501537 | 5.29895531 | 6.97307691 | 6.95681478 | 6.98282347 | 4.95462149 | 4.28542982 | 5.6479111 |
| ENSMUSG00 Gm4766 | 941 | 0 | 0 | 0 | 0 | 0.05114976 | 0 | 0 | 0.04058704 | 0 | 0 | 0 | 0 | 0 | 0 | 0 | 0.0406914 | 0 | 0.05845814 |
| ENSMUSG00 Uqcrq | 2021 | 118.619096 | 91.7464469 | 94.8800962 | 88.800076 | 103.599143 | 90.9897052 | 145.051637 | 108.737795 | 116.671352 | 129.390302 | 113.941267 | 107.626738 | 150.680814 | 151.971672 | 116.469317 | 116.2928 | 111.148157 | 129.289093 |
| ENSMUSG00 Olfr821 | 932 | 0 | 0 | 0 | 0 | 0 | 0 | 0 | 0 | 0 | 0 | 0 | 0 | 0 | 0 | 0 | 0 | 0 | 0 |
| ENSMUSG00 Olfr649 | 2019 | 0 | 0 | 0 | 0 | 0 | 0 | 0 | 0 | 0 | 0 | 0 | 0 | 0.02443384 | 0 | 0 | 0 | 0 | 0 |
| ENSMUSG00 Psg22 | 1866 | 0 | 0 | 0 | 0 | 0 | 0 | 0 | 0 | 0 | 0 | 0 | 0 | 0 | 0 | 0 | 0 | 0 | 0 |
| ENSMUSG00 4930503L19f | 2282 | 0.26076795 | 0.10218679 | 0.14838506 | 0.21961941 | 0.25310389 | 0.39086293 | 0.60125666 | 0.70292765 | 0.88437304 | 0.35125414 | 1.0049071 | 0.22013189 | 0 | 0.12211416 | 0.93886105 | 0.41948518 | 0.10077028 | 0.12052828 |
| ENSMUSG00 Syt16 | 3526 | 0 | 0 | 0 | 0 | 0 | 0 | 0 | 0 | 0 | 0 | 0 | 0 | 0 | 0 | 0 | 0 | 0 | 0 |
| ENSMUSG00 1700029I15R | 640 | 1.32828675 | 1.30128493 | 1.20933826 | 1.92210656 | 2.63221458 | 2.09050595 | 1.18281697 | 1.90962012 | 1.54449434 | 2.06284327 | 2.31409955 | 4.01175075 | 1.92702802 | 3.33816878 | 2.18883266 | 2.75213741 | 2.21573904 | 3.95377955 |
| ENSMUSG00 Rassf9 | 2116 | 0.44192527 | 0.34635334 | 0.13716502 | 0.25838021 | 0.25021321 | 0.07025435 | 0 | 0.45123349 | 0.01946433 | 0.13389765 | 0.67733873 | 0.23740121 | 0 | 0.021949 | 0.01947143 | 0.3981065 | 0.36225233 | 0.98787621 |
| ENSMUSG00 Olfr1030 | 988 | 0 | 0 | 0 | 0 | 0 | 0 | 0 | 0 | 0 | 0 | 0 | 0 | 0 | 0 | 0 | 0 | 0 | 0 |
| ENSMUSG00 H1fx | 1060 | 2.00496113 | 3.48842572 | 3.33138464 | 3.39558727 | 3.63259801 | 2.9451153 | 1.16049967 | 2.34198694 | 2.05932579 | 1.46789925 | 1.53240433 | 2.42218913 | 2.23389814 | 2.89180167 | 3.88693923 | 3.14271966 | 2.71176629 | 3.68457224 |
| ENSMUSG00 Sstr3 | 3925 | 0 | 0 | 0.13556913 | 0.02321582 | 0.09810328 | 0.0252498 | 0.01205419 | 0 | 0.13641394 | 0.14415502 | 0 | 0.0568822 | 0.03770592 | 0 | 0 | 0.06828898 | 0.16599925 | 0.12613553 |
| ENSMUSG00 Zfp367 | 5580 | 1.31019539 | 1.17610039 | 1.12698189 | 0.8736616 | 1.31974629 | 1.65175779 | 1.71275289 | 1.10881168 | 0.75287179 | 0.98864378 | 0.9589239 | 0.96026946 | 0.9548113 | 0.98215095 | 1.09280112 | 0.43231331 | 0.70058822 | 1.48859769 |
| ENSMUSG00 BC030307 | 4629 | 1.19370769 | 1.08668115 | 1.00320996 | 1.19094542 | 1.09178051 | 0.79215863 | 1.48203466 | 1.24585283 | 1.02321221 | 0.98803591 | 1.05272106 | 1.01285713 | 1.33214294 | 1.29429408 | 1.42411945 | 0.62039221 | 0.70376652 | 1.09328966 |
| ENSMUSG00 Klhl31 | 6250 | 0.47605797 | 0.3464541 | 0.23993271 | 0.33532929 | 0.2002288 | 0.29335218 | 0.01514006 | 0.0794402 | 0.05930858 | 0.09807346 | 0.02293198 | 0.12502708 | 0.09471728 | 0.26008689 | 0.08569924 | 0.03063249 | 0.11651194 | 0.22883788 |
| ENSMUSG00 Wdr96 | 6627 | 0 | 0.14577847 | 0.03649731 | 0.1306262 | 0.07263003 | 0.037387 | 0 | 0.01728945 | 0 | 0.02134482 | 0 | 0.09264709 | 0.11910528 | 0.24529094 | 0.05595503 | 0.03466782 | 0.00578336 | 0.09960907 |
| ENSMUSG00 Ubtd2 | 1009 | 5.30788125 | 6.801245 | 2.97240722 | 4.87670247 | 4.24553142 | 5.10751664 | 3.18856508 | 4.20154278 | 2.1634146 | 4.34590228 | 3.83525435 | 3.59565715 | 4.4980539 | 2.11737167 | 3.06255371 | 2.69438367 | 2.12712847 | 2.56236672 |
| ENSMUSG00 Pwwp2a | 4345 | 13.8814372 | 11.462089 | 10.943885 | 11.0206347 | 11.7421954 | 12.6248276 | 9.4298688 | 10.1612007 | 9.27052069 | 8.78988945 | 11.2262906 | 9.64731015 | 8.54935183 | 10.208079 | 6.47656218 | 10.381205 | 10.0821536 | 8.98882987 |
| ENSMUSG00 Mylk4 | 6649 | 0 | 0 | 0 | 0 | 0.01447794 | 0.00745266 | 0.03557879 | 0 | 0.04336075 | 0 | 0 | 0 | 0 | 0 | 0 | 0 | 0 | 0 |
| ENSMUSG00 Kctd21 | 2964 | 2.93979794 | 2.68616117 | 2.07268501 | 2.65926473 | 1.73755595 | 1.67181962 | 2.29859169 | 3.68523181 | 3.62674785 | 1.98847433 | 1.78914318 | 1.46883323 | 1.83080665 | 1.58260892 | 1.34836401 | 1.44687855 | 1.51288014 | 3.84172911 |
| ENSMUSG00 4921523A10f | 2473 | 0.0171877 | 0 | 0.03912133 | 0.14738712 | 0.11677782 | 0 | 0 | 0.07721877 | 0.01665447 | 0 | 0.01931862 | 0.02257002 | 0.05984462 | 0.05634139 | 0.03332111 | 0.2167685 | 0 | 0.02224388 |
| ENSMUSG00 Fbxo48 | 2265 | 0.09383041 | 0 | 0.02135697 | 0.02011525 | 0 | 0.02187759 | 0 | 0.16861988 | 0 | 0 | 0.0210927 | 0 | 0 | 0 | 0.12733373 | 0 | 0.05076331 | 0 |
| ENSMUSG00 Napepld | 6390 | 18.3590432 | 19.0858381 | 18.244164 | 22.5024501 | 17.3018824 | 19.6970177 | 14.164187 | 14.727086 | 16.9580161 | 13.8500579 | 12.769895 | 13.8534649 | 7.84369765 | 7.81336368 | 8.80771915 | 8.04761905 | 9.74651525 | 8.72914463 |
| ENSMUSG00 Wdr72 | 8548 | 11.2478601 | 12.521564 | 11.7538398 | 13.8687222 | 12.6129515 | 14.5388694 | 8.54041455 | 10.6516948 | 11.0723694 | 11.0871448 | 13.9501985 | 10.8262013 | 8.36818906 | 8.38905837 | 9.37494457 | 8.02275127 | 11.0701482 | 10.6504529 |
| ENSMUSG00 Sft2d3 | 2809 | 13.0284644 | 12.7013563 | 14.5516672 | 12.5378025 | 13.3994889 | 12.9659164 | 13.1377321 | 14.24907 | 16.2165491 | 12.7402577 | 15.2219964 | 15.3796094 | 14.1550464 | 14.1365948 | 16.0171232 | 12.486364 | 12.3752052 | 14.3348758 |
| ENSMUSG00 Olfr124 | 1121 | 0 | 0 | 0 | 0 | 0 | 0 | 0 | 0 | 0 | 0 | 0 | 0 | 0 | 0 | 0 | 0 | 0 | 0 |
| ENSMUSG00 Tst | 1079 | 1.65451102 | 2.13029629 | 1.3449545 | 1.26675749 | 1.87353178 | 1.60736393 | 0.8331241 | 1.20346773 | 1.03061717 | 1.3109556 | 1.50542038 | 1.75878826 | 1.60020121 | 2.53957475 | 1.33647308 | 1.80984335 | 2.09569186 | 3.31380161 |
| ENSMUSG00 Ucn3 | 1476 | 0 | 0 | 0 | 0 | 0 | 0 | 0 | 0 | 0 | 0 | 0 | 0 | 0 | 0 | 0 | 0 | 0 | 0 |
| ENSMUSG00 1110034G24 | 2111 | 81.8088727 | 75.9836976 | 61.435071 | 72.1077447 | 80.4402779 | 63.6838305 | 67.5735324 | 77.633159 | 61.2432368 | 75.1820309 | 83.6684138 | 64.9904778 | 109.343459 | 110.950987 | 87.2825002 | 132.556968 | 103.72247 | 108.246248 |
| ENSMUSG00 Olfr1426 | 1668 | 0 | 0 | 0 | 0 | 0 | 0 | 0 | 0 | 0 | 0 | 0 | 0 | 0 | 0 | 0 | 0 | 0 | 0 |
| ENSMUSG00 E130304I02R | 434 | 1.07732013 | 1.15136731 | 1.44897672 | 1.46971111 | 1.3308366 | 1.94100569 | 0.43606156 | 0.61600649 | 1.42349709 | 1.41234825 | 1.76128879 | 0.64303759 | 1.25034813 | 0.85611221 | 2.18349259 | 1.23518088 | 0.97140384 | 1.14074184 |
| ENSMUSG00 Spata21 | 2511 | 0.08463795 | 0.19900176 | 0.05779394 | 0.16330123 | 0.03833686 | 0.03946853 | 0.16957949 | 0.01521004 | 0.09841461 | 0.01877766 | 0.22831521 | 0.3334269 | 0.03929265 | 0.18496251 | 0.1312674 | 0.01524915 | 0.09158016 | 0 |
| ENSMUSG00 Fzd5 | 6910 | 4.78567684 | 3.08542001 | 2.14915685 | 3.27696658 | 2.9603571 | 3.20550968 | 2.73879473 | 3.04544338 | 2.95040887 | 2.72941842 | 3.04902412 | 4.37801579 | 2.84854341 | 1.57278096 | 1.98554531 | 2.53793027 | 2.15204098 | 2.00612082 |
| ENSMUSG00 Tubg2 | 1743 | 1.63387653 | 1.2231929 | 0.30528332 | 1.41152977 | 1.29787746 | 1.16561221 | 1.68295243 | 1.46809579 | 1.18148353 | 1.43372764 | 1.58975765 | 0.19213653 | 2.40574467 | 1.17242676 | 3.38027682 | 2.50437708 | 1.78108437 | 1.32552064 |
| ENSMUSG00 9030612E09f | 1858 | 0 | 0 | 0 | 0 | 0 | 0 | 0 | 0 | 0 | 0 | 0 | 0 | 0 | 0 | 0 | 0 | 0 | 0 |
| ENSMUSG00 Prrt3 | 4054 | 0.13630175 | 0.13969393 | 0.20284904 | 0.53944996 | 0.35618098 | 0.4155878 | 0.01167062 | 0.83846172 | 0.37590061 | 0.53501045 | 0.5774477 | 0.52318579 | 0.12168702 | 0.22912722 | 0.11179505 | 0.60448912 | 0.70904595 | 0.37993463 |
| ENSMUSG00 Gm4779 | 1417 | 0.08998979 | 0 | 0 | 0 | 0 | 0 | 0 | 0 | 0 | 0 | 0 | 0.27573016 | 0 | 0 | 0 | 0.02702231 | 0.02704749 | 0 |
| ENSMUSG00 Olfr711 | 1356 | 0 | 0 | 0 | 0 | 0 | 0 | 0 | 0 | 0 | 0 | 0 | 0 | 0 | 0 | 0 | 0 | 0 | 0 |
| ENSMUSG00 Acer1 | 2423 | 0 | 0 | 0 | 0 | 0 | 0 | 0 | 0 | 0 | 0 | 0 | 0 | 0 | 0 | 0 | 0 | 0 | 0 |
| ENSMUSG00 1700024P04f | 475 | 0 | 0 | 0 | 0 | 0 | 0.10432154 | 0 | 0 | 0 | 0 | 0 | 0 | 0 | 0 | 0 | 0 | 0 | 0 |
| ENSMUSG00 Prss22 | 1317 | 0 | 0.15176717 | 0.11019027 | 0.27675653 | 0.43855967 | 1.35451664 | 0 | 0.26099592 | 0 | 0 | 0.18137797 | 0.76285644 | 0 | 0.10579519 | 0.0908532 | 0.69777872 | 0.34921455 | 0.20884247 |
| ENSMUSG00 Olfr1443 | 4332 | 0 | 0 | 0 | 0 | 0 | 0 | 0 | 0 | 0 | 0 | 0 | 0 | 0 | 0 | 0 | 0 | 0 | 0 |
| ENSMUSG00 Cetn4 | 2672 | 0.07953813 | 0.31168501 | 0.1629348 | 0.27282062 | 0.162121 | 0.22254222 | 0.74368732 | 0.14293564 | 0.36993876 | 0.3352782 | 0.19867835 | 0.10444548 | 0.88620211 | 0.39978069 | 0.43175283 | 0.17196381 | 0.44465389 | 0.45291929 |
| ENSMUSG00 Ankrd34b | 4743 | 0 | 0 | 0 | 0 | 0 | 0 | 0 | 0 | 0 | 0 | 0 | 0 | 0 | 0 | 0 | 0 | 0 | 0 |
| ENSMUSG00 Tmem232 | 3643 | 0 | 0 | 0.01327849 | 0 | 0 | 0 | 0 | 0 | 0 | 0.07765694 | 0 | 0 | 0 | 0 | 0 | 0 | 0 | 0 |
| ENSMUSG00 Prkce | 8164 | 1.90034165 | 1.85252988 | 3.02778957 | 1.77467076 | 2.10473993 | 2.37323847 | 1.21700913 | 1.82447782 | 2.37110025 | 1.72685697 | 1.62097789 | 1.91430494 | 1.41397215 | 0.96711108 | 1.30205799 | 1.42112372 | 0.99993891 | 1.44193396 |
| ENSMUSG00 Megf8 | 10118 | 3.98669817 | 3.65131446 | 3.70044599 | 2.94494198 | 3.47265312 | 4.34406747 | 2.99737371 | 4.57116024 | 4.67306979 | 3.44846021 | 2.73863174 | 3.66845381 | 4.61724903 | 3.0249707 | 2.56542599 | 4.09094155 | 3.0568612 | 3.08807794 |
| ENSMUSG00 Lrfn4 | 4210 | 5.60341394 | 7.19273654 | 6.05530891 | 6.66641409 | 6.58527032 | 6.20291938 | 5.49546318 | 7.25746364 | 6.21221794 | 5.64464476 | 6.83147862 | 8.08730503 | 8.0969964 | 7.72229482 | 7.74119494 | 7.18517336 | 6.43626886 | 5.43557919 |
| ENSMUSG00 Gm9795 | 698 | 0 | 0 | 0 | 0 | 0 | 0 | 0 | 0 | 0 | 0 | 0 | 0 | 0 | 0 | 0 | 0 | 0 | 0 |
| ENSMUSG00 Prlhr | 1571 | 0 | 0 | 0 | 0 | 0 | 0 | 0 | 0 | 0 | 0 | 0 | 0 | 0 | 0 | 0 | 0 | 0 | 0 |
| ENSMUSG00 Kcng3 | 3354 | 0 | 0.08939059 | 0.07211319 | 0.02716818 | 0.02870121 | 0 | 0 | 0.06832272 | 0 | 0.07029026 | 0.08546504 | 0.06656609 | 0.14708383 | 0.11077898 | 0.07370582 | 0.0114164 | 0.12569746 | 0.16401046 |
| ENSMUSG00 Rpsa-ps2 | 509 | 0 | 0 | 0.09503641 | 0.35804357 | 0 | 0 | 0.27885665 | 0 | 0 | 0 | 0 | 0.10965749 | 0.19383857 | 0 | 0 | 0 | 0 | 0 |
| ENSMUSG00 Pcdhb7 | 3505 | 1.22482818 | 1.41615442 | 0.81427626 | 1.62485893 | 2.12851588 | 1.49859958 | 1.87630881 | 1.4383444 | 2.92594648 | 1.06274053 | 0.83146147 | 0.50958665 | 0.90078251 | 0.79504857 | 0.83461069 | 0.41513355 | 0.75449771 | 0.87889015 |
| ENSMUSG00 Fam164c | 4479 | 1.96440532 | 1.66601659 | 2.28961564 | 1.6173713 | 1.91281143 | 2.50031654 | 3.81332375 | 3.0015016 | 3.6138202 | 2.91599571 | 2.86927078 | 4.05003133 | 2.58830109 | 2.88266495 | 1.99614593 | 2.33385484 | 2.62696431 | 2.14927298 |
| ENSMUSG00 9930022D16f | 2704 | 0.01571937 | 0 | 0.0894481 | 0.0505485 | 0.03560054 | 0.01832572 | 0.03499459 | 0.01412441 | 0.0609268 | 0.01743739 | 0 | 0 | 0 | 0.06870427 | 0.1980844 | 0.01416073 | 0.04252178 | 0.18309244 |
| ENSMUSG00 Olfr907 | 932 | 0 | 0 | 0 | 0 | 0 | 0 | 0 | 0 | 0 | 0 | 0 | 0 | 0 | 0 | 0 | 0 | 0 | 0 |
| ENSMUSG00 E130308A19f | 11702 | 4.01369163 | 4.04525916 | 3.6046589 | 4.35676026 | 4.70954133 | 4.3765273 | 4.4272247 | 3.75657624 | 4.12850649 | 4.94796305 | 4.09896243 | 4.43587132 | 4.61195672 | 5.07622524 | 5.39753762 | 5.44484453 | 5.48922181 | 6.42133309 |
| ENSMUSG00 Gm9796 | 2249 | 0 | 0 | 0 | 0 | 0.04280296 | 0 | 0 | 0 | 0 | 0 | 0 | 0.12408996 | 0 | 0 | 0 | 0 | 0 | 0 |
| ENSMUSG00 Rnf216 | 4628 | 17.4686354 | 21.3784131 | 21.2287468 | 20.4079613 | 21.7571271 | 19.5191515 | 16.1627799 | 16.6204621 | 13.4292248 | 13.8864323 | 18.3956301 | 17.3790773 | 13.1750756 | 12.4439646 | 14.3422011 | 15.1242935 | 14.7574462 | 14.881461 |
| ENSMUSG00 Lingo2 | 11470 | 0 | 0 | 0 | 0 | 0 | 0 | 0 | 0 | 0 | 0 | 0 | 0 | 0 | 0 | 0 | 0 | 0 | 0 |
| ENSMUSG00 S1pr5 | 2497 | 0 | 0 | 0.07749064 | 0.14597051 | 0 | 0 | 0 | 0.04588595 | 0 | 0.05664882 | 0 | 0 | 0 | 0 | 0 | 0 | 0 | 0 |
| ENSMUSG00 Aqp12 | 1085 | 0 | 0 | 0 | 0 | 0 | 0 | 0 | 0 | 0 | 0 | 0 | 0 | 0 | 0 | 0 | 0 | 0 | 0 |
| ENSMUSG00 S1pr1 | 5095 | 0 | 0 | 0 | 0 | 0 | 0 | 1.09575979 | 0.11244083 | 0.67903186 | 0.00925431 | 0.21566713 | 0.0876399 | 0.30983736 | 0.34639359 | 0.08895331 | 0.1503086 | 0 | 0 |
| ENSMUSG00 Arhgef37 | 3584 | 3.98486025 | 4.80545935 | 3.53623464 | 4.67814295 | 3.80059554 | 4.68704707 | 5.98009028 | 5.94625015 | 2.19492871 | 7.0120884 | 6.27846134 | 5.79336676 | 5.39567845 | 6.12947917 | 4.80531539 | 2.95940241 | 5.05813014 | 5.47942274 |
| ENSMUSG00 Magi1 | 7856 | 6.56839151 | 6.33945734 | 6.2560472 | 7.40598951 | 6.37184326 | 7.44300224 | 5.546713 | 6.40267235 | 5.73025226 | 6.72809806 | 5.69212845 | 7.38193399 | 5.14292773 | 5.97104484 | 6.40365323 | 8.27615216 | 6.87882875 | 8.50764571 |
| ENSMUSG00 Suv420h1 | 11245 | 5.1293485 | 5.97232501 | 6.09562406 | 6.16259212 | 5.00362759 | 6.5614958 | 6.83709233 | 5.9470784 | 6.6110859 | 4.96874908 | 5.02603609 | 6.26406102 | 5.53201847 | 5.22883189 | 5.63521503 | 6.06792907 | 6.5984625 | 5.42019478 |
| ENSMUSG00 Slc25a26 | 1905 | 7.25153922 | 6.08550509 | 7.59248587 | 4.49631304 | 5.25534915 | 5.6445896 | 3.99860436 | 6.37542465 | 8.36702446 | 6.38576454 | 4.96558624 | 5.18602219 | 6.26683674 | 5.85122361 | 7.05076495 | 5.56771562 | 3.78233271 | 8.02757568 |
| ENSMUSG00 Poln | 2872 | 0 | 0.11599197 | 0.05052945 | 0 | 0 | 0.05176121 | 0.18121151 | 0.06649095 | 0 | 0 | 0 | 0.01943442 | 0.08588426 | 0.0808567 | 0.53079899 | 0.05332954 | 0.17348254 | 0.13407512 |
| ENSMUSG00 Dmd | 23967 | 2.93866886 | 3.07456595 | 2.91851817 | 3.11382278 | 2.64286776 | 2.79738176 | 2.23662808 | 1.98555236 | 1.95389779 | 1.77255325 | 2.12691954 | 2.34748562 | 1.35026235 | 0.89915536 | 1.1707387 | 1.3532 | 1.49518453 | 1.43679646 |
| ENSMUSG00 Gm5514 | 305 | 5.01700438 | 4.15045895 | 4.44064869 | 4.92955562 | 5.99676426 | 6.1737832 | 3.87808844 | 6.51149156 | 4.18617045 | 7.26584608 | 4.22925862 | 4.75805648 | 8.57243153 | 3.95916811 | 9.05083359 | 9.03909421 | 5.65469954 | 3.96786999 |
| ENSMUSG00 Ccdc73 | 4330 | 0.35339176 | 0.29235334 | 0.46921207 | 0.48402033 | 0.30012978 | 0.30698933 | 0.40428848 | 0.59978831 | 0.76095179 | 0.59891194 | 0.61787475 | 0.68319402 | 0.64940399 | 0.6221148 | 0.74219893 | 0.43331172 | 0.46026962 | 0.44464636 |
| ENSMUSG00 1810063B07f | 2292 | 13.2225962 | 9.69445918 | 10.3627415 | 10.157807 | 11.6129816 | 9.72893984 | 12.8602962 | 9.74806082 | 9.79347779 | 10.9030858 | 11.3809456 | 10.1549438 | 12.3760264 | 11.7528668 | 10.0666978 | 11.0427973 | 9.09664289 | 12.2642467 |
| ENSMUSG00 Krtap4-7 | 977 | 0 | 0 | 0 | 0 | 0 | 0 | 0 | 0 | 0 | 0 | 0 | 0 | 0 | 0 | 0 | 0 | 0 | 0 |
| ENSMUSG00 Rassf8 | 1130 | 0 | 0 | 0 | 0 | 0 | 0 | 0 | 0 | 0 | 0 | 0 | 0 | 0 | 0 | 0 | 0 | 0 | 0 |
| ENSMUSG00 Taar6 | 2001 | 0 | 0 | 0 | 0 | 0 | 0 | 0 | 0 | 0 | 0 | 0 | 0 | 0 | 0 | 0 | 0 | 0 | 0 |
| ENSMUSG00 Prrt2 | 2059 | 0 | 0.08089581 | 0.0704811 | 0.15489427 | 0.09350544 | 0.02406641 | 0.11489237 | 0.01854901 | 0 | 0 | 0.04640598 | 0.05421628 | 0 | 0.10005235 | 0 | 0 | 0 | 0 |
| ENSMUSG00 Olfr1445 | 1000 | 0 | 0 | 0 | 0 | 0 | 0 | 0 | 0 | 0 | 0 | 0 | 0 | 0 | 0 | 0 | 0 | 0 | 0 |
| ENSMUSG00 Rpl18a | 1286 | 106.329045 | 101.855599 | 106.978476 | 102.707206 | 104.834773 | 107.274347 | 116.184635 | 101.598918 | 103.414665 | 106.400732 | 103.611477 | 95.4855814 | 158.429874 | 154.934008 | 127.92987 | 102.664086 | 97.6337081 | 126.828928 |
| ENSMUSG00 Olfr620 | 941 | 0 | 0 | 0 | 0 | 0 | 0 | 0 | 0 | 0 | 0 | 0 | 0 | 0 | 0 | 0 | 0 | 0 | 0 |
| ENSMUSG00 Tubb2b | 1918 | 11.9892076 | 14.9369599 | 9.33170294 | 8.69413046 | 10.2136042 | 9.7658672 | 2.9601259 | 5.39631963 | 3.39284124 | 3.19582452 | 3.31286208 | 5.06356897 | 4.01239786 | 5.10933391 | 2.98593132 | 5.49005059 | 3.21717091 | 2.81068429 |
| ENSMUSG00 Pigw | 4399 | 1.26578269 | 0.89359434 | 1.96837052 | 1.21178499 | 1.17074695 | 0.92369269 | 1.75311813 | 1.34572002 | 1.76955023 | 0.54664375 | 2.13950144 | 1.44646182 | 1.15507785 | 1.82651219 | 0.73055725 | 0.90525646 | 0.85382521 | 1.21297644 |
| ENSMUSG00 Olfr1255 | 1860 | 0 | 0 | 0 | 0 | 0 | 0 | 0 | 0 | 0 | 0 | 0 | 0 | 0 | 0 | 0 | 0 | 0 | 0 |
| ENSMUSG00 Gm8946 | 1024 | 0.08301792 | 0 | 0.04723978 | 0 | 0.42303449 | 0 | 0.04620379 | 0 | 0.04022121 | 0 | 0 | 0 | 0 | 0 | 0.04023589 | 0 | 0.22456815 | 0 |
| ENSMUSG00 Olfr1161 | 1011 | 0 | 0 | 0 | 0 | 0 | 0 | 0 | 0 | 0 | 0 | 0 | 0 | 0 | 0 | 0 | 0 | 0 | 0 |
| ENSMUSG00 Bola3 | 3224 | 27.2381184 | 25.2016591 | 26.3323653 | 25.9036583 | 25.8574726 | 23.1932615 | 35.0295796 | 27.0095153 | 29.3057901 | 31.6483008 | 31.4449323 | 27.7173932 | 33.5714102 | 32.585771 | 29.342051 | 27.6134188 | 28.3048737 | 31.957834 |
| ENSMUSG00 AI467606 | 1916 | 0.19965897 | 0 | 0 | 0 | 0.12560523 | 0.38793894 | 0.19754772 | 0.33886787 | 0 | 0.07382678 | 0.3241516 | 0 | 0.51494694 | 1.04232555 | 0.17203155 | 0.19984659 | 0.08001315 | 0 |
| ENSMUSG00 Fam123c | 4348 | 0 | 0 | 0 | 0 | 0 | 0 | 0 | 0 | 0 | 0 | 0 | 0 | 0 | 0 | 0 | 0 | 0 | 0 |
| ENSMUSG00 2310047M10 | 1801 | 2.64329801 | 2.55257046 | 2.60534839 | 3.01041881 | 1.57678151 | 1.89646675 | 4.49220883 | 3.5838512 | 4.27644556 | 2.64420934 | 3.68723999 | 3.22311433 | 3.06783717 | 2.52721853 | 4.04923675 | 3.78441317 | 2.93671792 | 3.29871378 |
| ENSMUSG00 4933411O13 | 2224 | 0 | 0 | 0 | 0 | 0 | 0 | 0 | 0 | 0 | 0 | 0 | 0 | 0 | 0 | 0 | 0 | 0 | 0 |
| ENSMUSG00 Sox3 | 3187 | 0.01333705 | 0.04181097 | 0 | 0.0142959 | 0 | 0.01554839 | 0.17814627 | 0.01198381 | 0.09046301 | 0 | 0 | 0 | 0.09287465 | 0 | 0 | 0 | 0 | 0.1553442 |
| ENSMUSG00 Shroom2 | 10971 | 3.62636448 | 3.14879877 | 3.75224449 | 4.63459352 | 3.08858575 | 3.93856382 | 5.32596468 | 4.21575055 | 4.30222834 | 5.48393924 | 4.00628583 | 4.07513861 | 4.0154409 | 3.46288064 | 4.4069533 | 4.01020034 | 3.88118821 | 4.72824607 |
| ENSMUSG00 Cirbp | 2590 | 145.420218 | 152.930622 | 142.804638 | 150.474584 | 154.375247 | 153.785665 | 136.421269 | 147.505251 | 158.162581 | 154.759895 | 167.802624 | 157.49068 | 164.642893 | 155.381474 | 178.757484 | 174.495768 | 165.690945 | 179.002607 |
| ENSMUSG00 Lrrc3b | 3073 | 0 | 0 | 0 | 0 | 0 | 0 | 0 | 0.04971351 | 0 | 0.03068708 | 0 | 0 | 0 | 0 | 0 | 0 | 0.03741584 | 0 |
| ENSMUSG00 Olfr123 | 940 | 0 | 0 | 0 | 0 | 0 | 0 | 0 | 0 | 0 | 0 | 0 | 0 | 0 | 0 | 0 | 0 | 0 | 0 |
| ENSMUSG00 Olfr835 | 935 | 0 | 0 | 0 | 0 | 0 | 0 | 0 | 0 | 0 | 0 | 0 | 0 | 0 | 0 | 0 | 0 | 0 | 0 |
| ENSMUSG00 Dpy19l4 | 8944 | 2.31440303 | 2.72641308 | 3.10447299 | 3.31112241 | 3.26117461 | 2.5430126 | 3.67646599 | 2.56210213 | 2.97478636 | 2.67805873 | 2.49985248 | 3.28878066 | 4.10915456 | 3.98285052 | 3.8096696 | 3.81450483 | 3.8694819 | 3.19205349 |
| ENSMUSG00 Vcpip1 | 9746 | 3.7158229 | 4.20769267 | 3.28082327 | 3.45471083 | 4.14845228 | 3.71162482 | 3.37392898 | 3.97364006 | 3.592093 | 4.13645098 | 4.25984393 | 3.61948479 | 3.09273563 | 3.90766998 | 3.23829179 | 4.37674294 | 4.77407392 | 2.99146589 |
| ENSMUSG00 Nudt18 | 3713 | 27.0851728 | 26.2340163 | 26.6686282 | 25.7929747 | 24.487262 | 26.050885 | 21.1397076 | 22.7220622 | 26.2448953 | 22.616591 | 23.1347631 | 23.9918657 | 21.4440384 | 24.7793528 | 21.9712041 | 21.4604778 | 18.9412213 | 22.0154949 |
| ENSMUSG00 Asxl3 | 15518 | 0 | 0.00214673 | 0.00623451 | 0 | 0 | 0 | 0 | 0.00492234 | 0 | 0 | 0.02155076 | 0.064743 | 0.02861111 | 0.00299292 | 0.02389573 | 0.00740249 | 0.01975837 | 0.03544858 |
| ENSMUSG00 Hs6st1 | 3717 | 16.6841678 | 19.6453764 | 19.2088595 | 20.9235143 | 17.3777352 | 20.6369738 | 16.5218879 | 19.0294133 | 17.4851552 | 18.9896698 | 18.8811983 | 15.9323158 | 13.5904986 | 14.544234 | 16.4495854 | 15.8539803 | 15.6625356 | 17.6851984 |
| ENSMUSG00 Ppp1r2-ps9 | 1675 | 0 | 0 | 0 | 0 | 0 | 0 | 0 | 0 | 0 | 0 | 0 | 0 | 0 | 0 | 0 | 0 | 0 | 0 |
| ENSMUSG00 Olfr1152 | 932 | 0 | 0 | 0 | 0 | 0 | 0 | 0 | 0 | 0 | 0 | 0 | 0 | 0 | 0 | 0 | 0 | 0 | 0 |
| ENSMUSG00 BC106179 | 4126 | 0.23694112 | 0.67013335 | 0.33999815 | 0.34231517 | 0.41995862 | 0.9127503 | 0.64214979 | 0.58316077 | 0.44919855 | 0.15998784 | 0.19684302 | 0.22997243 | 0.23912708 | 0.24764177 | 0.25963171 | 0.18560643 | 0.23222429 | 0.25331387 |
| ENSMUSG00 Rln3 | 473 | 0 | 0 | 0 | 0 | 0 | 0 | 0 | 0 | 0 | 0 | 0 | 0 | 0 | 0 | 0 | 0 | 0 | 0 |
| ENSMUSG00 Krtap5-4 | 1001 | 0 | 0 | 0 | 0 | 0 | 0 | 0 | 0 | 0 | 0 | 0 | 0 | 0 | 0 | 0 | 0 | 0 | 0 |
| ENSMUSG00 1110012L19f | 1091 | 57.8163898 | 56.6105462 | 53.4281431 | 53.9549673 | 53.9112839 | 51.5966135 | 49.4376297 | 52.4401639 | 52.2098545 | 56.8315076 | 54.9127387 | 45.4813236 | 97.7142743 | 85.7789515 | 98.0755643 | 111.116465 | 83.1515534 | 90.757463 |
| ENSMUSG00 A730035I17R | 682 | 0 | 0 | 0 | 0 | 0 | 0 | 0 | 0 | 0 | 0 | 0 | 0 | 0 | 0 | 0 | 0 | 0 | 0 |
| ENSMUSG00 Kcng4 | 3990 | 0 | 0 | 0.02424738 | 0 | 0.08444197 | 0 | 0.17786721 | 0.04786015 | 0.16515896 | 0.46087154 | 0.01197367 | 0.1119111 | 0.04945556 | 0 | 0.15489307 | 0 | 0 | 0 |
| ENSMUSG00 Med26 | 3037 | 4.60461077 | 5.70388706 | 3.99794406 | 4.32057319 | 4.26324908 | 4.03013671 | 4.26857887 | 5.0177045 | 4.69230637 | 6.10149041 | 4.29455504 | 3.96976724 | 5.40913021 | 5.27599009 | 4.99248356 | 5.24494326 | 4.78289975 | 4.61880876 |
| ENSMUSG00 Zfp688 | 3370 | 10.8722438 | 10.5078951 | 10.9235183 | 10.5317666 | 11.3402811 | 9.66057744 | 9.08347278 | 10.3810803 | 10.7304928 | 10.7593148 | 9.03045951 | 11.9581331 | 12.1939131 | 10.8047966 | 12.0059133 | 13.8164328 | 10.8155218 | 14.1195482 |
| ENSMUSG00 Zfp574 | 4193 | 5.23078245 | 6.77698516 | 4.03785729 | 6.73690615 | 4.91306066 | 6.45260971 | 5.48389267 | 3.99867986 | 4.33180383 | 6.13982446 | 4.96777532 | 4.25972145 | 2.92956055 | 4.56354973 | 3.58658905 | 4.12767816 | 5.73112048 | 4.90660766 |
| ENSMUSG00 Morn2 | 639 | 13.9688372 | 10.0095081 | 15.0646832 | 8.69866574 | 10.9972783 | 13.105496 | 18.2142708 | 9.68258089 | 12.1174726 | 14.314924 | 13.9063259 | 9.60833002 | 11.7346658 | 11.1204153 | 9.15590131 | 9.64755523 | 9.17671898 | 8.6086239 |
| ENSMUSG00 Klhdc9 | 1554 | 3.22754876 | 2.93685103 | 3.08171139 | 2.34548491 | 3.68577794 | 4.49609745 | 4.59730664 | 3.46533381 | 4.47909985 | 4.73327523 | 3.56621312 | 4.59742909 | 4.47606199 | 6.03713362 | 5.09053972 | 3.17856393 | 4.31602443 | 3.1858556 |
| ENSMUSG00 Tas2r119 | 1007 | 0 | 0 | 0 | 0 | 0 | 0 | 0 | 0 | 0 | 0 | 0 | 0 | 0 | 0 | 0 | 0 | 0 | 0 |
| ENSMUSG00 Zfp691 | 2442 | 2.41941829 | 2.81017863 | 3.12981892 | 1.69781123 | 2.54259548 | 2.15093766 | 3.13867895 | 3.84739189 | 2.61421374 | 2.25906318 | 3.77582596 | 3.10848899 | 1.17168354 | 4.62158609 | 4.08303707 | 1.00352124 | 2.7936449 | 3.35641151 |
| ENSMUSG00 Cenph | 1775 | 0.7662905 | 0.48796352 | 1.14461311 | 0.48769569 | 1.30159568 | 0.97709616 | 0.879616 | 1.50617925 | 1.69386797 | 1.14224239 | 1.29194254 | 0.69179976 | 1.91769143 | 0.88963322 | 0.71957647 | 2.71890382 | 1.85693608 | 1.7664962 |
| ENSMUSG00 Lca5l | 6832 | 0.05599335 | 0.19504036 | 0.07080435 | 0.22006945 | 0.13385635 | 0.06527731 | 0.20082958 | 0.1956578 | 0.06631318 | 0.25535363 | 0.04195693 | 0.13888558 | 0.15885571 | 0.05438418 | 0.18092018 | 0.14011493 | 0.06731785 | 0.12882695 |
| ENSMUSG00 Gpr20 | 2142 | 0.31749898 | 0.74650743 | 0.40650026 | 0.44667691 | 0.31458774 | 0.27760635 | 0 | 0.3031143 | 0 | 0 | 0 | 0.05211546 | 0.04606155 | 0 | 0 | 0 | 0 | 0.07704357 |
| ENSMUSG00 Tmem86b | 1217 | 2.20035011 | 1.50551289 | 1.27194164 | 2.62060239 | 1.18648949 | 0.97721085 | 2.25483597 | 0.65903077 | 0.43995456 | 0.50366404 | 1.29545902 | 1.51348962 | 2.51321189 | 0.53427874 | 1.42191072 | 2.01363917 | 1.29118995 | 0.85881103 |
| ENSMUSG00 Dcaf12l1 | 3529 | 2.37277407 | 1.57643903 | 1.85050343 | 1.45888297 | 1.44573078 | 1.93773795 | 2.06465077 | 0.59523438 | 0.78194858 | 1.97741686 | 0.67689068 | 0.52193734 | 0.65701335 | 1.97410403 | 0.89698549 | 0.62931574 | 0.73850615 | 0.31175464 |
| ENSMUSG00 Rtn4rl1 | 3523 | 0.07239031 | 0.23639579 | 0.247154 | 0.42677107 | 0.50550133 | 0.53448875 | 1.15495044 | 1.05156487 | 0.67806356 | 0.48181247 | 1.73579185 | 1.26745757 | 0.53210697 | 0.59323983 | 1.13441695 | 0.39127501 | 1.1314036 | 0.62457118 |
| ENSMUSG00 Ush1g | 3303 | 0.20589852 | 0.11094212 | 0 | 0 | 0.13114966 | 0 | 0.27215892 | 0.09250355 | 0 | 0.18557649 | 0.23142578 | 0.01689847 | 0.20909683 | 0.59056969 | 0.23700562 | 0 | 0 | 0 |
| ENSMUSG00 E330021D16f | 1115 | 0 | 0 | 0 | 0 | 0 | 0 | 0 | 0 | 0 | 0 | 0 | 0 | 0 | 0 | 0 | 0 | 0 | 0 |
| ENSMUSG00 Insig1 | 2642 | 5.6630666 | 5.98925994 | 4.68721566 | 7.81194288 | 7.68799084 | 6.18940276 | 4.47697567 | 5.24747997 | 2.85281317 | 5.55029096 | 5.85885183 | 5.00693113 | 12.4356763 | 15.3641682 | 14.612361 | 20.1743094 | 19.8884758 | 16.8441966 |
| ENSMUSG00 Preb | 7654 | 13.5778881 | 13.7708384 | 12.9118268 | 11.5896725 | 12.249933 | 11.4332542 | 13.8711329 | 15.0544131 | 12.6615981 | 12.7887196 | 15.2799972 | 12.8782937 | 15.5652705 | 16.5048233 | 16.18671 | 15.7684864 | 15.1973231 | 16.0485152 |
| ENSMUSG00 Olfr734 | 1032 | 0 | 0 | 0 | 0 | 0 | 0 | 0 | 0 | 0 | 0 | 0 | 0 | 0 | 0 | 0 | 0 | 0 | 0 |
| ENSMUSG00 Lhfpl2 | 8342 | 32.4266291 | 34.9501858 | 33.4648338 | 34.8999129 | 34.9247823 | 35.260732 | 34.8691381 | 36.1596253 | 32.8623171 | 34.1732381 | 33.3714557 | 33.4746775 | 45.1332046 | 47.8025575 | 53.2972896 | 56.7014962 | 59.0285618 | 50.9338696 |
| ENSMUSG00 Sowahb | 3899 | 1.55892284 | 3.45175924 | 3.6103353 | 2.60582531 | 1.27150247 | 3.36790828 | 1.77164686 | 2.39008622 | 0.94013847 | 1.65674438 | 1.02926302 | 1.81805313 | 0.96158649 | 2.9064779 | 0.65516709 | 1.72842957 | 2.02493387 | 1.32620057 |
| ENSMUSG00 Fahd1 | 1406 | 19.5896678 | 18.7888514 | 22.6041319 | 17.6929802 | 16.6031173 | 19.1021206 | 22.3776184 | 19.557987 | 23.258957 | 20.3894933 | 21.5429045 | 22.2706876 | 14.9469394 | 14.50139 | 19.3113978 | 14.8151426 | 19.3539622 | 12.6372272 |
| ENSMUSG00 Adra2c | 3195 | 0.17294751 | 0.42748941 | 0.37850963 | 0.04278032 | 0.13558288 | 0.09305678 | 0.0592334 | 0.08367662 | 0.06445464 | 0.23612246 | 0 | 0.84921923 | 0.36341226 | 0.69636432 | 0.25167535 | 0.09596569 | 0 | 0 |
| ENSMUSG00 5430407P10f | 3638 | 6.72979148 | 8.03978041 | 6.75474256 | 9.02955277 | 7.55451578 | 8.88080877 | 4.96796134 | 4.60870386 | 7.30217226 | 5.98780234 | 7.47222413 | 6.13696125 | 4.63757991 | 5.10655166 | 8.21086532 | 5.35731698 | 6.22617962 | 6.47165961 |
| ENSMUSG00 Tlr9 | 3469 | 0 | 0 | 0 | 0 | 0 | 0 | 0 | 0 | 0 | 0 | 0 | 0 | 0.09954552 | 0 | 0 | 0.05518969 | 0 | 0 |
| ENSMUSG00 Fndc7 | 6412 | 0.16572511 | 0.22340213 | 0.30176875 | 0.54713045 | 0.11259807 | 0.18547499 | 0.14019665 | 0.23825579 | 0.20554718 | 0.10294913 | 0.12666474 | 0.17409751 | 0.18464847 | 0.18832599 | 0.14136529 | 0.49565197 | 0.31679565 | 0.0686327 |
| ENSMUSG00 6330549D23 | 5288 | 0.1286087 | 0.18899146 | 0.03659117 | 0.15508676 | 0.54612621 | 0.36546078 | 0.04473589 | 0.12278193 | 0.14798483 | 0.15158131 | 0.19876117 | 0.24276858 | 0.11294838 | 0.35131685 | 0.07012368 | 0.39825707 | 0.21018582 | 0.32248153 |
| ENSMUSG00 Cenpe | 7767 | 0.42138516 | 0.29165402 | 0.22421103 | 0.11145357 | 0.18599932 | 0.24881635 | 0.42031349 | 0.29503594 | 0.33937647 | 0.15783678 | 0.22143686 | 0.4239892 | 0.19689577 | 0.938808 | 0.48272713 | 0.27607493 | 0.6513546 | 0.26913172 |
| ENSMUSG00 4933402E13f | 1966 | 0 | 0 | 0 | 0 | 0 | 0 | 0 | 0 | 0 | 0 | 0 | 0 | 0 | 0 | 0 | 0 | 0 | 0 |
| ENSMUSG00 2310079G19 | 878 | 0 | 0 | 0 | 0 | 0 | 0 | 0 | 0 | 0 | 0 | 0 | 0 | 0 | 0 | 0 | 0 | 0 | 0 |
| ENSMUSG00 Zfp423 | 6289 | 0.03379327 | 0.02118804 | 0.09999299 | 0.02897824 | 0.09184021 | 0.0551549 | 0.06770776 | 0.06680179 | 0.01309795 | 0.05997863 | 0 | 0.00887513 | 0.19610398 | 0.50217808 | 0.22929789 | 0.47490338 | 0.14626032 | 0.2973938 |
| ENSMUSG00 Hsfy2 | 1398 | 0 | 0 | 0 | 0 | 0 | 0 | 0 | 0 | 0 | 0 | 0 | 0 | 0 | 0 | 0 | 0 | 0 | 0 |
| ENSMUSG00 Defb11 | 315 | 4.18304907 | 1.48057308 | 0.46070029 | 1.30174412 | 1.52799758 | 0.78655133 | 80.3564547 | 47.1645858 | 46.2857987 | 45.8035403 | 19.7166495 | 45.8928781 | 226.77021 | 499.384521 | 183.379623 | 81.0788417 | 121.427442 | 210.431662 |
| ENSMUSG00 A930104D05 | 935 | 0 | 0 | 0 | 0 | 0 | 0 | 0 | 0 | 0 | 0 | 0 | 0 | 0 | 0 | 0 | 0 | 0 | 0 |
| ENSMUSG00 Vmn1r70 | 896 | 0 | 0 | 0 | 0 | 0 | 0 | 0 | 0 | 0 | 0 | 0 | 0 | 0 | 0 | 0 | 0 | 0 | 0 |
| ENSMUSG00 Olfr167 | 938 | 0 | 0 | 0 | 0 | 0 | 0 | 0 | 0 | 0 | 0 | 0 | 0 | 0 | 0 | 0 | 0 | 0 | 0 |
| ENSMUSG00 9530056K15f | 2313 | 0.60642923 | 0.05760985 | 0.39736147 | 0.66972568 | 0.56185125 | 0.06427073 | 0.24546137 | 0.41280158 | 0.92503983 | 0.22423594 | 0.12392985 | 0.55501956 | 0.74648383 | 0.36143259 | 0.32063468 | 0.39730851 | 0.24854927 | 0.47565159 |
| ENSMUSG00 6430598A04f | 5273 | 3.44200838 | 3.07037106 | 3.19248787 | 2.71309841 | 2.77491083 | 3.48645252 | 1.88425234 | 1.72383686 | 2.49946616 | 2.23547806 | 1.54931119 | 1.69362906 | 3.12476017 | 0.99529336 | 1.33613997 | 3.40570735 | 1.7734907 | 2.37854662 |
| ENSMUSG00 Sh2d5 | 3825 | 0.23336175 | 0.78383281 | 0.65762708 | 0.33351876 | 0.88084566 | 0.37569393 | 0.37107983 | 0.73888569 | 0.72143701 | 0.49307925 | 1.17407741 | 0.52532388 | 0.06448617 | 0.20641817 | 0.19388967 | 0.24025479 | 0.59117687 | 0.0431444 |
| ENSMUSG00 1700030F18f | 5632 | 0.01509417 | 0.01774479 | 0.00858905 | 0.0323587 | 0 | 0.00879843 | 0.21001722 | 0.00815718 | 0.09506831 | 0.02511579 | 0.02544831 | 0.00910451 | 0.18394358 | 0.40407675 | 0.09510302 | 0.03399379 | 0.03402548 | 0.13674139 |
| ENSMUSG00 Tnfrsf26 | 3099 | 0 | 0 | 0 | 0 | 0 | 0 | 0 | 0 | 0 | 0 | 0.12333 | 0 | 0 | 0 | 0 | 0 | 0 | 0 |
| ENSMUSG00 Ifne | 663 | 0 | 0 | 0 | 0 | 0 | 0 | 0 | 0 | 0 | 0 | 0 | 0 | 0 | 0 | 0 | 0 | 0 | 0 |
| ENSMUSG00 Rbm15b | 3015 | 8.95216809 | 8.12105381 | 7.86170145 | 10.1246765 | 6.89651444 | 8.74363325 | 6.55943608 | 6.99243964 | 6.06527794 | 8.21032146 | 7.25735688 | 5.23908209 | 4.84319984 | 5.62258438 | 8.90992187 | 5.99441681 | 7.04237761 | 6.2945744 |
| ENSMUSG00 Wdr81 | 8168 | 1.81614917 | 1.79452417 | 2.13795843 | 2.20330711 | 1.73835914 | 3.07581243 | 1.91729882 | 1.80955677 | 1.51272707 | 2.41872486 | 1.48565615 | 1.56486126 | 1.8904126 | 1.53524775 | 2.09841421 | 1.0547731 | 1.86751545 | 1.7106162 |
| ENSMUSG00 Tmem88 | 1580 | 0.37662814 | 0.31626165 | 0.30616158 | 0.40370546 | 0.2132427 | 0.37634988 | 0.32939207 | 0.53179294 | 0.78202245 | 0.41779104 | 0.75593289 | 0.74185374 | 0.31222732 | 0.55850485 | 1.0691543 | 1.21172808 | 0.6306859 | 1.98450575 |
| ENSMUSG00 Pxt1 | 707 | 0 | 0 | 0 | 0 | 0.2042373 | 0 | 0 | 0 | 0 | 0 | 0 | 0 | 0 | 0 | 0 | 0 | 0 | 0 |
| ENSMUSG00 Olfr433 | 1003 | 0 | 0 | 0 | 0 | 0 | 0 | 0 | 0 | 0 | 0 | 0 | 0 | 0 | 0 | 0 | 0 | 0 | 0 |
| ENSMUSG00 Cxcr4 | 2608 | 2.16763359 | 1.08573467 | 0.55644398 | 2.06142762 | 0.27683228 | 0.8170121 | 0.29026183 | 0.10251028 | 0.31584751 | 0.01807926 | 0.27477929 | 0 | 0.37831225 | 0.26712474 | 0.45678 | 0.01468198 | 0.02939133 | 0.02109245 |
| ENSMUSG00 1700120B22f | 451 | 0 | 0.29545804 | 0 | 0 | 0.21344534 | 0.21974605 | 0 | 0.25405146 | 0 | 0.41818806 | 0 | 0.37127935 | 0 | 0 | 0 | 0 | 0 | 0 |
| ENSMUSG00 Olfr1033 | 1103 | 0 | 0 | 0 | 0 | 0 | 0 | 0.60052358 | 0 | 0 | 0 | 0 | 0 | 1.56538269 | 0 | 0 | 0 | 1.21615629 | 0 |
| ENSMUSG00 Epcam | 2030 | 558.534764 | 564.185864 | 567.757505 | 577.278631 | 566.46295 | 566.316955 | 446.370654 | 561.992735 | 586.491927 | 499.774228 | 524.159544 | 587.109282 | 418.081924 | 380.040756 | 443.880802 | 466.051393 | 438.449061 | 423.461729 |
| ENSMUSG00 Olfr1499 | 1051 | 0 | 0 | 0 | 0 | 0 | 0 | 0 | 0 | 0 | 0 | 0 | 0 | 0 | 0 | 0 | 0 | 0 | 0 |
| ENSMUSG00 Kcnk13 | 3405 | 0 | 0 | 0.22730587 | 0 | 0 | 0 | 0 | 0 | 0 | 0 | 0 | 0.13113812 | 0 | 0 | 0 | 0 | 0.06753532 | 0 |
| ENSMUSG00 Trim39 | 8694 | 4.59079368 | 4.09993062 | 4.30654618 | 4.72694525 | 3.4269221 | 3.91565769 | 3.62436671 | 4.64775267 | 4.77051085 | 3.62822872 | 6.08864205 | 3.94831292 | 3.13785947 | 5.17648041 | 3.80074164 | 3.66434154 | 3.56636464 | 3.00544349 |
| ENSMUSG00 Akr1e1 | 4350 | 10.8168345 | 3.65293116 | 5.47121309 | 9.8453751 | 5.08981262 | 7.65504298 | 10.104248 | 4.2055542 | 6.89282379 | 13.1371615 | 7.17173515 | 7.82702395 | 14.9696852 | 3.39522294 | 8.10770847 | 13.8374333 | 7.14543145 | 4.96978826 |
| ENSMUSG00 2410002F23f | 6409 | 30.7995065 | 34.5499778 | 32.5534462 | 31.0943997 | 35.9055589 | 32.2413636 | 23.0103948 | 30.0044853 | 32.0803693 | 27.9269885 | 32.1580856 | 27.9644395 | 22.5068694 | 28.6606905 | 24.9947963 | 21.4544502 | 24.2791021 | 21.7238335 |
| ENSMUSG00 1190002N15 | 4352 | 0.76181152 | 1.03337333 | 0.27788103 | 0.59673243 | 0.60828488 | 0.71733047 | 0.60880285 | 0.85125529 | 1.05048328 | 0.61755287 | 0.21955404 | 1.00037263 | 1.99504077 | 0.74703266 | 0.08520542 | 1.09979916 | 0.89827258 | 0.83423737 |
| ENSMUSG00 Vmn1r230 | 950 | 0 | 0 | 0 | 0 | 0 | 0 | 0 | 0 | 0 | 0 | 0 | 0 | 0 | 0 | 0 | 0 | 0 | 0 |
| ENSMUSG00 Olfr1390 | 935 | 0 | 0 | 0 | 0 | 0 | 0 | 0 | 0 | 0 | 0 | 0 | 0 | 0 | 0 | 0 | 0 | 0 | 0 |

| | | | | | | | | | | | | | | | | | | | |
|---|---|---|---|---|---|---|---|---|---|---|---|---|---|---|---|---|---|---|---|
| ENSMUSG00 Asphd1 | 1889 | 0 | 0.0176352 | 0 | 0.02411913 | 0 | 0.02623226 | 0.05009283 | 0.02021832 | 0 | 0 | 0 | 0 | 0.02611536 | 0 | 0 | 0 | 0.0202892 | 0 |
| ENSMUSG00 Jrk | 5740 | 0.79234387 | 0.9401897 | 1.07871287 | 1.34936891 | 0.8301499 | 0.89781956 | 1.27760718 | 0.81175489 | 1.08347803 | 1.32251973 | 0.89890166 | 1.04046618 | 0.6445808 | 0.58257392 | 0.76804294 | 0.61371705 | 0.79456957 | 1.15959964 |
| ENSMUSG00 Pcdhb17 | 3268 | 4.17508002 | 4.8827651 | 5.28437893 | 4.89349038 | 3.63787795 | 6.18651019 | 2.73625958 | 6.6848391 | 4.41104055 | 3.00102395 | 4.86813377 | 4.37234077 | 4.84563814 | 4.40565088 | 4.96738465 | 5.41317644 | 3.48314265 | 5.10029355 |
| ENSMUSG00 Gm9806 | 549 | 0 | 0 | 0 | 0 | 0 | 0 | 0 | 0 | 0 | 0 | 0 | 0 | 0 | 0 | 0 | 0 | 0 | 0.20039747 |
| ENSMUSG00 Olfr609 | 964 | 0 | 0 | 0 | 0 | 0 | 0 | 0 | 0 | 0 | 0 | 0 | 0 | 0 | 0 | 0 | 0 | 0 | 0 |
| ENSMUSG00 Rbp1 | 2624 | 10.1565341 | 9.12803771 | 6.91313791 | 8.23015816 | 7.79575745 | 7.59153921 | 2.45218153 | 1.71749371 | 0.64353931 | 2.91098092 | 2.18483042 | 1.63788339 | 5.26407654 | 11.8765178 | 9.31117479 | 8.34688545 | 7.65357453 | 7.50505343 |
| ENSMUSG00 Yod1 | 3823 | 4.09152623 | 4.29590815 | 3.68210759 | 4.33801207 | 4.60797386 | 3.14970292 | 3.83649764 | 5.8941976 | 3.2320049 | 3.99603135 | 5.48605984 | 6.16118483 | 3.61311453 | 3.97259131 | 5.17309621 | 3.83607184 | 6.75697727 | 3.22313364 |
| ENSMUSG00 1700067K01l | 2152 | 0.29627214 | 1.09907783 | 0.60691697 | 0.67748765 | 0.6933502 | 0.64473817 | 0.59360703 | 0.53242197 | 1.12918421 | 0.78876641 | 0.97681142 | 1.11527579 | 0.16046627 | 0.1726546 | 0.28718557 | 0.28468853 | 0.37400197 | 0.12780926 |
| ENSMUSG00 Kcnk6 | 4484 | 0.39813055 | 0.43832756 | 0.06472819 | 0.33530653 | 0.0429366 | 0.2983773 | 0.08441156 | 0.02555245 | 0 | 0.02103064 | 0.08523632 | 0.13692513 | 0.18703002 | 0.21751245 | 0.33078858 | 0.03415754 | 0.0940208 | 0.0368036 |
| ENSMUSG00 Gm6816 | 767 | 0.05541744 | 0 | 0 | 0 | 0 | 0 | 0 | 0 | 0 | 0 | 0 | 0 | 0.0727714 | 0.06431801 | 0 | 0 | 0.19969026 | 0 |
| ENSMUSG00 D230002A01l | 3310 | 0 | 0 | 0.0730718 | 0 | 0 | 0 | 0 | 0 | 0 | 0.01424493 | 0.01443352 | 0.05058821 | 0 | 0.09822012 | 0 | 0.03470448 | 0.06947365 | 0 |
| ENSMUSG00 B430212C06 | 5708 | 0.08935916 | 0.02918088 | 0.01694938 | 0 | 0.00843236 | 0.08681278 | 0 | 0.01338206 | 0.10823366 | 0 | 0.01673965 | 0.02933549 | 0.06049815 | 0 | 0.03609106 | 0.09391529 | 0 | 0 |
| ENSMUSG00 BC046404 | 2507 | 13.7501786 | 14.0985165 | 10.4002125 | 14.7750814 | 9.84909328 | 15.3184557 | 7.11483046 | 6.18512779 | 4.38643786 | 6.03724601 | 5.08811883 | 5.83314862 | 6.25749888 | 4.90932715 | 6.29445389 | 5.8344683 | 3.92894228 | 5.06864924 |
| ENSMUSG00 Olfr518 | 1001 | 0 | 0 | 0 | 0 | 0 | 0 | 0 | 0 | 0 | 0 | 0 | 0 | 0 | 0 | 0 | 0 | 0 | 0 |
| ENSMUSG00 Ngfrap1 | 1044 | 301.933319 | 267.268967 | 266.008082 | 250.673026 | 265.094407 | 274.723393 | 266.065841 | 234.53208 | 228.301118 | 246.09596 | 231.324152 | 220.536023 | 295.188206 | 295.391514 | 256.562562 | 306.985042 | 244.091521 | 265.19237 |
| ENSMUSG00 Hnrnpa1 | 1116 | 32.1835786 | 41.9396204 | 37.3637842 | 36.2120487 | 36.7890062 | 38.8074276 | 41.1654223 | 37.7817314 | 46.7592433 | 39.2922527 | 39.2559471 | 41.1115364 | 32.6227195 | 38.4953233 | 31.3072754 | 33.5557473 | 32.6597745 | 35.6376202 |
| ENSMUSG00 Gm13078 | 2102 | 0 | 0 | 0 | 0 | 0 | 0 | 0 | 0 | 0 | 0 | 0 | 0 | 0 | 0 | 0 | 0 | 0 | 0 |

[illegible]

| | | | | | | | | | | | | | | | | | | | | |
|---|---|---|---|---|---|---|---|---|---|---|---|---|---|---|---|---|---|---|---|---|
| ENSMUSG00 | A730018C14 | 3885 | 0 | 0 | 0 | 0 | 0 | 0 | 0 | 0 | 0 | 0 | 0 | 0 | 0 | 0 | 0 | 0 | 0 | 0 |
| ENSMUSG00 | Cdca4 | 3079 | 6.06033526 | 7.93061106 | 6.58282209 | 7.60583852 | 8.11317583 | 8.99642029 | 4.4408458 | 8.29838169 | 7.2768641 | 7.15147074 | 7.57199732 | 7.19675805 | 7.54638942 | 6.18450011 | 6.97174752 | 7.0761142 | 8.6760081 | 7.87886199 |
| ENSMUSG00 | BC051628 | 1139 | 0 | 0.32172242 | 0 | 0 | 0.04225805 | 0.39154926 | 0 | 0 | 0.32544218 | 0 | 0 | 0.29402456 | 0 | 0 | 0 | 0 | 0 | 0 |
| ENSMUSG00 | Diras2 | 4346 | 0 | 0 | 0 | 0.02096689 | 0 | 0 | 0.31570817 | 0 | 0.02843064 | 0 | 0.41772858 | 0 | 0.3972888 | 0 | 0.13272475 | 0.00881054 | 0 | 0 |
| ENSMUSG00 | Bri3 | 1642 | 6.39389676 | 6.12697567 | 7.60071305 | 5.82693015 | 6.80061284 | 7.06171721 | 6.85774518 | 6.62901016 | 6.87278034 | 8.04031481 | 6.19736063 | 7.24039959 | 7.96160662 | 6.98641266 | 7.00075158 | 6.71601397 | 5.92867199 | 8.04030793 |
| ENSMUSG00 | Bex4 | 834 | 65.2357618 | 55.6413209 | 55.9717707 | 57.9073225 | 47.9587873 | 45.1559682 | 81.6342266 | 73.9577097 | 67.3602008 | 84.5772807 | 61.2940115 | 58.8272991 | 95.7062856 | 106.698887 | 74.3011272 | 96.6447585 | 81.1561635 | 89.7690579 |
| ENSMUSG00 | Stx19 | 2810 | 0.18151677 | 0.6875971 | 0.74023552 | 0.6323419 | 0.59950795 | 0.52903274 | 0.87553712 | 0.99218697 | 0.36642808 | 0.70474361 | 0.91809529 | 0.19863225 | 1.33424403 | 1.38836418 | 1.05569823 | 0.57231513 | 0.98202611 | 1.40948601 |
| ENSMUSG00 | Foxl1 | 2355 | 1589.11602 | 1598.97649 | 1690.13623 | 1688.60258 | 1675.72672 | 1665.56101 | 1570.60013 | 1454.87916 | 1505.88745 | 1535.95168 | 1512.16365 | 1530.629 | 1488.58799 | 1437.24236 | 1537.80652 | 1799.3984 | 1659.52052 | 1606.17248 |
| ENSMUSG00 | Lonp2 | 5658 | 18.4880237 | 22.3734532 | 21.4338 | 22.3054247 | 23.4449596 | 23.9093253 | 19.00702 | 20.2032274 | 17.2811574 | 22.7170055 | 20.5859577 | 19.3154944 | 18.963754 | 18.6416618 | 22.5232259 | 21.7846351 | 23.3155026 | 23.051713 |
| ENSMUSG00 | Gimap6 | 3254 | 0.09143707 | 0 | 0 | 0.0280031 | 0 | 0.0304565 | 2.21005753 | 0.38732307 | 1.8985794 | 0.49266254 | 3.94943572 | 0.3087529 | 0.78834041 | 1.92684444 | 0.88632726 | 0.48245695 | 0 | 0.69310798 |
| ENSMUSG00 | Olfr770 | 935 | 0 | 0 | 0 | 0 | 0 | 0 | 0 | 0 | 0 | 0 | 0 | 0 | 0 | 0 | 0 | 0 | 0 | 0 |
| ENSMUSG00 | Gpr157 | 5529 | 0.38438394 | 0.28318069 | 0.37620941 | 0.23897093 | 0.25245538 | 0.43019194 | 0.61611736 | 0.93944054 | 0.2607213 | 0.79309376 | 0.50116614 | 0.67636994 | 0.47288689 | 1.29361358 | 0.76754571 | 0.34627064 | 0.54068569 | 0.29847589 |
| ENSMUSG00 | A4galt | 2168 | 18.9391144 | 17.394002 | 20.8398881 | 24.6718939 | 22.2232729 | 20.182225 | 35.4845092 | 32.2204354 | 39.3437611 | 45.3455795 | 40.5249763 | 42.093454 | 76.7056703 | 62.7680701 | 78.4691994 | 99.4354796 | 82.4863936 | 79.0375311 |
| ENSMUSG00 | Usp14 | 43693 | 4.13348804 | 4.15372366 | 4.00225007 | 3.80709434 | 4.14419465 | 3.77092072 | 3.87224818 | 3.54538219 | 3.74885618 | 3.77266058 | 3.79308656 | 3.53853902 | 3.81734402 | 3.6438407 | 3.72476474 | 3.55362216 | 3.6358799 | 3.47859295 |
| ENSMUSG00 | Cxcr5 | 2612 | 0.68346761 | 0.05101515 | 0.05555918 | 0.12210081 | 0.07370892 | 0.15176947 | 2.17362997 | 0.2339504 | 0.3626684 | 0.28882513 | 5.99930566 | 0.55559235 | 1.60536484 | 2.59603245 | 0.47321848 | 1.12878131 | 0.24944374 | 0.29484207 |
| ENSMUSG00 | Rell1 | 5351 | 0.38922699 | 0.24902182 | 0.58760596 | 0.93659407 | 0.5396964 | 0.28707433 | 1.7595259 | 2.20546671 | 1.30848583 | 2.00022606 | 1.24102396 | 1.05351933 | 1.72398963 | 2.23063548 | 1.30126387 | 1.91059469 | 1.66168957 | 1.12053684 |
| ENSMUSG00 | Klk9 | 1470 | 0 | 0 | 0 | 0 | 0 | 0 | 0 | 0 | 0 | 0 | 0 | 0 | 0 | 0 | 0 | 0 | 0 | 0 |
| ENSMUSG00 | Tnrc6b | 17020 | 4.53272001 | 4.15334674 | 4.06144388 | 4.1063832 | 4.65482646 | 3.5257556 | 3.4191889 | 3.8461679 | 3.55481737 | 3.7759347 | 3.92978507 | 4.43705003 | 2.92455373 | 2.96619994 | 3.47865075 | 3.94604732 | 3.49260203 | 3.94953751 |
| ENSMUSG00 | Serpinb6d | 1456 | 0 | 0 | 0 | 0 | 0 | 0 | 0 | 0 | 0 | 0 | 0 | 0 | 0 | 0 | 0 | 0 | 0 | 0 |
| ENSMUSG00 | Ang2 | 437 | 0 | 0 | 0 | 0 | 0 | 0 | 0 | 0 | 0 | 0 | 0 | 0 | 0 | 0 | 0 | 0 | 0 | 0 |
| ENSMUSG00 | Ripply2 | 498 | 0 | 0 | 0 | 0 | 0 | 0 | 0 | 0 | 0 | 0 | 0 | 0 | 0 | 0 | 0 | 0 | 0 | 0 |
| ENSMUSG00 | Ccr4 | 4199 | 0 | 0 | 0 | 0 | 0 | 0 | 0 | 0 | 0 | 0 | 0 | 0 | 0 | 0 | 0 | 0 | 0 | 0 |
| ENSMUSG00 | Sstr2 | 1886 | 0.38313255 | 0.07065301 | 0 | 0 | 0 | 0 | 0 | 0 | 0 | 0 | 0 | 0 | 0 | 0 | 0 | 0 | 0 | 0 |
| ENSMUSG00 | Gm8566 | 1120 | 0.18975525 | 0.02974366 | 0 | 0.08135901 | 0.08594986 | 0.17697405 | 0.38019117 | 0.03410036 | 0.03677367 | 0 | 0.04265621 | 0.04983541 | 0.13213906 | 0.20733968 | 0.11036131 | 0 | 0.06843982 | 0 |
| ENSMUSG00 | Tshz2 | 6370 | 6.47920344 | 6.90314292 | 6.50802441 | 4.97094597 | 5.80303256 | 5.95099548 | 6.91493 | 6.21750727 | 6.45276966 | 6.49895094 | 5.9849948 | 5.9846307 | 6.69891812 | 6.18282356 | 5.80185175 | 5.92692918 | 5.84220319 | 5.10367066 |
| ENSMUSG00 | Ankrd16 | 5870 | 7.7189979 | 8.49563929 | 8.78469904 | 8.55336812 | 9.5853865 | 7.40336411 | 8.94669056 | 9.45375821 | 8.66530613 | 8.50640456 | 9.85612857 | 11.0490289 | 11.1353987 | 14.637404 | 13.0202532 | 12.948357 | 14.3250248 | 11.8452318 |
| ENSMUSG00 | Pcdhb16 | 5267 | 0.98455126 | 0.82222826 | 1.34090287 | 1.66085447 | 1.26109644 | 1.37359011 | 1.57199902 | 0.97166925 | 0.57865999 | 0.75197628 | 1.82319473 | 1.03852951 | 0.17795831 | 0.56434813 | 0.69621008 | 1.19953488 | 2.29943228 | 0.48042888 |
| ENSMUSG00 | Npm2 | 988 | 0 | 0.10115251 | 0.04896106 | 0.18445767 | 0 | 0.15046377 | 1.2450705 | 0.38656278 | 0.87542189 | 0.09544677 | 0 | 0.33896151 | 0 | 0.56409823 | 0.04170198 | 0.62009081 | 0 | 0 |
| ENSMUSG00 | Trappc9 | 6350 | 11.218689 | 11.6831205 | 13.6969461 | 13.7615879 | 13.7498125 | 13.2270683 | 11.839346 | 11.7584483 | 10.5139121 | 12.9274605 | 11.4584664 | 12.7628885 | 11.12493 | 10.407864 | 10.1868414 | 9.96761796 | 12.3006288 | 12.8036944 |
| ENSMUSG00 | Gm9821 | 311 | 0 | 0 | 0 | 0 | 0 | 0 | 0 | 0 | 0 | 0 | 0 | 0 | 0 | 0 | 0 | 0 | 0 | 0 |
| ENSMUSG00 | Gm5607 | 5052 | 0 | 0 | 0 | 0 | 0 | 0 | 0 | 0 | 0 | 0 | 0 | 0 | 0 | 0 | 0 | 0 | 0 | 0 |
| ENSMUSG00 | 4930483J18R | 1329 | 0.09594848 | 0 | 0 | 0 | 0 | 0 | 0 | 0 | 0.06198121 | 0.106435 | 0 | 0 | 0 | 0.03494664 | 0 | 0 | 0 | 0.62087028 |
| ENSMUSG00 | B930007M17 | 2428 | 0.070025 | 0.06860151 | 0 | 0 | 0.21806061 | 0.04081774 | 0.13640393 | 0.17302983 | 0 | 0.31071304 | 0.37385676 | 0.7126382 | 0.264133 | 0.3634422 | 0.27150943 | 0.28386776 | 0.63140522 | 0.49843507 |
| ENSMUSG00 | Marcksl1 | 1603 | 123.64419 | 135.994747 | 126.803228 | 131.22604 | 129.172511 | 126.061165 | 95.5403254 | 101.639918 | 105.779716 | 91.7717992 | 96.891073 | 112.536632 | 78.4756014 | 85.355135 | 89.5227818 | 84.2011173 | 86.3835999 | 86.5801474 |
| ENSMUSG00 | Gp5 | 2194 | 0 | 0.0455509 | 0 | 0 | 0.04387596 | 0 | 0 | 0 | 0 | 0 | 0 | 0 | 0 | 0 | 0 | 0 | 0 | 0 |
| ENSMUSG00 | Kcna3 | 1967 | 0 | 0 | 0 | 0 | 0.02446971 | 0 | 0 | 0 | 0 | 0 | 0 | 0 | 0.32603707 | 0 | 0.29324951 | 0 | 0 | 0 |
| ENSMUSG00 | Olfr186 | 1004 | 0 | 0 | 0 | 0 | 0 | 0 | 0 | 0 | 0 | 0 | 0 | 0 | 0 | 0 | 0 | 0 | 0 | 0 |
| ENSMUSG00 | Stbd1 | 2337 | 1.40047007 | 0.69847318 | 0.49677566 | 0.42890157 | 0.7414417 | 1.46304605 | 0.02024505 | 0.47393225 | 0.07049468 | 0 | 0 | 0.35825201 | 0.63327237 | 0.4769611 | 0.14104084 | 0.16384513 | 0.04919935 | 0 |
| ENSMUSG00 | Rpl9-ps7 | 578 | 0.8824604 | 0.34580859 | 0.75322106 | 0.47295202 | 0.41636612 | 0.85731373 | 0.40927923 | 0.59469139 | 0.35628474 | 0.65260489 | 0.57858946 | 0.19313378 | 0.93884272 | 0.64282474 | 0.42769781 | 0.33123363 | 0.39785083 | 0.66620025 |
| ENSMUSG00 | Olfr1093 | 1629 | 0 | 0 | 0 | 0 | 0 | 0 | 0 | 0 | 0 | 0 | 0 | 0 | 0 | 0 | 0 | 0 | 0 | 0 |
| ENSMUSG00 | Kcna1 | 8968 | 0.04265684 | 0.02600248 | 0.03236409 | 0.03556281 | 0.03756952 | 0.02210202 | 0 | 0.10220982 | 0 | 0 | 0 | 0 | 0 | 0.06214641 | 0 | 0 | 0 | 0 |
| ENSMUSG00 | Synb | 4052 | 0.02097985 | 0 | 0 | 0 | 0 | 0 | 0 | 0 | 0 | 0 | 0.02358093 | 0 | 0 | 0 | 0.05084101 | 0 | 0.04729306 | 0.09503054 |
| ENSMUSG00 | 9230110F15R | 1132 | 10.8515865 | 20.0995643 | 6.92271371 | 10.3840543 | 4.29445609 | 19.3483288 | 0 | 0 | 0.03638385 | 0 | 0.04220403 | 0.04930712 | 0 | 0.04102835 | 0.07279427 | 0 | 0 | 0 |
| ENSMUSG00 | Palm3 | 3606 | 0 | 0 | 0 | 0 | 0 | 0 | 0.0918438 | 0.12709618 | 0.07995164 | 0.16998312 | 0 | 0.12382843 | 0 | 0 | 0 | 0 | 0 | 0 |
| ENSMUSG00 | 4933428G20 | 1422 | 0.08967337 | 0.02342679 | 0.17008977 | 0 | 0.23693633 | 0.06969442 | 0 | 0.08057469 | 0.46342071 | 0.39789623 | 0 | 0.11775456 | 0.1387677 | 0 | 0.11589748 | 0.13463645 | 0.02695239 | 0.07736865 |
| ENSMUSG00 | Ino80c | 3105 | 12.3066516 | 12.8638197 | 11.513056 | 12.2963463 | 16.3850059 | 13.1342672 | 15.770893 | 17.4172115 | 14.6971528 | 15.382822 | 16.6481498 | 17.3109446 | 19.875758 | 18.4430144 | 18.7496936 | 18.7815121 | 18.9471387 | 18.8855742 |
| ENSMUSG00 | C2cd4a | 1350 | 0 | 0 | 0 | 0 | 0 | 0 | 0 | 0 | 0 | 0 | 0 | 0 | 0 | 0 | 0 | 0 | 0 | 0 |
| ENSMUSG00 | Fam69c | 2774 | 0 | 0 | 0 | 0 | 0 | 0 | 0 | 0 | 0 | 0 | 0 | 0 | 0 | 0 | 0 | 0 | 0 | 0 |
| ENSMUSG00 | Cypt4 | 642 | 0 | 0 | 0 | 0 | 0 | 0 | 0 | 0 | 0 | 0 | 0 | 0 | 0 | 0 | 0 | 0 | 0 | 0 |
| ENSMUSG00 | Prrg1 | 10315 | 1.43812956 | 1.34349627 | 2.02592003 | 1.81095765 | 1.96913929 | 1.65255844 | 1.49987843 | 2.08456835 | 1.55722163 | 1.15191248 | 0.57895005 | 1.53675697 | 0.52607958 | 0.50428869 | 0.99059485 | 0.46772821 | 1.48251987 | 1.0185884 |
| ENSMUSG00 | Gigyf2 | 12332 | 5.14253346 | 5.25140013 | 5.44457186 | 5.08737902 | 4.87485993 | 5.67373174 | 4.93383921 | 4.30485236 | 4.11798361 | 5.40634888 | 5.15250525 | 5.28646561 | 4.70837388 | 3.83393456 | 3.87225132 | 5.17600085 | 4.12414823 | 4.545433 |
| ENSMUSG00 | Hes5 | 1439 | 0 | 0 | 0.23531252 | 0 | 0 | 0.34435534 | 0 | 0 | 0 | 0 | 0 | 0 | 0 | 0 | 0.02863207 | 0 | 0 | 0 |
| ENSMUSG00 | Catsper4 | 2389 | 0.14233629 | 0 | 0 | 0 | 0 | 0.08298816 | 0 | 0.18964072 | 0 | 0 | 0.29996835 | 0 | 0.04129922 | 0 | 0.05173908 | 0.17630669 | 0.04812846 | 0 |
| ENSMUSG00 | Tmem196 | 1164 | 0 | 0 | 0 | 0 | 0 | 0 | 0 | 0 | 0 | 0 | 0 | 0 | 0 | 0 | 0 | 0 | 0 | 0 |
| ENSMUSG00 | Timm8a1 | 1419 | 1.94703061 | 2.95801598 | 1.77267342 | 2.50441259 | 1.2889451 | 1.74604417 | 2.03387838 | 2.26086103 | 1.21905121 | 1.62817791 | 3.93916148 | 1.69138371 | 2.32927305 | 1.44012674 | 1.36467451 | 1.53810051 | 1.48551538 | 2.36473257 |
| ENSMUSG00 | Zfp473 | 13029 | 0.03588587 | 0.01022731 | 0.06311689 | 0.01748447 | 0.03694215 | 0.02281959 | 0 | 0.02931338 | 0.13276795 | 0.03618904 | 0.01466727 | 0 | 0 | 0.00712934 | 0.06640822 | 0.12637164 | 0.12354781 | 0.03799846 |
| ENSMUSG00 | Krt35 | 2690 | 0 | 0 | 0 | 0 | 0 | 0.01842109 | 0 | 0 | 0 | 0 | 0.05328062 | 0.08299727 | 0.09169501 | 0 | 0.12253251 | 0 | 0 | 0 |
| ENSMUSG00 | Neurod4 | 3342 | 0 | 0 | 0 | 0 | 0 | 0 | 0 | 0 | 0 | 0 | 0 | 0 | 0 | 0 | 0 | 0 | 0 | 0 |
| ENSMUSG00 | Tmem229a | 5156 | 0 | 0 | 0 | 0.01767302 | 0.02800539 | 0.04805347 | 0 | 0 | 0 | 0 | 0.0092659 | 0 | 0 | 0.00900777 | 0.07990992 | 0 | 0 | 0 |
| ENSMUSG00 | Rgmb | 5744 | 24.8193524 | 23.2563903 | 25.6015899 | 26.6986989 | 23.3620827 | 26.6915316 | 13.6567586 | 16.5961414 | 19.381468 | 21.4000493 | 17.2335853 | 21.5139062 | 13.8101885 | 17.7965593 | 17.674205 | 15.6322205 | 16.5342207 | 18.1863325 |
| ENSMUSG00 | Eno4 | 1813 | 0.93778656 | 1.15759092 | 0.64035561 | 1.1057286 | 0.66370551 | 0.1639914 | 1.06995027 | 0.50558062 | 0.29532526 | 0.57215415 | 0.05270266 | 0.2770772 | 1.19724455 | | 0.3863356 | 0.86592107 | 0.48621336 | 0.81922001 |
| ENSMUSG00 | Fcrl5 | 1779 | 0 | 0 | 0 | 0 | 0 | 0 | 0 | 0 | 0 | 0.15902429 | 1.20847281 | 0 | 0 | 0.44381646 | 0 | 0.04304734 | 0 | 0 |
| ENSMUSG00 | 4932418E24R | 3254 | 0.09143707 | 0.01023752 | 0.01486587 | 0.01400155 | 0 | 0 | 0.08723911 | 0.03521119 | 0.02531439 | 0.04347022 | 0.02936383 | 0.17152939 | 0 | 0.02854564 | 0.01266182 | 0 | 0 | 0 |
| ENSMUSG00 | Isg20l2 | 2967 | 4.0940658 | 4.44630305 | 4.9921084 | 3.0562104 | 4.78504504 | 4.58537507 | 4.91518027 | 4.61146654 | 5.68138728 | 4.97945163 | 5.09468081 | 6.16333942 | 4.44615147 | 6.50249177 | 5.85354062 | 4.61015225 | 4.311214 | 5.27951179 |
| ENSMUSG00 | Arxes2 | 1539 | 11.0750978 | 7.72755246 | 9.38667449 | 7.48989228 | 13.0415934 | 9.65940226 | 13.7726317 | 10.9336545 | 15.4951219 | 9.00734682 | 12.8517432 | 11.5693284 | 18.848062 | 28.125984 | 16.2503862 | 13.3855405 | 16.8595861 | 20.087796 |
| ENSMUSG00 | Zbtb33 | 5390 | 1.33272259 | 1.16193397 | 1.38210087 | 1.31864989 | 1.07158272 | 0.91934571 | 1.15867785 | 1.28961359 | 0.83289985 | 0.97975487 | 1.22318076 | 1.03554105 | 1.13490867 | 1.37005749 | 0.94786511 | 1.03718528 | 1.06659453 | 1.64312916 |
| ENSMUSG00 | Ldlrad3 | 6230 | 8.11214355 | 11.0953861 | 8.0907253 | 11.6571918 | 11.0402117 | 10.2843795 | 10.0397049 | 10.4645957 | 8.36952311 | 13.2445796 | 10.0381093 | 10.1328273 | 11.0462319 | 9.19935856 | 12.0364096 | 14.756942 | 12.8205462 | 13.0856334 |
| ENSMUSG00 | Fpr-rs4 | 971 | 0 | 0 | 0 | 0 | 0 | 0 | 0 | 0 | 0 | 0 | 0 | 0 | 0 | 0 | 0 | 0 | 0 | 0 |
| ENSMUSG00 | Cyb5r2 | 2727 | 2.83679576 | 2.418758 | 2.57211658 | 3.25794046 | 2.13566658 | 2.56213254 | 0.46844237 | 0.92434857 | 0.72495517 | 0.29393544 | 0.92851955 | 0.92105054 | 0.77787768 | 0.42578005 | 1.04250361 | 0.33699104 | 0.77299095 | 0.70602081 |
| ENSMUSG00 | Olfr1349 | 1053 | 0 | 0 | 0 | 0 | 0 | 0 | 0 | 0 | 0 | 0 | 0 | 0 | 0 | 0.4851709 | 0 | 0 | 0 | 0 |
| ENSMUSG00 | Pirt | 4198 | 0.10125101 | 0.38090017 | 0.1958909 | 0.90080197 | 0.26370516 | 0.55478287 | 0 | 0 | 0 | 0 | 0 | 0 | 0.01175129 | 0.09957046 | 0.0294437 | 0 | 0 | 0 |
| ENSMUSG00 | Arf1 | 3364 | 186.105451 | 176.754414 | 181.673954 | 187.742924 | 191.154132 | 182.405321 | 182.260703 | 183.605093 | 183.625257 | 184.355886 | 185.305487 | 174.432242 | 185.361306 | 171.68021 | 174.530967 | 186.75208 | 183.314541 | 171.699055 |
| ENSMUSG00 | H1fnt | 1325 | 0 | 0 | 0 | 0 | 0 | 0 | 0 | 0 | 0 | 0 | 0 | 0 | 0.03723164 | 0 | 0 | 0 | 0 | 0 |
| ENSMUSG00 | Odz4 | 20988 | 0.00810085 | 0.00634894 | 0.00230482 | 0.00217081 | 0 | 0 | 0 | 0.00181973 | 0 | 0.00449311 | 0.04097338 | 0 | 0.03055627 | 0 | 0 | 0.06750298 | 0 | 0.04193566 |
| ENSMUSG00 | Olfr731 | 980 | 0 | 0 | 0 | 0 | 0 | 0 | 0 | 0 | 0 | 0 | 0 | 0 | 0 | 0 | 0 | 0 | 0 | 0 |
| ENSMUSG00 | Gm4737 | 2080 | 0.22478699 | 0.33633211 | 0.13953903 | 0.24094783 | 0.532228 | 0.4049983 | 0.52316905 | 0.55085196 | 0.49503024 | 0.20401747 | 0.75796809 | 0.32201344 | 0.26088995 | 0.51356443 | 0.31693505 | 0.27613419 | 0.64491365 | 0.52893372 |
| ENSMUSG00 | Lmod1 | 4399 | 0.01932493 | 0.06058267 | 0 | 0.01035714 | 0.04376624 | 0.02252909 | 0.01075533 | 0 | 0.1310778 | 0.0107185 | 0 | 0 | 0.07850044 | 0.0422315 | 0 | 0 | 0 | 0 |
| ENSMUSG00 | Taf13 | 2839 | 13.3848635 | 10.6310258 | 10.1722429 | 10.9930664 | 9.68063699 | 10.1409434 | 12.2156371 | 14.0850459 | 13.2887807 | 13.9841114 | 13.6812403 | 12.0124586 | 15.6041077 | 14.3961948 | 13.5258366 | 13.4199205 | 13.391929 | 16.6635547 |
| ENSMUSG00 | Olfr19 | 1019 | 0 | 0 | 0 | 0 | 0 | 0 | 0 | 0 | 0 | 0 | 0 | 0 | 0 | 0 | 0 | 0 | 0 | 0 |
| ENSMUSG00 | 4632415L05R | 1076 | 7.07102835 | 9.93814038 | 5.97925608 | 8.8496824 | 11.0041387 | 7.96712366 | 7.87078952 | 9.37062661 | 7.50237648 | 7.01125699 | 9.63491263 | 10.3227852 | 8.1608562 | 6.51771115 | 8.65385839 | 6.04963128 | 6.41146231 | 6.90170019 |
| ENSMUSG00 | Kctd12 | 3803 | 0 | 0 | 0 | 0 | 0 | 0 | 0 | 0 | 0 | 0 | 0 | 0 | 0 | 0 | 0 | 0 | 0 | 0 |
| ENSMUSG00 | Tmem72 | 3859 | 0.66087343 | 0.50931867 | 0.77718551 | 0.80283779 | 1.15995566 | 0.3467022 | 0.01226035 | 0.17814544 | 0.46960526 | 0.26880422 | 0.64376726 | 0.49176795 | 0.20453762 | 0.27681109 | 0.06406047 | 0.02976725 | 0.00993167 | 0 |
| ENSMUSG00 | Rbm15 | 3817 | 2.02670737 | 1.71059137 | 3.06690971 | 2.06498839 | 1.7906034 | 2.67431572 | 2.57821253 | 2.55149662 | 3.11839221 | 2.33468245 | 1.57706177 | 2.80759948 | 2.58485288 | 2.84724006 | 2.21261607 | 2.31729901 | 2.46003216 | 1.78703936 |
| ENSMUSG00 | Arid4a | 5722 | 8.79519895 | 8.50579141 | 9.76431799 | 7.74744777 | 8.89119946 | 8.79859792 | 10.6499005 | 10.2122292 | 10.5377594 | 9.4186043 | 9.00059511 | 9.53997169 | 9.06114035 | 11.0712575 | 8.55425523 | 11.4162489 | 11.6144353 | 12.7572675 |
| ENSMUSG00 | Entpd1 | 3846 | 58.2871291 | 72.420465 | 67.466359 | 68.0572542 | 65.1645675 | 66.6629858 | 33.7192025 | 41.2112506 | 45.0524368 | 53.5134737 | 47.1414372 | 37.9360744 | 42.9056847 | 43.4492528 | 47.4792761 | 47.4003073 | 51.3309296 | 44.6966871 |
| ENSMUSG00 | Col6a3 | 12596 | 0.0539834 | 0.0026443 | 0 | 0 | 0 | 0 | 0.05257799 | 0.07275898 | 0 | 0 | 0.00379227 | 0 | 0.03132682 | 0 | 0.00327048 | 0 | 0 | 0 |
| ENSMUSG00 | Dmrt2 | 2672 | 244.892453 | 248.258879 | 238.663267 | 278.703319 | 276.02001 | 256.519875 | 307.142863 | 302.723387 | 309.099252 | 381.758351 | 421.857175 | 329.671702 | 367.361161 | 382.830695 | 394.945902 | 432.231026 | 444.453082 | 407.442078 |
| ENSMUSG00 | Nat8l | 6526 | 0 | 0 | 0 | 0 | 0 | 0 | 0 | 0 | 0 | 0 | 0 | 0 | 0 | 0 | 0 | 0 | 0 | 0 |
| ENSMUSG00 | Nwd1 | 8977 | 0.02840939 | 0 | 0.02155443 | 0.05075308 | 0.00536169 | 0.01103993 | 0.08432693 | 0.21697811 | 0 | 0.04201912 | 0.06918508 | 0.08082919 | 0 | 0 | 0.00917936 | 0.06398119 | 0.09392654 | 0.01838335 |
| ENSMUSG00 | Gm12663 | 1708 | 2.53836531 | 1.40429062 | 3.03042608 | 2.37408252 | 2.1698818 | 0.6672792 | 2.49305685 | 1.49817972 | 2.19436354 | 2.62255083 | 1.79016238 | 1.60126901 | 3.14823125 | 1.87625412 | 1.37499338 | 2.24183884 | 1.72782486 | 0.90178863 |
| ENSMUSG00 | Olfr49 | 1029 | 0.24784359 | 0 | 0 | 0.17710804 | 0 | 0 | 0 | 0.14846415 | 0 | 0 | 0 | 0.05424263 | 0 | 0 | 0.20020192 | 0 | 0.48420006 | 0 |
| ENSMUSG00 | Mll2 | 19750 | 4.36673429 | 5.69102304 | 5.98117272 | 4.9182859 | 4.80588119 | 4.88753038 | 4.15872459 | 5.91547134 | 5.93502906 | 4.94903331 | 5.01697539 | 5.16330205 | 5.77745441 | 4.67497953 | 4.65629735 | 6.07415053 | 5.12505064 | 4.72660527 |
| ENSMUSG00 | 1700014N06 | 1048 | 0 | 0 | 0 | 0 | 0 | 0 | 0 | 0 | 0 | 0 | 0 | 0 | 0 | 0 | 0 | 0 | 0 | 0 |
| ENSMUSG00 | Selplg | 4229 | 0 | 0.05514076 | 0.1715779 | 0.01077348 | 0 | 0 | 0.32444258 | 0.14449715 | 0.39930176 | 0 | 0.71171019 | 0.14518143 | 0.0699909 | 0.64795487 | 0.29227871 | 0 | 0 | 0 |
| ENSMUSG00 | Gm7204 | 872 | 4.82570232 | 5.95964621 | 3.77224778 | 4.44115684 | 5.24373022 | 4.37564276 | 4.17783977 | 4.99304343 | 4.15643737 | 3.73096161 | 5.25962846 | 3.90453603 | 5.14817027 | 3.94153604 | 5.81168736 | 4.52285844 | 5.31821322 | 5.55137774 |
| ENSMUSG00 | Mcmbp | 5644 | 10.9200045 | 12.2355829 | 13.130448 | 13.9169455 | 12.8260831 | 11.7560436 | 12.1886313 | 11.8014794 | 11.1431271 | 12.1385492 | 12.3669763 | 11.0266591 | 10.1478306 | 10.8128155 | 10.6288918 | 12.9376862 | 11.7953185 | 10.3215174 |
| ENSMUSG00 | Olfr1324 | 1151 | 0 | 0 | 0 | 0 | 0 | 0 | 0 | 0 | 0 | 0 | 0 | 0 | 0 | 0 | 0 | 0 | 0 | 0 |
| ENSMUSG00 | Tmem81 | 1738 | 1.02716766 | 1.6292267 | 1.47514218 | 1.65152232 | 1.82779457 | 1.11194281 | 1.17056685 | 1.07677084 | 2.4645556 | 1.05804225 | 2.85880074 | 0.9955613 | 2.01528546 | 0.64134528 | 0.45041977 | 0.99141389 | 1.65389659 | 2.08895342 |
| ENSMUSG00 | Asb8 | 3128 | 16.6460488 | 16.8481453 | 17.7070627 | 14.8422967 | 12.7869657 | 17.1882084 | 16.9859778 | 17.0693665 | 13.864898 | 16.3098021 | 16.8464767 | 17.8081941 | 15.1086882 | 22.1826939 | 17.1234088 | 15.0812111 | 17.6683249 | 17.269483 |
| ENSMUSG00 | Gm7075 | 338 | 0 | 0 | 0 | 0 | 0 | 0 | 0 | 0 | 0 | 0 | 0 | 0 | 0.14595242 | 0 | 0 | 0 | 0 | 0 |
| ENSMUSG00 | Bend7 | 4643 | 5.21816721 | 6.98831768 | 5.83441245 | 5.3872525 | 4.91374797 | 5.03745213 | 5.23771634 | 4.49951413 | 3.52165556 | 2.74191038 | 3.69399313 | 4.56815675 | 4.65375399 | 4.48135928 | 2.87514627 | 5.64091653 | 5.34900719 | 3.54247747 |
| ENSMUSG00 | Gm8181 | 1437 | 0.88737319 | 0.95047228 | 0.37029146 | 0.91946436 | 0.63639983 | 0.86208653 | 0.69141702 | 1.24916 | 0.74519792 | 0.45936663 | 0.53194108 | 0.77683594 | 0.34329796 | 0.74336396 | 0.68812619 | 0.71944774 | 0.64010517 | 0.80389091 |
| ENSMUSG00 | Muc6 | 8772 | 0.1695943 | 0.05316695 | 0.01654362 | 0.01558175 | 0.005487 | 0.09603243 | 0.01618081 | 0.12190917 | 0.04225703 | 0.03225082 | 0.02178521 | 0.05090348 | 0.03374276 | 0.08471334 | 0.0704541 | 0.09166698 | 0.05242905 | 0.01681296 |
| ENSMUSG00 | Olfr1153 | 956 | 0 | 0 | 0 | 0 | 0 | 0 | 0 | 0 | 0 | 0 | 0 | 0 | 0 | 0 | 0 | 0 | 0 | 0 |
| ENSMUSG00 | Efcab4a | 2752 | 3.21260051 | 3.12308383 | 2.58390587 | 3.60912342 | 2.69342887 | 3.45716746 | 2.78512136 | 3.30297666 | 3.11293433 | 3.75218169 | 4.25322123 | 2.41354064 | 4.91168072 | 3.27403814 | 3.51830147 | 5.63506395 | 2.82712146 | 4.71735072 |
| ENSMUSG00 | Dnajb8 | 988 | 0 | 0 | 0 | 0 | 0 | 0 | 0 | 0 | 0 | 0 | 0 | 0 | 0 | 0 | 0 | 0 | 0 | 0 |
| ENSMUSG00 | A630023P12R | 1434 | 0 | 0 | 0 | 0 | 0 | 0 | 0 | 0 | 0 | 0.13152218 | 0 | 0 | 0 | 0 | 0 | 0.16021175 | 0 | 0 |
| ENSMUSG00 | Gpr85 | 6082 | 0 | 0 | 0 | 0 | 0 | 0 | 0 | 0 | 0.14898115 | 0 | 0 | 0.13765783 | 0.0486668 | 0 | 0 | 0 | 0.00630159 | 0 |
| ENSMUSG00 | Naga | 2722 | 0 | 0 | 0 | 0 | 0 | 0 | 0 | 0 | 0 | 0 | 0 | 0 | 0 | 0 | 0 | 0 | 0 | 0 |
| ENSMUSG00 | Amigo2 | 2869 | 28.7861823 | 25.7074757 | 25.4934743 | 26.2027616 | 22.3799185 | 24.2323058 | 21.834084 | 21.7386522 | 20.5717243 | 27.6593355 | 26.0772342 | 23.3262389 | 18.2780858 | 15.3464604 | 18.5543431 | 24.8642041 | 21.0133374 | 16.0483173 |
| ENSMUSG00 | Mfap1b | 3355 | 4.244183 | 4.61713735 | 4.67154213 | 4.20981334 | 4.57647799 | 5.28759422 | 4.8229318 | 4.67871159 | 5.43833874 | 5.62154435 | 5.02669459 | 4.37541558 | 4.22004777 | 4.83129255 | 5.0473441 | 4.74780706 | 5.89459589 | 5.49271259 |
| ENSMUSG00 | Olfr1212 | 1869 | 0 | 0 | 0 | 0 | 0 | 0 | 0 | 0 | 0 | 0 | 0 | 0 | 0 | 0 | 0 | 0 | 0 | 0 |
| ENSMUSG00 | Fbxo43 | 4030 | 0.03164157 | 0 | 0 | 0 | 0.08360384 | 0 | 0.02348024 | 0 | 0 | 0 | 0 | 0 | 0 | 0.01152459 | 0.02044742 | 0 | 0 | 0 |
| ENSMUSG00 | H2-M10.4 | 1404 | 0 | 0 | 0 | 0 | 0 | 0 | 0 | 0 | 0 | 0 | 0 | 0 | 0 | 0 | 0 | 0 | 0 | 0 |
| ENSMUSG00 | Fbxo10 | 5752 | 6.25163055 | 7.39578684 | 6.22329841 | 8.74467889 | 5.81566185 | 9.32128959 | 5.06686548 | 6.28793551 | 4.1172195 | 5.84465428 | 5.66455523 | 4.43458933 | 4.02236947 | 6.22538098 | 6.48967483 | 5.44536107 | 4.27772646 | 3.08900234 |
| ENSMUSG00 | Rnf149 | 2831 | 96.2559815 | 103.362933 | 91.8068939 | 88.8528879 | 84.226616 | 93.9594044 | 61.7855789 | 61.3830558 | 55.8076561 | 58.3429578 | 56.5840465 | 56.9394679 | 71.2184903 | 67.6564523 | 77.6877094 | 76.6623676 | 77.0181219 | 71.972353 |
| ENSMUSG00 | Ovch2 | 1937 | 3.88405584 | 20.7582154 | 2.64718339 | 8.32659251 | 1.86365218 | 15.4516526 | 0.0488515 | 0.43378051 | 0.36147174 | 0.07302639 | 0.02466441 | 0.11526208 | 0 | 0.16784131 | 0.02127081 | 0.019768 | 0.03957284 | 0 |
| ENSMUSG00 | Gng7 | 6897 | 0.52384224 | 0.56028646 | 0.72241172 | 1.30797256 | 0.55131535 | 0.09867043 | 1.13188227 | 1.07428245 | 0.94352175 | 1.5245189 | 1.57933747 | 0.58267764 | 0.72957163 | 0.73400109 | 0.85425873 | 0.48300462 | 0.7390746 | 0.58223355 |
| ENSMUSG00 | D630032N06 | 445 | 0 | 0 | 0 | 0 | 0 | 0 | 0 | 0 | 0 | 0 | 0 | 0 | 0 | 0 | 0 | 0 | 0 | 0 |
| ENSMUSG00 | A930001N09R | 8674 | 7.10543062 | 9.20578827 | 9.63678354 | 9.80134987 | 9.3112022 | 9.11189878 | 7.72908301 | 7.95198049 | 9.40632784 | 8.02877435 | 11.5334059 | 9.94823776 | 8.21819466 | 9.01681383 | 9.12476238 | 9.02305759 | 11.9742062 | 10.0835231 |
| ENSMUSG00 | Bcl11b | 10299 | 2.22864308 | 2.36447477 | 2.90269364 | 3.35768838 | 2.45356442 | 3.20921189 | 1.88350309 | 2.31772517 | 2.19149535 | 1.84500761 | 1.81376918 | 2.03774047 | 1.2310227 | 1.44757278 | 1.72023197 | 1.13624028 | 1.59646387 | 2.16853067 |
| ENSMUSG00 | Gm4879 | 2620 | 0.01622335 | 0.02542969 | 0 | 0 | 0.01837096 | 0 | 0.05417482 | 0.07288626 | 0.01572004 | 0.01799645 | 0.03646943 | 0.02130369 | 0 | 0 | 0.03145157 | 0.01461474 | 0.05851343 | 0.02099584 |
| ENSMUSG00 | Dip2c | 8336 | 0.98410496 | 1.47462308 | 1.10256367 | 1.26801371 | 1.45504376 | 1.19482959 | 1.13514105 | 1.12707905 | 1.33895703 | 0.75228449 | 1.45571491 | 1.25879853 | 0.79300347 | 0.75215353 | 1.13185656 | 1.41017472 | 1.29194931 | 0.67309608 |
| ENSMUSG00 | Rbm33 | 5591 | 6.32522026 | 6.69713702 | 6.11698908 | 6.62513487 | 5.9917401 | 5.88499644 | 6.2113229 | 7.81472158 | 6.95404595 | 6.37559143 | 6.77616564 | 7.39749704 | 6.71464658 | 6.07237129 | 7.24398666 | 6.9992847 | 5.53198383 | 7.11350101 |
| ENSMUSG00 | Syngr2 | 1435 | 87.6169412 | 90.6528585 | 92.1288212 | 87.5343549 | 90.5282662 | 89.4711726 | 80.9755676 | 88.2816716 | 93.3943769 | 81.4869297 | 93.3526018 | 92.805694 | 95.5686467 | 85.4441747 | 87.9732176 | 88.2952055 | 86.0538875 | 87.8995691 |
| ENSMUSG00 | Sacs | 20700 | 0.11293646 | 0.04988888 | 0.15423444 | 0.06162847 | 0.00232521 | 0.07660326 | 0.07771165 | 0.02767565 | 0 | 0.02961155 | 0.0369275 | 0 | 0.06672916 | 0.00448735 | 0.00199041 | 0.00184979 | 0 | 0.00265744 |
| ENSMUSG00 | Zfp738 | 5555 | 2.24959888 | 1.84105464 | 2.21185899 | 1.76338875 | 2.78134068 | 2.29253871 | 1.74601245 | 2.42698795 | 1.92772171 | 1.81642673 | 2.69191035 | 1.75836921 | 1.78500727 | 1.72231899 | 1.85425544 | 1.88178863 | 2.07672644 | 1.92111012 |
| ENSMUSG00 | Dleu7 | 1312 | 0.12958895 | 0 | 0.11061021 | 0 | 0 | 0 | 0 | 0 | 0 | 0.07187607 | 0 | 0 | 0 | 0 | 0.15701812 | 0 | 0 | 0 |
| ENSMUSG00 | Tas2r126 | 929 | 0 | 0 | 0 | 0 | 0 | 0 | 0.61114332 | 0 | 0 | 0 | 0 | 0.12016289 | 0 | 0 | 0 | 0 | 0 | 0 |
| ENSMUSG00 | Frmd6 | 4678 | 0.93587711 | 1.15363165 | 0.92031727 | 0.64280225 | 1.09063358 | 1.09104993 | 0.4551241 | 0.22043499 | 0.4578236 | 0.70554708 | 0.63318671 | 0.38180872 | 0.44291161 | 0.88360919 | 1.49727758 | 1.34238128 | 0.95856707 | 2.36358069 |
| ENSMUSG00 | Olfr1417 | 947 | 0 | 0 | 0 | 0.14433277 | 0 | 0 | 0.09992118 | 0 | 0 | 0 | 0 | 0 | 0 | 0 | 0 | 0 | 0 | 0 |
| ENSMUSG00 | Krtap4-13 | 818 | 0 | 0 | 0 | 0 | 0 | 0 | 0 | 0 | 0 | 0 | 0 | 0 | 0 | 0 | 0 | 0 | 0 | 0 |
| ENSMUSG00 | Olfr148 | 1016 | 0 | 0 | 0 | 0 | 0 | 0 | 0 | 0 | 0 | 0 | 0 | 0 | 0 | 0 | 0 | 0 | 0 | 0 |
| ENSMUSG00 | Slitrk3 | 3729 | 0 | 0 | 0 | 0 | 0 | 0 | 0 | 0 | 0 | 0 | 0 | 0 | 0 | 0 | 0 | 0 | 0 | 0 |
| ENSMUSG00 | Ankrd46 | 2475 | 43.7245164 | 43.9326411 | 39.5197084 | 45.6899039 | 39.1278507 | 44.3872365 | 39.799999 | 31.7729925 | 34.1473658 | 38.8255084 | 39.7062988 | 38.1350648 | 31.6920106 | 25.5770873 | 34.3928947 | 38.1977817 | 36.003491 | 33.5388857 |
| ENSMUSG00 | Pskh1 | 3252 | 13.8939121 | 16.6974224 | 12.0487576 | 14.6266083 | 16.4731953 | 15.7099841 | 14.5924406 | 16.0544324 | 13.9694732 | 16.2243656 | 13.8241808 | 13.0957413 | 12.3329793 | 10.3970774 | 15.4062398 | 14.1058265 | 18.3381667 | 17.5582573 |
| ENSMUSG00 | Gm4884 | 1444 | 0 | 0 | 0 | 0 | 0 | 0 | 0 | 0 | 0 | 0 | 0 | 0 | 0 | 0 | 0 | 0 | 0 | 0 |
| ENSMUSG00 | Gm9823 | 325 | 0 | 0 | 0 | 0 | 0 | 0 | 0 | 0 | 0 | 0 | 0 | 0 | 0 | 0 | 0 | 0 | 0 | 0 |
| ENSMUSG00 | Ckap2l | 3128 | 1.86163974 | 1.61878514 | 1.80936798 | 1.34003072 | 1.08173361 | 1.18812501 | 0.89240667 | 0.45176435 | 0.77685564 | 0.13566379 | 0.33601314 | 0.60669198 | 0.53621649 | 0.59391416 | 0.73762376 | 0.89361072 | 0.35532692 | 1.10791999 |
| ENSMUSG00 | Mfsd6l | 2058 | 8.03426311 | 7.7373972 | 10.3422514 | 9.36458783 | 9.26153633 | 5.36941671 | 9.05791812 | 8.2583183 | 8.02516658 | 7.19403344 | 9.72677726 | 7.29563325 | 4.9859275 | 12.9537931 | 10.4905808 | 9.35868587 | 9.59088577 | 7.56442041 |
| ENSMUSG00 | Ric3 | 9489 | 0.398668 | 0.68107298 | 0.43841539 | 0.58097654 | 0.7811483 | 0.66320973 | 0.45871709 | 0.66813561 | 0.98694136 | 0.98385913 | 0.57899886 | 1.05878589 | 1.51806511 | 1.8648116 | 1.90181067 | 1.85218556 | 1.50655588 | 1.52464907 |
| ENSMUSG00 | Lhfp | 2225 | 0.0764138 | 0.29944175 | 0.36959551 | 0.45049122 | 0.21632325 | 0 | 0.14884888 | 0.27464199 | 0.22212952 | 0.67812247 | 0.34355026 | 0.30102829 | 0.62080615 | 0.98106612 | 0.53700904 | 1.29069463 | 0.17225302 | 0.37084791 |
| ENSMUSG00 | Gm8258 | 622 | 0.13667259 | 0 | 0 | 0.07324927 | 0.30953006 | 0 | 0 | 0.2456103 | 0.06621626 | 0 | 0.15361723 | 0 | 0.15862353 | 0 | 0.06624044 | 0 | 0.18485352 | 0.08843908 |
| ENSMUSG00 | Ppyr1 | 3625 | 0 | 0 | 0 | 0 | 0 | 0 | 0 | 0.08428668 | 0 | 0.28615599 | 0 | 0 | 0 | 0 | 0 | 0 | 0 | 0 |
| ENSMUSG00 | Pcdhb18 | 9970 | 0.32827468 | 0.13031122 | 0.23774353 | 0.34730585 | 0.26069447 | 0.3081514 | 0.36065833 | 0.1302449 | 0.21894537 | 0.146607 | 0.15333989 | 0.08397542 | 0.36615465 | 0.09316768 | 0.18183234 | 0.39173941 | 0.29984465 | 0.13241911 |
| ENSMUSG00 | Pou4f1 | 4587 | 0.11119732 | 0.05083721 | 0.03163736 | 0.03973058 | 0.04197246 | 0.05401432 | 0.04125806 | 0.03330491 | 0.01795793 | 0.17474645 | 0.06249177 | 0 | 0.09679251 | 0.01012516 | 0.06287571 | 0.03339054 | 0.07519675 | 0 |
| ENSMUSG00 | 2010305A19R | 3983 | 11.2159026 | 8.76523052 | 8.99944414 | 9.70016711 | 8.44695321 | 10.0399337 | 12.3538002 | 9.97240736 | 10.692156 | 10.3464009 | 10.7592613 | 9.58521547 | 9.83418085 | 12.3718746 | 9.18578499 | 9.20012837 | 8.59286546 | 9.75054716 |
| ENSMUSG00 | Arxes1 | 1579 | 6.59516664 | 5.67521757 | 5.32308802 | 4.15502874 | 4.87720569 | 5.80573509 | 8.83929087 | 5.80505166 | 6.2340578 | 5.94236222 | 6.92873052 | 8.48369792 | 6.15477372 | 8.47111917 | 8.37599711 | 5.2622304 | 7.52447881 | 4.70312185 |
| ENSMUSG00 | Olfr1496 | 1066 | 0 | 0 | 0 | 0 | 0 | 0 | 0 | 0 | 0 | 0 | 0 | 0 | 0 | 0 | 0 | 0 | 0 | 0 |
| ENSMUSG00 | Gm9825 | 1000 | 0.38254658 | 0.53300631 | 4.40199126 | 0.31892731 | 3.85055389 | 11.5953397 | 0.70969018 | 0.45830883 | 3.45966732 | 0.28290422 | 6.40184444 | 13.395759 | 0.44398726 | 0.3715527 | 3.62573692 | 0.5360685 | 4.75246082 | 11.8819671 |
| ENSMUSG00 | Omd | 1553 | 0.05473944 | 0.49336546 | 0.3737813 | 0 | 0.12397147 | 0.38289298 | 0.09139603 | 0.07377798 | 0.45085046 | 0 | 0.09228904 | 0.10782163 | 0 | 0.08971813 | 0 | 0.07403663 | 0 | 0 |
| ENSMUSG00 | Pdp2 | 3483 | 3.61226876 | 3.51970861 | 3.63877834 | 3.57110683 | 2.52889521 | 3.62789178 | 5.50147429 | 5.17565717 | 4.15057911 | 5.414953 | 4.27958284 | 3.87809083 | 5.73626945 | 5.08044711 | 6.31686214 | 5.19995903 | 5.51291265 | 5.62252139 |
| ENSMUSG00 | Fgfbp1 | 1152 | 94.4190499 | 111.62133 | 91.4981967 | 73.7203009 | 103.742479 | 85.5558916 | 7.43367612 | 13.1286383 | 21.8803365 | 17.6405843 | 15.9343629 | 15.5558018 | 12.3329793 | 3.50749618 | 3.43346299 | 4.18803518 | 3.6928984 | 1.28927594 |
| ENSMUSG00 | F2r | 3305 | 2.97086102 | 4.6769086 | 2.38574446 | 4.59056816 | 4.79134732 | 4.01819443 | 0.28630971 | 0.08089162 | 0.23677573 | 0.44226076 | 0.98296435 | 0.21954724 | 0.0597058 | 0.71668637 | 0.27426149 | 0.52135471 | 0.95090964 | 0 |
| ENSMUSG00 | Foxl2 | 2589 | 0 | 0 | 0 | 0.14078345 | 0 | 0 | 0 | 0 | 0 | 0 | 0 | 0 | 0 | 0 | 0 | 0 | 0 | 0 |
| ENSMUSG00 | Olfr1394 | 938 | 0 | 0 | 0 | 0 | 0 | 0 | 0 | 0 | 0 | 0 | 0 | 0 | 0 | 0 | 0 | 0 | 0 | 0 |
| ENSMUSG00 | Socs4 | 1921 | 11.0411675 | 9.64183716 | 8.86386397 | 8.91772653 | 11.95155 | 7.79017467 | 12.3638547 | 9.12561827 | 12.9284066 | 10.5052061 | 7.5604307 | 9.44304547 | 11.376387 | 7.44652729 | 9.17973239 | 12.4180367 | 9.75614742 | 8.5906986 |
| ENSMUSG00 | Scrt1 | 7172 | 0 | 0 | 0 | 0 | 0 | 0 | 0 | 0 | 0 | 0 | 0 | 0 | 0 | 0.025903 | 0 | 0 | 0 | 0 |
| ENSMUSG00 | Osr1 | 2007 | 0 | 0 | 0 | 0 | 0 | 0 | 0 | 0 | 0 | 0 | 0 | 0 | 0 | 0 | 0 | 0 | 0 | 0 |
| ENSMUSG00 | Fam171b | 5736 | 0 | 0 | 0 | 0 | 0 | 0 | 0 | 0.01331674 | 0 | 0 | 0 | 0 | 0 | 0 | 0 | 0 | 0 | 0 |
| ENSMUSG00 | Olfr843 | 1106 | 0 | 0 | 0 | 0 | 0 | 0 | 0 | 0 | 0 | 0 | 0 | 0 | 0 | 0 | 0 | 0 | 0 | 0 |
| ENSMUSG00 | Gm5065 | 1434 | 0 | 0 | 0 | 0 | 0 | 0 | 0 | 0 | 0 | 0 | 0 | 0 | 0 | 0 | 0 | 0 | 0 | 0 |
| ENSMUSG00 | Tprg | 2032 | 0.16734321 | 0.06557656 | 0 | 0.04484355 | 0 | 0.07315856 | 0.09313519 | 0.09397737 | 0.08107582 | 0 | 0 | 0.02746834 | 0 | 0.04571268 | 0.08110542 | 0.05653141 | 0.07544547 | 0 |
| ENSMUSG00 | Prss54 | 1432 | 0 | 0 | 0 | 0 | 0 | 0 | 0 | 0 | 0 | 0 | 0 | 0 | 0 | 0 | 0 | 0 | 0 | 0 |
| ENSMUSG00 | Gli2 | 6853 | 0 | 0 | 0 | 0.01994501 | 0 | 0 | 0 | 0 | 0 | 0 | 0 | 0 | 0 | 0 | 0 | 0 | 0 | 0 |
| ENSMUSG00 | B330016D10 | 1698 | 5.6823763 | 4.80663079 | 6.38143157 | 3.97116287 | 5.66924896 | 5.07784196 | 7.46748996 | 7.49002945 | 7.42230496 | 6.10904282 | 7.87808503 | 6.87012572 | 3.54446049 | 5.60720725 | 3.56691915 | 5.50230165 | 4.44657273 | 7.96951723 |
| ENSMUSG00 | Zfp407 | 10280 | 1.2569624 | 0.94624174 | 0.80465697 | 1.32960246 | 1.01133225 | 1.05564676 | 0.74558891 | 0.82477756 | 0.56090586 | 0.83018262 | 0.87835284 | 1.03704196 | 0.69582957 | 0.89906356 | 1.39075291 | 0.41717393 | 0.60024648 | 0.87757719 |
| ENSMUSG00 | Gm597 | 2858 | 0 | 0 | 0 | 0 | 0 | 0 | 0 | 0 | 0 | 0 | 0 | 0 | 0 | 0 | 0 | 0 | 0 | 0 |
| ENSMUSG00 | Mlf1 | 1847 | 0.25314398 | 1.49706607 | 1.07380333 | 0.81403057 | 0.36483321 | 0.05365754 | 0 | 0.18610266 | 1.20415368 | 0.5616218 | 0.02586625 | 0.60439266 | 0.24038292 | 0.70407929 | 0.28999471 | 1.14021841 | 1.07902947 | 0.77435667 |
| ENSMUSG00 | Ranbp3l | 3011 | 0.09881642 | 0.1438285 | 0.04819681 | 0.48420904 | 0.03197072 | 0.4278881 | 1.00564977 | 1.71237938 | 1.60040596 | 1.11182329 | 1.95161737 | 0.88978805 | 0.8847305 | 1.6041797 | 0.87575542 | 1.03006948 | 0.70008181 | 1.37020359 |
| ENSMUSG00 | Olfr686 | 1017 | 0 | 0 | 0 | 0 | 0 | 0 | 0 | 0 | 0 | 0 | 0 | 0 | 0 | 0 | 0 | 0 | 0 | 0 |
| ENSMUSG00 | 1810026J23R | 3144 | 22.9154814 | 23.7979518 | 23.8175016 | 26.5048187 | 25.9796038 | 25.7062686 | 21.1432296 | 26.2390551 | 23.4621659 | 27.6845414 | 25.2094962 | 28.3694112 | 22.0142112 | 23.0300039 | 20.0372707 | 24.296998 | 26.7333235 | 21.0833249 |
| ENSMUSG00 | Nupl2 | 3964 | 3.15250296 | 4.55489119 | 4.16129512 | 3.3791491 | 3.72767421 | 3.82521102 | 2.98389751 | 4.19114406 | 3.335235 | 4.09178656 | 3.13357447 | 4.06930537 | 3.90772503 | 3.05799869 | 3.9289072 | 4.85877284 | 4.04147127 | 3.91336228 |
| ENSMUSG00 | Cyp4f16 | 2922 | 14.2120318 | 14.8665346 | 12.9625169 | 12.6610431 | 13.9025816 | 14.6691015 | 12.6458598 | 9.94675504 | 10.1627234 | 11.795744 | 12.3443168 | 9.91387025 | 13.4895599 | 11.8891777 | 11.9994949 | 12.2917829 | 10.8473126 | 11.2390269 |
| ENSMUSG00 | 2210020M01 | 1810 | 62.8419067 | 36.3681099 | 47.6252105 | 43.5724684 | 45.4727014 | 40.7646521 | 99.4611842 | 74.8022467 | 96.4583648 | 95.9686137 | 99.0076626 | 77.3709931 | 99.0726626 | 77.8515807 | 83.199827 | 82.2084532 | 78.1771758 | 86.9508587 |
| ENSMUSG00 | Ccdc57 | 4103 | 3.00426543 | 3.39390692 | 3.17145495 | 3.15362822 | 2.39310564 | 4.09416931 | 3.65540317 | 3.14624226 | 2.77052848 | 2.65459723 | 2.51508434 | 3.53694182 | 4.36448598 | 4.39198292 | 3.94643224 | 3.28498509 | 3.48420681 | 2.37304701 |
| ENSMUSG00 | Msx1 | 1933 | 11.1705274 | 11.5293979 | 10.1601931 | 14.6134752 | 11.6034539 | 11.3050986 | 1.5664829 | 2.21290692 | 2.10939734 | 2.80513755 | 1.75479672 | 1.79025922 | 12.7859755 | 14.0077098 | 14.941692 | 21.0766716 | 16.6351594 | 11.8384834 |
| ENSMUSG00 | Sprr1b | 953 | 0 | 0 | 0 | 0 | 0 | 0 | 0 | 0 | 0 | 0 | 0 | 0 | 0 | 0 | 0 | 0 | 0 | 0 |
| ENSMUSG00 | Olfr1448 | 944 | 0 | 0 | 0 | 0 | 0 | 0 | 0 | 0 | 0 | 0 | 0 | 0 | 0 | 0 | 0 | 0 | 0 | 0 |
| ENSMUSG00 | 6530418L21R | 4778 | 0.25798453 | 0.14641498 | 0.23285709 | 0.03814236 | 0.11081021 | 0.04148408 | 0.27726141 | 0.10391403 | 0.38790147 | 0.2763122 | 0 | 0.03504541 | 0.28909453 | 0.09720403 | 0.13797089 | 0.31254368 | 0.02406423 | 0.28782496 |
| ENSMUSG00 | Olfr564 | 950 | 0 | 0 | 0 | 0 | 0 | 0 | 0 | 0 | 0 | 0 | 0 | 0 | 0 | 0 | 0 | 0 | 0 | 0 |
| ENSMUSG00 | Gm766 | 2286 | 0 | 0 | 0 | 0 | 0 | 0 | 0 | 0 | 0 | 0.14438098 | 0 | 0 | 0 | 0 | 0 | 0 | 0 | 0 |
| ENSMUSG00 | 4732415M23 | 515 | 0.66027458 | 0.90559324 | 0.56357511 | 0.88468047 | 1.02806051 | 0.6735323 | 2.02112415 | 1.5573601 | 2.31924069 | 0.91554764 | 2.04087201 | 1.30055913 | 1.34106183 | 3.06621159 | 2.80010574 | 1.1896111 | 1.71165986 | 1.06813799 |
| ENSMUSG00 | Cxcr1 | 1123 | 0 | 0 | 0 | 0 | 0 | 0 | 0 | 0 | 0 | 0 | 0 | 0 | 0 | 0 | 0 | 0 | 0 | 0 |
| ENSMUSG00 | Mypop | 2207 | 0.5585184 | 0.46792013 | 0.24110051 | 0.66060418 | 0.1962788 | 0.53886072 | 0.62168903 | 0.31149218 | 0.41055883 | 0.29909825 | 0.54117533 | 0.83457946 | 0.87174661 | 0.2314839 | 0.54138882 | 0.72868397 | 0.29521842 | 0.34894766 |
| ENSMUSG00 | Bdnf | 6208 | 0 | 0.00536612 | 0 | 0 | 0 | 0 | 0 | 0.03691276 | 0.00683443 | 0 | 0 | 0 | 0 | 0 | 0.00683685 | 0 | 0 | 0 |
| ENSMUSG00 | Zdhhc22 | 1016 | 0 | 0 | 0.04761174 | 0 | 0 | 0 | 0 | 0 | 0.08107582 | 0 | 0 | 0 | 0 | 0 | 0 | 0 | 0 | 0 |
| ENSMUSG00 | Gm7461 | 826 | 0.41167241 | 0.2016519 | 0.81989035 | 0.60674514 | 0.75752422 | 0.53992083 | 0.34367563 | 1.94198656 | 1.24656525 | 1.31291304 | 0.28919466 | 1.1487485 | 1.73199225 | 1.74305897 | 2.14487519 | 1.25163002 | 0.69599813 | 1.26534265 |
| ENSMUSG00 | Zbtb8b | 6016 | 2.53646247 | 2.76869134 | 2.91881508 | 3.15807106 | 2.08016957 | 2.78404653 | 2.32788447 | 2.55843387 | 2.93700387 | 3.16637037 | 2.04885959 | 3.25653218 | 4.11645985 | 1.85282264 | 2.93807638 | 2.94690014 | 3.01335413 | 3.32834356 |
| ENSMUSG00 | Fitm2 | 3965 | 6.81798032 | 7.31791447 | 5.80726367 | 6.83702935 | 5.40194351 | 6.01131522 | 7.07601982 | 4.96067774 | 7.27126382 | 5.88640558 | 9.21761494 | 6.54584694 | 7.51487466 | 5.96217919 | 8.19874593 | 7.10766382 | 6.64064716 | 5.20262674 |
| ENSMUSG00 | 8430408G22 | 1315 | 1.18081304 | 1.36798197 | 1.58179605 | 1.10870982 | 1.83011117 | 1.58267286 | 1.36721049 | 2.35253581 | 0.09396163 | 1.72109031 | 2.25250755 | 2.84383984 | 5.66472966 | 0.0706374 | 0.31331982 | 0.90368831 | 0.61205493 | 1.54778475 |
| ENSMUSG00 | Nrip1 | 15604 | 13.4892099 | 12.1795733 | 12.8777009 | 14.0911048 | 12.5357689 | 13.6774945 | 13.1653199 | 10.9358917 | 12.4398903 | 12.0354557 | 10.8445849 | 12.6733461 | 8.57080414 | 9.8787443 | 10.7809506 | 10.402069 | 11.5489777 | 12.1553063 |
| ENSMUSG00 | 1110034B05R | 2811 | 5.51917085 | 4.24262402 | 5.30026587 | 6.35358569 | 5.75322887 | 5.28844542 | 5.13353507 | 5.38035978 | 6.34427653 | 3.63967997 | 6.27141931 | 3.89180714 | 6.80924365 | 5.46886977 | 5.05675446 | 5.61214168 | 7.13079092 | 6.86879988 |
| ENSMUSG00 | Mmgt2 | 4612 | 12.0179422 | 10.7262897 | 11.2438041 | 10.639472 | 9.58046529 | 12.4633305 | 7.71446976 | 7.90844412 | 9.26964513 | 8.50593779 | 7.34441578 | 8.20533808 | 7.45540901 | 6.95855689 | 7.98659821 | 7.39742654 | 7.52897333 | 8.79048388 |
| ENSMUSG00 | Cd300e | 2084 | 0 | 0 | 0 | 0 | 0 | 0 | 0 | 0 | 0 | 0 | 0 | 0 | 0 | 0 | 0 | 0 | 0 | 0 |
| ENSMUSG00 | Defb15 | 343 | 41.6377235 | 39.6258334 | 23.4110963 | 52.069765 | 41.1156083 | 16.6138903 | 10.2074001 | 10.912115 | 8.84054114 | 24.8812749 | 20.196411 | 4.23092486 | 16.252206 | 26.8102895 | 4.68472501 | 22.9966913 | 23.4650797 | 16.8395225 |
| ENSMUSG00 | Olfr938 | 1118 | 0 | 0 | 0 | 0 | 0 | 0 | 0 | 0 | 0 | 0 | 0 | 0 | 0 | 0 | 0 | 0 | 0 | 0 |
| ENSMUSG00 | Duxbl | 4649 | 0 | 0 | 0 | 0 | 0 | 0 | 0 | 0 | 0 | 0 | 0 | 0 | 0 | 0 | 0 | 0 | 0 | 0 |
| ENSMUSG00 | Tmem136 | 3497 | 1.05746363 | 1.53370774 | 2.03343122 | 0.72960208 | 0.63313368 | 1.36032669 | 0.08117703 | 0.31672281 | 0.01177767 | 0.14831505 | 0.16394038 | 0.3671033 | 0.01410693 | 0 | 0.03534592 | 0.39418412 | 0.02191953 | 0 |
| ENSMUSG00 | Olfr453 | 953 | 0 | 0 | 0 | 0 | 0 | 0 | 0 | 0 | 0 | 0 | 0 | 0 | 0 | 0 | 0 | 0 | 0 | 0 |
| ENSMUSG00 | Adam26a | 2487 | 0 | 0 | 0 | 0 | 0 | 0 | 0 | 0 | 0 | 0 | 0 | 0 | 0 | 0 | 0 | 0 | 0 | 0 |
| ENSMUSG00 | Fbxl13 | 3737 | 0.04549658 | 0 | 0 | 0 | 0 | 0 | 0 | 0 | 0 | 0 | 0 | 0.1642955 | 0.01320094 | 0 | 0 | 0 | 0.0923031 | 0 |
| ENSMUSG00 | Cxcr6 | 1891 | 0.06743285 | 0.33471443 | 0.2046474 | 0 | 0.05090632 | 0.15722708 | 0 | 0 | 0.02178028 | 0 | 0.15158633 | 0.11806592 | 0.2608774 | 0 | 0 | 0.14174207 | 0 | 0.37816943 |
| ENSMUSG00 | Nkx1-2 | 3263 | 0 | 0 | 0 | 0 | 0 | 0 | 0 | 0 | 0 | 0 | 0 | 0 | 0.07559289 | 0 | 0.25840592 | 0 | 0 | 0 |
| ENSMUSG00 | Amica1 | 2446 | 0.55607753 | 0.66734743 | 0.41530831 | 0.52154916 | 0.72807898 | 0.40517362 | 0.09671439 | 0.53088376 | 0.30308965 | 0.52046974 | 0.25391433 | 0.15973411 | 1.75465119 | 0.72153529 | 0.65693405 | 0.50094008 | 0.09401381 | 0.58472476 |
| ENSMUSG00 | Phldb1 | 8540 | 0.19411029 | 0.41348557 | 0.33419652 | 0.38945623 | 0.44524847 | 0.51061365 | 0.35456809 | 0.34435773 | 0.09645554 | 0.35335422 | 0.08950812 | 0.23528851 | 0.27727541 | 0.21753671 | 0.22192876 | 0.4618188 | 0.1660507 | 0.46377701 |
| ENSMUSG00 | Gm9826 | 1298 | 0 | 0 | 0.03726774 | 0 | 0.03708161 | 0.11452866 | 0 | 0.20596827 | 0.03173075 | 0.07265131 | 0 | 0 | 0.0380061 | 0.07156254 | 0.06348468 | 0.0589994 | 0.02952719 | 0 |
| ENSMUSG00 | Nhlh2 | 3350 | 0 | 0 | 0 | 0 | 0 | 0 | 0 | 0 | 0 | 0 | 0 | 0 | 0 | 0 | 0 | 0 | 0 | 0 |
| ENSMUSG00 | 5930422O12 | 839 | 0 | 0 | 0 | 0 | 0 | 0 | 0 | 0 | 0 | 0 | 0 | 0 | 0.05535648 | 0 | 0 | 0 | 0 | 0 |
| ENSMUSG00 | Tob2 | 3992 | 17.2597421 | 20.4867624 | 19.339718 | 18.5120275 | 19.7133505 | 20.4318135 | 17.7896621 | 17.0296784 | 16.5179388 | 16.6303082 | 14.947626 | 19.2810117 | 19.6240192 | 18.5566932 | 25.1729947 | 21.054079 | 22.4274122 | 23.3568226 |
| ENSMUSG00 | Thnsl1 | 3444 | 4.31963171 | 5.30065797 | 5.77279935 | 6.56163705 | 6.49864823 | 5.89913495 | 3.6542313 | 5.24535608 | 6.39802147 | 3.75124643 | 3.88414297 | 3.37098078 | 3.79586471 | 1.73963045 | 3.18223399 | 5.20325327 | 5.69775378 | 3.68963521 |
| ENSMUSG00 | 4930555K19R | 170 | 0 | 0 | 0 | 0 | 0 | 0 | 0 | 0 | 0 | 0 | 0 | 0 | 0 | 0 | 0 | 0 | 0 | 0 |
| ENSMUSG00 | Sp8 | 1921 | 0.24339247 | 0 | 0 | 0.07115207 | 0 | 0.02579528 | 0 | 0 | 0.12864086 | 0 | 0 | 0 | 0 | 0 | 0 | 0 | 0 | 0 |
| ENSMUSG00 | E030010A14R | 2460 | 1.45139625 | 0.29792019 | 0.80622551 | 0.2592905 | 0.27392152 | 0.3222942 | 0.07693119 | 0.54338784 | 0 | 0.34500515 | 0 | 0.22689294 | 0.02005362 | 0.64191015 | 0.167486 | 0.15565288 | 0.12463836 | 0.51431279 |
| ENSMUSG00 | Cypt3 | 673 | 0 | 0 | 0 | 0 | 0 | 0 | 0 | 0 | 0 | 0 | 0 | 0 | 0 | 0 | 0 | 0 | 0 | 0 |
| ENSMUSG00 | Gm5593 | 2038 | 0 | 0 | 0 | 0 | 0 | 0 | 0 | 0 | 0 | 0 | 0 | 0 | 0 | 0 | 0 | 0 | 0 | 0 |
| ENSMUSG00 | Mlec | 11165 | 16.9335444 | 19.1493198 | 19.1934384 | 21.7460641 | 18.5802589 | 21.2013801 | 22.7431391 | 24.3897742 | 21.9194157 | 26.681428 | 19.9358291 | 22.0563102 | 17.899113 | 18.1200577 | 17.9456486 | 17.229916 | 21.4064297 | 17.0077417 |
| ENSMUSG00 | E130311K13R | 2392 | 2.62991892 | 2.66001789 | 3.90304823 | 2.49519097 | 2.53537725 | 2.75523142 | 3.52075955 | 2.42694196 | 3.63309248 | 3.86351916 | 2.17703615 | 1.9600818 | 2.94919071 | 3.70853708 | 2.4114623 | 1.07252119 | 4.00567484 | 1.58680116 |
| ENSMUSG00 | Gja3 | 1546 | 0.79731572 | 1.20667663 | 0.56321057 | 1.29669208 | 0.49813116 | 1.28208884 | 3.12153509 | 1.95161694 | 3.1702428 | 2.13489601 | 3.33744859 | 0.79427204 | 3.22284841 | 7.99102668 | 3.86431152 | 3.76465515 | 4.7102176 | 5.08816446 |
| ENSMUSG00 | Igf2 | 7654 | 0 | 0 | 0 | 0 | 0 | 0 | 0 | 0 | 0 | 0.01848081 | 0 | 0.05104646 | 0.03222623 | 0 | 0 | 0 | 0 | 0 |
| ENSMUSG00 | Gm5946 | 970 | 0.08763954 | 0.03434319 | 0.04986962 | 0 | 0 | 0.10217058 | 0.09755192 | 0.03937361 | 0 | 0 | 0 | 0.05754192 | 0 | 0 | 0.04247583 | 0 | 0.07902329 | 0.17013126 |
| ENSMUSG00 | Gm5763 | 3050 | 0.06968062 | 0 | 0.01586017 | 0 | 0.03156192 | 0 | 0 | 0 | 0.0540151 | 0 | 0 | 0.03660043 | 0.0161744 | 0 | 0.01350871 | 0 | 0.025132 | 0 |
| ENSMUSG00 | Morc2b | 6790 | 0 | 0 | 0 | 0 | 0 | 0 | 0 | 0 | 0.00606576 | 0.01388828 | 0 | 0 | 0 | 0 | 0 | 0 | 0.00564452 | 0 |
| ENSMUSG00 | Gm9828 | 2788 | 0.76228795 | 0.43015215 | 0.52051862 | 0.57196433 | 0.55244676 | 0.28437724 | 0.54304366 | 0.82193119 | 1.01902295 | 0.77795278 | 0.4798059 | 0.8007986 | 1.16782874 | 0.98285551 | 0.84235606 | 0.46695863 | 0.75607831 | 0.69057343 |
| ENSMUSG00 | Myof | 7091 | 16.1784614 | 19.2849289 | 16.1745368 | 14.6046639 | 19.3858093 | 21.8728044 | 8.48038569 | 10.8690267 | 8.35811538 | 8.31171613 | 10.9752373 | 12.6807266 | 5.49601109 | 6.87065966 | 7.18165605 | 6.43126688 | 9.23160557 | 9.97626727 |
| ENSMUSG00 | Nog | 1921 | 0.02212659 | 0 | 0 | 0.11858679 | 0 | 0 | 0 | 0 | 0 | 0 | 0 | 0 | 0 | 0 | 0 | 0.01993264 | 0.11970733 | 0 |
| ENSMUSG00 | Rtbdn | 3387 | 0 | 0 | 0 | 0.08071044 | 0 | 0 | 0 | 0 | 0 | 0.04176324 | 0 | 0 | 0 | 0 | 0 | 0 | 0 | 0 |
| ENSMUSG00 | Olfr1336 | 944 | 0 | 0 | 0 | 0 | 0 | 0 | 0 | 0 | 0 | 0 | 0 | 0 | 0 | 0 | 0 | 0 | 0 | 0 |
| ENSMUSG00 | Gm5377 | 1060 | 0 | 0 | 0 | 0.21491059 | 0 | 0 | 0.62488444 | 0.57648909 | 1.70962896 | 0.17792718 | 0.04507072 | 0 | 0.13961863 | 0 | 0 | 0.14449286 | 0 | 0 |
| ENSMUSG00 | Klf17 | 2397 | 0 | 0 | 0 | 0 | 0 | 0.02067281 | 0 | 0.06373367 | 0 | 0 | 0 | 0 | 0 | 0 | 0 | 0 | 0 | 0 |
| ENSMUSG00 | 4930542C16 | 1249 | 0 | 0 | 0 | 0 | 0 | 0 | 0 | 0 | 0 | 0 | 0 | 0 | 0 | 0 | 0 | 0 | 0 | 0 |
| ENSMUSG00 | A730049H05R | 3733 | 0.01138633 | 0.01784779 | 0.05183341 | 0.0610247 | 0 | 0.10619391 | 0.07604502 | 0 | 0.09929779 | 0.02526156 | 0 | 0.01495196 | 0.06607543 | 0 | 0.0662227 | 0 | 0 | 0 |
| ENSMUSG00 | Krtap16-10a | 167 | 0 | 0 | 0 | 0 | 0 | 0 | 0 | 0 | 0 | 0 | 0 | 0 | 0 | 0 | 0 | 0 | 0 | 0 |
| ENSMUSG00 | Ctxn1 | 1222 | 22.7482693 | 24.5893867 | 19.5948425 | 19.2012585 | 22.9631027 | 25.181872 | 13.396225 | 14.7518936 | 12.2683238 | 9.72338561 | 14.7390829 | 19.092428 | 11.5053981 | 9.91972734 | 8.76628849 | 9.1183034 | 7.46453246 | 6.79736098 |
| ENSMUSG00 | Exd1 | 10243 | 1.40674165 | 2.00989638 | 1.91737316 | 1.65911057 | 1.86550558 | 1.76092893 | 0.90994804 | 1.40196654 | 1.7812776 | 1.40397974 | 1.69775309 | 1.56935574 | 0.88135711 | 0.69373673 | 1.71354709 | 1.48781233 | 1.08883651 | 1.3801953 |
| ENSMUSG00 | Gm9830 | 1375 | 2.99854696 | 4.91819456 | 3.90506318 | 4.87088983 | 3.88555893 | 6.88332517 | 4.26674341 | 5.91635034 | 7.33868826 | 6.78970127 | 3.82199668 | 6.17016779 | 4.59235303 | 6.11373078 | 6.23267173 | 7.26825348 | 4.98938702 | 8.48140409 |
| ENSMUSG00 | Samd13 | 2922 | 0.49458453 | 0.44462795 | 0.54631297 | 0.31184835 | 0.57652888 | 0.37308697 | 0.61529151 | 0.39211912 | 0.50743141 | 0.37113832 | 0.5722531 | 0.76407478 | 0.27012686 | 0.93778274 | 0.32431067 | 0.20966794 | 0.23809628 | 0.47064602 |
| ENSMUSG00 | A730037C10R | 3314 | 0.11543349 | 0 | 0 | 0 | 0.02904763 | 0 | 0.05710643 | 0.16134388 | 0.02485607 | 0 | 0 | 0 | 0 | 0 | 0 | 0.13865036 | 0.12721462 | 0 |
| ENSMUSG00 | Lemd3 | 6499 | 4.656668 | 4.69015206 | 4.65201669 | 4.87227663 | 5.68784696 | 4.84929044 | 5.19791549 | 5.52405882 | 5.45013134 | 4.36029738 | 4.35186574 | 4.56041188 | 3.09700296 | 3.28731808 | 2.94160978 | 4.66038936 | 4.08680162 | 2.81859248 |
| ENSMUSG00 | A430010J10R | 405 | 0 | 0 | 0 | 0 | 0 | 0 | 0 | 0.18860446 | 0 | 0 | 0 | 0 | 0 | 0 | 0 | 0 | 0 | 0 |
| ENSMUSG00 | 5930416I19R | 3351 | 5.02299305 | 3.41976592 | 4.17187415 | 3.68458461 | 3.46159165 | 3.65249931 | 4.02390734 | 3.80670319 | 4.09284087 | 4.44632117 | 4.73329908 | 3.84763296 | 3.8717679 | 4.1717906 | 3.94679184 | 3.88505118 | 4.23179047 | 4.25167372 |
| ENSMUSG00 | Tpcn2 | 2985 | 3.27510569 | 3.39266996 | 2.25256976 | 2.716873 | 3.19267031 | 3.20391209 | 3.34438032 | 4.01755925 | 4.04276352 | 2.65370792 | 2.54479678 | 2.24384573 | 2.89215595 | 2.75397101 | 4.61015734 | 4.68210107 | 2.58076572 | 2.69056267 |
| ENSMUSG00 | Hmgb4 | 682 | 0 | 0 | 0 | 0 | 0 | 0 | 0 | 0 | 0 | 0 | 0 | 0 | 0 | 0 | 0 | 0 | 0 | 0 |
| ENSMUSG00 | Olfr435 | 941 | 0 | 0 | 0 | 0 | 0 | 0 | 0 | 0 | 0 | 0 | 0 | 0 | 0 | 0 | 0 | 0 | 0 | 0 |
| ENSMUSG00 | Mex3d | 4655 | 4.43770474 | 5.44599624 | 4.75941502 | 4.51206046 | 5.09753778 | 4.92866072 | 2.58161556 | 3.45413564 | 3.17636072 | 3.76800464 | 3.46894435 | 2.92567598 | 3.35944528 | 2.91335629 | 4.08032594 | 3.15866865 | 3.48271183 | 3.87604447 |
| ENSMUSG00 | Vmn1r26 | 1019 | 0 | 0 | 0 | 0 | 0 | 0 | 0 | 0 | 0 | 0 | 0 | 0 | 0 | 0 | 0 | 0 | 0 | 0 |
| ENSMUSG00 | 4732456N10 | 2525 | 0 | 0 | 0 | 0 | 0 | 0 | 0 | 0 | 0 | 0 | 0 | 0 | 0 | 0 | 0 | 0 | 0 | 0 |
| ENSMUSG00 | Ccdc6 | 6362 | 10.1753196 | 7.97477804 | 9.38273129 | 7.65557315 | 9.07864011 | 7.56298402 | 7.16160167 | 5.94317485 | 8.36419025 | 7.12226121 | 7.17900979 | 8.69432908 | 7.35093643 | 5.3583716 | 6.00990944 | 7.90849703 | 6.69896922 | 7.05555346 |
| ENSMUSG00 | Speer4b | 2646 | 0 | 0 | 0 | 0 | 0 | 0 | 0 | 0 | 0 | 0 | 0 | 0 | 0.07457584 | 0 | 0 | 0 | 0 | 0 |
| ENSMUSG00 | D4Bwg0951e | 3064 | 0.24970404 | 0.21744709 | 0.2841787 | 0.1933073 | 0.07854426 | 0.04851769 | 1.12722766 | 1.05951508 | 1.41141781 | 0.27699499 | 0.48336283 | 1.2387288 | 0.56351733 | 0.68210964 | 0.76647803 | 0.18745402 | 0.47532614 | 1.41831574 |
| ENSMUSG00 | Tprn | 4562 | 3.04673226 | 3.6949418 | 3.91272089 | 3.2258258 | 2.99637578 | 3.58448095 | 3.66097669 | 3.80919401 | 5.69677585 | 4.40293722 | 4.33556177 | 3.59706374 | 5.0391656 | 3.63448908 | 3.6396815 | 4.56599966 | 3.08324221 | 5.31762737 |
| ENSMUSG00 | Gm8666 | 291 | 0 | 0.1144773 | 0.16623206 | 0.15656716 | 0.827009 | 0.34056862 | 0.16258653 | 0 | 0.14153442 | 0.16202991 | 0.16417512 | 0.1918064 | 0 | 0.31920335 | 0 | 0.26316569 | 0.52682195 | 0.56710419 |
| ENSMUSG00 | Tbc1d12 | 4965 | 4.50306598 | 5.39447471 | 4.29662173 | 4.10187046 | 5.12825531 | 5.0999893 | 3.53534821 | 3.76154779 | 4.62052151 | 3.15287683 | 3.98365213 | 4.03581528 | 2.94103676 | 3.7030183 | 2.4213958 | 3.54756886 | 4.73964679 | 2.92495552 |
| ENSMUSG00 | C030019I05R | 2288 | 0.07430975 | 0 | 0.08456911 | 0.07965218 | 0.04207336 | 0 | 0 | 0 | 0 | 0 | 0 | 0 | 0.06488346 | 0.10149495 | 0.01800767 | 0.05020622 | 0.05025301 | 0 |
| ENSMUSG00 | Zfp454 | 2541 | 1.20439696 | 0.76038877 | 0.32363243 | 1.16547339 | 0.85239534 | 1.24808144 | 0.78202775 | 1.06716276 | 0.79423033 | 0.92779818 | 0.71446219 | 0.43902045 | 1.08720479 | 1.35256295 | 1.10259969 | 0.67810993 | 1.59881443 | 0.45462072 |
| ENSMUSG00 | Ggnbp1 | 6932 | 1.03626295 | 0.72565595 | 0.99091767 | 0.73612766 | 0.74989148 | 0.63620792 | 0.62792361 | 1.16803077 | 1.03382295 | 0.99307598 | 1.20609027 | 1.41713165 | 1.3236781 | 1.27299143 | 0.95098802 | 0.88932311 | 1.1113078 | 1.41252467 |
| ENSMUSG00 | Klhl11 | 2391 | 0.67553186 | 1.60225129 | 1.29481637 | 1.61969417 | 1.28834927 | 1.28493077 | 1.56323782 | 2.5557442 | 0.86128222 | 2.05088797 | 1.17888856 | 1.44733211 | 1.87754265 | 2.38921905 | 1.61980186 | 2.38615663 | 2.03573388 | 2.967869 |
| ENSMUSG00 | Olfr820 | 1060 | 0 | 0 | 0 | 0 | 0 | 0 | 0 | 0 | 0 | 0 | 0 | 0 | 0 | 0 | 0 | 0 | 0 | 0 |
| ENSMUSG00 | E130114P18R | 2035 | 0.18798358 | 0.67116888 | 0.52295708 | 0.49255183 | 0.28382461 | 0.21915214 | 0.41849053 | 0.15014212 | 0 | 0.57924697 | 0.42257949 | 0.32913413 | 0.36362593 | 0 | 0 | 0.26342433 | 0.37667122 | 0.24328352 |
| ENSMUSG00 | Prss50 | 1450 | 0 | 0 | 0 | 0 | 0.03319443 | 0 | 0 | 0 | 0 | 0 | 0 | 0 | 0 | 0 | 0 | 0 | 0 | 0 |
| ENSMUSG00 | Mcat | 1889 | 20.7238047 | 21.7442025 | 20.2303277 | 27.5440512 | 21.734532 | 25.2616835 | 27.0000359 | 25.0100592 | 28.2789364 | 26.2087022 | 25.038226 | 26.6520528 | 24.5745549 | 26.9715002 | 27.2641317 | 22.6216611 | 24.3064604 | 24.0246231 |
| ENSMUSG00 | Foxo3 | 8073 | 4.60696507 | 2.80599134 | 3.52330433 | 3.5893502 | 2.90949817 | 2.91558881 | 3.89144293 | 3.74212688 | 3.86713476 | 3.34079057 | 3.01219545 | 4.31425411 | 3.59921953 | 4.30900798 | 3.10300334 | 5.53039121 | 3.90716491 | 3.69998079 |
| ENSMUSG00 | Rpl29 | 829 | 39.2749877 | 35.241747 | 36.8199007 | 32.810547 | 33.5007478 | 36.1035598 | 42.6327758 | 33.9999915 | 42.5279342 | 37.4248043 | 37.5168854 | 34.7417152 | 52.5453353 | 51.6543409 | 47.5631954 | 38.5677409 | 41.6549982 | 42.9986745 |
| ENSMUSG00 | Hoxb3 | 2999 | 1.06298373 | 1.17744808 | 1.20974151 | 0.95710096 | 0.94691014 | 1.47055461 | 1.05700216 | 1.32444476 | 1.07120648 | 0.78610709 | 1.09919044 | 1.24696545 | 1.20081026 | 1.09954325 | 0.92047491 | 0.99588776 | 0.71566276 | 0.62364442 |
| ENSMUSG00 | Tmprss11f | 6795 | 0 | 0 | 0 | 0 | 0 | 0 | 0 | 0 | 0 | 0 | 0 | 0 | 0 | 0 | 0 | 0 | 0.00564037 | 0 |
| ENSMUSG00 | Skint10 | 3727 | 0 | 0 | 0 | 0 | 0 | 0 | 0 | 0 | 0 | 0 | 0 | 0 | 0 | 0 | 0 | 0 | 0 | 0 |
| ENSMUSG00 | H1f0 | 531 | 303.459741 | 354.521968 | 360.751752 | 390.143255 | 352.151645 | 389.982813 | 319.338307 | 321.434739 | 343.298529 | 405.087586 | 411.080575 | 389.553381 | 242.200204 | 265.544726 | 345.751663 | 376.199809 | 427.868716 | 429.402537 |
| ENSMUSG00 | Tmem53 | 1687 | 11.0861159 | 10.7027793 | 10.4947908 | 9.9116202 | 10.9274018 | 9.63443783 | 7.79863589 | 9.53112117 | 6.66503782 | 10.2294946 | 11.8092221 | 7.64280857 | 9.18211146 | 8.20411265 | 8.67015551 | 10.5997117 | 7.65617194 | 9.22796515 |
| ENSMUSG00 | Serpinb13 | 1751 | 0 | 0 | 0 | 0 | 0 | 0 | 0 | 0 | 0 | 0 | 0 | 0 | 0 | 0 | 0 | 0 | 0 | 0 |
| ENSMUSG00 | Pthlh | 1599 | 0.47848228 | 0.12500148 | 0.33277601 | 0.68384307 | 0.68222784 | 0.21692879 | 0.14794459 | 0.09554072 | 0.38636506 | 0.29487619 | 0.44817034 | 1.29154441 | 0 | 0.23236567 | 0.36073908 | 0 | 0.35953374 | 0 |
| ENSMUSG00 | P2ry6 | 1935 | 0.15376549 | 0.03443193 | 0 | 0 | 0 | 0 | 0 | 0 | 0.29799029 | 0 | 0 | 0 | 0 | 0.02400211 | 0 | 0 | 0 | 0 |
| ENSMUSG00 | Insc | 3795 | 0 | 0 | 0 | 0 | 0 | 0 | 0 | 0 | 0 | 0 | 0 | 0 | 0 | 0 | 0 | 0.01009916 | 0 | 0 |
| ENSMUSG00 | Dcun1d3 | 8440 | 1.67704071 | 1.70906199 | 1.74236413 | 1.19840662 | 1.28313778 | 1.74961074 | 1.42386498 | 1.64263532 | 1.25413916 | 1.35753802 | 0.78115453 | 1.52765617 | 1.22745292 | 2.00303884 | 1.23018864 | 1.07522203 | 1.87544557 | 1.53165167 |
| ENSMUSG00 | E030011O05R | 1796 | 0 | 0 | 0 | 0 | 0 | 0 | 0 | 0 | 0 | 0 | 0 | 0 | 0 | 0 | 0 | 0 | 0 | 0 |
| ENSMUSG00 | Ccdc37 | 3752 | 0.11328672 | 0.08878703 | 0 | 0.10928822 | 0.05131335 | 0.01320702 | 0 | 0 | 0.03293165 | 0.15080182 | 0 | 0.01487624 | 0 | 0 | 0.02196245 | 0.01020539 | 0.05107449 | 0 |
| ENSMUSG00 | Cyb561d1 | 8649 | 2.44740174 | 2.64222979 | 2.53361189 | 2.86040549 | 2.95502965 | 2.53235151 | 2.49445966 | 3.1705567 | 2.99529638 | 2.55133878 | 2.76187759 | 2.12962986 | 2.85188561 | 2.65272045 | 2.1341539 | 1.52294704 | 2.11372917 | 2.24513986 |
| ENSMUSG00 | Cep120 | 5457 | 6.76873703 | 8.62582363 | 9.14815527 | 8.18211901 | 8.44094758 | 8.98069516 | 7.72504983 | 6.08894816 | 7.62293951 | 6.51486719 | 7.42407271 | 8.6428871 | 6.15631953 | 6.32361314 | 6.64419446 | 7.31850898 | 9.11627879 | 7.92324682 |
| ENSMUSG00 | Ifnb1 | 769 | 0 | 0 | 0 | 0 | 0 | 0 | 0 | 0 | 0 | 0 | 0 | 0 | 0 | 0 | 0 | 0 | 0 | 0 |
| ENSMUSG00 | Slc35e4 | 4023 | 1.97575638 | 2.78228498 | 2.08019407 | 2.29900373 | 3.09872439 | 1.67516077 | 3.90450147 | 3.39867762 | 2.27278312 | 3.76221122 | 2.86198484 | 1.92850537 | 2.90620542 | 4.07525798 | 3.0314841 | 4.81606944 | 3.03904765 | 3.2406558 |
| ENSMUSG00 | Olfr202 | 923 | 0 | 0 | 0 | 0 | 0 | 0 | 0 | 0 | 0 | 0 | 0 | 0 | 0 | 0 | 0 | 0 | 0 | 0 |
| ENSMUSG00 | Lonrf2 | 9989 | 0 | 0 | 0 | 0 | 0.00963699 | 0.01488219 | 0 | 0.06882203 | 0 | 0.04248236 | 0 | 0.02793856 | 0.16791322 | 0.01394857 | 0.07424447 | 0 | 0.02685795 | 0.07709756 |
| ENSMUSG00 | Gm568 | 504 | 0 | 0 | 0 | 0 | 0 | 0 | 0 | 0 | 0 | 0 | 0 | 0 | 0 | 0 | 0 | 0 | 0 | 0 |
| ENSMUSG00 | Dact2 | 2804 | 31.7879294 | 25.9232294 | 31.6394631 | 31.1810428 | 30.5201713 | 29.4241446 | 30.8612303 | 36.7622304 | 42.3175292 | 42.1396799 | 31.2990723 | 42.1006157 | 26.4428929 | 19.9921589 | 31.386064 | 42.7969913 | 28.1159746 | 29.4075075 |
| ENSMUSG00 | Pkd1l3 | 6724 | 0.11378542 | 0.16844712 | 0.13668904 | 0.08131061 | 0.07158228 | 0.0221086 | 0.38700139 | 0.05680012 | 0.0245012 | 0.1051845 | 0.09236681 | 0.15771826 | 0.02934677 | 0.1243298 | 0.07353044 | 0.12528158 | 0.1139985 | 0.12271514 |
| ENSMUSG00 | 2310057N15R | 1025 | 0 | 0 | 0 | 0 | 0 | 0 | 0 | 0 | 0 | 0 | 0 | 0 | 0 | 0 | 0 | 0 | 0 | 0 |
| ENSMUSG00 | Vps37c | 2410 | 50.8298412 | 57.4751096 | 44.6602096 | 57.6602428 | 51.0677713 | 52.0406509 | 32.3532344 | 42.0116426 | 39.8535082 | 40.5769953 | 45.1185915 | 41.5259266 | 33.7954338 | 37.8104977 | 32.6193231 | 38.767254 | 45.6416897 | 39.0541832 |
| ENSMUSG00 | Slc39a9 | 5222 | 13.1536508 | 16.5926537 | 14.5065011 | 15.2160975 | 15.2451554 | 16.606144 | 14.1883675 | 17.1361162 | 16.152812 | 17.9140167 | 16.8886582 | 17.0910081 | 10.8450864 | 12.3981344 | 16.8767001 | 16.0216348 | 19.0237 | 11.8508704 |
| ENSMUSG00 | Vstm2a | 4810 | 0.7599678 | 0.60254091 | 0.62352576 | 0.97563151 | 0.75049777 | 0.60781939 | 0.40328895 | 0.90694459 | 1.08746102 | 0.84302713 | 0.64560755 | 0.84709841 | 2.66590012 | 3.08983534 | 3.00660002 | 2.14140819 | 2.14340413 | 2.19578971 |
| ENSMUSG00 | Gm12185 | 9403 | 0.06780577 | 0 | 0 | 0.00484537 | 0 | 0.0737784 | 0 | 0 | 0 | 0 | 0 | 0 | 0 | 0 | 0 | 0 | 0 | 0 |
| ENSMUSG00 | Slc25a47 | 1573 | 3.99921554 | 4.61679017 | 3.65953599 | 3.35987358 | 4.22263539 | 3.15020557 | 3.91013875 | 4.41895565 | 3.53476137 | 2.84762671 | 3.34090619 | 4.00964391 | 7.24454729 | 8.1491215 | 6.4696658 | 4.28426377 | 3.45984371 | 4.12655727 |
| ENSMUSG00 | Arhgap30 | 5526 | 0 | 0 | 0 | 0 | 0.01742017 | 0 | 0.35103508 | 0.17278503 | 0.4024741 | 0 | 0.79538476 | 0 | 0.08927238 | 0.33618594 | 0.02982378 | 0.00692917 | 0.04854942 | 0.0199092 |
| ENSMUSG00 | Phf3 | 9536 | 8.39316762 | 7.44090443 | 6.77716407 | 6.46913318 | 6.13258046 | 7.92968451 | 6.64342251 | 5.48294871 | 5.34267198 | 5.8987824 | 4.3787032 | 5.35563457 | 5.44223752 | 5.49380564 | 5.66434981 | 6.89439733 | 6.12111474 | 6.36273539 |
| ENSMUSG00 | Hdhd1a | 1043 | 0 | 0 | 0 | 0 | 0 | 0 | 0 | 0 | 0 | 0 | 0 | 0 | 0 | 0 | 0 | 0 | 0 | 0 |
| ENSMUSG00 | Hexim1 | 3400 | 9.86370114 | 6.80954749 | 7.88203995 | 7.26296648 | 6.61106128 | 7.60760204 | 9.54602875 | 6.09955133 | 9.25485237 | 9.01410503 | 9.90627816 | 7.68286179 | 10.5047965 | 9.37077763 | 9.19764145 | 11.0029187 | 7.82307354 | 10.7267758 |
| ENSMUSG00 | Cdk5r1 | 7396 | 0.00574705 | 0.00450418 | 0.0065405 | 0.04928182 | 0.02603133 | 0.01339987 | 0.00639706 | 0 | 0 | 0.00637516 | 0.01291913 | 0.03773368 | 0.00667008 | 0.03767773 | 0 | 0 | 0 | 0 |
| ENSMUSG00 | Zfp710 | 4796 | 11.0605629 | 11.3983443 | 10.2072587 | 10.9912695 | 10.447317 | 12.0885526 | 7.2212012 | 8.09876423 | 11.6019564 | 9.57564324 | 6.52472848 | 8.20476274 | 8.81514869 | 6.34296857 | 7.4482379 | 6.54676774 | 7.63968721 | 5.80371309 |
| ENSMUSG00 | Rimkla | 4152 | 0.10237277 | 0.05616336 | 0.01165066 | 0.30725174 | 0.11592467 | 0.32223598 | 0.18232246 | 0.00918856 | 0.40670692 | 0.31797199 | 0 | 0.26886157 | 0 | 0 | 0 | 0.4611104 | 0.06461563 | 0.370967 |
| ENSMUSG00 | Neurog1 | 1658 | 0 | 0 | 0 | 0 | 0 | 0 | 0 | 0 | 0 | 0 | 0 | 0 | 0 | 0 | 0 | 0 | 0 | 0 |
| ENSMUSG00 | 4930539E08R | 4580 | 0.06496424 | 0.10910337 | 0.05280953 | 0.61676523 | 0.09458238 | 0.49769121 | 0.14462391 | 0.15010114 | 0.2428026 | 0.12353896 | 0.21905549 | 0.02437365 | 0.42007528 | 0.25351576 | 0.20690738 | 0.48490289 | 0.21757286 | 0.14412866 |
| ENSMUSG00 | 2810453I06R | 2328 | 0 | 0 | 0 | 0 | 0 | 0 | 0 | 0 | 0 | 0 | 0 | 0 | 0 | 0 | 0 | 0 | 0 | 0 |
| ENSMUSG00 | Rnf24 | 12308 | 0.21756793 | 0.24088764 | 0.27511758 | 0.17028015 | 0.14078236 | 0.36637136 | 0.12300989 | 0.27927496 | 0.34132472 | 0.13025056 | 0.17855444 | 0.09523309 | 0.27255203 | 0.11320463 | 0.3213641 | 0.05599861 | 0.08719015 | 0.01340813 |
| ENSMUSG00 | Efna5 | 4171 | 0.02038129 | 0 | 0 | 0.02184658 | 0.01153966 | 0 | 0 | 0.05493992 | 0.03949798 | 0 | 0 | 0.02676368 | 0.01182736 | 0 | 0.1185372 | 0 | 0 | 0 |
| ENSMUSG00 | Gm5540 | 1847 | 0.02301309 | 0.03607244 | 0.15714195 | 0.09867037 | 0.13029757 | 0.18780137 | 0.05123192 | 0.10339037 | 0.11149571 | 0.05105653 | 0.23279622 | 0.06043927 | 0.10683685 | 0.12572844 | 0.28999471 | 0.06219373 | 0.22825623 | 0.02978295 |
| ENSMUSG00 | Olfr223 | 1101 | 0 | 0 | 0 | 0 | 0 | 0 | 0 | 0 | 0 | 0 | 0 | 0 | 0 | 0 | 0 | 0 | 0 | 0 |
| ENSMUSG00 | Fkrp | 2771 | 16.750506 | 15.8810297 | 16.1128721 | 14.1895277 | 12.784228 | 15.7545862 | 13.9837907 | 16.8426978 | 18.3117241 | 16.2500619 | 16.0169384 | 14.180522 | 11.0556191 | 12.4029681 | 15.6271509 | 12.9615986 | 13.9833584 | 12.8639124 |
| ENSMUSG00 | Zfp689 | 2963 | 1.31976922 | 1.85508861 | 2.08971039 | 2.02971915 | 2.03054015 | 1.93996527 | 3.0817911 | 2.34593899 | 1.86263689 | 2.51427982 | 2.30571011 | 2.01561792 | 1.36524375 | 1.70854051 | 1.96065453 | 2.0288982 | 2.03078927 | 2.78480122 |
| ENSMUSG00 | Cdca2 | 7365 | 0.09811106 | 0.01356941 | 0.01313606 | 0.04948925 | 0 | 0.03364069 | 0.15417574 | 0.0414853 | 0.07829073 | 0.09602995 | 0.07784107 | 0 | 0.08707603 | 0.00630605 | 0.22936372 | 0.04159197 | 0.01561153 | 0.23900766 |
| ENSMUSG00 | Ccdc125 | 2692 | 4.62630631 | 3.95992799 | 4.38452504 | 4.62041794 | 4.1659503 | 4.19688828 | 4.65745168 | 3.90152699 | 4.63577796 | 3.71320546 | 6.30018965 | 5.49448387 | 3.8116786 | 4.03711607 | 5.26498337 | 4.93567636 | 6.89076066 | 5.98724676 |
| ENSMUSG00 | Tada3 | 4275 | 10.5492378 | 9.36657283 | 10.4781027 | 8.08908366 | 8.09517032 | 9.43530412 | 11.4435813 | 11.6319317 | 12.331869 | 11.0514631 | 10.2702191 | 11.8290036 | 9.79714568 | 10.1796748 | 12.2014432 | 11.9842884 | 9.60174596 | 9.41910319 |
| ENSMUSG00 | Olfr722 | 1578 | 0 | 0 | 0 | 0 | 0 | 0 | 0 | 0 | 0 | 0 | 0 | 0 | 0 | 0 | 0 | 0 | 0 | 0 |
| ENSMUSG00 | Nr1h5 | 2824 | 0.73751899 | 0.16514891 | 0.29120043 | 0.20973569 | 0.30678988 | 0.71942708 | 0 | 0.28400866 | 0.33544259 | 0.23374995 | 0.43985443 | 0.69176636 | 0.20962571 | 0.04933862 | 0.04376936 | 0 | 0 | 0 |
| ENSMUSG00 | Atp13a5 | 5579 | 0 | 0.01791337 | 0 | 0 | 0 | 0 | 0 | 0 | 0 | 0 | 0 | 0 | 0 | 0 | 0.00738511 | 0 | 0 | 0 |
| ENSMUSG00 | Gm6206 | 3780 | 1.59675529 | 0.80197708 | 0.85741443 | 0.84372304 | 1.33699788 | 1.08806267 | 1.31424108 | 0.47487908 | 0.71912964 | 0.93552983 | 1.47874871 | 1.03362338 | 1.72270187 | 0.89693608 | 1.02458896 | 1.10414714 | 1.39921401 | 0.98958181 |
| ENSMUSG00 | Prex2 | 11014 | 0.01929598 | 0.2631398 | 0.1185841 | 0.2481989 | 0.10051155 | 0.17096458 | 0.03866117 | 0 | 0 | 0 | 0.07374018 | 0 | 0.14780763 | 0.00421682 | 0.1496334 | 0.01390616 | 0.04175736 | 0 |
| ENSMUSG00 | Rassf10 | 1237 | 2.86169003 | 4.28193224 | 3.32396935 | 2.68872776 | 2.49025312 | 3.80558584 | 2.79209827 | 5.40312888 | 2.330684 | 4.72650542 | 4.17113623 | 3.20364757 | 9.2921073 | 6.79577997 | 5.16268485 | 4.92175148 | 3.31520919 | 5.64766092 |
| ENSMUSG00 | Mrgpre | 3757 | 0 | 0.25713972 | 0.28326262 | 0.31530134 | 0 | 0.32973605 | 0.08815245 | 0.26430728 | 0 | 0 | 0 | 0.05942578 | 0.38078935 | 0.63046272 | 0.46059764 | 0.38728855 | 0.15301955 | 1.30311698 |
| ENSMUSG00 | Yjefn3 | 1006 | 1.85907331 | 1.82128149 | 2.21191093 | 2.08330819 | 2.05732875 | 0.83737224 | 2.3515248 | 3.07513379 | 1.71951656 | 2.34347432 | 2.56446099 | 2.55220724 | 1.96150765 | 3.09319469 | 2.29352597 | 1.82698723 | 1.52390843 | 1.03893939 |
| ENSMUSG00 | C1galt1c1 | 2397 | 22.5559382 | 20.471378 | 24.1161322 | 22.4288829 | 23.4133596 | 21.2723249 | 20.5475589 | 21.7172484 | 23.2651407 | 21.3820669 | 22.6218514 | 20.8872129 | 22.1448239 | 13.8537849 | 17.2403674 | 20.0318822 | 21.5855239 | 19.3231823 |
| ENSMUSG00 | Nrsn1 | 4226 | 0 | 0.00788284 | 0 | 0 | 0.05694738 | 0 | 0 | 0.04518741 | 0.21441158 | 0 | 0 | 0 | 0 | 0.07799632 | 0 | 0 | 0 | 0 |
| ENSMUSG00 | Krt31 | 1574 | 0 | 0.02116448 | 0 | 0 | 0 | 0 | 0 | 0 | 0 | 0 | 0 | 0.14184412 | 0.15670876 | 0.17704226 | 0 | 0.07298083 | 0.02434962 | 0 |
| ENSMUSG00 | Gphb5 | 1611 | 0 | 0 | 0 | 0 | 0 | 0 | 0 | 0.02370726 | 0 | 0 | 0 | 0 | 0 | 0 | 0 | 0 | 0 | 0 |
| ENSMUSG00 | Elfn1 | 3819 | 0 | 0 | 0 | 0 | 0 | 0 | 0 | 0 | 0 | 0 | 0 | 0 | 0 | 0 | 0 | 0 | 0 | 0 |
| ENSMUSG00 | Prss32 | 1827 | 0 | 0 | 0.05295406 | 0 | 0 | 0 | 0 | 0 | 0 | 0 | 0 | 0 | 0 | 0.05084191 | 0 | 0.06287456 | 0 | 0.36130776 |
| ENSMUSG00 | 4930557A04R | 551 | 0 | 0 | 0 | 0 | 0 | 0 | 0 | 0 | 0 | 0 | 0 | 0 | 0 | 0 | 0 | 0 | 0 | 0 |
| ENSMUSG00 | Olfr1366 | 2159 | 0 | 0 | 0 | 0 | 0 | 0 | 0 | 0 | 0 | 0 | 0 | 0 | 0 | 0 | 0 | 0 | 0 | 0 |
| ENSMUSG00 | Atxn7l2 | 4508 | 1.8669088 | 1.84004229 | 1.84566265 | 2.58731751 | 2.5090954 | 1.79172484 | 1.67924326 | 2.25358896 | 1.27908434 | 2.03957124 | 3.3277145 | 1.70864273 | 1.37884241 | 2.20475481 | 1.43492553 | 2.21691405 | 2.184973 | 3.68517086 |
| ENSMUSG00 | A930038C07R | 4714 | 0.34263825 | 0.1272024 | 0.3899436 | 0.35760684 | 0.25526052 | 0.08409458 | 0.2910623 | 0.43750312 | 0.50674966 | 0.19004314 | 0.22296332 | 0.3315313 | 1.03603305 | 0.54188053 | 0.56811649 | 0.38176889 | 0.56912193 | 0.29173264 |
| ENSMUSG00 | BB014433 | 2360 | 0 | 0 | 0 | 0 | 0 | 0 | 0 | 0 | 0 | 0 | 0 | 0 | 0 | 0 | 0 | 0 | 0 | 0 |
| ENSMUSG00 | Olfr982 | 965 | 0 | 0 | 0 | 0 | 0 | 0 | 0 | 0 | 0 | 0 | 0 | 0 | 0 | 0 | 0 | 0 | 0 | 0 |
| ENSMUSG00 | Olfr729 | 971 | 0 | 0 | 0 | 0 | 0 | 0 | 0 | 0 | 0 | 0 | 0 | 0 | 0 | 0 | 0 | 0 | 0 | 0 |
| ENSMUSG00 | Prss48 | 943 | 0 | 0.10597952 | 0.30778492 | 0.048315 | 0.15312383 | 0.05254797 | 0 | 0 | 0.1747042 | 0.25000373 | 0 | 0.05918946 | 0 | 0.39401135 | 0 | 0.0406051 | 0.04064294 | 0.11666831 |
| ENSMUSG00 | Olfr1467 | 926 | 0 | 0 | 0 | 0 | 0 | 0 | 0 | 0 | 0 | 0 | 0 | 0 | 0 | 0 | 0 | 0 | 0 | 0 |
| ENSMUSG00 | Olfr368 | 984 | 0 | 0 | 0 | 0 | 0 | 0 | 0 | 0 | 0 | 0 | 0 | 0 | 0 | 0 | 0 | 0 | 0 | 0 |
| ENSMUSG00 | Olfr873 | 1061 | 0 | 0 | 0 | 0 | 0 | 0 | 0 | 0 | 0 | 0 | 0 | 0 | 0 | 0 | 0 | 0 | 0 | 0 |
| ENSMUSG00 | Tmem121 | 1596 | 6.41838811 | 4.52938474 | 5.18287825 | 7.16530209 | 5.12683397 | 4.12939446 | 0.02964454 | 0.04786015 | 0.10322435 | 0.11817219 | 0.11973674 | 0.24480554 | 2.13277086 | 0.43650458 | 1.21332903 | 0.98365595 | 0.31218162 | 0.24126801 |
| ENSMUSG00 | Clec4a1 | 1552 | 0 | 0 | 0 | 0 | 0 | 0 | 0 | 0.39373611 | 0 | 0.03038061 | 0 | 0 | 0 | 0 | 0 | 0 | 0 | 0 |
| ENSMUSG00 | Mterfd3 | 1535 | 2.99059219 | 3.14682063 | 4.56948658 | 5.69884072 | 4.23310078 | 3.71241978 | 5.30148585 | 6.14561785 | 7.70067103 | 5.86696047 | 6.72273031 | 3.96345747 | 5.65629801 | 3.29798406 | 7.7571659 | 6.03669511 | 7.29073531 | 6.80894481 |
| ENSMUSG00 | Olfr398 | 2246 | 0 | 0 | 0 | 0 | 0 | 0 | 0 | 0 | 0 | 0 | 0 | 0 | 0 | 0 | 0 | 0 | 0 | 0 |
| ENSMUSG00 | Rapgef4 | 6796 | 0.17512433 | 0.45096914 | 0.14947677 | 0.41565402 | 0.25496605 | 0.33540697 | 0.37593947 | 0.27537194 | 0.4363492 | 0.3399624 | 0.2952543 | 0.32030766 | 0.17421513 | 0.21185502 | 0.19400379 | 0.30425144 | 0.30453503 | 0.38043379 |
| ENSMUSG00 | Armcx3 | 3658 | 7.25074627 | 6.65711473 | 6.10950547 | 6.28986533 | 6.46057259 | 6.73256112 | 3.47925386 | 2.24476941 | 2.04919242 | 2.70684737 | 3.10837619 | 3.08222084 | 1.76667063 | 1.9679698 | 1.79088228 | 2.177268 | 2.46218693 | 2.13539999 |
| ENSMUSG00 | Olfr812 | 1042 | 0 | 0 | 0 | 0 | 0 | 0 | 0 | 0 | 0 | 0 | 0 | 0 | 0 | 0 | 0 | 0 | 0 | 0 |
| ENSMUSG00 | Olfr1257 | 1482 | 0 | 0 | 0 | 0 | 0 | 0 | 0 | 0 | 0 | 0 | 0 | 0 | 0 | 0 | 0 | 0 | 0 | 0 |
| ENSMUSG00 | Gm9244 | 2127 | 0 | 0 | 0 | 0 | 0 | 0 | 0 | 0 | 0 | 0 | 0 | 0 | 0 | 0 | 0 | 0 | 0 | 0 |
| ENSMUSG00 | Olfr985 | 1115 | 0 | 0 | 0 | 0 | 0 | 0 | 0 | 0 | 0 | 0 | 0 | 0 | 0 | 0 | 0 | 0 | 0 | 0 |
| ENSMUSG00 | Acap2 | 6590 | 4.08926883 | 5.08034275 | 3.65554144 | 4.61832741 | 5.22950794 | 3.63186196 | 3.55383552 | 3.56423786 | 3.60616382 | 4.82954851 | 3.18258069 | 4.52284732 | 3.08418056 | 2.91773174 | 3.19483992 | 4.3461873 | 4.34442243 | 2.85479734 |
| ENSMUSG00 | Bmyc | 838 | 46.0050056 | 47.5046642 | 35.4431356 | 39.9610592 | 38.8271842 | 34.7696985 | 11.9893173 | 10.9837339 | 10.3703518 | 8.32733184 | 9.06469976 | 11.2563806 | 8.12387182 | 9.25556395 | 6.24414988 | 6.67115594 | 6.95178655 | 7.68026908 |
| ENSMUSG00 | Zadh2 | 3319 | 12.7297816 | 10.950397 | 10.2314608 | 10.5837798 | 9.61478319 | 10.2270631 | 10.5772847 | 11.2770577 | 11.9501701 | 10.1006779 | 10.9685201 | 8.1730678 | 10.9989813 | 10.5790088 | 10.4152171 | 11.8944309 | 10.1387432 | 11.0051361 |
| ENSMUSG00 | Sephs2 | 1358 | 61.5980754 | 70.6979095 | 70.6723742 | 87.4986772 | 72.268772 | 84.9475433 | 79.3654511 | 91.712389 | 83.6165125 | 97.7387553 | 104.943079 | 95.1496752 | 78.6477179 | 82.6622675 | 84.6482628 | 97.2773163 | 104.508304 | 111.0309 |
| ENSMUSG00 | Gpr137c | 4095 | 0.69544488 | 0.98433705 | 0.86233644 | 0.77882127 | 0.681722 | 0.8228537 | 1.24780692 | 1.45494866 | 0.95548693 | 1.05930762 | 0.62999945 | 0.9132233 | 0.77099944 | 1.75795691 | 0.58356294 | 0.90700584 | 0.8329769 | 0.69852834 |
| ENSMUSG00 | Il23r | 2477 | 0 | 0 | 0 | 0 | 0.01943154 | 0 | 0 | 0 | 0 | 0 | 0 | 0 | 0 | 0 | 0 | 0 | 0 | 0 |
| ENSMUSG00 | Olfr367-ps | 736 | 0 | 0 | 0 | 0 | 0 | 0 | 0 | 0 | 0 | 0 | 0 | 0 | 0 | 0 | 0 | 0 | 0 | 0 |
| ENSMUSG00 | Ankrd34a | 925 | 0 | 0 | 0 | 0 | 0 | 0.05357052 | 0 | 0.04128908 | 0 | 0.1529212 | 0.0516486 | 0 | 0 | 0 | 0 | 0.28976676 | 0.99441203 | 1.72460983 |
| ENSMUSG00 | Olfr147 | 1164 | 0 | 0 | 0 | 0 | 0 | 0 | 0 | 0 | 0 | 0 | 0 | 0 | 0 | 0 | 0 | 0 | 0 | 0 |
| ENSMUSG00 | Pcdh10 | 14114 | 0 | 0 | 0.00342734 | 0 | 0 | 0 | 0 | 0.01623597 | 0 | 0 | 0.03384934 | 0 | 0 | 0 | 0 | 0.00271295 | 0 | 0 |
| ENSMUSG00 | Ccr2 | 4749 | 0 | 0 | 0.03055814 | 0 | 0 | 0 | 0 | 0 | 0.03469068 | 0 | 0 | 0 | 0 | 0.01955952 | 0.13013757 | 0 | 0 | 0 |
| ENSMUSG00 | Dcaf5 | 5698 | 8.22801144 | 9.87460132 | 9.02440554 | 11.9299891 | 8.58231563 | 12.0968502 | 9.32469962 | 8.19749178 | 8.57992176 | 9.33415115 | 10.3213362 | 8.05203135 | 8.45863166 | 7.98792658 | 7.4839611 | 7.28448902 | 7.62086599 | 9.64445378 |
| ENSMUSG00 | Ntf3 | 1424 | 0 | 0 | 0.03397018 | 0 | 0 | 0 | 0 | 0.08046152 | 0.14461558 | 0 | 0 | 0 | 0.2425024 | 0 | 0 | 0 | 0 | 0 |
| ENSMUSG00 | Themis | 9376 | 0 | 0 | 0 | 0 | 0 | 0 | 0 | 0 | 0 | 0 | 0 | 0 | 0 | 0 | 0 | 0 | 0 | 0 |
| ENSMUSG00 | Oxtr | 6547 | 0.03246157 | 0.27985477 | 0.15516177 | 0.06959072 | 0.12497979 | 0.33302586 | 0.05781296 | 0.08750359 | 0.0314545 | 0 | 0 | 0 | 0 | 0 | 0 | 0 | 0 | 0 |
| ENSMUSG00 | Agtr1a | 2463 | 0 | 0 | 0 | 0 | 0 | 0 | 0.09604685 | 0 | 0.28427559 | 0.36372853 | 0 | 0 | 0.56081757 | 0.58455814 | 0 | 0 | 0.42014211 | 0.69235985 |
| ENSMUSG00 | Fam110b | 8443 | 0.19634038 | 0.19728115 | 0.10885906 | 0.33996752 | 0.18812667 | 0.38735999 | 0.10647174 | 0.05880626 | 0.20000558 | 0.40767516 | 0.13014616 | 0.15205025 | 0.33304741 | 0.36305931 | 0.09759933 | 0.2675762 | 0.14526134 | 0.54077412 |
| ENSMUSG00 | Frmd3 | 6532 | 0 | 0 | 0 | 0 | 0 | 0 | 0 | 0 | 0 | 0 | 0 | 0 | 0 | 0 | 0 | 0 | 0 | 0 |
| ENSMUSG00 | Catsperg2 | 4623 | 0 | 0 | 0 | 0 | 0 | 0 | 0 | 0 | 0 | 0 | 0 | 0 | 0 | 0 | 0 | 0 | 0 | 0 |
| ENSMUSG00 | Gm8186 | 230 | 18.8501215 | 18.9738658 | 17.6668546 | 13.6683133 | 17.5786156 | 14.0040334 | 18.3079497 | 19.2622551 | 15.2211037 | 18.4502752 | 17.6559629 | 17.9580827 | 23.3790391 | 17.1641193 | 15.7640736 | 17.6469756 | 13.9974302 | 14.8285418 |
| ENSMUSG00 | Ivl | 3593 | 0 | 0 | 0 | 0 | 0 | 0 | 0 | 0 | 0 | 0 | 0 | 0 | 0 | 0 | 0 | 0 | 0 | 0 |
| ENSMUSG00 | C5ar1 | 4912 | 0 | 0 | 0 | 0 | 0.05879307 | 0 | 0 | 0.06997794 | 0.20962192 | 0 | 0 | 0 | 0 | 0 | 0 | 0 | 0 | 0 |
| ENSMUSG00 | Flg2 | 7005 | 0 | 0 | 0 | 0 | 0 | 0 | 0 | 0 | 0 | 0 | 0 | 0 | 0 | 0 | 0 | 0 | 0 | 0 |
| ENSMUSG00 | Nrap | 5726 | 0 | 0 | 0 | 0 | 0 | 0 | 0 | 0 | 0 | 0 | 0 | 0 | 0 | 0 | 0 | 0 | 0 | 0 |
| ENSMUSG00 | Plcxd3 | 2266 | 0 | 0.04410357 | 0 | 0 | 0 | 0 | 0 | 0 | 0 | 0 | 0 | 0 | 0 | 0 | 0 | 0 | 0 | 0 |
| ENSMUSG00 | Olfr1258 | 1444 | 0 | 0 | 0 | 0 | 0 | 0 | 0 | 0 | 0 | 0 | 0 | 0 | 0 | 0 | 0 | 0 | 0 | 0 |
| ENSMUSG00 | Ugt3a2 | 2521 | 0 | 0 | 0 | 0 | 0 | 0.05896795 | 0 | 0 | 0 | 0 | 0 | 0 | 0 | 0 | 0 | 0 | 0 | 0 |
| ENSMUSG00 | Fam183b | 751 | 0 | 1.02023511 | 1.73912826 | 1.57734641 | 0 | 0 | 0 | 0 | 0 | 0 | 0 | 0.44593073 | 0 | 0 | 0 | 0 | 0 | 0 |
| ENSMUSG00 | 4930469G21R | 1189 | 0 | 0.19612301 | 0.08136843 | 0 | 0 | 0 | 0 | 0 | 0 | 0 | 0 | 0 | 0 | 0 | 0 | 0 | 0 | 0 |
| ENSMUSG00 | Zfp518a | 6454 | 3.4378218 | 4.20669489 | 4.41462804 | 4.57445874 | 4.03460965 | 3.60090363 | 5.32212657 | 5.23118744 | 3.22268213 | 5.53038153 | 5.45555383 | 5.46568001 | 5.21294818 | 3.38220035 | 5.34969071 | 4.60982367 | 6.01557774 | 4.83268725 |
| ENSMUSG00 | Olfr449 | 1036 | 0 | 0 | 0 | 0 | 0 | 0 | 0 | 0 | 0 | 0 | 0 | 0 | 0 | 0 | 0 | 0 | 0 | 0 |
| ENSMUSG00 | Myoz3 | 863 | 0 | 0 | 0 | 0 | 0 | 0 | 0 | 0 | 0 | 0 | 0 | 0 | 0 | 0.10763404 | 0 | 0 | 0 | 0 |
| ENSMUSG00 | Frmpd4 | 13397 | 0.05710929 | 0.07211121 | 0.04332928 | 0.04081007 | 0.03592739 | 0.07027707 | 0.15538985 | 0.16534742 | 0.39966015 | 0.21116983 | 0.07845406 | 0.27497453 | 0.62599283 | 0.58241447 | 0.3905798 | 0.25151702 | 0.15448384 | 0.44345626 |
| ENSMUSG00 | Purg | 3000 | 0.3542098 | 0.62184069 | 0.7578519 | 0.5163585 | 0.48131924 | 0.34686914 | 0.28387607 | 0.47103963 | 0.50796703 | 0.75441125 | 0.44589961 | 0.29768353 | 0.57553903 | 0.80503085 | 0.3433463 | 0.28079779 | 0.44714013 | 0.55009107 |
| ENSMUSG00 | Rgag4 | 4922 | 0.18998656 | 0.20304487 | 0.10810826 | 0.06479628 | 0.04889468 | 0.11074361 | 0.01922498 | 0.1707064 | 0.16735683 | 0.14369373 | 0.06794488 | 0.15876052 | 0.28063667 | 0.10379621 | 0.21764333 | 0.19448703 | 0.35040296 | 0.1452902 |
| ENSMUSG00 | 4930515G13 | 489 | 0 | 0 | 0 | 0 | 0 | 0 | 0 | 0 | 0 | 0 | 0 | 0 | 0 | 0 | 0 | 0 | 0 | 0 |
| ENSMUSG00 | Skint7 | 4009 | 0 | 0 | 0 | 0 | 0 | 0 | 0 | 0 | 0 | 0 | 0 | 0 | 0 | 0 | 0 | 0 | 0 | 0 |
| ENSMUSG00 | Olfr788 | 935 | 0 | 0 | 0 | 0 | 0 | 0 | 0 | 0 | 0 | 0 | 0 | 0 | 0 | 0 | 0 | 0 | 0 | 0 |
| ENSMUSG00 | Pdp1 | 6267 | 1.09874557 | 1.41395083 | 1.37394102 | 1.35221864 | 1.01378872 | 1.44696829 | 1.8873735 | 2.21219755 | 2.78651391 | 1.58748977 | 1.53227488 | 1.92375668 | 1.43264863 | 2.18621442 | 2.17611537 | 2.23621546 | 1.45550641 | 1.47463378 |
| ENSMUSG00 | Olfr844 | 929 | 0 | 0 | 0 | 0 | 0 | 0 | 0 | 0 | 0 | 0 | 0 | 0 | 0 | 0 | 0 | 0 | 0 | 0 |
| ENSMUSG00 | Gm9833 | 1764 | 0.69878124 | 1.03866734 | 1.206596 | 1.21392805 | 0.87314147 | 0.87082468 | 0.64370992 | 1.1908062 | 1.02732806 | 0.88207098 | 0.97499915 | 1.01252903 | 0.61525067 | 0.63189235 | 1.09777388 | 1.19386814 | 1.10807321 | 0.65487032 |
| ENSMUSG00 | Gm12497 | 236 | 0 | 0.14115633 | 0 | 0.19305527 | 0 | 0 | 0.40095491 | 0 | 0 | 0.59937335 | 0 | 0.47301409 | 0 | 0.9839849 | 0 | 0 | 0.16239956 | 0 |
| ENSMUSG00 | Tigd2 | 3005 | 11.9948051 | 10.6423888 | 12.8942422 | 10.9619418 | 11.9328729 | 12.23565 | 11.1787028 | 12.7350373 | 16.9406101 | 12.1917126 | 12.7187909 | 11.478895 | 15.1525323 | 12.0244592 | 13.0665833 | 15.2015623 | 15.8024232 | 14.5348522 |
| ENSMUSG00 | Apoo-ps | 596 | 0.85580891 | 0.44715294 | 0.48698185 | 0.61155764 | 1.21137392 | 0.74827953 | 0.87322058 | 0.89713697 | 0.76015381 | 0.23733575 | 0.72143394 | 0.84285397 | 1.2415751 | 0.70133689 | 0.55304102 | 0.83519781 | 0.96458801 | 1.29216023 |
| ENSMUSG00 | Gm7324 | 2775 | 0.24507489 | 0.16806505 | 0.19175093 | 0.22985752 | 0.26017256 | 0.19642525 | 0.08524807 | 0.0963412 | 0.22262981 | 0.06796498 | 0.17216201 | 0.30170628 | 0.07110907 | 0.11715626 | 0.17816889 | 0.12418575 | 0.15192406 | 0.0594693 |
| ENSMUSG00 | Gpr81 | 4315 | 0 | 0 | 0.01121055 | 0.01055876 | 0.17847295 | 0 | 0 | 0 | 0.12408452 | 0.06556297 | 0.13286199 | 0 | 0.01143266 | 0 | 0 | 0 | 0.19540638 | 0 |
| ENSMUSG00 | Hspe1-ps2 | 311 | 0.82003555 | 0.32134625 | 0.46662569 | 0.29299707 | 0.46429508 | 0.15933355 | 0.760654 | 0 | 1.32432527 | 0.30321996 | 0 | 0.89735792 | 1.58623528 | 0.89602741 | 0.52992355 | 0.61560462 | 0 | 0 |
| ENSMUSG00 | Rpl37-ps1 | 269 | 0.15801181 | 0 | 0.17982725 | 0 | 0 | 0 | 0 | 0.28395838 | 0.15310972 | 0.17528142 | 0 | 0 | 0 | 0 | 0.15316564 | 0 | 0 | 0 |
| ENSMUSG00 | Gm5316 | 269 | 0 | 0 | 0 | 0 | 0 | 0 | 0 | 0 | 0 | 0 | 0 | 0 | 0 | 0 | 0 | 0 | 0 | 0 |
| ENSMUSG00 | Lrp1b | 23227 | 0.04940973 | 0.0229477 | 0.08747097 | 0.07061599 | 0.24452435 | 0.07466938 | 0.13036602 | 0.07399398 | 0.2783951 | 0.15427965 | 0.23859711 | 0.19224407 | 0.02123904 | 0 | 0 | 0 | 0 | 0 |
| ENSMUSG00 | Kcnk3 | 3808 | 0.20091732 | 0.0524888 | 0.20325013 | 0.16750384 | 0.18959529 | 0.16916637 | 0.89456746 | 0.59174153 | 0.3244736 | 0.45813446 | 0.43910807 | 0.24917707 | 0.57001165 | 0.23173258 | 0.38951051 | 0.9150329 | 0.70452752 | 0.37558739 |
| ENSMUSG00 | Olfr509 | 1074 | 0 | 0 | 0 | 0 | 0 | 0 | 0 | 0 | 0 | 0 | 0 | 0 | 0 | 0 | 0 | 0 | 0 | 0 |
| ENSMUSG00 | Scn3b | 3429 | 0 | 0 | 0.01410718 | 0.01328698 | 0 | 0 | 0.04139342 | 0.02227612 | 0.25223588 | 0 | 0 | 0.08138767 | 0.07193339 | 0.14898949 | 0 | 0 | 0 | 0 |
| ENSMUSG00 | Mblac1 | 1086 | 6.84936078 | 5.98159703 | 6.05782701 | 5.16022878 | 4.25475568 | 4.42598081 | 8.19040851 | 6.99842365 | 10.1638916 | 5.99152583 | 8.00648475 | 8.42888459 | 4.95136186 | 8.12557698 | 7.20837534 | 5.21824116 | 5.36426992 | 5.57182481 |
| ENSMUSG00 | A230051G13 | 4880 | 1.86395649 | 1.24240712 | 1.85365782 | 1.53114985 | 1.84439953 | 0.87326539 | 2.18142475 | 1.55743608 | 1.90740831 | 1.43964237 | 1.68387149 | 1.64701955 | 1.41525992 | 1.71310158 | 2.27115134 | 1.42020491 | 2.01055984 | 2.09665858 |
| ENSMUSG00 | Lix1l | 2503 | 0.39057892 | 0.29280211 | 0.23191465 | 0 | 0.07691878 | 0.03959467 | 0 | 0.16784516 | 0.0822743 | 0 | 0 | 0.24529456 | 0.43360055 | 0.05566611 | 0.32921739 | 0.38244714 | 0.16843359 | 0.06593181 |
| ENSMUSG00 | Prss38 | 1238 | 0 | 0.08072591 | 0 | 0.07360427 | 0.03887878 | 0 | 0.11465108 | 0 | 0 | 0.07617238 | 0 | 0 | 0 | 0 | 0 | 0.12371763 | 0 | 0 |
| ENSMUSG00 | Zfp219 | 4172 | 18.7972315 | 19.0519093 | 18.8531544 | 20.5308637 | 17.6168378 | 18.5050716 | 15.27569 | 15.2971008 | 13.7518736 | 15.1781866 | 17.0395825 | 16.3085553 | 17.3820514 | 15.073177 | 17.05539 | 16.3276583 | 12.0435703 | 18.5649143 |
| ENSMUSG00 | Trappc1 | 841 | 94.815351 | 99.3445169 | 110.494116 | 99.3024902 | 114.349089 | 93.7436375 | 107.22707 | 106.629918 | 101.864391 | 104.168855 | 110.717472 | 93.7119091 | 111.627394 | 105.258538 | 97.1004563 | 102.897471 | 95.5650947 | 103.21566 |
| ENSMUSG00 | Prmt6 | 5938 | 7.19395451 | 8.50494234 | 7.16071627 | 8.42472662 | 6.04688701 | 7.14333781 | 8.03949024 | 7.36448312 | 5.43789632 | 7.38466723 | 6.32386618 | 7.21900116 | 6.43026338 | 8.41593769 | 6.00884934 | 6.86109907 | 6.97721909 | 7.15173971 |
| ENSMUSG00 | Syt12 | 5673 | 0.32967173 | 0.54611302 | 0.30697111 | 0.28109229 | 0.2460472 | 0.53282509 | 0.2668792 | 0.55204953 | 0.20328264 | 0.14129419 | 0.1347434 | 0.24597066 | 0.37392428 | 0.43390387 | 0.21788237 | 0.26998489 | 0.08782687 | 0.37816943 |
| ENSMUSG00 | Ccdc71 | 3106 | 18.3924522 | 19.1125491 | 15.153717 | 18.5412621 | 17.8054025 | 17.0227839 | 16.9539638 | 19.9692407 | 18.4318277 | 21.6474253 | 23.9951498 | 19.87512 | 14.3421511 | 16.3287004 | 18.5181494 | 16.1372843 | 15.8685183 | 15.4967702 |
| ENSMUSG00 | Fut4 | 3663 | 0.58019623 | 0.91853735 | 0.76594725 | 0.59703252 | 0.65700142 | 0.98753742 | 0.37457485 | 0.63601871 | 0.87702654 | 0.48914188 | 0.7303843 | 0.53331919 | 1.91240302 | 0.93826439 | 1.16979574 | 0.39722716 | 1.39159091 | 2.04238016 |
| ENSMUSG00 | Sorl1 | 11237 | 9.14256567 | 8.55278987 | 11.2657764 | 9.8404071 | 10.656957 | 10.5482014 | 5.25462519 | 9.60503063 | 7.16924667 | 8.23679189 | 8.76249795 | 9.00541036 | 7.76614414 | 6.05091608 | 7.92351716 | 7.94299242 | 8.31534323 | 9.67322194 |
| ENSMUSG00 | Fbxw13 | 1502 | 0 | 0 | 0 | 0 | 0 | 0 | 0.03149979 | 0.07628309 | 0.02742112 | 0 | 0.06361512 | 0.18580447 | 0 | 0.09276449 | 0.05486226 | 0 | 0 | 0 |
| ENSMUSG00 | Olfr348 | 1008 | 0 | 0 | 0 | 0 | 0 | 0 | 0 | 0 | 0 | 0 | 0 | 0 | 0 | 0 | 0 | 0 | 0 | 0 |
| ENSMUSG00 | Zfp2 | 5574 | 3.30189114 | 3.19143981 | 3.65361612 | 2.39676443 | 2.82367044 | 3.58266086 | 3.03874041 | 2.41186323 | 4.65509597 | 3.57817501 | 4.1826659 | 3.59487314 | 2.79671436 | 2.3996945 | 3.31149929 | 3.68205338 | 3.36919386 | 2.31918552 |
| ENSMUSG00 | Smcr8 | 10931 | 1.43096741 | 1.5176856 | 0.991279 | 1.71307193 | 1.77450967 | 1.49596579 | 1.83520043 | 1.36962965 | 1.95928901 | 1.72108047 | 1.75697862 | 1.13867832 | 1.69238578 | 1.83974841 | 1.07423323 | 1.49225128 | 1.83724998 | 1.87205084 |
| ENSMUSG00 | Setd6 | 4012 | 16.0506834 | 15.3196136 | 15.5658593 | 15.5693399 | 13.148701 | 17.958543 | 17.524088 | 14.9456809 | 16.8975586 | 18.0517149 | 12.8963809 | 15.3868701 | 12.2100184 | 14.0304671 | 12.6623924 | 11.4051037 | 12.3041551 | 17.1800625 |
| ENSMUSG00 | Olfr902 | 1055 | 0 | 0 | 0 | 0 | 0 | 0 | 0 | 0 | 0 | 0 | 0 | 0 | 0 | 0 | 0 | 0 | 0 | 0 |
| ENSMUSG00 | Odz2 | 12573 | 0 | 0 | 0 | 0 | 0.04593837 | 0 | 0.04139342 | 0 | 0.01637895 | 0.01875078 | 0.01139942 | 0.06658991 | 0.07062551 | 0.12559445 | 0.18023428 | 0.02740917 | 0.02133811 | 0.02625107 |
| ENSMUSG00 | Fam134a | 1694 | 52.0148937 | 55.2199569 | 55.8837066 | 56.3461558 | 53.4167748 | 52.4194206 | 62.6460088 | 61.0762799 | 53.2215366 | 56.0018978 | 65.542505 | 65.140239 | 58.1266274 | 56.5060591 | 71.1420018 | 61.278534 | 62.8967564 | 59.1331661 |
| ENSMUSG00 | Gm5105 | 1661 | 0 | 0 | 0.11649255 | 0.05485978 | 0 | 0 | 0 | 0 | 0 | 0 | 0 | 0 | 0 | 0.22324744 | 0 | 0 | 0 | 0 |
| ENSMUSG00 | Zg16 | 633 | 0 | 0 | 0 | 0 | 0 | 0 | 0 | 0 | 0 | 0 | 0 | 0 | 0 | 0 | 0 | 0 | 0 | 0 |
| ENSMUSG00 | Rd3 | 2357 | 0 | 0 | 0 | 0 | 0 | 0 | 0 | 0.16203819 | 0 | 0 | 0 | 0 | 0 | 0.05244152 | 0 | 0 | 0 | 0 |
| ENSMUSG00 | Dcaf7 | 6487 | 13.0588586 | 15.3649113 | 14.9736471 | 15.3040721 | 14.4907734 | 14.0247909 | 11.9102212 | 13.1880656 | 12.1521491 | 12.2764819 | 15.7678721 | 12.5535766 | 15.0801899 | 14.4479988 | 15.4593166 | 13.9598099 | 13.7305864 | 11.6513816 |
| ENSMUSG00 | 4933408B17R | 2726 | 0.03118502 | 0.04888172 | 0.05323573 | 0.05014055 | 0.15890951 | 0.03635564 | 0.32976555 | 0.25218754 | 0.10576141 | 0.25944994 | 0.14020531 | 0.06142589 | 0.23525859 | 0.42593624 | 0.21160007 | 0.42139333 | 0.1546549 | 0 |
| ENSMUSG00 | Olfr173 | 1163 | 0 | 0 | 0 | 0 | 0 | 0 | 0 | 0 | 0 | 0 | 0 | 0 | 0 | 0 | 0 | 0 | 0 | 0 |
| ENSMUSG00 | Olfr1183 | 923 | 0 | 0 | 0 | 0 | 0 | 0 | 0 | 0 | 0 | 0 | 0 | 0 | 0 | 0 | 0 | 0 | 0 | 0 |
| ENSMUSG00 | Krt8 | 1796 | 423.3241 | 484.353901 | 436.977815 | 414.36197 | 441.843555 | 481.732031 | 289.671613 | 374.247266 | 310.66466 | 259.932691 | 281.728611 | 313.730019 | 228.256254 | 251.020466 | 234.706636 | 290.910545 | 279.999633 | 321.509239 |
| ENSMUSG00 | Cox7b2 | 539 | 0 | 0.061805 | 0 | 0.4226442 | 0.17859712 | 0.09193457 | 0 | 0.21257367 | 0 | 0 | 0 | 0 | 0 | 0 | 0 | 0 | 0 | 0 |
| ENSMUSG00 | Gemin4 | 3731 | 0.20506384 | 0.35714708 | 0.27227128 | 0.13432632 | 0.37411573 | 0.19922032 | 0.55796244 | 0.46064275 | 0.39740407 | 0.24011347 | 0.2304876 | 0.49367914 | 0.29088775 | 0.32365218 | 0.07730123 | 0.20525654 | 0.11299632 | 0.29487594 |
| ENSMUSG00 | Ogfr | 2691 | 23.2348993 | 25.7738557 | 26.5146627 | 26.8524029 | 26.4180049 | 25.8720144 | 21.9245301 | 22.6798436 | 24.1517361 | 26.5627893 | 20.7717211 | 27.0885378 | 26.7466619 | 26.1301926 | 25.5232232 | 24.0899288 | 25.3087438 | 25.7772143 |
| ENSMUSG00 | Rarres1 | 1319 | 0.16112652 | 2.62664215 | 0.14669759 | 0.27633689 | 0.29192979 | 0.07513682 | 0 | 0 | 1.12412022 | 0 | 0.03622059 | 0.25389991 | 2.13185693 | 5.3521617 | 2.06163964 | 2.03210198 | 1.65625392 | 1.50138578 |
| ENSMUSG00 | Prokr1 | 4052 | 0.2517582 | 0.08221346 | 0.28651647 | 0.29234629 | 0 | 0.06114602 | 0.42034957 | 0.16966023 | 0 | 0.31418287 | 0.50698993 | 0.9366893 | 0 | 0.34386047 | 0.0101682 | 0 | 0 | 0 |
| ENSMUSG00 | Gm13060 | 1466 | 0 | 0.04544733 | 0 | 0 | 0 | 0 | 0.06454663 | 0.07815635 | 0 | 0 | 0 | 0 | 0 | 0.15840412 | 0.02810474 | 0 | 0 | 0.22513959 |
| ENSMUSG00 | 6030446N20R | 3319 | 4.26460489 | 5.53040214 | 4.64933901 | 5.18893485 | 4.59711353 | 5.53903711 | 4.96077501 | 7.5026955 | 4.97613523 | 5.41259956 | 6.75096581 | 5.44871186 | 5.67785249 | 6.11511485 | 7.47313548 | 7.706576 | 6.54745717 | 5.55229007 |

| | | | | | | | | | | | | | | | | | | | |
|---|---|---|---|---|---|---|---|---|---|---|---|---|---|---|---|---|---|---|---|
| ENSMUSG00 Gm5417 | 382 | 0.55635047 | 0.08720653 | 0.12683228 | 0.2385395 | 0.3779994 | 0.1297192 | 0.99084144 | 0.29994033 | 1.40163535 | 0.12343116 | 0.12506534 | 0.7305715 | 0 | 0.36474414 | 0.64714486 | 0.20047438 | 0.30099186 | 0.57601159 |
| ENSMUSG00 Tmem200a | 1475 | 0.31698775 | 0.06775504 | 0.16397807 | 0.15444422 | 0.13052725 | 0 | 0 | 0.18125208 | 0 | 0.22376605 | 0 | 0 | 0 | 0 | 0 | 0 | 0.10393572 | 0 |
| ENSMUSG00 Zfp260 | 4501 | 25.0724822 | 27.0292579 | 30.2643549 | 25.9437806 | 27.3114715 | 24.8259085 | 19.5410509 | 29.2997924 | 20.7625426 | 22.0197241 | 21.9291411 | 25.14867 | 17.0979318 | 17.9647092 | 22.3079742 | 18.0266157 | 21.9773767 | 23.5020023 |
| ENSMUSG00 Chchd10 | 935 | 395.139027 | 338.972059 | 335.614002 | 332.425074 | 339.600321 | 333.566743 | 411.645608 | 359.04943 | 369.092851 | 390.064909 | 392.470006 | 356.086019 | 404.574483 | 358.53842 | 337.456166 | 391.751819 | 315.833281 | 357.94161 |
| ENSMUSG00 Gm9837 | 730 | 0.05822627 | 0 | 0 | 0 | 0 | 0 | 0 | 0 | 0 | 0 | 0 | 0 | 0 | 0 | 0 | 0.15735866 | 0 | 0 |
| ENSMUSG00 Upk1b | 1761 | 0.5792869 | 1.79711809 | 0.90648864 | 1.86280249 | 0.10932862 | 3.09528716 | 0 | 0 | 0 | 0 | 0 | 0 | 0 | 0.21098961 | 0.42114026 | 0 | 0 | 0 |
| ENSMUSG00 Cyp20a1 | 2273 | 19.9155356 | 18.2905816 | 14.0459877 | 18.6012535 | 15.5851719 | 14.9334019 | 8.92966971 | 13.610139 | 17.0145791 | 14.2095168 | 17.6765245 | 16.9436019 | 14.0638286 | 12.9340551 | 13.9755388 | 16.3404704 | 15.1248079 | 17.3037885 |
| ENSMUSG00 Gm7103 | 311 | 0 | 0 | 0 | 0 | 0 | 0 | 0 | 0 | 0 | 0 | 0 | 0 | 0 | 0 | 0 | 0 | 0 | 0 |
| ENSMUSG00 Olfr1404 | 1033 | 0 | 0 | 0 | 0 | 0 | 0 | 0 | 0 | 0 | 0 | 0 | 0 | 0 | 0 | 0 | 0 | 0 | 0 |
| ENSMUSG00 Gm8396 | 822 | 0 | 0.20263318 | 0 | 0 | 0 | 0 | 0.057558 | 0 | 0.15031575 | 0 | 0 | 0 | 0 | 0 | 0.05012355 | 0 | 0 | 0 |
| ENSMUSG00 Aff4 | 10704 | 23.8773938 | 25.0406901 | 24.2319572 | 25.7557809 | 24.1738809 | 25.8318622 | 17.4903093 | 20.0060538 | 16.9032477 | 19.1571757 | 17.7995641 | 20.1487033 | 17.4025897 | 16.5747771 | 16.0202612 | 20.0503414 | 20.6061135 | 15.7668105 |
| ENSMUSG00 1700104B16I | 553 | 0 | 0 | 0 | 0 | 0 | 0 | 0 | 0.13812804 | 0 | 0.08526348 | 0 | 0 | 0 | 0 | 0 | 0 | 0 | 0 |
| ENSMUSG00 Gm8508 | 406 | 0.10469255 | 0 | 0 | 0.11221932 | 0.23710307 | 0.12205107 | 0.11653369 | 0 | 0.10144462 | 0.34840421 | 0.11767231 | 0.412431 | 0 | 0 | 0 | 0 | 0 | 0.13549041 |
| ENSMUSG00 Ctu2 | 2574 | 10.1556656 | 11.4278499 | 10.411397 | 10.655691 | 10.1163056 | 9.27910551 | 10.4036427 | 9.49616843 | 9.79259815 | 9.26894167 | 10.2825668 | 8.76050027 | 11.3842886 | 15.3370141 | 12.1171631 | 12.3321342 | 11.1971155 | 11.2625483 |
| ENSMUSG00 Tmem67 | 8243 | 0.91786016 | 1.2285721 | 1.60208344 | 1.63606322 | 0.99265159 | 1.34657434 | 1.09055065 | 1.76992572 | 1.66884584 | 1.47578326 | 1.27508078 | 1.71313389 | 1.69367131 | 2.59180883 | 2.12930521 | 2.25293517 | 2.20853949 | 1.75511283 |
| ENSMUSG00 Fam58b | 1222 | 31.931057 | 21.7542468 | 21.6928761 | 19.7978024 | 21.3482018 | 23.5193007 | 22.1850778 | 20.2838537 | 27.7048412 | 20.6043171 | 25.2949248 | 22.7008055 | 21.476743 | 16.9509517 | 18.7800873 | 20.4926819 | 20.699964 | 19.401739 |
| ENSMUSG00 Slc36a3 | 2681 | 0 | 0 | 0 | 0 | 0 | 0 | 0 | 0 | 0 | 0 | 0 | 0 | 0 | 0.0866171 | 0 | 0 | 0 | 0 |
| ENSMUSG00 Pls1 | 3873 | 4.12650302 | 6.21875097 | 7.91862077 | 4.52904778 | 5.7663859 | 5.5399777 | 6.85319207 | 6.54783248 | 10.4853872 | 8.37585434 | 6.73512195 | 5.37548209 | 13.2341498 | 12.0876943 | 16.2444554 | 29.3037336 | 21.0383959 | 17.2143138 |
| ENSMUSG00 Gm12669 | 1650 | 0.15456428 | 0.24227559 | 0.35180749 | 0.3037403 | 0.49590467 | 0.18019176 | 0.34409221 | 0.46293821 | 0.34946135 | 0.34291421 | 0.40536328 | 0.50741511 | 0.38867571 | 0.36592311 | 0.42450088 | 0.20885786 | 0.41810506 | 0.33338853 |
| ENSMUSG00 Olfr1449 | 947 | 0 | 0 | 0 | 0 | 0 | 0 | 0 | 0 | 0 | 0 | 0 | 0 | 0 | 0 | 0 | 0 | 0 | 0 |
| ENSMUSG00 Dtx3l | 5719 | 2.72764462 | 2.95325011 | 3.61173238 | 3.83193956 | 3.30754433 | 4.89548776 | 1.68767031 | 2.9918161 | 1.32511259 | 3.48745366 | 1.55379302 | 2.166651 | 1.98397289 | 2.3144894 | 2.19732026 | 1.66713783 | 2.48628364 | 2.54894445 |
| ENSMUSG00 2810046L04F | 5458 | 3.97171854 | 5.52366604 | 3.93511313 | 5.1838418 | 4.73558868 | 4.35768056 | 5.12308414 | 5.3181066 | 5.26716526 | 4.04296979 | 3.83390103 | 4.20304825 | 5.3055763 | 4.05896476 | 4.33303281 | 4.69635503 | 3.43377779 | 4.34388512 |
| ENSMUSG00 4933407P14F | 4439 | 0 | 0 | 0 | 0 | 0 | 0.03348912 | 0 | 0 | 0 | 0 | 0 | 0 | 0 | 0 | 0 | 0 | 0 | 0 |
| ENSMUSG00 4930429B21I | 1661 | 0.17913078 | 0.92257263 | 0.17473882 | 0.19200922 | 0.23182143 | 0.56682838 | 0.99695591 | 0.1149681 | 0.49592433 | 0.17032163 | 0.83412029 | 1.0081095 | 0.65340288 | 0.25165369 | 0.64493706 | 0.29968567 | 0.299965 | 0 |
| ENSMUSG00 Htr1b | 1272 | 0 | 0 | 0 | 0 | 0 | 0 | 0 | 0 | 0 | 0 | 0 | 0 | 0 | 0 | 0 | 0 | 0 | 0 |
| ENSMUSG00 Espnl | 5729 | 0 | 0 | 0.02533088 | 0 | 0.00840145 | 0.01729891 | 0 | 0.07333155 | 0.03594564 | 0 | 0 | 0.00974265 | 0 | 0.02432052 | 0 | 0 | 0.04682913 | 0 |
| ENSMUSG00 Spty2d1 | 5217 | 11.53677 | 15.9700112 | 13.1388331 | 14.9774183 | 14.8538235 | 16.1471434 | 8.8331511 | 11.1055922 | 8.25782931 | 8.81194857 | 11.9689229 | 12.143143 | 11.6876078 | 12.9708713 | 14.1603201 | 13.6075879 | 14.5091885 | 14.7091631 |
| ENSMUSG00 Rps23 | 569 | 134.089264 | 110.242847 | 122.931678 | 110.579617 | 121.387189 | 119.83227 | 133.539828 | 102.361008 | 118.709817 | 113.940627 | 102.01507 | 114.279872 | 161.607546 | 176.716079 | 137.507477 | 120.726449 | 124.745697 | 135.444215 |
| ENSMUSG00 Cdc42ep1 | 2539 | 31.9751423 | 38.3773988 | 27.3780083 | 39.2086971 | 34.2553706 | 34.817675 | 21.000941 | 32.9125547 | 21.671991 | 39.2396361 | 25.2704881 | 30.9085552 | 19.9736947 | 16.7923089 | 17.4769892 | 24.1295675 | 24.3482934 | 23.9622182 |
| ENSMUSG00 Tmem202 | 1250 | 16.3899959 | 13.0320042 | 15.7117227 | 14.6524319 | 14.7091159 | 11.8530139 | 14.2316538 | 11.2743972 | 13.4432787 | 12.4100651 | 15.6319664 | 11.5650053 | 10.8135563 | 9.17735167 | 7.54812504 | 5.60574492 | 8.30914117 | 10.1656829 |
| ENSMUSG00 Olfr429 | 942 | 0 | 0 | 0 | 0 | 0 | 0 | 0 | 0 | 0 | 0 | 0 | 0 | 0 | 0 | 0 | 0 | 0 | 0 |
| ENSMUSG00 Clrn2 | 941 | 0.27102131 | 0.10620476 | 0.10281303 | 0.14525306 | 0.51149759 | 0 | 0 | 0 | 0.65653319 | 0 | 0 | 0 | 0.41939993 | 0 | 0 | 0 | 0 | 0 |
| ENSMUSG00 Sall2 | 9915 | 3.99973063 | 3.28257162 | 2.9858397 | 3.67153549 | 3.41268203 | 2.88370421 | 3.93198663 | 3.88664993 | 3.65548501 | 4.35603068 | 3.43073832 | 4.08695623 | 4.0301415 | 3.48037891 | 4.42142768 | 4.40641281 | 3.63355714 | 3.11801492 |
| ENSMUSG00 Tceal1 | 2801 | 13.7333753 | 10.1092324 | 10.223895 | 9.10895567 | 10.6024266 | 11.2515311 | 11.6888161 | 10.8536781 | 12.3662477 | 11.9686362 | 10.8478663 | 11.6573233 | 10.5497388 | 11.9053391 | 10.4290979 | 11.4694108 | 9.24975962 | 10.5069161 |
| ENSMUSG00 Tecrl | 2338 | 0 | 0 | 0 | 0 | 0 | 0 | 0 | 0 | 0 | 0 | 0 | 0 | 0 | 0 | 0 | 0 | 0 | 0 |
| ENSMUSG00 Adamts16 | 7966 | 0 | 0 | 0 | 0 | 0.00604217 | 0 | 0 | 0.01917771 | 0.02585144 | 0 | 0 | 0 | 0 | 0 | 0 | 0 | 0 | 0 |
| ENSMUSG00 Hist1h1a | 746 | 0.17093234 | 0.08931071 | 0.45390712 | 0.06107379 | 0 | 0 | 0 | 0 | 0 | 0 | 0 | 0 | 0 | 0 | 0 | 0.51327892 | 0.10275147 | 0 |
| ENSMUSG00 4930474N05I | 2268 | 0 | 0 | 0 | 0 | 0 | 0 | 0 | 0 | 0 | 0 | 0 | 0 | 0 | 0 | 0 | 0 | 0.03379744 | 0 |
| ENSMUSG00 Krt82 | 2086 | 0 | 0 | 0 | 0 | 0 | 0 | 0 | 0 | 0 | 0 | 0 | 0 | 0 | 0 | 0 | 0 | 0 | 0 |
| ENSMUSG00 Clip1 | 11466 | 20.299873 | 20.6542268 | 19.7063283 | 19.8440481 | 19.2888705 | 21.9326812 | 18.028004 | 19.7490628 | 17.7770859 | 20.5240851 | 20.2333157 | 19.8368066 | 17.2184138 | 20.3339716 | 20.708579 | 20.6013219 | 22.5558915 | 23.4601894 |
| ENSMUSG00 Fzd9 | 2292 | 0 | 0.15987864 | 0 | 0 | 0 | 0 | 0 | 0 | 0 | 0.08228744 | 0.20844223 | 0.36528575 | 0 | 0.4457984 | 0.01797625 | 0 | 0.15049593 | 0 |
| ENSMUSG00 Polr1a | 6542 | 1.8517235 | 1.71605707 | 1.9688846 | 2.45146555 | 1.89820182 | 1.26495055 | 2.1913393 | 2.04331104 | 2.46791718 | 2.49375471 | 1.69425105 | 1.92820846 | 2.08880176 | 1.1146013 | 1.90199784 | 2.34707025 | 2.37855037 | 3.69978706 |
| ENSMUSG00 Tmie | 2796 | 0.31924488 | 0.67912553 | 0.84774785 | 0.58662289 | 0.2926476 | 0.63801803 | 0.35535274 | 0.80591979 | 0.29461027 | 0.69140874 | 1.12773507 | 0.01996268 | 0.45873743 | 0.8305452 | 0.50102035 | 0.62995992 | 0.35639618 | 0.29511323 |
| ENSMUSG00 Lingo1 | 6146 | 0.26972045 | 1.12199302 | 0.33057082 | 0.64494156 | 0.26626837 | 0.72563391 | 0.36951002 | 0.14914052 | 0.09381894 | 0.23015312 | 0.07773342 | 0.11806112 | 0.21672011 | 0.12090879 | 0.19441021 | 0.16196435 | 0.08730364 | 0 |
| ENSMUSG00 Defb20 | 688 | 453.903384 | 604.909866 | 303.600151 | 449.782868 | 266.544519 | 460.235418 | 8.32097987 | 6.9390266 | 4.72926562 | 4.93437592 | 1.94432971 | 6.32793849 | 2.58132125 | 6.27805251 | 2.93441314 | 1.28006391 | 1.94973894 | 2.5585631 |
| ENSMUSG00 Olfr95 | 973 | 0 | 0 | 0 | 0 | 0 | 0 | 0 | 0 | 0 | 0 | 0 | 0 | 0 | 0 | 0 | 0 | 0 | 0 |
| ENSMUSG00 Gm962 | 3090 | 3.87911315 | 3.06176762 | 3.20924716 | 3.61244526 | 3.55148175 | 3.92893842 | 5.08343346 | 3.75744024 | 4.78510005 | 4.51670167 | 3.3859922 | 4.46164034 | 7.23215486 | 4.40392156 | 3.96014955 | 5.97283908 | 4.40318298 | 8.02883726 |
| ENSMUSG00 Aknad1 | 5979 | 0.00710908 | 0 | 0 | 0 | 0.11270228 | 0 | 0 | 0 | 0 | 0.00788605 | 0 | 0 | 0 | 0 | 0 | 0 | 0 | 0 |
| ENSMUSG00 9330101J02F | 5244 | 0 | 0.02541029 | 0.04612274 | 0.05212934 | 0.01835695 | 0.07569532 | 0.036089 | 0.0218492 | 0.26703691 | 0.00899136 | 0 | 0.07450603 | 0 | 0 | 0.06285516 | 0.15333767 | 0.0073086 | 0.14685879 |
| ENSMUSG00 Olfr780 | 1024 | 0 | 0 | 0 | 0 | 0 | 0 | 0 | 0 | 0 | 0 | 0 | 0 | 0 | 0 | 0 | 0 | 0 | 0 |
| ENSMUSG00 Zfa | 3421 | 0.02484956 | 0 | 0 | 0 | 0 | 0 | 0 | 0 | 0 | 0 | 0 | 0 | 0.01442032 | 0 | 0 | 0.01119281 | 0 | 0 |
| [illegible] | | | | | | | | | | | | | | | | | | | |

| | | | | | | | | | | | | | | | | | | | |
|---|---|---|---|---|---|---|---|---|---|---|---|---|---|---|---|---|---|---|---|
| ENSMUSG00 Zfp366 | 59456 | 0.00357451 | 0 | 0 | 0 | 0 | 0 | 0.00079576 | 0 | 0 | 0 | 0 | 0 | 0 | 0 | 0 | 0 | 0 | 0 |
| ENSMUSG00 P2ry10 | 2978 | 0 | 0 | 0.09746178 | 0 | 0 | 0 | 0.54017162 | 0.87208978 | 0.44256834 | 0.14249709 | 1.42779426 | 0.09371334 | 0.44726722 | 1.91827493 | 0.34588277 | 0.16715174 | 0 | 0 |
| ENSMUSG00 Dcaf12l2 | 4735 | 0 | 0 | 0 | 0 | 0 | 0 | 0 | 0 | 0 | 0 | 0 | 0 | 0 | 0 | 0 | 0 | 0.00809425 | 0 |
| ENSMUSG00 4933403G14 | 7022 | 1.4769671 | 1.97353517 | 1.72221341 | 1.71291878 | 1.65877606 | 1.73596874 | 1.78551124 | 2.15382959 | 2.90335023 | 1.92040745 | 2.45610368 | 2.79793695 | 1.56665018 | 1.80564453 | 1.98908109 | 1.95215572 | 1.67015763 | 1.12807055 |
| ENSMUSG00 Sgms2 | 5866 | 3.90560687 | 6.18440876 | 5.5910763 | 5.72425668 | 6.44931674 | 6.75795889 | 6.27501947 | 10.5475097 | 6.93697197 | 7.37081399 | 8.59232547 | 9.11547984 | 8.16592084 | 10.1423246 | 8.29509675 | 7.94402816 | 8.33038332 | 9.09628938 |
| ENSMUSG00 Vmn1r231 | 1560 | 0 | 0 | 0 | 0 | 0 | 0 | 0 | 0 | 0 | 0 | 0 | 0 | 0 | 0 | 0 | 0 | 0 | 0 |
| ENSMUSG00 Hist2h2bb | 1889 | 0.02250142 | 0.31743361 | 0.28168811 | 0.0723574 | 0 | 0 | 0.15027849 | 0.16174654 | 0.52328024 | 0.17472468 | 0 | 0 | 0 | 0.1475196 | 0.37079219 | 0.06081092 | 0.0405784 | 0.40769057 |
| ENSMUSG00 Efcab5 | 5524 | 0.09233565 | 0.06030575 | 0.04378488 | 0 | 0.00871324 | 0.01794089 | 0.04282466 | 0 | 0 | 0.04267804 | 0 | 0 | 0 | 0.17656152 | 0 | 0.04852177 | 0 | 0 |
| ENSMUSG00 Zfp438 | 3204 | 3.31657116 | 4.09652944 | 4.89170532 | 3.89629405 | 4.26637526 | 4.37684881 | 4.63676067 | 2.70589118 | 3.40650021 | 4.51787325 | 3.9663355 | 4.42483717 | 4.03400822 | 4.07327982 | 4.46221595 | 4.01549441 | 4.44986968 | 5.27084762 |
| ENSMUSG00 Amigo1 | 2386 | 4.54267388 | 3.47649231 | 2.61533337 | 3.60898465 | 3.22762271 | 1.70298581 | 2.55797803 | 4.30584922 | 2.96901958 | 3.99180304 | 4.48515956 | 1.94161777 | 2.08823288 | 4.08770258 | 3.81623799 | 1.60480333 | 2.44157466 | 2.88186854 |
| ENSMUSG00 Gja1 | 1531 | 9.35613607 | 21.3890105 | 10.0475393 | 9.61216024 | 11.412076 | 20.9409658 | 0.67986867 | 1.1475183 | 0.05380342 | 1.0163117 | 0.21843547 | 0.87496793 | 3.86664276 | 1.75947555 | 4.06364137 | 1.42558107 | 1.3768428 | 5.35359693 |
| ENSMUSG00 Zfp169 | 9532 | 0.86954567 | 1.07291844 | 1.12154324 | 1.62035187 | 1.11089207 | 0.98772759 | 1.29548984 | 1.59068231 | 0.99812056 | 1.76097885 | 1.40337688 | 1.51660267 | 0.82806408 | 0.86242168 | 1.11519108 | 1.06452066 | 1.36305231 | 1.62742007 |
| ENSMUSG00 Insl6 | 703 | 102.604956 | 82.5477407 | 90.0036667 | 85.9373326 | 94.9629645 | 92.973052 | 100.278651 | 79.0468642 | 105.221881 | 95.1733256 | 99.8992731 | 92.2586059 | 98.4533143 | 85.6209634 | 95.4141294 | 89.980204 | 91.7541361 | 87.7172242 |
| ENSMUSG00 Olfr1392 | 1017 | 0 | 0 | 0 | 0 | 0 | 0 | 0.09304362 | 0.03755398 | 0.20249024 | 0 | 0 | 0 | 0 | 0.13700321 | 0 | 0 | 0 | 0 |
| ENSMUSG00 AY358078 | 1452 | 0 | 0 | 0 | 0 | 0 | 0 | 0.03258449 | 0 | 0 | 0 | 0 | 0 | 0 | 0 | 0 | 0 | 0 | 0 |
| ENSMUSG00 Kcna2 | 4858 | 0.01749904 | 0.03428663 | 0.0298725 | 0 | 0 | 0.02040047 | 0.02921738 | 0.01572351 | 0 | 0.01941157 | 0.01966857 | 0 | 0.01015478 | 0.05736199 | 0 | 0.00788197 | 0.0473359 | 0 |
| ENSMUSG00 Prkca | 10419 | 0.66905162 | 0.48919021 | 0.67320874 | 0.70840668 | 0.65136781 | 0.62779161 | 1.621137 | 1.24265519 | 1.27682547 | 1.3712125 | 1.09590317 | 0.94285024 | 2.12592675 | 2.20652877 | 2.55458346 | 2.56887749 | 2.95751442 | 2.98830544 |
| ENSMUSG00 Lin28a | 4066 | 0 | 0 | 0 | 0 | 0 | 0 | 0 | 0 | 0 | 0 | 0 | 0 | 0 | 0 | 0 | 0 | 0 | 0 |
| ENSMUSG00 Creg2 | 5795 | 0.09535242 | 0.18395386 | 0.07512714 | 0.06289704 | 0.12458652 | 0.16246798 | 0.02449319 | 0.00659058 | 0.07817975 | 0.01627289 | 0.1071742 | 0 | 0.08512842 | 0.07213059 | 0.13508707 | 0.03964515 | 0.13888736 | 0.03797005 |
| ENSMUSG00 D330012F22R | 2246 | 4.31486204 | 6.19981736 | 4.45829064 | 4.32079361 | 5.42180618 | 5.95691811 | 4.10773481 | 6.47876461 | 5.39128924 | 6.48689551 | 10.0612446 | 4.2495451 | 6.98466148 | 5.52118047 | 6.54895612 | 6.9216325 | 6.22844985 | 3.50236076 |
| ENSMUSG00 Gm9847 | 596 | 0 | 0.16768235 | 0.24349092 | 0 | 0 | 0 | 0.07938369 | 0 | 0 | 0 | 0.08015933 | 0 | 0 | 0 | 0 | 0 | 0.1929176 | 0 |
| ENSMUSG00 Gm5215 | 404 | 0 | 0 | 0 | 0 | 0 | 0 | 0 | 0 | 0 | 0 | 0.11825485 | 0 | 0 | 0 | 0 | 0 | 0 | 0 |
| ENSMUSG00 Apol10a | 3344 | 0 | 0 | 0 | 0 | 0 | 0 | 0 | 0 | 0 | 0 | 0 | 0 | 0 | 0 | 0 | 0 | 0 | 0 |
| ENSMUSG00 Sepn1 | 5412 | 24.7632705 | 26.5973791 | 25.080585 | 31.7462487 | 25.2132305 | 29.418502 | 18.3585783 | 19.4419935 | 15.4563587 | 22.5037447 | 20.7801649 | 27.5571786 | 23.4353953 | 19.5919903 | 24.4377299 | 25.8100768 | 22.57654 | 22.5342183 |
| ENSMUSG00 9130014G24R | 6059 | 0 | 0 | 0 | 0 | 0 | 0 | 0 | 0 | 0 | 0 | 0 | 0 | 0 | 0 | 0 | 0 | 0 | 0 |
| ENSMUSG00 Cetn1 | 1245 | 0 | 0 | 0 | 0 | 0 | 0 | 0 | 0 | 0 | 0 | 0 | 0 | 0 | 0 | 0 | 0 | 0 | 0 |
| ENSMUSG00 Fam160a1 | 5229 | 13.046626 | 11.4674335 | 10.407386 | 11.1963936 | 12.0398845 | 10.3578386 | 9.74483748 | 8.24616988 | 8.57757041 | 8.88189763 | 9.52027285 | 10.1909105 | 11.8683404 | 9.44158824 | 8.55706439 | 11.855516 | 10.6791766 | 9.95192483 |
| ENSMUSG00 Gm9848 | 238 | 0 | 0 | 0 | 0 | 0 | 0 | 0 | 0 | 0 | 0 | 0 | 0 | 0 | 0 | 0 | 0 | 0 | 0 |
| ENSMUSG00 Olfr161 | 941 | 0 | 0 | 0 | 0 | 0 | 0 | 0 | 0 | 0 | 0 | 0 | 0 | 0 | 0 | 0 | 0 | 0 | 0 |
| ENSMUSG00 Pddc1 | 3646 | 9.13989523 | 9.94087463 | 9.06174472 | 9.94695318 | 9.50493281 | 10.1524663 | 12.9765987 | 13.6491224 | 14.5271144 | 12.6605975 | 12.5399438 | 11.9867427 | 13.7874996 | 12.6619372 | 12.4870322 | 12.8020435 | 12.5091315 | 12.9300067 |
| ENSMUSG00 4930412M03R | 1109 | 0.76654961 | 0.51065753 | 0.34895243 | 0.94490894 | 1.51904177 | 1.07237656 | 0.12798741 | 0.86096489 | 1.41126023 | 0.2976149 | 0.81850695 | 1.96285919 | 1.55691353 | 1.21449823 | 1.07740768 | 0.20716289 | 1.65884784 | 0.89284393 |
| ENSMUSG00 Hs3st1 | 3690 | 0.1958233 | 0.06319519 | 0.03932807 | 0.12347166 | 0.09130717 | 0.1074314 | 0.42312152 | 0.15525367 | 0.82596265 | 0.03833391 | 0.33662572 | 0.25714533 | 0.7219305 | 0.99433141 | 0.71460693 | 0.45658177 | 0.02077306 | 0.05963047 |
| ENSMUSG00 Serpinb1b | 1689 | 15.3763544 | 16.1140524 | 16.9837202 | 17.0483008 | 29.0102417 | 19.5687823 | 6.41480371 | 9.83640915 | 8.51041682 | 11.9481948 | 10.2677975 | 9.35218029 | 10.5732114 | 9.54179889 | 11.0749002 | 11.4713128 | 14.7949944 | 13.5161511 |
| ENSMUSG00 Gm9726 | 680 | 0.12501522 | 0.04898955 | 0 | 0.13400307 | 0 | 0 | 0.27830988 | 0 | 0 | 0 | 0.14051458 | 0.08208186 | 0.07254694 | 0 | 0.06059052 | 0 | 0 | 0.16179149 |
| ENSMUSG00 Zfp11 | 3926 | 2.39267547 | 2.43525233 | 2.14391092 | 3.00568275 | 3.15076525 | 3.68553189 | 3.97686807 | 2.99624553 | 4.34315271 | 2.45001107 | 3.20041113 | 3.71061842 | 1.87225055 | 2.78002051 | 2.99094331 | 3.32580161 | 2.23553795 | 2.10172338 |
| ENSMUSG00 Ttc24 | 2635 | 0 | 0 | 0 | 0 | 0 | 0 | 0 | 0 | 0 | 0 | 0 | 0 | 0 | 0 | 0 | 0 | 0 | 0 |
| ENSMUSG00 Zfp455 | 2767 | 2.45783454 | 1.5651161 | 0.90907972 | 1.30080322 | 1.07848907 | 0.59097948 | 1.94927553 | 0.92478893 | 1.6671087 | 0.97130108 | 2.02011931 | 1.79529959 | 0.07131466 | 0.83924986 | 0.7742974 | 1.12090322 | 0.48479234 | 0.83497741 |
| ENSMUSG00 Rhox11 | 806 | 0 | 0 | 0 | 0 | 0 | 0 | 0 | 0 | 0 | 0 | 0 | 0 | 0 | 0 | 0 | 0 | 0 | 0 |
| ENSMUSG00 Olfml1 | 1682 | 0 | 0 | 0 | 0 | 0 | 0 | 0 | 0 | 0 | 0.05606505 | 0 | 0 | 0 | 0 | 0 | 0 | 0 | 0 |
| ENSMUSG00 Gprc5c | 5023 | 23.0338621 | 27.715426 | 28.6023064 | 26.9393394 | 25.7668212 | 30.907568 | 25.7332747 | 31.9727358 | 28.4525602 | 32.0470836 | 29.3421754 | 28.8245727 | 20.7325657 | 21.5993208 | 27.7165155 | 22.6862129 | 24.9811459 | 22.9542281 |
| ENSMUSG00 Olfr214 | 1007 | 0 | 0 | 0 | 0 | 0 | 0 | 0 | 0 | 0 | 0 | 0 | 0 | 0 | 0 | 0 | 0 | 0 | 0 |
| ENSMUSG00 P4ha3 | 3194 | 0 | 0 | 0.03029025 | 0.02852914 | 0 | 0 | 0 | 0 | 0 | 0.01476227 | 0.02991544 | 0 | 0 | 0 | 0 | 0 | 0 | 0 |
| ENSMUSG00 Spink14 | 563 | 0 | 0.29585164 | 0.08592101 | 0.16185096 | 0.25647561 | 0.17603103 | 0.08403673 | 0 | | 0.16749806 | 0.06485783 | 0.29741916 | 74.5674274 | 195.923104 | 101.284109 | 27.068671 | 46.0187863 | 78.5565219 |
| ENSMUSG00 Olfr523 | 1055 | 0 | 0 | 0 | 0 | 0 | 0 | 0 | 0 | 0 | 0 | 0 | 0 | 0 | 0 | 0 | 0 | 0 | 0 |
| ENSMUSG00 Olfr335-ps | 955 | 0 | 0 | 0 | 0 | 0 | 0 | 0 | 0 | 0 | 0 | 0 | 0 | 0 | 0 | 0 | 0 | 0 | 0 |
| ENSMUSG00 1700014D04R | 794 | 0 | 0 | 0 | 0 | 0 | 0 | 0 | 0 | 0 | 0 | 0 | 0 | 0 | 0 | 0 | 0 | 0 | 0 |
| ENSMUSG00 Gja10 | 1776 | 0 | 0 | 0 | 0 | 0 | 0.05580263 | 0 | 0 | 0 | 0 | 0 | 0 | 0 | 0 | 0 | 0 | 0 | 0 |
| ENSMUSG00 Fbll1 | 1503 | 0 | 0 | 0 | 0 | 0 | 0 | 0 | 0 | 0 | 0 | 0 | 0 | 0 | 0.03090092 | 0 | 0 | 0 | 0 |
| ENSMUSG00 1600021P15R | 4551 | 2.68984634 | 3.8575907 | 3.61393107 | 3.09346577 | 3.71221824 | 2.26477007 | 2.84853308 | 3.28130726 | 4.02724665 | 3.11851498 | 2.62442092 | 2.72271525 | 3.78308924 | 4.98016143 | 4.26410302 | 3.97966541 | 3.57914221 | 2.78006912 |
| ENSMUSG00 Lingo3 | 3431 | 0 | 0 | 0 | 0 | 0 | 0.0866559 | 0 | 0.05565783 | 0 | 0.16491065 | 0 | 0 | 0 | 0.12182943 | 0.04803446 | 0 | 0.03351177 | 0 |
| ENSMUSG00 4930579K19R | 651 | 4.76632542 | 4.60546925 | 4.53269639 | 3.49931216 | 6.50631226 | 6.85060833 | 7.41304647 | 4.45871364 | 8.35118292 | 7.09795533 | 8.58628286 | 7.54497436 | 7.5020888 | 9.77394773 | 4.93659195 | 6.29354069 | 5.29856639 | 7.43594684 |
| [illegible] | | | | | | | | | | | | | | | | | | | |

[illegible]

| | | | | | | | | | | | |
|---|---|---|---|---|---|---|---|---|---|---|---|
| ENSMUSG00 | Cyp3a44 | 1949 | 0 | 0 | 0 | 0 | 0 | 0 | 0 | 0 | [illegible] |
| ENSMUSG00 | 2900041M22 | 1782 | 0 | 0.09347052 | 0.16287384 | 0.12783682 | 0.18907041 | 0.13903685 | 0.13275162 | 0.30005254 | [illegible] |
| ENSMUSG00 | Fabp1 | 549 | 0 | 0 | 0 | 0 | 0 | 0 | 0 | 0 | [illegible] |
| ENSMUSG00 | Cadps | 6733 | 0.01262592 | 0.03463393 | 0 | 0 | 0.09293257 | 0.03679841 | 0.2108095 | 0.00567242 | [illegible] |
| ENSMUSG00 | A930005H10l | 308 | 9.52226346 | 9.40981103 | 10.8369273 | 9.31930452 | 10.0014387 | 11.4228704 | 17.9726735 | 17.6081855 | [illegible] |
| ENSMUSG00 | Atpif1 | 1439 | 179.000255 | 161.911315 | 172.921084 | 151.24747 | 156.50401 | 164.670724 | 205.295599 | 173.604242 | [illegible] |
| ENSMUSG00 | Olfr450 | 932 | 0 | 0 | 0 | 0 | 0 | 0 | 0 | 0 | [illegible] |
| ENSMUSG00 | Tmem120b | 2424 | 3.50702773 | 2.84478923 | 2.57433392 | 3.70277464 | 2.32319929 | 3.29125004 | 1.99088005 | 2.11129617 | [illegible] |
| ENSMUSG00 | Gimap4 | 1881 | 0 | 0.30107347 | 0.2314523 | 0 | 0.02558848 | 0 | 3.92385854 | 0.85278091 | [illegible] |
| ENSMUSG00 | Cpa1 | 1740 | 0 | 0 | 0 | 0 | 0 | 0 | 0 | 0 | [illegible] |
| ENSMUSG00 | Gm9945 | 931 | 0 | 0 | 0 | 0 | 0 | 0 | 0 | 0.16409195 | [illegible] |
| ENSMUSG00 | Aes | 1445 | 568.686898 | 571.391061 | 565.987043 | 583.528199 | 571.087773 | 598.542154 | 619.485042 | 549.44198 | [illegible] |
| ENSMUSG00 | Sytl5 | 2284 | 3.83365423 | 6.19876534 | 4.32057802 | 5.16651072 | 5.31052748 | 4.46929208 | 0.26929283 | 1.2541288 | [illegible] |
| ENSMUSG00 | Vapb | 7124 | 5.24453253 | 3.96069924 | 3.63955815 | 4.84773605 | 4.56050652 | 4.54210205 | 6.10336215 | 5.68131063 | [illegible] |
| ENSMUSG00 | 9430021M05 | 1198 | 2.0933267 | 1.50158288 | 2.14006436 | 1.48320595 | 2.61149836 | 2.10950702 | 1.30327079 | 0.38256163 | [illegible] |
| ENSMUSG00 | Vsnl1 | 1930 | 0 | 0 | 0.05012801 | 0 | 0 | 0.02567499 | 0 | 0 | [illegible] |
| ENSMUSG00 | Lclat1 | 4838 | 13.2663943 | 16.4223342 | 16.3578122 | 17.6198251 | 17.838061 | 19.6858937 | 14.5028323 | 18.8514793 | [illegible] |
| ENSMUSG00 | Thnsl2 | 2515 | 1.92667597 | 1.92062412 | 2.03880486 | 2.2463497 | 1.56931123 | 1.97028762 | 1.82478324 | 2.06527504 | [illegible] |
| ENSMUSG00 | Kcnn2 | 3778 | 1.00131304 | 0.78476643 | 0.92188835 | 0.51856138 | 0.67522286 | 0.80008384 | 0.76391567 | 0.54589458 | [illegible] |
| ENSMUSG00 | Tmem62 | 3333 | 2.28275623 | 2.01896329 | 2.74305348 | 2.85696318 | 1.92065582 | 3.58302094 | 2.83904464 | 4.04497991 | [illegible] |
| ENSMUSG00 | Gm9946 | 2098 | 0 | 0 | 0 | 0 | 0 | 0 | 0 | 0 | [illegible] |
| ENSMUSG00 | Gm9947 | 2919 | 0 | 0 | 0 | 0 | 0 | 0 | 0 | 0 | [illegible] |
| ENSMUSG00 | Tas2r130 | 1089 | 0 | 0 | 0 | 0 | 0 | 0 | 0 | 0 | [illegible] |
| [illegible] | | | | | | | | | | | |

| | | | | | | | | | | | | | | | | | | | |
|---|---|---|---|---|---|---|---|---|---|---|---|---|---|---|---|---|---|---|---|
| ENSMUSG00 | Gm13247 | 3398 | 0.03752664 | 0.06862574 | 0.05694353 | 0.02681639 | 0.05665912 | 0.18957785 | 0.02784737 | 0.10115704 | 0.13332892 | 0.01387602 | 0 | 0.29566861 | 0.07258964 | 0.0410042 | 0.08487666 | 0.12395429 | 0.02255815 | 0.04856602 |
| ENSMUSG00 | Usp39 | 2412 | 15.1728676 | 14.8195421 | 12.1134379 | 12.3913951 | 15.3455428 | 12.4703604 | 10.3962354 | 13.6808606 | 12.8238281 | 13.781611 | 16.2220941 | 11.6166926 | 13.7442158 | 13.8446513 | 12.5039548 | 15.4781684 | 15.2224678 | 15.1890817 |
| ENSMUSG00 | 6030405A18l | 5977 | 0 | 0 | 0 | 0 | 0 | 0 | 0 | 0 | 0 | 0 | 0 | 0 | 0 | 0 | 0 | 0 | 0 | 0 |
| ENSMUSG00 | Gm9994 | 2842 | 0 | 0 | 0 | 0 | 0 | 0 | 0 | 0 | 0 | 0 | 0 | 0 | 0 | 0 | 0 | 0 | 0 | 0 |
| ENSMUSG00 | Tyw1 | 7370 | 4.34856211 | 4.04093995 | 4.05628789 | 4.48192091 | 5.2572861 | 4.8208019 | 5.03299054 | 3.96434028 | 3.58775348 | 4.43997124 | 4.68026052 | 4.36982867 | 3.97600527 | 4.76414248 | 4.09220338 | 3.87581996 | 3.67142003 | 3.99319838 |
| ENSMUSG00 | 1810011O10l | 1330 | 14.4134093 | 14.6025694 | 10.111159 | 13.6683133 | 14.6567136 | 13.5618008 | 11.6325158 | 21.7380817 | 13.5946469 | 12.9398547 | 14.0810404 | 11.6247658 | 12.1660668 | 12.0126061 | 13.2898252 | 11.1129131 | 21.4973064 | 21.0937176 |
| ENSMUSG00 | Gm9996 | 912 | 0 | 0 | 0 | 0 | 0 | 0 | 0 | 0 | 0 | 0 | 0 | 0 | 0 | 0 | 0 | 0 | 0 | 0 |
| ENSMUSG00 | Myh1 | 6334 | 0.01342127 | 0 | 0 | 0 | 0 | 0 | 0 | 0 | 0 | 0 | 0 | 0 | 0 | 0.17598012 | 0 | 0 | 0 | 0 |
| ENSMUSG00 | 1810010K12l | 2529 | 0 | 0 | 0 | 0 | 0 | 0 | 0 | 0 | 0 | 0 | 0 | 0 | 0 | 0 | 0 | 0 | 0 | 0 |
| ENSMUSG00 | Gm16294 | 1456 | 0 | 0 | 0 | 0 | 0 | 0 | 0 | 0 | 0 | 0 | 0 | 0 | 0 | 0 | 0 | 0 | 0 | 0 |
| ENSMUSG00 | Usp34 | 17049 | 3.43801031 | 3.18103066 | 3.91550659 | 2.74451244 | 2.99254144 | 2.94718 | 2.77787504 | 2.87635901 | 2.75639712 | 2.62455378 | 3.02358966 | 3.28365943 | 3.30441959 | 2.27194374 | 3.23590142 | 3.24759331 | 2.92690707 | 3.10069515 |
| ENSMUSG00 | Krtap13-1 | 747 | 0 | 0 | 0 | 0 | 0 | 0 | 0 | 0 | 0 | 0 | 0 | 0 | 0 | 0 | 0 | 0 | 0 | 0 |
| ENSMUSG00 | Hsd3b4 | 1568 | 0 | 0 | 0 | 0 | 0 | 0 | 0 | 0 | 0 | 0 | 0 | 0 | 0 | 0 | 0 | 0 | 0 | 0 |
| ENSMUSG00 | D830013O20 | 679 | 0 | 0 | 0 | 0 | 0 | 0 | 0 | 0 | 0 | 0 | 0.07036076 | 0 | 0 | 0.13680144 | 0 | 0 | 0 | 0 |
| ENSMUSG00 | Six3os1 | 7534 | 0 | 0 | 0 | 0 | 0 | 0 | 0 | 0 | 0 | 0 | 0 | 0 | 0 | 0 | 0 | 0 | 0 | 0 |
| ENSMUSG00 | Fabp3-ps1 | 401 | 0 | 0.08307455 | 0.24126449 | 0 | 0 | 0 | 0.58993365 | 0.57145739 | 0 | 0.58791401 | 0.11913955 | 0.41757354 | 0 | 0 | 0.61648213 | 0 | 0 | 0.13717982 |
| ENSMUSG00 | Actr3b | 1831 | 2.20534775 | 2.6562985 | 2.32488841 | 3.01086093 | 1.57723398 | 2.62513116 | 6.56330991 | 6.2576301 | 4.09390818 | 5.48503539 | 3.73119556 | 4.42013712 | 11.3428384 | 12.6066147 | 12.6012405 | 16.7717461 | 13.8569135 | 12.9185778 |
| ENSMUSG00 | Sftpb | 1120 | 0 | 0 | 0 | 0 | 0 | 0 | 0.04224346 | 0 | 0 | 0 | 0 | 0 | 0 | 0.04404635 | 0 | 0 | 0 | 0 |
| ENSMUSG00 | Taar1 | 1113 | 0 | 0 | 0 | 0 | 0 | 0 | 0 | 0 | 0 | 0 | 0 | 0 | 0 | 0 | 0 | 0 | 0 | 0 |
| ENSMUSG00 | Gpr50 | 1238 | 0 | 0 | 0 | 0 | 0.03887878 | 0.04002644 | 0.19108513 | 0.1851005 | 0 | 0.60937904 | 0 | 0 | 0.19924038 | 1.57564755 | 0.23296518 | 0.58765876 | 0 | 0.6220739 |
| ENSMUSG00 | AI987944 | 1818 | 26.1156665 | 19.4966554 | 21.5259549 | 18.8960547 | 23.8012098 | 23.0319361 | 23.2919954 | 20.9869142 | 15.8583944 | 24.3274806 | 17.1338135 | 24.1929275 | 21.5725381 | 20.4374422 | 19.0596856 | 25.737691 | 28.3125505 | 25.8403615 |
| ENSMUSG00 | Lig1 | 4837 | 5.49219248 | 4.44217837 | 6.27045763 | 4.86034709 | 5.48287987 | 5.71644105 | 4.09841068 | 4.78490714 | 4.78536735 | 3.32404172 | 4.22734799 | 4.17723173 | 5.84395051 | 4.23441028 | 4.52305689 | 5.91339337 | 4.22326158 | 4.42392857 |
| ENSMUSG00 | Prss34 | 1211 | 0 | 0 | 0 | 0 | 0 | 0 | 0 | 0 | 0 | 0 | 0 | 0 | 0 | 0 | 0 | 0 | 0 | 0 |
| ENSMUSG00 | Gm12500 | 2033 | 0 | 0.19663292 | 0.3093241 | 0 | 0.16572723 | 0.21936773 | 0.44217457 | 0.28179343 | 0 | 0.30150474 | 0 | 0.24709344 | 0.14559346 | 0.34267649 | 0.18239744 | 0.0565036 | 0.22622507 | 0.24352285 |
| ENSMUSG00 | Gm12396 | 305 | 4.04147575 | 2.7305651 | 2.22042434 | 3.13698994 | 3.78743006 | 3.73676352 | 4.18833551 | 2.75486181 | 2.02556635 | 2.62807198 | 3.28942337 | 3.66004345 | 3.39662381 | 4.72054659 | 3.24208964 | 3.5152033 | 3.7697997 | 1.62321954 |
| ENSMUSG00 | Adap1 | 3444 | 0.08639263 | 0.07738187 | 0.08427444 | 0.06614553 | 0.29348734 | 0.15826947 | 0.1236394 | 0.16634322 | 0.23917837 | 0.10952544 | 0.11097551 | 0.19447966 | 0.2435083 | 0.33713768 | 0.46656814 | 0.56702119 | 0.33385276 | 0 |
| ENSMUSG00 | BC043934 | 1650 | 0.05152143 | 0.16151706 | 0.14658646 | 0.33135305 | 0.26253777 | 0.18019176 | 0 | 0.02314691 | 0.74884574 | 0.1714571 | 0 | 0.10148302 | 0.20928692 | 0 | 0.7491192 | 0.162445 | 0.83621011 | 0 |
| ENSMUSG00 | Uts2d | 830 | 0 | 0 | 0 | 0 | 0 | 0 | 0 | 0 | 0 | 0 | 0 | 0 | 0 | 0 | 0 | 0 | 0 | 0 |
| ENSMUSG00 | Nat3 | 1392 | 0 | 0 | 0 | 0 | 0 | 0 | 0 | 0 | 0.08876404 | 0 | 0.58345854 | 0 | 1.20494626 | 0.26692004 | 0 | 0 | 0 | 0.39518036 |
| ENSMUSG00 | Slit3 | 6030 | 0.09163637 | 0.17126032 | 0.12033217 | 0.15111457 | 0.32726515 | 0.12326551 | 1.05139287 | 0.62703944 | 0.83329269 | 0.6255483 | 0.64175317 | 0.50909808 | 0.62176214 | 0.17714992 | 0.3621364 | 0.1778005 | 0.12076279 | 0.20981915 |
| ENSMUSG00 | Tgoln1 | 5009 | 54.1221826 | 62.7484841 | 51.5604469 | 61.4696621 | 61.7287837 | 66.0241453 | 51.3459219 | 59.2138545 | 46.65448 | 55.311945 | 52.448492 | 54.0773429 | 48.0614407 | 53.9081501 | 48.8677268 | 57.6614198 | 67.4019464 | 57.5789073 |
| ENSMUSG00 | Cyct | 651 | 0 | 0 | 0.44583899 | 0.41991746 | 0.2218061 | 0.53282509 | 0 | 0 | 1.26533075 | 0 | 0 | 0.25721503 | 0.07577867 | 0 | 0 | 0 | 0.05887296 | 0 |
| ENSMUSG00 | 5730446D14l | 1677 | 0.02534596 | 0.0794583 | 0 | 0.05433637 | 0 | 0.02954844 | 0 | 0 | 0.07367892 | 0 | 0 | 0.09984913 | 0.05883353 | 0 | 0 | 0.0228328 | 0.11427041 | 0 |
| ENSMUSG00 | Prl2c3 | 845 | 0 | 0 | 0 | 0 | 0 | 0 | 0 | 0.04519811 | 0 | 0 | 0 | 0 | 0 | 0 | 0 | 0 | 0 | 0 |
| ENSMUSG00 | Stk30 | 2130 | 0.31928771 | 0.48483555 | 1.06739715 | 0.91977695 | 0.58752583 | 0.72119002 | 0.97735111 | 0.71722821 | 0.52208259 | 0.42059313 | 1.2784848 | 1.7294994 | 0.48637102 | 0.15263315 | 0.63833396 | 0.79097968 | 0.1619421 | 1.52372643 |
| ENSMUSG00 | Zbtb25 | 4499 | 2.05959733 | 2.45089308 | 2.96756932 | 2.75452413 | 2.4071311 | 1.92747908 | 2.75526159 | 2.37694436 | 2.92946989 | 3.1440789 | 2.12380344 | 3.20080928 | 1.72151834 | 2.81823424 | 2.9671714 | 3.37032241 | 2.18082507 | 2.76329365 |
| ENSMUSG00 | 5730596B20l | 3353 | 0.0633838 | 0.10928775 | 0 | 0 | 0.15790372 | 0.02955725 | 0 | 0 | 0 | 0 | 0 | 0 | 0 | 0 | 0.04915187 | 0 | 0 | 0 |
| ENSMUSG00 | Med12l | 10362 | 0.26663158 | 0.28612696 | 0.25209136 | 0.25062532 | 0.32050789 | 0.07651455 | 0.06392371 | 0.09951697 | 0 | 0.00455035 | 0.00461059 | 0 | 0.00476085 | 0.00448215 | 0.01590487 | 0.05173408 | 0.01109621 | 0 |
| ENSMUSG00 | Cd248 | 2559 | 0 | 0 | 0 | 0 | 0 | 0 | 0 | 0 | 0 | 0 | 0 | 0 | 0 | 0 | 0 | 0 | 0 | 0 |
| ENSMUSG00 | Chn1 | 7551 | 0.71489304 | 1.37204478 | 1.74890396 | 1.97304483 | 1.69554916 | 1.68653854 | 1.54137452 | 1.3201188 | 1.74542247 | 1.07401946 | 1.11354691 | 0.96093711 | 1.27396687 | 0.79344289 | 1.1349389 | 0.8874131 | 1.10649354 | 1.56627704 |
| ENSMUSG00 | Mettl7a2 | 2309 | 0 | 0.02885482 | 0 | 0 | 0 | 0.04292138 | 0 | 0 | 0 | 0 | 0 | 0.02417309 | 0.06409517 | 0.02011437 | 0.05353169 | 0.0994992 | 0 | 0 |
| ENSMUSG00 | Gpr116 | 8633 | 80.2689546 | 69.817004 | 66.0184101 | 99.5661604 | 84.3827945 | 89.1409652 | 72.1062107 | 67.8199385 | 59.6591428 | 81.3899896 | 75.7382233 | 75.3540539 | 51.5205104 | 40.5477918 | 46.4561502 | 64.0024864 | 77.4472682 | 54.0915449 |
| ENSMUSG00 | Foxk1 | 7954 | 3.81552624 | 3.69817521 | 3.70373145 | 3.08170001 | 3.3887198 | 3.11495685 | 2.75405712 | 2.91460753 | 3.14309971 | 3.0054572 | 3.5197543 | 2.36483257 | 2.98943728 | 2.80276112 | 2.96812818 | 4.43851396 | 3.08383581 | 3.45103649 |
| ENSMUSG00 | Cngb3 | 4690 | 0 | 0 | 0 | 0 | 0 | 0 | 0 | 0 | 0 | 0 | 0 | 0 | 0 | 0 | 0 | 0 | 0 | 0 |
| ENSMUSG00 | Tmem154 | 4826 | 0.31707135 | 0.60054327 | 0.02004705 | 0.11328896 | 0 | 0.27723245 | 0 | 0.13453602 | 0.5205921 | 0.05862085 | 0 | 0.04626246 | 0.57243833 | 0 | 0.02561224 | 0.15075042 | 0 | 0 |
| ENSMUSG00 | Cebpb | 1503 | 25.3956408 | 32.3155022 | 22.9154715 | 28.4339718 | 23.5055436 | 26.9028813 | 21.4370821 | 22.8951128 | 16.3321114 | 21.5832893 | 21.0425965 | 19.6821696 | 30.1636939 | 40.4184074 | 29.4140183 | 44.4813044 | 35.0368143 | 43.8828469 |
| ENSMUSG00 | 1700001K19l | 948 | 0.62771357 | 0.07028037 | 0.30616158 | 0.62478225 | 0.76158107 | 0.3136249 | 0.74861834 | 0.68488485 | 0 | 0.29842217 | 0 | 0.47101825 | 0.26018944 | 1.51874125 | 0 | 0.80781872 | 0.56600017 | 0.46421187 |
| ENSMUSG00 | Gm9999 | 441 | 0.09638362 | 0.07553944 | 0.10969055 | 0.20662605 | 0 | 0.11236448 | 0 | 0 | 0 | 0 | 0 | 0 | 0.22372752 | 0.10531539 | 0 | 0.08682677 | 0 | 0.1247372 |
| ENSMUSG00 | Rab31 | 3805 | 35.4563545 | 31.7982738 | 31.2362061 | 36.9038472 | 34.7865414 | 36.0087538 | 30.2154559 | 32.6417063 | 26.0757731 | 26.9025432 | 30.284678 | 26.2722343 | 31.8291345 | 38.1683735 | 41.2990103 | 34.9898665 | 40.1494401 | 35.5932775 |
| ENSMUSG00 | Ptafr | 3635 | 0.08185316 | 0 | 0.09315398 | 0 | 0.02648249 | 0 | 0.01301587 | 0.16810961 | 0.73648515 | 0 | 0 | 0 | 0.01357137 | 0 | 0.44205246 | 0 | 0.2214174 | 0 |
| ENSMUSG00 | Ccdc18 | 4628 | 0.4684019 | 0.74140625 | 0.55397517 | 0.40363069 | 0.42640641 | 0.53535797 | 1.22677646 | 1.32864667 | 1.38830952 | 0.88636802 | 1.18714784 | 1.21810327 | 0.9913285 | 1.3046092 | 0.45403616 | 0.57088416 | 1.01861161 | 0.84391672 |
| ENSMUSG00 | Pign | 4412 | 3.66091725 | 4.58316564 | 5.30661575 | 4.79155135 | 3.63280385 | 5.8515354 | 4.86853042 | 4.81300447 | 4.32215703 | 4.3175168 | 3.78994458 | 4.1241854 | 3.32084756 | 3.2527704 | 2.23190659 | 3.44546037 | 4.64745441 | 3.52846265 |
| ENSMUSG00 | Rlim | 8912 | 5.63269893 | 7.39746607 | 7.09970576 | 7.02944748 | 6.54576879 | 6.53882571 | 6.57769403 | 8.35244527 | 6.38224621 | 7.35936134 | 7.08690475 | 7.42162928 | 6.27166318 | 7.75978974 | 7.32307726 | 7.95715944 | 8.56664985 | 9.60437276 |
| ENSMUSG00 | Defb21 | 424 | 45.5126177 | 25.0632388 | 37.5351214 | 16.118294 | 23.6118871 | 30.6198495 | 12.3861023 | 14.6824566 | 2.33131221 | 0.66722693 | 1.69015183 | 1.44804785 | 0 | 1.42399325 | 0.09717348 | 0 | 0 | 0.12973846 |
| ENSMUSG00 | Zfp953 | 4964 | 0.68501492 | 0.57713717 | 0.59443702 | 0.6241239 | 1.03749312 | 0.63887489 | 1.0770211 | 0.27697955 | 0.57249589 | 0.69339269 | 0.79881578 | 0.7870863 | 0.05962762 | 0.75785074 | 0.19920172 | 0.50138789 | 1.12724403 | 0.01108161 |
| ENSMUSG00 | Ptprn2 | 5423 | 0.00783795 | 0.01842867 | 0 | 0 | 0.00887552 | 0.01827503 | 0 | 0.07746938 | 0 | 0.01738916 | 0.07047753 | 0.2367251 | 0.07277436 | 0.17984987 | 0 | 0 | 0 | 0 |
| ENSMUSG00 | Olfr324 | 1121 | 0 | 0 | 0 | 0 | 0 | 0 | 0 | 0 | 0 | 0 | 0 | 0 | 0 | 0 | 0 | 0 | 0 | 0 |
| ENSMUSG00 | Mpz | 1203 | 0 | 0 | 0.36189674 | 0 | 0.04000991 | 0 | 0 | 0 | 0 | 0 | 0 | 0 | 0 | 0 | 0 | 0 | 0 | 0 |
| [illegible] | [illegible] | [illegible] | [illegible] | [illegible] | [illegible] | [illegible] | [illegible] | [illegible] | [illegible] | [illegible] | [illegible] | [illegible] | [illegible] | [illegible] | [illegible] | [illegible] | [illegible] | [illegible] | [illegible] |

| | | | | | | | | | | | | | | | | | | | |
|---|---|---|---|---|---|---|---|---|---|---|---|---|---|---|---|---|---|---|---|
| ENSMUSG00 Sdhc | 3742 | 80.1941589 | 80.5937016 | 82.5530104 | 80.9189473 | 86.9641731 | 82.830665 | 89.5299518 | 79.4875548 | 82.1859201 | 88.7068281 | 85.0680247 | 79.9348305 | 80.9718428 | 68.9090254 | 69.8290399 | 68.0471778 | 79.8584011 | 72.326244 |
| ENSMUSG00 Olfr1503-ps1 | 796 | 0 | 0 | 0 | 0 | 0 | 0 | 0 | 0 | 0 | 0 | 0 | 0 | 0 | 0 | 0 | 0 | 0 | 0 |
| ENSMUSG00 Olfr605 | 1860 | 0 | 0 | 0 | 0 | 0 | 0 | 0 | 0 | 0 | 0 | 0 | 0 | 0 | 0 | 0 | 0 | 0 | 0 |
| ENSMUSG00 Olfr825 | 970 | 0 | 0 | 0 | 0 | 0 | 0 | 0 | 0 | 0 | 0 | 0 | 0 | 0 | 0 | 0 | 0 | 0 | 0 |
| ENSMUSG00 AA987161 | 5457 | 3.78550771 | 3.19881924 | 1.93245916 | 2.01213335 | 2.81364919 | 2.86946377 | 3.52005638 | 2.68753574 | 3.23031496 | 2.01321493 | 4.54374262 | 3.11962197 | 1.63626114 | 2.05113159 | 3.12579149 | 2.91196666 | 2.34579131 | 2.19754174 |
| ENSMUSG00 Nfam1 | 5283 | 0 | 0 | 0.00915645 | 0 | 0.00911072 | 0.05627795 | 0 | 0.18073255 | 0.3664142 | 0 | 0.40694172 | 0.13734689 | 0.13073005 | 0.06153864 | 0.11698341 | 0.10871836 | 0 | 0 |
| ENSMUSG00 1700084J12R | 1004 | 0 | 0 | 0 | 0 | 0.04794016 | 0 | 0 | 0.11412072 | 0 | 0 | 0 | 0 | 0 | 0 | 0 | 0.38138055 | 0 | 0 |
| ENSMUSG00 Olfr127 | 1094 | 0 | 0 | 0 | 0 | 0 | 0 | 0 | 0 | 0 | 0 | 0 | 0 | 0 | 0 | 0 | 0 | 0 | 0 |
| ENSMUSG00 Gm5771 | 819 | 0 | 0 | 0 | 0 | 0 | 0 | 0 | 0 | 0 | 0 | 0 | 0 | 0 | 0 | 0 | 0 | 0 | 0 |
| ENSMUSG00 H2-M10.3 | 1586 | 0 | 0 | 0 | 0 | 0 | 0 | 0 | 0 | 0 | 0 | 0 | 0 | 0 | 0 | 0 | 0 | 0 | 0 |
| ENSMUSG00 Tpm3-rs7 | 2105 | 0.94904669 | 0.72797774 | 0.80431048 | 0.51946084 | 0.9374864 | 0.80037669 | 0.5619083 | 0.67131539 | 0.76307559 | 0.53758522 | 0.52200667 | 0.68940961 | 0.7030677 | 0.48540139 | 0.43061009 | 0.50932874 | 0.40055987 | 0.26132592 |
| ENSMUSG00 Olfr884 | 1854 | 0 | 0 | 0 | 0 | 0 | 0 | 0 | 0 | 0 | 0 | 0 | 0 | 0 | 0.10020299 | 0 | 0 | 0 | 0 |
| ENSMUSG00 Vmn1r82 | 914 | 0 | 0 | 0 | 0 | 0 | 0 | 0 | 0 | 0 | 0 | 0 | 0 | 0 | 0 | 0 | 0 | 0 | 0 |
| ENSMUSG00 Gstm1 | 1773 | 1000.68164 | 1039.83954 | 1024.65669 | 1018.27521 | 1088.7642 | 972.888131 | 1064.2284 | 1114.98981 | 1041.71918 | 1033.7785 | 1067.35131 | 1051.84034 | 1012.90418 | 935.194791 | 941.45439 | 822.783734 | 879.170232 | 980.235599 |
| ENSMUSG00 Adamts17 | 7290 | 0.01166123 | 0.08682373 | 0.033178 | 0.09999681 | 0.01320492 | 0.00679736 | 0 | 0.06810716 | 0.06779673 | 0.07114647 | 0.03276746 | 0.03062588 | 0.1421084 | 0.05096745 | 0.20346447 | 0.07878726 | 0.01051476 | 0.03018332 |
| ENSMUSG00 Xlr3c | 1575 | 0.02698741 | 0.04230209 | 0.03071335 | 0.05785529 | 0 | 0.03146205 | 0 | 0.19399316 | 0.10460067 | 0 | 0 | 0 | 0 | 0 | 0 | 0 | 0 | 0 |
| ENSMUSG00 Chsy3 | 2303 | 0 | 0.043396 | 0 | 0 | 0 | 0 | 0 | 0 | 0 | 0 | 0 | 0 | 0 | 0 | 0.25046539 | 0 | 0.0665676 | 0 |
| ENSMUSG00 Sez6l | 6266 | 0.01356693 | 0.06911389 | 0 | 0.04362692 | 0.0537701 | 0 | 0 | 0 | 0 | 0 | 0 | 0 | 0 | 0 | 0 | 0 | 0 | 0 |
| ENSMUSG00 T2 | 1758 | 0.38685029 | 0.15159451 | 0.11006492 | 0.10366563 | 0.08213639 | 0.14093496 | 0.16147672 | 0.1303495 | 0.51541715 | 0.85826081 | 0.35328468 | 0.47624285 | 0.11224555 | 0.47553673 | 0.02343661 | 0.28315011 | 0.30521511 | 0.21903512 |
| ENSMUSG00 Gm5431 | 5381 | 0.11848683 | 0 | 0.06292784 | 0 | 0.05366875 | 0.0276265 | 0 | 0 | 0 | 0 | 0.00887845 | 0 | 0.08251018 | 0 | 0 | 0.10673836 | 0.07122523 | 0 |
| ENSMUSG00 Krtap6-1 | 593 | 0 | 0 | 0 | 0 | 0 | 0 | 0 | 0 | 0 | 0 | 0 | 0 | 0 | 0 | 0 | 0 | 0 | 0 |
| ENSMUSG00 BC017612 | 951 | 40.9855378 | 30.4755184 | 30.8756392 | 24.3375505 | 28.1402204 | 27.8766693 | 34.6263139 | 33.5739731 | 38.9777752 | 36.2926549 | 29.7902738 | 34.4519389 | 27.4412032 | 21.1953038 | 25.3448057 | 27.6207746 | 26.6389929 | 23.4844346 |
| ENSMUSG00 Gm5148 | 4833 | 0.83550418 | 0.47560308 | 0.68061247 | 0.88614487 | 0.91622946 | 0.18455394 | 0.930003 | 0.27658475 | 0.62210131 | 0.85852719 | 0.73150154 | 0.57744323 | 0.51036538 | 0.38439137 | 0.88660497 | 1.20425664 | 0.68992093 | 0.61462689 |
| ENSMUSG00 Gm5407 | 746 | 0 | 0 | 0 | 0 | 0 | 0 | 0 | 0 | 0 | 0 | 0 | 0 | 0 | 0 | 0 | 0 | 0 | 0 |
| ENSMUSG00 Mmel1 | 3539 | 0 | 0 | 0.0136687 | 0 | 0 | 0 | 0 | 0 | 0 | 0.02664634 | 0 | 0 | 0 | 0.19685259 | 0.13970576 | 0 | 0 | 0 |
| ENSMUSG00 Gm13242 | 4132 | 0 | 0 | 0 | 0 | 0 | 0.01199243 | 0 | 0 | 0 | 0 | 0 | 0 | 0 | 0 | 0 | 0 | 0 | 0 |
| ENSMUSG00 Olfr746 | 944 | 0 | 0 | 0 | 0 | 0 | 0 | 0 | 0 | 0 | 0 | 0 | 0 | 0 | 0 | 0 | 0 | 0 | 0 |
| ENSMUSG00 Hist1h2bm | 523 | 0.16254369 | 0 | 0 | 0.08711481 | 0 | 0 | 0 | 0.07302563 | 0 | 0.45077154 | 0.36539165 | 0 | 0 | 0.08880323 | 0.47267559 | 0.1464268 | 0.29312655 | 0 |
| ENSMUSG00 Zfp846 | 3089 | 2.24290829 | 2.55589379 | 1.51901342 | 3.0531357 | 2.39958441 | 2.39020955 | 2.49659005 | 2.16370036 | 1.74666027 | 3.34283069 | 2.75297592 | 1.60815603 | 1.64492958 | 1.84934372 | 1.97404671 | 2.045306 | 3.10183691 | 1.54930148 |
| ENSMUSG00 Olfr1126 | 944 | 0 | 0 | 0 | 0 | 0 | 0 | 0 | 0 | 0 | 0 | 0 | 0 | 0 | 0 | 0 | 0 | 0 | 0 |
| ENSMUSG00 Olfr66 | 935 | 0 | 0 | 0 | 0 | 0 | 0 | 0 | 0 | 0 | 0 | 0 | 0 | 0 | 0 | 0 | 0 | 0 | 0 |
| ENSMUSG00 Olfr344 | 929 | 0 | 0 | 0 | 0 | 0 | 0 | 0 | 0 | 0 | 0 | 0 | 0 | 0 | 0 | 0 | 0 | 0 | 0 |
| ENSMUSG00 Serpina3k | 1649 | 0 | 0 | 0 | 0 | 0 | 0 | 0 | 0 | 0 | 0 | 0 | 0 | 0 | 0 | 0 | 0 | 0 | 0 |
| ENSMUSG00 BC021614 | 679 | 0.12519934 | 0 | 0 | 0 | 0 | 0 | 0 | 0.16874405 | 0 | 0.27776556 | 0 | 0 | 0 | 0 | 0 | 0 | 0 | 0 |
| ENSMUSG00 Olfr318 | 920 | 0 | 0 | 0 | 0 | 0 | 0 | 0 | 0 | 0 | 0 | 0 | 0 | 0 | 0 | 0 | 0 | 0 | 0 |
| ENSMUSG00 Grlf1 | 7841 | 6.46169747 | 6.86990617 | 6.14462903 | 8.43120461 | 6.83214272 | 8.15872708 | 5.86505853 | 6.50259625 | 4.58561768 | 5.22560402 | 5.34353253 | 5.58085441 | 5.01435251 | 5.37237435 | 6.01655165 | 6.82696967 | 6.19790135 | 5.91412791 |
| ENSMUSG00 Usf2 | 3310 | 8.16715769 | 6.70283612 | 6.56184747 | 6.89609765 | 6.3254945 | 5.73374531 | 5.36019776 | 4.75385794 | 5.86067943 | 5.18515287 | 5.22493504 | 5.46352709 | 4.90338392 | 5.23372949 | 4.60561199 | 5.10155827 | 5.51157623 | 4.45390955 |
| ENSMUSG00 Cryzl1 | 4910 | 11.0807791 | 11.4254407 | 12.7682475 | 10.9309389 | 11.9790653 | 11.0408738 | 12.1317032 | 11.2321444 | 13.1780074 | 14.193226 | 12.9799989 | 13.4707862 | 10.1276115 | 10.4806617 | 11.6304188 | 12.0876663 | 13.1839339 | 11.3155189 |
| [illegible] | | | | | | | | | | | | | | | | | | | |

| | | | | | | | | | | | | | | | | | | | |
|---|---|---|---|---|---|---|---|---|---|---|---|---|---|---|---|---|---|---|---|
| ENSMUSG00 Akap17b | 6781 | 2.99623568 | 2.26474624 | 3.36710018 | 3.07726859 | 3.23671393 | 2.94495674 | 4.8352288 | 4.04396769 | 6.26209965 | 5.44447731 | 5.04451707 | 4.99632904 | 6.19831788 | 7.66419905 | 6.28869051 | 6.92856146 | 5.94026516 | 8.4529552 |
| ENSMUSG00 Rcan3 | 5288 | 13.6164463 | 12.5175342 | 12.7611715 | 15.1381912 | 12.1057977 | 14.3373076 | 14.2975909 | 11.4548317 | 13.5990273 | 17.7439296 | 10.4801346 | 13.2467292 | 10.1033408 | 10.7239468 | 9.87964691 | 10.6153613 | 11.8573793 | 10.7251114 |
| ENSMUSG00 Flot1 | 3006 | 47.2279734 | 44.4393566 | 39.2974588 | 43.5906732 | 40.7183905 | 45.3161892 | 32.8638967 | 33.5803458 | 34.0617692 | 38.0374103 | 36.5543594 | 39.7728374 | 35.858363 | 30.7155175 | 35.0884841 | 39.9975074 | 35.0623142 | 35.391754 |
| ENSMUSG00 Hoxa3 | 2083 | 0 | 0 | 0 | 0 | 0 | 0 | 0 | 0 | 0 | 0 | 0 | 0 | 0 | 0 | 0 | 0 | 0 | 0 |
| ENSMUSG00 Olfr1385 | 929 | 0 | 0 | 0 | 0 | 0 | 0 | 0 | 0 | 0 | 0 | 0 | 0 | 0 | 0 | 0 | 0 | 0 | 0 |
| ENSMUSG00 Olfr837 | 943 | 0 | 0 | 0 | 0 | 0 | 0 | 0 | 0 | 0 | 0 | 0 | 0 | 0 | 0 | 0 | 0 | 0 | 0 |
| ENSMUSG00 Olfr1354 | 932 | 0 | 0 | 0 | 0 | 0 | 0 | 0 | 0 | 0 | 0 | 0 | 0 | 0 | 0 | 0 | 0 | 0 | 0 |
| ENSMUSG00 Ndufs8 | 956 | 161.172869 | 140.395032 | 154.683977 | 137.874583 | 151.142296 | 142.282692 | 176.036819 | 145.21902 | 165.392295 | 154.324844 | 162.414869 | 145.552769 | 160.535141 | 177.517464 | 148.644525 | 137.862208 | 134.382581 | 159.388311 |
| ENSMUSG00 Myl3 | 1298 | 0 | 0 | 0 | 0 | 0 | 0 | 0 | 0 | 0 | 0 | 0 | 0 | 0 | 0 | 0 | 0 | 0.17716316 | 0 |
| ENSMUSG00 Kcnh7 | 15210 | 0.22076982 | 0.19273732 | 0.17174034 | 0.14677785 | 0.09809925 | 0.1303162 | 0.1150933 | 0.16070439 | 0.21392076 | 0.18289885 | 0.21673058 | 0.12843841 | 0.18162968 | 0.04885637 | 0.0758477 | 0.15860015 | 0.17134702 | 0.15189891 |
| ENSMUSG00 Fdps | 1223 | 60.8904893 | 60.2247008 | 54.1087404 | 50.4040008 | 46.8725438 | 50.4846329 | 59.6534349 | 51.4957249 | 59.3718948 | 41.9845592 | 36.4074091 | 40.8006561 | 121.373458 | 148.484368 | 97.0577944 | 84.5649473 | 74.8663314 | 113.166731 |
| ENSMUSG00 Gm9000 | 861 | 51.045705 | 51.9232335 | 59.0482932 | 8.41371202 | 22.1373307 | 38.0422264 | 46.7631705 | 52.6975308 | 60.6556353 | 7.99535735 | 19.1987638 | 37.599401 | 64.4580801 | 86.6308994 | 75.8950844 | 14.9426761 | 25.0612139 | 57.7563676 |
| ENSMUSG00 Olfr767 | 1085 | 0 | 0 | 0 | 0 | 0 | 0 | 0 | 0 | 0 | 0 | 0 | 0 | 0 | 0 | 0 | 0 | 0 | 0 |
| ENSMUSG00 Taar2 | 1018 | 0 | 0 | 0 | 0 | 0 | 0 | 0 | 0 | 0 | 0 | 0 | 0 | 0 | 0 | 0 | 0 | 0 | 0 |
| ENSMUSG00 Olfr682-ps1 | 894 | 0 | 0 | 0 | 0 | 0 | 0 | 0 | 0 | 0 | 0 | 0 | 0 | 0 | 0 | 0 | 0 | 0 | 0 |
| ENSMUSG00 4933400F03I | 941 | 0 | 0 | 0 | 0 | 0 | 0 | 0 | 0 | 0 | 0 | 0 | 0 | 0 | 0 | 0 | 0 | 0 | 0 |
| ENSMUSG00 Six1b | 4494 | 3.90623892 | 3.31349882 | 3.49830827 | 3.49767697 | 3.00958401 | 2.70147301 | 3.81112367 | 3.39941276 | 5.26974779 | 3.73512469 | 4.67756603 | 4.29733406 | 3.68836765 | 4.0098589 | 3.99730271 | 3.89381566 | 3.98272823 | 4.18627381 |
| ENSMUSG00 Rps26-ps1 | 412 | 1.23801484 | 0.88942193 | 1.8785837 | 0.5529253 | 1.28507563 | 0.36082088 | 1.03352939 | 2.31750015 | 1.19960726 | 1.02999109 | 2.2032141 | 0.81284946 | 0.47895065 | 4.05822123 | 1.10004154 | 0.74350694 | 3.44192472 | 2.67034499 |
| ENSMUSG00 Gm10071 | 568 | 0.22449917 | 0.05864946 | 0.08516467 | 0 | 0.42369651 | 0 | 0 | 0.13448029 | 0.07251147 | 0.0830118 | 0.33644337 | 0.19653402 | 0 | 0.16353552 | 0.1450759 | 0.06741304 | 0 | 0.19369404 |
| ENSMUSG00 Olfr781 | 935 | 0 | 0 | 0 | 0 | 0 | 0 | 0 | 0 | 0 | 0 | 0 | 0 | 0 | 0 | 0 | 0 | 0 | 0 |
| ENSMUSG00 Nrm | 1641 | 0.67345191 | 0.7511134 | 0.08843424 | 0.83292586 | 0.93858718 | 0.21137668 | 1.18209618 | 1.07059751 | 1.12942913 | 0.80452144 | 0.87339961 | 0.71427722 | 0.96198742 | 0.53774385 | 1.50645543 | 0.37333926 | 0.51381995 | 0.30169528 |
| ENSMUSG00 Eif4a1 | 3828 | 105.563403 | 101.157023 | 97.8717327 | 87.3015309 | 93.0072726 | 86.290105 | 83.9342745 | 91.3804111 | 91.2171579 | 87.0834566 | 75.1070272 | 83.1985817 | 89.6041852 | 82.0051167 | 83.1995891 | 92.2455542 | 81.8688949 | 79.8120634 |
| ENSMUSG00 Rgs3 | 9762 | 0.43976877 | 0.6756764 | 0.58472409 | 0.49472144 | 0.56208147 | 0.54821709 | 0.65913996 | 0.37167366 | 0.58645008 | 0.37674194 | 0.709626 | 0.49743522 | 0.49018602 | 0.88967879 | 0.85678302 | 0.60012937 | 0.73810119 | 0.9523191 |
| ENSMUSG00 Atl2 | 5091 | 9.65153869 | 11.6016031 | 11.2596019 | 8.9672297 | 9.89866904 | 9.94753363 | 8.64285826 | 10.9528398 | 9.12559218 | 9.1226562 | 12.0680803 | 10.722396 | 10.6977876 | 8.83997656 | 8.49766916 | 11.9286787 | 12.7679041 | 9.57337823 |
| ENSMUSG00 Gm8355 | 2023 | 0.12606577 | 0.13173661 | 0.2391178 | 0.1801722 | 0.1427541 | 0.07348403 | 0.16371169 | 0.13215364 | 0.20359128 | 0.25638049 | 0.40147024 | 0.4138581 | 0.04877105 | 0.27549632 | 0.30549844 | 0.17034872 | 0.13261694 | 0.35349401 |
| ENSMUSG00 Fam159a | 590 | 0 | 0 | 0 | 0.07722211 | 0 | 0 | 0 | 0 | 0 | 0 | 0 | 0.09460282 | 0 | 0 | 0 | 0 | 0 | 0 |
| ENSMUSG00 AU019823 | 6572 | 4.41091449 | 4.70395098 | 4.98309191 | 4.00010995 | 4.99482227 | 4.60692639 | 5.55052883 | 6.22399011 | 5.73427601 | 5.44543271 | 5.79376779 | 5.72424781 | 5.48716244 | 5.8302453 | 5.02167472 | 5.09803431 | 6.67736001 | 4.36088628 |
| ENSMUSG00 Olfr847 | 938 | 0 | 0 | 0 | 0 | 0 | 0 | 0 | 0 | 0 | 0 | 0 | 0 | 0 | 0 | 0 | 0 | 0 | 0 |
| ENSMUSG00 Dbp | 3079 | 8.29672322 | 9.27221512 | 8.75091147 | 8.30131402 | 8.363293 | 7.5640743 | 8.77412786 | 9.01782285 | 13.617367 | 12.7103227 | 11.6683237 | 11.5836987 | 9.30881583 | 10.1667148 | 8.33665779 | 7.62330053 | 10.0825919 | 10.7910037 |
| ENSMUSG00 Gm5670 | 371 | 0 | 0 | 0 | 0 | 0 | 0 | 0 | 0 | 0 | 0 | 0 | 0 | 0 | 0 | 0 | 0 | 0 | 0 |
| ENSMUSG00 Gm16380 | 457 | 0.83708224 | 0.65605262 | 0.21170035 | 0.69787158 | 1.26385795 | 0.65058294 | 0.62117303 | 0.25071599 | 0.72098934 | 0.41269762 | 0.41816156 | 1.22134929 | 0.64768382 | 1.4227948 | 0.63109604 | 0.83786887 | 0.41932491 | 0.60185018 |
| ENSMUSG00 Kprp | 2727 | 0 | 0 | 0 | 0 | 0 | 0 | 0 | 0 | 0 | 0 | 0 | 0 | 0 | 0 | 0 | 0 | 0 | 0 |
| ENSMUSG00 Sclt1 | 7366 | 2.68903231 | 2.65019767 | 2.63998903 | 3.10502909 | 2.72481838 | 2.89270641 | 3.23082779 | 2.76358274 | 4.12088815 | 3.14295348 | 3.54128799 | 3.70538406 | 3.89779743 | 3.68223555 | 3.65813434 | 5.23987677 | 5.026229 | 4.75710032 |
| ENSMUSG00 Rpl13-ps3 | 623 | 0 | 0.16041522 | 0.07764612 | 0.14626338 | 0.38629152 | 0.1590778 | 0.0759433 | 0 | 0 | 0.07568331 | 0 | 0 | 0.63347566 | 0.22364729 | 0.06613412 | 0 | 0 | 0.08829712 |
| ENSMUSG00 Zfp874b | 3767 | 1.74895203 | 2.13124702 | 1.48960168 | 1.13690952 | 1.59715701 | 1.30228846 | 1.80860785 | 1.51066312 | 2.897379 | 1.58963082 | 2.18138913 | 2.31145298 | 1.71555114 | 1.99733001 | 1.55312475 | 2.01262019 | 1.62787563 | 0.71554187 |
| ENSMUSG00 Zfp341 | 3330 | 0.49780837 | 0.29011229 | 0.45032416 | 0.23259392 | 0.24571853 | 0.35713682 | 0.42624035 | 0.82578167 | 0.27210311 | 0.21239056 | 0.27258985 | 0.28494482 | 0.19258706 | 0.71130584 | 0.37118519 | 0.32196307 | 0.25320677 | 0.36342353 |
| ENSMUSG00 Gm11567 | 1042 | 0 | 0 | 0 | 0 | 0 | 0 | 0 | 0 | 0 | 0 | 0 | 0 | 0 | 0 | 0 | 0 | 0 | 0 |
| ENSMUSG00 Suv420h2 | 3443 | 18.7156105 | 19.834863 | 20.0069437 | 21.7549686 | 19.3478311 | 16.9253601 | 17.3969943 | 19.800592 | 17.9196633 | 18.6521225 | 20.3699213 | 19.8426869 | 17.6666436 | 16.9562062 | 20.3554594 | 17.7495816 | 18.3115766 | 15.8013379 |
| ENSMUSG00 Kcng2 | 704 | 0.06037667 | 0.37855562 | 0.1374248 | 0.06471739 | 0.54695368 | 0.28154962 | 0.33602755 | 0 | 0.40952501 | 0.06697543 | 0.06786216 | 0 | 0 | 0.72568887 | 0.11704987 | 0.10878013 | 0.05444076 | 0 |
| ENSMUSG00 Hydin | 16424 | 0 | 0.02636798 | 0.0029453 | 0.0943178 | 0 | 0 | 0.00576141 | 0 | 0.03260014 | 0.00287084 | 0 | 0.04757789 | 0.02102554 | 0.02545036 | 0 | 0 | 0.00233355 | 0 |
| ENSMUSG00 Ntng1 | 6181 | 0.08939772 | 0 | 0.08608782 | 0.02211343 | 0 | 0 | 0 | 0 | 0 | 0 | 0 | 0 | 0 | 0 | 0 | 0 | 0 | 0 |
| ENSMUSG00 Olfr826 | 941 | 0 | 0 | 0 | 0 | 0 | 0 | 0 | 0 | 0 | 0 | 0 | 0 | 0 | 0 | 0 | 0 | 0 | 0 |
| ENSMUSG00 Olfr1393 | 935 | 0 | 0 | 0 | 0 | 0 | 0 | 0.20240718 | 0 | 0 | 0 | 0 | 0 | 0 | 0 | 0 | 0 | 0 | 0 |
| ENSMUSG00 Tnip2 | 3693 | 7.58486623 | 9.03859193 | 7.91161991 | 7.48864173 | 8.25006978 | 6.87002426 | 6.85412489 | 9.16286719 | 7.69528727 | 9.29480422 | 7.1668906 | 9.91472371 | 7.61418707 | 10.4005579 | 9.82901997 | 7.3201107 | 8.83175702 | 7.95420065 |
| ENSMUSG00 Olfr960 | 1066 | 0 | 0 | 0 | 0 | 0 | 0 | 0 | 0 | 0 | 0 | 0 | 0 | 0 | 0 | 0 | 0 | 0 | 0 |
| ENSMUSG00 Olfr1029 | 1060 | 0 | 0 | 0 | 0 | 0 | 0 | 0 | 0 | 0 | 0 | 0 | 0 | 0 | 0 | 0 | 0 | 0 | 0 |
| ENSMUSG00 Olfr603 | 938 | 0 | 0 | 0 | 0 | 0 | 0 | 0 | 0 | 0 | 0 | 0 | 0 | 0 | 0 | 0 | 0 | 0 | 0 |
| ENSMUSG00 Dph3b-ps | 248 | 0 | 0 | 0 | 0 | 0 | 0 | 0 | 0 | 0 | 0 | 0 | 0 | 0 | 0 | 0 | 0 | 0 | 0 |
| ENSMUSG00 Zfp422 | 5850 | 12.5989701 | 12.9607089 | 11.9238685 | 12.0795179 | 11.7902644 | 10.8168959 | 9.09050446 | 11.248805 | 12.3137121 | 10.3167351 | 10.0204913 | 11.5829426 | 9.18332614 | 9.58256641 | 7.47262407 | 9.17665497 | 9.77484359 | 10.4376254 |
| ENSMUSG00 Irak4 | 5187 | 1.00793072 | 1.50283733 | 1.36158385 | 0.87836985 | 0.57531893 | 1.95841727 | 0.90301816 | 1.25909019 | 1.40543923 | 0.70903313 | 0.60789421 | 1.70018791 | 1.1222607 | 1.69229468 | 0.80226666 | 1.04824875 | 0.65022443 | 1.45291067 |
| ENSMUSG00 Olfr1507 | 1603 | 0 | 0 | 0 | 0 | 0 | 0 | 0 | 0 | 0 | 0 | 0 | 0 | 0 | 0 | 0 | 0 | 0 | 0 |
| ENSMUSG00 Ube4a | 7175 | 7.15034808 | 8.89581956 | 8.38025062 | 6.87701896 | 7.33215227 | 8.32900302 | 7.25351175 | 7.57992767 | 7.38774157 | 7.34696672 | 8.58950473 | 7.1023972 | 6.83427537 | 7.54109572 | 7.10332051 | 6.81492762 | 7.75606733 | 7.22976831 |
| ENSMUSG00 Tsks | 2056 | 0.4341482 | 0.27544708 | 0.21175184 | 0.33240062 | 0.3745675 | 0.0964061 | 0.57530008 | 0.46440178 | 0.40064704 | 0.27519866 | 0.9759476 | 0.70584009 | 0.02399412 | 0.06776861 | 0.48095201 | 0.27935754 | 0.24233553 | 0.68888505 |
| ENSMUSG00 Ptp4a3 | 4611 | 26.1336313 | 22.844366 | 22.3561035 | 23.5957003 | 21.1484011 | 25.3083252 | 23.8769473 | 21.0054072 | 20.1779092 | 23.73385 | 21.9033316 | 21.0140946 | 36.8036858 | 27.8402641 | 33.2221611 | 41.6039781 | 27.1966262 | 35.2053511 |
| ENSMUSG00 Zfp930 | 1064 | 7.94974627 | 7.57680489 | 9.00184118 | 8.1787213 | 8.5497496 | 6.10094746 | 7.69275701 | 7.71744973 | 8.67084542 | 8.68565586 | 12.0335422 | 7.55399945 | 7.51106259 | 7.11502467 | 11.9654894 | 9.86055115 | 11.0584334 | 11.477464 |
| ENSMUSG00 Dsc3 | 5522 | 0 | 0.06032759 | 0 | 0 | 0 | 0 | 0 | 0 | 0 | 0 | 0 | 0 | 0 | 0 | 0 | 0 | 0 | 0 |
| ENSMUSG00 Tmem40 | 2384 | 0.57053928 | 1.00609412 | 0.5884364 | 0.30577882 | 0.74701392 | 1.18477593 | 1.19075534 | 1.39376636 | 0.76015381 | 1.40423655 | 1.36270655 | 1.1472179 | 3.45571736 | 2.80534756 | 2.3331418 | 2.28073249 | 3.4403639 | 2.67661761 |
| ENSMUSG00 Adamts14 | 5666 | 0 | 0 | 0.00853751 | 0 | 0 | 0.09620192 | 0 | 0 | 0 | 0 | 0 | 0 | 0 | 0 | 0 | 0 | 0 | 0.00970863 |
| ENSMUSG00 Mug1 | 5027 | 0 | 0 | 0 | 0 | 0 | 0 | 0 | 0 | 0 | 0 | 0 | 0 | 0 | 0.00923893 | 0.0163921 | 0.02285097 | 0.01524818 | 0 |
| ENSMUSG00 Olfr1265 | 1851 | 0 | 0 | 0 | 0 | 0 | 0 | 0 | 0 | 0 | 0 | 0 | 0 | 0 | 0 | 0 | 0 | 0 | 0 |
| ENSMUSG00 Rpl21-ps6 | 467 | 0 | 0 | 0 | 0.09756112 | 0.10306822 | 0 | 0 | 0.08178245 | 0 | 0 | 0.10230184 | 0 | 0 | 0 | 0.17645206 | 0.24597821 | 0.24620747 | 0 |
| ENSMUSG00 4930453N24 | 5938 | 15.2110979 | 12.2749432 | 13.7348892 | 16.5271959 | 14.4687578 | 11.9751049 | 12.8201584 | 11.6995588 | 14.1218838 | 13.7926527 | 13.484478 | 10.6029079 | 11.5645047 | 11.2395004 | 10.289981 | 14.9602912 | 12.7732808 | 11.1815412 |
| ENSMUSG00 Unc5c | 8029 | 0.10587913 | 0.174261 | 0.26509345 | 0.18726049 | 0.18583754 | 0.24686877 | 0.13553265 | 0.38530136 | 0.25135624 | 0.2466471 | 0.0892545 | 0.66736874 | 0.20275916 | 0.64208416 | 0.02565796 | 0.15260922 | 0.0859227 | 0.16443126 |
| ENSMUSG00 Grb2 | 4774 | 21.2614915 | 23.138994 | 25.6560073 | 23.0954603 | 21.9386397 | 24.3819378 | 20.3957883 | 20.9442207 | 20.2136587 | 22.7358439 | 22.4464248 | 21.0097917 | 21.5142547 | 19.5738783 | 21.0322982 | 20.9096478 | 21.0417311 | 19.3349981 |
| ENSMUSG00 Prh1 | 972 | 0 | 0 | 0 | 0 | 0 | 0 | 0 | 0 | 0 | 0 | 0 | 0 | 0 | 0 | 0 | 0 | 0 | 0 |
| ENSMUSG00 Gdap10 | 5001 | 1.6828684 | 2.85767479 | 2.73739434 | 2.29581747 | 2.17512792 | 1.65473036 | 2.59221636 | 4.18505029 | 3.09660666 | 3.08303939 | 4.93894292 | 2.52236348 | 2.73244173 | 4.92208885 | 5.61052981 | 2.58027088 | 4.94310368 | 3.35488453 |
| ENSMUSG00 9430015G10I | 3723 | 2.68297512 | 2.7380461 | 3.4172077 | 3.32866077 | 1.86166989 | 2.6752886 | 3.59642443 | 4.18546876 | 3.80557653 | 3.83740615 | 4.51699849 | 4.28774631 | 3.2596432 | 4.10424517 | 2.87735819 | 2.57121994 | 3.42805718 | 4.00415469 |
| ENSMUSG00 Olfr887 | 929 | 0 | 0 | 0 | 0 | 0 | 0 | 0 | 0 | 0 | 0 | 0 | 0 | 0 | 0 | 0 | 0 | 0 | 0 |
| ENSMUSG00 Serpinb12 | 2489 | 0 | 0.05353619 | 0 | 0.03660992 | 0 | 0 | 0.28513065 | 0.16878924 | 0.01654741 | 0.0568309 | 0 | 0.11212467 | 0.13873982 | 0.07463895 | 0.26485532 | 0 | 0.20017753 | 0.46411862 |
| ENSMUSG00 Olfr119 | 1233 | 0 | 0 | 0 | 0 | 0 | 0 | 0 | 0 | 0 | 0 | 0 | 0 | 0 | 0 | 0 | 0 | 0 | 0 |
| ENSMUSG00 Hspa2 | 2518 | 16.1546678 | 19.2891977 | 11.661133 | 16.031408 | 17.5859292 | 16.6684533 | 13.8104921 | 9.75286527 | 9.11075825 | 15.2612483 | 11.3840251 | 15.1398322 | 20.3361915 | 23.1113747 | 20.1753449 | 21.4111101 | 26.3017637 | 27.4390143 |
| ENSMUSG00 Gm14773 | 449 | 0 | 0 | 0 | 0 | 0 | 0 | 0 | 0 | 0 | 0 | 0 | 0 | 0 | 0 | 0 | 0 | 0 | 0 |
| ENSMUSG00 Ntm | 8123 | 0 | 0.00410106 | 0 | 0 | 0 | 0 | 0.02329813 | 0 | 0 | 0 | 0 | 0 | 0.00607312 | 0.0057176 | 0 | 0.0377108 | 0 | 0 |
| ENSMUSG00 Zfp74 | 6145 | 2.11661251 | 1.47454959 | 1.99949173 | 1.42355094 | 1.3393912 | 1.57246266 | 1.70156258 | 1.11252075 | 2.41287969 | 1.51925781 | 1.6171182 | 1.689457 | 1.05969293 | 0.74068683 | 1.89748419 | 0.81005353 | 0.87941544 | 1.95150289 |
| ENSMUSG00 Taok2 | 9636 | 5.81821577 | 6.4682882 | 5.87349839 | 6.71878829 | 6.12388319 | 5.76469638 | 4.36007253 | 4.97024415 | 5.60779459 | 5.54886857 | 3.83746553 | 4.54704184 | 4.36185072 | 4.16435155 | 4.2843461 | 5.9327394 | 3.97342991 | 4.59550964 |
| ENSMUSG00 Olfr996 | 1067 | 0 | 0 | 0 | 0 | 0 | 0 | 0 | 0 | 0 | 0 | 0 | 0 | 0 | 0 | 0 | 0 | 0 | 0 |
| ENSMUSG00 Nptx2 | 2546 | 0 | 0.13084405 | 0 | 0 | 0.09452459 | 0 | 0.26016398 | 0.25501604 | 0.01617695 | 0.05555857 | 0 | 0.30692038 | 0 | 0 | 0 | 0 | 0 | 0 |
| ENSMUSG00 Fcrl1 | 2031 | 0.14649741 | 0 | 0.07145278 | 0.13459688 | 0 | 0.04879639 | 0.48920052 | 0 | 0.14195254 | 0.2089396 | 0.14113725 | 0.02748186 | 0.89871046 | 0.82323345 | 0 | 0.0942654 | 0 | 0 |
| ENSMUSG00 Atxn7l3 | 6610 | 4.50773501 | 3.95622117 | 4.35434956 | 3.58422739 | 3.32772906 | 3.95072475 | 3.39277002 | 4.2583662 | 4.40527483 | 3.43822072 | 2.88384394 | 4.08695623 | 3.03006935 | 2.72621875 | 3.14154072 | 4.50682187 | 2.84693068 | 3.36212997 |
| ENSMUSG00 Olfr484 | 1097 | 0 | 0 | 0 | 0 | 0 | 0 | 0 | 0 | 0 | 0 | 0 | 0 | 0 | 0 | 0 | 0 | 0 | 0 |
| ENSMUSG00 Gm10072 | 802 | 0 | 0 | 0 | 0 | 0 | 0 | 0 | 0 | 0 | 0.0587914 | 0 | 0 | 0.06151112 | 0 | 0 | 0 | 0 | 0 |
| ENSMUSG00 Chpt1 | 5097 | 10.1738895 | 10.529149 | 11.3317628 | 10.9410866 | 9.53761878 | 11.1802322 | 10.6933895 | 10.0108004 | 12.4925159 | 12.8029377 | 10.7697523 | 10.9835412 | 8.23650414 | 4.92961571 | 7.77631877 | 7.84292606 | 7.95550758 | 5.87109163 |
| ENSMUSG00 Kif13b | 5492 | 1.88069151 | 3.46958767 | 2.95948765 | 3.60871345 | 3.22515439 | 3.37449424 | 1.58512981 | 2.71213346 | 2.69227224 | 2.6013589 | 1.93987905 | 1.94114923 | 1.66176342 | 1.91120971 | 2.70075749 | 4.10654912 | 2.91704153 | 3.65592206 |
| ENSMUSG00 Olfr121 | 1240 | 0 | 0 | 0 | 0 | 0 | 0 | 0 | 0 | 0 | 0 | 0 | 0 | 0 | 0 | 0 | 0 | 0 | 0 |
| ENSMUSG00 Gm10073 | 344 | 4.69533921 | 4.64831082 | 3.93738038 | 4.63557137 | 3.77779633 | 5.18575119 | 10.3152643 | 4.7740503 | 8.14152056 | 8.49809187 | 8.05508021 | 7.95048682 | 9.17803112 | 10.9359624 | 8.02472164 | 11.3536103 | 10.4728835 | 12.3130849 |
| ENSMUSG00 Vmn1r213 | 1154 | 0 | 0 | 0 | 0 | 0 | 0 | 0 | 0 | 0 | 0 | 0 | 0 | 0 | 0 | 0 | 0 | 0 | 0 |
| ENSMUSG00 4930473A02I | 1395 | 0.73127185 | 0.69252612 | 0.41611639 | 0.26128197 | 0.37953488 | 0.46178175 | 0.50873848 | 0.60231746 | 0.5609633 | 0.64219596 | 0.54795651 | 0.60016841 | 0.67190425 | 0.76574481 | 0.35442199 | 0.41172696 | 0.6868512 | 0.35489746 |
| ENSMUSG00 Olfr317 | 1127 | 0 | 0 | 0 | 0 | 0 | 0 | 0 | 0 | 0 | 0 | 0 | 0 | 0 | 0 | 0 | 0 | 0 | 0 |
| ENSMUSG00 H2afj | 1826 | 68.2503703 | 62.904085 | 58.1489042 | 51.5743037 | 56.7513864 | 60.4890707 | 71.927709 | 69.377984 | 66.6968933 | 69.7190026 | 60.2550544 | 65.1692183 | 81.6975454 | 85.8935833 | 74.7991006 | 74.505218 | 60.1340858 | 79.2902349 |
| ENSMUSG00 Ctf2 | 1751 | 0.26702281 | 0.13317548 | 0.16575739 | 0.02602001 | 0.13744124 | 0.39619547 | 0.02702038 | 0.19630589 | 0 | 0.10771149 | 0.08185316 | 0.22313514 | 0 | 0.05304864 | 0.1411818 | 0.10933926 | 0.10944117 | 0.09424747 |
| ENSMUSG00 Rpl3 | 1351 | 138.778928 | 126.988457 | 139.176841 | 118.674548 | 122.984012 | 136.040776 | 140.04693 | 134.874872 | 174.288165 | 123.687707 | 119.455078 | 147.616108 | 180.238596 | 157.517993 | 174.687278 | 131.622192 | 125.702311 | 172.519308 |
| ENSMUSG00 Dhps | 2573 | 27.3896548 | 25.5446328 | 23.4817487 | 26.1714821 | 24.0191955 | 24.0541641 | 21.33024 | 25.6347761 | 22.0739235 | 22.9614579 | 21.4458131 | 22.3219265 | 28.0308057 | 23.7545352 | 21.95388 | 27.5311402 | 20.4069284 | 23.23937 |
| ENSMUSG00 Tmem26 | 5393 | 0 | 0 | 0 | 0 | 0 | 0 | 0 | 0 | 0 | 0 | 0 | 0 | 0 | 0 | 0 | 0 | 0 | 0 |
| ENSMUSG00 Olfr235 | 1078 | 0 | 0 | 0 | 0 | 0 | 0 | 0 | 0 | 0 | 0 | 0 | 0 | 0 | 0 | 0 | 0 | 0 | 0 |
| ENSMUSG00 Olfr193 | 1006 | 0 | 0 | 0 | 0 | 0 | 0 | 0 | 0 | 0 | 0 | 0 | 0 | 0 | 0 | 0 | 0 | 0 | 0 |
| ENSMUSG00 Olfr497 | 944 | 0 | 0 | 0 | 0 | 0 | 0 | 0 | 0 | 0 | 0 | 0 | 0 | 0 | 0 | 0 | 0 | 0 | 0 |
| ENSMUSG00 Alox5ap | 877 | 0 | 0 | 0.55157959 | 0 | 0 | 0 | 0 | 0.91452731 | 0.2348148 | 0 | 0.05447544 | 0 | 0 | 0 | 0.65772153 | 0 | 0 | 0 |
| ENSMUSG00 Gm12222 | 519 | 0.4094911 | 0.44930686 | 0 | 0.35114485 | 0.37095895 | 0.09547733 | 0.09116123 | 0 | 0.07935745 | 0.4542457 | 0.6443636 | 0.10754463 | 0.47525932 | 0.44743822 | 0.15877286 | 0.22133299 | 0.22153929 | 0.10599057 |
| ENSMUSG00 Defa26 | 370 | 0 | 0 | 0 | 0 | 0 | 0 | 0 | 0 | 0 | 0 | 0 | 0 | 0 | 0 | 0 | 0 | 0 | 0 |
| ENSMUSG00 Psma3 | 3888 | 49.0755492 | 46.3450218 | 46.8058696 | 44.1431209 | 50.2116981 | 45.2066734 | 51.7421581 | 47.9566123 | 49.2903441 | 44.7131283 | 50.9206348 | 44.1873995 | 45.9695309 | 45.7034667 | 46.3623475 | 46.6026631 | 48.5584718 | 48.8120932 |
| ENSMUSG00 Hist1h2aa | 389 | 0 | 0 | 0 | 0 | 0 | 0 | 0 | 0 | 0 | 0 | 0 | 0 | 0 | 0 | 0 | 0 | 0 | 0 |
| ENSMUSG00 Olfr748 | 923 | 0 | 0 | 0 | 0 | 0 | 0 | 0 | 0 | 0 | 0 | 0 | 0 | 0 | 0 | 0 | 0 | 0 | 0 |
| ENSMUSG00 Gm10074 | 271 | 0 | 0 | 0 | 0 | 0 | 0 | 0.17458553 | 0 | 0 | 0 | 0 | 0 | 0.1820366 | 0 | 0 | 0 | 0 | 0 |
| ENSMUSG00 Olfr805 | 959 | 0 | 0 | 0 | 0 | 0 | 0 | 0 | 0 | 0 | 0 | 0 | 0 | 0 | 0 | 0 | 0 | 0 | 0 |
| ENSMUSG00 Rp2h | 5407 | 4.91321158 | 5.25539093 | 5.39471774 | 6.44612519 | 6.28465658 | 5.63619958 | 4.83889614 | 6.62557305 | 3.97620885 | 4.93569411 | 5.32796374 | 3.79880966 | 3.93232039 | 3.45302814 | 4.37390292 | 3.7886952 | 6.30856376 | 3.84565237 |
| ENSMUSG00 Hist1h4a | 538 | 0.0790059 | 0 | 0.53948175 | 0 | 0 | 0 | 0 | 0 | 0.6124389 | 0.17528142 | 0 | 0.10374658 | 0.36678005 | 0.6906184 | 0.61266254 | 0.85406559 | 0.21371541 | 0.30674223 |
| ENSMUSG00 Amd-ps3 | 1335 | 0.03183908 | 0.02495348 | 0.10870456 | 0.10238437 | 0.10816163 | 0.03711815 | 0.17720104 | 0.05721708 | 0 | 0.10595664 | 0.10735946 | 0.04180949 | 0.11085824 | 0.06957916 | 0.03086259 | 0.14341051 | 0.08612651 | 0.16482129 |
| ENSMUSG00 Prmt7 | 6063 | 10.2284433 | 11.2746262 | 9.61406962 | 11.557461 | 10.645705 | 10.8373618 | 11.2682679 | 10.3559805 | 10.1760516 | 10.957503 | 11.4729242 | 12.9343569 | 9.75572634 | 10.3413356 | 10.4719772 | 10.0036817 | 9.98139911 | 8.56483535 |
| ENSMUSG00 Olfr898 | 950 | 0 | 0 | 0 | 0 | 0 | 0 | 0 | 0 | 0 | 0 | 0 | 0 | 0 | 0 | 0 | 0 | 0 | 0 |
| ENSMUSG00 Olfr504 | 956 | 0 | 0 | 0 | 0 | 0 | 0 | 0 | 0 | 0 | 0 | 0 | 0 | 0 | 0 | 0 | 0 | 0 | 0 |
| ENSMUSG00 Olfr229 | 923 | 0 | 0 | 0 | 0 | 0 | 0 | 0 | 0 | 0 | 0 | 0 | 0 | 0 | 0 | 0 | 0 | 0 | 0 |
| ENSMUSG00 Olfr741 | 935 | 0 | 0 | 0 | 0 | 0 | 0 | 0 | 0 | 0 | 0 | 0 | 0 | 0 | 0 | 0 | 0 | 0 | 0 |
| ENSMUSG00 Olfr60 | 935 | 0 | 0 | 0 | 0 | 0 | 0 | 0 | 0 | 0 | 0 | 0 | 0 | 0 | 0 | 0 | 0 | 0 | 0 |
| ENSMUSG00 Olfr912 | 993 | 0 | 0 | 0 | 0 | 0 | 0 | 0 | 0 | 0 | 0 | 0 | 0 | 0 | 0 | 0 | 0 | 0 | 0 |
| ENSMUSG00 Sip1 | 1732 | 7.90223249 | 6.48178138 | 7.90398908 | 7.89163585 | 8.53146684 | 6.0081259 | 9.45160907 | 8.18093608 | 6.99124459 | 8.03086458 | 6.34424968 | 8.21766395 | 8.51630674 | 6.67700794 | 8.89687177 | 8.06932546 | 7.67853639 | 8.09891583 |
| ENSMUSG00 Gm5528 | 482 | 0 | 0 | 0 | 0 | 0 | 0.411226 | 0 | 0.1584747 | 0 | 0 | 0 | 0 | 0.10234838 | 0.19271406 | 0.08548041 | 0 | 0.07951514 | 0 |
| ENSMUSG00 Tpt1 | 2530 | 104.09568 | 91.2878639 | 92.6936265 | 83.1804205 | 93.010662 | 81.7130453 | 104.293601 | 93.1411553 | 98.668566 | 90.9840843 | 85.9571585 | 89.7242291 | 124.109744 | 120.38748 | 94.8775749 | 103.535986 | 94.3160176 | 108.561451 |
| ENSMUSG00 Gm10075 | 663 | 139.311836 | 145.310543 | 132.206391 | 126.924961 | 145.339534 | 140.511522 | 124.739914 | 131.225177 | 137.971722 | 146.572548 | 121.851365 | 122.912319 | 131.849408 | 112.502571 | 131.186251 | 138.319766 | 134.055328 | 128.271612 |
| ENSMUSG00 Olfr951 | 944 | 0 | 0 | 0 | 0 | 0 | 0 | 0 | 0 | 0 | 0 | 0 | 0 | 0 | 0 | 0 | 0 | 0 | 0 |
| ENSMUSG00 Atp8b4 | 7184 | 0 | 0 | 0 | 0 | 0 | 0 | 0 | 0 | 0 | 0 | 0 | 0 | 0 | 0 | 0 | 0 | 0 | 0 |
| ENSMUSG00 Olfr105-ps | 952 | 0 | 0 | 0 | 0 | 0 | 0 | 0 | 0 | 0 | 0 | 0 | 0 | 0 | 0 | 0 | 0 | 0 | 0 |
| ENSMUSG00 Gm10076 | 376 | 6.33055813 | 4.34130802 | 6.56130332 | 5.81630353 | 6.27251133 | 4.87619985 | 7.92739035 | 6.80556107 | 5.8055461 | 5.7684371 | 5.33656451 | 9.3520924 | 12.3329793 | 11.3639788 | 8.76628849 | 7.12857053 | 6.93135157 | 7.31504079 |
| ENSMUSG00 Serpinb6a | 5883 | 18.2722404 | 17.1802348 | 17.7361389 | 15.411606 | 16.2321497 | 17.0482293 | 15.4572444 | 12.7567688 | 16.2281733 | 15.260061 | 13.1314139 | 14.6488837 | 18.1378441 | 16.1208272 | 17.9499554 | 20.4112434 | 18.9318747 | 18.4579257 |
| ENSMUSG00 BC002059 | 1483 | 4.09861104 | 3.90859312 | 5.15375441 | 2.42705496 | 4.54380938 | 3.50845383 | 5.0088271 | 5.35672266 | 4.55467875 | 3.65632561 | 2.41613074 | 3.80133643 | 6.02095551 | 2.53673033 | 6.1955138 | 4.67336476 | 4.00578289 | 6.56548341 |
| ENSMUSG00 Vmn1r49 | 932 | 0 | 0 | 0 | 0 | 0 | 0 | 0 | 0 | 0 | 0 | 0 | 0 | 0 | 0 | 0 | 0 | 0 | 0 |
| ENSMUSG00 Pop5 | 2026 | 28.3227975 | 23.0362116 | 24.3777762 | 23.1853093 | 27.0356017 | 23.528988 | 37.9949302 | 36.7974183 | 31.7132106 | 32.9542921 | 34.0980207 | 34.9054514 | 43.0254185 | 46.4440874 | 40.4084361 | 38.4418043 | 34.6185209 | 34.8354807 |
| ENSMUSG00 Klk1b7-ps | 445 | 0 | 0.29944175 | 0 | 0 | 0 | 0.11135446 | 0 | 0 | 0 | 0.74169646 | 0 | 0 | 0 | 0 | 0 | 0 | 0 | 0 |
| ENSMUSG00 Zdhhc8 | 4856 | 4.963022 | 5.39207884 | 6.73404171 | 5.7608075 | 4.33147666 | 5.02058174 | 4.02391606 | 5.25381483 | 4.8938673 | 4.91314989 | 3.83695095 | 5.13789151 | 5.05916285 | 3.634422 | 4.76838435 | 3.7533267 | 3.7252908 | 3.5570139 |
| ENSMUSG00 Olfr1371 | 937 | 0 | 0 | 0 | 0 | 0 | 0 | 0 | 0 | 0 | 0 | 0 | 0 | 0 | 0 | 0 | 0 | 0 | 0 |
| ENSMUSG00 Kif27 | 4445 | 0.04781235 | 0.13490036 | 0.17412294 | 0.03074986 | 0 | 0 | 0 | 0.01718443 | 0.1204555 | 0.12729099 | 0 | 0.15068345 | 0.16647441 | 0.17762643 | 0 | 0.05168586 | 0 | 0.06187751 |
| ENSMUSG00 Klk1b22 | 850 | 0.30003654 | 0.43110804 | 0.05691004 | 0.53601229 | 0.22650317 | 0.174892 | 0.27830988 | 0.26959343 | 0 | 0 | 0.11241167 | 0.19699646 | 0 | 0.16392031 | 0.29083451 | 0 | 0 | 0.32358298 |
| ENSMUSG00 Myh13 | 6022 | 0 | 0 | 0.06426241 | 0 | 0.01598536 | 0.0493717 | 0 | 0 | 0 | 0 | 0 | 0 | 0 | 0.13882324 | 0.33525012 | 0 | 0.0954657 | 0.01826938 |
| ENSMUSG00 Slc35e3 | 3477 | 2.23711453 | 3.13296417 | 3.36680884 | 2.476571 | 3.28077823 | 2.82181227 | 2.99361213 | 2.83394875 | 3.28118058 | 3.58003614 | 2.77553685 | 3.80451885 | 2.35521952 | 2.4043531 | 2.18035268 | 1.64086871 | 2.61240505 | 3.51222942 |
| ENSMUSG00 Cxcl11 | 2841 | 0.40395627 | 0.48075631 | 0.54486201 | 1.04240334 | 0.06776758 | 1.41280233 | 0.01665353 | 0.04032989 | 0.07248595 | 0.43150943 | 0.01681625 | 0.53045508 | 0 | 0.06539118 | 0 | 0.08086718 | 0 | 0 |
| ENSMUSG00 Lrrc10 | 1427 | 0 | 0 | 0 | 0.09578356 | 0.03372945 | 0 | 0 | 0 | 0 | 0 | 0 | 0 | 0 | 0 | 0 | 0 | 0 | 0 |
| ENSMUSG00 Cxcl17 | 757 | 0 | 0 | 0.12780325 | 0.06018632 | 0.06358246 | 0 | 0.31250118 | 0.05045231 | 0 | 0 | 0 | 0.07373271 | 0.26067063 | 0 | 0 | 0 | 0.40503352 | 0 |
| ENSMUSG00 Gm11353 | 668 | 0 | 0.24934801 | 0 | 0 | 0 | 0 | 0 | 0 | 0 | 0 | 0.0715194 | 0 | 0 | 0.06952708 | 0 | 0 | 0 | 0 |
| ENSMUSG00 Spink7 | 254 | 0 | 0 | 0 | 0 | 0 | 0 | 0 | 0 | 0 | 0 | 0 | 0 | 0.19422015 | 0 | 0 | 0 | 0 | 0.21657129 |
| ENSMUSG00 Gm5868 | 1377 | 0 | 0 | 0 | 0 | 0 | 0.10795803 | 0 | 0 | 0 | 0 | 0 | 0 | 0 | 0 | 0 | 0 | 0 | 0 |
| ENSMUSG00 Olfr57 | 1079 | 0 | 0 | 0 | 0 | 0 | 0.09184937 | 0 | 0 | 0.11451302 | 0 | 0.13283121 | 0.20691627 | 0 | 0 | 0 | 0.10646137 | 0.1065606 | 0 |
| ENSMUSG00 Zfp462 | 10688 | 1.16921049 | 1.03167739 | 1.47999106 | 1.01881452 | 1.0537865 | 0.8530785 | 1.08897072 | 1.09703102 | 0.65895342 | 0.80290307 | 0.89846245 | 1.14367797 | 1.103137 | 0.74741608 | 1.09094689 | 1.45452719 | 1.31961801 | 0.9624535 |
| ENSMUSG00 Defa17 | 395 | 0 | 0 | 0 | 0 | 0 | 0 | 0 | 0 | 0 | 0 | 0 | 0 | 0 | 0 | 0 | 0 | 0 | 0 |
| ENSMUSG00 Gm6570 | 463 | 0.09180384 | 0.14390019 | 0.10447847 | 0 | 0 | 0.64215206 | 0.10218721 | 0.08248899 | 0.17791152 | 0 | 0 | 0.24110437 | 0.10654842 | 0.40124482 | 0.17797648 | 0 | 0.08277818 | 0 |
| ENSMUSG00 Pcnxl2 | 8540 | 0.16922435 | 0.17163552 | 0.30021043 | 0.0640202 | 0.07326874 | 0.26691168 | 0.3157872 | 0.35777426 | 0.28454384 | 0.27605798 | 0.23495881 | 0.21568113 | 0.31193484 | 0.30455139 | 0.23640237 | 0.24660227 | 0.37697997 | 0.16103368 |
| ENSMUSG00 Gm8203 | 1054 | 1.65342715 | 1.61191424 | 1.23917013 | 0.90776274 | 1.36997885 | 1.59847528 | 1.57110414 | 1.9567265 | 1.56305563 | 0.93943526 | 1.63178226 | 1.85346128 | 1.02969846 | 1.23380877 | 1.60271707 | 1.52581168 | 1.92722366 | 1.35696089 |
| ENSMUSG00 Arrb2 | 2465 | 6.70771338 | 5.16248503 | 6.08348658 | 5.69282014 | 4.37385432 | 6.67403957 | 4.721671 | 5.73273383 | 5.44697937 | 4.93504318 | 4.78718651 | 4.347508 | 4.32279681 | 3.59871022 | 6.4685607 | 4.75331678 | 4.91322914 | 4.5524778 |
| ENSMUSG00 Pyroxd2 | 3103 | 1.38350718 | 1.09504196 | 0.37414268 | 0.57263317 | 0.7755708 | 0.94218862 | 1.57048209 | 1.88315745 | 0.70347578 | 1.59549592 | 0.61585509 | 0.95334519 | 1.41493414 | 2.17028446 | 1.56680103 | 1.09824817 | 0.70402801 | 1.45367282 |
| ENSMUSG00 Casc4 | 4893 | 14.5853649 | 16.0062568 | 16.0256474 | 14.9728509 | 14.3421918 | 17.9759048 | 14.3591515 | 15.9388689 | 12.9881042 | 14.7243561 | 13.8647948 | 16.3351786 | 14.417462 | 11.4947455 | 15.7547744 | 12.8496173 | 12.5639445 | 11.9619231 |
| ENSMUSG00 Ccnd1 | 3325 | 0.01278351 | 0.03005675 | 0 | 0 | 0.02895153 | 0 | 0 | 0 | 0 | 0.05672265 | 0 | 0.05036 | 0 | 0 | 0.11515972 | 0 | 0 | 0 |
| ENSMUSG00 Alyref2 | 1272 | 5.14606691 | 3.84983919 | 3.72689149 | 4.22657487 | 6.54624434 | 5.14226481 | 8.0342285 | 6.30534945 | 7.96531673 | 5.5972926 | 7.39910914 | 8.42500567 | 6.12770671 | 5.84202357 | 6.89931714 | 5.11745539 | 6.29732394 | 5.23278452 |
| ENSMUSG00 Vmn1r228 | 1439 | 0 | 0 | 0 | 0 | 0 | 0 | 0 | 0 | 0 | 0 | 0 | 0 | 0 | 0 | 0 | 0 | 0 | 0 |
| ENSMUSG00 Olfr980 | 936 | 0 | 0 | 0 | 0 | 0 | 0 | 0 | 0 | 0 | 0 | 0 | 0 | 0 | 0 | 0 | 0 | 0 | 0 |
| ENSMUSG00 Tdpoz4 | 1112 | 0 | 0 | 0 | 0 | 0 | 0 | 0 | 0 | 0 | 0 | 0 | 0 | 0 | 0 | 0 | 0 | 0 | 0 |
| ENSMUSG00 Scrt2 | 3393 | 0.05010926 | 0.06872687 | 0.11405489 | 0.09399567 | 0.02837131 | 0.08762641 | 0.23705144 | 0.12381858 | 0.10924805 | 0.11117171 | 0.05632179 | 0.19740287 | 0.23262914 | 0.3832698 | 0.15786037 | 0.05642589 | 0.12425266 | 0.34046308 |
| ENSMUSG00 Pwwp2b | 2637 | 20.3257592 | 25.1267905 | 23.3887945 | 20.7676812 | 19.6214706 | 24.3347706 | 19.7360435 | 18.2344462 | 19.5702329 | 21.9035311 | 18.8599665 | 18.1818939 | 22.9729217 | 20.6594291 | 23.9834616 | 23.6539247 | 23.0364735 | 24.7822597 |
| ENSMUSG00 Gtf2i | 8067 | 38.8168627 | 36.3852623 | 43.4264419 | 35.2594025 | 39.1463433 | 36.4627528 | 33.3247355 | 35.4559194 | 37.6177325 | 34.4731434 | 36.7476903 | 38.7187861 | 31.5425845 | 26.4432495 | 30.3125368 | 29.984104 | 29.2946506 | 30.1537461 |
| ENSMUSG00 Hist1h2ao | 518 | 0 | 0.06431061 | 0 | 0 | 0 | 0 | 0 | 0.07373051 | 0 | 0 | 0 | 0 | 0 | 0.0896604 | 0 | 0 | 0 | 0 |
| ENSMUSG00 Gm1661 | 2376 | 0 | 0.07010289 | 0.2035923 | 0.03835105 | 0 | 0 | 0.07965097 | 0.25718789 | 0 | 0.11906743 | 0.08042922 | 0.04698288 | 0.02076259 | 0 | 0 | 0.11280903 | 0 | 0 |
| ENSMUSG00 Nrg2 | 3930 | 0.05407783 | 0.01695313 | 0.01230879 | 0.06955885 | 0.09797847 | 0.1008707 | 0.0240777 | 0.10689985 | 0.23056065 | 0.09598108 | 0.02431296 | 0.15622705 | 0.33892157 | 0.7327057 | 1.16370807 | 0.30203787 | 0.1267791 | 0.62987527 |
| ENSMUSG00 Ap2a1 | 4492 | 24.0723882 | 32.1189103 | 30.174228 | 32.3146677 | 29.3699027 | 32.0570445 | 22.0237782 | 24.5631902 | 24.1140998 | 27.3856155 | 25.6636186 | 21.9435575 | 29.1686492 | 30.4492069 | 34.1206117 | 26.1947989 | 30.8692213 | 29.0597975 |
| ENSMUSG00 Sp7 | 4305 | 0 | 0 | 0 | 0 | 0 | 0 | 0 | 0.01774328 | 0 | 0 | 0 | 0 | 0 | 0.05394203 | 0 | 0 | 0 | 0 |
| ENSMUSG00 Ppih | 2054 | 5.93913609 | 5.49808722 | 6.21743525 | 4.54723179 | 5.36621739 | 5.54874816 | 7.0945984 | 6.76827384 | 7.71996522 | 6.45051004 | 6.79176625 | 6.98375136 | 7.25328092 | 7.21307881 | 7.62248843 | 5.685801 | 8.02351883 | 6.5346748 |
| ENSMUSG00 2610008E11I | 4223 | 6.3410516 | 7.42302474 | 7.34251313 | 5.59938006 | 7.28304504 | 6.31289857 | 5.27688178 | 6.99998093 | 5.15928649 | 6.93359856 | 4.09531967 | 8.08884336 | 6.03945092 | 6.96166406 | 7.86371159 | 5.00507111 | 7.69611645 | 6.59119299 |
| ENSMUSG00 Olfr1447 | 1102 | 0 | 0 | 0 | 0 | 0 | 0 | 0 | 0 | 0 | 0 | 0 | 0 | 0 | 0 | 0 | 0 | 0 | 0 |
| ENSMUSG00 Zfp941 | 4711 | 1.36240322 | 2.22745949 | 0.8419931 | 1.74081681 | 1.72665997 | 1.65140717 | 1.37589408 | 1.64573502 | 2.04577472 | 2.04176257 | 1.78484243 | 1.55208062 | 1.94772588 | 1.47879709 | 1.67919735 | 1.78814129 | 2.14776956 | 1.03922957 |
| ENSMUSG00 Acnat2 | 2525 | 0.03366747 | 0 | 0 | 0 | 0 | 0 | 0 | 0 | 0 | 0 | 0 | 0.02210521 | 0.01953739 | 0 | 0.01631745 | 0 | 0.01517873 | 0 |
| ENSMUSG00 Gm2228 | 437 | 0 | 0 | 0 | 0 | 0 | 0 | 0 | 0 | 0 | 0 | 0 | 0 | 0 | 0.21255875 | 0 | 0 | 0 | 0 |
| ENSMUSG00 Tmc2 | 3201 | 0 | 0 | 0 | 0 | 0 | 0.01548039 | 0 | 0 | 0.01286677 | 0 | 0 | 0.01743695 | 0 | 0 | 0 | 0 | 0 | 0 |
| ENSMUSG00 Olfr384 | 938 | 0 | 0 | 0 | 0 | 0 | 0 | 0 | 0 | 0 | 0 | 0 | 0 | 0 | 0 | 0 | 0 | 0 | 0 |
| ENSMUSG00 Zfp937 | 2793 | 1.81099747 | 1.5863283 | 2.23422321 | 0.94612981 | 1.63714026 | 0.92257148 | 0.84698673 | 1.42213027 | 1.91702365 | 1.87387328 | 0.9921044 | 2.43806331 | 2.79070638 | 2.0619645 | 2.38977875 | 1.80965277 | 1.71528361 | 1.59532318 |
| ENSMUSG00 Gm15454 | 1133 | 0.1500624 | 0.38223091 | 0 | 0.36191474 | 0.12744552 | 0.17494345 | 0.08351753 | 0 | 0 | 0 | 0 | 0 | 0 | 0.16396853 | 0 | 0 | 0 | 0.09710345 |
| ENSMUSG00 Calm5 | 586 | 0 | 0 | 0 | 0 | 0 | 0 | 0 | 0 | 0 | 0 | 0 | 0 | 0 | 0 | 0 | 0 | 0 | 0 |
| ENSMUSG00 Caln1 | 3555 | 0.01195645 | 0 | 0 | 0 | 0 | 0.01393888 | 0.27948418 | 0.05371646 | 0.03475655 | 0.13263208 | 0 | 0 | 0.19427478 | 0.01306444 | 0.13907698 | 0.02154183 | 0.07546669 | 0.07736865 |
| ENSMUSG00 Hnrnpc | 3193 | 51.1180319 | 47.0637225 | 47.6160369 | 51.9107671 | 51.6894351 | 49.9872705 | 47.786843 | 47.0676804 | 48.0874822 | 46.043186 | 53.0716498 | 49.260425 | 35.5350485 | 32.6257714 | 42.2338529 | 38.5424404 | 42.0472966 | 41.0888567 |
| ENSMUSG00 3110018I06R | 1004 | 0 | 0 | 0 | 0 | 0 | 0 | 0 | 0 | 0 | 0 | 0 | 0 | 0 | 0 | 0 | 0 | 0 | 0 |
| ENSMUSG00 Bckdha | 1764 | 36.7703507 | 38.8461584 | 31.6182996 | 43.2881578 | 38.8820812 | 33.7374337 | 40.1514061 | 44.4495477 | 41.0230772 | 47.3378092 | 53.2457871 | 44.899334 | 54.2259567 | 45.1013162 | 46.4101426 | 48.3408065 | 46.9736133 | 48.0238233 |
| ENSMUSG00 Rpl36a-ps1 | 320 | 0 | 0 | 0.30233456 | 0 | 0.30082452 | 0.15485229 | 0 | 0 | 0 | 0.14734595 | 0 | 0.34884789 | 0 | 0.29027555 | 0.25750972 | 0 | 0.23953936 | 0 |
| ENSMUSG00 C030014I23F | 3077 | 2.34835227 | 2.19776325 | 2.51536069 | 2.25065932 | 1.72067325 | 1.61042358 | 3.04449478 | 2.22178747 | 2.81090943 | 2.39048089 | 1.75449149 | 2.43071134 | 2.85378007 | 2.9282265 | 3.82958888 | 3.60879953 | 2.65307158 | 2.19893407 |
| ENSMUSG00 Gm5580 | 2697 | 0.09456101 | 0.074111 | 0.01793605 | 0.03378646 | 0.12492527 | 0.11023967 | 0.07017083 | 0.01416107 | 0.16798356 | 0.12237854 | 0.10628467 | 0.04139093 | 0.0182914 | 0.12054454 | 0.06110724 | 0.08518489 | 0.099475 | 0.06118922 |
| ENSMUSG00 Zfp128 | 4084 | 1.08240409 | 2.27578293 | 1.42135741 | 1.13791051 | 1.16676309 | 1.09200441 | 1.54078998 | 1.29053661 | 2.01696943 | 1.38542713 | 2.02376783 | 1.50336016 | 0.5314898 | 1.39878128 | 1.67469595 | 1.56575207 | 1.56721146 | 0.60612385 |
| ENSMUSG00 Chst8 | 3870 | 2.37238192 | 4.01992806 | 3.06240696 | 3.29640631 | 4.80075518 | 3.36753719 | 0.12225498 | 0.10855722 | 0.01064251 | 0.20712195 | 0.18517426 | 0.11538121 | 1.6953863 | 3.85233903 | 0.93688292 | 0.51449912 | 0.75266113 | 1.13713916 |
| ENSMUSG00 Olfr1369-ps1 | 953 | 0 | 0 | 0 | 0 | 0 | 0 | 0 | 0 | 0 | 0 | 0 | 0 | 0 | 0 | 0 | 0 | 0 | 0 |
| ENSMUSG00 Cyp2a12 | 1647 | 0 | 0 | 0 | 0 | 0 | 0 | 0 | 0 | 0 | 0 | 0 | 0 | 0 | 0 | 0 | 0 | 0 | 0 |
| ENSMUSG00 Npr2 | 74 | 0 | 2.25087123 | 0 | 0 | 0 | 0 | 0 | 0 | 0 | 0 | 0 | 0 | 0 | 0 | 0 | 0 | 0 | 0 |
| ENSMUSG00 Tas2r124 | 929 | 0 | 0 | 0 | 0 | 0 | 0 | 0 | 0 | 0 | 0 | 0 | 0 | 0 | 0 | 0 | 0 | 0 | 0 |
| ENSMUSG00 Gm839 | 1840 | 0 | 0 | 0 | 0 | 0 | 0 | 0 | 0 | 0 | 0 | 0 | 0 | 0 | 0 | 0 | 0 | 0 | 0 |
| ENSMUSG00 Gm8603 | 1392 | 0 | 0.14359006 | 0.0695022 | 0 | 0 | 0 | 0.10196698 | 0.02743707 | 0 | 0 | 0 | 0.12029238 | 0 | 0.06673001 | 0 | 0 | 0 | 0.07903607 |
| ENSMUSG00 Rps16-ps2 | 934 | 2.18442018 | 1.81901243 | 2.12346332 | 2.19512526 | 1.95825814 | 1.6977382 | 2.68476658 | 2.53525583 | 2.11665177 | 2.22123227 | 2.04603677 | 2.15135316 | 3.16907391 | 3.08301222 | 2.02919868 | 1.80384018 | 2.29793643 | 2.65033169 |
| ENSMUSG00 Vmn1r62 | 1985 | 0 | 0 | 0 | 0 | 0 | 0 | 0 | 0 | 0 | 0 | 0 | 0 | 0 | 0 | 0 | 0 | 0 | 0 |
| ENSMUSG00 Olfr1034 | 1015 | 0 | 0 | 0 | 0 | 0 | 0 | 0 | 0 | 0 | 0 | 0 | 0 | 0 | 0 | 0 | 0 | 0 | 0 |
| ENSMUSG00 2610017I09R | 1173 | 73.7771001 | 57.0550763 | 58.9307544 | 63.0785473 | 65.9403251 | 60.6630225 | 72.8445849 | 59.2258994 | 57.3380906 | 67.7715991 | 60.563822 | 56.7673363 | 55.93474 | 46.5232759 | 49.2802923 | 45.7985696 | 49.6967584 | 50.3194983 |
| ENSMUSG00 Zfp868 | 3929 | 8.93593164 | 7.42735946 | 7.14091311 | 7.73459317 | 7.43600857 | 8.38701141 | 6.34608852 | 8.99159385 | 11.3213126 | 7.60843621 | 8.81568972 | 6.98938814 | 8.26174639 | 8.21548504 | 10.4550652 | 9.03420541 | 7.81353113 | 7.89644596 |
| ENSMUSG00 Sntb1 | 2861 | 5.25929127 | 3.31848125 | 3.77046392 | 6.01959971 | 4.96292118 | 4.43393911 | 6.71406768 | 7.19528308 | 6.40615153 | 5.90001809 | 5.5940619 | 6.92574632 | 4.79352429 | 3.78240907 | 9.66314016 | 6.10294197 | 7.76974911 | 8.26770915 |
| ENSMUSG00 Rps10-ps1 | 463 | 3.5803496 | 1.79875239 | 2.50748321 | 2.26329162 | 2.39100269 | 2.14050685 | 2.96342913 | 1.89724677 | 1.69015939 | 3.05512116 | 2.27008442 | 1.92883499 | 3.08990411 | 3.41058093 | 1.60178835 | 2.39833178 | 1.82111994 | 3.20787447 |
| ENSMUSG00 Trim5 | 4267 | 0.59768234 | 0.55430408 | 0.68019963 | 0.93962313 | 0.56400192 | 0.58065073 | 0.53222606 | 0.70710096 | 0.21235138 | 0.69615522 | 0.7165684 | 0.6017156 | 0.36996048 | 0.63129999 | 0.44416957 | 0.61020888 | 0.38622704 | 0.65747936 |
| ENSMUSG00 Sycp2 | 6210 | 0 | 0 | 0.00778962 | 0 | 0 | 0 | 0 | 0 | 0 | 0 | 0 | 0 | 0 | 0 | 0 | 0 | 0 | 0 |
| ENSMUSG00 Rnf14 | 3591 | 64.6514262 | 71.0508095 | 74.7088831 | 78.320336 | 74.1346894 | 79.8418592 | 79.5527584 | 79.5861173 | 76.8562635 | 76.7462714 | 79.2657206 | 74.7317476 | 63.1519977 | 74.4450479 | 78.662731 | 68.9039 | 80.3027173 | 71.8442524 |
| ENSMUSG00 Kng2 | 1878 | 0.24896535 | 0.05321549 | 0.05151601 | 0.09704163 | 0.17940547 | 0.23747316 | 0 | 0 | 0 | 0.05021374 | 0 | 0.148604 | 0.15760996 | 0.37095916 | 0.28520779 | 0.32622456 | 0.30612058 | 0.7322831 |
| ENSMUSG00 Dppa5a | 616 | 0 | 0 | 0 | 0 | 0 | 0 | 0 | 0 | 0 | 0 | 0 | 0 | 0 | 0 | 0 | 0 | 0 | 0 |
| ENSMUSG00 Thap6 | 2995 | 3.27836249 | 3.94860683 | 4.70006589 | 3.75745508 | 3.24629335 | 3.60684338 | 5.13409705 | 5.04981347 | 5.98201481 | 2.97545339 | 4.96093893 | 5.55361184 | 5.23791309 | 3.78375871 | 5.97044249 | 3.6436805 | 3.53190582 | 7.56719598 |
| ENSMUSG00 Gm10080 | 451 | 2.35616275 | 2.36366433 | 3.32500985 | 4.24293539 | 1.92100804 | 2.30733349 | 1.99321707 | 2.37114693 | 2.55703424 | 1.67275222 | 3.07200398 | 4.82683158 | 2.40643499 | 4.32517 | 1.91847599 | 3.2262596 | 2.88934389 | 2.073514 |
| ENSMUSG00 Krtap16-5 | 525 | 0 | 0 | 0 | 0 | 0 | 0 | 0 | 0 | 0 | 0 | 0 | 0 | 0 | 0 | 0 | 0 | 0 | 0 |
| ENSMUSG00 Gpr97 | 2424 | 0.19288652 | 0.16491532 | 0 | 0.03759162 | 0 | 0.18398292 | 0.07807373 | 0 | 0.13592909 | 0 | 0.01970914 | 0 | 0 | 0 | 0 | 0 | 0 | 0 |
| ENSMUSG00 Wtap | 6706 | 3.96782586 | 3.45249948 | 4.63104779 | 3.67559275 | 4.12702895 | 3.74638174 | 3.4570851 | 3.86205111 | 4.88882591 | 4.06398844 | 3.43386967 | 2.99636721 | 3.8547457 | 3.08195927 | 4.07346131 | 3.96838237 | 3.73775696 | 3.77336551 |
| ENSMUSG00 Irak2 | 5674 | 3.98532845 | 4.80847028 | 5.32842025 | 6.51216196 | 5.17456352 | 6.09584209 | 6.0120623 | 6.09166799 | 6.37323949 | 6.16598904 | 5.5824458 | 6.09899732 | 6.1034554 | 7.62881379 | 6.55710345 | 8.79320787 | 9.64574409 | 7.15486795 |
| ENSMUSG00 Olfr171 | 1010 | 0 | 0 | 0 | 0 | 0 | 0 | 0 | 0 | 0 | 0 | 0 | 0 | 0 | 0 | 0 | 0 | 0 | 0 |
| ENSMUSG00 Samd5 | 2493 | 0.35804601 | 0.76166665 | 1.28064701 | 0.32896061 | 0.73365949 | 0.63605595 | 2.14453779 | 1.65454451 | 2.11467068 | 0.8889222 | 1.07316393 | 1.54483783 | 0.73216243 | 1.50901367 | 1.14035594 | 1.73559512 | 1.27600587 | 1.3680564 |
| ENSMUSG00 Vmn1r218 | 896 | 0 | 0 | 0 | 0 | 0 | 0 | 0 | 0 | 0 | 0 | 0 | 0 | 0 | 0 | 0 | 0 | 0 | 0 |
| ENSMUSG00 4930522H14I | 1417 | 0 | 0 | 0 | 0 | 0 | 0 | 0 | 0 | 0 | 0 | 0 | 0 | 0 | 0.0655527 | 0 | 0 | 0.02704749 | 0 |
| ENSMUSG00 Rpl10l | 1020 | 0 | 0 | 0.04742503 | 0 | 0.04718816 | 0 | 0 | 0 | 0 | 0 | 0 | 0 | 0.04836462 | 0 | 0 | 0 | 0 | 0 |
| ENSMUSG00 Olfr715 | 1130 | 0 | 0 | 0 | 0 | 0 | 0 | 0 | 0 | 0 | 0 | 0 | 0 | 0 | 0 | 0 | 0 | 0 | 0 |
| ENSMUSG00 Nlrp9b | 3530 | 0 | 0.03774832 | 0 | 0 | 0 | 0 | 1.03203294 | 0.7681758 | 0.35002705 | 0.14692854 | 0.46015541 | 0 | 0.23757581 | 0.01315697 | 0.01167183 | 0.13016637 | 0.03257192 | 0 |
| ENSMUSG00 Xcr1 | 4451 | 0.04774789 | 0 | 0 | 0 | 0 | 0.07793061 | 0.03188902 | 0.07722571 | 0 | 0 | 0 | 0 | 0.11083333 | 0 | 0.12959375 | 0.06021888 | 0.02583215 | 0 |
| ENSMUSG00 Zfp266 | 7677 | 11.5052437 | 11.5642651 | 12.8353369 | 13.6321114 | 12.6709154 | 13.3354107 | 11.9560502 | 12.2930085 | 11.2663388 | 11.9703948 | 13.3486109 | 10.694912 | 9.47825687 | 10.3390576 | 11.1040796 | 11.8757244 | 10.8683533 | 11.9734554 |
| ENSMUSG00 0610040J01R | 1866 | 0.36446024 | 0.67839763 | 0.2592365 | 0.7569091 | 0.85120765 | 0.34522269 | 0.02535513 | 0 | 0.41936967 | 0.12634165 | 0 | 0 | 1.66554705 | 0.2986758 | 0.97152651 | 1.60057201 | 0.55456057 | 1.53294402 |
| ENSMUSG00 Tor3a | 4715 | 4.11980179 | 3.14405895 | 4.11405847 | 4.84116293 | 4.3385085 | 5.26530637 | 5.49890733 | 3.06187235 | 4.51610364 | 4.59006846 | 4.26580223 | 4.47472332 | 3.95492359 | 3.50670098 | 5.06827199 | 5.9689494 | 5.42989742 | 6.98843159 |
| ENSMUSG00 Olfr726 | 965 | 0 | 0 | 0 | 0 | 0 | 0 | 0 | 0 | 0 | 0 | 0 | 0 | 0 | 0 | 0 | 0 | 0 | 0 |
| ENSMUSG00 A930017M01 | 1672 | 0 | 0 | 0 | 0 | 0 | 0 | 0 | 0 | 0 | 0 | 0 | 0 | 0 | 0 | 0 | 0 | 0 | 0 |
| ENSMUSG00 Dcc | 8127 | 0 | 0 | 0 | 0 | 0 | 0 | 0 | 0 | 0 | 0 | 0 | 0 | 0 | 0 | 0 | 0 | 0 | 0 |
| ENSMUSG00 Tmem219 | 2138 | 9.24457757 | 8.33601422 | 7.67007801 | 6.43565734 | 8.66734828 | 7.09220602 | 9.11731699 | 9.21762378 | 7.18548661 | 8.07163582 | 6.86010863 | 7.88415814 | 11.3292663 | 10.07954 | 7.70842954 | 8.32793847 | 7.81583978 | 8.72221103 |
| ENSMUSG00 Tnfrsf19 | 5059 | 3.47838365 | 3.21342012 | 3.17454281 | 2.99897762 | 3.65342137 | 2.71320561 | 2.42221464 | 3.67655663 | 4.23344301 | 5.07016853 | 3.80575376 | 4.21458452 | 4.77814577 | 3.74563899 | 3.65675007 | 4.82890048 | 4.94703931 | 5.36064234 |
| ENSMUSG00 Olfr488 | 944 | 0 | 0 | 0 | 0 | 0 | 0 | 0 | 0 | 0 | 0 | 0 | 0 | 0 | 0 | 0 | 0 | 0 | 0 |
| ENSMUSG00 H2-Q7 | 3685 | 2.20311767 | 1.26561878 | 2.34975901 | 2.21314164 | 2.05066812 | 2.82390069 | 1.63058622 | 3.11965078 | 2.54831088 | 3.54429981 | 2.04842427 | 2.62038253 | 1.66001567 | 0.71840244 | 3.55552097 | 1.29886728 | 1.3832829 | 3.0900638 |
| ENSMUSG00 Olfr1418 | 1071 | 0 | 0 | 0 | 0 | 0 | 0 | 0 | 0 | 0 | 0 | 0 | 0 | 0 | 0 | 0 | 0 | 0 | 0 |
| ENSMUSG00 Gm10083 | 2230 | 0.03812123 | 0 | 0 | 0 | 0 | 0 | 0 | 0.01712664 | 0.03693858 | 0 | 0 | 0 | 0 | 0.20826945 | 0 | 0 | 0.05156004 | 0 |
| ENSMUSG00 Ces4a | 2419 | 0 | 0 | 0 | 0 | 0 | 0 | 0 | 0 | 0 | 0 | 0 | 0 | 0 | 0 | 0 | 0 | 0 | 0.04548087 |
| ENSMUSG00 Olfr823 | 947 | 0 | 0 | 0 | 0 | 0 | 0 | 0 | 0 | 0 | 0 | 0 | 0 | 0 | 0 | 0 | 0 | 0 | 0 |
| ENSMUSG00 Gm5591 | 3438 | 0 | 0 | 0 | 0 | 0 | 0 | 0 | 0 | 0 | 0 | 0 | 0.01623492 | 0 | 0 | 0 | 0 | 0 | 0 |
| ENSMUSG00 Rpl21-ps3 | 482 | 0 | 0.06911389 | 0 | 0.18904998 | 0.39943505 | 0.51403251 | 0 | 0.4754241 | 0 | 0.19564607 | 0.59470903 | 0.23160026 | 0 | 0.38542811 | 0.08548041 | 0.07944109 | 0.39757569 | 0.11412678 |
| ENSMUSG00 Fam78b | 5379 | 0.42671119 | 0.1114765 | 0.29677012 | 0.27104544 | 0.45635399 | 0.09212258 | 0.51015716 | 0.39761564 | 0.53598386 | 0.33309662 | 0.31974317 | 0.39431031 | 0.03668482 | 0.39717941 | 0.16851352 | 0.34880828 | 0.19237963 | 0.11249306 |
| ENSMUSG00 Mfap2 | 1201 | 0.60165528 | 1.24819337 | 1.24860903 | 1.32775733 | 0.80153079 | 1.15526773 | 1.30001532 | 0.82681304 | 0.3772287 | 1.45260285 | 0.67624837 | 1.06890944 | 0.82151403 | 0.54139652 | 1.26932354 | 1.02023267 | 0.7658877 | 0.6870413 |
| ENSMUSG00 Fhit | 1775 | 3.78355933 | 4.42920734 | 4.25142013 | 4.95396144 | 5.01656669 | 4.02005275 | 11.4616631 | 7.31572778 | 9.65272707 | 8.42071715 | 7.96697899 | 7.51546105 | 13.5905958 | 14.36496 | 13.6719529 | 12.70601 | 12.804222 | 13.4191229 |
| ENSMUSG00 Olfr881 | 947 | 0 | 0 | 0 | 0 | 0 | 0 | 0 | 0 | 0 | 0 | 0 | 0 | 0 | 0 | 0 | 0.16173435 | 0.44518402 | 0 |
| ENSMUSG00 H2-Eb1 | 1287 | 4.52463801 | 3.59789611 | 7.29173651 | 4.21271506 | 5.90896965 | 10.1646633 | 23.9320544 | 33.6223713 | 19.80921 | 12.0899239 | 40.3136013 | 7.97986163 | 33.6545636 | 27.5344512 | 38.4483828 | 21.5998298 | 11.6735885 | 17.3533002 |
| ENSMUSG00 Ifitm2 | 649 | 266.88535 | 246.689938 | 271.830917 | 236.791067 | 262.389439 | 259.903706 | 261.787103 | 254.223696 | 265.142162 | 259.946404 | 290.845703 | 246.139332 | 397.163741 | 334.697992 | 334.056375 | 425.326639 | 332.476197 | 347.006599 |
| ENSMUSG00 Olfr1454 | 923 | 0 | 0 | 0 | 0 | 0 | 0 | 0 | 0 | 0 | 0 | 0 | 0 | 0 | 0 | 0 | 0 | 0 | 0 |
| ENSMUSG00 Layn | 2236 | 0.15207576 | 0.02979686 | 0 | 0.22413752 | 0.08610362 | 0.19945197 | 0.33855226 | 0.17080681 | 0 | 0.08434831 | 0.06409878 | 0 | 0.1765006 | 0 | 0.22111747 | 0.11987221 | 0.1371245 | 0 |
| ENSMUSG00 Eno3 | 1729 | 6.93259667 | 6.85910373 | 5.76341657 | 6.50872062 | 4.34272995 | 5.18741745 | 10.0426565 | 10.404061 | 9.59986458 | 8.39931556 | 9.44999179 | 7.13433281 | 7.87484625 | 8.971848 | 11.8195325 | 12.2467935 | 5.85201969 | 6.90397696 |
| ENSMUSG00 Nr1h2 | 3640 | 70.588953 | 75.6587078 | 66.1415002 | 70.3943169 | 68.1648534 | 69.795842 | 67.2905876 | 67.6970825 | 65.1063768 | 70.1431478 | 73.591811 | 68.1288431 | 65.9340048 | 72.218004 | 62.9568829 | 66.3668797 | 68.5240495 | 66.0411529 |
| ENSMUSG00 Cyp2c70 | 1972 | 0 | 0 | 0 | 0 | 0 | 0 | 0 | 0 | 0 | 0 | 0 | 0 | 0 | 0 | 0 | 0 | 0 | 0 |
| ENSMUSG00 Angl4 | 707 | 0 | 0 | 0 | 0 | 0 | 0 | 0 | 0 | 0 | 0 | 0 | 0 | 0 | 0 | 0 | 0 | 0 | 0 |
| ENSMUSG00 Nkpd1 | 2749 | 0.2783169 | 0.31507284 | 0.36953224 | 0.28175255 | 0.21010661 | 0.18025731 | 0.41306086 | 0.08335919 | 0.55434743 | 0.39449479 | 0.34758064 | 0.22334387 | 0.14356324 | 1.2840126 | 1.37888074 | 0.86359318 | 0.68315335 | 0.86045529 |
| ENSMUSG00 Zfp735 | 4073 | 0 | 0 | 0 | 0 | 0 | 0 | 0 | 0 | 0 | 0 | 0 | 0 | 0 | 0 | 0 | 0 | 0 | 0 |
| ENSMUSG00 Rpl35a | 1977 | 14.9853858 | 9.8910819 | 12.650033 | 12.029775 | 12.0025485 | 10.4519423 | 14.7179047 | 10.4125973 | 13.1871849 | 11.7340142 | 13.000975 | 11.7164896 | 17.3422774 | 17.6896296 | 13.2336813 | 14.3323467 | 13.0856097 | 18.4198425 |
| ENSMUSG00 Hist1h4i | 536 | 0.47580421 | 0.12430184 | 0 | 0.1700039 | 0.08979837 | 0.9244913 | 1.67712854 | 0.49878137 | 0.84524566 | 0.35187092 | 0.53479431 | 1.24960439 | 1.74870602 | 0.95314358 | 0.76868574 | 0.5715016 | 0.78654714 | 1.23154717 |
| ENSMUSG00 Olfr1512 | 941 | 0 | 0 | 0 | 0 | 0 | 0.05265965 | 0 | 0 | 0 | 0 | 0 | 0 | 0 | 0 | 0 | 0 | 0.04072933 | 0 |
| ENSMUSG00 Gm7099 | 550 | 0.54097497 | 0.90853348 | 0.17590375 | 0.82838262 | 0.52507553 | 0.36038352 | 0.86023053 | 0.90272951 | 0.44930744 | 0 | 0.26059068 | 0.20296605 | 0.26908319 | 0.84443795 | 0.97385496 | 0.83543143 | 0.62715759 | 0.60009935 |
| ENSMUSG00 4921513D23 | 6447 | 1.58891693 | 2.35107288 | 2.31850797 | 1.46287206 | 1.44089674 | 1.32970729 | 1.08612944 | 1.68835655 | 1.61628486 | 1.96735523 | 1.28200214 | 1.75749643 | 2.92303279 | 1.23908532 | 2.00671453 | 1.81148368 | 2.25308927 | 2.94371674 |
| ENSMUSG00 Olfr175-ps1 | 1027 | 0 | 0 | 0 | 0 | 0 | 0 | 0 | 0 | 0 | 0 | 0 | 0 | 0 | 0 | 0 | 0 | 0 | 0 |
| ENSMUSG00 Atp8b2 | 10316 | 1.9241874 | 1.79223113 | 1.68341386 | 1.72245127 | 1.87096756 | 2.23361973 | 1.53183741 | 1.97329881 | 1.6209505 | 0.93698082 | 0.9447549 | 1.6664622 | 1.84588213 | 1.86838855 | 1.63751822 | 1.68514306 | 1.22974062 | 2.78884866 |
| ENSMUSG00 4930595M18 | 2640 | 0 | 0 | 0 | 0 | 0 | 0 | 0 | 0 | 0 | 0 | 0 | 0 | 0 | 0 | 0 | 0 | 0 | 0 |
| ENSMUSG00 Pla2g16 | 6349 | 6.41360193 | 6.75282641 | 5.9962149 | 6.12121134 | 5.34462217 | 5.47897605 | 2.31756862 | 2.46033904 | 2.607809 | 2.16853132 | 2.00159694 | 1.705503 | 5.14375953 | 4.99626897 | 3.98452596 | 7.11050971 | 5.24579493 | 4.67867658 |
| ENSMUSG00 Hist1h4c | 394 | 0.32364347 | 0.76095444 | 0.98220366 | 0.34691151 | 1.58810916 | 0.37730508 | 1.32091236 | 0.09693503 | 0.94080874 | 0.23934367 | 0.48502496 | 0.99164883 | 1.50249494 | 1.65029752 | 1.77773211 | 1.36057996 | 0.58364919 | 1.2565532 |
| ENSMUSG00 Mrps9 | 2496 | 16.2289394 | 14.6277773 | 17.1710528 | 14.6941669 | 14.3470157 | 14.9094964 | 14.4250596 | 14.2456437 | 12.8377842 | 13.9789745 | 13.5706913 | 13.8197434 | 16.1475066 | 12.076207 | 13.1065847 | 13.760687 | 14.495202 | 13.906549 |
| ENSMUSG00 Gm8894 | 455 | 0.28025391 | 0.87858183 | 0.10631545 | 0.10013416 | 0.31735334 | 0.43562843 | 0.93585519 | 0.25181804 | 0.36207926 | 0.62176752 | 1.36499881 | 0.73603072 | 0.1084218 | 0.81659934 | 0 | 0.16831036 | 0 | 0.48359654 |
| ENSMUSG00 Slc9a6 | 4969 | 7.20252731 | 8.31313923 | 8.13851306 | 7.28023127 | 7.76852541 | 6.19284515 | 7.31256539 | 10.8605081 | 9.39937792 | 6.6422806 | 6.28794986 | 10.5475764 | 8.4486741 | 10.4590328 | 8.54047144 | 6.48066026 | 7.64366276 | 7.87109577 |
| ENSMUSG00 Gm14511 | 733 | 0 | 0 | 0 | 0 | 0 | 0 | 0 | 0 | 0 | 0 | 0 | 0 | 0 | 0 | 0 | 0 | 0 | 0 |
| ENSMUSG00 Olfr292 | 1076 | 0 | 0 | 0 | 0 | 0 | 0 | 0 | 0 | 0 | 0 | 0 | 0 | 0 | 0 | 0 | 0 | 0 | 0 |
| ENSMUSG00 Krtap16-2 | 492 | 0 | 0 | 0 | 0 | 0 | 0 | 0 | 0 | 0 | 0 | 0 | 0 | 0 | 0 | 0 | 0 | 0 | 0 |
| ENSMUSG00 Vmn1r35 | 890 | 0 | 0 | 0 | 0 | 0 | 0 | 0 | 0 | 0 | 0 | 0 | 0 | 0 | 0 | 0 | 0 | 0 | 0 |
| ENSMUSG00 Fpr-rs3 | 1031 | 0 | 0 | 0 | 0 | 0 | 0 | 0 | 0 | 0 | 0 | 0 | 0 | 0 | 0 | 0 | 0 | 0 | 0 |
| ENSMUSG00 Cd302 | 1348 | 0.75676871 | 0.98851318 | 0.4306249 | 0.33798994 | 0.7141235 | 0.77196395 | 0.73706696 | 0.90664457 | 1.98599668 | 0.524674 | 1.48853728 | 0.745313 | 2.04939716 | 3.68658557 | 2.78141067 | 3.29503743 | 1.25100672 | 3.42786718 |
| ENSMUSG00 Cno | 1305 | 24.7539722 | 23.8168048 | 20.2019724 | 21.4363841 | 18.5519982 | 18.9097788 | 21.3541516 | 21.3643324 | 20.198751 | 21.5700918 | 23.8325655 | 24.4220255 | 25.7055201 | 28.7917752 | 23.9000596 | 25.4097058 | 26.1676096 | 26.7247308 |
| ENSMUSG00 1700019A02I | 618 | 0.20633581 | 0 | 0 | 0 | 0 | 0 | 1.37803919 | 0.61800004 | 0.1332897 | 0 | 1.08228061 | 0.09031661 | 0.23947533 | 0.30060898 | 0.13333837 | 0 | 0.06201666 | 0.178023 |
| ENSMUSG00 Plekhh1 | 6429 | 13.8444297 | 18.9648768 | 16.7490245 | 17.6532216 | 16.320982 | 17.5889467 | 12.0471077 | 13.1882304 | 11.2239502 | 14.3747672 | 10.4036307 | 11.9462361 | 7.34338853 | 11.3274738 | 11.6189798 | 8.71946836 | 8.76932381 | 11.4655783 |
| ENSMUSG00 Vmn1r47 | 932 | 0 | 0 | 0 | 0 | 0 | 0 | 0 | 0 | 0 | 0 | 0 | 0 | 0 | 0 | 0 | 0 | 0 | 0 |
| ENSMUSG00 Olfr487 | 944 | 0 | 0 | 0 | 0 | 0 | 0 | 0 | 0 | 0 | 0 | 0 | 0 | 0 | 0 | 0 | 0 | 0 | 0 |
| ENSMUSG00 Tmsb15a | 560 | 0 | 0 | 0 | 0 | 0 | 0 | 0 | 0 | 0 | 0 | 0 | 0 | 0 | 0 | 0 | 0 | 0 | 0 |
| ENSMUSG00 Gm12571 | 242 | 0.35128245 | 0.41296976 | 0.59967186 | 0.94134389 | 1.39224573 | 0.81905345 | 0 | 1.57819845 | 0 | 0.38967523 | 0.78966874 | 0.92257294 | 1.01925449 | 1.34342402 | 0.34050873 | 0.47467695 | 0.63349251 | 0.22731036 |
| ENSMUSG00 Gm10086 | 760 | 1.00670154 | 0.65749133 | 0.50919506 | 0.4196412 | 0.56998331 | 0.71721062 | 1.05830992 | 0.50253161 | 0.86708454 | 0.74448479 | 1.0057886 | 0.9547416 | 1.62276044 | 1.40554475 | 1.0300389 | 0.80611805 | 1.0590161 | 1.80951009 |
| ENSMUSG00 Ipmk | 3365 | 3.86525523 | 4.52421773 | 5.54893989 | 3.16828659 | 3.79047839 | 4.97736224 | 5.73657444 | 4.95990486 | 3.87997786 | 6.20735856 | 4.99758881 | 4.31265149 | 3.45983135 | 6.51459413 | 5.0935653 | 6.09918739 | 4.88617574 | 4.98596658 |
| ENSMUSG00 Rxfp3 | 4251 | 0 | 0 | 0 | 0 | 0 | 0 | 0 | 0 | 0 | 0 | 0 | 0 | 0 | 0 | 0 | 0 | 0 | 0 |
| ENSMUSG00 Prl7c1 | 1009 | 0 | 0 | 0 | 0 | 0 | 0 | 0 | 0 | 0 | 0 | 0 | 0 | 0 | 0 | 0 | 0 | 0 | 0 |
| ENSMUSG00 Nsa2 | 2680 | 6.16959458 | 6.50098645 | 5.10810041 | 4.79410989 | 5.11850681 | 5.10319197 | 5.20792548 | 7.08269552 | 5.33273171 | 5.06694125 | 6.45318469 | 5.99810105 | 6.51623086 | 7.76378774 | 5.7805168 | 6.38653041 | 6.44968653 | 6.99929306 |
| ENSMUSG00 Olfr1076 | 941 | 0 | 0 | 0 | 0 | 0 | 0 | 0 | 0 | 0 | 0 | 0 | 0 | 0 | 0 | 0 | 0 | 0 | 0 |
| ENSMUSG00 H3f3a | 4006 | 31.2687853 | 29.3130185 | 31.4440023 | 27.8643431 | 27.778584 | 27.6213814 | 29.7741547 | 29.5738473 | 32.1184911 | 30.9669247 | 29.4926291 | 32.3663965 | 28.7420606 | 28.0449944 | 27.6151217 | 27.5566204 | 26.8073599 | 27.7379919 |
| ENSMUSG00 Il28b | 697 | 0 | 0 | 0 | 0 | 0 | 0 | 0 | 0 | 0 | 0 | 0 | 0 | 0 | 0 | 0 | 0 | 0 | 0 |
| ENSMUSG00 Krtap2-4 | 829 | 0 | 0 | 0 | 0 | 0 | 0 | 0 | 0 | 0 | 0 | 0 | 0 | 0 | 0 | 0 | 0 | 0 | 0 |
| ENSMUSG00 Olfr503 | 971 | 0 | 0 | 0 | 0 | 0 | 0 | 0 | 0 | 0 | 0 | 0 | 0 | 0 | 0 | 0 | 0 | 0 | 0 |
| ENSMUSG00 Olfr325 | 2044 | 0 | 0 | 0 | 0 | 0 | 0 | 0 | 0 | 0 | 0 | 0 | 0 | 0 | 0 | 0 | 0 | 0 | 0 |
| ENSMUSG00 Tsga10 | 8336 | 0.85153124 | 0.54349252 | 0.9400806 | 0.56841994 | 0.56007637 | 0.74305323 | 0.47108354 | 0.8476001 | 0.40020487 | 0.55431489 | 0.72212629 | 0.72313958 | 0.52669633 | 0.2952899 | 0.54862916 | 0.35369203 | 0.50574528 | 0.56751238 |
| ENSMUSG00 Gm10087 | 661 | 0 | 0 | 0 | 0 | 0 | 0 | 0 | 0 | 0 | 0 | 0 | 0 | 0 | 0 | 0 | 0 | 0 | 0 |
| ENSMUSG00 Lrrtm1 | 3416 | 0 | 0 | 0.05664348 | 0 | 0.08454085 | 0 | 0.16620379 | 0.01118045 | 0 | 0.16563479 | 0.02797129 | 0.04901844 | 0.37547712 | 0.61182199 | 0.663374 | 0 | 0 | 0.22544716 |
| ENSMUSG00 Gm8994 | 1462 | 0 | 0 | 0 | 0 | 0 | 0 | 0.03236161 | 0 | 0 | 0 | 0 | 0.03817761 | 0 | 0 | 0.02818164 | 0.02619057 | 0 | 0 |
| ENSMUSG00 Olfr464 | 1083 | 0 | 0 | 0 | 0 | 0 | 0 | 0 | 0 | 0 | 0 | 0 | 0 | 0 | 0 | 0 | 0 | 0 | 0 |
| ENSMUSG00 Gmfg | 2893 | 0.42608023 | 0.20727 | 0.36785955 | 0.37796926 | 0.16637374 | 0.18841344 | 1.60271087 | 0.73929296 | 0.85419666 | 0.57043713 | 0.84221323 | 0.54021381 | 1.27891248 | 1.92647442 | 1.02541031 | 0.74119392 | 0.5961574 | 0.43733476 |
| ENSMUSG00 Tssk5 | 1506 | 0 | 0 | 0 | 0 | 0 | 0.06580706 | 0.06283224 | 0.07608048 | 0 | 0 | 0 | 0 | 0 | 0 | 0 | 0 | 0.02544907 | 0 |
| ENSMUSG00 Gm13363 | 521 | 0 | 0 | 0 | 0 | 0.18476746 | 0.09511081 | 0.09081128 | 0 | 0 | 0 | 0.45849288 | 0 | 0 | 0 | 0.07908168 | 0.1469889 | 0 | 0 |
| ENSMUSG00 Gm6280 | 683 | 0 | 0 | 0 | 0 | 0 | 0 | 0 | 0 | 0 | 0 | 0 | 0 | 0 | 0 | 0 | 0 | 0 | 0 |
| ENSMUSG00 Intu | 7412 | 1.09531687 | 1.14608581 | 1.26611777 | 0.90359869 | 1.42863238 | 1.03624848 | 0.85535604 | 1.10784761 | 1.339173 | 1.40586959 | 1.57273204 | 0.75304456 | 0.89851711 | 0.95244216 | 0.82825578 | 1.25534506 | 0.80148084 | 1.69955281 |
| ENSMUSG00 B2m | 856 | 2399.90381 | 1948.14272 | 2284.51909 | 2201.99295 | 2227.05736 | 2271.55288 | 3428.78743 | 2740.03717 | 3286.98227 | 3399.58774 | 2489.7674 | 2696.82242 | 4857.75308 | 4636.21592 | 4500.83679 | 4256.92249 | 3641.84891 | 4819.40182 |
| ENSMUSG00 Gstp1 | 746 | 208.993278 | 203.360483 | 208.5379 | 203.620002 | 222.852097 | 179.678478 | 178.786115 | 174.630408 | 188.707119 | 194.670464 | 182.710398 | 185.927509 | 241.237043 | 246.352887 | 227.437007 | 193.557481 | 178.838927 | 236.848862 |
| ENSMUSG00 Serpina6 | 1566 | 0 | 0 | 0 | 0 | 0 | 0 | 0 | 0 | 0 | 0 | 0 | 0 | 0 | 0 | 0 | 0 | 0 | 0 |
| ENSMUSG00 3110043A19I | 673 | 0.69473542 | 0.5444901 | 0.3593873 | 0.13539686 | 0.50062922 | 0.07362962 | 0.56240926 | 0 | 0.48958711 | 0 | 0 | 0 | 0 | 0.55208425 | 0.48976589 | 0.22758162 | 0.34169061 | 0 |
| ENSMUSG00 Gm10088 | 422 | 0 | 0 | 0 | 0 | 0 | 0 | 0 | 0 | 0 | 0 | 0 | 0 | 0 | 0 | 0 | 0 | 0 | 0 |
| ENSMUSG00 Gm4980 | 680 | 0 | 0 | 0 | 0 | 0.3539112 | 0.07287167 | 0 | 0 | 0 | 0.13867854 | 0 | 0 | 0.72546937 | 0 | 0 | 0 | 0 | 0 |
| ENSMUSG00 Vmn1r52 | 929 | 0 | 0 | 0 | 0 | 0 | 0 | 0 | 0 | 0 | 0 | 0 | 0 | 0 | 0 | 0 | 0 | 0 | 0 |
| ENSMUSG00 Taar7f | 1076 | 0 | 0.15479969 | 0.08991363 | 0 | 0.17892908 | 0.04605273 | 0.04397089 | 0.21296879 | 0.03827743 | 0.04382036 | 0.04440052 | 0.46885963 | 0.27508504 | 0.3453092 | 0.03829141 | 0.24910246 | 0.21371541 | 0 |
| ENSMUSG00 Olfr1193 | 956 | 0 | 0 | 0 | 0 | 0 | 0 | 0 | 0 | 0 | 0 | 0 | 0 | 0 | 0 | 0 | 0 | 0 | 0 |
| ENSMUSG00 Ctnna3 | 7204 | 0 | 0 | 0 | 0 | 0 | 0 | 0.05254045 | 0 | 0 | 0.00654507 | 0 | 0 | 0 | 0 | 0 | 0 | 0 | 0 |
| ENSMUSG00 Olfr671 | 941 | 0 | 0 | 0 | 0 | 0 | 0 | 0 | 0 | 0 | 0 | 0 | 0 | 0 | 0 | 0 | 0 | 0 | 0 |
| ENSMUSG00 Ube2s | 984 | 37.1056364 | 31.1462019 | 31.1183381 | 33.3836514 | 26.5116897 | 29.7618552 | 35.8691651 | 33.4183521 | 32.8571289 | 35.7942839 | 35.2000456 | 29.779698 | 43.3158298 | 38.6090076 | 35.6326464 | 36.2282068 | 33.8471057 | 35.7223772 |
| ENSMUSG00 Zbtb40 | 10078 | 0.65794875 | 0.509048 | 0.68158775 | 0.46564671 | 0.46804212 | 0.52611059 | 0.23473248 | 0.4320236 | 0.453632 | 0.44914343 | 0.51671666 | 0.42645426 | 0.26433057 | 0.64979324 | 0.33114964 | 0.39893965 | 0.60086872 | 0.34387515 |
| ENSMUSG00 Tcl1b2 | 1252 | 0 | 0 | 0 | 0.10917183 | 0 | 0 | 0 | 0 | 0 | 0 | 0 | 0 | 0 | 0 | 0.06581718 | 0.03058355 | 0 | 0 |
| ENSMUSG00 Olfr1420 | 1051 | 0 | 0.06339276 | 0 | 0 | 0 | 0.09429635 | 0 | 0 | 0 | 0 | 0 | 0 | 0 | 0 | 0 | 0 | 0 | 0.52339778 |
| ENSMUSG00 Gm6564 | 644 | 0 | 0 | 0 | 0 | 0 | 0 | 0 | 0 | 0 | 0 | 0 | 0 | 0 | 0 | 0 | 0.05945746 | 0 | 0 |
| ENSMUSG00 Kcnd2 | 5146 | 0.14867726 | 0.12299747 | 0.07520176 | 0.15936626 | 0.3647775 | 0.10592306 | 0 | 0 | 0 | 0 | 0 | 0 | 0 | 0.00902528 | 0.10408477 | 0 | 0 | 0 |
| ENSMUSG00 Gm12960 | 1307 | 0 | 0.02548806 | 0.03701112 | 0.06971851 | 0 | 0 | 0 | 0 | 0 | 0 | 0.03655314 | 0.04270517 | 0 | 0 | 0 | 0.11718625 | 0 | 0 |
| ENSMUSG00 Olfr566 | 950 | 0 | 0 | 0 | 0 | 0 | 0 | 0 | 0 | 0 | 0 | 0 | 0 | 0 | 0 | 0 | 0 | 0 | 0 |
| ENSMUSG00 Arr3 | 2307 | 0 | 0.05775968 | 0.10484077 | 0 | 0.02086343 | 0 | 0 | 0 | 0 | 0 | 0 | 0 | 0.19245221 | 0 | 0.01785937 | 0.08298788 | 0.01661305 | 0 |
| ENSMUSG00 Rpl21-ps11 | 467 | 0 | 0 | 0 | 0.09756112 | 0 | 0 | 0 | 0.08178245 | 0 | 0 | 0 | 0 | 0 | 0.09945201 | 0 | 0 | 0 | 0 |
| ENSMUSG00 Arl1 | 2316 | 64.2532907 | 63.9791682 | 63.954987 | 67.4169685 | 64.342157 | 64.8507493 | 60.4891374 | 67.0345923 | 70.5648077 | 65.310646 | 64.8963814 | 64.6121725 | 64.0292501 | 62.4267893 | 71.7113436 | 69.0421315 | 67.5674742 | 62.41966 |
| ENSMUSG00 Olfr944 | 935 | 0 | 0 | 0 | 0 | 0 | 0 | 0 | 0 | 0 | 0 | 0 | 0 | 0 | 0 | 0 | 0 | 0 | 0 |
| ENSMUSG00 Trim65 | 1777 | 0.02391963 | 0 | 0 | 0 | 0 | 0.02788561 | 0 | 0 | 0 | 0 | 0 | 0 | 0 | 0.02613623 | 0.04637204 | 0.04309579 | 0 | 0 |
| ENSMUSG00 Olfr51 | 923 | 0 | 0 | 0 | 0 | 0 | 0 | 0 | 0 | 0 | 0 | 0 | 0 | 0 | 0 | 0 | 0 | 0 | 0 |
| ENSMUSG00 Acyp2 | 859 | 25.4833128 | 15.7450932 | 17.795152 | 18.4047525 | 21.2923527 | 19.5557354 | 16.0830061 | 13.4273638 | 22.2953781 | 14.4361292 | 16.6294675 | 12.4756778 | 22.0529176 | 21.032305 | 15.4445879 | 21.4409571 | 21.4163242 | 13.3840551 |
| ENSMUSG00 Csmd1 | 22294 | 0 | 0 | 0 | 0 | 0 | 0 | 0 | 0 | 0 | 0 | 0 | 0.01752533 | 0 | 0 | 0.0092405 | 0 | 0 | 0 |
| ENSMUSG00 AI597468 | 3775 | 2.84869127 | 3.85635358 | 5.40758406 | 3.52419204 | 3.03454247 | 3.75419386 | 4.33647335 | 5.14965108 | 5.66246492 | 4.48400065 | 4.27759893 | 4.13996968 | 3.75053252 | 3.40795025 | 3.47075358 | 4.64558892 | 3.21839367 | 2.76867627 |
| ENSMUSG00 Rpl26 | 1398 | 42.6571974 | 30.9299976 | 40.553489 | 36.8268813 | 48.3040693 | 33.0706012 | 45.315935 | 34.9414039 | 40.2143019 | 38.6851621 | 45.2119242 | 26.9096971 | 40.1218812 | 47.9722906 | 41.6731045 | 42.4264312 | 45.0429941 | 46.3131034 |
| ENSMUSG00 Trmt61a | 2625 | 3.75664794 | 3.02036908 | 2.89319782 | 3.05475954 | 3.33714671 | 3.09586602 | 5.02866187 | 3.6955696 | 3.59303318 | 3.16134239 | 2.78459758 | 2.31767894 | 3.57069116 | 2.42393904 | 3.54725776 | 2.10051332 | 2.35067954 | 3.20624508 |
| ENSMUSG00 Gm10092 | 177 | 0 | 0 | 0 | 0 | 0 | 0 | 0 | 0 | 0 | 0 | 0 | 0 | 0 | 0 | 0 | 0 | 0 | 0 |
| ENSMUSG00 Slc4a4 | 10849 | 0.57984755 | 0.31934197 | 0.36116287 | 0.27297151 | 0.33717635 | 0.59834161 | 0.01308305 | 0 | 0 | 0.00869217 | 0 | 0 | 0.00909428 | 0.1541144 | 0.01519091 | 0.02117648 | 0 | 0 |
| ENSMUSG00 Dmkn | 2707 | 0 | 0.38149232 | 0.14295835 | 0.31978568 | 0 | 0 | 0 | 0 | 0 | 0.01741807 | 0.01764867 | 0 | 0 | 0 | 0 | 0 | 0 | 0 |
| ENSMUSG00 Etd | 476 | 0 | 0 | 0 | 0 | 0 | 0 | 0 | 0 | 0 | 0 | 0 | 0 | 0 | 0 | 0 | 0 | 0 | 0 |
| ENSMUSG00 Irx1 | 2369 | 0.14353795 | 0.14062007 | 0.02041939 | 0.32694713 | 0.0406348 | 0.18825437 | 0.61911906 | 0.46753046 | 0.34771225 | 0.29854814 | 0 | 0.35341281 | 0.45812671 | 0.76459241 | 0.34783922 | 0.22626472 | 0.19413911 | 0.60373017 |
| ENSMUSG00 Olfr530 | 1079 | 0 | 0 | 0 | 0 | 0 | 0 | 0 | 0 | 0 | 0 | 0 | 0 | 0 | 0 | 0 | 0 | 0 | 0 |
| ENSMUSG00 Hist1h4h | 469 | 1.17818185 | 1.20750363 | 1.13456041 | 0.19429017 | 0.92364033 | 1.05656148 | 2.32023799 | 3.01304667 | 2.54671419 | 2.31229462 | 1.32425258 | 2.49920877 | 6.73186078 | 4.55528362 | 3.60184177 | 3.59229579 | 2.04297958 | 7.27199279 |
| ENSMUSG00 Tdrd5 | 7409 | 0.01147393 | 0.03597019 | 0.07181925 | 0.07379303 | 0.07795696 | 0.04012909 | 0.15326013 | 0 | 0.06114883 | 0 | 0 | 0 | 0.21306807 | 0.01880581 | 0.13346435 | 0.19122047 | 0.03621057 | 0.10394486 |
| ENSMUSG00 Galnt13 | 8139 | 13.7505994 | 15.5779427 | 18.3414074 | 16.0546843 | 17.4987544 | 17.0046424 | 22.0606483 | 23.1528828 | 20.7273582 | 22.3674733 | 20.0104231 | 20.1962313 | 13.8740335 | 15.6297279 | 17.6115263 | 11.1969094 | 18.9347635 | 16.2344114 |
| ENSMUSG00 Gm11847 | 5253 | 0 | 0.06878365 | 0.05525246 | 0.02602001 | 0 | 0 | 0.05404075 | 0.02181177 | 0.03920285 | 0.04487979 | 0.02728439 | 0.02125097 | 0 | 0 | 0 | 0.02186785 | 0.02188823 | 0.01047194 |
| ENSMUSG00 Copz1 | 2457 | 108.312946 | 100.386109 | 103.066924 | 98.7990412 | 110.877773 | 99.0247952 | 115.229577 | 102.810154 | 114.457277 | 113.971521 | 113.127679 | 100.749883 | 122.898114 | 98.9559615 | 123.168672 | 109.884849 | 113.122631 | 113.331745 |
| ENSMUSG00 Olfr507 | 950 | 0 | 0 | 0 | 0 | 0 | 0 | 0 | 0 | 0 | 0 | 0 | 0 | 0 | 0 | 0 | 0 | 0 | 0 |
| ENSMUSG00 Mkx | 3497 | 0.01215475 | 0.04763068 | 0 | 0 | 0.02752755 | 0.09919049 | 0 | 0.08737181 | 0.21199808 | 0.21573098 | 0.05464679 | 0.01596101 | 0 | 0.07968674 | 0.09425578 | 0.05474779 | 0.16439647 | 0 |
| ENSMUSG00 Vmn1r222 | 1053 | 0 | 0 | 0 | 0 | 0 | 0 | 0 | 0 | 0 | 0 | 0 | 0 | 0 | 0 | 0 | 0 | 0 | 0 |
| ENSMUSG00 Rrs1 | 2046 | 15.5603015 | 14.149025 | 13.7838554 | 15.7437235 | 13.2680376 | 15.6941209 | 17.3433574 | 16.9495119 | 12.9034979 | 18.9201991 | 16.2285416 | 19.2326697 | 11.7181387 | 15.5494623 | 12.5457328 | 19.2201631 | 14.6111982 | 15.9703858 |
| ENSMUSG00 Vmn2r50 | 2532 | 0 | 0 | 0 | 0 | 0 | 0 | 0 | 0 | 0 | 0 | 0 | 0 | 0 | 0 | 0 | 0 | 0 | 0 |
| ENSMUSG00 Clasrp | 2132 | 28.4896325 | 25.9846825 | 27.6809132 | 32.332955 | 30.6129871 | 30.5172323 | 25.4094828 | 27.4619854 | 28.0114671 | 27.887916 | 21.1088231 | 26.4941138 | 27.2112264 | 25.7708046 | 28.0216961 | 27.550559 | 21.2304675 | 23.1440754 |
| ENSMUSG00 Rrp1 | 2486 | 141.381859 | 151.328445 | 133.737439 | 152.755955 | 148.268009 | 146.066948 | 132.840909 | 128.726524 | 136.482107 | 137.848476 | 139.635096 | 137.945065 | 136.010041 | 146.468884 | 139.283136 | 138.66868 | 139.121683 | 155.025664 |
| ENSMUSG00 Olfr920 | 1958 | 0.08683386 | 0.05104121 | 0.02470558 | 0.0930767 | 0 | 0.25307831 | 0.14496267 | 0.25357571 | 0.04206999 | 0.02408105 | 0 | 0 | 0.37792582 | 0.18976134 | 0.0841707 | 0 | 0.09787103 | 0.47760716 |
| ENSMUSG00 Haghl | 2023 | 25.6123626 | 25.8862431 | 26.2790459 | 26.1024471 | 29.0028744 | 24.9355822 | 26.1003211 | 27.9976929 | 26.8129715 | 27.45602 | 26.0719497 | 28.6113901 | 29.4089433 | 25.2767871 | 26.6598303 | 24.586999 | 21.8628505 | 27.8716433 |
| ENSMUSG00 Cdh3 | 4072 | 1.78496687 | 2.43792792 | 1.60373934 | 2.7077045 | 2.69500948 | 2.42165864 | 0.74361774 | 2.03530239 | 0.5461866 | 1.38950992 | 2.00626867 | 2.91963068 | 0.72689466 | 1.03792042 | 1.35584688 | 0.38553902 | 1.16710727 | 2.12093069 |
| ENSMUSG00 Gm10093 | 1976 | 5.52825417 | 4.63615684 | 5.1164311 | 5.46455846 | 4.38448697 | 4.28821733 | 4.908451 | 4.67740961 | 5.41927839 | 5.82225283 | 4.18272663 | 5.64935856 | 6.11655857 | 4.44227354 | 6.48465784 | 6.54970916 | 4.34468143 | 5.42853027 |
| ENSMUSG00 Gm5382 | 508 | 0 | 0 | 0 | 0 | 0 | 0 | 0 | 0 | 0 | 0 | 0 | 0 | 0 | 0 | 0 | 0 | 0 | 0 |
| ENSMUSG00 Mcpt4 | 985 | 0 | 0 | 0 | 0 | 0 | 0 | 0 | 0 | 0 | 0 | 0 | 0 | 0 | 0 | 0 | 0 | 0 | 0 |
| ENSMUSG00 Olfr1337 | 947 | 0 | 0 | 0 | 0 | 0 | 0 | 0 | 0 | 0 | 0 | 0 | 0 | 0 | 0 | 0 | 0 | 0 | 0 |
| ENSMUSG00 Zfp143 | 3001 | 4.60319301 | 9.35780401 | 6.93122894 | 6.96851694 | 8.06743006 | 7.00111931 | 5.40761375 | 7.92864602 | 6.21709152 | 6.31608887 | 7.81656268 | 6.47245937 | 6.72334361 | 3.7452413 | 4.09132411 | 5.97134431 | 7.76487455 | 6.69054447 |
| ENSMUSG00 Lsamp | 11077 | 0.08825666 | 0.09924397 | 0.07423942 | 0.0781493 | 0.02172606 | 0 | 0.30753028 | 0.21376988 | 0.10782784 | 0.22134482 | 0.25446624 | 0.08062206 | 0.32510878 | 0.31027015 | 0.07439118 | 0.19012218 | 0.40135871 | 0.04966065 |
| ENSMUSG00 Plac1 | 2282 | 0 | 0 | 0 | 0 | 0 | 0 | 0 | 0 | 0 | 0 | 0 | 0 | 0 | 0 | 0 | 0 | 0 | 0 |
| ENSMUSG00 Myl4 | 4314 | 0.85719757 | 1.10425217 | 1.05403613 | 0.9927534 | 1.03761449 | 0.76959507 | 0.23031207 | 0.48692214 | 0.30550962 | 0.53555504 | 0.27685998 | 0.46577743 | 0.48028292 | 0.39833826 | 0.42022913 | 0.82545816 | 0.43532419 | 0.47179809 |
| ENSMUSG00 Olfr38 | 953 | 0 | 0 | 0 | 0 | 0 | 0 | 0 | 0 | 0 | 0 | 0 | 0 | 0 | 0 | 0 | 0 | 0 | 0 |
| ENSMUSG00 Olfr890 | 944 | 0 | 0 | 0 | 0 | 0 | 0 | 0 | 0 | 0 | 0 | 0 | 0 | 0 | 0 | 0 | 0 | 0 | 0 |
| ENSMUSG00 Gapdhs | 1614 | 5.05637782 | 3.61199287 | 3.74640105 | 4.14961184 | 3.87679683 | 3.68421811 | 3.01933453 | 3.95175415 | 4.61881001 | 3.82697777 | 3.31523875 | 2.83573998 | 3.3621505 | 3.65452237 | 2.47617777 | 3.55860664 | 3.65690813 | 2.79476255 |
| ENSMUSG00 Retnla | 594 | 0 | 0 | 0 | 0 | 0 | 0.33368844 | 0 | 6.81547922 | 7.28044471 | 0 | 0 | 0 | 0 | 3.19069288 | 0.0644623 | 0.06452239 | 0 | 0 |
| ENSMUSG00 Gm10094 | 779 | 2.56449714 | 1.71054656 | 1.61452091 | 0.99427183 | 1.85360425 | 2.16276372 | 2.490141 | 2.40234624 | 1.53325925 | 1.39212603 | 2.39181435 | 2.14951204 | 1.45652644 | 2.26556524 | 2.27428357 | 2.75259282 | 1.91877481 | 1.83599073 |
| ENSMUSG00 Fam195b | 1278 | 173.64595 | 155.694771 | 150.684683 | 150.301536 | 151.287901 | 145.013477 | 167.889671 | 162.990112 | 167.968795 | 162.776919 | 160.969461 | 154.519416 | 184.859587 | 165.93404 | 156.907647 | 156.188526 | 144.278415 | 145.657916 |
| ENSMUSG00 Gm17113 | 169 | 0 | 0 | 1.43116954 | 0 | 0.28480428 | 0 | 0 | 0 | 0 | 0 | 0 | 0 | 0 | 0 | 0 | 0 | 0 | 0 |
| ENSMUSG00 Dnajc30 | 2287 | 7.24836845 | 6.30716362 | 5.54169871 | 7.31128258 | 7.26082801 | 6.93348262 | 7.81993556 | 5.77812472 | 7.13155236 | 6.55615375 | 8.73193382 | 7.98063912 | 8.28310286 | 7.27021569 | 7.1341566 | 6.04412299 | 7.00498124 | 7.33615109 |
| ENSMUSG00 Prcp | 2947 | 4.00965013 | 3.2216406 | 3.87382191 | 4.6380432 | 4.54043936 | 3.2284102 | 5.8920099 | 2.69562935 | 5.08717059 | 5.77584115 | 5.8847336 | 5.53042873 | 4.36906359 | 2.5846048 | 4.37600509 | 5.11927361 | 4.70787902 | 4.27454545 |
| ENSMUSG00 Cyp4f39 | 2431 | 0.01748465 | 0.93182921 | 0.49746535 | 0.58099234 | 0.41578379 | 0.63189418 | 0.09731115 | 0.01571057 | 0.01694221 | 0 | 0 | 0.09183984 | 0 | 0 | 0 | 0 | 0 | 0.49781997 |
| ENSMUSG00 Ppm1b | 4961 | 21.5310436 | 22.5353906 | 21.0128921 | 21.1687577 | 22.1692089 | 22.3441776 | 19.1882907 | 20.5165798 | 18.2479262 | 19.4552489 | 19.4046647 | 18.8114872 | 16.5367826 | 16.832588 | 17.7811996 | 20.8394759 | 20.6966639 | 18.6394494 |
| ENSMUSG00 Blnk | 5974 | 78.4502964 | 74.0757427 | 74.7789677 | 76.3494501 | 75.879889 | 71.3015229 | 64.6807246 | 64.5831351 | 69.2256118 | 66.9849378 | 67.7198441 | 68.9240279 | 59.4229121 | 55.4157106 | 58.4642767 | 70.1011672 | 65.9579274 | 61.5928883 |
| ENSMUSG00 Prpf40a | 10284 | 17.8882149 | 16.9285478 | 17.0887822 | 15.6832066 | 16.0252535 | 17.8956488 | 16.5898017 | 15.4752762 | 15.3067739 | 14.964984 | 16.5056814 | 16.0489026 | 15.9738705 | 16.3213664 | 14.6873691 | 16.5948305 | 16.9382565 | 15.1590634 |
| ENSMUSG00 Maml3 | 9769 | 2.42352165 | 2.45183447 | 1.97574354 | 2.63040526 | 1.76386822 | 2.36883269 | 1.21078613 | 1.72020238 | 1.39550995 | 1.98371778 | 1.24217826 | 2.19400291 | 2.17648238 | 1.32167635 | 1.72920851 | 2.23025444 | 1.77331213 | 1.9145354 |
| ENSMUSG00 Spink12 | 1003 | 19.0701189 | 15.4109501 | 22.7640143 | 12.9460595 | 14.4923639 | 9.28804976 | 15.4249711 | 13.8604531 | 4.35271253 | 2.49151273 | 2.1434428 | 6.90044084 | 2.75432439 | 3.88963444 | 1.35558459 | 0.80169766 | 0.4967516 | 3.78427554 |
| ENSMUSG00 Vmn1r229 | 920 | 0 | 0 | 0 | 0 | 0 | 0 | 0 | 0 | 0 | 0 | 0 | 0 | 0 | 0 | 0 | 0 | 0 | 0 |
| ENSMUSG00 Olfr854 | 947 | 0 | 0 | 0 | 0 | 0 | 0 | 0 | 0 | 0 | 0 | 0 | 0 | 0 | 0 | 0 | 0 | 0 | 0 |
| ENSMUSG00 Olfr151 | 2250 | 0 | 0 | 0 | 0 | 0 | 0 | 0 | 0 | 0 | 0 | 0 | 0 | 0 | 0 | 0 | 0 | 0 | 0 |
| ENSMUSG00 Rpl15-ps3 | 1296 | 0.0327972 | 0 | 0.18662627 | 0.03515513 | 0.11141649 | 0.03823513 | 0.07301339 | 0.05893889 | 0.03177972 | 0.03638172 | 0 | 0 | 0 | 0.07167297 | 0.09537397 | 0 | 0.1478638 | 0 |
| ENSMUSG00 Slc38a11 | 4669 | 0 | 0 | 0 | 0.03903281 | 0 | 0 | 0 | 0 | 0 | 0 | 0 | 0 | 0 | 0 | 0 | 0 | 0 | 0 |
| ENSMUSG00 Fnip2 | 12115 | 0.50872889 | 0.65443408 | 0.49910782 | 0.5565856 | 0.59593797 | 0.60943932 | 0.53893105 | 0.41297604 | 0.24817298 | 0.30357036 | 0.47321461 | 0.28103635 | 0.50899626 | 0.43703062 | 0.3672941 | 0.4330015 | 0.40176968 | 0.52216651 |
| ENSMUSG00 Tmprss11c | 1605 | 0.02648298 | 0 | 0 | 0 | 0 | 0 | 0 | 0 | 0 | 0 | 0 | 0 | 0 | 0 | 0 | 0 | 0 | 0 |
| ENSMUSG00 Sfmbt2 | 9143 | 0.11622327 | 0.2586914 | 0.32802788 | 0.16942749 | 0.37376863 | 0.14091338 | 0.14489282 | 0.33835553 | 0.26127288 | 0.12376866 | 0.18811096 | 0.09767588 | 0.02158238 | 0.09651511 | 0.07660795 | 0 | 0.13413994 | 0.05414874 |
| ENSMUSG00 Olfr1289 | 1013 | 0 | 0 | 0 | 0 | 0 | 0 | 0 | 0 | 0 | 0 | 0 | 0 | 0 | 0 | 0 | 0 | 0 | 0 |
| ENSMUSG00 Olfr601 | 989 | 0 | 0 | 0 | 0 | 0 | 0 | 0 | 0 | 0 | 0 | 0 | 0 | 0 | 0 | 0 | 0 | 0 | 0 |
| ENSMUSG00 Stk19 | 2797 | 7.87189173 | 6.46725118 | 7.281107 | 6.49941247 | 7.19311551 | 5.91727316 | 7.30749993 | 7.82418541 | 7.30372249 | 6.84418503 | 7.3276572 | 6.82479678 | 9.48894226 | 7.83754353 | 7.3653121 | 9.21329238 | 6.96094345 | 5.60514673 |
| ENSMUSG00 Vmn1r8 | 2140 | 0 | 0 | 0 | 0 | 0 | 0 | 0 | 0 | 0 | 0 | 0 | 0 | 0 | 0 | 0 | 0 | 0 | 0 |
| ENSMUSG00 Olfr47 | 1058 | 0 | 0 | 0 | 0 | 0 | 0 | 0 | 0 | 0 | 0 | 0 | 0 | 0 | 0 | 0 | 0 | 0 | 0 |
| ENSMUSG00 Vmn1r189 | 1136 | 0 | 0 | 0 | 0 | 0 | 0 | 0 | 0 | 0 | 0 | 0 | 0 | 0 | 0 | 0 | 0 | 0 | 0 |
| ENSMUSG00 Gm10096 | 898 | 0 | 0 | 0 | 0 | 0 | 0 | 0 | 0 | 0 | 0 | 0 | 0 | 0 | 0 | 0 | 0 | 0 | 0 |
| ENSMUSG00 H2-K1 | 2339 | 676.7391 | 606.88146 | 696.525066 | 714.796102 | 657.446272 | 733.969412 | 750.631125 | 744.939625 | 774.074063 | 861.069812 | 721.097535 | 826.305664 | 775.875691 | 689.19531 | 867.152712 | 826.399381 | 728.642057 | 913.188801 |
| ENSMUSG00 Olfr611 | 971 | 0 | 0 | 0 | 0 | 0 | 0 | 0 | 0 | 0 | 0 | 0 | 0 | 0 | 0 | 0 | 0 | 0 | 0 |
| ENSMUSG00 Rpl21-ps7 | 482 | 0.17637003 | 0 | 0 | 0.09452499 | 0 | 0 | 0 | 0 | 0.0854492 | 0 | 0 | 0 | 0 | 0 | 0.08548041 | 0.07944109 | 0 | 0 |
| ENSMUSG00 Exoc5 | 10131 | 7.8163205 | 9.26618655 | 8.59464561 | 8.58064086 | 8.03386467 | 8.53514165 | 7.61691633 | 6.99308128 | 5.47608693 | 7.27901489 | 6.27662321 | 8.50650311 | 5.83841366 | 8.82946524 | 7.46272382 | 8.13736825 | 7.23323262 | 9.39352035 |
| ENSMUSG00 Olfr387-ps1 | 1144 | 0 | 0 | 0 | 0 | 0 | 0 | 0 | 0 | 0 | 0 | 0 | 0 | 0 | 0 | 0 | 0 | 0 | 0 |
| ENSMUSG00 Tmprss11d | 2786 | 0 | 0 | 0 | 0 | 0 | 0 | 0 | 0 | 0 | 0 | 0 | 0 | 0 | 0 | 0 | 0 | 0 | 0 |
| ENSMUSG00 Gm10097 | 317 | 0 | 0 | 0 | 0 | 0 | 0 | 0 | 0 | 0 | 0.14874039 | 0 | 0 | 0.46686357 | 0 | 0 | 0 | 0 | 0.52059692 |
| ENSMUSG00 Gm14173 | 303 | 0 | 0 | 0 | 0.75183241 | 0 | 0 | 0 | 0.25209507 | 0 | 0 | 0 | 0 | 0 | 0.45984245 | 0 | 0 | 0 | 0 |
| ENSMUSG00 Mmgt1 | 4230 | 6.68225581 | 8.15102731 | 7.85641025 | 8.6598297 | 10.5708173 | 9.06709593 | 8.38877286 | 7.87323284 | 7.20520607 | 5.57336918 | 9.92770414 | 8.39214218 | 5.70299959 | 4.00758674 | 5.51302143 | 3.72948706 | 6.6232915 | 4.4475448 |
| ENSMUSG00 Tgif2lx1 | 1031 | 0 | 0 | 0 | 0 | 0 | 0 | 0 | 0 | 0 | 0 | 0 | 0 | 0 | 0 | 0 | 0 | 0 | 0 |
| ENSMUSG00 Exosc5 | 1024 | 31.7958641 | 30.7753886 | 26.9739119 | 32.5245346 | 32.9026822 | 27.4378906 | 36.7320115 | 33.1945685 | 30.3670111 | 31.1728771 | 34.0116648 | 28.0713537 | 38.4923847 | 32.2931545 | 31.9875361 | 36.4957352 | 32.300385 | 39.2154765 |
| ENSMUSG00 Taok3 | 10929 | 19.057129 | 20.6601516 | 19.5060983 | 19.6601597 | 18.2724267 | 21.4324981 | 16.8012176 | 17.3541468 | 17.9345437 | 18.2062373 | 18.1543968 | 16.7309096 | 19.8745019 | 18.9023059 | 18.4311837 | 19.1330412 | 21.6372268 | 18.0545032 |
| ENSMUSG00 Cyp3a59 | 1986 | 0 | 0 | 0 | 0 | 0 | 0 | 0 | 0 | 0 | 0 | 0 | 0 | 0 | 0 | 0 | 0 | 0 | 0 |
| ENSMUSG00 Olfr1261 | 1116 | 0 | 0 | 0 | 0 | 0 | 0 | 0 | 0 | 0 | 0 | 0 | 0 | 0 | 0 | 0 | 0 | 0 | 0 |
| ENSMUSG00 Vmn1r210 | 920 | 0 | 0 | 0 | 0 | 0 | 0 | 0 | 0 | 0 | 0 | 0 | 0 | 0 | 0 | 0 | 0 | 0 | 0 |
| ENSMUSG00 Agbl4 | 9508 | 0 | 0.04204404 | 0 | 0 | 0.05062255 | 0.0469052 | 0 | 0.01606748 | 0.06497662 | 0.04959056 | 0.06532125 | 0.03522234 | 0 | 0.00976948 | 0.05200028 | 0.03221759 | 0.06046429 | 0.00578556 |
| ENSMUSG00 Olfr342 | 1053 | 0 | 0 | 0 | 0 | 0 | 0 | 0 | 0 | 0 | 0 | 0 | 0 | 0 | 0 | 0 | 0 | 0 | 0 |
| ENSMUSG00 Slc38a10 | 6789 | 16.9544877 | 21.0996384 | 16.4380225 | 20.7504418 | 18.5324567 | 20.108673 | 16.0287467 | 17.9682624 | 12.1939751 | 19.6825885 | 14.1094111 | 14.9384385 | 10.1657611 | 13.545337 | 13.1148273 | 11.6129563 | 14.0117645 | 12.2755629 |
| ENSMUSG00 Rag1 | 6606 | 0 | 0 | 0 | 0 | 0 | 0 | 0 | 0 | 0 | 0 | 0 | 0 | 0 | 0 | 0 | 0 | 0 | 0 |
| ENSMUSG00 Ddhd2 | 4566 | 13.9077383 | 13.0231091 | 13.2534574 | 12.7223678 | 13.8935338 | 14.73776 | 14.7865074 | 14.9139408 | 13.9904262 | 12.5260191 | 15.1507096 | 12.8476262 | 12.4140107 | 12.3586435 | 13.5894751 | 14.0046681 | 15.3691304 | 13.9872039 |
| ENSMUSG00 Naca | 7025 | 28.4557783 | 27.0391634 | 28.6109635 | 24.9045424 | 27.385523 | 26.5574437 | 29.8760205 | 25.7044382 | 31.0496495 | 28.0487956 | 26.1283118 | 25.091226 | 33.0681848 | 30.2531166 | 27.8176483 | 27.171342 | 26.9457054 | 27.5946039 |
| ENSMUSG00 Dnaic1 | 2761 | 0 | 0.18098276 | 0 | 0.23102304 | 0 | 0.01794739 | 0 | 0 | 0 | 0 | 0 | 0 | 0.05360223 | 0.01682147 | 0.07461347 | 0.02773677 | 0 | 0.15938894 |
| ENSMUSG00 Gm11361 | 968 | 0.17564122 | 0.06882829 | 0.09994531 | 0.04706719 | 0.09944612 | 0 | 0.1466302 | 0.19727481 | 0.17019221 | 0.09741881 | 0.09870859 | 0 | 0.30577635 | 0.28787657 | 0.12769077 | 0 | 0 | 0 |
| ENSMUSG00 Gm17132 | 1157 | 0 | 0 | 0 | 0 | 0 | 0 | 0 | 0 | 0 | 0 | 0 | 0 | 0 | 0 | 0 | 0 | 0 | 0 |
| ENSMUSG00 Olfr116 | 965 | 0 | 0 | 0 | 0 | 0 | 0 | 0 | 0 | 0 | 0 | 0 | 0 | 0 | 0 | 0 | 0 | 0 | 0 |
| ENSMUSG00 Olfr1168 | 938 | 0 | 0 | 0 | 0 | 0 | 0 | 0 | 0 | 0 | 0 | 0 | 0 | 0 | 0 | 0 | 0 | 0 | 0 |
| ENSMUSG00 Olfr320 | 927 | 0 | 0 | 0 | 0 | 0 | 0 | 0 | 0 | 0 | 0 | 0 | 0 | 0 | 0 | 0 | 0 | 0 | 0 |
| ENSMUSG00 Cxcl12 | 8314 | 0.19427432 | 0.11619845 | 0.12800309 | 0 | 0.1215745 | 0.20264529 | 0.11381448 | 0.24346853 | 0 | 0.37430195 | 0.04022429 | 0.04028073 | 0.26107822 | 0.64241882 | 0.33203082 | 0.07829448 | 0.35956834 | 0.11247953 |
| ENSMUSG00 Gm600 | 2726 | 0.03118502 | 0 | 0.01774524 | 0.05014055 | 0.07062645 | 0.01817782 | 0.03471216 | 0.16812503 | 0.27195792 | 0.05188999 | 0 | 0 | 0 | 0.1511429 | 0 | 0.14059537 | 0 | 0 |
| ENSMUSG00 Phf5a | 1138 | 53.9718624 | 44.8758057 | 45.1854682 | 44.6402148 | 49.9083216 | 44.5016641 | 44.0698064 | 47.6566005 | 45.3485978 | 48.9737533 | 53.4004808 | 44.240534 | 51.5860998 | 60.5650489 | 54.8871342 | 47.7118465 | 44.9273111 | 39.347463 |
| ENSMUSG00 Olfr168 | 938 | 0 | 0 | 0 | 0 | 0 | 0 | 0 | 0 | 0 | 0 | 0 | 0 | 0 | 0 | 0 | 0 | 0 | 0 |
| ENSMUSG00 Olfr771 | 953 | 0 | 0 | 0 | 0 | 0 | 0 | 0 | 0 | 0 | 0 | 0 | 0 | 0 | 0 | 0 | 0 | 0 | 0 |
| ENSMUSG00 Zfp873 | 1891 | 5.91161359 | 6.00724322 | 9.41378063 | 6.55346592 | 7.61049454 | 7.07521844 | 7.40589792 | 6.82656373 | 8.58143163 | 8.17844033 | 9.60046759 | 6.64120787 | 3.54793271 | 4.78931626 | 6.23143574 | 5.48744294 | 7.70174131 | 6.31252044 |
| ENSMUSG00 Fiz1 | 2829 | 20.1482648 | 23.9395949 | 18.6893845 | 20.3567197 | 20.9099096 | 23.5765215 | 20.9052134 | 24.0575684 | 24.8371141 | 23.0336769 | 24.9598404 | 20.3019854 | 16.5137242 | 21.2437784 | 18.671048 | 16.0931538 | 17.7338716 | 23.4114403 |
| ENSMUSG00 Vmn1r219 | 938 | 0 | 0 | 0 | 0 | 0 | 0 | 0 | 0 | 0 | 0 | 0 | 0 | 0 | 0 | 0 | 0 | 0 | 0 |
| ENSMUSG00 Olfr1490 | 950 | 0 | 0 | 0 | 0 | 0 | 0 | 0 | 0 | 0 | 0 | 0 | 0 | 0 | 0 | 0 | 0 | 0 | 0 |
| ENSMUSG00 Csn1s2b | 2252 | 0 | 0 | 0 | 0 | 0 | 0 | 0.67229384 | 0.03391865 | 0.16459975 | 0.06281177 | 0.04242892 | 0 | 0 | 0 | 0.02062348 | 0 | 0 | 0 |
| ENSMUSG00 Gm7866 | 395 | 0 | 0.08433644 | 0 | 0 | 0.60926486 | 0 | 0 | 0.38675851 | 0.20853932 | 0.23873774 | 0.24189652 | 0 | 0.12489093 | 1.29337965 | 0.10430774 | 0.19387649 | 0.0970286 | 0.27852712 |
| ENSMUSG00 Mageb5 | 1355 | 0 | 0 | 0 | 0 | 0 | 0 | 0 | 0 | 0 | 0 | 0 | 0 | 0 | 0 | 0 | 0 | 0 | 0 |
| ENSMUSG00 Acvr2b | 1839 | 4.25282892 | 5.88727059 | 4.68215791 | 6.56534897 | 5.1298842 | 4.79629504 | 3.85910921 | 6.50039258 | 5.82299842 | 7.28156592 | 6.0270747 | 6.92004952 | 4.61396942 | 4.94999518 | 3.78633113 | 6.4546429 | 3.37621539 | 4.78600167 |
| ENSMUSG00 Gm13287 | 967 | 0 | 0 | 0 | 0.04711587 | 0 | 0 | 0 | 0 | 0.25555232 | 0.04875978 | 0 | 0 | 0 | 0 | 0.12782282 | 0.19798659 | 0 | 0.17065907 |
| ENSMUSG00 Krt79 | 2088 | 0.46820836 | 1.22849275 | 0.46334799 | 0.67643313 | 0.76070569 | 0.5903038 | 0.06797799 | 0.01829138 | 0 | 0.04516351 | 0 | 0.08019492 | 0.09450559 | 0 | 0 | 0 | 0 | 0.02634536 |
| ENSMUSG00 Zcchc14 | 9917 | 2.82024865 | 2.50594122 | 2.45355307 | 2.45791658 | 1.96080439 | 2.17858141 | 2.45222517 | 2.69969488 | 2.40465792 | 2.26791222 | 2.18231887 | 2.40327598 | 2.63647435 | 1.73749687 | 2.51771129 | 2.70663666 | 1.97100045 | 1.69736681 |
| ENSMUSG00 8430427H17I | 13426 | 2.01350305 | 2.42415445 | 2.65899489 | 2.28042766 | 1.99324814 | 2.03732377 | 2.41038823 | 2.12496087 | 2.71181885 | 2.77088521 | 2.4695234 | 1.79594565 | 2.12377835 | 2.34538144 | 2.47344362 | 3.1885073 | 2.60057027 | 2.67957365 |
| ENSMUSG00 Efcab4b | 1087 | 0 | 0 | 0 | 0 | 0 | 0 | 0 | 0 | 0 | 0 | 0 | 0 | 0.81690387 | 0 | 0 | 0 | 0 | 0 |
| ENSMUSG00 Hipk2 | 9601 | 2.87322771 | 4.55228801 | 3.27998836 | 2.99912301 | 3.45912169 | 3.66961708 | 2.20276716 | 3.59209867 | 2.86988637 | 2.77963733 | 2.55270644 | 3.08116875 | 2.38926586 | 2.44773547 | 2.49329278 | 2.22142155 | 2.97397054 | 2.76735742 |
| ENSMUSG00 Tmem151a | 4876 | 0.12204111 | 0.34160064 | 0.17857333 | 0.11212726 | 0.11845428 | 0.3353651 | 0.5336746 | 0.2428147 | 0.49836022 | 0.39646818 | 0.06818183 | 0.33196354 | 2.3674464 | 1.38113057 | 1.343529 | 2.63856524 | 2.41307899 | 2.35785548 |
| ENSMUSG00 Gm10100 | 344 | 0 | 0 | 0 | 0 | 0 | 0 | 0 | 0 | 0 | 0 | 0 | 0 | 0 | 0 | 0 | 0 | 0 | 0 |
| ENSMUSG00 Stx17 | 2428 | 13.0421566 | 15.7783478 | 16.9546435 | 16.2878857 | 15.5814217 | 15.0821541 | 15.0239191 | 14.9592153 | 15.4364618 | 17.0697974 | 14.6591203 | 20.8504143 | 13.470783 | 17.3687114 | 12.7100352 | 15.7073497 | 14.269758 | 20.2772449 |
| ENSMUSG00 Olfr871 | 1146 | 0 | 0 | 0 | 0 | 0 | 0 | 0 | 0 | 0 | 0 | 0 | 0 | 0 | 0 | 0 | 0 | 0 | 0 |
| ENSMUSG00 Nol10 | 3459 | 5.33311547 | 4.85391694 | 5.20235712 | 3.64857163 | 3.85445009 | 4.32637339 | 4.44539481 | 4.31720999 | 3.14346348 | 2.56268639 | 3.83967576 | 3.95340773 | 4.34987996 | 4.72631361 | 6.28922276 | 2.77853208 | 3.4016112 | 3.78495733 |
| ENSMUSG00 1810013D10I | 6728 | 18.2896083 | 14.4283255 | 14.7967785 | 17.0515323 | 16.1965926 | 14.2810271 | 16.4694254 | 14.5946294 | 15.4633084 | 15.5160013 | 17.0848023 | 13.3566018 | 14.7453159 | 14.7174189 | 14.5748667 | 12.3101343 | 13.6602943 | 14.725238 |

| | | | | | | | | | | | | | | | | | | | |
|---|---|---|---|---|---|---|---|---|---|---|---|---|---|---|---|---|---|---|---|
| ENSMUSG00 Obscn | 29009 | 0.15971127 | 0.11024295 | 0.12840022 | 0.07852915 | 0.11946287 | 0.1349466 | 0.03588124 | 0.0315977 | 0.08660683 | 0.0178792 | 0.04940704 | 0.10582445 | 0.00170057 | 0.00480307 | 0.02982635 | 0 | 0.03435085 | 0 |
| ENSMUSG00 Gm3724 | 353 | 0.48164505 | 0.37748322 | 1.23331947 | 0.77440889 | 0.81810635 | 0.84225609 | 0.40209076 | 0.75735642 | 0.58337841 | 1.20214258 | 1.35339826 | 0.6324721 | 0.97825332 | 0.13156965 | 0.35015487 | 0.75930383 | 0.5428654 | 0.31166633 |
| ENSMUSG00 Gm12569 | 793 | 0 | 0.04200869 | 0 | 0 | 0 | 0 | 0 | 0 | 0 | 0 | 0 | 0 | 0 | 0 | 0 | 0 | 0 | 0 |
| ENSMUSG00 Mrps36 | 1464 | 5.86478522 | 4.25513061 | 5.55106086 | 5.91297707 | 4.63565658 | 5.28020932 | 8.95192081 | 5.40015526 | 4.36059422 | 5.24970262 | 7.40772922 | 4.68943067 | 7.8513229 | 6.15447606 | 5.71305727 | 4.55093285 | 4.79078711 | 4.65924128 |
| ENSMUSG00 Rps7 | 954 | 60.3716075 | 60.4101751 | 52.8356589 | 49.2384033 | 54.8925922 | 58.8503639 | 66.3071821 | 52.324392 | 57.8942533 | 54.9597296 | 52.0316372 | 52.7732994 | 68.1545775 | 67.7187899 | 59.5133585 | 53.3419468 | 51.583821 | 58.4111374 |
| ENSMUSG00 Snrpa | 2081 | 55.5978324 | 49.8492804 | 54.9752039 | 48.3415598 | 50.7455264 | 49.2431778 | 49.2452007 | 47.699209 | 52.3094479 | 45.3380861 | 51.6088932 | 50.3172432 | 55.6376789 | 44.6363166 | 46.5076669 | 43.0378331 | 43.7409684 | 45.0963652 |
| ENSMUSG00 Hist1h4d | 392 | 8.24079943 | 5.09891238 | 5.42968198 | 5.69513054 | 6.99877461 | 5.30922145 | 8.32799706 | 5.94320548 | 7.14459967 | 13.7121943 | 9.62811663 | 10.2518564 | 19.3803961 | 13.1512594 | 13.7688873 | 13.5775368 | 12.2213957 | 11.9279951 |
| ENSMUSG00 Gm5161 | 857 | 0 | 0 | 0.05644519 | 0 | 0 | 0 | 0 | 0 | 0 | 0 | 0 | 0 | 0 | 0 | 0 | 0.04467982 | 0 | 0 |
| ENSMUSG00 Rpl27a-ps1 | 496 | 0 | 0 | 0 | 0.64299861 | 0 | 0 | 0.09538847 | 0.23100243 | 0.08303733 | 0 | 0 | 0 | 0 | 0.28091182 | 0 | 0.30879522 | 0.07727076 | 0 |
| ENSMUSG00 Olfr525 | 1049 | 0 | 0 | 0 | 0 | 0 | 0 | 0 | 0 | 0 | 0 | 0 | 0 | 0 | 0 | 0 | 0 | 0 | 0 |
| ENSMUSG00 Gm5959 | 415 | 0 | 0 | 0 | 0 | 0 | 0 | 0 | 0 | 0 | 0 | 0 | 0 | 0 | 0 | 0 | 0.09226652 | 0 | 0 |
| ENSMUSG00 Olfr197 | 922 | 0 | 0 | 0 | 0 | 0 | 0 | 0 | 0 | 0 | 0 | 0 | 0 | 0 | 0 | 0 | 0 | 0 | 0 |
| ENSMUSG00 Gm10101 | 2913 | 0.43774641 | 0.33164227 | 0.13284869 | 0.10948414 | 0.26437033 | 0.47630503 | 0.35732198 | 0.44577469 | 0 | 0.37228499 | 0.13120483 | 0.45986128 | 0.01693509 | 0.5739743 | 0.24044849 | 0.26289466 | 0.2631397 | 0.79312821 |
| ENSMUSG00 Sox21 | 6609 | 0.88110291 | 1.01314749 | 0.56358932 | 1.57178364 | 0.80838909 | 0.72728327 | 0.11454121 | 0.05200963 | 0.00623188 | 0.11414908 | 0.05060141 | 0.10979023 | 0 | 0 | 0 | 0.04055595 | 0.01159821 | 0.04161682 |
| ENSMUSG00 Cox5b | 671 | 282.966649 | 216.360049 | 237.325875 | 208.24996 | 252.925727 | 231.221474 | 333.304073 | 272.412575 | 266.822331 | 289.93115 | 294.624111 | 255.953493 | 347.161421 | 318.671503 | 286.445988 | 279.047795 | 260.915833 | 277.095053 |
| ENSMUSG00 Olfr153 | 1257 | 0 | 0 | 0 | 0 | 0 | 0 | 0 | 0 | 0 | 0 | 0 | 0 | 0 | 0 | 0 | 0 | 0 | 0 |
| ENSMUSG00 Zic2 | 5491 | 0 | 0 | 0 | 0 | 0 | 0 | 0 | 0 | 0 | 0 | 0 | 0 | 0 | 0 | 0 | 0 | 0 | 0 |
| ENSMUSG00 4921509C19I | 2680 | 0 | 0 | 0 | 0 | 0 | 0 | 0 | 0 | 0 | 0 | 0 | 0 | 0 | 0 | 0 | 0 | 0 | 0 |
| ENSMUSG00 Krt5 | 2415 | 2.71047499 | 1.36562175 | 1.24188774 | 2.22617111 | 2.88990846 | 1.29268001 | 0.78364686 | 1.5182073 | 0.69923277 | 5.2910313 | 0.03956518 | 1.82585397 | 4.59614136 | 1.8654561 | 4.74287062 | 1.07816203 | 0.92046593 | 1.73113545 |
| ENSMUSG00 Fam23a | 3632 | 0 | 0 | 0 | 0 | 0 | 0 | 0 | 0 | 0 | 0 | 0.0263078 | 0 | 0 | 0 | 0.01134404 | 0 | 0 | 0 |
| ENSMUSG00 Zfp955b | 1634 | 5.61879928 | 6.11619845 | 5.21036802 | 4.46130177 | 7.1579299 | 5.42835943 | 5.21192302 | 8.34436209 | 7.61219569 | 7.30056789 | 7.30951017 | 6.21692203 | 5.88722391 | 6.45214678 | 5.16910585 | 5.46004377 | 5.18366684 | 5.62210577 |
| ENSMUSG00 4930534B04I | 11877 | 0.54039586 | 0.47121043 | 0.69238866 | 0.5063617 | 0.51467158 | 0.36297784 | 0.71305629 | 0.36336966 | 0.57564719 | 0.64312654 | 0.75622566 | 0.841206 | 0.20352479 | 0.84074083 | 0.67645899 | 0.64156192 | 0.79059887 | 0.18063107 |
| ENSMUSG00 C1qtnf7 | 4529 | 0 | 0 | 0 | 0 | 0 | 0 | 0 | 0 | 0 | 0 | 0 | 0 | 0 | 0 | 0 | 0 | 0 | 0 |
| ENSMUSG00 Sec22c | 8129 | 0.28758576 | 0.54913616 | 0.48200959 | 0.7174085 | 0.57433837 | 0.26821507 | 0.7624506 | 0.38525981 | 0.22799769 | 0.58003079 | 0.65235827 | 0.39137566 | 0.78285365 | 0.33708958 | 0.58287353 | 0.54169269 | 0.4667614 | 0.31128293 |
| ENSMUSG00 Orm2 | 769 | 0.05527331 | 0.38987783 | 0.56614015 | 0 | 0 | 0.06443789 | 0 | 0 | 0.32135123 | 0 | 0 | 0 | 0 | 0 | 0.0535781 | 0 | 0.14951741 | 0.0715333 |
| ENSMUSG00 Zfp229 | 4519 | 2.26681731 | 1.99774161 | 1.8839879 | 1.4518235 | 2.52428987 | 2.05053356 | 2.2614602 | 1.74101237 | 1.87749994 | 1.52334647 | 1.01491392 | 1.86505091 | 1.33181985 | 2.3638283 | 1.69583744 | 1.54213112 | 2.24750358 | 2.15459435 |
| ENSMUSG00 Olfr301 | 1206 | 0 | 0 | 0 | 0 | 0 | 0 | 0 | 0 | 0 | 0 | 0 | 0 | 0 | 0 | 0 | 0 | 0 | 0 |
| ENSMUSG00 Wdr61 | 4664 | 22.9112377 | 21.1991145 | 21.147862 | 19.1563482 | 22.1258243 | 21.8440009 | 20.3391769 | 18.8423495 | 22.1562968 | 20.8761154 | 20.4457155 | 23.324341 | 18.753321 | 16.978381 | 18.7014781 | 20.69696 | 18.9330592 | 21.0176304 |
| ENSMUSG00 Olfr373 | 944 | 0 | 0 | 0 | 0.19305527 | 0 | 0.10498461 | 0 | 0 | 0 | 0.14984334 | 0 | 0 | 0 | 0 | 0 | 0 | 0 | 0 |
| ENSMUSG00 Dpp6 | 10758 | 0 | 0 | 0 | 0 | 0.00894812 | 0.00460613 | 0.17591626 | 0.04615182 | 0.24119265 | 0.0350628 | 0.00444088 | 0.10895417 | 0.33933462 | 0.29788455 | 0.23745087 | 0.02135561 | 0.16031636 | 0.04090657 |
| ENSMUSG00 Gpr114 | 3192 | 0 | 0 | 0 | 0 | 0 | 0 | 0 | 0 | 0 | 0 | 0 | 0 | 0 | 0 | 0 | 0 | 0 | 0 |
| ENSMUSG00 Ksr2 | 3870 | 0.25261474 | 0.04303992 | 0.17499468 | 0.12950168 | 0.13680909 | 0.35852107 | 0.37899045 | 0.2861963 | 0.53212553 | 0.30459111 | 0.25924396 | 0.37498895 | 0.16571445 | 0.1920169 | 0.31939191 | 0.08904792 | 0.15845497 | 0.08528544 |
| ENSMUSG00 Lyg2 | 2560 | 0 | 0 | 0 | 0 | 0 | 0 | 0 | 0 | 0 | 0 | 0 | 0 | 0.01927028 | 0 | 0 | 0.01495727 | 0 | 0 |
| ENSMUSG00 Dot1l | 8986 | 4.12942563 | 4.52649052 | 3.90282768 | 3.22468328 | 3.32627694 | 3.82701949 | 3.16962305 | 4.76023711 | 2.76379579 | 3.98781821 | 2.83906386 | 4.32313612 | 3.70016829 | 3.43704853 | 3.69099182 | 4.16739529 | 3.1860387 | 3.63625744 |
| ENSMUSG00 Pclo | 20404 | 0.49787968 | 0.1861238 | 0.45756182 | 0.41532809 | 0.27363768 | 0.26228657 | 0.74201418 | 0.31446401 | 0.79530912 | 0.77875843 | 0.23648651 | 0.44315513 | 0.67938976 | 0.18665042 | 0.42808909 | 0.74126592 | 0.30617459 | 0.45292736 |
| ENSMUSG00 Vmn1r78 | 941 | 0 | 0 | 0 | 0 | 0 | 0 | 0 | 0 | 0 | 0 | 0 | 0 | 0 | 0 | 0 | 0 | 0 | 0 |
| ENSMUSG00 Akap6 | 10252 | 0 | 0 | 0 | 0 | 0 | 0 | 0 | 0.03725361 | 0 | 0 | 0 | 0 | 0 | 0 | 0 | 0 | 0 | 0.08585112 |
| ENSMUSG00 Mdc1 | 7408 | 2.01968439 | 1.6548522 | 2.08303944 | 1.64826672 | 1.53985771 | 1.65220372 | 1.95433042 | 1.6497798 | 2.07934083 | 2.0112881 | 2.16044966 | 1.95897304 | 2.15094618 | 1.68021266 | 1.83538248 | 1.9279693 | 1.9297663 | 1.06929149 |
| ENSMUSG00 U2af1 | 965 | 54.2656755 | 41.3217973 | 43.6614973 | 44.0974253 | 43.8424465 | 42.8258858 | 54.8140674 | 48.9179372 | 41.1865158 | 46.5155124 | 45.6461261 | 43.2065284 | 45.8045574 | 45.0002298 | 42.3970415 | 45.5122556 | 42.0199193 | 52.7289365 |
| ENSMUSG00 Olfr845 | 941 | 0 | 0 | 0 | 0 | 0 | 0 | 0 | 0 | 0 | 0 | 0 | 0 | 0 | 0 | 0 | 0 | 0 | 0 |
| ENSMUSG00 Hist1h2ab | 505 | 0.08416867 | 0 | 0 | 0.18043978 | 0.09531074 | 0.29437267 | 0 | 0.22688556 | 0 | 0.09336773 | 0 | 0.22105213 | 0 | 0 | 0.08158724 | 0.60658388 | 0 | 0.10892892 |
| ENSMUSG00 Olfr12 | 1120 | 0 | 0 | 0 | 0 | 0 | 0 | 0 | 0 | 0 | 0 | 0 | 0 | 0 | 0 | 0 | 0 | 0 | 0 |
| ENSMUSG00 Vma21-ps | 475 | 0 | 0 | 0.10183901 | 0 | 0 | 0 | 0.09960564 | 0.16081012 | 0 | 0 | 0 | 0.35251997 | 0.10385667 | 0.09777703 | 0 | 0.08061181 | 0 | 0 |
| ENSMUSG00 Olfr68 | 947 | 0 | 0 | 0 | 0 | 0 | 0 | 0 | 0 | 0 | 0 | 0 | 0 | 0 | 0 | 0 | 0 | 0 | 0 |
| ENSMUSG00 1700029P11I | 1018 | 0.08350722 | 0 | 0 | 0 | 0 | 0 | 0 | 0 | 0 | 0.23158499 | 0 | 0 | 0.24229822 | 0 | 0 | 0 | 0.03764862 | 0.27018225 |
| ENSMUSG00 Olfr1457 | 953 | 0 | 0 | 0 | 0 | 0 | 0 | 0 | 0 | 0 | 0 | 0 | 0 | 0 | 0 | 0 | 0 | 0 | 0 |
| ENSMUSG00 Med9 | 1918 | 51.3031713 | 51.0288234 | 46.1288775 | 47.841472 | 49.1859074 | 49.6044048 | 50.2728048 | 47.6508962 | 48.5949349 | 48.1832004 | 50.5398283 | 45.7176256 | 50.5922217 | 45.3302042 | 49.5363858 | 36.5337912 | 43.9413592 | 46.6344148 |
| ENSMUSG00 Vmn1r46 | 929 | 0 | 0 | 0 | 0 | 0 | 0 | 0 | 0 | 0 | 0 | 0 | 0 | 0 | 0 | 0 | 0 | 0 | 0 |
| ENSMUSG00 Cd2ap | 5381 | 54.6619249 | 51.17965 | 58.0104797 | 54.4260162 | 51.2626007 | 56.1923026 | 45.9762122 | 48.1078059 | 48.3889895 | 46.1254975 | 47.3576712 | 51.5732158 | 40.8608726 | 36.3284081 | 42.9932608 | 46.9862257 | 45.1069389 | 44.9089399 |
| ENSMUSG00 Gdpd1 | 2326 | 38.4666361 | 45.6870733 | 42.8415617 | 43.5239211 | 41.5515489 | 44.9724078 | 21.4595341 | 23.3489236 | 23.125361 | 22.7442343 | 20.7243908 | 26.4200535 | 13.9130429 | 11.700875 | 13.3559644 | 13.7293064 | 18.7347376 | 16.5074619 |
| ENSMUSG00 Gm6404 | 407 | 0 | 0.08184986 | 0 | 0 | 0.11826025 | 0.12175119 | 0.11624737 | 0.37535531 | 0 | 0 | 0 | 0.13713922 | 0.12120864 | 0 | 0.10123232 | 0.09408012 | 0 | 0.13515751 |
| [illegible] | [illegible] | [illegible] | [illegible] | [illegible] | [illegible] | [illegible] | [illegible] | [illegible] | [illegible] | [illegible] | [illegible] | [illegible] | [illegible] | [illegible] | [illegible] | [illegible] | [illegible] | [illegible] | [illegible] |

| | | | | | | | | | | | | | | | | | | | |
|---|---|---|---|---|---|---|---|---|---|---|---|---|---|---|---|---|---|---|---|
| ENSMUSG00 Olfr177 | 929 | 0 | 0 | 0 | 0 | 0 | 0 | 0 | 0 | 0 | 0 | 0 | 0 | 0 | 0 | 0 | 0 | 0 | 0 |
| ENSMUSG00 Kcnma1 | 5370 | 79.9287277 | 91.2723673 | 90.4413864 | 96.5266297 | 90.7246427 | 103.34098 | 73.4359737 | 73.5683818 | 71.9651913 | 76.8986702 | 72.3118925 | 74.5144293 | 64.4438361 | 62.3232949 | 68.4544286 | 66.505865 | 73.3481105 | 65.2938634 |
| ENSMUSG00 Bbs5 | 2568 | 4.86624679 | 4.76083808 | 4.70926113 | 5.48222847 | 4.85442688 | 4.34165306 | 3.75848384 | 5.1607335 | 4.08978252 | 4.82890769 | 5.00446411 | 7.25951375 | 7.2614738 | 6.27573922 | 5.35876934 | 4.4732018 | 3.82069003 | 5.91219371 |
| ENSMUSG00 Clip2 | 4978 | 0.94778516 | 0.72943059 | 0.71909226 | 0.58575871 | 0.64781818 | 0.64703248 | 0.55125259 | 1.00506322 | 0.69499143 | 0.92823803 | 0.63341648 | 1.12124674 | 0.98108875 | 0.77437912 | 1.10080493 | 0.72304482 | 0.81610837 | 0.85088413 |
| ENSMUSG00 Csn2 | 1113 | 0 | 0 | 0 | 0 | 0 | 0 | 0 | 0 | 0 | 0 | 0 | 0 | 0 | 0 | 0 | 0 | 0 | 0 |
| ENSMUSG00 Numbl | 5769 | 2.28403615 | 1.85937804 | 1.2158367 | 1.61900054 | 1.48508969 | 1.19393829 | 1.58283013 | 1.15854921 | 0.78532098 | 1.14423617 | 0.87782035 | 1.27711344 | 1.03469613 | 0.90972125 | 0.76418209 | 1.6128649 | 1.74723801 | 1.38261752 |
| ENSMUSG00 Gm16416 | 482 | 0 | 0 | 0 | 0 | 0.39943505 | 0.1028065 | 0 | 0 | 0 | 0 | 0.09911817 | 0 | 0 | 0 | 0 | 0 | 0.07951514 | 0.11412678 |
| ENSMUSG00 Speer2 | 1251 | 0 | 0 | 0 | 0 | 0 | 0 | 0 | 0 | 0 | 0 | 0 | 0 | 0 | 0 | 0 | 0 | 0 | 0 |
| ENSMUSG00 Gm5449 | 356 | 0 | 0.09357555 | 0 | 0 | 0 | 0 | 0.13290078 | 0.10728203 | 0 | 0 | 0.67099661 | 0 | 0 | 0 | 0 | 0 | 0 | 0 |
| ENSMUSG00 Rps4y2 | 942 | 151.791308 | 137.813534 | 148.612523 | 129.524922 | 152.111186 | 125.670361 | 181.465721 | 158.161955 | 192.859576 | 161.073209 | 183.999522 | 179.534456 | 176.903627 | 167.829802 | 172.679132 | 169.624952 | 146.754727 | 154.165649 |
| ENSMUSG00 Hspb11 | 2063 | 6.18107262 | 4.39219933 | 4.17377043 | 4.81449717 | 4.59621375 | 4.37159356 | 7.95807057 | 5.40580781 | 4.89127308 | 6.23952593 | 6.22950258 | 4.84294891 | 6.40860583 | 6.52873744 | 5.03286092 | 6.19925491 | 5.4990712 | 4.18634501 |
| ENSMUSG00 Olfr811 | 959 | 0 | 0 | 0 | 0 | 0 | 0 | 0 | 0 | 0 | 0 | 0 | 0 | 0 | 0 | 0 | 0 | 0 | 0 |
| ENSMUSG00 Olfr955 | 944 | 0 | 0 | 0 | 0 | 0 | 0 | 0 | 0 | 0 | 0 | 0 | 0 | 0 | 0 | 0 | 0 | 0 | 0 |
| ENSMUSG00 Klk1b27 | 877 | 20.7436891 | 15.6118581 | 21.5667621 | 13.9748243 | 19.977218 | 18.080815 | 11.0594061 | 9.84205581 | 10.9423696 | 11.2365986 | 13.1285804 | 10.7558118 | 16.5377237 | 19.5414689 | 25.0873784 | 9.29977124 | 11.6246237 | 15.3047002 |
| ENSMUSG00 Pstk | 2181 | 10.855288 | 8.52296881 | 10.9123232 | 10.02719 | 10.0192083 | 8.13382789 | 9.56666273 | 11.942787 | 12.9734692 | 11.4147507 | 10.8211048 | 11.3883401 | 12.7570845 | 10.7113141 | 10.5412509 | 10.1651819 | 9.47174417 | 9.18079544 |
| ENSMUSG00 Gm10126 | 170 | 0 | 0 | 0 | 0 | 0.28312896 | 0.58297334 | 0 | 0 | 0 | 0 | 0 | 0 | 0 | 0 | 0 | 0 | 0 | 0 |
| ENSMUSG00 Olfr107 | 1032 | 0 | 0 | 0 | 0 | 0 | 0 | 0 | 0 | 0 | 0 | 0 | 0 | 0 | 0 | 0 | 0 | 0 | 0 |
| ENSMUSG00 Cd300lb | 2913 | 0 | 0 | 0 | 0 | 0 | 0 | 0 | 0 | 0 | 0 | 0 | 0 | 0 | 0 | 0 | 0 | 0 | 0 |
| ENSMUSG00 Nol7 | 1803 | 40.7842233 | 41.9413588 | 42.5247064 | 39.8501203 | 44.7149041 | 41.1702689 | 47.627572 | 40.4801337 | 47.9481401 | 40.2729246 | 43.0584068 | 44.0210051 | 41.752915 | 47.0365522 | 46.2974777 | 42.0283483 | 45.4686351 | 38.5033237 |
| ENSMUSG00 Gm15315 | 373 | 0 | 0 | 0 | 0 | 0 | 0 | 0 | 0 | 0 | 0 | 0 | 0 | 0 | 0 | 0 | 0 | 0 | 0 |
| ENSMUSG00 Olfr1351 | 959 | 0 | 0 | 0 | 0 | 0 | 0 | 0 | 0 | 0 | 0 | 0 | 0 | 0 | 0 | 0 | 0 | 0 | 0 |
| ENSMUSG00 Olfr930 | 2024 | 0 | 0 | 0 | 0 | 0 | 0 | 0 | 0 | 0 | 0 | 0 | 0 | 0 | 0 | 0 | 0 | 0 | 0 |
| ENSMUSG00 Olfr917 | 929 | 0 | 0 | 0 | 0 | 0 | 0 | 0 | 0 | 0 | 0 | 0 | 0 | 0 | 0 | 0 | 0 | 0 | 0 |
| ENSMUSG00 Ldha | 2895 | 633.261053 | 570.404203 | 610.007763 | 548.196151 | 586.145412 | 596.669686 | 926.119133 | 891.248404 | 889.059669 | 875.292951 | 925.629506 | 911.829342 | 930.677718 | 669.436573 | 791.809936 | 751.011737 | 779.751489 | 769.500447 |
| ENSMUSG00 Olfr535 | 990 | 0 | 0 | 0 | 0 | 0 | 0 | 0 | 0 | 0 | 0 | 0 | 0 | 0 | 0 | 0 | 0 | 0 | 0 |
| ENSMUSG00 Serpina11 | 1624 | 0 | 0 | 0 | 0 | 0 | 0 | 0 | 0 | 0 | 0 | 0 | 0 | 0 | 0 | 0 | 0 | 0 | 0 |
| ENSMUSG00 Gpr84 | 1548 | 0 | 0 | 0 | 0 | 0 | 0.03201081 | 0 | 0 | 0 | 0 | 0 | 0 | 0.03186816 | 0 | 0 | 0 | 0 | 0 |
| ENSMUSG00 Ptpmt1 | 2004 | 20.107239 | 17.6039695 | 18.2004201 | 16.0054767 | 18.18157 | 18.6935462 | 20.7760267 | 16.7711148 | 20.7165708 | 15.8815992 | 16.3779424 | 18.187439 | 19.4472129 | 17.4976477 | 17.6607468 | 17.502094 | 13.769927 | 19.0775096 |
| ENSMUSG00 1110038F14I | 1005 | 36.0342388 | 30.32965 | 33.8855377 | 32.0514013 | 29.7412185 | 29.7809463 | 34.2722689 | 32.9480725 | 34.4655321 | 39.5502914 | 29.2354224 | 35.5998405 | 36.6676042 | 37.8022522 | 31.1983921 | 36.5380025 | 33.4068021 | 35.8517064 |
| ENSMUSG00 Grm4 | 5599 | 0 | 0 | 0 | 0 | 0 | 0 | 0 | 0 | 0 | 0 | 0 | 0 | 0 | 0 | 0 | 0 | 0 | 0 |
| ENSMUSG00 Olfr133 | 620 | 0 | 0 | 0 | 0 | 0 | 0 | 0 | 0 | 0 | 0 | 0 | 0 | 0 | 0 | 0 | 0 | 0 | 0 |
| ENSMUSG00 Tgif2lx2 | 1031 | 0 | 0 | 0 | 0 | 0 | 0 | 0 | 0 | 0 | 0 | 0 | 0 | 0 | 0 | 0 | 0 | 0 | 0 |
| ENSMUSG00 Gm13248 | 1496 | 1.02148621 | 0.84505339 | 0.6781336 | 0.54746248 | 1.09244687 | 0.8600608 | 1.16860422 | 0.81585906 | 0.85232442 | 0.72394271 | 0.57406492 | 0.81972268 | 1.28434231 | 0.93012191 | 1.37521882 | 1.32917996 | 0.58845449 | 0.5141038 |
| ENSMUSG00 Krtap4-1 | 1010 | 0 | 0.09894919 | 0 | 0 | 0 | 0.04906211 | 0 | 0 | 0 | 0 | 0 | 0 | 0 | 0 | 0 | 0 | 0 | 0 |
| ENSMUSG00 Gm4744 | 735 | 0 | 0 | 0 | 0 | 0 | 0 | 0 | 0 | 0 | 0 | 0 | 0 | 0 | 0 | 0 | 0 | 0 | 0 |
| ENSMUSG00 Scoc | 2057 | 17.7294317 | 14.8183268 | 15.873667 | 16.0582193 | 17.6897104 | 13.75528 | 18.5616587 | 17.861493 | 18.4007817 | 16.412204 | 21.7855669 | 16.8233888 | 12.9984925 | 17.4532229 | 14.9022643 | 14.8545964 | 18.9116147 | 14.9757413 |
| ENSMUSG00 B230325K18I | 2622 | 0.03242195 | 0.05082059 | 0.05534729 | 0 | 0 | 0 | 0 | 0.05826453 | 0.10995637 | 0.05394817 | 0 | 0.1702995 | 0.03762923 | 0 | 0 | 0.29207176 | 0.0438516 | 0 |
| ENSMUSG00 Syt10 | 1838 | 0 | 0.18124534 | 0 | 0 | 0 | 0 | 0 | 0 | 0 | 0 | 0 | 0.15183804 | 0 | 0 | 0 | 0 | 0 | 0 |
| ENSMUSG00 Parp10 | 4264 | 6.90808794 | 7.4532132 | 6.00131275 | 8.46255795 | 7.13399994 | 8.99479732 | 5.53682617 | 6.18029026 | 4.91649543 | 6.04864791 | 6.63292106 | 7.02931773 | 4.08399784 | 4.46577763 | 5.48838737 | 4.46304688 | 5.18627423 | 5.67636186 |
| ENSMUSG00 Vmn1r202 | 908 | 0 | 0 | 0 | 0 | 0 | 0 | 0 | 0 | 0 | 0 | 0 | 0 | 0 | 0 | 0 | 0 | 0 | 0 |
| ENSMUSG00 Naa15 | 6070 | 9.75448274 | 8.50658749 | 9.96956944 | 6.94299275 | 7.5250734 | 7.9839495 | 7.06962105 | 7.64477247 | 8.41371986 | 7.74452243 | 6.98128306 | 6.71259204 | 6.76993198 | 8.19466517 | 6.50943857 | 7.43102727 | 7.15382115 | 5.66403488 |
| ENSMUSG00 Ptpla | 1120 | 4.2505176 | 3.77744426 | 4.10311195 | 4.47474543 | 5.97351553 | 2.9200718 | 6.75895413 | 6.41086755 | 4.8173514 | 6.06223328 | 6.01452602 | 6.82745158 | 5.99030424 | 8.00331149 | 8.4978209 | 5.53846286 | 4.79076711 | 9.47924785 |
| ENSMUSG00 Gm10128 | 610 | 0 | 0 | 0 | 0 | 0 | 0 | 0 | 0 | 0 | 0 | 0 | 0 | 0 | 0 | 0 | 0 | 0 | 0 |
| ENSMUSG00 Zfp35 | 923 | 22.703198 | 32.9519745 | 25.6804224 | 21.9166887 | 32.5398921 | 22.9778656 | 24.4508644 | 24.7029948 | 23.2482933 | 23.8052305 | 28.1059615 | 23.1003067 | 27.7926295 | 36.531319 | 25.6226361 | 23.7293905 | 24.8726456 | 32.7193928 |
| ENSMUSG00 Gm8995 | 1129 | 0 | 0 | 0 | 0 | 0 | 0 | 0.54478727 | 0.20297114 | 0 | 0 | 0.42316172 | 0 | 0.48064756 | 0.2468242 | 0 | 0 | 0 | 0 |
| ENSMUSG00 Olfr339 | 929 | 0 | 0 | 0 | 0 | 0 | 0 | 0 | 0 | 0 | 0 | 0 | 0 | 0 | 0 | 0 | 0 | 0 | 0 |
| ENSMUSG00 Tmem117 | 2773 | 174.189982 | 187.191315 | 178.858207 | 207.727473 | 203.185106 | 204.572555 | 151.885131 | 157.686554 | 132.693953 | 157.673448 | 171.924382 | 146.815522 | 87.9364108 | 97.6447997 | 106.131523 | 98.0946538 | 126.630196 | 109.145368 |
| ENSMUSG00 Luzp2 | 1026 | 0 | 0 | 0 | 0 | 0 | 0 | 0.09222744 | 0 | 0 | 0 | 0 | 0 | 0 | 0.09053428 | 0.20078731 | 0 | 0 | 0 |
| ENSMUSG00 Gm10129 | 359 | 0 | 0 | 0 | 0 | 0 | 0 | 0 | 0 | 0 | 0 | 0 | 0 | 0 | 0 | 0 | 0 | 0 | 0 |
| ENSMUSG00 Psg20 | 2157 | 0 | 0 | 0 | 0 | 0 | 0 | 0 | 0 | 0 | 0 | 0 | 0 | 0 | 0 | 0 | 0 | 0 | 0 |
| ENSMUSG00 Gm12657 | 452 | 0.47019 | 0.29480437 | 0.21404217 | 1.7135791 | 0.63891934 | 0.98666947 | 0.52337034 | 1.01395759 | 1.00232671 | 1.04315715 | 0.52848405 | 0.37045794 | 0.76398987 | 0.71926684 | 0.54692331 | 0.67770987 | 1.01751231 | 1.82552345 |
| ENSMUSG00 Rpl27 | 1284 | 44.756229 | 36.5559719 | 38.1638522 | 33.4965933 | 38.3480981 | 38.3609168 | 46.0966989 | 37.5677603 | 39.5506028 | 38.5945398 | 41.7100689 | 33.4285394 | 57.515483 | 55.7039677 | 43.1268622 | 45.686301 | 39.60981 | 45.4981864 |
| ENSMUSG00 Usp31 | 10182 | 0.68879926 | 0.93571869 | 1.46327316 | 1.24843168 | 0.59562192 | 0.8565391 | 1.14308771 | 1.06902717 | 1.73531873 | 0.84280377 | 0.68504655 | 0.8825694 | 0.90601734 | 0.72982263 | 0.70004606 | 0.91383005 | 1.08406733 | 0.54566095 |
| ENSMUSG00 1190007I07R | 1178 | 11.943305 | 6.41937775 | 5.54365586 | 11.0228333 | 9.19327914 | 6.01531486 | 13.6957755 | 8.68893365 | 9.40507024 | 13.4087314 | 11.4773457 | 9.33419824 | 15.3691117 | 12.6952429 | 10.1779735 | 14.1070659 | 9.43516647 | 14.0091104 |
| ENSMUSG00 Gm5299 | 275 | 0.15456428 | 0.1211378 | 0.35180749 | 0.16567652 | 0.52507553 | 0 | 1.20432274 | 0.83328878 | 0 | 0.1714571 | 0.34745424 | 0.20296605 | 0.17938879 | 0 | 0 | 0 | 0 | 0 |
| ENSMUSG00 Gm9396 | 566 | 0.07509748 | 0 | 0 | 0 | 0 | 0 | 0.16718261 | 0 | 0 | 0 | 0 | 0 | 0 | 0 | 0 | 0 | 0 | 0.09718923 |
| ENSMUSG00 Krr1 | 6066 | 7.19630989 | 5.00297505 | 6.72253315 | 5.89604679 | 5.97483325 | 5.43233891 | 5.60794859 | 5.80504369 | 4.73244337 | 5.63538738 | 6.49758337 | 5.76927472 | 5.87981803 | 5.42843026 | 5.28434067 | 6.32495691 | 6.83630947 | 4.94229528 |
| ENSMUSG00 Olfr874 | 1857 | 0 | 0 | 0 | 0 | 0 | 0 | 0 | 0 | 0 | 0 | 0 | 0 | 0 | 0 | 0 | 0 | 0 | 0 |
| ENSMUSG00 Slc39a4 | 2252 | 1.47220414 | 1.58280625 | 3.28647875 | 2.30637613 | 2.2655346 | 2.70647701 | 0.92440403 | 0.15263394 | 2.32268539 | 0.4396824 | 0.8485783 | 1.58623553 | 1.55531356 | 0.32997575 | 0.85989038 | 1.10519071 | 1.31044621 | 3.37089553 |
| ENSMUSG00 Mapk1 | 6724 | 42.0184273 | 42.5130793 | 46.9562809 | 43.8941769 | 47.6236895 | 44.0329541 | 40.0300164 | 42.2024911 | 37.431707 | 37.6981232 | 43.7889752 | 39.9110211 | 40.5792437 | 36.1730645 | 36.8264948 | 38.951183 | 43.7640255 | 37.4690227 |
| ENSMUSG00 Olfr969 | 935 | 0 | 0 | 0 | 0 | 0 | 0 | 0 | 0 | 0 | 0 | 0 | 0 | 0 | 0 | 0 | 0 | 0 | 0 |
| ENSMUSG00 Alg11 | 3127 | 10.2898683 | 10.099336 | 13.056367 | 9.4414956 | 10.1743529 | 10.37961 | 10.3340453 | 9.96642161 | 8.29789093 | 9.06222343 | 10.175287 | 7.78242049 | 6.51553624 | 6.22324036 | 10.7780213 | 6.89402365 | 11.4598937 | 7.95142828 |
| ENSMUSG00 3300002I08R | 2392 | 0.63971001 | 0.69633976 | 0.56624534 | 0.83807941 | 0.68414942 | 0.49718462 | 0.37581141 | 0.68656911 | 1.18807257 | 0.92645613 | 0.2197009 | 0.58335768 | 0.94869072 | 0.83490625 | 0.36171935 | 0.73635783 | 0.57681718 | 0.7589049 |
| ENSMUSG00 Olfr763 | 1043 | 0 | 0 | 0 | 0 | 0 | 0 | 0 | 0 | 0 | 0 | 0 | 0 | 0 | 0 | 0 | 0 | 0 | 0 |
| ENSMUSG00 Ifna13 | 818 | 0 | 0 | 0 | 0 | 0 | 0 | 0 | 0 | 0 | 0 | 0 | 0 | 0 | 0 | 0 | 0 | 0 | 0 |
| ENSMUSG00 Olfr1287 | 917 | 0 | 0 | 0 | 0 | 0 | 0 | 0 | 0 | 0 | 0 | 0 | 0 | 0 | 0 | 0 | 0 | 0 | 0 |
| ENSMUSG00 Olfr945 | 962 | 0 | 0 | 0 | 0 | 0 | 0 | 0 | 0 | 0 | 0 | 0 | 0 | 0 | 0 | 0 | 0 | 0 | 0 |
| ENSMUSG00 Bcl9l | 5295 | 4.54351834 | 4.44171923 | 3.78217971 | 4.23343415 | 3.29969562 | 4.94123576 | 4.52128716 | 4.39266725 | 4.48812457 | 4.79076078 | 3.06770272 | 5.50250725 | 6.92230681 | 6.21885892 | 5.29122909 | 3.89052819 | 3.82901059 | 4.83082807 |
| ENSMUSG00 Zfp947 | 2867 | 0.99331943 | 1.42918939 | 1.1304592 | 1.65272013 | 1.79634316 | 1.71109893 | 1.46872285 | 1.91827902 | 0.9912346 | 2.07219693 | 1.11647095 | 0.85660591 | 0.55061764 | 0.93957345 | 0.96285464 | 1.50918683 | 1.08281481 | 0.61398375 |
| ENSMUSG00 Olfr1387 | 935 | 0 | 0 | 0 | 0 | 0 | 0 | 0 | 0 | 0 | 0 | 0 | 0 | 0 | 0 | 0 | 0 | 0 | 0 |
| ENSMUSG00 BC023105 | 2038 | 0 | 0.22884226 | 0.6408662 | 0.29062491 | 0.14170341 | 1.14277649 | 0.58038124 | 0.39354291 | 0 | 0 | 0 | 0.16432482 | 0 | 0 | 0 | 0 | 0.01880584 | 1.32259383 |
| ENSMUSG00 Gm10130 | 1064 | 1.31829863 | 1.06450978 | 1.90948146 | 1.28461591 | 1.04044572 | 1.44373566 | 1.73420534 | 1.32811926 | 2.43867528 | 0.84197684 | 2.06548673 | 1.4163749 | 2.36459378 | 3.36108527 | 3.09786135 | 2.59109373 | 1.51288014 | 0.93060519 |
| ENSMUSG00 Olfr293 | 1010 | 0 | 0 | 0 | 0 | 0 | 0 | 0 | 0 | 0 | 0 | 0 | 0 | 0 | 0 | 0 | 0 | 0 | 0 |
| ENSMUSG00 Tmed5 | 7682 | 10.7618546 | 10.9062652 | 9.93686114 | 8.96156444 | 10.5511793 | 10.7465314 | 9.47239003 | 9.31194607 | 8.29413432 | 7.42675748 | 9.39080803 | 8.53001665 | 8.4767158 | 7.53310764 | 8.23280243 | 9.50037721 | 9.84849129 | 10.125342 |
| ENSMUSG00 Lrrc43 | 2180 | 0.01949779 | 0.29034174 | 0.04437939 | 0.02089956 | 0 | 0 | 0 | 0 | 0 | 0 | 0 | 0 | 0.14913239 | 0 | 0 | 0 | 0 | 0 |
| ENSMUSG00 Stk24 | 2665 | 3.82785825 | 3.53754186 | 4.84642874 | 4.13725056 | 3.44960504 | 3.55143419 | 3.60393014 | 4.42831232 | 4.899109 | 3.46774403 | 4.69682519 | 3.70708153 | 3.70220768 | 4.37428365 | 4.3907099 | 4.81326585 | 4.30002355 | 5.55251396 |
| ENSMUSG00 Gm10131 | 314 | 1.48903483 | 2.01574838 | 0.92433497 | 1.16079094 | 1.53286381 | 1.42030128 | 1.95880518 | 1.58121411 | 1.18050523 | 1.65177623 | 0.76074775 | 2.8441102 | 1.57108017 | 1.03537774 | 1.04972117 | 1.58527992 | 1.58675752 | 1.75188238 |
| ENSMUSG00 Cyp26b1 | 4595 | 0 | 0 | 0 | 0 | 0.02094969 | 0 | 0.03088967 | 0 | 0 | 0 | 0 | 0.11809599 | 0.01010753 | 0 | 0 | 0 | 0 | 0.20351574 |
| ENSMUSG00 Gm8017 | 557 | 0.68679818 | 0.71769251 | 1.3026983 | 0.73617486 | 0.60488953 | 0.71170892 | 1.35907157 | 0.27427219 | 0.5915478 | 0.59255821 | 1.11503494 | 0.80166122 | 1.23994047 | 1.58426869 | 1.25749812 | 1.3061428 | 0.61927589 | 0.79007694 |
| ENSMUSG00 Ddo | 8576 | 0.27755246 | 0.25637255 | 0.27074737 | 0.35594566 | 0.14030995 | 0.22534475 | 0.51306893 | 0.57448926 | 0.48505575 | 0.4893205 | 0.6016436 | 0.34494288 | 0.52346134 | 0.08664942 | 0.17775858 | 0.17412939 | 0.36645946 | 0.16677201 |
| ENSMUSG00 Wscd2 | 4481 | 2.46626775 | 2.64659459 | 2.80676588 | 2.43005793 | 2.36309378 | 2.57660945 | 2.29119646 | 2.38649245 | 2.41732953 | 3.48290176 | 3.51835222 | 2.60331923 | 2.87338326 | 1.87600532 | 2.3906281 | 1.54666368 | 1.4625746 | 3.92834505 |
| ENSMUSG00 Sorcs3 | 5607 | 1.63743856 | 1.7229783 | 0.79371585 | 0.94258626 | 1.02152647 | 1.90893356 | 1.34166505 | 1.36912304 | 2.16693814 | 0.99229231 | 0.85205918 | 0.78641651 | 1.46051333 | 1.77261186 | 0.84504707 | 2.5472439 | 1.1551889 | 0.96145755 |
| ENSMUSG00 Olfr46 | 956 | 0 | 0 | 0 | 0 | 0 | 0 | 0 | 0 | 0 | 0 | 0 | 0 | 0 | 0 | 0 | 0 | 0 | 0 |
| ENSMUSG00 B9d2 | 1018 | 21.5031097 | 20.6487586 | 17.1540712 | 17.946934 | 20.6144585 | 18.1076787 | 20.5424402 | 16.1323507 | 17.8420957 | 22.8805967 | 27.548036 | 17.9838284 | 18.6569628 | 21.8077346 | 16.1487434 | 19.1829173 | 17.1677715 | 17.8860652 |

| | | | | | | | | | | | | | | | | | | | |
|---|---|---|---|---|---|---|---|---|---|---|---|---|---|---|---|---|---|---|---|
| ENSMUSG00 U6 | 106 | 0 | 0 | 0 | 0 | 0 | 0 | 0 | 0 | 0 | 0 | 0 | 0 | 0 | 0 | 0 | 0 | 0 | 0 |
| ENSMUSG00 U6 | 103 | 0 | 0 | 0 | 0 | 0 | 0 | 0 | 0 | 0 | 0 | 0 | 0 | 0 | 0 | 0 | 0 | 0 | 0 |
| ENSMUSG00 SNORA65 | 128 | 0 | 1.8217989 | 0 | 0 | 0 | 0.77426146 | 0 | 0 | 0.96530896 | 1.47345948 | 0.74648373 | 0 | 0 | 0 | 0.96566147 | 0 | 0.59884839 | 0 |
| ENSMUSG00 U6 | 105 | 0 | 0 | 0 | 0 | 0 | 0 | 0 | 0 | 0 | 0 | 0 | 0 | 0 | 0 | 0 | 0 | 0 | 0 |
| ENSMUSG00 Y_RNA | 106 | 0 | 0 | 0 | 0 | 0 | 0 | 0 | 0 | 0 | 0 | 0 | 0 | 0 | 0.43815177 | 0 | 0 | 0.36156884 | 0.51895384 |
| ENSMUSG00 U6 | 104 | 0 | 0 | 0 | 0 | 0 | 0 | 0 | 0 | 0 | 0 | 0 | 0 | 0 | 0 | 0 | 0 | 0 | 0 |
| ENSMUSG00 SNORA36 | 86 | 0 | 0 | 0 | 0 | 0 | 0 | 0 | 0 | 0 | 0 | 0 | 0 | 0 | 0 | 0 | 0 | 0 | 0 |
| ENSMUSG00 U6 | 106 | 0 | 0 | 0 | 0 | 0 | 0 | 0 | 0 | 0 | 0 | 0 | 0 | 0 | 0 | 0 | 0 | 0 | 0 |
| ENSMUSG00 U6 | 106 | 0 | 0 | 0 | 0 | 0 | 0 | 0 | 0 | 0 | 0 | 0 | 0 | 0 | 0 | 0 | 0 | 0 | 0 |
| ENSMUSG00 U6 | 106 | 0 | 0 | 0 | 0 | 0 | 0 | 0 | 0 | 0 | 0 | 0 | 0 | 0 | 0 | 0 | 0 | 0 | 0 |
| ENSMUSG00 U6 | 106 | 0 | 0 | 0 | 0 | 0 | 0 | 0 | 0 | 0 | 0 | 0 | 0 | 0 | 0 | 0 | 0 | 0 | 0 |
| ENSMUSG00 SNORD38 | 64 | 0 | 0 | 0 | 0 | 0 | 0 | 0 | 0 | 0 | 0 | 0 | 0 | 0 | 0 | 0 | 0 | 0 | 0 |
| ENSMUSG00 Snord42a | 64 | 0 | 0 | 0 | 0 | 0 | 0 | 0 | 0 | 0 | 0 | 0 | 0 | 0 | 0 | 0 | 0 | 0 | 0 |
| ENSMUSG00 U1 | 159 | 0 | 0 | 0 | 0 | 0 | 0 | 0 | 0 | 0 | 0 | 0 | 0 | 0 | 0 | 0 | 0 | 0 | 0 |
| ENSMUSG00 SNORD38 | 69 | 0.61601704 | 0 | 0 | 0 | 0 | 0 | 4.11414599 | 0.55351308 | 0 | 0 | 0 | 0 | 0 | 0 | 1.194248 | 1.10987268 | 0.55545358 | 6.37786745 |
| ENSMUSG00 U1 | 158 | 0 | 0 | 0 | 0 | 0 | 0 | 0 | 0 | 0 | 0 | 0 | 0 | 0 | 0 | 0 | 0 | 0 | 0 |
| ENSMUSG00 SNORD113 | 74 | 0 | 0 | 0 | 0 | 0 | 0 | 0 | 0 | 0 | 0 | 0 | 0 | 0 | 0 | 0 | 0 | 0 | 0 |
| ENSMUSG00 U6 | 106 | 0 | 0 | 0 | 0 | 0 | 0 | 0 | 0 | 0 | 0 | 0 | 0 | 0 | 0 | 0 | 0 | 0 | 0 |
| ENSMUSG00 7SK | 318 | 0 | 0 | 0.15211802 | 0 | 0 | 0 | 0 | 0 | 0 | 0 | 0 | 0 | 0 | 0 | 0 | 0 | 0 | 0 |
| ENSMUSG00 n-R5s65 | 116 | 0 | 0 | 0 | 0 | 0 | 0 | 0 | 0 | 0 | 0 | 0 | 0 | 0 | 0 | 0 | 0 | 0 | 0 |
| ENSMUSG00 U6 | 106 | 0 | 0 | 0 | 0 | 0 | 0 | 0 | 0 | 0 | 0 | 0 | 0 | 0 | 0 | 0 | 0 | 0 | 0 |
| ENSMUSG00 U2 | 193 | 0 | 0 | 0 | 0 | 0 | 0 | 0 | 0 | 0 | 0 | 0 | 0 | 0 | 0 | 0 | 0 | 0 | 0 |
| ENSMUSG00 U1 | 163 | 0 | 0 | 0 | 0 | 0 | 0 | 0 | 0 | 0 | 0 | 0 | 0 | 0 | 0 | 0 | 0 | 0 | 0 |
| ENSMUSG00 U1 | 161 | 0 | 0 | 0 | 0 | 0 | 0 | 0 | 0 | 0 | 0 | 0 | 0 | 0 | 0 | 0 | 0 | 0 | 0 |
| ENSMUSG00 n-R5s173 | 118 | 0 | 0 | 0 | 0 | 0 | 0 | 0 | 0 | 0 | 0 | 0 | 0 | 0 | 0 | 0 | 0 | 0 | 0 |
| ENSMUSG00 U1 | 160 | 0 | 0 | 0 | 0 | 0 | 0 | 0 | 0 | 0 | 0 | 0 | 0 | 0 | 0 | 0 | 0 | 0 | 0 |
| ENSMUSG00 SNORA19 | 129 | 0 | 0 | 0 | 0 | 0 | 0 | 0 | 0 | 0 | 0 | 0 | 0 | 0 | 0 | 0 | 0 | 0 | 0 |
| ENSMUSG00 U6 | 106 | 0 | 0 | 0 | 0 | 0 | 0 | 0 | 0 | 0 | 0 | 0 | 0 | 0 | 0 | 0 | 0 | 0 | 0 |
| ENSMUSG00 SNORA55 | 129 | 0.65899498 | 0 | 0 | 1.05955917 | 1.11934706 | 0 | 0 | 1.48032567 | 0.95782595 | 0.36550933 | 0.37034852 | 0 | 0.76483593 | 0 | 1.59695953 | 0 | 0.29710308 | 0 |
| ENSMUSG00 U6 | 106 | 0 | 0 | 0 | 0 | 0 | 0 | 0 | 0 | 0 | 0 | 0 | 0 | 0 | 0 | 0 | 0 | 0 | 0 |
| ENSMUSG00 Snora7a | 139 | 2.14054843 | 0.71898333 | 0.69602202 | 0.98333189 | 2.42390982 | 1.42597794 | 2.72303188 | 1.37382742 | 0.29630587 | 1.69606846 | 2.7496379 | 5.22016988 | 3.19415292 | 5.01195188 | 4.149797 | 5.78491191 | 4.68738883 | 6.33198351 |
| ENSMUSG00 U6 | 106 | 0 | 0 | 0 | 0 | 0 | 0 | 0 | 0 | 0 | 0 | 0 | 0 | 0 | 0 | 0 | 0 | 0 | 0 |
| ENSMUSG00 U1 | 162 | 0 | 1.02817575 | 0.29860204 | 0.56248203 | 0 | 0 | 0.87616072 | 0 | 0 | 1.74632234 | 1.47453576 | 1.03362338 | 0.30451801 | 0.2866919 | 0 | 0.23636177 | 1.41949248 | 0 |
| ENSMUSG00 U6 | 110 | 0 | 0 | 0 | 0 | 0 | 0 | 0 | 0 | 0 | 0 | 0 | 0 | 0 | 0 | 0.3745596 | 0 | 0 | 0 |
| ENSMUSG00 SNORA38 | 144 | 0 | 0 | 0 | 0 | 0 | 0 | 0 | 0 | 0 | 0 | 0 | 0 | 0 | 0 | 0.28612192 | 0 | 0 | 0 |
| ENSMUSG00 U6 | 106 | 0.80198445 | 0.62854517 | 0 | 0 | 0 | 0 | 0 | 0 | 0 | 0 | 0.45070716 | 0 | 0 | 0 | 0 | 0 | 0 | 0 |
| ENSMUSG00 SNORA71 | 130 | 0 | 0 | 0 | 0 | 0 | 0 | 0 | 0 | 0 | 0 | 0 | 0 | 0 | 0 | 0 | 0 | 0 | 0 |
| ENSMUSG00 SNORA9 | 133 | 0 | 0 | 0 | 0 | 0 | 0 | 0 | 0 | 0 | 0 | 0 | 0 | 0 | 0 | 0 | 0 | 0 | 0 |
| ENSMUSG00 U6 | 106 | 0 | 0 | 0 | 0 | 0 | 0 | 0 | 0 | 0 | 0 | 0 | 0 | 0 | 0 | 0 | 0 | 0 | 0 |
| ENSMUSG00 U6 | 105 | 0 | 0 | 0 | 0 | 0 | 0 | 0 | 0 | 0 | 0 | 0 | 0 | 0 | 0 | 0 | 0 | 0 | 0 |
| ENSMUSG00 SNORD81 | 76 | 0 | 0 | 0 | 0 | 0 | 0 | 0 | 0 | 0 | 0 | 0 | 0 | 0 | 0 | 0 | 0 | 0 | 0 |
| ENSMUSG00 U6 | 106 | 0 | 0 | 0 | 0 | 0 | 0 | 0 | 0 | 0 | 0 | 0 | 0 | 0 | 0 | 0 | 0 | 0 | 0 |
| ENSMUSG00 U6 | 106 | 0 | 0 | 0 | 0 | 0 | 0 | 0 | 0 | 0 | 0 | 0 | 0 | 0 | 0 | 0 | 0 | 0 | 0 |
| ENSMUSG00 U6 | 106 | 0 | 0 | 0 | 0 | 0 | 0 | 0 | 0 | 0 | 0 | 0 | 0 | 0 | 0 | 0 | 0 | 0 | 0 |
| ENSMUSG00 U1 | 162 | 0 | 0 | 0 | 0 | 0 | 0 | 0 | 0 | 0 | 0 | 0 | 0 | 0 | 0 | 0 | 0 | 0 | 0 |
| ENSMUSG00 U1 | 159 | 0 | 0.20951506 | 0.30423604 | 0 | 0 | 0 | 0 | 0 | 0 | 0 | 0 | 0 | 0 | 0 | 1.81390498 | 0 | 0.48209178 | 0 |
| ENSMUSG00 SNORA30 | 129 | 0 | 0 | 0 | 0 | 0 | 0 | 0 | 0 | 0 | 0 | 0 | 0 | 0 | 0 | 0 | 0 | 0 | 0 |
| ENSMUSG00 U4 | 142 | 1.79599335 | 0.70379354 | 2.04395199 | 5.45449122 | 4.40644372 | 7.32822116 | 5.99738184 | 3.49648754 | 4.35068828 | 3.32047206 | 2.01866022 | 7.07522483 | 5.55852589 | 1.96242623 | 1.74091081 | 4.5840868 | 2.15922799 | 2.32432845 |
| ENSMUSG00 U1 | 155 | 0 | 0 | 0 | 0 | 0 | 0 | 0 | 0 | 0 | 0 | 0 | 0 | 0 | 0 | 0 | 0 | 0 | 0 |
| ENSMUSG00 SNORA71 | 132 | 0 | 0 | 0 | 0 | 0 | 0 | 0 | 0 | 0 | 0 | 0 | 0 | 0 | 0 | 0 | 0 | 0 | 0 |
| ENSMUSG00 U6 | 106 | 0 | 0 | 0 | 0 | 0 | 0 | 0 | 0 | 0 | 0 | 0 | 0 | 0 | 0 | 0 | 0 | 0 | 0 |
| ENSMUSG00 U4 | 150 | 0 | 0 | 0 | 0 | 0.64175898 | 0 | 0 | 0 | 0 | 0 | 0 | 0 | 0 | 0 | 0.54935408 | 0 | 0 | 0 |
| ENSMUSG00 n-R5s74 | 115 | 0 | 0 | 0 | 0 | 0 | 0 | 0 | 0 | 0 | 0 | 0 | 0 | 0 | 0 | 0 | 0 | 0 | 0 |
| ENSMUSG00 U6 | 106 | 0 | 0 | 0 | 0 | 0 | 0 | 0 | 0 | 0 | 0 | 0 | 0 | 0 | 0 | 0 | 0 | 0 | 0 |
| ENSMUSG00 U6 | 106 | 0.80198445 | 0.31427259 | 0.45635406 | 0 | 0 | 0 | 0 | 0 | 0 | 0 | 0 | 1.57968856 | 0.93079089 | 0.43815177 | 0 | 0 | 0 | 0.51895384 |
| ENSMUSG00 U6 | 105 | 0 | 0 | 0 | 0 | 0 | 0 | 0 | 0 | 0 | 0 | 0 | 0 | 0 | 0 | 0 | 0 | 0 | 0 |
| ENSMUSG00 U1 | 163 | 0.5215359 | 0.20437358 | 0.59354025 | 0 | 0.29528788 | 0.9120135 | 0 | 0 | 0 | 0 | 0 | 0 | 0 | 0 | 0 | 0 | 0 | 0 |
| ENSMUSG00 SNORA44 | 125 | 0 | 0.79950946 | 0 | 0 | 0 | 0 | 0 | 0 | 0.65898425 | 0 | 0 | 0 | 0 | 0 | 0 | 0 | 0 | 0 |
| ENSMUSG00 SNORA51 | 132 | 0 | 0 | 0 | 0 | 0 | 0 | 0 | 0 | 0 | 0 | 0 | 0 | 0 | 0 | 0 | 0 | 0 | 0 |
| ENSMUSG00 SNORA2 | 135 | 0.31485316 | 0 | 0.35832245 | 0 | 0 | 0 | 0 | 0 | 0 | 0 | 0 | 0.41344935 | 0 | 0 | 0.91559013 | 0.56726826 | 0.2838985 | 0.81494973 |
| ENSMUSG00 SNORA13 | 132 | 0.32200891 | 0 | 0 | 0 | 0 | 0 | 0 | 0 | 0 | 0 | 0 | 0 | 0 | 0 | 0 | 0 | 0.58070147 | 1.66694263 |
| ENSMUSG00 U6 | 104 | 0 | 0 | 0 | 0 | 0 | 0 | 0 | 0 | 0 | 0 | 0 | 0 | 0 | 0 | 0 | 0 | 0 | 0 |
| ENSMUSG00 Snora41 | 131 | 1.94680196 | 3.05156283 | 4.0618995 | 2.78235385 | 2.93935412 | 5.29571199 | 4.33398586 | 6.99708136 | 6.91681944 | 2.15957419 | 7.29388679 | 4.2607376 | 8.66132899 | 3.54535018 | 6.91934526 | 3.21524184 | 4.38850728 | 7.97842006 |
| ENSMUSG00 Snora44 | 128 | 0 | 0 | 0 | 0 | 0 | 0 | 0 | 0 | 0 | 0 | 0 | 0 | 0 | 0 | 0 | 0 | 0 | 0 |
| ENSMUSG00 U1 | 148 | 0 | 0 | 0 | 0 | 0 | 0 | 0 | 0 | 0 | 0 | 0 | 0 | 0 | 0 | 0 | 0 | 0 | 0 |
| ENSMUSG00 U4 | 140 | 0.3036084 | 0.23794924 | 0 | 0 | 0 | 0.3539481 | 0 | 0 | 0.2941894 | 0 | 0.3412497 | 0 | 0.35237084 | 0 | 0.29429683 | 0 | 0.54751853 | 0 |
| ENSMUSG00 U6 | 106 | 0 | 0 | 0 | 0 | 0 | 0 | 0 | 0 | 0 | 0 | 0 | 0 | 0 | 0 | 0 | 0 | 0 | 0 |
| ENSMUSG00 SNORA36 | 130 | 0 | 0 | 0 | 0 | 0 | 0 | 0 | 0 | 0 | 0 | 0 | 0 | 0 | 0 | 0 | 0 | 0 | 0 |
| ENSMUSG00 U6 | 106 | 0 | 0.31427259 | 0 | 0 | 0 | 0 | 0 | 0 | 0 | 0 | 0 | 0 | 0 | 0 | 0 | 0 | 0 | 0 |
| ENSMUSG00 SNORD51 | 72 | 2.36139867 | 2.31339543 | 2.01556376 | 4.42954598 | 3.3424947 | 4.12939446 | 7.88544649 | 4.24360027 | 6.29238435 | 0.65487088 | 1.32708218 | 0.77521754 | 9.59231725 | 12.2560786 | 4.57795065 | 3.72269794 | 2.6615484 | 8.40416908 |
| ENSMUSG00 U1 | 139 | 0 | 0 | 0 | 0 | 0 | 0 | 0 | 0 | 0 | 0 | 0 | 0 | 0 | 0 | 0 | 0 | 0 | 0 |
| ENSMUSG00 U6 | 106 | 0 | 0.62854517 | 0 | 0.85964235 | 0.45407475 | 0.46747862 | 1.78538411 | 0.72061137 | 0.77710407 | 0.44481796 | 0.90141431 | 1.05312571 | 0 | 0 | 0 | 0 | 0.36156884 | 0 |
| ENSMUSG00 U6 | 106 | 0 | 0 | 0 | 0 | 0 | 0 | 0 | 0 | 0 | 0 | 0 | 0 | 0 | 0 | 0 | 0 | 0 | 0 |
| ENSMUSG00 n-R5s176 | 108 | 0 | 0 | 0 | 0 | 0 | 0 | 0 | 0 | 0 | 0 | 0 | 0 | 0 | 0 | 0 | 0 | 0 | 0 |
| ENSMUSG00 n-R5s95 | 117 | 0 | 0 | 0 | 0 | 0 | 0 | 0 | 0 | 0 | 0 | 0 | 0 | 0 | 0 | 0 | 0 | 0 | 0 |
| ENSMUSG00 U6 | 105 | 0 | 0 | 0 | 0 | 0 | 0 | 0 | 0 | 0 | 0 | 0 | 0 | 0 | 0 | 0 | 0 | 0 | 0 |
| ENSMUSG00 SNORA32 | 121 | 0 | 0 | 0.39978124 | 0 | 0 | 0 | 0 | 0.63127938 | 0 | 0 | 0 | 0 | 0 | 0 | 0 | 0 | 0 | 0 |
| ENSMUSG00 AF357355 | 71 | 0 | 0 | 0 | 0 | 0 | 0 | 0 | 0 | 0 | 0 | 0 | 0 | 0 | 0 | 0 | 0 | 0 | 0 |
| ENSMUSG00 SNORA61 | 128 | 0.66414338 | 2.08205589 | 1.88959103 | 1.42378264 | 0.37603065 | 1.54852292 | 0 | 4.17729401 | 0.32176965 | 0.36836487 | 1.11972559 | 2.18029932 | 0.77081121 | 0.72568887 | 1.28754862 | 0.59829074 | 0.29942419 | 0.42975865 |
| ENSMUSG00 U6 | 106 | 0 | 0 | 0 | 0 | 0 | 0 | 0 | 0 | 0 | 0 | 0 | 0 | 0 | 0 | 0 | 0 | 0 | 0 |
| ENSMUSG00 U6 | 107 | 0 | 0 | 0 | 0 | 0 | 0 | 0 | 0 | 0 | 0 | 0 | 0 | 0 | 0 | 0 | 0 | 0 | 0 |
| ENSMUSG00 U6 | 106 | 0 | 0 | 0 | 0 | 0 | 0 | 0 | 0 | 0 | 0 | 0 | 0 | 0 | 0 | 0 | 0 | 0 | 0 |
| ENSMUSG00 U6 | 106 | 0 | 0.31427259 | 0.45635406 | 0 | 0 | 0 | 0.89269206 | 0 | 0 | 0 | 0 | 0 | 0 | 0 | 0 | 0 | 0 | 0 |
| ENSMUSG00 U6 | 108 | 0 | 0 | 0 | 0 | 0 | 0 | 0 | 0 | 0 | 0 | 0 | 0 | 0 | 0 | 0 | 0 | 0 | 0 |
| ENSMUSG00 n-R5s42 | 103 | 0 | 0 | 0 | 0 | 0 | 0 | 0 | 0 | 0 | 0 | 0 | 0 | 0 | 0 | 0 | 0 | 0 | 0 |
| ENSMUSG00 U6 | 106 | 0 | 0 | 0 | 0 | 0 | 0 | 0 | 0 | 0 | 0 | 0 | 0 | 0 | 0 | 0 | 0 | 0 | 0 |
| ENSMUSG00 SNORA1 | 132 | 2.89808018 | 0.25237041 | 1.09939842 | 0 | 0 | 2.25239698 | 1.07528816 | 2.89336382 | 0 | 0.3572023 | 1.44772601 | 0.84569186 | 1.12117994 | 2.81479318 | 0.624266 | 1.4504018 | 1.7421044 | 0 |
| ENSMUSG00 Snora20 | 127 | 1.67343213 | 2.09845003 | 0.38089394 | 0 | 0 | 0.390179 | 1.11762234 | 0.90218273 | 0 | 0.74253076 | 1.88090388 | 0 | 1.94220147 | 0 | 0 | 0 | 0.30178187 | 2.16571286 |
| ENSMUSG00 SNORA32 | 78 | 0 | 0 | 0 | 0 | 0 | 0 | 0 | 0 | 0 | 0 | 0 | 0 | 0 | 0 | 0 | 0 | 0 | 0 |
| ENSMUSG00 U6 | 106 | 0 | 0 | 0 | 0 | 0 | 0 | 0 | 0 | 0 | 0 | 0 | 0 | 0 | 0 | 0 | 0 | 0 | 0 |
| ENSMUSG00 U6 | 99 | 0 | 0 | 0 | 0 | 0 | 0 | 0 | 0 | 0 | 0 | 0 | 0.56379457 | 0 | 0.4691322 | 0 | 0 | 0 | 0 |
| ENSMUSG00 SNORA42 | 124 | 0 | 0 | 0 | 0.36742778 | 0 | 0 | 0 | 0 | 0 | 0 | 0 | 0 | 0 | 0 | 0 | 0 | 0 | 0 |
| ENSMUSG00 SNORD58 | 64 | 10.626294 | 13.5333633 | 18.1400739 | 8.54269581 | 13.5371035 | 4.64556877 | 7.39260608 | 14.3221509 | 16.0884827 | 6.63056765 | 10.4507722 | 13.0817959 | 16.1870354 | 21.770666 | 10.300389 | 10.7692333 | 18.5643001 | 11.1737248 |
| ENSMUSG00 U1 | 151 | 1.40745616 | 0.66184558 | 7.04779912 | 1.50864385 | 1.27501785 | 0.6563276 | 3.75994799 | 4.29980689 | 2.72758383 | 0.62451263 | 2.21473317 | 5.17496209 | 1.30680576 | 2.4606139 | 2.18286389 | 0.76074054 | 0.76144961 | 0.72859744 |
| ENSMUSG00 U6 | 106 | 0 | 0 | 0 | 0 | 0 | 0 | 0 | 0 | 0 | 0 | 0 | 0 | 0 | 0 | 0 | 0 | 0 | 0 |
| ENSMUSG00 U6 | 106 | 0 | 0 | 0 | 0 | 0 | 0 | 0 | 0 | 0 | 0 | 0 | 0 | 0 | 0 | 0 | 0 | 0 | 0 |
| ENSMUSG00 SNORA31 | 127 | 0 | 0 | 0 | 0 | 0 | 0 | 0.74508156 | 0 | 0.32430327 | 0 | 0 | 0.43949341 | 0 | 0 | 0 | 0 | 0 | 0 |
| ENSMUSG00 U6 | 106 | 0.80198445 | 0.31427259 | 0 | 0 | 0.45407475 | 0 | 1.78538411 | 0.36030568 | 0 | 0 | 4.0563644 | 0 | 2.32697723 | 1.3144553 | 0 | 2.16739287 | 0.36156884 | 0 |
| ENSMUSG00 SNORA55 | 130 | 0 | 0 | 0.37210408 | 0 | 0.37024557 | 0 | 0 | 0 | 0 | 0 | 0.73499906 | 0 | 0 | 0.35726221 | 0.95080514 | 0 | 0 | 0 |
| ENSMUSG00 U6 | 106 | 0 | 0 | 0 | 0 | 0 | 0 | 0 | 0 | 0 | 0 | 0 | 0 | 0 | 0 | 0 | 0 | 0 | 0 |
| ENSMUSG00 SNORA2 | 134 | 0 | 0 | 0 | 0 | 0 | 0 | 0 | 0 | 0 | 0 | 0 | 0 | 0 | 0.34659767 | 0 | 0 | 0 | 0 |
| ENSMUSG00 U6 | 109 | 0 | 0 | 0 | 0 | 0 | 0 | 0 | 0 | 0 | 0 | 0 | 0 | 0 | 0 | 0 | 0 | 0 | 0 |
| ENSMUSG00 U6 | 106 | 0.40099223 | 0 | 0 | 0 | 0 | 0 | 0 | 0.72061137 | 0 | 2.66890773 | 0 | 0 | 0 | 0.43815177 | 0.38869392 | 0 | 1.08470652 | 0 |
| ENSMUSG00 U4 | 85 | 0 | 0 | 0 | 0 | 0 | 0 | 0 | 0 | 0.48454724 | 0 | 0 | 0 | 0.5803755 | 0.54640103 | 0 | 0 | 0.45089761 | 1.94149788 |
| ENSMUSG00 SNORA4 | 105 | 0 | 0 | 0 | 0 | 0 | 0 | 1.35179083 | 0.36373717 | 0 | 0 | 0 | 0 | 0 | 0 | 0 | 0 | 0 | 0 |
| ENSMUSG00 Snora52 | 136 | 0 | 0 | 0 | 0 | 0 | 0 | 0 | 0 | 0 | 0 | 0 | 0 | 0 | 0 | 0 | 0 | 0 | 0 |
| ENSMUSG00 U6 | 106 | 0 | 0 | 0 | 0 | 0 | 0 | 0 | 0 | 0 | 0 | 0 | 0 | 0 | 0 | 0 | 0 | 0 | 0 |
| ENSMUSG00 U6 | 102 | 0 | 0.326597 | 0 | 0 | 0 | 0 | 0 | 0 | 0 | 0 | 0 | 0 | 0.48364625 | 0.91066838 | 0 | 0 | 0 | 0 |
| ENSMUSG00 U6 | 106 | 0 | 0 | 0 | 0 | 0 | 0 | 0 | 0 | 0 | 0 | 0 | 0 | 0 | 0 | 0 | 0 | 0 | 0 |
| ENSMUSG00 SNORA65 | 128 | 0 | 0 | 0 | 0 | 0 | 0 | 0 | 0 | 0 | 0 | 0 | 0 | 0 | 0 | 0 | 0 | 0 | 0 |
| ENSMUSG00 SNORA2 | 135 | 1.25941262 | 0.24676218 | 0 | 0 | 0.35653277 | 0 | 0 | 0 | 0.3050853 | 2.09558681 | 0 | 0 | 0 | 0 | 0 | 0 | 0 | 0 |
| ENSMUSG00 U6 | 106 | 0 | 0 | 0 | 0 | 0 | 0 | 0 | 0 | 0 | 0 | 0 | 0 | 0 | 0 | 0 | 0 | 0 | 0 |
| ENSMUSG00 U6 | 106 | 0 | 0 | 0 | 0 | 0 | 0 | 0 | 0 | 0 | 0 | 0 | 0 | 0 | 0 | 0 | 0 | 0 | 0 |
| ENSMUSG00 U1 | 157 | 0 | 0 | 0 | 0 | 0 | 0 | 0 | 0 | 0 | 0 | 0 | 0 | 0 | 0 | 0 | 0 | 0 | 0 |
| ENSMUSG00 U1 | 155 | 0 | 0 | 0 | 0 | 0 | 0 | 0 | 0 | 0 | 0 | 0 | 0 | 0 | 0 | 0 | 0 | 0 | 0 |
| ENSMUSG00 U6 | 106 | 0 | 0 | 0 | 0 | 0 | 0 | 0 | 0 | 0 | 0 | 0 | 0 | 0 | 0 | 0 | 0 | 0 | 0 |
| ENSMUSG00 SNORD113 | 71 | 0 | 0 | 0 | 0 | 0 | 0 | 0 | 0 | 0 | 0 | 0 | 0 | 0 | 0 | 0 | 0 | 0 | 0 |
| ENSMUSG00 U6 | 106 | 0 | 0 | 0 | 0 | 0 | 0 | 0 | 0 | 0 | 0 | 0 | 0 | 0 | 0 | 0 | 0 | 0 | 0 |
| ENSMUSG00 U2 | 190 | 0 | 0 | 0 | 0 | 0 | 0.26080386 | 0 | 0 | 0.21677114 | 0 | 0 | 0 | 0 | 0 | 0 | 0 | 0 | 0 |
| ENSMUSG00 U6 | 106 | 0 | 0 | 0 | 0.42982117 | 0 | 0 | 0 | 0 | 0 | 0 | 0 | 0 | 0 | 0 | 0 | 0 | 0 | 0 |
| ENSMUSG00 U6 | 102 | 0 | 0 | 0 | 0 | 0 | 0 | 0 | 0 | 0 | 0 | 0 | 0 | 0 | 0 | 0 | 0 | 0 | 0 |
| ENSMUSG00 U6 | 107 | 0 | 0 | 0 | 0 | 0 | 0 | 0 | 0 | 0 | 0 | 0 | 0 | 1.38313787 | 0.86811378 | 1.15518381 | 0 | 0 | 0 |
| ENSMUSG00 U6 | 100 | 0 | 0 | 0 | 0 | 0 | 0 | 0 | 0 | 0 | 0 | 0 | 0 | 0 | 0 | 0 | 0 | 0 | 0 |
| ENSMUSG00 U6 | 106 | 0 | 0 | 0 | 0 | 0 | 0 | 0 | 0 | 0 | 0 | 0 | 0 | 0 | 0 | 0 | 0 | 0 | 0 |
| ENSMUSG00 U6 | 106 | 0 | 0 | 0 | 0 | 0 | 0 | 0 | 0 | 0 | 0 | 0 | 0 | 0 | 0 | 0 | 0 | 0 | 0 |
| ENSMUSG00 U6 | 106 | 0 | 0 | 0 | 0 | 0 | 0 | 0 | 0 | 0 | 0 | 0 | 1.57968856 | 0 | 0 | 0 | 0 | 0 | 0 |
| ENSMUSG00 U6 | 106 | 0 | 0 | 0 | 0 | 0 | 0 | 0 | 0 | 0 | 0 | 0 | 0 | 0 | 0 | 0 | 0 | 0 | 0 |
| ENSMUSG00 SNORA67 | 140 | 0 | 0.23794924 | 2.41867652 | 0 | 0 | 0 | 0 | 0.27280287 | 0 | 0 | 0 | 0 | 0 | 0 | 0 | 0 | 0 | 0 |
| ENSMUSG00 U6 | 103 | 0 | 0 | 1.40893778 | 2.21170118 | 0 | 0 | 0 | 0 | 0 | 0 | 1.39150365 | 1.08379927 | 0 | 2.70548082 | 0 | 0.37175347 | 0 | 0.534069 |
| ENSMUSG00 7SK | 320 | 0 | 0 | 0 | 0 | 0 | 0.15485229 | 0 | 0 | 0 | 0 | 0 | 0 | 0 | 0.58055109 | 0 | 0 | 0.23953936 | 0 |
| ENSMUSG00 SNORD20 | 79 | 0 | 1.68672882 | 0 | 1.73016624 | 0.60926486 | 1.25449958 | 3.59336802 | 0.48344813 | 0 | 2.38737738 | 10.8854336 | 2.11958212 | 0 | 2.9394992 | 3.12923209 | 0.48469123 | 0.485143 | 2.78527123 |
| ENSMUSG00 SNORA67 | 143 | 0 | 0 | 0 | 0 | 0 | 0 | 0 | 0 | 0 | 0 | 0 | 0 | 0 | 0 | 0 | 0 | 0 | 0 |
| ENSMUSG00 U1 | 162 | 0 | 0 | 0 | 0 | 0 | 0 | 0 | 0 | 0 | 0 | 0 | 0 | 0 | 0 | 0 | 0 | 0 | 0 |
| ENSMUSG00 U6 | 106 | 0 | 0 | 0 | 0 | 1.81629901 | 0 | 0 | 0 | 0 | 0.44481796 | 0 | 0 | 0 | 0.43815177 | 0 | 0 | 1.08470652 | 0 |
| ENSMUSG00 U2 | 190 | 0 | 0 | 0 | 0 | 0 | 0 | 0 | 0 | 0 | 0 | 0 | 0 | 0 | 0 | 0 | 0 | 0 | 0 |
| ENSMUSG00 U6 | 106 | 0 | 0 | 0 | 0 | 0 | 0 | 0.89269206 | 0 | 0 | 0 | 0 | 0 | 0 | 0 | 0 | 0 | 0 | 0 |
| ENSMUSG00 n-R5s180 | 105 | 0 | 0 | 0 | 0 | 0 | 0 | 0 | 0 | 0 | 0 | 0 | 0 | 0 | 0 | 0 | 0 | 0 | 0 |
| ENSMUSG00 Y_RNA | 109 | 0 | 0 | 0 | 0 | 0 | 0 | 0 | 0 | 0 | 0 | 0 | 0 | 0 | 0 | 0 | 0 | 0 | 0 |
| ENSMUSG00 U1 | 159 | 0 | 0 | 0 | 0 | 0 | 0 | 0 | 0 | 0 | 0 | 0 | 0 | 0 | 0 | 0 | 0 | 0 | 0 |
| ENSMUSG00 U6 | 106 | 0 | 0 | 0 | 0 | 0 | 0 | 0 | 0 | 0 | 0 | 0 | 0 | 0 | 0 | 0 | 0 | 0 | 0 |
| ENSMUSG00 U6 | 106 | 0 | 0 | 0 | 0 | 0 | 0 | 0 | 0 | 0 | 0 | 0 | 0 | 0 | 0 | 0 | 0 | 0 | 0 |
| ENSMUSG00 U6 | 106 | 0 | 0 | 0 | 0 | 0 | 0 | 0 | 0 | 0 | 0 | 0 | 0 | 0 | 0 | 0 | 0 | 0 | 0 |
| ENSMUSG00 n-R5s62 | 95 | 0 | 0 | 0 | 0 | 0 | 0 | 0 | 0 | 0 | 0 | 0 | 0 | 0 | 0 | 0 | 0 | 0 | 0 |
| ENSMUSG00 U6 | 107 | 0 | 0 | 0 | 0 | 0 | 0 | 0 | 0 | 0 | 0 | 0 | 0 | 0 | 0 | 0 | 0 | 0 | 0 |
| ENSMUSG00 U8 | 134 | 0 | 0 | 0 | 0.34000779 | 1.07758038 | 0 | 1.41231877 | 0.28501793 | 0 | 1.05561276 | 0.35652954 | 0 | 0 | 0 | 0 | 0 | 0 | 0 |
| ENSMUSG00 U6 | 106 | 0 | 0 | 0 | 0 | 0 | 0 | 0 | 0 | 0 | 0 | 0 | 0 | 0 | 0 | 0 | 0 | 0 | 0 |
| ENSMUSG00 U6 | 106 | 0.80198445 | 1.57136293 | 0 | 0 | 0 | 0.93495724 | 1.33903808 | 0 | 0.38855204 | 0 | 0 | 0 | 0.46539545 | 3.50521414 | 0 | 0.36123215 | 2.53098187 | 0 |
| ENSMUSG00 SNORD113 | 74 | 0 | 0 | 0 | 0 | 0 | 0 | 0 | 0 | 0 | 0 | 0 | 0 | 0 | 0 | 0 | 0 | 0 | 0 |
| ENSMUSG00 SNORD25 | 71 | 0 | 0 | 0 | 0 | 0 | 0 | 0 | 0 | 0 | 0 | 0 | 0 | 0 | 0 | 0 | 0 | 0 | 0 |
| ENSMUSG00 SNORA44 | 128 | 0 | 0 | 0 | 0 | 0 | 0 | 0 | 0 | 0.64353931 | 0 | 0 | 0 | 0 | 0 | 0 | 0 | 0 | 0 |
| ENSMUSG00 U6 | 106 | 0 | 0 | 0 | 0 | 0 | 0 | 0 | 0 | 0 | 0 | 0 | 0 | 0 | 0 | 0 | 0 | 0 | 0 |
| ENSMUSG00 SNORA63 | 129 | 0 | 0 | 1.12496582 | 0 | 0 | 0 | 0 | 0 | 0.95782595 | 0 | 0 | 0 | 0 | 0 | 0 | 0 | 0 | 0 |
| ENSMUSG00 Snord96a | 80 | 0 | 0 | 0 | 0 | 0 | 0 | 0 | 0 | 0 | 0 | 0 | 0 | 0 | 0 | 0 | 0 | 0 | 0 |
| ENSMUSG00 SNORD113 | 71 | 0 | 0 | 0 | 0 | 0 | 0 | 0 | 0 | 0 | 0 | 0 | 0 | 0 | 0 | 0 | 0 | 0 | 0 |
| ENSMUSG00 SNORA71 | 131 | 0 | 0 | 0 | 0 | 0 | 0 | 0 | 0 | 0 | 0 | 0 | 0 | 0 | 0 | 0 | 0 | 0 | 0 |
| ENSMUSG00 U6 | 106 | 0 | 0 | 0 | 0 | 0 | 0 | 0 | 0 | 0 | 0 | 0 | 0 | 0 | 0 | 0 | 0 | 0 | 0 |
| ENSMUSG00 Snord45b | 70 | 3.036084 | 4.75898489 | 3.45525217 | 7.15959268 | 8.25118691 | 9.20265052 | 4.73126789 | 7.09287473 | 2.94189398 | 4.71507033 | 7.50749348 | 6.37893287 | 4.93319173 | 2.65394785 | 5.2973429 | 7.65812148 | 7.66525938 | 0 |
| ENSMUSG00 U1 | 162 | 0 | 0 | 0 | 0 | 0 | 0 | 0 | 0 | 0 | 0 | 0 | 0 | 0 | 0 | 0 | 0 | 0 | 0 |
| ENSMUSG00 U6 | 106 | 0 | 0 | 0 | 0 | 0 | 0 | 0 | 0 | 0 | 0 | 0 | 0 | 0 | 0 | 0 | 0 | 0 | 0 |
| ENSMUSG00 U6 | 106 | 0 | 0 | 0 | 0 | 0 | 0 | 0 | 0 | 0 | 0 | 0 | 0 | 0 | 0 | 0 | 0 | 0 | 1.55686151 |
| ENSMUSG00 U6 | 110 | 0 | 0 | 0 | 0 | 0 | 0 | 0 | 0 | 0 | 0 | 0 | 0 | 0 | 0 | 0 | 0 | 0 | 0 |
| ENSMUSG00 U1 | 161 | 0 | 0 | 0 | 0 | 0 | 0 | 1.17547028 | 0 | 0 | 0 | 0 | 0 | 0.30640942 | 0 | 0 | 0 | 0 | 0 |
| ENSMUSG00 U6 | 93 | 0 | 0 | 0 | 0 | 0 | 0 | 0 | 0 | 0 | 0 | 0 | 0 | 0 | 0 | 0 | 0 | 0 | 0 |
| ENSMUSG00 SNORA70 | 136 | 0 | 0 | 0 | 0 | 0 | 0 | 0 | 0 | 0 | 0 | 0 | 0 | 0 | 0 | 0 | 0 | 0 | 0 |
| ENSMUSG00 Snord14a | 86 | 9.39067842 | 3.87359235 | 5.0623462 | 5.29779585 | 6.71608237 | 5.18575119 | 2.75073715 | 5.77327013 | 6.22586866 | 6.03090391 | 13.8880693 | 7.78823199 | 8.03077723 | 5.40047528 | 11.4981086 | 8.4595528 | 9.35874692 | 7.67568931 |
| ENSMUSG00 U6 | 106 | 0 | 0 | 0 | 0 | 0.9081495 | 0 | 0 | 0 | 0 | 0 | 0 | 0 | 0 | 0 | 0 | 0 | 0 | 0 |
| ENSMUSG00 U6 | 106 | 0 | 0 | 0 | 0 | 0 | 0 | 0 | 0 | 0 | 0 | 0 | 0 | 0 | 0 | 0 | 0 | 0 | 0 |
| ENSMUSG00 n-R5s86 | 118 | 0 | 0 | 0 | 0 | 0 | 0 | 0 | 0 | 0 | 0 | 0 | 0 | 0 | 0 | 0 | 0 | 0 | 0 |
| ENSMUSG00 U6 | 102 | 0 | 0 | 0 | 0 | 0 | 0 | 0 | 0 | 0 | 0 | 0 | 0 | 0 | 0 | 0 | 0 | 0 | 0 |
| ENSMUSG00 U6 | 106 | 0 | 0 | 0 | 0 | 0 | 0 | 0 | 0 | 0 | 0 | 0 | 0 | 0 | 0 | 0 | 0 | 0 | 0 |
| ENSMUSG00 U6 | 106 | 0 | 0 | 0 | 0 | 0 | 0 | 0 | 0 | 0 | 0 | 0 | 0 | 0 | 0 | 0 | 0 | 0 | 0 |
| ENSMUSG00 n-R5s28 | 118 | 0 | 0 | 0 | 0 | 0 | 0 | 0 | 0 | 0 | 0 | 0 | 0 | 0 | 0 | 0 | 0 | 0 | 0 |
| ENSMUSG00 U6 | 106 | 0 | 0 | 0 | 0 | 0 | 0 | 0 | 0 | 0 | 0 | 0 | 0 | 0 | 0 | 0 | 0 | 0 | 0 |
| ENSMUSG00 SNORD46 | 103 | 0 | 0 | 0 | 0 | 0 | 0 | 0 | 0 | 0 | 0 | 0 | 0 | 0 | 0 | 0 | 0 | 0 | 0 |
| ENSMUSG00 U6 | 106 | 0 | 0 | 0.91270812 | 0 | 2.27037376 | 0 | 0.89269206 | 0.72061137 | 0.77710407 | 0 | 0 | 0 | 0 | 0 | 0 | 0 | 0 | 0 |
| ENSMUSG00 U6 | 106 | 0 | 0 | 0 | 0 | 0 | 0 | 0 | 0 | 0.38855204 | 0 | 0 | 0 | 0 | 0 | 0 | 0 | 0 | 0 |
| ENSMUSG00 U6 | 106 | 0 | 0 | 0 | 0 | 0 | 0 | 0 | 0 | 0 | 0 | 0 | 0 | 0 | 0 | 0 | 0 | 0 | 0 |
| ENSMUSG00 U6 | 106 | 0 | 0 | 0 | 0 | 0 | 0 | 0 | 0 | 0 | 0 | 0 | 0 | 0 | 0 | 0 | 0 | 0 | 0 |
| ENSMUSG00 U2 | 179 | 0 | 0 | 0 | 0 | 0 | 0 | 0 | 0 | 0 | 0 | 0 | 0 | 0 | 0 | 0 | 0 | 0 | 0 |
| ENSMUSG00 SNORA61 | 129 | 0 | 0 | 0 | 0 | 0 | 0 | 0 | 0 | 0.63855063 | 0 | 0 | 0 | 0 | 0 | 0 | 0 | 0 | 0 |
| ENSMUSG00 U1 | 170 | 0 | 0 | 0 | 0 | 0 | 0 | 0 | 0 | 0 | 0 | 0 | 0 | 0 | 0 | 0 | 0 | 0 | 0 |
| ENSMUSG00 U6 | 106 | 0 | 0 | 0 | 0 | 0 | 0 | 0 | 0 | 0 | 0 | 0 | 0 | 0 | 0 | 0 | 0 | 0 | 0 |
| ENSMUSG00 U2 | 191 | 0 | 0 | 0 | 0 | 0 | 0 | 0 | 0 | 0 | 0 | 0 | 0 | 0 | 0.24316276 | 0 | 0 | 0 | 0 |
| ENSMUSG00 Y_RNA | 110 | 0 | 0.90853348 | 2.6385562 | 1.24257394 | 2.18781471 | 0 | 3.4409221 | 0.69440732 | 2.24653722 | 0.85728551 | 2.60590683 | 0.50741511 | 3.13930383 | 0 | 0.3745596 | 2.08857859 | 0 | 4.00066231 |
| ENSMUSG00 SNORA24 | 127 | 0 | 0 | 0 | 0 | 0 | 0 | 0 | 0 | 0 | 0 | 0 | 0 | 0 | 0 | 0 | 0 | 0 | 0 |
| ENSMUSG00 Snord35b | 86 | 6.42520103 | 8.13454394 | 3.37489747 | 0 | 3.91771472 | 2.88097288 | 1.10029486 | 3.55278162 | 4.31021677 | 7.12743189 | 3.88865941 | 5.19215466 | 6.3098964 | 0 | 2.87452716 | 3.56191697 | 4.01089154 | 3.19820388 |
| ENSMUSG00 SNORD60 | 77 | 0 | 0 | 0 | 0 | 0 | 0 | 0 | 0 | 0 | 0 | 0 | 0 | 0 | 0 | 0 | 0 | 0 | 0 |
| ENSMUSG00 U1 | 152 | 0 | 0 | 0 | 0 | 0 | 0 | 0 | 0 | 0 | 0 | 0 | 0 | 0 | 0 | 0 | 0 | 0 | 0 |
| ENSMUSG00 n-R5s29 | 118 | 0.72042671 | 0 | 0.40994517 | 0 | 0 | 0 | 0 | 0 | 0 | 0.39958223 | 0 | 0 | 0 | 0 | 0 | 0 | 0 | 0.46617887 |
| ENSMUSG00 U2 | 190 | 0 | 0 | 0 | 0 | 0 | 0 | 0 | 0 | 0 | 0 | 0 | 0 | 0 | 0 | 0 | 0 | 0 | 0 |
| ENSMUSG00 U6 | 102 | 0 | 0 | 0 | 0 | 0 | 0 | 0 | 0 | 0 | 0 | 0 | 0 | 0 | 0 | 0 | 0 | 0 | 0 |
| ENSMUSG00 U3 | 213 | 0.19955482 | 0.15639856 | 0 | 0 | 0.22597147 | 0 | 0.22212525 | 0 | 0 | 0 | 0 | 0.26204536 | 0 | 0 | 0 | 0 | 0 | 0 |
| ENSMUSG00 U6 | 105 | 0 | 0 | 0 | 0 | 0 | 0 | 0 | 0 | 0 | 0 | 0 | 0 | 0 | 0 | 0 | 0 | 0 | 0 |
| ENSMUSG00 SNORD59 | 73 | 12.2275164 | 8.67047932 | 13.253022 | 6.86536285 | 9.23077988 | 8.8244594 | 5.83307 | 21.973711 | 23.1321527 | 8.39670058 | 18.3246416 | 15.2919624 | 10.1366953 | 8.27086488 | 8.46607313 | 6.29434642 | 14.7004974 | 9.04259289 |
| ENSMUSG00 U6 | 106 | 0 | 0 | 0 | 0 | 0 | 0 | 0 | 0 | 0 | 0 | 0 | 0 | 0 | 0 | 0 | 0 | 0 | 0 |
| ENSMUSG00 Snord95 | 67 | 0 | 0 | 0 | 0 | 0 | 0 | 0 | 0 | 0 | 0 | 0 | 0 | 0 | 0 | 0 | 0 | 0 | 0 |
| ENSMUSG00 U6 | 103 | 0 | 0 | 0 | 0 | 0 | 0 | 0 | 0 | 0 | 0 | 0 | 0 | 0 | 0 | 0 | 0 | 0 | 0 |
| ENSMUSG00 U6 | 105 | 0 | 0 | 0 | 0 | 0 | 0 | 0 | 0 | 0 | 0 | 0 | 0 | 0 | 0 | 0 | 0 | 0 | 0 |
| ENSMUSG00 SNORA71 | 131 | 0 | 0 | 0 | 0 | 0 | 0 | 0 | 0 | 0 | 0 | 0 | 0 | 0 | 0 | 0 | 0 | 0 | 0 |
| ENSMUSG00 SNORA2 | 136 | 0 | 0 | 0 | 0 | 0 | 0 | 0 | 0 | 0 | 0 | 0 | 0 | 0 | 0 | 0 | 0 | 0 | 0 |
| ENSMUSG00 Snord14e | 89 | 0 | 0 | 0 | 0 | 0 | 0 | 0 | 0 | 0 | 0 | 0 | 0 | 0 | 0 | 0 | 0 | 0 | 0 |
| ENSMUSG00 SNORA32 | 121 | 0 | 0.27531318 | 0 | 0.37653756 | 0.39778449 | 0 | 0.78202775 | 0.63127938 | 1.70192214 | 0 | 0 | 0.46128647 | 0 | 0.38383543 | 0 | 0.9493539 | 0 | 0 |
| ENSMUSG00 U6 | 105 | 0 | 0 | 0 | 0 | 0 | 0 | 0 | 0 | 0 | 0 | 0 | 0 | 0 | 0 | 0 | 0 | 0 | 0 |
| ENSMUSG00 U2 | 183 | 0 | 0 | 0 | 0 | 0 | 0 | 0 | 0 | 0 | 0 | 0 | 0 | 0 | 0 | 0 | 0 | 0 | 0 |
| ENSMUSG00 Telomerase-v | 422 | 0.20144633 | 0.07894051 | 0.11462922 | 0.21592912 | 0 | 0 | 0.33634606 | 0 | 0.39039351 | 0.67038915 | 0 | 0.26452921 | 0 | 0 | 0.09763402 | 0 | 0.54492365 | 0 |
| ENSMUSG00 snoZ40 | 73 | 2.32905074 | 4.10706915 | 1.9879533 | 4.36886727 | 1.97802426 | 2.71521828 | 1.29623778 | 3.13910157 | 2.25679538 | 5.16720036 | 2.61780594 | 3.05839247 | 2.02733907 | 6.36220375 | 3.95083413 | 5.24528869 | 5.25017766 | 2.26064822 |
| ENSMUSG00 n-R5s193 | 108 | 0 | 0 | 0 | 0 | 0.89133192 | 0 | 0 | 0.35363336 | 0 | 1.30974176 | 0 | 0 | 0.45677701 | 0 | 0 | 0 | 0 | 0 |
| ENSMUSG00 SNORA73 | 206 | 0 | 0 | 0 | 0 | 0 | 0 | 0 | 0 | 0 | 0 | 0 | 0 | 0 | 0 | 0 | 0 | 0 | 0 |
| ENSMUSG00 U6 | 106 | 0 | 0 | 0 | 0 | 0 | 0 | 0 | 0 | 0 | 0 | 0 | 0 | 0 | 0 | 0 | 0 | 0 | 0 |
| ENSMUSG00 U1 | 155 | 0 | 0 | 0.31208729 | 0 | 0 | 0 | 0 | 0 | 0 | 0 | 0 | 0 | 0 | 0 | 0 | 0 | 0 | 0 |
| ENSMUSG00 SNORA44 | 128 | 0 | 0 | 0 | 0 | 0 | 0 | 0 | 0 | 0 | 0 | 0 | 0 | 0 | 0 | 0 | 0 | 0 | 0 |
| ENSMUSG00 U2 | 179 | 0 | 0 | 0 | 0 | 0 | 0 | 0 | 0 | 0 | 0 | 0 | 0 | 0 | 0 | 0 | 0 | 0 | 0 |
| ENSMUSG00 U1 | 158 | 0 | 0 | 0 | 0 | 0 | 0 | 0 | 0 | 0 | 0 | 0 | 0 | 0 | 0 | 0 | 0 | 0 | 0 |
| ENSMUSG00 U3 | 213 | 5.78708969 | 1.56398564 | 2.95237509 | 1.92511455 | 3.38957209 | 1.62849359 | 3.7761293 | 1.25514937 | 2.90045886 | 1.54955363 | 2.2429558 | 0.78613609 | 1.62123672 | 0 | 2.90151802 | 1.07860866 | 1.079614 | 1.5495523 |
| ENSMUSG00 SNORA32 | 121 | 0.70256489 | 0.55062635 | 0 | 0 | 0 | 0 | 0.39101388 | 0.94691907 | 3.06345985 | 0.38967523 | 0 | 0.46128647 | 1.63080718 | 0 | 1.36203491 | 0 | 0.63349251 | 0.45462072 |
| ENSMUSG00 U6 | 106 | 0 | 0 | 0 | 0 | 0 | 0 | 0 | 0 | 0 | 0 | 0 | 0 | 0 | 0 | 0 | 0 | 0 | 0 |
| ENSMUSG00 SNORA32 | 132 | 0 | 0 | 0 | 0 | 0 | 0 | 0 | 0 | 0 | 0 | 0 | 0 | 0 | 0 | 0 | 0 | 0 | 0 |
| ENSMUSG00 SNORD24 | 82 | 0 | 0 | 0 | 0 | 0 | 0 | 0 | 0 | 0 | 0 | 0 | 0 | 0 | 0 | 0 | 0 | 0 | 0 |
| ENSMUSG00 U1 | 164 | 0 | 0 | 0 | 0 | 0 | 0 | 0 | 0 | 0 | 0 | 0 | 0.34033941 | 0 | 0 | 0 | 0 | 0 | 0 |
| [illegible] | [illegible] | [illegible] | [illegible] | [illegible] | [illegible] | [illegible] | [illegible] | [illegible] | [illegible] | [illegible] | [illegible] | [illegible] | [illegible] | [illegible] | [illegible] | [illegible] | [illegible] | [illegible] | [illegible] |

| | | | | | | | | | | | | | | | | | | | |
|---|---|---|---|---|---|---|---|---|---|---|---|---|---|---|---|---|---|---|---|
| ENSMUSG00 U6 | 106 | 0 | 0 | 0 | 0 | 0 | 0 | 0 | 0 | 0 | 0 | 0 | 0 | 0 | 0 | 0 | 0 | 0 | 0 |
| ENSMUSG00 U6 | 103 | 0 | 0 | 0 | 0 | 0 | 0 | 0 | 0 | 0 | 0 | 0 | 0 | 0 | 0 | 0 | 0 | 0 | 0.534069 |
| ENSMUSG00 SNORA46 | 130 | 0 | 0 | 0 | 0 | 0 | 0 | 0 | 0 | 0 | 0 | 0 | 0 | 0 | 0 | 0 | 0 | 0 | 0 |
| ENSMUSG00 U6 | 106 | 0 | 0 | 0 | 0 | 0 | 0 | 0 | 0 | 0 | 0 | 0 | 0 | 0 | 0 | 0 | 0 | 0 | 0 |
| ENSMUSG00 Vault | 142 | 0.59866445 | 1.40758708 | 1.70329332 | 2.24596697 | 0.67791442 | 0.69792583 | 1.99912728 | 0.26896058 | 1.16018354 | 2.65637765 | 1.34577348 | 0.78613609 | 11.1170518 | 13.7369836 | 8.41440226 | 5.3930433 | 3.77864899 | 5.03604498 |
| ENSMUSG00 SNORD36 | 67 | 0 | 0 | 0 | 0 | 0 | 0 | 0 | 0 | 0 | 0 | 0 | 0 | 0 | 0 | 0 | 0 | 0 | 0 |
| ENSMUSG00 Snora31 | 129 | 13.1798995 | 8.78014266 | 8.99972658 | 6.71054141 | 9.32789218 | 6.91433492 | 17.6047177 | 12.4347357 | 8.30115822 | 14.6203731 | 21.4802139 | 20.3359391 | 11.8549569 | 41.0436121 | 29.3840554 | 26.4175508 | 27.9276892 | 36.2463106 |
| ENSMUSG00 n-R5s92 | 118 | 0 | 0 | 0 | 0 | 0 | 0 | 0 | 0 | 0 | 0 | 0 | 0 | 0 | 0 | 0 | 0 | 0 | 0 |
| ENSMUSG00 U6 | 106 | 0 | 0 | 0 | 0 | 0 | 0 | 0 | 0 | 0 | 0 | 0 | 0 | 0 | 0 | 0 | 0 | 0 | 0 |
| ENSMUSG00 SNORA67 | 132 | 0 | 0 | 0 | 0 | 0 | 0 | 0 | 0 | 0 | 0 | 0 | 0 | 0 | 0 | 0 | 0 | 0 | 0 |
| ENSMUSG00 U6 | 106 | 0 | 0 | 0 | 0 | 0 | 0 | 0 | 0 | 0 | 0 | 0 | 0 | 0 | 0 | 0 | 0 | 0 | 0 |
| ENSMUSG00 U1 | 157 | 0 | 0 | 0 | 0 | 0.30657276 | 0 | 0 | 0 | 0 | 0 | 0 | 0 | 0 | 0 | 0 | 0 | 0.97646616 | 0 |
| ENSMUSG00 U6 | 106 | 0 | 0 | 0 | 0 | 0 | 0 | 0 | 0.36030568 | 0 | 0.44481796 | 0 | 0 | 0 | 0 | 0 | 0 | 0 | 0 |
| ENSMUSG00 U6 | 106 | 0 | 0 | 0 | 0 | 0 | 0 | 0 | 0 | 0 | 0 | 0 | 0 | 0 | 0 | 0 | 0 | 0 | 0 |
| ENSMUSG00 SNORA73 | 226 | 0 | 0 | 0 | 0 | 0 | 0.65777965 | 0 | 0 | 0 | 1.04315715 | 0 | 0 | 0 | 0 | 0 | 0 | 0 | 0 |
| ENSMUSG00 U6 | 106 | 0 | 0 | 0 | 0 | 0 | 0.46747862 | 0 | 0.72061137 | 0 | 0 | 0 | 0 | 0 | 0 | 0 | 0 | 0.36156884 | 0 |
| ENSMUSG00 SNORD79 | 83 | 0 | 0 | 0 | 0 | 0 | 0 | 0 | 0 | 0 | 0 | 0 | 0 | 0 | 0 | 0 | 0 | 0 | 0 |
| ENSMUSG00 n-R5s167 | 120 | 0 | 0 | 0 | 0 | 0 | 0 | 0 | 0 | 0 | 0 | 0 | 0 | 0 | 0 | 0 | 0 | 0 | 0 |
| ENSMUSG00 U6 | 107 | 0 | 0.62267092 | 1.35626721 | 0 | 0 | 0.92621932 | 1.32652371 | 2.14163004 | 0.38492071 | 0.44066078 | 0 | 0 | 0 | 1.30217067 | 0 | 0 | 1.07456907 | 0 |
| ENSMUSG00 U6 | 106 | 0 | 0 | 0 | 0 | 0 | 0 | 0 | 0 | 0 | 0 | 0 | 0 | 0 | 0 | 0 | 0 | 0 | 0 |
| ENSMUSG00 U6 | 101 | 0 | 0 | 0 | 0 | 0 | 0 | 0 | 0 | 0 | 0 | 0 | 0 | 0 | 0 | 0 | 0 | 0 | 0 |
| ENSMUSG00 U6 | 106 | 0 | 0 | 0 | 0 | 0 | 0 | 0 | 0 | 0 | 0 | 0 | 0 | 0 | 0 | 0 | 0 | 0 | 0 |
| ENSMUSG00 U6 | 106 | 0 | 0 | 0 | 0 | 0 | 0 | 0 | 0 | 0 | 0 | 0 | 0 | 0 | 0 | 0 | 0 | 0 | 0 |
| ENSMUSG00 Snora19 | 131 | 1.29786797 | 2.03437522 | 1.47705436 | 2.78235385 | 1.10225779 | 1.51306057 | 2.52815842 | 0.58309011 | 1.8864053 | 0.35992903 | 1.09408302 | 0.42607376 | 1.12973856 | 2.12721011 | 1.88709416 | 1.16917885 | 2.34053722 | 2.09958423 |
| ENSMUSG00 U6 | 106 | 0 | 0 | 0 | 0 | 0 | 0 | 0 | 0 | 0 | 0 | 0 | 0 | 0 | 0 | 0 | 0 | 0 | 0 |
| ENSMUSG00 U6 | 106 | 0 | 0 | 0 | 0 | 0 | 0 | 0 | 0 | 0 | 0 | 0 | 0 | 0 | 0 | 0 | 0 | 0 | 0 |
| ENSMUSG00 U6 | 106 | 0 | 0 | 0 | 0 | 0 | 0 | 0 | 0 | 0 | 0 | 0 | 0 | 0 | 0 | 0 | 0 | 0 | 0 |
| ENSMUSG00 U6 | 105 | 0 | 0 | 0 | 0 | 0 | 0 | 0 | 0 | 0 | 0 | 0 | 0 | 0 | 0 | 0 | 0 | 0 | 0 |
| ENSMUSG00 U1 | 163 | 0 | 0 | 0 | 0 | 0 | 0 | 0 | 0 | 0 | 0 | 0 | 0 | 0 | 0 | 0 | 0 | 0 | 0 |
| ENSMUSG00 U12 | 150 | 1.13347136 | 1.99877365 | 1.28996081 | 0.91122089 | 2.24615644 | 0.66070311 | 0.94625358 | 1.5276961 | 1.37288386 | 1.25735209 | 2.86649751 | 1.86052209 | 0.6577589 | 2.47701799 | 1.09870816 | 1.5316243 | 0 | 0.36672738 |
| ENSMUSG00 U1 | 141 | 0 | 0 | 0 | 0 | 0 | 0 | 0 | 0 | 0 | 0 | 0 | 0 | 0 | 0 | 0 | 0 | 0 | 0 |
| ENSMUSG00 SNORA73 | 209 | 0 | 0 | 0 | 0 | 0 | 0 | 0 | 0 | 0 | 0 | 0 | 0 | 0 | 0 | 0 | 0 | 0 | 0 |
| ENSMUSG00 SNORA42 | 135 | 0 | 0 | 0 | 0 | 0 | 0 | 0 | 0 | 0 | 0 | 0 | 0 | 0 | 0 | 0 | 0 | 0 | 0 |
| ENSMUSG00 U6 | 109 | 0 | 0 | 0 | 0 | 0 | 0 | 0 | 0 | 0 | 0 | 0 | 0 | 0 | 0 | 0 | 0 | 0 | 0 |
| ENSMUSG00 SNORA36 | 114 | 0 | 0 | 0 | 0 | 0 | 0 | 0 | 0 | 0 | 0 | 0 | 0 | 0 | 0 | 0 | 0 | 0 | 0 |
| ENSMUSG00 U1 | 139 | 0 | 0.47932222 | 0 | 0 | 0.34627283 | 0.71298897 | 0 | 0 | 0 | 0 | 0 | 0 | 0 | 0 | 0 | 0 | 0 | 0 |
| ENSMUSG00 U1 | 168 | 0.253007 | 1.38803726 | 0.86381304 | 0.54239338 | 0.28649955 | 1.76974048 | 0.28162309 | 1.36401437 | 0 | 0.28065895 | 1.13749901 | 0.66447217 | 2.05549655 | 1.38226451 | 0.24524736 | 2.5071231 | 2.05319448 | 4.58409223 |
| ENSMUSG00 U6 | 108 | 0 | 0 | 0 | 0 | 0 | 0 | 0 | 0 | 0 | 0 | 0 | 0 | 0 | 0 | 0 | 0 | 0 | 0 |
| ENSMUSG00 U4 | 130 | 0 | 0 | 0 | 0 | 0 | 0 | 0 | 0 | 0 | 0 | 0 | 0 | 0 | 0 | 0 | 0 | 0 | 0 |
| ENSMUSG00 U6 | 106 | 0.40099223 | 0 | 0 | 0 | 0 | 0 | 0 | 0 | 0 | 0 | 0 | 0 | 0 | 0 | 0 | 0.72246429 | 0 | 0.51895384 |
| ENSMUSG00 U6 | 106 | 0 | 0 | 0 | 0 | 0 | 0 | 0 | 0 | 0 | 0 | 0 | 0 | 0 | 0 | 0 | 0 | 0 | 0 |
| ENSMUSG00 Snord85 | 72 | 0 | 0.46267969 | 0 | 0 | 0 | 0 | 0 | 0 | 0 | 0 | 0 | 0 | 0 | 0 | 0 | 0 | 0 | 0 |
| ENSMUSG00 U6 | 103 | 0 | 0 | 0 | 0 | 0 | 0 | 0 | 2.59560016 | 1.99934543 | 0 | 0 | 0 | 0 | 0 | 0 | 0.37175347 | 0 | 0 |
| ENSMUSG00 U6 | 106 | 0 | 0 | 0 | 0 | 0 | 0 | 0 | 0 | 0 | 0 | 0 | 0 | 0 | 0 | 0 | 0 | 0 | 0 |
| ENSMUSG00 SNORA65 | 129 | 0 | 0 | 0 | 0 | 0 | 0 | 0 | 0 | 0 | 0 | 0.37034852 | 0 | 0 | 0 | 0 | 0 | 0 | 0 |
| ENSMUSG00 U6 | 106 | 0 | 0 | 0 | 0 | 0 | 0 | 0 | 0 | 0 | 0 | 0 | 0 | 0 | 0 | 0 | 0 | 0 | 0 |
| ENSMUSG00 SNORA20 | 112 | 0 | 0 | 0 | 0 | 0.85949864 | 0 | 0 | 0 | 0 | 0 | 0 | 0 | 0 | 0 | 0 | 0 | 0 | 0 |
| ENSMUSG00 U6 | 106 | 0 | 0 | 0 | 0 | 0 | 0 | 0 | 0 | 0 | 0 | 0 | 0 | 0 | 0 | 0 | 0 | 0 | 0 |
| ENSMUSG00 SNORA32 | 108 | 0 | 0 | 0 | 0 | 0 | 0 | 1.31424108 | 1.06090007 | 0 | 0.43658059 | 0 | 0 | 0 | 0 | 0 | 0 | 0 | 0 |
| ENSMUSG00 U6 | 106 | 0 | 0 | 0 | 0 | 0 | 0 | 0 | 0 | 0 | 0 | 0 | 0 | 0 | 0 | 0 | 0 | 0 | 0 |
| ENSMUSG00 SNORA4 | 138 | 4.00411079 | 1.20698892 | 0.70106566 | 1.65076248 | 1.74391028 | 0 | 1.0285365 | 2.49080885 | 2.3876241 | 1.36668705 | 3.46195351 | 0.80892265 | 3.21729895 | 3.70206494 | 3.88130599 | 1.94227719 | 2.22181431 | 5.58063402 |
| ENSMUSG00 U6 | 106 | 0 | 0 | 0 | 0 | 0 | 0 | 0 | 0 | 0 | 0 | 0 | 0 | 0 | 0 | 0 | 0 | 0 | 0 |
| ENSMUSG00 SNORA76 | 134 | 1.26881122 | 0.49720738 | 1.80498248 | 0 | 0 | 0 | 0 | 0 | 0.30736206 | 0 | 0.71305908 | 1.24960439 | 1.47259455 | 0.34659767 | 0.3074743 | 0.2857508 | 2.28813713 | 0 |
| ENSMUSG00 U1 | 158 | 0 | 0 | 0 | 0 | 0 | 0 | 0 | 0 | 0 | 0 | 0 | 0 | 0 | 0 | 0 | 0 | 0 | 0 |
| ENSMUSG00 U6 | 106 | 0 | 0 | 0 | 0 | 0 | 0 | 0 | 0 | 0 | 0 | 0 | 0 | 0 | 0 | 0 | 0 | 0 | 0 |
| ENSMUSG00 U1 | 163 | 0 | 0 | 0 | 0 | 0 | 0 | 0 | 0 | 0 | 0 | 0 | 0 | 0 | 0 | 0 | 0 | 0 | 0 |
| ENSMUSG00 U1 | 163 | 0 | 0 | 0 | 0 | 0 | 0 | 0 | 0 | 0 | 0 | 0 | 0 | 0 | 0 | 0 | 0 | 0 | 0 |
| ENSMUSG00 Snord32a | 82 | 0 | 0 | 0 | 0 | 0 | 0 | 0 | 0 | 0 | 0 | 0 | 0 | 0 | 0 | 0 | 0 | 0 | 0 |
| ENSMUSG00 U6 | 106 | 0 | 0 | 0 | 0 | 0 | 0 | 0 | 0 | 0 | 0 | 0 | 0 | 0 | 0 | 0 | 0 | 0 | 0 |
| ENSMUSG00 U6 | 106 | 0 | 0 | 0 | 0 | 0 | 0 | 0 | 0 | 0 | 0 | 0 | 0 | 0 | 0 | 0 | 0 | 0 | 0 |
| ENSMUSG00 U6 | 99 | 0 | 0 | 0 | 0 | 0 | 0 | 0 | 0 | 0 | 0 | 0 | 0 | 0 | 0 | 0 | 0 | 0 | 0 |
| ENSMUSG00 SNORA8 | 136 | 3.12538059 | 3.67421627 | 3.20118951 | 0.67001536 | 2.12346722 | 1.45743334 | 0 | 2.52743839 | 2.72557825 | 2.77357078 | 1.05385938 | 2.4624557 | 6.52922435 | 1.02450193 | 4.24133664 | 3.66013159 | 1.97267705 | 3.23582981 |
| ENSMUSG00 U6 | 106 | 0 | 0 | 0 | 0 | 0 | 0.46747862 | 0 | 0 | 0.38855204 | 0 | 0 | 0 | 0 | 0 | 0 | 0 | 0 | 0.51895384 |
| ENSMUSG00 SNORD81 | 75 | 0 | 0 | 0 | 0 | 0 | 0 | 0 | 0 | 0 | 0 | 0 | 0 | 0 | 0 | 0 | 0 | 0 | 0 |
| ENSMUSG00 SNORA75 | 130 | 0 | 0 | 0 | 0 | 0 | 0 | 0 | 0 | 0 | 0 | 0 | 0 | 0 | 0 | 0 | 0 | 0 | 0 |
| ENSMUSG00 U2 | 190 | 2.68453743 | 1.22731716 | 4.07356045 | 0.47958994 | 4.30654054 | 2.86884247 | 1.2450705 | 0.40202529 | 0.21677114 | 0.99264638 | 2.76591865 | 0.29376665 | 0.25964167 | 0 | 0.43370059 | 1.6122361 | 2.01717352 | 1.73712969 |
| ENSMUSG00 n-R5s60 | 118 | 0 | 0 | 0 | 0 | 0 | 0 | 0 | 0 | 0 | 0 | 0 | 0 | 0 | 0 | 0 | 0 | 0 | 0 |
| ENSMUSG00 U6 | 106 | 0 | 0 | 0 | 0 | 0 | 0 | 0 | 0 | 0 | 0 | 0 | 0 | 0 | 0 | 0 | 0 | 0 | 0 |
| ENSMUSG00 U6 | 102 | 0 | 0 | 0 | 0 | 0 | 0 | 0 | 0 | 0 | 0 | 0 | 0 | 0 | 0 | 0 | 0 | 0 | 0 |
| ENSMUSG00 n-R5s197 | 118 | 0.36021336 | 0 | 0 | 0 | 0.40789766 | 0 | 0 | 0 | 0 | 0.39958223 | 1.21461759 | 0.94602818 | 0 | 2.75515773 | 0 | 0 | 0.64959825 | 0 |
| ENSMUSG00 U6 | 106 | 0 | 0 | 0 | 0 | 0 | 0 | 0 | 0 | 0 | 0 | 0 | 0 | 0 | 0 | 0 | 0 | 0 | 0 |
| ENSMUSG00 U2 | 185 | 0 | 0 | 0 | 0 | 0 | 0 | 0 | 0 | 0 | 0 | 0 | 0 | 0 | 0 | 0 | 0 | 0 | 0 |
| ENSMUSG00 U6 | 96 | 0 | 0 | 0 | 0 | 0 | 0 | 0 | 0 | 0 | 0 | 0 | 0 | 0 | 0 | 0 | 0 | 0 | 0 |
| ENSMUSG00 U6 | 106 | 0 | 0 | 0 | 0 | 0 | 0 | 0 | 0 | 0 | 0 | 0 | 0 | 0 | 0 | 0 | 0 | 0 | 0 |
| ENSMUSG00 U6 | 103 | 0 | 0 | 0 | 0 | 0 | 0 | 0 | 0 | 0 | 0 | 0 | 0 | 0 | 0 | 0 | 0 | 0 | 0 |
| ENSMUSG00 U6 | 102 | 0 | 0 | 0 | 0 | 0 | 0 | 0 | 0 | 0 | 0 | 0 | 0 | 0 | 0 | 0 | 0 | 0 | 0 |
| ENSMUSG00 U6 | 106 | 0 | 0 | 0 | 0 | 0 | 0 | 0 | 0 | 0 | 0 | 0 | 0 | 0 | 0 | 0 | 0 | 0 | 0 |
| ENSMUSG00 U6 | 105 | 0.4048112 | 0.95179698 | 0 | 0 | 0 | 1.88772318 | 0 | 0 | 0 | 0 | 0 | 0 | 0 | 0 | 0 | 1.82336226 | 0 | 1.57168876 |
| ENSMUSG00 SNORA32 | 108 | 0 | 0 | 0 | 0 | 0 | 0 | 0 | 0 | 0 | 0 | 0 | 0 | 0 | 0 | 0 | 0 | 0 | 0 |
| ENSMUSG00 U6 | 106 | 0 | 0 | 0 | 0 | 0 | 0 | 0 | 0 | 0 | 0 | 0 | 0 | 0 | 0 | 0 | 0 | 0 | 0 |
| ENSMUSG00 U2 | 190 | 0.22371145 | 0 | 0 | 0 | 0.50665183 | 0 | 0 | 0.60303793 | 0 | 0 | 0 | 0.29376665 | 0 | 0 | 0.43370059 | 0 | 0.20171735 | 0 |
| ENSMUSG00 U6 | 106 | 0 | 0.94281776 | 0 | 0 | 0 | 0 | 0 | 0 | 0 | 0 | 0 | 0 | 0 | 0 | 0 | 0 | 1.08470652 | 0 |
| ENSMUSG00 U6 | 106 | 0 | 0 | 0 | 0 | 0 | 0 | 0 | 0 | 0 | 0 | 0 | 0 | 0 | 0 | 0 | 0 | 0 | 0 |
| ENSMUSG00 7SK | 320 | 0 | 0 | 0 | 0 | 0 | 0 | 0 | 0 | 0 | 0 | 0 | 0 | 0.46248672 | 0 | 0 | 0 | 0 | 0 |
| ENSMUSG00 SNORA55 | 130 | 0 | 0 | 0 | 0 | 0 | 0 | 0 | 0 | 0 | 0 | 0 | 0 | 0 | 0 | 0 | 0 | 0 | 0 |
| ENSMUSG00 U1 | 164 | 0 | 0 | 0 | 0 | 0 | 0 | 0 | 0 | 0 | 0 | 0 | 0 | 0 | 0 | 0 | 0 | 0 | 0 |
| ENSMUSG00 SNORD24 | 73 | 1.16452537 | 0.45634102 | 0.6626511 | 0 | 7.25275562 | 0.67880457 | 0.64811889 | 0 | 0 | 5.16720036 | 2.61780594 | 0.76459812 | 4.05467813 | 5.089763 | 2.82202438 | 3.67170208 | 1.05003553 | 3.76774704 |
| ENSMUSG00 Snora30 | 129 | 0 | 0.51647898 | 0 | 0 | 0.74623137 | 0.38412972 | 0.73352991 | 0.29606513 | 0.63855063 | 0 | 1.11104555 | 1.29803866 | 0.76483593 | 0 | 3.51331097 | 2.0777849 | 1.18841231 | 4.69069962 |
| ENSMUSG00 U6 | 105 | 0 | 0 | 0 | 0 | 0 | 0 | 0 | 0 | 0 | 0 | 0 | 0 | 0 | 0 | 0 | 0 | 0 | 0 |
| ENSMUSG00 U6 | 106 | 0 | 0 | 0 | 0 | 0 | 0 | 0 | 0 | 0 | 0 | 0 | 0 | 0 | 0 | 0 | 0 | 0 | 0 |
| ENSMUSG00 SNORA32 | 121 | 0 | 0 | 0 | 0 | 0 | 0 | 0 | 0 | 0 | 0 | 0 | 0 | 0 | 0 | 0 | 0 | 0 | 0 |
| ENSMUSG00 U6 | 106 | 0 | 0 | 0 | 0 | 0 | 0 | 0 | 0 | 0 | 0 | 0 | 0 | 0 | 0 | 0 | 0 | 0 | 0 |
| ENSMUSG00 SNORA32 | 99 | 4.29345212 | 2.01896329 | 5.37483671 | 9.20425138 | 6.80653466 | 3.50372864 | 7.64649356 | 6.94407316 | 3.32820329 | 6.19150649 | 11.5818081 | 6.76553486 | 5.97962634 | 5.62958635 | 5.41030532 | 10.4428929 | 9.6783578 | 2.22259017 |
| ENSMUSG00 SNORA16 | 132 | 0 | 0 | 0 | 0 | 0 | 0 | 0 | 0 | 0 | 0 | 0 | 0 | 0 | 0 | 0 | 0 | 0 | 0 |
| ENSMUSG00 U6 | 106 | 0 | 0 | 0 | 0 | 0 | 0 | 0 | 0 | 0 | 0 | 0 | 0 | 0 | 0 | 0 | 0 | 0 | 0 |
| ENSMUSG00 SNORA62 | 136 | 0 | 0 | 0 | 0 | 0 | 0 | 0 | 0 | 0 | 0 | 0 | 0 | 0 | 0 | 0 | 0 | 0 | 0 |
| ENSMUSG00 U6 | 106 | 0 | 0 | 0 | 0 | 0 | 0 | 0 | 0 | 0 | 0 | 0 | 0 | 0 | 0 | 0 | 0 | 0 | 0 |
| ENSMUSG00 Snord57 | 71 | 0 | 0 | 0 | 0 | 0 | 0 | 0 | 0 | 0 | 0 | 0 | 0 | 0 | 0 | 0 | 0 | 0 | 0 |
| ENSMUSG00 SNORD83 | 75 | 0 | 0.44417192 | 0 | 0 | 0 | 0 | 1.26167144 | 0 | 0 | 0 | 0 | 0 | 0 | 0 | 0.54935408 | 0 | 0 | 0.73345476 |
| ENSMUSG00 SNORA42 | 135 | 0 | 0 | 0 | 0 | 0 | 0 | 0 | 0 | 0 | 0 | 0 | 0 | 0 | 0 | 0 | 0 | 0 | 0 |
| ENSMUSG00 n-R5s217 | 115 | 0 | 0 | 0 | 0 | 0 | 0 | 0 | 0 | 0 | 0 | 0 | 0 | 0 | 0 | 0 | 0 | 0 | 0 |
| ENSMUSG00 U6 | 101 | 0 | 0 | 0 | 0 | 0 | 0 | 0 | 0 | 0 | 0 | 0 | 0 | 0 | 0 | 0 | 0 | 0 | 0 |
| ENSMUSG00 U1 | 163 | 0 | 0 | 0 | 0 | 0 | 0 | 0 | 0 | 0 | 0 | 0 | 0 | 0 | 0 | 0 | 0 | 0 | 0 |
| ENSMUSG00 SNORD28 | 75 | 0 | 0 | 0 | 0 | 0 | 0 | 0 | 0 | 0 | 0 | 0 | 0 | 0 | 0 | 0 | 0 | 0 | 0 |
| ENSMUSG00 SNORD27 | 71 | 0 | 0 | 0 | 0 | 0 | 0 | 0 | 0 | 0 | 0 | 0 | 0 | 0 | 0 | 0 | 0 | 0 | 0 |
| ENSMUSG00 SNORA18 | 130 | 8.17407231 | 4.35630155 | 4.09314488 | 3.85516529 | 2.96196453 | 1.52469949 | 1.45577474 | 2.6440894 | 0.95045806 | 1.81348859 | 2.57249776 | 0.8587025 | 3.79476287 | 4.64440874 | 2.85241541 | 3.53451761 | 1.76890601 | 1.26944092 |
| ENSMUSG00 SNORA51 | 131 | 0 | 0 | 0 | 0 | 0 | 0 | 0 | 0 | 0 | 0 | 0 | 0 | 0 | 0 | 0 | 0 | 0 | 0 |
| ENSMUSG00 U1 | 163 | 0 | 0 | 0 | 0 | 0 | 0 | 0 | 0 | 0 | 0 | 0 | 0 | 0 | 0 | 0 | 0 | 0 | 0 |
| ENSMUSG00 SNORA51 | 132 | 0 | 0 | 0 | 0 | 0 | 0 | 0 | 0 | 0 | 0 | 0 | 0 | 0 | 0 | 0 | 0 | 0 | 0 |
| ENSMUSG00 SNORD56 | 69 | 0 | 0 | 0 | 0 | 0 | 0 | 0 | 0 | 0 | 0 | 0 | 0 | 0 | 0 | 0 | 0 | 0 | 0 |
| ENSMUSG00 U6 | 106 | 0 | 0 | 0 | 0 | 0 | 0 | 0 | 0 | 0 | 0 | 0 | 0 | 0 | 0 | 0 | 0 | 0 | 0 |
| ENSMUSG00 U6 | 106 | 0 | 0 | 0 | 0 | 0 | 0 | 0 | 0 | 0 | 0 | 0 | 0 | 0 | 0 | 0 | 0 | 0 | 0 |
| ENSMUSG00 Y_RNA | 107 | 0 | 0 | 0 | 0 | 0 | 0 | 0 | 0 | 0 | 0 | 0 | 0 | 0 | 0 | 0 | 0 | 0 | 0 |
| ENSMUSG00 U1 | 165 | 0 | 0 | 0 | 0 | 0 | 0 | 0 | 0 | 0 | 0 | 0 | 0 | 0 | 0 | 0 | 0 | 0 | 0 |
| ENSMUSG00 SNORD36 | 70 | 0 | 0 | 0 | 0 | 0 | 0 | 0 | 0 | 0 | 0 | 0 | 0 | 0 | 0 | 0 | 0 | 0 | 0 |
| ENSMUSG00 U6 | 81 | 0 | 0 | 0 | 0 | 0 | 0 | 0 | 0 | 0 | 0 | 0 | 0 | 0 | 0 | 0 | 0 | 0 | 0 |
| ENSMUSG00 SNORA42 | 134 | 0 | 0 | 0 | 0 | 0 | 0 | 0 | 0 | 0 | 0 | 0 | 0 | 0 | 0 | 0 | 0 | 0 | 0 |
| ENSMUSG00 U1 | 161 | 0 | 0 | 0 | 0.28298785 | 0 | 0 | 0 | 0.23721969 | 0.25581687 | 0 | 0 | 0 | 0 | 0 | 0 | 0 | 0 | 0 |
| ENSMUSG00 U6 | 102 | 0 | 0 | 0 | 0 | 0.4718816 | 0 | 0 | 0 | 0 | 0 | 0 | 0 | 0 | 0.45533419 | 0 | 0 | 0 | 0 |
| ENSMUSG00 U6 | 106 | 0 | 0 | 0 | 0 | 0 | 0 | 0 | 0 | 0 | 0 | 0 | 0 | 0 | 0 | 0 | 0 | 0 | 0 |
| ENSMUSG00 U6 | 107 | 0 | 0 | 0 | 0 | 0 | 0 | 0 | 0 | 1.53968283 | 0 | 0 | 0 | 0 | 0.43405689 | 0 | 0 | 0 | 0 |
| ENSMUSG00 U1 | 160 | 0 | 0 | 0 | 0 | 0 | 0 | 0 | 0 | 0 | 0 | 0 | 0 | 0 | 0 | 0 | 0 | 0 | 0 |
| ENSMUSG00 SNORA22 | 134 | 2.22041964 | 3.23184795 | 3.24896846 | 3.74008573 | 3.5919346 | 1.10938956 | 1.76539847 | 2.56516136 | 7.37668939 | 3.16683828 | 5.34794311 | 0 | 2.20889182 | 3.46597667 | 1.84484579 | 2.28600641 | 4.29025712 | 0.82103144 |
| ENSMUSG00 SNORD31 | 66 | 0 | 0 | 0 | 0 | 0 | 0 | 0 | 0 | 0 | 0 | 0 | 0 | 0 | 0 | 0 | 0 | 0 | 0 |
| ENSMUSG00 U6 | 106 | 0.80198445 | 0.62854517 | 1.36906218 | 0 | 1.81629901 | 0.46747862 | 0 | 2.52213978 | 0.38855204 | 0.44481796 | 1.35212147 | 0.52656285 | 0 | 0 | 4.27563316 | 1.80616073 | 1.44627535 | 2.07581535 |
| ENSMUSG00 U6 | 106 | 0 | 0 | 0 | 0 | 0 | 0 | 0 | 0 | 0 | 0 | 0 | 0 | 0 | 0 | 0 | 0 | 0 | 0 |
| ENSMUSG00 U1 | 164 | 0 | 0 | 0 | 0 | 0 | 0 | 0 | 0 | 0 | 0 | 0 | 0 | 0 | 0 | 0 | 0 | 0 | 0 |
| ENSMUSG00 U6 | 91 | 0 | 0 | 0 | 0 | 0 | 0 | 0 | 0 | 0 | 0 | 0 | 0 | 0 | 0 | 0 | 0 | 0 | 0 |
| ENSMUSG00 n-R5s156 | 104 | 0 | 0 | 0 | 0 | 0 | 0 | 0 | 0 | 0 | 0 | 0 | 0 | 0 | 0 | 0 | 0 | 0 | 0 |
| ENSMUSG00 U6 | 106 | 0 | 0 | 0 | 0 | 0 | 0 | 0 | 0 | 0 | 0 | 0 | 0 | 0 | 0 | 0 | 0 | 0 | 0 |
| ENSMUSG00 U6 | 106 | 0 | 0 | 0 | 0 | 0 | 0 | 0 | 0 | 0 | 0 | 0 | 0 | 0 | 0 | 0 | 0 | 0 | 0 |
| ENSMUSG00 U6 | 104 | 1.22611085 | 0 | 0 | 0 | 0.46280696 | 0 | 0 | 0 | 0 | 0 | 0 | 0 | 2.37172679 | 0 | 0 | 0 | 0 | 0 |
| ENSMUSG00 U6 | 108 | 0 | 0.30845272 | 0 | 0 | 0.44566596 | 0 | 0 | 0 | 0 | 0 | 0.88472145 | 0 | 0 | 0 | 0 | 0 | 0 | 0 |
| ENSMUSG00 U6 | 106 | 0 | 0 | 0 | 0 | 0 | 0 | 0 | 0 | 0 | 0 | 0 | 0 | 0 | 0 | 0 | 0 | 0 | 0 |
| ENSMUSG00 U6 | 106 | 0 | 0 | 0 | 0 | 0 | 0 | 0 | 0 | 0 | 0 | 0 | 0 | 0 | 0 | 0 | 0 | 0 | 0 |
| ENSMUSG00 U6 | 108 | 0 | 0 | 0 | 0 | 0 | 0 | 0 | 0 | 0 | 0 | 0 | 0 | 0 | 0 | 0 | 0 | 0 | 0 |
| ENSMUSG00 U6 | 80 | 0 | 0 | 0 | 0 | 0 | 0 | 0 | 0 | 0 | 0 | 0 | 0 | 0 | 0 | 0 | 0 | 0 | 0 |
| ENSMUSG00 U6 | 106 | 0 | 0 | 0 | 0 | 0 | 0 | 0 | 0 | 0 | 0 | 0 | 0 | 0 | 0 | 0 | 0 | 0 | 0 |
| ENSMUSG00 U6 | 106 | 0 | 0 | 0 | 0 | 0 | 0 | 0 | 0 | 0 | 0 | 0 | 0 | 0 | 0 | 0 | 0 | 0 | 0 |
| ENSMUSG00 U6 | 106 | 0 | 0 | 0 | 0 | 0 | 0 | 0 | 0 | 0 | 0 | 0 | 0 | 0 | 0 | 0 | 0 | 0 | 0 |
| ENSMUSG00 U6 | 106 | 0 | 0 | 0 | 0 | 0 | 0 | 0 | 0 | 0 | 0 | 0 | 0 | 0 | 0 | 0 | 0 | 0 | 0 |
| ENSMUSG00 7SK | 309 | 0 | 0 | 0 | 0 | 0 | 0 | 0 | 0 | 0 | 0 | 0 | 0 | 0 | 0 | 0 | 0 | 0 | 0 |
| ENSMUSG00 SNORA73 | 199 | 0 | 0 | 0 | 0 | 0 | 0 | 0 | 0 | 0 | 0 | 0 | 0 | 0 | 0 | 0 | 0 | 0 | 0 |
| ENSMUSG00 U1 | 154 | 0 | 0 | 0 | 0 | 0 | 0 | 0 | 0 | 0 | 0 | 0 | 0 | 0 | 0 | 0 | 0 | 0 | 0 |
| ENSMUSG00 U6 | 106 | 0 | 0 | 0 | 0 | 0 | 0 | 0 | 0 | 0 | 0 | 0 | 0 | 0 | 0 | 0 | 0 | 0 | 0 |
| ENSMUSG00 U6 | 106 | 0 | 0.31427259 | 0 | 0 | 0 | 0.46747862 | 0 | 0 | 0 | 0 | 0 | 0 | 4.18855901 | 0 | 0 | 0 | 0 | 1.55686151 |
| ENSMUSG00 U6 | 106 | 0 | 0 | 0 | 0 | 0 | 0 | 0 | 0 | 0 | 0 | 0 | 0 | 0 | 0 | 0 | 0 | 0 | 0 |
| ENSMUSG00 U6 | 106 | 0 | 1.25709035 | 0 | 0 | 0 | 0 | 0.44634603 | 0.36030568 | 0 | 0.44481796 | 1.35212147 | 0 | 1.39618634 | 0 | 0.77738785 | 2.88985716 | 2.16941303 | 0.51895384 |
| ENSMUSG00 Snora34 | 137 | 1.24102704 | 1.45895687 | 1.41236585 | 0.66512473 | 1.05398373 | 2.5318915 | 0 | 2.50898994 | 3.30694652 | 1.72082859 | 1.3948893 | 0.4074136 | 0 | 3.05107143 | 0.60148257 | 0 | 2.23803194 | 8.43205286 |
| ENSMUSG00 U1 | 138 | 0 | 0 | 0 | 0 | 0 | 0 | 0 | 0 | 0 | 0 | 0 | 0 | 0 | 0 | 0 | 0 | 0 | 0 |
| ENSMUSG00 U6 | 106 | 0 | 0 | 0 | 0 | 0 | 0 | 0 | 0 | 0 | 0 | 0 | 0 | 0 | 0 | 0 | 0 | 0 | 0 |
| ENSMUSG00 U6 | 103 | 0 | 0 | 0 | 0 | 0 | 0 | 0 | 0 | 0 | 0 | 0 | 0 | 0 | 0 | 0 | 0 | 0 | 0 |
| ENSMUSG00 U6 | 105 | 0 | 0.31726566 | 0 | 0 | 0 | 0.4719308 | 0 | 0 | 4.31477784 | 0 | 0 | 0 | 0.46982778 | 0 | 0 | 0 | 0 | 0 |
| ENSMUSG00 U6 | 106 | 0 | 0 | 0 | 0 | 0 | 0 | 0 | 0 | 0 | 0.44481796 | 0 | 0 | 0 | 0 | 0 | 0 | 0 | 0 |
| ENSMUSG00 7SK | 326 | 0 | 0 | 0.14838506 | 0.69878902 | 0 | 0 | 0 | 0 | 0 | 0 | 0 | 0 | 0 | 0 | 0 | 0.2349117 | 0 | 0 |
| ENSMUSG00 U1 | 163 | 0 | 0 | 0 | 0 | 0 | 0 | 0 | 0 | 0 | 0 | 0 | 0 | 0 | 0 | 0 | 0 | 0 | 0 |
| ENSMUSG00 SNORA4 | 109 | 0 | 0 | 0 | 0 | 0 | 0 | 0 | 0 | 0 | 0 | 0 | 0 | 0 | 0 | 0 | 0 | 0 | 0 |
| ENSMUSG00 U6 | 106 | 0 | 0 | 0 | 0 | 0 | 0 | 0 | 0 | 0 | 0 | 0 | 0 | 0 | 0 | 0 | 0 | 0 | 0 |
| ENSMUSG00 U6 | 106 | 0 | 0 | 0 | 0 | 0 | 0 | 0 | 0 | 0 | 0 | 0 | 0 | 0 | 0 | 0 | 0 | 0 | 0 |
| ENSMUSG00 Snora73b | 204 | 0 | 0 | 0 | 0 | 0 | 0 | 0 | 0 | 0 | 0 | 0 | 0 | 0 | 0 | 0 | 0 | 0 | 0 |
| ENSMUSG00 SNORD16 | 98 | 1.73490514 | 2.71941994 | 2.96164472 | 1.39472585 | 5.40256286 | 1.01128028 | 1.93112975 | 1.55887357 | 1.68108228 | 6.25468513 | 0.97499915 | 0 | 1.51016073 | 1.42175778 | 0.84084806 | 0.39072048 | 3.91084682 | 2.24526966 |
| ENSMUSG00 SNORA32 | 108 | 1.18069933 | 0 | 0.89580612 | 0 | 0 | 0 | 0 | 0 | 1.90678314 | 0.43658059 | 0 | 0 | 1.82710805 | 0 | 0 | 0.70908532 | 0 | 0 |
| ENSMUSG00 Telomerase-v | 411 | 0.62051352 | 0.64842616 | 0.35309146 | 0 | 0.35132791 | 0.12056626 | 1.38139209 | 0.18585111 | 0.7014735 | 0.34416572 | 0.4649631 | 0.54321813 | 0.120029 | 0.56501323 | 0.40098838 | 0.09316449 | 0.83926198 | 0.93689476 |
| ENSMUSG00 SNORA71 | 130 | 0.32696289 | 1.53751819 | 0 | 0 | 0 | 1.14352462 | 0.36394368 | 1.76272626 | 0 | 0.36269772 | 0 | 1.28805375 | 1.51790515 | 1.78631105 | 2.53548036 | 2.35634507 | 0.29481767 | 0 |
| ENSMUSG00 U6 | 106 | 0 | 0 | 0 | 0 | 0 | 0 | 0 | 0 | 0 | 0 | 0 | 0 | 0 | 0 | 0 | 0 | 0 | 0 |
| ENSMUSG00 U6 | 106 | 0 | 0 | 0 | 0 | 0.45407475 | 0 | 0 | 0 | 0 | 0 | 0 | 0 | 0 | 0 | 0 | 0 | 0 | 0 |
| ENSMUSG00 U1 | 163 | 1.56460771 | 1.63498867 | 0 | 0.27951561 | 3.24816663 | 2.1280315 | 0.8707855 | 0.23430922 | 0.75803403 | 0.86780435 | 0.2930979 | 0 | 0 | 0 | 0.25277028 | 0.93964681 | 0 | 0.67495836 |
| ENSMUSG00 U1 | 159 | 0 | 0 | 0 | 0.5730949 | 0 | 0 | 0 | 0 | 0 | 0 | 0.30047144 | 0 | 0 | 0 | 0 | 0 | 0 | 0 |
| ENSMUSG00 U1 | 164 | 0 | 0 | 0 | 0 | 0 | 0 | 0 | 0 | 0 | 0 | 0 | 0 | 0 | 0 | 0.251229 | 0 | 0 | 0 |
| ENSMUSG00 U6 | 108 | 0 | 0 | 0 | 0 | 0 | 0 | 0 | 0 | 0 | 0 | 0 | 0 | 0 | 0 | 0 | 0 | 0 | 0 |
| ENSMUSG00 U1 | 146 | 0 | 0 | 0 | 0 | 0.32967071 | 0.67880457 | 0.97217833 | 0 | 1.41049712 | 0.96885007 | 0 | 0.38229906 | 1.68944922 | 3.49921206 | 0.28220244 | 0 | 1.05003553 | 0 |
| ENSMUSG00 SNORD29 | 64 | 0 | 0 | 0 | 0 | 0 | 0 | 0 | 0 | 0 | 0 | 0 | 0 | 0 | 0 | 0 | 0 | 0 | 0 |
| ENSMUSG00 SNORA25 | 132 | 0 | 0 | 0 | 0 | 0 | 0 | 0 | 0 | 0 | 0 | 0 | 0 | 0 | 0.70369829 | 0 | 0 | 0 | 0 |
| ENSMUSG00 U6 | 106 | 0 | 0 | 0 | 0 | 0 | 0 | 0 | 0 | 0 | 0 | 0 | 0 | 0 | 0 | 0 | 0 | 0 | 0 |
| ENSMUSG00 U6 | 106 | 0 | 0 | 0 | 0 | 0 | 0 | 0 | 0 | 0 | 0 | 0 | 0 | 0 | 0 | 0 | 0 | 0 | 0 |
| ENSMUSG00 U6 | 104 | 0 | 0 | 0 | 0 | 0 | 0 | 0 | 0 | 0 | 0 | 0 | 0 | 0 | 0 | 0 | 0 | 0 | 0 |
| ENSMUSG00 U6 | 101 | 0 | 0 | 0 | 0 | 0 | 0 | 0 | 0 | 0 | 0 | 0 | 0 | 0 | 0 | 0 | 0 | 0 | 0 |
| ENSMUSG00 n-R5s153 | 115 | 0 | 0 | 0 | 0 | 0 | 0 | 0 | 0 | 0 | 0 | 0 | 0 | 0 | 0 | 0 | 0 | 0 | 0 |
| ENSMUSG00 n-R5s85 | 112 | 0 | 0 | 0 | 0 | 0.42974932 | 0 | 0 | 0 | 0 | 0 | 0 | 0 | 0 | 0 | 0 | 0 | 0 | 0 |
| ENSMUSG00 U1 | 138 | 0 | 0 | 0 | 0 | 0 | 0 | 0 | 0 | 0 | 0 | 0 | 0 | 0 | 0 | 0 | 0 | 0 | 0 |
| ENSMUSG00 U1 | 140 | 0 | 0 | 0 | 0 | 0 | 0 | 0 | 0 | 0 | 0 | 0 | 0 | 0 | 0 | 0 | 0 | 0 | 0 |
| ENSMUSG00 U6 | 106 | 0 | 0 | 0 | 0 | 0 | 0 | 0 | 0 | 0 | 0 | 0 | 0 | 0 | 0 | 0 | 0 | 0 | 0 |
| ENSMUSG00 U6 | 106 | 0 | 0 | 0 | 0 | 0 | 0 | 0 | 0 | 0 | 0 | 0 | 0 | 0 | 0 | 0 | 0 | 0 | 0 |
| ENSMUSG00 SNORD26 | 74 | 0 | 0 | 0 | 0 | 0 | 0 | 0 | 0 | 0 | 0 | 0 | 0 | 0 | 0 | 0 | 0 | 0 | 0 |
| ENSMUSG00 U6 | 106 | 0 | 0 | 0 | 0 | 0 | 0 | 0 | 0 | 0 | 0 | 0 | 0 | 0 | 0 | 0 | 0 | 0 | 0 |
| ENSMUSG00 Mir25 | 83 | 0 | 0 | 0 | 0 | 0 | 0 | 0 | 0 | 0 | 0 | 0 | 0 | 0 | 0 | 0 | 0 | 0 | 0 |
| ENSMUSG00 Mir193 | 65 | 0 | 0.51250606 | 0 | 0 | 0 | 0 | 0 | 0 | 0 | 0 | 0 | 0 | 0 | 0 | 0 | 0 | 0 | 0 |
| ENSMUSG00 Mir99b | 69 | 0.61601704 | 0 | 0 | 0 | 0.69756411 | 0 | 0 | 0 | 0 | 0 | 0 | 0 | 0 | 0 | 0 | 0 | 0 | 0 |
| ENSMUSG00 Mir155 | 64 | 0 | 0 | 0 | 0 | 0 | 0 | 0 | 0 | 0 | 0 | 0 | 0 | 0 | 0 | 0 | 0 | 0 | 0 |
| ENSMUSG00 Mir188 | 67 | 0 | 0 | 0 | 0 | 0 | 0 | 0 | 0 | 0 | 0 | 0 | 0 | 0 | 0 | 0 | 0 | 0 | 0 |
| ENSMUSG00 Mir133a-1 | 86 | 0 | 0 | 0 | 0 | 2.23869412 | 0 | 0 | 0 | 0 | 0 | 0.55552277 | 0 | 0 | 0.54004753 | 0 | 0.44523962 | 0 | 0 |
| ENSMUSG00 Mir200a | 89 | 0 | 0 | 0 | 0.51192185 | 0 | 0 | 0.53160313 | 0 | 0.92553968 | 0 | 0 | 0 | 0 | 0 | 0 | 0 | 0 | 0 |
| ENSMUSG00 Mir144 | 65 | 0 | 0 | 0 | 0 | 0 | 0 | 0 | 0 | 0 | 0 | 0 | 0 | 0 | 0 | 0 | 0 | 0 | 0 |
| ENSMUSG00 Mir122a | 65 | 0 | 0 | 0 | 0 | 0 | 0 | 0 | 0 | 0 | 0 | 0 | 0 | 1.51790515 | 0 | 0 | 0 | 0 | 0.84629395 |
| ENSMUSG00 Mir18 | 95 | 0 | 0 | 0 | 0 | 0 | 0 | 0 | 0 | 0 | 0 | 0 | 0 | 0 | 0 | 0 | 0 | 0 | 0 |
| ENSMUSG00 Mir201 | 65 | 0 | 0 | 0 | 0 | 0 | 0 | 0 | 0 | 0 | 0 | 0 | 0 | 0 | 0 | 0 | 0 | 0 | 0 |
| ENSMUSG00 Mir30a | 70 | 0 | 0 | 0 | 0 | 0 | 0 | 0 | 0 | 0 | 0.67358148 | 0 | 0 | 0 | 0 | 0 | 0 | 0 | 0 |
| ENSMUSG00 Mirlet7i | 84 | 2.024056 | 0.79316415 | 0 | 0 | 0.57299909 | 0 | 0 | 2.27335729 | 0.98063133 | 0 | 0 | 1.99341652 | 0.58728473 | 0 | 1.96197885 | 0 | 0.91253088 | 3.27435159 |
| ENSMUSG00 Mir135a-1 | 89 | 0 | 0 | 0 | 0 | 0 | 0 | 0 | 0 | 0 | 0 | 0 | 0 | 0 | 0 | 0 | 0 | 0 | 0 |
| ENSMUSG00 Mir31 | 105 | 0.4048112 | 0 | 0 | 0 | 0 | 0 | 0 | 0 | 0 | 0.89810863 | 0 | 0 | 0 | 0 | 0 | 0 | 0 | 0 |
| ENSMUSG00 Mir30e | 91 | 0 | 0 | 0 | 0 | 0 | 0 | 0 | 0 | 0 | 0 | 0 | 0 | 0 | 0 | 0 | 0 | 0 | 0 |
| ENSMUSG00 Mir298 | 81 | 0 | 0 | 0 | 0 | 0 | 0 | 0 | 0 | 0 | 0 | 0 | 0 | 0 | 0 | 0 | 0 | 0 | 0 |
| ENSMUSG00 Mir195 | 93 | 0 | 0 | 0 | 0 | 0 | 0 | 0 | 0 | 0 | 0 | 0 | 0 | 0 | 0 | 0 | 0 | 0 | 0 |
| ENSMUSG00 Mir29b-2 | 80 | 0 | 0 | 0 | 0 | 0 | 0 | 0 | 0 | 0 | 0 | 0 | 0 | 0 | 0 | 0 | 0 | 0 | 0 |
| ENSMUSG00 Mir367 | 74 | 0 | 0.45017425 | 0 | 0 | 0 | 0 | 0 | 0 | 0 | 0 | 0 | 0 | 0 | 0 | 0 | 0 | 0 | 0 |
| ENSMUSG00 Mir138-1 | 98 | 0 | 0 | 0 | 0 | 0 | 0 | 0 | 0 | 0 | 0 | 0 | 0 | 0 | 0 | 0 | 0 | 0 | 0 |
| ENSMUSG00 Mir217 | 107 | 0 | 0 | 0 | 0 | 0 | 0 | 0 | 0 | 0 | 0 | 0 | 0 | 0 | 0 | 0 | 0 | 0 | 0 |
| ENSMUSG00 Mir19a | 81 | 0 | 0 | 0 | 0 | 0 | 0 | 0 | 0 | 0 | 0 | 0 | 0 | 0 | 0 | 0 | 0 | 0 | 0 |
| ENSMUSG00 Mir340 | 97 | 0 | 0 | 0 | 0 | 0.4962054 | 0 | 0 | 0 | 0.42460326 | 0 | 0 | 1.1508384 | 0 | 0 | 0 | 0 | 0 | 0 |
| ENSMUSG00 Mir322 | 94 | 0 | 0 | 0 | 0 | 0 | 0 | 0 | 0 | 0 | 0 | 0 | 0 | 0 | 0 | 0 | 0 | 0 | 0 |
| ENSMUSG00 Mir300 | 78 | 0 | 0 | 0 | 0 | 0 | 0 | 0 | 0 | 0 | 0 | 0 | 0 | 0 | 0 | 0 | 0 | 0 | 0 |
| ENSMUSG00 Mir142 | 63 | 0 | 0 | 0 | 0 | 0 | 0 | 0 | 0 | 0 | 0 | 0 | 0 | 0 | 0 | 0 | 0 | 0 | 0 |
| ENSMUSG00 Mirlet7a-1 | 93 | 0.91408981 | 0.35820316 | 0 | 0 | 0.51754757 | 0 | 0 | 0 | 0 | 0 | 0 | 0 | 0 | 0 | 0 | 0 | 0 | 0 |
| ENSMUSG00 Mir221 | 94 | 0 | 0 | 0 | 0 | 0 | 0 | 0 | 0.81260431 | 0 | 0 | 0.50824424 | 0 | 0 | 1.48225811 | 0 | 0 | 0 | 0 |
| ENSMUSG00 Mir181a-2 | 75 | 1.13347136 | 0 | 0 | 1.21496118 | 0 | 0 | 0.63083572 | 0 | 0 | 0.62867604 | 0 | 1.48841767 | 0 | 0 | 1.09870816 | 0 | 0 | 0.73345476 |
| ENSMUSG00 Mir9-2 | 86 | 0 | 0 | 0 | 0 | 0 | 0 | 0 | 0 | 0 | 0 | 0 | 0 | 0 | 0 | 0 | 0 | 0 | 0 |
| ENSMUSG00 Mir325 | 97 | 0 | 0 | 0 | 0 | 0 | 0 | 0 | 0 | 0 | 0 | 0 | 0 | 0 | 0 | 0 | 0 | 0 | 0 |
| ENSMUSG00 Mir134 | 70 | 0 | 0 | 0 | 0 | 0 | 0 | 0 | 0 | 0 | 0 | 0 | 0 | 0 | 0 | 0 | 0 | 0 | 0 |
| ENSMUSG00 Mir429 | 82 | 0 | 0 | 0 | 0 | 0 | 0 | 0 | 0 | 0 | 0 | 0 | 0 | 0 | 0 | 0 | 0 | 0 | 0 |
| ENSMUSG00 Mir382 | 79 | 0 | 0 | 0 | 0 | 0 | 0 | 0 | 0 | 0 | 0 | 0 | 0 | 0 | 0 | 0 | 0 | 0 | 0 |
| ENSMUSG00 Mir345 | 95 | 0 | 0 | 0 | 0 | 0 | 1.04321544 | 0 | 0.40202529 | 2.16771136 | 0 | 0 | 0.58753329 | 1.03856668 | 0 | 0.86740118 | 4.43364928 | 0 | 0 |
| ENSMUSG00 Mir26a-2 | 83 | 0 | 0 | 0 | 0 | 0 | 0 | 0 | 0 | 0 | 0 | 0 | 0 | 0 | 0 | 0 | 0 | 0 | 0 |
| ENSMUSG00 Mir186 | 70 | 3.036084 | 2.85539093 | 6.2194539 | 3.90523237 | 2.75039564 | 1.41579239 | 0 | 2.72802874 | 0 | 2.02074443 | 5.45999525 | 4.78419965 | 2.11422503 | 1.99046089 | 2.35437462 | 2.18803471 | 2.19007411 | 0 |
| ENSMUSG00 Mir208a | 82 | 0 | 0 | 0 | 0 | 0 | 0 | 0 | 0 | 0 | 0 | 0 | 0 | 0 | 0 | 0 | 0 | 0 | 0 |
| ENSMUSG00 Mir370 | 78 | 0 | 0 | 0 | 0 | 0 | 0 | 0 | 0 | 0 | 0 | 0 | 0 | 0 | 0 | 0 | 0 | 0 | 0 |
| ENSMUSG00 Mir7-1 | 107 | 1.98622318 | 6.84938012 | 4.52089069 | 1.27741246 | 4.94814169 | 1.38932898 | 1.76869828 | 3.21244506 | 3.46428637 | 4.40660778 | 4.91144433 | | 1.84418382 | 1.30217067 | 1.15518381 | 3.93641758 | 1.07456907 | 3.59872661 |
| ENSMUSG00 Mir135b | 96 | 0 | 0 | 0 | 0 | 0 | 0 | 0 | 0 | 0 | 0 | 0 | 0 | 0 | 0 | 0 | 0 | 0 | 0 |
| ENSMUSG00 Mir342 | 98 | 0 | 0 | 0 | 0 | 0 | 0.50564014 | 0 | 0 | 0 | 0 | 0 | 0 | 0 | 0 | 0 | 0 | 0 | 0 |
| ENSMUSG00 Mir30d | 81 | 0 | 0 | 0 | 0 | 0 | 0 | 0 | 0 | 0 | 0 | 0 | 0 | 0 | 0 | 0 | 0 | 0 | 0 |
| ENSMUSG00 Mir377 | 67 | 0 | 0 | 0 | 0 | 0 | 0 | 0 | 0 | 0 | 0 | 0 | 0 | 0 | 0 | 0 | 0 | 0 | 0 |
| ENSMUSG00 Mir140 | 69 | 0 | 0 | 0 | 1.32060998 | 0.69756411 | 1.43631112 | 0 | 0 | 0 | 0.68334353 | 0.6923907 | 3.23569058 | 0 | 0 | 0 | 1.66480902 | 0 | 1.59446686 |
| ENSMUSG00 Mirlet7g | 87 | 0 | 0 | 0 | 0 | 0 | 0 | 0 | 0 | 0 | 0 | 0 | 0 | 0 | 0 | 0 | 0 | 0 | 0 |
| ENSMUSG00 Mir128-2 | 83 | 0 | 0 | 0 | 0 | 0 | 0 | 0 | 0 | 0 | 0 | 0 | 0 | 0 | 0 | 0 | 0 | 0 | 0 |
| ENSMUSG00 Mir20a | 106 | 0 | 0 | 0 | 0 | 0 | 0 | 0 | 0 | 0 | 0 | 0 | 0 | 0 | 0 | 0 | 0 | 0 | 0 |
| ENSMUSG00 Mir196b | 84 | 0 | 0 | 0 | 0 | 0 | 0 | 0 | 0 | 0 | 0 | 0 | 0 | 0 | 0 | 0 | 0 | 0 | 0 |
| ENSMUSG00 Mir27a | 86 | 0 | 0 | 0 | 0.52977959 | 1.11934706 | 1.15238915 | 0 | 0.4440977 | 1.43673892 | 0 | 0 | 0.64901933 | 1.72088084 | 2.70023764 | 1.43726358 | 0.89047924 | 2.67392769 | 7.03604853 |
| ENSMUSG00 Mir143 | 62 | 0 | 0 | 0 | 0 | 0 | 0 | 0 | 0 | 0 | 0 | 0 | 0 | 0 | 0 | 0 | 0 | 0 | 0 |
| ENSMUSG00 Mir139 | 67 | 0 | 0 | 0 | 0 | 0 | 0 | 0 | 0 | 0 | 0 | 0 | 0 | 0 | 0 | 0 | 0 | 0 | 0 |
| ENSMUSG00 Mir302d | 65 | 0 | 0.51250606 | 0 | 0 | 0 | 0 | 0 | 0 | 0 | 0 | 0 | 0 | 0 | 0 | 0 | 0 | 0 | 0 |
| ENSMUSG00 Mir154 | 83 | 0 | 0 | 0 | 0 | 0 | 0 | 0 | 0 | 0 | 0 | 0 | 0 | 0 | 0 | 0 | 0 | 0 | 0 |
| ENSMUSG00 Mir365-1 | 86 | 0 | 0 | 0 | 0 | 0 | 0 | 0 | 0 | 0 | 0 | 0 | 0 | 0 | 0 | 0 | 0 | 0 | 0 |
| ENSMUSG00 Mir448 | 111 | 0 | 0 | 0 | 0 | 0 | 0 | 0 | 0 | 0 | 0 | 0 | 0 | 0 | 0 | 0 | 0 | 0 | 0 |
| ENSMUSG00 Mir101a | 82 | 0 | 0 | 0 | 0 | 0 | 0 | 0 | 0 | 0 | 0 | 0 | 0 | 0 | 0 | 0 | 0 | 0 | 0 |
| ENSMUSG00 Mir207 | 78 | 0 | 0 | 0 | 0 | 0 | 0 | 0 | 0 | 2.11212901 | 0 | 0 | 0 | 0 | 1.19087404 | 0.52822508 | 0.98181045 | 0 | 0.70524496 |
| ENSMUSG00 Mirlet7d | 102 | 0 | 0 | 0.4742503 | 0.44667691 | 0 | 0.48581111 | 0 | 0 | 0 | 0 | 0 | 0 | 0 | 0.91066838 | 0 | 0 | 1.12724403 | 0 |
| ENSMUSG00 Mir124a-3 | 86 | 0 | 0 | 0 | 0 | 0 | 0 | 0 | 0 | 0 | 0 | 0 | 0 | 0 | 0 | 0 | 0 | 0 | 0 |
| ENSMUSG00 Mir21 | 91 | 0 | 0 | 0 | 0 | 0 | 0 | 0 | 0 | 0 | 0 | 0 | 0 | 0 | 0 | 0 | 0 | 0 | 0 |
| ENSMUSG00 Mir106a | 64 | 0 | 0 | 0 | 0 | 0 | 0 | 0 | 0 | 0 | 0 | 0 | 0 | 0 | 0 | 0 | 0 | 0 | 0 |
| ENSMUSG00 Mir383 | 73 | 0 | 0 | 0 | 0 | 0 | 0 | 0 | 0 | 0 | 0 | 0 | 0 | 0 | 0 | 0 | 0 | 0 | 0 |
| ENSMUSG00 Mir181b-1 | 79 | 0 | 0 | 0 | 0 | 0 | 0 | 0 | 0 | 0 | 0 | 0 | 0 | 0 | 0 | 0 | 0 | 0 | 0 |
| ENSMUSG00 Mir133a-2 | 103 | 0 | 0 | 0 | 0 | 0 | 0 | 0 | 0 | 0 | 0 | 0 | 0 | 0 | 0 | 0 | 0 | 0 | 0 |
| ENSMUSG00 Mir200c | 68 | 0 | 0.4898955 | 0 | 0 | 1.41564481 | 0 | 0 | 0 | 0 | 0 | 0 | 0 | 0 | 0 | 0 | 0 | 1.69086604 | 0 |
| ENSMUSG00 Snord45c | 79 | 0 | 0 | 0 | 0 | 0 | 0 | 0 | 0 | 0 | 0 | 0 | 0 | 0 | 0 | 0 | 0 | 0 | 0 |
| ENSMUSG00 Mir185 | 79 | 0 | 0 | 0 | 0 | 0 | 0 | 0 | 0 | 0 | 0 | 0 | 0 | 0 | 0 | 0 | 0.48469123 | 0 | 0 |
| ENSMUSG00 Mir33 | 68 | 0 | 0 | 0 | 0 | 0 | 0 | 0 | 0 | 0 | 0 | 0 | 0 | 0 | 0 | 0 | 0 | 0 | 0 |
| ENSMUSG00 n-R5s89 | 103 | 0 | 0.32342616 | 0 | 0 | 0 | 0 | 0 | 0 | 0 | 0 | 2.78300729 | 0 | 0 | 0.45091347 | 1.60006042 | 3.34578123 | 1.48839688 | 2.13627599 |
| ENSMUSG00 Mir26b | 84 | 0 | 0 | 0 | 0 | 0 | 0 | 0 | 0 | 0 | 0 | 0 | 0.66447217 | 0 | 0 | 0 | 0.45584056 | 0 | 0.65487032 |
| ENSMUSG00 Mir129-1 | 72 | 0 | 0 | 0 | 0 | 0 | 0 | 0 | 0 | 0 | 0 | 0 | 0 | 0 | 0 | 0 | 0 | 0.53230968 | 0 |
| ENSMUSG00 Mir149 | 65 | 0 | 0 | 0 | 0 | 0 | 0 | 0 | 0 | 0 | 0 | 0 | 0 | 0 | 0 | 0 | 0 | 0 | 0 |
| ENSMUSG00 Mir222 | 78 | 0 | 0 | 0 | 0 | 0 | 0 | 0 | 0 | 0 | 0 | 0 | 0 | 0 | 0 | 0 | 0 | 0 | 0 |
| ENSMUSG00 Mir125b-2 | 70 | 0 | 0 | 0 | 0 | 0 | 0 | 0 | 0 | 0 | 0 | 0 | 0 | 0 | 0 | 0 | 0 | 0 | 0 |
| ENSMUSG00 Mir19b-2 | 83 | 0 | 0 | 0 | 0 | 0 | 0 | 0 | 0 | 0 | 0 | 0 | 0 | 0 | 0 | 0 | 0 | 0 | 0 |
| ENSMUSG00 Mir141 | 71 | 0 | 0 | 0 | 0.64170485 | 1.35582884 | 0 | 0 | 0.53792116 | 0.58009177 | 0.66409441 | 0 | 0 | 0 | 0 | 0 | 0 | 3.23884199 | 0 |
| ENSMUSG00 Mir27b | 96 | 0 | 0 | 0 | 0.47459421 | 0 | 0 | 0 | 0 | 0 | 0 | 0 | 0 | 0 | 0 | 0 | 0 | 0.39923226 | 0 |
| ENSMUSG00 Mir30b | 95 | 0 | 0 | 0.50919506 | 0 | 0 | 0 | 0 | 0 | 0 | 0 | 0 | 0 | 0 | 0 | 0 | 0 | 0 | 0 |
| ENSMUSG00 Mir411 | 95 | 0 | 0 | 0 | 0 | 0 | 0 | 0 | 0 | 0 | 0 | 0 | 0 | 0 | 0 | 0 | 0 | 0 | 0 |
| ENSMUSG00 Mir125a | 84 | 0 | 0 | 0 | 0 | 0 | 0 | 0 | 0 | 0 | 0 | 0 | 0 | 0 | 0 | 0 | 0 | 0 | 0 |
| ENSMUSG00 Mir133b | 118 | 0 | 0 | 0 | 0 | 0 | 0 | 0 | 0 | 0 | 0 | 0 | 0 | 0 | 0 | 0 | 0 | 0 | 0 |
| ENSMUSG00 Mir346 | 97 | 0 | 0 | 0 | 0 | 0 | 0 | 0 | 0 | 0 | 0 | 0 | 0 | 0 | 0 | 0 | 0 | 0 | 0 |
| ENSMUSG00 Mir302c | 67 | 0 | 0 | 0 | 0 | 0 | 0 | 0 | 0 | 0 | 0 | 0 | 0 | 0 | 0 | 0 | 0 | 0 | 0 |
| ENSMUSG00 Mir181c | 105 | 0 | 0 | 0 | 0 | 0 | 0 | 0 | 0 | 0 | 0 | 0 | 0 | 0 | 0 | 0 | 0 | 0 | 0 |
| ENSMUSG00 Mir130a | 63 | 0 | 0 | 0 | 1.44638236 | 5.34799152 | 3.14620531 | 0 | 0 | 0 | 0 | 0 | 1.7719258 | 2.34913892 | 0 | 0.65399295 | 2.43114968 | 0 | 2.61948127 |
| ENSMUSG00 Mir219-2 | 96 | 0 | 0 | 0 | 0 | 0 | 0 | 0 | 0 | 0 | 0 | 0 | 0 | 0.51387414 | 0 | 0 | 0 | 0 | 0 |
| ENSMUSG00 Mir450-1 | 90 | 0 | 0 | 0 | 0 | 0 | 0 | 0 | 0 | 0 | 0 | 0 | 0 | 0 | 0 | 0 | 0 | 0 | 0 |
| ENSMUSG00 Mir196a-2 | 109 | 0 | 0 | 0 | 0 | 0 | 0 | 0 | 0 | 0 | 0 | 0 | 0 | 0 | 0 | 0 | 0 | 0 | 0 |
| ENSMUSG00 Mir365-2 | 111 | 0 | 0 | 0 | 0 | 0 | 0 | 0 | 0 | 0 | 0 | 0 | 0 | 0 | 0 | 0 | 0 | 0 | 0 |
| ENSMUSG00 Mir30c-1 | 88 | 0 | 0 | 0 | 0 | 0 | 0 | 0 | 0 | 0 | 0 | 0 | 0 | 0 | 0 | 0 | 0 | 0 | 0 |
| ENSMUSG00 Mir7b | 110 | 0 | 0 | 0 | 0 | 0 | 0 | 0 | 0 | 0 | 0 | 0 | 0 | 0 | 0 | 0 | 0 | 0 | 0 |
| ENSMUSG00 Mir34b | 83 | 0 | 0 | 0 | 0 | 0 | 0 | 0 | 0 | 0 | 0 | 1.15120382 | 0 | 0 | 0 | 0 | 0.92266524 | 0 | 0 |
| ENSMUSG00 Mir34a | 101 | 0 | 0 | 0 | 0 | 0 | 0 | 0 | 0 | 0 | 0 | 0 | 0 | 0 | 0 | 0 | 0 | 0 | 0 |
| ENSMUSG00 Mir28 | 85 | 1.50018268 | 0 | 1.13820071 | 0.53601229 | 0 | 0 | 0 | 0 | 0.96909449 | 0 | 0 | 0 | 0 | 1.09280206 | 0.96944837 | 0 | 0 | 1.94149788 |
| ENSMUSG00 Mir150 | 64 | 0 | 0 | 0 | 0 | 0.75206131 | 0 | 1.47852122 | 0 | 0 | 1.47345948 | 0 | 0 | 0 | 0 | 0.64377431 | 1.79487222 | 1.79654517 | 4.29758646 |
| ENSMUSG00 Mir410 | 80 | 0 | 0 | 0 | 0 | 0 | 0 | 0 | 0 | 0 | 0 | 0 | 0 | 0 | 0 | 0 | 0 | 0 | 0 |
| ENSMUSG00 Mir379 | 122 | 0 | 0 | 0 | 0 | 0 | 0 | 0 | 0 | 0 | 0 | 0 | 0 | 0 | 0 | 0 | 0 | 0 | 0 |
| ENSMUSG00 Mir384 | 87 | 0 | 0 | 0 | 0 | 0 | 0 | 0 | 0 | 0 | 0 | 0 | 0 | 0 | 0 | 0 | 0 | 0 | 0 |
| ENSMUSG00 Mir10b | 67 | 1.90321684 | 0 | 0.72199299 | 0 | 0 | 1.47918608 | 0 | 0.57003586 | 1.22944823 | 0 | 0.71305908 | 0 | 0 | 2.77278134 | 1.22989719 | 0 | 0 | 2.46309433 |
| ENSMUSG00 Mir335 | 97 | 0 | 0 | 0 | 0 | 0 | 0 | 0 | 0 | 0 | 0 | 0 | 0 | 0 | 0 | 0 | 0 | 0 | 0 |
| ENSMUSG00 Mir202 | 71 | 0 | 0 | 0 | 0 | 0 | 0 | 0 | 0 | 0 | 0 | 0 | 0 | 0 | 0 | 0 | 0 | 0 | 0 |
| ENSMUSG00 Mir351 | 98 | 0 | 0 | 0 | 0 | 0 | 0 | 0 | 0 | 0 | 0 | 0 | 0 | 0 | 0 | 0 | 0 | 0 | 0 |
| ENSMUSG00 Mir148a | 98 | 0 | 0 | 0 | 0 | 0 | 0 | 0 | 0 | 0 | 0 | 0 | 0 | 0 | 0 | 0 | 0 | 0 | 0 |
| ENSMUSG00 Mir204 | 67 | 0 | 0 | 1.44398598 | 0 | 0 | 0 | 0 | 0 | 0 | 0 | 0 | 0 | 0 | 0 | 0 | 0 | 0 | 0 |
| ENSMUSG00 Mir17 | 83 | 0 | 0 | 0 | 0 | 0 | 0 | 0 | 0 | 0 | 0 | 0 | 0 | 0 | 0 | 0 | 0 | 0 | 0 |
| ENSMUSG00 Mir421 | 84 | 0 | 0 | 0 | 0 | 0 | 0 | 0 | 0 | 0 | 0 | 0 | 0 | 0 | 0 | 0 | 0 | 1.82506176 | 0 |
| ENSMUSG00 Mir361 | 69 | 0 | 0 | 0 | 0 | 0 | 0 | 0 | 0 | 0 | 0 | 2.07717211 | 0 | 0 | 0 | 0 | 0 | 0 | 0 |
| ENSMUSG00 Mir129-2 | 89 | 0 | 0 | 0 | 0 | 0 | 0 | 0 | 0 | 0 | 0 | 0 | 0 | 0 | 0 | 0 | 0 | 0 | 0 |
| ENSMUSG00 Mir138-2 | 70 | 0 | 0 | 0 | 0 | 0 | 0 | 0 | 0 | 0 | 0 | 0 | 0 | 0 | 0 | 0 | 0 | 0 | 0 |
| ENSMUSG00 Mir26a-1 | 89 | 1.43275874 | 0.37430218 | 0.54352281 | 2.04768739 | 0.54080813 | 0 | 1.06320627 | 0 | 2.3138492 | 0.52978318 | 0 | 0.62714228 | 0 | 1.04368736 | 2.77763298 | 1.72092618 | 0 | 0.61807985 |
| ENSMUSG00 Mir106b | 81 | 0 | 0 | 0 | 0 | 0.59422128 | 0 | 0 | 0 | 0 | 0.58210745 | 0 | 0 | 0 | 0 | 0.50866118 | 0 | 0 | 0 |
| ENSMUSG00 Mir152 | 72 | 0 | 0 | 0 | 0 | 0 | 0 | 0 | 0 | 0 | 0 | 0 | 0 | 0 | 0 | 0 | 0 | 0 | 0 |
| ENSMUSG00 Mir214 | 109 | 0 | 0 | 0 | 0 | 0 | 0 | 0 | 0 | 0 | 0 | 0 | 0 | 0 | 0 | 0 | 0 | 0 | 0 |
| ENSMUSG00 Mir423 | 108 | 1.18069933 | 0.30845272 | 1.79161224 | 0 | 0 | 1.83528643 | 0 | 0 | 0 | 0.87316117 | 0 | 0.51681169 | 0.45677701 | 0.43003785 | 0 | 0 | 0 | 0 |
| ENSMUSG00 Mir10a | 109 | 0 | 0 | 0 | 0 | 0 | 0 | 0 | 0 | 0 | 0 | 0 | 0 | 0 | 0 | 0 | 0 | 0 | 0 |
| ENSMUSG00 Mir128-1 | 81 | 0 | 0.8225406 | 0.59720408 | 0 | 0 | 0 | 0.58410715 | 0.47151114 | 0 | 0 | 0 | 0 | 0 | 0 | 0 | 1.89089419 | 0 | 0 |
| ENSMUSG00 Mir296 | 78 | 0 | 0 | 0 | 0 | 0 | 0 | 0 | 0 | 0 | 0 | 0 | 0 | 0 | 0 | 0 | 0 | 0 | 0 |
| ENSMUSG00 Mir192 | 88 | 0 | 0 | 0 | 0 | 0 | 0 | 0 | 0 | 0 | 0 | 0 | 0 | 0 | 0 | 0 | 0 | 0 | 0 |
| ENSMUSG00 Mir135a-2 | 99 | 0 | 0.67298776 | 0.48862152 | 0 | 0 | 0 | 0 | 0 | 0 | 0 | 0 | 0 | 0 | 0 | 0 | 0 | 0 | 0 |
| ENSMUSG00 Mir337 | 96 | 0 | 0 | 0 | 0 | 0 | 0 | 0 | 0 | 0 | 0 | 0 | 0 | 0 | 0 | 0 | 0 | 0 | 0 |
| ENSMUSG00 Mir93 | 87 | 0 | 0 | 0 | 0 | 0 | 0 | 0 | 0 | 0 | 0 | 0 | 0 | 0 | 0 | 0 | 0 | 0 | 0 |
| ENSMUSG00 Mir320 | 81 | 0 | 0 | 0 | 0 | 0 | 0 | 0 | 0 | 0 | 0 | 0 | 0.68908225 | 0 | 0 | 1.52598355 | 0.47272355 | 0 | 1.35824955 |
| ENSMUSG00 Mir22 | 94 | 0 | 0 | 0 | 0 | 0 | 0 | 0 | 0 | 0 | 0 | 0 | 0 | 0 | 0 | 0 | 0 | 0 | 0 |
| ENSMUSG00 Mir99a | 64 | 0 | 0 | 0 | 0 | 0 | 0 | 0 | 0 | 0 | 0 | 0 | 0 | 0 | 0 | 0 | 0 | 0 | 0 |
| ENSMUSG00 Mir187 | 60 | 0 | 0 | 0 | 0 | 0 | 0 | 0 | 0 | 0 | 0 | 0 | 0 | 0 | 0 | 0 | 0 | 0 | 0 |
| ENSMUSG00 Mir205 | 67 | 0 | 0 | 0 | 0 | 0 | 0 | 0 | 0 | 0 | 0 | 0 | 0 | 0 | 0 | 0 | 0 | 0 | 0 |
| ENSMUSG00 Mir324 | 88 | 0 | 0 | 0 | 0 | 0 | 0 | 0 | 0 | 0 | 0 | 0 | 0 | 0 | 0 | 0 | 0 | 0 | 0 |
| ENSMUSG00 Mir98 | 107 | 0 | 0 | 0 | 0 | 0 | 0 | 0 | 0 | 0 | 0 | 0 | 0 | 0 | 0 | 0 | 0 | 0 | 0 |
| ENSMUSG00 Mir132 | 100 | 0 | 0 | 0 | 0 | 0 | 0 | 0 | 0 | 0 | 0.47150703 | 0.47774958 | 0 | 0.49331917 | 0.92888175 | 2.88410891 | 0.38290607 | 1.53305188 | 1.10018213 |
| ENSMUSG00 Mir153 | 68 | 0 | 0 | 0 | 0 | 0 | 0 | 2.78309876 | 0 | 0 | 0 | 0 | 0 | 0 | 0 | 0 | 0 | 0 | 0 |
| ENSMUSG00 Mir216a | 71 | 0 | 0 | 0 | 0 | 0 | 0 | 0 | 0 | 0 | 0 | 0 | 0 | 0 | 0 | 0 | 0 | 0 | 0 |
| ENSMUSG00 Mir126 | 72 | 0 | 0 | 0 | 0 | 0 | 0 | 0 | 0 | 0 | 0 | 0 | 0 | 0 | 0 | 0 | 0 | 0 | 0 |
| ENSMUSG00 Mir24-2 | 106 | 0.40099223 | 0.62854517 | 1.36906218 | 1.71928469 | 0.45407475 | 0.46747862 | 0.89269206 | 1.44122273 | 1.55420814 | 0 | 2.25353578 | 2.10625142 | 0.93079089 | 3.94336591 | 3.10955139 | 5.05725003 | 1.80784419 | 5.70849221 |
| ENSMUSG00 Mir224 | 81 | 0 | 0 | 0 | 0 | 0 | 0 | 0 | 0 | 0 | 0 | 0 | 0 | 0 | 0 | 0 | 0 | 0 | 0 |
| ENSMUSG00 Mir330 | 97 | 0 | 0 | 0 | 0 | 0 | 0 | 0 | 0 | 0 | 0 | 0 | 0 | 0 | 0 | 0 | 0 | 0 | 0 |
| ENSMUSG00 Mir32 | 69 | 0 | 0 | 0 | 0 | 0 | 0 | 0 | 0 | 0 | 0 | 0 | 0 | 0 | 0 | 0 | 0 | 0 | 0 |
| ENSMUSG00 Mir196a-1 | 101 | 0 | 0 | 0 | 0 | 0 | 0 | 0 | 0 | 0 | 0 | 0 | 0 | 0 | 0 | 0 | 0 | 0 | 0 |
| ENSMUSG00 Mir199a-1 | 69 | 0.61601704 | 0.48279557 | 0 | 0.66030499 | 0.69756411 | 0.71815556 | 2.057073 | 0 | 0 | 0.68334353 | 2.07717211 | 10.5159944 | 0.71495532 | 3.36551358 | 0.597124 | 1.66480902 | 0 | 1.59446686 |
| ENSMUSG00 Mir29c | 87 | 0 | 0 | 0 | 0 | 0 | 0 | 0 | 0 | 0 | 0 | 0 | 0 | 0 | 0 | 0 | 0 | 0 | 0 |
| ENSMUSG00 Mir200b | 69 | 0 | 0 | 0 | 0 | 0 | 0 | 0 | 0 | 0 | 0 | 0 | 0 | 0 | 0 | 0 | 0 | 0 | 0 |
| ENSMUSG00 Mir210 | 109 | 0 | 0 | 0 | 0 | 0.44157728 | 0 | 0 | 0 | 0 | 0 | 0 | 0 | 0 | 0 | 0 | 0 | 0 | 0 |
| ENSMUSG00 Mir302a | 68 | 0 | 0.4898955 | 0 | 0 | 0 | 0 | 0 | 0 | 0 | 0 | 0 | 0 | 0 | 0 | 0 | 0 | 0 | 0 |
| ENSMUSG00 Mir103-1 | 85 | 0 | 0 | 0 | 0 | 0 | 0 | 0 | 0 | 0 | 0 | 0 | 0 | 0 | 0 | 0 | 0 | 0 | 0 |
| ENSMUSG00 Mir219-1 | 109 | 0 | 0 | 0.88758771 | 0 | 0 | 0 | 0 | 0 | 0 | 0 | 0 | 0.5120703 | 0.90517279 | 0 | 0 | 0 | 0 | 0 |
| ENSMUSG00 Mir101b | 96 | 0 | 0 | 0 | 0 | 0 | 0 | 0 | 0 | 0 | 0 | 0 | 0 | 0 | 0 | 0 | 0 | 0 | 0 |
| ENSMUSG00 Mirlet7c-1 | 93 | 0 | 0 | 0 | 0 | 0 | 0 | 0 | 0 | 0 | 0 | 0 | 0 | 0 | 0 | 0 | 0 | 0 | 0 |
| ENSMUSG00 Mir206 | 72 | 0 | 0 | 0 | 0 | 0 | 0 | 0 | 0 | 0 | 0 | 0 | 0 | 0 | 0 | 0 | 0 | 0 | 0 |
| ENSMUSG00 Mir148b | 96 | 0 | 0 | 0 | 0 | 0 | 0 | 0 | 0 | 0 | 0 | 0 | 0 | 0 | 0 | 0 | 0 | 0 | 0 |
| ENSMUSG00 Mir369 | 78 | 0 | 0 | 0 | 0 | 0 | 0 | 0 | 0 | 0 | 0 | 0 | 0 | 0 | 0 | 0 | 0 | 0 | 0 |
| ENSMUSG00 Mir215 | 111 | 0.38292951 | 2.70104548 | 0 | 0 | 0 | 0 | 0 | 0.3440757 | 0 | 1.69912444 | 0 | 0 | 0 | 0 | 0.37118519 | 0.68992085 | 0.69056391 | 0.49557754 |
| ENSMUSG00 Mir103-2 | 85 | 0 | 0 | 0 | 0 | 0 | 1.16594667 | 0 | 0 | 0 | 0 | 0 | 0 | 0 | 0 | 0 | 0 | 0 | 0 |
| ENSMUSG00 Mirlet7b | 84 | 0 | 0.79316415 | 0 | 0 | 0 | 0 | 1.12649236 | 0 | 0.98063133 | 0 | 0 | 0 | 0 | 0.5529058 | 0.49049471 | 0.45584056 | 0 | 3.27435159 |
| ENSMUSG00 Mir181a-1 | 86 | 0 | 0 | 0 | 0 | 0 | 0 | 0 | 0 | 0 | 0 | 0 | 0 | 0 | 0 | 0 | 0 | 0 | 0 |
| ENSMUSG00 Mir381 | 74 | 0 | 0 | 0 | 0 | 0 | 0 | 0 | 0 | 0 | 0 | 0 | 0 | 0 | 0 | 0 | 0 | 0 | 0 |
| ENSMUSG00 Mir30c-2 | 83 | 0 | 0 | 0 | 0 | 0 | 0 | 0 | 0 | 0 | 0 | 0 | 0 | 0 | 0 | 0 | 0 | 0 | 0 |
| ENSMUSG00 Mir344 | 94 | 0 | 0 | 0 | 0 | 0 | 0 | 0 | 0 | 0 | 0 | 0 | 0 | 0 | 0 | 0 | 0 | 0 | 0 |
| ENSMUSG00 Mir137 | 101 | 0 | 0 | 0 | 0 | 0 | 0 | 0 | 0 | 0 | 0 | 0 | 0 | 0 | 0 | 0 | 0 | 0 | 0 |
| ENSMUSG00 Mir412 | 79 | 0 | 0 | 0 | 0 | 0 | 0 | 0 | 0 | 0 | 0 | 0 | 0 | 0 | 0 | 0 | 0 | 0 | 0 |
| ENSMUSG00 Mir326 | 94 | 0 | 0 | 0 | 0 | 0 | 0 | 0 | 0 | 0 | 0 | 0 | 0 | 0 | 0 | 0 | 0 | 0 | 0 |
| ENSMUSG00 Mir130b | 81 | 0 | 0 | 0 | 0 | 0 | 0 | 0 | 0 | 0 | 0 | 0 | 0 | 0 | 0 | 0 | 0 | 0 | 0 |
| ENSMUSG00 Mir350 | 98 | 0 | 0 | 0 | 0 | 0 | 0 | 0 | 0 | 0 | 0 | 0 | 0 | 0 | 0 | 0 | 0 | 0 | 0 |
| ENSMUSG00 Mir203 | 75 | 0 | 0 | 0 | 0 | 0 | 0 | 0 | 0 | 0 | 0 | 0 | 0 | 0 | 0 | 0 | 0 | 0 | 0 |
| ENSMUSG00 Mir449a | 90 | 0 | 0 | 0 | 0 | 0 | 0 | 0 | 0 | 0 | 0 | 0 | 0 | 0 | 0 | 0 | 0 | 0 | 0 |
| ENSMUSG00 Mir329 | 96 | 0 | 0 | 0 | 0 | 0 | 0 | 0 | 0 | 0 | 0 | 0 | 0 | 0 | 0 | 0 | 0 | 0 | 0 |
| ENSMUSG00 Mir181b-2 | 88 | 0 | 1.89277808 | 2.74849604 | 0 | 0 | 0 | 0 | 0.43400457 | 0 | 0 | 0.54289726 | 0.63426889 | 0 | 0 | 0 | 0.87024108 | 1.3065783 | 0.62510349 |
| ENSMUSG00 Mir425 | 84 | 3.036084 | 6.34531318 | 2.30350145 | 3.79675369 | 2.29199637 | 1.76974048 | 0.56324618 | 5.91072894 | 1.96126266 | 5.61317896 | 0.56874951 | 2.65788869 | 6.46013202 | 8.29358703 | 8.82890483 | 7.74928959 | 4.56265439 | 2.61948127 |
| ENSMUSG00 Mir15b | 97 | 0 | 0 | 0 | 0 | 0 | 0 | 0 | 0 | 0 | 0 | 0 | 0 | 0 | 0 | 0 | 0 | 0 | 0 |
| ENSMUSG00 Mir194-1 | 66 | 0.64401782 | 1.00948164 | 0 | 0 | 0 | 0.75079899 | 0 | 1.73601829 | 0 | 0.71440459 | 0 | 0 | 0 | 0 | 0 | 0 | 0 | 1.66694263 |
| ENSMUSG00 Mir194-2 | 85 | 0 | 0 | 0 | 0 | 0 | 0 | 0 | 0 | 0 | 0 | 0 | 0 | 0 | 0 | 0 | 0 | 0 | 0 |
| ENSMUSG00 Mir218-2 | 109 | 0 | 0 | 0 | 0 | 0 | 0 | 0 | 0 | 0 | 0 | 0 | 0 | 0 | 0 | 0 | 0 | 0 | 0 |
| ENSMUSG00 Mir211 | 105 | 0 | 0 | 0 | 0 | 0 | 0 | 0 | 0 | 0 | 0 | 0 | 0 | 0 | 0 | 0 | 0 | 0 | 0 |
| ENSMUSG00 Mir96 | 105 | 0 | 0 | 0 | 0 | 0 | 0 | 0 | 0 | 0 | 0 | 0 | 0 | 0 | 0 | 0 | 0 | 0 | 0 |
| ENSMUSG00 Mir34c | 76 | 0 | 0 | 0 | 0 | 0 | 0 | 0 | 0 | 0 | 0 | 0 | 0 | 0 | 0 | 0 | 0 | 0 | 0 |
| ENSMUSG00 Mir301 | 85 | 0 | 0 | 0 | 0 | 0 | 0 | 0 | 0 | 0 | 0 | 0 | 0 | 0 | 0 | 0 | 0 | 0 | 0 |
| ENSMUSG00 Mir212 | 90 | 0 | 0 | 0 | 0 | 0 | 0 | 0 | 0 | 0 | 1.57169011 | 0 | 0 | 0 | 2.06418166 | 0.91559013 | 0.42545119 | 0 | 0.6112123 |
| ENSMUSG00 U6 | 106 | 0.80198445 | 2.19990811 | 0.45635406 | 0.42982117 | 3.17852326 | 1.40243586 | 1.33903808 | 0.72061137 | 0.38855204 | 0.88963591 | 0.45070716 | 4.73906569 | 1.86158178 | 0.87630354 | 3.49824531 | 1.80616073 | 1.08470652 | 0 |
| ENSMUSG00 Mir145 | 87 | 0 | 0 | 0 | 0 | 0 | 0 | 0 | 0 | 0 | 0 | 0 | 0 | 0 | 0 | 0 | 0 | 0 | 0 |
| ENSMUSG00 Mir339 | 95 | 0 | 0.70132409 | 0 | 0 | 0 | 0 | 1.9921128 | 0 | 0 | 0 | 0 | 0 | 0 | 0 | 0 | 0 | 0 | 0.57904323 |
| ENSMUSG00 Mir107 | 86 | 0 | 0 | 0 | 0 | 0 | 0 | 0 | 0 | 0 | 0 | 0 | 0 | 0 | 0 | 0 | 0 | 0 | 0.63964078 |
| ENSMUSG00 Mir184 | 68 | 0 | 0 | 0 | 0 | 0 | 0 | 0 | 0 | 0 | 0 | 0 | 0 | 0 | 0 | 0 | 0 | 0 | 0 |
| ENSMUSG00 Mir124a-1 | 84 | 0 | 0 | 0 | 0 | 0 | 0 | 0 | 0 | 0 | 0 | 0 | 0 | 0 | 0 | 0 | 0 | 0 | 0 |
| ENSMUSG00 Mir23b | 96 | 0 | 0 | 0 | 0.94918842 | 0.5013742 | 0 | 0 | 0 | 0 | 0 | 0.49765582 | 0 | 0 | 0 | 0 | 0 | 0.39923226 | 0 |
| ENSMUSG00 Mir338 | 97 | 0 | 0 | 0 | 0 | 0 | 0 | 0 | 0 | 0 | 0 | 0 | 0 | 0 | 0 | 0 | 0 | 0 | 0 |
| ENSMUSG00 Mir146 | 64 | 0 | 0 | 0 | 0 | 0 | 0 | 0.73926061 | 0 | 0 | 0 | 0 | 0 | 0 | 0 | 0 | 0 | 0 | 0 |
| ENSMUSG00 Mirlet7f-2 | 82 | 0 | 0 | 0 | 0 | 0 | 0 | 0 | 0 | 0 | 0 | 0 | 0 | 0 | 0 | 0 | 0 | 0 | 0 |
| ENSMUSG00 Mir218-1 | 109 | 0 | 0 | 0 | 0 | 0 | 0 | 0 | 0 | 0 | 0 | 0 | 0 | 0 | 0 | 0 | 0 | 0 | 0 |
| ENSMUSG00 Mir29b-1 | 80 | 0 | 0 | 0 | 0 | 0 | 0 | 0 | 0 | 0 | 0 | 0 | 0 | 1.23329793 | 0 | 0 | 0 | 0.47863259 | 0 |
| ENSMUSG00 Mir16-2 | 94 | 0 | 0 | 0 | 0 | 0 | 0 | 0 | 0 | 0 | 0 | 0 | 0 | 0 | 0 | 0 | 0 | 0 | 0 |
| ENSMUSG00 Mir331 | 95 | 0 | 0 | 0 | 0 | 0 | 0 | 0 | 0 | 0 | 0 | 0 | 0 | 0 | 0 | 0 | 0 | 0 | 0 |
| ENSMUSG00 Mirlet7c-2 | 94 | 0 | 0 | 0.51461203 | 0.48469196 | 2.04816696 | 0 | 2.01330549 | 0 | 0.87630885 | 0 | 1.01648848 | 0 | 0 | 5.4349464 | 0.87682885 | 0.40734689 | 0.40772656 | 1.17040653 |
| ENSMUSG00 Mir7-2 | 96 | 0 | 0 | 0 | 0 | 0 | 0 | 0 | 0 | 0 | 0 | 0 | 0 | 0 | 0 | 0 | 0 | 0 | 0 |
| ENSMUSG00 Mir29a | 87 | 0 | 0 | 0 | 0 | 0 | 0 | 0 | 0 | 0 | 0 | 0 | 0 | 0 | 0 | 0 | 0 | 0 | 0 |
| ENSMUSG00 Mir23a | 74 | 1.14878854 | 0 | 0.65369636 | 0 | 0.6504314 | 0.66963153 | 0 | 2.06445418 | 0.55657454 | 0.63717167 | 5.81046792 | 3.77132855 | 3.33323765 | 7.53147363 | 2.22711113 | 0 | 4.14338345 | 11.8938609 |
| ENSMUSG00 Mir151 | 67 | 0 | 0 | 0 | 0 | 0 | 0 | 0 | 0 | 0 | 0 | 0 | 0 | 0 | 0 | 0 | 0 | 0.57203428 | 0 |
| ENSMUSG00 Mir92-2 | 90 | 0 | 0 | 0 | 0 | 0 | 0 | 0 | 0 | 0 | 0 | 0 | 0 | 0 | 0 | 0 | 0 | 0 | 0 |
| ENSMUSG00 Mir375 | 74 | 0 | 0 | 0 | 0 | 0 | 0 | 0 | 0 | 0 | 0 | 0 | 0 | 0 | 0 | 0 | 0 | 0 | 0 |
| ENSMUSG00 Mir323 | 85 | 0 | 0 | 0 | 0 | 0 | 0 | 0 | 0 | 0 | 0 | 0 | 0 | 0 | 0 | 0 | 0 | 0 | 0 |
| ENSMUSG00 Mir183 | 69 | 0 | 0 | 0 | 0 | 0 | 0 | 0 | 0 | 0 | 0 | 0 | 0 | 0 | 0 | 0 | 0 | 0 | 0 |
| ENSMUSG00 U2 | 198 | 0 | 0 | 0 | 0 | 0 | 0 | 0 | 0 | 0 | 0 | 0 | 0 | 0 | 0 | 0 | 0 | 0 | 0 |
| ENSMUSG00 U2 | 189 | 0 | 0 | 0 | 0 | 0 | 0 | 0 | 0 | 0 | 0 | 0 | 0 | 0 | 0 | 0 | 0 | 0 | 0 |
| ENSMUSG00 U6 | 103 | 0 | 0.97027847 | 1.8785837 | 2.21170118 | 0.93460046 | 0 | 1.83738559 | 1.11240007 | 0.79973817 | 0.45777382 | 1.39150365 | 2.16759855 | 0 | 0 | 0 | 0 | 0.74419994 | 0 |
| ENSMUSG00 U6 | 106 | 0 | 0 | 0 | 0 | 0 | 0 | 0 | 0 | 0 | 0 | 0 | 0 | 0 | 0 | 0 | 0 | 0 | 0 |
| ENSMUSG00 U6 | 102 | 0 | 0 | 0 | 0 | 0 | 0 | 0 | 0 | 0 | 0 | 0 | 0 | 0 | 0 | 0 | 0 | 0 | 0 |
| ENSMUSG00 U6 | 106 | 0 | 0 | 0 | 0 | 0 | 0 | 0 | 0 | 0 | 0 | 0 | 0 | 0 | 0 | 0 | 0 | 0 | 0 |
| ENSMUSG00 Snord33 | 83 | 0 | 0 | 0 | 0 | 0 | 0 | 0 | 0 | 0 | 0 | 0 | 0 | 0 | 0 | 0 | 0 | 0 | 0 |
| ENSMUSG00 U3 | 213 | 0 | 0 | 0 | 0 | 0 | 0 | 0 | 0 | 0 | 0 | 0 | 0 | 0 | 0 | 0 | 0 | 0 | 0 |
| ENSMUSG00 U4 | 83 | 0 | 0 | 0 | 0 | 0 | 0 | 0 | 0 | 0 | 0 | 0 | 0 | 0 | 0 | 0 | 0 | 0 | 0 |
| ENSMUSG00 SNORA24 | 131 | 0 | 0 | 0 | 0 | 0 | 0 | 0 | 0 | 0 | 0 | 0 | 0 | 0 | 0 | 0 | 0 | 0 | 0 |
| ENSMUSG00 snoR38 | 75 | 0 | 0 | 0 | 0 | 0 | 0 | 0 | 0 | 0 | 0 | 0 | 0 | 0 | 0 | 0 | 0 | 0 | 0 |
| ENSMUSG00 U6 | 103 | 0 | 0 | 0 | 0 | 0 | 0 | 0 | 0 | 0 | 0 | 0 | 0 | 0 | 0 | 0 | 0 | 0 | 0 |
| ENSMUSG00 SNORA14 | 133 | 0 | 0 | 1.09113226 | 0.68512849 | 3.61894163 | 2.23546167 | 0 | 2.58444828 | 0.30967305 | 1.0635497 | 0.71842043 | 0.41966664 | 0.37091667 | 0 | 0.9293584 | 0.86369791 | 0.57633529 | 0.41360231 |
| ENSMUSG00 U6 | 106 | 0 | 0 | 0 | 0 | 0 | 0 | 0 | 0 | 0 | 0 | 0 | 0 | 0 | 0 | 0 | 0 | 0 | 0 |
| ENSMUSG00 Snord1c | 75 | 0 | 0 | 0 | 0 | 0 | 0 | 0 | 0 | 0 | 0 | 0 | 0 | 0 | 0 | 0 | 0 | 0 | 0 |
| ENSMUSG00 Snora69 | 130 | 0.98088868 | 2.05002426 | 0.37210408 | 0.35046957 | 0.37024557 | 0 | 0.36394368 | 0.29376771 | 0 | 0 | 0 | 0.42935125 | 3.41528658 | 0.71452442 | 0 | 0 | 0 | 3.3851758 |
| ENSMUSG00 U6 | 106 | 0.80198445 | 0 | 0 | 0 | 1.36222425 | 0 | 0 | 0 | 0.38855204 | 0 | 0 | 0 | 0 | 0 | 0.38869392 | 0 | 0 | 0 |
| ENSMUSG00 SNORA32 | 90 | 0 | 0 | 0 | 0 | 0 | 0 | 0 | 0 | 0 | 0 | 0 | 0 | 0 | 0 | 0 | 0 | 0 | 0 |
| ENSMUSG00 7SK | 317 | 0 | 0 | 0 | 0 | 0 | 0 | 0 | 0 | 0 | 0 | 0 | 0 | 0 | 0 | 0 | 0 | 0 | 0 |
| ENSMUSG00 SNORA2 | 135 | 0 | 0 | 0 | 0 | 0.35653277 | 0.36705729 | 0 | 0 | 0.3050853 | 1.04779341 | 0.35388858 | 0.41344935 | 0 | 0 | 0 | 0 | 0 | 0 |
| ENSMUSG00 U6 | 99 | 0 | 0 | 0 | 0 | 0 | 0 | 0 | 0 | 0 | 0 | 0 | 0 | 0 | 0 | 0 | 0 | 0 | 0 |
| ENSMUSG00 Snora74a | 195 | 0.21797526 | 0 | 0 | 0.46729276 | 0.49366076 | 0.50823316 | 1.45577474 | 1.76272626 | 2.32334191 | 0.24179848 | 0.73499936 | 0 | 1.26492096 | 0.47634961 | 0.63387009 | 0 | 0.19654511 | 0 |
| ENSMUSG00 SNORA61 | 126 | 0 | 0.26438805 | 0 | 0 | 0 | 0 | 0 | 0.3031143 | 1.63438555 | 0.37421193 | 0.37916634 | 0.8859629 | 0.39152315 | 2.21162321 | 0.98098943 | 0.30389371 | 0 | 0 |
| ENSMUSG00 SNORA38 | 133 | 1.27835116 | 0.50094578 | 1.09113226 | 1.02769273 | 0 | 0 | 0 | 0.28716092 | 0 | 0 | 0.35921021 | 1.67866654 | 0 | 1.04761099 | 0 | 2.30319443 | 0.86450294 | 1.65440922 |
| ENSMUSG00 U6 | 106 | 0 | 0 | 0 | 0 | 0 | 0 | 0 | 0 | 0 | 0 | 0 | 0 | 0 | 0 | 0 | 0 | 0 | 0 |
| ENSMUSG00 U6 | 103 | 0 | 0 | 0 | 0 | 0 | 0 | 0 | 0 | 0 | 0 | 0 | 0 | 0 | 0 | 0 | 0 | 0 | 0 |
| ENSMUSG00 U6 | 103 | 0 | 0 | 0 | 0 | 0 | 0 | 0 | 0 | 0.39986909 | 0 | 0 | 2.70949818 | 0 | 0 | 0 | 0 | 0 | 0.534069 |
| ENSMUSG00 U5 | 115 | 0 | 0 | 0 | 0 | 0 | 0 | 0 | 0 | 0 | 0 | 0 | 0 | 0 | 0 | 0 | 0 | 0 | 0 |
| ENSMUSG00 U6 | 106 | 0 | 0 | 0 | 0 | 0 | 0 | 0 | 0 | 0 | 0 | 0 | 0 | 0 | 0 | 0 | 0 | 0 | 0 |
| ENSMUSG00 n-R5s170 | 112 | 0 | 0 | 0 | 0 | 0 | 0 | 0 | 0 | 0 | 0 | 0 | 0 | 0 | 0 | 0 | 0 | 0 | 0 |
| ENSMUSG00 U6 | 106 | 0 | 0 | 0 | 0 | 0 | 0 | 0 | 0 | 0 | 0 | 0 | 0 | 0 | 0 | 0 | 0 | 0 | 0 |
| ENSMUSG00 n-R5s161 | 113 | 0 | 0 | 0 | 0 | 0 | 0 | 0 | 0 | 0 | 0 | 0 | 0 | 0 | 0 | 0 | 0 | 0 | 0 |
| ENSMUSG00 SNORA25 | 132 | 3.22008909 | 1.00948164 | 4.76405981 | 2.07095656 | 0.72927157 | 2.62779648 | 3.58429386 | 3.76137296 | 2.18413341 | 1.42880919 | 3.25738353 | 0.84569186 | 4.11099311 | 6.68513379 | 2.18493099 | 1.4504018 | 6.96841762 | 4.16735657 |
| ENSMUSG00 U6 | 106 | 0 | 0.62854517 | 0.45635406 | 0 | 0 | 0 | 0 | 0 | 0 | 0 | 0 | 0 | 1.86158178 | 0 | 0 | 0 | 0 | 0 |
| ENSMUSG00 U6 | 108 | 0 | 0 | 0 | 0 | 0 | 0 | 0 | 0 | 0 | 0 | 0 | 0 | 0 | 0 | 0 | 0 | 0 | 0 |
| ENSMUSG00 U6 | 106 | 0 | 0 | 0 | 0 | 0 | 0 | 0 | 0 | 0 | 0 | 0 | 0 | 0 | 0 | 0 | 0 | 0 | 0 |
| ENSMUSG00 U6 | 103 | 0 | 0 | 0.46964593 | 0 | 0.93460046 | 0 | 0 | 0 | 0 | 0.45777382 | 0 | 0 | 0 | 0 | 0 | 0 | 0 | 0 |
| ENSMUSG00 SNORA6 | 135 | 0.31485316 | 0 | 0.35832245 | 0.33748922 | 0 | 0 | 0.35046429 | 0.28290668 | 0 | 0 | 0.35388858 | 0 | 0 | 0 | 0 | 0 | 0 | 0.40747486 |
| ENSMUSG00 SNORA71 | 132 | 0 | 0 | 0 | 0 | 0 | 0 | 0 | 0 | 0 | 0 | 0 | 0.42284593 | 0 | 0 | 0 | 0 | 0 | 0 |
| ENSMUSG00 U6 | 106 | 0.40099223 | 2.5141807 | 0.45635406 | 0 | 0 | 0 | 0 | 0 | 0 | 0 | 0 | 0 | 1.39618634 | 0 | 0 | 0 | 0 | 0 |
| ENSMUSG00 U6 | 106 | 0 | 0 | 0 | 0 | 0 | 0 | 0 | 0 | 0 | 0 | 0 | 0 | 0 | 0 | 0 | 0 | 0 | 0 |
| ENSMUSG00 Snord42b | 68 | 0.62507612 | 1.46968651 | 0 | 0.67001536 | 0.70782241 | 0 | 0.69577469 | 0 | 0 | 0.69339269 | 0 | 0 | 3.62734686 | 0.68300129 | 0 | 2.25238867 | 0 | 0 |
| ENSMUSG00 U6 | 103 | 0 | 0 | 0.46964593 | 0.44234024 | 0.93460046 | 0 | 0 | 1.11240007 | 0 | 0 | 0 | 1.62569891 | 0 | 0 | 0 | 0 | 0 | 0 |
| ENSMUSG00 Snord38a | 70 | 0 | 0 | 0 | 0 | 0 | 0 | 0 | 0 | 0 | 0 | 0 | 0 | 0 | 0 | 0 | 0 | 0 | 0 |
| ENSMUSG00 U6 | 106 | 0 | 0 | 0 | 0 | 0 | 0 | 0 | 0 | 0 | 0 | 0 | 0 | 0 | 0 | 0 | 0 | 0 | 0 |
| ENSMUSG00 U6 | 105 | 0 | 0 | 0 | 0 | 0 | 0 | 0 | 0 | 0 | 0 | 0 | 0 | 0 | 0 | 0 | 0 | 0 | 0 |

| | | | | | | | | | | | | | | | | | | | | |
|---|---|---|---|---|---|---|---|---|---|---|---|---|---|---|---|---|---|---|---|---|
| ENSMUSG00 | U6 | 106 | 0 | 0 | 0 | 0 | 0 | 0 | 0 | 0 | 0 | 0 | 0 | 0 | 0 | 0.43815177 | 0 | 0 | 0 | 0 |
| ENSMUSG00 | Snora5c | 135 | 0 | 0 | 0 | 0 | 0 | 0 | 0 | 0 | 0 | 0 | 0 | 0 | 0 | 0 | 0 | 0 | 0 | 0 |
| ENSMUSG00 | snoU6-53 | 108 | 0 | 0 | 0 | 0 | 0 | 0 | 0 | 0 | 0 | 0 | 0 | 0 | 0 | 0 | 0 | 0 | 0 | 0 |
| ENSMUSG00 | n-R5s7 | 120 | 0 | 0 | 0 | 0 | 0 | 0 | 0 | 0 | 0 | 0 | 0 | 0 | 0 | 0 | 0 | 0 | 0 | 0 |
| ENSMUSG00 | U6 | 106 | 0 | 0 | 0 | 0 | 0 | 0 | 0 | 0 | 0 | 0 | 0 | 0 | 0 | 0 | 0 | 0 | 0 | 0 |
| ENSMUSG00 | n-R5s77 | 118 | 0.36021336 | 0 | 0 | 0 | 0 | 0 | 0 | 0 | 0 | 0 | 0 | 0 | 0 | 0 | 0.69833146 | 0 | 0 | 0 |
| ENSMUSG00 | SNORA36 | 136 | 0 | 0 | 0.35568772 | 0 | 0 | 0 | 0 | 0 | 0 | 0 | 0 | 0 | 0 | 0 | 0 | 0 | 1.69086604 | 0 |
| ENSMUSG00 | U1 | 150 | 0 | 0 | 0 | 0 | 0 | 0 | 0 | 0 | 0 | 0 | 0 | 0 | 0 | 0 | 0 | 0 | 0 | 0 |
| ENSMUSG00 | U6 | 106 | 0 | 0 | 0 | 0 | 0 | 0 | 0 | 0 | 0 | 0 | 0 | 0 | 0 | 0 | 0 | 0 | 0 | 0 |
| ENSMUSG00 | SNORA41 | 130 | 0 | 0 | 0 | 0 | 0 | 0 | 0 | 0 | 0 | 0 | 0 | 0 | 0 | 0 | 0 | 0 | 0 | 0 |
| ENSMUSG00 | U6 | 106 | 0 | 0 | 0 | 0 | 0 | 0 | 0 | 0 | 0 | 0 | 0 | 0 | 0 | 0 | 0 | 0 | 0 | 0 |
| ENSMUSG00 | Y_RNA | 111 | 0 | 0.60023233 | 0.87159514 | 0 | 0.43362093 | 0 | 0 | 0.3440757 | 0 | 0.42478111 | 1.72162013 | 0 | 0 | 0.4184152 | 1.11355556 | 0 | 0.69056391 | 0 |
| ENSMUSG00 | U6 | 106 | 0 | 0 | 0 | 0 | 0 | 0 | 0 | 0 | 0 | 0 | 0 | 0 | 0 | 0 | 0 | 0 | 0 | 0 |
| ENSMUSG00 | U6 | 106 | 0 | 0 | 0 | 0 | 0 | 0 | 0 | 0 | 0 | 0 | 0 | 0 | 0 | 0 | 0 | 0 | 0 | 0 |
| ENSMUSG00 | U6 | 105 | 0 | 0 | 0 | 0 | 0 | 0 | 0 | 0 | 0 | 0 | 0 | 0 | 0 | 0 | 0 | 0 | 0 | 0 |
| ENSMUSG00 | SNORA61 | 129 | 0 | 0 | 0 | 0 | 0 | 0 | 0 | 0 | 0 | 0 | 0 | 0 | 0 | 0 | 0 | 0 | 0 | 0 |
| ENSMUSG00 | U1 | 164 | 0.2591779 | 0.2031274 | 0 | 0 | 0 | 0 | 0 | 0 | 0 | 0 | 0.29131072 | 0 | 0 | 0 | 0 | 0 | 0 | 0 |
| ENSMUSG00 | SNORA44 | 128 | 0.33207169 | 0 | 0 | 0 | 0 | 0 | 0 | 0 | 0 | 0 | 0 | 0 | 0 | 0 | 0 | 0 | 0 | 0 |
| ENSMUSG00 | U6 | 106 | 0 | 0 | 0 | 0 | 0 | 0 | 0 | 0 | 0 | 0 | 0 | 0 | 0 | 0 | 0 | 0 | 0 | 0 |
| ENSMUSG00 | U6 | 102 | 0 | 0 | 0 | 0 | 0 | 0 | 0 | 0 | 0 | 0 | 0 | 0 | 0 | 0 | 0 | 0 | 0 | 0 |
| ENSMUSG00 | Snord7 | 96 | 3.542098 | 2.77607452 | 2.01556376 | 4.74594212 | 5.51511625 | 5.16174308 | 10.3496485 | 5.96756288 | 5.14831447 | 1.47345948 | 1.99062327 | 9.30261043 | 0.51387414 | 1.93517031 | 4.72101161 | 5.98290741 | 1.19769678 | 6.30312681 |
| ENSMUSG00 | SNORA51 | 118 | 0 | 0 | 0 | 0 | 0 | 0 | 0 | 0 | 0 | 0 | 0 | 0 | 0 | 0 | 0 | 0 | 0 | 0 |
| ENSMUSG00 | U1 | 161 | 0 | 0 | 0 | 0 | 0 | 0 | 0 | 0 | 0 | 0 | 0 | 0 | 0 | 0 | 0 | 0 | 0 | 0 |
| ENSMUSG00 | n-R5s12 | 118 | 0 | 0 | 0 | 0 | 0 | 0 | 0 | 0 | 0 | 0 | 0 | 0 | 0 | 0 | 0 | 0 | 0 | 0 |
| ENSMUSG00 | Y_RNA | 111 | 0 | 0 | 0 | 0 | 0 | 0 | 0 | 0 | 0 | 0 | 0 | 0 | 0 | 0 | 0 | 0 | 0 | 0 |
| ENSMUSG00 | U6 | 106 | 0 | 0 | 0 | 0 | 0 | 0 | 0 | 0 | 0 | 0 | 0 | 0 | 0 | 0 | 0 | 0 | 0 | 0 |
| ENSMUSG00 | U6 | 106 | 0 | 0 | 0 | 0 | 0 | 0 | 0 | 0 | 0 | 0 | 0 | 0 | 0 | 0 | 0 | 0 | 0 | 0 |
| ENSMUSG00 | SNORA70 | 137 | 0 | 0.24315981 | 0 | 0 | 0 | 0 | 0 | 0 | 0.90189451 | 0 | 1.04616697 | 0 | 0 | 0 | 0.90222385 | 0 | 0 | 0.40152633 |
| ENSMUSG00 | SNORA73 | 177 | 0 | 0 | 0 | 0 | 0 | 0 | 0 | 0 | 0 | 0 | 0 | 0 | 0 | 0 | 0 | 0 | 0 | 0 |
| ENSMUSG00 | U6 | 103 | 0 | 0 | 0 | 0 | 0 | 0 | 0 | 0 | 0 | 0 | 0 | 0 | 0 | 0 | 0 | 0 | 0 | 0 |
| ENSMUSG00 | SNORA36 | 130 | 0 | 0 | 0 | 0 | 0 | 0 | 0 | 0 | 0 | 0.36269772 | 0 | 0 | 1.13842886 | 0 | 0 | 0.29454313 | 0 | 0.42314697 |
| ENSMUSG00 | U6 | 104 | 0 | 0 | 0 | 0 | 0 | 0 | 0 | 0 | 0 | 0 | 0 | 0 | 0 | 0 | 0 | 0 | 0 | 0 |
| ENSMUSG00 | U6 | 106 | 0.40099223 | 0 | 0 | 0 | 0 | 0 | 0 | 0 | 0 | 0 | 0 | 0 | 1.39618634 | 0.43815177 | 0.38869392 | 0 | 1.44627535 | 0 |
| ENSMUSG00 | U6 | 106 | 0 | 0 | 0 | 0 | 0 | 0 | 0 | 0 | 0 | 0 | 0 | 0 | 0 | 0 | 0 | 0 | 0 | 0 |
| ENSMUSG00 | Snord49a | 71 | 4.78931561 | 2.34597847 | 5.45053863 | 5.1336388 | 0 | 4.18755495 | 10.6620122 | 2.6896058 | 5.80091771 | 3.32047206 | 3.3644337 | 2.35840828 | 2.08444721 | 9.81213114 | 15.6681973 | 9.1681736 | 3.23884199 | 17.0450753 |
| ENSMUSG00 | SNORD77 | 80 | 0 | 0 | 0 | 0 | 0 | 0 | 0 | 0 | 0 | 0 | 0 | 0 | 0 | 0 | 0 | 0 | 0 | 0 |
| ENSMUSG00 | U6 | 97 | 0 | 0 | 0 | 0 | 0 | 0 | 0 | 0 | 0 | 0 | 0 | 0 | 0 | 0 | 0 | 0 | 0 | 0 |
| ENSMUSG00 | SNORD45 | 80 | 2.1252588 | 4.16411178 | 9.67470607 | 5.69513054 | 6.01649046 | 3.71645502 | 1.18281697 | 5.72886036 | 7.72247171 | 3.53630274 | 4.18030887 | 4.18617469 | 2.46659586 | 2.32220437 | 2.57509724 | 4.30769333 | 4.3117084 | 1.37522767 |
| ENSMUSG00 | SNORA67 | 142 | 0 | 0 | 0 | 0 | 0 | 0 | 0 | 0 | 0 | 0 | 0 | 0 | 0 | 0 | 0 | 0 | 0 | 0 |
| ENSMUSG00 | U2 | 132 | 0 | 0 | 0 | 0 | 0 | 0 | 0 | 0 | 0 | 0 | 0 | 0 | 0 | 0 | 0 | 0 | 0 | 0 |
| ENSMUSG00 | U6 | 80 | 0 | 0 | 0 | 1.70853916 | 0 | 0 | 0 | 0 | 0 | 0 | 0 | 0 | 0 | 0 | 0 | 0 | 0 | 0 |
| ENSMUSG00 | U1 | 160 | 0 | 0 | 0 | 0 | 0 | 0 | 0 | 0 | 0 | 0 | 0 | 0 | 0 | 0 | 0 | 0 | 0 | 0 |
| ENSMUSG00 | U1 | 155 | 0 | 0 | 0 | 0 | 0 | 0 | 0 | 0 | 0 | 0 | 0 | 0 | 0 | 0 | 0 | 0 | 0 | 0 |
| ENSMUSG00 | U6 | 106 | 0 | 0 | 0 | 0 | 0 | 0 | 0 | 0 | 0 | 0 | 0 | 0 | 0 | 0 | 0 | 0 | 0 | 0 |
| ENSMUSG00 | AF357428 | 68 | 0 | 0 | 0 | 0 | 0 | 0 | 0 | 0 | 0 | 0 | 0 | 0 | 0 | 0 | 0 | 0 | 0 | 0 |
| ENSMUSG00 | SNORA5 | 135 | 0 | 0.74028654 | 0.71664489 | 0.33748922 | 0.71306554 | 0.73411457 | 0 | 0.84872005 | 0 | 0 | 0 | 0 | 0.73084322 | 0 | 0 | 0.56726826 | 0 | 0 |
| ENSMUSG00 | U2 | 149 | 0 | 0 | 0 | 0 | 0 | 0 | 0 | 0 | 0 | 0 | 0 | 0 | 0 | 0 | 0 | 0 | 0 | 0 |
| ENSMUSG00 | SNORD18 | 67 | 0 | 0.49720738 | 0 | 0 | 0 | 0.73959304 | 1.41231877 | 1.71010757 | 0 | 2.11122552 | 0 | 0 | 0 | 0 | 0 | 0.5715016 | 6.86441139 | 0 |
| ENSMUSG00 | SNORA44 | 128 | 0 | 0 | 0 | 0 | 0 | 0 | 0 | 0 | 0 | 0 | 0 | 0 | 0 | 0 | 0 | 0 | 0 | 0 |
| ENSMUSG00 | SNORD113 | 81 | 0 | 0 | 0 | 0 | 0 | 0 | 0 | 0 | 0 | 0 | 0 | 0 | 0 | 0 | 0 | 0 | 0 | 0 |
| ENSMUSG00 | n-R5s10 | 118 | 0 | 0 | 0 | 0 | 0 | 0 | 0 | 0 | 0 | 0 | 0 | 0 | 0 | 0 | 0 | 0 | 0 | 0 |
| ENSMUSG00 | U6 | 105 | 0 | 0 | 0 | 0 | 0 | 0 | 0 | 0 | 0 | 0 | 0 | 0 | 0 | 0 | 0 | 0 | 0 | 0 |
| ENSMUSG00 | U6 | 109 | 0 | 0 | 0 | 0 | 0 | 0 | 0 | 0 | 0 | 0 | 0 | 0 | 0 | 0 | 0 | 0 | 0 | 0 |
| ENSMUSG00 | U1 | 168 | 0 | 0.99145519 | 0.57587536 | 0 | 0 | 0 | 0 | 0.90934291 | 0 | 0 | 0.56874951 | 0 | 0 | 0 | 0 | 0 | 0 | 0 |
| ENSMUSG00 | U6 | 102 | 2.08358706 | 0 | 0.4742503 | 1.34003072 | 1.41564481 | 1.45743334 | 1.39154938 | 1.12330595 | 0.40378937 | 0.92452359 | 1.40514584 | 1.09442476 | 3.38552373 | 0.91066838 | 0.80787364 | 0.75079622 | 1.50299204 | 1.07860994 |
| ENSMUSG00 | U2 | 190 | 0.67113436 | 0.87665511 | 1.7821827 | 3.35712958 | 2.53325914 | 3.65125405 | 2.9881692 | 3.41721495 | 3.0347959 | 4.96323192 | 3.26881295 | 8.51923271 | 11.9435168 | 3.66663848 | 6.72235912 | 7.65812148 | 5.84980321 | 6.36947552 |
| ENSMUSG00 | U1 | 151 | 0 | 0 | 0 | 0 | 0 | 0 | 0 | 0 | 0 | 0 | 0 | 0 | 0 | 0 | 0 | 0 | 0 | 0 |
| ENSMUSG00 | U6 | 106 | 0 | 0 | 0 | 0 | 0 | 0 | 0 | 0 | 0 | 0 | 0 | 0 | 0 | 0 | 0 | 0 | 0 | 0 |
| ENSMUSG00 | U1 | 161 | 0 | 0 | 0 | 0 | 0 | 0 | 0 | 0 | 0 | 0 | 0 | 0 | 0 | 0 | 0 | 0 | 0 | 0 |
| ENSMUSG00 | n-R5s54 | 111 | 0 | 0 | 0 | 0 | 0 | 0 | 0 | 0 | 0 | 0 | 0 | 0 | 0 | 0 | 0 | 0 | 0 | 0 |
| ENSMUSG00 | U1 | 165 | 0 | 0 | 0 | 0 | 0 | 0 | 0 | 0 | 0 | 0 | 0 | 0 | 0 | 0 | 0 | 0 | 0 | 0 |
| ENSMUSG00 | U6 | 106 | 10.0248057 | 10.685268 | 12.3215596 | 13.3244564 | 5.90297177 | 15.894273 | 6.69519041 | 8.28703071 | 16.7077375 | 8.89635911 | 7.66202164 | 4.73906569 | 4.65395446 | 3.50521414 | 7.38518455 | 3.9735536 | 14.8243224 | 2.07581535 |
| ENSMUSG00 | U6 | 106 | 2.40595336 | 1.88563552 | 0.45635406 | 0.42982117 | 2.27037376 | 0.46747862 | 0 | 1.08091705 | 2.33131221 | 0.44481796 | 0.45070716 | 0 | 0 | 0 | 0.38869392 | 2.16739287 | 1.44627535 | 1.03790767 |
| ENSMUSG00 | SNORA55 | 130 | 0 | 0 | 0 | 0 | 0 | 0 | 1.45577474 | 0 | 0.95045806 | 0.72539543 | 0 | 0 | 0.37947629 | 0.35726221 | 0 | 1.17817254 | 0.58963534 | 0 |
| ENSMUSG00 | SNORD30 | 68 | 0 | 0 | 0 | 0 | 0 | 0 | 0 | 0 | 0 | 0 | 0 | 0 | 0 | 0 | 0 | 0 | 0 | 0 |
| ENSMUSG00 | U6 | 106 | 0 | 0 | 0 | 0 | 0 | 0 | 0 | 0 | 0 | 0 | 0 | 0 | 0 | 0 | 0 | 0.36123215 | 0 | 0 |
| ENSMUSG00 | U6 | 105 | 0 | 0 | 0 | 0 | 0 | 0 | 0 | 0 | 0 | 0 | 0 | 0 | 0 | 0 | 0 | 0 | 0 | 0 |
| ENSMUSG00 | U6 | 106 | 0 | 0 | 0 | 0 | 0 | 0 | 0 | 0 | 0 | 0 | 0 | 0 | 0 | 0 | 0 | 0 | 0 | 0 |
| ENSMUSG00 | U6 | 106 | 0 | 0 | 0 | 0 | 0 | 0 | 0 | 0 | 0 | 0 | 0 | 0 | 0 | 0 | 0 | 0 | 0 | 0 |
| ENSMUSG00 | SNORD103 | 82 | 0 | 0 | 0 | 0 | 0 | 0 | 0 | 0 | 0 | 0 | 0 | 0 | 0 | 0 | 0 | 0 | 0 | 0 |
| ENSMUSG00 | U6 | 106 | 0 | 0 | 0 | 0 | 0 | 0 | 0 | 0 | 0 | 0 | 0 | 0 | 0 | 0 | 0 | 0 | 0 | 0 |
| ENSMUSG00 | U1 | 156 | 0 | 0 | 0 | 0 | 0 | 0 | 0 | 0 | 0 | 0 | 0 | 0 | 0 | 0 | 0 | 0 | 0 | 0 |
| ENSMUSG00 | U1 | 134 | 0 | 0 | 0 | 0 | 0 | 0 | 0 | 0 | 0 | 0 | 0 | 0 | 0 | 0 | 0 | 0 | 0 | 0 |
| ENSMUSG00 | U1 | 161 | 0 | 0 | 0 | 0 | 0 | 0 | 0 | 0 | 0 | 0 | 0 | 0 | 0 | 0 | 0 | 0 | 0 | 0 |
| ENSMUSG00 | U6 | 106 | 0.40099223 | 0.31427259 | 0 | 0 | 0 | 0.93495724 | 0.44634603 | 3.24275115 | 0.77710407 | 0.44481796 | 0 | 0 | 5.58474535 | 5.25782121 | 3.10955139 | 0 | 0 | 1.55686151 |
| ENSMUSG00 | U6 | 106 | 0 | 0 | 0 | 0 | 0 | 0 | 0 | 0 | 0 | 0 | 0 | 0 | 1.39618634 | 0 | 0 | 0 | 0 | 0 |
| ENSMUSG00 | SNORA5 | 133 | 0 | 1.00189156 | 1.09113226 | 0.34256424 | 0.36189416 | 0.74515389 | 0.35573443 | 0 | 0.6193461 | 0 | 1.07763064 | 0.41966664 | 0.37091667 | 0 | 1.23914454 | 0.2878993 | 0 | 0.41360231 |
| ENSMUSG00 | U6 | 100 | 0 | 0 | 0 | 0 | 0 | 0 | 0 | 0 | 0 | 0 | 0 | 0 | 0 | 0 | 0 | 0 | 0 | 0 |
| ENSMUSG00 | U6 | 105 | 0 | 0 | 0 | 0 | 0 | 0 | 0 | 0 | 0 | 0 | 0 | 0 | 0 | 0 | 0 | 0 | 0 | 0 |
| ENSMUSG00 | U1 | 162 | 0 | 0 | 0 | 0 | 0 | 0 | 0 | 0 | 0 | 0 | 0 | 0 | 0 | 0 | 0 | 0 | 0 | 0 |
| ENSMUSG00 | U1 | 160 | 1.5939441 | 0.83282236 | 0.90700369 | 0.56951305 | 0.30082452 | 0 | 0 | 1.67091761 | 0 | 0 | 0 | 1.74423946 | 0 | 0 | 0.25750972 | 0 | 0 | 0 |
| ENSMUSG00 | n-R5s69 | 118 | 0 | 0 | 0 | 0 | 0 | 0 | 0 | 0 | 0 | 0 | 0 | 0 | 0 | 0 | 0 | 0 | 0 | 0 |
| ENSMUSG00 | U6 | 103 | 0 | 0 | 0 | 0 | 0 | 0 | 0 | 0 | 0 | 0 | 0 | 0 | 0 | 0 | 0 | 0 | 0 | 0 |
| ENSMUSG00 | 7SK | 323 | 0 | 0.20627179 | 0 | 0.14105586 | 0 | 0 | 0.58591553 | 0 | 0 | 0 | 0 | 0 | 0.45819118 | 0.28757949 | 0 | 0 | 0.47462906 | 0 |
| ENSMUSG00 | U4 | 82 | 0 | 0 | 0 | 0 | 0 | 0 | 0 | 0 | 0 | 0 | 0 | 0 | 0 | 0 | 0 | 0 | 0 | 0 |
| ENSMUSG00 | SNORA42 | 133 | 0 | 0 | 0 | 0 | 0 | 0 | 0 | 0 | 0 | 0 | 0 | 0 | 0 | 0 | 0 | 0 | 0 | 0 |
| ENSMUSG00 | n-R5s3 | 111 | 0 | 0 | 0 | 0 | 0 | 0 | 0 | 0 | 0 | 0 | 0 | 0 | 0 | 0 | 0 | 0 | 0 | 0 |
| ENSMUSG00 | U6 | 106 | 0 | 0 | 0 | 0 | 0 | 0 | 0 | 0 | 0 | 0 | 0 | 0 | 0 | 0 | 0 | 0 | 0 | 0 |
| ENSMUSG00 | U6 | 106 | 0 | 0 | 0 | 0 | 0 | 0 | 0 | 0 | 0 | 0 | 0 | 0 | 0 | 0 | 0 | 0 | 0 | 0 |
| ENSMUSG00 | SNORA3 | 129 | 0 | 0.51647898 | 0.37498861 | 0.35318639 | 0 | 0.76825944 | 0 | 0.59213027 | 0 | 0.73101866 | 0.74069703 | 0 | 0 | 1.44012674 | 0 | 0 | 0 | 1.70570874 |
| ENSMUSG00 | Snord35a | 89 | 2.38793124 | 1.12290655 | 0 | 1.02384369 | 1.08161626 | 0 | 2.12641254 | 2.5747687 | 13.4203254 | 2.11913273 | 4.29437829 | 5.01713821 | 13.3029889 | 10.9587172 | 4.6293883 | 3.44185235 | 3.87569295 | 16.688156 |
| ENSMUSG00 | U6 | 106 | 0 | 0 | 0 | 0 | 0 | 0 | 0 | 0 | 0 | 0 | 0 | 0 | 0 | 0 | 0 | 0 | 0 | 0 |
| ENSMUSG00 | U2 | 190 | 0 | 0 | 0 | 0 | 0.25332591 | 0 | 0 | 0 | 0.21677114 | 0 | 0.25144715 | 0.29376665 | 0.25964167 | 0 | 0.21685029 | 0 | 1.00858676 | 0.86856484 |
| ENSMUSG00 | U6 | 103 | 0 | 0 | 0 | 0 | 0 | 0 | 0 | 0 | 0 | 0 | 0 | 0 | 0 | 0 | 0 | 0 | 0 | 0 |
| ENSMUSG00 | Snord15a | 147 | 0.28915086 | 0.45323666 | 0 | 0 | 0.32742805 | 0 | 0.32185496 | 0.77943678 | 0 | 0 | 0 | 0 | 2.01354764 | 0 | 0 | 1.82336226 | 1.04289243 | 0 |
| ENSMUSG00 | n-R5s182 | 106 | 0 | 0 | 0 | 0 | 0 | 0 | 0 | 0 | 0 | 0 | 0 | 0 | 0 | 0 | 0 | 0 | 0 | 0 |
| ENSMUSG00 | 7SK | 320 | 0 | 0 | 0 | 0.14237826 | 0 | 0 | 0 | 0 | 0.25741572 | 0 | 0 | 0 | 0 | 0 | 0 | 0 | 0 | 0 |
| ENSMUSG00 | U6 | 106 | 0 | 0 | 0 | 0 | 0 | 0 | 0 | 0 | 0 | 0 | 0 | 0 | 0 | 0 | 0 | 0 | 0 | 0 |
| ENSMUSG00 | U6 | 106 | 0 | 0 | 0 | 0 | 0 | 0.46747862 | 0 | 0 | 0 | 0 | 0 | 0 | 0 | 0 | 0.38869392 | 0 | 0 | 0 |
| ENSMUSG00 | U6 | 106 | 0 | 0 | 0 | 0 | 0 | 0 | 0 | 0 | 0 | 0 | 0 | 0 | 0 | 0 | 0 | 0 | 0 | 0 |
| ENSMUSG00 | Snord16a | 98 | 0.86745257 | 1.01978248 | 3.45525217 | 0.46490862 | 0 | 4.04512111 | 4.82782438 | 4.28690231 | 2.10135285 | 6.73581475 | 6.33749449 | 2.84773789 | 4.02709529 | 8.0566274 | 2.94296828 | 3.90720484 | 3.51976196 | 4.49053932 |
| ENSMUSG00 | U6 | 106 | 0 | 0 | 0 | 0 | 0 | 0 | 0 | 0 | 0 | 0 | 0 | 0 | 0 | 0 | 0 | 0 | 1.08470652 | 0 |
| ENSMUSG00 | U12 | 144 | 0.8855245 | 0 | 0 | 0 | 0 | 0 | 0 | 0 | 0 | 0 | 0 | 0 | 0 | 0 | 0 | 0.265907 | 0 | 0.76401537 |
| ENSMUSG00 | U6 | 106 | 0 | 0 | 0 | 0 | 0 | 0 | 0 | 0 | 0 | 0 | 0 | 0 | 0 | 0 | 0 | 0 | 0 | 0 |
| ENSMUSG00 | SNORA71 | 132 | 0 | 0 | 0 | 0 | 0 | 0 | 0 | 0 | 0 | 0 | 0 | 0 | 0 | 0 | 0 | 0 | 0 | 0 |
| ENSMUSG00 | U6 | 109 | 0 | 0 | 0 | 0 | 0 | 0 | 0 | 0 | 0 | 0 | 0 | 0 | 0 | 0 | 0 | 0 | 0 | 0 |
| ENSMUSG00 | U6 | 107 | 0 | 0 | 0 | 0 | 0.44983106 | 0 | 0 | 0 | 0 | 0 | 0 | 0 | 0 | 0 | 0 | 0 | 0 | 0 |
| ENSMUSG00 | U1 | 163 | 0 | 0 | 0 | 0 | 0 | 0 | 0 | 0 | 0 | 0 | 0 | 0 | 0 | 0 | 0 | 0 | 0 | 0 |
| ENSMUSG00 | U6 | 106 | 0 | 0 | 0 | 0 | 0 | 0 | 0 | 0 | 0 | 0 | 0 | 0 | 0 | 0 | 0 | 0 | 0 | 0 |
| ENSMUSG00 | SNORA32 | 121 | 3.86410691 | 1.1012527 | 1.99890621 | 4.14191312 | 0.79556899 | 1.6381069 | 0.78202775 | 0.94691907 | 2.72307542 | 1.1690257 | 1.97417184 | 1.3838594 | 0.4077018 | 3.07068346 | 1.70254363 | 1.2658052 | 3.48420881 | 3.63696573 |
| ENSMUSG00 | U6 | 106 | 0 | 0 | 0 | 0 | 0 | 0 | 0 | 0 | 0 | 0 | 0 | 0 | 0 | 0 | 0 | 0 | 0 | 0 |
| ENSMUSG00 | U6 | 106 | 0 | 0 | 0 | 0 | 0 | 0 | 0 | 0 | 0 | 0 | 0 | 0 | 0 | 0 | 0 | 0 | 0 | 0 |
| ENSMUSG00 | SNORA2 | 135 | 0 | 0.24676218 | 1.43328979 | 1.01246765 | 0 | 0 | 0.35046429 | 0 | 1.22034121 | 1.74632234 | 0.35388858 | 3.30759482 | 0.73084322 | 1.72015138 | 0.91559013 | 0.85090239 | 0.2838985 | 0.81494973 |
| ENSMUSG00 | SNORD58 | 69 | 0 | 0 | 0 | 0 | 0 | 0 | 0.685691 | 0 | 0 | 0 | 0 | 0 | 0 | 0 | 0 | 0 | 0 | 0 |
| ENSMUSG00 | U6 | 106 | 18.0446502 | 9.11390502 | 20.0795786 | 13.7542775 | 10.897794 | 12.1544441 | 8.49057452 | 14.772533 | 7.38248867 | 13.7893566 | 11.2676789 | 10.5312571 | 9.77330437 | 6.57227652 | 12.8268995 | 11.5594286 | 8.67765213 | 11.9359383 |
| ENSMUSG00 | U1 | 155 | 0 | 0 | 0 | 0 | 0 | 0 | 0 | 0 | 0 | 0 | 0 | 0 | 0 | 0 | 0 | 0 | 0 | 0 |
| ENSMUSG00 | U6 | 104 | 0 | 0 | 0 | 0 | 0 | 0 | 0 | 0 | 0 | 0 | 0 | 0 | 0 | 0 | 0 | 0 | 0 | 0 |
| ENSMUSG00 | SNORA67 | 142 | 0 | 0 | 0 | 0 | 0 | 0 | 0 | 0 | 0 | 0 | 0 | 0 | 0 | 0 | 0 | 0 | 0 | 0 |
| ENSMUSG00 | SNORA71 | 130 | 0 | 0 | 0 | 0 | 0 | 0 | 0 | 0.58757542 | 0 | 0 | 0 | 0 | 0 | 0 | 0 | 0 | 0 | 0 |
| ENSMUSG00 | U1 | 163 | 0 | 0 | 0 | 0 | 0 | 0 | 0 | 0 | 0 | 0 | 0 | 0 | 0 | 0 | 0 | 0 | 0 | 0 |
| ENSMUSG00 | U1 | 158 | 0 | 0 | 0 | 0 | 0 | 0 | 0 | 0 | 0 | 0 | 0 | 0 | 0 | 0 | 0 | 0 | 0 | 0 |
| ENSMUSG00 | U6 | 103 | 0 | 0 | 0 | 0 | 0 | 0 | 0 | 0 | 0 | 0 | 0 | 0 | 0 | 0 | 0 | 0 | 0 | 0 |
| ENSMUSG00 | U3 | 211 | 7.85640694 | 4.1049064 | 8.94107907 | 3.2389368 | 5.24660779 | 7.04541236 | 4.26038341 | 4.34415952 | 9.5646411 | 1.78770439 | 4.98127529 | 6.34870096 | 1.16900278 | 3.74194069 | 2.92902056 | 3.81091353 | 2.17969461 | 2.60706667 |
| ENSMUSG00 | U6 | 105 | 0 | 0 | 0 | 0 | 0 | 0 | 0 | 0 | 0 | 0 | 0 | 0 | 0 | 0 | 0 | 0 | 0 | 0 |
| ENSMUSG00 | n-R5s183 | 114 | 0 | 0.58443674 | 0 | 0.79931657 | 0 | 0 | 0 | 0 | 0 | 0.82720532 | 0 | 0 | 0 | 0 | 0.72283431 | 0 | 0 | 0 |
| ENSMUSG00 | SNORA71 | 130 | 0 | 0 | 0 | 0 | 0 | 0 | 0 | 0 | 0 | 0 | 0 | 0 | 0.75895257 | 0.71452442 | 0 | 0 | 0 | 0 |
| ENSMUSG00 | U6 | 106 | 0 | 0 | 0 | 0 | 0 | 0 | 0 | 0 | 0 | 0 | 0 | 0 | 0 | 0 | 0 | 0 | 0 | 0 |
| ENSMUSG00 | U1 | 163 | 0 | 0 | 0 | 0 | 0 | 0 | 0 | 0 | 0 | 0 | 0 | 0 | 0.3026498 | 0 | 0 | 0 | 0 | 0 |
| ENSMUSG00 | Snord34 | 69 | 0 | 0 | 0 | 0 | 0 | 0 | 0 | 0 | 0 | 0 | 0 | 0 | 0 | 0 | 0 | 0 | 0 | 0 |
| ENSMUSG00 | U1 | 176 | 0 | 0 | 0 | 0 | 0 | 0 | 0 | 0 | 0 | 0 | 0 | 0 | 0 | 0 | 0 | 0 | 0 | 0 |
| ENSMUSG00 | n-R5s168 | 103 | 0 | 0 | 0 | 0 | 0 | 0 | 0 | 0 | 0 | 0 | 0 | 0 | 0 | 0 | 0 | 0 | 0 | 0 |
| ENSMUSG00 | U3 | 212 | 0 | 0 | 0.22817703 | 0 | 0 | 0.23373931 | 0 | 0 | 0 | 0 | 0 | 0 | 0 | 0 | 0 | 0 | 0 | 0 |
| ENSMUSG00 | SNORA63 | 129 | 0 | 0 | 0 | 0 | 0 | 0 | 0 | 0 | 0 | 0 | 0 | 0 | 0 | 0 | 0 | 0 | 0 | 0 |
| ENSMUSG00 | U6 | 106 | 0 | 0 | 0 | 0 | 0 | 0 | 0 | 0 | 0 | 0 | 0 | 0 | 0 | 0 | 0 | 0 | 0 | 0 |
| ENSMUSG00 | n-R5s185 | 118 | 0 | 0 | 0 | 0 | 0 | 0 | 0 | 0 | 0 | 0 | 0 | 0 | 0 | 0 | 0 | 0 | 0 | 0 |
| ENSMUSG00 | U1 | 163 | 0 | 0 | 0 | 0 | 0 | 0 | 0 | 0 | 0 | 0 | 0 | 0 | 0 | 0 | 0 | 0 | 0 | 0 |
| ENSMUSG00 | U6 | 106 | 0 | 0 | 0 | 0 | 0 | 0 | 0 | 0 | 0 | 0 | 0 | 0 | 0 | 0 | 0 | 0 | 0 | 0 |
| ENSMUSG00 | n-R5s151 | 118 | 0 | 0 | 0 | 0 | 0 | 0 | 0 | 0 | 0 | 0 | 0 | 0 | 0 | 0 | 0 | 0 | 0 | 0 |
| ENSMUSG00 | n-R5s46 | 115 | 0 | 0 | 0 | 0 | 0 | 0 | 0 | 0 | 0 | 0 | 0 | 0 | 0 | 0 | 0 | 0 | 0 | 0 |
| ENSMUSG00 | U6 | 107 | 0 | 0 | 0 | 0 | 0 | 0 | 0 | 0 | 0 | 0 | 0 | 0 | 0 | 0 | 0 | 0 | 0 | 0 |
| ENSMUSG00 | SNORA72 | 131 | 0 | 0 | 0 | 0 | 0 | 0 | 0 | 0 | 0 | 0 | 0 | 0 | 0 | 0 | 0 | 0 | 0 | 0 |
| ENSMUSG00 | U6 | 105 | 0 | 0 | 0 | 0 | 0 | 0 | 0 | 0 | 0 | 0 | 0 | 0 | 0 | 0 | 0 | 0 | 0 | 0 |
| ENSMUSG00 | SNORD63 | 69 | 0.61601704 | 1.44838671 | 0.70106566 | 1.32060998 | 3.48782056 | 0.71815556 | 0.685691 | 3.32107847 | 0.59690603 | 2.05003058 | 0.6923907 | 0 | 0 | 0 | 0 | 0 | 1.66636073 | 0 |
| ENSMUSG00 | U1 | 164 | 0 | 0 | 0 | 0 | 0 | 0 | 0 | 0 | 0 | 0 | 0 | 0 | 0 | 0 | 0 | 0 | 0 | 0 |
| ENSMUSG00 | U6 | 106 | 0 | 0 | 0 | 0 | 0 | 0 | 0 | 0 | 0 | 0 | 0 | 0 | 0 | 0 | 0 | 0 | 0 | 0 |
| ENSMUSG00 | U2 | 190 | 0 | 0 | 0 | 0 | 0 | 0 | 0 | 0 | 0 | 0 | 0 | 0 | 0 | 0 | 0 | 0 | 0 | 0 |
| ENSMUSG00 | U6 | 106 | 0 | 0 | 0 | 0 | 0 | 0 | 0 | 0 | 0 | 0 | 0 | 0 | 0 | 0 | 0 | 0 | 0 | 0 |
| ENSMUSG00 | 7SK | 337 | 0 | 0 | 0 | 0 | 0 | 0 | 0 | 0 | 0 | 0 | 0 | 0 | 0 | 0.27563257 | 0 | 0 | 0 | 0.48969531 |
| ENSMUSG00 | U6 | 106 | 0 | 0 | 0 | 0 | 0 | 0 | 0 | 0 | 0 | 0 | 0 | 0 | 0 | 0 | 0 | 0 | 0 | 0 |
| ENSMUSG00 | U6 | 106 | 0 | 0 | 0 | 0 | 0 | 0 | 0 | 0 | 0 | 0 | 0 | 0 | 0 | 0 | 0 | 0 | 0 | 0 |
| ENSMUSG00 | U6 | 106 | 0.40099223 | 0 | 0 | 0 | 0 | 0 | 0 | 0 | 0 | 0 | 0 | 0 | 0.46539545 | 0 | 0 | 0 | 0 | 0 |
| ENSMUSG00 | U1 | 155 | 0 | 0 | 0 | 0 | 0 | 0 | 0 | 0 | 0 | 0 | 0 | 0 | 0 | 0 | 0 | 0 | 0 | 0 |
| ENSMUSG00 | U6 | 105 | 0 | 0 | 0 | 0 | 0 | 0 | 0 | 1.81868583 | 0 | 0 | 0.4549996 | 0 | 0 | 0 | 0 | 0 | 0.36501235 | 0 |
| ENSMUSG00 | U6 | 106 | 0 | 0.62854517 | 0 | 0 | 0 | 0 | 0 | 0 | 0 | 0 | 0.90141431 | 0 | 0 | 0 | 0 | 0 | 0 | 0 |
| ENSMUSG00 | n-R5-8s1 | 147 | 26.6018789 | 19.7157945 | 19.4152265 | 22.9354917 | 26.8491003 | 24.2707267 | 23.8172669 | 15.3289234 | 20.7333481 | 12.1886172 | 34.4499701 | 22.0225063 | 30.2032147 | 32.858402 | 23.8240289 | 24.22467 | 28.4188188 | 29.1885056 |
| ENSMUSG00 | SNORA71 | 130 | 0 | 0 | 0 | 0 | 0 | 0 | 0 | 0 | 0 | 0 | 0 | 0 | 0 | 0 | 0 | 0 | 0 | 0 |
| ENSMUSG00 | U6 | 106 | 0 | 0 | 0 | 0 | 0 | 0 | 0 | 0 | 0 | 0 | 0 | 0 | 0 | 0 | 0 | 0 | 0 | 0 |
| ENSMUSG00 | U6 | 106 | 0 | 0 | 0 | 0 | 0 | 0 | 0 | 0 | 0 | 0 | 0 | 0 | 0 | 0 | 0 | 0 | 0 | 0 |
| ENSMUSG00 | U6 | 106 | 0 | 0 | 0 | 0 | 0 | 0 | 0 | 0 | 0 | 0 | 0 | 0 | 0 | 0 | 0.77738785 | 0 | 0 | 0 |
| ENSMUSG00 | U6 | 106 | 0 | 0 | 0 | 0 | 0 | 0 | 0 | 0 | 0 | 0 | 0 | 0 | 0 | 0 | 0 | 0 | 0 | 0 |
| ENSMUSG00 | U1 | 161 | 0 | 0.20691239 | 0 | 0 | 0.5979121 | 0 | 1.76320543 | 0 | 0.25581687 | 0.58572302 | 0 | 0 | 0 | 0.86541778 | 0 | 2.14046874 | 0 | 0 |
| ENSMUSG00 | U6 | 106 | 0 | 0 | 0 | 0 | 0 | 0 | 0 | 0 | 0 | 0 | 0 | 0 | 0 | 0 | 0 | 0 | 0 | 0 |
| ENSMUSG00 | U6 | 104 | 0 | 0 | 0 | 0 | 0 | 0 | 0 | 0 | 0 | 0 | 0 | 0 | 0 | 0 | 0 | 0 | 0 | 0 |
| ENSMUSG00 | U1 | 152 | 0 | 0 | 0 | 0 | 0 | 0 | 0 | 0.25126581 | 0 | 0 | 0 | 0 | 0 | 0 | 0 | 0 | 0 | 0 |
| ENSMUSG00 | U6 | 106 | 0 | 0 | 0 | 0 | 0 | 0 | 0 | 0 | 0 | 0 | 0 | 0 | 0 | 0 | 0 | 0 | 0 | 0 |
| ENSMUSG00 | Snora2b | 136 | 0.31253806 | 0.24494775 | 0 | 0 | 0.70782241 | 0 | 0 | 0 | 1.81705217 | 1.38678539 | 0 | 0 | 0 | 0 | 0 | 1.40774292 | 0.28181101 | 0.40447873 |
| ENSMUSG00 | SNORA21 | 136 | 0 | 0 | 0 | 0 | 0 | 0 | 0 | 0 | 0 | 0 | 0 | 0 | 0 | 0 | 0 | 0 | 0 | 0 |
| ENSMUSG00 | U1 | 163 | 4.69382312 | 1.43061509 | 4.45155187 | 3.0746717 | 3.24816683 | 1.824027 | 0.29026183 | 0.93723687 | 2.02142409 | 0 | 1.75858743 | 6.1636928 | 0.6052996 | 0.85479915 | 2.52770282 | 0.70473511 | 0 | 1.01243755 |
| ENSMUSG00 | U6 | 106 | 0 | 0 | 0 | 0 | 0 | 0 | 0 | 0 | 0 | 0 | 0 | 0 | 0 | 0 | 0 | 0 | 0 | 0 |
| ENSMUSG00 | U6 | 103 | 0 | 0 | 0.46964593 | 0 | 0 | 0 | 1.83738559 | 0 | 0.79973817 | 0 | 0 | 0 | 0 | 0 | 0 | 0 | 0 | 1.06813799 |
| ENSMUSG00 | U2 | 190 | 0.89484581 | 0.35066204 | 1.01839011 | 0.47958994 | 3.79988871 | 1.04321544 | 0.9960564 | 0.80405058 | 2.81802476 | 1.24080798 | 2.0115772 | 1.46883323 | 2.85605837 | 2.68886822 | 2.60220353 | 4.03059025 | 5.84980321 | 3.47425937 |
| ENSMUSG00 | U1 | 159 | 0 | 0 | 0 | 0 | 0 | 0 | 0 | 0 | 0 | 0 | 0 | 0 | 0 | 0 | 0 | 0 | 0 | 0 |
| ENSMUSG00 | U6 | 103 | 0 | 0 | 0 | 0 | 0 | 0 | 0 | 0 | 0 | 0 | 0 | 0 | 0 | 0 | 0 | 0 | 0 | 0 |
| ENSMUSG00 | Gm5908 | 446 | 0 | 0.07469259 | 0.21692166 | 0 | 0 | 0 | 0.10608224 | 0.17126638 | 0.09234645 | 0.3171572 | 0 | 0 | 0.11060968 | 0 | 0.18476034 | 0.17170676 | 0 | 0.37001641 |
| ENSMUSG00 | C330021F23I | 2868 | 0.25194839 | 0.092923 | 0.01686664 | 0 | 0.0167824 | 0.0172778 | 0.1649675 | 0.06658369 | 0.12924639 | 0.13152218 | 0 | 0.50599973 | 0.08600404 | 0.14574504 | 0.30168503 | 0.05340392 | 0.02672685 | 0 |
| ENSMUSG00 | Tacc1 | 7682 | 12.7371668 | 12.80131 | 13.1040506 | 12.6742973 | 13.1576464 | 13.171919 | 9.68179267 | 9.23239928 | 9.27527627 | 10.1212587 | 10.1370974 | 10.4045859 | 10.1656372 | 12.1763072 | 11.6385546 | 11.8181502 | 11.5697322 | 13.0970654 |
| ENSMUSG00 | Gm14850 | 401 | 0 | 0 | 0 | 0 | 0 | 0 | 0 | 0.0952429 | 0 | 0 | 0 | 0 | 0 | 0 | 0 | 0 | 0 | 0 |
| ENSMUSG00 | Gm15308 | 748 | 0 | 0 | 0 | 0 | 0 | 0 | 0 | 0 | 0 | 0 | 0 | 0 | 0 | 0 | 0 | 0 | 0 | 0 |
| ENSMUSG00 | Defa20 | 423 | 0 | 0 | 0 | 0 | 0 | 0 | 0 | 0 | 0 | 0 | 0 | 0 | 0 | 0 | 0 | 0 | 0 | 0 |
| ENSMUSG00 | Ifitm7 | 871 | 0 | 0 | 0 | 0 | 0 | 0 | 0 | 0 | 0 | 0 | 0 | 0.12816455 | 0 | 0 | 0 | 0 | 0 | 0 |
| ENSMUSG00 | Cpped1 | 7055 | 12.4171747 | 13.5518081 | 15.9965197 | 13.7813279 | 15.3640102 | 15.6630186 | 11.3738205 | 12.8408758 | 12.0436261 | 11.0675499 | 12.2975655 | 12.0729555 | 9.50277471 | 9.05840741 | 12.6495493 | 10.4369721 | 12.8750282 | 12.327342 |
| ENSMUSG00 | Cd209b | 2285 | 0 | 0 | 0 | 0 | 0 | 0 | 0.74540763 | 0.0501432 | 0.45061833 | 0 | 0 | 0 | 0 | 0 | 0 | 0 | 0 | 0 |
| ENSMUSG00 | Aurkaip1 | 1742 | 48.1904263 | 40.0251938 | 39.2653111 | 41.7424953 | 46.3083261 | 40.3362665 | 54.5100727 | 42.5772936 | 46.2698113 | 50.1822066 | 43.0577984 | 41.301027 | 50.7761927 | 54.2292042 | 47.3983456 | 44.840895 | 45.3667188 | 44.9356825 |
| ENSMUSG00 | Gm13154 | 2952 | 0.02879754 | 0.29340625 | 0.14748028 | 0.13890562 | 0.35870675 | 0.537157 | 0 | 0.07762683 | 0.0976645 | 0.04791738 | 0.17802322 | 0.18907745 | 0.06684542 | 0.45625967 | 0.22331467 | 0.11673966 | 0.28562958 | 0.3540559 |
| ENSMUSG00 | 2610305D13I | 5687 | 0.73246127 | 1.99748495 | 2.13500196 | 1.81058485 | 1.58267447 | 2.78826706 | 0.84026386 | 1.8871224 | 3.20830429 | 0.89542394 | 1.67174551 | 2.82660644 | 0.59854093 | 2.50718038 | 3.4413116 | 0.17505817 | 2.56092717 | 2.4181953 |
| ENSMUSG00 | Gm13237 | 1139 | 2.91080222 | 2.57377936 | 3.73737548 | 2.20005043 | 3.29612822 | 2.69733932 | 3.07387027 | 3.28608906 | 3.83298566 | 2.7321742 | 3.73307402 | 3.57729883 | 3.55155155 | 3.83296244 | 3.87055881 | 3.29453865 | 3.26396032 | 4.20174915 |
| ENSMUSG00 | Zfp600 | 4929 | 0 | 0.0135171 | 0 | 0 | 0 | 0.02010661 | 0 | 0 | 0.02506787 | 0 | 0 | 0.04529573 | 0 | 0.01884524 | 0.01671802 | 0 | 0.02332702 | 0 |
| ENSMUSG00 | Gm13051 | 1925 | 0 | 0 | 0 | 0.02366807 | 0 | 0 | 0 | 0 | 0 | 0 | 0 | 0 | 0 | 0.14476079 | 0 | 0 | 0 | 0 |
| ENSMUSG00 | Dhrs3 | 2920 | 26.9296492 | 26.3194681 | 22.0331491 | 25.0117651 | 21.7747504 | 22.8926841 | 12.9137689 | 10.3459556 | 13.1599381 | 15.4047161 | 13.0399459 | 10.2456148 | 25.7472061 | 21.3770046 | 25.9767344 | 33.5173947 | 26.1852611 | 32.0446885 |
| ENSMUSG00 | Gm2042 | 1889 | 0 | 0 | 0 | 0 | 0 | 0 | 0 | 0 | 0 | 0 | 0 | 0 | 0 | 0 | 0 | 0 | 0 | 0 |
| ENSMUSG00 | Oog2 | 2633 | 0 | 0 | 0 | 0 | 0 | 0 | 0 | 0 | 0 | 0 | 0 | 0 | 0 | 0 | 0 | 0 | 0 | 0 |
| ENSMUSG00 | Gm13023 | 1919 | 0 | 0 | 0 | 0 | 0 | 0 | 0 | 0 | 0 | 0 | 0 | 0 | 0 | 0 | 0 | 0 | 0 | 0 |
| ENSMUSG00 | Gm17372 | 2729 | 0 | 0.07324198 | 0.08862867 | 0 | 0 | 0 | 0.05201101 | 0.18193523 | 0 | 0.01727765 | 0.05251919 | 0.02045279 | 0.03615384 | 0 | 0.01509768 | 0 | 0.08426449 | 0 |
| ENSMUSG00 | Ubr4 | 17888 | 9.15781241 | 9.72122696 | 8.55082195 | 7.82443695 | 8.83907476 | 7.72322346 | 6.98528539 | 8.17524088 | 7.22974393 | 6.66615209 | 7.44881257 | 7.62597716 | 7.15378988 | 6.01841428 | 6.45847287 | 7.67182116 | 7.34044573 | 6.11656492 |
| ENSMUSG00 | Hnrnpr | 3264 | 30.6938419 | 28.6384746 | 29.8777687 | 30.2065257 | 27.2216701 | 29.1486668 | 24.7144968 | 23.7415394 | 25.1737435 | 27.5190226 | 24.7656954 | 29.2074607 | 20.3735982 | 19.0955776 | 25.9529408 | 22.4534996 | 23.2376659 | 26.2742639 |
| ENSMUSG00 | Gm10151 | 3199 | 1.03638754 | 0.83308269 | 1.19459484 | 1.182109 | 1.11339867 | 0.54215245 | 0.29579668 | 1.11031367 | 0.46349314 | 0.89909125 | 0.25388381 | 1.16900575 | 0.47805234 | 0.98724537 | 0.56669849 | 1.48422486 | 0.8146884 | 0.58465446 |
| ENSMUSG00 | Med18 | 2153 | 9.16042809 | 9.4229227 | 9.03212225 | 9.33229008 | 8.36104944 | 10.541176 | 8.26268801 | 9.61462244 | 9.46926396 | 11.3880008 | 7.85524166 | 8.84028841 | 11.0670302 | 8.82286658 | 8.99430156 | 7.04276848 | 7.9394001 | 11.1142413 |
| ENSMUSG00 | Phactr4 | 7298 | 5.15443694 | 4.0716774 | 5.26289163 | 5.2752922 | 4.61665478 | 4.85478275 | 3.72122194 | 3.95111864 | 4.63898547 | 4.10258928 | 3.92123871 | 4.48177285 | 3.92735598 | 3.11833418 | 3.52285159 | 3.5730205 | 3.94396396 | 3.33913871 |
| ENSMUSG00 | 2610044O15I | 387 | 6.37028478 | 10.5017393 | 9.24971898 | 9.53603254 | 5.8454791 | 5.76194576 | 6.47951417 | 5.82261432 | 7.02405695 | 10.1124247 | 8.51801585 | 8.50936458 | 9.05055847 | 6.48057033 | 7.66540575 | 5.04604904 | 7.03143948 | 6.25426536 |
| ENSMUSG00 | Cldn19 | 1962 | 0 | 0 | 0 | 0 | 0 | 0 | 0 | 0.05839817 | 0 | 0.02403196 | 0 | 0 | 0 | 0.16570266 | 0 | 0 | 0 | 0 |
| ENSMUSG00 | Gm12866 | 1669 | 0 | 0 | 0.11593417 | 0.21838727 | 0.05767756 | 0.1187603 | 0.25513128 | 0.13730043 | 0.4441925 | 0.33901045 | 0.0572498 | 0.33442578 | 0 | 0.72351484 | 0.44435471 | 0.9635743 | 0.36741806 | 0 |
| ENSMUSG00 | Olfr1335 | 2106 | 0 | 0 | 0 | 0 | 0 | 0 | 0 | 0 | 0 | 0 | 0 | 0 | 0 | 0 | 0 | 0 | 0 | 0 |
| ENSMUSG00 | Olfr1338 | 941 | 0.04517022 | 0 | 0 | 0.14525306 | 0 | 0.15797896 | 0.25139574 | 0.68997964 | 0.13130664 | 0.3006421 | 0 | 1.06767474 | 0 | 0.19742439 | 0 | 0 | 0 | 0 |
| ENSMUSG00 | Gm13611 | 700 | 0.12144336 | 0.04758985 | 0.4837353 | 0.19526162 | 0 | 0 | 0.13517908 | 0 | 0 | 0.06735815 | 0 | 0 | 0.2114225 | 0.19904609 | 0 | 0.10940174 | 0.10950371 | 0.07858444 |
| ENSMUSG00 | Cyp4a12a | 2415 | 0.36961023 | 0.22070655 | 0.06009134 | 0.01886586 | 0.0199304 | 0.04103746 | 5.21125159 | 6.46819569 | 5.54269881 | 2.38194029 | 0.77152107 | 3.8828287 | 23.756944 | 35.2898138 | 21.871799 | 4.02725229 | 6.3797811 | 13.4390778 |
| ENSMUSG00 | Cyp4a10 | 2318 | 0 | 0 | 0 | 0 | 0 | 0 | 0.20410992 | 0 | 0 | 0 | 0 | 0 | 0 | 0 | 0 | 0 | 0 | 0 |
| ENSMUSG00 | Insl5 | 953 | 0 | 0 | 0 | 0 | 0 | 0 | 0 | 0 | 0 | 0 | 0 | 0 | 0 | 0 | 0 | 0 | 0 | 0 |
| ENSMUSG00 | Gm10152 | 1343 | 0 | 0 | 0.03601901 | 0.06784966 | 0 | 0.11069114 | 0.24660369 | 0 | 0 | 0 | 0 | 0.33248347 | 0.22039576 | 0.48415281 | 0.46018119 | 0.14255624 | 0.11415129 | 1.43359548 |
| ENSMUSG00 | Cyp2j11 | 1908 | 0 | 0 | 0.025353 | 0 | 0 | 0 | 1.01667706 | 0.22018681 | 0 | 0.19769687 | 0 | 0 | 0.82736968 | 0.77893648 | 0.66941731 | 0.08027381 | 0 | 0.20181538 |
| ENSMUSG00 | Krtap5-1 | 564 | 0 | 0 | 0 | 0 | 0 | 0 | 0 | 0 | 0 | 0 | 0 | 0 | 0 | 0 | 0 | 0 | 0 | 0 |
| ENSMUSG00 | Gm10153 | 559 | 0 | 0 | 0 | 0 | 0 | 0 | 0 | 0 | 0 | 0 | 0 | 0 | 0 | 0 | 0 | 0 | 0 | 0 |
| ENSMUSG00 | Gm12686 | 760 | 0 | 0 | 0 | 0 | 0 | 0 | 0 | 0 | 0 | 0 | 0 | 0 | 0 | 0 | 0 | 0 | 0 | 0 |
| ENSMUSG00 | Muc5b | 15048 | 0.33330747 | 0.34313521 | 0.63006459 | 0.254328 | 0.54055656 | 0.29661 | 0.10375587 | 0.43400457 | 0.03558112 | 0.02193347 | 0 | 0 | 0.00983491 | 0 | 0 | 0.00254456 | 0.01018775 | 0 |
| ENSMUSG00 | Adamtsl1 | 10912 | 0 | 0.01221147 | 0 | 0 | 0 | 0.01816449 | 0.05203007 | 0.02450026 | 0.00377442 | 0.02592597 | 0 | 0 | 0.05877153 | 0.04681864 | 0 | 0.02105422 | 0.05970922 | 0.00504116 |
| ENSMUSG00 | Gm10154 | 407 | 0 | 0 | 0 | 0 | 0 | 0 | 0 | 0 | 0 | 0 | 0 | 0 | 0 | 0 | 0 | 0 | 0 | 0 |
| ENSMUSG00 | Olfr45 | 1034 | 0 | 0 | 0 | 0 | 0 | 0 | 0 | 0 | 0 | 0 | 0 | 0 | 0 | 0 | 0 | 0 | 0 | 0 |
| ENSMUSG00 | Olfr53 | 1001 | 0 | 0 | 0 | 0 | 0 | 0 | 0 | 0 | 0 | 0 | 0 | 0 | 0 | 0 | 0 | 0 | 0 | 0 |
| ENSMUSG00 | Olfr61 | 1095 | 0 | 0 | 0 | 0 | 0 | 0 | 0 | 0 | 0 | 0 | 0 | 0 | 0 | 0 | 0 | 0 | 0 | 0 |
| ENSMUSG00 | Olfr538 | 956 | 0 | 0 | 0 | 0 | 0 | 0 | 0 | 0 | 0 | 0 | 0 | 0 | 0 | 0 | 0 | 0 | 0 | 0 |
| ENSMUSG00 | Kndc1 | 8150 | 0.47459767 | 0.36378498 | 0.42141358 | 0.5981634 | 0.6673506 | 0.44992666 | 0.93464311 | 1.0778224 | 1.22296157 | 0.67110203 | 0.65653931 | 1.15740454 | 0.006053 | 0.35331698 | 0.18705001 | 0.06107704 | 0.0376209 | 0.0404975 |
| ENSMUSG00 | Gm11237 | 1858 | 0 | 0 | 0 | 0 | 0 | 0 | 0 | 0 | 0 | 0 | 0 | 0 | 0 | 0 | 0 | 0 | 0 | 0 |
| ENSMUSG00 | Gm11487 | 1317 | 0 | 0 | 0 | 0 | 0 | 0 | 0 | 0 | 0 | 0 | 0 | 0 | 0 | 0 | 0 | 0 | 0 | 0 |
| ENSMUSG00 | Gm11232 | 1677 | 0 | 0 | 0 | 0 | 0 | 0 | 0 | 0 | 0 | 0 | 0 | 0 | 0 | 0 | 0 | 0 | 0 | 0 |
| ENSMUSG00 | Prpf4 | 5543 | 4.4552602 | 4.53747702 | 6.23977525 | 5.52354714 | 4.94952129 | 5.39063654 | 6.07732769 | 5.7739912 | 4.77772499 | 5.25692488 | 6.67108386 | 5.80007607 | 6.33669946 | 4.37377117 | 5.52277756 | 4.49705672 | 4.90228117 | 3.2153979 |
| ENSMUSG00 | Cdc26 | 7175 | 6.01885141 | 5.29756269 | 4.63172339 | 4.84502813 | 5.21904343 | 5.11066521 | 5.45332202 | 4.65228707 | 7.00314275 | 5.07321852 | 4.84741042 | 5.51544317 | 6.13297146 | 6.00051136 | 5.37486499 | 6.0517838 | 5.08524525 | 5.08307146 |
| ENSMUSG00 | Slc31a1 | 4276 | 57.3263214 | 62.1072013 | 55.330902 | 61.7141181 | 56.2364571 | 60.8515912 | 51.7164316 | 60.6648263 | 50.5969989 | 57.5269455 | 53.9982166 | 46.286796 | 40.2985002 | 42.9032145 | 40.1416468 | 40.3860242 | 49.0820397 | 41.1539131 |
| ENSMUSG00 | Fkbp15 | 7485 | 8.73386249 | 8.80777791 | 9.48728692 | 8.93568645 | 9.15053138 | 8.26871934 | 8.00870597 | 9.55703002 | 9.1947452 | 10.4380381 | 8.38056386 | 8.74706376 | 8.89754019 | 8.93513505 | 8.27333848 | 7.30002629 | 9.24746723 | 9.28209777 |
| ENSMUSG00 | Slc31a2 | 2026 | 144.173529 | 125.737267 | 146.290533 | 143.99179 | 142.827801 | 135.548494 | 161.81123 | 133.729962 | 151.776173 | 155.020155 | 143.513523 | 153.479297 | 135.358405 | 129.956333 | 136.233575 | 146.35857 | 139.287524 | 141.459746 |
| ENSMUSG00 | Mup21 | 671 | 0 | 0 | 0 | 0 | 0 | 0 | 0 | 0 | 0 | 0 | 0 | 0 | 0 | 0 | 0 | 0 | 0 | 0 |
| ENSMUSG00 | Mup3 | 962 | 0 | 0 | 0 | 0 | 0 | 0 | 0 | 0 | 0 | 0 | 0 | 0 | 0 | 0 | 0 | 0 | 0 | 0 |
| ENSMUSG00 | AY512931 | 2776 | 0 | 0 | 0 | 0 | 0 | 0 | 0 | 0 | 0 | 0.01698512 | 0 | 0 | 0 | 0 | 0 | 0.02758689 | 0 | 0 |
| ENSMUSG00 | E230001N04 | 2054 | 1.17954968 | 1.47588772 | 1.27174812 | 1.1978074 | 1.33569603 | 1.18212461 | 1.49723667 | 1.91519837 | 2.96767494 | 2.11191076 | 2.37246629 | 1.5489254 | 4.53930495 | 3.45956445 | 2.36698325 | 1.1371602 | 2.1458248 | 2.89239704 |
| ENSMUSG00 | 2510046G10 | 347 | 5.51219862 | 4.41611854 | 4.87917453 | 4.33289471 | 4.29997013 | 2.9988686 | 9.13530112 | 10.4561332 | 8.1898259 | 6.11464451 | 8.81151972 | 7.72089857 | 5.97101016 | 6.95992088 | 6.88671539 | 6.06911645 | 6.51657498 | 6.65816277 |
| ENSMUSG00 | Gm12511 | 1601 | 0 | 0.04161511 | 0 | 0 | 0 | 0 | 0 | 0 | 0 | 0 | 0.14920349 | 0.209178 | 0 | 0 | 0 | 0 | 0 | 0 |
| ENSMUSG00 | 6030445D17 | 1727 | 0.19689717 | 0.01928946 | 0 | 0.36934257 | 0 | 0.02869296 | 0.05479175 | 0.04422977 | 0 | 0.13651043 | 0 | 0 | 0.08569528 | 0 | 0.0238573 | 0 | 0.02219241 | 0 |
| ENSMUSG00 | Gm10155 | 540 | 0 | 0 | 0 | 0 | 0 | 0.27529296 | 0 | 0 | 0 | 0.17463223 | 0 | 0 | 0.1827108 | 0 | 0 | 0.07090853 | 0 | 0 |
| ENSMUSG00 | Cacng3 | 1519 | 0 | 0 | 0 | 0 | 0 | 0 | 0 | 0 | 0 | 0 | 0 | 0 | 0 | 0 | 0 | 0 | 0 | 0 |
| ENSMUSG00 | Anks6 | 6347 | 0.12054406 | 0.41463977 | 0.56398948 | 0.79679784 | 0.5156721 | 0.43720704 | 0.27581048 | 0.8665048 | 0.37636961 | 1.35947356 | 0.5043205 | 0.87940228 | 0.57516337 | 0.29269946 | 0.85687811 | 0.40420209 | 0.90577352 | 0.95336407 |
| ENSMUSG00 | Spag8 | 1655 | 0.3595604 | 0.96617458 | 0.26305847 | 0.66070397 | 0.90156473 | 0.2095886 | 1.40079835 | 0.60000149 | 0.32351946 | 0.71224627 | 0.31753749 | 0.43843119 | 0 | 0.58932075 | 1.6679784 | 0.50899901 | 0.23157883 | 0.06647626 |
| ENSMUSG00 | Gpr139 | 1036 | 0 | 0 | 0 | 0 | 0 | 0 | 0 | 0 | 0 | 0 | 0 | 0 | 0 | 0 | 0 | 0 | 0 | 0 |
| ENSMUSG00 | Il11ra2 | 2666 | 0 | 0 | 0 | 0 | 0 | 0 | 0 | 0 | 0 | 0 | 0 | 0 | 0 | 0 | 0 | 0 | 0 | 0 |
| ENSMUSG00 | Ccl27b | 927 | 0 | 0 | 0 | 0 | 0 | 0 | 0 | 0 | 0 | 0 | 0 | 0 | 0 | 0 | 0 | 0 | 0 | 0 |
| ENSMUSG00 | Arid3c | 1937 | 0 | 0 | 0 | 0 | 0 | 0 | 0 | 0 | 0 | 0.07302639 | 0 | 0 | 0 | 0 | 0.06381243 | 0 | 0 | 0.1135965 |
| ENSMUSG00 | Ipo7 | 3092 | 13.4443927 | 12.0667663 | 12.6565932 | 12.9669208 | 9.72912429 | 10.6092851 | 8.47711 | 10.8697652 | 8.68486684 | 8.2650974 | 9.98144346 | 10.2172267 | 7.1796128 | 5.81302129 | 7.475444 | 7.82654071 | 11.614405 | 7.20526786 |
| ENSMUSG00 | Tmem42 | 1041 | 25.0702959 | 25.408682 | 23.8382527 | 25.2971024 | 27.8342152 | 16.7555833 | 31.8144815 | 30.5979477 | 33.115383 | 32.0226198 | 36.1180522 | 36.1380947 | 25.5900435 | 28.7765959 | 27.0719157 | 26.2995046 | 26.3608344 | 28.3236131 |
| ENSMUSG00 | C85492 | 4176 | 7.3691924 | 6.3817805 | 6.15094459 | 7.63714824 | 7.43416924 | 6.44327929 | 5.59685426 | 6.25565212 | 5.02009974 | 7.40681545 | 5.73162217 | 4.75823177 | 6.29643484 | 4.14838233 | 5.76190341 | 7.31702699 | 4.37778822 | 4.87389116 |
| ENSMUSG00 | Olfr519 | 959 | 0 | 0 | 0 | 0 | 0 | 0 | 0 | 0 | 0 | 0 | 0 | 0 | 0 | 0 | 0 | 0 | 0 | 0 |
| ENSMUSG00 | Olfr517 | 1054 | 0 | 0 | 0 | 0 | 0 | 0 | 0 | 0 | 0 | 0 | 0 | 0 | 0 | 0 | 0 | 0 | 0 | 0 |
| ENSMUSG00 | Olfr514 | 932 | 0 | 0 | 0 | 0 | 0 | 0 | 0 | 0 | 0 | 0 | 0 | 0 | 0 | 0 | 0 | 0 | 0 | 0 |
| ENSMUSG00 | Olfr467 | 926 | 0 | 0 | 0 | 0 | 0 | 0 | 0 | 0 | 0 | 0 | 0 | 0 | 0 | 0 | 0 | 0 | 0 | 0 |
| ENSMUSG00 | Olfr470 | 944 | 0 | 0 | 0 | 0 | 0 | 0 | 0 | 0 | 0 | 0 | 0 | 0 | 0 | 0 | 0 | 0 | 0 | 0 |
| ENSMUSG00 | Gm10156 | 1591 | 0.2137281 | 0.10469169 | 0.30404482 | 0.20045714 | 0.18151574 | 0.15572826 | 0.20816389 | 0.02400528 | 0.28475907 | 0.23708713 | 0.09008477 | 0.07016425 | 0.09302059 | 0.08757527 | 0.15537985 | 0.19253605 | 0.28907326 | 0.24202624 |
| ENSMUSG00 | Bhmt-ps1 | 1871 | 0 | 0 | 0 | 0 | 0 | 0 | 0 | 0 | 0 | 0 | 0 | 0 | 0 | 0 | 0 | 0 | 0 | 0 |
| ENSMUSG00 | Olfr206 | 1004 | 0 | 0 | 0 | 0 | 0 | 0 | 0 | 0 | 0 | 0 | 0 | 0 | 0 | 0 | 0 | 0 | 0 | 0 |
| ENSMUSG00 | Trim12a | 6000 | 3.77587647 | 4.03641235 | 3.2652133 | 3.46263937 | 3.56176235 | 4.04680658 | 3.020126 | 3.1572386 | 4.21475345 | 3.42628444 | 3.19295972 | 2.56752048 | 3.65878386 | 4.32704081 | 4.42916726 | 5.20114084 | 4.03703661 | 2.72295078 |
| ENSMUSG00 | Olfr640 | 944 | 0 | 0 | 0 | 0 | 0 | 0 | 0 | 0 | 0 | 0 | 0 | 0 | 0 | 0 | 0 | 0 | 0 | 0 |
| ENSMUSG00 | Olfr639 | 920 | 0 | 0 | 0 | 0 | 0 | 0 | 0 | 0 | 0 | 0 | 0 | 0 | 0 | 0 | 0 | 0 | 0 | 0 |
| ENSMUSG00 | Olfr612 | 971 | 0 | 0 | 0 | 0 | 0 | 0 | 0 | 0 | 0 | 0 | 0 | 0 | 0 | 0 | 0 | 0 | 0 | 0 |
| ENSMUSG00 | Olfr586 | 953 | 0 | 0 | 0 | 0 | 0 | 0 | 0 | 0 | 0 | 0 | 0 | 0 | 0 | 0 | 0 | 0 | 0 | 0 |
| ENSMUSG00 | Olfr575 | 954 | 0 | 0 | 0 | 0 | 0 | 0 | 0 | 0 | 0 | 0 | 0 | 0 | 0 | 0 | 0 | 0 | 0 | 0 |
| ENSMUSG00 | Gm10157 | 680 | 0.25003045 | 0.34292685 | 0.21341263 | 0 | 0.07078224 | 0.14574333 | 0.06957747 | 0.22466119 | 0.18170522 | 0.06933927 | 0.07025729 | 0.16416371 | 0.14509387 | 0.20490039 | 0.24236209 | 0.16892915 | 0.22544881 | 0.32358298 |
| ENSMUSG00 | Olfr559 | 1021 | 0 | 0 | 0 | 0 | 0 | 0 | 0 | 0.26184801 | 0 | 0 | 0 | 0 | 0 | 0 | 0 | 0 | 0 | 0 |
| ENSMUSG00 | Olfr33 | 917 | 0 | 0 | 0 | 0 | 0 | 0 | 0 | 0 | 0 | 0 | 0 | 0 | 0 | 0 | 0 | 0 | 0 | 0 |
| ENSMUSG00 | Vps37b | 2508 | 42.301802 | 44.8289545 | 43.7637721 | 51.7194152 | 39.1887512 | 47.7547676 | 52.5382021 | 55.4155312 | 55.3751719 | 62.6232028 | 59.2996195 | 62.1806066 | 54.60343 | 53.1292212 | 47.3949317 | 57.3290394 | 55.7931858 | 71.5030654 |
| ENSMUSG00 | Chrna10 | 2508 | 0 | 0 | 0 | 0 | 0 | 0 | 0 | 0 | 0 | 0 | 0 | 0 | 0 | 0.14814701 | 0 | 0 | 0 | 0 |
| ENSMUSG00 | Numa1 | 7308 | 25.7078377 | 24.8570351 | 24.3787236 | 24.632141 | 24.9682707 | 26.532546 | 24.4008603 | 23.2561837 | 21.4048148 | 29.0530401 | 17.5985479 | 24.1730394 | 18.8471145 | 23.2601751 | 19.2758784 | 20.9843846 | 23.8883802 | 22.3182815 |
| ENSMUSG00 | Rad54b | 1601 | 0.34513884 | 0.64503418 | 0.42300401 | 0.45532586 | 0.39082761 | 0.58807117 | 0.17731173 | 0.54867286 | 0.12862747 | 0.44176174 | 0.59681397 | 0 | 0 | 0.11603769 | 0.30881866 | 0.19133345 | 0.4548405 | 0.03435922 |
| ENSMUSG00 | Gm12918 | 1323 | 0 | 0.02517981 | 0 | 0 | 0 | 0 | 0 | 0 | 0 | 0.10691769 | 0 | 0 | 0 | 0 | 0 | 0 | 0 | 0 |
| ENSMUSG00 | Rtp3 | 2450 | 0 | 0 | 0 | 0 | 0 | 0 | 0 | 0 | 0 | 0 | 0 | 0 | 0 | 0 | 0 | 0 | 0 | 0 |
| ENSMUSG00 | Impad1 | 6538 | 14.2767462 | 13.0948514 | 13.1329178 | 13.5052469 | 13.3323514 | 15.7268159 | 12.2297992 | 13.0676686 | 12.8762983 | 13.212005 | 12.1373824 | 11.8409795 | 9.2129506 | 13.5894064 | 10.7761793 | 12.5448885 | 13.7641703 | 12.082147 |
| ENSMUSG00 | Wdr6 | 5246 | 31.8505236 | 42.8381213 | 37.981428 | 44.3364719 | 41.9479899 | 43.0634602 | 27.7869546 | 29.8127884 | 26.677808 | 36.8685064 | 33.386008 | 29.3229062 | 24.0265057 | 24.4526438 | 31.9732242 | 25.7144129 | 25.1027747 | 27.053659 |
| ENSMUSG00 | Tcl1b1 | 1033 | 0 | 0 | 0 | 0 | 0 | 0 | 0 | 0 | 0 | 0 | 0 | 0 | 0 | 0 | 0 | 0 | 0 | 0 |
| ENSMUSG00 | Serpina3c | 2208 | 0.30800852 | 0.24139778 | 0.35053283 | 0 | 0.34878206 | 0.17953889 | 0.42855687 | 0.25945926 | 0.31710633 | 0.19219037 | 0.51929303 | 0.05055767 | 0 | 0 | 0 | 0 | 0 | 0 |
| ENSMUSG00 | Rps13-ps1 | 455 | 1.02759766 | 0.65893637 | 0.42526181 | 0.20026833 | 0.63470669 | 0.54453553 | 0.31195173 | 0.41969673 | 0.45259907 | 0.5181396 | 0.41999963 | 0.49068714 | 0.21684359 | 0.51037459 | 0.63387009 | 0.42077591 | 0.33693448 | 0.48359654 |
| ENSMUSG00 | Serpina3f | 2283 | 2.5320648 | 4.1732316 | 3.8986989 | 4.19089764 | 4.44846075 | 4.73171087 | 3.85464664 | 8.33106281 | 5.03330613 | 5.10128064 | 3.70397181 | 4.47405443 | 3.75985703 | 6.08269038 | 3.80794056 | 4.69617612 | 3.76044262 | 6.04786938 |
| ENSMUSG00 | Serpina3b | 2065 | 0 | 0 | 0 | 0 | 0 | 0 | 0 | 0 | 0 | 0 | 0 | 0 | 0 | 0 | 0 | 0 | 0 | 0 |
| ENSMUSG00 | Serpina1a | 2531 | 0 | 0 | 0 | 0 | 0 | 0 | 0 | 0 | 0 | 0 | 0 | 0 | 0 | 0 | 0 | 0 | 0 | 0 |
| ENSMUSG00 | 4921517D21I | 3992 | 0 | 0 | 0 | 0 | 0 | 0 | 0 | 0 | 0 | 0 | 0.02393535 | 0 | 0 | 0 | 0 | 0 | 0 | 0 |
| ENSMUSG00 | Vmn2r65 | 2532 | 0 | 0 | 0 | 0 | 0 | 0 | 0 | 0 | 0 | 0 | 0 | 0 | 0 | 0 | 0 | 0 | 0 | 0 |
| ENSMUSG00 | Gm10160 | 204 | 0 | 0 | 0 | 0 | 0 | 0 | 0 | 0.18721766 | 0 | 0 | 0 | 0 | 0 | 0 | 0 | 0 | 0 | 0 |
| ENSMUSG00 | Iqcf5 | 542 | 0 | 0 | 0 | 0 | 0 | 0 | 0 | 0 | 0 | 0 | 0 | 0 | 0 | 0 | 0 | 0 | 0 | 0 |
| ENSMUSG00 | Iqcf1 | 631 | 0 | 0 | 0 | 0 | 0 | 0 | 0.22494142 | 0 | 0.06527182 | 0 | 0 | 0 | 0 | 0 | 0 | 0 | 0 | 0 |
| ENSMUSG00 | Rpl31-ps1 | 377 | 0.11274583 | 0.17672623 | 0.2566235 | 0 | 0.25534177 | 0.26287922 | 0.12549782 | 0.10130611 | 0.10924805 | 0.12506818 | 0 | 0 | 0 | 0 | 0 | 0 | 0.20332253 | 0.2918255 |
| ENSMUSG00 | Nrxn3 | 11016 | 0 | 0 | 0 | 0 | 0 | 0 | 0.03435924 | 0.01733497 | 0 | 0.0042802 | 0 | 0 | 0.02686924 | 0.02529634 | 0.02618109 | 0.03475908 | 0 | 0.09987129 |
| ENSMUSG00 | Akap13 | 15343 | 15.6246622 | 18.3554199 | 14.8844057 | 16.4183676 | 15.6256998 | 17.5596828 | 14.6936658 | 16.3294114 | 15.2526744 | 15.4946129 | 15.1579546 | 16.501326 | 14.6101957 | 14.1333145 | 14.9225736 | 19.7804441 | 17.9004265 | 17.101834 |
| ENSMUSG00 | Gm10263 | 209 | 0 | 0.15939184 | 0.2314523 | 0 | 0 | 0 | 0 | 0 | 0.19706467 | 0 | 0 | 0 | 0 | 0 | 0 | 0 | 0 | 0 |
| ENSMUSG00 | Msl2 | 4906 | 4.34928625 | 4.0809314 | 4.71311608 | 3.9004563 | 4.17941286 | 4.47449266 | 4.45545407 | 4.18824144 | 4.65929805 | 4.60358477 | 3.13565769 | 4.59631628 | 4.22327869 | 4.53459394 | 3.4012699 | 3.50437887 | 4.21073666 | 2.5340518 |
| ENSMUSG00 | Gm17627 | 1872 | 0 | 0.21354419 | 0.72353571 | 0.04867633 | 0.051423 | 0.07941143 | 0.379108 | 0.30602887 | 0.50603091 | 0.05037468 | 0.07656243 | 0.08944818 | 0 | 0.49619751 | 0 | 0.32727015 | 0.0409469 | 0.20569645 |
| ENSMUSG00 | Vmn2r68-ps | 2550 | 0 | 0 | 0 | 0 | 0 | 0 | 0 | 0 | 0 | 0 | 0 | 0 | 0 | 0 | 0 | 0 | 0 | 0 |
| ENSMUSG00 | 3830431G21I | 3925 | 52.8147117 | 59.1823723 | 54.9301464 | 58.8637085 | 59.5732192 | 56.1681813 | 36.4880202 | 30.4858081 | 29.8011987 | 34.0926614 | 32.2191886 | 30.2328914 | 26.494696 | 29.6295528 | 29.654623 | 28.8959947 | 27.819521 | 29.7259402 |
| ENSMUSG00 | Zfyve26 | 9326 | 1.63621683 | 2.33611761 | 1.78431208 | 1.93949545 | 1.74443386 | 2.17317907 | 1.68430296 | 2.50220436 | 1.87251584 | 1.83021173 | 1.83395187 | 2.31617643 | 2.07356976 | 2.32071035 | 2.06317034 | 2.41009935 | 2.06713782 | 2.73688886 |
| ENSMUSG00 | Rdh11 | 4147 | 32.3390503 | 36.4297023 | 30.2699087 | 30.015137 | 32.0802115 | 35.1182744 | 29.3094459 | 25.1791725 | 27.0637221 | 31.3011541 | 28.2594582 | 24.9804364 | 28.1692742 | 23.7876155 | 29.0209175 | 32.464378 | 30.1009764 | 27.7366873 |
| ENSMUSG00 | Mthfs | 3056 | 2.80956988 | 2.27827058 | 3.2291231 | 2.59411705 | 2.97674528 | 3.34026934 | 4.84583393 | 2.79944311 | 5.05397363 | 4.3663773 | 4.76811595 | 4.58433616 | 3.98723284 | 4.34653436 | 3.7480473 | 4.02201734 | 4.06339012 | 4.15808366 |
| ENSMUSG00 | Gm10163 | 1029 | 0.08261453 | 0.12949619 | 0.23505117 | 0.17710804 | 0.04677544 | 0.09631241 | 0.09195856 | 0.07423207 | 0.08005154 | 0.09164374 | 0.13928559 | 0 | 0 | 0.09027034 | 0.12012115 | 0.18605737 | 0 | 0.1603764 |
| ENSMUSG00 | Gm8226 | 851 | 0 | 0 | 0 | 0 | 0 | 0 | 0 | 0 | 0 | 0 | 0 | 0 | 0 | 0 | 0.04841546 | 0 | 0 | 0 |
| ENSMUSG00 | Hmgn3 | 2043 | 162.197921 | 134.132094 | 135.981001 | 135.880295 | 141.191688 | 141.017911 | 163.47538 | 149.161615 | 160.835058 | 161.992553 | 152.14091 | 147.17522 | 145.725952 | 116.189785 | 126.428073 | 151.119514 | 122.876772 | 124.34266 |
| ENSMUSG00 | Gm5068 | 305 | 0 | 0 | 0 | 0 | 0.15780959 | 0 | 0 | 0 | 0 | 0 | 0 | 0 | 0 | 0 | 0 | 0.12554298 | 0 | 0.54107318 |
| ENSMUSG00 | Omt2a | 1751 | 0 | 0 | 0 | 0 | 0 | 0 | 0 | 0 | 0 | 0 | 0 | 0 | 0 | 0 | 0 | 0 | 0 | 0 |
| ENSMUSG00 | Gm10268 | 603 | 0.35244756 | 0.11049053 | 0.08022144 | 0.68001559 | 0.07982077 | 0 | 0.23538646 | 0.19001195 | 0.3415134 | 0.15638708 | 0.07922879 | 0.27768986 | 0.16362162 | 0.0770217 | 0.20498287 | 0 | 0.57203428 | 0.09122572 |
| ENSMUSG00 | Gm16551 | 4217 | 0 | 0 | 0 | 0 | 0 | 0 | 0 | 0 | 0 | 0 | 0 | 0 | 0 | 0 | 0 | 0 | 0 | 0 |
| ENSMUSG00 | Gm5745 | 314 | 0 | 0 | 0 | 0 | 0 | 0 | 0 | 0 | 0 | 0 | 0 | 0.17775689 | 0 | 0 | 0 | 0 | 0 | 0 |
| ENSMUSG00 | Gm5786 | 706 | 0 | 0.18874161 | 0.06851775 | 0.06453406 | 0.13635106 | 0 | 0.33507563 | 0.16229066 | 0.11667568 | 0 | 0.27067965 | 0 | 0.06987524 | 0.06578483 | 0.05835914 | 0.21694395 | 0.05428654 | 0 |
| ENSMUSG00 | Rpl26-ps2 | 437 | 0 | 0.15246176 | 0 | 0 | 0.2202834 | 0 | 0 | 0 | 0 | 0 | 0 | 0 | 0 | 0 | 0.18856547 | 0 | 0 | 0 |
| ENSMUSG00 | Gm6265 | 230 | 0.73922045 | 1.88290272 | 3.36511516 | 0.39618299 | 1.8834231 | 1.07723334 | 2.2627803 | 2.49080885 | 2.14886169 | 2.46003669 | 1.8694549 | 1.45606076 | 1.50140618 | 1.81737733 | 2.86619519 | 2.16425172 | 1.66636073 | 1.6741902 |
| ENSMUSG00 | Izumo2 | 713 | 0 | 0 | 0 | 0 | 0 | 0 | 0 | 0 | 0 | 0 | 0 | 0 | 0 | 0 | 0 | 0 | 0 | 0 |
| ENSMUSG00 | Ankdd1a | 1429 | 0 | 0 | 0 | 0.06376633 | 0 | 0 | 0 | 0 | 0 | 0 | 0 | 0 | 0 | 0 | 0 | 0.18756771 | 0 | 0 |
| ENSMUSG00 | Klk1b5 | 857 | 4.41418981 | 3.84828066 | 6.20897122 | 3.66827545 | 5.56016388 | 4.45222927 | 5.68635464 | 4.76847965 | 6.34378072 | 3.85128265 | 3.7907785 | 4.03800593 | 5.41096876 | 4.33550407 | 3.31727813 | 2.81482878 | 3.53299586 | 3.20939946 |
| ENSMUSG00 | Klk1b4 | 834 | 1.98765212 | 1.7575148 | 2.37807523 | 2.56758883 | 2.71247052 | 2.73312439 | 1.30478611 | 4.25886502 | 4.64212528 | 1.58299723 | 1.66123956 | 1.87390714 | 1.83368038 | 1.78202733 | 1.33386332 | 1.19371198 | 1.1948246 | 2.83620095 |
| ENSMUSG00 | Klk1b3 | 883 | 4.38048813 | 4.64041448 | 5.20440021 | 6.08856538 | 5.0693872 | 4.88231916 | 4.76877511 | 3.20072229 | 3.40500074 | 3.68448304 | 3.08400072 | 3.35020398 | 1.89953022 | 2.62990302 | 2.70633096 | 1.30092664 | 2.43065983 | 1.86893907 |
| ENSMUSG00 | Klk1b21 | 893 | 2.0467218 | 2.87244441 | 3.46685996 | 2.60203053 | 2.1020661 | 2.10862696 | 0 | 0.9836789 | 1.38364555 | 0.47520306 | 1.60498181 | 1.500085 | 0.05524291 | 0.41607245 | 0.32296852 | 0.04287862 | 0 | 0.80080447 |
| ENSMUSG00 | Hmg1l1 | 1280 | 1.42790826 | 1.63961901 | 1.2849219 | 1.3525935 | 1.50412261 | 1.47109678 | 1.25674303 | 1.2531882 | 1.57667131 | 1.3997865 | 1.30634652 | 1.9186634 | 1.54162241 | 1.26995551 | 1.54505835 | 1.8547013 | 1.79654517 | 1.3322518 |
| ENSMUSG00 | Vmn2r57 | 2619 | 0 | 0 | 0 | 0 | 0 | 0 | 0 | 0 | 0 | 0 | 0 | 0 | 0 | 0 | 0 | 0 | 0 | 0 |
| ENSMUSG00 | Gm6254 | 1523 | 0 | 0.02187321 | 0.06352401 | 0.23932262 | 0.03160336 | 0.03253627 | 0.09319635 | 0 | 0.08112965 | 0.0309591 | 0 | 0.073297 | 0.03239128 | 0.15247566 | 0.02705289 | 0.12570784 | 0.10066 | 0 |
| ENSMUSG00 | Gm10166 | 1157 | 0.11021221 | 0 | 0.04180945 | 0.11813581 | 0.12480188 | 0.08565727 | 0.12267765 | 0.09902957 | 0 | 0.04075255 | 0.0412921 | 0 | 0.08527557 | 0 | 0.14244272 | 0.13237894 | 0 | 0.14263381 |
| ENSMUSG00 | Hmgb1 | 5623 | 47.1162657 | 43.5858017 | 46.0851862 | 42.6116899 | 46.8308295 | 42.6789772 | 42.5923494 | 45.1204213 | 45.9841625 | 42.22013 | 41.1902897 | 43.0305704 | 44.1907998 | 41.3313557 | 44.1471411 | 44.4737608 | 43.2678877 | 40.2565311 |
| ENSMUSG00 | Gm6969 | 1372 | 7.31138596 | 5.85160897 | 6.69895829 | 7.47174561 | 7.85827325 | 7.40401631 | 6.44859399 | 5.67875371 | 7.44479294 | 8.59160045 | 7.24285085 | 6.75320699 | 7.62271608 | 9.27527693 | 8.25832935 | 7.89813549 | 5.97800841 | 6.97637359 |
| ENSMUSG00 | Gm10167 | 1478 | 0.66144726 | 0.42824424 | 0.49093569 | 0.70900137 | 0.61874597 | 0.63701078 | 0.25609028 | 0.41344955 | 0.78025876 | 0.41472202 | 0.35556464 | 0.4154075 | 0.43390726 | 0.53420128 | 0.696914 | 0.54404787 | 0.49269259 | 0.40940472 |
| ENSMUSG00 | Lsm14a | 3092 | 34.8207021 | 39.6587204 | 40.1913323 | 38.5323839 | 36.8617061 | 40.2896417 | 36.7545455 | 36.6360497 | 35.1657185 | 38.3061341 | 40.2811503 | 34.6230404 | 30.5851505 | 30.627262 | 29.1289137 | 36.5197287 | 37.570183 | 36.2398287 |
| ENSMUSG00 | 4931406P16I | 9677 | 3.19766127 | 3.83492448 | 3.23923196 | 3.64883842 | 3.59609185 | 3.51278033 | 2.71349972 | 3.21262949 | 3.0005677 | 2.68472021 | 3.01648234 | 2.28984376 | 2.36540349 | 1.86217897 | 2.52054574 | 2.56009331 | 3.62390841 | 2.33065348 |
| ENSMUSG00 | Abpa | 1063 | 0 | 0 | 0 | 0 | 0 | 0 | 0 | 0 | 0 | 0 | 0 | 0 | 0 | 0 | 0 | 0 | 0 | 0 |
| ENSMUSG00 | Abpb | 1805 | 0 | 0 | 0 | 0 | 0 | 0 | 0 | 0 | 0 | 0 | 0 | 0 | 0 | 0 | 0 | 0 | 0 | 0 |
| ENSMUSG00 | Abpg | 576 | 0 | 0 | 0 | 0 | 0 | 0 | 0 | 0 | 0 | 0 | 0 | 0 | 0 | 0 | 0 | 0 | 0 | 0 |
| ENSMUSG00 | Gm7589 | 536 | 0.1586014 | 0 | 0 | 0.1700039 | 0 | 0 | 0.08826992 | 0.07125448 | 0.15368103 | 0.08796773 | 0 | 0 | 0 | 0 | 0 | 0.2143131 | 0 | 0.20525786 |
| ENSMUSG00 | Mfsd7b | 4077 | 3.22151077 | 3.2356895 | 2.59843099 | 3.35254189 | 3.96672219 | 3.51256807 | 5.4774551 | 6.48249754 | 4.38433845 | 4.41784858 | 5.46066486 | 5.571983 | 2.09330922 | 3.7820547 | 5.04281982 | 6.13288365 | 4.72850606 | 5.77480934 |
| ENSMUSG00 | 6030419C18I | 877 | 0 | 0 | 0 | 0 | 0 | 0 | 0 | 0 | 0 | 0.16129089 | 0 | 0 | 0 | 0 | 0 | 0.0873218 | 0 | 0 |
| ENSMUSG00 | Zfp932 | 2985 | 5.06929403 | 4.14039657 | 5.42885517 | 5.90690927 | 4.77288087 | 4.99677481 | 7.84582113 | 6.94756265 | 6.27801362 | 5.13366116 | 6.14592431 | 6.65674234 | 6.61064218 | 4.02982199 | 5.56255512 | 5.68265966 | 6.50969264 | 6.24726539 |
| ENSMUSG00 | Gm4883 | 527 | 0 | 0 | 0 | 0 | 0 | 0 | 0 | 0 | 0 | 0 | 0 | 0 | 0.09360895 | 0 | 0 | 0 | 0 | 0 |
| ENSMUSG00 | Tecpr1 | 8858 | 6.77069354 | 8.19095548 | 6.05624804 | 7.31404437 | 6.8845278 | 7.91008865 | 7.00236194 | 7.23060046 | 6.91401546 | 8.47946154 | 6.13771763 | 5.37488826 | 4.2882791 | 5.3165844 | 6.47931445 | 6.7909849 | 6.7410669 | 6.27220567 |
| ENSMUSG00 | Gm5121 | 678 | 0 | 0.19653625 | 0 | 0 | 0 | 0.07308663 | 0 | 0 | 0 | 0 | 0 | 0 | 0.07276094 | 0 | 0 | 0.16942747 | 0 | 0 |
| ENSMUSG00 | Rpl36-ps3 | 383 | 0.22195914 | 0 | 0 | 0.11895834 | 0.37701246 | 0.12938051 | 0 | 0.19943813 | 0.10753659 | 0 | 0 | 0.4371984 | 0.38641188 | 0 | 0.10757586 | 0.09997548 | 0.10006866 | 0.28725382 |
| ENSMUSG00 | Pgk1-rs7 | 1253 | 0 | 0.02658651 | 0 | 0 | 0 | 0 | 0 | 0 | 0 | 0 | 0 | 0.04454562 | 0 | 0 | 0 | 0.03055914 | 0 | 0.04390192 |
| ENSMUSG00 | Ttc32 | 997 | 16.4990001 | 13.7996242 | 19.5046732 | 12.5669882 | 14.917517 | 13.9165149 | 21.4022249 | 20.6859552 | 23.2990922 | 17.8765956 | 18.7362174 | 23.5691013 | 22.3156416 | 16.490679 | 16.5715385 | 22.2753784 | 21.4504249 | 17.6558818 |
| ENSMUSG00 | 1700001J11R | 2288 | 0.05573231 | 0 | 0.02114228 | 0 | 0 | 0 | 0 | 0 | 0 | 0 | 0 | 0.02439496 | 0 | 0 | 0 | 0 | 0 | 0 |
| ENSMUSG00 | Fbxl18 | 11703 | 1.46732386 | 1.29232282 | 1.12015951 | 1.02388744 | 1.22149717 | 1.29989654 | 0.9257971 | 1.19116696 | 0.88686678 | 1.28523236 | 0.9634188 | 0.74878741 | 0.74189673 | 1.27787714 | 0.97168499 | 0.8997622 | 0.82527786 | 1.72505701 |
| ENSMUSG00 | Wdr35 | 7304 | 3.85246804 | 4.46057099 | 4.91417847 | 4.22924261 | 4.88304428 | 5.51565887 | 4.57321349 | 3.3256254 | 3.59761734 | 4.73830999 | 4.41512828 | 5.09707653 | 3.53239221 | 1.97756177 | 4.0784125 | 4.46653854 | 4.08240129 | 4.26275389 |
| ENSMUSG00 | Gm5113 | 2999 | 2.14014057 | 1.34406809 | 1.69363811 | 1.94458609 | 2.11851081 | 1.07400056 | 1.68804823 | 2.8908554 | 2.44454812 | 2.27971056 | 2.21431118 | 1.45169112 | 1.6284961 | 1.68803119 | 1.62113491 | 1.87686538 | 1.23963014 | 1.3023163 |
| ENSMUSG00 | Lefty2 | 2607 | 0 | 0 | 0 | 0 | 0 | 0 | 0.70778461 | 1.17199547 | 0.11058993 | 0.2351205 | 0.34818727 | 1.43446467 | 1.2489093 | 0.74823616 | 0.28447564 | 0 | 0.26462345 | 0.84402158 |
| ENSMUSG00 | Gm16432 | 3403 | 0.31226253 | 0.42093872 | 0.42644899 | 0.91041758 | 0.25459143 | 0.26210673 | 0.61174195 | 1.47023353 | 1.71862628 | 1.20543967 | 0.7721489 | 0.45925317 | 0.20295235 | 0.12283185 | 0.07264453 | 0.9676733 | 0.90100022 | 0 |
| ENSMUSG00 | Olfr220 | 977 | 0 | 0 | 0 | 0 | 0 | 0 | 0 | 0 | 0 | 0 | 0 | 0 | 0 | 0 | 0 | 0 | 0 | 0 |
| ENSMUSG00 | Olfr417 | 929 | 0 | 0 | 0 | 0 | 0 | 0 | 0 | 0 | 0 | 0 | 0 | 0 | 0 | 0 | 0 | 0 | 0 | 0 |
| ENSMUSG00 | Pydc3 | 2749 | 0 | 0 | 0 | 0 | 0 | 0 | 0 | 0 | 0 | 0 | 0 | 0 | 0 | 0 | 0 | 0.06964461 | 0 | 0 |
| ENSMUSG00 | Olfr1407-ps1 | 238 | 0 | 0 | 0 | 0 | 0 | 0 | 0 | 0 | 0 | 0 | 0 | 0 | 0 | 0 | 0 | 0 | 0 | 0 |
| ENSMUSG00 | Pilrb2 | 1824 | 0 | 0 | 0 | 0 | 0 | 0 | 0 | 0.04187763 | 0 | 0 | 0 | 0 | 0 | 0 | 0.11294286 | 0 | 0 | 0 |
| ENSMUSG00 | Pilrb1 | 1236 | 0 | 0 | 0 | 0 | 0 | 0 | 0 | 0.09270001 | 0 | 0 | 0 | 0 | 0 | 0 | 0 | 0 | 0 | 0 |
| ENSMUSG00 | Zbtb16 | 5093 | 7.14400759 | 5.95877609 | 5.88878634 | 5.15279041 | 5.69871392 | 5.40964458 | 3.81808581 | 5.51926334 | 5.98429642 | 4.74932471 | 2.81415424 | 5.44676699 | 3.61295997 | 4.82405699 | 4.71637681 | 5.87929609 | 5.71169435 | 3.43469388 |
| ENSMUSG00 | Gm13083 | 2461 | 0 | 0 | 0 | 0 | 0 | 0 | 0 | 0 | 0 | 0 | 0 | 0 | 0 | 0 | 0 | 0 | 0 | 0 |
| ENSMUSG00 | F830001A07I | 433 | 0 | 0 | 0 | 0 | 0 | 0 | 0 | 0 | 0 | 0 | 0 | 0 | 0 | 0 | 0 | 0 | 0 | 0 |
| ENSMUSG00 | Gm7493 | 625 | 0 | 0 | 0 | 0 | 0.07701108 | 0 | 0 | 0.06110784 | 0 | 0 | 0 | 0 | 0 | 0 | 0 | 0 | 0 | 0 |
| ENSMUSG00 | Gm10172 | 556 | 0 | 0 | 0 | 0 | 0 | 0 | 0 | 0 | 0 | 0 | 0 | 0 | 0 | 0 | 0 | 0 | 0 | 0 |
| ENSMUSG00 | Cyp2b19 | 4139 | 0 | 0.02414561 | 0.01168725 | 0.01100774 | 0 | 0.01197215 | 0 | 0 | 0 | 0 | 0 | 0 | 0 | 0 | 0.10949918 | 0 | 0.0092598 | 0 |
| ENSMUSG00 | Fxyd6 | 1783 | 1.74025679 | 3.0080628 | 2.17043322 | 3.75629474 | 2.24057749 | 2.36230082 | 1.35330714 | 1.79930555 | 2.35615514 | 2.40645765 | 3.08139104 | 1.00173932 | 1.79841538 | 0.65120706 | 0.32351194 | 0.68721225 | 1.87009974 | 0.37022394 |
| ENSMUSG00 | Gm8062 | 1024 | 0 | 0.13012849 | 0 | 0.13347962 | 0.42303449 | 0.04839134 | 0 | 0 | 0.08044241 | 0 | 0 | 0 | 0 | 0.04535555 | 0 | 0 | 0.03742802 | 0.05371983 |
| ENSMUSG00 | Cldn9 | 1433 | 80.6500863 | 82.6661911 | 66.8385137 | 71.1235563 | 77.2865013 | 90.4605381 | 85.0175494 | 101.997435 | 111.143235 | 89.3329793 | 109.785721 | 113.578836 | 93.7065449 | 100.407385 | 90.5974198 | 175.180197 | 128.298148 | 117.426977 |
| ENSMUSG00 | Zfp575 | 2737 | 0.4503654 | 0.24342634 | 0.12371747 | 0.39951226 | 0.2110278 | 0.25346667 | 0 | 0.02790822 | 0.0601922 | 0.05168144 | 0.17455228 | 0.12235805 | 0.07209634 | 0.32241054 | 0.01505355 | 0.12590993 | 0 | 0.44216308 |
| ENSMUSG00 | Vmn1r183 | 917 | 0 | 0 | 0 | 0 | 0 | 0 | 0 | 0 | 0 | 0 | 0 | 0 | 0 | 0 | 0 | 0 | 0 | 0 |
| ENSMUSG00 | Gm10175 | 440 | 0 | 0.15142225 | 0 | 0 | 0 | 0.33785955 | 0.10752882 | 0 | 0 | 0 | 0 | 0 | 0 | 0 | 0 | 0 | 0 | 0 |
| ENSMUSG00 | Vkorc1l1 | 3221 | 2.96916008 | 4.59202888 | 5.43657808 | 4.46981993 | 4.30363055 | 5.01527201 | 5.66988329 | 5.86936951 | 5.17868329 | 5.56264118 | 5.39897078 | 5.44120399 | 3.47666725 | 3.82107518 | 3.58163168 | 4.61246683 | 4.79524919 | 5.08932437 |
| ENSMUSG00 | Olfr878 | 1064 | 0 | 0 | 0 | 0 | 0 | 0 | 0 | 0 | 0 | 0 | 0 | 0 | 0.04636458 | 0 | 0 | 0 | 0 | 0 |
| ENSMUSG00 | Olfr145 | 1050 | 0 | 0 | 0 | 0 | 0 | 0 | 0 | 0 | 0 | 0 | 0 | 0 | 0 | 0 | 0 | 0 | 0 | 0 |
| ENSMUSG00 | Olfr877 | 935 | 0 | 0 | 0 | 0 | 0 | 0 | 0 | 0 | 0 | 0 | 0 | 0 | 0 | 0 | 0 | 0 | 0 | 0 |
| ENSMUSG00 | Olfr876 | 1047 | 0 | 0 | 0 | 0 | 0 | 0 | 0 | 0 | 0 | 0 | 0 | 0 | 0 | 0 | 0 | 0 | 0 | 0 |
| ENSMUSG00 | Gm10176 | 2263 | 0 | 0 | 0 | 0 | 0 | 0 | 0 | 0 | 0 | 0 | 0 | 0 | 0 | 0 | 0 | 0 | 0 | 0 |
| ENSMUSG00 | Tnfsf18 | 2063 | 0 | 0 | 0 | 0 | 0 | 0 | 0 | 0 | 0 | 0 | 0 | 0 | 0 | 0.45025775 | 0 | 0 | 0 | 0 |
| ENSMUSG00 | Igfl3 | 571 | 0 | 0 | 0 | 0 | 0 | 0 | 0 | 0 | 0 | 0 | 0 | 0 | 0 | 0 | 0 | 0 | 0 | 0 |
| ENSMUSG00 | Psg16 | 5736 | 0.02223074 | 0.0464615 | 0.00843332 | 0.12708799 | 0.0671296 | 0.0518334 | 0.197961 | 0.09321716 | 0.0789839 | 0.03288055 | 0.09161864 | 0.18488452 | 0.01720081 | 0 | 0.01436595 | 0.0600794 | 0.21381477 | 0 |
| ENSMUSG00 | Gm3867 | 877 | 0.24233282 | 0.03798506 | 0.11031592 | 0.05195102 | 0 | 0.11300509 | 0 | 0 | 0 | 0 | 0 | 0 | 0.67500913 | 0 | 0 | 0 | 0 | 0 |
| ENSMUSG00 | Gm5614 | 317 | 0 | 0 | 0 | 0.43117708 | 0 | 0 | 0.44775406 | 0 | 0 | 0.29748078 | 0 | 0 | 0 | 0.29302263 | 0.12997336 | 0 | 0 | 0.69412122 |
| ENSMUSG00 | Obox3 | 2589 | 0 | 0 | 0 | 0 | 0 | 0 | 0 | 0 | 0 | 0 | 0 | 0 | 0 | 0 | 0 | 0 | 0 | 0 |
| ENSMUSG00 | Zfp648 | 1577 | 0.18867231 | 1.81668288 | 0.85888323 | 0.4333644 | 0.15260597 | 0.81697595 | 0.72004077 | 0.29062069 | 0.33952106 | 0.86707064 | 0 | 0.17696786 | 0.31282129 | 0 | 0.39189812 | 0.16996465 | 0.12151648 | 0.24417486 |
| ENSMUSG00 | Zbtb6 | 9497 | 1.49486457 | 1.51533856 | 1.52806772 | 1.35766827 | 1.59138928 | 1.32008441 | 1.29528235 | 1.89413726 | 1.43114143 | 1.28091834 | 1.29787715 | 1.83368292 | 1.57392555 | 1.60894017 | 0.99348808 | 1.17327227 | 1.4487881 | 0.62556423 |
| ENSMUSG00 | Rnasel | 4749 | 13.3986626 | 13.2648311 | 13.6085569 | 11.6373019 | 12.0304471 | 12.0308069 | 8.56788879 | 11.3475426 | 10.3985328 | 10.1271251 | 11.5488845 | 11.4122991 | 9.25557766 | 7.7749104 | 7.9817712 | 8.65146804 | 8.38513845 | 8.95389335 |
| ENSMUSG00 | Vmn1r84 | 956 | 0 | 0 | 0 | 0 | 0 | 0 | 0 | 0 | 0 | 0 | 0 | 0 | 0 | 0 | 0 | 0 | 0 | 0 |
| ENSMUSG00 | Vmn1r83 | 995 | 0 | 0 | 0 | 0 | 0 | 0 | 0 | 0 | 0 | 0 | 0 | 0 | 0 | 0 | 0 | 0 | 0 | 0 |
| ENSMUSG00 | Vmn1r81 | 920 | 0 | 0 | 0 | 0 | 0 | 0 | 0 | 0 | 0 | 0 | 0 | 0 | 0 | 0 | 0 | 0 | 0 | 0 |
| ENSMUSG00 | Gm10179 | 688 | 0.06178078 | 0 | 0 | 0 | 0 | 0 | 0.06876843 | 0 | 0.17959237 | 0.068533 | 0 | 0.08112742 | 0.2151101 | 0.06750594 | 0 | 0 | 0 | 0 |
| ENSMUSG00 | Gm10180 | 77 | 0 | 2.16317495 | 0.62822767 | 2.95850937 | 1.25017984 | 2.57416798 | 1.22890075 | 2.48002613 | 0 | 1.83704039 | 1.24090801 | 0 | 0.64067425 | 0 | 1.07017028 | 0 | 0.99548823 | 2.85761593 |
| ENSMUSG00 | Gm10181 | 68 | 0 | 0 | 0 | 0 | 0 | 0 | 0 | 0 | 0 | 0 | 0 | 0 | 0 | 0 | 0 | 0 | 0 | 0 |
| ENSMUSG00 | Vmn2r28 | 2553 | 0 | 0 | 0 | 0 | 0 | 0 | 0 | 0 | 0 | 0 | 0 | 0 | 0 | 0 | 0 | 0 | 0 | 0 |
| ENSMUSG00 | Gm10182 | 272 | 0 | 0.36742163 | 0 | 0.16750384 | 0 | 0.18217917 | 0.52183102 | 0.14041324 | 0 | 0.69339269 | 0 | 0.20520464 | 0 | 0 | 0 | 0 | 0 | 0 |
| ENSMUSG00 | Vmn2r37 | 3075 | 0 | 0 | 0 | 0 | 0 | 0 | 0 | 0 | 0 | 0 | 0 | 0 | 0 | 0 | 0 | 0 | 0 | 0 |
| ENSMUSG00 | Zfp810 | 1381 | 12.3114196 | 16.3307961 | 14.2563554 | 14.4502081 | 15.0913273 | 14.6038831 | 16.890044 | 12.3067481 | 10.8856468 | 13.4521195 | 13.1113029 | 12.9738072 | 11.3238362 | 14.3267062 | 11.9636668 | 15.1110652 | 12.5164083 | 14.5788074 |
| ENSMUSG00 | Vmn2r39 | 3199 | 0 | 0 | 0 | 0 | 0 | 0 | 0 | 0 | 0 | 0 | 0 | 0 | 0 | 0 | 0 | 0 | 0 | 0 |
| ENSMUSG00 | Zfp772 | 2564 | 4.9069938 | 2.26070343 | 5.24486796 | 5.75732386 | 5.80061015 | 5.99116514 | 5.05603511 | 4.91555881 | 5.92738858 | 4.26636629 | 7.19232994 | 3.78780238 | 4.38676955 | 3.60496981 | 3.77627365 | 4.68925535 | 3.96118123 | 2.38143949 |
| ENSMUSG00 | Ecsit | 1580 | 22.8667086 | 24.7105772 | 25.5644923 | 20.8773393 | 25.9242196 | 25.1213542 | 22.7280527 | 30.4814047 | 26.4584263 | 26.5297311 | 30.7815872 | 25.6469437 | 28.6000229 | 27.7488725 | 32.1789367 | 27.8697459 | 33.4748669 | 28.3053188 |
| ENSMUSG00 | Hmcn1 | 20289 | 0 | 0 | 0 | 0 | 0 | 0 | 0 | 0 | 0 | 0 | 0 | 0 | 0 | 0 | 0 | 0 | 0 | 0 |
| ENSMUSG00 | Olfr1348 | 955 | 0 | 0 | 0 | 0 | 0 | 0 | 0 | 0 | 0 | 0 | 0 | 0 | 0 | 0 | 0 | 0 | 0 | 0 |
| ENSMUSG00 | Olfr5 | 956 | 0 | 0 | 0 | 0 | 0 | 0 | 0 | 0 | 0 | 0 | 0 | 0 | 0 | 0 | 0 | 0 | 0 | 0 |
| ENSMUSG00 | Vmn1r65 | 2679 | 0 | 0 | 0 | 0 | 0 | 0 | 0 | 0 | 0 | 0 | 0 | 0 | 0 | 0 | 0 | 0 | 0 | 0 |
| ENSMUSG00 | Cdkn2d | 1269 | 0.83737541 | 0.42002073 | 0.49555232 | 0.75396527 | 0.15171607 | 0.31238918 | 0.55925152 | 0.66212203 | 0.61666178 | 0.7059601 | 0.79060215 | 0.92366345 | 1.0884899 | 1.39076069 | 1.03896752 | 1.08625837 | 0.60403935 | 0.56352907 |
| ENSMUSG00 | Oas1g | 2683 | 1.91693116 | 0.89397256 | 0.99163033 | 1.69813807 | 0.89697957 | 1.69916194 | 0.61719857 | 0.68327816 | 1.04386249 | 1.12472792 | 1.22865156 | 0.64490702 | 0.82740823 | 0.67510973 | 0.82925234 | 2.05510528 | 1.14278932 | 1.53771264 |
| ENSMUSG00 | Oas1e | 1709 | 0 | 0 | 0 | 0 | 0.05632759 | 0 | 0 | 0 | 0 | 0 | 0 | 0 | 0.02717618 | 0 | 0 | 0 | 0 | 0 |
| ENSMUSG00 | Nck2 | 4186 | 1.75666399 | 1.68713177 | 1.05159849 | 1.4911283 | 0.82787829 | 0.94701832 | 0.83639232 | 2.23534128 | 0.90519815 | 1.41925194 | 0.59347775 | 1.26671953 | 1.17849778 | 1.48674336 | 1.28939413 | 1.03364516 | 0.3570773 | 0.11827089 |
| ENSMUSG00 | Gm10184 | 1324 | 0 | 0 | 0 | 0 | 0 | 0 | 0 | 0 | 0 | 0 | 0 | 0 | 0 | 0 | 0.031119 | 0 | 0.02894735 | 0 |
| ENSMUSG00 | Zfp617 | 3019 | 7.25080015 | 7.76822707 | 8.01151546 | 7.19861482 | 7.254066 | 7.6487492 | 7.45970029 | 7.47655178 | 6.94400017 | 7.16865611 | 8.38712421 | 8.57849203 | 7.22249335 | 7.78426904 | 7.32866363 | 6.0498906 | 7.921699 | 7.57992328 |
| ENSMUSG00 | Ptprv | 288 | 3.83727284 | 1.85071635 | 1.67963647 | 4.42954598 | 4.67949258 | 6.3661498 | 0.32856027 | 1.5913501 | 0.85805241 | 2.12833036 | 2.32239381 | 3.29467453 | 0 | 0 | 0.57224383 | 0.265907 | 1.06461936 | 0.19100384 |
| ENSMUSG00 | Fbxl12 | 5116 | 5.10128579 | 4.39527045 | 4.70876038 | 4.87136264 | 4.35595791 | 4.4942276 | 5.27135008 | 5.83785353 | 4.74176266 | 5.18878928 | 5.01468974 | 5.54229019 | 4.33920304 | 4.07611322 | 4.92067057 | 6.41420065 | 4.19521624 | 5.39768795 |
| ENSMUSG00 | Vsig10 | 5321 | 6.86185796 | 8.07623258 | 6.94556985 | 8.34843417 | 6.30482067 | 7.76676937 | 4.62367845 | 6.34506366 | 7.6629676 | 6.56618514 | 5.83606897 | 7.91971909 | 4.98789165 | 5.05377309 | 5.37377932 | 6.24624079 | 6.08639746 | 6.73011247 |
| ENSMUSG00 | Olfr18 | 1154 | 0 | 0 | 0 | 0 | 0 | 0 | 0 | 0 | 0 | 0 | 0 | 0 | 0 | 0 | 0 | 0 | 0 | 0 |
| ENSMUSG00 | Olfr872 | 1122 | 0 | 0 | 0 | 0 | 0 | 0 | 0 | 0 | 0 | 0 | 0 | 0 | 0 | 0 | 0 | 0 | 0 | 0 |
| ENSMUSG00 | Olfr870 | 1810 | 0 | 0 | 0 | 0 | 0 | 0 | 0 | 0 | 0 | 0 | 0 | 0 | 0 | 0 | 0 | 0 | 0 | 0 |
| ENSMUSG00 | Suds3 | 3580 | 19.1629481 | 19.2247038 | 18.8224379 | 19.344354 | 18.2444191 | 20.7761601 | 22.4008354 | 22.6167299 | 21.8932792 | 23.8387634 | 19.7772314 | 22.4665837 | 18.8370757 | 23.7798917 | 21.3143244 | 17.1238163 | 18.7670387 | 21.6809636 |
| ENSMUSG00 | Gm10186 | 460 | 0 | 0 | 0 | 0 | 0 | 0 | 0 | 0 | 0 | 0 | 0 | 0 | 0 | 0 | 0 | 0 | 0 | 0 |
| ENSMUSG00 | Olfr859 | 929 | 0 | 0 | 0 | 0 | 0 | 0 | 0 | 0 | 0 | 0 | 0 | 0 | 0 | 0 | 0 | 0 | 0 | 0 |
| ENSMUSG00 | Olfr860 | 1217 | 0 | 0 | 0 | 0 | 0 | 0 | 0 | 0 | 0 | 0 | 0 | 0 | 0 | 0 | 0 | 0 | 0 | 0 |
| ENSMUSG00 | Olfr834 | 938 | 0 | 0 | 0 | 0 | 0 | 0 | 0 | 0 | 0 | 0 | 0 | 0 | 0 | 0 | 0 | 0 | 0 | 0 |
| ENSMUSG00 | Olfr833-ps1 | 265 | 0 | 0 | 0 | 0 | 0 | 0 | 0 | 0 | 0 | 0 | 0 | 0 | 0 | 0 | 0 | 0 | 0 | 0 |
| ENSMUSG00 | Olfr831-ps1 | 328 | 0 | 0 | 0 | 0 | 0 | 0 | 0 | 0 | 0 | 0 | 0 | 0 | 0 | 0 | 0 | 0 | 0 | 0 |
| ENSMUSG00 | Gm17644 | 3197 | 0 | 0 | 0 | 0 | 0 | 0 | 0 | 0 | 0 | 0 | 0 | 0 | 0 | 0 | 0.01288757 | 0 | 0 | 0 |
| ENSMUSG00 | Gm10188 | 719 | 0 | 0 | 0 | 0 | 0 | 0 | 0 | 0 | 0 | 0 | 0 | 0 | 0 | 0 | 0 | 0 | 0 | 0 |
| ENSMUSG00 | Gm10190 | 323 | 0 | 0 | 0 | 0 | 0 | 0 | 0 | 0 | 0 | 0 | 0 | 0 | 0 | 0 | 0 | 0 | 0 | 0 |
| ENSMUSG00 | 4921513D11I | 1344 | 0 | 0.04957276 | 0 | 0.03389959 | 0 | 0 | 0 | 0.02841697 | 0 | 0.03508237 | 0 | 0.04152951 | 0 | 0 | 0 | 0.02849004 | 0 | 0 |
| ENSMUSG00 | Myo1h | 5121 | 0.08300171 | 0 | 0.00944611 | 0.01779381 | 0.00939893 | 0.00967638 | 0.10162848 | 0.007458 | 0 | 0 | 0 | 0 | 0.02889978 | 0 | 0 | 0 | 0 | 0 |
| ENSMUSG00 | Gm6396 | 416 | 0 | 0 | 0 | 0 | 0 | 0.2382343 | 0.3411972 | 0 | 0 | 0 | 0 | 0 | 0.11858634 | 0 | 0 | 0 | 0 | 0.39670029 |
| ENSMUSG00 | Gm10191 | 377 | 0.11274583 | 0.17672623 | 0 | 0 | 0.12767089 | 0.26287922 | 0.12549782 | 0.30391832 | 0.32774416 | 0.37520454 | 0.12672403 | 0.29610431 | 0 | 0.3695816 | 0.10928795 | 0.2031332 | 0.20332253 | 0.43773825 |
| ENSMUSG00 | Gm16391 | 446 | 0 | 0 | 0 | 0 | 0 | 0 | 0 | 0 | 0.1846929 | 0 | 0 | 0 | 0 | 0 | 0 | 0 | 0.1718668 | 0 |
| ENSMUSG00 | Tmem211 | 1822 | 0 | 0.20112066 | 0 | 0 | 0 | 0 | 0 | 0 | 0.29386647 | 0 | 0 | 0 | 0 | 0 | 0 | 0 | 0 | 0 |
| ENSMUSG00 | Cryba4 | 1165 | 6.56732333 | 14.354569 | 4.31832374 | 6.53106816 | 4.79253489 | 10.0381503 | 0.4873409 | 0.19669907 | 0.14141293 | 0.28330895 | 0.32806838 | 0.43119396 | 30.1072903 | 57.407284 | 32.1478234 | 8.31547096 | 7.89554614 | 28.5197429 |
| ENSMUSG00 | Bub3 | 2047 | 66.8620747 | 65.4866079 | 68.3657173 | 66.6388699 | 71.3161331 | 66.691637 | 59.6782301 | 61.999684 | 59.8582728 | 59.0132397 | 60.0979571 | 59.1695104 | 61.0925307 | 51.844035 | 64.388753 | 52.5443655 | 57.3115754 | 59.604396 |
| ENSMUSG00 | Gm16519 | 527 | 0 | 0 | 0 | 0.08645359 | 0 | 0 | 0.26933214 | 0 | 0.07815278 | 0 | 0 | 0.10591207 | 0 | 0.0881292 | 0.15636264 | 0.0726577 | 0.07272542 | 0 |
| ENSMUSG00 | Gm10192 | 326 | 0 | 0 | 0 | 0 | 0 | 0 | 0 | 0 | 0 | 0 | 0 | 0 | 0 | 0 | 0 | 0 | 0 | 0 |
| ENSMUSG00 | Serpinb7 | 1948 | 0 | 0 | 0 | 0 | 0 | 0 | 0 | 0 | 0 | 0 | 0 | 0 | 0 | 0 | 0 | 0 | 0 | 0 |
| ENSMUSG00 | Serpinb5 | 2799 | 5.95285066 | 7.67660478 | 6.89569082 | 7.63420143 | 8.90758716 | 7.32934323 | 7.26847158 | 4.36640542 | 5.50330722 | 6.182318 | 5.8715919 | 5.00526306 | 6.46831498 | 4.19805434 | 6.02052031 | 4.88379666 | 4.55972021 | 6.28900113 |
| ENSMUSG00 | Vmn2r10 | 5081 | 0 | 0 | 0 | 0 | 0 | 0 | 0 | 0 | 0 | 0 | 0 | 0 | 0 | 0 | 0 | 0 | 0 | 0 |
| ENSMUSG00 | Gm10193 | 230 | 0 | 0.14483867 | 0 | 0 | 0.20926923 | 0.86178667 | 0.2057073 | 0.16605392 | 0 | 0 | 0 | 0 | 0 | 0.40386163 | 0.5374116 | 0 | 0.33327215 | 0.23917003 |
| ENSMUSG00 | Gm3608 | 868 | 0 | 0.15351564 | 0 | 0.15746905 | 0.2218061 | 0.28544201 | 0.05450769 | 0.30800325 | 0.0474499 | 0.16296326 | 0 | 0 | 0.11366801 | 0.26753507 | 0.14240169 | 0 | 0.17661888 | 0.06337455 |
| ENSMUSG00 | Gm626 | 3147 | 0 | 0.04234241 | 0 | 0 | 0 | 0 | 0 | 0 | 0 | 0 | 0 | 0 | 0 | 0 | 0 | 0 | 0 | 0 |
| ENSMUSG00 | Cntnap5b | 5100 | 0 | 0 | 0 | 0 | 0.01887526 | 0 | 0 | 0.13479671 | 0 | 0.01849047 | 0 | 0 | 0 | 0.20945373 | 0 | 0 | 0.10520944 | 0 |
| ENSMUSG00 | Zfp960 | 2444 | 0.74784066 | 1.19948228 | 0.77191804 | 1.08123591 | 0.94530783 | 0.60825778 | 1.54869653 | 0.78135029 | 0.67408373 | 0.59806539 | 0.87965349 | 0.75364847 | 0.94869072 | 1.02617869 | 0.69118813 | 1.09670316 | 1.01931641 | 0.49517199 |
| ENSMUSG00 | Rps12-ps3 | 468 | 0 | 0 | 0 | 0 | 0 | 0 | 0 | 0 | 0 | 0 | 0 | 0 | 0 | 0 | 0 | 0 | 0 | 0 |
| ENSMUSG00 | Olfr719-ps | 242 | 0 | 0 | 0 | 0 | 0 | 0 | 0 | 0 | 0 | 0 | 0 | 0 | 0 | 0 | 0 | 0 | 0 | 0 |
| ENSMUSG00 | Unc93a | 3173 | 0.12056306 | 0.08399091 | 0.03049072 | 0 | 0 | 0.04685099 | 0 | 0.31295382 | 0.15576369 | 0.20803966 | 0 | 0.29904389 | 0.07773703 | 0.04391184 | 0.16880562 | 0 | 0.18118325 | 0.27738598 |

| | | | | | | | | | | | | | | | | | | | |
|---|---|---|---|---|---|---|---|---|---|---|---|---|---|---|---|---|---|---|---|
| [illegible] | | | | | | | | | | | | | | | | | | | |
| ENSMUSG00 Gm6990 | 347 | 0 | 0 | 0 | 0 | 0 | 0 | 0 | 0 | 0 | 0 | 0 | 0 | 0 | 0 | 0 | 0 | 0 | 0 |
| ENSMUSG00 Olfr1416 | 938 | 0 | 0 | 0 | 0 | 0 | 0 | 0 | 0 | 0 | 0 | 0 | 0 | 0 | 0 | 0 | 0 | 0 | 0 |
| ENSMUSG00 Hes6 | 2566 | 17.8071178 | 19.6424041 | 17.6263643 | 20.4190183 | 17.388267 | 19.3692096 | 21.5174966 | 20.9864721 | 25.0393471 | 21.425456 | 19.307339 | 24.0142212 | 24.8197604 | 22.7695487 | 20.1993598 | 18.5782565 | 18.4014021 | 20.0013627 |
| ENSMUSG00 Asb18 | 1719 | 0.04945338 | 0.58137686 | 0.39396709 | 0.18553072 | 0 | 0.14413244 | 0 | 0.0222178 | 0 | 0 | 0 | 0 | 0.02869803 | 0.10807234 | 0.07190498 | 0 | 0 | 0 |
| ENSMUSG00 Gm10197 | 200 | 0 | 0 | 0 | 0 | 0 | 0 | 0 | 0 | 0 | 0 | 0 | 0 | 0 | 0 | 0 | 0 | 0 | 0 |
| ENSMUSG00 Gm6941 | 314 | 0.1353668 | 0 | 0.30811166 | 0 | 0 | 0 | 0 | 0.36489556 | 0 | 0 | 0 | 0 | 0.15710802 | 0 | 0.13121515 | 0.12194461 | 0.36617481 | 0 |
| ENSMUSG00 1700010H22 | 376 | 0 | 0 | 0 | 0 | 0 | 0 | 0 | 0 | 0 | 0 | 0 | 0 | 0 | 0 | 0 | 0 | 0 | 0 |
| ENSMUSG00 AY702103 | 2120 | 0 | 0 | 0 | 0 | 0 | 0 | 0 | 0 | 0 | 0 | 0 | 0 | 0 | 0 | 0 | 0 | 0 | 0 |
| ENSMUSG00 Gm6529 | 971 | 0.17509856 | 0.06861564 | 0.39854608 | 0.1876871 | 0.19827775 | 0 | 0.14617717 | 0.15733224 | 0.12724979 | 0.09711782 | 0.09840362 | 0 | 0.25402635 | 0.28698715 | 0.25459252 | 0.35490779 | 0.11841286 | 0.05665202 |
| ENSMUSG00 Gm7027 | 861 | 0.04936722 | 0 | 0.16854889 | 0 | 0 | 0 | 0.16485254 | 0.04435819 | 0.14350702 | 0.27381361 | 0.05548776 | 0.06482655 | 0.05729607 | 0.05394203 | 0.23926571 | 0.0889445 | 0.17805481 | 0 |
| ENSMUSG00 Gm10198 | 71 | 0 | 0 | 0 | 0 | 0 | 0 | 0 | 0 | 0 | 0 | 0 | 0 | 0 | 0 | 0 | 0 | 0 | 0 |
| ENSMUSG00 Rpl30-ps6 | 347 | 0 | 0 | 0 | 0 | 0 | 0 | 0 | 0 | 0 | 0 | 0 | 0 | 0 | 0 | 0 | 0 | 0 | 0 |
| ENSMUSG00 Slc22a7 | 2017 | 0 | 0 | 0 | 0 | 0 | 0 | 0 | 0 | 0 | 0 | 0 | 0 | 0 | 0 | 0 | 0 | 0 | 0 |
| ENSMUSG00 Rpl7a-ps11 | 1651 | 0 | 0 | 0 | 0 | 0 | 0 | 0 | 0 | 0 | 0 | 0 | 0 | 0 | 0 | 0 | 0 | 0.06964197 | 0 |
| ENSMUSG00 Polr1c | 2435 | 13.5108855 | 12.5590295 | 10.6084046 | 12.0123985 | 14.1134265 | 11.9048662 | 15.0584502 | 14.273136 | 15.3582572 | 12.8575224 | 13.9106553 | 13.845035 | 15.2756738 | 16.975457 | 15.6853562 | 12.4071003 | 14.0713386 | 16.5140275 |
| ENSMUSG00 Igj | 2001 | 0 | 0 | 0.09669871 | 0.02276914 | 0.14432361 | 0 | 0.07093355 | 0.13360661 | 0 | 0.42414426 | 0.23875541 | 0.02789388 | 0.14792179 | 0.69631315 | 0 | 0.21049309 | 0.134075 | 0 |
| ENSMUSG00 Xpo5 | 3677 | 8.27677619 | 7.68271262 | 10.3666962 | 8.94617188 | 8.27287891 | 8.75966191 | 9.39305293 | 11.456682 | 9.68896822 | 8.47609868 | 8.31546734 | 10.7320298 | 8.27788767 | 10.5342314 | 10.0398678 | 7.62271543 | 6.94188571 | 8.36282041 |
| ENSMUSG00 Gm7337 | 1277 | 0 | 0 | 0 | 0 | 0 | 0 | 0.1111496 | 0 | 0 | 0 | 0 | 0.08741686 | 0 | 0 | 0 | 0 | 0 | 0 |
| ENSMUSG00 Col4a4 | 9107 | 0.22403079 | 0.13168598 | 0.19122072 | 0.18010295 | 0.17969533 | 0.04352936 | 0.10390398 | 0.06709986 | 0.04974763 | 0.2329836 | 0.07344344 | 0.11644862 | 0.1191723 | 0.15299469 | 0.02714498 | 0.17659004 | 0.04629288 | 0.01812093 |
| ENSMUSG00 Gm5560 | 2155 | 0.80868316 | 0.88112992 | 0.76320187 | 0.90910669 | 1.04974497 | 0.98875524 | 1.09774197 | 1.16969771 | 1.26139677 | 1.37841963 | 1.04195965 | 1.37272859 | 1.28194309 | 1.25000328 | 0.95595257 | 1.26154669 | 1.17379842 | 1.42947099 |
| ENSMUSG00 Mrgpra4 | 1502 | 0 | 0 | 0 | 0 | 0 | 0 | 0 | 0 | 0 | 0 | 0 | 0 | 0 | 0 | 0 | 0 | 0 | 0 |
| ENSMUSG00 Olfr132 | 941 | 0 | 0 | 0 | 0 | 0 | 0 | 0 | 0 | 0 | 0 | 0 | 0 | 0 | 0 | 0 | 0 | 0 | 0 |
| ENSMUSG00 Olfr130 | 1848 | 0 | 0 | 0 | 0 | 0 | 0 | 0 | 0 | 0 | 0 | 0 | 0 | 0 | 0 | 0 | 0 | 0 | 0 |
| ENSMUSG00 Gm7335 | 620 | 0.34278368 | 0.05373047 | 0.78021823 | 0.22045667 | 0.07763213 | 0.15984753 | 0.68679695 | 0.06160065 | 0.19928959 | 0.60839617 | 0 | 0.54015157 | 0.07956761 | 0 | 0.19936237 | 0.74110853 | 0.06181661 | 0.53234619 |
| ENSMUSG00 Olfr91 | 938 | 0 | 0 | 0 | 0 | 0 | 0 | 0 | 0 | 0 | 0 | 0 | 0 | 0 | 0 | 0 | 0 | 0 | 0 |
| ENSMUSG00 Eif1ax | 5259 | 21.7092418 | 20.5679725 | 21.3031178 | 17.9506529 | 20.8031684 | 20.7482638 | 20.7279734 | 21.9321269 | 22.3984474 | 20.3521765 | 22.8109756 | 21.1630407 | 19.1736906 | 21.1864167 | 18.4658808 | 17.190364 | 18.3432762 | 17.3845095 |
| ENSMUSG00 Frat1 | 577 | 0.8839898 | 0.51961187 | 0.3353451 | 0.71065754 | 0.83417545 | 0.17175991 | 0.40998855 | 0.39714803 | 0.92794576 | 1.30747184 | 0.68239111 | 1.06407676 | 0.25649177 | 1.04640058 | 0.35703255 | 0.19908461 | 0.99635087 | 0 |
| ENSMUSG00 H2-M9 | 1437 | 0 | 0 | 0 | 0 | 0 | 0 | 0 | 0 | 0 | 0 | 0 | 0 | 0 | 0 | 0 | 0 | 0 | 0 |
| ENSMUSG00 H2-K2 | 1111 | 14.9590674 | 12.8334102 | 15.6310508 | 15.9115078 | 17.3725485 | 15.6552741 | 21.6761058 | 25.1636711 | 26.5432451 | 31.6177083 | 27.9941476 | 29.1386897 | 28.0183977 | 33.6521065 | 27.4058954 | 37.0498676 | 38.7057652 | 48.6714239 |
| ENSMUSG00 Lrrc66 | 3209 | 0.3708772 | 0 | 0 | 0 | 0.08999425 | 0.0617672 | 0.19166869 | 0 | 0.28236315 | 0.01469327 | 0.50618529 | 0.38265646 | 0.09223793 | 0.17367686 | 0.11555438 | 0.01193226 | 0 | 0 |
| ENSMUSG00 4921511M17 | 981 | 0 | 0 | 0 | 0 | 0 | 0 | 0 | 0 | 0 | 0 | 0 | 0 | 0 | 0 | 0 | 0 | 0 | 0 |
| ENSMUSG00 Gm15144 | 981 | 0 | 0 | 0 | 0 | 0 | 0 | 0 | 0 | 0 | 0 | 0 | 0 | 0 | 0 | 0 | 0 | 0 | 0 |
| ENSMUSG00 H2-T23 | 3276 | 40.0399796 | 36.5974073 | 37.7419852 | 35.3529227 | 38.4790928 | 39.4637002 | 31.7728613 | 32.258357 | 41.714548 | 40.9042426 | 34.0812204 | 32.661363 | 46.4406694 | 30.3956421 | 43.5282616 | 48.0385825 | 39.5781021 | 48.4100289 |
| ENSMUSG00 Usp51 | 2202 | 1.29330009 | 1.60361798 | 1.14233587 | 0.82763023 | 1.7268038 | 2.0478196 | 1.11728397 | 1.6997527 | 1.23447322 | 1.00639557 | 1.34516232 | 1.06460392 | 5.17514664 | 4.55582329 | 2.88148938 | 7.18166251 | 4.57757315 | 3.59732578 |
| ENSMUSG00 Nipal1 | 4199 | 0 | 0.14280355 | 0.046081 | 0 | 0 | 0 | 0 | 0.22738987 | 0 | 0.02245806 | 0 | 0 | 0 | 0.12166825 | 0 | 0.06383288 | 0.13691223 | 0 |
| ENSMUSG00 Cnga1 | 5965 | 0 | 0 | 0 | 0 | 0 | 0 | 0 | 0 | 0 | 0 | 0 | 0 | 0 | 0 | 0 | 0 | 0 | 0 |
| ENSMUSG00 Gm3695 | 1008 | 1.012028 | 0.5618246 | 0.76783382 | 0.45199449 | 0.81174871 | 0.54075404 | 0.46937181 | 0.71989647 | 0.65375422 | 0.60809439 | 0.71093688 | 0.66447217 | 0.19576158 | 1.05973612 | 0.36787103 | 0.53181399 | 0.5703318 | 0.32743516 |
| ENSMUSG00 Cyp2c54 | 1760 | 0 | 0 | 0 | 0 | 0 | 0 | 0 | 0 | 0 | 0 | 0 | 0 | 0 | 0 | 0 | 0 | 0 | 0 |
| ENSMUSG00 Cyp2c66 | 2123 | 0 | 0 | 0.04557092 | 0 | 0 | 0 | 0 | 0 | 0 | 0 | 0 | 0 | 0 | 0 | 0 | 0 | 0 | 0 |
| ENSMUSG00 Cyp2c65 | 1919 | 0 | 0 | 0 | 0 | 0 | 0 | 0 | 0 | 0 | 0 | 0 | 0 | 0 | 0 | 0 | 0 | 0 | 0 |
| ENSMUSG00 H2-Q10 | 1789 | 0.02375918 | 0.03724192 | 1.35197122 | 0.02546732 | 0.13452187 | 1.88350245 | 0.05289288 | 0 | 1.7496787 | 0.1054236 | 0.05340968 | 2.5271485 | 0.13787568 | 0.02596092 | 1.70425664 | 0.06421007 | 0.04284661 | 1.96790544 |
| ENSMUSG00 Lgi1 | 8140 | 0.19320535 | 0.25373458 | 0.12479658 | 0.10634642 | 0.27199859 | 0.21306458 | 0.06974842 | 0.37066336 | 0.56669407 | 0.13901927 | 0.28758882 | 0.27427844 | 0.39392809 | 0.24534346 | 0.25308081 | 0.15523219 | 0.34371249 | 0.27031502 |
| ENSMUSG00 Foxd3 | 2372 | 0 | 0 | 0 | 0 | 0 | 0 | 0 | 0 | 0 | 0 | 0 | 0 | 0 | 0 | 0 | 0 | 0 | 0 |
| ENSMUSG00 Vmn1r239-ps | 1894 | 0 | 0 | 0 | 0 | 0 | 0 | 0 | 0 | 0 | 0 | 0 | 0 | 0 | 0 | 0 | 0 | 0 | 0 |
| ENSMUSG00 Rplp0 | 1654 | 173.027418 | 167.934046 | 166.587442 | 154.863477 | 164.038485 | 161.930184 | 190.967016 | 162.583256 | 184.791495 | 173.52257 | 166.750203 | 170.112911 | 233.566048 | 239.268482 | 194.17541 | 181.753058 | 165.099677 | 214.149721 |
| ENSMUSG00 Capn6 | 7782 | 0.01092397 | 0.01284229 | 0 | 0.01170934 | 0 | 0 | 0 | 0 | 0 | 0 | 0 | 0 | 0 | 0 | 0.10059491 | 0.06396529 | 0.02954996 | 0 |
| ENSMUSG00 Ppp1r3c | 2717 | 1.43926249 | 1.79009295 | 2.24330689 | 1.71042566 | 1.36406261 | 1.98794551 | 2.2289374 | 2.67079737 | 3.04692295 | 3.1237124 | 2.81339468 | 1.91051035 | 2.16065446 | 3.17946274 | 4.56447123 | 1.5079481 | 4.71144025 | 3.27989521 |
| ENSMUSG00 Gm16223 | 2527 | 0.15138369 | 0.10546227 | 0.01914267 | 0 | 0.09523531 | 0.05882794 | 0.03744573 | 0.06045493 | 0.01629858 | 0 | 0 | 0.13234061 | 0.04417544 | 0.05856579 | 0.0918957 | 0 | 0.12133375 | 0 |
| ENSMUSG00 Rps28 | 1361 | 23.0171306 | 15.9588589 | 21.2544975 | 17.1397904 | 18.7435118 | 17.5127589 | 22.45701 | 16.865271 | 20.2452454 | 18.1535404 | 17.5514175 | 20.3823544 | 29.0337 | 28.7673517 | 22.8561166 | 23.463899 | 21.8243057 | 26.3930541 |
| ENSMUSG00 Gm10203 | 195 | 0 | 0 | 0 | 0 | 0 | 0 | 0 | 0 | 0 | 0 | 0 | 0 | 0 | 0 | 0 | 0 | 0 | 0 |
| ENSMUSG00 2010002M12 | 6495 | 0 | 0.07180608 | 0.11171716 | 0.00701479 | 0.06669551 | 0 | 0 | 0.01176056 | 0 | 0 | 0 | 0 | 0.01519074 | 0 | 0.00634358 | 0.0117908 | 0.02950446 | 0.00846945 |
| ENSMUSG00 Crygd | 1024 | 0 | 0 | 0 | 0 | 0 | 0 | 0 | 0 | 0 | 0 | 0 | 0 | 0 | 0 | 0 | 0 | 0 | 0 |
| ENSMUSG00 Gm7205 | 327 | 0 | 0 | 0 | 0 | 0 | 0 | 0 | 0 | 0 | 0 | 0 | 0 | 0.15086213 | 0 | 0 | 0 | 0 | 0 |
| ENSMUSG00 Gm7931 | 272 | 0 | 0 | 0 | 0 | 0 | 0 | 0 | 0 | 0 | 0 | 0 | 0 | 0 | 0 | 0 | 0 | 0.42271651 | 0 |
| ENSMUSG00 Gm10205 | 230 | 0 | 0 | 0 | 0 | 0 | 0 | 0 | 0 | 0 | 0 | 0 | 0 | 0 | 0 | 0 | 0 | 0 | 0 |
| ENSMUSG00 Bmpr2 | 3494 | 1.04620639 | 1.35387263 | 1.43985322 | 1.38221829 | 1.11582307 | 1.84369072 | 1.40827665 | 1.07122365 | 0.81335707 | 0.91764391 | 1.36734283 | 1.16615437 | 1.5389751 | 1.76790602 | 1.01411958 | 1.31507524 | 0.87753399 | 1.35397343 |
| ENSMUSG00 Tuba3b | 1770 | 0 | 0 | 0 | 0 | 0 | 0 | 0.26730327 | 0.02157763 | 0 | 0 | 0 | 0 | 0 | 0 | 0 | 0 | 0 | 0 |
| ENSMUSG00 H2-Eb2 | 2204 | 0 | 0 | 0 | 0 | 0 | 0 | 0 | 0.03465735 | 0 | 0.10838239 | 0 | 0 | 0 | 0.0632179 | 0 | 0 | 0 | 0 |
| ENSMUSG00 Rps25-ps1 | 816 | 0.36462774 | 0.24494775 | 0.29640644 | 0.27917307 | 0.4128964 | 0.24290556 | 0.40586857 | 0.18721766 | 0.3533157 | 0.2311309 | 0.4098342 | 0.68401547 | 0.84638093 | 0.45533419 | 0.25246051 | 0.32847335 | 0.32877951 | 0.87637057 |
| ENSMUSG00 Rps15a-ps2 | 392 | 0.54215786 | 0 | 0 | 0 | 0 | 0 | 0.36208683 | 0 | 0 | 0 | 0 | 0 | 0.12584673 | 0 | 0 | 0.09768012 | 0 | 0.14032935 |
| [illegible] | | | | | | | | | | | | | | | | | | | |

| | | | | | | | | | | | | | | | | | | | |
|---|---|---|---|---|---|---|---|---|---|---|---|---|---|---|---|---|---|---|---|
|ENSMUSG00 Gm10257|1024|1.20375987|0.74823883|0.42515798|0.75638453|0.75206131|0.91943549|0.50824167|0.70864809|0.92508776|1.24323143|0.51320756|1.47170204|0.57810841|0.54426665|0.40235894|0.97222245|0.82341654|0.26859915|
|ENSMUSG00 Slc7a14|9768|0|0|0|0|0|0|0|0|0|0|0|0|0|0|0|0|0|0|
|ENSMUSG00 Gm10258|2122|0|0|0|0|0|0|0|0|0|0|0|0|0|0|0|0|0|0|
|ENSMUSG00 Lcn11|646|0|0|0|0|0|0|0|0|0|0|0|0|0|0|0|0|0|0|
|ENSMUSG00 Gm10259|1324|0|0|0.0730718|0|0|0|0|0|0|0|0|0|0|0|0.031119|0|0.02894735|0|
|ENSMUSG00 Dytn|3374|0|0|0|0|0|0|0|0|0|0|0|0|0|0|0|0|0|0|
|ENSMUSG00 Cdk7|8373|7.67560327|7.66280118|7.14077195|6.63853745|6.78894086|6.50997336|7.80916306|7.51714788|7.12266509|7.42202638|7.2692341|6.70614553|8.71984206|6.70618675|8.6753067|8.43737856|9.07691947|7.3516291|
|ENSMUSG00 Pde7a|10891|3.22369621|3.08322444|2.73602926|2.80285554|2.49255394|1.95644802|1.99398766|2.43370923|2.94594581|2.03478382|2.93466598|1.95260008|2.53657825|1.17698725|2.05043093|2.15518707|2.05514254|1.38394043|
|ENSMUSG00 Gm8910|1675|0.17763357|0.21877125|0.1443986|0.05440125|0.11494191|0.32542094|0.08473913|0.15961004|0.12294482|0.11259869|0.17113418|0.33322784|0.08835567|0.11091125|0.27057738|0.22860064|0.18305097|0.09852377|
|ENSMUSG00 Zbtb10|5388|3.27387677|3.25833245|4.71345647|3.55998509|6.09242241|3.89027585|4.19737575|5.33757962|5.84010728|4.27926761|6.09157321|4.15406101|3.64404289|3.61174325|3.91521838|5.55739699|5.58391668|4.54325399|
|ENSMUSG00 Gm17352|425|0.10001218|0|0.11382007|0|0|0.23318933|0|0|0|0|0|0|0|0|0|0.09009555|0.18035904|0|
|ENSMUSG00 1700008P02f|784|0|0|0|0|0|0|0|0|0|0|0|0|0|0|0|0|0|0|
|ENSMUSG00 Rps24-ps2|398|0|0|0|0|0|0|0.11887608|0.09596081|0|0|0|0|0|0|0|0|0|0|
|ENSMUSG00 Nxph2|2577|0|0|0|0|0|0|0|0|0|0|0|0|0|0|0|0|0|0|
|ENSMUSG00 Fgfr1op|4173|6.52907209|7.55986361|7.33775335|6.71460395|7.68173045|6.94664489|6.24713302|8.1912773|6.63247989|5.22013537|7.65910549|6.99534904|8.19243198|9.92766018|9.89311263|9.46024832|10.6538472|12.2725805|
|ENSMUSG00 A130006I12R|405|0.94455947|0.8225406|0.83608571|0.78747484|0.59422128|0.36705729|0.46728572|0.75441783|0.2033902|0.93137192|0.94370288|0.13781645|0.1218072|0.11467676|0.40692895|0.28363413|0|0.27164991|
|ENSMUSG00 Gpr98|25128|0|0.00132573|0.00962542|0|0|0|0|0.00911948|0.01475162|0.00375284|0.0209139|0.00666376|0|0.036966|0.00163967|0|0|0|
|ENSMUSG00 Nr2f1|4942|0|0|0|0|0|0|0|0|0|0|0|0|0|0|0|0|0|0|
|ENSMUSG00 Gm17681|1421|0.11964863|0|0|0|0|0.03487173|0.03329534|0.18813991|0|0|0|0|0|0|0.3769319|0|0|0|
|ENSMUSG00 Zfp72|1626|2.35268502|3.29850921|4.61125289|3.78274353|4.29220722|5.48554246|5.15027317|4.90910953|3.39421471|2.78380536|5.49441404|3.77596733|1.85070538|4.6558341|2.38188576|2.14295527|2.94636354|3.9920508|
|ENSMUSG00 Gm13192|520|0.16348145|0|0.18605204|0.08761739|0|0.09529372|0.18197184|0.14689386|0.07920484|0.27202329|0.09187492|0|0.09486907|0.17863111|0|0.29454313|0.51593092|0.21157349|
|ENSMUSG00 Zdhhc11|1451|0|0|0|0|0|0|0|0|0|0|0|0|0|0|0|0|0|0.07582234|
|ENSMUSG00 Gm3511|313|4.20977782|2.23505041|2.62731635|3.78462349|3.99817896|2.69136253|6.19750746|5.00283865|2.50013993|3.61538939|1.67899215|2.31822241|6.30439837|5.93534663|3.94903092|3.67002627|3.30610229|4.56944656|
|ENSMUSG00 Gm6132|1029|0.33045812|0.29136642|0.51711257|0.04427701|0.23387718|0.38524963|0.18391712|0.44539245|0.48030922|0.18328748|0.18571412|0.32545576|0.47941611|0.63189235|0.92092885|0.63259507|0.40970774|0.26729401|
|ENSMUSG00 Zfp748|4599|3.24403496|3.70143269|3.91279698|2.58565614|5.19100547|4.48226509|3.20973164|3.3301051|3.68968134|2.77839543|4.633101|3.61667046|2.41349889|2.93873221|5.49174902|1.37376608|5.42518358|2.76301449|
|ENSMUSG00 Zfp874a|3435|3.93497699|2.99670577|2.54894003|3.05066672|3.37694137|3.13040559|4.28362246|3.16880195|5.33566216|4.1316919|2.99028124|3.49355676|3.18826365|2.21741785|4.46200256|4.40314117|2.75592295|4.61211724|
|ENSMUSG00 Zfp825|819|16.4519424|17.2869109|16.7151515|18.35793|15.1624375|18.7562239|20.1613247|13.7567261|10.2086236|17.0986067|13.3583217|12.8123865|16.1428008|12.2489901|11.9227946|14.6804038|13.8985472|11.7540826|
|ENSMUSG00 4930451E10f|896|0.42694931|0.03717957|0|0.15254814|0.10743733|0.16591317|0.31682597|0.38362904|0|0.21049421|0.4798824|0.24917707|0.27528972|0.36284443|0.2299194|0|0.04277488|0|
|ENSMUSG00 Gprin1|8285|0|0|0|0|0|0|0|0.09219651|0.15907888|0.09105748|0|0|0.05954365|0.10651028|0|0.00462168|0.06013782|0|
|ENSMUSG00 Gm7251|1007|0.37988737|0.29773192|0.09607454|0.54293201|0.23898671|0.14762483|0.70475689|0.37926914|0.53170279|0.60869825|0.61675716|0.27713834|0.39191195|0.32284867|0.6546424|0.45629324|0.3425389|0.16388016|
|ENSMUSG00 C78339|4357|6.59478976|6.26958303|6.6947989|6.46241115|5.19210561|5.1406554|6.17877308|6.61814639|6.8439379|6.47145296|6.29396974|6.32842261|5.95560902|7.07800643|6.17503236|6.67910526|6.078373|8.06766564|
|ENSMUSG00 Serpinb6e|1486|0.08581126|0|0|0|0|0|0|0|0|0|0|0|0|0|0|0|0|0|
|ENSMUSG00 Dusp22|1820|13.1018702|13.2702463|12.0402249|12.0411331|12.4296726|14.3757381|10.0864392|9.3172674|11.0886773|11.9690247|13.7287381|10.3964339|8.48400555|6.58383217|9.41749849|8.43655691|9.81321668|9.21855909|
|ENSMUSG00 Prl2b1|1184|0|0|0|0|0|0|0|0|0|0|0|0|0|0|0|0|0|0|
|ENSMUSG00 Prl8a1|1153|0|0|0|0|0|0|0|0|0|0|0|0|0|0|0|0|0|0|
|ENSMUSG00 Hist1h3a|533|0.71772342|0.25000296|0.45378546|0.25644115|0.27091139|0.74375585|0.71013402|1.07483309|1.46818724|1.41540573|0.80670662|0.6283189|0.27766558|0.697097|1.54602461|0.35919894|0.35953374|3.92185001|
|ENSMUSG00 Hist1h4b|490|0|0|0|0|0|0|0|0|0.25216234|0|0|0|0.60406429|0|0|0.39072048|0|0.22452697|
|ENSMUSG00 Hist1h3b|619|0|0|0|0|0|0|0|0|0|0|0|0|0.1593923|0|0|0|0|0|
|ENSMUSG00 Hist1h2bf|460|0.27720767|0|0.10515985|0|0|0|0|0|0|0|0.20771721|0|0|0.10096541|0.3582744|0.08324045|0|0.71751009|
|ENSMUSG00 Hist1h2ac|2481|0.35977779|0.17455366|0.40944947|0.31218773|0.3492038|0.43940352|0.40047008|0.73890984|1.12885251|1.59639624|1.19389255|1.43982362|2.22699506|1.16063419|2.3249568|2.62370143|2.8887616|2.66065812|
|ENSMUSG00 Hist1h2bn|380|0.44742291|0|0|0|0|0|0|0.20101264|0|0|0.12572357|0|0|0|0|0|0|0|
|ENSMUSG00 Hist1h2ae|703|1.02786343|0.52125439|0.89453186|0.97214177|0.7531311|0.7048753|1.21141994|0.97789935|1.17173587|1.40848473|0.81550427|2.22309894|1.47364191|1.05704893|1.875462|0.59914179|0.59970024|0.70424176|
|ENSMUSG00 Hist1h3e|835|0.20361761|0.19947841|0.4055266|0.32738475|0.40350116|1.00885805|0.56661891|0.73183047|2.07165708|1.97637678|1.02987934|1.67112762|3.24940772|2.1692448|4.29285672|2.10942268|2.24908808|2.96456264|
|ENSMUSG00 Hist1h4f|390|0|0.08541768|0|0|0.12341519|0|0.24262912|0.09792924|0|0|0|0|0|0.1190874|0.84516012|0|0.09827256|0.42314697|
|ENSMUSG00 Gm10264|428|0|0|0|0|0|0|0|0|0|0|0|0|0|0|0|0|0|0|
|ENSMUSG00 Vmn1r223|1085|0|0|0|0|0|0|0|0|0|0|0|0|0|0|0|0|0|0|
|ENSMUSG00 Vmn1r217|896|0|0|0|0|0|0|0|0|0|0|0|0|0|0|0|0|0|0|
|ENSMUSG00 Vmn1r204|908|0|0|0|0|0|0|0|0|0|0|0|0|0|0|0|0|0|0|
|ENSMUSG00 Vmn1r203|935|0|0|0|0|0|0|0|0|0|0|0|0|0|0|0|0|0|0|
|ENSMUSG00 Vmn1r201|902|0|0|0|0|0|0|0|0|0|0|0|0|0|0|0|0|0|0|
|ENSMUSG00 Vmn1r200|1062|0|0|0|0|0|0|0|0|0|0|0|0|0|0|0|0|0|0|
|ENSMUSG00 Vmn1r199|1103|0|0|0|0|0|0|0|0|0|0|0|0|0|0|0|0|0|0|
|ENSMUSG00 Vmn1r198|902|0|0|0|0|0|0|0|0|0|0|0|0|0|0|0|0|0|0|
|ENSMUSG00 Vmn1r197|896|0|0|0|0|0|0|0|0|0|0|0|0|0|0|0|0|0|0|
|ENSMUSG00 V1rh19|905|0|0|0|0|0|0|0|0|0|0|0|0|0|0|0|0|0|0|
|ENSMUSG00 Vmn1r195|1087|0|0|0|0|0|0|0|0|0|0|0|0|0|0|0|0|0|0|
|ENSMUSG00 Vmn1r194|890|0|0|0|0|0|0|0|0|0|0|0|0|0|0|0|0|0|0|
|ENSMUSG00 Vmn1r188|926|0|0|0|0|0|0|0|0|0|0|0|0|0|0|0|0|0|0|
|ENSMUSG00 Hist1h2bj|462|0.64401782|0|0|0.09861698|0|0|0|0|0|0.1020578|0|0.12081312|0|0.10052833|0|0.33152041|0.08295735|0|
|ENSMUSG00 Hist1h2ag|470|0.09043654|0.0708785|0|0|0|0|0|0|0|0|0|0.11875673|0|0|0|0|0.32618125|0|
|ENSMUSG00 Hist1h2ah|530|0|0|0.09127081|0|0|0|0|0|0|0|0|0.10531257|0|0|0|0|0|0|
|ENSMUSG00 Hist1h2br|2603|0|0|0|0|0|0|0|0|0|0|0|0|0.01895195|0|0|0|0|0.02113296|
|ENSMUSG00 Hist1h2ap|518|0|0|0|0|0|0|0|0|0.1590213|0|0|0|0|0|0|0|0|0|
|ENSMUSG00 Hist1h4n|429|0.29723899|0|0.33827644|0.2124058|0.22439125|0.11550754|0.33085789|0.08902658|0.96005864|0.2198168|0.22272708|0.78063864|0.11499281|0.32478383|1.53665477|0.0892555|0.26801606|1.41048992|
|ENSMUSG00 Hist1h4m|401|0|0|0.48252898|0|0.12002973|0|0.58993365|0.3809716|0.20541903|0.4703312|0.11913955|0.41757354|0.24604448|0.23164133|0.41098809|0.1909756|0.2867304|0.68589909|
|ENSMUSG00 Hist1h2bq|3928|0|0|0|0|0|0|0|0|0|0|0|0|0|0|0|0|0|0|
|ENSMUSG00 Hist1h2bp|1087|0|0.03064664|0|0|0|0.09117338|0.08705185|0|0.11367024|0.04337691|0.08790241|0|0|0.12818055|0.34113524|0.1409038|0.03525878|0|
|ENSMUSG00 Hist1h2an|392|0.10843157|0|0|0.11622715|0|0|0.48278244|0.0974296|0.31520293|0|0|0|0.37754018|0|0.21021202|0.19536024|0.19554233|0|
|ENSMUSG00 Hist1h3c|629|0|0|0.07690545|0|0.38260671|0|0.07521889|0|0|0|0.5316768|0|0|0.44302786|0|0|0.06093211|0|
|ENSMUSG00 Hist1h3h|487|0.08727962|0|0|0|0|0|0|0|0|0|0.19620106|0|0|0|0|0|0|0|
|ENSMUSG00 Hist1h2bl|380|0.22371145|0|0|0|0|0|0|0|0.21677114|0|0|0.14688332|0|0|0.32527544|0.30229427|0|0|
|ENSMUSG00 Gm7731|469|0|0|0|0|0|0|0|0|0|0|0|0|0|0|0|0|0|0|
|ENSMUSG00 Gm5096|1223|0|0|0|0|0|0|0|0|0|0|0|0|0|0|0|0|0|0|
|ENSMUSG00 Gm5152|923|0|0|0|0|0|0|0|0|0|0|0|0|0|0|0|0|0|0|
|ENSMUSG00 Rps2-ps10|706|0.48164505|0.23592701|0.1370355|0.12906811|0.13635106|0.63169207|0.13403025|0.10819377|0.29168921|0.2003571|0.27067965|0.31623605|0.34937618|0.26313931|0.35015487|0.05423599|0.5428654|0.23374975|
|ENSMUSG00 Ctxn3|1470|0|0|0|0|0|0|0|0|0|0|0|0|0|0|0|0|0|0|
|ENSMUSG00 Gm5507|1007|0.29546796|0.33081325|0.6244845|0.135733|0.81255482|0.44287448|0.65777309|0.68268445|0.77710407|0.6555212|0.52187144|0.83141503|0.8328129|0.64569734|0.2454909|0.38024436|0.53283829|0.60089392|
|ENSMUSG00 Prdm6|3389|0.0250842|0|0.07136844|0|0.15622637|0|0|0.13523424|0|0.18086726|0|0|0.10189537|0.06852182|0.25530619|0.18077596|0|0|
|ENSMUSG00 Gm4950|617|0.48223052|0.32395035|0.07840118|0.07384286|0.0780096|0|0.07668181|0.30950083|0.66752862|0.45851575|0.23229315|0.18092597|0.15990897|0.1505481|0.20033171|0|0.06211717|0.26746729|
|ENSMUSG00 Gm10267|254|0.66937285|0.39345938|0.19044697|1.07624514|0|0.1950895|4.47048935|6.46564293|4.05379092|3.15575573|1.5047231|5.27392087|1.55376117|9.6910891|3.73084955|0.60300169|1.96158212|3.24856929|
|ENSMUSG00 Olfr707|923|0|0|0|0|0|0|0|0|0|0|0|0|0|0|0|0|0|0|
|ENSMUSG00 Rps19-ps13|470|0|0|0|0|0|0|0|0.08126043|0|0|0|0.11875673|0|0|0|0|0|0|
|ENSMUSG00 Rpl17-ps3|581|0.87790381|0.57337167|0.24977727|0.70576489|0.57990269|0.85288698|0.89576501|0.26294253|0.56711209|0.32461758|0.82228844|0.48034133|0.76417772|0.31975275|0.28365959|0.85676058|0.46176261|0.18936009|
|ENSMUSG00 Olfr810|1005|0|0|0|0|0|0|0|0|0|0|0|0|0|0|0|0|0|0|
|ENSMUSG00 Olfr764|929|0|0|0|0|0|0|0|0|0|0|0|0|0|0|0|0|0|0|
|ENSMUSG00 Olfr9|1051|0|0|0|0|0|0|0|0|0|0|0|0|0|0|0|0|0|0|
|ENSMUSG00 Gm8444|356|0|0|0.1358807|0.12798046|0.4056061|0|0.26580157|0.32184609|0.11569246|0.1324458|0.13419932|0|0.1385728|0.39138276|0.11573471|0.10755789|0.64594882|0.15451996|
|ENSMUSG00 Dsg1a|5671|0|0|0|0|0|0.00873792|0|0|0|0|0|0|0|0|0|0|0|0.05820043|
|ENSMUSG00 Rdh16|6568|0|0.02536|0|0.06243139|0|0|0.0504246|0.09885366|0.26337297|0|0|0|0|0.00707127|0.0501846|0|0.09336491|0|
|ENSMUSG00 Gm6020|65|9.15496099|11.7876395|12.6515387|13.3178437|13.3288404|6.86114773|21.836621|14.6893855|13.3064128|18.8602813|30.8699732|9.44572751|14.4200989|17.1485861|4.43709063|14.7271567|24.1750488|27.0814064|
|ENSMUSG00 Zfp616|5195|0|0|0|0|0|0|0|0|0|0|0|0|0|0|0|0|0|0|
|ENSMUSG00 Epc2|4683|4.08441794|3.64215286|3.36745054|2.78250239|3.61785973|3.98919081|3.94019748|3.46610528|4.300706|2.92992839|4.35616213|4.70791944|6.12040229|4.61168068|3.95035204|4.90590742|4.23119699|5.03927115|
|ENSMUSG00 Lyz1|1260|0.13493707|0.0264388|0.07678338|0.28927647|0.34379945|0.19663783|0.30039796|0.3031143|0.45762795|0.82326625|0.41708297|0.35438516|0|0|0.6212933|0.24311497|0.21292387|0.21829011|
|ENSMUSG00 Lyz2|1309|0.12988595|0.8907191|0.07390914|0.10441798|0.51477993|0.18927706|3.72284639|2.50920291|17.5569716|0.0720408|1.49638907|0.1705597|1.13060162|0.2483641|1.32197506|0.38027341|0|0|
|ENSMUSG00 Gm10271|117|0|0|0|0|0|0|0|0|0|0|0|0|0|0|0|0|0|0|
|ENSMUSG00 Tmem19|3003|24.6283737|26.4351072|29.993844|31.8305265|29.2830585|25.4611615|19.8672288|24.5713358|21.8344766|21.8560702|25.995432|20.1664981|22.522164|20.8325627|23.420265|24.0352298|22.1559945|23.8867383|
|ENSMUSG00 Scyl2|11527|2.48902523|2.69057903|2.32488382|2.56915753|2.22558474|2.94900453|2.19591249|1.56056403|2.36535729|2.5197218|2.47018958|2.37266198|1.85310316|1.95414092|1.62275582|3.70382816|2.85942703|1.96614487|
|ENSMUSG00 Gm4925|1956|0.36942126|0.13624906|0.09892337|0.18634374|0.04921465|0.45600675|0.21769638|0.37098959|0.33690401|0.48211353|0.2930979|0.42803422|0.3026498|0.47488842|0.40021961|0.35236755|0.25472488|0|
|ENSMUSG00 I830134H01F|2644|0|0|0|0|0|0|0|0|0|0|0|0|0|0|0|0|0|0|
|ENSMUSG00 Dazap1|2931|16.6482232|18.2987921|17.8574411|17.0057259|16.323143|15.807508|16.6587119|15.9102434|15.7523317|19.09515|17.0985778|16.7770722|13.3133902|13.7858601|15.9970558|14.1744487|15.6914215|15.9903647|
|ENSMUSG00 C330046G03|2630|0.95353817|0.79798948|0.84607696|0.72759082|0.45752779|0.52755762|0.77355331|1.23435521|0.89263551|0.84261713|0.74478072|1.1672477|1.18171513|0.30020893|1.12795134|0.43677499|1.19496439|0.56473227|
|ENSMUSG00 Krtap10-4|634|0|0|0|0|0|0|0|0|0|0|0|0|0|0|0|0|0|0|
|ENSMUSG00 Krtap12-1|670|0|0|0|0|0|0|0|0|0|0|0|0|0|0|0|0|0|0|
|ENSMUSG00 Gm10272|314|0|0|0|0|0|0|0|0|0|0|0|0|0|0|0|0|0|0|
|ENSMUSG00 Gm11733|2308|0|0|0|0|0|0|0|0.01654783|0|0|0|0|0|0|0.03570325|0|0|0|
|ENSMUSG00 Ank3|10574|25.0512821|28.7856927|28.5968355|30.1097577|28.2901367|30.2499428|19.159532|21.6714975|21.0957225|21.6757678|19.8211408|23.8169349|13.6649504|16.0845705|16.2912526|16.8711878|19.0399128|16.9907076|
|ENSMUSG00 Cd300lh|848|0|0|0|0|0|0|0|0|0|0|0|0|0|0|0|0|0|0|
|ENSMUSG00 Gm11710|848|0|0|0|0|0|0|0|0|0|0|0|0|0|0|0|0|0|0|
|ENSMUSG00 Dppa5b|590|0|0|0|0|0|0|0|0|0|0|0|0|0|0|0|0|0|0|
|ENSMUSG00 Gm10273|629|0|0|0|0|0|0|0|0|0|0|0|0|0|0|0|0|0|0|
|ENSMUSG00 Strada|3939|8.72980132|8.73628325|9.05084841|9.1376555|9.2500295|9.88790266|9.57304014|8.35792101|8.96086418|10.0310458|9.14504156|10.0181958|9.74364855|8.2064191|10.1670252|9.75977909|8.75696045|9.35671528|
|ENSMUSG00 Pex11c|3739|10.9588098|10.9320463|9.13391614|9.72391371|9.10116877|10.0324737|11.4643721|12.4311724|10.4535981|14.3633728|13.4418979|12.942546|13.8139923|10.7321989|11.6805695|8.8173878|10.5374253|13.6382567|
|ENSMUSG00 Gm3699|199|1.7087508|0.33480296|1.4584984|1.37369983|1.45121378|0.49801742|1.18876078|0.57576486|2.06967416|0.71081462|0.7202255|1.40240358|3.71848623|2.80064849|1.44930096|2.50139646|1.15556674|3.04070439|
|ENSMUSG00 Marcks|4184|12.7190441|16.4414738|13.0761144|13.6770248|13.3444148|13.3474978|6.27594092|5.93333211|5.11878303|5.24021917|5.62931514|7.3504852|3.83194864|3.19691615|5.39637969|3.55084983|2.89462472|3.91795263|
|ENSMUSG00 Sult3a1|1082|0|0|0|0|0|0|0|0|0|0|0|0|0|0|0|0|0|0|
|ENSMUSG00 Nkain2|557|0.07631091|0|0|0.16359441|0|0.08896362|0|0|0|0|0|0|0|0|0|0|0|0|
|ENSMUSG00 Nkain2|217|0|0|0|0|0|0|0|0|0.37959922|0|0|0|0|0|0|0|0|0|
|ENSMUSG00 Pcgf1|1461|14.0811124|10.1694393|9.96607299|9.1059719|11.4646882|9.56459334|13.1154243|11.7635736|12.234735|11.0373309|14.420778|11.9577703|12.4933671|12.5885411|10.9137592|9.27780631|11.1752241|11.4837629|
|ENSMUSG00 Gm10275|490|1.56141463|1.01978248|0.78977192|1.02279895|1.76811148|1.1124083|1.64146029|1.48092989|1.17675759|1.25093703|0.77999932|1.13909515|1.7115155|2.65394785|1.0931025|1.87545832|2.03364024|1.23489831|
|ENSMUSG00 Taar7e|1214|0.070025|0.10976242|0|0|0.11894215|0|0|0.18875981|0.10177887|0|0|0.13792997|0.32508677|0.19128537|0.03393868|0.15770431|0.12628104|0|
|ENSMUSG00 Taar7d|1076|0.86906494|0.27863945|0.31469769|0.12702893|0.80518088|0.59868544|0.21985446|0.39044278|0.26794202|0.35056285|0.53280623|0.98559255|0.91695014|1.68338235|1.18703367|0.28468853|1.24667323|0.66460817|
|ENSMUSG00 Taar7b|1076|0|0|0|0|0|0|0|0|0|0|0|0|0|0|0|0|0|0|
|ENSMUSG00 Taar5|1121|0|0|0|0|0|0|0|0|0|0|0|0|0|0|0|0|0|0|
|ENSMUSG00 Taar4|1043|0|0|0|0|0|0|0|0|0|0|0|0|0|0|0.03950293|0|0|0|
|ENSMUSG00 Taar3|1106|0|0|0|0|0|0|0|0|0|0|0|0|0|0|0|0|0|0|
|ENSMUSG00 4930444G20I|1772|0|0|0|0|0|0|0|0|0|0|0|0|0|0|0|0|0|0|
|ENSMUSG00 4933406P04f|1788|1.30748584|0.98746275|2.48901834|1.68178352|1.02293798|0.94227793|1.29660026|1.26026384|3.31703483|1.42401453|3.66060923|1.09258847|0.88289785|5.35094072|2.39651108|1.24208905|2.29357593|1.96900606|
|ENSMUSG00 Gm11568|1134|0|0|0|0|0|0|0|0|0|0|0|0|0|0|0|0|0|0|
|ENSMUSG00 Gm11563|1017|0|0|0|0|0|0|0|0|0|0|0|0|0|0|0|0|0|0|
|ENSMUSG00 4930572O03|1164|0|0|0|0|0|0|0|0|0|0|0|0|0|0|0|0|0|0|
|ENSMUSG00 Krtap3-2|1030|0|0|0|0|0|0|0|0|0|0|0|0|0|0|0|0|0|0|
|ENSMUSG00 Krtap3-3|731|0|0|0|0|0|0|0|0|0|0|0|0|0|0|0|0|0|0|
|ENSMUSG00 Gm5595|3144|2.16311328|1.7376955|3.13874052|1.15931411|1.60745928|1.6233879|2.76893541|2.02866769|2.135306|1.40972204|1.42838616|2.94701017|1.67891703|3.58966706|2.89616535|2.43578927|3.34014165|0.8923233|
|ENSMUSG00 Arid1b|8111|17.0523786|21.0408035|18.6313536|21.2891577|19.7310623|21.254341|16.4961479|19.291613|16.4624194|15.2479712|17.2286714|17.4032582|15.8195434|14.0231254|13.6441104|15.0924759|16.3067502|14.4593041|
|ENSMUSG00 Ube2u|1896|0|0|0|0.14418052|0|0|0|0|0.13033768|0|0|0|0|0.07348748|0|0|0|0|
|ENSMUSG00 Zfp820|3690|0.18430429|0.37917115|0.47193688|0.12347166|0.19565823|0.22829173|0.05128746|0.04140098|0.33484972|0.11500172|0.50493859|0|0.05347633|0.35242126|0.49129227|0.08301487|0.27004979|0.81991893|
|ENSMUSG00 Psmb3|2350|49.6315758|44.9511436|41.6218206|45.037577|46.12472|42.2779706|42.2995483|38.0461336|39.4689504|42.0945427|43.7496641|45.6025839|46.1830715|47.1358078|45.0236577|47.7084674|40.5280203|43.7966122|
|ENSMUSG00 Zfp125|541|7.30680475|6.34238097|6.16963696|8.08476942|6.22779049|7.69395494|4.02288952|5.15350347|5.93816678|5.75221149|4.23881332|6.60296193|3.19152884|4.97921824|3.65559091|4.74209001|5.24241399|4.47393844|
|ENSMUSG00 Tmem100|1764|0.21686314|0.35881235|0.24680373|0.36159559|0.38199939|0.53373126|0.69735241|0.36806737|0|0.05345885|0.43333296|0.41133992|0|0.28961732|0.16349824|0.10853347|0.36935774|0|
|ENSMUSG00 Msi2|12668|8.08296251|6.79511515|8.18317616|7.05282665|6.44773243|6.63415189|6.92061841|7.29297611|8.48569673|6.58427487|6.49419628|7.20386867|6.59290622|5.8733366|5.91288511|6.91878752|6.73463348|6.21392735|
|ENSMUSG00 Gm11444|572|0|0|0|0|0|0|0.16542895|0|0|0|0|0|0|0|0|0|0|0|
|ENSMUSG00 Gm11428|530|0|0.31427259|0|0|0.09081495|0|2.23173014|1.72946728|6.13912216|0|1.98311148|1.05312571|0.65155362|2.27838919|0.69964906|0.14449286|0|0|
|ENSMUSG00 Slfn9|6294|0|0|0|0.02895522|0|0|0.04510265|0|0.01308755|0.00749137|0|0|0.02351378|0|0|0.04258568|0|0|
|ENSMUSG00 Gm11426|2217|0.01917238|0|0|0|0|0|0|0|0|0|0|0|0|0|0|0|0|0|
|ENSMUSG00 Gm10277|2294|0.7411539|1.32147924|0.99108802|0.29791441|1.23791783|0.73443459|0.57748693|1.48174534|0.95156292|1.13046586|0.3748689|1.09490184|1.16125699|1.2754915|1.23927958|0.60089881|0.26731506|1.19897791|
|ENSMUSG00 Fbp1|2096|0.20279187|0.01589356|0.16155282|0.06521142|0|0.04728314|0.18058274|0|0.07860022|0|0|0|0|0|0.255544|0|0|0|
|ENSMUSG00 Cacng7|2206|0|0|0|0|0|0|0|0|0|0|0|0|0|0|0|0|0|0|
|ENSMUSG00 Fam57a|6608|4.08456216|3.64989943|3.90179959|3.6956295|4.20280266|3.20203045|4.46062333|3.96488923|3.09771464|3.7888958|4.43190822|3.750326|4.70325483|3.66182953|3.75353157|4.61828301|4.88358686|2.94691643|
|ENSMUSG00 E130309D14|1380|1.4168392|1.20698892|1.36707803|0.82538124|1.18585899|0.61043223|1.5085202|1.49448531|2.62638651|2.05003058|2.00793304|1.73918369|2.50234363|2.79337627|2.80648279|1.22085995|1.52749734|3.26865707|
|ENSMUSG00 Olfr23|1917|0|0|0|0|0|0|0|0|0|0|0|0|0|0|0|0|0|0|
|ENSMUSG00 Olfr390|1029|0|0|0|0|0|0|0|0|0|0|0|0|0|0|0|0|0|0|
|ENSMUSG00 Olfr382|938|0|0|0|0|0|0|0|0|0|0|0|0|0|0|0|0|0|0|
|ENSMUSG00 Olfr1|1879|0|0|0|0|0|0|0|0|0|0|0|0|0|0|0|0|0|0|
|ENSMUSG00 Olfr377-ps1|1326|0|0|0|0|0|0|0|0|0|0|0|0|0|0|0|0|0|0|
|ENSMUSG00 Spata22|3439|0|0|0|0|0|0|0|0|0|0|0|0|0|0|0|0|0|0|
|ENSMUSG00 Nlrp1a|5565|0|0|0|0|0|0|0|0|0|0|0|0|0|0|0|0|0|0|
|ENSMUSG00 Ahnak|9814|11.6419312|9.60961927|10.8438727|9.86057246|9.72054949|11.613133|9.94559371|10.4645782|8.51512538|7.43726194|9.3904519|10.1348595|6.27833347|4.99744816|4.9119442|8.09198286|8.22058844|7.61742165|
|ENSMUSG00 Sat2|1393|3.14288093|3.51543105|2.98644911|2.55115683|2.66055859|3.02367721|5.50226417|4.96254475|5.02635153|5.44957877|6.65351181|4.8883782|5.56002226|6.10141277|9.40566746|5.22269591|2.94394384|4.65978076|
|ENSMUSG00 Sco1|3919|4.89151171|4.52218926|5.12248406|5.16180242|5.23199783|4.6151436|5.23952607|5.28203167|3.16334301|4.8967431|5.30291068|5.66844443|4.12882594|4.62189183|5.06740238|5.87207325|4.8213484|3.3968369|
|ENSMUSG00 Slc47a2|2404|0.05304306|0.08314366|0.42256412|0.34113927|0.36038878|0.24735142|0.1377657|0.17475725|0.20558993|0|0.03974622|0.11608915|0.16416611|0.4829876|0.32563626|0.19113448|0.54205245|0.25170556|
|ENSMUSG00 Gm10279|866|1.07980817|0.76935091|1.00545444|1.05221811|0.88927342|0.68664296|0.7648701|0.92614371|0.80851128|0.81669809|0.33100433|0.90233173|0.9114442|0.85808937|1.37972878|1.14960253|0.7081071|1.07985544|
|ENSMUSG00 Pabpn1l|1141|0.07450513|0.02919623|0|0|0.08436796|0|0|0|0|0|0|0|0|0|0|0|0|0|
|ENSMUSG00 4930438A08I|2055|0.10341892|0.06484262|0|0.19953742|0|0.09645301|0|0|0|0|0|0|0|0|0|0|0|0|
|ENSMUSG00 Irgm2|3472|4.12564641|3.05112338|3.09300799|3.24123789|4.46384776|6.39390111|4.45600403|3.33303512|3.03679379|3.82963661|2.76577381|3.87430146|3.22532984|3.39769533|2.83616701|3.93713907|2.95836624|3.16872735|
|ENSMUSG00 Gm10281|71|0|0|0|0|0|0|0|0|0|0|0|0|0|0|0|0|0|0|
|ENSMUSG00 1700024J04R|1819|0|0|0|0|0|0|0|0|0|0|0|0|0|0|0|0|0|0|
|ENSMUSG00 9930111J21R|7013|0.37577655|0.16150555|0.39316858|0.1754097|0.13040162|0.55820133|0.30358913|0.09258104|0.20555084|0.18152987|0.19755793|0.20693102|0.30951153|0.16556426|0.24675108|0.21839787|0.31697208|0.04706326|
|ENSMUSG00 9930111J21R|4891|0.04345244|0.05448848|0.05934189|0.15835979|0.118091|0.21275964|0.08706075|0.11713066|0.23578459|0.0192806|0.12698312|0.19400251|0.07060385|0.35134558|0.11793535|0.18789094|0.13321346|0.1799521|
|ENSMUSG00 Atxn1l|7522|2.220757|2.04164374|1.90355823|1.90191012|1.80446723|2.04877694|1.13218322|1.61462164|1.47837799|1.46053096|2.03878778|1.84766019|1.27887847|1.54360833|1.29816123|2.27035508|1.59480603|1.69663823|
|ENSMUSG00 Gm12166|1739|0.02444231|0.01915635|0.02781687|0.02619957|0|0|0.05441366|0.04392456|0.04736805|0.02711369|0.02747266|0.03209641|0|0|0.02369267|0|0.02203927|0.03163261|
|ENSMUSG00 Ccdc99|3448|1.86145057|0.85987459|0.81370788|1.75743007|1.24238434|1.69583021|1.55056053|0.80859785|0.63308739|1.03928464|0.96990925|1.63497155|1.58812147|0.76778219|1.23078894|0.54415306|1.83406003|1.14868204|
|ENSMUSG00 Fam196b|5367|0|0|0.00901314|0|0|0.00923286|0.13223219|0.14943925|0|0|0|0|0|0.09519004|0.10747564|0.20689913|0.18566866|0|
|ENSMUSG00 Anp32-ps|387|0.1098325|0|0|0|0|0|0|0|0|0|0|0|0|0|0.31939191|0|0.09903436|0|
|ENSMUSG00 Hba-a2|754|0|0.08836311|0|0|0|0|0|0|0|0|0|0|0|0|0|0.0507833|0|0|
|ENSMUSG00 Hba-a1|930|0|0|0|0|0|0|0|0.2053355|0|0|0|0|0.10609014|0|0.17721099|0|0|0|
|ENSMUSG00 B3gnt9-ps|4838|0|0.13771349|0.03999465|0.03766932|0.03979489|0|0|0|0.00851313|0|0.07899952|0|0.10196758|0.10559838|0.04258119|0.11871829|0.04753158|0|
|ENSMUSG00 Ces3a|2036|0|0|0|0|0|0|0|0|0|0|0|0|0|0|0|0|0|0|
|ENSMUSG00 4932416K20I|3552|0|0|0|0|0|0|0|0|0|0|0|0|0|0|0|0|0|0|
|ENSMUSG00 Gm12070|1375|0.09273857|0.07268268|0|0.0331353|0.07001007|0|0|0.13888146|0.23963064|0.13716568|0.1389817|0.08118642|0.17938879|0.16888759|0.11985907|0.11139086|0.05574734|0.16002649|
|ENSMUSG00 Gm5356|617|0.13778015|0|0.07840118|0|0.1560192|0|0.07668181|0.06190017|0|0.15283858|0.23229315|0|0.23986346|0.45164429|0.26710895|0|0|0.35662306|
|ENSMUSG00 4933402J07R|1298|0|0|0|0|0|0|0|0|0|0|0|0|0|0|0|0|0|0|
|ENSMUSG00 Gm10159|520|2.45222169|2.81878336|3.53498876|2.89137397|2.86940314|3.81174874|2.1836621|2.05651398|1.6633016|2.99225617|1.19437396|1.50272938|4.07937008|4.01919987|3.01088293|3.01966712|3.4641076|2.75045534|
|ENSMUSG00 Gm8054|467|0|0|0|0.09756112|0|0.10610864|0|0|0|0.1009651|0|0|0|0.09945201|0|0|0|0|
|ENSMUSG00 Gm7125|272|6.8758373|5.51132441|9.07003694|6.86765742|8.67082448|5.46537503|11.3063387|5.61652977|11.205155|8.49406051|8.2552318|11.9018692|9.97520386|10.074269|13.3299151|8.44645752|5.49531463|8.89853197|
|ENSMUSG00 Olfr372|1015|0|0|0|0|0|0|0|0|0|0|0|0|0|0|0|0|0|0|
|ENSMUSG00 Fcho1|4777|2.05541044|1.20643305|0.88099166|1.44017536|1.37030388|1.63896418|1.89171482|1.11131136|1.75885477|1.09560981|1.60016608|0.95810851|1.65231458|1.25419453|0.79349867|1.16226462|0.75417038|1.08244841|
|ENSMUSG00 Ell|3073|7.44151796|11.6210292|7.9022168|10.5562849|8.55191354|8.53022976|8.42174923|11.3222514|7.66634787|10.1267374|10.3385445|7.99182933|8.49221745|10.0958836|8.87583144|10.6286652|11.8982386|10.1855456|
|ENSMUSG00 Sabp4|1457|33.7532523|32.3068768|29.7478951|33.2717578|33.6956501|30.4730606|44.4226114|36.9866025|40.8472994|46.1474968|44.2008538|40.1474361|58.8123132|48.8029498|42.2478548|51.1943056|37.5371487|50.1387054|
|ENSMUSG00 Gm10283|351|0|0|0|0|0|0|0|0|0|0|0|0|0|0|0|0|0|0|
|ENSMUSG00 Gm10284|2235|0.03803595|0.04471529|0.02164364|0.10192627|0.06460661|0.04434249|0.08467594|0.01708832|0.1289958|0.08438605|0.08550328|0.19978761|0.02207245|0.06234106|0.0737388|0.05139679|0.06859293|0.12306288|
|ENSMUSG00 Sp140|3310|1.57949748|2.03299234|1.47605032|1.69305391|2.06487407|2.1857097|1.01486411|0.9346177|1.18209033|1.12534911|1.90522493|1.26470535|1.35625513|1.65235848|0.9460176|1.6773831|1.60947289|1.14671552|
|ENSMUSG00 Sp110|6090|0.46762673|0.224274|0.21446393|0.29177023|0.26871682|0.25223887|0.04661348|0.1442406|0.15554842|0.17033095|0.30594801|0.29328427|0.18631102|0.31267776|0.11501255|0.07544947|0.08810643|0.17162119|
|ENSMUSG00 Gm4885|1357|0.37587481|0.46642962|0.24953184|0.53719728|0.35469362|0.18258192|0.27892515|0.42217099|0.21245806|0.48644793|0.28165046|0.24678996|0.29082928|0.4449323|0.42507132|0.47969073|0.59311145|0.44591022|
|ENSMUSG00 Fam149a|8586|3.47031544|3.63547425|3.8029505|2.95037743|2.91504779|3.41086251|3.05829686|3.84326062|3.36265404|3.40477941|3.0325358|3.35440041|3.20605752|3.42407493|2.91760377|2.44389155|3.02200128|3.28670763|
|ENSMUSG00 Fat1|14749|3.8732766|3.261497|2.67958331|2.95008457|3.28624633|2.98008507|1.52052409|1.51226273|0.99412839|1.46736543|1.46735753|1.63106316|2.05034004|2.32708527|1.81299137|1.23057481|1.33306599|1.92078717|
|ENSMUSG00 Mfhas1|6661|7.15970688|6.57655846|8.0973557|8.20796475|4.73298454|8.01949359|8.86446829|6.41031465|8.02583658|8.61468336|5.15691265|6.6281623|6.88025089|6.23345055|9.37720443|6.30033114|4.66636042|9.86051245|
|ENSMUSG00 U6|104|0|0|0|0|0|0|0|0|0|0|0|0|0|0|0|0|0|0|
|ENSMUSG00 Snora33|128|2.6565735|0.78077096|1.13375462|0.35594566|0.37603065|1.93565366|3.69630304|1.79026886|2.25238758|9.20912173|3.73241863|3.48847891|4.62486724|1.81422216|1.60943578|2.69230833|3.59309033|0|
|ENSMUSG00 Mir451|71|0|0|0|0|0|0|0|0|0|0|0|0|0|0|0|0|0|0|
|ENSMUSG00 Mir465|73|0|0|0|0|0|0|0|0|0|0|0|0|0|0|0|0|0|0|
|ENSMUSG00 Mir433|123|0|0|0|0|0|0|0|0|0|0|0|0|0|0|0|0|0|0|
|ENSMUSG00 Mir470|74|0|0|0|0|0|0|0|0|0|0|0|0|0|0|0|0|0|0|
|ENSMUSG00 Mir484|66|0|0.50474082|0.73293228|0.69031885|0.72927157|0|0.71685877|0|1.87211435|0.71440459|0.72386301|0|0|1.40739659|0|0|0|0|
|ENSMUSG00 Mir490|83|0|0|0|0|0|0|0|0|0|0|0|0|0|0|0|0|0|0|
|ENSMUSG00 Mir127|96|0|0|0|0|0|0|0|0|0|0|0|0|0|0|0|0|0|0|
|ENSMUSG00 Mir491|85|0|0|0|0|0|0|0|2.24661191|0|0|0|0|0|0|0|0|0|0|
|ENSMUSG00 Mir463|74|0|0|0|0|0|0|0|0|0|0|0|0|0|0|0|0|0|0|
|ENSMUSG00 Mir431|90|0|0|0|0|0|0|0|0|0|0|0|0|0|0|0|0|0|0|
|ENSMUSG00 Mir505|89|0|0|0|0|0|0|0|0|0|0|0|0|0|0|0|0|0|0|
|ENSMUSG00 Mir486|127|0|0|0|0|0|0|0|0|0|0|0|0|0|0|0|0|0|0|
|ENSMUSG00 Mir466|72|0|0|0|0|0|0|0|0|0|0|0|0|0|0|0|0|0|0|
|ENSMUSG00 Mir362|64|0|0|0|0|0|0|0|0|0|0|0|0|0|0|0|0|0|0|
|ENSMUSG00 Mir341|95|0|0|0|0|0|0|0|0|0|0|0|0|0|0|0|0|0|0|
|ENSMUSG00 Mir455|81|1.04951052|0|0|0|0|0|0|0|0|0|0|0|0|0|0|0|0|0|
|ENSMUSG00 Mir488|108|0|0|0|0|0|0|0|0|0|0|0|0|0|0|0|0|0|0|
|ENSMUSG00 Mir471|66|0|0|0|0|0|0|0|0|0|0|0|0|0|0|0|0|0|0|
|ENSMUSG00 Mir495|79|0|0|0|0|0|0|0|0|0|0|0|0|0|0|0|0|0|0|
|ENSMUSG00 Mir363|74|0|0|0|0|0|0|0|0|0|0|0|0|0|0|0|0|0|0|
|ENSMUSG00 Mir489|106|0|0|0|0|0|0|0|0|0|0|0|0|0|0|0|0|0|0|
|ENSMUSG00 Mir500|91|0|0|0|0|0|0|0|0|0|0|0|0|0|0|0|0|0|0|
|ENSMUSG00 Mir497|83|0|0|0|0|0|0|0|0|0|0|0|0|0|0|0|0|0|0|
|ENSMUSG00 Mir504|78|0|0|0|0|0|0|0|0|0|0|0|0|0|0|0|0|0|0|
|ENSMUSG00 Gm10286|5075|0.02512621|0.10502587|0.00953173|0.01795509|0.00948412|0|0|0|0|0.01858156|0.00941378|0|0.04860287|0.06406081|0.0487112|0.00754495|0|0.02167847|
|ENSMUSG00 Mir199a-2|109|0|0|0|0|0|0|0|0|0|0|0|0|0|0|0|0|0|0|
|ENSMUSG00 Mir146b|108|0|0|0|0|0|0|0|0|0|0|0|0|0|0|0|0|0|0|
|ENSMUSG00 Mir485|78|0|0|0|0|0|0|0|0|0|0|0|0|0|0|0|0|0|0|
|ENSMUSG00 Mir136|81|0|0|0|0|0|0|0|0|0|0|0|0|0|0|0|0|0|0|
|ENSMUSG00 Mir328|96|0|0.34700931|1.00778188|0|0|0|0.98568081|0|0|0|0|0.58141315|0.51387414|0|0.42918287|0|0|0|
|ENSMUSG00 Mir434|93|0|0|0|0|0|0|0|0|0|0|0|0|0|0|0|0|0|0|
|ENSMUSG00 Mir496|78|0|0|0|0|0|0|0|0|0|0|0|0|0|0|0|0|0|0|
|ENSMUSG00 Mir452|84|0|0|0|0|0|0|0|0|0|0|0|0|0|0|0|0|0|0|
|ENSMUSG00 Mir532|95|0|0|0|0|0|0|0|0|0|0|0|0|0|0|0|0|0|0|
|ENSMUSG00 Mir483|72|0|0|0|0|0|0|0|0|0|0|0|0|0|0|0|0|0|0|
|ENSMUSG00 Mir494|84|0|0|0|0|0|0|0|0|0|0|0|0|0|0|0|0|0|0|
|ENSMUSG00 Mir1a-1|76|0|0|0|0|0|0|0|0|0|3.10201995|0|0|0.64910417|0|0|0|0|0|
|ENSMUSG00 U6|106|0|0|0.45635406|0|0|0|0|0|0|0|0|0|0|0|0|0|0|0|
|ENSMUSG00 U6|106|0|0|0|0|0|0|0|0|0|0|0|0|0|0|0|0|0|0|
|ENSMUSG00 n-R5s152|118|0|0|0|0|0|0|0|0|0|0|0|0|0|0|0|0|0|0|
|ENSMUSG00 U6|106|0|0|0|0|0|0|0|0|0|0|0.45070716|0|0|0|0|0|0|0|
|ENSMUSG00 U6|106|0|0|0|0|0|0|0|0|0|0|0|0|0|0|0|0|0|0|
|ENSMUSG00 U6|106|0|2.5141807|0.45635406|1.71928469|0|0|0.44634603|0.72061137|1.55420814|0.88963591|2.25353578|4.21250284|0.46539545|0|0.77738785|0.36123215|1.08470652|0|
|ENSMUSG00 SNORA21|132|4.18611582|8.32822355|3.66466139|5.1773914|5.104901|2.62779648|1.07528816|8.68009146|11.8567242|6.42964135|2.53352053|5.49699707|4.11099311|4.57403891|3.43346299|4.06112503|5.22631321|2.9171496|
|ENSMUSG00 SNORA57|147|2.60235772|0.67985498|4.93607453|1.54969539|0.98228416|1.68546713|2.25298471|1.29906131|2.80180379|1.9245185|1.62499859|1.13909515|7.7185993|6.9508158|4.76480578|4.42816548|1.04289243|7.1100206|
|ENSMUSG00 U3|213|0.39910963|0.62559426|1.36263466|0.42780323|0|0.93056777|0.66637576|0|0.58009177|0.88545922|0.22429558|0.26204536|0.23160525|0|0.19343453|0|1.079614|0|
|ENSMUSG00 SNORA71|131|0.64893399|2.03437522|0|0.34779423|1.83709632|0|0.72233098|0.58309011|0|0.35992903|2.55286038|0.42607376|1.50631808|0|0.94354708|0.87688414|0|0|
|ENSMUSG00 U6|106|0|0|0|0|0|0|0|0|0|0|0|0|0|0|0|0|0|0|
|ENSMUSG00 U6|103|0|0|0|0|0|0|0|0|0|0|0|0|0|0|0|0|0|0|
|ENSMUSG00 U6|106|0|0|0|0|0|0.46747862|0|0|0|0|0|0|0|0|0.38869392|0|0|0|
|ENSMUSG00 U6|101|0|0|0|0|0|0|0|0|0|0|0|0|0|0|0|0|0|0|
|ENSMUSG00 n-R5s71|118|0|0|0|0|0|0|0|0|0|0|0|0|0|0|0|0|0|0|
|ENSMUSG00 U6|104|0|0|0|0|0|0|0|0|0|0|0|0|0|0|0|0|0|0|
|ENSMUSG00 U3|213|0|0.31279713|0|1.06950808|0|1.86113553|0.22212525|0.35861411|0.58009177|1.54955363|0|1.04818146|2.31605245|0.87218943|2.90151802|1.43814488|1.97929233|1.29129359|
|ENSMUSG00 U6|106|0|0|0|0|0|0|0|0|0|0|0|0|0|0|0|0|0|0|
|ENSMUSG00 n-R5s58|118|0|0|0|0|0|0|0|0|0|0|0|0|0|0|0|0|0|0|
|ENSMUSG00 U1|163|0.26076795|0|0.29677012|0|0|1.824027|0|0|0.75803403|0.28926812|0.87929371|0.68485476|0|1.42466526|1.51662169|0.4698234|0.23513066|0|
|ENSMUSG00 n-R5s211|118|0|0|0|0|0|0|0|0|0|0|0|0|0|0|0|0|0|0|
|ENSMUSG00 n-R5s210|118|0|0|0|0|0|0|0|0|0|0|0|0|0|0|0|0|0|0|
|ENSMUSG00 U6|107|0|0|0|0|0|0|0|0|0|0|0|0|0|0|0|0|0|0|
|ENSMUSG00 n-R5s27|118|0|0|0|0|0|0|0|0|0|0|0|0|0|0|0|0|0|0|
|ENSMUSG00 U6|106|0|0.31427259|0|0|0.45407475|0.46747862|0|0|0|0|0.45070716|0|0|0|0|0|0|0|
|ENSMUSG00 n-R5s13|118|0|0|0|0|0|0|0|0|0|0|0|0|0|0|0|0|0|0|
|ENSMUSG00 U6|106|0|0|0|0|0|0|0|0|0|0|0|0|0|0|0|0|0|0|
|ENSMUSG00 n-R5s33|118|0.36021336|1.12925065|0.81989035|0|0.81579532|0|0|0|0|0|0|0|0.4180671|0|0|0.64899335|0|0|
|ENSMUSG00 U6|106|0|0|0|0|0|0|0|0|0|0|0|0|0|0|0|0|0|0|
|ENSMUSG00 U6|106|0|0|0|0|0|0|0|0|0|0|0|0|0|0|0|0|0|0|
|ENSMUSG00 U5|115|0.73922045|0.28967734|0|1.58473198|0|0.86178667|0.4114146|0.33210785|1.07443085|0|0.41543442|0.48535359|0.42897319|0|0|0|0|3.34838041|
|ENSMUSG00 Y_RNA|109|0|0|0|0|0|0|0|0|0|0|0|0|0|0|0|0|0|0|
|ENSMUSG00 U6|110|0|0|0|0|0|0|0|0|0|0|0|0|0|0|0|0|0|0|
|ENSMUSG00 U5|115|0|0|0.84127879|0|0|0.86178667|0.8228292|0|0|0.41000612|0|0.48535359|1.71589277|0|0|0|0.33327215|0|
|ENSMUSG00 U6|106|0|0|0|0|0|0|0|0|0|0|0|0|0|0|0|0|0|0|
|ENSMUSG00 U6|106|0|0|0|0|0|0|0|0|0|0|0|0|0|0|0|0|0|0|
|ENSMUSG00 U3|213|9.97774085|4.3791598|12.7179235|2.5668194|11.5245451|7.21190019|5.99738184|2.15168464|7.3478291|3.54183686|11.214779|10.2197692|5.55852589|5.66923132|2.51464895|3.95489842|2.87897066|4.39039819|
|ENSMUSG00 U1|163|0|0|0|0|0|0.608009|0|0.70292765|0.75803403|0.86780435|0|1.02728213|3.02649799|2.56439746|2.7804731|2.34911702|2.1161759|1.68739591|
|ENSMUSG00 U6|106|0|0|0|0|0|0|0|0|0|0|0|0|0|0|0|0|0|0|
|ENSMUSG00 U6|106|0|0|0|0|0|0|0|0|0|0|0|0|0|0|0|0|0|0|
|ENSMUSG00 U3|213|2.79376744|1.72038421|4.7692213|3.63632748|4.29345798|2.32641942|1.99912728|1.97237759|3.86727848|3.76320167|0.44859116|5.24090729|3.93728917|2.6165683|7.54394685|1.43814488|3.41877766|2.06606974|
|ENSMUSG00 U6|106|0|0|0|0|0|0|0|0|0|0|0|0|0|0|0|0|0|0|
|ENSMUSG00 U6|106|0|0|0|0|0|0|0|0|0|0|0|0|0|0|0|0|0|0|
|ENSMUSG00 U5|116|0|0|0|0|0|0|0|0|1.06516851|0|0|0|0|1.20114019|0|0|1.32159644|0|
|ENSMUSG00 U6|106|0|0|0|0|0|0|0|0|0|0|0|0|0|0|0|0|0|0|
|ENSMUSG00 Gm13268|873|0|0|0|0.46970149|0|0|0|0|0.14153442|0.05400997|0|0.12787093|0.0565085|0|0|0|0.13170549|0.06301158|
|ENSMUSG00 Slc22a14|2783|0|0|0|0|0|0|0|0|0|0|0|0|0|0|0|0|0|0|
|ENSMUSG00 3000002C10|5186|0.21309961|0.46892427|0.63428462|0.50955275|0.92811268|0.42042427|0.84845336|0.75854559|0.92919829|0.44550414|1.31735809|0.9471227|0.78002646|1.11945833|0.91364806|0.71619532|0.58383676|0.70007733|
|ENSMUSG00 Ndufa3|1061|53.9627447|50.3618118|50.881112|48.1375407|58.1121529|50.5335134|54.2245217|54.714846|54.9671125|54.2166428|57.2759163|52.2384099|62.629682|62.2464583|60.3072727|58.2840066|52.3780683|59.0531315|
|ENSMUSG00 Gmppb|2730|11.6149675|11.567994|10.7024221|9.66294677|11.9536426|10.4187799|11.4728914|11.8214579|8.62955568|13.2643737|11.6199899|10.4271018|9.10743088|15.1070878|11.6209517|12.945872|12.3542642|12.9765072|
|ENSMUSG00 BY080835|1140|0.93213105|0.52599307|0.25459753|0.23979497|0.54887281|0.17386924|0|0.26801686|0.43354227|0.2481616|0.25144715|0.14688332|0.51928334|0.08148086|0.83125946|0.60458854|0.20171735|0.53078963|
|ENSMUSG00 Tmem22|1578|11.7980146|8.6765523|7.5411207|9.03713997|8.72352989|7.85055982|4.76724712|5.71191823|5.14178936|7.70905034|6.6606406|7.8523936|7.97188777|3.47300527|3.15930815|2.71771104|4.17751779|3.52086171|
|ENSMUSG00 Rpl7a-ps10|865|0.04913893|0|0.05592316|0|0.05564384|0|0|0|0.04761447|0|0.05523117|0|0|0|0|0|0.04430786|0|
|ENSMUSG00 Ddx43|2190|0.01940876|0.07605684|0|0|0|0|0.19443567|0.06975781|0|0|0|0|0|0|0|0|0.05250178|0|
|ENSMUSG00 Trcg1|2687|0.12655058|0.12397802|0.12601962|0.11869271|0|0.22129989|0.14086395|0.21320656|0.0919684|0.22812026|0.08890018|0.16617987|0.22031374|0.36297947|0.46000993|0.17100383|0.11410881|0.04094463|
|ENSMUSG00 Scn2b|4768|0.05348806|0.02794706|0.02029091|0.03822235|0.0605687|0.09353494|0.02976888|0|0.25914335|0.03955596|0.06011949|0.02341261|0.04138584|0.04870395|0.00864127|0.04818449|0.00803823|0.25381719|
|ENSMUSG00 Mpzl3|6511|1.32522665|2.78843916|2.20656405|1.51147068|2.83128963|1.98637129|1.15538565|2.20555111|1.5497921|1.13694677|1.65829222|1.62020584|1.92448272|2.66780659|1.20231848|1.18794389|2.34867032|1.34333405|
|ENSMUSG00 Ccdc153|1079|0|0.46310789|0.62764544|1.09785649|0|0|0|0|0|0|0|0|0|0|0.22910967|0|0|0|
|ENSMUSG00 Gm4893|329|0|0|0|0|0|0|0|0|0|0|0|0|0|0|0|0|0|0|
|ENSMUSG00 Olfr895|950|0|0|0|0|0|0|0|0|0|0|0|0|0|0|0|0|0|0|
|ENSMUSG00 Olfr143|941|0|0|0|0|0|0|0|0|0|0|0|0|0|0|0|0|0|0|
|ENSMUSG00 Olfr894|948|0|0|0|0|0|0|0|0|0|0|0|0|0|0|0|0|0|0|
|ENSMUSG00 Olfr893|941|0|0|0|0|0|0|0|0|0|0|0|0|0|0|0|0|0|0|
|ENSMUSG00 A630095E13f|614|0|0|0|0|0|0|0|0|0|0|0|0|0|0|0.13420702|0.49890042|0.06242068|0.17918276|
|ENSMUSG00 4930581F22f|3403|0.23731953|0.72440616|0.54016872|0.72297866|0.52332682|0.6843898|0.61174195|0.9090757|0.19364803|0.33253508|0.71599262|0.72168356|1.14523111|1.2283185|0.85962694|0.41632456|0.57438764|1.06688247|
|ENSMUSG00 Eif3g|1064|158.954977|156.263774|140.84699|131.758772|154.664518|144.699571|142.738439|147.528923|158.668729|143.756467|153.69707|143.683364|154.950439|177.439112|149.626703|161.511509|143.867697|162.959309|
|ENSMUSG00 Mmp27|2344|0|0|0|0|0|0|0|0|0|0|0|0.02381214|0|0.01981403|0|0|0|0|
|ENSMUSG00 Fbxw22|1814|0|0|0|0|0|0|0|0|0|0|0|0|0|0|0|0|0|0|
|ENSMUSG00 Rnf213|16895|2.41269392|2.29512926|2.29627531|1.99287433|1.94863781|2.11761312|1.70544075|1.5733597|2.074562|2.0261267|2.07840157|2.8147348|2.37096874|1.58066589|2.19725374|2.06694489|1.70818003|1.71913633|
|ENSMUSG00 Cldn27|1816|0|0|0|0|0|0|0|0|0|0|0|0|0|0|0|0|0|0|
|ENSMUSG00 Qrich2|2737|0.09317905|0.08519922|0|0|0.05275695|0|0.13829062|0.41862334|0|0.08613574|0.10473137|0.02039301|0.34245759|0.4242244|0|0.30777982|0.21004547|0|
|ENSMUSG00 4933422H20I|1926|0|0.25944622|0|0|0|0|0|0.03965982|0.21384484|0|0|0|0|0|0.23531522|0|0|0|
|ENSMUSG00 Krtap31-1|1415|0|0|0|0|0|0|0|0|0|0|0|0|0|0|0|0|0|0|
|ENSMUSG00 Krtap9-1|611|0|0|0|0|0|0|0|0|0|0|0|0|0|0|0|0|0|0|
|ENSMUSG00 Fbxo47|2191|0.64019663|0.76022123|1.23638416|1.24767807|1.6256332|1.17605757|0.60463487|0.59267078|1.10908463|0.62408507|0.69776297|1.50302332|1.03572259|0.127186|0.99666018|0.20971579|0.80465982|0.1506411|
|ENSMUSG00 Gpr179|9531|0.02675806|0.17476075|0.01522617|0.06214391|0.16665129|0.04679201|0.05460492|0.15227272|0.07346247|0.01978836|0.05513845|0.11126824|0.06728727|0.0389836|0.0345832|0.17275166|0.0482547|0.0115432|
|ENSMUSG00 Gm10287|4052|0|0|0|0|0|0|0|0|0|0|0|0|0.06087354|0|0.03050461|0|0|0|
|ENSMUSG00 Gm10288|536|1.11020982|0.93226383|0.63174387|0.93502143|0.53879019|0.73959304|1.76539847|1.06881723|0.30736206|0.8796773|1.42611816|1.77027288|1.01240875|0.95314358|1.30676577|1.00012781|0.92955571|1.64206289|
|ENSMUSG00 Hsf5|4152|0|0|0.06990395|0|0|0.02386933|0|0.00919856|0.03967872|0.14762985|0|0.02688616|0.03564445|0.05592978|0|0.12911091|0.16615447|0|
|ENSMUSG00 Ccnd1|3830|0.4661142|0.73062222|0.65676856|0.22602085|0.40214662|0.78922108|0.87707577|1.00716257|0.73124884|0.35701577|0.52390294|0.99098304|0.79858456|1.46729362|0.79606139|0.88978174|0.66045316|0.51705688|
|ENSMUSG00 Evi2b|3937|0.26990841|0.56691997|0.29488563|0.18516045|0.1467064|0.25172839|0.96139556|0.48504448|0.24061211|0.59881513|0.75236155|0.26936693|0.40097063|0.46007605|0.19883911|0.29177501|0.45754025|0.04191702|
|ENSMUSG00 E030019B06I|1503|0|0|0|0|0|0|0|0|0|0|0|0|0|0.09270277|0|0|0|0|
|ENSMUSG00 Amy2a1|886|0|0|0|0|0|0|0|0|0|0|0|0|0.05567936|0|0|0|0|0|
|ENSMUSG00 Ppapdc1a|1081|0|0|0|0|0|0|0|0|0|0|0|0|0|0|0|0|0|0|
|ENSMUSG00 Prok1|434|0|0|0|0|0|0|0|0|0|0|0|0|0|0|0|0|0|0.25349819|
|ENSMUSG00 Itgad|4579|0|0|0|0|0|0|0|0|0|0|0|0|0|0|0|0.06689776|0|0|
|ENSMUSG00 Prss36|3418|1.23113295|1.3839763|1.44356352|1.47960091|1.07022417|1.47875331|1.93790961|2.89404102|2.38587774|2.06922337|1.63536283|2.05756977|1.90515304|2.71761775|2.26620612|1.60197685|1.4577|2.20487057|
|ENSMUSG00 Capza1|5642|7.15702184|7.78796659|7.26203123|8.72943795|7.87411654|8.07142186|7.02729971|6.56622303|5.6720884|6.96981328|6.43547828|6.68759091|5.80581231|6.40437788|6.53587248|6.14197088|7.55385363|6.06445664|
|ENSMUSG00 Olfr59|1352|0|0|0|0|0|0|0|0|0|0|0|0|0|0|0|0|0|0|
|ENSMUSG00 Olfr406-ps|1381|0|0|0|0|0|0|0|0|0|0|0|0|0|0|0|0|0|0|
|ENSMUSG00 Olfr43|1091|0|0|0|0|0|0|0|0|0|0|0|0|0|0|0|0|0|0|
|ENSMUSG00 Olfr403|941|0|0|0|0|0|0|0|0|0|0|0|0|0|0|0|0|0|0|
|ENSMUSG00 Olfr402|999|0|0|0|0|0|0|0|0|0|0|0|0|0|0|0|0|0|0|
|ENSMUSG00 Olfr401|947|0|0|0|0|0|0|0|0|0|0|0|0|0|0|0|0|0|0|
|ENSMUSG00 Olfr391-ps|2056|0|0|0|0|0|0|0|0|0|0|0|0|0|0|0|0|0|0|
|ENSMUSG00 Olfr389|2080|0|0|0|0|0|0|0|0|0|0|0|0|0|0|0|0|0|0|
|ENSMUSG00 Ampd1|2598|0|0|0|0|0|0|0.14568954|0|0.01585316|0.09074423|0|0|0|0|0|0.02947699|0|0.02117364|
|ENSMUSG00 Fbxo39|1704|0|0|0|0|0|0|0|0|0|0|0|0|0|0|0|0|0|0|
|ENSMUSG00 Nlrp1b|6928|0|0|0|0|0|0|0|0.06064036|0|0|0|0|0|0|0|0|0|0|
|ENSMUSG00 Hist2h3c1|2955|0|0|0|0|0|0|0|0|0|0|0|0|0.01669439|0|0.02788599|0|0|0|
|ENSMUSG00 1810027O10I|916|127.283513|103.357255|113.646111|104.352698|107.613733|106.083963|154.64428|111.825353|123.019986|127.605451|118.863679|106.878463|153.704467|159.461414|130.531567|123.775642|108.911955|127.373707|
|ENSMUSG00 Gm12302|248|0|0|0|0|0|0|0|0|0|0|0|0|0|0|0|0|0|0|
|ENSMUSG00 Hs3st3b1|4909|1.43733127|2.23262216|1.25146432|2.06969095|1.40209107|1.61508197|0.21203482|0.48236483|0.31043004|1.43113766|0.64231967|0.95508569|1.72847622|2.45040103|0.99038167|1.31041394|1.18671769|2.28597632|
|ENSMUSG00 Gm10290|1248|0.03405863|0.29362327|0|0.25555073|0.19283623|0.2382343|0.2653756|0.18361732|0.6270383|0.18890506|0.38281217|0.31306862|0.1976439|0.29771851|0.23109847|0.39886049|0.09213052|0.30854467|
|ENSMUSG00 Olfr485|940|0|0|0|0|0|0|0|0|0|0|0|0|0|0|0|0|0|0|
|ENSMUSG00 Olfr478|944|0|0|0|0|0|0|0|0|0|0|0|0|0|0|0|0|0|0|
|ENSMUSG00 Olfr2|1248|0|0|0|0|0|0|0|0|0|0|0|0|0|0|0|0|0|0|
|ENSMUSG00 Cyp3a57|1697|0|0|0|0.05369599|0|0|0|0|0|0|0|0|0.3488409|0.0547367|0|0|0|0|
|ENSMUSG00 Zfp496|1991|0.25618388|0.15058566|0.31584927|0.32036897|0.24174748|0.37332547|0.23763274|0.32610288|0|0.16577344|0.74385922|0.33640781|0.39643932|0.18661612|0.60012311|0.19231847|0.51974386|0.60783543|
|ENSMUSG00 Olfr658|2910|0|0|0|0|0|0|0.03251731|0.20999259|0.07076721|0|0|0|0|0.12768134|0.07079305|0.07894971|0.07902329|0|
|ENSMUSG00 Olfr558|2913|0|0|0|0|0|0|0|0|0|0|0|0|0|0|0|0|0|0|
|ENSMUSG00 Art5|2145|0.55484612|0.66781093|0.27062115|0.40357102|0.13463475|0.41582713|0.24262912|0.40952226|0.38402346|0.15387176|0.04454542|0|0|0.75782893|0.46099643|0.37487308|0.64323855|0.28209798|
|ENSMUSG00 Trpc2|8025|1.28707262|1.49025907|1.68779919|1.62373317|1.66137606|1.67954437|2.48207325|1.97981799|2.33518563|1.58050332|1.97648426|1.87791014|2.50809|1.76516469|1.29380587|1.86562336|1.59036223|1.33666988|
|ENSMUSG00 Rnf121|5709|6.41039702|5.8234837|6.36337735|6.32060731|6.49178161|6.70945228|6.9614031|6.45571349|6.08166628|6.48332494|5.6821154|6.30602599|7.15481302|5.85737308|5.71582278|5.62722361|6.17624683|7.18808786|
|ENSMUSG00 Il18bp|1393|6.16370823|4.6394124|5.00056595|4.93877796|7.49793786|8.50186886|9.78180297|9.56866363|6.7412244|10.4929778|8.47122379|7.65311669|12.7136815|12.1028024|13.3986395|14.3761577|14.5271247|9.91190652|
|ENSMUSG00 Gm6109|347|0.1224933|0.19200515|0.83642992|0.13129984|0.41612614|0.28560653|1.36347778|0.44025824|0.23738626|0.40764297|0.55071998|0.16085205|0.56866763|0.66922316|0.83115531|1.43451843|0.22090085|0|
|ENSMUSG00 Olfr311|926|0|0|0|0|0|0|0|0|0|0|0|0|0|0|0|0|0|0|
|ENSMUSG00 Serpinh1|2429|22.1187907|18.9536516|19.1980582|22.3397298|19.7560839|20.1352606|13.6932126|13.3964293|18.3126542|14.7527931|13.4533024|13.6264668|11.3733527|8.77638374|12.4673296|12.7214986|10.2561107|10.5307677|
|ENSMUSG00 Olfr312|926|0|0|0|0|0|0|0|0|0|0|0|0|0|0|0|0|0|0|
|ENSMUSG00 Olfr313|1073|0|0|0|0|0|0|0|0|0|0|0|0|0|0|0|0|0|0|
|ENSMUSG00 Gm10291|1250|0.03400414|0.10660126|0.34828942|0.03644884|0.30804431|0.03964219|0.30280115|0.15276961|0.09884764|0.3017645|0.30575973|0|0.55251747|0.3715527|0.32961245|0.18379492|0.06132208|0.35205828|
|ENSMUSG00 Vmn2r89|3224|0|0|0|0|0|0|0|0|0|0|0|0|0|0|0|0|0|0|
|ENSMUSG00 Vmn2r73|2550|0|0|0|0|0|0|0|0|0|0|0|0|0|0|0|0|0|0|
|ENSMUSG00 Olfr290|947|0|0|0|0|0|0|0|0.04032989|0|0|0|0|0|0|0|0|0|0|
|ENSMUSG00 Olfr291|1154|0|0|0|0|0.12512632|0|0|0.13238268|0|0|0|0|0|0.12073853|0.03570325|0|0|0|
|ENSMUSG00 9230112E08I|1359|0.06255361|0.17158959|0.81868374|0|0.10625149|0|0|0.33723976|0.27275838|0.03469515|0.21092697|0.04107113|0.43560192|0.06835039|0.51539842|0.05635115|0.42302756|0|
|ENSMUSG00 Mesdc1|3846|3.83497038|4.52140686|5.92405949|5.04654313|4.29257666|4.61255294|2.85401495|4.21049886|4.48703851|4.56059844|3.77628377|3.75877707|4.00196521|5.68777045|5.78492984|4.6096077|4.76338271|6.16456708|
|ENSMUSG00 Ccl26|279|0|0|0|0|0|0|0|0|0|0|0|0|0|0|0|0|0|0|
|ENSMUSG00 Gm9696|3045|0|0|0|0|0|0|0|0|0|0|0|0|0|0|0|0|0|0|
|ENSMUSG00 Adamtsl3|7878|0|0|0.00614033|0|0|0|0|0|0|0.00598511|0|0|0|0|0|0|0|0|
|ENSMUSG00 Fam194a|2181|0.05846654|0.01527414|0|0.02088998|0|0.11360095|0.6724865|0.91059373|1.01974867|0.25942615|0.30667099|0.89571215|0.79166305|0.27683317|0.05667339|0|0.42174742|0.47921734|
|ENSMUSG00 Cldn3|1856|159.600524|145.600322|148.48276|139.481602|151.475521|153.330468|142.269429|146.596269|160.352243|129.207153|142.707096|165.251652|168.993712|173.139354|137.612309|155.019194|134.410488|180.646824|
|ENSMUSG00 9030418K01f|4681|0.08172326|0.04981633|0.0620041|0.14599779|0.06169441|0.10585929|0.27289945|0.1223854|0.07918792|0.15109176|0.10206144|0.07154325|0.07377129|0|0.13202806|0.06544005|0.14737734|0|
|ENSMUSG00 4930527J03R|541|0|0|0|0|0.08896844|0|0|0|0|0|0|0|0|0|0|0|0|0|
|ENSMUSG00 Gm10293|1231|0.10358694|0.21649322|0.23577675|0.33310268|0.31279885|0.12076215|0.23060607|0.43435714|0.53532433|0.42133041|0.97024692|0.45341724|0.44082136|0.41501622|0.60245979|0.24884229|0.31134279|0.31280564|
|ENSMUSG00 Chchd2|2503|92.7879672|79.1231147|83.1414014|74.4121251|87.0143646|82.7924618|98.1222998|86.0435306|90.4030034|86.9547128|89.1748326|83.5339481|107.532937|103.891508|93.6623464|106.029644|91.3828765|105.161237|
|ENSMUSG00 Ctcfl|3206|0|0|0|0|0|0|0|0|0|0|0|0|0|0|0|0|0|0|
|ENSMUSG00 Tmem132b|1219|0.62764001|1.99494773|1.38890366|2.39204827|1.93475329|0.56910441|0.42693968|0.06266186|0|0.3481184|0.11757578|0.13736422|0|0|0.50699207|0.15705745|0.91178229|0.58664347|
|ENSMUSG00 BC094916|3937|0|0|0|0|0|0|0|0|0.0104614|0|0|0|0.02359364|0|0|0|0|0|
|ENSMUSG00 Fcrl6|1907|0|0|0|0|0|0|0|0|0|0|0|0|0|0|0|0|0|0|
|ENSMUSG00 Rgma|3777|0.47265486|0.51155623|0.19211092|0.44632212|0.59894107|0.44606644|0.28811004|0.1719012|0.19628204|0.26215641|0.29092508|0.29555553|0.13061138|0.11066899|0.29453058|0.06082702|0.142062|0.01456423|
|ENSMUSG00 Gm10295|2524|0.0168404|0|0|0|0|0|0|0|0|0|0|0|0|0|0|0|0|0|
|ENSMUSG00 Rpl35a-ps6|332|0|0|0|0.1457034|0|0|0|0|0|0|0|0|0|0|0|0|0|0|
|ENSMUSG00 Ndnl2|5945|7.8504093|6.97637566|7.9496954|8.81332229|6.11263286|7.55168656|8.56323676|8.39012238|8.65297867|7.90735932|7.40933755|7.98976095|7.34377574|9.20288224|9.36304491|8.27643911|6.82717383|8.8551245|
|ENSMUSG00 Gm6505|1861|0.04567993|0.07160214|0.0259933|0.12241011|0.05172695|0.07988082|0.05084651|0.20522516|0.11065695|0.17735353|0.1283583|0.29992296|0.13254142|0.12478261|0.08855788|0.04115057|0.08237785|0.11823559|
|ENSMUSG00 Fcrlb|1278|0|0.02606643|0|0|0|0|0.03702088|0.02988451|0|0|0|0|0|0|0|0|0|0.38738808|
|ENSMUSG00 Peg12|2639|0|0|0.05499075|0.05179353|0.0182387|0|0|0|0|0|0|0.20562754|0.10559474|0.07806282|0|0|0|0.08337871|
|ENSMUSG00 Mkrn3|2560|1.62715127|1.88686315|2.26750924|2.65179516|2.29378699|1.72273175|3.14185759|2.56605204|2.20412213|1.52871421|2.27677537|3.29225197|1.69578466|2.05007104|2.55900289|2.36324843|2.96429953|2.55706395|
|ENSMUSG00 Gm10297|215|0|0.77471847|0|0.42382367|0|0.23047783|0|0.17763908|0|0.2193056|0|0|0|0.21601901|0|0.17809585|0|0.25585631|
|ENSMUSG00 Wfdc10|715|76.6876602|84.9828239|74.0825395|71.8781231|59.845147|78.8685466|369.303582|294.802614|236.462444|145.804482|95.1490082|267.524861|1122.35286|1645.67966|765.59982|260.965217|355.764521|840.600699|
|ENSMUSG00 Wfdc16|1334|1.84805113|0.67424898|0.8702884|0.27322965|0.72161805|0.4828977|66.0746033|45.2353792|48.3802625|36.3350247|17.2978298|38.828287|171.552297|198.867037|104.239319|26.9526839|30.6552915|100.69883|
|ENSMUSG00 Wfdc6b|1153|6.37761965|4.91170166|6.62880988|3.83297598|5.84429255|4.16879025|11.0792917|11.0966651|10.1448139|5.11174146|4.72362989|10.5531782|102.856834|176.672825|83.7252779|23.9108737|34.9357659|76.1921452|
|ENSMUSG00 1700084C01|1730|0|0|0|0|0|0|0|0|0|0|0|0|0|0|0|0|0|0|
|ENSMUSG00 Wfdc8|3918|1.36693522|2.70380305|2.38287171|1.81406915|1.7444444|1.17621343|3.57441372|3.95766089|2.29164381|0.69799408|0.58529811|1.73800685|8.14643963|22.6411963|8.99115117|1.29003578|4.0889209|10.5721943|
|ENSMUSG00 Gm9343|395|0|0|0|0|0|0|0|0|0|0|0|0|0|0|0|0|0|0|
|ENSMUSG00 Top1|10716|14.2715214|11.7633438|12.779532|11.3902611|12.2395943|11.7315496|14.6582768|12.9981048|13.2368757|13.6268876|13.2009754|11.3964791|14.137581|13.4140025|12.4189087|12.6277535|14.6996156|12.6845374|
|ENSMUSG00 Mrgprb3|938|0|0|0|0|0|0|0|0|0|0|0|0|0|0|0|0|0|0|
|ENSMUSG00 Mrgprb1|4183|0|0|0|0|0|0.01184622|0|0|0|0|0|0|0|0|0|0|0|0|
|ENSMUSG00 Mrgprb4|965|0|0|0|0|0|0|0|0|0|0|0|0|0|0|0|0|0|0|
|ENSMUSG00 Mrgprb5|1031|0|0|0|0|0|0|0|0|0|0|0|0|0|0|0|0.07427858|0|0|
|ENSMUSG00 Mrgprx1|1182|0|0|0|0|0|0|0|0|0|0|0|0|0|0|0|0|0|0|
|ENSMUSG00 Kcnj11|3074|0.304201|0.01083699|0|0.11857136|0.07828875|0.12895962|0|0|0|0|0|0|0|0|0|0|0|0|
|ENSMUSG00 Spaca4|709|0|0|0|0|0.27154823|0|0|0|0|0|0|0|0|0|0|0.1620195|0|0|
|ENSMUSG00 Ntn5|1568|0.16264736|0.23370015|0.24680373|0.11622715|0.36835656|0.22121756|0.0301739|0.38971839|0|0.39091782|0.09140617|0.24917707|0.06292336|0|0.18393552|0.21978027|0.17109954|0.38590572|
|ENSMUSG00 Rasal2|14058|0.63494715|0.66113939|0.6056154|0.53151311|0.52041915|0.72964973|0.39040196|0.33959669|0.53907518|0.40918949|0.44179432|0.4049792|0.31231616|0.49225843|0.57444195|0.29961352|0.32988205|0.3247799|
|ENSMUSG00 1700039E15f|3443|0|0|0|0.0264659|0|0|0|0.05546384|0|0|0|0|0|0|0|0|0|0|
|ENSMUSG00 Slc17a7|2927|0.17426106|0.11381242|0.01652666|0|0|0|0|0|0|0|0.11425511|0|0.10112453|0|0|0|0.01309405|0|
|ENSMUSG00 Gm13072|377|1.8039332|0.88363115|0.89818226|1.32936734|0.76602531|1.70871495|1.63147169|1.21567329|0.87398442|0.12506818|1.39396431|1.48052155|0.52341557|1.23193866|0.54643973|0.71096619|0.71162885|2.48051675|
|ENSMUSG00 2310016G11I|879|0|0|0|0|0|0.05637399|0|0|0|0|0|0|0.22449109|0|0.3281125|0.04356156|0|0|
|ENSMUSG00 Mn1|3349|0.40614083|0.46751449|0.47665766|0.28569183|0.503021|0.48827716|0.49445917|0.4789731|0.7132929|0.87290045|0.44222864|0.88331744|0.04419103|0.12481242|0.36907933|0.30870302|0.18310563|0.03285106|
|ENSMUSG00 Gm572|1374|0|0|0|0|0|0|0|0|0|0|0|0|0|0|0|0|0|0|
|ENSMUSG00 Fv1|1379|2.80491009|3.74437897|3.89373595|2.70921366|4.15351626|2.73097009|2.74475295|3.5727483|5.34618297|3.04308382|3.25659615|2.99518421|2.89766882|2.35756789|6.06520366|2.22135503|2.3901824|5.78413377|
|ENSMUSG00 Gm4788|3528|0|0|0|0|0|0|0|0|0|0|0|0|0|0|0|0|0|0|
|ENSMUSG00 Vmn2r84|3028|0|0|0|0|0|0|0|0|0|0|0|0|0|0|0|0|0|0|
|ENSMUSG00 Vsig10l|3759|1.11944997|1.4977066|1.78875252|1.90292204|1.48531608|1.62143821|2.54247437|2.67214734|1.17237488|2.67174775|2.64357312|3.05879928|3.04469402|2.94059399|2.49905686|3.35131946|2.80386583|2.97069664|
|ENSMUSG00 Gm13251|4546|1.48665266|1.14316135|1.41523967|1.32293398|1.11171403|0.92652493|1.23849731|1.26019806|1.30463226|1.50392696|1.15601527|0.73667834|1.23709603|1.74701693|1.30510868|2.2826121|1.3742161|1.23425624|
|ENSMUSG00 Gm10298|263|0.16161664|0|0|0|0|0|0|0|0|0|0|0|0|0|0|0|0|0|
|ENSMUSG00 Gm13177|1220|0|0|0|0|0|0|0|0|0|0|0|0|0|0|0|0|0|0|
|ENSMUSG00 Gm13127|1388|1.316803|1.20003221|0.73187618|1.24734848|1.14434689|0.71401633|0.78399972|0.93554876|1.06823816|1.12101816|0.96375997|0.92489931|0.92408491|0.76960664|0.92020766|0.85519368|0.85599078|0.43595113|
|ENSMUSG00 Gm13040|2146|0|0|0|0|0|0|0|0|0|0|0|0|0|0|0|0|0|0|
|ENSMUSG00 Gm13089|1803|0|0|0|0|0|0|0|0|0|0|0|0|0|0|0|0|0|0|
|ENSMUSG00 BC080695|2091|0|0|0|0|0|0|0|0|0|0|0|0|0|0|0|0|0|0|
|ENSMUSG00 Gm13119|2090|0|0|0|0|0|0|0|0|0|0|0|0|0|0|0|0|0|0|
|ENSMUSG00 Vmn2r-ps60|1156|0|0|0|0|0|0|0|0|0|0|0|0|0|0|0|0|0|0|
|ENSMUSG00 Btbd8|1527|0.30619315|0.41450229|0.57021843|0.59673928|0.85105562|0.6165697|0.18590444|0.27512536|0.43155485|0.89546195|1.25147239|1.1696799|0.61382215|0.39539826|0.21585622|0.45136276|0.15059449|1.15277761|
|ENSMUSG00 Gm694|903|0.04707107|0|0.0535698|0.0504552|0|0|0.31436996|0|0.27366456|0|0.10581386|0|0.05463114|0|0.27376449|0|0.0424433|0.12183634|
|ENSMUSG00 Lrrc8b|2799|12.0879314|17.3765007|16.0035331|14.7312416|16.9381725|17.5975052|11.4436169|12.04855|11.1684764|13.8470447|12.1357612|12.2838329|7.79817772|8.03106048|12.2912823|11.8880092|12.802818|12.9710648|
|ENSMUSG00 Gm16410|773|0|0|0|0|0|0|0|0|0|0|0|0|0|0|0|0|0|0|
|ENSMUSG00 Sox13|4571|16.8402699|13.1473334|12.0854456|21.1807524|12.878001|15.393761|16.9336125|15.2652634|14.3085073|17.659594|13.3573391|16.570084|13.7386853|10.4857358|14.3407734|11.3925238|13.7676175|17.1850806|
|ENSMUSG00 Etnk2|4560|31.1052132|37.9518608|39.2080193|31.5929873|34.8639789|31.6007345|41.1080776|48.3267899|48.8818911|51.2143494|52.4791156|34.1748531|55.0332156|66.0809735|57.8719223|58.7794412|61.6582706|54.0802249|
|ENSMUSG00 Ren1|1878|0.38476464|0.07095398|0|0.04852081|0.17940547|0|0.20154496|0.06101023|0.02193105|0|0.05087855|0.1188832|0.3940249|0|0.1096953|0|0.08163216|0|
|ENSMUSG00 Rnf186|1253|12.6870996|13.5857055|11.9679125|9.30856133|11.5240041|8.58175833|9.74195624|8.16884585|7.0342493|6.51003325|11.9342075|6.14729564|10.94515|6.37540545|5.91881888|15.6462817|15.1714631|14.3559281|
|ENSMUSG00 Ldlrad2|1707|0|0.09757731|0|0|0|0|0|0|0|0|0|0|0|0|0|0|0|0|
|ENSMUSG00 Gm13004|792|0.21467261|0|0.12215538|0.23010628|0.42540842|0.12513317|0.41816762|0.14466819|0.05200318|0|0.24128767|0.35237161|0.43601442|0.29320762|0.05202217|0|0.14517537|0.27782377|
|ENSMUSG00 D5Ertd577e|2458|0|0|0|0|0|0|0|0|0|0|0|0|0|0|0|0|0|0|
|ENSMUSG00 Lactbl1|3204|0|0.01039728|0|0|0|0|0|0|0.01285472|0|0|0|0|0|0|0|0|0|

| | | | | | | | | | | | | | | | | | | | |
|---|---|---|---|---|---|---|---|---|---|---|---|---|---|---|---|---|---|---|---|
| ENSMUSG00 C87414 | 4644 | 0 | 0 | 0 | 0 | 0 | 0 | 0 | 0 | 0 | 0 | 0 | 0 | 0 | 0 | 0 | 0 | 0 | 0 |
| ENSMUSG00 Htr1d | 3268 | 0 | 0.06116198 | 0 | 0.01394157 | 0 | 0.09097809 | 0 | 0 | 0 | 0 | 0 | 0 | 0 | 0 | 0 | 0.16403565 | 0.02345551 | 0 |
| ENSMUSG00 5830473C10 | 1075 | 0 | 0 | 0 | 0 | 0 | 0 | 0 | 0 | 0 | 0 | 0 | 0 | 0 | 0 | 0 | 0 | 0 | 0 |
| ENSMUSG00 Runx3 | 4994 | 0 | 0 | 0 | 0 | 0 | 0 | 0.06631733 | 0.12236252 | 0.1979328 | 0 | 0.16263001 | 0.15647162 | 0.03951295 | 0.01859995 | 0.04125106 | 0.00766732 | 0 | 0.04406016 |
| ENSMUSG00 Gm14059 | 892 | 0.14295463 | 0.07469259 | 0.10846083 | 0.0510774 | 0.10791911 | 0.16665718 | 0 | 0.12844978 | 0.04617322 | 0.10571907 | 0.05355937 | 0 | 0.33182904 | 0.20826945 | 0 | 0.17170676 | 0.0859334 | 0.18500821 |
| ENSMUSG00 Cntnap5a | 5708 | 0 | 0 | 0 | 0 | 0 | 0 | 0 | 0 | 0 | 0 | 0 | 0 | 0 | 0 | 0 | 0 | 0 | 0 |
| ENSMUSG00 Utp3 | 1600 | 43.6740684 | 33.4378176 | 37.0662176 | 36.819019 | 35.2867165 | 38.0626935 | 31.0193751 | 33.2035198 | 30.3235722 | 31.9446015 | 36.8165774 | 33.0707801 | 32.5590654 | 29.0856097 | 31.9312058 | 31.0632553 | 31.8108264 | 36.5122946 |
| ENSMUSG00 Sars2 | 1844 | 6.38499662 | 5.61838943 | 7.03042632 | 7.56056376 | 6.60378345 | 6.71810379 | 8.44136191 | 6.64846051 | 5.38283639 | 8.59145135 | 9.27518174 | 5.47865235 | 5.93909199 | 6.34702279 | 5.18370985 | 5.7518971 | 5.65333664 | 4.14656499 |
| ENSMUSG00 Csn1s1 | 1381 | 0 | 0 | 0 | 0 | 0 | 0 | 0 | 0 | 0 | 0 | 0 | 0 | 0 | 0 | 0 | 0 | 0 | 0 |
| ENSMUSG00 Ugt2b36 | 2078 | 0 | 0 | 0 | 0 | 0 | 0 | 0 | 0 | 0 | 0 | 0 | 0 | 0 | 0 | 0 | 0 | 0 | 0 |
| ENSMUSG00 Eid2b | 2423 | 4.87678041 | 3.6571316 | 3.79321947 | 3.74190996 | 3.85373223 | 4.04929478 | 5.68220783 | 5.78481704 | 6.52728933 | 4.98166737 | 4.43638698 | 4.10036646 | 5.76183763 | 4.48531426 | 3.96201507 | 2.71811163 | 3.21099392 | 2.76975279 |
| ENSMUSG00 Gtsf1l | 823 | 0 | 0 | 0 | 0.05535971 | 0 | 0 | 0 | 0 | 0 | 0 | 0 | 0 | 0 | 0 | 0 | 0 | 0 | 0 |
| ENSMUSG00 1700049G17 | 7777 | 2.26818157 | 2.39448475 | 1.91578338 | 1.96843396 | 1.82575768 | 1.80956363 | 1.67909212 | 1.79249413 | 1.91712983 | 2.06742829 | 2.47567292 | 2.10286603 | 1.68097699 | 1.39147131 | 1.64234053 | 1.52138327 | 2.42958266 | 0.91952988 |
| ENSMUSG00 Gm10282 | 1150 | 0.22176614 | 0.37658054 | 0 | 0.0792366 | 0.75336924 | 0.30162533 | 0.57598044 | 0.46495099 | 0.42977234 | 0.16400245 | 0.08308688 | 0.14560608 | 0.30028124 | 0.52502012 | 0.28661952 | 0.29966562 | 0.16663607 | 0.28700404 |
| ENSMUSG00 Gm10300 | 2295 | 0.3518947 | 0.17418507 | 0.12646675 | 0.03970461 | 0.31458774 | 0.04318321 | 0.32984874 | 0.41603924 | 0.39481627 | 0.08217987 | 0.74941111 | 0.0972822 | 0.62336627 | 0.04047415 | 0.19748022 | 0.30031849 | 0.25049867 | 0.31159843 |
| ENSMUSG00 Pla2g4d | 3668 | 0 | 0 | 0 | 0 | 0 | 0 | 0 | 0 | 0 | 0 | 0 | 0 | 0 | 0 | 0 | 0 | 0 | 0 |
| ENSMUSG00 Tmem200b | 854 | 30.2115244 | 34.8342091 | 32.2867826 | 44.3873406 | 30.8855904 | 42.6478445 | 38.9470882 | 52.6375382 | 47.8419481 | 54.9907499 | 62.0962575 | 52.4824087 | 50.9493572 | 43.4529576 | 58.8111202 | 60.4399751 | 58.5216524 | 65.1864356 |
| ENSMUSG00 Gm12966 | 809 | 1.1033482 | 0.70002374 | 1.4948557 | 0.95740143 | 1.13041601 | 1.10253301 | 1.22814123 | 1.18023493 | 1.32367047 | 1.04908858 | 1.06297806 | 1.17288784 | 0.67076773 | 1.60745914 | 0.86579289 | 0.75729261 | 1.08962278 | 1.49592132 |
| ENSMUSG00 Fam82a2 | 3525 | 15.8806573 | 15.6121706 | 16.6459836 | 14.3081067 | 13.9002265 | 15.9412198 | 17.6499781 | 15.8837055 | 16.2058716 | 16.4659619 | 18.9202389 | 16.8001186 | 16.8638185 | 15.9556851 | 16.7027017 | 16.1852497 | 18.4618588 | 16.9318811 |
| ENSMUSG00 Rbm44 | 3835 | 0 | 0 | 0 | 0 | 0 | 0 | 0 | 0 | 0 | 0 | 0 | 0 | 0 | 0 | 0 | 0 | 0 | 0 |
| ENSMUSG00 Fryl | 15704 | 3.62690626 | 4.31260277 | 3.78264743 | 3.24358428 | 3.20286935 | 3.54353794 | 4.03109809 | 3.51183323 | 3.93663781 | 3.60295746 | 4.21346902 | 4.0944755 | 4.5926683 | 4.07243431 | 3.71506643 | 4.95212835 | 4.65167613 | 3.43281486 |
| ENSMUSG00 Gm12942 | 1906 | 14.7407772 | 14.9960458 | 13.679608 | 16.1591112 | 16.843648 | 14.897018 | 21.7697898 | 24.6667615 | 21.4576129 | 21.942641 | 22.2080867 | 24.2473078 | 17.3412301 | 20.151763 | 17.1420954 | 21.9377457 | 18.0571955 | 16.6816703 |
| ENSMUSG00 Dgkd | 5664 | 6.3487604 | 7.89887305 | 6.94344636 | 6.86150615 | 7.12121328 | 6.6577737 | 7.11694958 | 5.17188783 | 7.8388178 | 6.37666644 | 6.74787549 | 6.59263388 | 6.92423627 | 6.8304952 | 7.41977172 | 8.00425126 | 4.79078711 | 5.48731379 |
| ENSMUSG00 Ceacam20 | 2885 | 0 | 0 | 0 | 0 | 0 | 0.05152797 | 0.01639954 | 0.13238268 | 0.11420871 | 0 | 0.08279889 | 0 | 0.13679561 | 0 | 0 | 0 | 0 | 0.09533641 |
| ENSMUSG00 Rpl7a-ps8 | 812 | 0 | 0 | 0 | 0 | 0 | 0 | 0 | 0 | 0 | 0 | 0 | 0 | 0 | 0 | 0 | 0 | 0 | 0 |
| ENSMUSG00 Rbm47 | 9723 | 19.0514818 | 18.0149334 | 17.0449157 | 20.4914581 | 16.9993856 | 18.1025722 | 16.5689264 | 15.4725777 | 18.2994701 | 16.5946422 | 13.9497179 | 16.4180598 | 14.906631 | 10.9100376 | 13.6024884 | 13.8347119 | 12.8936868 | 12.3788877 |
| ENSMUSG00 BC016548 | 1174 | 0.03620543 | 0 | 0 | 0.15523354 | 0 | 0 | 0 | 0.22772301 | 0 | 0.04016244 | 0 | 0.47543154 | 0 | 0.19780276 | 0 | 0.16307754 | 0.29381318 | 0.32799297 |
| ENSMUSG00 Psg21 | 2154 | 0 | 0 | 0 | 0 | 0 | 0 | 0 | 0 | 0 | 0 | 0 | 0 | 0 | 0 | 0 | 0 | 0 | 0 |
| ENSMUSG00 Psg27 | 1936 | 0 | 0 | 0 | 0 | 0 | 0 | 0 | 0 | 0 | 0 | 0 | 0 | 0 | 0 | 0 | 0 | 0 | 0 |
| ENSMUSG00 Psg25 | 3060 | 0.01389058 | 0 | 0 | 0 | 0 | 0 | 0 | 0 | 0 | 0 | 0 | 0 | 0 | 0 | 0 | 0 | 0 | 0 |
| ENSMUSG00 Psg26 | 2052 | 0 | 0 | 0 | 0 | 0 | 0 | 0 | 0 | 0 | 0 | 0 | 0 | 0 | 0 | 0 | 0 | 0 | 0 |
| ENSMUSG00 Pnmal2 | 2396 | 0.07096023 | 0.22245672 | 0.12113572 | 0.17113915 | 0.30132673 | 0.37226595 | 0.19746527 | 0.07970034 | 0.12032788 | 0.11807355 | 0.11963679 | 0 | 0 | 0 | 0.03439195 | 0 | 0 | 0 |
| ENSMUSG00 Cited4 | 1492 | 47.1203493 | 42.7575016 | 45.6825096 | 40.5529939 | 36.7118828 | 52.2097166 | 41.7949804 | 42.006523 | 44.7475483 | 43.7376497 | 37.2721526 | 49.4933791 | 43.5126026 | 46.0394137 | 35.1814894 | 32.6702032 | 27.2548264 | 42.6947356 |
| ENSMUSG00 Zmynd12 | 1983 | 0.27865219 | 0.36958329 | 0.36591173 | 0.2757098 | 0.21845049 | 0.3498428 | 0.23859142 | 0.63557704 | 0.10384901 | 0.28532952 | 0.33729169 | 0.50664747 | 0.1492645 | 0.65579145 | 0.47787987 | 0.23171321 | 0.30923891 | 0.44384554 |
| ENSMUSG00 Gltscr1 | 5346 | 1.54246243 | 2.72310789 | 2.14450555 | 2.05391165 | 1.75565378 | 1.90017029 | 2.92938584 | 2.20038532 | 1.91063522 | 2.13439397 | 1.55496498 | 1.8688746 | 1.19961639 | 1.02514072 | 1.32560187 | 1.71899472 | 1.41949248 | 0.76144293 |
| ENSMUSG00 Sult2a6 | 982 | 0 | 0 | 0 | 0 | 0 | 0 | 0 | 0 | 0 | 0 | 0 | 0 | 0 | 0 | 0 | 0 | 0 | 0 |
| ENSMUSG00 Sult2a2 | 736 | 0 | 0 | 0 | 0 | 0 | 0 | 0 | 0 | 0 | 0 | 0 | 0 | 0 | 0 | 0 | 0 | 0 | 0 |
| ENSMUSG00 6330408A02 | 13611 | 0.40597112 | 0.55802953 | 0.5331004 | 0.46863171 | 0.43849523 | 0.58614283 | 0.82035062 | 0.4208993 | 0.75951917 | 0.57505082 | 0.39663289 | 0.48799235 | 0.17397194 | 0.34804915 | 0.56606355 | 0.32070599 | 0.61103566 | 0.49710676 |
| ENSMUSG00 Vmn1r87 | 887 | 0 | 0 | 0 | 0 | 0 | 0 | 0 | 0 | 0 | 0 | 0 | 0 | 0 | 0 | 0 | 0 | 0 | 0 |
| ENSMUSG00 Vmn1r86 | 950 | 0 | 0 | 0 | 0 | 0 | 0 | 0 | 0 | 0 | 0 | 0 | 0 | 0 | 0 | 0 | 0 | 0 | 0 |
| ENSMUSG00 Vmn1r85 | 926 | 0 | 0 | 0 | 0 | 0 | 0 | 0 | 0 | 0 | 0 | 0 | 0 | 0 | 0 | 0 | 0 | 0 | 0 |
| ENSMUSG00 Olfr1339 | 947 | 0 | 0 | 0 | 0 | 0 | 0 | 0 | 0 | 0 | 0 | 0 | 0 | 0 | 0 | 0 | 0 | 0 | 0 |
| ENSMUSG00 Olfr1340 | 950 | 0 | 0 | 0 | 0 | 0 | 0 | 0 | 0 | 0 | 0 | 0 | 0 | 0 | 0 | 0 | 0 | 0 | 0 |
| ENSMUSG00 Zscan18 | 3620 | 2.4892534 | 2.23619704 | 2.25832227 | 2.64304401 | 2.23374673 | 2.13542167 | 3.2935898 | 3.56603094 | 3.85696929 | 3.03483808 | 3.15420306 | 2.79078313 | 4.15641844 | 3.91310681 | 3.25515055 | 3.88194832 | 3.0703387 | 5.74404485 |
| ENSMUSG00 Gm1043 | 1576 | 0 | 0 | 0 | 0 | 0 | 0 | 0 | 0 | 0 | 0 | 0 | 0 | 0 | 0 | 0 | 0 | 0 | 0 |
| ENSMUSG00 Vmn1r72 | 920 | 0 | 0 | 0 | 0 | 0 | 0 | 0 | 0 | 0 | 0 | 0 | 0 | 0 | 0 | 0 | 0 | 0 | 0 |
| ENSMUSG00 Zscan4-ps2 | 2273 | 0 | 0 | 0 | 0 | 0 | 0 | 0 | 0 | 0 | 0 | 0 | 0 | 0 | 0 | 0 | 0 | 0 | 0 |
| ENSMUSG00 Zscan4f | 2487 | 0 | 0 | 0 | 0 | 0 | 0 | 0 | 0 | 0 | 0 | 0 | 0 | 0 | 0 | 0 | 0 | 0 | 0 |
| ENSMUSG00 Gm13015 | 203 | 0 | 0 | 0 | 0 | 0.94841229 | 0.24410214 | 0.46613477 | 0 | 0.20288924 | 0 | 0.47068925 | 0.274954 | 0 | 0 | 0 | 0 | 0 | 0 |
| ENSMUSG00 Aurkc | 941 | 0 | 0 | 0 | 0 | 0 | 0 | 0 | 0 | 0 | 0 | 0 | 0 | 0 | 0 | 0 | 0 | 0 | 0 |
| ENSMUSG00 Vmn2r34 | 4916 | 0 | 0 | 0 | 0 | 0 | 0 | 0 | 0 | 0 | 0 | 0 | 0 | 0 | 0 | 0 | 0 | 0 | 0 |
| ENSMUSG00 Vmn2r42 | 4107 | 0 | 0 | 0 | 0 | 0 | 0 | 0 | 0 | 0 | 0 | 0 | 0 | 0 | 0 | 0.0300961 | 0 | 0 | 0 |
| ENSMUSG00 Vmn2r-ps44 | 2556 | 0 | 0 | 0 | 0 | 0 | 0 | 0 | 0 | 0 | 0 | 0 | 0 | 0 | 0 | 0 | 0 | 0 | 0 |
| ENSMUSG00 Vmn2r30 | 4586 | 0 | 0 | 0 | 0 | 0.0104954 | 0 | 0 | 0 | 0 | 0 | 0 | 0 | 0 | 0 | 0 | 0 | 0 | 0 |
| ENSMUSG00 Gm10302 | 3450 | 0 | 0 | 0 | 0 | 0 | 0 | 0 | 0 | 0 | 0 | 0 | 0 | 0 | 0 | 0 | 0 | 0 | 0 |
| ENSMUSG00 Vmn2r32 | 2553 | 0 | 0 | 0 | 0 | 0 | 0 | 0 | 0 | 0 | 0 | 0 | 0 | 0 | 0 | 0 | 0 | 0 | 0 |
| ENSMUSG00 Gm4073 | 2556 | 0 | 0 | 0 | 0 | 0 | 0 | 0 | 0 | 0 | 0 | 0 | 0 | 0 | 0 | 0 | 0 | 0 | 0 |
| ENSMUSG00 Olfr1121 | 944 | 0 | 0 | 0 | 0 | 0 | 0 | 0 | 0 | 0 | 0 | 0 | 0 | 0 | 0 | 0 | 0 | 0 | 0 |
| ENSMUSG00 Olfr1120 | 1010 | 0 | 0 | 0 | 0 | 0 | 0 | 0 | 0 | 0 | 0 | 0 | 0 | 0 | 0 | 0 | 0 | 0 | 0 |
| ENSMUSG00 Olfr1116-ps | 1918 | 0 | 0 | 0 | 0 | 0 | 0 | 0 | 0 | 0 | 0 | 0 | 0 | 0 | 0 | 0 | 0 | 0 | 0 |
| ENSMUSG00 Olfr1115 | 1101 | 0 | 0 | 0 | 0 | 0 | 0 | 0 | 0 | 0 | 0 | 0 | 0 | 0 | 0 | 0 | 0 | 0 | 0 |
| ENSMUSG00 Olfr1113 | 1102 | 0 | 0 | 0 | 0 | 0 | 0 | 0 | 0 | 0 | 0 | 0 | 0 | 0 | 0 | 0 | 0 | 0 | 0 |
| ENSMUSG00 Gm1673 | 1015 | 1.54944977 | 1.83795278 | 1.38210087 | 0.9426423 | 1.42261843 | 0.97640854 | 3.63585119 | 3.72517028 | 2.63756012 | 2.18333306 | 1.41206774 | 2.41959523 | 7.24182825 | 7.18396229 | 4.70874924 | 2.37665839 | 1.85023502 | 4.17310465 |
| ENSMUSG00 Zfp804a | 5241 | 0 | 0 | 0 | 0 | 0 | 0 | 0 | 0 | 0 | 0 | 0 | 0 | 0 | 0 | 0 | 0 | 0 | 0 |
| ENSMUSG00 Gm12824 | 6610 | 1.85196531 | 2.9734656 | 1.41241927 | 1.27515782 | 2.36654693 | 2.18151974 | 0.91619106 | 1.50805099 | 0.93464105 | 1.14131808 | 1.16365633 | 0.67552996 | 1.86580625 | 2.04466406 | 2.33745589 | 2.08541886 | 1.82644229 | 2.33851119 |
| ENSMUSG00 Skint3 | 3833 | 0 | 0 | 0 | 0 | 0 | 0 | 0 | 0 | 0 | 0 | 0 | 0 | 0 | 0 | 0 | 0 | 0 | 0 |
| ENSMUSG00 Crygs | 1060 | 0 | 0 | 0 | 0 | 0 | 0 | 0 | 0 | 0 | 0 | 0 | 0.05265629 | 0.09307909 | 0 | 0.03886939 | 0 | 0.10847065 | 0 |
| ENSMUSG00 Ccnyl1 | 3270 | 1.68980822 | 2.4246082 | 1.58286475 | 1.35150498 | 1.32473184 | 1.48506663 | 0.89705997 | 2.24248968 | 1.58700336 | 1.64378599 | 1.89931028 | 2.3896614 | 1.35775919 | 1.3066838 | 1.61278262 | 0.9601926 | 2.05110152 | 2.69157709 |
| ENSMUSG00 Lilra5 | 2103 | 0 | 0 | 0 | 0 | 0 | 0 | 0 | 0.03632183 | 0 | 0 | 0 | 0 | 0 | 0 | 0 | 0 | 0 | 0 |
| ENSMUSG00 Olfr1100 | 968 | 0 | 0 | 0 | 0 | 0 | 0 | 0 | 0 | 0 | 0 | 0 | 0 | 0 | 0 | 0 | 0 | 0 | 0 |
| ENSMUSG00 Ldlrad1 | 1088 | 0 | 0 | 0 | 0 | 0 | 0 | 0 | 0 | 0 | 0.13001113 | 0 | 0.05130116 | 0 | 0.2134379 | 0 | 0 | 0 | 0 |
| ENSMUSG00 Gad1 | 7618 | 5.08298968 | 4.21112065 | 4.9402214 | 6.46514688 | 4.79550145 | 4.70939605 | 3.98723285 | 2.02041719 | 1.97876933 | 2.18485144 | 2.10089408 | 2.07348812 | 1.22390816 | 2.21917141 | 2.10929467 | 1.553137 | 1.54955362 | 2.28903739 |
| ENSMUSG00 4930578N16 | 2549 | 1.38404457 | 2.05183381 | 1.76490323 | 1.5014232 | 2.39809898 | 2.11896742 | 2.96980331 | 2.2175267 | 2.68221327 | 1.72028851 | 3.07379097 | 2.27729655 | 2.65142121 | 2.56909232 | 2.58620986 | 2.56873043 | 2.82673355 | 3.17235727 |
| ENSMUSG00 Gm10304 | 2305 | 0 | 0 | 0 | 0 | 0 | 0 | 0 | 0 | 0 | 0 | 0 | 0 | 0 | 0 | 0 | 0 | 0 | 0 |
| ENSMUSG00 Pgap1 | 2844 | 0.83695143 | 0.6559501 | 0.15308079 | 0.86508312 | 0.20308829 | 0.38331932 | 1.03142971 | 1.19519121 | 2.67915099 | 0.64658137 | 1.07510455 | 1.74669268 | 0.72853042 | 2.26994661 | 1.26038515 | 0.78089143 | 0.90290503 | 0.83171294 |
| ENSMUSG00 Gm12794 | 1449 | 0 | 0 | 0 | 0 | 0 | 0 | 0 | 0 | 0 | 0 | 0 | 0 | 0 | 0 | 0 | 0 | 0 | 0 |
| ENSMUSG00 Gm12689 | 791 | 0.429888 | 0.42114911 | 0.48923925 | 0.46079438 | 0.24339784 | 0.18793704 | 0.05981375 | 0.53112064 | 0.15620676 | 0.65569878 | 0.42278724 | 0.846761 | 0.74839824 | 0.70458792 | 0.41670347 | 0.14522354 | 0.33917077 | 0 |
| ENSMUSG00 Gm10305 | 2474 | 0.01718075 | 0 | 0.23463313 | 0 | 0 | 0.0600882 | 0.01912396 | 0.01543751 | 0 | 0 | 0 | 0 | 0 | 0 | 0 | 0 | 0 | 0 |
| ENSMUSG00 Rpl23a-ps1 | 467 | 0 | 0 | 0 | 0 | 0 | 0 | 0 | 0 | 0 | 0 | 0 | 0 | 0.21127159 | 0 | 0 | 0 | 0 | 0 |
| ENSMUSG00 Gm10306 | 532 | 0 | 0.12523644 | 0.09092769 | 0 | 0.18094708 | 0.09314424 | 0.26680082 | 0.14358046 | 0.07741826 | 0 | 0 | 0.31474998 | 0 | 0.08730092 | 0.15489307 | 0.07197483 | 0.21612573 | 0.31020173 |
| ENSMUSG00 Zfp352 | 1903 | 0 | 0.01750546 | 0.02541962 | 0 | 0 | 0.02603927 | 0.02486215 | 0 | 0 | 0 | 0 | 0 | 0 | 0.02440572 | 0 | 0.02012118 | 0.04027987 | 0 |
| ENSMUSG00 Ifna1 | 569 | 0 | 0 | 0 | 0 | 0 | 0 | 0 | 0 | 0 | 0 | 0 | 0 | 0 | 0 | 0 | 0 | 0 | 0 |
| ENSMUSG00 Ifna4 | 643 | 0 | 0 | 0 | 0 | 0 | 0 | 0 | 0 | 0 | 0 | 0 | 0 | 0 | 0 | 0 | 0 | 0 | 0 |
| ENSMUSG00 Ifna11 | 646 | 0 | 0 | 0 | 0 | 0 | 0 | 0 | 0 | 0 | 0 | 0 | 0 | 0 | 0 | 0 | 0 | 0 | 0 |
| ENSMUSG00 Ifna7 | 575 | 0 | 0 | 0 | 0 | 0 | 0 | 0 | 0 | 0 | 0 | 0 | 0 | 0 | 0 | 0 | 0 | 0 | 0 |
| ENSMUSG00 Gm13288 | 2849 | 0.10443532 | 0 | 0 | 0 | 0 | 0 | 0 | 0 | 0 | 0.03309983 | 0 | 0.1175479 | 0 | 0 | 0.04338528 | 0 | 0 | 0 |
| ENSMUSG00 Klhl9 | 7049 | 7.71231666 | 4.91493971 | 6.52620618 | 9.59179881 | 5.61960181 | 6.65717672 | 6.81936187 | 5.6186014 | 5.59748647 | 7.49835982 | 5.89646955 | 7.18976048 | 6.06763687 | 4.82955257 | 7.44071225 | 7.91994688 | 6.5734846 | 6.58642163 |
| ENSMUSG00 Speer4d | 1313 | 0 | 0 | 0 | 0 | 0 | 0 | 0 | 0 | 0 | 0 | 0 | 0 | 0 | 0 | 0 | 0 | 0 | 0 |
| ENSMUSG00 Rraga | 1608 | 65.3173445 | 49.5135679 | 50.8403397 | 52.4462022 | 45.3481744 | 51.6174308 | 44.7528512 | 42.4676713 | 47.538665 | 47.3852839 | 34.791341 | 50.7478226 | 40.7417824 | 41.5339538 | 46.0955218 | 45.1248141 | 38.5408098 | 41.7015554 |
| ENSMUSG00 Tgfbrap1 | 6619 | 7.59686104 | 9.45685575 | 9.47153578 | 9.60918838 | 8.81339953 | 9.39547978 | 7.79133102 | 8.03200244 | 7.39851446 | 7.18765064 | 9.06562137 | 8.31458578 | 6.17860091 | 8.81307958 | 8.95740126 | 7.93117129 | 8.6276148 | 8.84267858 |
| ENSMUSG00 Gm6455 | 1122 | 0 | 0 | 0 | 0 | 0 | 0 | 0 | 0 | 0 | 0 | 0 | 0 | 0 | 0 | 0 | 0 | 0 | 0 |
| ENSMUSG00 Il1rl2 | 1939 | 0.78916263 | 1.97575185 | 1.87107518 | 1.97376365 | 3.20217543 | 1.73779571 | 0 | 0.17727283 | 0.78592114 | 0.04863404 | 0 | 0.20150059 | 0 | 0.38324157 | 0 | 0.37520451 | 0.43485226 | 0.02836983 |
| ENSMUSG00 Olfr356 | 947 | 0 | 0 | 0 | 0 | 0 | 0 | 0 | 0 | 0 | 0 | 0 | 0 | 0 | 0 | 0 | 0 | 0 | 0 |
| ENSMUSG00 Gm428 | 1316 | 0 | 0 | 0 | 0 | 0 | 0 | 0 | 0 | 0 | 0 | 0 | 0 | 0 | 0 | 0 | 0 | 0 | 0 |
| ENSMUSG00 Rabepk | 4007 | 11.6154649 | 12.3956889 | 11.9153168 | 10.4834746 | 11.0269793 | 10.6599592 | 10.9573661 | 10.1223687 | 8.53127229 | 9.31953007 | 10.0152147 | 10.9346881 | 13.8872979 | 12.1470935 | 10.26183 | 14.6396832 | 13.0751305 | 12.3142423 |
| ENSMUSG00 Gm10307 | 671 | 0.88684421 | 0.89363949 | 0.4325502 | 0.40740129 | 1.14770608 | 0.51694357 | 0.35255349 | 0.45534906 | 0.42966559 | 0.56215444 | 0.21359892 | 1.08137647 | 1.17631994 | 0.4845135 | 0.52262892 | 0.4565199 | 0.51406359 | 0.65584628 |
| ENSMUSG00 Dnajc25 | 4535 | 1.80893031 | 1.68951834 | 1.82400743 | 1.33618939 | 2.02716591 | 1.85754459 | 2.11785531 | 2.35807556 | 2.4067093 | 2.41211977 | 2.27549968 | 2.11693362 | 3.12199785 | 2.40669472 | 1.74436576 | 2.11928169 | 2.91567198 | 4.03925745 |
| ENSMUSG00 Defb44-ps | 217 | 0.58762916 | 0.15351564 | 0.22291949 | 0.20995873 | 0 | 0 | 1.52621545 | 0.70400742 | 0.94899806 | 0.65185304 | 0 | 1.28607517 | 0.45467205 | 2.14028052 | 2.84803382 | 0 | 0.17661888 | 1.01399275 |
| ENSMUSG00 Actl7a | 1503 | 0 | 0 | 0 | 0 | 0 | 0 | 0 | 0 | 0 | 0 | 0 | 0 | 0 | 0 | 0 | 0 | 0 | 0 |
| ENSMUSG00 Actl7b | 1418 | 0 | 0 | 0 | 0 | 0 | 0 | 0.26692626 | 0 | 0 | 0.13300622 | 0 | 0 | 0 | 0 | 0 | 0.1620195 | 0 | 0 |
| ENSMUSG00 Olfr270 | 1882 | 0 | 0 | 0 | 0 | 0 | 0 | 0 | 0 | 0 | 0 | 0 | 0 | 0 | 0 | 0 | 0 | 0 | 0 |
| ENSMUSG00 4930552N02 | 1184 | 0 | 0 | 0 | 0 | 0 | 0 | 0 | 0 | 0 | 0 | 0 | 0 | 0 | 0 | 0 | 0 | 0 | 0 |
| ENSMUSG00 Acnat1 | 2921 | 1.2514362 | 1.18608045 | 1.40765152 | 1.73135088 | 1.40062085 | 1.17053701 | 1.02043779 | 1.03293385 | 0.90240911 | 1.38820968 | 0.81778429 | 0.80255318 | 1.09776605 | 1.43100577 | 1.66442437 | 0.68165409 | 1.46954647 | 1.50658286 |
| ENSMUSG00 Foxe1 | 2803 | 0 | 0 | 0 | 0.01625439 | 0 | 0 | 0 | 0.01362554 | 0.14693727 | 0 | 0 | 0 | 0 | 0 | 0 | 0 | 0 | 0 |
| ENSMUSG00 Gm6313 | 2587 | 0 | 0 | 0 | 0 | 0 | 0 | 0 | 0 | 0 | 0 | 0 | 0 | 0 | 0 | 0 | 0 | 0 | 0 |
| ENSMUSG00 1700055D18 | 1569 | 0.70435601 | 1.23145179 | 2.28147944 | 0.84210981 | 1.50316396 | 1.13696521 | 0.69355743 | 1.7769569 | 2.59876677 | 0.81138877 | 1.12662426 | 2.41903445 | 2.01226431 | 0.97683549 | 1.86444262 | 1.12260544 | 1.58776883 | 0.94661943 |
| ENSMUSG00 Ccin | 2029 | 0 | 0 | 0 | 0 | 0 | 0 | 0.04663645 | 0.09411632 | 0 | 0 | 0 | 0 | 0 | 0 | 0 | 0.03777851 | 0 | 0 |
| ENSMUSG00 Olfr155 | 1346 | 0 | 0 | 0 | 0 | 0 | 0 | 0 | 0 | 0 | 0 | 0 | 0 | 0 | 0 | 0 | 0 | 0 | 0 |
| ENSMUSG00 Hrct1 | 1441 | 0 | 0 | 0 | 0 | 0.03340175 | 0 | 0 | 0 | 0 | 0.32720821 | 0 | 0 | 0 | 0 | 0 | 0.13286123 | 0 | 0 |
| ENSMUSG00 Ccl21a | 866 | 0 | 0 | 0 | 0 | 0 | 0 | 0 | 0 | 0 | 0 | 0 | 0 | 0 | 0 | 0 | 0 | 0 | 0 |
| ENSMUSG00 Ccl19 | 1101 | 0 | 0 | 0 | 0 | 0 | 0 | 0 | 0 | 0 | 0 | 0 | 0 | 0 | 0 | 0 | 0 | 0 | 0 |
| ENSMUSG00 Ndufb6 | 837 | 111.874436 | 103.162511 | 101.486164 | 100.103657 | 107.707399 | 100.526334 | 122.83208 | 103.3522 | 108.15766 | 109.229646 | 114.728395 | 92.2925651 | 126.188333 | 117.247738 | 101.896321 | 110.297076 | 105.683504 | 109.229552 |
| ENSMUSG00 Gm136 | 1306 | 0 | 0 | 0 | 0 | 0 | 0 | 0 | 0 | 0 | 0 | 0 | 0 | 0 | 0 | 0.22083529 | 0 | 0 | 0 |
| ENSMUSG00 Sdr16c6 | 2172 | 0 | 0 | 0 | 0 | 0 | 0 | 0 | 0 | 0 | 0 | 0 | 0 | 0 | 0 | 0 | 0 | 0 | 0 |
| ENSMUSG00 Gm10308 | 740 | 0 | 0 | 0 | 0 | 0 | 0 | 0 | 0 | 0 | 0 | 0 | 0 | 0 | 0 | 0 | 0 | 0 | 0 |
| ENSMUSG00 Gm5499 | 887 | 0.09584031 | 0.48823858 | 0.27268055 | 0.82184522 | 0.92248332 | 0.78211755 | 1.06680223 | 1.03339082 | 0.97510353 | 0.74420501 | 1.02336439 | 0.81804241 | 1.3904148 | 0.52360865 | 1.30061281 | 1.33822867 | 0.77776025 | 0.80622141 |
| ENSMUSG00 Gm10309 | 2003 | 0.2970906 | 0.0498945 | 0.36225809 | 0.06823921 | 0.26432909 | 0.59374219 | 0.25982999 | 0.47669 | 0.08224966 | 0.51788091 | 0.40547893 | 0.72451684 | 1.23145076 | 1.36804851 | 1.58388408 | 1.68226333 | 0.86105011 | 1.6478015 |
| ENSMUSG00 1700106N22 | 3380 | 3.67204479 | 3.33128942 | 3.64948232 | 2.68244018 | 4.04422081 | 2.93221441 | 4.33932854 | 4.90399487 | 4.0211687 | 3.18057998 | 4.83403426 | 4.88799885 | 4.02828674 | 5.13907949 | 3.21810969 | 4.31618977 | 3.87798625 | 2.32730836 |
| ENSMUSG00 Gm9386 | 380 | 0 | 0.17533102 | 0.25459753 | 0.35969246 | 0 | 0 | 0 | 0 | 0 | 0.2481616 | 0 | 0 | 0 | 0.12222128 | 0.10842515 | 0 | 0.10085868 | 0 |
| ENSMUSG00 Gm15210 | 3198 | 0.21265879 | 0.17708543 | 0.10588327 | 0.09972711 | 0.16555696 | 0.15494914 | 0.26630026 | 0.13136849 | 0.19318253 | 0.16218191 | 0.17926814 | 0.12217312 | 0.10798106 | 0.07261427 | 0.23190369 | 0.09578638 | 0.15579795 | 0.15480987 |
| ENSMUSG00 Rasgrp3 | 4906 | 0 | 0 | 0 | 0 | 0 | 0.02020087 | 0 | 0.05449385 | 0 | 0 | 0.06816647 | 0 | 0.13072053 | 0 | 0 | 0.02341456 | 0 | 0 |
| ENSMUSG00 Ces1a | 1966 | 0 | 0.06777801 | 0 | 0 | 0 | 0 | 0 | 0 | 0 | 0 | 0 | 0 | 0 | 0 | 0 | 0 | 0.01949456 | 0 |
| ENSMUSG00 Rpl7a-ps5 | 806 | 0 | 0 | 0 | 0 | 0 | 0 | 0 | 0 | 0 | 0 | 0 | 0 | 0 | 0 | 0 | 0 | 0 | 0 |
| ENSMUSG00 Safb | 3173 | 23.7375266 | 20.7772511 | 20.4287837 | 19.4133413 | 17.0501992 | 20.6769049 | 22.456002 | 22.8456287 | 21.0410785 | 22.4534235 | 19.3930496 | 21.9533397 | 24.7825642 | 23.683118 | 21.2305528 | 23.471551 | 23.8437158 | 25.103431 |
| ENSMUSG00 Zfp827 | 8497 | 3.50666452 | 2.76398616 | 1.62820168 | 2.80969604 | 2.87194131 | 2.89840045 | 1.49783578 | 1.75747079 | 1.58018173 | 1.54264982 | 1.79922169 | 1.32034226 | 1.24824788 | 0.99480098 | 0.92130112 | 2.06391646 | 1.31257531 | 1.38542413 |
| ENSMUSG00 Olfr806 | 964 | 0 | 0 | 0 | 0 | 0 | 0 | 0 | 0 | 0 | 0 | 0 | 0 | 0 | 0 | 0 | 0 | 0 | 0 |
| ENSMUSG00 Treml2 | 3525 | 0 | 0 | 0 | 0 | 0 | 0 | 0 | 0 | 0 | 0 | 0 | 0 | 0 | 0 | 0 | 0 | 0 | 0 |
| ENSMUSG00 Ptges3 | 2247 | 40.9729467 | 41.0814552 | 41.4845541 | 38.9914946 | 42.005653 | 42.3194018 | 39.6272638 | 38.3963672 | 40.9482315 | 39.65947 | 36.357445 | 36.0181178 | 38.8376331 | 34.1664782 | 35.6456719 | 38.8529528 | 38.1045604 | 36.3055208 |
| ENSMUSG00 Gm88 | 1378 | 0.18507334 | 0.21757333 | 0.17552079 | 0 | 0.27943062 | 0.07191979 | 0.17167155 | 0.97005376 | 0.41844065 | 0.1026503 | 0.24268847 | 0.20252417 | 0.17899825 | 0.13481593 | 0.68768925 | 0.30565797 | 0.08343896 | 0.35927573 |
| ENSMUSG00 Yipf3 | 2821 | 41.5860637 | 40.8351607 | 37.210408 | 41.1680617 | 35.642534 | 39.0484675 | 31.8492653 | 37.7591671 | 37.7409228 | 38.4091868 | 43.0838335 | 35.4561033 | 40.133552 | 39.1012788 | 35.4616724 | 39.7022427 | 36.2748007 | 38.2002269 |
| ENSMUSG00 Jund | 4434 | 103.904737 | 113.665038 | 109.020453 | 125.524068 | 120.774872 | 135.191569 | 103.353768 | 102.759441 | 98.7120214 | 120.716009 | 118.564647 | 110.762746 | 113.728204 | 140.117852 | 115.799231 | 128.196677 | 144.013178 | 188.462639 |
| ENSMUSG00 Gm9104 | 515 | 0.24760297 | 0 | 0.09392919 | 0.08846805 | 0.09346005 | 0.0962189 | 0 | 0 | 0.63979054 | 0 | 0.37106764 | 0.10837993 | 0 | 0.36073078 | 0.32001208 | 0 | 0.14883999 | 0.3204414 |
| ENSMUSG00 Nr2c2ap | 1170 | 5.41305233 | 5.38131368 | 3.96911018 | 4.51716337 | 4.89546916 | 4.7858623 | 4.93345883 | 4.24360027 | 4.68188598 | 4.9568688 | 7.18666042 | 5.39074348 | 6.07162059 | 6.5498072 | 6.51477593 | 4.90905223 | 4.81535525 | 6.67631894 |
| ENSMUSG00 Gm10311 | 386 | 4.95526664 | 3.53841623 | 7.51920161 | 1.41640552 | 5.86062283 | 3.33774889 | 3.67715121 | 5.63981072 | 5.76184417 | 7.20697278 | 4.57946493 | 6.21780697 | 4.08969262 | 6.97864526 | 7.68526431 | 7.34068639 | 8.63831044 | 3.70527662 |
| ENSMUSG00 Olfr761 | 971 | 0 | 0 | 0 | 0 | 0 | 0 | 0 | 0 | 0 | 0 | 0 | 0 | 0 | 0 | 0 | 0 | 0 | 0 |

[illegible]

| | | | | | | | | | | | | | | | | | | | | |
|---|---|---|---|---|---|---|---|---|---|---|---|---|---|---|---|---|---|---|---|---|
| ENSMUSG00 | Gm17184 | 459 | 0 | 0 | 0 | 0 | 0 | 0 | 0 | 0 | 0 | 0 | 0 | 0 | 0 | 0 | 0 | 0 | 0 | 0 |
| ENSMUSG00 | Eddm3b | 1255 | 47.585476 | 53.5926163 | 68.3782015 | 44.2178104 | 58.5637031 | 60.371418 | 79.0932274 | 67.8942229 | 67.0798711 | 65.4098601 | 54.094196 | 63.4208246 | 73.8209886 | 89.5575231 | 73.1779029 | 52.0813282 | 60.8029141 | 79.6864988 |
| ENSMUSG00 | Gm16020 | 518 | 0.16411265 | 0.12862121 | 0.56031116 | 0 | 0.18583754 | 0 | 0.09133722 | 0.14746101 | 0 | 0.18204905 | 0 | 0.10775224 | 0 | 0 | 0.07953968 | 0.07392009 | 0.14797798 | 0 |
| ENSMUSG00 | Ptrh2 | 3129 | 11.6553023 | 9.16663532 | 11.2237721 | 10.3673581 | 8.96801262 | 9.13772045 | 8.64904198 | 8.48313189 | 10.3723153 | 9.52356806 | 10.855863 | 8.65151689 | 7.33120535 | 8.31214092 | 9.71772076 | 11.3807174 | 6.8225463 | 9.96809316 |
| ENSMUSG00 | Gm7489 | 1220 | 0 | 0 | 0 | 0 | 0 | 0 | 0 | 0 | 0 | 0 | 0 | 0 | 0 | 0 | 0 | 0 | 0 | 0 |
| ENSMUSG00 | Gm6616 | 335 | 0 | 0 | 0.1443986 | 0 | 0 | 0 | 0 | 0 | 0 | 0 | 0 | 0 | 0 | 0 | 0 | 0 | 0 | 0.32841258 |
| ENSMUSG00 | Gm10371 | 1407 | 0.06041958 | 0 | 0 | 0.06476339 | 0.0342089 | 0.07043743 | 0.06725327 | 0.08143369 | 0 | 0 | 0 | 0 | 0 | 0 | 0 | 0 | 0 | 0 |
| ENSMUSG00 | 5930412G12I | 3544 | 5.38510836 | 5.18869007 | 5.73275473 | 6.543615 | 4.31883514 | 4.82384116 | 2.72341606 | 3.07134162 | 5.31101515 | 5.21531481 | 5.23044128 | 4.07907918 | 3.09020475 | 3.5645575 | 3.08081611 | 3.27371728 | 3.03885142 | 2.96465558 |
| ENSMUSG00 | Gm10373 | 665 | 0 | 0.35066204 | 0.07274215 | 0.06851285 | 0.36189416 | 0.14903078 | 0.78261574 | 0 | 0.68128071 | 0.70903313 | 0 | 0.16786665 | 0.14836667 | 0.90792953 | 0.43370059 | 0.63337847 | 0.69160235 | 0.41360231 |
| ENSMUSG00 | Gm16439 | 563 | 0.83047413 | 2.07096145 | 1.03105216 | 1.861286 | 1.7953293 | 2.02435679 | 2.68917536 | 2.10295644 | 3.51146138 | 3.68495727 | 4.24289152 | 3.56902993 | 1.05147959 | 3.29975754 | 2.04910047 | 1.29222299 | 2.4507046 | 4.59223449 |
| ENSMUSG00 | 4930503E14I | 1283 | 0 | 0 | 0 | 0 | 0 | 0 | 0 | 0 | 0 | 0 | 0 | 0 | 0 | 0 | 0 | 0 | 0 | 0 |
| ENSMUSG00 | Ear2 | 717 | 0 | 0 | 0 | 0 | 0 | 0 | 0 | 0 | 0.22977136 | 0 | 0 | 0 | 0 | 0 | 0 | 0 | 0 | 0 |
| ENSMUSG00 | Ang6 | 1155 | 0 | 0 | 0 | 0 | 0 | 0 | 0 | 0 | 0 | 0 | 0 | 0 | 0 | 0 | 0 | 0 | 0 | 0 |
| ENSMUSG00 | Ear-ps2 | 465 | 0 | 0 | 0 | 0 | 0 | 0 | 0 | 0 | 0 | 0 | 0 | 0 | 0 | 0 | 0 | 0 | 0 | 0 |
| ENSMUSG00 | Ear-ps9 | 778 | 0 | 0 | 0 | 0 | 0 | 0 | 0 | 0 | 0 | 0 | 0 | 0 | 0 | 0 | 0 | 0 | 0 | 0 |
| ENSMUSG00 | Ear1 | 711 | 0 | 0 | 0 | 0 | 0 | 0 | 0 | 0 | 0 | 0 | 0 | 0 | 0 | 0 | 0 | 0 | 0 | 0 |
| ENSMUSG00 | Gm10375 | 1241 | 0 | 0 | 0 | 0 | 0 | 0 | 0 | 0 | 0 | 0 | 0 | 0 | 0 | 0 | 0 | 0 | 0 | 0 |
| ENSMUSG00 | 1700001F09I | 894 | 0 | 0 | 0 | 0 | 0 | 0 | 0 | 0 | 0 | 0 | 0 | 0 | 0 | 0 | 0 | 0 | 0 | 0 |
| ENSMUSG00 | Gm10376 | 895 | 0 | 0 | 0 | 0 | 0 | 0 | 0 | 0 | 0 | 0 | 0 | 0 | 0 | 0 | 0 | 0 | 0 | 0 |
| ENSMUSG00 | Gm10377 | 894 | 0 | 0 | 0 | 0 | 0 | 0 | 0 | 0 | 0 | 0 | 0 | 0 | 0 | 0 | 0 | 0 | 0 | 0 |
| ENSMUSG00 | Gm10378 | 1398 | 0 | 0 | 0 | 0 | 0 | 0 | 0 | 0 | 0 | 0 | 0 | 0 | 0 | 0 | 0 | 0 | 0 | 0 |
| ENSMUSG00 | Gm3633 | 1407 | 0 | 0 | 0 | 0 | 0 | 0 | 0 | 0 | 0 | 0 | 0 | 0 | 0 | 0 | 0 | 0 | 0 | 0 |
| ENSMUSG00 | Gm9611 | 1407 | 0 | 0 | 0 | 0 | 0 | 0 | 0 | 0 | 0 | 0 | 0 | 0 | 0 | 0 | 0 | 0 | 0 | 0 |
| ENSMUSG00 | Gm7995 | 1568 | 0 | 0 | 0 | 0 | 0 | 0 | 0 | 0 | 0 | 0 | 0 | 0 | 0 | 0 | 0 | 0 | 0 | 0 |
| ENSMUSG00 | Gm10382 | 1232 | 3.72610309 | 3.13660368 | 4.27980098 | 3.77209945 | 5.82114986 | 3.05682447 | 6.06769746 | 4.3090454 | 5.24860631 | 7.19507485 | 4.61462667 | 4.9835413 | 8.60906024 | 11.6110218 | 11.97253 | 10.1631726 | 10.2348634 | 15.8061881 |
| ENSMUSG00 | 1700049E17I | 2041 | 0 | 0 | 0 | 0 | 0 | 0 | 0 | 0 | 0 | 0 | 0 | 0 | 0 | 0 | 0 | 0 | 0 | 0 |
| ENSMUSG00 | Gm7853 | 474 | 0 | 0 | 0 | 0 | 0 | 0 | 0 | 0 | 0 | 0 | 0 | 0 | 0 | 0 | 0 | 0 | 0 | 0 |
| ENSMUSG00 | Gm10384 | 2044 | 0 | 0 | 0 | 0 | 0 | 0.07272906 | 0 | 0 | 0 | 0.04613572 | 0 | 0 | 0 | 0 | 0 | 0 | 0.0562519 | 0 |
| ENSMUSG00 | Slfn2 | 1662 | 0 | 0.02004386 | 0.2037393 | 0.41120076 | 0.14480121 | 0.23852098 | 0.79708484 | 1.49368601 | 1.33819004 | 0 | 2.70207346 | 0 | 1.45443077 | 0.27944698 | 1.04119455 | 0.50685521 | 0.09224139 | 0 |
| ENSMUSG00 | Slfn10-ps | 3616 | 0 | 0 | 0.06688818 | 0.05039939 | 0 | 0.1918524 | 0.05233703 | 0 | 0.25058168 | 0.07823679 | 0 | 0.10805023 | 0.0545707 | 0.01284405 | 0 | 0.02117843 | 0 | 0 |
| ENSMUSG00 | Zfp9 | 3962 | 0.52568239 | 0.95852346 | 0.39069989 | 0.51747779 | 0.86253573 | 0.72540599 | 0.48960622 | 1.05072485 | 0.37423386 | 0.85685276 | 0.56673979 | 0.9157037 | 0.34863546 | 1.54735475 | 0.1663869 | 1.07275553 | 0.38693889 | 0.30545188 |
| ENSMUSG00 | Gm5460 | 1898 | 0 | 0 | 0 | 0 | 0 | 0 | 0 | 0 | 0 | 0 | 0 | 0 | 0 | 0 | 0 | 0 | 0 | 0 |
| ENSMUSG00 | Gdf2 | 2986 | 0 | 0 | 0 | 0 | 0 | 0 | 0 | 0 | 0 | 0 | 0 | 0.07476981 | 0 | 0 | 0 | 0.06411689 | 0 | 0 |
| ENSMUSG00 | 1810012P15I | 8174 | 1.08680961 | 1.1941128 | 1.47949383 | 1.72233456 | 1.5251001 | 1.75198679 | 1.71330474 | 2.27079858 | 2.72091002 | 2.06507851 | 2.93406278 | 2.37629687 | 1.56915812 | 2.53414032 | 2.20272571 | 1.85503799 | 2.10996252 | 1.43344014 |
| ENSMUSG00 | Cux2 | 412 | 0 | 0 | 0 | 0 | 0 | 0 | 0 | 0 | 0 | 0 | 0 | 0 | 0 | 0 | 0 | 0 | 0 | 0 |
| ENSMUSG00 | Adam1a | 3091 | 0.46754319 | 0.2047703 | 0.23474699 | 0.3537577 | 0.20243125 | 0.16031295 | 0.13775934 | 0.38303606 | 0.35976575 | 0.09152514 | 0.35549144 | 0.45143693 | 0.38303656 | 0.55594689 | 0.42654474 | 0.21059215 | 0.21078843 | 0.07118616 |
| ENSMUSG00 | Zfp783 | 3693 | 0.86322453 | 0.28865763 | 0.96930443 | 0.90061095 | 0.33886543 | 0.63064676 | 0.40996635 | 0.37230612 | 0.79183391 | 1.09801259 | 0.78913416 | 1.2997961 | 0.60112003 | 0.81745618 | 0.58014647 | 0.74652687 | 0.78873506 | 0.31280564 |
| ENSMUSG00 | Gm17524 | 1308 | 0 | 0 | 0 | 0 | 0 | 0 | 0 | 0 | 0 | 0 | 0 | 0 | 0 | 0 | 0 | 0 | 0 | 0 |
| ENSMUSG00 | Gm6288 | 1298 | 0.45845336 | 0.69294926 | 0.7826226 | 0.70201917 | 0.85287692 | 0.7635244 | 2.07767542 | 0.58848078 | 1.11057631 | 0.47223355 | 0.73613187 | 1.07503202 | 0.72211589 | 0.71562538 | 0.92052783 | 0.35399637 | 1.12203335 | 0.55093867 |
| ENSMUSG00 | Gm5887 | 1275 | 0.20002436 | 0.10451104 | 0 | 0 | 0 | 0.03886489 | 0 | 0.17972895 | 0.58145669 | 0 | 0.14988222 | 0.26266194 | 0.0773834 | 0.21856041 | 0 | 0 | 0.06011968 | 0.17257759 |
| ENSMUSG00 | Spef2 | 7201 | 0.11215086 | 0.22668127 | 0.00671761 | 0.13286793 | 0 | 0.11698326 | 0.01971088 | 0 | 0.00571956 | 0.11131259 | 0.03980694 | 0.14727088 | 0.03425352 | 0.11609409 | 0.2231441 | 0.14356984 | 0.18628252 | 0.19097732 |
| ENSMUSG00 | Ugt3a1 | 2660 | 0 | 0 | 0 | 0 | 0 | 0 | 0 | 0 | 0 | 0 | 0 | 0 | 0 | 0 | 0 | 0 | 0 | 0 |
| ENSMUSG00 | Gm10390 | 8062 | 0.03163372 | 0.11156638 | 0.01200038 | 0.14128332 | 0.0835831 | 0.04917165 | 0.05868603 | 0.03316135 | 0.03065233 | 0.05848512 | 0.07111132 | 0.02769321 | 0.02447627 | 0.03456519 | 0.06132705 | 0.04749517 | 0.09507888 | 0.07505584 |
| ENSMUSG00 | Gm10391 | 5073 | 0 | 0 | 0 | 0 | 0 | 0 | 0 | 0 | 0 | 0 | 0 | 0 | 0 | 0 | 0 | 0 | 0 | 0 |
| ENSMUSG00 | Gm10392 | 1730 | 0 | 0 | 0 | 0 | 0 | 0 | 0 | 0 | 0 | 0 | 0 | 0 | 0.05703112 | 0 | 0 | 0 | 0 | 0 |
| ENSMUSG00 | Gm10393 | 446 | 0 | 0 | 0 | 0 | 0 | 0 | 0 | 0 | 0 | 0 | 0 | 0 | 0 | 0 | 0 | 0 | 0 | 0 |
| ENSMUSG00 | Gm10394 | 4758 | 0 | 0 | 0 | 0 | 0 | 0 | 0 | 0 | 0 | 0 | 0 | 0 | 0 | 0 | 0 | 0 | 0 | 0 |
| ENSMUSG00 | Gm10395 | 1617 | 0 | 0 | 0 | 0 | 0 | 0 | 0 | 0 | 0 | 0 | 0 | 0 | 0 | 0 | 0 | 0 | 0 | 0 |
| ENSMUSG00 | Gm10396 | 1205 | 0 | 0 | 0 | 0 | 0 | 0 | 0 | 0 | 0 | 0 | 0 | 0 | 0 | 0 | 0 | 0 | 0 | 0 |
| ENSMUSG00 | Cphx | 1706 | 0 | 0 | 0 | 0 | 0 | 0 | 0 | 0 | 0 | 0 | 0 | 0 | 0 | 0 | 0 | 0 | 0 | 0 |
| ENSMUSG00 | D6Ertd474e | 2623 | 0 | 0 | 0 | 0 | 0 | 0 | 0 | 0 | 0 | 0.10785521 | 0 | 0 | 0.11284464 | 0 | 0.34557157 | 0 | 0.02922325 | 0 |
| ENSMUSG00 | D14Ertd449e | 1272 | 0 | 0 | 0 | 0 | 0 | 0 | 0 | 0 | 0 | 0 | 0 | 0 | 0 | 0 | 0 | 0 | 0 | 0 |
| ENSMUSG00 | Plac9 | 1327 | 0 | 0 | 0 | 0 | 0 | 0 | 0 | 0 | 0 | 0 | 0 | 0 | 0 | 0 | 0 | 0 | 0 | 0 |
| ENSMUSG00 | Ssh2 | 1511 | 0 | 0 | 0 | 0 | 0 | 0 | 0 | 0 | 0 | 0 | 0 | 0 | 0 | 0 | 0 | 0 | 0 | 0 |
| ENSMUSG00 | Gm7457 | 4108 | 0 | 0 | 0 | 0 | 0 | 0 | 0 | 0 | 0 | 0 | 0 | 0 | 0 | 0 | 0 | 0 | 0 | 0 |
| ENSMUSG00 | Gm10397 | 1698 | 0 | 0 | 0 | 0 | 0 | 0 | 0 | 0 | 0 | 0 | 0 | 0 | 0 | 0 | 0 | 0 | 0 | 0 |
| ENSMUSG00 | Gm10398 | 395 | 0 | 0 | 0 | 0 | 0 | 0 | 0 | 0 | 0 | 0 | 0 | 0 | 0 | 0 | 0 | 0 | 0 | 0 |
| ENSMUSG00 | Gm10399 | 497 | 3.07884575 | 3.08328598 | 3.50391769 | 2.93350788 | 4.45486617 | 2.59229592 | 2.47510996 | 4.2265234 | 5.13795569 | 1.99228324 | 1.44190016 | 3.03223921 | 3.2755599 | 3.73795472 | 2.90151802 | 1.84904342 | 3.7015337 | 2.76705768 |
| ENSMUSG00 | Gm13826 | 1011 | 3.61567274 | 3.98700317 | 4.06701294 | 2.02793966 | 2.47562812 | 4.46023616 | 4.25860908 | 3.96656998 | 4.39974649 | 3.26463821 | 3.21335032 | 3.58854408 | 6.34337215 | 7.0745692 | 3.91231391 | 3.14354146 | 4.05629453 | 6.09398611 |
| ENSMUSG00 | Gm10401 | 606 | 0 | 0 | 0 | 0 | 0 | 0 | 0 | 0 | 0 | 0 | 0 | 0 | 0 | 0 | 0 | 0 | 0 | 0 |
| ENSMUSG00 | 1500011B03I | 1331 | 25.0049232 | 19.9977479 | 21.0430309 | 19.2718768 | 21.3357137 | 19.880661 | 29.1838538 | 24.4477287 | 27.9734112 | 25.4705903 | 27.925558 | 27.6771881 | 34.4693336 | 33.1494238 | 33.7722746 | 36.4782045 | 31.5018549 | 29.7983215 |
| ENSMUSG00 | 2700089E24I | 2508 | 41.6408363 | 41.3356168 | 37.7653 | 39.4571724 | 41.0886956 | 37.2633395 | 43.8227086 | 37.1416545 | 39.2158691 | 38.6342485 | 43.2222411 | 43.0635196 | 41.1296009 | 34.1293672 | 37.0452586 | 38.4280139 | 37.4399631 | 42.0025675 |
| ENSMUSG00 | Olfr31 | 953 | 0 | 0 | 0 | 0 | 0 | 0.15598972 | 0 | 0.08015195 | 0 | 0.19790432 | 0 | 0.11713675 | 0.51764866 | 0 | 0 | 0.20089511 | 0 | 0 |
| ENSMUSG00 | Olfr381 | 1998 | 0 | 0 | 0 | 0 | 0 | 0 | 0 | 0 | 0 | 0 | 0 | 0 | 0 | 0 | 0 | 0 | 0 | 0 |
| ENSMUSG00 | Olfr380 | 2212 | 0 | 0 | 0 | 0 | 0 | 0 | 0 | 0 | 0 | 0 | 0 | 0 | 0 | 0 | 0 | 0 | 0 | 0 |
| ENSMUSG00 | Rpl21-ps4 | 440 | 0 | 0 | 0 | 0 | 0 | 0 | 0 | 0 | 0 | 0 | 0 | 0 | 0 | 0 | 0 | 0 | 0 | 0 |
| ENSMUSG00 | Klra11 | 838 | 0 | 0 | 0 | 0 | 0 | 0 | 0 | 0 | 0 | 0 | 0 | 0 | 0 | 0 | 0 | 0 | 0 | 0 |
| ENSMUSG00 | Klra10 | 819 | 0 | 0 | 0 | 0 | 0 | 0 | 0 | 0 | 0 | 0 | 0 | 0 | 0 | 0 | 0 | 0 | 0 | 0 |
| ENSMUSG00 | Myo18b | 8236 | 0 | 0 | 0 | 0 | 0 | 0 | 0.07468004 | 0.03709801 | 0 | 0 | 0.07540972 | 0 | 0.07187749 | 0.13533974 | 0 | 0 | 0 | 0 |
| ENSMUSG00 | Klra14 | 877 | 0 | 0 | 0 | 0 | 0 | 0 | 0 | 0 | 0 | 0 | 0 | 0 | 0 | 0 | 0 | 0 | 0 | 0 |
| ENSMUSG00 | Gm6588 | 2312 | 0 | 0.05763477 | 0 | 0 | 0 | 0 | 0 | 0 | 0 | 0 | 0 | 0 | 0 | 0 | 0 | 0 | 0 | 0 |
| ENSMUSG00 | Gm10044 | 2622 | 0 | 0 | 0.16604187 | 0.12163513 | 0 | 0.03779766 | 0.1984895 | 0 | 0.17278859 | 0 | 0.05466242 | 0 | 0 | 0 | 0.12571032 | 0.10222512 | 0.0438516 | 0.18881844 |
| ENSMUSG00 | Gm5797 | 992 | 0 | 0 | 0 | 0 | 0 | 0 | 0 | 0 | 0 | 0 | 0 | 0 | 0 | 0 | 0 | 0 | 0 | 0 |
| ENSMUSG00 | Gm16434 | 2061 | 0 | 0 | 0 | 0 | 0 | 0 | 0 | 0 | 0 | 0 | 0 | 0 | 0 | 0 | 0 | 0 | 0 | 0 |
| ENSMUSG00 | Gm6676 | 2064 | 0 | 0 | 0 | 0 | 0 | 0 | 0 | 0 | 0 | 0 | 0 | 0 | 0 | 0 | 0 | 0 | 0 | 0 |
| ENSMUSG00 | Gm3715 | 2172 | 0 | 0 | 0 | 0 | 0 | 0 | 0 | 0 | 0 | 0 | 0 | 0 | 0 | 0 | 0 | 0 | 0 | 0 |
| ENSMUSG00 | Gm10406 | 2714 | 0 | 0 | 0 | 0 | 0 | 0 | 0 | 0 | 0 | 0 | 0 | 0 | 0 | 0 | 0 | 0 | 0 | 0 |
| ENSMUSG00 | Gm3591 | 1740 | 0 | 0 | 0 | 0 | 0 | 0 | 0 | 0 | 0 | 0 | 0 | 0 | 0 | 0 | 0 | 0 | 0 | 0 |
| ENSMUSG00 | Gm16440 | 1084 | 0 | 0 | 0 | 0 | 0 | 0 | 0 | 0 | 0 | 0 | 0 | 0 | 0 | 0 | 0 | 0 | 0 | 0 |
| ENSMUSG00 | Gm10408 | 1507 | 0 | 0 | 0 | 0 | 0 | 0 | 0 | 0 | 0 | 0 | 0 | 0 | 0 | 0 | 0 | 0 | 0 | 0 |
| ENSMUSG00 | Gm10409 | 2061 | 0 | 0 | 0 | 0 | 0 | 0 | 0 | 0 | 0 | 0 | 0 | 0 | 0 | 0 | 0 | 0 | 0 | 0 |
| ENSMUSG00 | Gm10410 | 1686 | 0 | 0 | 0 | 0 | 0 | 0 | 0 | 0 | 0 | 0 | 0 | 0 | 0 | 0 | 0 | 0 | 0 | 0 |
| ENSMUSG00 | Gm3483 | 606 | 0 | 0 | 0 | 0 | 0 | 0 | 0 | 0 | 0 | 0 | 0 | 0 | 0 | 0 | 0 | 0 | 0 | 0 |
| ENSMUSG00 | Gm10412 | 608 | 0 | 0 | 0 | 0 | 0 | 0 | 0 | 0 | 0 | 0 | 0 | 0 | 0 | 0 | 0 | 0 | 0 | 0 |
| ENSMUSG00 | Gm8165 | 1399 | 0 | 0 | 0 | 0 | 0 | 0 | 0 | 0 | 0 | 0 | 0 | 0 | 0 | 0 | 0.05890144 | 0 | 0 | 0 |
| ENSMUSG00 | Gm10413 | 1111 | 0 | 0 | 0 | 0 | 0 | 0 | 0 | 0 | 0 | 0 | 0 | 0 | 0 | 0 | 0 | 0 | 0 | 0 |
| ENSMUSG00 | 9230020A06I | 2053 | 0 | 0.03245289 | 0 | 0 | 0 | 0 | 0 | 0 | 0 | 0 | 0 | 0 | 0 | 0 | 0 | 0 | 0 | 0 |
| ENSMUSG00 | Gm1679 | 681 | 10.4234426 | 9.00084073 | 17.0479402 | 6.95792748 | 9.11750096 | 9.82322912 | 3.82114147 | 10.4314051 | 6.22938491 | 2.14636094 | 2.59570259 | 7.62240326 | 0.72440407 | 1.43219653 | 1.08902791 | 0.05622703 | 0.16883831 | 1.29243129 |
| ENSMUSG00 | Gm6712 | 468 | 4.63197431 | 6.47750721 | 4.03112753 | 3.01793242 | 5.5536835 | 2.85881155 | 5.05477339 | 7.99755435 | 4.75229028 | 7.45545308 | 6.94166063 | 4.17424827 | 5.27050398 | 4.56501714 | 5.28225076 | 5.23632238 | 6.38771615 | 3.05606148 |
| ENSMUSG00 | 4930522L14I | 2405 | 3.57008131 | 4.07234549 | 4.44513522 | 4.50874368 | 3.44228311 | 5.33644823 | 4.99684842 | 6.65389464 | 3.75045611 | 5.7835582 | 6.09850821 | 4.89692508 | 7.26133003 | 3.76573682 | 6.5442804 | 6.8142952 | 6.62941352 | 3.95699603 |
| ENSMUSG00 | 5430403G16I | 3337 | 3.32449833 | 2.63548219 | 2.13092867 | 2.45759304 | 2.85589478 | 2.25712182 | 2.90653257 | 2.66671554 | 2.28333995 | 2.10532058 | 3.3644337 | 3.29508107 | 3.29667892 | 3.3681358 | 3.27192458 | 2.86864005 | 3.14695995 | 2.52214364 |
| ENSMUSG00 | Gm10418 | 574 | 0 | 0.1741092 | 0 | 0 | 0.08385353 | 0 | 0 | 0.06653729 | 0 | 0 | 0 | 0 | 0 | 0 | 0.14355943 | 0.2668335 | 0 | 0 |
| ENSMUSG00 | Gm10419 | 1666 | 3.21467718 | 1.53967158 | 1.65503675 | 0.71103671 | 1.097847 | 1.75486872 | 0.68157521 | 1.28377823 | 0.86526294 | 0.90565576 | 0.6308818 | 1.70864273 | 0.97716283 | 0.55755207 | 0.86557891 | 0.29878625 | 0.48310458 | 0.33018672 |
| ENSMUSG00 | Acrbp | 3129 | 1.07315721 | 1.26693334 | 1.20585982 | 0.7717275 | 1.21521955 | 1.12439888 | 1.36086325 | 1.17177074 | 2.85633555 | 1.32606644 | 1.5573812 | 2.06922878 | 2.01805222 | 2.34521119 | 1.80396713 | 0.83213848 | 1.16362998 | 1.59981742 |
| ENSMUSG00 | Grcc10 | 1110 | 8.3478634 | 7.23279956 | 6.4933838 | 5.91062197 | 7.97862518 | 6.07132592 | 7.54445421 | 6.39980797 | 9.90702676 | 6.79649777 | 8.09161459 | 8.04550091 | 9.33306543 | 7.53147363 | 7.90624451 | 6.89920854 | 7.31997743 | 9.71331975 |
| ENSMUSG00 | Zfp951 | 1775 | 0.7662905 | 0.91962356 | 0.65406464 | 0.79571401 | 0.97619676 | 0.69792583 | 1.2261314 | 0.86067386 | 1.06736886 | 0.79691329 | 1.37268895 | 0.84902698 | 0.30571892 | 0.57564503 | 1.62485009 | 0.21572173 | 0.38866104 | 0.49585674 |
| ENSMUSG00 | Vmn2r27 | 2568 | 0 | 0 | 0 | 0 | 0 | 0 | 0 | 0 | 0 | 0 | 0 | 0 | 0 | 0 | 0 | 0 | 0 | 0 |
| ENSMUSG00 | Vmn2r25 | 2562 | 0.08295311 | 0 | 0 | 0 | 0.05636057 | 0 | 0.46167719 | 0.29814522 | 0.14468331 | 0.11042319 | 0.31700792 | 0.19607376 | 0.1540419 | 0.03625612 | 0.11257256 | 0.1942927 | 0.43382616 | 0.45089432 |
| ENSMUSG00 | Vmn2r24 | 2559 | 0 | 0 | 0 | 0 | 0 | 0 | 0 | 0 | 0 | 0 | 0.01866939 | 0 | 0.03855562 | 0 | 0.08050324 | 0 | 0 | 0 |
| ENSMUSG00 | Gm10420 | 1830 | 0 | 0.01820377 | 0.02643362 | 0.04979349 | 0 | 0 | 0 | 0 | 0.04501259 | 0 | 0 | 0.03050036 | 0 | 0 | 0 | 0.02092383 | 0 | 0 |
| ENSMUSG00 | Abcb5 | 4348 | 0.47901417 | 0.36009798 | 0.47839508 | 0.61823864 | 0.17711839 | 0.17095009 | 16.2134065 | 12.5170592 | 11.5754191 | 5.56308895 | 1.71409695 | 9.47377162 | 14.3865849 | 22.356825 | 11.4090785 | 2.91494734 | 5.56207299 | 9.57725248 |
| ENSMUSG00 | Cecr6 | 787 | 0 | 0 | 0 | 0 | 0 | 0 | 0 | 0 | 0 | 0 | 0 | 0 | 0 | 0 | 0 | 0 | 0 | 0 |
| ENSMUSG00 | 9330160F10I | 2765 | 1.3527868 | 1.45781562 | 1.29462613 | 0.82388869 | 0.83556323 | 0.7885426 | 1.59134869 | 1.51940841 | 2.30882819 | 1.34716295 | 0.69113864 | 1.81678468 | 1.30243398 | 1.64611955 | 2.19046247 | 0.99707911 | 1.80195971 | 1.77063671 |
| ENSMUSG00 | Ahnak2 | 16977 | 0.01502215 | 0.00981118 | 0.04843907 | 0.00536738 | 0.04536201 | 0.00583763 | 0 | 0.01349793 | 0.03396426 | 0.01110931 | 0.02532689 | 0.00986317 | 0 | 0 | 0 | 0 | 0.0158028 | 0 |
| ENSMUSG00 | E330014E10I | 2119 | 0 | 0 | 0 | 0 | 0 | 0 | 0 | 0 | 0 | 0 | 0 | 0 | 0 | 0 | 0 | 0 | 0 | 0 |
| ENSMUSG00 | Gm7982 | 1586 | 0 | 0 | 0 | 0 | 0 | 0 | 0 | 0 | 0 | 0 | 0 | 0 | 0 | 0 | 0 | 0 | 0 | 0 |
| ENSMUSG00 | Gm7978 | 1592 | 0 | 0 | 0 | 0 | 0 | 0 | 0 | 0 | 0 | 0 | 0 | 0 | 0 | 0 | 0 | 0 | 0 | 0 |
| ENSMUSG00 | Gm6468 | 1215 | 0 | 0 | 0 | 0 | 0 | 0 | 0 | 0 | 0 | 0 | 0 | 0 | 0 | 0 | 0 | 0 | 0 | 0 |
| ENSMUSG00 | Gm10424 | 1844 | 0 | 0 | 0 | 0 | 0 | 0 | 0 | 0 | 0 | 0 | 0 | 0 | 0 | 0 | 0 | 0 | 0 | 0 |
| ENSMUSG00 | Gm16513 | 1689 | 0 | 0 | 0 | 0 | 0 | 0 | 0 | 0 | 0 | 0 | 0 | 0 | 0 | 0 | 0 | 0 | 0 | 0 |
| ENSMUSG00 | Gm6367 | 2511 | 0 | 0 | 0 | 0 | 0 | 0 | 0 | 0 | 0 | 0 | 0 | 0 | 0 | 0 | 0 | 0 | 0 | 0 |
| ENSMUSG00 | Gm6351 | 1592 | 0 | 0 | 0 | 0 | 0 | 0 | 0 | 0 | 0 | 0 | 0 | 0 | 0 | 0 | 0 | 0 | 0 | 0 |
| ENSMUSG00 | BC061212 | 1893 | 0 | 0 | 0 | 0 | 0 | 0 | 0 | 0 | 0 | 0 | 0 | 0 | 0 | 0 | 0 | 0 | 0 | 0 |
| ENSMUSG00 | Gm16429 | 1844 | 0 | 0 | 0 | 0 | 0 | 0 | 0 | 0 | 0 | 0 | 0 | 0 | 0 | 0 | 0 | 0 | 0 | 0 |
| ENSMUSG00 | Gm7792 | 2063 | 0 | 0 | 0 | 0 | 0 | 0 | 0 | 0 | 0 | 0 | 0 | 0 | 0 | 0 | 0 | 0 | 0 | 0 |
| ENSMUSG00 | AW555464 | 6621 | 20.6009832 | 26.4903169 | 21.004969 | 24.3253726 | 23.8515091 | 25.4985898 | 18.3076701 | 20.0047201 | 17.6104857 | 21.805687 | 13.9551079 | 17.4924483 | 14.1044131 | 13.4962127 | 15.6380471 | 16.2045434 | 17.5162928 | 16.0931339 |
| ENSMUSG00 | Gm16427 | 1706 | 0 | 0 | 0 | 0 | 0 | 0 | 0 | 0 | 0 | 0 | 0 | 0 | 0 | 0 | 0 | 0 | 0 | 0 |
| ENSMUSG00 | 4932430I15R | 3304 | 0 | 0.05041298 | 0 | 0 | 0 | 0.0449934 | 0 | 0.01155944 | 0 | 0 | 0 | 0 | 0.01493097 | 0.05622771 | 0.03741061 | 0.01158917 | 0 | 0 |
| ENSMUSG00 | Gm7292 | 994 | 0 | 0.06702796 | 0.09733105 | 0.04583606 | 0.04842246 | 0.09970369 | 0 | 0.03842294 | 0.24861076 | 0.04743532 | 0.14419002 | 0.16845773 | 0 | 0.09344887 | 0.12435077 | 0.03852174 | 0.50124936 | 0.11068231 |
| ENSMUSG00 | Gm10425 | 3362 | 1.03671161 | 0.72333173 | 0.7338043 | 1.05703791 | 0.74445569 | 0.75169227 | 1.57615111 | 1.32912287 | 1.13930576 | 0.58903318 | 1.43523819 | 1.02931918 | 1.26191103 | 2.05834891 | 1.37256819 | 1.52615746 | 0.99178698 | 0.60539469 |
| ENSMUSG00 | Gm12298 | 727 | 0 | 0 | 0 | 0 | 0 | 0 | 0 | 0 | 0 | 0 | 0 | 0 | 0.06785683 | 0 | 0 | 0 | 0 | 0 |
| ENSMUSG00 | Mir546 | 120 | 0 | 0 | 0 | 0 | 0 | 0 | 0 | 0 | 0 | 0 | 0 | 0 | 0 | 0 | 0 | 0 | 0 | 0 |
| ENSMUSG00 | Gm10428 | 2510 | 0.03386867 | 0.22562518 | 0.19272323 | 0.1815181 | 0.03835213 | 0.0789685 | 0.01884967 | 0.0152161 | 0.18049867 | 0.05635542 | 0.26647387 | 0.15566121 | 0 | 0.03700724 | 0.16414963 | 0.21357311 | 0.0763472 | 0.02191598 |
| ENSMUSG00 | G530011O06 | 2378 | 1.10820896 | 0.42026359 | 0.54923689 | 0.76637585 | 0.38456962 | 0.41675974 | 0.21885596 | 0.44970028 | 0.20783776 | 0.35690187 | 0.08036158 | 0.72762218 | 0.24894155 | 0.07812294 | 0.10395683 | 0.14491819 | 0.29010654 | 0.2544576 |
| ENSMUSG00 | Tmprss11a | 2648 | 0 | 0 | 0 | 0 | 0 | 0 | 0 | 0 | 0 | 0 | 0 | 0 | 0 | 0 | 0 | 0 | 0 | 0 |
| ENSMUSG00 | A530017D24 | 4334 | 1.47110669 | 1.12990204 | 1.08265631 | 1.45072084 | 0.95508663 | 0.83464457 | 1.32091236 | 1.1896572 | 1.06434928 | 0.62011769 | 1.15744593 | 0.84998471 | 0.910603 | 0.80371633 | 1.09325771 | 1.26339568 | 1.03465084 | 1.32001548 |
| ENSMUSG00 | Serpina1e | 1809 | 0 | 0 | 0 | 0 | 0 | 0 | 0 | 0 | 0 | 0 | 0 | 0 | 0 | 0 | 0 | 0 | 0 | 0 |
| ENSMUSG00 | 2310040G07 | 937 | 1.63306973 | 2.27537378 | 1.13577126 | 1.06973637 | 1.79788402 | 1.79807144 | 2.92863968 | 1.30432964 | 1.23076034 | 1.55994856 | 1.47862732 | 2.38273906 | 2.21124923 | 0.79306873 | 1.75887112 | 1.3076835 | 1.47251514 | 2.75926149 |
| ENSMUSG00 | Gm10430 | 1843 | 0 | 0 | 0.02624717 | 0 | 0 | 0 | 0 | 0 | 0 | 0 | 0 | 0 | 0 | 0.52920555 | 0.0894228 | 0 | 0 | 0 |
| ENSMUSG00 | Rybp | 4453 | 4.3526522 | 5.16937119 | 5.01876735 | 5.11576963 | 5.09098047 | 5.33028506 | 3.66559044 | 5.02599322 | 3.61642211 | 4.74365934 | 4.95666535 | 5.27698045 | 3.92173787 | 4.19279657 | 3.91382397 | 4.5143878 | 4.68212565 | 4.76836182 |
| ENSMUSG00 | Gm6116 | 1428 | 0 | 0 | 0 | 0 | 0 | 0 | 0 | 0 | 0 | 0 | 0 | 0 | 0 | 0 | 0 | 0 | 0 | 0 |
| ENSMUSG00 | Gpr27 | 1139 | 0.03731798 | 0.05849499 | 0.29729123 | 0.60001375 | 0 | 0.43505473 | 0.41538787 | 0.13412608 | 0 | 0.37256921 | 0.12583396 | 0 | 0.12993481 | 0.53009055 | 0.21704068 | 0.33617741 | 0.20189465 | 0.2897758 |
| ENSMUSG00 | 1700123L14I | 1702 | 0 | 0 | 0 | 0 | 0 | 0 | 0 | 0 | 0 | 0 | 0 | 0 | 0 | 0 | 0 | 0 | 0 | 0 |
| ENSMUSG00 | Gm10431 | 389 | 0 | 0 | 0 | 0 | 0 | 0 | 0 | 0 | 0 | 0 | 0 | 0 | 0 | 0 | 0 | 0 | 0 | 0 |
| ENSMUSG00 | Gm10433 | 2012 | 0.44364249 | 0 | 0.52893522 | 0 | 0.07176728 | 0.29554314 | 0 | 0.32269923 | 0 | 0.02343474 | 0.18996007 | 0 | 0 | 0.04616708 | 0.22525702 | 0.01903112 | 0 | 0.35542663 |
| ENSMUSG00 | Gm5647 | 920 | 0 | 0 | 0 | 0 | 0 | 0 | 0 | 0 | 0 | 0 | 0 | 0 | 0 | 0 | 0 | 0 | 0 | 0 |
| ENSMUSG00 | Nbl1 | 5924 | 3.63776574 | 3.55959859 | 4.31964847 | 3.4301529 | 3.42058404 | 3.05313095 | 3.70578714 | 4.15835576 | 2.39860701 | 2.94492914 | 3.68554288 | 5.01248016 | 2.39831063 | 3.22223496 | 2.90024457 | 3.77476616 | 3.00839434 | 2.90645685 |
| ENSMUSG00 | 4933439C10 | 4836 | 8.88600764 | 8.63131854 | 9.18256842 | 9.0255336 | 9.1864693 | 8.38174857 | 12.7771627 | 11.1986821 | 13.4903724 | 9.34044122 | 9.82962855 | 12.2919108 | 11.3740876 | 9.97837196 | 9.43137353 | 8.55125227 | 9.60534984 | 8.48568933 |
| ENSMUSG00 | Ppp1r2-ps2 | 873 | 0 | 0 | 0 | 0.05218905 | 0 | 0 | 0 | 0 | 0 | 0 | 0.16417512 | 0 | 0 | 0.05320056 | 0 | 0 | 0 | 0 |
| ENSMUSG00 | Gm10434 | 469 | 0.18125875 | 1.06544438 | 0 | 0.5828705 | 0 | 0 | 0.40351965 | 0.24430108 | 0.08781773 | 0.10053455 | 0.10186558 | 0 | 0 | 0.39611162 | 0.2635494 | 0 | 0.08171918 | 0.58645103 |
| ENSMUSG00 | Mir540 | 83 | 0 | 0 | 0 | 0 | 0 | 0 | 0 | 0 | 0 | 0 | 0 | 0 | 0 | 0 | 0 | 0 | 0 | 0 |
| ENSMUSG00 | Gm10435 | 4148 | 0.05123575 | 0 | 0 | 0 | 0 | 0 | 0 | 0 | 0 | 0 | 0 | 0 | 0 | 0 | 0 | 0 | 0 | 0 |
| ENSMUSG00 | Gm5662 | 2838 | 0 | 0 | 0 | 0 | 0 | 0 | 0 | 0 | 0 | 0 | 0 | 0 | 0 | 0 | 0 | 0 | 0 | 0 |
| ENSMUSG00 | Gm8332 | 434 | 0 | 0 | 0 | 0 | 0 | 0 | 0 | 0 | 0 | 0 | 0 | 0 | 0 | 0 | 0 | 0 | 0 | 0 |
| ENSMUSG00 | Gm10436 | 2731 | 0 | 0 | 0 | 0 | 0 | 0 | 0 | 0 | 0 | 0 | 0 | 0 | 0 | 0 | 0 | 0 | 0 | 0 |
| ENSMUSG00 | Gm16368 | 434 | 0 | 0 | 0 | 0 | 0 | 0 | 0 | 0 | 0 | 0 | 0 | 0 | 0 | 0 | 0 | 0 | 0 | 0 |
| ENSMUSG00 | Gm16381 | 493 | 0 | 0 | 0 | 0 | 0 | 0 | 0 | 0 | 0 | 0 | 0 | 0 | 0 | 0 | 0 | 0 | 0 | 0 |
| ENSMUSG00 | Gm12258 | 2523 | 4.9698878 | 4.58167828 | 4.79325509 | 2.49204285 | 2.91882058 | 3.14246428 | 2.55034655 | 3.58763352 | 3.95051003 | 4.14881337 | 2.93504501 | 3.7608651 | 2.11171108 | 3.57120609 | 3.72332729 | 2.73179125 | 4.11669697 | 2.63816961 |
| ENSMUSG00 | Gm1965 | 2402 | 0 | 0 | 0 | 0 | 0 | 0 | 0 | 0 | 0 | 0 | 0 | 0 | 0 | 0 | 0 | 0 | 0 | 0 |
| ENSMUSG00 | 1700022K14I | 930 | 0.31993143 | 0.07164063 | 0.52014549 | 0.19596148 | 0.10350951 | 0.37297756 | 0.61048618 | 0.41067099 | 0.44286576 | 0.15209904 | 0.667822 | 0.48013473 | 0.10609014 | 0.29963927 | 0.39872473 | 0 | 0.45332179 | 0.05914958 |
| ENSMUSG00 | 4933437F05I | 1095 | 0 | 0 | 0 | 0.04160826 | 0 | 0 | 0 | 0 | 0 | 0 | 0 | 0 | 0.45051979 | 0.16965877 | 0 | 0.45459169 | 0.10500355 | 0.05023663 |
| ENSMUSG00 | Gm15091 | 2019 | 0 | 0 | 0 | 0 | 0 | 0 | 0 | 0 | 0 | 0 | 0 | 0 | 0 | 0 | 0 | 0 | 0 | 0 |
| ENSMUSG00 | Gm15097 | 2107 | 0.04034663 | 0 | 0.04591697 | 0 | 0 | 0.07055444 | 0.22454997 | 0.03625287 | 0.01954747 | 0.08951249 | 0.0453488 | 0.05298117 | 0.14048007 | 0.19838481 | 0.27376449 | 0.32711482 | 0.10913991 | 0 |
| ENSMUSG00 | Gm10439 | 1970 | 0 | 0 | 0 | 0 | 0 | 0 | 0 | 0 | 0 | 0 | 0 | 0 | 0 | 0 | 0 | 0 | 0 | 0 |
| ENSMUSG00 | Gm15085 | 2036 | 0 | 0 | 0 | 0 | 0 | 0 | 0 | 0 | 0 | 0 | 0 | 0 | 0 | 0 | 0 | 0 | 0 | 0 |
| ENSMUSG00 | Ott | 2288 | 0 | 0 | 0 | 0 | 0 | 0 | 0 | 0 | 0 | 0 | 0 | 0 | 0 | 0 | 0 | 0 | 0 | 0 |
| ENSMUSG00 | Gm15127 | 2019 | 0.06315777 | 0.0329994 | 0.02395915 | 0 | 0 | 0.02454321 | 0.1171686 | 0 | 0.14279624 | 0.07006048 | 0 | 0 | 0.1954707 | 0.29904563 | 0.14284839 | 0.13275594 | 0.17084531 | 0.1362286 |
| ENSMUSG00 | Gm15109 | 2020 | 0 | 0 | 0 | 0 | 0 | 0 | 0 | 0 | 0 | 0 | 0 | 0 | 0 | 0 | 0 | 0 | 0.07589366 | 0 |
| ENSMUSG00 | Gm15107 | 2021 | 0 | 0 | 0 | 0.06763144 | 0.02381589 | 0.04903784 | 0.02341053 | 0.0755911 | 0.04075855 | 0.1399823 | 0 | 0.02761784 | 0 | 0.04596149 | 0.3465742 | 0.2273564 | 0.13274818 | 0.05443751 |
| ENSMUSG00 | Gm15080 | 2022 | 0 | 0 | 0 | 0 | 0 | 0 | 0 | 0 | 0 | 0 | 0 | 0 | 0 | 0.09187752 | 0.02037663 | 0 | 0 | 0 |
| ENSMUSG00 | Gm15128 | 2105 | 0 | 0 | 0 | 0 | 0 | 0 | 0 | 0 | 0 | 0 | 0 | 0 | 0 | 0 | 0.01957319 | 0.01819031 | 0 | 0 |
| ENSMUSG00 | Gm15070 | 3581 | 0.16617494 | 0.13954019 | 0.4863019 | 0.30535187 | 0.36290476 | 0 | 0.01321214 | 0.31995869 | 0.37954622 | 0.21067055 | 0.0400237 | 0.31173227 | 0.09643212 | 0 | 0.51775203 | 0.84472437 | 0.55653936 | 0.93704426 |
| ENSMUSG00 | Gm10443 | 515 | 1.32054916 | 0.90559324 | 1.22107941 | 0.53080828 | 1.02806051 | 0.1924378 | 0.73495424 | 0.74160005 | 0.71976435 | 0.73243811 | 0.83490219 | 0.32513978 | 1.62843222 | 1.89383657 | 1.04003927 | 0.59480555 | 0.44651996 | 0.9613242 |
| ENSMUSG00 | Sod3 | 3568 | 0.07147731 | 0 | 0 | 0 | 0 | 0.0277762 | 0 | 0 | 0 | 0.10571907 | 0 | 0 | 0.04147863 | 0 | 0.01154752 | 0 | 0 | 0 |
| ENSMUSG00 | Nup62cl | 2316 | 0 | 0 | 0.12532003 | 0 | 0 | 0 | 0 | 0 | 0 | 0 | 0 | 0.12050014 | 0 | 0 | 0 | 0.01653308 | 0 | 0 |
| ENSMUSG00 | Ripply1 | 807 | 0 | 0 | 0 | 0 | 0 | 0 | 0 | 0 | 0 | 0 | 0 | 0 | 0 | 0 | 0 | 0 | 0 | 0 |
| ENSMUSG00 | Ptgr2 | 5971 | 25.221209 | 21.9928704 | 23.072821 | 26.1722295 | 25.2065861 | 26.2660194 | 26.8139517 | 25.5149042 | 24.8181349 | 28.8542404 | 25.3236884 | 24.0986063 | 22.4063238 | 20.3401924 | 21.597868 | 22.1047938 | 25.9124536 | 19.8165447 |
| ENSMUSG00 | Acot1 | 2025 | 5.54141554 | 6.21840692 | 4.84929712 | 4.04987061 | 7.41588157 | 5.75056414 | 2.03269287 | 2.75362506 | 1.66779965 | 2.84068434 | 2.14692406 | 3.25246824 | 3.8978305 | 4.54119966 | 3.96755723 | 3.57379002 | 2.17655513 | 4.75387342 |
| ENSMUSG00 | Gm10444 | 168 | 0 | 0 | 0 | 0 | 0 | 0 | 0 | 0 | 0 | 0 | 0 | 0 | 0 | 0 | 0 | 0 | 0 | 0 |
| ENSMUSG00 | Gm5878 | 4671 | 0 | 0.03565927 | 0 | 0 | 0.04121766 | 0 | 0 | 0 | 0 | 0 | 0.14319191 | 0 | 0 | 0 | 0 | 0.04098759 | 0 | 0.01177673 |
| ENSMUSG00 | Tmsb15l | 812 | 0.15703883 | 0.28718012 | 0.41701319 | 0.11221932 | 0.29637884 | 0.48820427 | 0.46613477 | 0.18813991 | 0.81155696 | 0.11613474 | 0.41185309 | 0.54990801 | 0.66828952 | 0.51477437 | 0.50740832 | 0.70733881 | 0.47199873 | 0.20323562 |
| ENSMUSG00 | BC065397 | 1285 | 2.24930114 | 2.74798972 | 2.10810716 | 2.48192459 | 2.4721455 | 1.5039351 | 3.16645166 | 2.22912855 | 3.65390101 | 3.33907704 | 3.30892709 | 1.99807041 | 2.80251359 | 3.54203935 | 2.34063314 | 1.63889759 | 2.20711749 | 1.49830252 |
| ENSMUSG00 | Gm16401 | 2071 | 0.20523986 | 0.35387913 | 0.25693329 | 0.06598862 | 0.2324091 | 0.1196348 | 0.09138132 | 0.18441527 | 0 | 0.22767119 | 0.06920564 | 0 | 0.28584404 | 0.47094439 | 0.15915618 | 0 | 0.4441483 | 0 |
| ENSMUSG00 | Gm10447 | 2306 | 0.25805397 | 0.89566324 | 0.67127189 | 0.55321303 | 0.91838883 | 0.62316968 | 0.77965386 | 1.10966651 | 1.89322232 | 0.8996665 | 0.55937722 | 1.01659056 | 1.02685691 | 1.04730813 | 1.03629239 | 0.64758616 | 0.89749351 | 0.59636933 |
| ENSMUSG00 | Bhlhb9 | 3195 | 15.2725953 | 14.0028848 | 16.7906871 | 14.6308706 | 18.062653 | 16.5796157 | 12.1724639 | 12.5514937 | 14.914804 | 13.9607403 | 13.6222182 | 13.9233437 | 7.3959275 | 13.0246956 | 11.6060721 | 14.5492324 | 11.8877497 | 12.4652874 |
| ENSMUSG00 | Gprasp2 | 4480 | 1.87857698 | 2.80333954 | 1.41449386 | 2.10516433 | 2.22395272 | 1.59276644 | 1.79534719 | 1.83289431 | 2.09609946 | 1.18929229 | 1.43964719 | 1.7317806 | 0.42945196 | 0.37321142 | 0.50582267 | 0.24786331 | 0.95815742 | 0.36836455 |
| ENSMUSG00 | Gm17728 | 398 | 0.10679692 | 0 | 0 | 0 | 0 | 0 | 0 | 0 | 0 | 0 | 0.48015034 | 0 | 0 | 0 | 0 | 0 | 0 | 0 |
| ENSMUSG00 | Armcx5 | 3603 | 5.54466632 | 6.49985141 | 6.35045236 | 6.15826494 | 8.12217863 | 6.53276393 | 4.2152012 | 6.06329564 | 5.7270176 | 6.29461234 | 6.53706759 | 7.23450303 | 4.94277606 | 6.93503164 | 5.64906151 | 8.76762451 | 8.08437015 | 6.79407507 |
| ENSMUSG00 | Adam4 | 3919 | 0.16268886 | 0.08500356 | 0.13577669 | 0.01162568 | 0.13509854 | 0 | 0.08450848 | 0.26312704 | 0.07356612 | 0.03609393 | 0.0487624 | 0.01424232 | 0.05035154 | 0.15406306 | 0 | 0.08793454 | 0.06845728 | 0.30880335 |
| ENSMUSG00 | Gm4787 | 2494 | 0.05112892 | 0.21371544 | 0.03879192 | 0 | 0 | 0.07947511 | 0.11382361 | 0.19907828 | 0.13211392 | 0.05671696 | 0.1340917 | 0.11189988 | 0 | 0 | 0.13216217 | 0 | 0.1229392 | 0 |
| ENSMUSG00 | Gm5830 | 316 | 0.5380402 | 0 | 0.45924238 | 0.14418052 | 0 | 0.3136249 | 0.59889467 | 0.12086203 | 0.39101123 | 0 | 0.30237315 | 0.52989553 | 0.46834099 | 0.44092488 | 0.13038467 | 0 | 0 | 1.74079452 |
| ENSMUSG00 | Oip5 | 2166 | 0.37285242 | 1.16887348 | 0.6699935 | 0.42069293 | 0.57776086 | 0.38891804 | 0.96110705 | 0.81110365 | 0.62749539 | 1.11019661 | 1.23517898 | 0.48961108 | 2.64196787 | 0.7075969 | 2.45383228 | 1.290496 | 1.928701 | 1.44760807 |
| ENSMUSG00 | 9230009I02R | 1145 | 0.03712242 | 0.17456538 | 0 | 0.07958261 | 0 | 0.12983249 | 0 | 0 | 0 | 0 | 0 | 0 | 0 | 0 | 0 | 0 | 0 | 0 |
| ENSMUSG00 | 4933414I15R | 3399 | 0 | 0 | 0 | 0 | 0 | 0 | 0 | 0 | 0 | 0 | 0 | 0 | 0 | 0 | 0 | 0 | 0 | 0 |
| ENSMUSG00 | Cpxcr1 | 1593 | 0 | 0 | 0 | 0 | 0 | 0 | 0 | 0 | 0 | 0 | 0 | 0 | 0 | 0 | 0 | 0 | 0 | 0 |
| ENSMUSG00 | Gm15401 | 1591 | 0.10686405 | 0.04187667 | 0.03040448 | 0.11454694 | 0.18151574 | 0.28031088 | 0.71370477 | 0 | 0.23298469 | 0.05927178 | 0.39036735 | 0.17541063 | 0 | 0.02919176 | 0.59562274 | 0.12033503 | 0.16862607 | 0.13830071 |
| ENSMUSG00 | Gm10451 | 1594 | 4.23985131 | 4.03349347 | 6.25153529 | 3.77293466 | 4.28778743 | 4.16566267 | 2.84944616 | 2.587691 | 2.68720053 | 2.21850862 | 2.30782422 | 2.10096597 | 2.22829237 | 2.94281859 | 2.40385489 | 4.29988628 | 2.38036599 | 2.2431568 |
| ENSMUSG00 | Cylc1 | 1918 | 0 | 0 | 0 | 0 | 0 | 0 | 0 | 0 | 0 | 0 | 0 | 0 | 0 | 0 | 0 | 0 | 0 | 0 |
| ENSMUSG00 | Vamp5 | 2775 | 0.30634361 | 0.5402091 | 0.38350186 | 0.18060234 | 0.60706931 | 0.58927575 | 0.971828 | 1.07351618 | 0.43041764 | 0.88354471 | 0.41318883 | 0.54307131 | 1.10219058 | 1.12135274 | 0.28210074 | 0.99348603 | 0.48339474 | 0.73345476 |
| ENSMUSG00 | Gm732 | 2077 | 0 | 0 | 0 | 0 | 0 | 0 | 0 | 0 | 0 | 0 | 0 | 0 | 0 | 0 | 0 | 0 | 0 | 0 |
| ENSMUSG00 | Fam46d | 3153 | 0.01348087 | 0.03169638 | 0.03068413 | 0.08670037 | 0.07632719 | 0.12572847 | 0.03001121 | 0.79946037 | 0 | 0.20935929 | 0.03030445 | 0.47796476 | 0.07823013 | 0.14730126 | 0.58803362 | 0.24288365 | 0.86304062 | 0.08723296 |
| ENSMUSG00 | Gpr174 | 5528 | 0.01537814 | 0 | 0 | 0 | 0 | 0 | 0 | 0 | 0 | 0.09382376 | 0 | 0 | 0 | 0.08401608 | 0 | 0.00692667 | 0 | 0 |
| ENSMUSG00 | Gm5127 | 2143 | 0 | 0 | 0 | 0 | 0 | 0 | 0 | 0 | 0 | 0 | 0 | 0 | 0 | 0 | 0 | 0 | 0 | 0 |
| ENSMUSG00 | Fnd3c2 | 3642 | 0 | 0 | 0 | 0 | 0 | 0 | 0 | 0 | 0 | 0 | 0 | 0 | 0 | 0 | 0 | 0 | 0 | 0 |
| ENSMUSG00 | Uprt | 2297 | 1.16579281 | 0.50759743 | 0.73708035 | 0.57521562 | 0.25145106 | 0.4314561 | 0.5561351 | 0.96437063 | 0.64550046 | 0.36948744 | 0.43677585 | 0.21869437 | 0.42953345 | 0.02021945 | 0.62779907 | 0.68346317 | 0.73415632 | 1.14951551 |
| ENSMUSG00 | Gm1070 | 2710 | 0 | 0.12292581 | 0 | 0.06724877 | 0 | 0 | 0.13966842 | 0.15502451 | 0 | 0 | 0.08814568 | 0 | 0.49149881 | 0.41131295 | 0 | 0.0565175 | 0.5657018 | 0.40597127 |
| ENSMUSG00 | Dmrtc1b | 1376 | 0 | 0 | 0 | 0 | 0 | 0 | 0 | 0 | 0 | 0 | 0 | 0 | 0 | 0 | 0 | 0 | 0 | 0 |
| ENSMUSG00 | Igkv4-71 | 287 | 0 | 0 | 0 | 0 | 0 | 0 | 0 | 0 | 0 | 0 | 0 | 0 | 0 | 0 | 0 | 0 | 0 | 0 |
| ENSMUSG00 | Atoh1 | 2087 | 0.02036664 | 0 | 0 | 0 | 0 | 0 | 0 | 0 | 0 | 0 | 0 | 0 | 0 | 0 | 0 | 0 | 0 | 0 |
| ENSMUSG00 | Gm5570 | 2521 | 0 | 0 | 0 | 0 | 0 | 0 | 0 | 0 | 0 | 0 | 0 | 0 | 0 | 0 | 0 | 0 | 0 | 0 |
| ENSMUSG00 | Gm14812 | 1787 | 0 | 0.09320899 | 0.08120906 | 0 | 0 | 0 | 0 | 0 | 0.04609571 | 0 | 0 | 0 | 0.11042399 | 0 | 0 | 0 | 0 | 0 |
| ENSMUSG00 | D130052B06 | 2291 | 0 | 0 | 0 | 0 | 0 | 0 | 0 | 0 | 0.25168539 | 0 | 0.08341328 | 0 | 0 | 0.22299649 | 0 | 0 | 0.03345814 | 0 |
| ENSMUSG00 | Gm10462 | 319 | 0 | 0 | 0 | 0 | 0 | 0 | 0 | 0 | 0 | 0 | 0 | 0 | 0 | 0 | 0 | 0 | 0 | 0 |
| ENSMUSG00 | Gm10463 | 2885 | 0 | 0 | 0.05030176 | 0 | 0.23356913 | 0 | 0.03279908 | 0 | 0.1284848 | 0.0163434 | 0 | 0.0386937 | 0.0341989 | 0.28977247 | 0.0285626 | 0 | 0.07970807 | 0 |
| ENSMUSG00 | Zxda | 4452 | 0.0286423 | 0 | 0 | 0 | 0 | 0 | 0 | 0 | 0 | 0 | 0 | 0 | 0 | 0 | 0 | 0 | 0 | 0 |
| ENSMUSG00 | Zxdb | 5622 | 0.49143302 | 0.41478168 | 0.64532458 | 0.8185104 | 0.30820869 | 0.3261208 | 0.47127535 | 0.48233023 | 0.78387712 | 0.44450147 | 0.4673822 | 0.49640397 | 0.50016352 | 0.3800121 | 0.90875008 | 0.41546194 | 0.38176319 | 0.27396923 |
| ENSMUSG00 | Hbq1b | 772 | 0 | 0 | 0 | 0 | 0 | 0 | 0 | 0 | 0 | 0 | 0 | 0 | 0 | 0 | 0 | 0 | 0 | 0 |
| ENSMUSG00 | 9130019P16I | 5283 | 0.0160913 | 0 | 0.0183129 | 0.00862409 | 0 | 0 | 0.06268952 | 0.12289813 | 0 | 0.07139989 | 0 | 0.01056515 | 0.00933786 | 0.0263737 | 0.02339668 | 0.02899156 | 0 | 0.0832998 |
| ENSMUSG00 | Gm8914 | 1034 | 0 | 0 | 0 | 0 | 0 | 0 | 0 | 0 | 0 | 0 | 0 | 0 | 0 | 0 | 0 | 0 | 0 | 0 |
| ENSMUSG00 | Mageb2 | 1142 | 0 | 0 | 0 | 0 | 0 | 0 | 0 | 0 | 0 | 0 | 0 | 0 | 0 | 0 | 0 | 0 | 0 | 0 |
| ENSMUSG00 | G930045G22 | 1742 | 0 | 0 | 0 | 0 | 0 | 0 | 0 | 0 | 0 | 0 | 0 | 0 | 0 | 0 | 0 | 0 | 0 | 0 |
| ENSMUSG00 | Gm8098 | 1497 | 0.05678714 | 0.48956825 | 0.71090025 | 0.82174228 | 0.99671986 | 0.09930408 | 0 | 0.43371466 | 0.11005081 | 0.56694232 | 0.3829656 | 0.44742014 | 0.13181541 | 0.15512387 | 0.74311423 | 0.53714279 | 1.12649102 | 0.73492461 |
| ENSMUSG00 | Gm7173 | 2816 | 0.09056501 | 0.04731945 | 0.2233153 | 0.24269022 | 0 | 0 | 0.35282893 | 0.16275171 | 0.40952501 | 0.13395086 | 0.27144863 | 0.43605986 | 0.33285029 | 0.09895757 | 0 | 0.48951061 | 0.78939106 | 0.64463797 |
| ENSMUSG00 | Srp54c | 2889 | 5.82625466 | 6.00762128 | 6.14506253 | 5.1569614 | 6.03106832 | 5.00844521 | 6.1576903 | 6.00185209 | 5.51719682 | 5.41849549 | 6.61473984 | 4.81069574 | 6.02774898 | 6.9127579 | 6.66013382 | 6.65347349 | 6.0361596 | 5.23624242 |
| ENSMUSG00 | Fam177a | 3832 | 0 | 0 | 0 | 0 | 0 | 0 | 0 | 0 | 0 | 0 | 0 | 0 | 0 | 0 | 0 | 0 | 0 | 0 |
| ENSMUSG00 | 4930428D18I | 1100 | 0 | 0 | 0 | 0 | 0 | 0 | 0 | 0 | 0 | 0 | 0 | 0 | 0 | 0 | 0 | 0 | 0 | 0 |
| ENSMUSG00 | Gm7932 | 378 | 0.89958045 | 1.762587 | 4.73497519 | 4.21861522 | 1.65533071 | 1.96637832 | 3.6298087 | 2.12180013 | 4.24940242 | 2.74422082 | 3.28610826 | 3.69151208 | 4.69827784 | 3.19456686 | 3.81495888 | 3.0389371 | 3.24455424 | 4.80238233 |
| ENSMUSG00 | 4930428E23I | 815 | 0 | 0 | 0 | 0.11180624 | 0 | 0 | 0 | 0.04686184 | 0 | 0 | 0 | 0 | 0 | 0 | 0 | 0 | 0 | 0 |
| ENSMUSG00 | Lrrc61 | 1847 | 2.34733511 | 3.51706246 | 4.24283266 | 5.20486213 | 4.09134381 | 3.2999384 | 5.17442401 | 5.16951846 | 5.61938385 | 2.62941117 | 4.08686705 | 3.20328112 | 0.88140404 | 3.394668 | 3.65839478 | 3.23407404 | 3.67285033 | 2.56133361 |
| ENSMUSG00 | 5730522E02I | 83 | 0 | 0 | 0 | 0 | 0 | 0 | 0 | 0 | 0 | 0 | 0 | 0 | 0 | 0 | 0 | 0 | 0 | 0 |
| ENSMUSG00 | Ccdc164 | 2469 | 0.08607772 | 0.26984928 | 0.23510829 | 0.11071943 | 0 | 0 | 0.01916269 | 0 | 0 | 0.01909709 | 0 | 0.04521317 | 0 | 0 | 0.15018793 | 0 | 0 | 0 |
| ENSMUSG00 | Gm10466 | 2463 | 0 | 0 | 0 | 0 | 0 | 0 | 0 | 0 | 0 | 0 | 0 | 0 | 0 | 0 | 0 | 0 | 0 | 0 |
| ENSMUSG00 | Olfr237-ps1 | 999 | 0 | 0 | 0 | 0 | 0 | 0 | 0 | 0 | 0 | 0 | 0 | 0 | 0 | 0 | 0 | 0 | 0 | 0 |
| ENSMUSG00 | Olfr441 | 999 | 0 | 0 | 0 | 0 | 0 | 0 | 0 | 0 | 0 | 0 | 0 | 0 | 0 | 0 | 0 | 0 | 0 | 0 |
| ENSMUSG00 | Olfr444 | 1049 | 0 | 0 | 0 | 0 | 0 | 0 | 0 | 0 | 0 | 0 | 0 | 0 | 0 | 0 | 0 | 0 | 0 | 0 |
| ENSMUSG00 | Olfr446 | 926 | 0 | 0 | 0 | 0 | 0 | 0 | 0 | 0 | 0 | 0 | 0 | 0 | 0 | 0 | 0 | 0 | 0 | 0 |
| ENSMUSG00 | Heatr5b | 1780 | 1.62379324 | 0.72988926 | 0.92398878 | 0.97265151 | 0.54080813 | 0.30622476 | 1.19610705 | 1.20155873 | 2.5915111 | 0.31786991 | 1.69091145 | 0.75257073 | 1.27486977 | 0 | 0.83328989 | 0.77441678 | 1.098113 | 1.54519963 |
| ENSMUSG00 | Gm10471 | 5635 | 0 | 0 | 0 | 0 | 0 | 0 | 0 | 0 | 0 | 0 | 0.01695651 | 0 | 0 | 0 | 0 | 0 | 0 | 0 |
| ENSMUSG00 | Gm7347 | 2990 | 0 | 0 | 0 | 0 | 0 | 0 | 0 | 0 | 0 | 0 | 0 | 0 | 0 | 0 | 0 | 0 | 0 | 0 |
| ENSMUSG00 | Speer4a | 2923 | 0 | 0 | 0 | 0 | 0 | 0 | 0 | 0 | 0 | 0 | 0 | 0 | 0 | 0 | 0 | 0 | 0 | 0 |
| ENSMUSG00 | Xlr3b | 3811 | 0.61343077 | 0.54195734 | 0.40618026 | 0.96836646 | 0.27785419 | 0.74114558 | 0.9931814 | 2.14462716 | 0.32421818 | 0.66810233 | 0.46383455 | 1.0545074 | 1.4756858 | 1.0846297 | 0.29190291 | 0.41194303 | 0.1609081 | 0.62067478 |
| ENSMUSG00 | Gm10472 | 533 | 0 | 0 | 0 | 0 | 0 | 0 | 0 | 0 | 0 | 0 | 0 | 0 | 0 | 0 | 0 | 0 | 0 | 0 |
| ENSMUSG00 | Gm1141 | 2294 | 0.01852885 | 0 | 0 | 0 | 0 | 0 | 0 | 0.03329765 | 0 | 0 | 0 | 0.14598691 | 0 | 0 | 0.12572402 | 0 | 0 | 0 |
| ENSMUSG00 | Vma21 | 7368 | 6.06623815 | 6.29728957 | 6.98473827 | 6.15277496 | 6.92299293 | 6.62270072 | 6.64674493 | 7.35821815 | 6.6479909 | 7.36729738 | 6.34511167 | 7.41301769 | 6.69642236 | 6.58303471 | 6.08711603 | 6.94978798 | 7.18083381 | 6.5845164 |
| ENSMUSG00 | BC023829 | 2839 | 17.1727499 | 20.6988184 | 22.2528463 | 20.4936434 | 19.3443201 | 21.2593271 | 19.6650092 | 14.7442314 | 18.2503124 | 14.9806039 | 13.3951627 | 17.3797273 | 27.5243948 | 33.2094038 | 30.0703148 | 33.0979748 | 32.1703295 | 32.8814562 |
| ENSMUSG00 | 4930567H17 | 839 | 0 | 0 | 0 | 0 | 0 | 0 | 0 | 0 | 0 | 0 | 0 | 0 | 0 | 0 | 0 | 0 | 0 | 0 |
| ENSMUSG00 | 6030469F06I | 1401 | 0 | 0 | 0 | 0 | 0 | 0 | 0 | 0 | 0 | 0 | 0 | 0 | 0 | 0 | 0 | 0 | 0 | 0 |
| ENSMUSG00 | 4930599N23 | 2423 | 1.59635618 | 1.74607411 | 2.27593168 | 1.67351752 | 2.2645643 | 2.12690231 | 2.34317849 | 2.41165397 | 1.81880198 | 1.7319078 | 1.49851294 | 3.455365 | 2.15814413 | 2.10848106 | 2.02351843 | 0.72693683 | 2.3568379 | 2.72434701 |
| ENSMUSG00 | Gm12000 | 627 | 0.33895675 | 0 | 0 | 0 | 0 | 0 | 0.98096463 | 1.76647475 | 0.26275289 | 0.37600242 | 0.07619611 | 0.26706059 | 2.20302023 | 4.29626327 | 2.36563957 | 0.79390414 | 1.46703529 | 2.54428085 |
| ENSMUSG00 | Gm10474 | 2739 | 0 | 0 | 0.03532204 | 0 | 0 | 0 | 0 | 0 | 0 | 0.01721457 | 0 | 0 | 0 | 0 | 0 | 0 | 0 | 0 |
| ENSMUSG00 | 5031425E22I | 2109 | 18.6627752 | 15.2111508 | 18.9686627 | 17.1312983 | 20.5171168 | 15.8596942 | 13.0339813 | 17.656516 | 17.5369802 | 11.0443089 | 17.3068128 | 13.7091101 | 16.4673636 | 17.1330014 | 16.2735401 | 11.6197196 | 11.8304501 | 18.2581198 |
| ENSMUSG00 | Gm10475 | 1792 | 0.04743881 | 0.07435914 | 0 | 0 | 0 | 0 | 0.05280433 | 0 | 0.04596709 | 0 | 0 | 0 | 0 | 0 | 0 | 0 | 0 | 0 |
| ENSMUSG00 | 9330158H04 | 8495 | 0 | 0 | 0 | 0 | 0 | 0 | 0 | 0 | 0 | 0 | 0 | 0 | 0 | 0 | 0 | 0 | 0 | 0 |
| ENSMUSG00 | 1810058I24R | 2847 | 54.5684645 | 45.6926069 | 45.3151407 | 43.7686885 | 49.9239798 | 43.9656496 | 53.5113545 | 43.5181431 | 43.211845 | 46.0576416 | 46.4492747 | 44.876028 | 56.6268724 | 56.9172536 | 46.237784 | 59.3390094 | 46.1342534 | 49.5603649 |
| ENSMUSG00 | 9030624G23 | 464 | 1.00766581 | 1.6512857 | 2.39782586 | 2.35660574 | 3.5269082 | 1.3883309 | 1.93737263 | 1.39929061 | 1.2426966 | 2.43882948 | 0.92666945 | 0.60146188 | 2.12637574 | 0.80076013 | 2.13111496 | 2.80577727 | 1.81719511 | 0.82987877 |
| ENSMUSG00 | 2410018L13I | 4513 | 1.31857404 | 0.94483724 | 0.68599733 | 0.99945566 | 1.04518691 | 1.03211987 | 0.70240405 | 0.42313763 | 0.5019407 | 0.44925332 | 0.39168479 | 1.01415562 | 0.87448557 | 0.8027119 | 0.66645548 | 0.94178095 | 1.1209996 | 0.24378066 |
| ENSMUSG00 | Gm20472 | 1234 | 0 | 0 | 0 | 0 | 0 | 0 | 0 | 0 | 0 | 0 | 0 | 0 | 0 | 0 | 0 | 0 | 0 | 0 |
| ENSMUSG00 | Gm10477 | 163 | 0 | 0 | 0 | 0 | 0 | 0 | 0 | 0 | 0 | 0 | 0 | 0 | 0 | 0 | 0 | 0 | 0 | 0 |
| ENSMUSG00 | [illegible] | 876 | 0.04852189 | 0.19014209 | 0.16566278 | 0 | 0.65934142 | 0 | 0.37806935 | 0 | 0 | 0 | 0.2181505 | 0 | 0.61946471 | 0.15905509 | 0.28220244 | 0.08742148 | 0.08750296 | 0.31397892 |
| ENSMUSG00 | Zfp449 | 3733 | 1.66240442 | 1.07979111 | 0.64791763 | 1.17167433 | 1.31515034 | 1.24777845 | 1.41950711 | 0.43993391 | 0.20962866 | 1.50306287 | 1.62534683 | 0.65788619 | 0.31716207 | 0.68428203 | 0.55185582 | 1.52834195 | 0.86241868 | 0.722059 |
| ENSMUSG00 | Gm773 | 943 | 0 | 0 | 0 | 0 | 0 | 0 | 0 | 0 | 0 | 0 | 0 | 0 | 0 | 0 | 0 | 0 | 0 | 0 |
| ENSMUSG00 | Gm10478 | 394 | 0.7551681 | 1.77556035 | 0.49110183 | 1.85019469 | 1.09946019 | 1.50922031 | 0.60041471 | 0.96935031 | 0.94080874 | 0.95737469 | 0.24251248 | 0 | 0.50083165 | 0.35363518 | 0.20914495 | 0.5831057 | 0.48637433 | 1.2565532 |
| ENSMUSG00 | Gm17680 | 407 | 0 | 0 | 0 | 0.1119436 | 0 | 0.24350238 | 0.23249474 | 0.37535531 | 0 | 0 | 0 | 0 | 0 | 0 | 0 | 0 | 0 | 0 |
| ENSMUSG00 | Gm10479 | 315 | 0.8096224 | 0.42302088 | 0.15356676 | 0 | 0.15279976 | 0 | 0 | 0.48498289 | 0 | 0.44905432 | 0 | 0.17719258 | 0.62643704 | 1.03209083 | 0.26159718 | 0 | 0.36501235 | 0 |
| ENSMUSG00 | Gm5784 | 407 | 0.41774129 | 0.98219836 | 0.35656165 | 0.44777439 | 0.35478076 | 0.73050713 | 1.51121579 | 0.56303296 | 0.91075833 | 0.57924697 | 0 | 0.6856961 | 0.48483457 | 0.91290589 | 1.41725254 | 1.1289614 | 0.84751025 | 1.08126008 |
| ENSMUSG00 | 5730507C01I | 1175 | 1.08523854 | 0.87889338 | 1.31740678 | 0.77550714 | 1.43371688 | 0.92779586 | 1.36904773 | 0.68258762 | 1.15672768 | 0.72230865 | 1.17912663 | 0.90255114 | 0.46183071 | 1.10675272 | 0.77143339 | 0.78210602 | 1.01116188 | 1.21722279 |
| ENSMUSG00 | Gm10480 | 656 | 0.77753371 | 1.57423738 | 1.10610207 | 1.31960342 | 1.24732119 | 1.35967867 | 2.5964275 | 1.57194339 | 1.63239239 | 1.36564537 | 2.11200274 | 1.53152733 | 1.42882077 | 1.76997284 | 1.004916 | 1.45924571 | 1.51903006 | 2.01252829 |
| ENSMUSG00 | Ccdc160 | 1534 | 31.9758625 | 28.0575354 | 25.9526828 | 26.0179106 | 25.1013944 | 26.8437559 | 29.8865619 | 24.7229828 | 28.7822327 | 29.3846625 | 26.6281548 | 21.7222624 | 25.7272059 | 27.7635125 | 27.1006323 | 26.4080369 | 24.5598109 | 18.6471548 |
| ENSMUSG00 | Speer4c | 1154 | 0 | 0 | 0 | 0 | 0 | 0 | 0 | 0 | 0 | 0 | 0 | 0 | 0 | 0 | 0 | 0 | 0 | 0 |
| ENSMUSG00 | Klf14 | 2969 | 0 | 0 | 0 | 0 | 0 | 0 | 0 | 0 | 0 | 0 | 0 | 0 | 0 | 0 | 0 | 0.02581765 | 0 | 0 |
| ENSMUSG00 | Gm10481 | 1231 | 0.17364491 | 0 | 0.27507288 | 0 | 0.03909986 | 0.12076215 | 0.19217173 | 0.34128061 | 0.06691554 | 0.22981659 | 0.19404938 | 0.13602517 | 0.12022401 | 0.15091499 | 0.13387995 | 0.06221057 | 0.18689567 | 0.17874608 |
| ENSMUSG00 | Gm10482 | 632 | 0 | 0 | 0 | 0 | 0.07615811 | 0 | 0 | 0.06043102 | 0 | 0 | 0 | 0 | 0 | 0 | 0 | 0 | 0 | 0 |
| ENSMUSG00 | Rpl15-ps2 | 686 | 0.0619609 | 0 | 0 | 0 | 0.07016315 | 0.28893722 | 0 | 0 | 0 | 0 | 0.0696428 | 0 | 0 | 0 | 0 | 0 | 0 | 0.0801882 |
| ENSMUSG00 | Gm10483 | 2745 | 0 | 0 | 0 | 0 | 0 | 0 | 0 | 0 | 0 | 0 | 0 | 0 | 0 | 0 | 0 | 0 | 0 | 0 |
| ENSMUSG00 | Gm9144 | 1263 | 0.16827069 | 0.39564007 | 0.61280799 | 0.21644202 | 0.22865522 | 0.15693661 | 0.03746055 | 0.27215489 | 0.03261007 | 0.03733231 | 0.34043913 | 0.13257877 | 0.19529658 | 0 | 0.29359779 | 0.27285468 | 0.15172722 | 0.26132592 |
| ENSMUSG00 | Gm8773 | 1113 | 0 | 0.17958434 | 0 | 0 | 0 | 0 | 0 | 0 | 0 | 0 | 0 | 0.05014884 | 0 | 0 | 0 | 0 | 0 | 0 |
| ENSMUSG00 | 2500004C02I | 2492 | 2.81434753 | 1.77793537 | 1.90232985 | 2.01112154 | 2.74266981 | 1.84927938 | 2.90483141 | 3.09585284 | 3.22286139 | 2.36510349 | 2.89487108 | 1.52305981 | 3.30595112 | 3.98837669 | 2.00055709 | 2.12042689 | 3.38354146 | 2.67098793 |
| ENSMUSG00 | 2410017P09I | 581 | 15.3633166 | 17.487836 | 17.4011495 | 12.9390229 | 18.2255133 | 17.7400492 | 19.9511297 | 15.5793449 | 12.9726891 | 17.691658 | 24.4219667 | 15.5630591 | 19.8686207 | 19.1851652 | 17.3741501 | 17.0693069 | 16.6894202 | 16.0009278 |
| ENSMUSG00 | Gm10485 | 1381 | 0.27700694 | 0.67542436 | 0 | 0.03299134 | 0.31367655 | 0 | 0.27407779 | 0.05531123 | 0 | 0 | 0.17297233 | 0.76792005 | 0 | 0.03363077 | 0.1491729 | 0.05545345 | 0.22202055 | 0.23899684 |
| ENSMUSG00 | Gm9 | 1659 | 0 | 0 | 0 | 0 | 0 | 0 | 0 | 0 | 0 | 0 | 0 | 0 | 0 | 0 | 0 | 0 | 0 | 0 |
| ENSMUSG00 | Gm14819 | 919 | 0 | 0 | 0 | 0 | 0 | 0 | 0 | 0 | 0 | 0 | 0 | 0 | 0 | 0 | 0 | 0 | 0 | 0 |
| ENSMUSG00 | Gm10486 | 915 | 0 | 0 | 0 | 0 | 0 | 0 | 0 | 0 | 0 | 0 | 0 | 0 | 0 | 0 | 0 | 0 | 0 | 0 |
| ENSMUSG00 | Gm10487 | 898 | 0 | 0 | 0 | 0 | 0 | 0 | 0 | 0 | 0 | 0 | 0 | 0 | 0 | 0 | 0 | 0 | 0 | 0 |
| ENSMUSG00 | Gm14632 | 855 | 0 | 0 | 0 | 0 | 0 | 0 | 0 | 0 | 0 | 0 | 0 | 0 | 0 | 0 | 0 | 0 | 0 | 0 |
| ENSMUSG00 | Gm10488 | 865 | 0 | 0 | 0 | 0 | 0 | 0 | 0 | 0 | 0 | 0 | 0 | 0 | 0 | 0 | 0 | 0 | 0 | 0 |
| ENSMUSG00 | Gm4836 | 898 | 0 | 0 | 0 | 0 | 0 | 0 | 0 | 0 | 0 | 0 | 0 | 0 | 0 | 0 | 0 | 0 | 0 | 0 |
| ENSMUSG00 | Gm5935 | 1399 | 0 | 0 | 0 | 0 | 0 | 0 | 0 | 0 | 0 | 0 | 0 | 0 | 0 | 0 | 0 | 0 | 0 | 0 |
| ENSMUSG00 | Gm14636 | 7006 | 2.19017536 | 1.72603206 | 1.17377964 | 1.43069223 | 1.73126531 | 1.2377574 | 2.22179152 | 1.27017267 | 0.9229636 | 1.37292941 | 2.78221283 | 0.81261741 | 1.04916581 | 1.68381362 | 1.43493857 | 1.85277133 | 2.62036772 | 1.97863187 |
| ENSMUSG00 | Gm10489 | 1464 | 0 | 0.20479238 | 0 | 0.0933628 | 0.032877 | 0 | 0 | 0.33914019 | 0 | 0.03220676 | 0 | 0 | 0 | 0.09517231 | 0.02814314 | 0 | 0.05235833 | 0.07514905 |
| ENSMUSG00 | Gm10490 | 358 | 0 | 0 | 0 | 0 | 0 | 0 | 0 | 0 | 0 | 0.13170588 | 0 | 0 | 0 | 0 | 0 | 0.106957 | 0 | 0 |
| ENSMUSG00 | Gm10491 | 112 | 0 | 0 | 0 | 0 | 0 | 0 | 0 | 0 | 0 | 0 | 0 | 0 | 0 | 0 | 0 | 0 | 0 | 0 |
| ENSMUSG00 | Nudt10 | 1773 | 0.69523413 | 0.82671593 | 0.08185031 | 0.30836578 | 0.24432449 | 0.16769115 | 1.3342549 | 0.75393913 | 2.18360546 | 0.71803101 | 0.4311333 | 0.88146562 | 1.2799031 | 0.86444156 | 0.48800489 | 0 | 0.19454973 | 0.71359811 |
| ENSMUSG00 | AU022751 | 3756 | 0 | 0 | 0.05151601 | 0 | 0 | 0 | 0 | 0 | 0 | 0 | 0 | 0 | 0 | 0 | 0 | 0 | 0 | 0 |
| ENSMUSG00 | Nudt11 | 3034 | 1.06473084 | 0.96622765 | 0.52614585 | 0.79589168 | 0.74561648 | 0.45730934 | 0.82649044 | 1.17069658 | 0.74662438 | 0.37297854 | 0.29918399 | 0.60709191 | 0.89428327 | 1.02562751 | 1.35799459 | 0.42909712 | 0.48002613 | 0.81588985 |
| ENSMUSG00 | Gm10492 | 1592 | 0.61408232 | 1.12995998 | 0.9115615 | 0.82994365 | 1.14887757 | 0.52914351 | 0.98072764 | 1.34345134 | 1.47464284 | 0.88851828 | 1.32041343 | 2.45420627 | 1.42541972 | 1.72123188 | 1.31969909 | 1.25069823 | 1.56482996 | 0.93294339 |
| ENSMUSG00 | Gm10493 | 610 | 0 | 0 | 0 | 0 | 0 | 0 | 0 | 0 | 0 | 0 | 0 | 0 | 0 | 0 | 0 | 0 | 0 | 0 |
| ENSMUSG00 | Gm6594 | 287 | 10.0709128 | 6.84829533 | 10.1129332 | 6.98496847 | 8.38535256 | 5.87035868 | 13.1882032 | 8.11754894 | 11.7675759 | 15.1145111 | 7.82377369 | 14.9749339 | 14.9542746 | 16.0207828 | 15.3608588 | 12.1409243 | 7.61184294 | 13.6085247 |
| ENSMUSG00 | C030034I22I | 1470 | 1.4168392 | 2.01690312 | 1.41500803 | 1.48770757 | 1.8663399 | 1.24724568 | 1.86675876 | 1.0912115 | 1.0366674 | 0.7698074 | 1.33249884 | 0.72142693 | 2.11422503 | 1.57973086 | 1.54155481 | 0.52096064 | 1.12110937 | 0.33679045 |
| ENSMUSG00 | Lrrc30 | 1758 | 0 | 0 | 0 | 0 | 0 | 0 | 0 | 0 | 0 | 0 | 0 | 0 | 0 | 0 | 0 | 0 | 0 | 0 |
| ENSMUSG00 | Gm10495 | 465 | 0.09140898 | 0.14328127 | 0 | 0.29394222 | 0.20701903 | 0 | 0.1017477 | 0 | 0.08857315 | 0 | 0 | 0 | 0 | 0 | 0.26581649 | 0.08234539 | 0 | 0 |
| ENSMUSG00 | AU016765 | 1456 | 0 | 0 | 0 | 0 | 0 | 0 | 0 | 0 | 0 | 0 | 0 | 0 | 0 | 0 | 0 | 0 | 0 | 0 |
| ENSMUSG00 | Arrdc5 | 1051 | 0 | 0.09508914 | 0.09205239 | 0.04335018 | 0.04579631 | 0 | 0 | 0 | 0 | 0.04486271 | 0 | 0 | 0 | 0.30933265 | 0.07840448 | 0 | 0 | 0 |
| ENSMUSG00 | Kcnh8 | 1603 | 0 | 0.12468956 | 0.18106125 | 0 | 0 | 0 | 0 | 0.26208136 | 0.15416038 | 0 | 0 | 0.0348195 | 0 | 0 | 0 | 0.04777368 | 0.23909106 | 0.10294905 |
| ENSMUSG00 | 9830107B12I | 3665 | 0 | 0 | 0 | 0 | 0 | 0 | 0 | 0 | 0 | 0 | 0 | 0 | 0 | 0 | 0 | 0 | 0 | 0 |
| ENSMUSG00 | A330017A19I | 1189 | 1.46569573 | 1.51294894 | 1.58668434 | 1.37947653 | 1.33587341 | 0.54178767 | 0.87542383 | 3.11578052 | 2.77117011 | 1.94313243 | 1.4866892 | 1.31441426 | 1.03725646 | 1.21090556 | 1.49004786 | 1.54579408 | 1.03148991 | 1.29542051 |
| ENSMUSG00 | B230354K17I | 4894 | 4.42074675 | 3.51235256 | 4.64559854 | 3.84485315 | 3.54055834 | 4.20195841 | 3.63497492 | 4.05023638 | 3.61034231 | 3.42020835 | 2.80167819 | 2.93106361 | 2.05633656 | 2.40097142 | 3.90631834 | 3.26260386 | 4.3150367 | 1.6298162 |
| ENSMUSG00 | Gm10497 | 790 | 0.37662814 | 0.08433644 | 0.06123232 | 0 | 0.12185297 | 0.06272496 | 0 | 0 | 0 | 0 | 0 | 0 | 0 | 0.05878998 | 0.1564616 | 0 | 0 | 0 |
| ENSMUSG00 | Gm6084 | 1046 | 0 | 0 | 0 | 0 | 0 | 0 | 0 | 0 | 0 | 0 | 0 | 0 | 0 | 0 | 0 | 0 | 0 | 0 |
| ENSMUSG00 | Trim40 | 2669 | 0 | 0 | 0 | 0 | 0 | 0 | 0 | 0 | 0 | 0 | 0 | 0 | 0 | 0 | 0 | 0 | 0 | 0 |
| ENSMUSG00 | Trim10 | 2230 | 0 | 0.05975407 | 0.04338433 | 0.18387865 | 0 | 0.08888383 | 0 | 0.06850655 | 0.57254798 | 0 | 0 | 0.17520612 | 0 | 0.31240418 | 0.12933224 | 0.0686827 | 0.03437336 | 0 |
| ENSMUSG00 | Gm8909 | 1767 | 0 | 0 | 0 | 0 | 0 | 0 | 0 | 0 | 0 | 0 | 0 | 0 | 0 | 0 | 0 | 0 | 0 | 0 |
| ENSMUSG00 | Gm10499 | 1704 | 0 | 0.05864946 | 1.50457577 | 0 | 0.05649287 | 2.87894403 | 0 | 0.13448029 | 1.5227409 | 0.1106824 | 0 | 2.29289694 | 0 | 0.02725592 | 1.13642789 | 0.06741304 | 0 | 1.58183464 |
| ENSMUSG00 | C920025E04 | 1809 | 0.82237765 | 0.51562246 | 4.70632468 | 0.42815796 | 0.37249692 | 6.05370598 | 0.68000534 | 0.29557415 | 4.07539321 | 0.99045148 | 0.18486717 | 6.6337023 | 0.59994593 | 0.46213022 | 4.78293352 | 0.61383506 | 0.78389883 | 3.43616863 |
| ENSMUSG00 | H2-Bl | 1226 | 0 | 0.16303211 | 0.07891277 | 0.11148706 | 0 | 0.08083643 | 0 | 0.12460816 | 0.03359422 | 0.03845897 | 0 | 0.22763321 | 0.04023811 | 0 | 0.10081947 | 0.18739286 | 0 | 0.04486877 |
| ENSMUSG00 | Gm6034 | 1630 | 0 | 0.04087472 | 0.05935402 | 0.05590312 | 0 | 0.03040045 | 0.02902618 | 0 | 0 | 0.02892681 | 0.08792937 | 0.06848548 | 0.18158988 | 0 | 0.07583108 | 0 | 0 | 0 |
| ENSMUSG00 | Dpcr1 | 2608 | 0 | 0.2255467 | 0 | 0.01746973 | 0 | 0 | 0 | 0.0585773 | 0 | 0 | 0.05495586 | 0 | 0 | 0 | 0 | 0 | 0.20573932 | 0 |
| ENSMUSG00 | H2-Q6 | 1650 | 7.52212812 | 7.79319828 | 7.82771673 | 12.1772246 | 9.42218869 | 9.73035496 | 11.9858787 | 11.8280713 | 8.3870723 | 9.25868355 | 6.83326679 | 7.67888206 | 7.74361611 | 2.64590558 | 11.2118173 | 5.31427218 | 6.73613703 | 12.0353258 |
| ENSMUSG00 | H2-D1 | 2664 | 361.772658 | 338.168393 | 353.159456 | 454.071219 | 377.430888 | 373.505589 | 466.360224 | 454.868072 | 453.561867 | 554.905724 | 515.983898 | 458.886878 | 484.541647 | 444.147737 | 523.015396 | 569.558412 | 503.392316 | 587.775609 |
| ENSMUSG00 | Lst1 | 402 | 2.11468537 | 1.07728265 | 2.7676398 | 2.60672642 | 2.51435422 | 3.94449621 | 1.53001201 | 1.04506574 | 2.76625852 | 3.87058012 | 2.37686361 | 3.19343343 | 3.31333773 | 2.19511856 | 4.61211447 | 2.95275828 | 5.24364759 | 4.10515722 |
| ENSMUSG00 | Ly6g6d | 1397 | 4.0162371 | 3.07613698 | 2.87401791 | 2.4133982 | 3.23865485 | 3.19237367 | 3.01419358 | 3.49937545 | 4.36335314 | 2.56510626 | 1.88090388 | 1.23857233 | 2.50720955 | 2.19420885 | 2.50689495 | 3.86469266 | 1.50890933 | 1.89007668 |
| ENSMUSG00 | AU023871 | 2480 | 0.61701062 | 0.63133308 | 0.58516367 | 0.60625583 | 0.29112051 | 0.49952352 | 0.49602002 | 0.47740503 | 0.94662556 | 0.66543331 | 0.42381012 | 0.47263263 | 0.69621657 | 0.58055109 | 0.34888414 | 0.26247594 | 0.43271626 | 0.06654327 |
| ENSMUSG00 | Gm10501 | 2198 | 0.11602869 | 0.28796405 | 0.30811166 | 0.16582728 | 0.30657276 | 0.29307804 | 0.60270929 | 0.39964752 | 0.28107267 | 0.38612951 | 0.3042991 | 0.40630146 | 0.44888005 | 0.23243174 | 0.05623506 | 0.45293712 | 0.33130102 | 0.20021513 |
| ENSMUSG00 | C4b | 5655 | 0 | 0.02945437 | 0.18819057 | 0.01611354 | 0.05957975 | 0 | 0.04183261 | 0 | 0.10196485 | 0 | 0 | 0 | 0 | 0.00821292 | 0.08014449 | 0.08125328 | 0 | 0 |
| ENSMUSG00 | Btnl5 | 1919 | 0 | 0 | 0 | 0 | 0 | 0 | 0 | 0 | 0 | 0 | 0 | 0 | 0 | 0 | 0 | 0 | 0 | 0 |
| ENSMUSG00 | H2-Ab1 | 1801 | 2.36008751 | 3.47741483 | 5.50614865 | 3.08631172 | 2.05783349 | 7.18115684 | 25.1668775 | 33.0817034 | 24.1722083 | 7.93262803 | 44.6447835 | 5.57846711 | 21.0639891 | 21.7650249 | 25.0274859 | 7.3774796 | 0.82994202 | 3.11545191 |
| ENSMUSG00 | H2-Ke6 | 1545 | 0.46769449 | 0.62529057 | 0.78274321 | 0.50131893 | 0.37384018 | 0.8659701 | 0.61246186 | 0.66744004 | 0.98834374 | 0.91554764 | 0.61844606 | 0.86703942 | 1.08562148 | 0.96194874 | 0.48001813 | 0.39653703 | 0.52093996 | 0.4984644 |
| ENSMUSG00 | Zfp414 | 2074 | 15.1452869 | 14.6326165 | 14.7872798 | 13.9275324 | 14.5509721 | 14.1920558 | 14.8736098 | 16.7206853 | 15.6683515 | 12.2309925 | 15.6178505 | 16.308723 | 14.7710321 | 14.9588478 | 14.3827996 | 14.0866603 | 14.7096106 | 18.2214351 |
| ENSMUSG00 | Cyp4f15 | 2412 | 5.70965051 | 6.47750721 | 6.63832444 | 9.52021822 | 9.2592092 | 6.57416034 | 8.88583895 | 7.07794522 | 5.19100364 | 9.03135361 | 9.26976806 | 4.92899508 | 15.6054116 | 19.5827682 | 19.1658979 | 14.2240399 | 15.2542475 | 15.6908231 |
| ENSMUSG00 | Gm4924 | 2644 | 0.98064135 | 1.0331533 | 1.17091753 | 0.9649843 | 1.11045664 | 1.16197787 | 0.98418961 | 1.30004395 | 1.05925986 | 1.58714546 | 0.83118309 | 1.75215582 | 1.0448515 | 0.75533122 | 1.35572442 | 0.89788868 | 1.31909721 | 0.27046838 |
| ENSMUSG00 | C130040N14 | 1108 | 0.84396559 | 1.20263156 | 0.74219316 | 0.90464167 | 0.99912838 | 0.71556294 | 0.85401948 | 0.79280258 | 1.33819004 | 1.82985581 | 1.12107303 | 1.30975382 | 0.84594443 | 0.92217502 | 1.59897735 | 1.10586592 | 1.0031251 | 1.43976904 |
| ENSMUSG00 | Gm10505 | 1585 | 0 | 0 | 0 | 0 | 0 | 0 | 0 | 0 | 0 | 0 | 0 | 0 | 0 | 0 | 0 | 0 | 0 | 0 |
| ENSMUSG00 | Arhgdig | 2958 | 17.9475878 | 16.9267343 | 15.9609755 | 15.6028864 | 13.7008383 | 15.2109135 | 11.7082086 | 11.0522977 | 12.7541746 | 12.337608 | 11.9033619 | 12.453799 | 6.90446712 | 4.27072068 | 7.00621454 | 8.12931083 | 5.1568175 | 6.28569238 |
| ENSMUSG00 | Wdr90 | 8098 | 1.04452087 | 1.44802899 | 1.27235885 | 1.84539671 | 1.86631564 | 1.6093318 | 2.17925775 | 1.90537547 | 1.84113592 | 2.03205673 | 2.0058639 | 1.82651897 | 2.28444912 | 1.4854308 | 2.01479577 | 1.51781736 | 1.23053065 | 1.44689303 |
| ENSMUSG00 | Nme3 | 927 | 0 | 0 | 0 | 0 | 0 | 0 | 0 | 0 | 0 | 0 | 0 | 0 | 0 | 0 | 0 | 0 | 0 | 0 |
| ENSMUSG00 | Eme2 | 6688 | 1.29650956 | 1.94756903 | 1.69972782 | 1.8257117 | 1.5760902 | 1.2669733 | 1.35825873 | 1.70746536 | 2.0322294 | 1.70611097 | 1.48582407 | 1.7525851 | 1.81454122 | 2.12498366 | 1.76190864 | 1.16795513 | 1.36388437 | 2.36881321 |
| ENSMUSG00 | D330041H03 | 282 | 0.30145515 | 0.47252332 | 0.34307468 | 0.64625595 | 0.34136116 | 0.35143783 | 0.67110183 | 0.13543405 | 0.29210295 | 0.16720108 | 0.33882949 | 0 | 0 | 0.73052404 | 0.40734689 | 0 | 0 | 0.58520326 |
| ENSMUSG00 | Gm10506 | 109 | 3.11964595 | 2.13936018 | 3.106557 | 1.67196493 | 1.78630913 | 1.3638367 | 0.86812255 | 2.8031121 | 3.02286354 | 2.1628763 | 3.0681166 | 5.63277329 | 2.71551838 | 2.98264781 | 6.04793481 | 1.05386993 | 4.57102624 | 1.00934141 |
| ENSMUSG00 | Gm16386 | 1420 | 1.61639402 | 2.36943825 | 1.97582025 | 3.30477998 | 2.20322186 | 2.26825893 | 1.9324897 | 3.73855207 | 4.03163781 | 2.55676349 | 3.02799033 | 2.16187426 | 2.91822609 | 3.20529617 | 2.26318406 | 3.74816509 | 3.31981304 | 3.25405983 |
| ENSMUSG00 | Vmn2r112 | 2568 | 0 | 0 | 0 | 0 | 0 | 0 | 0 | 0 | 0 | 0.03672173 | 0 | 0 | 0 | 0 | 0 | 0 | 0 | 0 |
| ENSMUSG00 | Vmn2r111 | 2568 | 0 | 0 | 0 | 0 | 0 | 0 | 0 | 0 | 0 | 0 | 0 | 0 | 0 | 0 | 0 | 0 | 0 | 0 |
| ENSMUSG00 | Gm10509 | 258 | 3.62447237 | 3.87359235 | 3.37489747 | 3.88505029 | 2.79836766 | 3.84129718 | 3.48426705 | 6.21736783 | 6.38550632 | 4.20335727 | 6.85144753 | 9.08627065 | 4.78022454 | 6.66058618 | 4.63118264 | 2.37461131 | 4.75364923 | 3.41141747 |
| ENSMUSG00 | Zfp97 | 2705 | 1.24137113 | 1.39162922 | 1.55582149 | 1.02743945 | 1.72598765 | 0.87930914 | 1.25933933 | 1.17189257 | 2.0402932 | 1.34218268 | 1.1126885 | 1.44439792 | 1.24013692 | 0.82414647 | 1.1119089 | 1.07581744 | 1.81359187 | 1.15915678 |
| ENSMUSG00 | Gm3435 | 2658 | 0.33581968 | 0.27572749 | 0.49137896 | 0.2228343 | 0.48892473 | 0.35421442 | 0.46280273 | 0.38795894 | 0.17044833 | 0.28382666 | 0.23366233 | 0.44098153 | 0.38975555 | 0.17473321 | 0.27901731 | 0.12965217 | 0.23070758 | 0.14486973 |
| ENSMUSG00 | Smok2b | 1940 | 0 | 0 | 0 | 0 | 0 | 0 | 0 | 0 | 0 | 0 | 0 | 0 | 0 | 0 | 0 | 0 | 0 | 0 |
| ENSMUSG00 | Smok2a | 1784 | 0 | 0 | 0 | 0 | 0 | 0 | 0 | 0 | 0 | 0 | 0 | 0 | 0 | 0 | 0 | 0 | 0 | 0 |
| ENSMUSG00 | Pnldc1 | 2133 | 0.03985483 | 0.09370716 | 0.3175009 | 0.10680039 | 0.04513073 | 0.18585179 | 0.08872514 | 0 | 0 | 0.02210535 | 0 | 0.05233536 | 0.3700472 | 0 | 0.03863249 | 0.05385458 | 0.0898413 | 0 |
| ENSMUSG00 | Park2 | 306 | 1.38905804 | 0.21773133 | 0.15808343 | 0.1488923 | 0.4718816 | 0.16193704 | 0.46384979 | 0.37443532 | 0.13459646 | 0.15408727 | 1.24901852 | 0.18240413 | 0.9672925 | 0 | 0.26929121 | 0 | 0.12524934 | 0.89884161 |
| ENSMUSG00 | 6530411M01 | 1030 | 0 | 0 | 0 | 0 | 0 | 0.27560784 | 0 | 0 | 0 | 0 | 0 | 0 | 0 | 0 | 0 | 0 | 0 | 0 |
| ENSMUSG00 | Sft2d1 | 3743 | 11.3218436 | 11.5611674 | 10.6879267 | 9.67700514 | 10.0815998 | 9.0950382 | 11.0476306 | 9.85676215 | 9.91425346 | 10.1406134 | 9.4579867 | 9.61824803 | 12.2440158 | 9.91419338 | 10.5232935 | 11.928092 | 11.7344206 | 11.1546652 |
| ENSMUSG00 | Rnaset2a | 1004 | 10.9226448 | 9.78815119 | 11.7561169 | 9.84735719 | 10.6906564 | 10.9568793 | 10.2730717 | 9.31985915 | 11.363212 | 13.4783385 | 11.8009858 | 12.4528968 | 11.3994072 | 15.1729688 | 15.1838403 | 14.0348043 | 13.9333649 | 15.8342947 |
| ENSMUSG00 | Gm1604b | 715 | 0 | 0 | 0 | 0 | 0 | 0 | 0 | 0 | 0 | 0 | 0 | 0 | 0 | 0 | 0 | 0 | 0 | 0 |
| ENSMUSG00 | Rsph3a | 2181 | 2.06581782 | 1.2983017 | 1.75218198 | 1.37873862 | 1.30205571 | 1.59041327 | 1.3666661 | 1.62856186 | 0.9253275 | 1.14579884 | 1.24858901 | 1.17722168 | 1.42499348 | 0.70273035 | 1.43572593 | 1.3518463 | 1.37067912 | 1.31154221 |
| ENSMUSG00 | A330023F24I | 6398 | 0.51819377 | 0.74977442 | 1.0433803 | 0.7762041 | 0.73725047 | 0.56538755 | 0.40672043 | 0.50143198 | 0.811113 | 0.55272003 | 0.39576005 | 0.7066378 | 0.42407869 | 0.93643127 | 0.90156577 | 1.11316864 | 0.61700666 | 0.72222022 |
| ENSMUSG00 | Gm10516 | 2596 | 0.60581337 | 0.46196618 | 0.37267743 | 0.68446869 | 1.46472341 | 0.22905732 | 1.73139619 | 1.35350579 | 1.99903736 | 1.25323518 | 0.5520989 | 0.60201793 | 1.29220739 | 0.69773475 | 0.41265841 | 1.59298367 | 1.3139601 | 0.88997784 |
| ENSMUSG00 | D730003I15I | 545 | 42.8171406 | 36.6747459 | 45.9770435 | 32.1017267 | 48.0436082 | 40.0967991 | 71.2728613 | 54.2402192 | 61.7419878 | 53.9853924 | 57.5052711 | 64.6232717 | 46.2543298 | 45.5919023 | 47.2494907 | 42.0142811 | 40.0140604 | 56.2203164 |
| ENSMUSG00 | Mosc2 | 6716 | 39.5558889 | 35.7384459 | 36.0280522 | 38.5667863 | 37.6469617 | 35.5560785 | 43.9875472 | 37.1858427 | 45.0684491 | 45.1287553 | 42.1267578 | 39.9336299 | 49.4494442 | 48.2282706 | 44.2076253 | 45.9475886 | 40.3407672 | 44.6559931 |
| ENSMUSG00 | Gm10517 | 401 | 0.74198562 | 0.66459639 | 0 | 0.22723713 | 1.80044602 | 0.74143741 | 0.94389384 | 1.14291479 | 0 | 0.1175828 | 0.23827909 | 1.11352943 | 0.36966671 | 0 | 0.20549404 | 0.0954878 | 0.2867304 | 1.09743854 |
| ENSMUSG00 | Gm16067 | 3131 | 0.12218032 | 0.34047033 | 0.30899732 | 0.36378988 | 0.3996902 | 0.34818273 | 0.15111044 | 0.43913334 | 0.13154429 | 0.15059311 | 0.41198463 | 0.33870891 | 0.04726789 | 0.34117343 | 0.22370695 | 0.30573784 | 0.20809551 | 0.22839936 |
| ENSMUSG00 | Gm10518 | 1300 | 0 | 0 | 0 | 0 | 0 | 0 | 0 | 0.23503017 | 0 | 0 | 0.03674997 | 0 | 0 | 0 | 0 | 0 | 0 | 0 |
| ENSMUSG00 | Ifi204 | 2676 | 0 | 0 | 0.01807681 | 0 | 0 | 0 | 0.03536075 | 0.04281659 | 0 | 0 | 0 | 0 | 0 | 0 | 0.10016228 | 0 | 0 | 0 |
| ENSMUSG00 | AI607873 | 3313 | 0.03848944 | 0 | 0 | 0 | 0.0435846 | 0 | 0.31418018 | 0.08069629 | 0 | 0 | 0.08652271 | 0 | 0 | 0 | 0 | 0 | 0 | 0 |
| ENSMUSG00 | Pydc4 | 4183 | 0 | 0 | 0 | 0 | 0 | 0 | 0 | 0 | 0 | 0 | 0 | 0 | 0 | 0 | 0 | 0.00915386 | 0 | 0 |
| ENSMUSG00 | Gm10521 | 2573 | 0 | 0 | 0 | 0 | 0 | 0 | 0 | 0 | 0 | 0 | 0 | 0 | 0 | 0 | 0 | 0 | 0 | 0 |
| ENSMUSG00 | Gm17286 | 1253 | 0 | 0.05317302 | 0.30884936 | 0.07272314 | 0 | 0.27683091 | 0.07551904 | 0 | 0.1314813 | 0.18815125 | 0.22877075 | 0.17818248 | 0.82679191 | 0.88959146 | 0.39458793 | 0.12223658 | 0.39763915 | 0.08780384 |
| ENSMUSG00 | Sh2d1b2 | 2765 | 0.06149031 | 0 | 0 | 0 | 0 | 0 | 0 | 0 | 0 | 0 | 0 | 0 | 0 | 0 | 0 | 0 | 0 | 0 |
| ENSMUSG00 | AA792892 | 1878 | 0 | 0 | 0 | 0 | 0 | 0 | 0 | 0 | 0 | 0 | 0 | 0 | 0 | 0 | 0 | 0 | 0 | 0 |
| ENSMUSG00 | Gm6502 | 1215 | 0 | 0 | 0 | 0 | 0 | 0 | 0 | 0 | 0 | 0 | 0 | 0 | 0 | 0 | 0 | 0 | 0 | 0 |
| ENSMUSG00 | Gm16522 | 1844 | 0 | 0 | 0 | 0 | 0 | 0 | 0 | 0 | 0 | 0 | 0 | 0 | 0 | 0 | 0 | 0 | 0 | 0 |
| ENSMUSG00 | Gm3106 | 1433 | 0 | 0 | 0 | 0 | 0 | 0 | 0 | 0 | 0 | 0 | 0 | 0 | 0 | 0 | 0 | 0 | 0 | 0 |
| ENSMUSG00 | Gm3176 | 1840 | 0 | 0 | 0 | 0 | 0 | 0 | 0 | 0 | 0 | 0 | 0 | 0 | 0 | 0 | 0 | 0 | 0 | 0 |
| ENSMUSG00 | Gm7647 | 1211 | 0 | 0 | 0 | 0 | 0 | 0 | 0 | 0 | 0 | 0 | 0 | 0 | 0 | 0 | 0 | 0 | 0 | 0 |
| ENSMUSG00 | Dok6 | 5138 | 0 | 0 | 0 | 0 | 0 | 0 | 0 | 0 | 0 | 0 | 0 | 0 | 0 | 0 | 0 | 0 | 0 | 0 |
| ENSMUSG00 | Gm10524 | 1982 | 5.64019238 | 6.47097088 | 6.07719932 | 7.74675678 | 6.16826872 | 7.07539031 | 4.17745651 | 6.60948235 | 6.62890945 | 4.21073384 | 4.91730148 | 8.22309459 | 3.21080592 | 4.12419747 | 4.84357342 | 2.85923809 | 3.15196085 | 4.80150124 |
| ENSMUSG00 | Gm10525 | 245 | 0 | 0 | 0 | 0 | 0 | 0 | 0 | 0 | 0 | 0 | 0 | 0 | 0 | 0 | 0 | 0 | 0 | 0 |
| ENSMUSG00 | Gm10526 | 245 | 0 | 0 | 0 | 0 | 0 | 0 | 0 | 0 | 0 | 0 | 0 | 0 | 0 | 0 | 0 | 0 | 0 | 0 |
| ENSMUSG00 | Gm10527 | 245 | 0 | 0 | 0 | 0 | 0 | 0 | 0 | 0 | 0 | 0 | 0 | 0 | 0 | 0 | 0 | 0 | 0 | 0 |
| ENSMUSG00 | Gm10528 | 245 | 0 | 0 | 0 | 0 | 0 | 0 | 0 | 0 | 0 | 0 | 0 | 0 | 0 | 0 | 0 | 0 | 0 | 0 |
| ENSMUSG00 | Gm10529 | 376 | 0 | 0 | 0 | 0 | 0 | 0 | 0 | 0 | 0 | 0 | 0 | 0 | 0 | 0 | 0 | 0 | 0 | 0 |
| ENSMUSG00 | Gm10530 | 1166 | 0 | 0 | 0 | 0 | 0 | 0 | 0 | 0 | 0 | 0 | 0 | 0 | 0 | 0 | 0 | 0 | 0 | 0 |
| ENSMUSG00 | F830208F22I | 2763 | 0 | 0 | 0.07003045 | 0.0164897 | 0.06968067 | 0 | 0 | 0 | 0 | 0.0853252 | 0 | 0.06060333 | 0 | 0 | 0 | 0 | 0 | 0.09954598 |
| ENSMUSG00 | Pappa2 | 9056 | 0 | 0 | 0 | 0 | 0 | 0 | 0 | 0 | 0 | 0 | 0 | 0 | 0.00544743 | 0 | 0 | 0 | 0 | 0 |
| ENSMUSG00 | Gm7276 | 1251 | 0 | 0.02662901 | 0.07733578 | 0.0364197 | 0.07694952 | 0 | 0.26473921 | 0.61058997 | 0.49384311 | 0.11307123 | 0.03818942 | 0.04461684 | 0.63094379 | 0 | 0 | 0.061216 | 0.03063653 | 0 |
| ENSMUSG00 | Gm10532 | 2297 | 0.05551394 | 0 | 0 | 0 | 0 | 0 | 0 | 0.01793057 | 0 | 0 | 0.06239655 | 0 | 0 | 0.02021945 | 0 | 0 | 0 | 0 |
| ENSMUSG00 | Gm5532 | 1749 | 9.13776226 | 8.55202373 | 8.85050298 | 6.38219317 | 8.53110139 | 6.60136474 | 0.05410255 | 0.2838772 | 0 | 0 | 0 | 0.1914774 | 0.0282057 | 0 | 0.02355721 | 0.02189286 | 0 | 0 |
| ENSMUSG00 | E330020D12I | 2120 | 0 | 0 | 0 | 0 | 0 | 0 | 0.11158651 | 0 | 0 | 0 | 0.33803037 | 0 | 0.02326977 | 0 | 0 | 0.10836964 | 0 | 0 |
| ENSMUSG00 | Cep76 | 3967 | 1.72506512 | 1.35199798 | 1.4998599 | 2.38888259 | 1.77142951 | 1.84870294 | 2.93393472 | 1.34785388 | 0.45682044 | 1.99680316 | 2.68561022 | 1.39292932 | 0.78344109 | 2.09566212 | 2.37841097 | 1.31271051 | 3.27517385 | 2.95360215 |
| ENSMUSG00 | Chmp1b | 2382 | 51.9626669 | 53.2698632 | 50.6480204 | 50.3045956 | 55.8104002 | 51.5498211 | 44.6908176 | 45.8565369 | 50.160404 | 46.3391252 | 47.1732587 | 50.168486 | 40.6750149 | 50.2851811 | 49.0371163 | 42.9041273 | 49.2674732 | 49.2125972 |
| ENSMUSG00 | F730048M01 | 2146 | 0 | 0 | 0 | 0 | 0 | 0 | 0 | 0 | 0 | 0 | 0 | 0 | 0 | 0 | 0 | 0 | 0 | 0 |
| ENSMUSG00 | Spink13 | 790 | 0.5380402 | 1.2228784 | 0.67355549 | 0.6920665 | 0.73111783 | 0.37634988 | 3.71314695 | 2.03048215 | 4.06651675 | 1.25337312 | 0.18142389 | 0.77718011 | 2.37292767 | 7.81906788 | 3.91154012 | 0.67856773 | 1.5039433 | 2.71563945 |
| ENSMUSG00 | Gm6960 | 637 | 0 | 0 | 0 | 0 | 0 | 0 | 0 | 0 | 0 | 0 | 0 | 0 | 0 | 0 | 0 | 0 | 0 | 0 |
| ENSMUSG00 | Gm4951 | 2642 | 0 | 0.16391659 | 0 | 0 | 0 | 0.18755766 | 0 | 0 | 0 | 0.08923297 | 0 | 0 | 0 | 0 | 0 | 0 | 0 | 0 |
| ENSMUSG00 | Ppp1r12b | 15492 | 2.91105033 | 5.1242336 | 5.35818346 | 3.42913618 | 4.25643787 | 3.9918546 | 1.35597918 | 2.22123383 | 2.72768946 | 1.62221307 | 1.68219356 | 1.35107626 | 1.29284524 | 1.28311835 | 2.14890635 | 2.07123218 | 1.90246077 | 1.9955537 |
| ENSMUSG00 | Gm8618 | 502 | 0.254015 | 0 | 0.09636161 | 0.1815181 | 0.09588033 | 0.09871062 | 0 | 0 | 0 | 0 | 0 | 0 | 0 | 0 | 0 | 0.07627611 | 0 | 0 |
| ENSMUSG00 | Csnk1g3 | 4381 | 12.1082994 | 11.8393463 | 14.5308299 | 12.7916194 | 11.2721647 | 11.9781659 | 11.1127018 | 15.2298852 | 13.622292 | 11.4082962 | 9.52009558 | 14.9317408 | 9.68396459 | 9.07467218 | 10.6742219 | 11.1699147 | 11.9064346 | 11.1499856 |
| ENSMUSG00 | Prr16 | 3817 | 0 | 0 | 0 | 0 | 0 | 0 | 0 | 0 | 0 | 0 | 0 | 0 | 0 | 0 | 0 | 0 | 0 | 0 |
| ENSMUSG00 | Tnfaip8 | 5038 | 0 | 0.01322465 | 0 | 0.06330435 | 0 | 0.00983579 | 0.01878233 | 0 | 0.01635034 | 0.03743605 | 0 | 0 | 0.03916786 | 0.07375004 | 0 | 0.02006319 | 0 | 0 |
| ENSMUSG00 | Gm949 | 2165 | 0 | 0 | 0 | 0 | 0 | 0 | 0.02185343 | 0.0529225 | 0 | 0 | 0.24273651 | 0 | 0 | 0 | 0 | 0 | 0 | 0 |
| ENSMUSG00 | Gm10540 | 275 | 0 | 0.24227559 | 0.17590375 | 0 | 0 | 0 | 0 | 0 | 0.14976915 | 0 | 0.17372712 | 0 | 0 | 0 | 0 | 0 | 0 | 0 |
| ENSMUSG00 | Gm10542 | 267 | 0 | 0 | 0 | 0 | 0 | 0 | 0 | 0 | 0 | 0 | 0 | 0 | 0 | 0 | 0.92587766 | 0.14341051 | 0 | 0.61807985 |
| ENSMUSG00 | Spink11 | 741 | 232.143654 | 187.10967 | 286.45486 | 117.069134 | 120.427242 | 122.109705 | 192.634753 | 327.547527 | 147.404372 | 51.2867299 | 92.1972883 | 187.106756 | 4.06106202 | 11.3446421 | 5.72707187 | 3.46217368 | 1.65511674 | 4.23146975 |
| ENSMUSG00 | Grxcr2 | 1117 | 0 | 0 | 0 | 0 | 0 | 0 | 0 | 0 | 0 | 0 | 0 | 0 | 0 | 0 | 0 | 0 | 0 | 0 |
| ENSMUSG00 | 9630014M24 | 3864 | 0 | 0 | 0 | 0 | 0 | 0 | 0 | 0 | 0 | 0 | 0 | 0 | 0 | 0 | 0 | 0 | 0 | 0 |
| ENSMUSG00 | Gm10543 | 1586 | 0 | 0 | 0 | 0 | 0 | 0 | 0 | 0 | 0 | 0 | 0 | 0 | 0 | 0 | 0 | 0 | 0 | 0 |
| ENSMUSG00 | 3222401L13I | 2830 | 2.98887987 | 1.25953346 | 1.76059139 | 1.86751984 | 1.42865074 | 1.80350938 | 0.96965914 | 2.99601178 | 2.38678042 | 2.28256055 | 2.24525423 | 2.97815019 | 2.09181275 | 0.80415558 | 1.77618015 | 2.11071899 | 1.66577191 | 2.5269201 |
| ENSMUSG00 | Pcdhb22 | 5324 | 0.74248335 | 0.86348223 | 0.49064062 | 0.57336401 | 0.46106839 | 0.41883415 | 0.15107354 | 0.43759139 | 0.26302433 | 0.5493157 | 0.50251647 | 0.35644863 | 0.3521061 | 0.25298244 | 0.13929902 | 0.74078373 | 0.29514992 | 0.15498434 |
| ENSMUSG00 | Gm10544 | 4057 | 0.02095399 | 0 | 0 | 0 | 0 | 0 | 0 | 0 | 0 | 0 | 0 | 0 | 0 | 0 | 0 | 0 | 0 | 0 |
| ENSMUSG00 | Gm10545 | 4966 | 0 | 0 | 0 | 0 | 0.00969229 | 0 | 0 | 0 | 0 | 0 | 0 | 0 | 0 | 0 | 0 | 0 | 0.03087096 | 0 |
| ENSMUSG00 | 2900060B14I | 77 | 0 | 0 | 0 | 0.59170187 | 1.25017984 | 1.28708399 | 0 | 0.49600523 | 2.13955926 | 1.22469359 | 0.62045401 | 4.34927241 | 2.562697 | 4.22218976 | 4.28068113 | 2.98368369 | 3.48426881 | 2.85761593 |
| ENSMUSG00 | 1700066B19I | 2833 | 0.09002155 | 0.24693638 | 0.1707502 | 0.17690487 | 0.13591789 | 0.08745629 | 0.16700557 | 0.04044377 | 0.17445753 | 0.36615442 | 0.0505912 | 0.17731767 | 0.29602633 | 0.06557584 | 0.07271718 | 0.64876426 | 0.28409892 | 0.2135899 |
| ENSMUSG00 | Ecscr | 1829 | 0.34859357 | 0.89247229 | 0.55540959 | 0.27401394 | 1.18421901 | 0.92115524 | 0.33628476 | 0.45039467 | 0.51792775 | 0.36091298 | 0.05224162 | 0.51878965 | 0.08091621 | 0.20314527 | 0.92359967 | 1.71669208 | 0.83819129 | 0.03007606 |
| ENSMUSG00 | Gm1614 | 3020 | 0.28149123 | 1.19132204 | 0.75283309 | 0.92027275 | 0.51000714 | 0.88604226 | 0.5013264 | 1.08759821 | 0.73644763 | 1.68618409 | 1.13900563 | 0.77624431 | 1.32314083 | 1.2918223 | 1.44614733 | 0.59591343 | 0.72337713 | 0.38251366 |
| ENSMUSG00 | Serpinb3c | 1651 | 0 | 0 | 0 | 0 | 0 | 0 | 0 | 0 | 0 | 0 | 0 | 0 | 0 | 0 | 0 | 0 | 0 | 0 |
| ENSMUSG00 | Serpinb3b | 2491 | 0 | 0 | 0 | 0 | 0 | 0 | 0 | 0 | 0 | 0 | 0 | 0 | 0 | 0 | 0 | 0 | 0 | 0 |
| ENSMUSG00 | Gm10548 | 2898 | 0.89469142 | 0.40232964 | 0.53414526 | 0.53453261 | 0.79721613 | 0.59846297 | 0.84895076 | 0.61940749 | 0.4405735 | 0.66707344 | 0.52753577 | 0.57780189 | 0.42556864 | 0.70515523 | 0.34121371 | 0.63421296 | 0.29095187 | 0.77825168 |
| ENSMUSG00 | Gm6086 | 4053 | 0 | 0 | 0 | 0 | 0 | 0 | 0 | 0 | 0 | 0 | 0 | 0 | 0 | 0 | 0 | 0 | 0 | 0 |
| ENSMUSG00 | D2hgdh | 3942 | 4.67966677 | 6.5831417 | 6.87193734 | 5.21259031 | 6.5934142 | 5.51842974 | 4.3808036 | 9.72733943 | 8.18088327 | 5.8131004 | 8.70178087 | 5.63537131 | 4.71794338 | 6.16191215 | 7.044609 | 5.92523351 | 7.85582138 | 5.12134504 |
| ENSMUSG00 | Gm10549 | 447 | 0 | 0.14905098 | 0 | 0 | 0 | 0 | 0 | 0 | 0 | 0 | 0 | 0 | 0 | 0.10390176 | 0.27652051 | 0 | 0 | 0 |
| ENSMUSG00 | Myeov2 | 1495 | 75.8838226 | 59.027329 | 68.9848607 | 57.0808268 | 70.7330009 | 65.4294957 | 102.378941 | 75.0308267 | 81.6291948 | 89.9805728 | 75.0977608 | 76.4618575 | 112.753954 | 105.780681 | 86.2338919 | 81.9598286 | 89.2400264 | 94.2697869 |
| ENSMUSG00 | Hdac4 | 2391 | 11.05739 | 13.0269996 | 10.0955214 | 11.8142399 | 10.3067942 | 9.69915488 | 10.3292423 | 12.5550934 | 10.1459046 | 10.1558395 | 12.7879437 | 10.574862 | 9.57340427 | 14.005097 | 13.8889394 | 15.2778082 | 12.9196969 | 13.3669138 |
| ENSMUSG00 | 5830472F04I | 350 | 0.36433008 | 0.1903594 | 0.55284035 | 0.26034882 | 0 | 0 | 0.54071633 | 0 | 0 | 0 | 0 | 0 | 0 | 1.19427653 | 0.47087492 | 0.54700868 | 0 | 0 |
| ENSMUSG00 | C130036L24I | 3784 | 0.41561615 | 0.42257371 | 0.4985644 | 0.33713246 | 0.58511324 | 0.49762259 | 1.11279821 | 0.56521526 | 0.92517279 | 0.39873745 | 0.51764622 | 0.5605167 | 0.48236811 | 0.45413088 | 0.38109262 | 0.38452513 | 0.52668273 | 0.94492387 |
| ENSMUSG00 | Gm10552 | 1053 | 0 | 0.03163618 | 0 | 0 | 0 | 0.09411725 | 0.17972528 | 0 | 0 | 0.13433249 | 0 | 0 | 0 | 0.13231934 | 0.07825557 | 0 | 0 | 0 |
| ENSMUSG00 | Gm10553 | 371 | 0 | 0 | 0 | 0 | 0 | 0 | 0 | 0 | 0 | 0 | 0 | 0 | 0 | 0 | 0 | 0 | 0 | 0 |
| ENSMUSG00 | Gm15433 | 890 | 0 | 0 | 0 | 0.05119218 | 0 | 0 | 0 | 0 | 0 | 0 | 0.05367973 | 0 | 0.11085824 | 0 | 0 | 0 | 0 | 0 |
| ENSMUSG00 | Fbxo36 | 2256 | 11.0784767 | 10.9123355 | 11.6859814 | 10.8449826 | 9.13141105 | 10.4113456 | 20.8880444 | 20.3489662 | 21.2139766 | 18.496619 | 19.4826958 | 16.749647 | 17.6247896 | 18.363531 | 18.4092058 | 17.5498617 | 17.1245156 | 16.5807591 |
| ENSMUSG00 | Rab18 | 2346 | 154.330388 | 155.232975 | 169.245498 | 171.504511 | 163.127419 | 158.775745 | 134.032423 | 142.057504 | 149.911193 | 140.406996 | 149.352747 | 149.127269 | 146.145279 | 134.2246 | 154.356354 | 157.340774 | 172.615368 | 144.111667 |
| ENSMUSG00 | Rpl27-ps3 | 474 | 0.5380402 | 0.63252331 | 1.3267092 | 0.57672208 | 0.30463243 | 1.04541632 | 1.29760512 | 0.56402282 | 1.21647937 | 0.99474058 | 0.50395526 | 0 | 1.04075775 | 1.56773291 | 0.78230802 | 0.32312749 | 0.64685733 | 0.69631781 |
| ENSMUSG00 | Wdfy1 | 9597 | 11.475556 | 9.65681689 | 9.01742688 | 7.06416942 | 7.05152492 | 9.29404193 | 9.54930281 | 6.95239419 | 8.17121246 | 7.0993817 | 7.92515723 | 8.72392351 | 9.01616994 | 8.35770959 | 7.79639736 | 9.6035732 | 9.24910947 | 8.91311461 |
| ENSMUSG00 | Gm10556 | 3896 | 0 | 0 | 0.0124162 | 0 | 0 | 0 | 0 | 0.00980298 | 0.02114298 | 0 | 0 | 0 | 0.01192097 | 0 | 0 | 0 | 0 | 0.01411938 |
| ENSMUSG00 | Gm10557 | 748 | 3.97775711 | 5.21070672 | 5.10896912 | 4.50737604 | 5.01910434 | 2.78237274 | 5.56619752 | 5.51441104 | 6.66252461 | 4.28642757 | 6.38702653 | 5.74572997 | 4.28686447 | 5.77446541 | 6.44462839 | 5.63097168 | 8.71052202 | 6.32457644 |
| ENSMUSG00 | Gm216 | 2092 | 0 | 0.74842544 | 0.18498482 | 0.13067221 | 0.11503806 | 0 | 0.6558641 | 0 | 0.15750102 | 0.18030862 | 0 | 0.48024949 | 0.07074367 | 0.11100403 | 0.49237041 | 0.43928039 | 0.16488369 | 0 |
| ENSMUSG00 | Apol7d | 2216 | 1.66875014 | 1.54838813 | 1.70267842 | 1.70648316 | 1.65073384 | 1.11806709 | 1.49453408 | 1.70624902 | 1.07798642 | 0.72343137 | 1.05639574 | 1.38531654 | 1.40248682 | 1.06888468 | 1.26430767 | 1.7279155 | 2.07543124 | 1.88659391 |
| ENSMUSG00 | Gm10558 | 1658 | 0.05127283 | 0.02009222 | 0 | 0 | 0.05806022 | 0 | 0.05707199 | 0 | 0.12420542 | 0 | 0.05762962 | 0 | 0 | 0 | 0 | 0 | 0 | 0 |
| ENSMUSG00 | Crygb | 620 | 0 | 0 | 0 | 0 | 0 | 0 | 0 | 0 | 0 | 0 | 0 | 0 | 0 | 0 | 0 | 0 | 0 | 0 |
| ENSMUSG00 | Nbeal1 | 16026 | 6.99136678 | 7.19222601 | 7.83587201 | 6.53024578 | 6.00072279 | 7.60946445 | 6.11705171 | 6.90875918 | 6.07286514 | 5.53416154 | 6.66571865 | 6.90643073 | 6.45507491 | 5.63380169 | 5.36293807 | 6.52512472 | 8.0928609 | 4.75056806 |
| ENSMUSG00 | 9930005F22I | 2899 | 0.02932403 | 0 | 0.05005884 | 0 | 0 | 0 | 0.09792207 | 0.17126638 | 0.14207146 | 0 | 0.09887884 | 0 | 0.17016874 | 0 | 0 | 0.02644105 | 0 | 0 |
| ENSMUSG00 | Gm6644 | 841 | 0.1516237 | 0.31688841 | 0.05751906 | 0.10834969 | 0.05723178 | 0.35352723 | 0.11251529 | 0.2724785 | 0 | 0.16819514 | 0.34084394 | 0.19910462 | 0.23463456 | 0.27612418 | 0.29394689 | 0.5008284 | 0.5012952 | 0.06540916 |
| ENSMUSG00 | Gm10561 | 1940 | 0.02190988 | 0.01717159 | 0 | 0.02348507 | 0.09924108 | 0 | 0 | 0 | 0 | 0 | 0 | 0 | 0 | 0 | 0 | 0 | 0 | 0 |
| ENSMUSG00 | Hspe1 | 1568 | 42.3696366 | 34.3964131 | 37.7918206 | 28.7952774 | 37.9100292 | 34.6363495 | 51.7180687 | 40.481998 | 44.8113494 | 42.1589834 | 39.7007467 | 40.1175075 | 54.5230948 | 67.9777937 | 54.8653372 | 41.4896314 | 45.341378 | 68.2351483 |
| ENSMUSG00 | Pgap1 | 2966 | 0.30094696 | 0.16847384 | 0.30987764 | 0.23041661 | 0.09736734 | 0.35084538 | 1.26018261 | 1.41644109 | 1.27753184 | 0.54050031 | 0.72483922 | 0.92210636 | 0.84825616 | 1.45627112 | 1.29188965 | 0.4905742 | 0.77531282 | 1.65064413 |
| ENSMUSG00 | Gm10562 | 686 | 0 | 0 | 0 | 0 | 0 | 0 | 0 | 0 | 0 | 0 | 0 | 0 | 0 | 0 | 0 | 0 | 0 | 0 |

| | | | | | | | | | | | | | | | | | | | | |
|---|---|---|---|---|---|---|---|---|---|---|---|---|---|---|---|---|---|---|---|---|
| ENSMUSG00 | Tmem88b | 3469 | 0 | 0 | 0 | 0 | 0 | 0 | 0 | 0 | 0 | 0.01359202 | 0.20657953 | 0.06435937 | 0.01422079 | 0 | 0 | 0 | 0.13257872 | 0 |
| ENSMUSG00 | 1500002C15I | 386 | 0.33035111 | 0.34521134 | 0.50128011 | 0.70820276 | 0.62347051 | 0.8986247 | 0.49028683 | 0.29683214 | 0.42680327 | 0.85506457 | 0.49507729 | 0.86760097 | 0.38340868 | 0.96257176 | 0.32021935 | 0.79358772 | 0.19858185 | 1.85263831 |
| ENSMUSG00 | Gm10563 | 250 | 0 | 0.13325158 | 0.19349412 | 0 | 0.38505539 | 0.5946328 | 0.37850143 | 0.61107844 | 0.65898425 | 0.94301407 | 0 | 0 | 0.78931068 | 0.55732905 | 0.82403112 | 0.15316243 | 0.15330519 | 0 |
| ENSMUSG00 | 2610002J02R | 1682 | 62.443692 | 58.20844 | 61.8617502 | 51.8995012 | 58.6053387 | 60.6299202 | 61.4614765 | 57.7881475 | 63.4448646 | 67.5583798 | 72.0884926 | 65.4722368 | 80.8022783 | 60.9682194 | 67.9262274 | 70.594039 | 67.3330603 | 69.2683044 |
| ENSMUSG00 | Gm10564 | 3111 | 0 | 0 | 0.01554919 | 0 | 0.09282917 | 0 | 0 | 0 | 0.06619498 | 0 | 0 | 0 | 0 | 0 | 0 | 0 | 0 | 0 |
| ENSMUSG00 | Gm833 | 881 | 0 | 0 | 0 | 0 | 0 | 0 | 0 | 0 | 0 | 0 | 0 | 0 | 0 | 0 | 0 | 0 | 0 | 0 |
| ENSMUSG00 | Klhl21 | 3982 | 8.4327195 | 10.741777 | 10.2651012 | 12.3799724 | 8.93256946 | 11.4362035 | 11.6796493 | 14.5307608 | 8.89512897 | 13.0250561 | 12.3936545 | 9.74180952 | 10.2330898 | 11.5585361 | 12.4577281 | 12.8853375 | 12.2717299 | 9.32474813 |
| ENSMUSG00 | Rpl31 | 1032 | 40.075132 | 33.2806143 | 40.123781 | 32.9346309 | 41.2759229 | 38.6050366 | 54.6021324 | 41.0050212 | 44.5389066 | 46.5567506 | 40.9235109 | 45.3231834 | 62.2385235 | 64.4006677 | 46.7110663 | 47.1211932 | 48.5763531 | 60.3927499 |
| ENSMUSG00 | Apitd1 | 1851 | 0.94149769 | 0.8638676 | 1.01921539 | 0.76304288 | 1.01412481 | 0.77635293 | 1.96816655 | 1.32053687 | 1.4018101 | 1.52838584 | 1.16146577 | 0.99509285 | 0.74624186 | 1.85675984 | 1.42458108 | 1.84109349 | 1.42869502 | 2.52608 |
| ENSMUSG00 | Gm13241 | 1534 | 0.08312616 | 0.02171636 | 0 | 0.05940162 | 0.03137674 | 0 | 0 | 0 | 0.0536982 | 0.03073709 | 0.03114404 | 0.0727714 | 0.16079504 | 0 | 0.08057671 | 0 | 0 | 0 |
| ENSMUSG00 | Vps13d | 2060 | 0 | 0 | 0 | 0 | 0 | 0 | 0 | 0 | 0 | 0 | 0 | 0 | 0 | 0 | 0 | 0 | 0 | 0 |
| ENSMUSG00 | Gm13125 | 1447 | 0 | 0 | 0 | 0 | 0 | 0 | 0 | 0 | 0 | 0 | 0 | 0 | 0 | 0 | 0 | 0 | 0 | 0 |
| ENSMUSG00 | 4931408C20I | 4316 | 0 | 0 | 0 | 0 | 0 | 0 | 0 | 0 | 0 | 0 | 0 | 0 | 0 | 0 | 0 | 0 | 0 | 0 |
| ENSMUSG00 | Gm13102 | 2289 | 0 | 0 | 0 | 0 | 0 | 0 | 0 | 0 | 0 | 0 | 0 | 0 | 0 | 0 | 0 | 0 | 0 | 0 |
| ENSMUSG00 | Pramel4 | 2527 | 0 | 0 | 0 | 0 | 0 | 0 | 0 | 0 | 0 | 0 | 0 | 0 | 0 | 0 | 0 | 0 | 0 | 0 |
| ENSMUSG00 | Lmbrd1 | 5374 | 51.5614519 | 60.5570643 | 58.0050297 | 65.6370683 | 60.2768601 | 62.7385187 | 50.5525498 | 54.1900016 | 54.4912778 | 63.3209202 | 60.629925 | 53.0425361 | 55.353826 | 58.5347607 | 60.3916367 | 67.1475036 | 69.6349019 | 65.9822668 |
| ENSMUSG00 | Gm13043 | 2109 | 0 | 0 | 0 | 0 | 0 | 0 | 0 | 0 | 0 | 0 | 0 | 0 | 0 | 0 | 0 | 0 | 0 | 0 |
| ENSMUSG00 | Gm10565 | 4264 | 0.39873523 | 0.50000592 | 0.30630519 | 0.76932345 | 0.14674367 | 0.72051348 | 0.14424597 | 0.17913885 | 0.11590952 | 0.11057857 | 0.01120426 | 0.44505922 | 0.41649836 | 0.26141137 | 0.27055431 | 0.51185849 | 0.26965031 | 0.3483222 |
| ENSMUSG00 | Gm6420 | 419 | 0.60866601 | 0.79505714 | 0.92359962 | 0.76116303 | 1.03385994 | 0.70958568 | 0.67750853 | 0.36460527 | 0.29489152 | 1.46290965 | 0.91217104 | 0.66605803 | 1.17737273 | 0.88676062 | 0.88499762 | 0.91385698 | 0.54882525 | 2.62573302 |
| ENSMUSG00 | 4933415F23I | 1780 | 0 | 0 | 0 | 0 | 0 | 0 | 0 | 0 | 0 | 0 | 0 | 0 | 0 | 0 | 0 | 0 | 0 | 0 |
| ENSMUSG00 | AI507597 | 940 | 0 | 0 | 0 | 0 | 0 | 0 | 0 | 0 | 0 | 0 | 0 | 0 | 0 | 0 | 0 | 0 | 0 | 0 |
| ENSMUSG00 | 6330545A04I | 1871 | 0.47707573 | 0.44512152 | 0.28439809 | 0.41396994 | 0.46305432 | 0.26484625 | 0.15172425 | 0.2857796 | 0.22013103 | 0.32761044 | 0.35748232 | 0.35798394 | 0.23729944 | 0.64540154 | 0.24223256 | 0.20465317 | 0.51210979 | 0.7350228 |
| ENSMUSG00 | Defb18 | 256 | 64.7539791 | 45.8052296 | 55.9318945 | 37.3742942 | 51.5161996 | 34.0675043 | 0 | 0 | 0.16088483 | 0 | 0 | 0 | 10.9840597 | 9.79679968 | 1.77037935 | 0.89743611 | 1.04798468 | 3.00831052 |
| ENSMUSG00 | Gm10566 | 998 | 0 | 0 | 0.04847047 | 0 | 0.04822838 | 0 | 0.14222248 | 0.03826894 | 0.04126905 | 0.09449039 | 0.0957414 | 0.05592752 | 0 | 0.18614865 | 0.04128412 | 0 | 0.11520931 | 0.11023869 |
| ENSMUSG00 | Gm10567 | 2638 | 0.0322253 | 0.21467748 | 0.14669759 | 0 | 0.36491223 | 0.18784205 | 0.2869609 | 0.05791115 | 0.12490225 | 0.14298924 | 0.12677207 | 0.29621656 | 0 | 0 | 0 | 0 | 0 | 0 |
| ENSMUSG00 | Gm16287 | 2321 | 0.43951927 | 0.22964511 | 0.50020023 | 0.0981496 | 0.4147516 | 0.51239363 | 0.22423071 | 0.52656479 | 0.55009995 | 0.20314823 | 0.61751347 | 0.43286597 | 1.19025738 | 0.90046701 | 0.86983035 | 0.87436544 | 1.27148852 | 1.06652727 |
| ENSMUSG00 | 4732440D04I | 680 | 0 | 0 | 0.28455018 | 0.20100461 | 0 | 0 | 0.20873241 | 0 | 0 | 0 | 0.07025729 | 0 | 0.07254694 | 0 | 0.18177157 | 0.05630972 | 0 | 0.08089575 |
| ENSMUSG00 | Gm7534 | 2330 | 0 | 0 | 0 | 0 | 0 | 0 | 0 | 0 | 0 | 0 | 0 | 0 | 0 | 0 | 0 | 0 | 0 | 0 |
| ENSMUSG00 | A330069E16I | 885 | 2.35339393 | 1.54330922 | 1.20250584 | 1.80184921 | 1.52281792 | 1.063844 | 1.9780442 | 0.86310514 | 1.1634609 | 1.91799471 | 2.26728617 | 1.9551249 | 1.83949522 | 2.0466886 | 1.90877265 | 1.55758403 | 1.77556856 | 1.61608675 |
| ENSMUSG00 | Gm10570 | 512 | 0 | 0 | 0 | 0 | 0 | 0 | 0 | 0 | 0 | 0 | 0 | 0 | 0 | 0 | 0.24141537 | 0 | 0 | 0 |
| ENSMUSG00 | Gm10571 | 1636 | 0.02598116 | 0.0203624 | 0 | 0.1113962 | 0 | 0.06057791 | 0.17351838 | 0.25679488 | 0 | 0.08646217 | 0 | 0 | 0 | 0.4542209 | 0 | 0.51491037 | 0.25769515 | 0.03362415 |
| ENSMUSG00 | 5730409E04I | 3047 | 2.0087776 | 0.81997606 | 1.42882105 | 0.67287397 | 0.63185984 | 1.04081882 | 1.00929574 | 1.98043964 | 1.75721925 | 0.86657019 | 0.42334226 | 1.37386764 | 1.14951301 | 1.02125168 | 0.44622624 | 0.49009967 | 1.53456128 | 0.86656945 |
| ENSMUSG00 | 1700029G01I | 2891 | 3.01403012 | 3.00749408 | 2.67719296 | 1.48140372 | 2.51398149 | 2.65675327 | 4.30412818 | 2.58931542 | 3.06298889 | 4.43617824 | 3.38770892 | 2.25888361 | 5.98945104 | 4.46608658 | 4.48927364 | 5.21843629 | 5.72707031 | 5.86053437 |
| ENSMUSG00 | 4933407E24I | 966 | 0 | 0 | 0 | 0 | 0 | 0 | 0 | 0 | 0 | 0 | 0 | 0 | 0 | 0 | 0 | 0 | 0 | 0 |
| ENSMUSG00 | 4933427I04R | 2181 | 0.03897769 | 0 | 0 | 0 | 0.04413748 | 0.02272019 | 0 | 0.15760276 | 0 | 0 | 0 | 0 | 0 | 0 | 0 | 0 | 0 | 0 |
| ENSMUSG00 | Gm12888 | 1177 | 0 | 0 | 0 | 0 | 0 | 0 | 0 | 0.03244894 | 0 | 0.36054064 | 0 | 0 | 1.08974499 | 0.43405689 | 0.7001114 | 0 | 0.29306429 | 0.09347342 |

[illegible]

| | | | | | | | | | | | | | | | | | | | | |
|---|---|---|---|---|---|---|---|---|---|---|---|---|---|---|---|---|---|---|---|---|
| ENSMUSG00 | Gm7381 | 299 | 0 | 0 | 0 | 0 | 0 | 0 | 0 | 0.63866894 | 0.13774754 | 0 | 0 | 0 | 0 | 0 | 0 | 0 | 0 | 0 |
| ENSMUSG00 | Bglap2 | 467 | 0 | 0 | 0 | 0 | 0 | 0 | 0 | 0 | 0 | 0 | 0 | 0 | 0 | 0 | 0 | 0 | 0 | 0 |
| ENSMUSG00 | Bglap-rs1 | 2542 | 19.0788379 | 25.4892129 | 21.9031996 | 16.6149048 | 21.0553498 | 16.0042464 | 9.17590508 | 10.8477241 | 5.52502041 | 5.71298844 | 5.35635844 | 4.04015811 | 33.0302609 | 15.9137687 | 14.6361153 | 7.56171712 | 7.29737518 | 16.3382673 |
| ENSMUSG00 | Clec4g | 2451 | 0 | 0 | 0 | 0 | 0 | 0 | 0 | 0 | 0 | 0 | 0 | 0 | 0 | 0 | 0 | 0 | 0 | 0 |
| ENSMUSG00 | A430078G23 | 3294 | 0 | 0 | 0 | 0 | 0 | 0 | 0.34471897 | 0 | 0 | 0.04294235 | 0.31907987 | 0 | 0.29952591 | 0 | 0.07504837 | 0.25573569 | 0 | 0 |
| ENSMUSG00 | Zfp558 | 2418 | 1.74028636 | 1.3225963 | 1.26035253 | 1.07401966 | 0.87584973 | 0.901704 | 1.0566107 | 0.7423668 | 1.0901311 | 1.50149055 | 2.09435302 | 1.36192063 | 2.34622435 | 0.97958993 | 1.12460823 | 1.28268784 | 2.04470318 | 1.02374268 |
| ENSMUSG00 | Gm5612 | 377 | 0.11274583 | 0 | 0 | 0 | 0.12767089 | 0.13143961 | 0 | 0 | 0 | 0.25013636 | 0 | 0 | 0 | 0 | 0 | 0 | 0 | 0 |
| ENSMUSG00 | Ubtfl1 | 2869 | 0 | 0 | 0 | 0 | 0 | 0 | 0 | 0 | 0 | 0 | 0 | 0 | 0 | 0 | 0 | 0 | 0 | 0 |
| ENSMUSG00 | Fat3 | 19888 | 0 | 0 | 0 | 0 | 0.00242015 | 0 | 0 | 0.00960187 | 0 | 0 | 0 | 0.03087149 | 0 | 0 | 0 | 0 | 0 | 0 |
| ENSMUSG00 | Gm10705 | 464 | 0.82445385 | 0.50256521 | 0.62551979 | 0.09819191 | 0.20746519 | 1.28153621 | 0.71376886 | 1.23466818 | 1.33146064 | 0.40647158 | 0.51481636 | 1.20292376 | 0.63791272 | 0.40038006 | 1.06555748 | 0.66018289 | 0.33039911 | 1.42264931 |
| ENSMUSG00 | Gm14340 | 1911 | 0 | 0 | 0 | 0 | 0 | 0 | 0 | 0 | 0 | 0 | 0 | 0 | 0 | 0 | 0 | 0 | 0 | 0 |
| ENSMUSG00 | Gm10706 | 1955 | 0.76096223 | 1.75510389 | 0.44538289 | 0.79236599 | 0.19695928 | 1.06456001 | 0 | 0.03907151 | 0 | 0.07235402 | 0.09774928 | 0.28550211 | 0.20186974 | 0.33259193 | 0 | 0.01958599 | 0.27445942 | 0 |
| ENSMUSG00 | Arfip1 | 5157 | 16.863601 | 19.3921482 | 18.2631886 | 18.6944289 | 18.0973046 | 18.3336466 | 15.6883229 | 17.5742817 | 20.0302078 | 17.4540804 | 16.1380604 | 15.1093009 | 16.1091621 | 17.6247972 | 19.9176806 | 18.555018 | 21.522776 | 19.6163947 |
| ENSMUSG00 | Gm10709 | 482 | 0.08818501 | 0 | 0 | 0 | 0.09985876 | 0 | 0.09815908 | 0 | 0 | 0 | 0 | 0 | 0 | 0 | 0 | 0.07944109 | 0.15903028 | 0 |
| ENSMUSG00 | Gm10710 | 3970 | 0 | 0 | 0 | 0 | 0 | 0 | 0 | 0 | 0 | 0 | 0 | 0 | 0 | 0 | 0 | 0 | 0 | 0 |
| ENSMUSG00 | Gm10711 | 67 | 0 | 0 | 0 | 0 | 0 | 0 | 0 | 0 | 0 | 0 | 0 | 0 | 0 | 0 | 0 | 0 | 0 | 0 |
| ENSMUSG00 | Etohi1 | 2043 | 0.81140571 | 0.94574051 | 1.04181857 | 1.15965458 | 0.65966415 | 0.75190149 | 0.57896083 | 0.99079654 | 1.24990895 | 1.43090729 | 0.74831066 | 0.6556906 | 0.94172529 | 1.02299752 | 1.16969664 | 1.01208654 | 0.69411306 | 1.07702607 |
| ENSMUSG00 | Gm14327 | 1448 | 0.14677202 | 0.06901843 | 0.06681427 | 0.09439443 | 0.03324028 | 0 | 0.39209402 | 0.05275194 | 0.19910608 | 0.29306376 | 0.09898127 | 0.03854673 | 0.57917306 | 1.25091119 | 0.73980694 | 0.21155032 | 0.44996343 | 0.56984572 |
| ENSMUSG00 | Gm14296 | 9862 | 0.00431 | 0.00675581 | 0 | 0.01847943 | 0.01464163 | 0 | 0.01918989 | 0.00774537 | 0.02505771 | 0.0095621 | 0.00484435 | 0 | 0.00500222 | 0.0094188 | 0.02924467 | 0.00388264 | 0.00388626 | 0 |
| ENSMUSG00 | C330013J21 | 859 | 0 | 0.19390509 | 0.16894132 | 0.21215853 | 0.33619504 | 0.34611921 | 0.16523636 | 0.17784588 | 0.1917882 | 0.4391218 | 0.22246779 | 0.12995498 | 0.40200631 | 0.75694671 | 0.04796456 | 0 | 0.71387747 | 0.64038541 |
| ENSMUSG00 | Gm10712 | 611 | 0 | 0.05452192 | 0 | 0 | 0 | 0 | 0.23230448 | 0.06250802 | 0 | 0 | 0 | 0 | 0 | 0.07601324 | 0 | 0.18800626 | 0.06272716 | 0 |
| ENSMUSG00 | Gm10713 | 335 | 0.12688112 | 0.89497328 | 1.15518878 | 0.8160187 | 1.43677384 | 1.77502329 | 0.84739126 | 3.30620797 | 0.98355859 | 1.26673531 | 0.99828271 | 2.83243661 | 0.44177836 | 2.21822507 | 1.35288691 | 0.34290096 | 0.80084799 | 0.82103144 |
| ENSMUSG00 | Gm10714 | 1003 | 0 | 0 | 0 | 0 | 0 | 0 | 0 | 0 | 0 | 0 | 0 | 0 | 0 | 0 | 0 | 0 | 0 | 0 |
| ENSMUSG00 | Gm10715 | 671 | 0 | 0 | 0 | 0 | 0 | 0 | 0 | 0 | 0 | 0 | 0 | 0 | 0 | 0 | 0 | 0 | 0 | 0 |
| ENSMUSG00 | Gm10717 | 681 | 0 | 0.04891761 | 0 | 0 | 0 | 0 | 0.2779012 | 0 | 0 | 0.06923745 | 0.21046237 | 0 | 0.07244041 | 0.2045995 | 0 | 0 | 0 | 0 |
| ENSMUSG00 | Gm10718 | 671 | 0 | 0.04964664 | 0 | 0 | 0.07173163 | 0 | 0 | 0 | 0.0613808 | 0 | 0 | 0 | 0 | 0.41529735 | 0 | 0 | 0 | 0 |
| ENSMUSG00 | Gm10719 | 671 | 0 | 0 | 0 | 0 | 0 | 0 | 0 | 0 | 0 | 0 | 0 | 0 | 0 | 0.06921623 | 0 | 0 | 0 | 0 |
| ENSMUSG00 | Gm10720 | 673 | 0.06315777 | 0.0989982 | 0.14375492 | 0.06769843 | 0 | 0.29451848 | 0 | 0 | 0 | 0 | 0 | 0 | 0 | 0 | 0.06122074 | 0 | 0 | 0 |
| ENSMUSG00 | Gm10721 | 641 | 0 | 0 | 0 | 0 | 0 | 0 | 0 | 0 | 0 | 0 | 0 | 0 | 0 | 0 | 0 | 0 | 0 | 0 |
| ENSMUSG00 | Gcnt7 | 3189 | 0.75973504 | 0.94015694 | 1.03148324 | 1.04294645 | 0.96595896 | 1.0410891 | 0.57861225 | 1.02996131 | 1.0648753 | 1.28633151 | 0.74905862 | 1.03265102 | 0.58783721 | 0.37865985 | 0.762274 | 0.80447498 | 0.66100543 | 0.62098709 |
| ENSMUSG00 | Cass4 | 5545 | 0 | 0 | 0 | 0 | 0 | 0 | 0 | 0 | 0.08170454 | 0 | 0.05169518 | 0 | 0 | 0 | 0 | 0 | 0 | 0 |
| ENSMUSG00 | 1700101G07 | 1152 | 0 | 0 | 0 | 0 | 0 | 0 | 0 | 0 | 0 | 0 | 0 | 0 | 0 | 0 | 0 | 0 | 0 | 0 |
| ENSMUSG00 | Kcng1 | 5893 | 0.00721282 | 0 | 0 | 0 | 0.04900586 | 0 | 0.04817174 | 0 | 0 | 0 | 0 | 0 | 0 | 0 | 0 | 0 | 0 | 0 |
| ENSMUSG00 | Mocs3 | 1972 | 1.33636963 | 2.17919034 | 2.33036784 | 2.17177392 | 1.61090617 | 1.68358679 | 1.51151053 | 2.16914253 | 1.27402508 | 2.31927901 | 1.69586567 | 2.34923935 | 1.17576071 | 2.44938392 | 2.94595303 | 1.88346294 | 1.94352418 | 1.95265592 |
| ENSMUSG00 | Fam65c | 4976 | 0 | 0 | 0 | 0.1098739 | 0 | 0 | 0 | 0 | 0 | 0.01895125 | 0.10561185 | 0 | 0 | 0 | 0 | 0 | 0.02310669 | 0 |
| ENSMUSG00 | 1500012F01 | 1233 | 11.5139731 | 8.96989528 | 8.04263887 | 10.2355306 | 10.422728 | 7.79661826 | 10.8209047 | 8.70402683 | 10.5889096 | 8.33645848 | 8.21434809 | 10.1853399 | 18.4444557 | 17.5154101 | 15.3712212 | 11.1486846 | 11.563165 | 13.473439 |
| ENSMUSG00 | Lekr1 | 3371 | 2.0804966 | 2.26302367 | 1.9372372 | 2.20304071 | 2.47013432 | 2.11675872 | 3.29827338 | 3.54619459 | 3.66536776 | 3.63666059 | 2.91950206 | 4.07316909 | 3.11708644 | 2.59017752 | 3.49559329 | 3.40764824 | 4.19531402 | 3.06784695 |
| ENSMUSG00 | 4931440P22 | 3378 | 1.57286767 | 1.31160892 | 1.67545975 | 1.7938481 | 1.1968862 | 2.31774183 | 2.01688152 | 2.18209996 | 2.20685594 | 2.48455452 | 2.84273732 | 2.89157518 | 2.45345237 | 2.08984644 | 2.8419072 | 2.1310344 | 2.87050122 | 2.47524696 |
| ENSMUSG00 | Arfgef2 | 8731 | 7.35600973 | 7.825535 | 6.37149925 | 8.76153859 | 7.10595002 | 7.16814827 | 5.03960502 | 6.19844625 | 6.34001571 | 7.64153535 | 6.42945553 | 6.46952818 | 7.18140727 | 6.97379436 | 7.52207996 | 7.73179943 | 6.83473191 | 6.14922554 |
| ENSMUSG00 | 4930449A18 | 475 | 0 | 0 | 0 | 0 | 0 | 0 | 0 | 0 | 0 | 0 | 0 | 0 | 0 | 0 | 0 | 0 | 0 | 0 |
| ENSMUSG00 | F630111L10 | 2976 | 0.04284796 | 0.12313234 | 0.03250909 | 0 | 0.16173361 | 0 | 0 | 0 | 0 | 0.01584365 | 0 | 0.20630789 | 0 | 0 | 0 | 0 | 0.07727076 | 0 |
| ENSMUSG00 | Gm410 | 2692 | 0 | 0 | 0 | 0 | 0 | 0 | 0 | 0 | 0 | 0 | 0.01774701 | 0.0207339 | 0 | 0 | 0 | 0 | 0.04271133 | 0 |
| ENSMUSG00 | F830104G03 | 522 | 0 | 0 | 0 | 0 | 0 | 0 | 0 | 0 | 0 | 0 | 0 | 0 | 0 | 0 | 0 | 0 | 0 | 0 |
| ENSMUSG00 | Gm14316 | 439 | 9.68227244 | 19.5020816 | 87.3808874 | 33.0032166 | 26.6424999 | 47.7467114 | 403.504943 | 528.951724 | 345.535165 | 145.640897 | 279.358356 | 517.470949 | 0.8989871 | 0.63477113 | 16.5181636 | 0.52333404 | 0.6984291 | 1.12774934 |
| ENSMUSG00 | Wfdc9 | 572 | 10.1804355 | 5.4744966 | 15.1378705 | 9.16000017 | 10.8549268 | 8.3165427 | 26.0550592 | 36.4563841 | 19.4411875 | 8.65528644 | 7.85112954 | 17.9546887 | 89.1769274 | 221.90855 | 108.838376 | 33.7385772 | 45.5627306 | 102.132293 |
| ENSMUSG00 | Wfdc6a | 862 | 2.66273725 | 2.85980762 | 2.46918252 | 1.95563647 | 2.28933744 | 3.10423157 | 42.921711 | 22.9508868 | 20.4976975 | 17.120847 | 9.47739896 | 15.6698264 | 80.1214434 | 76.2394242 | 42.1575085 | 19.234145 | 18.096059 | 43.586102 |
| ENSMUSG00 | Spint3 | 555 | 3.06343611 | 2.04078992 | 2.70194494 | 2.05229929 | 1.38758699 | 1.8749683 | 220.110517 | 156.554442 | 143.225181 | 69.239321 | 38.9086148 | 117.665451 | 492.963627 | 607.287824 | 260.497765 | 68.3711566 | 88.0468983 | 305.771341 |
| ENSMUSG00 | Ufm1 | 1607 | 1.37540084 | 1.43036073 | 1.08366341 | 1.21911942 | 0.8086882 | 0.18501332 | 1.47208086 | 0.95065096 | 0.46132996 | 0.76286141 | 1.2486299 | 0.27786266 | 0.33767958 | 0.2312089 | 0.05127761 | 0 | 0.28619512 | 0.17115466 |
| ENSMUSG00 | Gm10728 | 1827 | 0.11632506 | 0.01823366 | 0.02647703 | 0.02493763 | 0.02634479 | 0 | 0 | 0.14633104 | 0.0450865 | 0.07742316 | 0.05229881 | 0.274954 | 0.13500798 | 0 | 0.02255148 | 0.25149824 | 0.25173266 | 0.1505449 |
| ENSMUSG00 | Gm10729 | 1319 | 0.06445061 | 0 | 0 | 0 | 0 | 0 | 0 | 0 | 0 | 0 | 0 | 0 | 0 | 0 | 0 | 0 | 0 | 0 |
| ENSMUSG00 | Mgst2 | 3166 | 0 | 0.02104415 | 0 | 0 | 0.1368248 | 0 | 0.1942719 | 0.0120633 | 0.02601801 | 0.01489283 | 0.21126008 | 0 | 0.34279917 | 0.19070535 | 0.01301376 | 0.29026361 | 0 | 0 |
| ENSMUSG00 | Tox2 | 2322 | 18.8911893 | 22.2803293 | 26.1242063 | 21.7994489 | 24.5634494 | 21.0417723 | 43.0745061 | 44.4920106 | 32.9740451 | 46.6024393 | 35.8209314 | 32.8355707 | 44.9553561 | 63.6255995 | 60.3828143 | 51.0706336 | 56.367056 | 58.349453 |
| ENSMUSG00 | Zfp850 | 940 | 0 | 0.0708785 | 0 | 0 | 0 | 0.10543135 | 1.00665274 | 0.32504172 | 0.43815442 | 0.15048097 | 0.71154193 | 0.47502692 | 0 | 0 | 0 | 0.12220407 | 0 | 0 |
| ENSMUSG00 | 1700034I23R | 2344 | 0.07253443 | 0.01421199 | 0 | 0 | 0.0205341 | 0 | 0 | 0.08146844 | 0.10542623 | 0.04023098 | 0 | 0 | 0 | 0.13869821 | 0 | 0.13068467 | 0 | 0 |
| ENSMUSG00 | Gm10731 | 1805 | 0.02354857 | 0.07382359 | 0.02679974 | 0 | 0 | 0 | 0 | 0 | 0 | 0.02612227 | 0 | 0 | 0 | 0 | 0 | 0.0636409 | 0.02123341 | 0 |
| ENSMUSG00 | Mafb | 3388 | 1.68113742 | 2.88095573 | 1.15651002 | 2.00371771 | 1.32121278 | 2.01838171 | 0.58652081 | 0.50727807 | 1.30075478 | 1.1690257 | 1.17040188 | 0.42833743 | 2.65006167 | 2.41267986 | 2.11601851 | 1.74048215 | 1.50454471 | 1.93213805 |
| ENSMUSG00 | Gm826 | 1811 | 0 | 0 | 0 | 0.0503159 | 0 | 0 | 0 | 0 | 0 | 0 | 0 | 0.06164071 | 0 | 0 | 0 | 0 | 0 | 0 |
| ENSMUSG00 | Arhgap40 | 2191 | 0.19399898 | 1.30758051 | 0.88313154 | 0.83178538 | 0.48329636 | 0.24878141 | 0.38869385 | 0.48808182 | 0.13158631 | 0.6025649 | 0.50151714 | 0.43307452 | 0.02251571 | 0.84790666 | 0.13163436 | 0.10485789 | 0.15743344 | 0.50213699 |
| ENSMUSG00 | 4922505G16 | 4818 | 0.13233243 | 0.07605684 | 0 | 0.15130276 | 0.03996009 | 0.02056984 | 0 | 0.01585405 | 0.05983927 | 0.03914546 | 0.06941152 | 0.0579241 | 0 | 0.15423524 | 0.00855159 | 0.15894814 | 0 | 0 |
| ENSMUSG00 | Gm1332 | 635 | 0 | 0 | 0 | 0 | 0 | 0 | 0 | 0 | 0 | 0 | 0 | 0 | 0 | 0.0731403 | 0 | 0 | 0 | 0 |
| ENSMUSG00 | 4930518I15R | 1330 | 2.65257865 | 1.87854667 | 1.01839011 | 1.88410334 | 1.15806132 | 0.93144236 | 1.20949706 | 1.34965633 | 1.98190753 | 1.31171129 | 1.65236698 | 0.71343328 | 2.07713336 | 1.81585906 | 1.11523008 | 1.29554687 | 1.58492205 | 0.99264553 |
| ENSMUSG00 | Gm7120 | 1445 | 1.91199754 | 2.02874373 | 2.14249546 | 2.27016968 | 1.43229946 | 1.47457962 | 1.40792055 | 1.7708588 | 1.39663617 | 1.89255418 | 2.41354462 | 2.12447159 | 1.29730994 | 1.99275669 | 1.42565937 | 2.75586379 | 1.80359044 | 0.91364606 |
| ENSMUSG00 | 3110070M22 | 1122 | 1.36380244 | 1.48453183 | 1.59520555 | 1.25881673 | 1.67303478 | 2.11990304 | 1.77106285 | 1.39562255 | 2.75310934 | 1.80702339 | 1.74578725 | 1.98986319 | 0.65951761 | 1.61436667 | 2.53378552 | 1.46746535 | 1.94705786 | 1.66694263 |
| ENSMUSG00 | Sox2 | 4828 | 0 | 0.01379987 | 0.02003874 | 0 | 0.06978531 | 0 | 0.00979964 | 0 | 0 | 0 | 0 | 0 | 0 | 0 | 0 | 0 | 0 | 0 |
| ENSMUSG00 | BC089597 | 1627 | 0 | 0 | 0 | 0 | 0 | 0 | 0 | 0 | 0 | 0 | 0 | 0 | 0 | 0 | 0 | 0.07060346 | 0 | 0 |
| ENSMUSG00 | Cpne1 | 7739 | 8.1396396 | 8.94915455 | 10.5322908 | 9.21346872 | 9.74579718 | 9.75815557 | 8.79738273 | 7.99972662 | 7.91372903 | 7.03696372 | 8.76604743 | 7.79645061 | 6.82704269 | 6.97951591 | 7.8740278 | 7.80754341 | 8.97367231 | 7.34260334 |
| ENSMUSG00 | 6430550D23 | 4905 | 0.45928121 | 0.57049605 | 0.39448343 | 0.49230078 | 0.17663091 | 0.53543219 | 0.57874837 | 0.73971014 | 0.33587373 | 0.55754145 | 1.01296548 | 1.09241664 | 0.34195417 | 0.29353042 | 0.7139923 | 0.43716086 | 0.85169549 | 0.82990294 |
| ENSMUSG00 | Fam83c | 3968 | 0.08569592 | 0.04197693 | 0.03657273 | 0.06889271 | 0 | 0.01248809 | 0.02384712 | 0 | 0.031139 | 0.04753095 | 0.03612018 | 0.07033224 | 0.23621634 | 0.15216057 | 0 | 0 | 0.2801065 | 0 |
| ENSMUSG00 | BC029722 | 1613 | 22.1617192 | 22.2430174 | 24.0218213 | 20.4502146 | 23.8421618 | 21.4431544 | 19.4178509 | 18.9422951 | 19.9421381 | 18.7082766 | 18.3043548 | 19.031999 | 26.1798643 | 24.0714551 | 19.7706164 | 19.9880294 | 18.0107458 | 23.360966 |
| ENSMUSG00 | Gm10735 | 2942 | 0 | 0 | 0 | 0 | 0 | 0.03368643 | 0 | 0 | 0 | 0 | 0 | 0.03794403 | 0 | 0 | 0 | 0 | 0 | 0 |
| ENSMUSG00 | Gm6320 | 2347 | 0 | 0 | 0 | 0.13588722 | 0 | 0.04222645 | 0.08063516 | 0.0813643 | 0.10529148 | 0.0803591 | 0 | 0.11890853 | 0.16815311 | 0.03957741 | 0 | 0.03262941 | 0.01632991 | 0.11719026 |
| ENSMUSG00 | Myh7b | 6592 | 0.01289599 | 0 | 0.05136752 | 0.05529253 | 0 | 0.0225513 | 0.05024101 | 0 | 0 | 0.15020703 | 0.04348449 | 0.11007336 | 0.02245081 | 0.04931866 | 0.03125118 | 0.17425944 | 0.09302499 | 0.01668966 |
| ENSMUSG00 | Lrrc31 | 4361 | 0.13645321 | 0.02291646 | 0 | 0 | 0.0331107 | 0.0568135 | 0.04339617 | 0.07006173 | 0.01888856 | 0.07568331 | 0.02191009 | 0 | 0 | 0 | 0.22674555 | 0.04390118 | 0.25486416 | 0.07568325 |
| ENSMUSG00 | Gm1527 | 2105 | 0 | 0 | 0 | 0 | 0 | 0 | 0 | 0 | 0 | 0 | 0 | 0 | 0 | 0 | 0 | 0 | 0 | 0 |
| ENSMUSG00 | Eif2s2 | 7784 | 12.575722 | 11.0543687 | 11.0431351 | 10.5883773 | 12.0267975 | 10.9303756 | 10.3876372 | 10.1074446 | 10.778126 | 9.24963051 | 10.6241589 | 10.4044869 | 10.2669201 | 11.0919268 | 9.76578502 | 8.54947015 | 9.6652776 | 10.2258707 |
| ENSMUSG00 | Kif5a | 6690 | 0.33038403 | 0.44317602 | 0.34707466 | 0.36094699 | 0.25181126 | 0.22220957 | 0.21216448 | 0.34824164 | 0.17853647 | 0.60612264 | 0.33563872 | 0.10011778 | 0.26546323 | 0.18050019 | 0.27098183 | 0.37203131 | 0.27498688 | 0.39468417 |
| ENSMUSG00 | Gm10736 | 199 | 0.21359385 | 0 | 0 | 0 | 0.96747585 | 0 | 0.23775216 | 0 | 0 | 0.23693821 | 0 | 0.28048072 | 0 | 1.16693687 | 0.20704299 | 0 | 0.38518891 | 0 |
| ENSMUSG00 | Pde4d | 1854 | 0.66485982 | 0.66482043 | 0.70446889 | 0.61436144 | 0.46730023 | 0.42763956 | 0.51038489 | 0.22660001 | 1.17739231 | 0.50863758 | 0.64421465 | 0.15052768 | 0.07982511 | 0.25050748 | 0.02222306 | 0.04130594 | 0.06201666 | 0 |
| ENSMUSG00 | Gm10737 | 391 | 0 | 0.85199218 | 0.12371747 | 0.34957323 | 0.61549775 | 1.26733334 | 0 | 0 | 0.31600907 | 0 | 0.85530616 | 0.57100422 | 0 | 0.95026266 | 0 | 0 | 0 | 0.42206476 |
| ENSMUSG00 | A830092H15 | 1228 | 0.41536003 | 0.32553317 | 0 | 0.4081201 | 0.11758613 | 0 | 0 | 0 | 0 | 0 | 0 | 0 | 0.04017257 | 0 | 0.33551756 | 0.12472511 | 0 | 0 |
| ENSMUSG00 | Bpifb4 | 3166 | 0 | 0 | 0 | 0 | 0 | 0 | 0 | 0 | 0 | 0 | 0 | 0 | 0 | 0 | 0 | 0 | 0 | 0 |
| ENSMUSG00 | Tspyl3 | 4029 | 3.27044045 | 3.68765223 | 5.37883882 | 4.1614456 | 4.92190433 | 4.77201803 | 2.88878606 | 3.6211213 | 4.20145395 | 5.39500477 | 4.00792652 | 4.84871728 | 2.57128385 | 2.76658748 | 4.75520563 | 4.98946857 | 4.76484173 | 3.16756336 |
| ENSMUSG00 | Ttll9 | 3524 | 0.15680116 | 0.03781259 | 0.09608817 | 0.05171515 | 0.06829161 | 0.08436901 | 0.09398092 | 0.45518754 | 0.47918477 | 0.1471787 | 0.10845621 | 0.41180682 | 0.15398726 | 0.01317937 | 0 | 0.05432833 | 0 | 0 |
| ENSMUSG00 | Gm10742 | 3384 | 0 | 0 | 0 | 0 | 0 | 0 | 0 | 0 | 0 | 0 | 0 | 0 | 0 | 0 | 0 | 0 | 0 | 0 |
| ENSMUSG00 | Foxs1 | 1310 | 0 | 0.07628907 | 0.55389539 | 0.06955885 | 0 | 0 | 0 | 0 | 0 | 0.50390065 | 0 | 0 | 0 | 0 | 0 | 0 | 0 | 0 |
| ENSMUSG00 | Sirpb1c | 2841 | 0 | 0 | 0 | 0 | 0 | 0 | 0 | 0 | 0 | 0 | 0 | 0 | 0 | 0 | 0 | 0.05391145 | 0 | 0 |
| ENSMUSG00 | Defb25 | 291 | 102.976457 | 187.857249 | 63.334416 | 92.0614916 | 58.0560316 | 148.828485 | 0.32517305 | 0.26249074 | 0.14153442 | 0 | 0.32835023 | 0.1918064 | 0.33905098 | 0 | 0.1415861 | 0.39474853 | 0.13170549 | 0.37806946 |
| ENSMUSG00 | Defb28 | 394 | 32.903753 | 16.6564471 | 37.9376164 | 14.454646 | 10.8724396 | 23.3929148 | 835.537107 | 541.188281 | 512.009021 | 311.864804 | 166.242305 | 526.99052 | 1268.13114 | 2157.88189 | 921.806384 | 319.833475 | 426.453009 | 1082.31116 |
| ENSMUSG00 | Defb26 | 559 | 0.30415153 | 0.05959373 | 0 | 0.08150455 | 0 | 0 | 32.3317412 | 30.6769028 | 40.0076531 | 15.9418299 | 7.00813344 | 31.152928 | 8.20727783 | 12.711888 | 1.91635144 | 0.34249202 | 0.34281124 | 5.90437639 |
| ENSMUSG00 | Defb23 | 622 | 36.8332635 | 37.2761646 | 118.989552 | 32.8889211 | 23.8338143 | 54.4920736 | 444.298003 | 559.86869 | 334.193481 | 132.734536 | 214.06561 | 515.980804 | 1.03105293 | 3.58410964 | 8.213815 | 0.61560462 | 0.86264977 | 0.61907355 |
| ENSMUSG00 | Zcchc3 | 3088 | 8.30007161 | 8.94313125 | 9.03870693 | 8.27711978 | 6.18794485 | 8.16785667 | 9.59123608 | 8.52155611 | 8.66944147 | 8.68806676 | 8.19971761 | 10.7546371 | 8.96217798 | 10.4228473 | 9.16627879 | 8.71706509 | 10.4007244 | 11.1336437 |
| ENSMUSG00 | Gm9049 | 2821 | 0 | 0 | 0 | 0 | 0 | 0 | 0 | 0 | 0 | 0 | 0 | 0 | 0 | 0 | 0 | 0 | 0 | 0 |
| ENSMUSG00 | Gm9048 | 1244 | 0 | 0 | 0 | 0 | 0 | 0 | 0 | 0 | 0 | 0 | 0 | 0 | 0 | 0 | 0 | 0 | 0 | 0 |
| ENSMUSG00 | Gm9046 | 1244 | 0 | 0 | 0 | 0 | 0 | 0 | 0 | 0 | 0 | 0 | 0 | 0 | 0 | 0 | 0 | 0 | 0 | 0 |
| ENSMUSG00 | Gm9045 | 1244 | 0 | 0 | 0 | 0 | 0 | 0 | 0 | 0 | 0 | 0 | 0 | 0 | 0 | 0 | 0 | 0 | 0 | 0 |
| ENSMUSG00 | Gm9044 | 1244 | 0 | 0 | 0 | 0 | 0 | 0 | 0 | 0 | 0 | 0 | 0 | 0 | 0 | 0 | 0 | 0 | 0 | 0 |
| ENSMUSG00 | Gm9040 | 1244 | 0 | 0 | 0 | 0 | 0 | 0 | 0 | 0 | 0 | 0 | 0 | 0 | 0 | 0 | 0 | 0 | 0 | 0 |
| ENSMUSG00 | Il22 | 1100 | 0 | 0 | 0 | 0 | 0 | 0 | 0 | 0 | 0 | 0 | 0 | 0 | 0 | 0 | 0 | 0 | 0 | 0 |
| ENSMUSG00 | Csnk2a1 | 7150 | 4.63692829 | 3.62481562 | 3.96459983 | 3.81693225 | 3.40625921 | 3.61076562 | 4.81067379 | 3.92073054 | 4.11865158 | 3.68632771 | 4.08926917 | 3.53629303 | 4.55371544 | 3.67005726 | 3.19240027 | 3.16499986 | 3.7843868 | 3.34670789 |
| ENSMUSG00 | Rad21l | 1637 | 0 | 0 | 0 | 0 | 0 | 0 | 0 | 0 | 0 | 0 | 0 | 0 | 0 | 0 | 0 | 0 | 0 | 0 |
| ENSMUSG00 | Gm10748 | 2371 | 0 | 0 | 0 | 0 | 0 | 0 | 0 | 0 | 0 | 0 | 0 | 0 | 0 | 0 | 0 | 0 | 0 | 0 |
| ENSMUSG00 | Zfp345 | 4224 | 0 | 0 | 0 | 0 | 0 | 0 | 0 | 0.0723341 | 0 | 0 | 0.12441395 | 0.02642787 | 0 | 0 | 0 | 0.00906501 | 0 | 0 |
| ENSMUSG00 | Zfp950 | 1387 | 1.93066048 | 1.41705895 | 2.51109891 | 1.57673405 | 1.49219374 | 2.03641371 | 3.07003684 | 2.28548623 | 2.96946761 | 1.46177379 | 2.54891631 | 4.18516864 | 1.35155938 | 1.84169056 | 2.34673606 | 2.15332904 | 2.21060112 | 1.78472228 |
| ENSMUSG00 | 5830428H23 | 4686 | 2.61235397 | 2.218016 | 2.033629 | 2.22652137 | 1.86940036 | 1.90343407 | 3.94777155 | 2.47769747 | 3.76180724 | 1.74073232 | 2.1104175 | 2.82294324 | 2.68451534 | 2.5471896 | 2.83117819 | 2.84360465 | 2.06108127 | 2.93475815 |
| ENSMUSG00 | 4933416C03 | 2253 | 0 | 0 | 0 | 0 | 0 | 0 | 0 | 0 | 0 | 0 | 0 | 0 | 0 | 0 | 0 | 0 | 0 | 0 |
| ENSMUSG00 | Gm10749 | 988 | 0 | 0 | 0 | 0 | 0 | 0 | 0 | 0 | 0 | 0 | 0 | 0 | 0 | 0 | 0 | 0 | 0 | 0 |
| ENSMUSG00 | Tmem90b | 2456 | 0 | 0 | 0 | 0 | 0 | 0 | 0 | 0 | 0 | 0 | 0 | 0 | 0 | 0 | 0 | 0 | 0 | 0 |
| ENSMUSG00 | C530025M09 | 689 | 0 | 0 | 0 | 0 | 0 | 0 | 0 | 0 | 0 | 0 | 0 | 0 | 0 | 0 | 0 | 0 | 0 | 0 |
| ENSMUSG00 | B930041F14 | 2139 | 0 | 0 | 0 | 0 | 0 | 0 | 0 | 0 | 0 | 0 | 0 | 0 | 0 | 0 | 0 | 0 | 0 | 0 |
| ENSMUSG00 | Gm10750 | 1505 | 0 | 0 | 0 | 0 | 0 | 0 | 0 | 0 | 0 | 0 | 0 | 0 | 0 | 0 | 0 | 0 | 0 | 0 |
| ENSMUSG00 | Thbd | 3717 | 0 | 0 | 0.07808479 | 0.01225748 | 0 | 0 | 0.03818618 | 0 | 0 | 0 | 0.17994335 | 0 | 0 | 0 | 0 | 0 | 0 | 0 |
| ENSMUSG00 | Gm807 | 1721 | 0 | 0.29035062 | 0.22486243 | 0.4235774 | 0.05593483 | 0 | 0 | 0 | 0 | 0 | 0 | 0.38918533 | 0 | 0.26986686 | 0 | 0 | 0 | 0 |
| ENSMUSG00 | Pdzd8 | 7071 | 6.33580194 | 6.7275934 | 7.00530266 | 6.48202667 | 6.32365395 | 8.24827711 | 5.51345601 | 6.96764236 | 6.06934655 | 5.52125333 | 6.4389104 | 6.01492503 | 7.05339674 | 4.56493293 | 6.35707785 | 4.88989089 | 6.00017122 | 5.36010105 |
| ENSMUSG00 | Atxn7l3b | 4044 | 60.5835397 | 55.043709 | 53.5649528 | 54.2924512 | 59.1532296 | 50.5697654 | 49.8280662 | 55.6925314 | 56.0458448 | 53.0153926 | 51.2482121 | 58.3966539 | 53.2477297 | 53.1511465 | 56.3719606 | 49.24566 | 48.9977634 | 50.3025907 |
| ENSMUSG00 | Plk1s1 | 2398 | 16.6262949 | 15.3644958 | 13.777782 | 14.9146872 | 14.2308315 | 18.4117955 | 13.8899608 | 15.1145079 | 13.0189237 | 18.7383737 | 10.6587168 | 13.6862425 | 16.4988314 | 9.27719677 | 11.9584166 | 13.3490191 | 11.2677395 | 8.12061042 |
| ENSMUSG00 | Gm14092 | 614 | 0 | 0 | 0 | 0 | 0 | 0 | 0 | 0 | 0 | 0 | 0 | 0 | 0 | 0 | 0 | 0 | 0 | 0 |
| ENSMUSG00 | Gm561 | 469 | 28.0044763 | 20.8827098 | 26.8168825 | 22.6348045 | 24.3225286 | 29.2667531 | 38.6370065 | 31.5148395 | 33.5463732 | 30.2608991 | 28.1149009 | 29.7524854 | 32.39708 | 40.7004689 | 30.3960306 | 25.3093567 | 28.4382757 | 27.9150691 |
| ENSMUSG00 | Gm5535 | 1827 | 0 | 0.03646732 | 0 | 0 | 0 | 0 | 0 | 0 | 0 | 0 | 0 | 0 | 0 | 0 | 0 | 0 | 0 | 0 |
| ENSMUSG00 | Gm14105 | 3564 | 0 | 0 | 0 | 0 | 0 | 0 | 0 | 0 | 0 | 0 | 0 | 0 | 0 | 0 | 0 | 0 | 0 | 0 |
| ENSMUSG00 | Sel1l2 | 2716 | 0 | 0 | 0 | 0 | 0 | 0 | 0 | 0 | 0 | 0 | 0 | 0 | 0 | 0 | 0 | 0 | 0 | 0 |
| ENSMUSG00 | Ism1 | 2938 | 3.29856369 | 3.70773193 | 4.01740688 | 4.10948834 | 4.3741401 | 4.36833152 | 7.3754959 | 3.2368646 | 6.33638704 | 3.53068574 | 4.53683234 | 3.59059232 | 3.66043498 | 9.27933264 | 7.48864222 | 5.2913501 | 5.12669662 | 11.1216506 |
| ENSMUSG00 | Bhmt | 5199 | 0 | 0 | 0 | 0 | 0 | 0 | 0 | 0 | 0 | 0 | 0 | 0 | 0 | 0 | 0.0316996 | 0 | 0 | 0 |
| ENSMUSG00 | Ankrd5 | 7593 | 0 | 0 | 0 | 0.05400361 | 0 | 0 | 0 | 0.0201198 | 0 | 0 | 0 | 0 | 0 | 0 | 0 | 0 | 0 | 0 |
| ENSMUSG00 | Gm10754 | 314 | 0 | 0 | 0 | 0 | 0 | 0 | 0 | 0 | 0 | 0 | 0 | 0 | 0 | 0 | 0 | 0 | 0 | 0 |
| ENSMUSG00 | Gm6843 | 424 | 0 | 0 | 0 | 0 | 0 | 0 | 0 | 0 | 0 | 0 | 0 | 0 | 0.11634886 | 0.10953794 | 0 | 0.09039221 | 0 | 0 |
| ENSMUSG00 | Ube2n | 3939 | 22.8010756 | 21.9633375 | 22.9771199 | 21.4677071 | 21.5304518 | 20.1909463 | 24.2629123 | 22.3685891 | 26.934873 | 22.3843146 | 22.8383465 | 23.2813236 | 23.0691525 | 20.7400736 | 20.7524465 | 22.6205238 | 23.8675822 | 22.134916 |
| ENSMUSG00 | 4833422C13 | 5911 | 0 | 0 | 0 | 0 | 0 | 0 | 0.0320167 | 0 | 0 | 0 | 0.10507096 | 0.00944268 | 0 | 0 | 0 | 0 | 0 | 0 |
| ENSMUSG00 | AU019990 | 2430 | 0.01749184 | 0 | 0.03981361 | 0 | 0 | 0 | 0.08156829 | 0 | 0 | 0.01694918 | 0.01940358 | 0.01966048 | 0.09187763 | 0.0609036 | 0 | 0.03154428 | 0 | 0 |
| ENSMUSG00 | Plxnc1 | 7008 | 0 | 0.0095071 | 0.01380523 | 0.1365271 | 0.04120884 | 0 | 0.1755322 | 0.11989624 | 0.09991021 | 0.13456251 | 0.16361287 | 0.1752204 | 0.26749613 | 0.17230968 | 0.00587922 | 0.15298759 | 0.07656509 | 0.02354842 |
| ENSMUSG00 | 5830416P10 | 8653 | 0.03929752 | 0.0115496 | 0.01118075 | 0.03685743 | 0 | 0.03435992 | 0.23511443 | 0.05296531 | 0.05235776 | 0.00544906 | 0.08833923 | 0.02580176 | 0.45609076 | 0.5689441 | 0.47139189 | 0.06195175 | 0.07086799 | 0.19071688 |
| ENSMUSG00 | Hspa12b | 3744 | 0.13623454 | 0.04448837 | 0.07752168 | 0.01216908 | 0.24425923 | 0.03970572 | 0 | 0.03060289 | 0.18701142 | 0 | 0.06380203 | 0.04472409 | 0 | 0.19847901 | 0.27511723 | 0 | 0.05118362 | 0.10284822 |
| ENSMUSG00 | Arrdc3 | 6363 | 13.8811497 | 15.1093844 | 17.1128111 | 15.8314426 | 14.5311057 | 15.6687254 | 16.811876 | 26.2599026 | 21.858693 | 20.3926977 | 23.5909036 | 21.710477 | 28.189667 | 30.7729487 | 32.4212149 | 22.4760991 | 31.6946369 | 16.8321123 |
| ENSMUSG00 | Slc4a11 | 4363 | 0.56504703 | 1.16820372 | 1.02002402 | 0.81452245 | 0.35301892 | 0.85181183 | 0.16266106 | 1.05919796 | 1.17999415 | 0.92939731 | 0.56940129 | 1.48398278 | 0.31659264 | 0.41515457 | 1.52983086 | 0.31594359 | 0.22839416 | 0.37824277 |
| ENSMUSG00 | Itpa | 2086 | 21.5174812 | 21.159916 | 18.6444479 | 18.1719985 | 18.7359166 | 18.5763364 | 16.9427474 | 14.9950036 | 17.5723869 | 12.9291478 | 15.6883253 | 16.1881476 | 19.0611339 | 15.4516762 | 15.4258941 | 17.2179241 | 17.1788531 | 17.5892014 |
| ENSMUSG00 | Gm10760 | 710 | 0 | 0 | 0 | 0 | 0 | 0 | 0 | 0 | 0 | 0 | 0 | 0 | 0 | 0 | 0 | 0 | 0 | 0 |
| ENSMUSG00 | Gm4149 | 360 | 18.5369795 | 18.229556 | 17.6025902 | 13.7948718 | 18.0494714 | 14.7281736 | 23.130643 | 18.2474811 | 20.7076649 | 19.6461264 | 18.844567 | 18.6052209 | 30.5583821 | 29.2855773 | 22.3175094 | 26.1652484 | 24.2733214 | 23.3788704 |
| ENSMUSG00 | Gas2l3 | 7784 | 0.02184233 | 0.14550853 | 0 | 0.11121015 | 0.05565099 | 0.06365973 | 0.0060782 | 0.10303706 | 0 | 0 | 0 | 0.01434113 | 0 | 0 | 0.03705176 | 0 | 0.02954237 | 0 |
| ENSMUSG00 | AU015228 | 4300 | 0 | 0 | 0 | 0 | 0 | 0 | 0 | 0 | 0 | 0 | 0 | 0 | 0 | 0 | 0 | 0 | 0 | 0 |
| ENSMUSG00 | Gm14047 | 2997 | 0 | 0 | 0 | 0 | 0 | 0 | 0 | 0 | 0 | 0 | 0.06376371 | 0 | 0 | 0.04649058 | 0 | 0 | 0 | 0 |
| ENSMUSG00 | Gm10762 | 1132 | 2.81615566 | 2.23655474 | 2.09390723 | 3.01862043 | 3.35903001 | 3.76460697 | 3.21826526 | 3.71127585 | 2.80155628 | 3.37385774 | 2.15240537 | 2.36674188 | 4.96805527 | 3.81563615 | 3.89449336 | 4.05907499 | 4.06285833 | 4.42211017 |
| ENSMUSG00 | Gm10764 | 2334 | 0 | 0 | 0 | 0 | 0 | 0 | 0 | 0 | 0 | 0 | 0 | 0 | 0 | 0 | 0 | 0 | 0 | 0 |
| ENSMUSG00 | Hps6 | 2695 | 1.98725498 | 2.26206295 | 2.42316386 | 2.07940944 | 1.21446041 | 2.83158478 | 1.52734808 | 2.26745246 | 1.86447307 | 3.21919458 | 1.13454447 | 1.96752799 | 1.97693769 | 1.46484135 | 2.20149315 | 0.71040088 | 2.24695915 | 2.20444658 |
| ENSMUSG00 | Gm355 | 1508 | 0 | 0 | 0 | 0 | 0 | 0 | 0 | 0 | 0 | 0 | 0 | 0 | 0 | 0 | 0 | 0 | 0 | 0 |
| ENSMUSG00 | Gm14005 | 2036 | 0.12526083 | 0 | 0.11879551 | 0 | 0.04728087 | 0.07301483 | 0 | 0.03751709 | 0 | 0.09263399 | 0.11732554 | 0 | 0.02422982 | 0.20530294 | 0.16189216 | 0.56420345 | 0.15059449 | 0.29720048 |
| ENSMUSG00 | Papolb | 4238 | 0 | 0 | 0 | 0 | 0 | 0 | 0 | 0 | 0 | 0 | 0 | 0 | 0 | 0 | 0 | 0.15373927 | 0 | 0 |
| ENSMUSG00 | Pdzd7 | 5174 | 0.23002415 | 0.08370074 | 0.17763763 | 0.30820188 | 0.13023713 | 0.05746355 | 0.10973176 | 0 | 0 | 0.15492114 | 0.12003758 | 0.06472632 | 0.10488038 | 0.10771725 | 0.41408599 | 0.48103778 | 0.37037396 | 0.23390034 |
| ENSMUSG00 | Gm10766 | 323 | 2.50030447 | 4.5379794 | 5.39147707 | 1.97478211 | 5.36454877 | 5.36949125 | 3.22253541 | 3.90201015 | 4.97298488 | 3.21150301 | 4.88103291 | 3.28327427 | 2.5964167 | 2.5882154 | 3.5716519 | 1.06692095 | 1.89851626 | 2.21398882 |
| ENSMUSG00 | Zfp706 | 1376 | 0.43246545 | 1.16207771 | 0.52732773 | 0.72844693 | 0.59465313 | 0.75625538 | 0.37822636 | 0.72165877 | 0.71836946 | 0.75386299 | 0.90272451 | 0.24338225 | 0.93214379 | 1.08009506 | 0.41920188 | 0.52872205 | 0.27853413 | 0.67961832 |
| ENSMUSG00 | Itpripl1 | 5052 | 0.17668422 | 0.05275201 | 0.1915025 | 0.16233151 | 0.10480031 | 0.17655368 | 0.01873028 | 0.12095773 | 0.01630503 | 0.02799923 | 0.07565314 | 0 | 0.02929449 | 0.10112529 | 0.02446648 | 0.18948242 | 0.02275908 | 0.18510586 |
| ENSMUSG00 | Gm10767 | 2200 | 2.91740072 | 2.69531599 | 4.24367789 | 3.49991659 | 3.28172207 | 3.28849959 | 4.53771602 | 3.50675695 | 2.20909494 | 2.48612799 | 3.51797422 | 2.53707557 | 3.81201179 | 2.95553283 | 2.13498971 | 3.68982217 | 2.85705122 | 3.70061263 |
| ENSMUSG00 | Gm10768 | 1925 | 0 | 0 | 0 | 0 | 0 | 0 | 0 | 0 | 0 | 0.09797549 | 0 | 0 | 0.10250788 | 0 | 0 | 0.07963906 | 0 | 0 |
| ENSMUSG00 | Zfp640 | 807 | 0 | 0 | 0 | 0 | 0 | 0 | 0 | 0 | 0 | 0 | 0 | 0 | 0 | 0 | 0 | 0 | 0 | 0 |
| ENSMUSG00 | 2410141K09 | 1742 | 0 | 0 | 0 | 0 | 0 | 0 | 0 | 0 | 0 | 0 | 0 | 0 | 0 | 0 | 0 | 0 | 0 | 0 |
| ENSMUSG00 | Vmn2r-ps104 | 668 | 0 | 0 | 0 | 0 | 0 | 0 | 0 | 0 | 0 | 0 | 0 | 0 | 0 | 0 | 0 | 0 | 0 | 0 |
| ENSMUSG00 | Gm10770 | 712 | 0 | 0 | 0 | 0 | 0 | 0 | 0 | 0 | 0 | 0 | 0 | 0 | 0 | 0 | 0 | 0 | 0 | 0 |
| ENSMUSG00 | Gm16512 | 283 | 0 | 0 | 0 | 0 | 0 | 0 | 0 | 0 | 0 | 0 | 0 | 0 | 0 | 0 | 0 | 0 | 0 | 0 |
| ENSMUSG00 | Gm5792 | 548 | 0 | 0 | 0 | 0 | 0 | 0 | 0 | 0 | 0 | 0 | 0 | 0 | 0 | 0 | 0 | 0 | 0 | 0 |
| ENSMUSG00 | Gm17449 | 170 | 0 | 0 | 0 | 0 | 0 | 0 | 0 | 0 | 0 | 0 | 0 | 0 | 0 | 0 | 0 | 0 | 0 | 0 |
| ENSMUSG00 | Gm6917 | 536 | 0 | 0 | 0 | 0 | 0 | 0 | 0 | 0 | 0 | 0 | 0 | 0 | 0 | 0 | 0 | 0 | 0 | 0 |
| ENSMUSG00 | Gm6901 | 548 | 0 | 0 | 0 | 0 | 0 | 0 | 0 | 0 | 0 | 0 | 0 | 0 | 0 | 0 | 0 | 0 | 0 | 0 |
| ENSMUSG00 | Gm10774 | 203 | 0 | 0 | 0 | 0 | 0 | 0 | 0 | 0 | 0 | 0 | 0 | 0 | 0 | 0 | 0 | 0 | 0 | 0 |
| ENSMUSG00 | Gm10775 | 1201 | 0 | 0.02773763 | 0 | 0 | 0 | 0 | 0 | 0 | 0 | 0 | 0 | 0 | 0 | 0.03867118 | 0 | 0 | 0 | 0 |
| ENSMUSG00 | 4932441B19 | 3386 | 0 | 0 | 0 | 0 | 0 | 0 | 0 | 0 | 0 | 0 | 0 | 0 | 0 | 0 | 0 | 0 | 0 | 0 |
| ENSMUSG00 | Hpse2 | 1767 | 0 | 0 | 0 | 0 | 0 | 0 | 0 | 0 | 0 | 0 | 0 | 0 | 0 | 0 | 0 | 0 | 0 | 0 |
| ENSMUSG00 | Gm8290 | 467 | 0 | 0.07133382 | 0 | 0.29268337 | 0 | 0 | 0.20262389 | 0.16356489 | 0 | 0 | 0 | 0 | 0 | 0 | 0.08822603 | 0 | 0 | 0.23558504 |
| ENSMUSG00 | Gm8158 | 467 | 0 | 0 | 0 | 0 | 0 | 0 | 0 | 0 | 0 | 0 | 0 | 0 | 0 | 0 | 0 | 0 | 0 | 0 |
| ENSMUSG00 | Gm8137 | 467 | 0 | 0 | 0 | 0 | 0 | 0 | 0 | 0 | 0 | 0 | 0 | 0 | 0 | 0 | 0 | 0 | 0 | 0 |
| ENSMUSG00 | Gm8264 | 467 | 0 | 0 | 0 | 0 | 0 | 0 | 0 | 0 | 0 | 0 | 0 | 0 | 0 | 0 | 0 | 0 | 0 | 0 |
| ENSMUSG00 | Gm8112 | 467 | 0 | 0 | 0 | 0 | 0 | 0 | 0 | 0 | 0 | 0 | 0 | 0 | 0 | 0 | 0 | 0 | 0 | 0 |
| ENSMUSG00 | Gm10778 | 1589 | 0.10699856 | 0.18868222 | 0.03044275 | 0.11469111 | 0 | 0.09355456 | 0.05955026 | 0.12017748 | 0.15551862 | 0.08901958 | 0.06013211 | 0.21075769 | 0.15522944 | 0.116914 | 0.05185847 | 0.12048649 | 0.14471855 | 0.20771217 |
| ENSMUSG00 | 1810034E14 | 1753 | 3.34610057 | 1.93834296 | 2.45592938 | 2.2611585 | 3.2673696 | 3.75956279 | 2.64497578 | 1.98260617 | 2.53746649 | 1.61382898 | 1.77146167 | 3.37504862 | 1.04123271 | 1.48366737 | 2.44435928 | 1.66006056 | 1.42111198 | 1.56899905 |
| ENSMUSG00 | BC025920 | 2339 | 0 | 0.05696946 | 0 | 0.05843657 | 0.02057799 | 0.04237087 | 0.02022774 | 0.0489855 | 0.0176086 | 0.02015849 | 0.02042538 | 0.07158914 | 0.04218206 | 0 | 0.07046012 | 0 | 0 | 0.09407286 |
| ENSMUSG00 | Gm5665 | 3442 | 0.06174488 | 0.05807012 | 0.04216171 | 0.01323679 | 0 | 0 | 0.06872847 | 0.17753586 | 0.01196587 | 0 | 0.02776 | 0.03243211 | 0.08599405 | 0.01349334 | 0.02394047 | 0.04449809 | 0.03340467 | 0.07990864 |
| ENSMUSG00 | Zfp934 | 1487 | 1.20054969 | 1.09773491 | 1.91932636 | 1.40942034 | 1.26237056 | 1.7328461 | 3.91355717 | 1.90063065 | 1.606468 | 1.64884773 | 0.48192628 | 3.19053888 | 1.42654502 | 1.40550365 | 1.9395487 | 2.36902211 | 1.44335752 | 1.07280706 |
| ENSMUSG00 | Zfp808 | 2567 | 0.77824046 | 0.37634357 | 0.56533148 | 0.42597003 | 0.43125604 | 0.17373378 | 0.82940029 | 0.44634674 | 0.22462455 | 0.99187299 | 1.63778588 | 1.39158645 | 0.46122556 | 0.47041148 | 0.22470658 | 0.73090758 | 1.19443076 | 1.37147753 |
| ENSMUSG00 | Gm6478 | 743 | 0.0572075 | 0.04483566 | 0.06510569 | 0 | 0.12956103 | 0 | 0.0636779 | 0.10280593 | 0 | 0 | 0.12860016 | 0 | 0 | 0.06250887 | 0 | 0.61842165 | 0 | 0.07403648 |
| ENSMUSG00 | Cts3 | 1372 | 0 | 0 | 0 | 0 | 0 | 0 | 0 | 0 | 0 | 0 | 0 | 0 | 0 | 0 | 0 | 0 | 0 | 0 |
| ENSMUSG00 | Ctsm | 1196 | 0 | 0 | 0 | 0 | 0 | 0 | 0 | 0 | 0 | 0 | 0 | 0 | 0 | 0 | 0 | 0 | 0 | 0 |
| ENSMUSG00 | Ctsr2 | 1026 | 0 | 0 | 0 | 0 | 0 | 0 | 0 | 0 | 0 | 0 | 0 | 0 | 0 | 0 | 0 | 0 | 0 | 0 |
| ENSMUSG00 | AI606181 | 2795 | 0.63871821 | 0.58401854 | 0.76151533 | 0.61943459 | 0.56828389 | 1.11693103 | 0.66017692 | 0.46459452 | 0.55995978 | 0.62417747 | 0.39313919 | 0.39939651 | 0.37065126 | 0.66467388 | 0.19163514 | 0.28769329 | 0.23311164 | 0.57075638 |
| ENSMUSG00 | Ctla2b | 1511 | 0 | 0.02204692 | 0 | 0 | 0 | 0 | 0 | 0.37914364 | 0 | 0 | 0 | 0 | 0 | 0 | 0 | 0 | 0 | 0 |
| ENSMUSG00 | Spata5l1 | 2658 | 0.28784544 | 0.58905419 | 0.41858209 | 0.3599631 | 0.32594982 | 0.27964297 | 0.87220514 | 0.57475399 | 0.69728864 | 0.37252249 | 0.55719478 | 1.07095515 | 0.4825545 | 0.48925299 | 0.69754327 | 0.99399997 | 1.42750316 | 1.05547947 |
| ENSMUSG00 | Gm10780 | 283 | 0 | 0 | 0 | 0 | 0 | 0 | 0 | 0 | 0 | 0 | 0 | 0 | 0 | 0 | 0 | 0 | 0 | 0 |
| ENSMUSG00 | Mageb3 | 1608 | 0.10573427 | 0 | 0 | 0 | 0 | 0 | 0 | 0 | 0 | 0 | 0 | 0 | 0.06135811 | 0 | 0 | 0 | 0 | 0 |
| ENSMUSG00 | Cyp2c68 | 1715 | 0.09913744 | 0.07769771 | 0.22564912 | 0.23909586 | 0.14032631 | 0.05778744 | 0.13793784 | 0.28950509 | 0.21613915 | 0.35741058 | 0.22285695 | 0.16272788 | 0.40270953 | 0.35205431 | 0.28829077 | 0.64747966 | 0.24582464 | 0.19245169 |
| ENSMUSG00 | Serf2 | 5521 | 12.2026271 | 11.6936042 | 11.0047372 | 10.7692742 | 11.847724 | 11.69484 | 11.5603702 | 12.7976715 | 12.4432364 | 11.3670687 | 11.3531508 | 11.6969248 | 11.6874022 | 13.0221785 | 11.9403169 | 14.1344427 | 12.7592345 | 13.4708046 |
| ENSMUSG00 | Gm10782 | 427 | 0 | 0 | 0 | 0 | 0 | 0 | 0 | 0 | 0 | 0 | 0 | 0 | 0 | 0 | 0 | 0 | 0 | 0 |
| ENSMUSG00 | Grk6 | 3224 | 11.2986774 | 14.4142331 | 12.3034413 | 14.4710017 | 13.316897 | 15.6619837 | 10.8302596 | 12.0950505 | 14.3590706 | 12.9430435 | 13.751601 | 11.1839076 | 11.3995901 | 12.7778863 | 12.7668593 | 10.5584212 | 11.2458675 | 15.0490174 |
| ENSMUSG00 | EU599041 | 807 | 0 | 0 | 0 | 0 | 0 | 0 | 0 | 0 | 0 | 0 | 0 | 0 | 0 | 0 | 0 | 0 | 0 | 0 |
| ENSMUSG00 | Lcmt2 | 3379 | 5.19521684 | 4.87024852 | 4.46657043 | 6.37773719 | 5.4128828 | 4.89808021 | 6.86096853 | 5.62882995 | 4.59524014 | 7.2840092 | 8.14394084 | 7.86275685 | 4.05867801 | 5.77286674 | 4.84078653 | 5.16736223 | 7.02100556 | 5.95837025 |
| ENSMUSG00 | B3galt5 | 6152 | 0 | 0 | 0 | 0 | 0 | 0 | 0 | 0 | 0 | 0 | 0 | 0 | 0 | 0 | 0 | 0 | 0 | 0 |
| ENSMUSG00 | Eif4e1b | 2188 | 0 | 0 | 0 | 0 | 0 | 0 | 0 | 0 | 0 | 0 | 0 | 0 | 0 | 0 | 0 | 0 | 0 | 0 |
| ENSMUSG00 | Ifit3 | 1743 | 0.24386217 | 0.42047256 | 0.58281362 | 0.81032265 | 0.05522883 | 0.90974611 | 0.10857758 | 0 | 0.25992638 | 0.5410293 | 0 | 0.03202276 | 0 | 0 | 0.61459578 | 0.79085592 | 0.39579653 | 0.15780008 |
| ENSMUSG00 | Spnb5 | 10233 | 0.03738362 | 0.09115226 | 0.0519993 | 0 | 0.04703599 | 0.00484244 | 0.06472955 | 0.15675568 | 0.01207462 | 0.10136963 | 0 | 0.0709082 | 0 | 0 | 0.02818439 | 0.06735375 | 0 | 0.0376296 |
| ENSMUSG00 | Gm8765 | 3148 | 0 | 0 | 0 | 0 | 0 | 0 | 0 | 0 | 0 | 0 | 0 | 0 | 0 | 0 | 0 | 0 | 0 | 0 |
| ENSMUSG00 | Gm904 | 645 | 0 | 0 | 0 | 0 | 0 | 0 | 0 | 0 | 0 | 0 | 0 | 0 | 0 | 0 | 0 | 0 | 0 | 0 |
| ENSMUSG00 | Gm2058 | 3268 | 0 | 0 | 0 | 0 | 0 | 0 | 0 | 0 | 0 | 0 | 0 | 0 | 0 | 0 | 0 | 0 | 0 | 0 |
| ENSMUSG00 | Gm906 | 3148 | 0 | 0 | 0 | 0 | 0 | 0 | 0 | 0 | 0 | 0 | 0 | 0 | 0 | 0 | 0 | 0 | 0 | 0 |
| ENSMUSG00 | Ranbp6 | 4866 | 5.34590377 | 7.05828071 | 6.41202776 | 6.35757277 | 7.71535151 | 6.46649935 | 8.11880125 | 7.28371341 | 5.12927015 | 6.76349997 | 7.10831688 | 7.73114603 | 3.68012453 | 7.69295817 | 5.92706312 | 5.90962724 | 6.33258173 | 7.23506541 |
| ENSMUSG00 | Tes3-ps | 3258 | 0.53490246 | 0.54192247 | 0.44542846 | 0.55937439 | 0.45797717 | 0.82131603 | 0.66801204 | 0.04890611 | 0.41717465 | 0.41969625 | 0.30794172 | 0.4111651 | 0.46939516 | 0.44191734 | 0.139109 | 0.42310063 | 0.34114874 | 0.37145499 |
| ENSMUSG00 | Gm17362 | 625 | 0.40804969 | 0.53300631 | 0.54178354 | 0.65607904 | 0.77011078 | 0.79284374 | 0 | 0.61107844 | 0.32949213 | 0.603529 | 0.61151947 | 0.62513542 | 1.42075922 | 0.44586324 | 0.26368996 | 1.47035932 | 0.91983113 | 0.26404371 |
| ENSMUSG00 | Gm14207 | 1372 | 11.9274729 | 7.67264911 | 12.1286403 | 8.60080939 | 9.99824945 | 8.12635937 | 12.9661569 | 13.7793289 | 18.672021 | 15.3617816 | 14.3464161 | 14.9709649 | 13.8431401 | 10.3585209 | 9.78987407 | 10.7448133 | 9.16255494 | 10.1839017 |
| ENSMUSG00 | Gm815 | 606 | 0 | 0 | 0 | 0 | 0 | 0 | 0 | 0 | 0 | 0 | 0 | 0 | 0 | 0 | 0 | 0 | 0 | 0 |
| ENSMUSG00 | Chst14 | 4077 | 1.86618261 | 2.03456234 | 1.67296242 | 1.3521919 | 1.85349816 | 2.11483337 | 1.57824978 | 1.27401686 | 1.49512002 | 1.57284661 | 1.11322567 | 1.67022586 | 1.33100586 | 1.67458446 | 1.25312557 | 1.85019614 | 1.11867288 | 1.49767252 |
| ENSMUSG00 | 5430417L22 | 3052 | 3.84384947 | 5.97056132 | 5.81686948 | 4.38890794 | 5.22007429 | 5.50405527 | 3.86004491 | 5.20577962 | 3.54916568 | 4.49569287 | 4.88394071 | 5.41331459 | 5.25323497 | 4.1696199 | 6.9119255 | 4.99333609 | 5.83036044 | 4.95658727 |
| ENSMUSG00 | Gm10787 | 1545 | 0 | 0 | 0 | 0 | 0 | 0 | 0 | 0 | 0 | 0 | 0 | 0 | 0 | 0 | 0 | 0 | 0 | 0.178023 |
| ENSMUSG00 | Fam122a | 854 | 162.206521 | 150.883226 | 146.196817 | 136.042931 | 147.551963 | 142.797749 | 148.087577 | 145.927177 | 159.875058 | 152.825699 | 166.820757 | 152.349309 | 153.887854 | 150.589787 | 150.477345 | 124.691076 | 140.69431 | 138.424555 |
| ENSMUSG00 | Pak6 | 5366 | 0.03960602 | 0.16141171 | 0.32453356 | 0.10188828 | 0.17042612 | 0 | 0.1587082 | 0.09964473 | 0.29166746 | 0.12301712 | 0 | 0.10401726 | 0.26660932 | 0 | 0.02303479 | 0.11417252 | 0.00714243 | 0.01025142 |
| ENSMUSG00 | Ptar1 | 2057 | 2.43831345 | 2.47781858 | 2.9630845 | 2.39212095 | 1.47414253 | 2.48125015 | 1.81706448 | 3.17496393 | 2.74309804 | 2.95694736 | 2.1367507 | 2.27929784 | 2.80594765 | 3.02552635 | 3.64544636 | 2.08485563 | 2.70165924 | 1.12318059 |
| ENSMUSG00 | Krtap14 | 199 | 0 | 0 | 0 | 0 | 0 | 0 | 0 | 0 | 0 | 0 | 0 | 0 | 0 | 0 | 0 | 0 | 0 | 0 |
| ENSMUSG00 | Gm13981 | 446 | 0 | 0 | 0 | 0 | 0 | 0 | 0 | 0 | 0 | 0 | 0 | 0 | 0 | 0 | 0 | 0 | 0.1718668 | 0.1233388 |
| ENSMUSG00 | Gm17719 | 3187 | 0 | 0 | 0 | 0 | 0 | 0 | 0 | 0 | 0 | 0 | 0 | 0.1225948 | 0 | 0 | 0 | 0 | 0 | 0 |
| ENSMUSG00 | Grem1 | 4146 | 0 | 0 | 0 | 0 | 0 | 0 | 0 | 0 | 0 | 0 | 0 | 0 | 0 | 0 | 0 | 0 | 0 | 0 |
| ENSMUSG00 | Chrm5 | 1745 | 0 | 0 | 0 | 0 | 0 | 0 | 0 | 0 | 0 | 0 | 0 | 0 | 0 | 0 | 0 | 0 | 0 | 0 |
| ENSMUSG00 | Olfr1316 | 944 | 0 | 0 | 0 | 0 | 0 | 0 | 0 | 0 | 0 | 0 | 0 | 0 | 0 | 0 | 0 | 0 | 0 | 0 |
| ENSMUSG00 | Olfr1314 | 938 | 0 | 0.03551481 | 0 | 0 | 0 | 0 | 0 | 0 | 0 | 0 | 0 | 0 | 0 | 0 | 0 | 0 | 0 | 0 |
| ENSMUSG00 | Olfr1313 | 937 | 0 | 0 | 0 | 0 | 0 | 0 | 0 | 0 | 0 | 0 | 0 | 0 | 0 | 0 | 0 | 0 | 0 | 0 |
| ENSMUSG00 | Olfr1312 | 953 | 0 | 0 | 0 | 0 | 0 | 0 | 0 | 0 | 0 | 0 | 0 | 0 | 0 | 0 | 0 | 0 | 0 | 0 |
| ENSMUSG00 | Olfr1311 | 942 | 0 | 0 | 0 | 0 | 0 | 0 | 0 | 0 | 0 | 0 | 0 | 0 | 0 | 0 | 0 | 0 | 0 | 0 |
| ENSMUSG00 | Olfr1310 | 1008 | 0 | 0 | 0 | 0 | 0 | 0 | 0 | 0 | 0 | 0 | 0 | 0 | 0 | 0 | 0 | 0 | 0 | 0 |
| ENSMUSG00 | Olfr1309 | 963 | 0 | 0 | 0 | 0 | 0 | 0 | 0 | 0 | 0 | 0 | 0 | 0 | 0 | 0 | 0 | 0 | 0 | 0 |
| ENSMUSG00 | Olfr1308 | 962 | 0 | 0 | 0 | 0 | 0 | 0 | 0 | 0 | 0 | 0 | 0 | 0 | 0 | 0 | 0 | 0 | 0 | 0 |
| ENSMUSG00 | Olfr1307 | 938 | 0 | 0 | 0 | 0 | 0 | 0 | 0 | 0 | 0 | 0 | 0 | 0 | 0 | 0 | 0 | 0 | 0 | 0 |
| ENSMUSG00 | Olfr1306 | 938 | 0 | 0 | 0 | 0 | 0 | 0 | 0 | 0 | 0 | 0 | 0 | 0 | 0 | 0 | 0 | 0 | 0 | 0 |
| ENSMUSG00 | Olfr1305 | 938 | 0 | 0 | 0 | 0 | 0 | 0 | 0 | 0 | 0 | 0 | 0 | 0 | 0 | 0 | 0 | 0 | 0 | 0 |
| ENSMUSG00 | Olfr1303 | 938 | 0 | 0 | 0 | 0 | 0 | 0 | 0 | 0 | 0 | 0 | 0 | 0 | 0 | 0 | 0 | 0 | 0 | 0 |
| ENSMUSG00 | Olfr1299 | 994 | 0 | 0 | 0 | 0 | 0 | 0 | 0 | 0 | 0 | 0 | 0 | 0 | 0 | 0 | 0 | 0 | 0 | 0 |
| ENSMUSG00 | Olfr1297 | 938 | 0 | 0 | 0 | 0 | 0 | 0 | 0 | 0 | 0 | 0 | 0 | 0 | 0 | 0 | 0 | 0 | 0 | 0 |
| ENSMUSG00 | Olfr1295 | 1042 | 0 | 0 | 0 | 0 | 0 | 0 | 0 | 0 | 0 | 0 | 0 | 0 | 0 | 0 | 0 | 0 | 0 | 0 |
| ENSMUSG00 | Olfr1294 | 938 | 0 | 0 | 0 | 0 | 0 | 0 | 0 | 0 | 0 | 0 | 0 | 0 | 0 | 0 | 0 | 0 | 0 | 0 |
| ENSMUSG00 | Olfr1290 | 945 | 0 | 0 | 0 | 0 | 0 | 0 | 0 | 0 | 0 | 0 | 0 | 0 | 0 | 0 | 0 | 0 | 0 | 0 |
| ENSMUSG00 | Olfr1286 | 917 | 0 | 0 | 0 | 0 | 0 | 0 | 0 | 0 | 0 | 0 | 0 | 0 | 0 | 0 | 0 | 0 | 0 | 0 |
| ENSMUSG00 | Olfr1282 | 917 | 0 | 0 | 0 | 0 | 0 | 0 | 0 | 0 | 0 | 0 | 0 | 0 | 0 | 0 | 0 | 0 | 0 | 0 |
| ENSMUSG00 | Olfr1277 | 961 | 0 | 0 | 0 | 0 | 0 | 0 | 0 | 0 | 0 | 0 | 0 | 0 | 0 | 0 | 0 | 0 | 0 | 0 |
| ENSMUSG00 | Olfr1275 | 1019 | 0 | 0 | 0 | 0 | 0 | 0 | 0 | 0 | 0 | 0 | 0 | 0 | 0 | 0 | 0 | 0 | 0 | 0 |
| ENSMUSG00 | Ano3 | 7416 | 0 | 0.00898406 | 0 | 0 | 0.02596112 | 0.04677306 | 0.08931736 | 0.01545 | 0 | 0 | 0.01932644 | 0.05268469 | 0 | 0 | 0.04444612 | 0.02065297 | 0.09302499 | 0.1483525 |
| ENSMUSG00 | Gm10794 | 205 | 0 | 0 | 0 | 0 | 0 | 0 | 0 | 0 | 0 | 0 | 0 | 0 | 0 | 0 | 0 | 0 | 0 | 0 |
| ENSMUSG00 | Fibin | 2068 | 0.12332256 | 0 | 0 | 0 | 0.25602087 | 0 | 0.50332637 | 0.03693656 | 0 | 0.18240117 | 0.13861207 | 0 | 0.47709765 | 0 | 0.6973184 | 0.20367344 | 0.31506143 | 0.34580193 |
| ENSMUSG00 | Gm11382 | 167 | 0.76356604 | 1.19687045 | 0.57902372 | 0.81846187 | 0 | 0.89016887 | 0.28330945 | 1.37218212 | 1.23312921 | 1.4116977 | 2.00254317 | 2.00535315 | 0.5908014 | 2.22486646 | 1.72701132 | 2.29285074 | 0.22949878 | 1.31758339 |
| ENSMUSG00 | Actg-ps1 | 407 | 0.31330597 | 0.08184986 | 0.23770777 | 0.44777439 | 0.59130127 | 0.12175119 | 0.11624737 | 0.28151648 | 0.20239074 | 0 | 0.23476638 | 0.13713922 | 0.12120864 | 0.11411324 | 0.50616162 | 0 | 0 | 0.13515751 |
| ENSMUSG00 | Dcdc5 | 5648 | 0.04515422 | 0.04128723 | 0.00856472 | 0.00806676 | 0 | 0 | 0.06701513 | 0.03381055 | 0.28439697 | 0.00834821 | 0.05075243 | 0.01976475 | 0 | 0.14801586 | 0 | 0 | 0.00678582 | 0 |
| ENSMUSG00 | Olfr1452-ps1 | 919 | 0 | 0 | 0 | 0 | 0 | 0 | 0 | 0 | 0 | 0 | 0 | 0 | 0 | 0 | 0 | 0 | 0 | 0 |
| ENSMUSG00 | AI314831 | 5437 | 0 | 0 | 0 | 0 | 0 | 0 | 0 | 0 | 0 | 0 | 0 | 0 | 0 | 0 | 0 | 0 | 0 | 0 |
| ENSMUSG00 | Gabrr3 | 1674 | 0 | 0 | 0 | 0 | 0 | 0 | 0 | 0 | 0 | 0 | 0 | 0 | 0 | 0 | 0 | 0 | 0 | 0 |
| ENSMUSG00 | Qser1 | 8961 | 6.74504635 | 6.92578082 | 6.26194568 | 6.32495694 | 6.16621453 | 6.37588459 | 4.23971445 | 5.90722796 | 5.2028943 | 4.09365552 | 3.40145335 | 5.34425159 | 3.90317256 | 2.70548082 | 2.90585686 | 3.95682429 | 3.98617439 | 4.04541919 |
| ENSMUSG00 | Olfr201 | 926 | 0 | 0 | 0 | 0 | 0 | 0 | 0 | 0 | 0 | 0 | 0 | 0 | 0 | 0 | 0 | 0 | 0 | 0 |
| ENSMUSG00 | Olfr199 | 930 | 0 | 0 | 0 | 0 | 0 | 0 | 0 | 0 | 0 | 0 | 0 | 0 | 0 | 0 | 0 | 0 | 0 | 0 |
| ENSMUSG00 | Pin1-ps1 | 1560 | 0 | 0 | 0 | 0.0308538 | 0 | 0 | 0.03032864 | 0.02448231 | 0 | 0.15112405 | 0.06124995 | 0.14311708 | 0.06324605 | 0 | 0.23770128 | 0 | 0 | 0.31736023 |
| ENSMUSG00 | Gm10797 | 2221 | 0 | 0 | 0 | 0 | 0 | 0 | 0 | 0 | 0 | 0 | 0 | 0 | 0 | 0 | 0 | 0 | 0 | 0 |
| ENSMUSG00 | Nrbf2 | 3323 | 6.38281157 | 4.82199683 | 6.53617669 | 8.07567111 | 6.32971732 | 5.92008283 | 7.48915712 | 7.02243692 | 7.41183762 | 7.73311263 | 6.52718361 | 7.86088778 | 7.37826148 | 8.76329906 | 7.17896505 | 8.08907806 | 7.43919997 | 7.00236297 |
| ENSMUSG00 | Gm813 | 698 | 0 | 0 | 0 | 0 | 0 | 0 | 0 | 0 | 0 | 0 | 0 | 0 | 0 | 0 | 0 | 0 | 0 | 0 |
| ENSMUSG00 | Jmjd1c | 593 | 0 | 0 | 0 | 0 | 0 | 0 | 0 | 0 | 0 | 0 | 0 | 0 | 0 | 0 | 0 | 0 | 0 | 0 |
| ENSMUSG00 | Gm10799 | 310 | 0 | 0 | 0 | 0 | 0 | 0 | 0 | 0 | 0 | 0 | 0 | 0 | 0 | 0 | 0 | 0 | 0 | 0 |
| ENSMUSG00 | AW112010 | 771 | 3.36292573 | 2.63565052 | 5.20752661 | 3.66379345 | 3.68324708 | 2.6993707 | 1.77959493 | 0.94118761 | 5.34196054 | 0.67270783 | 3.77986053 | 1.52027097 | 3.51913807 | 4.63838486 | 6.25237619 | 7.05222601 | 5.61721342 | 7.20612163 |
| ENSMUSG00 | Fjx1 | 2424 | 1.80611928 | 1.56669552 | 0.99780384 | 1.86078523 | 2.64090175 | 2.00336959 | 0.9759216 | 0.74052925 | 0.59468979 | 0.46683865 | 0.17738227 | 0.66776164 | 0.79370659 | 0.51732276 | 0.45892822 | 0.66345112 | 0.53758007 | 0.11346763 |
| ENSMUSG00 | Gm10800 | 655 | 0.38936039 | 0.30515628 | 0.59082174 | 0.06955885 | 0.07348385 | 0.83218331 | 0.28893239 | 0.29154506 | 1.13184318 | 0.57588645 | 0.65644981 | 0 | 2.03352941 | 2.33993112 | 0.69193453 | 0.11691789 | 0.11702686 | 2.35153433 |
| ENSMUSG00 | Gm10801 | 507 | 0.08383664 | 0 | 0 | 0 | 0.4746738 | 0.19547429 | 0.09331889 | 0 | 0.40617866 | 0 | 0.09423069 | 0 | 0.19460322 | 0.64123087 | 0.16253079 | 0.22657164 | 0.07559427 | 0.65099535 |
| ENSMUSG00 | E530001K10 | 1227 | 0.62354781 | 0.95024556 | 0.70963614 | 0.59411305 | 0.98068304 | 0.80770552 | 1.6195049 | 1.58745927 | 3.08814951 | 2.11351971 | 1.90788343 | 2.72937225 | 1.48759653 | 2.30895626 | 2.61916981 | 2.71498194 | 1.811675 | 1.16563714 |
| ENSMUSG00 | Gm13889 | 745 | 0 | 0 | 0 | 0 | 0 | 0 | 0 | 0 | 0 | 0 | 0 | 0 | 0 | 0 | 0 | 0 | 0 | 0 |
| ENSMUSG00 | Accsl | 3201 | 0.09295103 | 0 | 0 | 0 | 0.03007305 | 0 | 0.05912237 | 0 | 0 | 0 | 0.17436946 | 0 | 0 | 0 | 0 | 0 | 0.15565194 | 0 |
| ENSMUSG00 | Gm10803 | 2481 | 0 | 0 | 0 | 0 | 0 | 0 | 0 | 0 | 0 | 0 | 0 | 0 | 0 | 0 | 0 | 0 | 0 | 0 |
| ENSMUSG00 | Gm10804 | 2202 | 0.03860597 | 0 | 0 | 0.02069076 | 0 | 0 | 0 | 0.10406649 | 0.11222484 | 0.21412672 | 0 | 0.12673856 | 0.02240323 | 0 | 0 | 0.13911211 | 0.10443133 | 0 |
| ENSMUSG00 | 4631405J19R | 4921 | 0.33686281 | 0.14892983 | 0.11796024 | 0.32404726 | 0.10759016 | 0.05034824 | 0.38457776 | 0.3958164 | 0.72815015 | 0.75694078 | 0.65046174 | 0.55577473 | 0.34084235 | 0.52852447 | 0.58608188 | 1.12047297 | 0.62306518 | 0.34653166 |
| ENSMUSG00 | Prdm11 | 17034 | 0.25452201 | 0.51629706 | 0.47992994 | 0.6232079 | 0.54817384 | 0.42472109 | 0.51940067 | 0.69281744 | 0.92605586 | 0.54530284 | 0.67592844 | 0.69466482 | 0.68637222 | 0.76887767 | 0.62404611 | 0.7912583 | 0.59624684 | 0.83317773 |
| ENSMUSG00 | 4930558J22R | 1097 | 0 | 0 | 0 | 0 | 0 | 0 | 0 | 0.13926127 | 0 | 0.12894449 | 0 | 0 | 0 | 0 | 0 | 0 | 0.13974949 | 0 |
| ENSMUSG00 | Hist1h2bb | 468 | 0.09082303 | 0.42708839 | 0.20672449 | 0.09735266 | 0.20569198 | 0.21176382 | 0.10109547 | 0 | 0.17601075 | 0 | 0.51041622 | 0.23852847 | 0 | 0 | 0.17607503 | 0 | 0 | 0.70524496 |
| ENSMUSG00 | Hist1h3i | 482 | 0 | 0 | 0 | 0 | 0 | 0 | 0 | 0 | 0.17089841 | 0 | 0.09911817 | 0 | 0 | 0 | 0.08548041 | 0 | 0 | 0.34238033 |
| ENSMUSG00 | Fam55c | 6424 | 0.21834851 | 0.01555708 | 0.08283139 | 0.11347707 | 0.00749252 | 0.10799164 | 0.78805365 | 0.63019842 | 0.36544698 | 0.25689206 | 0.23798236 | 0.61689166 | 2.0964529 | 1.51102339 | 0.99412222 | 1.57358661 | 1.19918831 | 1.30158534 |
| ENSMUSG00 | Gm10805 | 397 | 0 | 0 | 0 | 0 | 0 | 0 | 0 | 0 | 0 | 0 | 0 | 0 | 0 | 0 | 0 | 0 | 0 | 0.55424793 |
| ENSMUSG00 | Zfp408 | 6776 | 2.2331527 | 3.05302646 | 2.04173992 | 2.81730779 | 2.5927025 | 2.34015271 | 2.07377002 | 2.7167559 | 2.9297374 | 2.34500989 | 3.22213046 | 2.85832865 | 2.22051871 | 1.73411365 | 3.02201495 | 2.26036643 | 2.76021737 | 2.92256175 |
| ENSMUSG00 | 4930431P03 | 2980 | 0 | 0 | 0 | 0 | 0 | 0 | 0 | 0 | 0 | 0 | 0 | 0 | 0 | 0 | 0 | 0 | 0 | 0 |
| ENSMUSG00 | Tmem223 | 746 | 121.077076 | 104.180941 | 107.251768 | 107.978454 | 111.942209 | 108.803455 | 112.066359 | 91.4876985 | 104.291325 | 103.655702 | 117.195944 | 90.9062065 | 110.03795 | 101.106163 | 102.12021 | 94.1866817 | 86.7222375 | 93.2057787 |
| ENSMUSG00 | Slc22a29 | 2712 | 0 | 0 | 0 | 0.03359959 | 0 | 0 | 0 | 0 | 0 | 0 | 0.03523227 | 0.04116199 | 0 | 0.0171254 | 0.04557694 | 0 | 0.02826423 | 0.08113438 |
| ENSMUSG00 | Gm4981 | 1812 | 0 | 0 | 0 | 0 | 0 | 0 | 0 | 0 | 0 | 0 | 0.06160669 | 0 | 0 | 0 | 0 | 0 | 0 | 0 |
| ENSMUSG00 | Gm10807 | 2443 | 0 | 0 | 0 | 0 | 0 | 0 | 0 | 0 | 0 | 0 | 0 | 0 | 0 | 0 | 0 | 0 | 0 | 0 |
| ENSMUSG00 | Vdac3-ps1 | 851 | 1.2486832 | 0.62632939 | 0.45474529 | 0.80307364 | 0.79182953 | 0.81520361 | 0.77835194 | 0.94246821 | 0.72596679 | 0.88649971 | 0.89823659 | 0.5902949 | 1.27532571 | 0.38203127 | 0.77464735 | 0.80990709 | 0.7656252 | 0.96960823 |
| ENSMUSG00 | 1600012F09 | 3433 | 4.12298969 | 5.88045846 | 4.53722015 | 4.44595102 | 4.52857889 | 4.47461329 | 6.64279384 | 5.15091241 | 6.15458275 | 4.68348087 | 4.32799653 | 5.60920582 | 6.0497341 | 6.52084186 | 5.42473151 | 5.84453198 | 6.00627665 | 5.11153657 |
| ENSMUSG00 | Olfr1272 | 926 | 0 | 0 | 0 | 0 | 0 | 0 | 0 | 0 | 0 | 0 | 0 | 0 | 0 | 0 | 0 | 0 | 0 | 0 |
| ENSMUSG00 | Olfr1271 | 925 | 0 | 0 | 0 | 0 | 0 | 0 | 0 | 0 | 0 | 0 | 0 | 0 | 0 | 0 | 0 | 0 | 0 | 0 |
| ENSMUSG00 | Olfr142 | 917 | 0 | 0 | 0 | 0 | 0 | 0 | 0 | 0 | 0 | 0 | 0 | 0 | 0 | 0 | 0 | 0 | 0 | 0 |
| ENSMUSG00 | Olfr1506 | 1082 | 0 | 0 | 0 | 0 | 0 | 0 | 0 | 0 | 0 | 0 | 0 | 0 | 0 | 0 | 0 | 0 | 0 | 0 |
| ENSMUSG00 | Olfr1270 | 1017 | 0 | 0 | 0 | 0 | 0 | 0 | 0 | 0 | 0 | 0 | 0 | 0 | 0 | 0 | 0 | 0 | 0 | 0 |
| ENSMUSG00 | Olfr32 | 1007 | 0 | 0 | 0 | 0 | 0 | 0 | 0 | 0 | 0 | 0 | 0 | 0 | 0 | 0 | 0 | 0 | 0 | 0 |
| ENSMUSG00 | Olfr1269 | 1017 | 0 | 0 | 0 | 0 | 0 | 0 | 0 | 0 | 0 | 0 | 0 | 0 | 0 | 0 | 0 | 0 | 0 | 0 |
| ENSMUSG00 | Olfr140 | 2321 | 0 | 0 | 0 | 0 | 0 | 0 | 0 | 0 | 0 | 0 | 0 | 0 | 0 | 0 | 0 | 0 | 0 | 0 |
| ENSMUSG00 | Olfr1264 | 926 | 0 | 0 | 0 | 0 | 0 | 0 | 0 | 0 | 0 | 0 | 0 | 0 | 0 | 0 | 0 | 0 | 0 | 0 |
| ENSMUSG00 | 4932412D23 | 5084 | 0 | 0 | 0 | 0 | 0 | 0 | 0 | 0 | 0 | 0 | 0 | 0 | 0 | 0 | 0 | 0 | 0 | 0 |
| ENSMUSG00 | Olfr48 | 905 | 0 | 0 | 0 | 0 | 0 | 0 | 0 | 0 | 0 | 0 | 0 | 0 | 0 | 0 | 0 | 0 | 0 | 0 |
| ENSMUSG00 | Olfr1256 | 2063 | 0 | 0 | 0 | 0 | 0 | 0 | 0 | 0 | 0 | 0 | 0 | 0 | 0 | 0 | 0 | 0 | 0 | 0 |
| ENSMUSG00 | Olfr1254 | 2062 | 0 | 0 | 0 | 0 | 0 | 0 | 0 | 0 | 0 | 0 | 0 | 0 | 0 | 0 | 0 | 0 | 0 | 0 |
| ENSMUSG00 | Olfr1253 | 956 | 0 | 0 | 0 | 0 | 0 | 0 | 0 | 0 | 0 | 0 | 0 | 0 | 0 | 0 | 0 | 0 | 0 | 0 |
| ENSMUSG00 | Olfr1252 | 1098 | 0 | 0 | 0 | 0 | 0 | 0 | 0 | 0 | 0 | 0 | 0 | 0 | 0 | 0 | 0 | 0 | 0 | 0 |
| ENSMUSG00 | Olfr1251 | 1039 | 0 | 0 | 0 | 0 | 0 | 0 | 0 | 0 | 0 | 0 | 0 | 0 | 0 | 0 | 0 | 0 | 0 | 0 |
| ENSMUSG00 | Olfr1250 | 2799 | 0 | 0 | 0 | 0 | 0 | 0 | 0 | 0 | 0 | 0 | 0 | 0 | 0 | 0 | 0 | 0 | 0 | 0 |
| ENSMUSG00 | Olfr1249 | 956 | 0 | 0 | 0 | 0 | 0 | 0 | 0 | 0 | 0 | 0 | 0 | 0 | 0 | 0 | 0 | 0 | 0 | 0 |
| ENSMUSG00 | Olfr1248 | 1077 | 0 | 0 | 0 | 0 | 0 | 0 | 0 | 0 | 0 | 0 | 0 | 0 | 0 | 0 | 0 | 0 | 0 | 0 |
| ENSMUSG00 | Olfr1247 | 2717 | 0 | 0 | 0 | 0 | 0 | 0 | 0 | 0 | 0 | 0 | 0 | 0 | 0 | 0 | 0 | 0 | 0 | 0 |
| ENSMUSG00 | Olfr1246 | 1077 | 0 | 0 | 0 | 0 | 0 | 0 | 0 | 0 | 0 | 0 | 0 | 0 | 0 | 0 | 0 | 0 | 0 | 0 |
| ENSMUSG00 | Olfr1245 | 1061 | 0 | 0 | 0 | 0 | 0 | 0 | 0 | 0 | 0 | 0 | 0 | 0 | 0 | 0 | 0 | 0 | 0 | 0 |
| ENSMUSG00 | Olfr1243 | 1352 | 0 | 0 | 0 | 0 | 0 | 0 | 0 | 0 | 0 | 0 | 0 | 0 | 0 | 0 | 0 | 0 | 0 | 0 |
| ENSMUSG00 | Olfr1242 | 2257 | 0 | 0 | 0 | 0 | 0 | 0 | 0 | 0 | 0 | 0 | 0 | 0 | 0 | 0 | 0 | 0 | 0 | 0 |
| ENSMUSG00 | Olfr1241 | 944 | 0 | 0 | 0 | 0 | 0 | 0 | 0 | 0 | 0 | 0 | 0 | 0 | 0 | 0 | 0 | 0 | 0 | 0 |
| ENSMUSG00 | Olfr1240 | 1073 | 0 | 0 | 0 | 0 | 0 | 0 | 0 | 0 | 0 | 0 | 0 | 0 | 0 | 0 | 0 | 0 | 0 | 0 |
| ENSMUSG00 | Olfr1239 | 917 | 0 | 0 | 0 | 0 | 0 | 0 | 0 | 0 | 0 | 0 | 0 | 0 | 0 | 0 | 0 | 0 | 0 | 0 |
| ENSMUSG00 | Olfr1238 | 948 | 0 | 0 | 0 | 0 | 0 | 0 | 0 | 0 | 0 | 0 | 0 | 0 | 0 | 0 | 0 | 0 | 0 | 0 |
| ENSMUSG00 | Olfr1234 | 2332 | 0 | 0 | 0 | 0 | 0 | 0 | 0 | 0 | 0 | 0 | 0 | 0 | 0 | 0 | 0 | 0 | 0 | 0 |
| ENSMUSG00 | Olfr1233 | 1041 | 0 | 0 | 0 | 0 | 0 | 0 | 0 | 0 | 0 | 0 | 0 | 0 | 0 | 0 | 0 | 0 | 0 | 0 |
| ENSMUSG00 | Olfr1232 | 935 | 0 | 0 | 0 | 0 | 0 | 0 | 0 | 0 | 0 | 0 | 0 | 0 | 0 | 0 | 0 | 0 | 0 | 0 |
| ENSMUSG00 | Olfr1231 | 941 | 0 | 0 | 0 | 0 | 0 | 0 | 0 | 0 | 0 | 0 | 0 | 0 | 0 | 0 | 0 | 0 | 0 | 0 |
| ENSMUSG00 | Olfr1230 | 917 | 0 | 0 | 0 | 0 | 0 | 0 | 0 | 0 | 0 | 0 | 0 | 0 | 0 | 0 | 0 | 0 | 0 | 0 |
| ENSMUSG00 | Olfr1229 | 2841 | 0 | 0 | 0 | 0 | 0 | 0 | 0 | 0 | 0 | 0 | 0 | 0 | 0 | 0 | 0 | 0 | 0 | 0 |
| ENSMUSG00 | Olfr1228 | 1093 | 0 | 0 | 0 | 0 | 0 | 0 | 0 | 0 | 0 | 0 | 0 | 0 | 0 | 0 | 0 | 0 | 0 | 0 |
| ENSMUSG00 | Olfr1226 | 932 | 0 | 0 | 0 | 0 | 0 | 0 | 0 | 0 | 0 | 0 | 0 | 0 | 0 | 0 | 0 | 0 | 0 | 0 |
| ENSMUSG00 | Olfr1225 | 1026 | 0 | 0 | 0 | 0 | 0 | 0 | 0 | 0 | 0 | 0 | 0 | 0 | 0 | 0 | 0 | 0 | 0 | 0 |
| ENSMUSG00 | Olfr1224-ps1 | 935 | 0 | 0 | 0 | 0 | 0 | 0 | 0 | 0 | 0 | 0 | 0 | 0 | 0 | 0 | 0 | 0 | 0 | 0 |
| ENSMUSG00 | Olfr1223 | 2464 | 0 | 0 | 0 | 0 | 0 | 0 | 0 | 0 | 0 | 0 | 0 | 0 | 0 | 0 | 0 | 0 | 0 | 0 |
| ENSMUSG00 | Olfr1222 | 1587 | 0 | 0 | 0 | 0 | 0 | 0 | 0 | 0 | 0 | 0 | 0 | 0 | 0 | 0 | 0 | 0 | 0 | 0 |
| ENSMUSG00 | Olfr1221 | 935 | 0 | 0 | 0 | 0 | 0 | 0 | 0 | 0 | 0 | 0 | 0 | 0 | 0 | 0 | 0 | 0 | 0 | 0 |
| ENSMUSG00 | Olfr1220 | 1025 | 0 | 0 | 0 | 0 | 0 | 0 | 0 | 0 | 0 | 0 | 0 | 0 | 0 | 0 | 0 | 0 | 0 | 0 |
| ENSMUSG00 | Olfr1219 | 935 | 0 | 0 | 0 | 0 | 0 | 0 | 0 | 0 | 0 | 0 | 0 | 0 | 0 | 0 | 0 | 0 | 0 | 0 |
| ENSMUSG00 | Olfr1218 | 935 | 0 | 0 | 0 | 0 | 0 | 0 | 0 | 0 | 0 | 0 | 0 | 0 | 0 | 0 | 0 | 0 | 0 | 0 |
| ENSMUSG00 | Olfr1217 | 1104 | 0 | 0 | 0 | 0 | 0 | 0 | 0 | 0 | 0 | 0 | 0 | 0 | 0 | 0 | 0 | 0 | 0 | 0 |
| ENSMUSG00 | Olfr1216 | 935 | 0 | 0 | 0 | 0 | 0 | 0 | 0 | 0 | 0 | 0 | 0 | 0 | 0 | 0 | 0 | 0 | 0 | 0 |
| ENSMUSG00 | Olfr1215 | 942 | 0 | 0 | 0 | 0 | 0 | 0 | 0 | 0 | 0 | 0 | 0 | 0 | 0 | 0 | 0 | 0 | 0 | 0 |
| ENSMUSG00 | Olfr1214 | 935 | 0 | 0 | 0 | 0 | 0 | 0 | 0 | 0 | 0 | 0 | 0 | 0 | 0 | 0 | 0 | 0 | 0 | 0 |
| ENSMUSG00 | Gm13762 | 1443 | 0 | 0 | 0 | 0 | 0 | 0 | 0 | 0 | 0 | 0 | 0 | 0 | 0 | 0 | 0 | 0 | 0 | 0 |
| ENSMUSG00 | Olfr1211 | 935 | 0 | 0 | 0 | 0 | 0 | 0 | 0 | 0 | 0 | 0 | 0 | 0 | 0 | 0 | 0 | 0 | 0 | 0 |
| ENSMUSG00 | Olfr1209 | 932 | 0 | 0 | 0 | 0 | 0 | 0 | 0 | 0 | 0 | 0 | 0 | 0 | 0 | 0 | 0 | 0 | 0 | 0 |
| ENSMUSG00 | Olfr1208 | 1074 | 0 | 0 | 0 | 0 | 0 | 0 | 0 | 0 | 0 | 0 | 0 | 0 | 0 | 0 | 0 | 0 | 0 | 0 |
| ENSMUSG00 | Olfr1200 | 1010 | 0 | 0 | 0 | 0 | 0 | 0 | 0 | 0 | 0 | 0 | 0 | 0 | 0 | 0 | 0 | 0 | 0 | 0 |
| ENSMUSG00 | Gm13760 | 895 | 0 | 0 | 0 | 0 | 0 | 0 | 0 | 0 | 0 | 0 | 0 | 0 | 0 | 0 | 0 | 0 | 0 | 0 |
| ENSMUSG00 | Olfr1198 | 926 | 0 | 0 | 0 | 0 | 0 | 0 | 0 | 0 | 0 | 0 | 0 | 0 | 0 | 0 | 0 | 0 | 0 | 0 |
| ENSMUSG00 | Gpr137b-ps | 2992 | 0.79555142 | 0.54556545 | 1.50358901 | 1.02025066 | 0.56304055 | 1.10963675 | 1.78687591 | 1.67219409 | 1.00488491 | 1.27647292 | 1.19756748 | 1.34315766 | 0.98927642 | 1.35047981 | 0.936399 | 0.85744341 | 1.20410157 | 0.66187428 |
| ENSMUSG00 | Olfr1197 | 944 | 0 | 0 | 0 | 0 | 0 | 0 | 0 | 0 | 0 | 0 | 0 | 0 | 0 | 0 | 0 | 0 | 0 | 0 |
| ENSMUSG00 | Olfr1196 | 944 | 0 | 0 | 0 | 0 | 0 | 0 | 0 | 0 | 0 | 0 | 0 | 0 | 0 | 0 | 0 | 0 | 0 | 0 |
| ENSMUSG00 | Olfr1195 | 926 | 0 | 0 | 0 | 0 | 0 | 0 | 0 | 0 | 0 | 0 | 0 | 0 | 0 | 0 | 0 | 0 | 0 | 0 |
| ENSMUSG00 | Cd80 | 3090 | 0.04126716 | 0 | 0.03130973 | 0.01474467 | 0 | 0 | 0 | 0.25956002 | 0.06664485 | 0.12207302 | 0.61844606 | 0.03612664 | 0.15965022 | 0.0601218 | 0 | 0 | 0.02480686 | 0 |
| ENSMUSG00 | Olfr1182 | 917 | 0 | 0 | 0 | 0 | 0 | 0 | 0 | 0 | 0 | 0 | 0 | 0 | 0 | 0 | 0 | 0 | 0 | 0 |
| ENSMUSG00 | Olfr1181 | 935 | 0 | 0 | 0 | 0 | 0 | 0 | 0 | 0 | 0 | 0 | 0 | 0 | 0 | 0 | 0 | 0 | 0 | 0 |
| ENSMUSG00 | Olfr1180 | 1062 | 0 | 0 | 0 | 0 | 0 | 0 | 0 | 0 | 0 | 0 | 0 | 0 | 0 | 0 | 0 | 0 | 0 | 0 |
| ENSMUSG00 | Olfr1179 | 923 | 0 | 0 | 0 | 0 | 0 | 0 | 0 | 0 | 0 | 0 | 0 | 0 | 0 | 0 | 0 | 0 | 0 | 0 |
| ENSMUSG00 | Olfr1177-ps | 957 | 0 | 0 | 0 | 0 | 0 | 0 | 0 | 0 | 0 | 0 | 0 | 0 | 0 | 0 | 0 | 0 | 0 | 0 |
| ENSMUSG00 | Olfr1173 | 938 | 0 | 0 | 0 | 0 | 0 | 0 | 0 | 0 | 0 | 0 | 0 | 0 | 0 | 0 | 0 | 0 | 0 | 0 |
| ENSMUSG00 | Olfr1170 | 950 | 0 | 0 | 0 | 0 | 0 | 0 | 0 | 0 | 0 | 0 | 0 | 0 | 0 | 0 | 0 | 0 | 0 | 0 |
| ENSMUSG00 | Olfr1167 | 1099 | 0 | 0 | 0 | 0 | 0 | 0 | 0 | 0 | 0 | 0 | 0 | 0 | 0 | 0 | 0 | 0 | 0 | 0 |
| ENSMUSG00 | Olfr1166 | 1627 | 0 | 0 | 0 | 0 | 0 | 0 | 0 | 0 | 0 | 0 | 0 | 0 | 0 | 0 | 0 | 0 | 0 | 0 |
| ENSMUSG00 | Olfr1164 | 1111 | 0 | 0 | 0 | 0 | 0 | 0 | 0 | 0 | 0 | 0 | 0 | 0 | 0 | 0 | 0 | 0 | 0 | 0 |
| ENSMUSG00 | Olfr1163 | 1042 | 0 | 0 | 0 | 0 | 0 | 0 | 0 | 0 | 0 | 0 | 0 | 0 | 0 | 0 | 0 | 0 | 0 | 0 |
| ENSMUSG00 | Olfr1162 | 944 | 0 | 0 | 0 | 0 | 0 | 0 | 0 | 0 | 0 | 0 | 0 | 0 | 0 | 0 | 0 | 0 | 0 | 0 |
| ENSMUSG00 | Olfr73 | 941 | 0 | 0 | 0 | 0 | 0 | 0 | 0 | 0 | 0 | 0 | 0 | 0 | 0 | 0 | 0 | 0 | 0 | 0 |
| ENSMUSG00 | Olfr1160 | 959 | 0 | 0 | 0 | 0 | 0 | 0 | 0 | 0 | 0 | 0 | 0 | 0 | 0 | 0 | 0 | 0 | 0 | 0 |
| ENSMUSG00 | Olfr74 | 956 | 0 | 0 | 0 | 0 | 0 | 0 | 0 | 0 | 0 | 0 | 0 | 0 | 0 | 0 | 0 | 0 | 0 | 0 |
| ENSMUSG00 | Olfr1157 | 2048 | 0 | 0 | 0 | 0 | 0 | 0 | 0 | 0 | 0 | 0 | 0 | 0 | 0 | 0 | 0 | 0 | 0 | 0 |
| ENSMUSG00 | Olfr1156 | 1075 | 0 | 0 | 0 | 0 | 0 | 0 | 0 | 0 | 0 | 0 | 0 | 0 | 0 | 0 | 0 | 0 | 0 | 0 |
| ENSMUSG00 | Olfr1155 | 944 | 0 | 0 | 0 | 0 | 0 | 0 | 0 | 0 | 0 | 0 | 0 | 0 | 0 | 0 | 0 | 0 | 0 | 0 |
| ENSMUSG00 | Olfr1154 | 932 | 0 | 0 | 0 | 0 | 0 | 0 | 0 | 0 | 0 | 0 | 0 | 0 | 0 | 0 | 0 | 0 | 0 | 0 |
| ENSMUSG00 | Olfr1141 | 935 | 0 | 0 | 0 | 0 | 0 | 0 | 0 | 0 | 0 | 0 | 0 | 0 | 0 | 0 | 0 | 0 | 0 | 0 |
| ENSMUSG00 | Olfr1138 | 935 | 0 | 0 | 0 | 0 | 0 | 0 | 0 | 0 | 0 | 0 | 0 | 0 | 0 | 0 | 0 | 0 | 0 | 0 |
| ENSMUSG00 | Olfr1137 | 932 | 0 | 0 | 0 | 0 | 0 | 0 | 0 | 0 | 0 | 0 | 0 | 0 | 0 | 0 | 0 | 0 | 0 | 0 |
| ENSMUSG00 | Olfr1136 | 1056 | 0 | 0 | 0 | 0 | 0 | 0 | 0 | 0 | 0 | 0 | 0 | 0 | 0 | 0 | 0 | 0 | 0 | 0 |
| ENSMUSG00 | Olfr1135 | 932 | 0 | 0 | 0 | 0 | 0 | 0 | 0 | 0 | 0 | 0 | 0 | 0 | 0 | 0 | 0 | 0 | 0 | 0 |
| ENSMUSG00 | Olfr1134 | 2061 | 0 | 0 | 0 | 0 | 0 | 0 | 0 | 0 | 0 | 0 | 0 | 0 | 0 | 0 | 0 | 0 | 0 | 0 |
| ENSMUSG00 | Olfr1133 | 941 | 0 | 0 | 0 | 0 | 0 | 0 | 0 | 0 | 0 | 0 | 0 | 0 | 0 | 0 | 0 | 0 | 0 | 0 |
| ENSMUSG00 | Olfr1128 | 935 | 0 | 0 | 0 | 0 | 0 | 0 | 0 | 0 | 0 | 0 | 0 | 0 | 0 | 0 | 0 | 0 | 0 | 0 |
| ENSMUSG00 | Olfr1111 | 939 | 0 | 0 | 0 | 0 | 0 | 0 | 0 | 0 | 0 | 0 | 0 | 0 | 0 | 0 | 0 | 0 | 0 | 0 |
| ENSMUSG00 | Olfr1110 | 938 | 0 | 0 | 0 | 0 | 0 | 0 | 0 | 0 | 0 | 0 | 0 | 0 | 0 | 0 | 0 | 0 | 0 | 0 |
| ENSMUSG00 | Olfr259 | 938 | 0 | 0 | 0 | 0 | 0 | 0 | 0 | 0 | 0 | 0 | 0 | 0 | 0 | 0 | 0 | 0 | 0 | 0 |
| ENSMUSG00 | Olfr1109 | 938 | 0 | 0 | 0 | 0 | 0 | 0 | 0 | 0 | 0 | 0 | 0 | 0 | 0 | 0 | 0 | 0 | 0 | 0 |
| ENSMUSG00 | Gm10812 | 544 | 0 | 0 | 0 | 0 | 0 | 0 | 0 | 0 | 0 | 0 | 0 | 0 | 0 | 0 | 0 | 0 | 0 | 0 |
| ENSMUSG00 | Olfr1107 | 1074 | 0 | 0 | 0 | 0 | 0 | 0 | 0 | 0 | 0 | 0 | 0 | 0 | 0 | 0 | 0 | 0 | 0 | 0 |
| ENSMUSG00 | Olfr1106 | 938 | 0 | 0 | 0 | 0 | 0 | 0 | 0 | 0 | 0 | 0 | 0 | 0 | 0 | 0 | 0 | 0 | 0 | 0 |
| ENSMUSG00 | Olfr1105 | 938 | 0 | 0 | 0 | 0 | 0 | 0 | 0 | 0 | 0 | 0 | 0 | 0 | 0 | 0 | 0 | 0 | 0 | 0 |
| ENSMUSG00 | Olfr1104 | 932 | 0 | 0 | 0 | 0 | 0 | 0 | 0 | 0 | 0 | 0 | 0 | 0 | 0 | 0 | 0 | 0 | 0 | 0 |
| ENSMUSG00 | Olfr1101 | 932 | 0 | 0 | 0 | 0 | 0 | 0 | 0 | 0 | 0 | 0 | 0 | 0 | 0 | 0 | 0 | 0 | 0 | 0 |
| ENSMUSG00 | Olfr1099 | 1030 | 0 | 0 | 0 | 0 | 0 | 0 | 0 | 0 | 0 | 0 | 0 | 0 | 0 | 0 | 0 | 0 | 0 | 0 |
| ENSMUSG00 | Olfr1098 | 2155 | 0 | 0 | 0 | 0 | 0 | 0 | 0 | 0 | 0 | 0 | 0 | 0 | 0 | 0 | 0 | 0 | 0 | 0 |
| ENSMUSG00 | Olfr1097 | 1221 | 0 | 0 | 0 | 0 | 0 | 0 | 0 | 0 | 0 | 0 | 0 | 0 | 0 | 0 | 0 | 0 | 0 | 0 |
| ENSMUSG00 | Olfr1095 | 926 | 0 | 0 | 0 | 0 | 0 | 0 | 0 | 0 | 0 | 0 | 0 | 0 | 0 | 0 | 0 | 0 | 0 | 0 |
| ENSMUSG00 | Olfr1090 | 941 | 0 | 0 | 0 | 0 | 0 | 0 | 0 | 0 | 0 | 0 | 0 | 0 | 0 | 0 | 0 | 0 | 0 | 0 |
| ENSMUSG00 | Olfr1089 | 935 | 0 | 0 | 0 | 0 | 0 | 0 | 0 | 0 | 0 | 0 | 0 | 0 | 0 | 0 | 0 | 0 | 0 | 0 |
| ENSMUSG00 | Olfr1087 | 941 | 0 | 0 | 0 | 0 | 0 | 0 | 0 | 0 | 0 | 0 | 0 | 0 | 0 | 0 | 0 | 0 | 0 | 0 |
| ENSMUSG00 | Olfr1086 | 932 | 0 | 0 | 0 | 0 | 0 | 0 | 0 | 0 | 0 | 0 | 0 | 0 | 0 | 0 | 0 | 0 | 0 | 0 |
| ENSMUSG00 | Olfr1085 | 941 | 0 | 0 | 0 | 0 | 0 | 0 | 0 | 0 | 0 | 0 | 0 | 0 | 0 | 0 | 0 | 0 | 0 | 0 |
| ENSMUSG00 | Olfr1084 | 1929 | 0 | 0 | 0 | 0 | 0 | 0 | 0 | 0 | 0 | 0 | 0 | 0 | 0 | 0 | 0 | 0 | 0 | 0 |
| ENSMUSG00 | Olfr1080 | 1918 | 0 | 0 | 0 | 0 | 0 | 0 | 0 | 0 | 0 | 0 | 0 | 0 | 0 | 0 | 0 | 0 | 0 | 0 |
| ENSMUSG00 | Olfr1079 | 2080 | 0 | 0 | 0 | 0 | 0 | 0 | 0 | 0 | 0 | 0 | 0 | 0 | 0 | 0 | 0 | 0 | 0 | 0 |
| ENSMUSG00 | Olfr228 | 941 | 0 | 0 | 0 | 0 | 0 | 0 | 0 | 0 | 0 | 0 | 0 | 0 | 0 | 0 | 0 | 0 | 0 | 0 |
| ENSMUSG00 | Olfr1066 | 941 | 0 | 0 | 0 | 0 | 0 | 0 | 0 | 0 | 0 | 0 | 0 | 0 | 0 | 0 | 0 | 0 | 0 | 0 |
| ENSMUSG00 | Olfr1065 | 991 | 0 | 0 | 0 | 0 | 0 | 0 | 0 | 0 | 0 | 0 | 0 | 0 | 0 | 0 | 0 | 0 | 0 | 0 |
| ENSMUSG00 | F930017D23 | 1832 | 0 | 0 | 0 | 0.02486957 | 0.05254577 | 0 | 0 | 0 | 0 | 0 | 0 | 0 | 0 | 0.15210946 | 0 | 0 | 0 | 0.09008041 |
| ENSMUSG00 | Olfr1061 | 941 | 0 | 0 | 0 | 0 | 0 | 0 | 0 | 0 | 0 | 0 | 0 | 0 | 0 | 0 | 0 | 0 | 0 | 0 |
| ENSMUSG00 | Olfr1058 | 950 | 0 | 0 | 0 | 0 | 0 | 0 | 0 | 0 | 0 | 0 | 0 | 0 | 0 | 0 | 0 | 0 | 0 | 0 |
| ENSMUSG00 | Olfr1057 | 947 | 0 | 0 | 0 | 0 | 0 | 0 | 0 | 0 | 0 | 0 | 0 | 0 | 0 | 0 | 0 | 0 | 0 | 0 |
| ENSMUSG00 | Olfr1056 | 1078 | 0 | 0 | 0 | 0 | 0 | 0 | 0 | 0 | 0 | 0 | 0 | 0 | 0 | 0 | 0 | 0 | 0 | 0 |
| ENSMUSG00 | Olfr1055 | 1040 | 0 | 0 | 0 | 0 | 0 | 0 | 0 | 0 | 0 | 0 | 0 | 0 | 0 | 0 | 0 | 0 | 0 | 0 |
| ENSMUSG00 | Olfr1054 | 938 | 0 | 0 | 0 | 0 | 0 | 0 | 0 | 0 | 0 | 0 | 0 | 0 | 0 | 0 | 0 | 0 | 0 | 0 |
| ENSMUSG00 | Olfr1053 | 941 | 0 | 0 | 0 | 0 | 0 | 0 | 0 | 0 | 0 | 0 | 0 | 0 | 0 | 0 | 0 | 0 | 0 | 0 |
| ENSMUSG00 | Olfr1051 | 926 | 0 | 0 | 0 | 0 | 0 | 0 | 0 | 0 | 0 | 0 | 0 | 0 | 0 | 0 | 0 | 0 | 0 | 0 |
| ENSMUSG00 | Olfr1049 | 926 | 0 | 0 | 0 | 0 | 0 | 0 | 0 | 0 | 0 | 0 | 0 | 0 | 0 | 0 | 0 | 0 | 0 | 0 |
| ENSMUSG00 | Olfr1048 | 962 | 0 | 0 | 0 | 0 | 0 | 0 | 0 | 0 | 0 | 0 | 0 | 0 | 0 | 0 | 0 | 0 | 0 | 0 |
| ENSMUSG00 | Olfr1047 | 959 | 0 | 0 | 0 | 0 | 0 | 0 | 0 | 0 | 0 | 0 | 0 | 0 | 0 | 0 | 0 | 0 | 0 | 0 |
| ENSMUSG00 | Olfr1046 | 950 | 0 | 0 | 0 | 0 | 0 | 0 | 0 | 0 | 0 | 0 | 0 | 0 | 0 | 0 | 0 | 0 | 0 | 0 |
| ENSMUSG00 | Olfr1045 | 947 | 0 | 0 | 0 | 0 | 0 | 0 | 0 | 0 | 0 | 0 | 0 | 0 | 0 | 0 | 0 | 0 | 0 | 0 |
| ENSMUSG00 | Olfr52 | 959 | 0 | 0 | 0 | 0 | 0 | 0 | 0 | 0 | 0 | 0 | 0 | 0 | 0 | 0 | 0 | 0 | 0 | 0 |
| ENSMUSG00 | Olfr1044 | 944 | 0 | 0 | 0 | 0 | 0 | 0 | 0 | 0 | 0 | 0 | 0 | 0 | 0 | 0 | 0 | 0 | 0 | 0 |
| ENSMUSG00 | Olfr1043 | 944 | 0 | 0 | 0 | 0 | 0 | 0 | 0 | 0 | 0 | 0 | 0 | 0 | 0 | 0 | 0 | 0 | 0 | 0 |
| ENSMUSG00 | Olfr1042 | 1096 | 0 | 0 | 0 | 0 | 0 | 0 | 0 | 0 | 0 | 0 | 0 | 0 | 0 | 0 | 0 | 0 | 0 | 0 |
| ENSMUSG00 | Olfr1040 | 941 | 0 | 0 | 0 | 0 | 0 | 0 | 0 | 0 | 0 | 0 | 0 | 0 | 0 | 0 | 0 | 0 | 0 | 0 |
| ENSMUSG00 | Olfr1039 | 959 | 0 | 0 | 0 | 0 | 0 | 0 | 0 | 0 | 0 | 0 | 0 | 0 | 0 | 0 | 0 | 0 | 0 | 0 |
| ENSMUSG00 | Olfr1037 | 1124 | 0 | 0 | 0 | 0 | 0 | 0 | 0 | 0 | 0 | 0 | 0 | 0 | 0 | 0 | 0 | 0 | 0 | 0 |
| ENSMUSG00 | Olfr1024 | 983 | 0 | 0 | 0 | 0 | 0 | 0 | 0 | 0 | 0 | 0 | 0 | 0 | 0 | 0 | 0 | 0 | 0 | 0 |
| ENSMUSG00 | Olfr1019 | 1863 | 0 | 0 | 0 | 0 | 0 | 0 | 0 | 0 | 0 | 0 | 0 | 0 | 0 | 0 | 0 | 0 | 0 | 0 |
| ENSMUSG00 | Olfr1016 | 929 | 0 | 0 | 0 | 0 | 0 | 0 | 0 | 0 | 0 | 0 | 0 | 0 | 0 | 0 | 0 | 0 | 0 | 0 |
| ENSMUSG00 | Olfr1012 | 935 | 0 | 0 | 0 | 0 | 0 | 0 | 0 | 0 | 0 | 0 | 0 | 0 | 0 | 0 | 0 | 0 | 0 | 0 |
| ENSMUSG00 | Olfr1006 | 1947 | 0 | 0 | 0 | 0 | 0 | 0 | 0 | 0 | 0 | 0 | 0 | 0 | 0 | 0 | 0 | 0 | 0 | 0 |
| ENSMUSG00 | Olfr154 | 1078 | 0 | 0 | 0 | 0 | 0 | 0 | 0 | 0 | 0 | 0 | 0 | 0 | 0 | 0 | 0 | 0 | 0 | 0 |
| ENSMUSG00 | Olfr1002 | 956 | 0 | 0 | 0 | 0 | 0 | 0 | 0 | 0 | 0 | 0 | 0 | 0 | 0 | 0 | 0 | 0 | 0 | 0 |
| ENSMUSG00 | Olfr1000 | 944 | 0 | 0 | 0 | 0 | 0 | 0 | 0 | 0 | 0 | 0 | 0 | 0 | 0 | 0 | 0 | 0 | 0 | 0 |
| ENSMUSG00 | 4833423E24 | 2260 | 0 | 0 | 0 | 0 | 0 | 0 | 0 | 0 | 0 | 0 | 0 | 0 | 0 | 0 | 0.2370001 | 0 | 0 | 0 |
| ENSMUSG00 | Olfr995 | 1112 | 0 | 0 | 0 | 0 | 0 | 0 | 0 | 0 | 0 | 0 | 0 | 0 | 0 | 0 | 0 | 0 | 0 | 0 |
| ENSMUSG00 | Olfr994 | 944 | 0 | 0 | 0 | 0 | 0 | 0 | 0 | 0 | 0 | 0 | 0 | 0 | 0 | 0 | 0 | 0 | 0 | 0 |
| ENSMUSG00 | Olfr993 | 944 | 0 | 0 | 0 | 0 | 0 | 0 | 0 | 0 | 0 | 0 | 0 | 0 | 0 | 0 | 0 | 0 | 0 | 0 |
| ENSMUSG00 | Olfr992 | 929 | 0 | 0 | 0 | 0 | 0 | 0 | 0 | 0 | 0 | 0 | 0 | 0 | 0 | 0 | 0 | 0 | 0 | 0 |
| ENSMUSG00 | Olfr988 | 2208 | 0 | 0 | 0 | 0 | 0 | 0 | 0 | 0 | 0 | 0 | 0 | 0 | 0 | 0 | 0 | 0 | 0 | 0 |
| ENSMUSG00 | Olfr987 | 2488 | 0 | 0 | 0 | 0 | 0 | 0 | 0 | 0 | 0 | 0 | 0 | 0 | 0 | 0 | 0 | 0 | 0 | 0 |
| ENSMUSG00 | Lrrc55 | 4076 | 0 | 0 | 0 | 0 | 0 | 0 | 0 | 0 | 0 | 0 | 0 | 0 | 0 | 0 | 0 | 0 | 0 | 0 |
| ENSMUSG00 | Ccdc162 | 5527 | 0.16149967 | 0.39780189 | 0.09627444 | 0.23905741 | 0.36575734 | 0.20620823 | 0.52217721 | 0.6011831 | 0.69302442 | 0.45214172 | 0.39762043 | 0.32315926 | 0.6247936 | 0.78989401 | 0.30563846 | 0.11777462 | 0.15949065 | 0.05971678 |
| ENSMUSG00 | Znhit2-ps | 1280 | 14.6111543 | 12.7005409 | 11.2619625 | 11.06991 | 10.8672859 | 14.1689848 | 12.0499479 | 11.7262611 | 11.9698311 | 14.4030664 | 9.44301914 | 13.6922797 | 17.4974144 | 16.4368528 | 10.300389 | 8.97436111 | 12.1266799 | 14.1820353 |
| ENSMUSG00 | Gm10813 | 88 | 0 | 0 | 0 | 0 | 0 | 0 | 0 | 0 | 0 | 0 | 0 | 0 | 0 | 0 | 0 | 0 | 0 | 0 |
| ENSMUSG00 | Ccdc58 | 1219 | 19.7357913 | 17.2986563 | 18.2144794 | 16.8190894 | 19.5449567 | 14.5528127 | 20.3766665 | 22.495607 | 20.001983 | 22.0861785 | 29.864248 | 19.4141435 | 23.674464 | 21.0312849 | 20.8880735 | 21.9880436 | 17.795475 | 19.3592344 |
| ENSMUSG00 | Amd1 | 3195 | 7.82254883 | 8.54978819 | 6.51036559 | 6.95893259 | 6.83940324 | 7.78575031 | 7.0783914 | 7.63848048 | 6.12319092 | 6.00636502 | 5.77187292 | 7.47702773 | 7.00991876 | 9.21613502 | 7.13128651 | 7.09484807 | 7.78521649 | 8.2126272 |

| | | | | | | | | | | | | | | | | | | | |
|---|---|---|---|---|---|---|---|---|---|---|---|---|---|---|---|---|---|---|---|
| ENSMUSG00 | Gm13695 | 2492 | 0 | 0 | 0 | 0 | 0 | 0 | 0 | 0 | 0 | 0 | 0 | 0 | 0 | 0 | 0 | 0 | 0 | 0 |
| ENSMUSG00 | Gm13693 | 2492 | 0 | 0 | 0 | 0 | 0 | 0 | 0 | 0 | 0 | 0 | 0 | 0 | 0 | 0 | 0 | 0 | 0 | 0 |
| ENSMUSG00 | Gm13698 | 2492 | 0 | 0 | 0 | 0 | 0 | 0 | 0 | 0 | 0 | 0 | 0 | 0 | 0 | 0 | 0 | 0 | 0 | 0 |
| ENSMUSG00 | Gm8055 | 1007 | 0.16883883 | 0.19848795 | 0.14411181 | 0.36195467 | 0.3345814 | 0.44287448 | 0.46983792 | 0.2654884 | 0.53170279 | 0.51505237 | 0.42698573 | 0.44342135 | 0.34292296 | 0.46121239 | 0.5728121 | 0.19012218 | 0.53283829 | 0.2731336 |
| ENSMUSG00 | 4930547M16 | 2398 | 0.15952735 | 0.0833517 | 0.06051734 | 0.09499801 | 0.18064525 | 0.06199258 | 0.13811041 | 0.22297483 | 0.10305217 | 0.45223777 | 0.75706773 | 0.18620738 | 0.02057211 | 0.09683921 | 0.24054286 | 0.33532225 | 0.07991305 | 0 |
| ENSMUSG00 | Gm6043 | 494 | 0.08604287 | 0 | 0.48961063 | 0.2766865 | 0.19486609 | 0 | 0.19154931 | 0.38656278 | 0.33349405 | 0.2863403 | 0.29013133 | 0 | 0.09986218 | 0 | 0.25021188 | 0.3100454 | 0.15516719 | 0.3340634 |
| ENSMUSG00 | Fsip2 | 19266 | 0 | 0 | 0 | 0 | 0 | 0 | 0 | 0 | 0 | 0 | 0 | 0 | 0 | 0 | 0 | 0 | 0 | 0 |
| ENSMUSG00 | Heg1 | 12921 | 0.27961767 | 0.45634102 | 0.28078436 | 0.29619439 | 0.30173251 | 0.28762905 | 0.16477599 | 0.12414526 | 0.23906731 | 0.10217628 | 0.06655439 | 0.01727905 | 0.35888865 | 0.31631295 | 0.10841676 | 0.65492023 | 0.40636968 | 0.15326429 |
| ENSMUSG00 | Cerkl | 5266 | 0.04035812 | 0 | 0.05511606 | 0 | 0 | 0 | 0 | 0.06527376 | 0 | 0.0089538 | 0 | 0 | 0 | 0 | 0 | 0 | 0 | 0 |
| ENSMUSG00 | Nkain2 | 322 | 0 | 0 | 0 | 0 | 0 | 0 | 0 | 0 | 0.63954217 | 0 | 0 | 0 | 0 | 0 | 0 | 0 | 0 | 0 |
| ENSMUSG00 | Gm10818 | 1389 | 0.12240511 | 0.02398337 | 0 | 0.09840398 | 0.03465221 | 0 | 0 | 0 | 0.17791152 | 0.03394579 | 0 | 0.44202469 | 0 | 0.06687414 | 0 | 0.13783516 | 0 | 0 |
| ENSMUSG00 | Cenpw | 1507 | 5.41539071 | 4.17792768 | 3.72350997 | 5.32099788 | 4.98246854 | 5.32683686 | 8.91625801 | 7.57765648 | 9.56554779 | 8.51027955 | 7.79869926 | 7.92604631 | 11.4573132 | 10.3243326 | 11.1547676 | 9.96013145 | 12.0548538 | 10.9142156 |
| ENSMUSG00 | Dfnb59 | 1478 | 0.28758576 | 0 | 0 | 0 | 0.22795904 | 0.06705377 | 0 | 0.20672478 | 0.13933192 | 0 | 0 | 0 | 0 | 0 | 0 | 0 | 0 | 0 |
| ENSMUSG00 | Gm10819 | 560 | 0.8349231 | 0.41641118 | 0.0863813 | 0.56951305 | 0.85949864 | 0.26546107 | 0.33794771 | 1.36401437 | 0.36773675 | 0.50518611 | 0.51187456 | 0.59802496 | 0.52855626 | 0.91229457 | 0.36787103 | 1.02564127 | 0.95815742 | 0.88407493 |
| ENSMUSG00 | Bex6 | 698 | 0 | 0 | 0 | 0 | 0 | 0 | 0 | 0 | 0 | 0 | 0 | 0 | 0 | 0 | 0 | 0 | 0 | 0 |
| ENSMUSG00 | Pde11a | 6561 | 0.78389366 | 1.23888831 | 0.59720408 | 0.81247404 | 0.59422128 | 1.01960357 | 0.66343034 | 0.58793364 | 0.20087921 | 0.21559535 | 0.1383515 | 0.08507188 | 0.01503793 | 0.01415762 | 0.0188393 | 0.00583609 | 0 | 0 |
| ENSMUSG00 | Ttc30a1 | 2861 | 0.7576945 | 0.64040866 | 1.31881698 | 0.76439361 | 0.77387923 | 1.12580485 | 1.75293393 | 0.90775371 | 1.15166769 | 0.92290786 | 1.03531892 | 0.99496637 | 0.98284491 | 0.92531037 | 1.00807722 | 0.80301868 | 1.13867013 | 0.67295307 |
| ENSMUSG00 | Ttc30a2 | 2128 | 0 | 0 | 0 | 0 | 0 | 0 | 0 | 0 | 0 | 0 | 0 | 0 | 0 | 0 | 0 | 0.10796224 | 0 | 0 |
| ENSMUSG00 | Ttc30b | 2599 | 5.41331792 | 4.3579777 | 6.84934943 | 4.71562944 | 4.51873389 | 3.50815813 | 7.09963247 | 5.37838372 | 4.5798011 | 5.96867309 | 4.98153665 | 3.90859969 | 3.22678951 | 4.16370618 | 4.21685797 | 4.96496141 | 7.13733271 | 5.52419271 |
| ENSMUSG00 | Gm13653 | 997 | 0 | 0.03341313 | 0 | 0 | 0 | 0 | 0 | 0.03830732 | 0 | 0 | 0.14375614 | 0 | 0 | 0.09316768 | 0 | 0 | 0 | 0 |
| ENSMUSG00 | 6720416L17F | 2524 | 13.0849928 | 10.3739837 | 13.5882857 | 14.2423391 | 11.5753081 | 11.7010417 | 10.928404 | 10.5921877 | 13.2175427 | 10.1997956 | 9.91841452 | 13.0693568 | 10.2221049 | 7.4892011 | 8.2925477 | 8.8444628 | 8.15420818 | 7.95496195 |
| ENSMUSG00 | Mrpl23-ps1 | 1122 | 1.06073523 | 1.03917228 | 1.25029624 | 0.97456779 | 0.68637324 | 1.01578687 | 0.75902693 | 1.15734553 | 0.77087062 | 0.5463094 | 0.63870265 | 0.54721238 | 0.70348545 | 1.24182052 | 0.77115212 | 0.75079622 | 0.78565493 | 0.88249904 |
| ENSMUSG00 | Gm10822 | 2318 | 0.018337 | 0 | 0 | 0 | 0 | 0 | 0 | 0 | 0 | 0.02034111 | 0 | 0.0722377 | 0.04256421 | 0 | 0 | 0.09911288 | 0.13227367 | 0.09492512 |
| ENSMUSG00 | Wipf1 | 8308 | 0.01534852 | 0.08420448 | 0.15138563 | 0.10419594 | 0.20856394 | 0.13121812 | 0.27335202 | 0.18388253 | 0.26770244 | 0.30646822 | 0.14376191 | 0.25529552 | 0.20782584 | 0.13416684 | 0.18349273 | 0.17513759 | 0.23988534 | 0.19863664 |
| ENSMUSG00 | Gm1968 | 345 | 0 | 0 | 0 | 0 | 0 | 0 | 0 | 0 | 0 | 0.68334353 | 0 | 0 | 0 | 0 | 0.1194248 | 0 | 0 | 0 |
| ENSMUSG00 | Gm6913 | 347 | 0 | 0.09600258 | 0.27880997 | 0 | 0.27741743 | 0 | 0 | 0 | 0 | 0 | 0.13768 | 0 | 0 | 0 | 0 | 0 | 0.11045042 | 0 |
| ENSMUSG00 | Carns1 | 3958 | 0.49399649 | 0.7574938 | 0.92885 | 0.25324481 | 0.86340742 | 0.51330522 | 1.03997046 | 0.70440762 | 0.92612428 | 0.75050387 | 0.54317158 | 1.11405693 | 0.1495662 | 0.36376117 | 0.63499113 | 0.4643631 | 1.57836948 | 0.76440143 |
| ENSMUSG00 | Aldh3b2 | 2131 | 0 | 0.07816259 | 0.20429928 | 0.25656149 | 0.13551926 | 0.11626639 | 0.06660631 | 0.10753375 | 0 | 0.19913483 | 0.44838065 | 0.23573016 | 0.76393865 | 1.91791633 | 0.8120438 | 0.16171538 | 0.57552393 | 0.59371631 |
| ENSMUSG00 | H60b | 1206 | 0.28195805 | 0.0828679 | 0.16044289 | 0.03777864 | 0 | 0.24653101 | 0.510004 | 0.9183911 | 0.54642144 | 0.70374184 | 0.63383029 | 1.01819617 | 0.16362162 | 0.38510852 | 0.34163811 | 0 | 0.03177968 | 0 |
| ENSMUSG00 | 4933404M02 | 3669 | 4.12424166 | 2.59675327 | 2.71598453 | 3.21621981 | 2.68930072 | 3.25489474 | 2.28245957 | 1.75920305 | 3.69320351 | 2.12043228 | 2.16152715 | 2.63180966 | 3.02526067 | 3.65831051 | 3.34643327 | 3.06825799 | 2.35034527 | 3.77822156 |
| ENSMUSG00 | Sp5 | 2098 | 64.1021816 | 55.9872569 | 62.62274 | 68.8193277 | 55.1062348 | 62.2597739 | 54.5967568 | 46.8940079 | 52.3567386 | 58.0506513 | 62.4399124 | 55.0973485 | 72.0462319 | 58.4203749 | 73.8800063 | 58.7682344 | 57.1788891 | 66.9653282 |
| ENSMUSG00 | Kbtbd10 | 2523 | 0.01684708 | 0.22446262 | 0.11503812 | 0 | 0 | 0 | 0.48756625 | 0.21192772 | 0 | 0.13081844 | 0.0757431 | 0 | 0 | 0.09204139 | 0 | 0.03035324 | 0 | 0 |
| ENSMUSG00 | Gm10827 | 2584 | 0 | 0.01289199 | 0.05616122 | 0.1057919 | 0.09313453 | 0.01917675 | 0 | 0.05912137 | 0.17532959 | 0.14597741 | 0.01848876 | 0 | 0.0763652 | 0.10784231 | 0 | 0.01481835 | 0.08899295 | 0.02128835 |
| ENSMUSG00 | Gm13597 | 1623 | 0.78567793 | 0.73891817 | 0.83454026 | 0.61758655 | 0.41518603 | 0.67169448 | 0.34981648 | 0.61183146 | 0.32989815 | 0.52292832 | 0.35323444 | 0.41268512 | 0.42553718 | 0.42924295 | 0.25386048 | 0.51903473 | 0.18891582 | 0.23725431 |
| ENSMUSG00 | Gm1322 | 702 | 0 | 0 | 0 | 0 | 0 | 0 | 0 | 0 | 0 | 0 | 0 | 0 | 0 | 0 | 0 | 0 | 0 | 0 |
| ENSMUSG00 | Scn9a | 13206 | 0 | 0.03027069 | 0.01465199 | 0.12075091 | 0 | 0 | 0 | 0.0057841 | 0.00311877 | 0 | 0 | 0 | 0 | 0 | 0 | 0 | 0 | 0 |
| ENSMUSG00 | Scn2a1 | 9136 | 0.01395748 | 0.00364633 | 0 | 0 | 0.00526838 | 0 | 0.08285933 | 0 | 0.08114681 | 0.05677077 | 0 | 0.08553188 | 0.03239837 | 0 | 0 | 0 | 0.0293656 | 0.08429592 |
| ENSMUSG00 | Gm10828 | 1571 | 0 | 0 | 0 | 0 | 0 | 0 | 0 | 0 | 0 | 0 | 0 | 0 | 0 | 0 | 0 | 0 | 0 | 0 |
| ENSMUSG00 | Fign | 13526 | 0.04399471 | 0.05910908 | 0.02503436 | 0.0033684 | 0.02846779 | 0.11356903 | 0 | 0.00282363 | 0 | 0 | 0.05651333 | 0.07840438 | 0 | 0 | 0.0030461 | 0 | 0 | 0 |
| ENSMUSG00 | Gm13582 | 2651 | 0 | 0.02513232 | 0.14597821 | 0.06874545 | 0.10893683 | 0 | 0 | 0 | 0 | 0 | 0 | 0 | 0 | 0 | 0 | 0 | 0.07228649 | 0 |
| ENSMUSG00 | Zbtb2 | 3334 | 6.54023854 | 6.28488616 | 5.84719038 | 6.61414081 | 5.57256225 | 7.56518938 | 3.93089744 | 6.4952286 | 7.90622978 | 7.39646605 | 8.87003578 | 7.41641827 | 4.64613738 | 5.65575869 | 5.07913601 | 7.25844747 | 7.94345266 | 7.17725298 |
| ENSMUSG00 | A930003A15F | 744 | 0 | 0 | 0 | 0 | 0 | 0 | 0 | 0 | 0 | 0 | 0 | 0 | 0 | 0 | 0 | 0 | 0 | 0 |
| ENSMUSG00 | Rprm | 1459 | 1.04879118 | 2.80842083 | 0.66310528 | 0.53086892 | 1.484535 | 1.80006503 | 4.44265731 | 2.85330491 | 2.76646919 | 3.296348 | 0.75313505 | 2.71618372 | 3.68552363 | 6.62122699 | 2.06148977 | 2.12579794 | 0.8143353 | 2.71463721 |
| ENSMUSG00 | 4930588K23F | 1961 | 0 | 0 | 0 | 0 | 0 | 0 | 0 | 0 | 0 | 0 | 0 | 0 | 0 | 0 | 0 | 0 | 0 | 0 |
| ENSMUSG00 | Igll1 | 1327 | 0 | 0 | 0 | 0 | 0 | 0 | 0 | 0 | 0 | 0 | 0 | 0 | 0 | 0 | 0 | 0 | 0 | 0 |
| ENSMUSG00 | Rc3h2 | 14765 | 3.88635202 | 3.86036993 | 3.72179685 | 3.40357819 | 3.78144473 | 4.44682506 | 2.97366516 | 3.59290531 | 4.69188754 | 3.93747491 | 4.67880731 | 4.29060461 | 3.29436305 | 3.16757169 | 3.69739665 | 3.35318357 | 4.32971644 | 3.90074736 |
| ENSMUSG00 | Olfr362 | 953 | 0 | 0 | 0 | 0 | 0 | 0 | 0 | 0 | 0 | 0 | 0 | 0 | 0 | 0 | 0 | 0 | 0 | 0 |
| ENSMUSG00 | Olfr361 | 968 | 0 | 0 | 0 | 0 | 0 | 0 | 0 | 0 | 0 | 0 | 0 | 0 | 0 | 0 | 0 | 0 | 0 | 0 |
| ENSMUSG00 | Olfr358 | 1058 | 0 | 0 | 0 | 0 | 0 | 0 | 0 | 0 | 0 | 0 | 0 | 0 | 0 | 0 | 0 | 0 | 0 | 0 |
| ENSMUSG00 | Olfr355 | 932 | 0 | 0 | 0 | 0 | 0 | 0 | 0 | 0 | 0 | 0 | 0 | 0 | 0 | 0 | 0 | 0 | 0 | 0 |
| ENSMUSG00 | Olfr353 | 935 | 0 | 0 | 0 | 0 | 0 | 0 | 0 | 0 | 0.0880995 | 0 | 0 | 0 | 0 | 0 | 0 | 0 | 0.04099069 | 0 |
| ENSMUSG00 | Olfr351 | 932 | 0 | 0 | 0 | 0 | 0 | 0 | 0 | 0 | 0 | 0 | 0 | 0 | 0 | 0 | 0 | 0 | 0 | 0 |
| ENSMUSG00 | Olfr3 | 941 | 0 | 0 | 0 | 0 | 0 | 0 | 0 | 0 | 0 | 0 | 0 | 0 | 0 | 0 | 0 | 0 | 0 | 0 |
| ENSMUSG00 | Olfr341 | 941 | 0 | 0 | 0 | 0 | 0 | 0 | 0 | 0 | 0 | 0 | 0 | 0 | 0 | 0 | 0 | 0 | 0 | 0 |
| ENSMUSG00 | 2810410L24F | 3069 | 2.99156664 | 3.03929957 | 3.08934896 | 2.88003995 | 2.82298673 | 2.85768004 | 2.92910036 | 2.97425356 | 3.52950591 | 2.13553201 | 2.33504196 | 3.31001974 | 1.80031761 | 1.67979593 | 2.10773681 | 1.68433757 | 1.83576593 | 2.3839072 |
| ENSMUSG00 | Rpl36-ps8 | 314 | 0 | 0 | 0 | 0 | 0 | 0 | 0 | 0 | 0 | 0 | 0 | 0 | 0 | 0 | 0 | 0.12194461 | 0 | 0 |
| ENSMUSG00 | Gm13443 | 554 | 0.84396559 | 0.5411842 | 5.32632735 | 0.32896061 | 7.90614631 | 21.3774428 | 0.34160779 | 0.89621161 | 4.53497737 | 0.25532872 | 8.27869317 | 19.0418055 | 0.53428069 | 1.34134548 | 3.34669678 | 0.82939944 | 7.67909559 | 16.8800508 |
| ENSMUSG00 | Gm5561 | 651 | 0 | 0.15351564 | 0.0743065 | 0 | 0 | 0.07611787 | 0.14535385 | 0.11733457 | 0.12653307 | 0 | 0 | 0 | 0 | 0.07134268 | 0 | 0 | 0.17661888 | 0 |
| ENSMUSG00 | Hoxc4 | 2553 | 0.08324555 | 0.22182499 | 0.05684316 | 0.48184418 | 0.26394318 | 0.11645766 | 0.1667897 | 0.04487944 | 0.08066298 | 0.31396865 | 0.07485305 | 0.30607884 | 0.4637548 | 0.50937503 | 0.17752335 | 0.49494322 | 0 | 0.90496768 |
| ENSMUSG00 | A630010A05F | 1953 | 0.58762916 | 1.10872408 | 0.44583899 | 0.55988995 | 0.78864391 | 1.14176805 | 1.35663595 | 0.54756132 | 1.37077497 | 0.98985091 | 1.19865487 | 1.28607517 | 1.5155735 | 1.04635937 | 1.07592389 | 1.09793856 | 2.02130496 | 1.52098913 |
| ENSMUSG00 | Krt76 | 2303 | 0 | 0 | 0 | 0 | 0 | 0 | 0 | 0 | 0 | 0 | 0 | 0 | 0 | 0 | 0 | 0 | 0 | 0 |
| ENSMUSG00 | Ciita | 1618 | 0 | 0 | 0 | 0 | 0 | 0 | 0.43862187 | 0 | 0 | 0 | 0 | 0 | 0 | 0 | 0 | 0 | 0 | 0 |
| ENSMUSG00 | 9430097D07 | 1500 | 0 | 0.0222086 | 0 | 0 | 0 | 0 | 0 | 0 | 0.02745768 | 0 | 0 | 0 | 0 | 0 | 0 | 0 | 0 | 0.14669095 |
| ENSMUSG00 | 6030408B16F | 1152 | 0 | 0 | 0 | 0.07909904 | 0 | 0 | 0 | 0 | 0 | 0 | 0.41471318 | 0.09690219 | 0.17129138 | 0 | 0.25035668 | 0.06647675 | 0.33269355 | 0.09550192 |
| ENSMUSG00 | Gm11744 | 1344 | 0.253007 | 0.66923225 | 0.57587536 | 0.37289545 | 0.07162489 | 0.11060878 | 0.84486927 | 0.19891876 | 0.64353931 | 0.24557658 | 0.42656213 | 0.3737656 | 0.25693707 | 0.10366984 | 0.33721512 | 0.14245018 | 0.14258295 | 0.45022334 |
| ENSMUSG00 | Bin2 | 1465 | 0 | 0 | 0 | 0.18659813 | 0.03285456 | 0 | 0.2906581 | 0.23462909 | 0 | 0.06436056 | 0.88049412 | 0.07619886 | 0.33673664 | 0.34872694 | 0.05624786 | 0.05227387 | 0.10464518 | 0.15019551 |
| ENSMUSG00 | Fnbp1 | 8586 | 17.406033 | 18.2510895 | 17.3527223 | 18.8696801 | 18.0957197 | 18.6241173 | 14.7624816 | 16.6630258 | 18.2907273 | 15.0524199 | 16.2421505 | 17.7536192 | 9.18150522 | 9.15250359 | 12.4669977 | 12.3532474 | 14.261882 | 10.3918904 |
| ENSMUSG00 | Gm14488 | 1389 | 0.27541151 | 0.19186692 | 0.34826156 | 0.06560266 | 0.31186992 | 0.57080183 | 0.40874885 | 0.10998532 | 0.29651919 | 0.23762053 | 0.30955697 | 0.36165656 | 0.67480664 | 0.70217843 | 0.62291769 | 0.30323735 | 0.30351999 | 0.19801694 |
| ENSMUSG00 | Dolk | 2125 | 28.8435124 | 24.0479905 | 23.7656309 | 22.1480277 | 24.4170417 | 24.4149233 | 17.8340969 | 22.5919293 | 22.0759724 | 23.275806 | 26.2368831 | 24.2962296 | 23.2150199 | 25.9431208 | 22.1228119 | 21.586893 | 20.6871824 | 26.5855777 |
| ENSMUSG00 | 1110017F19F | 976 | 2.74367427 | 3.4814705 | 5.30324564 | 3.68783043 | 6.11512145 | 2.84318963 | 3.10247075 | 3.13052479 | 5.14831447 | 5.21749585 | 5.97186981 | 5.37568881 | 5.71158468 | 6.70964787 | 4.7280474 | 4.59016503 | 3.88760594 | 4.28349602 |
| ENSMUSG00 | Gm13547 | 3521 | 0.07243143 | 0.15137924 | 0.01373858 | 0.0258796 | 0.01366996 | 0.05629393 | 0 | 0.0325411 | 0 | 0 | 0 | 0.03170444 | 0 | 0.03957179 | 0.01170166 | 0.01087492 | 0 | 0 |
| ENSMUSG00 | Olfr288 | 4628 | 0 | 0 | 0 | 0 | 0 | 0 | 0 | 0 | 0 | 0 | 0 | 0 | 0 | 0 | 0 | 0 | 0.12422093 | 0 |
| ENSMUSG00 | Gm11691 | 656 | 0 | 0 | 0 | 0 | 0 | 0 | 0 | 0 | 0 | 0 | 0 | 0 | 0 | 0 | 0 | 0 | 0 | 0 |
| ENSMUSG00 | Gm11681 | 867 | 0 | 0.19211588 | 0 | 0 | 0 | 0 | 0 | 0 | 0 | 0 | 0 | 0 | 0 | 0 | 0 | 0.08832897 | 0 | 0.19034293 |
| ENSMUSG00 | 4930594M22 | 4562 | 0.44722675 | 0.30669477 | 0.15905369 | 0.34954769 | 0.23211362 | 0.20637921 | 0.76745687 | 0.17580895 | 0.28890147 | 0.20671067 | 0.09425134 | 0.22022839 | 0.61637862 | 0.38686438 | 0.14450348 | 0.63789701 | 0.44526386 | 0.33762713 |
| ENSMUSG00 | Muc19 | 308 | 0 | 0 | 0 | 0 | 0 | 0 | 0 | 0 | 0 | 0 | 0 | 0 | 0 | 0 | 0 | 0 | 0 | 0 |
| ENSMUSG00 | Gm10837 | 1535 | 0.08307201 | 0.10851106 | 0 | 0.05936292 | 0.0313563 | 0.06456382 | 0.15411296 | 0.2239294 | 0.10732643 | 0.12286828 | 0.09337125 | 0 | 0.16069028 | 0.06051347 | 0.10736562 | 0.14967013 | 0.12484136 | 0.10750965 |
| ENSMUSG00 | Dnlz | 2913 | 21.0701937 | 17.256834 | 18.980757 | 15.7813573 | 20.5382702 | 16.7387195 | 19.9613053 | 17.0312155 | 19.2571351 | 21.9486281 | 18.1226671 | 18.8926342 | 26.7913124 | 17.8410346 | 17.680036 | 19.9142706 | 20.2222857 | 17.5432407 |
| ENSMUSG00 | Gm17205 | 1206 | 0 | 0 | 0 | 0 | 0 | 0 | 0 | 0 | 0 | 0 | 0 | 0 | 0 | 0 | 0 | 0.03175009 | 0 | 0 |
| ENSMUSG00 | Alg10b | 3724 | 3.25294714 | 2.19163778 | 2.94865505 | 2.86285832 | 2.30061289 | 2.60803861 | 2.37579779 | 2.18447414 | 1.71426153 | 1.9245185 | 2.41184001 | 2.77279742 | 2.76862801 | 2.83104439 | 1.38297381 | 2.80701821 | 2.74784434 | 2.64410046 |
| [illegible] | | | | | | | | | | | | | | | | | | | | |

| | | | | | | | | | | | | | | | | | | | | |
|---|---|---|---|---|---|---|---|---|---|---|---|---|---|---|---|---|---|---|---|---|
| ENSMUSG00 | Mir669a-6 | 96 | 0 | 0 | 0 | 0 | 0 | 0 | 0 | 0 | 0 | 0 | 0 | 0 | 0 | 0 | 0 | 0 | 0 | 0 |
| ENSMUSG00 | Mir669a-5 | 96 | 0 | 0 | 0 | 0 | 0 | 0 | 0 | 0 | 0 | 0 | 0 | 0 | 0 | 0 | 0 | 0 | 0 | 0 |
| ENSMUSG00 | Mir669a-4 | 96 | 0 | 0 | 0 | 0 | 0 | 0 | 0 | 0 | 0 | 0 | 0 | 0 | 0 | 0 | 0 | 0 | 0 | 0 |
| ENSMUSG00 | Mir669a-1 | 96 | 0 | 0 | 0 | 0 | 0 | 0 | 0 | 0 | 0 | 0 | 0 | 0 | 0 | 0 | 0 | 0 | 0 | 0 |
| ENSMUSG00 | Mir669a-3 | 108 | 0 | 0 | 0 | 0 | 0 | 0 | 0 | 0 | 0 | 0 | 0 | 0 | 0 | 0 | 0 | 0 | 0 | 0 |
| ENSMUSG00 | AC101886.1 | 108 | 0 | 0 | 0 | 0.84372304 | 0 | 0 | 0 | 0 | 0 | 0 | 0 | 0 | 0 | 0 | 0 | 0 | 0 | 0 |
| ENSMUSG00 | AC124732.1 | 102 | 0 | 0 | 0 | 0 | 0 | 0 | 0 | 0 | 0 | 0 | 0 | 0 | 0 | 0 | 0 | 0 | 0 | 0 |
| ENSMUSG00 | AL645948.1 | 108 | 0 | 0 | 0 | 0 | 0 | 0 | 0 | 0.35363336 | 0 | 0 | 0.44236073 | 0 | 0 | 0 | 0 | 0 | 0.70974624 | 0 |
| ENSMUSG00 | AC122459.1 | 108 | 0 | 0 | 0 | 0 | 0 | 0 | 0 | 0 | 0 | 0 | 0 | 0 | 0 | 0 | 0 | 0 | 0 | 0 |
| ENSMUSG00 | Mir692-2b | 108 | 0 | 0 | 0 | 0 | 0 | 0 | 0 | 0 | 0 | 0 | 0 | 0 | 0 | 0 | 0 | 0 | 0 | 0 |
| ENSMUSG00 | Mir692-2a | 108 | 1.57426578 | 0.30845272 | 0.44790306 | 1.26558456 | 0.89133192 | 0.91764321 | 0 | 0 | 0 | 1.30974176 | 0 | 0 | 0.91355402 | 1.29011354 | 1.14448766 | 1.77271331 | 0 | 0.50934358 |
| ENSMUSG00 | AC151602.1 | 108 | 0 | 0 | 0 | 0 | 0 | 0 | 0 | 0 | 0 | 0 | 0 | 0 | 0 | 0 | 0 | 0 | 0 | 0 |
| ENSMUSG00 | AC123859.1 | 108 | 0 | 0 | 0 | 0 | 0 | 0 | 0 | 0 | 0 | 0 | 0 | 0 | 0 | 0 | 0 | 0 | 0 | 0 |
| ENSMUSG00 | AC154650.1 | 108 | 0 | 0 | 0 | 0 | 0 | 0 | 0 | 0 | 0 | 0 | 0 | 0 | 0 | 0 | 0 | 0 | 0 | 0 |
| ENSMUSG00 | AC115005.1 | 107 | 0 | 0 | 0 | 0 | 0 | 0 | 0 | 0 | 0 | 0 | 0 | 0 | 0 | 0 | 0 | 0 | 0 | 0 |
| ENSMUSG00 | Mir20b | 79 | 0 | 0 | 0 | 0 | 0 | 0 | 0 | 0 | 0 | 0 | 0 | 0 | 0 | 0 | 0 | 0 | 0 | 0 |
| ENSMUSG00 | Mir710 | 109 | 0 | 0 | 0 | 0 | 0 | 0 | 0 | 0 | 0 | 0 | 0 | 0 | 0 | 0 | 0 | 0 | 0 | 0 |
| ENSMUSG00 | Mir467b | 72 | 0 | 0 | 0 | 0 | 0 | 0 | 0 | 0 | 0 | 0 | 0 | 0 | 0 | 0 | 0 | 0 | 0 | 0 |
| ENSMUSG00 | Mir720 | 63 | 0 | 0 | 0 | 0 | 0 | 0 | 0 | 0 | 0 | 0 | 0 | 0 | 0 | 0 | 0 | 0 | 0 | 0 |
| ENSMUSG00 | Mir598 | 78 | 0 | 0 | 0 | 0 | 0 | 0 | 0 | 0 | 0 | 0 | 0 | 0 | 0 | 0 | 0 | 0 | 0 | 0 |
| ENSMUSG00 | Mir450b | 81 | 0 | 0 | 0 | 0 | 0 | 0 | 0 | 0 | 0 | 0 | 0 | 0 | 0 | 0 | 0 | 0 | 0 | 0 |
| ENSMUSG00 | Mir707 | 72 | 0 | 0 | 0 | 0 | 0 | 0 | 0 | 0 | 0 | 0 | 0 | 0 | 0 | 0 | 0 | 0 | 0 | 0 |
| ENSMUSG00 | Mir541 | 89 | 0 | 0 | 0 | 0 | 0 | 0 | 0 | 0 | 0 | 0 | 0 | 0 | 0 | 0 | 0 | 0 | 0 | 0 |
| ENSMUSG00 | AC162934.1 | 108 | 0 | 0 | 0 | 0 | 0 | 0 | 0 | 0 | 0 | 0 | 0 | 0 | 0 | 0 | 0 | 0 | 0 | 0 |
| ENSMUSG00 | AC122194.1 | 108 | 0 | 0 | 0 | 0 | 0 | 0 | 0 | 0 | 0 | 0 | 0 | 0 | 0 | 0 | 0 | 0 | 0 | 0 |
| ENSMUSG00 | AC122313.1 | 108 | 0 | 0 | 0 | 0 | 0 | 0 | 0 | 0 | 0 | 0 | 0 | 0 | 0 | 0 | 0 | 0 | 0 | 0 |
| ENSMUSG00 | Mir683-1 | 108 | 0 | 0 | 0 | 0 | 0 | 0 | 0 | 0 | 0 | 0 | 0 | 0 | 0 | 0 | 0 | 0 | 0 | 0 |
| ENSMUSG00 | Mir683-2 | 108 | 0 | 0 | 0 | 0 | 0 | 0 | 0 | 0 | 0 | 0 | 0 | 0 | 0 | 0 | 0 | 0 | 0 | 0 |
| ENSMUSG00 | Mir704 | 76 | 0 | 0.43832756 | 0.63649382 | 0 | 1.26662957 | 0.65200965 | 1.86760575 | 1.50759483 | 3.79349487 | 0 | 1.88585362 | 0.73441661 | 0.64910417 | 0 | 0 | 0 | 3.53005366 | 0.72380404 |
| ENSMUSG00 | Mir92-1 | 79 | 0 | 0 | 0 | 0 | 0 | 0 | 0 | 0 | 0 | 0 | 0 | 0 | 0 | 0 | 0 | 0 | 0 | 0 |
| ENSMUSG00 | Mir539 | 73 | 0 | 0 | 0 | 0 | 0 | 0 | 0 | 0 | 0 | 0 | 0 | 0 | 0 | 0 | 0 | 0 | 0 | 0 |
| ENSMUSG00 | Mir690 | 108 | 0.39356644 | 0 | 0 | 0 | 0 | 0 | 0 | 0 | 0 | 0 | 0 | 0 | 0 | 0 | 0 | 0 | 0 | 0 |
| ENSMUSG00 | Mir681 | 109 | 0 | 0 | 0 | 0 | 0 | 0 | 0 | 0 | 0 | 0 | 0 | 0 | 0 | 0 | 0 | 0 | 0 | 0 |
| ENSMUSG00 | Mir223 | 109 | 0 | 0 | 0 | 0 | 0 | 0 | 0 | 0 | 0 | 0 | 0 | 0 | 0 | 0 | 0 | 0 | 0 | 0 |
| ENSMUSG00 | Mir592 | 95 | 0 | 0 | 0 | 0 | 0 | 0 | 0 | 0 | 0 | 0 | 0 | 0 | 0 | 0 | 0 | 0 | 0 | 0 |
| ENSMUSG00 | Mir678 | 83 | 0 | 0 | 0 | 0 | 0 | 0 | 0 | 0 | 0 | 0 | 0 | 0 | 0 | 0 | 0 | 0 | 0 | 0 |
| ENSMUSG00 | Mir680-2 | 109 | 0 | 0 | 0 | 0.41799123 | 0 | 0 | 0 | 0 | 0 | 0 | 0 | 0 | 0 | 0 | 0 | 0 | 0 | 0 |
| ENSMUSG00 | Mir669c | 108 | 0 | 0 | 0 | 0 | 0 | 0 | 0 | 0 | 0 | 0 | 0 | 0 | 0 | 0 | 0 | 0 | 0 | 0 |
| ENSMUSG00 | Mir698 | 108 | 0 | 0.30845272 | 0 | 0 | 0.89133192 | 0 | 0 | 0.35363336 | 0 | 0 | 0 | 1.03362338 | 0 | 0 | 0 | 0 | 0 | 0 |
| ENSMUSG00 | Mir551b | 97 | 0 | 0 | 0 | 0 | 0 | 0 | 0 | 0 | 0 | 0 | 0 | 0 | 0 | 0 | 0 | 0 | 0 | 0 |
| ENSMUSG00 | Mir687 | 88 | 0 | 0 | 0 | 0 | 0 | 0.56309925 | 0 | 0 | 0.46802859 | 0 | 0 | 0 | 0 | 0 | 0 | 1.30536162 | 2.17763051 | 0 |
| ENSMUSG00 | Mir503 | 70 | 0 | 0 | 0 | 0 | 0 | 0 | 0 | 0 | 0 | 0 | 0 | 0 | 0 | 0 | 0 | 0 | 0 | 0 |
| ENSMUSG00 | Mir700 | 78 | 0 | 0 | 0 | 0.58411595 | 0 | 0 | 0 | 0.48964618 | 0 | 0 | 1.22499894 | 0 | 0 | 0 | 0 | 0 | 0.49136278 | 0.70524496 |
| ENSMUSG00 | Mir669b | 96 | 0 | 0 | 0 | 0 | 0 | 0 | 0 | 0 | 0 | 0 | 0 | 0 | 0 | 0 | 0 | 0 | 0 | 0 |
| ENSMUSG00 | Mir718 | 87 | 0 | 0 | 0 | 0 | 0 | 0 | 0 | 0 | 0 | 0 | 0 | 0 | 0 | 0 | 0 | 0 | 0 | 0 |
| ENSMUSG00 | Mir686 | 108 | 0 | 0 | 0 | 0 | 0 | 0 | 0 | 0 | 0 | 0 | 0 | 0 | 0 | 0 | 0 | 0 | 0 | 0 |
| ENSMUSG00 | AL672269.1 | 108 | 0 | 0 | 0 | 0 | 0 | 0 | 0 | 0 | 0 | 0 | 0 | 0 | 0 | 0 | 0 | 0 | 0 | 0 |
| ENSMUSG00 | AC163389.1 | 109 | 0 | 0 | 0 | 0 | 0 | 0 | 0 | 0 | 0 | 0 | 0.87660474 | 0 | 0 | 0 | 0.37799593 | 0 | 0 | 0 |
| ENSMUSG00 | AC122119.1 | 108 | 0.39356644 | 0 | 0 | 0 | 0 | 0.91764321 | 0.43808036 | 0 | 1.14406988 | 0 | 0 | 1.03362338 | 0 | 0.43003785 | 0.38149589 | 0.35454266 | 0 | 1.52803074 |
| ENSMUSG00 | CT009594.1 | 109 | 0 | 0 | 0 | 0.83598246 | 0.44157728 | 0 | 0 | 0 | 0 | 0 | 0 | 0 | 0 | 0 | 0 | 0 | 0 | 0 |
| ENSMUSG00 | mmu-mir-70 | 107 | 0 | 0 | 0 | 0 | 0 | 0 | 0 | 0 | 0 | 0 | 0 | 0 | 0 | 0 | 0 | 0 | 0 | 0 |
| ENSMUSG00 | Mir542 | 84 | 0 | 0 | 0 | 0 | 0 | 0 | 0 | 0 | 0 | 0 | 0 | 0 | 0 | 0 | 0 | 0 | 0 | 0 |
| ENSMUSG00 | Mir693 | 88 | 0 | 0 | 0 | 0 | 0 | 0 | 0 | 0 | 0 | 0 | 0 | 0 | 0 | 0 | 0 | 0 | 0 | 0 |
| ENSMUSG00 | Mir711 | 81 | 0 | 0 | 0 | 0 | 0 | 0 | 0 | 0 | 0 | 0 | 0 | 0 | 0 | 0 | 0 | 0 | 0 | 0 |
| ENSMUSG00 | Mir708 | 108 | 0 | 0 | 0 | 0 | 0 | 0 | 0 | 0 | 0 | 0 | 0 | 0 | 0 | 0 | 0 | 0 | 0 | 0 |
| ENSMUSG00 | Mir709 | 87 | 2.44282621 | 0 | 0 | 0 | 0 | 0 | 0 | 0 | 0 | 0 | 0 | 0 | 0 | 1.06768017 | 0 | 0.88024385 | 0 | 0 |
| ENSMUSG00 | Mir679 | 73 | 0 | 0 | 0 | 0 | 0 | 0 | 0 | 0 | 0 | 0 | 0 | 0 | 0 | 0 | 0 | 0 | 0 | 0 |
| ENSMUSG00 | Mir449c | 108 | 0 | 0 | 0 | 0 | 0 | 0 | 0 | 0 | 0 | 0 | 0 | 0 | 0 | 0 | 0 | 0 | 0 | 0 |
| ENSMUSG00 | Mir694 | 69 | 0 | 0 | 0 | 0 | 0 | 0 | 0 | 0 | 0 | 0 | 0 | 0 | 0 | 0 | 0 | 0 | 0 | 0 |
| ENSMUSG00 | mmu-mir-46 | 82 | 0 | 0 | 0 | 0 | 0 | 0 | 0 | 0 | 0 | 0 | 0 | 0 | 0 | 0 | 0 | 0 | 0 | 0 |
| ENSMUSG00 | Mir467a-9 | 82 | 0 | 0 | 0 | 0 | 0 | 0 | 0 | 0 | 0 | 0 | 0 | 0 | 0 | 0 | 0 | 0 | 0 | 0 |
| ENSMUSG00 | Mir467a-8 | 82 | 0 | 0 | 0 | 0 | 0 | 0 | 0 | 0 | 0 | 0 | 0 | 0 | 0 | 0 | 0 | 0 | 0 | 0 |
| ENSMUSG00 | Mir467a-7 | 82 | 0 | 0 | 0 | 0 | 0 | 0 | 0 | 0 | 0 | 0 | 0 | 0 | 0 | 0 | 0 | 0 | 0 | 0 |
| ENSMUSG00 | Mir467a-5 | 82 | 0 | 0 | 0 | 0 | 0 | 0 | 0 | 0 | 0 | 0 | 0 | 0 | 0 | 0 | 0 | 0 | 0 | 0 |
| ENSMUSG00 | Mir467a-4 | 82 | 0 | 0 | 0 | 0 | 0 | 0 | 0 | 0 | 0 | 0 | 0 | 0 | 0 | 0 | 0 | 0 | 0 | 0 |
| ENSMUSG00 | Mir467a-2 | 82 | 0 | 0 | 0 | 0 | 0 | 0 | 0 | 0 | 0 | 0 | 0 | 0 | 0 | 0 | 0 | 0 | 0 | 0 |
| ENSMUSG00 | Mir467a-1 | 72 | 0 | 0 | 0 | 0 | 0 | 0 | 0 | 0 | 0 | 0 | 0 | 0 | 0 | 0 | 0 | 0 | 0 | 0 |
| ENSMUSG00 | Mir701 | 108 | 0 | 0 | 1.79161224 | 0 | 0 | 0 | 0 | 0 | 0 | 0 | 0 | 0 | 0 | 0 | 0 | 0 | 0 | 0 |
| ENSMUSG00 | AC102703.1 | 109 | 0 | 0 | 0 | 0 | 0 | 0.45461223 | 0 | 0 | 0 | 0 | 0 | 0 | 0 | 0 | 0 | 0 | 0 | 0 |
| ENSMUSG00 | AC153658.1 | 108 | 0 | 0 | 0 | 0 | 0 | 0 | 0 | 0 | 0 | 0 | 0 | 0 | 0 | 0 | 0 | 0 | 0 | 0 |
| ENSMUSG00 | AC102628.1 | 109 | 0 | 0 | 0 | 0 | 0 | 0 | 0 | 0 | 0 | 0 | 0 | 0 | 0 | 0 | 0 | 0 | 0 | 0 |
| ENSMUSG00 | AC102561.1 | 109 | 0 | 0 | 0 | 0 | 0 | 0 | 0 | 0 | 0 | 0 | 0 | 0 | 0 | 0 | 0 | 0 | 0 | 0 |
| ENSMUSG00 | AC138317.1 | 109 | 0 | 0 | 0 | 0 | 0 | 0 | 0 | 0 | 0 | 0 | 0 | 0 | 0 | 0 | 0 | 0 | 0 | 0 |
| ENSMUSG00 | AC112681.1 | 109 | 0 | 0 | 0 | 0 | 0 | 0 | 0 | 0 | 0 | 0 | 0 | 0 | 0 | 0 | 0 | 0 | 0 | 0 |
| ENSMUSG00 | Mir680-1 | 108 | 0 | 0 | 0 | 0 | 0 | 0 | 0 | 0 | 0 | 0 | 0 | 0 | 0 | 0 | 0 | 0 | 0 | 0 |
| ENSMUSG00 | Mir717 | 108 | 0 | 0 | 0 | 0 | 0 | 0 | 0 | 0 | 0 | 0 | 0 | 0 | 0 | 0 | 0 | 0 | 0 | 0 |
| ENSMUSG00 | Mir695 | 108 | 0 | 0.30845272 | 0 | 0 | 0.44566596 | 0 | 0.87616072 | 0.35363336 | 1.14406988 | 0 | 0.88472145 | 0 | 1.37033104 | 0 | 0.38149589 | 0.70908532 | 0 | 0 |
| ENSMUSG00 | Mir706 | 83 | 0 | 0 | 0 | 0 | 0 | 0 | 0 | 0 | 0 | 0 | 0 | 0 | 0 | 0 | 0 | 0 | 0 | 0 |
| ENSMUSG00 | Mir692-1 | 108 | 0 | 0 | 0 | 0 | 0 | 0.45882161 | 0 | 0 | 0 | 0.43658059 | 0 | 0 | 0.45677701 | 0.43003785 | 0 | 0.70908532 | 0 | 0 |
| ENSMUSG00 | Mir487b | 81 | 0 | 0 | 0 | 0 | 0 | 0 | 0 | 0 | 0 | 0 | 0 | 0 | 0 | 0 | 0 | 0 | 0 | 0 |
| ENSMUSG00 | Mir297b | 108 | 0 | 0 | 0 | 0 | 0 | 0 | 0 | 0 | 0 | 0 | 0 | 0 | 0 | 0 | 0 | 0 | 0 | 0 |
| ENSMUSG00 | Mir297-1 | 75 | 0 | 0 | 0 | 0 | 0 | 0 | 0 | 0 | 0 | 0 | 0 | 0 | 0 | 0 | 0 | 0 | 0 | 0 |
| ENSMUSG00 | AL590969.1 | 95 | 0 | 0 | 0 | 0 | 0 | 0 | 0 | 0 | 0 | 0 | 0 | 0 | 0 | 0 | 0 | 0 | 0 | 0 |
| ENSMUSG00 | AC159474.1 | 96 | 0 | 0 | 0 | 0 | 0 | 0 | 0 | 0 | 0 | 0 | 0 | 0 | 0 | 0 | 0 | 0 | 0 | 0 |
| ENSMUSG00 | AC155261.1 | 81 | 0 | 0 | 0 | 0 | 0 | 0 | 0 | 0 | 0 | 0 | 0 | 0 | 1.21807203 | 0 | 0.50866118 | 0 | 0 | 0 |
| ENSMUSG00 | AL928935.1 | 95 | 0 | 0 | 0 | 0 | 0 | 0 | 0 | 0 | 0 | 0 | 0 | 0 | 0 | 0 | 0 | 0 | 0 | 0 |
| ENSMUSG00 | AL929404.1 | 95 | 0 | 0 | 0 | 0 | 0 | 0 | 0 | 0 | 0 | 0 | 0 | 0 | 0 | 0 | 0 | 0 | 0 | 0 |
| ENSMUSG00 | AL596117.1 | 92 | 0 | 0 | 0 | 0 | 0 | 0 | 0 | 0 | 0 | 0 | 0 | 0 | 0 | 0 | 0 | 0 | 0 | 0 |
| ENSMUSG00 | AL928544.1 | 95 | 0 | 0 | 0 | 0 | 0 | 0 | 0 | 0 | 0 | 0 | 0 | 0 | 0 | 0 | 0 | 0 | 0 | 0 |
| ENSMUSG00 | AL928544.2 | 95 | 0 | 0 | 0 | 0 | 0 | 0 | 0 | 0 | 0 | 0 | 0 | 0 | 0 | 0 | 0 | 0 | 0 | 0 |
| ENSMUSG00 | Mir682 | 95 | 0 | 0 | 0 | 0 | 0 | 0 | 0 | 0 | 0 | 0 | 0 | 0 | 0 | 0 | 0 | 0 | 0 | 0 |
| ENSMUSG00 | AL590614.1 | 93 | 0 | 0 | 0 | 0 | 0 | 0 | 0 | 0 | 0 | 0 | 0 | 0 | 0 | 0 | 0 | 0 | 0 | 0 |
| ENSMUSG00 | AC122372.1 | 93 | 0 | 0 | 0 | 0 | 0 | 0 | 0 | 0 | 0 | 0 | 0 | 0 | 0 | 0 | 0 | 0 | 0 | 0 |
| ENSMUSG00 | Mir702 | 108 | 0 | 0 | 0 | 0 | 0 | 0 | 0 | 0 | 0 | 0 | 0 | 0 | 0 | 0 | 0 | 0 | 0 | 0 |
| ENSMUSG00 | Mir543 | 79 | 0 | 0 | 0 | 0 | 0 | 0 | 0 | 0 | 0 | 0 | 0 | 0 | 0 | 0 | 0 | 0 | 0 | 0 |
| ENSMUSG00 | CT030240.1 | 107 | 0 | 0 | 0 | 0 | 0 | 0 | 0 | 0 | 0 | 0 | 0 | 0 | 0 | 0 | 0 | 0 | 0 | 0 |
| ENSMUSG00 | AC122357.1 | 106 | 0 | 0 | 0 | 0 | 0 | 0 | 0 | 0 | 0 | 0 | 0 | 0 | 0 | 0 | 0 | 0 | 0 | 0 |
| ENSMUSG00 | CT573034.1 | 107 | 0 | 0 | 0 | 0 | 0 | 0 | 0 | 0 | 0 | 0 | 0 | 0 | 0 | 0 | 0 | 0 | 0 | 0 |
| ENSMUSG00 | AC159290.1 | 107 | 0 | 0 | 0 | 0 | 0 | 0 | 0 | 0 | 0 | 0 | 0 | 0 | 0 | 0 | 0 | 0 | 0 | 0 |
| ENSMUSG00 | Mir713 | 107 | 0 | 0 | 0 | 0 | 0 | 0 | 0 | 0 | 0 | 0 | 0 | 0 | 0 | 0 | 0 | 0 | 0 | 0 |
| ENSMUSG00 | Mir501 | 108 | 0 | 0 | 0 | 0 | 0 | 0 | 0 | 0 | 0 | 0 | 0 | 0 | 0 | 0 | 0 | 0 | 0 | 0 |
| ENSMUSG00 | Mir302b | 73 | 0 | 0.45634102 | 0 | 0 | 0 | 0 | 0 | 0 | 0 | 0 | 0 | 0 | 0 | 0 | 0 | 0 | 0 | 0 |
| ENSMUSG00 | Mir680-3 | 86 | 0 | 0 | 0 | 0 | 0 | 0 | 0 | 0 | 0 | 0 | 0 | 0 | 0 | 0 | 0 | 0 | 0 | 0 |
| ENSMUSG00 | Mir92b | 82 | 0 | 0 | 0 | 0 | 0 | 0 | 0 | 0 | 0 | 0 | 0 | 0 | 0 | 0 | 0 | 0 | 0 | 0 |
| ENSMUSG00 | Mir19b-1 | 86 | 0 | 0 | 0 | 0 | 0 | 0 | 0 | 0 | 0 | 0 | 0 | 0 | 0 | 0 | 0 | 0 | 0 | 0 |
| ENSMUSG00 | mmu-mir-68 | 56 | 2400.78342 | 2302.75381 | 2081.78943 | 1907.86873 | 2315.48933 | 2165.27748 | 3361.73481 | 2602.53942 | 2754.34824 | 2585.71089 | 2428.84476 | 2655.23081 | 5495.22321 | 7852.3682 | 3476.38128 | 2672.13739 | 3064.05056 | 4334.91407 |
| ENSMUSG00 | Mir374 | 94 | 0 | 0 | 0 | 0 | 0 | 0 | 0 | 0 | 0 | 0 | 0 | 0 | 0 | 0 | 0 | 0 | 0 | 0 |
| ENSMUSG00 | Mir719 | 109 | 0 | 0 | 0 | 0 | 0 | 0 | 0 | 0 | 0 | 0 | 0 | 0 | 0 | 0 | 0 | 0 | 0 | 0 |
| ENSMUSG00 | Mir697 | 108 | 0 | 0.30845272 | 0 | 0.84372304 | 0.44566596 | 0 | 1.31424108 | 0 | 0.76271325 | 0 | 1.32708218 | 0 | 0 | 0.43003785 | 0.76299178 | 0 | 0 | 1.01868716 |
| ENSMUSG00 | Mir666 | 98 | 0 | 0 | 0 | 0 | 0 | 0 | 0 | 0 | 0 | 0 | 0 | 0 | 0 | 0 | 0 | 0 | 0 | 0 |
| ENSMUSG00 | AC134908.1 | 91 | 0 | 0 | 0 | 0 | 0 | 0 | 0 | 0 | 0 | 0 | 0 | 0 | 0 | 0 | 0 | 0 | 0 | 0 |
| ENSMUSG00 | Mir675 | 83 | 0 | 0 | 0 | 0 | 0 | 0 | 0 | 0 | 0 | 0 | 0 | 0 | 0 | 0 | 0 | 0 | 0 | 0 |
| ENSMUSG00 | mmu-mir-21 | 56 | 3.795105 | 4.16411178 | 3.45525217 | 0.81359008 | 5.15699182 | 4.42435121 | 5.0692156 | 5.45605749 | 12.5030494 | 3.36790738 | 4.26562129 | 2.99012478 | 3.52370838 | 19.9046089 | 5.88593656 | 4.78632593 | 4.79078711 | 12.7699712 |
| ENSMUSG00 | AC156273.1 | 110 | 0 | 0 | 0 | 0 | 0 | 0 | 0 | 0 | 0 | 0 | 0 | 0 | 0 | 0 | 0 | 0 | 0 | 0 |
| ENSMUSG00 | AC134452.1 | 97 | 3.94377922 | 4.80804659 | 5.48565808 | 4.69701488 | 4.46584859 | 3.57597046 | 5.85311492 | 3.14988886 | 3.39682604 | 2.43044862 | 7.38788018 | 7.48044962 | 6.1029176 | 6.22446532 | 5.0970997 | 5.52647942 | 7.11209633 | 6.23814612 |
| ENSMUSG00 | Mir672 | 99 | 0 | 0 | 0 | 0 | 0 | 0 | 0 | 0 | 0 | 0 | 0 | 0 | 0 | 0 | 0 | 0 | 0 | 0 |
| ENSMUSG00 | AC109299.1 | 163 | 0 | 0 | 0 | 0 | 0 | 0 | 0 | 0 | 0 | 0 | 0 | 0 | 0 | 0 | 0 | 0 | 0 | 0 |
| ENSMUSG00 | Mir301b | 96 | 0 | 0 | 0 | 0 | 0 | 0 | 0 | 0 | 0 | 0 | 0 | 0 | 0 | 0 | 0 | 0 | 0 | 0 |
| ENSMUSG00 | AC122184.1 | 105 | 0 | 0 | 0 | 0 | 0 | 0 | 0 | 0 | 0 | 0 | 0 | 0 | 0 | 0 | 0 | 0 | 0 | 0 |
| ENSMUSG00 | AL627124.1 | 95 | 0 | 0 | 0 | 1.43876982 | 1.01330366 | 0 | 0 | 0 | 0 | 0 | 0 | 0 | 0 | 0 | 0 | 0 | 0 | 0.57904323 |
| ENSMUSG00 | CR974423.1 | 102 | 0 | 0 | 0 | 0 | 0 | 0 | 0 | 0 | 0 | 0 | 0 | 0 | 0 | 0 | 0 | 0 | 0 | 0 |
| ENSMUSG00 | AC157912.1 | 108 | 0 | 0 | 0 | 0 | 0 | 0 | 0 | 0 | 0 | 0 | 0 | 0 | 0 | 0 | 0 | 0 | 0 | 0 |
| ENSMUSG00 | AC133509.1 | 115 | 0 | 0 | 0 | 0 | 0 | 0 | 0 | 0 | 0 | 0 | 0 | 0 | 0 | 0 | 0 | 0 | 0 | 0 |
| ENSMUSG00 | CT573086.1 | 109 | 0 | 0 | 0 | 0 | 0 | 0 | 0 | 0 | 0 | 0 | 0 | 0 | 0 | 0 | 0 | 0 | 0 | 0 |
| ENSMUSG00 | AC133509.2 | 92 | 0 | 0 | 0 | 0 | 0 | 0 | 0 | 0 | 0 | 0 | 0 | 0 | 0 | 0 | 0 | 0 | 0 | 0 |
| ENSMUSG00 | Mir147 | 78 | 0 | 0 | 0 | 0 | 0 | 0 | 0 | 0 | 0 | 0 | 0 | 0 | 0 | 0 | 0 | 0 | 0 | 0 |
| ENSMUSG00 | Mir499 | 78 | 0 | 0 | 0 | 0 | 0 | 0 | 0 | 0 | 0 | 0 | 0 | 0 | 0 | 0 | 0 | 0 | 0 | 0 |
| ENSMUSG00 | Mir665 | 93 | 0 | 0 | 0 | 0 | 0 | 0 | 0 | 0 | 0 | 0 | 0 | 0 | 0 | 0 | 0 | 0 | 0 | 0 |
| ENSMUSG00 | Mir343 | 74 | 0 | 0 | 0 | 0 | 0 | 0 | 0 | 0 | 0 | 0 | 0 | 0 | 0 | 0 | 0 | 0 | 0 | 0 |
| ENSMUSG00 | Mir673 | 90 | 0 | 0 | 0 | 0 | 0 | 0 | 0 | 0 | 0 | 0 | 0 | 0 | 0 | 0 | 0 | 0 | 0 | 0 |
| ENSMUSG00 | Mir216b | 85 | 0 | 0 | 0 | 0 | 0 | 0 | 0 | 0 | 0 | 0 | 0 | 0 | 0 | 0 | 0 | 0 | 0 | 0 |
| ENSMUSG00 | AC167419.1 | 98 | 0 | 0 | 0 | 0 | 0 | 0 | 0 | 0 | 0 | 0 | 0 | 0 | 0 | 0 | 0.42042404 | 0 | 0 | 0 |
| ENSMUSG00 | AL603804.1 | 98 | 0 | 0 | 0 | 0 | 0 | 0 | 0 | 0 | 0 | 0 | 0 | 0 | 0 | 0 | 0 | 0 | 0 | 0 |
| ENSMUSG00 | AC144778.1 | 109 | 0 | 0 | 0 | 0 | 0 | 0 | 0 | 0 | 0.75571589 | 0.86515052 | 0 | 0 | 0 | 0 | 0 | 0 | 0 | 0 |
| ENSMUSG00 | AC151412.1 | 94 | 0 | 0 | 0 | 0 | 0 | 0 | 0 | 0 | 0 | 0 | 0 | 0 | 0 | 0 | 0 | 0 | 0 | 0 |
| ENSMUSG00 | Mir599 | 87 | 0 | 0 | 0 | 0 | 0 | 0.56957165 | 0 | 0 | 0 | 0 | 0 | 0 | 0 | 0 | 0 | 0 | 0 | 0 |
| ENSMUSG00 | Mir670 | 99 | 0 | 0 | 0 | 0 | 0 | 0 | 0 | 0 | 0 | 0 | 0 | 0 | 0 | 0 | 0 | 0 | 0 | 0 |
| ENSMUSG00 | BX255277.1 | 100 | 0 | 0 | 0 | 0 | 0 | 0 | 0 | 0 | 0 | 0 | 0 | 0 | 0 | 0 | 0 | 0 | 0 | 0 |
| ENSMUSG00 | Mir181d | 71 | 0 | 0 | 0 | 0 | 0 | 0 | 0 | 0 | 0 | 0 | 0 | 0 | 0 | 0 | 0 | 0 | 0 | 0 |
| ENSMUSG00 | AC137692.1 | 112 | 0 | 0 | 0 | 0 | 0 | 0 | 0 | 0 | 0 | 0 | 0 | 0 | 0 | 0 | 0 | 0 | 0 | 0 |
| ENSMUSG00 | AC151983.1 | 100 | 0 | 0 | 0 | 0 | 0 | 0 | 0 | 0 | 0 | 0 | 0 | 0 | 0 | 0 | 0 | 0 | 0 | 0 |
| ENSMUSG00 | AL607145.1 | 89 | 0 | 0 | 0.54352281 | 1.53576554 | 0 | 0 | 5.31603134 | 0 | 0 | 0 | 0 | 1.88142683 | 0 | 2.08737471 | 0 | 0 | 0 | 0 |
| ENSMUSG00 | Mir653 | 84 | 0 | 0 | 0 | 0 | 0 | 0 | 0 | 0 | 0 | 0 | 0 | 0 | 0 | 0 | 0 | 0 | 0 | 0 |
| ENSMUSG00 | AC171202.1 | 103 | 0 | 0 | 0 | 0 | 0 | 0 | 0 | 0 | 0 | 0 | 0 | 0 | 0 | 0 | 0 | 0 | 0 | 0 |
| ENSMUSG00 | Mir182 | 74 | 0 | 0 | 0 | 0 | 0 | 0 | 0 | 0 | 0 | 0 | 0 | 0 | 0 | 0 | 0 | 0 | 0 | 0 |
| ENSMUSG00 | AL731552.1 | 99 | 0 | 0 | 0 | 0 | 0 | 0 | 0 | 0 | 0 | 0 | 0 | 0 | 0 | 0 | 0 | 0 | 0 | 0 |
| ENSMUSG00 | AC099576.1 | 105 | 0 | 0 | 0 | 0 | 0 | 0 | 0 | 0 | 0 | 0 | 0 | 0 | 0 | 0 | 0 | 0 | 0 | 0 |
| ENSMUSG00 | AC132253.1 | 103 | 0 | 0 | 0 | 0 | 0 | 0 | 0 | 0 | 0 | 0 | 0 | 0 | 0 | 0 | 0 | 0 | 0 | 0 |
| ENSMUSG00 | AL671993.1 | 108 | 0 | 0 | 0 | 0 | 0 | 0 | 0 | 0 | 0 | 0 | 0 | 0 | 0 | 0 | 0 | 0 | 0 | 0 |
| ENSMUSG00 | Mir568 | 82 | 0 | 0 | 0 | 0 | 0 | 0 | 0 | 0 | 0 | 0 | 0 | 0 | 0 | 0 | 0 | 0 | 0 | 0 |
| ENSMUSG00 | Mir674 | 99 | 1.28803564 | 0.33649388 | 0.48862152 | 0 | 0 | 0.50053266 | 0 | 0.77156368 | 0 | 0 | 0.96515068 | 0 | 0 | 0.4691322 | 0.41617733 | 0 | 0.77426862 | 0 |
| ENSMUSG00 | AC153972.1 | 82 | 0 | 0 | 0 | 0 | 0 | 0 | 0 | 0 | 0 | 0 | 0 | 0 | 0 | 0 | 0 | 0 | 0 | 0 |
| ENSMUSG00 | Mir190 | 84 | 0 | 0 | 0 | 0 | 0 | 0 | 0 | 0 | 0 | 0 | 0 | 0 | 2.34913892 | 0 | 0 | 0 | 0 | 0 |
| ENSMUSG00 | AC163625.1 | 81 | 0.52475526 | 0 | 0 | 0 | 0 | 0 | 0 | 0 | 0 | 0 | 0 | 0 | 0 | 0 | 0 | 0 | 0 | 0.67912477 |
| ENSMUSG00 | AC114901.1 | 105 | 0 | 0 | 0 | 0 | 0 | 0 | 0 | 0 | 0 | 0 | 0 | 0 | 0 | 0 | 0 | 0 | 0 | 0 |
| ENSMUSG00 | Mir671 | 97 | 0 | 0 | 0 | 0 | 0 | 0 | 0 | 0 | 0 | 0 | 0 | 0 | 0 | 0 | 0 | 0 | 0 | 0 |
| ENSMUSG00 | AC099601.1 | 95 | 0 | 0 | 0 | 0 | 0 | 0 | 0 | 0 | 0 | 0 | 0 | 0 | 0 | 0 | 0 | 0 | 0 | 0 |
| ENSMUSG00 | AC116121.1 | 108 | 0 | 0 | 0.44790306 | 0.42186152 | 0 | 0 | 0 | 0 | 0.38135663 | 0 | 0 | 1.03362338 | 0 | 0 | 0 | 0 | 0 | 0 |
| ENSMUSG00 | AL663045.1 | 98 | 0 | 0 | 0 | 0 | 0 | 0 | 0 | 0 | 0 | 0 | 0 | 0 | 0 | 0 | 0 | 0 | 0 | 0 |
| ENSMUSG00 | AC113593.1 | 85 | 0 | 0 | 0 | 0 | 0 | 0 | 0 | 0 | 0 | 0 | 0 | 0 | 0 | 0 | 0 | 0 | 0 | 0 |
| ENSMUSG00 | Mir667 | 91 | 0 | 0 | 0 | 0 | 0 | 0 | 0 | 0 | 0 | 0 | 0 | 0 | 0 | 0 | 0 | 0 | 0 | 0 |
| ENSMUSG00 | Mir676 | 88 | 0 | 0 | 0 | 0 | 0 | 0 | 0 | 0 | 0 | 0 | 0 | 0 | 0 | 0 | 0 | 0 | 0 | 0 |
| ENSMUSG00 | Mir450-2 | 68 | 0 | 0 | 0 | 0 | 0 | 0 | 0 | 0 | 0 | 0 | 0 | 0 | 0 | 0 | 0 | 0 | 0 | 0 |
| ENSMUSG00 | CT010465.1 | 109 | 0 | 0 | 0 | 0 | 0 | 0 | 0 | 0 | 0 | 0 | 0 | 0 | 0 | 0 | 0 | 0 | 0 | 0 |
| ENSMUSG00 | AC103388.1 | 68 | 0 | 0 | 0 | 0 | 0 | 0 | 0 | 0 | 0 | 0 | 0 | 0 | 0 | 0 | 0 | 0 | 0 | 0 |
| ENSMUSG00 | AC154309.1 | 67 | 0.63440561 | 0 | 0 | 0 | 0 | 0 | 0 | 0 | 0 | 0 | 0 | 0 | 0 | 0 | 0 | 0 | 0 | 0 |
| ENSMUSG00 | BX255946.1 | 108 | 0 | 0 | 0 | 0 | 0 | 0 | 0 | 0 | 0 | 0 | 0 | 0 | 0 | 0 | 0 | 0 | 0 | 0 |
| ENSMUSG00 | Hus1b | 1186 | 0 | 0 | 0 | 0 | 0.04058341 | 0 | 0.07978529 | 0 | 0 | 0.03975607 | 0 | 0.09412422 | 0.33276167 | 0.31328221 | 0 | 0 | 0 | 0 |
| ENSMUSG00 | Sox4 | 5703 | 99.7228222 | 104.296287 | 88.1970838 | 94.6691873 | 94.3058241 | 99.2184227 | 68.4677432 | 79.1104418 | 76.1044193 | 87.8939376 | 82.1715882 | 84.0769349 | 70.7410871 | 72.9195789 | 78.0828364 | 85.7728405 | 81.6727137 | 80.4543151 |
| ENSMUSG00 | Ywhaq | 4139 | 54.8284936 | 49.635327 | 48.7942714 | 49.0504986 | 54.9464458 | 50.929531 | 48.7186851 | 50.1973107 | 49.0874806 | 47.8797791 | 48.3174622 | 48.9516442 | 44.3141021 | 42.987992 | 41.4205543 | 44.3038702 | 48.5861512 | 45.9184498 |
| ENSMUSG00 | BC100451 | 3542 | 0 | 0 | 0 | 0 | 0 | 0.01399004 | 0 | 0 | 0 | 0 | 0 | 0 | 0 | 0 | 0 | 0 | 0 | 0 |
| ENSMUSG00 | Wfdc3 | 2524 | 0.0168404 | 0.03959536 | 0.05749627 | 0 | 0 | 0.07853048 | 0.05623536 | 0.13618527 | 0 | 0 | 0 | 0.06634191 | 9.59666061 | 3.73540006 | 3.64023255 | 0.56131239 | 1.98920162 | 5.53578174 |
| ENSMUSG00 | Acsf2 | 3415 | 0.17425255 | 0.33166571 | 0.25497029 | 0.80048687 | 0.32416815 | 0.36275793 | 0.18010683 | 0.53681854 | 0.34975372 | 0.13806941 | 0.13989739 | 0.16344264 | 0.56338061 | 0.34000064 | 0.21716779 | 0.3812242 | 0.63970686 | 0.3060536 |
| ENSMUSG00 | Oxct2a | 1759 | 0.24164398 | 0.2272625 | 0.1925041 | 0.51803348 | 0.30099554 | 0.11268387 | 0.26897487 | 0.13027539 | 0.16390313 | 0.21444322 | 0.16296177 | 0.06346295 | 0.19631803 | 0.95053277 | 0.18738627 | 0.30475753 | 0.19609816 | 0 |
| ENSMUSG00 | 2700094K13F | 996 | 29.1476257 | 26.3225379 | 30.9376896 | 27.1262041 | 31.5563716 | 29.2540234 | 38.1921625 | 30.9833947 | 36.9272676 | 34.1795259 | 34.9198492 | 33.9601321 | 46.0134048 | 40.0556938 | 44.3040827 | 41.4814914 | 35.0554784 | 40.4283796 |
| ENSMUSG00 | Oxct2b | 1685 | 0 | 0 | 0 | 0 | 0 | 0 | 0 | 0 | 0.02444304 | 0 | 0 | 0 | 0 | 0 | 0 | 0 | 0.02274558 | 0 |
| ENSMUSG00 | Mog | 1706 | 0 | 0 | 0.05670988 | 0 | 0 | 0.02904615 | 0 | 0.04477421 | 0.19313724 | 0 | 0 | 0 | 0 | 0 | 0 | 0 | 0 | 0.25795595 |
| ENSMUSG00 | Ifna5 | 569 | 0 | 0 | 0 | 0 | 0 | 0 | 0 | 0 | 0 | 0 | 0 | 0 | 0 | 0 | 0 | 0 | 0 | 0 |
| ENSMUSG00 | Ass1 | 2291 | 0.50093398 | 0.31989685 | 0.61232317 | 0.33807846 | 0.33614613 | 0.54073258 | 0.53694005 | 0.2000475 | 0.46741572 | 0.32929343 | 0.35450646 | 0.24363013 | 2.36862108 | 2.98004402 | 1.6725206 | 1.06966341 | 1.53907434 | 1.41664657 |
| ENSMUSG00 | Mir761 | 75 | 0 | 0 | 0 | 0 | 0.64175898 | 0.66070311 | 0 | 1.01846406 | 0 | 0 | 0 | 0 | 0.6577589 | 0 | 0 | 0 | 0 | 0 |
| ENSMUSG00 | AC119211.1 | 95 | 0 | 0 | 0 | 0 | 0 | 0 | 0 | 0 | 0 | 0 | 0 | 0 | 0 | 0 | 0 | 0 | 0 | 0 |
| ENSMUSG00 | Mir763 | 119 | 0 | 0 | 0 | 0 | 0 | 0 | 0 | 0 | 0 | 0 | 0 | 0 | 0 | 0 | 0 | 0 | 0 | 0 |
| ENSMUSG00 | Mir804 | 94 | 0 | 0 | 0 | 0 | 0 | 0 | 0 | 0 | 0 | 0 | 0 | 0 | 0 | 0 | 0 | 0 | 0 | 0 |
| ENSMUSG00 | Mir770 | 93 | 0 | 0 | 0 | 0 | 0 | 0 | 0 | 0 | 0 | 0 | 0 | 0 | 0 | 0 | 0 | 0 | 0 | 0 |
| ENSMUSG00 | Mir759 | 97 | 0 | 0 | 0 | 0 | 0 | 0 | 0 | 0 | 0 | 0 | 0 | 0 | 0 | 0 | 0 | 0 | 0 | 0 |
| ENSMUSG00 | Mir762 | 75 | 0 | 0 | 0 | 0 | 0 | 0 | 0 | 0 | 0 | 0 | 0 | 0 | 0 | 0 | 0 | 0 | 0 | 0 |
| ENSMUSG00 | Mir767 | 106 | 0 | 0 | 0 | 0 | 0 | 0 | 0 | 0 | 0 | 0 | 0 | 0 | 0 | 0 | 0 | 0 | 0 | 0 |
| ENSMUSG00 | Mir760 | 118 | 0.36021336 | 1.12925065 | 1.63978069 | 1.93055273 | 0 | 3.77944578 | 0 | 2.26565099 | 3.83942096 | 0 | 0.80974506 | 3.78411272 | 2.92646967 | 0.78718792 | 1.04749718 | 0 | 2.92319214 | 0.93235774 |
| ENSMUSG00 | Mir802 | 96 | 0 | 0 | 0 | 0 | 0 | 0 | 0 | 0 | 0 | 0 | 0 | 0 | 0 | 0 | 0 | 0 | 0 | 0 |
| ENSMUSG00 | Mir764 | 107 | 0 | 0 | 0 | 0 | 0 | 0 | 0 | 0 | 0 | 0 | 0 | 0 | 0 | 0 | 0 | 0 | 0 | 0 |
| ENSMUSG00 | Mir758 | 80 | 0 | 0 | 0 | 0 | 0 | 0 | 0 | 0 | 0 | 0 | 0 | 0 | 0 | 0 | 0 | 0 | 0 | 0 |
| ENSMUSG00 | Mir744 | 99 | 0 | 0 | 0.48862152 | 0.46021257 | 0 | 0 | 1.43371754 | 1.15734553 | 0 | 0 | 0.48257534 | 0 | 0 | 0.4691322 | 0.41617733 | 0 | 0.38713431 | 2.22259017 |
| ENSMUSG00 | Trbv1 | 348 | 0 | 0 | 0 | 0 | 0 | 0 | 0 | 0 | 0 | 0 | 0 | 0 | 0 | 0 | 0 | 0 | 0 | 0 |
| ENSMUSG00 | Trbv2 | 342 | 0 | 0 | 0 | 0 | 0 | 0 | 0 | 0 | 0 | 0 | 0 | 0 | 0 | 0 | 0 | 0 | 0 | 0 |
| ENSMUSG00 | Trbv3 | 345 | 0 | 0 | 0 | 0 | 0 | 0 | 0 | 0 | 0 | 0 | 0 | 0 | 0 | 0 | 0 | 0 | 0 | 0 |
| ENSMUSG00 | Trbv4 | 337 | 0 | 0 | 0 | 0 | 0 | 0 | 0 | 0 | 0 | 0 | 0 | 0 | 0 | 0 | 0 | 0 | 0 | 0 |
| ENSMUSG00 | Trbv5 | 339 | 0 | 0 | 0 | 0 | 0 | 0 | 0 | 0 | 0 | 0 | 0 | 0 | 0 | 0 | 0 | 0 | 0 | 0 |
| ENSMUSG00 | Trbv12-1 | 370 | 0 | 0 | 0 | 0 | 0 | 0 | 0 | 0 | 0 | 0 | 0 | 0 | 0 | 0 | 0 | 0 | 0 | 0 |
| ENSMUSG00 | Trbv13-1 | 332 | 0 | 0 | 0 | 0 | 0 | 0 | 0 | 0 | 0 | 0 | 0 | 0 | 0 | 0 | 0 | 0 | 0 | 0 |
| ENSMUSG00 | Trbv12-2 | 370 | 0 | 0 | 0 | 0 | 0 | 0 | 0 | 0 | 0 | 0 | 0 | 0 | 0 | 0 | 0 | 0 | 0 | 0 |
| ENSMUSG00 | Trbv13-2 | 317 | 0 | 0 | 0 | 0 | 0 | 0 | 0 | 0 | 0 | 0 | 0 | 0 | 0 | 0 | 0 | 0 | 0 | 0 |
| ENSMUSG00 | Trbv13-3 | 338 | 0 | 0 | 0 | 0 | 0 | 0 | 0 | 0 | 0 | 0 | 0 | 0 | 0 | 0 | 0 | 0 | 0 | 0 |
| ENSMUSG00 | Trbv14 | 337 | 0 | 0 | 0 | 0 | 0 | 0 | 0 | 0 | 0 | 0 | 0 | 0 | 0 | 0 | 0 | 0 | 0 | 0 |
| ENSMUSG00 | Trbv15 | 345 | 0 | 0 | 0 | 0 | 0 | 0 | 0 | 0 | 0 | 0 | 0 | 0 | 0 | 0 | 0 | 0 | 0 | 0 |
| ENSMUSG00 | Trbv16 | 340 | 0 | 0 | 0 | 0 | 0 | 0 | 0 | 0 | 0 | 0 | 0 | 0 | 0 | 0 | 0 | 0 | 0 | 0 |
| ENSMUSG00 | Trbv17 | 337 | 0 | 0 | 0 | 0 | 0 | 0 | 0 | 0 | 0 | 0 | 0 | 0 | 0 | 0 | 0 | 0 | 0 | 0 |
| ENSMUSG00 | Trbv19 | 350 | 0 | 0 | 0 | 0 | 0 | 0 | 0 | 0 | 0 | 0 | 0 | 0 | 0 | 0 | 0 | 0 | 0 | 0 |
| ENSMUSG00 | Trbv20 | 333 | 0 | 0 | 0 | 0 | 0 | 0 | 0 | 0 | 0 | 0 | 0 | 0 | 0 | 0 | 0 | 0 | 0 | 0 |
| ENSMUSG00 | Trbv21 | 289 | 0 | 0 | 0 | 0 | 0 | 0 | 0 | 0 | 0 | 0 | 0 | 0 | 0 | 0 | 0 | 0 | 0 | 0 |
| ENSMUSG00 | Trbv23 | 342 | 0 | 0 | 0 | 0 | 0 | 0 | 0 | 0 | 0 | 0 | 0 | 0 | 0 | 0 | 0 | 0 | 0 | 0 |
| ENSMUSG00 | Trbv26 | 342 | 0 | 0 | 0 | 0 | 0 | 0 | 0 | 0 | 0 | 0 | 0 | 0 | 0 | 0 | 0 | 0 | 0 | 0 |
| ENSMUSG00 | Trbv29 | 342 | 0 | 0 | 0 | 0 | 0 | 0 | 0 | 0 | 0 | 0 | 0 | 0 | 0 | 0 | 0 | 0 | 0 | 0 |
| ENSMUSG00 | Trbv30 | 330 | 0 | 0 | 0 | 0 | 0 | 0 | 0 | 0 | 0 | 0 | 0 | 0 | 0 | 0 | 0 | 0 | 0 | 0 |
| ENSMUSG00 | Trbd1 | 11 | 0 | 0 | 0 | 0 | 0 | 0 | 0 | 0 | 0 | 0 | 0 | 0 | 0 | 0 | 0 | 0 | 0 | 0 |
| ENSMUSG00 | Trbj1-1 | 47 | 0 | 0 | 0 | 0 | 0 | 0 | 0 | 0 | 0 | 0 | 0 | 0 | 0 | 0 | 0 | 0 | 0 | 0 |
| ENSMUSG00 | Trbj1-2 | 47 | 0 | 0 | 0 | 0 | 0 | 0 | 0 | 0 | 0 | 0 | 0 | 0 | 0 | 0 | 0 | 0 | 0 | 0 |
| ENSMUSG00 | Trbj1-3 | 53 | 0 | 0 | 0 | 0 | 0 | 0 | 0 | 0 | 0 | 0 | 0 | 0 | 0 | 0 | 0 | 0 | 0 | 0 |
| ENSMUSG00 | Trbj1-4 | 50 | 0 | 0 | 0 | 0 | 0 | 0 | 0 | 0 | 0 | 0 | 0 | 0 | 0 | 0 | 0 | 0 | 0 | 0 |
| ENSMUSG00 | Trbj1-5 | 49 | 0 | 0 | 0 | 0 | 0 | 0 | 0 | 0 | 0 | 0 | 0 | 0 | 0 | 0 | 0 | 0 | 0 | 0 |
| ENSMUSG00 | Trbj1-6 | 52 | 0 | 0 | 0 | 0 | 0 | 0 | 0 | 0 | 0 | 0 | 0 | 0 | 0 | 0 | 0 | 0 | 0 | 0 |
| ENSMUSG00 | Trbj1-7 | 44 | 0 | 0 | 0 | 0 | 0 | 0 | 0 | 0 | 0 | 0 | 0 | 0 | 0 | 0 | 0 | 0 | 0 | 0 |
| ENSMUSG00 | Trbc1 | 509 | 0 | 0 | 0 | 0 | 0 | 0 | 0 | 0.15006838 | 0 | 0 | 0 | 0 | 0 | 0 | 0.08094608 | 0 | 0 | 0 |
| ENSMUSG00 | Trbd2 | 13 | 0 | 0 | 0 | 0 | 0 | 0 | 0 | 0 | 0 | 0 | 0 | 0 | 0 | 0 | 0 | 0 | 0 | 0 |
| ENSMUSG00 | Trbj2-1 | 49 | 0 | 0 | 0 | 0 | 0 | 0 | 0 | 0 | 0 | 0 | 0 | 0 | 0 | 0 | 0 | 0 | 0 | 0 |
| ENSMUSG00 | Trbj2-2 | 50 | 0 | 0 | 0 | 0 | 0 | 0 | 0 | 0 | 0 | 0 | 0 | 0 | 0 | 0 | 0 | 0 | 0 | 0 |
| ENSMUSG00 | Trbj2-3 | 48 | 0 | 0 | 0 | 0 | 0 | 0 | 0 | 0 | 0 | 0 | 0 | 0 | 0 | 0 | 0 | 0 | 0 | 0 |
| ENSMUSG00 | Trbj2-4 | 48 | 0 | 0 | 0 | 0 | 0 | 0 | 0 | 0 | 0 | 0 | 0 | 0 | 0 | 0 | 0 | 0 | 0 | 0 |
| ENSMUSG00 | Trbj2-5 | 48 | 0 | 0 | 0 | 0 | 0 | 0 | 0 | 0 | 0 | 0 | 0 | 0 | 0 | 0 | 0 | 0 | 0 | 0 |
| ENSMUSG00 | Trbj2-7 | 46 | 0 | 0 | 0 | 0 | 0 | 0 | 0 | 0 | 0 | 0 | 0 | 0 | 0 | 0 | 0 | 0 | 0 | 0 |
| ENSMUSG00 | Trbc2 | 517 | 0 | 0 | 0 | 0 | 0 | 0 | 0 | 0 | 0 | 0 | 0 | 0 | 0 | 0 | 0 | 0 | 0 | 0 |
| ENSMUSG00 | Trbv31 | 332 | 0 | 0 | 0 | 0 | 0 | 0 | 0 | 0 | 0 | 0 | 0 | 0 | 0 | 0 | 0 | 0 | 0 | 0 |
| ENSMUSG00 | AC158671.1 | 357 | 0 | 0 | 0 | 0 | 0 | 0 | 0 | 0 | 0 | 0 | 0 | 0 | 0 | 0 | 0 | 0 | 0 | 0 |
| ENSMUSG00 | Igkv2-137 | 360 | 0 | 0 | 0 | 0 | 0 | 0 | 0 | 0.10609001 | 0.80084892 | 0 | 0 | 0 | 0 | 0 | 0 | 0 | 0 | 0.6112123 |
| ENSMUSG00 | Igkv1-135 | 360 | 0 | 0 | 0 | 0.12655846 | 0.13369979 | 0 | 5.12554022 | 0 | 1.37288386 | 0 | 0.53083287 | 0 | 0.68516552 | 0 | 0 | 2.23361877 | 0 | 0 |
| ENSMUSG00 | Igkv1-133 | 360 | 0 | 0 | 0 | 0 | 0 | 0 | 0 | 0 | 0 | 0 | 2.52145614 | 0 | 0 | 0 | 0 | 0 | 0 | 0 |
| ENSMUSG00 | Igkv1-132 | 360 | 0 | 0 | 0 | 0 | 0 | 0 | 0 | 0 | 0 | 0 | 0 | 0 | 0 | 0 | 0 | 0 | 0 | 0 |
| ENSMUSG00 | Igkv1-131 | 360 | 0 | 0 | 0 | 0 | 0 | 0 | 0 | 0 | 0 | 0 | 0 | 0 | 0 | 0 | 0 | 0 | 0 | 0 |
| ENSMUSG00 | Igkv14-130 | 348 | 0 | 0 | 0 | 0 | 0 | 0 | 0 | 0 | 0 | 0 | 0 | 0 | 0 | 0 | 0 | 0 | 0 | 0 |
| ENSMUSG00 | Igkv9-129 | 348 | 0 | 0 | 0 | 0 | 0 | 0 | 0 | 0 | 0 | 0 | 0 | 0 | 0 | 0 | 0 | 0 | 0 | 0 |
| ENSMUSG00 | Igkv17-127 | 339 | 0 | 0 | 0 | 0 | 0 | 0 | 0 | 0 | 0 | 0 | 6.34180865 | 0 | 0 | 0 | 0 | 0 | 0 | 0 |
| ENSMUSG00 | Igkv14-126 | 351 | 0 | 0 | 0 | 0 | 0 | 0 | 6.47010994 | 0 | 0 | 0 | 3.8111078 | 0 | 0 | 0 | 0 | 0 | 0 | 0 |
| ENSMUSG00 | Igkv11-125 | 348 | 0 | 0 | 0 | 0 | 0 | 0 | 0 | 0 | 0 | 0 | 0 | 0 | 0 | 0.26692004 | 0 | 0 | 0 | 0 |
| ENSMUSG00 | Igkv9-124 | 286 | 0 | 0 | 0 | 0 | 0 | 0 | 0 | 0 | 0 | 1.15403819 | 0 | 0 | 0 | 0 | 0 | 0 | 0 | 0 |
| ENSMUSG00 | Igkv9-123 | 351 | 0 | 0 | 0 | 0 | 0 | 0 | 0 | 0 | 0 | 0 | 0 | 0 | 0 | 0 | 0 | 0 | 0 | 0 |
| ENSMUSG00 | Igkv1-122 | 357 | 0 | 0 | 0 | 0 | 0 | 0 | 0 | 0 | 0 | 0 | 0 | 0 | 0 | 0 | 0 | 3.1104415 | 0 | 0 |
| ENSMUSG00 | Igkv17-121 | 348 | 0 | 0 | 0 | 0 | 0 | 0 | 0 | 0 | 0 | 0 | 0 | 2.56623736 | 0 | 0 | 0.9471622 | 0 | 0 | 0 |
| ENSMUSG00 | Igkv9-120 | 351 | 0.12109737 | 0.09490853 | 0 | 0 | 0 | 0.14117588 | 5.79614015 | 3.15549763 | 0 | 0 | 5.44443971 | 0 | 0 | 0 | 3.5215095 | 0 | 0 | 0 |
| ENSMUSG00 | Igkv1-117 | 357 | 0 | 0 | 0 | 0 | 0 | 0 | 1.98792769 | 0 | 0 | 0 | 12.5794008 | 7.34828051 | 8.15289389 | 0 | 0 | 4.18300753 | 0 | 0 |
| ENSMUSG00 | Igkv2-116 | 362 | 0 | 0 | 0 | 0 | 0 | 0 | 0 | 0 | 0 | 0 | 0 | 0 | 0 | 0 | 0 | 0.21155032 | 0 | 0 |
| ENSMUSG00 | Igkv2-112 | 356 | 0 | 0 | 0 | 0 | 0 | 0 | 0 | 0 | 0 | 0 | 0 | 0 | 0 | 0 | 0 | 0 | 0 | 0 |
| ENSMUSG00 | Igkv14-111 | 351 | 0 | 0 | 0 | 0 | 0 | 0 | 0 | 0 | 0.704043 | 0 | 0 | 0 | 0 | 0 | 0 | 0 | 0 | 0 |
| ENSMUSG00 | Igkv1-110 | 357 | 0 | 0 | 0 | 0 | 0 | 0 | 5.43366901 | 1.28377823 | 0 | 1.05659839 | 6.69117066 | 0 | 6.90923211 | 5.20381932 | 0 | 1.60884905 | 0 | 0 |
| ENSMUSG00 | Igkv2-109 | 362 | 0 | 0 | 0 | 0 | 0 | 0 | 0.26139602 | 0.94953487 | 0 | 0 | 1.05580019 | 0 | 0 | 0.12829858 | 0 | 0 | 0 | 0 |
| ENSMUSG00 | Igkv16-104 | 346 | 0 | 0.09628004 | 0 | 0 | 0 | 0 | 11.8965407 | 0 | 0 | 0 | 3.17579204 | 0 | 0 | 3.08732373 | 1.54803534 | 0 | 0 | 0 |
| ENSMUSG00 | Igkv15-103 | 345 | 0 | 0 | 0 | 0 | 0 | 0 | 10.0110886 | 0 | 0 | 0 | 0 | 0 | 0 | 0 | 0 | 0 | 0 | 0 |
| ENSMUSG00 | Igkv14-100 | 348 | 0 | 0 | 0 | 0 | 0 | 0 | 0 | 0 | 0 | 0 | 0 | 0 | 0 | 0 | 0 | 0 | 0 | 0 |
| ENSMUSG00 | Igkv1-99 | 357 | 0 | 0 | 0 | 0 | 0 | 0 | 0 | 0 | 0 | 0 | 0 | 0 | 0 | 0 | 0 | 0 | 0 | 0 |
| ENSMUSG00 | Igkv12-98 | 342 | 0 | 0 | 0 | 0 | 0 | 0 | 0 | 0 | 0 | 0 | 0 | 0 | 0 | 0 | 0 | 0 | 0 | 0 |
| ENSMUSG00 | Igkv10-96 | 345 | 0 | 0 | 0 | 0 | 0 | 0 | 0.2742764 | 1.21772877 | 0 | 0 | 5.26216934 | 0 | 4.71870513 | 3.09627249 | 0 | 0 | 0 | 0 |
| ENSMUSG00 | Igkv10-95 | 286 | 0 | 0 | 0 | 0 | 0 | 0 | 0 | 0 | 0 | 0 | 0 | 0 | 0 | 0 | 0 | 0 | 0 | 0 |
| ENSMUSG00 | Igkv10-94 | 345 | 0 | 0 | 0 | 0 | 0 | 0 | 0 | 0 | 0 | 1.09334964 | 0 | 0 | 0.57196426 | 0 | 0 | 0 | 0 | 0 |
| ENSMUSG00 | Igkv19-93 | 342 | 3.35567179 | 0 | 0 | 0 | 0 | 0 | 0 | 0 | 6.02142043 | 0 | 0 | 0 | 0 | 11.6789226 | 0 | 0 | 0 | 0 |
| ENSMUSG00 | Igkv4-92 | 349 | 0 | 0 | 0 | 0 | 0 | 0 | 0 | 0 | 0 | 0.54040921 | 0 | 0.95958159 | 0 | 0 | 0 | 0 | 0 | 0 |
| ENSMUSG00 | Igkv4-91 | 343 | 0 | 0 | 0 | 0 | 0 | 0 | 0 | 0 | 3.48224186 | 0 | 0 | 0 | 0 | 4.46838159 | 0 | 0 | 0 | 0 |
| ENSMUSG00 | Igkv4-90 | 324 | 0 | 0 | 0 | 0 | 0 | 0 | 0 | 0 | 0 | 0 | 0 | 0 | 1.97936705 | 0 | 0 | 0 | 0 | 0 |
| ENSMUSG00 | Igkv12-89 | 333 | 0 | 0 | 0 | 0 | 0 | 0 | 0 | 0 | 0 | 0 | 0 | 0 | 0 | 0 | 0 | 0 | 0 | 0 |
| ENSMUSG00 | Igkv1-88 | 360 | 0 | 0 | 0 | 0 | 0 | 0 | 0 | 0 | 0 | 0 | 0 | 0 | 0 | 1.54813625 | 0 | 0 | 0 | 0 |
| ENSMUSG00 | Igkv4-86 | 338 | 0 | 0 | 0 | 0 | 0 | 0 | 0 | 0 | 0 | 0 | 0 | 0 | 0 | 0 | 0 | 0 | 0 | 0 |
| ENSMUSG00 | Igkv13-84 | 345 | 0 | 0 | 0 | 0 | 0 | 0 | 0 | 0 | 0 | 0 | 0 | 0 | 0 | 0 | 0 | 0 | 0 | 0 |
| ENSMUSG00 | Igkv4-81 | 289 | 0 | 0 | 0 | 0 | 0 | 0 | 0 | 0 | 0 | 0 | 0 | 0 | 0 | 0 | 0 | 0 | 0 | 0 |
| ENSMUSG00 | Igkv4-80 | 344 | 0 | 0 | 0 | 0 | 0 | 0 | 0 | 0 | 0 | 0 | 0 | 0 | 0 | 0 | 0 | 0 | 0 | 0 |
| ENSMUSG00 | Igkv4-79 | 350 | 0 | 0 | 0 | 0 | 0 | 0 | 0 | 0 | 0 | 0 | 0 | 0 | 0.28189667 | 0 | 0 | 0 | 0 | 0 |
| ENSMUSG00 | Igkv4-78 | 354 | 0 | 0 | 0 | 0 | 0 | 0 | 0 | 0 | 0 | 0 | 0 | 0 | 0.1393557 | 0 | 0 | 0 | 0 | 0 |
| ENSMUSG00 | Igkv4-74 | 356 | 0 | 0 | 0 | 0 | 0 | 0 | 0 | 0 | 0 | 0 | 0 | 0 | 0 | 0 | 0 | 0 | 0 | 3.55395914 |
| ENSMUSG00 | Igkv4-73 | 354 | 0 | 0 | 0 | 0.25740703 | 0 | 0 | 0 | 0 | 0 | 1.33194077 | 0 | 0 | 0.4180671 | 0.26239597 | 0 | 0 | 0 | 0 |
| ENSMUSG00 | Igkv4-72 | 350 | 0 | 0 | 0 | 0 | 0 | 0 | 4.32573064 | 0 | 0 | 0 | 0.27299976 | 0 | 0 | 0 | 1.41262477 | 0 | 0 | 0 |
| ENSMUSG00 | Igkv4-70 | 346 | 0 | 0 | 0 | 0 | 0 | 0 | 0.13674185 | 0 | 0 | 0 | 0 | 0 | 0 | 0 | 0 | 0 | 0 | 0 |
| ENSMUSG00 | Igkv4-69 | 346 | 0 | 0 | 0 | 0 | 0 | 0 | 0 | 0 | 0 | 0 | 0 | 0 | 0 | 0 | 0 | 0 | 0 | 0 |
| ENSMUSG00 | Igkv4-68 | 350 | 0 | 0 | 0 | 0 | 0 | 0 | 0 | 0 | 0 | 0 | 0 | 0 | 0 | 0 | 0 | 0 | 0 | 0 |
| ENSMUSG00 | Igkv4-63 | 285 | 0 | 0 | 0 | 0 | 0 | 0 | 0 | 0 | 0 | 0 | 0 | 0 | 0 | 0 | 0 | 0 | 0 | 0 |
| ENSMUSG00 | Igkv4-62 | 278 | 0 | 0 | 0 | 0 | 0 | 0 | 0 | 0 | 0 | 0 | 0 | 0 | 0 | 0 | 0 | 0 | 0 | 0 |
| ENSMUSG00 | Igkv4-61 | 283 | 0 | 0 | 0 | 0 | 0 | 0 | 0 | 0 | 0 | 0 | 0 | 0 | 0 | 0 | 0 | 0.270605 | 0 | 0 |
| ENSMUSG00 | Igkv4-59 | 350 | 0 | 0 | 0 | 0 | 0 | 0 | 1.21661174 | 0 | 0 | 0 | 0 | 0 | 3.24181171 | 0.13269739 | 0 | 0 | 0 | 0 |
| ENSMUSG00 | Igkv4-58 | 291 | 0 | 0 | 0 | 0 | 0 | 0 | 0 | 0 | 0 | 0 | 0 | 0 | 0 | 0 | 0 | 0 | 0 | 0 |
| ENSMUSG00 | Igkv4-57-1 | 356 | 0 | 0 | 0 | 0 | 0 | 0 | 0 | 0 | 0 | 0 | 2.95238507 | 0 | 0 | 0 | 0 | 0 | 0 | 0 |
| ENSMUSG00 | Igkv4-57 | 348 | 0 | 0 | 0 | 0 | 0 | 0 | 0 | 0 | 0.94681645 | 8.40041265 | 0 | 0 | 3.11868443 | 0 | 0 | 0 | 0 | 0 |
| ENSMUSG00 | Igkv4-56 | 285 | 0 | 0 | 0 | 0 | 0 | 0 | 0 | 0 | 0 | 0 | 0 | 0 | 0 | 0 | 0 | 0.26870602 | 0 | 0 |
| ENSMUSG00 | Igkv4-55 | 350 | 0 | 0 | 0 | 0 | 0 | 0 | 0 | 0 | 0 | 0 | 0 | 0 | 0 | 0 | 0 | 0 | 0 | 0 |
| ENSMUSG00 | Igkv4-50 | 350 | 0.12144336 | 0 | 0 | 0 | 0 | 0 | 0 | 0 | 0 | 0 | 2.45699786 | 0 | 0 | 0 | 2.23665589 | 0 | 0 | 0 |
| ENSMUSG00 | Igkv5-48 | 345 | 0 | 0 | 0 | 0 | 0 | 0 | 0 | 0 | 0 | 0 | 0 | 0 | 0 | 0 | 0 | 0 | 0 | 0 |
| ENSMUSG00 | Igkv12-46 | 345 | 0 | 0 | 0 | 0 | 0 | 0 | 0 | 0 | 0 | 0 | 0 | 0.48535359 | 0 | 0 | 0 | 0 | 0 | 0 |
| ENSMUSG00 | Igkv5-45 | 345 | 0 | 0 | 0 | 0 | 0 | 0 | 0 | 0 | 0 | 0 | 0 | 0 | 0 | 0 | 0 | 0 | 0 | 0 |
| ENSMUSG00 | Igkv12-44 | 345 | 0 | 0 | 0 | 0 | 0 | 0 | 0 | 0 | 0 | 0 | 0 | 0 | 0 | 0 | 0 | 0 | 0 | 0 |
| ENSMUSG00 | Igkv5-43 | 345 | 0 | 0 | 0 | 0 | 0 | 0 | 0 | 0 | 0.59690603 | 0 | 0 | 0 | 0 | 0.40386163 | 0 | 0 | 0 | 0 |
| ENSMUSG00 | Igkv12-41 | 345 | 0 | 0 | 0 | 0 | 0 | 0 | 0 | 0 | 0 | 0 | 0 | 0 | 0 | 0 | 0 | 0 | 0 | 0 |
| ENSMUSG00 | Igkv5-39 | 286 | 0 | 0 | 0 | 0 | 0 | 0 | 0 | 0 | 0 | 0 | 0 | 2.14675625 | 0 | 0 | 2.44904353 | 0 | 0 | 0 |
| ENSMUSG00 | Igkv12-38 | 342 | 0 | 0 | 0 | 0 | 0 | 0 | 0 | 0 | 0 | 0 | 0 | 0 | 0 | 0 | 0 | 0 | 0 | 0 |
| ENSMUSG00 | Igkv5-37 | 345 | 0 | 0 | 0 | 0 | 0 | 0 | 0 | 0 | 0 | 0 | 0 | 0 | 0 | 0 | 0 | 0 | 0 | 0 |
| ENSMUSG00 | Igkv18-36 | 342 | 0 | 0 | 0 | 0 | 0 | 0 | 0 | 0 | 0 | 0 | 0 | 0 | 0 | 0.48188954 | 0 | 0 | 0 | 0 |
| ENSMUSG00 | Igkv1-35 | 357 | 0 | 0 | 0 | 0 | 0 | 0 | 0 | 0 | 0 | 0 | 0 | 0 | 0 | 0 | 0 | 0 | 0 | 0 |
| ENSMUSG00 | Igkv7-33 | 360 | 0 | 0 | 0 | 0 | 0 | 0 | 0 | 0 | 0 | 0 | 0 | 0 | 0 | 0 | 0 | 0 | 0 | 0 |
| ENSMUSG00 | Igkv6-32 | 345 | 0 | 0 | 0.14021313 | 0 | 0.13951282 | 0.14363111 | 0.2742764 | 0 | 0 | 0 | 0 | 0 | 0 | 0 | 0 | 0 | 0.22218143 | 0.15944669 |
| ENSMUSG00 | Igkv8-30 | 363 | 0 | 0 | 0 | 0 | 0 | 0 | 0.26067592 | 0 | 0 | 0 | 0 | 0 | 0 | 0 | 0 | 0 | 0 | 0 |
| ENSMUSG00 | Igkv6-29 | 342 | 0 | 0 | 0 | 0 | 0 | 0 | 0 | 0 | 0 | 0 | 0 | 0 | 0 | 0 | 0 | 0 | 0 | 0 |
| ENSMUSG00 | Igkv8-28 | 304 | 0 | 0 | 0 | 0 | 0 | 0 | 0 | 0 | 0 | 0 | 0 | 0 | 0 | 0.1527766 | 3.65934871 | 0 | 0 | 0 |
| ENSMUSG00 | Igkv8-27 | 299 | 0 | 0 | 0 | 0 | 0 | 0 | 0 | 0 | 3.7191837 | 0 | 7.66955855 | 0 | 0 | 3.26195931 | 0 | 0 | 0 | 0 |
| ENSMUSG00 | Igkv8-26 | 369 | 0 | 0 | 0 | 0 | 0 | 0 | 0 | 0 | 0 | 0 | 0 | 0 | 0 | 0 | 0 | 0 | 0 | 0 |
| ENSMUSG00 | Igkv6-25 | 345 | 0 | 0 | 0 | 0 | 0 | 0 | 0 | 0 | 0 | 0 | 0 | 0 | 0 | 0 | 0 | 0 | 0 | 0 |
| ENSMUSG00 | Igkv8-24 | 363 | 0 | 0 | 0 | 0 | 0 | 0 | 0 | 0 | 0 | 0 | 0 | 0 | 0 | 0 | 2.49706399 | 0 | 0 | 0 |
| ENSMUSG00 | Igkv6-23 | 345 | 0 | 0 | 0 | 0 | 0 | 0 | 1.6456584 | 0 | 0 | 0 | 2.90804095 | 1.77962982 | 0 | 0 | 0 | 0 | 0 | 0 |
| ENSMUSG00 | Igkv8-21 | 360 | 0 | 0 | 0 | 0 | 0 | 0 | 0 | 0 | 0 | 0.13097418 | 0.39812465 | 0 | 0 | 0 | 0 | 0 | 0.10646194 | 0 |
| ENSMUSG00 | Igkv6-20 | 372 | 0 | 0 | 0 | 0 | 0 | 0 | 0 | 0 | 0 | 0.2534984 | 0 | 0 | 0 | 0 | 0 | 0 | 0 | 0 |
| ENSMUSG00 | Igkv8-19 | 304 | 0 | 0 | 0 | 0 | 0 | 0 | 0 | 0 | 0 | 0 | 0 | 0 | 0 | 5.04162791 | 0 | 0 | 0 | 0 |
| ENSMUSG00 | Igkv8-18 | 368 | 0 | 0 | 0 | 0 | 0 | 0 | 0 | 0 | 0 | 0 | 0 | 0 | 0 | 0 | 0 | 0 | 0 | 0 |
| ENSMUSG00 | Igkv6-17 | 345 | 0 | 0 | 0 | 0 | 0 | 0 | 0 | 1.54983662 | 0 | 0 | 0 | 0 | 0 | 0 | 0 | 0 | 0 | 0.31889337 |
| ENSMUSG00 | Igkv8-16 | 360 | 0 | 0 | 0 | 0 | 0 | 0 | 0 | 0 | 0 | 0 | 2.25603971 | 0 | 0 | 0 | 1.25893643 | 0 | 0 | 0 |
| ENSMUSG00 | Igkv6-15 | 345 | 0 | 0 | 0 | 0 | 0 | 0 | 0 | 5.86723863 | 12.1768829 | 0 | 12.7399889 | 0 | 0 | 0 | 2.26907119 | 0 | 0 | 0 |
| ENSMUSG00 | Igkv6-14 | 345 | 0 | 0 | 0 | 2.24503697 | 0 | 0 | 0 | 0 | 0 | 0 | 0.13847814 | 0 | 0 | 0 | 0 | 0 | 0 | 0 |
| ENSMUSG00 | Igkv6-13 | 286 | 0 | 0 | 0 | 0 | 0 | 0 | 0 | 0 | 0 | 0 | 0 | 0 | 0 | 0 | 0 | 0 | 0 | 0 |
| ENSMUSG00 | Igkv3-12 | 357 | 0 | 0 | 0 | 0 | 0 | 0 | 0 | 0 | 0 | 0 | 0 | 0 | 0 | 0 | 0 | 0.75079622 | 0 | 0 |
| ENSMUSG00 | Igkv3-10 | 354 | 0 | 0 | 0 | 0 | 0 | 0 | 0 | 0 | 0 | 0 | 0 | 0 | 0 | 0 | 0 | 0 | 0 | 0 |
| ENSMUSG00 | Igkv3-9 | 357 | 0 | 0 | 0 | 0 | 0 | 0 | 0 | 0 | 0 | 0 | 0 | 0 | 0 | 0 | 0 | 0 | 0 | 0 |
| ENSMUSG00 | Igkv3-7 | 298 | 0 | 0 | 0 | 0 | 0 | 0 | 0 | 0 | 0 | 0 | 0 | 0 | 0 | 0 | 0 | 0 | 0 | 0 |
| ENSMUSG00 | Igkv3-5 | 357 | 0 | 0 | 0 | 0 | 0 | 0 | 0 | 0 | 0 | 0 | 0 | 0 | 0 | 0 | 0 | 0 | 0 | 0 |
| ENSMUSG00 | Igkv3-4 | 357 | 0 | 0 | 0 | 0 | 0 | 0 | 0 | 0 | 0 | 0 | 0 | 0 | 0 | 0 | 0 | 0 | 0 | 0 |
| ENSMUSG00 | Igkv3-3 | 357 | 0 | 0 | 0 | 0 | 0 | 0 | 0 | 0 | 0 | 0 | 0 | 0 | 0 | 0 | 0 | 0 | 0 | 0 |
| ENSMUSG00 | Igkv3-2 | 357 | 0 | 0 | 0 | 0 | 0 | 0 | 0 | 0 | 0 | 0 | 0 | 0 | 0 | 0 | 0 | 0 | 0 | 0 |
| ENSMUSG00 | Igkv3-1 | 357 | 0 | 0 | 0 | 0 | 0 | 0 | 0 | 0 | 0 | 0 | 0 | 0 | 0 | 0 | 0 | 0 | 0 | 0 |
| ENSMUSG00 | Igkj1 | 37 | 0 | 0 | 0 | 0 | 0 | 0 | 0 | 0 | 5.56574537 | 0 | 0 | 0 | 0 | 0 | 0 | 0 | 0 | 0 |
| ENSMUSG00 | Igkj2 | 42 | 0 | 0 | 0 | 0 | 0 | 0 | 1.12649236 | 0 | 0 | 0 | 0 | 0 | 0 | 0 | 0 | 0 | 0 | 0 |
| ENSMUSG00 | Igkj3 | 35 | 0 | 0 | 0 | 0 | 0 | 0 | 0 | 0 | 0 | 0 | 0 | 0 | 0 | 0 | 0 | 0 | 0 | 0 |
| ENSMUSG00 | Igkj4 | 37 | 0 | 0 | 0 | 0 | 0 | 1.33926307 | 0 | 0 | 0 | 0 | 0 | 0 | 0 | 0 | 0 | 0 | 0 | 0 |
| ENSMUSG00 | Igkj5 | 41 | 0 | 0 | 0 | 0 | 0 | 0 | 0 | 0 | 0 | 0 | 0 | 0 | 0 | 1.13278262 | 0 | 0 | 0 | 0 |
| ENSMUSG00 | Igkc | 322 | 9.90027392 | 4.03479154 | 5.55844914 | 7.92365988 | 3.43799455 | 4.46282383 | 133.268943 | 25.8569681 | 57.17507 | 27.0896898 | 156.974864 | 19.9341652 | 94.9869215 | 56.6848644 | 44.1445242 | 33.6529251 | 0.23805153 | 11.7876657 |
| ENSMUSG00 | Ighe | 1163 | 0 | 0 | 0 | 0 | 0 | 0 | 0 | 0 | 0 | 0 | 0 | 0 | 0 | 0 | 0 | 0 | 0 | 0 |
| ENSMUSG00 | Ighg2c | 1206 | 0 | 0 | 0 | 0 | 0 | 0 | 0 | 0 | 0 | 0 | 0 | 0 | 0 | 0 | 0 | 0 | 0 | 0 |
| ENSMUSG00 | Ighg2b | 1006 | 0 | 0 | 0 | 0 | 0 | 0 | 0 | 0 | 0 | 0 | 0 | 0 | 0.39230153 | 0 | 0 | 0 | 0 | 0 |
| ENSMUSG00 | Ighg1 | 970 | 0 | 0.13737276 | 0 | 0 | 0.14886162 | 0.30651175 | 0 | 0 | 0 | 0 | 0 | 0 | 0 | 0 | 0 | 0 | 0 | 0 |
| ENSMUSG00 | Ighg3 | 988 | 0 | 0 | 0 | 0.09222883 | 0 | 0 | 0.14366198 | 0 | 0 | 0 | 0.09671044 | 0 | 1.44800162 | 0 | 0 | 4.65068106 | 0 | 0 |
| ENSMUSG00 | Ighm | 1363 | 2.49480123 | 1.14872049 | 0.95824308 | 2.64073551 | 1.97754052 | 2.94480662 | 43.6679018 | 8.96666821 | 25.4733916 | 8.99426474 | 56.3975849 | 4.87312894 | 50.3091453 | 25.7606237 | 13.3610328 | 13.3441222 | 0.05623815 | 2.58296613 |
| ENSMUSG00 | Ighj4 | 53 | 0 | 0 | 0 | 0 | 0 | 0 | 0.89269206 | 0 | 0.77710407 | 0 | 1.80282862 | 0 | 0 | 0 | 1.55477569 | 0 | 0 | 0 |
| ENSMUSG00 | Ighj3 | 47 | 0 | 0 | 0 | 0 | 0 | 0 | 0 | 0 | 0 | 0 | 5.08244239 | 0 | 0 | 0 | 0 | 0 | 0 | 1.17040653 |
| ENSMUSG00 | Ighj2 | 53 | 0 | 0 | 0 | 0 | 0 | 0 | 0 | 0 | 0 | 0 | 0.90141431 | 0 | 0 | 0 | 0.77738785 | 0 | 0 | 0 |
| ENSMUSG00 | Ighj1 | 52 | 0 | 0 | 0 | 0 | 0 | 0 | 0.90985921 | 0 | 0 | 0 | 0 | 0 | 0 | 0.89315553 | 0 | 1.47271567 | 0 | 0 |
| ENSMUSG00 | Ighd4-1 | 10 | 0 | 0 | 0 | 0 | 0 | 0 | 0 | 0 | 0 | 0 | 0 | 0 | 0 | 0 | 0 | 0 | 0 | 0 |
| ENSMUSG00 | Ighd3-2 | 15 | 0 | 0 | 0 | 0 | 0 | 0 | 0 | 0 | 0 | 0 | 0 | 0 | 0 | 0 | 0 | 0 | 0 | 0 |
| ENSMUSG00 | Ighd2-8 | 16 | 0 | 0 | 0 | 0 | 0 | 0 | 0 | 0 | 0 | 0 | 0 | 0 | 0 | 0 | 0 | 0 | 0 | 0 |
| ENSMUSG00 | Ighd2-7 | 16 | 0 | 0 | 0 | 0 | 0 | 0 | 0 | 0 | 0 | 0 | 0 | 0 | 0 | 0 | 0 | 0 | 0 | 0 |
| ENSMUSG00 | Ighd2-6 | 16 | 0 | 0 | 0 | 0 | 0 | 0 | 0 | 0 | 0 | 0 | 0 | 0 | 0 | 0 | 0 | 0 | 0 | 0 |
| ENSMUSG00 | Ighd2-5 | 16 | 0 | 0 | 0 | 0 | 0 | 0 | 0 | 0 | 0 | 0 | 0 | 0 | 0 | 0 | 0 | 0 | 0 | 0 |
| ENSMUSG00 | Ighd2-4 | 16 | 0 | 0 | 0 | 0 | 0 | 0 | 0 | 0 | 0 | 0 | 0 | 0 | 0 | 0 | 0 | 0 | 0 | 0 |
| ENSMUSG00 | Ighd2-3 | 16 | 0 | 0 | 0 | 0 | 0 | 0 | 0 | 0 | 0 | 0 | 0 | 0 | 0 | 0 | 0 | 0 | 0 | 0 |
| ENSMUSG00 | Ighd1-1 | 22 | 0 | 0 | 0 | 0 | 0 | 0 | 0 | 0 | 0 | 0 | 2.17158902 | 0 | 2.24235988 | 6.33328464 | 0 | 0 | 0 | 0 |
| ENSMUSG00 | Ighd3-1 | 16 | 0 | 0 | 0 | 0 | 0 | 0 | 0 | 0 | 0 | 0 | 0 | 0 | 0 | 0 | 0 | 0 | 0 | 0 |
| ENSMUSG00 | Gm16968 | 22 | 0 | 0 | 0 | 0 | 0 | 0 | 0 | 0 | 0 | 0 | 0 | 0 | 0 | 0 | 0 | 0 | 0 | 0 |
| ENSMUSG00 | Ighv5-2 | 348 | 0 | 0 | 0 | 0 | 0 | 0 | 0.67977987 | 0 | 0 | 0 | 0 | 0 | 0 | 0 | 0.23679055 | 0 | 0 | 0 |
| ENSMUSG00 | Ighv2-2 | 348 | 0 | 0 | 0 | 0 | 0 | 0 | 0.40786792 | 0 | 0 | 0 | 0 | 0.16038984 | 0 | 0 | 0 | 0 | 0 | 0 |
| ENSMUSG00 | Ighv5-4 | 351 | 0 | 0 | 0 | 0 | 0 | 0 | 0 | 0 | 0 | 0 | 0 | 0 | 0 | 0 | 0 | 0.10909005 | 0 | 0 |
| ENSMUSG00 | Ighv2-3 | 345 | 0 | 0 | 0 | 0 | 0 | 0 | 0 | 0 | 0 | 0 | 0 | 0 | 0 | 0 | 0 | 0.11098727 | 0 | 0 |
| ENSMUSG00 | Ighv5-6 | 351 | 0 | 0 | 0 | 0 | 0 | 0 | 1.07835166 | 0.32643079 | 0 | 0 | 0 | 0 | 0 | 0 | 0 | 0 | 0 | 0 |
| ENSMUSG00 | Ighv2-4 | 348 | 0 | 0 | 0 | 0 | 0 | 0 | 0 | 0 | 0 | 0 | 0 | 0 | 0 | 0 | 0 | 0 | 0 | 0 |
| ENSMUSG00 | Ighv5-9 | 351 | 0 | 0 | 0 | 0 | 0 | 0 | 0 | 0 | 0 | 0 | 0 | 0 | 0 | 0 | 0 | 1.52726069 | 0 | 0 |
| ENSMUSG00 | Ighv2-5 | 345 | 0 | 0 | 0 | 0 | 0 | 0 | 0 | 0 | 0 | 0 | 0 | 0 | 0 | 0 | 0 | 0 | 0 | 0 |
| ENSMUSG00 | Ighv5-12 | 348 | 0 | 0 | 0 | 0 | 0 | 0 | 0.40786792 | 0 | 0 | 0 | 0 | 0 | 0.14175838 | 0 | 0.11839528 | 0 | 0 | 0 |
| ENSMUSG00 | Ighv2-6 | 343 | 0 | 0 | 0 | 0 | 0 | 0 | 0 | 0 | 0 | 0 | 0 | 0 | 0 | 0.40621651 | 0 | 0.78144097 | 0 | 0 |
| ENSMUSG00 | Ighv5-9-1 | 348 | 0 | 0 | 0 | 0 | 0 | 0 | 0.13595597 | 0 | 0 | 0 | 0 | 0 | 0 | 0 | 0.11839528 | 0 | 0 | 0 |
| ENSMUSG00 | Ighv2-9-1 | 345 | 0 | 0 | 0 | 0 | 0 | 0 | 1.5085202 | 0.55351308 | 3.22329254 | 0 | 0 | 0 | 0 | 0 | 0 | 0 | 0 | 0 |
| ENSMUSG00 | Ighv2-6-8 | 340 | 0 | 0 | 0 | 0 | 0 | 0 | 0 | 0 | 0 | 0 | 0 | 0 | 0 | 0 | 0 | 0 | 0 | 0 |
| ENSMUSG00 | Ighv2-7 | 348 | 0 | 0 | 0 | 0 | 0 | 0 | 0.5438239 | 0 | 0 | 0 | 0 | 0 | 0 | 0 | 1.30234803 | 0 | 0 | 0 |
| ENSMUSG00 | Ighv5-15 | 348 | 0 | 0 | 0 | 0 | 0 | 0 | 0 | 0 | 0 | 0 | 0 | 0 | 0 | 0 | 0 | 0 | 0 | 0 |
| ENSMUSG00 | Ighv5-16 | 348 | 0 | 0 | 0 | 0 | 0 | 0 | 0 | 0 | 0 | 0 | 0 | 0 | 4.81978502 | 0 | 0 | 0 | 0 | 0 |
| ENSMUSG00 | Ighv5-17 | 346 | 0 | 0 | 0 | 0 | 0 | 0 | 0 | 0 | 2.26168728 | 0 | 1.51885706 | 0 | 0 | 1.34231466 | 0.95263713 | 0.1106665 | 0 | 0 |
| ENSMUSG00 | Ighv2-9 | 345 | 0 | 0 | 0 | 0 | 0 | 0 | 2.6056258 | 0 | 0 | 0 | 0.41543442 | 0.16178453 | 0 | 0 | 0 | 0.44394907 | 0 | 0 |
| ENSMUSG00 | Ighv7-1 | 355 | 0 | 0 | 0 | 0 | 0 | 0 | 2.53222789 | 0 | 0 | 0 | 0 | 0 | 0 | 0 | 0 | 0 | 0 | 0 |
| ENSMUSG00 | Ighv7-2 | 352 | 0 | 0 | 0 | 0 | 0 | 0 | 0 | 0 | 0 | 0 | 0 | 0 | 0 | 0 | 0 | 0 | 0 | 0 |
| ENSMUSG00 | Ighv14-1 | 349 | 0 | 0 | 0 | 0 | 0 | 0 | 0 | 0 | 0 | 0 | 0 | 0 | 0 | 0 | 0 | 0 | 0 | 0 |
| ENSMUSG00 | Ighv4-1 | 348 | 0 | 0 | 0 | 0 | 0 | 0 | 0 | 0 | 0 | 0 | 0 | 0 | 0 | 0 | 0 | 0 | 0 | 0 |
| ENSMUSG00 | Ighv11-1 | 351 | 0 | 0 | 0 | 0 | 0 | 0 | 3.50464288 | 0 | 0 | 0 | 0 | 0 | 0 | 0 | 0 | 0 | 0 | 0 |
| ENSMUSG00 | Ighv14-2 | 349 | 0 | 0 | 0 | 0 | 0 | 0 | 0.67783208 | 0.21886764 | 1.29814233 | 0 | 0 | 0.4797908 | 0 | 1.06462091 | 0.3541681 | 0 | 0 | 0 |
| ENSMUSG00 | Ighv11-2 | 351 | 0 | 0 | 0 | 0 | 0 | 0 | 0.26958791 | 0 | 0 | 0 | 2.44999787 | 0 | 0 | 0 | 0 | 0 | 0 | 0 |
| ENSMUSG00 | Ighv14-3 | 349 | 0 | 0 | 0 | 0 | 0 | 0 | 0.81339849 | 0.43773527 | 0 | 0 | 2.46403797 | 0 | 0 | 0 | 0 | 0.10971521 | 0 | 0 |
| ENSMUSG00 | Ighv16-1 | 298 | 0 | 0 | 0 | 0 | 0 | 0 | 0 | 0 | 0 | 0 | 0 | 0 | 0 | 0 | 0 | 0 | 0 | 0 |
| ENSMUSG00 | Ighv9-1 | 293 | 0 | 0 | 0 | 0.15549844 | 0 | 0 | 0 | 0 | 0 | 0 | 0 | 0 | 2.52552478 | 0 | 0 | 0 | 0 | 0 |
| ENSMUSG00 | Ighv9-2 | 346 | 0 | 0 | 0 | 0 | 0 | 0 | 0 | 0 | 0 | 0 | 0 | 0 | 0 | 0 | 0 | 0 | 0 | 0 |
| ENSMUSG00 | Ighv9-3 | 349 | 0 | 0 | 0 | 0 | 0 | 0 | 0 | 0 | 0 | 0 | 0 | 0 | 1.97893078 | 0 | 0 | 0 | 0 | 0 |
| ENSMUSG00 | Ighv7-3 | 352 | 0 | 0 | 0 | 0 | 0 | 0 | 0 | 0 | 1.17007147 | 0 | 0 | 0 | 1.12117994 | 4.88190691 | 0.936399 | 5.98290741 | 0 | 0 |
| ENSMUSG00 | Ighv14-4 | 331 | 0 | 0 | 0 | 0 | 0 | 0 | 1.28644746 | 0 | 0 | 0 | 0 | 0 | 0 | 0 | 0 | 0 | 0 | 0 |
| ENSMUSG00 | Ighv3-3 | 295 | 0 | 0 | 0 | 0 | 0 | 0 | 0 | 0 | 0 | 0 | 1.13364308 | 0 | 0 | 0 | 0 | 0 | 0 | 0 |
| ENSMUSG00 | Ighv7-4 | 358 | 0 | 0 | 0 | 0 | 0 | 0 | 0 | 0 | 0 | 0 | 0 | 0 | 0 | 0 | 0 | 0 | 0 | 0 |
| ENSMUSG00 | Ighv3-1 | 295 | 0 | 0 | 0 | 0 | 0 | 0 | 0 | 0 | 1.67538369 | 0 | 0 | 0 | 0 | 0 | 0 | 0 | 0 | 0 |
| ENSMUSG00 | Ighv3-5 | 352 | 0 | 0 | 0 | 0 | 0 | 0 | 0 | 0 | 0 | 0 | 0 | 0 | 0 | 0 | 0 | 0 | 0 | 0 |
| ENSMUSG00 | Ighv13-1 | 299 | 0 | 0 | 0 | 0 | 0 | 0 | 0 | 0 | 0 | 0 | 0 | 0 | 0 | 0 | 0 | 0 | 0 | 0 |
| ENSMUSG00 | Ighv3-6 | 348 | 0 | 0 | 0 | 0 | 0 | 0 | 0 | 0 | 0 | 0 | 0 | 0 | 0 | 0 | 0 | 0 | 0 | 0 |
| ENSMUSG00 | Ighv9-4 | 293 | 0 | 0 | 0 | 0 | 0 | 0 | 0 | 0 | 0 | 0 | 0 | 0 | 0 | 0 | 0 | 0 | 0 | 0 |
| ENSMUSG00 | Ighv3-8 | 345 | 0 | 0 | 0 | 0 | 0 | 0 | 0 | 0 | 0 | 0 | 0 | 0 | 0 | 0 | 0 | 0 | 0 | 0 |
| ENSMUSG00 | Ighv12-3 | 346 | 0 | 0 | 0 | 0 | 0 | 0 | 0 | 0 | 0 | 0 | 0 | 0 | 0 | 0 | 0 | 0 | 0 | 0 |
| ENSMUSG00 | Ighv6-3 | 354 | 0 | 0 | 0 | 0 | 0 | 0 | 0 | 0 | 3.60672878 | 1.46513485 | 3.37393775 | 0 | 0 | 0 | 2.56054867 | 0 | 0 | 0 |
| ENSMUSG00 | Ighv6-6 | 355 | 0 | 0 | 0 | 0 | 0 | 0 | 2.13240243 | 0 | 0 | 0 | 0 | 1.10059053 | 0 | 0 | 0 | 0 | 0 | 2.32432845 |
| ENSMUSG00 | Ighv10-1 | 357 | 0 | 0 | 0 | 0 | 0 | 0 | 0 | 0.42792608 | 0.23073678 | 0.2641496 | 0 | 0 | 0 | 5.59410576 | 0 | 0 | 0 | 0 |
| ENSMUSG00 | Ighv1-4 | 349 | 0 | 0 | 0 | 0 | 0 | 0 | 0 | 0 | 0 | 0 | 0 | 0 | 0.1413522 | 0 | 0 | 0 | 0 | 0 |
| ENSMUSG00 | Ighv1-5 | 293 | 0 | 0 | 0 | 0 | 0 | 0 | 0.32295344 | 0 | 0 | 0 | 0 | 0 | 0 | 0 | 0 | 0 | 0 | 0 |
| ENSMUSG00 | Ighv10-3 | 357 | 0 | 0 | 0 | 0 | 0 | 0 | 0 | 0 | 0 | 0 | 3.0779385 | 0.93807836 | 0 | 0 | 0 | 0 | 0 | 0 |
| ENSMUSG00 | Ighv1-7 | 293 | 0 | 0 | 0 | 0 | 0 | 0 | 0 | 0 | 0 | 0 | 1.79359912 | 0 | 0.33673664 | 0 | 0.14061964 | 0 | 0 | 0 |
| ENSMUSG00 | Ighv15-2 | 349 | 0 | 0 | 0 | 0 | 0 | 0 | 0 | 0 | 0 | 0 | 0 | 0 | 0 | 0 | 0 | 0 | 0 | 0 |
| ENSMUSG00 | Ighv1-9 | 349 | 0 | 0 | 0 | 0 | 0 | 0 | 0 | 0 | 0 | 0 | 0 | 0 | 1.27216979 | 0 | 0 | 0 | 0 | 0 |
| ENSMUSG00 | Ighv1-12 | 293 | 0 | 0 | 0 | 0 | 0 | 0 | 0 | 0 | 0 | 0 | 0 | 0 | 0 | 0.47553673 | 0 | 0 | 0 | 0 |
| ENSMUSG00 | Ighv1-14 | 293 | 0 | 0 | 0 | 0 | 0 | 0 | 0 | 0 | 0 | 0 | 0 | 0 | 0 | 0 | 0 | 0 | 0 | 0 |
| ENSMUSG00 | Ighv1-15 | 337 | 0 | 0 | 0 | 0 | 0 | 0 | 0 | 0 | 0 | 0 | 0 | 0 | 0.87831307 | 0 | 0.12225981 | 0 | 0 | 0 |
| ENSMUSG00 | Ighv1-30 | 291 | 0 | 0 | 0 | 0 | 0 | 0 | 0 | 0 | 0 | 0 | 0 | 0 | 0 | 0 | 0 | 0 | 0 | 0 |
| ENSMUSG00 | Ighv1-18 | 304 | 0 | 0 | 0 | 0 | 0 | 0 | 0 | 0 | 0 | 0 | 1.57154469 | 0 | 0.16227604 | 0 | 0 | 0 | 0 | 0 |
| ENSMUSG00 | Ighv1-19 | 346 | 0 | 0 | 0 | 0 | 0 | 0 | 0 | 0 | 0 | 0 | 2.89963621 | 0 | 0 | 0 | 0 | 0 | 0 | 0 |
| ENSMUSG00 | Ighv1-20 | 346 | 0 | 0 | 0 | 0 | 0 | 0 | 0 | 0 | 0 | 0 | 0 | 0 | 0 | 0 | 0 | 0 | 0 | 0 |
| ENSMUSG00 | Ighv1-22 | 349 | 0 | 0 | 0 | 0 | 0 | 0 | 0 | 0.10943382 | 0 | 0 | 3.01160197 | 0 | 2.54433957 | 0 | 0.11805603 | 0 | 0 | 0 |
| ENSMUSG00 | Ighv1-23 | 337 | 0 | 0 | 0 | 0 | 0 | 0 | 0 | 0 | 0 | 0 | 0 | 0 | 0.14638551 | 0 | 0.24451962 | 0 | 0 | 0 |
| ENSMUSG00 | Ighv1-24 | 293 | 0 | 0 | 0 | 0 | 0 | 0 | 0 | 0 | 0 | 0 | 0 | 0 | 0 | 0 | 0 | 0 | 0 | 0 |
| ENSMUSG00 | Ighv1-26 | 349 | 0 | 0 | 0 | 0 | 0 | 0 | 0 | 0 | 0.23602588 | 0.67551151 | 0 | 0 | 0 | 0.9315433 | 0 | 0 | 0 | 0 |
| ENSMUSG00 | Ighv1-31 | 293 | 0 | 0 | 0 | 0 | 0 | 0 | 0 | 0 | 0 | 0 | 0 | 0 | 0 | 0 | 0 | 0 | 0 | 0 |
| ENSMUSG00 | Ighv1-34 | 349 | 0 | 0 | 0 | 0 | 0 | 0 | 0 | 0 | 0 | 0 | 0 | 0 | 0 | 0 | 0 | 0 | 0 | 0 |
| ENSMUSG00 | Ighv1-36 | 346 | 0 | 0 | 0 | 0 | 0 | 0 | 0 | 0 | 0 | 0 | 0 | 0 | 0 | 0 | 0 | 0 | 0 | 0 |
| ENSMUSG00 | Ighv1-37 | 346 | 0 | 0 | 0 | 0 | 0 | 0 | 0 | 0 | 0 | 0 | 0 | 0 | 0 | 0 | 0 | 0 | 0 | 0 |
| ENSMUSG00 | Ighv1-39 | 349 | 0 | 0 | 0 | 0 | 0 | 0 | 0 | 0 | 0 | 0 | 0 | 0 | 0 | 0 | 0 | 0 | 0 | 0 |
| ENSMUSG00 | Ighv1-42 | 293 | 0 | 0 | 0 | 0 | 0 | 0 | 0 | 0 | 0 | 0 | 0 | 0 | 0 | 0 | 0 | 0 | 0 | 0 |
| ENSMUSG00 | Ighv1-43 | 293 | 0 | 0 | 0 | 0 | 0 | 0 | 0 | 0 | 0 | 0 | 0 | 0 | 0 | 0 | 0 | 0 | 0 | 0 |
| ENSMUSG00 | Ighv1-47 | 293 | 0 | 0 | 0 | 0 | 0 | 0 | 0 | 0 | 0 | 0 | 0 | 0 | 0 | 0 | 0 | 0 | 0 | 0 |
| ENSMUSG00 | Ighv1-49 | 293 | 0 | 0 | 0 | 0 | 0 | 0 | 0 | 0 | 0 | 0 | 0 | 0 | 0 | 0 | 0 | 0 | 0 | 0 |
| ENSMUSG00 | Ighv8-5 | 334 | 0 | 0 | 0 | 0 | 0 | 0 | 0 | 0 | 0 | 0 | 0 | 0 | 0 | 0 | 0 | 0 | 0 | 0 |
| ENSMUSG00 | Ighv1-50 | 349 | 0 | 0 | 0 | 0 | 0 | 0 | 0 | 0 | 0 | 0 | 0 | 0 | 0 | 0 | 0 | 0 | 0 | 0 |
| ENSMUSG00 | Ighv1-52 | 346 | 0 | 0 | 0 | 0 | 0 | 0 | 0 | 0 | 0 | 0 | 0.41423374 | 0 | 0 | 0 | 0 | 0 | 0 | 0 |
| ENSMUSG00 | Ighv1-53 | 349 | 0 | 0 | 0 | 1.04437924 | 0 | 0 | 0 | 0 | 0 | 0 | 0 | 0 | 0 | 0 | 0 | 0 | 0 | 0 |
| ENSMUSG00 | Ighv8-6 | 300 | 0 | 0 | 0 | 0 | 0 | 0 | 0 | 0 | 0 | 0 | 0 | 0 | 0 | 0 | 0 | 0 | 0 | 0 |
| ENSMUSG00 | Ighv1-54 | 349 | 0 | 0 | 0 | 0 | 0 | 0 | 0 | 0 | 0 | 0 | 1.23201899 | 0 | 0 | 0 | 0 | 0 | 0 | 0 |

| | | | | | | | | | | | | | | | | | | | |
|---|---|---|---|---|---|---|---|---|---|---|---|---|---|---|---|---|---|---|---|
| ENSMUSG00 Ighv1-55 | 349 | 0 | 0 | 0 | 0 | 0 | 0 | 0 | 0 | 0 | 0 | 0 | 0 | 0 | 0 | 0.82639224 | 0 | 0 | 0 |
| ENSMUSG00 Ighv1-56 | 293 | 0 | 0 | 0 | 0 | 0 | 0 | 0 | 0 | 0 | 0 | 0 | 0 | 0 | 0 | 0 | 0 | 0 | 0 |
| ENSMUSG00 Ighv8-8 | 334 | 0 | 0 | 0 | 0 | 0 | 0 | 0 | 0 | 0 | 0 | 0 | 0 | 1.47700351 | 0 | 0 | 0 | 0 | 0 |
| ENSMUSG00 Ighv1-58 | 343 | 0 | 0 | 0 | 0 | 0 | 0 | 0 | 0 | 0 | 0 | 0 | 0 | 0 | 0 | 0 | 0 | 0 | 0 |
| ENSMUSG00 Ighv1-59 | 346 | 0 | 0 | 0 | 0 | 0 | 0 | 0 | 0 | 0 | 0 | 1.65693498 | 0 | 0 | 0 | 0 | 0 | 0 | 0 |
| ENSMUSG00 Ighv1-61 | 346 | 0 | 0 | 0 | 0 | 0 | 0 | 0 | 0 | 0 | 0 | 0 | 0 | 0 | 0 | 0 | 0 | 0 | 0 |
| ENSMUSG00 Ighv1-62-3 | 349 | 0 | 0 | 0 | 0 | 0 | 0 | 0 | 0 | 0 | 0 | 0 | 0 | 0 | 0 | 0 | 0 | 0 | 0 |
| ENSMUSG00 Ighv8-9 | 300 | 0 | 0 | 0 | 0 | 0 | 0 | 0 | 0 | 0 | 0 | 0 | 0 | 0 | 0 | 0 | 0 | 0 | 0 |
| ENSMUSG00 Ighv1-63 | 349 | 0 | 0 | 0 | 0 | 0 | 0 | 0 | 0 | 0 | 0 | 0 | 0 | 0 | 0 | 0 | 0.76800645 | 0 | 0 |
| ENSMUSG00 Ighv1-64 | 349 | 0 | 0 | 0 | 0 | 0 | 0 | 0.13556642 | 0 | 0 | 0 | 2.60092897 | 0 | 0 | 0.79846569 | 0 | 0 | 0 | 0 |
| ENSMUSG00 Ighv8-11 | 334 | 0 | 0 | 0 | 0 | 0 | 0 | 0 | 0 | 0 | 0 | 0 | 0 | 0.14770035 | 0 | 0 | 0 | 0 | 0 |
| ENSMUSG00 Ighv1-66 | 349 | 0 | 0 | 0 | 0 | 0 | 0 | 0 | 0 | 0 | 0 | 0 | 0 | 0 | 0 | 0 | 0 | 0 | 0 |
| ENSMUSG00 Ighv1-67 | 293 | 0 | 0 | 0 | 0 | 0 | 0 | 0 | 0 | 0 | 0 | 0.32610893 | 0 | 0 | 0 | 0 | 0 | 0 | 0 |
| ENSMUSG00 Ighv1-69 | 346 | 0 | 0 | 0 | 0 | 0 | 0 | 0 | 0 | 0 | 0.13627371 | 0 | 0 | 0 | 0 | 0 | 0 | 0 | 0 |
| ENSMUSG00 Ighv8-12 | 313 | 0 | 0 | 0 | 0 | 0 | 0 | 0 | 0 | 0 | 0 | 0.76317825 | 0.1783248 | 0 | 0 | 0 | 0 | 0 | 0 |
| ENSMUSG00 Ighv1-72 | 349 | 0 | 0 | 0 | 0 | 0 | 0 | 0 | 0 | 0.47205176 | 0 | 0.136891 | 0 | 0 | 0 | 0 | 0 | 0 | 0 |
| ENSMUSG00 Ighv8-13 | 302 | 0 | 0 | 0 | 0 | 0 | 0 | 0 | 0 | 0 | 0 | 0 | 0 | 0 | 0 | 0 | 0 | 0 | 0 |
| ENSMUSG00 Gm16710 | 346 | 0.12284733 | 0 | 0 | 0 | 0 | 0 | 0 | 0 | 0 | 0 | 2.62348038 | 0 | 0 | 0 | 0 | 0 | 0 | 0 |
| ENSMUSG00 Ighv1-75 | 349 | 0 | 0 | 0 | 0 | 0 | 0 | 0 | 0 | 0 | 0 | 0 | 0 | 0 | 0 | 0 | 0 | 0 | 0 |
| ENSMUSG00 Ighv1-80 | 349 | 0 | 0 | 0 | 0 | 0 | 0 | 0 | 0 | 0 | 0 | 0 | 0 | 0 | 0 | 0 | 0 | 0 | 0 |
| ENSMUSG00 Ighv1-81 | 349 | 0 | 0 | 0 | 0 | 0 | 0 | 0 | 0 | 0 | 0 | 0 | 0 | 0 | 0 | 0 | 0 | 0 | 0 |
| ENSMUSG00 Ighv1-82 | 293 | 0 | 0 | 0 | 0 | 0 | 0 | 4.35987144 | 0 | 0 | 0 | 0 | 0 | 2.02041982 | 0 | 0 | 0 | 0 | 0 |
| ENSMUSG00 Ighv1-83 | 290 | 0 | 0 | 0 | 0 | 0 | 0 | 0 | 0 | 0 | 0 | 0 | 0 | 0 | 0 | 0 | 0.13203658 | 0 | 0 |
| ENSMUSG00 Ighv1-84 | 349 | 0 | 0 | 0 | 0 | 0 | 0 | 0 | 0 | 0 | 0 | 0 | 0 | 0 | 0 | 0 | 0 | 0 | 0 |
| ENSMUSG00 Ighv1-85 | 349 | 0 | 0 | 0 | 0 | 0 | 0 | 0.40669925 | 0 | 0 | 0 | 0 | 0 | 0 | 0 | 0 | 0 | 0 | 0 |
| ENSMUSG00 Tcrg-V7 | 348 | 0 | 0 | 0 | 0 | 0 | 0 | 0 | 0 | 0 | 0 | 0 | 0 | 0 | 0 | 0 | 0 | 0 | 0 |
| ENSMUSG00 Tcrg-V4 | 412 | 0 | 0 | 0 | 0 | 0 | 0 | 0 | 0 | 0 | 0 | 0 | 0 | 0 | 0 | 0 | 0 | 0 | 0 |
| ENSMUSG00 Tcrg-V6 | 401 | 0 | 0 | 0 | 0 | 0 | 0 | 0 | 0 | 0 | 0 | 0 | 0 | 0 | 0 | 0 | 0 | 0 | 0 |
| ENSMUSG00 Tcrg-V5 | 292 | 0 | 0 | 0 | 0 | 0 | 0 | 0 | 0 | 0 | 0 | 0 | 0 | 0 | 0 | 0 | 0 | 0 | 0 |
| ENSMUSG00 Trgj1 | 59 | 0 | 0 | 0 | 0 | 0 | 0 | 0 | 0 | 0 | 0 | 0 | 0 | 0 | 0 | 0 | 0 | 0 | 0 |
| ENSMUSG00 Gm17004 | 515 | 0 | 0 | 0 | 0 | 0 | 0 | 0 | 0 | 0 | 0 | 0 | 0 | 0 | 0 | 0 | 0 | 0 | 0 |
| ENSMUSG00 Tcrg-V3 | 348 | 0 | 0 | 0 | 0 | 0 | 0 | 0 | 0 | 0 | 0 | 0 | 0 | 0 | 0 | 0 | 0 | 0 | 0 |
| ENSMUSG00 Trgj3 | 59 | 0 | 0 | 0 | 0 | 0 | 0 | 0 | 0 | 0 | 0 | 0 | 0 | 0 | 0 | 0 | 0 | 0 | 0 |
| ENSMUSG00 Gm16603 | 500 | 0 | 0 | 0 | 0 | 0 | 0 | 0 | 0 | 0 | 0 | 0 | 0 | 0 | 0 | 0 | 0 | 0 | 0 |
| ENSMUSG00 Trgj2 | 59 | 0 | 0 | 0 | 0 | 0 | 0 | 0 | 0 | 0 | 0 | 0 | 0 | 0 | 0 | 0 | 0 | 0 | 0 |
| ENSMUSG00 Trgv2 | 348 | 0 | 0 | 0 | 0 | 0 | 0 | 0 | 0 | 0 | 0 | 0 | 0 | 0 | 0 | 0 | 0 | 0 | 0 |
| ENSMUSG00 Gm16602 | 345 | 0 | 0 | 0 | 0 | 0 | 0 | 0 | 0 | 0 | 0 | 0 | 0 | 0 | 0 | 0 | 0 | 0 | 0 |
| ENSMUSG00 Trgj4 | 60 | 0 | 0 | 0 | 0 | 0 | 0 | 0 | 0 | 0 | 0 | 0 | 0 | 0 | 0 | 0 | 0 | 0 | 0 |
| ENSMUSG00 Tcrg-C | 638 | 0 | 0 | 0 | 0 | 0 | 0 | 0 | 0 | 0 | 0 | 0 | 0.17497073 | 0 | 0.14559275 | 0.06457924 | 0 | 0 | 0 |
| ENSMUSG00 Trav1 | 367 | 0 | 0 | 0 | 0 | 0 | 0.13502107 | 0 | 0 | 0 | 0 | 0 | 0 | 0 | 0 | 0 | 0 | 0 | 0 |
| ENSMUSG00 Trav2 | 323 | 0 | 0 | 0 | 0 | 0 | 0 | 0 | 0 | 0 | 0 | 0 | 0 | 0 | 0 | 0 | 0 | 0 | 0 |
| ENSMUSG00 Trav3-1 | 341 | 0 | 0 | 0 | 0 | 0 | 0 | 0 | 0 | 0 | 0 | 0 | 0 | 0 | 0 | 0 | 0 | 0 | 0 |
| ENSMUSG00 Trav5-1 | 341 | 0 | 0.09769177 | 0 | 0.1336101 | 0 | 0 | 0.41624058 | 0 | 0 | 0 | 0 | 0 | 0 | 0 | 0 | 0 | 0 | 0 |
| ENSMUSG00 Trav6-1 | 336 | 0 | 0 | 0 | 0 | 0 | 0 | 0 | 0 | 0 | 0 | 0 | 0 | 0 | 0 | 0 | 0 | 0 | 0 |
| ENSMUSG00 Trav6-2 | 336 | 0 | 0 | 0 | 0 | 0 | 0 | 0 | 0 | 0 | 0 | 0 | 0 | 0 | 0 | 0 | 0 | 0 | 0 |
| ENSMUSG00 Trav6-4 | 374 | 0 | 0 | 0 | 0 | 0 | 0 | 0 | 0 | 0 | 0 | 0 | 0 | 0 | 0 | 0 | 0 | 0 | 0 |
| ENSMUSG00 Trav7d-3 | 399 | 0 | 0 | 0 | 0 | 0 | 0 | 0 | 0 | 0 | 0 | 0 | 0 | 0 | 0 | 0 | 0 | 0 | 0 |
| ENSMUSG00 Trav6d-4 | 374 | 0 | 0 | 0 | 0 | 0 | 0 | 0 | 0 | 0 | 0 | 0 | 0 | 0 | 0 | 0 | 0 | 0 | 0 |
| ENSMUSG00 Trav7d-4 | 447 | 0 | 0 | 0 | 0 | 0 | 0 | 0 | 0 | 0 | 0 | 0 | 0 | 0 | 0 | 0 | 0 | 0 | 0 |
| ENSMUSG00 Trav8d-1 | 397 | 0 | 0 | 0 | 0 | 0 | 0 | 0 | 0 | 0 | 0 | 0 | 0 | 0 | 0 | 0.31134677 | 0 | 0 | 0 |
| ENSMUSG00 Trav9d-1 | 442 | 0 | 0 | 0 | 0 | 0 | 0 | 0 | 0 | 0 | 0 | 0 | 0 | 0 | 0 | 0 | 0 | 0 | 0 |
| ENSMUSG00 Trav6d-5 | 471 | 0 | 0 | 0 | 0 | 0 | 0 | 0 | 0 | 0 | 0 | 0 | 0 | 0 | 0 | 0 | 0 | 0 | 0 |
| ENSMUSG00 Trav10d | 403 | 0 | 0 | 0 | 0 | 0 | 0 | 0 | 0 | 0 | 0 | 0 | 0 | 0 | 0 | 0 | 0 | 0 | 0 |
| ENSMUSG00 Trav6d-6 | 402 | 0 | 0 | 0 | 0 | 0 | 0 | 0 | 0 | 0 | 0 | 0 | 0 | 0 | 0 | 0 | 0 | 0 | 0 |
| ENSMUSG00 Trav11d | 357 | 0 | 0 | 0 | 0 | 0 | 0 | 0 | 0 | 0 | 0 | 0 | 0 | 0 | 0 | 0 | 0 | 0 | 0 |
| ENSMUSG00 Trav7d-5 | 355 | 0 | 0 | 0 | 0 | 0 | 0 | 0 | 0 | 0 | 0 | 0 | 0 | 0 | 0 | 0 | 0 | 0 | 0 |
| ENSMUSG00 Trav12d-1 | 432 | 0 | 0 | 0 | 0 | 0 | 0 | 0 | 0 | 0 | 0 | 0 | 0 | 0 | 0 | 0 | 0 | 0 | 0 |
| ENSMUSG00 Trav13d-1 | 393 | 0 | 0 | 0 | 0 | 0 | 0 | 0 | 0 | 0 | 0 | 0 | 0 | 0 | 0 | 0 | 0 | 0 | 0 |
| ENSMUSG00 Trav14d-1 | 412 | 0 | 0 | 0 | 0 | 0 | 0 | 0 | 0 | 0 | 0 | 0 | 0 | 0 | 0 | 0 | 0 | 0 | 0 |
| ENSMUSG00 Trav15d-1-dv | 415 | 0 | 0 | 0 | 0 | 0 | 0 | 0 | 0 | 0 | 0 | 0 | 0 | 0 | 0 | 0 | 0 | 0 | 0 |
| ENSMUSG00 Trav9d-2 | 335 | 0 | 0 | 0 | 0 | 0 | 0 | 0 | 0 | 0 | 0 | 0 | 0 | 0 | 0 | 0 | 0 | 0 | 0 |
| ENSMUSG00 Trav4d-3 | 329 | 0 | 0 | 0 | 0 | 0 | 0 | 0 | 0 | 0 | 0 | 0 | 0 | 0 | 0 | 0 | 0 | 0 | 0 |
| ENSMUSG00 Trav12d-2 | 339 | 0 | 0 | 0 | 0 | 0 | 0 | 0 | 0 | 0 | 0 | 0 | 0 | 0 | 0 | 0 | 0 | 0 | 0 |
| ENSMUSG00 Trav9d-3 | 338 | 0 | 0 | 0 | 0 | 0 | 0 | 0 | 0 | 0 | 0 | 0 | 0 | 0 | 0 | 0 | 0 | 0 | 0 |
| ENSMUSG00 Trav12d-3 | 345 | 0 | 0 | 0 | 0 | 0 | 0 | 0 | 0 | 0 | 0 | 0 | 0 | 0 | 0 | 0 | 0 | 0 | 0 |
| ENSMUSG00 Trav13d-2 | 263 | 0 | 0 | 0 | 0 | 0 | 0 | 0 | 0 | 0 | 0 | 0 | 0 | 0 | 0 | 0 | 0 | 0 | 0 |
| ENSMUSG00 Trav14d-2 | 275 | 0 | 0 | 0 | 0 | 0 | 0 | 0 | 0 | 0 | 0 | 0 | 0 | 0 | 0 | 0 | 0 | 0 | 0 |
| ENSMUSG00 Trav15d-2-dv | 346 | 0 | 0 | 0 | 0 | 0 | 0 | 0 | 0 | 0 | 0 | 0 | 0 | 0 | 0 | 0 | 0 | 0 | 0 |
| ENSMUSG00 Trav3d-3 | 275 | 0 | 0 | 0 | 0 | 0 | 0 | 0 | 0 | 0 | 0 | 0 | 0 | 0 | 0 | 0 | 0 | 0 | 0 |
| ENSMUSG00 Trav4d-4 | 329 | 0 | 0 | 0 | 0 | 0 | 0 | 0 | 0 | 0 | 0 | 0 | 0 | 0 | 0 | 0 | 0 | 0 | 0 |
| ENSMUSG00 Trav5d-4 | 341 | 0 | 0 | 0 | 0 | 0 | 0 | 0 | 0 | 0 | 0 | 0 | 0 | 0 | 0 | 0 | 0 | 0 | 0 |
| ENSMUSG00 Trav6d-7 | 315 | 0 | 0 | 0 | 0 | 0 | 0 | 0 | 0 | 0 | 0 | 0 | 0 | 0 | 0 | 0 | 0 | 0 | 0 |
| ENSMUSG00 C920008G01 | 1408 | 0 | 0 | 0 | 0 | 0 | 0 | 0 | 0 | 0 | 0 | 0 | 0 | 0 | 0 | 0 | 0 | 0 | 0 |
| ENSMUSG00 Trav8d-2 | 332 | 0 | 0 | 0 | 0 | 0 | 0 | 0 | 0 | 0 | 0 | 0 | 0 | 0 | 0 | 0 | 0 | 0 | 0 |
| ENSMUSG00 Trav16d-dv11 | 351 | 0 | 0 | 0 | 0 | 0 | 0 | 0 | 0 | 0 | 0 | 0 | 0 | 0 | 0 | 0 | 0 | 0 | 0 |
| ENSMUSG00 Trav13d-4 | 263 | 0 | 0 | 0 | 0 | 0 | 0 | 0 | 0 | 0 | 0 | 0 | 0 | 0 | 0 | 0 | 0 | 0 | 0 |
| ENSMUSG00 Trav14d-3-dv | 326 | 0 | 0 | 0 | 0 | 0 | 0 | 0 | 0 | 0 | 0 | 0 | 0 | 0 | 0 | 0 | 0 | 0 | 0 |
| ENSMUSG00 Trav7n-4 | 339 | 0 | 0 | 0 | 0 | 0 | 0 | 0 | 0 | 0 | 0 | 0 | 0 | 0 | 0 | 0 | 0 | 0 | 0 |
| ENSMUSG00 Trav6n-5 | 336 | 0 | 0 | 0 | 0 | 0 | 0 | 0 | 0 | 0 | 0 | 0 | 0 | 0 | 0 | 0 | 0 | 0 | 0 |
| ENSMUSG00 Trav10n | 341 | 0 | 0 | 0 | 0 | 0 | 0 | 0 | 0 | 0 | 0 | 0 | 0 | 0 | 0 | 0 | 0 | 0 | 0 |
| ENSMUSG00 Trav6n-6 | 286 | 0 | 0 | 0 | 0 | 0 | 0 | 0 | 0 | 0 | 0 | 0 | 0 | 0 | 0 | 0 | 0 | 0 | 0 |
| ENSMUSG00 Trav14n-1 | 338 | 0 | 0 | 0 | 0 | 0 | 0 | 0 | 0 | 0 | 0 | 0 | 0 | 0 | 0 | 0 | 0 | 0 | 0 |
| ENSMUSG00 Trav15n-1 | 347 | 0 | 0 | 0 | 0 | 0 | 0 | 0 | 0 | 0 | 0 | 0 | 0 | 0 | 0 | 0 | 0 | 0 | 0 |
| ENSMUSG00 Trav9n-2 | 335 | 0 | 0 | 0 | 0 | 0 | 0 | 0 | 0 | 0 | 0 | 0 | 0 | 0 | 0 | 0 | 0 | 0 | 0 |
| ENSMUSG00 Trav4n-3 | 329 | 0 | 0 | 0 | 0 | 0 | 0 | 0 | 0 | 0 | 0 | 0 | 0 | 0 | 0 | 0 | 0 | 0 | 0 |
| ENSMUSG00 Trav12n-2 | 339 | 0 | 0 | 0 | 0 | 0 | 0 | 0 | 0 | 0 | 0 | 0 | 0 | 0 | 0 | 0 | 0 | 0 | 0 |
| ENSMUSG00 Trav9n-3 | 338 | 0 | 0 | 0 | 0 | 0 | 0 | 0 | 0 | 0 | 0 | 0 | 0 | 0 | 0 | 0 | 0 | 0 | 0 |
| ENSMUSG00 Trav12n-3 | 345 | 0 | 0 | 0 | 0 | 0 | 0 | 0 | 0 | 0 | 0 | 0 | 0 | 0 | 0 | 0 | 0 | 0 | 0 |
| ENSMUSG00 Trav13n-2 | 263 | 0 | 0 | 0 | 0 | 0 | 0 | 0 | 0 | 0 | 0 | 0 | 0 | 0 | 0 | 0 | 0 | 0 | 0 |
| ENSMUSG00 Trav14n-2 | 275 | 0 | 0 | 0 | 0 | 0 | 0 | 0 | 0 | 0 | 0 | 0 | 0 | 0 | 0 | 0 | 0 | 0 | 0 |
| ENSMUSG00 Trav15n-2 | 346 | 0 | 0 | 0 | 0 | 0 | 0 | 0 | 0 | 0 | 0 | 0 | 0 | 0 | 0 | 0 | 0 | 0 | 0 |
| ENSMUSG00 Trav3n-3 | 275 | 0 | 0 | 0 | 0 | 0 | 0 | 0 | 0 | 0 | 0 | 0 | 0 | 0 | 0 | 0 | 0 | 0 | 0 |
| ENSMUSG00 Trav9n-4 | 338 | 0 | 0 | 0 | 0 | 0 | 0 | 0 | 0 | 0 | 0 | 0 | 0 | 0 | 0 | 0 | 0 | 0 | 0 |
| ENSMUSG00 Trav4n-4 | 329 | 0 | 0 | 0 | 0 | 0 | 0 | 0 | 0 | 0 | 0 | 0 | 0 | 0 | 0 | 0 | 0 | 0 | 0 |
| ENSMUSG00 Trav5n-4 | 341 | 0 | 0 | 0 | 0 | 0 | 0 | 0 | 0 | 0 | 0 | 0 | 0 | 0 | 0 | 0 | 0 | 0 | 0 |
| ENSMUSG00 Trav6n-7 | 315 | 0 | 0 | 0 | 0 | 0 | 0 | 0 | 0 | 0 | 0 | 0 | 0 | 0 | 0 | 0 | 0 | 0 | 0 |
| ENSMUSG00 Trav7n-6 | 1408 | 0 | 0 | 0 | 0 | 0 | 0 | 0 | 0 | 0 | 0 | 0 | 0 | 0 | 0 | 0 | 0 | 0 | 0 |
| ENSMUSG00 Trav8n-2 | 331 | 0 | 0 | 0 | 0 | 0 | 0 | 0 | 0 | 0 | 0 | 0 | 0 | 0 | 0 | 0 | 0 | 0 | 0 |
| ENSMUSG00 Trav16 | 351 | 0 | 0 | 0 | 0 | 0 | 0 | 0 | 0 | 0 | 0 | 0 | 0 | 0 | 0 | 0 | 0 | 0 | 0 |
| ENSMUSG00 Trav13n-4 | 266 | 0 | 0 | 0 | 0 | 0 | 0 | 0 | 0 | 0 | 0 | 0 | 0 | 0 | 0 | 0 | 0 | 0 | 0 |
| ENSMUSG00 Trav14n-3 | 279 | 0 | 0 | 0 | 0 | 0 | 0 | 0 | 0 | 0 | 0 | 0 | 0 | 0 | 0 | 0 | 0 | 0 | 0 |
| ENSMUSG00 Trav7-2 | 399 | 0 | 0 | 0 | 0 | 0 | 0 | 0 | 0 | 0 | 0 | 0 | 0 | 0 | 0 | 0 | 0 | 0 | 0 |
| ENSMUSG00 Trav4-2 | 367 | 0 | 0 | 0 | 0 | 0 | 0 | 0 | 0 | 0 | 0 | 0 | 0 | 0 | 0 | 0 | 0 | 0 | 0 |
| ENSMUSG00 Trav6-3 | 384 | 0 | 0 | 0 | 0 | 0 | 0 | 0 | 0 | 0 | 0 | 0 | 0 | 0 | 0 | 0 | 0 | 0 | 0 |
| ENSMUSG00 Trav7-3 | 399 | 0 | 0 | 0 | 0 | 0 | 0 | 0 | 0 | 0 | 0 | 0 | 0 | 0 | 0 | 0 | 0 | 0 | 0 |
| ENSMUSG00 Trav6d-3 | 343 | 0 | 0 | 0 | 0 | 0 | 0 | 0 | 0 | 0 | 0 | 0 | 0 | 0 | 0 | 0 | 0 | 0 | 0 |
| ENSMUSG00 Trav7-4 | 447 | 0 | 0 | 0 | 0 | 0 | 0 | 0 | 0 | 0 | 0 | 0 | 0 | 0 | 0 | 0 | 0 | 0 | 0 |
| ENSMUSG00 Trav8-1 | 325 | 0 | 0 | 0 | 0 | 0 | 0 | 0 | 0 | 0 | 0 | 0 | 0 | 0 | 0 | 0 | 0 | 0 | 0 |
| ENSMUSG00 Trav9-1 | 395 | 0 | 0 | 0 | 0 | 0 | 0 | 0 | 0 | 0 | 0 | 0 | 0 | 0 | 0 | 0 | 0 | 0 | 0 |
| ENSMUSG00 Trav6-5 | 371 | 0 | 0 | 0 | 0 | 0 | 0 | 0 | 0 | 0 | 0 | 0 | 0 | 0 | 0 | 0 | 0 | 0 | 0 |
| ENSMUSG00 Trav10 | 341 | 0 | 0 | 0 | 0 | 0 | 0 | 0 | 0 | 0 | 0 | 0 | 0 | 0 | 0 | 0 | 0 | 0 | 0 |
| ENSMUSG00 Trav6-6 | 342 | 0 | 0 | 0 | 0 | 0 | 0 | 0 | 0 | 0 | 0 | 0 | 0 | 0 | 0 | 0 | 0 | 0 | 0 |
| ENSMUSG00 Trav11 | 341 | 0 | 0 | 0 | 0 | 0 | 0 | 0 | 0 | 0 | 0 | 0 | 0 | 0 | 0 | 0 | 0 | 0 | 0 |
| ENSMUSG00 Trav12-1 | 342 | 0 | 0 | 0 | 0 | 0 | 0 | 0 | 0 | 0 | 0 | 0 | 0 | 0 | 0 | 0 | 0 | 0 | 0 |
| ENSMUSG00 Trav13-1 | 325 | 0 | 0 | 0 | 0 | 0 | 0 | 0 | 0 | 0 | 0 | 0 | 0 | 0 | 0 | 0 | 0 | 0 | 0 |
| ENSMUSG00 Trav14-1 | 355 | 0 | 0 | 0 | 0 | 0 | 0 | 0 | 0 | 0 | 0 | 0 | 0 | 0 | 0 | 0 | 0 | 0 | 0 |
| ENSMUSG00 Trav15-1-dv6 | 347 | 0 | 0 | 0 | 0 | 0 | 0 | 0 | 0 | 0 | 0 | 0 | 0 | 0 | 0 | 0 | 0 | 0 | 0 |
| ENSMUSG00 Trav9-2 | 338 | 0 | 0 | 0 | 0 | 0 | 0 | 0 | 0 | 0 | 0 | 0 | 0 | 0 | 0 | 0 | 0 | 0 | 0 |
| ENSMUSG00 Trav4-3 | 329 | 0 | 0 | 0 | 0 | 0 | 0 | 0 | 0 | 0 | 0 | 0 | 0 | 0 | 0.28233488 | 2.12895578 | 0 | 0.46597321 | 1.00320559 |
| ENSMUSG00 Trav12-2 | 422 | 0 | 0 | 0 | 0 | 0 | 0 | 0 | 0 | 0 | 0 | 0 | 0 | 0 | 0 | 0 | 0 | 0 | 0 |
| ENSMUSG00 Trav12-3 | 438 | 0 | 0 | 0 | 0 | 0 | 0 | 0 | 0 | 0 | 0 | 0 | 0 | 0 | 0 | 0 | 0 | 0 | 0 |
| ENSMUSG00 Trav13-2 | 387 | 0 | 0 | 0 | 0 | 0 | 0 | 0 | 0 | 0 | 0 | 0 | 0 | 0 | 0 | 0 | 0 | 0 | 0 |
| ENSMUSG00 Trav14-2 | 384 | 0 | 0 | 0 | 0 | 0 | 0 | 0 | 0 | 0 | 0 | 0 | 0 | 0 | 0.24189629 | 0 | 0 | 0 | 0 |
| ENSMUSG00 Trav15-2-dv6 | 384 | 0 | 0 | 0 | 0 | 0 | 0 | 0 | 0 | 0 | 0 | 0 | 0 | 0 | 0 | 0 | 0 | 0 | 0 |
| ENSMUSG00 Trav3-3 | 416 | 0 | 0 | 0 | 0 | 0 | 0 | 0 | 0 | 0 | 0 | 0 | 0 | 0 | 0 | 0 | 0 | 0 | 0 |
| ENSMUSG00 Trav9-4 | 392 | 0 | 0 | 0 | 0 | 0 | 0 | 0 | 0 | 0 | 0 | 0 | 0 | 0 | 0 | 0 | 0 | 0 | 0 |
| ENSMUSG00 Trav4-4-dv10 | 358 | 0 | 0.09305278 | 0 | 0 | 0 | 0 | 0 | 0 | 0 | 0 | 0 | 0 | 0.2755973 | 0.12973209 | 0.69052887 | 0 | 0.21411339 | 0 |
| ENSMUSG00 Trav5-4 | 340 | 0 | 0 | 0 | 0 | 0 | 0 | 0 | 0 | 0 | 0 | 0 | 0 | 0 | 0 | 0 | 0 | 0 | 0 |
| ENSMUSG00 Trav6-7-dv9 | 375 | 0 | 0 | 0 | 0 | 0 | 0 | 0 | 0 | 0 | 0 | 0 | 0 | 0 | 0 | 0 | 0 | 0 | 0 |
| ENSMUSG00 Trav7-6 | 378 | 0 | 0 | 0 | 0 | 0 | 0 | 0 | 0 | 0 | 0 | 0 | 0 | 0 | 0 | 0 | 0 | 0 | 0 |
| ENSMUSG00 Gm13896 | 419 | 0 | 0 | 0 | 0 | 0 | 0 | 0 | 0 | 0 | 0 | 0 | 0 | 0 | 0 | 0 | 0 | 0 | 0 |
| ENSMUSG00 Trav13-4-dv7 | 382 | 0 | 0 | 0 | 0 | 0 | 0 | 0 | 0 | 0 | 0 | 0 | 0 | 0 | 0 | 0 | 0 | 0 | 0 |
| ENSMUSG00 Trav14-3 | 962 | 0 | 0 | 0 | 0 | 0 | 0 | 0 | 0 | 0 | 0 | 0 | 0 | 0 | 0 | 0 | 0 | 0 | 0 |
| ENSMUSG00 Trav3-4 | 455 | 0 | 0 | 0 | 0 | 0 | 0 | 0 | 0 | 0 | 0 | 0 | 0 | 0 | 0 | 0 | 0 | 0 | 0 |
| ENSMUSG00 Trav17 | 451 | 0 | 0 | 0 | 0 | 0 | 0 | 0 | 0 | 0 | 0 | 0 | 0 | 0 | 0 | 0 | 0 | 0 | 0 |
| ENSMUSG00 Trav18 | 325 | 0 | 0 | 0 | 0 | 0 | 0 | 0 | 0 | 0 | 0 | 0 | 0 | 0 | 0 | 0 | 0 | 0 | 0 |
| ENSMUSG00 Trav19 | 443 | 0 | 0 | 0 | 0 | 0 | 0 | 0 | 0 | 0 | 0 | 0 | 0 | 0 | 0 | 0 | 0 | 0 | 0 |
| ENSMUSG00 Trav21-dv12 | 357 | 0 | 0 | 0 | 0 | 0 | 0 | 0 | 0 | 0 | 0 | 0 | 0 | 0 | 0 | 0 | 0 | 0 | 0 |
| ENSMUSG00 Trdv1 | 414 | 0 | 0 | 0 | 0 | 0 | 0 | 0 | 0 | 0 | 0 | 0 | 0 | 0 | 0 | 0 | 0 | 0 | 0 |
| ENSMUSG00 Trdv2-1 | 343 | 0 | 0 | 0 | 0 | 0 | 0 | 0 | 0 | 0 | 0 | 0 | 0 | 0 | 0 | 0 | 0 | 0 | 0 |
| ENSMUSG00 Trdv2-2 | 343 | 0 | 0 | 0 | 0 | 0 | 0 | 0 | 0 | 0 | 0 | 0 | 0 | 0 | 0 | 0 | 0 | 0 | 0 |
| ENSMUSG00 Trdv4 | 343 | 0 | 0 | 0 | 0 | 0 | 0 | 0 | 0 | 0 | 0 | 0 | 0 | 0 | 0 | 0 | 0 | 0 | 0 |
| ENSMUSG00 Trdd1 | 10 | 0 | 0 | 0 | 0 | 0 | 0 | 0 | 0 | 0 | 0 | 0 | 0 | 0 | 0 | 0 | 0 | 0 | 0 |
| ENSMUSG00 Trdd2 | 15 | 0 | 0 | 0 | 0 | 0 | 0 | 0 | 0 | 0 | 0 | 0 | 0 | 0 | 0 | 0 | 0 | 0 | 0 |
| ENSMUSG00 Trdj1 | 50 | 0 | 0 | 0 | 0 | 0 | 0 | 0 | 0 | 0 | 0 | 0 | 0 | 0 | 0 | 0 | 0 | 0 | 0 |
| ENSMUSG00 Trdj2 | 58 | 0 | 0 | 0 | 0 | 0 | 0 | 0 | 0 | 0 | 0 | 0 | 0 | 0 | 0 | 0 | 0 | 0 | 0 |
| ENSMUSG00 Trdv5 | 343 | 0 | 0 | 0 | 0 | 0 | 0 | 0 | 0 | 0 | 0 | 0 | 0 | 0 | 0 | 0 | 0 | 0 | 0 |
| ENSMUSG00 Traj59 | 63 | 0 | 0 | 0 | 0 | 0 | 0 | 0 | 0 | 0 | 0 | 0 | 0 | 0 | 0 | 0 | 0 | 0 | 0 |
| ENSMUSG00 Traj58 | 62 | 0 | 0 | 0 | 0 | 0 | 0 | 0 | 0 | 0 | 0 | 0 | 0 | 0 | 0 | 0 | 0 | 0 | 0 |
| ENSMUSG00 Traj57 | 64 | 0 | 0 | 0 | 0 | 0 | 0 | 0 | 0 | 0 | 0 | 0 | 0 | 0 | 0 | 0 | 0 | 0 | 0 |
| ENSMUSG00 Traj56 | 62 | 0 | 0 | 0 | 0 | 0 | 0 | 0 | 0 | 0 | 0 | 0 | 0 | 0 | 0 | 0 | 0 | 0 | 0 |
| ENSMUSG00 Traj53 | 65 | 0 | 0 | 0 | 0 | 0 | 0 | 0 | 0 | 0 | 0 | 0 | 0 | 0 | 0 | 0 | 0 | 0 | 0 |
| ENSMUSG00 Traj52 | 65 | 0 | 0 | 0 | 0 | 0 | 0 | 0 | 0 | 0 | 0 | 0 | 0 | 0 | 0 | 0 | 0 | 0 | 0 |
| ENSMUSG00 Traj51 | 42 | 0 | 0 | 0 | 0 | 0 | 0 | 0 | 0 | 0 | 0 | 0 | 0 | 0 | 0 | 0 | 0 | 0 | 0 |
| ENSMUSG00 Traj50 | 62 | 0 | 0 | 0 | 0 | 0 | 0 | 0 | 0 | 0 | 0 | 0 | 0 | 0 | 0 | 0 | 0 | 0 | 0 |
| ENSMUSG00 Traj49 | 58 | 0 | 0 | 0 | 0 | 0 | 0 | 0 | 0 | 0 | 0 | 0 | 0 | 0 | 0 | 0 | 0 | 0 | 0 |
| ENSMUSG00 Traj48 | 59 | 0 | 0 | 0 | 0 | 0 | 0 | 0 | 0 | 0 | 0 | 0 | 0 | 0 | 0 | 0 | 0 | 0 | 0 |
| ENSMUSG00 Traj46 | 62 | 0 | 0 | 0 | 0 | 0 | 0 | 0 | 0 | 0 | 0 | 0 | 0 | 0 | 0 | 0 | 0 | 0 | 0 |
| ENSMUSG00 Traj45 | 62 | 0 | 0 | 0 | 0 | 0 | 0 | 0 | 0 | 0 | 0 | 0 | 0 | 0 | 0 | 0 | 0 | 0 | 0 |
| ENSMUSG00 Traj44 | 60 | 0 | 0 | 0 | 0 | 0 | 0 | 0 | 0 | 0 | 0 | 0 | 0 | 0 | 0 | 0 | 0 | 0 | 0 |
| ENSMUSG00 Traj43 | 56 | 0 | 0 | 0 | 0 | 0 | 0 | 0 | 0 | 0 | 0 | 0 | 0 | 0 | 0 | 0 | 0 | 0 | 0 |
| ENSMUSG00 Traj42 | 63 | 0 | 0 | 0 | 0 | 0 | 0 | 0 | 0 | 0 | 0 | 0 | 0 | 0 | 0 | 0 | 0 | 0 | 0 |
| ENSMUSG00 Traj41 | 54 | 0 | 0 | 0 | 0 | 0 | 0 | 0 | 0 | 0 | 0 | 0 | 0 | 0 | 0 | 0 | 0 | 0 | 0 |
| ENSMUSG00 Traj40 | 60 | 0 | 0 | 0 | 0 | 0 | 0 | 0 | 0 | 0 | 0 | 0 | 0 | 0 | 0 | 0 | 0 | 0 | 0 |
| ENSMUSG00 Traj39 | 62 | 0 | 0 | 0 | 0 | 0 | 0 | 0 | 0 | 0 | 0 | 0 | 0 | 0 | 0 | 0 | 0 | 0 | 0 |
| ENSMUSG00 Traj38 | 61 | 0 | 0 | 0 | 0 | 0 | 0 | 0 | 0 | 0 | 0 | 0 | 0 | 0 | 0 | 0 | 0 | 0 | 0 |
| ENSMUSG00 Traj37 | 59 | 0 | 0 | 0 | 0 | 0 | 0 | 0 | 0 | 0 | 0 | 0 | 0 | 0 | 0 | 0 | 0 | 0 | 0 |
| ENSMUSG00 Traj35 | 64 | 0 | 0 | 0 | 0 | 0 | 0 | 0 | 0 | 0 | 0 | 0 | 0 | 0 | 0 | 0 | 0 | 0 | 0 |
| ENSMUSG00 Traj34 | 57 | 0 | 0 | 0 | 0 | 0 | 0 | 0 | 0 | 0 | 0 | 0 | 0 | 0 | 0 | 0 | 0 | 0 | 0 |
| ENSMUSG00 Traj33 | 56 | 0 | 0 | 0 | 0 | 0 | 0 | 0 | 0 | 0 | 0 | 0 | 0 | 0 | 0 | 0 | 0 | 0 | 0 |
| ENSMUSG00 Traj32 | 65 | 0 | 0 | 0 | 0 | 0 | 0 | 0 | 0 | 0 | 0 | 0 | 0 | 0 | 0 | 0 | 0 | 0 | 0 |
| ENSMUSG00 Traj31 | 56 | 0 | 0 | 0 | 0 | 0 | 0 | 0 | 0 | 0 | 0 | 0 | 0 | 0 | 0 | 0 | 0 | 0 | 0 |
| ENSMUSG00 Traj30 | 58 | 0 | 0 | 0 | 0 | 0 | 0 | 0 | 0 | 0 | 0 | 0 | 0 | 0 | 0 | 0 | 0 | 0 | 0 |
| ENSMUSG00 Traj29 | 59 | 0 | 0 | 0 | 0 | 0 | 0 | 0 | 0 | 0 | 0 | 0 | 0 | 0 | 0 | 0 | 0 | 0 | 0 |
| ENSMUSG00 Traj28 | 64 | 0 | 0 | 0 | 0 | 0 | 0 | 0 | 0 | 0 | 0 | 0 | 0 | 0 | 0 | 0 | 0 | 0 | 0 |
| ENSMUSG00 Traj27 | 58 | 0 | 0.57436025 | 0 | 0 | 0 | 0 | 0 | 0.6584897 | 0 | 0 | 0 | 0 | 2.55165089 | 0 | 0 | 0 | 0 | 0 |
| ENSMUSG00 Traj26 | 59 | 0 | 0 | 0 | 0 | 0 | 0 | 0 | 0 | 0 | 0 | 0 | 0 | 0 | 0 | 0 | 0 | 0 | 0 |
| ENSMUSG00 Traj25 | 56 | 0 | 0 | 0 | 0 | 0 | 0 | 1.68973853 | 1.36401437 | 3.67736748 | 0 | 0.85312426 | 0.99670826 | 0 | 0 | 0 | 0 | 0 | 0.98230548 |
| ENSMUSG00 Traj24 | 62 | 0 | 0 | 0 | 0 | 0 | 0.79923764 | 0 | 0 | 0 | 0 | 0 | 0 | 0 | 0 | 0 | 0.61759044 | 0 | 0 |
| ENSMUSG00 Traj23 | 59 | 0 | 0 | 0 | 0 | 0 | 0 | 0 | 0 | 0 | 0 | 0 | 0 | 0 | 0 | 0 | 0 | 0 | 0 |
| ENSMUSG00 Traj22 | 59 | 0 | 0 | 0 | 0 | 0 | 0 | 0 | 1.29465771 | 0 | 0 | 0 | 0 | 0 | 0 | 0 | 0 | 0 | 0 |
| ENSMUSG00 Traj21 | 56 | 0 | 0 | 0 | 0 | 0 | 0 | 0 | 0.68200719 | 0 | 0 | 0 | 0 | 0 | 0 | 0 | 0 | 0 | 4.91152739 |
| ENSMUSG00 Traj20 | 57 | 0 | 0 | 0 | 0 | 0 | 0 | 0 | 0 | 0 | 0 | 0 | 0 | 0 | 0.81480855 | 0 | 0 | 0 | 0 |
| ENSMUSG00 Traj19 | 60 | 0 | 0 | 0 | 0 | 0 | 0 | 0 | 0 | 0 | 0 | 0 | 0 | 0 | 0 | 0 | 0 | 0 | 0.91681845 |
| ENSMUSG00 Traj18 | 65 | 0 | 0 | 0 | 0 | 0.74049113 | 0 | 0 | 0 | 0 | 0 | 0 | 0.8587025 | 0 | 0 | 0 | 0 | 0 | 0 |
| ENSMUSG00 Traj17 | 62 | 0 | 0 | 0 | 0 | 0 | 0 | 0 | 0 | 0 | 0 | 0 | 0 | 0 | 0 | 0 | 0 | 0 | 0 |
| ENSMUSG00 Traj16 | 60 | 0 | 0 | 0 | 0.75935074 | 0 | 0 | 0 | 0 | 0 | 0 | 0 | 0 | 0 | 0 | 0 | 0 | 0 | 0 |
| ENSMUSG00 Traj15 | 59 | 0 | 0 | 0 | 0 | 0 | 0 | 0 | 0 | 0 | 0 | 0 | 0 | 0 | 0 | 0 | 0 | 0 | 0 |
| ENSMUSG00 Traj14 | 32 | 0 | 0 | 0 | 0 | 0 | 0 | 0 | 0 | 0 | 0 | 0 | 0 | 0 | 0 | 0 | 0 | 0 | 0 |
| ENSMUSG00 Traj13 | 56 | 0 | 0 | 0 | 0 | 0 | 0 | 0 | 0 | 0 | 0 | 0 | 0 | 0 | 0 | 0 | 0 | 0 | 0 |
| ENSMUSG00 Traj12 | 57 | 0 | 0 | 0 | 0 | 0 | 0 | 0 | 0 | 0 | 0 | 0 | 0 | 0 | 0 | 0 | 0 | 0 | 0 |
| ENSMUSG00 Traj11 | 58 | 0 | 0 | 0 | 0 | 0 | 0 | 0 | 0 | 0 | 0 | 0 | 0 | 0 | 0 | 0.71037165 | 0 | 0.66079822 | 0 |
| ENSMUSG00 Traj9 | 57 | 0 | 0 | 0 | 0 | 0 | 0.8693462 | 0 | 0 | 0 | 0 | 0 | 1.9584443 | 0 | 0 | 0 | 0 | 3.36195587 | 0 |
| ENSMUSG00 Traj8 | 36 | 0 | 0 | 0 | 0 | 0 | 0 | 0 | 0 | 0 | 0 | 0 | 0 | 0 | 1.29011354 | 0 | 0 | 0 | 0 |
| ENSMUSG00 Traj7 | 58 | 0 | 0 | 0 | 0 | 0 | 0 | 0 | 0 | 0 | 0 | 0 | 0 | 0 | 0 | 0.71037165 | 0 | 0 | 0 |
| ENSMUSG00 Traj6 | 61 | 0 | 0 | 0 | 0 | 0 | 0 | 0 | 0 | 0 | 0 | 0 | 0 | 0 | 0 | 0 | 0 | 0 | 0 |
| ENSMUSG00 Traj5 | 61 | 0 | 0 | 0 | 0 | 0 | 0 | 0 | 0 | 0 | 0 | 0 | 0 | 0 | 0 | 0 | 0 | 0 | 0 |
| ENSMUSG00 Traj4 | 62 | 0 | 0 | 0 | 0 | 0 | 0 | 0 | 0 | 0 | 0 | 0 | 0 | 0 | 0 | 0 | 0 | 0 | 0 |
| ENSMUSG00 Traj3 | 65 | 0.65392579 | 0 | 0 | 0 | 0 | 0 | 0 | 0 | 0 | 0 | 0 | 0 | 0 | 0 | 0 | 0 | 0 | 0 |
| ENSMUSG00 Traj2 | 65 | 0 | 0 | 0 | 0 | 0 | 0 | 0 | 0 | 0 | 0 | 0 | 0 | 0 | 1.42904884 | 0 | 0 | 0 | 0 |
| ENSMUSG00 Traj1 | 26 | 0 | 0 | 0 | 0 | 0 | 0 | 0 | 0 | 0 | 0 | 0 | 0 | 0 | 0 | 0 | 0 | 0 | 0 |
| ENSMUSG00 Trac | 410 | 6.84229663 | 2.92503461 | 4.83735304 | 1.66686748 | 2.70008352 | 2.53806684 | 5.76983889 | 6.33434967 | 6.73047941 | 6.44009605 | 3.72877725 | 5.71770202 | 7.58027022 | 3.39834786 | 9.34571878 | 8.59203873 | 5.70220515 | 7.78177607 |
| ENSMUSG00 Iglc3 | 579 | 0.36705679 | 0 | 0 | 0.0786892 | 0.0831294 | 0 | 3.59543674 | 0 | 8.03812829 | 1.30295553 | 3.96061832 | 0.38560043 | 1.10762509 | 2.48664371 | 5.47930881 | 0.33066155 | 0 | 1.99514895 |
| ENSMUSG00 Iglv1 | 349 | 0 | 0 | 0 | 0 | 0 | 0 | 6.64275435 | 0 | 7.31680223 | 0 | 4.92807595 | 1.75923292 | 0 | 1.19769853 | 2.83334482 | 7.13148848 | 0 | 0 |
| ENSMUSG00 Iglc2 | 604 | 0.42223685 | 0 | 0.08008863 | 0 | 0 | 0.08204095 | 7.91155724 | 0.56909209 | 5.59154684 | 2.4980505 | 15.2658394 | 0.7392803 | 1.79685791 | 5.07501617 | 4.70680026 | 3.86709777 | 0.06345413 | 0 |
| ENSMUSG00 Iglv3 | 364 | 0 | 0 | 0 | 0 | 0 | 0 | 1.0398391 | 0.31477255 | 0.33944931 | 0 | 1.18124897 | 0 | 0.27105449 | 0 | 0 | 0 | 0 | 0 |
| ENSMUSG00 Iglv2 | 346 | 0 | 0 | 0 | 0 | 0 | 0.14321599 | 2.87157878 | 0 | 2.02361494 | 1.36273709 | 6.6277399 | 0 | 0 | 2.28193493 | 2.26251318 | 4.09466033 | 0 | 0 |
| ENSMUSG00 AC134447.1 | 109 | 0 | 0 | 0 | 0 | 0 | 0 | 0 | 0 | 0 | 0 | 0 | 0 | 0 | 0 | 0 | 0 | 0 | 0 |
| ENSMUSG00 AC158299.1 | 65 | 0 | 0 | 0 | 0 | 0 | 0 | 0 | 0 | 0 | 0 | 0 | 0 | 0 | 0 | 0 | 0 | 0 | 0 |
| ENSMUSG00 Mir582 | 80 | 0 | 0 | 0 | 0 | 0 | 0 | 0 | 0 | 0 | 0 | 0 | 0 | 0 | 0 | 0 | 0 | 0 | 0 |
| ENSMUSG00 Mir467e | 86 | 0 | 0 | 0 | 0 | 0 | 0 | 0 | 0 | 0 | 0 | 0 | 0 | 0 | 0 | 0 | 0 | 0 | 0 |
| ENSMUSG00 AC121540.1 | 109 | 0 | 0 | 0 | 0 | 0 | 0 | 0 | 0 | 0 | 0 | 0 | 0 | 0 | 0 | 0 | 0 | 0 | 0 |
| ENSMUSG00 AL929011.1 | 109 | 0 | 0 | 0 | 0 | 0 | 0 | 0 | 0 | 0 | 0 | 0 | 0 | 0 | 0 | 0 | 0 | 0 | 0 |
| ENSMUSG00 AC127431.1 | 109 | 0 | 0 | 0 | 0 | 0 | 0 | 0 | 0 | 0 | 0 | 0 | 0 | 0 | 0 | 0 | 0 | 0 | 0 |
| ENSMUSG00 AC087183.1 | 110 | 0 | 0 | 0 | 0 | 0 | 0 | 0 | 0 | 0 | 0 | 0 | 0 | 0 | 0 | 0 | 0 | 0 | 0 |
| ENSMUSG00 Mir466b-1 | 89 | 0 | 0 | 0 | 0 | 0 | 0 | 0 | 0 | 0 | 0 | 0 | 0 | 0 | 0 | 0 | 0 | 0 | 0 |
| ENSMUSG00 Mir297a-3 | 101 | 0 | 0 | 0 | 0 | 0 | 0 | 0 | 0 | 0 | 0 | 0 | 0 | 0 | 0 | 0 | 0 | 0 | 0 |
| ENSMUSG00 AC159715.1 | 109 | 0 | 0 | 0 | 0 | 0 | 0 | 0 | 0 | 0 | 0 | 0 | 0 | 0 | 0 | 0 | 0 | 0 | 0 |
| ENSMUSG00 AC154424.1 | 109 | 0 | 0 | 0 | 0 | 0 | 0 | 0 | 0 | 0 | 0 | 0 | 0 | 0 | 0 | 0 | 0 | 0 | 0 |
| ENSMUSG00 Mir669l | 97 | 0 | 0 | 0 | 0 | 0 | 0 | 0 | 0 | 0 | 0 | 0 | 0 | 0 | 0 | 0 | 0 | 0 | 0 |
| ENSMUSG00 AC158749.1 | 109 | 0 | 0 | 0 | 0 | 0 | 0 | 0 | 0 | 0.75571589 | 0 | 0 | 0 | 0 | 0 | 0 | 0 | 0 | 0 |
| ENSMUSG00 AC132266.1 | 109 | 0 | 0 | 0 | 0 | 0 | 0 | 0 | 0 | 0 | 0 | 0 | 0 | 0 | 0 | 0 | 0 | 0 | 0 |
| ENSMUSG00 Mir466b-2 | 81 | 0 | 0 | 0 | 0 | 0 | 0 | 0 | 0 | 0 | 0 | 0 | 0 | 0 | 0 | 0 | 0 | 0 | 0 |
| ENSMUSG00 Mir193b | 78 | 0 | 0 | 0 | 0 | 0 | 0 | 0 | 0 | 0 | 0 | 0 | 0 | 0 | 0 | 0 | 0 | 0 | 0 |
| ENSMUSG00 AL691478.1 | 109 | 0 | 0 | 0 | 0 | 0 | 0 | 0 | 0 | 0 | 0 | 0 | 0 | 0 | 0 | 0 | 0 | 0 | 0 |
| ENSMUSG00 AC098884.1 | 109 | 0 | 0 | 0 | 0 | 0 | 0 | 0 | 0 | 0 | 0 | 0 | 0 | 0 | 0 | 0 | 0 | 0 | 0 |
| ENSMUSG00 Mir297c | 97 | 0 | 0 | 0 | 0 | 0 | 0 | 0 | 0 | 0 | 0 | 0 | 0 | 0 | 0 | 0 | 0 | 0 | 0 |
| ENSMUSG00 AL683818.1 | 109 | 0 | 0 | 0 | 0 | 0 | 0 | 0 | 0 | 0 | 0 | 0 | 0 | 0 | 0 | 0 | 0 | 0 | 0 |
| ENSMUSG00 mmu-mir-466 | 89 | 0 | 0 | 0 | 0 | 0 | 0 | 0 | 0 | 0 | 0 | 0 | 0 | 0 | 0 | 0 | 0 | 0 | 0 |
| ENSMUSG00 AC117684.1 | 109 | 0 | 0 | 0 | 0 | 0 | 0 | 0 | 0 | 0 | 0 | 0 | 0 | 0 | 0 | 0 | 0 | 0 | 0 |
| ENSMUSG00 mmu-mir-297 | 89 | 0 | 0 | 0 | 0 | 0 | 0 | 0 | 0 | 0 | 0 | 0 | 0 | 0 | 0 | 0 | 0 | 0 | 0 |
| ENSMUSG00 AC174081.1 | 109 | 0 | 0 | 0 | 0 | 0 | 0 | 0 | 0 | 0 | 0 | 0 | 0 | 0 | 0 | 0 | 0 | 0 | 0 |
| ENSMUSG00 Mir467a-6 | 82 | 0 | 0 | 0 | 0 | 0 | 0 | 0 | 0 | 0 | 0 | 0 | 0 | 0 | 0 | 0 | 0 | 0 | 0 |
| ENSMUSG00 BX813309.1 | 109 | 0 | 0 | 0 | 0 | 0 | 0 | 0 | 0 | 0 | 0 | 0 | 0 | 0 | 0 | 0 | 0 | 0 | 0 |
| ENSMUSG00 AC155315.1 | 75 | 0 | 0 | 0 | 0 | 0 | 0 | 0 | 0 | 0 | 0 | 0 | 0 | 0 | 0 | 0 | 0 | 0 | 0 |
| ENSMUSG00 Mir466b-5 | 87 | 0 | 0 | 0 | 0 | 0 | 0 | 0 | 0 | 0 | 0 | 0 | 0 | 0 | 0 | 0 | 0 | 0 | 0 |
| ENSMUSG00 AC137969.1 | 109 | 0 | 0 | 0 | 0 | 0 | 0 | 0 | 0 | 0 | 0 | 0 | 0 | 0 | 0 | 0 | 0 | 0 | 0 |
| ENSMUSG00 Mir466b-3 | 80 | 0 | 0 | 0 | 0 | 0 | 0 | 0 | 0 | 0 | 0 | 0 | 0 | 0 | 0 | 0 | 0 | 0 | 0 |
| ENSMUSG00 AC122327.1 | 109 | 0 | 0 | 0 | 0 | 0 | 0 | 0 | 0 | 0 | 0 | 0 | 0 | 0 | 0 | 0 | 0 | 0 | 0 |
| ENSMUSG00 AC161167.1 | 109 | 0 | 0 | 0 | 0 | 0 | 0 | 0 | 0 | 0 | 0 | 0 | 0 | 0 | 0 | 0 | 0 | 0 | 0 |
| ENSMUSG00 AC122386.1 | 109 | 0 | 0 | 0 | 0 | 0 | 0 | 0 | 0 | 0 | 0 | 0 | 0 | 0 | 0 | 0 | 0 | 0 | 0 |
| ENSMUSG00 AC132082.1 | 95 | 0 | 0 | 0 | 0 | 0 | 0 | 1.4940846 | 0 | 0 | 0 | 0 | 0 | 0.51928334 | 0 | 0 | 0 | 0.4034347 | 0 |
| ENSMUSG00 Mir544 | 77 | 0 | 0 | 0 | 0 | 0 | 0 | 0 | 0 | 0 | 0 | 0 | 0 | 0 | 0 | 0 | 0 | 0 | 0 |
| ENSMUSG00 AC137969.2 | 109 | 0 | 0 | 0 | 0 | 0 | 0 | 0 | 0 | 0 | 0 | 0 | 0 | 0 | 0 | 0 | 0 | 0 | 0 |
| ENSMUSG00 AC122386.2 | 109 | 0 | 0 | 0 | 0 | 0 | 0 | 0 | 0 | 0 | 0 | 0 | 0 | 0 | 0 | 0 | 0 | 0 | 0 |
| ENSMUSG00 AL731707.1 | 66 | 0 | 0 | 0.73293228 | 0 | 0 | 0 | 1.43371754 | 0 | 0 | 0 | 0 | 0.84569186 | 0 | 0 | 0 | 0 | 1.7421044 | 0.83347131 |
| ENSMUSG00 AC134254.1 | 85 | 0 | 0 | 0 | 0 | 0 | 0 | 0 | 0 | 0 | 0 | 0 | 0 | 0 | 0 | 0 | 0 | 0 | 0 |
| ENSMUSG00 AC118240.1 | 109 | 0 | 0 | 0 | 0 | 0 | 0 | 0 | 0 | 0 | 0 | 0 | 0 | 0 | 0 | 0 | 0 | 0 | 0 |
| ENSMUSG00 AC102463.1 | 109 | 0 | 0 | 0 | 0 | 0 | 0 | 0 | 0 | 0 | 0 | 0 | 0 | 0 | 0 | 0 | 0 | 0 | 0 |
| ENSMUSG00 AC124506.1 | 109 | 0 | 0 | 0 | 0 | 0 | 0 | 0 | 0 | 0 | 0 | 0 | 0 | 0 | 0 | 0 | 0 | 0 | 0 |
| ENSMUSG00 AC154322.1 | 109 | 0 | 0 | 0 | 0 | 0 | 0 | 0 | 0 | 0 | 0 | 0 | 0 | 0 | 0 | 0 | 0 | 0 | 0 |
| ENSMUSG00 AC115121.1 | 108 | 0 | 0 | 0 | 0 | 0 | 0 | 0 | 0 | 0 | 0 | 0 | 0 | 0 | 0 | 0 | 0 | 0 | 0 |
| ENSMUSG00 Mir467d | 84 | 0 | 0 | 0 | 0 | 0 | 0 | 0 | 0 | 0 | 0 | 0 | 0 | 0 | 0 | 0 | 0 | 0 | 0 |
| ENSMUSG00 AC123045.1 | 109 | 0 | 0 | 0 | 0 | 0 | 0 | 0 | 0 | 0 | 0 | 0 | 0 | 0 | 0 | 0 | 0 | 0 | 0 |
| ENSMUSG00 Mir493 | 82 | 0 | 0 | 0 | 0 | 0 | 0 | 0 | 0 | 0 | 0 | 0 | 0 | 0 | 0 | 0 | 0 | 0 | 0 |
| ENSMUSG00 AL606907.1 | 109 | 0 | 0 | 0 | 0 | 0 | 0 | 0 | 0 | 0 | 0 | 0 | 0 | 0 | 0 | 0 | 0 | 0 | 0 |
| ENSMUSG00 mmu-mir-466 | 89 | 0 | 0 | 0 | 0 | 0 | 0 | 0 | 0 | 0 | 0 | 0 | 0 | 0 | 0 | 0 | 0 | 0 | 0 |
| ENSMUSG00 AC115121.2 | 108 | 0 | 0 | 0 | 0 | 0 | 0 | 0 | 0 | 0 | 0 | 0 | 0 | 0 | 0 | 0 | 0 | 0 | 0 |
| ENSMUSG00 Mir467a-10 | 82 | 0 | 0 | 0 | 0 | 0 | 0 | 0 | 0 | 0 | 0 | 0 | 0 | 0 | 0 | 0 | 0 | 0 | 0 |
| ENSMUSG00 AC167229.1 | 109 | 0 | 0 | 0 | 0 | 0 | 0 | 0 | 0 | 0 | 0 | 0 | 0 | 0 | 0 | 0 | 0 | 0 | 0 |
| ENSMUSG00 Mir466b-4 | 89 | 0 | 0 | 0 | 0 | 0 | 0 | 0 | 0 | 0 | 0 | 0 | 0 | 0 | 0 | 0 | 0 | 0 | 0 |
| ENSMUSG00 AC126045.1 | 109 | 0 | 0 | 0 | 0 | 0 | 0 | 0 | 0 | 0 | 0 | 0 | 0 | 0 | 0 | 0 | 0 | 0 | 0 |
| ENSMUSG00 AL833773.1 | 109 | 0 | 0 | 0 | 0 | 0 | 0 | 0 | 0 | 0 | 0 | 0 | 0 | 0 | 0 | 0 | 0 | 0 | 0 |
| ENSMUSG00 Mir669f | 120 | 0 | 0 | 0 | 0 | 0 | 0 | 0 | 0 | 0 | 0 | 0 | 0 | 0 | 0 | 0 | 0 | 0 | 0 |
| ENSMUSG00 AC111074.1 | 108 | 0 | 0 | 0 | 0 | 0 | 0 | 0 | 0 | 0 | 0 | 0 | 0 | 0 | 0 | 0 | 0 | 0 | 0 |
| ENSMUSG00 AL671906.1 | 109 | 0 | 0 | 0 | 0 | 0 | 0 | 0 | 0 | 0 | 0 | 0 | 0 | 0 | 0 | 0 | 0 | 0 | 0 |
| ENSMUSG00 Mir574 | 77 | 0 | 0 | 0 | 0 | 0 | 0 | 0 | 0 | 0 | 0 | 0 | 0 | 0 | 0 | 0 | 0 | 0 | 0 |
| ENSMUSG00 AC167229.2 | 109 | 0 | 0 | 0 | 0 | 0 | 0 | 0 | 0 | 0 | 0 | 0 | 0 | 0 | 0 | 0 | 0 | 0 | 0 |
| ENSMUSG00 BX537299.1 | 109 | 0 | 0 | 0 | 0 | 0 | 0 | 0 | 0 | 0 | 0 | 0 | 0 | 0 | 0 | 0 | 0 | 0 | 0 |
| ENSMUSG00 AC102227.1 | 109 | 0 | 0 | 0 | 0 | 0 | 0 | 0 | 0 | 0 | 0 | 0 | 0 | 0 | 0 | 0 | 0 | 0 | 0 |
| ENSMUSG00 Mir467c | 96 | 0 | 0 | 0 | 0 | 0 | 0 | 0 | 0 | 0 | 0 | 0 | 0 | 0 | 0 | 0 | 0 | 0 | 0 |
| ENSMUSG00 AC131665.1 | 109 | 0 | 0 | 0 | 0 | 0 | 0 | 0 | 0 | 0 | 0 | 0 | 0 | 0 | 0 | 0 | 0 | 0 | 0 |
| ENSMUSG00 AC122806.1 | 109 | 0 | 0 | 0 | 0 | 0 | 0 | 0 | 0 | 0 | 0 | 0 | 0 | 0 | 0 | 0 | 0 | 0 | 0 |
| ENSMUSG00 AC154006.1 | 109 | 0 | 0 | 0 | 0 | 0 | 0 | 0 | 0 | 0 | 0 | 0 | 0 | 0 | 0 | 0 | 0 | 0 | 0 |
| ENSMUSG00 Mir467a-3 | 82 | 0 | 0 | 0 | 0 | 0 | 0 | 0 | 0 | 0 | 0 | 0 | 0 | 0 | 0 | 0 | 0 | 0 | 0 |
| ENSMUSG00 AC121967.1 | 81 | 0 | 0 | 0 | 0 | 0 | 0 | 0 | 0 | 0 | 0 | 0 | 0 | 0 | 0 | 0 | 0 | 0 | 0 |
| ENSMUSG00 Mir669p-2 | 86 | 0 | 0 | 0 | 0 | 0 | 0 | 0 | 0 | 0 | 0 | 0 | 0 | 0 | 0 | 0 | 0 | 0 | 0 |
| ENSMUSG00 AC124426.1 | 109 | 0 | 0 | 0 | 0 | 0 | 0 | 0 | 0 | 0 | 0 | 0 | 0 | 0 | 0 | 0 | 0 | 0 | 0 |
| ENSMUSG00 AC123824.1 | 109 | 0 | 0 | 0 | 0 | 0 | 0 | 0 | 0 | 0 | 0 | 0 | 0 | 0 | 0 | 0 | 0 | 0 | 0 |
| ENSMUSG00 AC110903.1 | 95 | 0 | 0 | 0 | 0 | 0 | 0 | 0 | 0 | 0 | 0 | 0 | 0 | 0 | 0.97777026 | 0 | 0 | 0 | 0 |
| ENSMUSG00 Mir466b-6 | 89 | 0 | 0 | 0 | 0 | 0 | 0 | 0 | 0 | 0 | 0 | 0 | 0 | 0 | 0 | 0 | 0 | 0 | 0 |
| ENSMUSG00 AC143329.1 | 109 | 0 | 0 | 0 | 0 | 0 | 0 | 0 | 0 | 0 | 0 | 0 | 0 | 0 | 0 | 0 | 0 | 0 | 0 |
| ENSMUSG00 AL591544.1 | 65 | 0 | 0 | 0 | 0 | 0 | 0 | 0 | 0 | 0 | 0 | 0 | 0 | 0 | 0 | 0 | 0 | 0 | 0 |
| ENSMUSG00 AC124080.1 | 109 | 0 | 0 | 0 | 0 | 0 | 0 | 0 | 0 | 0 | 0 | 0 | 0 | 0 | 0 | 0 | 0 | 0 | 0 |
| ENSMUSG00 AC020968.1 | 109 | 0 | 0 | 0 | 0 | 0 | 0 | 0 | 0 | 0 | 0 | 0 | 0 | 0 | 0 | 0 | 0 | 0 | 0 |
| ENSMUSG00 Mir466e | 85 | 0 | 0 | 0 | 0 | 0 | 0 | 0 | 0 | 0 | 0 | 0 | 0 | 0 | 0 | 0 | 0 | 0 | 0 |
| ENSMUSG00 Mir669p-1 | 86 | 0 | 0 | 0 | 0 | 0 | 0 | 0 | 0 | 0 | 0 | 0 | 0 | 0 | 0 | 0 | 0 | 0 | 0 |
| ENSMUSG00 AC126798.1 | 109 | 0 | 0 | 0 | 0 | 0 | 0 | 0 | 0 | 0.37785794 | 0 | 0 | 0 | 0 | 0 | 0 | 0 | 0 | 0 |
| ENSMUSG00 AC087801.1 | 109 | 0 | 0 | 0 | 0 | 0 | 0 | 0 | 0 | 0 | 0 | 0 | 0 | 0 | 0 | 0 | 0 | 0 | 0 |
| ENSMUSG00 AC108942.1 | 109 | 0 | 0 | 0 | 0 | 0 | 0 | 0 | 0 | 0 | 0 | 0 | 0 | 0 | 0 | 0 | 0 | 0 | 0 |
| ENSMUSG00 AC154654.1 | 109 | 0.38995574 | 0 | 0 | 0 | 0 | 0 | 0 | 0 | 0 | 0 | 0 | 0 | 0 | 0 | 0 | 0 | 0 | 0 |
| ENSMUSG00 AC164303.1 | 109 | 0 | 0 | 0 | 0 | 0 | 0 | 0 | 0 | 0 | 0 | 0 | 0 | 0 | 0 | 0 | 0 | 0 | 0 |
| ENSMUSG00 AC159378.1 | 79 | 0 | 0 | 0 | 0 | 0 | 0 | 0 | 0 | 0 | 0 | 0 | 0 | 0 | 0.58789984 | 0 | 0 | 0 | 0 |
| ENSMUSG00 AC138311.1 | 109 | 0 | 0 | 0 | 0 | 0 | 0 | 0 | 0 | 0 | 0 | 0 | 0 | 0 | 0 | 0 | 0 | 0 | 0 |
| ENSMUSG00 AC102528.1 | 109 | 0 | 0 | 0 | 0 | 0 | 0 | 0 | 0 | 0 | 0 | 0 | 0 | 0 | 0 | 0 | 0 | 0 | 0 |
| ENSMUSG00 AC163215.1 | 109 | 0 | 0 | 0 | 0 | 0 | 0 | 0 | 0 | 0 | 0 | 0 | 0 | 0 | 0 | 0 | 0 | 0 | 0 |
| ENSMUSG00 Mir669o | 95 | 0 | 0 | 0 | 0 | 0 | 0 | 0 | 0 | 0 | 0 | 0 | 0 | 0 | 0 | 0 | 0 | 0 | 0 |
| ENSMUSG00 Mir466b-7 | 87 | 0 | 0 | 0 | 0 | 0 | 0 | 0 | 0 | 0 | 0 | 0 | 0 | 0 | 0 | 0 | 0 | 0 | 0 |
| ENSMUSG00 AC155640.1 | 66 | 0 | 0 | 0 | 0 | 0 | 0 | 0 | 0 | 0 | 0 | 0 | 0 | 0 | 0 | 0 | 0 | 0 | 0 |
| ENSMUSG00 AC122806.2 | 109 | 0 | 0 | 0 | 0 | 0 | 0 | 0 | 0 | 0 | 0 | 0 | 0 | 0 | 0 | 0 | 0 | 0 | 0 |
| ENSMUSG00 Mir742 | 64 | 0 | 0 | 0 | 0 | 0 | 0 | 0 | 0 | 0 | 0 | 0 | 0 | 0 | 0 | 0 | 0 | 0 | 0 |
| ENSMUSG00 AC122520.1 | 109 | 0 | 0 | 0 | 0 | 0 | 0 | 0 | 0 | 0 | 0 | 0 | 0 | 0 | 0 | 0 | 0 | 0 | 0 |
| ENSMUSG00 CR974486.1 | 108 | 0 | 0 | 0 | 0 | 0 | 0 | 0 | 0 | 0 | 0 | 0 | 0 | 0 | 0 | 0 | 0 | 0 | 0 |
| ENSMUSG00 AL772216.1 | 103 | 0 | 0 | 0 | 0 | 0 | 0 | 0 | 0 | 0 | 0 | 0 | 0 | 0 | 0 | 0 | 0 | 0 | 0 |
| ENSMUSG00 AC154375.1 | 64 | 0 | 0 | 0 | 0 | 0 | 0 | 0 | 0 | 0 | 0 | 0 | 0 | 0 | 0 | 0 | 0 | 0 | 0 |
| ENSMUSG00 Mir466b-8 | 85 | 0 | 0 | 0 | 0 | 0 | 0 | 0 | 0 | 0 | 0 | 0 | 0 | 0 | 0 | 0 | 0 | 0 | 0 |
| ENSMUSG00 AL606904.1 | 109 | 0 | 0 | 0 | 0 | 0 | 0 | 0 | 0 | 0 | 0 | 0 | 0 | 0 | 0 | 0 | 0 | 0 | 0 |
| ENSMUSG00 AL772216.2 | 99 | 0 | 0 | 0 | 0 | 0 | 0 | 0 | 0 | 0 | 0 | 0 | 0 | 0 | 0 | 0 | 0 | 0 | 0 |
| ENSMUSG00 Mir344c | 91 | 0 | 0 | 0 | 0 | 0 | 0 | 0 | 0 | 0 | 0 | 0 | 0 | 0 | 0 | 0 | 0 | 0 | 0 |
| ENSMUSG00 AC138311.2 | 109 | 0 | 0 | 0 | 0 | 0 | 0 | 0 | 0 | 0 | 0 | 0 | 0 | 0 | 0 | 0 | 0 | 0 | 0 |
| ENSMUSG00 AC131111.1 | 109 | 0 | 0 | 0 | 0 | 0 | 0 | 0 | 0 | 0 | 0 | 0 | 0 | 0 | 0 | 0 | 0 | 0 | 0 |
| ENSMUSG00 Mir18b | 82 | 0 | 0 | 0 | 0 | 0 | 0 | 0 | 0 | 0 | 0 | 0 | 0 | 0 | 0 | 0 | 0 | 0 | 0 |
| ENSMUSG00 Mir466p | 88 | 0 | 0 | 0 | 0 | 0 | 0 | 0 | 0 | 0 | 0 | 0 | 0 | 0 | 0 | 0 | 0 | 0 | 0 |
| ENSMUSG00 AC117684.2 | 109 | 0 | 0 | 0 | 0 | 0 | 0 | 0 | 0 | 0 | 0 | 0 | 0 | 0 | 0 | 0 | 0 | 0 | 0 |
| ENSMUSG00 AC122520.2 | 109 | 0 | 0 | 0 | 0 | 0 | 0.45461223 | 0 | 0 | 0 | 0 | 0 | 0 | 0 | 0 | 1.13398778 | 0 | 0 | 0 |
| ENSMUSG00 Mir741 | 70 | 0 | 0 | 0 | 0 | 0 | 0 | 0 | 0 | 0 | 0 | 0 | 0 | 0 | 0 | 0 | 0 | 0 | 0 |
| ENSMUSG00 AC164603.1 | 109 | 0 | 0 | 0 | 0 | 0 | 0 | 0 | 0 | 0 | 0 | 0 | 0 | 0 | 0 | 0 | 0 | 0 | 0 |
| ENSMUSG00 Mir743 | 61 | 0 | 0 | 0 | 0 | 0 | 0 | 0 | 0 | 0 | 0 | 0 | 0 | 0 | 0 | 0 | 0 | 0 | 0 |
| ENSMUSG00 AC161197.1 | 109 | 0 | 0 | 0 | 0 | 0 | 0 | 0 | 0 | 0 | 0 | 0 | 0 | 0 | 0 | 0 | 0 | 0 | 0 |
| ENSMUSG00 AL772216.3 | 103 | 0 | 0 | 0 | 0 | 0 | 0 | 0 | 0 | 0 | 0 | 0 | 0 | 0 | 0 | 0 | 0 | 0 | 0 |
| ENSMUSG00 AC125134.1 | 109 | 0 | 0 | 0 | 0 | 0 | 0 | 0 | 0 | 1.13357383 | 0 | 0 | 0 | 0 | 0 | 0 | 0 | 0 | 0 |
| ENSMUSG00 AC122775.1 | 109 | 0 | 0 | 0 | 0 | 0 | 0 | 0 | 0 | 0 | 0 | 0 | 0 | 0 | 0 | 0.75599185 | 0 | 0 | 0 |
| ENSMUSG00 AC134445.1 | 109 | 0 | 0 | 0 | 0 | 0 | 0 | 0 | 0 | 0 | 0 | 0 | 0 | 0 | 0 | 0 | 0 | 0 | 0 |
| ENSMUSG00 Mir466c-1 | 89 | 0 | 0 | 0 | 0 | 0 | 0 | 0 | 0 | 0 | 0 | 0 | 0 | 0 | 0 | 0 | 0 | 0 | 0 |
| ENSMUSG00 AC114404.1 | 109 | 0 | 0 | 0 | 0 | 0 | 0 | 0 | 0 | 0 | 0 | 0 | 0 | 0 | 0 | 0 | 0 | 0 | 0 |
| ENSMUSG00 Mir466c-2 | 89 | 0 | 0 | 0 | 0 | 0 | 0 | 0 | 0 | 0 | 0 | 0 | 0 | 0 | 0 | 0 | 0 | 0 | 0 |
| ENSMUSG00 SNORA17 | 130 | 0 | 0 | 0 | 0 | 0 | 0 | 0 | 0 | 0 | 0 | 0 | 0 | 0 | 0 | 0 | 0 | 0 | 0 |
| ENSMUSG00 SNORA17 | 125 | 0 | 0 | 0 | 0 | 0 | 0 | 0 | 0 | 0 | 0 | 0 | 0 | 0 | 0 | 0 | 0 | 0 | 0 |
| ENSMUSG00 SNORA17 | 119 | 0 | 0 | 0 | 0 | 0 | 0 | 0 | 0 | 0 | 0 | 0 | 0 | 0 | 0 | 0 | 0 | 0 | 0 |
| ENSMUSG00 SNORA17 | 130 | 0 | 0 | 0 | 0 | 0 | 0 | 0 | 0 | 0 | 0 | 0 | 0 | 0 | 0 | 0 | 0 | 0 | 0 |
| ENSMUSG00 U6 | 103 | 0 | 0 | 0 | 0 | 0 | 0 | 0.4593464 | 0 | 0.39986909 | 1.37332145 | 0 | 0 | 0 | 0 | 0 | 0 | 0 | 0 |
| ENSMUSG00 SNORA17 | 131 | 0 | 0 | 0 | 0 | 0 | 0 | 0 | 0 | 0 | 0 | 0 | 0 | 0 | 0 | 0 | 0 | 0 | 0 |
| ENSMUSG00 SNORA17 | 132 | 0 | 0.50474082 | 0 | 0 | 0 | 0 | 0 | 0.28933638 | 0 | 0.71440459 | 0 | 0 | 0 | 0 | 0.624266 | 0 | 0 | 0 |
| ENSMUSG00 SNORA17 | 143 | 0 | 0 | 0 | 0 | 0 | 0 | 0 | 0 | 0 | 0 | 0 | 0 | 0 | 0 | 0 | 0 | 0 | 0 |
| ENSMUSG00 SNORA17 | 130 | 0 | 0 | 0 | 0 | 0 | 0 | 0 | 0 | 0 | 0 | 0 | 0 | 0 | 0 | 0 | 0 | 0 | 0 |
| ENSMUSG00 U6 | 102 | 0 | 0 | 0 | 0 | 0 | 0 | 0 | 0 | 0.40378937 | 0 | 0 | 0 | 0 | 0.91066838 | 0.80787364 | 0 | 0 | 0 |
| ENSMUSG00 n-R5s90 | 117 | 0 | 0 | 0 | 0 | 0 | 0 | 0 | 0 | 0 | 0 | 0 | 0 | 0 | 0 | 0 | 0 | 0 | 0 |
| ENSMUSG00 SNORA17 | 128 | 0 | 0 | 0 | 0 | 0 | 0 | 1.84815152 | 0 | 0 | 0 | 0 | 0 | 0 | 0 | 0 | 0 | 0 | 0 |
| ENSMUSG00 SNORA17 | 125 | 0 | 0 | 0 | 0 | 0 | 0 | 0 | 0 | 0 | 0 | 0 | 0 | 0 | 0 | 0 | 0 | 0 | 0 |
| ENSMUSG00 SNORA17 | 132 | 0 | 0 | 0 | 0 | 0 | 0 | 0 | 0 | 0 | 0 | 0 | 0 | 0 | 0 | 0 | 0 | 0 | 0 |
| ENSMUSG00 U6 | 106 | 0 | 0 | 0 | 0 | 0 | 0 | 0 | 0 | 0 | 0 | 0 | 0 | 0 | 0 | 0 | 0 | 0 | 0 |
| ENSMUSG00 SNORA17 | 132 | 0 | 0 | 0 | 0 | 0 | 0 | 0 | 0 | 0 | 0 | 0 | 0 | 0 | 0 | 0 | 0 | 0 | 0 |
| ENSMUSG00 SNORA17 | 131 | 0 | 0 | 0 | 0 | 0 | 0 | 0 | 0 | 0 | 0 | 0 | 0 | 0 | 0 | 0 | 0 | 0 | 0 |
| ENSMUSG00 SNORA17 | 128 | 0 | 0 | 0 | 0 | 0 | 0 | 0 | 0 | 0 | 0 | 0 | 0 | 0 | 0 | 0 | 0 | 0 | 0 |
| ENSMUSG00 SNORA63 | 245 | 4.33726286 | 3.67121691 | 2.76420173 | 2.60348825 | 3.73267979 | 1.8203045 | 4.24848545 | 6.54726898 | 8.74162784 | 2.50187405 | 4.87499576 | 7.97366608 | 6.64470722 | 7.96184355 | 5.21325809 | 10.6275972 | 6.10092073 | 9.20560561 |
| ENSMUSG00 SNORA17 | 132 | 0 | 0 | 0 | 0 | 0 | 0 | 0 | 0 | 0 | 0 | 0 | 0 | 0 | 0 | 0 | 0 | 0 | 0 |
| ENSMUSG00 SNORA43 | 131 | 1.29786797 | 0 | 0 | 0 | 0 | 0 | 0 | 2.0408154 | 0 | 1.0797871 | 0 | 0 | 0 | 0 | 0 | 0 | 0.5851343 | 0 |
| ENSMUSG00 SNORA17 | 127 | 0 | 0 | 0 | 0 | 0 | 0 | 0 | 0 | 0 | 0 | 0 | 0 | 0 | 0 | 0 | 0 | 0 | 0 |
| ENSMUSG00 SNORA17 | 131 | 0 | 0 | 0 | 0 | 0 | 0 | 0 | 0 | 0 | 0 | 0 | 0 | 0 | 0 | 0 | 0 | 0 | 0 |
| ENSMUSG00 SNORA17 | 122 | 0 | 0 | 0 | 0 | 0 | 0 | 0 | 0 | 0 | 0 | 0 | 0 | 0 | 0 | 0 | 0 | 0 | 0 |
| ENSMUSG00 SNORA17 | 128 | 0 | 0 | 0 | 0 | 0 | 0 | 0 | 0 | 0 | 0 | 0 | 0 | 0 | 0 | 0 | 0 | 0 | 0 |
| ENSMUSG00 SNORA17 | 128 | 0 | 0 | 0 | 0 | 0 | 0 | 0 | 0 | 0 | 0 | 0 | 0 | 0 | 0 | 0 | 0 | 0 | 0 |
| ENSMUSG00 SNORA17 | 131 | 0 | 0 | 0 | 0 | 0 | 0 | 0 | 0 | 0 | 0 | 0 | 0 | 0 | 0 | 0 | 0 | 0 | 0 |
| ENSMUSG00 SNORA17 | 135 | 0 | 0 | 0 | 0 | 0 | 0 | 0 | 0 | 0 | 0 | 0 | 0 | 0 | 0 | 0 | 0 | 0 | 0 |
| ENSMUSG00 SNORA17 | 130 | 0 | 0 | 0 | 0 | 0 | 0 | 0 | 0 | 0 | 0 | 0 | 0 | 0 | 0 | 0 | 0 | 0 | 0 |
| ENSMUSG00 SNORA17 | 131 | 0 | 0 | 0 | 0 | 0 | 0 | 0 | 0 | 0 | 0 | 0 | 0 | 0 | 0 | 0 | 0 | 0 | 0 |
| ENSMUSG00 SNORA17 | 131 | 0 | 0 | 0 | 0 | 0 | 0 | 0 | 0 | 0 | 0 | 0 | 0 | 0 | 0 | 0 | 0 | 0 | 0 |
| ENSMUSG00 SNORA17 | 131 | 0 | 0 | 0 | 0 | 0 | 0 | 0 | 0 | 0 | 0 | 0 | 0 | 0 | 0 | 0 | 0 | 0 | 0 |
| ENSMUSG00 SNORA17 | 128 | 0 | 0 | 0 | 0 | 0 | 0 | 0 | 0 | 0 | 0 | 0 | 0 | 0 | 0 | 0 | 0 | 0 | 0 |
| ENSMUSG00 SNORD102 | 70 | 0 | 0 | 0 | 0 | 0 | 0 | 0 | 0 | 0 | 0 | 0 | 0 | 0 | 0 | 0 | 0 | 0 | 0 |
| ENSMUSG00 SNORA17 | 131 | 0 | 0 | 0 | 0 | 0 | 0 | 0 | 0 | 0 | 0 | 0 | 0 | 0 | 0 | 0 | 0 | 0 | 0 |
| ENSMUSG00 SNORA17 | 129 | 0 | 0.25823949 | 0 | 0 | 0 | 0 | 0 | 0 | 0 | 0 | 0 | 0 | 0 | 0 | 0 | 0 | 0 | 0 |
| ENSMUSG00 Snord64 | 66 | 0 | 0 | 0 | 0 | 0 | 0 | 0.71685877 | 0 | 0 | 0 | 0 | 0 | 0 | 0 | 0 | 0 | 0 | 0 |
| ENSMUSG00 Snora17 | 131 | 0 | 0 | 0 | 0 | 0 | 0 | 0 | 0 | 0 | 0 | 0 | 0 | 0 | 0 | 0 | 0 | 0 | 0 |
| ENSMUSG00 U6 | 106 | 0 | 0 | 0 | 0 | 0 | 0 | 0 | 0 | 0 | 0 | 0 | 0 | 0 | 0 | 0 | 0 | 0 | 0 |
| ENSMUSG00 U6 | 106 | 0 | 0 | 0 | 0 | 0 | 0 | 0 | 0 | 0 | 0 | 0 | 0 | 0 | 0 | 0 | 0 | 0 | 0 |
| ENSMUSG00 SNORA17 | 130 | 0 | 0 | 0 | 0 | 0 | 0 | 0 | 0 | 0 | 0 | 0 | 0 | 0 | 0 | 0 | 0 | 0 | 0 |
| ENSMUSG00 SNORA17 | 130 | 0 | 0 | 0 | 0 | 0 | 0 | 0 | 0 | 0 | 0 | 0 | 0 | 0 | 0 | 0 | 0 | 0 | 0 |
| ENSMUSG00 SNORA17 | 131 | 0 | 0 | 0 | 0 | 0 | 0 | 0 | 0 | 0 | 0 | 0 | 0 | 0 | 0 | 0 | 0 | 0 | 0 |
| ENSMUSG00 SNORA17 | 128 | 1.66035844 | 2.08205589 | 1.13375462 | 2.13567395 | 1.88015327 | 1.16139219 | 0.3696303 | 0.29837814 | 0 | 0 | 1.11972559 | 0.43605986 | 2.31243362 | 1.45137773 | 4.82830733 | 2.69230833 | 1.49712097 | |
| ENSMUSG00 SNORA17 | 132 | 0 | 0 | 0 | 0 | 0 | 0 | 0 | 0 | 0 | 0 | 0 | 0 | 0 | 0 | 0 | 0 | 0 | 0 |
| ENSMUSG00 SNORA61 | 125 | 0 | 0 | 0 | 0 | 0 | 0 | 0 | 0 | 0 | 0 | 0 | 0 | 0 | 0 | 0 | 0 | 0 | 0 |
| ENSMUSG00 SNORA17 | 128 | 0 | 0 | 0 | 0 | 0 | 0 | 0 | 0 | 0 | 0 | 0 | 0 | 0 | 0 | 0 | 0 | 0 | 0 |
| ENSMUSG00 SNORA17 | 132 | 0 | 0 | 0 | 0 | 0 | 0 | 0 | 0 | 0.31201906 | 0 | 0 | 0 | 0 | 0 | 0 | 0 | 0 | 0 |
| ENSMUSG00 U6 | 103 | 0 | 0 | 0 | 0 | 0 | 0 | 0 | 0 | 0 | 0 | 0 | 0 | 0 | 0 | 0 | 0 | 0 | 0 |
| ENSMUSG00 SNORA17 | 126 | 0 | 0 | 0 | 0 | 0 | 0 | 0 | 0 | 0 | 0 | 0 | 0 | 0 | 0 | 0 | 0 | 0 | 0 |
| ENSMUSG00 SNORA17 | 131 | 0 | 0 | 0 | 0 | 0 | 0 | 0 | 0 | 0 | 0 | 0 | 0 | 0 | 0 | 0 | 0 | 0 | 0 |
| ENSMUSG00 Snord69 | 79 | 5.38040203 | 8.0119619 | 2.44929268 | 2.8836104 | 5.48338371 | 6.89974771 | 7.18673604 | 6.76827384 | 5.73483131 | 10.7431982 | 4.83797048 | 2.11958212 | 2.4978186 | 8.23059777 | 3.65077078 | 7.27036849 | 7.76228798 | 4.17790684 |
| ENSMUSG00 SNORA17 | 128 | 0 | 0 | 0 | 0 | 0 | 0 | 0 | 0 | 0 | 0 | 0 | 0 | 0 | 0 | 0 | 0 | 0 | 0 |
| ENSMUSG00 SNORA17 | 126 | 0 | 0 | 0 | 0 | 0 | 0 | 0 | 0 | 0 | 0 | 0 | 0 | 0 | 0 | 0 | 0 | 0 | 0 |
| ENSMUSG00 SNORA17 | 131 | 0 | 0 | 0 | 0 | 0 | 0 | 0 | 0 | 0 | 0 | 0 | 0 | 0 | 0 | 0 | 0 | 0 | 0 |
| ENSMUSG00 SNORA17 | 131 | 0 | 0 | 0 | 0 | 0 | 0 | 0 | 0 | 0 | 0 | 0 | 0 | 0 | 0 | 0 | 0 | 0 | 0 |
| ENSMUSG00 SNORD78 | 65 | 0 | 0 | 0 | 0 | 0 | 0 | 0 | 0 | 0 | 0 | 0 | 0 | 0 | 0 | 0 | 0 | 0 | 0 |
| ENSMUSG00 Snord67 | 110 | 0 | 0 | 0 | 0 | 0 | 0 | 0 | 0 | 0 | 0 | 0.86863561 | 0 | 0 | 0 | 0 | 0 | 0 | 0 |
| ENSMUSG00 SNORA66 | 131 | 0.32446699 | 0 | 0 | 0 | 0 | 0.37826514 | 0 | 0 | 0 | 0 | 0 | 0 | 0 | 0 | 0 | 0 | 0 | 0.41991685 |
| ENSMUSG00 SNORA17 | 127 | 0 | 0 | 0 | 0 | 0 | 0 | 0 | 0 | 0 | 0 | 0 | 0 | 0 | 0 | 0 | 0 | 0 | 0 |
| ENSMUSG00 SNORA17 | 126 | 0 | 0 | 0 | 0 | 0 | 0 | 0 | 0 | 0 | 0 | 0 | 0 | 0 | 0 | 0 | 0 | 0 | 0 |
| ENSMUSG00 SNORD115 | 78 | 0 | 0 | 0 | 0 | 0 | 0 | 0 | 0 | 0 | 0 | 0 | 0 | 0 | 0 | 0 | 0 | 0 | 0 |
| ENSMUSG00 SNORA17 | 133 | 0 | 0 | 0 | 0 | 0 | 0 | 0 | 0 | 0 | 0 | 0 | 0 | 0 | 0 | 0 | 0 | 0 | 0 |
| ENSMUSG00 SNORA17 | 131 | 0 | 0 | 0 | 0 | 0 | 0 | 0 | 0 | 0 | 0 | 0 | 0 | 0 | 0 | 0 | 0 | 0 | 0 |
| ENSMUSG00 U11 | 144 | 0 | 0 | 0 | 0 | 0 | 0 | 0 | 0 | 0.85805241 | 0 | 0.33177054 | 0 | 0 | 0 | 0 | 0.53181399 | 0 | 0 |
| ENSMUSG00 SNORA17 | 128 | 0 | 0 | 0 | 0 | 0 | 0 | 0 | 0 | 0 | 0 | 0 | 0 | 0 | 0 | 0 | 0 | 0 | 0 |
| ENSMUSG00 SNORA17 | 130 | 0 | 0 | 0 | 0 | 0 | 0 | 0 | 0 | 0 | 0 | 0 | 0 | 0 | 0 | 0 | 0 | 0 | 0 |
| ENSMUSG00 SNORA17 | 131 | 0 | 0 | 0 | 0 | 0 | 0 | 0 | 0 | 0 | 0 | 0 | 0 | 0 | 0 | 0 | 0 | 0 | 0.83983369 |
| ENSMUSG00 SNORA17 | 131 | 0 | 0 | 0 | 0 | 0 | 0 | 0 | 0 | 0 | 0 | 0 | 0 | 0 | 0 | 0 | 0 | 0 | 0 |
| ENSMUSG00 Snord66 | 74 | 1.72318281 | 0 | 1.30739271 | 0 | 1.9512942 | 0 | 1.27872105 | 1.03222709 | 0 | 0.63717167 | 0 | 0 | 1.99994259 | 0 | 0 | 0 | 0.51792293 | 0.74336631 |
| ENSMUSG00 SNORA17 | 123 | 0 | 0 | 0 | 0 | 0 | 0 | 0 | 0 | 0 | 0 | 0 | 0 | 0 | 0 | 0 | 0 | 0 | 0 |
| ENSMUSG00 SNORA17 | 128 | 0 | 0 | 0 | 0 | 0 | 0 | 0 | 0 | 0 | 0 | 0 | 0 | 0 | 0 | 0 | 0 | 0 | 0 |
| ENSMUSG00 SNORA17 | 130 | 0 | 0 | 0 | 0 | 0 | 0 | 0 | 0 | 0 | 0 | 0 | 0 | 0 | 0 | 0 | 0 | 0 | 0 |
| ENSMUSG00 SNORA17 | 129 | 0 | 0 | 0 | 0 | 0 | 0 | 0 | 0 | 0 | 0 | 0 | 0 | 0 | 0 | 0 | 0 | 0 | 0 |
| ENSMUSG00 SNORA17 | 131 | 0 | 0 | 0 | 0 | 0 | 0 | 0 | 0 | 0 | 0 | 0 | 0 | 0 | 0 | 0 | 0 | 0 | 0 |
| ENSMUSG00 SNORA17 | 134 | 0 | 0 | 0 | 0 | 0 | 0 | 0 | 0 | 0 | 0 | 0 | 0 | 0 | 0 | 0 | 0 | 0 | 0 |
| ENSMUSG00 U6 | 106 | 0 | 0 | 0 | 0 | 0 | 0 | 0 | 0 | 0 | 0 | 0 | 0 | 0 | 0 | 0 | 0 | 0 | 0 |
| ENSMUSG00 U6 | 103 | 0 | 0 | 0 | 0 | 0 | 0 | 0 | 0 | 0 | 0 | 0 | 0 | 0 | 0 | 0 | 0 | 0 | 0 |
| ENSMUSG00 SNORA65 | 120 | 0 | 0 | 0 | 0 | 0 | 0 | 0 | 0 | 0 | 0 | 0 | 0 | 0 | 0 | 0 | 0 | 0 | 0 |
| ENSMUSG00 SNORA17 | 128 | 0 | 0 | 0 | 0 | 0 | 0 | 0 | 0 | 0 | 0 | 0 | 0 | 0 | 0 | 0 | 0 | 0 | 0 |
| ENSMUSG00 SNORA17 | 122 | 0 | 0 | 0 | 0 | 0 | 0 | 0 | 0 | 0 | 0 | 0 | 0 | 0 | 0 | 0 | 0 | 0 | 0 |
| ENSMUSG00 U6 | 104 | 0 | 0 | 0 | 0 | 0 | 0 | 0 | 0 | 0 | 0 | 0 | 0 | 0 | 0 | 0 | 0 | 0 | 0 |
| ENSMUSG00 SNORA49 | 132 | 0 | 0 | 0 | 0 | 0 | 0 | 0 | 0 | 0 | 0 | 0 | 0 | 0 | 0 | 0 | 0 | 0 | 0 |
| ENSMUSG00 SNORA17 | 127 | 0 | 0 | 0 | 0 | 0 | 0 | 0 | 0 | 0 | 0 | 0 | 0 | 0 | 0 | 0 | 0 | 0 | 0 |
| ENSMUSG00 SNORA17 | 131 | 0 | 0 | 0 | 0 | 0 | 0 | 0 | 0 | 0 | 0 | 0 | 0 | 0 | 0 | 0 | 0 | 0 | 0 |
| ENSMUSG00 SNORA17 | 130 | 0 | 0.7687591 | 0 | 0.70093914 | 0 | 0 | 0 | 0 | 0 | 0 | 0 | 0 | 0 | 0 | 1.58467523 | 0 | 0.88445301 | 0 |
| ENSMUSG00 SNORA17 | 126 | 0 | 0 | 0 | 0 | 0 | 0 | 0 | 0 | 0 | 0 | 0 | 0 | 0 | 0 | 0 | 0 | 0 | 0 |
| ENSMUSG00 SNORA17 | 127 | 0 | 0 | 0 | 0 | 0 | 0 | 0 | 0 | 0 | 0 | 0 | 0 | 0 | 0 | 0 | 0 | 0 | 0 |
| ENSMUSG00 SNORA17 | 127 | 0 | 0 | 0 | 0 | 0 | 0 | 0 | 0 | 0 | 0 | 0 | 0 | 0 | 0 | 0 | 0 | 0 | 0 |
| ENSMUSG00 snoU2-30 | 65 | 0 | 0 | 0 | 0 | 0 | 0 | 0 | 0 | 0 | 0 | 0 | 0 | 0 | 0 | 0 | 0 | 0 | 0 |
| ENSMUSG00 n-R5s165 | 99 | 0 | 0 | 0 | 0 | 0 | 0 | 0 | 0 | 0 | 0 | 0 | 0 | 0 | 0 | 0 | 0 | 0 | 0 |
| ENSMUSG00 SNORA17 | 125 | 0 | 0 | 0 | 0 | 0 | 0 | 0 | 0 | 0 | 0 | 0 | 0 | 0 | 0 | 0 | 0 | 0 | 0 |
| ENSMUSG00 SNORA17 | 131 | 0 | 0 | 0 | 0 | 0 | 0 | 0 | 0 | 0 | 0 | 0 | 0 | 0 | 0 | 0 | 0 | 0 | 0 |
| ENSMUSG00 U6 | 103 | 0 | 0 | 0 | 0 | 0 | 0 | 0 | 0.37080002 | 0 | 1.37332145 | 0 | 0 | 0 | 0 | 0 | 0 | 0 | 0 |
| ENSMUSG00 SNORA17 | 132 | 0 | 0 | 0 | 0 | 0 | 0 | 0 | 0 | 0 | 0 | 0 | 0 | 0 | 0 | 0 | 0 | 0 | 0 |
| ENSMUSG00 SNORA17 | 131 | 0 | 0 | 0 | 0 | 0 | 0 | 0 | 0 | 0 | 0 | 0 | 0 | 0 | 0 | 0 | 0 | 0 | 0 |
| ENSMUSG00 SNORA17 | 131 | 0 | 0 | 0 | 0 | 0 | 0 | 0 | 0 | 0 | 0 | 0 | 0 | 0 | 0 | 0 | 0 | 0 | 0 |
| ENSMUSG00 SNORA17 | 136 | 0 | 0.24494775 | 1.06706317 | 0 | 0 | 0 | 0.69577469 | 0.28082649 | 0 | 0 | 0.70257292 | 0 | 0 | 0 | 0.30295262 | 0 | 0 | 0 |
| ENSMUSG00 SNORA17 | 131 | 0 | 0 | 0 | 0 | 0 | 0 | 0 | 0 | 0 | 0 | 0 | 0 | 0 | 0 | 0 | 0 | 0 | 0 |
| ENSMUSG00 U6 | 106 | 0 | 0 | 0 | 0 | 0 | 0 | 0 | 0 | 0 | 0 | 0 | 0 | 0 | 0 | 0 | 0 | 0 | 0 |
| ENSMUSG00 SNORA26 | 121 | 1.05384734 | 0.27531318 | 0.39978124 | 0.37653756 | 0.79656899 | 0 | 0 | 1.26255876 | 0.68076886 | 0 | 1.57933747 | 0.46128647 | 0 | 0 | 0 | 0 | 0 | 0.90924143 |
| ENSMUSG00 SNORA17 | 131 | 0 | 0 | 0 | 0 | 0 | 0 | 0 | 0 | 0 | 0 | 0 | 0 | 0 | 0 | 0 | 0 | 0 | 0 |
| ENSMUSG00 SNORA69 | 131 | 0 | 0 | 0 | 0 | 0 | 0 | 0 | 0 | 0 | 0 | 1.09408302 | 0 | 0 | 0 | 0 | 0 | 0 | 0 |
| ENSMUSG00 U6 | 106 | 0 | 0 | 0 | 0 | 0 | 0 | 0 | 0 | 0 | 0 | 0 | 0 | 0 | 0 | 0 | 0 | 0 | 0 |
| ENSMUSG00 SNORA17 | 127 | 0 | 0 | 0 | 0 | 0 | 0 | 0 | 0 | 0 | 0 | 0 | 0 | 0 | 0 | 0 | 0 | 0 | 0 |

| | | | | | | | | | | | | | | | | | | | | |
|---|---|---|---|---|---|---|---|---|---|---|---|---|---|---|---|---|---|---|---|---|
| ENSMUSG00 | SNORA17 | 128 | 0 | 0 | 0 | 0 | 0 | 0 | 0 | 0 | 0 | 0 | 0 | 0 | 0 | 0 | 0 | 0 | 0 | 0 |
| ENSMUSG00 | SNORA17 | 133 | 0 | 0 | 0 | 0 | 0 | 0 | 0 | 0 | 0 | 0 | 0 | 0 | 0 | 0 | 0 | 0 | 0 | 0 |
| ENSMUSG00 | SNORA17 | 131 | 0 | 0 | 0 | 0 | 0 | 0 | 0 | 0 | 0 | 0 | 0 | 0 | 0 | 0 | 0 | 0 | 0 | 0 |
| ENSMUSG00 | SNORA17 | 131 | 0 | 0 | 0 | 0 | 0 | 0 | 0 | 0 | 0 | 0 | 0 | 0 | 0 | 0 | 0 | 0 | 0 | 0 |
| ENSMUSG00 | n-R5s158 | 117 | 0 | 0 | 0 | 0 | 1.23415189 | 0 | 0 | 0 | 0 | 0 | 0 | 0 | 0 | 0 | 0 | 0 | 0 | 0 |
| ENSMUSG00 | SNORA42 | 114 | 0 | 0 | 0 | 0 | 0 | 0 | 0 | 0 | 0 | 0 | 0 | 0 | 0 | 0 | 0 | 0 | 0 | 0 |
| ENSMUSG00 | SNORA17 | 131 | 0 | 0 | 0 | 0 | 0 | 0 | 0 | 0 | 0 | 0 | 0 | 0 | 0 | 0 | 0 | 0 | 0 | 0 |
| ENSMUSG00 | SNORA17 | 128 | 0 | 0 | 0 | 0 | 0 | 0 | 0 | 0 | 0 | 0 | 0 | 0 | 0 | 0 | 0 | 0 | 0 | 0 |
| ENSMUSG00 | SNORA17 | 131 | 0 | 0 | 0 | 0 | 0 | 0 | 0 | 0 | 0 | 0 | 0 | 0 | 0 | 0 | 0 | 0 | 0 | 0 |
| ENSMUSG00 | SNORA17 | 130 | 0 | 0 | 0 | 0 | 0 | 0 | 0 | 0 | 0 | 0 | 0 | 0 | 0 | 0 | 0 | 0 | 0 | 0 |
| ENSMUSG00 | SNORA17 | 133 | 0 | 0 | 0 | 0 | 0 | 0 | 0 | 0 | 0 | 0 | 0 | 0 | 0 | 0 | 0 | 0 | 0 | 0 |
| ENSMUSG00 | SNORA17 | 131 | 0 | 0 | 0 | 0 | 0 | 0 | 0 | 0 | 0 | 0 | 0 | 0 | 0 | 0 | 0 | 0 | 0 | 0 |
| ENSMUSG00 | SNORA17 | 132 | 0 | 0 | 0 | 0 | 0 | 0 | 0 | 0 | 0 | 0 | 0 | 0 | 0 | 0 | 0 | 0 | 0 | 0 |
| ENSMUSG00 | SNORA17 | 136 | 0.62507612 | 0 | 0 | 0 | 0 | 0 | 0 | 0.28082649 | 0 | 0 | 0 | 0 | 0 | 0 | 0 | 0 | 0 | 0 |
| ENSMUSG00 | SNORD12 | 82 | 0 | 0 | 0 | 0 | 0 | 0 | 0 | 0 | 0 | 0 | 0 | 0 | 0 | 0 | 0 | 0 | 0 | 0 |
| ENSMUSG00 | SNORA17 | 131 | 0 | 0 | 0 | 0 | 0 | 0 | 0 | 0 | 0 | 0 | 0 | 0 | 0 | 0 | 0 | 0 | 0 | 0 |
| ENSMUSG00 | SNORA17 | 132 | 0 | 0 | 0 | 0 | 0 | 0 | 0 | 0 | 0 | 0 | 0 | 0 | 0 | 0 | 0 | 0 | 0 | 0 |
| ENSMUSG00 | SNORD115 | 78 | 0 | 0 | 0 | 0 | 0 | 0 | 0 | 0 | 0 | 0 | 0 | 0 | 0 | 0 | 0 | 0 | 0 | 0 |
| ENSMUSG00 | SNORA17 | 129 | 0 | 0 | 0 | 0 | 0 | 0 | 0 | 0 | 0 | 0 | 0 | 0 | 0 | 0 | 0 | 0 | 0 | 0 |
| ENSMUSG00 | U6 | 105 | 0 | 0 | 0 | 0 | 0 | 0 | 0 | 0 | 0 | 0 | 0 | 0 | 0 | 0 | 0 | 0 | 0 | 0 |
| ENSMUSG00 | SNORA17 | 131 | 0 | 0 | 0 | 0 | 0 | 0 | 0 | 0 | 0 | 0 | 0 | 0 | 0 | 0 | 0 | 0 | 0 | 0 |
| ENSMUSG00 | U11 | 133 | 0 | 0 | 0 | 0 | 0 | 0 | 0 | 0 | 0 | 0 | 0 | 0 | 0 | 0 | 0 | 0 | 0 | 0 |
| ENSMUSG00 | SNORA17 | 129 | 0 | 0 | 0 | 0 | 0 | 0 | 0 | 0 | 0 | 0 | 0 | 0 | 0 | 0 | 0 | 0 | 0 | 0 |
| ENSMUSG00 | SNORA17 | 129 | 0 | 0 | 0 | 0 | 0 | 0 | 0 | 0 | 0 | 0 | 0 | 0 | 0 | 0 | 0 | 0 | 0 | 0 |
| ENSMUSG00 | SNORA17 | 132 | 0 | 0 | 0 | 0 | 0 | 0 | 0 | 0 | 0 | 0 | 0 | 0 | 0 | 0 | 0 | 0 | 0 | 0 |
| ENSMUSG00 | SNORA17 | 131 | 0 | 0 | 0 | 0 | 0 | 0 | 0 | 0 | 0 | 0 | 0 | 0 | 0 | 0 | 0 | 0 | 0 | 0 |
| ENSMUSG00 | n-R5s8 | 117 | 0 | 0 | 0 | 0 | 0 | 0 | 0 | 0 | 0 | 0 | 0 | 0 | 1.68656127 | 1.19087404 | 0 | 0 | 0 | 0 |
| ENSMUSG00 | SNORA17 | 131 | 0 | 0 | 0 | 0 | 0 | 0 | 0 | 0 | 0 | 0 | 0 | 0 | 0 | 0 | 0 | 0 | 0 | 0 |
| ENSMUSG00 | SNORA17 | 132 | 0 | 0 | 0 | 0 | 0 | 0 | 0 | 0 | 0 | 0 | 0 | 0 | 0 | 0 | 0 | 0 | 0 | 0 |
| ENSMUSG00 | SNORA17 | 130 | 0 | 0 | 0 | 0 | 0 | 0 | 0 | 0 | 0 | 0 | 0 | 0 | 0 | 0 | 0.63387009 | 0 | 0 | 0 |
| ENSMUSG00 | SNORA24 | 116 | 0 | 0 | 0 | 0 | 0 | 0 | 0 | 0 | 0 | 0 | 0 | 0 | 0 | 0 | 0 | 0 | 0 | 0 |
| ENSMUSG00 | n-R5s171 | 78 | 0 | 0 | 0 | 0 | 0 | 0 | 0 | 0 | 0 | 0 | 0 | 0 | 0 | 0 | 0 | 0 | 0 | 0 |
| ENSMUSG00 | U11 | 124 | 0 | 0 | 0 | 0 | 0 | 0 | 0 | 0 | 0 | 0 | 0 | 0 | 0 | 0 | 0 | 0 | 0 | 0 |
| ENSMUSG00 | SNORA60 | 127 | 0 | 0 | 0 | 0 | 0 | 0 | 0 | 0 | 0 | 0 | 0 | 0 | 0 | 0 | 0 | 0 | 0 | 0 |
| ENSMUSG00 | SNORA17 | 131 | 0 | 0 | 0 | 0 | 0 | 0 | 0 | 0 | 0 | 0 | 0 | 0 | 0 | 0 | 0 | 0 | 0 | 0 |
| ENSMUSG00 | U11 | 133 | 0.95876337 | 0.75141867 | 1.45484302 | 0.34256424 | 2.17136498 | 0.74515389 | 4.98028199 | 1.72296552 | 3.09673051 | 2.83613253 | 1.79605107 | 3.35733309 | 1.11275001 | 1.39681466 | 0.9293584 | 1.15159722 | 1.72900588 | 0.82720461 |
| ENSMUSG00 | SNORA17 | 131 | 0 | 0 | 0 | 0 | 0 | 0 | 0 | 0 | 0 | 0 | 0 | 0 | 0 | 0 | 0 | 0 | 0 | 0 |
| ENSMUSG00 | SNORA17 | 132 | 0 | 0 | 0 | 0 | 0 | 0 | 0 | 0 | 0 | 0 | 0 | 0 | 0 | 0 | 0 | 0 | 0 | 0 |
| ENSMUSG00 | U1 | 165 | 0 | 0 | 0 | 0 | 0 | 0 | 0 | 0 | 0 | 0 | 0 | 0 | 0 | 0 | 0 | 0 | 0 | 0 |
| ENSMUSG00 | SNORA17 | 129 | 0 | 0 | 0 | 0 | 0 | 0 | 0 | 0 | 0 | 0 | 0 | 0 | 0 | 0.36003169 | 0 | 0 | 0 | 0 |
| ENSMUSG00 | SNORA17 | 131 | 0 | 0 | 0 | 0 | 0 | 0 | 0 | 0 | 0 | 0 | 0 | 0 | 0 | 0 | 0 | 0 | 0 | 0 |
| ENSMUSG00 | SNORA17 | 129 | 0 | 0 | 0 | 0 | 0 | 0 | 0 | 0 | 0 | 0 | 0 | 0 | 0 | 0 | 0 | 0 | 0 | 0 |
| ENSMUSG00 | SNORA17 | 127 | 0 | 0 | 0 | 0 | 0 | 0 | 0 | 0 | 0 | 0 | 0 | 0 | 0 | 0 | 0 | 0 | 0 | 0 |
| ENSMUSG00 | U6 | 105 | 0 | 0 | 0 | 0 | 0 | 0 | 0 | 0 | 0 | 0 | 0 | 0 | 0 | 0 | 0 | 0 | 0 | 0 |
| ENSMUSG00 | SNORA17 | 131 | 0 | 0 | 0 | 0 | 0 | 0 | 0 | 0 | 0 | 0 | 0 | 0 | 0 | 0 | 0 | 0 | 0 | 0 |
| ENSMUSG00 | SNORA24 | 82 | 0 | 1.21876442 | 0 | 0 | 0 | 0 | 0 | 0 | 0 | 0.57500858 | 0 | 0 | 0 | 0 | 0 | 0 | 0 | 0 |
| ENSMUSG00 | U1 | 163 | 0 | 0 | 0 | 0 | 0 | 0 | 1.741571 | 0 | 0 | 0 | 0 | 0 | 0 | 0 | 0 | 0 | 0 | 0 |
| ENSMUSG00 | SNORA17 | 125 | 0 | 0 | 0 | 0 | 0 | 0 | 0 | 0 | 0 | 0 | 0 | 0 | 0 | 0 | 0 | 0 | 0 | 0 |
| ENSMUSG00 | SNORA17 | 133 | 0 | 0 | 0 | 0 | 0 | 0 | 0 | 0 | 0 | 0 | 0 | 0 | 0 | 0 | 0 | 0 | 0 | 0 |
| ENSMUSG00 | SNORA17 | 131 | 0 | 0 | 0 | 0 | 0 | 0 | 0 | 0 | 0 | 0 | 0 | 0 | 0 | 0 | 0 | 0 | 0 | 0 |
| ENSMUSG00 | SNORA17 | 124 | 0 | 0 | 0 | 0 | 0 | 0 | 0 | 0 | 0 | 0 | 0 | 0 | 0 | 0 | 0 | 0 | 0 | 0 |
| ENSMUSG00 | Snord70 | 86 | 1.4827387 | 0.38735924 | 0 | 0 | 0 | 0 | 0 | 0.8881954 | 0 | 0 | 0 | 0 | 0.57362695 | 0 | 0 | 0 | 0 | 0 |
| ENSMUSG00 | U1 | 157 | 0 | 0 | 0 | 0 | 0 | 0 | 0.30135464 | 0 | 0 | 0 | 0 | 0 | 0 | 0 | 1.04972117 | 0 | 0 | 0 |
| ENSMUSG00 | SNORA17 | 128 | 0 | 0 | 0 | 0 | 0 | 0 | 0 | 0 | 0 | 0 | 0 | 0 | 0 | 0 | 0 | 0 | 0 | 0 |
| ENSMUSG00 | U6 | 106 | 0 | 0 | 0 | 0 | 0 | 0 | 0 | 0 | 0 | 0 | 0 | 0 | 0 | 0 | 0 | 0 | 0 | 0 |
| ENSMUSG00 | n-R5s31 | 109 | 0.77991149 | 0 | 0 | 0 | 0 | 0 | 0.43406127 | 0 | 0 | 0 | 0 | 0 | 0 | 1.27827763 | 0 | 0 | 0 | 0 |
| ENSMUSG00 | SNORA17 | 131 | 0 | 0 | 0 | 0 | 0 | 0 | 0 | 0 | 0 | 0 | 0 | 0 | 0 | 0 | 0 | 0 | 0 | 0 |
| ENSMUSG00 | SNORA17 | 131 | 0 | 0 | 0 | 0 | 0 | 0 | 0 | 0 | 0 | 0 | 0 | 0 | 0 | 0 | 0 | 0 | 0 | 0 |
| ENSMUSG00 | SNORA17 | 131 | 0 | 0 | 0 | 0 | 0 | 0 | 0 | 0 | 0 | 0 | 0 | 0 | 0 | 0 | 0 | 0 | 0 | 0 |
| ENSMUSG00 | SNORA42 | 133 | 0 | 0 | 0 | 0 | 0 | 0 | 0 | 0 | 0 | 0 | 0 | 0 | 0 | 0 | 0 | 0 | 0 | 0 |
| ENSMUSG00 | SNORA17 | 131 | 0 | 0 | 0 | 0 | 0 | 0 | 0 | 0 | 0 | 0 | 0 | 0 | 0 | 0 | 0 | 0 | 0 | 0 |
| ENSMUSG00 | SNORA17 | 130 | 0 | 0 | 0 | 0 | 0 | 0 | 0 | 0 | 0 | 0 | 0 | 0 | 0 | 0 | 0 | 0 | 0 | 0 |
| ENSMUSG00 | SNORD70 | 83 | 0 | 0.80272034 | 0 | 0 | 0.57990269 | 0 | 0 | 1.38044828 | 0 | 1.13616152 | 0 | 0 | 0.59436045 | 1.11913464 | 0 | 0 | 1.38528784 | 0 |
| ENSMUSG00 | U1 | 157 | 0 | 0 | 0 | 0 | 0 | 0 | 0 | 0 | 0 | 0 | 0 | 0 | 0 | 0 | 0 | 0 | 0 | 0 |
| ENSMUSG00 | SNORA24 | 121 | 0 | 1.1012527 | 0.79956249 | 0 | 0.79556899 | 0 | 0 | 0 | 0 | 0 | 0 | 0 | 0 | 0 | 0 | 0.3164513 | 0 | 0 |
| ENSMUSG00 | SNORA17 | 130 | 0 | 0 | 0 | 0 | 0 | 0 | 0 | 0 | 0 | 0 | 0 | 0 | 0 | 0 | 0 | 0 | 0 | 0 |
| ENSMUSG00 | SNORA17 | 130 | 0 | 0 | 0 | 0 | 0 | 0 | 0 | 0 | 0 | 0 | 0 | 0 | 0 | 0 | 0 | 0 | 0 | 0 |
| ENSMUSG00 | U6 | 107 | 0 | 0 | 0 | 0 | 0 | 0 | 0 | 0 | 0 | 0 | 2.2324747 | 0 | 0 | 0.43405689 | 0 | 0.35785614 | 0 | 0 |
| ENSMUSG00 | SNORA17 | 130 | 0 | 0 | 0 | 0 | 0 | 0 | 0 | 0 | 0 | 0 | 0 | 0 | 0 | 0 | 0 | 0 | 0 | 0 |
| ENSMUSG00 | U1 | 154 | 0 | 0 | 0 | 0 | 0 | 0 | 0 | 0 | 0 | 0 | 0 | 0 | 0 | 0 | 0 | 0 | 0 | 0 |
| ENSMUSG00 | SNORA17 | 119 | 0 | 0 | 0 | 0 | 0 | 0 | 0 | 0 | 0 | 0 | 0 | 0 | 0 | 0 | 0 | 0 | 0 | 0 |
| ENSMUSG00 | SNORA17 | 131 | 0 | 0 | 0 | 0 | 0 | 0 | 0 | 0 | 0 | 0 | 0 | 0 | 0 | 0 | 0 | 0 | 0 | 0 |
| ENSMUSG00 | SNORA17 | 128 | 0 | 0 | 0 | 0 | 0 | 0 | 0 | 0 | 0 | 0 | 0 | 0 | 0 | 0 | 0 | 0 | 0 | 0 |
| ENSMUSG00 | SNORA17 | 128 | 0 | 0 | 0 | 0 | 0 | 0 | 0 | 0 | 0 | 0 | 0 | 0 | 0 | 0 | 0 | 0 | 0 | 0 |
| ENSMUSG00 | SNORA68 | 131 | 0 | 1.27148451 | 0 | 0 | 0 | 0 | 0 | 0 | 0 | 0.71985806 | 0 | 0.42607376 | 0 | 0 | 0 | 0 | 0 | 0 |
| ENSMUSG00 | SNORA17 | 129 | 0 | 0 | 0 | 0 | 0 | 0 | 0 | 0 | 0 | 0 | 0 | 0 | 0 | 0 | 0 | 0 | 0 | 0 |
| ENSMUSG00 | n-R5s200 | 112 | 0 | 0.59487311 | 0 | 0 | 0 | 0.44243512 | 1.2673039 | 0 | 0 | 0 | 0 | 0 | 0.44046355 | 0 | 0 | 2.39316296 | 0 | 0 |
| ENSMUSG00 | SNORA17 | 129 | 0 | 0 | 0 | 0 | 0 | 0 | 0 | 0 | 0 | 0 | 0 | 0 | 0 | 0 | 0 | 0 | 0 | 0 |
| ENSMUSG00 | U1 | 154 | 0 | 0 | 0 | 0 | 0 | 0 | 0 | 0 | 0 | 0 | 0 | 0 | 0 | 0 | 0 | 0 | 0 | 0 |
| ENSMUSG00 | SNORA17 | 131 | 0 | 1.27148451 | 1.10779077 | 0 | 0 | 0 | 0 | 0 | 0 | 0 | 0 | 0 | 0 | 0 | 0 | 0.29229471 | 0 | 0 |
| ENSMUSG00 | U1 | 170 | 0.50006089 | 0 | 0.85365054 | 0 | 0 | 0 | 0.27830988 | 0 | 0.24227362 | 0.27735708 | 0 | 0 | 0 | 0.54640103 | 0 | 0 | 1.12724403 | 0 |
| ENSMUSG00 | SNORA12 | 145 | 0 | 0 | 1.66805277 | 0.3142141 | 0 | 0 | 0.65258867 | 0.26339588 | 1.42022468 | 0 | 0 | 0 | 1.36088048 | 0 | 0.56829732 | 0 | 0 | 0 |
| ENSMUSG00 | SNORA17 | 125 | 0 | 0 | 0 | 0 | 0 | 0 | 0 | 0 | 0 | 0 | 0 | 0 | 0 | 0 | 0 | 0 | 0 | 0 |
| ENSMUSG00 | SNORA24 | 126 | 0 | 0 | 0 | 0 | 0 | 0 | 0 | 0 | 0 | 0 | 0 | 0 | 0 | 0 | 0 | 0 | 0 | 0 |
| ENSMUSG00 | snoU2_19 | 75 | 0 | 0 | 0.6449804 | 0 | 0 | 0.66070311 | 0 | 0 | 0 | 0 | 0 | 0 | 0 | 0 | 1.64806224 | 0.51054143 | 0 | 0 |
| ENSMUSG00 | U6 | 96 | 0 | 0 | 0 | 0 | 0 | 0 | 0 | 0 | 0 | 0 | 0 | 0 | 0 | 0 | 0 | 0 | 0 | 0 |
| ENSMUSG00 | SNORA17 | 124 | 0 | 0 | 0 | 0 | 0 | 0 | 0 | 0 | 0 | 0 | 0 | 0 | 0 | 0 | 0 | 0 | 0 | 0 |
| ENSMUSG00 | SNORA17 | 127 | 0 | 0 | 0 | 0 | 0 | 0 | 0 | 0 | 0 | 0 | 0 | 0 | 0 | 0 | 0 | 0 | 0 | 0 |
| ENSMUSG00 | SNORA17 | 131 | 0 | 0 | 0 | 0 | 0 | 0 | 0 | 0 | 0 | 0 | 0 | 0 | 0 | 0 | 0 | 0 | 0 | 0 |
| ENSMUSG00 | SNORA17 | 130 | 0 | 0 | 0 | 0 | 0 | 0 | 0 | 0 | 0 | 0 | 0 | 0 | 0 | 0 | 0 | 0 | 0 | 0 |
| ENSMUSG00 | n-R5s82 | 108 | 0 | 0 | 0 | 0 | 0 | 0 | 0 | 0 | 0 | 0 | 0 | 0 | 0 | 0 | 0 | 0 | 0 | 0 |
| ENSMUSG00 | SNORA17 | 128 | 0 | 0 | 0 | 0 | 0 | 0 | 0 | 0 | 0 | 0 | 0 | 0 | 0 | 0 | 0 | 0 | 0 | 0 |
| ENSMUSG00 | U3 | 189 | 0 | 0 | 0 | 0 | 0 | 0 | 0 | 0 | 0 | 0 | 0 | 0 | 0 | 0 | 0 | 0 | 0 | 0 |
| ENSMUSG00 | SNORA17 | 126 | 0 | 0 | 0 | 0 | 0 | 0 | 0 | 0 | 0 | 0 | 0 | 0 | 0 | 0 | 0 | 0 | 0 | 0 |
| ENSMUSG00 | SNORA17 | 131 | 0 | 0 | 0 | 0 | 0 | 0 | 0 | 0 | 0 | 0 | 0 | 0 | 0 | 0 | 0 | 0 | 0 | 0 |
| ENSMUSG00 | U1 | 163 | 0 | 0 | 0 | 0 | 0 | 0 | 0 | 0 | 0 | 0.28926812 | 0 | 0 | 0 | 0 | 0 | 0 | 0 | 0 |
| ENSMUSG00 | SNORA17 | 128 | 0 | 0 | 0 | 0 | 0 | 0 | 0 | 0 | 0 | 0 | 0 | 0 | 0 | 0 | 0 | 0 | 0 | 0 |
| ENSMUSG00 | SNORA17 | 131 | 0 | 0 | 0 | 0 | 0 | 0 | 0 | 0 | 0 | 0 | 0 | 0 | 0 | 0 | 0 | 0 | 0 | 0 |
| ENSMUSG00 | U6 | 106 | 0 | 0 | 0 | 0 | 0 | 0 | 0 | 0 | 0 | 0 | 0 | 0 | 0 | 0 | 0 | 0 | 0 | 0 |
| ENSMUSG00 | SNORA17 | 131 | 0 | 0 | 0 | 0 | 0 | 0 | 0 | 0 | 0 | 0 | 0 | 0 | 0 | 0 | 0 | 0 | 0 | 0 |
| ENSMUSG00 | SNORA17 | 130 | 0 | 0 | 0 | 0 | 0 | 0 | 0 | 0 | 0 | 0 | 0 | 0 | 0 | 0 | 0 | 0 | 0 | 0 |
| ENSMUSG00 | SNORD30 | 68 | 0 | 0 | 0 | 0 | 0 | 0 | 0 | 0 | 0 | 0 | 0 | 0 | 0 | 0 | 0 | 0 | 0 | 0 |
| ENSMUSG00 | SNORA17 | 130 | 0 | 0 | 0 | 0 | 0 | 0 | 0 | 0 | 0 | 0 | 0 | 0 | 0 | 0 | 0 | 0 | 0 | 0 |
| ENSMUSG00 | SNORA17 | 130 | 0.65392579 | 0 | 0 | 0 | 0 | 0.38117487 | 0 | 0.29378771 | 0 | 0.36269772 | 0 | 0 | 0 | 0 | 0 | 1.17817254 | 0.29481767 | 0 |
| ENSMUSG00 | SNORA17 | 124 | 0 | 0 | 0 | 0 | 0 | 0 | 0 | 0 | 0 | 0 | 0 | 0 | 0 | 0 | 0 | 0 | 0 | 0 |
| ENSMUSG00 | SNORA17 | 124 | 0 | 0 | 0 | 0 | 0 | 0 | 0 | 0 | 0 | 0 | 0 | 0 | 0 | 0 | 0 | 0 | 0 | 0 |
| ENSMUSG00 | U2 | 148 | 0 | 0 | 0 | 0 | 0 | 0 | 0 | 0 | 0 | 0 | 0 | 0 | 0 | 0 | 0 | 0 | 0 | 0 |
| ENSMUSG00 | AC161183.1 | 65 | 0 | 0 | 0 | 1.40187829 | 2.2214734 | 0 | 0 | 0.58757542 | 0 | 0.72539543 | 0 | 0 | 0 | 1.42904884 | 0.63387009 | 1.17817254 | 1.17927067 | 0 |
| ENSMUSG00 | U6 | 106 | 0 | 0 | 0 | 0 | 0 | 0.93495724 | 0 | 0 | 0 | 0 | 0 | 0 | 0 | 0 | 0 | 0 | 0 | 0 |
| ENSMUSG00 | SNORA17 | 135 | 0 | 0 | 0 | 0 | 0 | 0 | 0 | 0 | 0 | 0 | 0 | 0 | 0 | 0 | 0 | 0 | 0 | 0 |
| ENSMUSG00 | SNORA17 | 130 | 0 | 0 | 0 | 0 | 0 | 0 | 0 | 0 | 0 | 0 | 0 | 0 | 0 | 0 | 0 | 0 | 0 | 0 |
| ENSMUSG00 | SNORA17 | 133 | 0 | 0 | 0 | 0 | 0 | 0 | 0 | 0 | 0 | 0 | 0 | 0 | 0 | 0 | 0 | 0 | 0 | 0 |
| ENSMUSG00 | SNORA66 | 131 | 1.62233496 | 0.5085938 | 4.0618995 | 1.73897116 | 0.73483853 | 0 | 2.52815842 | 0.29154506 | 0.94320265 | 0 | 2.91755472 | 0 | 2.25947713 | 3.54535018 | 1.57257847 | 2.3383577 | 2.04797006 | 3.35933476 |
| ENSMUSG00 | SNORA17 | 126 | 0 | 0 | 0 | 0 | 0 | 0 | 0 | 0 | 0 | 0 | 0 | 0 | 0 | 0 | 0 | 0 | 0 | 0 |
| ENSMUSG00 | SNORA42 | 141 | 0 | 0 | 0 | 0 | 0 | 0 | 0 | 0 | 0 | 0 | 0 | 0 | 0 | 0 | 0 | 0 | 0 | 0 |
| ENSMUSG00 | SNORA17 | 130 | 0 | 0 | 0 | 0 | 0 | 0 | 0 | 0 | 0 | 0 | 0 | 0 | 0 | 0 | 0 | 0 | 0 | 0 |
| ENSMUSG00 | U6 | 105 | 0 | 0 | 0 | 0 | 0 | 0 | 0 | 0 | 0 | 0 | 0 | 0 | 0 | 0 | 0 | 0 | 0 | 0 |
| ENSMUSG00 | U6 | 108 | 0 | 0 | 0 | 0 | 0 | 0 | 0 | 0 | 0 | 0 | 0 | 0 | 0 | 0 | 0 | 0 | 0 | 0 |
| ENSMUSG00 | SNORA17 | 127 | 0 | 0 | 0 | 0 | 0 | 0 | 0 | 0 | 0 | 0 | 0 | 0 | 0 | 0 | 0 | 0 | 0 | 0 |
| ENSMUSG00 | n-R5s178 | 113 | 0 | 0.58960875 | 0 | 1.20958525 | 2.12973114 | 0 | 0 | 0 | 0.72896488 | 0 | 0 | 2.46971958 | 0 | 0 | 0 | 0 | 0 | 0 |
| ENSMUSG00 | SNORA17 | 129 | 0 | 0 | 0 | 0 | 0 | 0 | 0 | 0 | 0 | 0 | 0 | 0 | 0 | 0 | 0 | 0.89047924 | 0 | 0 |
| ENSMUSG00 | SNORA17 | 129 | 0 | 0 | 0 | 0 | 0 | 0 | 0 | 0 | 0 | 0 | 0 | 0 | 0 | 0 | 0 | 0 | 0 | 0 |
| ENSMUSG00 | SNORD101 | 73 | 6.40488954 | 2.7380461 | 3.9759066 | 3.74474337 | 0.65934142 | 6.7880457 | 1.94435667 | 2.61591797 | 0.56419885 | 0.64590004 | 1.96335446 | 0 | 1.35155938 | 1.27244075 | 5.64404875 | 4.72075982 | 13.1254441 | 3.01419763 |
| ENSMUSG00 | SNORA17 | 131 | 0 | 0 | 0 | 0 | 0 | 0 | 0 | 0 | 0 | 0 | 0 | 0 | 0 | 0 | 0 | 0 | 0 | 0 |
| ENSMUSG00 | SNORA17 | 131 | 0 | 0 | 0 | 0 | 0 | 0 | 0 | 0 | 0 | 0 | 0 | 0 | 0 | 0 | 0 | 0 | 0 | 0 |
| ENSMUSG00 | U12 | 134 | 0 | 0 | 0 | 0 | 0 | 0 | 0 | 0 | 0 | 0 | 0 | 0 | 0 | 0 | 0 | 0 | 0 | 0 |
| ENSMUSG00 | SNORA17 | 131 | 0 | 0 | 0 | 0 | 0 | 0 | 0 | 0 | 0 | 0 | 0 | 0 | 0 | 0 | 0 | 0 | 0 | 0 |
| ENSMUSG00 | SNORA17 | 131 | 0 | 0 | 0 | 0 | 0 | 0 | 0 | 0 | 0 | 0 | 0 | 0 | 0 | 0 | 0 | 0 | 0 | 0 |
| ENSMUSG00 | SNORA17 | 131 | 5.51593887 | 3.56015683 | 3.32337231 | 3.47794232 | 4.40903118 | 1.89132571 | 5.41748232 | 5.83090113 | 13.2048371 | 3.59929032 | 4.37633207 | 12.356139 | 3.76579521 | 7.7997704 | 2.51612555 | 4.09212598 | 2.63310437 | 2.09958423 |
| ENSMUSG00 | SNORA17 | 133 | 0 | 0 | 0 | 0 | 0 | 0 | 0.35573443 | 0 | 0 | 0 | 0 | 0 | 0 | 0 | 0 | 0 | 0 | 0 |
| ENSMUSG00 | SNORA42 | 129 | 0.32949749 | 0 | 1.49995443 | 1.76593195 | 0.74623137 | 0.76825944 | 0 | 0 | 1.59637658 | 0.73101866 | 0.37034852 | 1.29803866 | 0 | 2.5202218 | 0.63878381 | 0 | 0.89130923 | 0 |
| ENSMUSG00 | Rab11b | 3701 | 40.1622806 | 43.097038 | 39.7470699 | 45.7333909 | 42.4096739 | 43.380399 | 41.4449352 | 37.7383452 | 34.5539399 | 40.8953681 | 41.3077188 | 38.5476448 | 33.3233702 | 34.484829 | 36.113982 | 34.6591556 | 36.5865461 | 36.69751 |
| ENSMUSG00 | SNORA17 | 130 | 0 | 0 | 0 | 0 | 0 | 0 | 0 | 0 | 0 | 0 | 0 | 0 | 0 | 0 | 0 | 0 | 0 | 0 |
| ENSMUSG00 | SNORA17 | 125 | 0 | 0 | 0 | 0 | 0 | 0 | 0 | 0 | 0 | 0 | 0 | 0 | 0 | 0 | 0 | 0 | 0 | 0 |
| ENSMUSG00 | U6 | 102 | 0 | 0 | 0 | 0 | 0 | 0 | 0 | 0 | 0 | 0 | 0 | 0 | 0 | 0 | 0 | 0 | 0 | 0 |
| ENSMUSG00 | SNORA17 | 129 | 0 | 0 | 0 | 0 | 0 | 0 | 0 | 0 | 0 | 0 | 0 | 0 | 0 | 0 | 0 | 0 | 0 | 0 |
| ENSMUSG00 | SNORA17 | 130 | 0 | 0 | 0 | 0 | 0 | 0 | 0 | 0 | 0 | 0 | 0 | 0 | 0 | 0 | 0 | 0 | 0 | 0 |
| ENSMUSG00 | Snord65 | 71 | 4.78931561 | 7.50713109 | 4.7692213 | 1.92511455 | 3.38957209 | 1.39585165 | 8.66288487 | 2.6896058 | 6.96110126 | 12.6177938 | 6.72886739 | 1.57227219 | 12.5066833 | 21.5866885 | 12.1863757 | 19.9542602 | 10.79614 | 17.8198515 |
| ENSMUSG00 | SNORA17 | 131 | 0 | 0 | 0 | 0 | 0 | 0 | 0 | 0 | 0 | 0 | 0 | 0 | 0 | 0 | 0 | 0 | 0 | 0 |
| ENSMUSG00 | SNORA17 | 132 | 0 | 0.25237041 | 1.46586456 | 0 | 0.72927157 | 0 | 0 | 0.57867276 | 0 | 2.14321378 | 0 | 0 | 0.37372665 | 0 | 1.248532 | 0 | 0 | 0 |
| ENSMUSG00 | U6 | 102 | 0 | 0 | 0 | 0 | 0 | 0 | 0 | 0 | 0 | 0 | 0 | 0 | 0 | 0 | 0 | 0 | 0 | 0 |
| ENSMUSG00 | SNORA17 | 134 | 0 | 0 | 0 | 0 | 0 | 0 | 0 | 0 | 0 | 0 | 0 | 0 | 0 | 0 | 0 | 0 | 0 | 0 |
| ENSMUSG00 | SNORA64 | 136 | 0 | 0 | 0 | 0 | 0 | 0 | 0 | 0 | 0 | 0 | 0 | 0 | 0 | 0 | 0 | 0 | 0 | 0 |
| ENSMUSG00 | SNORA17 | 131 | 0 | 0 | 0 | 0 | 0 | 0 | 0 | 0 | 0 | 0 | 0 | 0 | 0 | 0 | 0 | 0 | 0 | 0 |
| ENSMUSG00 | SNORA17 | 127 | 0 | 0 | 0 | 0 | 0 | 0 | 0 | 0 | 0 | 0 | 0 | 0 | 0 | 0 | 0 | 0 | 0 | 0 |
| ENSMUSG00 | SNORA17 | 131 | 0 | 0 | 0 | 0 | 0 | 0 | 0 | 0 | 0 | 0 | 0 | 0 | 0 | 0 | 0 | 0 | 0 | 0 |
| ENSMUSG00 | SNORA17 | 133 | 0 | 0 | 0 | 0 | 0 | 0 | 0 | 0 | 0 | 0 | 0 | 0 | 0 | 0 | 0 | 0 | 0 | 0 |
| ENSMUSG00 | SNORA17 | 129 | 0 | 0 | 0 | 0 | 0 | 0 | 0 | 0 | 0 | 0 | 0 | 0 | 0 | 0 | 0 | 0 | 0 | 0 |
| ENSMUSG00 | SNORA17 | 130 | 0 | 0 | 0 | 0 | 0 | 0 | 0 | 0 | 0 | 0 | 0 | 0.42935125 | 0 | 0 | 0 | 0 | 0 | 0 |
| ENSMUSG00 | U1 | 155 | 0 | 0 | 0 | 0 | 0 | 0 | 0 | 0 | 0 | 0 | 0 | 0 | 0 | 0 | 0 | 0 | 0 | 0 |
| ENSMUSG00 | SNORA17 | 130 | 0 | 0 | 0 | 0 | 0 | 0 | 0 | 0 | 0 | 0 | 0 | 0 | 0 | 0 | 0 | 0 | 0 | 0 |
| ENSMUSG00 | SNORA17 | 131 | 0 | 0 | 0 | 0 | 0 | 0 | 0 | 0 | 0 | 0 | 0 | 0 | 0 | 0 | 0 | 0 | 0 | 0 |
| ENSMUSG00 | SNORA17 | 127 | 0 | 0 | 0 | 0 | 0 | 0 | 0 | 0 | 0 | 0 | 0 | 0 | 0 | 0 | 0 | 0 | 0 | 0 |
| ENSMUSG00 | SNORA17 | 139 | 0 | 0.95864444 | 0.69602202 | 0.98333189 | 0 | 0 | 0 | 1.09906194 | 0 | 0 | 0 | 0 | 0 | 1.00239038 | 0 | 0 | 0 | 0 |
| ENSMUSG00 | SNORA17 | 131 | 0 | 0 | 0 | 0 | 0 | 0 | 0 | 0 | 0 | 0 | 0 | 0 | 0 | 0 | 0 | 0 | 0 | 0 |
| ENSMUSG00 | SNORA17 | 131 | 0 | 0 | 0 | 0 | 0 | 0 | 0 | 0 | 0 | 0 | 0 | 0 | 0 | 0 | 0 | 0 | 0 | 0 |
| ENSMUSG00 | SNORA17 | 127 | 0 | 0 | 0 | 0 | 0 | 0 | 0 | 0 | 0 | 0 | 0 | 0 | 0 | 0 | 0 | 0 | 0 | 0 |
| ENSMUSG00 | SNORA42 | 114 | 0 | 0 | 0 | 0 | 0 | 0 | 0 | 0 | 0 | 0 | 0 | 0 | 0 | 0 | 0 | 0 | 0 | 0 |
| ENSMUSG00 | SNORA17 | 134 | 0 | 0 | 0 | 0 | 0 | 0 | 0 | 0 | 0 | 0 | 0 | 0 | 0 | 0 | 0 | 0 | 0 | 0 |
| ENSMUSG00 | SNORA17 | 128 | 0 | 0 | 0 | 0 | 0 | 0 | 0 | 0 | 0 | 0 | 0 | 0 | 0 | 0 | 0 | 0 | 0 | 0 |
| ENSMUSG00 | SNORA17 | 129 | 0 | 0 | 0 | 0 | 0 | 0 | 0 | 0 | 0 | 0 | 0 | 0 | 0 | 0 | 0 | 0 | 0 | 0 |
| ENSMUSG00 | SNORA17 | 123 | 0 | 0 | 0 | 0 | 0 | 0 | 0 | 0 | 0 | 0 | 0 | 0 | 0 | 0 | 0 | 0 | 0 | 0 |
| ENSMUSG00 | SNORA17 | 131 | 0 | 0 | 0 | 0 | 0 | 0 | 0 | 0 | 0 | 0 | 0 | 0 | 0 | 0 | 0 | 0 | 0 | 0 |
| ENSMUSG00 | Snord91a | 92 | 5.08214061 | 4.34516012 | 3.15479546 | 1.98091497 | 1.04634617 | 2.69308335 | 2.057073 | 1.66053924 | 3.58143615 | 4.10006115 | 8.82798146 | 1.21338397 | 0.53621649 | 0.50482704 | 4.47842999 | 3.74582029 | 0.41659018 | 7.17510088 |
| ENSMUSG00 | SNORA17 | 129 | 0 | 0 | 0 | 0 | 0 | 0 | 0 | 0 | 0 | 0 | 0 | 0 | 0 | 0 | 0 | 0 | 0 | 0 |
| ENSMUSG00 | SNORA17 | 133 | 0 | 0 | 0 | 0 | 0 | 0 | 0 | 0 | 0 | 0 | 0 | 0 | 0 | 0 | 0 | 0 | 0 | 0 |
| ENSMUSG00 | SNORA17 | 131 | 0 | 0 | 0 | 0 | 0 | 0 | 0 | 0 | 0 | 0 | 0 | 0 | 0 | 0 | 0 | 0 | 0 | 0 |
| ENSMUSG00 | SNORA26 | 119 | 0 | 0 | 0 | 0 | 0 | 0 | 0 | 0 | 0 | 0 | 0 | 0 | 0 | 0 | 0 | 0 | 0 | 0 |
| ENSMUSG00 | SNORA17 | 131 | 0 | 0 | 0 | 0 | 0 | 0 | 0 | 0 | 0 | 0 | 0 | 0 | 0 | 0 | 0 | 0 | 0 | 0 |
| ENSMUSG00 | SNORA17 | 131 | 0 | 0.5085938 | 0 | 0 | 0 | 0 | 0 | 0 | 0 | 0 | 0 | 0 | 0 | 0 | 0 | 0 | 0 | 0 |
| ENSMUSG00 | SNORA17 | 128 | 0 | 0 | 0 | 0 | 0 | 0 | 0 | 0 | 0 | 0 | 0 | 0 | 0 | 0 | 0 | 0 | 0 | 0 |
| ENSMUSG00 | SNORA17 | 131 | 0 | 0 | 0 | 0 | 0 | 0 | 0 | 0 | 0 | 0 | 0 | 0 | 0 | 0 | 0 | 0 | 0 | 0 |
| ENSMUSG00 | SNORA17 | 132 | 0 | 0 | 0 | 0 | 0 | 0 | 0 | 0 | 0 | 0 | 0 | 0 | 0 | 0 | 0 | 0 | 0 | 0 |
| ENSMUSG00 | U6 | 106 | 0 | 0 | 0 | 0 | 0 | 0 | 0 | 0 | 0 | 0 | 0 | 0 | 0 | 0 | 0 | 0 | 0 | 0 |
| ENSMUSG00 | SNORA26 | 121 | 0 | 0 | 0.39978124 | 0 | 0 | 0 | 0 | 0 | 0 | 0 | 0.39483437 | 0 | 0 | 0.34050873 | 0 | 0 | 0 | 0 |
| ENSMUSG00 | Scarna9 | 69 | 0 | 0.48279557 | 0 | 0 | 0 | 0.71815556 | 0 | 0 | 0 | 0 | 0 | 0 | 0 | 0 | 0 | 0 | 0 | 0 |
| ENSMUSG00 | SNORA17 | 128 | 0 | 0 | 0 | 0 | 0 | 0 | 0 | 0 | 0 | 0 | 0 | 0 | 0 | 0 | 0 | 0 | 0 | 0 |
| ENSMUSG00 | AC120367.1 | 109 | 0 | 0 | 0 | 0 | 0 | 0 | 0 | 0 | 0 | 0 | 0 | 0 | 0 | 0 | 0 | 0 | 0 | 0 |
| ENSMUSG00 | U6 | 106 | 0 | 0 | 0 | 0 | 0 | 0 | 0 | 0 | 0 | 0 | 0 | 0 | 0 | 0 | 0 | 0 | 0 | 0 |
| ENSMUSG00 | SNORA17 | 131 | 0 | 0 | 0 | 0 | 0 | 0 | 0 | 0 | 0 | 0 | 0 | 0 | 0 | 0 | 0 | 0 | 0 | 0 |
| ENSMUSG00 | SNORA66 | 128 | 0 | 0 | 0 | 0 | 0 | 0 | 0 | 0 | 0 | 0 | 0 | 0 | 0 | 0 | 0 | 0 | 0 | 0 |
| ENSMUSG00 | SNORD12 | 92 | 0 | 0 | 0 | 0 | 0 | 0 | 0 | 0 | 0 | 0 | 0 | 0 | 0 | 0 | 0 | 0 | 0 | 0 |
| ENSMUSG00 | n-R5s219 | 131 | 0 | 0 | 0 | 0 | 0 | 0 | 0 | 0 | 0 | 0 | 0.72938868 | 0 | 0 | 0 | 0 | 0 | 0 | 0 |
| ENSMUSG00 | SNORA32 | 118 | 0 | 0 | 0 | 0 | 0 | 0 | 0 | 0 | 0 | 0 | 0 | 0 | 0 | 0 | 0 | 0 | 0 | 0 |
| ENSMUSG00 | U6 | 106 | 0 | 0 | 0 | 0 | 0 | 0 | 0 | 0 | 0 | 0 | 0 | 0 | 0 | 0 | 0 | 0 | 0 | 0 |
| ENSMUSG00 | SNORA17 | 131 | 0 | 0 | 0 | 0 | 0 | 0 | 0 | 0 | 0 | 0 | 0 | 0 | 0 | 0 | 0 | 0 | 0 | 0 |
| ENSMUSG00 | SNORA17 | 131 | 0.64893399 | 0 | 2.21558154 | 0 | 0 | 0 | 0 | 0.29154506 | 0.62880177 | 0.71985806 | 0.36469434 | 0.42607376 | 0 | 0.70907004 | 0.94354708 | 0.29229471 | 0.29256715 | 0 |
| ENSMUSG00 | SNORA17 | 128 | 0 | 0 | 0 | 0 | 0 | 0 | 0 | 0 | 0 | 0 | 0 | 0 | 0 | 0 | 0 | 0 | 0 | 0 |
| ENSMUSG00 | SNORA17 | 128 | 0 | 0 | 0 | 0 | 0 | 0 | 0 | 0 | 0 | 0 | 0 | 0 | 0 | 0 | 0 | 0 | 0 | 0 |
| ENSMUSG00 | n-R5s67 | 113 | 0.376152 | 0.58960875 | 0.42808434 | 0.40319508 | 1.70378491 | 1.3155593 | 0.83739255 | 2.7038869 | 2.18689464 | 0.83452572 | 2.11393622 | 0.49394392 | 0 | 0.41100962 | 0 | 0 | 2.71336615 | 1.46041876 |
| ENSMUSG00 | SNORA65 | 113 | 0 | 0.29480437 | 0 | 0 | 0 | 0 | 0 | 0.67597172 | 0 | 0 | 0 | 0 | 0 | 0 | 0 | 0 | 0 | 0 |
| ENSMUSG00 | SNORD41 | 69 | 0 | 0 | 0 | 0 | 0 | 0 | 0 | 0 | 0 | 0 | 0 | 0 | 0 | 0 | 0 | 0 | 0 | 0 |
| ENSMUSG00 | SNORA17 | 132 | 0 | 0 | 0 | 0 | 0 | 0 | 0 | 0 | 0 | 0 | 0 | 0 | 0 | 0 | 0 | 0 | 0 | 0 |
| ENSMUSG00 | SNORA17 | 131 | 0 | 0 | 0 | 0 | 0 | 0 | 0 | 0 | 0 | 0 | 0 | 0 | 0 | 0 | 0 | 0 | 0 | 0 |
| ENSMUSG00 | U6 | 93 | 0 | 0 | 0 | 0 | 0 | 0 | 0 | 0 | 0 | 0 | 0 | 0 | 0 | 0 | 0 | 0 | 0 | 0 |
| ENSMUSG00 | SNORA17 | 126 | 0 | 0 | 0 | 0 | 0 | 0 | 0 | 0 | 0 | 0 | 0 | 0 | 0 | 0 | 0 | 0 | 0 | 0 |
| ENSMUSG00 | SNORA17 | 131 | 0 | 0 | 0 | 0 | 0 | 0 | 0 | 0 | 0 | 0 | 0 | 0 | 0 | 0 | 0 | 0 | 0 | 0 |
| ENSMUSG00 | SNORA17 | 129 | 0 | 0 | 0 | 0 | 0 | 0 | 0 | 0 | 0 | 0 | 0 | 0 | 0 | 0 | 0 | 0 | 0 | 0 |
| ENSMUSG00 | SNORA17 | 131 | 0.64893399 | 0.5085938 | 0 | 0 | 0 | 0 | 0 | 0 | 0 | 0 | 0 | 0 | 0 | 0 | 0.31451569 | 0 | 0 | 0 |
| ENSMUSG00 | U6 | 106 | 0 | 0 | 0 | 0 | 0 | 0 | 0 | 0 | 0 | 0 | 0 | 0 | 0 | 0 | 0 | 0 | 0 | 0 |
| ENSMUSG00 | SNORA17 | 129 | 0 | 0 | 0 | 0 | 0 | 0 | 0 | 0 | 0 | 0 | 0 | 0 | 0 | 0 | 0 | 0 | 0 | 0 |
| ENSMUSG00 | SNORA17 | 126 | 0 | 0 | 0 | 0 | 0 | 0 | 0 | 0 | 0 | 0 | 0 | 0 | 0 | 0 | 0 | 0 | 0 | 0 |
| ENSMUSG00 | SNORA17 | 130 | 1.30785157 | 0 | 0 | 0.35046957 | 0 | 0.76234975 | 0 | 0 | 0 | 1.45079087 | 1.10249604 | 1.28805375 | 5.6921443 | 0.35726221 | 3.80322054 | 0 | 0.88445301 | 0 |
| ENSMUSG00 | SNORA17 | 135 | 0 | 0 | 0 | 0 | 0 | 0 | 0 | 0 | 0 | 0 | 0 | 0 | 0 | 0 | 0 | 0 | 0 | 0 |
| ENSMUSG00 | U6 | 95 | 0 | 0 | 0 | 0 | 0 | 0 | 0 | 0 | 0 | 0 | 0 | 0 | 0 | 0 | 0 | 0 | 0 | 0 |
| ENSMUSG00 | Snord71 | 84 | 1.012028 | 0.39658207 | 0 | 0 | 0 | 0 | 0 | 0 | 0 | 0 | 0 | 0 | 0 | 0.5529058 | 0.49049471 | 0 | 0.91253088 | 0 |
| ENSMUSG00 | CT009748.1 | 109 | 0 | 0 | 0 | 0 | 0 | 0 | 0 | 0 | 0 | 0 | 0 | 0 | 0 | 0 | 0 | 0 | 0 | 0 |
| ENSMUSG00 | SNORA17 | 129 | 0.65899498 | 0 | 0 | 0 | 1.49246275 | 0.76825944 | 0 | 0 | 0 | 0 | 0 | 0 | 0 | 0 | 0 | 0 | 0 | 0 |
| ENSMUSG00 | SNORA17 | 129 | 0 | 0 | 0 | 0 | 0 | 0 | 0 | 0 | 0 | 0 | 0 | 0 | 0 | 0 | 0 | 0 | 0 | 0 |
| ENSMUSG00 | SNORA17 | 124 | 0 | 0 | 0 | 0 | 0 | 0 | 0 | 0 | 0 | 0 | 0 | 0 | 0 | 0 | 0 | 0 | 0 | 0 |
| ENSMUSG00 | SNORA17 | 122 | 0 | 0 | 0 | 0 | 0 | 0 | 0 | 0 | 0 | 0 | 0 | 0 | 0 | 0 | 0 | 0 | 0 | 0 |
| ENSMUSG00 | SNORD86 | 81 | 0 | 0.4112703 | 1.19440816 | 0 | 1.18844256 | 1.22352429 | 1.16821429 | 0.47151114 | 1.01695101 | 0.58210745 | 1.1796286 | 2.75632902 | 0 | 0 | 0 | 0 | 0 | 0 |
| ENSMUSG00 | SNORA17 | 129 | 0 | 0 | 0 | 0 | 0 | 0 | 0 | 0 | 0 | 0 | 0 | 0 | 0 | 0 | 0 | 0 | 0 | 0 |
| ENSMUSG00 | SNORA17 | 131 | 0 | 0 | 0 | 0 | 0 | 0 | 0 | 0 | 0 | 0 | 0 | 0 | 0 | 0 | 0 | 0 | 0 | 0 |
| ENSMUSG00 | SNORA17 | 132 | 0 | 0.50474082 | 0 | 0 | 0 | 0 | 0.35842939 | 0 | 0 | 0 | 0 | 0 | 0 | 0 | 0 | 0 | 0 | 0 |
| ENSMUSG00 | U6 | 106 | 0 | 0 | 0 | 0 | 0 | 0 | 0 | 0 | 0 | 0 | 0 | 0 | 0 | 0 | 0 | 0 | 0 | 0 |
| ENSMUSG00 | Snora68 | 130 | 5.88533207 | 3.84379549 | 4.09314488 | 5.95798272 | 3.3322101 | 4.95527336 | 3.63943684 | 7.34469277 | 7.9204838 | 3.62697717 | 4.77749585 | 1.717405 | 11.3842886 | 7.50256642 | 10.1419215 | 7.65812148 | 7.96007705 | 13.1175562 |
| ENSMUSG00 | SNORA27 | 128 | 1.32828675 | 0 | 0.37791821 | 0 | 0 | 0 | 0 | 0 | 0 | 0 | 0 | 0 | 0.3854056 | 0.36284443 | 0 | 0 | 0 | 0 |
| ENSMUSG00 | SNORD12 | 88 | 0.48301336 | 0.75711123 | 0 | 0.51773914 | 2.73476839 | 0.56309925 | 0 | 4.34004573 | 5.14831447 | 0.53580345 | 1.62869177 | 1.26853779 | 0 | 1.05554744 | 4.21379549 | 0.43512054 | 2.61315661 | 0.62510349 |
| ENSMUSG00 | SNORA17 | 131 | 0 | 0 | 0 | 0 | 0 | 0 | 0 | 0 | 0 | 0 | 0 | 0 | 0 | 0 | 0 | 0 | 0 | 0 |
| ENSMUSG00 | SNORA58 | 135 | 1.57426578 | 3.94819487 | 2.14993468 | 2.36242452 | 3.20879491 | 1.10117186 | 1.75232144 | 4.24360027 | 3.96610893 | 1.74632234 | 1.06166574 | 6.20174029 | 2.19252966 | 1.72015138 | 0.61039342 | 0 | 2.83898496 | 3.25979892 |
| ENSMUSG00 | SNORA17 | 131 | 0 | 0 | 0 | 0 | 0 | 0 | 0 | 0 | 0 | 0 | 0 | 0 | 0 | 0 | 0 | 0 | 0 | 0 |
| ENSMUSG00 | Snora36b | 131 | 0 | 0 | 0 | 0 | 0 | 0 | 0 | 0 | 0 | 0 | 0 | 0 | 0 | 0 | 0 | 0 | 0 | 0 |
| ENSMUSG00 | SNORA17 | 122 | 0 | 0 | 0 | 0 | 0 | 0 | 0 | 0 | 0 | 0 | 0 | 0 | 0 | 0 | 0 | 0 | 0 | 0 |
| ENSMUSG00 | SNORA17 | 131 | 0 | 0 | 0 | 0 | 0 | 0 | 0 | 0 | 0 | 0 | 0 | 0 | 0 | 0 | 0 | 0 | 0 | 0 |
| ENSMUSG00 | U6 | 107 | 0 | 0 | 0 | 0 | 0 | 0 | 0 | 0 | 0 | 0 | 0 | 0 | 0 | 0 | 0 | 0 | 0 | 0 |
| ENSMUSG00 | SNORA3 | 129 | 2.9654774 | 3.87359235 | 1.87494304 | 3.17867751 | 2.23869412 | 1.92064859 | 5.86823925 | 3.25671648 | 0.95782595 | 3.65509328 | 6.68627328 | 1.73071822 | 1.52967185 | 6.84060202 | 4.15209478 | 3.56191697 | 5.34785538 | 6.82283494 |
| ENSMUSG00 | SNORA17 | 131 | 0 | 0 | 0 | 0 | 0 | 0 | 0 | 0 | 0 | 0 | 0 | 0 | 0 | 0 | 0 | 0 | 0 | 0 |
| ENSMUSG00 | U11 | 101 | 0 | 0 | 0 | 0 | 0 | 0 | 0 | 0 | 0 | 0 | 0 | 0 | 0 | 0 | 0 | 0 | 0 | 0 |
| ENSMUSG00 | SNORA17 | 129 | 0 | 0 | 0 | 0 | 0 | 0 | 0 | 0.59213027 | 0.31927532 | 0 | 0 | 0 | 0 | 0 | 0 | 0 | 0 | 0 |
| ENSMUSG00 | SNORA17 | 135 | 0 | 0 | 0 | 0 | 0 | 0 | 0 | 0 | 0 | 0 | 0 | 0 | 0 | 0 | 0 | 0 | 0 | 0 |
| ENSMUSG00 | SNORA17 | 131 | 0 | 0 | 0 | 0 | 0 | 0 | 0 | 0 | 0 | 0 | 0 | 0 | 0 | 0 | 0 | 0 | 0 | 0 |
| ENSMUSG00 | U6 | 105 | 0 | 0 | 0 | 0 | 0 | 0 | 0 | 0 | 0 | 0 | 0 | 0 | 0 | 0 | 0 | 0 | 0 | 0 |
| ENSMUSG00 | SNORA17 | 131 | 0 | 0 | 0 | 0 | 0 | 0 | 0 | 0 | 0 | 0 | 0 | 0 | 0 | 0 | 0 | 0 | 0 | 0 |
| ENSMUSG00 | SNORA17 | 130 | 0 | 0 | 0 | 0 | 0 | 0 | 0 | 0 | 0 | 0 | 0 | 0 | 0 | 0 | 0 | 0 | 0 | 0 |
| ENSMUSG00 | SNORA17 | 126 | 0 | 0 | 0 | 0 | 0 | 0 | 0 | 0 | 0 | 0 | 0 | 0 | 0 | 0 | 0 | 0 | 0 | 0 |
| ENSMUSG00 | Y_RNA | 109 | 0 | 0 | 0 | 0 | 0 | 0 | 0 | 0 | 0 | 0 | 0.43830237 | 0 | 0 | 0 | 0 | 0 | 0 | 0 |
| ENSMUSG00 | SNORA17 | 131 | 0 | 0 | 0 | 0 | 0 | 0 | 0 | 0 | 0 | 0 | 0 | 0 | 0 | 0 | 0 | 0 | 0 | 0 |
| ENSMUSG00 | SNORA17 | 124 | 0 | 0 | 0 | 0 | 0 | 0 | 0 | 0 | 0 | 0 | 0 | 0 | 0 | 0 | 0 | 0 | 0 | 0 |
| ENSMUSG00 | SNORA17 | 128 | 0 | 0 | 0 | 0 | 0 | 0 | 0 | 0 | 0 | 0 | 0 | 0 | 0 | 0 | 0 | 0 | 0 | 0 |
| ENSMUSG00 | SNORA17 | 118 | 0 | 0 | 0 | 0.38611055 | 0 | 0 | 0 | 0 | 0.69807654 | 0 | 0 | 0 | 0 | 0 | 0.34916573 | 0 | 0 | 0 |
| ENSMUSG00 | SNORA17 | 131 | 0 | 0.2542969 | 1.10779077 | 0 | 0.36741926 | 0 | 0 | 0.29154506 | 0.62880177 | 0 | 0 | 0.42607376 | 0 | 1.41814007 | 0.62903139 | 0 | 0.29256715 | 0 |
| ENSMUSG00 | SNORA17 | 127 | 0 | 0 | 0 | 0 | 0 | 0 | 0 | 0 | 0 | 0 | 0 | 0 | 0 | 0 | 0 | 0 | 0 | 0 |
| ENSMUSG00 | SNORA17 | 118 | 0 | 0 | 0 | 0 | 0 | 0 | 0 | 0 | 0 | 0 | 0 | 0 | 0 | 0 | 0 | 0 | 0 | 0 |
| ENSMUSG00 | U6 | 106 | 0 | 0 | 0 | 0 | 0 | 0 | 0 | 0 | 0 | 0 | 0 | 0 | 0 | 0 | 0 | 0 | 0 | 0 |
| ENSMUSG00 | U6 | 73 | 0 | 0 | 0 | 0 | 0 | 0 | 0 | 0 | 0 | 0 | 0 | 0 | 0 | 0 | 0 | 0 | 0 | 0 |
| ENSMUSG00 | U6 | 103 | 0 | 0 | 0 | 0 | 0 | 0 | 0 | 0 | 0 | 0 | 0 | 0 | 0 | 0 | 0 | 0 | 0 | 0 |
| ENSMUSG00 | SNORA17 | 127 | 0 | 0 | 0 | 0 | 0 | 0 | 0 | 0 | 0 | 0 | 0 | 0 | 0 | 0.36570148 | 0 | 0 | 0 | 0 |
| ENSMUSG00 | SNORA17 | 130 | 0 | 0 | 0 | 0 | 0 | 0 | 0 | 0 | 0 | 0 | 0 | 0 | 0 | 0 | 0 | 0 | 0 | 0 |
| ENSMUSG00 | SNORA17 | 133 | 0.31958779 | 0.75141867 | 0.72742151 | 0 | 0 | 0.74515389 | 0.35573443 | 0.28716092 | 0.30967305 | 0.35451657 | 0.71842043 | 1.67866654 | 0 | 0 | 0.61957227 | 3.16689234 | 0 | 0.82720461 |
| ENSMUSG00 | SNORA3 | 130 | 0 | 0 | 0 | 0 | 0 | 0 | 0 | 0.29378771 | 0 | 0 | 0 | 0 | 0 | 0 | 0 | 0 | 0 | 0 |
| ENSMUSG00 | Snord1b | 74 | 0 | 0 | 0 | 0 | 0 | 0 | 0 | 0 | 0 | 0 | 0 | 0 | 0 | 0 | 0 | 0 | 0 | 0 |
| ENSMUSG00 | SNORA17 | 131 | 0 | 0 | 0 | 0 | 0 | 0 | 0 | 0 | 0 | 0 | 0 | 0 | 0 | 0 | 0 | 0 | 0 | 0 |
| ENSMUSG00 | SNORA17 | 130 | 0 | 0 | 0 | 0 | 0 | 0 | 0 | 0 | 0 | 0 | 0 | 0 | 0 | 0 | 0 | 0 | 0 | 0 |
| ENSMUSG00 | SNORA17 | 133 | 0 | 0 | 0 | 0 | 0 | 0 | 0 | 0 | 0 | 0 | 0 | 0 | 0 | 0 | 0 | 0 | 0 | 0 |
| ENSMUSG00 | SNORA17 | 131 | 0 | 0 | 0 | 0 | 0 | 0 | 0 | 0 | 0 | 0 | 0 | 0 | 0 | 0 | 0 | 0 | 0 | 0 |
| ENSMUSG00 | SNORA74 | 196 | 0 | 0 | 0 | 0 | 0 | 0 | 0 | 0 | 1.47094699 | 0 | 0 | 0.85432137 | 0.50338691 | 0 | 1.89190818 | 0.78144097 | 0.78216932 | 0 |
| ENSMUSG00 | SNORA17 | 128 | 0 | 0 | 0 | 0 | 0 | 0 | 0 | 0 | 0 | 0 | 0 | 0 | 0 | 0 | 0 | 0 | 0 | 0 |
| ENSMUSG00 | SNORA17 | 131 | 0 | 0 | 0 | 0 | 0 | 0 | 0 | 0 | 0 | 0 | 0 | 0 | 0 | 0 | 0 | 0 | 0 | 0 |
| ENSMUSG00 | SNORA40 | 122 | 2.43882158 | 0.81916953 | 0 | 2.2407071 | 0.39452396 | 0 | 0 | 0 | 1.68797196 | 0.39648117 | 0.78319604 | 0 | 0.80871996 | 0 | 0.33771767 | 1.56928719 | 1.2565999 | 3.15626022 |
| ENSMUSG00 | SNORD115 | 68 | 0 | 0 | 0 | 0 | 0 | 0 | 0 | 0 | 0 | 0 | 0 | 0 | 0 | 0 | 0 | 0 | 0 | 0 |
| ENSMUSG00 | SNORA17 | 130 | 0 | 0 | 0 | 0 | 0 | 0 | 0 | 0 | 0 | 0 | 0 | 0 | 0 | 0 | 0 | 0 | 0 | 0 |
| ENSMUSG00 | SNORA17 | 131 | 0 | 0 | 0 | 0 | 0 | 0 | 0 | 1.74927034 | 0 | 0 | 0 | 0 | 0 | 0 | 0 | 0 | 0 | 0 |
| ENSMUSG00 | SNORA17 | 119 | 0 | 0 | 0 | 0 | 0 | 0 | 0 | 0 | 0 | 0 | 0 | 0 | 0 | 0 | 0 | 0 | 0 | 0 |
| ENSMUSG00 | SNORA17 | 130 | 0 | 0 | 0 | 0 | 0 | 0 | 0 | 0 | 0 | 0 | 0 | 0 | 0 | 0 | 0 | 0 | 0 | 0 |
| ENSMUSG00 | n-R5s9 | 116 | 0 | 0 | 0 | 0 | 0 | 0 | 0 | 0 | 0 | 0 | 0 | 0 | 0 | 0 | 0 | 0 | 0 | 0 |
| ENSMUSG00 | SNORA63 | 101 | 0 | 0 | 0 | 0 | 0 | 0 | 0 | 0 | 0 | 0.46683865 | 0 | 0 | 0 | 0 | 0 | 0 | 0.37946829 | 0 |
| ENSMUSG00 | SNORA61 | 128 | 0 | 0 | 0 | 0 | 0 | 0 | 0 | 0 | 0 | 0 | 0 | 0 | 0 | 0 | 0 | 0 | 0 | 0 |
| ENSMUSG00 | SNORA17 | 128 | 0 | 0 | 0 | 0 | 0 | 0 | 0 | 0 | 0 | 0 | 0 | 0 | 0 | 0 | 0 | 0 | 0 | 0 |
| ENSMUSG00 | Snord4a | 70 | 0 | 0 | 0 | 0 | 0 | 0 | 0 | 0 | 0 | 0 | 0 | 0 | 0 | 0 | 0 | 0 | 0 | 0 |
| ENSMUSG00 | SNORA17 | 131 | 0 | 0 | 0 | 0 | 0 | 0 | 0 | 0 | 0 | 0 | 0 | 0 | 0 | 0 | 0 | 0 | 0 | 0 |
| ENSMUSG00 | SNORA17 | 130 | 0 | 0 | 0 | 0 | 0 | 0 | 0 | 0 | 0 | 0 | 0 | 0 | 0 | 0 | 0 | 0 | 0 | 0 |
| ENSMUSG00 | SNORA17 | 120 | 0 | 0 | 0 | 0 | 0 | 0 | 0 | 0 | 0 | 0 | 0 | 0 | 0 | 0 | 0 | 0 | 0 | 0 |
| ENSMUSG00 | SNORA17 | 135 | 0 | 0.49352436 | 0 | 0 | 0.35653277 | 0 | 0 | 0.28290668 | 0 | 0 | 0 | 0 | 0.36542161 | 0.68806055 | 0.30519671 | 0.28363413 | 0 | 0 |
| ENSMUSG00 | SNORA17 | 133 | 0 | 0 | 0 | 0 | 0 | 0 | 0 | 0 | 0 | 0 | 0 | 0 | 0 | 0 | 0 | 0 | 0 | 0 |
| ENSMUSG00 | U6 | 106 | 0 | 0.31427259 | 0 | 0 | 0 | 0 | 0 | 0 | 0.38855204 | 0 | 0 | 0 | 0 | 0 | 0 | 0 | 0 | 0 |
| ENSMUSG00 | SNORA17 | 134 | 0 | 0 | 0 | 0 | 0 | 0 | 0 | 0 | 0 | 0 | 0 | 0 | 0 | 0 | 0 | 0 | 0 | 0 |
| ENSMUSG00 | SNORD91 | 90 | 1.4168392 | 0.74028654 | 2.14993468 | 1.01246765 | 1.60439746 | 0.55058593 | 1.05139287 | 2.54616016 | 4.11865158 | 4.71507033 | 0.53083287 | 4.3412182 | 3.28879449 | 0 | 0 | 1.70180477 | 0.85169549 | 1.22242459 |
| ENSMUSG00 | SNORA33 | 120 | 0 | 0 | 0 | 0.37967537 | 0 | 0 | 0 | 0.63654004 | 0 | 0 | 0 | 0 | 0 | 0.38703406 | 1.7167315 | 0 | 0 | 0 |
| ENSMUSG00 | SNORA17 | 131 | 0 | 0 | 0 | 0 | 0 | 0 | 0 | 0 | 0 | 0 | 0 | 0 | 0 | 0 | 0 | 0 | 0 | 0 |
| ENSMUSG00 | SNORA17 | 133 | 0 | 0 | 0 | 0 | 0 | 0 | 0 | 0 | 0 | 0 | 0 | 0 | 0 | 0 | 0 | 0 | 0 | 0 |
| ENSMUSG00 | SNORA17 | 132 | 0 | 0 | 0 | 0 | 0 | 0 | 0 | 0 | 0 | 0 | 0 | 0 | 0 | 0 | 0 | 0 | 0 | 0 |
| ENSMUSG00 | SNORA17 | 128 | 0 | 0 | 0 | 0 | 0 | 0 | 0 | 0 | 0 | 0 | 0 | 0 | 0 | 0 | 0 | 0 | 0 | 0 |
| ENSMUSG00 | SNORA17 | 131 | 0 | 0 | 0 | 0 | 0 | 0 | 0 | 0 | 0 | 0 | 0 | 0 | 0 | 0 | 0 | 0 | 0 | 0 |
| ENSMUSG00 | SNORD115 | 68 | 0 | 0 | 0 | 0 | 0 | 0 | 0 | 0 | 0 | 0 | 0 | 0 | 0 | 0 | 0 | 0 | 0 | 0 |
| ENSMUSG00 | SNORA17 | 131 | 0 | 0 | 0 | 0 | 0 | 0 | 0 | 0 | 0 | 0 | 0 | 0 | 0 | 0 | 0 | 0 | 0 | 0 |
| ENSMUSG00 | SNORA26 | 120 | 0 | 0 | 0 | 0 | 0 | 0 | 0 | 0 | 0 | 0 | 0 | 0 | 0 | 0 | 0 | 0 | 0 | 0 |
| ENSMUSG00 | SNORD86 | 87 | 4.39708718 | 0.38290683 | 1.11203518 | 0 | 0 | 0 | 0.5438239 | 0 | 3.31385759 | 0 | 0 | 0 | 0 | 0 | 0 | 0.44012192 | 1.76212859 | 2.52915433 |
| ENSMUSG00 | SNORA40 | 126 | 0 | 0 | 0 | 0 | 0 | 0 | 0.37549745 | 0 | 0 | 0 | 0 | 0 | 0 | 0 | 0 | 0 | 0 | 0 |
| ENSMUSG00 | U1 | 166 | 0 | 0 | 0 | 0 | 0 | 0 | 0 | 0.46014943 | 0 | 0 | 0 | 0 | 0 | 0 | 0 | 0 | 0 | 0 |
| ENSMUSG00 | SNORA17 | 125 | 0 | 0 | 0 | 0 | 0 | 0 | 0 | 0 | 0 | 0 | 0 | 0 | 0 | 0 | 0 | 0 | 0 | 0 |
| ENSMUSG00 | SNORA17 | 130 | 0 | 0 | 0 | 0 | 0 | 0 | 0 | 0 | 0 | 0 | 0 | 0 | 0 | 0 | 0 | 0 | 0 | 0 |
| ENSMUSG00 | SNORA17 | 132 | 0 | 0 | 0 | 0 | 0 | 0 | 0 | 0 | 0 | 0 | 0 | 0 | 0 | 0 | 0 | 0 | 0 | 0 |
| ENSMUSG00 | SNORA17 | 130 | 0 | 0 | 0 | 0 | 0 | 0 | 0 | 0 | 0 | 0 | 0 | 0 | 0 | 0 | 0 | 0 | 0 | 0 |
| ENSMUSG00 | n-R5s51 | 122 | 0 | 0 | 0 | 0 | 0 | 0 | 0 | 0 | 0 | 0 | 0 | 0 | 0 | 0 | 0 | 0 | 0 | 0 |
| ENSMUSG00 | SNORA17 | 131 | 0 | 0 | 0 | 0 | 0 | 0 | 0 | 0 | 0 | 0 | 0 | 0 | 0 | 0 | 0 | 0 | 0 | 0 |
| ENSMUSG00 | SNORA17 | 126 | 0 | 0 | 0 | 0 | 0 | 0 | 0 | 0 | 0 | 0 | 0 | 0 | 0 | 0 | 0 | 0 | 0 | 0 |
| ENSMUSG00 | U1 | 184 | 0 | 0 | 0 | 0 | 0 | 0 | 0 | 0 | 0 | 0 | 0 | 0 | 0 | 0 | 0.2239215 | 0 | 0 | 0 |
| ENSMUSG00 | SNORA24 | 113 | 0 | 0.29480437 | 0 | 0 | 0 | 0 | 0 | 0 | 0 | 0 | 0 | 0 | 0 | 0 | 0 | 0 | 0.67834154 | 0 |
| ENSMUSG00 | U6 | 107 | 0 | 0 | 0 | 0 | 0 | 0 | 0 | 0 | 0 | 0 | 0 | 0 | 0 | 0 | 0 | 0 | 0 | 0 |
| ENSMUSG00 | SNORA17 | 127 | 0 | 0 | 0 | 0 | 0 | 0 | 0 | 0 | 0 | 0 | 0 | 0 | 0 | 0 | 0 | 0 | 0 | 0 |
| ENSMUSG00 | SNORA17 | 130 | 0 | 0 | 0 | 0 | 0 | 0 | 0 | 0 | 0 | 0 | 0 | 0 | 0 | 0 | 0 | 0 | 0 | 0 |
| ENSMUSG00 | SNORA17 | 131 | 0 | 0 | 0 | 0 | 0 | 0 | 0 | 0 | 0 | 0 | 0 | 0 | 0 | 0 | 0 | 0 | 0 | 0 |
| ENSMUSG00 | SNORA17 | 130 | 0 | 0 | 0 | 0 | 0 | 0 | 0 | 0 | 0 | 0 | 0 | 0 | 0 | 0 | 0 | 0 | 0 | 0 |
| ENSMUSG00 | SNORA17 | 131 | 0 | 0 | 0 | 0 | 0 | 0 | 0 | 0 | 0 | 0 | 0 | 0 | 0 | 0 | 0 | 0 | 0 | 0 |
| ENSMUSG00 | SNORA17 | 130 | 0 | 0 | 0 | 0 | 0 | 0 | 0 | 0 | 0 | 0 | 0 | 0 | 1.13842886 | 0.35726221 | 0 | 0 | 0 | 0 |
| ENSMUSG00 | U6 | 103 | 0.82534322 | 0 | 0 | 0 | 0 | 0 | 0 | 0 | 0 | 0 | 0.9276691 | 0 | 0 | 0 | 0 | 0.37175347 | 0 | 0 |
| ENSMUSG00 | SNORA17 | 130 | 0 | 0 | 0 | 0 | 0 | 0 | 0 | 0 | 0 | 0 | 0 | 0 | 0 | 0 | 0 | 0 | 0 | 0 |
| ENSMUSG00 | SNORA17 | 125 | 0 | 0 | 0 | 0 | 0 | 0 | 0 | 0 | 0 | 0 | 0 | 0 | 0 | 0 | 0 | 0 | 0 | 0 |
| ENSMUSG00 | SNORD115 | 87 | 0 | 0 | 0 | 0 | 0 | 0 | 0 | 0 | 0 | 0 | 0 | 0 | 0 | 0 | 0.9471622 | 0 | 0 | 0 |
| ENSMUSG00 | SNORA17 | 131 | 0 | 0 | 0 | 0 | 0 | 0 | 0 | 0 | 0 | 0 | 0 | 0 | 0 | 0 | 0 | 0 | 0 | 0 |
| ENSMUSG00 | SNORA17 | 127 | 0 | 0 | 0 | 0 | 0 | 0 | 0 | 0 | 0 | 0 | 0 | 0 | 0 | 0 | 0 | 0 | 0 | 0 |
| ENSMUSG00 | SNORD86 | 85 | 0 | 0 | 0 | 0 | 0 | 0 | 0 | 0 | 0 | 0 | 0 | 0 | 0 | 0 | 0 | 0 | 0 | 0 |
| ENSMUSG00 | SNORA17 | 131 | 0.97340098 | 0.5085938 | 0.36926359 | 0.34779423 | 0 | 0.75653028 | 0.36116549 | 0 | 0 | 0.71985806 | 2.18816604 | 0.42607376 | 0.75315904 | 0 | 0.62903139 | 0.29229471 | 0.29256715 | 0.83983369 |
| ENSMUSG00 | SNORA17 | 130 | 0 | 0 | 0 | 0 | 0 | 0 | 0 | 0 | 0 | 0 | 0 | 0 | 0 | 0 | 0 | 0 | 0 | 0 |
| ENSMUSG00 | SNORA17 | 134 | 1.26881122 | 0.24860369 | 0 | 1.70003897 | 0 | 1.10938956 | 0 | 0 | 0.92208617 | 0 | 0 | 0 | 0 | 0.34659767 | 0 | 1.14300321 | 0 | 0 |
| ENSMUSG00 | SNORA17 | 131 | 0 | 0 | 0 | 0 | 0 | 0 | 0 | 0 | 0 | 0 | 0 | 0 | 0 | 0 | 0 | 0 | 0 | 0 |
| ENSMUSG00 | SNORA17 | 131 | 0 | 0 | 0 | 0 | 0 | 0 | 0 | 0 | 0 | 0 | 0 | 0 | 0 | 0 | 0 | 0 | 0 | 0 |
| ENSMUSG00 | SNORA17 | 132 | 0 | 0 | 0 | 0 | 0 | 0 | 0 | 0 | 0 | 0 | 0 | 0 | 0 | 0 | 0 | 0 | 0 | 0 |
| ENSMUSG00 | SNORA17 | 131 | 0 | 0 | 0 | 0 | 0 | 0 | 0 | 0 | 0 | 0 | 0 | 0 | 0 | 0 | 0 | 0 | 0 | 0 |
| ENSMUSG00 | n-R5s172 | 113 | 0 | 0 | 0 | 0 | 0 | 0 | 0 | 0 | 0 | 0 | 0 | 0 | 0 | 0 | 0 | 0 | 0 | 0 |
| ENSMUSG00 | SNORA17 | 127 | 0 | 0 | 0.38089394 | 0 | 0 | 0 | 0 | 0 | 0 | 0 | 0 | 0 | 0 | 0 | 0 | 0 | 0 | 0 |
| ENSMUSG00 | Snord12 | 89 | 0 | 0 | 0 | 0 | 0 | 0 | 0 | 0 | 0 | 0 | 0 | 0 | 0 | 0 | 0 | 0 | 0 | 0 |
| ENSMUSG00 | SNORA17 | 126 | 0 | 0 | 0 | 0 | 0 | 0 | 0 | 0 | 0 | 0 | 0 | 0 | 0 | 0 | 0 | 0 | 0 | 0 |
| ENSMUSG00 | SNORA17 | 131 | 0 | 0 | 0 | 0 | 0 | 0 | 0 | 0 | 0 | 0 | 0 | 0 | 0 | 0 | 0 | 0 | 0 | 0 |
| ENSMUSG00 | SNORD105 | 84 | 0 | 0 | 0 | 0 | 0 | 0 | 0 | 0 | 0 | 0 | 0 | 0 | 0 | 0 | 0 | 0 | 0 | 0 |
| ENSMUSG00 | SNORA17 | 130 | 0 | 0 | 0 | 0 | 0 | 0 | 0 | 0 | 0 | 0 | 0 | 0 | 0 | 0 | 0 | 0 | 0 | 0 |
| ENSMUSG00 | Snord89 | 109 | 5.06942466 | 1.22249153 | 3.55035085 | 1.2539737 | 2.20788641 | 0.90922447 | 0 | 3.50389013 | 0.75571589 | 0.86515052 | 1.75320949 | 0 | 3.62069118 | 0 | 0.37799593 | 3.16160979 | 1.05485221 | 2.52335352 |
| ENSMUSG00 | n-R5s202 | 113 | 0 | 0 | 0 | 0 | 0 | 0 | 0 | 0 | 0 | 0 | 0 | 0 | 0 | 0 | 0 | 0 | 0 | 0 |
| ENSMUSG00 | n-R5s72 | 118 | 0 | 0 | 0 | 0 | 0 | 0 | 0 | 0 | 0 | 0 | 0 | 0 | 0 | 0 | 0 | 0 | 0 | 0 |
| ENSMUSG00 | SNORA17 | 129 | 0 | 0 | 0 | 0 | 0 | 0 | 0 | 0 | 0 | 0 | 0 | 0 | 0 | 0 | 0 | 0 | 0 | 0 |
| ENSMUSG00 | SNORA17 | 131 | 0 | 0 | 0 | 0 | 0 | 0 | 0 | 0 | 0 | 0 | 0 | 0 | 0 | 0 | 0 | 0 | 0 | 0 |
| ENSMUSG00 | Snora64 | 133 | 0 | 0 | 0 | 0 | 0 | 0 | 0 | 0 | 0 | 0 | 0 | 0 | 0 | 0 | 0 | 0 | 0 | 0 |
| ENSMUSG00 | SNORA17 | 130 | 0 | 0 | 0 | 0 | 0 | 0 | 0 | 0 | 0 | 0 | 0 | 0 | 0 | 0 | 0 | 0 | 0 | 0 |
| ENSMUSG00 | SNORA40 | 124 | 2.74226942 | 1.07460949 | 1.17032735 | 2.57299444 | 3.88160675 | 3.59656937 | 3.0524309 | 0.30800325 | 0 | 3.42222846 | 0.77056385 | 2.70075787 | 0.39783804 | 3.74549092 | 0.33227061 | 1.5439761 | 0.30908304 | 0 |
| ENSMUSG00 | SNORA17 | 131 | 0 | 0 | 0 | 0 | 0 | 0 | 0 | 0 | 0 | 0 | 0 | 0 | 0 | 0 | 0 | 0 | 0 | 0 |
| ENSMUSG00 | U1 | 158 | 0 | 0 | 0 | 0 | 0 | 0 | 0 | 0 | 0 | 0 | 0 | 0 | 0 | 0 | 0 | 0 | 0 | 0 |
| ENSMUSG00 | Snord17 | 237 | 0.17934673 | 1.12448588 | 1.42875406 | 0.57672208 | 1.421618 | 1.04541632 | 0.79852623 | 0.16114938 | 0.86891383 | 1.59158492 | 2.62056734 | 1.884073 | 0.62445465 | 0.58789984 | 1.73846227 | 0.64625498 | 0.80857166 | 1.39263561 |
| ENSMUSG00 | SNORA17 | 129 | 0 | 0 | 0 | 0 | 0 | 0 | 0 | 0 | 0 | 0 | 0 | 0 | 0 | 0 | 0 | 0 | 0 | 0 |
| ENSMUSG00 | SNORA17 | 128 | 0 | 0.52051397 | 0 | 0 | 0 | 0 | 0 | 0 | 0 | 0 | 0 | 0 | 0 | 0 | 0 | 0 | 0.29942419 | 0 |
| ENSMUSG00 | SNORA17 | 125 | 0 | 0 | 0 | 0 | 0 | 0 | 0 | 0 | 0 | 0 | 0 | 0 | 0 | 0 | 0 | 0 | 0 | 0 |
| ENSMUSG00 | SNORA17 | 116 | 0 | 0 | 0 | 0 | 0 | 0 | 0 | 0 | 0 | 0 | 0 | 0 | 0 | 0 | 0 | 0 | 0 | 0 |
| ENSMUSG00 | SNORA17 | 129 | 0 | 0 | 0 | 0 | 0 | 0 | 0 | 0 | 0 | 0 | 0 | 0 | 0 | 0 | 0 | 0 | 0 | 0 |
| ENSMUSG00 | SNORA17 | 130 | 0 | 0 | 0 | 0 | 0 | 0 | 0 | 0 | 0 | 0 | 0 | 0 | 0 | 0 | 0 | 0 | 0 | 0 |
| ENSMUSG00 | SNORA17 | 126 | 0 | 0 | 0 | 0 | 0 | 0 | 0 | 0 | 0 | 0 | 0 | 0 | 0 | 0 | 0 | 0 | 0 | 0 |
| ENSMUSG00 | SNORA17 | 127 | 0 | 0 | 0 | 0 | 0 | 0 | 0 | 0 | 0 | 1.48506152 | 0 | 0 | 0 | 0 | 1.2976868 | 0 | 0.60356373 | 0.43314257 |
| ENSMUSG00 | Snord83b | 93 | 2.74226942 | 0.71640633 | 1.04029098 | 1.95961481 | 3.1052854 | 0 | 6.61360028 | 0.82134199 | 2.2143288 | 3.04198086 | 2.56854615 | 1.20033683 | 4.77405651 | 0 | 7.97449469 | 2.05863481 | 3.29688576 | 0.59149577 |
| ENSMUSG00 | SNORA17 | 129 | 0 | 0 | 0 | 0 | 0 | 0 | 0 | 0 | 0 | 0 | 0 | 0 | 0 | 0 | 0 | 0 | 0 | 0 |
| ENSMUSG00 | SNORA17 | 123 | 0 | 0 | 0 | 0 | 0 | 0 | 0 | 0 | 0 | 0 | 0 | 0 | 0 | 0 | 0 | 0 | 0 | 0 |
| ENSMUSG00 | Snord72 | 80 | 7.96972051 | 12.0759242 | 4.83735304 | 10.820748 | 9.62638473 | 11.1493651 | 8.8711273 | 14.7995559 | 24.7119095 | 10.6089082 | 9.5549917 | 7.67465361 | 8.01643656 | 27.2859013 | 14.935564 | 13.4017126 | 16.2886762 | 17.8779597 |
| ENSMUSG00 | SNORA17 | 125 | 0 | 0 | 0 | 0 | 0 | 0 | 0 | 0 | 0 | 0 | 0 | 0 | 0 | 0 | 0 | 0 | 0 | 0 |
| ENSMUSG00 | SNORA17 | 131 | 0 | 0 | 0 | 0 | 0 | 0 | 0 | 0 | 0 | 0 | 0 | 0 | 0 | 0 | 0 | 0 | 0 | 0 |
| ENSMUSG00 | n-R5s16 | 118 | 0 | 0 | 0 | 0 | 0 | 0 | 0 | 0 | 0 | 0 | 0 | 0 | 0 | 0 | 0 | 0 | 0 | 0 |
| ENSMUSG00 | SNORA17 | 130 | 0 | 0 | 0 | 0 | 0 | 0 | 0 | 0 | 0 | 0 | 0 | 0 | 0 | 0 | 0 | 0 | 0 | 0 |
| ENSMUSG00 | SNORA17 | 128 | 0 | 0 | 0 | 0 | 0 | 0 | 0 | 0 | 0 | 0 | 0 | 0 | 0 | 0 | 0 | 0 | 0 | 0 |
| ENSMUSG00 | SNORA17 | 133 | 0 | 0 | 0 | 0 | 0 | 0 | 0 | 0 | 0 | 0 | 0 | 0 | 0 | 0 | 0 | 0 | 0 | 0 |
| ENSMUSG00 | SNORA17 | 131 | 0 | 0 | 0 | 0 | 0 | 0 | 0 | 0 | 0 | 0 | 0 | 0 | 0 | 0 | 0 | 0 | 0 | 0 |
| ENSMUSG00 | SNORA17 | 131 | 0 | 0 | 0 | 0 | 0 | 0 | 0 | 0 | 0 | 0 | 0 | 0 | 0 | 0 | 0 | 0 | 0 | 0 |
| ENSMUSG00 | SNORA17 | 136 | 0 | 0 | 0 | 0 | 0 | 0 | 0 | 0 | 0 | 0 | 0 | 0 | 0 | 0 | 0 | 0 | 0 | 0 |
| ENSMUSG00 | SNORD95 | 67 | 0 | 0 | 0 | 0 | 0 | 0 | 0 | 0 | 0 | 0 | 0 | 0 | 0 | 0 | 0 | 0 | 0 | 0 |
| ENSMUSG00 | Snord90 | 110 | 0 | 0 | 0 | 0 | 0 | 0.4504794 | 0 | 0.34720366 | 0 | 0 | 0.4343178 | 0 | 0 | 0.84443795 | 0 | 0 | 1.04526264 | 0 |
| ENSMUSG00 | SNORA42 | 111 | 0 | 0 | 0 | 0 | 0 | 0 | 0 | 0 | 0 | 0 | 0 | 0 | 0 | 0 | 0 | 0 | 0 | 0 |
| ENSMUSG00 | SNORA17 | 129 | 0 | 0.77471847 | 0 | 2.11911834 | 0 | 0.76825944 | 0 | 2.66458621 | 2.87347784 | 0 | 1.11104555 | 0 | 0 | 0.72006337 | 1.59695953 | 0.89047924 | 0.29710308 | 0.42642718 |
| ENSMUSG00 | SNORA17 | 131 | 0 | 0 | 0 | 0 | 0 | 0 | 0 | 0 | 0 | 0 | 0 | 0 | 0 | 0 | 0 | 0 | 0 | 0 |
| ENSMUSG00 | SNORA17 | 128 | 0 | 0 | 0 | 0 | 0 | 0.77426146 | 0 | 0 | 0 | 0 | 0 | 0 | 0 | 0 | 0 | 0 | 0 | 0 |
| ENSMUSG00 | SNORA17 | 131 | 0 | 0 | 0 | 0 | 0 | 0 | 0 | 0 | 0 | 0 | 0 | 0 | 0 | 0 | 0 | 0 | 0 | 0 |
| ENSMUSG00 | Snora35 | 127 | 0 | 0 | 0 | 0 | 0 | 0 | 0 | 0 | 0 | 0 | 0 | 0 | 0 | 0 | 0 | 0 | 0 | 0 |
| ENSMUSG00 | SNORA17 | 125 | 0 | 0 | 0 | 0 | 0 | 0 | 0 | 0 | 0 | 0 | 0 | 0 | 0 | 0 | 0 | 0 | 0 | 0 |
| ENSMUSG00 | SNORA29 | 120 | 0 | 0 | 0 | 0 | 0 | 0 | 0 | 0 | 0 | 0 | 0 | 0 | 0 | 0 | 0 | 0 | 0 | 0 |
| ENSMUSG00 | SNORA17 | 131 | 0 | 0 | 0 | 0 | 0 | 0 | 0 | 0 | 0 | 0 | 0 | 0 | 0 | 0 | 0 | 0 | 0 | 0 |
| ENSMUSG00 | SNORD116 | 88 | 0 | 0 | 0 | 0 | 0 | 0 | 0 | 0 | 0 | 0 | 0 | 0 | 0 | 0 | 0 | 0 | 0 | 0 |
| ENSMUSG00 | SNORA17 | 131 | 0 | 0 | 0 | 0 | 0 | 0 | 0 | 0 | 0 | 0 | 0 | 0 | 0 | 0 | 0 | 0 | 0 | 0 |
| ENSMUSG00 | SNORA17 | 129 | 0 | 0 | 0 | 0 | 0 | 0 | 0 | 0 | 0 | 0 | 0 | 0 | 0 | 0 | 0 | 0 | 0 | 0 |
| ENSMUSG00 | SNORA17 | 135 | 0 | 0 | 0 | 0 | 0 | 0 | 0 | 0 | 0 | 0 | 0 | 0 | 0 | 0 | 0 | 0 | 0 | 0 |
| ENSMUSG00 | SNORA17 | 129 | 0 | 1.29119745 | 2.99990886 | 0 | 1.11934706 | 2.30477831 | 0 | 2.36852108 | 5.42768037 | 0 | 1.48139406 | 0 | 0.38241796 | 0 | 0.95817572 | 0 | 0.29710308 | 1.27928155 |
| ENSMUSG00 | SNORA17 | 125 | 0 | 0 | 0 | 0 | 0 | 0 | 0 | 0 | 0 | 0 | 0 | 0 | 0 | 0 | 0 | 0 | 0 | 0 |
| ENSMUSG00 | U2 | 188 | 0 | 0 | 0 | 0 | 0 | 0 | 0 | 0 | 0 | 0 | 0 | 0 | 0 | 0 | 0 | 0 | 0 | 0 |
| ENSMUSG00 | SNORA17 | 133 | 0 | 0 | 0 | 0 | 0 | 0 | 0 | 0 | 0 | 0 | 0 | 0 | 0 | 0 | 0 | 0 | 0 | 0 |
| ENSMUSG00 | SNORA17 | 131 | 2.59573594 | 0.5085938 | 0 | 1.04338269 | 1.10225779 | 1.51306057 | 2.16699293 | 2.33236045 | 2.51520707 | 1.0797871 | 0 | 1.27822128 | 5.2721133 | 2.48174513 | 0.94354708 | 2.63065242 | 0 | 1.25975054 |
| ENSMUSG00 | SNORA17 | 130 | 0 | 0 | 0 | 0 | 0 | 0 | 0 | 0 | 0 | 0 | 0 | 0 | 0 | 0 | 0 | 0 | 0 | 0 |
| ENSMUSG00 | SNORA17 | 131 | 0 | 0 | 0 | 1.39117693 | 0 | 0 | 0 | 0.29154506 | 0 | 0 | 0 | 0 | 0 | 0 | 0 | 0 | 0 | 0 |
| ENSMUSG00 | n-R5s26 | 121 | 0 | 0 | 0 | 0 | 0 | 0 | 0.39101388 | 0 | 1.02115328 | 0 | 0 | 0 | 0 | 0 | 0 | 0 | 0 | 0 |
| ENSMUSG00 | U6 | 73 | 0 | 0 | 0 | 0 | 0 | 0 | 0 | 0 | 0 | 0 | 0 | 0 | 0 | 0 | 0 | 0 | 0 | 0 |
| ENSMUSG00 | SNORA17 | 131 | 0 | 0 | 0 | 0 | 0 | 0 | 0 | 0 | 0 | 0 | 0 | 0 | 0 | 0 | 0 | 0 | 0 | 0 |
| ENSMUSG00 | U6 | 104 | 0 | 0 | 0 | 0 | 0 | 0 | 0 | 0 | 0 | 0 | 0 | 0 | 0 | 0 | 0 | 0 | 0 | 0 |
| ENSMUSG00 | U2 | 175 | 0 | 0 | 0 | 0 | 0 | 0 | 0 | 0 | 0 | 0 | 0 | 0 | 0 | 0 | 0 | 0 | 0 | 0 |
| ENSMUSG00 | SNORA17 | 124 | 0 | 0.26865237 | 0 | 0 | 0 | 0 | 0 | 0 | 0.66429864 | 2.28148564 | 0 | 0 | 0 | 0 | 0.66454122 | 0.30879522 | 0 | 0 |
| ENSMUSG00 | SNORA17 | 132 | 0 | 0 | 0 | 0 | 0 | 0 | 0 | 0 | 0 | 0 | 0 | 0 | 0 | 0 | 0 | 0 | 0 | 0 |
| ENSMUSG00 | Snord19 | 75 | 2.26694272 | 3.99754731 | 3.22490202 | 1.82244177 | 2.56703593 | 4.6249218 | 0.63083572 | 1.01846406 | 4.94238189 | 4.4007323 | 1.91099834 | 0 | 0 | 1.8577635 | 6.59224894 | 3.06324859 | 1.53305188 | 4.40072854 |
| ENSMUSG00 | SNORA17 | 131 | 0 | 0 | 0 | 0 | 0 | 0 | 0 | 0 | 1.57200442 | 0 | 0 | 0 | 0 | 0.70907004 | 0 | 0 | 0 | 0 |
| ENSMUSG00 | SNORA17 | 122 | 0 | 0 | 0 | 0 | 0 | 0 | 0 | 0 | 0 | 0 | 0 | 0 | 0 | 0 | 0 | 0 | 0 | 0 |
| ENSMUSG00 | n-R5s52 | 112 | 0 | 0 | 0 | 0.40679504 | 0 | 0 | 0 | 0 | 0.7354735 | 0.42098842 | 2.13281065 | 0 | 0 | 1.65871741 | 0 | 2.05128254 | 0.34219908 | 0 |
| ENSMUSG00 | SNORA17 | 129 | 0 | 0 | 0 | 0.70637278 | 0 | 0 | 0 | 1.18426054 | 0 | 0 | 0 | 0 | 0 | 0 | 0 | 0 | 0 | 0 |
| ENSMUSG00 | CR974466.1 | 88 | 0 | 0 | 0 | 0 | 0 | 0 | 0 | 0 | 0 | 0 | 0 | 0 | 0 | 0 | 0 | 0 | 0 | 0 |
| ENSMUSG00 | AC127299.1 | 83 | 0.51211055 | 0 | 0 | 0 | 0 | 0 | 0 | 1.84059771 | 0 | 0 | 0 | 0 | 0 | 0 | 0 | 0 | 0 | 0 |
| ENSMUSG00 | AC158665.1 | 92 | 0 | 0 | 0 | 0 | 0 | 0 | 0 | 0 | 0 | 0 | 0 | 0 | 0 | 0 | 0 | 0 | 0 | 0 |
| ENSMUSG00 | Mir290 | 82 | 0 | 0 | 0 | 0 | 0 | 0 | 0 | 0 | 0 | 0 | 0 | 0 | 0 | 0 | 0 | 0 | 0 | 0 |
| ENSMUSG00 | AC166933.1 | 93 | 0 | 0 | 0 | 0 | 0 | 0 | 0 | 0 | 0 | 0 | 0 | 0 | 0 | 0 | 0 | 0 | 0 | 0 |
| ENSMUSG00 | AC123803.1 | 84 | 0 | 0 | 0 | 0 | 0 | 0 | 0 | 0 | 0 | 0 | 0 | 0 | 0 | 0 | 0 | 0 | 0 | 0 |
| ENSMUSG00 | AC100600.1 | 93 | 0 | 0 | 0 | 0 | 0 | 0 | 0 | 0 | 0 | 0 | 0 | 0 | 0 | 0 | 0 | 0 | 0 | 0 |
| ENSMUSG00 | Mir465b-1 | 78 | 0 | 0 | 0 | 0 | 0 | 0 | 0 | 0 | 0 | 0 | 0 | 0 | 0 | 0 | 0 | 0 | 0 | 0 |
| ENSMUSG00 | AL593843.1 | 77 | 0 | 0 | 0 | 0 | 0 | 0 | 0 | 0 | 0 | 0 | 0 | 0 | 0 | 0 | 0 | 0 | 0 | 0 |
| ENSMUSG00 | AC107454.1 | 91 | 0.46708985 | 0 | 0 | 0 | 0 | 0 | 0 | 0 | 0 | 0 | 0 | 0 | 0 | 0 | 0 | 0 | 0 | 0 |
| ENSMUSG00 | CR974568.1 | 62 | 5.48453884 | 1.61191424 | 4.68130939 | 1.46971111 | 1.5526427 | 0 | 2.28932317 | 0 | 0 | 3.04198086 | 2.31169154 | 0 | 1.59135217 | 0 | 6.64541224 | 0.61759044 | 1.23633216 | 0 |
| ENSMUSG00 | AC116151.1 | 92 | 0 | 0 | 0 | 0 | 0 | 0 | 0 | 0 | 0 | 0 | 0 | 0 | 0 | 0 | 0 | 0 | 0 | 0 |
| ENSMUSG00 | Mir669d | 120 | 0 | 0 | 0 | 0 | 0 | 0 | 0 | 0 | 0 | 0 | 0 | 0 | 0 | 0 | 0 | 0 | 0 | 0 |
| ENSMUSG00 | AL596212.1 | 88 | 0 | 0 | 0 | 0 | 0 | 0 | 0 | 0 | 0 | 0 | 0 | 0 | 0 | 0 | 0 | 0 | 0 | 0 |
| ENSMUSG00 | AC157101.1 | 92 | 0 | 0 | 0 | 0 | 0 | 0 | 0 | 0 | 0 | 0 | 0 | 0 | 0 | 0 | 0 | 0 | 0 | 0 |
| ENSMUSG00 | AC154473.1 | 76 | 0 | 0 | 0 | 0 | 0 | 0 | 0 | 0 | 0 | 0 | 0 | 0 | 0 | 0 | 0 | 0 | 0 | 0 |
| ENSMUSG00 | CT030170.1 | 90 | 0 | 0 | 0 | 0 | 0 | 0 | 0 | 0 | 0 | 0 | 0 | 0 | 0 | 0 | 0 | 0 | 0 | 0 |
| ENSMUSG00 | AC155640.2 | 90 | 0 | 0 | 0 | 0 | 0 | 0 | 0 | 0 | 0 | 0 | 0 | 0 | 0 | 0 | 0 | 0 | 0 | 0 |
| ENSMUSG00 | AC126023.1 | 72 | 0 | 0 | 0.67185459 | 0 | 1.33699788 | 0 | 0 | 0 | 0 | 0 | 0 | 0 | 0 | 0 | 0 | 0.53181399 | 0 | 0 |
| ENSMUSG00 | AC131591.1 | 93 | 0.91408981 | 0 | 0 | 0 | 0 | 0 | 0 | 0 | 0 | 0 | 0 | 0 | 0 | 0 | 0 | 0 | 0 | 0 |
| ENSMUSG00 | Mir105 | 79 | 0 | 0 | 0 | 0 | 0 | 0 | 0 | 0 | 0 | 0 | 0 | 0 | 0 | 0 | 0 | 0 | 0 | 0 |
| ENSMUSG00 | Mir876 | 80 | 0 | 0 | 0 | 0 | 0 | 0 | 0 | 0 | 0 | 0 | 0 | 0 | 0 | 0 | 0 | 0 | 0 | 0 |
| ENSMUSG00 | AC158626.1 | 92 | 0 | 0 | 0 | 0 | 0 | 0 | 0 | 0 | 0 | 0 | 0 | 0 | 0 | 0 | 0 | 0 | 0 | 0 |
| ENSMUSG00 | AC154495.1 | 87 | 0 | 0 | 0 | 0.52369016 | 0 | 0 | 0 | 0 | 0.94681645 | 0 | 0 | 0 | 0 | 0 | 0 | 0 | 0 | 0 |
| ENSMUSG00 | Mir465b-2 | 78 | 0 | 0 | 0 | 0 | 0 | 0 | 0 | 0 | 0 | 0 | 0 | 0 | 0 | 0 | 0 | 0 | 0 | 0 |
| ENSMUSG00 | AC156022.1 | 109 | 0 | 0 | 0 | 0 | 0 | 0 | 0 | 0 | 0 | 0 | 0 | 0 | 0 | 0 | 0 | 0 | 0 | 0 |
| ENSMUSG00 | U6 | 106 | 0 | 0 | 0 | 0 | 0 | 0 | 0 | 0 | 0 | 0 | 0 | 0 | 0 | 0 | 0 | 0 | 0 | 0 |
| ENSMUSG00 | AL662925.1 | 93 | 0 | 0 | 0 | 0 | 0 | 0 | 0 | 0 | 0 | 0 | 0 | 0 | 0 | 0 | 0 | 0 | 0 | 0 |
| ENSMUSG00 | AC102463.2 | 90 | 0 | 0 | 0 | 0 | 0 | 0 | 0 | 0 | 0 | 0 | 0 | 0 | 0 | 0 | 0 | 0 | 0 | 0 |
| ENSMUSG00 | AC147251.1 | 93 | 0 | 0 | 0 | 0 | 0 | 0 | 0 | 0 | 0 | 0 | 0 | 0 | 0 | 0 | 0 | 0 | 0 | 0 |
| ENSMUSG00 | AC122209.1 | 88 | 0 | 0 | 0 | 0 | 0 | 0 | 0 | 0 | 0 | 0 | 0 | 0 | 0 | 0 | 0 | 0 | 0 | 0 |
| ENSMUSG00 | AC154247.1 | 81 | 0 | 0 | 0 | 0 | 0 | 0 | 0 | 0 | 0 | 0 | 0 | 0 | 0 | 0 | 0 | 0 | 0 | 0 |
| ENSMUSG00 | Mir344e | 65 | 0 | 0 | 0 | 0 | 0 | 0 | 0 | 0 | 0 | 0 | 0 | 0 | 0 | 0 | 0 | 0 | 0 | 0 |
| ENSMUSG00 | AC124717.1 | 91 | 0 | 0 | 0 | 0 | 0 | 0 | 0 | 0 | 0 | 0 | 0 | 0 | 0 | 0 | 0 | 0 | 0 | 0 |
| ENSMUSG00 | Mir878 | 77 | 0 | 0 | 0 | 0 | 0 | 0 | 0 | 0 | 0 | 0 | 0 | 0 | 0 | 0 | 0 | 0 | 0 | 0 |
| ENSMUSG00 | AL671913.1 | 80 | 0 | 0 | 0 | 0 | 0 | 0 | 0 | 0 | 0 | 0 | 0 | 0 | 0 | 0 | 0 | 0 | 0 | 0 |
| ENSMUSG00 | AC118561.1 | 90 | 0 | 0 | 0 | 0 | 0 | 0 | 0 | 0 | 0 | 0 | 0 | 0 | 0 | 0 | 0 | 0 | 0 | 0 |
| ENSMUSG00 | AC163663.1 | 96 | 0 | 0 | 0 | 0 | 0 | 0 | 0 | 0 | 0 | 0 | 0 | 0 | 0 | 0 | 0 | 0 | 0 | 0 |

| | | | | | | | | | | | | | | | | | | | |
|---|---|---|---|---|---|---|---|---|---|---|---|---|---|---|---|---|---|---|---|
| ENSMUSG00 AC152063.1 | 86 | 0 | 0 | 0 | 0 | 0 | 0 | 0 | 0 | 0 | 0 | 0 | 0 | 0 | 0 | 0 | 0 | 0 | 0 |
| ENSMUSG00 Mir882 | 76 | 0 | 0 | 0 | 0 | 0 | 0 | 0 | 0 | 0 | 0 | 0 | 0 | 0 | 0 | 0 | 0 | 0 | 0 |
| ENSMUSG00 AL683882.1 | 86 | 0 | 0 | 0 | 0 | 0 | 0 | 0 | 0 | 0 | 0 | 0 | 0 | 0 | 0 | 0 | 0 | 0 | 0 |
| ENSMUSG00 AC121892.1 | 90 | 0 | 0 | 0 | 0 | 0 | 0 | 0 | 0 | 0 | 0 | 0 | 0 | 0 | 0 | 0 | 0 | 0 | 0 |
| ENSMUSG00 Mir295 | 68 | 0 | 0 | 0 | 0 | 0 | 0 | 0 | 0 | 0 | 0 | 0 | 0 | 0 | 0 | 0 | 0 | 0 | 0 |
| ENSMUSG00 Mir875 | 77 | 0 | 0 | 0 | 0 | 0 | 0 | 0 | 0 | 0 | 0 | 0 | 0 | 0 | 0 | 0 | 0 | 0 | 0 |
| ENSMUSG00 AC124987.1 | 83 | 0 | 0 | 0 | 0 | 0 | 0 | 0 | 0 | 0 | 0 | 0 | 0 | 0.59436045 | 0 | 0 | 0 | 0 | 0 |
| ENSMUSG00 Mir465c-2 | 80 | 0 | 0 | 0 | 0 | 0 | 0 | 0 | 0 | 0 | 0 | 0 | 0 | 0 | 0 | 0 | 0 | 0 | 0 |
| ENSMUSG00 Mir879 | 75 | 0 | 0 | 0 | 0 | 0 | 0 | 0 | 0 | 0 | 0 | 0 | 0 | 0 | 0 | 0 | 0 | 0 | 0 |
| ENSMUSG00 Mir511 | 78 | 0 | 0 | 0 | 0 | 0 | 0 | 0 | 0 | 0 | 0 | 0 | 0 | 0 | 0 | 0 | 0 | 0 | 0 |
| ENSMUSG00 AC126435.1 | 91 | 0 | 0 | 0 | 0 | 0 | 0 | 0 | 0 | 0 | 0 | 0 | 0 | 0 | 0 | 0 | 0 | 0 | 0 |
| ENSMUSG00 Mir294 | 83 | 0 | 0 | 0 | 0 | 0 | 0 | 0 | 0 | 0 | 0 | 0 | 0 | 0 | 0 | 0 | 0 | 0 | 0 |
| ENSMUSG00 AC153556.1 | 80 | 0 | 0 | 0 | 0 | 0 | 0 | 0 | 0 | 0 | 0 | 0 | 0 | 0 | 0 | 0 | 0 | 0 | 0 |
| ENSMUSG00 n-R5s194 | 118 | 0.36021336 | 0 | 0 | 0 | 0 | 0 | 0 | 0 | 0 | 0 | 0 | 0 | 0 | 0 | 0 | 0 | 0 | 0 |
| ENSMUSG00 AC104196.1 | 93 | 0 | 0 | 0 | 0 | 0 | 0 | 0 | 0 | 0 | 0 | 0 | 0 | 0 | 0 | 0 | 0 | 0 | 0 |
| ENSMUSG00 Mir872 | 80 | 0 | 0 | 0 | 0 | 0 | 0 | 0 | 0 | 0 | 0 | 0 | 0 | 0 | 0 | 0 | 0 | 0 | 0 |
| ENSMUSG00 AC125010.1 | 92 | 0 | 0 | 0 | 0 | 0 | 0 | 0 | 0 | 0 | 0 | 0.51929303 | 0.60669198 | 0 | 0 | 0 | 0 | 0 | 0 |
| ENSMUSG00 Mir453 | 81 | 0 | 0 | 0 | 0 | 0 | 0 | 0 | 0 | 0 | 0 | 0 | 0 | 0 | 0 | 0 | 0 | 0 | 0 |
| ENSMUSG00 AL583883.1 | 70 | 0 | 0 | 0 | 0 | 0 | 0 | 0 | 0 | 0 | 0 | 0 | 0 | 0 | 0 | 0 | 0 | 0 | 0 |
| ENSMUSG00 Mir208b | 76 | 0 | 0 | 0 | 0 | 0 | 0 | 0 | 0 | 0 | 0 | 0 | 0 | 0 | 0 | 0 | 0 | 0 | 0 |
| ENSMUSG00 Mir877 | 84 | 0 | 0 | 0 | 0 | 0 | 0 | 0 | 0 | 0 | 0 | 0 | 0 | 0 | 0 | 0 | 0 | 0 | 0 |
| ENSMUSG00 AC128668.1 | 90 | 0 | 0 | 0 | 0 | 0 | 0 | 0 | 0 | 0 | 0 | 0 | 0 | 0 | 0 | 0 | 0 | 0 | 0 |
| ENSMUSG00 AC084389.1 | 74 | 0 | 0 | 0 | 0 | 0 | 0 | 0 | 0 | 0 | 0 | 0 | 0 | 0 | 0 | 0 | 0 | 0 | 0 |
| ENSMUSG00 Mir466h | 80 | 0 | 0 | 0 | 0 | 0 | 0 | 0 | 0 | 0 | 0 | 0 | 0 | 0 | 0 | 0 | 0 | 0 | 0 |
| ENSMUSG00 Mir871 | 76 | 0 | 0 | 0 | 0 | 0 | 0 | 0 | 0 | 0 | 0 | 0 | 0 | 0 | 0 | 0 | 0 | 0 | 0 |
| ENSMUSG00 CT485999.1 | 93 | 0 | 0 | 0 | 0 | 0 | 0 | 0 | 0 | 0.88573152 | 0 | 0 | 0 | 0 | 0 | 0 | 0 | 0 | 0 |
| ENSMUSG00 Mir465c-1 | 80 | 0 | 0 | 0 | 0 | 0 | 0 | 0 | 0 | 0 | 0 | 0 | 0 | 0 | 0 | 0 | 0 | 0 | 0 |
| ENSMUSG00 Mir297a-4 | 97 | 0 | 0 | 0 | 0 | 0 | 0 | 0 | 0 | 0 | 0 | 0 | 0 | 0 | 0 | 0 | 0 | 0 | 0 |
| ENSMUSG00 AC160978.1 | 92 | 0 | 0 | 0 | 0 | 0 | 0 | 0 | 0 | 0 | 0 | 0 | 0 | 0 | 0 | 0 | 0 | 0 | 0 |
| ENSMUSG00 Mir743b | 76 | 0 | 0 | 0 | 0 | 0 | 0 | 0 | 0 | 0 | 0 | 0 | 0 | 0 | 0 | 0 | 0 | 0 | 0 |
| ENSMUSG00 AC157352.1 | 89 | 0 | 0 | 0 | 0 | 0 | 0 | 0 | 0 | 0 | 0 | 0 | 0 | 0 | 0 | 0 | 0 | 0 | 0 |
| ENSMUSG00 Mir874 | 75 | 0 | 0 | 0 | 0 | 0 | 0 | 0 | 0 | 0 | 0 | 0 | 0 | 0 | 0 | 0 | 0 | 0 | 0 |
| ENSMUSG00 AC122270.1 | 100 | 0 | 0 | 0 | 0 | 0.48131924 | 0.49552734 | 0 | 0 | 0 | 0 | 0 | 0 | 0 | 0 | 0 | 0 | 0.76652594 | 0 |
| ENSMUSG00 AC161218.1 | 95 | 0 | 0 | 0 | 0 | 0 | 0 | 0 | 0 | 0 | 0 | 0 | 0 | 0 | 0 | 0 | 0 | 0 | 0 |
| ENSMUSG00 AL732528.1 | 94 | 0 | 0 | 0 | 0 | 0 | 0 | 0 | 0 | 0 | 0 | 0 | 0 | 0 | 0 | 0 | 0 | 0 | 0 |
| ENSMUSG00 AL845457.1 | 85 | 0 | 0 | 0 | 0 | 0.56625793 | 0 | 0 | 0 | 0.96909449 | 0 | 0.56205834 | 0 | 0 | 0.54640103 | 0 | 0 | 0 | 3.88299577 |
| ENSMUSG00 AC154816.1 | 96 | 0 | 0 | 0 | 0 | 0 | 0 | 0 | 0 | 0 | 0 | 0 | 0 | 0 | 0 | 0 | 0 | 0 | 0 |
| ENSMUSG00 Mir466f-1 | 93 | 0 | 0 | 0 | 0 | 0 | 0 | 0 | 0 | 0 | 0 | 0 | 0 | 0 | 0 | 0 | 0 | 0 | 0 |
| ENSMUSG00 AC162467.1 | 91 | 0 | 0 | 0 | 0 | 0 | 0 | 0 | 0 | 0 | 0 | 0 | 0 | 0 | 0 | 0 | 0 | 0 | 0 |
| ENSMUSG00 AC142100.1 | 91 | 0 | 0 | 0 | 0.50067082 | 0 | 0 | 0.51991955 | 2.09848365 | 0 | 0.5181396 | 0 | 0 | 0 | 0 | 0 | 0 | 0 | 0 |
| ENSMUSG00 AC168608.1 | 109 | 0 | 0 | 0 | 0 | 0 | 0 | 0 | 0 | 0 | 0 | 0 | 0 | 0 | 0 | 0 | 0 | 0 | 0 |
| ENSMUSG00 AL844871.1 | 89 | 0 | 0 | 0 | 0 | 0 | 0 | 0 | 0 | 0 | 0 | 0 | 0 | 0 | 0 | 0 | 0 | 0 | 0 |
| ENSMUSG00 Mir654 | 83 | 0 | 0 | 0 | 0 | 0 | 0 | 0 | 0 | 0 | 0 | 0 | 0 | 0 | 0 | 0 | 0 | 0 | 0 |
| ENSMUSG00 Mir466f-3 | 93 | 0 | 0 | 0 | 0 | 0 | 0 | 0 | 0 | 0 | 0 | 0 | 0 | 0 | 0 | 0 | 0 | 0 | 0 |
| ENSMUSG00 AL672259.1 | 86 | 0 | 0 | 0 | 0 | 0 | 0 | 0 | 0 | 0 | 0 | 0 | 0 | 0 | 0 | 0 | 0 | 0 | 0 |
| ENSMUSG00 Mir466n | 93 | 0 | 0 | 0 | 0 | 0 | 0 | 0 | 0 | 0 | 0 | 0 | 0 | 0 | 0 | 0 | 0 | 0 | 0 |
| ENSMUSG00 AC113244.1 | 83 | 0 | 0 | 0 | 0.54892825 | 0 | 0 | 0 | 0 | 0 | 0.56808076 | 0.57560191 | 0 | 0.59436045 | 0 | 0 | 0 | 0 | 0 |
| ENSMUSG00 AC122735.1 | 92 | 0 | 0 | 0 | 0 | 0 | 0 | 0 | 0 | 0 | 0 | 0 | 0 | 0 | 0 | 0 | 0 | 0 | 0 |
| ENSMUSG00 Mir190b | 79 | 0 | 0 | 0 | 0 | 0 | 0 | 0 | 0 | 0 | 0 | 0 | 0 | 0 | 0 | 0 | 0 | 0 | 0 |
| ENSMUSG00 Mir466f-2 | 93 | 0 | 0 | 0 | 0 | 0 | 0 | 0 | 0 | 0 | 0 | 0 | 0 | 0 | 0 | 0 | 0 | 0 | 0 |
| ENSMUSG00 AL589870.1 | 88 | 0 | 0 | 0 | 0 | 0 | 0 | 0 | 0 | 0.46802859 | 0 | 0 | 0 | 0 | 0.52777372 | 0 | 0 | 0 | 0 |
| ENSMUSG00 Mir880 | 77 | 0 | 0 | 0 | 0 | 0 | 0 | 0 | 0 | 0 | 0 | 0 | 0 | 0 | 0 | 0 | 0 | 0 | 0 |
| ENSMUSG00 CT025680.1 | 91 | 0 | 0 | 0 | 0 | 0 | 0 | 0 | 0 | 0 | 0 | 0 | 0 | 0 | 0 | 0 | 0 | 0 | 0 |
| ENSMUSG00 AC141869.1 | 94 | 0 | 0 | 0 | 0 | 0 | 0 | 0 | 0 | 0 | 0 | 0 | 0 | 0 | 0 | 0 | 0 | 0 | 0 |
| ENSMUSG00 Mir291a | 81 | 0 | 0 | 0 | 0 | 0 | 0 | 0 | 0 | 0 | 0 | 0 | 0 | 0 | 0 | 0 | 0 | 0 | 0 |
| ENSMUSG00 AL954388.1 | 84 | 0 | 0 | 0.57587536 | 0 | 0 | 0 | 0 | 1.81868583 | 0 | 0 | 0 | 0 | 0 | 0 | 0.49049471 | 0 | 1.36879632 | 0 |
| ENSMUSG00 AL732472.1 | 91 | 0 | 0 | 0 | 0 | 0 | 0 | 0 | 0 | 0 | 0 | 0 | 0 | 0 | 0 | 0 | 0 | 0 | 0 |
| ENSMUSG00 AL603708.1 | 78 | 0 | 0 | 0 | 0 | 0 | 0 | 0 | 0 | 0 | 0 | 0 | 0 | 0 | 0 | 0 | 0 | 0 | 0 |
| ENSMUSG00 Mir883a | 75 | 0 | 0 | 0 | 0 | 0 | 0 | 0 | 0 | 0 | 0 | 0 | 0 | 0 | 0 | 0 | 0 | 0 | 0 |
| ENSMUSG00 AC124406.1 | 86 | 4.94246233 | 0 | 0 | 0 | 0 | 0 | 0 | 0.4440977 | 0 | 0.54826399 | 0 | 0 | 0 | 0 | 0 | 0.44523962 | 0 | 0 |
| ENSMUSG00 AL805935.1 | 91 | 0 | 0 | 0 | 0 | 0 | 0 | 0 | 0 | 0 | 0 | 0 | 0 | 0 | 0 | 0 | 0 | 0 | 0 |
| ENSMUSG00 AC168268.1 | 109 | 0 | 0 | 0 | 0 | 0 | 0 | 0 | 0 | 0 | 0 | 0 | 0 | 0 | 0 | 0 | 0 | 0 | 0 |
| ENSMUSG00 AC159288.1 | 109 | 0 | 0 | 0 | 0 | 0 | 0 | 0 | 0 | 0 | 0 | 0 | 0 | 0 | 0 | 0 | 0 | 0 | 0 |
| ENSMUSG00 Mir883b | 77 | 0 | 0 | 0 | 0 | 0 | 0 | 0 | 0 | 0 | 0 | 0 | 0 | 0 | 0 | 0 | 0 | 0 | 0 |
| ENSMUSG00 Mir466g | 79 | 0 | 0 | 0 | 0 | 0 | 0 | 0 | 0 | 0 | 0 | 0 | 0 | 0 | 0 | 0 | 0 | 0 | 0 |
| ENSMUSG00 Mir4660 | 83 | 0 | 0 | 0 | 0 | 0 | 0 | 0 | 0 | 0 | 0 | 0 | 0 | 0 | 0 | 0 | 0 | 0 | 0 |
| ENSMUSG00 Mir449b | 79 | 0 | 0 | 0 | 0 | 0 | 0 | 0 | 0 | 0 | 0 | 0 | 0 | 0 | 0 | 0 | 0 | 0 | 0 |
| ENSMUSG00 Mir466d | 95 | 0 | 0 | 0 | 0 | 0 | 0 | 0 | 0 | 0 | 0 | 0 | 0 | 0 | 0 | 0 | 0 | 0 | 0 |
| ENSMUSG00 Mir291b | 78 | 0 | 0 | 0 | 0 | 0 | 0 | 0 | 0 | 0 | 0 | 0 | 0 | 0 | 0 | 0 | 0 | 0 | 0 |
| ENSMUSG00 BX119956.1 | 87 | 0 | 0 | 0 | 0 | 0 | 0 | 0 | 0 | 0 | 0 | 0 | 0 | 0 | 0 | 0 | 0 | 0 | 0 |
| ENSMUSG00 Mir293 | 79 | 0 | 0 | 0 | 0 | 0 | 0 | 0 | 0 | 0 | 0 | 0 | 0 | 0 | 0 | 0 | 0 | 0 | 0 |
| ENSMUSG00 AC114558.1 | 95 | 0 | 0 | 0 | 0 | 0 | 0 | 0 | 0 | 0 | 0 | 0 | 0 | 0 | 0 | 0 | 0 | 0 | 0 |
| ENSMUSG00 Mir881 | 77 | 0 | 0 | 0 | 0 | 0 | 0 | 0 | 0 | 0 | 0 | 0 | 0 | 0 | 0 | 0 | 0 | 0 | 0 |
| ENSMUSG00 Mir292 | 81 | 0 | 0 | 0 | 0 | 0 | 0 | 0 | 0 | 0 | 0 | 0 | 0 | 0 | 0 | 0 | 0 | 0 | 0 |
| ENSMUSG00 AC153982.1 | 96 | 0 | 0 | 0 | 0 | 0 | 0 | 0 | 0 | 0 | 0 | 0 | 0 | 0 | 0 | 0 | 0 | 0 | 0 |
| ENSMUSG00 AC157475.1 | 94 | 0 | 0 | 0 | 0 | 0 | 0 | 0 | 0 | 0 | 0 | 0 | 0 | 0 | 0 | 0 | 0 | 0 | 0 |
| ENSMUSG00 AL626775.1 | 81 | 0.52475526 | 1.2338109 | 1.79161224 | 0.56248203 | 0 | 3.05881071 | 0 | 0.94302228 | 0.5084755 | 0 | 0.5898143 | 2.06724676 | 0.60903602 | 0.57338379 | 1.52598355 | 0 | 0.94632832 | 2.7164991 |
| ENSMUSG00 AL590992.1 | 77 | 0 | 0 | 0 | 0 | 0 | 0 | 0 | 0 | 0 | 0 | 0 | 0 | 0 | 0 | 0 | 0 | 0 | 0 |
| ENSMUSG00 Mir873 | 76 | 0 | 0 | 0 | 0 | 0 | 0 | 0 | 0 | 0 | 0 | 0 | 0 | 0 | 0 | 0 | 0 | 0 | 0 |
| ENSMUSG00 Mir466l | 120 | 0 | 0 | 0 | 0 | 0 | 0 | 0 | 0 | 0 | 0 | 0 | 0 | 0 | 0 | 0 | 0 | 0 | 0 |
| ENSMUSG00 AL731851.1 | 91 | 0 | 0 | 0 | 0 | 0 | 0 | 0 | 0 | 0 | 0 | 0 | 0 | 0 | 0 | 0 | 0 | 0 | 0 |
| ENSMUSG00 AC167132.1 | 86 | 0 | 1.54943694 | 1.68744873 | 0 | 0 | 2.30477831 | 2.20058972 | 0.4440977 | 0 | 0 | 0 | 1.29803866 | 0 | 0 | 0.95817572 | 1.33571886 | 0 | 0 |
| ENSMUSG00 AL713967.1 | 89 | 0 | 0 | 0 | 0 | 0 | 0 | 0 | 0 | 0 | 0 | 0 | 0 | 0 | 0 | 0 | 0 | 0 | 0 |
| ENSMUSG00 AC121600.1 | 80 | 0 | 0 | 0 | 0 | 0 | 0 | 0 | 0 | 0 | 0 | 0 | 0 | 0 | 0 | 0 | 0 | 0 | 0 |
| ENSMUSG00 Olfr585 | 956 | 0 | 0 | 0 | 0 | 0 | 0 | 0 | 0 | 0 | 0 | 0 | 0 | 0 | 0 | 0 | 0 | 0 | 0 |
| ENSMUSG00 Gm15887 | 494 | 0 | 0 | 0 | 0 | 0 | 0 | 0 | 0 | 0 | 0 | 0 | 0 | 0 | 0 | 0 | 0 | 0 | 0 |
| ENSMUSG00 Gm7336 | 597 | 0.35598975 | 0.66960591 | 0.48616613 | 0.38158329 | 0.24186896 | 0.41501452 | 0.79250718 | 0.51179099 | 0.75888052 | 0.55285582 | 0.96030067 | 0.65445501 | 0.66106422 | 0.62236633 | 0.41408599 | 0.83379882 | 0.44938707 | 0.55285534 |
| ENSMUSG00 Gm10912 | 1660 | 0 | 0 | 0 | 0 | 0 | 0.02985104 | 0 | 0 | 0.07443346 | 0 | 0 | 0.03362389 | 0 | 0 | 0 | 0 | 0 | 0 |
| ENSMUSG00 Golgb1 | 497 | 0 | 0 | 0 | 0 | 0 | 0 | 0 | 0 | 0 | 0 | 0 | 0 | 0 | 0 | 0 | 0 | 0 | 0 |
| ENSMUSG00 Gm10913 | 926 | 0 | 0 | 0 | 0.14760598 | 0.05197832 | 0 | 0 | 0 | 0 | 0 | 0 | 0.12055219 | 0 | 0 | 0 | 0.12405164 | 0.08277818 | 0.17821525 |
| ENSMUSG00 Gm6551 | 1248 | 0 | 0 | 0 | 0 | 0 | 0 | 0 | 0 | 0 | 0 | 0 | 0 | 0 | 0 | 0 | 0 | 0 | 0 |
| ENSMUSG00 Gm4478 | 386 | 0.33035111 | 0.17260567 | 0 | 0 | 0 | 0 | 0.12257171 | 0 | 0 | 0 | 0 | 0.14460016 | 0.38340868 | 0 | 0 | 0 | 0 | 0.28502128 |
| ENSMUSG00 Gm16496 | 4170 | 0 | 0 | 0 | 0 | 0 | 0.01188315 | 0 | 0 | 0 | 0 | 0 | 0 | 0 | 0 | 0 | 0 | 0 | 0 |
| ENSMUSG00 Olfr828 | 938 | 0 | 0 | 0 | 0 | 0 | 0 | 0 | 0 | 0 | 0 | 0 | 0 | 0 | 0 | 0 | 0 | 0 | 0 |
| ENSMUSG00 Gm16485 | 3982 | 0 | 0 | 0 | 0 | 0 | 0 | 0 | 0 | 0 | 0 | 0 | 0 | 0 | 0 | 0 | 0 | 0 | 0 |
| ENSMUSG00 Olfr483 | 1064 | 0 | 0 | 0 | 0 | 0 | 0 | 0 | 0 | 0 | 0 | 0 | 0 | 0 | 0 | 0 | 0 | 0 | 0 |
| ENSMUSG00 Gm14958 | 398 | 0 | 0 | 0 | 0 | 0 | 0 | 0 | 0 | 0 | 0 | 0 | 0 | 0 | 0 | 0 | 0 | 0 | 0 |
| ENSMUSG00 Gm7671 | 398 | 0 | 0 | 0 | 0 | 0 | 0 | 0 | 0 | 0 | 0 | 0 | 0 | 0 | 0 | 0 | 0 | 0 | 0 |
| ENSMUSG00 F630028O10R | 3307 | 0 | 0 | 0 | 0 | 0 | 0 | 0 | 0 | 0 | 0 | 0 | 0 | 0 | 0 | 0 | 0 | 0 | 0 |
| ENSMUSG00 Gm5071 | 1034 | 0 | 0 | 0 | 0 | 0 | 0 | 0 | 0 | 0 | 0 | 0 | 0 | 0 | 0 | 0 | 0 | 0 | 0 |
| ENSMUSG00 Gm10916 | 482 | 0 | 0 | 0 | 0 | 0 | 0 | 0 | 0 | 0.17089841 | 0.09782304 | 0 | 0 | 0 | 0 | 0 | 0 | 0 | 0 |
| ENSMUSG00 Rpl23a-ps3 | 470 | 0.18087309 | 0 | 0 | 0.09693839 | 0.10240835 | 0.2108627 | 0 | 0 | 0.08763088 | 0 | 0.2032977 | 0.11875673 | 0.31488458 | 0.19763441 | 0 | 0.08146938 | 0.24463594 | 0 |
| ENSMUSG00 Fam170b | 1692 | 0 | 0 | 0 | 0 | 0 | 0 | 0 | 0 | 0 | 0 | 0 | 0 | 0 | 0 | 0 | 0 | 0 | 0 |
| ENSMUSG00 Gm2178 | 356 | 0 | 0.09357555 | 0 | 0 | 0 | 0 | 0 | 0.10728203 | 0.11569246 | 0 | 0 | 0 | 0 | 0 | 0 | 0.10755789 | 0.53829069 | 1.39067966 |
| ENSMUSG00 1700007I08R | 1500 | 0 | 0.0222086 | 0 | 0 | 0 | 0 | 0 | 0.0509232 | 0 | 0 | 0 | 0 | 0 | 0 | 0 | 0 | 0 | 0 |
| ENSMUSG00 Gm11555 | 1127 | 0 | 0 | 0 | 0 | 0 | 0 | 0 | 0 | 0 | 0 | 0 | 0 | 0 | 0 | 0 | 0 | 0 | 0 |
| ENSMUSG00 Krtap1-3 | 878 | 0 | 0 | 0 | 0 | 0 | 0 | 0 | 0 | 0 | 0 | 0 | 0 | 0 | 0 | 0 | 0 | 0 | 0 |
| ENSMUSG00 Gm11939 | 299 | 0 | 0 | 0 | 0 | 0 | 0 | 0 | 0 | 0 | 0 | 0 | 0 | 0 | 0 | 0 | 0 | 0 | 0 |
| ENSMUSG00 Gm12355 | 1310 | 0.55159389 | 0.55945319 | 0.51696903 | 0.38257365 | 0.44090312 | 0.2647856 | 0.28893239 | 0.46647209 | 0.31440088 | 0.21595742 | 0.54704151 | 0.42607376 | 0.33892157 | 0.70907004 | 0.47177354 | 0.52613048 | 0.55587759 | 0.46190853 |
| ENSMUSG00 Eid1 | 968 | 0 | 0 | 0 | 0 | 0 | 0 | 0 | 0 | 0 | 0 | 0 | 0 | 0 | 0 | 0 | 0 | 0 | 0 |
| ENSMUSG00 Gm1337 | 4860 | 0 | 0 | 0 | 0 | 0 | 0.03058811 | 0 | 0 | 0 | 0 | 0 | 0 | 0 | 0 | 0 | 0 | 0 | 0.01131875 |
| ENSMUSG00 AK157302 | 1923 | 0.0884143 | 0.05197019 | 0.07546573 | 0.21323422 | 0.0500592 | 0.02576845 | 0.09841431 | 0.03972169 | 0.08567138 | 0.22067412 | 0.07453192 | 0.11610122 | 0.15392174 | 0.09660757 | 0.064277 | 0.07964765 | 0.05979141 | 0.02860588 |
| ENSMUSG00 Gm2399 | 227 | 0 | 0 | 0 | 0 | 0 | 0 | 0.6252777 | 0 | 0.1814384 | 0 | 0 | 0 | 0 | 0 | 0 | 0 | 0 | 0 |
| ENSMUSG00 Gm17344 | 1638 | 6.072168 | 6.99611454 | 7.26488917 | 7.81602775 | 8.16891012 | 8.07727709 | 7.27887368 | 7.2980598 | 4.6768571 | 5.67075003 | 6.12499468 | 7.49660914 | 5.93308163 | 3.96958012 | 5.70986153 | 8.2285066 | 5.21780477 | 5.13821327 |
| ENSMUSG00 Capns2 | 1000 | 0 | 0 | 0 | 0 | 0 | 0 | 0 | 0 | 0 | 0 | 0 | 0 | 0 | 0 | 0 | 0 | 0 | 0 |
| ENSMUSG00 Gm12226 | 875 | 0 | 0.11421564 | 0 | 0.05206976 | 0.11001583 | 0.16989509 | 0.10814327 | 0.04364846 | 0.09414061 | 0 | 0.21839981 | 0.51031463 | 0.05637933 | 0.05307896 | 0.28252495 | 0.04376069 | 0 | 0 |
| ENSMUSG00 Psme2b-ps | 836 | 9.81279782 | 10.4003175 | 7.00143203 | 7.35734568 | 7.71492556 | 9.42450316 | 8.37592881 | 8.77146084 | 7.38992508 | 9.58806167 | 7.02909078 | 7.8782873 | 7.61222168 | 7.27772183 | 8.57544345 | 10.0764756 | 7.70193526 | 5.59303119 |
| ENSMUSG00 Gm12184 | 2220 | 0.53610132 | 0.42016263 | 0.41400769 | 0.43098285 | 0.34689675 | 0.62498943 | 0.68198456 | 0.39568705 | 0.81630932 | 0.382303 | 0.45192528 | 0.50284381 | 0.37776693 | 0.58578128 | 0.6866926 | 0.60368075 | 0.29348966 | 0.66902968 |
| ENSMUSG00 4931440F15R | 3273 | 0.06493305 | 0 | 0.22169354 | 0.01392027 | 0.23529202 | 0.06055941 | 0.5203961 | 0.21004071 | 0 | 0.38896089 | 0.467094 | 0.01705336 | 0.79883642 | 0.46827219 | 0.23917799 | 0.25737652 | 0.23419674 | 0.18487631 |
| ENSMUSG00 Gm16503 | 3282 | 0 | 0.08120145 | 0 | 0 | 0.01466542 | 0 | 0 | 0 | 0 | 0.12929809 | 0.04366998 | 0.20407921 | 0 | 0 | 0 | 0 | 0.01167773 | 0 |
| ENSMUSG00 BC007180 | 5630 | 0 | 0 | 0 | 0.16185096 | 0 | 0 | 0 | 0 | 0 | 0 | 0 | 0 | 0 | 0 | 0 | 0 | 0 | 0 |
| ENSMUSG00 Gm2574 | 1231 | 0.03452898 | 0.02706165 | 0 | 0 | 0 | 0 | 0.03843435 | 0.03102551 | 0.10037331 | 0.03830276 | 0.03880988 | 0.04534172 | 0.12022401 | 0 | 0.10040997 | 0.06221057 | 0 | 0 |
| ENSMUSG00 Gm4392 | 338 | 0 | 0 | 0.14311695 | 0 | 0.56960856 | 0 | 0.13997834 | 0 | 0 | 0.27899824 | 0 | 0 | 0 | 0 | 0 | 0 | 0 | 0 |
| ENSMUSG00 Gm16499 | 65 | 0 | 0 | 0 | 0 | 0 | 0 | 0 | 0 | 0 | 0 | 0 | 0 | 0 | 0 | 0 | 0 | 0 | 0 |
| ENSMUSG00 Lenep | 2471 | 0 | 0 | 0 | 0 | 0 | 0 | 0 | 0.01545625 | 0 | 0 | 0 | 0 | 0 | 0 | 0 | 0 | 0 | 0 |
| ENSMUSG00 Gm6525 | 320 | 2.6565735 | 1.97795309 | 1.20933826 | 1.70853916 | 1.65453488 | 1.85822751 | 1.47852122 | 1.43221509 | 1.80191006 | 0.58938379 | 2.83663816 | 1.74423946 | 0.77081121 | 0.87082664 | 1.93132293 | 5.26495852 | 4.67101744 | 4.9852003 |
| ENSMUSG00 Gm4877 | 809 | 0 | 0 | 0 | 0 | 0 | 0 | 0 | 0 | 0 | 0 | 0 | 0 | 0 | 0 | 0 | 0 | 0 | 0 |
| ENSMUSG00 Rnf148 | 1238 | 0 | 0 | 0 | 0 | 0 | 0 | 0.38217027 | 0 | 0 | 0 | 0 | 0 | 0 | 0 | 0 | 0 | 0 | 0.0888677 |
| ENSMUSG00 Gm6083 | 356 | 0 | 0 | 0 | 0 | 0 | 0 | 0 | 0 | 0 | 0.1324458 | 0 | 0 | 0 | 0 | 0 | 0 | 0 | 0 |
| ENSMUSG00 Gm15610 | 642 | 1.5227711 | 1.97179125 | 2.4111417 | 1.63224925 | 1.94926794 | 3.08739773 | 3.8321796 | 2.85550672 | 3.14351912 | 2.93773852 | 2.15805887 | 2.52126825 | 3.99573162 | 3.3277695 | 2.82378265 | 2.98213453 | 2.86551753 | 2.48483504 |
| ENSMUSG00 B020018G12 | 1337 | 0.69941202 | 0.69765223 | 0.18090325 | 0.20446243 | 0.10799983 | 0.07412526 | 0.28309756 | 0.3713547 | 0.21563621 | 0.56425673 | 0.07146591 | 0.2922286 | 0.25828229 | 0.24316276 | 0.21571495 | 0.20047438 | 0.51598605 | 0.16457474 |
| ENSMUSG00 Chml | 1865 | 0.31907371 | 0.16075928 | 0.2593755 | 0.29315417 | 0.23227202 | 0.18598881 | 0.07610619 | 0.163828 | 0 | 0.20225503 | 0.256166 | 0.17956782 | 0.02645143 | 0.32373895 | 0 | 0.24637388 | 0.18495264 | 0 |
| ENSMUSG00 Gm13821 | 513 | 0 | 0 | 0 | 0 | 0 | 0 | 0 | 0 | 0 | 0 | 0 | 0 | 0 | 0 | 0 | 0 | 0 | 0 |
| ENSMUSG00 Gm13820 | 137 | 0 | 0 | 0 | 0 | 0 | 0 | 0 | 0 | 0 | 0 | 0 | 0 | 0 | 0 | 0 | 0 | 0 | 0 |
| ENSMUSG00 Dnm3os | 8528 | 0.05482609 | 0 | 0 | 0 | 0 | 0 | 0 | 0 | 0.13522777 | 0.00552893 | 0 | 0 | 0 | 0 | 0 | 0 | 0 | 0 |
| ENSMUSG00 Gm6559 | 1250 | 0.13601656 | 0.05330063 | 0.11609647 | 0.14579534 | 0.11551662 | 0.03964219 | 0.11355043 | 0.09166177 | 0.06589843 | 0.07544113 | 0.19109983 | 0.04465253 | 0 | 0 | 0.13184498 | 0.15316243 | 0.06132208 | 0.13202186 |
| ENSMUSG00 Gm7290 | 1228 | 0 | 0.02712776 | 0 | 0 | 0 | 0.08070478 | 0 | 0.0311013 | 0 | 0 | 0 | 0.04545249 | 0.04017257 | 0.03782092 | 0 | 0 | 0 | 0.04479569 |
| ENSMUSG00 Gm2000 | 444 | 9.66897022 | 9.07851397 | 8.71595142 | 8.31181214 | 6.50431401 | 9.93286777 | 11.9347298 | 8.68791136 | 6.02955749 | 8.38942693 | 7.74729056 | 7.41694615 | 12.777411 | 14.9583435 | 8.72285192 | 10.2625727 | 8.20044641 | 11.8938609 |
| ENSMUSG00 Gm17374 | 100 | 0 | 0.33312894 | 0 | 0 | 0 | 0 | 0 | 0 | 0 | 0 | 0 | 0 | 0 | 0 | 0 | 0 | 0 | 0 |
| ENSMUSG00 Olfr364-ps1 | 923 | 0 | 0 | 0 | 0 | 0 | 0 | 0 | 0 | 0 | 0 | 0 | 0 | 0 | 0 | 0 | 0 | 0 | 0 |
| ENSMUSG00 Tmem203 | 1675 | 5.76040296 | 6.28470124 | 5.66042504 | 4.406501 | 5.89077275 | 5.11798383 | 6.94860837 | 6.08798295 | 7.3029225 | 7.37521448 | 6.7598001 | 6.03142384 | 8.39378891 | 6.57149177 | 5.70672297 | 5.94361667 | 6.31525847 | 7.12655293 |
| ENSMUSG00 Nrarp | 2572 | 0.59494026 | 0.56989399 | 0.80673321 | 0.3719992 | 0.50527292 | 0.8862464 | 0.82778793 | 0.40093113 | 0.64053679 | 0.89828323 | 0.01857502 | 1.01995962 | 0.80557563 | 1.95021829 | 2.43492865 | 0.49128695 | 0.90898294 | 0.44914123 |
| ENSMUSG00 Gm14458 | 855 | 0 | 0 | 0 | 0 | 0 | 0 | 0 | 0 | 0 | 0 | 0 | 0 | 0 | 0 | 0 | 0 | 0 | 0 |
| ENSMUSG00 Gm5634 | 841 | 0 | 0 | 0 | 0 | 0 | 0 | 0 | 0 | 0 | 0 | 0 | 0 | 0 | 0 | 0 | 0 | 0 | 0 |
| ENSMUSG00 Gm14374 | 1959 | 0 | 0 | 0 | 0 | 0 | 0 | 0 | 0 | 0 | 0 | 0 | 0 | 0 | 0 | 0 | 0 | 0 | 0 |
| ENSMUSG00 Gm6924 | 869 | 0 | 0 | 0 | 0 | 0 | 0 | 0 | 0 | 0 | 0 | 0 | 0 | 0 | 0 | 0 | 0 | 0 | 0 |
| ENSMUSG00 Gm3763 | 1959 | 0 | 0 | 0 | 0 | 0 | 0 | 0 | 0 | 0 | 0 | 0 | 0 | 0 | 0 | 0 | 0 | 0 | 0 |
| ENSMUSG00 Gm3757 | 1451 | 0 | 0 | 0 | 0 | 0 | 0 | 0 | 0 | 0 | 0 | 0 | 0 | 0 | 0 | 0 | 0 | 0 | 0 |
| ENSMUSG00 Gm3750 | 1490 | 0 | 0 | 0 | 0 | 0 | 0 | 0 | 0 | 0 | 0 | 0 | 0 | 0 | 0 | 0 | 0 | 0 | 0 |
| ENSMUSG00 Gm6925 | 3455 | 0 | 0 | 0 | 0 | 0 | 0 | 0 | 0 | 0 | 0 | 0 | 0 | 0 | 0 | 0 | 0 | 0 | 0 |
| ENSMUSG00 Gm10921 | 1959 | 0 | 0 | 0 | 0 | 0 | 0 | 0 | 0 | 0 | 0 | 0 | 0 | 0 | 0 | 0 | 0 | 0 | 0 |
| ENSMUSG00 Gm10922 | 1959 | 0 | 0 | 0 | 0 | 0 | 0 | 0 | 0 | 0 | 0 | 0 | 0 | 0 | 0 | 0 | 0 | 0 | 0 |
| ENSMUSG00 Gm14347 | 1959 | 0 | 0 | 0 | 0 | 0 | 0 | 0 | 0 | 0 | 0 | 0 | 0 | 0 | 0 | 0 | 0 | 0 | 0 |
| ENSMUSG00 Gm3706 | 1942 | 0 | 0 | 0 | 0 | 0 | 0 | 0 | 0 | 0 | 0 | 0 | 0 | 0 | 0 | 0 | 0 | 0 | 0 |
| ENSMUSG00 Gm3701 | 1959 | 0 | 0 | 0 | 0 | 0 | 0 | 0 | 0 | 0 | 0 | 0 | 0 | 0 | 0 | 0 | 0 | 0 | 0 |
| ENSMUSG00 Gm14351 | 1959 | 0 | 0 | 0 | 0 | 0 | 0 | 0 | 0 | 0 | 0 | 0 | 0 | 0 | 0 | 0 | 0 | 0 | 0 |
| ENSMUSG00 Gm14345 | 1959 | 0 | 0 | 0 | 0 | 0 | 0 | 0 | 0 | 0 | 0 | 0 | 0 | 0 | 0 | 0 | 0 | 0 | 0 |
| ENSMUSG00 Gm14346 | 1959 | 0 | 0 | 0 | 0 | 0 | 0 | 0 | 0 | 0 | 0 | 0 | 0 | 0 | 0 | 0 | 0 | 0 | 0 |
| ENSMUSG00 Gm6919 | 482 | 0 | 0 | 0 | 0 | 0 | 0 | 0 | 0 | 0 | 0 | 0 | 0 | 0 | 0 | 0 | 0 | 0 | 0 |
| ENSMUSG00 Gm16508 | 2740 | 0 | 0 | 0 | 0 | 0 | 0 | 0 | 0 | 0 | 0 | 0 | 0 | 0 | 0 | 0.03007413 | 0.04192402 | 0 | 0 |
| ENSMUSG00 Klhdc7a | 9877 | 1.97098012 | 1.97307311 | 1.66518177 | 1.35156282 | 1.79818567 | 1.69574 | 0.79038089 | 0.88936444 | 1.13839413 | 0.68265167 | 0.51755802 | 0.73463968 | 0.70923684 | 1.1050279 | 0.55897626 | 1.03121409 | 0.70622517 | 0.36201194 |
| ENSMUSG00 Fam43b | 2366 | 1.58091948 | 0.83071038 | 1.10404507 | 1.92565699 | 0.85441285 | 1.92681804 | 0.51991955 | 0.45198109 | 0.88779049 | 0.4782827 | 0 | 1.1795364 | 0 | 0 | 0.01741401 | 0.06473475 | 0 | 0 |
| ENSMUSG00 Pou3f1 | 3849 | 0 | 0 | 0 | 0 | 0 | 0 | 0 | 0 | 0 | 0 | 0 | 0 | 0 | 0 | 0 | 0 | 0 | 0 |
| ENSMUSG00 Gm12854 | 601 | 1.34375764 | 0.72057841 | 1.28781445 | 0.83389599 | 2.08224628 | 1.07185613 | 1.33829541 | 1.14386563 | 0.82235972 | 0.86299124 | 1.03340176 | 0.46435659 | 0.49249834 | 1.08189222 | 1.30254503 | 1.01938389 | 0.63770877 | 1.00682225 |
| ENSMUSG00 Gm3550 | 644 | 0.13200365 | 0 | 0.15022836 | 0.28298785 | 0.22421704 | 0.38472619 | 0.14693379 | 0.05930497 | 0 | 0 | 0 | 0.26001085 | 0.15320471 | 0.1442363 | 0 | 0.23782986 | 0 | 0 |
| ENSMUSG00 Airn | 3669 | 1.1700798 | 1.76143404 | 1.50302057 | 0.8692486 | 1.75788437 | 1.93132758 | 0.94135338 | 1.40528055 | 2.503296 | 1.24655716 | 1.43233727 | 1.47563894 | 1.93616683 | 1.13926625 | 1.87535019 | 2.36902913 | 1.99518199 | 1.79915312 |
| ENSMUSG00 Hmga1-rs1 | 1625 | 0 | 0 | 0 | 0 | 0 | 0 | 0 | 0 | 0 | 0 | 0 | 0 | 0 | 0 | 0 | 0 | 0 | 0 |
| ENSMUSG00 Gm16487 | 4035 | 0 | 0.0495359 | 0 | 0.01129146 | 0 | 0 | 0 | 0 | 0 | 0 | 0 | 0 | 0.09780801 | 0 | 0 | 0 | 0.00949846 | 0 |
| ENSMUSG00 Krtap17-1 | 760 | 0 | 0 | 0 | 0 | 0 | 0 | 0.31126762 | 0 | 0 | 0 | 0 | 0 | 0 | 0 | 0 | 0 | 0 | 0 |
| ENSMUSG00 Gm11570 | 1887 | 0 | 0 | 0 | 0 | 0 | 0.02626006 | 0 | 0 | 0 | 0 | 0 | 0 | 0 | 0 | 0 | 0 | 0 | 0 |
| ENSMUSG00 Krtap29-1 | 1028 | 0 | 0 | 0 | 0 | 0 | 0 | 0 | 0 | 0 | 0 | 0 | 0 | 0 | 0 | 0 | 0 | 0 | 0 |
| ENSMUSG00 Krtap9-5 | 1076 | 0 | 0 | 0 | 0 | 0 | 0 | 0 | 0 | 0 | 0 | 0 | 0 | 0 | 0 | 0 | 0 | 0 | 0 |
| ENSMUSG00 Gm11565 | 920 | 0 | 0 | 0 | 0 | 0 | 0 | 0 | 0 | 0 | 0 | 0 | 0 | 0 | 0 | 0 | 0 | 0 | 0 |
| ENSMUSG00 2300003K06R | 918 | 0 | 0 | 0 | 0 | 0 | 0 | 0 | 0 | 0 | 0 | 0 | 0 | 0 | 0 | 0 | 0 | 0 | 0 |
| ENSMUSG00 Gm11564 | 690 | 0 | 0 | 0 | 0 | 0 | 0 | 0 | 0 | 0 | 0 | 0 | 0 | 0 | 0 | 0 | 0 | 0 | 0 |
| ENSMUSG00 Gm11554 | 889 | 0 | 0 | 0 | 0 | 0 | 0 | 0 | 0 | 0 | 0 | 0 | 0 | 0 | 0 | 0 | 0 | 0 | 0 |
| ENSMUSG00 Gm11569 | 868 | 0 | 0 | 0 | 0 | 0 | 0 | 0 | 0 | 0 | 0 | 0 | 0 | 0 | 0 | 0 | 0 | 0 | 0 |
| ENSMUSG00 Gm11596 | 1078 | 0 | 0 | 0 | 0 | 0 | 0 | 0 | 0 | 0 | 0 | 0 | 0 | 0 | 0 | 0 | 0 | 0 | 0 |
| ENSMUSG00 Krtap4-9 | 1058 | 0 | 0 | 0 | 0 | 0 | 0 | 0 | 0 | 0 | 0 | 0 | 0 | 0 | 0 | 0 | 0 | 0 | 0 |
| ENSMUSG00 Gm14188 | 167 | 0 | 0 | 0 | 0 | 0 | 0 | 0 | 0 | 0 | 0 | 0 | 0 | 0 | 0 | 0 | 0 | 0 | 0 |
| ENSMUSG00 Gm14194 | 140 | 0 | 0 | 0 | 0 | 0 | 0 | 0 | 0 | 0 | 0 | 0 | 0 | 0 | 0 | 0 | 0 | 0 | 0 |
| ENSMUSG00 Gm14181 | 122 | 0 | 0 | 0 | 0 | 0 | 0 | 0 | 0 | 0 | 0 | 0 | 0 | 0 | 0 | 0 | 0 | 0 | 0 |
| ENSMUSG00 Gm14178 | 122 | 0 | 0 | 0 | 0 | 0 | 0 | 0 | 0 | 0 | 0 | 0 | 0 | 0 | 0 | 0 | 0 | 0 | 0 |
| ENSMUSG00 Gm14179 | 128 | 0 | 0 | 0 | 0 | 0 | 0 | 0 | 0 | 0 | 0 | 0 | 0 | 0 | 0 | 0 | 0 | 0 | 0 |
| ENSMUSG00 Gm14191 | 152 | 0 | 0 | 0 | 0 | 0 | 0 | 0 | 0 | 0 | 0 | 0 | 0 | 0 | 0 | 0 | 0 | 0 | 0 |
| ENSMUSG00 Gm14180 | 77 | 0 | 0 | 0 | 0 | 0 | 0 | 0 | 0 | 0 | 0 | 0 | 0 | 0 | 0 | 0 | 0 | 0 | 0 |
| ENSMUSG00 Gm14193 | 152 | 0 | 0 | 0 | 0 | 0 | 0 | 0 | 0 | 0 | 0 | 0 | 0 | 0 | 0 | 0 | 0 | 0 | 0 |
| ENSMUSG00 Gm14185 | 137 | 0 | 0 | 0 | 0 | 0 | 0 | 0 | 0 | 0 | 0 | 0 | 0 | 0 | 0 | 0 | 0 | 0 | 0 |
| ENSMUSG00 Gm14196 | 149 | 0 | 0 | 0 | 0 | 0 | 0 | 0 | 0 | 0 | 0 | 0 | 0 | 0 | 0 | 0 | 0 | 0 | 0 |
| ENSMUSG00 Gm14183 | 152 | 0 | 0 | 0 | 0 | 0 | 0 | 0 | 0 | 0 | 0 | 0 | 0 | 0 | 0 | 0 | 0 | 0 | 0 |
| ENSMUSG00 Gm14184 | 137 | 0 | 0 | 0 | 0 | 0 | 0 | 0 | 0 | 0 | 0 | 0 | 0 | 0 | 0 | 0 | 0 | 0 | 0 |
| ENSMUSG00 Gm14192 | 149 | 0 | 0 | 0 | 0 | 0 | 0 | 0 | 0 | 0 | 0 | 0 | 0 | 0 | 0 | 0 | 0 | 0 | 0 |
| ENSMUSG00 Gm14190 | 152 | 0 | 0 | 0 | 0 | 0 | 0 | 0 | 0 | 0 | 0 | 0 | 0 | 0 | 0 | 0 | 0 | 0 | 0 |
| ENSMUSG00 Gm14177 | 152 | 0 | 0 | 0 | 0 | 0 | 0 | 0 | 0 | 0 | 0 | 0 | 0 | 0 | 0 | 0 | 0 | 0 | 0 |
| ENSMUSG00 Gm14187 | 137 | 0 | 0 | 0 | 0 | 0 | 0 | 0 | 0 | 0 | 0 | 0 | 0 | 0 | 0 | 0 | 0 | 0 | 0 |
| ENSMUSG00 Gm14189 | 152 | 0 | 0 | 0 | 0 | 0 | 0 | 0 | 0 | 0 | 0 | 0 | 0 | 0 | 0 | 0 | 0 | 0 | 0 |
| ENSMUSG00 Tas2r122 | 929 | 0 | 0 | 0 | 0 | 0 | 0 | 0 | 0 | 0 | 0 | 0 | 0 | 0 | 0 | 0 | 0 | 0 | 0 |
| ENSMUSG00 Gm6134 | 170 | 0 | 0 | 0 | 0 | 0 | 0 | 0 | 0 | 0 | 0 | 0 | 0 | 0 | 0.27320051 | 0 | 0 | 0 | 0 |
| ENSMUSG00 Cdc73 | 1446 | 0 | 0 | 0 | 0 | 0 | 0 | 0 | 0 | 0 | 0 | 0 | 0 | 0 | 0 | 0 | 0 | 0 | 0 |
| ENSMUSG00 Gm10923 | 461 | 0.27660635 | 0.5058357 | 0.94438563 | 0 | 0 | 0 | 0.10263054 | 0.99416232 | 0.26802505 | 1.53418774 | 0.31089995 | 0.72645114 | 0 | 0 | 0.35749723 | 1.57813783 | 0.16627461 | 0 |
| ENSMUSG00 Gm3940 | 431 | 0 | 0.1545842 | 0 | 0.21142016 | 0.22334999 | 0.11497154 | 0 | 0 | 0 | 0 | 0 | 0.1295027 | 0 | 0.32327671 | 0.19119051 | 0.26652395 | 0 | 0.12763134 |
| ENSMUSG00 Gm3150 | 451 | 0.9424651 | 1.32956119 | 1.60687573 | 0.50511136 | 1.92100804 | 1.53822233 | 0.83924929 | 1.60899256 | 1.46116242 | 0.7318291 | 0.31779352 | 0.37127935 | 1.42198431 | 1.85364429 | 1.461696 | 2.20744078 | 1.35969124 | 1.58562835 |
| ENSMUSG00 Gm2606 | 1252 | 0.33949821 | 0.07982323 | 0.57955508 | 0.43668733 | 0.07688806 | 0.39578861 | 0.26452776 | 0.45757671 | 0.65793156 | 0.52724429 | 0.3052713 | 0.891624 | 0.31521992 | 0.25967142 | 0.82271477 | 0.24466842 | 0.30612058 | 0.65905479 |
| ENSMUSG00 Foxd1 | 2460 | 0 | 0 | 0 | 0 | 0.01956582 | 0 | 0 | 0 | 0 | 0 | 0 | 0 | 0 | 0 | 0 | 0 | 0 | 0 |
| ENSMUSG00 Gm3436 | 1912 | 0.04446148 | 0.26134593 | 0.20239971 | 0.16680299 | 0.125868 | 0.0518334 | 0.14847075 | 0.09987553 | 0.06462319 | 0.12330205 | 0.27485593 | 0.3794998 | 0.18060848 | 0.14574504 | 0.30168503 | 0.12015881 | 0.28063188 | 0.23016363 |
| ENSMUSG00 Gm16493 | 3757 | 0.49779818 | 0.24827283 | 0.45064508 | 0.32742832 | 0.43558302 | 0.39568326 | 0.5289147 | 0.25414162 | 0.52620515 | 0.15060113 | 0.81384012 | 0.29712889 | 0.43331202 | 0.4326705 | 0.49349748 | 0.20383608 | 0.67328602 | 0.5271035 |
| ENSMUSG00 AI593442 | 5613 | 0.01514526 | 0.00593495 | 0.05170874 | 0 | 0.00857508 | 0 | 0 | 0.09525987 | 0 | 0 | 0 | 0 | 0 | 0 | 0 | 0 | 0.04096878 | 0 |
| ENSMUSG00 Gm7293 | 3449 | 0.07394348 | 0.03863484 | 0.12622841 | 0.05283972 | 0.02791065 | 0.02873455 | 0.10974237 | 0.13288166 | 0.14329898 | 0.16405 | 0.19392561 | 0.12946515 | 0.04290976 | 0.14812553 | 0.05972971 | 0.07771361 | 0.11112293 | 0.07974646 |
| ENSMUSG00 Sox4 | 404 | 0 | 0 | 0 | 0 | 0 | 0 | 0 | 0 | 0 | 0 | 0 | 0 | 0 | 0 | 0 | 0 | 0 | 0 |
| ENSMUSG00 Gm6344 | 482 | 0 | 0 | 0 | 0.18904998 | 0 | 0 | 0 | 0 | 0 | 0 | 0 | 0 | 0.10234838 | 0 | 0 | 0 | 0 | 0 |
| ENSMUSG00 Gm14762 | 3198 | 0 | 0.04166716 | 0 | 0 | 0 | 0 | 0.02958892 | 0.34633511 | 0 | 0.02948762 | 0 | 0 | 0.01542587 | 0 | 0 | 0.11973298 | 0 | 0.15480987 |
| ENSMUSG00 Fam47a-ps | 2638 | 0 | 0 | 0 | 0 | 0 | 0 | 0 | 0 | 0 | 0 | 0 | 0 | 0 | 0 | 0 | 0 | 0 | 0 |
| ENSMUSG00 F8a | 3693 | 7.94166571 | 8.81307816 | 7.87232379 | 9.78334908 | 8.57590192 | 8.96323478 | 7.78936063 | 9.55585698 | 7.8848813 | 8.61812204 | 10.0000116 | 7.82900439 | 6.41194701 | 9.78432169 | 6.98407094 | 7.88000586 | 8.45814567 | 7.6264995 |
| ENSMUSG00 Gm362 | 1814 | 0 | 0 | 0 | 0 | 0 | 0 | 0 | 0 | 0 | 0 | 0 | 0 | 0 | 0 | 0 | 0 | 0 | 0 |
| ENSMUSG00 Gm2964 | 1959 | 0 | 0 | 0 | 0 | 0 | 0 | 0 | 0 | 0 | 0 | 0 | 0 | 0 | 0 | 0 | 0 | 0 | 0 |
| ENSMUSG00 Gm2933 | 1700 | 0 | 0 | 0 | 0 | 0 | 0 | 0 | 0 | 0 | 0 | 0 | 0 | 0 | 0 | 0 | 0 | 0 | 0 |
| ENSMUSG00 Gm2927 | 1959 | 0 | 0 | 0 | 0 | 0 | 0 | 0 | 0 | 0 | 0 | 0 | 0 | 0 | 0 | 0 | 0 | 0 | 0 |
| ENSMUSG00 Gm2913 | 1959 | 0 | 0 | 0 | 0 | 0 | 0 | 0 | 0 | 0 | 0 | 0 | 0 | 0 | 0 | 0 | 0 | 0 | 0 |
| ENSMUSG00 Gm2882 | 777 | 0 | 0 | 0 | 0 | 0 | 0 | 0 | 0 | 0 | 0 | 0 | 0 | 0 | 0 | 0 | 0 | 0 | 0 |
| ENSMUSG00 Gm2825 | 1959 | 0 | 0 | 0 | 0 | 0 | 0 | 0 | 0 | 0 | 0 | 0 | 0 | 0 | 0 | 0 | 0 | 0 | 0 |
| ENSMUSG00 Gm5926 | 777 | 0 | 0 | 0 | 0 | 0 | 0 | 0 | 0 | 0 | 0 | 0 | 0 | 0 | 0 | 0 | 0 | 0 | 0 |
| ENSMUSG00 Gmcl1l | 1959 | 0 | 0 | 0 | 0 | 0 | 0 | 0 | 0 | 0 | 0 | 0 | 0 | 0 | 0 | 0 | 0 | 0 | 0 |
| ENSMUSG00 Gm2799 | 1496 | 0 | 0 | 0 | 0 | 0 | 0 | 0 | 0 | 0 | 0 | 0 | 0 | 0 | 0 | 0 | 0 | 0 | 0 |
| ENSMUSG00 Gm2784 | 1959 | 0 | 0 | 0 | 0 | 0 | 0 | 0 | 0 | 0 | 0 | 0 | 0 | 0 | 0 | 0 | 0 | 0 | 0 |
| ENSMUSG00 Gm2777 | 1959 | 0 | 0 | 0 | 0 | 0 | 0 | 0 | 0 | 0 | 0 | 0 | 0 | 0 | 0 | 0 | 0 | 0 | 0 |
| ENSMUSG00 Gm5132 | 508 | 0 | 0 | 0 | 0 | 0 | 0 | 0 | 0 | 0 | 0 | 0 | 0 | 0 | 0 | 0 | 0 | 0 | 0 |
| ENSMUSG00 1700012L04R | 499 | 0.08518071 | 0 | 0 | 0 | 0 | 0 | 0 | 0 | 0.16507622 | 0 | 0 | 0 | 0 | 0 | 0.08256825 | 0.07673468 | 0 | 0 |
| ENSMUSG00 Sf3b5 | 732 | 102.488573 | 89.2439693 | 99.1260868 | 87.8855118 | 87.7815821 | 86.3111138 | 106.712067 | 82.7502052 | 85.7489754 | 88.8262565 | 87.0000268 | 94.093617 | 118.073113 | 118.457802 | 97.0938305 | 102.317525 | 81.7313517 | 113.700517 |
| ENSMUSG00 AW011738 | 1847 | 0.02301309 | 0.32465192 | 0.05238065 | 0 | 0 | 0.0804863 | 0.17931172 | 0.51695185 | 0.13379485 | 0.12764132 | 0.28452872 | 0.2719767 | 0.16025528 | 0.77951636 | 0.51306756 | 0 | 0.58101587 | 0.38717834 |
| ENSMUSG00 1190007F08R | 6103 | 8.42024542 | 9.02281077 | 8.79806959 | 8.83897698 | 8.12319865 | 10.084302 | 9.86100731 | 9.48708538 | 7.99030553 | 8.5679387 | 8.07861005 | 8.18532165 | 9.73220193 | 13.3708441 | 9.61347134 | 9.19150251 | 9.3507875 | 11.24879 |
| ENSMUSG00 Ifnab | 669 | 0 | 0 | 0 | 0 | 0 | 0 | 0 | 0 | 0 | 0 | 0 | 0 | 0 | 0 | 0 | 0 | 0 | 0 |
| ENSMUSG00 Ifna2 | 1031 | 0 | 0 | 0 | 0 | 0 | 0 | 0 | 0 | 0 | 0 | 0 | 0 | 0 | 0 | 0 | 0 | 0 | 0 |
| ENSMUSG00 Gm13280 | 1009 | 0 | 0 | 0 | 0 | 0 | 0 | 0 | 0 | 0 | 0 | 0 | 0 | 0 | 0 | 0 | 0 | 0 | 0 |
| ENSMUSG00 Ptprd | 371 | 0 | 0 | 0 | 0 | 0 | 0 | 0 | 0 | 0 | 0 | 0 | 0 | 0 | 0 | 0 | 0 | 0 | 0 |
| ENSMUSG00 Gm10929 | 926 | 0 | 0 | 0 | 0 | 0 | 0 | 0 | 0 | 0 | 0 | 0 | 0 | 0 | 0 | 0 | 0 | 0 | 0 |
| ENSMUSG00 Gm2500 | 490 | 0 | 0 | 0 | 0 | 0 | 0 | 0 | 0 | 0 | 0 | 0 | 0 | 0 | 0 | 0 | 0 | 0 | 0 |
| ENSMUSG00 Gm10930 | 4967 | 0 | 0 | 0 | 0 | 0 | 0 | 0 | 0 | 0 | 0 | 0 | 0 | 0 | 0 | 0 | 0 | 0 | 0 |
| ENSMUSG00 Gm10931 | 2384 | 0 | 0 | 0 | 0 | 0 | 0 | 0 | 0 | 0 | 0 | 0 | 0 | 0 | 0 | 0 | 0 | 0 | 0 |
| ENSMUSG00 Mos | 2619 | 0 | 0.0381591 | 0 | 0 | 0 | 0.13244335 | 0 | 0 | 0 | 0 | 0 | 0.02131182 | 0 | 0 | 0 | 0 | 0 | 0 |
| ENSMUSG00 Vmn1r3 | 1010 | 0 | 0 | 0 | 0 | 0 | 0 | 0 | 0 | 0 | 0 | 0 | 0 | 0 | 0 | 0 | 0 | 0 | 0 |
| ENSMUSG00 Vmn1r2 | 1010 | 0 | 0 | 0 | 0 | 0 | 0 | 0 | 0 | 0 | 0 | 0 | 0 | 0 | 0 | 0 | 0 | 0 | 0 |
| ENSMUSG00 Gm2001 | 493 | 0 | 0 | 0 | 0 | 0 | 0 | 0 | 0 | 0 | 0 | 0 | 0 | 0 | 0 | 0 | 0 | 0 | 0 |
| ENSMUSG00 Gm6316 | 1019 | 0.25027582 | 0.19615051 | 0.18988628 | 0.40240373 | 0.37787575 | 0.04862879 | 0.371444 | 0.07496055 | 0.36376707 | 0.18508618 | 0.42195744 | 0.16432482 | 0.04841209 | 0.63809345 | 0.28303326 | 0.33818986 | 0.18805838 | 0.4858508 |
| ENSMUSG00 2010109K11R | 809 | 0 | 0 | 0 | 0 | 0 | 0 | 0 | 0 | 0 | 0 | 0 | 0 | 0 | 0 | 0 | 0 | 0 | 0 |
| ENSMUSG00 Gm10476 | 1482 | 0 | 0 | 0 | 0 | 0 | 0 | 0 | 0 | 0 | 0 | 0 | 0 | 0 | 0 | 0 | 0 | 0 | 0 |
| ENSMUSG00 Gm9257 | 335 | 0 | 0 | 0 | 0 | 0 | 0 | 0 | 0 | 0 | 0 | 0 | 0 | 0 | 0 | 0 | 0 | 0 | 0 |
| ENSMUSG00 Gm4294 | 682 | 0.81021597 | 0.19538354 | 0.49650251 | 0.4008303 | 0.42344801 | 0.36328984 | 0.34686715 | 0.16800177 | 0.18117236 | 0.27654371 | 0 | 0.08184115 | 0.21700257 | 0.27239934 | 0.18123852 | 0.44915669 | 0.44957533 | 0.24197554 |
| ENSMUSG00 Gm6096 | 455 | 0.56050782 | 0.07321515 | 0.31894635 | 0.10013416 | 0.2115689 | 0.32672132 | 0.41593564 | 0 | 0.09051981 | 0 | 0 | 0 | 0.21684359 | 0.10207492 | 0.45276435 | 0 | 1.09503705 | 0 |
| ENSMUSG00 Gm5329 | 467 | 0 | 0 | 0 | 0 | 0 | 0 | 0 | 0 | 0 | 0 | 0 | 0 | 0 | 0 | 0 | 0 | 0 | 0 |
| ENSMUSG00 Gm6666 | 467 | 0 | 0 | 0 | 0 | 0 | 0 | 0 | 0 | 0 | 0 | 0 | 0 | 0 | 0 | 0 | 0 | 0 | 0 |
| ENSMUSG00 Gm10935 | 2827 | 0 | 0 | 0 | 0 | 0 | 0 | 0 | 0 | 0 | 0 | 0 | 0 | 0 | 0 | 0 | 0 | 0 | 0 |
| ENSMUSG00 Gm8660 | 917 | 0 | 0 | 0 | 0 | 0 | 0 | 0 | 0 | 0 | 0 | 0 | 0 | 0 | 0 | 0 | 0 | 0 | 0 |
| ENSMUSG00 Gm4179 | 920 | 0 | 0 | 0 | 0 | 0 | 0 | 0 | 0 | 0 | 0 | 0 | 0 | 0 | 0 | 0 | 0 | 0 | 0 |
| ENSMUSG00 Vmn1r137 | 923 | 0 | 0 | 0 | 0 | 0 | 0 | 0 | 0 | 0 | 0 | 0 | 0 | 0 | 0 | 0 | 0 | 0 | 0 |
| ENSMUSG00 Gm5725 | 923 | 0 | 0 | 0 | 0 | 0 | 0 | 0 | 0 | 0 | 0 | 0 | 0 | 0 | 0 | 0 | 0 | 0 | 0 |
| ENSMUSG00 Gm4177 | 917 | 0 | 0 | 0 | 0 | 0 | 0 | 0 | 0 | 0 | 0 | 0 | 0 | 0 | 0 | 0 | 0 | 0 | 0 |
| ENSMUSG00 Vmn1r121 | 944 | 0 | 0 | 0 | 0 | 0 | 0 | 0 | 0 | 0 | 0 | 0 | 0 | 0 | 0 | 0 | 0 | 0 | 0 |
| ENSMUSG00 Vmn1r127 | 917 | 0 | 0 | 0 | 0 | 0 | 0 | 0 | 0 | 0 | 0 | 0 | 0 | 0 | 0 | 0 | 0 | 0 | 0 |
| ENSMUSG00 Vmn1r124 | 923 | 0 | 0 | 0 | 0 | 0 | 0 | 0 | 0 | 0 | 0 | 0 | 0 | 0 | 0 | 0 | 0 | 0 | 0 |
| ENSMUSG00 Gm5728 | 917 | 0 | 0 | 0 | 0 | 0 | 0 | 0 | 0 | 0 | 0 | 0 | 0 | 0 | 0 | 0 | 0 | 0 | 0 |
| ENSMUSG00 Olfr141 | 922 | 0 | 0 | 0 | 0 | 0 | 0 | 0 | 0 | 0 | 0 | 0 | 0 | 0 | 0 | 0 | 0 | 0 | 0 |
| ENSMUSG00 Sarnp | 1051 | 49.339976 | 39.9691338 | 43.5868061 | 37.0644081 | 40.8961064 | 41.8204326 | 42.4508623 | 40.6634617 | 41.029764 | 45.1767442 | 39.1836482 | 38.7151456 | 36.5647608 | 37.0315369 | 41.3583649 | 42.8082433 | 42.228213 | 40.09227 |
| ENSMUSG00 Ctdsp2 | 5057 | 11.8597594 | 12.4108152 | 12.1675164 | 11.6853222 | 10.0794359 | 11.9839812 | 13.3414831 | 13.2695375 | 17.8689372 | 15.1139589 | 12.4420843 | 17.0967889 | 14.4864341 | 13.5373907 | 13.589914 | 17.3394287 | 12.0049148 | 12.5638755 |
| ENSMUSG00 Gm10936 | 136 | 0 | 0 | 0 | 0 | 0 | 0 | 0 | 0 | 0 | 0 | 0 | 0 | 0 | 0 | 0 | 0 | 0 | 0 |
| ENSMUSG00 AU041133 | 1491 | 3.6775102 | 4.26744654 | 5.54786968 | 5.98924526 | 7.06967893 | 4.65283883 | 4.72809467 | 7.24912802 | 6.82298417 | 5.97684971 | 4.64612272 | 7.07522483 | 5.95556345 | 9.03339057 | 5.99647058 | 3.95489842 | 5.21813432 | 6.78851485 |
| ENSMUSG00 Gm4767 | 1494 | 0 | 0 | 0.06475707 | 0 | 0.06443363 | 0 | 0 | 0 | 0.08270385 | 0 | 0 | 0 | 0 | 0 | 0 | 0 | 0 | 0 |
| ENSMUSG00 Gm3055 | 729 | 0 | 0 | 0 | 0 | 0 | 0 | 0 | 0 | 0 | 0 | 0 | 0 | 0 | 0 | 0 | 0 | 0 | 0 |
| ENSMUSG00 2210404O07 | 1054 | 27.8662966 | 28.9196378 | 28.500913 | 28.789047 | 28.7695559 | 31.5463797 | 56.3353056 | 37.6488673 | 33.0586265 | 41.5588267 | 39.9333382 | 36.8574015 | 67.9132941 | 63.5852164 | 44.6024433 | 80.8316902 | 65.6710552 | 59.706279 |
| ENSMUSG00 Dohh | 3063 | 15.1675342 | 16.6836369 | 13.6608239 | 19.0395484 | 18.11822 | 16.7440677 | 18.0724239 | 15.7856942 | 19.201549 | 22.5670686 | 18.5921484 | 18.4229954 | 21.0663231 | 20.7732288 | 22.3696335 | 18.0139619 | 16.2664662 | 18.2465728 |
| ENSMUSG00 Scamp4 | 1795 | 63.0355312 | 59.220861 | 58.6680644 | 59.8766037 | 59.8766493 | 63.0244517 | 55.0355842 | 55.235363 | 66.7932855 | 59.6541767 | 63.9039974 | 58.0545081 | 50.0739573 | 48.902131 | 54.7670821 | 49.4471465 | 44.5609926 | 57.8898065 |
| ENSMUSG00 Ccdc105 | 1711 | 0 | 0 | 0 | 0 | 0 | 0 | 0 | 0 | 0.24071605 | 0 | 0 | 0 | 0 | 0 | 0 | 0 | 0 | 0 |
| ENSMUSG00 Gm10941 | 706 | 1.143907 | 0.70778104 | 0.27407099 | 0.64534057 | 1.158984 | 0.14037602 | 0.60313613 | 1.02784086 | 0.9917433 | 1.13535688 | 1.15038852 | 0.6324721 | 0.69875237 | 1.44726618 | 0.70030973 | 0.54235988 | 0.32571924 | 1.16874873 |
| ENSMUSG00 Gm10942 | 70 | 0 | 0 | 0 | 0 | 0 | 0 | 1.35179083 | 0 | 0 | 0 | 0 | 0 | 0 | 0 | 0 | 0 | 0 | 0 |
| ENSMUSG00 Gm10943 | 57 | 0 | 0 | 0 | 0 | 0 | 0.8693462 | 0 | 0 | 0 | 0 | 0 | 0 | 0 | 0 | 0 | 0 | 0 | 0 |
| ENSMUSG00 Ppil6 | 1203 | 1.05997945 | 1.46765037 | 0.92484721 | 1.43916848 | 1.24030726 | 1.56525675 | 1.33718295 | 1.80961508 | 1.91724429 | 1.01905094 | 0.79426365 | 1.34551473 | 1.76331874 | 0.96517222 | 0.65073114 | 0.54109753 | 0.79647334 | 1.64615781 |
| ENSMUSG00 Raet1d | 1264 | 0.23539259 | 0.31626165 | 0.34443178 | 0.14418052 | 0.22847432 | 0.3136249 | 2.24585501 | 0.90646525 | 1.36853929 | 1.19368869 | 0.49135638 | 1.10394902 | 0.31222732 | 0.77161854 | 0.58673102 | 0.39381163 | 0.78835737 | 1.47967534 |
| ENSMUSG00 3110003A17R | 804 | 53.6072742 | 51.046624 | 53.246983 | 44.541021 | 63.5772425 | 46.6559942 | 55.9042848 | 54.7234423 | 53.4297711 | 46.8574775 | 59.4215901 | 48.1791913 | 57.247113 | 58.9216034 | 49.5546076 | 55.1499047 | 63.0191101 | 52.5460124 |
| ENSMUSG00 Mrpl48-ps | 635 | 0.73631014 | 0.52461251 | 0.53325152 | 0.14349935 | 0.22739491 | 0.62428641 | 0.22352447 | 0.12029103 | 0.51888524 | 0.37126538 | 0.45141693 | 0.52739209 | 0.15537612 | 0.21942089 | 0.19465302 | 0.24120068 | 0.42249461 | 0.51977109 |
| ENSMUSG00 Gm10944 | 49 | 0 | 0 | 0 | 0 | 0 | 0 | 0 | 0 | 0 | 0 | 0 | 0 | 0 | 0 | 0 | 0 | 0 | 0 |
| ENSMUSG00 Klhl17 | 3002 | 12.3466068 | 10.6419272 | 9.6521468 | 12.3388171 | 13.0671278 | 13.7169626 | 11.3947591 | 12.1243703 | 12.3751623 | 10.224886 | 11.6811524 | 13.1265349 | 10.7471932 | 6.9000876 | 10.1425549 | 8.43107645 | 11.9370712 | 11.9839959 |
| ENSMUSG00 Plekhn1 | 2194 | 7.90433538 | 6.66561557 | 6.74671845 | 9.34479031 | 6.4717035 | 8.08563292 | 4.37760885 | 3.46412401 | 4.0923703 | 6.12486346 | 2.70013439 | 4.27394317 | 5.48632079 | 2.83660333 | 4.84503255 | 7.6965168 | 5.50263606 | 6.01740456 |
| ENSMUSG00 2310042D19R | 3838 | 1.68337331 | 0.7290993 | 0.7562308 | 0.72413333 | 0.94056651 | 0.64555411 | 0.01232743 | 0.3781426 | 0.11804369 | 0.68797274 | 0.19916606 | 0.78531677 | 0.35989934 | 0.31462904 | 0.52602299 | 0.53874226 | 0.28959422 | 0.11466203 |
| ENSMUSG00 E230028L10R | 1316 | 0.0968963 | 0.17719625 | 0 | 0.48469196 | 0.07314882 | 0.26357837 | 0.53927826 | 1.04477697 | 0 | 0 | 0 | 0.25412212 | 0.84826235 | 0.1874313 | 0 | 0 | 0.14561663 | 0.91960513 |
| ENSMUSG00 Gm10945 | 58 | 0 | 0 | 0 | 0 | 0 | 0 | 0 | 0 | 0 | 0 | 0 | 0 | 0 | 0 | 0 | 0 | 0 | 0 |
| ENSMUSG00 Gm17106 | 457 | 0 | 0.07289474 | 0 | 0 | 0 | 0.21686098 | 0.51764419 | 0.25071599 | 0 | 0 | 0.20908078 | 0.24426986 | 0 | 0.1016282 | 0.18031315 | 0.6702951 | 0.25159495 | 0 |
| ENSMUSG00 C030017K20R | 3614 | 0.08232879 | 0.13826602 | 0.26770078 | 0.0630341 | 0.02663637 | 0.04113398 | 0.13091499 | 0.10567903 | 0.19373845 | 0.03914004 | 0.34370474 | 0.09266574 | 0 | 0.01285116 | 0.01140054 | 0 | 0 | 0 |
| ENSMUSG00 Gm13090 | 742 | 0.0572846 | 0 | 0 | 0 | 0 | 0 | 0 | 0 | 0 | 0 | 0 | 0 | 0 | 0 | 0 | 0 | 0 | 0 |
| ENSMUSG00 1700045H11R | 589 | 0 | 0 | 0 | 0 | 0 | 0 | 0 | 0 | 0 | 0 | 0 | 0 | 0.08375538 | 0 | 0 | 0 | 0.19521034 | 0 |
| ENSMUSG00 A930039A15R | 457 | 3.72036552 | 3.35315784 | 5.61005932 | 3.58905382 | 4.00221685 | 3.25291468 | 3.41645165 | 3.42645186 | 3.96544134 | 3.71427859 | 4.70431757 | 2.32056365 | 5.39736513 | 6.7074612 | 5.77002096 | 5.11100011 | 6.54146862 | 9.02775274 |
| ENSMUSG00 Gm13201 | 597 | 0 | 0.11160099 | 0 | 0.07631666 | 0.24186896 | 0 | 0.23775216 | 0 | 0.27595655 | 0 | 0 | 0 | 0.24789908 | 0 | 0 | 0.19241511 | 0.19259446 | 0.18428511 |
| ENSMUSG00 Gm13157 | 4939 | 1.11017771 | 1.08592346 | 1.74336676 | 0.69185631 | 1.3643388 | 1.04342666 | 1.4848077 | 0.8892744 | 0.85892106 | 0.95466093 | 0.98664624 | 0.81367234 | 2.0475892 | 0.77109034 | 0.64234052 | 0.89931372 | 1.08639027 | 0.42323265 |
| ENSMUSG00 Gm13152 | 1484 | 0.31506532 | 0.22448042 | 0.16298359 | 0.42982117 | 0.25947129 | 0.16695665 | 0.19129115 | 0.30883344 | 0.22202973 | 0.06354542 | 0.12877347 | 0.07522326 | 0.29918279 | 0.21907588 | 0.02776385 | 0.36123215 | 0.23243711 | 0.25947692 |
| ENSMUSG00 Gm13145 | 1423 | 0 | 0.04682065 | 0.0679881 | 0 | 0.03382426 | 0.10446817 | 0 | 0.02683936 | 0 | 0 | 0 | 0 | 0 | 0 | 0 | 0 | 0 | 0 |
| ENSMUSG00 Gm13151 | 1531 | 0 | 0.08703565 | 0.18957621 | 0.05951802 | 0.09431468 | 0.12946501 | 0 | 0.07483815 | 0.05380342 | 0 | 0 | 0.036457 | 0 | 0.03033579 | 0.0807346 | 0 | 0.10013402 | 0.21558108 |
| ENSMUSG00 Rex2 | 746 | 0 | 0 | 0 | 0 | 0 | 0 | 0 | 0 | 0 | 0 | 0 | 0 | 0 | 0 | 0 | 0 | 0 | 0 |
| ENSMUSG00 Gm13235 | 1410 | 0 | 0 | 0 | 0 | 0 | 0.14057513 | 0 | 0 | 0 | 0 | 0.0677659 | 0.07917115 | 0 | 0.03293907 | 0 | 0 | 0.08154531 | 0 |
| ENSMUSG00 Gm13212 | 2159 | 0.3740613 | 0.37031471 | 0.35848841 | 0.12661708 | 0.28981705 | 0.68855118 | 0.43828327 | 0.05306957 | 0.01907686 | 0 | 0 | 0.3877883 | 0.25134372 | 0.19360666 | 0.47709074 | 0.05320603 | 0.19527062 | 0.35670565 |
| ENSMUSG00 Gm13225 | 3638 | 0.35050997 | 0.91569253 | 1.02384877 | 0.16280747 | 1.13780798 | 1.04880717 | 0.40315917 | 0.44092383 | 1.17740452 | 0.10368489 | 0.81419665 | 1.25807706 | 0.33900438 | 1.12344137 | 0.57759191 | 0.11577699 | 0.96921916 | 2.28322571 |
| ENSMUSG00 Gm438 | 1428 | 0 | 0 | 0 | 0 | 0 | 0 | 0 | 0 | 0 | 0 | 0 | 0 | 0 | 0 | 0 | 0 | 0 | 0 |
| ENSMUSG00 Gm436 | 1220 | 0 | 0 | 0 | 0 | 0 | 0 | 0 | 0 | 0 | 0 | 0 | 0 | 0 | 0 | 0 | 0 | 0 | 0 |
| ENSMUSG00 Gm13124 | 1217 | 0 | 0 | 0 | 0 | 0 | 0 | 0 | 0 | 0 | 0 | 0 | 0 | 0 | 0 | 0 | 0 | 0 | 0 |
| ENSMUSG00 Aadacl3 | 3149 | 0 | 0 | 0 | 0 | 0 | 0 | 0 | 0 | 0 | 0 | 0 | 0 | 0 | 0 | 0 | 0 | 0 | 0 |
| ENSMUSG00 Gm13128 | 1742 | 0 | 0 | 0 | 0 | 0 | 0 | 0 | 0 | 0 | 0 | 0 | 0 | 0 | 0 | 0 | 0 | 0 | 0 |
| ENSMUSG00 Pramef17 | 1750 | 0 | 0 | 0 | 0 | 0.02750396 | 0 | 0 | 0 | 0 | 0 | 0 | 0 | 0 | 0 | 0 | 0 | 0 | 0 |
| ENSMUSG00 Gm13101 | 1675 | 0 | 0 | 0 | 0 | 0 | 0 | 0 | 0 | 0 | 0 | 0 | 0 | 0 | 0 | 0 | 0 | 0 | 0 |
| ENSMUSG00 Gm13109 | 2888 | 0 | 0 | 0 | 0 | 0 | 0 | 0 | 0 | 0 | 0 | 0 | 0 | 0 | 0 | 0 | 0 | 0 | 0 |
| ENSMUSG00 Pramef6 | 2081 | 0 | 0 | 0 | 0 | 0 | 0 | 0 | 0 | 0 | 0 | 0 | 0 | 0 | 0 | 0 | 0 | 0 | 0 |
| ENSMUSG00 Gm13088 | 1815 | 0 | 0 | 0 | 0 | 0 | 0 | 0 | 0 | 0 | 0 | 0 | 0 | 0 | 0 | 0 | 0 | 0 | 0 |
| ENSMUSG00 Gm13057 | 2122 | 0 | 0 | 0 | 0 | 0 | 0 | 0 | 0 | 0 | 0 | 0 | 0 | 0 | 0 | 0 | 0 | 0 | 0 |
| ENSMUSG00 Ddi2 | 9543 | 4.16455408 | 4.14709403 | 4.01465326 | 4.02472602 | 3.29352889 | 4.01385969 | 3.68863388 | 3.24973601 | 3.83250823 | 3.68094665 | 3.28912792 | 3.93628219 | 2.78632541 | 3.13423371 | 3.14311356 | 3.24204242 | 3.53824453 | 3.52777673 |
| ENSMUSG00 C230096C10R | 6783 | 8.30928253 | 11.7918707 | 8.81463711 | 10.3440968 | 9.99111728 | 11.9005459 | 7.74943038 | 10.912115 | 6.43634184 | 8.73779065 | 9.09294875 | 10.1954306 | 6.2183089 | 7.0525446 | 6.99144786 | 5.00154477 | 9.67904269 | 7.62325819 |
| ENSMUSG00 Gm13030 | 652 | 0 | 0 | 0 | 0 | 0 | 0 | 0 | 0 | 0 | 0 | 0 | 0 | 0 | 0 | 0 | 0 | 0 | 0 |
| ENSMUSG00 2310026L22R | 398 | 1.81554772 | 2.42732144 | 1.823123 | 2.63292467 | 2.90242756 | 2.2410784 | 2.85302587 | 2.78286349 | 1.86270674 | 1.65856745 | 3.24101477 | 2.24384573 | 1.11554587 | 1.28363056 | 1.24225797 | 0.48103778 | 2.98521408 | 0.13821384 |
| ENSMUSG00 Gm13011 | 950 | 0 | 0 | 0 | 0 | 0 | 0 | 0 | 0 | 0 | 0 | 0 | 0 | 0 | 0 | 0 | 0 | 0 | 0 |
| ENSMUSG00 2610002D18R | 1272 | 0.10024806 | 0 | 0 | 0.10745529 | 0 | 0 | 0 | 0.06005095 | 0.19427692 | 0.14827265 | 0 | 0 | 0.03878295 | 0.47466442 | 0.12956464 | 0.15051339 | 0.39169958 | 0 |
| ENSMUSG00 Gm5776 | 715 | 2.08067295 | 1.21137797 | 0.5412423 | 2.03909569 | 1.07707801 | 1.59400402 | 1.85280421 | 0.85465516 | 0.86405278 | 0.98917559 | 1.93772559 | 1.32708568 | 1.37991377 | 1.94870297 | 1.4982384 | 1.8208121 | 1.60809637 | 1.46178046 |
| ENSMUSG00 Gm10948 | 260 | 0 | 0 | 0 | 0 | 0 | 0 | 0 | 0 | 0 | 0 | 0 | 0 | 0 | 0.35726221 | 0 | 0 | 0 | 0 |
| ENSMUSG00 Gm17068 | 979 | 0.30391852 | 0.10208241 | 0.24705582 | 0.04653835 | 0 | 0.30369397 | 0.19331023 | 0.39011647 | 0 | 0.09632422 | 0.19519901 | 0.1710388 | 0.15117033 | 0.09488067 | 0.04208535 | 0.11733588 | 0.39148414 | 0 |
| ENSMUSG00 Gm831 | 4509 | 0 | 0 | 0 | 0 | 0 | 0 | 0 | 0 | 0 | 0 | 0 | 0 | 0 | 0 | 0 | 0 | 0 | 0 |
| ENSMUSG00 Nkain1 | 2545 | 0 | 0.11780591 | 0.13305097 | 0.01790218 | 0 | 0.09735311 | 0.11154266 | 0.07503419 | 0 | 0 | 0 | 0.13158899 | 0 | 0.1459932 | 0 | 0 | 0 | 0 |
| ENSMUSG00 2210404O09 | 1740 | 5.3497894 | 3.96308569 | 4.42033984 | 4.34662837 | 3.62372529 | 3.64525856 | 4.56812072 | 4.14848509 | 5.2074905 | 5.41962106 | 5.98559825 | 3.43234247 | 7.65495268 | 3.81695661 | 4.07279748 | 4.33520095 | 5.52867846 | 4.48924894 |
| ENSMUSG00 Dear1 | 518 | 0 | 0 | 0 | 0 | 0 | 0 | 0.09133722 | 0 | 0 | 0 | 0 | 0 | 0 | 0 | 0 | 0 | 0 | 0 |
| ENSMUSG00 Dcdc2b | 1391 | 1.58897854 | 2.70622361 | 1.84313236 | 2.71859574 | 2.38756487 | 1.17558606 | 1.42856399 | 2.416198 | 1.98382211 | 1.86433406 | 3.40023069 | 2.12669311 | 2.12790441 | 1.66944958 | 2.48808821 | 2.17466426 | 3.00328279 | 3.95464462 |
| ENSMUSG00 1700125D06 | 730 | 3.78470745 | 1.87099817 | 1.85542308 | 1.93478407 | 3.09890467 | 1.96853325 | 4.083149 | 5.17951759 | 5.02136973 | 5.23179036 | 4.842941 | 4.58758871 | 8.51482408 | 7.57102246 | 5.64404875 | 5.71736467 | 6.98273629 | 4.52129644 |
| ENSMUSG00 Ebf1 | 451 | 0 | 0 | 0 | 0 | 0 | 0 | 0 | 0 | 0 | 0 | 0 | 0 | 0 | 0 | 0 | 0 | 0 | 0 |
| ENSMUSG00 Gm9631 | 2015 | 0.04218876 | 0.06612981 | 0 | 0.06783282 | 0.11943405 | 0 | 0.02348024 | 0.09477023 | 0.08175983 | 0.09359941 | 0.14225794 | 0.08310024 | 0.12241171 | 0.06914752 | 0.10223711 | 0.13301948 | 0.01902049 | 0.05459961 |
| ENSMUSG00 Bnip3 | 2259 | 85.5936458 | 75.0314324 | 98.3316739 | 78.1132912 | 84.609061 | 81.7762686 | 174.464195 | 161.83164 | 181.392052 | 158.087396 | 148.421717 | 156.155373 | 148.563538 | 119.183875 | 148.993143 | 131.584765 | 141.530752 | 128.695498 |
| ENSMUSG00 1110065P20R | 1402 | 12.0663766 | 11.0726168 | 10.592492 | 11.6989843 | 12.2218009 | 10.567951 | 14.0048229 | 11.8772378 | 12.3677055 | 14.0914013 | 13.2897531 | 13.8941985 | 19.9157384 | 16.9279092 | 12.6367112 | 13.3825946 | 10.6340439 | 12.4378651 |
| ENSMUSG00 1810043H04R | 777 | 27.5162208 | 18.1784648 | 22.1634193 | 20.8748157 | 26.1411477 | 19.3236529 | 23.9303511 | 20.6936955 | 28.5708262 | 24.2125233 | 24.0411953 | 22.771641 | 27.0468427 | 29.5281585 | 24.9754605 | 25.1821112 | 21.9993931 | 25.416048 |
| ENSMUSG00 Gm10955 | 202 | 0 | 0 | 0 | 0 | 0 | 0 | 0 | 0 | 0 | 0 | 0 | 0 | 0 | 0 | 0 | 0 | 0 | 0 |
| ENSMUSG00 Gm12887 | 356 | 0 | 0 | 0 | 0 | 0 | 0 | 0 | 0 | 0 | 0 | 0 | 0 | 0 | 0 | 0 | 0 | 0 | 0 |
| ENSMUSG00 Gm12886 | 1145 | 0 | 0 | 0 | 0 | 0 | 0 | 0 | 0 | 0 | 0 | 0 | 0 | 0 | 0 | 0 | 0 | 0 | 0 |
| ENSMUSG00 Tmco2 | 717 | 0 | 0 | 0 | 0 | 0 | 0 | 0 | 0 | 0 | 0 | 0 | 0 | 0 | 0 | 0 | 0 | 0 | 0 |
| ENSMUSG00 Ube2d3 | 2666 | 152.466991 | 149.333233 | 160.579799 | 162.932857 | 159.163931 | 160.405135 | 152.213371 | 162.382176 | 153.221254 | 161.950109 | 159.632157 | 156.978934 | 149.031981 | 139.924572 | 159.845346 | 164.867923 | 162.017017 | 162.840161 |
| ENSMUSG00 E430018J23R | 4890 | 1.37337788 | 2.05054829 | 2.0576062 | 1.77958271 | 1.68314089 | 1.23628497 | 2.40917322 | 2.21812726 | 2.20672129 | 1.92845412 | 3.10683779 | 1.86052209 | 0.86759609 | 2.00402913 | 1.8283717 | 1.94976713 | 2.24941661 | 1.78863967 |
| ENSMUSG00 Hivep3 | 2972 | 0.22883002 | 0.81825077 | 0.30925204 | 0.39858249 | 0.3886831 | 0.13338555 | 0.65269846 | 0.15420889 | 0.01385818 | 0.52354415 | 0.46617557 | 0.30048809 | 0.7303514 | 0.35942598 | 0.47135024 | 0.73437571 | 0.67058124 | 0.37018241 |
| ENSMUSG00 AU022252 | 9537 | 1.99222121 | 2.10279567 | 2.358571 | 1.82970326 | 1.85219838 | 1.85491516 | 1.90500878 | 1.81010442 | 1.96496431 | 2.21984542 | 2.02882019 | 2.25322744 | 1.851822 | 1.90412479 | 1.79719485 | 1.49757749 | 1.51102943 | 1.43622393 |
| ENSMUSG00 Gm11735 | 1572 | 0 | 0 | 0 | 0 | 0 | 0 | 0 | 0 | 0 | 0 | 0 | 0 | 0 | 0 | 0 | 0 | 0 | 0 |
| ENSMUSG00 Gm2464 | 2549 | 0 | 0 | 0 | 0 | 0 | 0 | 0 | 0 | 0 | 0 | 0 | 0 | 0 | 0 | 0 | 0 | 0 | 0 |
| ENSMUSG00 Ccdc24 | 5446 | 0.03121937 | 0.07952031 | 0.05329438 | 0.09202561 | 0.16792261 | 0.06369246 | 0.06950081 | 0.19636197 | 0.09831522 | 0.12986789 | 0.05263492 | 0.04099571 | 0.11775889 | 0.06822488 | 0.0529583 | 0.2109288 | 0 | 0.06060497 |
| ENSMUSG00 Gm10959 | 68 | 0 | 1.46968651 | 0 | 0 | 0 | 0 | 0.69577469 | 0 | 0 | 0 | 0.70257292 | 0 | 0.72546937 | 3.41500643 | 0.60590523 | 0.56309717 | 0 | 0 |
| ENSMUSG00 Hs3st4 | 3187 | 0 | 0 | 0 | 0 | 0 | 0 | 0 | 0 | 0 | 0 | 0 | 0 | 0 | 0 | 0 | 0 | 0 | 0 |
| ENSMUSG00 Gm4609 | 1001 | 0.12738814 | 0.19967769 | 0.14497562 | 0.13654659 | 0.19233536 | 0.29701938 | 0.18906165 | 0.15261699 | 0.12343611 | 0.1884144 | 0.28636339 | 0 | 0.1478479 | 0.13919307 | 0.28812277 | 0.03825236 | 0.22972805 | 0.65944983 |
| ENSMUSG00 1700042G07 | 383 | 0 | 0 | 0 | 0 | 0 | 0.25876101 | 0 | 0 | 0 | 0 | 1.12264915 | 0 | 0 | 0 | 0 | 0 | 0 | 0 |
| ENSMUSG00 Dnahc7c | 607 | 0 | 0 | 0 | 0 | 0 | 0 | 0 | 0 | 0 | 0 | 0 | 0 | 0 | 0 | 0 | 0 | 0 | 0 |
| ENSMUSG00 Cyp4a12b | 2383 | 0 | 0.01397939 | 0 | 0.0191192 | 0.10099019 | 0 | 0.21839676 | 0.20835133 | 0.15555126 | 0 | 0.04009648 | 0.0702673 | 0.35192723 | 0.91601851 | 0.70888115 | 0 | 0.01608321 | 0.53093137 |
| ENSMUSG00 Skint5 | 4530 | 0 | 0 | 0 | 0 | 0 | 0 | 0 | 0 | 0 | 0 | 0 | 0 | 0 | 0 | 0 | 0 | 0 | 0 |
| ENSMUSG00 Skint8 | 4834 | 0 | 0 | 0 | 0 | 0 | 0 | 0 | 0 | 0 | 0.00975397 | 0 | 0 | 0 | 0 | 0 | 0 | 0 | 0 |
| ENSMUSG00 1700092K14R | 392 | 0 | 0.50989124 | 2.09783167 | 1.16227154 | 0 | 0.50564014 | 0.36208683 | 1.75373276 | 0.10506764 | 0 | 0.97499915 | 0.14238689 | 0.50338691 | 0 | 0.52553005 | 0.97680121 | 0.09777117 | 0 |
| ENSMUSG00 Gm12525 | 552 | 0 | 0 | 0 | 0 | 0 | 0 | 0 | 0 | 0 | 0.17083588 | 0 | 0 | 0 | 0 | 0 | 0.13873408 | 0 | 0.19930836 |
| ENSMUSG00 Gm10961 | 141 | 4.22037209 | 4.72523322 | 8.2337924 | 2.90815177 | 8.87539018 | 5.2715674 | 6.37546737 | 9.20951547 | 6.13416192 | 10.7008688 | 5.42127188 | 7.91711526 | 4.5483328 | 2.30573483 | 3.79872501 | 4.07346887 | 8.69816667 | 7.41257466 |
| ENSMUSG00 E030025P04R | 1075 | 0 | 0 | 0 | 0 | 0 | 0 | 0 | 0 | 0 | 0.43861119 | 0 | 0 | 0 | 0 | 0 | 0 | 0 | 0 |
| ENSMUSG00 Gm4070 | 9598 | 0 | 0 | 0 | 0 | 0 | 0 | 0 | 0 | 0 | 0 | 0 | 0 | 0 | 0 | 0 | 0 | 0 | 0 |
| ENSMUSG00 1810010H24R | 1247 | 0.68171894 | 1.68300909 | 1.3965093 | 0.9499496 | 1.11934706 | 1.51002717 | 3.18706097 | 2.32768451 | 3.26981962 | 2.26867858 | 2.79676982 | 4.96835489 | 4.43077364 | 2.04845614 | 3.17189203 | 1.84237085 | 2.30511008 | 2.99969467 |
| ENSMUSG00 Gm5901 | 999 | 0.21273862 | 0.36680865 | 0.43579757 | 0.41045986 | 0.24090052 | 0.24801168 | 0.71040058 | 0.07646127 | 0.49473292 | 1.13274963 | 0 | 0.11174307 | 0.24690649 | 0.51139636 | 0.1237284 | 0.45994724 | 0.46037594 | 0.33038503 |
| ENSMUSG00 1700024P16R | 3359 | 293.62611 | 267.187461 | 264.232669 | 248.490126 | 259.502571 | 268.165999 | 245.310395 | 269.222895 | 248.921541 | 255.924865 | 259.298978 | 274.159755 | 233.867654 | 252.034184 | 255.869144 | 265.286995 | 267.200125 | 253.395626 |
| ENSMUSG00 Trim30c | 1535 | 0 | 0 | 0 | 0 | 0 | 0 | 0 | 0 | 0 | 0 | 0 | 0 | 0.16069028 | 0 | 0.13420702 | 0 | 0 | 0 |
| ENSMUSG00 Gm10963 | 61 | 0 | 0 | 0 | 0 | 0 | 0 | 0 | 0 | 0 | 0 | 0 | 0 | 0 | 0 | 0 | 0 | 0 | 0 |
| ENSMUSG00 4930456L15R | 1181 | 0 | 0 | 0 | 0 | 0 | 0 | 0 | 0 | 0 | 0 | 0 | 0.09452271 | 0.16708524 | 0.23595641 | 0.10466102 | 0.06484438 | 0 | 0 |
| ENSMUSG00 Smarcd2 | 3848 | 24.2460658 | 21.3140191 | 23.105652 | 23.6448559 | 19.0501351 | 23.4628588 | 20.5456046 | 17.9349457 | 23.2798004 | 20.37724 | 18.3625685 | 18.1749026 | 18.6533107 | 19.4683765 | 19.4015643 | 19.9015631 | 19.2727091 | 22.5868993 |
| ENSMUSG00 Gm10964 | 118 | 0 | 0.28231266 | 0 | 0 | 0 | 0 | 0 | 0 | 0 | 0 | 0 | 0 | 0 | 0 | 0 | 0 | 0 | 0 |
| ENSMUSG00 Hbb-bh2 | 441 | 0 | 0 | 0 | 0 | 0 | 0 | 0 | 0 | 0 | 0 | 0 | 0 | 0 | 0 | 0 | 0 | 0 | 0 |
| ENSMUSG00 Ccdc47 | 6145 | 26.6582504 | 27.9459674 | 29.2209186 | 28.115131 | 28.573679 | 29.6832567 | 22.0972154 | 27.8005884 | 22.4598884 | 27.492428 | 24.7932209 | 27.1312261 | 19.339396 | 20.1421469 | 22.8368591 | 25.0742724 | 25.7026313 | 24.7697638 |
| ENSMUSG00 Olfr613 | 5136 | 0 | 0 | 0 | 0 | 0 | 0 | 0 | 0 | 0 | 0 | 0 | 0 | 0 | 0 | 0 | 0 | 0 | 0 |
| ENSMUSG00 Gm12789 | 1296 | 0 | 0 | 0 | 0 | 0 | 0 | 0 | 0 | 0 | 0 | 0 | 0 | 0 | 0 | 0 | 0 | 0 | 0 |
| ENSMUSG00 Gm12790 | 947 | 0.22442015 | 0 | 0 | 0 | 0 | 0 | 0 | 0 | 0 | 0 | 0 | 0 | 0 | 0 | 0 | 0 | 0 | 0 |
| ENSMUSG00 10-Mar | 4234 | 0.88343304 | 0.75532306 | 0.68550114 | 1.11911871 | 1.023116 | 0.76072926 | 0.42462962 | 0.84791824 | 0.46692318 | 0.42317589 | 0.7560043 | 0.36911633 | 0 | 0 | 0 | 0.09043601 | 0.06336421 | 0 |
| ENSMUSG00 Tomt | 3654 | 0.16285508 | 0.1549861 | 0.11914663 | 0.06234407 | 0.06586196 | 0.05424492 | 0.23306738 | 0.10452217 | 0.090173 | 0.16775018 | 0 | 0.10692656 | 0.16200958 | 0.06355239 | 0.14658463 | 0.1571864 | 0.19928835 | 0.03010898 |
| ENSMUSG00 Gm10967 | 90 | 0 | 0 | 0 | 0 | 0 | 0 | 0 | 0 | 0 | 0 | 0 | 0 | 0 | 0 | 0 | 0 | 0 | 0 |
| ENSMUSG00 Lrrc37a | 9636 | 0 | 0 | 0 | 0 | 0 | 0 | 0 | 0 | 0 | 0 | 0 | 0 | 0 | 0 | 0 | 0 | 0 | 0 |
| ENSMUSG00 1700028N14 | 744 | 0.68556736 | 0.44775395 | 0.9102546 | 0.79609352 | 0.58224101 | 0.33301568 | 1.01747697 | 0.51333874 | 0.71965686 | 0.69712061 | 0.19264096 | 0.30008421 | 0.06630634 | 0.81152303 | 0.49640592 | 0.61759044 | 0.41211072 | 0.07393697 |
| ENSMUSG00 Gm3200 | 1001 | 0.25477628 | 0.43263499 | 0.53157726 | 0.50067082 | 0.2404192 | 0.4950323 | 0.23632707 | 0.53415947 | 0.24687222 | 0.5652432 | 0.71590847 | 0.83639854 | 1.08421796 | 0.74236304 | 1.15249107 | 0.7650471 | 0.49774412 | 1.09908305 |
| ENSMUSG00 Gm12695 | 1893 | 0 | 0 | 0 | 0 | 0 | 0 | 0 | 0 | 0 | 0 | 0 | 0 | 0 | 0 | 0 | 0 | 0 | 0 |
| ENSMUSG00 Gm11627 | 1508 | 0 | 0 | 0 | 0 | 0 | 0 | 0 | 0 | 0 | 0 | 0 | 0 | 0 | 0 | 0 | 0 | 0 | 0 |
| ENSMUSG00 Ifna6 | 569 | 0 | 0 | 0 | 0 | 0 | 0 | 0 | 0 | 0 | 0 | 0 | 0 | 0 | 0 | 0 | 0 | 0 | 0 |
| ENSMUSG00 Gm13283 | 3016 | 0.01409323 | 0.01104539 | 0.01603897 | 0.07553224 | 0 | 0.0328599 | 0.07843614 | 0 | 0 | 0.06253409 | 0 | 0.0925326 | 0.01635674 | 0 | 0.09562695 | 0.1015666 | 0 | 0.10943456 |
| ENSMUSG00 Ifna9 | 572 | 0 | 0 | 0 | 0 | 0 | 0 | 0 | 0 | 0 | 0 | 0 | 0 | 0 | 0 | 0 | 0 | 0 | 0 |
| ENSMUSG00 Fam63a | 1124 | 0 | 0 | 0 | 0 | 0 | 0 | 0 | 0 | 0 | 0 | 0 | 0 | 0 | 0 | 0 | 0 | 0 | 0 |
| ENSMUSG00 Gm17546 | 730 | 4.13406507 | 6.29750603 | 5.10241348 | 3.30785664 | 4.61538994 | 5.56619747 | 10.1106547 | 8.26630079 | 7.78594408 | 7.81539054 | 10.7330044 | 8.71641854 | 5.74412735 | 10.6248803 | 11.6831809 | 8.86453788 | 7.35024872 | 10.6250466 |
| ENSMUSG00 G6pc | 2409 | 0 | 0 | 0 | 0 | 0 | 0 | 0.21603963 | 0.06341619 | 0 | 0.05871819 | 0 | 0 | 0 | 0 | 0 | 0 | 0 | 0 |
| ENSMUSG00 Aoc2 | 4135 | 0.08223492 | 0.26585865 | 0.11698556 | 0.02203678 | 0.19788215 | 0.21570718 | 0.09153602 | 0.42487316 | 0.37849761 | 0.05701415 | 0.15019939 | 0.05399339 | 0.28632794 | 0.13478332 | 0.0797128 | 0.52782699 | 0.31513763 | 0.02660658 |
| ENSMUSG00 Psme3 | 2937 | 59.235167 | 66.5350485 | 69.0273972 | 66.8445829 | 71.0589108 | 67.6562689 | 65.9671162 | 64.4472408 | 69.7380126 | 71.0391086 | 71.670571 | 58.0581713 | 61.9462414 | 65.594169 | 71.3767506 | 71.9529664 | 72.1113778 | 56.2265367 |
| ENSMUSG00 Cntd1 | 4250 | 0.61007429 | 0.32920978 | 0.52357233 | 0.52529204 | 0.64553404 | 0.53633547 | 0.82379723 | 1.3210078 | 1.02724016 | 0.90973122 | 0.67447 | 0.94558299 | 0.99824586 | 1.4862108 | 1.54142291 | 1.30638543 | 1.44287235 | 0.88014571 |
| ENSMUSG00 Gm10972 | 79 | 0 | 0 | 0 | 0 | 0 | 0 | 0 | 0 | 0 | 0 | 0 | 0 | 0 | 0 | 0 | 0 | 0 | 0 |
| ENSMUSG00 Vps25 | 1714 | 0.32238465 | 0.68025163 | 0.84667789 | 0.69112436 | 0.67395926 | 0.57821159 | 0.63488426 | 1.42608737 | 0.64879576 | 0.79776569 | 0.75258103 | 0.68385584 | 0.77710721 | 0.37935661 | 1.08172113 | 0.24573902 | 0.98387227 | 0.51350391 |
| ENSMUSG00 Crnn | 1625 | 0 | 0 | 0 | 0 | 0 | 0 | 0 | 0 | 0 | 0 | 0 | 0 | 0 | 0 | 0 | 0 | 0 | 0 |
| ENSMUSG00 Lce1d | 676 | 0 | 0 | 0 | 0 | 0 | 0 | 0 | 0 | 0 | 0 | 0 | 0 | 0 | 0 | 0 | 0 | 0 | 0 |
| ENSMUSG00 Gm11756 | 1745 | 0 | 0 | 0 | 0 | 0 | 0 | 0 | 0 | 0 | 0 | 0 | 0 | 0 | 0 | 0 | 0 | 0 | 0 |
| ENSMUSG00 Gm13871 | 1731 | 0 | 0 | 0 | 0 | 0 | 0 | 0 | 0 | 0 | 0 | 0 | 0 | 0 | 0 | 0 | 0 | 0 | 0 |
| ENSMUSG00 Gm11757 | 1734 | 0 | 0 | 0 | 0 | 0 | 0 | 0 | 0 | 0 | 0 | 0 | 0 | 0 | 0 | 0 | 0 | 0 | 0 |
| ENSMUSG00 Gm11758 | 1739 | 0 | 0 | 0 | 0 | 0 | 0 | 0 | 0 | 0 | 0 | 0 | 0 | 0 | 0 | 0 | 0 | 0 | 0 |
| ENSMUSG00 Sprr2a1 | 3539 | 0 | 0 | 0 | 0 | 0 | 0 | 0 | 0 | 0 | 0 | 0 | 0 | 0 | 0 | 0 | 0 | 0 | 0 |
| ENSMUSG00 Gm10973 | 76 | 0 | 0.43832756 | 0 | 0.59948743 | 0.63331479 | 0 | 0 | 0.50253161 | 0.54192784 | 0.62040399 | 0.62861787 | 0 | 1.29820835 | 0 | 0.54212574 | 0 | 0 | 0 |
| ENSMUSG00 1700094D03 | 3093 | 1.97890247 | 1.64787353 | 2.09571712 | 2.07698262 | 1.68064913 | 1.9865952 | 1.91208693 | 2.09916211 | 2.2903591 | 1.60065433 | 2.40960023 | 1.64216789 | 2.52002681 | 1.48657118 | 1.46529943 | 2.26549666 | 1.56130404 | 2.08084885 |
| ENSMUSG00 Gm11595 | 1935 | 0 | 0 | 0 | 0 | 0 | 0 | 0 | 0 | 0 | 0 | 0 | 0 | 0 | 0 | 0 | 0 | 0 | 0 |
| ENSMUSG00 Fam174b | 2675 | 0.85804841 | 1.30760893 | 0.47017263 | 1.48179845 | 1.16956076 | 0.79654861 | 4.22718888 | 5.79667865 | 3.55666734 | 7.20921033 | 7.28679741 | 8.53405832 | 2.76627574 | 3.69816472 | 1.89450145 | 7.24300835 | 7.13513864 | 3.94831719 |
| ENSMUSG00 Chd2 | 9502 | 8.4589863 | 9.19242356 | 8.20649663 | 9.11029091 | 8.7986899 | 8.67251062 | 7.99664937 | 9.69882835 | 7.48569909 | 9.96903419 | 8.9496344 | 7.75968115 | 11.3924188 | 10.1471191 | 10.4499842 | 9.8607784 | 12.6046809 | 10.4379519 |
| ENSMUSG00 Mup20 | 921 | 0 | 0 | 0 | 0 | 0 | 0 | 0 | 0 | 0 | 0 | 0 | 0 | 0 | 0 | 0 | 0 | 0 | 0 |
| ENSMUSG00 Mup19 | 1354 | 0 | 0 | 0 | 0 | 0 | 0 | 0 | 0 | 0 | 0 | 0 | 0 | 0 | 0 | 0 | 0 | 0 | 0 |
| ENSMUSG00 Mup18 | 923 | 0 | 0.03609198 | 0 | 0 | 0 | 0 | 0 | 0 | 0 | 0 | 0 | 0 | 0 | 0 | 0 | 0 | 0 | 0 |
| ENSMUSG00 Mup16 | 1031 | 0 | 0.03231125 | 0 | 0 | 0 | 0 | 0 | 0 | 0 | 0 | 0 | 0 | 0 | 0 | 0 | 0 | 0 | 0 |
| ENSMUSG00 Casc3 | 4015 | 15.3293885 | 16.0383125 | 14.7229026 | 15.2059276 | 15.8241941 | 15.4150347 | 14.7417587 | 14.0498576 | 12.2995348 | 14.8791883 | 16.4921774 | 13.8044715 | 12.8029534 | 10.9545581 | 14.5719077 | 16.7849238 | 15.149149 | 16.2218636 |
| ENSMUSG00 Gm10974 | 123 | 0 | 1.08334615 | 0 | 0 | 0 | 0 | 0 | 0 | 0 | 0 | 0 | 0 | 0 | 0 | 0 | 0 | 0 | 0 |
| ENSMUSG00 Mup-ps8 | 539 | 0 | 0 | 0 | 0 | 0 | 0 | 0 | 0 | 0 | 0 | 0 | 0 | 0 | 0 | 0 | 0 | 0 | 0 |
| ENSMUSG00 Mup-ps7 | 638 | 0 | 0 | 0 | 0 | 0 | 0 | 0 | 0 | 0 | 0 | 0 | 0 | 0 | 0 | 0 | 0 | 0 | 0 |
| ENSMUSG00 Mup10 | 1384 | 0 | 0 | 0 | 0 | 0 | 0 | 0 | 0 | 0 | 0 | 0 | 0 | 0 | 0 | 0 | 0 | 0 | 0 |
| ENSMUSG00 Tm2d3 | 1482 | 42.1036427 | 39.6293067 | 36.6555159 | 37.4141639 | 33.3545787 | 36.4122314 | 35.1812228 | 32.2651065 | 40.6584835 | 35.5380132 | 36.234179 | 33.5948522 | 35.2846372 | 39.4241983 | 28.7187633 | 35.112642 | 32.2489151 | 34.334294 |
| ENSMUSG00 Mup1 | 1694 | 0 | 0 | 0 | 0 | 0 | 0 | 0 | 0 | 0 | 0 | 0 | 0 | 0 | 0 | 0 | 0 | 0 | 0 |
| ENSMUSG00 5830417I10R | 9873 | 1.06338281 | 0.92451464 | 0.78883201 | 0.83064803 | 0.9408955 | 1.10418327 | 0.87695941 | 0.78141044 | 0.86352768 | 1.11274322 | 1.10328072 | 0.9723786 | 0.69953088 | 0.54097742 | 1.10588598 | 0.92691737 | 0.80355955 | 0.65188549 |
| ENSMUSG00 Mup-ps4 | 623 | 0 | 0 | 0 | 0 | 0 | 0 | 0 | 0 | 0 | 0 | 0 | 0 | 0 | 0 | 0 | 0 | 0 | 0 |
| ENSMUSG00 Mup9 | 1591 | 0 | 0 | 0 | 0 | 0 | 0 | 0 | 0 | 0 | 0 | 0 | 0 | 0 | 0 | 0 | 0 | 0 | 0 |
| ENSMUSG00 Mup8 | 1229 | 0 | 0 | 0 | 0 | 0 | 0 | 0 | 0 | 0 | 0 | 0 | 0 | 0 | 0 | 0 | 0 | 0 | 0 |
| ENSMUSG00 Mup2 | 1510 | 0 | 0 | 0.03203545 | 0 | 0 | 0 | 0 | 0 | 0 | 0 | 0 | 0 | 0 | 0 | 0 | 0 | 0 | 0 |
| ENSMUSG00 Mup6 | 992 | 0.72841531 | 1.0074464 | 0.34134548 | 0.45928472 | 0.14556025 | 0.64938058 | 0 | 0 | 0 | 0 | 0 | 0 | 0 | 0 | 0 | 0 | 0 | 0 |
| ENSMUSG00 A230006K03R | 2292 | 0 | 0 | 0 | 0 | 0 | 0 | 0 | 0 | 0 | 0 | 0 | 0 | 0 | 0 | 0 | 0 | 0 | 0 |
| ENSMUSG00 Gm6821 | 560 | 0 | 0 | 0 | 0 | 0 | 0 | 0 | 0 | 0 | 0 | 0 | 0 | 0 | 0 | 0 | 0 | 0 | 0.09823055 |
| ENSMUSG00 Cisd3 | 729 | 23.730599 | 17.5475328 | 19.5086666 | 15.1870148 | 16.9022942 | 19.98423 | 30.7629764 | 24.4661892 | 26.3277316 | 22.3141188 | 25.7552245 | 23.5053613 | 26.729914 | 28.3506432 | 30.2370815 | 28.0482639 | 22.2387155 | 24.1466586 |
| ENSMUSG00 Gm12537 | 2253 | 0.16979431 | 0.16264618 | 0.23617791 | 0.12133434 | 0.29908874 | 0.24193523 | 0.23099843 | 0.16951799 | 0.12796521 | 0.27206353 | 0.27566554 | 0.12386965 | 0.19706492 | 0.2679863 | 0.16458677 | 0.2719262 | 0.37424702 | 0.17091156 |
| ENSMUSG00 Mrgpra3 | 1293 | 0 | 0 | 0 | 0 | 0 | 0 | 0 | 0 | 0 | 0 | 0 | 0 | 0 | 0 | 0 | 0 | 0 | 0 |
| ENSMUSG00 D030028A08 | 6217 | 3.17233419 | 2.82921154 | 3.16680503 | 3.0926107 | 2.60130712 | 2.6780953 | 3.08974548 | 2.80745181 | 2.81554917 | 2.96540534 | 2.55127653 | 2.99862173 | 2.06310093 | 1.45674715 | 2.2598891 | 2.06942964 | 1.87408626 | 1.9465986 |
| ENSMUSG00 Gm10979 | 46 | 0 | 0 | 0 | 0 | 0 | 0 | 0 | 0 | 0 | 0 | 0 | 0 | 0 | 0 | 0 | 0 | 0 | 0 |
| ENSMUSG00 Gm53 | 3931 | 18.8034854 | 19.6775732 | 17.8924226 | 17.2925663 | 19.578465 | 18.4294318 | 14.2263003 | 13.7671417 | 12.0594453 | 12.0545553 | 14.5111423 | 10.6065378 | 21.4219239 | 21.5502456 | 16.7699032 | 17.2897044 | 18.1150495 | 21.0184885 |
| ENSMUSG00 Gm10982 | 64 | 0 | 0 | 0 | 0 | 0 | 0 | 0 | 0 | 0 | 0 | 0 | 0 | 3.08324483 | 0 | 0 | 0 | 0 | 0 |
| ENSMUSG00 Tomm5 | 1290 | 15.3216332 | 12.7570308 | 13.7995808 | 13.4917201 | 13.32023 | 12.5994547 | 15.7342165 | 13.7670288 | 16.155331 | 14.10866 | 16.4434741 | 13.0669226 | 20.2681521 | 17.9295779 | 17.087467 | 15.7021173 | 12.359488 | 15.5219495 |
| ENSMUSG00 Tmem8b | 3405 | 2.95851005 | 7.1224044 | 5.228035 | 5.63324513 | 6.2479619 | 5.13718501 | 6.53067815 | 5.85504671 | 8.22520726 | 6.67449015 | 6.72076508 | 6.13070714 | 6.92530292 | 4.80126836 | 5.7476473 | 4.83552458 | 4.25472547 | 5.23434672 |
| ENSMUSG00 Msmp | 740 | 0 | 0 | 0 | 0 | 0 | 0 | 0 | 0 | 0 | 0 | 0 | 0 | 0 | 0 | 0 | 0 | 0 | 0 |
| ENSMUSG00 Gm12429 | 4142 | 0 | 0 | 0 | 0 | 0 | 0 | 0 | 0 | 0 | 0 | 0 | 0 | 0 | 0 | 0 | 0 | 0 | 0 |
| ENSMUSG00 Gm12394 | 3344 | 0 | 0 | 0 | 0 | 0 | 0 | 0 | 0 | 0 | 0 | 0 | 0 | 0 | 0 | 0 | 0 | 0 | 0 |
| ENSMUSG00 Gm2163 | 907 | 0 | 0 | 0 | 0 | 0 | 0 | 0 | 0 | 0 | 0 | 0 | 0 | 0 | 0 | 0 | 0 | 0 | 0 |
| ENSMUSG00 Gm2542 | 3923 | 0 | 0 | 0 | 0 | 0 | 0 | 0 | 0 | 0 | 0 | 0 | 0 | 0 | 0 | 0 | 0 | 0 | 0 |
| ENSMUSG00 Gm7819 | 3538 | 0 | 0 | 0 | 0 | 0 | 0 | 0 | 0 | 0 | 0 | 0 | 0 | 0 | 0 | 0 | 0 | 0 | 0 |

| | | | | | | | | | | | | | | | | | | | | |
|---|---|---|---|---|---|---|---|---|---|---|---|---|---|---|---|---|---|---|---|---|
| ENSMUSG00 | Gm7819 | 907 | 0 | 0 | 0 | 0 | 0 | 0 | 0 | 0 | 0 | 0 | 0 | 0 | 0 | 0 | 0 | 0 | 0 | 0 |
| ENSMUSG00 | Ccl21c | 864 | 0 | 0 | 0 | 0 | 0 | 0 | 0 | 0 | 0 | 0 | 0 | 0 | 0 | 0 | 0 | 0 | 0 | 0 |
| ENSMUSG00 | Gm2506 | 927 | 0 | 0 | 0 | 0 | 0 | 0 | 0 | 0 | 0 | 0 | 0 | 0 | 0 | 0 | 0 | 0 | 0 | 0 |
| ENSMUSG00 | Gm2002 | 1939 | 0 | 0 | 0 | 0.09398875 | 0 | 0 | 0 | 0 | 0 | 0 | 0 | 0 | 0 | 0 | 0 | 0 | 0 | 0 |
| ENSMUSG00 | Gm2023 | 716 | 0 | 0 | 0 | 0 | 0 | 0 | 0 | 0 | 0 | 0 | 0 | 0 | 0 | 0 | 0 | 0 | 0 | 0 |
| ENSMUSG00 | Gm2128 | 1569 | 0 | 0 | 0 | 0 | 0 | 0 | 0 | 0 | 0 | 0 | 0 | 0 | 0 | 0 | 0 | 0 | 0 | 0 |
| ENSMUSG00 | Gm1987 | 864 | 0 | 0 | 0 | 0 | 0 | 0 | 0 | 0 | 0 | 0 | 0 | 0 | 0 | 0 | 0 | 0 | 0 | 0 |
| ENSMUSG00 | Gm10985 | 79 | 0 | 0 | 0 | 0 | 0 | 0 | 0 | 0 | 0 | 0 | 0 | 0 | 0 | 0 | 0 | 0 | 0 | 0 |
| ENSMUSG00 | Gm3893 | 907 | 0 | 0 | 0 | 0 | 0 | 0 | 0 | 0 | 0 | 0 | 0 | 0 | 0 | 0 | 0 | 0 | 0 | 0 |
| ENSMUSG00 | Gm13298 | 4126 | 0 | 0 | 0 | 0 | 0 | 0 | 0 | 0 | 0 | 0 | 0 | 0 | 0 | 0 | 0 | 0 | 0 | 0 |
| ENSMUSG00 | Gm13308 | 1516 | 0 | 0.06592261 | 0 | 0 | 0.06349858 | 0 | 0 | 0 | 0.02716789 | 0.09330614 | 0.03151382 | 0 | 0.03254084 | 0.06127188 | 0.05435561 | 0.02525766 | 0 | 0.07257138 |
| ENSMUSG00 | Gm12776 | 1067 | 0 | 0 | 0 | 0 | 0 | 0 | 0 | 0 | 0 | 0 | 0 | 0 | 0 | 0 | 0 | 0 | 0.17959839 | 0 |
| ENSMUSG00 | Gm12769 | 404 | 0 | 0 | 0 | 0 | 0 | 0 | 0 | 0 | 0 | 0 | 0 | 0 | 0 | 0 | 0 | 0 | 0 | 0 |
| ENSMUSG00 | Gm4362 | 336 | 0 | 0 | 0 | 0 | 0 | 0 | 0 | 0 | 0 | 0 | 0 | 0 | 0 | 0 | 0 | 0 | 0 | 0 |
| ENSMUSG00 | Gm5326 | 415 | 0 | 0 | 0 | 0 | 0 | 0 | 0 | 0 | 0 | 0 | 0 | 0 | 0 | 0 | 0 | 0 | 0 | 0 |
| ENSMUSG00 | Gm4684 | 517 | 0 | 0 | 0 | 0 | 0 | 0 | 0 | 0 | 0 | 0 | 0 | 0 | 0 | 0 | 0 | 0 | 0 | 0 |
| ENSMUSG00 | Gm6662 | 416 | 0 | 0 | 0 | 0 | 0 | 0 | 0 | 0 | 0 | 0 | 0 | 0 | 0 | 0 | 0 | 0 | 0 | 0 |
| ENSMUSG00 | Haus5 | 5172 | 4.12559907 | 4.09004018 | 4.39589313 | 4.72171689 | 3.15481866 | 3.64076542 | 4.07994099 | 4.08360374 | 4.18076581 | 4.38505187 | 3.97200931 | 3.5289485 | 4.33991152 | 4.86711047 | 3.87160792 | 4.67157256 | 3.58660628 | 4.08420282 |
| ENSMUSG00 | Slfn1 | 1862 | 0 | 0 | 0 | 0 | 0 | 0 | 0 | 0 | 0 | 0 | 0 | 0 | 0 | 0 | 0 | 0 | 0 | 0 |
| ENSMUSG00 | U2af1l4 | 2388 | 17.5324951 | 19.0419182 | 16.2257947 | 18.8502143 | 18.8859349 | 17.4513605 | 20.5655614 | 19.7999138 | 17.8681869 | 19.4881676 | 21.4267088 | 20.9659336 | 19.8112683 | 15.4035667 | 16.0803392 | 16.178904 | 15.600151 | 18.1981551 |
| ENSMUSG00 | Fut9 | 622 | 0 | 0 | 0 | 0 | 0 | 0 | 0 | 0 | 0 | 0 | 0 | 0 | 0 | 0 | 0 | 0 | 0 | 0 |
| ENSMUSG00 | Gm10988 | 868 | 0 | 0 | 0 | 0 | 0 | 0.11417681 | 0 | 0 | 0 | 0.48888978 | 0 | 0 | 0 | 0 | 0 | 0 | 0 | 0.25349819 |
| ENSMUSG00 | Zfp566 | 1688 | 19.1625823 | 17.0314145 | 19.1430795 | 15.3579587 | 16.338621 | 17.2906162 | 20.6852826 | 17.5350189 | 19.9344688 | 15.9496751 | 17.29295 | 16.66534 | 9.3812473 | 8.30930947 | 11.8625333 | 8.89213158 | 11.7158585 | 8.99438001 |
| ENSMUSG00 | Gm10989 | 61 | 2.78722466 | 0 | 0 | 0 | 0 | 0 | 0 | 0 | 0 | 0 | 0.78319604 | 1.83002172 | 0.80871996 | 0 | 0 | 0.62771488 | 0 | 0 |
| ENSMUSG00 | Evi2a | 3133 | 0.04070078 | 0.22329102 | 0.10808002 | 0.10179614 | 0.07681443 | 0.07908192 | 0.36243354 | 1.30436867 | 0.55213331 | 0.24079517 | 0.71670062 | 0.49883133 | 0.51961483 | 0.57814217 | 0.71014491 | 0.40331632 | 0.0122331 | 0.08778983 |
| ENSMUSG00 | Gm12353 | 549 | 0.23226872 | 0.18203767 | 0.44056039 | 0.08298915 | 0.35068797 | 0 | 0 | 0.62610496 | 0.60016781 | 0.08588471 | 0.08702178 | 0.30500362 | 0.44928886 | 0.33839044 | 0.07504837 | 0.41847658 | 0.34905553 | 0.30059621 |
| ENSMUSG00 | Rad54b | 3587 | 0.04739914 | 0.26932644 | 0.06742895 | 0 | 0.01341843 | 0.09670174 | 0.01319004 | 0.09582705 | 0 | 0.18402839 | 0.2264216 | 0 | 0.0550119 | 0.25895783 | 0.18378168 | 0.18147207 | 0.39533677 | 0.04600706 |
| ENSMUSG00 | Hps3 | 61 | 0 | 0 | 0 | 0 | 0.78904793 | 0 | 0 | 0 | 0 | 0 | 0 | 0 | 0 | 0 | 0 | 0 | 0 | 0 |
| ENSMUSG00 | 9530053A07f | 10869 | 0 | 0 | 0 | 0 | 0 | 0.03191362 | 0.00870599 | 0.00702777 | 0.00378936 | 0 | 0 | 0.01027062 | 0 | 0.00427308 | 0 | 0 | 0 | 0 |
| ENSMUSG00 | Zfp59 | 3603 | 2.61897005 | 1.85842124 | 2.17499637 | 1.84621495 | 2.03054466 | 2.29678227 | 2.7444768 | 2.88324548 | 2.76634383 | 1.9368038 | 2.50609691 | 2.78845941 | 1.80733086 | 1.16013541 | 2.10410388 | 1.66850551 | 1.92535657 | 1.70996946 |
| ENSMUSG00 | Gm5150 | 909 | 0 | 0 | 0 | 0 | 0 | 0 | 0 | 0.12604753 | 0 | 0 | 0 | 0 | 0 | 0 | 0 | 0 | 0 | 0 |
| ENSMUSG00 | Sirpb1b | 1490 | 0 | 0 | 0 | 0 | 0 | 0 | 0.03175348 | 0.07689745 | 0 | 0.0949343 | 0 | 0.18730088 | 0.03310867 | 0 | 0 | 0 | 0 | 0 |
| ENSMUSG00 | Gm9733 | 615 | 0 | 0 | 0 | 0 | 0 | 0 | 0 | 0 | 0 | 0 | 0 | 0 | 0 | 0 | 0 | 0 | 0 | 0 |
| ENSMUSG00 | 1810022K09f | 524 | 76.979794 | 68.5330152 | 73.5757704 | 63.9071901 | 81.0159479 | 71.8703769 | 75.1224215 | 68.8046333 | 74.6702099 | 67.2167468 | 74.671166 | 71.0477995 | 85.2952616 | 75.7818601 | 70.4515154 | 78.6272778 | 83.3816383 | 84.4032859 |
| ENSMUSG00 | BC024978 | 2589 | 0.41044009 | 0.90069625 | 1.19579218 | 0.63352553 | 0.89236475 | 1.20580232 | 0.85890147 | 1.15064017 | 1.11357903 | 0.4006626 | 1.23635466 | 0.77611582 | 0.81934046 | 1.50687653 | 1.59140811 | 0.8282248 | 1.02144244 | 0.67991171 |
| ENSMUSG00 | Cyp2t4 | 1608 | 0.21146854 | 0.39362251 | 0.3609965 | 0.14166991 | 0 | 0.06163275 | 0.02942331 | 0.23751494 | 0.23052154 | 0.05864515 | 0 | 0.03471123 | 0.12271621 | 0.23106511 | 0.07686857 | 0.19050053 | 0.19067809 | 0.41051572 |
| ENSMUSG00 | Dph1 | 3226 | 4.9804577 | 4.98764039 | 4.64841736 | 5.29615363 | 3.77476029 | 5.8369618 | 5.23578003 | 4.90131885 | 6.7793056 | 6.32865918 | 7.68605191 | 5.96912697 | 5.38277585 | 6.1618318 | 6.87118322 | 6.65872001 | 6.3916763 | 6.46263219 |
| ENSMUSG00 | Gm5890 | 1334 | 0 | 0 | 0 | 0 | 0 | 0 | 0 | 0 | 0 | 0 | 0 | 0 | 0 | 0 | 0 | 0 | 0 | 0 |
| ENSMUSG00 | Gm6155 | 1833 | 0 | 0 | 0 | 0 | 0 | 0 | 0 | 0 | 0 | 0 | 0 | 0 | 0 | 0 | 0 | 0 | 0 | 0 |
| ENSMUSG00 | Dact3 | 2248 | 2.77947548 | 2.69704037 | 2.88347556 | 2.77662948 | 2.11968881 | 3.01989524 | 2.56768097 | 2.02175084 | 2.03367582 | 2.68473755 | 1.67892425 | 2.88016764 | 1.448357 | 1.11564979 | 0.91640471 | 1.26045594 | 1.85834803 | 2.07997067 |
| ENSMUSG00 | Ceacam15 | 1388 | 0 | 0 | 0 | 0 | 0 | 0 | 0 | 0 | 0 | 0 | 0 | 0 | 0.07108345 | 0 | 0 | 0 | 0 | 0 |
| ENSMUSG00 | Zfp541 | 4545 | 0 | 0 | 0 | 0 | 0 | 0 | 0.01040983 | 0.01680634 | 0.0453097 | 0.12449031 | 0 | 0 | 0 | 0.01021872 | 0.05439149 | 0 | 0 | 0 |
| ENSMUSG00 | Sult2a1 | 1009 | 0 | 0 | 0 | 0 | 0 | 0 | 0 | 0 | 0 | 0 | 0 | 0 | 0 | 0 | 0 | 0 | 0 | 0 |
| ENSMUSG00 | Sult2a5 | 1162 | 0 | 0 | 0 | 0 | 0 | 0 | 0 | 0 | 0.24811154 | 0 | 0.08222884 | 0.09606827 | 0 | 0 | 0 | 0 | 0.06596609 | 0 |
| ENSMUSG00 | 9230107M04 | 525 | 2.59079168 | 1.64978143 | 0.18428012 | 0.26034882 | 0.27503956 | 0.09438616 | 201.056356 | 114.649955 | 115.165343 | 57.20952 | 34.5799699 | 70.3808926 | 102.98625 | 215.58903 | 53.8386997 | 12.5447323 | 21.6087312 | 70.0973189 |
| ENSMUSG00 | Gm10991 | 312 | 0 | 0 | 0 | 0 | 0 | 0 | 0 | 0 | 0 | 0 | 0 | 0 | 0 | 0 | 0 | 0 | 0 | 0 |
| ENSMUSG00 | Vmn2r49 | 2553 | 0 | 0 | 0 | 0 | 0 | 0 | 0 | 0 | 0 | 0 | 0 | 0 | 0 | 0 | 0 | 0 | 0 | 0 |
| ENSMUSG00 | Vmn2r38 | 2696 | 0 | 0 | 0 | 0 | 0 | 0 | 0 | 0 | 0 | 0 | 0 | 0 | 0 | 0 | 0 | 0 | 0 | 0 |
| ENSMUSG00 | Vmn1r58 | 3579 | 0 | 0 | 0 | 0 | 0 | 0 | 0 | 0 | 0 | 0 | 0 | 0 | 0 | 0 | 0 | 0 | 0 | 0 |
| ENSMUSG00 | Gp6 | 1346 | 0 | 0 | 0 | 0 | 0 | 0 | 0 | 0 | 0 | 0 | 0 | 0 | 0 | 0 | 0 | 0 | 0 | 0 |
| ENSMUSG00 | Eif5a | 2967 | 227.181355 | 209.241354 | 220.004186 | 210.959766 | 211.621484 | 217.484225 | 224.444205 | 214.09371 | 209.250266 | 211.470984 | 216.235227 | 209.605026 | 230.065972 | 224.753693 | 207.757492 | 217.109164 | 198.413185 | 211.526424 |
| ENSMUSG00 | Leng1 | 2218 | 6.40068927 | 6.65356724 | 6.67371519 | 5.54620468 | 5.51195158 | 5.85338873 | 5.18349007 | 4.92471916 | 6.35067105 | 5.90978156 | 5.55723142 | 5.51110465 | 8.58526604 | 7.8314196 | 7.87622169 | 8.57999634 | 7.11922197 | 5.33226237 |
| ENSMUSG00 | Gm3144 | 554 | 0 | 0 | 0 | 0 | 0 | 0 | 0.1708039 | 0 | 0 | 0 | 0 | 0 | 0 | 0 | 0 | 0 | 0 | 0 |
| ENSMUSG00 | Cacng6 | 779 | 0 | 0 | 0 | 0 | 0 | 0 | 0 | 0 | 0 | 0 | 0 | 0 | 0 | 0 | 0 | 0 | 0 | 0 |
| ENSMUSG00 | Prkcc | 2735 | 0.15541198 | 0.09744174 | 0.10612109 | 0.08329259 | 0 | 0 | 0 | 0.13964315 | 0 | 0.25859618 | 0.19214791 | 0.06122376 | 0.46896887 | 0.08490692 | | | | |
| ENSMUSG00 | Nlrp12 | 4341 | 0 | 0 | 0 | 0 | 0 | 0 | 0 | 0 | 0 | 0 | 0 | 0 | 0 | 0 | 0 | 0 | 0 | 0 |
| ENSMUSG00 | Gm10995 | 90 | 0 | 0 | 0 | 0 | 0 | 0 | 0 | 0 | 0 | 0 | 0 | 0 | 0 | 0 | 0 | 0 | 0 | 0 |
| ENSMUSG00 | Gm17414 | 3516 | 0.15715793 | 0.37898628 | 0.20637172 | 0.47945354 | 0.56126532 | 0.26777643 | 0.45751737 | 0.51053553 | 0.11714026 | 0.17433423 | 0.38046042 | 0.28574571 | 0.64541189 | 0.47553673 | 0.28123929 | 0.37027322 | 0.43602158 | 0.34419805 |
| ENSMUSG00 | Gm17382 | 1512 | 0.05622378 | 0.33048506 | 0 | 0 | 0 | 0.03277297 | 0 | 0.05051905 | 0.05447952 | 0 | 0.12638878 | 0.07383024 | 0.03262693 | 0 | 0 | 0.07597343 | 0.17743656 | 0.10914505 |
| ENSMUSG00 | Gm10999 | 200 | 0 | 0 | 0 | 0 | 0 | 0 | 0 | 0 | 0 | 0 | 0 | 0 | 0 | 0 | 0 | 0 | 0 | 0 |
| ENSMUSG00 | Gm6746 | 482 | 0 | 0 | 0 | 0 | 0 | 0.3084195 | 0 | 0 | 0 | 0.19564607 | 0 | 0.11580013 | 0 | 0 | 0 | 0 | 0 | 0.11412678 |
| ENSMUSG00 | Gm2348 | 299 | 0 | 0.55707181 | 0 | 0 | 0 | 0 | 0 | 0 | 0 | 0 | 0 | 0 | 0 | 0 | 0 | 0 | 0 | 0 |
| ENSMUSG00 | Hist3h2a | 1989 | 9.29600381 | 4.5388609 | 6.68814522 | 6.18475715 | 5.73517642 | 7.59858408 | 10.6566517 | 5.76054335 | 8.80053756 | 8.41553527 | 7.90244411 | 10.9161854 | 9.69772733 | 7.5655527 | 10.5437868 | 9.77960209 | 9.32625827 | 11.2009493 |
| ENSMUSG00 | Igtp | 2196 | 9.73593057 | 8.88950638 | 8.48078743 | 10.0831819 | 10.739819 | 14.6898131 | 9.11350783 | 8.15675625 | 8.77745418 | 9.51173112 | 12.4876258 | 9.96345161 | 11.2322216 | 8.67125493 | 10.3941994 | 12.9553376 | 10.9777963 | 11.7733516 |
| ENSMUSG00 | Gm11003 | 64 | 0 | 0 | 0 | 0 | 0 | 0 | 0 | 0 | 0 | 0 | 0 | 0 | 0 | 0 | 0 | 0 | 0 | 0 |
| ENSMUSG00 | Gm17491 | 2307 | 2.26620574 | 1.84830969 | 2.13875167 | 1.36268403 | 2.12806945 | 1.56798854 | 1.86625652 | 3.26133649 | 5.28444242 | 2.41169614 | 2.67142161 | 2.85489735 | 2.45911161 | 1.73133572 | 2.23242067 | 1.82573334 | 1.91050028 | 2.93286525 |
| ENSMUSG00 | Gm11006 | 49 | 0 | 0 | 0 | 0 | 0 | 0 | 0 | 0 | 0 | 0 | 0 | 0 | 2.01354764 | 0 | 0 | 0 | 0 | 0 |
| ENSMUSG00 | Zfp931 | 2596 | 2.97994696 | 3.55457307 | 2.12426135 | 2.98358149 | 2.39176354 | 3.26406681 | 4.48340486 | 3.60444476 | 5.12451641 | 4.12296211 | 3.80948244 | 2.68758006 | 3.81961301 | 3.13086105 | 3.60275547 | 3.30396612 | 2.84937415 | 3.45396163 |
| ENSMUSG00 | Gm14326 | 2857 | 1.74067399 | 1.60909325 | 1.76088455 | 1.75418792 | 1.63416052 | 1.59667781 | 2.38467825 | 2.29929759 | 1.96057618 | 2.14546427 | 2.90964045 | 1.36755211 | 2.22744744 | 2.0970554 | 2.71119794 | 2.92171943 | 3.39396329 | 2.23348141 |
| ENSMUSG00 | Gm14325 | 1973 | 1.89582133 | 2.11054829 | 1.78979611 | 2.40159585 | 1.80525208 | 1.83342603 | 2.66178782 | 2.14868559 | 2.79725956 | 2.36589945 | 2.61515231 | 1.92370251 | 2.65037163 | 3.03663825 | 2.35974446 | 2.61986581 | 3.51599581 | 2.23047569 |
| ENSMUSG00 | Gm14322 | 1248 | 0.57899679 | 0.13346512 | 0.19380421 | 0.36507247 | 0.77134493 | 0.39705716 | 0.75821601 | 0.58145484 | 0.42902621 | 1.09564935 | 0.42109338 | 0.80503359 | 0.63246048 | 0.63265183 | 0.79233761 | 0.70567626 | 1.01343574 | 0.96971182 |
| ENSMUSG00 | Gm14406 | 1616 | 0 | 0 | 0 | 0 | 0 | 0 | 0 | 0 | 0 | 0 | 0.02956371 | 0 | 0 | 0 | 0 | 0 | 0 | 0 |
| ENSMUSG00 | Gm14420 | 2169 | 1.05822015 | 0.92151851 | 0.64676458 | 0.50413327 | 0.7988701 | 1.0509109 | 1.00340398 | 1.07410629 | 1.55707436 | 0.56519976 | 0.68281407 | 1.26093475 | 0.95525151 | 1.92714056 | 2.3174688 | 1.11217532 | 1.16622204 | 1.19199079 |
| ENSMUSG00 | Gm14418 | 2542 | 0.16721155 | 0.13104994 | 0.1522377 | 0.23300298 | 0.113608 | 0.15594881 | 0.27918575 | 0.28546642 | 0.35645293 | 0.05564599 | 0.0939712 | 0.21957381 | 0.21347407 | 0.20097756 | 0.27554148 | 0.06025273 | 0.27138999 | 0.77904321 |
| ENSMUSG00 | Gm14412 | 1577 | 0.6199233 | 0.29573907 | 1.19630164 | 0.54892825 | 0.21364836 | 0.91124241 | 0.87004926 | 0.62967816 | 0.96632916 | 1.07636355 | 1.03002447 | 0.95562644 | 0.84461748 | 1.26638919 | 0.33964504 | 0.9469459 | 2.13869 | 0.38370334 |
| ENSMUSG00 | Gm14409 | 1047 | 0 | 0 | 0 | 0 | 0 | 0 | 0 | 0 | 0 | 0 | 0 | 0 | 0 | 0 | 0 | 0 | 0.03660582 | 0 |
| ENSMUSG00 | Gm14410 | 2392 | 0.56863112 | 0.22282872 | 0.4651301 | 0.59046504 | 0.52317308 | 0.31074039 | 0.51426825 | 0.38320136 | 0.25827665 | 0.43366031 | 0.2197009 | 0.28001169 | 0.1856134 | 0.34949564 | 0.60286558 | 0.27213224 | 0.60886258 | 0.66691643 |
| ENSMUSG00 | Gm14401 | 1324 | 0.03210361 | 0.05032159 | 0 | 0 | 0 | 0.07485307 | 0.03573465 | 0.08653868 | 0 | 0 | 0 | 0 | 0 | 0.03507862 | 0 | 0 | 0 | 0 |
| ENSMUSG00 | Gm14419 | 2743 | 0.03099174 | 0.03643408 | 0 | 0.01660993 | 0 | 0 | 0.01724852 | 0.01392359 | 0.01501514 | 0 | 0 | 0 | 0 | 0 | 0 | 0 | 0.0279448 | 0.04010872 |
| ENSMUSG00 | Gm14408 | 404 | 0 | 0 | 0 | 0 | 0 | 0 | 0 | 0 | 0 | 0 | 0 | 0 | 0 | 0 | 0 | 0 | 0 | 0 |
| ENSMUSG00 | Gm14295 | 3522 | 1.09823141 | 0.87018349 | 0.76914188 | 0.8667206 | 1.06595402 | 0.80196076 | 1.54484897 | 1.18199088 | 0.97060784 | 1.19148569 | 1.11230738 | 1.14103569 | 1.69482169 | 1.4637404 | 1.46229259 | 1.4676979 | 1.58876756 | 1.29635317 |
| ENSMUSG00 | Gm14305 | 2564 | 0 | 0 | 0.01886643 | 0 | 0 | 0 | 0 | 0 | 0 | 0 | 0 | 0 | 0.05772065 | 0 | 0 | 0 | 0 | 0 |
| ENSMUSG00 | Gm14306 | 2798 | 0.01519127 | 0 | 0 | 0 | 0.05160678 | 0.01771005 | 0.03381893 | 0.04094968 | 0.04415995 | 0.01685157 | 0.03414936 | 0 | 0 | 0.09959418 | 0 | 0 | 0.01369775 | 0 |
| ENSMUSG00 | Gm14308 | 3323 | 0 | 0 | 0 | 0 | 0 | 0 | 0 | 0 | 0 | 0 | 0 | 0 | 0 | 0 | 0 | 0 | 0 | 0 |
| ENSMUSG00 | Gm14434 | 2844 | 0 | 0 | 0 | 0 | 0 | 0 | 0 | 0 | 0 | 0 | 0 | 0 | 0 | 0 | 0 | 0 | 0 | 0 |
| ENSMUSG00 | Gm14431 | 1753 | 0 | 0 | 0 | 0 | 0 | 0 | 0 | 0 | 0 | 0 | 0 | 0 | 0 | 0 | 0 | 0 | 0 | 0 |
| ENSMUSG00 | Gm14430 | 1497 | 0 | 0 | 0 | 0 | 0 | 0 | 0 | 0 | 0 | 0 | 0 | 0 | 0 | 0 | 0 | 0 | 0 | 0 |
| ENSMUSG00 | Gm14436 | 2035 | 0 | 0 | 0 | 0 | 0 | 0 | 0 | 0 | 0 | 0 | 0 | 0 | 0 | 0 | 0 | 0 | 0 | 0 |
| ENSMUSG00 | Gm14432 | 1497 | 0 | 0 | 0 | 0 | 0 | 0 | 0 | 0 | 0 | 0 | 0 | 0 | 0 | 0 | 0.110091 | 0 | 0 | 0 |
| ENSMUSG00 | Gm2026 | 3093 | 0 | 0 | 0 | 0 | 0 | 0 | 0 | 0 | 0 | 0 | 0 | 0 | 0 | 0 | 0 | 0 | 0 | 0 |
| ENSMUSG00 | Gm6710 | 3151 | 0 | 0 | 0 | 0.01445923 | 0 | 0 | 0 | 0 | 0 | 0.01496373 | 0 | 0 | 0 | 0 | 0 | 0 | 0 | 0 |
| ENSMUSG00 | Gm2007 | 1497 | 0 | 0 | 0 | 0 | 0 | 0 | 0 | 0 | 0 | 0 | 0 | 0 | 0 | 0 | 0 | 0 | 0 | 0 |
| ENSMUSG00 | Gm14288 | 3574 | 0 | 0 | 0 | 0 | 0 | 0 | 0 | 0 | 0 | 0 | 0 | 0 | 0 | 0 | 0 | 0 | 0 | 0 |
| ENSMUSG00 | Gm11007 | 1497 | 0 | 0 | 0 | 0 | 0 | 0 | 0 | 0 | 0 | 0 | 0 | 0 | 0 | 0 | 0 | 0 | 0 | 0 |
| ENSMUSG00 | Gm11008 | 1026 | 0 | 0 | 0 | 0 | 0 | 0 | 0 | 0 | 0 | 0 | 0 | 0 | 0 | 0 | 0 | 0 | 0 | 0 |
| ENSMUSG00 | Gm2004 | 1749 | 0 | 0 | 0 | 0 | 0.02751968 | 0 | 0 | 0 | 0 | 0 | 0 | 0 | 0 | 0 | 0 | 0 | 0 | 0 |
| ENSMUSG00 | 2210418O10 | 5784 | 0 | 0 | 0 | 0 | 0 | 0 | 0 | 0 | 0 | 0 | 0 | 0 | 0 | 0 | 0 | 0 | 0 | 0 |
| ENSMUSG00 | Gm11009 | 414 | 0 | 0 | 0 | 0 | 0 | 0 | 0 | 0 | 0 | 0 | 0 | 0 | 0 | 0 | 0 | 0 | 0 | 0 |
| ENSMUSG00 | Gm8923 | 1296 | 0 | 0 | 0 | 0 | 0 | 0 | 0 | 0 | 0 | 0 | 0 | 0 | 0 | 0 | 0 | 0 | 0 | 0 |
| ENSMUSG00 | Gm4724 | 2625 | 0.06476979 | 0.06345313 | 0.01842801 | 0.01735659 | 0.05500791 | 0.07550893 | 0.23431041 | 0.05819795 | 0.15690101 | 0.17962173 | 0.07279994 | 0 | 0.09396556 | 0.14154389 | 0 | 0.04376069 | 0.14600494 | 0.06286755 |
| ENSMUSG00 | Gm4723 | 3148 | 0 | 0 | 0 | 0 | 0 | 0 | 0 | 0 | 0 | 0 | 0 | 0 | 0 | 0 | 0 | 0 | 0 | 0 |
| ENSMUSG00 | Gm4631 | 5702 | 0 | 0 | 0 | 0 | 0 | 0 | 0 | 0 | 0 | 0 | 0 | 0 | 0 | 0 | 0 | 0 | 0 | 0 |
| ENSMUSG00 | Gm14440 | 3379 | 0 | 0 | 0 | 0 | 0 | 0 | 0 | 0 | 0 | 0 | 0 | 0 | 0 | 0 | 0 | 0 | 0 | 0 |
| ENSMUSG00 | Gm14443 | 5759 | 0.22880022 | 0.50325088 | 0.33598562 | 0.36391874 | 0.32594982 | 0.42161555 | 0.59151118 | 0.60349169 | 0.47201077 | 0.23743191 | 0.26546252 | 0.48459509 | 0.35977436 | 0.51613502 | 0.35056021 | 0.57844814 | 0.27951111 | 0.19103701 |
| ENSMUSG00 | Gm14391 | 2792 | 0 | 0 | 0 | 0 | 0 | 0 | 0 | 0 | 0 | 0 | 0 | 0 | 0 | 0 | 0 | 0 | 0 | 0 |
| ENSMUSG00 | Gm14393 | 933 | 0.27334518 | 0.49987194 | 0.3629311 | 0.24416422 | 0.7222368 | 0.3186671 | 0.20284107 | 0.57309071 | 0.22072088 | 0.10107332 | 0 | 0.41876703 | 0.31724706 | 0.54757231 | 0.13248089 | 0.57456431 | 0.49294273 | 0.11791877 |
| ENSMUSG00 | Gm14444 | 2006 | 0 | 0 | 0 | 0 | 0 | 0 | 0 | 0 | 0 | 0 | 0 | 0 | 0 | 0 | 0 | 0 | 0 | 0 |
| ENSMUSG00 | Fam188b | 3179 | 0 | 0.16766477 | 0 | 0.11465503 | 0 | 0.01558752 | 0.35718915 | 0.0360419 | 0 | 0.20764701 | 0.04508489 | 0.3160371 | 0 | 0 | 0 | 0 | 0 | 0 |
| ENSMUSG00 | Mon1b | 2104 | 7.69699243 | 8.47072168 | 8.43777835 | 8.05546982 | 10.1342406 | 8.50215628 | 7.26330575 | 7.35167442 | 7.67353335 | 9.43462266 | 6.9709659 | 7.8523936 | 7.71397376 | 5.16535953 | 6.12901891 | 6.67901755 | 8.48861899 | 4.99369743 |
| ENSMUSG00 | Gm17407 | 1217 | 1.71138342 | 1.36864808 | 2.18614969 | 0.74874354 | 0.94919159 | 1.5065334 | 0.77752965 | 0.75317803 | 1.52291965 | 1.04607148 | 1.21694635 | 1.60521626 | 1.25660595 | 0.4579532 | 0.9817955 | 0.44048357 | 1.4171597 | 0.27120348 |
| ENSMUSG00 | Gm11011 | 252 | 0 | 0 | 0 | 0 | 0 | 0 | 0 | 0 | 0 | 0 | 0 | 0 | 0 | 0 | 0 | 0 | 0 | 0 |
| ENSMUSG00 | Hsp25-ps1 | 592 | 0.64619355 | 0.73153315 | 0.08171204 | 0.46176734 | 0.40651963 | 0.66963153 | 0.71928059 | 0.51611355 | 0.62614635 | 0.07964646 | 0.24210283 | 0.09428321 | 0.58331659 | 0.7845285 | 0.69597223 | 0.3234004 | 0.5826633 | 1.02212867 |
| ENSMUSG00 | Gm11281 | 1048 | 0 | 0 | 0 | 0 | 0 | 0.04728314 | 0 | 0 | 0 | 0 | 0 | 0 | 0 | 0 | 0 | 0 | 0 | 0 |
| ENSMUSG00 | Dpm1 | 3646 | 3.32253844 | 3.22530216 | 3.38322826 | 2.81163878 | 3.22111612 | 2.85410698 | 2.69913253 | 3.66630303 | 3.09520168 | 2.74162071 | 3.17102028 | 3.15360024 | 3.63968342 | 2.81517919 | 2.93812521 | 3.74924488 | 3.52147818 | 3.60591786 |
| ENSMUSG00 | Ifi47 | 2313 | 33.0228281 | 29.9715231 | 27.1460624 | 35.9682088 | 35.7711962 | 33.1636972 | 21.9892477 | 31.818746 | 35.3103592 | 25.0532703 | 29.5159601 | 31.9739528 | 25.4444346 | 25.6215545 | 24.0832268 | 30.7583003 | 28.7157253 | 27.4926621 |
| ENSMUSG00 | Tgtp2 | 2935 | 0.36205431 | 0.14755285 | 0.18129773 | 0.24837367 | 0.06559717 | 0.92858615 | 0 | 0.13012744 | 0.12629596 | 0.16064976 | 0.11394368 | 0.13312083 | 0 | 0.06329688 | 0.07019004 | 0.07827722 | 0.16975873 | 0.29987929 |
| ENSMUSG00 | Tgtp1 | 3614 | 0.39988267 | 0.29496752 | 0.36139605 | 0.49166595 | 0.42618195 | 2.61886334 | 0.13091499 | 0.90883968 | 0.54702622 | 0.45663382 | 0.23794943 | 0.20077576 | 0.19110317 | 1.06664617 | 0.70683355 | 1.48331075 | 1.03928531 | 1.55255365 |
| ENSMUSG00 | Ube2v1 | 3900 | 0.18527897 | 0.07687591 | 0.21085898 | 0.10514087 | 0.12341519 | 0.10164663 | 0.12131456 | 0.13710093 | 0.12672774 | 0.19343878 | 0.06124995 | 0.18605221 | 0.16443972 | 0.23817481 | 0.08451601 | 0.24545261 | 0.29481767 | 0.12694409 |
| ENSMUSG00 | Gm12169 | 1058 | 0 | 0 | 0 | 0 | 0 | 0 | 0 | 0 | 0 | 0 | 0 | 0 | 0 | 0 | 0 | 0 | 0 | 0 |
| ENSMUSG00 | D030018L15l | 4301 | 0.03953051 | 0 | 0 | 0 | 0.01119087 | 0 | 0.04400156 | 0 | 0 | 0 | 0 | 0 | 0 | 0 | 0 | 0 | 0 | 0 |
| ENSMUSG00 | Cdc20b | 1548 | 0 | 0 | 0 | 0 | 0 | 0 | 0 | 0 | 0 | 0 | 0 | 0 | 0 | 0.03000264 | 0 | 0 | 0 | 0 |
| ENSMUSG00 | Gm17594 | 2195 | 0.01936454 | 0 | 0 | 0 | 0 | 0.04515055 | 0 | 0.12179809 | 0 | 0.08592383 | 0 | 0 | 0 | 0.14811326 | 0 | 0 | 0 | 0 |
| ENSMUSG00 | Pdf | 1837 | 0.37021384 | 0.34455361 | 0.23699607 | 0.198415 | 0.20961099 | 0.21579851 | 0.5151081 | 0.31185957 | 0.40356956 | 0.4106757 | 0.52014108 | 0.33422552 | 0.13427305 | 0.27810831 | 0.04485744 | 0.06253229 | 0.12518115 | 0.23956062 |
| ENSMUSG00 | CN725425 | 1917 | 0 | 0 | 0 | 0 | 0 | 0 | 0 | 0 | 0 | 0 | 0 | 0 | 0 | 0 | 0 | 0 | 0 | 0 |
| ENSMUSG00 | Ipo11 | 1613 | 0.52703256 | 0.14456929 | 0.08996937 | 0.4801846 | 0.26856002 | 0.21504596 | 0 | 0.11838934 | 0.05106822 | 0.26308514 | 0.11847479 | 0.17301817 | 0.12233581 | 0 | 0.0766303 | 0.33234253 | 0.16632615 | 0.2387252 |
| ENSMUSG00 | 9230113P08f | 2035 | 38.7663915 | 30.7755485 | 36.0840389 | 26.0604696 | 15.9414823 | 23.2544769 | 1.37171895 | 0.11260659 | 0.85004111 | 0.13901927 | 0.02347664 | 0.19199491 | 0.02424173 | 0.20540383 | 0.30369697 | 0.11289614 | 0.09416781 | 0.35140953 |
| ENSMUSG00 | 1700025C18l | 699 | 0 | 0 | 0 | 0 | 0 | 0 | 0 | 0 | 0 | 0 | 0 | 0 | 0 | 0 | 0 | 0 | 0 | 0 |
| ENSMUSG00 | 2410002O22l | 1607 | 0 | 0 | 0 | 0 | 0 | 0 | 0 | 0 | 0 | 0 | 0 | 0 | 0 | 0 | 0 | 0 | 0 | 0 |
| ENSMUSG00 | Cpt1b | 3353 | 0 | 0 | 0.01442694 | 0 | 0 | 0.02955725 | 0 | 0 | 0 | 0.05624897 | 0 | 0 | 0 | 0.0138515 | 0 | 0 | 0 | 0 |
| ENSMUSG00 | Syce3 | 549 | 0 | 0 | 0 | 0 | 0 | 0 | 0 | 0 | 0 | 0 | 0 | 0 | 0 | 0 | 0 | 0 | 0 | 0 |
| ENSMUSG00 | Gm11016 | 46 | 0 | 0 | 0 | 0 | 0 | 0 | 0 | 0 | 0 | 0 | 0 | 0 | 0 | 0 | 0 | 0 | 0 | 0 |
| ENSMUSG00 | Wfdc11 | 632 | 6.72550254 | 1.84485965 | 3.06161585 | 4.18123508 | 1.29468782 | 1.25449958 | 7.93535438 | 3.32370591 | 5.0179774 | 3.20803835 | 0.75593289 | 3.97421648 | 8.74236509 | 7.05479808 | 7.23634922 | 2.36286976 | 3.69921537 | 8.18173423 |
| ENSMUSG00 | Taf9 | 983 | 21.3606887 | 14.063938 | 21.8000752 | 16.8710276 | 19.8795127 | 16.5343811 | 22.140216 | 21.6410663 | 26.2286458 | 19.7620445 | 19.2946679 | 17.7156528 | 20.4253208 | 26.3167413 | 16.3045832 | 18.2299128 | 17.1941983 | 12.9268603 |
| ENSMUSG00 | Naip6 | 7485 | 0.039751 | 0 | 0 | 0 | 0 | 0.03310136 | 0.031605 | 0.02551263 | 0 | 0 | 0 | 0 | 0.00659077 | 0.11789414 | 0 | 0 | 0 | 0 |
| ENSMUSG00 | Naip2 | 5301 | 0 | 0 | 0 | 0 | 0 | 0 | 0 | 0 | 0 | 0 | 0 | 0 | 0 | 0.07885244 | 0 | 0.00722328 | 0 | 0 |
| ENSMUSG00 | R3hdml | 1089 | 0 | 0 | 0 | 0 | 0 | 0 | 0 | 0 | 0 | 0 | 0 | 0 | 0 | 0 | 0 | 0 | 0 | 0 |
| ENSMUSG00 | Gm11019 | 2722 | 0 | 0 | 0 | 0 | 0 | 0.03640906 | 0 | 0 | 0 | 0 | 0 | 0 | 0 | 0 | 0 | 0.04220126 | 0 | 0 |
| ENSMUSG00 | Gm11020 | 183 | 0 | 0 | 0 | 0 | 0 | 0 | 0 | 0 | 0 | 0 | 0 | 0 | 0 | 0 | 0 | 0 | 0 | 0 |
| ENSMUSG00 | Arhgap8 | 1998 | 17.1254588 | 16.7231396 | 20.4340639 | 20.0897298 | 15.2730929 | 19.196104 | 14.4684919 | 15.8465974 | 15.6871564 | 20.1063059 | 15.9728089 | 16.0071945 | 18.4686057 | 13.3660412 | 16.7033335 | 22.0008095 | 15.6719642 | 18.2813047 |
| ENSMUSG00 | Gm14222 | 573 | 0 | 0 | 0 | 0 | 0 | 0 | 0 | 0 | 0 | 0 | 0 | 0 | 0 | 0 | 0 | 0 | 0 | 0 |
| ENSMUSG00 | Gm14221 | 337 | 0 | 0 | 0 | 0 | 0 | 0 | 0 | 0 | 0 | 0 | 0.14176546 | 0 | 0 | 0.41344885 | 0 | 0 | 0 | 0 |
| ENSMUSG00 | 1700060C20l | 594 | 0 | 0 | 0 | 0.07670209 | 0 | 0 | 0 | 0 | 0 | 0 | 0 | 0 | 0 | 0 | 0 | 0 | 0 | 0 |
| ENSMUSG00 | Atp6ap1l | 1158 | 0 | 0 | 0 | 0 | 0 | 0 | 0 | 0 | 0 | 0 | 0 | 0 | 0 | 0 | 0 | 0 | 0 | 0 |
| ENSMUSG00 | Tob2 | 859 | 1.33601834 | 0.50415323 | 1.40784431 | 0.63647559 | 0.44826006 | 0.9806711 | 0.9363394 | 0.75584498 | 0.8630469 | 0.6586827 | 0.38931864 | 0.71475237 | 1.60802524 | 0.81101433 | 1.24707853 | 0.75778851 | 1.07081621 | 1.34480936 |
| ENSMUSG00 | 4932414J04R | 3485 | 0 | 0 | 0 | 0 | 0 | 0 | 0 | 0 | 0 | 0 | 0 | 0 | 0 | 0 | 0 | 0 | 0 | 0 |
| ENSMUSG00 | Hsbp1l1 | 1246 | 18.0459375 | 22.1373005 | 19.9162288 | 20.5865714 | 21.2846629 | 16.9417853 | 31.9341597 | 26.2381191 | 31.4352941 | 32.9222406 | 27.2232909 | 27.7734437 | 23.5573762 | 18.1899797 | 17.3271391 | 18.3463023 | 24.8844095 | 23.1779944 |
| ENSMUSG00 | Ces1b | 1951 | 0 | 0 | 0.07438267 | 0.30358461 | 0 | 0 | 0 | 0 | 0 | 0 | 0 | 0 | 0 | 0 | 0 | 0 | 0 | 0.16917204 |
| ENSMUSG00 | Gm12033 | 1250 | 0.06800828 | 0.07995095 | 0.11609647 | 0.18224418 | 0.07701108 | 0 | 0.03785014 | 0 | 0.03294921 | 0.03772056 | 0.1146599 | 0.22326265 | 0.1183966 | 0.26008689 | 0.03296124 | 0.12252994 | 0.09198311 | 0 |
| ENSMUSG00 | Fam81b | 1522 | 0 | 0 | 0 | 0.05986997 | 0 | 0 | 0 | 0 | 0 | 0 | 0 | 0 | 0 | 0 | 0 | 0 | 0 | 0 |
| ENSMUSG00 | Gm3272 | 1228 | 0.03461334 | 0 | 0.11817638 | 0.03710183 | 0 | 0.04035239 | 0.03852824 | 0.12440522 | 0.26831606 | 0.07679268 | 0 | 0.09090499 | 0.12051771 | 0.07564184 | 0.20131053 | 0.03118128 | 0.1560517 | 0.17918276 |
| ENSMUSG00 | Wdr92 | 3890 | 6.63255575 | 6.90236317 | 7.51095433 | 6.04357298 | 6.74341861 | 6.07626065 | 7.9178802 | 6.86285071 | 7.42204307 | 8.50894439 | 5.96879944 | 6.68640071 | 8.40798487 | 6.16070671 | 6.69392887 | 6.81159391 | 8.43375582 | 5.11909939 |
| ENSMUSG00 | Ighv1-33 | 289 | 0 | 0 | 0 | 0 | 0 | 0 | 0 | 0 | 0 | 0 | 0 | 0 | 0 | 0 | 0 | 0 | 0 | 0 |
| ENSMUSG00 | Gm15557 | 1378 | 0 | 0 | 0 | 0 | 0 | 0 | 0 | 0 | 0 | 0 | 0 | 0 | 0 | 0 | 0 | 0 | 0 | 0 |
| ENSMUSG00 | Sec61g | 2858 | 0.81797924 | 1.25884975 | 1.25249869 | 1.30720981 | 0.58943923 | 0.814898 | 1.52301136 | 1.36305985 | 1.09523275 | 1.36931713 | 1.77191938 | 1.34754399 | 1.50170637 | 1.33254554 | 1.78761124 | 1.74170013 | 1.94447622 | 1.23183444 |
| ENSMUSG00 | Gm2035 | 434 | 0 | 0 | 0 | 0 | 0 | 0 | 0 | 0 | 0 | 0 | 0 | 0 | 0 | 0 | 0 | 0 | 0 | 0 |
| ENSMUSG00 | Gm11025 | 1016 | 0 | 0 | 0 | 0 | 0 | 0 | 0.41910838 | 0 | 0.08107582 | 0 | 0 | 0 | 0 | 0 | 0 | 0 | 0 | 0 |
| ENSMUSG00 | Gm11027 | 196 | 0 | 0 | 0 | 0 | 0.49114208 | 0 | 0 | 0 | 0 | 0 | 0 | 0 | 0 | 0 | 0 | 0 | 0 | 0 |
| ENSMUSG00 | Ighv1-2 | 122 | 0 | 0 | 0 | 0 | 0 | 0 | 0.38780884 | 0 | 0 | 0 | 0 | 0 | 0 | 0 | 0 | 0 | 0 | 0 |
| ENSMUSG00 | Ighv3-2 | 158 | 0 | 0 | 0 | 0 | 0 | 0 | 0 | 0 | 0 | 0 | 0 | 0 | 0 | 0 | 0 | 0 | 0 | 0 |
| ENSMUSG00 | Gm11029 | 160 | 0 | 0 | 0 | 0 | 0 | 0 | 0 | 0 | 0 | 0 | 0 | 0 | 0 | 0 | 0 | 0 | 0 | 0 |
| ENSMUSG00 | Zfp429 | 2029 | 3.07947801 | 4.69565685 | 3.74304794 | 4.37871052 | 5.0527845 | 3.24815848 | 3.63764313 | 4.31052743 | 6.17087274 | 4.48501022 | 7.81729237 | 5.72186191 | 2.43134141 | 4.23467529 | 4.71106997 | 4.24612453 | 6.29012167 | 2.54847513 |
| ENSMUSG00 | Zfp456 | 4825 | 0.58141795 | 0.66280577 | 0.50128011 | 0.85928602 | 0.61848275 | 0.49295984 | 0.79426466 | 0.75197476 | 1.10115244 | 0.42020316 | 0.64360048 | 0.47428853 | 0.43964196 | 0.85668887 | 0.74290888 | 0.34124272 | 0.92936305 | 0.53584 |
| ENSMUSG00 | Vmn2r-ps106 | 668 | 0 | 0 | 0 | 0 | 0 | 0 | 0 | 0 | 0 | 0 | 0 | 0 | 0 | 0 | 0 | 0 | 0 | 0 |
| ENSMUSG00 | Vmn2r-ps105 | 668 | 0 | 0 | 0 | 0 | 0 | 0 | 0 | 0 | 0 | 0 | 0 | 0 | 0 | 0 | 0 | 0 | 0 | 0 |
| ENSMUSG00 | Bpifa6 | 1561 | 0 | 0 | 0 | 0 | 0 | 0 | 0 | 0 | 0 | 0 | 0 | 0 | 0 | 0 | 0 | 0 | 0 | 0 |
| ENSMUSG00 | 7530422B04l | 716 | 1.24666019 | 2.46589859 | 2.49974947 | 1.27265487 | 1.4116905 | 2.28385504 | 2.1806123 | 3.6272114 | 3.6239532 | 3.02923513 | 2.8691665 | 3.74183213 | 0.55119461 | 1.75138318 | 0.86316109 | 1.33696255 | 1.44526539 | 0.84511197 |
| ENSMUSG00 | 4930404H24l | 329 | 0 | 0 | 0.44109602 | 0.69241709 | 0.43889292 | 0 | 0.28761507 | 0.23217266 | 0.62593489 | 0.14331521 | 0.29042528 | 0.16965247 | 0.29989007 | 0 | 0.12523269 | 0.34915447 | 0 | 0 |
| ENSMUSG00 | C030006K11l | 1755 | 12.836321 | 11.1612432 | 13.6969428 | 11.3967513 | 11.0525158 | 13.0728864 | 18.73636 | 19.6076094 | 27.1056557 | 22.8096564 | 19.6544274 | 21.4993663 | 21.8690779 | 17.9160383 | 23.9462034 | 18.8725786 | 20.5717218 | 17.6154518 |
| ENSMUSG00 | Samd1 | 2066 | 3.92956855 | 2.67664107 | 3.72284188 | 3.28586428 | 2.67917291 | 3.57374506 | 3.02288171 | 4.54759487 | 5.02371828 | 2.67019956 | 2.45118374 | 4.26857439 | 3.51006381 | 2.76506425 | 2.37317771 | 2.61325055 | 2.26321792 | 4.81928767 |
| ENSMUSG00 | Gm11031 | 58 | 0 | 0 | 0 | 0 | 0 | 0 | 0 | 0 | 0 | 0 | 0 | 0 | 0 | 0 | 0 | 0 | 0 | 0 |
| ENSMUSG00 | Gm14147 | 2677 | 0 | 0 | 0 | 0 | 0 | 0 | 0 | 0 | 0 | 0 | 0 | 0 | 0 | 0 | 0 | 0 | 0 | 0 |
| ENSMUSG00 | Gm14151 | 2712 | 0 | 0 | 0 | 0 | 0 | 0 | 0 | 0 | 0 | 0 | 0 | 0 | 0 | 0 | 0 | 0 | 0 | 0 |
| ENSMUSG00 | Tcl1b4 | 1707 | 0 | 0 | 0 | 0 | 0 | 0 | 0 | 0 | 0 | 0 | 0 | 0 | 0 | 0 | 0 | 0 | 0 | 0 |
| ENSMUSG00 | Gm14124 | 3014 | 0.02820516 | 0 | 0 | 0 | 0 | 0 | 0.01569764 | 0 | 0 | 0.03128779 | 0.09510609 | 0 | 0.03273518 | 0.12327561 | 0 | 0.03811275 | 0 | 0.0182512 |
| ENSMUSG00 | Gm14139 | 1724 | 0 | 0 | 0 | 0 | 0 | 0 | 0 | 0 | 0 | 0 | 0 | 0 | 0 | 0 | 0 | 0 | 0 | 0 |
| ENSMUSG00 | Gm11032 | 100 | 0 | 0 | 0 | 0 | 0 | 0 | 0 | 0 | 0 | 0 | 0 | 0 | 0 | 0 | 0 | 0 | 0 | 0 |
| ENSMUSG00 | Gm11033 | 3710 | 0 | 0 | 0 | 0 | 0 | 0 | 0 | 0 | 0 | 0 | 0 | 0 | 0 | 0 | 0 | 0 | 0 | 0 |
| ENSMUSG00 | Serpina3m | 1654 | 0 | 0 | 0 | 0 | 0 | 0 | 0 | 0 | 0 | 0 | 0 | 0 | 0 | 0 | 0 | 0 | 0 | 0 |
| ENSMUSG00 | Serpina3j | 2151 | 0 | 0 | 0 | 0 | 0 | 0 | 0 | 0 | 0 | 0 | 0 | 0 | 0 | 0 | 0 | 0 | 0 | 0 |
| ENSMUSG00 | Serpina3i | 2131 | 5.76442791 | 5.12746565 | 6.56027699 | 6.88439994 | 4.1785105 | 5.09246769 | 4.8844624 | 6.39825793 | 5.4309765 | 6.77058433 | 5.44782492 | 4.74079537 | 3.68079533 | 1.41664274 | 4.00221589 | 4.59990403 | 5.1797154 | 4.28508293 |
| ENSMUSG00 | Serpina1c | 1356 | 0 | 0 | 0 | 0 | 0 | 0 | 0 | 0 | 0 | 0 | 0 | 0 | 0 | 0 | 0 | 0 | 0 | 0 |
| ENSMUSG00 | Gm11034 | 88 | 0 | 0 | 0 | 0 | 0 | 0 | 0 | 0 | 0 | 0 | 0 | 0 | 0 | 0 | 0 | 0 | 0 | 0 |
| ENSMUSG00 | Ifi27l2a | 459 | 0.46301935 | 0 | 0 | 0 | 0.10486258 | 0 | 0.3092332 | 0 | 0.17946194 | 0.41089937 | 0 | 0 | 0.10747694 | 0 | 0.08976374 | 0.41710901 | 0.16699912 | 0 |
| ENSMUSG00 | Ly6c1 | 836 | 0.25421756 | 0.67741531 | 4.57118289 | 3.70592227 | 0.92118514 | 0.71128326 | 0.05659411 | 0.27410815 | 0.39412934 | 0.4512029 | 0.51432372 | 0.46735603 | 14.2802918 | 6.72217054 | 11.4832088 | 1.00764756 | 2.15470808 | 11.844066 |
| ENSMUSG00 | Insl3 | 621 | 0.27378535 | 0.05364395 | 0.07789618 | 0 | 0.07750712 | 0 | 0 | 0 | 0.92852048 | 0.15185412 | 0 | 0 | 0.07943948 | 0 | 0.19904133 | 0 | 0.18515119 | 0.08858149 |
| ENSMUSG00 | Slc45a4 | 10298 | 5.72073936 | 5.4993125 | 4.67388451 | 7.14518223 | 4.70195331 | 6.64520539 | 4.13951483 | 3.85706919 | 3.56352549 | 4.38174626 | 3.53974493 | 2.86178577 | 2.94132814 | 4.06802941 | 4.79311167 | 3.75543926 | 3.8445392 | 3.64305795 |
| ENSMUSG00 | Flrt3 | 733 | 0 | 0 | 0 | 0 | 0 | 0 | 0 | 0 | 0 | 0 | 0 | 0 | 0 | 0 | 0 | 0 | 0 | 0 |
| ENSMUSG00 | Col22a1 | 9137 | 0 | 0 | 0.08470794 | 0 | 0.03160682 | 0 | 0.01553442 | 0.07523949 | 0 | 0.00516041 | 0 | 0 | 0 | 0.01524924 | 0.00901862 | 0.04609792 | 0 | 0 |
| ENSMUSG00 | Gm11035 | 112 | 0.3795105 | 0 | 0 | 0 | 0 | 0 | 0 | 0 | 0 | 0 | 0 | 0 | 0 | 0 | 0 | 0 | 0 | 0 |
| ENSMUSG00 | Gsdmc | 2390 | 0 | 0 | 0 | 0 | 0 | 0 | 0 | 0 | 0 | 0 | 0 | 0 | 0 | 0 | 0 | 0 | 0 | 0 |
| ENSMUSG00 | Gm2016 | 1840 | 0 | 0 | 0 | 0 | 0 | 0 | 0 | 0 | 0 | 0 | 0 | 0 | 0 | 0 | 0 | 0 | 0 | 0 |
| ENSMUSG00 | Gm2046 | 433 | 0 | 0 | 0 | 0 | 0 | 0 | 0 | 0 | 0 | 0 | 0 | 0 | 0 | 0 | 0 | 0 | 0 | 0 |
| ENSMUSG00 | BB287469 | 1849 | 0 | 0 | 0 | 0 | 0 | 0 | 0 | 0 | 0 | 0 | 0 | 0 | 0 | 0 | 0 | 0 | 0 | 0 |
| ENSMUSG00 | Mef2b | 1396 | 0 | 0 | 0 | 0 | 0 | 0 | 0 | 0 | 0 | 0 | 0 | 0 | 0 | 0 | 0 | 0 | 0 | 0 |
| ENSMUSG00 | Gm8300 | 1858 | 0 | 0 | 0 | 0 | 0 | 0 | 0 | 0 | 0 | 0 | 0 | 0 | 0 | 0 | 0 | 0 | 0 | 0 |
| ENSMUSG00 | Alkbh1 | 6126 | 0.23590858 | 0.18489037 | 0.17372146 | 0.23055703 | 0.20428175 | 0.26693441 | 0.26259077 | 0.28055144 | 0.18825048 | 0.2232077 | 0.22616288 | 0.38267349 | 0.09663451 | 0.13646647 | 0.26902746 | 0.23751928 | 0.13763933 | 0.1436738 |
| ENSMUSG00 | Prnp | 2399 | 49.9112467 | 62.293307 | 60.7340865 | 60.8683398 | 67.9544916 | 64.1562278 | 52.1839718 | 53.8896549 | 55.6593098 | 64.4661553 | 58.3296596 | 53.6519041 | 65.9061254 | 55.9690941 | 69.7111777 | 72.1280762 | 76.1253876 | 79.9798934 |
| ENSMUSG00 | D130040H23 | 1216 | 0.62918846 | 1.26019172 | 0.75583641 | 0.5245515 | 0.91039 | 1.2225181 | 0.38908453 | 1.00506322 | 0.37257539 | 2.05508822 | 2.00371948 | 0.91802077 | 0.56796615 | 0.53471811 | 1.45696291 | 0.97615858 | 1.22921511 | 0.90475504 |
| ENSMUSG00 | Gm11037 | 58 | 0 | 0 | 0 | 0 | 0 | 0 | 0 | 0 | 0 | 0 | 0 | 0 | 0 | 0 | 0 | 0 | 0 | 0 |
| ENSMUSG00 | Trim61 | 2959 | 0 | 0 | 0 | 0 | 0 | 0 | 0 | 0 | 0 | 0 | 0 | 0 | 0 | 0 | 0 | 0.16822504 | 0 | 0 |
| ENSMUSG00 | Fastkd5 | 3013 | 1.66465674 | 1.27148451 | 1.57338399 | 1.87506455 | 1.50162656 | 1.5623995 | 1.69590751 | 1.82532557 | 0.94320265 | 1.00154166 | 1.60148384 | 1.18559655 | 0.85139718 | 1.51062747 | 1.55890387 | 1.4106397 | 1.65364042 | 1.36929406 |
| ENSMUSG00 | C230021G24 | 3677 | 0 | 0 | 0 | 0 | 0 | 0 | 0 | 0.04154735 | 0 | 0 | 0 | 0 | 0 | 0 | 0 | 0 | 0 | 0.31416678 |
| ENSMUSG00 | Gm17566 | 826 | 0 | 0 | 0 | 0.05515865 | 0 | 0 | 0 | 0 | 0 | 0 | 0 | 0 | 0 | 0 | 0 | 0 | 0 | 0 |
| ENSMUSG00 | 4933413L06f | 1198 | 0 | 0 | 0 | 0 | 0 | 0 | 0 | 0 | 0 | 0 | 0 | 0 | 0 | 0 | 0 | 0 | 0 | 0 |
| ENSMUSG00 | Serpinb1c | 1346 | 0.12631553 | 0.22274595 | 0.57501968 | 0.57543667 | 0.14303692 | 0.773111 | 0.07030116 | 0.48237061 | 1.101571 | 0.38533264 | 0.35494026 | 1.57577651 | 0.21990453 | 0.79362111 | 1.59173916 | 1.2232512 | 1.33828823 | 0.32694863 |
| ENSMUSG00 | Gm14025 | 1857 | 0 | 0 | 0.18234503 | 0 | 0 | 0.08005288 | 0.35669225 | 0.06170017 | 0 | 0 | 0.05145391 | 0 | 0 | 0 | 0 | 0.18557645 | 0.0825553 | 0 |
| ENSMUSG00 | Gm14010 | 372 | 0 | 0 | 0 | 0 | 0 | 0 | 0 | 0 | 0 | 0 | 0 | 0 | 0 | 0.37454909 | 0 | 0 | 0 | 0 |
| ENSMUSG00 | Slc8a3 | 4893 | 0 | 0 | 0.41522344 | 0 | 0 | 0.07089089 | 0 | 0 | 0 | 0.02890908 | 0 | 0.12547972 | 0 | 0.18034696 | 0 | 0 | 0 | 0 |
| ENSMUSG00 | Kcnip3 | 3793 | 1.09820914 | 0.24591644 | 0.48462804 | 0.73272441 | 0.68524226 | 0.52257035 | 1.25984196 | 1.00691807 | 1.1510073 | 0.41022231 | 0.76832915 | 0.51504039 | 4.13592135 | 8.52230024 | 2.63959348 | 1.1104579 | 1.54598561 | 3.91575503 |
| ENSMUSG00 | Cyp4v3 | 6220 | 70.5093901 | 77.3533973 | 68.2984475 | 77.5636493 | 71.5246576 | 78.3682058 | 38.4282402 | 41.6432272 | 42.2261111 | 46.4229754 | 45.3170828 | 44.9666053 | 41.3055667 | 41.1052574 | 37.4390987 | 48.5404243 | 45.7327614 | 37.4627936 |
| ENSMUSG00 | Adam34 | 2333 | 0 | 0 | 0 | 0 | 0 | 0 | 0 | 0 | 0 | 0 | 0 | 0 | 0 | 0 | 0 | 0 | 0 | 0 |
| ENSMUSG00 | Gm11042 | 199 | 0 | 0 | 0 | 0.45789994 | 0.72560689 | 0 | 0.47550431 | 0 | 0 | 0 | 0 | 0 | 0 | 2.10048636 | 0 | 0 | 0 | 0 |
| ENSMUSG00 | Gm11043 | 46 | 0 | 0 | 0 | 0 | 0 | 0 | 0 | 0 | 0 | 0 | 0 | 0 | 0 | 0 | 0 | 0 | 0 | 0 |
| ENSMUSG00 | Gm11044 | 318 | 0 | 0 | 0 | 0 | 0 | 0 | 0 | 0 | 0 | 0 | 0 | 0 | 0 | 0 | 0 | 0 | 0 | 0 |
| ENSMUSG00 | BC005561 | 5179 | 17.1941193 | 17.6052117 | 16.1867789 | 15.1840824 | 14.9999855 | 14.6581942 | 16.9737319 | 15.8108729 | 19.6747326 | 14.9218001 | 20.4881604 | 21.9102611 | 15.5835135 | 20.1326465 | 13.1106708 | 14.4467748 | 18.29361 | 20.4146559 |
| ENSMUSG00 | Hmgn2-ps1 | 1433 | 2.49158045 | 1.4645585 | 2.05916633 | 2.32097435 | 1.78017582 | 1.76356554 | 3.16958631 | 2.29207719 | 2.90288771 | 3.19163867 | 1.66695598 | 3.19391789 | 2.68519857 | 3.66237395 | 3.99652217 | 2.67265913 | 2.08614875 | 3.14776466 |
| ENSMUSG00 | 8430423G03l | 636 | 0 | 0 | 0 | 0 | 0.3027165 | 0.0779131 | 1.33903808 | 0.18015284 | 0 | 0 | 0 | 0.08776048 | 0 | 0 | 0.06478232 | 0 | 0 | 0 |
| ENSMUSG00 | Gm3985 | 2511 | 0 | 0 | 0 | 0 | 0 | 0 | 0 | 0 | 0 | 0 | 0 | 0 | 0 | 0 | 0 | 0 | 0 | 0 |
| ENSMUSG00 | Gm14085 | 5151 | 0 | 0.01293453 | 0 | 0 | 0 | 0 | 0.06429601 | 0.02224368 | 0.03198332 | 0.05492219 | 0.05564934 | 0.05417944 | 0 | 0 | 0.0159975 | 0 | 0 | 0 |
| ENSMUSG00 | Gm3086 | 538 | 0.0790059 | 0 | 0.08991363 | 0 | 0.26839363 | 0 | 0 | 0 | 0 | 0 | 0 | 0 | 0 | 0 | 0.61266254 | 0 | 0 | 0 |
| ENSMUSG00 | Gm3415 | 918 | 0 | 0 | 0 | 0 | 0 | 0 | 0 | 0 | 0 | 0 | 0 | 0 | 0 | 0 | 0 | 0 | 0 | 0 |
| ENSMUSG00 | Gm3409 | 918 | 0 | 0 | 0 | 0 | 0 | 0 | 0 | 0 | 0 | 0 | 0 | 0 | 0 | 0 | 0 | 0 | 0 | 0 |
| ENSMUSG00 | Gm3404 | 1125 | 0 | 0 | 0 | 0 | 0 | 0 | 0 | 0 | 0 | 0 | 0 | 0 | 0 | 0 | 0 | 0 | 0 | 0 |
| ENSMUSG00 | Prl2c2 | 1042 | 0 | 0 | 0 | 0 | 0 | 0 | 0 | 1.31950718 | 0 | 0 | 0 | 0 | 0 | 0 | 0 | 0 | 0 | 0 |
| ENSMUSG00 | 4930449I24R | 1134 | 0 | 0 | 0 | 0 | 0 | 0 | 0 | 0 | 0 | 0 | 0 | 0 | 0 | 0 | 0 | 0 | 0 | 0 |
| ENSMUSG00 | Gm6370 | 972 | 0 | 0 | 0 | 0 | 0 | 0 | 0 | 0 | 0 | 0 | 0 | 0 | 0 | 0 | 0 | 0 | 0 | 0 |
| ENSMUSG00 | Gm6309 | 2458 | 0.13834069 | 0 | 0 | 0.05560746 | 0.01958174 | 0.04031956 | 0 | 0.031076 | 0 | 0 | 0 | 0.13624653 | 0 | 0.13226551 | | | | |
| ENSMUSG00 | Trav12n-1 | 342 | 0 | 0 | 0 | 0 | 0 | 0 | 0 | 0 | 0 | 0 | 0 | 0 | 0 | 0 | 0 | 0 | 0 | 0 |
| ENSMUSG00 | Gm5456 | 1691 | 0 | 0 | 0 | 0 | 0 | 0 | 0 | 0 | 0 | 0 | 0 | 0 | 0 | 0 | 0 | 0 | 0 | 0 |
| ENSMUSG00 | Gm2904 | 1694 | 0.0250916 | 0.07866091 | 0.0285558 | 0 | 0.02841318 | 0 | 0 | 0.02254569 | 0 | 0 | 0.05640491 | 0.06589807 | 0 | 0 | 0.02432205 | 0.02260366 | 0.04524947 | 0.06494582 |
| ENSMUSG00 | Dab2 | 1373 | 0 | 0 | 0 | 0 | 0 | 0 | 0 | 0 | 0 | 0 | 0 | 0 | 0 | 0 | 0 | 0 | 0 | 0 |
| ENSMUSG00 | Tgm7 | 2139 | 0 | 0 | 0 | 0 | 0 | 0 | 0 | 0 | 0 | 0 | 0 | 0 | 0 | 0 | 0 | 0 | 0 | 0 |
| ENSMUSG00 | Prps1l3 | 1718 | 20.5845788 | 21.7561509 | 21.4273903 | 24.2125922 | 20.9841739 | 21.8055102 | 22.3895273 | 26.1655749 | 22.3193517 | 22.0933156 | 18.7151031 | 26.2509053 | 21.9093439 | 25.3036471 | 29.9059023 | 24.2046564 | 25.6326631 | 19.7879092 |
| ENSMUSG00 | C7 | 2658 | 0 | 0 | 0 | 0 | 0 | 0 | 0 | 0 | 0 | 0 | 0 | 0 | 0 | 0 | 0 | 0 | 0 | 0 |
| ENSMUSG00 | Srp54b | 3510 | 0 | 0 | 0 | 0 | 0 | 0 | 0.0134794 | 0.01088103 | 0.03520215 | 0.01343325 | 0.0408333 | 0 | 0.01405468 | 0.05292773 | 0 | 0.02181801 | 0.05459586 | 0.01567211 |
| ENSMUSG00 | Prss2 | 7870 | 1.5068544 | 2.03602314 | 1.56122957 | 2.02043259 | 1.90203663 | 1.9644794 | 2.12215701 | 2.9360106 | 2.34454372 | 2.01903266 | 2.50712298 | 1.78723596 | 2.15631252 | 1.36322544 | 2.33492934 | 1.98021311 | 3.16545019 | 1.64957423 |
| ENSMUSG00 | Capn3 | 4267 | 0.16934333 | 0.60114667 | 0.71420961 | 0.38439128 | 0.31584107 | 0.49905963 | 7.98339087 | 7.24107184 | 4.91303879 | 3.51392633 | 1.05245963 | 4.00271684 | 8.40503957 | 11.9185085 | 2.93538153 | 1.0319709 | 1.21257326 | 5.65947922 |
| ENSMUSG00 | Kdelr2 | 1841 | 30.0606948 | 31.0872093 | 31.5833696 | 27.4455177 | 27.4778119 | 29.3386636 | 33.2807818 | 30.9522349 | 26.5329754 | 32.475335 | 28.6753553 | 30.560667 | 34.8351398 | 31.6606897 | 30.3696422 | 32.1341599 | 27.8547448 | 29.0732541 |
| ENSMUSG00 | C86695 | 1882 | 0 | 0 | 0 | 0 | 0 | 0 | 0 | 0 | 0 | 0 | 0.30462248 | 0 | 0 | 0 | 0 | 0 | 0 | 0 |
| ENSMUSG00 | Gm7861 | 504 | 0 | 0 | 0 | 0 | 0 | 0 | 0 | 0 | 0 | 0 | 0 | 0 | 0 | 0 | 0 | 0 | 0 | 0 |
| ENSMUSG00 | Gm7849 | 486 | 0 | 0 | 0 | 0 | 0 | 0 | 0 | 0 | 0 | 0 | 0 | 0 | 0 | 0 | 0 | 0 | 0 | 0 |
| ENSMUSG00 | Gm11048 | 93 | 0 | 0 | 0 | 0 | 0 | 0 | 0 | 0 | 0 | 0 | 0 | 0 | 0 | 0 | 0 | 0 | 0 | 0 |
| ENSMUSG00 | Gm15293 | 416 | 0 | 0 | 0 | 0 | 0 | 0 | 0 | 0 | 0 | 0 | 0 | 0 | 0 | 0 | 0 | 0 | 0 | 0 |
| ENSMUSG00 | AY761185 | 485 | 43.64449 | 45.1956379 | 87.7705293 | 9.58191036 | 18.3595997 | 20.0254346 | 178.520005 | 223.799603 | 176.889716 | 45.4979982 | 34.279764 | 170.554251 | 37.5329432 | 72.2037977 | 40.6918459 | 5.2896303 | 7.58623609 | 26.3136346 |
| ENSMUSG00 | Gm11051 | 64 | 0 | 0 | 0 | 0 | 0 | 0 | 0 | 0 | 0 | 0 | 0 | 0 | 3.85405604 | 3.62844433 | 1.28754862 | 1.79487222 | 0 | 0.85951729 |
| ENSMUSG00 | Gm1977 | 1496 | 0 | 0 | 0 | 0 | 0 | 0 | 0 | 0 | 0 | 0 | 0 | 0 | 0 | 0 | 0 | 0 | 0 | 0 |
| ENSMUSG00 | Gm3090 | 475 | 0 | 0 | 0 | 0 | 0 | 0 | 0 | 0 | 0 | 0 | 0 | 0 | 0 | 0 | 0 | 0 | 0 | 0 |
| ENSMUSG00 | Rpl27-ps1 | 410 | 0.31101348 | 0 | 0 | 0 | 0 | 0 | 0 | 0 | 0.10045492 | 0 | 0 | 0 | 0 | 0 | 0 | 0.7471338 | 0 | 0 |
| ENSMUSG00 | Gm8818 | 1046 | 0.36572331 | 0.12739156 | 0.73993928 | 0.30490183 | 0.32210656 | 0.23686775 | 0.99510415 | 0.36512813 | 0 | 0.09015431 | 0.68510935 | 0.5869716 | 0.51878689 | 1.50965485 | 0.39389633 | 2.34282875 | 0.80609802 | 0.36812978 |
| ENSMUSG00 | Gm4204 | 779 | 1.5823493 | 2.22371053 | 2.35968441 | 1.28670472 | 2.03896467 | 2.73526 | 2.24720041 | 1.56887917 | 1.7976143 | 1.08948993 | 1.28790004 | 0.93145522 | 1.8998171 | 2.98100689 | 1.85116105 | 1.08137787 | 1.77117675 | 2.542141 |
| ENSMUSG00 | Gm6374 | 1572 | 0 | 0 | 0 | 0.05796571 | 0.03061827 | 0 | 0 | 0 | 0.02620007 | 0 | 0 | 0.03550615 | 0 | 0 | 0 | 0 | 0 | 0 |
| ENSMUSG00 | Gm11052 | 1831 | 0.27857024 | 0.03638765 | 0.31703024 | 0 | 0 | 0.10825283 | 0 | 0.1877289 | 0.044988 | 0.15450804 | 0.31310732 | 0.15241852 | 0.16165565 | 0 | 0 | 0.16729921 | 0.35584219 | 0 |
| ENSMUSG00 | A130010J15R | 1508 | 5.8627829 | 7.95268032 | 7.76286097 | 8.7617393 | 9.6072341 | 9.33221242 | 6.49451229 | 11.4729166 | 11.4983575 | 8.16069864 | 12.7357648 | 9.58637706 | 9.84675537 | 9.23953993 | 8.98893361 | 9.57265185 | 15.6812501 | 6.49311737 |
| ENSMUSG00 | Gm15353 | 958 | 0 | 0.03477338 | 0 | 0 | 0 | 0 | 0 | 0 | 0 | 0 | 0 | 0 | 0 | 0 | 0 | 0 | 0.12001972 | 0 |
| ENSMUSG00 | Gm11053 | 102 | 1.25015224 | 0 | 0.4742503 | 1.34003072 | 1.88752642 | 0.48581111 | 0.92769959 | 0.37443532 | 0 | 0 | 0 | 0 | 0 | 1.21181047 | 0 | 0 | 1.12724463 | 0 |
| ENSMUSG00 | Smok3c | 2328 | 0 | 0 | 0 | 0 | 0 | 0 | 0 | 0 | 0 | 0 | 0 | 0 | 0 | 0 | 0 | 0 | 0 | 0 |
| ENSMUSG00 | Fam155a | 1935 | 0 | 0 | 0 | 0 | 0 | 0 | 0 | 0 | 0 | 0.07310187 | 0 | 0.17307182 | 0 | 0 | 0.08517117 | 0 | 0 | 0.25585631 |
| ENSMUSG00 | Gm17608 | 1534 | 0.19396104 | 0.10858179 | 0.03153424 | 0.29700811 | 0.34514417 | 0.22612069 | 0.64769639 | 1.3693495 | 0 | 0.06147419 | 0.03114404 | 0.0363857 | 0 | 0.99912313 | 0.08057671 | 0.02496128 | 0.02498455 | 0 |
| ENSMUSG00 | Gm17607 | 82 | 0.51835581 | 0 | 0 | 0 | 0 | 0 | 0 | 0.93152201 | 0 | 0 | 0.58262144 | 0 | 0 | 0 | 0 | 0 | 0 | 0 |
| ENSMUSG00 | Trim43b | 2255 | 0 | 0 | 0 | 0 | 0 | 0 | 0 | 0 | 0 | 0 | 0 | 0 | 0 | 0 | 0 | 0 | 0 | 0 |
| ENSMUSG00 | Gm15498 | 721 | 0 | 0 | 0 | 0 | 0 | 0 | 0 | 0 | 0 | 0 | 0.06626208 | 0 | 0 | 0 | 0 | 0 | 0 | 0 |
| ENSMUSG00 | Tlr5 | 3520 | 0.47093803 | 0.06624723 | 0.04122744 | 0.12943479 | 0.51960599 | 0.15485229 | 0.53764408 | 0.34720366 | 0 | 0.10716069 | 0 | 0 | 0.32233923 | 0.60693978 | 0.33944464 | 0.65268081 | 0.25042751 | 0 |
| ENSMUSG00 | Sap25 | 1593 | 0.05336494 | 0.14638434 | 0.12146524 | 0.02860078 | 0.06042928 | 0.03110655 | 0.02970036 | 0.11987571 | 0.10341875 | 0.02959868 | 0 | 0.03503808 | 0 | 0.05831022 | 0.20691302 | 0.02403679 | 0.02405919 | 0.06906354 |
| ENSMUSG00 | Cd209g | 1080 | 0 | 0 | 0 | 0 | 0.0445666 | 0 | 0 | 0.07072667 | 0.6101706 | 0 | 0 | 0 | 0.31974391 | 0 | 0 | 0 | 0 | 0 |
| ENSMUSG00 | Gm15130 | 653 | 0 | 0 | 0 | 0 | 0 | 0 | 0 | 0 | 0 | 0 | 0 | 0 | 0 | 0 | 0 | 0 | 0 | 0 |
| ENSMUSG00 | Gm13941 | 726 | 0 | 0 | 0 | 0 | 0 | 0 | 0 | 0 | 0 | 0 | 0 | 0 | 0 | 0 | 0 | 0 | 0 | 0 |
| ENSMUSG00 | Gm13942 | 322 | 0 | 0 | 0 | 0 | 0 | 0 | 0 | 0 | 0 | 0 | 0 | 0 | 0 | 0 | 0 | 0 | 0 | 0 |
| ENSMUSG00 | Gm17290 | 1070 | 0 | 0 | 0 | 0 | 0 | 0 | 0 | 0 | 0 | 0 | 0 | 0 | 0 | 0 | 0 | 0 | 0 | 0 |
| ENSMUSG00 | Zan | 16867 | 0.00756006 | 0 | 0 | 0.00810358 | 0 | 0 | 0 | 0 | 0.10255728 | 0 | 0 | 0 | 0.06141995 | 0.0578245 | 0 | 0 | 0 | 0 |
| ENSMUSG00 | Gm3054 | 240 | 0 | 0 | 0 | 0 | 0 | 0 | 0 | 0 | 0 | 0 | 0 | 0 | 0 | 0 | 0 | 0 | 0 | 0 |
| ENSMUSG00 | Gm11060 | 70 | 0 | 0 | 0 | 0 | 0 | 0 | 0 | 0 | 0 | 0 | 0 | 0 | 0 | 0 | 0 | 0 | 0 | 0 |
| ENSMUSG00 | 4930417G10l | 2996 | 0.05674923 | 0.22238247 | 0.03229208 | 0.28893853 | 0.0160654 | 0.06615852 | 0.61588601 | 0.26770376 | 0 | 0.2045925 | 0 | 0.14904049 | 0.21405705 | 0.18602438 | 0 | 0.08946404 | 0.01279249 | 0 |
| ENSMUSG00 | 1700012B15l | 771 | 11.6875452 | 12.7029713 | 12.8619633 | 13.7687722 | 12.8601508 | 12.1471682 | 13.807202 | 10.3530637 | 13.0878033 | 12.1087409 | 11.3395816 | 11.5830169 | 13.1807717 | 12.5899018 | 18.1158592 | 12.9621901 | 12.6263027 | 13.9841568 |
| ENSMUSG00 | Gm11062 | 658 | 0 | 0 | 0 | 0 | 0 | 0 | 0 | 0 | 0 | 0 | 0 | 0 | 0 | 0 | 0 | 0 | 0 | 0 |
| ENSMUSG00 | C030005K15l | 1881 | 0 | 0 | 0 | 0 | 0 | 0 | 0 | 0 | 0 | 0 | 0 | 0 | 0 | 0 | 0 | 0 | 0 | 0 |
| ENSMUSG00 | Mphosph8 | 3572 | 10.1146136 | 7.70337364 | 9.72345879 | 7.24486931 | 8.30046612 | 8.60097839 | 9.72214623 | 9.0134925 | 8.98216567 | 7.68244941 | 9.26876994 | 7.68793561 | 10.6342599 | 12.3521509 | 9.00851488 | 11.277077 | 8.99144367 | 10.6260592 |
| ENSMUSG00 | Gzmc | 929 | 0 | 0 | 0 | 0 | 0 | 0 | 0 | 0 | 0 | 0.20301702 | 0 | 0 | 0.47791954 | 0.34995545 | 0 | 0.04121702 | 0 | 0 |
| ENSMUSG00 | Gm7694 | 811 | 1.10062725 | 1.47874746 | 1.55081602 | 1.23593462 | 1.1276283 | 1.03871328 | 0.87508037 | 0.70639462 | 0.45706368 | 1.22091833 | 1.06035666 | 1.10117213 | 1.39905561 | 0.45814143 | 1.06687136 | 0.23607033 | 0.89790338 | 0.27131495 |
| ENSMUSG00 | Psme2 | 1879 | 116.001353 | 101.020155 | 109.052834 | 98.2266049 | 107.918996 | 110.550856 | 94.9275144 | 93.5195548 | 93.6834318 | 96.0077651 | 90.8970072 | 92.3230864 | 104.754843 | 93.8521553 | 96.4585654 | 101.911297 | 84.6074931 | 83.4943972 |
| ENSMUSG00 | Zfp664 | 5840 | 17.3587063 | 16.6450386 | 15.2078428 | 16.4924739 | 15.7417764 | 15.2052224 | 14.6393854 | 13.105749 | 12.9765735 | 14.0886947 | 14.0788876 | 13.5047143 | 11.0321034 | 10.4022031 | 11.8242821 | 12.5559098 | 12.1410358 | 13.4131794 |
| ENSMUSG00 | Gm8778 | 1723 | 0.1480157 | 0.23201087 | 0.19652624 | 0 | 0 | 0.02875957 | 0.16475686 | 0.22166223 | 0.11951978 | 0 | 0.16636666 | 0.03239446 | 0.11452564 | 0.24259825 | 0.11956342 | 0.04444644 | 0 | 0.06385271 |
| ENSMUSG00 | Gm6565 | 599 | 0 | 0 | 0 | 0.15212369 | 0 | 0 | 0.07898611 | 0.06376027 | 0 | 0 | 0 | 0.09318141 | 0 | 0 | 0.1375678 | 0.06392422 | 0.31991901 | 0 |
| ENSMUSG00 | Gm9531 | 439 | 0.3872909 | 0.98648662 | 0.33057082 | 0.1037837 | 0.32891975 | 0.67725832 | 0 | 0.26099592 | 0.6567326 | 0.10740479 | 0.21765357 | 0.76285644 | 0.67424033 | 0.10579519 | 0.0938532 | 0.34888936 | 0.34921455 | 0.25061096 |
| ENSMUSG00 | Gm336 | 569 | 0.07470154 | 0 | 0 | 0 | 0 | 0 | 0 | 0 | 0 | 0 | 0 | 0 | 0 | 0 | 0 | 0 | 0 | 0 |
| ENSMUSG00 | Ccr5 | 2310 | 0 | 0 | 0 | 0 | 0 | 0 | 0 | 0.13226806 | 0.57054914 | 0 | 0.1447726 | 0.16913837 | 0.23491389 | 1.46771358 | | | | |
| ENSMUSG00 | Gm4694 | 1269 | 0 | 0 | 0 | 0 | 0 | 0 | 0 | 0 | 0 | 0 | 0 | 0 | 0 | 0 | 0 | 0 | 0 | 0 |
| ENSMUSG00 | Gm11065 | 226 | 0 | 0 | 0 | 0 | 0 | 0 | 0 | 0 | 0 | 0 | 0 | 0 | 0 | 0 | 0 | 0 | 0 | 0 |
| ENSMUSG00 | Gm11067 | 52 | 0 | 1.28126516 | 0 | 0 | 0 | 0 | 1.81971842 | 0 | 0 | 0 | 0 | 1.07337813 | 0 | 0 | 0 | 1.47271567 | 0 | 0 |
| ENSMUSG00 | Ccdc13 | 4012 | 0.97469496 | 1.73539255 | 1.19366388 | 1.0674821 | 1.82354247 | 0.93868588 | 1.54485567 | 1.86583023 | 1.76572301 | 1.70410069 | 1.8338344 | 1.48860316 | 1.29108956 | 1.63225731 | 1.54043703 | 1.87062788 | 1.24187901 | 1.64533719 |
| ENSMUSG00 | Trav9n-1 | 334 | 0 | 0 | 0 | 0 | 0 | 0 | 0 | 0 | 0 | 0 | 0 | 0 | 0 | 0 | 0 | 0 | 0 | 0 |
| ENSMUSG00 | Olfr1082 | 1929 | 0 | 0 | 0 | 0 | 0 | 0 | 0 | 0 | 0 | 0 | 0 | 0 | 0 | 0 | 0 | 0 | 0 | 0 |
| ENSMUSG00 | C730034F03l | 1279 | 1.62842348 | 1.53671678 | 1.21028379 | 1.38927344 | 1.3171363 | 0.73612349 | 2.03455617 | 0.98541773 | 3.4456272 | 2.02759084 | 1.38207464 | 3.84032706 | 3.00851411 | 1.92457907 | 2.64153838 | 2.3052203 | 2.15753978 | 3.65580459 |
| ENSMUSG00 | Xirp1 | 5836 | 0 | 0.01141634 | 0 | 0 | 0 | 0 | 0 | 0 | 0 | 0 | 0 | 0 | 0 | 0 | 0 | 0 | 0 | 0 |
| ENSMUSG00 | Gm5622 | 1576 | 0 | 0 | 0 | 0 | 0 | 0 | 0 | 0 | 0 | 0 | 0 | 0 | 0 | 0 | 0 | 0 | 0 | 0 |
| ENSMUSG00 | Slc35f4 | 1309 | 0 | 0 | 0 | 0 | 0 | 0 | 0 | 0 | 0 | 0 | 0 | 0 | 0 | 0 | 0 | 0 | 0 | 0 |
| ENSMUSG00 | Gm13691 | 2492 | 0 | 0 | 0 | 0 | 0 | 0 | 0 | 0 | 0 | 0 | 0 | 0 | 0 | 0 | 0 | 0 | 0 | 0 |
| ENSMUSG00 | Gm13694 | 2492 | 0 | 0 | 0 | 0 | 0 | 0 | 0 | 0 | 0 | 0 | 0 | 0 | 0 | 0 | 0 | 0 | 0 | 0 |
| ENSMUSG00 | Gm13696 | 2492 | 0 | 0 | 0 | 0 | 0 | 0 | 0 | 0 | 0 | 0 | 0 | 0 | 0 | 0 | 0 | 0 | 0 | 0 |
| ENSMUSG00 | Gm13697 | 2492 | 0 | 0 | 0 | 0 | 0 | 0 | 0 | 0 | 0 | 0 | 0 | 0 | 0 | 0 | 0 | 0 | 0 | 0 |
| ENSMUSG00 | Tor1aip2 | 3228 | 28.5474664 | 31.414018 | 32.2640058 | 30.176429 | 29.210173 | 28.3377776 | 27.1886677 | 30.9514637 | 30.902646 | 28.1180619 | 24.4942863 | 27.855958 | 26.927769 | 30.0275125 | 30.5948356 | 38.4448137 | 31.1787037 | 29.9244087 |
| ENSMUSG00 | Itprip | 3762 | 18.1567831 | 25.1662003 | 17.4103562 | 26.946657 | 25.1022951 | 25.7642602 | 13.2681755 | 18.9033103 | 7.68552208 | 18.499319 | 17.3092154 | 14.9108827 | 11.7232148 | 13.5554513 | 13.5914755 | 24.1428285 | 19.8151641 | 18.0731674 |
| ENSMUSG00 | Gm11074 | 146 | 0.29113134 | 0 | 0 | 0.6241239 | 0 | 0.33940228 | 0 | 0.78477539 | 0.28209942 | 0 | 0 | 0.38229906 | 0 | 0.63622038 | 0.56440488 | 0.26226443 | 1.31254441 | 2.26064822 |
| ENSMUSG00 | Calhm1 | 1045 | 0 | 0 | 0 | 0 | 0 | 0 | 0 | 0 | 0 | 0 | 0 | 0.16023635 | 0 | 0 | 0 | 0 | 0 | 0 |
| ENSMUSG00 | Trim71 | 5186 | 0 | 0 | 0 | 0 | 0 | 0 | 0 | 0 | 0 | 0 | 0 | 0 | 0 | 0 | 0 | 0 | 0 | 0 |
| ENSMUSG00 | Glb1 | 2919 | 1.93668668 | 2.54497273 | 3.64582963 | 3.60555027 | 4.60048191 | 2.76707625 | 1.91260573 | 2.28971237 | 3.16059593 | 4.13517644 | 2.02949464 | 1.75917813 | 2.21393668 | 2.38664375 | 2.8088762 | 3.89596108 | 3.25622529 | 3.27906289 |
| ENSMUSG00 | Gm15217 | 1030 | 0 | 0 | 0 | 0 | 0 | 0 | 0 | 0 | 0 | 0 | 0 | 0 | 0 | 0 | 0 | 0 | 0 | 0 |
| ENSMUSG00 | Slco1a6 | 3790 | 1.88413445 | 1.08113087 | 1.97833699 | 1.25022391 | 1.23187245 | 0.4053126 | 1.78514857 | 1.73326998 | 2.01042359 | 2.18958411 | 1.52526912 | 2.2826986 | 0.83304557 | 2.2548053 | 1.326277 | 0.84865726 | 1.68878406 | 1.34982768 |
| ENSMUSG00 | Gm6614 | 1939 | 0 | 0.0343609 | 0.02494767 | 0.02349719 | 0 | 0 | 0 | 0 | 0.06372334 | 0.07295106 | 0.02463897 | 0 | 0 | 0 | 0.12749321 | 0 | 0 | 0.02836983 |
| ENSMUSG00 | Gm11077 | 82 | 0 | 0 | 0 | 0 | 0 | 0 | 0 | 0 | 0 | 0 | 0 | 0 | 0 | 0 | 0 | 0 | 0 | 0 |
| ENSMUSG00 | Gm8257 | 2855 | 0 | 0 | 0 | 0 | 0 | 0 | 0 | 0 | 0 | 0 | 0 | 0 | 0 | 0 | 0 | 0 | 0 | 0 |
| ENSMUSG00 | Gm6490 | 458 | 0 | 0 | 0 | 0 | 0 | 0 | 0 | 0 | 0 | 0 | 0 | 0 | 0 | 0 | 0 | 0 | 0 | 0 |
| ENSMUSG00 | Gm6930 | 1741 | 0 | 0 | 0 | 0 | 0 | 0 | 0 | 0 | 0 | 0 | 0 | 0 | 0 | 0 | 0 | 0 | 0 | 0 |
| ENSMUSG00 | Gm3676 | 894 | 0 | 0 | 0 | 0 | 0 | 0 | 0 | 0 | 0 | 0 | 0 | 0 | 0 | 0 | 0 | 0 | 0 | 0 |
| ENSMUSG00 | Gm3543 | 894 | 0 | 0 | 0 | 0 | 0 | 0 | 0 | 0 | 0 | 0 | 0 | 0 | 0 | 0 | 0 | 0 | 0 | 0 |
| ENSMUSG00 | 1700049E17f | 1776 | 0 | 0 | 0 | 0 | 0 | 0 | 0 | 0 | 0 | 0 | 0 | 0 | 0 | 0 | 0 | 0 | 0 | 0 |
| ENSMUSG00 | Gm7152 | 601 | 0 | 0 | 0 | 0 | 0 | 0 | 0 | 0 | 0 | 0 | 0 | 0 | 0 | 0 | 0 | 0 | 0 | 0 |
| ENSMUSG00 | Gm17165 | 490 | 0 | 0 | 0 | 0 | 0 | 0 | 0 | 0 | 0 | 0 | 0 | 0 | 0 | 0 | 0 | 0 | 0 | 0 |
| ENSMUSG00 | Gm11078 | 69 | 1.23203409 | 1.44838671 | 2.10319697 | 0.66030499 | 2.79025644 | 0.71815556 | 0 | 1.66053924 | 2.3876241 | 0.68334353 | 0 | 1.61784529 | 0 | 0.67310272 | 0.597124 | 3.32961803 | 0 | 0 |
| ENSMUSG00 | Gm4736 | 1313 | 0 | 0 | 0 | 0 | 0 | 0 | 0 | 0 | 0 | 0 | 0 | 0 | 0 | 0 | 0 | 0 | 0 | 0 |
| ENSMUSG00 | Hoxd3 | 5006 | 34.0907475 | 32.5409614 | 35.1833847 | 37.0149355 | 32.1520481 | 32.6656054 | 35.8483802 | 35.2322242 | 32.259754 | 36.5639293 | 33.3928445 | 34.9767746 | 24.478722 | 22.0808885 | 28.8229822 | 23.6046373 | 21.3757533 | 25.8123036 |
| ENSMUSG00 | Tmem233 | 2474 | 0 | 0 | 0 | 0 | 0 | 0 | 0 | 0 | 0 | 0 | 0 | 0 | 0 | 0 | 0 | 0 | 0 | 0 |
| ENSMUSG00 | Gm11081 | 31 | 0 | 0 | 0 | 0 | 0 | 0 | 0 | 0 | 0 | 0 | 0 | 0 | 0 | 0 | 0 | 0 | 0 | 0 |
| ENSMUSG00 | Gm11082 | 70 | 0 | 0 | 0 | 0 | 0 | 0 | 0 | 0 | 0 | 0 | 0 | 0 | 0 | 0 | 0 | 0 | 0 | 0 |
| ENSMUSG00 | 2310009B15l | 568 | 10.4766279 | 10.0290579 | 10.2197599 | 10.58813 | 11.0161093 | 10.0326837 | 16.4095031 | 12.5739071 | 11.7468584 | 11.704664 | 9.84096856 | 14.8383188 | 14.6779824 | 17.2529973 | 12.6216034 | 11.8646953 | 13.6976026 | 14.9144409 |
| ENSMUSG00 | Fbxw28 | 1826 | 0 | 0 | 0 | 0 | 0 | 0 | 0 | 0 | 0 | 0 | 0 | 0 | 0 | 0 | 0 | 0 | 0 | 0 |
| ENSMUSG00 | Gm11084 | 67 | 0 | 0 | 0 | 0 | 0 | 1.47918608 | 0 | 0 | 0 | 0 | 0 | 0 | 0 | 0 | 0 | 0 | 0.57203428 | 0 |
| ENSMUSG00 | Clec7a | 2264 | 0 | 0.0588567 | 0 | 0 | 0 | 0.10943625 | 0 | 0.67477743 | 0 | 0 | 0 | 0 | 0.10894858 | 0 | 1.2738997 | 0 | 0 | 0 |
| ENSMUSG00 | Klrb1-ps1 | 544 | 0 | 0 | 0 | 0 | 0 | 0 | 0 | 0 | 0 | 0 | 0 | 0.10260232 | 0 | 0 | 0 | 0 | 0 | 0 |
| ENSMUSG00 | Gm2223 | 2461 | 0.03454301 | 0.10829059 | 0.03931209 | 0 | 0.01955787 | 0.08054081 | 0.03844996 | 0.03103812 | 0.05020705 | 0.03831833 | 0 | 0.0907203 | 0.10022738 | 0.01887204 | 0.01674179 | 0.04667689 | 0 | 0.04470468 |
| ENSMUSG00 | Klrb1b | 3155 | 0 | 0 | 0 | 0 | 0 | 0 | 0 | 0 | 0 | 0 | 0 | 0 | 0 | 0 | 0 | 0 | 0 | 0 |
| ENSMUSG00 | Klrb1 | 639 | 0 | 0 | 0 | 0 | 0 | 0 | 0 | 0 | 0 | 0 | 0 | 0 | 0 | 0 | 0 | 0 | 0 | 0 |
| ENSMUSG00 | 4933413G19l | 1871 | 0 | 0 | 0 | 0 | 0 | 0 | 0 | 0 | 0 | 0 | 0 | 0 | 0 | 0 | 0 | 0 | 0 | 0 |
| ENSMUSG00 | Gm4461 | 947 | 0 | 0 | 0 | 0 | 0 | 0 | 0 | 0 | 0 | 0 | 0 | 0 | 0 | 0 | 0 | 0 | 0 | 0 |
| ENSMUSG00 | Olfr239 | 947 | 0 | 0 | 0 | 0 | 0 | 0 | 0 | 0 | 0 | 0 | 0 | 0 | 0 | 0 | 0 | 0 | 0 | 0 |
| ENSMUSG00 | Olfr55 | 947 | 0 | 0 | 0 | 0 | 0 | 0 | 0 | 0 | 0 | 0 | 0 | 0 | 0 | 0 | 0 | 0 | 0 | 0 |
| ENSMUSG00 | Gm3222 | 989 | 0.30084553 | 0.97681894 | 0.19564623 | 0.36854232 | 0.58400716 | 1.00207752 | 1.3873283 | 0.61787506 | 0.45809067 | 0.52442643 | 1.2076582 | 0.39505525 | 0.14964181 | 0.65744916 | 0.37493832 | 0.46459787 | 0.4650309 | 0.88993499 |
| ENSMUSG00 | Rab9 | 2336 | 34.4626727 | 34.7674812 | 31.8072528 | 35.4385349 | 34.5742157 | 34.6402457 | 34.7958829 | 34.4974183 | 35.1566406 | 34.3134399 | 36.6901864 | 36.1272611 | 37.8647807 | 37.4773565 | 34.5697986 | 47.2895558 | 39.9833842 | 39.7968281 |
| ENSMUSG00 | Trappc2 | 2438 | 7.9675412 | 7.48788599 | 8.59136122 | 6.97057815 | 7.91669458 | 7.82518557 | 9.29564118 | 8.41235443 | 8.36231555 | 7.63926488 | 7.48565797 | 6.47901252 | 7.00116627 | 7.10567867 | 7.70627953 | 9.54909323 | 7.59294561 | 6.58845741 |
| ENSMUSG00 | Gm11091 | 40 | 10.626294 | 4.99693413 | 1.20933826 | 1.13902611 | 3.60989428 | 0 | 4.73126789 | 4.7740503 | 3.08898868 | 7.07260549 | 0 | 11.1631325 | 1.23329793 | 0 | 3.09011669 | 14.3589778 | 0.95815742 | 1.37522767 |
| ENSMUSG00 | Gm11092 | 108 | 0.39356644 | 0 | 0 | 0 | 0 | 0 | 0 | 0 | 0.38135663 | 0 | 0 | 1.55043507 | 0.45677701 | 0 | 0 | 0 | 0 | 1.52803074 |
| ENSMUSG00 | Cdhr4 | 2250 | 0 | 0 | 0.04299869 | 0 | 0 | 0 | 0 | 0 | 0 | 0.12573521 | 0 | 0 | 0 | 0 | 0 | 0 | 0 | 0 |
| ENSMUSG00 | 4932414N04 | 4321 | 0 | 0 | 0 | 0 | 0 | 0 | 0 | 0 | 0 | 0 | 0 | 0.0129173 | 0 | 0 | 0 | 0 | 0 | 0 |
| ENSMUSG00 | Gm15262 | 370 | 0 | 0 | 0 | 0 | 0 | 0 | 0 | 0 | 0 | 0 | 0 | 0 | 0 | 0 | 0 | 0 | 0 | 0 |
| ENSMUSG00 | Lemd1 | 978 | 0 | 0 | 0 | 0 | 0 | 0 | 0 | 0 | 0 | 0 | 0 | 0 | 0 | 0 | 0 | 0 | 0 | 0 |
| ENSMUSG00 | Gm11096 | 71 | 0 | 0 | 0 | 0 | 0 | 0 | 0 | 0 | 0 | 0 | 0 | 0 | 0 | 0 | 0 | 0 | 0 | 0 |
| ENSMUSG00 | Nat6 | 2620 | 21.7068418 | 20.1784592 | 19.4601912 | 19.3895284 | 19.4548501 | 20.7289298 | 15.2953584 | 14.9562614 | 14.588201 | 16.17881 | 16.5024189 | 17.2559873 | 15.4962473 | 17.4076694 | 17.3612663 | 19.5545163 | 15.3890322 | 15.9568401 |
| ENSMUSG00 | Gm14446 | 4404 | 0 | 0 | 0 | 0 | 0.02185828 | 0 | 0 | 0 | 0 | 0 | 0 | 0 | 0 | 0 | 0.06548839 | 0 | 0 | 0 |
| ENSMUSG00 | Gm5097 | 2215 | 0 | 0 | 0 | 0 | 0 | 0 | 0 | 0 | 0 | 0.021287 | 0 | 0 | 0 | 0 | 0 | 0 | 0 | 0 |
| ENSMUSG00 | Gm5077 | 4386 | 0 | 0.03038112 | 0 | 0 | 0 | 0.04519173 | 0 | 0 | 0 | 0 | 0 | 0.01272587 | 0.01124759 | 0 | 0 | 0 | 0 | 0 |
| ENSMUSG00 | Lipo4 | 2706 | 0 | 0 | 0 | 0 | 0 | 0 | 0 | 0 | 0 | 0 | 0 | 0 | 0 | 0 | 0 | 0 | 0 | 0 |
| ENSMUSG00 | 1700013D24l | 532 | 0 | 0.50094578 | 0.72742151 | 0.08564106 | 0.27142062 | 0.27943271 | 0.08893361 | 0.14358046 | 0.23225479 | 0.35451657 | 0.08980255 | 0.52458329 | 0.37091667 | 0.17460183 | 0.15489307 | 0.57579861 | 0.79246103 | |
| ENSMUSG00 | Magea5 | 2754 | 0 | 0 | 0 | 0 | 0 | 0 | 0 | 0 | 0 | 0 | 0 | 0 | 0 | 0 | 0 | 0 | 0 | 0 |
| ENSMUSG00 | Magea8 | 1642 | 0 | 0 | 0 | 0 | 0 | 0 | 0 | 0 | 0 | 0 | 0 | 0 | 0 | 0 | 0 | 0 | 0 | 0 |
| ENSMUSG00 | Gm15140 | 981 | 0 | 0 | 0 | 0 | 0 | 0 | 0 | 0 | 0 | 0 | 0 | 0 | 0 | 0 | 0 | 0 | 0 | 0 |
| ENSMUSG00 | Gm11099 | 111 | 0 | 0 | 0 | 0 | 0 | 0 | 0 | 0 | 0 | 0 | 0 | 1.00568761 | 0 | 0 | 0 | 0 | 0 | 0 |
| ENSMUSG00 | Gm2237 | 1963 | 0.3031444 | 0.40728959 | 0.2710692 | 0.20888915 | 0.19615659 | 0.53011075 | 0.02410223 | 0.33075438 | 0.35668404 | 0.38431546 | 0.51109227 | 0.65397873 | 0.22617792 | 0.61515348 | 0.25186891 | 0.64370354 | 0.52715742 | 0.30825276 |
| ENSMUSG00 | Ccrl1 | 2106 | 164.773152 | 182.604013 | 158.971132 | 149.230809 | 172.41569 | 158.328748 | 76.8325555 | 87.1026157 | 69.0157712 | 68.1961264 | 85.7272403 | 69.9948551 | 70.0859907 | 88.9406479 | 77.5708305 | 93.1992657 | 86.4798494 | 84.3159528 |
| ENSMUSG00 | Mettl4-ps1 | 2135 | 0 | 0.03120646 | 0 | 0 | 0 | 0.02320971 | 0 | 0 | 0 | 0 | 0 | 0 | 0 | 0 | 0 | 0 | 0.03590285 | 0 |
| ENSMUSG00 | Gm11100 | 2128 | 0.09987118 | 0.01565456 | 0 | 0.04282053 | 0 | 0.02328606 | 0 | 0 | 0 | 0 | 0 | 0.07868749 | 0.04636458 | 0.5238055 | 0.03872327 | 0.03598741 | 0.03602096 | 0.07755043 |
| ENSMUSG00 | Gm9006 | 840 | 0 | 0 | 0 | 0 | 0 | 0 | 0 | 0 | 0 | 0 | 0 | 0 | 0 | 0 | 0 | 0 | 0 | 0 |
| ENSMUSG00 | Gm3839 | 1001 | 0.25477628 | 0.29951653 | 0.14497562 | 0.18206212 | 0.09616768 | 0.14850969 | 0.47265414 | 0.22892549 | 0.4114537 | 0.75365759 | 0.42954508 | 0.27879951 | 0.29569581 | 0.18559076 | 0.37044356 | 0.30601884 | 0.38288009 | 0.27477076 |
| ENSMUSG00 | Gm11102 | 58 | 2.19854359 | 0.57436025 | 1.66805277 | 2.35660574 | 6.63888602 | 1.70871495 | 0 | 1.97546909 | 2.13033702 | 0.81294316 | 0 | 2.88701703 | 2.55365089 | 2.40228038 | 3.55185827 | 0 | 0 | 1.89686575 |
| ENSMUSG00 | Gbp6 | 4303 | 2.0645089 | 2.60123924 | 3.19267549 | 2.17058194 | 2.50558933 | 3.00563873 | 1.902183 | 3.27515605 | 1.6175973 | 1.86279794 | 2.05400124 | 2.19215593 | 0.4815107 | 0.88506046 | 1.66606338 | 1.31699045 | 1.4696349 | 0.81816938 |
| ENSMUSG00 | Gbp4 | 4527 | 4.32844845 | 4.56976084 | 5.34278003 | 5.76683861 | 6.84713029 | 8.52696697 | 4.2640983 | 5.67781905 | 5.71352593 | 4.83276481 | 5.17113533 | 8.84025405 | 2.32111738 | 1.53890283 | 2.12970269 | 5.21875519 | 3.20020769 | 6.42805775 |
| ENSMUSG00 | Gm3558 | 2043 | 0 | 0 | 0 | 0 | 0 | 0 | 0 | 0 | 0 | 0 | 0 | 0 | 0 | 0 | 0 | 0 | 0 | 0 |
| ENSMUSG00 | Gm3618 | 382 | 0 | 0 | 0 | 0 | 0 | 0 | 0 | 0 | 0 | 0 | 0 | 0 | 0 | 0 | 0 | 0 | 0 | 0 |
| ENSMUSG00 | Gm3358 | 2662 | 0 | 0 | 0 | 0 | 0 | 0 | 0 | 0 | 0 | 0 | 0 | 0 | 0 | 0 | 0 | 0 | 0 | 0 |
| ENSMUSG00 | Gm3476 | 1992 | 0 | 0 | 0 | 0 | 0 | 0 | 0 | 0 | 0 | 0 | 0 | 0 | 0 | 0 | 0 | 0 | 0 | 0 |
| ENSMUSG00 | Gm3468 | 1954 | 0.08701162 | 0.13638851 | 0.04951231 | 0.04663362 | 0.19706008 | 0.05071928 | 0 | 0 | 0.02107805 | 0 | 0.14669895 | 0 | 0 | 0.07130617 | 0 | 0 | 0.19614277 | 0.0563041 |
| ENSMUSG00 | Gm10021 | 2059 | 0 | 0 | 0 | 0 | 0 | 0 | 0 | 0 | 0 | 0 | 0 | 0 | 0 | 0 | 0 | 0 | 0 | 0 |
| ENSMUSG00 | Gm8334 | 1659 | 0 | 0.06024032 | 0 | 0 | 0 | 0 | 0 | 0.02302134 | 0 | 0 | 0 | 0 | 0 | 0 | 0.1986814 | 0 | 0.11551024 | 0 |
| ENSMUSG00 | Gm3194 | 2134 | 0 | 0 | 0 | 0 | 0 | 0 | 0 | 0 | 0 | 0 | 0 | 0 | 0 | 0 | 0 | 0 | 0 | 0 |
| ENSMUSG00 | Gm3383 | 3201 | 0 | 0 | 0 | 0 | 0 | 0 | 0 | 0 | 0 | 0 | 0 | 0 | 0 | 0 | 0 | 0 | 0 | 0 |
| ENSMUSG00 | Gm3029 | 979 | 0 | 0 | 0 | 0 | 0 | 0 | 0 | 0 | 0 | 0 | 0 | 0 | 0 | 0 | 0 | 0 | 0 | 0 |
| ENSMUSG00 | Gm8279 | 1950 | 0 | 0 | 0 | 0 | 0 | 0 | 0 | 0 | 0 | 0 | 0 | 0 | 0 | 0 | 0 | 0 | 0 | 0 |
| ENSMUSG00 | Gm3242 | 482 | 0 | 0 | 0 | 0 | 0 | 0 | 0 | 0 | 0 | 0 | 0 | 0 | 0 | 0 | 0 | 0 | 0 | 0 |
| ENSMUSG00 | Gm8271 | 1467 | 0 | 0 | 0 | 0 | 0 | 0 | 0 | 0 | 0 | 0 | 0 | 0 | 0 | 0 | 0 | 0 | 0 | 0 |
| ENSMUSG00 | 4930555G01l | 2060 | 0 | 0 | 0 | 0 | 0 | 0 | 0 | 0 | 0 | 0 | 0 | 0 | 0 | 0 | 0 | 0 | 0 | 0 |
| ENSMUSG00 | Gm3443 | 475 | 0 | 0 | 0 | 0 | 0 | 0 | 0 | 0 | 0 | 0 | 0 | 0 | 0 | 0 | 0 | 0 | 0 | 0 |
| ENSMUSG00 | Gm3298 | 870 | 0 | 0 | 0 | 0 | 0 | 0 | 0 | 0 | 0 | 0 | 0 | 0 | 0 | 0.10676802 | 0 | 0 | 0 | 0 |
| ENSMUSG00 | Gm3173 | 596 | 0 | 0 | 0 | 0 | 0 | 0 | 0 | 0 | 0 | 0 | 0 | 0 | 0 | 0 | 0 | 0 | 0 | 0 |
| ENSMUSG00 | Luzp4 | 2107 | 0 | 0 | 0 | 0 | 0 | 0 | 0 | 0 | 0.03909494 | 0 | 0 | 0 | 0 | 0 | 0 | 0 | 0 | 0 |
| ENSMUSG00 | 2610042L04f | 1879 | 0 | 0 | 0 | 0.04849499 | 0 | 0 | 0 | 0 | 0 | 0 | 0 | 0 | 0 | 0 | 0 | 0 | 0 | 0 |
| ENSMUSG00 | Gm3149 | 1946 | 0 | 0 | 0 | 0 | 0 | 0 | 0 | 0 | 0 | 0 | 0 | 0 | 0 | 0 | 0 | 0 | 0 | 0 |
| ENSMUSG00 | Gm11105 | 477 | 0 | 0 | 0 | 0 | 0 | 0 | 0 | 0 | 0 | 0 | 0 | 0 | 0 | 0 | 0 | 0 | 0 | 0 |
| ENSMUSG00 | Gm2974 | 2173 | 0 | 0 | 0 | 0 | 0 | 0 | 0 | 0 | 0 | 0 | 0 | 0 | 0 | 0 | 0 | 0 | 0 | 0 |
| ENSMUSG00 | Gm17170 | 315 | 0 | 0 | 0 | 0 | 0 | 0 | 0 | 0 | 0 | 0 | 0 | 0 | 0 | 0 | 0 | 0 | 0 | 0 |
| ENSMUSG00 | Gm3099 | 2060 | 0 | 0 | 0 | 0 | 0 | 0 | 0 | 0 | 0 | 0 | 0 | 0 | 0 | 0 | 0 | 0 | 0 | 0 |
| ENSMUSG00 | Gm15114 | 1967 | 0.15126397 | 0.01693589 | 0 | 0 | 0 | 0 | 0 | 0.01941657 | 0 | 0 | 0 | 0 | 0.07523932 | 0.18889309 | 0.3141959 | 0.077866 | 0.09742323 | 0.25169393 |
| ENSMUSG00 | Gm5796 | 2067 | 0 | 0 | 0 | 0 | 0 | 0 | 0 | 0 | 0 | 0 | 0 | 0 | 0 | 0 | 0 | 0 | 0 | 0 |
| ENSMUSG00 | Gm2916 | 382 | 0 | 0 | 0 | 0 | 0 | 0 | 0 | 0 | 0 | 0 | 0 | 0 | 0 | 0 | 0 | 0 | 0 | 0 |
| ENSMUSG00 | Gm3033 | 499 | 0 | 0 | 0 | 0 | 0 | 0 | 0 | 0 | 0 | 0 | 0 | 0 | 0 | 0 | 0 | 0 | 0 | 0 |
| ENSMUSG00 | Gm3002 | 1222 | 0 | 0 | 0 | 0 | 0 | 0 | 0 | 0 | 0 | 0 | 0 | 0.13702699 | 0 | 0 | 0.16858247 | 0 | 0 | 0 |
| ENSMUSG00 | Gm3020 | 1932 | 0 | 0 | 0 | 0 | 0 | 0 | 0 | 0 | 0 | 0 | 0 | 0 | 0 | 0 | 0 | 0 | 0 | 0 |
| ENSMUSG00 | Gm3012 | 1111 | 0 | 0 | 0 | 0 | 0 | 0 | 0 | 0 | 0 | 0 | 0 | 0 | 0 | 0 | 0 | 0 | 0 | 0 |
| ENSMUSG00 | Gm11109 | 1698 | 0 | 0 | 0 | 0 | 0 | 0 | 0 | 0 | 0 | 0 | 0 | 0 | 0 | 0 | 0 | 0 | 0 | 0 |
| ENSMUSG00 | AC195488.1 | 561 | 0 | 0 | 0 | 0 | 0 | 0 | 0 | 0 | 0 | 0 | 0 | 0 | 0 | 0 | 0 | 0 | 0 | 0 |
| ENSMUSG00 | Gm1973 | 810 | 0 | 0 | 0 | 0 | 0 | 0 | 0 | 0 | 0 | 0 | 0 | 0 | 0 | 0 | 0 | 0 | 0 | 0 |
| ENSMUSG00 | 1700110I01R | 3678 | 0 | 0 | 0 | 0 | 0 | 0 | 0 | 0 | 0 | 0 | 0 | 0 | 0 | 0 | 0 | 0 | 0 | 0 |
| ENSMUSG00 | D830030K20l | 603 | 0 | 0 | 0 | 0 | 0 | 0 | 0 | 0 | 0 | 0 | 0 | 0 | 0 | 0 | 0 | 0 | 0 | 0 |
| ENSMUSG00 | Gm5795 | 1934 | 0 | 0 | 0 | 0 | 0 | 0 | 0 | 0 | 0 | 0 | 0 | 0 | 0 | 0 | 0 | 0 | 0 | 0 |
| ENSMUSG00 | Gm2897 | 2013 | 0 | 0.01654888 | 0 | 0 | 0 | 0.04923272 | 0 | 0 | 0 | 0.09369241 | 0 | 0.0277276 | 0 | 0 | 0 | 0 | 0 | 0 |
| ENSMUSG00 | Gm11110 | 1008 | 0.3795105 | 0.49572759 | 0.43190652 | 0.85878953 | 0.14324977 | 0.24579729 | 1.12649236 | 1.89446441 | 1.2666488 | 1.30974176 | 0.18958317 | 0.8859629 | 0 | 0 | 0.16349824 | 0.03798671 | 0.68439816 | 0 |
| ENSMUSG00 | Cntf | 1071 | 0 | 0.03110448 | 0 | 0.17016263 | 0.17976442 | 0.09253545 | 0.04417617 | 0 | 0.07691226 | 0 | 0.17843122 | 0.26057732 | 0.18424619 | 0 | 0 | 0.0715044 | 0 | 0 |
| ENSMUSG00 | Gm11111 | 96 | 0 | 0 | 0 | 0 | 0 | 0 | 0 | 0 | 0 | 0 | 0 | 0 | 0 | 0 | 0 | 0 | 0 | 0 |
| ENSMUSG00 | Atg4a | 2668 | 79.1953636 | 75.1537895 | 77.800157 | 73.1572391 | 76.2028657 | 74.8305711 | 119.168367 | 103.382882 | 109.280114 | 102.289457 | 99.7758878 | 98.5560669 | 121.369829 | 105.178102 | 99.2358314 | 88.4358032 | 95.4566128 | 111.68828 |
| ENSMUSG00 | Ms4a6c | 1840 | 0 | 0.0543145 | 0.07886989 | 0 | 0 | 0 | 0.02571341 | 1.10010724 | 0.49244747 | 0 | 0 | 0 | 0 | 0 | 0.1791372 | 0.08324045 | 0 | 0 |
| ENSMUSG00 | Gm17418 | 3536 | 1.44248335 | 1.40373904 | 2.18884753 | 2.21620465 | 1.25230118 | 1.23321283 | 0.78943667 | 1.77136708 | 1.87529102 | 0.69339269 | 0.94577124 | 2.36774587 | 1.3532794 | 0.89315553 | 1.65458737 | 2.16575834 | 1.506605 | 0.63783184 |
| ENSMUSG00 | 4930402F06f | 2135 | 0 | 0 | 0 | 0 | 0 | 0 | 0 | 0 | 0 | 0 | 0 | 0 | 0 | 0 | 0 | 0 | 0 | 0 |
| ENSMUSG00 | Gm7971 | 1844 | 0 | 0 | 0 | 0 | 0 | 0 | 0 | 0 | 0 | 0 | 0 | 0 | 0 | 0 | 0 | 0 | 0 | 0 |
| ENSMUSG00 | Gm3286 | 1282 | 0 | 0 | 0 | 0 | 0 | 0 | 0 | 0 | 0 | 0 | 0 | 0 | 0 | 0 | 0 | 0 | 0 | 0 |
| ENSMUSG00 | Gm3259 | 2888 | 0 | 0 | 0 | 0 | 0 | 0 | 0 | 0 | 0 | 0 | 0 | 0 | 0 | 0 | 0 | 0 | 0 | 0 |
| ENSMUSG00 | Arpc4 | 2409 | 63.1666792 | 59.2966751 | 60.9839816 | 62.7717336 | 61.2588119 | 63.0876853 | 63.1032119 | 58.7075409 | 60.762506 | 63.0241862 | 66.3772462 | 55.1900823 | 59.9805669 | 60.5180533 | 61.4517187 | 65.0097901 | 60.6156875 | 64.6910749 |
| ENSMUSG00 | Gm2382 | 1190 | 11.6085565 | 9.54596381 | 10.3251065 | 12.0219898 | 10.5566656 | 13.6165915 | 16.7781097 | 14.4745996 | 14.2595332 | 14.3433232 | 12.0441072 | 11.9135952 | 12.7268055 | 14.2064267 | 10.7331784 | 12.6133766 | 10.9503705 | 12.6659624 |
| ENSMUSG00 | Tceal7 | 1212 | 0.98196776 | 0.32983064 | 0.23947292 | 0.15036648 | 0.87368178 | 0.73593169 | 0.50747923 | 1.00838026 | 1.63114914 | 0.85587085 | 0.03941828 | 0.27631516 | 0.93616675 | 0.61312327 | 1.93769694 | 1.42168097 | 1.42300607 | 2.58706195 |
| ENSMUSG00 | Heatr7b1 | 8657 | 0.08837856 | 0.39250493 | 0.12851926 | 0.17893907 | 0.25575471 | 0.2289603 | 0.03279151 | 0.14999904 | 0.04757597 | 0.06535849 | 0.1769682 | 0.0322373 | 0.0854775 | 0.13948784 | 0.02855601 | 0.04865388 | 0.06646605 | 0 |
| ENSMUSG00 | Gm15023 | 3693 | 0 | 0 | 0 | 0 | 0 | 0 | 0 | 0 | 0 | 0 | 0 | 0 | 0 | 0 | 0 | 0 | 0 | 0 |
| ENSMUSG00 | Gm11114 | 55 | 0 | 0 | 0 | 0 | 0 | 0 | 0 | 0 | 0 | 0 | 0 | 0 | 0 | 0 | 0 | 0 | 0 | 0 |
| ENSMUSG00 | Neu2 | 2584 | 0.72377235 | 1.32787465 | 0.78625707 | 0.91686312 | 1.11761433 | 1.16978202 | 1.0619719 | 0.7833581 | 1.78517406 | 2.71882925 | 1.47910088 | 0.32400733 | 7.48378931 | 4.49342951 | 5.46909198 | 7.0387146 | 6.03668841 | 6.42908291 |
| ENSMUSG00 | Rpl36a | 1243 | 23.0478589 | 18.7066775 | 20.1588807 | 16.384382 | 22.9236515 | 20.0523129 | 19.4884084 | 19.3880981 | 24.5859973 | 18.8906277 | 18.9101203 | 18.051405 | 32.3455451 | 31.647741 | 22.6724571 | 21.5326907 | 21.6760955 | 26.5973235 |
| ENSMUSG00 | Kcnj13 | 1287 | 0.75961076 | 0.56945118 | 0.33827644 | 0.28320773 | 0.86016647 | 0.46203015 | 0.25733392 | 0.59351053 | 0 | 0.29308906 | 0.29696944 | 0.39031932 | 0.22998563 | 0.21652255 | 0.09604092 | 0.41652564 | 0.11911825 | 0.98306873 |
| ENSMUSG00 | Tmem179b | 843 | 84.5061388 | 77.2558816 | 76.9500643 | 68.9090528 | 83.5314405 | 70.7140433 | 96.3090831 | 94.2351091 | 97.7630107 | 75.7319718 | 78.7748426 | 84.617339 | 109.958093 | 86.0565415 | 85.7754811 | 93.5689813 | 91.3375213 | 89.7242251 |
| ENSMUSG00 | Gm11115 | 52 | 0 | 0 | 0 | 0 | 0 | 0 | 0 | 0 | 0 | 0 | 0 | 0 | 0 | 0 | 0 | 0 | 0 | 0 |
| ENSMUSG00 | Gm11116 | 52 | 0 | 0 | 0 | 0 | 0 | 0 | 0 | 0 | 0 | 0 | 0 | 0 | 0 | 0 | 0 | 0 | 0 | 0 |
| ENSMUSG00 | Alpi | 2324 | 0 | 0 | 0 | 0 | 0 | 0 | 0 | 0 | 0 | 0.12173159 | 0 | 0 | 0 | 0.07993819 | 0 | 0 | 0 | 0 |
| ENSMUSG00 | St6galnac4 | 4117 | 0.27875632 | 1.06808405 | 0.27024319 | 0.1217322 | 0.40918565 | 0.97492626 | 0.4941572 | 0.417454 | 0.61024471 | 0.09162148 | 0.39454666 | 0.39316352 | 0.53921211 | 1.0378567 | 0.57043689 | 0.23251523 | 0.13963917 | 1.42967559 |
| ENSMUSG00 | Prickle4 | 3549 | 0.02395333 | 0.07509246 | 0.08178112 | 0.11553942 | 0.1627453 | 0.1116996 | 0.21330033 | 0.118376 | 0.11605104 | 0.30556951 | 0.13461527 | 0.14154437 | 0.11120184 | 0.14395181 | 0.13931211 | 0.18341514 | 0.10799182 | 0.15499889 |
| ENSMUSG00 | B3gnt7 | 2591 | 2.11623609 | 5.23286297 | 3.13653149 | 2.84866429 | 2.52641514 | 3.19386588 | 0.87649888 | 1.06130952 | 2.30491887 | 0.70971726 | 2.2495349 | 0.66780607 | 3.37003063 | 5.0907452 | 5.08857502 | 3.85713953 | 3.1359224 | 3.18462602 |
| ENSMUSG00 | 3110007F17f | 5920 | 0.33027671 | 0.28698608 | 0.30233456 | 0.29245265 | 0.39025884 | 0.30970458 | 0.21578418 | 0.12257697 | 0.306116 | 0.31858583 | 0.47613557 | 0.16028146 | 0.46665327 | 0.17259627 | 0.11135556 | 0.34927243 | 0.27838358 | 0.16725742 |
| ENSMUSG00 | Tmprss11g | 4063 | 0.10461525 | 0 | 0 | 0 | 0 | 0 | 0 | 0 | 0 | 0 | 0 | 0 | 0 | 0 | 0 | 0 | 0 | 0 |
| ENSMUSG00 | Gm16026 | 3306 | 0 | 0.0100704 | 0 | 0 | 0.05820063 | 0 | 0.02860501 | 0 | 0 | 0 | 0.04332675 | 0 | 0.04473874 | 0.01403993 | 0 | 0 | 0 | 0 |

| | | | | | | | | | | | | | | | | | | | |
|---|---|---|---|---|---|---|---|---|---|---|---|---|---|---|---|---|---|---|---|
| ENSMUSG00 Gm7609 | 2268 | 0 | 0 | 0 | 0 | 0 | 0 | 0 | 0 | 0 | 0 | 0 | 0 | 0 | 0 | 0 | 0.01688298 | 0 | 0 |
| ENSMUSG00 4933403O08 | 1186 | 0 | 0 | 0 | 0 | 0 | 0 | 0 | 0 | 0 | 0 | 0 | 0 | 0 | 0 | 0 | 0 | 0 | 0 |
| ENSMUSG00 Gm15737 | 4892 | 0 | 0.05447734 | 0 | 0 | 0.00983891 | 0 | 0 | 0.03122846 | 0.01683831 | 0 | 0 | 0.01140958 | 0 | 0 | 0.00842223 | 0.00782719 | 0 | 0 |
| ENSMUSG00 Gm17682 | 1791 | 0.0474653 | 0 | 0.10803692 | 0.05087777 | 0 | 0 | 0.13208453 | 0.0852985 | 0 | 0 | 0.02667502 | 0 | 0.05508868 | 0.10372772 | 0.09201911 | 0 | 0.04279877 | 0 |
| ENSMUSG00 Col4a3 | 10370 | 0.11476807 | 0.14455933 | 0.20058455 | 0.18013528 | 0.25063875 | 0.10034787 | 0.13231125 | 0 | 0.12709436 | 0.24098238 | 0.06910553 | 0.11303075 | 0.0999007 | 0.03135088 | 0.15495281 | 0.25477839 | 0.01478353 | 0.05304639 |
| ENSMUSG00 Prdm12 | 2466 | 0 | 0 | 0 | 0 | 0 | 0 | 0 | 0 | 0 | 0 | 0 | 0 | 0 | 0 | 0 | 0 | 0 | 0 |
| ENSMUSG00 Gm14966 | 1040 | 3.1878882 | 4.35630155 | 3.16288468 | 5.56370445 | 3.65617497 | 3.04939899 | 5.23169046 | 4.40681566 | 4.91069996 | 4.6243959 | 3.30749713 | 4.88387048 | 3.93706647 | 5.58222204 | 4.63517504 | 3.35042815 | 4.68023049 | 4.65461672 |
| ENSMUSG00 Pigb | 2181 | 6.02205383 | 5.63615679 | 5.25654594 | 6.16254382 | 6.17924742 | 4.95300134 | 6.76825118 | 5.90134783 | 5.62750192 | 6.76669882 | 5.45436252 | 6.06525082 | 4.93092983 | 6.13291938 | 6.21518198 | 7.46148929 | 7.97805538 | 6.12893761 |
| ENSMUSG00 Utp14b | 3750 | 5.44066253 | 3.44677412 | 4.99214833 | 5.28508114 | 3.28580599 | 4.08314525 | 1.79157344 | 4.1349641 | 2.75675079 | 3.20624782 | 2.22949806 | 2.00936385 | 2.40739756 | 1.95684422 | 2.74677039 | 4.25791554 | 3.69976519 | 6.14635086 |
| ENSMUSG00 Gm7325 | 941 | 0 | 0.03540159 | 0 | 0 | 0 | 0 | 0.30167489 | 0.24352223 | 0 | 0.05010702 | 0 | 0 | 0.68152489 | 0.24678049 | 0.52541835 | 0 | 0.65166924 | 0 |
| ENSMUSG00 Gm550 | 838 | 0.10144433 | 0 | 0 | 0 | 0 | 0 | 0 | 0 | 0 | 0 | 0 | 0 | 0 | 0 | 0 | 0 | 0 | 0 |
| ENSMUSG00 1700011M02 | 417 | 0 | 0 | 0 | 0 | 0 | 0 | 0 | 0 | 0 | 0 | 0 | 0 | 0 | 0 | 0 | 0 | 0 | 0 |
| ENSMUSG00 Rab7 | 4013 | 109.901248 | 105.077153 | 112.70683 | 109.207995 | 112.419766 | 114.009566 | 102.666037 | 106.030789 | 99.9540187 | 107.284593 | 105.073956 | 100.824256 | 100.802821 | 96.9157209 | 101.109624 | 106.217553 | 104.406448 | 101.95807 |
| ENSMUSG00 Sssca1 | 3014 | 8.64488152 | 8.02427379 | 7.73591295 | 7.58846857 | 7.21819161 | 7.61211534 | 7.81742339 | 6.8553715 | 6.81886907 | 8.07225046 | 8.76561116 | 7.59270792 | 8.60935252 | 6.93425326 | 8.12001466 | 6.91111162 | 7.55335778 | 7.24572507 |
| ENSMUSG00 Gm9112 | 417 | 0 | 0 | 0 | 0 | 0 | 0 | 0 | 0 | 0 | 0 | 0 | 0 | 0 | 0 | 0 | 0 | 0 | 0 |
| ENSMUSG00 Pin4 | 646 | 17.3705363 | 14.2327536 | 13.403811 | 13.541363 | 13.9329253 | 12.2731229 | 14.0619727 | 15.8445261 | 14.9827108 | 12.5540572 | 13.5337731 | 11.1458521 | 17.7930909 | 13.0848867 | 11.7354277 | 13.3957853 | 12.7556561 | 15.3276149 |
| ENSMUSG00 Nhsl2 | 15927 | 0.06404999 | 0.02300759 | 0.00607441 | 0.02288493 | 0.09368303 | 0.02177869 | 0.11288264 | 0.11989829 | 0.0232736 | 0.08585235 | 0.06899128 | 0.0770983 | 0.07123966 | 0.21578844 | 0.1008891 | 0.13703551 | 0.28876472 | 0.13469926 |
| [illegible] | | | | | | | | | | | | | | | | | | | |

| | | | | | | | | | | | | | | | | | | | |
|---|---|---|---|---|---|---|---|---|---|---|---|---|---|---|---|---|---|---|---|
| ENSMUSG00 AC108410.1 | 120 | 0 | 0 | 0 | 0 | 0 | 0 | 0 | 0 | 0 | 0 | 0 | 0 | 0 | 0 | 0 | 0 | 0 | 0 |
| ENSMUSG00 U6atac | 102 | 0 | 0 | 0 | 0 | 0 | 0 | 0 | 0 | 0 | 0 | 0 | 0 | 0 | 0 | 0 | 0 | 0 | 0 |
| ENSMUSG00 Mir466f-4 | 120 | 0 | 0 | 0 | 0 | 0 | 0 | 0 | 0 | 0 | 0 | 0 | 0 | 0 | 0 | 0 | 0 | 0 | 0 |
| ENSMUSG00 SNORA11 | 127 | 0 | 1.57383752 | 0 | 0.35874838 | 1.5159661 | 0 | 0 | 0 | 0.97290982 | 0.74253076 | 0 | 0.43949341 | 0 | 0.36570148 | 0.3244217 | 0.60300169 | 3.01781865 | 0 |
| ENSMUSG00 AL929465.1 | 120 | 0 | 0.5552149 | 0 | 0 | 0 | 0 | 0 | 0 | 0 | 0 | 0 | 0 | 0 | 0 | 0 | 0 | 0 | 0 |
| ENSMUSG00 AC163028.1 | 89 | 0 | 0 | 0 | 0 | 0 | 0 | 0 | 0 | 0 | 0 | 0 | 0 | 0 | 0 | 0 | 0 | 0 | 0 |
| ENSMUSG00 Mir1187 | 121 | 0 | 0 | 0 | 0 | 0 | 0 | 0 | 0 | 0 | 0 | 0 | 0 | 0 | 0 | 0 | 0 | 0 | 0 |
| ENSMUSG00 AC131713.1 | 120 | 0 | 0 | 0 | 0 | 0 | 0 | 0 | 0 | 0 | 0 | 0 | 0 | 0 | 0 | 0 | 0 | 0 | 0 |
| ENSMUSG00 AC111132.1 | 120 | 0 | 0 | 0 | 0 | 0 | 0 | 0 | 0 | 0 | 0 | 0 | 0 | 0 | 0 | 0 | 0 | 0 | 0 |
| ENSMUSG00 AL590988.1 | 62 | 0 | 0 | 0 | 0 | 0 | 0 | 0 | 0 | 0 | 0 | 0 | 0 | 0 | 0 | 0 | 0 | 0 | 0 |
| ENSMUSG00 Mir669h | 124 | 0 | 0 | 0 | 0 | 0 | 0 | 0 | 0 | 0 | 0 | 0 | 0 | 0 | 0 | 0 | 0 | 0 | 0 |
| ENSMUSG00 U6atac | 125 | 0 | 0 | 0 | 0 | 0 | 0 | 0 | 0 | 0 | 0 | 0 | 0 | 0 | 0 | 0 | 0 | 0 | 0 |
| ENSMUSG00 AC161111.1 | 115 | 0 | 0 | 0 | 0 | 0 | 0 | 0 | 0 | 0 | 0 | 0 | 0 | 0 | 0 | 0 | 0 | 0 | 0 |
| ENSMUSG00 AL606976.1 | 92 | 0 | 0 | 0 | 0 | 0 | 0 | 0 | 0 | 0 | 0 | 0 | 0 | 0 | 0 | 0 | 0 | 0 | 0 |
| ENSMUSG00 AL772216.26 | 123 | 0 | 0 | 0 | 0 | 0 | 0 | 0 | 0 | 0 | 0 | 0 | 0 | 0 | 0 | 0 | 0 | 0 | 0 |
| ENSMUSG00 AL772216.27 | 125 | 0 | 0 | 0 | 0 | 0 | 0 | 0 | 0 | 0 | 0 | 0 | 0 | 0 | 0 | 0 | 0 | 0 | 0 |
| ENSMUSG00 AL591376.1 | 96 | 0 | 0 | 0 | 0 | 0 | 0 | 0 | 0 | 0 | 0 | 0 | 0 | 0 | 0 | 0 | 0 | 0 | 0 |
| ENSMUSG00 AC119834.1 | 113 | 1.128456 | 0.88441312 | 1.71233736 | 1.61278033 | 2.98162359 | 1.3155593 | 1.25608882 | 1.68992931 | 0 | 0 | 0.42278724 | 5.927327 | 0.87313128 | 0 | 2.18769323 | 0 | 2.03502461 | 1.94722502 |
| ENSMUSG00 AL929160.1 | 89 | 0 | 0 | 0 | 0 | 0 | 0 | 0 | 0 | 0 | 1.05956637 | 0 | 0 | 0.55429121 | 0 | 0 | 0 | 0 | 0.61807985 |
| ENSMUSG00 AC138376.1 | 90 | 0 | 0 | 0 | 0 | 0 | 0 | 0 | 0 | 0 | 0 | 0 | 0 | 0 | 0 | 0 | 0 | 0 | 0 |
| ENSMUSG00 AL772216.28 | 126 | 0 | 0 | 0 | 0 | 0 | 0 | 0 | 0 | 0 | 0 | 0 | 0 | 0 | 0 | 0 | 0 | 0 | 0 |
| ENSMUSG00 AC141438.1 | 120 | 0 | 0 | 0 | 0 | 0 | 0 | 0 | 0 | 0 | 0 | 0 | 0 | 0 | 0 | 0 | 0 | 0 | 0 |
| ENSMUSG00 SNORD23 | 108 | 0 | 0 | 0 | 0 | 0 | 0 | 0 | 0 | 0 | 0 | 0 | 0 | 0 | 0 | 0 | 0 | 0 | 0 |
| ENSMUSG00 AC158296.1 | 120 | 0 | 0 | 0 | 0 | 0 | 0 | 0 | 0 | 0 | 0 | 0 | 0.93026104 | 0 | 0 | 0 | 0 | 0 | 0 |
| ENSMUSG00 Snord110 | 67 | 0 | 1.98882951 | 0 | 2.04004676 | 0 | 0 | 0.70615939 | 1.14007171 | 0 | 2.81496736 | 0 | 0 | 0 | 4.85236734 | 0.6149486 | 0 | 2.28813713 | 1.64206289 |
| ENSMUSG00 Mir669j | 120 | 0 | 0 | 0 | 0 | 0 | 0 | 0 | 0 | 0 | 0 | 0 | 0 | 0 | 0 | 0 | 0 | 0 | 0 |
| ENSMUSG00 AL772216.29 | 125 | 0 | 0 | 0 | 0 | 0 | 0 | 0 | 0 | 0 | 0 | 0 | 0 | 0 | 0 | 0 | 0 | 0 | 0 |
| ENSMUSG00 AC121604.1 | 57 | 0 | 0 | 0 | 0 | 0 | 0 | 0 | 0 | 0 | 0 | 0 | 0 | 0 | 0 | 0 | 0 | 0 | 0 |
| ENSMUSG00 AC102689.1 | 86 | 0 | 0 | 0 | 0 | 0 | 0 | 0 | 0 | 0 | 0 | 0 | 0 | 0 | 0 | 0 | 0 | 0 | 0 |
| ENSMUSG00 Snord99 | 73 | 0 | 0 | 0 | 0 | 0 | 0 | 0 | 0 | 0 | 0 | 0 | 0 | 0 | 0 | 0 | 0 | 0 | 0 |
| ENSMUSG00 U3 | 124 | 0 | 0 | 0 | 0 | 0 | 0 | 0 | 0 | 0 | 0 | 0 | 0 | 0 | 0 | 0 | 0 | 0 | 0 |
| ENSMUSG00 mmu-mir-119 | 120 | 0 | 0 | 0 | 0 | 0 | 0 | 0 | 0 | 0 | 0 | 0 | 0 | 0 | 0 | 0 | 0 | 0 | 0 |
| ENSMUSG00 SNORA11 | 128 | 0 | 0 | 0 | 0 | 0 | 0 | 0 | 0 | 0 | 0 | 0 | 0 | 0 | 0 | 0 | 0 | 0 | 0 |
| ENSMUSG00 AL772216.30 | 100 | 0 | 0 | 0 | 0 | 0 | 0 | 0 | 0 | 0 | 0 | 0 | 0 | 0 | 0 | 0 | 0 | 0 | 0 |
| ENSMUSG00 AC163659.2 | 99 | 0 | 0 | 0 | 0 | 0 | 0 | 0 | 0 | 0 | 0 | 0 | 0 | 0 | 0 | 0 | 0 | 0 | 0 |
| ENSMUSG00 Mir1192 | 120 | 0.7084196 | 0.5552149 | 0.80622551 | 0.75935074 | 0 | 1.23881834 | 0.78854465 | 7.63848048 | 2.05932579 | 3.14338022 | 0.39812465 | 1.39539156 | 1.64439724 | 0.38703406 | 1.0300389 | 1.27635358 | 1.59692904 | 0 |
| ENSMUSG00 AL772216.31 | 107 | 0 | 0 | 0 | 0 | 0 | 0 | 0 | 0 | 0 | 0 | 0 | 0 | 0 | 0 | 0 | 0 | 0 | 0 |
| ENSMUSG00 AL691457.1 | 143 | 0 | 0 | 0 | 0 | 0 | 0 | 0 | 0 | 0 | 0 | 0 | 0 | 0 | 0 | 0 | 0 | 0 | 0 |
| ENSMUSG00 AC098726.1 | 88 | 0 | 0 | 0 | 0 | 1.64086103 | 1.12619849 | 0.53764408 | 0.43400457 | 0 | 0 | 0 | 0 | 0.56058997 | 0 | 0 | 0 | 0 | 0 |
| ENSMUSG00 AC109145.1 | 120 | 0 | 0 | 0 | 0 | 0 | 0 | 0 | 0 | 0 | 0 | 0 | 0 | 0 | 0 | 0 | 0 | 0 | 0 |
| ENSMUSG00 AC139348.1 | 105 | 0 | 0 | 0 | 0 | 0 | 0 | 0 | 0 | 0 | 0 | 0 | 0 | 0 | 0 | 0 | 0 | 0 | 0 |
| ENSMUSG00 AC161265.2 | 122 | 0 | 0 | 0 | 0 | 0 | 0 | 0 | 0 | 0 | 0 | 0 | 0 | 0 | 0 | 0 | 0 | 0 | 0 |
| ENSMUSG00 AC122490.1 | 81 | 0 | 0 | 0 | 0 | 0 | 0 | 0 | 0 | 0 | 0 | 0 | 0 | 0 | 0 | 0 | 0 | 0 | 0 |
| ENSMUSG00 AL772216.32 | 125 | 0 | 0 | 0 | 0 | 0 | 0 | 0 | 0 | 0 | 0 | 0 | 0 | 0 | 0 | 0 | 0 | 0 | 0 |
| ENSMUSG00 AL772216.33 | 120 | 0 | 0 | 0 | 0 | 0 | 0 | 0 | 0 | 0 | 0 | 0 | 0 | 0 | 0 | 0 | 0 | 0 | 0 |
| ENSMUSG00 Mir1198 | 120 | 0 | 0.27760745 | 0 | 1.51870148 | 1.20329809 | 0 | 0 | 0 | 1.02966289 | 0.78584505 | 0 | 1.39539156 | 0.41109931 | 0 | 0 | 0.3190884 | 0.31938581 | 0.45840922 |
| ENSMUSG00 AC131332.1 | 90 | 0 | 0 | 0 | 0 | 0 | 0 | 0 | 0 | 0 | 0 | 0 | 0 | 0 | 0 | 0 | 0 | 0 | 0 |
| ENSMUSG00 AL772216.34 | 122 | 0 | 0 | 0 | 0 | 0 | 0 | 0 | 0 | 0 | 0 | 0 | 0 | 0 | 0 | 0 | 0 | 0 | 0 |
| ENSMUSG00 AC119939.1 | 93 | 0 | 0 | 0 | 0 | 0 | 0 | 0 | 0 | 0 | 0 | 0 | 0 | 0 | 0 | 0 | 0 | 0 | 0 |
| ENSMUSG00 AC103351.1 | 87 | 0 | 0 | 0 | 0 | 0 | 0 | 0 | 0 | 0 | 0 | 0 | 0 | 0 | 0 | 0 | 0 | 0 | 0 |
| ENSMUSG00 Mir669k | 109 | 0 | 0 | 0 | 0 | 0 | 0 | 0 | 0 | 0 | 0 | 0 | 0 | 0 | 0 | 0 | 0 | 0 | 0 |
| ENSMUSG00 Mir669e | 118 | 0 | 0 | 0 | 0 | 0 | 0 | 0 | 0 | 0 | 0 | 0 | 0 | 0 | 0 | 0 | 0 | 0 | 0 |
| ENSMUSG00 Mir1196 | 118 | 0 | 0 | 0 | 0 | 0 | 0 | 0 | 0 | 0 | 0 | 0 | 0 | 0 | 0 | 0 | 0 | 0 | 0 |
| ENSMUSG00 AC155833.1 | 92 | 0 | 0 | 0 | 0 | 0 | 0 | 0 | 0 | 0 | 0 | 0 | 0 | 0 | 0 | 0 | 0 | 0 | 0 |
| ENSMUSG00 Mir669i | 126 | 0 | 0 | 0 | 0 | 0 | 0 | 0 | 0 | 0 | 0 | 0 | 0 | 0 | 0 | 0 | 0 | 0 | 0 |
| ENSMUSG00 AL844480.1 | 86 | 0 | 0 | 0 | 0 | 0 | 0 | 0 | 0 | 0 | 0 | 0 | 0 | 0 | 0 | 0 | 0 | 0 | 0 |
| ENSMUSG00 AC132253.2 | 120 | 0 | 0 | 0 | 0 | 0 | 0 | 0 | 0 | 0 | 0 | 0 | 0 | 0 | 0 | 0 | 0 | 0 | 0 |
| ENSMUSG00 AC108410.2 | 120 | 0 | 0 | 0 | 0 | 0 | 0 | 0 | 0 | 0 | 0 | 0 | 0 | 0 | 0 | 0 | 0 | 0 | 0 |
| ENSMUSG00 AC101970.1 | 89 | 0 | 0 | 0 | 0 | 0 | 0 | 0 | 0 | 0 | 0 | 0 | 0 | 0 | 0 | 0 | 0 | 0 | 0 |
| ENSMUSG00 Mir1b | 121 | 0 | 0 | 0.79956249 | 0 | 0 | 0 | 0 | 0.31563969 | 1.02115328 | 0.38967523 | 1.97417184 | 0 | 1.22310539 | 0.76767087 | 0 | 0 | 0 | 0 |
| ENSMUSG00 AL772302.1 | 65 | 0 | 0 | 0 | 0 | 0 | 0 | 0 | 0 | 0 | 0 | 0 | 0 | 0 | 0 | 0 | 0 | 0 | 0 |
| ENSMUSG00 U3 | 192 | 0 | 0 | 0 | 0 | 0.2506871 | 0 | 0 | 0 | 0 | 0 | 0 | 0 | 0 | 0 | 0 | 0 | 0 | 0 |
| ENSMUSG00 Mir1199 | 118 | 0 | 0.56462533 | 0.81989035 | 0 | 0 | 0 | 0.40095491 | 0 | 0 | 0.79916446 | 0 | 0 | 0 | 0.78718792 | 0 | 0.32449667 | 0 | 0 |
| ENSMUSG00 AC123619.1 | 113 | 0 | 0 | 0 | 0 | 0 | 0 | 0 | 0 | 0 | 0 | 0 | 0 | 0 | 0 | 0 | 0 | 0 | 0 |
| ENSMUSG00 AC122402.1 | 100 | 0 | 0 | 0 | 0 | 0 | 0 | 0 | 0 | 0 | 0 | 0 | 0 | 0 | 0 | 0 | 0 | 0 | 0 |
| ENSMUSG00 Mir1224 | 84 | 0 | 0 | 0 | 0 | 0 | 0 | 0 | 0 | 0 | 0 | 0 | 0 | 0 | 0 | 0 | 0 | 0 | 0 |
| ENSMUSG00 AC138299.1 | 88 | 0 | 0 | 0 | 0 | 0 | 0 | 0 | 0 | 0 | 0 | 0 | 0 | 0 | 0 | 0 | 0 | 0 | 0 |
| ENSMUSG00 U6atac | 122 | 0 | 0 | 0 | 0 | 0 | 0 | 0 | 0 | 0 | 0 | 0 | 0 | 0 | 0 | 0 | 0 | 0 | 0 |
| ENSMUSG00 AL772216.35 | 126 | 0 | 0 | 0 | 0 | 0 | 0 | 0 | 0 | 0 | 0 | 0 | 0 | 0 | 0 | 0 | 0 | 0 | 0 |
| ENSMUSG00 AC161513.1 | 120 | 0 | 0 | 0 | 0 | 0 | 0 | 0 | 0 | 0 | 0 | 0 | 0 | 0 | 0 | 0 | 0 | 0 | 0 |
| ENSMUSG00 AC165239.1 | 123 | 0 | 0 | 0 | 0 | 0 | 0 | 0 | 0 | 0 | 0 | 0 | 0 | 0 | 0 | 0 | 0 | 0 | 0 |
| ENSMUSG00 AC112790.1 | 97 | 0 | 0 | 0 | 0.46970149 | 0 | 2.04341169 | 0 | 0 | 0 | 0 | 0 | 0 | 0 | 0 | 0 | 0 | 0 | 0 |
| ENSMUSG00 AC161049.1 | 65 | 0 | 0 | 0 | 0 | 1.48098227 | 0 | 1.45577474 | 0 | 0 | 2.1761863 | 0 | 0 | 0 | 0 | 0 | 0 | 0 | 0 |
| ENSMUSG00 AL627312.1 | 116 | 0 | 0.28718012 | 0 | 0 | 0 | 0 | 0 | 0 | 0 | 0 | 0 | 0 | 1.7011006 | 0 | 0 | 0 | 0 | 1.42264931 |
| ENSMUSG00 mmu-mir-119 | 120 | 0 | 0 | 0 | 0 | 0 | 0 | 0 | 0 | 0 | 0 | 0 | 0 | 0 | 0 | 0 | 0 | 0 | 0 |
| ENSMUSG00 SNORD88 | 92 | 0 | 0 | 0 | 0 | 0 | 0 | 0 | 0 | 0 | 0 | 0 | 0 | 0 | 0 | 0 | 0 | 0 | 0 |
| ENSMUSG00 AL772216.36 | 121 | 0 | 0 | 0 | 0 | 0 | 0 | 0 | 0 | 0 | 0 | 0 | 0 | 0 | 0 | 0 | 0 | 0 | 0 |
| ENSMUSG00 AL772216.37 | 119 | 0 | 0 | 0 | 0 | 0 | 0 | 0 | 0 | 0 | 0 | 0 | 0 | 0 | 0 | 0 | 0 | 0 | 0 |
| ENSMUSG00 AL772216.38 | 89 | 0 | 0 | 0 | 0 | 0 | 0 | 0 | 0 | 0 | 0 | 0 | 0 | 0 | 0 | 0 | 0 | 0 | 0 |
| ENSMUSG00 Mir1197 | 119 | 0 | 0 | 0 | 0 | 0 | 0 | 0 | 0 | 0 | 0 | 0 | 0 | 0 | 0 | 0 | 0 | 0 | 0 |
| ENSMUSG00 SNORA79 | 128 | 0 | 0 | 0 | 0 | 0 | 0 | 0 | 0 | 0 | 0 | 0 | 0 | 0.3854056 | 0.36284443 | 0 | 0 | 0 | 0 |
| ENSMUSG00 Gm11621 | 588 | 0 | 0 | 0 | 0 | 0 | 0 | 0 | 0 | 0 | 0 | 0 | 0 | 0 | 0 | 0 | 0 | 0 | 0 |
| ENSMUSG00 Gm5394 | 974 | 0 | 0 | 0 | 0 | 0 | 0 | 0 | 0 | 0 | 0 | 0 | 0 | 0 | 0 | 0 | 0 | 0 | 0 |
| ENSMUSG00 Gm12825 | 561 | 0 | 0 | 0 | 0 | 0 | 0 | 0 | 0 | 0 | 0 | 0 | 0 | 0 | 0 | 0 | 0 | 0 | 0 |
| ENSMUSG00 Gm11299 | 580 | 0 | 0 | 0 | 0 | 0 | 0 | 0 | 0 | 0 | 0 | 0 | 0 | 0 | 0 | 0 | 0 | 0 | 0 |
| ENSMUSG00 Gm11333 | 147 | 0 | 0 | 0 | 0 | 0 | 0 | 0 | 0 | 0 | 0 | 0 | 0 | 0 | 0 | 0 | 0 | 0 | 0 |
| ENSMUSG00 Olfr1296-ps1 | 938 | 0 | 0 | 0 | 0 | 0 | 0 | 0 | 0 | 0 | 0 | 0 | 0 | 0 | 0 | 0 | 0 | 0 | 0 |
| ENSMUSG00 Gm15981 | 463 | 0 | 0 | 0 | 0 | 0 | 0 | 0 | 0 | 0 | 0 | 0 | 0 | 0 | 0 | 0 | 0 | 0.08277818 | 0 |
| ENSMUSG00 Gm12340 | 461 | 0.18440423 | 0.21678673 | 0.20986347 | 0.49415449 | 0.10440764 | 0 | 0 | 0 | 0.08934168 | 0.10227918 | 0 | 0.60537595 | 0.21402133 | 0 | 0.08937431 | 0.08305989 | 0 | 0 |
| ENSMUSG00 Gm7438 | 631 | 0 | 0 | 0 | 0 | 0 | 0 | 0 | 0 | 0 | 0 | 0 | 0 | 0 | 0 | 0 | 0 | 0 | 0 |
| ENSMUSG00 Gm13183 | 788 | 0 | 0 | 0 | 0 | 0 | 0 | 0 | 0 | 0 | 0 | 0 | 0 | 0 | 0 | 0 | 0 | 0 | 0 |
| ENSMUSG00 Gm6031 | 1053 | 0 | 0 | 0.04593878 | 0 | 0 | 0 | 0 | 0 | 0 | 0 | 0 | 0 | 0 | 0 | 0 | 0 | 0 | 0 |
| ENSMUSG00 Gm15439 | 383 | 0 | 0 | 0 | 0 | 0 | 0 | 0 | 0 | 0 | 0 | 0 | 0 | 0 | 0 | 0 | 0 | 0 | 0 |
| ENSMUSG00 Gm12890 | 372 | 0 | 0 | 0 | 0 | 0 | 0 | 0 | 0 | 0 | 0 | 0 | 0 | 0 | 0 | 0 | 0 | 0 | 0 |
| ENSMUSG00 Gm13823 | 344 | 0 | 0 | 0 | 0 | 0 | 0 | 0 | 0 | 0 | 0 | 0 | 0 | 0 | 0 | 0 | 0 | 0 | 0 |
| ENSMUSG00 Ccnd3-ps | 880 | 0 | 0.03785556 | 0 | 0 | 0.10939074 | 0 | 0 | 0 | 0 | 0.05358034 | 0 | 0 | 0 | 0 | 0 | 0 | 0 | 0 |
| ENSMUSG00 Gm14324 | 3048 | 0 | 0 | 0 | 0 | 0 | 0 | 0 | 0 | 0 | 0 | 0 | 0 | 0 | 0 | 0 | 0 | 0 | 0 |
| ENSMUSG00 Gm8107 | 2008 | 0 | 0.03318017 | 0 | 0 | 0 | 0 | 0 | 0 | 0 | 0 | 0 | 0 | 0 | 0 | 0 | 0 | 0 | 0 |
| ENSMUSG00 Gm11686 | 332 | 0 | 0 | 0 | 0 | 0 | 0 | 0 | 0.11503736 | 0 | 0 | 0 | 0 | 0 | 0 | 0 | 0 | 0 | 0 |
| ENSMUSG00 Hist3h2bb-ps | 464 | 0.09160598 | 0.14359006 | 0.31275989 | 0.19638381 | 0.51866297 | 0.32038405 | 0 | 0.57617848 | 0.26629213 | 0.60970737 | 0 | 0.84204663 | 0.21263757 | 0.50047508 | 0.44398228 | 0 | 0 | 0.71132466 |
| ENSMUSG00 Olfr1174-ps | 938 | 0 | 0 | 0 | 0 | 0 | 0 | 0 | 0 | 0 | 0 | 0 | 0 | 0 | 0 | 0 | 0 | 0 | 0 |
| ENSMUSG00 Gm14602 | 339 | 0 | 0 | 0 | 0 | 0 | 0 | 0 | 0 | 0 | 0 | 0 | 0 | 0 | 0 | 0 | 0 | 0 | 0 |
| ENSMUSG00 Gm5406 | 765 | 0.05556232 | 0.08709253 | 0 | 0.05955692 | 0.06291755 | 0 | 0 | 0.04992471 | 0 | 0.36980944 | 0 | 0.36480825 | 0 | 0 | 0.10771649 | 0.05005308 | 0.1502992 | 0.07190733 |
| ENSMUSG00 Gm13503 | 445 | 0 | 0.07486044 | 0.21740913 | 0 | 0.10816163 | 0 | 0 | 0.25747687 | 0 | 0 | 0 | 0 | 0 | 0 | 0 | 0.08604631 | 0 | 0 |
| ENSMUSG00 B230307C23 | 1295 | 3.38072056 | 3.54994548 | 2.76420173 | 2.84976416 | 3.08490322 | 3.67340728 | 3.1420003 | 2.09394639 | 4.19816222 | 3.93225942 | 3.5785104 | 4.26698888 | 3.27609644 | 3.37123105 | 3.05432383 | 4.28736531 | 3.16672878 | 6.41418928 |
| ENSMUSG00 Gm8019 | 586 | 0 | 0 | 0.08254869 | 0 | 0 | 0 | 0 | 0.06517475 | 0 | 0 | 0 | 0 | 0.16836832 | 0 | 0 | 0 | 0 | 0 |
| ENSMUSG00 Gm15532 | 883 | 0 | 0 | 0 | 0 | 0 | 0 | 0 | 0 | 0 | 0 | 0 | 0 | 0 | 0 | 0 | 0 | 0 | 0 |
| ENSMUSG00 Gm16077 | 749 | 0 | 0 | 0 | 0 | 0 | 0 | 0 | 0 | 0 | 0 | 0 | 0 | 0 | 0 | 0 | 0 | 0 | 0 |
| ENSMUSG00 Gm14944 | 131 | 0 | 0 | 0 | 0 | 0 | 0 | 0 | 0 | 0 | 0 | 0 | 0 | 0 | 0 | 0 | 0 | 0 | 0 |
| ENSMUSG00 Defa-ps7 | 172 | 0 | 0 | 0 | 0 | 0 | 0 | 0 | 0 | 0 | 0 | 0 | 0 | 0 | 0 | 0 | 0 | 0 | 0 |
| ENSMUSG00 Gm12564 | 320 | 0 | 0 | 0 | 0 | 0 | 0 | 0 | 0 | 0 | 0 | 0 | 0 | 0 | 0 | 0 | 0 | 0 | 0.85951729 |
| ENSMUSG00 Gm6121 | 1105 | 0 | 0 | 0 | 0 | 0 | 0 | 0 | 0 | 0 | 0 | 0 | 0 | 0 | 0 | 0 | 0 | 0 | 0 |
| ENSMUSG00 Gm15366 | 4236 | 0 | 0 | 0 | 0 | 0 | 0 | 0 | 0 | 0.00972297 | 0 | 0 | 0 | 0 | 0 | 0 | 0.00903933 | 0 | 0 |
| ENSMUSG00 C920021L13I | 2249 | 2.02225604 | 1.76266537 | 2.62408657 | 2.30945267 | 1.84052709 | 2.51178818 | 2.48239045 | 2.93787711 | 3.44289238 | 3.52215124 | 3.25014168 | 3.42488281 | 3.1367115 | 2.68462933 | 2.61975211 | 1.92389446 | 3.37421378 | 2.59269245 |
| ENSMUSG00 Gm13616 | 221 | 0 | 0 | 0 | 0 | 0 | 0 | 0 | 0 | 0 | 0 | 0 | 0 | 0 | 0 | 0 | 0 | 0 | 0 |
| ENSMUSG00 Gm8084 | 748 | 0 | 0 | 0 | 0 | 0 | 0 | 0 | 0 | 0 | 0 | 0 | 0 | 0 | 0 | 0 | 0 | 0 | 0 |
| ENSMUSG00 Gm11313 | 431 | 0 | 0 | 0 | 0 | 0 | 0 | 0 | 0 | 0 | 0 | 0 | 0 | 0 | 0 | 0 | 0 | 0 | 0 |
| ENSMUSG00 Rps2-ps9 | 816 | 1.09388321 | 0.44907088 | 1.89700119 | 0.8375192 | 1.59260042 | 0.72871667 | 1.97136162 | 1.73176334 | 1.81705217 | 1.73348174 | 0.87821615 | 0.68401547 | 1.2695714 | 0.56916774 | 1.51476308 | 1.54851721 | 2.11358255 | 2.02239363 |
| ENSMUSG00 Gm13222 | 137 | 0 | 0 | 0 | 0 | 0 | 0 | 0 | 0 | 0 | 0 | 0 | 0 | 0 | 0 | 0 | 0 | 0 | 0 |
| ENSMUSG00 Gm14867 | 861 | 0 | 0 | 0 | 0 | 0 | 0 | 0 | 0 | 0 | 0 | 0 | 0 | 0 | 0 | 0 | 0 | 0 | 0 |
| ENSMUSG00 Gm12006 | 402 | 0 | 0 | 0 | 0 | 0 | 0 | 0 | 0 | 0 | 0 | 0 | 0 | 0 | 0 | 0 | 0 | 0 | 0 |
| ENSMUSG00 Gm12075 | 396 | 0 | 0 | 0 | 0 | 0 | 0 | 0 | 0 | 0 | 0 | 0 | 0 | 0 | 0 | 0 | 0 | 0 | 0 |
| ENSMUSG00 Gm15617 | 770 | 0.27600764 | 0.086527 | 0.62822767 | 0.4733615 | 0 | 0.321771 | 0 | 0 | 0 | 0 | 0 | 0 | 0 | 0.060317 | 0 | 0 | 0 | 0 |
| ENSMUSG00 Gm6982 | 1559 | 0 | 0 | 0 | 0 | 0 | 0 | 0 | 0 | 0 | 0 | 0 | 0 | 0 | 0 | 0 | 0 | 0 | 0 |
| ENSMUSG00 Mup-ps22 | 242 | 0 | 0 | 0 | 0 | 0 | 0 | 0 | 0 | 0 | 0 | 0 | 0 | 0 | 0 | 0 | 0 | 0 | 0 |
| ENSMUSG00 Gm13795 | 294 | 0 | 0 | 0 | 0 | 0 | 0 | 0 | 0 | 0 | 0 | 0 | 0 | 0 | 0 | 0 | 0 | 0 | 0 |
| ENSMUSG00 Gm15013 | 800 | 0.05313147 | 0.33312894 | 0 | 0.22780522 | 0 | 0.24776367 | 0 | 0 | 0.41186516 | 0.05893838 | 0.0597187 | 0 | 0.0616649 | 0 | 0.10300389 | 0.14358978 | 0.19163148 | 0.06876138 |
| ENSMUSG00 Gm11398 | 2417 | 0.29896069 | 0.38591685 | 0.64044393 | 0.30160393 | 0.27879476 | 0.51254379 | 0.41107417 | 1.31153057 | 0.68161383 | 0.33163507 | 0.59298686 | 0.92371804 | 0.71436372 | 2.20979315 | 0.9205147 | 0.74458359 | 1.10998791 | 0.97864774 |
| ENSMUSG00 Gm14035 | 1474 | 0 | 0.02260034 | 0 | 0 | 0 | 0 | 0 | 0.02591072 | 0 | 0 | 0 | 0 | 0 | 0 | 0 | 0 | 0 | 0 |
| ENSMUSG00 Gm7150 | 972 | 0 | 0.06854505 | 0 | 0 | 0 | 0.05098018 | 0.14602679 | 0.27504817 | 0.04237296 | 0 | 0.14745358 | 0.11484704 | 0.20301201 | 0 | 0.08477686 | 0 | 0.07886069 | 0.05659373 |
| ENSMUSG00 Gm12710 | 215 | 0 | 0 | 0 | 0 | 0.22386941 | 0 | 0 | 0 | 0 | 0 | 0 | 0 | 0 | 0 | 0 | 0.17809585 | 0 | 0 |
| ENSMUSG00 Rpsa-ps12 | 1733 | 0 | 0.0768907 | 0.08373952 | 0.02629027 | 0 | 0.02859361 | 0.08190308 | 0.06611495 | 0.07129807 | 0 | 0.02756778 | 0 | 0.05693239 | 0 | 0.0475494 | 0.04418997 | 0 | 0.03174213 |
| ENSMUSG00 Gm14016 | 485 | 0 | 0 | 0 | 0 | 0 | 0 | 0 | 0 | 0 | 0 | 0 | 0 | 0 | 0 | 0.25485498 | 0 | 0 | 0 |
| ENSMUSG00 Gm15661 | 555 | 0 | 0 | 0 | 0 | 0 | 0 | 0 | 0 | 0 | 0 | 0 | 0 | 0 | 0 | 0 | 0 | 0 | 0 |
| ENSMUSG00 Gm12134 | 519 | 0 | 0 | 0 | 0 | 0 | 0 | 0 | 0 | 0 | 0 | 0 | 0 | 0 | 0 | 0 | 0 | 0 | 0 |
| ENSMUSG00 Gm15553 | 429 | 0 | 0 | 0 | 0 | 0 | 0 | 0 | 0 | 0 | 0 | 0 | 0 | 0 | 0 | 0 | 0 | 0 | 0 |
| ENSMUSG00 Olfr634-ps1 | 954 | 0 | 0 | 0 | 0 | 0 | 0 | 0 | 0 | 0 | 0 | 0 | 0 | 0 | 0 | 0 | 0 | 0 | 0 |
| ENSMUSG00 Olfr1069-ps1 | 927 | 0 | 0 | 0 | 0 | 0 | 0 | 0 | 0 | 0 | 0 | 0 | 0 | 0 | 0 | 0 | 0 | 0 | 0 |
| ENSMUSG00 Gm15613 | 325 | 0.52314063 | 0.41000485 | 0.89304979 | 0.56075131 | 0 | 0.76234975 | 0 | 0 | 0.12672774 | 0.29015817 | 0.73499936 | 0 | 0 | 0.14290488 | 1.77483625 | 0.94253803 | 0.47170827 | 0 |
| ENSMUSG00 Rpl36-ps7 | 314 | 0 | 0 | 0 | 0 | 0 | 0 | 0.15067732 | 0 | 0 | 0 | 0 | 0 | 0 | 0 | 0 | 0 | 0 | 0 |
| ENSMUSG00 Gm15062 | 366 | 0 | 0 | 0 | 0 | 0 | 0 | 0 | 0 | 0 | 0 | 0 | 0 | 0 | 0 | 0 | 0 | 0 | 0 |
| ENSMUSG00 Gm12633 | 1704 | 0 | 0 | 0 | 0 | 0 | 0 | 0 | 0 | 0 | 0 | 0 | 0 | 0 | 0 | 0 | 0 | 0 | 0 |
| ENSMUSG00 Gm15029 | 198 | 0 | 0 | 0 | 0 | 0 | 0 | 0 | 0 | 0 | 0 | 0 | 0 | 0 | 0 | 0 | 0 | 0 | 0 |
| ENSMUSG00 Gm5392 | 787 | 0 | 0 | 0 | 0 | 0 | 0 | 0 | 0 | 0 | 0 | 0 | 0 | 0 | 0 | 0 | 0 | 0 | 0 |
| ENSMUSG00 Gm15573 | 539 | 0 | 0.185415 | 0 | 0 | 0.08929856 | 0 | 0 | 0.21257367 | 0 | 0 | 0 | 0.1035541 | 0 | 0 | 0 | 0 | 0 | 0 |
| ENSMUSG00 Gm7571 | 923 | 0 | 0 | 0 | 0 | 0 | 0 | 0 | 0 | 0.08924489 | 0 | 0 | 0 | 0.05344736 | 0 | 0 | 0 | 0 | 0 |
| ENSMUSG00 Gm7386 | 631 | 0 | 0 | 0 | 0 | 0 | 0 | 0 | 0 | 0 | 0 | 0 | 0 | 0 | 0 | 0 | 0 | 0 | 0 |
| ENSMUSG00 Gm11879 | 701 | 0 | 0 | 0 | 0 | 0 | 0 | 0 | 0 | 0 | 0 | 0 | 0 | 0 | 0 | 0 | 0 | 0 | 0 |
| ENSMUSG00 Gm12341 | 502 | 0.08467167 | 0.13272069 | 0 | 0.27227716 | 0.19176065 | 0 | 0.09424836 | 0 | 0 | 0 | 0.09516924 | 0 | 0.09827075 | 0 | 0.16414963 | 0.07627611 | 0 | 0 |
| ENSMUSG00 Gm14606 | 1154 | 0 | 0 | 0 | 0 | 0 | 0 | 0 | 0 | 0 | 0 | 0 | 0 | 0 | 0 | 0 | 0 | 0 | 0 |
| ENSMUSG00 Gm8908 | 1023 | 0 | 0 | 0 | 0 | 0 | 0 | 0 | 0 | 0 | 0 | 0 | 0 | 0 | 0 | 0 | 0 | 0 | 0 |
| ENSMUSG00 Gm13593 | 690 | 0 | 0 | 0.07010657 | 0 | 0 | 0 | 0 | 0 | 0 | 0 | 0 | 0 | 0 | 0 | 0 | 0 | 0 | 0 |
| ENSMUSG00 Gm15595 | 960 | 0 | 0 | 0 | 0 | 0 | 0 | 0 | 0 | 0 | 0 | 0 | 0 | 0 | 0 | 0 | 0 | 0 | 0 |
| ENSMUSG00 Gm12219 | 442 | 0.28849667 | 0.30147416 | 0.98498139 | 0.41231714 | 0.54447877 | 0 | 0.85633808 | 0 | 0.46591081 | 0.4267032 | 0.64852885 | 0.50511912 | 0.11161067 | 0 | 0.27964857 | 0.259891 | 0.08671108 | 0.37336498 |
| ENSMUSG00 Gm11278 | 996 | 0 | 0 | 0 | 0 | 0 | 0 | 0 | 0 | 0 | 0 | 0 | 0 | 0 | 0 | 0 | 0 | 0 | 0 |
| ENSMUSG00 Gm11925 | 534 | 0 | 0 | 0 | 0 | 0 | 0 | 0 | 0 | 0 | 0 | 0 | 0 | 0 | 0 | 0 | 0 | 0 | 0 |
| ENSMUSG00 Gm12543 | 542 | 0 | 0.12292581 | 0.08925006 | 0.08406097 | 0 | 0 | 0.08729277 | 0 | 0.37994941 | 0.08699392 | 0.35258272 | 0.92682835 | 0.27305489 | 0.25707059 | 0.60814105 | 0.42388126 | 0.49498907 | 0 |
| ENSMUSG00 Gm12955 | 956 | 0 | 0.24392287 | 0 | 0.28594798 | 0.0503472 | 0 | 0 | 0 | 0.47390342 | 0 | 0 | 0.35030751 | 0 | 0 | 0 | 0 | 0 | 0 |
| ENSMUSG00 Gm9143 | 744 | 0 | 0 | 0 | 0.06123796 | 0 | 0 | 0.06359231 | 0.05133387 | 0 | 0.0633746 | 0 | 0 | 0.06630634 | 0 | 0 | 0 | 0 | 0.22181091 |
| ENSMUSG00 Gm6368 | 2146 | 0.63381437 | 1.73860399 | 1.73567653 | 0.33969092 | 0.89714676 | 0.62345005 | 0.55117287 | 1.24579132 | 1.36264801 | 0.17577149 | 0.69013221 | 0.80628404 | 0.32182984 | 1.62316242 | 1.15195403 | 0.35685561 | 0.58936058 | 0.92279955 |
| ENSMUSG00 Gm12174 | 812 | 0.10469255 | 0.08205146 | 0.05957331 | 0.11221932 | 0.1778273 | 0.24410214 | 0.40786792 | 0.04703498 | 0.05072231 | 0.23226947 | 0.05883616 | 0.137477 | 0.18226078 | 0.40038006 | 0.1522225 | 0.09431184 | 0.09439975 | 0.20323562 |
| ENSMUSG00 Gm15365 | 299 | 0 | 0 | 0 | 0 | 0 | 0 | 0 | 0 | 0.13774754 | 0 | 0 | 0 | 0 | 0 | 0 | 0 | 0 | 0 |
| ENSMUSG00 Gm7199 | 1484 | 0 | 0 | 0 | 0 | 0 | 0 | 0 | 0 | 0 | 0 | 0 | 0 | 0 | 0 | 0 | 0 | 0 | 0 |
| ENSMUSG00 Gm8731 | 411 | 0 | 0 | 0 | 0 | 0.1171093 | 0 | 0 | 0 | 0.400842 | 0.11472191 | 0 | 0 | 0 | 0 | 0 | 0 | 0 | 0.93689476 |
| ENSMUSG00 Gm11252 | 938 | 0.04531469 | 0 | 0 | 0 | 0 | 0 | 0 | 0 | 0 | 0.05026727 | 0.10186558 | 0.29752485 | 0 | 0 | 0 | 0 | 0 | 0 |
| ENSMUSG00 Olfr388-ps1 | 940 | 0 | 0 | 0 | 0 | 0 | 0 | 0 | 0 | 0 | 0 | 0 | 0 | 0 | 0 | 0 | 0 | 0 | 0 |
| ENSMUSG00 Gm13495 | 536 | 0 | 0.06215092 | 0 | 0 | 0 | 0 | 0.08826992 | 0 | 0 | 0 | 0 | 0.1041337 | 0 | 0 | 0 | 0 | 0 | 0.20525786 |
| ENSMUSG00 Gm8250 | 773 | 1.04475853 | 1.20667663 | 0.81352638 | 0.35364329 | 0.24906558 | 0.57693998 | 0.24482628 | 0.39526419 | 0.53281392 | 0.4269792 | 0.55624143 | 0.50544584 | 0.19145634 | 0.54074617 | 0.31980509 | 0.34674547 | 0.59497485 | 0.78279453 |
| ENSMUSG00 Gm13765 | 227 | 0 | 0 | 0 | 0 | 0 | 0 | 0 | 0 | 0 | 0 | 0 | 0 | 0 | 0 | 0 | 0 | 0 | 0 |
| ENSMUSG00 Gm11591 | 701 | 0 | 0 | 0 | 0 | 0 | 0 | 0 | 0 | 0 | 0 | 0 | 0 | 0 | 0 | 0 | 0 | 0 | 0 |
| ENSMUSG00 Gm11320 | 434 | 0 | 0 | 0 | 0 | 0 | 0 | 0 | 0 | 0 | 0 | 0 | 0 | 0 | 0 | 0 | 0 | 0 | 0 |
| ENSMUSG00 Gm7841 | 1628 | 0 | 0 | 0 | 0 | 0 | 0 | 0 | 0 | 0 | 0 | 0 | 0.06856961 | 0 | 0 | 0 | 0 | 0 | 0 |
| ENSMUSG00 Gm14007 | 209 | 0 | 0 | 0 | 0 | 0 | 0 | 0 | 0 | 0 | 0 | 0 | 0 | 0 | 0 | 0 | 0 | 0 | 0 |
| ENSMUSG00 Gm7638 | 966 | 0 | 0.0689708 | 0 | 0.04716464 | 0.04982601 | 0.05129683 | 0 | 0 | 0.08527229 | 0.0976205 | 0 | 0.05778019 | 0.05106824 | 0 | 0.12795514 | 0.07927662 | 0 | 0 |
| ENSMUSG00 Gm11442 | 457 | 0 | 0.14578947 | 0.31755053 | 0.09969594 | 0 | 0.21686098 | 0.41411535 | 0.25071599 | 0 | 0.41269762 | 0 | 0.73280957 | 0 | 0.2032564 | 0 | 0.08378689 | 0.33545993 | 0.24074007 |
| ENSMUSG00 Gm14367 | 1496 | 0 | 0 | 0 | 0 | 0 | 0 | 0 | 0 | 0 | 0 | 0 | 0 | 0 | 0 | 0 | 0 | 0 | 0 |
| ENSMUSG00 Cbx3-ps3 | 548 | 0 | 0 | 0 | 0 | 0 | 0 | 0 | 0 | 0 | 0 | 0 | 0 | 0 | 0 | 0 | 0 | 0 | 0 |
| ENSMUSG00 Gm11328 | 891 | 0 | 0 | 0 | 0 | 0 | 0 | 0 | 0 | 0 | 0 | 0 | 0 | 0 | 0 | 0 | 0 | 0 | 0 |
| ENSMUSG00 Gm6783 | 1436 | 0.0295997 | 0 | 0 | 0.03172775 | 0 | 0 | 0 | 0.05319276 | 0.02868142 | 0 | 0 | 0 | 0 | 0 | 0 | 0 | 0 | 0.03830718 |
| ENSMUSG00 Gm9159 | 565 | 0 | 0 | 0 | 0 | 0 | 0 | 0 | 0 | 0 | 0.08345257 | 0 | 0 | 0 | 0 | 0 | 0 | 0 | 0 |
| ENSMUSG00 Gm15760 | 282 | 0.90436545 | 1.77196246 | 1.54383608 | 0.16156399 | 0 | 0.35143783 | 0.83887729 | 0.2708681 | 2.77497801 | 0.6688043 | 0.33882949 | 0.79171153 | 0.17493588 | 0.49408604 | 0.58441923 | 0.40734689 | 0 | 0.39013551 |
| ENSMUSG00 Gm5135 | 572 | 0 | 0 | 0 | 0 | 0 | 0 | 0.49628684 | 0 | 0 | 0 | 0 | 0 | 0 | 0 | 0 | 0 | 0 | 0 |
| ENSMUSG00 Gm15166 | 384 | 0 | 0 | 0 | 0 | 0 | 0 | 0 | 0 | 0 | 0 | 0 | 0 | 0 | 0 | 0 | 0 | 0 | 0 |
| ENSMUSG00 Gm11582 | 235 | 0 | 0 | 0 | 0 | 0 | 0 | 0 | 0 | 0 | 0 | 0 | 0 | 0 | 0 | 0 | 0 | 0 | 0 |
| ENSMUSG00 Gm13310 | 1391 | 0 | 0 | 0 | 0 | 0 | 0 | 0 | 0 | 0 | 0 | 0 | 0 | 0 | 0 | 0 | 0.0275274 | 0 | 0 |
| ENSMUSG00 Gm13804 | 171 | 0 | 0 | 0 | 0 | 0 | 0 | 0 | 0 | 0 | 0 | 0 | 0 | 0 | 0 | 0 | 0 | 0 | 0 |
| ENSMUSG00 Gm12683 | 414 | 0 | 0 | 0 | 0 | 0 | 0 | 0 | 0 | 0 | 0 | 0 | 0 | 0 | 0 | 0 | 0 | 0 | 0 |
| ENSMUSG00 Gm11219 | 357 | 0 | 0 | 0 | 0 | 0 | 0 | 0 | 0 | 0 | 0 | 0 | 0 | 0 | 0 | 0 | 0 | 0 | 0 |
| ENSMUSG00 Gm13005 | 583 | 0 | 0 | 0 | 0 | 0.16511809 | 0 | 0 | 0 | 0 | 0 | 0 | 0 | 0 | 0 | 0 | 0 | 0 | 0 |
| ENSMUSG00 Gm13117 | 690 | 0 | 0.14483867 | 0 | 0 | 0 | 0 | 0.1371382 | 0 | 0 | 0 | 0.06923907 | 0 | 0 | 0 | 0 | 0 | 0 | 0 |
| ENSMUSG00 Gm12608 | 1712 | 0 | 0 | 0 | 0 | 0 | 0 | 0 | 0 | 0 | 0 | 0 | 0 | 0 | 0 | 0 | 0 | 0 | 0 |
| ENSMUSG00 Olfr1273-ps | 924 | 0 | 0 | 0 | 0 | 0 | 0 | 0 | 0 | 0 | 0 | 0 | 0 | 0 | 0 | 0 | 0 | 0 | 0 |
| ENSMUSG00 Gm11737 | 467 | 0.54610505 | 0.57067056 | 0 | 0.68292786 | 0.41226487 | 0.74276046 | 0.40524779 | 0.49069468 | 2.02845795 | 1.11061614 | 1.02301838 | 0.47807848 | 0.31690739 | 0.09945201 | 0.17645206 | 1.55786197 | 0.82069158 | 0.35337756 |
| ENSMUSG00 Gm14513 | 122 | 0 | 0 | 0 | 0.37345118 | 0 | 0 | 0.77561769 | 1.25220991 | 0 | 0 | 0 | 0 | 0 | 0 | 0 | 0 | 0 | 1.35268295 |
| ENSMUSG00 Gm13526 | 188 | 0 | 0 | 0 | 0 | 0 | 0 | 0 | 0 | 0 | 0 | 0 | 0 | 0 | 0 | 0 | 0 | 0 | 0 |
| ENSMUSG00 Gm14828 | 1154 | 0 | 0.02886733 | 0 | 0.03948097 | 0 | 0 | 0 | 0 | 0 | 0.0408585 | 0 | 0.04836713 | 0 | 0.04024618 | 0 | 0 | 0 | 0 |
| ENSMUSG00 Gm12314 | 537 | 0 | 0 | 0 | 0 | 0 | 0 | 0 | 0 | 0 | 0 | 0 | 0 | 0 | 0 | 0 | 0 | 0 | 0 |
| ENSMUSG00 Hmgb1-rs18 | 626 | 0.06789964 | 0 | 0 | 0 | 0 | 0 | 0 | 0 | 0 | 0 | 0 | 0.0891624 | 0 | 0.07419183 | 0 | 0 | 0 | 0 |
| ENSMUSG00 Gm12447 | 469 | 0 | 0.21308888 | 0 | 0 | 0 | 0 | 0.16286739 | 0.52690639 | 0.10053455 | 0.10186558 | 0 | 0.10518532 | 0.69319533 | 0 | 0 | 0 | 0.32687673 | 0.11729021 |
| ENSMUSG00 Gm13751 | 751 | 0 | 0 | 0 | 0 | 0 | 0 | 0 | 0 | 0 | 0 | 0 | 0 | 0 | 0 | 0 | 0 | 0 | 0 |
| ENSMUSG00 Gm9434 | 1543 | 0 | 0.02158969 | 0.03135031 | 0 | 0.03119373 | 0 | 0.03066279 | 0 | 0 | 0 | 0 | 0 | 0 | 0.03009986 | 0.05340448 | 0 | 0.04967764 | 0.0713015 |
| ENSMUSG00 Gm2005 | 535 | 0 | 0 | 0 | 0 | 0 | 0 | 0 | 0 | 0 | 0 | 0 | 0 | 0 | 0 | 0 | 0 | 0 | 0 |
| ENSMUSG00 Gm13197 | 224 | 0 | 0 | 0 | 0 | 0 | 0 | 0 | 0 | 0 | 0 | 0 | 0 | 0 | 0 | 0 | 0 | 0 | 0 |
| ENSMUSG00 Gm14694 | 279 | 0 | 0 | 0 | 0 | 0 | 0 | 0 | 0 | 0 | 0 | 0 | 0 | 0 | 0 | 0 | 0 | 0 | 0 |
| ENSMUSG00 Gm11477 | 313 | 0.13579928 | 0 | 0 | 0.14556244 | 0.15377611 | 0.31683089 | 0 | 0.12202045 | 0 | 0.6025649 | 0.6105426 | 0 | 0 | 0.39490309 | 0 | 0 | 0 | 0.17574794 |
| ENSMUSG00 Gm9001 | 398 | 2.02914157 | 0.83700739 | 1.70158147 | 1.6026498 | 0.72560689 | 1.74306098 | 3.09077802 | 2.49498106 | 3.10451124 | 2.48785118 | 1.20037584 | 4.90841254 | 1.48739449 | 2.21718005 | 3.20916641 | 2.79001913 | 2.88891685 | 2.21142138 |
| ENSMUSG00 Gm14892 | 265 | 0 | 0.25141807 | 0 | 0 | 0 | 0 | 0 | 0.14412227 | 0 | 0 | 0.21062514 | 0 | 0 | 0 | 0 | 0 | 0 | 0 |
| ENSMUSG00 Gm12019 | 531 | 0 | 0 | 0 | 0 | 0 | 0 | 0 | 0 | 0 | 0 | 0 | 0 | 0 | 0 | 0 | 0 | 0 | 0 |
| ENSMUSG00 Gm12086 | 227 | 0 | 0 | 0 | 0 | 0 | 0 | 0 | 0 | 0 | 0 | 0 | 0 | 0 | 0 | 0 | 0 | 0 | 0 |
| ENSMUSG00 Gm14669 | 249 | 0 | 0 | 0 | 0 | 0 | 0 | 0 | 0 | 0 | 0 | 0 | 0 | 0 | 0 | 0 | 0 | 0 | 0 |
| ENSMUSG00 Gm8545 | 800 | 0.15939441 | 0.04164112 | 0 | 0.05695131 | 0 | 0.06194092 | 0 | 0.0477405 | 0 | 0 | 0 | 0 | 0 | 0.11611022 | 0 | 0.04786326 | 0.09581574 | 0 |
| ENSMUSG00 Gm12671 | 2253 | 0.0565981 | 0.02957203 | 0.19323647 | 0.10111195 | 0.02136348 | 0.17595289 | 0.20999857 | 0.11866259 | 0.2924919 | 0.08371186 | 0.10602521 | 0.1734175 | 0.13137661 | 0.04122866 | 0.10972452 | 0.05098616 | 0.08505614 | 0.12207969 |
| ENSMUSG00 Gm12529 | 267 | 1.11436791 | 1.24767394 | 1.44939417 | 1.19448431 | 1.98296315 | 1.4847261 | 1.41760836 | 3.14693952 | 0.77128307 | 1.58934955 | 3.93651343 | 3.34475881 | 0.73905494 | 2.78316629 | 1.23450355 | 2.0077472 | 1.43544183 | 1.85423955 |
| ENSMUSG00 M6pr-ps | 3450 | 0.01232034 | 0.04827956 | 0 | 0.0264122 | 0.04185385 | 0.01436311 | 0.04114146 | 0.06642157 | 0.02387624 | 0.01366687 | 0.04154344 | 0 | 0.04289732 | 0.02692411 | 0.03582744 | 0.03329618 | 0.03332721 | 0.06377867 |
| ENSMUSG00 Gm11976 | 293 | 0 | 0 | 0 | 0 | 0 | 0 | 0 | 0 | 0 | 0 | 0 | 0 | 0 | 0 | 0 | 0 | 0 | 0 |
| ENSMUSG00 Gm15216 | 276 | 0 | 0 | 0 | 0 | 0 | 0 | 0 | 0 | 0 | 0 | 0 | 0 | 0 | 0 | 0 | 0 | 0 | 0 |
| ENSMUSG00 Gm12137 | 345 | 0 | 0 | 0 | 0 | 0 | 0 | 0 | 0 | 0 | 0 | 0 | 0 | 0 | 0 | 0 | 0 | 0 | 0 |
| ENSMUSG00 Gm6444 | 457 | 0 | 0.29157894 | 0.74095123 | 0.09969594 | 0.31596449 | 0.21686098 | 0.51764419 | 0.083572 | 0.09012367 | 0.30952322 | 0.10454039 | 0.24426986 | 0.32384191 | 0.4065128 | 0.27046973 | 0.50272132 | 0.08386498 | 0.60185018 |
| ENSMUSG00 Gm14361 | 867 | 0 | 0 | 0 | 0 | 0 | 0 | 0 | 0 | 0 | 0 | 0 | 0 | 0 | 0 | 0 | 0 | 0 | 0 |
| ENSMUSG00 Gm12573 | 285 | 0 | 0 | 0 | 0 | 0 | 0 | 0 | 0 | 0 | 0 | 0 | 0 | 0 | 0 | 0 | 0 | 0 | 0 |
| ENSMUSG00 Gm11625 | 295 | 0 | 0 | 0 | 0 | 0 | 0 | 0.48114589 | 0 | 0 | 0.15983289 | 0 | 0 | 0 | 0.15743758 | 0 | 0.12979867 | 0 | 0 |
| ENSMUSG00 Gm11349 | 1115 | 0.07624247 | 0.08963111 | 0.17353733 | 0 | 0 | 0 | 0.0424329 | 0.03425328 | 0.22163147 | 0.21143813 | 0.08569499 | 0.05005889 | 0.08848774 | 0.08330778 | 0.1476827 | 0.13736541 | 0 | 0 |
| ENSMUSG00 Gm14880 | 291 | 0 | 0 | 0 | 0 | 0 | 0 | 0 | 0 | 0 | 0 | 0 | 0 | 0 | 0 | 0 | 0 | 0 | 0 |
| ENSMUSG00 Gm15400 | 606 | 0 | 0 | 0 | 0 | 0 | 0 | 0 | 0 | 0 | 0 | 0 | 0 | 0 | 0 | 0 | 0 | 0 | 0 |
| ENSMUSG00 Gm14726 | 895 | 0 | 0 | 0 | 0 | 0 | 0 | 0 | 0 | 0 | 0 | 0 | 0 | 0 | 0 | 0 | 0 | 0 | 0 |
| ENSMUSG00 Gm14107 | 329 | 0.12919506 | 0 | 0 | 0 | 0 | 0 | 0 | 0 | 0.25037396 | 0 | 0 | 0 | 0 | 0 | 0 | 0 | 0 | 0.16720093 |
| ENSMUSG00 Gm9115 | 1347 | 0.15777719 | 0 | 0.03591205 | 0 | 0 | 0 | 0 | 0.02835368 | 0.15288239 | 0.03500423 | 0 | 0 | 0 | 0 | 0 | 0.02842658 | 0 | 0.04083824 |
| ENSMUSG00 Gm13379 | 620 | 0 | 0 | 0 | 0.14697111 | 0.15526427 | 0 | 0.38155386 | 0 | 0 | 0.22814856 | 0 | 0 | 0 | 0.14981964 | 0.26581649 | 0.06175904 | 0 | 0 |
| ENSMUSG00 Gm4773 | 701 | 0 | 0 | 0 | 0 | 0 | 0 | 0 | 0 | 0 | 0 | 0 | 0 | 0 | 0 | 0 | 0 | 0 | 0 |
| ENSMUSG00 Gm9385 | 671 | 1.58365037 | 1.63833906 | 2.09065929 | 1.56170495 | 1.07597445 | 0.66464173 | 1.83327817 | 1.19529128 | 1.78004316 | 1.12430887 | 0.85439568 | 1.24774208 | 1.7644799 | 1.86883809 | 1.71928996 | 1.0271698 | 1.14236354 | 1.88555805 |
| ENSMUSG00 Gm15702 | 443 | 0 | 0 | 0 | 0 | 0 | 0 | 0 | 0 | 0 | 0 | 0 | 0 | 0 | 0 | 0 | 0 | 0 | 0 |
| ENSMUSG00 Gm12439 | 364 | 0 | 0 | 0 | 0 | 0 | 0 | 0 | 0 | 0 | 0 | 0 | 0 | 0 | 0 | 0 | 0 | 0 | 0 |
| ENSMUSG00 Gm12094 | 471 | 0 | 0 | 0 | 0.09673258 | 0 | 0 | 0 | 0 | 0.69955865 | 0.2002153 | 0 | 0 | 0.10473868 | 0 | 0 | 0.08129641 | 0 | 0 |
| ENSMUSG00 Gm14918 | 267 | 0 | 0 | 0 | 0 | 0 | 0 | 0 | 0 | 0 | 0 | 0 | 0 | 0 | 0 | 0 | 0 | 0 | 0 |
| ENSMUSG00 Gm15256 | 882 | 0 | 0 | 0 | 0 | 0 | 0 | 0 | 0 | 0 | 0 | 0 | 0 | 0 | 0 | 0 | 0 | 0 | 0 |
| ENSMUSG00 Gm15084 | 1501 | 0 | 0 | 0 | 0 | 0 | 0 | 0 | 0 | 0 | 0 | 0 | 0 | 0 | 0 | 0 | 0 | 0 | 0 |
| ENSMUSG00 Rpl35-ps1 | 371 | 0 | 0 | 0 | 0 | 0 | 0 | 0 | 0 | 0 | 0 | 0 | 0 | 0 | 0 | 0 | 0 | 0 | 0 |
| ENSMUSG00 Gm14433 | 987 | 0 | 0 | 0 | 0 | 0 | 0 | 0 | 0 | 0 | 0 | 0 | 0 | 0 | 0 | 0 | 0 | 0 | 0 |
| ENSMUSG00 Gm12859 | 425 | 0 | 0 | 0 | 0 | 0 | 0 | 0 | 0 | 0.09690945 | 0 | 0 | 0 | 0 | 0 | 0 | 0 | 0 | 0 |
| ENSMUSG00 Gm14310 | 201 | 0 | 0 | 0 | 0 | 0 | 0 | 0 | 0 | 0 | 0 | 0 | 0 | 0 | 0 | 0 | 0 | 0 | 0 |
| ENSMUSG00 Rpl10-ps1 | 644 | 0 | 0 | 0.07511418 | 0 | 0.22421704 | 0 | 0.07346689 | 0 | 0.06395422 | 0 | 0 | 0 | 0.07660236 | 0.07211815 | 0.06397757 | 0 | 0.05951288 | 0.08541787 |
| ENSMUSG00 Gm13815 | 411 | 0 | 0 | 0 | 0 | 0.1171093 | 0 | 0 | 0 | 0.3006315 | 0 | 0 | 0.27160906 | 0 | 0 | 0 | 0 | 0.09325133 | 0 |
| ENSMUSG00 Gm11325 | 468 | 0 | 0 | 0 | 0 | 0 | 0 | 0 | 0 | 0 | 0 | 0 | 0 | 0 | 0 | 0 | 0 | 0 | 0 |
| ENSMUSG00 Gm14523 | 557 | 0.22893273 | 0 | 0 | 0 | 0 | 0 | 0.16988395 | 0.27427219 | 0 | 0 | 0 | 0 | 0 | 0 | 0 | 0.13748872 | 0.13761687 | 0 |
| ENSMUSG00 Gm14787 | 387 | 0 | 0 | 0 | 0 | 0 | 0 | 0 | 0 | 0 | 0 | 0 | 0 | 0 | 0 | 0 | 0 | 0 | 0 |
| ENSMUSG00 Rps12-ps6 | 398 | 0 | 0 | 0 | 0 | 0 | 0 | 0 | 0 | 0 | 0 | 0 | 0 | 0 | 0 | 0 | 0 | 0 | 0 |
| ENSMUSG00 Gm13637 | 341 | 0 | 0 | 0 | 0 | 0 | 0 | 0 | 0 | 0 | 0 | 0 | 0.16368229 | 0 | 0 | 0 | 0 | 0 | 0 |
| ENSMUSG00 Gm5687 | 639 | 0.06651827 | 0.46919569 | 0.07570193 | 0.14260108 | 0.15064765 | 0 | 0.296167 | 0 | 0.12890928 | 0 | 0.29906077 | 0 | 0 | 0 | 0.38686907 | 0.11984541 | 0 | 0.25825872 |
| ENSMUSG00 Gm14079 | 445 | 0 | 0 | 0 | 0 | 0 | 0 | 0 | 0 | 0 | 0 | 0 | 0 | 0 | 0 | 0 | 0 | 0 | 0 |
| ENSMUSG00 Gm15745 | 114 | 0 | 0 | 0 | 0 | 0 | 0 | 0.4150235 | 0 | 0 | 0 | 0 | 0 | 0 | 0 | 0.36141716 | 0 | 0 | 0 |
| ENSMUSG00 Gm11856 | 274 | 0 | 0 | 0 | 0 | 0 | 0 | 0 | 0 | 0.15031575 | 0 | 0 | 0 | 0 | 0 | 0 | 0 | 0 | 0 |
| ENSMUSG00 Gm14111 | 998 | 0.25554214 | 0.36717619 | 0.82399801 | 0.63913289 | 0.28937028 | 0.49652038 | 0.8059274 | 0.65057198 | 0.61903581 | 0.51969713 | 0.6223191 | 0.61520269 | 0.98861558 | 0.79113175 | 0.33027299 | 0.57551013 | 0.80646517 | 0.27559673 |
| ENSMUSG00 Gm12875 | 341 | 0 | 0 | 0 | 0 | 0 | 0 | 0 | 0 | 0 | 0 | 0 | 0 | 0 | 0 | 0 | 0 | 0 | 0 |
| ENSMUSG00 Gm11506 | 469 | 0 | 0 | 0 | 0 | 0 | 0 | 0 | 0 | 0 | 0 | 0 | 0 | 0 | 0 | 0 | 0 | 0 | 0 |
| ENSMUSG00 Rpl30-ps3 | 780 | 0 | 0 | 0 | 0 | 0.06170759 | 0 | 0 | 0 | 0 | 0 | 0 | 0 | 0 | 0 | 0 | 0 | 0 | 0 |
| ENSMUSG00 Gm15010 | 406 | 0 | 0 | 0 | 0 | 0 | 0 | 0 | 0.56441974 | 0 | 0 | 0 | 0 | 0 | 0 | 0 | 0 | 0 | 0 |
| ENSMUSG00 Gm7332 | 700 | 1.2144336 | 0.99938683 | 0.96747061 | 0.97630809 | 1.37519782 | 0.77868581 | 1.01384312 | 0.76384805 | 1.05908183 | 0.67358148 | 1.16024899 | 0.71762995 | 0.98663835 | 1.06157914 | 0.41201556 | 1.36752169 | 0.27375926 | 0.94301326 |
| ENSMUSG00 Gm14332 | 897 | 0 | 0 | 0 | 0 | 0 | 0 | 0 | 0 | 0 | 0 | 0 | 0 | 0 | 0 | 0 | 0 | 0 | 0 |
| ENSMUSG00 Rpl22-ps1 | 385 | 0 | 0.34610799 | 0 | 0 | 0.37505395 | 0 | 0 | 0.09920105 | 0.10697796 | 0.24493872 | 0.1240908 | 0 | 0.12813485 | 0.12063399 | 0.10701703 | 0 | 0 | 0.1428808 |
| ENSMUSG00 Gm15618 | 451 | 0 | 0 | 0 | 0 | 0 | 0 | 0 | 0 | 0 | 0 | 0 | 0 | 0 | 0 | 0 | 0 | 0 | 0 |
| ENSMUSG00 Gm14278 | 866 | 0 | 0 | 0 | 0 | 0 | 0 | 0 | 0 | 0 | 0 | 0 | 0 | 0 | 0 | 0 | 0 | 0 | 0 |
| ENSMUSG00 Gm12674 | 205 | 0 | 0 | 0 | 0 | 0 | 0 | 0 | 0 | 0 | 0 | 0 | 0 | 0 | 0 | 0 | 0 | 0 | 0 |
| ENSMUSG00 Gm4167 | 641 | 0.06631073 | 0.93546349 | 0.45279436 | 0.7818588 | 0.90106565 | 1.08227499 | 0.29524293 | 0.29791266 | 0.57828181 | 0.29423216 | 0.67078725 | 0.95783508 | 0.76960869 | 1.01437944 | 0.32138499 | 0.71682884 | 1.25561971 | 1.45889987 |
| ENSMUSG00 Gm14847 | 518 | 0 | 0 | 0 | 0 | 0 | 0 | 0 | 0 | 0 | 0 | 0 | 0 | 0 | 0 | 0 | 0 | 0 | 0 |
| ENSMUSG00 Gm14831 | 408 | 0 | 0 | 0 | 0 | 0 | 0 | 0 | 0 | 0 | 0 | 0 | 0 | 0 | 0 | 0 | 0 | 0 | 0 |
| ENSMUSG00 Gm14387 | 2182 | 0 | 0.04580141 | 0 | 0 | 0 | 0.02270978 | 0 | 0 | 0 | 0.04321788 | 0 | 0 | 0 | 0 | 0 | 0 | 0.01756476 | 0 |
| ENSMUSG00 Gm7974 | 404 | 0 | 0 | 0 | 0 | 0 | 0 | 0 | 0 | 0 | 0 | 0 | 0 | 0 | 0 | 0 | 0 | 0 | 0 |
| ENSMUSG00 Gm16458 | 497 | 0 | 0 | 0 | 0 | 0 | 0 | 0 | 0 | 0 | 0 | 0 | 0 | 0 | 0 | 0 | 0 | 0 | 0 |
| ENSMUSG00 Olfr587-ps1 | 1423 | 0 | 0 | 0 | 0 | 0 | 0 | 0 | 0 | 0 | 0 | 0 | 0 | 0 | 0 | 0 | 0 | 0 | 0 |
| ENSMUSG00 Gm9266 | 483 | 0 | 0 | 0 | 0 | 0 | 0 | 0 | 0 | 0 | 0 | 0 | 0 | 0 | 0 | 0 | 0 | 0 | 0.11389049 |
| ENSMUSG00 Gm15920 | 401 | 84.5863603 | 81.8284309 | 90.8360807 | 38.5166933 | 105.746196 | 53.0127748 | 96.1591854 | 102.005145 | 91.5141784 | 43.8583848 | 113.420849 | 47.8817654 | 91.0364558 | 97.0577188 | 107.576132 | 39.436461 | 102.840637 | 64.2001545 |
| ENSMUSG00 Defa-ps15 | 474 | 0 | 0 | 0 | 0 | 0 | 0 | 0 | 0 | 0 | 0 | 0 | 0 | 0 | 0 | 0 | 0 | 0 | 0 |
| ENSMUSG00 Gm11645 | 648 | 0 | 0 | 0 | 0.21093076 | 0.07427766 | 0.07647027 | 0 | 0 | 0 | 0 | 0 | 0 | 0 | 0 | 0 | 0 | 0 | 0 |
| ENSMUSG00 Gm14567 | 349 | 0.24358267 | 0.28635726 | 0.55442442 | 0.91383183 | 0.13791382 | 0.85190946 | 0.94896491 | 0.76603672 | 0.47205176 | 1.08081841 | 0.95823699 | 0.63972106 | 0.56540879 | 1.73000899 | 1.77084051 | 0.76800645 | 0.76872229 | 2.52190747 |
| ENSMUSG00 Gm14500 | 112 | 0 | 0 | 0 | 0 | 0 | 0 | 0 | 0 | 0 | 0 | 0 | 0 | 0 | 0 | 0 | 0 | 0 | 0 |
| ENSMUSG00 Gm14836 | 470 | 0 | 0 | 0 | 0 | 0 | 0 | 0 | 0 | 0 | 0 | 0 | 0 | 0 | 0 | 0 | 0 | 0 | 0 |
| ENSMUSG00 Gm9109 | 1404 | 0.03027434 | 0 | 0 | 0 | 0 | 0 | 0.03369849 | 0 | 0 | 0.03402775 | 0 | 0 | 0.38650363 | 0 | 0 | 0 | 0 | 0 |
| ENSMUSG00 Gm3657 | 631 | 0 | 0 | 0 | 0 | 0 | 0 | 0 | 0 | 0 | 0 | 0 | 0 | 0 | 0 | 0 | 0 | 0 | 0 |
| ENSMUSG00 Gm6215 | 1383 | 0 | 0 | 0 | 0 | 0 | 0 | 0 | 0 | 0 | 0 | 0 | 0 | 0 | 0 | 0 | 0 | 0 | 0 |
| ENSMUSG00 Gm11226 | 737 | 0.11534647 | 0.04520067 | 0.26254291 | 0.1854588 | 0.26123161 | 0.06723573 | 0.06419631 | 0.25910721 | 0.11176802 | 0.57578878 | 0.06482355 | 0.1514672 | 0.20080835 | 0 | 0.33542651 | 0.25977346 | 0.26001558 | 0.29855689 |
| ENSMUSG00 Gm16383 | 230 | 0 | 0 | 0 | 0 | 0 | 0 | 0 | 0 | 0 | 0 | 0 | 0 | 0 | 0 | 0 | 0 | 0 | 0 |
| ENSMUSG00 Gm11966 | 86 | 1.97698493 | 0.77471847 | 1.12496582 | 0.52977959 | 1.11934706 | 0.57619458 | 2.20058972 | 1.33229311 | 0.95782595 | 2.19305597 | 1.11104555 | 1.29803866 | 1.14725389 | 1.62014258 | 1.91635144 | 1.33571886 | 1.78261846 | 1.27928155 |
| ENSMUSG00 Gm13295 | 255 | 0 | 0 | 0 | 0 | 0 | 0 | 0 | 0 | 0 | 0 | 0 | 0 | 0 | 0 | 0 | 0 | 0 | 0 |
| ENSMUSG00 Gm12095 | 815 | 0 | 0 | 0 | 0 | 0 | 0 | 0 | 0 | 0 | 0 | 0 | 0 | 0 | 0 | 0 | 0 | 0 | 0 |
| ENSMUSG00 4930455H04 | 1338 | 0.22237387 | 0.09959012 | 0.14461444 | 0.30646442 | 0.57556859 | 0.14813971 | 0.31824672 | 0.37107715 | 0.21547505 | 0.3171572 | 0.24994373 | 0.45887316 | 0.29495915 | 0.03471158 | 0.09238017 | 0.25756014 | 0.20051127 | 0.49335522 |
| ENSMUSG00 Gm13044 | 252 | 0 | 0 | 0 | 0 | 0 | 0 | 0 | 0 | 0 | 0 | 0 | 0 | 0 | 0 | 0 | 0 | 0 | 0 |
| ENSMUSG00 Gm14821 | 434 | 0 | 0 | 0 | 0 | 0 | 0 | 0 | 0 | 0 | 0 | 0 | 0 | 0 | 0 | 0 | 0 | 0 | 0 |
| ENSMUSG00 Gm11918 | 178 | 0 | 0 | 0 | 0 | 0 | 0 | 0 | 0 | 0 | 0 | 0 | 0 | 0 | 0 | 0 | 0 | 0 | 0 |
| ENSMUSG00 Gm12269 | 716 | 0 | 0.09305278 | 0 | 0 | 0 | 0 | 0 | 0 | 0 | 0 | 0 | 0 | 0 | 0 | 0.05754407 | 0 | 0 | 0 |
| ENSMUSG00 Gm12719 | 240 | 0 | 0 | 0 | 0 | 0 | 0 | 0 | 0 | 0 | 0 | 0 | 0 | 0 | 0 | 0 | 0 | 0 | 0 |
| ENSMUSG00 Gm7129 | 996 | 0.04267588 | 0.06689336 | 0.0971356 | 0.04574402 | 0.1933009 | 0 | 0.42752421 | 0.30676628 | 0.12405577 | 0.18936025 | 0.14390048 | 0.05603982 | 0.09906007 | 0.13989183 | 0.24820214 | 0.19222192 | 0.15392087 | 0.11046005 |
| ENSMUSG00 Gm6446 | 817 | 0 | 0 | 0 | 0 | 0 | 0 | 0 | 0 | 0 | 0 | 0 | 0 | 0 | 0 | 0 | 0 | 0 | 0 |
| ENSMUSG00 Gm12463 | 848 | 0 | 0 | 0 | 0 | 0 | 0 | 0 | 0.04503821 | 0 | 0 | 0 | 0.13164071 | 0.05817443 | 0 | 0 | 0 | 0 | 0 |
| ENSMUSG00 Gm15925 | 1824 | 0 | 0 | 0 | 0 | 0 | 0 | 0 | 0 | 0 | 0 | 0 | 0 | 0 | 0 | 0 | 0 | 0 | 0 |
| ENSMUSG00 Gm12202 | 806 | 0 | 0 | 0 | 0 | 0 | 0 | 0.17610178 | 0 | 0 | 0 | 0 | 0 | 0 | 0 | 0 | 0 | 0 | 0 |
| ENSMUSG00 Gm11854 | 445 | 0 | 0 | 0 | 0 | 0 | 0 | 0 | 0 | 0 | 0 | 0 | 0 | 0 | 0 | 0 | 0 | 0 | 0 |
| ENSMUSG00 Gm6228 | 1383 | 0 | 0 | 0 | 0 | 0 | 0 | 0 | 0 | 0 | 0 | 0 | 0 | 0 | 0 | 0 | 0 | 0 | 0 |
| ENSMUSG00 Rpl38-ps2 | 502 | 0.67737332 | 0.59724312 | 0.57816969 | 0.36303621 | 1.15056391 | 0.49355312 | 0.47124182 | 0.68472435 | 0.73840367 | 0.56355422 | 0.28550772 | 0.22237316 | 0.78616601 | 1.20273533 | 1.14904738 | 0.07627611 | 0.53443043 | 1.09579894 |
| ENSMUSG00 Gm2214 | 329 | 0 | 0.101255 | 0 | 0 | 0 | 0 | 0 | 0 | 0 | 0.14331521 | 0 | 0 | 0 | 0 | 0 | 0 | 0 | 0.16720093 |
| ENSMUSG00 Gm5400 | 942 | 0 | 0 | 0 | 0 | 0 | 0 | 0.05022577 | 0.04054395 | 0 | 0 | 0 | 0 | 0 | 0 | 0 | 0 | 0 | 0 |
| ENSMUSG00 Olfr1078-ps1 | 630 | 0 | 0 | 0 | 0 | 0 | 0 | 0 | 0 | 0 | 0 | 0 | 0 | 0 | 0 | 0 | 0 | 0 | 0 |
| ENSMUSG00 Gm13682 | 701 | 0 | 0 | 0 | 0 | 0 | 0 | 0 | 0 | 0 | 0 | 0 | 0 | 0 | 0 | 0 | 0 | 0 | 0 |
| ENSMUSG00 Gm11430 | 215 | 0 | 0 | 0 | 0 | 0 | 0 | 0 | 0 | 0 | 0 | 0 | 0 | 0 | 0 | 0 | 0 | 0 | 0 |
| ENSMUSG00 Hmgb1-ps6 | 615 | 0.55291286 | 0.21666923 | 0.15731229 | 0.296332 | 0.62610632 | 0.1611471 | 0.15386237 | 0.1863044 | 0 | 0.07666781 | 0.07768286 | 0.27227152 | 0 | 0 | 0.0669944 | 0 | 0 | 1.78891404 |
| ENSMUSG00 Gm12891 | 239 | 0 | 0 | 0 | 0 | 0 | 0 | 0 | 0 | 0 | 0 | 0 | 0 | 0 | 0 | 0 | 0 | 0 | 0 |
| ENSMUSG00 Gm13367 | 801 | 0 | 0 | 0 | 0 | 0 | 0 | 0 | 0 | 0 | 0 | 0 | 0 | 0 | 0 | 0 | 0 | 0 | 0 |
| ENSMUSG00 Gm10224 | 1300 | 0.06539258 | 0.07687591 | 0.03721041 | 0.21028174 | 0.14809823 | 0.15246995 | 0.47312679 | 0.05875754 | 0.03168194 | 0.10880932 | 0.18374984 | 0.08587025 | 0.15179051 | 0.35726221 | 0.0316935 | 0.05890863 | 0 | 0 |
| ENSMUSG00 Rhox3g-ps | 641 | 0 | 0 | 0 | 0 | 0 | 0 | 0 | 0 | 0 | 0 | 0 | 0 | 0 | 0 | 0 | 0 | 0 | 0 |
| ENSMUSG00 Gm12931 | 323 | 0 | 0 | 0 | 0 | 0 | 0 | 0 | 0 | 0 | 0 | 0 | 0 | 0 | 0 | 0 | 0 | 0 | 0 |
| ENSMUSG00 Got2-ps1 | 1292 | 0.06579749 | 0.1031359 | 0.11232244 | 0.03526397 | 0.22352287 | 0.11506053 | 0.10985916 | 0.08868205 | 0.03187811 | 0.03649435 | 0.03697752 | 0.08640195 | 0.0381826 | 0.03594744 | 0.127559 | 0.14818347 | 0 | 0 |
| ENSMUSG00 Olfr337-ps1 | 452 | 0 | 0 | 0 | 0 | 0 | 0 | 0 | 0 | 0 | 0 | 0 | 0 | 0 | 0 | 0 | 0 | 0 | 0 |
| ENSMUSG00 Gm13510 | 428 | 0 | 0 | 0 | 0 | 0 | 0 | 0 | 0 | 0 | 0.55082597 | 0 | 0 | 0 | 0.43405689 | 0 | 0.08946404 | 0 | 0 |
| ENSMUSG00 Gm6062 | 630 | 0 | 0 | 0 | 0 | 0 | 0 | 0 | 0 | 0 | 0 | 0 | 0 | 0 | 0 | 0 | 0 | 0 | 0 |
| ENSMUSG00 Gm9208 | 947 | 0 | 0 | 0 | 0 | 0 | 0 | 0 | 0 | 0 | 0 | 0 | 0 | 0 | 0 | 0 | 0 | 0 | 0 |
| ENSMUSG00 Gm15145 | 308 | 0 | 0 | 0 | 0 | 0 | 0 | 0 | 0 | 0 | 0 | 0 | 0 | 0 | 0 | 0 | 0 | 0 | 0 |
| ENSMUSG00 Gm16056 | 134 | 0 | 0 | 0 | 0 | 0.35919346 | 0 | 0 | 0 | 0 | 0 | 0 | 0 | 0 | 0 | 0 | 0 | 0 | 0 |
| ENSMUSG00 Gm14556 | 898 | 0.04733316 | 0.11129029 | 0 | 0.05073613 | 0 | 0 | 0 | 0.04253051 | 0 | 0.05250635 | 0.05320151 | 0 | 0.2746766 | 0.15515842 | 0 | 0.1705595 | 0.04267962 | 0.06125736 |
| ENSMUSG00 Gm13130 | 1164 | 0 | 0 | 0 | 0 | 0 | 0 | 0 | 0 | 0 | 0.08101495 | 0 | 0 | 0 | 0.11970126 | 0 | 0 | 0 | 0 |
| ENSMUSG00 Gm12967 | 288 | 0 | 0 | 0 | 0.15819807 | 0.66849894 | 0 | 0.16428014 | 0 | 0 | 0.32743544 | 0 | 0.38760877 | 0.17129138 | 0 | 0 | 0.39886049 | 0 | 0 |
| ENSMUSG00 Gm12985 | 182 | 0 | 0 | 0 | 0 | 0.26446112 | 0 | 0 | 0 | 0 | 0.2590698 | 0 | 0 | 0 | 0 | 0 | 0 | 0 | 0.30224784 |
| ENSMUSG00 Gm12779 | 191 | 0 | 0 | 0 | 0 | 0 | 0 | 0 | 0 | 0 | 0 | 0 | 0 | 0 | 0 | 0 | 0 | 0 | 0 |
| ENSMUSG00 Gm14481 | 628 | 0 | 0 | 0 | 0 | 0 | 0 | 0 | 0 | 0 | 0 | 0 | 0 | 0 | 0 | 0 | 0 | 0 | 0 |
| ENSMUSG00 Gm13683 | 1074 | 0 | 0 | 0 | 0 | 0 | 0 | 0 | 0 | 0 | 0 | 0 | 0 | 0 | 0 | 0 | 0 | 0 | 0 |
| ENSMUSG00 Gm7278 | 1400 | 0 | 0.11897462 | 0.03455252 | 0.0325436 | 0 | 0 | 0.06758954 | 0.30008316 | 0 | 0.06735815 | 0.03412497 | 0.11960499 | 0.03523708 | 0.03317435 | 0.20600778 | 0 | 0 | 0 |
| ENSMUSG00 Gm12208 | 162 | 0.78713289 | 0.61690545 | 0 | 0.56248203 | 0 | 0.30588107 | 0 | 0 | 0 | 0.29105372 | 0 | 1.03362338 | 0.60903602 | 0 | 0.25433059 | 0.23636177 | 0.23658208 | 0 |
| ENSMUSG00 Gm5379 | 825 | 0 | 0 | 0 | 0 | 0 | 0 | 0 | 0 | 0 | 0 | 0 | 0 | 0 | 0 | 0 | 0 | 0 | 0 |
| ENSMUSG00 Gm15025 | 417 | 0 | 0 | 0 | 0 | 0 | 0 | 0 | 0 | 0 | 0 | 0 | 0 | 0 | 0 | 0 | 0 | 0 | 0 |
| ENSMUSG00 Gm14919 | 389 | 0 | 0 | 0 | 0 | 0 | 0 | 0 | 0 | 0 | 0 | 0 | 0 | 0 | 0 | 0 | 0 | 0 | 0 |
| ENSMUSG00 Gm14263 | 745 | 0 | 0 | 0 | 0 | 0 | 0.06651374 | 0.06350695 | 0 | 0 | 0 | 0 | 0 | 0 | 0 | 0 | 0 | 0 | 0 |
| ENSMUSG00 Gm16179 | 372 | 0.11426123 | 0 | 0 | 0 | 0 | 0 | 0.50873848 | 0 | 0 | 0 | 0.12842731 | 0.30008421 | 0 | 0.1248497 | 0 | 0 | 0 | 0 |
| ENSMUSG00 Gm15739 | 871 | 0.1464013 | 0.26772705 | 0.16661377 | 0.20923557 | 0 | 0.11378354 | 0.32591972 | 0 | 0 | 0.05413399 | 0.43880559 | 0.12816455 | 0 | 0 | 0.04730374 | 0.13188499 | 0.13200791 | 0 |
| ENSMUSG00 Gm11902 | 269 | 0 | 0 | 0 | 0 | 0 | 0 | 0 | 0 | 0.15310972 | 0 | 0 | 0 | 0 | 0 | 0.15316564 | 0 | 0 | 0 |
| ENSMUSG00 Gm2003 | 628 | 0 | 0 | 0 | 0 | 0 | 0 | 0 | 0 | 0 | 0 | 0 | 0 | 0 | 0 | 0 | 0 | 0 | 0 |
| ENSMUSG00 Olfr343-ps1 | 302 | 0 | 0 | 0 | 0 | 0 | 0 | 0 | 0 | 0 | 0 | 0 | 0 | 0 | 0 | 0 | 0 | 0 | 0 |
| ENSMUSG00 Gm11798 | 419 | 0 | 0 | 0 | 0 | 0 | 0 | 0 | 0 | 0 | 0 | 0 | 0 | 0 | 0 | 0 | 0 | 0 | 0 |
| ENSMUSG00 Gm11453 | 1006 | 0.16900666 | 0.09934263 | 0.09617004 | 0.18115723 | 0.19137942 | 0.04925719 | 0.28218298 | 0.18982307 | 0.16376348 | 0.3280864 | 0 | 0.38837936 | 0.34326384 | 0.23083542 | 0.36860239 | 0.2283734 | 0.07619542 | 0.10936204 |
| ENSMUSG00 Ifna-ps1 | 553 | 0 | 0 | 0 | 0 | 0 | 0 | 0 | 0 | 0 | 0 | 0 | 0 | 0 | 0 | 0 | 0 | 0 | 0 |
| ENSMUSG00 Gm6461 | 632 | 0 | 0 | 0 | 0 | 0.07615811 | 0 | 0 | 0 | 0 | 0 | 0 | 0 | 0 | 0 | 0 | 0 | 0 | 0 |
| ENSMUSG00 Gm6263 | 1341 | 0.15848313 | 0.07452549 | 0.07214546 | 0.37372967 | 0.14357024 | 0.18476038 | 0.10584492 | 0.19936377 | 0.03071329 | 0.21096512 | 0.03562637 | 0.33297934 | 0.25751187 | 0.03463392 | 0.122898 | 0.08566131 | 0.2000627 | 0 |
| ENSMUSG00 Gm12706 | 697 | 0 | 0 | 0 | 0 | 0 | 0 | 0 | 0 | 0 | 0.47353647 | 0 | 0 | 0 | 0 | 0 | 0 | 0 | 0 |
| ENSMUSG00 Gm1862 | 809 | 0.05254039 | 0.04117787 | 0 | 0.05631773 | 0.35697348 | 0 | 0.05848292 | 0 | 0.10182081 | 0 | 0 | 0.1379868 | 0 | 0 | 0 | 0 | 0.0473749 | 0.06799642 |
| ENSMUSG00 Gm12372 | 767 | 0 | 0.04343272 | 0 | 0 | 0 | 0 | 0 | 0 | 0.0536982 | 0 | 0 | 0 | 0 | 0 | 0 | 0 | 0.0499691 | 0 |
| ENSMUSG00 Gm12173 | 272 | 0 | 0 | 0 | 0.50251152 | 0 | 0.18217917 | 0 | 0 | 0 | 0 | 0 | 0.41040928 | 0.18136734 | 2.56125482 | 0 | 0 | 0 | 0.80895745 |
| ENSMUSG00 Gm16128 | 625 | 0.06800828 | 0 | 0.30959059 | 0 | 0 | 0 | 0.07570029 | 0 | 0 | 0 | 0 | 0.26791518 | 0 | 0 | 0 | 0 | 0 | 0 |
| ENSMUSG00 Gm16061 | 856 | 0.2482779 | 0.1167508 | 0 | 0.31935311 | 0.44983106 | 0.17366612 | 0.05527182 | 0.04461729 | 0.04811509 | 0.16524779 | 0.27905934 | 0 | 0.05763074 | 0 | 0.09626532 | 0.22366009 | 0.08954742 | 0.12852595 |
| ENSMUSG00 Gm9009 | 325 | 0 | 0.10250121 | 0.14884163 | 0 | 0.14809823 | 0 | 0 | 0.23503017 | 0 | 0 | 0.14699987 | 0 | 0.30358103 | 0.14290488 | 0 | 0 | 0.23585413 | 0 |
| ENSMUSG00 Gm11490 | 271 | 0.62738267 | 0 | 0 | 0 | 0 | 0 | 0 | 0 | 0 | 0 | 0 | 0 | 0 | 0 | 0 | 0 | 0 | 0 |
| ENSMUSG00 Gm13772 | 316 | 0 | 0 | 0 | 0 | 0 | 0 | 0.14972367 | 0.12086203 | 0 | 0.14921109 | 0 | 0.52989553 | 0 | 0 | 0 | 0 | 0 | 0 |
| ENSMUSG00 Gm12297 | 973 | 0 | 0 | 0 | 0 | 0 | 0 | 0 | 0 | 0 | 0 | 0 | 0 | 0 | 0 | 0.08468973 | 0 | 0 | 0 |
| ENSMUSG00 Gm13675 | 872 | 0 | 0 | 0 | 0 | 0.05519716 | 0 | 0 | 0 | 0.04723224 | 0 | 0.0547878 | 0 | 0 | 0.10652314 | 0 | 0 | 0.04395218 | 0.12616768 |
| ENSMUSG00 1700071K01f | 818 | 0 | 0 | 0 | 0.1113962 | 0 | 0 | 0 | 0.14006994 | 0 | 0 | 0 | 0 | 0 | 0 | 0 | 0 | 0 | 0 |
| ENSMUSG00 Gm12161 | 360 | 0 | 0 | 0 | 0 | 0 | 0 | 0.13142411 | 0 | 0 | 0 | 0 | 0.15504351 | 0 | 0 | 0 | 0 | 0 | 0 |
| ENSMUSG00 Gm13549 | 795 | 0.05346563 | 0 | 0 | 0.22923796 | 0 | 0 | 0 | 0 | 0 | 0 | 0.06009429 | 0 | 0 | 0 | 0 | 0 | 0 | 0 |
| ENSMUSG00 Gm5419 | 525 | 0 | 0 | 0 | 0 | 0 | 0 | 0 | 0 | 0 | 0 | 0 | 0 | 0 | 0 | 0 | 0 | 0 | 0 |
| ENSMUSG00 Gm14469 | 201 | 0 | 0 | 0 | 0 | 0 | 0 | 0 | 0 | 0 | 0 | 0 | 0 | 0 | 0 | 0 | 0 | 0 | 0 |
| ENSMUSG00 Gm13559 | 862 | 0.0986199 | 0 | 0 | 0 | 0 | 0 | 0 | 0 | 0 | 0 | 0 | 0 | 0 | 0 | 0 | 0 | 0 | 0 |
| ENSMUSG00 Olfr1081-ps1 | 944 | 0 | 0 | 0 | 0 | 0 | 0 | 0 | 0 | 0 | 0 | 0 | 0 | 0 | 0 | 0 | 0 | 0 | 0 |
| ENSMUSG00 Gm12073 | 523 | 0 | 0 | 0 | 0 | 0 | 0 | 0 | 0 | 0 | 0 | 0 | 0 | 0 | 0 | 0 | 0 | 0 | 0 |
| ENSMUSG00 Gm14048 | 1267 | 0 | 0 | 0 | 0 | 0 | 0 | 0 | 0 | 0 | 0 | 0 | 0 | 0 | 0 | 0 | 0 | 0 | 0 |
| ENSMUSG00 Olfr118 | 965 | 0 | 0 | 0 | 0 | 0 | 0 | 0 | 0 | 0 | 0 | 0 | 0 | 0 | 0 | 0 | 0 | 0 | 0 |
| ENSMUSG00 Psmb5-ps | 797 | 0.05333146 | 0 | 0 | 0 | 0.12078274 | 0 | 0 | 0 | 0 | 0.05916023 | 0 | 0.0700322 | 0 | 0 | 0 | 0 | 0 | 0 |
| ENSMUSG00 Gm12903 | 543 | 0 | 0 | 0 | 0 | 0 | 0 | 0 | 0 | 0 | 0 | 0 | 0 | 0 | 0 | 0 | 0 | 0 | 0 |
| ENSMUSG00 Gm13464 | 932 | 0.18242565 | 0.10723035 | 0.72664101 | 0.14665572 | 0.67136803 | 0.26584085 | 0.71070548 | 0.20489486 | 0.39772386 | 0.55649972 | 0.3588248 | 0.41921635 | 0.37051869 | 0.24916356 | 0.26524607 | 0.20542171 | 0.37010373 | 0.17706794 |
| ENSMUSG00 Gm15165 | 501 | 0 | 0 | 0 | 0 | 0 | 0 | 0 | 0 | 0 | 0 | 0 | 0 | 0 | 0 | 0 | 0 | 0 | 0 |
| ENSMUSG00 Gm4986 | 1123 | 0 | 0 | 0 | 0 | 0 | 0.08825064 | 0.04213061 | 0 | 0 | 0.04198638 | 0 | 0.04970228 | 0 | 0 | 0 | 0 | 0 | 0 |
| ENSMUSG00 Gm11885 | 1427 | 0 | 0 | 0 | 0 | 0 | 0 | 0 | 0 | 0 | 0 | 0 | 0 | 0 | 0 | 0 | 0 | 0 | 0 |
| ENSMUSG00 Gm5384 | 462 | 0 | 0 | 0 | 0 | 0 | 0 | 0 | 0.08266754 | 0 | 0 | 0.103409 | 0 | 0 | 0 | 0.08918086 | 0 | 0 | 0 |
| ENSMUSG00 Gm11336 | 333 | 0.12764317 | 0 | 1.16212686 | 0.13681995 | 0.14454031 | 0.14880701 | 0 | 0.5734595 | 0.49473292 | 0 | 0.28693669 | 0.50284381 | 0.59257558 | 0.69735867 | 0.49491358 | 0 | 0 | 0.66077005 |
| ENSMUSG00 Gm11200 | 447 | 0 | 0 | 0 | 0 | 0 | 0 | 0 | 0 | 0 | 0 | 0 | 0 | 0 | 0 | 0 | 0 | 0 | 0 |
| ENSMUSG00 Gm14678 | 481 | 0 | 0 | 0 | 0 | 0 | 0 | 0 | 0 | 0 | 0 | 0 | 0 | 0 | 0 | 0 | 0 | 0 | 0 |
| ENSMUSG00 Gm14240 | 545 | 0 | 0 | 0 | 0 | 0 | 0 | 0 | 0.14015561 | 0 | 0 | 0 | 0 | 0 | 0 | 0 | 0 | 0 | 0 |
| ENSMUSG00 Gm14301 | 323 | 0 | 0 | 0 | 0 | 0 | 0 | 0 | 0 | 0 | 0 | 0 | 0 | 0 | 0 | 0 | 0 | 0 | 0 |
| ENSMUSG00 Gm9157 | 224 | 0 | 0 | 0.86381304 | 0 | 0.21487466 | 0 | 1.47852122 | 0 | 0.36773675 | 0.21049421 | 0 | 0.7475312 | 0.44046355 | 0 | 1.1036131 | 0.34188042 | 1.02659724 | 0.49115274 |
| ENSMUSG00 Gm13198 | 825 | 0 | 0 | 0 | 0 | 0.05834173 | 0 | 0 | 0 | 0 | 0 | 0.2895452 | 0 | 0 | 0 | 0.14982384 | 0.13923857 | 0 | 0 |
| ENSMUSG00 Gm12832 | 312 | 0 | 0 | 0 | 0 | 0 | 0 | 0 | 0 | 0 | 0 | 0 | 0 | 0 | 0 | 0 | 0 | 0 | 0 |
| ENSMUSG00 Gm11543 | 367 | 0 | 0 | 0 | 0 | 0 | 0 | 0 | 0 | 0 | 0 | 0 | 0 | 0 | 0 | 0 | 0 | 0 | 0 |
| ENSMUSG00 Gm13314 | 318 | 0 | 0 | 0 | 0 | 0 | 0 | 0 | 0 | 0 | 0 | 0 | 0 | 0 | 0 | 0 | 0 | 0 | 0 |
| ENSMUSG00 Gm14065 | 584 | 0 | 0 | 0 | 0 | 0 | 0 | 0 | 0 | 0 | 0 | 0 | 0 | 0 | 0 | 0 | 0 | 0 | 0 |
| ENSMUSG00 Gm13880 | 949 | 0.13436831 | 0.14041262 | 0.05097316 | 0.28805718 | 0.05071857 | 0 | 0.2991318 | 0.16097957 | 0.21699956 | 0.14905386 | 0.05034242 | 0.2940762 | 0.20793221 | 0 | 0 | 0.12104512 | 0.28270187 | 0.23186136 |
| ENSMUSG00 Gm11913 | 334 | 0 | 0 | 0 | 0 | 0 | 0 | 0 | 0 | 0 | 0 | 0 | 0 | 0 | 0 | 0 | 0 | 0 | 0 |
| ENSMUSG00 Gm15224 | 387 | 0 | 0 | 0 | 0 | 0 | 0 | 0 | 0 | 0 | 0 | 0 | 0 | 0 | 0 | 0 | 0 | 0 | 0 |
| ENSMUSG00 Gm14497 | 579 | 0 | 0 | 0 | 0 | 0 | 0 | 0 | 0 | 0 | 0.08143472 | 0 | 0 | 0 | 0.08021431 | 0 | 0 | 0 | 0 |
| ENSMUSG00 Gm13714 | 220 | 0 | 0 | 0 | 0 | 0 | 0 | 0 | 0 | 0 | 0 | 0 | 0 | 0 | 0 | 0 | 0 | 0 | 0 |
| ENSMUSG00 Gm13805 | 103 | 0 | 0 | 0 | 0 | 0 | 0 | 0 | 0 | 0 | 0 | 0 | 0 | 0 | 0 | 0 | 0 | 0 | 0 |
| ENSMUSG00 Olfr1397-ps1 | 961 | 0 | 0 | 0 | 0 | 0 | 0 | 0 | 0.07948471 | 0 | 0 | 0 | 0 | 0 | 0 | 0 | 0 | 0 | 0 |
| ENSMUSG00 Gm15000 | 1242 | 0 | 0 | 0 | 0 | 0 | 0 | 0 | 0 | 0 | 0 | 0 | 0 | 0 | 0 | 0 | 0 | 0 | 0 |
| ENSMUSG00 Gm15012 | 105 | 0 | 0 | 0 | 0 | 0 | 0 | 0 | 0 | 0 | 0 | 0 | 0 | 0 | 0 | 0 | 0 | 0 | 0 |
| ENSMUSG00 Gm12567 | 285 | 0 | 0 | 0 | 0 | 0 | 0 | 0 | 0 | 0 | 0 | 0 | 0 | 0 | 0 | 0 | 0 | 0 | 0 |
| ENSMUSG00 Gm14509 | 1285 | 0 | 0 | 0 | 0 | 0.0749135 | 0 | 0 | 0 | 0 | 0 | 0 | 0 | 0 | 0 | 0 | 0 | 0 | 0 |
| ENSMUSG00 Gm11964 | 443 | 0.38379391 | 0.15039681 | 0.10919533 | 0.51423301 | 0.32594982 | 0.55928593 | 0.42720252 | 0.25863929 | 0.09297182 | 0.21286999 | 0.75490905 | 0.25198945 | 1.00222857 | 0.94355934 | 0.18601154 | 0.8643478 | 0.08651534 | 0 |
| ENSMUSG00 Gm15936 | 459 | 0 | 0 | 0 | 0 | 0 | 0 | 0 | 0 | 0 | 0 | 0 | 0 | 0 | 0 | 0 | 0 | 0 | 0 |
| ENSMUSG00 Gm15766 | 812 | 0 | 0 | 0 | 0 | 0 | 0 | 0 | 0.04703498 | 0 | 0 | 0 | 0 | 0 | 0 | 0 | 0 | 0 | 0 |
| ENSMUSG00 Gm14627 | 566 | 0 | 0 | 0 | 0 | 0 | 0 | 0 | 0 | 0.0727677 | 0 | 0 | 0 | 0 | 0 | 0 | 0 | 0 | 0 |
| ENSMUSG00 Gm11943 | 326 | 0 | 0 | 0 | 0 | 0 | 0 | 0 | 0 | 0 | 0 | 0 | 0 | 0 | 0 | 0 | 0 | 0 | 0 |
| ENSMUSG00 Gm11855 | 1867 | 0 | 0 | 0 | 0 | 0 | 0.02654137 | 0 | 0 | 0 | 0 | 0 | 0 | 0 | 0 | 0 | 0 | 0 | 0 |
| ENSMUSG00 Olfr1077-ps1 | 941 | 0 | 0 | 0 | 0 | 0 | 0 | 0 | 0 | 0 | 0 | 0 | 0 | 0 | 0 | 0 | 0 | 0 | 0 |
| ENSMUSG00 Gm15077 | 361 | 0 | 0 | 0 | 0 | 0 | 0 | 0 | 0 | 0.34227021 | 0 | 0 | 0 | 0 | 0 | 0 | 0 | 0 | 0 |
| ENSMUSG00 Gm12704 | 463 | 0 | 0 | 0 | 0 | 0.10395664 | 0 | 0.30656163 | 0.08248899 | 0 | 0.10183737 | 0 | 0.12055219 | 2.55716202 | 2.5077801 | 0.53392945 | 0 | 0.24833454 | 1.66334232 |
| ENSMUSG00 Gm15042 | 882 | 0 | 0 | 0 | 0 | 0 | 0 | 0 | 0 | 0 | 0 | 0 | 0 | 0 | 0 | 0 | 0 | 0 | 0 |
| ENSMUSG00 Gm14797 | 1479 | 0 | 0 | 0 | 0 | 0 | 0 | 0 | 0 | 0 | 0 | 0 | 0 | 0 | 0 | 0 | 0 | 0 | 0 |
| ENSMUSG00 Gm5944 | 1000 | 0.12751553 | 0.09993868 | 0.04837353 | 0.13668313 | 0.04813192 | 0.04955273 | 0.09462536 | 0.19096201 | 0.16474606 | 0.0471507 | 0 | 0 | 0.14799575 | 0.23222044 | 0.04120156 | 0.07658121 | 0.22995778 | 0 |
| ENSMUSG00 Olfr1300-ps1 | 933 | 0 | 0 | 0 | 0 | 0 | 0 | 0 | 0 | 0 | 0 | 0 | 0 | 0 | 0 | 0 | 0 | 0 | 0 |
| ENSMUSG00 Gm14628 | 634 | 0 | 0 | 0 | 0 | 0 | 0 | 0 | 0 | 0 | 0 | 0 | 0 | 0 | 0 | 0 | 0 | 0 | 0 |
| ENSMUSG00 Gm13929 | 2245 | 0 | 0 | 0 | 0 | 0 | 0 | 0 | 0 | 0 | 0 | 0 | 0 | 0 | 0 | 0 | 0 | 0 | 0 |
| ENSMUSG00 Olfr626-ps1 | 284 | 0 | 0 | 0 | 0 | 0 | 0 | 0 | 0 | 0 | 0 | 0 | 0 | 0 | 0 | 0 | 0 | 0 | 0 |
| ENSMUSG00 Gm14693 | 446 | 0 | 0 | 0 | 0 | 0 | 0 | 0 | 0 | 0 | 0 | 0 | 0 | 0 | 0 | 0 | 0 | 0 | 0 |
| ENSMUSG00 Nlk-ps1 | 1228 | 0 | 0 | 0 | 0 | 0 | 0 | 0 | 0 | 0 | 0 | 0 | 0 | 0 | 0 | 0 | 0 | 0 | 0 |
| ENSMUSG00 Psmb7-ps2 | 819 | 0 | 0 | 0 | 0 | 0 | 0 | 0 | 0 | 0 | 0.05757107 | 0.05833328 | 0 | 0 | 0 | 0 | 0.09350576 | 0 | 0 |
| ENSMUSG00 Gm11512 | 662 | 0.12841443 | 0.20128637 | 0.0730718 | 0 | 0 | 0.22455922 | 0 | 0 | 0.55993753 | 0.42734776 | 0 | 0.08431369 | 0.37259756 | 0.14031446 | 0.871332 | 0 | 0.23157883 | 0.08309533 |
| ENSMUSG00 AU015836 | 4188 | 0 | 0 | 0 | 0 | 0 | 0 | 0 | 0 | 0 | 0 | 0 | 0 | 0 | 0 | 0 | 0 | 0 | 0 |
| ENSMUSG00 Gm12797 | 272 | 0 | 0 | 0 | 0 | 0 | 0 | 0 | 0 | 0 | 0 | 0 | 0 | 0 | 0 | 0 | 0 | 0 | 0 |
| ENSMUSG00 Gm12846 | 1371 | 0.55805483 | 0.4859649 | 0.70566784 | 0.29908782 | 0.56171465 | 0.25300447 | 0.48313458 | 0.94714929 | 1.20164889 | 0.17195734 | 0.17423398 | 0.77352122 | 0.50375408 | 0.23713247 | 0.15026096 | 0.61443717 | 0.44727991 | 0.72222022 |
| ENSMUSG00 Olfr997-ps1 | 921 | 0 | 0 | 0 | 0 | 0 | 0 | 0 | 0 | 0 | 0 | 0 | 0 | 0 | 0 | 0 | 0 | 0 | 0 |
| ENSMUSG00 Gm11760 | 795 | 0.05346563 | 0 | 0 | 0 | 0 | 0 | 0 | 0 | 0 | 0 | 0 | 0 | 0 | 0 | 0 | 0 | 0 | 0 |
| ENSMUSG00 Rps24-ps3 | 401 | 2.1199589 | 1.41226734 | 1.08569021 | 0.68171139 | 1.20029735 | 0.74143741 | 2.00577442 | 1.61912928 | 1.23251419 | 1.29341081 | 1.42967457 | 1.25272061 | 1.72231133 | 1.62148934 | 0.71922916 | 1.05036579 | 1.0513448 | 1.92051744 |
| ENSMUSG00 Gm12461 | 1238 | 0.03433374 | 0.05381728 | 0.03907393 | 0.03680214 | 0.03887878 | 0.12007932 | 0.07643405 | 0.15425041 | 0.23288014 | 0.19043095 | 0.07718087 | 0.0901707 | 0 | 0.07503084 | 0.06656148 | 0.03092941 | 0.06191647 | 0.04463385 |
| ENSMUSG00 Gm15427 | 1193 | 0.85509155 | 1.22863985 | 1.58136436 | 1.14570941 | 1.45243022 | 0.87226103 | 0.99146435 | 1.3765912 | 0.96665754 | 1.30425248 | 0.76087528 | 0.70178955 | 1.73674814 | 1.75187253 | 1.13969098 | 0.89868985 | 0.83527554 | 0.92219793 |
| ENSMUSG00 Gm6305 | 1011 | 0 | 0.13180176 | 0 | 0 | 0.04760823 | 0 | 0.0467979 | 0 | 0 | 0.18655075 | 0 | 0 | 0 | 0.09187752 | 0.04075327 | 0.03787399 | 0 | 0.92498003 |
| ENSMUSG00 Mup-ps17 | 216 | 0 | 0 | 0 | 0 | 0 | 0 | 0 | 0 | 0 | 0 | 0 | 0 | 0 | 0 | 0 | 0 | 0 | 0 |
| ENSMUSG00 Gm12025 | 143 | 0 | 0 | 0 | 0.3186087 | 0 | 0 | 0 | 0 | 0 | 0 | 0 | 0 | 0 | 0 | 0 | 0 | 0 | 0.38467907 |
| ENSMUSG00 Gm6987 | 426 | 0 | 0 | 0 | 0 | 0 | 0 | 0 | 0 | 0 | 0 | 0 | 0 | 0 | 0 | 0 | 0 | 0.08996783 | 0 |
| ENSMUSG00 Gm13998 | 334 | 0 | 0 | 0.28966186 | 0 | 0 | 0 | 0 | 0 | 0 | 0.28233954 | 0.1430388 | 0 | 0 | 0 | 0 | 0 | 0 | 0.16469792 |
| ENSMUSG00 Olfr1017-ps1 | 686 | 0 | 0 | 0 | 0 | 0 | 0 | 0 | 0 | 0 | 0 | 0 | 0 | 0 | 0 | 0 | 0 | 0 | 0 |
| ENSMUSG00 Hist2h3c2 | 1010 | 0.252506 | 0.26386451 | 0.38315668 | 0.22554972 | 0.09531074 | 0.04906211 | 1.45217133 | 1.09661353 | 1.06024694 | 0.32678705 | 0.56762327 | 0.99473458 | 1.61183492 | 2.48314923 | 2.0804746 | 1.40272522 | 1.44197949 | 2.45090079 |
| ENSMUSG00 Gm11945 | 475 | 0 | 0.07013241 | 0.10183901 | 0 | 0 | 0 | 0.59763384 | 0 | 0.08670845 | 0.29779392 | 0.10057886 | 0 | 0.20771334 | 0.09777703 | 0.17348024 | 0 | 0.08068694 | 0.11580865 |
| ENSMUSG00 Gm11318 | 960 | 0 | 0 | 0 | 0 | 0 | 0 | 0 | 0 | 0 | 0 | 0 | 0 | 0 | 0 | 0 | 0 | 0 | 0 |
| ENSMUSG00 Gm14712 | 771 | 0 | 0 | 0 | 0 | 0 | 0 | 0 | 0 | 0 | 0 | 0 | 0 | 0 | 0 | 0 | 0 | 0 | 0 |
| ENSMUSG00 Gm13640 | 263 | 0 | 0 | 0 | 0 | 0 | 0 | 0 | 0 | 0.15660272 | 0 | 0 | 0.21222685 | 0 | 0 | 0 | 0 | 0 | 0 |
| ENSMUSG00 Gm11825 | 735 | 0 | 0 | 0 | 0 | 0 | 0 | 0 | 0 | 0 | 0 | 0 | 0 | 0 | 0 | 0 | 0 | 0 | 0 |
| ENSMUSG00 Gm6071 | 438 | 0.09704378 | 0 | 0 | 0 | 0 | 0 | 0.10801981 | 0 | 0.09403314 | 0 | 0 | 0 | 0 | 0 | 0 | 0 | 0.08750296 | 0 |
| ENSMUSG00 Gm13576 | 777 | 0 | 0 | 0 | 0 | 0 | 0 | 0 | 0 | 0 | 0 | 0 | 0 | 0 | 0 | 0 | 0 | 0 | 0 |

| | | | | | | | | | | | | | | | | | | | |
|---|---|---|---|---|---|---|---|---|---|---|---|---|---|---|---|---|---|---|---|
| ENSMUSG00 Gm12817 | 608 | 0 | 0 | 0 | 0 | 0 | 0 | 0 | 0 | 0 | 0 | 0 | 0 | 0 | 0 | 0 | 0 | 0 | 0 |
| ENSMUSG00 Gm12912 | 1380 | 0.12320341 | 0.26553756 | 0.2103197 | 0.16507625 | 0.10463462 | 0.21544667 | 0.23999185 | 0.30443219 | 0.14922651 | 0.30750459 | 0.34619535 | 0.08089226 | 0.17873883 | 0.20193081 | 0.2388496 | 0.1664809 | 0.22218143 | 0.39861672 |
| ENSMUSG00 Gm12804 | 998 | 0.12777107 | 0.06675931 | 0.14541141 | 0.04565235 | 0 | 0.04965204 | 0 | 0 | 0 | 0 | 0 | 0 | 0 | 0.13961149 | 0.04128412 | 0 | 0.0384031 | 0 |
| ENSMUSG00 Gm13928 | 1731 | 0 | 0 | 0 | 0 | 0 | 0 | 0 | 0 | 0 | 0 | 0 | 0 | 0 | 0 | 0 | 0 | 0 | 0 |
| ENSMUSG00 Gm13181 | 1068 | 0 | 0.03119185 | 0 | 0.04266015 | 0 | 0.04639769 | 0.08860052 | 0.07152135 | 0 | 0.0882972 | 0 | 0.31357114 | 0.04619093 | 0.17394789 | 0.03857824 | 0.03585263 | 0.03588605 | 0.10301331 |
| ENSMUSG00 Gm9836 | 215 | 0 | 0 | 0.22499316 | 0 | 0.22386941 | 0 | 0 | 0.35527816 | 0 | 0 | 0 | 0 | 0 | 0 | 0.38327029 | 0 | 0.89130923 | 0.51171262 |
| ENSMUSG00 Gm14977 | 810 | 0 | 0 | 0 | 0 | 0 | 0 | 0 | 0 | 0 | 0 | 0 | 0 | 0 | 0 | 0 | 0 | 0 | 0 |
| ENSMUSG00 Gm13758 | 741 | 0 | 0 | 0 | 0 | 0 | 0 | 0 | 0 | 0 | 0 | 0 | 0 | 0 | 0 | 0 | 0 | 0 | 0.22270893 |
| ENSMUSG00 Gm7698 | 398 | 0 | 0 | 0 | 0 | 0 | 0 | 0 | 0 | 0 | 0 | 0 | 0 | 0 | 0 | 0 | 0 | 0 | 0 |
| ENSMUSG00 Gm13160 | 1732 | 0 | 0 | 0 | 0 | 0 | 0 | 0 | 0 | 0 | 0 | 0 | 0 | 0 | 0 | 0 | 0 | 0 | 0 |
| ENSMUSG00 Rpsa-ps4 | 1917 | 0.02217276 | 0 | 0 | 0 | 0 | 0 | 0 | 0 | 0.02148488 | 0 | 0 | 0 | 0 | 0 | 0 | 0 | 0 | 0 |
| ENSMUSG00 Gm13899 | 1254 | 0 | 0.15939184 | 0 | 0 | 0.15353086 | 0 | 0 | 0.06091292 | 0 | 0 | 0.07619611 | 0 | 0 | 0 | 0 | 0 | 0 | 0 |
| ENSMUSG00 Gm12568 | 623 | 1.09162571 | 0.48124566 | 0.54352281 | 0.36565846 | 0.30903322 | 0.63631119 | 0.53160313 | 0.12260803 | 0.39665986 | 0.75683312 | 0.15337065 | 0.26877526 | 0.47510675 | 0 | 0.33067059 | 0.1229233 | 0.06151894 | 0 |
| ENSMUSG00 Gm13059 | 396 | 0 | 0 | 0 | 0 | 0 | 0 | 0 | 0 | 0 | 0 | 0 | 0 | 0 | 0 | 0 | 0 | 0 | 0 |
| ENSMUSG00 Gm11297 | 885 | 0 | 0 | 0 | 0 | 0 | 0 | 0 | 0 | 0 | 0 | 0 | 0 | 0 | 0 | 0 | 0 | 0 | 0 |
| ENSMUSG00 Gm14856 | 612 | 0 | 0 | 0 | 0 | 0 | 0 | 0 | 0 | 0 | 0 | 0 | 0 | 0 | 0 | 0 | 0 | 0 | 0 |
| ENSMUSG00 Gm14937 | 207 | 0 | 0 | 0 | 0 | 0 | 0 | 0 | 0 | 0 | 0 | 0 | 0 | 0 | 0 | 0 | 0 | 0 | 0 |
| ENSMUSG00 Gm6079 | 403 | 0 | 0 | 0 | 0 | 0 | 0 | 0 | 0 | 0 | 0 | 0 | 0 | 0 | 0 | 0 | 0 | 0 | 0 |
| ENSMUSG00 Gm5638 | 1257 | 0 | 0 | 0 | 0 | 0 | 0 | 0 | 0 | 0 | 0 | 0 | 0 | 0 | 0 | 0 | 0 | 0 | 0 |
| ENSMUSG00 Gm13463 | 232 | 0 | 0.14359006 | 0 | 0 | 0 | 0 | 0 | 0 | 0 | 0.20323579 | 0 | 0 | 0 | 0 | 0 | 0 | 0 | 0 |
| ENSMUSG00 Gm9506 | 319 | 0 | 0 | 0 | 0 | 0 | 0 | 0 | 0 | 0 | 0 | 0 | 0 | 0 | 0.2911855 | 0 | 0 | 0 | 0 |
| ENSMUSG00 Rps15a-ps7 | 391 | 0 | 0 | 0.12371747 | 0.11652441 | 0.12309955 | 0 | 0 | 0 | 0.10533636 | 0.12059003 | 0 | 0 | 0 | 0 | 0.10537482 | 0.09792994 | 0 | 0 |
| ENSMUSG00 Gm11679 | 778 | 0 | 0 | 0 | 0 | 0 | 0 | 0 | 0 | 0 | 0 | 0 | 0 | 0 | 0 | 0 | 0 | 0 | 0 |
| ENSMUSG00 Gm14624 | 190 | 0 | 0 | 0 | 0 | 0 | 0 | 0 | 0 | 0 | 0 | 0 | 0 | 0 | 0 | 0 | 0 | 0 | 0 |
| ENSMUSG00 Gm14128 | 1009 | 0 | 0 | 0 | 0 | 0 | 0 | 0 | 0 | 0 | 0 | 0 | 0 | 0 | 0 | 0 | 0 | 0 | 0 |
| ENSMUSG00 Gm12565 | 648 | 0.19678322 | 0.05140879 | 0.14930102 | 0.07031025 | 0.07427766 | 0 | 0.29205357 | 0.23575557 | 0.25423775 | 0.21829029 | 0.36863394 | 0.08613528 | 0.45677701 | 0 | 0.06358265 | 0.05909044 | 0 | 0 |
| ENSMUSG00 Rpl19-ps11 | 717 | 1.48204937 | 0.51107648 | 0.74213226 | 0.50835196 | 1.00694401 | 0.69111204 | 1.121779 | 0.47940254 | 0.91908543 | 0.72337202 | 0.46642219 | 0.46707667 | 1.23845817 | 1.0364092 | 0.80449342 | 0.48063524 | 0.80180537 | 0.61376967 |
| ENSMUSG00 Gm14946 | 203 | 0 | 0 | 0 | 0 | 0 | 0 | 0 | 0 | 0 | 0 | 0 | 0 | 0 | 0 | 0 | 0 | 0 | 0 |
| ENSMUSG00 Gm12691 | 455 | 0 | 0 | 0.2126309 | 0 | 0.10578445 | 0 | 0.83187128 | 0 | 0 | 0 | 0 | 0.36801536 | 0 | 0 | 0.09055287 | 0 | 0 | 0.72539481 |
| ENSMUSG00 Gm12135 | 465 | 0 | 0 | 0 | 0 | 0 | 0 | 0 | 0 | 0 | 0 | 0 | 0 | 0 | 0 | 0 | 0 | 0 | 0 |
| ENSMUSG00 Gm12220 | 171 | 0 | 0 | 0 | 0 | 0 | 0 | 0.830047 | 0 | 0 | 0 | 0 | 0 | 1.73094447 | 0 | 0 | 0.22392168 | 0 | 0 |
| ENSMUSG00 Gm5270 | 998 | 0.12777107 | 0.23365757 | 0 | 0.1826094 | 0.19291352 | 0.04965204 | 0.18962998 | 0.11480682 | 0.20634527 | 0.18898077 | 0.0957414 | 0.27963759 | 0.14829234 | 0.04653716 | 0.20642062 | 0.03836734 | 0.0384031 | 0.33071607 |
| ENSMUSG00 Gm14630 | 245 | 0 | 0.27194199 | 0.19744298 | 0 | 0 | 0 | 0.19311298 | 0.31177471 | 0.16810823 | 0 | 0 | 0 | 0 | 0.1895677 | 0 | 0.15628819 | 0.15643386 | 0 |
| ENSMUSG00 Gm12024 | 1252 | 0 | 0 | 0 | 0 | 0 | 0 | 0 | 0 | 0 | 0 | 0 | 0 | 0 | 0 | 0 | 0 | 0 | 0 |
| ENSMUSG00 Gm11717 | 332 | 0 | 0.20068009 | 0 | 0 | 0 | 0.14925522 | 0 | 0 | 0 | 0.14202019 | 0.14390048 | 0 | 0 | 0 | 0.49640429 | 0.46133262 | 0 | 0 |
| ENSMUSG00 Gm15989 | 497 | 0.08552349 | 0 | 0.77864838 | 0 | 0.38737967 | 0.19940738 | 0.38078615 | 0.3842294 | 0.33148101 | 0.47435315 | 0.09612668 | 0.11230516 | 0 | 0.18689774 | 0.16580103 | 0.07704348 | 0.07711529 | 0.33204692 |
| ENSMUSG00 Gm14087 | 139 | 0 | 0 | 0 | 0 | 0 | 0 | 0 | 0 | 0 | 0 | 0 | 0 | 0 | 0 | 0 | 0 | 0 | 0 |
| ENSMUSG00 Gm7155 | 1670 | 0.12726101 | 0 | 0 | 0 | 0 | 0 | 0 | 0 | 0 | 0 | 0 | 0 | 0 | 0 | 0 | 0 | 0 | 0 |
| ENSMUSG00 Gm12235 | 169 | 0.25150992 | 0 | 0 | 0.26959198 | 0 | 0 | 0 | 0 | 0 | 0 | 0 | 0 | 0.58380967 | 0 | 0 | 0 | 0 | 0.65099535 |
| ENSMUSG00 Gm4234 | 2005 | 0 | 0 | 0 | 0 | 0 | 0 | 0 | 0 | 0 | 0 | 0 | 0 | 0 | 0.02316413 | 0 | 0 | 0 | 0 |
| ENSMUSG00 Gm12745 | 213 | 0 | 0 | 0 | 0 | 0 | 0 | 0 | 0 | 0 | 0 | 0 | 0 | 0 | 0 | 0 | 0 | 0 | 0 |
| ENSMUSG00 Gm6913 | 2278 | 0.39183876 | 0.83355354 | 0.65828773 | 0.68001559 | 0.57048373 | 0.2392801 | 0.41538787 | 0.78799074 | 0.6870446 | 0.82793158 | 0.69208676 | 0.53904503 | 0.34649283 | 0.75435963 | 0.32556102 | 0.45383951 | 0.43743798 | 0.36221975 |
| ENSMUSG00 Gm12140 | 543 | 0 | 0 | 0 | 0 | 0 | 0 | 0 | 0 | 0 | 0 | 0 | 0 | 0 | 0 | 0 | 0 | 0 | 0 |
| ENSMUSG00 Gm7308 | 446 | 0.6671216 | 0.07469259 | 0.21692166 | 0.20430962 | 0.10791911 | 1.22215262 | 0.42432896 | 0.59943232 | 0.73877158 | 0.74003346 | 0.42847496 | 1.37661948 | 0.11060968 | 0.20826945 | 0.73904136 | 0.34341352 | 0.51560041 | 0.1233388 |
| ENSMUSG00 Gm15390 | 737 | 0 | 0 | 0 | 0 | 0 | 0 | 0 | 0 | 0 | 0 | 0 | 0 | 0 | 0 | 0 | 0 | 0 | 0 |
| ENSMUSG00 Gm15913 | 131 | 0 | 0 | 0 | 0 | 0 | 0 | 0 | 0 | 0 | 0 | 0 | 0 | 0 | 0 | 0 | 0 | 0 | 0 |
| ENSMUSG00 Gm13329 | 264 | 0 | 0 | 0 | 0 | 0 | 0 | 0 | 0 | 0 | 0 | 0 | 0 | 0 | 0 | 0 | 0 | 0 | 0 |
| ENSMUSG00 Gm11233 | 632 | 0 | 0 | 0 | 0 | 0 | 0 | 0 | 0 | 0 | 0 | 0 | 0 | 0 | 0 | 0 | 0 | 0 | 0 |
| ENSMUSG00 Gm6758 | 1054 | 0 | 0 | 0 | 0 | 0 | 0 | 0 | 0 | 0 | 0.04473501 | 0 | 0 | 0.1872179 | 0.0440646 | 0 | 0 | 0 | 0.0521908 |
| ENSMUSG00 Gm14601 | 235 | 0 | 0 | 0 | 0 | 0 | 0 | 0 | 0 | 0 | 0 | 0 | 0 | 0 | 0 | 0 | 0 | 0 | 0 |
| ENSMUSG00 Gm12458 | 494 | 0 | 0 | 0 | 0 | 0 | 0 | 0 | 0 | 0 | 0 | 0 | 0 | 0 | 0 | 0 | 0 | 0 | 0 |
| ENSMUSG00 Gm12791 | 654 | 0.51994099 | 0.81499435 | 0.36982821 | 0.27866082 | 0.22078864 | 0.60614965 | 0.795779 | 0.29199084 | 0.18892897 | 0.36047938 | 0.14610079 | 0.8534505 | 0 | 0.07101542 | 0.62999321 | 0.46838663 | 0.4688232 | 0.75700606 |
| ENSMUSG00 Gm13678 | 339 | 0 | 0 | 0 | 0 | 0 | 0 | 0 | 0 | 0 | 0 | 0 | 0 | 0 | 0 | 0 | 0 | 0 | 0 |
| ENSMUSG00 Gm11469 | 495 | 0 | 0 | 0 | 0 | 0 | 0 | 0 | 0 | 0 | 0 | 0 | 0 | 0 | 0 | 0 | 0 | 0 | 0 |
| ENSMUSG00 Gm14906 | 389 | 0 | 0 | 0 | 0 | 0 | 0 | 0 | 0 | 0 | 0 | 0 | 0 | 0 | 0 | 0 | 0 | 0 | 0 |
| ENSMUSG00 Rpl26-ps5 | 896 | 0 | 0 | 0 | 0 | 0 | 0 | 0 | 0 | 0 | 0 | 0 | 0 | 0 | 0 | 0 | 0 | 0 | 0 |
| ENSMUSG00 Gm15784 | 1461 | 0 | 0.02280143 | 0.03310988 | 0.09355451 | 0 | 0.033917 | 0.06476753 | 0 | 0.0845719 | 0.0645458 | 0 | 0.03820374 | 0.06753171 | 0 | 0 | 0 | 0.05246584 | 0.03765168 |
| ENSMUSG00 Gm12677 | 424 | 0 | 0 | 0 | 0 | 0 | 0 | 0 | 0 | 0 | 0 | 0 | 0 | 0 | 0 | 0 | 0 | 0 | 0 |
| [illegible] | [illegible] | [illegible] | [illegible] | [illegible] | [illegible] | [illegible] | [illegible] | [illegible] | [illegible] | [illegible] | [illegible] | [illegible] | [illegible] | [illegible] | [illegible] | [illegible] | [illegible] | [illegible] | [illegible] |

| | | | | | | | | | | | | | | | | | | | |
|---|---|---|---|---|---|---|---|---|---|---|---|---|---|---|---|---|---|---|---|
| ENSMUSG00 Gm16200 | 330 | 0.25760713 | 0 | 0 | 0 | 0.14585431 | 0 | 0 | 0.23146911 | 0 | 0 | 0 | 0.16913837 | 0.14949066 | 0 | 0.1248532 | 0.58016072 | 0.46456117 | 0 |
| ENSMUSG00 Mup-ps9 | 539 | 0 | 0 | 0 | 0 | 0 | 0 | 0 | 0 | 0 | 0 | 0 | 0 | 0 | 0 | 0 | 0 | 0 | 0 |
| ENSMUSG00 Gm13170 | 642 | 0.19862232 | 0 | 0 | 0 | 0 | 0 | 0 | 0 | 0 | 0 | 0 | 0.08694028 | 0 | 0 | 0 | 0 | 0 | 0 |
| ENSMUSG00 Gm13058 | 1432 | 0 | 0 | 0 | 0 | 0 | 0 | 0 | 0 | 0 | 0 | 0 | 0 | 0 | 0 | 0 | 0 | 0 | 0 |
| ENSMUSG00 Gm11658 | 1007 | 0.88640387 | 0.36389457 | 0.48037269 | 0.40719901 | 0.28678405 | 0.1968331 | 0.42285413 | 0.87231902 | 0.08180043 | 0.46822943 | 0.85397145 | 0.49884902 | 0.68584592 | 0.73793982 | 0.85921815 | 0.95061091 | 1.25597597 | 0.10925344 |
| ENSMUSG00 Gm13325 | 805 | 0 | 0 | 0 | 0 | 0 | 0 | 0 | 0 | 0 | 0 | 0 | 0 | 0 | 0 | 0 | 0 | 0 | 0 |
| ENSMUSG00 Gm7219 | 296 | 0.14359857 | 0.22508712 | 0 | 0 | 0 | 0 | 0.15984013 | 0.25805677 | 0 | 0.15929292 | 0 | 0 | 0 | 0.1569057 | 0 | 0 | 0 | 0.18584158 |
| ENSMUSG00 Gm12924 | 317 | 1.07268583 | 0.73561596 | 0.76298944 | 0.28745138 | 0 | 0.46895331 | 0 | 0.60240382 | 1.16933326 | 0.29748078 | 0.75354824 | 0.17607465 | 1.86745428 | 0.43953395 | 0.64986681 | 0.84553392 | 0.36270943 | 2.25589397 |
| ENSMUSG00 Gm14884 | 195 | 0 | 0.17083535 | 0 | 0 | 0 | 0 | 0 | 0 | 0 | 0 | 0 | 0 | 0 | 0 | 0 | 0 | 0 | 0 |
| ENSMUSG00 Gm12646 | 237 | 0 | 0 | 0 | 0 | 0 | 0 | 0 | 0 | 0 | 0 | 0 | 0 | 0 | 0 | 0 | 0 | 0 | 0 |
| ENSMUSG00 Gm11310 | 1105 | 0 | 0 | 0 | 0 | 0 | 0 | 0 | 0 | 0 | 0 | 0 | 0 | 0 | 0 | 0 | 0 | 0 | 0 |
| ENSMUSG00 Gm15781 | 322 | 0.39601096 | 0 | 0 | 0.28298785 | 0 | 0.30778095 | 0 | 0.11860995 | 0 | 0.43929227 | 0.14836944 | 0 | 0.45961414 | 0 | 0 | 0.23782986 | 0 | 0 |
| ENSMUSG00 Gm15111 | 876 | 0 | 0 | 0 | 0 | 0 | 0 | 0 | 0 | 0 | 0 | 0 | 0 | 0 | 0 | 0 | 0 | 0 | 0 |
| ENSMUSG00 Slc48a1 | 2874 | 97.2561021 | 112.758467 | 108.798365 | 97.7168675 | 107.953507 | 122.899055 | 95.7283326 | 111.786531 | 99.4839222 | 100.076301 | 117.991181 | 107.572357 | 97.1704884 | 98.3179637 | 102.330699 | 91.1966624 | 93.46889 | 124.966756 |
| ENSMUSG00 Gm15453 | 464 | 1.55730171 | 1.22051552 | 0.93827968 | 1.47287859 | 1.24479113 | 0.74756279 | 1.32557075 | 0.90542333 | 2.21910106 | 1.32103263 | 1.132596 | 0.72175426 | 0.95686909 | 1.10104518 | 0.79916811 | 1.48541149 | 1.486796 | 1.06698698 |
| ENSMUSG00 Gm15193 | 863 | 0 | 0 | 0 | 0 | 0 | 0 | 0 | 0 | 0 | 0 | 0 | 0 | 0 | 0 | 0 | 0 | 0 | 0 |
| ENSMUSG00 Gm15235 | 193 | 0 | 0.17260567 | 0 | 0 | 0 | 0 | 0.73543024 | 0 | 0 | 0 | 0 | 0 | 0 | 0 | 0 | 0 | 0 | 0 |
| ENSMUSG00 Gm13636 | 400 | 0 | 0 | 0 | 0 | 0 | 0 | 0 | 0 | 0 | 0 | 0 | 0 | 0 | 0 | 0 | 0 | 0 | 0 |
| ENSMUSG00 Gm11990 | 896 | 0 | 0 | 0 | 0 | 0 | 0 | 0 | 0 | 0 | 0 | 0 | 0 | 0 | 0 | 0 | 0 | 0 | 0 |
| ENSMUSG00 Gm12538 | 485 | 0 | 0 | 0 | 0 | 0 | 0 | 0.09755192 | 0.07874722 | 0 | 0 | 0.19701014 | 0 | 0 | 0.095761 | 0 | 0 | 0.07902329 | 0.11342084 |
| ENSMUSG00 Gm13940 | 300 | 0.42505176 | 1.77668769 | 1.12871571 | 0.3037403 | 0.80219873 | 2.97316401 | 0.78854465 | 0.76384805 | 1.9220374 | 0.94301407 | 3.18499723 | 0.74420883 | 0.98663835 | 0.6192545 | 0.82403112 | 0.63817679 | 1.27754323 | 0.55009107 |
| ENSMUSG00 Gm12512 | 216 | 0 | 0 | 0 | 0 | 0 | 0 | 0 | 0 | 0 | 0 | 0 | 0 | 0 | 0 | 0 | 0 | 0 | 0 |
| ENSMUSG00 Gm12733 | 766 | 0.05548979 | 0 | 0 | 0 | 0 | 0 | 0 | 0 | 0 | 0 | 0 | 0 | 0.12880396 | 0 | 0 | 0 | 0 | 0 |
| ENSMUSG00 Gm13077 | 483 | 0 | 0 | 0.20030447 | 0.09432928 | 0 | 0 | 0.09795586 | 0 | 0.08527229 | 0 | 0 | 0 | 0 | 0 | 0 | 0 | 0.07935051 | 0 |
| ENSMUSG00 Gm8346 | 1768 | 0 | 0 | 0 | 0 | 0 | 0 | 0 | 0 | 0 | 0 | 0 | 0 | 0 | 0 | 0 | 0 | 0 | 0 |
| ENSMUSG00 Gm14927 | 453 | 0 | 0 | 0 | 0 | 0 | 0 | 0.104443 | 0 | 0 | 0.20817088 | 0 | 0 | 0 | 0 | 0 | 0 | 0.08460551 | 0 |
| ENSMUSG00 Gm11553 | 239 | 0 | 0 | 0 | 0 | 0 | 0 | 0 | 0 | 0 | 0 | 0 | 0 | 0 | 0 | 0 | 0 | 0 | 0 |
| ENSMUSG00 Gm11692 | 900 | 0 | 0 | 0 | 0 | 0 | 0 | 0 | 0 | 0 | 0 | 0 | 0 | 0 | 0 | 0 | 0 | 0 | 0 |
| ENSMUSG00 Gm15877 | 963 | 0 | 0 | 0 | 0.04731157 | 0 | 0 | 0 | 0 | 0.17107587 | 0 | 0 | 0 | 0 | 0 | 0 | 0 | 0 | 0 |
| ENSMUSG00 Gm14654 | 446 | 0 | 0 | 0 | 0 | 0 | 0 | 0 | 0 | 0 | 0 | 0 | 0 | 0 | 0 | 0 | 0 | 0 | 0 |
| ENSMUSG00 Gm12328 | 196 | 31.0114294 | 25.1546344 | 19.9911018 | 28.1269712 | 28.2406695 | 24.7763668 | 28.4841638 | 23.3831035 | 30.8898868 | 27.1838238 | 27.0562265 | 23.6362245 | 33.726923 | 41.9418544 | 36.7871035 | 29.6947568 | 30.5046037 | 42.0988062 |
| ENSMUSG00 Gm12699 | 940 | 0.36174618 | 0.24807474 | 0.10292241 | 0 | 0.40963339 | 0.15814702 | 0.15099791 | 0.20315108 | 0.70104708 | 0.15048097 | 0.30494654 | 0.05937836 | 0.36736534 | 0.44467743 | 0.17532577 | 0.20367344 | 0.32618125 | 0.29260163 |
| ENSMUSG00 Gm7350 | 628 | 0 | 0 | 0 | 0 | 0 | 0 | 0 | 0 | 0 | 0 | 0 | 0 | 0 | 0 | 0 | 0 | 0 | 0 |
| ENSMUSG00 Gm6427 | 978 | 0 | 0 | 0 | 0 | 0 | 0 | 0 | 0 | 0 | 0 | 0 | 0 | 0 | 0 | 0 | 0 | 0 | 0 |
| ENSMUSG00 Gm15778 | 607 | 0 | 0 | 0 | 0 | 0 | 0 | 0 | 0 | 0 | 0 | 0 | 0 | 0 | 0.06787736 | 0 | 0 | 0 | 0 |
| ENSMUSG00 Gm5697 | 2483 | 0.27389562 | 0.241495 | 0.175337 | 0.11009515 | 0.27138418 | 0.199568 | 0.30487429 | 0.19996022 | 0.31516061 | 0.22787291 | 0.32709396 | 0.17963298 | 0.29801803 | 0.3366869 | 0.26549533 | 0.2313166 | 0.26240316 | 0.24369721 |
| ENSMUSG00 Gm8320 | 1761 | 0 | 0 | 0 | 0 | 0 | 0 | 0 | 0 | 0 | 0 | 0 | 0 | 0 | 0 | 0 | 0 | 0 | 0 |
| ENSMUSG00 Gm12411 | 482 | 0 | 0 | 0 | 0 | 0 | 0 | 0 | 0 | 0 | 0 | 0 | 0 | 0.10234838 | 0 | 0 | 0 | 0 | 0 |
| ENSMUSG00 Gm9218 | 2692 | 0 | 0 | 0 | 0 | 0 | 0 | 0 | 0 | 0 | 0 | 0 | 0 | 0 | 0 | 0 | 0 | 0 | 0 |
| ENSMUSG00 Gm12425 | 362 | 0 | 0 | 0 | 0 | 0 | 0 | 0 | 0 | 0 | 0 | 0 | 0 | 0 | 0 | 0 | 0 | 0 | 0 |
| ENSMUSG00 Gm11575 | 227 | 0.56174241 | 1.32077554 | 0.21309925 | 0.40141889 | 0.63610472 | 0.21829398 | 0 | 1.68248469 | 1.08863037 | 0.20771235 | 0.63138712 | 0 | 1.52124855 | 0.61379851 | 0.54451395 | 1.01208654 | 1.18186819 | 0.24233087 |
| [illegible] | | | | | | | | | | | | | | | | | | | |

| | | | | | | | | | | | | | | | | | | | |
|---|---|---|---|---|---|---|---|---|---|---|---|---|---|---|---|---|---|---|---|
| ENSMUSG00 Gm14115 | 481 | 0 | 0 | 0 | 0 | 0 | 0 | 0 | 0 | 0 | 0.09802641 | 0 | 0 | 0 | 0 | 0 | 0 | 0 | 0 |
| ENSMUSG00 Gm16397 | 493 | 0 | 0 | 0 | 0 | 0 | 0.10051264 | 0 | 0 | 0 | 0 | 0 | 0 | 0 | 0 | 0 | 0 | 0 | 0 |
| ENSMUSG00 Gm13600 | 742 | 0 | 0.04489608 | 0 | 0 | 0.06486782 | 0 | 0 | 0 | 0 | 0 | 0 | 0 | 0.06648506 | 0 | 0 | 0.15481378 | 0 | 0 |
| ENSMUSG00 Gm11960 | 904 | 0.47019 | 0.44220656 | 0.16053163 | 0.40319508 | 0.26621639 | 0.16444491 | 0.78505551 | 0.21124116 | 0.50116336 | 0.46942072 | 0.21139362 | 0.30871495 | 0.38199493 | 0.41100962 | 0.63807719 | 0.42356867 | 0.59354885 | 0.73020938 |
| ENSMUSG00 Gm12482 | 629 | 0.27030319 | 0.26480838 | 0.23071636 | 0.50703865 | 0.45912805 | 0.3909009 | 0.5265322 | 0.66791165 | 0.19643807 | 0.29984549 | 0.37976915 | 0.26621143 | 0.31371649 | 0.14767595 | 0.65503269 | 0.12175074 | 0.18279633 | 0.26236458 |
| ENSMUSG00 Gm14823 | 237 | 0 | 0 | 0 | 0 | 0 | 0 | 0 | 0 | 0 | 0 | 0 | 0 | 0 | 0 | 0 | 0 | 0 | 0 |
| ENSMUSG00 Flg | 3161 | 0 | 0 | 0 | 0 | 0 | 0 | 0 | 0 | 0 | 0 | 0 | 0 | 0 | 0 | 0 | 0 | 0 | 0 |
| ENSMUSG00 Gm14895 | 1360 | 0 | 0 | 0 | 0 | 0 | 0 | 0 | 0 | 0 | 0 | 0 | 0 | 0 | 0 | 0 | 0 | 0 | 0 |
| ENSMUSG00 Gm8076 | 649 | 0 | 0 | 0 | 0 | 0 | 0 | 0 | 0 | 0 | 0 | 0 | 0 | 0 | 0 | 0 | 0 | 0 | 0 |
| ENSMUSG00 Gm13702 | 824 | 0 | 0 | 0 | 0 | 0 | 0 | 0 | 0 | 0 | 0 | 0 | 0 | 0 | 0 | 0 | 0 | 0 | 0 |
| ENSMUSG00 Gm6168 | 343 | 0 | 0 | 0 | 0 | 0 | 0 | 0 | 0 | 0 | 0 | 0 | 0 | 0 | 0 | 0 | 0 | 0 | 0 |
| ENSMUSG00 Gm13249 | 344 | 0 | 0 | 0 | 0 | 0 | 0 | 0 | 0 | 0 | 0 | 0 | 0 | 0 | 0 | 0 | 0 | 0 | 0 |
| ENSMUSG00 Dnajb3 | 1016 | 1.71526793 | 2.6886391 | 2.85670455 | 2.69061285 | 2.08445339 | 2.09721215 | 3.44600221 | 4.47332272 | 3.24303274 | 4.26955187 | 5.12546307 | 3.95544066 | 4.3699533 | 5.5769475 | 5.92069602 | 5.35164001 | 4.82850985 | 3.84414033 |
| ENSMUSG00 Gng2-ps1 | 216 | 0 | 0 | 0 | 0 | 0 | 0 | 0 | 0 | 0 | 0 | 0 | 0 | 0 | 0 | 0 | 0 | 0 | 0 |
| ENSMUSG00 Gm14841 | 757 | 0 | 0 | 0 | 0 | 0 | 0 | 0 | 0 | 0 | 0 | 0 | 0 | 0 | 0 | 0 | 0 | 0 | 0 |
| ENSMUSG00 Gm12639 | 492 | 0 | 0 | 0 | 0 | 0 | 0 | 0 | 0 | 0 | 0 | 0 | 0 | 0 | 0 | 0 | 0 | 0 | 0 |
| ENSMUSG00 Gm15746 | 1026 | 0 | 0 | 0 | 0 | 0 | 0 | 0 | 0 | 0 | 0 | 0 | 0 | 0.04808179 | 0.04526714 | 0.16062985 | 0 | 0.07471013 | 0.96507205 |
| ENSMUSG00 Gm13300 | 2993 | 0 | 0 | 0 | 0 | 0 | 0 | 0 | 0 | 0 | 0 | 0 | 0 | 0 | 0 | 0 | 0 | 0 | 0 |
| ENSMUSG00 Gm13319 | 306 | 0 | 0 | 0 | 0 | 0 | 0 | 0 | 0 | 0 | 0 | 0 | 0 | 0 | 0 | 0 | 0 | 0 | 0 |
| ENSMUSG00 Gm13859 | 1420 | 0 | 0 | 0.03406587 | 0 | 0 | 0 | 0.19991273 | 0 | 0.02900459 | 0 | 0 | 0 | 0 | 0 | 0 | 0 | 0 | 0 |
| ENSMUSG00 Gm13408 | 797 | 0.15999439 | 0.6269679 | 0.60694517 | 0.34299406 | 0.24156549 | 0.31087035 | 0.35618077 | 0.47920204 | 0.72347706 | 0.17748069 | 0.4196044 | 0.0700322 | 0.37138206 | 0.05827364 | 0.20678322 | 0.72065133 | 0.19235281 | 0.13804042 |
| ENSMUSG00 Gm8475 | 525 | 0 | 0 | 0 | 0 | 0 | 0 | 0 | 0 | 0 | 0 | 0 | 0 | 0 | 0 | 0 | 0 | 0 | 0 |
| ENSMUSG00 Gm14517 | 336 | 0 | 0 | 0 | 0 | 0 | 0 | 0 | 0 | 0 | 0 | 0 | 0 | 0 | 0.13822645 | 0 | 0 | 0 | 0 |
| ENSMUSG00 Gm11719 | 350 | 1.33587696 | 0.1903594 | 0 | 0.13017441 | 0.13751978 | 0.14157924 | 0.54071633 | 0 | 0.11767576 | 0 | 0 | 0 | 1.55043169 | 0 | 0 | 0 | 0 | 0 |
| ENSMUSG00 Gm7153 | 454 | 1.12348483 | 0.51363493 | 0.74584738 | 0.10035472 | 0.63610472 | 0.65488194 | 0.52106475 | 0.75711811 | 0.72575358 | 0.4154247 | 1.05231186 | 1.10647789 | 0.43464244 | 1.73909578 | 0.63526628 | 0.92774599 | 0 | 0.60582717 |
| ENSMUSG00 Gm11886 | 302 | 0.28149123 | 0 | 0 | 0 | 0 | 0 | 0 | 0 | 0 | 0 | 0 | 0 | 0 | 0 | 0 | 0 | 0 | 0 |
| ENSMUSG00 Gm13245 | 259 | 0.16411265 | 0.51448485 | 0.18677039 | 0 | 0 | 0.57396989 | 0.36534887 | 1.03222709 | 0.1590213 | 0.72819619 | 0.3689186 | 1.07752244 | 0 | 0.3586416 | 0.79539683 | 0.73920092 | 0.14797798 | 0 |
| ENSMUSG00 Gm13461 | 1525 | 0.25085022 | 0.30582329 | 0.50752556 | 0.26888485 | 0.37874301 | 0.35742955 | 0.18614824 | 0.22539778 | 0.29708306 | 0.24734795 | 0.18796705 | 0.36600434 | 0.16174399 | 0.18273084 | 0.21613931 | 0.47706331 | 0.27645198 | 0.50500164 |
| ENSMUSG00 Gm12488 | 498 | 0 | 0 | 0 | 0 | 0 | 0 | 0 | 0 | 0 | 0 | 0 | 0 | 0 | 0 | 0 | 0 | 0 | 0 |
| ENSMUSG00 Gm15607 | 357 | 0 | 0 | 0 | 0 | 0 | 0 | 0 | 0 | 0 | 0 | 0 | 0 | 0 | 0 | 0 | 0 | 0 | 0 |
| ENSMUSG00 Gm6568 | 696 | 0 | 0.04786335 | 0 | 0 | 0.41493038 | 0 | 0 | 0.10974828 | 0 | 0 | 0 | 0.16038984 | 0 | 0 | 0 | 0.05501524 | 0.33039911 | 0 |
| ENSMUSG00 Gm16422 | 401 | 0 | 0 | 0 | 0 | 0 | 0 | 0 | 0 | 0 | 0 | 0 | 0 | 0 | 0 | 0 | 0 | 0 | 0 |
| ENSMUSG00 Gm13519 | 271 | 0 | 0 | 0 | 0 | 0 | 0 | 0 | 0 | 0 | 0 | 0 | 0 | 0 | 0 | 0 | 0 | 0 | 0 |
| ENSMUSG00 Gm7189 | 428 | 0 | 0 | 0 | 0.10645104 | 0 | 0 | 0 | 0 | 0.19246035 | 0.11016519 | 0 | 0 | 0 | 0.10851422 | 0 | 0 | 0.08954742 | 0 |
| ENSMUSG00 Gm13649 | 605 | 0 | 0 | 0 | 0 | 0 | 0 | 0 | 0.06312794 | 0 | 0.15587009 | 0 | 0.09225729 | 0 | 0 | 0 | 0 | 0 | 0 |
| ENSMUSG00 Gm15206 | 453 | 0 | 0 | 0 | 0 | 0 | 0 | 0 | 0.08430994 | 0 | 0 | 0 | 0 | 0 | 0 | 0 | 0 | 0 | 0 |
| ENSMUSG00 Gm12390 | 396 | 0 | 0 | 0 | 0 | 0 | 0 | 0 | 0 | 0.20801271 | 0 | 0 | 0 | 0 | 0 | 0 | 0 | 0 | 0 |
| ENSMUSG00 Gm12954 | 609 | 0 | 0 | 0 | 0 | 0 | 0 | 0 | 0 | 0 | 0 | 0 | 0 | 0 | 0 | 0 | 0 | 0 | 0 |
| ENSMUSG00 Olfr1171-ps1 | 841 | 0 | 0 | 0 | 0 | 0 | 0 | 0 | 0 | 0 | 0 | 0 | 0 | 0 | 0 | 0 | 0 | 0 | 0 |
| ENSMUSG00 Gm15043 | 1542 | 0 | 0 | 0 | 0 | 0 | 0 | 0 | 0 | 0 | 0 | 0.03098246 | 0 | 0 | 0 | 0 | 0 | 0 | 0 |
| ENSMUSG00 Gm5257 | 789 | 0.05387221 | 0 | 0 | 0 | 0 | 0 | 0 | 0.04840609 | 0.05220091 | 0 | 0.06055128 | 0 | 0.06252461 | 0 | 0 | 0.09706111 | 0 | 0 |
| [illegible] | | | | | | | | | | | | | | | | | | | |

| | | | | | | | | | | | | | | | | | | | |
|---|---|---|---|---|---|---|---|---|---|---|---|---|---|---|---|---|---|---|---|
| ENSMUSG00 Gm6517 | 520 | 2.61570314 | 1.60158145 | 1.30236428 | 0.96379132 | 3.14708732 | 3.23998643 | 1.54676066 | 3.23166482 | 0.79204838 | 0.36269772 | 0.55124952 | 1.93208063 | 1.61277422 | 1.6969955 | 1.66390899 | 2.72452399 | 2.1374281 | 0.95208069 |
| ENSMUSG00 Gm15004 | 629 | 0 | 0 | 0 | 0 | 0 | 0 | 0 | 0 | 0 | 0 | 0 | 0 | 0 | 0 | 0 | 0 | 0 | 0 |
| ENSMUSG00 Gm6973 | 720 | 0 | 0 | 0 | 0 | 0 | 0 | 0 | 0 | 0 | 0 | 0 | 0 | 0 | 0 | 0 | 0 | 0 | 0 |
| ENSMUSG00 Gm15148 | 155 | 0 | 0 | 0.31208729 | 0 | 0.31052854 | 0 | 0 | 0 | 0 | 0 | 0 | 0 | 0 | 0 | 0 | 0 | 0.24726643 | 0 |
| ENSMUSG00 Gm13292 | 1997 | 0.12770709 | 0.16681469 | 0.1211155 | 0.06844423 | 0.09640846 | 0.09925435 | 0.18953502 | 0.11474933 | 0.16499355 | 0.23610768 | 0.23923364 | 0.05589951 | 0.19762411 | 0.02325693 | 0.06189518 | 0.09587032 | 0.26868711 | 0.16527523 |
| ENSMUSG00 Mup-ps13 | 539 | 0 | 0 | 0 | 0 | 0 | 0 | 0 | 0 | 0 | 0 | 0 | 0 | 0 | 0 | 0 | 0 | 0 | 0 |
| ENSMUSG00 Gm14610 | 965 | 0 | 0 | 0 | 0 | 0 | 0 | 0 | 0 | 0 | 0 | 0 | 0 | 0 | 0 | 0 | 0 | 0 | 0 |
| ENSMUSG00 Olfr1327-ps1 | 866 | 0 | 0 | 0 | 0 | 0 | 0 | 0 | 0 | 0 | 0 | 0 | 0 | 0 | 0 | 0 | 0 | 0 | 0 |
| ENSMUSG00 Gm13742 | 456 | 0 | 0 | 0 | 0 | 0 | 0 | 0 | 0 | 0 | 0 | 0 | 0 | 0 | 0 | 0 | 0 | 0 | 0 |
| ENSMUSG00 Gm13171 | 390 | 0.10898763 | 0.51250606 | 0 | 0 | 0 | 0.25411658 | 0 | 0 | 0.10560645 | 0 | 0.61249947 | 0.14311708 | 0 | 0 | 0 | 0 | 0.39309022 | 0 |
| ENSMUSG00 Olfr1315-ps1 | 944 | 0 | 0 | 0 | 0 | 0 | 0 | 0 | 0 | 0 | 0 | 0 | 0 | 0 | 0 | 0 | 0 | 0 | 0 |
| ENSMUSG00 Gm7110 | 596 | 6.48988426 | 1.17377647 | 6.2496004 | 6.72713406 | 0.64606609 | 5.98623627 | 5.00117244 | 1.15346182 | 6.01212562 | 6.40806538 | 1.5230272 | 7.49203525 | 9.02211239 | 1.87023171 | 5.66867044 | 7.70951827 | 1.28611735 | 4.61485795 |
| ENSMUSG00 Gm11231 | 1705 | 0 | 0 | 0 | 0 | 0 | 0 | 0 | 0 | 0 | 0 | 0 | 0 | 0 | 0 | 0 | 0 | 0 | 0 |
| ENSMUSG00 Gm9025 | 1566 | 0 | 0.0212726 | 0 | 0.0290939 | 0 | 0 | 0 | 0 | 0 | 0 | 0 | 0 | 0 | 0 | 0 | 0 | 0 | 0 |
| ENSMUSG00 Gm5123 | 836 | 0 | 0 | 0 | 0 | 0 | 0 | 0 | 0 | 0 | 0 | 0 | 0 | 0 | 0 | 0 | 0 | 0 | 0 |
| ENSMUSG00 Gm13813 | 786 | 0 | 0 | 0 | 0 | 0 | 0 | 0 | 0 | 0 | 0 | 0 | 0 | 0 | 0 | 0 | 0 | 0 | 0 |
| ENSMUSG00 Gm11688 | 307 | 0 | 0 | 0.31513701 | 0.14840731 | 0 | 0 | 0.30822592 | 0 | 0.67079016 | 0 | 0.15561876 | 0 | 0.16069028 | 0.15128367 | 0.80524213 | 0 | 0.12484136 | 0 |
| ENSMUSG00 Gm15136 | 532 | 3.1958779 | 2.50472889 | 2.63690297 | 2.14102652 | 2.4427856 | 0.93144236 | 1.86760575 | 1.86654598 | 2.32254788 | 1.77258283 | 1.61644596 | 1.67866654 | 7.7892501 | 5.23805497 | 2.78807521 | 2.80701821 | 7.70848453 | 2.58501441 |
| ENSMUSG00 Gm15682 | 482 | 0 | 0.06911389 | 0 | 0 | 0 | 0.205613 | 0.49079542 | 0 | 0 | 0 | 0 | 0 | 0.4093935 | 0 | 0 | 0 | 0 | 0 |
| ENSMUSG00 Gm4914 | 458 | 0 | 0 | 0 | 0 | 0 | 0 | 0 | 0 | 0 | 0 | 0 | 0 | 0 | 0 | 0 | 0 | 0 | 0 |
| ENSMUSG00 Gm6664 | 463 | 0 | 0 | 0 | 0 | 0 | 0 | 0 | 0 | 0 | 0 | 0 | 0 | 0 | 0 | 0 | 0 | 0 | 0 |
| ENSMUSG00 Gm12686 | 396 | 0 | 0 | 0 | 0 | 0 | 0 | 0 | 0 | 0 | 0 | 0 | 0 | 0 | 0 | 0 | 0 | 0 | 0 |
| ENSMUSG00 Gm12604 | 377 | 0 | 0 | 0 | 0 | 0 | 0 | 0 | 0 | 0 | 0 | 0 | 0 | 0 | 0 | 0 | 0 | 0 | 0 |
| ENSMUSG00 Gm14709 | 303 | 0 | 0 | 0 | 0 | 0 | 0 | 0 | 0 | 0 | 0 | 0 | 0 | 0 | 0 | 0 | 0 | 0 | 0 |
| ENSMUSG00 Gm13863 | 600 | 0 | 0 | 0.24186765 | 0 | 0.40109936 | 0 | 0 | 0 | 0 | 0 | 0 | 0 | 0 | 0 | 0 | 0 | 0 | 0 |
| ENSMUSG00 Gm15364 | 584 | 0 | 0 | 0 | 0 | 0 | 0 | 0 | 0 | 0 | 0 | 0 | 0 | 0 | 0 | 0 | 0 | 0 | 0 |
| ENSMUSG00 Gm11676 | 489 | 0 | 0 | 0 | 0 | 0 | 0 | 0 | 0 | 0 | 0 | 0 | 0 | 0 | 0 | 0 | 0 | 0 | 0 |
| ENSMUSG00 Gm15244 | 129 | 0 | 0 | 0 | 0 | 0 | 0 | 0 | 0 | 0 | 0 | 0 | 0 | 0 | 0 | 0 | 0 | 0 | 0 |
| ENSMUSG00 Mageb7-ps | 1040 | 0 | 0.03203163 | 0.18605204 | 0 | 0 | 0.38117487 | 0.86436625 | 0.51412849 | 2.05932579 | 0.04533721 | 0 | 0.96604031 | 0.18973814 | 0 | 0 | 0 | 0 | 1.05786744 |
| ENSMUSG00 Gm11894 | 327 | 0 | 0 | 0 | 0 | 0 | 0 | 0 | 0 | 0 | 0 | 0 | 0 | 0 | 0 | 0 | 0 | 0 | 0 |
| ENSMUSG00 Gm15069 | 190 | 0 | 0 | 0 | 0 | 0 | 0 | 0 | 0 | 0 | 0 | 0 | 0 | 0 | 0 | 0 | 0 | 0 | 0 |
| ENSMUSG00 Gm11502 | 1210 | 0 | 0.02753132 | 0 | 0 | 0.03977845 | 0 | 0 | 0.03156397 | 0 | 0 | 0 | 0 | 0 | 0 | 0 | 0 | 0 | 0 |
| ENSMUSG00 Gm12183 | 1847 | 0.48327488 | 0.68537628 | 0.60237748 | 0.96203613 | 0.59936884 | 0.85852056 | 0.33300749 | 0.62034222 | 0.62437598 | 1.02113055 | 0.9311849 | 1.2692246 | 0.5876027 | 0.65378791 | 0.53537485 | 0.70486229 | 0.58101587 | 0.77435667 |
| ENSMUSG00 Gm13665 | 1063 | 0.47983266 | 0.31338565 | 0.45506614 | 0.42860813 | 0.58863124 | 0.69923895 | 0.53410362 | 0.25150218 | 0.5036921 | 0.17742504 | 0.22471758 | 0.05250768 | 0.0928164 | 0.56798978 | 0.19379848 | 0.57634028 | 0 | 0.2069957 |
| ENSMUSG00 Gm11598 | 356 | 5.49224185 | 5.42738164 | 6.79403516 | 4.73527708 | 6.21929351 | 5.84610902 | 3.85412272 | 5.36410146 | 5.2061607 | 5.03294024 | 5.90477015 | 4.86035264 | 7.89864968 | 9.39318621 | 4.05071476 | 6.8837047 | 5.05993246 | 4.94463881 |
| ENSMUSG00 Gm13776 | 1260 | 0.06746853 | 0.21151044 | 0 | 0 | 0 | 0 | 0 | 0 | 0.39225253 | 0 | 0 | 0.08859629 | 0.27406621 | 0 | 0.09809894 | 0.27350434 | 0 | 0 |
| ENSMUSG00 Gm9430 | 1526 | 0 | 0 | 0 | 0 | 0 | 0 | 0 | 0 | 0 | 0 | 0 | 0 | 0 | 0 | 0 | 0 | 0 | 0 |
| ENSMUSG00 Gm11988 | 1067 | 0 | 0 | 0 | 0 | 0 | 0.09288235 | 0 | 0 | 0 | 0 | 0 | 0 | 0 | 0 | 0 | 0 | 0 | 0 |
| ENSMUSG00 Gm13685 | 140 | 0.3036084 | 0 | 0 | 0 | 0 | 0 | 0 | 0 | 0 | 0 | 0 | 0 | 0 | 0 | 0 | 0 | 0 | 0 |
| ENSMUSG00 Gm8844 | 2613 | 0 | 0 | 0 | 0 | 0 | 0 | 0.05431995 | 0 | 0.12609725 | 0.01804466 | 0.1645521 | 0.02136076 | 0 | 0.26661359 | 0 | 0 | 0.02933509 | 0 |
| ENSMUSG00 Gm15147 | 482 | 0 | 0 | 0 | 0 | 0 | 0 | 0 | 0 | 0 | 0 | 0 | 0 | 0 | 0 | 0 | 0 | 0 | 0 |
| ENSMUSG00 Gm15765 | 909 | 0 | 0 | 0.10643241 | 0 | 0 | 0 | 0 | 0.3781426 | 0 | 0 | 0 | 0 | 0 | 0 | 0 | 0.12637164 | 0.12648943 | 0 |
| ENSMUSG00 Rab9b-ps1 | 605 | 0 | 0 | 0.07995625 | 0 | 0 | 0 | 0.07820278 | 0.06312794 | 0 | 0 | 0.15793375 | 0 | 0.08154036 | 0 | 0 | 0.06329026 | 0 | 0 |
| ENSMUSG00 Rps6-ps3 | 749 | 0.62424157 | 0.26685897 | 0.45208907 | 0.36497499 | 0.51409264 | 0.99237784 | 0.56851016 | 0.40792953 | 0.6048754 | 0.3147577 | 0.31892496 | 0.37260122 | 0.52690966 | 0.31004064 | 0.88014005 | 0.76683459 | 0.35818969 | 0.6609906 |
| ENSMUSG00 Gm14060 | 446 | 0 | 0 | 0 | 0 | 0 | 0 | 0 | 0 | 0 | 0 | 0 | 0 | 0 | 0 | 0 | 0 | 0 | 0 |
| ENSMUSG00 Gm6743 | 1065 | 0 | 0 | 0.04542116 | 0 | 0.04519429 | 0 | 0 | 0 | 0 | 0 | 0.08971823 | 0 | 0.04632105 | 0 | 0 | 0.03595362 | 0.07197427 | 0.10330349 |
| ENSMUSG00 Gm12980 | 335 | 0 | 0 | 0 | 0 | 0 | 0 | 0 | 0 | 0 | 0 | 0 | 0 | 0 | 0 | 0 | 0 | 0 | 0 |
| ENSMUSG00 Gm13079 | 3575 | 0 | 0 | 0 | 0 | 0 | 0 | 0 | 0 | 0 | 0 | 0 | 0 | 0 | 0 | 0 | 0 | 0 | 0 |
| ENSMUSG00 Defa-ps17 | 170 | 0 | 0 | 0 | 0 | 0 | 0 | 0 | 0 | 0 | 0 | 0 | 0 | 0 | 0 | 0 | 0 | 0 | 0 |
| ENSMUSG00 Gm13501 | 978 | 0 | 0.20437358 | 0 | 0.1397578 | 0.04921465 | 0.05066742 | 0 | 0 | 0.042113 | 0.09642271 | 0 | 0 | 0.05044163 | 0.04748884 | 0.08425676 | 0 | 0 | 0 |
| ENSMUSG00 Kif22-ps | 1983 | 0 | 0 | 0 | 0 | 0 | 0 | 0 | 0 | 0 | 0 | 0 | 0 | 0 | 0 | 0 | 0 | 0 | 0 |
| ENSMUSG00 Gm8213 | 440 | 0 | 0 | 0.10993984 | 0 | 0.21878147 | 0.4504794 | 0 | 0 | 0 | 0.10716069 | 0.10857945 | 0 | 0.22423599 | 0.21110949 | 0 | 0.43512054 | 0.26131566 | 0.1250207 |
| ENSMUSG00 Gm7206 | 379 | 0.11215086 | 0 | 0 | 0.12021384 | 0.25399432 | 0.130746 | 0 | 0 | 0.10867155 | 0 | 0 | 0.73635439 | 0.26032674 | 0.24508753 | 0 | 0.20206125 | 0.40449918 | 0 |
| ENSMUSG00 Gm12242 | 737 | 0 | 0 | 0 | 0 | 0 | 0 | 0 | 0 | 0 | 0 | 0 | 0 | 0 | 0 | 0 | 0 | 0 | 0 |
| ENSMUSG00 Gm11619 | 236 | 0 | 0 | 0 | 0 | 0 | 0 | 0 | 0 | 0 | 0 | 0 | 0 | 0 | 1.18078188 | 0 | 0 | 0 | 0 |
| ENSMUSG00 Gm4746 | 2405 | 0 | 0 | 0 | 0 | 0 | 0 | 0 | 0 | 0 | 0 | 0 | 0 | 0 | 0 | 0 | 0 | 0 | 0 |
| ENSMUSG00 Gm11687 | 885 | 0.09605689 | 0.11292507 | 0.10931871 | 0 | 0.10877271 | 0.05599179 | 0.05346065 | 0.08631051 | 0.23269218 | 0.05327763 | 0.053983 | 0 | 0 | 0.10495839 | 0.23277715 | 0 | 0.12991965 | 0 |
| ENSMUSG00 Gm12334 | 399 | 0.10652926 | 0.16698193 | 0.12123692 | 0.11418808 | 0 | 0 | 0.71146886 | 0.95720307 | 0.10322435 | 0 | 0 | 0 | 0.37091667 | 0 | 0.10326204 | 0.09596643 | 0.09605588 | 0 |
| ENSMUSG00 H2afb2-ps | 335 | 0 | 0 | 0 | 0 | 0 | 0 | 0 | 0 | 0 | 0 | 0 | 0 | 0 | 0 | 0 | 0 | 0 | 0 |
| ENSMUSG00 Gm13486 | 281 | 0 | 0 | 0 | 0 | 0 | 0 | 0 | 0 | 0 | 0 | 0 | 0 | 0 | 0 | 0 | 0 | 1.22753264 | 0 |
| ENSMUSG00 Gm16177 | 717 | 0.53353777 | 0.23230749 | 0 | 0.12708799 | 0.0671296 | 0.20733361 | 0.065987 | 0.15980085 | 0 | 0.13152218 | 0.06663174 | 0.15569222 | 0.41281939 | 0 | 0.11492763 | 0.16021175 | 0.05345369 | 0 |
| ENSMUSG00 Gm15693 | 636 | 0 | 0 | 0 | 0 | 0 | 0 | 0 | 0 | 0 | 0 | 0 | 0 | 0 | 0 | 0 | 0 | 0 | 0 |
| ENSMUSG00 Gm11919 | 720 | 0 | 0 | 0 | 0 | 0 | 0 | 0 | 0 | 0 | 0 | 0 | 0 | 0 | 0 | 0 | 0 | 0 | 0 |
| ENSMUSG00 Gm11628 | 170 | 1.50018268 | 0.5878746 | 0.28455018 | 0.80401843 | 0 | 0.87446 | 1.1132395 | 1.12330595 | 0.24227382 | 0.27735708 | 0.28102917 | 0.98498228 | 0 | 0 | 0.24236209 | 1.12619434 | 0.22544881 | 0 |
| ENSMUSG00 1700119I11R | 331 | 0 | 0 | 0 | 0 | 0 | 0 | 0 | 0 | 0 | 0 | 0 | 0 | 0 | 0 | 0 | 0 | 0 | 0 |
| ENSMUSG00 Gm14356 | 868 | 0 | 0 | 0 | 0 | 0 | 0 | 0 | 0 | 0 | 0 | 0 | 0 | 0 | 0 | 0 | 0 | 0 | 0 |
| ENSMUSG00 Gm11880 | 398 | 0 | 0 | 0 | 0 | 0 | 0 | 0 | 0 | 0 | 0 | 0 | 0 | 0 | 0 | 0 | 0 | 0 | 0 |
| ENSMUSG00 Gm5909 | 1280 | 0.03320717 | 0.15615419 | 0.15116728 | 0.03559457 | 0.1128092 | 0.03871307 | 0.22177818 | 0.08951344 | 0.12870786 | 0.11050946 | 0.14929675 | 0.04360599 | 0.03854056 | 0.07256889 | 0.19313229 | 0.08974361 | 0.02994242 | 0.12892759 |
| ENSMUSG00 Gm14871 | 193 | 0 | 0 | 0 | 0 | 0 | 0 | 0 | 0 | 0 | 0 | 0 | 0 | 0 | 0 | 0 | 0 | 0 | 0 |
| ENSMUSG00 Gm14214 | 775 | 0 | 0 | 0 | 0 | 0 | 0 | 0 | 0 | 0 | 0 | 0 | 0 | 0 | 0 | 0 | 0 | 0 | 0 |
| ENSMUSG00 Gm13687 | 281 | 0 | 0.11855122 | 0 | 0 | 0 | 0 | 0 | 0 | 0 | 0.33559219 | 0 | 0 | 0 | 0 | 0.14662475 | 0 | 0 | 0 |
| ENSMUSG00 Gm12570 | 915 | 0 | 0 | 0 | 0 | 0 | 0 | 0 | 0.04174033 | 0 | 0 | 0 | 0 | 0 | 0 | 0 | 0 | 0 | 0 |
| ENSMUSG00 Gm6798 | 359 | 0 | 0 | 0 | 0 | 0 | 0 | 0 | 0 | 0 | 0 | 0 | 0 | 0 | 0 | 0 | 0 | 0 | 0 |
| ENSMUSG00 Gm40 | 658 | 0 | 0 | 0 | 0 | 0 | 0.07530811 | 0 | 0 | 0 | 0.21497281 | 0 | 0 | 0.07497252 | 0.21175116 | 0 | 0.11638482 | 0.05824665 | 0 |
| ENSMUSG00 Gm12937 | 446 | 0 | 0 | 0 | 0 | 0 | 0 | 0 | 0 | 0 | 0.52859533 | 0 | 0.5005889 | 0 | 0 | 0 | 0 | 0.0859334 | 0.1233388 |
| ENSMUSG00 Gm13615 | 1610 | 0 | 0 | 0 | 0 | 0 | 0 | 0 | 0 | 0 | 0 | 0 | 0 | 0 | 0 | 0 | 0 | 0 | 0 |
| ENSMUSG00 Gm15195 | 262 | 0 | 0 | 0.1846318 | 0 | 0.18370963 | 0.18913257 | 0.18058274 | 0 | 0 | 0.35992903 | 0 | 0 | 0 | 0 | 0 | 0.43844207 | 0 | 0.20995842 |
| ENSMUSG00 Gm11835 | 662 | 0 | 0 | 0 | 0 | 0 | 0 | 0 | 0 | 0 | 0 | 0 | 0 | 0 | 0 | 0 | 0 | 0 | 0 |
| ENSMUSG00 Gm12636 | 623 | 0 | 0 | 0 | 0 | 0 | 0 | 0 | 0 | 0 | 0 | 0 | 0 | 0 | 0 | 0 | 0 | 0 | 0 |
| ENSMUSG00 Gm12229 | 467 | 0.36407003 | 1.14134113 | 1.03583577 | 0.19512225 | 0.82452974 | 0.10610864 | 0.70918362 | 1.88099626 | 0.70555059 | 0.80772083 | 1.43222574 | 0.83663734 | 0.63381478 | 0 | 0.52935618 | 1.31188377 | 0.41034579 | 0 |
| ENSMUSG00 Gm15003 | 445 | 0 | 0 | 0 | 0 | 0 | 0 | 0 | 0 | 0 | 0 | 0 | 0 | 0 | 0 | 0 | 0 | 0 | 0 |
| ENSMUSG00 Gm14816 | 293 | 0 | 0 | 0 | 0 | 0 | 0 | 0 | 0 | 0 | 0 | 0 | 0 | 0 | 0 | 0 | 0 | 0 | 0 |
| ENSMUSG00 Gm9378 | 470 | 0 | 0.85054198 | 0.61753443 | 0.29081518 | 0 | 0 | 0.10066527 | 0.08126043 | 0.78867796 | 0 | 0.2032977 | 0 | 0 | 0 | 0.17532577 | 0.08146938 | 0 | 0 |
| ENSMUSG00 Gm13591 | 605 | 0 | 0.16518791 | 0.07995625 | 0.22592253 | 0 | 0.24571603 | 0 | 0 | 0 | 0.07793505 | 0.63173499 | 0 | 0 | 0.61413669 | 0.27240698 | 0 | 0 | 0 |
| ENSMUSG00 Gm15673 | 937 | 0 | 0 | 0 | 0 | 0 | 0 | 0 | 0.16304121 | 0 | 0 | 0 | 0 | 0 | 0 | 0 | 0 | 0.0409032 | 0 |
| ENSMUSG00 Gm11793 | 980 | 0.08674526 | 0.03399275 | 0 | 0 | 0 | 0 | 0 | 0.03897184 | 0 | 0.04811296 | 0.14624987 | 0.11390952 | 0.05033869 | 0 | 0 | 0.03907205 | 0.07821693 | 0 |
| ENSMUSG00 Gm4915 | 1568 | 0 | 0 | 0 | 0 | 0 | 0 | 0 | 0 | 0 | 0 | 0 | 0 | 0 | 0 | 0 | 0 | 0 | 0 |
| ENSMUSG00 Gm13937 | 987 | 0.04306502 | 0 | 0.09802134 | 0.13848342 | 0.04876588 | 0.0502054 | 0 | 0 | 0 | 0.04777174 | 0.04840421 | 0.05655082 | 0.19992672 | 0 | 0.04174423 | 0.07758988 | 0 | 0.05573364 |
| ENSMUSG00 Gm15374 | 843 | 0 | 0 | 0 | 0 | 0 | 0 | 0 | 0 | 0 | 0 | 0 | 0 | 0 | 0 | 0 | 0 | 0 | 0 |
| ENSMUSG00 Gm11452 | 689 | 0 | 0 | 0 | 0 | 0 | 0 | 0 | 0 | 0 | 0 | 0 | 0 | 0.0715993 | 0 | 0 | 0 | 0 | 0 |
| ENSMUSG00 Gm12902 | 321 | 0.26482976 | 0.31133546 | 0 | 0 | 0.14994369 | 0.46310966 | 0.44217457 | 0.23795889 | 0.64153451 | 0.73443463 | 1.19065317 | 0.34776114 | 0 | 1.30217067 | 0.38506127 | 0.35785614 | 0.83577594 | 0.34273587 |
| ENSMUSG00 Gm13342 | 206 | 0.20633581 | 0 | 0 | 0.22117012 | 0 | 0 | 0.4593464 | 0.37080002 | 0.19993454 | 0.22888691 | 0.69575182 | 0 | 0 | 0.90182694 | 0.20000755 | 0.18587673 | 0.18604998 | 0.2670345 |
| ENSMUSG00 Gm5757 | 527 | 0 | 0 | 0 | 0 | 0 | 0 | 0 | 0 | 0 | 0 | 0 | 0 | 0 | 0 | 0 | 0 | 0 | 0 |
| ENSMUSG00 Gm6221 | 1383 | 0 | 0 | 0 | 0 | 0 | 0 | 0 | 0 | 0 | 0 | 0 | 0 | 0 | 0 | 0 | 0 | 0 | 0 |
| ENSMUSG00 Vamp7-ps | 654 | 0.71491886 | 0.71312006 | 0.59172514 | 1.3236389 | 0.58876971 | 1.06076188 | 1.08515319 | 0.58398169 | 0.56678691 | 0.64886289 | 1.02270553 | 0.59741535 | 0.4525864 | 0.63913882 | 0.88199049 | 0.46838663 | 1.34786671 | 0.75700606 |
| ENSMUSG00 Gm13349 | 410 | 0 | 0.08125096 | 0.35395266 | 0.222249 | 0 | 0.36258098 | 0.80777745 | 0.1863044 | 0 | 0 | 0 | 0 | 0.36096525 | 0.45311305 | 0 | 0.7471338 | 0.09347877 | 0 |
| ENSMUSG00 Gm8402 | 1769 | 0 | 0 | 0 | 0.05151051 | 0 | 0 | 0 | 0 | 0 | 0 | 0 | 0 | 0 | 0 | 0 | 0 | 0 | 0 |
| ENSMUSG00 Gm12667 | 1331 | 0 | 0 | 0 | 0 | 0 | 0 | 0 | 0 | 0 | 0 | 0 | 0 | 0 | 0 | 0 | 0 | 0 | 0 |
| ENSMUSG00 Gm6272 | 170 | 0.25003045 | 0.1959582 | 0 | 0 | 0 | 0 | 0 | 0 | 0 | 0 | 0.28102917 | 0 | 0 | 0.27320051 | 0 | 0 | 0 | 0 |
| ENSMUSG00 Gm12627 | 381 | 0 | 0 | 0 | 0 | 0 | 0 | 0 | 0 | 0 | 0 | 0 | 0 | 0 | 0 | 0 | 0 | 0 | 0 |
| ENSMUSG00 Gm14067 | 313 | 0 | 0 | 0 | 0 | 0 | 0 | 0 | 0 | 0 | 0 | 0 | 0 | 0 | 0 | 0 | 0 | 0 | 0 |
| ENSMUSG00 Gm8200 | 221 | 0 | 0 | 0 | 0 | 0 | 0 | 0 | 0 | 0 | 0 | 0 | 0 | 0 | 0 | 0 | 0 | 0 | 0 |
| ENSMUSG00 Gm15182 | 1675 | 0 | 0.0198883 | 0 | 0 | 0.02873548 | 0 | 0 | 0 | 0 | 0 | 0 | 0 | 0 | 0 | 0 | 0 | 0.09152549 | 0 |
| ENSMUSG00 Gm11199 | 506 | 0.50401394 | 0.06583576 | 0 | 0 | 0 | 0 | 0 | 0 | 0 | 0 | 0.09441691 | 0 | 0 | 0 | 0 | 0 | 0 | 0 |
| ENSMUSG00 Gm12168 | 367 | 0.81072543 | 0.6353958 | 1.58169582 | 0.49657814 | 1.1803469 | 1.21518965 | 1.41809119 | 1.2487979 | 1.23447322 | 0.77085618 | 1.30176999 | 0.45625882 | 0.40325818 | 1.26550647 | 0.78586074 | 1.25200896 | 0.83545061 | 0.14988857 |
| ENSMUSG00 Gm11556 | 287 | 0 | 0 | 0 | 0 | 0 | 0 | 0 | 0 | 0 | 0 | 0 | 0 | 0 | 0 | 0 | 0 | 0 | 0 |
| ENSMUSG00 Rpl31-ps6 | 377 | 0 | 0 | 0 | 0 | 0 | 0 | 0 | 0.10130611 | 0 | 0.62534089 | 0 | 0.14805216 | 0.26170778 | 0.12319387 | 0.32786384 | 0.71096619 | 0.10166126 | 0 |
| ENSMUSG00 Gm11253 | 200 | 0 | 0 | 0 | 0 | 0 | 0 | 0 | 0 | 0 | 0 | 0 | 0 | 0 | 0 | 0 | 0 | 0 | 0 |
| ENSMUSG00 Gm13712 | 2292 | 0 | 0 | 0 | 0.01987829 | 0 | 0 | 0 | 0 | 0 | 0 | 0.02084422 | 0.04870477 | 0 | 0 | 0 | 0 | 0.03344354 | 0 |
| ENSMUSG00 Gm7860 | 878 | 0.09682272 | 0.07588359 | 0.22038055 | 0.15567565 | 0.27409979 | 0.05643819 | 0.05388688 | 0.13049796 | 0.04690947 | 0 | 0 | 0.06357137 | 0.05618669 | 0.21159038 | 0.0938532 | 0.04361117 | 0.30556273 | 0.06265274 |
| ENSMUSG00 Gm13456 | 1364 | 0.84137812 | 0.63499652 | 0.92207609 | 0.86846565 | 0.91747069 | 0.6539217 | 0.76310772 | 1.09201151 | 0.90586178 | 0.76049521 | 0.49035881 | 0.61380861 | 0.86801027 | 0.81719802 | 0.936399 | 0.44915669 | 0.95534758 | 0.8469144 |
| ENSMUSG00 Gm14981 | 942 | 0 | 0 | 0 | 0 | 0 | 0 | 0 | 0 | 0 | 0 | 0 | 0 | 0 | 0 | 0 | 0 | 0 | 0 |
| ENSMUSG00 Gm14704 | 873 | 0 | 0 | 0 | 0 | 0 | 0 | 0 | 0 | 0 | 0 | 0 | 0 | 0 | 0 | 0 | 0 | 0 | 0 |
| ENSMUSG00 Gm15207 | 441 | 0 | 0 | 0 | 0 | 0 | 0 | 0 | 0 | 0 | 0 | 0 | 0 | 0 | 0 | 0 | 0 | 0 | 0 |
| ENSMUSG00 Gm11311 | 543 | 0 | 0 | 0 | 0 | 0 | 0 | 0 | 0 | 0 | 0 | 0 | 0 | 0 | 0 | 0 | 0 | 0 | 0 |
| ENSMUSG00 Gm9095 | 880 | 0.19320535 | 0 | 0 | 0 | 0 | 0.05630992 | 0 | 0.04340046 | 0 | 0 | 0 | 0 | 0 | 0 | 0.14045985 | 0 | 0 | 0 |
| ENSMUSG00 Gm13311 | 379 | 0 | 0 | 0 | 0 | 0 | 0 | 0 | 0 | 0 | 0 | 0 | 0 | 0 | 0 | 0 | 0 | 0 | 0.58057105 |
| ENSMUSG00 Gm14336 | 474 | 0.08967337 | 0 | 0 | 0 | 0 | 0 | 0 | 0.80574689 | 0 | 0 | 0.10079105 | 0 | 0.10407577 | 0.09798331 | 0 | 0 | 0 | 0.11605297 |
| ENSMUSG00 Gm11667 | 485 | 0 | 0 | 0 | 0 | 0 | 0 | 0 | 0 | 0 | 0 | 0 | 0 | 0 | 0 | 0 | 0 | 0 | 0 |
| ENSMUSG00 Gm12196 | 261 | 0 | 0 | 0 | 0 | 0 | 0 | 0.18127463 | 0.29266209 | 0 | 0.18065404 | 0 | 0 | 0 | 0 | 0 | 0.14670731 | 0 | 0 |
| ENSMUSG00 Gm14898 | 644 | 0 | 0 | 0 | 0 | 0 | 0 | 0 | 0 | 0 | 0 | 0 | 0 | 0 | 0 | 0 | 0 | 0 | 0 |
| ENSMUSG00 Olfr1073-ps1 | 593 | 0 | 0 | 0 | 0 | 0 | 0 | 0 | 0 | 0 | 0 | 0 | 0 | 0 | 0 | 0 | 0 | 0 | 0 |
| ENSMUSG00 Gm12377 | 251 | 0 | 0 | 0 | 0 | 0 | 0 | 0 | 0 | 0 | 0 | 0 | 0 | 0 | 0 | 0 | 0 | 0 | 0 |
| ENSMUSG00 Gm14813 | 394 | 0 | 0.42275246 | 0.12277546 | 0.11563717 | 0 | 0 | 0.12008294 | 0.29080509 | 0.31360291 | 0.23934367 | 0.12125624 | 0.14166412 | 0.25041582 | 0 | 0.10457248 | 0.68028998 | 0.19454973 | 0.13961702 |
| ENSMUSG00 Gm12344 | 328 | 0.90712266 | 0.50781851 | 0.5899211 | 0.27781125 | 1.4674367 | 0.15107541 | 0.86547583 | 1.28084276 | 0.50227458 | 0.28750429 | 0.14565536 | 0.85084852 | 0 | 0.28319565 | 0.502458 | 0 | 0.11684847 | 0.16771069 |
| ENSMUSG00 Gm15065 | 272 | 0 | 0 | 0 | 0 | 0 | 0 | 0 | 0 | 0 | 0 | 0 | 0 | 0 | 0 | 0 | 0 | 0 | 0 |
| ENSMUSG00 Gm11850 | 478 | 0 | 0 | 0 | 0 | 0 | 0 | 0 | 0 | 0 | 0 | 0 | 0 | 0 | 0 | 0 | 0.08010587 | 0.08018054 | 0 |
| ENSMUSG00 Gm11404 | 492 | 0 | 0 | 0 | 0 | 0 | 0 | 0 | 0 | 0 | 0 | 0 | 0 | 0 | 0 | 0 | 0 | 0 | 0 |
| ENSMUSG00 Gm15806 | 954 | 0 | 0 | 0 | 0 | 0 | 0.15582621 | 0 | 0 | 0 | 0 | 0 | 0 | 0 | 0 | 0 | 0 | 0 | 0 |
| ENSMUSG00 Gm11406 | 483 | 0 | 0 | 0 | 0 | 0 | 0 | 0 | 0 | 0 | 0 | 0 | 0 | 0 | 0 | 0 | 0 | 0 | 0 |
| ENSMUSG00 Gm5387 | 856 | 0 | 0 | 0 | 0 | 0 | 0 | 0 | 0 | 0 | 0 | 0 | 0 | 0 | 0 | 0 | 0 | 0 | 0 |
| ENSMUSG00 Gm13317 | 487 | 0 | 0 | 0 | 0 | 0 | 0 | 0 | 0 | 0 | 0 | 0 | 0 | 0 | 0 | 0 | 0 | 0 | 0 |
| ENSMUSG00 Snrp1c-ps2 | 364 | 0 | 0 | 0 | 0 | 0 | 0.13613388 | 0 | 0 | 0 | 0 | 0 | 0 | 0 | 0 | 0 | 0 | 0 | 0 |
| ENSMUSG00 Gm12812 | 432 | 0 | 0 | 0 | 0 | 0 | 0 | 0 | 0 | 0 | 0 | 0 | 0 | 0 | 0 | 0 | 0 | 0 | 0 |
| ENSMUSG00 Gm15157 | 701 | 0 | 0 | 0 | 0 | 0 | 0 | 0 | 0 | 0 | 0 | 0 | 0 | 0.49261544 | 0.92755667 | 0.1763262 | 0 | 0.10934749 | 0.54930634 |
| ENSMUSG00 Gm15580 | 555 | 0 | 0 | 0 | 0 | 0 | 0 | 0 | 0 | 0 | 0 | 0 | 0 | 0 | 0 | 0 | 0 | 0 | 0 |
| ENSMUSG00 Gm11726 | 423 | 0 | 0 | 0 | 0 | 0 | 0 | 0 | 0 | 0 | 0 | 0 | 0 | 0 | 0 | 0 | 0 | 0 | 0 |
| ENSMUSG00 Gm15026 | 457 | 0 | 0 | 0 | 0 | 0 | 0.10843049 | 0 | 0 | 0 | 0 | 0 | 0 | 0 | 0 | 0 | 0 | 0 | 0 |
| ENSMUSG00 Gm15586 | 527 | 0 | 0 | 0 | 0 | 0 | 0 | 0 | 0 | 0 | 0 | 0 | 0 | 0 | 0 | 0 | 0 | 0 | 0 |
| ENSMUSG00 Gm5255 | 640 | 0 | 0 | 0 | 0 | 0 | 0 | 0 | 0 | 0 | 0 | 0 | 0 | 0 | 0 | 0 | 0 | 0 | 0 |
| ENSMUSG00 Gm11485 | 1174 | 0 | 0 | 0 | 0 | 0 | 0 | 0 | 0 | 0 | 0 | 0 | 0 | 0 | 0 | 0 | 0 | 0 | 0 |
| ENSMUSG00 Gm14896 | 1350 | 0 | 0 | 0 | 0 | 0 | 0 | 0 | 0 | 0 | 0 | 0 | 0 | 0 | 0 | 0 | 0 | 0 | 0 |
| ENSMUSG00 Gm14683 | 4287 | 0 | 0 | 0 | 0 | 0 | 0 | 0.04414526 | 0 | 0 | 0 | 0 | 0 | 0 | 0 | 0 | 0 | 0 | 0 |
| ENSMUSG00 Gm12634 | 991 | 0 | 0 | 0 | 0.04597482 | 0 | 0 | 0 | 0.03853926 | 0 | 0 | 0.09641768 | 0 | 0.14933981 | 0 | 0.08315148 | 0 | 0.03867437 | 0.11101737 |
| ENSMUSG00 Gm15711 | 534 | 0.07959771 | 0.0623837 | 0 | 0 | 0.8112122 | 0 | 0 | 0 | 0 | 0 | 0 | 0 | 0 | 0 | 0 | 0 | 0.35886046 | 0 |
| ENSMUSG00 Gm8737 | 641 | 0 | 0 | 0 | 0 | 0 | 0 | 0.07381073 | 0 | 0 | 0 | 0 | 0 | 0 | 0 | 0 | 0 | 0 | 0 |
| ENSMUSG00 Gm11903 | 726 | 0 | 0 | 0 | 0 | 0 | 0 | 0 | 0 | 0 | 0 | 0 | 0 | 0 | 0 | 0 | 0 | 0 | 0 |
| ENSMUSG00 Olfr1268-ps1 | 923 | 0 | 0 | 0 | 0 | 0 | 0 | 0 | 0 | 0 | 0 | 0 | 0 | 0 | 0 | 0 | 0 | 0 | 0 |
| ENSMUSG00 Gm13050 | 1344 | 0.253007 | 0.09914552 | 0.3599221 | 0.13559835 | 0.10743733 | 0.22121756 | 0.42243463 | 0.11366786 | 0.33709202 | 0.31574132 | 0.24882791 | 0.29070658 | 0.62399003 | 0.48379258 | 0.12262368 | 0.17094021 | 0.22813272 | 0.24557637 |
| ENSMUSG00 Gm12058 | 436 | 0.09748894 | 0 | 0 | 0 | 0 | 0 | 0 | 0 | 0.18892897 | 0 | 0 | 0 | 0 | 0 | 0 | 0 | 0.26371305 | 0 |
| ENSMUSG00 Olfr1003-ps1 | 724 | 0 | 0 | 0 | 0 | 0 | 0 | 0 | 0 | 0 | 0 | 0 | 0 | 0 | 0 | 0 | 0 | 0 | 0 |
| ENSMUSG00 Gm13527 | 690 | 0.12320341 | 0.14483867 | 0 | 0 | 0 | 0 | 0 | 0 | 0 | 0 | 0.13847814 | 0 | 0.07149553 | 0 | 0 | 0 | 0 | 0 |
| ENSMUSG00 Gm11979 | 742 | 0 | 0 | 0.13038687 | 0 | 0 | 0 | 0 | 0 | 0 | 0 | 0 | 0.07522326 | 0 | 0 | 0.0555277 | 0 | 0 | 0 |
| ENSMUSG00 Gm13182 | 768 | 0.05534528 | 0 | 0.06298637 | 0.11864855 | 0 | 0 | 0.06160505 | 0 | 0.10725655 | 0.12278829 | 0 | 0 | 0.12846853 | 0 | 0 | 0 | 0 | 0 |
| ENSMUSG00 Gm13761 | 799 | 0.05319797 | 0 | 0 | 0.05702258 | 0.0602402 | 0.06201844 | 0 | 0.04780025 | 0.05154758 | 0.05901214 | 0 | 0 | 0.06174207 | 0 | 0 | 0 | 0 | 0 |
| ENSMUSG00 Olfr336-ps1 | 938 | 0 | 0 | 0 | 0 | 0 | 0 | 0 | 0 | 0 | 0 | 0 | 0 | 0 | 0 | 0 | 0 | 0 | 0 |
| ENSMUSG00 Gm8569 | 646 | 0 | 0 | 0 | 0 | 0 | 0 | 0 | 0 | 0 | 0 | 0 | 0 | 0 | 0 | 0 | 0 | 0 | 0 |
| ENSMUSG00 Gm15387 | 647 | 0.06569579 | 0 | 0 | 0 | 0 | 0 | 0.07312624 | 0 | 0 | 0.14575179 | 0 | 0 | 0 | 0.07178375 | 0 | 0.11836355 | 0 | 0 |
| ENSMUSG00 Gm15436 | 342 | 0 | 0 | 0 | 0 | 0 | 0 | 0 | 0 | 0 | 0 | 0 | 0 | 0 | 0 | 0 | 0 | 0 | 0 |
| ENSMUSG00 Gm12129 | 1146 | 0.07418006 | 0.02906884 | 0.08442152 | 0 | 0 | 0 | 0 | 0 | 0 | 0 | 0 | 0 | 0 | 0.04052713 | 0 | 0 | 0 | 0 |
| ENSMUSG00 Gm15443 | 819 | 0.83038195 | 0.85417677 | 1.06315451 | 0.611931 | 0.35261483 | 0.24201579 | 0.98207026 | 0.55959564 | 0.80462058 | 0.11514213 | 0.87499924 | 0.34075496 | 0.78304631 | 0.73720774 | 0.8049144 | 0.70129318 | 0.51476101 | 0.33583093 |
| ENSMUSG00 Gm12433 | 254 | 0 | 0 | 0 | 0 | 0 | 0 | 0 | 0 | 0 | 0 | 0 | 0 | 0 | 0 | 0 | 0 | 0 | 0 |
| ENSMUSG00 Gm14943 | 542 | 0 | 0 | 0 | 0 | 0 | 0 | 0 | 0 | 0 | 0 | 0 | 0 | 0 | 0 | 0 | 0 | 0 | 0 |
| ENSMUSG00 Gm11558 | 458 | 0 | 0 | 0 | 0 | 0 | 0 | 0 | 0 | 0 | 0 | 0 | 0 | 0 | 0 | 0 | 0 | 0 | 0 |
| ENSMUSG00 Gm12950 | 416 | 0 | 0 | 0 | 0 | 0.23140348 | 0 | 0 | 0.18361732 | 0 | 0 | 0 | 0 | 0 | 0 | 0 | 0.09204473 | 0 | 0 |
| ENSMUSG00 Gm11331 | 287 | 0 | 0 | 0 | 0 | 0 | 0 | 0 | 0 | 0 | 0 | 0 | 0 | 0 | 0 | 0 | 0 | 0 | 0 |
| ENSMUSG00 Gm15074 | 935 | 0 | 0 | 0 | 0 | 0 | 0 | 0 | 0 | 0 | 0 | 0 | 0 | 0 | 0 | 0 | 0 | 0 | 0 |
| ENSMUSG00 Gm14793 | 310 | 0 | 0 | 0 | 0 | 0 | 0 | 0 | 0 | 0 | 0 | 0 | 0 | 0 | 0 | 0 | 0 | 0 | 0 |
| ENSMUSG00 Gm14227 | 628 | 0 | 0.05304601 | 0.61622332 | 0.36274717 | 0.61314552 | 0.23671688 | 0.15067732 | 0.18244778 | 0.13116725 | 0.75080738 | 0.22822432 | 0.35551377 | 0.31421603 | 0.07395555 | 0.19682272 | 0.12194461 | 0.06102914 | 1.05112943 |
| ENSMUSG00 Gm12645 | 274 | 0 | 0 | 0 | 0 | 0 | 0 | 0 | 0 | 0 | 0 | 0 | 0 | 0 | 0.33900794 | 0 | 0 | 0 | 0 |
| ENSMUSG00 Gm15212 | 465 | 0 | 0 | 0 | 0 | 0 | 0 | 0 | 0 | 0 | 0 | 0 | 0 | 0 | 0 | 0 | 0 | 0 | 0 |
| ENSMUSG00 Gm13141 | 455 | 0.09341797 | 0 | 0 | 0 | 0 | 0 | 0 | 0 | 0 | 0 | 0 | 0 | 0 | 0 | 0 | 0 | 0 | 0 |
| ENSMUSG00 Gm9673 | 780 | 0 | 0.04270884 | 0 | 0 | 0 | 0 | 0.06065728 | 0 | 0.05280323 | 0.06044962 | 0 | 0 | 0.06324605 | 0 | 0.05282251 | 0.04909052 | 0 | 0 |
| ENSMUSG00 Gm12700 | 456 | 0 | 0.14610919 | 0.1060823 | 0 | 0 | 0 | 0 | 0 | 0 | 0 | 0 | 0.12240277 | 0 | 0 | 0 | 0 | 0 | 0 |
| ENSMUSG00 Gm15021 | 741 | 0 | 0 | 0 | 0 | 0 | 0 | 0 | 0 | 0 | 0 | 0 | 0 | 0 | 0 | 0 | 0 | 0 | 0 |
| ENSMUSG00 Gm8482 | 470 | 0.99480199 | 0.35439249 | 0.72045684 | 0.09693839 | 0.2048167 | 0.42172539 | 0 | 0.65008345 | 0.35052354 | 0.40128258 | 0.71154193 | 0.8312971 | 0.52480763 | 1.18580649 | 0 | 0.16293875 | 0.24463594 | 0 |
| ENSMUSG00 Gm16148 | 892 | 0.14295463 | 0.48550182 | 0.27115208 | 0 | 0.37771689 | 0.11110478 | 0.15912336 | 0.29971616 | 0.13851967 | 0 | 0.05355937 | 0.43801529 | 0.38713388 | 0.20826945 | 0 | 0.25756014 | 0.2578002 | 0.18500821 |
| ENSMUSG00 Gm14638 | 213 | 0 | 0 | 0 | 0 | 0 | 0 | 0 | 0 | 0 | 0 | 0 | 0 | 0 | 0 | 0 | 0 | 0 | 0 |
| ENSMUSG00 Gm12045 | 572 | 0 | 0 | 0 | 0 | 0 | 0 | 0 | 0 | 0 | 0 | 0.08352265 | 0 | 0.17248922 | 0 | 0 | 0 | 0.06700402 | 0 |
| ENSMUSG00 Gm7670 | 640 | 0 | 0 | 0 | 0.07118913 | 0 | 0 | 0 | 0 | 0 | 0 | 0 | 0 | 0 | 0 | 0 | 0 | 0 | 0 |
| ENSMUSG00 Gm15464 | 430 | 0.09884925 | 0.38735924 | 0.22499316 | 0.10595592 | 0 | 0.23047783 | 0.11002949 | 0.26645862 | 0.28734778 | 0.32895839 | 0.11110455 | 0 | 0 | 0.32402852 | 0.38327029 | 0 | 0 | 0.51171262 |
| ENSMUSG00 Gm13829 | 137 | 1.24102704 | 0.48631962 | 0 | 0 | 0 | 1.44679514 | 1.03604406 | 0.55755332 | 0 | 0.68833143 | 1.04616697 | 0 | 3.60086987 | 1.35603175 | 1.8044477 | 0.55898697 | 2.23803194 | 0 |
| ENSMUSG00 Gm7746 | 502 | 0 | 0.06636035 | 0 | 0 | 0 | 0 | 0 | 0 | 0 | 0 | 0 | 0 | 0 | 0 | 0 | 0 | 0 | 0 |
| ENSMUSG00 4930503H13 | 723 | 0 | 0 | 0 | 0 | 0 | 0 | 0 | 0 | 0 | 0 | 0 | 0 | 0 | 0 | 0 | 0 | 0 | 0 |
| ENSMUSG00 Gm12518 | 778 | 0 | 0 | 0 | 0 | 0 | 0 | 0 | 0 | 0 | 0 | 0 | 0 | 0 | 0 | 0 | 0 | 0 | 0 |
| ENSMUSG00 Gm12149 | 651 | 0 | 0.05117188 | 0 | 0 | 0 | 0 | 0 | 0 | 0 | 0 | 0 | 0 | 0 | 0 | 0 | 0 | 0 | 0 |
| ENSMUSG00 Gm13239 | 636 | 0 | 0 | 0 | 0 | 0 | 0 | 0 | 0 | 0 | 0 | 0 | 0 | 0 | 0 | 0 | 0 | 0 | 0 |
| ENSMUSG00 Gm6025 | 200 | 0 | 0 | 0 | 0 | 0 | 0 | 0 | 0.19096201 | 0 | 0 | 0 | 0 | 0 | 0 | 0 | 0 | 0 | 0 |
| ENSMUSG00 Smt3h2-ps | 284 | 0.14966611 | 0.11729892 | 0 | 0.16042621 | 0.1694786 | 0.34896291 | 0 | 0 | 0.14502294 | 0.1660236 | 0 | 0 | 0.17370393 | 0 | 0 | 0 | 0.13495175 | 0 |
| ENSMUSG00 Gm14852 | 641 | 0 | 0 | 0 | 0 | 0 | 0 | 0 | 0 | 0 | 0 | 0 | 0 | 0 | 0 | 0 | 0 | 0 | 0 |
| ENSMUSG00 Gm16219 | 273 | 0 | 0 | 0 | 0 | 0 | 0 | 0 | 0 | 0 | 0 | 0 | 0 | 0 | 0 | 0 | 0 | 0 | 0 |
| ENSMUSG00 Gm8244 | 263 | 0 | 0 | 0 | 0 | 0 | 0 | 0 | 0 | 0 | 0 | 0 | 0 | 0 | 0 | 0 | 0 | 0 | 0 |
| ENSMUSG00 Rpl7a-ps13 | 562 | 0 | 0 | 0 | 0 | 0 | 0 | 0 | 0 | 0 | 0 | 0 | 0 | 0 | 0 | 0 | 0 | 0 | 0 |
| ENSMUSG00 Gm12632 | 494 | 0.2581286 | 0 | 0.09792213 | 0.09222883 | 0.09743304 | 0 | 0.09577465 | 0 | 0 | 0.09544677 | 0 | 0.45194868 | 0 | 0.09401637 | 0.08340396 | 0.07751135 | 0 | 0.3340634 |
| ENSMUSG00 Gm15493 | 527 | 0 | 0 | 0 | 0 | 0 | 0 | 0 | 0 | 0 | 0 | 0 | 0 | 0 | 0 | 0 | 0 | 0 | 0 |
| ENSMUSG00 Gm12535 | 483 | 0 | 0 | 0 | 0 | 0 | 0 | 0 | 0 | 0 | 0 | 0 | 0 | 0 | 0 | 0 | 0.15855324 | 0 | 0 |
| ENSMUSG00 Gm15204 | 2280 | 0.11185573 | 0 | 0.12729876 | 0.21981206 | 0.06333148 | 0 | 0.12450705 | 0.2177637 | 0.01806426 | 0.10340067 | 0 | 0 | 0 | 0 | 0.01807086 | 0.13435301 | 0.18490757 | 0 |
| ENSMUSG00 Gm12270 | 455 | 0 | 0 | 0.10631545 | 0 | 0.10578445 | 0 | 0 | 0 | 0.18103963 | 0.10362792 | 0.10499991 | 0 | 0 | 0.30622475 | 0 | 0.16831036 | 0 | 0.12089914 |
| ENSMUSG00 Gm12566 | 358 | 0 | 0 | 0 | 0 | 0 | 0 | 0 | 0 | 0 | 0 | 0 | 0 | 0 | 0 | 0 | 0 | 0 | 0 |
| ENSMUSG00 Gm5639 | 2402 | 0 | 0 | 0 | 0 | 0 | 0 | 0 | 0 | 0 | 0 | 0 | 0 | 0 | 0 | 0 | 0 | 0 | 0 |
| ENSMUSG00 Gm14834 | 793 | 0 | 0 | 0 | 0 | 0 | 0 | 0 | 0 | 0 | 0 | 0 | 0 | 0 | 0 | 0 | 0 | 0 | 0 |
| ENSMUSG00 Gm13365 | 966 | 0.04400122 | 0.0344854 | 0.20030447 | 0 | 0 | 0 | 0.04897793 | 0.03953665 | 0.04263614 | 0.24405126 | 0 | 0.17334057 | 0.05106824 | 0 | 0 | 0.07927662 | 0.03967526 | 0.11389049 |
| ENSMUSG00 Gm15779 | 411 | 0 | 0.08105327 | 0.11769715 | 0.22170824 | 0.1171093 | 0 | 0.11511601 | 0 | 0.200421 | 0.11472191 | 0 | 0 | 0.120029 | 0 | 0.20049419 | 0.37265798 | 0.37300532 | 0.53536844 |
| ENSMUSG00 Gm12900 | 572 | 0 | 0 | 0 | 0 | 0 | 0 | 0 | 0 | 0 | 0 | 0 | 0 | 0 | 0 | 0 | 0 | 0 | 0 |
| ENSMUSG00 Gm14750 | 512 | 0 | 0 | 0 | 0 | 0 | 0 | 0 | 0 | 0 | 0 | 0 | 0 | 0 | 0 | 0 | 0 | 0 | 0 |
| ENSMUSG00 Gm14923 | 153 | 0 | 0 | 0 | 0 | 0 | 0 | 0 | 0 | 0 | 0 | 0 | 0 | 0 | 0 | 0 | 0 | 0 | 0 |
| ENSMUSG00 Gm11248 | 280 | 0 | 0 | 0 | 0 | 0 | 0 | 0 | 0 | 0 | 0 | 0 | 0 | 0 | 0 | 0 | 0 | 0 | 0 |
| ENSMUSG00 Gm12815 | 134 | 0 | 0 | 0 | 0 | 0 | 0 | 0 | 0 | 0 | 0 | 0 | 0 | 0 | 0 | 0 | 0 | 0 | 0 |
| ENSMUSG00 Gm2012 | 1311 | 0 | 0 | 0 | 0 | 0 | 0 | 0 | 0 | 0 | 0 | 0 | 0 | 0 | 0 | 0 | 0 | 0 | 0 |
| ENSMUSG00 Gm11402 | 308 | 0 | 0 | 0 | 0 | 0.15627248 | 0 | 0 | 0 | 0 | 0 | 0 | 0 | 0 | 0 | 0 | 0.12432015 | 0 | 0 |
| ENSMUSG00 Rpl30-ps5 | 360 | 0 | 0 | 0 | 0 | 0 | 0 | 0 | 0 | 0.22881398 | 0 | 0 | 0 | 0 | 0 | 0 | 0 | 0.10646194 | 0 |
| ENSMUSG00 Gm14362 | 272 | 0 | 0 | 0 | 0 | 0 | 0 | 0 | 0 | 0 | 0 | 0 | 0 | 0 | 0 | 0 | 0 | 0 | 0 |
| ENSMUSG00 Mup-ps19 | 513 | 0 | 0 | 0 | 0 | 0 | 0 | 0 | 0 | 0 | 0 | 0 | 0 | 0 | 0 | 0 | 0 | 0 | 0 |
| ENSMUSG00 Gm15481 | 538 | 0 | 0 | 0 | 0 | 0 | 0 | 0 | 0 | 0 | 0 | 0 | 0 | 0 | 0 | 0 | 0 | 0 | 0 |
| ENSMUSG00 Gm12732 | 179 | 0 | 0 | 0 | 0 | 0 | 0 | 0 | 0 | 0 | 0 | 0 | 0 | 0 | 0 | 0 | 0 | 0.21411339 | 0.30731345 |
| ENSMUSG00 Gm14056 | 641 | 0 | 0 | 0.07546573 | 0 | 0 | 0 | 0 | 0 | 0 | 0 | 0 | 0 | 0 | 0.14491135 | 0 | 0 | 0 | 0 |
| ENSMUSG00 Gm11366 | 428 | 0 | 0 | 0 | 0 | 0.22491553 | 0 | 0 | 0 | 0 | 0 | 0 | 0.52164171 | 0.23052298 | 0.65108534 | 0 | 0.08946404 | 0.62683196 | 0 |
| ENSMUSG00 Gm12754 | 453 | 0 | 0.0735384 | 0.10678484 | 0 | 0 | 0 | 0.104443 | 0 | 0.09091946 | 0 | 0.10546348 | 0 | 0 | 0 | 0 | 0.08452673 | 0 | 0 |
| ENSMUSG00 Gm15358 | 519 | 0 | 0 | 0 | 0 | 0 | 0 | 0 | 0 | 0 | 0 | 0 | 0 | 0 | 0 | 0 | 0 | 0 | 0 |
| ENSMUSG00 Gm14001 | 246 | 0.34557054 | 0 | 0 | 0 | 0 | 0 | 1.34629574 | 0.465761 | 0 | 0.95834763 | 0 | 0 | 0 | 0 | 0 | 0.15565288 | 0 | 0 |
| ENSMUSG00 Gm14725 | 827 | 0 | 0 | 0 | 0 | 0 | 0 | 0 | 0 | 0 | 0 | 0 | 0 | 0 | 0 | 0 | 0 | 0 | 0 |
| ENSMUSG00 Gm15897 | 1485 | 0 | 0 | 0 | 0 | 0 | 0 | 0 | 0 | 0 | 0 | 0 | 0 | 0 | 0 | 0 | 0 | 0 | 0 |
| ENSMUSG00 Gm7857 | 357 | 0.23812424 | 0 | 0 | 0 | 0 | 0 | 0 | 0 | 0 | 0 | 0.26764683 | 0 | 0 | 0 | 0.11541052 | 0 | 0.10735657 | 0 |
| ENSMUSG00 Gm11323 | 289 | 0 | 0 | 0 | 0 | 0 | 0 | 0 | 0 | 0 | 0 | 0 | 0 | 0 | 0 | 0 | 0 | 0 | 0 |
| ENSMUSG00 Gm14736 | 335 | 0 | 0 | 0 | 0 | 0 | 0 | 0 | 0 | 0 | 0 | 0 | 0 | 0 | 0 | 0 | 0 | 0 | 0 |
| ENSMUSG00 Fau-ps2 | 370 | 0 | 0 | 0 | 1.10824162 | 0 | 0 | 0 | 0.41289084 | 0 | 0 | 0.12912151 | 0.15085314 | 0 | 0.37657368 | 0.22271111 | 0 | 0.7250921 | 0 |
| ENSMUSG00 Gm11394 | 515 | 0 | 0 | 0 | 0 | 0 | 0 | 0 | 0 | 0 | 0 | 0 | 0 | 0 | 0 | 0 | 0 | 0 | 0 |
| ENSMUSG00 Gm14612 | 880 | 0 | 0 | 0 | 0 | 0 | 0 | 0 | 0 | 0 | 0 | 0 | 0 | 0 | 0 | 0 | 0 | 0 | 0 |
| ENSMUSG00 Gm11590 | 1452 | 0 | 0 | 0 | 0 | 0 | 0 | 0 | 0 | 0 | 0 | 0 | 0 | 0 | 0 | 0 | 0 | 0 | 0 |
| ENSMUSG00 Gm15516 | 192 | 0.22138113 | 0 | 0 | 0 | 0 | 0 | 0 | 0 | 0 | 0.24557658 | 0 | 0 | 0.51387414 | 0 | 0 | 0 | 0 | 0 |
| ENSMUSG00 Trav13-3 | 495 | 0 | 0 | 0 | 0 | 0 | 0 | 0 | 0 | 0 | 0 | 0 | 0 | 0 | 0 | 0 | 0 | 0 | 0 |
| ENSMUSG00 Gm8761 | 954 | 0 | 0 | 0 | 0 | 0 | 0 | 0 | 0 | 0 | 0 | 0 | 0 | 0 | 0 | 0 | 0 | 0 | 0 |
| ENSMUSG00 Gm11470 | 1473 | 0.0288562 | 0 | 0.09852043 | 0 | 0.03267612 | 0 | 0.16059972 | 0 | 0 | 0 | 0 | 0 | 0 | 0 | 0 | 0 | 0.02601921 | 0 |
| ENSMUSG00 Gm13677 | 594 | 0.14311507 | 0 | 0 | 0 | 0 | 0 | 0 | 0 | 0 | 0 | 0 | 0 | 0 | 0 | 0.06936289 | 0 | 0 | 0 |
| ENSMUSG00 Gm11348 | 317 | 0 | 0 | 0 | 0 | 0 | 0 | 0 | 0 | 0 | 0 | 0 | 0 | 0 | 0 | 0 | 0 | 0 | 0 |
| ENSMUSG00 Gm13258 | 289 | 0 | 0 | 0 | 0 | 0 | 0 | 1.30969353 | 0.39646092 | 0 | 0 | 0 | 0 | 0 | 0 | 0 | 0 | 0.66308472 | 0 |
| ENSMUSG00 Gm5393 | 271 | 0 | 0 | 0 | 0 | 0 | 0 | 0 | 0 | 0 | 0 | 0 | 0 | 0 | 0 | 0 | 0 | 0 | 0 |
| ENSMUSG00 Gm14050 | 345 | 0 | 0 | 0 | 0 | 0 | 0 | 0 | 0 | 0 | 0 | 0 | 0 | 0 | 0 | 0 | 0 | 0 | 0 |
| ENSMUSG00 Gm14006 | 952 | 0.08929659 | 0 | 0 | 0 | 0 | 0 | 0.04969819 | 0.04011807 | 0 | 0.04952805 | 0 | 0 | 0 | 0 | 0.04327895 | 0 | 0.04025872 | 0.05778268 |
| ENSMUSG00 Gm11876 | 268 | 0 | 0 | 0 | 0 | 0 | 0 | 0 | 0 | 0 | 0 | 0 | 0 | 0 | 0 | 0 | 0 | 0 | 0 |
| ENSMUSG00 Gm11857 | 323 | 0 | 0 | 0 | 0 | 0 | 0.15341404 | 0 | 0 | 0.12751243 | 0 | 0 | 0 | 0 | 0 | 0.127559 | 0 | 0 | 0 |
| ENSMUSG00 Gm11914 | 506 | 1.00802789 | 0.92170063 | 0.7647989 | 0.72033272 | 0.38048952 | 0.78344243 | 1.21554313 | 0.83026962 | 1.62792552 | 1.67729774 | 0.6609184 | 0.88246107 | 0.87744517 | 1.83573468 | 1.95422399 | 1.74048215 | 0.83318037 | 1.19585015 |
| ENSMUSG00 Gm6382 | 941 | 1.30993635 | 1.23905558 | 1.64500847 | 1.16202451 | 1.6367923 | 1.00053341 | 1.45809531 | 1.33937224 | 1.00668423 | 0.70149824 | 1.11694908 | 0.88972895 | 1.04849984 | 1.48068291 | 1.09462157 | 1.627656 | 1.3440678 | 0.46766509 |
| ENSMUSG00 Gm11843 | 424 | 0.10024806 | 0 | 0 | 0 | 0 | 0 | 0 | 0 | 0 | 0 | 0 | 0.13164071 | 0.11634886 | 0 | 0 | 0 | 0 | 0 |
| ENSMUSG00 Gm15371 | 783 | 0 | 0 | 0 | 0 | 0 | 0 | 0 | 0 | 0 | 0 | 0 | 0 | 0 | 0 | 0 | 0 | 0 | 0 |
| ENSMUSG00 Gm12818 | 1348 | 0.03153203 | 0.02471283 | 0.03588541 | 0 | 0 | 0.03676019 | 0.03509843 | 0 | 0.06110759 | 0 | 0 | 0.08281256 | 0 | 0.03445407 | 0.12225981 | 0 | 0 | 0 |
| ENSMUSG00 Gm13469 | 349 | 0 | 0 | 0 | 0 | 0 | 0 | 0 | 0 | 0 | 0 | 0 | 0 | 0 | 0 | 0 | 0 | 0 | 0 |
| ENSMUSG00 Gm14885 | 495 | 0 | 0 | 0 | 0 | 0 | 0 | 0 | 0 | 0 | 0 | 0 | 0 | 0 | 0 | 0 | 0 | 0 | 0 |
| ENSMUSG00 Gm12675 | 172 | 0 | 0 | 0 | 0 | 0 | 0 | 0 | 0 | 0 | 0 | 0 | 0 | 0 | 0 | 0 | 0 | 0 | 0 |
| ENSMUSG00 Gm15349 | 203 | 0.2093851 | 0 | 0 | 0.22443864 | 0 | 0 | 0.46613477 | 0.18813991 | 0 | 1.39361685 | 0.47068925 | 0.54990801 | 0.24301437 | 0 | 0.20296333 | 0.18862368 | 0 | 0.54196164 |
| ENSMUSG00 Gm12697 | 666 | 0 | 0 | 0 | 0 | 0 | 0 | 0 | 0 | 0 | 0 | 0 | 0 | 0 | 0 | 0 | 0 | 0 | 0 |
| ENSMUSG00 Gm12774 | 415 | 0.61453267 | 0.80272034 | 0.69937634 | 0.87828519 | 0 | 0.11940418 | 0.45602582 | 0.46014943 | 0.79395693 | 0 | 0.11512038 | 0.40348672 | 0.23774418 | 0 | 0.89352772 | 0.64586567 | 0.09235252 | 0.79531239 |
| ENSMUSG00 Gm14689 | 228 | 0 | 0 | 0 | 0 | 0 | 0 | 0 | 0 | 0 | 0 | 0 | 0 | 0 | 0 | 0 | 0 | 0 | 0 |
| ENSMUSG00 Gm12961 | 1049 | 0.08103942 | 0.06351362 | 0.04611395 | 0 | 0 | 0 | 0 | 0 | 0 | 0 | 0 | 0 | 0 | 0 | 0 | 0.25551406 | 0 | 0 |
| ENSMUSG00 Gm11474 | 276 | 0 | 0 | 0 | 0 | 0 | 0 | 0 | 0 | 0 | 0 | 0 | 0 | 0 | 0 | 0 | 0 | 0 | 0 |
| ENSMUSG00 Gm12124 | 678 | 0 | 0 | 0 | 0 | 0 | 0 | 0 | 0 | 0.06074707 | 0 | 0 | 0 | 0 | 0 | 0 | 0.05647582 | 0 | 0 |
| ENSMUSG00 Gm14156 | 482 | 0 | 0.13822778 | 0.10036002 | 0 | 0 | 0 | 0 | 0 | 0 | 0 | 0 | 0 | 0 | 0.09635703 | 0 | 0 | 0 | 0 |
| ENSMUSG00 Gm14972 | 363 | 0 | 0 | 0 | 0 | 0 | 0 | 0 | 0 | 0 | 0 | 0 | 0 | 0 | 0 | 0 | 0 | 0 | 0 |
| ENSMUSG00 Dynll1-ps1 | 341 | 1.12183749 | 1.75845189 | 1.4185786 | 0.9352707 | 1.27034403 | 1.74379121 | 2.21994974 | 1.23201298 | 2.05328671 | 2.62716528 | 2.52184532 | 0.49104688 | 1.30201541 | 2.04299505 | 2.77899057 | 1.90891591 | 2.69745198 | 1.93580434 |
| ENSMUSG00 Gm15652 | 304 | 0 | 0 | 0 | 0 | 0.47498609 | 0.16300241 | 0 | 0 | 0 | 0 | 0 | 0 | 0 | 0 | 0 | 0 | 0 | 0 |
| ENSMUSG00 Gm15190 | 275 | 0.30912855 | 0.96910238 | 0 | 0.16567652 | 0.52507553 | 0.18019176 | 0.17204611 | 0 | 0.44930744 | 0.34291421 | 0.17372712 | 0.40593209 | 0.53816637 | 0.16888759 | 0.59929536 | 0.83543143 | 0 | 0 |
| ENSMUSG00 Gm16187 | 196 | 0 | 0 | 0 | 0 | 0 | 0.25282007 | 0 | 0 | 0 | 0 | 0 | 0 | 0 | 0 | 0 | 0 | 0 | 0 |
| ENSMUSG00 Gm15040 | 1238 | 0 | 0 | 0 | 0 | 0 | 0 | 0 | 0 | 0 | 0 | 0 | 0 | 0 | 0 | 0 | 0 | 0 | 0 |
| ENSMUSG00 Gm12913 | 544 | 0 | 0 | 0 | 0 | 0 | 0 | 0 | 0 | 0 | 0 | 0 | 0 | 0.09068367 | 0 | 0 | 0 | 0 | 0 |
| ENSMUSG00 Gm15316 | 420 | 0.8096224 | 1.26906264 | 0.92140958 | 1.08478677 | 1.60439746 | 0.94386159 | 1.915037 | 1.18214579 | 0.19612627 | 1.01037221 | 2.16124812 | 1.19604991 | 3.17133754 | 1.54813625 | 1.47148414 | 1.91453037 | 1.91631485 | 0.91681845 |
| ENSMUSG00 Gm12736 | 235 | 0 | 0 | 0 | 0 | 0 | 0 | 0 | 0 | 0 | 0 | 0 | 0 | 0 | 0.98817207 | 0 | 0 | 0.16309063 | 0 |
| ENSMUSG00 Gm13792 | 422 | 0 | 0 | 0 | 0 | 0 | 0 | 0 | 0 | 0 | 0 | 0 | 0 | 0 | 0 | 0 | 0 | 0 | 0 |
| ENSMUSG00 Gm14460 | 638 | 0 | 0.05221457 | 0 | 0 | 0 | 0 | 0 | 0 | 0 | 0 | 0 | 0 | 0 | 0 | 0.06457924 | 0 | 0 | 0 |
| ENSMUSG00 Gm11907 | 1006 | 0.04225167 | 0.06622842 | 0.09617004 | 0 | 0.09568971 | 0 | 0 | 0.11389384 | 0 | 0 | 0.04749002 | 0.11096553 | 0.19615076 | 0 | 0.04095582 | 0 | 0 | 0.16404306 |
| ENSMUSG00 Gm12515 | 360 | 0 | 0 | 0 | 0 | 0 | 0 | 0 | 0 | 0 | 0 | 0 | 0 | 0 | 0 | 0 | 0 | 0 | 0 |
| ENSMUSG00 Gm15616 | 374 | 0 | 0.17814382 | 0.38802297 | 0.24364195 | 0 | 0 | 0 | 0 | 0 | 0 | 0.12774053 | 0 | 0 | 0 | 0 | 0 | 0 | 0 |
| ENSMUSG00 Gm11663 | 289 | 0.4412302 | 1.26796483 | 0.16738246 | 1.41885605 | 1.49891804 | 0.51438824 | 0.32742338 | 0.26430728 | 0.28502779 | 0.81575611 | 0.49593383 | 0.19313378 | 0.51209603 | 0.16070618 | 0.57026375 | 0 | 0.26523389 | 0.95171465 |
| ENSMUSG00 Gm6881 | 832 | 0 | 0 | 0 | 0 | 0 | 0 | 0 | 0 | 0 | 0 | 0 | 0 | 0 | 0 | 0 | 0 | 0 | 0.06611671 |
| ENSMUSG00 Gm15172 | 1045 | 0.04067481 | 0 | 0 | 0 | 0 | 0 | 0.04527529 | 0 | 0 | 0 | 0 | 0 | 0 | 0 | 0 | 0 | 0 | 0 |
| ENSMUSG00 Gm12362 | 341 | 0.49859444 | 0.09769177 | 0 | 0.2672202 | 0 | 0.29063187 | 0.27749372 | 0 | 0.12078157 | 0.27654371 | 0.42030755 | 0 | 0 | 0.13619967 | 0.24165135 | 0.22457834 | 0.11239383 | 0 |
| ENSMUSG00 Defa-ps14 | 416 | 0 | 0 | 0 | 0.43808696 | 0 | 0 | 0 | 0 | 0 | 0 | 0 | 0 | 0 | 0 | 0 | 0 | 0 | 0 |
| ENSMUSG00 Gm12987 | 488 | 0.95810848 | 0.7509054 | 0.49563043 | 0.28008839 | 0.19726198 | 0.30462746 | 0.8725699 | 0.31305248 | 0.33759439 | 0.19324059 | 0.19579901 | 0.34312907 | 0.80871996 | 0 | 0.50657651 | 0.70617923 | 0.62829995 | 0 |
| ENSMUSG00 Olfr990-ps1 | 895 | 0 | 0 | 0 | 0 | 0 | 0 | 0 | 0 | 0 | 0 | 0 | 0 | 0 | 0 | 0 | 0 | 0 | 0 |
| ENSMUSG00 Gm14721 | 524 | 0 | 0 | 0 | 0 | 0 | 0 | 0 | 0 | 0 | 0 | 0 | 0 | 0 | 0 | 0 | 0 | 0 | 0 |
| ENSMUSG00 Gm15120 | 300 | 0 | 0 | 0 | 0 | 0 | 0 | 0 | 0 | 0.13728839 | 0 | 0 | 0 | 0 | 0 | 0 | 0 | 0 | 0 |
| ENSMUSG00 Gm11633 | 443 | 0 | 0 | 0.10919533 | 0 | 0.21729988 | 0 | 0.10680063 | 0 | 0.18594364 | 0.106435 | 0 | 0.12599472 | 0 | 0.10483993 | 0.09300577 | 0.08643478 | 0 | 0 |
| ENSMUSG00 Rpl31-ps5 | 374 | 0 | 0 | 0 | 0 | 0 | 0 | 0 | 0 | 0 | 0 | 0 | 0 | 0 | 0 | 0 | 0 | 0 | 0 |
| ENSMUSG00 Gm13087 | 255 | 0 | 0 | 0 | 0 | 0 | 0 | 0 | 0 | 0 | 0 | 0 | 0 | 0 | 0 | 0 | 0 | 0 | 0 |
| ENSMUSG00 Gm14928 | 476 | 0 | 0 | 0 | 0 | 0 | 0 | 0 | 0 | 0 | 0 | 0 | 0 | 0 | 0 | 0 | 0 | 0 | 0 |
| ENSMUSG00 Gm12489 | 147 | 0.57830171 | 0.45323666 | 0 | 0.92981723 | 0 | 0 | 0 | 0 | 0 | 0 | 2.59999774 | 0.37969838 | 0 | 0.31594617 | 0 | 0.78144097 | 0 | 0 |
| ENSMUSG00 Gm12087 | 257 | 0 | 0 | 0 | 0 | 0 | 0 | 0 | 0 | 0 | 0 | 0 | 0 | 0 | 0 | 0 | 0 | 0 | 0 |
| ENSMUSG00 Gm11393 | 286 | 0 | 0 | 0 | 0 | 0 | 0 | 0 | 0 | 0 | 0 | 0 | 0 | 0 | 0 | 0 | 0 | 0 | 0 |
| ENSMUSG00 Gm14564 | 293 | 0 | 0 | 0 | 0 | 0 | 0 | 0 | 0 | 0 | 0 | 0 | 0 | 0 | 0 | 0 | 0 | 0 | 0 |
| ENSMUSG00 Gm14416 | 2435 | 0 | 0 | 0 | 0 | 0 | 0 | 0 | 0 | 0 | 0 | 0 | 0 | 0 | 0 | 0 | 0 | 0 | 0 |
| ENSMUSG00 Gm12451 | 629 | 0 | 0 | 0 | 0 | 0 | 0 | 0 | 0 | 0 | 0 | 0 | 0 | 0 | 0 | 0 | 0 | 0 | 0 |
| ENSMUSG00 Gm11424 | 828 | 0 | 0 | 0 | 0 | 0 | 0 | 0 | 0 | 0 | 0 | 0 | 0 | 0 | 0 | 0 | 0 | 0 | 0 |
| ENSMUSG00 Gm13260 | 868 | 0 | 0 | 0 | 0 | 0 | 0 | 0 | 0 | 0 | 0 | 0 | 0 | 0 | 0 | 0 | 0.39702243 | 0 | 0 |
| ENSMUSG00 Fam48b1 | 1558 | 0 | 0 | 0 | 0 | 0 | 0 | 0 | 0 | 0 | 0 | 0 | 0 | 0 | 0 | 0 | 0 | 0 | 0 |
| ENSMUSG00 Gm14845 | 305 | 0 | 0 | 0 | 0 | 0 | 0 | 0 | 0.25044198 | 0 | 0 | 0.31327842 | 0 | 0 | 0 | 0 | 0 | 0.12565999 | 0 |
| ENSMUSG00 Gm11363 | 508 | 0 | 0 | 0 | 0 | 0 | 0 | 0 | 0 | 0 | 0 | 0 | 0 | 0 | 0 | 0 | 0 | 0 | 0 |
| ENSMUSG00 Gm14467 | 272 | 0 | 0 | 0 | 0 | 0 | 0 | 0 | 0 | 0 | 0 | 0 | 0 | 0 | 0 | 0 | 0 | 0 | 0 |
| ENSMUSG00 Rpl23a-ps6 | 467 | 0 | 0 | 0 | 0 | 0 | 0 | 0 | 0 | 0 | 0 | 0 | 0 | 0 | 0 | 0 | 0 | 0 | 0 |
| ENSMUSG00 Gm15153 | 273 | 0 | 0 | 0 | 0 | 0 | 0 | 0 | 0.13989691 | 0 | 0 | 0 | 0 | 0 | 0 | 0 | 0 | 0 | 0 |
| ENSMUSG00 Gm15474 | 133 | 0 | 0 | 0 | 0 | 0 | 0 | 0 | 0 | 0 | 0 | 0 | 0 | 0 | 0 | 0 | 0 | 0 | 0 |
| ENSMUSG00 Gm16394 | 673 | 0 | 0 | 0 | 0 | 0 | 0 | 0 | 0 | 0 | 0 | 0 | 0 | 0 | 0 | 0 | 0 | 0 | 0 |
| ENSMUSG00 Gm5381 | 1247 | 0 | 0 | 0.03879192 | 0 | 0 | 0 | 0 | 0 | 0 | 0 | 0 | 0 | 0 | 0 | 0 | 0 | 0 | 0 |
| ENSMUSG00 Gm12026 | 1023 | 0 | 0 | 0 | 0.1336101 | 0.04704978 | 0 | 0 | 0.03733373 | 0 | 0 | 0 | 0 | 0 | 0 | 0 | 0 | 0 | 0 |
| ENSMUSG00 Gm15422 | 540 | 0 | 0.06169054 | 0 | 0 | 0 | 0.09176432 | 0 | 0 | 0 | 0 | 0 | 0 | 0 | 0 | 0 | 0 | 0 | 0 |
| ENSMUSG00 Olfr602-ps1 | 959 | 0 | 0 | 0 | 0 | 0 | 0 | 0 | 0 | 0 | 0 | 0 | 0 | 0 | 0 | 0 | 0 | 0 | 0 |
| ENSMUSG00 Gm14127 | 676 | 0 | 0.04927943 | 0 | 0 | 0 | 0 | 0 | 0 | 0 | 0 | 0 | 0 | 0 | 0.13740854 | 0 | 0 | 0 | 0 |
| ENSMUSG00 Gm9703 | 342 | 0.86998898 | 0.68184286 | 0.28288614 | 1.99829142 | 0.14073662 | 0.57956414 | 0.830047 | 1.11673691 | 1.08385568 | 0.55147021 | 0.55877144 | 1.14242584 | 1.44245372 | 0.13580143 | 1.56614101 | 0.5598042 | 1.34478235 | 0.96507205 |
| ENSMUSG00 Gm13354 | 368 | 0 | 0 | 0 | 0 | 0 | 0 | 0 | 0 | 0 | 0 | 0 | 0 | 0 | 0 | 0 | 0 | 0 | 0 |
| ENSMUSG00 Gm12988 | 477 | 0.62376569 | 0.13967671 | 0.10141201 | 0.66861071 | 0.5045275 | 0.20776827 | 0.19837601 | 0.40033965 | 0.94979386 | 0.2965453 | 0.30047144 | 0.3510419 | 0.10342121 | 0.19473412 | 0.51825856 | 0.16054762 | 0.88383494 | 0.4612923 |
| ENSMUSG00 Gm15625 | 1092 | 0 | 0 | 0 | 0 | 0 | 0 | 0 | 0 | 0 | 0 | 0 | 0 | 0 | 0 | 0 | 0 | 0 | 0 |
| ENSMUSG00 Gm15181 | 176 | 0 | 0 | 0 | 0 | 0 | 0 | 0 | 0 | 0 | 0 | 0 | 0 | 0 | 0 | 0 | 0 | 0 | 0 |
| ENSMUSG00 Gm9153 | 400 | 0 | 0 | 0 | 0 | 0 | 0 | 0 | 0 | 0 | 0 | 0 | 0 | 0 | 0 | 0 | 0 | 0 | 0 |
| ENSMUSG00 Gm14265 | 304 | 0 | 0 | 0 | 0 | 0 | 0 | 0 | 0 | 0 | 0 | 0 | 0 | 0 | 0 | 0 | 0 | 0 | 0 |
| ENSMUSG00 Gm12616 | 381 | 0 | 0 | 0 | 0 | 0 | 0 | 0 | 0 | 0 | 0 | 0 | 0 | 0 | 0 | 0 | 0 | 0 | 0 |
| ENSMUSG00 Rpl9-ps2 | 576 | 0 | 0 | 0 | 0 | 0 | 0 | 0 | 0 | 0 | 0 | 0 | 0 | 0 | 0 | 0 | 0 | 0 | 0 |
| ENSMUSG00 Gm12156 | 552 | 0 | 0 | 0 | 0 | 0 | 0 | 0 | 0 | 0 | 0 | 0 | 0 | 0 | 0 | 0 | 0 | 0 | 0 |
| ENSMUSG00 Gm6620 | 713 | 0 | 0 | 0 | 0 | 0 | 0 | 0 | 0 | 0 | 0 | 0 | 0 | 0 | 0 | 0 | 0 | 0 | 0 |
| ENSMUSG00 Gm14899 | 547 | 0 | 0 | 0 | 0 | 0 | 0 | 0 | 0 | 0 | 0 | 0 | 0 | 0 | 0 | 0 | 0 | 0 | 0 |
| ENSMUSG00 Gm12614 | 381 | 0 | 0 | 0 | 0 | 0 | 0 | 0 | 0 | 0 | 0 | 0 | 0 | 0 | 0 | 0 | 0 | 0 | 0 |
| ENSMUSG00 Olfr1021-ps1 | 948 | 0 | 0 | 0 | 0 | 0 | 0 | 0 | 0 | 0 | 0 | 0 | 0 | 0 | 0 | 0 | 0 | 0 | 0 |
| ENSMUSG00 Gm8692 | 437 | 0.68086094 | 0.68607791 | 1.43902951 | 0.72981078 | 0.66085021 | 0.45357193 | 1.407471 | 1.31095203 | 0.65973824 | 0.75527442 | 0.32797454 | 0.51089851 | 0.90310146 | 0.74395563 | 0.84854463 | 1.22670138 | 0.70162557 | 0.62939481 |
| ENSMUSG00 Gm14888 | 173 | 0 | 0 | 0 | 0 | 0 | 0 | 0 | 0 | 0 | 0 | 0 | 0 | 0 | 0 | 0 | 0 | 0 | 0 |
| ENSMUSG00 Gm6923 | 405 | 0 | 0 | 0 | 0 | 0 | 0 | 0 | 0 | 0 | 0 | 0 | 0 | 0 | 0 | 0 | 0 | 0 | 0 |
| ENSMUSG00 Gm13024 | 717 | 0 | 0.092923 | 0 | 0 | 0 | 0 | 0 | 0 | 0 | 0 | 0 | 0 | 0 | 0 | 0 | 0 | 0 | 0 |
| ENSMUSG00 Gm14826 | 900 | 0 | 0 | 0 | 0 | 0 | 0 | 0 | 0 | 0 | 0 | 0 | 0 | 0 | 0 | 0 | 0 | 0 | 0 |
| ENSMUSG00 Gm15546 | 825 | 0 | 0 | 0 | 0 | 0 | 0 | 0 | 0 | 0 | 0 | 0 | 0 | 0 | 0 | 0 | 0 | 0 | 0 |
| ENSMUSG00 Gm12366 | 872 | 0.09748894 | 0.11460858 | 0.11094846 | 0.36574233 | 0.16559148 | 0.73874488 | 0.70534957 | 0.13139588 | 0.14169673 | 0.10814381 | 0.21915119 | 0.19202636 | 0 | 0 | 0.18899796 | 0.26346748 | 0.39556958 | 0 |
| ENSMUSG00 Gm12457 | 167 | 0 | 0 | 0 | 0 | 0 | 0 | 0 | 0 | 0 | 0 | 0 | 0 | 0 | 0 | 0 | 0 | 0 | 0 |
| ENSMUSG00 Gm7415 | 631 | 0 | 0 | 0 | 0 | 0 | 0 | 0 | 0 | 0 | 0 | 0 | 0 | 0 | 0 | 0 | 0 | 0 | 0 |
| ENSMUSG00 Gm14411 | 911 | 0.09331542 | 0 | 0 | 0.25006062 | 0.15850249 | 0 | 0.05193488 | 0 | 0.04521023 | 0.51757084 | 0 | 0 | 0.32490835 | 0 | 0.27136041 | 0.0840628 | 0.25242347 | 0 |
| ENSMUSG00 1700064H15I | 392 | 0.32529471 | 0.33992749 | 0.24680373 | 0.11622715 | 0 | 0 | 0.84486927 | 0.1948592 | 0.42027057 | 0 | 0 | 1.56625584 | 0.75508037 | 0.59239907 | 0.52553005 | 1.36752169 | 1.36879632 | 0.98230548 |
| ENSMUSG00 Gm11622 | 690 | 0 | 0 | 0 | 0 | 0 | 0 | 0 | 0 | 0 | 0 | 0 | 0 | 0 | 0 | 0 | 0 | 0 | 0 |
| ENSMUSG00 Gm12727 | 358 | 0.23745908 | 0.18610555 | 0 | 0.25453097 | 0.13444671 | 0 | 0.39647496 | 0.32004806 | 0.11504613 | 0 | 0.53379842 | 0.46772902 | 0 | 0.25946418 | 0.23017629 | 0.64174202 | 0.32117009 | 0 |
| ENSMUSG00 Gm15793 | 655 | 0 | 0.10171876 | 0 | 0.06955885 | 0.07348385 | 0 | 0 | 0.05830901 | 0.06288018 | 0 | 0 | 0 | 0 | 0.070907 | 0 | 0.11691789 | 0 | 0.16796674 |
| ENSMUSG00 Gm14722 | 688 | 0 | 0 | 0 | 0 | 0 | 0 | 0 | 0 | 0 | 0 | 0 | 0 | 0 | 0 | 0 | 0 | 0 | 0 |
| ENSMUSG00 Gm12803 | 819 | 0 | 0 | 0 | 0 | 0 | 0 | 0 | 0 | 0 | 0 | 0 | 0 | 0 | 0 | 0 | 0 | 0 | 0 |
| ENSMUSG00 Gm381 | 2036 | 0 | 0 | 0 | 0 | 0 | 0 | 0 | 0 | 0 | 0 | 0 | 0 | 0 | 0 | 0 | 0 | 0 | 0 |
| ENSMUSG00 Gm15058 | 457 | 0 | 0 | 0 | 0 | 0 | 0 | 0 | 0 | 0 | 0 | 0 | 0 | 0 | 0 | 0 | 0 | 0 | 0 |
| ENSMUSG00 Gm8805 | 312 | 0 | 0.32031629 | 0 | 0.14602899 | 0.30853797 | 0 | 0 | 0 | 0 | 0 | 0 | 0 | 0 | 0 | 0.13205627 | 0 | 0 | 0.35262248 |
| ENSMUSG00 Itpa-ps3 | 394 | 0 | 0 | 0 | 0 | 0 | 0 | 0 | 0 | 0 | 0 | 0 | 0 | 0 | 0 | 0 | 0 | 0 | 0 |
| ENSMUSG00 Gm7061 | 1415 | 0.06007799 | 0 | 0 | 0 | 0 | 0 | 0.20061913 | 0 | 0.14553539 | 0 | 0 | 0.0394457 | 0 | 0 | 0.02911771 | 0 | 0.10834289 | 0 |
| ENSMUSG00 Gm15861 | 338 | 0 | 0 | 0 | 0 | 0 | 0 | 0 | 0.11299527 | 0 | 0 | 0 | 0 | 0 | 0 | 0 | 0 | 0 | 0 |
| ENSMUSG00 Gm15191 | 999 | 0.25528634 | 0.1667312 | 0.14526586 | 0.04560665 | 0.09636021 | 0.09920467 | 0.23680019 | 0.1146919 | 0.45350518 | 0.37758321 | 0.28693669 | 0 | 0.69133818 | 0.37192462 | 0.20621399 | 0.19164468 | 0.38364662 | 0.22025668 |
| ENSMUSG00 Gm13162 | 452 | 0 | 0 | 0 | 0 | 0 | 0 | 0 | 0 | 0 | 0 | 0 | 0 | 0 | 0 | 0 | 0 | 0 | 0 |
| ENSMUSG00 Gm8329 | 453 | 0 | 0 | 0 | 0 | 0 | 0 | 0 | 0 | 0 | 0 | 0 | 0 | 0 | 0 | 0 | 0 | 0 | 0 |
| ENSMUSG00 Gm15052 | 268 | 0 | 0 | 0 | 0 | 0 | 0 | 0 | 0 | 0 | 0 | 0 | 0 | 0 | 0 | 0 | 0 | 0 | 0 |
| ENSMUSG00 Gm5319 | 872 | 0 | 0.03820286 | 0 | 0 | 0 | 0 | 0 | 0 | 0 | 0 | 0.0547878 | 0.12801757 | 0 | 0 | 0.04724949 | 0 | 0 | 0 |
| ENSMUSG00 Gm9126 | 412 | 0 | 0 | 0 | 0 | 0 | 0 | 0 | 0 | 0 | 0 | 0 | 0 | 0 | 0 | 0 | 0 | 0 | 0 |
| ENSMUSG00 Gm15957 | 973 | 0 | 0 | 0 | 0 | 0 | 0 | 0 | 0 | 0 | 0 | 0 | 0 | 0 | 0.04773288 | 0 | 0 | 0 | 0 |
| ENSMUSG00 Gm11296 | 455 | 0 | 0 | 0 | 0 | 0 | 0 | 0 | 0 | 0 | 0 | 0 | 0 | 0 | 0 | 0 | 0 | 0 | 0 |
| ENSMUSG00 Gm14489 | 900 | 0 | 0 | 0 | 0 | 0 | 0 | 0 | 0 | 0 | 0 | 0 | 0 | 0 | 0 | 0 | 0 | 0 | 0 |
| ENSMUSG00 Gm13396 | 270 | 0 | 0 | 0 | 0 | 0 | 0 | 0 | 0 | 0 | 0 | 0 | 0 | 0 | 0 | 0 | 0 | 0 | 0 |
| ENSMUSG00 Gm13284 | 545 | 0.46794689 | 0 | 0.08875877 | 0.08359825 | 0.08831546 | 0.09092245 | 0.08681225 | 0 | 0.30228635 | 0.1730301 | 0 | 0.20482812 | 0 | 0 | 0.22679756 | 0.21077399 | 0 | 0.10093414 |
| ENSMUSG00 Gm12155 | 159 | 0 | 0 | 0 | 0 | 0 | 0 | 0 | 0 | 0 | 0 | 0 | 0 | 0 | 0 | 0 | 0 | 0 | 0 |
| ENSMUSG00 Gm4875 | 785 | 3.89856392 | 4.11637037 | 5.17627586 | 4.29492647 | 3.00441307 | 4.22934159 | 2.47110807 | 4.08683032 | 3.04308015 | 3.36361705 | 4.56448648 | 2.48859642 | 1.38255055 | 2.07075549 | 3.09667745 | 2.58522572 | 2.63645864 | 2.52271063 |
| ENSMUSG00 Gm12176 | 874 | 0.09726585 | 0.34303896 | 0.22138915 | 0.41703473 | 0.93620446 | 0.17008947 | 0.0541335 | 0.30588881 | 0.09424832 | 0.64737808 | 0.38263697 | 1.14952166 | 0.16933152 | 1.48791127 | 0.23570684 | 0.13143229 | 0.26310959 | 0.50351585 |
| ENSMUSG00 Gm9308 | 213 | 0.19955482 | 0 | 0 | 0 | 0.22597147 | 0 | 0 | 0.53792116 | 0 | 0 | 0 | 0 | 0.23160525 | 0 | 0 | 0.17976811 | 0.35987133 | 0 |
| ENSMUSG00 Gm6806 | 1005 | 0 | 0 | 0 | 0 | 0 | 0 | 0 | 0 | 0 | 0 | 0 | 0 | 0 | 0 | 0 | 0 | 0 | 0 |
| ENSMUSG00 Gm13193 | 224 | 0 | 0 | 0 | 0 | 0 | 0 | 0 | 0 | 0 | 0 | 0 | 0 | 0 | 0 | 0 | 0 | 0 | 0 |
| ENSMUSG00 Gm13385 | 219 | 0 | 0 | 0 | 0 | 0 | 0 | 0 | 0 | 0 | 0 | 0 | 1.01946416 | 0 | 0 | 0 | 0 | 0 | 0 |
| ENSMUSG00 Gm12015 | 1261 | 0 | 0 | 0 | 0 | 0 | 0 | 0 | 0 | 0 | 0 | 0 | 0 | 0 | 0 | 0 | 0 | 0 | 0 |
| ENSMUSG00 Gm11561 | 416 | 0 | 0 | 0 | 0 | 0 | 0 | 0 | 0 | 0 | 0 | 0 | 0 | 0 | 0 | 0 | 0 | 0 | 0 |
| ENSMUSG00 Gm14554 | 652 | 0 | 0 | 0 | 0 | 0 | 0 | 0 | 0 | 0 | 0 | 0 | 0 | 0 | 0 | 0 | 0 | 0 | 0 |
| ENSMUSG00 Gm13560 | 328 | 0 | 0 | 0 | 0 | 0 | 0 | 0 | 0 | 0 | 0 | 0.14565536 | 0.51050911 | 0 | 0.14159783 | 0 | 0 | 0 | 0 |
| ENSMUSG00 Gm15465 | 124 | 0 | 0 | 0 | 0 | 0 | 0 | 0 | 0 | 0 | 0 | 0 | 0 | 0 | 0 | 0 | 0 | 0 | 0 |
| ENSMUSG00 Gm14423 | 120 | 0 | 0 | 0 | 0 | 0 | 0 | 0 | 0 | 0 | 0 | 0 | 0 | 0 | 0 | 0 | 0 | 0 | 0 |
| ENSMUSG00 Rpl17-ps1 | 1129 | 0 | 0 | 0 | 0 | 0 | 0 | 0 | 0 | 0 | 0 | 0 | 0.09687628 | 0 | 0 | 0 | 0 | 0 | 0 |
| ENSMUSG00 Gm11395 | 497 | 0 | 0 | 0 | 0 | 0 | 0 | 0 | 0 | 0 | 0 | 0 | 0 | 0 | 0 | 0 | 0 | 0 | 0 |
| ENSMUSG00 Rpl13-ps1 | 667 | 0.70098491 | 0.44949932 | 0.4351442 | 0.88799637 | 0.72161805 | 0.22287586 | 0.28373421 | 0.51533976 | 0.1852467 | 0.7775978 | 0.35813312 | 0.41840827 | 0.44376537 | 0.69631315 | 0.06177145 | 0.22962883 | 0.28730357 | 1.23708636 |
| ENSMUSG00 Gm14782 | 194 | 0 | 0 | 0 | 0 | 0 | 0 | 0 | 0 | 0 | 0 | 0 | 0 | 0 | 0 | 0 | 0 | 0 | 0 |
| ENSMUSG00 Gm11391 | 232 | 0 | 0 | 0 | 0 | 0 | 0 | 0 | 0 | 0 | 0.20323579 | 0 | 0 | 0 | 0 | 0 | 0 | 0 | 0 |
| ENSMUSG00 Gm14150 | 1649 | 0.5928557 | 0.26262439 | 0.2933507 | 0.30392449 | 0.58377106 | 0.30050172 | 0.37299262 | 0.4863799 | 0.49953324 | 0.4289027 | 0.11588832 | 0.3046337 | 0.35899515 | 0.45064002 | 0.44974409 | 0.39474853 | 0.4648429 | 0.30023163 |
| ENSMUSG00 Gm8682 | 497 | 0 | 0 | 0 | 0.09167212 | 0.19368983 | 0 | 0 | 0 | 0 | 0 | 0 | 0 | 0 | 0 | 0 | 0 | 0 | 0 |
| ENSMUSG00 Gm14860 | 428 | 0 | 0 | 0 | 0 | 0 | 0 | 0 | 0 | 0 | 0 | 0 | 0 | 0 | 0 | 0 | 0 | 0 | 0 |
| ENSMUSG00 Mup-ps6 | 539 | 0 | 0 | 0 | 0 | 0 | 0 | 0 | 0 | 0 | 0 | 0 | 0 | 0 | 0 | 0 | 0 | 0 | 0 |
| ENSMUSG00 Gm12956 | 413 | 0 | 0 | 0 | 0 | 0 | 0 | 0 | 0 | 0 | 0 | 0.11567787 | 0 | 0 | 0 | 0 | 0 | 0 | 0 |
| ENSMUSG00 Gm5678 | 831 | 0.05114943 | 0 | 0.17463368 | 0.1644803 | 0 | 0.05963024 | 0.11386926 | 0.09191914 | 0.14868778 | 0.05673972 | 0.11498185 | 0 | 0.65300974 | 0 | 0.09916139 | 0 | 0.04612069 | 0 |
| ENSMUSG00 Gm11953 | 1002 | 1.10292872 | 0.39895682 | 0.04827698 | 0.13641031 | 0.43232267 | 0.54399209 | 0.70827364 | 0.60985872 | 0.41104307 | 0.4705659 | 0.52447559 | 0.50133829 | 0.73850176 | 0.23175692 | 0.61678976 | 0.38214179 | 0.26774858 | 0.54899308 |
| ENSMUSG00 Gm16428 | 714 | 0 | 0 | 0 | 0 | 0 | 0 | 0 | 0 | 0 | 0 | 0 | 0 | 0 | 0 | 0 | 0 | 0 | 0 |
| ENSMUSG00 Gm12175 | 344 | 0.12356156 | 0 | 0.14062073 | 0 | 0 | 0 | 0 | 0 | 0 | 0.137066 | 0 | 0 | 0 | 0 | 0.77916934 | 0 | 0 | 0 |
| ENSMUSG00 Trav4d-2 | 356 | 0 | 0 | 0 | 0 | 0 | 0 | 0 | 0 | 0 | 0 | 0 | 0 | 0 | 0 | 0 | 0 | 0 | 0 |
| ENSMUSG00 Gm13803 | 805 | 0 | 0.16552991 | 0 | 0.05659757 | 0.05979121 | 0.36933714 | 0.23509406 | 0.09488796 | 0.20465349 | 0.0585723 | 0 | 0.06933623 | 0.30640942 | 0.05769452 | 0 | 0.09513194 | 0.23805153 | 0.20500288 |
| ENSMUSG00 Gm12880 | 491 | 0 | 0 | 0 | 0 | 0 | 0 | 0 | 0 | 0 | 0 | 0 | 0 | 0 | 0 | 0 | 0 | 0 | 0 |
| ENSMUSG00 Gm14731 | 375 | 0 | 0 | 0 | 0 | 0 | 0 | 0 | 0 | 0 | 0 | 0 | 0 | 0 | 0.3715527 | 0 | 0 | 0 | 0 |
| ENSMUSG00 Gm15828 | 521 | 0 | 0 | 0 | 0 | 0 | 0 | 0 | 0 | 0 | 0 | 0 | 0 | 0 | 0 | 0 | 0 | 0 | 0 |
| ENSMUSG00 Gm14716 | 717 | 0 | 0 | 0 | 0 | 0 | 0 | 0 | 0 | 0 | 0 | 0 | 0 | 0 | 0 | 0 | 0 | 0 | 0 |
| ENSMUSG00 Gm15741 | 753 | 0 | 0 | 0 | 0 | 0 | 0 | 0 | 0 | 0 | 0 | 0 | 0 | 0 | 0 | 0 | 0 | 0 | 0 |
| ENSMUSG00 Gm6097 | 344 | 0 | 0 | 0.84372437 | 0.26488979 | 0.41975515 | 0.14404864 | 0.68768429 | 0.55512213 | 0.59864122 | 0.41119799 | 0.27776139 | 0.64901933 | 0.43022021 | 0.13501188 | 0.23954393 | 0.55654953 | 0.44565462 | 0.63964078 |
| ENSMUSG00 Gm5397 | 1451 | 0 | 0 | 0 | 0 | 0 | 0 | 0 | 0 | 0 | 0 | 0 | 0 | 0 | 0 | 0 | 0 | 0 | 0 |
| ENSMUSG00 Gm16106 | 293 | 0 | 0 | 0 | 0 | 0 | 0 | 0 | 0.1303495 | 0 | 0 | 0 | 0 | 0 | 0 | 0 | 0 | 0 | 0 |
| ENSMUSG00 Gm15780 | 934 | 0 | 0 | 0 | 0 | 0.05153311 | 0 | 0 | 0.5315859 | 0 | 0.20193021 | 0.20460368 | 0.23903924 | 0.26408949 | 0 | 0.08822603 | 0.28697457 | 0.24620747 | 0 |
| ENSMUSG00 Gm12591 | 1174 | 0 | 0 | 0 | 0 | 0 | 0 | 0 | 0 | 0 | 0 | 0 | 0 | 0 | 0 | 0.07019004 | 0 | 0 | 0 |
| ENSMUSG00 Gm14786 | 942 | 0 | 0 | 0 | 0 | 0 | 0 | 0 | 0 | 0 | 0 | 0 | 0 | 0 | 0 | 0 | 0 | 0 | 0 |
| ENSMUSG00 Gm12982 | 189 | 0 | 0 | 0 | 0 | 0 | 0 | 0 | 0 | 0 | 0 | 0 | 0 | 0 | 0 | 0 | 0 | 0.20278464 | 0 |
| ENSMUSG00 Defa-ps1 | 280 | 0 | 0 | 0 | 0 | 0 | 0 | 0 | 0 | 0 | 0 | 0 | 0 | 0 | 0 | 0 | 0 | 0 | 0 |
| ENSMUSG00 Gm13118 | 548 | 0 | 0 | 0 | 0 | 0 | 0 | 0 | 0 | 0 | 0 | 0 | 0 | 0 | 0 | 0 | 0 | 0 | 0 |
| ENSMUSG00 Gm13612 | 389 | 0 | 0.08563726 | 0.12435355 | 0.23424701 | 0 | 0 | 0.36487927 | 0.09818098 | 0 | 0 | 0.1228148 | 0 | 0.25363454 | 0 | 0 | 0.29530031 | 0.09852518 | 0 |
| ENSMUSG00 Gm15373 | 213 | 0 | 0 | 0 | 0 | 0 | 0 | 0 | 0 | 0 | 0 | 0 | 0 | 0 | 0 | 0 | 0 | 0 | 0 |
| ENSMUSG00 Gm14519 | 311 | 0.13667259 | 0 | 0 | 0 | 0.77382514 | 0.3186671 | 0.1521308 | 0.49122061 | 0 | 0.30321996 | 0.15361723 | 0.17947158 | 0.47587058 | 0.2986758 | 0.39744266 | 0.86184647 | 0.24647136 | 0.35375631 |
| ENSMUSG00 Gm14588 | 171 | 0 | 0 | 0 | 0 | 0 | 0 | 0 | 0 | 0 | 0 | 0 | 0 | 0 | 0 | 0 | 0 | 0 | 0 |
| ENSMUSG00 Gm16421 | 1556 | 0 | 0 | 0 | 0 | 0 | 0 | 0 | 0 | 0 | 0 | 0 | 0 | 0 | 0 | 0 | 0 | 0.0246313 | 0.0353529 |
| ENSMUSG00 Gm12994 | 446 | 0.09530309 | 0.14938518 | 0.10846083 | 0 | 0 | 0.11110478 | 0 | 0.51379914 | 0.1846929 | 0 | 0.10711874 | 0.12514723 | 0 | 0.20826945 | 0.18476034 | 0 | 0.3437336 | 0 |
| ENSMUSG00 Gm14748 | 408 | 0 | 0 | 0 | 0 | 0 | 0 | 0 | 0 | 0 | 0 | 0 | 0 | 0 | 0 | 0 | 0 | 0 | 0 |
| ENSMUSG00 4930448K20I | 997 | 0 | 0 | 0.04851909 | 0 | 0 | 0 | 0 | 0 | 0 | 0 | 0 | 0.05598361 | 0 | 0 | 0.20682766 | 0 | 0 | 0 |
| ENSMUSG00 Gm13537 | 266 | 0 | 0 | 0 | 0 | 0.36189416 | 0 | 0 | 0 | 0 | 0 | 0 | 0 | 0 | 0.17460183 | 0 | 0 | 0 | 0 |
| ENSMUSG00 Gm13879 | 241 | 0 | 0 | 0 | 0 | 0 | 0 | 0 | 0 | 0 | 0 | 0 | 0 | 0 | 0 | 0 | 0 | 0 | 0 |
| ENSMUSG00 Gm13484 | 411 | 0 | 0 | 0 | 0 | 0 | 0 | 0 | 0 | 0 | 0 | 0 | 0 | 0 | 0 | 0 | 0 | 0 | 0 |
| ENSMUSG00 Gm5759 | 810 | 0 | 0 | 0 | 0 | 0 | 0 | 0 | 0 | 0 | 0 | 0 | 0 | 0 | 0 | 0 | 0 | 0 | 0 |
| ENSMUSG00 Gm16066 | 1083 | 0.03924762 | 0.06151966 | 0 | 0.12620788 | 0.04444314 | 0 | 0.08737337 | 0.21159226 | 0.11409007 | 0.21768561 | 0.22056768 | 0.20615203 | 0.13665351 | 0.30019262 | 0.22826347 | 0.74247715 | 0.17694505 | 0.25396633 |
| ENSMUSG00 Gm14053 | 469 | 0.09062937 | 0 | 0 | 0 | 0 | 0 | 0 | 0.16286739 | 0 | 0 | 0 | 0.11900994 | 0 | 0 | 0 | 0.08164309 | 0 | 0 |
| ENSMUSG00 Gm14299 | 396 | 0 | 0.08412347 | 0 | 0 | 0 | 0 | 0 | 0 | 0 | 0 | 0 | 0 | 0.37372665 | 0 | 0 | 0 | 0 | 0 |
| ENSMUSG00 Gm11931 | 446 | 0 | 0 | 0 | 0 | 0 | 0 | 0 | 0 | 0 | 0 | 0 | 0 | 0 | 0 | 0 | 0 | 0 | 0 |
| ENSMUSG00 Gm12021 | 1903 | 0.02233588 | 0 | 0 | 0 | 0 | 0 | 0 | 0 | 0 | 0 | 0 | 0 | 0 | 0 | 0 | 0 | 0 | 0 |
| ENSMUSG00 Gm7134 | 2405 | 0 | 0 | 0 | 0 | 0 | 0 | 0 | 0 | 0 | 0 | 0 | 0 | 0 | 0 | 0 | 0 | 0 | 0 |
| ENSMUSG00 Gm14537 | 597 | 0.1423959 | 0 | 0 | 0 | 0 | 0 | 0.15850144 | 0.38384324 | 0.20696742 | 0 | 0.16005011 | 0 | 0 | 0.07779579 | 0.06901433 | 0 | 0 | 0 |
| ENSMUSG00 Gm15106 | 819 | 0 | 0 | 0 | 0 | 0 | 0 | 0 | 0 | 0 | 0 | 0 | 0 | 0 | 0 | 0 | 0 | 0 | 0 |
| ENSMUSG00 Gm12293 | 589 | 0 | 0 | 0 | 0 | 0 | 0 | 0 | 0 | 0 | 0 | 0 | 0 | 0 | 0 | 0 | 0 | 0 | 0 |
| ENSMUSG00 Gm11746 | 820 | 0 | 0 | 0 | 0 | 0 | 0 | 0 | 0 | 0 | 0 | 0 | 0 | 0 | 0 | 0 | 0 | 0 | 0 |
| ENSMUSG00 Defa-ps9 | 274 | 0 | 0 | 0 | 0 | 0 | 0 | 0 | 0 | 0 | 0 | 0 | 0 | 0 | 0 | 0 | 0.41924023 | 0 | 0 |
| ENSMUSG00 Gm13204 | 820 | 1.19221835 | 1.9093976 | 0.88488165 | 1.16680723 | 1.99571391 | 2.53806684 | 1.32706295 | 1.53701132 | 2.10955325 | 1.09251629 | 1.16524289 | 1.42942551 | 1.32353924 | 3.2850696 | 2.31130679 | 1.82113864 | 1.16848466 | 0.67084276 |
| ENSMUSG00 Gm11362 | 1478 | 0.4888958 | 0 | 0 | 0 | 0.06513116 | 0 | 0 | 0 | 0 | 0 | 0.06464812 | 0.03776432 | 0.20026489 | 0 | 0 | 0.05181408 | 0 | 0.03721861 |
| ENSMUSG00 Gm13806 | 324 | 0 | 0 | 0 | 0 | 0 | 0 | 0 | 0 | 0 | 0 | 0 | 0.17227056 | 0 | 0 | 0 | 0 | 0 | 0 |
| ENSMUSG00 Gm14674 | 341 | 0 | 0 | 0 | 0 | 0 | 0 | 0 | 0 | 0 | 0 | 0 | 0 | 0 | 0.68099835 | 0 | 0 | 0 | 0 |
| ENSMUSG00 Gm13358 | 568 | 0 | 0 | 0 | 0 | 0 | 0 | 0 | 0 | 0 | 0 | 0 | 0 | 0 | 0 | 0 | 0 | 0 | 0 |
| ENSMUSG00 Gm14745 | 1095 | 0 | 0 | 0 | 0 | 0 | 0 | 0 | 0 | 0 | 0 | 0 | 0 | 0 | 0 | 0 | 0 | 0 | 0 |
| ENSMUSG00 Gm15730 | 414 | 0 | 0 | 0 | 0 | 0 | 0 | 0 | 0 | 0 | 0 | 0 | 0 | 0 | 0 | 0 | 0.18497878 | 0.27772679 | 0 |
| ENSMUSG00 Mup-ps5 | 98 | 0 | 0 | 0 | 0 | 0 | 0 | 0 | 0 | 0 | 0 | 0 | 0 | 0 | 0 | 0 | 0 | 0 | 0 |
| ENSMUSG00 Gm11293 | 431 | 0 | 0 | 0 | 0 | 0 | 0 | 0 | 0 | 0 | 0 | 0 | 0 | 0 | 0 | 0 | 0 | 0 | 0 |
| ENSMUSG00 Gm12165 | 341 | 0 | 0.29307532 | 0 | 0 | 0.14114934 | 0.14531593 | 0.27749372 | 0.44800472 | 0.12078157 | 0.13827186 | 0.28020504 | 0 | 0 | 0.13619967 | 0 | 0.11228917 | 0.11239383 | 0 |
| ENSMUSG00 Gm13263 | 795 | 0 | 0 | 0 | 0 | 0 | 0 | 0 | 0 | 0 | 0 | 0 | 0 | 0 | 0 | 0 | 0 | 0 | 0 |
| ENSMUSG00 Gm15773 | 1447 | 0.05874938 | 0.02302204 | 0.06686044 | 0.09445966 | 0.0665265 | 0 | 0.03269708 | 0.07918259 | 0 | 0.06517029 | 0.03301656 | 0 | 0.03409255 | 0 | 0 | 0.07938619 | 0.05297346 | 0.03801597 |
| ENSMUSG00 Gm15367 | 1015 | 0 | 0 | 0 | 0 | 0 | 0 | 0 | 0 | 0 | 0 | 0 | 0 | 0 | 0 | 0 | 0 | 0 | 0 |

| | | | | | | | | | | | | | | | | | | | | |
|---|---|---|---|---|---|---|---|---|---|---|---|---|---|---|---|---|---|---|---|---|
| ENSMUSG00 | Rpl15-ps1 | 563 | 0 | 0.05917033 | 0 | 0 | 0 | 0.08801551 | 0 | 0 | 0 | 0 | 0 | 0 | 0 | 0 | 0.14636432 | 0.06801174 | 0.06807513 | 0 |
| ENSMUSG00 | Gm12598 | 829 | 0 | 0 | 0 | 0 | 0 | 0.23909642 | 0 | 0 | 0 | 0 | 0 | 0 | 0 | 0 | 0 | 0 | 0 | 0 |
| ENSMUSG00 | Gm11889 | 1078 | 0 | 0 | 0 | 0.04226442 | 0 | 0 | 0 | 0 | 0 | 0 | 0 | 0 | 0 | 0 | 0 | 0 | 0 | 0 |
| ENSMUSG00 | Gm13009 | 265 | 0 | 0 | 0 | 0 | 0 | 0 | 0 | 0 | 0 | 0 | 0 | 0 | 0 | 0 | 0 | 0 | 0 | 0 |
| ENSMUSG00 | Gm12583 | 852 | 0 | 0.03909964 | 0 | 0 | 0 | 0 | 0 | 0.04482676 | 0 | 0 | 0 | 0 | 0 | 0 | 0 | 0 | 0 | 0 |
| ENSMUSG00 | Gm13525 | 477 | 0.17821877 | 0.55870682 | 0.20282403 | 0 | 0 | 0.8310731 | 0 | 0.72061137 | 0.34537959 | 0.69193904 | 0.60094287 | 0.70208381 | 0.41368484 | 0 | 0 | 0.48164286 | 0.08034863 | 0 |
| ENSMUSG00 | Gm14789 | 227 | 0 | 0 | 0 | 0 | 0 | 0 | 0 | 0 | 0 | 0 | 0 | 0 | 0 | 0 | 0 | 0 | 0 | 0 |
| ENSMUSG00 | Olfr1185-ps1 | 911 | 0 | 0 | 0 | 0 | 0 | 0 | 0 | 0 | 0 | 0 | 0 | 0 | 0 | 0 | 0 | 0 | 0 | 0 |
| ENSMUSG00 | Olfr1293-ps | 1825 | 0 | 0 | 0 | 0 | 0 | 0 | 0 | 0 | 0 | 0 | 0 | 0 | 0 | 0 | 0 | 0 | 0 | 0 |
| ENSMUSG00 | Gm13339 | 509 | 0.91857944 | 0.52358183 | 0 | 0.35804357 | 0.37824694 | 0.09735311 | 1.48723549 | 0.37517095 | 0.56641574 | 0.27790198 | 0.657023 | 1.0965749 | 0.19383857 | 0.36498301 | 0.89040691 | 0.52658989 | 0.37648622 | 0.75651031 |
| ENSMUSG00 | Olfr475-ps1 | 931 | 0 | 0 | 0 | 0 | 0 | 0 | 0 | 0 | 0 | 0 | 0 | 0 | 0 | 0 | 0 | 0 | 0 | 0 |
| ENSMUSG00 | Gm5386 | 2449 | 0 | 0 | 0 | 0 | 0 | 0 | 0 | 0 | 0 | 0 | 0 | 0 | 0 | 0 | 0 | 0 | 0 | 0 |
| ENSMUSG00 | Gm14607 | 490 | 0 | 0 | 0 | 0 | 0 | 0 | 0 | 0 | 0 | 0 | 0 | 0 | 0 | 0 | 0 | 0 | 0 | 0 |
| ENSMUSG00 | Gm12934 | 342 | 0.24856828 | 0.09740612 | 0.56577228 | 0 | 0.84441971 | 0.4346731 | 0 | 0.67004215 | 0 | 0.13786755 | 0 | 0 | 0.72122686 | 1.76541853 | 0 | 0.11196084 | 0.22413039 | 0 |
| ENSMUSG00 | Gm12831 | 169 | 0 | 0 | 0 | 0 | 0 | 0 | 0 | 0 | 0 | 0 | 0 | 0 | 0 | 0 | 0 | 0 | 0 | 0 |
| ENSMUSG00 | Gm15014 | 257 | 0 | 0 | 0 | 0 | 0 | 0 | 0 | 0 | 0 | 0 | 0 | 0 | 0 | 0 | 0 | 0 | 0 | 0 |
| ENSMUSG00 | Gm12881 | 201 | 0.21146854 | 0 | 0 | 0 | 0 | 0 | 0 | 0 | 0 | 0 | 0 | 0 | 0 | 0 | 0 | 0.19050053 | 0 | 0 |
| ENSMUSG00 | Olfr1027-ps1 | 1341 | 0 | 0 | 0 | 0 | 0 | 0 | 0 | 0 | 0 | 0 | 0 | 0 | 0 | 0 | 0 | 0 | 0 | 0 |
| ENSMUSG00 | Gm11383 | 806 | 0.05273595 | 0.04133113 | 0 | 0 | 0.17915108 | 0 | 0.05870059 | 0 | 0.0510999 | 0 | 0.35564485 | 0 | 0.18361756 | 0.11524587 | 0.10223711 | 0 | 0.09510247 | 0 |
| ENSMUSG00 | Gm6480 | 1885 | 0 | 0 | 0 | 0 | 0 | 0 | 0 | 0 | 0 | 0 | 0 | 0 | 0 | 0 | 0 | 0 | 0 | 0 |
| ENSMUSG00 | Rpsa-ps9 | 884 | 0.09616556 | 0 | 0.21888475 | 0.05153964 | 0 | 0 | 0.05352113 | 0.12961223 | 0 | 0 | 0 | 0 | 0.22322135 | 0.10507712 | 0.09321619 | 0 | 0 | 0.18668249 |
| ENSMUSG00 | Gm5844 | 2190 | 0.23290507 | 0.15211367 | 0.2208837 | 0.2080413 | 0.26373657 | 0.24889501 | 0.15122774 | 0.22671289 | 0.01880663 | 0.08612001 | 0.10907525 | 0.15291962 | 0.29283787 | 0.16965877 | 0.18813496 | 0.27974873 | 0.21000711 | 0.1758282 |
| ENSMUSG00 | Gm11982 | 313 | 0.13579928 | 0 | 0 | 0 | 0 | 0 | 0 | 0.12202045 | 0.65793156 | 0 | 0 | 0 | 0.47282988 | 0.445151 | 0 | 0 | 0 | 0 |
| ENSMUSG00 | Gm12041 | 601 | 0 | 0 | 0 | 0 | 0 | 0 | 0 | 0 | 0 | 0 | 0 | 0 | 0 | 0 | 0 | 0 | 0 | 0 |
| ENSMUSG00 | Gm12379 | 389 | 0 | 0 | 0 | 0 | 0 | 0 | 0 | 0 | 0 | 0 | 0 | 0 | 0 | 0 | 0 | 0 | 0 | 0 |
| ENSMUSG00 | Gm11557 | 1007 | 0.12662912 | 0.16540662 | 0.04803727 | 0.09048867 | 0.28678405 | 0.04920828 | 0.18793517 | 0.15170766 | 0.16360086 | 0.09364589 | 0.42698573 | 0.27713834 | 0.58786793 | 0.36896991 | 0.20457575 | 0.03802444 | 0.41865866 | 0.16388016 |
| ENSMUSG00 | Gm14323 | 626 | 0 | 0 | 0 | 0 | 0.07688806 | 0 | 0 | 0 | 0 | 0 | 0 | 0.0891624 | 0 | 0 | 0 | 0.0611671 | 0.24489647 | 0 |
| ENSMUSG00 | Ccl19-ps1 | 323 | 0 | 0 | 0 | 0 | 0 | 0 | 0 | 0 | 0 | 0 | 0 | 0 | 0 | 0 | 0 | 0 | 0 | 0 |
| ENSMUSG00 | Gm14083 | 619 | 0 | 0 | 0 | 0 | 0 | 0 | 0 | 0 | 0 | 0 | 0 | 0 | 0 | 0 | 0 | 0 | 0 | 0 |
| ENSMUSG00 | Gm12615 | 381 | 0 | 0 | 0 | 0 | 0 | 0 | 0 | 0 | 0 | 0 | 0 | 0 | 0 | 0 | 0 | 0 | 0 | 0 |
| ENSMUSG00 | Gm7823 | 1622 | 0 | 0 | 0 | 0 | 0 | 0 | 0 | 0 | 0 | 0 | 0 | 0 | 0 | 0 | 0.0508034 | 0 | 0 | 0 |
| ENSMUSG00 | Smt3h1-ps | 279 | 0 | 0.11940105 | 0 | 0 | 0 | 0 | 0 | 0 | 0 | 0 | 0 | 0 | 0 | 0 | 0 | 0 | 0.13737024 | 0 |
| ENSMUSG00 | Gm14832 | 321 | 0 | 0 | 0 | 0 | 0 | 0 | 0 | 0 | 0 | 0 | 0 | 0 | 0 | 0 | 0 | 0 | 0 | 0 |
| ENSMUSG00 | Gm13736 | 512 | 31.878882 | 14.4442627 | 20.5020627 | 16.1065411 | 30.4584829 | 28.8412395 | 35.8541395 | 19.9167411 | 29.6832506 | 19.8917029 | 27.9931397 | 36.6290286 | 37.6733978 | 20.5007104 | 27.6018236 | 18.2478676 | 31.5892525 | 35.0253297 |
| ENSMUSG00 | Gm12398 | 306 | 0 | 0 | 0 | 0 | 0 | 0 | 0 | 0 | 0 | 0 | 0 | 0 | 0 | 0 | 0 | 0 | 0 | 0.17976832 |
| ENSMUSG00 | Gm14758 | 137 | 0 | 0 | 0 | 0 | 0 | 0 | 0 | 0 | 0 | 0 | 0 | 0 | 0 | 0 | 0 | 0 | 0 | 0 |
| ENSMUSG00 | Gm14119 | 234 | 0 | 0 | 0 | 0 | 0.20569198 | 0 | 0 | 0 | 0 | 0 | 0 | 0 | 0 | 0 | 0.17607503 | 0 | 0.16378759 | 0 |
| ENSMUSG00 | Gm8770 | 1108 | 0 | 0 | 0 | 0 | 0 | 0 | 0 | 0 | 0 | 0 | 0 | 0 | 0 | 0 | 0 | 0 | 0 | 0.04964721 |
| ENSMUSG00 | Gm14739 | 801 | 0 | 0 | 0 | 0 | 0 | 0 | 0 | 0 | 0 | 0 | 0 | 0 | 0 | 0 | 0 | 0 | 0 | 0 |
| ENSMUSG00 | Gm14527 | 631 | 0 | 0 | 0 | 0 | 0 | 0 | 0 | 0 | 0 | 0 | 0 | 0 | 0 | 0 | 0 | 0 | 0 | 0 |
| ENSMUSG00 | Gm15240 | 719 | 0 | 0 | 0 | 0 | 0 | 0 | 0 | 0 | 0 | 0 | 0 | 0 | 0.61750661 | 0 | 0 | 0 | 0 | 0 |
| ENSMUSG00 | Gm11440 | 526 | 0.08080832 | 0 | 0.09196489 | 0 | 0 | 0 | 0 | 0.29043652 | 0 | 0 | 0 | 0 | 0 | 0 | 0 | 0 | 0 | 0 |
| ENSMUSG00 | Defa-ps16 | 140 | 0 | 0 | 0 | 0 | 0 | 0 | 0 | 0 | 0 | 0 | 0 | 0 | 0 | 0 | 0 | 0 | 0 | 0 |
| ENSMUSG00 | Gm12641 | 983 | 0 | 0 | 0 | 0 | 0 | 0 | 0.09626181 | 0 | 0.0418988 | 0 | 0 | 0 | 0 | 0 | 0 | 0 | 0.03898911 | 0 |
| ENSMUSG00 | Gm13864 | 784 | 0 | 0 | 0 | 0 | 0 | 0.06320502 | 0 | 0.0487148 | 0.15760146 | 0 | 0 | 0.07119345 | 0.06292336 | 0 | 0.15765901 | 0.14652018 | 0 | 0 |
| ENSMUSG00 | Gm6939 | 858 | 0.84217715 | 0.54356704 | 0.56379406 | 0.1062029 | 0.22439125 | 0.34652261 | 0 | 0.62318605 | 0.48002932 | 0.4396336 | 0.38977239 | 0.26021288 | 0.28748204 | 0 | 0.91238877 | 0.31239423 | 0.22334672 | 1.08992403 |
| ENSMUSG00 | Gm11462 | 561 | 0.1515336 | 0.05938127 | 0 | 0.08121398 | 0 | 0 | 0 | 0 | 0.07341625 | 0 | 0 | 0 | 0.08793568 | 0 | 0.14688612 | 0 | 0 | 0 |
| ENSMUSG00 | Gm13135 | 3120 | 0 | 0 | 0 | 0 | 0 | 0 | 0 | 0 | 0 | 0 | 0 | 0 | 0 | 0 | 0 | 0 | 0 | 0 |
| ENSMUSG00 | Gm8172 | 1545 | 0 | 0 | 0 | 0 | 0 | 0.03207297 | 0 | 0 | 0 | 0 | 0 | 0 | 0 | 0 | 0.02666767 | 0 | 0 | 0 |
| ENSMUSG00 | Gm5863 | 1007 | 0.46430679 | 0.1323253 | 0.14411181 | 0.54293201 | 0.57356811 | 0.24604138 | 0.51682172 | 0.22756148 | 0.49080257 | 0.42140648 | 0.33210001 | 0.49884902 | 0.24494497 | 0.32284867 | 0.7364727 | 0.26617105 | 0.22835927 | 0.21850688 |
| ENSMUSG00 | Gm12785 | 171 | 0 | 0 | 0 | 0 | 0 | 0 | 0 | 0 | 0 | 0 | 0 | 0 | 0 | 0 | 0 | 0 | 0 | 0 |
| ENSMUSG00 | Gm6067 | 490 | 0.08674526 | 0 | 0 | 0 | 0 | 0 | 0 | 0.31177471 | 0 | 0 | 0 | 0 | 0.10067738 | 0 | 0 | 0 | 0 | 0 |
| ENSMUSG00 | Gm13462 | 960 | 0 | 0 | 0 | 0.04745942 | 0 | 0 | 0 | 0 | 0 | 0 | 0 | 0 | 0 | 0 | 0 | 0 | 0 | 0 |
| ENSMUSG00 | Gm13586 | 463 | 0.27541151 | 0.64755086 | 0.6268708 | 0.49201992 | 0.62373963 | 0.10702534 | 0.40874885 | 0.65991192 | 0.26686727 | 0.61102423 | 1.13504221 | 0.36165656 | 0.10654842 | 1.10342324 | 0.80089417 | 1.57132082 | 0.4138909 | 0.11881017 |
| ENSMUSG00 | Cyp2j8 | 1503 | 0 | 0.0664928 | 0 | 0 | 0 | 0 | 0 | 0.22869702 | 0 | 0.09411318 | 0.1271456 | 0 | 0 | 0 | 0 | 0 | 0 | 0 |
| ENSMUSG00 | Gm14811 | 116 | 0 | 0 | 0 | 0 | 0 | 0 | 0 | 0 | 0 | 0 | 0 | 0 | 0 | 0 | 0 | 0 | 0 | 0 |
| ENSMUSG00 | Gm13887 | 343 | 0 | 0 | 0 | 0 | 0 | 0 | 0 | 0 | 0 | 0 | 0 | 0 | 0 | 0 | 0 | 0 | 0 | 0 |
| ENSMUSG00 | Gm7658 | 1123 | 0.227098 | 0.17798519 | 0.25845163 | 0.16228333 | 0.25716077 | 0.2206266 | 0.04213061 | 0.44212042 | 0.25672806 | 0.20993189 | 0.12762678 | 0.14910685 | 0 | 0.16542863 | 0.11006649 | 0.34096712 | 0.34128492 | 0.34288847 |
| ENSMUSG00 | Gm16166 | 786 | 0.32446699 | 0.12714845 | 0.1846318 | 0.05796571 | 0 | 0 | 0.36116549 | 0 | 0.83840236 | 0.17996452 | 0 | 0.14202459 | 0 | 0 | 0 | 0.63330521 | 0 | 0 |
| ENSMUSG00 | Gm15095 | 1429 | 0 | 0 | 0 | 0 | 0 | 0 | 0 | 0 | 0 | 0 | 0 | 0 | 0 | 0 | 0 | 0 | 0 | 0 |
| ENSMUSG00 | Gm2810 | 989 | 0 | 0 | 0 | 0.09213558 | 0 | 0 | 0 | 0 | 0 | 0 | 0 | 0 | 0 | 0.09392131 | 0 | 0 | 0 | 0.05562094 |
| ENSMUSG00 | Gm14881 | 370 | 0 | 0 | 0 | 0 | 0 | 0 | 0 | 0 | 0 | 0 | 0 | 0 | 0 | 0 | 0 | 0 | 0 | 0 |
| ENSMUSG00 | Gm14103 | 365 | 0 | 0 | 0 | 0 | 0 | 0 | 0 | 0 | 0 | 0 | 0 | 0 | 0 | 0 | 0 | 0 | 0 | 0 |
| ENSMUSG00 | Gm12850 | 435 | 0.09771305 | 0 | 0.44481407 | 0 | 0 | 0.34174299 | 0 | 0.17559725 | 0 | 0 | 0 | 0 | 0 | 0 | 0 | 0 | 0 | 0 |
| ENSMUSG00 | Gm13355 | 316 | 0 | 0 | 0 | 0 | 0 | 0 | 0 | 0 | 0 | 0 | 0 | 0 | 0 | 0 | 0 | 0 | 0 | 0 |
| ENSMUSG00 | Gm13137 | 619 | 0 | 0 | 0 | 0 | 0 | 0 | 0 | 0 | 0 | 0 | 0 | 0 | 0 | 0 | 0 | 0 | 0 | 0 |
| ENSMUSG00 | Gm11540 | 777 | 0 | 0 | 0 | 0 | 0 | 0 | 0 | 0 | 0 | 0 | 0 | 0 | 0.12698048 | 0 | 0 | 0 | 0 | 0 |
| ENSMUSG00 | Gm12177 | 1995 | 0 | 0 | 0 | 0.02283762 | 0.07237883 | 0 | 0 | 0 | 0.08257948 | 0 | 0 | 0 | 0.12363889 | 0.11640122 | 0 | 0 | 0 | 0 |
| ENSMUSG00 | Gm12587 | 426 | 0.49888704 | 0.07819928 | 0.11355289 | 0.21390162 | 0.22597147 | 0 | 0.44425051 | 0 | 0.19336392 | 0.33204721 | 0.11214779 | 0.13102268 | 0.11580262 | 0.21804736 | 0 | 0.08988405 | 0.44983917 | 0.38738808 |
| ENSMUSG00 | Gm11920 | 562 | 0 | 0 | 0 | 0 | 0 | 0 | 0 | 0 | 0 | 0 | 0 | 0 | 0 | 0 | 0 | 0 | 0 | 0 |
| ENSMUSG00 | Gm7834 | 436 | 0 | 0 | 0 | 0 | 0 | 0 | 0 | 0 | 0 | 0 | 0 | 0 | 0 | 0 | 0 | 0 | 0 | 0 |
| ENSMUSG00 | Gm13359 | 694 | 0 | 0 | 0 | 0.13129984 | 0 | 0 | 0 | 0 | 0 | 0 | 0 | 0 | 0 | 0 | 0 | 0 | 0 | 0 |
| ENSMUSG00 | Gm12507 | 414 | 0 | 0 | 0 | 0 | 0 | 0 | 0 | 0 | 0 | 0 | 0 | 0 | 0 | 0 | 0 | 0 | 0 | 0 |
| ENSMUSG00 | Gm13217 | 1513 | 0 | 0.06605333 | 0 | 0 | 0.03181224 | 0 | 0 | 0 | 0 | 0.09349115 | 0 | 0.18445361 | 0.0978161 | 0.06139337 | 0 | 0.07592321 | 0.02533133 | 0 |
| ENSMUSG00 | Gm15504 | 264 | 0 | 0 | 0 | 0 | 0 | 0 | 0 | 0 | 0 | 0 | 0 | 0 | 0 | 0 | 0 | 0 | 0 | 0 |
| ENSMUSG00 | Gm8036 | 1716 | 0 | 0 | 0.11275881 | 0.05310145 | 0 | 0.08663065 | 0.11028596 | 0.06676993 | 0.09600586 | 0.0549542 | 0.05568177 | 0 | 0.05749641 | 0.08119596 | 0.02401023 | 0.11156937 | 0.08933869 | 0.03205659 |
| ENSMUSG00 | Naip3 | 4854 | 0 | 0 | 0 | 0 | 0 | 0 | 0.06823007 | 0.00786823 | 0 | 0 | 0 | 0 | 0 | 0 | 0 | 0 | 0.02368745 | 0 |
| ENSMUSG00 | Gm15303 | 2623 | 0 | 0 | 0 | 0 | 0 | 0 | 0 | 0 | 0 | 0 | 0 | 0 | 0 | 0 | 0 | 0 | 0 | 0 |
| ENSMUSG00 | Gm12731 | 267 | 0 | 0 | 0 | 0 | 0 | 0 | 0 | 0 | 0 | 0 | 0 | 0 | 0 | 0 | 0 | 0 | 0 | 0 |
| ENSMUSG00 | Gm2566 | 397 | 0 | 0 | 0 | 0 | 0 | 0 | 0 | 0 | 0 | 0 | 0 | 0 | 0 | 0 | 0 | 0 | 0 | 0 |
| ENSMUSG00 | Gm6884 | 614 | 6.64576042 | 13.6181375 | 8.7450519 | 7.56877284 | 11.1314872 | 7.90906823 | 3.85282402 | 10.8232541 | 6.90913864 | 3.91642649 | 10.7376943 | 3.7271045 | 6.42761137 | 7.56418361 | 7.24717921 | 14.5928373 | 14.5440182 | 7.43608446 |
| ENSMUSG00 | Gm15131 | 594 | 0 | 0 | 0 | 0 | 0 | 0 | 0 | 0.12859395 | 0 | 0.23813486 | 0 | 0 | 0 | 0.0781887 | 0 | 0 | 0.06452239 | 0 |
| ENSMUSG00 | Gm10601 | 2092 | 0 | 0.07961973 | 0 | 0 | 0 | 0.02368678 | 0.04523201 | 0.01825641 | 0 | 0.15777004 | 0.09134791 | 0 | 0.40088078 | 0 | 0 | 0.1098201 | 0.10992246 | 0.02629498 |
| ENSMUSG00 | Gm12465 | 299 | 0 | 0 | 0 | 0 | 0 | 0 | 0 | 0 | 0 | 0 | 0 | 0 | 0 | 0 | 0 | 0 | 0 | 0 |
| ENSMUSG00 | Rpl13-ps5 | 629 | 0 | 0 | 0.30762181 | 0 | 0 | 0 | 0 | 0 | 0 | 0 | 0 | 0 | 0 | 0 | 0 | 0 | 0 | 0 |
| ENSMUSG00 | Gm6754 | 912 | 0 | 0 | 0 | 0 | 0 | 0 | 0 | 0 | 0 | 0 | 0 | 0 | 0 | 0 | 0 | 0 | 0 | 0 |
| ENSMUSG00 | Gm14614 | 383 | 0 | 0 | 0 | 0 | 0 | 0 | 0 | 0 | 0 | 0 | 0 | 0 | 0 | 0 | 0 | 0 | 0 | 0 |
| ENSMUSG00 | Gm16082 | 376 | 0 | 0 | 0 | 0 | 0 | 0 | 0.12583159 | 0 | 0 | 0 | 0 | 0 | 0 | 0 | 0 | 0 | 0 | 0 |
| ENSMUSG00 | Gm14581 | 197 | 0 | 0 | 0 | 0 | 0 | 0 | 0 | 0 | 0 | 0 | 0 | 0 | 0 | 0 | 0 | 0 | 0 | 0 |
| ENSMUSG00 | Gm12630 | 783 | 0.05428503 | 0.12763561 | 0.30889866 | 0.17456339 | 0.24588467 | 0 | 0.36254926 | 0.43899313 | 0.05260091 | 0.48174409 | 0.30507636 | 0.85541245 | 0.75604471 | 0.83041791 | 0.78930184 | 0.19560974 | 0.2936881 | 0.42152572 |
| ENSMUSG00 | Gm15214 | 469 | 0 | 0 | 0 | 0 | 0 | 0 | 0 | 0 | 0 | 0 | 0 | 0 | 0 | 0 | 0 | 0 | 0 | 0 |
| ENSMUSG00 | Gm13517 | 1494 | 0 | 0 | 0 | 0 | 0 | 0 | 0 | 0 | 0 | 0 | 0 | 0 | 0 | 0 | 0 | 0 | 0.02565348 | 0 |
| ENSMUSG00 | Gm13810 | 1346 | 0 | 0 | 0 | 0 | 0 | 0 | 0 | 0 | 0 | 0 | 0 | 0 | 0 | 0 | 0 | 0 | 0 | 0 |
| ENSMUSG00 | Gm13672 | 458 | 0 | 0 | 0 | 0 | 0 | 0 | 0 | 0 | 0 | 0 | 0 | 0 | 0 | 0 | 0 | 0 | 0 | 0 |
| ENSMUSG00 | Gm15129 | 278 | 0.15289632 | 0 | 0 | 0 | 0 | 0 | 0 | 0 | 0 | 0 | 0 | 0 | 0 | 0 | 0 | 0 | 0 | 0 |
| ENSMUSG00 | Gm14363 | 407 | 0 | 0 | 0 | 0 | 0 | 0 | 0 | 0 | 0 | 0 | 0 | 0 | 0 | 0 | 0 | 0 | 0 | 0 |
| ENSMUSG00 | Gm15056 | 371 | 0.34370762 | 0.08979217 | 0.5215475 | 0.12280605 | 0 | 0.26713064 | 1.4028018 | 0.5147224 | 1.22116354 | 1.27090844 | 0 | 0.75223265 | 1.32970127 | 1.25186219 | 0.77738785 | 0.30962755 | 0.20661076 | 0.14827252 |
| ENSMUSG00 | Gm11911 | 1022 | 0 | 0 | 0 | 0 | 0 | 0 | 0 | 0 | 0 | 0 | 0 | 0 | 0 | 0 | 0 | 0 | 0 | 0 |
| ENSMUSG00 | Rpsa-ps11 | 888 | 0.04786619 | 0 | 0.0544747 | 0.15392245 | 0.05420262 | 0 | 0 | 0 | 0.04638121 | 0 | 0.05380063 | 0 | 0.16666188 | 0 | 0.04639815 | 0 | 0 | 0 |
| ENSMUSG00 | Gm14996 | 511 | 0 | 0 | 0 | 0 | 0 | 0 | 0 | 0 | 0 | 0 | 0 | 0 | 0 | 0 | 0 | 0 | 0 | 0 |
| ENSMUSG00 | Olfr1274-ps | 1963 | 0 | 0 | 0 | 0 | 0 | 0 | 0 | 0 | 0 | 0 | 0 | 0 | 0 | 0 | 0 | 0 | 0 | 0 |
| ENSMUSG00 | Gm11526 | 174 | 2.19854359 | 0.57436025 | 1.39004398 | 0.78553525 | 0 | 1.99350078 | 0.5438239 | 1.75597252 | 0.94681645 | 1.62588632 | 2.19654982 | 1.60389835 | 0.56703353 | 3.73688059 | 1.65753386 | 1.7604877 | 1.76212859 | 0.63228858 |
| ENSMUSG00 | Olfr1149-ps1 | 444 | 0 | 0 | 0 | 0 | 0 | 0 | 0 | 0 | 0 | 0 | 0 | 0 | 0 | 0 | 0 | 0 | 0 | 0 |
| ENSMUSG00 | Gm12318 | 272 | 0 | 0 | 0 | 0 | 0 | 0 | 0 | 0 | 0 | 0 | 0 | 0 | 0 | 0 | 0 | 0 | 0 | 0 |
| ENSMUSG00 | Gm10599 | 2993 | 0 | 0 | 0 | 0 | 0 | 0 | 0 | 0 | 0 | 0 | 0 | 0 | 0 | 0 | 0 | 0 | 0 | 0 |
| ENSMUSG00 | Gm14042 | 832 | 0.20435181 | 0 | 0 | 0 | 0 | 0 | 0 | 0 | 0.14850907 | 0.22668607 | 0 | 0.06708613 | 0 | 0 | 0 | 0.36817892 | 0 | 0 |
| ENSMUSG00 | Gm11389 | 427 | 0 | 0 | 0 | 0 | 0 | 0 | 0 | 0 | 0 | 0 | 0 | 0 | 0 | 0 | 0 | 0 | 0 | 0 |
| ENSMUSG00 | Gm11654 | 427 | 0 | 0 | 0 | 0 | 0 | 0 | 0 | 0.2683307 | 0 | 0 | 0 | 0 | 0 | 0 | 0.09649076 | 0.08967355 | 0 | 0 |
| ENSMUSG00 | Gm6903 | 2324 | 0.03657933 | 0.02866858 | 0.04162954 | 0.01960458 | 0.02071081 | 0 | 0 | 0.08216954 | 0 | 0.10144299 | 0 | 0.02401707 | 0 | 0.01998455 | 0.01772872 | 0.03295233 | 0.03298304 | 0.04734002 |
| ENSMUSG00 | Gm13557 | 380 | 0 | 0 | 0 | 0 | 0 | 0 | 0 | 0 | 0 | 0 | 0 | 0 | 0 | 0.36666385 | 0 | 0 | 0 | 0 |
| ENSMUSG00 | Gm14052 | 1030 | 0.16506864 | 0.03234262 | 0.04696459 | 0.13270207 | 0.32711016 | 0.0962189 | 0.13780392 | 0.14832001 | 0.11996073 | 0.09155476 | 0.23191727 | 0.05418996 | 0.62263585 | 0 | 0.72002719 | 0.44610416 | 0.33488997 | 0.4806621 |
| ENSMUSG00 | Gm11806 | 499 | 0 | 0 | 0 | 0 | 0 | 0.09930408 | 0 | 0.07653788 | 0 | 0 | 0 | 0 | 0 | 0 | 0.08256825 | 0 | 0 | 0.11023869 |
| ENSMUSG00 | Gm11566 | 202 | 0 | 0 | 0 | 0 | 0 | 0 | 0 | 0 | 0 | 0 | 0 | 0 | 0 | 0 | 0 | 0 | 0 | 0 |
| ENSMUSG00 | Gm11848 | 972 | 0 | 0 | 0 | 0 | 0 | 0 | 0 | 0 | 0 | 0 | 0 | 0 | 0 | 0 | 0 | 0 | 0 | 0 |
| ENSMUSG00 | Gm15126 | 831 | 0 | 0 | 0 | 0 | 0 | 0 | 0 | 0 | 0 | 0 | 0 | 0 | 0 | 0 | 0 | 0 | 0 | 0 |
| ENSMUSG00 | Gm6238 | 464 | 0 | 0.43077018 | 0 | 0 | 0 | 0 | 0 | 0.08231121 | 0 | 0.30485368 | 0.51481636 | 0 | 0 | 0 | 0.44398228 | 0.74270575 | 0.90859756 | 0 |
| ENSMUSG00 | Gm13975 | 781 | 0 | 0 | 0 | 0 | 0 | 0 | 0 | 0 | 0 | 0 | 0 | 0 | 0 | 0 | 0 | 0 | 0 | 0 |
| ENSMUSG00 | Gm11221 | 1280 | 0.29886452 | 0.36435978 | 0.11337546 | 0.07118913 | 0.30082452 | 0.15485229 | 0.11088909 | 0.14918907 | 0.19306179 | 0.18418243 | 0.22394512 | 0.21802993 | 0.30832448 | 0.21770666 | 0.06437743 | 0.5683762 | 0.32936681 | 0.30083105 |
| ENSMUSG00 | Gm13238 | 779 | 0 | 0.08552733 | 0.12419392 | 0 | 0 | 0 | 0.12147029 | 0 | 0 | 0 | 0 | 0.0716504 | 0.25330895 | 0 | 0 | 0 | 0 | 0 |
| ENSMUSG00 | Gm15678 | 958 | 0 | 0 | 0 | 0 | 0 | 0 | 0 | 0 | 0 | 0 | 0 | 0 | 0 | 0 | 0 | 0 | 0 | 0 |
| ENSMUSG00 | Gm13250 | 752 | 0 | 0 | 0 | 0 | 0 | 0 | 0 | 0 | 0 | 0 | 0 | 0 | 0 | 0 | 0 | 0 | 0 | 0 |
| ENSMUSG00 | Gm13413 | 601 | 0 | 0 | 0 | 0 | 0 | 0 | 0 | 0.06354809 | 0 | 0 | 0 | 0 | 0.65666446 | 0 | 0.068555 | 0.06371149 | 0 | 0 |
| ENSMUSG00 | Gm12392 | 937 | 0 | 0 | 0 | 0 | 0 | 0 | 0 | 0 | 0 | 0 | 0 | 0 | 0 | 0 | 0 | 0 | 0 | 0 |
| ENSMUSG00 | Gm15715 | 282 | 0 | 0.35439249 | 0 | 0 | 0 | 0.17571891 | 0.83887729 | 0.94803836 | 0 | 0 | 0 | 0 | 0 | 0.82347673 | 0 | 0 | 0.27181771 | 0.78027102 |
| ENSMUSG00 | Gm13623 | 624 | 0 | 0 | 0 | 0.07301449 | 0.46280696 | 0 | 0 | 0.06120577 | 0.1980121 | 0.07556202 | 0 | 0 | 0.07905756 | 0.07442963 | 0.06602813 | 0.18408946 | 0 | 0 |
| ENSMUSG00 | Gm12053 | 250 | 0 | 0 | 0 | 0 | 0 | 0 | 0 | 0 | 0 | 0 | 0 | 0 | 0 | 0 | 0 | 0 | 0 | 0 |
| ENSMUSG00 | Gm1335 | 384 | 0 | 0 | 0 | 0 | 0 | 0.12904358 | 0 | 0 | 0.10725655 | 0 | 0.12441395 | 0 | 0 | 0 | 0 | 0.19943025 | 0.19961613 | 0 |
| ENSMUSG00 | Gm14687 | 361 | 0 | 0 | 0 | 0 | 0 | 0 | 0 | 0 | 0.11409007 | 0 | 0 | 0 | 0 | 0 | 0 | 0 | 0 | 0 |
| ENSMUSG00 | Gm12816 | 1211 | 0.07019847 | 0 | 0 | 0 | 0 | 0.04091886 | 0.0781382 | 0 | 0.06802067 | 0 | 0 | 0 | 0 | 0.03835185 | 0.03402275 | 0.094857 | 0 | 0 |
| ENSMUSG00 | 2810453I06R | 3117 | 0.12272909 | 0.16031229 | 0.09311555 | 0.20463735 | 0.13897572 | 0.19077087 | 0.1973259 | 0.2695646 | 0.23784321 | 0.28741205 | 0.24523559 | 0.34023022 | 0.22157422 | 0.13410227 | 0.44942345 | 0.34396439 | 0.23362196 | 0.28236949 |
| ENSMUSG00 | Gm12978 | 521 | 0 | 0 | 0 | 0 | 0 | 0 | 0 | 0 | 0 | 0 | 0 | 0 | 0 | 0 | 0 | 0 | 0 | 0 |
| ENSMUSG00 | Gm11764 | 622 | 0.0683363 | 0 | 0 | 0 | 0 | 0 | 0 | 0 | 0.59594637 | 0.15160998 | 0.46085169 | 0 | 0.87242941 | 0.4480137 | 0.59616399 | 0.06156046 | 0.3080892 | 0.61907355 |
| ENSMUSG00 | Gm16518 | 506 | 0 | 0 | 0 | 0 | 0 | 0 | 0 | 0 | 0 | 0 | 0 | 0 | 0 | 0.18357347 | 0 | 0 | 0 | 0 |
| ENSMUSG00 | Gm11703 | 1022 | 0 | 0.03259579 | 0.09466444 | 0 | 0 | 0 | 0.09258841 | 0 | 0.04029992 | 0.04613572 | 0 | 0.1092283 | 0 | 0 | 0.04031463 | 0 | 0.03750127 | 0.05382496 |
| ENSMUSG00 | Gm12163 | 388 | 0 | 0 | 0 | 0 | 0 | 0 | 0 | 0 | 0 | 0 | 0 | 0 | 0 | 0 | 0 | 0 | 0 | 0 |
| ENSMUSG00 | Gm13555 | 318 | 0 | 0 | 0 | 0 | 0 | 0 | 0 | 0 | 0 | 0 | 0 | 0 | 0 | 0 | 0 | 0 | 0 | 0 |
| ENSMUSG00 | Gm6222 | 395 | 0 | 0.08433644 | 0 | 0.34603325 | 0.48741189 | 0.25089992 | 0 | 0.58013776 | 0.72988762 | 0.35810661 | 0.84664483 | 0.14130547 | 1.2489093 | 0.23515994 | 0 | 0 | 0 | 0.27852712 |
| ENSMUSG00 | Gm15331 | 384 | 0 | 0 | 0 | 0 | 0 | 0 | 0 | 0 | 0 | 0 | 0 | 0 | 0 | 0 | 0 | 0 | 0 | 0 |
| ENSMUSG00 | Gm14370 | 867 | 0 | 0 | 0 | 0 | 0 | 0 | 0 | 0 | 0 | 0 | 0 | 0 | 0 | 0 | 0 | 0 | 0 | 0 |
| ENSMUSG00 | Rps15a-ps6 | 392 | 2.06019986 | 2.12454683 | 2.22123354 | 1.85963446 | 2.33292487 | 0.88487024 | 3.8622595 | 2.72802874 | 3.04696163 | 2.40564813 | 3.41249703 | 2.84773789 | 2.01354764 | 2.72503574 | 3.67871035 | 2.63736326 | 4.39970245 | 4.06955126 |
| ENSMUSG00 | Gm11804 | 467 | 0 | 0 | 0 | 0 | 0 | 0 | 0 | 0 | 0 | 0 | 0 | 0 | 0 | 0 | 0 | 0 | 0 | 0 |
| ENSMUSG00 | Gm14982 | 441 | 0 | 0 | 0 | 0 | 0 | 0 | 0 | 0 | 0 | 0 | 0 | 0 | 0 | 0 | 0 | 0 | 0 | 0 |
| ENSMUSG00 | Gm11829 | 281 | 0 | 0 | 0 | 0 | 0 | 0 | 0 | 0 | 0 | 0 | 0 | 0 | 0 | 0 | 0 | 0 | 0 | 0 |
| ENSMUSG00 | Gm6373 | 359 | 0 | 0 | 0 | 0 | 0 | 0 | 0 | 0 | 0 | 0 | 0 | 0 | 0 | 0 | 0 | 0 | 0 | 0 |
| ENSMUSG00 | Gm13140 | 3149 | 0 | 0.04231552 | 0.03072311 | 0 | 0 | 0 | 0 | 0.07277053 | 0 | 0.01497323 | 0.01517147 | 0.10634931 | 0 | 0.01474884 | 0 | 0.01215961 | 0 | 0 |
| ENSMUSG00 | Gm14604 | 441 | 0 | 0 | 0 | 0 | 0 | 0 | 0 | 0 | 0 | 0 | 0 | 0 | 0 | 0 | 0 | 0 | 0 | 0 |
| ENSMUSG00 | Gm13720 | 195 | 0 | 0 | 0 | 0 | 0 | 0 | 0 | 0 | 0 | 0 | 0 | 0 | 0 | 0 | 0 | 0 | 0 | 0 |
| ENSMUSG00 | Gm14561 | 795 | 0 | 0 | 0 | 0 | 0 | 0 | 0 | 0 | 0 | 0 | 0 | 0 | 0 | 0 | 0 | 0 | 0 | 0 |
| ENSMUSG00 | Gm14910 | 1804 | 0 | 0 | 0 | 0 | 0 | 0 | 0 | 0 | 0 | 0 | 0 | 0 | 0 | 0 | 0 | 0 | 0 | 0 |
| ENSMUSG00 | Gm8087 | 566 | 0 | 0 | 0 | 0 | 0 | 0 | 0 | 0 | 0 | 0 | 0 | 0 | 0 | 0 | 0 | 0 | 0 | 0 |
| ENSMUSG00 | Gm9083 | 818 | 0.05196232 | 0.08144962 | 0.11827269 | 0.0556981 | 0 | 0 | 0 | 0 | 0.15105079 | 0 | 0 | 0 | 0 | 0.17033284 | 0 | 0.04681003 | 0.18741465 | 0 |
| ENSMUSG00 | Olfr334-ps1 | 905 | 0 | 0 | 0 | 0 | 0 | 0 | 0 | 0 | 0 | 0 | 0 | 0 | 0 | 0 | 0 | 0 | 0 | 0 |
| ENSMUSG00 | Gm15959 | 367 | 0 | 0 | 0 | 0 | 0 | 0 | 0 | 0 | 0 | 0 | 0 | 0 | 0 | 0 | 0 | 0 | 0 | 0 |
| ENSMUSG00 | Gm14900 | 732 | 0 | 0 | 0 | 0 | 0 | 0 | 0 | 0 | 0 | 0 | 0 | 0 | 0 | 0 | 0 | 0 | 0 | 0 |
| ENSMUSG00 | Gm13233 | 1441 | 0 | 0 | 0 | 0 | 0 | 0 | 0 | 0 | 0 | 0 | 0 | 0 | 0 | 0 | 0 | 0 | 0 | 0 |
| ENSMUSG00 | Gm11606 | 616 | 0 | 0 | 0 | 0 | 0 | 0 | 0 | 0 | 0 | 0 | 0 | 0 | 0 | 0 | 0 | 0 | 0 | 0 |
| ENSMUSG00 | Gm9835 | 425 | 0 | 0 | 0 | 0 | 0 | 0 | 0 | 0.17972895 | 0 | 0 | 0 | 0 | 0 | 0 | 0 | 0 | 0 | 0 |
| ENSMUSG00 | Gm11388 | 158 | 0 | 0 | 0 | 0 | 0 | 0 | 0 | 0 | 0 | 0 | 0 | 0 | 0 | 0 | 0 | 0 | 0 | 0 |
| ENSMUSG00 | Gm13433 | 239 | 0 | 0 | 0 | 0.57189595 | 0.60416641 | 0 | 0 | 0.15980085 | 0 | 0 | 0.59968567 | 0.70061501 | 0 | 0.38865345 | 0.51717434 | 0 | 0.6414443 | 0 |
| ENSMUSG00 | Gm12416 | 1007 | 0.37988737 | 0.29773192 | 0.76859631 | 0.54293201 | 0.76475748 | 0.29524965 | 0.46983792 | 0.68268445 | 1.30880686 | 0.6555212 | 1.09118574 | 0.66513203 | 0.48988994 | 0.46121239 | 0.53189695 | 1.02665978 | 0.87537719 | 0.71014736 |
| ENSMUSG00 | Gm16100 | 230 | 0 | 0 | 0 | 0 | 0 | 0 | 0 | 0 | 0 | 0 | 0 | 0 | 0 | 0 | 0 | 0 | 0 | 0 |
| ENSMUSG00 | Gm14699 | 227 | 0 | 0 | 0 | 0 | 0 | 0 | 0 | 0 | 0 | 0 | 0 | 0 | 0 | 0 | 0 | 0 | 0 | 0 |
| ENSMUSG00 | Gm14082 | 800 | 0 | 0 | 0 | 0.11390261 | 0 | 0 | 0 | 0 | 0 | 0 | 0 | 0 | 0 | 0 | 0 | 0 | 0 | 0 |
| ENSMUSG00 | Gm14595 | 694 | 0 | 0 | 0 | 0 | 0 | 0 | 0 | 0 | 0 | 0 | 0 | 0 | 0 | 0 | 0 | 0 | 0 | 0 |
| ENSMUSG00 | Gm14822 | 290 | 0 | 0 | 0 | 0 | 0 | 0 | 0 | 0 | 0 | 0 | 0 | 0 | 0 | 0 | 0 | 0 | 0 | 0 |
| ENSMUSG00 | Gm11242 | 2456 | 0 | 0 | 0 | 0 | 0 | 0 | 0 | 0 | 0 | 0 | 0 | 0 | 0 | 0 | 0 | 0 | 0 | 0 |
| ENSMUSG00 | Gm14625 | 750 | 0 | 0 | 0 | 0 | 0 | 0 | 0 | 0 | 0 | 0 | 0 | 0 | 0 | 0 | 0 | 0 | 0 | 0 |
| ENSMUSG00 | Gm12285 | 232 | 0 | 0 | 0 | 0 | 0 | 0 | 0 | 0 | 0 | 0 | 0 | 0 | 0 | 0 | 0 | 0 | 0 | 0 |
| ENSMUSG00 | Gm15001 | 790 | 0 | 0 | 0 | 0 | 0 | 0 | 0 | 0 | 0 | 0 | 0 | 0 | 0 | 0 | 0 | 0 | 0 | 0 |
| ENSMUSG00 | Gm12986 | 966 | 0 | 0 | 0 | 0 | 0 | 0 | 0 | 0 | 0 | 0 | 0 | 0 | 0 | 0 | 0 | 0 | 0 | 0 |
| ENSMUSG00 | Gm14140 | 994 | 0 | 0 | 0 | 0 | 0 | 0 | 0.28558961 | 0 | 0.04143513 | 0 | 0 | 0 | 0 | 0 | 0 | 0 | 0.07711529 | 0 |
| ENSMUSG00 | Gm2092 | 631 | 0 | 0 | 0 | 0 | 0 | 0 | 0 | 0 | 0 | 0 | 0 | 0 | 0 | 0 | 0 | 0 | 0 | 0 |
| ENSMUSG00 | Gm13356 | 334 | 0 | 0 | 0 | 0 | 0 | 0 | 0 | 0 | 0 | 0 | 0 | 0 | 0 | 0 | 0 | 0 | 0 | 0 |
| ENSMUSG00 | Gm14795 | 470 | 0 | 0 | 0 | 0 | 0 | 0 | 0 | 0 | 0 | 0 | 0 | 0 | 0 | 0 | 0 | 0 | 0 | 0 |
| ENSMUSG00 | Olfr268-ps1 | 1611 | 0 | 0 | 0 | 0 | 0 | 0 | 0 | 0 | 0 | 0 | 0 | 0 | 0 | 0 | 0 | 0 | 0 | 0 |
| ENSMUSG00 | Gm14533 | 92 | 0 | 0 | 0 | 0 | 0 | 0 | 0 | 0 | 0 | 0 | 0 | 0 | 0 | 0 | 0 | 0 | 0 | 0 |
| ENSMUSG00 | Gm12191 | 347 | 0 | 0 | 0 | 0 | 0.13870871 | 0 | 0.68173889 | 0.33019368 | 0 | 0 | 0 | 0.32170411 | 0.14216691 | 0 | 0.11873647 | 0.22069514 | 0.11045042 | 0 |
| ENSMUSG00 | Gm14980 | 735 | 0 | 0 | 0 | 0 | 0 | 0 | 0 | 0 | 0 | 0 | 0 | 0 | 0 | 0 | 0 | 0 | 0 | 0 |
| ENSMUSG00 | Gm14882 | 217 | 0 | 0 | 0 | 0 | 0 | 0 | 0 | 0 | 0 | 0 | 0 | 0 | 0 | 0 | 0 | 0 | 0 | 0 |
| ENSMUSG00 | Gm7785 | 891 | 0.66787033 | 0.78515239 | 0.5429128 | 0.66475149 | 0.59422128 | 1.33475377 | 0.69030845 | 0.77156368 | 0.78582578 | 1.05837718 | 0.91153119 | 0.87701378 | 1.66100732 | 0.67763539 | 1.06356429 | 0.85949736 | 1.50552233 | 0.86434062 |
| ENSMUSG00 | Gm13195 | 260 | 0 | 0.25625303 | 0 | 0 | 0.55536835 | 0 | 0 | 0.44068157 | 0 | 0 | 0.18374984 | 0 | 0 | 0 | 0 | 0.4418147 | 0 | 0 |
| ENSMUSG00 | Gm12748 | 503 | 0 | 0 | 0 | 0 | 0 | 0 | 0 | 0 | 0 | 0 | 0 | 0 | 0 | 0 | 0 | 0 | 0 | 0 |
| ENSMUSG00 | Gm14751 | 221 | 0 | 0 | 0 | 0 | 0 | 0 | 0 | 0 | 0 | 0 | 0 | 0 | 0 | 0 | 0 | 0 | 0 | 0 |
| ENSMUSG00 | Gm11800 | 577 | 0 | 0 | 0 | 0 | 0 | 0 | 0 | 0 | 0 | 0 | 0 | 0 | 0 | 0 | 0 | 0 | 0 | 0 |
| ENSMUSG00 | Gm16431 | 427 | 0 | 0 | 0 | 0 | 0 | 0 | 0 | 0 | 0 | 0 | 0 | 0 | 0 | 0 | 0 | 0 | 0 | 0 |
| ENSMUSG00 | Gm13350 | 478 | 0.08892296 | 0 | 0 | 0 | 0 | 0 | 0 | 0 | 0 | 0 | 0 | 0 | 0 | 0 | 0 | 0 | 0 | 0 |
| ENSMUSG00 | Gm13668 | 995 | 0.08543754 | 0.26784237 | 0.19446645 | 0.04578999 | 0.04837379 | 0.04980174 | 0.38040345 | 0.1535373 | 0.33114787 | 0.28432585 | 0.4321353 | 0.11219229 | 0.14873945 | 0.28006485 | 0.37267739 | 0.42331325 | 0.07703778 | 0.27642767 |
| ENSMUSG00 | Gm11845 | 326 | 0 | 0 | 0 | 0 | 0 | 0 | 0 | 0 | 0 | 0 | 0 | 0 | 0 | 0 | 0 | 0 | 0 | 0 |
| ENSMUSG00 | Gm14670 | 336 | 0 | 0 | 0 | 0 | 0 | 0 | 0 | 0 | 0 | 0 | 0 | 0 | 0 | 0 | 0 | 0 | 0 | 0 |
| ENSMUSG00 | Gm11980 | 985 | 0 | 0 | 0 | 0 | 0 | 0 | 0 | 0.03877401 | 0 | 0 | 0 | 0 | 0 | 0 | 0 | 0 | 0 | 0 |
| ENSMUSG00 | Gm11408 | 518 | 0 | 0 | 0 | 0.17591137 | 0 | 0 | 0.45668609 | 0.14746101 | 0.1590213 | 0.09102452 | 0.09222965 | 0.21550449 | 0.38094145 | 0.1793208 | 0 | 0 | 0.14797798 | 0.21239037 |
| ENSMUSG00 | Gm14126 | 394 | 0 | 0.16910099 | 0 | 0 | 0 | 0 | 0 | 0 | 0 | 0 | 0 | 0 | 0 | 0.23575679 | 0 | 0 | 0 | 0 |
| ENSMUSG00 | Gm13172 | 255 | 0 | 0 | 0 | 0.17867076 | 0 | 0 | 0 | 0 | 0 | 0 | 0 | 0 | 0 | 0.36426735 | 0.16157473 | 0 | 0 | 0 |
| ENSMUSG00 | Amy2b | 1564 | 0 | 0 | 0 | 0 | 0 | 0 | 0 | 0 | 0 | 0 | 0 | 0 | 0 | 0 | 0.02634371 | 0 | 0 | 0 |
| ENSMUSG00 | Gm11484 | 337 | 0 | 0 | 0 | 0 | 0 | 0 | 0 | 0 | 0 | 0 | 0 | 0 | 0 | 0 | 0 | 0 | 0 | 0 |
| ENSMUSG00 | Gm13223 | 746 | 0 | 0 | 0 | 0 | 0 | 0 | 0 | 0 | 0 | 0 | 0 | 0 | 0 | 0 | 0 | 0 | 0 | 0 |
| ENSMUSG00 | Gm15382 | 536 | 0 | 0 | 0 | 0 | 0 | 0 | 0 | 0 | 0 | 0 | 0 | 0 | 0 | 0 | 0 | 0 | 0 | 0 |
| ENSMUSG00 | Gm14667 | 1349 | 0 | 0 | 0 | 0 | 0 | 0 | 0.07014482 | 0 | 0 | 0 | 0 | 0 | 0 | 0 | 0 | 0 | 0 | 0 |
| ENSMUSG00 | Gm12765 | 597 | 0 | 0 | 0 | 0 | 0 | 0 | 0 | 0 | 0 | 0 | 0 | 0 | 0 | 0 | 0 | 0 | 0 | 0 |
| ENSMUSG00 | Gm15531 | 251 | 0 | 0 | 0.38544646 | 0.1815181 | 0 | 0.19742125 | 0 | 0.60864386 | 0.32817941 | 0.37570281 | 0.38067696 | 0 | 0.1965415 | 0 | 0.16414963 | 0 | 0 | 0.43831958 |
| ENSMUSG00 | Gm11249 | 858 | 0.34677883 | 0.2329573 | 0.11275881 | 0.37171015 | 0.22439125 | 0.34652261 | 0.38600088 | 0.71221263 | 0.24001466 | 0.93422139 | 0.83522655 | 0.19515966 | 1.32241736 | 0.81195957 | 0.57624554 | 0.62478847 | 0.58070147 | 0.25645271 |
| ENSMUSG00 | Gm14153 | 1138 | 0 | 0 | 0 | 0 | 0 | 0 | 0 | 0 | 0 | 0 | 0 | 0 | 0 | 0 | 0 | 0 | 0 | 0 |
| ENSMUSG00 | Gm4760 | 1029 | 0 | 0 | 0 | 0 | 0 | 0.09631241 | 0 | 0 | 0 | 0 | 0 | 0 | 0 | 0 | 0 | 0 | 0 | 0 |
| ENSMUSG00 | Gm11736 | 504 | 0 | 0.19829104 | 0.09597923 | 0.63279228 | 0.47749924 | 0.49159458 | 0.37549745 | 1.89446441 | 0 | 0.65487088 | 0.47395792 | 0 | 1.6639734 | 0.46075484 | 0 | 0.60778742 | 0.60835392 | 0 |
| ENSMUSG00 | Gm12438 | 335 | 0 | 0 | 0 | 0 | 0 | 0 | 0 | 0 | 0 | 0 | 0 | 0 | 0 | 0 | 0 | 0 | 0 | 0 |
| ENSMUSG00 | Gm15385 | 661 | 0 | 0 | 0 | 0 | 0 | 0 | 0 | 0 | 0 | 0 | 0.07227679 | 0 | 0 | 0 | 0 | 0.1158566 | 0 | 0.08322104 |
| ENSMUSG00 | Gm4908 | 1211 | 0.03509924 | 0.05501717 | 0.15978045 | 0.03762266 | 0 | 0.04091886 | 0.0781382 | 0 | 0.03401034 | 0.07787069 | 0.03945083 | 0.13827167 | 0.24441908 | 0 | 0.06804551 | 0 | 0.03164847 | 0.04542453 |
| ENSMUSG00 | Gm11863 | 230 | 0 | 0 | 0 | 0 | 0 | 0 | 0 | 0 | 0 | 0 | 0 | 0 | 0 | 0 | 0 | 0 | 0 | 0 |
| ENSMUSG00 | Gm11356 | 340 | 0 | 0 | 0 | 0 | 0 | 0 | 0 | 0 | 0 | 0 | 0 | 0 | 0 | 0 | 0 | 0 | 0 | 0 |
| ENSMUSG00 | Gm14494 | 1215 | 0.27986947 | 0.08225406 | 0.11944082 | 0.26249161 | 0.23768851 | 0.285489 | 0 | 0.18860446 | 0.13559347 | 0.07761433 | 0 | 0.09187763 | 0.1218072 | 0.03822559 | 0.27128596 | 0.06302981 | 0.18926566 | 0.40747486 |
| ENSMUSG00 | Gm14578 | 390 | 0 | 0 | 0 | 0 | 0 | 0 | 0 | 0 | 0 | 0 | 0 | 0 | 0 | 0 | 0 | 0 | 0 | 0 |
| ENSMUSG00 | Gm4732 | 467 | 0 | 0 | 0 | 0 | 0 | 0 | 0 | 0 | 0 | 0 | 0 | 0 | 0 | 0 | 0 | 0 | 0 | 0 |
| ENSMUSG00 | Gm6044 | 498 | 0.4267588 | 0.06689336 | 0.48567802 | 0.09148804 | 0.09665045 | 0.49751741 | 0.09500538 | 0.07669157 | 0 | 0.18936025 | 0.57560191 | 0 | 0.29718022 | 0.18652244 | 0.1654681 | 1.23022032 | 0.15392087 | 0.22092011 |
| ENSMUSG00 | Gm13430 | 287 | 0 | 0 | 0 | 0 | 0 | 0 | 0 | 0.26614915 | 0 | 0 | 0 | 0 | 0 | 0 | 0 | 0.2668335 | 0 | 0 |
| ENSMUSG00 | Gm16424 | 398 | 0 | 0 | 0 | 0 | 0 | 0 | 0 | 0 | 0 | 0 | 0 | 0 | 0 | 0 | 0 | 0 | 0 | 0 |
| ENSMUSG00 | Gm6429 | 474 | 0 | 0 | 0 | 0 | 0 | 0 | 0 | 0 | 0 | 0 | 0 | 0 | 0 | 0 | 0 | 0 | 0 | 0 |
| ENSMUSG00 | Gm12917 | 638 | 0 | 0.57436025 | 0.07582058 | 0 | 0.30176755 | 0.54368203 | 0.14831561 | 0.4789016 | 1.35566901 | 0.51732747 | 0.89858856 | 0.87485365 | 1.00519581 | 0 | 0.51663393 | 0.48013301 | 0.72087079 | 0.43110585 |
| ENSMUSG00 | Gm12393 | 847 | 0 | 0 | 0 | 0 | 0 | 0 | 0 | 0 | 0 | 0 | 0 | 0 | 0 | 0 | 0 | 0 | 0 | 0 |
| ENSMUSG00 | Gm15309 | 239 | 0 | 0 | 0 | 0 | 0 | 0 | 0 | 0 | 0 | 0 | 0 | 0 | 0 | 0 | 0 | 0 | 0 | 0 |
| ENSMUSG00 | Gm5303 | 1251 | 0 | 0 | 0 | 0 | 0 | 0 | 0 | 0.061059 | 0 | 0 | 0 | 0 | 0 | 0 | 0 | 0 | 0 | 0 |
| ENSMUSG00 | Gm12876 | 440 | 0 | 0 | 0 | 0 | 0 | 0 | 0 | 0 | 0 | 0.42864276 | 0 | 0 | 0 | 0 | 0 | 0.17404822 | 0 | 0 |
| ENSMUSG00 | Gm14073 | 380 | 0 | 0 | 0 | 0 | 0 | 0 | 0 | 0 | 0 | 0 | 0 | 0 | 0 | 0 | 0 | 0 | 0 | 0 |
| ENSMUSG00 | Gm14421 | 1955 | 0 | 0 | 0.02474349 | 0 | 0 | 0 | 0 | 0.03907151 | 0 | 0 | 0 | 0 | 0 | 0 | 0.04214993 | 0 | 0 | 0 |
| ENSMUSG00 | Gm14620 | 382 | 0 | 0.17441306 | 0 | 0 | 0 | 0 | 0 | 0 | 0 | 0 | 0 | 0 | 0 | 0 | 0 | 0.10023719 | 0 | 0.28800579 |
| ENSMUSG00 | Rpl10-ps4 | 285 | 0 | 0 | 0 | 0 | 0 | 0 | 0 | 0 | 0 | 0 | 0 | 0 | 0 | 0 | 0 | 0 | 0 | 0 |
| ENSMUSG00 | Gm14790 | 1291 | 0 | 0 | 0 | 0 | 0 | 0 | 0 | 0 | 0 | 0 | 0 | 0 | 0 | 0 | 0 | 0 | 0 | 0 |
| ENSMUSG00 | Gm13386 | 123 | 0 | 0 | 0 | 0 | 0 | 0 | 0 | 0 | 0 | 0 | 0 | 0 | 0 | 0 | 0 | 0 | 0 | 0 |
| ENSMUSG00 | Gm13410 | 1413 | 0.03008151 | 0.023576 | 0 | 0 | 0.1703182 | 0 | 0 | 0 | 0 | 0 | 0 | 0 | 0 | 0 | 0.02915892 | 0.08129641 | 0 | 0 |
| ENSMUSG00 | Gm15171 | 333 | 0 | 0 | 0 | 0 | 0 | 0 | 0 | 0 | 0 | 0 | 0 | 0 | 0 | 0 | 0 | 0 | 0 | 0 |
| ENSMUSG00 | Gm11609 | 791 | 0 | 0 | 0 | 0 | 0 | 0 | 0 | 0 | 0 | 0 | 0 | 0 | 0 | 0 | 0 | 0 | 0 | 0 |
| ENSMUSG00 | Gm14724 | 536 | 0 | 0.06215092 | 0.09024912 | 0 | 0 | 0 | 0 | 0 | 0 | 0 | 0 | 0 | 0 | 0 | 0 | 0 | 0 | 0 |
| ENSMUSG00 | Gm15854 | 802 | 0 | 0 | 0 | 0 | 0 | 0 | 0 | 0 | 0 | 0 | 0 | 0 | 0 | 0 | 0 | 0 | 0 | 0 |
| ENSMUSG00 | Gm6341 | 872 | 0.09748894 | 0.03820286 | 0.05547423 | 0.15674671 | 0.11039432 | 0.05682653 | 0.05425766 | 0 | 0.14169673 | 0.16221572 | 0 | 0.19202636 | 0 | 0.1597847 | 0.18899796 | 0.13173374 | 0.43952175 | 0 |
| ENSMUSG00 | Gm15578 | 474 | 0 | 0 | 0 | 0 | 0 | 0 | 0 | 0 | 0 | 0 | 0 | 0 | 0 | 0.09798331 | 0 | 0 | 0 | 0.11605297 |
| ENSMUSG00 | Gm6826 | 527 | 0 | 0 | 0 | 0 | 0 | 0 | 0 | 0 | 0 | 0 | 0 | 0 | 0 | 0 | 0 | 0 | 0 | 0 |
| ENSMUSG00 | Gm11327 | 925 | 0 | 0 | 0 | 0 | 0 | 0 | 0 | 0 | 0 | 0 | 0 | 0 | 0 | 0 | 0 | 0 | 0 | 0 |
| ENSMUSG00 | Gm16203 | 495 | 0 | 0 | 0 | 0 | 0 | 0 | 0 | 0 | 0 | 0 | 0 | 0 | 0 | 0 | 0 | 0 | 0 | 0 |
| ENSMUSG00 | Gm14109 | 308 | 0 | 0 | 0 | 0 | 0 | 0 | 0 | 0 | 0 | 0 | 0 | 0.18121968 | 0 | 0 | 0 | 0 | 0 | 0 |
| ENSMUSG00 | Gm15189 | 1964 | 0.02164215 | 0 | 0 | 0 | 0 | 0 | 0 | 0 | 0 | 0 | 0 | 0 | 0 | 0 | 0 | 0 | 0 | 0 |
| ENSMUSG00 | Gm12723 | 450 | 0 | 0 | 0 | 0 | 0 | 0 | 0 | 0 | 0 | 0 | 0 | 0 | 0 | 0 | 0 | 0 | 0 | 0 |
| ENSMUSG00 | Gm12210 | 152 | 0 | 0 | 0 | 0 | 0 | 0 | 0 | 0 | 0 | 0 | 0 | 0 | 0 | 0 | 0 | 0 | 0 | 0 |
| ENSMUSG00 | Gm14560 | 374 | 0 | 0 | 0 | 0 | 0 | 0 | 0 | 0 | 0 | 0 | 0 | 0 | 0 | 0 | 0 | 0 | 0 | 0 |
| ENSMUSG00 | Gm14422 | 150 | 0 | 0 | 0 | 0 | 0 | 0 | 0 | 0 | 0 | 0 | 0 | 0 | 0 | 0 | 0 | 0 | 0 | 0 |
| ENSMUSG00 | Gm14844 | 272 | 0 | 0 | 0 | 0 | 0 | 0 | 0 | 0 | 0 | 0 | 0 | 0 | 0 | 0 | 0 | 0 | 0 | 0 |
| ENSMUSG00 | Gm11504 | 172 | 0 | 0 | 0 | 0 | 0 | 0 | 0 | 0 | 0 | 0 | 0 | 0 | 0 | 0 | 0 | 0 | 0 | 0 |
| ENSMUSG00 | Gm15170 | 2021 | 0 | 0 | 0 | 0 | 0 | 0 | 0 | 0 | 0 | 0 | 0 | 0 | 0 | 0 | 0 | 0 | 0 | 0 |
| ENSMUSG00 | Gm11792 | 167 | 0 | 0 | 0 | 0 | 0 | 0 | 0 | 0 | 0 | 0 | 0 | 0 | 0 | 0 | 0 | 0 | 0 | 0 |
| ENSMUSG00 | Cyp4a29-ps | 1518 | 0 | 0 | 0 | 0 | 0 | 0 | 0 | 0 | 0 | 0 | 0 | 0 | 0 | 0 | 0.081426 | 0 | 0 | 0 |
| ENSMUSG00 | Gm12418 | 941 | 0.22585109 | 0.10620476 | 0 | 0.14525306 | 0.05114976 | 0 | 0.05027915 | 0.08117408 | 0.2188444 | 0 | 0.05077041 | 0 | 0.05242499 | 0 | 0.17513945 | 0.3255312 | 0 | 0.05845814 |
| ENSMUSG00 | Gm5073 | 819 | 0 | 0 | 0 | 0 | 0 | 0 | 0 | 0 | 0 | 0 | 0 | 0 | 0 | 0 | 0 | 0 | 0 | 0 |
| ENSMUSG00 | Gm15752 | 393 | 0 | 0 | 0 | 0 | 0 | 0 | 0 | 0 | 0 | 0 | 0 | 0 | 0 | 0 | 0 | 0 | 0 | 0 |
| ENSMUSG00 | Gm12288 | 1281 | 0 | 0 | 0 | 0 | 0 | 0 | 0 | 0 | 0 | 0 | 0.2237703 | 0 | 0 | 0 | 0 | 0 | 0 | 0 |
| ENSMUSG00 | Gm13424 | 404 | 0 | 0 | 0 | 0 | 0 | 0 | 0 | 0 | 0 | 0 | 0 | 0 | 0 | 0 | 0 | 0 | 0 | 0 |
| ENSMUSG00 | Gm14334 | 692 | 0 | 0 | 0 | 0 | 0 | 0 | 0 | 0 | 0 | 0 | 0 | 0 | 0 | 0 | 0 | 0 | 0 | 0 |
| ENSMUSG00 | Gm12579 | 782 | 0 | 0 | 0 | 0 | 0 | 0 | 0 | 0 | 0 | 0 | 0 | 0 | 0 | 0 | 0 | 0 | 0 | 0 |
| ENSMUSG00 | Gm15921 | 482 | 0.08818501 | 0 | 0 | 0.18904998 | 0.19971753 | 0.61683901 | 0 | 0.1584747 | 0 | 0 | 0 | 0 | 0 | 0.77085622 | 0 | 0.15888219 | 0.23854542 | 0.91301422 |
| ENSMUSG00 | Olfr1103-ps1 | 937 | 0 | 0 | 0 | 0 | 0 | 0 | 0 | 0 | 0 | 0 | 0 | 0 | 0 | 0 | 0 | 0 | 0 | 0 |
| ENSMUSG00 | Gm15102 | 1501 | 0 | 0 | 0 | 0 | 0 | 0 | 0 | 0 | 0 | 0 | 0 | 0 | 0 | 0 | 0 | 0 | 0 | 0 |
| ENSMUSG00 | Gm11380 | 999 | 0 | 0 | 0 | 0 | 0.14454031 | 0 | 0 | 0 | 0 | 0 | 0 | 0.05587153 | 0 | 0 | 0.0824856 | 0.11498681 | 0 | 0 |
| ENSMUSG00 | Gm11852 | 221 | 0 | 0 | 0 | 0 | 0 | 0 | 0 | 0 | 0 | 0 | 0 | 0 | 0 | 0 | 0 | 0 | 0 | 0 |
| ENSMUSG00 | Gm4913 | 1289 | 0 | 0 | 0 | 0 | 0 | 0 | 0 | 0 | 0 | 0 | 0 | 0 | 0 | 0 | 0 | 0 | 0 | 0 |
| ENSMUSG00 | Apc-ps1 | 469 | 0.63440561 | 0.49720738 | 0.41256742 | 0 | 0.71838692 | 0.10565615 | 0.40351965 | 0.16286739 | 0.35127092 | 0.2010691 | 0.10186558 | 0.11900994 | 0.63111195 | 0 | 0.0878498 | 0.5715016 | 0.40859592 | 0.23458041 |
| ENSMUSG00 | Gm12459 | 354 | 0 | 0 | 0 | 0 | 0 | 0 | 0 | 0 | 0 | 0 | 0 | 0 | 0 | 0 | 0 | 0 | 0 | 0 |
| ENSMUSG00 | Gm11892 | 1201 | 0 | 0 | 0 | 0 | 0 | 0 | 0 | 0 | 0 | 0 | 0 | 0 | 0 | 0 | 0 | 0 | 0 | 0 |
| ENSMUSG00 | Gm11866 | 334 | 0 | 0 | 0 | 0 | 0 | 0 | 0 | 0 | 0 | 0 | 0 | 0 | 0 | 0 | 0 | 0 | 0 | 0 |
| ENSMUSG00 | Gm12201 | 1232 | 0 | 0 | 0 | 0 | 0 | 0 | 0 | 0 | 0 | 0 | 0 | 0 | 0 | 0 | 0 | 0 | 0 | 0 |
| ENSMUSG00 | Gm13511 | 362 | 0 | 0.18404914 | 0.66814268 | 0.25171848 | 0 | 0.54754402 | 1.43767809 | 0 | 0.79642434 | 0.52100225 | 1.05580019 | 0.15418691 | 1.63531217 | 0 | 0.11381645 | 0.74042611 | 0.42349499 | 0.91175315 |
| ENSMUSG00 | Gm14655 | 577 | 0 | 0 | 0 | 0 | 0 | 0 | 0 | 0 | 0 | 0 | 0 | 0 | 0 | 0 | 0 | 0 | 0 | 0 |
| ENSMUSG00 | Gm12180 | 660 | 0 | 0 | 0 | 0.06903189 | 0 | 0 | 0 | 0.05786728 | 0 | 0 | 0 | 0 | 0 | 0 | 0 | 0 | 0.05807015 | 0 |
| ENSMUSG00 | Gm15967 | 1115 | 0 | 0 | 0 | 0 | 0.04316764 | 0 | 0 | 0 | 0 | 0.04228763 | 0 | 0 | 0 | 0 | 0 | 0 | 0.03437336 | 0 |
| ENSMUSG00 | Gm11427 | 1610 | 0 | 0 | 0 | 0 | 0 | 0 | 0 | 0 | 0 | 0 | 0 | 0 | 0 | 0 | 0 | 0 | 0 | 0 |
| ENSMUSG00 | Gm4917 | 649 | 0 | 0 | 0 | 0 | 0 | 0 | 0 | 0 | 0 | 0 | 0 | 0 | 0 | 0 | 0 | 0 | 0 | 0 |
| ENSMUSG00 | Gm11429 | 143 | 0 | 0 | 0 | 0 | 0 | 0 | 0 | 0 | 0 | 0 | 0 | 0 | 0 | 0 | 0 | 0 | 0 | 0 |
| ENSMUSG00 | Gm11344 | 334 | 0 | 0 | 0 | 0 | 0 | 0 | 0 | 0 | 0 | 0 | 0 | 0 | 0 | 0 | 0 | 0 | 0 | 0 |
| ENSMUSG00 | Gm8648 | 220 | 0 | 0 | 0 | 0 | 0 | 0.2252397 | 0 | 0.34720366 | 0 | 0 | 0 | 0 | 0 | 0 | 0 | 0 | 0 | 0 |
| ENSMUSG00 | Gm11287 | 602 | 0 | 0 | 0 | 0 | 0 | 0 | 0 | 0 | 0 | 0 | 0 | 0 | 0 | 0 | 0 | 0 | 0 | 0 |
| ENSMUSG00 | Gm14201 | 288 | 0 | 0 | 0 | 0 | 0 | 0 | 0 | 0 | 0 | 0 | 0 | 0 | 0 | 0 | 0 | 0 | 0 | 0 |
| ENSMUSG00 | Gm6580 | 1356 | 0.031346 | 0.07370109 | 0 | 0.03359959 | 0.07099104 | 0 | 0 | 0.02816549 | 0 | 0.03477191 | 0 | 0 | 0 | 0 | 0 | 0.02823791 | 0 | 0 |
| ENSMUSG00 | Gm14405 | 2327 | 0 | 0 | 0 | 0.01957931 | 0 | 0 | 0 | 0 | 0 | 0 | 0 | 0 | 0 | 0 | 0 | 0 | 0 | 0 |
| ENSMUSG00 | Gm12321 | 803 | 0 | 0 | 0 | 0.05673854 | 0 | 0.06170951 | 0 | 0 | 0 | 0.05871819 | 0 | 0 | 0 | 0 | 0 | 0 | 0 | 0 |
| ENSMUSG00 | Gm13409 | 542 | 0 | 0 | 0 | 0 | 0 | 0 | 0 | 0 | 0 | 0 | 0 | 0 | 0 | 0 | 0 | 0 | 0 | 0 |
| ENSMUSG00 | Gm13064 | 556 | 0 | 0 | 0 | 0 | 0 | 0 | 0 | 0 | 0 | 0.76323081 | 0 | 0 | 0 | 0.08353253 | 0 | 0 | 0 | 0 |
| ENSMUSG00 | Gm7190 | 1451 | 0 | 0 | 0 | 0 | 0 | 0.03415075 | 0 | 0 | 0.02838492 | 0 | 0 | 0.03846703 | 0 | 0 | 0 | 0 | 0.02641371 | 0 |
| ENSMUSG00 | Gm12206 | 600 | 0 | 0 | 0 | 0 | 0 | 0.08258789 | 0 | 0 | 0 | 0 | 0 | 0 | 0 | 0 | 0 | 0 | 0 | 0 |
| ENSMUSG00 | D11Bhm181e | 281 | 0 | 0 | 0 | 0 | 0 | 0 | 0 | 0 | 0 | 0 | 0 | 0 | 0.7022337 | 0 | 0 | 0 | 0 | 0 |
| ENSMUSG00 | Gm7820 | 1051 | 0.0404426 | 0 | 0.04602619 | 0 | 0.04579631 | 0 | 0.09003364 | 0.03633911 | 0 | 0 | 0 | 0 | 0 | 0 | 0 | 0.03643255 | 0.03646651 | 0 |
| ENSMUSG00 | Gm12187 | 1374 | 0 | 0 | 0 | 0 | 0 | 0 | 0 | 0 | 0 | 0 | 0 | 0 | 0 | 0 | 0 | 0 | 0 | 0 |
| ENSMUSG00 | Gm12693 | 482 | 0.61729509 | 0.48379722 | 0.10036002 | 0.18904998 | 0.19971753 | 0.205613 | 0 | 0.1584747 | 0.93994123 | 0.29346911 | 0.29735451 | 0.34740039 | 0.51174188 | 0.86721325 | 0.51288244 | 0.31776438 | 0.23854542 | 0.68476066 |
| ENSMUSG00 | Gm15007 | 709 | 0 | 0 | 0 | 0 | 0 | 0 | 0 | 0 | 0 | 0 | 0 | 0 | 0 | 0 | 0 | 0 | 0 | 0 |
| ENSMUSG00 | Gm13903 | 192 | 0 | 0 | 0 | 0 | 0 | 0 | 0 | 0 | 0 | 0 | 0 | 0 | 0 | 0 | 0 | 0 | 0 | 0 |
| ENSMUSG00 | Gm13828 | 362 | 0 | 0 | 0 | 0 | 0 | 0 | 0 | 0 | 0 | 0 | 0 | 0.30837383 | 0 | 0 | 0 | 0 | 0 | 0 |
| ENSMUSG00 | Gm15575 | 1580 | 0 | 0.02108411 | 0 | 0 | 0 | 0 | 0 | 0 | 0 | 0 | 0 | 0 | 0 | 0 | 0.05215387 | 0 | 0.02425715 | 0 |
| ENSMUSG00 | Gm14969 | 305 | 0 | 0 | 0 | 0 | 0 | 0 | 0 | 0 | 0 | 0 | 0 | 0 | 0 | 0 | 0 | 0.12554298 | 0 | 0 |
| ENSMUSG00 | Olfr1146-ps1 | 921 | 0 | 0 | 0 | 0 | 0 | 0 | 0 | 0 | 0 | 0 | 0 | 0 | 0 | 0 | 0 | 0 | 0 | 0 |
| ENSMUSG00 | Gm11805 | 599 | 0.42576136 | 0.22245672 | 0 | 0.07606184 | 0.24106139 | 0.08272577 | 0 | 0.25504108 | 0.68758791 | 0.0787157 | 0.23927358 | 0.09318141 | 0 | 0 | 0.0687839 | 0 | 0.0639838 | 0 |
| ENSMUSG00 | Olfr404-ps1 | 1009 | 0 | 0 | 0 | 0 | 0 | 0 | 0 | 0 | 0 | 0 | 0 | 0 | 0 | 0 | 0 | 0 | 0 | 0 |
| ENSMUSG00 | Gm15027 | 137 | 0 | 0 | 0 | 0 | 0 | 0 | 0 | 0 | 0 | 0 | 0 | 0 | 0 | 0 | 0 | 0 | 0 | 0 |
| ENSMUSG00 | Gm12836 | 1012 | 0 | 0 | 0 | 0 | 0 | 0 | 0 | 0 | 0 | 0 | 0 | 0 | 0 | 0 | 0 | 0 | 0 | 0.16307047 |
| ENSMUSG00 | Gm15571 | 636 | 0 | 0 | 0 | 0 | 0.15135825 | 0 | 0 | 0 | 0 | 0 | 0 | 0 | 0 | 0 | 0 | 0 | 0 | 0 |
| ENSMUSG00 | Gm7689 | 374 | 0 | 0 | 0 | 0 | 0 | 0 | 0 | 0 | 0 | 0 | 0 | 0 | 0 | 0 | 0 | 0 | 0 | 0 |
| ENSMUSG00 | 4930595D18 | 814 | 0.52217661 | 0.49109918 | 0.05942694 | 0.27985899 | 0.29565064 | 0.12175119 | 0.46498947 | 0.28151648 | 0.10119537 | 0.23169879 | 0 | 0.41141766 | 0.06060432 | 0.11411324 | 0.4049293 | 0.18816023 | 0.37667122 | 0.20273627 |
| ENSMUSG00 | Gm13567 | 246 | 0 | 0 | 0 | 0 | 0.97829113 | 0 | 0 | 0 | 0 | 0.19166953 | 0 | 0.22689294 | 0 | 0 | 0.167486 | 0 | 0.15579795 | 0 |
| ENSMUSG00 | Gm15272 | 232 | 0 | 0 | 0 | 0.19638381 | 0 | 0 | 0 | 0 | 0 | 0 | 0 | 0 | 0 | 0 | 0 | 0 | 0 | 0 |
| ENSMUSG00 | Gm13581 | 276 | 0 | 0 | 0 | 0 | 0 | 0 | 0 | 0 | 0 | 0 | 0 | 0 | 0 | 0 | 0 | 0 | 0 | 0 |
| ENSMUSG00 | Gm9432 | 758 | 0 | 0 | 0.19145197 | 0 | 0 | 0 | 0 | 0 | 0.05433577 | 0.12440819 | 0 | 0.22090632 | 0 | 0.24508753 | 0 | 0.05051531 | 0 | 0 |
| ENSMUSG00 | Gm15066 | 2607 | 0.01630425 | 0.0255565 | 0 | 0.01747643 | 0 | 0 | 0.01814832 | 0 | 0 | 0 | 0 | 0 | 0 | 0.03563029 | 0.0158042 | 0 | 0.05880521 | 0 |
| ENSMUSG00 | Gm12080 | 1880 | 0 | 0 | 0 | 0 | 0 | 0 | 0 | 0 | 0 | 0 | 0 | 0 | 0 | 0 | 0 | 0 | 0 | 0 |
| ENSMUSG00 | Gm12388 | 1886 | 0 | 0 | 0 | 0 | 0 | 0 | 0 | 0 | 0 | 0 | 0 | 0 | 0 | 0 | 0 | 0 | 0 | 0 |
| ENSMUSG00 | Gm11421 | 293 | 0 | 0 | 0 | 0 | 0 | 0 | 0 | 0 | 0 | 0 | 0 | 0 | 0 | 0 | 0 | 0 | 0 | 0 |
| ENSMUSG00 | Gm13788 | 600 | 0 | 0 | 0 | 0 | 0 | 0 | 0 | 0 | 0 | 0 | 0 | 0 | 0 | 0 | 0 | 0 | 0 | 0 |
| ENSMUSG00 | Gm7821 | 787 | 0 | 0 | 0 | 0 | 0 | 0 | 0 | 0 | 0 | 0 | 0 | 0 | 0 | 0 | 0 | 0 | 0 | 0 |
| ENSMUSG00 | Gm11338 | 1157 | 0 | 0 | 0 | 0 | 0 | 0 | 0 | 0 | 0 | 0 | 0 | 0 | 0 | 0 | 0 | 0 | 0 | 0 |
| ENSMUSG00 | Gm14780 | 2736 | 0 | 0.02435153 | 0 | 0 | 0.03518415 | 0.01811138 | 0.01729265 | 0 | 0 | 0 | 0 | 0 | 0 | 0 | 0 | 0.01399511 | 0 | 0 |
| ENSMUSG00 | Gm11522 | 175 | 0.48577344 | 2.28431275 | 1.10568069 | 1.56209295 | 0.82511869 | 0.84947543 | 0.27035817 | 0.4364846 | 4.00097582 | 0.26943259 | 0 | 1.91367986 | 1.97327669 | 1.32697393 | 0.94174985 | 3.28205206 | 2.84709634 | 0.31433775 |
| ENSMUSG00 | Gm11969 | 355 | 0 | 0 | 0.81758079 | 0.12834097 | 0.13558288 | 0.4187555 | 0.79965091 | 0.75308962 | 0.23203671 | 0 | 0.67288674 | 0.78613609 | 1.11170518 | 0.78497049 | 0 | 1.40219126 | 0.539807 | 1.08468661 |
| ENSMUSG00 | Gm14869 | 883 | 0 | 0 | 0 | 0 | 0 | 0 | 0 | 0 | 0.13993154 | 0 | 0 | 0 | 0 | 0 | 0 | 0 | 0 | 0 |
| ENSMUSG00 | Gm13737 | 231 | 0 | 0 | 0 | 0 | 0 | 0 | 0 | 0 | 0 | 0 | 0 | 0 | 0 | 0 | 0 | 0 | 0 | 0 |
| ENSMUSG00 | Gm12274 | 824 | 0 | 0 | 0 | 0 | 0 | 0.12027363 | 0.0574183 | 0 | 0 | 0.05722173 | 0 | 0 | 0 | 0 | 0 | 0 | 0 | 0 |
| ENSMUSG00 | Gm14875 | 318 | 0 | 0 | 0 | 0 | 0 | 0 | 0 | 0 | 0 | 0 | 0 | 0 | 0 | 0 | 0 | 0 | 0 | 0 |
| ENSMUSG00 | Gm11912 | 263 | 0 | 0 | 0 | 0 | 0 | 0 | 0 | 0 | 0 | 0 | 0 | 0 | 0 | 0 | 0 | 0 | 0 | 0 |
| ENSMUSG00 | Gm14081 | 566 | 0 | 0.0588567 | 0 | 0.16099309 | 0.08503873 | 0.175098 | 0 | 0 | 0 | 0 | 0 | 0 | 0.08205669 | 0 | 0 | 0.20295375 | 0 | 0.19437847 |
| ENSMUSG00 | Hbb-bh3 | 298 | 0 | 0 | 0 | 0 | 0 | 0 | 0 | 0 | 0 | 0 | 0 | 0 | 0 | 0 | 0 | 0 | 0 | 0 |
| ENSMUSG00 | Gm16425 | 2679 | 0.14279454 | 0.31087061 | 0.21667875 | 0.18707409 | 0.32339478 | 0.18496728 | 0.28256919 | 0.25661189 | 0.18448607 | 0.31680204 | 0.17833131 | 0.16667611 | 0.07365721 | 0.22537258 | 0.24607126 | 0.17151448 | 0.22889912 | 0.10266724 |
| ENSMUSG00 | Gm11410 | 1142 | 0.03721994 | 0 | 0 | 0.07979167 | 0 | 0.08678237 | 0.04142967 | 0 | 0.18032625 | 0 | 0.20917232 | 0 | 0.08639565 | 0.08133816 | 0 | 0.03352943 | 0.06712136 | 0 |
| ENSMUSG00 | Gm12919 | 1593 | 0.08004741 | 0.02091205 | 0.03036631 | 0.02860078 | 0 | 0.03110655 | 0 | 0 | 0 | 0 | 0 | 0 | 0.03096793 | 0 | 0 | 0 | 0 | 0 |
| ENSMUSG00 | Gm14528 | 230 | 0 | 0 | 0 | 0 | 0 | 0 | 0 | 0 | 0 | 0 | 0 | 0 | 0 | 0 | 0 | 0 | 0 | 0 |
| ENSMUSG00 | Gm12427 | 300 | 0 | 0 | 0 | 0 | 0 | 0 | 0 | 0 | 0 | 0 | 0 | 0 | 0 | 0 | 0 | 0 | 0.51101729 | 0 |
| ENSMUSG00 | Gm15016 | 719 | 0 | 0 | 0 | 0 | 0 | 0 | 0 | 0 | 0 | 0 | 0 | 0 | 0 | 0 | 0 | 0 | 0 | 0 |
| ENSMUSG00 | Gm11255 | 1179 | 0 | 0 | 0 | 0 | 0 | 0 | 0 | 0 | 0 | 0 | 0 | 0 | 0 | 0 | 0 | 0 | 0 | 0 |
| ENSMUSG00 | Gm15215 | 488 | 0 | 0 | 0 | 0 | 0 | 0 | 0 | 0 | 0 | 0 | 0 | 0 | 0 | 0 | 0 | 0 | 0 | 0 |
| ENSMUSG00 | Gm7831 | 551 | 0 | 0 | 0 | 0 | 0 | 0 | 0 | 0 | 0 | 0.34229186 | 0 | 0 | 0.17906322 | 0 | 0 | 0.06949294 | 0.34778854 | 0.29950512 |
| ENSMUSG00 | Gm11546 | 351 | 0 | 0 | 0 | 0 | 0 | 0 | 0 | 0 | 0 | 0 | 0 | 0 | 0 | 0 | 0 | 0 | 0 | 0 |
| ENSMUSG00 | Gm11779 | 367 | 0 | 0 | 0 | 0 | 0 | 0 | 0 | 0 | 0 | 0 | 0 | 0 | 0 | 0.12655065 | 0 | 0 | 0.10443133 | 0 |
| ENSMUSG00 | Gm11862 | 658 | 0 | 0 | 0 | 0 | 0 | 0 | 0 | 0 | 0 | 0 | 0 | 0 | 0 | 0 | 0 | 0 | 0 | 0 |
| ENSMUSG00 | Gm14158 | 307 | 0 | 0 | 0 | 0 | 0 | 0 | 0 | 0 | 0 | 0 | 0 | 0 | 0 | 0 | 0 | 0 | 0 | 0 |
| ENSMUSG00 | Gm15096 | 877 | 0 | 0 | 0 | 0 | 0 | 0 | 0 | 0 | 0 | 0 | 0 | 0 | 0 | 0 | 0 | 0 | 0 | 0 |
| ENSMUSG00 | Gm11796 | 971 | 0 | 0 | 0 | 0 | 0 | 0 | 0 | 0 | 0 | 0.04855891 | 0 | 0 | 0 | 0 | 0 | 0 | 0 | 0 |
| ENSMUSG00 | Gm13406 | 255 | 0 | 0.2612776 | 0 | 0 | 0.56625793 | 0.58297334 | 0 | 0.59909651 | 0.48454724 | 0 | 0 | 0 | 0.5803755 | 0 | 0.16157473 | 0 | 0 | 0 |
| ENSMUSG00 | Gm12262 | 304 | 0 | 0 | 0 | 0 | 0 | 0 | 0 | 0 | 0 | 0 | 0 | 0 | 0 | 0 | 0 | 0 | 0 | 0.18095101 |
| ENSMUSG00 | Gm13246 | 314 | 0 | 0 | 0 | 0 | 0 | 0 | 0 | 0 | 0 | 0 | 0 | 0 | 0 | 0 | 0 | 0 | 0 | 0 |
| ENSMUSG00 | Gm7222 | 905 | 0 | 0 | 0.10690283 | 0 | 0 | 0.1095088 | 0 | 0.1688062 | 0.09101992 | 0 | 0.05279001 | 0 | 0.05451041 | 0 | 0 | 0 | 0.084699 | 0.12156709 |
| ENSMUSG00 | Gm11698 | 454 | 0 | 0.22012926 | 0.10654963 | 0.10035472 | 0 | 0 | 0 | 0 | 0 | 0 | 0 | 0 | 0 | 0 | 0 | 0.59038381 | 0 | 0.24233087 |
| ENSMUSG00 | Hspe1-ps4 | 284 | 0 | 0 | 0 | 0 | 0 | 0 | 0 | 0 | 0 | 0 | 0 | 0 | 0 | 0 | 0 | 0 | 0 | 0 |
| ENSMUSG00 | Mup-ps1 | 538 | 0 | 0 | 0 | 0 | 0 | 0 | 0 | 0 | 0 | 0 | 0 | 0 | 0 | 0 | 0 | 0 | 0 | 0 |
| ENSMUSG00 | Gm13453 | 356 | 13.0142252 | 14.410634 | 11.8216212 | 6.27104262 | 20.550709 | 10.4394804 | 14.4861854 | 13.7320997 | 13.1889404 | 11.2578926 | 16.2381179 | 9.09356301 | 18.153037 | 17.873146 | 16.4343285 | 10.3255571 | 17.1176438 | 17.7697957 |
| ENSMUSG00 | Gm13259 | 455 | 0 | 0 | 0 | 0 | 0 | 0 | 0 | 0 | 0 | 0 | 0 | 0 | 0 | 0 | 0 | 0 | 0 | 0 |
| ENSMUSG00 | Gm15659 | 340 | 0 | 0 | 0 | 0 | 0 | 0 | 0 | 0 | 0 | 0 | 0.14051458 | 0 | 0 | 0 | 0 | 0 | 0 | 0 |
| ENSMUSG00 | Gm14668 | 1722 | 0 | 0 | 0 | 0 | 0 | 0 | 0.02747542 | 0 | 0 | 0 | 0 | 0.12965311 | 0 | 0 | 0 | 0 | 0.0445137 | 0 |
| ENSMUSG00 | Gm15180 | 491 | 0 | 0 | 0 | 0 | 0 | 0 | 0 | 0 | 0 | 0 | 0 | 0 | 0 | 0 | 0 | 0 | 0 | 0 |
| ENSMUSG00 | Gm11185 | 1006 | 0.33801333 | 0.16557104 | 0.28851012 | 0.27173585 | 0.28706913 | 0.14777157 | 0.37624397 | 0.45557538 | 0.53223132 | 0.89052024 | 0.5223902 | 0.22193106 | 0.39230153 | 0.36933668 | 0.53242567 | 0.2283734 | 0.3428794 | 0.10936204 |
| ENSMUSG00 | Gm11839 | 1952 | 0.06532558 | 0.44371683 | 0 | 0.16338489 | 0.41918171 | 0.15231373 | 0.460523 | 0.31305248 | 0.52749124 | 0.02415507 | 0.29369852 | 0.45750543 | 0.15163499 | 0.26172385 | 0.10553677 | 0.45117007 | 0.5890312 | 0.33817074 |
| ENSMUSG00 | Gm6574 | 361 | 0 | 0 | 0 | 0 | 0 | 0 | 0 | 0 | 0 | 0 | 0 | 0 | 0 | 0 | 0 | 0 | 0 | 0 |
| ENSMUSG00 | Gm14623 | 597 | 0 | 0 | 0 | 0 | 0 | 0 | 0 | 0 | 0 | 0 | 0 | 0 | 0 | 0 | 0 | 0 | 0 | 0 |
| ENSMUSG00 | Gm5403 | 2794 | 0.21298227 | 0.38153637 | 0.24238705 | 0.24460117 | 0.37899152 | 0.26603114 | 0.32173976 | 0.24604964 | 0.17689269 | 0.27001119 | 0.20518951 | 0.11986184 | 0.14125102 | 0.19947353 | 0.35391458 | 0.13704584 | 0.23319508 | 0.31501278 |
| ENSMUSG00 | Gm15300 | 277 | 0 | 0 | 0 | 0 | 0 | 0 | 0 | 0 | 0 | 0 | 0 | 0 | 0 | 0 | 0 | 0 | 0 | 0 |
| ENSMUSG00 | Gm13210 | 272 | 0 | 0 | 0 | 0 | 0 | 0 | 0 | 0 | 0 | 0 | 0 | 0 | 0 | 0 | 0 | 0 | 0 | 0 |
| ENSMUSG00 | Gm7108 | 3405 | 0 | 0 | 0 | 0 | 0 | 0.02910586 | 0 | 0 | 0 | 0.02769498 | 0 | 0 | 0 | 0 | 0 | 0 | 0 | 0 |
| ENSMUSG00 | Gm15116 | 662 | 0 | 0 | 0 | 0 | 0 | 0 | 0 | 0 | 0 | 0 | 0 | 0 | 0 | 0 | 0 | 0 | 0 | 0 |
| ENSMUSG00 | Gm12738 | 535 | 0 | 0 | 0 | 0 | 0 | 0 | 0 | 0.07138767 | 0 | 0 | 0 | 0 | 0 | 0 | 0 | 0 | 0 | 0 |
| ENSMUSG00 | Kars-ps1 | 1766 | 0 | 0 | 0 | 0 | 0 | 0 | 0 | 0 | 0 | 0 | 0 | 0 | 0 | 0 | 0 | 0 | 0 | 0 |
| ENSMUSG00 | Gm15568 | 1212 | 0 | 0 | 0 | 0 | 0 | 0 | 0 | 0 | 0 | 0 | 0 | 0 | 0 | 0 | 0 | 0 | 0 | 0 |
| ENSMUSG00 | Gm5939 | 4090 | 0 | 0 | 0 | 0.03341886 | 0 | 0.01211558 | 0.04627157 | 0.03735198 | 0.01007005 | 0.03458487 | 0 | 0 | 0.01206159 | 0.02271105 | 0.01007373 | 0.00936201 | 0.03748293 | 0.02689932 |
| ENSMUSG00 | Gm12625 | 381 | 0 | 0 | 0 | 0 | 0 | 0 | 0 | 0 | 0 | 0 | 0 | 0 | 0 | 0 | 0 | 0 | 0 | 0 |
| ENSMUSG00 | Gm13460 | 352 | 0 | 0 | 0 | 0 | 0 | 0 | 0 | 0 | 0.11700715 | 0 | 0 | 0 | 0 | 0 | 0 | 0 | 0 | 0 |
| ENSMUSG00 | Gm7816 | 2371 | 0.01792711 | 0.04215044 | 0 | 0.05764788 | 0 | 0.04179902 | 0.07981894 | 0 | 0 | 0 | 0.04029942 | 0.04708196 | 0.10403188 | 0.0195884 | 0.05213187 | 0.03229912 | 0.09698768 | 0.09280322 |
| ENSMUSG00 | Gm14470 | 465 | 0.09140898 | 0 | 0.31208729 | 0.19596148 | 0.20701903 | 0.21313004 | 0.20349539 | 0.0821342 | 0.08857315 | 0.10139936 | 0.20548369 | 0.36010105 | 0.10609014 | 0.09987976 | 0.0886055 | 0.08234539 | 0 | 0.23659831 |
| ENSMUSG00 | Gm14874 | 435 | 0 | 0 | 0 | 0 | 0 | 0 | 0 | 0.3511945 | 0.09468165 | 0 | 0.10982749 | 0 | 0 | 0 | 0 | 0 | 0 | 0.88520402 |
| ENSMUSG00 | Gm11461 | 385 | 0 | 0 | 0 | 0 | 0 | 0 | 0 | 0 | 0 | 0 | 0 | 0 | 0 | 0 | 0 | 0 | 0 | 0 |
| ENSMUSG00 | Gm12572 | 414 | 0 | 0 | 0 | 0 | 0 | 0 | 0 | 0 | 0 | 0 | 0 | 0 | 0 | 0 | 0 | 0 | 0 | 0 |
| ENSMUSG00 | Olfr1127-ps1 | 855 | 0 | 0 | 0 | 0 | 0 | 0 | 0 | 0 | 0 | 0 | 0 | 0 | 0 | 0 | 0 | 0 | 0 | 0 |
| ENSMUSG00 | Gm11511 | 539 | 0.70973392 | 0.12361 | 0.44873405 | 0 | 1.16088128 | 0.36773828 | 0.43889313 | 0 | 1.14619246 | 0 | 0.62045401 | 1.13909515 | 0.54914936 | 0 | 0.15288147 | 0.92352114 | 0.0711063 | 0 |
| ENSMUSG00 | Gm13254 | 653 | 0 | 0 | 0 | 0 | 0 | 0 | 0 | 0 | 0 | 0 | 0 | 0 | 0 | 0 | 0 | 0 | 0 | 0 |
| ENSMUSG00 | Gm13498 | 2165 | 0.01963288 | 0.06154807 | 0.02234343 | 0.06313309 | 0.02223184 | 0.0228881 | 0.02185343 | 0.01764083 | 0.01902379 | 0.02177862 | 0 | 0.02578091 | 0.0227861 | 0.02145223 | 0.01903074 | 0.07074477 | 0.05310803 | 0.15245018 |
| ENSMUSG00 | Gm12384 | 918 | 0 | 0 | 0 | 0 | 0 | 0 | 0 | 0 | 0 | 0 | 0 | 0 | 0 | 0 | 0 | 0 | 0 | 0 |
| ENSMUSG00 | Gm12550 | 328 | 0 | 0 | 0 | 0 | 0 | 0 | 0 | 0 | 0 | 0 | 0 | 0 | 0 | 0 | 0 | 0 | 0 | 0 |
| ENSMUSG00 | Gm14080 | 785 | 0.21658688 | 0 | 0 | 0 | 0 | 0 | 0.36162557 | 0 | 0 | 0.06006459 | 0 | 0 | 0 | 0 | 0 | 0 | 0.29293985 | 0 |
| ENSMUSG00 | Gm4993 | 1173 | 0.0362363 | 0 | 0.04123916 | 0 | 0 | 0.16897778 | 0.08066953 | 0 | 0.03511212 | 0 | 0.16291546 | 0 | 0 | 0.19797139 | 0 | 0 | 0.03267374 | 0 |
| ENSMUSG00 | Gm13551 | 429 | 0 | 0 | 0 | 0 | 0 | 0 | 0 | 0 | 0 | 0 | 0 | 0 | 0 | 0 | 0 | 0 | 0 | 0 |
| ENSMUSG00 | Gm15376 | 782 | 0 | 0 | 0 | 0 | 0 | 0 | 0 | 0 | 0 | 0 | 0 | 0 | 0 | 0 | 0 | 0 | 0 | 0 |
| ENSMUSG00 | Gm14589 | 726 | 0 | 0 | 0 | 0 | 0 | 0 | 0 | 0 | 0 | 0 | 0 | 0 | 0 | 0 | 0 | 0 | 0 | 0 |
| ENSMUSG00 | Gm13638 | 1156 | 0 | 0.02881738 | 0.08369123 | 0 | 0 | 0 | 0 | 0 | 0 | 0 | 0 | 0 | 0 | 0 | 0 | 0 | 0 | 0 |
| ENSMUSG00 | Gm11838 | 230 | 0 | 0 | 0 | 0.1980915 | 0 | 0 | 0 | 0 | 0 | 0 | 0 | 0 | 0.2144866 | 0 | 0 | 0 | 0 | 0 |
| ENSMUSG00 | Gm13332 | 1397 | 0.03042604 | 0 | 0 | 0 | 0.03445377 | 0.03547082 | 0 | 0.02733887 | 0 | 0 | 0 | 0 | 0.03531275 | 0 | 0.05898576 | 0 | 0 | 0.0393766 |
| ENSMUSG00 | Ctsf | 1961 | 93.8538563 | 102.350937 | 106.663511 | 110.28979 | 105.124441 | 120.71056 | 83.7441655 | 93.7963335 | 96.0458626 | 91.2958798 | 98.1567097 | 92.361971 | 90.2364035 | 111.05371 | 117.364554 | 142.69646 | 109.936471 | 120.313136 |
| ENSMUSG00 | Gm15361 | 443 | 0.19189696 | 0 | 0.54597664 | 0.4113864 | 0 | 0 | 0.42720252 | 0 | 0.27891546 | 0.106435 | 0.32353245 | 0.12599472 | 0.22271746 | 0.20967985 | 0.09300577 | 0.08643478 | 0 | 0.37252217 |
| ENSMUSG00 | Gm13688 | 607 | 0 | 0 | 0 | 0 | 0.15858953 | 0.08163548 | 0 | 0.18875981 | 0 | 0 | 0 | 0.09195332 | 0.89398862 | 0.15302829 | 0 | 0 | 0.69454574 | 0.18124912 |
| ENSMUSG00 | Gm15815 | 254 | 0 | 0 | 0 | 0.17937419 | 0 | 0 | 0 | 0 | 0 | 0 | 0 | 0 | 0 | 0 | 0 | 0 | 0.4526728 | 0 |
| ENSMUSG00 | Flg | 931 | 0 | 0 | 0 | 0 | 0 | 0 | 0 | 0 | 0 | 0 | 0 | 0 | 0 | 0 | 0 | 0 | 0 | 0 |
| ENSMUSG00 | Gm13502 | 3158 | 0.04037857 | 0.01054873 | 0.01531777 | 0 | 0.01524127 | 0.01569118 | 0.01498185 | 0.01209386 | 0 | 0.10451391 | 0.03025647 | 0.01767437 | 0.06248501 | 0.04412041 | 0.0521869 | 0.06062477 | 0.04854502 | 0.03483794 |
| ENSMUSG00 | Csnk2a1-ps | 1174 | 0 | 0 | 0 | 0 | 0 | 0 | 0 | 0 | 0 | 0 | 0 | 0 | 0 | 0 | 0 | 0 | 0 | 0 |
| ENSMUSG00 | Gm8812 | 411 | 0 | 0 | 0 | 0 | 0 | 0 | 0 | 0 | 0 | 0 | 0 | 0 | 0 | 0.11300265 | 0 | 0.09316449 | 0 | 0 |
| ENSMUSG00 | Gm15030 | 483 | 0 | 0 | 0 | 0 | 0 | 0 | 0 | 0 | 0 | 0 | 0 | 0 | 0 | 0 | 0 | 0 | 0 | 0 |
| ENSMUSG00 | Gm13679 | 483 | 0 | 0 | 0 | 0 | 0 | 0 | 0 | 0 | 0 | 0 | 0 | 0 | 0 | 0.09615753 | 0 | 0 | 0.07935051 | 0 |
| ENSMUSG00 | Gm12010 | 2015 | 0 | 0 | 0 | 0 | 0 | 0 | 0 | 0 | 0 | 0 | 0 | 0 | 0 | 0 | 0 | 0 | 0 | 0 |
| ENSMUSG00 | Gm14313 | 288 | 0 | 0 | 0 | 0 | 0 | 0 | 0 | 0 | 0 | 0 | 0 | 0 | 0 | 0 | 0 | 0 | 0 | 0 |
| ENSMUSG00 | Gm14957 | 1452 | 0 | 0 | 0 | 0 | 0 | 0 | 0 | 0 | 0 | 0 | 0 | 0 | 0 | 0 | 0 | 0 | 0 | 0 |
| ENSMUSG00 | Defa-ps8 | 277 | 0 | 0 | 0 | 0 | 0 | 0 | 0 | 0 | 0 | 0 | 0 | 0 | 0 | 0 | 0 | 0 | 0 | 0 |
| ENSMUSG00 | Gm12638 | 297 | 0 | 0 | 0 | 0 | 0 | 0 | 0 | 0 | 0 | 0 | 0 | 0 | 0 | 0 | 0 | 0 | 0 | 0 |
| ENSMUSG00 | Gm11316 | 1039 | 0 | 0 | 0 | 0 | 0 | 0 | 0 | 0 | 0 | 0 | 0 | 0 | 0 | 0 | 0 | 0 | 0 | 0 |
| ENSMUSG00 | Gm12085 | 939 | 0 | 0 | 0 | 0 | 0 | 0 | 0 | 0 | 0 | 0 | 0 | 0 | 0 | 0 | 0 | 0 | 0 | 0 |
| ENSMUSG00 | Gm12332 | 921 | 0.04615111 | 0 | 0.10504567 | 0 | 0 | 0.05380319 | 0.05137099 | 0 | 0.04471934 | 0 | 0.10374584 | 0 | 0.05356343 | 0.05042789 | 0.22367837 | 0.04157504 | 0 | 0 |
| ENSMUSG00 | Gm12309 | 181 | 0 | 0 | 0 | 0 | 0 | 0 | 0 | 0 | 0.22754981 | 0 | 0 | 0 | 0 | 0 | 0 | 0 | 0 | 0 |
| ENSMUSG00 | Gm7905 | 1383 | 0 | 0 | 0 | 0 | 0 | 0 | 0 | 0 | 0 | 0 | 0 | 0 | 0 | 0 | 0 | 0 | 0 | 0 |
| ENSMUSG00 | Gm12110 | 578 | 0 | 0 | 0 | 0 | 0 | 0 | 0 | 0.06607682 | 0 | 0 | 0 | 0 | 0 | 0.08035309 | 0.14256594 | 0 | 0 | 0 |
| ENSMUSG00 | Mup-ps15 | 533 | 0 | 0 | 0 | 0 | 0 | 0 | 0 | 0 | 0 | 0 | 0 | 0 | 0 | 0 | 0 | 0 | 0 | 0 |
| ENSMUSG00 | Gm13315 | 988 | 0.08604287 | 0.0337175 | 0.19584425 | 0.13834325 | 0.09743304 | 0 | 0.04788733 | 0.19328139 | 0 | 0 | 0.04835522 | 0 | 0 | 0 | 0 | 0.03875568 | 0.0775836 | 0.11135447 |
| ENSMUSG00 | Gm13868 | 293 | 3.6267215 | 2.04652592 | 1.98116848 | 1.86598134 | 0.82136389 | 0.84560979 | 3.06805768 | 2.86768892 | 1.26511482 | 1.93108682 | 4.07636164 | 2.47646285 | 0.50510496 | 9.19371013 | 0.84371787 | 2.48300867 | 1.70048416 | 3.75488783 |
| ENSMUSG00 | AA414768 | 1256 | 75.7377261 | 62.1964466 | 60.7750246 | 60.6150518 | 58.2488248 | 56.3780703 | 35.1454852 | 30.8032672 | 36.3661194 | 32.772742 | 26.7022459 | 43.3282413 | 49.4497483 | 37.902221 | 47.3030602 | 53.1983359 | 46.1380262 | 35.9135888 |
| ENSMUSG00 | Gm15926 | 548 | 0 | 0 | 0 | 0 | 0 | 0 | 0 | 0 | 0 | 0 | 0 | 0 | 0 | 0 | 0 | 0 | 0 | 0 |
| ENSMUSG00 | Gm5760 | 989 | 0 | 0 | 0.04891156 | 0 | 0 | 0 | 0 | 0.03861719 | 0.04164461 | 0.04767513 | 0 | 0.05643646 | 0 | 0 | 0.08331963 | 0.07743298 | 0.03875258 | 0 |
| ENSMUSG00 | Gm5643 | 965 | 0.08809363 | 0.03452113 | 0.15038403 | 0.04721352 | 0.04987764 | 0.10269997 | 0.14708605 | 0.03957762 | 0.08536065 | 0.04886083 | 0.04950773 | 0 | 0.05112116 | 0.19251435 | 0.08539183 | 0.07935877 | 0.07943274 | 0.05700426 |
| ENSMUSG00 | Olfr1374-ps1 | 935 | 0 | 0 | 0 | 0 | 0 | 0 | 0 | 0 | 0 | 0 | 0 | 0 | 0 | 0 | 0 | 0 | 0 | 0 |
| ENSMUSG00 | Gm15090 | 1195 | 0 | 0 | 0 | 0 | 0 | 0 | 0 | 0 | 0 | 0 | 0 | 0 | 0 | 0 | 0 | 0 | 0 | 0 |
| ENSMUSG00 | Gm11742 | 232 | 0 | 0 | 0 | 0 | 0 | 0 | 0 | 0 | 0 | 0 | 0 | 0 | 0 | 0 | 0 | 0 | 0 | 0 |
| ENSMUSG00 | Gm11871 | 1060 | 0 | 0 | 0 | 0 | 0 | 0 | 0 | 0 | 0 | 0 | 0 | 0 | 0 | 0 | 0 | 0 | 0 | 0 |
| ENSMUSG00 | Gm15185 | 745 | 0 | 0.04471529 | 0 | 0 | 0 | 0 | 0 | 0 | 0 | 0 | 0 | 0.29968141 | 0 | 0 | 0 | 0 | 0 | 0 |
| ENSMUSG00 | Gm14402 | 504 | 0 | 0 | 0 | 0 | 0 | 0 | 0 | 0 | 0 | 0 | 0 | 0 | 0 | 0 | 0 | 0 | 0 | 0 |
| ENSMUSG00 | Gm15192 | 292 | 0 | 0 | 0 | 0 | 0 | 0 | 0 | 0 | 0 | 0 | 0 | 0 | 0 | 0 | 0.13113222 | 0 | 0 | 0 |
| ENSMUSG00 | Gm11284 | 1885 | 0.02254917 | 0.05301787 | 0 | 0 | 0.17873924 | 0.07886377 | 0.02509956 | 0.10130611 | 0.10924805 | 0 | 0.15206883 | 0.08883129 | 0.02617078 | 0.19711019 | 0.04371518 | 0 | 0.12199352 | 0.0583651 |
| ENSMUSG00 | Gm13935 | 796 | 0 | 0 | 0 | 0 | 0 | 0 | 0 | 0.09596081 | 0 | 0 | 0 | 0 | 0.06197477 | 0 | 0.05176075 | 0.04810378 | 0 | 0.06910692 |
| ENSMUSG00 | Gm14931 | 252 | 0 | 0 | 0 | 0 | 0 | 0 | 0 | 0 | 0 | 0 | 0 | 0 | 0 | 0 | 0 | 0 | 0 | 0 |
| ENSMUSG00 | Gm15005 | 511 | 0 | 0 | 0 | 0 | 0 | 0 | 0 | 0 | 0 | 0 | 0 | 0 | 0 | 0 | 0 | 0 | 0 | 0 |
| ENSMUSG00 | Gm14997 | 635 | 0 | 0 | 0 | 0 | 0 | 0 | 0 | 0 | 0 | 0 | 0 | 0 | 0 | 0 | 0 | 0 | 0 | 0 |
| ENSMUSG00 | Gm14121 | 1379 | 0.18493913 | 0.12078642 | 0.31570832 | 0.29735272 | 0.0349035 | 0.17966908 | 0.03430941 | 0.30465296 | 0.14933472 | 0.34191953 | 0.24251248 | 0.12142639 | 0.28618951 | 0.13471817 | 0.1195114 | 0 | 0.25013537 | 0.11967173 |
| ENSMUSG00 | Rpl38-ps1 | 429 | 0 | 0.07765243 | 0 | 0 | 0 | 0 | 0 | 0.08902658 | 0.28801759 | 0.1099084 | 0.11136354 | 0.13010644 | 0 | 0 | 0.19208185 | 0 | 0.08933869 | 0.12822636 |
| ENSMUSG00 | Vcp-rs | 2405 | 0.65392579 | 1.24663637 | 1.40796138 | 1.13665807 | 1.26083625 | 0.86536998 | 0.6885421 | 0.98458584 | 0.87339389 | 1.11750108 | 0.95351268 | 1.04436791 | 1.02561159 | 0.83039325 | 1.74742566 | 0.85974753 | 1.17927067 | 1.18938609 |
| ENSMUSG00 | Gm11826 | 770 | 0.05520153 | 0.173054 | 0.12564553 | 0 | 0.25003597 | 0 | 0.12289008 | 0 | 0.32093389 | 0.12246936 | 0.0620454 | 0 | 0.06406743 | 0.12063399 | 0 | 0.24864031 | 0.19909765 | 0 |
| ENSMUSG00 | Gm14220 | 448 | 0 | 0 | 0 | 0 | 0 | 0 | 0 | 0 | 0 | 0 | 0 | 0 | 0 | 0 | 0 | 0 | 0 | 0 |
| ENSMUSG00 | Gm11859 | 454 | 0 | 0 | 0 | 0 | 0 | 0 | 0 | 0 | 0 | 0 | 0 | 0 | 0 | 0 | 0 | 0 | 0 | 0 |
| ENSMUSG00 | Gm12400 | 1000 | 0 | 0 | 0 | 0 | 0 | 0 | 0 | 0 | 0 | 0 | 0 | 0.05581566 | 0 | 0 | 0 | 0 | 0 | 0 |
| ENSMUSG00 | Gm7599 | 795 | 0 | 0 | 0 | 0 | 0.0605433 | 0 | 0.0595128 | 0.14412227 | 0 | 0.11861812 | 0 | 0.14041676 | 0.18615818 | 0 | 0.88103956 | 0 | 0 | 0 |

| | | | | | | | | | | | | | | | | | | | |
|---|---|---|---|---|---|---|---|---|---|---|---|---|---|---|---|---|---|---|---|
| ENSMUSG00 Magea7-ps | 1201 | 0 | 0 | 0 | 0 | 0 | 0 | 0 | 0 | 0 | 0 | 0 | 0 | 0 | 0.50272534 | 0 | 0 | 0 | 0 |
| ENSMUSG00 Gm13579 | 515 | 0 | 0 | 0 | 0 | 0 | 0 | 0 | 0 | 0 | 0 | 0 | 0 | 0 | 0 | 0 | 0 | 0 | 0 |
| ENSMUSG00 Gm13397 | 877 | 0 | 0 | 0 | 0 | 0 | 0 | 0 | 0 | 0 | 0 | 0 | 0 | 0 | 0 | 0 | 0 | 0 | 0 |
| ENSMUSG00 Olfr1332-ps1 | 953 | 0 | 0 | 0 | 0 | 0 | 0 | 0 | 0 | 0 | 0 | 0 | 0 | 0 | 0 | 0 | 0 | 0 | 0 |
| ENSMUSG00 Gm11539 | 556 | 3.05792633 | 2.81601804 | 1.47904679 | 1.88471946 | 2.42390982 | 3.20845038 | 3.82926358 | 2.47288936 | 1.55560581 | 1.61126504 | 1.89037605 | 1.40543035 | 2.75052057 | 2.00478075 | 2.37131257 | 2.13490797 | 2.68835536 | 2.27555657 |
| ENSMUSG00 Gm12475 | 1388 | 0 | 0 | 0 | 0 | 0.03467718 | 0 | 0 | 0.05503228 | 0 | 0 | 0 | 0 | 0 | 0 | 0 | 0 | 0 | 0 |
| ENSMUSG00 Gm11693 | 563 | 0.07549765 | 0 | 0 | 0 | 0 | 0 | 0.25211019 | 0 | 0 | 0 | 0 | 0 | 0.43811649 | 0 | 0 | 0 | 0.13615026 | 0 |
| ENSMUSG00 Gm13156 | 320 | 0 | 0 | 0 | 0 | 0.15041226 | 0 | 0 | 0 | 0 | 0.2946919 | 0.14929675 | 0 | 0 | 0 | 0 | 0 | 0 | 0 |
| ENSMUSG00 Gm14238 | 322 | 0 | 0 | 0 | 0.14149393 | 0.29895605 | 0.15389048 | 0.29386757 | 0 | 0.3837253 | 0 | 0 | 0 | 0.15320471 | 0.28847259 | 0 | 0 | 0.11902577 | 0 |
| ENSMUSG00 Gm14237 | 429 | 0 | 0 | 0 | 0 | 0 | 0 | 0 | 0 | 0 | 0 | 0 | 0 | 0 | 0 | 0 | 0 | 0 | 0 |
| ENSMUSG00 Gm14686 | 842 | 0 | 0 | 0 | 0 | 0 | 0 | 0 | 0 | 0 | 0 | 0 | 0 | 0 | 0 | 0 | 0 | 0 | 0 |
| ENSMUSG00 Gm7363 | 400 | 0.10626294 | 0 | 0 | 0 | 0.12032981 | 0 | 0 | 0 | 0 | 0 | 0 | 0 | 0 | 0.23222044 | 0 | 0 | 0 | 0 |
| ENSMUSG00 Gm6491 | 988 | 0 | 0 | 0 | 0 | 0 | 0 | 0 | 0 | 0 | 0 | 0 | 0 | 0 | 0 | 0 | 0 | 0 | 0 |
| ENSMUSG00 Gm8260 | 352 | 0 | 0 | 0 | 0 | 0 | 0 | 0 | 0 | 0 | 0 | 0 | 0 | 0 | 0 | 0 | 0 | 0 | 0 |
| ENSMUSG00 Gm13913 | 259 | 0 | 0 | 0 | 0 | 0 | 0 | 0 | 0 | 0 | 0 | 0 | 0 | 0 | 0 | 0 | 0 | 0 | 0 |
| ENSMUSG00 Gm11289 | 765 | 0 | 0 | 0 | 0 | 0 | 0 | 0 | 0 | 0 | 0 | 0 | 0 | 0 | 0 | 0 | 0 | 0 | 0 |
| ENSMUSG00 Gm7643 | 398 | 0 | 0 | 0 | 0 | 0 | 0 | 0 | 0 | 0 | 0 | 0 | 0 | 0 | 0 | 0 | 0 | 0 | 0 |
| ENSMUSG00 Gm9060 | 804 | 0 | 0 | 0 | 0 | 0 | 0 | 0 | 0 | 0 | 0 | 0 | 0 | 0 | 0 | 0 | 0 | 0 | 0 |
| ENSMUSG00 Gm15783 | 631 | 0 | 0 | 0 | 0 | 0 | 0 | 0 | 0 | 0 | 0 | 0 | 0 | 0 | 0 | 0 | 0 | 0 | 0 |
| ENSMUSG00 Gm14907 | 853 | 0 | 0 | 0 | 0 | 0 | 0 | 0 | 0 | 0 | 0 | 0 | 0 | 0 | 0 | 0 | 0 | 0 | 0 |
| ENSMUSG00 Gm12540 | 318 | 0 | 0.10475753 | 0 | 0 | 0 | 0.15582621 | 0 | 0.12010189 | 0 | 0.14827265 | 0 | 0 | 0 | 0 | 0 | 0 | 0 | 0 |
| ENSMUSG00 Gm11581 | 761 | 0 | 0 | 0 | 0 | 0 | 0 | 0 | 0 | 0 | 0 | 0 | 0 | 0 | 0 | 0 | 0 | 0 | 0 |
| ENSMUSG00 Gm11708 | 831 | 0 | 0 | 0 | 0 | 0 | 0 | 0 | 0 | 0 | 0 | 0 | 0 | 0 | 0 | 0 | 0 | 0 | 0 |
| ENSMUSG00 Olfr1004-ps1 | 453 | 0 | 0 | 0 | 0 | 0 | 0 | 0 | 0 | 0 | 0 | 0 | 0 | 0 | 0 | 0 | 0 | 0 | 0 |
| ENSMUSG00 Gm12656 | 576 | 0 | 0.11566977 | 0 | 0 | 0 | 0.08602905 | 0 | 0 | 0 | 0 | 0 | 0 | 0 | 0 | 0 | 0.1329535 | 0 | 0 |
| ENSMUSG00 Gm13105 | 2033 | 0 | 0 | 0 | 0 | 0.02367532 | 0 | 0 | 0 | 0 | 0 | 0 | 0 | 0 | 0 | 0 | 0 | 0 | 0 |
| ENSMUSG00 Olfr360 | 1423 | 0 | 0 | 0 | 0 | 0 | 0 | 0 | 0 | 0 | 0 | 0 | 0 | 0 | 0 | 0 | 0 | 0 | 0 |
| ENSMUSG00 Gm8606 | 887 | 0.28752092 | 0 | 0 | 0 | 0 | 0 | 0 | 0.12917385 | 0.0464335 | 0 | 0 | 0 | 0.11123318 | 0 | 0.04645046 | 0.04316867 | 0 | 0 |
| ENSMUSG00 Gm12237 | 469 | 0 | 0 | 0 | 0 | 0 | 0 | 0 | 0 | 0 | 0 | 0 | 0 | 0 | 0 | 0 | 0 | 0 | 0 |
| ENSMUSG00 Gm14535 | 1125 | 0.11334714 | 0.41456046 | 0.25799216 | 0.20249353 | 0.21391966 | 0.22023437 | 0.46261286 | 0.40738563 | 0.29288189 | 0.29338215 | 0.67946608 | 0.19845569 | 0.35080475 | 0.6192545 | 0.40285966 | 0.34036095 | 0.47694947 | 0.19558794 |
| ENSMUSG00 Gm12895 | 225 | 0 | 0 | 0 | 0 | 0 | 0 | 0 | 0 | 0 | 0 | 0 | 0 | 0 | 0 | 0.36623605 | 0 | 0 | 0 |
| ENSMUSG00 Gm14887 | 661 | 0 | 0 | 0 | 0 | 0 | 0 | 0 | 0 | 0 | 0 | 0 | 0 | 0 | 0 | 0 | 0 | 0 | 0 |
| ENSMUSG00 Gm8806 | 1406 | 0.18138766 | 0.07108014 | 0.2408355 | 0.06480945 | 0.13693293 | 0.24670635 | 0.06730111 | 0.29880258 | 0.11717359 | 0.03353535 | 0.03397934 | 0.27768737 | 0 | 0.03303278 | 0.11721638 | 0.10893487 | 0.10903641 | 0.15649817 |
| ENSMUSG00 Gm13214 | 407 | 0 | 0 | 0 | 0 | 0 | 0 | 0 | 0 | 0 | 0 | 0 | 0 | 0 | 0 | 0 | 0 | 0.65917464 | 0 |
| ENSMUSG00 Gm15332 | 356 | 0 | 0 | 0 | 0 | 0 | 0 | 0 | 0 | 0 | 0 | 0 | 0 | 0 | 0 | 0 | 0 | 0 | 0 |
| ENSMUSG00 Gm11422 | 1102 | 0 | 0 | 0 | 0 | 0 | 0 | 0 | 0 | 0 | 0.04278648 | 0 | 0 | 0 | 0 | 0 | 0 | 0 | 0 |
| ENSMUSG00 Gm11235 | 332 | 0 | 0 | 0 | 0 | 0 | 0 | 0 | 0 | 0 | 0 | 0 | 0 | 0 | 0 | 0 | 0 | 0 | 0 |
| ENSMUSG00 Gm13392 | 149 | 0 | 0 | 0 | 0 | 0 | 0 | 0.31753476 | 0 | 0 | 0 | 0.32063731 | 0 | 0 | 0 | 0 | 0 | 0 | 0 |
| ENSMUSG00 Gm11449 | 475 | 0 | 0 | 0 | 0 | 0 | 0 | 0 | 0 | 0 | 0 | 0 | 0 | 0 | 0 | 0 | 0 | 0 | 0 |
| ENSMUSG00 Gm15057 | 533 | 0 | 0 | 0 | 0 | 0 | 0 | 0 | 0 | 0 | 0 | 0 | 0 | 0 | 0 | 0 | 0 | 0 | 0 |
| ENSMUSG00 Gm11882 | 505 | 0 | 0 | 0 | 0 | 0 | 0 | 0 | 0 | 0 | 0 | 0 | 0 | 0 | 0 | 0 | 0 | 0 | 0 |
| ENSMUSG00 Gm12653 | 560 | 0 | 0 | 0 | 0 | 0 | 0 | 0 | 0 | 0 | 0 | 0 | 0 | 0 | 0 | 0 | 0 | 0 | 0 |
| ENSMUSG00 Gm13196 | 378 | 0 | 0 | 0 | 0 | 0 | 0 | 0 | 0 | 0 | 0 | 0.25277756 | 0 | 0.13050772 | 0 | 0 | 0 | 0.10139232 | 0 |
| ENSMUSG00 Gm4919 | 602 | 0 | 0 | 0 | 0 | 0 | 0 | 0 | 0.25377012 | 0 | 0.07832343 | 0 | 0 | 0 | 0 | 0 | 0 | 0 | 0 |
| ENSMUSG00 Gm3244 | 389 | 0.54633902 | 0.59946082 | 0.2487071 | 1.05411157 | 0.61866226 | 0.50953968 | 0.4865057 | 0.78544786 | 0.42351173 | 0.96968027 | 0 | 0.57393998 | 0.50726907 | 0.23878708 | 0.31774979 | 1.27963469 | 1.08377703 | 0.28282317 |
| ENSMUSG00 Gm12767 | 337 | 0 | 0 | 0 | 0 | 0 | 0 | 0 | 0 | 0 | 0 | 0 | 0 | 0 | 0 | 0 | 0 | 0 | 0 |
| ENSMUSG00 Gm6433 | 874 | 0 | 0.03811544 | 0 | 0 | 0 | 0 | 0 | 0.1310952 | 0.04712416 | 0 | 0 | 0 | 0 | 0.10627938 | 0 | 0.04381076 | 0.13155479 | 0 |
| ENSMUSG00 Olfr595-ps1 | 926 | 0 | 0 | 0 | 0 | 0 | 0 | 0 | 0 | 0 | 0 | 0 | 0 | 0 | 0 | 0 | 0 | 0 | 0 |
| ENSMUSG00 Gm12813 | 896 | 0 | 0 | 0 | 0 | 0 | 0 | 0 | 0 | 0 | 0 | 0 | 0 | 0 | 0 | 0 | 0 | 0 | 0 |
| ENSMUSG00 Gm11411 | 718 | 0 | 0 | 0 | 0 | 0 | 0 | 0 | 0 | 0 | 0 | 0 | 0 | 0 | 0 | 0 | 0 | 0 | 0 |
| ENSMUSG00 Gm15426 | 555 | 0 | 0 | 0 | 0 | 0 | 0 | 0 | 0 | 0 | 0 | 0 | 0 | 0 | 0 | 0 | 0 | 0 | 0 |
| ENSMUSG00 Gm11860 | 1009 | 0 | 0 | 0 | 0 | 0 | 0 | 0 | 0 | 0 | 0 | 0 | 0 | 0 | 0 | 0 | 0 | 0 | 0 |
| ENSMUSG00 Gm11550 | 698 | 0.06089567 | 0 | 0 | 0 | 0 | 0 | 0.13556642 | 0 | 0.05900647 | 0 | 0 | 0 | 0 | 0 | 0 | 0 | 0 | 0 |
| ENSMUSG00 Gm13681 | 209 | 0 | 0 | 0 | 0 | 0 | 0 | 0 | 0 | 0 | 0 | 0 | 0 | 0 | 0 | 0 | 0 | 0 | 0 |
| ENSMUSG00 Gm13121 | 631 | 0.26944644 | 0.10558762 | 0.0766617 | 0.07220451 | 0 | 0 | 0.22494142 | 0.06052679 | 0 | 0 | 0 | 0 | 0.07818053 | 0 | 0.06529565 | 0.06068242 | 0 | 0.17435533 |
| ENSMUSG00 Gm14148 | 1039 | 0.36818728 | 0.32062458 | 0.09311555 | 0.13155258 | 0.09265048 | 0.14307815 | 0.45536746 | 0.25731166 | 0.03964053 | 0.13614255 | 0.22990837 | 0.48348505 | 0.28488114 | 0.08940152 | 0.31724008 | 0.07370866 | 0.14755071 | 0.21177712 |
| ENSMUSG00 Gm6335 | 1193 | 0.14251526 | 0 | 0 | 0 | 0 | 0.08307248 | 0 | 0 | 0 | 0.0395228 | 0.08009213 | 0 | 0.04135115 | 0.1167915 | 0.10360827 | 0 | 0.06425196 | 0 |
| ENSMUSG00 Gm15194 | 433 | 0 | 0 | 0 | 0 | 0 | 0 | 0 | 0 | 0 | 0 | 0 | 0 | 0 | 0 | 0 | 0 | 0 | 0 |
| ENSMUSG00 Gm12618 | 482 | 0 | 0.06911389 | 0 | 0.09452499 | 0 | 0 | 0.09815908 | 0 | 0 | 0.09782304 | 0.39647268 | 0 | 0 | 0.09635703 | 0.17096081 | 0 | 0 | 0 |
| ENSMUSG00 Gm11203 | 740 | 0 | 0.04501742 | 0 | 0 | 0 | 0 | 0 | 0 | 0 | 0 | 0 | 0 | 0 | 0 | 0 | 0 | 0 | 0 |
| ENSMUSG00 Gm15542 | 1193 | 0.42754578 | 0.39093086 | 0.81095608 | 0.2673322 | 0.32276227 | 0.58150735 | 0.35692717 | 0.92839872 | 0.72499315 | 0.43475083 | 0.48055281 | 0.23392985 | 0.37216031 | 0.233583 | 0.58711354 | 0.2888646 | 0.09637795 | 0.23054948 |
| ENSMUSG00 Gm5466 | 1889 | 0 | 0.0176352 | 0 | 0 | 0 | 0.02623226 | 0 | 0 | 0 | 0 | 0 | 0 | 0 | 0 | 0 | 0 | 0 | 0.14560378 |
| ENSMUSG00 Gm13458 | 366 | 0 | 0 | 0 | 0 | 0 | 0 | 0 | 0 | 0 | 0 | 0 | 0 | 0 | 0 | 0 | 0 | 0 | 0 |
| ENSMUSG00 Gm15033 | 672 | 0 | 0 | 0 | 0 | 0 | 0 | 0 | 0 | 0 | 0 | 0 | 0 | 0 | 0 | 0 | 0 | 0 | 0 |
| ENSMUSG00 Gm11900 | 351 | 0 | 0 | 0 | 0 | 0 | 0 | 0 | 0 | 0 | 0 | 0 | 0 | 0 | 0 | 0 | 0 | 0 | 0 |
| ENSMUSG00 Gm13506 | 1081 | 0 | 0 | 0 | 0 | 0 | 0 | 0 | 0 | 0 | 0 | 0 | 0 | 0 | 0 | 0 | 0 | 0 | 0 |
| ENSMUSG00 Gm14394 | 294 | 0 | 0 | 0 | 0 | 0 | 0 | 0 | 0 | 0 | 0 | 0 | 0 | 0 | 0 | 0 | 0 | 0 | 0 |
| ENSMUSG00 Gm13041 | 1521 | 0 | 0 | 0 | 0 | 0 | 0 | 0 | 0 | 0 | 0 | 0 | 0 | 0 | 0 | 0 | 0 | 0 | 0 |
| ENSMUSG00 Gm13019 | 274 | 0 | 0 | 0 | 0 | 0 | 0 | 0.17267401 | 0 | 0 | 0 | 0 | 0 | 0.36008699 | 0.16950397 | 0.45111193 | 0 | 0 | 0 |
| ENSMUSG00 Gm15725 | 1089 | 0 | 0 | 0 | 0 | 0 | 0 | 0 | 0 | 0 | 0 | 0 | 0 | 0 | 0 | 0 | 0 | 0 | 0 |
| ENSMUSG00 Gm8410 | 604 | 0 | 0 | 0 | 0 | 0 | 0 | 0 | 0 | 0 | 0 | 0 | 0 | 0 | 0 | 0 | 0 | 0 | 0 |
| ENSMUSG00 Gm13176 | 476 | 0 | 0.06998507 | 0 | 0 | 0 | 0.52051191 | 0 | 0 | 0 | 0 | 0 | 0 | 0 | 0.08655789 | 0 | 0 | 0 | 0 |
| ENSMUSG00 Gm11340 | 411 | 0 | 0 | 0 | 0 | 0 | 0 | 0 | 0 | 0 | 0 | 0 | 0 | 0 | 0 | 0 | 0 | 0 | 0 |
| ENSMUSG00 Gm14300 | 945 | 0 | 0 | 0 | 0 | 0 | 0 | 0 | 0 | 0 | 0 | 0 | 0 | 0 | 0 | 0 | 0 | 0 | 0 |
| ENSMUSG00 Rpl30-ps10 | 779 | 0 | 0 | 0.12419392 | 0 | 0 | 0 | 0 | 0 | 0 | 0 | 0 | 0 | 0.12665447 | 0.05962014 | 0 | 0 | 0.04919935 | 0 |
| ENSMUSG00 Gm12268 | 441 | 0 | 0.90647331 | 0 | 0 | 0 | 0 | 0 | 0.43302044 | 0 | 0.21383539 | 0 | 0 | 0 | 0 | 0 | 0 | 0 | 0 |
| ENSMUSG00 Gm11306 | 1093 | 0 | 0 | 0 | 0 | 0 | 0 | 0 | 0 | 0 | 0 | 0 | 0 | 0 | 0 | 0 | 0 | 0 | 0 |
| ENSMUSG00 Gm14840 | 593 | 0.07167821 | 0 | 0.326297 | 0 | 0 | 0 | 0.07978529 | 0 | 0.0694545 | 0.07951215 | 0 | 0 | 0.16638083 | 0 | 0.06947986 | 0.12914201 | 0.06463119 | 0 |
| ENSMUSG00 Gm14690 | 374 | 0 | 0 | 0 | 0 | 0 | 0 | 0 | 0 | 0 | 0 | 0 | 0 | 0 | 0 | 0 | 0 | 0 | 0 |
| ENSMUSG00 Gm15378 | 206 | 0 | 0 | 0 | 0 | 0 | 0 | 0 | 0 | 0 | 0 | 0 | 0 | 0 | 0 | 0 | 0 | 0 | 0 |
| ENSMUSG00 Olfr269-ps1 | 301 | 0 | 0 | 0 | 0 | 0 | 0 | 0 | 0 | 0 | 0 | 0 | 0 | 0 | 0 | 0 | 0 | 0 | 0 |
| ENSMUSG00 Olfr1035-ps1 | 924 | 0 | 0 | 0 | 0 | 0 | 0 | 0 | 0 | 0 | 0 | 0 | 0 | 0 | 0 | 0 | 0 | 0 | 0 |
| ENSMUSG00 Gm14485 | 492 | 0 | 0 | 0 | 0 | 0 | 0 | 0 | 0 | 0 | 0 | 0 | 0 | 0 | 0 | 0 | 0 | 0 | 0 |
| ENSMUSG00 Gm15665 | 846 | 0 | 0 | 0 | 0 | 0 | 0 | 0 | 0 | 0 | 0 | 0 | 0 | 0 | 0 | 0 | 0 | 0 | 0 |
| ENSMUSG00 Gm14776 | 885 | 0 | 0 | 0 | 0 | 0 | 0 | 0 | 0 | 0 | 0 | 0 | 0 | 0 | 0 | 0 | 0.04326622 | 0 | 0 |
| ENSMUSG00 Gm15604 | 1455 | 0 | 0.04579092 | 0 | 0 | 0 | 0 | 0 | 0.1049963 | 0 | 0 | 0 | 0.03836128 | 0 | 0 | 0 | 0 | 0 | 0 |
| ENSMUSG00 Gm13221 | 429 | 0 | 0 | 0 | 0 | 0 | 0 | 0 | 0 | 0 | 0 | 0 | 0 | 0 | 0 | 0 | 0 | 0 | 0 |
| ENSMUSG00 Rpl35a-ps4 | 332 | 0.25605528 | 0 | 0.43711021 | 0 | 0.72487837 | 0 | 0.28501614 | 0.57518678 | 0.24811154 | 0.56808076 | 0.57560191 | 0.5043584 | 1.04013079 | 0.27978366 | 0.24820214 | 0.46133262 | 0.46176261 | 0.16569008 |
| ENSMUSG00 Gm13992 | 155 | 0.27422694 | 0 | 0 | 0 | 0 | 0 | 0 | 0 | 0 | 0 | 0 | 0 | 0 | 0 | 0 | 0 | 0.24726643 | 0 |
| ENSMUSG00 Defa-ps10 | 170 | 0 | 0 | 0 | 0 | 0 | 0 | 0 | 0 | 0 | 0 | 0 | 0 | 0 | 0 | 0 | 0 | 0 | 0 |
| ENSMUSG00 Gm12007 | 596 | 0 | 0 | 0 | 0 | 0 | 0 | 0 | 0 | 0 | 0 | 0 | 0 | 0 | 0 | 0 | 0 | 0 | 0 |
| ENSMUSG00 Gm14591 | 516 | 0 | 0 | 0 | 0 | 0 | 0 | 0 | 0 | 0 | 0 | 0 | 0 | 0 | 0 | 0 | 0 | 0 | 0 |
| ENSMUSG00 Gm15198 | 185 | 0 | 0.1800697 | 0 | 0.49255183 | 0 | 0.26785261 | 0.51148842 | 0 | 0 | 0.25486867 | 0 | 0 | 0 | 0.25104912 | 0.44542223 | 0.41395251 | 0.41433834 | 0 |
| ENSMUSG00 Gm13006 | 213 | 0 | 0 | 0 | 0 | 0 | 0 | 0 | 0 | 0 | 0 | 0 | 0 | 0 | 0 | 0 | 0 | 0 | 0 |
| ENSMUSG00 Gm13437 | 1978 | 0 | 0 | 0 | 0.0230339 | 0 | 0.02505194 | 0 | 0 | 0.0208223 | 0 | 0.02415316 | 0 | 0.0249403 | 0 | 0 | 0 | 0 | 0 |
| ENSMUSG00 Gm14878 | 292 | 0 | 0 | 0 | 0 | 0 | 0 | 0 | 0 | 0 | 0 | 0 | 0 | 0 | 0 | 0 | 0 | 0 | 0 |
| ENSMUSG00 Gm15405 | 341 | 0 | 0 | 0 | 0 | 0 | 0 | 0 | 0 | 0 | 0 | 0 | 0 | 0 | 0 | 0 | 0 | 0 | 0 |
| ENSMUSG00 Gm12760 | 480 | 0 | 0 | 0 | 0 | 0 | 0 | 0.09856808 | 0 | 0 | 0 | 0.09953116 | 0 | 0 | 0.19351703 | 0.08583657 | 0 | 0.07984645 | 0 |
| ENSMUSG00 Gm15824 | 957 | 0 | 0 | 0 | 0 | 0 | 0 | 0 | 0 | 0 | 0 | 0 | 0 | 0 | 0 | 0.08610565 | 0 | 0.08009675 | 0 |
| ENSMUSG00 Gm11471 | 197 | 0 | 0 | 0 | 0.23127434 | 0 | 0 | 0 | 0 | 0 | 0 | 0 | 0 | 0 | 0 | 0 | 0 | 0 | 0 |
| ENSMUSG00 Gm15187 | 263 | 0 | 0 | 0 | 0 | 0 | 0 | 0 | 0 | 0 | 0 | 0 | 0 | 0 | 0 | 0 | 0 | 0 | 0 |
| ENSMUSG00 Gm5402 | 1617 | 0 | 0 | 0 | 0 | 0 | 0 | 0 | 0.0236193 | 0 | 0 | 0 | 0 | 0 | 0 | 0 | 0 | 0 | 0 |
| ENSMUSG00 Gm14491 | 198 | 0 | 0.16824694 | 0 | 0.23010628 | 0 | 0 | 0.47790585 | 0 | 0.20801271 | 0 | 0.24128767 | 0 | 0 | 0.70369829 | 0 | 0.19338691 | 0.38713431 | 0 |
| ENSMUSG00 Rpl7a-ps12 | 792 | 0.05366815 | 0 | 0.06107769 | 0 | 0 | 0.12513317 | 0.05973823 | 0 | 0.05200318 | 0 | 0.06032192 | 0.07047432 | 0.31143887 | 0 | 0 | 0 | 0.09678358 | 0.06945594 |
| ENSMUSG00 Gm15329 | 154 | 0 | 0 | 0 | 0 | 0 | 0 | 0 | 0 | 0 | 0 | 0 | 0 | 0 | 0 | 0 | 0 | 0 | 0 |
| ENSMUSG00 Gm11187 | 445 | 0 | 0 | 0 | 0 | 0 | 0 | 0 | 0 | 0 | 0 | 0 | 0 | 0 | 0 | 0 | 0 | 0 | 0 |
| ENSMUSG00 Gm15519 | 1565 | 0.02715986 | 0.10643097 | 0.0309096 | 0 | 0.09226567 | 0.03166309 | 0.15115872 | 0.29284909 | 0.10526905 | 0.06025649 | 0.3052713 | 0.10699488 | 0.09456598 | 0 | 0.07898062 | 0.02446684 | 0.07346894 | 0.45694466 |
| ENSMUSG00 Gm14768 | 676 | 0 | 0 | 0 | 0 | 0 | 0 | 0 | 0 | 0 | 0 | 0 | 0 | 0 | 0 | 0 | 0 | 0 | 0 |
| ENSMUSG00 Gm15038 | 295 | 0 | 0 | 0 | 0 | 0 | 0 | 0 | 0 | 0 | 0 | 0 | 0 | 0 | 0 | 0 | 0 | 0 | 0 |
| ENSMUSG00 Gm12826 | 378 | 0 | 0 | 0 | 0 | 0 | 0 | 0 | 0 | 0 | 0 | 0 | 0 | 0 | 0 | 0 | 0 | 0 | 0 |
| ENSMUSG00 Gm11904 | 247 | 0 | 0 | 0 | 0 | 0 | 0 | 0 | 0 | 0 | 0 | 0 | 0 | 0 | 0 | 0 | 0 | 0 | 0 |
| ENSMUSG00 Gm8749 | 330 | 0 | 0 | 0 | 0 | 0 | 0 | 0 | 0 | 0 | 0 | 0 | 0 | 0 | 0 | 0 | 0 | 0 | 0 |
| ENSMUSG00 Gm13487 | 863 | 0 | 0 | 0 | 0 | 0 | 0 | 0 | 0 | 0 | 0 | 0 | 0 | 0 | 0 | 0.04774224 | 0 | 0 | 0 |
| ENSMUSG00 Gm14619 | 167 | 0 | 0 | 0 | 0 | 0 | 0 | 0 | 0 | 0 | 0 | 0 | 0 | 0 | 0 | 0 | 0 | 0 | 0 |
| ENSMUSG00 Gm11445 | 531 | 0 | 0 | 0 | 0 | 0 | 0 | 0 | 0 | 0.07756406 | 0 | 0 | 0 | 0 | 0 | 0 | 0 | 0 | 0 |
| ENSMUSG00 Gm14905 | 637 | 0 | 0 | 0 | 0 | 0 | 0 | 0 | 0 | 0 | 0 | 0 | 0 | 0 | 0 | 0 | 0 | 0 | 0 |
| ENSMUSG00 Gm13338 | 360 | 0 | 0 | 0.13437092 | 0 | 0 | 0 | 0 | 0 | 0 | 0 | 0 | 0.31008701 | 0 | 0 | 0 | 0 | 0 | 0 |
| ENSMUSG00 Gm13226 | 403 | 0.4218876 | 0.41331134 | 0 | 1.01749231 | 0.47773622 | 0.24591927 | 0 | 0 | 0.61319875 | 0.11699926 | 0.11854828 | 0.41550121 | 0.61205853 | 0.4609835 | 0.10223711 | 0.19002783 | 0.09510247 | 0.13649902 |
| ENSMUSG00 Gm15037 | 258 | 0 | 0 | 0 | 0 | 0 | 0 | 0 | 0 | 0 | 0 | 0 | 0 | 0 | 0 | 0 | 0 | 0 | 0 |
| ENSMUSG00 Gm13108 | 1453 | 0 | 0 | 0 | 0 | 0 | 0 | 0 | 0 | 0 | 0 | 0 | 0 | 0 | 0 | 0 | 0 | 0 | 0 |
| ENSMUSG00 Gm12839 | 330 | 0 | 0 | 0 | 0 | 0 | 0 | 0 | 0 | 0.24961525 | 0 | 0 | 0 | 0 | 0 | 0 | 0 | 0 | 0 |
| ENSMUSG00 Gm6510 | 485 | 0 | 0.13737276 | 0 | 0 | 0.19848216 | 0.10217058 | 0.29265575 | 0.07874722 | 0.08492065 | 0.09721794 | 0.19701014 | 0.23016768 | 0.50857647 | 0.095761 | 0.16990332 | 0.07894971 | 0.15804659 | 0.45368335 |
| ENSMUSG00 Gm12182 | 1007 | 0.16883883 | 0.09924397 | 0 | 0.271466 | 0.14339203 | 0 | 0.23491896 | 0.03792691 | 0.36810193 | 0.18729177 | 0.28465715 | 0.22171068 | 0.14696698 | 0.32284867 | 0.2454909 | 0.19012218 | 0.19029939 | 0.10925344 |
| ENSMUSG00 Gm12935 | 244 | 0 | 0 | 0 | 0 | 0.39452396 | 0 | 0 | 0 | 0 | 0 | 0 | 0 | 0.20217999 | 0 | 0 | 0 | 0 | 0 |
| ENSMUSG00 Gm12501 | 528 | 0 | 0 | 0.09161653 | 0.08628986 | 0 | 0.18769975 | 0 | 0 | 0 | 0.3572023 | 0 | 0 | 0.09343166 | 0.08796229 | 0 | 0 | 0 | 0.10418391 |
| ENSMUSG00 Gm4909 | 793 | 0 | 0 | 0 | 0 | 0 | 0 | 0 | 0 | 0 | 0 | 0 | 0 | 0 | 0 | 0 | 0 | 0 | 0 |
| ENSMUSG00 Gm11652 | 1266 | 0.94008288 | 0.47364305 | 0.61135583 | 0.64778736 | 0.34217007 | 0.43055298 | 0.11211535 | 0.27150997 | 1.13864775 | 0.85660835 | 0.52831708 | 0.44088201 | 0.27276732 | 0.18342847 | 0.39053607 | 0.21171742 | 0.27246183 | 0.17380444 |
| ENSMUSG00 Gm15225 | 1013 | 0 | 0 | 0 | 0 | 0 | 0 | 0 | 0 | 0 | 0 | 0 | 0 | 0 | 0 | 0 | 0 | 0 | 0 |
| ENSMUSG00 Gm15840 | 942 | 0 | 0 | 0 | 0 | 0 | 0 | 0 | 0 | 0 | 0 | 0 | 0 | 0 | 0 | 0 | 0 | 0 | 0 |
| ENSMUSG00 Gm9431 | 1050 | 0.04048112 | 0 | 0.04607003 | 0 | 0 | 0.09438616 | 0 | 0 | 0 | 0 | 0 | 0 | 0 | 0 | 0 | 0 | 0 | 0 |
| ENSMUSG00 Gm14281 | 1293 | 0 | 0 | 0 | 0 | 0 | 0 | 0 | 0 | 0 | 0 | 0 | 0 | 0 | 0 | 0 | 0 | 0 | 0 |
| ENSMUSG00 Nsa2-ps2 | 772 | 0 | 0 | 0 | 0 | 0 | 0 | 0 | 0 | 0 | 0 | 0 | 0 | 0 | 0 | 0 | 0 | 0 | 0 |
| ENSMUSG00 Gm12118 | 531 | 0 | 0 | 0 | 0.08580234 | 0 | 0.09331965 | 0 | 0 | 0.15512812 | 0 | 0 | 0 | 0 | 0.08746532 | 0 | 0 | 0 | 0 |
| ENSMUSG00 Gm13491 | 234 | 0 | 0 | 0 | 0 | 0 | 0 | 0 | 0 | 0 | 0 | 0 | 0 | 0 | 0 | 0 | 0 | 0 | 0 |
| ENSMUSG00 Gm13207 | 772 | 0.11011704 | 0.2589085 | 0 | 0 | 0 | 0 | 0 | 0 | 0.5868545 | 0 | 0 | 0.14460016 | 0 | 0 | 0 | 0 | 0.09929092 | 0 |
| ENSMUSG00 Gm15267 | 463 | 0 | 0 | 0 | 0 | 0.20791328 | 0.21405069 | 0 | 0 | 0 | 0 | 0 | 0.36165656 | 0 | 0.1003112 | 0.53392945 | 0 | 0 | 0.11881017 |
| ENSMUSG00 Gm9440 | 532 | 0 | 0 | 0 | 0 | 0 | 0 | 0 | 0 | 0 | 0 | 0 | 0 | 0 | 0 | 0 | 0 | 0 | 0 |
| ENSMUSG00 Gm6555 | 1013 | 0.7552746 | 0.6248223 | 0.57303294 | 0.44976352 | 0.23757119 | 0.04891681 | 0.46705507 | 0.79174773 | 0.93513313 | 0.55854732 | 0.33013298 | 0.82649056 | 0.24349416 | 0.45848063 | 0.36605528 | 0.6425867 | 0.52968229 | 0.21721266 |
| ENSMUSG00 Gm15375 | 848 | 0 | 0 | 0 | 0 | 0 | 0 | 0 | 0 | 0 | 0 | 0 | 0 | 0 | 0 | 0 | 0 | 0 | 0 |
| ENSMUSG00 Gm11883 | 326 | 0 | 0 | 0 | 0 | 0 | 0 | 0 | 0 | 0 | 0 | 0 | 0 | 0 | 0 | 0 | 0 | 0 | 0 |
| ENSMUSG00 Rps8-ps2 | 1310 | 0 | 0 | 0 | 0 | 0 | 0 | 0 | 0 | 0 | 0 | 0 | 0 | 0 | 0 | 0 | 0 | 0 | 0 |
| ENSMUSG00 Gm14706 | 254 | 0.33468643 | 0 | 0 | 0.35874838 | 0 | 0.1950895 | 0 | 0.15036379 | 0.48645491 | 0 | 0.18809039 | 0 | 0.19422015 | 0.91425369 | 0.3244217 | 0.15075042 | 0.15089093 | 0 |
| ENSMUSG00 Gm14044 | 951 | 0.04469524 | 0.03502933 | 0.10173193 | 0.09581713 | 0.10122381 | 0 | 0 | 0.04016025 | 0.04330864 | 0.04958013 | 0.05023655 | 0 | 0 | 0 | 0 | 0 | 0 | 0 |
| ENSMUSG00 Gm14574 | 278 | 0 | 0 | 0 | 0 | 0 | 0 | 0 | 0 | 0 | 0 | 0 | 0 | 0 | 0 | 0 | 0 | 0 | 0 |
| ENSMUSG00 Gm12308 | 388 | 0 | 0 | 0 | 0 | 0 | 0 | 0 | 0 | 0 | 0 | 0 | 0 | 0 | 0 | 0.31856873 | 0 | 0 | 0 |
| ENSMUSG00 Gm14521 | 349 | 0 | 0 | 0 | 0 | 0 | 0 | 0 | 0 | 0 | 0 | 0 | 0 | 0 | 0 | 0 | 0 | 0 | 0 |
| ENSMUSG00 Gm13888 | 421 | 0 | 0 | 0 | 0 | 0 | 0 | 0 | 0 | 0 | 0 | 0 | 0 | 0 | 0 | 0 | 0 | 0 | 0 |
| ENSMUSG00 Gm8534 | 765 | 0 | 0 | 0 | 0 | 0 | 0 | 0 | 0 | 0 | 0 | 0 | 0 | 0 | 0 | 0 | 0 | 0 | 0 |
| ENSMUSG00 Gm11764 | 471 | 0 | 0 | 0 | 0 | 0 | 0 | 0 | 0 | 0 | 0 | 0 | 0 | 0 | 0 | 0 | 0 | 0 | 0 |
| ENSMUSG00 Gm14940 | 492 | 0 | 0 | 0 | 0 | 0 | 0 | 0 | 0 | 0 | 0 | 0 | 0 | 0 | 0 | 0 | 0 | 0 | 0 |
| ENSMUSG00 Gm12423 | 134 | 0 | 0 | 0 | 0 | 0 | 0 | 0 | 0 | 0 | 0 | 0 | 0 | 0 | 0 | 0 | 0 | 0 | 0 |
| ENSMUSG00 Gm14515 | 215 | 0 | 0 | 0 | 0 | 0 | 0 | 0 | 0 | 0 | 0 | 0 | 0 | 0 | 0 | 0 | 0 | 0 | 0 |
| ENSMUSG00 Gm15044 | 597 | 0 | 0 | 0 | 0 | 0 | 0 | 0 | 0 | 0 | 0 | 0 | 0 | 0 | 0 | 0 | 0 | 0 | 0 |
| ENSMUSG00 Gm12873 | 418 | 0 | 0 | 0 | 0 | 0 | 0 | 0 | 0 | 0 | 0 | 0 | 0 | 0.5900947 | 0 | 0 | 1.00764756 | 0 | 0 |
| ENSMUSG00 Gm13877 | 416 | 0 | 0 | 0 | 0 | 0 | 0 | 0 | 0 | 0 | 0 | 0 | 0 | 0 | 0 | 0 | 0 | 0 | 0 |
| ENSMUSG00 Gm11263 | 748 | 0.1136502 | 0.13360786 | 0.12934099 | 0 | 0.12869498 | 0.13249394 | 0.12650449 | 0.20423745 | 0 | 0 | 0 | 0.29847948 | 0.06595176 | 0.12418205 | 0 | 0.15357195 | 0.05123836 | 0.14708317 |
| ENSMUSG00 Gm14172 | 643 | 0 | 0 | 0 | 0 | 0 | 0 | 0 | 0 | 0 | 0 | 0 | 0 | 0 | 0 | 0 | 0 | 0 | 0 |
| ENSMUSG00 Gm14609 | 901 | 0 | 0 | 0 | 0 | 0 | 0 | 0 | 0 | 0 | 0 | 0 | 0 | 0 | 0 | 0 | 0 | 0 | 0 |
| ENSMUSG00 Gm15470 | 1370 | 4.18846625 | 1.96959448 | 2.25978536 | 1.92886173 | 2.67009212 | 1.80849393 | 2.17569253 | 1.53327163 | 1.05221026 | 1.78966173 | 2.02258948 | 2.48522293 | 1.69240884 | 1.93234524 | 1.20296514 | 2.79493485 | 2.21005654 | 3.05160008 |
| ENSMUSG00 Gm12255 | 209 | 0 | 0 | 0 | 0 | 0 | 0 | 0 | 0 | 0 | 0 | 0 | 0 | 0 | 0 | 0 | 0 | 0 | 0 |
| ENSMUSG00 Gm14861 | 454 | 0 | 0 | 0 | 0 | 0 | 0 | 0 | 0 | 0 | 0 | 0 | 0 | 0 | 0 | 0 | 0 | 0 | 0 |
| ENSMUSG00 Gm13407 | 359 | 0 | 0.27838073 | 0.13474521 | 1.0152879 | 0 | 0.2760598 | 0 | 0 | 0.45890268 | 0 | 0 | 0.15547538 | 0 | 0.51748287 | 0.3443027 | 0 | 0 | 0 |
| ENSMUSG00 Gm12450 | 467 | 0 | 0 | 0 | 0 | 0 | 0 | 0 | 0 | 0 | 0 | 0 | 0 | 0 | 0 | 0 | 0 | 0 | 0 |
| ENSMUSG00 Gm7541 | 534 | 0 | 0 | 0 | 0.08532031 | 0 | 0 | 0 | 0 | 0 | 0 | 0 | 0 | 0.36952747 | 0 | 0 | 0.28682103 | 0 | 0 |
| ENSMUSG00 Gm13993 | 1235 | 0 | 0 | 0 | 0 | 0 | 0 | 0 | 0 | 0 | 0.03817871 | 0 | 0 | 0 | 0 | 0 | 0 | 0 | 0 |
| ENSMUSG00 Gm12842 | 475 | 0 | 0 | 0.40735605 | 0 | 0.10133037 | 0 | 0 | 0.08040506 | 0 | 0 | 0.20115772 | 0.11750666 | 0 | 0 | 0 | 0 | 0 | 0 |
| ENSMUSG00 Gm6027 | 755 | 0 | 0 | 0.0640709 | 0 | 0 | 0 | 0 | 0 | 0 | 0 | 0 | 0 | 0 | 0 | 0 | 0 | 0 | 0 |
| ENSMUSG00 Defa-ps4 | 182 | 0 | 0 | 0 | 0 | 0 | 0 | 0 | 0 | 0 | 0 | 0 | 0 | 0 | 0 | 0 | 0 | 0 | 0 |
| ENSMUSG00 Gm13478 | 412 | 0 | 0 | 0 | 0 | 0.11682506 | 0 | 0 | 0 | 0 | 0 | 0 | 0.13547491 | 0 | 0 | 0 | 0 | 0 | 0 |
| ENSMUSG00 Gm12749 | 1158 | 0.36705679 | 0.28767612 | 0.04177334 | 0.35410138 | 0.1246941 | 0.21395826 | 0.16342894 | 0.09894405 | 0.10670082 | 0.32573888 | 0.04125644 | 0.19280022 | 0.21300482 | 0 | 0 | 0.09919846 | 0.03309697 | 0.33252482 |
| ENSMUSG00 Olfr647-ps1 | 876 | 0 | 0 | 0 | 0 | 0 | 0 | 0 | 0 | 0 | 0 | 0 | 0 | 0 | 0 | 0 | 0 | 0 | 0 |
| ENSMUSG00 Gm11881 | 571 | 0 | 0 | 0 | 0 | 0 | 0 | 0 | 0 | 0 | 0 | 0 | 0 | 0 | 0 | 0 | 0 | 0 | 0 |
| ENSMUSG00 Gm14577 | 255 | 0 | 0 | 0 | 0 | 0 | 0 | 0 | 0 | 0 | 0 | 0 | 0 | 0 | 0.18213368 | 0 | 0 | 0 | 0 |
| ENSMUSG00 Gm15196 | 481 | 0 | 0.34628768 | 0 | 0 | 0 | 0 | 0 | 0 | 0 | 0 | 0 | 0 | 0 | 0 | 0 | 0 | 0 | 0 |
| ENSMUSG00 Gm14998 | 494 | 0 | 0 | 0 | 0 | 0 | 0 | 0 | 0 | 0 | 0 | 0 | 0 | 0 | 0 | 0 | 0 | 0 | 0 |
| ENSMUSG00 Gm9429 | 867 | 0 | 0 | 0 | 0 | 0 | 0.17146275 | 0 | 0 | 0 | 0 | 0 | 0 | 0 | 0 | 0 | 0 | 0.04420565 | 0 |
| ENSMUSG00 Gm13674 | 822 | 0 | 0 | 0 | 0 | 0 | 0 | 0 | 0 | 0 | 0 | 0 | 0 | 0 | 0 | 0 | 0 | 0 | 0 |
| ENSMUSG00 Gm6418 | 3575 | 0 | 0 | 0 | 0 | 0 | 0 | 0 | 0 | 0 | 0 | 0 | 0 | 0 | 0 | 0 | 0 | 0 | 0 |
| ENSMUSG00 Gm12157 | 747 | 0 | 0 | 0 | 0 | 0 | 0 | 0 | 0 | 0 | 0 | 0 | 0 | 0 | 0 | 0 | 0 | 0 | 0 |
| ENSMUSG00 Gm15321 | 287 | 0 | 0 | 0 | 0 | 0 | 0 | 0 | 0.13307457 | 0 | 0 | 0 | 0 | 0 | 0 | 0 | 0 | 0 | 0 |
| ENSMUSG00 Mup-ps21 | 443 | 0 | 0 | 0 | 0 | 0 | 0 | 0 | 0 | 0 | 0 | 0 | 0 | 0 | 0 | 0 | 0 | 0 | 0 |
| ENSMUSG00 Gm15045 | 882 | 0 | 0 | 0 | 0 | 0 | 0 | 0 | 0 | 0 | 0 | 0 | 0 | 0 | 0 | 0 | 0 | 0 | 0 |
| ENSMUSG00 Gm12001 | 911 | 0 | 0 | 0 | 0 | 0 | 0 | 0 | 0 | 0 | 0 | 0 | 0 | 0 | 0 | 0 | 0 | 0 | 0 |
| ENSMUSG00 Gm13432 | 240 | 0 | 0 | 0 | 0 | 0 | 0 | 0 | 0 | 0 | 0 | 0 | 0 | 0 | 0 | 0 | 0 | 0 | 0 |
| ENSMUSG00 Gm15035 | 1560 | 0 | 0 | 0 | 0 | 0.0308538 | 0 | 0 | 0 | 0 | 0 | 0 | 0 | 0 | 0 | 0 | 0 | 0 | 0 |
| ENSMUSG00 Gm13867 | 393 | 0 | 0 | 0 | 0 | 0 | 0 | 0 | 0 | 0 | 0 | 0 | 0 | 0 | 0 | 0 | 0 | 0 | 0 |
| ENSMUSG00 Gm13450 | 27 | 0 | 0 | 0 | 0 | 0 | 0 | 0 | 0 | 0 | 0 | 0 | 0 | 0 | 3.44030277 | 0 | 0 | 0 | 2.03737432 |
| ENSMUSG00 H2-M6-ps | 1029 | 0.16522906 | 0.32374047 | 0.37608187 | 0.22138506 | 0 | 0.09631241 | 0.6896892 | 0.63097263 | 0.16010307 | 0.32075308 | 0.13928559 | 0.21697051 | 0.33559127 | 0.76729785 | 0.04004038 | 0.70701802 | 0.03724616 | 0.26729401 |
| ENSMUSG00 Gm14381 | 413 | 0 | 0 | 0 | 0 | 0 | 0 | 0 | 0 | 0 | 0 | 0 | 0 | 0 | 0 | 0 | 0 | 0 | 0 |
| ENSMUSG00 Gm15808 | 603 | 0.07048951 | 0 | 0 | 0 | 0 | 0 | 0 | 0 | 0 | 0 | 0 | 0 | 0 | 0.0770217 | 0 | 0 | 0.06355936 | 0 |
| ENSMUSG00 Gm14942 | 335 | 0 | 0 | 0 | 0 | 0.14367738 | 0 | 0 | 0 | 0 | 0 | 0 | 0 | 0 | 0 | 0 | 0.11430032 | 0.22881371 | 0 |
| ENSMUSG00 Gm16016 | 549 | 0.92907489 | 0.66747147 | 1.49790531 | 3.81750098 | 1.1397359 | 1.08311986 | 0.51707846 | 0.97394104 | 1.05029366 | 1.03061646 | 1.30532673 | 1.83002172 | 1.79715546 | 0.67678087 | 1.12572557 | 3.20832047 | 0.55848884 | 1.10218611 |
| ENSMUSG00 Gm12466 | 619 | 0 | 0 | 0 | 0 | 0 | 0 | 0 | 0 | 0.06653718 | 0 | 0 | 0 | 0 | 0.07503084 | 0 | 0 | 0 | 0 |
| ENSMUSG00 Gm12585 | 289 | 0 | 0 | 0 | 0 | 0 | 0 | 0 | 0 | 0 | 0 | 0 | 0 | 0 | 0 | 0 | 0 | 0 | 0 |
| ENSMUSG00 Olfr1379-ps1 | 823 | 0 | 0 | 0 | 0 | 0 | 0 | 0 | 0 | 0 | 0 | 0 | 0 | 0 | 0 | 0 | 0 | 0 | 0 |
| ENSMUSG00 Gm14663 | 1152 | 0 | 0 | 0 | 0 | 0 | 0 | 0 | 0 | 0 | 0 | 0 | 0 | 0 | 0.04031605 | 0 | 0 | 0 | 0 |
| ENSMUSG00 Oat-rs1 | 1310 | 0 | 0.10171876 | 0 | 0 | 0.07348385 | 0.03782651 | 0 | 0.26239055 | 0.06288018 | 0 | 0 | 0 | 0 | 0.070907 | 0.09435471 | 0 | 0 | 0 |
| ENSMUSG00 Gm13320 | 329 | 0 | 0.101255 | 0.14703201 | 0 | 0.58519056 | 0 | 0.14380753 | 0 | 0 | 0 | 0.43563792 | 0.16965247 | 0 | 0.14116744 | 0 | 0 | 0 | 0.33440186 |
| ENSMUSG00 Tpt1-ps1 | 435 | 0 | 0.07658137 | 0 | 0.10473803 | 0 | 0 | 0 | 0 | 0 | 0 | 0 | 0 | 0.22681341 | 0 | 0 | 0.08802438 | 0 | 0 |
| ENSMUSG00 Gm13321 | 398 | 0 | 0 | 0 | 0 | 0 | 0 | 0 | 0 | 0 | 0 | 0.12003758 | 0 | 0 | 0 | 0 | 0 | 0 | 0 |
| ENSMUSG00 Gm11514 | 431 | 0 | 0 | 0 | 0 | 0 | 0 | 0 | 0 | 0 | 0 | 0 | 0 | 0 | 0 | 0 | 0 | 0 | 0 |
| ENSMUSG00 Gm11227 | 470 | 0 | 0 | 0 | 0 | 0 | 0 | 0 | 0 | 0 | 0 | 0 | 0 | 0 | 0 | 0 | 0 | 0 | 0 |
| ENSMUSG00 Gm13729 | 405 | 0 | 0 | 0 | 0 | 0 | 0 | 0 | 0 | 0 | 0 | 0 | 0 | 0 | 0 | 0 | 0 | 0 | 0 |
| ENSMUSG00 Gm14516 | 251 | 0 | 0 | 0 | 0 | 0 | 0 | 0 | 0 | 0 | 0 | 0 | 0 | 0 | 0 | 0 | 0 | 0 | 0 |
| ENSMUSG00 Gm14814 | 304 | 0 | 0 | 0 | 0 | 0 | 0 | 0 | 0 | 0 | 0 | 0 | 0 | 0 | 0 | 0 | 0 | 0 | 0 |
| ENSMUSG00 Gm11989 | 671 | 0 | 0 | 0 | 0 | 0 | 0 | 0 | 0 | 0 | 0 | 0 | 0 | 0 | 0 | 0.06140321 | 0 | 0 | 0 |
| ENSMUSG00 Gm6645 | 2724 | 0.18724747 | 0.19567045 | 0.19534098 | 0.23416102 | 0.21203491 | 0.14552932 | 0.10421295 | 0.19628988 | 0.0907192 | 0.20771235 | 0.12276972 | 0.06147099 | 0.10866061 | 0.05114988 | 0.07562694 | 0.18273785 | 0.21104789 | 0.04038848 |
| ENSMUSG00 Gm12035 | 622 | 0.27334518 | 0.10711542 | 0 | 0 | 0 | 0 | 0.2281962 | 0 | 0 | 0 | 0 | 0 | 0 | 0 | 0 | 0 | 0 | 0.08843908 |
| ENSMUSG00 Gm9043 | 311 | 0 | 0 | 0 | 0 | 0 | 0 | 0 | 0 | 0 | 0 | 0 | 0 | 0 | 0 | 0 | 0 | 0 | 0 |
| ENSMUSG00 Gm6877 | 1100 | 0.11592321 | 0.0605689 | 0 | 0.04141913 | 0.13126888 | 0 | 0.04301153 | 0.17360183 | 0.11232686 | 0.1714571 | 0 | 0.05074151 | 0.0448472 | 0 | 0.07491192 | 0 | 0.03484209 | 0.05000828 |
| ENSMUSG00 Gm12896 | 225 | 0 | 0 | 0 | 0 | 0 | 0 | 0 | 0 | 0 | 0 | 0 | 0 | 0 | 0 | 0 | 0 | 0 | 0 |
| ENSMUSG00 Gm14939 | 678 | 0 | 0 | 0 | 0 | 0 | 0 | 0 | 0 | 0 | 0 | 0 | 0 | 0 | 0 | 0 | 0 | 0 | 0 |
| ENSMUSG00 Gm5942 | 438 | 0 | 0 | 0 | 0 | 0 | 0 | 0 | 0 | 0 | 0 | 0 | 0 | 0 | 0 | 0 | 0 | 0 | 0 |
| ENSMUSG00 Gm13234 | 746 | 0 | 0 | 0 | 0 | 0 | 0 | 0 | 0 | 0 | 0 | 0 | 0 | 0 | 0 | 0 | 0 | 0 | 0 |
| ENSMUSG00 Gm13340 | 1544 | 2.11975295 | 1.9849652 | 2.19310047 | 1.7705069 | 2.93031141 | 2.56749915 | 3.98358048 | 3.04252947 | 4.29470793 | 4.24478481 | 3.34177171 | 3.43425385 | 4.0577419 | 3.91044777 | 4.72323536 | 2.95115433 | 4.89007804 | 3.34900002 |
| ENSMUSG00 Gm11929 | 944 | 0 | 0 | 0 | 0 | 0 | 0 | 0 | 0 | 0.04362978 | 0 | 0 | 0 | 0 | 0 | 0 | 0 | 0 | 0 |
| ENSMUSG00 Gm11941 | 609 | 0 | 0 | 0 | 0 | 0 | 0 | 0 | 0 | 0 | 0 | 0 | 0 | 0 | 0 | 0 | 0 | 0 | 0 |
| ENSMUSG00 Gm14711 | 215 | 0 | 0 | 0.22499316 | 0 | 0 | 0 | 0 | 0 | 0 | 0 | 0 | 0 | 0 | 0 | 0 | 0 | 0 | 0 |
| ENSMUSG00 Gm11451 | 1246 | 0.27290643 | 0.18715109 | 0.58234587 | 0.43879015 | 0.61806644 | 0.59654174 | 0.7974047 | 0.30652008 | 0.39665986 | 0.41625821 | 0.38342663 | 0.49275465 | 0.51469898 | 0.82004007 | 0.99201178 | 0.58388567 | 0.43063255 | 0.52978273 |
| ENSMUSG00 Gm12770 | 461 | 0 | 0 | 0 | 0 | 0 | 0 | 0 | 0 | 0 | 0 | 0 | 0 | 0 | 0 | 0 | 0 | 0 | 0 |
| ENSMUSG00 Gm13666 | 798 | 0.10652926 | 0.16698193 | 0 | 0 | 0 | 0 | 0.05928907 | 0.52646169 | 0 | 0 | 0 | 0 | 0 | 0.11640122 | 0 | 0.04798322 | 0 | 0 |
| ENSMUSG00 Gm14671 | 261 | 0 | 0 | 0 | 0 | 0 | 0 | 0 | 0 | 0 | 0 | 0 | 0 | 0 | 0 | 0 | 0 | 0 | 0 |
| ENSMUSG00 Gm11849 | 260 | 0 | 0 | 0 | 0 | 0 | 0 | 0 | 0 | 0 | 0 | 0 | 0 | 0 | 0 | 0 | 0 | 0 | 0 |
| ENSMUSG00 Gm7312 | 468 | 0 | 0.0711814 | 0 | 0 | 0 | 0 | 0 | 0 | 0 | 0 | 0 | 0 | 0 | 0 | 0 | 0 | 0 | 0 |
| ENSMUSG00 Gm15709 | 212 | 0 | 0 | 0 | 0 | 0 | 0 | 0 | 0 | 0 | 0 | 0 | 0 | 0 | 0 | 0 | 0 | 0 | 0 |
| ENSMUSG00 Gm12533 | 137 | 0 | 0 | 0 | 0 | 0 | 0 | 0 | 0 | 0 | 0 | 0 | 0 | 0 | 0 | 0 | 0 | 0 | 0 |
| ENSMUSG00 Gm16411 | 1099 | 2.82336474 | 2.78870452 | 4.09348346 | 1.98992732 | 1.9708249 | 1.93882397 | 10.2891085 | 9.73054838 | 7.38284222 | 4.59065082 | 4.60795778 | 7.51657695 | 18.763186 | 23.0318723 | 17.5453395 | 5.3307215 | 6.34703006 | 13.2141985 |
| ENSMUSG00 Gm12432 | 185 | 0 | 0 | 0 | 0 | 0 | 0 | 0 | 0 | 0 | 0 | 0 | 0 | 0 | 0 | 0 | 0 | 0 | 0 |
| ENSMUSG00 Gm11247 | 307 | 0 | 0 | 0 | 0 | 0 | 0 | 0 | 0 | 0 | 0 | 0 | 0 | 0 | 0 | 0 | 0 | 0 | 0 |
| ENSMUSG00 Gm15538 | 312 | 0 | 0 | 0 | 0 | 0.15426899 | 0 | 0 | 0 | 0 | 0 | 0 | 0 | 0 | 0.29771851 | 0 | 0 | 0 | 0 |
| ENSMUSG00 Rpsa-ps3 | 874 | 0 | 0 | 0 | 0 | 0 | 0 | 0 | 0 | 0 | 0 | 0 | 0 | 0 | 0 | 0 | 0 | 0 | 0 |
| ENSMUSG00 Gm14186 | 136 | 0 | 0 | 0 | 0 | 0 | 0 | 0 | 0 | 0 | 0 | 0 | 0 | 0 | 0 | 0 | 0 | 0 | 0 |
| ENSMUSG00 Gm15393 | 218 | 0 | 0 | 0 | 0 | 0 | 0 | 0 | 0 | 0 | 0 | 0 | 0 | 0 | 0 | 0 | 0 | 0 | 0 |
| ENSMUSG00 Gm15394 | 1307 | 0 | 0 | 0 | 0 | 0 | 0 | 0 | 0 | 0 | 0 | 0 | 0 | 0 | 0 | 0 | 0 | 0 | 0 |
| ENSMUSG00 Gm15885 | 387 | 0 | 0.17215966 | 0 | 0 | 0 | 0 | 0 | 0 | 0 | 0 | 0 | 0 | 0 | 0 | 0 | 0 | 0 | 0 |
| ENSMUSG00 Gm8688 | 964 | 0 | 0 | 0 | 0 | 0 | 0 | 0 | 0 | 0 | 0 | 0 | 0 | 0 | 0 | 0 | 0 | 0 | 0 |
| ENSMUSG00 Gm14741 | 1008 | 0 | 0 | 0 | 0 | 0 | 0 | 0 | 0 | 0 | 0 | 0 | 0 | 0 | 0 | 0 | 0 | 0 | 0 |
| ENSMUSG00 Gm8051 | 465 | 0 | 0 | 0.1040291 | 0 | 0 | 0 | 0 | 0 | 0 | 0 | 0 | 0 | 0 | 0 | 0 | 0 | 0.08242214 | 0 |
| ENSMUSG00 Gm14043 | 227 | 0 | 0 | 0 | 0 | 0 | 0 | 0 | 0 | 0 | 0 | 0 | 0 | 0 | 0 | 0 | 0 | 0 | 0 |
| ENSMUSG00 Gm13916 | 627 | 0 | 0 | 0 | 0 | 0 | 0 | 0 | 0 | 0 | 0 | 0 | 0 | 0 | 0 | 0 | 0 | 0 | 0 |
| ENSMUSG00 Gm13614 | 502 | 0 | 0 | 0 | 0 | 0 | 0 | 0 | 0 | 0 | 0 | 0 | 0 | 0 | 0 | 0 | 0 | 0 | 0 |
| ENSMUSG00 Gm14365 | 586 | 0 | 0 | 0 | 0 | 0 | 0 | 0 | 0 | 0 | 0 | 0 | 0 | 0 | 0 | 0 | 0 | 0 | 0 |
| ENSMUSG00 Gm13722 | 3065 | 0 | 0 | 0 | 0 | 0 | 0 | 0 | 0.02492163 | 0 | 0 | 0 | 0 | 0 | 0 | 0 | 0 | 0 | 0 |
| ENSMUSG00 Gm9200 | 1353 | 0 | 0.0246215 | 0 | 0.13469636 | 0.03557422 | 0.25637039 | 0.06993744 | 0 | 0.12176354 | 0 | 0 | 0.12375978 | 0.03646114 | 0.06865349 | 0.030452 | 0.16980314 | 0 | 0 |
| ENSMUSG00 Rpl21-ps15 | 1022 | 0.04159019 | 0 | 0 | 0 | 0.14128745 | 0 | 0 | 0 | 0 | 0.09227143 | 0 | 0.21845661 | 0.04826998 | 0.04544431 | 0.04031463 | 0 | 0 | 0 |
| ENSMUSG00 Gm14805 | 339 | 0 | 0 | 0 | 0 | 0 | 0 | 0 | 0 | 0 | 0 | 0 | 0 | 0 | 0 | 0 | 0 | 0 | 0 |
| ENSMUSG00 Gm14579 | 282 | 0 | 0 | 0 | 0 | 0 | 0 | 0 | 0 | 0 | 0 | 0 | 0 | 0 | 0 | 0 | 0 | 0 | 0 |
| ENSMUSG00 Gm12821 | 599 | 0 | 0.05561418 | 0 | 0 | 0 | 0 | 0 | 0.06376027 | 0 | 0 | 0 | 0 | 0.08235712 | 0 | 0 | 0 | 0 | 0.0918349 |
| ENSMUSG00 Gm12028 | 434 | 0 | 0 | 0 | 0 | 0 | 0 | 0 | 0 | 0 | 0 | 0 | 0 | 0 | 0 | 0 | 0 | 0 | 0 |
| ENSMUSG00 Gm14911 | 988 | 0 | 0 | 0 | 0 | 0 | 0 | 0.04788733 | 0 | 0 | 0.04772338 | 0 | 0 | 0 | 0 | 0 | 0 | 0.0387918 | 0.05567723 |
| ENSMUSG00 Gm14810 | 485 | 0 | 0 | 0 | 0 | 0 | 0 | 0 | 0 | 0 | 0 | 0 | 0 | 0 | 0 | 0 | 0.31579882 | 0 | 0 |
| ENSMUSG00 Gm7398 | 631 | 0 | 0 | 0 | 0 | 0 | 0 | 0 | 0 | 0 | 0 | 0 | 0 | 0 | 0 | 0 | 0 | 0 | 0 |
| ENSMUSG00 Gm14893 | 280 | 0 | 0 | 0 | 0 | 0 | 0 | 0 | 0 | 0 | 0 | 0 | 0 | 0 | 0 | 0 | 0 | 0 | 0 |
| ENSMUSG00 Gm15916 | 1169 | 0.21816172 | 0.05699383 | 0.08276053 | 0.50666687 | 0.57643022 | 0 | 0 | 0 | 0.21139358 | 0 | 0.12260468 | 0.19098601 | 0 | 0.4767571 | 0 | 0.06551002 | 0 | 0 |
| ENSMUSG00 Gm12640 | 309 | 0 | 0 | 0 | 0 | 0 | 0 | 0 | 0 | 0 | 0 | 0 | 0 | 0 | 0 | 0 | 0 | 0 | 0 |
| ENSMUSG00 Gm14863 | 306 | 0.41671741 | 0 | 0 | 0 | 0 | 0 | 0 | 0.37443532 | 0 | 0 | 0.62450926 | 0 | 0.16121542 | 0 | 0.13464561 | 0 | 0.12524904 | 0.35953665 |
| ENSMUSG00 Gm16007 | 285 | 0 | 0 | 0 | 0 | 0.16888394 | 0 | 0.3320188 | 0 | 0 | 0 | 0 | 0 | 0 | 0 | 0 | 0 | 0 | 0 |
| ENSMUSG00 Gm9329 | 535 | 0.47669356 | 0.43586964 | 0.6329247 | 0.17032166 | 0.62976349 | 0.46310966 | 0.35373966 | 0.28555067 | 0 | 0.88132156 | 0.08929899 | 0.41731337 | 0.27662757 | 0.26043413 | 0.15402451 | 0.21471369 | 0.28655175 | 0.5141038 |
| ENSMUSG00 Gm8832 | 394 | 0 | 0 | 0 | 0 | 0 | 0 | 0 | 0 | 0 | 0 | 0 | 0 | 0.37562374 | 0.23575679 | 0 | 0.09718428 | 0 | 0 |
| ENSMUSG00 Pou2f3-rs1 | 1045 | 0 | 0 | 0 | 0 | 0 | 0 | 0 | 0 | 0 | 0 | 0 | 0 | 0 | 0 | 0 | 0 | 0 | 0 |
| ENSMUSG00 Gm12795 | 537 | 0 | 0.18610555 | 0.36032425 | 0.33937463 | 0 | 0.09227697 | 0.44052774 | 0.42673075 | 0 | 0 | 0.1793281 | 0 | 0.18373154 | 0 | 0.07672543 | 0.21391401 | 0 | 0 |
| ENSMUSG00 Gm12878 | 331 | 0 | 0 | 0 | 0 | 0 | 0 | 0 | 0 | 0 | 0 | 0 | 0 | 0 | 0 | 0 | 0 | 0 | 0 |
| ENSMUSG00 H2afb3-ps | 347 | 0 | 0 | 0 | 0 | 0 | 0 | 0 | 0 | 0 | 0 | 0 | 0 | 0 | 0 | 0 | 0 | 0 | 0 |
| ENSMUSG00 Gm5390 | 1229 | 0 | 0 | 0 | 0 | 0 | 0 | 0 | 0 | 0 | 0 | 0 | 0 | 0 | 0 | 0 | 0 | 0 | 0 |
| ENSMUSG00 Gm11771 | 724 | 0 | 0 | 0.06681427 | 0 | 0 | 0 | 0 | 0 | 0.11377491 | 0.19537584 | 0.06598751 | 0.23128037 | 0 | 0 | 0.05690823 | 0.05288758 | 0.10587375 | 0.07597943 |
| ENSMUSG00 Gm14414 | 677 | 0 | 0 | 0.07145278 | 0 | 0.0710959 | 0.14638917 | 0.20965737 | 0.05641418 | 0.30418402 | 0.06964653 | 0.07056862 | 0.16489117 | 0 | 0.06860279 | 0.06085902 | 0.11311849 | 0 | 0 |
| ENSMUSG00 Gm15529 | 993 | 0 | 0 | 0 | 0 | 0 | 0 | 0 | 0 | 0 | 0 | 0 | 0 | 0 | 0 | 0 | 0 | 0 | 0 |
| ENSMUSG00 Gm14586 | 407 | 3.86410691 | 4.17434301 | 5.46727862 | 3.47025153 | 2.6017256 | 4.1395404 | 5.92861579 | 4.31658602 | 4.04781482 | 3.82302999 | 2.58243019 | 4.25131582 | 7.15130988 | 6.16211479 | 4.15052529 | 4.32768536 | 5.46173273 | 4.46019784 |
| ENSMUSG00 Gm15570 | 624 | 0 | 0 | 0 | 0 | 0 | 0 | 0 | 0 | 0 | 0 | 0 | 0 | 0 | 0 | 0 | 0 | 0 | 0 |
| ENSMUSG00 Gm7224 | 2064 | 0.2883103 | 0.48419904 | 0.32811503 | 0.55185373 | 0.39643542 | 0.16805675 | 0.29799652 | 0.49960992 | 0.37913944 | 0.52541966 | 0.48608243 | 0.59493439 | 0.50192358 | 0.31502772 | 0.35931589 | 0.53799788 | 0.38994779 | 0.47973058 |
| ENSMUSG00 Gm12469 | 206 | 0 | 0 | 0 | 0 | 0 | 0 | 0 | 0 | 0 | 0 | 0 | 0 | 0 | 0.22545673 | 0 | 0 | 0 | 0 |
| ENSMUSG00 Gm15188 | 698 | 0 | 0 | 0 | 0 | 0 | 0 | 0 | 0 | 0 | 0 | 0 | 0 | 0 | 0 | 0 | 0 | 0 | 0 |
| ENSMUSG00 Gm4459 | 467 | 0.09101751 | 0.21400146 | 0.10358358 | 0.19512225 | 0.30919865 | 0.74276046 | 0.10131195 | 0 | 0.17638765 | 0.20193021 | 0.30690551 | 0 | 0.31690739 | 0 | 0.35290412 | 0.16398547 | 0.08206916 | 0.47117008 |
| ENSMUSG00 Gm12502 | 443 | 0 | 0 | 0 | 0.1028496 | 0 | 0 | 0 | 0 | 0 | 0 | 0 | 0.12599472 | 0 | 0.10483993 | 0 | 0.34573912 | 0 | 0.12417406 |
| ENSMUSG00 Gm2030 | 1120 | 0 | 0 | 0 | 0 | 0 | 0 | 0 | 0 | 0 | 0 | 0 | 0 | 0 | 0 | 0 | 0 | 0 | 0 |
| ENSMUSG00 Gm11262 | 831 | 0 | 0 | 0 | 0 | 0 | 0 | 0 | 0 | 0 | 0 | 0 | 0 | 0 | 0 | 0 | 0 | 0 | 0 |
| ENSMUSG00 Gm15602 | 436 | 0.09748894 | 0 | 0 | 0 | 0 | 0.34095918 | 0 | 0.08759725 | 0 | 0 | 0 | 0 | 0 | 0.10652314 | 0.37799593 | 0 | 0 | 0 |
| ENSMUSG00 Gm13366 | 975 | 0 | 0 | 0 | 0 | 0 | 0 | 0 | 0 | 0 | 0 | 0 | 0 | 0 | 0 | 0 | 0 | 0 | 0 |
| ENSMUSG00 Gm11786 | 154 | 0 | 0 | 0 | 0 | 0 | 0 | 0 | 0 | 0 | 0 | 0 | 0 | 0 | 0 | 0 | 0 | 0 | 0 |
| ENSMUSG00 Gm13312 | 465 | 0 | 0 | 0 | 0 | 0 | 0 | 0 | 0 | 0 | 0 | 0 | 0 | 0 | 0 | 0 | 0 | 0.49453286 | 0 |
| ENSMUSG00 Gm15068 | 190 | 0 | 0 | 0 | 0 | 0 | 0 | 0 | 0 | 0 | 0 | 0 | 0 | 0 | 0 | 0 | 0 | 0 | 0 |
| ENSMUSG00 Gm11480 | 658 | 0 | 0 | 0 | 0 | 0 | 0 | 0 | 0 | 0 | 0 | 0 | 0 | 0 | 0 | 0 | 0 | 0 | 0 |
| ENSMUSG00 Gm15175 | 131 | 0 | 0 | 0 | 0 | 0 | 0 | 0 | 0 | 0 | 0 | 0 | 0 | 0 | 0 | 0 | 0 | 0 | 0 |
| ENSMUSG00 Gm14754 | 725 | 0 | 0 | 0 | 0 | 0 | 0 | 0 | 0 | 0 | 0 | 0 | 0 | 0 | 0 | 0 | 0 | 0 | 0 |
| ENSMUSG00 Gm11392 | 706 | 0 | 0 | 0 | 0 | 0 | 0 | 0 | 0 | 0 | 0 | 0 | 0 | 0 | 0 | 0 | 0 | 0 | 0 |
| ENSMUSG00 Defa-ps6 | 280 | 0 | 0 | 0 | 0 | 0 | 0 | 0 | 0 | 0 | 0 | 0 | 0 | 0 | 0 | 0 | 0 | 0 | 0 |
| ENSMUSG00 Gm8876 | 1121 | 0 | 0 | 0 | 0 | 0 | 0 | 0 | 0 | 0 | 0 | 0 | 0 | 0 | 0 | 0 | 0 | 0 | 0 |
| ENSMUSG00 Gm9225 | 834 | 0 | 0 | 0 | 0 | 0 | 0 | 0 | 0 | 0 | 0 | 0 | 0 | 0 | 0 | 0 | 0 | 0 | 0 |
| ENSMUSG00 Gm13148 | 744 | 0 | 0 | 0 | 0 | 0.06469345 | 0 | 0 | 0 | 0 | 0 | 0 | 0 | 0 | 0 | 0 | 0 | 0 | 0 |
| ENSMUSG00 Gm13876 | 260 | 0 | 0 | 0 | 0 | 0 | 0 | 0 | 0 | 0 | 0 | 0 | 0 | 0 | 0 | 0 | 0 | 0 | 0 |
| ENSMUSG00 Gm12787 | 394 | 0 | 0 | 0 | 0 | 0 | 0 | 0 | 0 | 0 | 0 | 0 | 0 | 0 | 0 | 0 | 0 | 0 | 0 |
| ENSMUSG00 Gm14086 | 171 | 0 | 0 | 0 | 0 | 0 | 0 | 0 | 0 | 0 | 0 | 0 | 0 | 0 | 0 | 0 | 0 | 0 | 0 |
| ENSMUSG00 Gm6088 | 2538 | 0 | 0.01312565 | 0 | 0.03590311 | 0.11378705 | 0 | 0 | 0 | 0.01622794 | 0 | 0 | 0 | 0.31099711 | 0 | 0.01623387 | 0.03017384 | 0 | 0 |
| ENSMUSG00 Gm14229 | 849 | 0 | 0 | 0 | 0 | 0 | 0 | 0 | 0 | 0 | 0 | 0 | 0 | 0 | 0 | 0 | 0 | 0 | 0 |
| ENSMUSG00 Rasl2-9-ps | 1005 | 0.38064337 | 0.19888295 | 0.19253146 | 0.22667186 | 0.38313969 | 0.59167443 | 1.64770524 | 0.87405498 | 0.32785286 | 0.37532898 | 0.76059635 | 0 | 0.73629727 | 0.18485209 | 1.14790404 | 0.19050053 | 0.91525485 | 0.43788344 |
| ENSMUSG00 Gm13357 | 536 | 0.0793007 | 0.31075461 | 0 | 0.25500585 | 0.17959673 | 0 | 0 | 0.49878137 | 0.30736206 | 0 | 0 | 0.1041337 | 0.55222295 | 0 | 0 | 0 | 0.21451286 | 0 |
| ENSMUSG00 Gm11524 | 806 | 0 | 0 | 0 | 0 | 0 | 0 | 0 | 0 | 0 | 0 | 0.05927414 | 0 | 0 | 0 | 0 | 0 | 0 | 0 |
| ENSMUSG00 Gm11384 | 292 | 0 | 0 | 0 | 0 | 0 | 0 | 0 | 0 | 0 | 0 | 0 | 0 | 0 | 0 | 0 | 0 | 0 | 0 |
| ENSMUSG00 Gm11875 | 798 | 0 | 0 | 0 | 0 | 0 | 0 | 0 | 0 | 0 | 0 | 0 | 0 | 0 | 0 | 0 | 0 | 0 | 0 |
| ENSMUSG00 Gm12099 | 749 | 0 | 0 | 0 | 0 | 0 | 0 | 0 | 0 | 0 | 0 | 0 | 0 | 0 | 0 | 0 | 0 | 0 | 0 |
| ENSMUSG00 Gm12245 | 356 | 0.11939656 | 0.09357555 | 0.40764211 | 0.25596092 | 0.13520203 | 0.13919307 | 0.13290078 | 0 | 0.46276984 | 0.26489159 | 0.26839864 | 0 | 0.4157184 | 0.13046092 | 0 | 0.10755789 | 0.32297441 | 0.61807985 |
| ENSMUSG00 Gm15798 | 407 | 0 | 0 | 0.11885388 | 0 | 0 | 0 | 0.12175119 | 0 | 0 | 0 | 0 | 0 | 0 | 0 | 0 | 0.09408012 | 0 | 0 |
| ENSMUSG00 Gm11258 | 1217 | 0 | 0 | 0 | 0 | 0 | 0 | 0 | 0.03138242 | 0 | 0 | 0 | 0 | 0 | 0 | 0 | 0 | 0 | 0 |
| ENSMUSG00 Gm11853 | 1382 | 0 | 0 | 0 | 0 | 0 | 0 | 0 | 0 | 0 | 0 | 0 | 0 | 0 | 0 | 0 | 0 | 0 | 0 |
| ENSMUSG00 Gm11473 | 358 | 0 | 0 | 0 | 0 | 0 | 0 | 0 | 0 | 0 | 0 | 0 | 0 | 0 | 0 | 0 | 0 | 0 | 0 |
| ENSMUSG00 Gm12152 | 683 | 0 | 0 | 0 | 0 | 0 | 0 | 0 | 0.0559186 | 0 | 0 | 0 | 0 | 0 | 0 | 0 | 0 | 0.05611464 | 0 |
| ENSMUSG00 Gm14903 | 535 | 0 | 0 | 0 | 0 | 0 | 0 | 0 | 0 | 0 | 0 | 0 | 0 | 0 | 0 | 0 | 0 | 0 | 0 |
| ENSMUSG00 Gm11433 | 226 | 0 | 0 | 0 | 0 | 0 | 0 | 0 | 0 | 0 | 0 | 0 | 0 | 0 | 0 | 0 | 0 | 0 | 0 |
| ENSMUSG00 Gm14575 | 559 | 0 | 0 | 0 | 0 | 0 | 0 | 0 | 0 | 0 | 0 | 0 | 0 | 0 | 0 | 0 | 0 | 0 | 0 |
| ENSMUSG00 Gm12358 | 518 | 0 | 0 | 0 | 0 | 0 | 0 | 0 | 0 | 0 | 0 | 0 | 0 | 0 | 0 | 0 | 0 | 0 | 0 |
| ENSMUSG00 Gm11334 | 155 | 0 | 0 | 0 | 0 | 0 | 0 | 0 | 0 | 0 | 0 | 0 | 0 | 0 | 0 | 0 | 0 | 0 | 0 |
| ENSMUSG00 Gm5648 | 1586 | 0 | 0 | 0.03050033 | 0.02872701 | 0 | 0 | 0 | 0 | 0 | 0 | 0 | 0 | 0 | 0 | 0 | 0 | 0 | 0 |
| ENSMUSG00 Gm10169 | 347 | 2.32737275 | 2.20805927 | 2.0910748 | 3.01989631 | 2.91288299 | 1.57083593 | 5.59025889 | 2.86167857 | 4.62903203 | 3.94054869 | 5.36951983 | 4.66470955 | 4.83367489 | 3.47996044 | 2.84967534 | 3.64146987 | 1.98810762 | 1.90233222 |
| ENSMUSG00 Gm6829 | 359 | 0 | 0.09279358 | 0.53898084 | 0 | 0 | 0 | 0.13179019 | 0.10638552 | 0 | 0.39401702 | 0.13307788 | 0 | 0 | 0 | 0 | 0.10665907 | 0.64055092 | 0.30645742 |
| ENSMUSG00 Gm14304 | 444 | 0 | 0 | 0 | 0 | 0 | 0 | 0 | 0 | 0 | 0 | 0 | 0 | 0 | 0 | 0 | 0 | 0 | 0 |
| ENSMUSG00 Gm9183 | 1568 | 0 | 0 | 0 | 0 | 0 | 0 | 0 | 0 | 0 | 0 | 0 | 0 | 0 | 0 | 0 | 0 | 0 | 0 |
| ENSMUSG00 Gm11308 | 1918 | 0 | 0 | 0 | 0 | 0 | 0 | 0 | 0 | 0 | 0 | 0 | 0 | 0 | 0 | 0 | 0 | 0 | 0 |
| ENSMUSG00 Zfp133-ps | 3107 | 0.49249641 | 0.43959725 | 0.45150704 | 0.36659997 | 0.44925194 | 0.27112857 | 0.791844 | 0.44252542 | 0.60977783 | 1.4872124 | 0.76882778 | 0.8622954 | 0.39694172 | 1.47987276 | 0.74260931 | 0.64084699 | 1.30755953 | 0.10622937 |
| ENSMUSG00 Gm15143 | 712 | 0 | 0 | 0 | 0 | 0 | 0 | 0 | 0 | 0 | 0 | 0 | 0 | 0 | 0 | 0 | 0 | 0 | 0 |
| ENSMUSG00 Gm11367 | 122 | 0 | 0 | 0 | 0 | 0 | 0 | 0 | 0 | 0 | 0 | 0 | 0 | 0 | 0 | 0.33771767 | 0 | 0 | 0 |
| ENSMUSG00 Gm11699 | 400 | 0 | 0 | 0 | 0 | 0 | 0 | 0 | 0 | 0 | 0 | 0 | 0 | 0 | 0 | 0 | 0 | 0 | 0 |
| ENSMUSG00 Gm12989 | 730 | 0.29113134 | 0.27380461 | 0.19879533 | 0.24964956 | 0.06593414 | 0.27152183 | 0.06481189 | 0.10463672 | 0.16925965 | 0.25836002 | 0.06544515 | 0.30583925 | 0.13515594 | 0.06362204 | 0.39508341 | 0.15735866 | 0.05250178 | 0.30141976 |
| ENSMUSG00 Gm12892 | 1145 | 0.66820364 | 0.02909423 | 0.5069715 | 1.98956525 | 0.50443937 | 0.77899494 | 1.15699128 | 0.16677905 | 1.33091798 | 2.01780302 | 0.79277224 | 0.97494607 | 0.90477752 | 0.04056252 | 0.89959729 | 3.14350838 | 0.60250947 | 0.67260043 |
| ENSMUSG00 Gm9119 | 491 | 0 | 0 | 0 | 0 | 0 | 0 | 0 | 0 | 0 | 0 | 0 | 0 | 0 | 0 | 0 | 0 | 0 | 0 |
| ENSMUSG00 Gm6640 | 1586 | 0 | 0.02100435 | 0 | 0 | 0 | 0.03124384 | 0 | 0 | 0.0519376 | 0 | 0 | 0 | 0 | 0 | 0 | 0 | 0 | 0 |
| ENSMUSG00 Pcna-ps1 | 736 | 0 | 0 | 0 | 0 | 0 | 0 | 0 | 0 | 0 | 0 | 0 | 0 | 0 | 0 | 0 | 0 | 0 | 0 |
| ENSMUSG00 Gm6539 | 1062 | 0 | 0 | 0 | 0 | 0 | 0 | 0 | 0 | 0 | 0 | 0 | 0 | 0 | 0 | 0 | 0 | 0 | 0 |
| ENSMUSG00 Gm14034 | 550 | 0.07728214 | 0 | 0 | 0 | 0 | 0.09009588 | 0 | 0 | 0 | 0 | 0 | 0 | 0 | 0 | 0 | 0 | 0 | 0 |
| ENSMUSG00 Gm12412 | 327 | 0 | 0 | 0 | 0 | 0 | 0 | 0 | 0 | 0 | 0 | 0 | 0 | 0 | 0 | 0 | 0 | 0 | 0 |
| ENSMUSG00 Gm13504 | 545 | 0 | 0 | 0 | 0 | 0.08831546 | 0.09092245 | 0.08681225 | 0 | 0 | 0 | 0 | 0 | 0 | 0 | 0.07559919 | 0 | 0.07032348 | 0.10093414 |
| ENSMUSG00 Gm15154 | 365 | 0 | 0 | 0 | 0 | 0 | 0 | 0 | 0 | 0 | 0 | 0 | 0 | 0 | 0 | 0 | 0 | 0 | 0 |
| ENSMUSG00 Gm13743 | 723 | 0.35274005 | 0.04607593 | 0 | 0 | 0 | 0.13707533 | 0 | 0 | 0 | 0.06521536 | 0 | 0 | 0 | 0 | 0 | 0 | 0 | 0 |
| ENSMUSG00 Gm14094 | 609 | 0 | 0 | 0 | 0.07481288 | 0 | 0 | 0.15537826 | 0.0627133 | 0 | 0 | 0 | 0.09165133 | 0.08100479 | 0 | 0 | 0 | 0 | 0 |
| ENSMUSG00 Defa-ps5 | 277 | 0 | 0 | 0 | 0 | 0 | 0 | 0 | 0 | 0 | 0 | 0 | 0 | 0 | 0 | 0 | 0 | 0 | 0 |
| ENSMUSG00 Gm14465 | 566 | 0 | 0 | 0 | 0 | 0 | 0 | 0 | 0 | 0 | 0 | 0 | 0 | 0 | 0 | 0 | 0 | 0 | 0 |
| ENSMUSG00 Gm9575 | 1338 | 0.12707078 | 0.09959012 | 0.07230722 | 0.10215481 | 0.25181126 | 0.18517464 | 0.24752523 | 0.19981077 | 0.09234645 | 0.10571907 | 0.10711874 | 0 | 0.47930861 | 1.1107704 | 0.03079339 | 0.20032455 | 0.1718668 | 0.1233388 |
| ENSMUSG00 Gm11971 | 287 | 0.44430498 | 0.1160728 | 0 | 0.63499713 | 0 | 0.34531522 | 0.16485254 | 0.13307457 | 0 | 0 | 0.16646327 | 0 | 0.51566464 | 0 | 0 | 1.067334 | 0.40062331 | 0.19166936 |
| ENSMUSG00 1700045I19R | 1935 | 0 | 0 | 0.09999696 | 0 | 0 | 0 | 0 | 0 | 0 | 0 | 0 | 0 | 0 | 0 | 0 | 0 | 0 | 0 |
| ENSMUSG00 Gm12563 | 101 | 0.84168665 | 1.31932254 | 1.91578338 | 1.35329835 | 0.9531074 | 0 | 2.81065419 | 0 | 0.40778728 | 0.46683865 | 0.47301939 | 1.10526065 | 1.46530447 | 1.37952735 | 3.26348958 | 0.75822985 | 2.27680972 | 1.08928924 |
| ENSMUSG00 Gm13914 | 990 | 0 | 0 | 0 | 0 | 0 | 0 | 0 | 0 | 0 | 0 | 0 | 0 | 0 | 0 | 0 | 0 | 0 | 0 |
| ENSMUSG00 Gm12687 | 705 | 0 | 0 | 0 | 0 | 0 | 0 | 0 | 0 | 0 | 0 | 0 | 0 | 0 | 0 | 0 | 0 | 0 | 0 |
| ENSMUSG00 Gm14932 | 451 | 0 | 0 | 0 | 0 | 0 | 0 | 0 | 0 | 0 | 0 | 0 | 0 | 0 | 0 | 0 | 0 | 0 | 0 |
| ENSMUSG00 Gm2986 | 381 | 0 | 0.17487084 | 0.25392929 | 0 | 0 | 0.13005967 | 0 | 0.20048505 | 0 | 0.12375513 | 0 | 0 | 0.1294801 | 0 | 0.10814057 | 0 | 0 | 0 |
| ENSMUSG00 Gm4912 | 1043 | 0 | 0 | 0 | 0 | 0 | 0 | 0 | 0 | 0 | 0 | 0 | 0 | 0 | 0 | 0 | 0 | 0 | 0 |
| ENSMUSG00 Gm14428 | 819 | 0 | 0 | 0 | 0 | 0 | 0 | 0 | 0 | 0 | 0 | 0 | 0 | 0 | 0 | 0.15092145 | 0 | 0 | 0 |
| ENSMUSG00 Gm13153 | 636 | 0 | 0 | 0 | 0 | 0 | 0 | 0 | 0 | 0 | 0 | 0 | 0 | 0 | 0 | 0 | 0 | 0 | 0 |
| ENSMUSG00 Gm8624 | 1098 | 0 | 0.03033961 | 0 | 0 | 0.043836 | 0 | 0.08617974 | 0.13913443 | 0 | 0.08588471 | 0 | 0.10166787 | 0 | 0 | 0 | 0.03487305 | 0 | 0 |
| ENSMUSG00 Olfr1117-ps1 | 920 | 0 | 0 | 0 | 0 | 0 | 0 | 0 | 0 | 0 | 0 | 0 | 0 | 0 | 0 | 0 | 0 | 0 | 0 |
| ENSMUSG00 Gm13592 | 1009 | 0 | 0 | 0 | 0 | 0.0477026 | 0 | 0 | 0 | 0.04081914 | 0 | 0 | 0.0553178 | 0 | 0 | 0.04083405 | 0 | 0.07596887 | 0 |
| ENSMUSG00 Gm13123 | 491 | 0 | 0 | 0 | 0 | 0 | 0 | 0.19271967 | 0 | 0 | 0 | 0 | 0 | 0 | 0 | 0 | 0.07798494 | 0 | 0 |
| ENSMUSG00 Gm14688 | 302 | 0 | 0 | 0 | 0 | 0 | 0 | 0 | 0 | 0 | 0 | 0 | 0 | 0 | 0 | 0 | 0 | 0 | 0 |
| ENSMUSG00 Gm15078 | 781 | 0 | 0 | 0 | 0 | 0 | 0 | 0 | 0 | 0 | 0 | 0 | 0 | 0 | 0 | 0 | 0 | 0 | 0 |
| ENSMUSG00 Gm3931 | 454 | 0 | 0 | 0 | 0 | 0 | 0 | 0 | 0 | 0 | 0 | 0 | 0 | 0 | 0 | 0 | 0 | 0 | 0 |
| ENSMUSG00 Gm5883 | 494 | 0.5162572 | 0.33717504 | 0.19584425 | 1.01451718 | 0.48716522 | 0.60185506 | 0.28732396 | 0.30925022 | 0.50024108 | 0.85902091 | 1.06381486 | 1.01688454 | 0.29958654 | 1.12819645 | 0.66723167 | 0.54257946 | 0.69825237 | 0.44541787 |
| ENSMUSG00 Gm14499 | 515 | 0 | 0 | 0 | 0 | 0 | 0 | 0 | 0 | 0 | 0 | 0 | 0 | 0 | 0 | 0 | 0 | 0 | 0 |
| ENSMUSG00 Gm14202 | 310 | 0 | 0 | 0 | 0 | 0 | 0 | 0 | 0 | 0 | 0 | 0 | 0 | 0 | 0 | 0 | 0 | 0 | 0 |
| ENSMUSG00 Gm13436 | 1181 | 1.04373421 | 0.90263558 | 1.06495494 | 1.00303739 | 0.69283887 | 0.46154113 | 0.96147696 | 0.45274651 | 1.08110245 | 0.51901706 | 0.64724753 | 0.661659 | 0.71011227 | 0.98315172 | 0.87217519 | 0.81055477 | 0.35697652 | 1.0247251 |
| ENSMUSG00 Gm13865 | 864 | 0 | 0 | 0 | 0 | 0 | 0 | 0 | 0 | 0 | 0 | 0 | 0 | 0 | 0 | 0.04768699 | 0 | 0 | 0 |
| ENSMUSG00 Ccnb2-ps | 2917 | 0.10200077 | 0.18272414 | 0.14924984 | 0.2811446 | 0.18150537 | 0.08493784 | 0.21085527 | 0.17020954 | 0.74833231 | 0.06465643 | 0.04913434 | 0.13394228 | 0.35514922 | 0.38620295 | 0.14124633 | 0.02625342 | 0.15766732 | 0.5657433 |
| ENSMUSG00 Gm15161 | 386 | 0 | 0 | 0 | 0 | 0 | 0 | 0 | 0 | 0 | 0 | 0 | 0.28920032 | 0 | 0 | 0 | 0 | 0 | 0 |
| ENSMUSG00 Gm12901 | 530 | 0 | 0 | 0.09127081 | 0 | 0 | 0.09349572 | 0 | 0 | 0 | 0 | 0 | 0 | 0 | 0 | 0 | 0 | 0 | 0 |
| ENSMUSG00 Gm7700 | 969 | 0 | 0 | 0 | 0 | 0 | 0.05113801 | 0 | 0 | 0 | 0 | 0 | 0 | 0 | 0 | 0 | 0 | 0 | 0 |
| ENSMUSG00 Gpr144 | 2644 | 0 | 0 | 0 | 0 | 0 | 0 | 0 | 0 | 0 | 0 | 0 | 0 | 0 | 0 | 0 | 0 | 0 | 0 |
| ENSMUSG00 Gm11671 | 423 | 0 | 0.15750777 | 0.22871646 | 0.53854662 | 0 | 0 | 0.33555091 | 0 | 0 | 0.33440215 | 0 | 0 | 0.69974351 | 0 | 0 | 0 | 0 | 0 |
| ENSMUSG00 Gm12651 | 446 | 0 | 0 | 0 | 0 | 0 | 0 | 0 | 0 | 0 | 0 | 0 | 0 | 0 | 0 | 0 | 0 | 0 | 0 |
| ENSMUSG00 Gm11865 | 857 | 0 | 0.03887152 | 0 | 0 | 0 | 0 | 0 | 0.08913046 | 0 | 0.05501832 | 0.05574674 | 0.13025826 | 0 | 0 | 0.14422948 | 0.08935964 | 0.04472147 | 0 |
| ENSMUSG00 Gm13309 | 320 | 0 | 0 | 0 | 0 | 0 | 0 | 0 | 0 | 0 | 0 | 0 | 0 | 0 | 0 | 0 | 0 | 0 | 0 |
| ENSMUSG00 Gm11887 | 229 | 0 | 0 | 0 | 0 | 0 | 0 | 0 | 0 | 0 | 0 | 0 | 0 | 0 | 0 | 0 | 0 | 0 | 0 |
| ENSMUSG00 Gm13854 | 1677 | 0 | 0 | 0 | 0 | 0 | 0 | 0 | 0 | 0 | 0 | 0 | 0 | 0.11766707 | 0 | 0 | 0 | 0 | 0 |
| ENSMUSG00 Sh3d2c-ps1 | 903 | 0 | 0 | 0 | 0 | 0 | 0 | 0 | 0 | 0 | 0 | 0 | 0 | 0 | 0 | 0 | 0 | 0 | 0.30459085 |
| ENSMUSG00 Gm2759 | 631 | 0 | 0 | 0 | 0 | 0 | 0 | 0 | 0 | 0 | 0 | 0 | 0 | 0 | 0 | 0 | 0 | 0 | 0 |
| ENSMUSG00 Gm16202 | 543 | 0 | 0 | 0 | 0 | 0 | 0 | 0 | 0 | 0 | 0 | 0 | 0 | 0 | 0 | 0 | 0 | 0 | 0 |
| ENSMUSG00 Gm14197 | 1048 | 0.04055837 | 0 | 0 | 0 | 0 | 0 | 0 | 0 | 0 | 0 | 0 | 0 | 0 | 0 | 0 | 0 | 0.29256715 | 0.47240645 |
| ENSMUSG00 Gm13035 | 341 | 0 | 0 | 0 | 0 | 0 | 0 | 0 | 0 | 0 | 0 | 0 | 0 | 0 | 0 | 0 | 0 | 0 | 0 |
| ENSMUSG00 Gm16209 | 626 | 0.40739785 | 0.58537035 | 0.61819208 | 0.50946855 | 1.15332085 | 0.39578861 | 0.45347616 | 0.48808182 | 0.46055209 | 0.15064122 | 0.83949608 | 1.0699488 | 0 | 0.74191833 | 0.26326873 | 0.79517236 | 0.24489647 | 0.17574794 |
| ENSMUSG00 Gm12495 | 446 | 0 | 0 | 0 | 0 | 0 | 0 | 0 | 0 | 0 | 0 | 0 | 0 | 0 | 0 | 0 | 0 | 0 | 0 |
| ENSMUSG00 Gm6039 | 1249 | 0 | 0 | 0.03872981 | 0 | 0 | 0.03967393 | 0 | 0 | 0 | 0 | 0 | 0.04468828 | 0 | 0 | 0 | 0 | 0 | 0 |
| ENSMUSG00 Prdx6-ps2 | 693 | 0.06133503 | 0.04807055 | 0 | 0.19723396 | 0 | 0 | 0 | 0 | 0.1188644 | 0 | 0.06893933 | 0 | 0.07118603 | 0 | 0 | 0.1105068 | 0 | 0.15875644 |
| ENSMUSG00 Gm14592 | 559 | 0 | 0 | 0 | 0 | 0 | 0 | 0 | 0 | 0 | 0 | 0 | 0 | 0 | 0 | 0 | 0 | 0 | 0 |
| ENSMUSG00 Gm13796 | 312 | 0 | 0 | 0 | 0 | 0 | 0 | 0 | 0 | 0 | 0 | 0 | 0 | 0 | 0 | 0 | 0 | 0 | 0 |
| ENSMUSG00 Gm11456 | 255 | 0 | 0 | 0 | 0 | 0 | 0 | 1.29877942 | 0 | 0.16151575 | 0 | 0 | 0 | 0 | 0.91068838 | 0.48472419 | 0 | 0 | 0 |
| ENSMUSG00 Gm15297 | 288 | 0 | 0 | 0 | 0 | 0 | 0 | 0 | 0 | 0 | 0 | 0 | 0 | 0 | 0 | 0 | 0 | 0 | 0 |
| ENSMUSG00 Gm11812 | 916 | 0 | 0 | 0 | 0 | 0 | 0 | 0 | 0 | 0 | 0 | 0 | 0 | 0 | 0 | 0 | 0 | 0 | 0 |
| ENSMUSG00 Gm12421 | 306 | 0.83343482 | 0 | 0 | 0.1488923 | 0 | 0 | 0 | 0 | 0 | 0 | 0 | 0 | 0.16121542 | 0 | 0 | 0 | 0 | 0 |
| ENSMUSG00 Gm14624 | 736 | 0 | 0 | 0 | 0 | 0 | 0 | 0 | 0 | 0 | 0 | 0 | 0 | 0 | 0 | 0 | 0 | 0 | 0 |
| ENSMUSG00 Gm13543 | 795 | 0 | 1.25709035 | 0.24338883 | 0.51578541 | 0.2421732 | 0.31165241 | 0.71415364 | 0.38432606 | 0.31084163 | 0.17792718 | 0.60094287 | 1.33395923 | 0.2482109 | 1.22682495 | 0.77738785 | 1.01145001 | 0.62671932 | 0.41516307 |
| ENSMUSG00 Gm14727 | 524 | 0 | 0 | 0 | 0 | 0 | 0 | 0 | 0 | 0 | 0 | 0 | 0 | 0 | 0 | 0 | 0 | 0 | 0 |
| ENSMUSG00 Gm11662 | 1191 | 0 | 0 | 0 | 0 | 0 | 0 | 0 | 0 | 0 | 0 | 0 | 0 | 0 | 0 | 0 | 0 | 0 | 0 |
| ENSMUSG00 Gm13100 | 1436 | 0 | 0 | 0 | 0 | 0 | 0 | 0 | 0 | 0 | 0 | 0 | 0 | 0 | 0 | 0 | 0 | 0 | 0 |
| ENSMUSG00 Gm14108 | 493 | 0 | 0 | 0 | 0 | 0 | 0 | 0.09596892 | 0.23240813 | 0 | 0 | 0 | 0.11321635 | 0 | 0.28262122 | 0.16714627 | 0 | 0.07774097 | 0 |
| ENSMUSG00 Gm14534 | 356 | 2.26853468 | 1.029331 | 0.81528422 | 1.53576554 | 1.89282846 | 0.41757922 | 0.39870235 | 2.03835856 | 2.66092658 | 2.78136171 | 1.34199322 | 1.09749898 | 2.07859202 | 2.08737471 | 0.69440825 | 1.72092618 | 1.7225302 | 6.18079851 |
| ENSMUSG00 Gm12014 | 1617 | 0 | 0 | 0 | 0 | 0 | 0 | 0 | 0 | 0 | 0 | 0 | 0 | 0 | 0 | 0 | 0 | 0 | 0 |
| ENSMUSG00 Gm12814 | 842 | 0 | 0 | 0 | 0 | 0 | 0 | 0 | 0 | 0 | 0 | 0 | 0 | 0 | 0 | 0 | 0 | 0 | 0 |
| ENSMUSG00 Gm7803 | 1580 | 0.02690201 | 0.14758877 | 0.06123232 | 0.17301662 | 0.03046324 | 0.15681245 | 0.05986947 | 0.14503444 | 0.07820225 | 0.02984222 | 0.03023732 | 0.07065274 | 0.15611366 | 0.14697496 | 0.0782308 | 0.14540737 | 0.02425715 | 0 |
| ENSMUSG00 Gm14559 | 495 | 0 | 0 | 0 | 0 | 0 | 0 | 0 | 0 | 0 | 0 | 0 | 0 | 0 | 0 | 0 | 0 | 0 | 0 |
| ENSMUSG00 Gm13468 | 444 | 0 | 0 | 0 | 0 | 0 | 0 | 0 | 0 | 0 | 0 | 0 | 0 | 0 | 0 | 0 | 0 | 0 | 0 |
| ENSMUSG00 Rps15a-ps4 | 391 | 0.10870889 | 0.34079687 | 0.24743494 | 0.11652441 | 0.86169684 | 0.50693334 | 0 | 0.48839389 | 1.26403629 | 0 | 0.36655978 | 0.14275106 | 0 | 0 | 0.10537482 | 0 | 0.88219098 | 0 |
| ENSMUSG00 Gm14524 | 682 | 0.12464861 | 0.14653766 | 0.14185786 | 0 | 0.35287334 | 0 | 0 | 0.72800767 | 0.06039079 | 0 | 0 | 0.16368229 | 0 | 0.13619967 | 0.42288987 | 0.05614459 | 0 | 0.96790217 |
| ENSMUSG00 Pgam1-ps1 | 748 | 0.28412551 | 0.13360786 | 0 | 0.18273146 | 0 | 0 | 0.06325224 | 0.40847489 | 0.44049749 | 0.1891071 | 0.12774053 | 0.29847948 | 0.26380704 | 0.49672821 | 0 | 0.20476261 | 0 | 0.29416635 |
| ENSMUSG00 Gm2318 | 631 | 0 | 0 | 0 | 0 | 0 | 0 | 0 | 0 | 0 | 0 | 0 | 0 | 0 | 0 | 0 | 0 | 0 | 0 |
| ENSMUSG00 Gm9050 | 447 | 0 | 0 | 0 | 0 | 0 | 0 | 0 | 0 | 0 | 0.10548256 | 0 | 0 | 0 | 0 | 0 | 0 | 0.08574116 | 0.12306288 |
| ENSMUSG00 Gm11516 | 1581 | 0.02688499 | 0 | 0 | 0 | 0.03044397 | 0 | 0 | 0 | 0.02605093 | 0.02982334 | 0 | 0.03530402 | 0 | 0 | 0.02606044 | 0 | 0.07272542 | 0 |
| ENSMUSG00 Gm15335 | 409 | 0 | 0 | 0.47309076 | 0 | 0 | 0.24231166 | 0 | 0 | 0 | 0 | 0 | 0 | 0 | 0 | 0 | 0 | 0 | 0 |
| ENSMUSG00 Gm11405 | 1555 | 0.05466904 | 0.10711542 | 0 | 0 | 0 | 0 | 0 | 0 | 0.02648651 | 0.030322 | 0 | 0 | 0 | 0 | 0 | 0.02462418 | 0 | 0 |
| ENSMUSG00 Gm15362 | 387 | 0.1098325 | 0 | 0 | 0 | 0 | 0 | 0 | 0 | 0 | 0 | 0 | 0 | 0 | 0 | 0 | 0 | 0 | 0 |
| ENSMUSG00 Gm11689 | 301 | 0.14121321 | 0 | 0 | 0 | 0 | 0 | 0 | 0 | 0 | 0 | 0.15872079 | 0 | 0 | 0 | 0.13688225 | 0 | 0 | 0 |
| ENSMUSG00 Gm15988 | 366 | 0.11613436 | 0 | 0.26433623 | 0 | 0 | 0 | 0 | 0 | 0.22506293 | 0 | 0 | 0.15250181 | 0 | 0 | 0 | 0 | 0 | 0 |
| ENSMUSG00 Gm15566 | 589 | 0 | 0 | 0.08212823 | 0.07735322 | 0 | 0 | 0 | 0 | 0 | 0 | 0 | 0 | 0 | 0 | 0.13990342 | 0 | 0 | 0 |
| ENSMUSG00 Gm13394 | 2249 | 20.146962 | 11.3462325 | 24.7567512 | 34.6823068 | 22.7925739 | 20.4909036 | 32.0396665 | 14.7912772 | 34.5021768 | 57.2139925 | 34.0096525 | 28.5158721 | 40.1411332 | 18.6891503 | 23.9991277 | 48.471925 | 21.7619747 | 23.603285 |
| ENSMUSG00 Gm7180 | 512 | 0.41508961 | 0.78077096 | 0.37791821 | 0.71189132 | 1.22209962 | 1.06460951 | 0.55444546 | 0.74594536 | 0.5630969 | 0.5525473 | 0.09331047 | 1.30817959 | 1.05986541 | 1.54208884 | 0.88518968 | 0.44871806 | 1.34740688 | 0.75207763 |
| ENSMUSG00 Gm14837 | 205 | 0 | 0 | 0 | 0 | 0 | 0 | 0 | 0 | 0 | 0 | 0 | 0 | 0 | 0 | 0 | 0 | 0 | 0 |
| ENSMUSG00 Gm13473 | 365 | 0 | 0 | 0 | 0 | 0 | 0 | 0 | 0 | 0 | 0 | 0 | 0 | 0 | 0 | 0 | 0 | 0 | 0 |
| ENSMUSG00 Gm13361 | 1430 | 0 | 0 | 0 | 0 | 0 | 0.03465226 | 0 | 0 | 0 | 0 | 0 | 0 | 0 | 0 | 0 | 0 | 0 | 0 |
| ENSMUSG00 Gm15094 | 1130 | 0 | 0 | 0 | 0 | 0 | 0 | 0 | 0 | 0 | 0 | 0 | 0 | 0 | 0 | 0 | 0 | 0 | 0 |
| ENSMUSG00 Gm14775 | 309 | 0 | 0 | 0 | 0 | 0 | 0 | 0 | 0 | 0 | 0 | 0 | 0 | 0 | 0 | 0 | 0 | 0 | 0 |
| ENSMUSG00 Gm12728 | 492 | 0.08639263 | 0.2031274 | 0.5899211 | 0.09260375 | 0.48914557 | 0.40286775 | 0.19232796 | 0.15525367 | 0.33484972 | 0.67084334 | 0.48551787 | 0 | 0.60160875 | 0.28319565 | 0.251229 | 0.31130575 | 0.07789898 | 0.22361425 |
| ENSMUSG00 Gm14842 | 268 | 0 | 0 | 0 | 0 | 0 | 0 | 0 | 0 | 0 | 0 | 0 | 0 | 0 | 0 | 0 | 0 | 0 | 0 |
| ENSMUSG00 Gm11660 | 411 | 0 | 0 | 0 | 0 | 0 | 0 | 0.69069604 | 0 | 0.5010525 | 0 | 0 | 0 | 0 | 0 | 0 | 0 | 0.46625665 | 0 |
| ENSMUSG00 Gm12739 | 381 | 0.44624857 | 0.69948334 | 0.38089394 | 0.59791397 | 0.63165254 | 0 | 0.12418026 | 0 | 1.08101091 | 0 | 0.75236155 | 1.31848022 | 0.2589602 | 0.60950246 | 1.51396793 | 0 | 0.80475164 | 0.28876172 |
| ENSMUSG00 Gm14729 | 688 | 0 | 0 | 0 | 0 | 0 | 0 | 0 | 0 | 0 | 0 | 0 | 0 | 0 | 0 | 0 | 0 | 0 | 0 |
| ENSMUSG00 Gm11888 | 808 | 0 | 0 | 0 | 0 | 0 | 0 | 0 | 0 | 0 | 0 | 0 | 0 | 0 | 0 | 0 | 0 | 0 | 0 |
| ENSMUSG00 Rps18-ps2 | 397 | 0 | 0 | 0 | 0 | 0 | 0 | 0 | 0 | 0 | 0 | 0 | 0 | 0 | 0 | 0 | 0 | 0 | 0 |
| ENSMUSG00 Gm11644 | 149 | 0 | 0 | 0 | 0 | 0 | 0 | 0 | 0 | 0 | 0 | 0 | 0 | 0 | 0 | 0 | 0 | 0 | 0 |
| ENSMUSG00 Gm11499 | 321 | 0 | 0 | 0 | 0 | 0 | 0 | 0 | 0 | 0 | 0 | 0 | 0 | 0 | 0 | 0 | 0.11928538 | 0 | 0 |
| ENSMUSG00 Gm6092 | 2028 | 0 | 0 | 0 | 0 | 0 | 0 | 0 | 0 | 0 | 0 | 0 | 0 | 0 | 0 | 0 | 0 | 0 | 0 |
| ENSMUSG00 Gm12721 | 267 | 0.79597708 | 0.74860436 | 0.18117427 | 0.51192185 | 1.44215502 | 0 | 0.70880418 | 0.14304271 | 0 | 0.52976318 | 0.17893243 | 0 | 0 | 0.34789579 | 0.30862589 | 0.43023154 | 0.43063255 | 0 |
| ENSMUSG00 Rpl12-ps2 | 320 | 0.26565735 | 0.10410279 | 0 | 0.28475653 | 0.15041226 | 0.15485229 | 0 | 0.35805377 | 0 | 0 | 0 | 0 | 0 | 0 | 0.12875486 | 0 | 0.11976968 | 0.17190346 |
| ENSMUSG00 Gm14274 | 614 | 0 | 0 | 0 | 0 | 0 | 0 | 0 | 0.06220261 | 0.06707902 | 0 | 0 | 0.18180998 | 0 | 0 | 0.06710351 | 0 | 0 | 0 |
| ENSMUSG00 Gm16123 | 1515 | 0 | 0 | 0 | 0 | 0 | 0 | 0 | 0 | 0 | 0 | 0 | 0 | 0 | 0 | 0 | 0 | 0 | 0 |
| ENSMUSG00 Gm15670 | 446 | 0 | 0 | 0 | 0 | 0 | 0 | 0 | 0 | 0 | 0 | 0 | 0 | 0 | 0 | 0 | 0 | 0 | 0 |
| ENSMUSG00 Gm13369 | 1570 | 0.02707336 | 0 | 0 | 0.02901977 | 0.06131455 | 0.0631245 | 0 | 0.02432637 | 0.02623345 | 0 | 0.21300937 | 0.35551377 | 0.0314216 | 0.17749333 | 0 | 0.12194461 | 0.04882331 | 0.10511294 |

| | | | | | | | | | | | | | | | | | | | | |
|---|---|---|---|---|---|---|---|---|---|---|---|---|---|---|---|---|---|---|---|---|
| ENSMUSG00 | Gm13800 | 165 | 0 | 0 | 0 | 0 | 0 | 0 | 0 | 0 | 0 | 0 | 0 | 0 | 0 | 0 | 0 | 0 | 0.46456117 | 0 |
| ENSMUSG00 | Gm14584 | 636 | 0.46782426 | 0.7333027 | 0.60847208 | 0.64473176 | 0.52975388 | 1.01287034 | 0.89269206 | 0.66056042 | 0.51806938 | 0.59309061 | 0.15023572 | 0.70208381 | 0.23269772 | 1.3874806 | 0.51825856 | 0.4214375 | 0.60261473 | 0.51895384 |
| ENSMUSG00 | Gm11322 | 434 | 0 | 0 | 0 | 0 | 0 | 0 | 0 | 0 | 0 | 0 | 0 | 0 | 0 | 0 | 0 | 0 | 0 | 0 |
| ENSMUSG00 | Gm12673 | 178 | 0 | 0 | 0 | 0 | 0 | 0 | 0 | 0 | 0 | 0 | 0 | 0 | 0 | 0 | 0 | 0 | 0 | 0 |
| ENSMUSG00 | Gm12455 | 415 | 0 | 0 | 0 | 0 | 0 | 0 | 0 | 0 | 0 | 0 | 0 | 0 | 0 | 0 | 0 | 0 | 0 | 0 |
| ENSMUSG00 | Gm13676 | 590 | 0 | 0 | 0 | 0 | 0 | 0 | 0 | 0.06473289 | 0.06980765 | 0 | 0 | 0 | 0 | 0.07871879 | 0 | 0 | 0 | 0 |
| ENSMUSG00 | Gm14848 | 416 | 0 | 0 | 0 | 0 | 0 | 0 | 0 | 0 | 0 | 0 | 0 | 0 | 0 | 0 | 0 | 0 | 0 | 0 |
| ENSMUSG00 | Gm12131 | 357 | 0 | 0 | 0 | 0 | 0 | 0 | 0 | 0 | 0 | 0 | 0 | 0 | 0 | 0 | 0 | 0 | 0 | 0 |
| ENSMUSG00 | Gm13466 | 998 | 0.12777107 | 0.03337965 | 0 | 0.0913047 | 0 | 0 | 0.04740749 | 0.15307576 | 0.08253811 | 0.04724519 | 0.0478707 | 0.16778255 | 0.04943078 | 0.09307432 | 0 | 0 | 0 | 0 |
| ENSMUSG00 | Gm3828 | 256 | 0.16603584 | 0 | 0 | 0 | 0 | 0 | 0 | 0 | 0.48265448 | 0.92091217 | 0 | 0 | 0 | 0 | 0 | 0 | 0.29942419 | 0 |
| ENSMUSG00 | Gm15110 | 879 | 0 | 0 | 0 | 0 | 0 | 0 | 0 | 0 | 0 | 0 | 0 | 0 | 0 | 0 | 0 | 0 | 0 | 0 |
| ENSMUSG00 | Gm14713 | 684 | 0 | 0 | 0 | 0 | 0 | 0 | 0 | 0 | 0 | 0 | 0 | 0 | 0 | 0 | 0 | 0 | 0 | 0 |
| ENSMUSG00 | Gm13071 | 375 | 0 | 0.71067508 | 0.12899608 | | 0.25670359 | 0.52856249 | 0 | 0.20369281 | 0.65898425 | 0.25147042 | 0.12739989 | 0 | 0 | 0.4954036 | 0.32961245 | 0 | 0.20440692 | 0.14669095 |
| ENSMUSG00 | Gm5054 | 772 | 0 | 0.08630283 | 0.12532003 | 0 | 0 | 0 | 0 | 0.04947202 | 0 | 0.06107604 | 0 | 0.07230008 | 0 | 0.30080367 | 0 | 0 | 0.19858185 | 0 |
| ENSMUSG00 | Gm15502 | 207 | 0.41067803 | 0.64372742 | 0 | 0.66030499 | 0 | 0 | 0 | 0 | 0.39793735 | 0 | 0 | 0 | 0 | 0 | 0 | 0 | 0 | 0 |
| ENSMUSG00 | Gm5133 | 1250 | 0 | 0 | 0 | 0 | 0 | 0 | 0 | 0 | 0 | 0 | 0 | 0 | 0 | 0 | 0 | 0 | 0 | 0 |
| ENSMUSG00 | Gm13435 | 1022 | 0 | 0 | 0 | 0 | 0 | 0 | 0 | 0 | 0 | 0 | 0 | 0 | 0 | 0 | 0 | 0 | 0 | 0 |
| ENSMUSG00 | Gm13033 | 1051 | 0.40442603 | 0.4437493 | 1.05860247 | 0.69360296 | 0.82433361 | 0.42433359 | 2.07077377 | 0.69044305 | 0.43106724 | 0.53835246 | 0.40911002 | 1.00903672 | 1.50201841 | 1.72342475 | 0.66643811 | 0.36432547 | 1.85979176 | 0.68041711 |
| ENSMUSG00 | Gm14400 | 2132 | 0 | 0 | 0 | 0.0213701 | 0 | 0 | 0 | 0 | 0 | 0 | 0 | 0 | 0 | 0 | 0 | 0 | 0 | 0 |
| ENSMUSG00 | Gm14908 | 1284 | 0 | 0 | 0 | 0 | 0 | 0 | 0 | 0 | 0 | 0 | 0 | 0 | 0 | 0 | 0 | 0 | 0 | 0 |
| ENSMUSG00 | Gm11977 | 1460 | 0.11645254 | 0.18253641 | 0.298193 | 0.09361858 | 0.06593414 | 0.03394023 | 0.16202972 | 0 | 0.31030937 | 0.19377001 | 0.19633545 | 0.26760934 | 0 | 0 | 0.02822024 | 0.07867933 | 0.23625799 | 0 |
| ENSMUSG00 | Gm6415 | 347 | 0.61246651 | 0.38401031 | 0 | 0.26259968 | 0.27741743 | 0.4284098 | 0.95443445 | 0 | 0.47477252 | 0 | 0 | 0.16085205 | 0.14216691 | 0.26768926 | 0.23747294 | 0.33104272 | 0.44180169 | 0 |
| ENSMUSG00 | Gm14631 | 593 | 0 | 0 | 0 | 0 | 0 | 0 | 0 | 0 | 0 | 0 | 0 | 0 | 0 | 0 | 0 | 0 | 0 | 0 |
| ENSMUSG00 | Hmgb1-ps5 | 556 | 1.29961869 | 0.95864444 | 0.69602202 | 1.55694216 | 1.73136416 | 0.89123622 | 1.27642119 | 1.23644468 | 1.77783521 | 1.35685477 | 1.37481895 | 1.30504247 | 1.59707646 | 1.25298797 | 2.22310553 | 1.03301998 | 2.20583004 | 1.08830967 |
| ENSMUSG00 | Amd-ps1 | 993 | 0 | 0.06709546 | 0 | 0.09176444 | 0 | 0.04990205 | 0 | 0 | 0 | 0 | 0 | 0 | 0 | 0 | 0.082984 | 0 | 0 | 0 |
| ENSMUSG00 | Gm5947 | 712 | 0 | 0 | 0 | 0 | 0 | 0 | 0 | 0 | 0 | 0 | 0 | 0 | 0 | 0 | 0 | 0 | 0 | 0 |
| ENSMUSG00 | Olfr1376-ps1 | 897 | 0 | 0 | 0 | 0 | 0 | 0 | 0 | 0 | 0 | 0 | 0 | 0 | 0 | 0 | 0 | 0 | 0 | 0 |
| ENSMUSG00 | Gm13039 | 793 | 0.16080142 | 0.25205216 | 0 | 0.05745403 | 0 | 0 | 0 | 0.04816192 | 0 | 0 | 0.18073755 | 0 | 0 | 0.05856758 | 0 | 0 | 0.04833077 | 0 |
| ENSMUSG00 | Gm15712 | 1155 | 1.14083156 | 0.51916199 | 0.67010951 | 0 | 0.37505395 | 0.5148336 | 0.04096336 | 0.62827329 | 1.10543895 | 0.73481615 | 0.74454481 | 0.62822824 | 1.11050203 | 1.00528328 | 0.14268937 | 0.29836837 | 0.29864647 | 2.42897354 |
| ENSMUSG00 | Gm15006 | 1121 | 0 | 0 | 0 | 0 | 0 | 0 | 0 | 0 | 0 | 0 | 0 | 0 | 0 | 0 | 0 | 0 | 0 | 0 |
| ENSMUSG00 | Gm2199 | 873 | 0.1460659 | 0.1526364 | 0.05541069 | 0.20875622 | 0.38593753 | 0.22704574 | 0.21678203 | 0.13124537 | 0.09435628 | 0.10801994 | 0 | 0.12787093 | 0.56508496 | 0.15960167 | 0.04719537 | 0.13158284 | 0.26341098 | 0.18903473 |
| ENSMUSG00 | Gm8839 | 1644 | 0 | 0 | 0 | 0 | 0 | 0 | 0 | 0 | 0 | 0 | 0 | 0 | 0 | 0 | 0 | 0 | 0 | 0 |
| ENSMUSG00 | Gm14248 | 543 | 0 | 0 | 0 | 0 | 0 | 0 | 0 | 0 | 0 | 0 | 0 | 0 | 0 | 0 | 0 | 0 | 0 | 0 |
| ENSMUSG00 | Gm13771 | 237 | 0.17934673 | 0.42168221 | 0.20410772 | 0.96120347 | 0.20308829 | 0.62724979 | 1.79668401 | 0.32229875 | 1.91161044 | 0 | 1.00791052 | 0.47101825 | 0.8326062 | 0 | 0.17384623 | 0 | 0.64685733 | 0 |
| ENSMUSG00 | Gm13841 | 1133 | 0.0375156 | 0.02940238 | 0.04269508 | 0.0804255 | 0 | 0 | 0.04175876 | 0.03370909 | 0 | 0 | 0 | 0.09852721 | 0 | 0.08198427 | 0 | 0.06759154 | 0.03382727 | 0.04855173 |
| ENSMUSG00 | Gm12577 | 573 | 0 | 0.17441306 | 0.08442152 | 0.15902633 | 0.2519996 | 0.34591786 | 0 | 0 | 0 | 0 | 0 | 0 | 0 | 0.48632552 | 0 | 0.06682479 | 0.06688708 | 0 |
| ENSMUSG00 | Gm11898 | 384 | 0 | 0 | 0 | 0 | 0 | 0 | 0 | 0 | 0 | 0 | 0 | 0 | 0 | 0 | 0 | 0 | 0 | 0 |
| ENSMUSG00 | Ldha-ps2 | 1056 | 0.2817578 | 0.37855562 | 0.2748496 | 0.17257971 | 0.18231789 | 0.28154962 | 0.26882204 | 0.21700229 | 0.35102144 | 0.17860115 | 0.49765582 | 0.10571148 | 0.37372665 | 0.26388686 | 0.1560665 | 0.14504018 | 0.14517537 | 0 |
| ENSMUSG00 | Gm14427 | 634 | 0 | 0 | 0 | 0 | 0 | 0 | 0 | 0 | 0 | 0 | 0.45212894 | 0 | 0 | 0 | 0 | 0 | 0 | 0 |
| ENSMUSG00 | Gm11957 | 778 | 0.0546339 | 0.08563726 | 0 | 0.11712351 | 0 | 0.06369246 | 0 | 0 | 0 | 0 | 0 | 0.28696999 | 0 | 0 | 0 | 0 | 0.04926259 | 0 |
| ENSMUSG00 | Gm14765 | 325 | 0 | 0 | 0 | 0 | 0 | 0 | 0 | 0 | 0 | 0 | 0 | 0 | 0 | 0 | 0 | 0 | 0 | 0 |
| ENSMUSG00 | Gm15459 | 3549 | 0 | 0 | 0.02726037 | 0 | 0 | 0 | 0 | 0 | 0 | 0 | 0 | 0 | 0 | 0 | 0 | 0 | 0 | 0 |
| ENSMUSG00 | Gm1947 | 909 | 0 | 0.03664785 | 0 | 0 | 0 | 0 | 0 | 0 | 0 | 0 | 0 | 0 | 0 | 0 | 0 | 0 | 0 | 0 |
| ENSMUSG00 | Gm12485 | 290 | 0.14656957 | 0 | 0 | 0.47132115 | 0 | 0 | 0 | 0 | 0 | 0 | 0 | 0 | 0 | 0 | 0 | 0 | 0 | 0 |
| ENSMUSG00 | Gm7071 | 461 | 0 | 0 | 0 | 0 | 0 | 0 | 0 | 0 | 0 | 0 | 0 | 0 | 0 | 0 | 0 | 0 | 0 | 0 |
| ENSMUSG00 | 2210015D19I | 3185 | 7.20652906 | 6.69395677 | 5.31577257 | 5.70764731 | 5.63679985 | 5.95877456 | 7.45713181 | 6.70315634 | 8.37954858 | 5.46267174 | 8.47499263 | 6.50160465 | 5.73086637 | 5.62870258 | 4.70874924 | 6.4198381 | 5.57145227 | 6.0967707 |
| ENSMUSG00 | Gm14571 | 569 | 0 | 0 | 0 | 0 | 0 | 0 | 0 | 0 | 0 | 0 | 0 | 0 | 0 | 0 | 0 | 0 | 0 | 0 |
| ENSMUSG00 | Gm13797 | 119 | 0 | 0 | 0 | 0 | 0 | 0 | 0 | 0 | 0 | 0 | 0 | 0 | 0 | 0 | 0 | 0 | 0 | 0 |
| ENSMUSG00 | Xlr3d-ps | 650 | 0 | 0 | 0 | 0 | 0 | 0 | 0 | 0 | 0 | 0 | 0 | 0 | 0 | 0 | 0 | 0 | 0 | 0 |
| ENSMUSG00 | Gm7309 | 580 | 0 | 0 | 0 | 0 | 0 | 0 | 0 | 0 | 0 | 0 | 0 | 0 | 0 | 0 | 0 | 0 | 0 | 0 |
| ENSMUSG00 | Gm13477 | 2686 | 0 | 0.02480484 | 0.0180095 | 0.1187369 | 0.14335644 | 0 | 0.01761455 | 0 | 0.12267019 | 0.1930967 | 0 | 0.14546152 | 0.05509894 | 0.01729117 | 0 | 0.04276687 | 0.11415129 | 0.02047994 |
| ENSMUSG00 | Gm12365 | 567 | 0 | 0 | 0 | 0 | 0 | 0 | 0 | 0 | 0 | 0 | 0 | 0 | 0 | 0 | 0 | 0 | 0 | 0 |
| ENSMUSG00 | Gm13007 | 364 | 0.23354492 | 0.09151894 | 0 | 0 | 0 | 0.40840165 | 0 | 0 | 0 | 0 | 0.26249977 | 0 | 0 | 0 | 0.22638218 | 0 | 0 | 0.15112392 |
| ENSMUSG00 | Gm15696 | 444 | 0.86159141 | 0.75029041 | 0.10894939 | 0.61568979 | 0.86724187 | 0.33481577 | 0.42624035 | 0.08601892 | 0.74209938 | 0.10619528 | 1.07601258 | 0.50284381 | 0 | 0.4184152 | 0.64957408 | 0.34496043 | 0.60424342 | 0.61947192 |
| ENSMUSG00 | Gm15118 | 350 | 1.09299024 | 0.66625788 | 1.52031095 | 0.91122089 | 0.82511869 | 0.42473772 | 1.08143266 | 1.0912115 | 0.5883788 | 1.61659554 | 0.81899929 | 0.47841997 | 0.84569001 | 1.32697393 | 2.35437462 | 0.98461562 | 1.09503705 | 0.62867551 |
| ENSMUSG00 | Olfr1175-ps | 1882 | 0 | 0 | 0 | 0 | 0 | 0 | 0 | 0 | 0 | 0 | 0 | 0 | 0 | 0 | 0 | 0 | 0 | 0 |
| ENSMUSG00 | Gm13886 | 1182 | 0.43152463 | 0.78913793 | 0.73665275 | 0.92509735 | 0.48864897 | 0.75461016 | 0.44030412 | 0.90472696 | 0.45298198 | 0.39890612 | 0.56586245 | 0.70832059 | 0.12520791 | 0.11787839 | 0.27885994 | 0.32394761 | 0.61607415 | 0.74462412 |
| ENSMUSG00 | Gm12584 | 1104 | 0 | 0 | 0 | 0 | 0 | 0 | 0 | 0 | 0 | 0 | 0 | 0 | 0 | 0 | 0 | 0 | 0 | 0 |
| ENSMUSG00 | Gm14979 | 1034 | 0 | 0 | 0 | 0 | 0 | 0 | 0 | 0 | 0 | 0 | 0 | 0 | 0 | 0 | 0 | 0 | 0 | 0 |
| ENSMUSG00 | Gm12003 | 1104 | 0.07700213 | 0.18104834 | 0.30671623 | 0.04126906 | 0.08719551 | 0.04488472 | 0.04285569 | 0.03459457 | 0.07461325 | 0 | 0 | 0 | 0 | 0.04206892 | 0 | 0 | 0 | 0 |
| ENSMUSG00 | Gm13313 | 1071 | 0 | 0.03110448 | 0 | 0 | 0 | 0 | 0 | 0 | 0 | 0 | 0 | 0.20846186 | 0 | 0 | 0 | 0 | 0 | 0 |
| ENSMUSG00 | Gm9599 | 480 | 0 | 0 | 0 | 0 | 0 | 0 | 0 | 0 | 0 | 0 | 0 | 0 | 0 | 0 | 0 | 0 | 0 | 0 |
| ENSMUSG00 | Gm13341 | 671 | 1.90038045 | 1.34045923 | 2.23484268 | 1.69750538 | 2.58233868 | 1.9939252 | 2.11532097 | 2.21982667 | 3.9283711 | 3.02158009 | 2.42078776 | 2.57866697 | 3.52895981 | 3.80689241 | 4.05261205 | 2.91031442 | 2.68455433 | 3.93507768 |
| ENSMUSG00 | Gm13719 | 2109 | 0 | 0.01579559 | 0.02293671 | 0 | 0 | 0.02349584 | 0.0224337 | 0 | 0 | 0.0223569 | 0 | 0 | 0 | 0 | 0 | 0.01815581 | 0.0545182 | 0.05216606 |
| ENSMUSG00 | Gm13472 | 313 | 0 | 0 | 0 | 0 | 0 | 0 | 0 | 0 | 0 | 0 | 0 | 0 | 0 | 0 | 0 | 0 | 0 | 0 |
| ENSMUSG00 | Gm12893 | 273 | 0 | 0 | 0 | 0 | 0 | 0 | 0 | 0 | 0 | 0 | 0 | 0 | 0 | 0 | 0 | 0 | 0 | 0 |
| ENSMUSG00 | Gm12477 | 570 | 0 | 0 | 0 | 0 | 0 | 0 | 0 | 0 | 0 | 0 | 0 | 0 | 0 | 0 | 0 | 0 | 0 | 0 |
| ENSMUSG00 | Gm12442 | 337 | 0 | 0 | 0 | 0 | 0 | 0 | 0 | 0 | 0 | 0 | 0 | 0 | 0 | 0 | 0 | 0 | 0 | 0 |
| ENSMUSG00 | Gm11761 | 444 | 1.3402533 | 0.82531945 | 0.98054453 | 0.10261496 | 0.10840523 | 1.22765781 | 0 | 0.68815139 | 1.66972361 | 1.27434333 | 0.43040503 | 0.2514219 | 1.44440298 | 0.4184152 | 0.18559259 | 0.34496043 | 0.43160244 | 0 |
| ENSMUSG00 | Gm16479 | 931 | 0 | 0 | 0 | 0 | 0 | 0 | 0 | 0 | 0 | 0 | 0 | 0 | 0 | 0 | 0 | 0 | 0 | 0 |
| ENSMUSG00 | Gm11928 | 436 | 0 | 0 | 0 | 0 | 0 | 0 | 0 | 0 | 0 | 0 | 0 | 0 | 0 | 0 | 0 | 0 | 0 | 0 |
| ENSMUSG00 | Gm11283 | 516 | 0 | 0 | 0 | 0 | 0 | 0 | 0 | 0 | 0 | 0 | 0 | 0 | 0 | 0 | 0 | 0 | 0 | 0 |
| ENSMUSG00 | Gm7494 | 440 | 0.19320535 | 0 | 0 | 0 | 0 | 0 | 0 | 0 | 0 | 0 | 0 | 0 | 0 | 0 | 0 | 0 | 0 | 0 |
| ENSMUSG00 | Gm13831 | 301 | 0.56485284 | 0.9960666 | 0.96425642 | 0.75682798 | 0.63962689 | 0.65850809 | 2.04340474 | 0.50754023 | 0.27366456 | 0.78323427 | 0.79360396 | 1.29803866 | 1.14725389 | 0.30859859 | 2.73764491 | 0.89047924 | 0.89130923 | 0.73101803 |
| ENSMUSG00 | Gm11439 | 408 | 0 | 0.326597 | 0 | 0 | 0 | 0 | 0.2319249 | 0 | 0 | 0 | 0 | 0 | 0 | 0 | 0 | 0 | 0 | 0 |
| ENSMUSG00 | Gm14740 | 1149 | 0 | 0 | 0 | 0 | 0 | 0 | 0 | 0 | 0 | 0 | 0 | 0 | 0 | 0 | 0 | 0 | 0 | 0 |
| ENSMUSG00 | Gm12034 | 398 | 0.4271877 | 0.08370074 | 0.12154153 | 0 | 0.72560689 | 0.12450436 | 0.11887608 | 0.19192162 | 0.10348371 | 0 | 0.12003758 | 0.14024036 | 0 | 0.11669369 | 0 | 0 | 0 | 0.27642767 |
| ENSMUSG00 | Gm14038 | 2386 | 0.12470085 | 0.13961817 | 0.10136951 | 0.09547578 | 0.06051793 | 0.14537684 | 0.07931715 | 0.17607562 | 0.13809393 | 0.15809121 | 0.10011517 | 0.14035791 | 0.14472901 | 0.09732625 | 0.17268045 | 0.06419213 | 0.12850393 | 0.16138464 |
| ENSMUSG00 | Hspe1-ps6 | 282 | 0 | 0 | 0 | 0 | 0 | 0 | 0 | 0 | 0 | 0 | 0 | 0 | 0.17493588 | 0 | 0 | 0 | 0 | 0 |
| ENSMUSG00 | Gm12605 | 711 | 0 | 0 | 0 | 0 | 0 | 0 | 0 | 0 | 0 | 0 | 0 | 0 | 0 | 0 | 0 | 0 | 0 | 0 |
| ENSMUSG00 | Gm11369 | 317 | 0 | 0 | 0 | 0 | 0 | 0 | 0 | 0 | 0 | 0 | 0 | 0 | 0 | 0 | 0 | 0 | 0 | 0 |
| ENSMUSG00 | Gm14941 | 443 | 0 | 0 | 0 | 0 | 0 | 0 | 0 | 0 | 0 | 0 | 0.10784415 | 0 | 0 | 0 | 0 | 0 | 0 | 0 |
| ENSMUSG00 | Gm15388 | 725 | 0 | 0 | 0 | 0 | 0 | 0 | 0 | 0 | 0 | 0 | 0 | 0 | 0 | 0 | 0 | 0 | 0 | 0 |
| ENSMUSG00 | Gm13213 | 473 | 0 | 0 | 0.10226962 | 0 | 0 | 0 | 0 | 0 | 0 | 0 | 0 | 0.11800351 | 0 | 0 | 0.34842753 | 0 | 0 | 0 |
| ENSMUSG00 | Olfr409-ps1 | 973 | 0 | 0 | 0 | 0 | 0 | 0 | 0 | 0 | 0 | 0 | 0 | 0 | 0 | 0 | 0 | 0 | 0 | 0 |
| ENSMUSG00 | Gm11930 | 189 | 0 | 0.3525174 | 0.25594461 | 0.24106373 | 0.25466626 | 0.52436755 | 0 | 0 | 0 | 0 | 0 | 0 | 0 | 0 | 0 | 0.20259581 | 0 | 0.58210695 |
| ENSMUSG00 | Gm12441 | 587 | 0 | 0 | 0 | 0 | 0 | 0 | 0 | 0 | 0 | 0 | 0 | 0 | 0 | 0 | 0 | 0 | 0 | 0 |
| ENSMUSG00 | E530001F21I | 482 | 0 | 0 | 0 | 0 | 0 | 0 | 0 | 0 | 0 | 0 | 0 | 0 | 0 | 0 | 0 | 0 | 0 | 0 |
| ENSMUSG00 | Gm15703 | 879 | 0 | 0 | 0 | 0 | 0 | 0 | 0 | 0 | 0 | 0 | 0 | 0 | 0 | 0 | 0 | 0 | 0 | 0 |
| ENSMUSG00 | Gm1848 | 894 | 0.09508988 | 0.03726275 | 0.05410909 | 0 | 0.05383884 | 0.16628434 | 0.10584492 | 0.08544162 | 0.04606993 | 0.10548256 | 0 | 0 | 0.11036223 | 0 | 0.13826025 | 0.12849197 | 0 | 0.12306288 |
| ENSMUSG00 | Gm13347 | 1352 | 0 | 0 | 0 | 0 | 0 | 0 | 0 | 0 | 0 | 0 | 0 | 0 | 0 | 0 | 0 | 0 | 0 | 0 |
| ENSMUSG00 | Gm11489 | 497 | 0 | 0 | 0.09733105 | 0 | 0 | 0 | 0 | 0 | 0 | 0 | 0 | 0 | 0.49629695 | 0.46724434 | 0.49740309 | 0 | 0 | 0 |
| ENSMUSG00 | Gm12074 | 218 | 0 | 0 | 0 | 0 | 0 | 0 | 0 | 0 | 0 | 0 | 0 | 0 | 0 | 0 | 0 | 0 | 0 | 0 |
| ENSMUSG00 | Scgb2b1-ps | 327 | 0 | 0 | 0 | 0 | 0 | 0 | 0 | 0 | 0 | 0 | 0 | 0 | 0 | 0 | 0 | 0 | 0 | 0 |
| ENSMUSG00 | Gm14843 | 211 | 1.00723166 | 0 | 0 | 0 | 0 | 0 | 0.89692282 | 0 | 0 | 0.89385219 | 0 | 0 | 0 | 0 | 0.39053607 | 0 | 0 | 0 |
| ENSMUSG00 | Gm7340 | 2694 | 0 | 0 | 0 | 0 | 0 | 0 | 0 | 0 | 0 | 0 | 0 | 0 | 0 | 0 | 0 | 0 | 0 | 0 |
| ENSMUSG00 | Gm12346 | 2255 | 0.37698604 | 0.31023094 | 0.51484023 | 0.30306681 | 0.46957974 | 0.37356828 | 0.54551204 | 0.23711469 | 0.21917436 | 0.39727865 | 0.27542105 | 0.54454305 | 0.26252018 | 0.70026562 | 0.3471528 | 0.30564565 | 0.47589193 | 0.24394282 |
| ENSMUSG00 | Gm13294 | 476 | 0.08929659 | 0 | 0 | 0 | 0 | 0 | 0 | 0 | 0 | 0 | 0 | 0 | 0 | 0 | 0 | 0 | 0 | 0 |
| ENSMUSG00 | Gm12885 | 327 | 0 | 0 | 0 | 0 | 0 | 0 | 0 | 0 | 0 | 0 | 0 | 0 | 0 | 0 | 0 | 0 | 0 | 0 |
| ENSMUSG00 | Gm15975 | 1405 | 0.09075838 | 0.26081269 | 0.10328868 | 0.3242779 | 0.44534876 | 0.17634425 | 0.16837252 | 0.21746564 | 0.32245671 | 0.36915141 | 0.13601412 | 0.27808515 | 0.17555842 | 0.06611258 | 0.29324951 | 0.32703722 | 0.38189904 | 0.23491433 |
| ENSMUSG00 | Gm5405 | 608 | 0.55927863 | 0.21916378 | 0.39780864 | 0.59948743 | 0 | 0.57050845 | 0.31126762 | 0.18844935 | 0.40644588 | 0 | 0.2357317 | 0.36720831 | 0 | 0.38194151 | 0 | 0.12595595 | 0.06303667 | 0 |
| ENSMUSG00 | Gm12805 | 1047 | 0 | 0 | 0 | 0 | 0 | 0 | 0 | 0 | 0 | 0 | 0 | 0 | 0 | 0 | 0 | 0 | 0 | 0 |
| ENSMUSG00 | Gm15355 | 284 | 0 | 0 | 0 | 0 | 0 | 0 | 0 | 0 | 0 | 0 | 0 | 0 | 0 | 0 | 0 | 0 | 0 | 0 |
| ENSMUSG00 | Gm15186 | 562 | 0.22689596 | 0 | 0.34429559 | 0.48641684 | 0.08564399 | 0.3526885 | 0.33674505 | 0.06795801 | 1.17256984 | 0 | 0 | 0.89384513 | 0 | 0.57848507 | 0.29324951 | 0 | 0 | 0 |
| ENSMUSG00 | Pik-ps1 | 1787 | 0 | 0.0186418 | 0.54139374 | 0.17847079 | 0 | 0 | 0.23828434 | 0.27784064 | 0.04609571 | 0.07915619 | 0.05346946 | 0.40604567 | 0 | 0 | 0 | 0 | 0.02144728 | 0.03078294 |
| ENSMUSG00 | Gm6447 | 834 | 0 | 0 | 0 | 0 | 0 | 0 | 0 | 0 | 0 | 0 | 0 | 0 | 0 | 0 | 0 | 0 | 0 | 0 |
| ENSMUSG00 | Gm15842 | 617 | 0.13778015 | 0.10798345 | 0 | 0 | 0 | 0 | 0.30672725 | 0 | 0 | 0.22925788 | 0 | 0 | 0 | 0 | 0 | 0 | 0 | 0 |
| ENSMUSG00 | Gm13604 | 481 | 0.26510505 | 0.06925758 | 0 | 0.37888602 | 0.20013274 | 0.41208094 | 0.19672632 | 0 | 0.42813426 | 0.19605282 | 0.19864848 | 0.58020439 | 0.20512232 | 0.09655735 | 0.08565812 | 0.23881876 | 0.3187218 | 0.45745619 |
| ENSMUSG00 | Gm4342 | 366 | 0.5806718 | 0.09101884 | 0 | 0 | 0.26301598 | 0.13538998 | 0 | | 0.22506293 | 0.25765412 | 0 | 0.15250181 | 0 | 0 | 0.11257256 | 0.31385744 | 0.20943332 | 0.15029811 |
| ENSMUSG00 | Gm5391 | 612 | 0.0694529 | 0.21773133 | 0.07904172 | 0 | 0 | 0.16193704 | 0.0773083 | 0.31202943 | 0.13459646 | 0.07704363 | 0.23419097 | 0.09120206 | 0 | 0 | 0.0673228 | 0 | 0.12524934 | 0.08968416 |
| ENSMUSG00 | Gm14219 | 287 | 0 | 0 | 0 | 0 | 0 | 0 | 0 | 0 | 0 | 0 | 0 | 0 | 0 | 0 | 0 | 0 | 0 | 0 |
| ENSMUSG00 | Rps18-ps1 | 968 | 0.04391031 | 0 | 0 | 0.09413439 | 0 | 0.05119084 | 0.1466302 | 0.03945496 | 0.12764416 | 0 | 0 | 0.05766081 | 0.10192545 | 0.14393829 | 0.04256359 | 0.15822565 | 0.19796641 | 0.17048277 |
| ENSMUSG00 | Gm13932 | 609 | 0 | 0.16410293 | 0.07943108 | 0 | 0 | 0.08136738 | 0.07768913 | 0.25085322 | 0.06762975 | 0.07742316 | 0 | 0.09165133 | 0 | 0 | 0 | 0 | 0 | 0.09032694 |
| ENSMUSG00 | Gm15476 | 999 | 0 | 0 | 0 | 0 | 0 | 0 | 0 | 0 | 0 | 0 | 0 | 0 | 0 | 0 | 0 | 0 | 0 | 0 |
| ENSMUSG00 | Gm12093 | 1403 | 0 | 0 | 0 | 0 | 0 | 0 | 0 | 0 | 0 | 0 | 0 | 0 | 0 | 0 | 0 | 0 | 0 | 0 |
| ENSMUSG00 | Gm14352 | 448 | 0 | 0 | 0 | 0 | 0 | 0 | 0 | 0 | 0 | 0 | 0 | 0 | 0 | 0 | 0 | 0 | 0 | 0 |
| ENSMUSG00 | Tpi-rs8 | 707 | 0 | 0 | 0 | 0 | 0 | 0 | 0 | 0.05402037 | 0 | 0 | 0 | 0 | 0 | 0 | 0 | 0 | 0 | 0 |
| ENSMUSG00 | Gm14987 | 675 | 0 | 0 | 0 | 0 | 0 | 0 | 0 | 0 | 0 | 0 | 0 | 0 | 0 | 0 | 0 | 0 | 0 | 0 |
| ENSMUSG00 | Gm15750 | 596 | 1.06976114 | 1.34145883 | 1.54210919 | 1.4524494 | 1.61516522 | 2.2448386 | 4.20733554 | 2.88365454 | 3.86987396 | 3.32270057 | 2.48493912 | 7.67933613 | 5.04907207 | 4.20802134 | 4.07867751 | 4.30448103 | 3.79404617 | 5.26093806 |
| ENSMUSG00 | Gm3835 | 402 | 0 | 0 | 0.12033217 | 0 | 0 | 0 | 0 | 0.09500598 | 1.02454019 | 0 | 0.23768636 | 0.13884493 | 0.12271621 | 0 | 0.10249143 | 0.66675187 | 0 | 0 |
| ENSMUSG00 | Gm649 | 1053 | 0 | 0 | 0 | 0 | 0 | 0 | 0 | 0 | 0 | 0 | 0 | 0 | 0 | 0 | 0 | 0 | 0 | 0 |
| ENSMUSG00 | Gm12304 | 384 | 0 | 0 | 0.25194547 | 0.23729711 | 0 | 0 | 0 | 0 | 0 | 0.24557658 | 0 | 0 | 0.77081121 | 0 | 0 | 0 | 0 | 0 |
| ENSMUSG00 | Gm12012 | 1065 | 0 | 0 | 0 | 0 | 0 | 0 | 0 | 0 | 0 | 0 | 0 | 0 | 0 | 0 | 0 | 0 | 0 | 0 |
| ENSMUSG00 | Gm14783 | 662 | 0 | 0 | 0 | 0 | 0 | 0 | 0 | 0 | 0 | 0 | 0 | 0 | 0 | 0 | 0 | 0 | 0 | 0 |
| ENSMUSG00 | Gm14749 | 331 | 0 | 0 | 0 | 0 | 0 | 0 | 0 | 0 | 0 | 0 | 0 | 0 | 0 | 0 | 0 | 0 | 0 | 0 |
| ENSMUSG00 | Gm4991 | 1551 | 0 | 0.02147833 | 0.03118861 | 0 | 0 | 0 | 0 | 0.04924875 | 0 | 0 | 0 | 0 | 0 | 0 | 0 | 0 | 0 | 0 |
| ENSMUSG00 | Gm13301 | 2214 | 0 | 0 | 0 | 0 | 0.04347961 | 0 | 0 | 0 | 0 | 0 | 0 | 0 | 0 | 0 | 0 | 0 | 0 | 0 |
| ENSMUSG00 | Gm13042 | 1953 | 0.02176404 | 0 | 0 | 0 | 0 | 0 | 0 | 0 | 0 | 0.02446234 | 0 | 0 | 0 | 0 | 0 | 0 | 0 | 0.02816647 |
| ENSMUSG00 | Gm14518 | 541 | 0 | 0 | 0 | 0 | 0 | 0 | 0 | 0 | 0 | 0 | 0 | 0 | 0 | 0 | 0 | 0 | 0 | 0 |
| ENSMUSG00 | Gm11438 | 188 | 0 | 0.17719625 | 0 | 0 | 0 | 0 | 0 | 0 | 0 | 0 | 0 | 0 | 0 | 0 | 0 | 0 | 0 | 0 |
| ENSMUSG00 | Gm15515 | 821 | 0 | 0 | 0 | 0 | 0 | 0 | 0 | 0 | 0 | 0 | 0 | 0 | 0 | 0 | 0 | 0 | 0 | 0 |
| ENSMUSG00 | Gm12368 | 433 | 0 | 0 | 0 | 0 | 0 | 0 | 0 | 0 | 0 | 0 | 0 | 0 | 0 | 0 | 0 | 0 | 0 | 0 |
| ENSMUSG00 | Gm12294 | 489 | 0.60845855 | 0.06812453 | 0.09892337 | 0.18634374 | 0.09842929 | 0 | 0.09675394 | 0.07810307 | 0 | 0 | 0.2930979 | 0 | 0 | 0 | 0 | 0 | 0 | 0.11249306 |
| ENSMUSG00 | Gm14829 | 1137 | 0 | 0 | 0 | 0.04007128 | 0 | 0.043582 | 0 | 0 | 0 | 0 | 0.16807373 | 0 | 0 | 0 | 0.03623708 | 0 | 0 | 0 |
| ENSMUSG00 | Cct3-ps1 | 1788 | 0 | 0 | 0.02705455 | 0.02548157 | 0.05383884 | 0 | 0 | 0.0213604 | 0 | 0 | 0 | 0 | 0 | 0.02597544 | 0 | 0 | 0 | 0 |
| ENSMUSG00 | Gm12448 | 521 | 0 | 0 | 0 | 0 | 0 | 0 | 0 | 0 | 0 | 0 | 0 | 0 | 0 | 0 | 0 | 0 | 0 | 0 |
| ENSMUSG00 | Gm14451 | 1539 | 0 | 0 | 0 | 0 | 0 | 0 | 0 | 0 | 0 | 0 | 0 | 0.07253497 | 0 | 0 | 0 | 0 | 0 | 0 |
| ENSMUSG00 | Gm11298 | 2033 | 0 | 0 | 0 | 0 | 0 | 0 | 0 | 0 | 0 | 0 | 0 | 0 | 0 | 0 | 0 | 0 | 0 | 0 |
| ENSMUSG00 | Gm12889 | 240 | 0 | 0 | 0 | 0 | 0 | 0 | 0 | 0 | 0 | 0 | 0 | 0 | 0 | 0 | 0 | 0 | 0 | 0 |
| ENSMUSG00 | Gm12759 | 302 | 0 | 0 | 0 | 0 | 0 | 0 | 0 | 0 | 0 | 0 | 0 | 0 | 0 | 0 | 0 | 0 | 0 | 0 |
| ENSMUSG00 | Gm11483 | 246 | 0 | 0 | 0 | 0 | 0 | 0.20143388 | 0.19232796 | 0 | 0 | 0 | 0 | 0 | 0 | 0.37759421 | | 0 | 0.15579795 | 0 |
| ENSMUSG00 | Gm14733 | 132 | 0 | 0 | 0 | 0 | 0 | 0 | 0 | 0 | 0 | 0 | 0 | 0 | 0 | 0 | 0 | 0 | 0 | 0 |
| ENSMUSG00 | Gm15242 | 282 | 0 | 0 | 0 | 0 | 0 | 0 | 0 | 0 | 0 | 0 | 0 | 0 | 0 | 0 | 0 | 0 | 0 | 0 |
| ENSMUSG00 | Gm11359 | 564 | 0 | 0 | 0 | 0 | 0 | 0 | 0 | 0 | 0 | 0 | 0 | 0 | 0 | 0 | 0 | 0 | 0 | 0 |
| ENSMUSG00 | Olfr122 | 1256 | 0 | 0 | 0 | 0 | 0 | 0 | 0 | 0 | 0 | 0 | 0 | 0 | 0 | 0 | 0 | 0 | 0 | 0 |
| ENSMUSG00 | Gm7158 | 902 | 0 | 0 | 0 | 0 | 0 | 0 | 0 | 0 | 0 | 0 | 0 | 0 | 0 | 0 | 0 | 0 | 0 | 0 |
| ENSMUSG00 | Gm12897 | 247 | 0 | 0 | 0 | 0 | 0 | 0 | 0 | 0 | 0 | 0 | 0 | 0 | 0 | 0 | 0 | 0 | 0 | 0 |
| ENSMUSG00 | Gm14466 | 450 | 0 | 0 | 0 | 0 | 0 | 0 | 0 | 0 | 0 | 0 | 0 | 0 | 0 | 0 | 0 | 0 | 0 | 0 |
| ENSMUSG00 | Gm13786 | 332 | 0 | 0.10034004 | 0 | 0.41169618 | 0 | 0 | 0.57003228 | 0.57518678 | 0 | 0.14202019 | 0.14390048 | 0 | 0 | 0 | 0 | 0 | 0 | 0 |
| ENSMUSG00 | Gm14675 | 446 | 0 | 0 | 0 | 0 | 0 | 0 | 0 | 0 | 0 | 0 | 0 | 0 | 0 | 0 | 0 | 0 | 0 | 0 |
| ENSMUSG00 | Gm14049 | 635 | 0.13387457 | 0.05246125 | 0 | 0.21524903 | 0.15159661 | 0 | 0.37254078 | 0 | 0.06486065 | 0.14850615 | 0.30094462 | 0 | 0.15537612 | 0 | 0 | 0 | 0.12071275 | 0 |
| ENSMUSG00 | Gm12179 | 1271 | 0 | 0 | 0 | 0 | 0 | 0 | 0 | 0 | 0 | 0 | 0 | 0 | 0 | 0 | 0 | 0 | 0 | 0 |
| ENSMUSG00 | Gm14785 | 537 | 0 | 0 | 0 | 0 | 0 | 0 | 0 | 0 | 0 | 0 | 0 | 0 | 0 | 0 | 0 | 0 | 0 | 0 |
| ENSMUSG00 | Gm12361 | 1041 | 0 | 0 | 0 | 0 | 0 | 0 | 0 | 0 | 0 | 0 | 0 | 0 | 0 | 0 | 0 | 0 | 0 | 0 |
| ENSMUSG00 | Gm12742 | 408 | 0 | 0 | 0 | 0 | 0 | 0 | 0 | 0 | 0 | 0 | 0 | 0 | 0 | 0 | 0 | 0 | 0 | 0 |
| ENSMUSG00 | Gm12601 | 843 | 0 | 0 | 0 | 0 | 0 | 0 | 0 | 0 | 0 | 0 | 0 | 0 | 0 | 0 | 0 | 0 | 0 | 0 |
| ENSMUSG00 | Gm7365 | 630 | 0 | 0 | 0 | 0 | 0 | 0 | 0 | 0 | 0 | 0 | 0 | 0 | 0 | 0 | 0 | 0 | 0 | 0 |
| ENSMUSG00 | Gm4918 | 1083 | 0 | 0 | 0 | 0 | 0 | 0 | 0 | 0 | 0 | 0 | 0 | 0 | 0 | 0 | 0 | 0 | 0 | 0 |
| ENSMUSG00 | Gm13725 | 206 | 0 | 0 | 0 | 0 | 0 | 0 | 0 | 0 | 0 | 0 | 0 | 0 | 0 | 0 | 0 | 0 | 0 | 0 |
| ENSMUSG00 | Gm15707 | 995 | 0 | 0.0334803 | 0.04861661 | 0 | 0 | 0 | 0 | 0.03838432 | 0.12418045 | 0 | 0 | 0.11219229 | 0 | 0 | 0.0414086 | 0.03848302 | 0 | 0.05528553 |
| ENSMUSG00 | Gm15203 | 454 | 0 | 0 | 0 | 0 | 0 | 0 | 0 | 0 | 0 | 0 | 0 | 0 | 0 | 0 | 0 | 0 | 0 | 0 |
| ENSMUSG00 | Gm14122 | 971 | 0.04377464 | 0.03430782 | 0 | 0.14076533 | 0.09913887 | 0 | 0 | 0 | 0 | 0.09711782 | 0 | 0 | 0 | 0.09566238 | 0 | 0.0394342 | 0.03947095 | 0 |
| ENSMUSG00 | Gm13482 | 253 | 0 | 0 | 0 | 0 | 0 | 0 | 0 | 0 | 0 | 0 | 0 | 0 | 0 | 0 | 0.325704 | 0 | 0 | 0 |
| ENSMUSG00 | Gm15115 | 263 | 0 | 0 | 0 | 0 | 0 | 0 | 0 | 0 | 0 | 0 | 0 | 0 | 0 | 0 | 0 | 0 | 0 | 0 |
| ENSMUSG00 | Olfr1050-ps1 | 946 | 0 | 0 | 0 | 0 | 0 | 0 | 0 | 0 | 0 | 0 | 0 | 0 | 0 | 0 | 0 | 0 | 0 | 0 |
| ENSMUSG00 | Gm15081 | 843 | 0 | 0 | 0 | 0 | 0 | 0 | 0 | 0 | 0 | 0 | 0 | 0 | 0 | 0 | 0 | 0 | 0 | 0 |
| ENSMUSG00 | Gm6943 | 458 | 0 | 0 | 0.10561906 | 0 | 0 | 0 | 0 | 0 | 0 | 0 | 0 | 0 | 0 | 0.1014063 | 0 | 0 | 0 | 0 |
| ENSMUSG00 | Gm14802 | 1884 | 0 | 0 | 0 | 0 | 0 | 0 | 0 | 0 | 0 | 0 | 0 | 0 | 0 | 0 | 0 | 0 | 0 | 0 |
| ENSMUSG00 | Gm13459 | 645 | 0 | 0 | 0 | 0 | 0 | 0 | 0 | 0 | 0 | 0 | 0 | 0 | 0 | 0 | 0 | 0 | 0 | 0 |
| ENSMUSG00 | Gm16053 | 187 | 0.45460081 | 0.35628764 | 0.25868198 | 0.97456779 | 0 | 0 | 0 | 0.61271234 | 1.76198998 | 0 | 0 | 0.29847948 | 1.84664931 | 0 | 0.22032918 | 0.20476261 | 0 | 0 |
| ENSMUSG00 | Gm14215 | 220 | 0 | 0 | 0 | 0 | 0 | 0 | 0 | 0 | 0 | 0 | 0 | 0 | 0 | 0 | 0 | 0 | 0 | 0 |
| ENSMUSG00 | Gm13664 | 344 | 0 | 0 | 0 | 0 | 0 | 0 | 0 | 0 | 0 | 0 | 0 | 0 | 0 | 0 | 0 | 0 | 0 | 0 |
| ENSMUSG00 | Mageb8-ps | 1032 | 0 | 0 | 0 | 0 | 0 | 0 | 0 | 0 | 0 | 0 | 0 | 0 | 0 | 0 | 0 | 0 | 0 | 0 |
| ENSMUSG00 | Gm5060 | 598 | 0 | 0 | 0 | 0 | 0 | 0 | 0 | 0.06386689 | 0.20662132 | 0.15769466 | 0 | 0 | 0 | 0 | 0.06889892 | 0.12806223 | 0.19227239 | 0 |
| ENSMUSG00 | Gm12527 | 563 | 21.2903368 | 25.1473891 | 28.0102503 | 23.0637613 | 43.3443789 | 25.1724366 | 13.4458768 | 17.7055365 | 12.5827366 | 11.2223699 | 28.0879418 | 11.5002076 | 13.6692346 | 16.6637756 | 11.2700526 | 12.8542181 | 17.7676083 | 19.0528878 |
| ENSMUSG00 | Gm15363 | 885 | 0 | 0 | 0 | 0 | 0 | 0 | 0 | 0 | 0 | 0 | 0 | 0 | 0 | 0 | 0 | 0 | 0 | 0 |
| ENSMUSG00 | Gm14650 | 888 | 0 | 0 | 0 | 0 | 0 | 0.05580263 | 0 | 0 | 0 | 0 | 0 | 0 | 0 | 0 | 0 | 0 | 0 | 0.06194719 |
| ENSMUSG00 | Gm12375 | 919 | 0 | 0 | 0 | 0 | 0 | 0 | 0 | 0 | 0 | 0 | 0 | 0 | 0 | 0 | 0 | 0 | 0 | 0 |
| ENSMUSG00 | Gm12468 | 403 | 0 | 0.08266227 | 0 | 0.3391641 | 0 | 0.12295964 | 0 | 0.09477023 | 0.10219979 | 0 | 0.11854828 | 0 | 0 | 0.4609835 | 0.20447422 | 0 | 0.28530742 | 0.13649902 |
| ENSMUSG00 | Gm12213 | 352 | 0 | 0 | 0 | 0 | 0 | 0 | 0 | 0 | 0 | 0 | 0 | 0 | 0 | 0 | 0 | 0 | 0 | 0 |
| ENSMUSG00 | Gm15311 | 280 | 0 | 0 | 0 | 0 | 0 | 0 | 0 | 0 | 0 | 0 | 0 | 0 | 0 | 0 | 0 | 0 | 0 | 0 |
| ENSMUSG00 | Gm15633 | 491 | 0.69254869 | 0.40708221 | 0.88668386 | 0.74233881 | 0 | 0 | 0 | 1.08898907 | 0 | 0 | | 0 | 0.10047234 | 0 | 0.58739489 | 0 | 0.46834579 | 0.22406968 |
| ENSMUSG00 | Gm12607 | 487 | 0 | 0 | 0 | 0 | 0 | 0 | 0 | 0 | 0 | 0 | 0 | 0 | 0 | 0 | 0.33841114 | 0 | 0 | 0 |
| ENSMUSG00 | Gm12101 | 389 | 0 | 0 | 0 | 0 | 0 | 0 | 0 | 0 | 0 | 0 | 0 | 0 | 0 | 0 | 0 | 0 | 0 | 0 |
| ENSMUSG00 | Gm11618 | 265 | 0 | 0 | 0 | 0 | 0 | 0 | 0 | 0 | 0 | 0 | 0 | 0 | 0 | 0 | 0 | 0 | 0 | 0 |
| ENSMUSG00 | Gm11478 | 598 | 7.03681008 | 4.84652474 | 6.55226749 | 4.26658609 | 5.47319533 | 4.72325387 | 6.80416453 | 4.21521498 | 7.23174608 | 5.36161843 | 4.31412669 | 5.69357093 | 6.92956698 | 8.23256399 | 7.09658906 | 6.59520495 | 5.25544539 | 5.33533142 |
| ENSMUSG00 | Gm14398 | 1072 | 0.03965035 | 0 | 0 | 0 | 0.04489918 | 0 | 0 | 0.07125448 | 0 | 0 | 0 | 0.15620055 | 0 | 0.08664942 | 0 | 0.10715655 | 0 | 0 |
| ENSMUSG00 | Gm16465 | 1756 | 0 | 0 | 0 | 0 | 0.02740998 | 0 | 0.08083032 | 0.02174966 | 0 | 0 | 0 | 0 | 0 | 0 | 0 | 0 | 0 | 0 |
| ENSMUSG00 | Gm12808 | 403 | 1.05471901 | 0.66129815 | 1.92053719 | 2.03498461 | 0.35830216 | 2.21327346 | 3.16983209 | 1.32678321 | 0.10219979 | 3.04198086 | 2.01532083 | 3.32400968 | 4.40682139 | 1.95917987 | 1.12460823 | 1.61523654 | 2.75797174 | 0.27299805 |
| ENSMUSG00 | Gm11649 | 374 | 0 | 0 | 0 | 0 | 0 | 0 | 0 | 0 | 0 | 0 | 0 | 0 | 0 | 0 | 0 | 0 | 0 | 0 |
| ENSMUSG00 | Gm12920 | 1185 | 0.03586935 | 0 | 0 | 0 | 0 | 0 | 0 | 0 | 0 | 0 | 0 | 0 | 0 | 0 | 0 | 0 | 0 | 0 |
| ENSMUSG00 | Prdx2-rs3 | 592 | 0 | 0 | 0 | 0.07696122 | 0 | 0 | 0.07992007 | 0 | 0 | 0 | 0 | 0 | 0 | 0.1569057 | 0.06959722 | 0 | 0.12948073 | 0 |
| ENSMUSG00 | Gm7790 | 1309 | 0 | 0 | 0 | 0 | 0 | 0 | 0 | 0 | 0 | 0 | 0 | 0 | 0 | 0 | 0 | 0 | 0 | 0 |
| ENSMUSG00 | Gm14653 | 1532 | 0 | 0 | 0 | 0 | 0 | 0 | 0 | 0 | 0 | 0 | 0 | 0 | 0.03220099 | 0 | 0 | 0 | 0 | 0 |
| ENSMUSG00 | Gm15113 | 834 | 0 | 0 | 0 | 0 | 0 | 0 | 0 | 0 | 0 | 0 | 0 | 0 | 0 | 0 | 0 | 0 | 0 | 0 |
| ENSMUSG00 | Gm13146 | 779 | 0 | 0 | 0 | 0 | 0 | 0 | 0 | 0 | 0 | 0 | 0 | 0 | 0 | 0 | 0 | 0 | 0.04919935 | 0 |
| ENSMUSG00 | Gm15269 | 203 | 0 | 0 | 0 | 0 | 0 | 0 | 0 | 0 | 0 | 0 | 0 | 0 | 0 | 0 | 0 | 0 | 0 | 0 |
| ENSMUSG00 | Gm12047 | 943 | 0 | 0 | 0 | 0 | 0 | 0 | 0 | 0 | 0 | 0 | 0 | 0 | 0 | 0 | 0 | 0.12181529 | 0.20321472 | 0 |
| ENSMUSG00 | Gm13701 | 300 | 0 | 0 | 0 | 0 | 0 | 0 | 0 | 0 | 0 | 0 | 0 | 0 | 0 | 0 | 0 | 0 | 0 | 0 |
| ENSMUSG00 | Gm14245 | 1842 | 0.02307556 | 0.01808518 | 0 | 0 | 0 | 0 | 0 | 0 | 0 | 0 | 0 | 0.03030166 | 0 | 0 | 0 | 0 | 0 | 0 |
| ENSMUSG00 | Gm9003 | 480 | 0.8855245 | 0 | 0.60466913 | 0.37967537 | 0 | 0.10323486 | 0.29570424 | 0.07956751 | 0.60063669 | 0 | 0 | 0.23256526 | 0 | 0 | 0.08583657 | 0 | 0.23953936 | 0 |
| ENSMUSG00 | Gm14992 | 511 | 0 | 0 | 0 | 0 | 0 | 0 | 0 | 0 | 0 | 0 | 0 | 0 | 0 | 0 | 0 | 0 | 0 | 0 |
| ENSMUSG00 | Gm7590 | 800 | 0 | 0 | 0 | 0.11390261 | 0 | 0 | 0 | 0.0477405 | 0 | 0 | 0 | 0 | 0 | 0 | 0 | 0 | 0.09581574 | 0 |
| ENSMUSG00 | Gm13302 | 1848 | 0 | 0 | 0 | 0 | 0 | 0 | 0 | 0 | 0 | 0 | 0 | 0 | 0 | 0 | 0 | 0 | 0 | 0 |
| ENSMUSG00 | Gm12969 | 1003 | 0.29664629 | 0.16606627 | 0.04822884 | 0.04542477 | 0.19195184 | 0 | 0.37736932 | 0.22846901 | 0.24637995 | 0.1880387 | 0.19052825 | 0.22259487 | 0.29510619 | 0.18522069 | 0.16431328 | 0.26723255 | 0.38211662 | 0.05484457 |
| ENSMUSG00 | Gm11956 | 511 | 0 | 0.06519157 | 0.09466444 | 0 | 0 | 0 | 0 | 0 | 0 | 0 | 0 | 0 | 0 | 0 | 0.2418878 | 0 | 0 | 0.10764992 |
| ENSMUSG00 | Gm14270 | 845 | 0.6036238 | 0.47308252 | 0.6297146 | 0.70093914 | 0.62656942 | 0.41049602 | 0.67189603 | 0.4971792 | 0.58489727 | 0.50219684 | 0.62192254 | 0 | 0.40866677 | 0.43970734 | 0.43883314 | 0.6344006 | 0.58963534 | 0.52079628 |
| ENSMUSG00 | Gm7209 | 332 | 0 | 0 | 0 | 0 | 0 | 0 | 0 | 0 | 0 | 0 | 0 | 0 | 0 | 0 | 0 | 0 | 0 | 0 |
| ENSMUSG00 | Gm14175 | 240 | 0 | 0 | 0 | 0 | 0 | 0 | 0 | 0 | 0 | 0 | 0 | 0 | 0 | 0 | 0 | 0 | 0 | 0 |
| ENSMUSG00 | Olfr1096-ps1 | 947 | 0 | 0 | 0 | 0 | 0 | 0 | 0 | 0 | 0 | 0 | 0 | 0 | 0 | 0 | 0 | 0 | 0 | 0 |
| ENSMUSG00 | Rpl21-ps9 | 481 | 0 | 0 | 0 | 0 | 0.10006637 | 0 | 0 | 0 | 0 | 0 | 0 | 0 | 0 | 0 | 0 | 0 | 0 | 0 |
| ENSMUSG00 | Gm11895 | 466 | 0 | 0 | 0 | 0 | 0.30986217 | 0.10633634 | 0 | 0 | 0.08838308 | 0 | 0 | 0 | 0.10586248 | 0 | 0 | 0 | 0.16449054 | 0 |
| ENSMUSG00 | Gm8754 | 519 | 0 | 0 | 0 | 0 | 0 | 0 | 0 | 0 | 0 | 0 | 0 | 0 | 0 | 0 | 0 | 0 | 0 | 0 |
| ENSMUSG00 | Gm12282 | 4151 | 0.02047949 | 0.07222743 | 0.04661386 | 0.06585552 | 0 | 0.03581262 | 0.0113979 | 0.01840154 | 0.00992207 | 0.04543551 | 0.03452779 | 0.04033895 | 0.27333994 | 0.0223773 | 0.00992569 | 0.04612215 | 0.05539816 | 0.1987802 |
| ENSMUSG00 | Gm5832 | 1581 | 0 | 0 | 0 | 0 | 0 | 0.03134265 | 0 | 0 | 0 | 0 | 0 | 0 | 0 | 0 | 0 | 0 | 0 | 0 |
| ENSMUSG00 | Gm16459 | 685 | 0.06205135 | 0.09726392 | 0.07061829 | 0 | 0 | 0 | 0 | 0.05575533 | 0.0601263 | 0.06883314 | 0 | 0.08148272 | 0 | 0 | 0 | 0 | 0 | 0 |
| ENSMUSG00 | Gm15937 | 1234 | 0 | 0 | 0 | 0 | 0 | 0 | 0 | 0.03095008 | 0 | 0 | 0.03871553 | 0.13569448 | 0 | 0 | 0 | 0 | 0 | 0 |
| ENSMUSG00 | Gm6276 | 578 | 0 | 0 | 0 | 0 | 0 | 0 | 0 | 0 | 0 | 0 | 0 | 0 | 0 | 0.24105928 | 0 | 0.06624673 | 0 | 0 |
| ENSMUSG00 | Gm11259 | 714 | 0 | 0 | 0 | 0 | 0 | 0 | 0 | 0 | 0 | 0 | 0 | 0 | 0 | 0 | 0 | 0 | 0 | 0 |
| ENSMUSG00 | Gm12376 | 606 | 0 | 0 | 0 | 0 | 0 | 0 | 0 | 0 | 0 | 0 | 0 | 0 | 0 | 0 | 0 | 0 | 0 | 0 |
| ENSMUSG00 | Gm13983 | 864 | 0 | 0 | 0 | 0 | 0 | 0 | 0 | 0 | 0 | 0 | 0 | 0 | 0 | 0 | 0 | 0 | 0 | 0 |
| ENSMUSG00 | Gm8855 | 877 | 0 | 0 | 0 | 0 | 0 | 0 | 0 | 0 | 0 | 0 | 0 | 0 | 0 | 0 | 0 | 0 | 0 | 0 |
| ENSMUSG00 | Gm16423 | 946 | 0 | 0 | 0 | 0 | 0 | 0 | 0 | 0 | 0 | 0 | 0 | 0 | 0 | 0 | 0 | 0 | 0 | 0 |
| ENSMUSG00 | Gm7416 | 1485 | 0 | 0 | 0 | 0 | 0.03241207 | 0 | 0 | 0 | 0 | 0 | 0 | 0 | 0 | 0 | 0 | 0.02578492 | 0 | 0 |
| ENSMUSG00 | Gm12991 | 77 | 0 | 0 | 0.62822767 | 0 | 0 | 0 | 0.61445038 | 0 | 0 | 0 | 0 | 0 | 0 | 0 | 0 | 0 | 0 | 0 |
| ENSMUSG00 | Gm14447 | 1446 | 0.47032007 | 0.85240462 | 0.76942683 | 0.97675821 | 0.33286254 | 0.65110784 | 0.94887115 | 0.23771205 | 0.8829751 | 0.68476125 | 0.42951207 | 0.57900065 | 0.64820638 | 0.64238019 | 0.37041509 | 0.26480365 | 0.58311102 | 0.38042259 |
| ENSMUSG00 | Gm12617 | 380 | 0 | 0 | 0 | 0 | 0 | 0 | 0 | 0 | 0 | 0 | 0 | 0 | 0 | 0 | 0 | 0 | 0 | 0 |
| ENSMUSG00 | Ccl19-ps3 | 320 | 0 | 0 | 0 | 0 | 0 | 0 | 0 | 0 | 0 | 0 | 0 | 0 | 0 | 0 | 0 | 0 | 0 | 0 |
| ENSMUSG00 | Gm11295 | 801 | 0 | 0.04158913 | 0.24156569 | 0.11376041 | 0 | 0.43304511 | 0.47253612 | 0.52448992 | 0.15425661 | 0.17659439 | 0.17893243 | 0.2787299 | 0 | 0.17394789 | 0.36006353 | 0.23901752 | 0.33493643 | 0.41205323 |
| ENSMUSG00 | Gm14551 | 255 | 0 | 0 | 0 | 0 | 0 | 0 | 0 | 0 | 0 | 0 | 0 | 0 | 0 | 0 | 0 | 0 | 0 | 0 |
| ENSMUSG00 | Gm14763 | 326 | 0 | 0 | 0 | 0 | 0 | 0 | 0 | 0 | 0 | 0 | 0 | 0 | 0 | 0 | 0 | 0 | 0 | 0 |
| ENSMUSG00 | Gm15312 | 270 | 0 | 0 | 0 | 0 | 0 | 0 | 0 | 0 | 0 | 0 | 0 | 0 | 0 | 0 | 0 | 0 | 0 | 0 |
| ENSMUSG00 | Gm13732 | 534 | 0 | 0 | 0 | 0 | 0 | 0 | 0 | 0 | 0 | 0 | 0 | 0 | 0 | 0 | 0 | 0 | 0 | 0 |
| ENSMUSG00 | Gm14647 | 679 | 0 | 0 | 0 | 0 | 0 | 0 | 0 | 0 | 0 | 0 | 0 | 0 | 0 | 0 | 0 | 0 | 0 | 0 |
| ENSMUSG00 | Gm11997 | 312 | 0 | 0 | 0 | 0 | 0 | 0 | 0 | 0 | 0 | 0 | 0 | 0 | 0 | 0 | 0 | 0 | 0 | 0 |
| ENSMUSG00 | Gm11833 | 852 | 0 | 0 | 0 | 0 | 0 | 0 | 0 | 0 | 0 | 0 | 0 | 0 | 0 | 0 | 0 | 0 | 0 | 0 |
| ENSMUSG00 | Gm14984 | 1262 | 0 | 0.07919072 | 0.03833085 | 0 | 0.1525576 | 0 | 0 | 0.06052679 | 0.03263591 | 0 | 0 | 0.26536765 | 0.03909027 | 0 | 0.03264783 | 0 | 0 | 0 |
| ENSMUSG00 | Gm12042 | 300 | 0 | 0.44417192 | 0 | 0 | 0 | 0.16517578 | 0 | 0 | 0.27457677 | 0 | 0 | 0 | 0 | 0 | 0 | 0 | 0 | 0 |
| ENSMUSG00 | Gm14439 | 886 | 0.09594848 | 0.45119044 | 0.21839066 | 0.46280971 | 0.5432497 | 0.61521453 | 0 | 0.21553274 | 0.41837318 | 0.106435 | 0.2156883 | 0.69297098 | 0.27839682 | 0.78629945 | 0.09300577 | 0.73469563 | 0.17303069 | 0.37252217 |
| ENSMUSG00 | Gm15146 | 446 | 0 | 0 | 0 | 0 | 0 | 0 | 0 | 0 | 0 | 0 | 0 | 0 | 0 | 0 | 0 | 0 | 0 | 0 |
| ENSMUSG00 | Gm11745 | 447 | 0 | 0.22357647 | 0.21643638 | 0.40770509 | 0.43071073 | 0.66513736 | 0 | 1.02529039 | 0 | 0.10548256 | 0.53439551 | 0 | 0.44144892 | 0.62341057 | 0.18434701 | 0.25698394 | 1.28611735 | 0.49225151 |
| ENSMUSG00 | Gm14904 | 334 | 0 | 0 | 0 | 0 | 0 | 0 | 0 | 0 | 0 | 0 | 0 | 0 | 0 | 0 | 0 | 0 | 0 | 0 |
| ENSMUSG00 | Gm14945 | 1117 | 0 | 0 | 0 | 0 | 0 | 0 | 0 | 0 | 0 | 0 | 0 | 0 | 0 | 0 | 0 | 0 | 0 | 0 |
| ENSMUSG00 | Gm12004 | 717 | 0.05928197 | 0 | 0 | 0 | 0 | 0 | 0 | 0 | 0 | 0 | 0 | 0 | 0 | 0 | 0 | 0 | 0 | 0 |
| ENSMUSG00 | Rhox3b | 644 | 0 | 0 | 0 | 0 | 0 | 0 | 0 | 0 | 0 | 0 | 0 | 0 | 0 | 0 | 0 | 0 | 0 | 0 |
| ENSMUSG00 | Gm8916 | 948 | 0 | 0 | 0 | 0 | 0 | 0 | 0 | 0 | 0 | 0 | 0 | 0 | 0 | 0 | 0 | 0 | 0 | 0 |
| ENSMUSG00 | Gm15263 | 640 | 0 | 0 | 0 | 0 | 0 | 0 | 0 | 0 | 0.19306179 | 0 | 0 | 0 | 0 | 0 | 0 | 0 | 0 | 0 |
| ENSMUSG00 | Gm13046 | 500 | 0 | 0 | 0 | 0 | 0 | 0 | 0 | 0 | 0 | 0 | 0 | 0 | 0 | 0 | 0 | 0 | 0 | 0 |
| ENSMUSG00 | Gm16412 | 478 | 0 | 0 | 0.10119985 | 0 | 0 | 0 | 0 | 0 | 0 | 0 | 0 | 0 | 0 | 0 | 0 | 0 | 0 | 0 |
| ENSMUSG00 | Gm8996 | 782 | 0 | 0 | 0 | 0 | 0.06154977 | 0 | 0 | 0 | 0 | 0 | 0 | 0 | 0 | 0 | 0.05268741 | 0 | 0 | 0 |
| ENSMUSG00 | Gm14611 | 711 | 0 | 0 | 0 | 0 | 0 | 0 | 0 | 0 | 0 | 0 | 0 | 0 | 0 | 0 | 0 | 0 | 0 | 0 |
| ENSMUSG00 | Gm5934 | 2580 | 0 | 0 | 0 | 0 | 0 | 0 | 0 | 0 | 0 | 0 | 0 | 0 | 0 | 0 | 0 | 0 | 0 | 0 |
| ENSMUSG00 | Gm12079 | 243 | 0 | 0 | 0 | 0.18749401 | 0 | 0 | 0 | 0 | 0 | 0 | 0 | 0 | 0 | 0 | 0 | 0 | 0 | 0.22637492 |
| ENSMUSG00 | Gm15159 | 763 | 0 | 0.04366041 | 0.06339912 | 0.05971303 | 0.06308247 | 0.0649446 | 0.06200875 | 0.50055573 | 0.05397971 | 0.06179647 | 0 | 0.0731529 | 0 | 0.18261109 | 0.05399942 | 0.15055285 | 0.05023106 | 0.07209581 |
| ENSMUSG00 | Xlr4d-ps | 639 | 0 | 0 | 0 | 0 | 0 | 0 | 0 | 0 | 0 | 0 | 0 | 0 | 0 | 0 | 0 | 0 | 0 | 0 |
| ENSMUSG00 | Gm14269 | 305 | 0.8361674 | 0.32766781 | 0.63440696 | 0.14938047 | 0.31561917 | 0.16246798 | 1.86148245 | 0.87654694 | 0.67518878 | 0.61836988 | 0.93983525 | 0.36600434 | 0.64697596 | 0.1522757 | 0.40526121 | 1.0043438 | 0.87961993 | 0.72143091 |
| ENSMUSG00 | Gm11573 | 393 | 0 | 0 | 0 | 0 | 0 | 0 | 0 | 0 | 0 | 0 | 0 | 0 | 0 | 0.11817834 | 0 | 0 | 0 | 0 |
| ENSMUSG00 | Gm12306 | 170 | 0 | 0 | 0 | 0 | 0 | 0 | 0 | 0 | 0 | 0.83207123 | 0 | 0 | 2.61168974 | 0 | 0 | 0 | 0 | 0 |
| ENSMUSG00 | Gm6591 | 430 | 0 | 0 | 0 | 0 | 0 | 0 | 0 | 0 | 0 | 0 | 0 | 0 | 0 | 0 | 0 | 0 | 0 | 0 |
| ENSMUSG00 | Gm13378 | 335 | 0 | 0 | 0 | 0 | 0 | 0 | 0 | 0 | 0 | 0 | 0 | 0 | 0 | 0 | 0 | 0 | 0 | 0 |
| ENSMUSG00 | Gm11285 | 321 | 0 | 0 | 0 | 0 | 0 | 0 | 0 | 0 | 0 | 0.14688693 | 0 | 0 | 0 | 0 | 0 | 0 | 0 | 0 |
| ENSMUSG00 | Gm11910 | 430 | 0 | 0 | 0 | 0 | 0 | 0 | 0 | 0 | 0 | 0 | 0 | 0 | 0 | 0 | 0 | 0 | 0 | 0 |
| ENSMUSG00 | Gm8310 | 1192 | 0 | 0 | 0 | 0 | 0 | 0 | 0 | 0 | 0 | 0 | 0 | 0 | 0 | 0 | 0 | 0 | 0 | 0 |
| ENSMUSG00 | Gm13382 | 131 | 0 | 0 | 0 | 0 | 0 | 0 | 0 | 0 | 0 | 0 | 0 | 0 | 0 | 0 | 0 | 0 | 0 | 0 |
| ENSMUSG00 | Gm14600 | 629 | 0 | 0 | 0 | 0 | 0 | 0 | 0 | 0 | 0 | 0 | 0 | 0 | 0 | 0 | 0 | 0.06087537 | 0 | 0 |
| ENSMUSG00 | Gm11952 | 820 | 0 | 0 | 0 | 0.05556225 | 0 | 0 | 0 | 0 | 0 | 0 | 0 | 0 | 0 | 0 | 0 | 0 | 0 | 0 |
| ENSMUSG00 | Olfr349-ps1 | 581 | 0 | 0 | 0 | 0 | 0 | 0 | 0 | 0 | 0 | 0 | 0 | 0 | 0 | 0 | 0 | 0 | 0 | 0 |
| ENSMUSG00 | Gm11631 | 155 | 0 | 0 | 0 | 0 | 0 | 0 | 0 | 0 | 0 | 0 | 0 | 0 | 0 | 0 | 0 | 0 | 0 | 0 |
| ENSMUSG00 | Gm15227 | 398 | 0 | 0 | 0 | 0 | 0 | 0 | 0 | 0 | 0 | 0 | 0 | 0 | 0 | 0 | 0 | 0 | 0 | 0 |
| ENSMUSG00 | Gm12057 | 2019 | 0.02105259 | 0.0329994 | 0 | 0 | 0 | 0.04908641 | 0 | 0.01891649 | 1.01997315 | 0 | 0.02366268 | 0.22116162 | 0 | 0 | 0 | 0.01896513 | 0.07593125 | 0 |
| ENSMUSG00 | Gm12872 | 317 | 0 | 0 | 0 | 0 | 0 | 0 | 0 | 0 | 0 | 0 | 0 | 0 | 0 | 0 | 0 | 0 | 0 | 0 |
| ENSMUSG00 | Gm15782 | 444 | 0.09573238 | 0 | 0 | 0 | 0 | 0 | 0 | 0 | 0 | 0 | 0 | 0.12571095 | 0 | 0 | 0 | 0 | 0 | 0 |
| ENSMUSG00 | Gm8383 | 1769 | 0 | 0 | 0 | 0 | 0 | 0 | 0 | 0 | 0 | 0 | 0 | 0 | 0 | 0 | 0 | 0 | 0 | 0 |
| ENSMUSG00 | Gm16140 | 1280 | 0 | 0 | 0 | 0 | 0 | 0 | 0.07392606 | 0 | 0 | 0 | 0 | 0 | 0 | 0 | 0 | 0 | 0 | 0 |
| ENSMUSG00 | Gm5761 | 833 | 0 | 0 | 0 | 0 | 0 | 0 | 0 | 0 | 0 | 0 | 0.05735289 | 0.0670056 | 0.11844398 | 0 | 0.04946165 | 0 | 0.04600996 | 0 |
| ENSMUSG00 | Gm13650 | 432 | 0 | 0.07711318 | 0.11197576 | 0.10546538 | 0.11141649 | 0.2294108 | 0 | 0 | 0 | 0 | 0 | 0 | 0 | 0 | 0 | 0 | 0.35487312 | 0 |
| ENSMUSG00 | Gm12941 | 219 | 0 | 0 | 0 | 0 | 0 | 0 | 0.64811889 | 0 | 0 | 0 | 0 | 0 | 0 | 0 | 0 | 0 | 0 | 0 |
| ENSMUSG00 | Gm14099 | 604 | 0 | 0 | 0 | 0 | 0 | 0 | 0 | 0 | 0 | 0 | 0 | 0 | 0 | 0 | 0 | 0 | 0 | 0 |
| ENSMUSG00 | Gm11245 | 1036 | 0 | 0 | 0 | 0 | 0 | 0 | 0 | 0 | 0 | 0 | 0 | 0 | 0 | 0 | 0 | 0 | 0 | 0 |
| ENSMUSG00 | Gm11995 | 227 | 0 | 0 | 0 | 0 | 0 | 0 | 0 | 0 | 0 | 0 | 0 | 0 | 0 | 0 | 0 | 0 | 0 | 0 |
| ENSMUSG00 | Gm5843 | 472 | 0 | 0 | 0 | 0 | 0 | 0 | 0 | 0 | 0 | 0 | 0 | 0 | 0 | 0 | 0 | 0 | 0 | 0 |
| ENSMUSG00 | Gm16418 | 170 | 0.75009134 | 0.78383281 | 1.42275089 | 0.80401843 | 0.28312896 | 0.58297334 | 0.55661975 | 2.02195072 | 1.93818898 | 0.27735708 | 1.68617501 | 0.98498228 | 1.16075099 | 2.18560411 | 1.21181047 | 1.12619434 | 1.57814164 | 0.97074894 |
| ENSMUSG00 | Gm13961 | 4771 | 0.00890907 | 0.00698237 | 0.01013908 | 0.01909916 | 0.04035374 | 0 | 0 | 0 | 0.03453072 | 0 | 0.02002723 | 0 | 0.02067991 | 0.01946933 | 0.01727366 | 0 | 0.05623225 | 0.01152989 |
| ENSMUSG00 | Gm11834 | 583 | 0 | 0 | 0 | 0 | 0 | 0 | 0 | 0 | 0 | 0 | 0 | 0 | 0 | 0 | 0 | 0 | 0 | 0 |
| ENSMUSG00 | Gm11332 | 594 | 0 | 0 | 0 | 0 | 0 | 0 | 0 | 0 | 0 | 0 | 0 | 0 | 0 | 0 | 0 | 0 | 0 | 0 |
| ENSMUSG00 | Gm14922 | 162 | 0 | 0 | 0 | 0 | 0 | 0 | 0 | 0 | 0 | 0 | 0 | 0 | 0 | 0 | 0 | 0 | 0 | 0 |
| ENSMUSG00 | Gm13422 | 503 | 0 | 0.13245684 | 0 | 0 | 0 | 0 | 0.18812198 | 0.07592923 | 0.16376348 | 0 | 0.09498004 | 0.3328966 | 0 | 0.46167085 | 0 | 0.07612447 | 0 | 0 |
| ENSMUSG00 | Gm12929 | 369 | 0.23038036 | 0.36111538 | 0.52437431 | 0.49388666 | 0.39131645 | 0 | 1.28218642 | 0.62101467 | 0 | 0.38333905 | 0 | 0.15126196 | 1.33690833 | 0.12586474 | 0 | 0.6226115 | 0.62319182 | |
| ENSMUSG00 | Mup-ps11 | 539 | 0 | 0 | 0 | 0 | 0 | 0 | 0 | 0 | 0 | 0 | 0 | 0 | 0 | 0 | 0 | 0 | 0 | 0 |
| ENSMUSG00 | Gm17014 | 167 | 0 | 0 | 0 | 0 | 0 | 0 | 0 | 0 | 0 | 0 | 0 | 0 | 0 | 0 | 0 | 0 | 0 | 0 |
| ENSMUSG00 | Gm15437 | 456 | 0 | 0 | 0 | 0 | 0 | 0 | 0 | 0 | 0 | 0 | 0 | 0 | 0 | 0 | 0 | 0 | 0 | 0 |
| ENSMUSG00 | Gm14629 | 143 | 0 | 0 | 0 | 0 | 0 | 0 | 0 | 0 | 0 | 0 | 0 | 0 | 0 | 0 | 0 | 0 | 0 | 0 |
| ENSMUSG00 | Gm13578 | 589 | 0.21649495 | 0.16967518 | 0 | 0.07735322 | 0.24515411 | 0.42065139 | 0.48196277 | 0.19452837 | 0.20977852 | 0.24015638 | 0.16222397 | 0.37905374 | 0.08375538 | 0.31540976 | 0.06995171 | 0.06500952 | 0.39042068 | 0 |
| ENSMUSG00 | Gm13252 | 320 | 0 | 0.31230838 | 0 | 0.14237826 | 0 | 0.30970458 | 0.29570424 | 0 | 0.25741572 | 0 | 0.29859349 | 0.17442395 | 0.30832448 | 0.29027555 | 0.25750972 | 0.59829074 | 0.11976968 | 0 |
| ENSMUSG00 | Gm6136 | 1817 | 0.18714442 | 0.09167004 | 0.53245493 | 0.20059898 | 0.10595911 | 0.2727173 | 0.15623339 | 0.27325329 | 0.1360039 | 0.23354779 | 0.39439977 | 0.21503007 | 0.38010283 | 0.15336518 | 0.18140476 | 0.18966179 | 0.14765222 | 0.36329625 |
| ENSMUSG00 | Gm13700 | 1127 | 0 | 0 | 0 | 0 | 0 | 0 | 0 | 0 | 0 | 0 | 0 | 0 | 0 | 0 | 0 | 0 | 0 | 0 |
| ENSMUSG00 | Gm15794 | 517 | 0 | 0 | 0 | 0 | 0 | 0 | 0 | 0 | 0 | 0 | 0 | 0.10796866 | 0 | 0 | 0 | 0 | 0 | 0 |
| ENSMUSG00 | Gm4992 | 1297 | 0.09831575 | 0 | 0 | 0 | 0 | 0 | 0 | 0 | 0 | 0 | 0 | 0 | 0.07607081 | 0 | 0 | 0 | 0.08864988 | 0 |
| ENSMUSG00 | Gm15710 | 732 | 0.81294053 | 0.31856593 | 0.33042029 | 0.99586982 | 0.52603195 | 0.81233989 | 0.45244365 | 0.62610496 | 0.45012586 | 0.25765412 | 0.13053267 | 0.53375634 | 0.87611328 | 0.82482669 | 0.73172162 | 0.62771488 | 0.41886663 | 0.30059621 |
| ENSMUSG00 | Gm15112 | 858 | 0 | 0 | 0 | 0 | 0 | 0 | 0 | 0 | 0 | 0 | 0 | 0 | 0 | 0 | 0 | 0 | 0 | 0 |
| ENSMUSG00 | Gm14277 | 359 | 0.23679764 | 0.18558715 | 0.13474521 | 0.25382197 | 0.26814442 | 0.96620929 | 0 | 0.42554209 | 0.34417701 | 0 | 0.26615576 | 0.15547538 | 0.13741481 | 0.12937072 | 0 | 0.31997722 | 0.10675849 | 0.30645742 |
| ENSMUSG00 | Gm8421 | 304 | 0 | 0 | 0 | 0 | 0 | 0 | 0 | 0 | 0 | 0 | 0 | 0 | 0 | 0 | 0 | 0 | 0 | 0 |
| ENSMUSG00 | Gm13566 | 1141 | 0 | 0 | 0 | 0 | 0 | 0 | 0 | 0 | 0 | 0 | 0 | 0 | 0 | 0 | 0 | 0 | 0 | 0 |
| ENSMUSG00 | Gm12460 | 356 | 0 | 0 | 0 | 0 | 0 | 0 | 0 | 0 | 0 | 0 | 0 | 0 | 0 | 0 | 0 | 0 | 0 | 0 |
| ENSMUSG00 | Olfr1063-ps1 | 941 | 0 | 0 | 0 | 0 | 0 | 0 | 0 | 0 | 0 | 0 | 0 | 0 | 0 | 0 | 0 | 0 | 0 | 0 |
| ENSMUSG00 | Gm14989 | 206 | 0 | 0 | 0 | 0.22117012 | 0 | 0 | 0 | 0 | 0 | 0 | 0 | 0 | 0 | 0 | 0 | 0 | 0 | 0 |
| ENSMUSG00 | Gm12351 | 291 | 0 | 0 | 0 | 0 | 0 | 0 | 0 | 0 | 0 | 0 | 0 | 0.1918064 | 0 | 0 | 0 | 0.13158284 | 0 | 0 |
| ENSMUSG00 | Gm11624 | 1040 | 0 | 0 | 0 | 0 | 0 | 0 | 0 | 0 | 0 | 0 | 0 | 0 | 0 | 0 | 0 | 0 | 0 | 0 |
| ENSMUSG00 | Gm13106 | 1440 | 0 | 0 | 0 | 0 | 0 | 0 | 0 | 0 | 0 | 0 | 0 | 0 | 0 | 0 | 0 | 0 | 0 | 0 |
| ENSMUSG00 | Gm13588 | 134 | 3.17202806 | 1.74022582 | 2.52697547 | 3.06007014 | 2.51435422 | 3.32816867 | 1.76539847 | 1.9951255 | 1.53681029 | 0.70374184 | 0.35652954 | 2.49920877 | 0.36814864 | 2.77278134 | 1.53737149 | 2.00025561 | 1.14406856 | 0.82103144 |
| ENSMUSG00 | Gm8757 | 371 | 0 | 0.08979217 | 0.65193437 | 0 | 0 | 0 | 0.38258231 | 0 | 0.22202973 | 0.12709084 | 0.77264084 | 0.30089306 | 0.26594025 | 0 | 0.33316622 | 0.10320918 | 0.41322153 | 0.74136262 |
| ENSMUSG00 | Gm15844 | 575 | 0 | 0 | 0 | 0 | 0 | 0 | 0 | 0 | 0 | 0 | 0 | 0 | 0 | 0 | 0 | 0 | 0 | 0 |
| ENSMUSG00 | Gm13169 | 1457 | 0.02917308 | 0.02286403 | 0 | 0.0625409 | 0.03303495 | 0 | 0 | 0.05242608 | 0 | 0 | 0 | 0.03830862 | 0 | 0 | 0.0565567 | 0 | 0 | 0 |
| ENSMUSG00 | Esrp2 | 5231 | 9.35260134 | 12.2845676 | 10.1722201 | 11.8627686 | 10.351446 | 12.0590002 | 10.7088916 | 10.3384519 | 9.50336925 | 9.67170801 | 9.23350851 | 9.94459903 | 4.80964974 | 5.753349 | 5.05666199 | 4.07720671 | 4.63784094 | 5.24747548 |
| ENSMUSG00 | Hmgb1-ps1 | 633 | 0.26859511 | 0.05262701 | 0.15283896 | 0.21592912 | 0 | 0 | 0.07474357 | 0.06033555 | 0.13013117 | 0.07448768 | 0.52831708 | 0 | 0.15586704 | 0.29348554 | 0.45562542 | 0.06049069 | 0.48437658 | 0 |
| ENSMUSG00 | Gm11704 | 213 | 0 | 0 | 0 | 0 | 0 | 0 | 0 | 0 | 0.19336392 | 0 | 0 | 0 | 0 | 0 | 0 | 0 | 0 | 0 |
| ENSMUSG00 | Rpl3-ps2 | 1213 | 0 | 0.02746323 | 0 | 0.15024252 | 0 | 0 | 0 | 0.0314859 | 0.06790852 | 0.03887115 | 0 | 0.04601456 | 0 | 0 | 0.03396666 | 0.03156687 | 0 | 0.04534963 |
| ENSMUSG00 | Gm15982 | 568 | 0 | 0.29324731 | 0 | 0 | 0 | 0.08724073 | 0 | 0.06724015 | 0 | 0.1660236 | 0.25233253 | 0 | 0 | 0 | 0.50776565 | 0.26965216 | 0.47233112 | 0.38738808 |
| ENSMUSG00 | Gm12856 | 1292 | 0.03289874 | 0 | 0 | 0 | 0 | 0 | 0 | 0 | 0 | 0 | 0 | 0 | 0 | 0 | 0 | 0 | 0 | 0 |
| ENSMUSG00 | Gm13930 | 366 | 0 | 0 | 0 | 0 | 0 | 0 | 0 | 0 | 0 | 0 | 0 | 0 | 0 | 0 | 0 | 0 | 0 | 0 |
| ENSMUSG00 | Pom121l12 | 1624 | 0 | 0 | 0 | 0 | 0 | 0 | 0 | 0 | 0 | 0 | 0 | 0 | 0 | 0 | 0 | 0 | 0 | 0 |
| ENSMUSG00 | Vmn1r207-ps | 2010 | 0 | 0 | 0 | 0 | 0 | 0 | 0 | 0 | 0 | 0 | 0 | 0 | 0 | 0 | 0 | 0 | 0 | 0 |
| ENSMUSG00 | Gm9085 | 1071 | 0.03968737 | 0.06220895 | 0.0451667 | 0.04254066 | 0.04494111 | 0.1850709 | 0 | 0 | 0.61529809 | 0.30817453 | 0.9813717 | 0.15634639 | 0.09212309 | 0.13009548 | 0 | 0.25026541 | 0.07157105 | 0.05136238 |
| ENSMUSG00 | Gm14915 | 203 | 0 | 0 | 0 | 0 | 0 | 0 | 0 | 0 | 0 | 0 | 0 | 0 | 0 | 0 | 0 | 0 | 0 | 0 |
| ENSMUSG00 | Gm12322 | 388 | 0 | 0 | 0 | 0 | 0.2481027 | 0.38313969 | 0 | 0 | 0 | 0.24304486 | 0 | 0 | 0 | 0 | 0.42475831 | 0 | 0.49389558 | 0 |
| ENSMUSG00 | Gm14000 | 232 | 0 | 0 | 0 | 0 | 0 | 0.21358937 | 0.61180188 | 0 | 0 | 0.20323579 | 0.20592655 | 0.24058475 | 0 | 0 | 0 | 0 | 0 | 0 |
| ENSMUSG00 | Olfr1372-ps1 | 905 | 0.98630795 | 0.2576688 | 0.74831981 | 0.35240587 | 0.26592223 | 0.1095088 | 0.0522792 | 0.21100775 | 0 | 0.05210022 | 0 | 0.24669906 | 0.27255203 | 0.56451377 | 0.22763291 | 0.50772076 | 0.2964465 | 0.72940252 |
| ENSMUSG00 | Gm9103 | 403 | 0 | 0 | 0.12003357 | 0 | 0 | 0 | 0 | 0 | 0 | 0 | 0 | 0 | 0.24482341 | 0 | 0 | 0.09501391 | 0 | 0 |
| ENSMUSG00 | Gm13400 | 379 | 0.89720688 | 0 | 0.25526929 | 0.12021384 | 0.12699716 | 0 | 0 | 0.10077151 | 0 | 0 | 0.6302765 | 0 | 0 | 0 | 1.08711229 | 0 | 0 | 0 |
| ENSMUSG00 | Gm12516 | 384 | 0 | 0 | 0 | 0 | 0 | 0.38713073 | 0 | 0 | 0 | 0 | 0 | 0.14535329 | 0 | 0 | 0 | 0 | 0 | 0 |
| ENSMUSG00 | Gm12263 | 1481 | 0.91841029 | 0.83226002 | 1.0125452 | 0.76909258 | 0.9099891 | 0.90339217 | 1.08617899 | 0.59312981 | 0.83429809 | 0.89143801 | 0.80646453 | 0.52762949 | 0.53295792 | 0.50175922 | 0.50076165 | 0.46538213 | 0.4658159 | 0.85429403 |
| ENSMUSG00 | Gm12104 | 978 | 0 | 0.13624906 | 0.64300194 | 0.04658593 | 0.14764394 | 0 | 0 | 0 | 0 | 0 | 0 | 0 | 0.05044163 | 0 | 0.71618246 | 0 | 0.23513066 | 0 |
| ENSMUSG00 | Gm15075 | 476 | 0 | 0 | 0 | 0 | 0 | 0 | 0 | 0 | 0 | 0 | 0 | 0 | 0 | 0 | 0 | 0 | 0 | 0 |
| ENSMUSG00 | Gm12380 | 543 | 0 | 0 | 0 | 0 | 0 | 0 | 0 | 0 | 0 | 0 | 0 | 0 | 0 | 0.08553239 | 0 | 0 | 0.0705825 | 0 |
| ENSMUSG00 | Gm12944 | 462 | 0 | 0.07210583 | 0 | 0 | 0 | 0 | 0.1024084 | 0 | 0 | 0 | 0.103409 | 0 | 0.21355808 | 0 | 0.08918086 | 0 | 0 | 0 |
| ENSMUSG00 | Gm14395 | 562 | 0 | 0 | 0 | 0 | 0 | 0 | 0 | 0 | 0 | 0 | 0 | 0 | 0 | 0 | 0 | 0 | 0 | 0 |
| ENSMUSG00 | Gm11577 | 735 | 0.28915086 | 0 | 0.46070029 | 0 | 0.39291366 | 0.33709343 | 0 | 0 | 0 | 0.51320493 | 0.06499694 | 0.15187935 | 0 | 0 | 0 | 0 | 0.26072311 | 0 |
| ENSMUSG00 | Gm11617 | 638 | 0 | 0 | 0 | 0 | 0 | 0 | 0 | 0 | 0 | 0 | 0 | 0 | 0 | 0 | 0 | 0 | 0 | 0 |
| ENSMUSG00 | Gm12186 | 1026 | 0 | 0 | 0 | 0 | 0 | 0 | 0.04611372 | 0 | 0 | 0 | 0 | 0.16320369 | 0 | 0 | 0 | 0 | 0 | 0 |
| ENSMUSG00 | Gm15644 | 386 | 0.22023407 | 0.51781701 | 0.50128011 | 0 | 0 | 0.12837496 | 0 | 0.39577619 | 0.42680327 | 0.24430416 | 0.74261593 | 0.43380049 | 0.12780289 | 0 | 0.10673978 | 1.09118311 | 0 | 0 |
| ENSMUSG00 | Gm7113 | 957 | 0 | 0 | 0 | 0 | 0 | 0 | 0 | 0 | 0 | 0 | 0 | 0 | 0 | 0 | 0 | 0 | 0 | 0 |
| ENSMUSG00 | Gm14734 | 446 | 0 | 0 | 0 | 0 | 0 | 0 | 0 | 0 | 0 | 0 | 0 | 0 | 0 | 0 | 0 | 0 | 0 | 0 |
| ENSMUSG00 | Gm15576 | 526 | 0 | 0 | 0 | 0 | 0 | 0 | 0 | 0 | 0 | 0 | 0 | 0 | 0 | 0 | 0 | 0 | 0 | 0 |
| ENSMUSG00 | Gm14652 | 234 | 0 | 0 | 0 | 0 | 0 | 0 | 0 | 0 | 0 | 0 | 0 | 0 | 0 | 0 | 0 | 0 | 0 | 0 |
| ENSMUSG00 | Gm12696 | 904 | 8.08726801 | 7.18585661 | 7.54498648 | 9.07188936 | 6.49567996 | 9.04447018 | 6.123433 | 8.49189478 | 7.56301064 | 8.55388864 | 6.76459589 | 8.21181762 | 4.31108569 | 4.21284863 | 5.0590406 | 4.40511413 | 5.21475057 | 5.29401801 |
| ENSMUSG00 | Gm13345 | 227 | 0 | 0 | 0 | 0 | 0 | 0 | 0 | 0 | 0 | 0 | 0 | 0 | 0 | 0 | 0 | 0 | 0 | 0 |
| ENSMUSG00 | Gm15061 | 1045 | 0 | 0 | 0 | 0 | 0 | 0 | 0 | 0 | 0 | 0 | 0 | 0 | 0 | 0 | 0 | 0 | 0 | 0 |
| ENSMUSG00 | Gm11251 | 2797 | 0 | 0 | 0 | 0 | 0 | 0 | 0 | 0 | 0 | 0 | 0 | 0 | 0 | 0 | 0 | 0 | 0 | 0 |
| ENSMUSG00 | Gm11901 | 436 | 0 | 0 | 0 | 0 | 0 | 0 | 0 | 0 | 0 | 0 | 0 | 0 | 0 | 0 | 0 | 0 | 0 | 0 |
| ENSMUSG00 | Gm13902 | 604 | 0 | 0 | 0 | 0 | 0 | 0 | 0 | 0 | 0 | 0 | 0 | 0.09241004 | 0.08167536 | 0.07689418 | 0 | 0 | 0 | 0 |
| ENSMUSG00 | Gm6451 | 716 | 0.11872954 | 0.69789583 | 0.20268239 | 0 | 0.4705635 | 0.41524637 | 0.85902909 | 0.05334134 | 0.23009227 | 0.32926469 | 0.0667248 | 0.54568385 | 1.17128854 | 0.06486604 | 0.11508814 | 0.37434951 | 0.53528348 | 0.61462689 |
| ENSMUSG00 | Gm13924 | 485 | 0 | 0 | 0 | 0 | 0 | 0 | 0 | 0 | 0 | 0 | 0 | 0 | 0 | 0 | 0 | 0 | 0 | 0 |
| ENSMUSG00 | Atp5l2 | 695 | 0.48926821 | 0.33552555 | 0.34801101 | 0.91777643 | 0.69254566 | 0.21389669 | 0.88498536 | 0.38467168 | 0.71113409 | 1.15332655 | 0.75615042 | 0.32124122 | 0.49686823 | 0.5346082 | 1.00780784 | 0.33056639 | 0.71689476 | 0.39574897 |
| ENSMUSG00 | Gm12240 | 228 | 0 | 1.89941941 | 0 | 0.19982914 | 0 | 0.21733655 | 0 | 0.16751054 | 0 | 0 | 0.20953929 | 0.73441661 | 5.1928334 | 11.4073197 | 3.07204583 | 0.33588252 | 1.34478235 | 2.65394813 |
| ENSMUSG00 | Gm11954 | 348 | 0.48856524 | 0 | 0 | 0.26184508 | 0.13831013 | 0.28478583 | 0.13595597 | 0 | 0.11835206 | 0 | 0 | 0.48116951 | 0.28351677 | 0 | 0 | 0.11003048 | 0.11013304 | 0.15807215 |
| ENSMUSG00 | Gm13457 | 265 | 0 | 0 | 0 | 0 | 0 | 0 | 0 | 0 | 0 | 0 | 0 | 0 | 0 | 0 | 0 | 0 | 0 | 0 |
| ENSMUSG00 | Gm16420 | 855 | 0 | 0 | 0 | 0 | 0 | 0 | 0 | 0 | 0 | 0 | 0 | 0 | 0 | 0 | 0 | 0 | 0 | 0 |
| ENSMUSG00 | Gm5287 | 597 | 0 | 0 | 0 | 0 | 0 | 0 | 0.07925072 | 0 | 0 | 0 | 0 | 0 | 0 | 0 | 0 | 0 | 0 | 0 |
| ENSMUSG00 | Sycn | 555 | 0 | 0 | 0 | 0 | 0 | 0 | 0 | 0 | 0 | 0 | 0 | 0 | 0 | 0 | 0 | 0 | 0 | 0 |
| ENSMUSG00 | Gm11372 | 569 | 0 | 0 | 0 | 0 | 0 | 0 | 0 | 0 | 0 | 0 | 0 | 0 | 0 | 0 | 0 | 0 | 0 | 0 |
| ENSMUSG00 | Gm13508 | 774 | 0 | 0 | 0 | 0 | 0 | 0 | 0 | 0 | 0.10642511 | 0 | 0 | 0 | 0 | 0 | 0 | 0 | 0.04951718 | 0 |
| ENSMUSG00 | Gm11759 | 463 | 0 | 0 | 0 | 0 | 0 | 0 | 0.10218721 | 0.08248899 | 0 | 0 | 0 | 0 | 0 | 0 | 0 | 0 | 0.16555636 | 0 |
| ENSMUSG00 | Gm11890 | 416 | 0 | 0 | 0 | 0 | 0 | 0 | 0 | 0 | 0 | 0 | 0 | 0 | 0 | 0 | 0 | 0 | 0 | 0 |
| ENSMUSG00 | Gm15237 | 468 | 0 | 0 | 0 | 0 | 0 | 0.21176382 | 0.10109547 | 0.0816077 | 0.08800538 | 0 | 0 | 0.23852847 | 0 | 0.89315553 | 0 | 0.16363507 | 0.0818938 | 0 |
| ENSMUSG00 | Gm11858 | 837 | 0 | 0 | 0 | 0 | 0 | 0 | 0 | 0 | 0 | 0 | 0 | 0 | 0 | 0 | 0 | 0 | 0 | 0 |
| ENSMUSG00 | Taar7c-ps | 1054 | 0 | 0 | 0 | 0 | 0 | 0 | 0 | 0 | 0 | 0 | 0 | 0 | 0 | 0 | 0 | 0 | 0.14545084 | 0 |
| ENSMUSG00 | Gm12009 | 1099 | 0 | 0.09093602 | 0.0880319 | 0.12437046 | 0 | 0 | 0 | 0.03475196 | 0 | 0.08580656 | 0 | 0.05078768 | 0.044888 | 0 | 0 | 0.03484132 | 0.06974758 | 0 |
| ENSMUSG00 | Gm13474 | 808 | 0 | 0 | 0 | 0 | 0 | 0 | 0 | 0 | 0 | 0 | 0 | 0 | 0 | 0 | 0 | 0 | 0 | 0 |
| ENSMUSG00 | Rplp1-ps1 | 344 | 0 | 0.09683981 | 0 | 0 | 0 | 0 | 0 | 0 | 0 | 0 | 0 | 0 | 0.14340674 | 0 | 0 | 0 | 0 | 0 |
| ENSMUSG00 | Rpl36-ps10 | 215 | 0 | 0 | 0 | 0.21191183 | 0 | 0 | 0 | 0.35527816 | 0.19156519 | 0 | 0 | 0 | 0 | 0.21601901 | 0.76654057 | 0 | 0 | 0 |
| ENSMUSG00 | Gm7567 | 401 | 0 | 0 | 0 | 0 | 0 | 0 | 0 | 0 | 0 | 0 | 0 | 0 | 0 | 0 | 0 | 0 | 0 | 0 |
| ENSMUSG00 | Gm12851 | 420 | 0 | 0 | 0 | 0 | 0 | 0 | 0 | 0 | 0 | 0 | 0 | 0 | 0.11745695 | 0.22116232 | 0.09809894 | 0 | 0 | 0.65487032 |
| ENSMUSG00 | Gm8089 | 962 | 0 | 0 | 0 | 0 | 0 | 0 | 0 | 0 | 0 | 0 | 0 | 0 | 0 | 0 | 0 | 0 | 0 | 0 |
| ENSMUSG00 | Gm11869 | 327 | 0.12998525 | 0.61124577 | 1.18345028 | 0.41799123 | 0.44157728 | 0.60614965 | 0.28937418 | 0.58398169 | 1.00762118 | 2.4512598 | 1.31490711 | 0.1706901 | 0 | 0.71015424 | 0 | 0.35128998 | 0.586029 | 1.17756498 |
| ENSMUSG00 | Gm15167 | 185 | 0 | 0 | 0 | 0 | 0 | 0 | 0 | 0 | 0 | 0 | 0 | 0 | 0 | 0 | 0 | 0 | 0 | 0 |
| ENSMUSG00 | Gm13282 | 544 | 0 | 0 | 0 | 0 | 0 | 0 | 0 | 0 | 0 | 0 | 0.08782161 | 0 | 0 | 0 | 0 | 0 | 0 | 0 |
| ENSMUSG00 | Gm6823 | 1838 | 0 | 0 | 0 | 0 | 0 | 0 | 0 | 0 | 0 | 0 | 0 | 0 | 0 | 0 | 0 | 0 | 0 | 0 |
| ENSMUSG00 | Gm12407 | 320 | 0 | 0 | 0 | 0 | 0 | 0 | 0 | 0 | 0 | 0 | 0 | 0 | 0 | 0 | 0 | 0.2393163 | 0 | 0 |
| ENSMUSG00 | Gm15372 | 833 | 0 | 0 | 0 | 0 | 0 | 0 | 0 | 0 | 0 | 0 | 0 | 0 | 0 | 0 | 0 | 0 | 0 | 0 |
| ENSMUSG00 | Gm12874 | 911 | 0 | 0 | 0 | 0 | 0 | 0 | 0 | 0 | 0 | 0 | 0 | 0 | 0 | 0 | 0 | 0 | 0 | 0 |
| ENSMUSG00 | Gm9089 | 525 | 0 | 0 | 0 | 0 | 0.09167985 | 0 | 0 | 0 | 0 | 0.17962173 | 0 | 0.2126311 | 0 | 0 | 0.07847915 | 0 | 0 | 0 |
| ENSMUSG00 | Gm13711 | 451 | 0 | 0 | 0 | 0 | 0 | 0 | 0 | 0 | 0 | 0 | 0 | 0 | 0 | 0 | 0 | 0 | 0.0849807 | 0 |
| ENSMUSG00 | Gm12422 | 301 | 0 | 0 | 0 | 0 | 0 | 0 | 0 | 0 | 0 | 0 | 0 | 0 | 0 | 0 | 0 | 0 | 0 | 0 |
| ENSMUSG00 | Gm11584 | 324 | 0 | 0 | 0 | 0 | 0 | 0 | 0 | 0 | 0 | 0 | 0 | 0 | 0 | 0 | 0 | 0 | 0 | 0 |
| ENSMUSG00 | Gm11926 | 429 | 0.89171698 | 0.2329573 | 1.35310574 | 0 | 0.22439125 | 0.34652261 | 0.44114386 | 0.08902658 | 0 | 0.4396336 | 0.89090832 | 0.91074508 | 0.34497844 | 0 | 0.96040923 | 0.17851099 | 0.62537081 | 0.51290542 |
| ENSMUSG00 | Gm6398 | 983 | 0 | 0.03388901 | 0 | 0 | 0 | 0 | 0 | 0 | 0 | 0.04796613 | 0.34020825 | 0 | 0 | 0 | 0.0419141 | 0 | 0 | 0 |
| ENSMUSG00 | Gm15593 | 1007 | 0 | 0.09924397 | 0 | 0.04524433 | 0 | 0.04920828 | 0.09396758 | 0 | 0 | 0.23411471 | 0 | 0.11085534 | 0 | 0.04612124 | 0.04091515 | 0.03802444 | 0 | 0.10925344 |
| ENSMUSG00 | Gm6584 | 397 | 0 | 0 | 0 | 0 | 0 | 0 | 0 | 0 | 0 | 0 | 0 | 0 | 0 | 0 | 0 | 0 | 0 | 0 |
| ENSMUSG00 | Gm8061 | 348 | 0 | 0.09572671 | 0 | 0 | 0 | 0 | 0 | 0 | 0 | 0.13549053 | 0 | 0 | 0.42527515 | 0.66730011 | 0.11839528 | 0 | 0 | 0 |
| ENSMUSG00 | Gm12062 | 258 | 0 | 0 | 0 | 0 | 0 | 0 | 0 | 0 | 0 | 0 | 0 | 0 | 0 | 0 | 0 | 0 | 0 | 0 |
| ENSMUSG00 | Gm14070 | 427 | 0 | 0 | 0 | 0 | 0 | 0 | 0 | 0 | 0 | 0 | 0 | 0 | 0 | 0 | 0 | 0 | 0 | 0 |
| ENSMUSG00 | Gm12792 | 1018 | 0 | 0 | 0 | 0 | 0 | 0 | 0 | 0 | 0 | 0 | 0 | 0.05482875 | 0 | 0 | 0.04047304 | 0 | 0 | 0 |
| ENSMUSG00 | Gm13811 | 255 | 0 | 0 | 0 | 0 | 0 | 0 | 0 | 0 | 0 | 0 | 0 | 0 | 0 | 0 | 0 | 0 | 0 | 0 |
| ENSMUSG00 | Gm15168 | 314 | 0 | 0 | 0 | 0 | 0 | 0 | 0 | 0 | 0 | 0 | 0 | 0 | 0 | 0 | 0 | 0 | 0 | 0 |
| ENSMUSG00 | Gm11949 | 371 | 0 | 0 | 0 | 0 | 0 | 0 | 0 | 0 | 0 | 0 | 0 | 0 | 0.13297013 | 0 | 0 | 0 | 0 | 0 |
| ENSMUSG00 | Gm12225 | 297 | 0 | 0 | 0 | 0 | 0 | 0 | 0 | 0 | 0 | 0 | 0 | 0 | 0 | 0 | 0 | 0 | 0 | 0 |
| ENSMUSG00 | Hspe1-ps5 | 296 | 0 | 0 | 0 | 0 | 0 | 0 | 0 | 0 | 0 | 0 | 0 | 0 | 0 | 0 | 0 | 0 | 0 | 0 |
| ENSMUSG00 | Gm14825 | 700 | 0 | 0 | 0 | 0 | 0 | 0 | 0 | 0 | 0 | 0 | 0 | 0 | 0 | 0 | 0 | 0 | 0 | 0 |
| ENSMUSG00 | Gm11814 | 928 | 0.13740897 | 0.32307764 | 0.62551979 | 0.04909595 | 0.41493038 | 0.26698671 | 0.30590094 | 0.16462242 | 0.6213483 | 0.35566263 | 0.20592655 | 0.48116951 | 0.47843454 | 0.30028505 | 0.39958405 | 0.4126143 | 0.61949833 | 0.59277055 |
| ENSMUSG00 | Gm7973 | 487 | 0.08727962 | 0 | 0 | 0 | 0 | 0.10175099 | 0 | 0 | 0 | 0 | 0 | 0 | 0.10129757 | 0 | 0 | 0 | 0 | 0 |
| ENSMUSG00 | Gm15488 | 677 | 0 | 0.19682655 | 0 | 0 | 0 | 0 | 0.13977158 | 0 | 0 | 0 | 0 | 0 | 0 | 0 | 0.06085902 | 0 | 0 | 0 |
| ENSMUSG00 | Gm8145 | 744 | 0.05713061 | 0 | 0 | 0 | 0 | 0 | 0 | 0 | 0 | 0 | 0 | 0 | 0 | 0 | 0 | 0.05146587 | 0.05151384 | 0 |
| ENSMUSG00 | Gm14876 | 325 | 0 | 0 | 0 | 0 | 0 | 0 | 0 | 0 | 0 | 0 | 0 | 0 | 0 | 0 | 0 | 0 | 0 | 0 |
| ENSMUSG00 | Gm12387 | 954 | 0 | 0 | 0 | 0 | 0 | 0 | 0 | 0 | 0 | 0 | 0 | 0 | 0 | 0 | 0 | 0 | 0 | 0 |
| ENSMUSG00 | Hmgb1-ps4 | 634 | 0.13408573 | 0.052544 | 0.07629894 | 0.07186285 | 0.07591786 | 0.15631777 | 0.07462568 | 0.06024038 | 0.19488888 | 0.14874039 | 0.07535482 | 0 | 0 | 0.07325566 | 0.19496004 | 0.12079056 | 0.42316101 | 0 |
| ENSMUSG00 | Gm13973 | 893 | 0 | 0 | 0 | 0 | 0 | 0 | 0 | 0 | 0 | 0 | 0 | 0 | 0 | 0 | 0 | 0 | 0 | 0 |
| ENSMUSG00 | Gm12244 | 290 | 0 | 0 | 0 | 0 | 0 | 0 | 0 | 0.6584897 | 0 | 0.4877659 | 0 | 0 | 0.51033018 | 0 | 0.14207433 | 0 | 0 | 0 |
| ENSMUSG00 | Gm12337 | 399 | 0.10652926 | 0.25047289 | 0.12123692 | 0.11418808 | 0.12063139 | 0 | 0 | 0 | 0.2064487 | 0.11817219 | 0.23947348 | 0.13988888 | 0 | 0.23280244 | 0 | 0.2878993 | 0.67239117 | 0 |
| ENSMUSG00 | Gm13426 | 794 | 0 | 0 | 0 | 0 | 0 | 0 | 0 | 0 | 0 | 0 | 0 | 0 | 0 | 0 | 0 | 0 | 0.19307958 | 0 |
| ENSMUSG00 | Gm8080 | 1148 | 0 | 0 | 0 | 0 | 0 | 0 | 0 | 0 | 0 | 0 | 0 | 0 | 0 | 0 | 0 | 0 | 0 | 0 |
| ENSMUSG00 | Gm7958 | 547 | 0 | 0 | 0 | 0 | 0 | 0 | 0 | 0 | 0 | 0 | 0 | 0 | 0 | 0 | 0 | 0 | 0 | 0 |
| ENSMUSG00 | Gm14388 | 476 | 0 | 0.13997014 | 0.20325013 | 0.28714944 | 0.10111749 | 0.10410238 | 0.29818915 | 0.08023614 | 0.17305259 | 0.1981122 | 0 | 0 | 0.20727696 | 0.09757161 | 0.08655789 | 0.40221226 | 0.16103486 | 0 |
| ENSMUSG00 | Gm11975 | 480 | 0 | 0.06940186 | 0 | 0 | 0.10027484 | 0.10323486 | 0 | 0 | 0 | 0.49115316 | 0 | 0.11628263 | 0 | 0 | 0 | 0 | 0 | 0 |
| ENSMUSG00 | 4933405O20I | 976 | 0 | 0 | 0 | 0 | 0 | 0 | 0 | 0 | 0 | 0 | 0 | 0 | 0 | 0 | 0 | 0 | 0 | 0 |
| ENSMUSG00 | Gm15421 | 398 | 0 | 0 | 0 | 0 | 0.12093448 | 0 | 0 | 0 | 0 | 0 | 0 | 0 | 0 | 0 | 0 | 0 | 0 | 0.69106918 |
| ENSMUSG00 | Olfr637-ps1 | 461 | 0 | 0 | 0 | 0 | 0 | 0 | 0 | 0 | 0 | 0 | 0 | 0 | 0 | 0 | 0 | 0 | 0 | 0 |
| ENSMUSG00 | Gm11781 | 471 | 0 | 0 | 0 | 0 | 0 | 0 | 0 | 0 | 0 | 0 | 0 | 0 | 0 | 0 | 0 | 0 | 0 | 0 |
| ENSMUSG00 | Gm13076 | 203 | 0 | 0 | 0 | 0 | 0 | 0 | 0 | 0 | 0 | 0 | 0 | 0 | 0 | 0 | 0 | 0 | 0 | 0 |
| ENSMUSG00 | Gm14463 | 497 | 0 | 0 | 0 | 0 | 0 | 0 | 0 | 0 | 0 | 0 | 0 | 0 | 0 | 0 | 0 | 0 | 0 | 0 |
| ENSMUSG00 | Gm15383 | 1442 | 0 | 0 | 0 | 0 | 0 | 0 | 0 | 0 | 0 | 0 | 0 | 0 | 0 | 0 | 0 | 0 | 0 | 0 |
| ENSMUSG00 | Gm13416 | 1611 | 0 | 0.08271358 | 0 | 0 | 0 | 0.03075899 | 0 | 0 | 0 | 0 | 0 | 0.03464659 | 0 | 0.02882935 | 0.02557514 | 0 | 0.02379038 | 0.03414594 |
| ENSMUSG00 | Mup-ps3 | 539 | 0 | 0 | 0 | 0 | 0 | 0 | 0 | 0 | 0 | 0 | 0 | 0 | 0 | 0 | 0 | 0 | 0 | 0 |
| ENSMUSG00 | Gm13784 | 517 | 0 | 0 | 0 | 0 | 0 | 0 | 0 | 0 | 0 | 0 | 0 | 0 | 0 | 0 | 0 | 0 | 0 | 0 |
| ENSMUSG00 | Gm14020 | 661 | 0 | 0 | 0 | 0.1378549 | 0 | 0 | 0 | 0 | 0 | 0 | 0 | 0 | 0 | 0 | 0 | 0 | 0 | 0 |
| ENSMUSG00 | Gm14994 | 1091 | 0 | 0 | 0 | 0 | 0 | 0 | 0 | 0 | 0 | 0 | 0 | 0 | 0 | 0 | 0 | 0 | 0 | 0 |
| ENSMUSG00 | Gm15162 | 282 | 0 | 0 | 0 | 0 | 0 | 0 | 0 | 0 | 0 | 0 | 0 | 0 | 0 | 0 | 0 | 0 | 0 | 0 |
| ENSMUSG00 | Gm14566 | 320 | 0 | 0 | 0 | 0 | 0 | 0 | 0 | 0 | 0 | 0 | 0 | 0 | 0 | 0 | 0 | 0 | 0 | 0 |
| ENSMUSG00 | Gm11409 | 402 | 0 | 0 | 0 | 0 | 0 | 0 | 0 | 0 | 0 | 0 | 0 | 0 | 0 | 0 | 0 | 0 | 0 | 0 |
| ENSMUSG00 | Gm12267 | 760 | 0.05592786 | 0 | 0 | 0 | 0.12666296 | 0 | 0 | 0.20101264 | 0 | 0 | 0.44003251 | 0.07344166 | 0.32455209 | 0.18333192 | 0.05421257 | 0.35267665 | 0 | 0 |
| ENSMUSG00 | Gm13550 | 524 | 0 | 0 | 0 | 0 | 0 | 0 | 0 | 0 | 0 | 0.62987581 | 0 | 0 | 0 | 0 | 0 | 0 | 0 | 0.10497921 |
| ENSMUSG00 | Gm12310 | 491 | 0 | 0 | 0 | 0 | 0 | 0 | 0 | 0 | 0 | 0 | 0 | 0 | 0 | 0 | 0.33565422 | 0 | 0.15611526 | 0 |

| | | | | | | | | | | | | | | | | | | | | |
|---|---|---|---|---|---|---|---|---|---|---|---|---|---|---|---|---|---|---|---|---|
| ENSMUSG00 | Gm12036 | 529 | 0.80350049 | 0 | 0.45721673 | 0 | 0 | 0.09367246 | 0.26831387 | 0.36098679 | 0 | 0.44565882 | 0 | 0 | 0 | 0.70236805 | 0 | 0.144766 | 0 | 0.20797394 |
| ENSMUSG00 | Gm11256 | 828 | 0 | 0 | 0 | 0 | 0 | 0 | 0 | 0 | 0 | 0 | 0 | 0 | 0 | 0 | 0 | 0 | 0 | 0 |
| ENSMUSG00 | Defa-ps11 | 297 | 0 | 0 | 0.32574768 | 0 | 0.16206035 | 0 | 0 | 0 | 0 | 0 | 0 | 0 | 0 | 0 | 0.13872578 | 0 | 0 | 0 |
| ENSMUSG00 | Gm14976 | 1393 | 0 | 0 | 0 | 0 | 0 | 0 | 0 | 0 | 0 | 0 | 0 | 0 | 0 | 0 | 0 | 0 | 0 | 0 |
| ENSMUSG00 | Gm11597 | 957 | 0 | 0 | 0 | 0 | 0 | 0 | 0 | 0 | 0 | 0 | 0 | 0 | 0 | 0 | 0 | 0 | 0 | 0 |
| ENSMUSG00 | Gm12054 | 377 | 0 | 0.08836311 | 0 | 0 | 0 | 0 | 0 | 0 | 0 | 0 | 0 | 0 | 0 | 0 | 0 | 0 | 0 | 0 |
| ENSMUSG00 | Gm12040 | 536 | 0.47580421 | 0.43505645 | 0.09024912 | 0.59501364 | 0.17959673 | 0.46224565 | 0.17653985 | 0.14250896 | 0.61472412 | 0.52780638 | 0.08913239 | 1.24960439 | 0.09203716 | 0.34659767 | 0.23060572 | 0.7858147 | 0.42902571 | 0.20525786 |
| ENSMUSG00 | Gm12844 | 581 | 1.90212492 | 1.20408051 | 1.74844086 | 2.43096794 | 1.73970808 | 2.13221745 | 2.7687282 | 0.72309196 | 1.27600221 | 1.78539668 | 1.31566151 | 1.24888746 | 3.8208886 | 0.79938188 | 1.41829797 | 1.44990252 | 0.72562696 | 1.98828097 |
| ENSMUSG00 | Gm13635 | 295 | 0 | 0 | 0 | 0 | 0 | 0 | 0.16038196 | 0 | 0 | 0 | 0 | 0 | 0 | 0 | 0 | 0 | 0 | 0 |
| ENSMUSG00 | Olfr588-ps1 | 941 | 0 | 0 | 0 | 0 | 0 | 0 | 0 | 0 | 0 | 0 | 0 | 0 | 0 | 0 | 0 | 0 | 0 | 0 |
| ENSMUSG00 | Gm14142 | 294 | 0 | 0 | 0 | 0 | 0 | 0 | 0 | 0 | 0 | 0 | 0 | 0 | 0 | 0 | 0 | 0.26048032 | 0.13036155 | 0 |
| ENSMUSG00 | Gm14541 | 156 | 0.54493815 | 0.85417677 | 0.31008673 | 2.62852179 | 0.61707594 | 3.49410301 | 1.81971842 | 0.48964618 | 0.79204838 | 1.51124049 | 0.9187492 | 0.71558542 | 1.89738143 | 0.29771851 | 2.64112538 | 2.2090735 | 0.98272556 | 1.41048992 |
| ENSMUSG00 | Gm7408 | 631 | 0 | 0 | 0 | 0 | 0 | 0 | 0 | 0 | 0 | 0 | 0 | 0 | 0 | 0 | 0 | 0 | 0 | 0 |
| ENSMUSG00 | Gm12417 | 402 | 0 | 0 | 0 | 0 | 0 | 0 | 0 | 0 | 0 | 0 | 0 | 0 | 0 | 0 | 0 | 0 | 0 | 0 |
| ENSMUSG00 | Olfr1291-ps1 | 1834 | 0 | 0 | 0 | 0 | 0 | 0 | 0 | 0 | 0 | 0 | 0 | 0 | 0 | 0 | 0 | 0 | 0 | 0 |
| ENSMUSG00 | Gm14784 | 662 | 0 | 0 | 0 | 0 | 0 | 0 | 0 | 0 | 0 | 0 | 0 | 0 | 0 | 0 | 0 | 0 | 0 | 0 |
| ENSMUSG00 | Gm15177 | 576 | 0 | 0 | 0 | 0 | 0 | 0 | 0 | 0 | 0 | 0 | 0 | 0 | 0 | 0 | 0 | 0 | 0 | 0 |
| ENSMUSG00 | Gm14846 | 215 | 0 | 0 | 0 | 0 | 0 | 0 | 0 | 0 | 0 | 0 | 0 | 0 | 0 | 0 | 0 | 0 | 0 | 0 |
| ENSMUSG00 | Gm13224 | 93 | 0 | 0 | 0 | 0 | 0 | 0 | 0 | 0 | 0 | 0 | 0 | 0 | 0 | 0 | 0 | 0 | 1.64844288 | 0 |
| ENSMUSG00 | Gm12628 | 380 | 0 | 0 | 0 | 0 | 0 | 0 | 0 | 0 | 0 | 0.37224239 | 0.12572357 | 0 | 0 | 0 | 0 | 0 | 0 | 0 |
| ENSMUSG00 | Gm12504 | 2535 | 0.83836639 | 0.26282362 | 0.66787912 | 0.35945597 | 0.24683038 | 0.29321144 | 0.57857714 | 0.46704713 | 0.19496576 | 0.79979497 | 0.16961524 | 0.61650436 | 0.64219064 | 0.20153253 | 0.3900739 | 0.36251463 | 0.21166397 | 0.56419597 |
| ENSMUSG00 | Gm5396 | 531 | 0 | 0.06273615 | 0 | 0 | 0.09064392 | 0.09331965 | 0 | 0 | 0 | 0 | 0 | 0.21022848 | 0.0929038 | 0 | 0.23277715 | 0 | 0 | 0.1035953 |
| ENSMUSG00 | Gm8317 | 470 | 0.45218272 | 0.35439249 | 0.30876722 | 0.19387678 | 0.40963339 | 0.2108627 | 0.10066527 | 0.24378129 | 0.43815442 | 0 | 0.2032977 | 0.23751346 | 0.20992305 | 0.09881721 | 0.17532577 | 0.08146938 | 0.08154531 | 0.58520326 |
| ENSMUSG00 | Gm11702 | 68 | 0.62507612 | 0.97979101 | 0 | 0 | 0 | 0.72871667 | 1.39154938 | 0 | 0 | 0.69339269 | 0 | 0 | 0 | 0 | 1.21181047 | 0 | 0 | 0 |
| ENSMUSG00 | Gm12189 | 911 | 0 | 0 | 0 | 0 | 0 | 0 | 0 | 0 | 0 | 0 | 0 | 0 | 0 | 0 | 0 | 0 | 0 | 0 |
| ENSMUSG00 | Gm13232 | 631 | 0 | 0 | 0 | 0 | 0 | 0 | 0 | 0 | 0 | 0 | 0 | 0 | 0 | 0 | 0 | 0 | 0 | 0 |
| ENSMUSG00 | Gm11972 | 388 | 0 | 0.08585797 | 0.12467405 | 0 | 0 | 0 | 0 | 0.09843403 | 0.10615081 | 0.12152243 | 0 | 0.1438548 | 0.12714412 | 0 | 0.10618958 | 0 | 0.19755823 | 0 |
| ENSMUSG00 | Gm14770 | 311 | 0 | 0 | 0 | 0 | 0 | 0 | 0 | 0 | 0 | 0 | 0 | 0 | 0 | 0 | 0 | 0 | 0 | 0 |
| ENSMUSG00 | Gm15054 | 280 | 0 | 0 | 0 | 0 | 0 | 0 | 0 | 0 | 0 | 0 | 0 | 0 | 0 | 0 | 0 | 0 | 0 | 0 |
| ENSMUSG00 | Gm14914 | 2030 | 0 | 0 | 0 | 0 | 0 | 0 | 0 | 0 | 0 | 0 | 0 | 0 | 0 | 0 | 0 | 0 | 0 | 0 |
| ENSMUSG00 | Tpt1-ps6 | 522 | 0.16285508 | 0 | 0.0926696 | 0.26184508 | 0.46103375 | 0.18965722 | 0.27191195 | 0.07316552 | 0.07890137 | 0 | 0 | 0.21385311 | 0 | 0.17794669 | 0.39465092 | 0.29341462 | 0 | 0.10538143 |
| ENSMUSG00 | Gm13451 | 591 | 0 | 0 | 0 | 0 | 0 | 0 | 0 | 0 | 0 | 0 | 0 | 0 | 0 | 0 | 0 | 0 | 0 | 0 |
| ENSMUSG00 | Gm12658 | 682 | 0 | 0 | 0 | 0 | 0 | 0 | 0 | 0 | 0 | 0 | 0 | 0 | 0 | 0 | 0 | 0 | 0 | 0 |
| ENSMUSG00 | Gm14576 | 1110 | 0 | 0 | 0 | 0 | 0 | 0 | 0 | 0 | 0 | 0 | 0 | 0 | 0 | 0 | 0 | 0 | 0 | 0 |
| ENSMUSG00 | Gm5228 | 356 | 0.11939656 | 0 | 0.95116492 | 1.663746 | 0.4056061 | 1.39193072 | 0.26580157 | 0.64369218 | 1.27261706 | 0.26489159 | 0.26839864 | 0.47035671 | 0.2771456 | 0.13046092 | 0.46293883 | 0.43023154 | 0.32297441 | 0.77259981 |
| ENSMUSG00 | Gm6977 | 510 | 4.7505785 | 6.53194004 | 6.54465411 | 6.70015358 | 5.19069765 | 6.89851781 | 10.1119255 | 6.29051334 | 8.56033465 | 10.9093784 | 9.64866808 | 9.74038033 | 10.833676 | 10.5637532 | 10.2599953 | 11.2619434 | 6.98891296 | 8.08957452 |
| ENSMUSG00 | Gm7375 | 631 | 0 | 0 | 0 | 0 | 0 | 0 | 0 | 0 | 0 | 0 | 0 | 0 | 0 | 0 | 0 | 0 | 0 | 0 |
| ENSMUSG00 | Gm6654 | 758 | 0 | 0 | 0 | 0 | 0 | 0 | 0 | 0 | 0 | 0 | 0 | 0 | 0 | 0 | 0 | 0 | 0 | 0 |
| ENSMUSG00 | Gm13535 | 1881 | 0 | 0 | 0 | 0 | 0 | 0 | 0 | 0 | 0 | 0 | 0 | 0 | 0 | 0 | 0 | 0 | 0 | 0.05848922 |
| ENSMUSG00 | Gm12143 | 331 | 0 | 0 | 0 | 0 | 0 | 0 | 0 | 0 | 0 | 0 | 0 | 0 | 0 | 0 | 0 | 0 | 0 | 0 |
| ENSMUSG00 | Gm13533 | 176 | 0 | 0 | 0 | 0 | 0 | 0 | 0 | 0 | 0 | 0 | 0 | 0 | 0 | 0 | 0 | 0 | 0 | 0 |
| ENSMUSG00 | Olfr1235-ps1 | 290 | 0 | 0 | 0 | 0 | 0 | 0 | 0 | 0 | 0 | 0 | 0 | 0 | 0 | 0 | 0 | 0 | 0 | 0 |
| ENSMUSG00 | Gm12777 | 304 | 0 | 0 | 0 | 0 | 0 | 0 | 0 | 0 | 0 | 0 | 0 | 0 | 0 | 0 | 0 | 0 | 0 | 0 |
| ENSMUSG00 | Gm13755 | 303 | 0 | 0 | 0 | 0 | 0 | 0 | 0 | 0 | 0 | 0 | 0 | 0 | 0 | 0 | 0 | 0 | 0 | 0 |
| ENSMUSG00 | Gm14331 | 407 | 0 | 0 | 0 | 0 | 0 | 0 | 0 | 0 | 0 | 0 | 0 | 0 | 0 | 0 | 0 | 0 | 0 | 0 |
| ENSMUSG00 | Gm6630 | 908 | 0 | 0 | 0 | 0 | 0 | 0 | 0 | 0 | 0 | 0 | 0 | 0 | 0.16299092 | 0 | 0 | 0 | 0 | 0 |
| ENSMUSG00 | Gm6954 | 224 | 0 | 0 | 0 | 0 | 0 | 0 | 0.21121732 | 0.1705018 | 0 | 0.21049421 | 0 | 1.24588533 | 0 | 0 | 0.18393552 | 0 | 0.17109954 | 0.24557637 |
| ENSMUSG00 | Gm7856 | 1492 | 0 | 0 | 0 | 0 | 0.03226 | 0 | 0 | 0 | 0 | 0.12640939 | 0 | 0 | 0 | 0 | 0 | 0 | 0 | 0 |
| ENSMUSG00 | Gm12194 | 1011 | 0 | 0 | 0 | 0 | 0 | 0 | 0 | 0 | 0 | 0 | 0 | 0 | 0 | 0 | 0.04075327 | 0 | 0 | 0 |
| ENSMUSG00 | Gm8216 | 600 | 0 | 0.05552149 | 0.08062255 | 0 | 0.24065962 | 0 | 0 | 0 | 0 | 0 | 0 | 0 | 0 | 0 | 0 | 0.12763536 | 0 | 0 |
| ENSMUSG00 | Gm6142 | 324 | 1.04951052 | 0.4112703 | 0.44790306 | 0.56248203 | 0.29711064 | 0.45882161 | 0.29205357 | 0 | 0.12711888 | 0.29105372 | 0.29490715 | 0 | 0.30451801 | 0 | 0.25433059 | 0.11818089 | 0.35487312 | 1.01868716 |
| ENSMUSG00 | Gm14645 | 416 | 0 | 0 | 0 | 0 | 0 | 0 | 0 | 0 | 0 | 0 | 0 | 0 | 0 | 0 | 0 | 0 | 0 | 0 |
| ENSMUSG00 | Gm11743 | 1733 | 0 | 0 | 0 | 0 | 0 | 0 | 0.05460205 | 0 | 0.38025635 | 0 | 0 | 0 | 0 | 0 | 0 | 0 | 0 | 0 |
| ENSMUSG00 | Gm12151 | 289 | 0.14707673 | 0 | 0 | 0 | 0 | 0 | 0 | 0 | 0 | 0 | 0 | 0 | 0 | 0 | 0 | 0 | 0 | 0 |
| ENSMUSG00 | Gm14593 | 293 | 0 | 0 | 0 | 0 | 0 | 0 | 0 | 0 | 0 | 0 | 0 | 0 | 0 | 0 | 0 | 0 | 0 | 0 |
| ENSMUSG00 | Mup-ps20 | 385 | 0 | 0 | 0 | 0 | 0 | 0 | 0 | 0 | 0 | 0 | 0 | 0 | 0 | 0 | 0 | 0 | 0 | 0 |
| ENSMUSG00 | Gm11950 | 837 | 0 | 0 | 0 | 0 | 0 | 0 | 0 | 0 | 0 | 0 | 0 | 0 | 0 | 0 | 0 | 0 | 0 | 0 |
| ENSMUSG00 | Gm11257 | 314 | 0 | 0 | 0 | 0 | 0 | 0 | 0 | 0 | 0 | 0 | 0 | 0 | 0.31421603 | 0.14791111 | 0 | 0.12194461 | 0 | 0.17518824 |
| ENSMUSG00 | Gm13122 | 452 | 0 | 0 | 0 | 0 | 0 | 0 | 0 | 0 | 0 | 0 | 0 | 0 | 0 | 0 | 0 | 0 | 0 | 0 |
| ENSMUSG00 | Gm13809 | 594 | 0 | 0.11216463 | 0 | 0 | 0.08103017 | 0 | 0 | 0 | 0 | 0 | 0 | 0.09396576 | 0 | 0 | 0 | 0 | 0 | 0 |
| ENSMUSG00 | Rps15a-ps3 | 394 | 0 | 0.08455049 | 0 | 0.11563717 | 0.12216224 | 0 | 0 | 0.19387006 | 0 | 0 | 0 | 0 | 0 | 0 | 0 | 0 | 0 | 0.13961702 |
| ENSMUSG00 | Vmn1r-ps128 | 1738 | 0 | 0 | 0 | 0 | 0 | 0 | 0 | 0 | 0 | 0 | 0 | 0 | 0 | 0 | 0 | 0 | 0 | 0 |
| ENSMUSG00 | Gm8668 | 1446 | 0 | 0 | 0 | 0 | 0 | 0 | 0 | 0 | 0 | 0.16303839 | 0 | 0 | 0 | 0 | 0 | 0 | 0 | 0 |
| ENSMUSG00 | Gm11987 | 378 | 0 | 0 | 0 | 0 | 0 | 0 | 0 | 0 | 0 | 0 | 0 | 0 | 0 | 0 | 0 | 0 | 0 | 0 |
| ENSMUSG00 | Tpt1-ps3 | 1204 | 2.85956749 | 1.77078507 | 1.80798079 | 1.24876617 | 1.67902059 | 2.14015128 | 1.96481225 | 1.49089943 | 2.22352452 | 1.9972474 | 1.74592872 | 1.947058 | 2.70424131 | 3.08598587 | 1.77946919 | 2.2261981 | 1.75078599 | 2.83269486 |
| ENSMUSG00 | Gm14125 | 1434 | 0 | 0.11615375 | 0.33733285 | 0 | 0.0671296 | 0.10366681 | 0.197961 | 0 | 0.34465704 | 0 | 0 | 0.11676917 | 0 | 0.19432673 | 0.22985526 | 0.32042349 | 0.08018054 | 0 |
| ENSMUSG00 | Gm14778 | 446 | 0 | 0 | 0 | 0 | 0.10791911 | 0 | 0 | 0 | 0 | 0 | 0 | 0 | 0 | 0 | 0.09238017 | 0 | 0 | 0 |
| ENSMUSG00 | Gm14438 | 1310 | 0 | 0 | 0 | 0 | 0 | 0 | 0 | 0 | 0 | 0 | 0 | 0 | 0 | 0 | 0 | 0 | 0 | 0 |
| ENSMUSG00 | Gm6500 | 450 | 0 | 0.07402865 | 0 | 0 | 0 | 0.11011719 | 0 | 0 | 0 | 0 | 0 | 0 | 0 | 0 | 0 | 0 | 0 | 0 |
| ENSMUSG00 | Gm15550 | 1073 | 0.31690718 | 0.186279 | 0.85656764 | 0.80676593 | 0.53828806 | 0.04618149 | 0 | 0.64069268 | 0.42222896 | 0.61520023 | 0.84596851 | 0.46816492 | 0.32182984 | 0.25970599 | 0 | 1.21330909 | 0.03571882 | 1.2303994 |
| ENSMUSG00 | Gm12820 | 341 | 0 | 0 | 0 | 0 | 0 | 0 | 0 | 0 | 0 | 0 | 0 | 0 | 0 | 0 | 0 | 0 | 0 | 0 |
| ENSMUSG00 | Gm14531 | 283 | 0 | 0 | 0 | 0 | 0 | 0 | 0 | 0 | 0 | 0 | 0 | 0 | 0 | 0 | 0 | 0 | 0 | 0 |
| ENSMUSG00 | Gm14046 | 508 | 0.16734321 | 0.13115313 | 0.28567045 | 0.35874838 | 0.28424364 | 0.48772376 | 0.37254078 | 0.45109137 | 0.64860655 | 0.27844903 | 0.18809039 | 0.43949341 | 0.48555037 | 0.36570148 | 0.08110542 | 0 | 0.2263364 | 0.10828564 |
| ENSMUSG00 | Gm6733 | 1919 | 1.08533279 | 2.13521938 | 1.8905757 | 1.59071911 | 1.47982465 | 1.65261852 | 1.13412362 | 1.65188609 | 1.80284905 | 1.22852275 | 1.79249453 | 1.71606258 | 0.82262707 | 1.93617873 | 1.99674033 | 0.6983696 | 2.77611009 | 2.14991298 |
| ENSMUSG00 | Gm15019 | 429 | 0 | 0 | 0 | 0 | 0 | 0 | 0 | 0 | 0 | 0 | 0 | 0 | 0 | 0 | 0 | 0 | 0 | 0 |
| ENSMUSG00 | Gm8163 | 1059 | 0 | 0 | 0 | 0 | 0 | 0 | 0.04467675 | 0 | 0.03889189 | 0 | 0 | 0 | 0.13975047 | 0 | 0.0389061 | 0 | 0 | 0 |
| ENSMUSG00 | Gm8840 | 385 | 0 | 0 | 0 | 0 | 0 | 0 | 0 | 0 | 0 | 0 | 0 | 0 | 0 | 0 | 0 | 0 | 0 | 0 |
| ENSMUSG00 | Gm11599 | 877 | 0.24233282 | 0.11395517 | 0.16547388 | 0 | 0 | 0 | 0.10789665 | 0.04354892 | 0 | 0 | 0 | 0 | 0 | 0 | 0 | 0.0436609 | 0 | 0 |
| ENSMUSG00 | Gm13494 | 1095 | 0 | 0 | 0 | 0 | 0 | 0 | 0 | 0 | 0 | 0 | 0 | 0 | 0.04505198 | 0 | 0.03762699 | 0 | 0 | 0 |
| ENSMUSG00 | Gm12619 | 383 | 0 | 0 | 0 | 0 | 0 | 0 | 0 | 0 | 0 | 0 | 0 | 0 | 0 | 0 | 0 | 0 | 0 | 0 |
| ENSMUSG00 | Olfr1267-ps1 | 929 | 0 | 0 | 0 | 0 | 0 | 0 | 0 | 0 | 0 | 0 | 0 | 0 | 0 | 0 | 0 | 0 | 0 | 0 |
| ENSMUSG00 | Gm14986 | 1001 | 0 | 0 | 0 | 0 | 0 | 0 | 0 | 0 | 0 | 0 | 0 | 0 | 0 | 0 | 0 | 0 | 0 | 0 |
| ENSMUSG00 | Gm13531 | 542 | 0 | 0.0614629 | 0 | 0 | 0 | 0 | 0 | 0 | 0 | 0 | 0 | 0 | 0 | 0 | 0 | 0 | 0 | 0 |
| ENSMUSG00 | Gm6141 | 693 | 0 | 0 | 0 | 0 | 0.06945444 | 0 | 0.06827226 | 0 | 0.0594322 | 0 | 0 | 0 | 0 | 0 | 0 | 0 | 0 | 0 |
| ENSMUSG00 | Gm14766 | 366 | 0.81294053 | 0.18203767 | 0.52867246 | 0 | 0 | 0 | 0 | 0.62610496 | 0 | 0.12882706 | 0.65266337 | 0 | 0.94350661 | 0.25379283 | 0 | 0 | 0.41886663 | 0.75149053 |
| ENSMUSG00 | Gm3928 | 1306 | 0 | 0 | 0 | 0 | 0 | 0 | 0 | 0 | 0 | 0 | 0 | 0 | 0 | 0 | 0 | 0 | 0 | 0 |
| ENSMUSG00 | Gm11224 | 1110 | 0 | 0.06002323 | 0.13073927 | 0 | 0.17344837 | 0.0892842 | 0 | 0 | 0 | 0 | 0.0430405 | 0 | 0 | 0 | 0.07423704 | 0.06899209 | 0.06905639 | 0.09911551 |
| ENSMUSG00 | Gm16198 | 669 | 0 | 0 | 0 | 0 | 0 | 0 | 0 | 0 | 0 | 0.07047938 | 0 | 0 | 0 | 0.06942315 | 0.05723559 | 0 | 0 | 0 |
| ENSMUSG00 | Gm15305 | 286 | 0 | 0 | 0 | 0 | 0 | 0 | 0 | 0 | 0 | 0 | 0 | 0 | 0 | 0 | 0 | 0 | 0 | 0 |
| ENSMUSG00 | Rpl17-ps4 | 1361 | 0 | 0.08607983 | 0 | 0.1177288 | 0 | 0.04268108 | 0.04075166 | 0.06579225 | 0 | 0.04061215 | 0.04114984 | 0.04807551 | 0 | 0 | 0.06596143 | 0 | 0 | 0 |
| ENSMUSG00 | Cyp4a30b-ps | 1514 | 0 | 0.24203556 | 0.03195081 | 0 | 0 | 0.09818904 | 1.50000567 | 2.42171112 | 2.3123209 | 1.27686845 | 0.06311091 | 0.73732711 | 0.81459573 | 2.48478935 | 0.81641128 | 0.05058204 | 0.43034812 | 2.10735019 |
| ENSMUSG00 | Akt2-ps | 1448 | 0.20548082 | 0.13803685 | 0.03340713 | 0.03146481 | 0 | 0.0342215 | 0.13069801 | 0.02637597 | 0.11377491 | 0.06512528 | 0.06598751 | 0.03854673 | 0.10220701 | 0.03207465 | 0.19917879 | 0.13221895 | 0.29115281 | 0.18994857 |
| ENSMUSG00 | Gm14608 | 1552 | 0 | 0 | 0 | 0 | 0 | 0 | 0 | 0 | 0 | 0 | 0 | 0 | 0 | 0 | 0 | 0.07408434 | 0 | 0 |
| ENSMUSG00 | Rpl3-ps1 | 1209 | 2.84774132 | 4.13311343 | 1.36038051 | 3.35395612 | 2.42849243 | 3.03300437 | 3.95250668 | 3.41172825 | 1.32859728 | 5.18696736 | 2.88467491 | 3.37017648 | 3.14190044 | 4.41775852 | 1.29500341 | 3.51551482 | 4.50151709 | 3.86747235 |
| ENSMUSG00 | 1700029I01R | 2410 | 0.31746605 | 0.17969611 | 0.4817281 | 0.39700495 | 0.57918082 | 0.1439291 | 0.53005906 | 0.14262723 | 0.35888665 | 0.46955057 | 0.39647268 | 0.27792031 | 0.3275148 | 0.59741357 | 0.39320987 | 0.8420756 | 0.44528478 | 0.38803104 |
| ENSMUSG00 | Gm13085 | 169 | 0 | 0 | 0 | 0 | 0 | 0 | 0 | 0 | 0 | 0 | 0 | 0 | 0 | 0 | 0 | 0 | 0 | 0 |
| ENSMUSG00 | Gm12370 | 602 | 0 | 0 | 0 | 0 | 0 | 0 | 0 | 0 | 0 | 0 | 0 | 0 | 0 | 0 | 0 | 0 | 0 | 0 |

| | | | | | | | | | | | | | | | | | | | | |
|---|---|---|---|---|---|---|---|---|---|---|---|---|---|---|---|---|---|---|---|---|
| ENSMUSG00 | Gm13605 | 656 | 3.62849064 | 5.3828762 | 7.96393488 | 4.16716869 | 4.10882275 | 3.92796059 | 10.674202 | 5.0069308 | 6.15285364 | 5.46258147 | 5.53490373 | 4.67966683 | 3.3840492 | 9.06226095 | 7.16002648 | 5.01980524 | 4.79078711 | 6.12144023 |
| ENSMUSG00 | Gm12061 | 476 | 0.35718635 | 0.20995522 | 0.20325013 | 0 | 0.40446995 | 0.41640953 | 0 | 0.08023614 | 0.08652629 | 0.0990561 | 0.60220536 | 0.70355877 | 0 | 0.48785806 | 0.25967367 | 0.24132736 | 0 | 0.34669605 |
| ENSMUSG00 | Gm15139 | 6636 | 0.11529433 | 0.14558076 | 0.1895286 | 0.02746236 | 0.10154414 | 0 | 0.04990789 | 0.09208536 | 0.11171749 | 0.19894812 | 0.01439872 | 0.26915329 | 0.01486797 | 0.06998808 | 0.08071432 | 0.19041441 | 0.04042858 | 0.04973699 |
| ENSMUSG00 | Gm15473 | 1177 | 0.57781038 | 0.53776125 | 0.36989106 | 0.38709468 | 0.98144959 | 0.63151317 | 1.48731446 | 0.77877456 | 1.04978375 | 0.2804205 | 0.44649494 | 0.85359552 | 1.29931133 | 1.02595265 | 0.77012254 | 1.39635982 | 0.91175558 | 2.0096785 |
| ENSMUSG00 | Gm12159 | 1444 | 0 | 0 | 0 | 0 | 0 | 0 | 0 | 0 | 0 | 0 | 0 | 0 | 0 | 0 | 0 | 0 | 0 | 0 |
| ENSMUSG00 | Gm14110 | 756 | 0 | 0 | 0 | 0 | 0 | 0 | 0 | 0 | 0 | 0 | 0 | 0 | 0 | 0 | 0 | 0 | 0 | 0 |
| ENSMUSG00 | Gm16741 | 1815 | 0.04683766 | 0 | 0.02665208 | 0 | 0.0795569 | 0 | 0.26067592 | 0.02104265 | 0 | 0.33771854 | 0 | 0.12300972 | 0.19026084 | 0 | 0.22700582 | 0.14767727 | 0 | 0.0606161 |
| ENSMUSG00 | Gm14951 | 568 | 0 | 0.11729892 | 0 | 0 | 0 | 0 | 0 | 0 | 0 | 0 | 0 | 0 | 0 | 0 | 0 | 0 | 0 | 0 |
| ENSMUSG00 | Gm16742 | 2093 | 0 | 0 | 0 | 0 | 0 | 0 | 0 | 0 | 0 | 0 | 0 | 0 | 0 | 0 | 0 | 0 | 0 | 0 |
| ENSMUSG00 | Gm16743 | 1153 | 0 | 0 | 0.33563594 | 0 | 0 | 0 | 0 | 0 | 0.1428847 | 0.24536359 | 0 | 0 | 0.08557141 | 0 | 0.21440532 | 0 | 0.132962 | 0 |
| ENSMUSG00 | 4921504A21 | 2152 | 0.41478099 | 0.41795917 | 0.26974088 | 0.29640085 | 0.33549203 | 0.36842181 | 0.43970891 | 0.0709896 | 0.09569358 | 0.02191018 | 0 | 0 | 0.29800879 | 0.41005467 | 0.21060275 | 0.4804119 | 0.37400197 | 0 |
| ENSMUSG00 | Gm15350 | 615 | 0 | 0.37917115 | 0.62924918 | 0.296332 | 0.70436961 | 0.08057355 | 0.15386237 | 0.06210147 | 0.13393989 | 0 | 0.07768286 | 0.27227152 | 0.320858 | 0.37759421 | 0.2679776 | 0.06226115 | 0 | 0.1788914 |
| ENSMUSG00 | D930015M05 | 1186 | 0.35839103 | 0.14044222 | 0.24472275 | 0.34574148 | 0.08116682 | 0.04178139 | 0.23935588 | 0.322027 | 0.34727248 | 0.23853644 | 0.24169456 | 0.18824844 | 0.9566898 | 0.27412193 | 0.2431795 | 0.12914201 | 0.90483669 | 0.6029686 |
| ENSMUSG00 | Gm11213 | 479 | 0 | 0 | 0 | 0 | 0 | 0 | 0 | 0 | 0 | 0 | 0 | 0 | 0 | 0 | 0 | 0 | 0 | 0 |
| ENSMUSG00 | Gm15419 | 740 | 0 | 0.04501742 | 0.06536964 | 0.30784489 | 1.04069024 | 0 | 0 | 0.51611355 | 0.55657454 | 0 | 0.38736453 | 0.07542657 | 0 | 0.3138114 | 0.16703333 | 0.67267283 | 0.05179229 | 0.37168315 |
| ENSMUSG00 | Gm15086 | 571 | 0 | 0 | 0 | 0 | 0 | 0 | 0 | 0 | 0 | 0 | 0 | 0 | 0 | 0 | 0 | 0 | 0 | 0 |
| ENSMUSG00 | Gm13972 | 3162 | 0.0134425 | 0.03160616 | 0.13768557 | 0.07204466 | 0.13699789 | 0.01567133 | 0.07481448 | 0.03623568 | 0.09117824 | 0.07455835 | 0.13598186 | 0.03530402 | 0.12481193 | 0 | 0 | 0.06054808 | 0.0121209 | 0.17396934 |
| ENSMUSG00 | Ubl5 | 4321 | 40.1148132 | 32.3646216 | 41.0632051 | 34.4264776 | 39.1204156 | 35.0917299 | 46.5571652 | 34.5243054 | 38.8798421 | 38.4647602 | 38.9519044 | 34.5925352 | 57.061079 | 52.7534323 | 46.5221919 | 43.5277629 | 39.6035445 | 49.9677722 |
| ENSMUSG00 | Gm12436 | 2955 | 0 | 0 | 0 | 0 | 0 | 0 | 0 | 0 | 0 | 0 | 0 | 0 | 0 | 0 | 0 | 0 | 0 | 0 |
| ENSMUSG00 | Gm11342 | 813 | 0 | 0 | 0 | 0 | 0 | 0 | 0 | 0 | 0 | 0 | 0 | 0 | 0 | 0 | 0 | 0 | 0 | 0 |
| ENSMUSG00 | Gm12974 | 331 | 0 | 0 | 0.1461436 | 0 | 0 | 0.14970614 | 0.28587721 | 0.1153849 | 0 | 0.14244925 | 1.01034655 | 0 | 0.5961561 | 0.84188678 | 0.124476 | 0.11568159 | 0 | 0 |
| ENSMUSG00 | Gm15879 | 1020 | 0.04167174 | 0.2612776 | 0.14227509 | 0.35734152 | 0.2359408 | 0.24290556 | 0.37107983 | 0.18721766 | 0.12113681 | 0.4622618 | 0.04683819 | 0.21888495 | 0.29018775 | 0.2276671 | 0.40393682 | 0.30031849 | 0.1127244 | 0.75502696 |
| ENSMUSG00 | Gm13667 | 614 | 0 | 0 | 0 | 0 | 0 | 0 | 0 | 0 | 0 | 0 | 0 | 0 | 0 | 0 | 0 | 0 | 0 | 0 |
| ENSMUSG00 | 1700056N10 | 1041 | 0.24498661 | 1.2480335 | 1.48698652 | 0.91909888 | 1.06343347 | 0.99962287 | 0.36359407 | 0.88051648 | 0.55390127 | 0.72469861 | 0.59661331 | 2.14469405 | 0.90039042 | 0.53537853 | 0.07915765 | 0.772433 | 0.51543531 | 0.6869533 |
| ENSMUSG00 | Gm2568 | 1164 | 0 | 0 | 0 | 0 | 0 | 0 | 0 | 0 | 0.17691802 | 0 | 0 | 0 | 0 | 0 | 0 | 0 | 0 | 0 |
| ENSMUSG00 | Gm13682 | 710 | 0 | 0 | 0 | 0 | 0 | 0 | 0 | 0 | 0 | 0 | 0 | 0 | 0 | 0 | 0 | 0 | 0 | 0 |
| ENSMUSG00 | Gm14617 | 924 | 0.27600764 | 0.54079374 | 0.99469381 | 0.29585094 | 0.72927157 | 0.214514 | 1.17769655 | 1.07467799 | 1.60466945 | 1.42880919 | 1.60283952 | 1.57057059 | 0.58728473 | 1.40739659 | 1.02557985 | 0.37296046 | 1.07844558 | 2.32181295 |
| ENSMUSG00 | A430104N18 | 267 | 0.15919542 | 0 | 0 | 0 | 0 | 0 | 0 | 0 | 0 | 0 | 0.89466214 | 0 | 0 | 1.04368736 | 0 | 0 | 0 | 0 |
| ENSMUSG00 | Gm14321 | 976 | 0.13065116 | 0.64850921 | 0.39650435 | 0.84026516 | 0.69041694 | 0.40616995 | 0.8725699 | 0.50871028 | 0.42199299 | 0.8212725 | 0.19579901 | 0.45750543 | 0.75817496 | 0.57103386 | 0.67543534 | 0.27462526 | 0.70683744 | 1.35268295 |
| ENSMUSG00 | 4930533B01 | 1023 | 0.20774768 | 0.19538354 | 0.04728595 | 0.0445367 | 0.23524889 | 0.14531593 | 0 | 0.29866981 | 0.28182367 | 0.04609062 | 0 | 0 | 0.04822279 | 0.04539989 | 0.08055045 | 0.48658641 | 0 | 0 |
| ENSMUSG00 | Gm11602 | 1572 | 1.78456846 | 2.09794945 | 0.86161504 | 2.55049103 | 1.77585978 | 1.41849428 | 2.19709005 | 2.25947419 | 1.91260538 | 1.82963925 | 2.06660126 | 2.16587495 | 2.10256899 | 1.80221968 | 1.75604596 | 1.14482096 | 1.36531338 | 2.76445256 |
| ENSMUSG00 | Gm12205 | 573 | 1.11270094 | 0.81392761 | 0.59095063 | 0.63610533 | 0.67199893 | 1.03775358 | 0.74313108 | 1.59968178 | 1.0063023 | 2.79777297 | 2.08442227 | 0.97409533 | 0.77484687 | 0.89159679 | 1.29428971 | 0.20047438 | 0.60198372 | 0.48000966 |
| ENSMUSG00 | Gm16929 | 2288 | 0.01857744 | 0.04367949 | 0.04228455 | 0 | 0.02103668 | 0.06497299 | 0.04135724 | 0.05007745 | 0 | 0.0824313 | 0 | 0.12197479 | 0 | 0.04059798 | 0.03601535 | 0 | 0 | 0 |
| ENSMUSG00 | Gm14089 | 316 | 0 | 0 | 0 | 0 | 0 | 0 | 0 | 0 | 0 | 0 | 0 | 0 | 0 | 0 | 0 | 0 | 0 | 0 |
| ENSMUSG00 | 5830444B04 | 5401 | 0.13378782 | 0.31456353 | 0.21495366 | 0.32899106 | 0.17823338 | 0.22019359 | 0.40295931 | 0.43135281 | 0.52617471 | 0.27062985 | 0.29190402 | 0.258358 | 0.42015704 | 0.06879332 | 0.03814253 | 0.50335736 | 0.36190356 | 0.37684446 |
| ENSMUSG00 | 1700095J12 | 735 | 0 | 0 | 0 | 0 | 0.26194244 | 0 | 0.25748397 | 0.05196245 | 0 | 0.12830123 | 0 | 0 | 0 | 0.37913541 | 0.05605654 | 0 | 0.46930159 | 0 |
| ENSMUSG00 | Gm12273 | 1081 | 0 | 0 | 0 | 0 | 0 | 0 | 0 | 0.07066124 | 0 | 0 | 0 | 0 | 0 | 0 | 0 | 0 | 0.28363587 | 0 |
| ENSMUSG00 | Gm15232 | 2314 | 0.0183687 | 0.01439624 | 0 | 0 | 0 | 0 | 0 | 0.16504928 | 0 | 0.12225766 | 0 | 0 | 0 | 0 | 0 | 0 | 0 | 0 |
| ENSMUSG00 | Gm13073 | 484 | 0 | 0 | 0 | 0 | 0.39778449 | 0 | 0 | 0 | 0 | 0 | 0 | 0 | 0.10192545 | 0 | 0 | 0 | 0 | 0 |
| ENSMUSG00 | 9430091E24 | 1917 | 0.66518272 | 0.59083902 | 0.85795515 | 0.85560647 | 0.27618736 | 0.43943478 | 0.78977868 | 0.51799815 | 1.0957289 | 0.83626704 | 1.02179097 | 0.72790379 | 0.77201748 | 0.94487189 | 0.92418722 | 0.67912397 | 0.75972837 | 0.68868991 |
| ENSMUSG00 | Gm12631 | 635 | 0 | 0 | 0 | 0 | 0 | 0 | 0 | 0 | 0 | 0 | 0 | 0 | 0 | 0 | 0 | 0 | 0 | 0 |
| ENSMUSG00 | Gm16761 | 1785 | 0.02381242 | 0 | 0 | 0 | 0.13482332 | 0 | 0 | 0 | 0 | 0 | 0 | 0 | 0 | 0 | 0 | 0 | 0.04294263 | 0 |
| ENSMUSG00 | 1700092E19 | 1000 | 0 | 0 | 0 | 0 | 0 | 0 | 0 | 0 | 0 | 0 | 0 | 0 | 0 | 0 | 0 | 0 | 0 | 0 |
| ENSMUSG00 | Gm17445 | 3900 | 0.01089876 | 0 | 0.01240347 | 0.0584116 | 0 | 0 | 0 | 0 | 0 | 0.01208992 | 0 | 0.01431171 | 0 | 0.04763496 | 0 | 0 | 0.02948177 | 0 |
| ENSMUSG00 | 9430076G02 | 1771 | 0 | 0 | 0 | 0 | 0 | 0 | 0 | 0 | 0 | 0 | 0 | 0 | 0 | 0 | 0 | 0 | 0 | 0 |
| ENSMUSG00 | Gm15525 | 502 | 0 | 0 | 0 | 0 | 0 | 0 | 0 | 0 | 0 | 0 | 0 | 0 | 0 | 0 | 0 | 0 | 0 | 0 |
| ENSMUSG00 | Gm11463 | 824 | 0 | 0.08085654 | 0.11741148 | 0.27646265 | 0.35047517 | 0 | 0.2296732 | 0.09270001 | 0.64978726 | 0 | 0 | 0 | 0.35921299 | 0 | 0.50001888 | 0.83644531 | 0.13953749 | 0.33379312 |
| ENSMUSG00 | Gm6526 | 877 | 0.48466563 | 0.68373101 | 0.55157959 | 0.31170612 | 0.82323701 | 0.45202037 | 0.43158658 | 0.4354892 | 0.42266664 | 0.80645445 | 0.65370525 | 0.06364386 | 0.61875837 | 0.58253701 | 0.46980109 | 0.26196539 | 0.48071752 | 0.06272418 |
| ENSMUSG00 | Gm14640 | 435 | 0 | 0 | 0 | 0 | 0 | 0 | 0 | 0 | 0 | 0 | 0 | 0 | 0 | 0 | 0 | 0 | 0 | 0 |
| ENSMUSG00 | Gm11967 | 478 | 0.08892296 | 0 | 0 | 0 | 0 | 0 | 0 | 0 | 0 | 0 | 0 | 0 | 0 | 0 | 0 | 0 | 0 | 0 |
| ENSMUSG00 | Gm15880 | 2879 | 0 | 0.08099696 | 0.28563738 | 0.06330121 | 0.03343656 | 0.13769429 | 0.21363836 | 0 | 0 | 0.11464221 | 0.01659429 | 0.15509736 | | 0 | 0 | 0 | 0 | 0.07642807 |
| ENSMUSG00 | Gm11768 | 1040 | 0.04087036 | 0 | 0 | 0 | 0 | 0 | 0 | 0 | 0 | 0 | 0.04593746 | 0 | | 0 | 0.15846752 | 0.22090735 | 0 | 0 |
| ENSMUSG00 | Gm17216 | 726 | 0.35128245 | 0.59651188 | 0.39978124 | 0.37653756 | 0 | 0.13650891 | 0.06516898 | 0.21042646 | 0.17019221 | 0 | 0 | 0.23064323 | | 0 | 0.05675145 | 0.10548377 | 0 | 0 |
| ENSMUSG00 | Gm16344 | 1565 | 0 | 0 | 0 | 0 | 0 | 0 | 0 | 0 | 0 | 0 | 0 | 0 | 0 | 0 | 0 | 0 | 0 | 0 |
| ENSMUSG00 | 2410152P15 | 869 | 0 | 0 | 0 | 0 | 0 | 0 | 0 | 0 | 0 | 0 | 0 | 0 | 0 | 0 | 0 | 0 | 0 | 0 |
| ENSMUSG00 | AI847159 | 1228 | 0 | 0 | 0 | 0 | 0 | 0 | 0 | 0 | 0 | 0 | 0 | 0 | 0 | 0 | 0 | 0 | 0 | 0 |
| ENSMUSG00 | B230344G16 | 2021 | 0.14722228 | 0.08241686 | 0.02393544 | 0.04508762 | 0.09526358 | 0.04903784 | 0.09364212 | 0.05669332 | 0.22417203 | 0.04666077 | 0.09455707 | 0.08285353 | 0 | 0.06894224 | 0 | 0 | 0.01896403 | 0.32662508 |
| ENSMUSG00 | Gm12367 | 2694 | 0 | 0 | 0 | 0.08456022 | 0 | 0 | 0 | 0 | 0 | 0 | 0 | 0 | 0 | 0 | 0 | 0 | 0 | 0 |
| ENSMUSG00 | Gm14266 | 1276 | 0 | 0 | 0 | 0 | 0 | 0 | 0 | 0 | 0 | 0 | 0 | 0 | 0 | 0 | 0 | 0 | 0 | 0 |
| ENSMUSG00 | Gm14539 | 349 | 5.11523608 | 5.5362403 | 5.26703196 | 8.22448651 | 6.89569107 | 5.96336622 | 11.1164461 | 4.48678653 | 6.0186599 | 6.34980818 | 8.21345991 | 6.23728035 | 9.7533017 | 6.78695833 | 6.25696981 | 6.36348203 | 6.80868312 | 6.14714946 |
| ENSMUSG00 | Gm14710 | 1062 | 0 | 0 | 0 | 0 | 0 | 0 | 0 | 0 | 0 | 0 | 0 | 0 | 0 | 0 | 0 | 0 | 0 | 0 |
| ENSMUSG00 | 4930563E18 | 1550 | 0 | 0 | 0 | 0 | 0 | 0 | 0 | 0.1232013 | 0 | 0 | 0 | 0 | 0.15913522 | 0 | 0 | 0 | 0 | 0 |
| ENSMUSG00 | Gm12649 | 578 | 0 | 0 | 0 | 0 | 0 | 0 | 0 | 0 | 0 | 0 | 0 | 0 | 0 | 0 | 0 | 0 | 0 | 0 |
| ENSMUSG00 | 4930565N06 | 2413 | 0.03523015 | 0.05522237 | 0.02004705 | 0.22657792 | 0.15957538 | 0.1642859 | 0.01960741 | 0 | 0.32430327 | 0.09770142 | 0.13859292 | 0.04626246 | 0.12266536 | 0.11548468 | 0.25612239 | 0.12694772 | 0.20648233 | 0.06839093 |
| ENSMUSG00 | Gm12352 | 758 | 0.95328231 | 1.49424591 | 1.7230677 | 0.66117611 | 1.33347018 | 0.58835699 | 0.74901339 | 1.10848661 | 1.57573741 | 1.30628597 | 0.56724885 | 1.61997965 | 1.43179707 | 1.83815649 | 1.41324598 | 0.90927564 | 1.11237273 | 1.66914176 |
| ENSMUSG00 | Gm16274 | 591 | 0 | 0 | 0 | 0 | 0 | 0 | 0 | 0 | 0 | 0 | 0 | 0 | 0 | 0 | 0 | 0 | 0 | 0 |
| ENSMUSG00 | 1700108N11 | 915 | 0 | 0.1092226 | 0.15860174 | 0.09958698 | 0.10520639 | 0.16246798 | 0 | 0 | 0 | 0.41224659 | 0.10442614 | 0 | 0.16174399 | 0 | 0.31520316 | 0 | 0.4607533 | 0 |
| ENSMUSG00 | Gm10241 | 572 | 0.29723899 | 0.17471798 | 0.33827644 | 0 | 0 | 0 | 0.16542895 | 0 | 0 | 0 | 0 | 0 | 0 | 0 | 0 | 0 | 0 | 0 |
| ENSMUSG00 | Gm14097 | 468 | 0.81740723 | 1.99307914 | 1.24034693 | 1.26558456 | 1.74838184 | 1.27058291 | 1.31424108 | 1.79536934 | 2.46415052 | 1.00749366 | 0.71458271 | 0.47705695 | 0.5270504 | 0.19847901 | 1.05645015 | 1.55453321 | 0.65515037 | 0.70524496 |
| ENSMUSG00 | Gm11468 | 380 | 0 | 0 | 0 | 0 | 0 | 0 | 0 | 0 | 0 | 0 | 0 | 0 | 0 | 0 | 0 | 0 | 0 | 0 |
| ENSMUSG00 | Gm16712 | 1957 | 0.13031735 | 0.37449345 | 0.44492772 | 0.09312426 | 0.31973174 | 0.22788687 | 0.50769865 | 0.52692635 | 0.6103265 | 0.28912031 | 0.39059751 | 0.45633654 | 0.30249515 | 0.47464576 | 0.10526713 | 0.4891493 | 0.76378415 | 0.22487116 |
| ENSMUSG00 | Pabpc1l2b-ps | 1291 | 0 | 0 | 0 | 0 | 0 | 0 | 0 | 0 | 0 | 0 | 0 | 0 | 0 | 0 | 0 | 0 | 0 | 0 |
| ENSMUSG00 | B230312C02 | 1696 | 0.32580618 | 0.17677833 | 0.08556639 | 0.08059147 | 0.31217639 | 0.29217414 | 0.05579325 | 0 | 0 | 0.05560224 | 0.76056832 | 0.16455089 | 0.17452329 | 0.52030522 | 0 | 0 | 0.09039221 | 0 |
| ENSMUSG00 | Hoxb3os | 2997 | 4.16967693 | 2.58989134 | 2.32425371 | 3.10125227 | 2.28052491 | 2.3478439 | 3.07840253 | 2.68888786 | 2.85845688 | 2.28123189 | 3.15630356 | 3.42678743 | 3.27562614 | 2.88241583 | 3.0932099 | 3.88399888 | 2.40418546 | 2.60637076 |
| ENSMUSG00 | Gm5151 | 5566 | 0.87820612 | 0.74813363 | 1.07767118 | 0.8267455 | 0.48425938 | 0.97040028 | 0.47601689 | 0.89888694 | 1.33933873 | 0.83017767 | 0.6609184 | 1.16324414 | 0.46088029 | 0.47562217 | 0.49595836 | 0.81864576 | 0.44757623 | 0.56333437 |
| ENSMUSG00 | A730011C13 | 3297 | 1.54704917 | 0.87904816 | 0.27876769 | 0.88441214 | 0.56934942 | 0.69136359 | 0.66011017 | 0.45177546 | 0.67457442 | 0.71505464 | 1.34761029 | 1.26969205 | 0.97257344 | 1.18328885 | 1.19968134 | 1.38203891 | 0.91834318 | 1.65177482 |
| ENSMUSG00 | Gm15507 | 1629 | 0 | 0 | 0 | 0 | 0 | 0 | 0 | 0 | 0 | 0 | 0 | 0 | 0 | 0 | 0 | 0 | 0 | 0 |
| ENSMUSG00 | 4933408N05 | 1581 | 0 | 0 | 0 | 0.05763573 | 0 | 0.03134265 | 0 | 0 | 0 | 0 | 0 | 0 | 0 | 0.0293764 | 0.07818132 | 0 | 0 | 0 |
| ENSMUSG00 | Gm16105 | 695 | 0 | 0 | 0 | 0 | 0 | 0 | 0 | 0 | 0 | 0 | 0 | 0 | 0 | 0 | 0 | 0 | 0 | 0 |
| ENSMUSG00 | Gm12092 | 597 | 0 | 0 | 0 | 0 | 0 | 0 | 0 | 0 | 0 | 0 | 0 | 0 | 0 | 0 | 0 | 0 | 0 | 0 |
| ENSMUSG00 | Gm11267 | 1067 | 0 | 0 | 0 | 0 | 0 | 0 | 0 | 0 | 0 | 0 | 0 | 0 | 0 | 0 | 0 | 0 | 0 | 0 |
| ENSMUSG00 | Gm16359 | 288 | 0 | 0 | 0 | 0 | 0 | 0 | 0 | 0 | 0 | 0 | 0 | 0 | 0 | 0 | 0 | 0 | 0 | 0 |
| ENSMUSG00 | Gm11791 | 334 | 0 | 0 | 0 | 0 | 0 | 0 | 0 | 0 | 0 | 0 | 0 | 0 | 0 | 0 | 0 | 0 | 0 | 0 |
| ENSMUSG00 | Gm12678 | 690 | 0 | 0 | 0 | 0 | 0 | 0 | 0 | 0 | 0 | 0 | 0 | 0 | 0 | 0 | 0 | 0 | 0 | 0 |
| ENSMUSG00 | Gm9850 | 226 | 0 | 0 | 0 | 0 | 0 | 0 | 0 | 0 | 0 | 0 | 0 | 0 | 0 | 0 | 0 | 0 | 0 | 0 |
| ENSMUSG00 | Gm11528 | 911 | 0 | 0 | 0 | 0 | 0 | 0 | 0 | 0 | 0 | 0 | 0 | 0 | 0 | 0 | 0 | 0 | 0 | 0 |
| ENSMUSG00 | Gm2449 | 582 | 0 | 0.3434319 | 0.24934809 | 0 | 0 | 0 | 0.16258653 | 0.32811342 | 0 | 0.16202991 | 0 | 0.1918064 | 0.16952549 | 0 | 0.07079305 | 0 | 0.32926372 | 0.37806946 |
| ENSMUSG00 | Gm1980 | 2304 | 0 | 0 | 0 | 0 | 0 | 0 | 0 | 0 | 0 | 0 | 0 | 0 | 0 | 0 | 0 | 0 | 0 | 0 |
| ENSMUSG00 | 1700080N15 | 1585 | 0 | 0 | 0 | 0 | 0 | 0 | 0 | 0 | 0 | 0 | 0 | 0 | 0 | 0 | 0 | 0 | 0 | 0 |
| ENSMUSG00 | 2610204G07 | 990 | 0 | 0 | 0 | 0 | 0.0486181 | 0 | 0.23895292 | 0 | 0.08320508 | 0 | 0.1447726 | 0 | 0 | 0.09382644 | 0.33294187 | 0 | 0 | 0.38895328 |
| ENSMUSG00 | 1700095J07 | 1974 | 0 | 0 | 0 | 0 | 0.09753176 | 0 | 0 | 0 | 0 | 0 | 0 | 0 | 0 | 0 | 0 | 0 | 0 | 0 |
| ENSMUSG00 | Gm16278 | 2069 | 0.06163148 | 0.22541349 | 0.0467603 | 0.04404161 | 0.04652675 | 0.19160071 | 0.11433707 | 0.18459354 | 0.01990648 | 0.06836738 | 0.20781761 | 0.21581697 | 0.1907469 | 0.0673428 | 0.17922378 | 0.07402727 | 0.01852407 | 0.02658729 |
| ENSMUSG00 | Gm12523 | 597 | 0.9967713 | 0.89280789 | 0.64822151 | 0.38158329 | 0.72560689 | 1.16204065 | 0.8717579 | 0.70371261 | 0.62090225 | 0.78979402 | 0.64020045 | 1.40240358 | 0.66106422 | 0.62236633 | 1.03521497 | 1.21862905 | 0.77037783 | 1.19785325 |
| ENSMUSG00 | 1700027A07 | 456 | 0 | 0 | 0 | 0 | 0 | 0 | 0.10375587 | 0 | 0.18064261 | 0 | 0 | 0 | 0 | 0 | 0.33588252 | 0 | 0 | 0.24126801 |
| ENSMUSG00 | Gm15723 | 598 | 0 | 0 | 0 | 0 | 0 | 0 | 0 | 0 | 0 | 0 | 0 | 0 | 0 | 0 | 0 | 0 | 0 | 0 |
| ENSMUSG00 | A930006K02 | 774 | 6.69978227 | 5.42302929 | 4.68735759 | 2.82549112 | 5.28580557 | 3.20108098 | 6.11274922 | 7.54966094 | 5.69374314 | 7.79753232 | 7.40697031 | 5.1200414 | 6.37363272 | 4.0803591 | 7.45247781 | 4.45239621 | 7.52661128 | 4.19320064 |
| ENSMUSG00 | Enox | 3390 | 0.564228 | 0.21618987 | 0.22831165 | 0.30911623 | 0.42594623 | 0.30696384 | 0.72574021 | 0.63090694 | 0.5831719 | 0.18081391 | 0.29595107 | 0.74091588 | 0.40746126 | 0.45211059 | 0.25523088 | 0.38403559 | 0.32786508 | 0.29208375 |
| ENSMUSG00 | Gm16831 | 1335 | 0 | 0 | 0 | 0 | 0 | 0 | 0 | 0 | 0 | 0 | 0 | 0 | 0 | 0 | 0 | 0 | 0 | 0 |
| ENSMUSG00 | Gm13063 | 667 | 0.31862951 | 0 | 0 | 0 | 0 | 0.07429195 | 0 | 0.05725997 | 0 | 0 | 0 | 0 | 0 | 0.30885724 | 0 | 0 | 0 | 0.16494485 |
| ENSMUSG00 | Gm13954 | 541 | 0 | 0 | 0 | 0 | 0 | 0 | 0 | 0 | 0 | 0.4357736 | 0 | 0 | 0 | 0 | 0 | 0 | 0 | 0 |
| ENSMUSG00 | Gm2800 | 639 | 0 | 0 | 0 | 0 | 0 | 0 | 0 | 0 | 0 | 0 | 0 | 0 | 0 | 0 | 0 | 0 | 0 | 0 |
| ENSMUSG00 | Gm15252 | 882 | 0 | 0 | 0 | 0 | 0 | 0 | 0 | 0 | 0 | 0 | 0 | 0 | 0 | 0 | 0 | 0 | 0 | 0 |
| ENSMUSG00 | C030047K22 | 648 | 0 | 0 | 0 | 0 | 0 | 0 | 0 | 0 | 0 | 0 | 0 | 0 | 0 | 0 | 0 | 0 | 0 | 0 |
| ENSMUSG00 | Gm12906 | 1475 | 0.08645121 | 0.09034005 | 0.39354737 | 0.33977728 | 0.06526363 | 0.13438029 | 0.09622918 | 0.18125208 | 0.08376918 | 0.19179947 | 0.12955921 | 0.03784113 | 0.06689074 | 0.15743758 | 0.11173303 | 0.0778792 | 0.10393572 | 0.07458862 |
| ENSMUSG00 | 4930570D08 | 194 | 0 | 0 | 0 | 0 | 0 | 0 | 0 | 0 | 0 | 0 | 0 | 0 | 0 | 0 | 0 | 0 | 0 | 0 |
| ENSMUSG00 | Gm14965 | 1229 | 0 | 0 | 0 | 0 | 0 | 0 | 0.07699378 | 0 | 0 | 0 | 0 | 0 | 0 | 0 | 0 | 0 | 0 | 0 |
| ENSMUSG00 | 5430416N02 | 3686 | 1.63747558 | 1.23816237 | 1.28611122 | 1.50798899 | 1.44944209 | 0.80660988 | 1.79700897 | 1.45060671 | 1.84367203 | 1.63735486 | 1.82752825 | 1.60511672 | 1.69971609 | 1.65061732 | 1.98965734 | 1.92180205 | 1.44529443 | 1.58192222 |
| ENSMUSG00 | Gm15817 | 584 | 0.14556567 | 0 | 0.24849416 | 0.07801549 | 0.24725303 | 0 | 0.32405944 | 0 | 0 | 0.08073751 | 0.16361287 | 0 | 0 | 0.56440488 | 0 | 0 | 0.59064499 | 0.28258103 |
| ENSMUSG00 | Gm14872 | 2941 | 0.27459991 | 0.15857889 | 0.21382383 | 0.34081706 | 0.26185338 | 0.32012987 | 0.59522921 | 0.36361349 | 0.53216171 | 0.40080503 | 0.09746676 | 0.41752621 | 0.43612032 | 0.47375812 | 0.1821218 | 0.42964639 | 0.23457101 | 0.24315484 |
| ENSMUSG00 | Gm15585 | 666 | 0.8935022 | 0.90034849 | 1.5252915 | 0.75250974 | 0.50589109 | 0.37201752 | 0.28416023 | 0.40142165 | 0.92762423 | 0.70796852 | 1.07601258 | 0.92188031 | 1.8517987 | 0.69735867 | 0.68050618 | 0.80490766 | 0.7481109 | 1.3215401 |

| | | | | | | | | | | | | | | | | | | | | |
|---|---|---|---|---|---|---|---|---|---|---|---|---|---|---|---|---|---|---|---|---|
| ENSMUSG00 | B430219N15 | 833 | 0.15307987 | 0.27994029 | 0.29035733 | 0 | 0.46225137 | 0.35692245 | 0.11359587 | 0.32094456 | 0.29666158 | 0.45282788 | 0 | 0.26802239 | 0.11844398 | 0 | 0 | 0.45967116 | 0 | 0.39622406 |
| ENSMUSG00 | 4931440J10 | 884 | 0.04808278 | 0.11305281 | 0.10944238 | 0 | 0.16334363 | 0.11211026 | 0.05352113 | 0 | 0 | 0.0533379 | 0.54044071 | 0.25255956 | 0 | 0.36776992 | 0 | 0.30320617 | 0.34684432 | 0 |
| ENSMUSG00 | Gm13091 | 466 | 1.27697954 | 2.43055451 | 1.55708789 | 1.5643277 | 1.54931085 | 2.55207212 | 1.72599902 | 0.9015374 | 0.97221389 | 1.41654473 | 1.53782055 | 0.7186566 | 0.84689987 | 1.19598508 | 1.76830712 | 1.64337371 | 1.56266017 | 3.77744939 |
| ENSMUSG00 | 0610005C13 | 3317 | 0.03844303 | 0.15064619 | 0.11666815 | 0.09614932 | 0.02902136 | 0.08963413 | 0.22821913 | 0.24179694 | 0.17383516 | 0.28429728 | 0.28806125 | 0.16827152 | 0.28257655 | 0.09801285 | 0.06210666 | 0.11543747 | 0.21953562 | 0.06633597 |
| ENSMUSG00 | 6030471H07 | 490 | 0 | 0 | 0 | 0 | 0 | 0 | 0 | 0 | 0 | 0 | 0 | 0 | 0 | 0 | 0 | 0 | 0 | 0 |
| ENSMUSG00 | BB218582 | 1472 | 11.492568 | 9.25609629 | 10.4173975 | 9.93552665 | 11.215523 | 9.05549275 | 9.86752203 | 8.84756061 | 7.72247171 | 11.3392316 | 10.4183164 | 11.9442484 | 12.9362229 | 9.33930018 | 7.13749779 | 11.0033471 | 10.310607 | 7.39932278 |
| ENSMUSG00 | Gm16617 | 710 | 1.13746246 | 1.78294363 | 1.77142506 | 1.60426212 | 0.33895721 | 1.81460715 | 1.06620122 | 0.59171328 | 1.2762019 | 0.19922832 | 0.40373204 | 0.78613609 | 0.27792629 | 0.39248525 | 0.17409108 | 0.05393043 | 0 | 0 |
| ENSMUSG00 | Gm12521 | 692 | 0 | 0 | 0 | 0 | 0 | 0 | 0 | 0 | 0 | 0 | 0 | 0 | 0 | 0 | 0 | 0.1106665 | 0 | 0 |
| ENSMUSG00 | Gm11309 | 537 | 0 | 0 | 0 | 0 | 0 | 0 | 0 | 0 | 0 | 0 | 0 | 0 | 0 | 0 | 0 | 0 | 0 | 0 |
| ENSMUSG00 | Gm13974 | 1445 | 0 | 0 | 0 | 0 | 0 | 0 | 0 | 0 | 0 | 0 | 0 | 0 | 0 | 0 | 0 | 0 | 0 | 0 |
| ENSMUSG00 | Gm11210 | 418 | 0 | 0 | 0 | 0 | 0 | 0 | 0 | 0.09136938 | 0 | 0 | 0 | 0 | 0 | 0 | 0 | 0 | 0 | 0 |
| ENSMUSG00 | Gm13425 | 671 | 0 | 0 | 0 | 0 | 0 | 0 | 0 | 0 | 0 | 0 | 0 | 0 | 0 | 0 | 0 | 0 | 0 | 0 |
| ENSMUSG00 | Gm14637 | 442 | 0 | 0 | 0 | 0 | 0 | 0 | 0 | 0 | 0 | 0 | 0 | 0 | 0 | 0 | 0 | 0 | 0 | 0 |
| ENSMUSG00 | Gm7855 | 686 | 0 | 0 | 0 | 0 | 0 | 0 | 0 | 0 | 0 | 0 | 0 | 0 | 0 | 0 | 0 | 0 | 0 | 0 |
| ENSMUSG00 | 6330418K02 | 1263 | 1.2452031 | 1.23967223 | 1.68522196 | 1.01006274 | 0.87651167 | 1.09855625 | 1.23619826 | 1.63292932 | 1.6305034 | 0.74664613 | 1.21045025 | 1.19320894 | 0.62494907 | 1.2502763 | 1.85945264 | 1.03078436 | 1.63865402 | 1.87283578 |
| ENSMUSG00 | Gm14376 | 295 | 0 | 0 | 0 | 0 | 0 | 0 | 0 | 0 | 0.13961531 | 0 | 0 | 0 | 0 | 0 | 0 | 0 | 0 | 0 |
| ENSMUSG00 | Gm11672 | 643 | 0 | 0.05180854 | 0.37615498 | 0 | 0.82340771 | 0.23119471 | 0.07358115 | 0.35638323 | 1.15296623 | 0.14665849 | 0.07430009 | 0.43402537 | 0.76721489 | 0.07223031 | 0 | 0.23819974 | 0.23842175 | 0.08555071 |
| ENSMUSG00 | Gm16013 | 351 | 0 | 0 | 0 | 0 | 0 | 0 | 0 | 0 | 0 | 0 | 0 | 0 | 0 | 0 | 0 | 0 | 0 | 0 |
| ENSMUSG00 | Gm11962 | 254 | 0 | 0.13115313 | 0.38089394 | 0.17937419 | 0.18949576 | 0.1950895 | 0.18627039 | 0.60145516 | 0.16215164 | 0.37126538 | 0 | 0 | 0 | 0.18285074 | 0 | 0 | 0.30178187 | 0.21657129 |
| ENSMUSG00 | B230208H11 | 1541 | 0.55165705 | 0.9079439 | 0.31391 | 0.6208838 | 0.68715271 | 0.90037413 | 0.52194389 | 0.32219418 | 0.3741799 | 1.34628874 | 0.58904881 | 1.4850371 | 0.28811632 | 0.96444568 | 1.01600203 | 0.86967635 | 1.31816595 | 0.35697019 |
| ENSMUSG00 | Gm15614 | 1533 | 0 | 0.0869221 | 0 | 0.02972019 | 0 | 0 | 0 | 0.0249135 | 0 | 0.03075714 | 0.03116436 | 0.07281887 | 0 | 0.21207346 | 0 | 0 | 0.02500085 | 0 |
| ENSMUSG00 | Gm12576 | 3034 | 0.12608655 | 0.23057705 | 0.39859534 | 0.18020189 | 0.11104926 | 0.19598972 | 0.09356496 | 0.17623389 | 0.20362483 | 0.12432618 | 0 | 0.11038035 | 0.06503878 | 0.03061575 | 0.21727913 | 0.11358453 | 0.01263227 | 0.1269162 |
| ENSMUSG00 | 2610206C17 | 1123 | 1.40043768 | 1.89850866 | 1.76608615 | 0.85198747 | 0.85720256 | 1.76501277 | 1.09539595 | 1.66645389 | 2.27387709 | 1.63746877 | 0.89338747 | 2.08749584 | 2.50393525 | 0.95121461 | 2.12795213 | 1.26157834 | 2.35486597 | 1.0286654 |
| ENSMUSG00 | Gm15406 | 356 | 0 | 0 | 0 | 0.38394138 | 0.67601016 | 0.55677229 | 1.19610705 | 0.42912812 | 0.23138492 | 0.66222898 | 0 | 0.31357114 | 0.4157184 | 0 | 0 | 0.10755789 | 0 | 0.15451996 |
| ENSMUSG00 | 4930479D17 | 737 | 0 | 0 | 0 | 0 | 0 | 0 | 0 | 0 | 0 | 0.38385919 | 0 | 0.1514672 | 0 | 0.4411243 | 0.16771325 | 0 | 0.36402182 | 0 |
| ENSMUSG00 | Gm12150 | 465 | 0 | 0.57312506 | 0.52014549 | 0.58788444 | 0.20701903 | 0.31969506 | 0 | 0 | 0 | 0.10139936 | 0 | 0 | 0.9548113 | 0 | 0 | 0 | 0 | 0 |
| ENSMUSG00 | Gm12119 | 697 | 0 | 0 | 0 | 0 | 0 | 0 | 0 | 0.4383633 | 0 | 0.2029442 | 0 | 0 | 0 | 0.99651408 | 0.41378894 | 0 | 0 | 0 |
| ENSMUSG00 | Snhg3 | 981 | 0.64992624 | 0.37353908 | 0.19724171 | 0.09288694 | 0.04906414 | 0.15153741 | 0.14468709 | 0.38932113 | 0.37785794 | 0.33644742 | 0.14610079 | 0.0568967 | 0.20114951 | 0.18937446 | 0.12599864 | 0.15612888 | 0.1172058 | 0.05607452 |
| ENSMUSG00 | Gm11537 | 789 | 0 | 0 | 0 | 0 | 0 | 0 | 0 | 0 | 0 | 0 | 0 | 0 | 0 | 0 | 0 | 0 | 0 | 0 |
| ENSMUSG00 | 4930480K23 | 4203 | 1.03153175 | 0.60237449 | 0.81715933 | 1.0081316 | 1.39711984 | 0.64844167 | 1.83487192 | 0.76330283 | 1.07792451 | 1.17792621 | 0.9775509 | 0.88975717 | 1.11504453 | 0.38675781 | 0.83324584 | 1.02946434 | 0.44682097 | 0.61513871 |
| ENSMUSG00 | B930086A06 | 4951 | 0 | 0 | 0 | 0 | 0 | 0 | 0 | 0 | 0 | 0 | 0 | 0 | 0 | 0 | 0 | 0 | 0 | 0 |
| ENSMUSG00 | Gm11713 | 521 | 0.08158383 | 0.19182089 | 0.2785424 | 0 | 0.18476746 | 0.09511081 | 0.27243385 | 0.14661191 | 0.39526407 | 0 | 0.09169858 | 0 | 0.94686981 | 0.17828824 | 0.07908168 | 0 | 0.2942518 | 0.1055837 |
| ENSMUSG00 | Gm15893 | 1556 | 0.08195085 | 0.21409315 | 0.03108839 | 0 | 0 | 0 | 0.12162642 | 0.02454525 | 0.18528638 | 0.09090753 | 0.33774071 | 0.32284124 | 0 | 0.02984839 | 0 | 0 | 0 | 0.10605869 |
| ENSMUSG00 | 4930545L23 | 2496 | 0.28949839 | 0.02669302 | 0.07752168 | 0.09126812 | 0.30853797 | 0.05955857 | 0.1137324 | 0.16831588 | 0.01650101 | 0.09445253 | 0.05742183 | 0.26834453 | 0.17787951 | 0.11164444 | 0.16507034 | 0.06136315 | 0.04606526 | 0.19835014 |
| ENSMUSG00 | Gm11587 | 499 | 0 | 0 | 0 | 0 | 0 | 0 | 0 | 0 | 0 | 0 | 0 | 0 | 0 | 0 | 0 | 0 | 0 | 0 |
| ENSMUSG00 | Gm12847 | 409 | 0.41569854 | 0.08144962 | 0 | 0.33418859 | 0 | 0 | 0 | 0 | 0 | 0 | 0 | 0 | 0 | 0 | 0 | 0.5617204 | 0 | 0.53798637 |
| ENSMUSG00 | Gm15234 | 1056 | 5.43390034 | 5.83606575 | 5.86345822 | 3.71046384 | 5.69743415 | 4.73941865 | 5.0180114 | 6.00372992 | 10.3356313 | 5.49198532 | 3.9360051 | 4.43988225 | 4.29785643 | 3.82635947 | 4.56494511 | 3.62600449 | 3.23015192 | 4.58409223 |
| ENSMUSG00 | Gm12326 | 736 | 0 | 0 | 0 | 0 | 0 | 0 | 0 | 0 | 0 | 0 | 0 | 0 | 0 | 0 | 0 | 0 | 0 | 0 |
| ENSMUSG00 | 8430437L04 | 1656 | 0 | 0 | 0 | 0 | 0 | 0 | 0 | 0 | 0 | 0 | 0 | 0 | 0 | 0 | 0 | 0 | 0 | 0 |
| ENSMUSG00 | Gm16833 | 200 | 0 | 0 | 0 | 0 | 0 | 0 | 0 | 0 | 0 | 0 | 0 | 0 | 0 | 0.69666131 | 0 | 0 | 0 | 0 |
| ENSMUSG00 | Gm15881 | 941 | 0 | 0 | 0 | 0 | 0 | 0 | 0 | 0 | 0 | 0 | 0 | 0 | 0 | 0 | 0 | 0 | 0 | 0 |
| ENSMUSG00 | 5330430P22 | 642 | 1.72139342 | 0.88211714 | 2.71253441 | 0.78064095 | 1.87429609 | 0.92621932 | 3.09522199 | 1.90367115 | 2.05291044 | 1.3954258 | 2.00922723 | 2.69514882 | 1.99786581 | 2.02559883 | 1.34771445 | 2.14713686 | 2.14913814 | 2.31346711 |
| ENSMUSG00 | 3010003L10 | 868 | 0 | 0 | 0 | 0 | 0 | 0 | 0 | 0 | 0 | 0 | 0 | 0 | 0 | 0 | 0 | 0 | 0 | 0 |
| ENSMUSG00 | Gm13264 | 299 | 0 | 0 | 0 | 0 | 0 | 0 | 0 | 0 | 0 | 0 | 0 | 0 | 0 | 0 | 0 | 0 | 0 | 0 |
| ENSMUSG00 | Gm4890 | 2505 | 0.84840671 | 0.55853954 | 0.96553953 | 0.38194887 | 0.57643022 | 0.53410132 | 1.13323782 | 1.60087914 | 1.56196367 | 0.84701862 | 0.80101727 | 1.18093019 | 2.0874983 | 1.87259594 | 1.25002724 | 1.52856716 | 0.74969603 | 1.95441537 |
| ENSMUSG00 | 1700013G23 | 524 | 0.97340098 | 1.65292987 | 0.73852718 | 1.13033125 | 0.64298371 | 1.22936171 | 2.34757567 | 3.20699562 | 4.48021259 | 1.88962742 | 2.37051321 | 1.70429504 | 3.20092593 | 3.45671643 | 2.64435201 | 1.38839989 | 1.90168649 | 1.36472975 |
| ENSMUSG00 | Gm13814 | 3075 | 0.16587386 | 0.11916808 | 0.36181828 | 0.296332 | 0.26609519 | 0.19337652 | 0.13847613 | 0.33534792 | 0.46878961 | 0.18400274 | 0.34180458 | 0 | 0.4331583 | 0.19634899 | 0.41536528 | 0.16187899 | 0.12463836 | 0 |
| ENSMUSG00 | Gm11574 | 943 | 0.27044651 | 0.60055059 | 0.10259497 | 0.19326 | 0.10208255 | 0.26273984 | 0.60207015 | 0.68851627 | 0.2620563 | 0.30000447 | 0.60795281 | 0.23675785 | 1.15090369 | 1.0835312 | 1.048608 | 0.48726118 | 0.73157301 | 0.23333661 |
| ENSMUSG00 | 1700125G02 | 516 | 0 | 0 | 0 | 0 | 0 | 0 | 0 | 0 | 0 | 0 | 0 | 0 | 0 | 0 | 0 | 0 | 0 | 0 |
| ENSMUSG00 | Gm15581 | 5635 | 0.01508613 | 0 | 0 | 0 | 0 | 0 | 0 | 0 | 0.00730905 | 0 | 0 | 0 | 0 | 0 | 0 | 0 | 0 | 0 |
| ENSMUSG00 | 1700081H22 | 1158 | 0 | 0 | 0 | 0 | 0 | 0 | 0 | 0 | 0 | 0 | 0 | 0 | 0 | 0 | 0 | 0 | 0 | 0 |
| ENSMUSG00 | 9330111N05 | 5464 | 0 | 0.01829039 | 0 | 0 | 0 | 0 | 0 | 0 | 0.0075378 | 0.04314669 | 0.00874359 | 0 | 0 | 0 | 0.00754055 | 0.08409357 | 0.02805732 | 0 |
| ENSMUSG00 | Gm14235 | 364 | 0.35031738 | 0.27455682 | 0 | 0 | 0.13223056 | 0 | 0 | 0 | 0.56574884 | 0.6476745 | 0 | 0.30667946 | 0.40658174 | 0.12759365 | 0 | 0.63116386 | 0.10529202 | 0.30224784 |
| ENSMUSG00 | 5033428I22 | 2035 | 0 | 0 | 0 | 0 | 0 | 0 | 0 | 0 | 0 | 0 | 0 | 0 | 0 | 0 | 0 | 0 | 0 | 0 |
| ENSMUSG00 | Gm15777 | 872 | 0.04874447 | 0.30562288 | 0.66569078 | 0.31349342 | 0.38638012 | 0.79557141 | 0.05425766 | 0.74457665 | 0.70848364 | 0.27035954 | 0.16436339 | 0.64008787 | 0.1697199 | 0.42609254 | 0.37799593 | 0 | 0.48347393 | 0.18925151 |
| ENSMUSG00 | 9530052E02 | 1844 | 0.43796006 | 0.18065561 | 0.02623293 | 0.17295407 | 0.10440764 | 0.26872415 | 0.33354925 | 0.10355857 | 0.26802505 | 0.20455837 | 0.33680828 | 0.24215038 | 0.240774 | 0 | 0.71499446 | 0.20764971 | 0.0831373 | 0.32814543 |
| ENSMUSG00 | E130006D01 | 1249 | 0 | 0 | 0 | 0 | 0 | 0 | 0 | 0 | 0 | 0 | 0 | 0 | 0 | 0 | 0 | 0 | 0 | 0 |
| ENSMUSG00 | Gm1078 | 3878 | 0.05480296 | 0.30065789 | 0.34926737 | 0.05874297 | 0.02482307 | 0.06388955 | 0.04880111 | 0.03939392 | 0.14868778 | 0.49849893 | 0.13551432 | 0.0143929 | 0 | 0 | 0 | 0.07899042 | 0.11859607 | 0.24114358 |
| [illegible] | [illegible] | [illegible] | [illegible] | [illegible] | [illegible] | [illegible] | [illegible] | [illegible] | [illegible] | [illegible] | [illegible] | [illegible] | [illegible] | [illegible] | [illegible] | [illegible] | [illegible] | [illegible] | [illegible] | [illegible] |

| | | | | | | | | | | | | | | | | | | | | |
|---|---|---|---|---|---|---|---|---|---|---|---|---|---|---|---|---|---|---|---|---|
| ENSMUSG00 | Gm12434 | 651 | 0 | 0 | 0 | 0 | 0 | 0 | 0 | 0 | 0 | 0 | 0 | 0 | 0 | 0 | 0 | 0 | 0 | 0 |
| ENSMUSG00 | 1700034P13R | 2844 | 0.01494556 | 0.05856697 | 0 | 0 | 0 | 0.06969442 | 0.06654385 | 0 | 0.27515605 | 0.06631604 | 0 | 0.03925152 | 0 | 0.0326611 | 0 | 0.10770916 | 0 | 0 |
| ENSMUSG00 | Gm16220 | 403 | 0 | 0 | 0 | 0 | 0 | 0 | 0 | 0 | 0 | 0 | 0 | 0 | 0 | 0 | 0 | 0 | 0 | 0 |
| ENSMUSG00 | Gm15718 | 309 | 1.51312924 | 0.97027847 | 1.09584049 | 0 | 0.62306697 | 0.80182417 | 0.61246186 | 0.61800004 | 2.26592482 | 0.76295636 | 0.61844606 | 1.4450657 | 2.07545283 | 0.75152245 | 0.80003021 | 2.85010994 | 2.23259982 | 0.178023 |
| ENSMUSG00 | 4930556N08 | 2015 | 0.04218876 | 0 | 0.04801343 | 0.06783282 | 0.26275492 | 0.2705112 | 0.09392095 | 0 | 0.18395962 | 0.09359941 | 0 | 0.30470089 | 0 | 0.0921967 | 0.10223711 | 0.13301948 | 0 | 0 |
| ENSMUSG00 | Gm12059 | 3103 | 0.12328282 | 0.02147141 | 0.03117856 | 0.10278031 | 0.06204566 | 0.0319386 | 0.39643237 | 0.1969315 | 0.46455948 | 0.28870879 | 0.18475653 | 0.12591352 | 0.1748795 | 0.44902437 | 0.61078684 | 0.04935947 | 0.34583832 | 0.12409402 |
| ENSMUSG00 | Gm9855 | 2434 | 1.76377271 | 4.5575981 | 4.76978114 | 1.83442167 | 4.52843489 | 3.84776773 | 2.33258894 | 3.57759562 | 4.17956836 | 1.76282416 | 3.9256334 | 3.43974913 | 0.97285622 | 3.49189318 | 2.84382144 | 0.83377247 | 3.27521354 | 2.5764331 |
| ENSMUSG00 | Gm12992 | 5867 | 0.55060395 | 0.75517555 | 0.89870714 | 0.51253263 | 0.50863802 | 0.55743658 | 0.66126464 | 0.55983409 | 1.20744515 | 0.92420843 | 1.03416051 | 0.65643101 | 1.13513019 | 0.87077716 | 1.02529865 | 1.01812421 | 1.08439838 | 0.87196982 |
| ENSMUSG00 | Gm13447 | 710 | 0 | 0 | 0 | 0 | 0 | 0 | 0 | 0 | 0 | 0 | 0 | 0 | 0 | 0 | 0 | 0 | 0 | 0 |
| ENSMUSG00 | Gm14817 | 641 | 0 | 0.10394039 | 0 | 0 | 0 | 0.23191607 | 0 | 0 | 0.1927606 | 0.07355804 | 0 | 0 | 0.07696087 | 0 | 0 | 0.23894295 | 0 | 0 |
| ENSMUSG00 | A830052D11 | 1127 | 0.41486862 | 0.82764955 | 0.55799103 | 0.48512203 | 0.25624804 | 0.61556191 | 1.17547028 | 0.10166567 | 1.53490121 | 0.37653623 | 0.46630394 | 0.24762938 | 1.88222932 | 1.64841481 | 0.43870335 | 0.16987847 | 0.7141546 | 1.02501441 |
| ENSMUSG00 | C330046G13 | 1330 | 0.41546413 | 0.30056747 | 0.18185538 | 0.10276927 | 0.36189416 | 0.18628847 | 0.07114689 | 0 | 0 | 0 | 0.07184204 | 0 | 0.22255 | 0.34920366 | 0 | 0 | 0.08645029 | 0.37224208 |
| ENSMUSG00 | Gm16193 | 775 | 0.05484539 | 0.17193752 | 0 | 0 | 0 | 0 | 0 | 0 | 0 | 0 | 0 | 0 | 0 | 0 | 0 | 0.09881447 | 0.14835986 | 0 |
| ENSMUSG00 | Gm16122 | 1156 | 0.40446102 | 0 | 0 | 0 | 0.04163661 | 0.08573137 | 0 | 0 | 0.07125695 | 0 | 0.04132782 | 0 | 0 | 0.28123582 | 0.14256594 | 0.29811027 | 0.03315424 | 0 |
| ENSMUSG00 | Gm12845 | 240 | 0 | 0 | 0 | 0 | 0 | 0 | 0 | 0 | 0 | 0 | 0 | 0 | 0 | 0 | 0 | 0 | 0 | 0 |
| ENSMUSG00 | Gm12022 | 1338 | 0 | 0 | 0 | 0 | 0 | 0 | 0 | 0 | 0 | 0 | 0 | 0 | 0 | 0 | 0 | 0 | 0 | 0 |
| ENSMUSG00 | 4930555O08 | 666 | 0 | 0 | 0 | 0 | 0 | 0 | 0 | 0 | 0 | 0 | 0 | 0 | 0 | 0 | 0 | 0 | 0 | 0 |
| ENSMUSG00 | 4930594O21 | 1152 | 0 | 0 | 0 | 0 | 0 | 0 | 0 | 0 | 0 | 0 | 0 | 0 | 0 | 0 | 0 | 0 | 0 | 0 |
| ENSMUSG00 | Gm16349 | 392 | 0.97588414 | 0.42490937 | 0 | 0.11622715 | 0 | 0 | 0 | 0 | 0 | 0.24056481 | 0.60937447 | 0.14238689 | 0.37754018 | 0.23695963 | 0.10510601 | 1.07448133 | 0 | 0 |
| ENSMUSG00 | Gm16328 | 195 | 0 | 0 | 0 | 0 | 0 | 0 | 0 | 0 | 0 | 0 | 0 | 0 | 0 | 0 | 0 | 0 | 0 | 0 |
| ENSMUSG00 | Gm13327 | 2739 | 0 | 0 | 0 | 0 | 0 | 0 | 0 | 0 | 0 | 0 | 0 | 0 | 0 | 0 | 0 | 0 | 0 | 0 |
| ENSMUSG00 | Gm13982 | 752 | 0 | 0 | 0 | 0 | 0 | 0 | 0 | 0 | 0 | 0 | 0 | 0 | 0 | 0 | 0 | 0 | 0 | 0 |
| ENSMUSG00 | Gm14267 | 601 | 0 | 0 | 0 | 0 | 0 | 0 | 0 | 0 | 0 | 0 | 0 | 0 | 0 | 0 | 0 | 0 | 0 | 0 |
| ENSMUSG00 | 9130409I20R | 1499 | 0.48204669 | 0.51113847 | 0.38724641 | 0.42552009 | 0.25687484 | 0.36362913 | 2.52502623 | 3.49056646 | 4.12139918 | 1.0065527 | 1.11549269 | 2.04793959 | 2.83025009 | 6.16569272 | 3.16089321 | 1.17502865 | 1.48293877 | 2.09174055 |
| ENSMUSG00 | 4930469K13R | 693 | 38.0890538 | 26.0542405 | 22.4067868 | 29.1906258 | 26.6705032 | 28.4588571 | 72.8465057 | 45.5222574 | 38.7497955 | 61.8470264 | 40.4673891 | 42.5262191 | 145.219497 | 84.1087008 | 70.0961535 | 256.210024 | 61.1119164 | 85.2522087 |
| ENSMUSG00 | Gm15253 | 272 | 0 | 0 | 0 | 0 | 0 | 0 | 0 | 0 | 0 | 0 | 0 | 0 | 0 | 0 | 0 | 0 | 0 | 0 |
| ENSMUSG00 | Gm9498 | 290 | 0 | 0 | 0 | 0 | 0 | 0 | 0 | 0 | 0 | 0 | 0 | 0 | 0 | 0 | 0 | 0 | 0 | 0 |
| ENSMUSG00 | Gm16153 | 4116 | 1.31150567 | 0.93075385 | 1.21051351 | 0.6088089 | 0.81857013 | 0.72234306 | 1.26443019 | 1.46608347 | 1.31084392 | 0.68732804 | 0.82410643 | 1.01704925 | 0.85096359 | 1.02682506 | 1.27128222 | 0.9488926 | 1.02426935 | 0.49449391 |
| ENSMUSG00 | Gm15395 | 699 | 0 | 0 | 0 | 0.06518032 | 0 | 0 | 0 | 0 | 0 | 0 | 0 | 0 | 0 | 0 | 0 | 0 | 0 | 0 |
| ENSMUSG00 | Gm16838 | 813 | 0 | 0 | 0 | 0 | 0 | 0 | 0 | 0 | 0 | 0 | 0 | 0 | 0 | 0.1142536 | 0 | 0 | 0 | 0 |
| ENSMUSG00 | Gm12629 | 708 | 0 | 0 | 0 | 0 | 0 | 0 | 0 | 0 | 0 | 0 | 0 | 0 | 0 | 0 | 0 | 0 | 0 | 0 |
| ENSMUSG00 | Gm14216 | 896 | 0 | 0 | 0 | 0 | 0 | 0 | 0 | 0 | 0 | 0 | 0 | 0 | 0 | 0 | 0 | 0 | 0 | 0 |
| ENSMUSG00 | Gm2061 | 2758 | 0 | 0 | 0 | 0 | 0 | 0 | 0 | 0 | 0 | 0 | 0 | 0 | 0 | 0 | 0 | 0 | 0 | 0 |
| ENSMUSG00 | Gm13371 | 467 | 0 | 0 | 0 | 0 | 0 | 0 | 0 | 0 | 0 | 0 | 0 | 0 | 0 | 0 | 0 | 0 | 0 | 0 |
| ENSMUSG00 | Gm14967 | 333 | 0 | 0 | 0 | 0 | 0 | 0 | 0 | 0 | 0 | 0 | 0 | 0 | 0 | 0 | 0 | 0 | 0 | 0 |
| ENSMUSG00 | Gm15243 | 380 | 0 | 0 | 0 | 0 | 0 | 0 | 0.2490141 | 0 | 0.3251567 | 0 | 0 | 0 | 0 | 0 | 0 | 0 | 0 | 0 |
| ENSMUSG00 | 2700086A05R | 2944 | 0.77964657 | 1.06365899 | 0.72297396 | 0.8511744 | 1.1117428 | 1.09406511 | 1.14103268 | 1.10270184 | 1.32904857 | 1.5054912 | 1.54165117 | 1.0806701 | 2.21189303 | 1.60913618 | 2.08526896 | 1.44370157 | 1.22373366 | 0.63661023 |
| ENSMUSG00 | Gm16022 | 528 | 0 | 0 | 0.36646614 | 0 | 0 | 0 | 0 | 0.21700229 | 0.23401429 | 0 | 0 | 0 | 0 | 0 | 0 | 0 | 0 | 0 |
| ENSMUSG00 | Gm14298 | 236 | 0 | 0 | 0 | 0 | 0 | 0 | 0 | 0 | 0 | 0 | 0 | 0 | 0 | 0 | 0 | 0 | 0 | 0 |
| ENSMUSG00 | Hbb-bh0 | 128 | 0 | 0 | 0 | 0 | 0 | 0 | 0 | 0 | 0 | 0 | 0 | 0 | 0 | 0 | 0 | 0 | 0 | 0 |
| ENSMUSG00 | Gm12968 | 1452 | 0 | 0 | 0 | 0 | 0 | 0 | 0 | 0 | 0 | 0 | 0 | 0 | 0 | 0 | 0 | 0.21096753 | 0 | 0 |
| ENSMUSG00 | Gm16722 | 4624 | 0 | 0.00720435 | 0.0104614 | 0.01970633 | 0.01040915 | 0.01071642 | 0.01023198 | 0 | 0.01781424 | 0.01019695 | 0 | 0 | 0 | 0.02008827 | 0.01782074 | 0.01656168 | 0.04973135 | 0.08327503 |
| ENSMUSG00 | Gm16723 | 1112 | 0 | 0 | 0 | 0 | 0 | 0 | 0 | 0 | 0 | 0 | 0 | 0 | 0 | 0 | 0.03705176 | 0 | 0.06893219 | 0 |
| ENSMUSG00 | 4921531C22 | 1964 | 5.88666389 | 5.20725994 | 4.63046014 | 4.96439078 | 5.63663057 | 3.30519761 | 8.5760253 | 6.22279469 | 6.87839978 | 7.058201 | 7.80843262 | 7.84374892 | 7.10841782 | 7.18890151 | 9.50321019 | 7.25259977 | 9.79623271 | 9.27088306 |
| ENSMUSG00 | Gm16046 | 1152 | 0 | 0.08675233 | 0 | 0 | 0 | 0 | 0.04107003 | 0 | 0 | 0 | 0 | 0 | 0 | 0 | 0 | 0 | 0 | 0 |
| ENSMUSG00 | Gm9428 | 767 | 0 | 0 | 0 | 0 | 0 | 0 | 0 | 0 | 0 | 0 | 0 | 0 | 0 | 0 | 0 | 0 | 0 | 0 |
| ENSMUSG00 | Gm12212 | 376 | 0 | 0.26579437 | 0.90057104 | 0.12117299 | 0 | 0 | 0.62915796 | 0 | 0 | 0.25080161 | 0 | 0 | 0 | 0 | 0 | 0 | 0.61158984 | 0.58520326 |
| ENSMUSG00 | Gm16063 | 1697 | 0.12523623 | 0.03926092 | 0.17103193 | 0.02684799 | 0.51053307 | 0.0292002 | 0.25092169 | 0.06751751 | 0.26697211 | 0 | 0.25337338 | 0.42758021 | 0.78489203 | 3.09262338 | 1.69963383 | 1.26356748 | 0.8808047 | 2.20425413 |
| ENSMUSG00 | Gm14211 | 406 | 0 | 0 | 0 | 0.11221932 | 0 | 0 | 0 | 0 | 0 | 0 | 0 | 0 | 0 | 0 | 0.10148166 | 0 | 0 | 0 |
| ENSMUSG00 | Gm12676 | 806 | 0.26367975 | 0.08266227 | 0.36010072 | 0 | 0.35830216 | 0.30739909 | 1.0566107 | 0.14215534 | 0.0510999 | 0.93599411 | 0.35564485 | 0 | 0.67326438 | 0 | 0.05111856 | 0.38005566 | 1.09367845 | 0.81899414 |
| ENSMUSG00 | Gm15163 | 2521 | 0.08430221 | 0.17178406 | 0.07675292 | 0.16265347 | 0.15273915 | 0.1179359 | 0.13137198 | 0.25754496 | 0.16337372 | 0.26184444 | 0.13265558 | 0.06642086 | 0.15654714 | 0.01842288 | 0.08171669 | 0.12150927 | 0.07601408 | 0.10910176 |
| ENSMUSG00 | Gm15124 | 951 | 0 | 0 | 0 | 0 | 0 | 0 | 0 | 0 | 0 | 0 | 0 | 0 | 0 | 0 | 0 | 0 | 0 | 0 |
| ENSMUSG00 | Gm15736 | 418 | 0 | 0 | 0 | 0 | 0 | 0 | 0 | 0 | 0 | 0 | 0 | 0 | 0 | 0 | 0 | 0 | 0 | 0 |
| ENSMUSG00 | Gm13008 | 505 | 0 | 0 | 0 | 0 | 0 | 0 | 0 | 0 | 0 | 0 | 0 | 0 | 0 | 0 | 0 | 0 | 0 | 0 |
| ENSMUSG00 | Tsix | 4302 | 0 | 0.01548717 | 0 | 0 | 0 | 0 | 0 | 0 | 0 | 0 | 0 | 0 | 0 | 0 | 0 | 0 | 0 | 0 |
| ENSMUSG00 | Gm13388 | 422 | 0 | 0 | 0 | 0.21592912 | 0 | 0.23484708 | 0 | 0.09050332 | 0.68318865 | 0 | 0 | 0 | 0 | 0 | 0.29290206 | 0.09073604 | 0 | 0.26070667 |
| ENSMUSG00 | 4930546G22 | 515 | 0 | 0 | 0 | 0 | 0 | 0 | 0 | 0 | 0 | 0 | 0 | 0 | 0 | 0 | 0 | 0 | 0 | 0 |
| ENSMUSG00 | Rpl9-ps3 | 1153 | 0 | 0 | 0 | 0 | 0.04174495 | 0 | 0 | 0.16562187 | 0.21432706 | 0 | 0.20717675 | 0 | 0 | 0.20140541 | 0.03573422 | 0.09962864 | 0.0997215 | 0 |
| ENSMUSG00 | 4933424L21R | 1182 | 0 | 0 | 0 | 0 | 0 | 0 | 0 | 0 | 0 | 0 | 0 | 0 | 0.08347194 | 0.11787839 | 0 | 0.06478952 | 0.03242496 | 0 |
| ENSMUSG00 | Gm7854 | 350 | 1.70020704 | 2.66503154 | 3.17883199 | 4.1655812 | 3.16295498 | 2.40684706 | 2.56840257 | 1.85505955 | 4.58935461 | 4.17620515 | 6.14249466 | 4.14630636 | 1.26853502 | 4.24631656 | 2.70753082 | 2.73504339 | 3.9421334 | 8.01561269 |
| ENSMUSG00 | Gm12796 | 341 | 0 | 0 | 0 | 0 | 0 | 0 | 0 | 0 | 0 | 0 | 0 | 0 | 0 | 0.13619967 | 0 | 0 | 0 | 0 |
| ENSMUSG00 | Gm4916 | 1705 | 0 | 0 | 0 | 0 | 0 | 0 | 0 | 0 | 0 | 0 | 0 | 0 | 0 | 0 | 0 | 0 | 0 | 0 |
| ENSMUSG00 | Gm15915 | 902 | 0.04712326 | 0.25852579 | 0.32177515 | 0.15153341 | 0.26680667 | 0.05493651 | 0 | 0 | 0 | 0.10454701 | 0 | 0 | 0 | 0 | 0.045678 | 0 | 0 | 0 |
| ENSMUSG00 | Gm13096 | 249 | 0 | 0 | 0 | 0 | 0 | 0 | 0 | 0.30676628 | 0 | 0 | 0 | 0 | 0 | 0 | 0 | 0 | 0.30784174 | 0 |
| ENSMUSG00 | Gm15873 | 759 | 0.05600155 | 0 | 0 | 0 | 0 | 0 | 0 | 0 | 0 | 0 | 0 | 0 | 0 | 0 | 0 | 0.10089752 | 0 | 0 |
| ENSMUSG00 | 9630001P10R | 5003 | 0.00849594 | 0.03329292 | 0 | 0.00910674 | 0.05772367 | 0.0099046 | 0.08511176 | 0.229017 | 0.23873855 | 0.05654692 | 0.05729557 | 0.12272083 | 0.16762794 | 0.12996547 | 0.07411833 | 0 | 0.05362464 | 0.05497612 |
| ENSMUSG00 | 4930471M09 | 1109 | 0.11498244 | 0.15019339 | 0.26171432 | 0 | 0.21700597 | 0.04468236 | 0.63993705 | 0 | 0.03713843 | 0.04251641 | 0 | 0 | 0.17793297 | 0.167517 | 0.26006392 | 0 | 0 | 0 |
| ENSMUSG00 | Gm14339 | 421 | 0 | 0 | 0 | 0 | 0 | 0 | 0 | 0 | 0 | 0 | 0 | 0 | 0 | 0 | 0 | 0 | 0 | 0 |
| ENSMUSG00 | Gm13307 | 825 | 0 | 0.16151706 | 0 | 0.16567652 | 0 | 0 | 0.0573487 | 0 | 0.0998461 | 0 | 0 | 0.20296605 | 0 | 0 | 0 | 0.04641286 | 0 | 0 |
| ENSMUSG00 | Gm16229 | 1230 | 0 | 0 | 0.11798422 | 0 | 0 | 0 | 0 | 0 | 0 | 0 | 0 | 0 | 0 | 0 | 0 | 0 | 0 | 0.04472285 |
| ENSMUSG00 | Gm4279 | 2327 | 0 | 0.02863162 | 0.02078794 | 0 | 0 | 0.04258937 | 0.20332049 | 0 | 0.07079762 | 0 | 0.14371496 | 0 | 0 | 0 | 0.03541174 | 0 | 0 | 0 |
| ENSMUSG00 | Gm11641 | 921 | 0.04615111 | 0.36170352 | 0.63027401 | 0.24734552 | 0.10452101 | 0.26901593 | 0.05137099 | 0 | 0.49191278 | 0.61434141 | 0.20749168 | 0.36361995 | 0.53563428 | 0.70599047 | 0.84997781 | 0.7067756 | 0.2912965 | 0.47782069 |
| ENSMUSG00 | Gm11684 | 755 | 0 | 0 | 0 | 0 | 0 | 0 | 0 | 0 | 0 | 0 | 0 | 0 | 0 | 0 | 0 | 0 | 0 | 0 |
| ENSMUSG00 | 4930550G17 | 665 | 0.31958779 | 0.30056747 | 0.72742151 | 0 | 0.07237883 | 0.37257694 | 0.85376263 | 0.86148276 | 2.53931902 | 0.07090331 | 0 | 0 | 0.89020001 | 0.13968147 | 0.6815295 | 0 | 0.28816765 | 0 |
| ENSMUSG00 | Gm16682 | 1273 | 0.10016931 | 0.10467524 | 0.37999631 | 0.03579029 | 0.11342951 | 0.2335557 | 0.14866513 | 0.36002265 | 0.25883121 | 0.03703904 | 0.11258828 | 0.1315373 | 0.34877239 | 0 | 0.06473143 | 0.27071129 | 0.06021413 | 0 |
| ENSMUSG00 | Gm13442 | 661 | 0 | 0 | 0 | 0 | 0 | 0 | 0 | 0.05777973 | 0 | 0 | 0 | 0.08444124 | 0 | 0 | 0 | 0 | 0 | 0 |
| ENSMUSG00 | Gm12335 | 613 | 0 | 0.05434404 | 0.47347664 | 0.22297412 | 0.47111181 | 0 | 0.07718218 | 0 | 0.06718844 | 0 | 0.07793631 | 0.81947955 | 0.24142863 | 0 | 0 | 0.31232143 | 0.06252251 | 0.17947506 |
| ENSMUSG00 | Gm15862 | 359 | 0 | 0 | 0 | 0 | 0 | 0 | 0 | 0 | 0 | 0 | 0 | 0 | 0 | 0 | 0 | 0 | 0 | 0 |
| ENSMUSG00 | 4930505K14R | 512 | 0 | 0 | 0 | 0 | 0 | 0 | 0 | 0 | 0 | 0 | 0 | 0 | 0 | 0 | 0 | 0 | 0 | 0 |
| ENSMUSG00 | 5430405H02 | 1479 | 2.78769579 | 2.29744098 | 2.94362254 | 2.89569856 | 3.41707369 | 3.01537933 | 3.10299517 | 3.22789067 | 3.28601005 | 3.25177264 | 3.23022032 | 4.07578875 | 4.03594455 | 3.14023579 | 3.90007967 | 4.99667832 | 3.16146601 | 4.24005285 |
| ENSMUSG00 | Gm13536 | 764 | 0 | 0 | 0 | 0 | 0 | 0 | 0 | 0 | 0 | 0 | 0 | 0 | 0 | 0 | 0 | 0 | 0 | 0 |
| ENSMUSG00 | 8430419K02R | 2581 | 0 | 0 | 0 | 0 | 0 | 0 | 0 | 0 | 0 | 0 | 0 | 0 | 0 | 0 | 0 | 0 | 0.16334338 | 0 |
| ENSMUSG00 | Gm12259 | 326 | 0.13038398 | 0.51093396 | 0.14838506 | 0.27951561 | 0 | 0 | 0 | 0.35146383 | 0.50535602 | 0.28926812 | 0 | 0 | 0 | 0.14246653 | 0.75831084 | 1.05710266 | 0.35269598 | 0 |
| ENSMUSG00 | Gm14372 | 2190 | 0 | 0 | 0 | 0 | 0 | 0 | 0 | 0 | 0 | 0 | 0 | 0 | 0 | 0 | 0 | 0 | 0 | 0 |
| ENSMUSG00 | 4930455B14R | 982 | 0 | 0 | 0 | 0 | 0 | 0 | 0 | 0 | 0 | 0 | 0 | 0 | 0 | 0 | 0 | 0 | 0 | 0 |
| ENSMUSG00 | Gm11194 | 3622 | 0.34032305 | 0.47826353 | 0.60099637 | 0.74215947 | 0.41195186 | 0.68405209 | 0 | 0.09490106 | 0.2842802 | 0.05207146 | 0.39570645 | 0.04623053 | 0 | 0.08975942 | 0 | 0.03171503 | 0.02116306 | 0.07593747 |
| ENSMUSG00 | Gm12280 | 1622 | 0 | 0 | 0.02982338 | 0 | 0 | 0 | 0 | 0 | 0 | 0 | 0 | 0 | 0 | 0.08590152 | 0 | 0 | 0.09451615 | 0 |
| ENSMUSG00 | Gm12843 | 1012 | 0.46201278 | 0.65835759 | 0.23899966 | 0 | 0.23780595 | 0.63654697 | 0.32726161 | 0.15095811 | 0.77326462 | 0.65228246 | 0.6609184 | 0.44123053 | 0.43872258 | 0.09178673 | 0.244278 | 0.22701941 | 0.26510284 | 0 |
| ENSMUSG00 | Gm14009 | 652 | 0 | 0 | 0 | 0 | 0 | 0 | 0 | 0 | 0 | 0 | 0 | 0 | 0 | 0 | 0 | 0 | 0 | 0 |
| ENSMUSG00 | Gm15412 | 1093 | 0 | 0 | 0 | 0 | 0 | 0 | 0 | 0 | 0 | 0 | 0 | 0 | 0 | 0 | 0 | 0 | 0 | 0 |
| ENSMUSG00 | Gm16131 | 657 | 0 | 0 | 0 | 0 | 0 | 0 | 0 | 0 | 0 | 0 | 0 | 0 | 0 | 0 | 0 | 0 | 0 | 0 |
| ENSMUSG00 | Gm15886 | 1452 | 0 | 0.04588553 | 0 | 0.03137813 | 0.06629742 | 0 | 0 | 0.18412315 | 0.34038443 | 0.09741881 | 0 | 0 | 0.10192545 | 0 | 0.19863009 | 0 | 0.02639552 | 0 |
| ENSMUSG00 | Gm13496 | 1163 | 0 | 0 | 0 | 0 | 0 | 0.12782304 | 0 | 0 | 0 | 0 | 0 | 0 | 0 | 0 | 0 | 0 | 0 | 0 |
| ENSMUSG00 | Gm15279 | 287 | 0 | 0 | 0 | 0 | 0 | 0 | 0 | 0 | 0 | 0 | 0 | 0 | 0 | 0 | 0 | 0 | 0 | 0 |
| ENSMUSG00 | 1700019B21R | 774 | 0 | 0 | 0 | 0 | 0 | 0 | 0 | 0 | 0 | 0 | 0 | 0 | 0.06373633 | 0 | 0 | 0 | 0 | 0 |
| ENSMUSG00 | 1700061E18R | 586 | 0 | 0 | 0 | 0 | 0 | 0 | 0 | 0 | 0 | 0 | 0 | 0 | 0 | 0 | 0 | 0 | 0 | 0 |
| ENSMUSG00 | Gm14302 | 379 | 0.22430172 | 0.61527773 | 1.40398109 | 0.24042767 | 0.12699716 | 0.261492 | 0.99868452 | 0.50385755 | 0.54335773 | 0.87085732 | 0 | 0.44181263 | 1.82228718 | 0.24508753 | 0.54355615 | 1.41442877 | 0.20224959 | 0.58057105 |
| ENSMUSG00 | 4930455G09 | 628 | 0.74451741 | 0.95482818 | 1.15541872 | 0.36274717 | 0.53650233 | 1.57811253 | 0 | 0.85142298 | 1.50842335 | 1.35145328 | 0.38037387 | 0.53327066 | 1.72818818 | 0.14791111 | 0.52486058 | 2.31694758 | 0.42720395 | 0.61315883 |

| | | | | | | | | | | | | | | | | | | | | |
|---|---|---|---|---|---|---|---|---|---|---|---|---|---|---|---|---|---|---|---|---|
| ENSMUSG00 | 4930596I21R | 451 | 0.56547906 | 0.44318706 | 0.21451676 | 0 | 0.42689068 | 0 | 0.41962465 | 0.16936764 | 0.54793591 | 0.10454701 | 0.21186234 | 0.24751957 | 0.87506727 | 1.23576286 | 0.365424 | 0.59431098 | 0.16996141 | 0 |
| ENSMUSG00 | Gm14642 | 467 | 0 | 0 | 0 | 0 | 0 | 0 | 0 | 0 | 0 | 0 | 0 | 0 | 0 | 0 | 0 | 0 | 0 | 0 |
| ENSMUSG00 | Gm15733 | 241 | 0 | 0 | 0 | 0 | 0 | 0 | 0 | 0 | 0 | 0 | 0 | 0 | 0 | 0 | 0 | 0 | 0 | 0 |
| ENSMUSG00 | Gm16608 | 2022 | 0.02102135 | 0.04942566 | 0.09569442 | 0.06759799 | 0.04760823 | 0.02450679 | 0 | 0.01888843 | 0 | 0.11659422 | 0.02362758 | 0.05520837 | 0 | 0.04593876 | 0.12225981 | 0.11362198 | 0.09477324 | 0.05441059 |
| ENSMUSG00 | Gm12065 | 654 | 0 | 0 | 0 | 0 | 0 | 0 | 0 | 0 | 0 | 0 | 0 | 0 | 0 | 0 | 0 | 0 | 0 | 0 |
| ENSMUSG00 | Gm14169 | 2515 | 0 | 0 | 0 | 0 | 0 | 0 | 0 | 0 | 0 | 0 | 0 | 0 | 0 | 0 | 0 | 0 | 0 | 0 |
| ENSMUSG00 | Gm2415 | 2052 | 0.18642621 | 0.09740612 | 0.14144307 | 0.19982914 | 0.18764883 | 0.04829701 | 0.16139803 | 0.29779651 | 0.14049981 | 0.34466888 | 0.162975 | 0.38080861 | 0.07212269 | 0.06790071 | 0.04015746 | 0.13062098 | 0.0560326 | 0.16084534 |
| ENSMUSG00 | Gm16986 | 1091 | 0 | 0 | 0 | 0 | 0 | 0 | 0 | 0 | 0 | 0 | 0 | 0 | 0 | 0 | 0 | 0 | 0 | 0 |
| ENSMUSG00 | Gm16068 | 501 | 0 | 0.26597121 | 0.09655395 | 0.18188042 | 0 | 0.09890765 | 0.37774594 | 0.30492936 | 0.16441723 | 0.28233954 | 0.1907184 | 0.22281702 | 0 | 0.74162215 | 0.2467159 | 0.53499851 | 0.07649959 | 0.10979862 |
| ENSMUSG00 | Gm14507 | 345 | 0 | 0 | 0 | 0 | 0 | 0 | 0 | 0 | 0 | 0 | 0 | 0 | 0 | 0 | 0 | 0 | 0 | 0 |
| ENSMUSG00 | Gm16060 | 956 | 0 | 0.13938449 | 0 | 0 | 0.1006944 | 0 | 0 | 0 | 0 | 0 | 0 | 0 | 0 | 0.14574504 | 0 | 0 | 0 | 0 |
| ENSMUSG00 | A530076I17R | 1738 | 0 | 0 | 0 | 0 | 0 | 0 | 0 | 0 | 0 | 0 | 0 | 0 | 0 | 0 | 0 | 0 | 0 | 0 |
| ENSMUSG00 | Gm15942 | 807 | 0.0526706 | 0 | 0.05994242 | 0.0564573 | 0.17892908 | 0.12280727 | 0.46902284 | 0.14197919 | 0.25518287 | 0.17528142 | 0 | 0.06916439 | 0.36678005 | 0.46041227 | 0.20422085 | 0.14234427 | 0.04749231 | 0 |
| ENSMUSG00 | 5730457N03 | 3073 | 0 | 0 | 0 | 0 | 0 | 0 | 0 | 0 | 0 | 0 | 0 | 0 | 0 | 0 | 0 | 0 | 0 | 0 |
| ENSMUSG00 | Gm11934 | 1534 | 0.30479592 | 0.17373087 | 0.06306849 | 0 | 0 | 0.25842364 | 0 | 0.64732885 | 0 | 0.27663385 | 0 | 0 | 0 | 0.09082938 | 0 | 0.17472898 | 0.37476822 | 0 |
| ENSMUSG00 | 1700091E21 | 1705 | 0 | 0 | 0 | 0 | 0 | 0 | 0 | 0 | 0 | 0 | 0 | 0 | 0 | 0 | 0 | 0 | 0 | 0 |
| ENSMUSG00 | Gm13558 | 551 | 0 | 0 | 0 | 0 | 0.26206129 | 0.44966183 | 0.08586693 | 0.20794411 | 0 | 0.08557296 | 0.43352957 | 0 | 0.26859483 | 0.08429054 | 0 | 0 | 0.34778854 | 0 |
| ENSMUSG00 | Gm16211 | 1618 | 0 | 0 | 0 | 0 | 0 | 0 | 0 | 0 | 0 | 0 | 0 | 0 | 0 | 0 | 0 | 0 | 0 | 0 |
| ENSMUSG00 | BC048594 | 393 | 0 | 0.08476563 | 0 | 0.69558846 | 0 | 0.50435352 | 0.96310797 | 0.38872674 | 0.20960059 | 0 | 1.09408302 | 1.13619669 | 0.37657952 | 1.29996173 | 0 | 0.97431571 | 0.09752238 | 0.69986141 |
| ENSMUSG00 | Gm13631 | 651 | 0 | 0 | 0 | 0.06998624 | 0 | 0 | 0 | 0 | 0 | 0 | 0 | 0 | 0 | 0 | 0 | 0 | 0 | 0 |
| ENSMUSG00 | Gm16331 | 497 | 0 | 0 | 0 | 0 | 0 | 0 | 0 | 0 | 0 | 0.47435315 | 0 | 0.11230516 | 0 | 0.18689774 | 0.16580103 | 0 | 0 | 0 |
| ENSMUSG00 | Gm16159 | 1151 | 0.22157346 | 0 | 0 | 0 | 0 | 0.12915569 | 0.04110572 | 0.09954579 | 0 | 0 | 0.0830147 | 0.33945234 | 0.21430025 | 0 | 0.21477788 | 0.03326725 | 0 | 0.04779245 |
| ENSMUSG00 | 1700001J04R | 898 | 0 | 0 | 0 | 0 | 0 | 0 | 0 | 0.08506103 | 0 | 0 | 0 | 0.06215553 | 0 | 0 | 0 | 0 | 0 | 0 |
| ENSMUSG00 | Gm12718 | 779 | 0 | 0 | 0 | 0 | 0 | 0 | 0 | 0 | 0 | 0 | 0 | 0 | 0 | 0 | 0 | 0 | 0 | 0 |
| ENSMUSG00 | Gm16248 | 242 | 0 | 0 | 0 | 0 | 0 | 0 | 0 | 0 | 0 | 0 | 0 | 0 | 0 | 0 | 0 | 0 | 0 | 0 |
| ENSMUSG00 | 4930471C06 | 679 | 0 | 0 | 0 | 0 | 0 | 0 | 0 | 0 | 0 | 0 | 0 | 0 | 0 | 0 | 0 | 0 | 0 | 0 |
| ENSMUSG00 | Gm11211 | 356 | 0 | 0 | 0 | 0 | 0 | 0 | 0 | 0 | 0 | 0 | 0 | 0 | 0 | 0 | 0 | 0 | 0 | 0 |
| ENSMUSG00 | Gm13825 | 440 | 0 | 0 | 0 | 0 | 0 | 0 | 0 | 0 | 0 | 0 | 0 | 0 | 0 | 0 | 0 | 0 | 0 | 0 |
| ENSMUSG00 | 9030622O22 | 1670 | 0 | 0 | 0 | 0.08184619 | 0 | 0 | 0 | 0 | 0 | 0 | 0 | 0 | 0.02954007 | 0.27810831 | 0 | 0.02292851 | 0.09179951 | 0 |
| ENSMUSG00 | Smek3-ps | 2444 | 0 | 0 | 0 | 0 | 0 | 0 | 0 | 0 | 0 | 0 | 0 | 0 | 0 | 0 | 0 | 0 | 0 | 0 |
| ENSMUSG00 | Gm16685 | 1589 | 0 | 0.06289407 | 0.09132825 | 0.08601833 | 0 | 0 | 0 | 0 | 0 | 0.11869277 | 0 | 0 | 0 | 0 | 0 | 0 | 0 | 0 |
| ENSMUSG00 | Gm11379 | 168 | 0.506014 | 0.59487311 | 1.15175072 | 0 | 0 | 0 | 0 | 0 | 0 | 0 | 0 | 0 | 0 | 0.2764529 | 0 | 0.22792028 | 0.45626544 | 0 |
| ENSMUSG00 | Gm15479 | 1131 | 0 | 0 | 0 | 0 | 0 | 0 | 0 | 0 | 0 | 0 | 0 | 0 | 0 | 0 | 0 | 0 | 0 | 0 |
| ENSMUSG00 | Gm15729 | 358 | 0 | 0 | 0 | 0 | 0 | 0 | 0 | 0 | 0 | 0 | 0 | 0.15590967 | 0 | 0 | 0 | 0.21391401 | 0 | 0 |
| ENSMUSG00 | Gm13150 | 2505 | 0 | 0.02659712 | 0.01931079 | 0.01818804 | 0.07685736 | 0.05934459 | 0 | 0.01524647 | 0 | 0 | 0 | 0.08912681 | 0 | 0.11124332 | 0 | 0 | 0.12239935 | 0.04391945 |
| ENSMUSG00 | Gm15271 | 2107 | 0 | 0 | 0 | 0 | 0 | 0 | 0 | 0 | 0 | 0 | 0.0226744 | 0.02649058 | 0 | 0 | 0 | 0 | 0 | 0 |
| ENSMUSG00 | Gm15598 | 761 | 0 | 0 | 0 | 0 | 0 | 0 | 0 | 0 | 0 | 0 | 0 | 0 | 0 | 0 | 0 | 0 | 0 | 0 |
| ENSMUSG00 | BC024582 | 2178 | 0 | 0.03059035 | 0 | 0 | 0.08839655 | 0.04550297 | 0 | 0 | 0.26474344 | 0 | 0 | 0.07688108 | 0.56625249 | 0 | 0 | 0.08790314 | 0.01759701 | 0.05051341 |
| ENSMUSG00 | Gm5698 | 1139 | 0 | 0 | 0 | 0 | 0 | 0 | 0 | 0 | 0 | 0 | 0 | 0 | 0 | 0 | 0 | 0 | 0 | 0 |
| ENSMUSG00 | Gm11454 | 711 | 0.05978224 | 0.42168221 | 0.13607182 | 0 | 0.20308829 | 0.76663863 | 0.93161393 | 0.48344813 | 0.92684142 | 0.59684435 | 0.40316421 | 0 | 0.06938385 | 1.24112189 | 0.34769245 | 0.48469123 | 0.21561911 | 0.07736865 |
| ENSMUSG00 | Gm12923 | 663 | 0.06411037 | 0 | 0 | 0 | 0.36298585 | 0 | 0 | 0 | 0 | 0 | 0 | 0 | 0 | 0 | 0 | 0 | 0.28903693 | 0 |
| ENSMUSG00 | Gm16196 | 462 | 0.73602036 | 0.86526998 | 0.20940922 | 1.38063771 | 0.93763488 | 1.39434099 | 0 | 1.40534814 | 0.26744491 | 0.91852019 | 0.93068101 | 0.84569186 | 0.53389521 | 2.51320819 | 0.98098943 | 0.0828801 | 1.41027499 | 1.42880797 |
| ENSMUSG00 | 9430041J12R | 2831 | 0.03002838 | 0.01176718 | 0 | 0 | 0.01700174 | 0.10502169 | 0.03342471 | 0.41821423 | 0.05819359 | 0 | 0.10125389 | 0.01971588 | 0 | 0.01640554 | 0.02910742 | 0 | 0 | 0 |
| ENSMUSG00 | Gm15354 | 1770 | 0 | 0 | 0 | 0 | 0 | 0 | 0 | 0 | 0 | 0 | 0 | 0 | 0 | 0 | 0 | 0 | 0 | 0 |
| ENSMUSG00 | Gm14643 | 1840 | 0 | 0 | 0 | 0 | 0 | 0 | 0 | 0 | 0 | 0 | 0 | 0 | 0 | 0 | 0 | 0 | 0 | 0 |
| ENSMUSG00 | Gm5918 | 707 | 19.6593954 | 21.5332287 | 22.9894006 | 20.5572463 | 19.8110181 | 17.7324492 | 26.9688962 | 19.069191 | 30.9918336 | 24.4756833 | 21.1507242 | 29.9999318 | 25.3986109 | 19.6418418 | 26.6906826 | 17.330968 | 18.3228972 | 16.9617896 |
| ENSMUSG00 | Gm13025 | 1159 | 0 | 0 | 0 | 0 | 0 | 0 | 0 | 0 | 0 | 0 | 0 | 0 | 0 | 0 | 0 | 0.1321505 | 0 | 0 |
| ENSMUSG00 | 4933438A12R | 1010 | 0 | 0 | 0 | 0 | 0 | 0 | 0 | 0 | 0 | 0 | 0 | 0 | 0 | 0 | 0 | 0 | 0 | 0 |
| ENSMUSG00 | Gm15389 | 663 | 0 | 0 | 0 | 0 | 0 | 0 | 0 | 0 | 0 | 0 | 0 | 0 | 0 | 0 | 0 | 0 | 0 | 0 |
| ENSMUSG00 | Gm16342 | 1014 | 0 | 0 | 0 | 0 | 0 | 0 | 0 | 0 | 0 | 0 | 0 | 0 | 0 | 0 | 0 | 0 | 0 | 0 |
| ENSMUSG00 | Gm14206 | 743 | 2.63154522 | 0.94154883 | 1.0416911 | 1.83961148 | 2.33209859 | 2.0674761 | 1.91033697 | 2.21032746 | 3.15966541 | 1.07881824 | 0.96450118 | 1.87805056 | 2.05826304 | 1.31268618 | 0.66543563 | 0.97916762 | 0.82533075 | 1.40669317 |
| ENSMUSG00 | Gm13029 | 591 | 0 | 0 | 0 | 0 | 0 | 0 | 0 | 0 | 0 | 0 | 0 | 0 | 0 | 0 | 0 | 0 | 0 | 0 |
| ENSMUSG00 | Gm15690 | 430 | 0.19769849 | 0.07747185 | 0.44998633 | 0 | 0.22386941 | 0.23047783 | 0 | 0.08881954 | 0.67047816 | 0.2193056 | 0.55552277 | 0.7788232 | 0.45890156 | 0.21601901 | 0 | 0.17809585 | 0.62391646 | 0 |
| ENSMUSG00 | Gm14342 | 2115 | 0 | 0 | 0.02287165 | 0 | 0 | 0 | 0 | 0.03611575 | 0 | 0 | 0 | 0 | 0.20992305 | 0 | 0.01948064 | 0 | 0 | 0.0780271 |
| ENSMUSG00 | Gm13827 | 796 | 0.05339846 | 0.04185037 | 0.1823123 | 0.05723749 | 0.06046724 | 0 | 0.41606627 | 0.23990203 | 0.05174185 | 0.17770366 | 0.24007517 | 0.3506009 | 0.30987385 | 0.58346843 | 0.20704299 | 0.24051889 | 0.14444584 | 0 |
| ENSMUSG00 | Gm15221 | 411 | 0 | 0 | 0 | 0 | 0 | 0 | 0 | 0 | 0.400842 | 0 | 0 | 0 | 0 | 0 | 0 | 0 | 0 | 0 |
| ENSMUSG00 | Gm12144 | 1290 | 0 | 0.07747185 | 0 | 0 | 0 | 0 | 0.0366765 | 0 | 0 | 0.03655093 | 0 | 0 | 0 | 0 | 0 | 0 | 0 | 0 |
| ENSMUSG00 | C630020P15R | 4601 | 0 | 0.00724036 | 0 | 0 | 0 | 0.02153998 | 0.06169878 | 0 | 0 | 0.01024793 | 0.0103836 | 0 | 0 | 0.06056608 | 0 | 0 | 0 | 0 |
| ENSMUSG00 | 2700068H02R | 1870 | 0.02273004 | 0 | 0.12934099 | 0 | 0 | 0 | 0 | 0 | 0 | 0 | 0.20438485 | 0 | 0 | 0.02483641 | 0 | 0 | 0 | 0 |
| ENSMUSG00 | Kis2 | 2210 | 0 | 0 | 0 | 0 | 0 | 0 | 0 | 0 | 0 | 0 | 0 | 0 | 0 | 0 | 0 | 0 | 0 | 0 |
| ENSMUSG00 | Gm6425 | 789 | 0.21548885 | 0 | 0.2452397 | 0.11549061 | 0.18301112 | 0 | 0.05996537 | 0 | 0.05220091 | 0.05976008 | 0.06055128 | 0 | 0.06252461 | 0.0588645 | 0.26109985 | 0 | 0 | 0.13944007 |
| ENSMUSG00 | Gm15802 | 207 | 0 | 0 | 0 | 0 | 0 | 0 | 0 | 0 | 0 | 0 | 0 | 0 | 0 | 0 | 0 | 0 | 0 | 0 |
| ENSMUSG00 | Gm15534 | 843 | 0 | 0 | 0 | 0.10809263 | 0 | 0 | 0 | 0 | 0 | 0 | 0 | 0 | 0 | 0 | 0 | 0 | 0 | 0 |
| ENSMUSG00 | Gm14616 | 311 | 0 | 0 | 0 | 0 | 0 | 0 | 0 | 0.12280515 | 0 | 0 | 0 | 0 | 0 | 1.04536531 | 0.26496177 | 0 | 0 | 0 |
| ENSMUSG00 | Gm16950 | 4925 | 0 | 0 | 0 | 0 | 0 | 0 | 0 | 0.0077548 | 0 | 0 | 0 | 0 | 0.02003327 | 0 | 0 | 0 | 0 | 0 |
| ENSMUSG00 | Gm14317 | 1461 | 0 | 0 | 0 | 0 | 0 | 0 | 0 | 0 | 0 | 0.09681869 | 0 | 0 | 0 | 0 | 0 | 0 | 0.23609628 | 0.75303363 |
| ENSMUSG00 | Rmst | 5137 | 2.29198633 | 2.76904922 | 2.693173 | 2.1463447 | 2.52980716 | 2.13181898 | 2.17360176 | 2.34195187 | 2.88634333 | 2.43234113 | 2.08323743 | 3.12924096 | 2.19914523 | 2.47725909 | 2.40616445 | 2.10199558 | 3.05894135 | 2.90197935 |
| ENSMUSG00 | C230034O21 | 2832 | 0.01500889 | 0 | 0 | 0 | 0 | 0 | 0 | 0 | 0 | 0 | 0 | 0 | 0 | 0.06559899 | 0 | 0.02704139 | 0 | 0 |
| ENSMUSG00 | Gm12764 | 615 | 0.13822821 | 0.10833462 | 0 | 0.074083 | 0.15652658 | 0.3222942 | 0.30772474 | 0 | 0.26787978 | 0.15333562 | 0.15536572 | 0.09075717 | 0.5615015 | 0.22655652 | 0.0669944 | 0.4980892 | 0.43623427 | 0.26833711 |
| ENSMUSG00 | 1700036O09 | 347 | 0 | 0 | 0 | 0 | 0 | 0 | 0 | 0 | 0 | 0 | 0 | 0 | 0 | 0 | 0 | 0 | 0 | 0 |
| ENSMUSG00 | 1700024P03R | 1330 | 0 | 0.27552018 | 0 | 0 | 0 | 0 | 0 | 0.02871609 | 0 | 0.07090331 | 0 | 0 | 0 | 0 | 0.06195723 | 0.02878993 | 0.11526706 | 0 |
| ENSMUSG00 | Gm12860 | 953 | 1.1150361 | 0.83893962 | 1.16746191 | 1.14739251 | 2.22225041 | 1.09192802 | 0.89362877 | 1.36258309 | 2.5066295 | 1.53375845 | 1.05275354 | 2.28416667 | 1.70824058 | 4.0937076 | 1.51317362 | 0.80358043 | 1.28692707 | 1.38532903 |
| ENSMUSG00 | Gm15169 | 4549 | 0 | 0 | 0 | 0 | 0 | 0 | 0 | 0 | 0 | 0 | 0 | 0 | 0 | 0 | 0 | 0 | 0 | 0 |
| ENSMUSG00 | Gm15462 | 4576 | 0.08359847 | 0 | 0 | 0 | 0 | 0 | 0 | 0 | 0 | 0 | 0 | 0 | 0 | 0 | 0.04501918 | 0.05020622 | 0 | 0 |
| ENSMUSG00 | 4930579M01 | 929 | 0 | 0 | 0 | 0 | 0 | 0 | 0 | 0 | 0 | 0 | 0 | 0 | 0 | 0 | 0 | 0 | 0 | 0 |
| ENSMUSG00 | 1110035M17 | 535 | 0.07944893 | 0.87173929 | 0.45208907 | 0.42580415 | 0.26989864 | 1.01884125 | 0.08843491 | 0.71387668 | 0 | 0.17626431 | 0.08929899 | 0.93895507 | 0.36883676 | 2.34390721 | 0.92414705 | 0.35785614 | 0.07163794 | 0.92538684 |
| ENSMUSG00 | Gm13609 | 587 | 0 | 0.22700439 | 0 | 0 | 0 | 0 | 0 | 0 | 0 | 0 | 0 | 0 | 0 | 0 | 0.14038009 | 0 | 0 | 0.18742455 |
| ENSMUSG00 | Gm11508 | 438 | 0 | 0 | 0 | 0 | 0 | 0 | 0 | 0 | 0 | 0 | 0 | 0.12743302 | 0.11262995 | 0 | 0.09406748 | 0 | 0.26250888 | 0 |
| ENSMUSG00 | Gm16329 | 368 | 0 | 0 | 0 | 0 | 0 | 0 | 0 | 0 | 0 | 0 | 0 | 0 | 0 | 0 | 0 | 0 | 0 | 0 |
| ENSMUSG00 | Gm6152 | 339 | 0 | 0 | 0 | 0 | 0 | 0 | 0 | 0 | 0 | 0 | 0 | 0 | 0 | 0 | 0 | 0 | 0 | 0 |
| ENSMUSG00 | Gm13571 | 3093 | 0.06871189 | 0.01077042 | 0 | 0.04419112 | 0 | 0.09612557 | 0 | 0 | 0 | 0.06097731 | 0 | 0.0721832 | 0 | 0 | 0.02664181 | 0 | 0.0123913 | 0 |
| ENSMUSG00 | Gm13976 | 477 | 0 | 0 | 0 | 0 | 0 | 0 | 0 | 0 | 0 | 0 | 0 | 0 | 0 | 0 | 0 | 0 | 0 | 0 |
| ENSMUSG00 | Gm16598 | 1357 | 0.31322901 | 0.34368498 | 0.35647406 | 0.63792177 | 0.74485659 | 0 | 0.59271595 | 0.84434198 | 0.48561846 | 0.5559405 | 0.4928883 | 0.78150154 | 0.32718294 | 1.06099242 | 0.72869369 | 1.58015771 | 1.52514372 | 0.85128315 |
| ENSMUSG00 | 8430403D17 | 3612 | 0.38833633 | 0.38735924 | 0.2142792 | 0.42886919 | 0.70625469 | 0.23322161 | 0.34056746 | 0.14803257 | 0.4447049 | 0.41772495 | 0.10581386 | 0.30905682 | 0.15023563 | 0.1028662 | 0.11406854 | 0.14841321 | 0.15916236 | 0.12183634 |
| ENSMUSG00 | Gm13659 | 652 | 0.19557596 | 0.56202736 | 0.29677012 | 0.55903122 | 0.07382197 | 0.38000563 | 0.29026183 | 0.23430922 | 0.18950851 | 0.21695109 | 0 | 0.17121369 | 0.1513249 | 0.14246653 | 0.18957771 | 0.4698234 | 0.52904398 | 0 |
| ENSMUSG00 | Gm6270 | 556 | 0 | 0 | 0 | 0 | 0 | 0 | 0 | 0 | 0 | 0 | 0 | 0 | 0 | 0 | 0 | 0 | 0 | 0 |
| ENSMUSG00 | 1700019E08 | 827 | 0 | 0 | 0 | 0 | 0 | 0 | 0 | 0 | 0 | 0 | 0 | 0 | 0 | 0 | 0 | 0 | 0 | 0 |
| ENSMUSG00 | Gm13985 | 618 | 0 | 0 | 0 | 0 | 0 | 0 | 0 | 0 | 0 | 0 | 0 | 0 | 0 | 0 | 0 | 0 | 0 | 0 |
| ENSMUSG00 | Gm15668 | 673 | 0 | 0 | 0 | 0 | 0 | 0 | 0 | 0 | 0 | 0 | 0 | 0 | 0 | 0 | 0 | 0 | 0 | 0 |
| ENSMUSG00 | Gm12679 | 634 | 0 | 0 | 0 | 0 | 0 | 0.07815889 | 0 | 0 | 0 | 0 | 0 | 0 | 0 | 0 | 0 | 0 | 0 | 0 |
| ENSMUSG00 | Gm13783 | 730 | 0 | 0 | 0 | 0 | 0 | 0 | 0 | 0 | 0 | 0 | 0 | 0 | 0 | 0.12724408 | 0 | 0.05245289 | 0.10500355 | 0 |
| ENSMUSG00 | Gm14717 | 645 | 0 | 0.15494369 | 0.07499772 | 0 | 0 | 0.15365189 | 0.07335299 | 0.05921303 | 0 | 0 | 0 | 0 | 0 | 0 | 0 | 0.11873057 | 0.05942062 | 0 |
| ENSMUSG00 | Gm2716 | 1565 | 0 | 0 | 0 | 0 | 0 | 0 | 0 | 0 | 0 | 0 | 0 | 0 | 0 | 0 | 0 | 0 | 0 | 0 |
| ENSMUSG00 | 4933405E24 | 1075 | 0 | 0 | 0.2699918 | 0 | 0 | 0 | 0 | 0 | 0 | 0 | 0 | 0 | 0 | 0 | 0 | 0 | 0 | 0 |
| ENSMUSG00 | A230087F16 | 3382 | 0.01256806 | 0 | 0 | 0 | 0.07115896 | 0 | 0 | 0 | 0 | 0.01394166 | 0 | 0.13202995 | 0.07293305 | 0 | 0.14619121 | 0.11321883 | 0.13598923 | 0 |
| ENSMUSG00 | Gm12462 | 3054 | 0 | 0 | 0 | 0 | 0 | 0 | 0 | 0 | 0 | 0 | 0 | 0 | 0.14537893 | 0 | 0 | 0 | 0 | 0 |
| ENSMUSG00 | Gm13845 | 2939 | 0.78097295 | 0.70275585 | 0.54315294 | 0.44956457 | 0.75334076 | 0.37092893 | 1.33615255 | 0.88366225 | 1.59757155 | 1.3155351 | 1.04035296 | 1.67124134 | 1.76245366 | 1.50125495 | 0.85515308 | 1.10741804 | 1.68223624 | 0.5615084 |
| ENSMUSG00 | A330040F15 | 1104 | 0.26950746 | 1.08629003 | 0.43816604 | 0.45395968 | 0.3051843 | 0.71815556 | 1.67137181 | 2.42161972 | 3.43220965 | 0.98230632 | 1.21168373 | 1.92119128 | 2.41297421 | 0.67310272 | 1.45548975 | 1.49139141 | 1.45806564 | 0.24913545 |
| ENSMUSG00 | Gm14062 | 4131 | 0 | 0 | 0 | 0 | 0 | 0 | 0 | 0 | 0 | 0 | 0 | 0 | 0 | 0 | 0 | 0 | 0 | 0 |
| ENSMUSG00 | Gm13610 | 949 | 0 | 0 | 0 | 0 | 0 | 0.05221574 | 0.0498553 | 0 | 0.08679982 | 0 | 0.10068484 | 0.05881524 | 0.05198305 | 0.04894003 | 0 | 0 | 0 | 0 |
| ENSMUSG00 | Gm9530 | 3198 | 0.0531647 | 0.01041679 | 0 | 0.01424673 | 0.03010127 | 0.04648474 | 0.04438338 | 0 | 0.09015185 | 0.10320667 | 0.17926814 | 0.05235991 | 0.01542587 | 0.02904571 | 0 | 0.0239466 | 0.2277047 | 0.24081535 |
| ENSMUSG00 | Gm16755 | 929 | 0.41178319 | 0.35858874 | 0.78105808 | 0.49043105 | 0 | 0.32003918 | 0.10185722 | 0.04111131 | 1.01968769 | 0.55829681 | 0.35998354 | 0.60081445 | 1.27445211 | 0.44994272 | 0.70960699 | 0.28851911 | 0.70134235 | 0.17763974 |
| ENSMUSG00 | 2410137F16 | 3651 | 0.01164206 | 0 | 0 | 0 | 0 | 0 | 0 | 0 | 0 | 0 | 0 | 0.03057555 | 0 | 0.03816277 | 0 | 0 | 0 | 0 |
| ENSMUSG00 | Gm16599 | 1177 | 0.03611315 | 0.02830322 | 0 | 0.03870947 | 0.1226812 | 0 | 0.04019769 | 0.12979576 | 0 | 0 | 0 | 0.14226592 | 0.12573981 | 0.03945972 | 0 | 0 | 0 | 0 |
| ENSMUSG00 | 4930412B13 | 1270 | 0 | 0 | 0 | 0 | 0 | 0 | 0 | 0 | 0 | 0 | 0 | 0 | 0 | 0 | 0 | 0 | 0 | 0 |
| ENSMUSG00 | Gm11665 | 283 | 0 | 0.94170726 | 0.34186241 | 0.16099309 | 0.17007747 | 0.175098 | 0 | 0 | 0 | 0 | 0 | 1.38059943 | 0 | 0 | 0 | 0.4059075 | 0 | 0 |
| ENSMUSG00 | Gm362 | 264 | 0 | 0 | 0 | 0 | 0 | 0 | 0 | 0 | 0 | 0 | 0 | 0 | 0 | 0 | 0 | 0 | 0 | 0 |
| ENSMUSG00 | 2700069I18R | 1134 | 0 | 0 | 0 | 0 | 0 | 0 | 0 | 0 | 0.03631968 | 0 | 0 | 0 | 0 | 0 | 0 | 0 | 0 | 0 |
| ENSMUSG00 | Gm8661 | 991 | 0.08578239 | 0.36976976 | 0 | 0.13792445 | 0 | 0 | 0.04774236 | 0 | 0 | 0.04757891 | 0.04820884 | 0.1689677 | 0.09955987 | 0.04686588 | 0 | 0.03863835 | 0.15469746 | 0 |
| ENSMUSG00 | Gm12660 | 662 | 0 | 0 | 0 | 0 | 0 | 0 | 0 | 0 | 0 | 0 | 0 | 0 | 0 | 0 | 0 | 0 | 0 | 0 |
| ENSMUSG00 | Gm11820 | 660 | 0 | 0 | 0 | 0 | 0 | 0 | 0 | 0 | 0 | 0 | 0 | 0 | 0 | 0 | 0 | 0 | 0 | 0 |
| ENSMUSG00 | Gm514 | 1374 | 0.18561212 | 0 | 0 | 0.03315942 | 0.07006102 | 0 | 0 | 0 | 0.26978067 | 0.17158189 | 0.13908285 | 0 | 0 | 0.0338021 | 0.05997315 | 0 | 0.1952577 | 0.12010722 |
| ENSMUSG00 | Gm4887 | 2101 | 0.06069278 | 0.06342293 | 0.11512025 | 0.17348327 | 0.06872716 | 0.14151185 | 0.13511474 | 0.05453461 | 0.09801646 | 0 | 0.09095661 | 0.10626495 | 0.07044062 | 0.02210571 | 0.1176627 | 0.01822494 | 0.18241931 | 0.07854703 |
| ENSMUSG00 | Gm15255 | 272 | 0 | 0 | 0 | 0 | 0 | 0 | 0 | 0 | 0 | 0 | 0 | 0 | 0 | 0 | 0 | 0 | 0 | 0 |
| ENSMUSG00 | 4732447D17 | 3483 | 0 | 0 | 0 | 0 | 0 | 0 | 0 | 0 | 0 | 0 | 0 | 0 | 0 | 0 | 0.07097598 | 0 | 0 | 0 |
| ENSMUSG00 | Gm7546 | 962 | 0 | 0 | 0 | 0 | 0 | 0 | 0 | 0 | 0 | 0 | 0 | 0 | 0 | 0 | 0 | 0 | 0 | 0 |
| ENSMUSG00 | Gm11816 | 980 | 0 | 0 | 0 | 0 | 0 | 0 | 0 | 0 | 0 | 0 | 0 | 0 | 0.05033869 | 0 | 0 | 0 | 0 | 0 |
| ENSMUSG00 | 1700112M01 | 822 | 0 | 0 | 0 | 0 | 0.05855465 | 0 | 0 | 0 | 0 | 0 | 0 | 0 | 0 | 0 | 0 | 0 | 0 | 0 |
| ENSMUSG00 | Gm16284 | 399 | 0 | 0 | 0 | 0 | 0 | 0 | 0 | 0 | 0 | 0 | 0 | 0 | 0 | 0 | 0 | 0 | 0 | 0 |
| ENSMUSG00 | 5830418P13R | 2381 | 0 | 0 | 0 | 0.01913526 | 0.020215 | 0 | 0 | 0 | 0 | 0 | 0.02006508 | 0 | 0 | 0 | 0 | 0.03216347 | 0 | 0 |
| ENSMUSG00 | Gm15591 | 625 | 0.13601656 | 0.10660126 | 0 | 0.29159068 | 0 | 0.31713749 | 0.15140057 | 0.06110784 | 0.06589843 | 0.15088225 | 0 | 0.17861012 | 0.07893107 | 0.07431054 | 0 | 0.4288548 | 0.2452883 | 0.26404371 |
| ENSMUSG00 | Gm14258 | 2029 | 0 | 0 | 0 | 0 | 0 | 0 | 0 | 0 | 0 | 0 | 0 | 0 | 0 | 0 | 0 | 0 | 0 | 0 |
| ENSMUSG00 | Gm14329 | 589 | 0 | 0 | 0 | 0 | 0 | 0 | 0 | 0 | 0 | 0 | 0 | 0 | 0 | 0 | 0 | 0 | 0 | 0 |
| ENSMUSG00 | Gm7846 | 545 | 1.09187608 | 1.71148814 | 0.53255263 | 1.58836668 | 1.05978548 | 1.3638367 | 2.77799216 | 2.10233408 | 1.20914542 | 0.95166557 | 1.31490711 | 1.74103902 | 1.99138015 | 1.53393316 | 1.88997963 | 1.89696587 | 1.26582265 | 1.61494625 |
| ENSMUSG00 | 4930483K19R | 1233 | 0 | 0 | 0 | 0 | 0 | 0 | 0.11511601 | 0 | 0 | 0.07648127 | 0 | 0 | 0 | 0.0753351 | 0.0668314 | 0 | 0 | 0 |
| ENSMUSG00 | Gm16718 | 701 | 0 | 0 | 0.27602585 | 0 | 0 | 0 | 0.13498625 | 0.16344823 | 0 | 0.20178618 | 0 | 0 | 0 | 0.06625405 | 0.2351016 | 0 | 0 | 0 |
| ENSMUSG00 | Gm12121 | 498 | 657.549952 | 492.134463 | 582.036534 | 681.677392 | 608.414577 | 582.194868 | 1225.56939 | 871.906472 | 956.139174 | 1068.65459 | 1098.24844 | 1057.1352 | 827.349745 | 610.301421 | 833.049129 | 753.356167 | 987.248467 | 770.017034 |
| ENSMUSG00 | 4930558G05 | 491 | 0.17313717 | 0 | 0 | 0 | 0 | 0 | 0 | 0 | 0 | 0 | 0 | 0 | 0 | 0 | 0 | 0 | 0 | 0 |
| ENSMUSG00 | Gm16170 | 3751 | 0.52125782 | 0.5950317 | 1.2509284 | 0.57087952 | 0.47477504 | 0.62089536 | 0.58021414 | 1.1200118 | 0.58194757 | 0.61593827 | 1.12082014 | 0.83329168 | 0.67073521 | 0.6314712 | 0.32952457 | 0.91872958 | 0.78675683 | 0.42529035 |
| ENSMUSG00 | Gm15277 | 290 | 0 | 0 | 0 | 0 | 0 | 0 | 0 | 0 | 0 | 0 | 0 | 0 | 0 | 0 | 0 | 0 | 0 | 0 |
| ENSMUSG00 | Gm15787 | 3091 | 0.23377159 | 0.4095406 | 0.20344739 | 0.38323751 | 0.51386395 | 0.25650072 | 0.56634394 | 0.29654405 | 0.31979178 | 0.51864248 | 1.22103582 | 0.37920703 | 0.36707671 | 0.51086994 | 0.85308948 | 0.65655199 | 0.21078843 | 0.03559308 |
| ENSMUSG00 | Gm16127 | 232 | 0 | 0 | 0 | 0 | 0 | 0 | 0 | 0 | 0 | 0 | 0 | 0 | 0 | 0 | 0 | 0 | 0 | 0 |
| ENSMUSG00 | Gm12724 | 660 | 0 | 0 | 0 | 0 | 0 | 0 | 0 | 0 | 0 | 0 | 0 | 0 | 0 | 0 | 0 | 0 | 0 | 0 |
| ENSMUSG00 | Gm14095 | 513 | 0 | 0 | 0 | 0 | 0 | 0 | 0 | 0 | 0 | 0 | 0 | 0 | 0 | 0 | 0 | 0 | 0 | 0 |
| ENSMUSG00 | Gm12694 | 743 | 0 | 0 | 0 | 0.06132038 | 0 | 0 | 0 | 0 | 0 | 0 | 0 | 0 | 0 | 0.1875266 | 0 | 0 | 0 | 0 |
| ENSMUSG00 | C030044B11 | 1777 | 25.4983217 | 28.6637115 | 26.568692 | 26.8187126 | 28.2507577 | 25.2922506 | 29.3940335 | 31.6586431 | 39.193246 | 32.1855954 | 31.6169675 | 37.8491128 | 27.2338834 | 30.1089413 | 26.803038 | 29.8869288 | 28.448163 | 27.9224616 |
| ENSMUSG00 | Gm13773 | 579 | 0 | 0 | 0 | 0 | 0 | 0 | 0 | 0 | 0 | 0 | 0 | 0 | 0 | 0 | 0 | 0 | 0 | 0 |
| ENSMUSG00 | 3100003L05R | 660 | 0 | 0 | 0 | 0 | 0 | 0 | 0 | 0 | 0 | 0 | 0 | 0 | 0 | 0 | 0 | 0 | 0 | 0 |
| ENSMUSG00 | Gm11534 | 670 | 0 | 0 | 0 | 0 | 0 | 0 | 0 | 0 | 0 | 0 | 0 | 0 | 0 | 0 | 0 | 0 | 0 | 0 |
| ENSMUSG00 | Gm12052 | 931 | 0 | 0 | 0 | 0.14681325 | 0.1550975 | 0.05322528 | 0.10163841 | 0 | 0.6193461 | 0 | 0 | 0 | 0 | 0.34920366 | 0.3540413 | 0 | 0 | 0 |
| ENSMUSG00 | 4930548K13 | 558 | 0 | 0 | 0 | 0 | 0 | 0 | 0 | 0 | 0 | 0 | 0 | 0 | 0 | 0 | 0 | 0 | 0 | 0 |
| ENSMUSG00 | Gm15239 | 601 | 0.07072409 | 0 | 0 | 0 | 0 | 0 | 0 | 0 | 0 | 0.1569075 | 0 | 0 | 0.08208306 | 0 | 0 | 0 | 0 | 0 |
| ENSMUSG00 | 5830403M04 | 1551 | 0 | 0.08591333 | 0.15594304 | 0.02937527 | 0.58962382 | 0.06389779 | 0.39656017 | 0.51711183 | 0.05310963 | 0.12160078 | 0.43123754 | 0.03598689 | 0.19083914 | 0.26950147 | 0.13282255 | 0.2468769 | 0.27181771 | 0.95760534 |
| ENSMUSG00 | Gm16259 | 552 | 0 | 0 | 0 | 0 | 0 | 0 | 0 | 0 | 0 | 0 | 0 | 0 | 0 | 0 | 0 | 0 | 0 | 0 |
| ENSMUSG00 | Gm3765 | 1446 | 0 | 0 | 0 | 0 | 0 | 0 | 0 | 0 | 0 | 0 | 0 | 0 | 0 | 0 | 0 | 0 | 0 | 0 |
| ENSMUSG00 | A930031H19 | 1043 | 0 | 0 | 0 | 0 | 0 | 0 | 0 | 0 | 0 | 0 | 0 | 0 | 0 | 0 | 0 | 0 | 0 | 0 |
| ENSMUSG00 | 4930519D14 | 1133 | 0 | 0 | 0 | 0 | 0 | 0 | 0 | 0 | 0 | 0 | 0 | 0 | 0 | 0 | 0 | 0 | 0 | 0 |
| ENSMUSG00 | Gm15850 | 2112 | 0 | 0 | 0 | 0 | 0 | 0 | 0 | 0 | 0 | 0 | 0 | 0 | 0 | 0 | 0 | 0 | 0 | 0 |
| ENSMUSG00 | Marcksl1-ps4 | 290 | 0.87941744 | 1.26359254 | 0.16680528 | 0.47132115 | 1.16180505 | 0.1708715 | 3.58923771 | 1.58037527 | 2.27235949 | 1.13812042 | 0 | 0.3849356 | 1.53099054 | 3.04288848 | 1.70489197 | 0.66018289 | 0.39647893 | 1.89686575 |
| ENSMUSG00 | Igf2as | 5671 | 0 | 0 | 0.00852998 | 0 | 0 | 0 | 0 | 0 | 0 | 0 | 0 | 0.01968459 | 0 | 0 | 0 | 0 | 0 | 0 |
| ENSMUSG00 | Gm12496 | 666 | 0 | 0 | 0 | 0 | 0 | 0 | 0 | 0 | 0 | 0 | 0 | 0 | 0 | 0 | 0 | 0 | 0 | 0 |
| ENSMUSG00 | Gm11670 | 508 | 0.16734321 | 0.06557656 | 0.28567045 | 0 | 0 | 0.29263425 | 0.55881117 | 0.60145516 | 0.97290982 | 0.64971441 | 0.28213558 | 0.43949341 | 0.77688059 | 1.00567906 | 0.72994882 | 0.45225127 | 0.52811826 | 0.32485693 |
| ENSMUSG00 | Gm16154 | 2910 | 0 | 0.02289546 | 0.04986962 | 0 | 0 | 0 | 0 | 0 | 0 | 0.06481196 | 0 | 0 | 0 | 0 | 0 | 0 | 0 | 0 |
| ENSMUSG00 | BC039966 | 1368 | 0 | 0 | 0 | 0 | 0 | 0 | 0 | 0 | 0 | 0 | 0 | 0 | 0 | 0 | 0 | 0 | 0 | 0 |
| ENSMUSG00 | 4930564K09R | 2806 | 0.01514796 | 0.09497618 | 0 | 0.06494803 | 0 | 0.03531913 | 0 | 0.32666346 | 0.01467802 | 0 | 0 | 0 | 0.05274261 | 0.04965512 | 0.04405013 | 0.12281378 | 0.01365869 | 0 |
| ENSMUSG00 | Gm16677 | 2171 | 0.01957862 | 0.15344493 | 0.02228168 | 0.06295861 | 0 | 0 | 0 | 0 | 0 | 0 | 0 | 0 | 0 | 0 | 0.15182517 | 0 | 0.01765375 | 0 |
| ENSMUSG00 | 1700121C08 | 673 | 0.31578883 | 0 | 0 | 0 | 0 | 0 | 0 | 0.28374742 | 0 | 0 | 0 | 0 | 0 | 0 | 0 | 0 | 0.05694844 | 0 |
| ENSMUSG00 | 2500002B13 | 2615 | 0.16254369 | 0.33121807 | 0.22198178 | 0.08711481 | 0.31290352 | 0.22739304 | 0.25329924 | 0.21907688 | 0.77175498 | 0.28849379 | 0.47500915 | 0.38419959 | 0.45275947 | 0.24864903 | 0.2363378 | 0.19035484 | 0.5276278 | 0.44175573 |
| ENSMUSG00 | 4930558K02R | 1603 | 0 | 0 | 0 | 0 | 0.06005231 | 0 | 0 | 0 | 0 | 0.05882808 | 0 | 0.06963901 | 0.03077475 | 0.08691969 | 0 | 0 | 0 | 0.0686327 |
| ENSMUSG00 | Olfr75-ps1 | 1731 | 0 | 0 | 0 | 0 | 0 | 0 | 0 | 0 | 0 | 0 | 0 | 0 | 0 | 0 | 0 | 0 | 0 | 0 |
| ENSMUSG00 | Gm15634 | 489 | 0.08692265 | 0 | 0.29677012 | 0 | 0.49214646 | 0.10133483 | 1.741571 | 0 | 0.58958203 | 0.38569082 | 0.0976993 | 0.34242738 | 0.80706613 | 0 | 0 | 0 | 0.07837689 | 0.89994449 |
| ENSMUSG00 | 4930540M05 | 2065 | 0 | 0.0322643 | 0 | 0.08825384 | 0 | 0.02399648 | 0 | 0 | 0.75791167 | 0.02283327 | 0.06940672 | 0.02702938 | 0.02388955 | 0 | 0.01995233 | 0.07417067 | 0 | 0 |
| ENSMUSG00 | 9530082P21R | 4763 | 0.34803734 | 0.60848663 | 0.34530759 | 0.67915896 | 0.6972712 | 0.5513951 | 0.59600267 | 1.20278404 | 0.83012923 | 0.61376099 | 0.3811565 | 0.85545735 | 0.88037224 | 0.7508282 | 0.5363209 | 0.35372386 | 0.74834046 | 0.53126578 |
| ENSMUSG00 | Gm15686 | 362 | 0 | 0 | 0 | 0 | 0 | 0 | 0 | 0 | 0 | 0 | 0 | 0 | 0.27255203 | 0 | 0 | 0 | 0 | 0 |
| ENSMUSG00 | 2810433D01 | 1201 | 0.10617446 | 0.08321289 | 0.12083313 | 0 | 0.04007654 | 0.04125956 | 0.15757762 | 0.19080301 | 0.51440278 | 0.70667166 | 0.31823453 | 0 | 0 | 0.46405416 | 0.54889666 | 0.35070498 | 0 | 0 |
| ENSMUSG00 | 2900093L17R | 412 | 0 | 0 | 0 | 0.11058506 | 0 | 0.12027363 | 0 | 0 | 0 | 0 | 0 | 0 | 0 | 0 | 0 | 0 | 0 | 0 |
| ENSMUSG00 | D630044L22 | 2957 | 0.07187213 | 0.06759465 | 0.01635899 | 0.15407861 | 0 | 0.01675777 | 0.04800069 | 0.01291593 | 0.02785696 | 0.07972726 | 0.09693938 | 0.15100619 | 0.0166831 | 0.01570649 | 0.06966783 | 0 | 0.06480605 | 0.22323614 |
| ENSMUSG00 | Gm15138 | 377 | 0 | 0.26508934 | 0 | 0.36255473 | 0.12767089 | 0 | 0 | 0 | 0 | 0 | 0 | 0.59220862 | 0 | 0 | 0 | 0 | 0 | 0 |
| ENSMUSG00 | Gm15972 | 415 | 1.02422111 | 0.88299238 | 0.46625089 | 1.64678474 | 3.01549401 | 1.43285013 | 1.71009683 | 1.47247817 | 3.27507234 | 1.36339383 | 1.84192611 | 1.07596458 | 3.20954643 | 5.93141357 | 2.97842573 | 2.67572919 | 3.04763325 | 3.31380161 |
| ENSMUSG00 | Gm15265 | 1137 | 0.7476724 | 0.61527773 | 0.68071811 | 0.52092663 | 1.35463637 | 0.479402 | 0.91546081 | 0.67181007 | 1.08671546 | 0.24881638 | 0.6302765 | 0.6872641 | 1.08469475 | 0.24508753 | 0.57979322 | 0.87559876 | 0.40449918 | 0.58057105 |
| ENSMUSG00 | Gm16933 | 2031 | 0.0627846 | 0.08201106 | 0.04763518 | 0.02243281 | 0.02369863 | 0.04879639 | 0 | 0 | 0 | 0 | 0.02352287 | 0.02748186 | 0 | 0 | 0 | 0 | 0.01887065 | 0 |
| ENSMUSG00 | Snhg12 | 1658 | 1.61509414 | 1.44663956 | 1.05032997 | 1.97852545 | 1.6256862 | 1.5840138 | 2.31141556 | 1.70460662 | 2.33506181 | 1.87692787 | 1.64244429 | 1.11092694 | 1.90424771 | 4.56597482 | 1.86376158 | 1.91683982 | 1.80304654 | 3.8154688 |
| ENSMUSG00 | Gm15513 | 1567 | 0.02712519 | 0 | 0 | 0 | 0 | 0 | 0 | 0 | 0 | 0 | 0 | 0 | 0 | 0 | 0 | 0 | 0 | 0 |
| ENSMUSG00 | Gm16052 | 503 | 0.25351 | 0.13245684 | 0.28851012 | 0.81520755 | 0.57413825 | 0.29554314 | 0.18812198 | 0.37964615 | 0.65505393 | 0.28121692 | 0.28494011 | 0.88772426 | 0.29422615 | 0.16496834 | 0.16382328 | 0.15224894 | 0.38097711 | 0 |
| ENSMUSG00 | Gm11269 | 592 | 0 | 0 | 0 | 0 | 0 | 0 | 0 | 0 | 0 | 0 | 0 | 0 | 0 | 0 | 0.06468008 | 0 | 0 | 0 |
| ENSMUSG00 | Gm16765 | 1094 | 0.54394192 | 0.45675815 | 0.75169106 | 0.70798698 | 0.21998137 | 0.86060506 | 0.51896906 | 0 | 0.11294291 | 0.25859618 | 0.52403976 | 0.71427722 | 0.9018632 | 0.63680193 | 0.15064554 | 0.73501166 | 0.21019907 | 0.45254293 |
| ENSMUSG00 | 4933433G15 | 1810 | 0.14090114 | 0.11042948 | 0.13362854 | 0 | 0.37229112 | 0.30114921 | 0.39209402 | 0.04220155 | 0.06826494 | 0.07815034 | 0.36953007 | 0.40088597 | 0.05451041 | 0.35923604 | 0.4780291 | 0.10577516 | 0.14822325 | 0.45587658 |
| ENSMUSG00 | D030055H07 | 5114 | 0.0249346 | 0.03908435 | 0.00945904 | 0.02672725 | 0.04705898 | 0.08720661 | 0 | 0.01493641 | 0.04832208 | 0.03687971 | 0.18683988 | 0.03274286 | 0.05787867 | 0.18163507 | 0.0402831 | 0.02246223 | 0.02997755 | 0 |
| ENSMUSG00 | Gm15632 | 244 | 0 | 0 | 0 | 0 | 0 | 0 | 0 | 0 | 0.33759439 | 0 | 0 | 0 | 0 | 0 | 0 | 0 | 0 | 0 |
| ENSMUSG00 | Gm11716 | 468 | 0 | 0 | 0 | 0 | 0 | 0 | 0 | 0 | 0 | 0 | 0 | 0 | 0.10541008 | 0 | 0.08803751 | 0.08181754 | 0 | 0.23508165 |
| ENSMUSG00 | 4732463B04R | 2929 | 0 | 0.02274694 | 0.0660615 | 0.07777577 | 0.04929866 | 0.01691797 | 0.06461274 | 0.0521576 | 0.07030815 | 0 | 0.17942115 | 0 | 0.1684258 | 0.28541945 | 0.3798267 | 0 | 0.17010647 | 0.01878085 |
| ENSMUSG00 | Gm13790 | 471 | 0 | 0 | 0 | 0 | 0 | 0 | 0 | 0 | 0 | 0 | 0 | 0 | 0 | 0 | 0 | 0.08129641 | 0 | 0 |
| ENSMUSG00 | Gm16619 | 1207 | 0.45780223 | 0 | 0 | 0 | 0 | 0 | 0 | 0 | 0 | 0 | 0 | 0 | 0 | 0 | 0 | 0.38068541 | 0.12701341 | 0.22787534 |
| ENSMUSG00 | Gm4262 | 1983 | 0.04286957 | 0 | 0.19515292 | 0.04595163 | 0.77671284 | 0.02498877 | 0.23859142 | 0.67409687 | 0 | 0.02377746 | 0.21683037 | 0.05629416 | 0 | 0 | 0.5817668 | 0 | 0.13529202 | 1.19283489 |
| ENSMUSG00 | Gm11754 | 658 | 0.71057285 | 0.101255 | 0.66164403 | 0.7616588 | 0.14629764 | 0.22592432 | 0.14380753 | 0.23217266 | 0.68852838 | 0 | 0.29042528 | 0.2544787 | 0 | 0.28233488 | 0.43831442 | 0.29096206 | 0 | 0 |
| ENSMUSG00 | G630016G05 | 3176 | 0 | 0 | 0 | 0 | 0 | 0 | 0 | 0 | 0 | 0 | 0 | 0 | 0 | 0 | 0 | 0 | 0 | 0 |
| ENSMUSG00 | A930007I19R | 5070 | 0.23474259 | 0.38766484 | 0.47705651 | 0.29655118 | 0.49366076 | 0.30298516 | 0.51325391 | 0.36158487 | 0.4549201 | 0.27899824 | 0.41461502 | 0.55045032 | 0.71030177 | 0.33894107 | 0.41445352 | 0.3776194 | 0.23434225 | 0.35804744 |
| ENSMUSG00 | Gm12278 | 405 | 0 | 0.16450812 | 0.11944082 | 0.67497843 | 0 | 0 | 0.11682143 | 0 | 0.7118657 | 0 | 0 | 0 | 0.24361441 | 0 | 0 | 0 | 0.18926566 | 0 |
| ENSMUSG00 | Gm12122 | 452 | 0 | 0 | 0 | 0 | 0 | 0 | 0 | 0 | 0 | 0.10431572 | 0 | 0 | 0.32742423 | 0.10275241 | 0.45576942 | 0 | 0 | 0 |
| ENSMUSG00 | Gm15336 | 1568 | 0.05421579 | 0.0637364 | 0.15425233 | 0 | 0.03069638 | 0.03160251 | 0 | 0.0487148 | 0.02626691 | 0.0300706 | 0.18281234 | 0 | 0.12584673 | 0 | 0 | 0.02442003 | 0.14665675 | 0.07016468 |
| ENSMUSG00 | Gm15940 | 815 | 0.36507513 | 0.57224604 | 0 | 0.22361249 | 0.05905758 | 0.3648054 | 0 | 0.18744737 | 0 | 0.17356087 | 0 | 0.13697095 | 0.24211984 | 0.51287949 | 0.20221623 | 0.09396468 | 0.04702613 | 0 |
| ENSMUSG00 | 4930555J06R | 746 | 0.28488724 | 0.04465535 | 0.12968775 | 0 | 0.12904001 | 0 | 0.19026546 | 0.35837375 | 0 | 0 | 0.0640415 | 0.44891954 | 0.66128575 | 0 | 0.1656899 | 0 | 0.10275147 | 0 |
| ENSMUSG00 | 2210013O21 | 1529 | 33.3035977 | 26.9509811 | 30.1504084 | 27.503495 | 27.6073885 | 27.482484 | 36.6990433 | 30.5988835 | 34.2906701 | 35.1857112 | 29.1836568 | 31.3210193 | 43.4921023 | 36.5416855 | 38.9919237 | 43.7249186 | 34.4410281 | 37.6321293 |
| ENSMUSG00 | Gm16766 | 2613 | 0 | 0 | 0 | 0 | 0 | 0 | 0 | 0.02923261 | 0 | 0 | 0 | 0 | 0 | 0 | 0 | 0 | 0 | 0 |
| ENSMUSG00 | Gm16767 | 2322 | 0.03661083 | 0 | 0 | 0 | 0 | 0 | 0 | 0 | 0 | 0 | 0.16459934 | 0 | 0 | 0 | 0 | 0 | 0 | 0 |
| ENSMUSG00 | Gm16087 | 961 | 0 | 0 | 0 | 0 | 0 | 0 | 0 | 0.23845413 | 0 | 0.19625683 | 0 | 0 | 0 | 0 | 0.04287363 | 0 | 0.03988168 | 0 |
| ENSMUSG00 | Gm12840 | 625 | 0.06800828 | 0 | 0 | 0.07289767 | 0 | 0.23785312 | 0 | 0.06110784 | 0 | 0 | 0 | 0 | 0 | 0.14862108 | 0 | 0.36758983 | 0 | 0.08801457 |
| ENSMUSG00 | Gm11413 | 1458 | 1.54511271 | 1.39374935 | 1.99068026 | 1.09371506 | 1.28747944 | 0.84966964 | 1.26556549 | 0.75065684 | 0.87570781 | 0.51742884 | 1.21239607 | 0.22969408 | 0.50753001 | 1.01934897 | 0.50866118 | 0.02626242 | 0.49945106 | 0.4150207 |
| ENSMUSG00 | E130218I03R | 2090 | 0 | 0 | 0.04629046 | 0.04359909 | 0 | 0 | 0.02263765 | 0.01827388 | 0.03941293 | 0 | 0.02285883 | 0.08011818 | 0.07081136 | 0.15555436 | 0.05914099 | 0.05496259 | 0.03667588 | 0.05264029 |
| ENSMUSG00 | Gm15564 | 933 | 110.932587 | 101.009837 | 82.2816642 | 88.1432851 | 93.8391952 | 85.8276714 | 147.161194 | 113.308221 | 139.054153 | 121.843886 | 94.8842423 | 107.922246 | 229.475371 | 267.513961 | 131.4652 | 121.766594 | 150.10106 | 142.386916 |
| ENSMUSG00 | Gm16938 | 1314 | 2.29670281 | 2.2563528 | 1.9143254 | 2.94725173 | 3.04029655 | 1.92327961 | 3.34861426 | 1.65674805 | 1.50453026 | 2.47595017 | 3.01774852 | 2.76104876 | 3.34135513 | 2.79230054 | 2.69660107 | 3.05975173 | 4.92933347 | 3.39097233 |
| ENSMUSG00 | Gm13200 | 256 | 0 | 0.13012849 | 0 | 0 | 0 | 0 | 0 | 0 | 0 | 0 | 0 | 0 | 0.1927028 | 0 | 0 | 0 | 0.1497121 | 0.42975865 |
| ENSMUSG00 | 1700003F17R | 571 | 0 | 0 | 0 | 0 | 0 | 0 | 0 | 0 | 0 | 0 | 0 | 0 | 0 | 0 | 0 | 0 | 0 | 0 |
| ENSMUSG00 | 2700033N17 | 719 | 0.05911707 | 0.13899678 | 0 | 0 | 0 | 0 | 0 | 0 | 0.05728305 | 0 | 0 | 0.77629572 | 0.06861185 | 0 | 0 | 0 | 0 | 0 |
| ENSMUSG00 | Gm14233 | 437 | 0 | 0 | 0 | 0 | 0 | 0 | 0 | 0 | 0 | 0 | 0 | 0 | 0 | 0 | 0 | 0 | 0 | 0 |
| ENSMUSG00 | 1700007J10R | 1291 | 0.52678762 | 0.87733416 | 0.97421517 | 0.88228204 | 0.89478402 | 0.61413148 | 0.54972129 | 0.85792383 | 0.79757002 | 0.98611076 | 1.40623425 | 0.77821993 | 0.42033392 | 0.71950561 | 0.63828901 | 0.71183159 | 0.74218236 | 0.72436469 |
| ENSMUSG00 | Gm16310 | 958 | 0.6655299 | 0.52160064 | 0.70692007 | 0.99872853 | 1.1555681 | 0.20690077 | 0.83957781 | 0.91693659 | 1.16078907 | 0.7382678 | 0.24934738 | 1.22351661 | 0.72092572 | 0.19392103 | 0.17203155 | 1.07917161 | 0.48007888 | 1.55036105 |
| ENSMUSG00 | 4930480G23 | 1010 | 0 | 0 | 0 | 0 | 0 | 0 | 0 | 0 | 0 | 0 | 0 | 0 | 0 | 0 | 0 | 0 | 0 | 0 |
| ENSMUSG00 | 1700120E14 | 1483 | 0 | 0 | 0 | 0 | 0 | 0 | 0 | 0.07726042 | 0 | 0.06358827 | 0 | 0.03763699 | 0.19958969 | 0 | 0 | 0 | 0 | 0 |
| ENSMUSG00 | Gm12107 | 886 | 0.2398712 | 0 | 0 | 0 | 0 | 0 | 0 | 0 | 0 | 0 | 0 | 0 | 0 | 0 | 0 | 0 | 0 | 0 |
| ENSMUSG00 | Gm12979 | 402 | 0 | 0 | 0 | 0 | 0 | 0.24653101 | 0 | 0 | 0 | 0.70374184 | 0.5942159 | 0 | 0.24543242 | 1.27085811 | 0.20498287 | 0.8572524 | 0.66737333 | 0.27367715 |
| ENSMUSG00 | Gm11535 | 776 | 0 | 0 | 0 | 0 | 0 | 0 | 0 | 0 | 0 | 0 | 0 | 0 | 0 | 0 | 0 | 0 | 0 | 0 |
| ENSMUSG00 | Gm13723 | 561 | 0 | 0 | 0 | 0 | 0 | 0 | 0 | 0 | 0 | 0 | 0 | 0 | 0 | 0 | 0 | 0 | 0 | 0 |
| ENSMUSG00 | Gm11916 | 648 | 0 | 0 | 0 | 0 | 0 | 0 | 0 | 0 | 0 | 0 | 0 | 0 | 0 | 0 | 0 | 0 | 0 | 0 |
| ENSMUSG00 | 1810059C17 | 484 | 0.17564122 | 0.82593953 | 0.09994531 | 0.75307511 | 0 | 0 | 0.78202775 | 0.31563969 | 0.51057664 | 0.29225643 | 0.09870859 | 0.23064323 | 1.01925449 | 0.28787657 | 0 | 0.23733848 | 0.15837313 | 0 |
| ENSMUSG00 | Gm15932 | 509 | 0.58455055 | 0.58902956 | 0.38014562 | 0.98461982 | 1.32386431 | 0.29205933 | 0.65066553 | 1.05047865 | 0 | 0.83370595 | 0.46930215 | 1.53520486 | 1.06611216 | 0.82121176 | 0.32378433 | 0.82749839 | 0.22589173 | 0.2161458 |
| ENSMUSG00 | Gm12932 | 618 | 0.55022882 | 0.48513924 | 1.40893778 | 1.32702071 | 1.47978406 | 1.04237142 | 0.84213506 | 0.92700006 | 2.46585936 | 1.75479964 | 0.38652879 | 0.81284946 | 2.31492816 | 2.40487184 | 0.60002266 | 0.74350694 | 0.68218328 | 2.31429899 |
| ENSMUSG00 | A530083I20R | 3182 | 0.37402418 | 0.24079088 | 0.39525826 | 0.47250612 | 0.24202099 | 0.28031088 | 0.47580318 | 0.39608714 | 0.33653344 | 0.3556307 | 0.52549452 | 0.36836232 | 0.48060636 | 0.78817747 | 0.54382946 | 0.34897159 | 0.49383349 | 0.55320283 |
| ENSMUSG00 | Gm11345 | 615 | 0 | 0 | 0 | 0 | 0 | 0 | 0 | 0 | 0 | 0 | 0 | 0 | 0 | 0 | 0 | 0 | 0 | 0 |
| ENSMUSG00 | Gm1989 | 628 | 0 | 0 | 0 | 0 | 0 | 0 | 0 | 0 | 0 | 0 | 0 | 0 | 0 | 0 | 0 | 0 | 0 | 0 |
| ENSMUSG00 | Gm12510 | 360 | 0 | 0 | 0 | 0 | 0 | 0 | 0 | 0 | 0 | 0.13097418 | 0 | 0 | 0 | 0 | 0.11444877 | 0 | 0 | 0 |
| ENSMUSG00 | Gm11626 | 331 | 0.25682886 | 0.80514548 | 0.1461436 | 0.13764666 | 0 | 0 | 0.42881582 | 0.57692451 | 0.12443056 | 0.14244925 | 0.43300567 | 0.50588214 | 1.1923122 | 0.56125785 | 0.124476 | 0.11568159 | 0.46315767 | 0.16619065 |
| ENSMUSG00 | 1700017J07R | 471 | 0 | 0 | 0 | 0 | 0 | 0 | 0 | 0 | 0 | 0 | 0 | 0 | 0 | 0 | 0 | 0 | 0 | 0 |
| ENSMUSG00 | Gm13497 | 2035 | 0.14620945 | 0 | 0 | 0 | 0 | 0 | 0 | 0 | 0 | 0 | 0 | 0 | 0 | 0 | 0 | 0 | 0 | 0 |
| ENSMUSG00 | B230369F24 | 2558 | 0.9305277 | 0.40371373 | 0.49167779 | 0.51652474 | 0.28224349 | 0.30994673 | 0.57337492 | 0.07465286 | 0.98216476 | 0.18432644 | 0.354857 | 0.74188136 | 0.84855526 | 0.01815641 | 0.49931518 | 0.70354126 | 0.26969247 | 0.73116092 |
| ENSMUSG00 | 6230400D17 | 1551 | 1.06879553 | 0.64434998 | 1.40348734 | 0.55813014 | 1.30337898 | 0.51118229 | 0.76261571 | 1.47746238 | 1.14185696 | 1.15520743 | 0.70846167 | 1.07960663 | 1.78116529 | 1.25767355 | 1.72669319 | 0.96281992 | 1.72974905 | 1.52507517 |
| ENSMUSG00 | 4930401G09 | 635 | 0 | 0 | 0 | 0 | 0 | 0 | 0 | 0 | 0 | 0 | 0 | 0 | 0 | 0 | 0 | 0 | 0 | 0 |
| ENSMUSG00 | Gm13481 | 1726 | 0 | 0 | 0 | 0.18477828 | 0.22309119 | 0 | 0.0548235 | 0.0221277 | 0.04772482 | 0.10927162 | 0.11071833 | 0.25870527 | 0.02858164 | 0.24217659 | 0.07161336 | 0.13310756 | 0.48851595 | 0.47806292 |
| ENSMUSG00 | Gm13938 | 488 | 0 | 0 | 0 | 0 | 0 | 0 | 0 | 0 | 0 | 0 | 0 | 0 | 0 | 0 | 0 | 0 | 0 | 0 |
| ENSMUSG00 | 4933428C19 | 811 | 0 | 0 | 0 | 0 | 0 | 0 | 0 | 0 | 0 | 0 | 0 | 0 | 0 | 0 | 0 | 0 | 0 | 0 |
| ENSMUSG00 | Gm13441 | 1164 | 0 | 0 | 0 | 0 | 0 | 0 | 0 | 0 | 0 | 0 | 0 | 0 | 0 | 0 | 0 | 0 | 0 | 0 |
| ENSMUSG00 | 1700112J16R | 2212 | 1.19137473 | 0.90360473 | 0.78727265 | 0.67970817 | 1.26204863 | 1.07528536 | 0.9197311 | 1.26041834 | 1.21027284 | 1.23632043 | 0.51835398 | 1.31212227 | 1.20430539 | 1.4697496 | 1.52736328 | 1.21172808 | 0.485143 | 1.14395068 |
| ENSMUSG00 | Gm13270 | 645 | 0.32949749 | 0.15494369 | 0.67497949 | 0.56509822 | 0.07462314 | 0.15365189 | 0.44011794 | 0.6513433 | 1.40481139 | 0.58481492 | 0.37034852 | 0.43267955 | 0.22945078 | 0.50404436 | 1.08593248 | 0.3561917 | 0.11884123 | 1.87627961 |
| ENSMUSG00 | 9630013K17 | 1068 | 0.31839083 | 0.65502882 | 0.27176141 | 0.21330077 | 0 | 0.27838614 | 0.17720104 | 0.14304271 | 0 | 0 | 0 | 0.05226186 | 0.32333654 | 0.13046092 | 0.15431294 | 0.03585263 | 0.21531627 | 0 |
| ENSMUSG00 | Gm16214 | 1384 | 0.06142366 | 0.26477011 | 0.03495197 | 0.09875949 | 0.34777401 | 0.25062799 | 0.30766915 | 0.22076533 | 0.2083133 | 0.17034214 | 0.06903896 | 0.08065847 | 0.17822224 | 0.13423147 | 0.47631857 | 0.05533325 | 0.22153929 | 0.23847879 |
| ENSMUSG00 | Gm6569 | 807 | 0 | 0 | 0 | 0 | 0 | 0 | 0 | 0 | 0 | 0 | 0 | 0 | 0 | 0 | 0 | 0 | 0 | 0 |
| ENSMUSG00 | A330102I10R | 1825 | 0.02329051 | 0.12777548 | 0.15903626 | 0.09985982 | 0.05274731 | 0 | 0.05184951 | 0.31391016 | 0.24824749 | 0.025836 | 0.10471224 | 0 | 0.02703119 | 0 | 0.15803337 | 0.06294346 | 0.12600426 | 0 |
| ENSMUSG00 | Gm11751 | 455 | 0 | 0 | 0 | 0 | 0 | 0 | 0 | 0 | 0 | 0 | 0 | 0 | 0 | 0 | 0 | 0 | 0 | 0 |
| ENSMUSG00 | 4930593C16 | 580 | 0 | 0 | 0 | 0 | 0 | 0 | 0 | 0 | 0 | 0 | 0 | 0 | 0 | 0 | 0 | 0 | 0 | 0.37937315 |
| ENSMUSG00 | 1700061J23R | 534 | 0 | 0 | 0 | 0 | 0 | 0 | 0 | 0 | 0 | 0 | 0 | 0 | 0 | 0 | 0 | 0 | 0 | 0 |
| ENSMUSG00 | Gm6182 | 1206 | 0 | 0 | 0 | 0 | 0 | 0 | 0 | 0 | 0 | 0 | 0 | 0 | 0 | 0 | 0 | 0 | 0 | 0 |
| ENSMUSG00 | Gm13830 | 2648 | 0.59391673 | 0.62901991 | 1.11434492 | 0.75705663 | 0.78159846 | 0.65496438 | 0.44668315 | 0.60577073 | 1.64870494 | 0.89030784 | 0.54125708 | 0.86421532 | 0.96875366 | 0.99974055 | 0.809094 | 0.30366418 | 0.3763156 | 0.43625047 |
| ENSMUSG00 | E330017L17 | 1604 | 0 | 0 | 0 | 0 | 0 | 0 | 0 | 0 | 0 | 0.0293957 | 0 | 0 | 0 | 0 | 0 | 0 | 0 | 0 |
| ENSMUSG00 | B230206F22 | 8653 | 0.36841421 | 0.21559252 | 0.43045901 | 0.33698218 | 0.55068305 | 0.30923929 | 0.62332664 | 0.27806788 | 0.47597961 | 0.19071705 | 0.52451416 | 0.2386663 | 0.21094198 | 0.32741123 | 0.3618766 | 0.31860901 | 0.73082619 | 0.23521749 |
| ENSMUSG00 | Gm12478 | 351 | 0 | 0 | 0 | 0 | 0 | 0 | 0 | 0 | 0 | 0 | 0 | 0 | 0 | 0 | 0 | 0 | 0 | 0 |
| ENSMUSG00 | Gm16246 | 1343 | 0.18989654 | 0.22324352 | 0.07203802 | 0.16962414 | 0.03583911 | 0.03689705 | 0 | 0.08531438 | 0.15333774 | 0 | 0 | 0.16624174 | 0.11019788 | 0.10374703 | 0.09203624 | 0.114045 | 0 | 0.36863884 |
| ENSMUSG00 | Gm11978 | 1868 | 0 | 0 | 0 | 0 | 0 | 0.07958148 | 0 | 0 | 0 | 0.07572383 | 0 | 0 | 0 | 0 | 0 | 0 | 0 | 0 |
| ENSMUSG00 | Gm12127 | 704 | 0 | 0 | 0 | 0 | 0 | 0 | 0 | 0 | 0 | 0 | 0 | 0 | 0 | 0 | 0 | 0 | 0 | 0 |
| ENSMUSG00 | 1700071G01 | 361 | 0 | 0 | 0 | 0 | 0 | 0 | 0 | 0 | 0 | 0 | 0 | 0 | 0 | 0.25730796 | 0 | 0 | 0 | 0 |
| ENSMUSG00 | 4930529C04 | 3533 | 0 | 0 | 0 | 0 | 0 | 0 | 0 | 0 | 0 | 0 | 0 | 0 | 0 | 0 | 0 | 0 | 0 | 0 |
| ENSMUSG00 | Gm13753 | 453 | 0 | 0 | 0 | 0 | 0 | 0 | 0 | 0 | 0 | 0 | 0 | 0 | 0 | 0 | 0 | 0 | 0 | 0 |
| ENSMUSG00 | 1700026D11 | 3169 | 0 | 0 | 0 | 0.0143771 | 0 | 0 | 0 | 0 | 0 | 0 | 0 | 0 | 0 | 0 | 0 | 0 | 0 | 0.19094357 |
| ENSMUSG00 | Gm11195 | 439 | 0 | 0 | 0 | 0 | 0 | 0 | 0 | 0 | 0 | 0 | 0 | 0 | 0 | 0 | 0 | 0 | 0 | 0 |
| ENSMUSG00 | AV064505 | 518 | 0 | 0 | 0 | 0 | 0 | 0 | 0 | 0 | 0 | 0 | 0 | 0 | 0 | 0 | 0 | 0 | 0 | 0 |
| ENSMUSG00 | Gm13705 | 846 | 2.46188372 | 2.52012439 | 2.11562721 | 2.53116913 | 2.78778281 | 1.05431348 | 1.62182942 | 1.17376178 | 1.46051474 | 2.17361398 | 1.80709063 | 3.89258167 | 2.56572619 | 2.96961777 | 1.94806411 | 2.12725597 | 2.80878299 | 0 |
| ENSMUSG00 | Gm16630 | 4422 | 0 | 0 | 0 | 0.02060653 | 0 | 0 | 0 | 0 | 0 | 0 | 0 | 0 | 0 | 0 | 0 | 0.00865912 | 0.01733437 | 0 |
| ENSMUSG00 | Gm12357 | 1051 | 0 | 0 | 0 | 0 | 0 | 0 | 0 | 0 | 0 | 0 | 0 | 0 | 0 | 0 | 0 | 0 | 0 | 0 |
| ENSMUSG00 | Gm12701 | 657 | 0 | 0 | 0 | 0 | 0 | 0 | 0 | 0 | 0 | 0 | 0 | 0.16991069 | 0 | 0 | 0 | 0 | 0 | 0 |
| ENSMUSG00 | Gm12108 | 2587 | 0 | 0 | 0 | 0 | 0 | 0 | 0 | 0 | 0 | 0 | 0 | 0 | 0 | 0 | 0 | 0 | 0 | 0 |
| ENSMUSG00 | Gm15563 | 408 | 0.10417935 | 0.24494775 | 0.4742503 | 0 | 0.70782241 | 0.12145278 | 0.34788735 | 0.09360883 | 0 | 0.2311309 | 0 | 0.54721238 | 0 | 0.34150064 | 0 | 0 | 0 | 0.13482624 |
| ENSMUSG00 | Gm16631 | 3670 | 0 | 0 | 0 | 0 | 0 | 0 | 0 | 0 | 0 | 0 | 0.0130177 | 0 | 0 | 0 | 0.01122658 | 0 | 0 | 0 |
| ENSMUSG00 | Gm15998 | 1841 | 1.33910929 | 0.45237499 | 0.81454614 | 0.74243961 | 0.60132224 | 1.42655887 | 0.2826939 | 1.72187366 | 1.76737357 | 0.43539487 | 0.88231862 | 1.33399737 | 0.80388784 | 0.35318698 | 0.69377959 | 0.97754402 | 1.31154628 | 1.49400073 |
| ENSMUSG00 | 1810019D21 | 1732 | 16.0007937 | 18.8875647 | 16.7017154 | 15.0204136 | 17.3408316 | 14.619773 | 23.2192708 | 19.8900387 | 18.7146581 | 16.0617292 | 20.2464316 | 20.0124287 | 8.40237621 | 11.6646524 | 9.51537088 | 6.36702941 | 6.61637574 | 9.68693854 |
| ENSMUSG00 | 1700042O10 | 1071 | 0 | 0 | 0 | 0 | 0 | 0 | 0 | 0 | 0 | 0.08804987 | 0 | 0 | 0 | 0 | 0 | 0 | 0 | 0 |
| ENSMUSG00 | A330050B17 | 8235 | 0.18581498 | 0.07686035 | 0.17035002 | 0.24896746 | 0.21041278 | 0.06017333 | 0.15512354 | 0.15768569 | 0.04001119 | 0.14314118 | 0.23205809 | 0.14233502 | 0.16773451 | 0.24815299 | 0.24015479 | 0.22318751 | 0.16289258 | 0.06011924 |
| ENSMUSG00 | Gm16794 | 3057 | 0.80644429 | 0.78460202 | 1.18678926 | 0.50673062 | 0.83447561 | 0.7132222 | 0.94408682 | 1.03695433 | 0.75447984 | 0.30847696 | 0.90642708 | 0.63904095 | 0.9036923 | 0.74444236 | 0.835622 | 0.57617532 | 0.4011912 | 0.52183974 |
| ENSMUSG00 | Gm11612 | 159 | 2.13862521 | 0.62854517 | 2.12965228 | 2.00583214 | 0.605433 | 0.62330482 | 1.19025607 | 2.40203789 | 0.77710407 | 1.77927182 | 3.90612868 | 2.45729332 | 0.62052726 | 0 | 0.25912928 | 0 | 0 | 0.69193845 |
| ENSMUSG00 | A630014C17 | 2982 | 0.49888704 | 0.32396845 | 0.61642996 | 0.56531142 | 0.64563278 | 0.79762951 | 0.79330448 | 0.47388293 | 0.64915032 | 0.64828264 | 0.28838003 | 0.65511341 | 0.84370482 | 0.73201613 | 0.51121984 | 1.20701445 | 0.88682578 | 0.7563291 |
| ENSMUSG00 | 4930555B11 | 1529 | 0 | 0 | 0 | 0 | 0 | 0 | 0 | 0 | 0 | 0 | 0 | 0 | 0 | 0 | 0 | 0 | 0 | 0 |
| ENSMUSG00 | 4933405D12 | 1376 | 0 | 0.0484199 | 0 | 0 | 0 | 0 | 0 | 0.02775611 | 0.11972824 | 0.1027995 | 0.38192191 | 0.44620079 | 0 | 0.16876485 | 0.08982897 | 0 | 0 | 0.43975303 |
| ENSMUSG00 | Gm15639 | 668 | 0.19089151 | 0 | 0 | 0 | 0 | 0 | 0 | 0 | 0 | 0 | 0 | 0 | 0 | 0 | 0 | 0 | 0 | 0 |
| ENSMUSG00 | Gm16144 | 825 | 0.56673568 | 1.89782549 | 1.23132623 | 0.49702957 | 0.87512588 | 0.54057528 | 0.45878961 | 0.78699496 | 0.14976915 | 0.34291421 | 1.27399889 | 0.33827674 | 0.05979626 | 0.05629586 | 0.64923664 | 0.37130286 | 0.789754 | 0.53342164 |
| ENSMUSG00 | Gm16789 | 908 | 0.1404356 | 0.22012926 | 0.10654963 | 0.20070945 | 0.53008727 | 0.21829398 | 0.15631942 | 0.04206212 | 0 | 0 | 0 | 0.06147099 | 0.16299092 | 0.15344963 | 0 | 0.33736218 | 0.16883831 | 0.12116543 |
| ENSMUSG00 | Gm15559 | 2258 | 0.35766091 | 0.41309169 | 1.11400513 | 0.46408504 | 0.57553673 | 0.57058063 | 0.37716042 | 0.47359902 | 0.47424686 | 0.33410596 | 0.44431981 | 0.42022421 | 0.50249517 | 0.63762919 | 0.49266697 | 0.5087326 | 0.56012746 | 0.21925685 |
| ENSMUSG00 | Gm15322 | 737 | 0 | 0 | 0 | 0 | 0 | 0 | 0.06419631 | 0 | 0 | 0 | 0 | 0 | 0 | 0 | 0 | 0 | 0 | 0.07463922 |
| ENSMUSG00 | 1500012K07R | 2338 | 0.01818014 | 0 | 0.04138027 | 0 | 0 | 0 | 0.08094556 | 0.21236152 | 0 | 0 | 0.02043411 | 0.02387325 | 0.0422001 | 0.13905415 | 0.03524513 | 0.04913252 | 0.01639277 | 0.04705655 |
| ENSMUSG00 | Gm6787 | 1450 | 0 | 0.04594882 | 0 | 0.15710705 | 0 | 0.0341743 | 0.48944151 | 0.34241464 | 0.14202247 | 0.16258863 | 0.13179299 | 0.07698712 | 0.30619811 | 0.28827365 | 0.05682973 | 0.23766584 | 0.39647893 | 0.7587463 |
| ENSMUSG00 | 9330198N18 | 1243 | 0 | 0.0268004 | 0 | 0 | 0.11616715 | 0 | 0 | 0 | 0 | 0 | 0 | 0 | 0.03968779 | 0.03736451 | 0.13258747 | 0 | 0 | 0 |
| ENSMUSG00 | Gm14123 | 2655 | 0 | 0.06273615 | 0.10931871 | 0.06864187 | 0.07251514 | 0 | 0 | 0.14385086 | 0.10858968 | 0.03551842 | 0.10796601 | 0.58863976 | 0.03716152 | 0 | 0.09311086 | 0.01442207 | 0.0866131 | 0 |
| ENSMUSG00 | Gm11588 | 655 | 0 | 0 | 0 | 0 | 0 | 0 | 0 | 0 | 0 | 0 | 0 | 0 | 0 | 0 | 0 | 0 | 0 | 0 |
| ENSMUSG00 | Gm13898 | 781 | 0 | 0 | 0 | 0 | 0 | 0 | 0 | 0 | 0 | 0 | 0 | 0 | 0 | 0 | 0 | 0 | 0 | 0 |
| ENSMUSG00 | Gm16158 | 802 | 0 | 0 | 0 | 0 | 0 | 0 | 0 | 0 | 0 | 0 | 0 | 0 | 0 | 0 | 0 | 0 | 0 | 0 |
| ENSMUSG00 | Gm14236 | 870 | 0 | 0 | 0 | 0 | 0 | 0 | 0 | 0 | 0 | 0 | 0 | 0 | 0 | 0.05338401 | 0 | 0 | 0 | 0 |
| ENSMUSG00 | Gm16626 | 1565 | 0.43455771 | 0.12771717 | 0.43273446 | 0.11644995 | 0.24604178 | 0.53827251 | 0 | 0.07321227 | 0.2105381 | 0.06025649 | 0.48843408 | 0.03566496 | 0 | 0.11870693 | 0.07898062 | 0.46486999 | 0.26938611 | 0.21089753 |
| ENSMUSG00 | Gm12415 | 510 | 0 | 0 | 0 | 0 | 0 | 0 | 0 | 0 | 0 | 0 | 0 | 0 | 0 | 0 | 0 | 0 | 0 | 0 |
| ENSMUSG00 | Gm15645 | 2203 | 0 | 0 | 0 | 0 | 0 | 0 | 0.04295295 | 0 | 0 | 0 | 0 | 0 | 0 | 0 | 0 | 0 | 0 | 0.02497009 |
| ENSMUSG00 | Gm4487 | 441 | 0 | 0 | 0 | 0 | 0 | 0 | 0 | 0 | 0 | 0 | 0 | 0 | 0 | 0 | 0 | 0 | 0 | 0 |
| ENSMUSG00 | Gm14002 | 2836 | 0 | 0 | 0 | 0 | 0.03394353 | 0 | 0 | 0 | 0 | 0.06650311 | 0.06738358 | 0 | 0 | 0 | 0 | 0 | 0 | 0 |
| ENSMUSG00 | Gm12996 | 634 | 0.80451437 | 0.78815996 | 0.83928838 | 0.57490277 | 1.21468577 | 0.78158886 | 1.34326218 | 0.96384612 | 2.20874059 | 1.78488467 | 1.35638683 | 2.64111968 | 4.51301451 | 2.05115843 | 1.23474694 | 0.42276696 | 1.20903145 | 1.38824244 |
| ENSMUSG00 | Gm13362 | 837 | 0.55861044 | 0.55720492 | 0.4045576 | 0.59877119 | 0.11501057 | 0.11840558 | 1.13052996 | 0.13689033 | 0 | 0.39433085 | 0 | 0.06668538 | 0 | 0 | 1.13218134 | 0 | 0.04579008 | 0.52577402 |
| ENSMUSG00 | 4933408J17R | 1855 | 0 | 0 | 0 | 0 | 0 | 0.02671306 | 0 | 0 | 0 | 0 | 0.02575469 | 0 | 0.05318805 | 0 | 0 | 0 | 0 | 0 |
| ENSMUSG00 | Gm8865 | 429 | 0.09907966 | 0.07765243 | 0 | 0 | 0 | 0 | 0 | 0 | 0 | 0 | 0 | 0 | 0 | 0 | 0 | 0 | 0.26801606 | 0 |
| ENSMUSG00 | Gm14091 | 872 | 0.1462334 | 0 | 0 | 0 | 0 | 0.11365306 | 0 | 0.04379863 | 0 | 0.05407191 | 0 | 0 | 0.3960131 | 0.10652314 | 0 | 0.04391125 | 0 | 0 |
| ENSMUSG00 | 4930421P07R | 398 | 0 | 0 | 0 | 0 | 0.12093448 | 0 | 0 | 0 | 0 | 0 | 0 | 0 | 0 | 0 | 0 | 0 | 0 | 0 |
| ENSMUSG00 | Gm11364 | 295 | 0 | 0 | 0 | 0 | 0 | 0 | 0 | 0 | 0 | 0 | 0 | 0 | 0 | 0 | 0 | 0 | 0 | 0 |
| ENSMUSG00 | Gm15569 | 591 | 0.07192077 | 0.056367 | 0.49110183 | 0.30836578 | 0.32576598 | 0.08384557 | 0.56038706 | 0.51698683 | 0.20906861 | 0.55846857 | 0.32334997 | 0.56665647 | 0 | 0 | 0.76686483 | 0.06478952 | 0.19454973 | 0 |
| ENSMUSG00 | F730016J06R | 3794 | 0 | 0 | 0 | 0 | 0 | 0 | 0 | 0 | 0 | 0 | 0 | 0 | 0 | 0 | 0 | 0 | 0 | 0 |
| ENSMUSG00 | Gm15295 | 486 | 0 | 0 | 0 | 0 | 0 | 0 | 0 | 0 | 0 | 0 | 0 | 0 | 0 | 0 | 0 | 0 | 0 | 0 |
| ENSMUSG00 | Hoxa11as | 3375 | 0.21410015 | 0.26650315 | 0.0286658 | 0.04049871 | 0.37079408 | 0.24959895 | 0 | 0 | 0 | 0 | 0 | 0 | 0 | 0 | 0 | 0 | 0 | 0 |
| ENSMUSG00 | Gm15199 | 539 | 0 | 0 | 0 | 0 | 0 | 0 | 0 | 0 | 0 | 0 | 0 | 0 | 0 | 0 | 0 | 0 | 0 | 0 |
| ENSMUSG00 | Gt(ROSA)26S | 1851 | 29.7605122 | 30.613308 | 31.2820723 | 24.5158294 | 30.0857027 | 28.1628718 | 23.3879531 | 24.3473986 | 24.4760493 | 15.0036544 | 21.5000218 | 25.8724141 | 19.0291674 | 18.7181465 | 16.5162369 | 15.7216973 | 19.7118502 | 14.3243595 |
| ENSMUSG00 | 4930551O13 | 1063 | 0 | 0.03133856 | 0.04550661 | 0 | 0 | 0.18646372 | 0.04450863 | 0.10778665 | 0 | 0.17742504 | 0.04494352 | 0 | 0.1392246 | 0 | 0.42635665 | 0.50429775 | 0.28843873 | 0.31049355 |
| ENSMUSG00 | Gm14929 | 1848 | 0 | 0 | 0 | 0 | 0 | 0 | 0 | 0 | 0 | 0 | 0 | 0 | 0 | 0 | 0 | 0 | 0 | 0 |
| ENSMUSG00 | B430119L08 | 1992 | 0 | 0 | 0 | 0 | 0 | 0 | 0 | 0 | 0 | 0 | 0 | 0 | 0 | 0 | 0 | 0 | 0 | 0 |
| ENSMUSG00 | 1700006D18 | 321 | 0.39724464 | 0.20755697 | 0 | 0.14193472 | 0 | 0.46310966 | 0.58956609 | 0 | 0 | 0.14688693 | 0.29766329 | 0 | 0 | 0 | 0 | 0 | 0.47758625 | 0.85683967 |
| ENSMUSG00 | Gm15200 | 408 | 0 | 0 | 0 | 0 | 0 | 0 | 0 | 0 | 0 | 0 | 0 | 0 | 0 | 0 | 0 | 0 | 0 | 0 |
| ENSMUSG00 | D930007P13 | 1317 | 0 | 0 | 0 | 0 | 0 | 0 | 0 | 0 | 0 | 0 | 0 | 0 | 0 | 0 | 0 | 0 | 0 | 0 |
| ENSMUSG00 | Gm13690 | 571 | 0 | 0.35004793 | 0.08471722 | 0.07979167 | 0 | 0 | 0 | 0 | 0.0721305 | 0.08257566 | 0.08366893 | 0 | 0 | 0 | 0 | 0.13411771 | 0.40272816 | 0.09633819 |
| ENSMUSG00 | Gm13021 | 277 | 0 | 0 | 0 | 0 | 0 | 0 | 0 | 0 | 0 | 0 | 0 | 0 | 0 | 0 | 0 | 0 | 0 | 0 |
| ENSMUSG00 | 2510003D18 | 494 | 0 | 0 | 0 | 0 | 0 | 0 | 0 | 0 | 0.08337351 | 0 | 0 | 0.11298717 | 0 | 0 | 0 | 0 | 0 | 0 |
| ENSMUSG00 | Gm12069 | 803 | 0 | 0 | 0 | 0 | 0 | 0 | 0 | 0 | 0 | 0 | 0 | 0 | 0 | 0 | 0 | 0 | 0 | 0 |
| ENSMUSG00 | Gm15046 | 406 | 0 | 0 | 0 | 0 | 0 | 0 | 0 | 0 | 0 | 0 | 0 | 0 | 0 | 0 | 0 | 0 | 0 | 0 |
| ENSMUSG00 | Gm11722 | 532 | 0.55927863 | 0.93927333 | 1.09113226 | 0.34256424 | 0.27142062 | 0.65200965 | 0.35573443 | 1.14864368 | 0.15483653 | 0.97492056 | 0.98782809 | 0.73441661 | 2.5964167 | 0.34920366 | 0.61957227 | 0 | 1.08062867 | 0.10340058 |
| ENSMUSG00 | 4933421A08 | 1636 | 0.02598116 | 0.18326164 | 0 | 0 | 0 | 0 | 0 | 0 | 0 | 0 | 0 | 0 | 0 | 0 | 0.07555298 | 0 | 0 | 0 |
| ENSMUSG00 | Gm15961 | 264 | 0 | 0.25237041 | 0 | 0 | 0 | 0 | 0 | 0 | 0 | 0 | 0 | 0 | 0 | 0 | 0 | 0 | 0 | 0 |
| ENSMUSG00 | Gm13191 | 414 | 0 | 0 | 0 | 0 | 0 | 0 | 0 | 0 | 0 | 0 | 0 | 0 | 0 | 0 | 0 | 0 | 0 | 0 |
| ENSMUSG00 | 4933426D04 | 984 | 2.89415325 | 3.08076562 | 2.90044542 | 2.36139559 | 3.86424997 | 2.56828192 | 2.30793556 | 1.74660377 | 1.84167347 | 2.63545597 | 2.86455544 | 1.41808086 | 1.65442405 | 1.27438045 | 1.2142735 | 1.47870232 | 2.37591881 | 2.12433542 |
| ENSMUSG00 | Gm13522 | 451 | 0 | 0 | 0 | 0 | 0 | 0 | 0 | 0 | 0 | 0 | 0 | 0 | 0 | 0 | 0 | 0 | 0 | 0 |
| ENSMUSG00 | 9330162012R | 3834 | 2.98223588 | 4.05767387 | 3.91126615 | 4.21861522 | 3.28914032 | 3.34745905 | 3.87485164 | 5.38917311 | 3.72762675 | 3.24668379 | 3.48904235 | 4.29463236 | 3.77001872 | 4.1671273 | 3.60003162 | 2.40689525 | 2.75901355 | 5.42343306 |
| ENSMUSG00 | 9030204H09 | 650 | 0 | 0 | 0 | 0 | 0 | 0 | 0 | 0 | 0 | 0 | 0.36749968 | 0 | 0 | 0 | 0 | 0 | 0 | 0 |
| ENSMUSG00 | Gm14077 | 3122 | 0 | 0.08536296 | 0 | 0 | 0 | 0 | 0.07577303 | 0 | 0 | 0 | 0 | 0 | 0 | 0 | 0 | 0 | 0 | 0 |
| ENSMUSG00 | 4933431K23R | 4433 | 0 | 0 | 0 | 0 | 0.02171528 | 0 | 0 | 0.04307738 | 0 | 0 | 0 | 0 | 0.02225667 | 0.11524587 | 0 | 0 | 0 | 0 |
| ENSMUSG00 | Gm14947 | 302 | 0 | 0 | 0 | 0 | 0 | 0 | 0 | 0 | 0 | 0 | 0 | 0 | 0 | 0 | 0 | 0 | 0 | 0 |
| ENSMUSG00 | Gm11457 | 781 | 1.03405678 | 0.554504 | 0.74325527 | 0.87505207 | 0.3697715 | 0.44413462 | 0.8481146 | 0.73352886 | 1.58206847 | 0.60372219 | 0.36702913 | 0.50026842 | 0.88431094 | 0.773077 | 0.10550975 | 1.12763633 | 0.63795373 | 0.35217098 |
| ENSMUSG00 | 4930500J02R | 1626 | 0.05228189 | 0.22536398 | 0.11900007 | 0.08406097 | 0.02960143 | 0 | 0.2618783 | 0.09395425 | 0 | 0.08699392 | 0.08814568 | 0.03432698 | 0.03033943 | 0 | 0.20271368 | 0.23548959 | 0.2828509 | 0 |
| ENSMUSG00 | Gm11815 | 706 | 0 | 0 | 0 | 0 | 0 | 0 | 0 | 0 | 0 | 0 | 0 | 0 | 0.13975047 | 0 | 0 | 0 | 0 | 0 |
| ENSMUSG00 | Gm16861 | 1644 | 1.21517231 | 1.31711565 | 1.11812296 | 0.80369239 | 1.72736222 | 1.23580418 | 2.84912118 | 1.41711469 | 1.65347326 | 1.43402382 | 1.74361162 | 2.00311684 | 3.0007249 | 2.31655424 | 2.53123914 | 1.21113843 | 2.02821644 | 2.47607901 |
| ENSMUSG00 | D930032P07R | 1460 | 0.11645254 | 0.15971906 | 0.43072322 | 0.06241239 | 0.09890121 | 0.10182069 | 0.06481189 | 0.36622852 | 0.11283977 | 0.25836002 | 0.1308903 | 0.30583925 | 0.67577969 | 1.08157464 | 0.50796439 | 0.44584954 | 0.28875977 | 0.18838735 |
| ENSMUSG00 | Gm2639 | 1296 | 0 | 0 | 0 | 0 | 0 | 0 | 0 | 0 | 0 | 0 | 0 | 0 | 0 | 0 | 0 | 0 | 0 | 0 |
| ENSMUSG00 | 1700030C12 | 1319 | 0 | 0 | 0 | 0.06908422 | 0.18245612 | 0.03756841 | 0.03587011 | 0.05791115 | 0 | 0.07149462 | 0 | 0 | 0.112203 | 0.03521159 | 0 | 0 | 0.05811417 | 0.08341032 |
| ENSMUSG00 | Gm12236 | 248 | 0 | 0 | 0 | 0 | 0 | 0 | 0 | 0 | 0 | 0 | 0 | 0 | 0 | 0 | 0 | 0 | 0 | 0 |
| ENSMUSG00 | Gm16899 | 1190 | 0.50006089 | 0.05598806 | 0.20325013 | 0.15314637 | 0.24268197 | 0 | 1.35179083 | 0.16047228 | 0.06922103 | 0.03962244 | 0.08029405 | 0 | 0.16582157 | 0 | 0.13849262 | 0.03217698 | 0 | 0 |
| ENSMUSG00 | Gm13990 | 2697 | 0 | 0 | 0 | 0 | 0 | 0 | 0 | 0 | 0 | 0 | 0 | 0 | 0 | 0 | 0 | 0 | 0 | 0 |
| ENSMUSG00 | Gm16898 | 5179 | 0 | 0 | 0 | 0 | 0 | 0 | 0 | 0 | 0 | 0 | 0 | 0 | 0 | 0 | 0 | 0 | 0 | 0 |
| ENSMUSG00 | Gm15835 | 354 | 0 | 0 | 0 | 0 | 0 | 0.27995895 | 0.13365164 | 0 | 0.34903827 | 0.66597038 | 0 | 1.89205636 | 1.25420129 | 0.39359396 | 0 | 0.21633112 | 0.32479913 | 0.31078591 |
| ENSMUSG00 | Gm16049 | 614 | 0.27690668 | 1.19362162 | 0.47270551 | 0.22261097 | 0.86229831 | 0.56493344 | 0.15411296 | 0.24881044 | 1.40865933 | 0.84471944 | 0.31123751 | 0.27271496 | 0.08034514 | 0.52949285 | 0.26841404 | 0.81071319 | 0.24968272 | 0 |
| ENSMUSG00 | 4930571N24 | 1054 | 0 | 0 | 0 | 0 | 0 | 0 | 0 | 0 | 0 | 0 | 0 | 0 | 0 | 0 | 0 | 0 | 0 | 0 |
| ENSMUSG00 | 2900053O20 | 933 | 2.36899159 | 2.03519289 | 2.22943387 | 3.3694663 | 2.73418216 | 3.0804486 | 4.20895214 | 2.08768759 | 2.64865053 | 4.69990933 | 3.4307848 | 4.18767029 | 4.81158036 | 4.52991635 | 4.90179282 | 4.51443388 | 2.75226355 | 2.47629419 |
| ENSMUSG00 | Gm16897 | 2486 | 0 | 0.0268004 | 0 | 0.18327049 | 0.01936119 | 0.01993272 | 0.01903165 | 0.03072599 | 0 | 0.07586597 | 0.0192176 | 0.08980798 | 0.03968779 | 0 | 0.08286717 | 0.03080499 | 0.2312528 | 0 |
| ENSMUSG00 | Gm12580 | 1749 | 0 | 0 | 0 | 0 | 0 | 0 | 0 | 0 | 0 | 0 | 0 | 0 | 0 | 0 | 0 | 0 | 0 | 0 |
| ENSMUSG00 | A530020G20 | 4167 | 1.30565456 | 1.21515717 | 1.16087186 | 1.08244382 | 1.4438422 | 1.11782024 | 0.79478942 | 0.81572446 | 1.6506235 | 1.459669 | 1.31848338 | 1.44662624 | 1.84683924 | 1.0476948 | 1.11729681 | 1.92969224 | 1.87630539 | 0.71286099 |
| ENSMUSG00 | Gm16172 | 680 | 0.87510657 | 0.34292685 | 0.85365054 | 0.60301382 | 0.77860465 | 0.36435834 | 1.18281697 | 0.33699179 | 0.78738927 | 0.9014105 | 0.35128646 | 1.31330971 | 1.52348568 | 0.06830013 | 0 | 0.50678745 | 0.73270862 | 1.21343618 |
| ENSMUSG00 | Gm16265 | 107 | 0.39724464 | 0 | 0 | 0 | 0 | 0 | 0 | 0 | 0 | 0 | 0.89298988 | 0 | 0 | 0 | 0 | 0 | 0 | 0 |
| ENSMUSG00 | 9130204K15 | 1964 | 0 | 0 | 0.04926021 | 0 | 0.02450709 | 0 | 0 | 0.01944623 | 0 | 0 | 0 | 0 | 0 | 0 | 0 | 0 | 0 | 0 |
| ENSMUSG00 | Gm12139 | 372 | 0 | 0.08955079 | 0 | 0 | 0 | 0 | 0 | 0 | 0 | 0 | 0 | 0 | 0 | 0 | 0 | 0 | 0 | 0 |
| ENSMUSG00 | 9530059O14 | 6795 | 0 | 0 | 0.00711899 | 0.05364067 | 0 | 0 | 0 | 0 | 0 | 0 | 0 | 0 | 0 | 0 | 0 | 0 | 0 | 0.00809553 |
| ENSMUSG00 | Gm15506 | 1129 | 0.33883666 | 1.00322268 | 0.85692702 | 0.40355221 | 0.81001466 | 0.87781636 | 2.2629625 | 1.58994058 | 1.7145848 | 1.33642383 | 0.88863962 | 1.87864941 | 2.09737115 | 1.31639575 | 1.09481548 | 1.15312724 | 1.32393763 | 1.41298857 |
| ENSMUSG00 | Gm14102 | 462 | 0 | 0 | 0 | 0 | 0.10418165 | 0.321771 | 0.1024084 | 0 | 0.17829661 | 0.510289 | 0 | 0 | 0 | 0 | 0 | 0 | 0 | 1.07160598 |
| ENSMUSG00 | Gm16014 | 536 | 0 | 0 | 0 | 0 | 0 | 0 | 0 | 0.14250896 | 0 | 0.17593546 | 0 | 0 | 0 | 0 | 0 | 0 | 0 | 0 |
| ENSMUSG00 | Gm15287 | 749 | 0 | 0 | 0 | 0 | 0 | 0 | 0.12633559 | 0 | 0 | 0 | 0 | 0 | 0 | 0 | 0 | 0.15336692 | 0 | 0 |
| ENSMUSG00 | Gm11707 | 488 | 0 | 0 | 0 | 0 | 0 | 0 | 0 | 0 | 0 | 0 | 0 | 0 | 0 | 0 | 0 | 0 | 0 | 0 |
| ENSMUSG00 | 1700012C08 | 361 | 0 | 0 | 0 | 0 | 0 | 0 | 0 | 0 | 0 | 0 | 0 | 0 | 0 | 0 | 0 | 0 | 0 | 0 |
| ENSMUSG00 | 8030443G20 | 897 | 0 | 0 | 0 | 0 | 0 | 0 | 0 | 0 | 0.04591585 | 0 | 0 | 0 | 0.16498969 | 0 | 0 | 0 | 0 | 0 |
| ENSMUSG00 | A630071L07R | 3786 | 0.56134675 | 0.10558762 | 0.15332339 | 0.19254535 | 0.0762788 | 0.05235365 | 0.19994793 | 0.0100878 | 0.07615045 | 0 | 0.0252377 | 0.13268383 | 0.09121062 | 0 | 0.06529565 | 0.04045495 | 0.03036949 | 0 |
| ENSMUSG00 | Gm15458 | 708 | 0 | 0 | 0 | 0 | 0 | 0 | 0 | 0 | 0 | 0 | 0.26991502 | 0 | 0 | 0 | 0 | 0 | 0.05413319 | 0 |
| ENSMUSG00 | 2310043D23 | 2014 | 0.46430679 | 0.23156927 | 0.50439133 | 0.18097734 | 0.47797342 | 0.14762483 | 0.30539465 | 0.36030568 | 0 | 0.58528678 | 0.49815001 | 0.41570752 | 0.04898899 | 0.09224248 | 0.1636606 | 0.74147651 | 0.01902994 | 0.16388016 |
| ENSMUSG00 | Gm11638 | 1611 | 0.29022777 | 0.10339197 | 0.21018914 | 0.16968732 | 0.26889343 | 0.15379495 | 0.35242219 | 0 | 0.02556581 | 0.05853594 | 0.11862187 | 0.27717275 | 0 | 0 | 0 | 0 | 0 | 0 |
| ENSMUSG00 | 1600020E01 | 2301 | 14.7779838 | 17.1559234 | 15.3466654 | 15.3256186 | 16.0648924 | 15.3116009 | 20.3356104 | 25.2956198 | 23.1439222 | 19.6307578 | 20.8249819 | 27.4105601 | 18.3091949 | 19.9822783 | 22.221265 | 19.7360541 | 20.2541404 | 20.4162439 |
| ENSMUSG00 | Gm16362 | 213 | 0 | 0 | 0 | 0 | 0 | 0 | 0 | 0 | 0 | 0 | 0 | 0 | 0 | 0 | 0 | 0 | 0 | 0 |
| ENSMUSG00 | Gm12292 | 2431 | 0 | 0 | 0 | 0 | 0 | 0 | 0 | 0.04713172 | 0 | 0 | 0 | 0 | 0 | 0 | 0 | 0.04725291 | 0 | 0.13576908 |
| ENSMUSG00 | Gm13291 | 653 | 0 | 0 | 0 | 0 | 0 | 0 | 0 | 0 | 0 | 0 | 0 | 0 | 0 | 0 | 0 | 0 | 0 | 0 |
| ENSMUSG00 | Gm16652 | 1879 | 0.02262117 | 0.01772905 | 0.05148859 | 0 | 0 | 0.10548746 | 0.17625798 | 0.20325919 | 0.04383876 | 0 | 0.02542574 | 0 | 0.23628912 | 0.0494349 | 0 | 0 | 0 | 0 |
| ENSMUSG00 | Gm12204 | 331 | 0.38524329 | 0.10064318 | 0 | 0.55058664 | 0.58165467 | 0.14970614 | 0 | 0.1153849 | 0 | 0.56979702 | 0 | 1.01176428 | 0.74519513 | 0.14031446 | 0.871332 | 1.27249753 | 1.27368358 | 0 |
| ENSMUSG00 | 2210417A02R | 548 | 0 | 0 | 0 | 0.08314059 | 0.0904247 | 0 | 0 | 0 | 0 | 0 | 0 | 0 | 0 | 0 | 0 | 0 | 0 | 0 |
| ENSMUSG00 | Gm13778 | 740 | 1.55086453 | 0.63024394 | 0.84960526 | 1.35451753 | 0.19512942 | 0.33481577 | 0.63936053 | 0.77417032 | 0.89051926 | 0.31858583 | 2.0013834 | 0.37713286 | 0.46665327 | 0.18828684 | 1.11355556 | 0.46569658 | 0.41433834 | 0.9663762 |
| ENSMUSG00 | Gm14204 | 7395 | 0.01149565 | 0 | 0 | 0 | 0 | 0 | 0 | 0 | 0 | 0 | 0 | 0 | 0 | 0 | 0 | 0 | 0 | 0 |
| ENSMUSG00 | Gm15259 | 290 | 0 | 0 | 0 | 0 | 0 | 0 | 0 | 0 | 0 | 0 | 0 | 0 | 0 | 0 | 0 | 0 | 0 | 0 |
| ENSMUSG00 | Gm12681 | 697 | 0 | 0 | 0 | 0 | 0 | 0 | 0 | 0 | 0 | 0 | 0 | 0 | 0 | 0.33317136 | 0 | 0 | 0 | 0 |
| ENSMUSG00 | Gm16217 | 614 | 0 | 0 | 0 | 0 | 0 | 0 | 0 | 0 | 0 | 0 | 0 | 0 | 0 | 0 | 0 | 0 | 0 | 0 |
| ENSMUSG00 | Gm15875 | 271 | 0 | 0.36877743 | 0.35700022 | 0.5043658 | 0 | 0.36570283 | 0 | 0 | 0 | 0.17398783 | 0 | 0 | 0 | 0.1713804 | 0.91221157 | 0 | 0 | 0 |
| ENSMUSG00 | 4930597A21R | 694 | 0 | 0 | 0 | 0 | 0 | 0 | 0.06817389 | 0.05503228 | 0 | 0.33970247 | 0 | 0 | 0 | 0 | 0 | 0 | 0 | 0 |
| ENSMUSG00 | B130055M24 | 2916 | 0.90374517 | 0.8225406 | 1.04510714 | 0.84372304 | 0.75928275 | 1.25751107 | 0.69768354 | 0.77275437 | 1.38418331 | 0.6629557 | 0.96025638 | 0.97619986 | 1.50567237 | 0.81229371 | 1.1868761 | 0.99797194 | 0.98575867 | 0.69798935 |
| ENSMUSG00 | Xist | 16001 | 0 | 0.00624578 | 0 | 0 | 0 | 0 | 0 | 0 | 0 | 0 | 0 | 0 | 0 | 0 | 0 | 0 | 0 | 0 |
| ENSMUSG00 | 4930511F01R | 493 | 0.0862174 | 0 | 0.09812075 | 0.18483182 | 0.09763068 | 0 | 1.05565815 | 0.15493875 | 0 | 0.09564037 | 0.96906609 | 0.11321635 | 0 | 0.09420707 | 0.50143881 | 0 | 0.93289161 | 0 |
| ENSMUSG00 | Gm16058 | 425 | 0 | 0 | 0 | 0 | 0 | 0 | 0 | 0 | 0 | 0 | 0 | 0 | 0 | 0 | 0 | 0 | 0 | 0 |
| ENSMUSG00 | Gm15911 | 806 | 0.1054719 | 0.08266227 | 0.12003357 | 0.05652735 | 0.41801919 | 0.12295964 | 0.29350297 | 0.23692557 | 0.10219979 | 0.23399853 | 0.11854828 | 0.27700081 | 0.06120585 | 0 | 0.05111856 | 0.04750696 | 0.09510247 | 0.06824951 |
| ENSMUSG00 | Gm11206 | 981 | 0 | 0 | 0 | 0 | 0 | 0 | 0 | 0 | 0 | 0 | 0 | 0 | 0 | 0 | 0 | 0 | 0 | 0 |
| ENSMUSG00 | Gm11291 | 191 | 0 | 0 | 0 | 0 | 0 | 0 | 0 | 0 | 0 | 0 | 0 | 0 | 0 | 0 | 0 | 0 | 0 | 0 |
| ENSMUSG00 | Nkx2-2as | 4339 | 0 | 0 | 0 | 0 | 0 | 0 | 0 | 0 | 0 | 0 | 0 | 0 | 0 | 0 | 0 | 0 | 0 | 0 |
| ENSMUSG00 | Gm16661 | 1252 | 0.10184946 | 0.23946969 | 0.11591102 | 0.03639061 | 0.11533209 | 0.11873658 | 0.83137295 | 0.15252557 | 0.2960692 | 0 | 0.11447674 | 0.4012308 | 0.11820747 | 0.07419183 | 0.42781168 | 0.24466842 | 0 | 0.21968493 |
| ENSMUSG00 | Gm12600 | 706 | 0 | 0 | 0 | 0 | 0 | 0 | 0 | 0 | 0 | 0 | 0 | 0 | 0 | 0 | 0 | 0 | 0 | 0 |
| ENSMUSG00 | 9130208D14 | 7292 | 0 | 0 | 0.05307025 | 0 | 0.0132013 | 0 | 0 | 0 | 0 | 0 | 0 | 0 | 0 | 0 | 0.05650241 | 0 | 0 | 0 |
| ENSMUSG00 | Gm11747 | 1645 | 0 | 0 | 0 | 0 | 0 | 0 | 0 | 0 | 0 | 0 | 0 | 0 | 0 | 0 | 0 | 0.04655393 | 0 | 0 |
| ENSMUSG00 | Gm16292 | 606 | 22.5852585 | 19.2950922 | 20.8341443 | 14.8110986 | 14.9320159 | 16.5175779 | 11.2426168 | 16.0080367 | 13.660874 | 13.4605143 | 9.38155125 | 16.3946996 | 14.3274215 | 9.42677021 | 14.4137456 | 13.6481373 | 10.3721332 | 11.437537 |
| ENSMUSG00 | 4930461G14 | 777 | 0 | 0 | 0 | 0 | 0 | 0 | 0 | 0 | 0 | 0 | 0 | 0 | 0 | 0 | 0 | 0 | 0 | 0 |
| ENSMUSG00 | Gm12534 | 467 | 0 | 0 | 0 | 0 | 0 | 0 | 0 | 0 | 0 | 0 | 0 | 0 | 0 | 0 | 0 | 0 | 0 | 0 |
| ENSMUSG00 | Gm14503 | 1163 | 0 | 0 | 0 | 0 | 0 | 0 | 0 | 0 | 0 | 0 | 0 | 0 | 0 | 0 | 0 | 0 | 0 | 0 |
| ENSMUSG00 | Gm13393 | 394 | 0 | 0 | 0 | 0 | 0 | 0 | 0 | 0 | 0 | 0 | 0 | 0 | 0 | 0 | 0 | 0 | 0 | 0 |
| ENSMUSG00 | 9330185C12 | 1006 | 0 | 0 | 0 | 0 | 0 | 0 | 0 | 0 | 0 | 0 | 0 | 0 | 0 | 0 | 0 | 0 | 0 | 0 |
| ENSMUSG00 | Olfr1343-ps1 | 967 | 0 | 0 | 0 | 0 | 0 | 0 | 0 | 0 | 0 | 0 | 0 | 0 | 0 | 0 | 0 | 0 | 0 | 0 |
| ENSMUSG00 | Gm4473 | 655 | 0.1946802 | 0.15257814 | 0.14770544 | 0.06955885 | 0 | 0.07565303 | 0 | 0.11661802 | 0 | 0 | 0 | 0 | 0.15063181 | 0 | 0 | 0 | 0.23405372 | 0 |
| ENSMUSG00 | Gm13499 | 600 | 0 | 0 | 0 | 0 | 0 | 0 | 0 | 0 | 0 | 0 | 0 | 0 | 0 | 0 | 0 | 0 | 0 | 0 |
| ENSMUSG00 | Pabpc1l2a-ps | 1114 | 0 | 0 | 0 | 0 | 0 | 0 | 0 | 0 | 0 | 0 | 0 | 0 | 0 | 0 | 0 | 0 | 0 | 0 |
| ENSMUSG00 | Gm15281 | 350 | 0 | 0 | 0 | 0 | 0 | 0 | 0 | 0 | 0 | 0 | 0 | 0 | 0 | 0 | 0 | 0 | 0 | 0 |
| ENSMUSG00 | Gm12762 | 454 | 0 | 0 | 0 | 0 | 0 | 0 | 0 | 0 | 0.0907192 | 0.20771235 | 0 | 0 | 0.10866061 | 0 | 0.81677093 | 1.09642708 | 0 | 0 |
| ENSMUSG00 | Gm15856 | 1160 | 0 | 0 | 0 | 0 | 0 | 0 | 0 | 0 | 0 | 0 | 0 | 0 | 0 | 0 | 0 | 0 | 0 | 0 |
| ENSMUSG00 | Gm15731 | 2189 | 0 | 0 | 0 | 0 | 0 | 0.06791147 | 0 | 0 | 0 | 0 | 0 | 0 | 0 | 0 | 0 | 0 | 0 | 0 |
| ENSMUSG00 | 4930471I20R | 625 | 0 | 0 | 0 | 0 | 0 | 0 | 0 | 0 | 0 | 0 | 0 | 0 | 0 | 0 | 0 | 0 | 0 | 0 |
| ENSMUSG00 | Gm12910 | 722 | 0 | 0 | 0 | 0 | 0 | 0 | 0 | 0 | 0 | 0 | 0 | 0 | 0 | 0 | 0 | 0 | 0 | 0 |
| ENSMUSG00 | Gm11351 | 521 | 0 | 0 | 0 | 0 | 0 | 0 | 0 | 0 | 0 | 0 | 0 | 0 | 0 | 0 | 0 | 0 | 0 | 0 |
| ENSMUSG00 | Gm16266 | 556 | 0.30579263 | 0.71898333 | 0.1740055 | 0 | 0.60597746 | 0.26737086 | 0.25528424 | 0.34345686 | 0.6666882 | 0.16960685 | 0 | 0.20077576 | 0.08872647 | 0.66826025 | 0.22231055 | 0.137736 | 0.06893219 | 0.19787448 |
| ENSMUSG00 | Gm16758 | 3842 | 0 | 0 | 0 | 0 | 0 | 0 | 0 | 0 | 0 | 0 | 0 | 0 | 0 | 0 | 0 | 0 | 0 | 0 |
| ENSMUSG00 | Gm12264 | 498 | 2.81660805 | 2.60884111 | 2.03984767 | 1.82976082 | 3.47941617 | 1.29354525 | 3.32518828 | 2.60751342 | 3.47356157 | 3.12444419 | 1.43900477 | 3.81070789 | 2.87274217 | 2.33153049 | 2.73022358 | 4.0751048 | 2.30881307 | 3.75564182 |
| ENSMUSG00 | Nespas | 2676 | 0.12707078 | 0.06224382 | 0.09038403 | 0.0340516 | 0.05395956 | 0.03703493 | 0.10608224 | 0.11417759 | 0.10773752 | 0.08809922 | 0.01785312 | 0.20857871 | 0.35026399 | 0.38182733 | 0.26174382 | 0.21463345 | 0.15754457 | 0.57558109 |
| ENSMUSG00 | Gm6634 | 1800 | 0 | 0 | 0 | 0 | 0 | 0 | 0 | 0 | 0 | 0 | 0 | 0 | 0 | 0 | 0 | 0 | 0 | 0 |
| ENSMUSG00 | Gm16759 | 1705 | 0.12464861 | 0.11723013 | 0.14185786 | 0.18705414 | 0.11291947 | 0.08718956 | 0.63823555 | 0.26880283 | 0 | 0.33185246 | 0.28020504 | 0.06547292 | 0.28933676 | 0.46307888 | 0.19332108 | 0.22457834 | 0.3371815 | 0.38716087 |
| ENSMUSG00 | Scml1 | 574 | 0.07405083 | 0 | 0 | 0 | 0.08385353 | 0 | 0.41213135 | 0.19961186 | 0 | 0.08214408 | 0 | 0 | 0.32365218 | 0 | 0 | 0 | 0 | 0 |
| ENSMUSG00 | Has2as | 1620 | 0 | 0 | 0 | 0 | 0 | 0 | 0 | 0 | 0 | 0 | 0 | 0 | 0 | 0 | 0 | 0 | 0 | 0 |
| ENSMUSG00 | Gm13782 | 349 | 0 | 0.19090484 | 0 | 0.78328443 | 0 | 0 | 3.38916038 | 2.95471308 | 2.1242329 | 4.32327365 | 0.68445499 | 3.03867504 | 0.4240566 | 4.25848366 | 1.29861637 | 0 | 0.10981747 | 1.26095374 |
| ENSMUSG00 | 4933406C10 | 2559 | 0.23254102 | 0.18225108 | 0.22683953 | 0.12462966 | 0.16927992 | 0.1161846 | 0.4437297 | 0.32834422 | 0.48284309 | 0.42378514 | 0.42939588 | 0.41441875 | 0.40483402 | 0.1633438 | 0.38641553 | 0.20948359 | 0.0748853 | 0.30094861 |
| ENSMUSG00 | Gm13704 | 3239 | 0.13122932 | 0.15427398 | 0.37336779 | 0.14066392 | 0.04458036 | 0 | 0.01460719 | 0.05895709 | 0 | 0.04367154 | 0 | 0 | 0.0456918 | 0.05735608 | 0 | 0.05910869 | 0.02366551 | 0 |
| ENSMUSG00 | Gm12 | 900 | 0 | 0 | 0 | 0 | 0 | 0 | 0 | 0 | 0 | 0 | 0 | 0 | 0 | 0 | 0 | 0 | 0 | 0 |
| ENSMUSG00 | Gm13709 | 612 | 1.11124643 | 0.87092534 | 1.18562574 | 0.81890766 | 1.57293868 | 0.80968519 | 0.54115809 | 0.81127652 | 0.9421752 | 1.77200355 | 0.46838195 | 1.7328392 | 1.37033104 | 1.51778063 | 0.33661402 | 0.43796446 | 0.81412069 | 0.08968416 |
| ENSMUSG00 | Gm11755 | 885 | 0.14408534 | 0 | 0.21863743 | 0 | 0 | 0.05599179 | 0 | 0 | 0 | 0 | 0 | 0 | 0 | 0 | 0.09311086 | 0 | 0 | 0 |
| ENSMUSG00 | Gm12972 | 1002 | 0 | 0 | 0 | 0 | 0 | 0 | 0 | 0 | 0 | 0 | 0 | 0 | 0 | 0 | 0 | 0 | 0 | 0 |
| ENSMUSG00 | Gm13648 | 1534 | 1.63481446 | 1.41156331 | 2.14432859 | 1.00982758 | 1.38057669 | 1.25981526 | 1.2337074 | 1.44404129 | 0.85917112 | 0.89137575 | 1.43262587 | 1.52819937 | 1.15772426 | 0.5146998 | 0.94006158 | 0.84868361 | 0.74953645 | 0.43031896 |
| ENSMUSG00 | 1810007I06R | 172 | 0 | 0 | 0 | 0 | 0 | 0 | 0 | 0 | 0 | 0 | 0 | 0 | 0 | 0 | 0 | 0 | 0 | 0 |
| ENSMUSG00 | A730090H04 | 1660 | 0.20484422 | 0 | 0 | 0 | 0 | 0 | 0 | 0 | 0 | 0 | 0 | 0 | 0 | 0 | 0 | 0.2998662 | 0 | 0.26510413 |
| ENSMUSG00 | 9230116N13 | 2094 | 0.52776245 | 0.85907177 | 0.57752543 | 0.71801073 | 0.89643984 | 0.66259625 | 1.89792981 | 0.63836394 | 1.00310998 | 0.87816496 | 0.75290049 | 0.9062715 | 1.76690248 | 1.41949455 | 1.84954453 | 1.84687266 | 1.40932419 | 0.3415083 |
| ENSMUSG00 | 9530034E10R | 499 | 29.6428883 | 21.2962189 | 18.2248972 | 21.3652993 | 19.3878089 | 19.3642947 | 18.9629976 | 21.2009929 | 26.082042 | 31.8432605 | 24.6055397 | 27.7400487 | 76.1233994 | 40.1150334 | 38.7245084 | 100.75264 | 57.6814609 | 44.8671472 |
| ENSMUSG00 | Gm13446 | 1068 | 0 | 0.0623837 | 0 | 0 | 0 | 0.18559076 | 0 | 0 | 0 | 0 | 0 | 0 | 0 | 0 | 0 | 0 | 0.07177209 | 0 |
| ENSMUSG00 | Gm15444 | 386 | 0 | 0 | 0 | 0.11803379 | 0 | 0 | 0 | 0 | 0 | 0 | 0 | 0 | 0 | 0 | 0 | 0 | 0 | 0 |
| ENSMUSG00 | Gm15999 | 1509 | 0 | 0 | 0 | 0 | 0 | 0 | 0 | 0 | 0 | 0 | 0 | 0 | 0 | 0.24622445 | 0 | 0 | 0 | 0 |
| ENSMUSG00 | Gm14223 | 513 | 0 | 0 | 0 | 0 | 0 | 0 | 0 | 0 | 0 | 0 | 0 | 0 | 0 | 0 | 0 | 0 | 0 | 0 |
| ENSMUSG00 | Gm14426 | 247 | 0.17208573 | 0.26974003 | 0.58753276 | 0.18445767 | 0.58459826 | 0 | 0 | 0.46387533 | 0 | 0 | 0.58026265 | 0.45194868 | 0 | 0 | 0 | 0.46506811 | 1.08617036 | 0 |
| ENSMUSG00 | Gm13372 | 686 | 0 | 0 | 0 | 0 | 0 | 0 | 0 | 0 | 0 | 0 | 0 | 0 | 0 | 0 | 0 | 0 | 0 | 0 |
| ENSMUSG00 | Gm15540 | 672 | 0 | 0 | 0 | 0.13559835 | 0 | 0 | 0 | 0 | 0 | 0 | 0 | 0 | 0.22023177 | 0 | 0 | 0 | 0 | 0 |
| ENSMUSG00 | Gm14651 | 441 | 0 | 0 | 0 | 0 | 0 | 0 | 0 | 0 | 0 | 0 | 0 | 0 | 0 | 0 | 0 | 0 | 0 | 0 |
| ENSMUSG00 | Gm14136 | 765 | 0 | 0 | 0 | 0 | 0 | 0 | 0 | 0 | 0 | 0 | 0 | 0 | 0 | 0 | 0 | 0 | 0 | 0 |
| ENSMUSG00 | Cd101 | 3681 | 0 | 0.09049958 | 0 | 0 | 0 | 0 | 0 | 0 | 0 | 0 | 0 | 0 | 0.01340177 | 0.01261725 | 0 | 0 | 0.0416477 | 0 |
| ENSMUSG00 | Gm13133 | 732 | 0.23226872 | 0.09101884 | 0.06608406 | 0 | 0 | 0.06769499 | 0 | 0 | 0.33759439 | 0 | 0 | 0 | 0 | 0 | 0 | 0 | 0.47122496 | 0.52604337 |

| | | | | | | | | | | | | | | | | | | | | |
|---|---|---|---|---|---|---|---|---|---|---|---|---|---|---|---|---|---|---|---|---|
| ENSMUSG00 | 2810429I04R | 7212 | 0.10019246 | 0 | 0 | 0.01263479 | 0 | 0.04122524 | 0 | 0 | 0 | 0 | 0.00662437 | 0 | 0 | 0 | 0.00571292 | 0 | 0.01062848 | 0 |
| ENSMUSG00 | Gm16247 | 384 | 4.87038475 | 6.59317698 | 3.52723659 | 3.08486238 | 5.26442915 | 2.19374081 | 6.89976568 | 5.27134721 | 5.57734068 | 4.91153159 | 3.98124654 | 3.19777234 | 8.60739182 | 12.6995551 | 6.65233454 | 4.38746543 | 5.48944357 | 11.6034834 |
| ENSMUSG00 | 9130020K20I | 1784 | 1.02450817 | 2.1847582 | 1.49133642 | 0.86831587 | 0.80939334 | 1.02771925 | 1.61775416 | 1.79829697 | 1.98544863 | 1.08362042 | 1.31220458 | 1.34533267 | 0.60835324 | 0.39050522 | 1.03927691 | 1.26633735 | 1.03120081 | 0.64752872 |
| ENSMUSG00 | 4930542H20I | 863 | 0 | 0 | 0.05605276 | 0 | 0 | 0.05741916 | 0 | 0 | 0 | 0.05463581 | 0 | 0.06467632 | 0 | 0 | 0 | 0 | 0 | 0 |
| ENSMUSG00 | Gm16961 | 1892 | 0.22465738 | 0.17607238 | 0.02556741 | 0.04816178 | 0.05087941 | 0.10476265 | 0.10002681 | 0 | 0.04353754 | 0.17444763 | 0.15150621 | 0.08850264 | 0.15644371 | 0.07364284 | 0.06533016 | 0.04047633 | 0.04051406 | 0 |
| ENSMUSG00 | Gm16962 | 1435 | 0.17772199 | 0.1160728 | 0.23596844 | 0.15874928 | 0.20124846 | 0.20718913 | 0.06594102 | 0.21291932 | 0.11480562 | 0.03285763 | 0.39951185 | 0.31116746 | 0.41253171 | 0.42074783 | 0.14355943 | 0.1067334 | 0.34720687 | 0.6900097 |
| ENSMUSG00 | Gm15384 | 484 | 0 | 0 | 0 | 0 | 0 | 0 | 0.39101388 | 0.31563969 | 0 | 0 | 0 | 0 | 0 | 0 | 0 | 0 | 0 | 0 |
| ENSMUSG00 | Gm16960 | 2650 | 0 | 0 | 0 | 0 | 0 | 0 | 0 | 0 | 0 | 0 | 0 | 0 | 0 | 0 | 0 | 0 | 0 | 0 |
| ENSMUSG00 | Gm14104 | 353 | 0 | 0.09437081 | 0 | 0.38720434 | 0 | 0 | 0 | 0 | 0.46670273 | 0 | 0 | 0.47435407 | 0 | 0 | 0 | 0.10847198 | 0 | 0 |
| ENSMUSG00 | Gm13660 | 720 | 0 | 0 | 0 | 0 | 0 | 0 | 0 | 0 | 0 | 0 | 0 | 0 | 0 | 0 | 0 | 0 | 0 | 0 |
| ENSMUSG00 | Gm7076 | 455 | 0 | 0 | 0 | 0 | 0 | 0 | 0 | 0 | 0 | 0 | 0 | 0 | 0 | 0 | 0 | 0 | 0 | 0 |
| ENSMUSG00 | Gm13583 | 1926 | 1.50070196 | 1.81612352 | 2.00928475 | 1.23010089 | 2.29913654 | 1.56942718 | 0.9580449 | 1.70537207 | 1.62522077 | 1.7136808 | 1.36429009 | 2.31840758 | 0.81963725 | 0.55462825 | 1.11239922 | 1.57059085 | 1.19396564 | 1.17101421 |
| ENSMUSG00 | A930012L18I | 2359 | 0.25225624 | 0.14121617 | 0.12303569 | 0.11588227 | 0.46928115 | 0.1890524 | 0.24067492 | 0.27523139 | 0.6110759 | 0.05996274 | 0.36453974 | 0.26026803 | 0.60645426 | 0.15750432 | 0.64623042 | 0.5681099 | 0.55239258 | 0.34978237 |
| ENSMUSG00 | Gm15280 | 639 | 0 | 0 | 0 | 0 | 0 | 0 | 0 | 0 | 0 | 0 | 0 | 0 | 0 | 0 | 0 | 0 | 0 | 0 |
| ENSMUSG00 | Gm16272 | 386 | 0 | 0 | 0 | 0.11803379 | 0.1246941 | 0.12837496 | 0 | 0.29683214 | 0.42680327 | 0.24430416 | 0.99015458 | 0 | 0 | 0.24064294 | 0.32021935 | 0 | 0 | 0 |
| ENSMUSG00 | Gm15500 | 1911 | 0.28915086 | 0.26148269 | 0.10125281 | 0.16689027 | 0.25186773 | 0.23337237 | 0.24758074 | 0.25981226 | 0.38794206 | 0.12336657 | 0.27499976 | 0.29207568 | 0.43885013 | 0.34024972 | 0.34496331 | 0.28051727 | 0.30083436 | 0.11514203 |
| ENSMUSG00 | Gm12002 | 2883 | 0 | 0 | 0.06711555 | 0.04741004 | 0.06678033 | 0 | 0 | 0 | 0.04285798 | 0.08177368 | 0 | 0.01936027 | 0.01711131 | 0 | 0 | 0.07968909 | 0 | 0 |
| ENSMUSG00 | Gm16126 | 585 | 0 | 0 | 0 | 0 | 0 | 0 | 0 | 0 | 0 | 0 | 0 | 0 | 0 | 0 | 0 | 0 | 0 | 0 |
| ENSMUSG00 | Gm11192 | 149 | 0 | 0 | 0 | 0 | 0 | 0 | 0 | 0.25632485 | 0.82925871 | 0.31644767 | 0 | 0 | 0.33108669 | 0 | 0.82956153 | 0 | 0 | 0.73837727 |
| ENSMUSG00 | Gm11837 | 1005 | 0.46523078 | 0.29832443 | 0.2887972 | 0.27200623 | 0.52681707 | 0.39444962 | 0.56492751 | 0.26601673 | 0.73766894 | 0.60990959 | 0.04753727 | 0.11107595 | 0.29451891 | 0.55455627 | 0.12298972 | 0.07620021 | 0.30508495 | 0.27367715 |
| ENSMUSG00 | Gm11729 | 710 | 0 | 0.09383914 | 0 | 0 | 0 | 0 | 0 | 0 | 0 | 0 | 0 | 0 | 0 | 0 | 0 | 0 | 0 | 0 |
| ENSMUSG00 | Gm12915 | 1495 | 0 | 0.06684862 | 0.09707063 | 0 | 0 | 0 | 0 | 0 | 0 | 0 | 0 | 0 | 0 | 0 | 0 | 0 | 0 | 0 |
| ENSMUSG00 | 6030455H03I | 857 | 0 | 0 | 0 | 0 | 0 | 0 | 0 | 0 | 0 | 0 | 0 | 0 | 0.17269049 | 0 | 0 | 0 | 0 | 0 |
| ENSMUSG00 | Gm12682 | 1076 | 0 | 0 | 0 | 0 | 0 | 0 | 0 | 0 | 0 | 0 | 0 | 0 | 0 | 0 | 0 | 0 | 0 | 0 |
| ENSMUSG00 | Gm16548 | 1223 | 0 | 0.05447734 | 0 | 0 | 0 | 0 | 0 | 0 | 0.20205977 | 0 | 0 | 0 | 0.16134724 | 0 | 0 | 0 | 0.03133794 | 0 |
| ENSMUSG00 | 4930505H01I | 1784 | 0.02382577 | 0.03734629 | 0.08134562 | 0 | 0 | 0.0277762 | 0.07956168 | 0 | 0 | 0 | 0 | 0 | 0 | 0 | 0.13857026 | 0 | 0 | 0 |
| ENSMUSG00 | Gm12214 | 415 | 0 | 0 | 0 | 0 | 0 | 0 | 0 | 0 | 0 | 0 | 0 | 0 | 0 | 0 | 0 | 0 | 0 | 0 |
| ENSMUSG00 | Susd5 | 3869 | 0.14281915 | 0.38745935 | 0.07501711 | 0.17663884 | 0.24880808 | 0.40984427 | 0.22011585 | 0 | 0 | 0 | 0.14817769 | 0.01442638 | 2.07834131 | 1.84864034 | 1.10751145 | 3.15706998 | 2.16941303 | 1.74880334 |
| ENSMUSG00 | F420014N23I | 2393 | 1.17231158 | 0.80741658 | 0.99051525 | 1.19947589 | 0.90511348 | 0.80758738 | 1.34444721 | 0.94164302 | 0.48191494 | 0.827551 | 1.35758344 | 0.76971035 | 0.92767918 | 0.40757452 | 1.0502695 | 1.20008172 | 0.59259214 | 1.35626297 |
| ENSMUSG00 | Gm13718 | 1087 | 0 | 0 | 0.04450187 | 0 | 0 | 0.13676007 | 0 | 0.0351356 | 0 | 0 | 0 | 0 | 0.04538355 | 0 | 0 | 0.0704519 | 0.24681148 | 0 |
| ENSMUSG00 | C030005K06I | 2686 | 0.23737068 | 0.06201209 | 0.03601901 | 0.01696241 | 0.14335644 | 0.11069114 | 0.49320738 | 0.02843813 | 0.03066755 | 0.03510849 | 0.03557331 | 0.16624174 | 0.05509894 | 0.31124109 | 0.12271498 | 0.32787936 | 0.08561347 | 0.08191974 |
| ENSMUSG00 | Gm16257 | 481 | 0 | 0 | 0 | 0 | 0 | 0 | 0 | 0 | 0 | 0 | 0 | 0 | 0 | 0 | 0 | 0 | 0.07968045 | 0 |
| ENSMUSG00 | Gm15609 | 2958 | 0 | 0 | 0 | 0 | 0 | 0 | 0 | 0 | 0 | 0 | 0 | 0 | 0 | 0 | 0 | 0 | 0 | 0 |
| ENSMUSG00 | Gm13620 | 509 | 0.66805778 | 2.42156598 | 1.04540046 | 1.61119607 | 2.26948167 | 2.23912156 | 2.04494879 | 2.40109406 | 1.61833068 | 2.13058188 | 3.37897545 | 3.83801216 | 1.93838575 | 3.7410758 | 3.07595113 | 3.61090207 | 1.12945865 | 3.35025994 |
| ENSMUSG00 | Gm15510 | 648 | 1.24629374 | 0.4112703 | 1.11975765 | 1.19527431 | 1.78266384 | 1.98822696 | 1.97136162 | 0.53045003 | 1.08051044 | 1.96461264 | 0.95844824 | 2.06724676 | 1.29420153 | 2.22186221 | 0.69940913 | 0.64999488 | 0.76889176 | 0.76401537 |
| ENSMUSG00 | Gm14290 | 2066 | 0.26745755 | 0.17736778 | 0.468282 | 0.1102639 | 0.60572605 | 0.69556112 | 1.03052786 | 0.42518163 | 0.3987078 | 0.15975553 | 0.2774925 | 0.35121182 | 0.07163396 | 0.04496039 | 0.13959869 | 1.05642044 | 0.92754833 | 0.61239567 |
| ENSMUSG00 | Gm13205 | 560 | 1.5939441 | 1.36820816 | 1.29571956 | 2.1153342 | 1.63304741 | 2.5661237 | 1.01384312 | 1.70501796 | 2.57415724 | 2.27333748 | 2.81531005 | 2.19275817 | 3.25943025 | 2.15633263 | 1.0300389 | 3.14529989 | 1.57411577 | 3.92922191 |
| ENSMUSG00 | 4930511M06I | 4153 | 0 | 0.01604281 | 0 | 0 | 0 | 0.08352255 | 0.13670892 | 0.02758902 | 0.04958646 | 0 | 0 | 0.08063905 | 0.1069076 | 0.03354979 | 0.10913005 | 0.01843997 | 0.03691432 | 0 |
| ENSMUSG00 | 4930562D21I | 1748 | 0 | 0 | 0 | 0 | 0 | 0 | 0 | 0 | 0 | 0 | 0 | 0 | 0 | 0 | 0 | 0 | 0 | 0 |
| ENSMUSG00 | 5830432E09I | 4491 | 0 | 0 | 0 | 0.01014497 | 0 | 0.04413514 | 0 | 0.04252104 | 0 | 0.06299359 | 0.01063793 | 0 | 0 | 0 | 0 | 0 | 0.01706805 | 0 |
| ENSMUSG00 | Gm15408 | 1375 | 1.32925278 | 0.38764095 | 0.73879574 | 1.3585475 | 1.6802417 | 0.64869033 | 0.89463975 | 0.77773619 | 1.67741446 | 0.78870267 | 0.59067221 | 0.56830493 | 1.29159929 | 2.73597897 | 0.38954198 | 1.36453801 | 1.1709416 | 1.0401722 |
| ENSMUSG00 | Gm12064 | 529 | 0 | 0 | 0 | 0 | 0 | 0 | 0 | 0 | 0 | 0 | 0 | 0 | 0 | 0 | 0 | 0 | 0 | 0 |
| ENSMUSG00 | 1700007M16 | 433 | 0 | 0 | 0 | 0 | 0 | 0 | 0 | 0 | 0 | 0 | 0 | 0 | 0 | 0 | 0 | 0 | 0 | 0 |
| ENSMUSG00 | Gm14330 | 382 | 0 | 0 | 0 | 0 | 0 | 0 | 0 | 0 | 0 | 0 | 0 | 0 | 0 | 0 | 0 | 0 | 0 | 0 |
| ENSMUSG00 | Gm16862 | 5206 | 0.30209211 | 0.01279788 | 0 | 0.12252298 | 0.05547283 | 0.03807356 | 0.02726432 | 0.04401737 | 0.25316337 | 0.21736782 | 0.16518426 | 0.26803526 | 0.19899544 | 0.18734649 | 0.05539971 | 0.06619583 | 0.54478409 | 0.03169945 |
| ENSMUSG00 | Gm16863 | 2426 | 0 | 0 | 0 | 0 | 0 | 0 | 0 | 0 | 0 | 0 | 0 | 0 | 0 | 0 | 0 | 0 | 0 | 0 |
| ENSMUSG00 | Gm11635 | 566 | 0 | 0 | 0 | 0 | 0 | 0 | 0 | 0 | 0 | 0 | 0 | 0 | 0 | 0 | 0 | 0 | 0 | 0 |
| ENSMUSG00 | Gm13256 | 518 | 0.49233795 | 0 | 0.37354077 | 0 | 0.09291877 | 0 | 0.36534887 | 0.07373051 | 0 | 0.18204905 | 0.1844593 | 0.10775224 | 0 | 0.2689812 | 0.15907937 | 0.36960046 | 0.51792293 | 0.10619519 |
| ENSMUSG00 | Gm14333 | 263 | 0 | 0 | 0 | 0 | 0 | 0 | 0 | 0 | 0 | 0 | 0 | 0 | 0 | 0 | 0 | 0 | 0 | 0 |
| ENSMUSG00 | Gm14157 | 1279 | 0 | 0.28650652 | 0 | 0 | 0 | 0.07748668 | 0 | 0 | 0 | 0 | 0.37353369 | 0.26184048 | 0.50141902 | 0 | 0.51542212 | 0 | 0.14982915 | 0 |
| ENSMUSG00 | Gm13348 | 684 | 0 | 0 | 0 | 0 | 0 | 0 | 0 | 0 | 0 | 0 | 0 | 0 | 0 | 0 | 0 | 0 | 0 | 0 |
| ENSMUSG00 | Gm16701 | 2122 | 0.28043 | 0.17268701 | 0.27355437 | 0.12882482 | 0.34023509 | 0.0700557 | 0.84725815 | 0.32396948 | 0.75696235 | 0.86657749 | 0.27016942 | 0.8417065 | 0 | 0.91925149 | 0.85432067 | 0.27066876 | 0.25285964 | 0.10369294 |
| ENSMUSG00 | Gm16130 | 2794 | 0 | 0 | 0 | 0 | 0.03465377 | 0 | 0 | 0 | 0 | 0 | 0 | 0 | 0 | 0 | 0 | 0 | 0 | 0 |
| ENSMUSG00 | Gm11831 | 627 | 2.10153183 | 1.32826532 | 1.92876915 | 0.94464685 | 0.15353086 | 0.47418884 | 1.28279991 | 1.82738767 | 1.05101157 | 0.60160387 | 2.43827539 | 1.78040391 | 0.23603788 | 0.81480855 | 0.85425873 | 0.97711279 | 0.79464411 | 1.22827352 |
| ENSMUSG00 | A630072L19I | 3401 | 0.03749354 | 0.00979503 | 0 | 0 | 0 | 0 | 0 | 0 | 0.02422024 | 0 | 0.11237861 | 0 | 0 | 0 | 0 | 0 | 0 | 0 |
| ENSMUSG00 | Gm16702 | 2381 | 0 | 0 | 0.04063295 | 0 | 0.020215 | 0.0624352 | 0.05961278 | 0.20852635 | 0 | 0 | 0 | 0 | 0 | 0.05851838 | 0 | 0 | 0.01609672 | 0 |
| ENSMUSG00 | Gm16157 | 1503 | 0 | 0 | 0 | 0 | 0 | 0.03296922 | 0 | 0 | 0 | 0 | 0 | 0 | 0 | 0 | 0 | 0 | 0 | 0 |
| ENSMUSG00 | 2810403D21I | 7266 | 0.85408143 | 0.65103647 | 0.29293082 | 0.52671728 | 0.47694722 | 0.69562054 | 1.05486567 | 0.662296 | 0.87293193 | 1.02529743 | 0.93366971 | 0.7912212 | 0.88262445 | 0.48579007 | 0.75984152 | 0.83790347 | 0.58022195 | 0.46181606 |
| ENSMUSG00 | Gm14397 | 571 | 0 | 0 | 0 | 0 | 0 | 0 | 0 | 0 | 0 | 0 | 0 | 0 | 0.1727913 | 0 | 0 | 0 | 0 | 0 |
| ENSMUSG00 | Gm12784 | 655 | 0 | 0 | 0.07385272 | 0 | 0 | 0 | 0 | 0 | 0 | 0 | 0 | 0 | 0 | 0 | 0 | 0 | 0 | 0 |
| ENSMUSG00 | 1700112J05R | 789 | 0 | 0 | 0 | 0 | 0 | 0 | 0 | 0 | 0 | 0 | 0 | 0 | 0 | 0 | 0 | 0 | 0 | 0 |
| ENSMUSG00 | Gm16081 | 682 | 0 | 0 | 0 | 0 | 0 | 0 | 0 | 0 | 0 | 0 | 0 | 0 | 0 | 0 | 0 | 0 | 0 | 0 |
| ENSMUSG00 | 4930448N21I | 2302 | 0 | 0 | 0 | 0.03958388 | 0 | 0 | 0 | 0 | 0.05367487 | 0.04096499 | 0 | 0.07273979 | 0.15001017 | 0.06052661 | 0 | 0.14970264 | 0 | 0 |
| ENSMUSG00 | 4932415G12I | 4132 | 8.4351995 | 9.61817106 | 7.78518821 | 9.31729973 | 6.81441804 | 8.79045346 | 10.2823779 | 9.55734368 | 12.2004587 | 8.86643186 | 8.49820776 | 8.69924654 | 8.78709853 | 11.8021035 | 10.180733 | 7.1818056 | 14.163663 | 9.43888109 |
| ENSMUSG00 | Gm12949 | 360 | 0 | 0 | 0.13437092 | 0 | 0 | 0 | 0 | 0 | 0 | 0 | 0 | 0 | 0 | 0 | 0 | 0 | 0 | 0 |
| ENSMUSG00 | Gm14040 | 694 | 0 | 0 | 0 | 0 | 0 | 0 | 0 | 0 | 0 | 0.06794049 | 0 | 0 | 0 | 0 | 0 | 0 | 0 | 0 |
| ENSMUSG00 | 4930405A21I | 1133 | 0.33764041 | 0.06820713 | 0.17078034 | 0.12063825 | 0.25489103 | 0.13120759 | 0.37582887 | 0.40450912 | 0.65433123 | 0.29131061 | 0.04216678 | 0.0492636 | 0 | 0.08198427 | 0.29092008 | 0.50693655 | 0.10148181 | 0.14565518 |
| ENSMUSG00 | Gm17481 | 1509 | 0 | 0.04415228 | 0 | 0.21135011 | 0 | 0 | 0 | 0.1771682 | 0.08188174 | 0.49994119 | 0 | 0.07397702 | 0.13076718 | 0 | 0 | 0.20299858 | 0.05079695 | 0.18227007 |
| ENSMUSG00 | Gm13663 | 1042 | 0 | 0 | 0 | 0 | 0 | 0 | 0 | 0 | 0 | 0.09050039 | 0 | 0 | 0.14203047 | 0 | 0 | 0.03674722 | 0.11034442 | 0.15837555 |
| ENSMUSG00 | Gm15254 | 272 | 0 | 0 | 0 | 0 | 0 | 0 | 0 | 0 | 0 | 0 | 0 | 0 | 0 | 0 | 0 | 0 | 0 | 0 |
| ENSMUSG00 | Gm15687 | 267 | 0 | 0 | 0 | 0 | 0 | 0 | 0 | 0 | 0 | 0 | 0 | 0.20904743 | 0 | 0 | 0 | 0 | 0 | 0 |
| ENSMUSG00 | Gm13470 | 418 | 6.71134358 | 5.73810618 | 8.67946119 | 7.73883767 | 7.02403671 | 10.3136072 | 24.2222806 | 25.948905 | 21.0859196 | 23.4625509 | 15.7725939 | 20.029544 | 78.9546714 | 55.2217977 | 54.5082785 | 60.0924365 | 52.9049601 | 75.4072205 |
| ENSMUSG00 | Gm15743 | 240 | 1.771049 | 0.41641118 | 0.80622551 | 0.37967537 | 1.80494714 | 0.41293945 | 0.39427232 | 0.95481006 | 0 | 0.19646126 | 0.19906233 | 2.79078313 | 1.23329793 | 0.58055109 | 0.3433463 | 0.47863259 | 0.47907871 | 2.0628415 |
| ENSMUSG00 | 5133400J02R | 1133 | 0 | 0 | 0 | 0 | 0 | 0 | 0 | 0.16854547 | 0 | 0 | 0 | 0 | 0 | 0 | 0 | 0 | 0 | 0 |
| ENSMUSG00 | Gm15511 | 554 | 0 | 0 | 0.08731684 | 0.1644803 | 0.26064219 | 0.08944537 | 0 | 0.13787871 | 0.07434389 | 0.25532872 | 0 | 0 | 0 | 0 | 0 | 0 | 0.48426729 | 0.09929442 |
| ENSMUSG00 | Gm15286 | 1093 | 0 | 0 | 0 | 0 | 0 | 0 | 0 | 0 | 0 | 0 | 0 | 0 | 0 | 0 | 0 | 0 | 0 | 0 |
| ENSMUSG00 | A430048G15I | 622 | 0 | 0 | 0 | 0 | 0 | 0 | 0 | 0 | 0 | 0 | 0 | 0 | 0 | 0 | 0 | 0 | 0 | 0 |
| ENSMUSG00 | 1300002E11I | 4033 | 12.4469658 | 8.47483994 | 9.97936456 | 9.4556395 | 9.46407525 | 8.61305882 | 10.4057392 | 10.7768296 | 14.3075399 | 10.6507044 | 11.0049433 | 11.403944 | 11.9751914 | 9.43161606 | 9.73594911 | 8.95314723 | 10.1303825 | 11.3755505 |
| ENSMUSG00 | Gm14029 | 1346 | 0.06315777 | 0.0989982 | 0 | 0 | 0.03575923 | 0 | 0 | 0 | 0 | 0 | 0 | 0 | 0 | 0 | 0 | 0 | 0 | 0 |
| ENSMUSG00 | 9130015L21I | 602 | 0 | 0 | 0 | 0 | 0 | 0 | 0 | 0 | 0 | 0 | 0 | 0 | 0 | 0 | 0 | 0 | 0 | 0 |
| ENSMUSG00 | Gm13165 | 631 | 0.13472322 | 0 | 0 | 0 | 0 | 0 | 0 | 0 | 0 | 0 | 0 | 0 | 0 | 0 | 0 | 0 | 0 | 0.08717767 |
| ENSMUSG00 | C330018A13I | 1597 | 0 | 0 | 0 | 0 | 0 | 0 | 0 | 0 | 0 | 0 | 0 | 0 | 0 | 0.20357458 | 0 | 0 | 0 | 0 |
| ENSMUSG00 | Gm15701 | 570 | 0 | 0 | 0 | 0 | 0 | 0 | 0 | 0 | 0 | 0.2481616 | 0 | 0 | 0 | 0 | 0 | 0 | 0.06723912 | 0 |
| ENSMUSG00 | 4921530L18I | 799 | 2.39390854 | 2.12635495 | 2.54278883 | 1.65365493 | 4.09633393 | 2.66679292 | 2.78309876 | 3.53721875 | 3.86606844 | 4.60294725 | 4.18554079 | 4.19141396 | 8.33518002 | 5.63839359 | 4.22844504 | 3.78592989 | 4.17320129 | 6.19626984 |
| ENSMUSG00 | Gm14260 | 258 | 0 | 0 | 0 | 0 | 0 | 0 | 0 | 0 | 0 | 0 | 0 | 0 | 0 | 0 | 0 | 0 | 0 | 0 |
| ENSMUSG00 | Gm14146 | 505 | 0 | 0 | 0 | 0 | 0 | 0 | 0 | 0 | 0 | 0 | 0 | 0 | 0 | 0 | 0 | 0 | 0 | 0 |
| ENSMUSG00 | Gm12594 | 497 | 0 | 0 | 0 | 0 | 0 | 0 | 0 | 0 | 0 | 0 | 0 | 0 | 0 | 0 | 0 | 0 | 0 | 0 |
| ENSMUSG00 | Gm4725 | 586 | 0 | 0 | 0 | 0 | 0 | 0 | 0 | 0 | 0 | 0 | 0 | 0 | 0 | 0 | 0 | 0 | 0 | 0 |
| ENSMUSG00 | BC046251 | 5199 | 0 | 0 | 0 | 0 | 0 | 0 | 0 | 0 | 0 | 0 | 0 | 0 | 0 | 0 | 0 | 0 | 0 | 0 |
| ENSMUSG00 | 9530036O11I | 3652 | 0.01163888 | 0 | 0 | 0 | 0 | 0 | 0.01295528 | 0.01045794 | 0 | 0.05164371 | 0 | 0.04585076 | 0.08104915 | 0.19076159 | 0.10153724 | 0.0314545 | 0 | 0.10543914 |
| ENSMUSG00 | B230209K01I | 2750 | 0 | 0.01211378 | 0 | 0 | 0 | 0 | 0 | 0 | 0 | 0.08572855 | 0 | 0.04059321 | 0 | 0.05066828 | 0.07491192 | 0 | 0.01393684 | 0.06000993 |
| ENSMUSG00 | Gm13067 | 402 | 0.31720281 | 0.0828679 | 0 | 0 | 0.11973115 | 0 | 0 | 0.09500598 | 0.71717814 | 0 | 0 | 0 | 0 | 0.46213022 | 0 | 0 | 0 | 0 |
| ENSMUSG00 | Gm12249 | 717 | 0.17784592 | 0.32523049 | 0.53973256 | 0.25417598 | 0 | 0.82933445 | 0.791844 | 0.2130678 | 0.11488568 | 0.52608874 | 0 | 0.6227689 | 0.2064097 | 0.58298018 | 1.0918125 | 1.01467439 | 0.37417584 | 0.23016363 |
| ENSMUSG00 | Gm13434 | 455 | 0 | 0 | 0 | 0 | 0 | 0 | 0 | 0 | 0 | 0 | 0 | 0 | 0 | 0 | 0 | 0 | 0 | 0 |
| ENSMUSG00 | Gm15273 | 867 | 0 | 0 | 0 | 0 | 0 | 0 | 0 | 0 | 0 | 0 | 0 | 0 | 0 | 0 | 0 | 0 | 0 | 0 |
| ENSMUSG00 | AA645442 | 283 | 1.20155975 | 0 | 0 | 0 | 0 | 0 | 0 | 0 | 0 | 0.33322052 | 0 | 0 | 0 | 0.16411338 | 0.43676561 | 0 | 0 | 0 |
| ENSMUSG00 | Gm13194 | 310 | 0.54845388 | 0 | 0 | 0.29394222 | 0.15526427 | 0 | 0 | 0.73920779 | 0.13285973 | 0 | 0 | 0 | 0 | 0.29963927 | 0.53163298 | 0.24703618 | 0.24726643 | 0 |
| ENSMUSG00 | Gm13618 | 380 | 0 | 0.08766551 | 0 | 0.11989749 | 0 | 0.26080386 | 0 | 0 | 0.10838557 | 0 | 0.12572357 | 0 | 0 | 0 | 0.21685029 | 0 | 0 | 0 |
| ENSMUSG00 | Gm14297 | 290 | 0 | 0 | 0 | 0 | 0 | 0 | 0 | 0 | 0 | 0 | 0 | 0 | 0 | 0 | 0 | 0 | 0 | 0 |
| ENSMUSG00 | Gm16736 | 1739 | 0.14665386 | 0.05746905 | 0 | 0 | 0 | 0.05698992 | 0 | 0.02196228 | 0 | 0.02711369 | 0 | 0 | 0.02836798 | 0.05341471 | 0.04738534 | 0.088075 | 0.04407855 | 0.09489783 |
| ENSMUSG00 | Gm12283 | 710 | 0 | 0 | 0.06813173 | 0 | 0 | 0 | 0 | 0 | 0 | 0 | 0 | 0 | 0 | 0 | 0 | 0 | 0.1079614 | 0 |
| ENSMUSG00 | 9430029A11I | 2041 | 0.1666053 | 0.13057479 | 0 | 0.40181225 | 0.21224268 | 0.24278654 | 0.48680365 | 0.33682667 | 0.46413026 | 0.25411942 | 0.6085982 | 0.21877771 | 0 | 0.31857777 | 0.16149556 | 0.05628213 | 0 | 0.35037648 |
| ENSMUSG00 | Gm14705 | 1821 | 0 | 0 | 0 | 0 | 0 | 0 | 0.25981702 | 0.23070644 | 0.29402784 | 0 | 0 | 0 | 0 | 0.30605659 | 0.06787736 | 0 | 0.29465577 | 1.38957656 |
| ENSMUSG00 | Gm12284 | 1008 | 3.4155945 | 1.81766784 | 2.92736642 | 2.89276472 | 1.43249773 | 2.06469723 | 2.39379626 | 1.32612508 | 2.45157832 | 1.82428316 | 2.51197698 | 3.32236087 | 3.86629114 | 4.05464255 | 2.86121916 | 5.47008677 | 4.10638895 | 3.54721422 |
| ENSMUSG00 | Gm16737 | 1179 | 0.32446699 | 0.67812507 | 0.16411715 | 0.23186282 | 0.48989235 | 0.08405892 | 0 | 0.19436337 | 0.48906804 | 0.07998423 | 0.12156478 | 0.52075682 | 0 | 0.07878556 | 0.13978475 | 0.25981752 | 0 | 0.55988913 |
| ENSMUSG00 | Gm15551 | 521 | 0 | 0.12788059 | 0.09284747 | 0 | 0 | 0.28533244 | 0 | 0 | 0 | 0 | 0 | 0 | 0 | 0 | 0 | 0 | 0 | 0 |
| ENSMUSG00 | Gm11592 | 634 | 0.40225719 | 0.10508799 | 0.07629894 | 0.07186285 | 0 | 0.23447666 | 0.82088244 | 0.30120191 | 0.19488888 | 0.52059136 | 0.15070965 | 0.61626126 | 0.23343179 | 0.07325566 | 0.38992009 | 0.6039528 | 0.24180629 | 0.43382576 |
| ENSMUSG00 | Gm16178 | 674 | 0.44144842 | 0.09885132 | 0 | 0 | 0 | 0 | 1.26354335 | 0.11333057 | 0.79439867 | 0.34978267 | 0 | 0.08281256 | 0 | 0 | 0.12225981 | 0 | 0 | 0 |
| ENSMUSG00 | Gm16023 | 2766 | 0.87592011 | 1.27663297 | 1.04931736 | 0.93889354 | 0.53943949 | 1.02115178 | 0.87235959 | 0.5523124 | 0.92319739 | 0.6136751 | 1.03633316 | 0.72645114 | 0.49938311 | 0.94029967 | 0.95332595 | 0.53988926 | 0.4018303 | 0.69606606 |
| ENSMUSG00 | Gm12867 | 455 | 0.09341797 | 0.43929091 | 0.74420816 | 0.40053665 | 1.48098227 | 0.98016396 | 0.31195173 | 0.41969673 | 0.09051981 | 0.62176752 | 0.41999963 | 0.73603072 | 0.86737437 | 0.30622475 | 0.09055287 | 0 | 0.50540172 | 0 |
| ENSMUSG00 | Gm12590 | 401 | 0 | 0 | 0 | 0 | 0 | 0 | 0 | 0 | 0 | 0 | 0 | 0 | 0 | 0 | 0 | 0 | 0 | 0 |
| ENSMUSG00 | Gm15015 | 630 | 0 | 0 | 0 | 0 | 0 | 0 | 0 | 0 | 0 | 0 | 0 | 0 | 0 | 0 | 0 | 0 | 0 | 0 |
| ENSMUSG00 | F630206G17I | 681 | 0 | 0 | 0 | 0 | 0 | 0 | 0 | 0 | 0.12095893 | 0.06923745 | 0 | 0 | 0 | 0 | 0 | 0 | 0 | 0 |
| ENSMUSG00 | 4930547E08I | 537 | 0 | 0 | 0 | 0 | 0.08963114 | 0 | 0.08810555 | 0.07112179 | 0 | 0 | 0 | 0.10393978 | 0.09186577 | 0 | 0.07672543 | 0 | 0.21411339 | 0.10243782 |
| ENSMUSG00 | Gm11560 | 947 | 0.13465209 | 0.17588645 | 0.56188895 | 0.43299831 | 0.20330274 | 0.31395607 | 0.29976354 | 0.2823092 | 0.04349157 | 0.39831639 | 0.20179497 | 0.11787891 | 0.20837135 | 0.14713016 | 0.1740298 | 0.16173435 | 0.28329892 | 0.29043879 |
| ENSMUSG00 | Gm16876 | 756 | 0.16867133 | 0.22032337 | 1.02377842 | 0.18079779 | 0.12733313 | 0 | 0 | 0.25259525 | 0.16343855 | 0.37421193 | 0 | 0.07383024 | 0 | 0.49147182 | 0 | 0 | 0.10139232 | 0 |
| ENSMUSG00 | 4933406J10R | 959 | 0.22161197 | 0.17368558 | 0.05044164 | 0.09501782 | 0 | 0 | 0 | 0.27877666 | 0 | 0.14749959 | 0.1992699 | 0 | 0.051441 | 0 | 0 | 0 | 0.11989457 | 0 |
| ENSMUSG00 | Gm15432 | 194 | 0 | 0 | 0 | 0 | 0 | 0 | 0 | 0 | 0 | 0 | 0 | 0 | 0 | 0 | 0 | 0 | 0 | 0 |
| ENSMUSG00 | A730081D07I | 1799 | 0.04725423 | 0.44441882 | 0.08066737 | 0.20260609 | 0 | 0.19281219 | 0 | 0 | 0.02289412 | 0.15725638 | 0 | 0 | 0 | 0 | 0 | 0.17027507 | 0.04260845 | 0 |
| ENSMUSG00 | Gm16237 | 1317 | 0.12909697 | 0 | 0 | 0 | 0 | 0.03762546 | 0 | 0 | 0 | 0 | 0 | 0 | 0 | 0 | 0 | 0 | 0 | 0 |
| ENSMUSG00 | Gm15247 | 401 | 0.10599795 | 0.08307455 | 0.24126449 | 0 | 0 | 0 | 0.11798673 | 0 | 0 | 0 | 0.23827909 | 0.41757354 | 0 | 0 | 0 | 0 | 0 | 0 |
| ENSMUSG00 | Gm11234 | 843 | 0 | 0 | 0 | 0 | 0 | 0 | 0 | 0 | 0 | 0 | 0 | 0 | 0 | 0 | 0 | 0 | 0 | 0 |
| ENSMUSG00 | 4933427G23I | 2236 | 0.03801894 | 0.02979686 | 0 | 0 | 0.12915543 | 0.11080665 | 0.1269571 | 0.05124204 | 0 | 0.02108708 | 0 | 0 | 0.13237545 | 0 | 0.27639684 | 0.32536742 | 0.05142169 | 0.09840627 |
| ENSMUSG00 | 4930511O05I | 388 | 0 | 0 | 0 | 0 | 0 | 0 | 0 | 0 | 0 | 0 | 0 | 0 | 0 | 0 | 0 | 0 | 0 | 0 |
| ENSMUSG00 | Gm16510 | 248 | 0 | 0 | 0 | 0 | 0.19408034 | 0 | 0.19077693 | 0 | 0.16607466 | 0.1901238 | 0 | 0 | 0 | 0 | 0 | 0 | 0.15454152 | 0.22181091 |
| ENSMUSG00 | Gm15747 | 977 | 0.26103486 | 0.17048564 | 0.7426847 | 0.65287065 | 0.78824031 | 0.35503494 | 1.16223572 | 0.27364055 | 1.13821487 | 0.82043189 | 1.12469196 | 0.57129644 | 1.00986525 | 0.71306173 | 0.92777301 | 0.70545643 | 0.23537132 | 0.28152051 |
| ENSMUSG00 | Gm13595 | 492 | 0 | 0 | 0 | 0.09260375 | 0 | 0 | 0 | 0 | 0 | 0 | 0.19420715 | 0 | 0.20053625 | 0 | 0.251229 | 0 | 0 | 0 |
| ENSMUSG00 | Gm12347 | 466 | 0 | 0 | 0 | 0 | 0 | 0 | 0 | 0 | 0 | 0 | 0 | 0 | 0 | 0 | 0 | 0 | 0 | 0 |
| ENSMUSG00 | 4930507D10I | 1149 | 0 | 0 | 0 | 0 | 0 | 0 | 0 | 0 | 0 | 0 | 0 | 0 | 0 | 0 | 0 | 0 | 0 | 0 |
| ENSMUSG00 | Gm14582 | 1185 | 0 | 0 | 0 | 0 | 0.12185297 | 0 | 0 | 0 | 0 | 0 | 0 | 0 | 0 | 0 | 0 | 0 | 0 | 0 |
| ENSMUSG00 | Gm16894 | 840 | 0 | 0 | 0 | 0 | 0 | 0 | 0 | 0 | 0 | 0 | 0 | 0 | 0.35237084 | 0 | 0 | 0.13675217 | 0 | 0 |
| ENSMUSG00 | Gm15848 | 1144 | 0.07430975 | 0.08735899 | 0.04228455 | 0.3186087 | 0.04207336 | 0 | 0.37221513 | 0.06676993 | 0.28801759 | 0.28850955 | 1.08579451 | 0.3415294 | 0.34497844 | 0 | 0.28812277 | 0 | 0.03350201 | 0.33659418 |
| ENSMUSG00 | Gm12088 | 518 | 0 | 0 | 0 | 0 | 0 | 0 | 0 | 0 | 0 | 0 | 0 | 0 | 0 | 0 | 0 | 0 | 0 | 0 |
| ENSMUSG00 | Gm15577 | 2291 | 0 | 0 | 0 | 0 | 0 | 0 | 0 | 0 | 0 | 0 | 0 | 0 | 0 | 0 | 0 | 0 | 0 | 0 |
| ENSMUSG00 | Gm16263 | 497 | 0 | 0 | 0 | 0 | 0 | 0 | 0 | 0 | 0 | 0 | 0 | 0 | 0 | 0 | 0 | 0 | 0 | 0 |
| ENSMUSG00 | Gm13465 | 976 | 0 | 0 | 0 | 0 | 0 | 0 | 0 | 0 | 0 | 0 | 0 | 0 | 0 | 0 | 0 | 0 | 0 | 0 |
| ENSMUSG00 | Gm15482 | 4527 | 0.01877852 | 0 | 0 | 0.02012858 | 0 | 0 | 0 | 0 | 0 | 0 | 0 | 0 | 0 | 0 | 0 | 0 | 0 | 0 |
| ENSMUSG00 | AI427809 | 1221 | 0.48736484 | 0.16369973 | 0.11885388 | 0.1119436 | 0.19710042 | 0.08116746 | 0 | 0 | 0.47224506 | 0.03861646 | 0 | 0 | 0 | 0.26626422 | 0.20246465 | 0 | 0 | 0 |
| ENSMUSG00 | 4930407G08I | 695 | 0 | 0 | 0 | 0 | 0 | 0 | 0 | 0 | 0 | 0 | 0 | 0 | 0 | 0 | 0 | 0 | 0 | 0 |
| ENSMUSG00 | 0610009E02I | 2076 | 0.30711832 | 0.40116684 | 0.16310921 | 0.39503796 | 0.02318493 | 0.16708533 | 0.66091893 | 0.22076533 | 0.4166266 | 0.31797199 | 0.39122076 | 0.53772315 | 0.54654822 | 0.29083484 | 0.15877286 | 0.44266598 | 0.36923215 | 0.26497643 |
| ENSMUSG00 | B230112J18I | 599 | 0 | 0 | 0 | 0 | 0 | 0 | 0 | 0 | 0 | 0 | 0.3987893 | 0 | 0 | 0.46521623 | 0.1375678 | 0.3196211 | 0.0639838 | 0 |
| ENSMUSG00 | Gm11629 | 382 | 0.11127009 | 0.17441306 | 0.12663228 | 0.35780925 | 1.0079984 | 0.38915759 | 0.24771036 | 0.19996022 | 0.64690862 | 0.6171558 | 0 | 0.1461143 | 0.25828229 | 0.24316276 | 0.4314299 | 0 | 0 | 0 |
| ENSMUSG00 | Gm15655 | 438 | 0 | 0 | 0 | 0 | 0 | 0 | 0 | 0 | 0 | 0 | 0 | 0 | 0 | 0 | 0 | 0 | 0 | 0 |
| ENSMUSG00 | Gm16069 | 543 | 0 | 0 | 0 | 0 | 0 | 0 | 0 | 0.07033592 | 0 | 0 | 0 | 0.10279128 | 0 | 0.34212956 | 0.37908818 | 0.42310063 | 0.0705825 | 0 |
| ENSMUSG00 | Gm16275 | 710 | 0.11973289 | 0.37535655 | 0.06813173 | 0.32085242 | 0.61012298 | 0.13958517 | 0.93292606 | 0 | 0.69611013 | 0.33204721 | 0.47102072 | 0.23584083 | 0.34740787 | 0.19624262 | 0.34818216 | 0 | 0.3778649 | 0 |
| ENSMUSG00 | Gm13851 | 437 | 0.29179755 | 0 | 0 | 0 | 0 | 0 | 0 | 0.216534 | 0 | 0 | 0 | 0 | 0 | 0 | 0 | 0 | 0 | 0 |
| ENSMUSG00 | Gm13750 | 464 | 0.09160598 | 0.07179503 | 0 | 0 | 0 | 0 | 0 | 0 | 0 | 0 | 0 | 0 | 0 | 0 | 0.08879646 | 0 | 0 | 0 |
| ENSMUSG00 | Gm15499 | 419 | 0 | 0 | 0 | 0 | 0 | 0 | 0.11291809 | 0 | 0 | 0 | 0 | 0 | 0 | 0 | 0 | 0 | 0 | 0 |
| ENSMUSG00 | Gm16601 | 1716 | 0.04953983 | 0.01941311 | 0 | 0 | 0 | 0 | 0.41357237 | 0 | 0 | 0 | 0.13920442 | 0.03252661 | 0 | 0.24358787 | 0 | 0 | 0 | 0 |
| ENSMUSG00 | Gm12833 | 476 | 0 | 0 | 0 | 0 | 0 | 0 | 0 | 0 | 0 | 0 | 0.10036756 | 0 | 0 | 0.19514322 | 0 | 0 | 0 | 0 |
| ENSMUSG00 | A630052C17I | 2201 | 2.76158118 | 2.37624916 | 4.17581589 | 2.91872206 | 2.05561146 | 2.2964011 | 3.8907746 | 3.29693615 | 3.19986106 | 2.37788644 | 2.80007708 | 3.29669974 | 3.04822388 | 4.0514606 | 3.72517475 | 3.02706301 | 3.65675709 | 1.94943677 |
| ENSMUSG00 | Gm9458 | 728 | 0 | 0 | 0 | 0 | 0 | 0 | 0 | 0 | 0 | 0 | 0 | 0 | 0 | 0 | 0 | 0 | 0 | 0 |
| ENSMUSG00 | 4931428L18I | 1542 | 0 | 0 | 0 | 0 | 0 | 0 | 0 | 0 | 0 | 0 | 0 | 0 | 0 | 0 | 0 | 0 | 0 | 0 |
| ENSMUSG00 | 2410133F24I | 2151 | 0.51377712 | 0.60399948 | 0.58471027 | 0.48717063 | 0.17901227 | 0.62200084 | 0.74785267 | 0.78124858 | 0.68931407 | 0.81105347 | 0.62189625 | 0.90820464 | 1.16965494 | 0.47502088 | 0.80449342 | 0.51623785 | 0.97998435 | 0.35803231 |
| ENSMUSG00 | Gm15589 | 269 | 0 | 0 | 0 | 0.16937191 | 0 | 0 | 0 | 0 | 0 | 0 | 0 | 0 | 0 | 0 | 0 | 0 | 0 | 0 |
| ENSMUSG00 | Gm11607 | 460 | 0 | 0 | 0 | 0 | 0 | 0 | 0 | 0 | 0 | 0 | 0 | 0 | 0 | 0.30289622 | 0 | 0 | 0 | 0 |
| ENSMUSG00 | Gm7518 | 204 | 0 | 0 | 0 | 0 | 0 | 0 | 0 | 0 | 0 | 0 | 0 | 0 | 0 | 0 | 0 | 0 | 0 | 0 |
| ENSMUSG00 | Gm6117 | 1331 | 0 | 0 | 0 | 0 | 0 | 0 | 0 | 0 | 0 | 0 | 0 | 0 | 0 | 0 | 0 | 0 | 0 | 0 |
| ENSMUSG00 | Gm11685 | 1330 | 0 | 0 | 0 | 0.1370257 | 0.07237883 | 0.14903078 | 0 | 0 | 0 | 0 | 0 | 0.20983332 | 0 | 0 | 0.03097861 | 0 | 0.02881676 | 0 |
| ENSMUSG00 | Astx1c | 1146 | 0 | 0 | 0 | 0 | 0 | 0 | 0 | 0 | 0 | 0 | 0 | 0 | 0 | 0 | 0 | 0 | 0 | 0 |
| ENSMUSG00 | Gm13977 | 4046 | 0.04202192 | 0 | 0.01195589 | 0.06756457 | 0.02379235 | 0 | 0.01169369 | 0.00943955 | 0 | 0.01165366 | 0 | 0.01379527 | 0.01219276 | 0.03443704 | 0.05091641 | 0.00946382 | 0.00947264 | 0 |
| ENSMUSG00 | Gm14902 | 1004 | 0 | 0 | 0 | 0 | 0 | 0 | 0 | 0 | 0 | 0 | 0 | 0 | 0 | 0 | 0 | 0 | 0 | 0 |
| ENSMUSG00 | Gm13763 | 588 | 0 | 0 | 0 | 0 | 0 | 0 | 0 | 0 | 0 | 0 | 0 | 0 | 0 | 0 | 0 | 0 | 0 | 0 |
| ENSMUSG00 | Gm12154 | 441 | 0 | 0 | 0 | 0 | 0 | 0 | 0 | 0 | 0 | 0 | 0 | 0 | 0 | 0 | 0 | 0 | 0 | 0 |
| ENSMUSG00 | Gm4409 | 1915 | 0 | 0 | 0 | 0 | 0 | 0 | 0 | 0 | 0 | 0 | 0 | 0 | 0 | 0 | 0 | 0 | 0 | 0 |
| ENSMUSG00 | Gm17029 | 5966 | 0.32060559 | 0.45228703 | 0.43784288 | 0.49639086 | 0.29850506 | 0.28239909 | 0.72959545 | 0.23686203 | 0.3865982 | 0.41887148 | 0.32832271 | 0.22453502 | 0.30594719 | 0.17126549 | 0.24861817 | 0.31448873 | 0.10921003 | 0.35037648 |
| ENSMUSG00 | Gm15816 | 1682 | 0.12635308 | 0.03961105 | 0.43139296 | 0.08126227 | 0.14307944 | 0.08838181 | 0.02812882 | 0 | 0.04897326 | 0 | 0 | 0.23228873 | 0 | 0.05522484 | 0.04899115 | 0 | 0.15050302 | 0.88302371 |
| ENSMUSG00 | Gm16201 | 2946 | 0.34627435 | 0.47492925 | 0.78816343 | 0.52582332 | 0.91493134 | 0.52143067 | 0.43361926 | 0.36299636 | 1.30018532 | 0.75223457 | 0.5027236 | 0.87152766 | 0.68656097 | 0.37836324 | 0.92304911 | 0.31193978 | 0.35125034 | 0.74689894 |
| ENSMUSG00 | Hnrnpa1l2-ps | 964 | 0 | 0 | 0 | 0 | 0 | 0 | 0 | 0 | 0 | 0 | 0 | 0 | 0 | 0 | 0 | 0 | 0.11927271 | 0 |
| ENSMUSG00 | Gm15997 | 1486 | 0 | 0 | 0 | 0 | 0 | 0 | 0 | 0 | 0 | 0 | 0 | 0 | 0 | 0 | 0 | 0 | 0 | 0 |
| ENSMUSG00 | 2410057H14I | 586 | 0 | 0 | 0 | 0 | 0 | 0 | 0 | 0 | 0 | 0 | 0 | 0 | 0 | 0 | 0 | 0 | 0 | 0 |
| ENSMUSG00 | Gm15222 | 342 | 0.49713656 | 0.09740612 | 0 | 0.53287771 | 0.28147324 | 0.28978207 | 0.13834117 | 0 | 0 | 0 | 0 | 0 | 0 | 0 | 0.12047239 | 0.44784336 | 0 | 0 |
| ENSMUSG00 | Gm13261 | 1016 | 0 | 0 | 0 | 0 | 0 | 0 | 0 | 0 | 0 | 0 | 0 | 0 | 0 | 0 | 0 | 0 | 0 | 0 |
| ENSMUSG00 | Gm12037 | 306 | 0.1389058 | 0.54432834 | 0 | 0 | 0 | 0 | 0.77308299 | 0.37443532 | 0.26919291 | 0.15408727 | 1.40514584 | 0.7296165 | 0 | 0.91066838 | 0.26929121 | 0.50053082 | 0.25049867 | 0.71907329 |
| ENSMUSG00 | Gm11770 | 546 | 0 | 0.06101263 | 0 | 0 | 0 | 0 | 0 | 0 | 0 | 0 | 0 | 0 | 0 | 0 | 0 | 0 | 0 | 0 |
| ENSMUSG00 | Astx5 | 1146 | 0 | 0 | 0 | 0 | 0 | 0 | 0 | 0 | 0 | 0 | 0 | 0 | 0 | 0 | 0 | 0 | 0 | 0 |
| ENSMUSG00 | Gm11674 | 3003 | 0 | 0 | 0 | 0 | 0 | 0 | 0 | 0 | 0 | 0 | 0 | 0 | 0 | 0 | 0 | 0 | 0 | 0 |
| ENSMUSG00 | Gm15751 | 1860 | 0.41134041 | 0.96714854 | 1.04029098 | 0.68586518 | 0.5951797 | 0.10656502 | 0.81398157 | 0.90347619 | 0.3764359 | 0.40559745 | 0.53939469 | 0.81022736 | 0.84872116 | 0.42448897 | 0.64238985 | 1.0293174 | 1.0302768 | 0.79851929 |
| ENSMUSG00 | Gm16098 | 794 | 0 | 0.04195579 | 0 | 0 | 0.1212391 | 0 | 0.05958776 | 0 | 0.10374437 | 0.23753503 | 0.06016997 | 0 | 0 | 0.11698763 | 0.05189113 | 0 | 0.53096885 | 0 |
| ENSMUSG00 | Gm11216 | 351 | 0 | 0 | 0 | 0 | 0 | 0 | 0 | 0 | 0 | 0 | 0 | 0 | 0 | 0 | 0 | 0 | 0 | 0 |
| ENSMUSG00 | Gm14051 | 374 | 0.34095061 | 0.26721573 | 0.38802297 | 0.97456779 | 1.15825485 | 1.05995152 | 2.15057631 | 0.51059362 | 1.10124374 | 0.1260714 | 0.89418371 | 0 | 0 | 1.98691283 | 2.09312717 | 1.12619434 | 0.40990692 | 0.88249904 |
| ENSMUSG00 | Gm2309 | 1937 | 0 | 0 | 0 | 0 | 0 | 0 | 0 | 0 | 0 | 0 | 0 | 0 | 0 | 0 | 0 | 0 | 0 | 0 |
| ENSMUSG00 | Gm13912 | 1857 | 0 | 0 | 0 | 0 | 0 | 0 | 0 | 0 | 0 | 0 | 0 | 0 | 0 | 0 | 0 | 0 | 0 | 0 |
| ENSMUSG00 | 1700025D23I | 844 | 0 | 0.03947025 | 0 | 0 | 0 | 0 | 0 | 0 | 0 | 0 | 0 | 0.1322646 | 0 | 0 | 0 | 0 | 0 | 0 |
| ENSMUSG00 | Gm11686 | 3003 | 0.07077119 | 0.05546602 | 0 | 0.09103106 | 0 | 0.04950323 | 0 | 0.20348932 | 0 | 0 | 0.07954539 | 0.13010644 | 0.01642754 | 0 | 0.10976105 | 0 | 0 | 0.09159625 |
| ENSMUSG00 | Gm14546 | 1645 | 0 | 0 | 0 | 0 | 0 | 0 | 0 | 0 | 0 | 0 | 0 | 0 | 0 | 0 | 0 | 0 | 0 | 0 |
| ENSMUSG00 | B230378P21I | 1975 | 0 | 0 | 0.04898585 | 0.11534442 | 0.17059416 | 0.10035997 | 0 | 0 | 0.25024718 | 0.21486396 | 0.4596072 | 0.08478328 | 0.1748473 | 0.23515994 | 0.14603083 | 0.09693825 | 0.07762288 | 0.2228217 |
| ENSMUSG00 | 4930527A07I | 557 | 0 | 0 | 0 | 0 | 0 | 0 | 0 | 0 | 0 | 0 | 0 | 0 | 0 | 0 | 0 | 0 | 0 | 0 |
| ENSMUSG00 | Gm11827 | 1364 | 1.58926978 | 0.73268829 | 0.95754056 | 0.5344404 | 1.37620603 | 0.69025069 | 1.28340845 | 2.15602272 | 0.51332168 | 0.89876707 | 0.70051259 | 0.69564975 | 0.54250642 | 2.38349422 | 1.14784393 | 0.44915669 | 1.34872599 | 0.68559737 |
| ENSMUSG00 | Gm15969 | 234 | 0 | 0 | 0 | 0 | 0 | 0 | 0.20219094 | 0 | 0 | 0 | 0 | 0 | 0.84328064 | 0 | 0 | 0 | 0 | 0 |
| ENSMUSG00 | Gm13070 | 765 | 0 | 0 | 0 | 0.11911384 | 0 | 0 | 0.18553992 | 0 | 0 | 0.18490472 | 0 | 0 | 0 | 0 | 0.05385824 | 0.05005308 | 0.05009973 | 0 |
| ENSMUSG00 | Gm16827 | 1787 | 0.21407196 | 0.33555238 | 0.29776656 | 0.25495828 | 0.29627933 | 0.38821392 | 0.23828434 | 0.25646829 | 0.29962211 | 0.15831238 | 0.18714309 | 0.31234282 | 0.13802999 | 0.36385967 | 0.13833762 | 0.08570925 | 0.17157827 | 0.18469762 |
| ENSMUSG00 | Gm15587 | 3852 | 0 | 0 | 0 | 0 | 0 | 0 | 0 | 0 | 0 | 0 | 0 | 0 | 0 | 0 | 0 | 0 | 0 | 0 |
| ENSMUSG00 | Gm15351 | 844 | 1.00723166 | 1.14463736 | 1.66212367 | 0.59380508 | 1.14056691 | 0.93938831 | 0.22423071 | 0.40726495 | 0.4391927 | 0 | 0.39623781 | 0.1983969 | 0.87675208 | 0.71537101 | 0.09763402 | 0.13610405 | 0.22705152 | 1.43388667 |
| ENSMUSG00 | 1700080G11I | 434 | 0 | 0 | 0 | 0 | 0 | 0 | 0 | 0 | 0 | 0 | 0 | 0 | 0 | 0 | 0 | 0 | 0 | 0 |
| ENSMUSG00 | Gm11705 | 108 | 0.39356644 | 0 | 0 | 0 | 0 | 0 | 0 | 0.35363336 | 0 | 0 | 0 | 0 | 0 | 0 | 0 | 0 | 0 | 0 |
| ENSMUSG00 | Gm16192 | 935 | 0.59098106 | 1.14012044 | 0.82778234 | 0.82838262 | 0.66921391 | 0.95395637 | 1.01203591 | 1.38881463 | 1.32149248 | 0.35299992 | 1.83946364 | 0.29847948 | 0.31656845 | 0.49672821 | 0.83725087 | 0.36857269 | 1.43467422 | 1.11783212 |
| ENSMUSG00 | Gm11915 | 356 | 0 | 0 | 0 | 0 | 0 | 0 | 0 | 0 | 0 | 0 | 0 | 0 | 0 | 0 | 0 | 0 | 0 | 0 |
| ENSMUSG00 | 5730588L14I | 4081 | 0.01041538 | 0 | 0 | 0 | 0 | 0 | 0 | 0.05615154 | 0.06055356 | 0 | 0 | 0.04103087 | 0 | 0 | 0.04038378 | 0.05629592 | 0 | 0.17523117 |
| ENSMUSG00 | Gm15100 | 571 | 0 | 0 | 0 | 0 | 0 | 0 | 0 | 0 | 0 | 0 | 0 | 0 | 0 | 0 | 0 | 0 | 0 | 0 |
| ENSMUSG00 | B230104C08 | 976 | 0 | 0 | 0 | 0 | 0 | 0 | 0 | 0 | 0 | 0 | 0 | 0 | 0 | 0 | 0 | 0 | 0 | 0 |
| ENSMUSG00 | Gm2165 | 2553 | 0 | 0 | 0 | 0 | 0 | 0 | 0 | 0 | 0 | 0 | 0 | 0 | 0 | 0 | 0 | 0 | 0 | 0 |
| ENSMUSG00 | Gm13562 | 1339 | 0.73011131 | 1.8410412 | 1.481191 | 0.74857578 | 0.71892343 | 1.59131258 | 1.48404221 | 1.62581549 | 1.07657061 | 1.69024179 | 0.60655287 | 1.41727597 | 0.77368952 | 1.1446265 | 1.53851964 | 1.42982104 | 1.88912292 | 1.68437145 |
| ENSMUSG00 | Gm16707 | 1191 | 0.67808425 | 0.41955786 | 0.64985431 | 0.76508891 | 0.44454338 | 0.45766589 | 0.15890068 | 0.64135017 | 1.10660664 | 0.71260509 | 1.40396603 | 1.0310198 | 0.53846761 | 0.66292988 | 0.44972311 | 0.80374911 | 1.12871972 | 0.04618733 |
| ENSMUSG00 | 1700092C02I | 356 | 0 | 0 | 0 | 0 | 0 | 0 | 0 | 0 | 0 | 0 | 0 | 0 | 0 | 0 | 0 | 0 | 0 | 0 |
| ENSMUSG00 | E130102H24I | 739 | 7.36219558 | 8.06902309 | 6.67672543 | 5.17879259 | 6.38285321 | 6.8394842 | 7.49064064 | 9.45765851 | 6.79939773 | 4.33863034 | 6.8527004 | 8.08156414 | 8.47788024 | 5.6562488 | 7.6939306 | 8.39388146 | 6.37907242 | 5.58279162 |
| ENSMUSG00 | Gm14262 | 324 | 0 | 0 | 0 | 0 | 0 | 0 | 0 | 0 | 0 | 0 | 0 | 0 | 0 | 0 | 0 | 0 | 0 | 0 |
| ENSMUSG00 | Isoc2a | 1082 | 1.29636858 | 0.49261211 | 0.44707514 | 0.58951444 | 0.40035796 | 0.45797351 | 1.7490824 | 1.55311063 | 0.64710792 | 1.26374343 | 1.05970333 | 2.0634256 | 1.64135769 | 1.54527463 | 1.67547917 | 1.13243941 | 1.70024238 | 0.86428356 |
| ENSMUSG00 | Gm6081 | 488 | 0 | 0.20479238 | 0.19825217 | 0.18672559 | 0 | 0 | 0 | 0.54784184 | 0.25319579 | 0 | 0 | 0.45750343 | 1.41525992 | 0 | 0 | 0.47078616 | 0.47122496 | 0 |
| ENSMUSG00 | Gm15908 | 1087 | 0.0782064 | 0.24517309 | 0 | 0.16765794 | 0.53135518 | 0.41028022 | 0 | 0.14054242 | 0 | 0.04337691 | 0 | 0 | 0 | 0.29908796 | 0.18951958 | 0.4931633 | 0.03525878 | 0.20242542 |
| ENSMUSG00 | Gm16706 | 3574 | 0.29732216 | 0.28894788 | 0.35190593 | 0.20396662 | 0.22894312 | 0.22183652 | 0.22504632 | 0.28852682 | 0.34571782 | 0.40897364 | 0.18714309 | 0.43727995 | 0.027606 | 0.77969928 | 0.23056271 | 0.10713656 | 0.11796006 | 0.13852321 |
| ENSMUSG00 | 1700029M20 | 1579 | 0 | 0 | 0 | 0 | 0 | 0 | 0 | 0 | 0 | 0 | 0 | 0 | 0 | 0 | 0 | 0 | 0 | 0 |
| ENSMUSG00 | Gm12228 | 925 | 0 | 0 | 0 | 0 | 0 | 0 | 0 | 0 | 0 | 0 | 0 | 0 | 0 | 0 | 0 | 0 | 0 | 0 |
| ENSMUSG00 | Gm12911 | 699 | 0 | 0 | 0 | 0 | 0 | 0 | 0 | 0 | 0 | 0 | 0 | 0 | 0 | 0 | 0 | 0 | 0 | 0 |
| ENSMUSG00 | Gm12147 | 180 | 0 | 0 | 0 | 0 | 0 | 0 | 0 | 0 | 0 | 0 | 0 | 0 | 0 | 0 | 0 | 0 | 0 | 0 |
| ENSMUSG00 | Gm12364 | 545 | 0.23397345 | 0 | 0 | 0 | 0 | 0 | 0 | 0 | 0 | 0.08651505 | 0 | 0 | 0 | 0 | 0 | 0 | 0 | 0 |
| ENSMUSG00 | Gm13764 | 414 | 0 | 0 | 0 | 0 | 0 | 0 | 0 | 0 | 0 | 0 | 0 | 0 | 0 | 0 | 0 | 0 | 0 | 0 |
| ENSMUSG00 | Gm11642 | 438 | 3.00835721 | 1.36902305 | 1.9879533 | 5.82515635 | 4.06593876 | 1.24447504 | 3.67267371 | 3.40069336 | 3.19712679 | 2.69125019 | 6.76266536 | 4.46015569 | 2.92837865 | 2.5448815 | 1.22287723 | 2.88490878 | 2.36257995 | 1.13032411 |
| ENSMUSG00 | Gm11579 | 843 | 0 | 0 | 0 | 0 | 0 | 0.17634425 | 0 | 0 | 0 | 0 | 0 | 0 | 0 | 0 | 0 | 0 | 0 | 0 |
| ENSMUSG00 | 4932702P03I | 4270 | 0.01990875 | 0.1092226 | 0 | 0.0960303 | 0.06763268 | 0.04641942 | 0.03324076 | 0.0626105 | 0.05787332 | 0.01104232 | 0.01118851 | 0.10457267 | 0 | 0.03263051 | 0.01929815 | 0.00896736 | 0.14361142 | 0 |
| ENSMUSG00 | Gm12965 | 345 | 0 | 0 | 0.14021313 | 0 | 0.13951282 | 0 | 0 | 0.22140523 | 0 | 0 | 0 | 0.32356906 | 0 | 0.13462054 | 0 | 0.77691087 | 0.11109072 | 0 |
| ENSMUSG00 | Gm16222 | 1295 | 0 | 0 | 0 | 0 | 0 | 0 | 0 | 0 | 0 | 0 | 0 | 0 | 0 | 0 | 0 | 0 | 0 | 0 |
| ENSMUSG00 | 4930502E09I | 752 | 0.05652284 | 0.26579437 | 0.12865301 | 0.66645145 | 0.19201565 | 0 | 0 | 0.40630215 | 0.16430791 | 0.25080161 | 0.31765265 | 0.07422296 | 0.8528124 | 0.37056453 | 0.16436791 | 0.05091836 | 0 | 0.14630082 |
| ENSMUSG00 | 4930556L07I | 1488 | 0 | 0 | 0 | 0 | 0 | 0 | 0 | 0 | 0 | 0 | 0 | 0 | 0 | 0 | 0 | 0 | 0 | 0 |
| ENSMUSG00 | Gm15943 | 3819 | 0 | 0.02616881 | 0 | 0 | 0.02520656 | 0.03892595 | 0.02477752 | 0 | 0.05392317 | 0.18519522 | 0.05003923 | 0.01461526 | 0 | 0 | 0.03236572 | 0.01002634 | 0.06021413 | 0.01440406 |
| ENSMUSG00 | 2700038G22I | 1257 | 0.304333 | 0.50353619 | 0.61573308 | 0.72491717 | 0.61265774 | 1.34032851 | 1.16682024 | 1.6711075 | 0.39318869 | 0.52514705 | 1.86234922 | 0.26642321 | 0.94189818 | 0.6281221 | 0.16388845 | 0.36554279 | 0.48784467 | 0.65643325 |
| ENSMUSG00 | Gm13847 | 1187 | 0 | 0 | 0 | 0 | 0 | 0 | 0 | 0 | 0 | 0 | 0 | 0 | 0 | 0 | 0 | 0.0322583 | 0.03228837 | 0 |
| ENSMUSG00 | 4932443L11I | 2817 | 0.54319714 | 0.20103628 | 0.37778405 | 0.54990256 | 0.15377611 | 0.10554363 | 0.23513578 | 0.20336742 | 0.19006912 | 0.21759288 | 0.06783807 | 0.03962773 | 0.5954154 | 0.21433196 | 0.1462604 | 0.33981725 | 0.17686967 | 0.58582648 |
| ENSMUSG00 | Gm13054 | 817 | 0 | 0 | 0 | 0 | 0 | 0 | 0 | 0.04674713 | 0 | 0 | 0.05847608 | 0 | 0 | 0 | 0.09373466 | 0 | 0 | 0 |
| ENSMUSG00 | C330013F16I | 4018 | 0.43372629 | 0.42283664 | 0.27690174 | 0.18142775 | 0.2874978 | 0.3576479 | 0.11775181 | 0.25664382 | 0.13325652 | 0.17602303 | 0.41615817 | 0.23615387 | 0.47883145 | 0.16182609 | 0.03076273 | 0.40025025 | 0.31477546 | 0.23274137 |
| ENSMUSG00 | Gm11981 | 659 | 0 | 0 | 0 | 0 | 0 | 0 | 0 | 0 | 0 | 0 | 0 | 0 | 0 | 0 | 0 | 0 | 0 | 0 |
| ENSMUSG00 | 2610037D02I | 4983 | 0.0682403 | 0.02005593 | 0.03883085 | 0.05485978 | 0.02897768 | 0.00994436 | 0.03797927 | 0 | 0.00826541 | 0.10408544 | 0.08628831 | 0.01120122 | 0 | 0 | 0.04961054 | 0.03842124 | 0 | 0.01103936 |
| ENSMUSG00 | Gm13110 | 3412 | 0.03737266 | 0.0195269 | 0.01417747 | 0.10682543 | 0.22570656 | 0.01452308 | 0 | 0.02238711 | 0 | 0.01381908 | 0.11201632 | 0 | 0 | 0.06805992 | 0.02415097 | 0.15711269 | 0.07862957 | 0 |
| ENSMUSG00 | Gm8978 | 2217 | 0 | 0 | 0 | 0 | 0 | 0 | 0 | 0 | 0 | 0 | 0 | 0 | 0 | 0 | 0 | 0 | 0 | 0 |
| ENSMUSG00 | Gm13657 | 1935 | 0.79079397 | 0.44761512 | 0.69997873 | 0.8711931 | 0.19899503 | 0.25608648 | 0.34231396 | 0.33554049 | 1.06425105 | 0.24367289 | 0.39503842 | 0.20191713 | 0 | 0.52804647 | 0.36197749 | 0.07915371 | 0.39613744 | 0.28428479 |
| ENSMUSG00 | Gm15677 | 764 | 0 | 0.39242938 | 0.37989684 | 0 | 0 | 0.1945788 | 0.12385518 | 0.29994033 | 0.21563621 | 0.55544022 | 0 | 0.21917145 | 0.7102763 | 0.48632552 | 1.18643224 | 1.10260911 | 0.70231434 | 0.36000724 |
| ENSMUSG00 | 3110082J24R | 498 | 0 | 0 | 0 | 0 | 0 | 0 | 0 | 0 | 0 | 0 | 0 | 0 | 0 | 0 | 0 | 0 | 0 | 0 |
| ENSMUSG00 | Gm16726 | 889 | 0 | 0 | 0.10882684 | 0 | 0.05414165 | 0.11147972 | 0.05322011 | 0 | 0.04632904 | 0.05303791 | 0 | 0 | 0.16647441 | 0.10448614 | 0.13903787 | 0.21535775 | 0.17244678 | 0 |
| ENSMUSG00 | Gm11482 | 715 | 0 | 0 | 0 | 0 | 0 | 0 | 0 | 0 | 0 | 0 | 0 | 0 | 0 | 0 | 0 | 0 | 0 | 0 |
| ENSMUSG00 | 1810035I16R | 319 | 0 | 0 | 0 | 0 | 0 | 0 | 0 | 0 | 0 | 0 | 0 | 0 | 0 | 0 | 0 | 0 | 0 | 0 |
| ENSMUSG00 | Gm13613 | 1439 | 0 | 0 | 0 | 0 | 0 | 0 | 0 | 0 | 0 | 0 | 0 | 0 | 0 | 0 | 0 | 0 | 0 | 0 |
| ENSMUSG00 | Gm11465 | 433 | 0.39265752 | 0.92322109 | 1.78747456 | 1.15743993 | 2.00086519 | 1.02996444 | 0 | 1.4112666 | 0.66583282 | 0.87114463 | 1.21368255 | 0.7734272 | 0.34179157 | 1.17987289 | 1.23699821 | 0.17686193 | 1.77026776 | 0.7622509 |
| ENSMUSG00 | Gm13377 | 674 | 0 | 0 | 0 | 0 | 0 | 0 | 0 | 0 | 0 | 0 | 0 | 0 | 0 | 0 | 0 | 0 | 0 | 0 |
| ENSMUSG00 | 5330413P13I | 5850 | 0 | 0 | 0 | 0 | 0 | 0 | 0 | 0 | 0 | 0 | 0 | 0 | 0 | 0 | 0 | 0 | 0 | 0 |
| ENSMUSG00 | 1700028P15I | 3524 | 0 | 0 | 0.01372688 | 0 | 0 | 0 | 0 | 0 | 0 | 0 | 0 | 0 | 0 | 0 | 0 | 0 | 0 | 0 |
| ENSMUSG00 | 4930448D08 | 1245 | 0 | 0 | 0 | 0 | 0 | 0 | 0 | 0 | 0 | 0 | 0 | 0 | 0 | 0 | 0 | 0 | 0 | 0 |
| ENSMUSG00 | Gm15675 | 3678 | 1.64103725 | 0.2988922 | 0.53923729 | 0.49549804 | 0.19629659 | 0.4850186 | 0.09004588 | 0.23883232 | 0.12317881 | 0.39740941 | 0.07793631 | 0.47044196 | 0.64380969 | 0.37882616 | 0.78415142 | 0.49971429 | 0.38555546 | 0.77772527 |
| ENSMUSG00 | Gm11739 | 720 | 0.3542098 | 1.75818053 | 1.07496734 | 0.56951305 | 0.93589852 | 0.55058593 | 1.24852903 | 1.27308008 | 0.45762795 | 0.78584505 | 1.19437396 | 1.0077828 | 1.30181448 | 0.38703406 | 0.45779507 | 0.1063628 | 0.37261678 | 1.22242459 |
| ENSMUSG00 | 4930461C15I | 1300 | 0 | 0 | 0 | 0 | 0 | 0 | 0 | 0.02937877 | 0 | 0.29015817 | 0 | 0 | 0.64510969 | 0.25008355 | 0 | 0.17672588 | 0 | 0.0423147 |
| ENSMUSG00 | Gm15579 | 755 | 0 | 0 | 0 | 0 | 0 | 0 | 0 | 0.20234385 | 0 | 0 | 0 | 0.44356818 | 0 | 0 | 0.16371479 | 0 | 0 | 0 |
| ENSMUSG00 | Gm13467 | 279 | 0 | 0 | 0 | 0 | 0 | 0 | 0 | 0 | 0 | 0 | 0 | 0 | 0 | 0 | 0 | 0 | 0 | 0 |
| ENSMUSG00 | Gm14280 | 1441 | 0 | 0 | 0 | 0 | 0 | 0 | 0 | 0 | 0.14290949 | 0 | 0 | 0 | 0 | 0.06446091 | 0 | 0 | 0 | 0 |
| ENSMUSG00 | 2810425M01 | 1847 | 0 | 0.16232596 | 0.1047613 | 0.17267315 | 0.20847612 | 0.24145891 | 0.10246384 | 0.02067807 | 0.22299142 | 0.12764132 | 0.07759874 | 0.21153743 | 0.21367371 | 0.07543707 | 0.53537485 | 0.24877493 | 0.04150113 | 0.20848064 |
| ENSMUSG00 | Gm12440 | 1065 | 0 | 0 | 0 | 0 | 0 | 0 | 0.22212525 | 0 | 0 | 0 | 0 | 0 | 0 | 0 | 0 | 0 | 0 | 0 |
| ENSMUSG00 | Gm14547 | 367 | 0 | 0.09077083 | 0 | 0 | 0.13114966 | 0 | 0 | 0 | 0 | 0 | 0 | 0 | 0 | 0 | 0 | 0 | 0 | 0 |
| ENSMUSG00 | Gm11775 | 770 | 0.44161222 | 0.77874298 | 0.12564553 | 1.00589319 | 0.87512588 | 1.73756339 | 0 | 0.04960052 | 0.64186778 | 0 | 0 | 0.07248787 | 0 | 0.060317 | 0 | 0.09945612 | 0 | 0.1428808 |
| ENSMUSG00 | Gm13748 | 374 | 0 | 0.08907191 | 0 | 0 | 0 | 0 | 0 | 0 | 0 | 0.1260714 | 0 | 0 | 0 | 0 | 0 | 0 | 0 | 0 |
| ENSMUSG00 | Gm16618 | 952 | 0.80366929 | 0.62986565 | 0.45731279 | 0.76573184 | 0.40446995 | 0.31230714 | 0.44728373 | 0.84247946 | 0.25957888 | 0.5943366 | 0.95349182 | 0.58629898 | 0.36273469 | 0 | 0.60590523 | 1.93061886 | 0.24155229 | 0 |
| ENSMUSG00 | Gm16212 | 664 | 0 | 0 | 0 | 0 | 0 | 0 | 0 | 0 | 0 | 0 | 0 | 0 | 0 | 0 | 0 | 0 | 0 | 0 |
| ENSMUSG00 | Gm11973 | 1213 | 0.14016546 | 0.02746323 | 0.03987925 | 0.11268189 | 0 | 0 | 0.23402809 | 0 | 0.27163407 | 0.11661345 | 0 | 0.32210193 | 0 | 0.07657723 | 0 | 0 | 0.03159629 | 0 |
| ENSMUSG00 | Gm14250 | 563 | 0 | 0 | 0 | 0 | 0 | 0 | 0 | 0 | 0 | 0 | 0 | 0 | 0 | 0 | 0 | 0 | 0 | 0 |
| ENSMUSG00 | 2410006H16I | 457 | 20.7410378 | 15.2349998 | 18.1003801 | 16.6492219 | 17.9046543 | 17.3488783 | 26.0892672 | 19.4722752 | 20.9988144 | 26.722171 | 31.7802787 | 20.3965332 | 35.8385045 | 30.8949728 | 42.1902782 | 31.922804 | 29.7720687 | 49.351715 |
| ENSMUSG00 | Gm15681 | 2703 | 0 | 0 | 0 | 0 | 0.01780685 | 0 | 0 | 0 | 0 | 0 | 0 | 0 | 0 | 0 | 0 | 0 | 0 | 0 |
| ENSMUSG00 | E030013I19R | 3315 | 0 | 0 | 0 | 0 | 0 | 0 | 0 | 0 | 0 | 0 | 0 | 0 | 0 | 0 | 0 | 0 | 0 | 0 |
| ENSMUSG00 | B230206H07 | 4996 | 0.02552352 | 0.03333957 | 0.03872981 | 0.03647802 | 0.1156091 | 0.08926633 | 0.03788045 | 0.04586758 | 0.08243896 | 0.05662614 | 0.07650113 | 0.10054863 | 0.09874283 | 0 | 0.03298763 | 0.02299276 | 0.01534279 | 0.01101063 |
| ENSMUSG00 | Gm13010 | 824 | 0 | 0 | 0 | 0 | 0.05841253 | 0 | 0.1722549 | 0 | 0 | 0 | 0 | 0 | 0 | 0.05636418 | 0 | 0.51116102 | 0 | 0 |
| ENSMUSG00 | Gm13642 | 766 | 0 | 0.04348942 | 0 | 0 | 0 | 0 | 0 | 0 | 0.0537683 | 0 | 0.0623694 | 0 | 0.06440198 | 0.06063197 | 0 | 0.24993869 | 0 | 0 |
| ENSMUSG00 | Gm16065 | 1342 | 19.2255155 | 14.2485852 | 19.5008047 | 18.7404594 | 17.1438595 | 16.4683451 | 25.1370641 | 18.0432065 | 32.4090616 | 24.24291 | 27.0202634 | 22.7089059 | 30.7681183 | 29.2092472 | 25.4823334 | 37.86262 | 25.7031797 | 26.8487071 |
| ENSMUSG00 | Lce6a | 1228 | 0 | 0 | 0 | 0 | 0 | 0 | 0 | 0 | 0 | 0 | 0 | 0 | 0 | 0.68077653 | 0 | 0.18708766 | 0.46815509 | 1.25427931 |
| ENSMUSG00 | Gm13840 | 254 | 0.16734321 | 0.91807189 | 0.38089394 | 0.71749676 | 0.37899152 | 0.58526851 | 1.86270389 | 1.20291031 | 0 | 0.55689807 | 0 | 0.2197467 | 1.55376117 | 0.54855221 | 1.2976868 | 0.15075042 | 0.60356373 | 0 |
| ENSMUSG00 | Gm13257 | 663 | 0 | 0.10049139 | 0 | 0 | 0 | 0.14948034 | 0.07136151 | 0 | 0 | 0.0711172 | 0 | 0 | 0 | 0 | 0 | 0 | 0 | 0 |
| ENSMUSG00 | A430108G06I | 2444 | 0 | 0 | 0 | 0.09320999 | 0 | 0 | 0 | 0 | 0.06740837 | 0.19292432 | 0 | 0.02283783 | 0 | 0.07601324 | 0 | 0.14100469 | 0.29795403 | 0 |
| ENSMUSG00 | Gm11374 | 671 | 0 | 0 | 0 | 0 | 0 | 0 | 0 | 0 | 0 | 0 | 0 | 0 | 0 | 0 | 0 | 0 | 0 | 0 |
| ENSMUSG00 | Gm14720 | 409 | 0 | 0 | 0 | 0 | 0 | 0 | 0.11567892 | 0 | 0 | 0 | 0 | 0 | 0.24123187 | 0 | 0.3022119 | 0 | 0.09370733 | 0 |
| ENSMUSG00 | Gm13781 | 490 | 0 | 0 | 0 | 0 | 0 | 0 | 0 | 0 | 0 | 0 | 0 | 0 | 0 | 0 | 0 | 0 | 0 | 0 |
| ENSMUSG00 | Gm13844 | 578 | 0.07353837 | 0 | 0.08369123 | 0.23647601 | 0 | 0.42865686 | 0.16371169 | 0.79292185 | 0 | 0 | 0 | 0.09656689 | 0.08534934 | 0 | 0 | 0.06624673 | 0 | 0.19034293 |
| ENSMUSG00 | Gm16080 | 687 | 0 | 0 | 0.07041271 | 0 | 0.14012205 | 0 | 0 | 0 | 0 | 0 | 0 | 0 | 0 | 0 | 0 | 0 | 0 | 0 |
| ENSMUSG00 | Gm16913 | 914 | 0 | 0 | 0 | 0 | 0 | 0 | 0 | 0 | 0 | 0 | 0 | 0 | 0 | 0 | 0 | 0 | 0 | 0 |
| ENSMUSG00 | Trav20 | 319 | 0 | 0 | 0 | 0 | 0 | 0 | 0 | 0 | 0 | 0 | 0 | 0 | 0 | 0 | 0 | 0 | 0 | 0 |
| ENSMUSG00 | 2810008D09I | 954 | 119.985785 | 89.5676873 | 99.6880091 | 88.1133405 | 95.1538868 | 91.9894037 | 114.165395 | 96.0014476 | 139.403836 | 95.3393151 | 106.867674 | 112.567437 | 129.379934 | 119.810167 | 130.471594 | 118.484144 | 101.962413 | 114.977106 |
| ENSMUSG00 | Gm1720 | 1494 | 0.14225293 | 0 | 0 | 0 | 0 | 0 | 0 | 0 | 0 | 0 | 0 | 0 | 0 | 0 | 0 | 0 | 0 | 0 |
| ENSMUSG00 | Gm15325 | 695 | 0.85621937 | 0.14379667 | 0 | 0 | 0 | 0.28519559 | 1.97419811 | 0.98915575 | 0.82965643 | 0.94979834 | 0.61866853 | 1.44558551 | 0.21294353 | 0.13365205 | 0.17784844 | 0 | 0 | 0 |
| ENSMUSG00 | Gm13546 | 1344 | 0 | 0.02478638 | 0 | 0 | 0.03581244 | 0 | 0 | 0 | 0 | 0.03508237 | 0 | 0 | 0 | 0 | 0 | 0.05696007 | 0 | 0 |
| ENSMUSG00 | Gm15543 | 359 | 0 | 0 | 0 | 0 | 0 | 0 | 0 | 0 | 0 | 0 | 0 | 0 | 0 | 0 | 0 | 0 | 0 | 0 |
| ENSMUSG00 | A630077J23R | 925 | 0 | 0 | 0 | 0 | 0 | 0 | 0 | 0 | 0 | 0 | 0 | 0 | 0 | 0 | 0 | 0 | 0 | 0 |
| ENSMUSG00 | Gm16978 | 1929 | 0.06610447 | 0.10361709 | 0.10030799 | 0.07085699 | 0.02495175 | 0.15412981 | 0.22074345 | 0.0593972 | 0.21351226 | 0.14665849 | 0.04953339 | 0.17361015 | 0.05114766 | 0.09630708 | 0.06407707 | 0.09924989 | 0.0993424 | 0.05703381 |
| ENSMUSG00 | 4930512H18I | 1490 | 0.02852696 | 0.02235765 | 0.09739637 | 0.03057748 | 0.16151652 | 0.13302747 | 0.06350695 | 0 | 0.24877761 | 0.0949343 | 0.38476477 | 0 | 0.09932601 | 0.46755793 | 0.41478076 | 0.07709518 | 0.8745598 | 0.11075659 |
| ENSMUSG00 | Gm4577 | 714 | 1.84546282 | 1.39970144 | 1.55825098 | 1.4038417 | 2.561643 | 2.2208508 | 3.24694855 | 3.3164271 | 3.74947272 | 3.56601957 | 2.74337997 | 3.12692788 | 6.70195515 | 4.81353287 | 3.86625244 | 3.2713264 | 4.24058467 | 3.38991694 |
| ENSMUSG00 | Gm15883 | 1686 | 0 | 0 | 0 | 0 | 0 | 0 | 0 | 0 | 0 | 0 | 0 | 0 | 0 | 0 | 0.04887492 | 0 | 0 | 0.13050796 |
| ENSMUSG00 | Gm7809 | 2738 | 0.915926 | 0.79084665 | 0.56535901 | 0.94849508 | 0.72074831 | 0.66963153 | 0.38016031 | 0.59980763 | 0.37606388 | 0.39607968 | 0.54091443 | 0.69310903 | 0.68466503 | 0.32229279 | 0.3762012 | 0.61533482 | 0.44793335 | 0.86391219 |
| ENSMUSG00 | Gm15288 | 647 | 0.06569579 | 0 | 0 | 0 | 0 | 0 | 0 | 0 | 0.2546307 | 0 | 0 | 0 | 0 | 0 | 0 | 0 | 0 | 0 |
| ENSMUSG00 | Gm16834 | 2246 | 0 | 0 | 0 | 0 | 0 | 0 | 0 | 0 | 0 | 0 | 0 | 0 | 0 | 0 | 0 | 0 | 0 | 0 |
| ENSMUSG00 | Gm14225 | 537 | 0.07915303 | 0.43424629 | 0.7206485 | 0.08484366 | 0.44815571 | 0 | 0.52863328 | 0.42673075 | 0.46018453 | 0 | 1.06759684 | 0.20787956 | 0.64306037 | 0.77839253 | 0.99743059 | 0.49913268 | 0.92782469 | 1.3316916 |
| ENSMUSG00 | Gm15672 | 1352 | 0.94316219 | 0.44351486 | 0.67980553 | 0.67397995 | 0.53400803 | 0.65972574 | 0.66489712 | 0.64972282 | 0.70065818 | 0.66262083 | 0.56538412 | 0.45412152 | 0.54732157 | 1.47714183 | 0.85328666 | 0.48146474 | 0.42521779 | 0.65099535 |
| ENSMUSG00 | Gm13175 | 464 | 0 | 0 | 0 | 0 | 0 | 0 | 0 | 0 | 0 | 0 | 0 | 0 | 0 | 0 | 0 | 0 | 0 | 0 |
| ENSMUSG00 | Gm8975 | 1996 | 0 | 0 | 0 | 0 | 0 | 0 | 0 | 0 | 0 | 0 | 0 | 0 | 0 | 0 | 0 | 0 | 0 | 0 |
| ENSMUSG00 | Gm12828 | 286 | 0 | 0 | 0 | 0 | 0 | 0 | 0 | 0 | 0 | 0 | 0 | 0 | 0 | 0 | 0 | 0 | 0 | 0 |
| ENSMUSG00 | A230072C01I | 1575 | 5.96421835 | 5.11855264 | 6.94121769 | 4.68627885 | 6.63150948 | 7.55089274 | 7.32971026 | 6.13503353 | 7.58354893 | 6.13707566 | 5.18699549 | 8.0091046 | 4.00919709 | 3.06678418 | 5.28426304 | 5.37284078 | 3.45545026 | 3.28308319 |
| ENSMUSG00 | Miat | 8753 | 0 | 0 | 0 | 0 | 0 | 0 | 0 | 0 | 0 | 0 | 0 | 0 | 0 | 0 | 0 | 0 | 0 | 0 |
| ENSMUSG00 | Gm14012 | 673 | 0 | 0 | 0 | 0 | 0 | 0 | 0 | 0 | 0 | 0 | 0 | 0 | 0 | 0 | 0 | 0 | 0 | 0 |
| ENSMUSG00 | C430042M11 | 1615 | 0 | 0.12376307 | 0 | 0 | 0 | 0.09204842 | 0.14647888 | 0 | 0.02550249 | 0.02919548 | 0.02958202 | 0 | 0.03054608 | 0.02875795 | 0 | 0 | 0 | 0.03406137 |
| ENSMUSG00 | Gm13594 | 474 | 5.20105529 | 6.46579381 | 5.00063921 | 5.57498011 | 6.09264857 | 4.28620691 | 5.78931514 | 5.6402282 | 6.77752791 | 6.26686562 | 7.35774677 | 3.65039143 | 5.41194029 | 2.9394992 | 3.91154012 | 4.20065735 | 6.46857332 | 5.68659542 |
| ENSMUSG00 | Gm13352 | 261 | 0.32571016 | 0.25527122 | 0.1853392 | 0.17456339 | 0.1844135 | 0.18985722 | 0.5438239 | 0.29266209 | 0.15780274 | 0 | 0 | 0 | 0.37802235 | 0.17794669 | 0 | 0.14670731 | 0.2936881 | 0.21076286 |
| ENSMUSG00 | Gm15992 | 229 | 0 | 0.14547115 | 0 | 0.7958261 | 0 | 0 | 0 | 0 | 0 | 0 | 0.20862427 | 0 | 0 | 0 | 0.17991946 | 0 | 0 | 0.24021444 |
| ENSMUSG00 | Gm16225 | 274 | 0 | 0 | 0 | 0 | 0 | 0 | 0 | 0 | 0 | 0 | 0 | 0 | 0 | 0 | 0 | 0 | 0 | 0 |
| ENSMUSG00 | Gm13846 | 459 | 0 | 0 | 0 | 0 | 0 | 0 | 0 | 0 | 0 | 0 | 0 | 0 | 0 | 0 | 0 | 0 | 0 | 0 |
| ENSMUSG00 | Gm15135 | 607 | 0 | 0 | 0 | 0 | 0 | 0 | 0 | 0 | 0 | 0 | 0 | 0 | 0 | 0 | 0 | 0 | 0 | 0 |
| ENSMUSG00 | 1700060C16I | 640 | 0 | 0 | 0 | 0 | 0 | 0 | 0 | 0 | 0 | 0 | 0 | 0 | 0 | 0 | 0 | 0 | 0 | 0 |
| ENSMUSG00 | A330033J07R | 4452 | 0.0572846 | 0 | 0.02173115 | 0.01023384 | 0.0108113 | 0 | 0.09564558 | 0.03431483 | 0.11101487 | 0.05295452 | 0.02146225 | 0.03761163 | 2.6483217 | 2.27421632 | 2.36918201 | 2.19319517 | 1.84227932 | 1.81633843 |
| ENSMUSG00 | Gm8619 | 495 | 5.83909489 | 5.31660332 | 5.2771124 | 4.6941682 | 4.66733805 | 5.80617888 | 5.16138316 | 9.7988588 | 6.65640659 | 5.23896703 | 6.46650953 | 3.83380309 | 7.1755516 | 6.9431565 | 9.32237224 | 6.11102623 | 8.05239369 | 6.44551149 |
| ENSMUSG00 | 1700021J08R | 573 | 0 | 0.05813769 | 0.08442152 | 0.15902633 | 0.08399987 | 0 | 0.16514024 | 0.06665341 | 0.86254483 | 0.16457488 | 0.16675378 | 0.48704767 | 0.68875277 | 0 | 0.21571495 | 0.13364959 | 0.40132248 | 0.67201352 |
| ENSMUSG00 | Gm15963 | 874 | 0 | 0 | 0.05534729 | 0.15638802 | 0 | 0 | 0 | 0 | 0 | 0 | 0 | 0 | 0 | 0 | 0 | 0 | 0 | 0 |
| ENSMUSG00 | 4933430M04 | 740 | 0 | 0 | 0 | 0 | 0 | 0 | 0 | 0 | 0 | 0 | 0 | 0 | 0 | 0 | 0 | 0 | 0 | 0 |
| ENSMUSG00 | Gm11365 | 772 | 0 | 0 | 0.31330007 | 0 | 0 | 0.19256244 | 0.18385756 | 0 | 0 | 0 | 0 | 0.21690024 | 0 | 0 | 0.04959923 | 0 | 0 | 0.49878724 |
| ENSMUSG00 | Gm15708 | 1762 | 0.55483487 | 1.5692226 | 1.01578923 | 1.75831499 | 0.84681591 | 0.89993614 | 0.34907198 | 1.10545546 | 0.79474548 | 0.8027929 | 0.62362318 | 1.48884003 | 0.8119328 | 0.89619692 | 1.6368382 | 0.82579063 | 0.50028651 | 1.46732578 |
| ENSMUSG00 | 9330136K24I | 2475 | 0.44651902 | 0.37687315 | 0.58634582 | 0.58907209 | 0.66120622 | 0.74078834 | 0.82199806 | 1.11105171 | 0.53251253 | 0.51437131 | 1.06166574 | 0.76676062 | 0.73748725 | 1.10715198 | 0.89894304 | 0.47959953 | 0.51101729 | 0.40006623 |
| ENSMUSG00 | Gm12835 | 1999 | 0 | 0 | 0 | 0 | 0 | 0 | 0 | 0 | 0 | 0 | 0 | 0 | 0 | 0 | 0 | 0 | 0 | 0 |
| ENSMUSG00 | 4632428C04 | 2427 | 0.17513464 | 0.05490382 | 0.09965705 | 0.31913381 | 0.01983186 | 0.22459006 | 0.01949431 | 0.09441879 | 0.0509104 | 0 | 0.01968478 | 0.0689934 | 0 | 0 | 0 | 0.33131552 | 0.0473749 | 0 |
| ENSMUSG00 | 1700062I23R | 792 | 0 | 0 | 0 | 0 | 0 | 0 | 0 | 0 | 0 | 0 | 0 | 0 | 0 | 0 | 0 | 0.09678358 | 0 | 0 |
| ENSMUSG00 | 1600002D24 | 2206 | 0.30800852 | 0.46770821 | 0.30671623 | 0.18571078 | 0.52317308 | 0.20198125 | 0.29998981 | 0.1037837 | 0.24249307 | 0.32031728 | 0.28128372 | 0.30334599 | 0.15639648 | 0.44172366 | 0.80238537 | 0.43354401 | 0.50337981 | 0.74740634 |
| ENSMUSG00 | Gm16040 | 564 | 0.52754651 | 0.41345791 | 0.08576867 | 0.24234598 | 0 | 0.52715674 | 0 | 0 | 0.94933458 | 0.16720108 | 1.10119585 | 0.69274759 | 0.08746794 | 0 | 0 | 0 | 0.06795443 | 0 |
| ENSMUSG00 | Gm11728 | 997 | 3.15484757 | 1.97137488 | 2.42595438 | 2.60479391 | 1.15864209 | 0.8946331 | 0.4270954 | 0.49799522 | 1.28062386 | 0.23646291 | 0.28751229 | 0 | 0.64324466 | 1.02484446 | 0.37192979 | 0.3072466 | 0.07688324 | 1.76558818 |
| ENSMUSG00 | Gm16258 | 2694 | 0 | 0 | 0 | 0 | 0 | 0 | 0 | 0 | 0 | 0 | 0 | 0 | 0 | 0 | 0 | 0 | 0 | 0 |
| ENSMUSG00 | Gm13404 | 1958 | 1.32421641 | 0.66353569 | 1.43292378 | 1.27980462 | 1.13078064 | 1.26539156 | 0.3382929 | 0.85825623 | 1.19899459 | 0.21672948 | 0.26839864 | 1.53934922 | 0.98260714 | 0.83020585 | 2.29365148 | 1.66225824 | 3.38676357 | 2.64088664 |
| ENSMUSG00 | Gm13716 | 806 | 0 | 0 | 0 | 0 | 0 | 0 | 0 | 0 | 0 | 0 | 0 | 0 | 0 | 0 | 0 | 0 | 0 | 0 |
| ENSMUSG00 | Gm15276 | 497 | 0 | 0 | 0 | 0 | 0 | 0 | 0 | 0 | 0 | 0 | 0 | 0 | 0 | 0 | 0 | 0 | 0 | 0 |
| ENSMUSG00 | Gm15298 | 1210 | 0 | 0 | 0 | 0 | 0 | 0 | 0 | 0 | 0.03403844 | 0 | 0 | 0 | 0 | 0 | 0 | 0 | 0 | 0 |
| ENSMUSG00 | Gm13598 | 622 | 0 | 0 | 0 | 0 | 0 | 0 | 0 | 0 | 0 | 0 | 0 | 0 | 0 | 0 | 0 | 0 | 0 | 0 |
| ENSMUSG00 | Gm4926 | 1605 | 0 | 0 | 0 | 0.02838694 | 0 | 0.2778658 | 0.0294783 | 0 | 0.23095242 | 0 | 0 | 0.10432834 | 0.36883676 | 0.86811378 | 0 | 0 | 0 | 0.99393402 |
| ENSMUSG00 | Gm15755 | 656 | 0 | 0 | 0 | 0 | 0 | 0.22661311 | 0 | 0 | 0.12556865 | 0 | 0 | 0 | 0 | 0 | 0 | 0 | 0.11684847 | 0 |
| ENSMUSG00 | Gm12027 | 947 | 0 | 0 | 0 | 0 | 0 | 0 | 0 | 0 | 0 | 0 | 0 | 0 | 0 | 0 | 0 | 0 | 0 | 0 |
| ENSMUSG00 | 4930505O20I | 3414 | 0 | 0 | 0 | 0 | 0 | 0 | 0 | 0 | 0 | 0 | 0 | 0 | 0 | 0 | 0 | 0 | 0 | 0 |
| ENSMUSG00 | Gm14019 | 401 | 0.42399178 | 0.08307455 | 0.48252898 | 0.11361856 | 0.60014867 | 0.1235729 | 0.11798673 | 0.76194319 | 1.43793322 | 0.1175828 | 0.23827909 | 1.67029414 | 0.36906671 | 0 | 0 | 0.0954878 | 0 | 0 |
| ENSMUSG00 | Gm16124 | 514 | 33.6568223 | 23.5264214 | 38.1153303 | 25.5283673 | 33.8983587 | 27.5721436 | 42.6182302 | 41.3131824 | 47.3564802 | 32.7486402 | 43.3134838 | 45.3969474 | 43.1894217 | 27.9206673 | 38.3158438 | 34.4168495 | 33.4050214 | 32.3205257 |
| ENSMUSG00 | Gm16365 | 1401 | 0.75847923 | 0.71333821 | 1.55375365 | 1.07317235 | 1.06501758 | 1.13182546 | 1.58722049 | 1.17221506 | 0.82314236 | 1.51447655 | 1.63682941 | 1.63343481 | 1.37326536 | 1.35917743 | 1.3822078 | 1.47586924 | 0.71126604 | 0.82454764 |
| ENSMUSG00 | Gm15903 | 308 | 0.96602673 | 1.29790497 | 2.04173992 | 1.18340375 | 1.25017984 | 1.76974048 | 2.6114141 | 2.60402744 | 0.66861227 | 1.07160689 | 0.93068101 | 3.44317399 | 1.2813485 | 1.20633993 | 2.00656928 | 0.99456123 | 2.11541249 | 1.60740896 |
| ENSMUSG00 | Gm11630 | 380 | 0 | 0.35066204 | 0 | 0.11989749 | 0.63331479 | 0 | 1.86760575 | 0 | 0 | 0 | 0.12572357 | 0 | 0 | 0.85554898 | 0.10842515 | 0 | 0.60515206 | 1.01332565 |
| ENSMUSG00 | 4930429F24I | 1517 | 0.0840577 | 0.04391944 | 0.41453915 | 0.0600673 | 0.09518508 | 0.06532991 | 0.03118832 | 0.02517627 | 0.27149977 | 0.09324463 | 0.34642356 | 0.33114104 | 0 | 0.18369448 | 0.29875881 | 0.12620503 | 0 | 0.32635594 |
| ENSMUSG00 | Gm11497 | 2152 | 0 | 0 | 0 | 0 | 0 | 0 | 0 | 0 | 0 | 0 | 0 | 0 | 0 | 0 | 0 | 0 | 0 | 0 |
| ENSMUSG00 | Gm12207 | 492 | 1.55506742 | 0.47396394 | 0.39328073 | 2.68550871 | 0.58697468 | 0.90645244 | 0.96163982 | 1.31965618 | 1.42311132 | 2.01253002 | 0.77682859 | 2.83616172 | 1.50402187 | 2.737558 | 2.00983199 | 1.08957013 | 2.88226217 | 4.91951361 |
| ENSMUSG00 | Gm13189 | 250 | 0 | 0 | 0 | 0 | 0 | 0 | 0 | 0 | 0 | 0 | 0 | 0 | 0 | 0 | 0 | 0 | 0 | 0 |
| ENSMUSG00 | Gm13835 | 1346 | 0.66315654 | 1.65821985 | 1.29379427 | 0.71083353 | 1.35885074 | 0.55222214 | 1.19511968 | 0.79449277 | 1.34636456 | 1.47127009 | 1.34877297 | 0.95375946 | 0.98957041 | 0.65560004 | 0.88770068 | 1.05256499 | 1.05354605 | 1.02171446 |
| ENSMUSG00 | 4930406D18I | 1036 | 0 | 0 | 0 | 0 | 0 | 0 | 0 | 0 | 0 | 0 | 0 | 0 | 0 | 0 | 0 | 0 | 0 | 0 |
| ENSMUSG00 | Gm11766 | 3016 | 0.39461039 | 0.44181557 | 0.41701319 | 0.60425788 | 0.41493038 | 0.85435748 | 0.10981059 | 0.56984685 | 0.58720828 | 0.64097441 | 0.49105561 | 0.88831293 | 0.78512335 | 0.30798466 | 0.10928795 | 0.1523499 | 0.29227614 | 0.16415184 |
| ENSMUSG00 | Gm6286 | 761 | 0 | 0 | 0 | 0 | 0 | 0 | 0 | 0 | 0 | 0 | 0 | 0 | 0 | 0 | 0 | 0 | 0.05036307 | 0 |
| ENSMUSG00 | Gm6226 | 2278 | 0.07463595 | 0.07311873 | 0.08494035 | 0.2400055 | 0.14790319 | 0 | 0 | 0 | 0.43392291 | 0 | 0.0838893 | 0.07350614 | 0.0433116 | 0.10194049 | 0.28938757 | 0.05042661 | 0.1345963 | 0 |
| ENSMUSG00 | Gm11823 | 2275 | 0 | 0 | 0 | 0 | 0 | 0 | 0 | 0 | 0 | 0 | 0 | 0 | 0 | 0 | 0 | 0 | 0 | 0 |
| ENSMUSG00 | Gm14210 | 501 | 0 | 0 | 0 | 0 | 0 | 0 | 0 | 0 | 0 | 0 | 0 | 0 | 0 | 0 | 0 | 0 | 0.07649959 | 0.10979862 |
| ENSMUSG00 | Gm11788 | 559 | 0 | 0.29796864 | 0 | 0 | 0.17220724 | 0.4432266 | 0 | 0.27329089 | 0.44207351 | 0 | 0.34186017 | 0.29954738 | 0.1765006 | 0.41542118 | 0.07370582 | 0.68498403 | 0.89130923 | 0.39362509 |
| ENSMUSG00 | Gm14873 | 1377 | 0.06173591 | 0.09676948 | 0.31616687 | 0.23161025 | 0.13981677 | 0.25190206 | 0.48102942 | 0.2218876 | 0.17946194 | 0.13696646 | 0.06938992 | 0 | 0.39408213 | 0.23609921 | 0.05984249 | 0.13903634 | 0.08349956 | 0 |
| ENSMUSG00 | Gm14532 | 277 | 0 | 0 | 0.34926737 | 0.1644803 | 0.34752291 | 0 | 0 | 0 | 0 | 0 | 0 | 0 | 0 | 0.50300456 | 0 | 0 | 0.27672417 | 0 |
| ENSMUSG00 | Gm11665 | 725 | 0.35176697 | 0.50543702 | 0.66722111 | 0.18852846 | 1.19499948 | 0.82018318 | 0 | 0 | 0.11361797 | 0.19510636 | 0.13179299 | 0.23096136 | 0 | 0.19218243 | 0.05682973 | 0.15844389 | 0.31718315 | 0 |
| ENSMUSG00 | 4933426K07I | 1546 | 0 | 0 | 0 | 0 | 0 | 0 | 0 | 0 | 0 | 0 | 0 | 0 | 0 | 0 | 0 | 0 | 0 | 0 |
| ENSMUSG00 | Gm16801 | 791 | 0 | 0 | 0 | 0 | 0 | 0 | 0 | 0 | 0 | 0 | 0 | 0 | 0 | 0 | 0 | 0 | 0 | 0 |
| ENSMUSG00 | Gm15956 | 500 | 0.08501035 | 0.26650315 | 0.77397649 | 0 | 0.38505539 | 0.19821093 | 0.56775215 | 0.38192402 | 1.23559547 | 0.47150703 | 0.28664975 | 1.00468193 | 0.19732767 | 0.83599357 | 0.32961245 | 0.07658121 | 0.22995778 | 0.33005464 |
| ENSMUSG00 | Gm9697 | 996 | 0 | 0 | 0 | 0 | 0 | 0 | 0 | 0 | 0 | 0 | 0 | 0 | 0 | 0 | 0 | 0 | 0 | 0 |
| ENSMUSG00 | Gm15063 | 893 | 0.14279454 | 0.03730447 | 0.05416969 | 0 | 0.10779826 | 0 | 0.15894517 | 0 | 0 | 0.15840102 | 0.26749697 | 0.12500708 | 0.11048582 | 0.57209962 | 0.13841508 | 0.08575724 | 0.04291859 | 0.36960206 |
| ENSMUSG00 | 4930481A15I | 2465 | 1.62088704 | 0.75680409 | 1.64842862 | 1.75590232 | 0.85914996 | 1.22625426 | 2.59116091 | 1.42543652 | 2.32248507 | 2.1614724 | 0.73649023 | 0.9283741 | 3.16204581 | 1.54499601 | 2.02246988 | 2.15918638 | 1.39933741 | 0.40168922 |
| ENSMUSG00 | Gm13530 | 388 | 0 | 0 | 0 | 0 | 0 | 0 | 0 | 0 | 0 | 0 | 0 | 0 | 0 | 0 | 0 | 0 | 0 | 0 |
| ENSMUSG00 | Gm12746 | 613 | 0 | 0 | 0 | 0 | 0 | 0 | 0 | 0 | 0 | 0 | 0 | 0 | 0 | 0 | 0 | 0 | 0 | 0 |
| ENSMUSG00 | Gm2020 | 576 | 0 | 0 | 0 | 0 | 0 | 0 | 0 | 0 | 0 | 0 | 0 | 0 | 0 | 0 | 0 | 0 | 0 | 0 |
| ENSMUSG00 | Gm15489 | 969 | 0.61410987 | 1.30638801 | 0.19968434 | 0.14105586 | 0.54638923 | 0.8693462 | 0.53708923 | 0.31531395 | 0.80757874 | 0.68122791 | 0.93676389 | 1.20962736 | 0.05691013 | 1.53375727 | 0.51023599 | 0.31612473 | 0 | 0.62445838 |
| ENSMUSG00 | 4732414G09I | 2411 | 0 | 0.05526818 | 0.04012736 | 0.01889716 | 0 | 0.06165832 | 0 | 0 | 0 | 0 | 0 | 0 | 0.4092237 | 0.75127309 | 0 | 0.20646118 | 0 | 0.09126355 |
| ENSMUSG00 | Gm15859 | 708 | 0.72042671 | 0.28231266 | 0.54659356 | 0.12870352 | 0.54386354 | 0.48992816 | 0.33412909 | 0.91704921 | 0.63990349 | 0.93235854 | 0.47235128 | 0.15767136 | 0.2787114 | 0.91838591 | 0.40736002 | 0.64899335 | 0.32479913 | 0.69926831 |
| ENSMUSG00 | 4930562A09I | 732 | 0 | 0 | 0 | 0 | 0 | 0 | 0.06463481 | 0 | 0.05626573 | 0 | 0 | 0 | 0 | 0 | 0 | 0 | 0.05235833 | 0 |
| ENSMUSG00 | Gm15527 | 3563 | 0 | 0 | 0 | 0 | 0 | 0 | 0 | 0 | 0 | 0 | 0 | 0 | 0 | 0 | 0 | 0 | 0 | 0 |
| ENSMUSG00 | 4930522O17I | 444 | 0 | 0 | 0 | 0 | 0 | 0 | 0 | 0 | 0 | 0 | 0 | 0 | 0 | 0 | 0 | 0 | 0 | 0 |
| ENSMUSG00 | Gm14697 | 644 | 0 | 0 | 0 | 0 | 0 | 0 | 0 | 0 | 0 | 0 | 0 | 0 | 0 | 0 | 0 | 0 | 0 | 0 |
| ENSMUSG00 | Gm13066 | 882 | 0 | 0 | 0 | 0 | 0 | 0 | 0 | 0 | 0 | 0 | 0 | 0 | 0 | 0 | 0.09342756 | 0.26048032 | 0 | 0 |
| ENSMUSG00 | Gm16907 | 2314 | 1.39602134 | 1.22368021 | 1.19156924 | 1.31918322 | 1.53922314 | 0.92081571 | 1.18588391 | 1.27087942 | 0.76535012 | 0.65204084 | 1.28005507 | 1.44725141 | 1.40704691 | 0.98347462 | 1.44223251 | 2.01877878 | 0.97720463 | 0.73694136 |
| ENSMUSG00 | 1700051A21I | 2479 | 0 | 0 | 0 | 0 | 0 | 0 | 0 | 0 | 0 | 0 | 0 | 0 | 0 | 0 | 0 | 0 | 0 | 0 |
| ENSMUSG00 | Gm12596 | 670 | 0.06344056 | 0.14916221 | 0.0721993 | 0.27200623 | 0.28735477 | 0 | 0.07061594 | 0.7980502 | 0.79914135 | 0.56299347 | 0.42783545 | 0.33322784 | 0.29451891 | 0.48523673 | 0.86092803 | 0.17145048 | 0.400424 | 0.90313459 |
| ENSMUSG00 | 4933437N03I | 1101 | 0 | 0 | 0.08787199 | 0 | 0.0874331 | 0 | 0 | 0.13875532 | 0 | 0.17130137 | 0.130177 | 0.15208627 | 0 | 0 | 0 | 0.06955605 | 0.03481044 | 0 |
| ENSMUSG00 | 4930583P06I | 703 | 0 | 0 | 0 | 0 | 0 | 0 | 0 | 0 | 0 | 0 | 0 | 0 | 0 | 0 | 0 | 0 | 0 | 0 |
| ENSMUSG00 | 1700021L23I | 724 | 0 | 0 | 0 | 0.06292962 | 0 | 0 | 0 | 0 | 0 | 0 | 0 | 0 | 0 | 0 | 0 | 0 | 0 | 0 |
| ENSMUSG00 | 1700125G22I | 858 | 0 | 0.15530487 | 0 | 0.05310145 | 0.44878251 | 0 | 0.38600088 | 0.89026579 | 0 | 0 | 0 | 0.06505322 | 0 | 0.16239191 | 0.19208185 | 0 | 0.08933869 | 0 |
| ENSMUSG00 | Gm13869 | 1052 | 0.16161664 | 0.0633325 | 0 | 0 | 0.09150556 | 0.14131008 | 0.35979224 | 0.14521826 | 0.11745204 | 0.17928024 | 0.04541346 | 0.15917014 | 0.14068037 | 0 | 0.43081475 | 0.10919375 | 0.29145473 | 1.46412071 |
| ENSMUSG00 | E330010L02I | 962 | 0 | 0 | 0 | 0 | 0 | 0 | 0 | 0 | 0 | 0 | 0 | 0 | 0 | 0 | 0 | 0 | 0 | 0 |
| ENSMUSG00 | Gm13017 | 398 | 0 | 0 | 0 | 0 | 0 | 0 | 0 | 0 | 0 | 0 | 0 | 0 | 0 | 0 | 0 | 0 | 0 | 0 |
| ENSMUSG00 | Gm12946 | 517 | 0 | 0 | 0 | 0 | 0.186197 | 0 | 0 | 0 | 0 | 0 | 0 | 0 | 0 | 0 | 0.07969353 | 0.07406307 | 0.1482642 | 0 |
| ENSMUSG00 | Gm12248 | 942 | 0.27073361 | 0.95482818 | 0.87298303 | 1.01569207 | 0.45985914 | 0.26301876 | 0.10045155 | 1.37849435 | 1.0930604 | 0.90096885 | 0.55788168 | 0.11850459 | 0.78554008 | 0.69025183 | 0.56859897 | 0.40648203 | 0.85440789 | 2.86140789 |
| ENSMUSG00 | Gm14135 | 587 | 0 | 0 | 0 | 0 | 0 | 0 | 0 | 0 | 0 | 0 | 0 | 0 | 0 | 0 | 0 | 0 | 0 | 0 |
| ENSMUSG00 | Gm15948 | 400 | 5.73819876 | 4.08082954 | 0.96747061 | 3.07537049 | 2.16593657 | 1.85822751 | 2.83876074 | 1.90962012 | 3.19195497 | 2.35753516 | 1.55268615 | 3.06986144 | 1.97327669 | 2.55442481 | 0.72102723 | 2.584616 | 1.24560465 | 3.16302364 |
| ENSMUSG00 | Gm16314 | 862 | 1.28205867 | 0.27052234 | 1.34682683 | 0.31713024 | 0.44669999 | 0.51737193 | 0.60375808 | 0.22153366 | 0.04778018 | 0.38289434 | 0.55423386 | 0 | 0.57229602 | 0.43103561 | 0.09559526 | 0.22210329 | 1.06708947 | 1.40394472 |
| ENSMUSG00 | Gm16866 | 2570 | 0 | 0 | 0 | 0 | 0 | 0 | 0 | 0 | 0 | 0 | 0 | 0 | 0.15356239 | 0 | 0 | 0 | 0 | 0 |
| ENSMUSG00 | 4930566F21I | 1723 | 0.02466928 | 0 | 0 | 0 | 0 | 0 | 0 | 0 | 0 | 0 | 0 | 0 | 0 | 0 | 0 | 0 | 0 | 0 |
| ENSMUSG00 | 4933431E20I | 8844 | 5.05121438 | 5.40148013 | 6.27368151 | 6.43954154 | 5.3879019 | 6.2921438 | 5.2266494 | 5.72626929 | 6.41734721 | 6.09376457 | 6.45534548 | 8.04038378 | 6.50953725 | 5.62958635 | 5.14320643 | 6.00509639 | 5.49499598 | 6.16395576 |
| ENSMUSG00 | 4930443O20I | 1440 | 0.05903497 | 0 | 0 | 0.03163961 | 0 | 0 | 0 | 0.07956751 | 0.17161048 | 0.06548709 | 0.19906233 | 0 | 0.06851655 | 0.29027555 | 0.17167315 | 0 | 0.31938581 | 0.11460231 |
| ENSMUSG00 | Gm14259 | 1236 | 0 | 0 | 0 | 0 | 0 | 0 | 0 | 0 | 0 | 0 | 0 | 0 | 0 | 0 | 0 | 0 | 0 | 0 |
| ENSMUSG00 | 4930527B05I | 599 | 0 | 0 | 0 | 0 | 0 | 0 | 0 | 0 | 0 | 0 | 0 | 0 | 0 | 0.54275227 | 0 | 0 | 0 | 0 |
| ENSMUSG00 | Gm15092 | 572 | 0 | 0 | 0 | 0 | 0 | 0 | 0 | 0 | 0 | 0 | 0 | 0 | 0 | 0 | 0 | 0 | 0 | 0 |
| ENSMUSG00 | Gm15208 | 491 | 0 | 0 | 0 | 0 | 0 | 0 | 0 | 0 | 0 | 0 | 0 | 0 | 0 | 0 | 0 | 0 | 0 | 0 |
| ENSMUSG00 | Gm12316 | 404 | 0 | 0 | 0 | 0 | 0 | 0 | 0 | 0 | 0 | 0 | 0 | 0 | 0 | 0 | 0 | 0 | 0 | 0 |
| ENSMUSG00 | 9430065F17I | 1449 | 0.90935849 | 0.87363008 | 0.6342975 | 0.31443095 | 0.73078145 | 0.71815556 | 1.40403395 | 1.87140136 | 1.19381205 | 0.52064269 | 0.85724563 | 2.0800868 | 1.46395614 | 1.41031045 | 1.27955143 | 0.89846836 | 1.34895869 | 2.3157733 |
| ENSMUSG00 | 4933435F18I | 2600 | 0 | 0.03843795 | 0 | 0 | 0 | 0 | 0 | 0 | 0 | 0 | 0 | 0 | 0 | 0 | 0 | 0 | 0 | 0 |
| ENSMUSG00 | Gm13791 | 606 | 0 | 0 | 0 | 0 | 0 | 0 | 0 | 0 | 0 | 0 | 0 | 0 | 0 | 0 | 0 | 0 | 0 | 0 |
| ENSMUSG00 | Gm12171 | 1384 | 0 | 0 | 0 | 0 | 0 | 0 | 0 | 0 | 0 | 0 | 0 | 0 | 0 | 0 | 0 | 0 | 0 | 0 |
| ENSMUSG00 | Gm8327 | 700 | 0 | 0 | 0 | 0 | 0 | 0 | 0 | 0 | 0 | 0 | 0 | 0 | 0 | 0 | 0 | 0 | 0.05475185 | 0 |
| ENSMUSG00 | Gm16972 | 588 | 0.79516486 | 0.73650957 | 1.15175072 | 0.15496954 | 0.49114208 | 0.84273356 | 0.56324618 | 1.4939205 | 1.96126266 | 1.04244752 | 1.46249873 | 1.32894435 | 1.84575201 | 0.94783852 | 2.03204953 | 0.78144097 | 2.15096564 | 1.02908193 |
| ENSMUSG00 | Gm6762 | 1315 | 0.61414323 | 0.43066099 | 0.4046455 | 0.41576618 | 0.58563557 | 0.41450956 | 0.39577146 | 0.43565478 | 0.62641089 | 0.57369677 | 0.3269769 | 0.50934445 | 0.41266243 | 0.52978046 | 0.12532793 | 0.58236665 | 0.29145473 | 0.16732808 |
| ENSMUSG00 | Gm16973 | 3109 | 1.99606166 | 1.69296793 | 1.19805784 | 2.18353027 | 2.33770359 | 1.60978646 | 2.55661951 | 1.68297174 | 1.70892909 | 1.42559219 | 1.39836643 | 2.85451603 | 3.09415371 | 2.59931528 | 2.06736659 | 2.62331919 | 3.30377857 | 1.87551152 |
| ENSMUSG00 | Gm12103 | 789 | 0.4309777 | 0.04222167 | 0 | 0.11549061 | 0.36602223 | 0.12560896 | 0.23986149 | 0.04840609 | 0 | 0 | 0 | 0.21222685 | 0 | 0.0588645 | 0 | 0.19412222 | 0.63148525 | 0 |
| ENSMUSG00 | Gm12484 | 579 | 0 | 0 | 0 | 0 | 0 | 0 | 0 | 0 | 0 | 0 | 0 | 0 | 0 | 0 | 0 | 0 | 0 | 0 |
| ENSMUSG00 | 1700086O06I | 1927 | 5.22767344 | 5.96416632 | 5.57287169 | 5.5089379 | 5.09543977 | 4.65440829 | 3.68287589 | 5.23237895 | 7.92952639 | 5.75008576 | 5.77662964 | 5.41646544 | 5.55528077 | 6.33876232 | 6.7350753 | 5.722727 | 4.73360595 | 6.73697415 |
| ENSMUSG00 | Gm16441 | 923 | 0 | 0 | 0 | 0 | 0 | 0 | 0 | 0 | 0 | 0 | 0 | 0 | 0 | 0 | 0 | 0 | 0 | 0 |
| ENSMUSG00 | 4833428L15I | 1254 | 2.2710102 | 4.25044902 | 3.04745526 | 3.81491998 | 4.33724671 | 1.81772388 | 0.79231759 | 3.65477535 | 4.07266982 | 2.29361475 | 2.85735398 | 5.11866124 | 1.337548 | 0.96295556 | 0.19713663 | 0.30534775 | 0.06112647 | 2.06174483 |
| ENSMUSG00 | Gm15334 | 323 | 0.26318994 | 0.61881537 | 0.59905301 | 0.98739105 | 0.14901524 | 0 | 0 | 1.06418459 | 0.3825373 | 0.14597741 | 0.29582018 | 0 | 0.15273039 | 0.86273467 | 0 | 0.59273386 | 0.23731453 | 0.17030683 |
| ENSMUSG00 | Gm15941 | 3358 | 0.27847346 | 0.38689782 | 0.34573101 | 0.46130897 | 0.30100369 | 0.32464566 | 0.36632807 | 0.13648268 | 0.1594475 | 0.42123916 | 0.73981472 | 0.23270377 | 0.47010761 | 0.15213965 | 0.30674178 | 0.7981961 | 0.18261488 | 0.21295962 |
| ENSMUSG00 | Gm15623 | 3849 | 0.12147491 | 0.09520443 | 0.2136529 | 0.07102267 | 0.37515139 | 0.02574837 | 0.0860454 | 0.08930414 | 0.19261036 | 0.12250118 | 0 | 0.1740161 | 0 | 0 | 0.03211345 | 0.06963737 | 0.24893672 | 0.10004254 |
| ENSMUSG00 | Gm13402 | 422 | 0 | 0 | 0 | 0 | 0.22811338 | 0 | 0 | 0.45251662 | 0 | 0 | 0 | 0 | 0 | 0 | 0 | 0 | 0 | 0 |
| ENSMUSG00 | Gm13544 | 2936 | 0 | 0 | 0 | 0 | 0 | 0 | 0 | 0 | 0 | 0 | 0 | 0 | 0 | 0.01581883 | 0 | 0 | 0 | 0 |
| ENSMUSG00 | 4933416E14I | 2427 | 0 | 0 | 0 | 0 | 0 | 0 | 0 | 0 | 0 | 0 | 0 | 0 | 0 | 0 | 0 | 0 | 0 | 0 |
| ENSMUSG00 | Gm12981 | 620 | 1.50824818 | 1.02087902 | 0.85824005 | 0.88182666 | 0.85395348 | 0.47954258 | 0.38155386 | 0.2464026 | 1.32859728 | 0.98864378 | 0.38528192 | 0.18005052 | 0.47740565 | 0.37454909 | 0.99681184 | 0.80286757 | 0.74179929 | 0.70979493 |
| ENSMUSG00 | Gm16033 | 594 | 0 | 0 | 0 | 0 | 0 | 0 | 0 | 0 | 0 | 0 | 0 | 0 | 0 | 0 | 0 | 0 | 0 | 0 |
| ENSMUSG00 | 1700040N02I | 1166 | 0 | 0 | 0 | 0 | 0 | 0 | 0 | 0 | 0 | 0 | 0 | 0 | 0 | 0 | 0 | 0 | 0 | 0 |
| ENSMUSG00 | Gm13395 | 1837 | 0 | 0 | 0 | 0 | 0 | 0.02697481 | 0 | 0 | 0 | 0 | 0 | 0 | 0 | 0 | 0.02242872 | 0 | 0 | 0 |
| ENSMUSG00 | Gm15475 | 618 | 0 | 0 | 0 | 0 | 0 | 0 | 0 | 0 | 0 | 0 | 0 | 0 | 0 | 0 | 0.33334592 | 0 | 0 | 0 |
| ENSMUSG00 | Gm16277 | 660 | 0 | 0 | 0 | 0 | 0 | 0 | 0 | 0 | 0 | 0 | 0 | 0 | 0 | 0 | 0 | 0 | 0 | 0 |
| ENSMUSG00 | 1700067G03I | 381 | 0 | 0 | 0 | 0 | 0 | 0 | 0 | 0 | 0 | 0 | 0 | 0 | 0 | 0 | 0 | 0 | 0 | 0 |
| ENSMUSG00 | Gm14154 | 575 | 0.29568818 | 0.11587094 | 0 | 0.1584732 | 0 | 0.517072 | 0 | 0.46495099 | 0.42977234 | 0.08200122 | 0.24926065 | 0 | 0.25738392 | 0.48463396 | 0.07165488 | 1.06547777 | 0.26661772 | 0 |
| ENSMUSG00 | Gm13889 | 1380 | 0 | 0 | 0 | 0.132061 | 0.03487821 | 0.07181556 | 0 | 0 | 0.0298453 | 0.03416718 | 0.06923907 | 0.1213384 | 0 | 0 | 0 | 0 | 0 | 0 |
| ENSMUSG00 | Gm2231 | 611 | 0 | 0 | 0 | 0 | 0 | 0 | 0 | 0 | 0 | 0 | 0 | 0 | 0 | 0 | 0 | 0 | 0 | 0 |
| ENSMUSG00 | Gm6530 | 984 | 0.17278527 | 0.1015637 | 0.49160092 | 0.41671687 | 0.3424019 | 1.00716938 | 0.38465593 | 0.27169392 | 0.75341187 | 1.00626501 | 0.24275894 | 0.73740205 | 1.05281531 | 0.33039493 | 1.2980165 | 0.50587185 | 0.3505454 | 1.17397484 |
| ENSMUSG00 | Nphs1as | 1336 | 0 | 0 | 0 | 0 | 0 | 0 | 0 | 0 | 0 | 0 | 0 | 0 | 0 | 0 | 0 | 0 | 0 | 0 |
| ENSMUSG00 | 1700041M05 | 304 | 0 | 0 | 0 | 0.29974371 | 0 | 0 | 0 | 0 | 0 | 0 | 0 | 0 | 0 | 0 | 0 | 0 | 0 | 0 |
| ENSMUSG00 | Gm12930 | 674 | 0 | 0 | 0 | 0 | 0 | 0 | 0.14039371 | 0 | 0 | 0 | 0 | 0 | 0 | 0 | 0 | 0 | 0 | 0 |
| ENSMUSG00 | Gm13113 | 637 | 0 | 0 | 0 | 0 | 0 | 0 | 0 | 0 | 0 | 0 | 0 | 0 | 0 | 0 | 0 | 0 | 0 | 0 |
| ENSMUSG00 | 2610027K06I | 2568 | 2.49933083 | 1.18048029 | 2.52416397 | 1.54354005 | 0.97463397 | 1.15777415 | 1.12386036 | 1.62109496 | 1.21891558 | 1.3954258 | 0.89298988 | 1.36930948 | 2.28601953 | 1.4468563 | 2.16596964 | 2.60936772 | 1.62677818 | 1.41378545 |
| ENSMUSG00 | Gm16364 | 1311 | 1.06992434 | 1.19428377 | 0.81176023 | 0.55604631 | 1.46855602 | 0.07559532 | 0.505246 | 1.6022747 | 1.79071808 | 0.28772359 | 1.67631433 | 0.63862314 | 1.35465219 | 1.52333773 | 0.37713095 | 1.05145833 | 1.11090716 | 1.0070317 |
| ENSMUSG00 | Gm14041 | 274 | 0 | 0 | 0 | 0 | 0 | 0 | 0 | 0 | 0 | 0 | 0 | 0 | 0 | 0 | 0 | 0 | 0 | 0 |
| ENSMUSG00 | Gm15979 | 588 | 0.07228771 | 0.22661833 | 0.16453582 | 0 | 0.16371403 | 0 | 0 | 0.1948592 | 0.63040585 | 0.40094135 | 0.40624965 | 0.0949246 | 0 | 0 | 0.07007067 | 0.52096064 | 0.26072311 | 0.65487032 |

| | | | | | | | | | | | | | | | | | | | |
|---|---|---|---|---|---|---|---|---|---|---|---|---|---|---|---|---|---|---|---|
| ENSMUSG00 | 4930417H01 | 1878 | 0 | 0 | 0.07727401 | 0.09704163 | 0.02562935 | 0.02638591 | 0.22673808 | 0.02033674 | 0 | 0.05021374 | 0.2035142 | 0 | 0 | 0.19784489 | 0.06581718 | 0 | 0.02040804 | 0 |
| ENSMUSG00 | 2900072N19 | 804 | 0 | 0 | 0 | 0.05666797 | 0 | 0.12326551 | 0 | 0 | 0 | 0 | 0 | 0 | 0 | 0 | 0 | 0 | 0 | 0 |
| ENSMUSG00 | 4930593A02 | 2016 | 0 | 0.01652425 | 0.04798961 | 0 | 0 | 0 | 0 | 0 | 0.02042982 | 0.04677649 | 0 | 0.08305902 | 0 | 0.11518871 | 0 | 0 | 0.01901106 | 0.05457253 |
| ENSMUSG00 | Gm16283 | 520 | 1.47133302 | 1.53751819 | 1.3953903 | 0.61332175 | 1.85122783 | 0.66705603 | 2.72957763 | 1.90962012 | 2.05932579 | 3.26427946 | 1.74562348 | 1.61006719 | 3.60502472 | 3.84056876 | 3.09011669 | 3.01906712 | 3.39040319 | 2.11573487 |
| ENSMUSG00 | Gm5753 | 1060 | 0 | 0 | 0 | 0 | 0 | 0 | 0 | 0 | 0 | 0 | 0 | 0 | 0 | 0 | 0 | 0 | 0 | 0 |
| ENSMUSG00 | 9130024F11 | 1219 | 0 | 0 | 0 | 0 | 0 | 0 | 0 | 0 | 0 | 0 | 0 | 0 | 0 | 0 | 0 | 0 | 0 | 0 |
| ENSMUSG00 | 1110020A21 | 851 | 4.14562821 | 4.22772336 | 3.18321704 | 4.71136534 | 5.71248448 | 4.42539101 | 6.44920183 | 5.7894476 | 8.08243024 | 3.98924869 | 4.37890336 | 3.27941613 | 5.8549044 | 2.89252248 | 4.84154593 | 4.31950448 | 3.15257436 | 2.71490303 |
| ENSMUSG00 | B230119M05 | 976 | 0.04355039 | 0.30718857 | 0 | 0.46681398 | 0 | 0.50771243 | 0 | 0.03913156 | 0.0421993 | 0.24155073 | 0.39159802 | 0.45750543 | 0.10108999 | 0.23793078 | 0 | 0.35308962 | 0.07853749 | 0 |
| ENSMUSG00 | A230103J11 | 3416 | 0.14931561 | 0.05851211 | 0.1840913 | 0.08002525 | 0.19726198 | 0.11604856 | 0.31855726 | 0.36895471 | 0.40993605 | 0.11042319 | 0.12587079 | 0.11437636 | 0.02888286 | 0.23113275 | 0.04824538 | 0.30264824 | 0.14585535 | 0.53141116 |
| ENSMUSG00 | Gm13206 | 1086 | 0 | 0 | 0 | 0 | 0 | 0 | 0.043566 | 0 | 0 | 0 | 0 | 0 | 0 | 0 | 0 | 0 | 0 | 0 |
| ENSMUSG00 | Gm13387 | 2878 | 0 | 0 | 0 | 0 | 0 | 0 | 0 | 0 | 0 | 0.11468204 | 0.13280044 | 0.01939391 | 0.25711562 | 0 | 0.21474056 | 0 | 0 | 0 |
| ENSMUSG00 | Gm14133 | 1433 | 0 | 0 | 0 | 0 | 0 | 0 | 0 | 0 | 0 | 0 | 0 | 0 | 0 | 0 | 0 | 0 | 0 | 0 |
| ENSMUSG00 | Gm16143 | 535 | 0 | 0 | 0 | 0 | 0 | 0.18524386 | 0 | 0 | 0.07698414 | 0 | 0.17859798 | 0 | 0.36883676 | 0 | 0.07701225 | 0 | 0 | 0 |
| ENSMUSG00 | Gm14241 | 262 | 0 | 0 | 0 | 0 | 0 | 0 | 0 | 0 | 0 | 0 | 0 | 0 | 0 | 0 | 0 | 0 | 0 | 0 |
| ENSMUSG00 | Gm13874 | 1673 | 0.1270328 | 0 | 0 | 0 | 0 | 0 | 0 | 0 | 0 | 0 | 0 | 0 | 0 | 0 | 0 | 0.27464871 | 0 | 0.23016363 |
| ENSMUSG00 | Gm14155 | 585 | 0 | 0 | 0 | 0 | 0 | 0 | 0 | 0 | 0 | 0 | 0 | 0 | 0 | 0 | 0 | 0 | 0 | 0 |
| ENSMUSG00 | Cbfa2t2-ps1 | 1258 | 0 | 0.07944251 | 0 | 0 | 0.07652134 | 0 | 0 | 0 | 0 | 0.03748069 | 0.03797691 | 0.04436857 | 0 | 0.03691899 | 0 | 0 | 0.09139816 | 0 |
| ENSMUSG00 | BC052486 | 2376 | 0 | 0 | 0 | 0.11505314 | 0 | 0 | 0 | 0 | 0 | 0 | 0 | 0.02349144 | 0 | 0 | 0 | 0 | 0 | 0 |
| ENSMUSG00 | Gm11748 | 1496 | 0 | 0.24494775 | 0.12934099 | 0 | 0 | 0.03312349 | 0 | 0.02552968 | 0.35790421 | 0 | 0 | 0 | 0 | 0 | 0 | 0 | 0.20495346 | 0 |
| ENSMUSG00 | 4930432L08 | 1602 | 0 | 0 | 0 | 0 | 0 | 0 | 0 | 0 | 0 | 0 | 0 | 0 | 0 | 0 | 0 | 0 | 0 | 0 |
| ENSMUSG00 | 2900079G21 | 6792 | 0 | 0 | 0 | 0 | 0 | 0 | 0 | 0 | 0 | 0 | 0 | 0 | 0 | 0 | 0 | 0 | 0 | 0 |
| ENSMUSG00 | Gm14526 | 492 | 0.60474844 | 0.67709135 | 0.5899211 | 0.55562249 | 0.78263291 | 0.50358469 | 0.57698389 | 0.465761 | 0.66969944 | 0.47917381 | 0 | 0.79412528 | 0.4010725 | 0 | 1.256145 | 0.38913219 | 0.38949489 | 0.22361425 |
| ENSMUSG00 | Gm14033 | 2359 | 0 | 0.01412162 | 0 | 0.03862742 | 0 | 0 | 0 | 0.04857024 | 0.06983725 | 0 | 0.18226987 | 0.09464292 | 0.02091222 | 0.25594453 | 0.01746569 | 0 | 0.01624684 | 0 |
| ENSMUSG00 | Gm16295 | 641 | 0.33155364 | 1.24728465 | 0.83012299 | 0.85293687 | 1.42668729 | 1.00496964 | 1.4024039 | 0.9533205 | 2.05611311 | 1.91250902 | 1.788766 | 2.87350525 | 0.46176522 | 0.94192377 | 0.83560098 | 0.65709311 | 0.71749698 | 0.34327056 |
| ENSMUSG00 | Gm11611 | 1324 | 0.77048657 | 0.22644717 | 0.40189489 | 0.37852831 | 0.436241 | 0.22455922 | 0.39308117 | 0.28846225 | 0.93322921 | 0.39173545 | 0.32475425 | 0.3794116 | 0.07451951 | 0.24555031 | 0.186714 | 0.23136319 | 0.08684206 | 0.41547664 |
| ENSMUSG00 | Gm13599 | 674 | 0 | 0 | 0 | 0 | 0 | 0 | 0 | 0 | 0 | 0 | 0 | 0 | 0 | 0 | 0 | 0 | 0 | 0 |
| ENSMUSG00 | 1700042G15 | 660 | 0 | 0 | 0 | 0 | 0 | 0 | 0 | 0 | 0 | 0 | 0 | 0 | 0 | 0 | 0 | 0 | 0 | 0 |
| ENSMUSG00 | 4930515B02 | 588 | 0 | 0 | 0 | 0 | 0 | 0 | 0 | 0 | 0 | 0 | 0 | 0 | 0 | 0 | 0 | 0 | 0 | 0 |
| ENSMUSG00 | AU022754 | 2971 | 0.62949436 | 0.62791049 | 0.73268558 | 0.35271088 | 0.85863075 | 0.7839039 | 0.44589532 | 0.35994186 | 0.24953123 | 0.12696251 | 0.57889549 | 0.01878683 | 1.01287343 | 0.51587172 | 0.59632006 | 0.82483974 | 1.34161389 | 0.64803727 |
| ENSMUSG00 | 1700110K17 | 1024 | 1.78488532 | 0.61811034 | 1.18099439 | 1.42378264 | 0.42303449 | 0.96782683 | 0.18481515 | 0.1118918 | 0.6033181 | 0.13813683 | 0.32658663 | 0.54507483 | 0.96351401 | 0 | 0.44259484 | 0.7852566 | 0 | 0 |
| ENSMUSG00 | Gm11932 | 477 | 0 | 0.34919176 | 0 | 0.09551582 | 0.201811 | 0.10388414 | 0 | 0.48040758 | 0.25903469 | 0 | 0 | 0.70208381 | 0.51710605 | 0 | 0 | 0.40136905 | 0.24104589 | 0.23064615 |
| ENSMUSG00 | A030003K02 | 2427 | 0.05254039 | 0.42550462 | 0.61787369 | 0.46931442 | 0.53546021 | 0.40834556 | 0.7212893 | 0.45635751 | 0.42425336 | 0.85481291 | 0.7480216 | 0.66693622 | 0.24391554 | 0.97595734 | 0.49231361 | 0.22087701 | 0.01579163 | 0.86128803 |
| ENSMUSG00 | Gm12730 | 289 | 0 | 0 | 0 | 0 | 0 | 0 | 0 | 0 | 0 | 0 | 0 | 0 | 0 | 0 | 0 | 0 | 0 | 0 |
| ENSMUSG00 | Gm8093 | 2974 | 0 | 0 | 0 | 0 | 0 | 0 | 0 | 0 | 0 | 0 | 0 | 0 | 0 | 0 | 0 | 0 | 0 | 0 |
| ENSMUSG00 | Gm15082 | 1958 | 0 | 0 | 0 | 0 | 0 | 0 | 0 | 0 | 0 | 0 | 0 | 0 | 0 | 0 | 0 | 0 | 0 | 0 |
| ENSMUSG00 | Gm12405 | 658 | 0 | 0.101255 | 0 | 0 | 0 | 0 | 0 | 0 | 0 | 0.0716576 | 0 | 0 | 0 | 0 | 0 | 0 | 0 | 0 |
| ENSMUSG00 | Gm11948 | 537 | 0.15830606 | 0.31017592 | 1.35121593 | 0.25453097 | 0.98594257 | 0.73821577 | 0.08810555 | 0.56897434 | 0.46018453 | 1.22925483 | 0.88966403 | 0.51969891 | 0.55119461 | 0.95136864 | 0.53707801 | 0.49913268 | 0.57096904 | 0.30731345 |
| ENSMUSG00 | Gm14004 | 586 | 0 | 0 | 0 | 0 | 0 | 0.42280489 | 0 | 0 | 0.07028416 | 0 | 0 | 0.09524857 | 0 | 0.47553673 | 0.07030982 | 0 | 0.32701619 | 0.37548878 |
| ENSMUSG00 | Gm11730 | 825 | 0 | 0.16151706 | 0 | 0.05522551 | 0 | 0 | 0 | 0 | 0 | 0 | 0 | 0 | 0 | 0.73184623 | 0.29964768 | 0 | 0.32519282 | 0.40006623 |
| ENSMUSG00 | Gm12301 | 521 | 0 | 0 | 0 | 0 | 0 | 0 | 0 | 0 | 0 | 0.09050039 | 0 | 0 | 0 | 0 | 0 | 0 | 0 | 0 |
| ENSMUSG00 | Gm12339 | 875 | 0.14573203 | 0.34264691 | 0.27642017 | 0.20827906 | 0.22003165 | 0.33979017 | 0 | 0.17459384 | 0.0470703 | 0.32331911 | 0.38219967 | 0.12757866 | 0.22551734 | 0 | 1.03592483 | 0.17504278 | 0.4818163 | 0.18860265 |
| ENSMUSG00 | 2810442I21R | 3297 | 0 | 0 | 0 | 0.04145682 | 0 | 0.01502964 | 0 | 0.02316797 | 0.12492119 | 0.04290328 | 0 | 0.11850459 | 0 | 0 | 0.09997344 | 0.1277515 | 0 | 0 |
| ENSMUSG00 | A130030D18 | 651 | 0 | 0 | 0 | 0 | 0 | 0 | 0 | 0 | 0 | 0 | 0 | 0 | 0 | 0 | 0 | 0 | 0 | 0 |
| ENSMUSG00 | Gm13587 | 363 | 0 | 0 | 0 | 0 | 0 | 0 | 0 | 0 | 0 | 0 | 0 | 0 | 0 | 0 | 0 | 0 | 0 | 0 |
| ENSMUSG00 | Gm15857 | 786 | 0 | 0 | 0 | 0.05796571 | 0 | 0 | 0 | 0 | 0 | 0.05998817 | 0.30391195 | 0 | 0 | 0 | 0 | 0.19486314 | 0 | 0 |
| ENSMUSG00 | Gm11721 | 2350 | 0.34365887 | 0.1559327 | 0.37052066 | 0.11632607 | 0.04096334 | 0.14760389 | 0.08053222 | 0.37379798 | 0.15773559 | 0.24076955 | 0.24395723 | 0.3087675 | 0.10496153 | 0.05929032 | 0.03506515 | 0.16293875 | 0.08154531 | 0.42134635 |
| ENSMUSG00 | Astx6 | 1146 | 0 | 0 | 0 | 0 | 0 | 0 | 0 | 0 | 0 | 0 | 0 | 0 | 0 | 0 | 0 | 0 | 0 | 0 |
| ENSMUSG00 | Gm15518 | 2105 | 0.04038497 | 0.11077922 | 0.1608621 | 0 | 0.11432761 | 0.11770246 | 0.04495266 | 0.03628732 | 0 | 0.02239938 | 0.09078377 | 0.02651575 | 0 | 0.1103185 | 0 | 0.05457094 | 0.07282907 | 0 |
| ENSMUSG00 | Gm11532 | 5270 | 0 | 0 | 0 | 0 | 0 | 0 | 0 | 0 | 0.02344583 | 0 | 0 | 0.06354724 | 0 | 0 | 0 | 0.00726577 | 0 | 0 |
| ENSMUSG00 | Astx4a | 1145 | 0 | 0 | 0 | 0 | 0 | 0 | 0 | 0 | 0 | 0 | 0 | 0 | 0 | 0 | 0 | 0 | 0 | 0 |
| ENSMUSG00 | Gm13793 | 436 | 0 | 0 | 0 | 0 | 0 | 0 | 0 | 0 | 0 | 0 | 0 | 0 | 0 | 0 | 0 | 0 | 0 | 0 |
| ENSMUSG00 | Gm12505 | 753 | 0 | 0 | 0 | 0 | 0 | 0 | 0 | 0 | 0 | 0 | 0 | 0 | 0 | 0.06167873 | 0 | 0 | 0 | 0 |
| ENSMUSG00 | Gm13116 | 2190 | 0 | 0 | 0 | 0 | 0 | 0 | 0 | 0 | 0 | 0 | 0 | 0 | 0 | 0 | 0 | 0 | 0 | 0 |
| ENSMUSG00 | Rpl29-ps3 | 394 | 1.40245505 | 1.94466134 | 1.96440732 | 0.46254867 | 0 | 0.25153672 | 1.20082941 | 0.67854522 | 0.20906861 | 0.83770285 | 0.84879368 | 1.27497706 | 0.37562374 | 0.35363518 | 0.83657982 | 0.09718428 | 1.65367271 | 0.83770213 |
| ENSMUSG00 | Gm14249 | 1011 | 0 | 0 | 0 | 0 | 0 | 0 | 0 | 0 | 0 | 0 | 0 | 0 | 0 | 0 | 0 | 0 | 0 | 0 |
| ENSMUSG00 | A230065H16 | 653 | 0.13018431 | 0 | 0.14815783 | 0.48840323 | 0 | 0 | 0 | 0 | 0 | 0.21661885 | 0 | 0 | 0 | 0 | 0.75714957 | 0.35182794 | 0 | 0 |
| ENSMUSG00 | Gm13229 | 680 | 0 | 0 | 0 | 0 | 0 | 0 | 0 | 0 | 0 | 0 | 0 | 0 | 0 | 0 | 0 | 0 | 0 | 0 |
| ENSMUSG00 | Gm12480 | 772 | 0 | 0 | 0 | 0 | 0 | 0 | 0 | 0 | 0 | 0 | 0 | 0 | 0 | 0 | 0 | 0 | 0 | 0 |
| ENSMUSG00 | 6720483E21 | 1116 | 0.03808708 | 0 | 0 | 0.36742778 | 0 | 0.26641255 | 0 | 0 | 0 | 0.59149628 | 0.34247282 | 0 | 0.4862465 | 0.16646626 | 0.29535166 | 0.92638566 | 0.61816608 | 0.64078709 |
| ENSMUSG00 | Gm13389 | 637 | 0 | 0 | 0 | 0 | 0 | 0 | 0 | 0 | 0 | 0 | 0 | 0 | 0.15488828 | 0 | 0 | 0 | 0 | 0 |
| ENSMUSG00 | Gm12199 | 515 | 0 | 0 | 0 | 0 | 0 | 0 | 0 | 0 | 0 | 0 | 0 | 0 | 0 | 0 | 0 | 0 | 0 | 0 |
| ENSMUSG00 | 6430590A07 | 1029 | 5.57648082 | 3.1079085 | 2.53855261 | 4.33914708 | 3.32105596 | 1.49284231 | 7.58658117 | 4.45392448 | 3.92252531 | 8.2937583 | 8.54284972 | 3.36304284 | 7.43094964 | 4.33297608 | 6.68674425 | 6.69806543 | 2.75621572 | 2.51256367 |
| ENSMUSG00 | Gm15423 | 1427 | 0 | 0 | 0.37288636 | 0 | 0 | 0 | 0.06631069 | 0 | 0 | 0 | 0.46871017 | 0 | 0.03457037 | 0 | 0 | 0 | 0 | 0 |
| ENSMUSG00 | Gm15083 | 1253 | 0 | 0 | 0 | 0 | 0 | 0 | 0 | 0.03048077 | 0 | 0 | 0 | 0 | 0.55119461 | 0.29653049 | 0.49323491 | 0 | 0.12235051 | 0.30731345 |
| ENSMUSG00 | Gm13016 | 1717 | 0 | 0 | 0 | 0 | 0 | 0 | 0 | 0 | 0 | 0 | 0 | 0 | 0 | 0 | 0 | 0 | 0 | 0 |
| ENSMUSG00 | Gm12068 | 372 | 0 | 0 | 0 | 0 | 0 | 0 | 0 | 0 | 0 | 0 | 0 | 0 | 0 | 0 | 0 | 0 | 0 | 0 |
| ENSMUSG00 | Gm13269 | 1060 | 0 | 0 | 0 | 0 | 0 | 0 | 0.0446346 | 0 | 0 | 0 | 0 | 0 | 0 | 0 | 0 | 0.03612321 | 0 | 0 |
| ENSMUSG00 | Gm16638 | 1861 | 6.46371027 | 4.74364157 | 5.1986599 | 3.89264162 | 5.4054659 | 4.10054861 | 5.1100744 | 3.48882773 | 6.0861321 | 4.10446745 | 4.10746553 | 4.70879045 | 3.18099413 | 3.91817395 | 4.67142842 | 3.9916055 | 2.43014671 | 2.45338842 |
| ENSMUSG00 | Gm12160 | 340 | 0 | 0 | 0 | 0 | 0 | 0 | 0 | 0 | 0 | 0 | 0 | 0 | 0 | 0 | 0 | 0 | 0 | 0 |
| ENSMUSG00 | Nctc1 | 3435 | 0 | 0 | 0 | 0 | 0 | 0 | 0 | 0 | 0 | 0 | 0 | 0 | 0 | 0 | 0 | 0 | 0 | 0 |
| ENSMUSG00 | 4930509E16 | 1176 | 0 | 0 | 0 | 0 | 0 | 0 | 0.16092748 | 0.29228879 | 0 | 0 | 0 | 0 | 0 | 0 | 0 | 0 | 0 | 0 |
| ENSMUSG00 | 4930511A02 | 1931 | 0 | 0 | 0 | 0 | 0 | 0 | 0 | 0 | 0 | 0 | 0 | 0 | 0 | 0 | 0 | 0 | 0 | 0 |
| ENSMUSG00 | Gm15486 | 657 | 0 | 0 | 0 | 0 | 0 | 0 | 0 | 0 | 0 | 0 | 0 | 0 | 0 | 0 | 0 | 0 | 0 | 0 |
| ENSMUSG00 | Gm13093 | 2482 | 0 | 0 | 0 | 0 | 0 | 0 | 0 | 0.18465303 | 0 | 0 | 0 | 0 | 0 | 0 | 0 | 0 | 0 | 0 |
| ENSMUSG00 | Emx2os | 5019 | 2.20190193 | 2.8474261 | 2.34205377 | 2.46006037 | 1.98511819 | 2.42876518 | 2.705467 | 2.29047881 | 1.9366443 | 2.20769382 | 2.01798988 | 2.50219647 | 1.78888403 | 1.90625264 | 2.49557143 | 1.56397181 | 1.64179196 | 2.27971592 |
| ENSMUSG00 | Gm16799 | 1754 | 0.02423328 | 0 | 0 | 0 | 0 | 0 | 0 | 0 | 0 | 0 | 0 | 0 | 0 | 0 | 0 | 0 | 0 | 0 |
| ENSMUSG00 | 4930586N03 | 877 | 0 | 0 | 0 | 0 | 0 | 0 | 0 | 0 | 0 | 0 | 0 | 0 | 0 | 0 | 0 | 0 | 0 | 0 |
| ENSMUSG00 | Gm11884 | 636 | 0 | 0 | 0 | 0 | 0 | 0 | 0 | 0 | 0 | 0 | 0 | 0 | 0 | 0 | 0 | 0 | 0 | 0 |
| ENSMUSG00 | Gm11782 | 2802 | 0 | 0 | 0 | 0 | 0 | 0 | 0 | 0 | 0 | 0 | 0 | 0 | 0 | 0 | 0 | 0 | 0 | 0 |
| ENSMUSG00 | Gm14641 | 450 | 0 | 0 | 0 | 0 | 0 | 0 | 0 | 0 | 0 | 0 | 0 | 0 | 0 | 0 | 0 | 0 | 0 | 0 |
| ENSMUSG00 | Gm16953 | 2429 | 0.31498278 | 0.12343188 | 0.27880997 | 0.24384256 | 1.18893183 | 0.65281493 | 0.56486937 | 0.47170526 | 0.2204301 | 0.15529256 | 0.15734857 | 0.41361957 | 0.40619117 | 0.4015339 | 0.93292943 | 0.40986241 | 0.3471299 | 0.11323406 |
| ENSMUSG00 | Gm13661 | 316 | 0 | 0 | 0 | 0 | 0 | 0 | 0 | 0 | 0 | 0 | 0 | 0 | 0 | 0 | 0 | 0 | 0 | 0 |
| ENSMUSG00 | Gm13344 | 1555 | 0 | 0 | 0 | 0 | 0 | 0 | 0 | 0 | 0 | 0 | 0 | 0 | 0 | 0 | 0 | 0 | 0.07394141 | 0 |
| ENSMUSG00 | 1700085B03 | 1285 | 0.03307796 | 0 | 0 | 0 | 0 | 0 | 0 | 0 | 0 | 0 | 0 | 0 | 0 | 0 | 0 | 0 | 0 | 0 |
| ENSMUSG00 | Gm16952 | 2478 | 3.31053227 | 3.67006462 | 4.00184573 | 3.03372571 | 3.8653159 | 2.01970383 | 2.34845016 | 2.01904952 | 2.24381744 | 1.5031903 | 2.1785998 | 2.47769285 | 1.65236043 | 1.49940557 | 3.05935685 | 1.06620335 | 1.65492888 | 1.99790945 |
| ENSMUSG00 | Gm13964 | 731 | 0.40702631 | 0.04557167 | 0.26469784 | 0.12465402 | 0 | 0 | 0 | 0 | 0.28171351 | 0.45151152 | 0 | 0.38177608 | 0 | 0.19060501 | 0 | 0 | 0.26214977 | 0 |
| ENSMUSG00 | AI662270 | 1485 | 0 | 0.31406096 | 0 | 0 | 0 | 0 | 0 | 0.61725095 | 0.36055536 | 0.12700526 | 0.12868676 | 0 | 0.16610073 | 0 | 0 | 0 | 0 | 0 |
| ENSMUSG00 | Gm16093 | 390 | 0 | 0.34167071 | 0.24806939 | 0.11682319 | 0 | 0 | 0 | 0 | 0 | 0 | 0 | 0.71558542 | 0 | 0 | 0 | 0.19636209 | 0 | 0 |
| ENSMUSG00 | 4930474H06 | 1588 | 0 | 0 | 0 | 0 | 0 | 0 | 0 | 0 | 0 | 0 | 0 | 0 | 0 | 0 | 0 | 0 | 0 | 0 |
| ENSMUSG00 | Gm11264 | 2963 | 0 | 0 | 0 | 0 | 0 | 0 | 0 | 0 | 0 | 0 | 0 | 0 | 0 | 0 | 0 | 0 | 0 | 0 |
| ENSMUSG00 | Gm11399 | 321 | 0 | 0.83022789 | 0.30139271 | 0 | 0.59977475 | 0 | 0.14739152 | 0 | 0.1283069 | 0 | 0 | 0.34776114 | 0 | 0 | 0.12835376 | 0.11928538 | 0.23879313 | 0.5141038 |
| ENSMUSG00 | Gm7237 | 1392 | 0.03053533 | 0 | 0.0695022 | 0.06546127 | 0 | 0 | 0.10196698 | 0.10974828 | 0 | 0.10161789 | 0 | 0 | 0.10631879 | 0 | 0 | 0 | 0.08259978 | 0.03951804 |
| ENSMUSG00 | Gm11714 | 874 | 0 | 0.03811544 | 0 | 0.15638802 | 0.1101417 | 0.22678597 | 0 | 1.00506322 | 0 | 0 | 0 | 0 | 0 | 0.26569844 | 0.18856547 | 0.26286458 | 0 | 0 |
| ENSMUSG00 | Gm16099 | 707 | 0 | 0 | 0 | 0 | 0 | 0 | 0 | 0 | 0 | 0 | 0 | 0 | 0 | 0 | 0 | 0 | 0 | 0 |
| ENSMUSG00 | Gm12360 | 3705 | 0 | 0 | 0 | 0 | 0 | 0 | 0 | 0 | 0 | 0 | 0 | 0 | 0 | 0 | 0 | 0 | 0 | 0 |
| ENSMUSG00 | Gm12324 | 3751 | 0 | 0 | 0 | 0 | 0 | 0 | 0 | 0 | 0 | 0 | 0 | 0 | 0 | 0 | 0 | 0 | 0 | 0 |
| ENSMUSG00 | Gm11523 | 782 | 0 | 0 | 0 | 0 | 0 | 0 | 0 | 0 | 0 | 0 | 0 | 0 | 0 | 0 | 0 | 0 | 0 | 0 |
| ENSMUSG00 | Gm16793 | 1608 | 1.13664339 | 1.88524464 | 1.11307253 | 1.13335931 | 0.86805086 | 1.10938956 | 2.70694432 | 1.73385906 | 1.38312926 | 2.02325779 | 1.12901021 | 2.77689864 | 2.45432424 | 3.00384645 | 3.15161155 | 2.90513315 | 1.85911142 | 3.45517399 |
| ENSMUSG00 | Atg4a-ps | 3250 | 0.53621914 | 0.29725352 | 0.31256743 | 0.36448835 | 0.42948486 | 0.24395192 | 0.65509863 | 0.29378771 | 0.43087432 | 0.4932689 | 0.48509958 | 0.36065505 | 0.62234111 | 0.42871465 | 0.35496725 | 0.30632486 | 0.46095203 | 0.57547989 |
| [illegible] | | | | | | | | | | | | | | | | | | | | |

| | | | | | | | | | | | | | | | | | | | |
|---|---|---|---|---|---|---|---|---|---|---|---|---|---|---|---|---|---|---|---|
| ENSMUSG00 Gm12862 | 470 | 0 | 0 | 0 | 0 | 0 | 0 | 0 | 0.16252086 | 0 | 0 | 0 | 0 | 0 | 0 | 0 | 0 | 0 | 0 |
| ENSMUSG00 Gm13905 | 576 | 0 | 0 | 0 | 0 | 0 | 0 | 0 | 0 | 0 | 0 | 0 | 0 | 0 | 0 | 0 | 0 | 0 | 0 |
| ENSMUSG00 A330041J22R | 321 | 0.13241488 | 0 | 0.30139271 | 0.14193472 | 0.29988737 | 0.15436989 | 0.29478305 | 0.35693834 | 1.15476212 | 0.73443463 | 0.14883165 | 0.34776114 | 0.92209191 | 0.72342815 | 1.54024508 | 2.14713686 | 1.67155189 | 1.0282076 |
| ENSMUSG00 Gm13601 | 400 | 0 | 0 | 0 | 0 | 0 | 0 | 0 | 0 | 0 | 0 | 0 | 0 | 0 | 0 | 0 | 0 | 0 | 0 |
| ENSMUSG00 Gm7023 | 867 | 0 | 0 | 0 | 0 | 0 | 0 | 0 | 0 | 0 | 0 | 0 | 0 | 0 | 0 | 0 | 0 | 0 | 0 |
| ENSMUSG00 Gm16142 | 2847 | 0 | 0 | 0 | 0 | 0 | 0 | 0 | 0 | 0 | 0 | 0 | 0 | 0 | 0 | 0 | 0 | 0 | 0 |
| ENSMUSG00 A630031M04 | 3562 | 0 | 0 | 0 | 0 | 0 | 0 | 0 | 0 | 0 | 0 | 0 | 0 | 0 | 0 | 0 | 0 | 0 | 0 |
| ENSMUSG00 Gm16836 | 2826 | 0.99268989 | 1.0727082 | 1.36938515 | 1.24140142 | 1.73724565 | 0.68384876 | 1.35609589 | 1.62175806 | 1.20965351 | 0.76749199 | 0.89599179 | 1.24429821 | 1.04738678 | 1.26546169 | 1.297372 | 0.79941466 | 1.2205827 | 1.65455558 |
| ENSMUSG00 A630075F10 | 1910 | 0 | 0 | 0 | 0 | 0 | 0.05188768 | 0.02477104 | 0 | 0 | 0 | 0 | 0 | 0 | 0 | 0 | 0 | 0 | 0 |
| ENSMUSG00 Gm13883 | 1814 | 0.02343174 | 0.11018598 | 0.05333355 | 0 | 0.02653359 | 0 | 0 | 0 | 0.11352402 | 0 | 0 | 0.21538569 | 0 | 0 | 0 | 0 | 0 | 0 |
| ENSMUSG00 Gm12100 | 1931 | 0 | 0 | 0 | 0 | 0 | 0 | 0 | 0 | 0 | 0 | 0 | 0 | 0 | 0 | 0 | 0 | 0 | 0 |
| ENSMUSG00 4933406I18R | 1642 | 0 | 0 | 0 | 0 | 0 | 0.03017828 | 0 | 0.06977905 | 0 | 0 | 0 | 0 | 0.30043799 | 0.02828507 | 0 | 0 | 0 | 0 |
| ENSMUSG00 1810058N05 | 469 | 0 | 0.21308888 | 0 | 0 | 0 | 0 | 0 | 0 | 0 | 0.10053455 | 0.40746233 | 0.35702982 | 0.21037065 | 0 | 0 | 0.32657234 | 0.40859592 | 0 |
| ENSMUSG00 Gm13822 | 344 | 0 | 0.09683981 | 0.14062073 | 0.1324449 | 0.55967353 | 0 | 0 | 0.33307328 | 0 | 0.274132 | 0.41664208 | 0.16225483 | 0.71703368 | 0 | 0 | 0.33392972 | 0 | 0 |
| ENSMUSG00 4930506C21 | 1915 | 0.26635097 | 0.36531111 | 0.10104132 | 0.30929168 | 0.67862242 | 0.18113271 | 0.54353991 | 0.25926957 | 0.23658051 | 0.64016621 | 0.52390294 | 0.49549152 | 0.15456475 | 0.41229738 | 0.17212138 | 0.67983324 | 0.58039823 | 0.43088073 |
| ENSMUSG00 Gm16835 | 2802 | 0.06067834 | 0.08322279 | 0 | 0.06504075 | 0.13742162 | 0.07073909 | 0 | 0.12267367 | 0.45566809 | 0.03365503 | 0 | 0.09959968 | 0.0528179 | 0.11602734 | 0.22056507 | 0 | 0 | 0.17668878 |
| ENSMUSG00 Gm15910 | 2968 | 0.40099223 | 0.54997703 | 0.45635406 | 0.72148554 | 0.42164084 | 0.25043497 | 0.03188186 | 0 | 0.18039916 | 0 | 0 | 0 | 0.36566785 | 0.01564828 | 0.374812 | 0.23222066 | 0.51652691 | 0.14827252 |
| ENSMUSG00 Gm15608 | 3923 | 0.20586244 | 0.03396675 | 0.14796899 | 0.15097975 | 0.03680749 | 0.1389447 | 0.03618099 | 0.02920653 | 0.27296697 | 0.07211425 | 0.01217817 | 0.11382241 | 0.0503002 | 0.01183892 | 0.17854358 | 0.08784488 | 0.00976964 | 0 |
| ENSMUSG00 Gm12720 | 469 | 0 | 0 | 0 | 0 | 0 | 0 | 0 | 0 | 0 | 0 | 0 | 0 | 0 | 0 | 0 | 0 | 0 | 0 |
| ENSMUSG00 D830044I16 | 2332 | 0.58326142 | 1.01424335 | 1.53500911 | 1.75835934 | 1.21774592 | 1.12619849 | 1.82596102 | 1.91617113 | 3.26736939 | 1.88036681 | 3.11397671 | 2.72855297 | 3.34238548 | 3.34588623 | 2.61485003 | 2.82417859 | 2.9254206 | 1.9814601 |
| ENSMUSG00 2900089D17 | 2893 | 0.91093015 | 0.40302499 | 0.63539376 | 0.29922566 | 0.5157586 | 0.2911844 | 1.06302251 | 0.5280664 | 0.56946444 | 0.4563497 | 0.41284962 | 0.42445371 | 0.35809549 | 0.40134884 | 0.48422153 | 0.48971741 | 0.2517109 | 0.49437842 |
| ENSMUSG00 Gm13383 | 1169 | 0 | 0.08549075 | 0 | 0 | 0 | 0 | 0 | 0.09801301 | 0 | 0 | 0 | 0 | 0 | 0 | 0.17622565 | 0 | 0 | 0 |
| ENSMUSG00 Gm11723 | 463 | 0 | 0 | 0 | 0 | 0 | 0 | 0 | 0 | 0 | 0 | 0 | 0 | 0 | 0 | 0 | 0 | 0 | 0 |
| ENSMUSG00 4933401P06 | 1488 | 0 | 0 | 0 | 0 | 0 | 0 | 0.57233079 | 0 | 0 | 0 | 0 | 0 | 0 | 0 | 0 | 0 | 0 | 0 |
| ENSMUSG00 Gm16990 | 4324 | 0 | 0 | 0 | 0 | 0 | 0 | 0 | 0 | 0 | 0 | 0 | 0 | 0 | 0 | 0 | 0 | 0 | 0 |
| ENSMUSG00 Gm15656 | 613 | 0.0693396 | 0 | 0.07891277 | 0 | 0 | 0 | 0 | 0.12460816 | 0 | 0 | 0.46761786 | 0.27315985 | 0 | 0 | 0 | 0.12492857 | 0 | 0 |
| ENSMUSG00 A330076H08 | 4519 | 0 | 0 | 0 | 0.02016422 | 0 | 0 | 0.01046972 | 0 | 0 | 0.04173552 | 0.07400414 | 0 | 0 | 0 | 0.03646962 | 0 | 0.06784916 | 0 |
| ENSMUSG00 Gm14552 | 2400 | 0 | 0 | 0 | 0 | 0 | 0 | 0 | 0 | 0 | 0 | 0 | 0 | 0 | 0 | 0 | 0 | 0 | 0 |
| ENSMUSG00 Gm15700 | 488 | 0.87100771 | 0.06826413 | 0.29737826 | 0.0933628 | 0.09863099 | 0 | 0.19390442 | 0 | 0.42199299 | 0.09662029 | 0.09789951 | 0 | 0.30326998 | 0.57103386 | 0 | 0 | 0.15707499 | 0 |
| ENSMUSG00 4930550C17 | 1406 | 0 | 0 | 0 | 0 | 0 | 0 | 0 | 0 | 0 | 0 | 0 | 0 | 0 | 0 | 0 | 0 | 0 | 0 |
| ENSMUSG00 Gm16728 | 1375 | 0 | 0 | 0 | 0 | 0 | 0 | 0 | 0 | 0 | 0 | 0 | 0 | 0 | 0 | 0 | 0 | 0 | 0 |
| ENSMUSG00 Gm14152 | 1235 | 0 | 0 | 0 | 0 | 0 | 0 | 0 | 0 | 0 | 0 | 0 | 0 | 0 | 0 | 0 | 0 | 0 | 0 |
| ENSMUSG00 Gm13523 | 486 | 0.08745921 | 0.06854505 | 0.99534013 | 0.28124101 | 1.28747944 | 1.01960357 | 0 | 0.47151114 | 0.76271325 | 0.87316117 | 0.49151192 | 0.91877634 | 1.11656603 | 0.47781983 | 1.44120669 | 0.23636177 | 1.10404971 | 0.9054997 |
| ENSMUSG00 2810001G20 | 1249 | 9.42668836 | 6.32118169 | 9.25642414 | 7.76981781 | 11.9462741 | 5.59402357 | 10.7959275 | 10.9164833 | 11.0797993 | 8.53167249 | 12.4696849 | 8.89296786 | 9.4003173 | 9.51936459 | 10.9848824 | 9.28907449 | 11.0774965 | 6.38616531 |
| ENSMUSG00 2210416O15 | 1501 | 6.03171385 | 8.1229309 | 9.41044028 | 6.34394288 | 6.50951399 | 7.49398436 | 5.7683013 | 3.7658065 | 5.98178577 | 2.04183592 | 4.77431297 | 4.64820641 | 3.05654118 | 2.81573081 | 2.77238984 | 0.71428182 | 2.09377505 | 3.40829242 |
| ENSMUSG00 Gm16727 | 2492 | 0.42641629 | 0.34756631 | 0.77646116 | 0.27424385 | 0.4056061 | 0.17896252 | 0.93030548 | 0.78162621 | 0.64457228 | 0.54870401 | 0.74768193 | 1.11989692 | 0.71266012 | 0.20501002 | 1.02507884 | 1.44434875 | 2.5837953 | 2.38402228 |
| ENSMUSG00 Gm12426 | 368 | 0 | 0 | 0 | 0 | 0 | 0 | 0 | 0 | 0 | 0 | 0 | 0 | 0 | 0 | 0 | 0 | 0 | 0 |
| ENSMUSG00 Gm12116 | 2766 | 0 | 0 | 0 | 0 | 0 | 0 | 0.08552545 | 0 | 0.05956112 | 0 | 0.06908888 | 0 | 0.05350533 | 0 | 0.11916574 | 0.02768663 | 0.04156865 | 0 |
| ENSMUSG00 Gm16091 | 961 | 0.66345228 | 0.90128538 | 1.10740652 | 0.94820071 | 0.85144922 | 1.28909297 | 0.59079308 | 1.19227063 | 1.07144942 | 0.6868989 | 0.84513454 | 1.2196971 | 0.82134306 | 1.25655179 | 1.11471431 | 0.75704635 | 0.79763365 | 0.85862289 |
| ENSMUSG00 Gm11459 | 419 | 0 | 0 | 0 | 0 | 0.11487333 | 0 | 0 | 0 | 0 | 0 | 0 | 0 | 0 | 0 | 0 | 0.0913857 | 0 | 0 |
| ENSMUSG00 Gm13658 | 926 | 0 | 0 | 0 | 0 | 0 | 0 | 0 | 0 | 0 | 0 | 0 | 0 | 0 | 0 | 0 | 0 | 0 | 0 |
| ENSMUSG00 Gm15241 | 469 | 0 | 0 | 0 | 0.19429017 | 0 | 0.10565615 | 0 | 0 | 0 | 0 | 0 | 0 | 0 | 0 | 0 | 0 | 0 | 0 |
| ENSMUSG00 Gm16141 | 788 | 0 | 0.16910099 | 0 | 0 | 0 | 0 | 0.06004147 | 0.04846752 | 0 | 0 | 0 | 0 | 0 | 0.35363518 | 0 | 0 | 0.09727487 | 0 |
| ENSMUSG00 Gm14174 | 159 | 0 | 0 | 0 | 0 | 0 | 0 | 0 | 0 | 0 | 0 | 0 | 0 | 0 | 0 | 0 | 0 | 0 | 0 |
| ENSMUSG00 Gm2863 | 1907 | 0 | 0 | 0 | 0 | 0 | 0 | 0 | 0 | 0 | 0 | 0 | 0 | 0 | 0 | 0 | 0 | 0 | 0 |
| ENSMUSG00 1700112K13 | 699 | 0 | 0 | 0 | 0 | 0 | 0 | 0 | 0 | 0 | 0 | 0 | 0 | 0 | 0 | 0 | 0 | 0 | 0 |
| ENSMUSG00 A330009N23 | 1389 | 0.30601279 | 0.40771721 | 0.55721849 | 0.45921859 | 0.24256549 | 0.67782717 | 0 | 0.21997064 | 0.20756343 | 0.16972895 | 0.10318566 | 0.04018406 | 0.24861297 | 0.13374827 | 0.5042667 | 0.35837142 | 0.05518545 | 0.43563727 |
| ENSMUSG00 Gm11985 | 717 | 0 | 0 | 0 | 0 | 0 | 0 | 0 | 0.1065339 | 0 | 0 | 0 | 0 | 0 | 0 | 0 | 0 | 0 | 0 |
| ENSMUSG00 Gm13236 | 937 | 0 | 0 | 0 | 0 | 0 | 0 | 0 | 0 | 0 | 0 | 0 | 0 | 0 | 0 | 0 | 0 | 0 | 0 |
| ENSMUSG00 Gm16076 | 478 | 0 | 0 | 0 | 0 | 0 | 0 | 0 | 0 | 0 | 0 | 0 | 0 | 0 | 0 | 0 | 0 | 0 | 0 |
| ENSMUSG00 Gm13554 | 475 | 0 | 0 | 0 | 0 | 0 | 0 | 0 | 0 | 0 | 0 | 0 | 0 | 0 | 0 | 0 | 0 | 0 | 0 |
| ENSMUSG00 Gm2301 | 1489 | 1.25602938 | 1.45422305 | 1.33197767 | 1.13214146 | 1.55159996 | 1.0649345 | 0.47662202 | 0.84644008 | 1.10642084 | 0.69665243 | 1.02672846 | 0.33736801 | 2.31916334 | 3.21272062 | 1.93694353 | 2.64871226 | 1.80177353 | 1.92107021 |
| ENSMUSG00 4930577H14 | 788 | 0 | 0 | 0 | 0 | 0 | 0.06288418 | 0 | 0 | 0 | 0 | 0 | 0 | 0 | 0 | 0 | 0 | 0 | 0 |
| ENSMUSG00 Gm13561 | 1821 | 0.04668333 | 0.05488121 | 0.05312853 | 0 | 0 | 0 | 0.0259817 | 0 | 0 | 0.07767826 | 0 | 0 | 0.08127169 | 0.17853301 | 0.20363207 | 0.02102724 | 0 | 0.18124912 |
| ENSMUSG00 AV039307 | 1327 | 0.99296191 | 0.77822134 | 0.65615942 | 0.72101125 | 0.76169585 | 0.70949656 | 1.10526982 | 0.89221136 | 0.18622388 | 1.27914893 | 1.5480987 | 1.09360002 | 0.63198387 | 0.66498693 | 1.4903351 | 0.57710034 | 1.12639456 | 1.16070459 |
| ENSMUSG00 Gm14389 | 225 | 0 | 0 | 0 | 0 | 0 | 0 | 0 | 0 | 0 | 0 | 0 | 0 | 0 | 0 | 0 | 0 | 0 | 0 |
| ENSMUSG00 AI646519 | 349 | 0 | 0 | 0 | 0 | 0 | 0 | 0 | 0 | 0 | 0 | 0 | 0 | 0 | 0 | 0 | 0 | 0 | 0 |
| ENSMUSG00 Gm371 | 2425 | 0 | 0 | 0 | 0 | 0 | 0 | 0 | 0 | 0 | 0 | 0 | 0 | 0 | 0 | 0 | 0 | 0 | 0 |
| ENSMUSG00 Gm12319 | 355 | 5.62744584 | 6.28722229 | 7.90328102 | 4.74861589 | 8.54172166 | 6.28133243 | 5.99738184 | 8.7143228 | 9.86156011 | 9.29732177 | 7.40175413 | 10.062542 | 9.03260457 | 7.7188765 | 7.66000757 | 7.22667802 | 5.93787698 | 7.59280628 |
| ENSMUSG00 Gm14285 | 378 | 0 | 0 | 0 | 0 | 0 | 0 | 0 | 0 | 0 | 0 | 0 | 0 | 0 | 0 | 0 | 0 | 0 | 0 |
| ENSMUSG00 Gm13062 | 559 | 0.45622729 | 0.11918746 | 0.17307166 | 0 | 0.17220724 | 0 | 0 | 0.06832272 | 0 | 0.67478645 | 0 | 0 | 0.4412515 | 1.49551623 | 0 | 0.20549521 | 0 | 0.29521882 |
| ENSMUSG00 Gm15738 | 578 | 0.95599877 | 0.92215624 | 1.42275089 | 0.39412668 | 0.08327322 | 1.28597059 | 0.81855846 | 0.52861457 | 0.49879863 | 1.14205856 | 0.49593383 | 0.86910201 | 0.6827947 | 0.24105928 | 0.85539562 | 1.12619434 | 0.99462708 | 0.76137172 |
| ENSMUSG00 Gm13656 | 343 | 0 | 0 | 0 | 0 | 0 | 0 | 0 | 0 | 0 | 0 | 0 | 0 | 0 | 0 | 0.36036346 | 0 | 0 | 1.44338764 |
| ENSMUSG00 9830144P21 | 4878 | 0.11327743 | 0.20487635 | 0.31733353 | 0.2521829 | 0.13814 | 0.20316824 | 0.15518714 | 0.11744281 | 0.03377328 | 0.12565788 | 0.17629136 | 0.1830772 | 0.06067887 | 0.1999438 | 0.11824965 | 0.21979028 | 0.1807103 | 0.10149282 |
| ENSMUSG00 C130060C02 | 1139 | 0.93294943 | 0.61419735 | 0.76446317 | 0.96002201 | 0.67612887 | 1.21815324 | 0.37384909 | 0.33531521 | 0.39776266 | 0.786535 | 0.37750187 | 0.7350614 | 0.43311604 | 0.97862871 | 0.90433617 | 0.63873709 | 0.1345963 | 0 |
| ENSMUSG00 Gm15533 | 463 | 0.64262685 | 0.35975048 | 0.41791387 | 0.5904239 | 0.20791328 | 0.21405069 | 1.43062096 | 1.97973576 | 3.46927455 | 1.01837372 | 0.41274262 | 0.84386531 | 1.27858101 | 2.00622408 | 1.86875307 | 0.49620658 | 0.74500361 | 1.42572199 |
| ENSMUSG00 Gm15606 | 591 | 0 | 0 | 0 | 0 | 0 | 0 | 0 | 0 | 0 | 0 | 0 | 0 | 0 | 0 | 0 | 0 | 0 | 0 |
| ENSMUSG00 Gm15938 | 633 | 0 | 0 | 0 | 0.07197637 | 0 | 0.07828236 | 0.14948714 | 0.18100665 | 0.13013117 | 0.14897537 | 0.15094774 | 0 | 0.07793352 | 0.36685693 | 0.06508935 | 0 | 0 | 0 |
| ENSMUSG00 Gm11418 | 622 | 0 | 0 | 0 | 0 | 0 | 0 | 0 | 0 | 0 | 0 | 0 | 0 | 0 | 0 | 0 | 0 | 0 | 0 |
| ENSMUSG00 D930049A15 | 3466 | 0.42922134 | 0.27872877 | 0.62804641 | 0.52580547 | 0.13886879 | 0.52898186 | 0.35491334 | 0.46280465 | 0.52285248 | 0.57135878 | 0.57892333 | 0.49921683 | 0.48392533 | 0.57619612 | 0.41605726 | 0.60761206 | 0.38702282 | 0.36503446 |
| ENSMUSG00 Gm14246 | 678 | 0.752304 | 0.34393844 | 1.56964258 | 0.20159754 | 0.35495519 | 0.73086628 | 0.48847899 | 0 | 0.66821781 | 0.6954381 | 0 | 0 | 0.80037034 | 0 | 0 | 0 | 0 | 0 |
| ENSMUSG00 Gm13799 | 834 | 0 | 0 | 0 | 0 | 0 | 0 | 0 | 0 | 0 | 0 | 0 | 0 | 0 | 0 | 0 | 0 | 0 | 0 |
| ENSMUSG00 Gm16341 | 373 | 0 | 0 | 0 | 0.12214757 | 0 | 0 | 0 | 0 | 0 | 0.25281878 | 0.128083 | 0 | 0 | 0 | 0 | 0 | 0.41100586 | 0 |
| ENSMUSG00 Gm13584 | 600 | 0 | 0 | 0 | 0 | 0 | 0 | 0 | 0 | 0 | 0 | 0 | 0 | 0 | 0 | 0 | 0 | 0 | 0 |
| ENSMUSG00 Gm11412 | 2798 | 0 | 0 | 0 | 0 | 0 | 0 | 0 | 0 | 0 | 0 | 0 | 0 | 0 | 0 | 0 | 0 | 0 | 0 |
| [illegible] | | | | | | | | | | | | | | | | | | | |

| | | | | | | | | | | | | | | | | | | | |
|---|---|---|---|---|---|---|---|---|---|---|---|---|---|---|---|---|---|---|---|
| ENSMUSG00 5S_rRNA | 134 | 0 | 0 | 0 | 0 | 0 | 0 | 0 | 0 | 0 | 0 | 0 | 0 | 0 | 0 | 0 | 0 | 0 | 0 |
| ENSMUSG00 7SK | 321 | 0 | 0 | 0 | 0 | 0 | 0 | 0 | 0 | 0 | 0 | 0 | 0 | 0 | 0 | 0 | 0 | 0 | 0 |
| ENSMUSG00 U7 | 63 | 0 | 0 | 0 | 0 | 0 | 0 | 0 | 0 | 0 | 0 | 0 | 0 | 0 | 0 | 0 | 0 | 0 | 0 |
| ENSMUSG00 AC124515.1 | 109 | 0 | 0 | 0 | 0 | 0 | 0 | 0 | 0 | 0 | 0 | 0 | 0 | 0 | 0 | 0 | 0 | 0 | 0 |
| ENSMUSG00 U1 | 113 | 0 | 0 | 0 | 0 | 0 | 0 | 0 | 0 | 0 | 0 | 0 | 0 | 0 | 0 | 0 | 0 | 0 | 0 |
| ENSMUSG00 Mir1948 | 84 | 0 | 0 | 0 | 0 | 0 | 0 | 0 | 0 | 0 | 0 | 0 | 0 | 0 | 0 | 0 | 0 | 0 | 0 |
| ENSMUSG00 U6 | 94 | 0 | 0 | 0 | 0 | 0 | 0 | 0 | 0 | 0 | 0 | 0 | 0 | 0 | 0 | 0 | 0 | 0 | 0 |
| ENSMUSG00 SNORA17 | 133 | 0 | 0 | 0 | 0 | 0 | 0 | 0 | 0 | 0 | 0 | 0 | 0 | 0 | 0 | 0 | 0 | 0 | 0 |
| ENSMUSG00 7SK | 83 | 0 | 0 | 0 | 0 | 0 | 0 | 0 | 0 | 0 | 0 | 0 | 0 | 0 | 0 | 0 | 0 | 0 | 0 |
| ENSMUSG00 5S_rRNA | 119 | 0 | 0 | 0 | 0 | 0 | 0 | 0 | 0 | 0 | 0 | 0 | 0 | 0 | 0 | 0 | 0 | 0 | 0 |
| ENSMUSG00 7SK | 268 | 0 | 0 | 0 | 0 | 0 | 0 | 0 | 0 | 0 | 0 | 0 | 0 | 0 | 0 | 0 | 0 | 0 | 0 |
| ENSMUSG00 5S_rRNA | 110 | 0 | 0 | 0 | 0 | 0 | 0 | 0 | 0 | 0 | 0 | 0 | 0 | 0 | 0 | 0 | 0 | 0 | 0 |
| ENSMUSG00 7SK | 296 | 0 | 0 | 0 | 0 | 0 | 0 | 0 | 0 | 0 | 0 | 0 | 0 | 0 | 0 | 0 | 0 | 0 | 0 |
| ENSMUSG00 7SK | 284 | 0 | 0 | 0 | 0 | 0 | 0 | 0 | 0 | 0 | 0 | 0 | 0 | 0 | 0 | 0 | 0 | 0 | 0 |
| ENSMUSG00 SNORA64 | 132 | 0.32200891 | 0 | 0 | 0 | 0 | 0 | 0 | 0 | 0 | 0 | 0 | 0 | 0 | 0 | 0 | 0 | 0 | 0 |
| ENSMUSG00 SNORA38 | 150 | 0 | 0 | 0 | 0 | 0 | 0 | 0 | 0 | 0 | 0 | 0 | 0 | 0 | 0 | 0 | 0 | 0 | 0 |
| ENSMUSG00 7SK | 285 | 0 | 0 | 0 | 0 | 0 | 0 | 0 | 0 | 0 | 0 | 0 | 0 | 0 | 0 | 0 | 0 | 0 | 0 |
| ENSMUSG00 U6 | 102 | 0 | 0 | 0 | 0 | 0 | 0 | 0 | 0 | 0 | 0 | 0 | 0 | 0 | 0 | 0 | 0 | 0 | 0 |
| ENSMUSG00 SNORA70 | 133 | 0 | 0 | 0 | 0 | 0 | 0 | 0 | 0 | 0 | 0 | 0 | 0 | 0 | 0 | 0 | 0 | 0 | 0 |
| ENSMUSG00 SCARNA6 | 271 | 0.78422834 | 1.47510971 | 1.78500112 | 1.68121935 | 0.7104343 | 3.29132548 | 1.92044084 | 2.95955886 | 2.12771668 | 1.91386618 | 1.41033088 | 2.2655804 | 2.54851233 | 2.57070594 | 1.9764584 | 4.38010638 | 2.2628072 | 2.63881324 |
| ENSMUSG00 U7 | 51 | 0 | 0 | 0 | 0 | 0 | 0 | 0 | 0 | 0 | 0 | 0 | 0 | 0 | 0 | 0 | 0 | 0 | 0 |
| ENSMUSG00 U7 | 65 | 0 | 0 | 0 | 0 | 0 | 0 | 0 | 0 | 0 | 0 | 0 | 0 | 0 | 0 | 0 | 0 | 0 | 0 |
| ENSMUSG00 SNORA48 | 143 | 0 | 0 | 0 | 0 | 0 | 0 | 0 | 0 | 0 | 0 | 0 | 0 | 0 | 0 | 0 | 0 | 0 | 0 |
| ENSMUSG00 7SK | 306 | 0 | 0 | 0 | 0 | 0 | 0 | 0 | 0 | 0 | 0 | 0 | 0 | 0 | 0 | 0 | 0 | 0 | 0 |
| ENSMUSG00 SNORA32 | 69 | 0 | 0 | 1.40213131 | 0 | 0 | 0 | 0 | 0 | 0 | 0 | 0 | 0 | 2.14486597 | 0 | 0 | 0 | 0 | 0 |
| ENSMUSG00 U6 | 105 | 0 | 0 | 0 | 0 | 0 | 0 | 0 | 0 | 0 | 0 | 0 | 0 | 0 | 0 | 0 | 0 | 0 | 0 |
| ENSMUSG00 U6 | 108 | 0 | 0 | 0 | 0 | 0 | 0 | 0 | 0 | 0 | 0 | 0 | 0 | 0 | 0 | 0 | 0 | 0 | 0 |
| ENSMUSG00 7SK | 258 | 0 | 0 | 0 | 0 | 0 | 0 | 0 | 0 | 0 | 0 | 0 | 0 | 0 | 0 | 0 | 0 | 0 | 0 |
| ENSMUSG00 5S_rRNA | 92 | 0 | 0 | 0 | 0 | 0 | 0 | 0 | 0 | 0 | 0 | 0 | 0 | 0 | 0 | 0 | 0 | 0 | 0 |
| ENSMUSG00 7SK | 277 | 0 | 0 | 0 | 0.1644803 | 0 | 0 | 0 | 0 | 0 | 0 | 0 | 0 | 0 | 0 | 0 | 0 | 0.13836208 | 0 |
| ENSMUSG00 snoU13 | 101 | 0 | 0.32983064 | 0.95789169 | 0.45109945 | 0 | 0 | 0 | 1.13442779 | 0 | 0.93367729 | 0 | 0 | 0 | 0 | 0 | 0 | 0.75893657 | 0 |
| ENSMUSG00 SNORA17 | 124 | 0 | 0 | 0 | 0 | 0 | 0 | 0 | 0 | 0 | 0 | 0 | 0 | 0 | 0 | 0 | 0 | 0 | 0 |
| ENSMUSG00 U3 | 88 | 0 | 0 | 0 | 0 | 0 | 0 | 0 | 0 | 0 | 0 | 0 | 0 | 0 | 0 | 0 | 0 | 0 | 0 |
| ENSMUSG00 U6 | 102 | 0 | 0 | 0 | 0 | 0 | 0 | 0 | 0 | 0 | 0 | 0 | 0 | 0 | 0 | 0 | 0 | 0 | 0 |
| ENSMUSG00 U6 | 99 | 0 | 0 | 0 | 0 | 0 | 0 | 0 | 0 | 0 | 0 | 0 | 0 | 0 | 0 | 0 | 0 | 0 | 0 |
| ENSMUSG00 U6 | 100 | 0 | 0 | 0 | 0 | 0 | 0 | 0 | 0 | 0 | 0 | 0 | 0 | 0 | 0 | 0 | 0 | 0 | 0 |
| ENSMUSG00 SCARNA4 | 126 | 0 | 0 | 0 | 0 | 0 | 0 | 0 | 0 | 0 | 0 | 0 | 0 | 0 | 0 | 0 | 0 | 0 | 0 |
| ENSMUSG00 7SK | 87 | 0.48856524 | 0.76581366 | 0 | 1.04738033 | 0 | 0 | 0.5438239 | 2.63395879 | 0 | 0.54196211 | 0.54913745 | 0.64155934 | 0 | 0 | 0.4735811 | 0.44012192 | 0.44053215 | 0 |
| ENSMUSG00 SNORA17 | 116 | 0 | 0 | 0 | 0 | 0 | 0 | 0.40786792 | 0 | 0 | 0.81294316 | 0 | 0 | 0 | 0 | 0 | 0 | 0 | 0.47421644 |
| ENSMUSG00 Mir1982 | 73 | 0 | 0 | 0 | 0 | 0 | 0 | 0 | 0 | 0 | 0 | 0 | 0 | 0.67577969 | 0.63622038 | 0 | 0 | 0 | 0 |
| ENSMUSG00 SNORD113 | 75 | 7.36756384 | 1.33251577 | 1.28996081 | 1.21496118 | 3.20879491 | 1.32140623 | 0 | 0.50923203 | 0 | 0 | 0 | 0 | 0 | 0 | 0 | 0 | 0 | 0 |
| ENSMUSG00 U6 | 96 | 0 | 0 | 0 | 0 | 0 | 0 | 0 | 0 | 0 | 0 | 0 | 0 | 0 | 0 | 0 | 0 | 0 | 0 |
| ENSMUSG00 U7 | 61 | 0 | 0 | 0 | 0 | 0 | 0 | 0 | 0 | 0 | 0 | 0 | 0 | 0 | 0 | 0 | 0 | 0 | 0 |
| ENSMUSG00 U1 | 129 | 0 | 0 | 0 | 0 | 0 | 0 | 0 | 0 | 0 | 0 | 0 | 0 | 0 | 0 | 0 | 0 | 0 | 0 |
| ENSMUSG00 U6 | 102 | 0 | 0 | 0 | 0 | 0 | 0 | 0 | 0 | 0 | 0 | 0 | 0 | 0 | 0 | 0 | 0 | 0 | 0 |
| ENSMUSG00 U6 | 105 | 0 | 0 | 0 | 0 | 0 | 0 | 0 | 0 | 0 | 0 | 0 | 0 | 0 | 0 | 0 | 0 | 0 | 0 |
| ENSMUSG00 U6 | 98 | 0.86745257 | 0 | 0.98721491 | 1.39472585 | 0 | 0 | 0.96556488 | 0 | 0 | 0 | 0 | 0.56954758 | 0 | 0 | 0.84084808 | 1.95360242 | 0 | 0 |
| ENSMUSG00 U6 | 73 | 0 | 0 | 0 | 0 | 0 | 0 | 0 | 0 | 0 | 0 | 0 | 0 | 0 | 0 | 0 | 0 | 0 | 0 |
| ENSMUSG00 5S_rRNA | 108 | 2.36139867 | 0.92535817 | 0.89580612 | 2.95303065 | 0.89133192 | 0.91764321 | 0 | 1.76816678 | 2.66949639 | 0.43658059 | 1.32708218 | 0.51681169 | 0 | 0 | 0 | 0.35454266 | 1.7743656 | 0 |
| ENSMUSG00 SNORA70 | 122 | 0 | 0 | 0 | 0.37345118 | 0 | 0 | 0.38780884 | 0 | 2.02556635 | 0 | 0.78319604 | 0.45750543 | 0 | 0.38068924 | 3.03945904 | 0 | 0 | 0 |
| ENSMUSG00 snoR38 | 71 | 1.79599335 | 0.46919569 | 0 | 0 | 0 | 0 | 2.66550304 | 1.61376348 | 0.58009177 | 0 | 2.01866022 | 0 | 1.38963147 | 0 | 0 | 2.15721732 | 0.539807 | 0 |
| ENSMUSG00 SCARNA20 | 128 | 0.66414338 | 0.78077096 | 0 | 1.06783698 | 0 | 0 | 0.3696303 | 2.98378144 | 0.32176965 | 0.36836487 | 1.49296745 | 0.43605986 | 0.77081121 | 0 | 0 | 1.79487222 | 0.89827258 | 0 |
| ENSMUSG00 SNORA17 | 133 | 0 | 0 | 0 | 0 | 0 | 0 | 0 | 0 | 0 | 0 | 0 | 0 | 0 | 0 | 0 | 0 | 0 | 0 |
| ENSMUSG00 5S_rRNA | 99 | 0 | 0 | 0 | 0 | 0 | 0 | 0 | 0 | 0 | 0 | 0 | 0 | 0 | 0 | 0 | 0 | 0 | 0 |
| ENSMUSG00 SNORA17 | 113 | 0 | 0 | 0 | 0 | 0 | 0 | 0 | 0 | 0 | 0 | 0 | 0 | 0 | 0 | 0 | 0 | 0 | 0 |
| ENSMUSG00 U3 | 130 | 0 | 0 | 0 | 0 | 0 | 0 | 0 | 0 | 0 | 0 | 0 | 0 | 0 | 0 | 0 | 0 | 0 | 0 |
| ENSMUSG00 U7 | 61 | 0 | 0 | 0 | 0 | 0 | 0 | 0 | 0 | 0 | 0 | 0 | 0 | 0 | 0 | 0 | 0 | 0 | 0 |
| ENSMUSG00 7SK | 273 | 0 | 0 | 1.06315451 | 0 | 0 | 0 | 0 | 0 | 0 | 0 | 0.3499997 | 0 | 0 | 0 | 0 | 0 | 0 | 0 |
| ENSMUSG00 Mir1932 | 89 | 4.29827623 | 3.74302182 | 0.54352281 | 3.07153108 | 2.70404066 | 1.67031686 | 0 | 4.29128117 | 0.46276984 | 0.52978318 | 1.61039186 | 6.27142276 | 0.55429121 | 1.04368736 | 0 | 0.86046309 | 3.87569295 | 0 |
| ENSMUSG00 U6 | 101 | 0 | 0.98949191 | 0 | 0 | 0 | 0.49062112 | 1.4053271 | 1.51257039 | 0 | 0 | 0 | 0 | 0 | 0 | 0 | 0 | 0 | 0 |
| ENSMUSG00 7SK | 269 | 0 | 0 | 0 | 0 | 0 | 0 | 0 | 0 | 0 | 0 | 0 | 0 | 0 | 0 | 0 | 0 | 0 | 0 |
| ENSMUSG00 U6 | 105 | 0 | 0 | 0 | 0 | 0 | 0 | 0 | 0 | 0 | 0 | 0 | 0 | 0 | 0 | 0 | 0 | 0 | 0 |
| ENSMUSG00 SNORA17 | 118 | 0 | 0 | 0 | 0 | 0 | 0 | 0 | 0 | 0 | 0 | 0 | 0 | 0 | 0 | 0 | 0 | 0 | 0 |
| ENSMUSG00 SNORA17 | 136 | 0 | 0 | 0 | 0 | 0 | 0 | 0 | 0 | 0 | 0 | 0 | 0 | 0 | 0 | 0 | 0 | 0 | 0 |
| ENSMUSG00 U1 | 147 | 0 | 0 | 0 | 0 | 0 | 0 | 0 | 0 | 0 | 0 | 0 | 0 | 0 | 0 | 0 | 0 | 0 | 0 |
| ENSMUSG00 7SK | 287 | 0.14810166 | 0 | 0 | 0 | 0.50312115 | 0 | 0.16485254 | 0 | 0.28701405 | 0 | 0 | 0 | 0.17188821 | 0.64730435 | 0.28711886 | 0 | 0 | 0 |
| ENSMUSG00 U7 | 61 | 0 | 0 | 0 | 0 | 0 | 0 | 0 | 0 | 0 | 0 | 0 | 0 | 0 | 0 | 0 | 0 | 0 | 0 |
| ENSMUSG00 SNORA5 | 91 | 0 | 0 | 0 | 0 | 0 | 0 | 0 | 0 | 0 | 0 | 0 | 0 | 0 | 0 | 0 | 0 | 0 | 0 |
| ENSMUSG00 RNaseP_nuc | 289 | 0 | 0 | 0 | 0 | 0 | 0 | 0 | 0 | 0 | 0.16315122 | 0 | 0 | 0 | 0 | 0 | 0 | 0 | 0 |
| ENSMUSG00 7SK | 202 | 0 | 0 | 0 | 0 | 0 | 0 | 0 | 0 | 0 | 0 | 0 | 0 | 0 | 0 | 0 | 0 | 0 | 0 |
| ENSMUSG00 SNORA17 | 101 | 0 | 0 | 0 | 0 | 0 | 0 | 0 | 0 | 0 | 0 | 0 | 0 | 0 | 0 | 0 | 0 | 0 | 0 |
| ENSMUSG00 7SK | 276 | 0 | 0 | 0 | 0 | 0 | 0 | 0 | 0 | 0 | 0 | 0 | 0 | 0 | 0 | 0 | 0 | 0 | 0 |
| ENSMUSG00 U7 | 58 | 0 | 0 | 0 | 0 | 0 | 0 | 0 | 0 | 0 | 0 | 0 | 0 | 0 | 0 | 0 | 0 | 0 | 0 |
| ENSMUSG00 5S_rRNA | 107 | 0 | 0 | 0 | 0 | 0 | 0 | 0 | 0 | 0 | 0 | 0 | 0 | 0 | 0 | 0 | 0 | 0 | 0 |
| ENSMUSG00 7SK | 322 | 0 | 0 | 0 | 0 | 0 | 0 | 0 | 0 | 0 | 0 | 0 | 0 | 0 | 0 | 0 | 0 | 0 | 0 |
| ENSMUSG00 U6 | 106 | 0 | 0 | 0 | 0 | 0 | 0 | 0 | 0 | 0 | 0 | 0 | 0 | 0 | 0 | 0 | 0 | 0 | 0 |
| ENSMUSG00 AL671963.1 | 90 | 0 | 0 | 0.53748367 | 0 | 0 | 0 | 0 | 0 | 0 | 0 | 0 | 0 | 0 | 1.03209083 | 0.45779507 | 0.42545119 | 0 | 0 |
| ENSMUSG00 SNORA70 | 129 | 0 | 0 | 0 | 0 | 0.74623137 | 0.38412972 | 0.36676495 | 1.48032567 | 2.55420253 | 1.09652798 | 0 | 0 | 0 | 0 | 1.59695953 | 1.18730566 | 0.29710308 | 0 |
| ENSMUSG00 snoU13 | 109 | 0 | 0 | 0 | 0 | 0 | 0 | 0 | 0 | 0 | 0 | 0 | 0 | 0 | 0 | 0 | 0 | 0 | 0 |
| ENSMUSG00 SNORA17 | 120 | 0 | 0 | 0 | 0 | 0 | 0 | 0 | 0 | 0 | 0 | 0 | 0 | 0 | 0 | 0 | 0 | 0 | 0 |
| ENSMUSG00 SNORA17 | 107 | 0 | 0 | 0 | 0 | 0 | 0 | 0 | 0 | 0 | 0 | 0 | 0 | 0 | 0 | 0 | 0 | 0 | 0 |
| ENSMUSG00 SNORA17 | 133 | 0 | 0 | 0 | 0 | 0 | 0 | 0 | 0 | 0 | 0 | 0 | 0 | 0 | 0 | 0 | 0 | 0 | 0 |
| ENSMUSG00 U3 | 89 | 0 | 0 | 0 | 0 | 0 | 0 | 0 | 0 | 0 | 0 | 0 | 0 | 0 | 0 | 0 | 0 | 0 | 0 |
| ENSMUSG00 SNORA17 | 131 | 0 | 0 | 0 | 0 | 0 | 0 | 0 | 0 | 0 | 0 | 0 | 0 | 0 | 0 | 0 | 0 | 0 | 0 |
| ENSMUSG00 U6 | 104 | 0 | 0 | 0 | 0 | 0 | 0 | 0 | 0 | 0 | 0 | 0 | 0 | 0 | 0 | 0 | 0 | 0 | 0 |
| ENSMUSG00 SNORA17 | 77 | 0 | 0 | 0 | 0 | 0 | 0 | 0 | 0 | 0 | 0 | 0 | 0 | 0 | 0 | 0 | 0 | 0 | 0 |
| ENSMUSG00 SNORA17 | 130 | 0 | 0 | 0 | 0 | 0 | 0 | 0 | 0 | 0 | 0 | 0 | 0 | 0 | 0 | 0 | 0 | 0 | 0 |
| ENSMUSG00 SNORA17 | 128 | 0 | 0 | 0 | 0 | 0 | 0 | 0 | 0 | 0 | 0 | 0 | 0 | 0 | 0 | 0 | 0 | 0 | 0 |
| ENSMUSG00 U6 | 107 | 0 | 0 | 0 | 0 | 0 | 0 | 0 | 0 | 0 | 0 | 0 | 0 | 0 | 0 | 0 | 0 | 0 | 0 |
| ENSMUSG00 7SK | 274 | 0 | 0 | 0 | 0 | 0 | 0 | 0 | 0 | 0 | 0 | 0 | 0 | 0 | 0 | 0 | 0 | 0 | 0 |
| ENSMUSG00 U7 | 59 | 0 | 0 | 0 | 0 | 0 | 0 | 0 | 0 | 0 | 0 | 0 | 0 | 0 | 0 | 0 | 0 | 0 | 0 |
| ENSMUSG00 U1 | 150 | 0 | 0 | 0 | 0 | 0 | 0 | 0 | 0 | 0 | 0 | 0 | 0 | 0 | 0 | 0 | 0 | 0 | 0 |
| ENSMUSG00 5S_rRNA | 100 | 0 | 0 | 0 | 0 | 0 | 0 | 0 | 0 | 0 | 0 | 0 | 0 | 0 | 0 | 0 | 0 | 0 | 0 |
| ENSMUSG00 Mir1968 | 67 | 0 | 0 | 0 | 0 | 0 | 0.73959304 | 0 | 0 | 0 | 0 | 1.66613918 | 0 | 0 | 0 | 0 | 0 | 0 | 0 |
| ENSMUSG00 SNORA17 | 67 | 0 | 0 | 0 | 0 | 0 | 0 | 0 | 0 | 0 | 0 | 0 | 0 | 0 | 0 | 0 | 0 | 0 | 0 |
| ENSMUSG00 SNORA48 | 143 | 0 | 0 | 0 | 0 | 0 | 0 | 0.99257368 | 0 | 0.86405278 | 0 | 0 | 0 | 0 | 0 | 0 | 0 | 0 | 0 |
| ENSMUSG00 U6 | 107 | 0 | 0 | 0 | 0 | 0 | 0 | 0 | 0 | 0 | 0 | 0 | 0 | 0 | 0 | 0 | 0 | 0 | 0 |
| ENSMUSG00 SNORA70 | 126 | 0 | 0 | 0 | 0 | 0 | 0 | 0 | 0 | 0 | 0 | 0 | 0 | 0 | 0 | 0 | 0 | 0 | 0 |
| ENSMUSG00 SNORA74 | 140 | 0 | 0.71384773 | 1.72762608 | 0.97630809 | 1.71899727 | 0.70789619 | 2.02768624 | 2.72802874 | 0.5883788 | 2.02074443 | 5.11874555 | 1.59473322 | 2.11422503 | 1.32697393 | 0 | 1.36752169 | 0.54751853 | 2.75045534 |
| ENSMUSG00 SNORA48 | 142 | 0 | 0 | 0 | 0 | 0 | 0 | 0 | 0 | 0 | 0 | 0 | 0 | 0 | 0 | 0 | 0 | 0 | 0 |
| ENSMUSG00 5S_rRNA | 130 | 0 | 0 | 0 | 0 | 0 | 0 | 0 | 0 | 0 | 0 | 0 | 0 | 0 | 0 | 0 | 0 | 0 | 0 |
| ENSMUSG00 U6 | 102 | 0 | 0 | 0 | 0 | 0 | 0 | 0 | 0 | 0 | 0 | 0 | 0 | 0 | 0 | 0 | 0 | 0 | 0 |
| ENSMUSG00 U1 | 157 | 0 | 0 | 0 | 0 | 0 | 0 | 0 | 0 | 0 | 0 | 0 | 0 | 0 | 0 | 0 | 0 | 0 | 0 |
| ENSMUSG00 U7 | 63 | 0 | 0 | 0 | 0 | 0 | 0 | 0 | 0 | 0 | 0 | 0 | 0 | 0 | 0 | 0 | 0 | 0 | 0 |
| ENSMUSG00 U6 | 103 | 0.41267161 | 0 | 0.93929185 | 0 | 0 | 0 | 0 | 0 | 0 | 0 | 0 | 0 | 0 | 0 | 0 | 0 | 0 | 1.06813799 |
| ENSMUSG00 U6 | 102 | 0 | 0 | 0 | 0 | 0 | 0 | 0 | 0 | 0 | 0 | 0 | 0 | 0 | 0 | 0 | 0 | 0 | 0 |
| ENSMUSG00 U6 | 100 | 0 | 0 | 0 | 0 | 0 | 0 | 0 | 0 | 0 | 0 | 0 | 0 | 0 | 0 | 0 | 0 | 0 | 0 |
| ENSMUSG00 7SK | 266 | 0.15979389 | 0 | 0 | 0 | 0 | 0.37257694 | 0 | 0 | 0.15483653 | 0.17725828 | 0 | 0 | 0 | 0 | 0 | 0 | 0 | 0 |
| ENSMUSG00 7SK | 302 | 0 | 0 | 0 | 0 | 0 | 0 | 0 | 0 | 0 | 0 | 0 | 0 | 0 | 0 | 0 | 0 | 0 | 0 |
| ENSMUSG00 U6 | 103 | 0.82534322 | 0.32342616 | 0 | 0 | 0 | 0 | 0.4593464 | 0 | 0 | 0 | 0 | 0 | 0 | 0 | 0 | 0 | 0.37209997 | 0 |
| ENSMUSG00 RNaseP_nuc | 355 | 0 | 0 | 0 | 0 | 0 | 0 | 0 | 0 | 0 | 0 | 0 | 0 | 0 | 0 | 0 | 0 | 0 | 0 |
| ENSMUSG00 5S_rRNA | 116 | 0 | 0 | 0 | 0 | 0 | 0 | 0 | 0 | 0 | 0.40647158 | 0 | 0 | 0 | 0 | 0 | 0 | 0 | 0 |
| ENSMUSG00 U7 | 61 | 0 | 0 | 0 | 0 | 0 | 0 | 0 | 0 | 0 | 0 | 0 | 0 | 0 | 0 | 0 | 0 | 0 | 0 |
| ENSMUSG00 U6 | 103 | 0 | 0 | 0 | 0 | 0 | 0 | 0 | 0 | 0 | 0 | 0 | 0 | 0 | 0 | 0 | 0 | 0 | 0 |
| ENSMUSG00 SCARNA4 | 123 | 0.69114107 | 1.35418269 | 1.96640367 | 2.59290496 | 3.52184807 | 1.61147101 | 3.07724741 | 0.31050734 | 1.00454916 | 1.91669525 | 5.04938586 | 3.630287 | 2.40643499 | 0.37759421 | 1.339888 | 1.86783451 | 5.29713047 | 0.44722851 |
| ENSMUSG00 U3 | 197 | 0 | 0 | 0 | 0 | 0 | 0 | 0 | 0 | 0 | 0 | 0 | 0 | 0 | 0 | 0 | 0 | 0 | 0 |
| ENSMUSG00 7SK | 231 | 0 | 0 | 0 | 0 | 0 | 0 | 0 | 0 | 0 | 0 | 0 | 0 | 0 | 0 | 0 | 0 | 0 | 0 |
| ENSMUSG00 SNORA17 | 121 | 0 | 0 | 0 | 0 | 0 | 0 | 0 | 0 | 0 | 0 | 0 | 0 | 0 | 0 | 0 | 0 | 0 | 0 |
| ENSMUSG00 U6 | 106 | 0 | 0 | 0 | 0 | 0 | 0 | 0 | 0 | 0 | 0 | 0 | 0 | 0 | 0 | 0 | 0 | 0 | 0 |
| ENSMUSG00 7SK | 267 | 0 | 0 | 0.18117427 | 0 | 0 | 0 | 0 | 0 | 0 | 0 | 0 | 0 | 0 | 0 | 0.92587766 | 0 | 0 | 0 |
| ENSMUSG00 RNase_MRP | 270 | 0 | 0 | 0 | 0 | 0 | 0 | 0 | 0 | 0 | 0.17463223 | 0 | 0 | 0 | 0 | 0 | 0 | 0 | 0 |
| ENSMUSG00 SNORA17 | 113 | 0 | 0 | 0 | 0 | 0 | 0 | 0 | 0 | 0 | 0 | 0 | 0 | 0 | 0 | 0 | 0 | 0 | 0 |
| ENSMUSG00 5S_rRNA | 110 | 0 | 0 | 0 | 0 | 0 | 0 | 0 | 0 | 0 | 0 | 0 | 0 | 0 | 0 | 0 | 0 | 0 | 0 |
| ENSMUSG00 5S_rRNA | 119 | 0 | 0 | 0 | 0 | 0 | 0 | 0 | 0 | 0 | 0 | 0 | 0 | 0 | 0 | 0 | 0 | 0 | 0 |
| ENSMUSG00 SNORA17 | 131 | 0 | 0 | 0 | 0 | 0 | 0 | 0 | 0 | 0 | 0 | 0 | 0 | 0 | 0 | 0 | 0 | 0 | 0 |
| ENSMUSG00 RNaseP_nuc | 153 | 0 | 0 | 0 | 0 | 0 | 0 | 0 | 0 | 0 | 0 | 0 | 0 | 0 | 0 | 0 | 0 | 0 | 0 |
| ENSMUSG00 SNORA17 | 118 | 0 | 0 | 0 | 0 | 0 | 0 | 0 | 0 | 0 | 0 | 0 | 0 | 0 | 0 | 0 | 0 | 0 | 0 |
| ENSMUSG00 SNORA17 | 127 | 0 | 0 | 0 | 0 | 0 | 0 | 0 | 0 | 0 | 0 | 0 | 0 | 0 | 0 | 0 | 0 | 0 | 0 |
| ENSMUSG00 U4 | 106 | 0 | 0 | 0 | 0 | 0 | 0 | 0 | 0 | 0 | 0 | 0 | 0 | 0 | 0 | 0 | 0 | 0 | 0 |
| ENSMUSG00 7SK | 102 | 0 | 0 | 0 | 0 | 0 | 0 | 0 | 0 | 0 | 0 | 0 | 0 | 0 | 0 | 0 | 0 | 0 | 0 |
| ENSMUSG00 SNORA17 | 130 | 0 | 0 | 0 | 0 | 0 | 0 | 0 | 0 | 0 | 0 | 0 | 0 | 0 | 0 | 0 | 0 | 0 | 0 |
| ENSMUSG00 7SK | 297 | 0 | 0 | 0 | 0 | 0 | 0 | 0 | 0 | 0 | 0 | 0 | 0 | 0 | 0 | 0 | 0 | 0 | 0 |
| ENSMUSG00 U3 | 121 | 0 | 0 | 0 | 0 | 0 | 0 | 0 | 0 | 0 | 0 | 0 | 0 | 0 | 0 | 0 | 0 | 0 | 0 |
| ENSMUSG00 U7 | 64 | 0 | 0 | 0 | 0 | 0 | 0 | 0 | 0 | 0 | 0 | 0 | 0 | 0 | 0 | 0 | 0 | 0 | 0 |
| ENSMUSG00 7SK | 275 | 0 | 0 | 0 | 0 | 0 | 0 | 0 | 0 | 0.2995383 | 0 | 0 | 0 | 0 | 0 | 0 | 0 | 0 | 0 |
| ENSMUSG00 SNORA48 | 143 | 0 | 0 | 0 | 0 | 0 | 0 | 0 | 0 | 0 | 0 | 0 | 0 | 0 | 0 | 0 | 0 | 0 | 0 |
| ENSMUSG00 SNORA17 | 94 | 0 | 0 | 0 | 0 | 0 | 0 | 0 | 0 | 0 | 0 | 0 | 0 | 0 | 0 | 0 | 0 | 0 | 0 |
| ENSMUSG00 SNORA17 | 130 | 0 | 0 | 0 | 0 | 0 | 0 | 0 | 0 | 0 | 0 | 0 | 0 | 0 | 0 | 0 | 0 | 0 | 0 |
| ENSMUSG00 U6 | 110 | 1.15923207 | 1.51422246 | 0 | 0 | 0.87512588 | 0 | 0 | 1.04161097 | 0 | 0.42864276 | 1.73727122 | 0.50741511 | 0 | 0 | 0.3745596 | 0 | 0 | 0 |
| ENSMUSG00 U7 | 61 | 0 | 0 | 0 | 0 | 0 | 0 | 0 | 0 | 0 | 0 | 0 | 0 | 0 | 0 | 0 | 0 | 0 | 0 |
| ENSMUSG00 Snora47 | 137 | 1.5512838 | 0.97263925 | 0 | 0 | 0 | 0 | 0 | 1.3938833 | 0.90189451 | 0.68833143 | 0 | 0 | 0 | 0 | 0.30074128 | 0 | 0 | 0 |
| ENSMUSG00 U6 | 73 | 0 | 0 | 0 | 0 | 0 | 0 | 0 | 0 | 0 | 0 | 0 | 0 | 0 | 0 | 0 | 0 | 0 | 0 |
| ENSMUSG00 SNORA17 | 105 | 0 | 0 | 0 | 0 | 0 | 0 | 0 | 0 | 0 | 0 | 0 | 0 | 0 | 0 | 0 | 0 | 0 | 0 |
| ENSMUSG00 7SK | 263 | 0 | 0 | 0 | 0 | 0 | 0 | 0 | 0 | 0 | 0 | 0 | 0 | 0 | 0 | 0 | 0 | 0 | 0 |
| ENSMUSG00 Mir1960 | 77 | 0 | 0 | 0 | 0 | 0.62508992 | 0 | 0 | 0 | 0 | 0 | 0 | 2.1746362 | 0 | 0 | 0 | 0 | 0 | 0 |
| ENSMUSG00 SNORA72 | 119 | 0 | 0 | 0 | 0 | 0 | 0 | 0 | 0 | 0 | 0 | 0 | 0 | 0 | 0 | 0 | 0 | 0 | 0 |
| ENSMUSG00 7SK | 265 | 0 | 0 | 0 | 0 | 0 | 0 | 0 | 0 | 0 | 0 | 0 | 0 | 0 | 0 | 0 | 0 | 0 | 0 |
| ENSMUSG00 AC141566.1 | 104 | 0 | 0 | 0 | 0 | 0 | 0 | 0 | 0 | 0 | 0 | 0 | 0 | 0 | 0 | 0 | 0 | 0 | 0 |
| ENSMUSG00 7SK | 311 | 0.13667259 | 0.21423083 | 0 | 0 | 0 | 0 | 0 | 0 | 0 | 0 | 0 | 0 | 0.95174117 | 0 | 0 | 0 | 0 | 0.70751263 |
| ENSMUSG00 U6 | 94 | 10.4002026 | 9.92298977 | 6.1753443 | 8.23976334 | 8.7047096 | 11.5974483 | 5.03326372 | 10.563856 | 6.57231635 | 6.52084194 | 9.1483963 | 8.90675467 | 12.0705755 | 12.3521509 | 9.64291734 | 8.96163152 | 6.11589844 | 8.77804895 |
| ENSMUSG00 U1 | 170 | 0 | 0 | 0 | 0 | 0 | 0 | 0 | 0 | 0 | 0 | 0 | 0 | 0 | 0 | 0 | 0 | 0 | 0 |
| ENSMUSG00 7SK | 277 | 0 | 0 | 0 | 0 | 0 | 0 | 0 | 0 | 0 | 0 | 0 | 0 | 0 | 0 | 0 | 0 | 0 | 0 |
| ENSMUSG00 7SK | 278 | 0 | 0 | 0 | 0 | 0 | 0 | 0 | 0 | 0 | 0 | 0 | 0 | 0 | 0 | 0 | 0 | 0 | 0 |
| ENSMUSG00 U6 | 108 | 0 | 0 | 0 | 0 | 0 | 0 | 0 | 0 | 0 | 0 | 0 | 0 | 0 | 0 | 0 | 0 | 0 | 0 |
| ENSMUSG00 SNORA17 | 136 | 0 | 0 | 0.71137545 | 0 | 0 | 0 | 0 | 0 | 0 | 0.34669635 | 0 | 0 | 0 | 0 | 0.30295262 | 0 | 0 | 0 |
| ENSMUSG00 SNORA17 | 111 | 0 | 0 | 0 | 0 | 0 | 0 | 0 | 0 | 0 | 0 | 0 | 0 | 0 | 0 | 0 | 0 | 0 | 0 |
| ENSMUSG00 U7 | 64 | 0 | 0 | 0 | 0 | 0 | 0 | 3.69630304 | 0 | 0.64353931 | 0 | 0.74648373 | 0 | 0.77081121 | 0 | 0 | 0 | 0 | 1.71903459 |
| ENSMUSG00 U6 | 104 | 0 | 0 | 0 | 0 | 0 | 0 | 0 | 0 | 0 | 0 | 0 | 0 | 0 | 0 | 0 | 0 | 0 | 0 |
| ENSMUSG00 U7 | 62 | 0 | 0 | 0 | 0 | 0 | 0 | 0 | 0 | 0 | 0 | 0 | 0 | 0 | 0 | 0 | 0 | 0 | 0 |
| ENSMUSG00 U7 | 59 | 0 | 0 | 0 | 0 | 0 | 0 | 0 | 0 | 0 | 0 | 0 | 0 | 0 | 0 | 0 | 0 | 0 | 0 |
| ENSMUSG00 SCARNA15 | 127 | 0 | 0 | 0 | 0 | 0 | 0 | 0 | 0 | 0 | 0 | 0 | 0 | 0 | 0 | 0 | 0 | 0 | 0 |
| ENSMUSG00 5S_rRNA | 121 | 0 | 0 | 0 | 0 | 0 | 0 | 0 | 0 | 0 | 0 | 0 | 0 | 0 | 0 | 0 | 0 | 0 | 0 |
| ENSMUSG00 7SK | 307 | 0 | 0 | 0 | 0 | 0 | 0 | 0 | 0 | 0 | 0 | 0 | 0 | 0 | 0 | 0 | 0 | 0 | 0 |
| ENSMUSG00 SNORA17 | 115 | 0.36961023 | 0 | 0 | 0 | 0 | 0 | 0 | 0 | 0 | 0 | 0 | 0 | 0 | 0 | 0 | 0 | 0 | 0 |
| ENSMUSG00 SNORA81 | 139 | 0 | 0 | 0 | 0 | 0 | 0 | 0 | 0 | 0 | 0 | 0 | 0 | 0 | 0 | 0 | 0 | 0 | 0 |
| ENSMUSG00 7SK | 297 | 0 | 0 | 0 | 0 | 0 | 0 | 0 | 0 | 0 | 0 | 0 | 0 | 0 | 0 | 0 | 0 | 0 | 0 |
| ENSMUSG00 snoU13 | 103 | 0 | 0 | 0 | 0 | 0 | 0 | 0 | 0 | 0 | 0 | 0 | 0 | 0 | 0 | 0 | 0 | 0 | 0 |
| ENSMUSG00 U6 | 102 | 0 | 0 | 0 | 0 | 0 | 0 | 0 | 0 | 0 | 0 | 0 | 0 | 0 | 0 | 0 | 0 | 0 | 0 |
| ENSMUSG00 U7 | 62 | 0 | 0 | 0 | 0 | 0 | 0 | 0 | 0 | 0 | 0 | 0 | 0 | 0 | 0 | 0 | 0 | 0 | 0 |
| ENSMUSG00 Mir1962 | 120 | 0 | 0 | 0 | 0 | 0 | 0 | 0 | 0 | 0 | 0 | 0 | 0 | 0 | 0 | 0 | 0 | 0 | 0 |
| ENSMUSG00 U1 | 165 | 0 | 0 | 0 | 0 | 0 | 0 | 0 | 0 | 0 | 0 | 0 | 0 | 0 | 0 | 0 | 0 | 0 | 0 |
| ENSMUSG00 SNORA22 | 120 | 0 | 0 | 0 | 0 | 0 | 0 | 0 | 0 | 0 | 0 | 0 | 0 | 0 | 0 | 0 | 0 | 0 | 0 |
| ENSMUSG00 SNORA17 | 109 | 0 | 0 | 0 | 0 | 0 | 0 | 0 | 0 | 0 | 0 | 0 | 0 | 0 | 0 | 0 | 0 | 0 | 0 |
| ENSMUSG00 SCARNA3 | 135 | 0 | 0 | 0 | 0 | 0 | 0 | 0 | 0 | 0 | 0 | 0 | 0 | 0 | 0 | 0 | 0 | 0 | 0 |
| ENSMUSG00 SNORA17 | 123 | 0 | 0 | 0 | 0 | 0 | 0 | 0 | 0 | 0 | 0 | 0 | 0 | 0 | 0 | 0 | 0 | 0 | 0 |
| ENSMUSG00 Mir1983 | 131 | 0 | 0 | 0 | 0 | 0 | 0 | 0 | 0 | 0 | 0 | 0 | 0 | 0 | 0 | 0 | 0 | 0 | 0 |
| ENSMUSG00 U6 | 103 | 0 | 0 | 0 | 0 | 0 | 0 | 0 | 0 | 0 | 0 | 0 | 0 | 0 | 0 | 0 | 0 | 0 | 0 |
| ENSMUSG00 7SK | 328 | 0 | 0 | 0 | 0 | 0 | 0 | 0 | 0 | 0 | 0 | 0 | 0 | 0 | 0 | 0 | 0 | 0 | 0.16771069 |
| ENSMUSG00 SCARNA11 | 130 | 1.96177736 | 0 | 0 | 0.35046957 | 0 | 3.04939899 | 0 | 0 | 2.21773546 | 0 | 0.36749968 | 0 | 0.37947629 | 0.35726221 | 0 | 0 | 1.17927067 | 0 |
| ENSMUSG00 U2 | 171 | 0 | 0 | 0 | 0 | 0 | 0 | 0 | 0 | 0.24085682 | 0 | 0 | 0 | 0 | 0 | 0 | 0 | 0 | 0 |
| ENSMUSG00 7SK | 239 | 0 | 0.83630697 | 1.01199854 | 0.95315992 | 0.2013888 | 0.41466723 | 0 | 1.59800847 | 1.20629962 | 0.19728328 | 0 | 0.93415335 | 0 | 0.38865345 | 0.34478289 | 0.32042349 | 0.80180537 | 0.69049088 |
| ENSMUSG00 U6 | 106 | 0 | 0 | 0 | 0 | 0 | 0 | 0 | 0 | 0 | 0 | 0 | 0 | 0 | 0 | 0 | 0 | 0 | 0 |
| ENSMUSG00 SNORA17 | 129 | 0 | 0 | 0 | 0 | 0 | 0 | 0 | 0 | 0 | 0 | 0 | 0 | 0 | 0 | 0 | 0 | 0 | 0 |
| ENSMUSG00 7SK | 287 | 0 | 0 | 0 | 0 | 0 | 0 | 0 | 0 | 0 | 0 | 0 | 0 | 0 | 0 | 0 | 0 | 0 | 0 |
| ENSMUSG00 RNaseP_nuc | 231 | 0 | 0 | 0 | 0 | 0 | 0 | 0 | 0 | 0 | 0 | 0 | 0 | 0 | 0 | 0 | 0 | 0 | 0 |
| ENSMUSG00 Scarna3b | 138 | 0 | 0 | 0 | 0 | 0 | 0 | 0 | 0 | 0 | 0 | 0.34619535 | 0 | 0 | 0 | 0 | 0 | 0.55545358 | 0 |
| ENSMUSG00 SNORA17 | 119 | 0 | 0 | 0 | 0 | 0 | 0 | 0 | 0 | 0 | 0 | 0 | 0 | 0 | 0 | 0 | 0 | 0 | 0 |
| ENSMUSG00 U6 | 100 | 0 | 0 | 0 | 0 | 0 | 0 | 0 | 0 | 0 | 0 | 0 | 0 | 0 | 0 | 0 | 0 | 0 | 0 |
| ENSMUSG00 7SK | 329 | 0 | 0 | 0 | 0 | 0 | 0 | 0 | 0 | 0 | 0 | 0 | 0 | 0 | 0 | 0 | 0 | 0 | 0 |
| ENSMUSG00 7SK | 223 | 0 | 0 | 0 | 0 | 0 | 0 | 0 | 0 | 0 | 0 | 0 | 0 | 0 | 0 | 0 | 0 | 0 | 0 |
| ENSMUSG00 SNORA17 | 96 | 0 | 0 | 0 | 0 | 0 | 0 | 0 | 0 | 0 | 0 | 0 | 0 | 0 | 0 | 0 | 0 | 0 | 0 |
| ENSMUSG00 7SK | 244 | 0 | 0 | 0 | 0 | 0 | 0 | 0 | 0 | 0 | 0 | 0 | 0 | 0 | 0 | 0 | 0.62771488 | 0 | 0 |
| ENSMUSG00 5S_rRNA | 133 | 0 | 0 | 0 | 0 | 0 | 0 | 0 | 0 | 0 | 0 | 0 | 0 | 0 | 0 | 0 | 0 | 0 | 0 |
| ENSMUSG00 U6 | 105 | 0 | 0 | 0 | 0 | 0 | 0 | 0 | 0 | 0 | 0 | 0 | 0 | 0 | 0 | 0 | 0 | 0 | 0 |
| ENSMUSG00 U1 | 163 | 0 | 0 | 0.89031037 | 0 | 0 | 0 | 0 | 0 | 0 | 0 | 0 | 0 | 0 | 0 | 0 | 0 | 0 | 0 |
| ENSMUSG00 SNORD124 | 103 | 0 | 0 | 0 | 3.53872189 | 0 | 0 | 0 | 0 | 0.39986909 | 0 | 0 | 0.54189964 | 0 | 0 | 0.80003021 | 0 | 0 | 0 |
| ENSMUSG00 7SK | 329 | 0 | 0 | 0 | 0 | 0 | 0 | 0.57523014 | 0.11608633 | 0 | 0.14331521 | 0 | 1.18756729 | 0 | 0 | 0 | 0 | 0.1164933 | 0 |
| ENSMUSG00 U6 | 94 | 0 | 0 | 0 | 0.48469196 | 0 | 0 | 0 | 2.03151077 | 0 | 1.50480968 | 2.03297696 | 0 | 0 | 0 | 0 | 1.62938755 | 0 | 0 |
| ENSMUSG00 U6 | 100 | 0 | 0 | 0 | 0 | 0 | 0 | 0 | 0 | 0 | 0 | 0 | 0 | 0 | 0 | 0 | 0 | 0 | 0 |
| ENSMUSG00 U7 | 56 | 0 | 0 | 0 | 0 | 0 | 0 | 0 | 0 | 0 | 0 | 0 | 0 | 0 | 0 | 0 | 0 | 0 | 0 |
| ENSMUSG00 U1 | 136 | 0 | 0 | 0 | 0 | 0 | 0 | 0 | 0 | 0 | 0 | 0 | 0.82081857 | 0 | 0 | 0 | 0 | 0 | 0 |
| ENSMUSG00 7SK | 321 | 0.79448927 | 0.51889243 | 0.75348178 | 0.99354302 | 0.14994369 | 1.23495909 | 0 | 0.95183557 | 0.64153451 | 1.17509541 | 0.89298988 | 0.69552228 | 0.30736397 | 0.28937126 | 0.8984763 | 0.47714153 | 0.47758625 | 0.17136793 |
| ENSMUSG00 7SK | 298 | 0 | 0 | 0 | 0 | 0 | 0 | 0 | 0 | 0 | 0 | 0 | 0 | 0 | 0 | 0 | 0 | 0 | 0 |
| ENSMUSG00 U6 | 100 | 0 | 0 | 0 | 0 | 0 | 0 | 0 | 0 | 0 | 0 | 0 | 0 | 0 | 0 | 0 | 0 | 0 | 0 |
| ENSMUSG00 U7 | 58 | 0 | 0 | 0 | 0 | 0 | 0 | 0 | 0 | 0 | 0 | 0 | 0 | 0 | 0 | 0 | 0 | 0 | 0 |
| ENSMUSG00 7SK | 312 | 0 | 0 | 0 | 0 | 0 | 0 | 0 | 0 | 0 | 0 | 0 | 0 | 0 | 0 | 0 | 0 | 0 | 0 |
| ENSMUSG00 5S_rRNA | 113 | 0 | 0.88441312 | 0 | 0 | 0 | 0 | 0 | 0 | 0 | 0 | 0 | 0 | 0 | 0 | 0 | 0 | 0 | 0 |
| ENSMUSG00 U6 | 95 | 0 | 0.35066204 | 1.01839011 | 0 | 0 | 0 | 0 | 0 | 0 | 0 | 0 | 0 | 0 | 0 | 0 | 0 | 0 | 0 |
| ENSMUSG00 U6 | 106 | 0 | 0 | 0 | 0 | 0 | 0 | 0 | 0 | 0 | 0 | 0 | 0 | 0 | 0 | 0 | 0.36123215 | 0 | 0 |
| ENSMUSG00 Scarna2 | 413 | 4.32256027 | 3.95237728 | 6.09061399 | 2.75793247 | 5.24439846 | 5.27922585 | 3.78043197 | 2.58931542 | 1.89477918 | 2.73999244 | 1.50381225 | 3.51381895 | 1.91116387 | 2.24910835 | 3.69118055 | 2.41054671 | 1.39199626 | 3.72943096 |
| ENSMUSG00 7SK | 318 | 0 | 0 | 0 | 0 | 0 | 0 | 0 | 0 | 0 | 0 | 0 | 0 | 0 | 0 | 0 | 0 | 0 | 0 |
| ENSMUSG00 U12 | 171 | 0 | 0 | 0 | 0 | 0 | 0 | 0 | 0 | 0 | 0 | 0 | 0 | 0 | 0 | 0 | 0.44784336 | 0 | 0 |
| ENSMUSG00 SNORA61 | 118 | 0 | 0.28231266 | 1.22983552 | 0.38611055 | 0 | 0.41993842 | 0.40095491 | 0.32366443 | 0 | 0 | 0 | 1.41904227 | 0 | 0 | 0.69833146 | 0.32449667 | 0 | 0.93235774 |
| ENSMUSG00 U3 | 88 | 0 | 0 | 0 | 0 | 0 | 0 | 0 | 0 | 0 | 0 | 0 | 0 | 0 | 0 | 0 | 0 | 0 | 0 |
| ENSMUSG00 SNORA17 | 130 | 0 | 0 | 0 | 0 | 0 | 0 | 0 | 0 | 0 | 0 | 0 | 0 | 0 | 0 | 0 | 0 | 0 | 0 |
| ENSMUSG00 SNORA17 | 123 | 0 | 0 | 0 | 0 | 0 | 0 | 0 | 0 | 0 | 0 | 0 | 0 | 0 | 0 | 0 | 0 | 0 | 0 |
| ENSMUSG00 5S_rRNA | 108 | 0 | 0 | 0 | 0 | 0 | 0 | 0 | 0 | 0 | 0 | 0 | 0 | 0 | 0 | 0 | 0 | 0 | 0 |
| ENSMUSG00 5S_rRNA | 107 | 0 | 0 | 0 | 0 | 0 | 0 | 0 | 0 | 0 | 0 | 0 | 0 | 0 | 0 | 0 | 0 | 0 | 0 |
| ENSMUSG00 U6 | 106 | 0 | 0 | 0 | 0 | 0 | 0 | 0 | 0 | 0 | 0 | 0 | 0 | 0 | 0 | 0 | 0 | 0 | 0 |
| ENSMUSG00 SNORA70 | 133 | 0 | 0 | 0 | 0 | 0 | 0 | 0 | 0 | 0 | 0 | 0 | 0 | 0 | 0 | 0 | 0 | 0 | 0 |
| ENSMUSG00 SNORA17 | 113 | 0 | 0 | 0 | 0 | 0 | 0 | 0 | 0 | 0 | 0 | 0 | 0 | 0 | 0 | 0 | 0 | 0 | 0 |
| ENSMUSG00 SNORA17 | 127 | 0 | 0 | 0 | 0 | 0 | 0 | 0 | 0 | 0 | 0 | 0 | 0 | 0 | 0 | 0 | 0 | 0 | 0 |
| ENSMUSG00 U1 | 148 | 0 | 0 | 0 | 0 | 0.6504314 | 0 | 0 | 0 | 0 | 0 | 0 | 0 | 0 | 0 | 0 | 0 | 0 | 0 |
| ENSMUSG00 U6 | 62 | 0 | 0 | 0 | 0 | 0 | 0 | 0 | 0 | 0 | 0 | 0 | 0 | 0 | 0 | 0.66454122 | 0 | 0 | 0 |
| ENSMUSG00 7SK | 342 | 0 | 0 | 0 | 0 | 0 | 0 | 0 | 0 | 0 | 0 | 0 | 0 | 0 | 0 | 0 | 0 | 0 | 0 |
| ENSMUSG00 AC117688.1 | 103 | 0 | 0 | 0 | 0 | 0 | 0 | 0 | 0 | 0 | 0 | 0 | 0 | 0 | 0 | 0 | 0 | 0 | 0 |
| ENSMUSG00 5S_rRNA | 112 | 0 | 0 | 0 | 0 | 0 | 0 | 0 | 0 | 0 | 0 | 0 | 0 | 0 | 0 | 0 | 0 | 0 | 0 |
| ENSMUSG00 5S_rRNA | 113 | 0 | 0 | 0 | 0 | 0 | 0 | 0 | 0 | 0 | 0 | 0 | 0 | 0 | 0 | 0 | 0 | 0 | 0 |
| ENSMUSG00 SCARNA11 | 135 | 0 | 0 | 0 | 0 | 0.35653277 | 0 | 0 | 0 | 0 | 0.34926447 | 1.41555433 | 0.41344935 | 1.46168644 | 1.72015138 | 0 | 0.85090239 | 0 | 0 |
| ENSMUSG00 5S_rRNA | 114 | 0 | 0 | 0 | 0 | 0 | 0 | 0 | 0 | 0 | 0 | 0 | 0 | 0 | 0 | 0 | 0 | 0 | 0 |
| ENSMUSG00 U6 | 104 | 0 | 0 | 0 | 0 | 0 | 0 | 0 | 0 | 0 | 0 | 0 | 0 | 0 | 0 | 0 | 0 | 0 | 0 |
| ENSMUSG00 U6 | 101 | 0 | 0 | 0 | 0 | 0 | 0 | 0 | 0 | 0 | 0 | 0 | 0 | 0 | 0 | 0 | 0 | 0 | 0 |
| ENSMUSG00 U1 | 160 | 0 | 0 | 0 | 0 | 0 | 0 | 0 | 0 | 0 | 0 | 0 | 0 | 0 | 0 | 0 | 0 | 0 | 0 |
| ENSMUSG00 7SK | 324 | 0 | 0 | 0 | 0 | 0 | 0 | 0 | 0 | 0 | 0 | 0 | 0 | 0 | 0 | 0 | 0 | 0 | 0 |
| ENSMUSG00 U7 | 62 | 0 | 0 | 0 | 0 | 0 | 0 | 0 | 0 | 0 | 0 | 0 | 0 | 0 | 0 | 0 | 0 | 0 | 0 |
| ENSMUSG00 U6 | 105 | 0 | 0 | 0 | 0 | 0 | 0 | 0 | 0 | 0 | 0 | 0 | 0 | 0 | 0 | 0 | 0 | 0 | 0 |
| ENSMUSG00 U6 | 105 | 0.4048112 | 0 | 3.22490202 | 4.33914708 | 2.29199637 | 2.83158478 | 4.05537248 | 3.63737166 | 1.96126266 | 1.34716295 | 1.36499881 | 3.72104417 | 2.34913892 | 2.65394785 | 0.78479154 | 1.45868981 | 0.7300247 | 1.57168876 |
| ENSMUSG00 U6 | 102 | 0 | 0 | 0 | 0 | 0 | 0 | 0 | 0 | 0 | 0 | 0 | 0 | 0 | 0 | 0 | 0 | 0 | 0 |
| ENSMUSG00 snoU83B | 92 | 3.23408948 | 2.89677341 | 1.57739773 | 0.49522874 | 1.56951925 | 1.07723334 | 2.057073 | 2.07567404 | 2.2383976 | 4.10006115 | 5.19293027 | 1.82007595 | 5.36216492 | 5.04827037 | 4.92627299 | 4.5782248 | 4.58249202 | 2.39170029 |
| ENSMUSG00 7SK | 249 | 0 | 0 | 0 | 0 | 0 | 0 | 0 | 0 | 0 | 0 | 0.1918673 | 0 | 0 | 0 | 0 | 0 | 0 | 0 |
| ENSMUSG00 U6 | 94 | 0 | 0 | 0 | 0 | 0 | 0 | 0 | 0 | 0 | 0 | 0 | 0 | 0 | 0 | 0 | 0 | 0 | 0 |
| ENSMUSG00 SNORA17 | 131 | 0 | 0 | 0 | 0 | 0 | 0 | 0 | 0 | 0 | 0 | 0 | 0 | 0 | 0 | 0 | 0 | 0 | 0 |
| ENSMUSG00 7SK | 268 | 0 | 0 | 0 | 0 | 0 | 0 | 0 | 0 | 0 | 0 | 0 | 0 | 0 | 0 | 0 | 0 | 0 | 0 |
| ENSMUSG00 SNORA17 | 104 | 0 | 0.32031629 | 0 | 0 | 0 | 0 | 0 | 0 | 0 | 0 | 0 | 0 | 0 | 0 | 0 | 0 | 0 | 0 |
| ENSMUSG00 SNORA40 | 126 | 0 | 0 | 0 | 0 | 0 | 0 | 0 | 0 | 0 | 0 | 0 | 0 | 0.39152315 | 0 | 0 | 0 | 0 | 0 |
| ENSMUSG00 7SK | 315 | 1.6192448 | 0.84604176 | 1.2285341 | 1.5910206 | 0.45839927 | 0.94386159 | 3.30437758 | 0.60622861 | 3.39952194 | 1.49684772 | 1.51666535 | 1.06315548 | 1.09626483 | 1.03209083 | 0.52319436 | 1.45868981 | 0.48668314 | 0.34926417 |
| ENSMUSG00 U6 | 79 | 0 | 0 | 0 | 0 | 0 | 0 | 0 | 0 | 0 | 0 | 0 | 0 | 0 | 0 | 0 | 0 | 0 | 0 |
| ENSMUSG00 SNORA70 | 95 | 0 | 0 | 0 | 0 | 0 | 0 | 0 | 0 | 0 | 0 | 0 | 0 | 0 | 0 | 0 | 0 | 0 | 0 |
| ENSMUSG00 7SK | 310 | 0 | 0 | 0 | 0 | 0 | 0 | 0 | 0 | 0 | 0 | 0 | 0 | 0 | 0 | 0 | 0 | 0 | 0 |
| ENSMUSG00 Y_RNA | 102 | 0.83343482 | 0 | 0 | 0 | 0 | 0 | 0 | 0 | 0 | 0 | 0 | 0 | 0 | 0 | 0 | 0 | 0 | 1.6179149 |
| ENSMUSG00 U1 | 169 | 0 | 0 | 0 | 0 | 0 | 0 | 0 | 0 | 0 | 0 | 0 | 0 | 0 | 0 | 0 | 0 | 0 | 0 |
| ENSMUSG00 SNORA17 | 131 | 0 | 0 | 0 | 0 | 0 | 0 | 0 | 0 | 0 | 0 | 0 | 0 | 0 | 0 | 0 | 0 | 0 | 0 |
| ENSMUSG00 SNORA63 | 113 | 0 | 0 | 0 | 0 | 0 | 0 | 0 | 0 | 0 | 0 | 0 | 0 | 0 | 0 | 0 | 0 | 0 | 0 |
| ENSMUSG00 U6 | 106 | 0 | 0 | 0 | 0 | 0 | 0 | 0 | 0 | 0 | 0 | 0 | 0 | 0 | 0 | 0 | 0 | 0 | 0 |
| ENSMUSG00 U6 | 95 | 0 | 0 | 0 | 0 | 0 | 0 | 0 | 0 | 0 | 0 | 0 | 0 | 0 | 0 | 0 | 0 | 0 | 0 |
| ENSMUSG00 U7 | 57 | 0 | 0 | 0 | 0 | 0 | 0 | 0 | 0 | 0 | 0 | 0 | 0 | 0 | 0 | 0 | 0 | 0 | 0 |
| ENSMUSG00 U6 | 73 | 0 | 0 | 0 | 0 | 0 | 0 | 0 | 0 | 0 | 0 | 0 | 0 | 0 | 0 | 0 | 0 | 0 | 0 |
| ENSMUSG00 7SK | 288 | 9.74076951 | 6.70884675 | 10.0778188 | 7.27711125 | 7.1863636 | 8.25878893 | 7.88546649 | 5.43711284 | 8.29450665 | 7.03986194 | 8.4601489 | 11.628263 | 7.02294656 | 2.74149127 | 8.01141365 | 13.5612568 | 5.98848389 | 7.44914987 |
| ENSMUSG00 SNORA70 | 135 | 0 | 0 | 0 | 0 | 0 | 0 | 0 | 0 | 0 | 0 | 0 | 0 | 0 | 0 | 0 | 0 | 0 | 0 |
| ENSMUSG00 SNORA17 | 118 | 0 | 0 | 0 | 0 | 0 | 0 | 0 | 0 | 0 | 0 | 0 | 0 | 0 | 0 | 0 | 0 | 0 | 0 |
| ENSMUSG00 SCARNA17 | 134 | 0 | 0 | 0 | 0.34000779 | 0 | 0.73959304 | 0 | 1.42508964 | 0 | 0.35187092 | 0 | 0 | 0.36814864 | 0 | 0.3074743 | 0 | 0 | 0 |
| ENSMUSG00 U1 | 152 | 0 | 0 | 0 | 0 | 0 | 0 | 0 | 0 | 0 | 0 | 0 | 0 | 0 | 0 | 0 | 0 | 0 | 0 |
| ENSMUSG00 SNORA40 | 120 | 0 | 0 | 0 | 0 | 0 | 0 | 0 | 0 | 0 | 0 | 0 | 0 | 0 | 0 | 0 | 0 | 0 | 0 |
| ENSMUSG00 SNORA17 | 139 | 0 | 0 | 0 | 0 | 0 | 0 | 0 | 0 | 0 | 0 | 0 | 0 | 0 | 0 | 0 | 0 | 0 | 0 |
| ENSMUSG00 mmu-mir-21 | 55 | 3.09128553 | 0 | 3.51807493 | 0 | 2.62537765 | 5.40575275 | 3.4409221 | 0 | 4.49307445 | 0.85728551 | 4.34317804 | 2.02966046 | 1.7938879 | 2.53331386 | 1.4982384 | 1.39238572 | 3.48426881 | 7.00115904 |
| ENSMUSG00 U7 | 56 | 0 | 0 | 0 | 0 | 0 | 0 | 0 | 0 | 2.20642049 | 0 | 0 | 0 | 0 | 0 | 0 | 0 | 0 | 0 |
| ENSMUSG00 Mir1969 | 93 | 0 | 0 | 0 | 0 | 0 | 0 | 0 | 0 | 0 | 0 | 0 | 0 | 0 | 0 | 0 | 0 | 0 | 0 |
| ENSMUSG00 5S_rRNA | 97 | 0 | 0 | 0 | 0 | 0 | 0 | 0 | 0 | 0 | 0 | 0 | 0 | 0 | 0 | 0 | 0.39474853 | 0.79023293 | 0 |
| ENSMUSG00 U4 | 128 | 0 | 0 | 0.37791821 | 0.35594566 | 1.12809196 | 0 | 0 | 0 | 0 | 0 | 0 | 0 | 0 | 0 | 0 | 0 | 0 | 0 |
| ENSMUSG00 U3 | 89 | 0 | 0 | 0 | 0 | 0 | 0 | 0 | 0 | 0 | 0 | 0 | 0 | 0 | 0 | 0 | 0 | 0 | 0 |
| ENSMUSG00 Snord13 | 103 | 11.5548051 | 6.14509699 | 12.2107941 | 3.09638165 | 22.430411 | 10.1029845 | 6.89019596 | 4.44960028 | 7.19764353 | 1.37332145 | 22.2640583 | 7.04469528 | 4.31055588 | 0.90182694 | 2.00007553 | 2.60227429 | 9.67459922 | 3.20441398 |
| ENSMUSG00 U6 | 99 | 0 | 0 | 0 | 0 | 0 | 0 | 0 | 0 | 0 | 0 | 0 | 0 | 0 | 0 | 0 | 0 | 0 | 0 |
| ENSMUSG00 SCARNA14 | 134 | 0 | 0 | 0 | 0 | 0 | 0 | 0 | 0 | 0 | 0 | 0 | 0 | 0 | 0 | 0 | 0 | 0 | 0 |
| ENSMUSG00 SNORA48 | 143 | 0 | 0 | 0 | 0 | 0 | 0 | 0 | 0 | 0 | 0 | 0 | 0 | 0 | 0 | 0 | 0 | 0 | 0 |
| ENSMUSG00 snoMe28S-C | 78 | 0 | 0 | 0 | 0 | 0 | 0 | 0 | 0 | 0 | 0 | 0 | 0 | 0 | 0 | 0 | 0 | 0 | 0 |
| ENSMUSG00 SNORA17 | 127 | 0 | 0 | 0 | 0 | 0 | 0 | 0 | 0 | 0 | 0 | 0 | 0 | 0 | 0 | 0 | 0 | 0 | 0 |
| ENSMUSG00 U1 | 161 | 0 | 0 | 0 | 0 | 0 | 0 | 0 | 0 | 0 | 0 | 0 | 0 | 0 | 0 | 0 | 0 | 0 | 0 |
| ENSMUSG00 7SK | 95 | 0 | 0 | 0 | 0.47958994 | 0 | 0 | 0 | 0 | 0 | 0 | 0 | 0 | 0 | 0 | 0 | 0 | 0 | 0 |
| ENSMUSG00 SNORA48 | 142 | 0 | 0 | 0 | 0 | 0 | 0 | 0 | 0 | 0 | 0 | 0 | 0 | 0 | 0 | 0 | 0 | 0 | 0 |
| ENSMUSG00 SNORA17 | 116 | 0 | 0 | 0 | 0 | 0 | 0 | 0 | 0 | 0 | 0 | 0 | 0 | 0 | 0 | 0 | 0 | 0 | 0 |
| ENSMUSG00 7SK | 207 | 0 | 0.16093186 | 0 | 0 | 0 | 0 | 0 | 0 | 0.19896868 | 0 | 0 | 0 | 0 | 0 | 0 | 0 | 0.37030239 | 0 |
| ENSMUSG00 snoU13 | 103 | 0.82534322 | 1.94055694 | 4.22681333 | 0.44234024 | 0 | 0.962189 | 1.37803919 | 2.22480014 | 2.7990836 | 0.91554764 | 1.85533819 | 0.54189964 | 0 | 0 | 0.40001511 | 0 | 0 | 1.06813799 |
| ENSMUSG00 U6 | 99 | 0 | 0 | 0 | 0 | 0 | 0 | 0 | 0 | 0 | 0 | 0 | 0 | 0 | 0 | 0 | 0 | 0 | 0 |
| ENSMUSG00 U7 | 61 | 0 | 0 | 0 | 0 | 0 | 0 | 0 | 0 | 0 | 0 | 0 | 0 | 0 | 0 | 0 | 0 | 0 | 0 |
| ENSMUSG00 SNORA48 | 131 | 0 | 0 | 0 | 0 | 0 | 0 | 0 | 0 | 0 | 0 | 0 | 0 | 0 | 0 | 0 | 0 | 0 | 0 |
| ENSMUSG00 U6 | 85 | 0 | 0 | 0 | 0 | 0 | 0 | 2.22647901 | 0.89864476 | 0 | 1.10942831 | 0 | 0 | 0 | 1.09280206 | 0.48472419 | 0 | 0 | 0 |
| ENSMUSG00 5S_rRNA | 106 | 0 | 0 | 0 | 0 | 0 | 0 | 0 | 0 | 0 | 0 | 0 | 0 | 0 | 0 | 0 | 0 | 0 | 0 |
| ENSMUSG00 U6 | 102 | 0 | 0 | 0 | 0 | 0 | 0 | 0 | 0 | 0 | 0 | 0 | 0 | 0 | 0 | 0 | 0 | 0 | 0 |
| ENSMUSG00 SNORA5 | 91 | 0 | 0 | 0 | 0 | 0 | 0 | 0 | 0 | 0 | 0 | 0 | 0 | 0 | 0 | 0 | 0 | 0 | 0 |
| ENSMUSG00 SNORA48 | 143 | 0.29723899 | 0.4659146 | 0.33827644 | 0.6372174 | 0 | 0 | 0.33085789 | 1.33539869 | 0 | 0 | 0 | 0 | 0 | 0 | 0 | 1.07106594 | 0 | 0 |
| ENSMUSG00 U6 | 106 | 0 | 0 | 0 | 0 | 0 | 0 | 0 | 0 | 0 | 0 | 0 | 0 | 0 | 0 | 0 | 0 | 0 | 0 |
| ENSMUSG00 U6 | 105 | 0 | 0 | 0 | 0 | 0 | 0 | 0 | 0 | 0 | 0 | 0 | 0 | 0 | 0 | 0 | 0 | 0 | 0 |
| ENSMUSG00 U1 | 172 | 0.74136935 | 0 | 0 | 0.26488979 | 0 | 0 | 0 | 0.22204885 | 0 | 0 | 0 | 0 | 0 | 0 | 0.47908786 | 0 | 0 | 0 |
| ENSMUSG00 7SK | 286 | 0 | 0 | 0 | 0 | 0 | 0 | 0 | 0 | 0 | 0 | 0 | 0 | 0 | 0 | 0 | 0 | 0 | 0 |
| ENSMUSG00 SNORA25 | 128 | 0 | 0 | 0 | 0 | 0 | 0 | 0 | 0 | 0 | 0 | 0 | 0 | 0 | 0 | 0 | 0 | 0 | 0 |
| ENSMUSG00 U6 | 101 | 0 | 0 | 0 | 0 | 0 | 0 | 0 | 0 | 0 | 0 | 0 | 0 | 0 | 0 | 0 | 0 | 0 | 0 |
| ENSMUSG00 SNORA17 | 122 | 0 | 0 | 0 | 0 | 0 | 0 | 0 | 0 | 0 | 0 | 0 | 0 | 0 | 0 | 0 | 0 | 0 | 0 |
| ENSMUSG00 U6 | 83 | 0 | 0 | 0 | 0 | 0 | 0 | 0 | 0 | 0 | 0 | 0 | 0 | 0 | 0 | 0 | 0 | 0 | 0 |
| ENSMUSG00 U6 | 98 | 0 | 0 | 0 | 0 | 0 | 0 | 0 | 0 | 0 | 0 | 0 | 0 | 0 | 0 | 0 | 0 | 0 | 0 |
| ENSMUSG00 U6 | 104 | 0 | 0 | 0 | 0 | 0 | 0 | 0 | 0 | 0 | 0 | 0 | 0 | 0 | 0 | 0 | 0 | 0 | 0 |
| ENSMUSG00 U6 | 103 | 0 | 0 | 0 | 0 | 0 | 0 | 0 | 0 | 0 | 0 | 0 | 0 | 0 | 0 | 0 | 0 | 0 | 0 |
| ENSMUSG00 7SK | 285 | 0 | 0 | 0 | 0 | 0 | 0 | 0 | 0 | 0 | 0 | 0 | 0 | 0 | 0 | 0 | 0 | 0 | 0 |
| ENSMUSG00 SCARNA4 | 126 | 0.67468533 | 0.26438805 | 1.15175072 | 0 | 0.76399879 | 1.17982699 | 0.7509949 | 1.81868583 | 0.65375422 | 0.74842386 | 1.13749901 | 0 | 0 | 0 | 0.98098943 | 1.21557484 | 0.91253088 | 2.18290106 |
| ENSMUSG00 U6 | 54 | 0 | 0 | 0 | 0 | 0 | 0 | 0 | 0 | 0 | 0 | 0 | 0 | 0 | 0 | 0 | 0 | 0 | 0 |
| ENSMUSG00 U6 | 106 | 0.40099223 | 0 | 0 | 0 | 0 | 0 | 0 | 0 | 0 | 0 | 0 | 0 | 0 | 0 | 0 | 0 | 0 | 0 |
| ENSMUSG00 AF357425 | 72 | 0 | 0 | 0 | 0 | 0 | 0 | 0 | 0 | 0 | 0 | 0 | 0 | 0 | 0 | 0 | 0 | 0 | 0 |
| ENSMUSG00 7SK | 260 | 0 | 0 | 0 | 0 | 0 | 0 | 0 | 0 | 0 | 0 | 0 | 0 | 0 | 0 | 0 | 0 | 0 | 0 |
| ENSMUSG00 U7 | 62 | 0 | 0 | 0 | 0 | 0 | 0 | 0 | 0 | 0 | 0 | 0 | 0 | 0 | 0 | 0 | 0 | 0 | 0 |
| ENSMUSG00 SNORA17 | 126 | 0 | 0 | 0 | 0 | 0 | 0 | 0 | 0 | 0 | 0 | 0 | 0 | 0 | 0 | 0 | 0 | 0 | 0 |
| ENSMUSG00 7SK | 291 | 0 | 0.1144773 | 0 | 0 | 0.827009 | 0 | 0 | 0.13124537 | 0.56613767 | 1.45826917 | 0.98505069 | 0 | 0 | 1.2768134 | 0 | 0 | 0 | 0.37806946 |
| ENSMUSG00 7SK | 268 | 0 | 0 | 0 | 0 | 0 | 0 | 0 | 0 | 0 | 0 | 0 | 0 | 0 | 0 | 0 | 0 | 0 | 0 |
| ENSMUSG00 5S_rRNA | 93 | 0 | 0 | 0 | 0 | 0 | 0 | 0 | 0 | 0 | 0 | 0 | 0 | 0 | 0 | 0 | 0 | 0 | 0 |
| ENSMUSG00 7SK | 307 | 0.27690668 | 0.32553317 | 1.10297952 | 0.89044386 | 1.5678151 | 0.64563822 | 1.07879072 | 0.49762088 | 0 | 0.46075606 | 0.31123751 | 0.7272399 | 0.96414171 | 0 | 0.40262107 | 0.74835063 | 0.62420679 | 2.32937585 |
| ENSMUSG00 SNORA70 | 103 | 0.41267161 | 0 | 0 | 0 | 0 | 0 | 0 | 0 | 0 | 0 | 0 | 0 | 0 | 0 | 0 | 0 | 0 | 0 |
| ENSMUSG00 U6 | 99 | 0 | 0 | 0 | 0 | 0 | 0 | 0 | 0 | 0 | 0 | 0 | 0 | 0 | 0 | 0 | 0 | 0 | 0 |
| ENSMUSG00 SNORD26 | 79 | 0 | 0 | 0 | 0 | 0 | 0 | 0 | 0 | 0 | 0 | 0 | 0 | 0 | 0 | 0 | 0 | 0 | 0 |
| ENSMUSG00 SNORA17 | 125 | 0 | 0 | 0 | 0 | 0 | 0 | 0 | 0 | 0 | 0 | 0 | 0 | 0 | 0 | 0 | 0 | 0 | 0 |
| ENSMUSG00 snoU109 | 133 | 0.31958779 | 0 | 0 | 0 | 3.25704747 | 0 | 1.06720328 | 0 | 0.6193461 | 1.41806626 | 0 | 0 | 0.37091667 | 0 | 0 | 0 | 0.57633529 | 0 |
| ENSMUSG00 U3 | 221 | 0 | 0 | 0 | 0 | 0.21779151 | 0 | 0 | 0.3456326 | 0 | 0 | 0 | 0 | 0 | 0.21015424 | 0 | 0.17326067 | 0 | 0 |
| ENSMUSG00 SCARNA17 | 138 | 0 | 0 | 0 | 0 | 0 | 0 | 0 | 0 | 0 | 0 | 0 | 0 | 0 | 0 | 0 | 0 | 0 | 0 |
| ENSMUSG00 SNORA17 | 130 | 0 | 0 | 0 | 0 | 0 | 0 | 0 | 0 | 0 | 0 | 0 | 0 | 0 | 0 | 0 | 0 | 0 | 0 |
| ENSMUSG00 U6 | 105 | 0 | 0 | 0 | 0 | 0 | 0 | 0 | 0 | 0 | 0 | 0 | 0 | 0 | 0 | 0 | 0 | 0 | 0 |
| ENSMUSG00 SNORA17 | 115 | 0 | 0 | 0 | 0 | 0 | 0 | 0 | 0 | 0 | 0 | 0 | 0 | 0 | 0 | 0 | 0 | 0 | 0 |
| ENSMUSG00 SNORA18 | 124 | 0 | 0 | 0 | 0 | 0 | 0 | 0 | 0 | 0 | 0 | 0 | 0 | 0 | 0 | 0 | 0 | 0 | 0 |
| ENSMUSG00 U6 | 103 | 0 | 0 | 0 | 0 | 0 | 0 | 0 | 0 | 0 | 0 | 0 | 0 | 0 | 0 | 0 | 0 | 0 | 0 |
| ENSMUSG00 7SK | 265 | 0 | 0 | 0 | 0 | 0 | 0 | 0 | 0 | 0 | 0 | 0 | 0 | 0 | 0 | 0 | 0 | 0 | 0 |
| ENSMUSG00 U6 | 100 | 0 | 0 | 0 | 0.45561044 | 0 | 0 | 0 | 0 | 0 | 0 | 0 | 0 | 0 | 0 | 0 | 0 | 0 | 0.55009107 |
| ENSMUSG00 SNORA17 | 109 | 0 | 0 | 0 | 0 | 0 | 0 | 0 | 0 | 0 | 0 | 0 | 0 | 0 | 0 | 0 | 0 | 0 | 0 |
| ENSMUSG00 U6 | 106 | 0 | 0 | 0 | 0 | 0 | 0 | 0 | 0 | 0 | 0 | 0 | 0 | 0 | 0 | 0 | 0 | 0 | 0 |
| ENSMUSG00 snoU109 | 134 | 3.17202806 | 2.73464057 | 1.08298949 | 1.70003897 | 0.71838692 | 3.32816867 | 0.35307969 | 1.42508964 | 1.84417235 | 0.70374184 | 0.71305908 | 3.74881316 | 0.73629727 | 1.38639067 | 0.3074743 | 0 | 3.43220569 | 0 |
| ENSMUSG00 5S_rRNA | 111 | 0 | 0 | 0 | 0 | 0 | 0 | 0 | 0 | 0 | 0 | 0 | 0 | 0 | 0 | 0 | 0 | 0 | 0 |
| ENSMUSG00 SNORA17 | 128 | 0 | 0 | 0.37791821 | 0 | 0.37603065 | 0 | 0 | 0.29837814 | 0 | 0 | 0 | 0 | 0 | 0 | 0 | 0 | 0 | 0 |
| ENSMUSG00 7SK | 274 | 0 | 0 | 0 | 0 | 0.17566395 | 0 | 0 | 0.13938833 | 0 | 0 | 0 | 0 | 0 | 0 | 0 | 0 | 0 | 0 |
| ENSMUSG00 U6 | 106 | 0 | 0 | 0 | 0 | 0 | 0 | 0 | 0 | 0 | 0 | 0 | 0 | 0 | 0 | 0 | 0 | 0 | 0 |
| ENSMUSG00 U7 | 59 | 0 | 0 | 0 | 0 | 0 | 0 | 0 | 0 | 0 | 0 | 0 | 0 | 0 | 0 | 0 | 0 | 0 | 0 |
| ENSMUSG00 RNaseP_nuc | 295 | 0 | 1.01632559 | 0.16397807 | 0 | 0 | 0 | 0.16038196 | 0.38839731 | 2.65269085 | 1.11883025 | 0.64779605 | 0.56761691 | 1.33781471 | 0.94462551 | 0 | 0 | 0 | 0 |
| ENSMUSG00 U1 | 164 | 0 | 0 | 0 | 0 | 0 | 0 | 0 | 0 | 0 | 0 | 0 | 0 | 0 | 0 | 0 | 0 | 0 | 0 |
| ENSMUSG00 7SK | 269 | 0 | 0 | 0 | 0 | 0 | 0 | 0 | 0 | 0 | 0 | 0 | 0 | 0 | 0 | 0 | 0 | 0 | 0 |
| ENSMUSG00 U1 | 104 | 0 | 0.32031629 | 0 | 0 | 0 | 0 | 0 | 0 | 0 | 0 | 0 | 0 | 0 | 0 | 0 | 0 | 0 | 0 |
| ENSMUSG00 U7 | 62 | 0 | 0 | 0 | 0 | 0 | 0 | 0 | 0 | 0 | 0 | 0 | 0 | 0.79567609 | 0 | 0 | 0 | 0 | 0 |
| ENSMUSG00 HAR1F | 131 | 0 | 0 | 0 | 0 | 0 | 0 | 0 | 0 | 0 | 0 | 0 | 0 | 0 | 0 | 0 | 0 | 0 | 0.83983369 |
| ENSMUSG00 SNORA25 | 133 | 0 | 0 | 0 | 0 | 0 | 1.49030778 | 0 | 0.28716092 | 0 | 0.35451657 | 0 | 0 | 0 | 1.04761099 | 0.30978613 | 0 | 0 | 1.24080692 |
| ENSMUSG00 snoZ178 | 107 | 0 | 0 | 0 | 0 | 0 | 0 | 0 | 0.35693834 | 0 | 0 | 0 | 0 | 0 | 0 | 0 | 0 | 0 | 0 |
| ENSMUSG00 U6 | 103 | 0 | 0 | 0 | 0 | 0 | 0 | 0 | 0 | 0 | 0 | 0 | 0 | 0 | 0 | 0 | 0 | 0 | 0 |
| ENSMUSG00 7SK | 306 | 0 | 0.10886567 | 0 | 0 | 0 | 0.48581111 | 0 | 0 | 0 | 0.15408727 | 0 | 0.18240413 | 0 | 0 | 0 | 0 | 0 | 0 |
| ENSMUSG00 U2 | 81 | 0 | 0 | 0 | 0 | 0 | 0 | 0 | 0 | 0 | 0 | 0 | 0 | 0 | 0 | 0 | 0 | 0 | 0 |
| ENSMUSG00 SNORA75 | 128 | 0 | 0 | 0 | 0 | 0 | 0 | 0 | 0 | 0 | 0 | 0 | 0 | 0 | 0 | 0 | 0 | 0 | 0 |
| ENSMUSG00 U6 | 104 | 0 | 0 | 0 | 0 | 0 | 0 | 0 | 0 | 0 | 0 | 0 | 0 | 0 | 0 | 0 | 0 | 0 | 0 |
| ENSMUSG00 5S_rRNA | 113 | 0 | 0 | 1.28425302 | 0.40319508 | 0.42594623 | 0.43851977 | 0 | 0 | 0 | 0 | 0 | 0 | 0 | 0 | 0 | 0 | 0.33917077 | 0 |
| ENSMUSG00 U7 | 61 | 0 | 0 | 0 | 0 | 0 | 0 | 0 | 0 | 0 | 0 | 0 | 0 | 0 | 0 | 0 | 0 | 0 | 0 |
| ENSMUSG00 U6 | 106 | 0 | 0 | 0 | 0 | 0 | 0 | 0 | 0 | 0 | 0 | 0 | 0 | 0 | 0 | 0 | 0 | 0 | 0 |
| ENSMUSG00 SNORA17 | 131 | 0 | 0 | 0 | 0 | 0 | 0 | 0 | 0 | 0 | 0 | 0 | 0 | 0 | 0 | 0 | 0 | 0 | 0 |
| ENSMUSG00 U6 | 96 | 0 | 0 | 0 | 0 | 0 | 0 | 0 | 0 | 0 | 0 | 0 | 0 | 0 | 0 | 0 | 0 | 0 | 0 |
| ENSMUSG00 U6 | 98 | 0 | 0 | 0 | 0 | 0 | 0 | 0 | 0 | 0 | 0 | 0 | 0 | 0 | 0 | 0 | 0 | 0 | 0 |
| ENSMUSG00 7SK | 231 | 0 | 0 | 0 | 0 | 0 | 0 | 0 | 0 | 0 | 0 | 0 | 0 | 0 | 0 | 0 | 0 | 0 | 0 |
| ENSMUSG00 snoU54 | 67 | 0 | 0 | 0 | 0 | 0 | 0 | 0 | 0 | 0 | 0 | 0 | 0 | 0 | 0 | 0 | 0 | 0 | 0 |
| ENSMUSG00 SNORA17 | 98 | 0 | 0 | 0 | 0 | 0 | 0 | 0 | 0 | 0 | 0 | 0 | 0 | 0 | 0 | 0 | 0 | 0 | 0 |
| ENSMUSG00 U1 | 156 | 0 | 0 | 0 | 0 | 0 | 0 | 0 | 0 | 0 | 0 | 0 | 0 | 0 | 0 | 0 | 0 | 0 | 0 |
| ENSMUSG00 AC164075.1 | 59 | 0 | 0 | 0 | 0 | 0 | 0 | 0 | 0 | 0 | 0 | 0 | 0 | 0 | 0 | 0 | 0 | 0 | 0 |
| ENSMUSG00 7SK | 281 | 0 | 0 | 0 | 0 | 0 | 0 | 0 | 0 | 0 | 0 | 0 | 0 | 0 | 0 | 0 | 0 | 0 | 0 |
| ENSMUSG00 7SK | 267 | 0.47758625 | 0.49906958 | 0 | 0.34128123 | 0.72107751 | 0 | 0 | 0.14304271 | 0 | 0.52978318 | 0 | 0.62714228 | 0 | 0 | 0 | 0 | 0 | 0.82410647 |
| ENSMUSG00 SNORA17 | 137 | 0 | 0 | 0 | 0 | 0 | 0 | 0 | 0 | 0 | 0 | 0 | 0 | 0 | 0 | 0.30074128 | 0 | 0 | 0 |
| ENSMUSG00 5S_rRNA | 92 | 0.46201278 | 0 | 0 | 0.49522874 | 0 | 0 | 0 | 1.66053924 | 0.89535904 | 0 | 2.07717211 | 0 | 0.53621649 | 0.50482704 | 1.343529 | 2.49721353 | 0 | 0 |
| ENSMUSG00 U6 | 97 | 0 | 0 | 0 | 0 | 0 | 0 | 0 | 0 | 0 | 0 | 0 | 0 | 0 | 0 | 0 | 0 | 0 | 0 |
| ENSMUSG00 7SK | 267 | 0 | 0 | 0 | 0 | 0 | 0 | 0 | 0 | 0 | 0 | 0 | 0 | 0 | 0 | 0 | 0 | 0 | 0 |
| ENSMUSG00 SNORA17 | 86 | 0 | 0 | 0 | 0 | 0 | 0 | 0 | 0 | 0 | 0 | 0 | 0 | 0 | 0 | 0 | 0 | 0 | 0 |
| ENSMUSG00 U1 | 129 | 0 | 0 | 0 | 0 | 0 | 0 | 0 | 0 | 0 | 0 | 0 | 0 | 0 | 0 | 0 | 0 | 0 | 0 |
| ENSMUSG00 U7 | 61 | 0 | 0 | 0 | 0 | 0 | 0 | 0 | 0 | 0 | 0 | 0 | 0 | 0 | 0 | 0 | 0 | 0 | 0 |
| ENSMUSG00 7SK | 241 | 0 | 0 | 0 | 0 | 0 | 0 | 0 | 0 | 0 | 0 | 0 | 0 | 0 | 0 | 0 | 0 | 0 | 0 |
| ENSMUSG00 7SK | 247 | 0 | 0 | 0 | 0 | 0 | 0 | 0 | 0 | 0 | 0 | 0 | 0 | 0 | 0.37606548 | 0 | 0 | 0.15516719 | 0 |
| ENSMUSG00 7SK | 305 | 0 | 0 | 0 | 0 | 0 | 0 | 0 | 0 | 0 | 0 | 0 | 0 | 0 | 0 | 0 | 0 | 0 | 0 |
| ENSMUSG00 7SK | 174 | 0 | 0 | 0 | 0 | 0 | 0 | 0 | 0 | 0 | 0 | 0 | 0 | 0 | 0 | 0 | 0 | 0 | 0 |
| ENSMUSG00 7SK | 255 | 0 | 0 | 0 | 0 | 0 | 0 | 0 | 0 | 0 | 0 | 0 | 0 | 0 | 0 | 0 | 0 | 0 | 0 |
| ENSMUSG00 U6 | 102 | 0 | 0 | 0 | 0 | 0 | 0 | 0 | 0 | 0 | 0 | 0 | 0 | 0 | 0 | 0 | 0 | 0 | 0 |
| ENSMUSG00 5S_rRNA | 114 | 0.37285242 | 0.29221837 | 0 | 0 | 0 | 0.4346731 | 0.4150235 | 1.34008429 | 0 | 0.41360266 | 0 | 0.97922215 | 0 | 0.81480855 | 0 | 1.00764756 | 0.33619559 | 0 |
| ENSMUSG00 SNORA17 | 133 | 0 | 0 | 0 | 0 | 0 | 0 | 0 | 0 | 0 | 0 | 0 | 0 | 0 | 0 | 0 | 0 | 0 | 0 |
| ENSMUSG00 Mir1956 | 64 | 1.99243013 | 0 | 0.75583641 | 0.71189132 | 0 | 0.77426146 | 3.69630304 | 0.59675629 | 0.64353931 | 2.21018922 | 1.49296745 | 0.87211973 | 3.08324483 | 0 | 1.28754862 | 0 | 1.19769678 | 2.57855188 |
| ENSMUSG00 SNORA17 | 123 | 0 | 0 | 0 | 0 | 0 | 0 | 0 | 0 | 0 | 0 | 0 | 0 | 0 | 0 | 0 | 0 | 0 | 0 |
| ENSMUSG00 RNase_MRP | 239 | 0 | 0 | 0 | 0 | 0 | 0 | 0 | 0 | 0 | 0 | 0 | 0 | 0 | 0 | 0 | 0 | 0 | 0 |
| ENSMUSG00 snoU13 | 102 | 0 | 0 | 0 | 0 | 0 | 0 | 0 | 0 | 0 | 0 | 0 | 0 | 0 | 0 | 0 | 0 | 0 | 0 |
| ENSMUSG00 SNORA40 | 105 | 0 | 0.31726566 | 1.84280116 | 0 | 0 | 0 | 0.45059694 | 0 | 1.56901012 | 0 | 0.4549996 | 0.53157774 | 0 | 0 | 1.56958308 | 0 | 0 | 0 |
| ENSMUSG00 U1 | 162 | 1.04951052 | 1.6450812 | 2.38881631 | 1.68744609 | 1.4855532 | 0 | 2.33642859 | 0.94302228 | 0.76271325 | 0 | 2.06435006 | 0 | 1.82710805 | 2.29353518 | 0.76299178 | 0.23636177 | 2.12923872 | 2.7164991 |
| ENSMUSG00 SNORA17 | 95 | 0 | 0 | 0 | 0 | 0 | 0 | 0 | 0 | 0 | 1.48896958 | 0 | 0 | 0 | 0 | 0 | 0 | 0 | 0 |
| ENSMUSG00 U6 | 92 | 0 | 0 | 0 | 0 | 0 | 0 | 0 | 0 | 0 | 0 | 0 | 0 | 0 | 0 | 0 | 0 | 0 | 0 |
| ENSMUSG00 snoU13 | 97 | 1.75279076 | 0.6868638 | 0 | 0 | 0 | 0 | 0 | 0 | 0 | 0 | 0 | 0 | 0.50857647 | 3.35163517 | 0.42475831 | 0 | 0 | 0 |
| ENSMUSG00 RNaseP_nuc | 240 | 0 | 0 | 0 | 0 | 0 | 0 | 0 | 0 | 0 | 0 | 0 | 0 | 0 | 0 | 0 | 0 | 0 | 0 |
| ENSMUSG00 5S_rRNA | 96 | 0 | 0 | 0 | 0 | 0 | 0 | 0 | 0 | 0 | 0 | 0 | 0 | 0 | 0 | 0 | 0 | 0 | 0 |
| ENSMUSG00 U6 | 103 | 0 | 0 | 0.46964593 | 3.09638165 | 0.93460046 | 3.36766151 | 2.29673199 | 4.8204003 | 6.79777445 | 3.66219054 | 1.85533819 | 2.70949818 | 3.35265457 | 0.90182694 | 7.2002719 | 2.97402776 | 0.74419994 | 1.06813799 |
| ENSMUSG00 U1 | 145 | 0 | 0 | 0 | 0 | 0 | 0 | 0 | 0 | 0 | 0 | 0 | 0 | 0 | 0 | 0 | 0 | 0 | 0 |
| ENSMUSG00 U1 | 139 | 0 | 0 | 0 | 0 | 0 | 0 | 0 | 0 | 0 | 0 | 0 | 0 | 0 | 0 | 0 | 0 | 0 | 0 |
| ENSMUSG00 U7 | 60 | 0 | 0 | 0 | 0 | 0 | 0 | 0 | 0 | 0 | 0 | 0 | 0 | 0 | 0 | 0 | 0 | 0 | 0 |
| ENSMUSG00 7SK | 284 | 0 | 0 | 0 | 0 | 0 | 0 | 0 | 0 | 0 | 0 | 0 | 0 | 0 | 0 | 0 | 0 | 0 | 0 |
| ENSMUSG00 U7 | 63 | 0 | 0 | 0 | 0 | 0 | 0 | 0 | 0 | 0 | 0 | 0 | 0 | 0 | 0 | 0 | 0 | 0 | 0 |
| ENSMUSG00 SNORA17 | 116 | 0 | 0 | 0 | 0 | 0 | 0 | 0 | 0 | 0 | 0 | 0 | 0 | 0 | 0 | 0 | 0 | 0 | 0 |
| ENSMUSG00 SNORA17 | 125 | 0 | 0 | 0 | 0 | 0 | 0 | 0 | 0 | 0 | 0 | 0 | 0 | 0 | 0 | 0 | 0 | 0 | 0 |
| ENSMUSG00 AC090489.1 | 68 | 0 | 0 | 0 | 0 | 2.83128963 | 0 | 0 | 0 | 0 | 0 | 0 | 0 | 0 | 0 | 1.21181047 | 0 | 3.38173208 | 0 |
| ENSMUSG00 U1 | 164 | 0 | 0 | 0 | 0 | 0 | 0 | 0 | 0 | 0 | 0 | 0 | 0 | 0 | 0 | 0 | 0 | 0 | 0 |
| ENSMUSG00 7SK | 307 | 0 | 0 | 0 | 0 | 0 | 0 | 0 | 0 | 0 | 0 | 0 | 0 | 0 | 0 | 0 | 0 | 0 | 0 |
| ENSMUSG00 SNORA13 | 76 | 0 | 0 | 0 | 0 | 0 | 0 | 0 | 0 | 0 | 0 | 0 | 0 | 0 | 0 | 0 | 0 | 0 | 0 |
| ENSMUSG00 SNORA40 | 118 | 0 | 0 | 0 | 0 | 0 | 0 | 0 | 0 | 0 | 0 | 0 | 0 | 0 | 0.78718792 | 0 | 0.32449667 | 0 | 0 |
| ENSMUSG00 U6 | 106 | 1.20297668 | 0 | 0 | 0 | 0 | 0.46747862 | 0 | 0.36030568 | 0 | 0 | 0 | 2.10625142 | 0.93079089 | 0.43815177 | 1.55477569 | 0.72246429 | 1.44627535 | 0 |
| ENSMUSG00 U2 | 83 | 0 | 0 | 0 | 0 | 0 | 0 | 0 | 0 | 0 | 0 | 0 | 0 | 0 | 0 | 0 | 0 | 0 | 0 |
| ENSMUSG00 7SK | 301 | 0 | 0 | 0 | 0 | 0 | 0 | 0 | 0 | 0 | 0 | 0 | 0 | 0 | 0 | 0 | 0 | 0 | 0 |
| ENSMUSG00 5S_rRNA | 108 | 0 | 0 | 0 | 0 | 0 | 0 | 0 | 0 | 0 | 0 | 0 | 0 | 0 | 0 | 0 | 0 | 0 | 0 |
| ENSMUSG00 U6 | 113 | 0 | 0 | 0 | 0 | 0 | 0 | 0 | 0 | 0 | 0 | 0 | 0 | 0 | 0 | 0 | 0 | 0 | 0 |
| ENSMUSG00 7SK | 270 | 0 | 0 | 0 | 0 | 0 | 0 | 0 | 0 | 0 | 0 | 0 | 0 | 0 | 0 | 0 | 0 | 0 | 0 |
| ENSMUSG00 U7 | 61 | 0 | 0 | 0 | 0 | 0 | 0 | 0 | 0 | 0 | 0 | 0 | 0 | 0 | 0 | 0 | 0 | 0 | 0 |
| ENSMUSG00 U6 | 102 | 0 | 0 | 0 | 0 | 0 | 0 | 0 | 0 | 0 | 0 | 0 | 0 | 0 | 0 | 0 | 0 | 0 | 0 |
| ENSMUSG00 U4 | 84 | 0 | 0 | 0 | 0 | 0 | 0 | 0 | 0 | 0 | 0 | 0 | 0 | 0 | 0 | 0 | 0 | 0 | 0 |
| ENSMUSG00 SNORA31 | 134 | 0.31720281 | 0 | 0.3609965 | 0.34000779 | 0 | 0 | 0 | 0.85505379 | 2.15153441 | 0 | 0.71305908 | 0 | 0 | 0 | 0 | 2.28600641 | 0.57203428 | 3.6946415 |
| ENSMUSG00 SNORA79 | 144 | 0 | 0 | 0 | 0 | 0 | 0 | 0 | 0 | 0 | 0 | 0 | 0 | 1.02774828 | 0 | 0 | 0 | 0 | 0 |
| ENSMUSG00 Mir1967 | 81 | 0 | 0 | 0 | 0.56248203 | 0 | 0 | 0 | 0 | 0 | 0 | 0 | 0 | 0 | 0 | 0.50866118 | 0 | 0 | 0 |
| ENSMUSG00 SNORA61 | 131 | 0 | 0 | 0 | 0 | 0 | 0 | 0 | 0 | 0 | 0 | 0 | 0 | 0 | 0 | 0 | 0 | 0 | 0 |
| ENSMUSG00 7SK | 254 | 0 | 0 | 0 | 0 | 0 | 0 | 0 | 0 | 0 | 0 | 0 | 0 | 0 | 0 | 0 | 0 | 0 | 0 |
| ENSMUSG00 5S_rRNA | 97 | 0 | 0 | 0 | 0 | 0 | 0 | 0 | 0 | 0 | 0 | 0 | 0 | 0 | 0 | 0 | 0 | 0 | 0 |
| ENSMUSG00 SNORA17 | 131 | 0 | 0 | 0 | 0 | 0 | 0 | 0 | 0 | 0 | 0 | 0 | 0 | 0 | 0 | 0 | 0 | 0 | 0 |
| ENSMUSG00 SNORA70 | 133 | 0 | 0 | 0 | 0 | 0 | 0 | 0 | 0 | 0 | 0 | 0 | 0 | 0 | 0 | 0 | 0 | 0 | 0 |
| ENSMUSG00 SNORA70 | 102 | 0 | 0 | 0 | 0 | 0 | 0 | 0 | 0 | 0 | 0 | 0 | 0 | 0 | 0 | 0 | 0 | 0 | 0 |
| ENSMUSG00 7SK | 336 | 0 | 0 | 0 | 0 | 0 | 0 | 0 | 0 | 0 | 0 | 0 | 0 | 0 | 0 | 0 | 0 | 0 | 0 |
| ENSMUSG00 7SK | 294 | 0 | 0 | 0 | 0 | 0 | 0 | 0 | 0 | 0 | 0 | 0 | 0 | 0 | 0 | 0 | 0 | 0 | 0 |
| ENSMUSG00 SNORA48 | 146 | 0 | 0 | 0 | 0 | 0 | 0 | 0 | 0 | 0 | 0 | 0 | 0 | 0 | 0 | 0 | 0 | 0 | 0 |
| ENSMUSG00 SNORA17 | 118 | 0 | 0 | 0 | 0 | 0 | 0 | 0 | 0 | 0 | 0 | 0 | 0 | 0 | 0 | 0 | 0 | 0 | 0 |
| ENSMUSG00 U6 | 61 | 0 | 0 | 0 | 0 | 0 | 0 | 0 | 0 | 0 | 0 | 0 | 0 | 0 | 0 | 0 | 0 | 0 | 0 |
| ENSMUSG00 5S_rRNA | 99 | 0 | 0 | 0.97724304 | 0 | 0 | 0 | 0 | 0 | 0 | 0 | 0 | 0 | 0 | 0 | 0 | 0 | 0 | 0 |
| ENSMUSG00 U6 | 101 | 0 | 0 | 0 | 0 | 0 | 0 | 0 | 0 | 0 | 0 | 0 | 0 | 0 | 0 | 0 | 0 | 0 | 0 |
| ENSMUSG00 U7 | 59 | 0 | 0 | 0 | 0 | 0 | 0 | 0 | 0 | 0 | 0 | 0 | 0 | 0 | 0 | 0 | 0 | 0 | 0 |
| ENSMUSG00 U6 | 104 | 0 | 0 | 0 | 0 | 0 | 0 | 0 | 0 | 0 | 0 | 0 | 0 | 0 | 0 | 0 | 0 | 0 | 0 |
| ENSMUSG00 SNORA17 | 135 | 0 | 0 | 0 | 0 | 0 | 0 | 0 | 0 | 0 | 0 | 0 | 0 | 0 | 0 | 0 | 0 | 0 | 0 |
| ENSMUSG00 U3 | 89 | 0 | 0 | 0 | 0 | 0 | 0 | 0 | 0 | 0 | 0 | 0 | 0 | 0 | 0 | 0 | 0 | 0 | 0 |
| ENSMUSG00 SNORA48 | 143 | 0 | 0 | 0 | 0 | 0 | 0 | 0 | 0 | 0 | 0 | 0 | 0 | 0 | 0 | 0 | 0 | 0 | 0 |
| ENSMUSG00 U6 | 96 | 0 | 0 | 0 | 0 | 0 | 0 | 0 | 0 | 0 | 0 | 0 | 0 | 0 | 0 | 0 | 0 | 0 | 0 |

| | | | | | | | | | | | | | | | | | | | |
|---|---|---|---|---|---|---|---|---|---|---|---|---|---|---|---|---|---|---|---|
| ENSMUSG00 U6 | 107 | 0 | 0 | 0.45208907 | 0 | 0 | 0 | 0 | 0.35693834 | 0 | 0 | 0 | 0 | 0 | 0 | 0.38506127 | 0 | 0.35818969 | 0 |
| ENSMUSG00 SNORA17 | 131 | 0 | 0 | 0 | 0 | 0 | 0 | 0 | 0 | 0 | 0 | 0 | 0 | 0 | 0 | 0 | 0 | 0 | 0 |
| ENSMUSG00 7SK | 287 | 0.14810166 | 0 | 0 | 0.47624785 | 0.50312115 | 0 | 0 | 0.39922372 | 0 | 0 | 0 | 0 | 0 | 0.16182609 | 0 | 0 | 0 | 0 |
| ENSMUSG00 SNORA17 | 134 | 0 | 0 | 0 | 0 | 0 | 0 | 0 | 0 | 0 | 0 | 0 | 0 | 0 | 0 | 0 | 0 | 0 | 0 |
| ENSMUSG00 U6 | 102 | 0 | 0 | 0 | 0 | 0 | 0 | 0 | 0 | 0 | 0 | 0 | 0 | 0 | 0 | 0 | 0 | 0 | 0 |
| ENSMUSG00 7SK | 294 | 0.72287714 | 0.33992749 | 0 | 0.15496954 | 0.32742805 | 0.16854671 | 0.48278244 | 0.25981226 | 0 | 0.16037654 | 0 | 1.13909515 | 0.67118255 | 0 | 0 | 0 | 0 | 0 |
| ENSMUSG00 SNORA17 | 109 | 0 | 0 | 0 | 0 | 0 | 0 | 0 | 0 | 0 | 0 | 0 | 0 | 0 | 0 | 0 | 0 | 0 | 0 |
| ENSMUSG00 SNORA17 | 128 | 0 | 0 | 0 | 0 | 0 | 0 | 0 | 0 | 0 | 0 | 0 | 0 | 0 | 0 | 0 | 0 | 0 | 0 |
| ENSMUSG00 SCARNA24 | 137 | 0 | 0 | 0 | 0 | 0 | 0 | 0 | 0 | 0 | 0 | 0 | 0 | 0 | 0 | 0 | 0 | 0 | 0 |
| ENSMUSG00 AL732470.1 | 91 | 0 | 0 | 0 | 0 | 0 | 0 | 0 | 0 | 0 | 0 | 0 | 0 | 0 | 0 | 0 | 0 | 0 | 0 |
| ENSMUSG00 SNORA40 | 121 | 0 | 0 | 0 | 0 | 0 | 0 | 0 | 0 | 0 | 0 | 0 | 0 | 0 | 0 | 0 | 0 | 0 | 0 |
| ENSMUSG00 5S_rRNA | 79 | 0 | 0 | 0 | 0 | 0 | 0 | 0 | 0 | 0 | 0 | 0 | 0 | 0 | 0 | 0 | 0 | 0 | 0 |
| ENSMUSG00 U6 | 102 | 0 | 0 | 0 | 0 | 0 | 0 | 0 | 0 | 0 | 0 | 0 | 0 | 0 | 0 | 0 | 0 | 0 | 0 |
| ENSMUSG00 U6 | 99 | 0 | 0 | 0 | 0 | 0 | 0 | 0 | 0 | 0 | 0 | 0 | 0 | 0 | 0.93826439 | 0 | 0 | 0 | 0 |
| ENSMUSG00 SNORA70 | 117 | 0 | 0 | 0 | 0 | 0 | 0 | 0 | 0 | 0 | 0 | 0 | 0 | 0 | 0 | 0 | 0 | 0 | 0 |
| ENSMUSG00 U7 | 55 | 0 | 0 | 0 | 0 | 0 | 0 | 0 | 0 | 0 | 0 | 0 | 0 | 0 | 0 | 0 | 0 | 0 | 0 |
| ENSMUSG00 7SK | 246 | 0 | 0 | 0 | 0 | 0 | 0 | 0 | 0 | 0 | 0 | 0 | 0 | 0 | 0 | 0 | 0.77826438 | 0 | 0 |
| ENSMUSG00 U6 | 103 | 0 | 0 | 0 | 0 | 0 | 0 | 0 | 0 | 0 | 0 | 0 | 0 | 0 | 0 | 0 | 0 | 0 | 0 |
| ENSMUSG00 7SK | 235 | 0 | 0 | 0 | 0 | 0 | 0 | 0 | 0 | 0 | 0 | 0 | 0 | 0 | 0 | 0 | 0 | 0 | 0 |
| ENSMUSG00 U6 | 102 | 0 | 0 | 0 | 0 | 0 | 0 | 0 | 0 | 0 | 0 | 0 | 0 | 0 | 0 | 0 | 0 | 0 | 0 |
| ENSMUSG00 Mir1941 | 82 | 2.07342322 | 2.84378365 | 0.5899211 | 6.6674699 | 0.58697468 | 1.20860326 | 2.30793556 | 2.79456603 | 2.51137291 | 3.45005146 | 0.58262144 | 3.40339406 | 7.82091371 | 8.49586964 | 2.51228999 | 3.73566901 | 2.80436319 | 5.36674212 |
| ENSMUSG00 5S_rRNA | 86 | 0 | 0 | 0 | 0 | 0 | 0 | 0 | 0 | 0 | 0 | 0 | 0 | 0 | 0 | 0 | 0 | 0 | 0 |
| ENSMUSG00 7SK | 107 | 0 | 0 | 0 | 0 | 0 | 0 | 0 | 0 | 0 | 0 | 0 | 0 | 0 | 0 | 0 | 0 | 0 | 0 |
| ENSMUSG00 7SK | 342 | 0 | 0 | 0 | 0 | 0 | 0 | 0 | 0 | 0 | 0 | 0 | 0 | 0 | 0 | 0 | 0 | 0 | 0 |
| ENSMUSG00 Mir1970 | 103 | 0 | 0 | 0 | 0 | 0 | 1.924378 | 0 | 0 | 0 | 0 | 0 | 0 | 0 | 0 | 0 | 0 | 0 | 0 |
| ENSMUSG00 SNORA17 | 84 | 0 | 0 | 0 | 0 | 0 | 0 | 0 | 0 | 0 | 0 | 0 | 0 | 0 | 0 | 0 | 0 | 0 | 0 |
| ENSMUSG00 snoU13 | 99 | 0 | 0 | 0 | 0 | 0 | 0 | 0 | 0 | 0 | 0 | 0 | 0 | 0 | 0 | 0 | 0 | 0 | 0 |
| ENSMUSG00 U6 | 103 | 0 | 0 | 0 | 0 | 0 | 0 | 0 | 0 | 0 | 0 | 0 | 0 | 0 | 0 | 0 | 0 | 0 | 0 |
| ENSMUSG00 SNORA17 | 101 | 0 | 0 | 0 | 0 | 0 | 0 | 0 | 0 | 0 | 0 | 0 | 0 | 0 | 0 | 0 | 0 | 0 | 0 |
| ENSMUSG00 7SK | 296 | 0 | 0 | 0 | 0 | 0 | 0 | 0 | 0 | 0 | 0 | 0 | 0 | 0 | 0 | 0 | 0 | 0 | 0 |
| ENSMUSG00 U6 | 102 | 0 | 0 | 0 | 0 | 0 | 0 | 0 | 0 | 0 | 0 | 0 | 0 | 0 | 0 | 0 | 0 | 0 | 0 |
| ENSMUSG00 SNORA17 | 116 | 0 | 0 | 0 | 0 | 0 | 0.42717874 | 0 | 0 | 0 | 0 | 0 | 0 | 0 | 0 | 0 | 0 | 0 | 0 |
| ENSMUSG00 U1 | 147 | 0 | 0 | 0 | 0 | 0 | 0 | 0 | 0 | 0 | 0.64150617 | 0 | 0 | 0 | 0 | 0 | 0 | 0 | 0 |
| ENSMUSG00 7SK | 292 | 0 | 0 | 0 | 0 | 0 | 0 | 0 | 0 | 0 | 0 | 0 | 0 | 0 | 0 | 0 | 0 | 0 | 0 |
| ENSMUSG00 5S_rRNA | 113 | 0 | 0.58960875 | 1.28425302 | 1.61278033 | 0 | 0 | 0.83739255 | 0.33798586 | 0 | 0 | 0 | 0 | 0 | 0 | 0 | 0 | 0 | 0 |
| ENSMUSG00 SNORA17 | 115 | 0 | 0 | 0 | 0 | 0 | 0 | 0 | 0 | 0 | 0 | 0 | 0 | 0 | 0 | 0 | 0 | 0 | 0 |
| ENSMUSG00 SNORA70 | 132 | 0 | 0 | 0 | 0 | 0 | 0 | 0 | 0 | 0 | 0 | 0 | 0 | 0 | 0 | 0 | 0 | 0 | 0 |
| ENSMUSG00 Mir1946b | 132 | 0 | 0 | 0 | 0 | 0.36463579 | 0 | 0 | 0 | 0 | 0 | 0 | 0 | 0 | 0 | 0 | 0 | 0 | 0.83347131 |
| ENSMUSG00 SNORA17 | 115 | 0 | 0.57935468 | 0.84127879 | 0 | 0 | 0 | 0 | 0.66421569 | 0 | 0 | 0 | 0 | 0 | 0.40386163 | 0 | 0.66592361 | 0 | 0 |
| ENSMUSG00 U6 | 63 | 0 | 0 | 0 | 0 | 0 | 0 | 0 | 0 | 0 | 0 | 0 | 0 | 0 | 0 | 0 | 0 | 0 | 0 |
| ENSMUSG00 SNORA17 | 76 | 0 | 0 | 0 | 0 | 0 | 0 | 0 | 0 | 0 | 0 | 0 | 0 | 0 | 0 | 0 | 0 | 0 | 0 |
| ENSMUSG00 U6 | 100 | 0 | 0 | 0 | 0 | 0 | 0 | 0 | 0 | 0 | 0 | 0 | 0 | 0 | 0 | 0 | 0 | 0 | 0 |
| ENSMUSG00 7SK | 257 | 0 | 0 | 0 | 0 | 0 | 0 | 0 | 0 | 0 | 0 | 0 | 0 | 0 | 0 | 0 | 0 | 0 | 0 |
| ENSMUSG00 SNORA17 | 116 | 0 | 0 | 0 | 0 | 0 | 0 | 0 | 0 | 0 | 0 | 0 | 0 | 0 | 0 | 0 | 0 | 0 | 0 |
| ENSMUSG00 U6 | 103 | 1.23801484 | 0 | 1.40893778 | 0 | 0 | 0 | 0 | 0 | 0 | 0 | 0 | 0 | 0 | 0 | 0 | 0.37175347 | 0.74419994 | 0 |
| ENSMUSG00 U7 | 61 | 0 | 0 | 0 | 0 | 0 | 0 | 0 | 0 | 0 | 0 | 0 | 0 | 0 | 0 | 1.35087068 | 0 | 0 | 0 |
| ENSMUSG00 7SK | 96 | 0 | 0 | 0 | 0 | 0 | 0 | 0 | 0 | 0 | 0 | 0 | 0 | 0 | 0 | 0 | 0 | 0 | 0 |
| ENSMUSG00 U6 | 106 | 0 | 0 | 0 | 0 | 0 | 0 | 0 | 0 | 0 | 0 | 0 | 0 | 0 | 0 | 0 | 0 | 0 | 0 |
| ENSMUSG00 U6 | 106 | 0.40099223 | 0 | 0.45635406 | 0 | 0.9081495 | 0 | 0 | 0 | 0 | 1.33445387 | 0 | 0 | 0 | 0.43815177 | 1.55477569 | 0 | 0.72313768 | 2.07581535 |
| ENSMUSG00 Mir1953 | 88 | 0 | 0 | 0 | 0 | 0 | 0 | 0 | 0 | 0 | 0 | 0 | 0 | 0 | 0 | 0.936399 | 0 | 0 | 0 |
| ENSMUSG00 SNORA40 | 90 | 1.4168392 | 2.22085961 | 1.61245101 | 3.54363678 | 3.20879491 | 1.10117186 | 0.52569643 | 0.42436003 | 0.45762795 | 1.04779341 | 1.59249862 | 1.24034806 | 0.54813241 | 1.03209083 | 0 | 0 | 3.40678195 | 0 |
| ENSMUSG00 U6 | 73 | 0 | 0 | 0 | 0 | 1.31868284 | 0 | 0.64811889 | 0 | 0 | 0 | 0 | 0 | 0 | 0 | 1.69321463 | 1.57358661 | 0 | 0 |
| ENSMUSG00 SNORA17 | 108 | 0 | 0 | 0 | 0 | 0 | 0 | 0 | 0 | 0 | 0 | 0 | 0 | 0 | 0 | 0 | 0 | 0 | 0 |
| ENSMUSG00 5S_rRNA | 93 | 0 | 0 | 0 | 0 | 0 | 0 | 0 | 0 | 0 | 0 | 0 | 0 | 0 | 0 | 0 | 0 | 0 | 0 |
| ENSMUSG00 U2 | 90 | 0 | 0 | 0 | 0 | 0 | 0 | 0 | 0 | 0 | 0 | 0 | 0 | 0 | 0 | 0 | 0 | 0 | 0 |
| ENSMUSG00 U6 | 104 | 0.40870362 | 0 | 0 | 0 | 0.46280696 | 0 | 0 | 0.36723464 | 0 | 0 | 0 | 0 | 0 | 0 | 0 | 0 | 0 | 0 |
| ENSMUSG00 7SK | 331 | 0 | 0 | 0 | 0 | 0 | 0 | 0 | 0 | 0 | 0 | 0 | 0 | 0 | 0 | 0 | 0 | 0 | 0 |
| ENSMUSG00 U7 | 60 | 0 | 0 | 0 | 0 | 0 | 0 | 0 | 0 | 0 | 0 | 0 | 0 | 0 | 0 | 0 | 0 | 0 | 0 |
| ENSMUSG00 U6 | 72 | 0 | 0 | 0 | 0 | 0 | 0 | 0 | 0 | 0 | 0 | 0 | 0 | 0 | 0 | 0 | 0 | 0 | 0 |
| ENSMUSG00 SNORA17 | 131 | 0 | 0 | 0 | 0 | 0 | 0 | 0 | 0 | 0 | 0 | 0 | 0 | 0 | 0 | 0 | 0 | 0 | 0 |
| ENSMUSG00 SNORA17 | 139 | 0 | 0 | 0 | 0 | 0.69254566 | 0 | 0 | 1.09906194 | 0 | 0 | 0 | 0 | 2.48434116 | 0 | 0 | 0 | 0 | 0 |
| ENSMUSG00 SCARNA13 | 263 | 0 | 0 | 0 | 0 | 0 | 0 | 0 | 0 | 0 | 0 | 0 | 0 | 0 | 0 | 0 | 0 | 0 | 0 |
| ENSMUSG00 U1 | 151 | 0 | 0 | 0 | 0 | 0 | 0 | 0 | 0 | 0 | 0 | 0 | 0 | 0 | 0 | 0 | 0 | 0 | 0 |
| ENSMUSG00 SNORA17 | 129 | 0 | 0 | 0 | 0 | 0 | 0 | 0 | 0 | 0 | 0 | 0 | 0 | 0 | 0 | 0 | 0 | 0 | 0 |
| ENSMUSG00 SNORA17 | 98 | 0 | 0 | 0 | 0 | 0 | 0 | 0 | 0 | 0 | 0 | 0 | 0 | 0 | 0 | 0 | 0 | 0 | 0 |
| ENSMUSG00 U6 | 109 | 0 | 0 | 0 | 0 | 0 | 0 | 0 | 0 | 0 | 0 | 0 | 0 | 0 | 0 | 0 | 0.35128998 | 0 | 0 |
| ENSMUSG00 SNORA17 | 125 | 0 | 0 | 0 | 0 | 0 | 0 | 0 | 0 | 0 | 0 | 0 | 0 | 0 | 0 | 0 | 0 | 0 | 0 |
| ENSMUSG00 7SK | 301 | 0 | 0 | 0 | 0 | 0 | 0 | 0 | 0 | 0 | 0.15664685 | 0 | 0 | 0 | 0 | 0.68441123 | 0 | 0.50931956 | 0.54826352 |
| ENSMUSG00 SNORA17 | 108 | 0 | 0 | 0 | 0 | 0 | 0 | 0 | 0 | 0 | 0 | 0 | 0 | 0 | 0 | 0 | 0 | 0 | 0 |
| ENSMUSG00 U6 | 99 | 0 | 0 | 0 | 0 | 0 | 0 | 0 | 0 | 0 | 0 | 0 | 0 | 0 | 0 | 0 | 0 | 0 | 0 |
| ENSMUSG00 U6 | 103 | 0 | 0 | 0 | 0 | 0 | 0 | 0 | 0 | 0 | 0 | 0 | 0 | 0 | 0 | 0 | 0 | 0 | 0 |
| ENSMUSG00 SNORA9 | 114 | 0 | 0 | 0 | 0 | 0 | 0 | 0 | 0 | 0 | 0 | 0 | 0 | 0 | 0 | 0 | 0 | 0 | 0 |
| ENSMUSG00 U8 | 116 | 0 | 0 | 0.41701319 | 0 | 0 | 0 | 0 | 0 | 0 | 0 | 0 | 0 | 0 | 0 | 0 | 0 | 0 | 0 |
| ENSMUSG00 U6 | 73 | 0 | 0 | 0 | 0 | 0 | 0 | 0 | 0 | 0 | 0 | 0 | 0 | 0 | 0 | 0 | 0 | 0 | 0 |
| ENSMUSG00 SNORA17 | 130 | 0 | 0 | 0 | 0 | 0 | 0 | 0 | 0 | 0 | 0 | 0 | 0 | 0 | 0 | 0 | 0 | 0 | 0 |
| ENSMUSG00 SNORA61 | 127 | 0 | 0 | 0 | 0 | 0 | 0 | 0 | 0 | 0 | 0 | 0 | 0 | 0 | 0 | 0 | 0 | 0 | 0 |
| ENSMUSG00 5S_rRNA | 106 | 0 | 0 | 0 | 0 | 0 | 0 | 0 | 0 | 0 | 0 | 0 | 0 | 0 | 0 | 0 | 0 | 0 | 0 |
| ENSMUSG00 Snord2 | 68 | 0 | 0 | 0 | 0 | 0 | 0 | 0 | 0 | 0 | 0 | 0 | 0 | 0 | 0 | 0 | 0 | 0 | 0 |
| ENSMUSG00 SNORA17 | 115 | 0 | 0 | 0 | 0 | 0 | 0 | 0 | 0 | 0 | 0 | 0 | 0 | 0 | 0 | 0 | 0 | 0 | 0 |
| ENSMUSG00 U7 | 61 | 0 | 0 | 0 | 0 | 0 | 0 | 0 | 0 | 0 | 0 | 0 | 0 | 0 | 0 | 0 | 0 | 0 | 0 |
| ENSMUSG00 7SK | 262 | 0 | 0 | 0 | 0 | 0 | 0 | 0 | 0 | 0 | 0 | 0 | 0 | 0.18828976 | 0 | 0 | 0 | 0 | 0 |
| ENSMUSG00 U6 | 103 | 0 | 0 | 0 | 0 | 0 | 0 | 0 | 0 | 0 | 0 | 0 | 0 | 0 | 0 | 0 | 0 | 0 | 0 |
| ENSMUSG00 SCARNA16 | 192 | 0 | 0 | 0 | 0 | 0 | 0 | 0 | 0 | 0 | 0 | 0 | 0 | 0 | 0 | 0 | 0 | 0 | 0 |
| ENSMUSG00 SNORA17 | 138 | 0 | 0 | 0 | 0 | 0 | 0 | 0 | 0 | 0 | 0 | 0 | 0 | 0 | 0 | 0 | 0 | 0 | 0 |
| ENSMUSG00 7SK | 100 | 0 | 0 | 0 | 0 | 0 | 0 | 0 | 0 | 0 | 0 | 0 | 0 | 0 | 0 | 0 | 0 | 0 | 0 |
| ENSMUSG00 U1 | 111 | 0 | 0 | 0 | 0 | 0 | 0 | 0 | 0 | 0 | 0 | 0 | 0 | 0 | 0 | 0 | 0 | 0 | 0 |
| ENSMUSG00 SNORA17 | 88 | 0 | 0 | 0 | 0 | 0 | 0 | 0 | 0 | 0 | 0 | 0 | 0 | 0 | 0 | 0 | 0 | 0 | 0 |
| ENSMUSG00 U6 | 104 | 0 | 0 | 0 | 0 | 0 | 0 | 0 | 0 | 0 | 0 | 0 | 0 | 0 | 0 | 0 | 0 | 0 | 0 |
| ENSMUSG00 SNORA48 | 142 | 0 | 0 | 0 | 0 | 0 | 0 | 0 | 0 | 0 | 0 | 0 | 0 | 0 | 0 | 0 | 0 | 0 | 0 |
| ENSMUSG00 5S_rRNA | 96 | 0 | 0 | 0 | 0 | 0 | 0 | 0 | 0 | 0 | 0 | 0 | 0.58141315 | 0 | 0 | 0 | 0 | 0 | 0 |
| ENSMUSG00 SNORA17 | 137 | 0 | 0 | 0 | 0 | 0 | 0 | 0 | 0 | 0 | 0 | 0 | 0 | 0 | 0 | 0 | 0 | 0 | 0 |
| ENSMUSG00 SNORA17 | 135 | 0 | 0 | 0 | 0 | 0 | 0 | 0 | 0 | 0 | 0 | 0 | 0 | 0 | 0 | 0 | 0 | 0 | 0 |
| ENSMUSG00 U7 | 57 | 0 | 0 | 0 | 0 | 0 | 0 | 0 | 0 | 0 | 0 | 0 | 0 | 0 | 0 | 0 | 0 | 0 | 0 |
| ENSMUSG00 SNORA48 | 143 | 0 | 0 | 0 | 0 | 0 | 0 | 0 | 0 | 0 | 0 | 0 | 0 | 0 | 0 | 0 | 0 | 0 | 0 |
| ENSMUSG00 U6 | 108 | 0 | 0 | 0 | 0 | 0 | 0 | 0 | 0 | 0 | 0 | 0 | 0 | 0 | 0 | 0 | 0 | 0 | 0 |
| ENSMUSG00 U1 | 183 | 0 | 0 | 0 | 0 | 0 | 0 | 0 | 0 | 0 | 0 | 0 | 0 | 0 | 0 | 0 | 0 | 0 | 0 |
| ENSMUSG00 Mir1945 | 77 | 0 | 0 | 0 | 0 | 0 | 0 | 0 | 0 | 0 | 0 | 0 | 0 | 1.2813485 | 0.60316997 | 0 | 0 | 0 | 0 |
| ENSMUSG00 U6 | 100 | 0 | 0 | 0 | 0 | 0 | 0 | 0 | 0.76384805 | 0 | 0 | 0 | 0.55815663 | 0 | 0 | 0 | 0 | 0 | 0 |
| ENSMUSG00 7SK | 294 | 0 | 0 | 0 | 0 | 0 | 0 | 0 | 0 | 0 | 0 | 0 | 0 | 0 | 0 | 0 | 0 | 0 | 0 |
| ENSMUSG00 U6 | 105 | 0 | 0 | 0 | 0 | 0 | 0 | 0 | 0 | 0 | 0 | 0 | 0 | 0 | 0 | 0 | 0 | 0 | 0 |
| ENSMUSG00 U6 | 89 | 0 | 0 | 0 | 0 | 0 | 0 | 0 | 0 | 0 | 0 | 0 | 0 | 1.10858241 | 0 | 0.46293883 | 0 | 0 | 0 |
| ENSMUSG00 U1 | 145 | 0.58627829 | 0.2297441 | 1.33444222 | 0.6284282 | 0.6638886 | 0.34174299 | 0 | 1.31697939 | 1.13617975 | 0.32517726 | 1.97689483 | 0 | 0.68044024 | 0.32030405 | 0.28414866 | 0.52814631 | 0.26431929 | 0.37937315 |
| ENSMUSG00 SNORA17 | 122 | 0 | 0 | 0 | 0 | 0 | 0 | 0 | 0 | 0 | 0 | 0 | 0 | 0 | 0 | 0 | 0 | 0 | 0 |
| ENSMUSG00 U7 | 62 | 0 | 0 | 0 | 0 | 0 | 0 | 0 | 0 | 0 | 0 | 0 | 0 | 0 | 0 | 0 | 0 | 0 | 0 |
| ENSMUSG00 Mir1957 | 65 | 0.65392579 | 0 | 0 | 0 | 0.74049113 | 0 | 0 | 0 | 0 | 0 | 0 | 0 | 2.27685772 | 0 | 0 | 0 | 0 | 0 |
| ENSMUSG00 U6 | 102 | 0 | 0 | 0 | 0 | 0 | 0 | 0 | 0 | 0 | 0 | 0 | 0 | 0 | 0 | 0 | 0 | 0 | 0 |
| ENSMUSG00 7SK | 277 | 0 | 0 | 0 | 0 | 0 | 0 | 0 | 0 | 0 | 0 | 0 | 0 | 0 | 0 | 0 | 0 | 0 | 0 |
| ENSMUSG00 U4 | 84 | 3.542098 | 1.98291037 | 6.33462898 | 3.79675369 | 1.14599818 | 4.71930796 | 2.25298471 | 4.54671457 | 4.41284097 | 5.05186106 | 0.56874951 | 3.98683304 | 4.11099311 | 4.97615222 | 6.86692598 | 2.73504339 | 2.2813272 | 7.85844382 |
| ENSMUSG00 SNORA17 | 97 | 0 | 0 | 0 | 0 | 0 | 0 | 0 | 0 | 0 | 0 | 0 | 0 | 0 | 0 | 0 | 0 | 0 | 0 |
| ENSMUSG00 Mir1981 | 81 | 0 | 0 | 0 | 0 | 0 | 0 | 0.58410715 | 0 | 0 | 0 | 0 | 0 | 0 | 0 | 0 | 0 | 0 | 0 |
| ENSMUSG00 snoU13 | 104 | 0 | 0 | 0 | 0.87617393 | 0 | 0.95293718 | 0 | 0 | 0 | 0 | 0 | 0 | 0.47434536 | 0 | 0 | 0 | 0 | 1.05786744 |
| ENSMUSG00 7SK | 332 | 0 | 0.7023803 | 0.43711021 | 0.41169618 | 1.30478106 | 0 | 0.57003228 | 0 | 0.49622308 | 0.28404038 | 1.00730334 | 0.67247786 | 0 | 0.41967549 | 0.99280858 | 0 | 0.80808457 | 0 |
| ENSMUSG00 U7 | 63 | 0 | 0 | 0 | 0 | 0 | 0 | 0 | 0 | 0 | 0 | 0 | 0 | 0 | 0 | 0 | 0 | 0 | 0 |
| ENSMUSG00 5S_rRNA | 119 | 0 | 0 | 0 | 0 | 0 | 0 | 0 | 0 | 0 | 0 | 0 | 0 | 0 | 0 | 0 | 0 | 0 | 0 |
| ENSMUSG00 7SK | 307 | 0 | 0 | 0 | 0 | 0 | 0 | 0 | 0 | 0 | 0 | 0 | 0 | 0 | 0 | 0 | 0 | 0 | 0 |
| ENSMUSG00 U6 | 101 | 0 | 0 | 0 | 0 | 0 | 0 | 0 | 0 | 0 | 0 | 0.47301939 | 0 | 0 | 0 | 0 | 0 | 0.37946829 | 0 |
| ENSMUSG00 U6 | 99 | 0 | 0 | 0 | 0 | 0 | 0 | 0 | 0 | 0 | 0 | 0 | 0 | 0 | 0 | 0 | 0 | 0 | 0 |
| ENSMUSG00 7SK | 324 | 0 | 0 | 0 | 0 | 0 | 0 | 0 | 0 | 0 | 0 | 0 | 0 | 0 | 0 | 0 | 0 | 0 | 0 |
| ENSMUSG00 7SK | 314 | 0 | 0 | 0 | 0 | 0 | 0 | 0 | 0 | 0 | 0 | 0 | 0 | 0 | 0 | 0 | 0 | 0 | 0 |
| ENSMUSG00 7SK | 287 | 0 | 0 | 0 | 0 | 0 | 0 | 0 | 0 | 0 | 0 | 0 | 0 | 0 | 0 | 0 | 0 | 0 | 0 |
| ENSMUSG00 7SK | 290 | 0 | 0 | 0 | 0 | 0 | 0 | 0 | 0 | 0 | 0 | 0 | 0 | 0 | 0 | 0 | 0 | 0 | 0 |
| ENSMUSG00 U6 | 101 | 0 | 0.32983064 | 0.47894585 | 0 | 0 | 0 | 0 | 0 | 0 | 0 | 0 | 0 | 0 | 0 | 0 | 0 | 0 | 0 |
| ENSMUSG00 7SK | 285 | 0 | 0 | 0 | 0.15986331 | 0 | 0.34773848 | 0.1660094 | 0 | 1.01159863 | 0 | 0 | 0 | 0 | 0 | 0 | 0 | 0 | 0 |
| ENSMUSG00 SNORA70 | 108 | 0 | 0 | 0 | 0 | 0 | 0 | 0 | 0 | 0 | 0 | 0 | 0 | 0 | 0 | 0 | 0 | 0 | 0 |
| ENSMUSG00 SNORA17 | 113 | 0 | 0 | 0 | 0 | 0 | 0 | 0 | 0 | 0 | 0 | 0 | 0 | 0 | 0 | 0 | 0 | 0 | 0 |
| ENSMUSG00 SNORA17 | 126 | 0 | 0 | 0 | 0 | 0 | 0 | 0 | 0 | 0 | 0 | 0 | 0 | 0 | 0 | 0 | 0 | 0 | 0 |
| ENSMUSG00 5S_rRNA | 117 | 0 | 0 | 0 | 0 | 0 | 0 | 0 | 0 | 0 | 0 | 0 | 0 | 0 | 0 | 0 | 0 | 0 | 0 |
| ENSMUSG00 7SK | 238 | 0 | 0 | 0 | 0 | 0 | 0 | 0 | 0 | 0 | 0 | 0 | 0 | 0 | 0 | 0 | 0 | 0 | 0 |
| ENSMUSG00 SNORA17 | 128 | 0 | 0 | 0 | 0 | 0 | 0 | 0 | 0 | 0 | 0 | 0 | 0 | 0 | 0 | 0 | 0 | 0 | 0 |
| ENSMUSG00 U6 | 100 | 0 | 0 | 0 | 0 | 0.91122089 | 0 | 0 | 0 | 0 | 0 | 1.43324875 | 0 | 0 | 0 | 0 | 0 | 0 | 0 |
| ENSMUSG00 U7 | 59 | 0 | 0 | 0 | 0 | 0 | 0 | 0 | 3.23664427 | 0 | 0 | 0 | 0 | 0 | 0 | 0 | 0 | 0 | 0 |
| ENSMUSG00 SNORA66 | 90 | 0 | 0 | 0 | 0 | 0 | 0 | 0 | 0 | 0 | 0 | 0 | 0 | 0 | 0 | 0 | 0 | 0 | 0 |
| ENSMUSG00 SNORD113 | 72 | 0 | 0 | 0 | 0 | 0 | 0 | 0 | 0 | 0 | 0 | 0 | 0 | 0 | 0 | 0 | 0 | 0 | 0 |
| ENSMUSG00 SNORA5 | 91 | 0 | 0 | 0 | 0 | 0 | 0 | 0 | 0 | 0 | 0 | 0 | 0 | 0 | 0 | 0 | 0 | 0 | 0 |
| ENSMUSG00 U6 | 105 | 0 | 0 | 0 | 0 | 0 | 0 | 0 | 0 | 0 | 0 | 0 | 0 | 0 | 0 | 0 | 0 | 0 | 0 |
| ENSMUSG00 5S_rRNA | 106 | 0 | 0 | 0 | 0 | 0 | 0 | 0 | 0.72061137 | 0 | 0 | 0 | 0 | 0.46539545 | 0 | 0 | 0 | 0 | 0 |
| ENSMUSG00 U7 | 65 | 0 | 0.51250606 | 0 | 0 | 0 | 0 | 0 | 0 | 0 | 0.72539543 | 0 | 0 | 0 | 0 | 0 | 0 | 0 | 0 |
| ENSMUSG00 SNORA17 | 141 | 0 | 0 | 0 | 0 | 0 | 0 | 0 | 0 | 0 | 0 | 0 | 0 | 0 | 0 | 0 | 0 | 0 | 0 |
| ENSMUSG00 5_8S_rRNA | 142 | 0 | 0 | 0 | 0 | 0 | 0 | 0 | 0 | 0 | 0 | 0 | 0 | 0 | 0 | 0 | 0 | 0 | 0 |
| ENSMUSG00 U6 | 62 | 0 | 0 | 0 | 0 | 0 | 0 | 0 | 0 | 0 | 0 | 0 | 0 | 0 | 0 | 0 | 0 | 0 | 0 |
| ENSMUSG00 U6 | 106 | 0 | 0 | 0 | 0 | 0 | 0 | 0 | 0 | 0 | 0 | 0 | 0 | 0 | 0 | 0 | 0 | 0 | 0 |
| ENSMUSG00 SNORA17 | 49 | 0 | 0 | 0 | 0 | 0 | 0 | 0 | 0 | 0 | 0 | 0 | 0 | 0 | 0 | 0 | 0 | 0 | 0 |
| ENSMUSG00 5S_rRNA | 93 | 0.4570449 | 0 | 0 | 1.46971111 | 0 | 1.06565018 | 0 | 0 | 0.88573152 | 0.50699681 | 0.51370923 | 1.80050524 | 0 | 0 | 0 | 0 | 1.23633216 | 0 |
| ENSMUSG00 7SK | 215 | 0 | 0 | 0 | 0 | 0 | 0 | 0 | 0 | 0 | 0 | 0 | 0 | 0 | 0 | 0 | 0 | 0 | 0 |
| ENSMUSG00 U6atac | 122 | 0.34840308 | 0 | 0 | 0 | 0 | 0 | 0 | 0.31305248 | 0 | 0 | 0 | 0 | 0 | 0 | 0 | 0 | 0 | 0 |
| ENSMUSG00 U1 | 144 | 0 | 0 | 0 | 0 | 0 | 0 | 0 | 0 | 0 | 0 | 0 | 0 | 0 | 0 | 0 | 0 | 0 | 0 |
| ENSMUSG00 SCARNA15 | 126 | 0 | 0.5287761 | 1.15175072 | 0.72319118 | 0.76399879 | 0.39327566 | 0 | 0 | 0.32687711 | 0.74842386 | 1.13749901 | 3.98683304 | 0.78304631 | 0.73720774 | 0 | 0 | 0 | 2.61948127 |
| ENSMUSG00 5S_rRNA | 107 | 0 | 0 | 0 | 0 | 0 | 0 | 0 | 0 | 0 | 0 | 0 | 0 | 0 | 0 | 0 | 0 | 0 | 0 |
| ENSMUSG00 Mir1951 | 88 | 0 | 0 | 0 | 0.51773914 | 0 | 0.56309925 | 0 | 0 | 0 | 0 | 0 | 0 | 0 | 0 | 0.4681995 | 0 | 0 | 0 |
| ENSMUSG00 5S_rRNA | 126 | 0 | 0 | 0 | 0 | 0 | 0 | 0 | 0 | 0 | 0 | 0 | 0 | 0 | 0 | 0 | 0 | 0 | 0 |
| ENSMUSG00 mmu-mir-21( | 56 | 0 | 0 | 0 | 1.62718015 | 0 | 0 | 0 | 0 | 0.7354735 | 1.68395369 | 0.85312426 | 0 | 0 | 0 | 0 | 0 | 0 | 0 |
| ENSMUSG00 7SK | 273 | 0 | 0 | 0 | 0 | 0 | 0 | 0 | 0 | 0 | 0 | 0 | 0 | 0 | 0 | 0 | 0 | 0 | 0 |
| ENSMUSG00 U7 | 64 | 0 | 0 | 0 | 0 | 0 | 0 | 0 | 0 | 0 | 0 | 0 | 0 | 0 | 0 | 0 | 0 | 0 | 0 |
| ENSMUSG00 U7 | 61 | 0 | 0 | 0 | 0 | 0 | 0 | 0 | 0 | 0 | 0 | 0 | 0 | 0 | 0 | 0 | 0 | 0 | 0 |
| ENSMUSG00 U6 | 102 | 0 | 0 | 0 | 0 | 0 | 0 | 0 | 0 | 0 | 0 | 0 | 0 | 0 | 0 | 0 | 0 | 0 | 0 |
| ENSMUSG00 U6 | 108 | 0 | 0 | 0 | 0 | 0 | 0 | 0 | 1.41453342 | 0 | 0 | 0 | 0 | 0 | 0 | 0 | 0 | 0 | 0 |
| ENSMUSG00 U6 | 106 | 0 | 0 | 0 | 0 | 0 | 0 | 0 | 0 | 0 | 0 | 0 | 0 | 0 | 0 | 0 | 0 | 0 | 0 |
| ENSMUSG00 7SK | 305 | 0 | 0 | 0 | 0 | 0 | 0 | 0 | 0 | 0 | 0 | 0 | 0 | 0 | 0 | 0 | 0 | 0 | 0 |
| ENSMUSG00 5S_rRNA | 117 | 0 | 0 | 0 | 0 | 0 | 0 | 0 | 0 | 0 | 0 | 0 | 0 | 0 | 0 | 0 | 0 | 0 | 0 |
| ENSMUSG00 Mir1965 | 125 | 0 | 0 | 0 | 0 | 0 | 0 | 0 | 0 | 0 | 0 | 0 | 0 | 0 | 0 | 0 | 0 | 0 | 0 |
| ENSMUSG00 U6 | 106 | 0 | 0 | 0 | 0 | 0 | 0 | 0 | 0 | 0 | 0 | 0 | 0 | 0 | 0 | 0 | 0 | 0 | 0 |
| ENSMUSG00 SNORA17 | 133 | 0 | 0 | 0 | 0 | 0 | 0 | 0 | 0 | 0 | 0 | 0 | 0 | 0 | 0 | 0 | 0 | 0 | 0 |
| ENSMUSG00 U6 | 113 | 0 | 0 | 0 | 0 | 1.27783868 | 0 | 0 | 0 | 0.72896488 | 0 | 0 | 0 | 0 | 0 | 0 | 0 | 0 | 0 |
| ENSMUSG00 U6atac | 118 | 0 | 0.84693799 | 0.40994517 | 0 | 0.81579532 | 0 | 1.20286472 | 0.32366443 | 1.39615308 | 0 | 0 | 0 | 0 | 0.78718792 | 0.34916573 | 0 | 0 | 0.46617887 |
| ENSMUSG00 SNORA42 | 128 | 0 | 0 | 0 | 0 | 0 | 0 | 0 | 0 | 0 | 0 | 0 | 0 | 0 | 0 | 0 | 0 | 0 | 0 |
| ENSMUSG00 U6 | 107 | 1.98622318 | 0.62267092 | 0 | 2.12902076 | 0 | 0 | 0.44217457 | 1.7846917 | 0.76984142 | 3.52528622 | 3.12546457 | 2.60820853 | 0 | 0.86811378 | 1.92530635 | 0.71571229 | 5.37284536 | 0 |
| ENSMUSG00 SNORA28 | 145 | 2.63825231 | 0 | 1.33444222 | 0.3142141 | 0 | 0 | 0 | 0.79018764 | 0 | 1.30070906 | 0 | 0 | 0 | 0 | 0.56829732 | 0 | 0.26431929 | 0 |
| ENSMUSG00 U6 | 105 | 1.6192448 | 0 | 0.92140058 | 0 | 0 | 2.35965398 | 0 | 0 | 0 | 0 | 0 | 0 | 0 | 0 | 0.39239577 | 0 | 0 | 0 |
| ENSMUSG00 SNORA40 | 108 | 0 | 0 | 0 | 0 | 0 | 0 | 0 | 0 | 0 | 0 | 0 | 0 | 0 | 0 | 0 | 0 | 0 | 0 |
| ENSMUSG00 7SK | 272 | 0 | 0 | 0 | 0 | 0 | 0 | 0 | 0 | 0 | 0 | 0 | 0 | 0 | 0 | 0 | 0 | 0 | 0 |
| ENSMUSG00 SNORA17 | 129 | 0 | 0 | 0 | 0 | 0 | 0 | 0 | 0 | 0 | 0 | 0 | 0 | 0 | 0 | 0 | 0 | 0 | 0 |
| ENSMUSG00 5S_rRNA | 109 | 0 | 0 | 0 | 0.41799123 | 1.76630913 | 0 | 0.43406127 | 0 | 0.75571589 | 0 | 0 | 0 | 0 | 0 | 0 | 0 | 0 | 0 |
| ENSMUSG00 7SK | 280 | 0 | 0.11897462 | 1.03657565 | 0.48815405 | 0 | 0.3539481 | 0.16897385 | 1.22761293 | 0.1470947 | 1.17876758 | 0.51187456 | 0.79736661 | 0.35237084 | 0.33174348 | 0.29429683 | 1.36752169 | 0 | 0.1964611 |
| ENSMUSG00 SNORA17 | 128 | 0 | 0 | 0 | 0 | 0 | 0 | 0 | 0 | 0 | 0 | 0 | 0 | 0 | 0 | 0 | 0 | 0 | 0 |
| ENSMUSG00 SNORD125 | 96 | 0 | 0.69401863 | 0.50389094 | 0.47459421 | 0 | 0 | 0 | 0 | 0 | 0 | 1.99062327 | 0 | 1.02774828 | 1.45137773 | 0 | 0 | 0.39923226 | 0 |
| ENSMUSG00 SNORA48 | 142 | 0 | 0 | 0 | 0 | 0 | 0 | 0 | 0 | 0 | 0 | 0 | 0 | 0 | 0 | 0 | 0 | 0 | 0 |
| ENSMUSG00 7SK | 215 | 0 | 0 | 0 | 0 | 0 | 0 | 0 | 0.17763908 | 0 | 0 | 0 | 0 | 0 | 0 | 0 | 0 | 0 | 0 |
| ENSMUSG00 SNORA17 | 134 | 0 | 0 | 0 | 0 | 0 | 0 | 0 | 0 | 0 | 0 | 0 | 0 | 0 | 0 | 0 | 0 | 0 | 0 |
| ENSMUSG00 Mir1955 | 97 | 0.43819769 | 0 | 0 | 0 | 0 | 0 | 0 | 0 | 0.42460326 | 0 | 1.97010138 | 1.72625761 | 0.50857647 | 0 | 0 | 0 | 0 | 1.70131258 |
| ENSMUSG00 U7 | 61 | 0 | 0 | 0 | 0 | 0 | 0 | 0 | 0 | 0 | 0 | 0 | 0 | 0 | 0 | 0 | 0 | 0 | 0 |
| ENSMUSG00 AC127677.1 | 103 | 0 | 0 | 0 | 0 | 0 | 0 | 0 | 0 | 0 | 0 | 0 | 0 | 0 | 0 | 0 | 0 | 0.37209997 | 0 |
| ENSMUSG00 U6 | 105 | 0 | 0 | 0 | 0 | 0 | 0 | 0 | 0 | 0 | 0 | 0 | 0 | 0 | 0 | 0 | 0 | 0 | 0 |
| ENSMUSG00 U6 | 63 | 0 | 0 | 0 | 0 | 0 | 0 | 0 | 0 | 0 | 0 | 0 | 0 | 0 | 0 | 0 | 0 | 0 | 0 |
| ENSMUSG00 5S_rRNA | 109 | 0 | 0 | 0 | 0 | 0 | 0 | 0 | 0 | 0 | 0 | 0 | 0 | 0 | 0 | 0 | 0 | 0 | 0 |
| ENSMUSG00 U6 | 93 | 0 | 0 | 0 | 0 | 0 | 0 | 0 | 0 | 0 | 0 | 0 | 0 | 0 | 0 | 0 | 0 | 0 | 0 |
| ENSMUSG00 U7 | 60 | 0 | 0 | 0 | 0 | 0 | 0 | 0 | 0 | 0 | 0 | 0 | 0 | 0 | 0 | 0 | 0 | 0 | 0 |
| ENSMUSG00 SNORA17 | 126 | 0 | 0 | 0 | 0 | 0 | 0 | 0 | 0 | 0 | 0 | 0 | 0 | 0 | 0 | 0 | 0 | 0 | 0 |
| ENSMUSG00 U6 | 97 | 0 | 0 | 0 | 0 | 0 | 0 | 0 | 0 | 0 | 0 | 0.98505069 | 0 | 0 | 0 | 0 | 0 | 0 | 0 |
| ENSMUSG00 SNORA40 | 120 | 0 | 0 | 0 | 0 | 0 | 0 | 0 | 0 | 0 | 0 | 0 | 0 | 0 | 0 | 0 | 0 | 0 | 0 |
| ENSMUSG00 U7 | 64 | 0 | 0 | 0 | 0 | 0 | 0 | 0 | 0 | 0 | 0 | 0 | 0 | 0 | 0 | 0 | 0 | 0 | 0 |
| ENSMUSG00 U6 | 102 | 0 | 0 | 0 | 0 | 0 | 0 | 0 | 0 | 0 | 0 | 0 | 0 | 0 | 0 | 0 | 0 | 0 | 1.07860994 |
| ENSMUSG00 SNORA17 | 118 | 0 | 0 | 0 | 0 | 0 | 0 | 0 | 0 | 0 | 0 | 0 | 0 | 0 | 0 | 0 | 0 | 0 | 0 |
| ENSMUSG00 U7 | 59 | 0 | 0 | 0 | 0 | 0 | 0 | 0 | 0 | 0 | 0 | 0 | 0 | 0 | 0 | 0 | 0 | 0 | 0 |
| ENSMUSG00 U6 | 106 | 0 | 0 | 0 | 0 | 0 | 0 | 0 | 0 | 0 | 0 | 0 | 0 | 0 | 0 | 0 | 0 | 0 | 0 |
| ENSMUSG00 U6 | 107 | 0 | 0 | 0 | 0 | 0 | 0 | 0 | 0 | 0 | 0 | 0 | 0 | 0 | 0 | 0 | 0 | 0 | 0 |
| ENSMUSG00 U6 | 107 | 0 | 0 | 0 | 0 | 0 | 0 | 0 | 0 | 0 | 0 | 0 | 0 | 0 | 0 | 0 | 0 | 0 | 0 |
| ENSMUSG00 5S_rRNA | 108 | 0 | 0 | 0 | 0 | 0.44566596 | 0 | 0 | 0 | 0 | 0 | 0 | 0 | 0 | 0 | 0 | 0 | 0.70974624 | 0 |
| ENSMUSG00 7SK | 245 | 0 | 0 | 0 | 0 | 0 | 0 | 0 | 0 | 0 | 0 | 0 | 0 | 0 | 0 | 0 | 0 | 0 | 0 |
| ENSMUSG00 U6 | 109 | 0.38995574 | 1.8337373 | 0.88758771 | 0.83598246 | 0 | 0.90922447 | 2.17030637 | 0 | 0 | 0 | 0 | 0 | 0 | 0 | 0 | 0 | 0 | 0 |
| ENSMUSG00 U3 | 88 | 0 | 0 | 0 | 0 | 0 | 0 | 0 | 0 | 0 | 0 | 0 | 0 | 0 | 0 | 0 | 0 | 0 | 0 |
| ENSMUSG00 U6 | 104 | 0 | 0 | 0 | 0 | 0 | 0 | 0 | 0 | 0 | 0 | 0 | 0 | 0 | 0 | 0 | 0 | 0 | 0 |
| ENSMUSG00 U7 | 60 | 0 | 0 | 0 | 0 | 0 | 0 | 0 | 0 | 0 | 0 | 0 | 0 | 0 | 0 | 0 | 0 | 0 | 0 |
| ENSMUSG00 5S_rRNA | 125 | 0 | 0 | 0 | 0 | 0 | 0 | 0 | 0 | 0 | 0 | 0 | 0 | 1.18396601 | 0 | 0 | 0 | 0 | 0 |
| ENSMUSG00 U7 | 56 | 0 | 0 | 0 | 0 | 0 | 0 | 0 | 0 | 0 | 0 | 0 | 0 | 0 | 0 | 0 | 0 | 0 | 0 |
| ENSMUSG00 U6 | 102 | 0 | 0 | 0 | 0 | 0 | 0 | 0 | 0 | 0 | 0 | 0 | 0 | 0 | 0 | 0 | 0 | 0 | 0 |
| ENSMUSG00 SNORA17 | 104 | 0.40870362 | 0 | 0 | 0 | 0.92561392 | 0 | 0 | 0.36723464 | 0 | 0 | 1.8374984 | 0 | 0 | 0 | 2.37701284 | 0 | 0 | 1.05786744 |
| ENSMUSG00 SNORA31 | 79 | 1.07608041 | 0.84336441 | 0 | 0 | 0.60926486 | 0.62724979 | 0.59889467 | 0.48344813 | 2.0853932 | 0 | 0.60474631 | 0 | 0 | 0 | 1.56461605 | 0 | 1.455429 | 0 |
| ENSMUSG00 5S_rRNA | 110 | 0 | 0 | 0 | 0 | 0 | 0 | 0 | 0 | 0 | 0 | 0 | 0 | 0 | 0 | 0 | 0 | 0 | 0 |
| ENSMUSG00 U6atac | 88 | 0 | 0 | 0 | 0 | 0 | 0 | 0 | 0 | 0 | 0 | 0 | 0 | 0 | 0 | 0 | 0 | 0 | 0 |
| ENSMUSG00 7SK | 282 | 0 | 0 | 0 | 0 | 0 | 0 | 0 | 0 | 0 | 0 | 0 | 0 | 0.34987175 | 0.32939069 | 0.43831442 | 0 | 0 | 0 |
| ENSMUSG00 RNaseP_nuc | 326 | 0 | 0 | 0 | 0 | 0 | 0 | 0 | 0 | 0 | 0 | 0 | 0 | 0 | 0 | 0 | 0 | 0 | 0 |
| ENSMUSG00 SNORA2 | 100 | 0 | 0 | 0 | 0 | 0 | 0 | 0 | 0 | 0 | 0 | 0 | 0 | 0 | 0 | 0 | 0 | 0 | 0 |
| ENSMUSG00 SNORA17 | 115 | 0 | 0 | 0 | 0 | 0 | 0 | 0 | 0 | 0 | 0 | 0 | 0 | 0 | 0 | 0 | 0 | 0 | 0 |
| ENSMUSG00 U7 | 61 | 0 | 0 | 0 | 0 | 0 | 0 | 0 | 0 | 0 | 0 | 0 | 0 | 0 | 0 | 0 | 0 | 0 | 0 |
| ENSMUSG00 5S_rRNA | 120 | 0 | 0 | 0 | 0 | 0 | 0 | 0 | 0 | 0 | 0 | 0 | 0 | 0 | 0 | 0 | 0 | 0 | 0 |
| ENSMUSG00 U3 | 88 | 0 | 0 | 0 | 0 | 0 | 0 | 0 | 0 | 0 | 0 | 0 | 0 | 0 | 0 | 0 | 0 | 0 | 0 |
| ENSMUSG00 5S_rRNA | 103 | 0 | 0 | 0 | 0 | 0 | 0 | 0 | 0 | 0 | 0 | 0 | 0 | 0 | 0 | 0 | 0 | 0 | 0 |
| ENSMUSG00 7SK | 322 | 0 | 0 | 0 | 0 | 0 | 0 | 0 | 0 | 0 | 0 | 0 | 0 | 0 | 0 | 0 | 0 | 0 | 0 |
| ENSMUSG00 SNORA17 | 87 | 0 | 0 | 0 | 0 | 0 | 0 | 0 | 0 | 0 | 0 | 0 | 0 | 0 | 0 | 0 | 0 | 0 | 0 |
| ENSMUSG00 SNORA70 | 128 | 0 | 0 | 0 | 0 | 0 | 0 | 0 | 0 | 0 | 0 | 0 | 0 | 0 | 0 | 0 | 0 | 0 | 0 |
| ENSMUSG00 7SK | 309 | 0 | 0 | 0 | 0 | 0 | 0 | 0 | 0 | 0 | 0 | 0 | 0 | 0 | 0 | 0 | 0 | 0 | 0 |
| ENSMUSG00 SNORA70 | 130 | 0 | 0 | 0 | 0 | 0 | 0 | 0 | 0 | 0 | 0 | 0 | 0 | 0 | 0 | 0 | 0 | 0 | 0 |
| ENSMUSG00 SNORA70 | 88 | 0 | 0 | 0 | 0 | 0 | 0 | 0 | 0.43400457 | 0 | 0.53580345 | 0 | 0 | 0 | 2.11109488 | 0 | 0 | 1.3065783 | 0 |
| ENSMUSG00 Scarna17 | 78 | 4.35950523 | 5.97923742 | 3.10086733 | 1.75234786 | 1.23415189 | 1.27058291 | 1.81971842 | 3.42752329 | 3.16819352 | 1.81348859 | 6.12499468 | 5.00909792 | 1.26492096 | 0 | 0.52822508 | 0.98181045 | 0.98272556 | 2.82097983 |
| ENSMUSG00 SNORA68 | 135 | 0 | 0 | 0 | 0 | 0 | 0 | 0 | 0 | 0 | 0 | 0 | 0 | 0 | 0 | 0 | 0 | 0 | 0 |
| ENSMUSG00 U1 | 161 | 0 | 0 | 0 | 0 | 0 | 0 | 0 | 0 | 0 | 0 | 0 | 0 | 0 | 0 | 0 | 0 | 0 | 0 |
| ENSMUSG00 SNORA17 | 114 | 0 | 0 | 0 | 0 | 0 | 0 | 0 | 0 | 0 | 0 | 0 | 0 | 0 | 0 | 0 | 0 | 0 | 0 |
| ENSMUSG00 U6 | 103 | 0 | 0 | 0 | 0 | 0 | 0 | 0 | 0 | 0 | 0 | 0 | 0 | 0 | 0 | 0 | 0 | 0 | 0 |
| ENSMUSG00 SNORA70 | 132 | 4.50812473 | 0.75711123 | 3.66466139 | 4.14191312 | 1.82317893 | 1.87699748 | 1.79214693 | 3.1827002 | 1.87211435 | 2.50041608 | 2.89545203 | 0 | 0 | 0.35184915 | 1.248532 | 4.35120539 | 1.45175367 | 2.08367828 |
| ENSMUSG00 U6 | 100 | 0 | 0 | 0 | 0 | 0 | 0 | 0 | 0 | 0 | 0 | 0 | 0 | 0 | 0 | 0 | 0 | 0 | 0 |
| ENSMUSG00 SNORA17 | 125 | 0 | 0 | 0 | 0 | 0 | 0 | 0 | 0 | 0 | 0 | 0 | 0 | 0 | 0 | 0 | 0 | 0 | 0 |
| ENSMUSG00 U7 | 64 | 0 | 0 | 0 | 0 | 0 | 0 | 0 | 0 | 0 | 0 | 0 | 0 | 0 | 0 | 0 | 0 | 0 | 0 |
| ENSMUSG00 AC152987.1 | 111 | 0 | 0 | 0 | 0 | 0 | 0 | 0 | 0 | 0 | 0 | 0.86081006 | 0 | 0 | 0 | 0 | 0 | 0 | 0 |
| ENSMUSG00 5S_rRNA | 112 | 0 | 0 | 0 | 0 | 0 | 0 | 0 | 0 | 0 | 0 | 0 | 0 | 0 | 0 | 0 | 0 | 0 | 0 |
| ENSMUSG00 5S_rRNA | 78 | 0 | 0 | 0 | 0 | 0 | 0 | 0 | 0 | 0 | 0 | 0 | 0 | 0 | 0 | 0 | 0 | 0 | 0 |
| ENSMUSG00 U1 | 143 | 0 | 0 | 0 | 0 | 0 | 0 | 0 | 0 | 0 | 0 | 0 | 0 | 0 | 0 | 0 | 0 | 0 | 0 |
| ENSMUSG00 5S_rRNA | 110 | 0 | 0 | 0 | 0 | 0 | 0 | 0 | 0 | 0 | 0 | 0 | 0 | 0 | 0 | 0 | 0 | 0 | 0 |
| ENSMUSG00 SNORA17 | 127 | 0 | 0 | 0 | 0 | 0 | 0 | 0 | 0 | 0 | 0 | 0 | 0 | 0 | 0 | 0 | 0 | 0 | 0 |
| ENSMUSG00 U6 | 93 | 0 | 0.35820316 | 3.12087293 | 0.4899037 | 0 | 0.53282509 | 0.50873848 | 1.64268397 | 0.88573152 | 0 | 3.59596462 | 1.20033683 | 0.53045072 | 0.99879758 | 0 | 0 | 0.41211072 | 0 |
| ENSMUSG00 U6 | 90 | 0 | 0 | 0 | 0 | 0 | 0 | 0 | 0 | 0 | 0 | 0 | 0 | 0 | 0 | 0 | 0 | 0 | 0 |
| ENSMUSG00 5S_rRNA | 102 | 0 | 0 | 0 | 0 | 0 | 0 | 0 | 0 | 0 | 0 | 0 | 0 | 0 | 0 | 0 | 0 | 0 | 0 |
| ENSMUSG00 U6 | 105 | 0 | 0 | 0 | 0.43391471 | 0 | 0 | 0.45059694 | 0 | 0.78450506 | 0 | 0.4549996 | 0 | 0.93965557 | 0 | 0 | 0 | 0.36501235 | 0 |
| ENSMUSG00 7SK | 314 | 0 | 0 | 0 | 0 | 0 | 0 | 0 | 0 | 0 | 0 | 0 | 0 | 0 | 0 | 0 | 0 | 0 | 0 |
| ENSMUSG00 U1 | 170 | 0 | 0 | 0 | 0 | 0 | 0 | 0 | 0 | 0 | 0 | 0 | 0 | 0 | 0 | 0 | 0 | 0 | 0 |
| ENSMUSG00 5S_rRNA | 93 | 0 | 0 | 0 | 0 | 0 | 0 | 0 | 0 | 0 | 0 | 0 | 0 | 0 | 0 | 0 | 0 | 0 | 0 |
| ENSMUSG00 SCARNA23 | 128 | 3.65278856 | 6.50642465 | 3.02334565 | 2.49161961 | 2.25618392 | 5.80696097 | 5.54445456 | 2.38702515 | 5.79185378 | 4.42037843 | 3.73241863 | 1.30817959 | 0.77081121 | 0.72568887 | 4.82830733 | 3.88888981 | 4.49136292 | 5.15710376 |
| ENSMUSG00 SNORA19 | 128 | 0 | 0.28025699 | 0 | 1.06783698 | 0 | 0 | 0 | 0.29837814 | 0 | 0 | 0 | 0 | 0 | 0 | 0 | 0 | 0 | 0 |
| ENSMUSG00 7SK | 178 | 0 | 0 | 0 | 0 | 0 | 0 | 0 | 0 | 0 | 0 | 0 | 0 | 0 | 0 | 0 | 0 | 0 | 0 |
| ENSMUSG00 SNORA17 | 115 | 0 | 0 | 0 | 0 | 0 | 0 | 0 | 0 | 0 | 0 | 0 | 0 | 0 | 0 | 0 | 0 | 0 | 0 |
| ENSMUSG00 SNORA17 | 126 | 0 | 0 | 0 | 0 | 0 | 0 | 0 | 0 | 0 | 0 | 0 | 0 | 0 | 0 | 0 | 0 | 0 | 0 |
| ENSMUSG00 SNORA70 | 109 | 0 | 0 | 0 | 0 | 0 | 0 | 0 | 0 | 0 | 0 | 0 | 0 | 0 | 0 | 0 | 0 | 0 | 0 |
| ENSMUSG00 SNORA17 | 120 | 0 | 0 | 0 | 0 | 0 | 0 | 0 | 0 | 0 | 0 | 0 | 0 | 0 | 0 | 0 | 0 | 0 | 0 |
| ENSMUSG00 U6 | 90 | 0 | 0 | 0 | 0 | 0 | 0 | 0 | 0 | 0 | 0 | 0 | 1.24034806 | 0.54813241 | 0 | 0 | 0 | 0 | 0 |
| ENSMUSG00 U6 | 111 | 0 | 0 | 0 | 0 | 0 | 0 | 0 | 0 | 0 | 0 | 0 | 0 | 0 | 0 | 0 | 0 | 0 | 0 |
| ENSMUSG00 U1 | 155 | 1.91958859 | 0.2149219 | 3.74504751 | 0 | 0.93158582 | 0 | 1.83145854 | 0.73920779 | 0 | 0.60839617 | 1.84935323 | 0 | 2.54616347 | 1.19855709 | 0.79744947 | 0 | 0 | 0 |
| ENSMUSG00 U1 | 164 | 0 | 0 | 0 | 0 | 0 | 0 | 0 | 0 | 0 | 0 | 0 | 0 | 0 | 0 | 0 | 0 | 0 | 0 |
| ENSMUSG00 U6 | 104 | 0 | 0 | 0 | 0 | 0 | 0 | 0 | 0 | 0 | 0 | 0 | 0 | 0 | 0 | 0 | 0 | 0 | 0 |
| ENSMUSG00 SNORA48 | 143 | 0 | 0 | 0 | 0 | 0 | 0 | 0 | 0 | 0 | 0 | 0 | 0 | 0 | 0 | 0 | 0 | 0 | 0 |
| ENSMUSG00 SNORA32 | 69 | 0 | 0 | 1.40213131 | 0 | 0 | 0 | 0 | 0 | 5.37215423 | 0.68334353 | 0 | 1.61784529 | 0 | 1.34620543 | 0 | 0 | 0 | 0 |
| ENSMUSG00 U7 | 59 | 0 | 0 | 0 | 0 | 0 | 0 | 0 | 0 | 0 | 0 | 0 | 0 | 0 | 0 | 0 | 0 | 0 | 0 |
| ENSMUSG00 SNORA17 | 129 | 0 | 0 | 0 | 0 | 0 | 0 | 0 | 0 | 0 | 0 | 0 | 0 | 0 | 0 | 0 | 0 | 0 | 0 |
| ENSMUSG00 SNORA71 | 133 | 0 | 0 | 0 | 0 | 0 | 0 | 0 | 0 | 0 | 0 | 0 | 0 | 0 | 0 | 0 | 0 | 0 | 0 |
| ENSMUSG00 7SK | 312 | 0 | 0 | 0 | 0 | 0 | 0 | 0 | 0 | 0 | 0 | 0 | 0 | 0 | 0 | 0 | 0 | 0 | 0 |
| ENSMUSG00 U1 | 159 | 0 | 0 | 0 | 0 | 0 | 0 | 0 | 0 | 0 | 0 | 0 | 0 | 0 | 0 | 0 | 0 | 0 | 0 |
| ENSMUSG00 SNORA17 | 129 | 0 | 0 | 0 | 0 | 0 | 0 | 0 | 0 | 0 | 0 | 0 | 0 | 0 | 0 | 0 | 0 | 0 | 0 |
| ENSMUSG00 snoU109 | 157 | 0 | 0 | 0 | 0 | 0 | 0 | 0 | 0 | 0 | 0 | 0 | 0 | 0 | 0 | 0 | 0 | 0 | 0 |
| ENSMUSG00 U6 | 102 | 0 | 0 | 0 | 0 | 0 | 0 | 0 | 0 | 0 | 0 | 0 | 0 | 0 | 0 | 0 | 0 | 0 | 0 |
| ENSMUSG00 5S_rRNA | 89 | 0 | 0 | 0 | 0 | 0 | 0 | 0 | 0 | 0 | 0 | 0 | 0 | 0 | 0 | 0 | 0 | 0 | 0 |
| ENSMUSG00 U7 | 62 | 0 | 0 | 0 | 0 | 0 | 0 | 0 | 0 | 0 | 0 | 0 | 0 | 0 | 0 | 0 | 0 | 0 | 0 |
| ENSMUSG00 SNORA17 | 137 | 0 | 0 | 0 | 0 | 0 | 0 | 0 | 0 | 0 | 0 | 0 | 0 | 0 | 0 | 0 | 0 | 0 | 0 |
| ENSMUSG00 SNORA17 | 136 | 0 | 0 | 0 | 0 | 0 | 0 | 0 | 0 | 0 | 0 | 0 | 0 | 0 | 0 | 0 | 0 | 0 | 0 |
| ENSMUSG00 U3 | 88 | 0 | 0 | 0 | 0 | 0 | 0 | 0 | 0 | 0 | 0 | 0 | 0 | 0 | 0 | 0 | 0 | 0 | 0 |
| ENSMUSG00 AC109196.1 | 75 | 0 | 0 | 0 | 0 | 0 | 0 | 0 | 0 | 0 | 0 | 0 | 0 | 0.6577589 | 0 | 0 | 0 | 0 | 0 |
| ENSMUSG00 7SK | 92 | 0 | 0 | 0 | 0 | 0 | 0 | 0 | 0 | 0 | 0 | 0 | 0 | 0 | 0 | 0 | 0 | 0 | 0 |
| ENSMUSG00 SNORA17 | 126 | 0 | 0 | 0 | 0 | 0 | 0 | 0 | 0 | 0 | 0 | 0 | 1.32894435 | 0 | 0 | 0 | 0 | 0 | 0 |
| ENSMUSG00 U3 | 89 | 0 | 0 | 0 | 0 | 0 | 0 | 0 | 0 | 0 | 0 | 0 | 0 | 0 | 0 | 0 | 0 | 0 | 0 |
| ENSMUSG00 SNORA17 | 113 | 0 | 0 | 0 | 0 | 0 | 0 | 0 | 0 | 0 | 0 | 0 | 0 | 0 | 0 | 0 | 0 | 0 | 0 |
| ENSMUSG00 7SK | 270 | 0 | 0 | 0 | 0 | 0 | 0 | 0 | 0 | 0 | 0 | 0 | 0 | 0 | 0 | 0 | 0 | 0 | 0 |
| ENSMUSG00 U6 | 86 | 0 | 0 | 0 | 0 | 0 | 0 | 0 | 0 | 0 | 0 | 0 | 0 | 0 | 0 | 0 | 0 | 0 | 0 |
| ENSMUSG00 7SK | 275 | 0 | 0 | 0 | 0 | 0 | 0 | 0 | 0 | 0 | 0 | 0 | 0 | 0 | 0 | 0 | 0 | 0 | 0 |
| ENSMUSG00 U6 | 109 | 0 | 0 | 0 | 0 | 0 | 0 | 0 | 0 | 0 | 0 | 0 | 0 | 0 | 0 | 0 | 0 | 0 | 0 |
| ENSMUSG00 SNORA17 | 122 | 0 | 0 | 0 | 0 | 0 | 0 | 0 | 0 | 0 | 0 | 0 | 0 | 0 | 0 | 0 | 0 | 0 | 0 |
| ENSMUSG00 Mir1934 | 82 | 0 | 0.40625481 | 0 | 1.66686748 | 1.17394936 | 0.60430163 | 2.88491945 | 0.93152201 | 3.01364749 | 1.72502573 | 1.16524289 | 0 | 4.21126123 | 0 | 0.502458 | 0.46695863 | 0.46739386 | 0 |
| ENSMUSG00 SNORA61 | 133 | 0 | 0 | 0 | 0 | 0 | 0 | 0 | 0 | 0 | 0 | 0 | 0 | 0 | 0 | 0 | 0 | 0 | 0 |
| ENSMUSG00 U6 | 103 | 0 | 0 | 0 | 0 | 0 | 0 | 0 | 0 | 0 | 0 | 0 | 0 | 0 | 0 | 0 | 0 | 0 | 0 |
| ENSMUSG00 U7 | 59 | 0 | 0 | 0 | 0 | 0 | 0 | 0 | 0 | 0 | 0 | 0 | 0 | 0 | 0 | 0 | 0 | 0 | 0 |
| ENSMUSG00 SNORA17 | 125 | 0 | 0.53300631 | 0 | 0 | 0 | 0 | 0 | 0 | 0 | 0 | 0 | 0 | 0 | 0 | 0 | 0 | 0 | 0 |
| ENSMUSG00 snR78 | 68 | 0 | 0 | 0 | 0 | 0 | 0 | 0 | 0 | 0 | 0 | 0 | 0 | 0 | 0 | 0 | 0 | 0 | 0 |
| ENSMUSG00 U6 | 105 | 0 | 0.31726566 | 0 | 0 | 0 | 0 | 0 | 0 | 0 | 0 | 0 | 0 | 0 | 1.76929857 | 0 | 0 | 0 | 0 |
| ENSMUSG00 SNORA17 | 113 | 0 | 0 | 0 | 0 | 0 | 0 | 0 | 0 | 0 | 0 | 0 | 0 | 0 | 0 | 0 | 0 | 0 | 0 |
| ENSMUSG00 U6 | 96 | 0 | 0 | 0 | 0 | 0 | 0 | 0 | 0 | 0 | 0 | 0 | 0 | 0 | 0 | 0 | 0 | 0 | 0 |
| ENSMUSG00 U6 | 102 | 0 | 0 | 0 | 0 | 0 | 0 | 0 | 0 | 0 | 0 | 0 | 0 | 0 | 0 | 0 | 0 | 0 | 0 |
| ENSMUSG00 U3 | 89 | 0 | 0 | 0 | 0 | 0 | 0 | 0 | 0 | 0 | 0 | 0 | 0 | 0 | 0 | 0 | 0 | 0 | 0 |
| ENSMUSG00 U6 | 101 | 0 | 0.65966127 | 0 | 0 | 0 | 0 | 0 | 0 | 0 | 0 | 0 | 0 | 0 | 0 | 0 | 0 | 0 | 0 |
| ENSMUSG00 SNORA17 | 124 | 0 | 0 | 0 | 0 | 0 | 0 | 0 | 0 | 0 | 0 | 0 | 0 | 0 | 0 | 0 | 0 | 0 | 0 |
| ENSMUSG00 SNORA17 | 122 | 0 | 0 | 0 | 0 | 0 | 0 | 0 | 0 | 0 | 0 | 0 | 0 | 0 | 0 | 0 | 0 | 0 | 0 |
| ENSMUSG00 7SK | 274 | 0 | 0 | 0 | 0 | 0 | 0 | 0 | 0 | 0 | 0 | 0 | 0 | 0 | 0 | 0 | 0 | 0 | 0 |
| ENSMUSG00 U6 | 99 | 0 | 0 | 0 | 0 | 0 | 0 | 0 | 0 | 0 | 0 | 0 | 0 | 0 | 0 | 0 | 0 | 0 | 0 |
| ENSMUSG00 7SK | 282 | 0 | 0.35439249 | 0 | 0.48469196 | 0.17068058 | 0 | 0 | 0.13543405 | 0 | 0.16720108 | 0.84707373 | 0 | 0 | 0.32939069 | 0.43831442 | 0.67891148 | 0.13590885 | 1.56054203 |
| ENSMUSG00 SNORA17 | 122 | 0 | 0 | 0 | 0 | 0 | 0 | 0 | 0 | 0 | 0 | 0 | 0 | 0 | 0 | 0 | 0 | 0 | 0 |
| ENSMUSG00 SNORA81 | 160 | 0 | 0 | 0 | 0 | 0 | 0 | 0 | 0 | 0 | 0 | 0 | 0 | 0 | 0 | 0 | 0 | 0 | 0 |
| ENSMUSG00 5S_rRNA | 107 | 0 | 0 | 0 | 0 | 0 | 0 | 0 | 0 | 0 | 0 | 0 | 0 | 0 | 0 | 0 | 0 | 0 | 0 |
| ENSMUSG00 SNORA48 | 143 | 0 | 0 | 0 | 0 | 0 | 0 | 0 | 0 | 0 | 0 | 0 | 0 | 0 | 0 | 0 | 0 | 0 | 0 |
| ENSMUSG00 7SK | 310 | 0.68556736 | 0.2149219 | 0.78021823 | 0.88182666 | 0 | 0.47954258 | 0.76310772 | 0.61600649 | 0.13285973 | 0 | 0.15411277 | 1.26035367 | 0.63654087 | 0 | 0 | 0.12351809 | 0.74179929 | 1.24214112 |
| ENSMUSG00 U6 | 105 | 0 | 0 | 0 | 0 | 0 | 0 | 0 | 0 | 0 | 0 | 0 | 0 | 0 | 0 | 0 | 0 | 0 | 0 |
| ENSMUSG00 SCARNA4 | 143 | 0 | 1.39774381 | 2.02965862 | 1.59304351 | 0 | 1.03956784 | 1.65428947 | 0.53415947 | 0.28801759 | 1.31890079 | 1.67045309 | 3.51287387 | 0.68995688 | 2.2734868 | 0.28812277 | 1.07106594 | 2.1441285 | 4.23146975 |
| ENSMUSG00 snoU13 | 108 | 0 | 0 | 0 | 0 | 0 | 0 | 0 | 0 | 0 | 0 | 0 | 0 | 0 | 0 | 0 | 1.41817064 | 0 | 0 |
| ENSMUSG00 U6 | 115 | 0 | 0 | 0 | 0 | 0 | 0 | 0 | 0 | 0 | 0 | 0 | 0 | 0 | 0 | 0 | 0 | 0 | 0 |
| ENSMUSG00 SNORA5 | 91 | 0 | 0 | 0 | 0 | 0 | 0 | 0 | 0 | 0 | 0 | 0 | 0 | 0 | 0 | 0 | 0 | 0 | 0 |
| ENSMUSG00 SNORA17 | 114 | 0 | 0 | 0 | 0 | 0 | 0 | 0 | 0 | 0 | 0 | 0 | 0 | 0 | 0 | 0 | 0 | 0 | 0 |
| ENSMUSG00 SNORA17 | 134 | 0 | 0 | 0 | 0 | 0 | 0 | 0 | 0 | 0 | 0 | 0 | 0 | 0 | 0 | 0 | 0 | 0 | 0 |
| ENSMUSG00 AC134903.1 | 100 | 0 | 0 | 0 | 0 | 0 | 0 | 0 | 0 | 0 | 0 | 0 | 0 | 0 | 0 | 0 | 0 | 0 | 0 |
| ENSMUSG00 SNORA55 | 93 | 0 | 0 | 0 | 0 | 0 | 0 | 0 | 0 | 0 | 0 | 0 | 0 | 0 | 0 | 0 | 0 | 0 | 0 |
| ENSMUSG00 7SK | 254 | 0 | 0 | 0 | 0 | 0 | 0 | 0 | 0 | 0 | 0 | 0 | 0 | 0 | 0 | 0 | 0 | 0 | 0 |
| ENSMUSG00 U6 | 100 | 0 | 0 | 0 | 0 | 0 | 0 | 0 | 0 | 0 | 0 | 0 | 0.55815663 | 1.97327669 | 0 | 1.23604668 | 1.5316243 | 0 | 0 |
| ENSMUSG00 Scarna13 | 274 | 0.46538514 | 1.45895887 | 1.23582012 | 0.33256237 | 1.58097559 | 1.80849393 | 0.86337005 | 1.11510664 | 0.3006315 | 1.54874573 | 0.17436116 | 2.44448157 | 0.54013048 | 0.33900794 | 0.15037064 | 1.39746742 | 1.53864696 | 0 |
| ENSMUSG00 SNORA18 | 126 | 0 | 0 | 0 | 0 | 0 | 2.75292964 | 0 | 0 | 0 | 0 | 0 | 0 | 0 | 0 | 0 | 0 | 0 | 0 |
| ENSMUSG00 U6 | 94 | 0 | 0 | 0 | 0 | 0 | 0 | 0 | 0 | 0 | 0 | 0 | 0 | 0 | 0 | 0 | 0 | 0 | 0 |
| ENSMUSG00 SNORA17 | 116 | 0 | 0 | 0 | 0 | 0 | 0 | 0 | 0 | 0 | 0 | 0 | 0 | 0 | 0 | 0 | 0 | 0 | 0 |
| ENSMUSG00 U6 | 116 | 0.73284786 | 0.28718012 | 0 | 0 | 0 | 0 | 0 | 0.32924485 | 1.06516851 | 0 | 0 | 0 | 0 | 0 | 0 | 0 | 0 | 0 |
| ENSMUSG00 7SK | 298 | 0 | 0 | 0 | 0 | 0 | 0 | 0 | 0 | 0 | 0 | 0 | 0 | 0 | 0 | 0 | 0 | 0 | 0 |
| ENSMUSG00 Mir1935 | 59 | 1.44085342 | 0 | 0 | 0 | 0 | 0 | 0 | 0 | 0 | 0 | 0 | 0 | 0 | 0 | 0 | 0 | 0 | 0 |
| ENSMUSG00 SNORA17 | 133 | 0 | 0 | 0 | 0 | 0 | 0 | 0 | 0 | 0 | 0 | 0 | 0 | 0 | 0 | 0 | 0 | 0 | 0 |
| ENSMUSG00 7SK | 298 | 0 | 0 | 0 | 0 | 0 | 0 | 0 | 0 | 0 | 0 | 0 | 0 | 0 | 0 | 0 | 0 | 0 | 0 |
| ENSMUSG00 U6atac | 103 | 0 | 0 | 0 | 0 | 0 | 0 | 0 | 0 | 0 | 0 | 0 | 0 | 0 | 0 | 0 | 0 | 0 | 0 |
| ENSMUSG00 5S_rRNA | 118 | 0 | 0 | 0 | 0 | 0 | 0 | 0 | 0 | 0 | 0 | 0 | 0 | 0 | 0 | 0 | 0 | 0 | 0 |
| ENSMUSG00 SNORA17 | 71 | 0 | 0 | 0 | 0 | 0 | 0 | 0 | 0 | 0 | 0 | 0 | 0 | 0 | 0 | 0 | 0 | 0 | 0 |
| ENSMUSG00 U6 | 105 | 0 | 0 | 0 | 0 | 0 | 0 | 0 | 0 | 0 | 0 | 0 | 0 | 0 | 0 | 0 | 0 | 0 | 0 |
| ENSMUSG00 7SK | 259 | 0 | 0 | 0 | 0 | 0 | 0 | 0 | 0 | 0 | 0 | 0 | 0 | 0 | 0 | 0 | 0 | 0 | 0 |
| ENSMUSG00 U6 | 93 | 0 | 0 | 0 | 0 | 0 | 0 | 0 | 0 | 0 | 0 | 0 | 0 | 0 | 0 | 0 | 0 | 0 | 0 |
| ENSMUSG00 SNORA84 | 132 | 0 | 0 | 0 | 0 | 0 | 0 | 0 | 0.28933638 | 0 | 0 | 0 | 0 | 0 | 0 | 0 | 0 | 0 | 0 |
| ENSMUSG00 SNORA17 | 135 | 0 | 0 | 0 | 0 | 0 | 0 | 0 | 0 | 0 | 0 | 0 | 0 | 0 | 0 | 0 | 0 | 0 | 0 |
| ENSMUSG00 AL806518.1 | 102 | 0 | 0 | 0 | 0 | 0 | 0 | 0 | 0 | 0 | 0 | 0 | 0 | 0 | 0 | 0 | 0 | 0 | 0 |
| ENSMUSG00 SNORA40 | 129 | 0 | 0 | 0 | 0 | 0 | 0 | 0 | 0 | 0 | 0 | 0 | 0 | 0 | 0 | 0 | 0 | 0 | 0 |
| ENSMUSG00 U6 | 106 | 0 | 0 | 0 | 0 | 0 | 0 | 0 | 0 | 0 | 0 | 0 | 0 | 0 | 0 | 0 | 0 | 0 | 0 |
| ENSMUSG00 U6 | 103 | 0 | 0 | 0 | 0 | 0 | 0 | 0 | 0 | 0 | 0 | 0 | 0 | 0 | 0 | 0 | 0 | 0 | 0 |
| ENSMUSG00 7SK | 326 | 0 | 0 | 0 | 0 | 0 | 0 | 0 | 0 | 0 | 0 | 0 | 0 | 0 | 0 | 0 | 0 | 0 | 0 |
| ENSMUSG00 5S_rRNA | 100 | 0 | 0 | 0 | 0 | 0 | 0 | 0 | 0 | 0 | 0 | 0 | 0 | 0 | 0 | 0 | 0 | 0 | 0 |
| ENSMUSG00 SCARNA3 | 130 | 0 | 0 | 0 | 0 | 0 | 0 | 0 | 0 | 0 | 0 | 0 | 0 | 0 | 0 | 0 | 0 | 0 | 0 |
| ENSMUSG00 SNORA17 | 128 | 0 | 0 | 0 | 0 | 0 | 0 | 0 | 0 | 0 | 0 | 0 | 0 | 0 | 0 | 0 | 0 | 0 | 0 |
| ENSMUSG00 7SK | 300 | 0 | 0 | 0 | 0 | 0 | 0 | 0 | 0 | 0 | 0 | 0 | 0 | 0 | 0 | 0 | 0 | 0 | 0 |
| ENSMUSG00 7SK | 247 | 0 | 0 | 0 | 0 | 0 | 0 | 0 | 0 | 0 | 0 | 0 | 0 | 0.39944872 | 0.56409823 | 0 | 0 | 0 | 0.22270893 |
| ENSMUSG00 SNORD113 | 67 | 0 | 0 | 0 | 0 | 0 | 0 | 0 | 0 | 0 | 0 | 0 | 0 | 0 | 0 | 0 | 0 | 0 | 0 |
| ENSMUSG00 U6 | 101 | 0 | 0.98949191 | 0 | 0 | 0 | 0 | 0 | 0 | 0 | 0 | 0 | 0.55263032 | 0 | 0 | 0 | 0 | 0 | 0 |
| ENSMUSG00 SNORA70 | 122 | 0 | 0 | 0 | 0 | 0 | 0 | 0 | 0 | 0 | 0 | 0 | 0 | 0 | 0 | 0 | 0 | 0 | 0 |
| ENSMUSG00 SNORA17 | 125 | 0 | 0 | 0 | 0 | 0 | 0 | 0 | 0 | 0 | 0 | 0 | 0 | 0 | 0 | 0 | 0 | 0 | 0 |
| ENSMUSG00 Y_RNA | 109 | 0 | 0 | 0 | 0.41799123 | 0 | 0 | 0 | 0 | 0 | 0 | 0 | 0 | 0 | 0 | 0 | 0 | 0 | 0 |
| ENSMUSG00 U3 | 89 | 0 | 0 | 0 | 0 | 0 | 0 | 0 | 0 | 0 | 0 | 0 | 0 | 0 | 0 | 0 | 0 | 0 | 0 |
| ENSMUSG00 U7 | 62 | 0 | 0 | 0 | 0 | 0 | 0 | 0 | 0 | 0 | 0 | 0 | 0 | 0 | 0 | 0 | 0 | 0 | 0 |
| ENSMUSG00 SNORA17 | 111 | 0 | 0 | 0 | 0 | 0 | 0 | 0 | 0 | 0 | 0 | 0 | 0 | 0 | 0 | 0 | 0 | 0 | 0 |
| ENSMUSG00 U6 | 103 | 0 | 0 | 0 | 0 | 0 | 0 | 0 | 0 | 0 | 0 | 0 | 0 | 0 | 0 | 0 | 0 | 0 | 0 |
| ENSMUSG00 U7 | 63 | 0 | 0 | 0 | 0 | 0 | 0 | 0 | 0 | 0 | 0 | 0 | 0 | 0 | 0 | 0 | 0 | 0 | 0 |
| ENSMUSG00 U6 | 105 | 0 | 0 | 0 | 0 | 0 | 0 | 0 | 0 | 0 | 0 | 0 | 0 | 0 | 0.44232464 | 0 | 0 | 0 | 0 |
| ENSMUSG00 AL772216.39 | 95 | 0 | 0 | 0 | 0 | 0 | 0 | 0 | 0 | 0 | 0 | 0 | 0 | 0 | 0 | 0 | 0 | 0 | 0 |
| ENSMUSG00 snoU89 | 270 | 0 | 0.24676218 | 0 | 0.84372304 | 0 | 0.18352864 | 0 | 0.70726671 | 1.06779856 | 0 | 0.70777716 | 0.20672468 | 0 | 0 | 0 | 0.28363413 | 0.56779699 | 0 |
| ENSMUSG00 SNORA48 | 136 | 0 | 0 | 0 | 0 | 0 | 0 | 0 | 0 | 0 | 0 | 0 | 0 | 0 | 0 | 0 | 0 | 0 | 0 |
| ENSMUSG00 U6 | 106 | 0 | 0 | 0 | 0 | 0 | 0 | 0 | 0 | 0 | 0 | 0 | 0 | 0 | 0 | 0 | 0 | 0 | 0 |
| ENSMUSG00 U7 | 57 | 0 | 0 | 0 | 0 | 0 | 0 | 0 | 0 | 0 | 0 | 0 | 0 | 0 | 0 | 0 | 0 | 0 | 0 |
| ENSMUSG00 SNORA17 | 96 | 0 | 0 | 0 | 0 | 0 | 0 | 0 | 0 | 0 | 0 | 0 | 0 | 0 | 0 | 0 | 0 | 0 | 0 |
| ENSMUSG00 7SK | 282 | 0 | 0 | 0 | 0 | 0 | 0 | 0 | 0 | 0 | 0 | 0 | 0 | 0 | 0 | 0 | 0 | 0 | 0 |
| ENSMUSG00 U6atac | 114 | 0 | 0 | 0 | 0.79931657 | 0 | 0 | 0 | 0 | 0 | 0 | 0 | 0 | 0 | 0 | 0.36141716 | 0 | 0 | 0 |
| ENSMUSG00 U6 | 104 | 0.40870362 | 0 | 0.4651301 | 0 | 0.92561392 | 0.47646859 | 0 | 0 | 0 | 0 | 0 | 0 | 0 | 0 | 0 | 0 | 0 | 0 |
| ENSMUSG00 SNORA17 | 128 | 0 | 0 | 0 | 0 | 0 | 0 | 0 | 0 | 0 | 0 | 0 | 0 | 0 | 0 | 0 | 0 | 0 | 0 |
| ENSMUSG00 SNORA17 | 124 | 0 | 0 | 0 | 0 | 0 | 0 | 0 | 0 | 0 | 0 | 0 | 0 | 0 | 0 | 0 | 0 | 0 | 0 |
| ENSMUSG00 7SK | 254 | 0.33468643 | 0.13115313 | 0.76178788 | 0 | 0.75798305 | 0 | 0 | 0.15036379 | 0 | 0 | 0 | 0 | 0 | 0 | 0 | 0.30150085 | 0.15089093 | 0 |
| ENSMUSG00 SNORA26 | 136 | 0 | 0 | 0 | 0 | 0 | 0 | 0 | 0 | 0 | 0 | 0 | 0 | 0 | 0 | 0 | 0 | 0 | 0 |
| ENSMUSG00 SNORA4 | 124 | 0 | 0 | 0 | 0 | 0 | 0 | 0 | 0 | 0 | 0 | 0 | 0 | 0 | 0 | 0 | 0 | 0 | 0 |
| ENSMUSG00 SNORA17 | 130 | 0 | 0 | 0 | 0 | 0 | 0 | 0 | 0 | 0 | 0 | 0 | 0 | 0 | 0 | 0 | 0 | 0 | 0 |
| ENSMUSG00 SNORA17 | 116 | 0 | 0 | 0 | 0 | 0 | 0 | 0 | 0 | 1.06516851 | 0 | 0 | 0 | 0 | 0 | 0 | 0.33009144 | 0 | 0 |
| ENSMUSG00 U1 | 124 | 0 | 0 | 0 | 0 | 0 | 0 | 0 | 0 | 0 | 0 | 0 | 0 | 0 | 0 | 0 | 0 | 0 | 0 |
| ENSMUSG00 U6 | 106 | 0 | 0 | 0 | 0 | 0 | 0 | 0 | 0 | 0 | 0 | 0 | 0 | 1.86158178 | 0 | 1.16608177 | 0 | 0 | 0 |
| ENSMUSG00 SNORA48 | 143 | 0.89171698 | 0 | 0 | 0 | 0.33658688 | 0 | 0.33085789 | 2.1366379 | 0 | 0 | 0 | 0 | 0 | 0 | 0 | 0.80329946 | 0.26801606 | 0 |
| ENSMUSG00 SNORA40 | 120 | 0 | 0 | 0 | 0 | 0 | 0 | 0 | 0 | 0 | 0 | 0 | 0 | 0 | 0 | 0 | 0 | 0 | 0 |
| ENSMUSG00 SNORA17 | 131 | 0 | 0 | 0 | 0 | 0 | 0 | 0 | 0 | 0 | 0 | 0 | 0 | 0 | 0 | 0 | 0 | 0 | 0 |
| ENSMUSG00 Mir1947 | 84 | 0 | 0 | 0 | 0 | 0 | 0 | 0 | 0 | 0 | 0 | 0 | 0 | 0 | 0 | 0 | 0 | 0 | 0 |
| ENSMUSG00 5S_rRNA | 129 | 0 | 0 | 0 | 0 | 0 | 0 | 0 | 0 | 0 | 0 | 1.11104555 | 0 | 0 | 0 | 0 | 0 | 0 | 0 |
| ENSMUSG00 U6 | 70 | 0 | 0 | 0 | 0 | 0 | 0 | 0 | 0 | 0 | 0 | 0 | 0 | 0 | 0 | 0 | 0 | 0 | 0 |
| ENSMUSG00 U7 | 61 | 0 | 0 | 0 | 0 | 0 | 0 | 0 | 0 | 0 | 0 | 0 | 0 | 0 | 0 | 0 | 0 | 0 | 0 |
| ENSMUSG00 7SK | 292 | 0 | 0 | 0 | 0 | 0 | 0 | 0 | 0 | 0 | 0.16147501 | 0 | 0.57344859 | 0 | 0 | 0 | 0 | 0 | 0 |
| ENSMUSG00 7SK | 310 | 0 | 0 | 0 | 0 | 0 | 0 | 0 | 0 | 0 | 0 | 0 | 0 | 0 | 0 | 0 | 0 | 0 | 0 |
| ENSMUSG00 SNORD19 | 59 | 0 | 0 | 0 | 0 | 0 | 0 | 0 | 0 | 0 | 0 | 0 | 0 | 0 | 0 | 0 | 0 | 0 | 0 |
| ENSMUSG00 5S_rRNA | 114 | 0 | 0 | 0 | 0 | 0 | 0 | 0 | 0 | 0 | 0 | 0 | 0 | 0 | 0 | 0 | 0 | 0 | 0 |
| ENSMUSG00 SNORA17 | 122 | 0 | 0 | 0 | 0 | 0 | 0 | 0 | 0 | 0 | 0 | 0 | 0 | 0 | 0 | 0 | 0 | 0 | 0 |
| ENSMUSG00 snoU13 | 99 | 0 | 1.00948164 | 0 | 0 | 0 | 0 | 0.9558117 | 0 | 0 | 0 | 0 | 0 | 0 | 0 | 0 | 0 | 0 | 0 |
| ENSMUSG00 SNORA17 | 111 | 0 | 0 | 0 | 0 | 0 | 0 | 0 | 0 | 0 | 0 | 0 | 0.50284381 | 0 | 0 | 0 | 0 | 0 | 0 |
| ENSMUSG00 AC154485.1 | 103 | 0 | 0 | 0 | 0 | 0 | 0 | 0 | 0 | 0 | 0 | 0 | 0 | 0 | 0 | 0 | 0 | 0 | 0 |
| ENSMUSG00 U6 | 91 | 0 | 0 | 0 | 0 | 0 | 0 | 0 | 0 | 0 | 0 | 0 | 0 | 0 | 0 | 0 | 0 | 0 | 0 |
| ENSMUSG00 RNaseP_nuc | 328 | 0 | 0 | 0 | 0 | 0 | 0 | 0 | 0 | 0 | 0 | 0 | 0 | 0 | 0 | 0 | 0 | 0 | 0 |
| ENSMUSG00 SNORA70 | 128 | 0.66414338 | 0 | 1.51167282 | 1.77972829 | 1.88015327 | 1.16139219 | 0.3696303 | 0.89513443 | 2.57415724 | 0.36836487 | 1.49296745 | 3.92453878 | 2.31243362 | 0 | 0.64377431 | 0.89743611 | 1.79654517 | 0 |
| ENSMUSG00 SNORA70 | 108 | 0 | 0 | 0 | 0 | 0 | 0 | 0 | 0 | 0 | 0 | 0 | 0 | 0 | 0 | 0 | 0 | 0 | 0 |
| ENSMUSG00 SNORA17 | 125 | 0 | 0 | 0 | 0 | 0 | 0 | 0 | 0 | 0 | 0 | 0 | 0 | 0 | 0 | 0 | 0 | 0 | 0 |
| ENSMUSG00 SNORA17 | 113 | 0 | 0 | 0 | 0 | 0 | 0 | 0 | 0 | 0 | 0 | 0 | 0 | 0 | 0 | 0 | 0 | 0 | 0 |
| ENSMUSG00 U6 | 102 | 0 | 0 | 0 | 0 | 0 | 0 | 0 | 0 | 0 | 0 | 0 | 0 | 0 | 0 | 0 | 0 | 0 | 0 |
| ENSMUSG00 7SK | 257 | 0 | 0 | 0 | 0 | 0 | 0 | 0 | 0 | 0 | 0 | 0 | 0 | 0 | 0 | 0 | 0 | 0 | 0 |
| ENSMUSG00 7SK | 202 | 0 | 0 | 0 | 0 | 0 | 0 | 0 | 0 | 0 | 0 | 0 | 0 | 0 | 0 | 0 | 0 | 0 | 0 |
| ENSMUSG00 SNORA17 | 131 | 0 | 0 | 0 | 0 | 0 | 0 | 0 | 0 | 0 | 0 | 0 | 0 | 0 | 0 | 0 | 0 | 0 | 0 |
| ENSMUSG00 SNORA17 | 116 | 0 | 0 | 0 | 0 | 0 | 0 | 0 | 0 | 0 | 0 | 0 | 0 | 0 | 0 | 0 | 0 | 0 | 0 |
| ENSMUSG00 5S_rRNA | 132 | 0 | 0 | 0 | 0 | 0 | 0 | 0 | 0 | 0 | 0 | 0 | 0 | 0 | 0 | 0.312133 | 1.16032144 | 0 | 0 |
| ENSMUSG00 U7 | 57 | 0 | 0 | 0 | 0 | 0 | 0 | 0 | 0 | 0 | 0 | 0 | 0 | 0 | 0 | 0 | 0 | 0 | 0 |
| ENSMUSG00 SNORA17 | 132 | 0 | 0 | 0 | 0 | 0 | 0 | 0 | 0 | 0 | 0 | 0 | 0 | 0 | 0 | 0 | 0 | 0 | 0 |
| ENSMUSG00 U6 | 106 | 0 | 0.94281776 | 0 | 0 | 0 | 0 | 0 | 0 | 0 | 0 | 0 | 0 | 0 | 0 | 0 | 0 | 0 | 0 |
| ENSMUSG00 SNORA17 | 135 | 0 | 0 | 0 | 0 | 0 | 0 | 0 | 0 | 0 | 0 | 0 | 0 | 0 | 0 | 0 | 0 | 0 | 0 |
| ENSMUSG00 SNORA76 | 123 | 0 | 0 | 0 | 0 | 0 | 0.80573551 | 1.5386237 | 0.62101467 | 0.33484972 | 0 | 0 | 0.45378587 | 0 | 0 | 0 | 0.6226115 | 0 | 1.78891404 |
| ENSMUSG00 AL772376.1 | 67 | 0 | 0 | 0 | 0 | 0 | 0 | 0 | 0 | 0 | 0 | 0 | 0 | 0 | 0 | 0.6149486 | 1.14300321 | 0 | 0 |
| ENSMUSG00 SNORA44 | 128 | 0 | 0 | 0 | 0 | 0 | 0 | 0 | 0 | 0 | 0 | 0 | 0 | 0 | 0 | 0 | 0 | 0 | 0 |
| ENSMUSG00 SNORA81 | 177 | 0 | 0 | 0 | 0 | 0 | 0 | 0 | 0 | 0.23269218 | 0 | 0 | 0 | 0 | 0 | 0 | 0 | 0 | 0 |
| ENSMUSG00 SNORA17 | 150 | 0 | 0 | 0 | 0 | 0 | 0 | 0 | 0 | 0 | 0 | 0 | 0 | 0 | 0 | 0 | 0 | 0 | 0 |
| ENSMUSG00 SNORA17 | 133 | 0 | 0 | 0 | 0 | 0 | 0 | 0 | 0 | 0 | 0 | 0 | 0 | 0 | 0 | 0 | 0 | 0 | 0 |
| ENSMUSG00 7SK | 304 | 0 | 0 | 0 | 0 | 0 | 0 | 0 | 0 | 0 | 0 | 0 | 0 | 0 | 0 | 0 | 0 | 0 | 0 |
| ENSMUSG00 SNORA17 | 122 | 0 | 0 | 0 | 0 | 0 | 0 | 0 | 0 | 0 | 0 | 0 | 0 | 0 | 0 | 0 | 0 | 0 | 0 |
| ENSMUSG00 SNORA17 | 144 | 0 | 0 | 0 | 0 | 0 | 0 | 0 | 0 | 1.14406988 | 0 | 0 | 0 | 1.02774828 | 0 | 0 | 0 | 0 | 0 |
| ENSMUSG00 Mir1943 | 72 | 0 | 0 | 0 | 0 | 0 | 0 | 0 | 0 | 0.57203494 | 0 | 0 | 3.10087014 | 0 | 0 | 0 | 0 | 0 | 0 |
| ENSMUSG00 7SK | 266 | 0 | 0 | 0 | 0 | 0 | 0 | 0 | 0.14358046 | 0.46450958 | 0 | 0 | 0 | 0 | 0 | 0 | 0.43184896 | 0 | 0 |
| ENSMUSG00 SCARNA16 | 166 | 0 | 0 | 0 | 0 | 0 | 0 | 0 | 0 | 0 | 0 | 0 | 0 | 0 | 0 | 0 | 0 | 0 | 0 |
| ENSMUSG00 U1 | 153 | 0 | 0 | 0 | 0 | 0 | 0 | 0 | 0 | 0 | 0 | 0 | 0 | 0 | 0 | 0 | 0 | 0 | 0 |
| ENSMUSG00 SNORA17 | 130 | 0 | 0 | 0 | 0 | 0 | 0 | 0 | 0 | 0 | 0 | 0 | 0 | 0 | 0 | 0 | 0 | 0 | 0 |
| ENSMUSG00 SNORA17 | 109 | 0 | 0 | 0 | 0 | 0 | 0 | 0 | 0 | 0 | 0 | 0 | 0 | 0 | 0 | 0 | 0 | 0 | 0 |
| ENSMUSG00 SNORA17 | 131 | 0 | 0 | 0 | 0 | 0 | 0 | 0 | 0 | 0 | 0 | 0 | 0 | 0 | 0 | 0 | 0 | 0 | 0 |
| ENSMUSG00 7SK | 285 | 0 | 0 | 0 | 0 | 0 | 0 | 0 | 0 | 0 | 0 | 0 | 0 | 0 | 0 | 0 | 0 | 0 | 0 |
| ENSMUSG00 SNORD86 | 65 | 0 | 0 | 0 | 0 | 0 | 0 | 0 | 0 | 0 | 0 | 0 | 0 | 0 | 0 | 0 | 0 | 0 | 0 |
| ENSMUSG00 U6 | 102 | 0 | 0 | 0 | 0 | 0 | 0 | 0 | 0 | 0 | 0 | 0 | 0 | 0 | 0 | 1.21181047 | 0 | 0 | 0 |
| ENSMUSG00 SNORA17 | 120 | 0 | 0 | 0 | 0 | 0 | 0 | 0 | 0 | 0 | 0 | 0 | 0 | 0 | 0 | 0 | 0 | 0 | 0 |
| ENSMUSG00 AC107678.1 | 59 | 0 | 0 | 0 | 0 | 0 | 0 | 0 | 0 | 0 | 0 | 0 | 0 | 0 | 0 | 0 | 0 | 0 | 0 |
| ENSMUSG00 U1 | 149 | 0 | 0 | 0 | 0 | 0 | 0 | 0 | 0 | 0 | 0 | 0 | 0 | 0 | 0 | 0 | 0 | 0 | 0 |
| ENSMUSG00 U6 | 107 | 0 | 0 | 0 | 0 | 0 | 0 | 0 | 0 | 0 | 0 | 0 | 0 | 0 | 0 | 0 | 0 | 0 | 0 |
| ENSMUSG00 U6 | 102 | 0 | 0 | 0 | 0 | 0 | 0 | 0 | 0 | 0 | 0 | 0 | 0 | 0 | 0 | 0 | 0 | 0 | 0 |
| ENSMUSG00 U6 | 104 | 0 | 1.60158145 | 0 | 0 | 0 | 0 | 0 | 0 | 0 | 0 | 0 | 0 | 0.94869072 | 0 | 0 | 0.73635783 | 0 | 0.52893372 |
| ENSMUSG00 SNORA17 | 131 | 0 | 0 | 0 | 0 | 0 | 0 | 0 | 0 | 0 | 0 | 0 | 0 | 0 | 0 | 0 | 0 | 0 | 0 |
| ENSMUSG00 U6 | 100 | 0 | 0 | 0 | 0 | 0 | 0 | 0 | 0 | 0 | 0 | 0 | 0 | 0 | 0 | 0 | 0 | 0 | 0 |
| ENSMUSG00 7SK | 276 | 0 | 0 | 0 | 0 | 0 | 0 | 0 | 0 | 0 | 0 | 0 | 0 | 0 | 0 | 0 | 0 | 0 | 0 |
| ENSMUSG00 snoU13 | 87 | 0 | 0 | 0 | 0 | 0 | 0 | 0 | 0 | 0 | 0 | 0 | 0 | 0 | 0 | 0 | 0 | 0 | 0 |

| | | | | | | | | | | | | | | | | | | | |
|---|---|---|---|---|---|---|---|---|---|---|---|---|---|---|---|---|---|---|---|
| ENSMUSG00 U6 | 105 | 0 | 0 | 0 | 0 | 0 | 0 | 0 | 0 | 0 | 0 | 0 | 0 | 0 | 0 | 0 | 0 | 0 | 0 |
| ENSMUSG00 SNORA17 | 130 | 0 | 0 | 0 | 0 | 0 | 0 | 0 | 0 | 0 | 0 | 0 | 0 | 0 | 0 | 0 | 0 | 0 | 0 |
| ENSMUSG00 SNORA17 | 59 | 0 | 0 | 0 | 0 | 0 | 0 | 0 | 0 | 0 | 0 | 0 | 0 | 0 | 0 | 0 | 0 | 0 | 0 |
| ENSMUSG00 SNORD70 | 67 | 0 | 0 | 0 | 0 | 0 | 0 | 0 | 0 | 0 | 0 | 0 | 0 | 0 | 0 | 0 | 0 | 0 | 0 |
| ENSMUSG00 5S_rRNA | 104 | 0 | 0 | 0 | 0 | 0 | 0 | 0 | 0 | 0 | 0 | 0 | 0 | 0 | 0 | 0 | 0 | 0 | 0 |
| ENSMUSG00 U6 | 105 | 0 | 0 | 0 | 0 | 0 | 0 | 0 | 0 | 0 | 0 | 0 | 0 | 0 | 0 | 0 | 0 | 0 | 0 |
| ENSMUSG00 SNORD5 | 70 | 10.9299024 | 3.80718791 | 3.45525217 | 3.90523237 | 4.12559346 | 1.41579239 | 4.05537248 | 6.54726898 | 4.11865158 | 6.73581475 | 9.5549917 | 0.79736661 | 4.93319173 | 3.31743481 | 4.12015559 | 2.18803471 | 1.09503705 | 3.92922191 |
| ENSMUSG00 SNORA70 | 136 | 0 | 0 | 0 | 0 | 0 | 0 | 0 | 0 | 0 | 0 | 0 | 0 | 0 | 0 | 0 | 0 | 0 | 0 |
| ENSMUSG00 U6 | 98 | 0 | 0 | 0 | 0 | 0 | 0 | 0 | 0 | 0 | 0 | 0 | 0 | 0 | 0 | 0 | 0 | 0 | 0 |
| ENSMUSG00 U6 | 103 | 0 | 0 | 0 | 0 | 0 | 0 | 0 | 0 | 0 | 0 | 0 | 0 | 0 | 0 | 0 | 0 | 0 | 0 |
| ENSMUSG00 U3 | 89 | 0 | 0 | 0 | 0 | 0 | 0 | 0 | 0 | 0 | 0 | 0 | 0 | 0 | 0 | 0 | 0 | 0 | 0 |
| ENSMUSG00 U6 | 102 | 0 | 0 | 0 | 0 | 0 | 0 | 0 | 0 | 0 | 0 | 0 | 0 | 0 | 0 | 0 | 0 | 0 | 0 |
| ENSMUSG00 5S_rRNA | 137 | 0 | 0 | 0 | 0 | 0 | 0 | 0 | 0 | 0 | 0 | 0 | 0 | 0 | 0 | 0 | 0 | 0 | 0 |
| ENSMUSG00 U6 | 85 | 0 | 0 | 0 | 0 | 0 | 0 | 0 | 0 | 0 | 0 | 0 | 0 | 0 | 0 | 0 | 0 | 0 | 0 |
| ENSMUSG00 7SK | 305 | 0 | 0 | 0 | 0 | 0 | 0 | 0 | 0 | 0 | 0 | 0 | 0 | 0 | 0 | 0 | 0 | 0 | 0 |
| ENSMUSG00 7SK | 82 | 0 | 0 | 0 | 0 | 0 | 0 | 0 | 0 | 0 | 0 | 0 | 0 | 0 | 0 | 0 | 0 | 0 | 0 |
| ENSMUSG00 SNORA17 | 135 | 0 | 0 | 0 | 0 | 0 | 0 | 0 | 0 | 0 | 0 | 0 | 0 | 0 | 0 | 0 | 0 | 0 | 0 |
| ENSMUSG00 U2 | 171 | 0.49713656 | 0.58443674 | 1.41443071 | 1.06575542 | 0.56294648 | 0.57956414 | 1.38341166 | 2.45682121 | 5.5397068 | 1.10294043 | 1.67631433 | 1.9584443 | 0.86547223 | 0.27160285 | 1.20472386 | 0.22392168 | 0.89652156 | 1.28676273 |
| ENSMUSG00 7SK | 302 | 0 | 0 | 0 | 0 | 0 | 0 | 0 | 0 | 0 | 0 | 0 | 0 | 0 | 0 | 0 | 0 | 0 | 0 |
| ENSMUSG00 U6 | 105 | 0 | 0 | 0 | 0 | 0 | 0 | 0 | 0 | 0 | 0 | 0 | 0 | 0 | 0 | 0 | 0 | 0 | 0 |
| ENSMUSG00 U3 | 88 | 0 | 0 | 0 | 0 | 0 | 0 | 0 | 0 | 0 | 0 | 0 | 0 | 0 | 0 | 0 | 0 | 0 | 0 |
| ENSMUSG00 U6 | 106 | 0 | 0 | 0 | 0 | 0 | 0 | 0 | 0 | 0 | 0 | 0 | 0 | 0 | 0 | 0 | 0 | 0 | 0 |
| ENSMUSG00 5S_rRNA | 131 | 0 | 0 | 0 | 0 | 0.73483853 | 0 | 0 | 0 | 0 | 0 | 0 | 0 | 0 | 0 | 0 | 0 | 0 | 0 |
| ENSMUSG00 snoU97 | 167 | 0 | 0 | 0 | 0 | 0 | 0 | 0 | 0 | 0 | 0 | 0 | 0 | 0 | 0 | 0 | 0 | 0 | 0 |
| ENSMUSG00 5S_rRNA | 106 | 0 | 0 | 0 | 0 | 0 | 0 | 0 | 0 | 0 | 0 | 0 | 0 | 0 | 0 | 0 | 0 | 0 | 0 |
| ENSMUSG00 5S_rRNA | 136 | 0 | 0 | 0 | 0 | 0 | 0 | 0 | 0 | 0 | 0.69339269 | 0 | 0 | 0 | 1.02450193 | 0 | 0 | 0 | 0 |
| ENSMUSG00 U6atac | 107 | 0 | 0 | 0 | 0 | 0 | 0 | 0 | 0 | 0 | 0 | 0 | 0 | 0 | 0 | 0 | 0 | 0 | 0 |
| ENSMUSG00 snoU13 | 101 | 2.10421664 | 3.95796763 | 5.74735014 | 3.60879559 | 2.8593222 | 0.98124225 | 2.34221183 | 5.29399637 | 8.1557457 | 6.0689024 | 4.73019391 | 4.42104258 | 3.41904377 | 3.21889715 | 2.85555338 | 1.5164597 | 1.51787314 | 1.08928924 |
| ENSMUSG00 SNORA17 | 113 | 0 | 0 | 0 | 0 | 0 | 0 | 0 | 0 | 0 | 0 | 0 | 0 | 0 | 0 | 0 | 0 | 0 | 0 |
| ENSMUSG00 RNaseP_nuc | 432 | 0 | 0 | 0 | 0 | 0 | 0 | 0 | 0 | 0 | 0 | 0 | 0 | 0 | 0 | 0 | 0 | 0 | 0 |
| ENSMUSG00 SNORA61 | 127 | 0 | 0 | 1.52357576 | 0 | 0 | 0 | 0 | 0 | 0 | 0 | 0 | 0 | 0 | 0 | 0 | 0 | 0 | 0 |
| ENSMUSG00 SNORA25 | 125 | 0 | 0 | 0 | 0 | 0 | 0 | 0 | 0 | 0 | 0 | 0 | 0 | 0 | 0 | 0.32961245 | 0 | 0 | 0 |
| ENSMUSG00 U6 | 105 | 0 | 0 | 0 | 0 | 0 | 0 | 0 | 0 | 0 | 0 | 0 | 0 | 0 | 0 | 0 | 0 | 0 | 0 |
| ENSMUSG00 Scarna8 | 130 | 0 | 0 | 0 | 0 | 1.1107367 | 0 | 0 | 0 | 0 | 0.72539543 | 0.73499936 | 0 | 0 | 0 | 0.95080514 | 0 | 0 | 0 |
| ENSMUSG00 U6 | 109 | 0 | 0 | 0 | 0 | 0 | 0 | 0 | 0 | 0 | 0 | 0 | 0 | 0 | 0 | 0 | 0 | 0 | 0 |
| ENSMUSG00 U6 | 110 | 0 | 0 | 0 | 0 | 0 | 0 | 0 | 0 | 0 | 0 | 0 | 0 | 0 | 0 | 0 | 0 | 0 | 0 |
| ENSMUSG00 U6 | 101 | 0 | 0 | 0 | 0 | 0 | 0 | 0 | 0 | 0 | 0.46683865 | 0 | 0 | 0 | 0 | 0 | 0 | 0 | 0 |
| ENSMUSG00 U6 | 104 | 0 | 0 | 0 | 0 | 0 | 0 | 0 | 0 | 0 | 0 | 0 | 0 | 0 | 0 | 0 | 0 | 0 | 0 |
| ENSMUSG00 U6 | 87 | 0 | 0 | 0 | 0 | 0 | 0 | 0 | 0 | 0 | 0 | 0 | 0 | 0 | 0 | 0 | 0 | 0 | 0 |
| ENSMUSG00 snoU13 | 90 | 0 | 0 | 0 | 0 | 1.0695983 | 0.55058593 | 0 | 0.84872005 | 0 | 1.57169011 | 0 | 0 | 0 | 0 | 0.45779507 | 0 | 1.27754323 | 0 |
| ENSMUSG00 SNORA25 | 116 | 0 | 0 | 0 | 0 | 0 | 0 | 0 | 0 | 0 | 0 | 0 | 0 | 0 | 0 | 0 | 0 | 0 | 0 |
| ENSMUSG00 U6 | 103 | 0 | 0 | 0 | 0 | 0 | 0 | 0 | 0 | 0 | 0 | 0 | 0 | 0 | 0 | 0 | 0 | 0 | 0 |
| ENSMUSG00 U6 | 109 | 0 | 0.30562288 | 0 | 0 | 0 | 0 | 0 | 0 | 0 | 0.86515052 | 0 | 0 | 0 | 0 | 0 | 0 | 0 | 0 |
| ENSMUSG00 U7 | 59 | 0 | 0 | 0 | 0 | 0 | 0 | 0 | 0 | 0 | 0 | 0 | 0 | 0 | 0 | 0 | 0 | 0 | 0 |
| ENSMUSG00 5S_rRNA | 47 | 0 | 0 | 0 | 0 | 0 | 0 | 0 | 0 | 0 | 0 | 0 | 0 | 0 | 0 | 0 | 0 | 0 | 0 |
| ENSMUSG00 U6 | 107 | 0 | 0 | 0 | 0 | 0 | 0 | 0 | 0 | 0 | 0 | 0 | 0 | 0 | 0 | 0 | 0 | 0 | 0 |
| ENSMUSG00 AL683804.1 | 105 | 0 | 0 | 0 | 0 | 0 | 0 | 1.35179083 | 0 | 0 | 0 | 0 | 0 | 2.34913892 | 0 | 0 | 0 | 0 | 0 |
| ENSMUSG00 U6 | 103 | 0 | 0 | 0 | 0 | 0 | 0 | 0 | 0 | 0 | 0 | 0 | 0 | 0 | 0 | 0 | 0 | 0 | 0 |
| ENSMUSG00 SNORA17 | 115 | 0 | 0 | 0 | 0 | 0 | 0 | 0 | 0 | 0 | 0 | 0 | 0 | 0 | 0 | 0 | 0 | 0 | 0 |
| ENSMUSG00 7SK | 267 | 0 | 0 | 0 | 0 | 0 | 0 | 0 | 0 | 0 | 0 | 0 | 0 | 0 | 0 | 0 | 0 | 0 | 0 |
| ENSMUSG00 5S_rRNA | 99 | 0 | 0 | 0.48862152 | 1.38063771 | 0 | 0 | 0 | 0 | 0 | 0 | 0 | 0 | 0 | 0 | 0 | 0 | 0 | 0 |
| ENSMUSG00 SNORA72 | 89 | 0 | 0 | 0 | 0 | 0 | 0 | 0 | 0 | 0 | 0 | 0 | 0 | 0 | 0 | 0 | 0 | 0 | 0 |
| ENSMUSG00 Mir669m-2 | 97 | 0 | 0 | 0 | 0 | 0 | 0 | 0 | 0 | 0 | 0 | 0 | 0 | 0 | 0 | 0 | 0 | 0 | 0 |
| ENSMUSG00 U6 | 104 | 0 | 0 | 0 | 0 | 0 | 0 | 0 | 0 | 0 | 0 | 0 | 0 | 0 | 0 | 0 | 0 | 0 | 0 |
| ENSMUSG00 U6 | 105 | 0 | 0 | 0 | 0 | 0 | 0 | 0.90119388 | 1.45494866 | 0 | 0.44905432 | 0 | 0.53157774 | 0.46982778 | 1.32697393 | 0 | 0 | 0.36501235 | 0 |
| ENSMUSG00 Mir1952 | 78 | 0 | 0 | 0 | 0 | 0 | 0 | 0 | 0 | 0 | 0 | 0 | 0 | 0 | 0 | 0 | 0 | 0 | 0 |
| ENSMUSG00 SNORA17 | 132 | 0.32200891 | 2.52370411 | 0 | 0 | 0 | 0.3753995 | 0 | 0 | 0 | 0.71440459 | 0 | 0.42284593 | 0.37372665 | 0 | 0 | 0.29008036 | 0.58070147 | 0.41673566 |
| ENSMUSG00 Mir1946a | 133 | 0 | 0 | 0 | 0 | 0 | 0 | 0 | 0 | 0 | 0 | 0 | 0 | 0 | 0 | 0 | 0 | 0 | 0 |
| ENSMUSG00 AC113291.1 | 104 | 0 | 0 | 0 | 0 | 0 | 0 | 0 | 0 | 0 | 0 | 0 | 0 | 0 | 0 | 0 | 0 | 0 | 0 |
| ENSMUSG00 U7 | 61 | 0 | 0 | 0 | 0 | 0 | 0 | 0 | 0 | 0 | 0 | 0 | 0 | 0 | 0 | 0 | 0 | 0 | 0 |
| ENSMUSG00 5S_rRNA | 94 | 0 | 0 | 0 | 0 | 0 | 0 | 0 | 0 | 0 | 0 | 0 | 0 | 0 | 0 | 0 | 0 | 0 | 0 |
| ENSMUSG00 U1 | 123 | 0 | 0 | 0 | 0 | 0 | 0 | 0 | 0 | 0 | 0 | 0 | 0 | 0 | 0 | 0 | 0 | 0 | 0 |
| ENSMUSG00 SCARNA1 | 164 | 0 | 0 | 0.29496055 | 0 | 0 | 0 | 1.15396778 | 0 | 0 | 0 | 0 | 0 | 0.60160875 | 0 | 0 | 0 | 0.7010908 | 0 |
| ENSMUSG00 U2 | 152 | 0 | 0 | 0 | 0 | 0 | 0 | 0 | 0 | 0 | 0 | 0 | 0 | 0 | 0 | 0 | 0 | 0 | 0 |
| ENSMUSG00 U6 | 97 | 0 | 0 | 0 | 0 | 0 | 0 | 0 | 0 | 0 | 0 | 0 | 0 | 0 | 0 | 0 | 0 | 0 | 0 |
| ENSMUSG00 U6 | 66 | 1.93205346 | 1.00948164 | 0 | 0 | 0 | 0 | 0 | 0 | 1.24807624 | 0 | 0 | 1.69138371 | 0.74745329 | 2.11109488 | 0.624266 | 0.58016072 | 0.58070147 | 0 |
| ENSMUSG00 7SK | 283 | 0 | 0 | 0 | 0 | 0 | 0 | 0 | 0 | 0 | 0 | 0 | 0 | 0 | 0 | 0 | 0 | 0 | 0 |
| ENSMUSG00 U6 | 91 | 0 | 0 | 0 | 0 | 0 | 0 | 0 | 0 | 0 | 0 | 0 | 0 | 0 | 0 | 0 | 0 | 0 | 0 |
| ENSMUSG00 SNORA25 | 125 | 0 | 0 | 0 | 0 | 0 | 0 | 0 | 0 | 0 | 0 | 0 | 0 | 0 | 0 | 0 | 0 | 0 | 0 |
| ENSMUSG00 U6 | 98 | 0 | 0 | 0.49360745 | 0 | 0 | 0 | 0 | 0 | 0 | 0 | 0.97499915 | 0 | 1.51016073 | 0 | 0 | 1.56288193 | 0.39108466 | 0 |
| ENSMUSG00 SNORA17 | 106 | 0 | 0 | 0 | 0 | 0 | 0 | 0 | 0 | 0 | 0 | 0 | 0 | 1.86158178 | 0 | 0 | 0 | 0 | 0 |
| ENSMUSG00 U6 | 103 | 0 | 0 | 0 | 0 | 0 | 0 | 0 | 0 | 0 | 0 | 0 | 0 | 0 | 0 | 0 | 0 | 0 | 0 |
| ENSMUSG00 SNORD19B | 76 | 0 | 0 | 0 | 0 | 0 | 0 | 0 | 0 | 0 | 0 | 0 | 0 | 0 | 0 | 0 | 0 | 0 | 0 |
| ENSMUSG00 SNORA70 | 134 | 0 | 0 | 0 | 0 | 0 | 0 | 0.35307969 | 0 | 0 | 0.70374184 | 0 | 0.4165348 | 0 | 0 | 0 | 0 | 0 | 2.05257861 |
| ENSMUSG00 U6 | 107 | 0 | 0 | 0 | 0 | 0 | 0 | 0 | 0 | 0 | 0 | 0 | 0 | 0 | 0 | 0 | 0 | 0 | 0 |
| ENSMUSG00 5S_rRNA | 110 | 0 | 0 | 0 | 0 | 0 | 0 | 0 | 0 | 0 | 0 | 0 | 0 | 0 | 0 | 0 | 0 | 0 | 0 |
| ENSMUSG00 U6 | 98 | 0 | 0 | 0 | 0 | 0 | 0 | 0 | 0 | 0 | 0 | 0 | 0 | 0 | 0 | 0 | 0 | 0 | 0 |
| ENSMUSG00 U3 | 89 | 0 | 0 | 0 | 0 | 0 | 0 | 0 | 0 | 0 | 0 | 0 | 0 | 0 | 0 | 0 | 0 | 0 | 0 |
| ENSMUSG00 SNORA68 | 128 | 0 | 0 | 0 | 0 | 0 | 0 | 0 | 0 | 0 | 0 | 0 | 0 | 0 | 0 | 0 | 0 | 0 | 0 |
| ENSMUSG00 U7 | 63 | 0 | 0 | 0 | 0 | 0 | 0 | 0 | 0 | 0 | 0 | 0 | 0 | 0 | 0 | 0 | 0 | 0 | 0 |
| ENSMUSG00 U6 | 105 | 0 | 0 | 0 | 0 | 0 | 0 | 0 | 0 | 0 | 0 | 0 | 0 | 0 | 0 | 0 | 0 | 0 | 0 |
| ENSMUSG00 SNORA48 | 142 | 0.29933223 | 0.70379354 | 0.34065866 | 0.32085242 | 0.67791442 | 0 | 2.66550304 | 0.26896058 | 0 | 0 | 0.67288674 | 0 | 0.69481574 | 0 | 0 | 0 | 0.2699035 | 0 |
| ENSMUSG00 AL591146.1 | 77 | 0 | 0 | 0 | 0 | 0 | 0 | 0 | 0 | 0 | 0 | 0 | 0 | 0 | 0 | 0 | 0 | 0 | 0 |
| ENSMUSG00 SNORA36 | 117 | 0 | 0 | 0 | 0 | 0 | 0 | 0 | 0 | 0 | 0 | 0 | 0 | 0 | 0 | 0 | 0 | 0 | 0 |
| ENSMUSG00 U6 | 98 | 0.86745257 | 0.67985498 | 0 | 3.25436031 | 0.49114208 | 2.52820069 | 0 | 1.16915518 | 0 | 4.33016663 | 3.89999661 | 1.70864273 | 2.01354764 | 0.47391926 | 1.68169616 | 2.73504339 | 2.73759264 | 0.56131742 |
| ENSMUSG00 7SK | 280 | 0 | 0 | 0 | 0 | 0 | 0 | 0 | 0 | 0 | 0 | 0 | 0 | 0 | 0 | 0 | 0 | 0 | 0 |
| ENSMUSG00 SNORA17 | 126 | 0 | 0 | 0 | 0 | 0 | 0 | 0 | 0 | 0 | 0 | 0 | 0 | 0 | 0 | 0 | 0 | 0 | 0 |
| ENSMUSG00 U6 | 107 | 0 | 0 | 0 | 0 | 0 | 0 | 0 | 0 | 0 | 0 | 0 | 0 | 0 | 0 | 0 | 0 | 0 | 0 |
| ENSMUSG00 U6 | 104 | 0 | 0 | 0 | 0 | 0 | 0 | 0 | 0 | 0 | 0 | 0 | 0 | 0 | 0 | 0 | 0 | 0 | 0 |
| ENSMUSG00 U1 | 163 | 0 | 0 | 0 | 0 | 0 | 0 | 0 | 0 | 0 | 0 | 0 | 0 | 0 | 0 | 0 | 0 | 0 | 0 |
| ENSMUSG00 SNORA31 | 125 | 0 | 0 | 0 | 0 | 0 | 0 | 0 | 0 | 0 | 0 | 0 | 0 | 0 | 0 | 0 | 0 | 0 | 0 |
| ENSMUSG00 SNORA17 | 130 | 0 | 0 | 0 | 0 | 0 | 0 | 0 | 0 | 0 | 0 | 0 | 0 | 0.75895257 | 0 | 0.63387009 | 0.29454313 | 0 | 0 |
| ENSMUSG00 SNORA17 | 127 | 0 | 0 | 0 | 0 | 0 | 0 | 0 | 0 | 0 | 0 | 0 | 0 | 0 | 0 | 0 | 0 | 0 | 0 |
| ENSMUSG00 U6 | 99 | 0 | 0 | 0.97724304 | 0 | 0 | 0 | 0.47790585 | 0.38578184 | 0 | 0 | 0.96515068 | 1.12758914 | 0 | 0 | 0 | 0 | 0.38713431 | 0.55564754 |
| ENSMUSG00 SNORA17 | 127 | 0 | 0 | 0 | 0 | 0 | 0 | 0 | 0 | 0 | 0 | 0 | 0 | 0 | 0 | 0 | 0 | 0 | 0 |
| ENSMUSG00 U7 | 64 | 1.99243013 | 1.04102794 | 3.02334565 | 1.42378264 | 5.26442915 | 0 | 0.73926061 | 2.38702515 | 3.21769654 | 0.73672974 | 0 | 0.87211973 | 0 | 3.62844433 | 7.08151742 | 0.59829074 | 0.59884839 | 3.43806917 |
| ENSMUSG00 SNORA17 | 101 | 0 | 0 | 0 | 0 | 0 | 0 | 0 | 0 | 0 | 0 | 0 | 0 | 0 | 0 | 0 | 0 | 0 | 0 |
| ENSMUSG00 7SK | 327 | 0 | 0 | 0 | 0 | 0 | 0 | 0 | 0 | 0 | 0 | 0 | 0 | 0 | 0 | 0 | 0 | 0 | 0 |
| ENSMUSG00 U7 | 58 | 0 | 0 | 0 | 0 | 0 | 0 | 0 | 0 | 0 | 0 | 0 | 0 | 0 | 0 | 0 | 0 | 0 | 0 |
| ENSMUSG00 U7 | 47 | 0 | 0 | 0 | 0 | 0 | 0 | 0 | 0 | 0 | 0 | 0 | 0 | 0 | 0 | 0 | 0 | 0 | 0 |
| ENSMUSG00 SNORA70 | 134 | 0 | 0 | 0 | 0 | 0 | 0 | 0 | 0 | 0 | 0 | 0 | 0 | 0 | 0 | 0 | 0 | 0 | 0 |
| ENSMUSG00 ACA64 | 140 | 0 | 0 | 0 | 0 | 0 | 0 | 0 | 0 | 0 | 0 | 0 | 0 | 0 | 0 | 0 | 0 | 0 | 0 |
| ENSMUSG00 U6 | 103 | 0 | 0 | 0 | 0 | 0 | 0 | 0 | 0 | 0 | 0 | 0 | 0 | 0 | 0 | 0 | 0 | 0 | 0 |
| ENSMUSG00 SNORA17 | 125 | 0 | 0 | 0 | 0 | 0 | 0 | 0 | 0 | 0 | 0 | 0 | 0 | 0 | 0 | 0 | 0 | 0 | 0 |
| ENSMUSG00 SNORA17 | 129 | 0 | 0 | 0 | 0 | 0 | 0 | 0 | 0 | 0 | 0 | 0 | 0 | 0 | 0 | 0 | 0 | 0 | 0 |
| ENSMUSG00 5S_rRNA | 105 | 0 | 0 | 0 | 0 | 0 | 0 | 0 | 0 | 0 | 0 | 0 | 0 | 0 | 0 | 0 | 0 | 0 | 0 |
| ENSMUSG00 U6 | 102 | 0 | 0 | 0 | 0 | 0 | 0 | 0 | 0 | 0 | 0 | 0 | 0 | 0 | 0 | 0 | 0 | 0 | 0 |
| ENSMUSG00 RNaseP_nuc | 218 | 0 | 0 | 0 | 0 | 0 | 0 | 0 | 0 | 0 | 0 | 0 | 0 | 0 | 0 | 0 | 0 | 0 | 0 |
| ENSMUSG00 SNORA61 | 135 | 0 | 0 | 0 | 0 | 0 | 0 | 0 | 0 | 0 | 0 | 0 | 0 | 0 | 0 | 0 | 0 | 0 | 0 |
| ENSMUSG00 7SK | 220 | 0 | 0 | 0 | 0 | 0 | 0 | 0 | 0 | 0 | 0 | 0 | 0 | 0 | 0 | 0 | 0 | 0 | 0 |
| ENSMUSG00 U1 | 147 | 0 | 0.67985498 | 0 | 0 | 0 | 0 | 0 | 0 | 0 | 0 | 0.97499915 | 0 | 0 | 0 | 0 | 0 | 0 | 0 |
| ENSMUSG00 SNORA70 | 134 | 0 | 0 | 0 | 0 | 0 | 0 | 1.05923908 | 0 | 0 | 0.70374184 | 0 | 0 | 0 | 0 | 0 | 0 | 0.57203428 | 0 |
| ENSMUSG00 SNORA17 | 133 | 0 | 0 | 0 | 0 | 0 | 0 | 0 | 0 | 0 | 0 | 0 | 0 | 0 | 0 | 0 | 0 | 0 | 0 |
| ENSMUSG00 5S_rRNA | 130 | 0 | 0 | 0 | 0 | 0 | 0 | 0 | 0 | 0 | 0 | 0 | 0 | 0 | 0 | 0 | 0 | 0 | 0 |
| ENSMUSG00 7SK | 252 | 0 | 0 | 0 | 0.18079779 | 0 | 0.39327566 | 0 | 0.3031143 | 0 | 0 | 0 | 0.8859629 | 0 | 0 | 0 | 0.15194685 | 0 | 0 |
| ENSMUSG00 7SK | 287 | 0 | 0 | 0 | 0 | 0 | 0 | 0 | 0 | 0 | 0 | 0 | 0 | 0 | 0 | 0 | 0 | 0 | 0 |
| ENSMUSG00 SNORA25 | 71 | 0 | 0 | 0 | 0 | 0 | 0 | 0 | 0 | 0 | 0 | 0 | 0 | 0 | 0 | 0 | 0 | 0 | 0 |
| ENSMUSG00 U6 | 103 | 0 | 0 | 0 | 0 | 0 | 0 | 0 | 0 | 0 | 0 | 0 | 0 | 0 | 0 | 0 | 0 | 0 | 0 |
| ENSMUSG00 SNORA40 | 124 | 0 | 0 | 0 | 0 | 0 | 0 | 0 | 0 | 0 | 0 | 0 | 0 | 0 | 0 | 0 | 0 | 0 | 0 |
| ENSMUSG00 U7 | 63 | 0 | 0 | 0 | 0 | 0 | 0 | 0 | 0 | 0 | 0 | 0 | 0 | 0 | 0 | 0 | 0 | 0 | 0 |
| ENSMUSG00 U6 | 102 | 0 | 0 | 0 | 0 | 0 | 0 | 0 | 0 | 0 | 0 | 0 | 0 | 0 | 0 | 0 | 0 | 0 | 0 |
| ENSMUSG00 U6 | 103 | 0 | 0 | 0 | 0 | 0 | 0 | 0 | 0 | 0 | 0 | 0 | 0 | 0 | 0 | 0 | 0 | 0 | 0 |
| ENSMUSG00 U7 | 61 | 0 | 0 | 0 | 0 | 0 | 0 | 0 | 0 | 0 | 0 | 0 | 0 | 0 | 0 | 0 | 0 | 0 | 0 |
| ENSMUSG00 U6 | 97 | 0 | 0 | 0 | 0 | 0 | 0 | 0 | 0 | 0 | 0 | 0 | 0 | 0 | 0 | 0 | 0 | 0 | 0 |
| ENSMUSG00 U3 | 200 | 0 | 0 | 0 | 0 | 0 | 0 | 0 | 0 | 0 | 0 | 0 | 0 | 0 | 0 | 0 | 0 | 0 | 0 |
| ENSMUSG00 U1 | 175 | 0 | 0 | 0 | 0 | 0 | 0 | 0 | 0 | 0 | 0 | 0 | 0 | 0 | 0 | 0 | 0 | 0 | 0 |
| ENSMUSG00 7SK | 303 | 0 | 0 | 0 | 0 | 0 | 0 | 0 | 0 | 0 | 0 | 0 | 0 | 0 | 0 | 0 | 0 | 0 | 0 |
| ENSMUSG00 U6 | 98 | 2.16863143 | 0.67985498 | 1.48082236 | 0 | 0 | 0 | 0 | 0.38971839 | 2.10135285 | 0 | 0 | 1.13909515 | 0 | 0 | 0 | 1.17216145 | 0 | 1.68395225 |
| ENSMUSG00 7SK | 331 | 0 | 0 | 0 | 0 | 0 | 0 | 0 | 0 | 0 | 0 | 0 | 0 | 0 | 0 | 0 | 0 | 0 | 0 |
| ENSMUSG00 7SK | 310 | 0 | 0 | 0 | 0 | 0 | 0 | 0 | 0 | 0 | 0 | 0 | 0 | 0 | 0 | 0 | 0 | 0 | 0 |
| ENSMUSG00 U7 | 60 | 0 | 0 | 0 | 0 | 0 | 0 | 0 | 0 | 0 | 0 | 0 | 0 | 0 | 0 | 0 | 0 | 0 | 0 |
| ENSMUSG00 AC099599.1 | 117 | 0 | 0 | 0 | 0 | 0 | 0 | 0 | 0 | 0 | 0 | 0 | 0 | 0 | 0 | 0 | 0 | 0 | 0 |
| ENSMUSG00 U7 | 61 | 0 | 0 | 0 | 0 | 0 | 0 | 0 | 0 | 0 | 0 | 0 | 0 | 0 | 0 | 0 | 0 | 0 | 0 |
| ENSMUSG00 SNORA64 | 135 | 1.25941262 | 0.24676218 | 0 | 0 | 0.71306554 | 0.73411457 | 0 | 0 | 0.3050853 | 0 | 1.41555433 | 0.41344935 | 0.73084322 | 0 | 0 | 0.56726826 | 0.56779699 | 0 |
| ENSMUSG00 U6 | 101 | 0 | 0.32983064 | 0 | 0 | 0 | 0 | 0.93688473 | 0 | 0 | 0 | 0 | 0 | 0 | 0 | 0 | 1.13734477 | 0.37946829 | 0 |
| ENSMUSG00 U6 | 103 | 0 | 0 | 0 | 0 | 0 | 0 | 0 | 0 | 0 | 0 | 0 | 0 | 0 | 0 | 0 | 0 | 0 | 0 |
| ENSMUSG00 SNORA4 | 123 | 0 | 0 | 0.78656147 | 0 | 0 | 0.40286775 | 0.38465593 | 0.31050734 | 0 | 0 | 1.94207148 | 1.36135762 | 1.20321749 | 0 | 0 | 0 | 0.31159591 | 1.34168553 |
| ENSMUSG00 5S_rRNA | 102 | 0 | 0 | 0 | 0 | 0 | 0 | 0 | 0 | 0 | 0 | 0 | 0 | 0 | 0 | 0 | 0 | 0 | 0 |
| ENSMUSG00 U6 | 103 | 0 | 0 | 0 | 0 | 0 | 0 | 0 | 0 | 0 | 0 | 0 | 0 | 0 | 0 | 0 | 0 | 0 | 0.534069 |
| ENSMUSG00 7SK | 299 | 0 | 0 | 0 | 0 | 0 | 0 | 0 | 0 | 0 | 0 | 0 | 0 | 0 | 0.31066279 | 0 | 0 | 0 | 0 |
| ENSMUSG00 U3 | 89 | 0 | 0 | 0 | 0 | 0 | 0 | 0 | 0 | 0 | 0 | 0 | 0 | 0 | 0 | 0 | 0 | 0 | 0 |
| ENSMUSG00 U7 | 61 | 0 | 0 | 0 | 0 | 0 | 0 | 0 | 0 | 0 | 0 | 0 | 0 | 0 | 0 | 0 | 0 | 0 | 0 |
| ENSMUSG00 U6 | 104 | 0 | 0 | 0 | 0 | 0 | 0 | 0 | 0 | 0 | 0 | 0 | 0 | 0 | 0 | 0 | 0 | 0 | 0 |
| ENSMUSG00 SNORA17 | 134 | 0 | 0 | 0 | 0 | 0 | 0 | 0 | 0 | 0 | 0 | 0 | 0 | 0 | 0 | 0 | 0 | 0 | 0 |
| ENSMUSG00 U6 | 99 | 0 | 0 | 0 | 0 | 0 | 0 | 0 | 0 | 0 | 0 | 0 | 0 | 0 | 0 | 0 | 0 | 0 | 0 |
| ENSMUSG00 U6 | 103 | 0 | 0 | 0 | 0 | 0 | 0 | 0 | 0.74160005 | 0 | 0 | 0 | 0 | 0 | 0 | 0 | 0 | 0 | 0 |
| ENSMUSG00 7SK | 275 | 0 | 0 | 0 | 0 | 0 | 0 | 0 | 0 | 0 | 0 | 0 | 0 | 0 | 0 | 0 | 0 | 0 | 0 |
| ENSMUSG00 U7 | 61 | 0 | 0 | 0 | 0 | 0 | 0 | 0 | 0 | 0 | 0 | 0 | 0 | 0 | 0 | 0 | 0 | 0 | 0 |
| ENSMUSG00 7SK | 295 | 0 | 0 | 0 | 0 | 0 | 0 | 0 | 0 | 0 | 0 | 0 | 0 | 0 | 0 | 0 | 0 | 0 | 0 |
| ENSMUSG00 U1 | 177 | 1.68099566 | 0 | 0.54659356 | 0 | 0.54386354 | 0.27995895 | 0 | 0 | 0.46538436 | 0 | 0.26991502 | 0.31534273 | 0.83613419 | 0.52479195 | 0 | 0 | 1.08266375 | 0 |
| ENSMUSG00 U7 | 64 | 0 | 0 | 0 | 0 | 0 | 0 | 0 | 0 | 0 | 0 | 0 | 0 | 0 | 0 | 0 | 0 | 0 | 0 |
| ENSMUSG00 SNORA17 | 131 | 0 | 0 | 0 | 0 | 0 | 0 | 0 | 0 | 0 | 0 | 0 | 0 | 0 | 0 | 0 | 0 | 0 | 0 |
| ENSMUSG00 AC112941.1 | 84 | 0 | 0 | 0.57587536 | 0 | 0 | 0 | 0 | 0 | 0 | 0 | 0 | 0 | 0 | 0 | 0 | 0 | 0 | 0 |
| ENSMUSG00 5S_rRNA | 127 | 0 | 0 | 0 | 0 | 0 | 0 | 0 | 0.30072758 | 0 | 0 | 0 | 0 | 0 | 0 | 0 | 0 | 0 | 0 |
| ENSMUSG00 7SK | 289 | 0 | 0 | 0 | 0 | 0 | 0 | 0 | 0 | 0 | 0 | 0 | 0 | 0 | 0 | 0.42769781 | 0 | 0 | 0 |
| ENSMUSG00 U7 | 65 | 0 | 0 | 0 | 0 | 0 | 0 | 0 | 0 | 0 | 0 | 0 | 0 | 0 | 0 | 0 | 0 | 0 | 0 |
| ENSMUSG00 U6 | 87 | 0 | 1.91453415 | 0 | 0 | 0 | 0 | 0 | 0 | 0 | 1.62588632 | 0.54913745 | 0 | 0 | 0 | 0 | 0 | 0 | 0 |
| ENSMUSG00 Snord11 | 85 | 0 | 0 | 0 | 0 | 0 | 0 | 0 | 0 | 0 | 0 | 0 | 0 | 0 | 0 | 0 | 0 | 0 | 0 |
| ENSMUSG00 5S_rRNA | 36 | 0 | 0 | 0 | 0 | 0 | 0 | 0 | 0 | 0 | 0 | 0 | 0 | 0 | 0 | 0 | 0 | 0 | 0 |
| ENSMUSG00 5S_rRNA | 103 | 0 | 0 | 0 | 0 | 0 | 0 | 0 | 0 | 0 | 0 | 0 | 0 | 0 | 0 | 0 | 0 | 0 | 0 |
| ENSMUSG00 U6 | 94 | 0 | 0 | 0 | 0 | 0 | 0 | 0 | 0 | 0 | 0 | 0 | 0 | 0 | 0 | 0 | 0 | 0 | 0 |
| ENSMUSG00 SNORA17 | 127 | 0 | 0 | 0 | 0 | 0 | 0 | 0 | 0 | 0 | 0 | 0 | 0 | 0 | 0 | 0 | 0 | 0 | 0 |
| ENSMUSG00 SNORA17 | 137 | 0 | 0 | 0 | 0 | 0 | 0 | 0 | 0 | 0 | 0 | 0 | 0 | 0 | 0 | 0 | 0 | 0 | 0 |
| ENSMUSG00 SNORA5 | 91 | 0 | 0 | 0 | 0 | 0 | 0 | 0 | 0 | 0 | 0 | 0 | 0 | 0 | 0 | 0 | 0 | 0 | 0 |
| ENSMUSG00 U6 | 98 | 0.43372629 | 0 | 0 | 0.46490862 | 0 | 0 | 0 | 0 | 0.84054114 | 1.44338888 | 0 | 0 | 1.00677382 | 0 | 0 | 0 | 0 | 0 |
| ENSMUSG00 5S_rRNA | 116 | 1.83211966 | 0.57436025 | 2.50207916 | 0.39276762 | 4.14930376 | 2.13589369 | 1.63147169 | 6.25565212 | 1.77528085 | 2.84530106 | 1.23555927 | 1.92467802 | 1.7011006 | 0.40038006 | 4.97260157 | 1.65045722 | 1.98239467 | 2.84529862 |
| ENSMUSG00 5S_rRNA | 107 | 0 | 0 | 0 | 0 | 0 | 0 | 0 | 0 | 0 | 0 | 0 | 0 | 0 | 0 | 0 | 0 | 0 | 0 |
| ENSMUSG00 U6 | 66 | 0 | 0 | 0 | 0 | 0 | 0 | 0 | 0 | 0 | 0 | 0 | 0 | 0 | 0 | 0 | 0 | 0 | 0 |
| ENSMUSG00 U6 | 107 | 0 | 0 | 0 | 0 | 0 | 0 | 0 | 0 | 0 | 0 | 0 | 0 | 0 | 0 | 0 | 0 | 0 | 0 |
| ENSMUSG00 U7 | 63 | 0 | 0 | 0 | 0 | 0 | 0 | 0 | 0 | 0 | 0 | 0 | 0 | 0 | 0 | 0 | 0 | 0 | 0 |
| ENSMUSG00 RNaseP_nuc | 398 | 0 | 0 | 0 | 0 | 0 | 0 | 0 | 0 | 0 | 0 | 0 | 0 | 0 | 0 | 0 | 0 | 0 | 0 |
| ENSMUSG00 SNORA74 | 124 | 0 | 0 | 0 | 0 | 0 | 0 | 0 | 0 | 0 | 0 | 0 | 0 | 0 | 0 | 0 | 0 | 0 | 0 |
| ENSMUSG00 RNaseP_nuc | 301 | 0 | 0 | 0 | 0 | 0 | 0 | 0 | 0 | 0 | 0 | 0 | 0 | 0 | 0 | 0 | 0 | 0 | 0 |
| ENSMUSG00 SNORA17 | 125 | 0 | 0 | 0 | 0 | 0 | 0 | 0 | 0 | 0 | 0 | 0 | 0 | 0 | 0 | 0 | 0 | 0 | 0 |
| ENSMUSG00 U6 | 100 | 0 | 0 | 0 | 0 | 0 | 0 | 0 | 0 | 0 | 0 | 0 | 0 | 0 | 0 | 0 | 0 | 0 | 0 |
| ENSMUSG00 SNORA17 | 142 | 0 | 0 | 0 | 0 | 0 | 0 | 0 | 0 | 0 | 0 | 0 | 0 | 0 | 0 | 0 | 0 | 0 | 0 |
| ENSMUSG00 U6 | 102 | 0 | 0 | 0 | 0 | 0 | 0 | 0 | 0 | 0 | 0 | 0 | 0 | 0 | 0 | 0 | 0 | 0 | 0 |
| ENSMUSG00 7SK | 317 | 0 | 0 | 0 | 0 | 0 | 0 | 0 | 0 | 0 | 0 | 0 | 0 | 0 | 0 | 0 | 0 | 0 | 0 |
| ENSMUSG00 SNORA17 | 129 | 0 | 0 | 0 | 0 | 0 | 0 | 0 | 0 | 0 | 0 | 0 | 0 | 0 | 0 | 0 | 0 | 0 | 0 |
| ENSMUSG00 5S_rRNA | 99 | 0 | 0 | 0 | 0 | 0 | 0 | 0 | 0 | 0 | 0 | 0 | 0 | 0 | 0 | 0 | 0 | 0 | 0 |
| ENSMUSG00 SNORA17 | 108 | 0 | 0 | 0 | 0 | 0 | 0 | 0 | 0 | 0 | 0 | 0 | 0 | 0 | 0 | 0 | 0 | 0 | 0 |
| ENSMUSG00 U6 | 112 | 0 | 0 | 0 | 0 | 0 | 0 | 0 | 0 | 0 | 0 | 0 | 0 | 0 | 0 | 0 | 0 | 0 | 0 |
| ENSMUSG00 U6 | 104 | 0 | 0 | 0 | 0.43808696 | 0 | 0 | 0 | 0 | 0.39602419 | 0 | 0 | 1.07337813 | 0 | 0 | 0 | 0 | 0.36852209 | 0 |
| ENSMUSG00 SNORA40 | 75 | 1.13347136 | 0.44417192 | 0 | 1.21496118 | 0 | 0 | 0 | 0 | 0 | 0 | 0 | 0 | 0 | 0 | 0 | 0 | 0 | 0 |
| ENSMUSG00 SNORA17 | 123 | 0 | 0 | 0 | 0 | 0 | 0 | 0 | 0 | 0 | 0 | 0 | 0 | 0 | 0 | 0 | 0 | 0 | 0 |
| ENSMUSG00 7SK | 306 | 0.1389058 | 0 | 0.15808343 | 0 | 0 | 0.64774815 | 0 | 0 | 0 | 0 | 0 | 0.18240413 | 0.16121542 | 0.45533419 | 0 | 0 | 0 | 0 |
| ENSMUSG00 SNORA17 | 91 | 0 | 0 | 0 | 0 | 0 | 0 | 0 | 0 | 0 | 0 | 0 | 0 | 0 | 0 | 0 | 0 | 0 | 0 |
| ENSMUSG00 SNORA18 | 128 | 0 | 0 | 0 | 0 | 0 | 0 | 0 | 0 | 0 | 0 | 0 | 0 | 0 | 0 | 0 | 0 | 0 | 0 |
| ENSMUSG00 5S_rRNA | 109 | 0 | 0 | 0 | 0 | 0 | 0 | 0 | 0 | 0 | 0 | 0 | 0 | 0 | 0 | 0 | 0 | 0 | 0 |
| ENSMUSG00 SNORA17 | 136 | 0 | 0 | 0 | 0 | 0 | 0 | 0 | 0 | 0 | 0 | 0 | 0 | 0 | 0 | 0 | 0 | 0 | 0 |
| ENSMUSG00 AC147047.1 | 94 | 0 | 0 | 0 | 0 | 0 | 0 | 0 | 0 | 0 | 0 | 0 | 0 | 0 | 0 | 0 | 0 | 0 | 0 |
| ENSMUSG00 U6atac | 101 | 0 | 0 | 0 | 0 | 0 | 0 | 0 | 0 | 0 | 0 | 0 | 0 | 0 | 0 | 0 | 0 | 0 | 0 |
| ENSMUSG00 U6 | 93 | 0 | 0 | 0 | 0 | 0 | 0 | 0 | 0 | 0 | 0 | 0 | 0 | 0 | 0 | 0 | 0 | 0 | 0 |
| ENSMUSG00 U6 | 98 | 0 | 0 | 0 | 0 | 0 | 0 | 0 | 0 | 0 | 0 | 0 | 0 | 0 | 0 | 0 | 0 | 0 | 0 |
| ENSMUSG00 Mir1964 | 82 | 0 | 0 | 0 | 0 | 0 | 0 | 0 | 0 | 0 | 0 | 0 | 0 | 1.20321749 | 0.56639131 | 0 | 0.93391725 | 0.46739386 | 0 |
| ENSMUSG00 7SK | 236 | 0 | 0 | 0 | 0 | 0 | 0 | 0 | 0 | 0 | 0 | 0 | 0 | 0 | 0 | 0 | 0 | 0 | 0 |
| ENSMUSG00 U7 | 65 | 0 | 0 | 0 | 0 | 0 | 0 | 0 | 0 | 1.26727741 | 0.72539543 | 0 | 2.5761075 | 0 | 0 | 0 | 0 | 0 | 0 |
| ENSMUSG00 U6 | 100 | 0 | 0 | 0 | 0 | 0 | 0 | 0 | 0 | 0 | 0 | 0 | 0 | 0 | 0 | 0 | 0 | 0 | 0 |
| ENSMUSG00 SNORA17 | 116 | 0 | 0 | 0 | 0 | 0 | 0 | 0 | 0 | 0 | 0.81294316 | 0 | 0 | 0 | 0 | 0 | 0 | 0 | 0 |
| ENSMUSG00 7SK | 346 | 0 | 0 | 0 | 0 | 0 | 0 | 0 | 0 | 0 | 0.40882113 | 0 | 0.32263389 | 0.71288898 | 0 | 0.47631857 | 0 | 0 | 0 |
| ENSMUSG00 7SK | 303 | 0 | 0 | 0 | 0 | 0 | 0 | 0 | 0 | 0 | 0 | 0 | 0 | 0 | 0 | 0 | 0 | 0 | 0 |
| ENSMUSG00 U7 | 62 | 0 | 0 | 0 | 0 | 0 | 0 | 0 | 0 | 0 | 0 | 0 | 0 | 0 | 0 | 0 | 0 | 0 | 0 |
| ENSMUSG00 7SK | 264 | 0 | 0 | 0 | 0 | 0 | 0 | 0 | 0 | 0 | 0 | 0 | 0 | 0 | 0 | 0 | 0 | 0 | 0 |
| ENSMUSG00 7SK | 311 | 0 | 0 | 0 | 0 | 0 | 0 | 0 | 0 | 0 | 0 | 0 | 0 | 0 | 0 | 0 | 0 | 0 | 0 |
| ENSMUSG00 U6atac | 94 | 0 | 0 | 0 | 0 | 0 | 0 | 0 | 0 | 0 | 0 | 0 | 0 | 0 | 0 | 0 | 0 | 0 | 0 |
| ENSMUSG00 Mir1950 | 73 | 0 | 0 | 0 | 0 | 0 | 0 | 0 | 0 | 0 | 0 | 0 | 0 | 0 | 0.63622038 | 0 | 0 | 0 | 0 |
| ENSMUSG00 SNORA17 | 86 | 0 | 0 | 0 | 0 | 0 | 0 | 0 | 0 | 0 | 0 | 0 | 0 | 0 | 0 | 0 | 0 | 0 | 0 |
| ENSMUSG00 U6 | 102 | 0 | 0 | 0 | 0 | 0 | 0 | 0 | 0 | 0 | 0 | 0 | 0 | 0 | 0 | 0 | 0 | 0 | 0 |
| ENSMUSG00 SNORA17 | 138 | 0 | 0 | 0 | 0.99045749 | 0.34878206 | 0 | 3.42845499 | 1.10702616 | 0.29845301 | 0 | 0 | 0 | 1.42991065 | 1.68275679 | 1.49281 | 0.27746817 | 0 | 0.79723343 |
| ENSMUSG00 U7 | 62 | 0 | 0 | 0 | 0 | 0 | 0 | 0 | 0 | 0 | 0 | 0 | 0 | 0 | 0 | 0 | 0 | 0 | 0 |
| ENSMUSG00 SNORA17 | 137 | 0 | 0 | 0 | 0 | 0 | 0 | 0 | 0 | 0 | 0 | 0 | 0 | 0 | 0 | 0 | 0 | 0 | 0 |
| ENSMUSG00 7SK | 288 | 0 | 0 | 0 | 0 | 0 | 0 | 0 | 0 | 0 | 0 | 0 | 0 | 0 | 0 | 0.57224383 | 0 | 0 | 0 |
| ENSMUSG00 SNORA48 | 150 | 0 | 0 | 0 | 0 | 0 | 0 | 0 | 0 | 0 | 0 | 0 | 0 | 0 | 0 | 0 | 0 | 0 | 0 |
| ENSMUSG00 U6 | 96 | 0 | 0 | 0 | 0 | 0 | 0 | 0 | 0 | 0 | 0 | 0 | 0 | 0 | 0 | 0 | 0 | 0 | 0 |
| ENSMUSG00 U6 | 106 | 0 | 0 | 0 | 0 | 0 | 0 | 0 | 0 | 0 | 0 | 0 | 0 | 0 | 0 | 0 | 0 | 0 | 0 |
| ENSMUSG00 SNORA17 | 131 | 0 | 0 | 0 | 0 | 0 | 0 | 0 | 0 | 0 | 0 | 0 | 0 | 0 | 0 | 0 | 0 | 0 | 0 |
| ENSMUSG00 U6 | 98 | 0 | 0 | 0 | 0 | 0 | 0 | 0 | 0 | 0 | 0 | 0 | 0 | 0 | 0 | 0 | 0 | 0 | 0 |
| ENSMUSG00 7SK | 311 | 0 | 0 | 0 | 0 | 0 | 0 | 0 | 0 | 0 | 0 | 0 | 0 | 0 | 0 | 0 | 0 | 0 | 0 |
| ENSMUSG00 Y_RNA | 106 | 0 | 0 | 0 | 0 | 1.36222425 | 0 | 0 | 0.36030568 | 1.16565611 | 0 | 0 | 0 | 0 | 0 | 0 | 0 | 0 | 0 |
| ENSMUSG00 SNORA17 | 106 | 0 | 0 | 0 | 0 | 0 | 0 | 0 | 0 | 0.38855204 | 0 | 0 | 0 | 0 | 0 | 0 | 0 | 0 | 0 |
| ENSMUSG00 snoU13 | 104 | 0.81740723 | 0 | 0 | 0 | 0 | 0.47646859 | 0 | 1.46893855 | 0 | 1.36011644 | 1.3781238 | 0 | 0.94869072 | 0 | 0 | 0.36817892 | 0 | 0 |
| ENSMUSG00 SNORA40 | 121 | 0 | 0 | 0 | 0 | 0 | 0 | 0 | 0 | 0 | 0 | 0 | 0 | 0 | 0 | 0 | 0 | 0 | 0 |
| ENSMUSG00 SCARNA14 | 141 | 0 | 0 | 0 | 0 | 0 | 0 | 0 | 0 | 0 | 0 | 0 | 0 | 0 | 0 | 0 | 0 | 0 | 0 |
| ENSMUSG00 snoU13 | 101 | 0 | 0 | 0 | 0 | 0 | 0 | 0 | 0 | 0 | 0 | 0 | 0 | 0 | 0 | 0 | 0 | 0 | 0 |
| ENSMUSG00 SNORA25 | 127 | 0 | 0 | 0 | 0.35874838 | 0 | 0.78035801 | 0 | 0.60145516 | 0.32430327 | 0.37126538 | 0.75236155 | 0 | 1.16532088 | 2.19420885 | 0.6488434 | 0.90450254 | 0.30178187 | 0.43314257 |
| ENSMUSG00 SNORA17 | 129 | 0 | 0 | 0 | 0 | 0 | 0 | 0 | 0 | 0 | 0 | 0 | 0 | 0 | 0 | 0 | 0 | 0 | 0 |
| ENSMUSG00 SNORA17 | 131 | 0 | 0 | 0 | 0 | 0 | 0 | 0 | 0.29154506 | 0 | 0 | 0 | 0 | 0.75315904 | 0 | 0.62903139 | 0 | 0.29256715 | 0 |
| ENSMUSG00 U3 | 89 | 0 | 0 | 0 | 0 | 0 | 0 | 0 | 0 | 0 | 0 | 0 | 0 | 0 | 0 | 0 | 0 | 0 | 0 |
| ENSMUSG00 U1 | 156 | 0 | 0 | 0 | 0 | 0 | 0 | 0 | 0 | 0 | 0 | 0 | 0 | 0 | 0 | 0 | 0 | 0 | 0 |
| ENSMUSG00 U2 | 81 | 0 | 0 | 0 | 0 | 0 | 0 | 0 | 0 | 0 | 0 | 0 | 0 | 0 | 0 | 0 | 0 | 0 | 0 |
| ENSMUSG00 SNORA70 | 90 | 0 | 0 | 0 | 0 | 0 | 0 | 0 | 0 | 0 | 0 | 0 | 0 | 0 | 0 | 0 | 0 | 0 | 0 |
| ENSMUSG00 U1 | 154 | 0 | 0 | 0 | 0 | 0 | 0 | 0 | 0 | 0 | 0 | 0.310227 | 0 | 0 | 0 | 0 | 0 | 0 | 0 |
| ENSMUSG00 7SK | 242 | 0 | 0.55062635 | 0.39978124 | 0.37653756 | 0.19889225 | 0 | 0 | 0.15781984 | 0 | 0 | 0 | 0 | 0 | 0 | 0 | 0 | 0 | 0 |
| ENSMUSG00 AC154701.1 | 104 | 0 | 0 | 0 | 0 | 0 | 0 | 0 | 0 | 0 | 0 | 0 | 0 | 0 | 0 | 0 | 0 | 0 | 0 |
| ENSMUSG00 SNORA70 | 123 | 0 | 0 | 0 | 0 | 0 | 0 | 0 | 0 | 0 | 0 | 0 | 0 | 0 | 0 | 0 | 0 | 0 | 0 |
| ENSMUSG00 U6 | 75 | 0 | 0 | 0 | 0 | 0 | 0 | 0 | 0 | 0 | 0 | 0 | 0 | 0 | 0 | 0 | 0 | 0 | 0.73345476 |
| ENSMUSG00 SNORA48 | 143 | 0 | 0 | 0 | 0 | 0 | 0 | 0 | 0 | 0 | 0 | 0 | 0 | 0 | 0 | 0 | 0 | 0 | 0 |
| ENSMUSG00 SNORA17 | 114 | 0 | 0 | 0 | 0 | 0 | 0 | 0 | 0 | 0 | 0 | 0 | 0 | 0 | 0 | 0 | 0 | 0 | 0 |
| ENSMUSG00 SNORA17 | 126 | 0 | 0 | 0 | 0 | 0 | 0 | 0 | 0 | 0 | 0 | 0 | 0 | 0 | 0 | 0 | 0 | 0 | 0 |
| ENSMUSG00 U6 | 97 | 0 | 0 | 0 | 0 | 0 | 0 | 0 | 0 | 0 | 0 | 0 | 1.1508384 | 0 | 1.43641507 | 0 | 0 | 0 | 0 |
| ENSMUSG00 SNORA17 | 123 | 0 | 0 | 0 | 0 | 0 | 0 | 0 | 0 | 0 | 0 | 0 | 0 | 0 | 0 | 0 | 0 | 0 | 0 |
| ENSMUSG00 U6 | 55 | 0 | 0 | 0 | 0 | 0 | 0 | 0 | 0 | 1.49769148 | 0 | 0 | 0 | 0.89694395 | 0 | 0.7491192 | 0 | 0 | 0 |
| ENSMUSG00 U7 | 60 | 0 | 0 | 0 | 0 | 0 | 0 | 0 | 0 | 0 | 0 | 0 | 0 | 0 | 0 | 0 | 0 | 0 | 0 |
| ENSMUSG00 SNORA70 | 128 | 0 | 0 | 0 | 0 | 0 | 0 | 0 | 0 | 0 | 0 | 0 | 0 | 0 | 0 | 0 | 0 | 0 | 0 |
| ENSMUSG00 Mir1927 | 108 | 0 | 0 | 0 | 0 | 0 | 0 | 0 | 0 | 0 | 0 | 0 | 0 | 0 | 0 | 0 | 0 | 0 | 0 |
| ENSMUSG00 U7 | 62 | 0 | 0 | 0 | 0 | 0 | 0 | 0 | 0 | 0 | 0 | 0 | 0 | 0 | 0 | 0 | 0 | 0 | 0 |
| ENSMUSG00 U6 | 106 | 0 | 0 | 0 | 0 | 0 | 0 | 0 | 0 | 0 | 0 | 0 | 0 | 0 | 0 | 0 | 0 | 0 | 0 |
| ENSMUSG00 5S_rRNA | 109 | 0 | 0 | 0 | 0 | 0 | 0 | 0 | 0 | 0.37785794 | 0 | 0 | 0 | 0 | 0 | 0 | 0 | 0 | 0 |
| ENSMUSG00 5S_rRNA | 72 | 0 | 0 | 0 | 0 | 0 | 0 | 0 | 0 | 0 | 0 | 0 | 0 | 0 | 0 | 0 | 0 | 0 | 0 |
| ENSMUSG00 SCARNA20 | 138 | 0 | 0 | 0 | 0 | 0 | 0 | 0 | 0 | 0 | 0 | 0 | 0 | 0 | 0 | 0 | 0 | 0 | 0 |
| ENSMUSG00 SCARNA21 | 143 | 0 | 0 | 0 | 0 | 0 | 0 | 0 | 0 | 0 | 0 | 0 | 0 | 0 | 0 | 0 | 0 | 0 | 0 |
| ENSMUSG00 U6 | 106 | 0 | 0 | 0 | 0 | 0 | 0 | 0 | 0 | 0 | 0 | 0 | 0 | 0 | 0 | 0 | 0 | 0 | 0 |
| ENSMUSG00 AC163703.1 | 56 | 0 | 0 | 0.86381304 | 2.44077023 | 0 | 0 | 0 | 0 | 0 | 0 | 0 | 0 | 0 | 0 | 0.73574207 | 0 | 2.05319448 | 0 |
| ENSMUSG00 Mir1931 | 118 | 0 | 0 | 0 | 1.54444218 | 0.40789766 | 0.41993842 | 0 | 0 | 1.39615308 | 0 | 2.02436265 | 0 | 0.4180671 | 0 | 0 | 0 | 0 | 0 |
| ENSMUSG00 SCARNA3 | 87 | 0 | 0 | 0 | 0 | 0 | 0 | 0 | 0 | 0 | 0 | 0 | 0 | 0 | 0 | 0 | 0 | 0 | 0 |
| ENSMUSG00 7SK | 294 | 0 | 0 | 0 | 0 | 0 | 0 | 0 | 0 | 0 | 0 | 0 | 0 | 0 | 0 | 0 | 0 | 0 | 0 |
| ENSMUSG00 U6 | 102 | 0 | 0 | 0 | 0 | 0 | 0 | 0 | 0 | 0 | 0 | 0 | 0 | 0 | 0 | 0 | 0 | 0 | 0 |
| ENSMUSG00 Mir1963 | 58 | 0 | 0 | 0 | 0 | 0 | 0 | 0 | 0 | 0 | 0 | 0 | 0 | 0 | 0 | 0 | 0 | 0 | 0 |
| ENSMUSG00 SNORA17 | 112 | 0 | 0 | 0 | 0 | 0 | 0 | 0 | 0 | 0 | 0 | 0 | 0 | 0 | 0 | 0 | 0 | 0 | 0 |
| ENSMUSG00 Y_RNA | 81 | 0 | 0 | 0 | 0 | 0 | 0 | 0 | 0 | 0 | 0 | 0 | 0 | 0 | 0 | 0 | 0 | 0 | 0 |
| ENSMUSG00 U6 | 72 | 0 | 0.46267909 | 0 | 0 | 0 | 1.37646482 | 0 | 0 | 0 | 0 | 0 | 0 | 0 | 0 | 0 | 0 | 0 | 0 |
| ENSMUSG00 U6 | 90 | 0 | 0 | 0 | 0 | 0 | 0 | 0 | 0 | 0 | 0 | 0 | 0 | 0 | 0 | 0 | 0.42545119 | 0 | 0 |
| ENSMUSG00 7SK | 191 | 0 | 0 | 0.25326456 | 0 | 0 | 0 | 0 | 0 | 0 | 0.24686232 | 0 | 0 | 0 | 0.72948828 | 0 | 0 | 0 | 0 |
| ENSMUSG00 SNORD79 | 83 | 0 | 0 | 0 | 0 | 0 | 0 | 0 | 0 | 0 | 0 | 0 | 0 | 0 | 0 | 0 | 0 | 0 | 0 |
| ENSMUSG00 5S_rRNA | 83 | 0 | 0 | 0 | 0 | 0 | 0 | 0 | 0 | 0 | 0 | 0 | 0 | 0 | 0 | 0 | 0 | 0 | 0 |
| ENSMUSG00 7SK | 248 | 0 | 0 | 0 | 0 | 0 | 0 | 0 | 0 | 0 | 0 | 0 | 0 | 0 | 0 | 0 | 0 | 0 | 0 |
| ENSMUSG00 U6 | 103 | 0 | 0 | 0 | 0 | 0 | 0 | 0 | 0 | 0 | 0 | 0 | 0 | 0 | 0 | 0 | 0 | 0 | 0 |
| ENSMUSG00 RNaseP_nuc | 211 | 0 | 0 | 0 | 0 | 0 | 0 | 0 | 0 | 0 | 0 | 0 | 0 | 0 | 0 | 0 | 0 | 0 | 0 |
| ENSMUSG00 U7 | 57 | 0 | 0 | 0 | 0 | 0 | 0 | 0 | 0 | 0 | 0 | 0 | 0 | 0 | 0 | 0 | 0 | 0 | 0 |
| ENSMUSG00 SNORA17 | 124 | 0 | 0 | 0 | 0 | 0 | 0 | 0 | 0 | 0 | 0 | 0 | 0 | 0 | 0 | 0 | 0 | 0 | 0 |
| ENSMUSG00 snoU13 | 95 | 0 | 0 | 0 | 0 | 0 | 0 | 0 | 0 | 0 | 0 | 0 | 0 | 0 | 0 | 0 | 0 | 0 | 0 |
| ENSMUSG00 7SK | 310 | 0 | 0 | 0 | 0 | 0 | 0 | 0 | 0 | 0 | 0 | 0 | 0 | 0 | 0 | 0 | 0 | 0 | 0 |
| ENSMUSG00 7SK | 316 | 0 | 0 | 0 | 0 | 0 | 0 | 0 | 0 | 0 | 0 | 0 | 0 | 0 | 0 | 0 | 0 | 0 | 0 |
| ENSMUSG00 SCARNA17 | 139 | 5.19847477 | 6.71051107 | 5.22016515 | 1.63888649 | 6.92545664 | 6.06040626 | 3.40378985 | 3.5719513 | 4.44458803 | 4.409778 | 6.53039001 | 6.42482447 | 1.7745294 | 4.67782175 | 1.48207036 | 1.65283197 | 1.65437253 | 3.56174072 |
| ENSMUSG00 U3 | 89 | 0 | 0 | 0 | 0 | 0 | 0 | 0 | 0 | 0 | 0 | 0 | 0 | 0 | 0 | 0 | 0 | 0 | 0 |
| ENSMUSG00 SNORA17 | 115 | 0 | 0 | 0 | 0 | 0 | 0 | 0 | 0 | 0 | 0 | 0 | 0 | 0 | 0 | 0 | 0 | 0 | 0 |
| ENSMUSG00 SNORD63 | 66 | 0.64401782 | 0 | 2.93172911 | 2.76127541 | 0 | 0 | 3.58429386 | 4.05070935 | 3.12019059 | 0.71440459 | 0 | 0 | 0 | 0 | 0.624266 | 2.32064287 | 0.58070147 | 0 |
| ENSMUSG00 U7 | 55 | 0 | 0 | 0 | 0 | 0 | 0 | 0 | 0 | 0 | 0 | 0 | 0 | 0 | 0 | 0 | 0 | 0 | 0 |
| ENSMUSG00 7SK | 301 | 0 | 0 | 0 | 0 | 0 | 0 | 0 | 0 | 0 | 0 | 0 | 0 | 0 | 0 | 0 | 0 | 0 | 0 |
| ENSMUSG00 U1 | 145 | 0.29313915 | 0 | 0.33361055 | 0 | 0.3319443 | 0 | 0 | 0 | 0 | 0.32517726 | 2.96534225 | 0 | 0.68044024 | 0 | 0.56829732 | 2.9048047 | 0 | 0 |
| ENSMUSG00 U7 | 61 | 0 | 0 | 0 | 0 | 0 | 0 | 0 | 0 | 0 | 0 | 0 | 0 | 0 | 0 | 0 | 0 | 0 | 0 |
| ENSMUSG00 AC102477.1 | 104 | 0 | 0 | 0 | 0 | 0 | 0 | 0 | 0 | 0 | 0 | 0 | 0 | 0 | 0 | 0 | 0 | 0 | 0 |
| ENSMUSG00 AC151572.1 | 104 | 0 | 0.32031629 | 0 | 0 | 0 | 0 | 0 | 0 | 3.16819352 | 0 | 0 | 0 | 0 | 0 | 0 | 0 | 0 | 0 |
| ENSMUSG00 U7 | 63 | 0 | 0 | 0 | 0 | 0 | 0 | 0 | 0 | 0 | 0 | 0 | 0 | 0 | 0 | 0 | 0 | 0 | 0 |
| ENSMUSG00 SNORD75 | 74 | 0 | 0 | 0 | 0 | 0 | 0 | 0 | 0 | 0 | 0 | 0 | 0 | 0 | 0 | 0 | 0 | 0 | 0 |
| ENSMUSG00 7SK | 289 | 0 | 0 | 0 | 0 | 0 | 0 | 0 | 0 | 0 | 0 | 0 | 0 | 0 | 0 | 0 | 0 | 0 | 0 |
| ENSMUSG00 U6 | 108 | 0 | 0.30845272 | 0.44790306 | 0.84372304 | 0 | 0 | 0.43808036 | 2.47543349 | 0 | 1.30974176 | 0 | 0 | 0 | 0 | 1.90747944 | 0.70908532 | 0 | 0 |
| ENSMUSG00 7SK | 295 | 0 | 0 | 0 | 0 | 0 | 0 | 0 | 0 | 0 | 0 | 0 | 0 | 0 | 0 | 0 | 0 | 0 | 0 |
| ENSMUSG00 SNORA17 | 124 | 0 | 0 | 0 | 0 | 0 | 0 | 0 | 0 | 0 | 0 | 0 | 0 | 0 | 0 | 0 | 0 | 0 | 0 |
| ENSMUSG00 U6 | 103 | 0 | 0 | 0 | 0 | 0 | 0 | 0.91869279 | 0 | 0 | 0.45777382 | 0 | 0.54189964 | 0 | 0 | 0 | 0.74350694 | 0 | 0 |
| ENSMUSG00 Snora78 | 127 | 2.6774914 | 3.93459381 | 2.28536364 | 1.43499352 | 4.92688982 | 1.17053701 | 5.96865246 | 4.81164125 | 1.94581964 | 7.05404222 | 8.65215783 | 6.59240109 | 12.4300894 | 9.50823836 | 10.3814944 | 7.2360203 | 3.62138238 | 18.6251306 |
| ENSMUSG00 7SK | 352 | 0 | 0 | 0.96197362 | 0 | 0 | 0 | 0 | 0 | 0 | 0 | 0 | 0 | 0 | 0 | 0 | 0 | 0 | 0 |
| ENSMUSG00 U6 | 117 | 0 | 0 | 0 | 0 | 0 | 0 | 0 | 0 | 0 | 0 | 0 | 0 | 0 | 0 | 0 | 0 | 0 | 0 |
| ENSMUSG00 SNORA17 | 127 | 0 | 0 | 0 | 0 | 0 | 0 | 0 | 0 | 0 | 0 | 0 | 0 | 0 | 0 | 0 | 0 | 0 | 0 |
| ENSMUSG00 5S_rRNA | 92 | 0 | 0 | 0 | 0 | 0 | 0 | 0 | 0 | 0 | 0 | 0 | 0 | 0 | 0 | 0 | 0 | 0 | 0 |
| ENSMUSG00 Mir1929 | 92 | 0 | 0 | 0 | 0 | 0 | 0 | 0 | 0.41513481 | 0 | 0 | 0 | 0 | 0 | 0.50482704 | 0 | 1.66480902 | 0 | 0 |
| ENSMUSG00 Mir669n | 53 | 1.60396891 | 0 | 0 | 0 | 0 | 0 | 0 | 0 | 0 | 0 | 1.80282862 | 0 | 0 | 0 | 0 | 0 | 0 | 0 |
| ENSMUSG00 SNORA17 | 115 | 0 | 0 | 0 | 0 | 0 | 0 | 0 | 0 | 0 | 0 | 0 | 0 | 0 | 0 | 0 | 0 | 0 | 0 |
| ENSMUSG00 AC161364.1 | 104 | 0 | 0 | 0 | 0 | 0 | 0 | 0 | 0 | 0 | 0 | 0 | 0 | 0 | 0 | 0 | 0 | 0 | 0 |
| ENSMUSG00 SNORA55 | 57 | 0 | 0 | 0 | 0 | 0 | 0 | 0 | 0 | 0 | 0 | 0 | 0 | 0 | 0 | 0 | 0 | 0 | 0 |
| ENSMUSG00 SNORD121A | 76 | 1.6778359 | 0.43832756 | 0 | 0 | 0 | 0.65200965 | 3.11267625 | 0 | 1.62578352 | 2.48161596 | 0 | 0 | 0 | 0 | 0 | 0.50382378 | 0.50429338 | 3.61902018 |
| ENSMUSG00 U1 | 163 | 0 | 0 | 0 | 0 | 0 | 0 | 0 | 0 | 0 | 0 | 0 | 0 | 0 | 0 | 0 | 0 | 0 | 0 |
| ENSMUSG00 SNORA48 | 143 | 0 | 0 | 0 | 0 | 0 | 0 | 0 | 0 | 0 | 0 | 0 | 0 | 0 | 0 | 0 | 0 | 0 | 0 |
| ENSMUSG00 Mir1961 | 111 | 0 | 0 | 0 | 0 | 0 | 0 | 0 | 0 | 0 | 0 | 0 | 0 | 0 | 0 | 0 | 0 | 0 | 0 |
| ENSMUSG00 U6 | 105 | 0 | 0 | 0 | 0 | 0 | 0 | 0 | 0 | 0 | 0 | 0 | 0 | 0 | 0 | 0 | 0 | 0 | 0 |
| ENSMUSG00 snoR9_plant | 91 | 0 | 0 | 0 | 0 | 0 | 0 | 0 | 0 | 0 | 0 | 0 | 0 | 0 | 0 | 0 | 0 | 0 | 0 |
| ENSMUSG00 U6 | 102 | 0 | 0 | 0 | 0 | 0 | 0 | 0 | 0 | 0 | 0 | 0 | 0 | 0 | 0 | 0 | 0 | 0 | 0 |
| ENSMUSG00 SNORA17 | 138 | 1.23203409 | 0 | 0 | 0 | 0 | 0.35907778 | 0.3428455 | 2.21405231 | 0 | 0 | 0 | 0.40446132 | 0.35747766 | 0 | 0 | 0 | 0 | 0 |
| ENSMUSG00 7SK | 278 | 0 | 0 | 0 | 0 | 0 | 0 | 0 | 0 | 0 | 0 | 0 | 0 | 0 | 0 | 0 | 0 | 0 | 0 |
| ENSMUSG00 U1 | 148 | 0 | 0 | 0 | 0 | 0 | 0 | 0 | 0 | 0 | 0 | 0 | 0 | 0 | 0 | 0 | 0 | 0 | 0 |
| ENSMUSG00 SNORA17 | 107 | 0 | 0 | 0 | 0 | 0 | 0 | 0 | 0 | 0 | 0 | 0 | 0 | 0 | 0 | 0 | 0 | 0 | 0 |
| ENSMUSG00 7SK | 84 | 0 | 0 | 0 | 0 | 0 | 0 | 0 | 0 | 0 | 0 | 0 | 0 | 0 | 0 | 0 | 0 | 0 | 0 |
| ENSMUSG00 U6 | 101 | 0 | 0 | 0 | 0 | 0 | 0 | 0 | 0 | 0 | 0 | 0 | 0 | 0 | 0 | 0 | 0 | 0 | 0 |
| ENSMUSG00 SNORA17 | 130 | 0 | 0 | 0 | 0 | 0 | 0 | 0 | 0 | 0 | 0 | 0 | 0 | 0 | 0 | 0 | 0 | 0 | 0 |
| ENSMUSG00 SNORA17 | 72 | 0 | 0 | 0 | 0 | 0 | 0 | 0 | 0 | 0 | 0 | 0 | 0 | 0 | 0 | 0 | 0 | 0 | 0 |
| ENSMUSG00 7SK | 270 | 0 | 0 | 0 | 0 | 0 | 0 | 0 | 0 | 0 | 0 | 0 | 0 | 0 | 0 | 0 | 0.28363413 | 0 | 0 |
| ENSMUSG00 Scarna6 | 264 | 0 | 0 | 0 | 0.34515943 | 0 | 0 | 0 | 0 | 0.15600953 | 0 | 0 | 0 | 0 | 0 | 0.1560665 | 0 | 0 | 0 |
| ENSMUSG00 7SK | 321 | 0 | 0.20755697 | 0 | 0 | 0.44983106 | 0 | 0.88434914 | 0 | 0 | 0 | 0.44649494 | 0.52164171 | 0.30736397 | 0.14468563 | 0.12835376 | 0.47714153 | 0.83577594 | 0 |
| ENSMUSG00 SNORA17 | 131 | 0 | 0 | 0 | 0 | 0 | 0 | 0 | 0 | 0 | 0 | 0 | 0 | 0 | 0 | 0 | 0 | 0 | 0 |
| ENSMUSG00 7SK | 293 | 0 | 0 | 0 | 0 | 0 | 0 | 0 | 0 | 0 | 0 | 0 | 0 | 0 | 0 | 0 | 0 | 0 | 0 |
| ENSMUSG00 U6 | 103 | 0.82534322 | 1.94055694 | 0.93929185 | 0 | 1.86920092 | 2.4054725 | 0.4593464 | 1.48320069 | 2.39921451 | 0.45777382 | 0 | 0.54189964 | 0.47895065 | 0 | 0.40001511 | 1.48701388 | 0.37209997 | 0 |
| ENSMUSG00 SNORA38 | 132 | 0 | 0 | 0 | 0 | 0 | 0 | 0 | 0 | 0 | 0 | 0 | 0 | 0 | 0 | 0 | 0 | 0 | 0 |
| ENSMUSG00 U6 | 107 | 0 | 0 | 0 | 0 | 0 | 0 | 0 | 0 | 0 | 0 | 0 | 0 | 0 | 0 | 0 | 0 | 0 | 0 |
| ENSMUSG00 SNORA17 | 130 | 0 | 0 | 0 | 0 | 0 | 0 | 0 | 0 | 0 | 0 | 0 | 0 | 0 | 0 | 0 | 0 | 0 | 0 |
| ENSMUSG00 U6 | 101 | 0 | 0 | 0 | 0 | 0 | 0 | 0 | 0 | 0 | 0 | 0 | 0 | 0 | 0 | 0 | 0 | 0 | 0 |
| ENSMUSG00 SNORA18 | 126 | 0 | 0 | 0 | 0 | 0 | 0 | 0 | 0 | 0 | 0 | 0 | 0 | 0 | 0 | 0 | 0 | 0 | 0 |
| ENSMUSG00 U6 | 109 | 0.77991149 | 2.44498306 | 0.88758771 | 0.83598246 | 0.88315456 | 0 | 0.43406127 | 1.40155605 | 0.75571589 | 0 | 0 | 0.5120703 | 1.35775919 | 1.27827763 | 2.26797555 | 0.70257995 | 1.05485221 | 0 |
| ENSMUSG00 U6 | 97 | 0 | 0 | 0 | 0 | 0 | 0 | 0 | 0 | 0 | 0 | 0 | 0 | 0 | 0 | 0 | 0 | 0 | 0 |
| ENSMUSG00 Mir2139 | 94 | 0 | 0 | 0 | 0 | 0 | 0 | 0 | 0 | 0 | 0 | 0 | 0 | 0 | 0 | 0 | 0 | 0 | 0 |
| ENSMUSG00 U3 | 85 | 0 | 0 | 0.56910036 | 0.53601229 | 0.56625793 | 0 | 0 | 0.44932238 | 0 | 1.66414247 | 2.81029168 | 1.31330971 | 1.16075099 | 0 | 1.45417256 | 0 | 0.90179522 | 1.94149788 |
| ENSMUSG00 U6 | 102 | 5.00060894 | 12.7372831 | 2.37125149 | 9.38021501 | 7.07822407 | 8.25878893 | 8.34929628 | 10.1097536 | 6.46062992 | 12.4810685 | 7.96249308 | 9.30261043 | 4.83646248 | 7.28534704 | 4.84724187 | 3.00318489 | 6.38771615 | 2.15721987 |
| ENSMUSG00 SNORA52 | 106 | 0 | 0 | 0 | 0 | 0 | 0 | 0 | 0 | 0 | 0 | 0 | 0 | 0 | 0 | 0 | 0 | 0 | 0 |
| ENSMUSG00 7SK | 287 | 0 | 0 | 0 | 0 | 0 | 0 | 0 | 0 | 0.28701405 | 0 | 0 | 0 | 0 | 0 | 0 | 0 | 0 | 0 |
| ENSMUSG00 SNORA17 | 84 | 1.012028 | 0.39658207 | 1.15175072 | 0 | 1.71899727 | 0 | 0 | 1.36401437 | 1.96126266 | 0 | 2.84374753 | 0 | 0 | 0 | 0 | 0 | 0 | 0 |
| ENSMUSG00 7SK | 356 | 0 | 0 | 0 | 0 | 0 | 0 | 0 | 0 | 0 | 0 | 0 | 0 | 0 | 0 | 0 | 0 | 0 | 0 |
| ENSMUSG00 SNORA17 | 123 | 0 | 0 | 0 | 0 | 0 | 0 | 0 | 0 | 0 | 0 | 0 | 0 | 0 | 0 | 0 | 0 | 0 | 0 |
| ENSMUSG00 SNORA17 | 126 | 0 | 0 | 0 | 0 | 0 | 0 | 0 | 0 | 0 | 0 | 0 | 0 | 0 | 0 | 0 | 0 | 0 | 0 |
| ENSMUSG00 7SK | 90 | 0 | 0 | 0 | 0 | 0 | 0 | 0 | 0 | 0 | 0 | 0 | 0 | 0 | 0 | 0 | 0 | 0 | 0 |
| ENSMUSG00 U6 | 53 | 0 | 0 | 0 | 0 | 0 | 0 | 0 | 0 | 0 | 0 | 0 | 0 | 0 | 0 | 0 | 0 | 2.16941303 | 0 |
| ENSMUSG00 SNORA17 | 75 | 0 | 0 | 0 | 0 | 0 | 0 | 0 | 0 | 0 | 0 | 0 | 0 | 0 | 0 | 0 | 0 | 0 | 0 |
| ENSMUSG00 SNORD121A | 80 | 0 | 0.41641118 | 0 | 0 | 0 | 0 | 0 | 1.43221509 | 0 | 0 | 0 | 0 | 0 | 0 | 0 | 0 | 0 | 0 |
| ENSMUSG00 SNORA17 | 129 | 0 | 0 | 0 | 0 | 0 | 0 | 0 | 0 | 0 | 0 | 0 | 0 | 0 | 0 | 0 | 0 | 0 | 0 |
| ENSMUSG00 SNORA17 | 109 | 0 | 0 | 0 | 0 | 0 | 0 | 0 | 0 | 0 | 0 | 0 | 0 | 0 | 0 | 0 | 0 | 0 | 0 |
| ENSMUSG00 SNORA17 | 124 | 0 | 0 | 0 | 0 | 0 | 0 | 0 | 0 | 0 | 0 | 0 | 0 | 0 | 0 | 0 | 0 | 0 | 0 |
| ENSMUSG00 7SK | 300 | 0 | 0 | 0 | 0 | 0 | 0 | 0 | 0 | 0 | 0 | 0 | 0 | 0 | 0 | 0 | 0 | 0 | 0 |
| ENSMUSG00 U8 | 133 | 0 | 0 | 0 | 0 | 0 | 0 | 1.42293771 | 0 | 0 | 0 | 0 | 0 | 0 | 0 | 0 | 1.43949652 | 0 | 0 |
| ENSMUSG00 U6 | 84 | 0 | 0 | 0 | 0 | 0 | 0 | 0 | 0 | 0 | 0 | 0 | 0 | 0 | 0 | 0 | 0 | 0 | 0 |
| ENSMUSG00 7SK | 323 | 0 | 0 | 0 | 0 | 0 | 0 | 0 | 0 | 0 | 0 | 0 | 0 | 0 | 0 | 0 | 0 | 0 | 0 |
| ENSMUSG00 SNORD123 | 70 | 0 | 0 | 0 | 0 | 0 | 0.70789619 | 0.67589541 | 0.54560575 | 0 | 0 | 0 | 0 | 2.8189667 | 0.66348696 | 0 | 0 | 0 | 0 |
| ENSMUSG00 7SK | 303 | 0 | 0 | 0 | 0 | 0 | 0 | 0.15614746 | 0 | 0 | 0 | 0 | 1.10526065 | 0 | 0 | 0 | 0 | 0 | 0 |
| ENSMUSG00 SNORA17 | 129 | 0 | 0 | 0 | 0 | 0 | 0 | 0 | 0 | 0 | 0 | 0 | 0 | 0 | 0 | 0 | 0 | 0 | 0 |
| ENSMUSG00 U6 | 99 | 0 | 0 | 0 | 0 | 0 | 0 | 0 | 0 | 0 | 0 | 0 | 0 | 0 | 0 | 0 | 0 | 0 | 0 |
| ENSMUSG00 SNORA17 | 84 | 0.506014 | 0 | 0 | 0 | 0 | 0 | 0 | 0 | 0 | 0 | 0 | 0 | 0 | 0 | 0 | 0 | 0 | 0 |
| ENSMUSG00 AL772326.1 | 89 | 0 | 0 | 0 | 0 | 0 | 0 | 0 | 0 | 0 | 0 | 0 | 0 | 0 | 0 | 0 | 0 | 0 | 0 |
| ENSMUSG00 Mir1936 | 92 | 0 | 0 | 0 | 0 | 0 | 0 | 0 | 0 | 0 | 0 | 0 | 0 | 0 | 0 | 0 | 0 | 0 | 0 |
| ENSMUSG00 5S_rRNA | 93 | 0 | 0 | 0 | 0 | 0 | 0 | 0 | 0 | 0 | 0 | 0 | 0 | 0 | 0 | 0 | 0 | 0 | 0 |
| ENSMUSG00 SNORA17 | 124 | 0 | 0 | 0 | 0 | 0 | 0 | 0 | 0 | 0 | 0 | 0 | 0 | 0 | 0 | 0 | 0 | 0 | 0 |
| ENSMUSG00 SNORA17 | 123 | 0 | 0 | 0 | 0 | 0 | 0 | 0 | 0 | 0 | 0 | 0 | 0 | 0 | 0 | 0 | 0 | 0 | 0 |
| ENSMUSG00 U7 | 63 | 0 | 0 | 0 | 0 | 0 | 0 | 0 | 0 | 0 | 0 | 0 | 0 | 0 | 0 | 0 | 0 | 0 | 0 |
| ENSMUSG00 SNORA17 | 110 | 0 | 0 | 0 | 0 | 0 | 0 | 0 | 0 | 0 | 0 | 0 | 0 | 0 | 0 | 0 | 0 | 0 | 0 |
| ENSMUSG00 SNORA17 | 100 | 0 | 0 | 0 | 0 | 0 | 0 | 0 | 0 | 0 | 0 | 0 | 0 | 0 | 0 | 0 | 0 | 0 | 0 |
| ENSMUSG00 SNORA17 | 129 | 0 | 0 | 0 | 0 | 0 | 0 | 0.36676495 | 0 | 0 | 0 | 0 | 0 | 0 | 0 | 0 | 0 | 0 | 0 |
| ENSMUSG00 SNORA17 | 141 | 0 | 0 | 0 | 0 | 0 | 0 | 0 | 0.2708681 | 0 | 0 | 0 | 0 | 0 | 0 | 0.58441923 | 0 | 0 | 0 |
| ENSMUSG00 SNORA73 | 193 | 0 | 0.34521134 | 0 | 0 | 0 | 0 | 0.24514341 | 0.79155238 | 0 | 1.22152081 | 0.24753864 | 0.28920032 | 0 | 0 | 0.21347956 | 0.19839693 | 0 | 0 |
| ENSMUSG00 7SK | 85 | 0 | 0 | 0 | 0 | 0 | 0 | 0 | 0 | 0 | 0 | 0 | 0 | 0 | 0 | 0 | 0 | 0 | 0 |
| ENSMUSG00 U3 | 89 | 0 | 0 | 0 | 0 | 0 | 0 | 0 | 0 | 0 | 0 | 0 | 0 | 0 | 0 | 0 | 0 | 0 | 0 |
| ENSMUSG00 7SK | 259 | 0 | 0 | 0 | 0 | 0 | 0 | 0 | 0 | 0 | 0 | 0 | 0 | 0 | 0 | 0 | 0 | 0 | 0 |
| ENSMUSG00 U1 | 163 | 0 | 0 | 0 | 0 | 0 | 0 | 0 | 0 | 0 | 0 | 0 | 0 | 0 | 0 | 0 | 0 | 0 | 0 |
| ENSMUSG00 7SK | 327 | 0 | 0 | 0 | 0 | 0 | 0.15153741 | 0 | 0.11679634 | 0 | 0.14419175 | 0.43830237 | 0 | 0.30172426 | 0 | 0 | 0.11709666 | 1.17205801 | 0.84111784 |
| ENSMUSG00 7SK | 104 | 0 | 0 | 0 | 0 | 0 | 0 | 0 | 0 | 0 | 0 | 0 | 0 | 0 | 0 | 0 | 0 | 0 | 0 |
| ENSMUSG00 7SK | 274 | 0 | 0 | 0 | 0 | 0 | 0 | 0 | 0 | 0 | 0 | 0 | 0 | 0 | 0 | 0 | 0 | 0 | 0 |
| ENSMUSG00 U6 | 101 | 0 | 0 | 1.91578338 | 0 | 0 | 2.94372675 | 0.93688473 | 0 | 0 | 0.46683865 | 0 | 0 | 0 | 0 | 0 | 0 | 0.37946829 | 0 |
| ENSMUSG00 7SK | 317 | 0 | 0 | 0 | 0 | 0 | 0 | 0 | 0 | 0 | 0 | 0 | 0 | 0 | 0 | 0 | 0 | 0 | 0 |
| ENSMUSG00 SNORA17 | 142 | 0 | 0 | 0 | 0 | 0 | 0 | 0 | 0 | 0 | 0 | 0 | 0 | 0 | 0 | 0 | 0 | 0 | 0 |
| ENSMUSG00 SNORD22 | 124 | 0 | 0 | 0 | 0 | 0 | 0 | 0 | 0 | 0 | 0 | 0 | 0 | 0 | 0 | 0 | 0 | 0 | 0 |
| ENSMUSG00 U7 | 62 | 0 | 0 | 0 | 0 | 0 | 0 | 0 | 0 | 0 | 0 | 0 | 0 | 0 | 0 | 0 | 0 | 0 | 0 |
| ENSMUSG00 U1 | 140 | 0 | 0 | 0 | 0 | 0 | 0 | 0 | 0.27280287 | 0 | 0 | 0 | 0 | 0 | 0 | 0 | 0.27350434 | 0 | 0 |
| ENSMUSG00 5S_rRNA | 117 | 0 | 0 | 0 | 0 | 0 | 0 | 0 | 0 | 0 | 0 | 0 | 0 | 0 | 0 | 0 | 0 | 0 | 0 |
| ENSMUSG00 Mir2137 | 65 | 0 | 0 | 0 | 0 | 0 | 0 | 0 | 0 | 0 | 0 | 0 | 0 | 0 | 2.85809769 | 0 | 0 | 0 | 0 |
| ENSMUSG00 7SK | 307 | 0 | 0 | 0 | 0 | 0 | 0 | 0 | 0 | 0 | 0 | 0 | 0 | 0 | 0 | 0 | 0 | 0 | 0 |
| ENSMUSG00 SNORA17 | 129 | 0 | 0 | 0 | 0 | 0 | 0 | 0 | 0 | 0 | 0 | 0 | 0 | 0 | 0 | 0 | 0 | 0 | 0 |
| ENSMUSG00 U7 | 65 | 0 | 0 | 0 | 0 | 0 | 0 | 0 | 0 | 0 | 0 | 0 | 0 | 0 | 0 | 0 | 0 | 0 | 0 |
| ENSMUSG00 5S_rRNA | 105 | 0 | 0 | 0 | 0 | 0 | 0 | 0 | 0.72747433 | 0 | 0 | 0 | 0 | 0 | 0 | 0 | 0 | 0 | 0 |
| ENSMUSG00 SNORA17 | 127 | 0 | 0 | 0 | 0 | 0 | 0 | 0 | 0 | 0 | 0 | 0 | 0 | 0 | 0 | 0 | 0 | 0 | 0 |
| ENSMUSG00 SNORA17 | 123 | 0 | 3.520875 | 0 | 0 | 3.52184807 | 7.25161955 | 0 | 2.48405869 | 0 | 0.7666781 | 3.49572867 | 3.630287 | 0 | 2.26556524 | 0 | 0.6226115 | 3.1159591 | 7.15565616 |
| ENSMUSG00 7SK | 275 | 0 | 0 | 0 | 0 | 0 | 0 | 0 | 0 | 0 | 0 | 0 | 0 | 0 | 0 | 0 | 0 | 0 | 0 |
| ENSMUSG00 snoMe18S-Ut | 81 | 0 | 0 | 0 | 0 | 0 | 0 | 0 | 0 | 0 | 0 | 0 | 0 | 0 | 0 | 0 | 0 | 0 | 0 |
| ENSMUSG00 U7 | 58 | 0 | 0 | 0 | 0 | 0 | 0 | 0 | 0 | 0 | 0 | 0 | 0 | 0 | 0 | 0 | 0 | 0 | 0 |
| ENSMUSG00 U6 | 108 | 0 | 0 | 0 | 0 | 0 | 0 | 0 | 0 | 1.90678314 | 0 | 0 | 0 | 0 | 0 | 0 | 0 | 0 | 0 |
| ENSMUSG00 U6 | 119 | 0 | 0 | 0 | 0 | 0 | 0 | 0 | 0 | 0 | 0 | 0 | 0 | 0 | 0 | 0 | 0 | 0 | 0 |
| ENSMUSG00 Mir1938 | 99 | 3.00541649 | 0.33649388 | 0 | 3.22148798 | 0 | 2.50266331 | 0.47790585 | 0.77156368 | 0.83205082 | 0.47626973 | 0.48257534 | 0 | 3.48811536 | 2.81479318 | 2.91324133 | 0.77354762 | 3.48420681 | 6.11212297 |
| ENSMUSG00 U6 | 104 | 0 | 0 | 0 | 0 | 0 | 0 | 0 | 0 | 0 | 0 | 0 | 0 | 0 | 0 | 0 | 0 | 0 | 0 |
| ENSMUSG00 SNORA17 | 127 | 0 | 0 | 0 | 0 | 0 | 0 | 0 | 0 | 0 | 0 | 0 | 0 | 0 | 0 | 0 | 0 | 0 | 0 |
| ENSMUSG00 Snord107 | 69 | 0 | 0 | 0 | 0 | 0 | 0 | 0 | 0 | 0 | 0 | 0 | 0 | 0 | 0 | 0 | 0 | 0 | 0 |
| ENSMUSG00 7SK | 267 | 0 | 0 | 0 | 0 | 0 | 0 | 0 | 0 | 0 | 0 | 0 | 0 | 0 | 0 | 0 | 0 | 0 | 0 |
| ENSMUSG00 5S_rRNA | 107 | 0 | 0 | 0 | 0 | 0 | 0 | 0 | 0 | 0 | 0 | 0 | 0 | 0 | 0 | 0 | 0 | 0 | 0 |
| ENSMUSG00 AC113092.1 | 89 | 0 | 0 | 0 | 0 | 0 | 0 | 0 | 0 | 0 | 0 | 0 | 0 | 0 | 0 | 0 | 0 | 0 | 0 |
| ENSMUSG00 SNORA17 | 149 | 0 | 0 | 0 | 0 | 0 | 0 | 0 | 0 | 0 | 0 | 0 | 0 | 0 | 0 | 0 | 0 | 0 | 0 |
| ENSMUSG00 U6 | 103 | 0 | 0 | 0 | 0 | 0 | 0 | 0 | 0 | 0 | 0 | 0 | 0 | 0 | 0 | 0 | 0 | 0 | 0 |
| ENSMUSG00 7SK | 258 | 0 | 0 | 0 | 0 | 0 | 0 | 0 | 0 | 0 | 0 | 0 | 0 | 0 | 0 | 0 | 0 | 0 | 0 |
| ENSMUSG00 U6 | 74 | 0 | 0 | 0 | 0 | 0 | 0 | 0 | 0 | 0 | 0 | 0 | 0 | 0 | 0 | 0 | 0 | 0 | 0 |
| ENSMUSG00 SNORA17 | 125 | 0 | 0 | 0 | 0 | 0 | 0 | 0 | 0 | 0 | 0 | 0 | 0 | 0 | 0 | 0 | 0 | 0 | 0 |
| ENSMUSG00 SNORA17 | 106 | 0 | 0 | 0 | 0 | 0 | 0 | 0 | 0 | 0 | 0 | 0 | 0 | 0 | 0 | 0 | 0 | 0 | 0 |
| ENSMUSG00 SNORA84 | 132 | 0 | 0 | 0 | 0 | 0 | 0 | 0 | 0 | 0 | 0 | 0 | 0 | 0 | 0 | 0 | 0 | 0 | 0 |
| ENSMUSG00 SNORA17 | 110 | 0 | 0 | 0 | 0 | 0 | 0 | 0 | 0 | 0 | 0 | 0 | 0 | 0 | 0 | 0 | 0 | 0 | 0 |
| ENSMUSG00 SCARNA20 | 136 | 0.93761418 | 0.4898955 | 0 | 0.67001536 | 1.06173361 | 0.72871667 | 0 | 0 | 0 | 1.04008904 | 0 | 1.64163713 | 0.36273469 | 0 | 0.90885785 | 1.40774292 | 0.28181101 | 0 |
| ENSMUSG00 U1 | 168 | 0 | 0 | 0 | 0 | 0 | 0 | 0 | 0 | 0 | 0 | 0 | 0 | 0 | 0 | 0 | 0 | 0 | 0 |
| ENSMUSG00 7SK | 252 | 0 | 0 | 0 | 0 | 0 | 0 | 0 | 0 | 0 | 0 | 0 | 1.32894435 | 0 | 0 | 0 | 0 | 0 | 0 |
| ENSMUSG00 5S_rRNA | 108 | 0.39356644 | 0.61690545 | 0 | 0 | 0 | 0 | 0 | 0.35363336 | 0 | 0 | 0 | 0 | 0 | 0 | 0 | 1.06362798 | 0 | 0 |
| ENSMUSG00 U2 | 179 | 0 | 0 | 0 | 0 | 0 | 0 | 0 | 0 | 0 | 0 | 0 | 0 | 0 | 0 | 0 | 0 | 0 | 0 |
| ENSMUSG00 U6 | 110 | 0 | 0 | 0 | 0 | 0 | 0 | 0 | 0 | 0 | 0 | 0 | 0 | 0 | 0 | 0 | 0 | 0 | 0 |
| ENSMUSG00 7SK | 291 | 0 | 0 | 0 | 0 | 0 | 0 | 0 | 0 | 0 | 0 | 0 | 0 | 0 | 0 | 0 | 0 | 0 | 0 |
| ENSMUSG00 U1 | 157 | 2.16586884 | 0.84873616 | 3.38922824 | 3.19217508 | 2.75915486 | 2.52498005 | 4.218965 | 2.67590081 | 3.67268293 | 0.90096885 | 3.65158918 | 3.19962397 | 1.57108017 | 5.32479983 | 4.72374526 | 5.1216736 | 0.24411654 | 3.50376476 |
| ENSMUSG00 U6 | 105 | 0 | 0 | 0 | 0 | 0 | 0 | 0 | 0 | 0 | 0 | 0 | 0 | 0 | 0 | 0 | 0 | 0 | 0 |
| ENSMUSG00 7SK | 264 | 0 | 0 | 0 | 0 | 0 | 0 | 0 | 0 | 0 | 0 | 0 | 0 | 0 | 0 | 0 | 0 | 0 | 0 |
| ENSMUSG00 U6 | 101 | 0 | 0 | 0 | 0 | 0 | 0 | 0 | 0 | 0 | 0 | 0 | 0 | 0 | 0 | 0 | 0 | 0 | 0 |
| ENSMUSG00 U1 | 164 | 0 | 0 | 0 | 0 | 0 | 0 | 0 | 0 | 0 | 0 | 0 | 0 | 0 | 0 | 0 | 0 | 0 | 0 |
| ENSMUSG00 5S_rRNA | 84 | 0 | 0 | 0 | 0 | 0 | 0 | 0 | 0 | 0 | 0 | 0 | 0 | 0 | 0 | 0 | 0 | 0 | 0 |
| ENSMUSG00 U7 | 65 | 0 | 0 | 0 | 0 | 0 | 0 | 0 | 0 | 0 | 0 | 0 | 0 | 0 | 0 | 0 | 0 | 0 | 0 |
| ENSMUSG00 U7 | 62 | 0 | 0 | 0 | 0 | 0 | 0 | 0 | 0 | 0 | 0 | 0 | 0 | 0 | 0 | 0 | 0 | 0 | 0 |
| ENSMUSG00 Y_RNA | 115 | 0 | 0 | 0 | 0 | 0 | 0 | 0 | 0 | 0 | 0 | 0 | 0 | 0 | 0 | 0 | 0 | 0 | 0 |
| ENSMUSG00 5S_rRNA | 91 | 0.46708985 | 0 | 0 | 0.50067082 | 0 | 0 | 0 | 2.51818038 | 0 | 0 | 0 | 0 | 0 | 1.02074917 | 0 | 0 | 0.8423362 | 0 |
| ENSMUSG00 Mir2136 | 74 | 0 | 0 | 0 | 0 | 0 | 0 | 0 | 0 | 0 | 0 | 0 | 0 | 0 | 0 | 0 | 0 | 0 | 0 |
| ENSMUSG00 U6 | 103 | 0 | 0 | 1.40893778 | 0 | 0 | 0 | 0 | 0.37080002 | 0 | 0 | 0.9276691 | 0 | 0 | 0 | 0.40001511 | 0 | 0.37209997 | 0 |
| ENSMUSG00 RNaseP_nuc | 269 | 0 | 0 | 0 | 0 | 0 | 0 | 0 | 0 | 0 | 0 | 0 | 0 | 0 | 0 | 0 | 0 | 0 | 0 |
| ENSMUSG00 SNORA17 | 118 | 0 | 0 | 0 | 0 | 0 | 0 | 0 | 0 | 0 | 0 | 0 | 0 | 0 | 0 | 0 | 0 | 0 | 0 |
| ENSMUSG00 7SK | 289 | 0 | 0 | 0 | 0 | 0 | 0 | 0 | 0 | 0 | 0 | 0 | 0 | 0 | 0 | 0 | 0 | 0 | 0 |
| ENSMUSG00 U3 | 88 | 0 | 0 | 0 | 0 | 0 | 0 | 0 | 0 | 0 | 0 | 0 | 0 | 0 | 0 | 0 | 0 | 0 | 0 |
| ENSMUSG00 snoU13 | 104 | 0 | 0 | 0 | 0 | 0 | 0 | 0 | 0 | 0 | 0 | 0 | 0 | 0 | 0 | 0 | 0 | 0 | 0 |
| ENSMUSG00 U6 | 102 | 0 | 0 | 0 | 0 | 0 | 0 | 0 | 0 | 0 | 0 | 0 | 0 | 0 | 0 | 0 | 0 | 0 | 0 |
| ENSMUSG00 snoZ159 | 86 | 0 | 0 | 0 | 0 | 0 | 0 | 0 | 0 | 0 | 0 | 0 | 0 | 0 | 0 | 0 | 0 | 0 | 0 |

| | | | | | | | | | | | | | | | | | | | |
|---|---|---|---|---|---|---|---|---|---|---|---|---|---|---|---|---|---|---|---|
| ENSMUSG00 SNORA17 | 122 | 0 | 0 | 0 | 0 | 0 | 0 | 0 | 0 | 0 | 0 | 0 | 0 | 0 | 0 | 0 | 0 | 0 | 0 |
| ENSMUSG00 U4 | 140 | 0 | 0 | 0 | 0 | 0 | 0 | 0 | 0 | 0 | 0 | 0 | 0 | 0 | 0 | 0 | 0 | 0 | 0 |
| ENSMUSG00 snoU90 | 342 | 0 | 0 | 0 | 0 | 0.28147324 | 0.28978207 | 0 | 0.22334738 | 0 | 0 | 0 | 0 | 0 | 0 | 0 | 0 | 0.33619559 | 0 |
| ENSMUSG00 Y_RNA | 109 | 0 | 0 | 0 | 0 | 0 | 0 | 0 | 0 | 0 | 0 | 0 | 0 | 0 | 0 | 0 | 0 | 0 | 0 |
| ENSMUSG00 snoU13 | 102 | 0 | 0 | 0 | 0 | 0 | 0 | 0 | 0 | 0 | 0 | 0 | 0 | 0 | 0 | 0 | 0 | 0 | 0 |
| ENSMUSG00 U7 | 61 | 0 | 0 | 0 | 0 | 0 | 0 | 0 | 0 | 0 | 0 | 0 | 0 | 0 | 0 | 0 | 0 | 0 | 0 |
| ENSMUSG00 7SK | 254 | 0 | 0 | 0 | 0 | 0 | 0 | 0 | 0 | 0 | 0 | 0 | 0 | 0 | 0 | 0 | 0 | 0 | 0 |
| ENSMUSG00 U6 | 98 | 0 | 0 | 0 | 0 | 0 | 0 | 0 | 0 | 0 | 0 | 0 | 0 | 0 | 0 | 0 | 0 | 0 | 0 |
| ENSMUSG00 U6atac | 119 | 0 | 0 | 0 | 0 | 0 | 0 | 0 | 0 | 0 | 0 | 0 | 0 | 0 | 0 | 0 | 0 | 0 | 0 |
| ENSMUSG00 Mir1930 | 83 | 0 | 0 | 0 | 0 | 0 | 0 | 0 | 0 | 0 | 0 | 0 | 0 | 0 | 0 | 0 | 0 | 0 | 0 |
| ENSMUSG00 SNORA17 | 80 | 0 | 0 | 0 | 0 | 0 | 0 | 0 | 0 | 0 | 0 | 0 | 0 | 0 | 0 | 0 | 0 | 0 | 0 |
| ENSMUSG00 U6 | 104 | 0 | 0 | 0 | 0 | 0 | 0 | 0 | 0 | 0 | 0 | 0 | 0 | 0 | 0 | 0 | 0 | 0 | 0 |
| ENSMUSG00 SNORA17 | 133 | 0 | 0 | 0 | 0 | 0 | 0 | 0 | 0 | 0 | 0 | 0 | 0 | 0 | 0 | 0 | 0 | 0 | 0 |
| ENSMUSG00 5S_rRNA | 115 | 1.84805113 | 0 | 0.42063939 | 0 | 0 | 1.29268001 | 1.6456584 | 1.32843139 | 0 | 1.23001835 | 0 | 0.97070718 | 0 | 0 | 2.14964639 | 0 | 0.66654429 | 0 |
| ENSMUSG00 U3 | 88 | 0 | 0 | 0 | 0 | 0 | 0 | 0 | 0 | 0 | 0 | 0 | 0 | 0 | 0 | 0 | 0 | 0 | 0 |
| ENSMUSG00 U6 | 106 | 0 | 0 | 0 | 0.42982117 | 0 | 2.33739309 | 0 | 0 | 0 | 0 | 0 | 0 | 0 | 0 | 0 | 0 | 0 | 0 |
| ENSMUSG00 Mir1966 | 107 | 0 | 0 | 0 | 0 | 0 | 0 | 0 | 0 | 0 | 0 | 0 | 0 | 0 | 0 | 0 | 0 | 0 | 0 |
| ENSMUSG00 U6 | 111 | 0 | 0 | 0 | 0 | 0 | 0 | 0 | 0 | 0 | 0 | 0 | 0 | 0 | 0 | 0 | 0 | 0 | 0 |
| ENSMUSG00 5S_rRNA | 90 | 0 | 0 | 0 | 0 | 0 | 0 | 0 | 0 | 0 | 0 | 0 | 0 | 0 | 0 | 0 | 0 | 0 | 0 |
| ENSMUSG00 U6 | 106 | 0 | 0 | 1.36906218 | 0.42982117 | 0.9081495 | 0 | 0 | 1.44122273 | 0.38855204 | 0.44481796 | 0.45070716 | 0.52656285 | 0.46539545 | 0.43815177 | 0.38869392 | 0 | 0 | 0 |
| ENSMUSG00 U6 | 104 | 0 | 0 | 0 | 0 | 0 | 0 | 0 | 0 | 0 | 0 | 0 | 0 | 0 | 0 | 0 | 0 | 0 | 0 |
| ENSMUSG00 SNORA17 | 128 | 0 | 0 | 0 | 0 | 0 | 0 | 0 | 0 | 0 | 0 | 0 | 0 | 0 | 0 | 0 | 0 | 0 | 0 |
| ENSMUSG00 SCARNA3 | 87 | 0 | 0 | 0 | 0 | 0 | 0 | 0 | 0 | 0 | 0 | 0 | 0 | 0 | 0 | 0 | 0 | 0 | 0 |
| ENSMUSG00 5S_rRNA | 105 | 0 | 0 | 0 | 0 | 0 | 0 | 0 | 0 | 0 | 0 | 0 | 0 | 0 | 0 | 0 | 0 | 0 | 0 |
| ENSMUSG00 SNORA17 | 122 | 0 | 0 | 0 | 0 | 0 | 0 | 0 | 0 | 0 | 0 | 0 | 0 | 0 | 0 | 0 | 0 | 0 | 0 |
| ENSMUSG00 U6 | 103 | 0 | 0 | 0 | 0 | 0 | 0 | 0 | 0 | 0 | 0 | 0 | 0 | 0 | 0 | 0 | 0 | 0 | 0 |
| ENSMUSG00 SNORA17 | 139 | 0 | 0 | 0 | 0 | 0 | 0 | 0 | 0 | 0 | 0 | 0 | 0 | 0 | 0 | 0 | 0 | 0 | 0 |
| ENSMUSG00 SNORA17 | 117 | 0 | 0 | 0 | 0 | 0 | 0 | 0 | 0 | 0 | 0 | 0 | 0 | 0 | 0 | 0 | 0 | 0 | 0 |
| ENSMUSG00 7SK | 273 | 0 | 0 | 0 | 0 | 0 | 0 | 0 | 0 | 0 | 0 | 0 | 0 | 0 | 0 | 0 | 0 | 0 | 0 |
| ENSMUSG00 SNORA50 | 139 | 0 | 0 | 0 | 0 | 0 | 0 | 0 | 0 | 0 | 0 | 0 | 0 | 0 | 0 | 0 | 0 | 0 | 0 |
| ENSMUSG00 U7 | 62 | 0 | 0 | 0 | 0 | 0 | 0 | 0 | 0 | 0 | 0 | 0 | 0 | 0 | 0 | 0 | 0 | 0 | 0 |
| ENSMUSG00 SNORD112 | 76 | 0 | 0 | 0 | 0 | 0 | 0 | 0 | 0 | 0 | 0 | 0 | 0 | 0 | 0 | 0 | 0 | 0 | 0 |
| ENSMUSG00 U6 | 97 | 0 | 0 | 0 | 0 | 0 | 0 | 0 | 0 | 0 | 0 | 0 | 0 | 0 | 0 | 0 | 0 | 0.79023293 | 0 |
| ENSMUSG00 U6 | 96 | 0 | 0 | 0 | 0 | 0 | 0 | 0 | 0 | 0 | 0 | 0 | 0 | 0 | 0 | 0 | 0 | 0.39923226 | 0 |
| ENSMUSG00 U6 | 89 | 0 | 0 | 0 | 0 | 0 | 0 | 0 | 0 | 0 | 0 | 0 | 0 | 0 | 0 | 0 | 0 | 0 | 0 |
| ENSMUSG00 7SK | 257 | 3.96935496 | 2.33319882 | 2.4469101 | 0.35456066 | 1.1237025 | 2.89218289 | 2.20915232 | 2.37773711 | 3.04491751 | 2.5685208 | 2.04484258 | 2.6061788 | 1.15171791 | 1.98787923 | 2.56507741 | 4.6187114 | 1.93868428 | 0.85617287 |
| ENSMUSG00 SNORA17 | 128 | 0 | 0 | 0 | 0 | 0 | 0 | 0 | 0 | 0 | 0 | 0 | 0 | 0 | 0 | 0 | 0 | 0 | 0 |
| ENSMUSG00 SNORA17 | 117 | 0 | 0 | 0 | 0 | 0 | 0 | 0 | 0 | 0 | 0 | 0 | 0 | 0 | 0 | 0 | 0 | 0 | 0 |
| ENSMUSG00 snoU13 | 101 | 0 | 0 | 0 | 0 | 0 | 0 | 0 | 0 | 0 | 0 | 0 | 0 | 0 | 0 | 0 | 0 | 0 | 0 |
| ENSMUSG00 snoU85 | 312 | 0 | 0 | 0 | 0 | 0 | 0 | 0 | 0 | 0 | 0 | 0 | 0 | 0 | 0 | 0 | 0 | 0 | 0 |
| ENSMUSG00 5S_rRNA | 92 | 0 | 0 | 0 | 0 | 0 | 0 | 0 | 0 | 0 | 0 | 0 | 0 | 0 | 0 | 0 | 0 | 0 | 0 |

[illegible]

| | | | | | | | | | | | | | | | | | | | |
|---|---|---|---|---|---|---|---|---|---|---|---|---|---|---|---|---|---|---|---|
| ENSMUSG00 Gm16168 | 1776 | 0 | 0 | 0 | 0.05130748 | 0 | 0 | 0 | 0.10752366 | 0 | 0 | 0.05380063 | 0 | 0 | 0 | 0 | 0 | 0 | 0 |
| ENSMUSG00 Pira2 | 4736 | 0 | 0 | 0 | 0 | 0 | 0 | 0 | 0 | 0.04348239 | 0 | 0.01008762 | 0.0235708 | 0.01041637 | 0 | 0.03479861 | 0 | 0 | 0 |
| ENSMUSG00 Ugt1a5 | 2198 | 0 | 0 | 0 | 0 | 0 | 0 | 0 | 0 | 0 | 0 | 0 | 0 | 0 | 0 | 0 | 0 | 0 | 0 |
| ENSMUSG00 Gm8292 | 622 | 3.82683257 | 2.19586602 | 3.26637986 | 2.56372436 | 2.7857705 | 1.51366871 | 3.57507381 | 3.8683623 | 3.17838064 | 2.50156464 | 1.84340676 | 3.05101693 | 1.82417057 | 2.01606167 | 2.51713686 | 2.58553941 | 2.40309579 | 2.29941604 |
| ENSMUSG00 Gm20459 | 8529 | 0 | 0 | 0 | 0 | 0 | 0 | 0 | 0 | 0 | 0 | 0 | 0 | 0 | 0 | 0 | 0 | 0 | 0 |
| ENSMUSG00 Gm15509 | 2159 | 0 | 0 | 0 | 0 | 0 | 0 | 0 | 0.15920872 | 0 | 0 | 0 | 0 | 0 | 0 | 0 | 0 | 0 | 0 |
| ENSMUSG00 Gm16583 | 552 | 0 | 0 | 0 | 0 | 0 | 0 | 0 | 0 | 0 | 0 | 0 | 0 | 0 | 0 | 0 | 0 | 0 | 0 |
| ENSMUSG00 Gm16137 | 662 | 1.09152265 | 2.26447166 | 1.68065136 | 0.75705663 | 1.45413667 | 0.5239715 | 1.78673259 | 1.55769617 | 2.98633347 | 0.35612314 | 1.9485255 | 1.51764641 | 1.71394879 | 2.17487418 | 1.369236 | 1.96658709 | 1.04210475 | 0.66476262 |
| ENSMUSG00 Gm16215 | 429 | 0 | 0 | 0 | 0 | 0 | 0 | 0 | 0 | 0 | 0 | 0 | 0 | 0 | 0 | 0 | 0 | 0 | 0 |
| ENSMUSG00 Gm14435 | 1504 | 0 | 0 | 0 | 0 | 0 | 0 | 0 | 0 | 0 | 0 | 0 | 0 | 0 | 0 | 0 | 0 | 0 | 0 |
| ENSMUSG00 4933413C19I | 2215 | 0.09594848 | 0 | 0 | 0.16455456 | 0 | 0 | 0 | 0 | 0 | 0.042574 | 0 | 0.20159156 | 0 | 0 | 0.26041615 | 0.01728696 | 0.13842455 | 0 |
| ENSMUSG00 Gm757 | 2019 | 0 | 0 | 0 | 0 | 0 | 0 | 0 | 0 | 0 | 0 | 0 | 0 | 0 | 0.02300351 | 0 | 0 | 0 | 0 |
| ENSMUSG00 Gm8783 | 521 | 0 | 0 | 0 | 0.08744922 | 0 | 0.09511081 | 0 | 0 | 0 | 0 | 0 | 0 | 0 | 0 | 0.15816336 | 0 | 0 | 0 |
| ENSMUSG00 C230030N03 | 2671 | 0 | 0 | 0 | 0 | 0 | 0 | 0 | 0 | 0 | 0 | 0 | 0 | 0 | 0 | 0 | 0 | 0 | 0 |
| ENSMUSG00 Gm16553 | 713 | 0 | 0 | 0 | 0 | 0 | 0 | 0 | 0 | 0 | 0 | 0 | 0 | 0 | 0 | 0 | 0 | 0 | 0 |
| ENSMUSG00 A830011K09I | 2637 | 0 | 0 | 0 | 0 | 0 | 0 | 0 | 0 | 0 | 0 | 0 | 0 | 0 | 0 | 0 | 0 | 0 | 0 |
| ENSMUSG00 Gm16165 | 794 | 0 | 0.04195579 | 0 | 0 | 0.06061955 | 0 | 0 | 0.04810126 | 0 | 0 | 0.06016997 | 0 | 0 | 0 | 0 | 0 | 0.0482699 | 0 |
| ENSMUSG00 Gm16268 | 1154 | 0 | 0 | 0 | 0 | 0 | 0 | 0 | 0 | 0 | 0 | 0.20699722 | 0 | 0 | 0.1609847 | 0 | 0 | 0 | 0.0476682 |
| ENSMUSG00 Ugt1a1 | 2219 | 0 | 0 | 0 | 0 | 0 | 0 | 0 | 0.05163461 | 0 | 0 | 0 | 0 | 0.0444632 | 0 | 0 | 0 | 0.01727188 | 0 |
| ENSMUSG00 Gm16567 | 573 | 0 | 0 | 0 | 0 | 0 | 0 | 0 | 0 | 0 | 0 | 0 | 0 | 0 | 0 | 0 | 0 | 0 | 0 |
| ENSMUSG00 Gm16273 | 267 | 0.31839083 | 0 | 0 | 0.17064062 | 0.18026938 | 0 | 0.17720104 | 0 | 0.30851323 | 0 | 0 | 0 | 0 | 0 | 0 | 0.14341051 | 0 | 0 |
| ENSMUSG00 Gm14555 | 230 | 0 | 0 | 0 | 0 | 0 | 0 | 0 | 0 | 0 | 0 | 0 | 0 | 0 | 0 | 0 | 0 | 0 | 0 |
| ENSMUSG00 Gm16582 | 1926 | 0 | 0 | 0 | 0 | 0 | 0 | 0 | 0 | 0 | 0 | 0 | 0 | 0 | 0 | 0 | 0 | 0 | 0 |
| ENSMUSG00 Gm16236 | 378 | 0 | 0 | 0 | 0 | 0 | 0 | 0 | 0 | 0 | 0 | 0 | 0 | 0 | 0 | 0 | 0 | 0 | 0 |
| ENSMUSG00 Gm16030 | 602 | 0 | 0 | 0 | 0 | 0 | 0 | 0 | 0 | 0 | 0 | 0 | 0 | 0 | 0 | 0 | 0 | 0 | 0 |
| ENSMUSG00 Gm16550 | 125 | 0 | 0 | 0.77397649 | 0 | 0.77011078 | 0 | 0 | 0 | 0 | 0.37720563 | 0 | 0.4465253 | 0 | 0 | 0 | 0 | 0 | 0 |
| ENSMUSG00 4921518J05R | 1704 | 0 | 0 | 0.05677644 | 0 | 0 | 0 | 0 | 0 | 0 | 0 | 0 | 0 | 0 | 0 | 0 | 0.02247101 | 0.02249196 | 0 |
| ENSMUSG00 Gm16318 | 631 | 0 | 0 | 0 | 0 | 0 | 0 | 0 | 0 | 0 | 0 | 0 | 0 | 0 | 0 | 0 | 0 | 0 | 0 |
| ENSMUSG00 Vmn1r190-ps | 1906 | 0 | 0 | 0 | 0 | 0 | 0 | 0 | 0 | 0 | 0 | 0 | 0 | 0 | 0 | 0 | 0 | 0 | 0 |
| ENSMUSG00 Gm7420 | 845 | 0.20120793 | 0 | 0.34348069 | 0 | 0 | 0 | 0 | 0 | 0 | 0.16739895 | 0.33923047 | 0.06605404 | 0.1751429 | 0.05496342 | 0 | 0.6344006 | 0.18142626 | 0 |
| ENSMUSG00 Gm2389 | 187 | 0 | 0 | 0 | 0 | 0 | 0 | 0 | 0 | 0 | 0 | 0.25548106 | 0 | 0 | 0 | 0 | 0 | 0 | 0 |
| ENSMUSG00 Gm16245 | 894 | 0 | 0 | 0 | 0 | 0 | 0 | 0 | 0.04272081 | 0 | 0 | 0 | 0 | 0 | 0 | 0 | 0 | 0 | 0 |
| ENSMUSG00 Gm15954 | 464 | 0.5496359 | 0.64615528 | 1.14678628 | 0.88372715 | 0.10373259 | 0 | 0.30590094 | 0.82311212 | 0.71011234 | 0 | 0.82370618 | 0.12029238 | 0 | 0 | 0.53277874 | 0.57766003 | 0.49559867 | 0.11855411 |
| ENSMUSG00 Gm11565 | 502 | 0 | 0 | 0 | 0 | 0 | 0 | 0 | 0 | 0 | 0 | 0 | 0 | 0 | 0 | 0 | 0 | 0 | 0 |
| ENSMUSG00 Gm7568 | 1010 | 0 | 0 | 0.14368375 | 0.09021989 | 0.04765537 | 0.04906211 | 0.14053271 | 0 | 0 | 0 | 0 | 0.11052606 | 0.09768696 | 0.13795273 | 0 | 0 | 0.2656278 | 0 |
| ENSMUSG00 2010320O07I | 523 | 0.32508739 | 0.63695782 | 1.10990892 | 0.43557404 | 0.18406089 | 1.32645941 | 0.09046401 | 0 | 1.26000813 | 0 | 0.7307833 | 0.42688843 | 0.66027423 | 0.08880323 | 0 | 0.87856078 | 0.36640819 | 0.52589968 |
| ENSMUSG00 Fbxo24 | 3114 | 0.05459881 | 0.23535121 | 0.13980789 | 0.17557243 | 0.18547947 | 0.14321599 | 0.24309662 | 0.12264741 | 0.33065604 | 0.24226437 | 0.4909437 | 0.05377231 | 0.07920989 | 0.25354833 | 0.48954964 | 0.08607394 | 0.39384762 | 0.30030662 |
| ENSMUSG00 Gpx2-ps1 | 302 | 0 | 0 | 0 | 0 | 0 | 0 | 0 | 0 | 0 | 0 | 0 | 0 | 0 | 0 | 0 | 0 | 0 | 0 |
| ENSMUSG00 Gm16238 | 587 | 0.21723259 | 0.45400878 | 0.49444835 | 0.93140125 | 0.49197878 | 0.42208461 | 0.40300408 | 0.45544603 | 1.05246633 | 0.64259902 | 0.89527179 | 0.19017262 | 0.58828521 | 0.23736331 | 0.14038009 | 0 | 0.06529182 | 0.09371228 |
| ENSMUSG00 Flt3l | 3926 | 0.66042174 | 0.45820079 | 0.62838768 | 0.25530896 | 0.63750892 | 0.47962401 | 0.71101581 | 0.53504384 | 0.35668404 | 0.67255206 | 0.76663841 | 0.55446022 | 0.77905728 | 0.5086846 | 0.50373782 | 0.87777755 | 0.56620612 | 0.68656297 |
| ENSMUSG00 Gm15852 | 913 | 0 | 0 | 0 | 0 | 0 | 0 | 0 | 0 | 0 | 0 | 0 | 0 | 0 | 0 | 0 | 0 | 0 | 0 |
| ENSMUSG00 Gm16332 | 3406 | 0 | 0 | 0 | 0 | 0 | 0 | 0 | 0 | 0 | 0 | 0 | 0 | 0 | 0 | 0 | 0 | 0 | 0 |
| ENSMUSG00 G6pd2 | 1634 | 0.15607776 | 0.18348595 | 0.29604364 | 0.11153254 | 0.0883695 | 0.27293427 | 0.11582051 | 0.30385632 | 0.02520595 | 0.057712 | 0.11695216 | 0.23911239 | 0.24152713 | 0 | 0.0504303 | 0.18746931 | 0 | 0.26932243 |
| ENSMUSG00 Gm5822 | 1377 | 0.06173591 | 0.02419237 | 0.21077791 | 0.13234871 | 0.06990839 | 0.10795803 | 0.06871849 | 0.24962355 | 0.02991032 | 0.17120807 | 0.41633951 | 0.04053425 | 0.39408213 | 0.16864229 | 0.65826741 | 0.05561454 | 0.25049867 | 0 |
| ENSMUSG00 Gm16239 | 2631 | 0 | 0.05064674 | 0 | 0 | 0 | 0 | 0 | 0.01451631 | 0 | 0.01792121 | 0 | 0 | 0 | 0 | 0 | 0 | 0 | 0 |
| ENSMUSG00 Gm15716 | 398 | 0 | 0.25110222 | 0 | 0 | 0 | 0 | 0.59438039 | 0 | 0 | 0 | 0 | 0 | 0.37184862 | 0.70016212 | 0.20704299 | 0.09620756 | 0 | 0 |
| ENSMUSG00 Tmsb15b2 | 447 | 0 | 0.14905098 | 0 | 0.10192627 | 0.32303304 | 0.33256868 | 0.84675935 | 0 | 0.09213986 | 0.10548256 | 0.85503281 | 0.62433627 | 0.55181116 | 0.83121409 | 0.27652051 | 0.68529051 | 0.51444694 | 0 |
| ENSMUSG00 1810020O05I | 1042 | 0 | 0 | 0 | 0.13117383 | 0 | 0 | 0 | 0 | 0.03952641 | 0 | 0 | 0 | 0 | 0 | 0 | 0 | 0 | 0 |
| ENSMUSG00 A130049A11I | 4356 | 0.17564122 | 0.50474082 | 0.43309635 | 0.28240317 | 0.53037932 | 0.44365395 | 0.29326041 | 0.76279592 | 0.4538459 | 0.11906743 | 0.27419053 | 0.28189729 | 0.1812008 | 0.30920077 | 0.30267442 | 0.11427408 | 0.30794775 | 0.63141766 |
| ENSMUSG00 Gm6485 | 438 | 2.32905074 | 2.05353458 | 2.31927885 | 2.18443363 | 2.85714615 | 3.95969332 | 1.08019815 | 3.0519043 | 4.1374582 | 3.87540027 | 2.61780594 | 4.20528965 | 1.57681927 | 1.27244075 | 2.35168698 | 2.62264434 | 2.88759771 | 2.26064822 |
| ENSMUSG00 Ier3ip1 | 2061 | 37.9267437 | 34.7837692 | 34.6899941 | 31.1919619 | 35.2640489 | 34.1170931 | 42.0327584 | 39.7119449 | 41.2864345 | 38.4572209 | 36.3469844 | 39.0790884 | 39.6139365 | 40.6526607 | 40.2020033 | 42.4150688 | 40.873945 | 39.1282632 |
| ENSMUSG00 Gm9719 | 542 | 0 | 0 | 0 | 0 | 0 | 0 | 0 | 0 | 0 | 0 | 0 | 0 | 0 | 0 | 0 | 0 | 0 | 0 |
| ENSMUSG00 Gm16006 | 564 | 0 | 0.11813083 | 0.6003807 | 0.48469196 | 0.08534029 | 0.79073511 | 0.67110183 | 1.15118943 | 0.29210295 | 0.25080161 | 0 | 0.29689182 | 0.61227557 | 0.08234767 | 0.58441923 | 0.20367344 | 0 | 0.39013551 |
| ENSMUSG00 Gm15882 | 853 | 0 | 0 | 0 | 0 | 0 | 0 | 0 | 0 | 0 | 0 | 0 | 0 | 0 | 0 | 0 | 0 | 0 | 0 |
| ENSMUSG00 Gm16125 | 338 | 0.25150992 | 1.1827063 | 2.28987126 | 1.75234786 | 0.28480428 | 0.43981716 | 1.3997834 | 2.03391492 | 2.43707194 | 1.53449034 | 2.2615365 | 0.82567548 | 0.72976209 | 0.27481709 | 0.97518475 | 1.92585895 | 1.36069693 | 0.97649302 |
| ENSMUSG00 Gm16540 | 424 | 0 | 0.94281776 | 0 | 0 | 0.11351869 | 0 | 0 | 0 | 0.77710407 | 0 | 0.11267679 | 0.26328143 | 0 | 0 | 0.87456133 | 0.27092411 | 0 | 0.25947692 |
| ENSMUSG00 Gm16227 | 561 | 0 | 0 | 0 | 0 | 0 | 0 | 0 | 0 | 0 | 0 | 0 | 0 | 0 | 0 | 0 | 0 | 0.3415891 | 0 |
| ENSMUSG00 Rpl30-ps2 | 290 | 0 | 0 | 0 | 0 | 0.16597215 | 0 | 0 | 0 | 0 | 0 | 0.82370618 | 0.1924678 | 0 | 0 | 0 | 0 | 0 | 0 |
| ENSMUSG00 Gm16111 | 284 | 0 | 0 | 0 | 0 | 0 | 0 | 0 | 0 | 0 | 0.66409441 | 0 | 0 | 0 | 0 | 0 | 0 | 0 | 0 |
| ENSMUSG00 Gm16282 | 448 | 0 | 0 | 0 | 0 | 0 | 0 | 0 | 0 | 0 | 0 | 0 | 0 | 0 | 0 | 0 | 0 | 0 | 0 |
| ENSMUSG00 Vmn2r-ps12 | 597 | 0 | 0 | 0 | 0 | 0 | 0 | 0 | 0 | 0 | 0 | 0 | 0 | 0 | 0 | 0 | 0 | 0 | 0 |
| ENSMUSG00 Gm16555 | 1197 | 0 | 0.11132128 | 0 | 0 | 0 | 0 | 0 | 0 | 0 | 0 | 0 | 0 | 0 | 0 | 0 | 0 | 0 | 0 |
| ENSMUSG00 Gm6507 | 650 | 0 | 0 | 0 | 0 | 0 | 0 | 0 | 0 | 0 | 0 | 0 | 0 | 0 | 0 | 0 | 0 | 0 | 0 |
| ENSMUSG00 Gm15446 | 3775 | 3.8395404 | 2.6209085 | 3.74174063 | 3.37936223 | 3.36604711 | 3.18975212 | 4.22367491 | 3.39937674 | 4.29867211 | 3.77205626 | 3.25249386 | 3.72597271 | 2.95338101 | 2.50982618 | 2.97960921 | 3.60083858 | 3.72602675 | 2.15664842 |
| ENSMUSG00 Gm16578 | 781 | 0 | 0 | 0 | 0 | 0 | 0 | 0 | 0 | 0 | 0 | 0 | 0 | 0 | 0 | 0 | 0 | 0 | 0 |
| ENSMUSG00 Gm16213 | 637 | 0 | 0 | 0 | 0 | 0 | 0 | 0 | 0 | 0 | 0 | 0 | 0 | 0 | 0 | 0 | 0 | 0 | 0 |
| ENSMUSG00 Gm13741 | 537 | 0 | 0 | 0 | 0 | 0 | 0 | 0 | 0 | 0 | 0 | 0 | 0 | 0 | 0 | 0 | 0 | 0 | 0 |
| ENSMUSG00 Gimap1 | 1738 | 0.68477844 | 0.11500424 | 0.1948301 | 0.07864392 | 0.24924471 | 0.08553406 | 1.06167691 | 0.35159864 | 0.16588355 | 0.4611979 | 0.68721172 | 0.89921666 | 0.56768604 | 0.58789984 | 0.40300716 | 0.24234562 | 0.11025977 | 0.18990486 |
| ENSMUSG00 Gm6493 | 1784 | 0.09530309 | 0.07469259 | 0.02711521 | 0.0510774 | 0.08093933 | 0 | 0.02652056 | 0.04281659 | 0.43864563 | 0 | 0.02677969 | 0.12514723 | 0.16591452 | 0.13016841 | 0.30023555 | 0.12878007 | 0.0429667 | 0.18500821 |
| ENSMUSG00 Gm16302 | 426 | 0 | 0 | 0 | 0 | 0 | 0 | 0.44425051 | 0 | 0 | 0 | 0 | 0 | 0 | 0 | 0 | 0 | 0 | 0 |
| ENSMUSG00 Gm16350 | 444 | 0 | 0.15005808 | 0 | 0 | 0.21681047 | 0.11160526 | 0.21312018 | 0 | 0.18552485 | 0 | 0 | 0 | 0 | 0.1046038 | 0.27838889 | 0.51744064 | 0.34528195 | 0 |
| ENSMUSG00 Gm16088 | 500 | 0 | 0 | 0 | 0 | 0 | 0 | 0 | 0 | 0 | 0 | 0 | 0 | 0 | 0 | 0 | 0 | 0 | 0 |
| ENSMUSG00 Gm15996 | 760 | 0.22371145 | 0.04383276 | 0.06364938 | 0 | 0 | 0 | 0.56028172 | 0.25126581 | 0.10838557 | 0 | 0 | 0.14688332 | 0 | 0 | 0.05421257 | 0 | 0.10085868 | 0 |
| ENSMUSG00 Gm15740 | 1291 | 0 | 0.02580395 | 0 | 0 | 0 | 0 | 0.07329617 | 0 | 0 | 0 | 0 | 0 | 0 | 0 | 0 | 0 | 0 | 0 |
| ENSMUSG00 Gm17018 | 651 | 0 | 0 | 0 | 0 | 0 | 0 | 0 | 0 | 0 | 0 | 0 | 0 | 0 | 0 | 0 | 0 | 0 | 0 |
| ENSMUSG00 A430072P03I | 1226 | 0 | 0 | 0 | 0 | 0 | 0 | 0 | 0 | 0 | 0 | 0 | 0 | 0 | 0.07576523 | 0 | 0 | 0 | 0 |
| ENSMUSG00 4732440D04 | 411 | 4.75727031 | 3.56634391 | 4.59018901 | 1.44110359 | 1.87374885 | 4.21981916 | 1.38139209 | 6.31893762 | 2.80589402 | 3.78582289 | 2.09233395 | 3.93833142 | 1.56037695 | 3.72908731 | 5.91457858 | 3.07442833 | 3.54355057 | 3.21221061 |
| ENSMUSG00 Gm11503 | 301 | 0 | 0 | 0 | 0 | 0 | 0 | 0 | 0 | 0 | 0 | 0 | 0 | 0 | 0 | 0 | 0 | 0 | 0 |
| ENSMUSG00 Gm16243 | 492 | 0.17278527 | 0.27083654 | 0.29496055 | 0 | 0.58697468 | 0 | 0 | 0.31050734 | 0.92083673 | 0.57500858 | 0.29131072 | 0 | 0 | 0.47199276 | 0.418715 | 0.07782644 | 1.09058568 | 0 |
| ENSMUSG00 Gm15917 | 484 | 0 | 0.06882829 | 0.39978124 | 0.09413439 | 0.09944612 | 0 | 0.48876734 | 0.07896992 | 0 | 0.09741881 | 0.98708592 | 0.80725132 | 0.30577635 | 0.57575315 | 1.53228927 | 0.6329026 | 0.23755969 | 0.45462072 |
| ENSMUSG00 Galnt4 | 5088 | 1.01918858 | 1.25709035 | 1.14088515 | 1.25364509 | 1.77845944 | 1.65565344 | 1.65519985 | 2.08677041 | 2.01561368 | 1.80707294 | 2.58217641 | 3.11549689 | 1.18288009 | 1.48789038 | 0.63162763 | 1.64059599 | 1.70238682 | 1.6109192 |
| ENSMUSG00 Gm15900 | 341 | 0 | 0 | 0 | 0 | 0 | 0 | 0 | 0.11200118 | 0 | 0 | 0 | 0 | 0 | 0 | 0 | 0 | 0 | 0 |
| ENSMUSG00 Krtap4-9 | 688 | 0 | 0 | 0 | 0 | 0 | 0 | 0 | 0 | 0 | 0 | 0 | 0 | 0 | 0 | 0 | 0 | 0 | 0 |
| ENSMUSG00 Gm16573 | 1296 | 0.45916085 | 0.74542742 | 0.7465051 | 0.28124101 | 1.00274841 | 0.72646754 | 0.10952009 | 0.85461394 | 0.85805241 | 0.61848916 | 0.77413127 | 0.90442046 | 0.79935977 | 0.07167297 | 1.49419223 | 1.56589675 | 0.65060072 | 0.80646067 |
| ENSMUSG00 Gm15894 | 2064 | 0.16474874 | 0.06455987 | 0.09374715 | 0.0441483 | 0 | 0 | 0.18338248 | 0.03700814 | 0.01995471 | 0 | 0.09258713 | 0.08112742 | 0 | 0 | 0 | 0 | 0 | 0 |
| ENSMUSG00 Gm16232 | 366 | 0 | 0 | 0 | 0 | 0 | 0 | 0 | 0 | 0 | 0 | 0 | 0 | 0 | 0 | 0 | 0 | 0 | 0 |
| ENSMUSG00 Gm16027 | 757 | 0 | 0 | 0 | 0.3009316 | 0 | 0 | 0.43750165 | 0 | 0.54407551 | 0 | 0 | 0.36866356 | 0 | 0 | 0 | 0 | 0 | 0 |
| ENSMUSG00 Gm8550 | 652 | 0 | 0 | 0 | 0 | 0 | 0 | 0 | 0 | 0 | 0 | 0 | 0 | 0 | 0 | 0 | 0 | 0 | 0 |
| ENSMUSG00 Gm16167 | 1064 | 0.39948474 | 0 | 0.31824691 | 0.17128212 | 0.18094708 | 0.04657212 | 0 | 0 | 0 | 0.79766227 | 0.26940766 | 0.62949995 | 0 | 0 | 0 | 0.14394965 | 0.07204191 | 0.98230548 |
| ENSMUSG00 Gm16557 | 190 | 0 | 0 | 0 | 0 | 0 | 0 | 0 | 0 | 0 | 0 | 0 | 0 | 0 | 0 | 0 | 0 | 0 | 0 |
| ENSMUSG00 Tmem229a | 893 | 0 | 0 | 0 | 0 | 0 | 0 | 0 | 0 | 0 | 0 | 0 | 0 | 0 | 0 | 0 | 0 | 0 | 0 |
| ENSMUSG00 Gm16297 | 458 | 0 | 0 | 0 | 0.09947826 | 0 | 0 | 0 | 0 | 0 | 0.20589827 | 0 | 0 | 0.10771161 | 0 | 0 | 0 | 0 | 0 |
| ENSMUSG00 Gm16317 | 736 | 0 | 0 | 0 | 0 | 0 | 0 | 0 | 0 | 0 | 0 | 0 | 0 | 0 | 0 | 0 | 0 | 0 | 0 |
| ENSMUSG00 Gm16339 | 927 | 0 | 0 | 0 | 0 | 0 | 0 | 0 | 0 | 0 | 0 | 0 | 0 | 0 | 0 | 0 | 0 | 0 | 0 |
| ENSMUSG00 Gm8194 | 858 | 0 | 0 | 0 | 0 | 0 | 0 | 0 | 0 | 0 | 0 | 0 | 0 | 0 | 0 | 0 | 0 | 0 | 0 |
| ENSMUSG00 Palm2 | 6583 | 0.36158132 | 0.36435187 | 0.264537 | 0.28376163 | 0.40213517 | 0.29356777 | 0.1006194 | 0.11023176 | 0.36913329 | 0.03581247 | 0.03628662 | 0.03391503 | 0.3821856 | 0.07760671 | 0.12517562 | 0.50022668 | 0.13972826 | 0.02506871 |
| ENSMUSG00 Gm15629 | 473 | 0 | 0 | 0 | 0 | 0 | 0 | 0 | 0 | 0 | 0 | 0 | 0 | 0 | 0 | 0 | 0 | 0 | 0 |
| ENSMUSG00 Gm16333 | 316 | 0 | 0 | 0 | 0 | 0 | 0 | 0 | 0 | 0 | 0 | 0 | 0 | 0 | 0 | 0 | 0.72703685 | 0 | 0 |
| ENSMUSG00 Olfr1062 | 955 | 0 | 0 | 0 | 0 | 0 | 0 | 0 | 0 | 0 | 0 | 0 | 0 | 0 | 0 | 0 | 0 | 0 | 0 |
| ENSMUSG00 3110047P20I | 10362 | 0 | 0 | 0 | 0 | 0 | 0 | 0 | 0 | 0 | 0 | 0 | 0 | 0 | 0 | 0 | 0 | 0 | 0 |
| ENSMUSG00 1810009N23 | 1384 | 0 | 0 | 0 | 0 | 0.0347774 | 0.035804 | 0 | 0.05519133 | 0 | 0 | 0 | 0 | 0 | 0 | 0 | 0 | 0 | 0 |
| ENSMUSG00 Dlx6as1 | 8889 | 0.02390886 | 0.00374765 | 0 | 0 | 0 | 0 | 0 | 0 | 0 | 0 | 0 | 0 | 0.02219909 | 0 | 0 | 0 | 0 | 0 |
| ENSMUSG00 Gm11554 | 867 | 0 | 0 | 0 | 0 | 0 | 0 | 0 | 0 | 0 | 0 | 0 | 0 | 0 | 0 | 0 | 0 | 0 | 0 |
| ENSMUSG00 1110002E22I | 9645 | 0.00881393 | 0.08980148 | 0 | 0.0141714 | 0.00499035 | 0 | 0 | 0 | 0 | 0 | 0 | 0 | 0.05114766 | 0.10593778 | 0 | 0.03572996 | 0 | 0 |
| ENSMUSG00 Gm2976 | 512 | 0 | 0 | 0 | 0 | 0 | 0 | 0 | 0 | 0 | 0 | 0 | 0 | 0 | 0 | 0.16094358 | 0 | 0.52399234 | 0 |
| ENSMUSG00 D030054H15 | 4878 | 0 | 0 | 0 | 0 | 0 | 0 | 0 | 0 | 0 | 0 | 0 | 0 | 0 | 0 | 0 | 0 | 0.00785697 | 0 |
| ENSMUSG00 Gm16577 | 469 | 0 | 0 | 0 | 0 | 0.1026267 | 0 | 0.40351965 | 0 | 0 | 0.70374184 | 0 | 0 | 0 | 0.69319533 | 0 | 0 | 0 | 0 |
| ENSMUSG00 Cdk5r2 | 3353 | 0 | 0.00993525 | 0 | 0 | 0 | 0.10345038 | 0 | 0 | 0 | 0 | 0 | 0 | 0 | 0 | 0 | 0 | 0 | 0 |
| ENSMUSG00 Gm13538 | 545 | 0 | 0 | 0 | 0 | 0 | 0 | 0 | 0 | 0 | 0 | 0 | 0 | 0 | 0 | 0 | 0 | 0 | 0 |
| ENSMUSG00 Gm6921 | 434 | 0.29381458 | 0.15351564 | 0.22291949 | 0.41991746 | 0 | 0 | 0 | 0 | 0 | 0.21728435 | 0.11008055 | 0 | 0 | 0 | 0 | 0 | 0.35323776 | 0 |
| ENSMUSG00 Gm16251 | 336 | 0.1265035 | 0.19829104 | 0.57587536 | 0.40679504 | 0.28649955 | 0 | 0.14081154 | 0.56833932 | 0.49031566 | 0.28065895 | 0.56874951 | 0.16611804 | 0.73410591 | 1.79694386 | 1.22623678 | 0.22792028 | 0.68439816 | 0.32743516 |
| ENSMUSG00 Lime1 | 1822 | 0.18663085 | 0.25597174 | 0.07964906 | 0.15003637 | 0.02641708 | 0.13598445 | 0.07790233 | 0.14673261 | 0.22605113 | 0.15527125 | 0.07866349 | 0.24507426 | 0.16245417 | 0.30588861 | 0.11306684 | 0.21015701 | 0.23138818 | 0.24153285 |
| ENSMUSG00 Gm16147 | 278 | 0 | 0 | 0 | 0 | 0 | 0 | 0 | 0 | 0 | 0 | 0 | 0 | 0 | 0 | 0 | 0 | 0 | 0 |
| ENSMUSG00 Gm15849 | 332 | 0 | 0 | 0 | 0 | 0 | 0 | 0 | 0 | 0 | 0 | 0 | 0 | 0 | 0 | 0 | 0 | 0 | 0 |
| ENSMUSG00 Gm15872 | 637 | 0 | 0 | 0 | 0.21457321 | 0 | 0 | 0 | 0 | 0 | 0 | 0 | 0.0876227 | 0 | 0 | 0.06468062 | 0 | 0 | 0 |
| ENSMUSG00 Gm16587 | 861 | 0 | 0 | 0 | 0 | 0 | 0 | 0.05495085 | 0 | 0.04783567 | 0.05476272 | 0 | 0 | 0.05729607 | 0 | 0 | 0 | 0 | 0 |
| ENSMUSG00 Rnf8 | 8382 | 10.6896816 | 10.679044 | 11.1382622 | 11.5451752 | 10.1810905 | 10.4757151 | 11.5431196 | 11.5643423 | 10.7855407 | 12.4880173 | 11.9180909 | 10.8474963 | 9.74632009 | 9.95151288 | 11.2761118 | 11.7539648 | 11.9752531 | 8.99098977 |
| ENSMUSG00 Srpx | 3186 | 0 | 0 | 0 | 0 | 0 | 0 | 0 | 0.03596271 | 0 | 0.10359539 | 0 | 0 | 0 | 0 | 0 | 0 | 0.0120296 | 0 |
| ENSMUSG00 AI480526 | 4686 | 3.81875354 | 3.49764062 | 3.62336943 | 3.06268224 | 3.18414348 | 2.8551511 | 3.91748174 | 3.51278819 | 3.81454286 | 1.93191102 | 2.56920391 | 3.01352169 | 2.61082277 | 3.58787017 | 3.02461272 | 2.85177592 | 2.00382901 | 2.23041619 |
| ENSMUSG00 Gm15939 | 2142 | 0 | 0 | 0 | 0 | 0 | 0 | 0 | 0 | 0 | 0 | 0 | 0 | 0 | 0 | 0 | 0 | 0 | 0 |
| ENSMUSG00 Gm15934 | 523 | 0 | 0 | 0 | 0 | 0 | 0 | 0.09046401 | 0.65723063 | 0 | 0 | 0.09134791 | 0 | 0.28297467 | 0.88803226 | 0 | 0 | 0.21984492 | 0.63107962 |
| ENSMUSG00 Magea9-ps | 1629 | 0 | 0 | 0 | 0 | 0 | 0 | 0 | 0 | 0 | 0 | 0 | 0 | 0 | 0 | 0 | 0 | 0 | 0 |
| ENSMUSG00 Mup-ps16 | 590 | 0 | 0 | 0 | 0 | 0 | 0 | 0 | 0 | 0 | 0 | 0 | 0 | 0 | 0 | 0 | 0 | 0 | 0 |
| ENSMUSG00 Ang-ps3 | 432 | 0 | 0 | 0 | 0 | 0 | 0 | 0 | 0 | 0 | 0 | 0 | 0 | 0 | 0 | 0 | 0 | 0 | 0 |
| ENSMUSG00 Gm14399 | 1788 | 0.64185668 | 0.67072941 | 0.56814549 | 0.50963137 | 0.43071073 | 0.44342491 | 1.03198796 | 0.89713697 | 0.73711885 | 1.42401453 | 0.50767573 | 1.6232743 | 2.20724462 | 0.62341057 | 0.64521452 | 0.96368978 | 1.60764668 | 0.92297159 |
| ENSMUSG00 Rpl29-ps1 | 506 | 0 | 0 | 0 | 0 | 0 | 0 | 0 | 0 | 0 | 0 | 0 | 0 | 0 | 0 | 0 | 0 | 0 | 0 |
| ENSMUSG00 Gm13757 | 286 | 0 | 0 | 0 | 0 | 0 | 0 | 0 | 0 | 0 | 0 | 0 | 0 | 0 | 0 | 0 | 0 | 0 | 0 |
| ENSMUSG00 Alms1-ps2 | 2411 | 0 | 0 | 0 | 0.03779431 | 0 | 0 | 0 | 0 | 0 | 0 | 0 | 0 | 0 | 0 | 0 | 0 | 0 | 0 |
| ENSMUSG00 Gm8837 | 1884 | 0.02256113 | 0 | 0 | 0 | 0 | 0 | 0 | 0 | 0 | 0 | 0 | 0 | 0 | 0 | 0 | 0 | 0 | 0 |
| ENSMUSG00 Ttbk2 | 11388 | 0.91818347 | 1.2549378 | 1.07893192 | 0.80015884 | 1.22569879 | 1.19661062 | 1.53305045 | 0.78477539 | 0.98734798 | 0.86120006 | 0.75933153 | 1.1272921 | 0.70610314 | 1.14193401 | 0.82851741 | 0.64557399 | 0.77069916 | 0.83566697 |
| ENSMUSG00 Snhg9 | 181 | 16.9081363 | 15.0920294 | 21.9150801 | 18.8788858 | 21.2737784 | 14.2361445 | 18.0363251 | 15.1925579 | 36.1804199 | 17.4535752 | 26.3950047 | 19.7359249 | 29.4356191 | 24.8899253 | 28.6817461 | 29.4054941 | 21.5982447 | 27.3525945 |
| ENSMUSG00 Gm4985 | 1232 | 0 | 0.02703969 | 0 | 0 | 0 | 0 | 0 | 0.03100033 | 0 | 0 | 0 | 0 | 0 | 0 | 0 | 0 | 0 | 0 |
| ENSMUSG00 Gm16094 | 425 | 0 | 0 | 0 | 0 | 0 | 0.11659467 | 0 | 0 | 0 | 0 | 0 | 0 | 0 | 0 | 0 | 0 | 0 | 0 |
| ENSMUSG00 Gm16241 | 2192 | 0 | 0.04559246 | 0 | 0 | 0.04391599 | 0.02260617 | 0 | 0 | 0 | 0 | 0 | 0 | 0 | 0 | 0 | 0.05240503 | 0 | 0 |
| ENSMUSG00 Gm15890 | 714 | 1.42874541 | 2.5661193 | 2.30350145 | 1.53146368 | 1.55046813 | 2.91486668 | 2.18672045 | 1.44425051 | 2.13431524 | 1.78300979 | 2.0742629 | 1.09442476 | 1.3127541 | 1.75628902 | 1.26951573 | 1.12619434 | 1.34195717 | 2.15721987 |
| ENSMUSG00 Gm15838 | 179 | 0 | 0 | 0 | 0 | 0 | 0 | 0 | 0 | 0 | 0.26341175 | 0 | 0 | 0 | 0 | 0 | 0.64174202 | 0 | 0 |
| ENSMUSG00 Gm11492 | 1970 | 0 | 0 | 0 | 0 | 0 | 0 | 0 | 0 | 0 | 0 | 0 | 0 | 0 | 0 | 0 | 0 | 0 | 0 |
| ENSMUSG00 Gm15717 | 344 | 1.23561558 | 0.19367962 | 0.28124146 | 0.1324449 | 0.27983677 | 0.14404864 | 0.68768429 | 0 | 0.11972824 | 0.41119799 | 0 | 0.16225483 | 1.14725389 | 0 | 1.07794768 | 0.77916934 | 0.22282731 | 0.47973058 |
| ENSMUSG00 Ear-ps10 | 459 | 0 | 0 | 0 | 0 | 0 | 0 | 0 | 0 | 0 | 0 | 0 | 0 | 0 | 0 | 0 | 0 | 0 | 0 |
| ENSMUSG00 Mtcp1 | 1455 | 7.27408167 | 8.37973834 | 8.34484957 | 6.5445074 | 9.36174185 | 8.17364678 | 5.88563222 | 9.02968139 | 11.1812191 | 7.45337577 | 7.48638525 | 8.36275907 | 7.32350112 | 8.33120743 | 7.27752568 | 8.21076942 | 6.61161548 | 9.9054199 |
| ENSMUSG00 Gm16584 | 188 | 0 | 0 | 0 | 0 | 0 | 0 | 0 | 0 | 0 | 0 | 0 | 0 | 0 | 0 | 0 | 0 | 0 | 0 |
| ENSMUSG00 Shprh | 14133 | 1.61503428 | 1.6287561 | 1.80378196 | 1.71180319 | 1.6210851 | 1.96696268 | 1.7106615 | 1.74301965 | 1.59698653 | 1.9249951 | 2.00794773 | 1.65871211 | 1.06810774 | 2.0013054 | 1.83953738 | 1.47386191 | 1.41015173 | 2.30810163 |
| ENSMUSG00 Nhlrc4 | 2478 | 0 | 0.3360865 | 0.66372076 | 0.75383487 | 0.19423698 | 0.17997361 | 0 | 0.12330073 | 0.04986261 | 0.17124953 | 0 | 0 | 0.37825118 | 0.20616827 | 0.18289633 | 0.91168113 | 0.4330655 | 0.02219899 |
| ENSMUSG00 Gm3076 | 450 | 0.18891189 | 0 | 0.10749673 | 0.10124677 | 0.32087949 | 0.22023437 | 0.21027857 | 0 | 0.27457677 | 0.20955868 | 0.4246663 | 0 | 0.32887945 | 0.41283633 | 0 | 0 | 0.08516955 | 0.24448492 |
| ENSMUSG00 Usp49 | 8574 | 0.1288937 | 0.24477634 | 0.12412149 | 0.18067127 | 0.23016199 | 0.09247069 | 0.17658103 | 0.3340833 | 0.12489496 | 0.12648311 | 0.34546856 | 0.36455296 | 0.02876832 | 0.04333481 | 0.12494057 | 0.18310181 | 0.12963175 | 0.11548448 |
| ENSMUSG00 Gm15680 | 463 | 0 | 0 | 0 | 0 | 0 | 0 | 0 | 0 | 0 | 0 | 0 | 0 | 0 | 0 | 0 | 0 | 0 | 0 |
| ENSMUSG00 Gm16541 | 857 | 0.09919528 | 0 | 0 | 0 | 0.11232654 | 0 | 0 | 0 | 0 | 0.16505497 | 0.11149349 | 0 | 0 | 0.1083876 | 0 | 0 | 0 | 0.06418799 |
| ENSMUSG00 Gm16163 | 1117 | 0.26637084 | 0 | 0.30314656 | 0.16315504 | 0.0861807 | 0.08872468 | 0.2117846 | 0.10257583 | 0 | 0.08442382 | 0 | 0.04996926 | 0.04416465 | 0 | 0.07377181 | 0.13711945 | 0 | 0.04924719 |
| ENSMUSG00 Gm8185 | 668 | 0 | 0 | 0.07241546 | 0 | 0.36026889 | 0.37090369 | 0 | 0.11434851 | 0.18496938 | 0 | 0 | 0 | 0.2954007 | 0 | 0.12335795 | 0.28660634 | 0.28687348 | 0.24704689 |
| ENSMUSG00 A730090N16I | 623 | 0.06822661 | 0 | 0.23293835 | 0.36565846 | 0.23177491 | 0.31815559 | 0 | 0.79695222 | 0.06610998 | 0.45409987 | 0.15337065 | 0.71673403 | 0.55429121 | 0.96913826 | 0.33067059 | 0.61461649 | 0.55367042 | 0.70637697 |
| ENSMUSG00 Abhd12b | 3390 | 0.4764592 | 0.61908918 | 1.0987498 | 0.34943574 | 0.14198208 | 0.23387721 | 0.11165234 | 0.13519434 | 0 | 0.13908762 | 0.05637163 | 0.13171838 | 0.66940065 | 0.41100962 | 0.40107709 | 0.30496944 | 0 | 0 |
| ENSMUSG00 Kcne1l | 2693 | 0.09470147 | 0.11133162 | 0.10777615 | 0.20301988 | 0.08936488 | 0.09200285 | 0 | 0.17018523 | 0 | 0.03501723 | 0 | 0 | 0.09159287 | 0.43115566 | 0.16829451 | 0.02843714 | 0.28463644 | 0.36768062 |
| ENSMUSG00 Ugt1a7c | 2792 | 0 | 0 | 0 | 0 | 0 | 0 | 0 | 0.04103768 | 0 | 0 | 0 | 0 | 0.0706761 | 0 | 0 | 0.15085841 | 0.02745437 | 0.15761922 |
| ENSMUSG00 Pou3f1 | 3006 | 0 | 0 | 0 | 0 | 0 | 0 | 0 | 0 | 0 | 0 | 0 | 0 | 0 | 0 | 0 | 0 | 0 | 0 |
| ENSMUSG00 4930519F09I | 1393 | 0 | 0 | 0 | 0.03270714 | 0.03455271 | 0 | 0.03396459 | 0 | 0.05913355 | 0 | 0 | 0 | 0 | 0.03334105 | 0.02957757 | 0 | 0 | 0 |
| ENSMUSG00 Gm11567 | 556 | 0 | 0 | 0 | 0 | 0 | 0 | 0 | 0 | 0 | 0 | 0 | 0 | 0 | 0 | 0 | 0 | 0 | 0 |
| ENSMUSG00 Gm16038 | 191 | 0 | 0 | 0 | 0 | 0 | 0 | 0 | 0 | 0 | 0 | 0 | 0 | 0 | 0 | 0 | 0 | 0 | 0 |
| ENSMUSG00 Olfr287 | 3193 | 0 | 0 | 0 | 0 | 0 | 0 | 0 | 0 | 0 | 0 | 0 | 0 | 0 | 0 | 0 | 0 | 0 | 0 |
| ENSMUSG00 Gm16162 | 294 | 0 | 0 | 0 | 0 | 0 | 0 | 0 | 0 | 0 | 0 | 0 | 0 | 0 | 0 | 0 | 0 | 0 | 0.18710581 |
| ENSMUSG00 Cypt2-ps | 695 | 0 | 0 | 0 | 0 | 0 | 0 | 0 | 0 | 0 | 0 | 0 | 0 | 0 | 0 | 0 | 0 | 0 | 0 |
| ENSMUSG00 Gm15809 | 659 | 0.25799803 | 0 | 0 | 0 | 0 | 0 | 0 | 0 | 0 | 0 | 0 | 0 | 0 | 0 | 0 | 0 | 0 | 0 |
| ENSMUSG00 Gm10177 | 664 | 1.21626257 | 1.15391049 | 1.31133064 | 0.68616031 | 2.24712294 | 1.41792461 | 2.13762104 | 1.26541092 | 1.36461347 | 1.84626248 | 1.22315406 | 1.68119466 | 2.89750719 | 2.16832336 | 1.92356662 | 2.01833021 | 2.42425372 | 1.90543593 |
| ENSMUSG00 Uba52 | 1758 | 2.68377391 | 1.76228621 | 1.92613602 | 2.38430949 | 2.10816731 | 1.94490251 | 3.4179239 | 2.49836535 | 2.694226 | 2.22611398 | 2.44581699 | 1.80972285 | 4.29339212 | 5.01955438 | 2.8358295 | 3.28889745 | 3.44457048 | 4.22424881 |
| ENSMUSG00 Gm3687 | 320 | 0 | 0 | 0 | 0 | 0 | 0 | 0 | 0 | 0 | 0 | 0 | 0 | 0 | 0 | 0 | 0 | 0 | 0 |
| ENSMUSG00 Gm614 | 768 | 0 | 0 | 0 | 0 | 0 | 0 | 0 | 0 | 0 | 0 | 0 | 0 | 0 | 0 | 0 | 0 | 0 | 0 |
| ENSMUSG00 Gm15795 | 302 | 0 | 0 | 0 | 0 | 0 | 0 | 0 | 0 | 0 | 0 | 0 | 0 | 0 | 0 | 0 | 0 | 0 | 0 |
| ENSMUSG00 Ugt1a6b | 2376 | 0 | 0 | 0 | 0 | 0 | 0 | 0 | 0 | 0 | 0 | 0 | 0 | 0 | 0 | 0 | 0 | 0 | 0 |
| ENSMUSG00 4930405H06 | 789 | 0 | 0 | 0 | 0 | 0 | 0 | 0 | 0 | 0 | 0 | 0 | 0 | 0 | 0 | 0 | 0 | 0 | 0 |
| ENSMUSG00 Gm3329 | 555 | 0 | 0 | 0 | 0 | 0 | 0 | 0 | 0 | 0.29683975 | 0 | 0 | 0 | 0 | 0 | 0 | 0 | 0 | 0 |
| ENSMUSG00 Acad11 | 4054 | 21.546161 | 30.4614945 | 25.5947762 | 26.7926812 | 30.5959465 | 23.3218095 | 20.1434839 | 23.2037215 | 18.866147 | 23.1915398 | 26.5625941 | 23.2129273 | 23.8993304 | 19.4529009 | 25.4587808 | 23.7450881 | 30.5551533 | 26.9346514 |
| ENSMUSG00 Gm16043 | 614 | 0 | 0 | 0 | 0 | 0 | 0 | 0 | 0 | 0 | 0 | 0 | 0 | 0 | 0 | 0 | 0 | 0 | 0 |
| ENSMUSG00 Gm16285 | 346 | 0 | 0 | 0 | 0 | 0 | 0 | 0 | 0 | 0 | 0 | 0 | 0 | 0 | 0 | 0 | 0 | 0 | 0 |
| ENSMUSG00 Gm16534 | 2045 | 0 | 0 | 0 | 0 | 0 | 0 | 0 | 0 | 0.02014011 | 0 | 0 | 0 | 0 | 0 | 0 | 0 | 0 | 0 |
| ENSMUSG00 Gm8580 | 463 | 0 | 0 | 0 | 0.49201992 | 0 | 0 | 0.20437442 | 0 | 0 | 0 | 0 | 0 | 0 | 0 | 0 | 0 | 0 | 0 |
| ENSMUSG00 4930480K15I | 2196 | 0 | 0 | 0 | 0 | 0 | 0 | 0 | 0 | 0 | 0 | 0 | 0 | 0 | 0 | 0 | 0 | 0 | 0 |
| ENSMUSG00 Gm6808 | 806 | 0 | 0 | 0 | 0 | 0 | 0 | 0 | 0 | 0 | 0 | 0 | 0 | 0 | 0 | 0 | 0 | 0 | 0 |
| ENSMUSG00 Gm16231 | 317 | 0 | 0 | 0 | 0 | 0 | 0 | 0 | 0 | 0 | 0 | 0 | 0 | 0 | 0 | 0 | 0 | 0 | 0 |
| ENSMUSG00 BC035044 | 2611 | 0 | 0 | 0 | 0 | 0 | 0 | 0.12684364 | 0.1609025 | 0.22083923 | 0.2347603 | 0.23786843 | 0 | 0 | 0.17787854 | 0 | 0 | 0.22018171 | 0 |
| ENSMUSG00 Ugt1a10 | 1917 | 0 | 0 | 0 | 0 | 0 | 0 | 0 | 0 | 0 | 0 | 0 | 0 | 0.02573392 | 0.07268245 | 0 | 0 | 0 | 0 |
| ENSMUSG00 Ear10 | 1146 | 0 | 0 | 0 | 0 | 0 | 0 | 0 | 0 | 0 | 0 | 0 | 0 | 0 | 0 | 0 | 0 | 0 | 0 |
| ENSMUSG00 D630014O11 | 3017 | 0.01406856 | 0 | 0 | 0.01510144 | 0.03190714 | 0 | 0.12545623 | 0.13924973 | 0.16381776 | 0.26568179 | 0.06334101 | 0.16650347 | 0 | 0.06157652 | 0 | 0 | 0 | 0.05469914 |
| ENSMUSG00 Ugt1a2 | 2206 | 0 | 0 | 0 | 0 | 0 | 0 | 0 | 0 | 0.09335112 | 0 | 0 | 0 | 0 | 0 | 0 | 0 | 0 | 0 |
| ENSMUSG00 Fbxw10 | 4683 | 0.07261187 | 0.00711358 | 0 | 0.00972903 | 0 | 0 | 0.08082456 | 0 | 0.0263847 | 0 | 0 | 0 | 0.05267128 | 0 | 0.01759622 | 0.08994164 | 0 | 0.04698621 |
| ENSMUSG00 Gm10612 | 1678 | 0 | 0 | 0 | 0 | 0 | 0 | 0 | 0 | 0 | 0 | 0 | 0 | 0 | 0 | 0 | 0 | 0 | 0 |
| ENSMUSG00 Ugt1a9 | 2185 | 0 | 0 | 0 | 0 | 0 | 0 | 0 | 0 | 0 | 0 | 0 | 0 | 0 | 0 | 0 | 0 | 0 | 0 |
| ENSMUSG00 Cd200r2 | 1744 | 0 | 0 | 0 | 0 | 0 | 0 | 0 | 0 | 0 | 0 | 0 | 0 | 0 | 0 | 0 | 0 | 0 | 0 |
| ENSMUSG00 Gm16307 | 209 | 0 | 0 | 0 | 0 | 0 | 0 | 0 | 0 | 0.98532334 | 0 | 0 | 0 | 0 | 0 | 0 | 0 | 0 | 0 |
| ENSMUSG00 Ang-ps1 | 435 | 0 | 0 | 0 | 0 | 0 | 0 | 0 | 0 | 0 | 0 | 0 | 0 | 0 | 0 | 0 | 0 | 0 | 0 |
| ENSMUSG00 Gm15523 | 435 | 0 | 0 | 0 | 0 | 0 | 0 | 0 | 0 | 0 | 0 | 0 | 0 | 0 | 0 | 0 | 0 | 0 | 0 |
| ENSMUSG00 Gm7592 | 1761 | 0.04827391 | 0 | 0 | 0.02587226 | 0.02733215 | 0 | 0 | 0 | 0.07016442 | 0 | 0.02712945 | 0 | 0 | 0.1054948 | 0 | 0 | 0 | 0 |
| ENSMUSG00 AK157302 | 389 | 0 | 0 | 0 | 0 | 0 | 0 | 0 | 0 | 0 | 0 | 0 | 0 | 0 | 0 | 0 | 0 | 0 | 0 |
| ENSMUSG00 Gm7418 | 800 | 0 | 0.04164112 | 0.06046691 | 0 | 0 | 0 | 0 | 0 | 0 | 0 | 0.0597187 | 0 | 0.18499469 | 0.23222044 | 0 | 0 | 0.04790787 | 0 |
| ENSMUSG00 Gm2666 | 890 | 0.04775862 | 0 | 0 | 0 | 0 | 0 | 0 | 0 | 0 | 0 | 0 | 0 | 0 | 0 | 0 | 0 | 0 | 0 |
| ENSMUSG00 9230105E05I | 2247 | 0.52965951 | 0.96365738 | 0.0430561 | 0.50690971 | 0.25704632 | 0.46310966 | 0.02105593 | 0 | 0 | 0 | 0 | 0 | 0 | 0 | 0 | 0 | 0 | 0 |
| ENSMUSG00 Gm16556 | 798 | 0 | 0 | 0 | 0 | 0 | 0 | 0 | 0 | 0 | 0 | 0 | 0 | 0 | 0.46560489 | 0 | 0 | 0 | 0 |
| ENSMUSG00 Gm15995 | 572 | 0 | 0 | 0 | 0 | 0 | 0 | 0 | 0 | 0 | 0 | 0 | 0 | 0 | 0 | 0 | 0 | 0 | 0 |
| ENSMUSG00 Gm16161 | 288 | 0.73793708 | 0 | 1.17574553 | 0 | 0.16712473 | 0 | 0 | 0.39783753 | 0 | 0.49115316 | 0 | 0 | 0.17129138 | 0 | 0 | 0.66476749 | 0 | 0 |
| ENSMUSG00 Dnaja1-ps | 641 | 0 | 0.15591058 | 0 | 0 | 0 | 0 | 0 | 0 | 0.12850707 | 0 | 0 | 0 | 0 | 0 | 0 | 0 | 0 | 0 |
| ENSMUSG00 Gm16029 | 2425 | 0 | 0 | 0 | 0 | 0 | 0 | 0 | 0 | 0 | 0 | 0 | 0 | 0 | 0 | 0 | 0 | 0 | 0 |
| ENSMUSG00 1700025N21 | 684 | 0 | 0 | 0 | 0 | 0 | 0 | 0 | 0 | 0 | 0 | 0 | 0 | 0.14424537 | 0 | 0 | 0 | 0 | 0 |
| ENSMUSG00 4930503B20I | 2571 | 0 | 0 | 0 | 0 | 0 | 0 | 0 | 0.04456523 | 0 | 0 | 0 | 0 | 0 | 0 | 0 | 0 | 0 | 0 |
| ENSMUSG00 AU015336 | 846 | 0 | 0 | 0 | 0 | 0 | 0 | 0 | 0 | 0 | 0 | 0 | 0 | 0 | 0 | 0 | 0 | 0 | 0 |
| ENSMUSG00 Tepp | 1568 | 0.759021 | 0.14871828 | 0.27765419 | 0.29056768 | 0.55253484 | 0.31602509 | 0.33191293 | 0.36536099 | 0.36773675 | 0.63148263 | 0.18281234 | 0.21358034 | 0.15730841 | 0.20733968 | 0.13138251 | 0.09768012 | 0.17109954 | 0.28065871 |
| ENSMUSG00 4930524O07I | 1339 | 0.28569573 | 0.62197338 | 0.50577254 | 0.74857578 | 0.2516232 | 0.33306542 | 1.16603316 | 1.39763086 | 0.36910992 | 0.63384067 | 0.82063035 | 0.91706092 | 1.36316724 | 2.25456735 | 0.70771903 | 1.25824251 | 0.62970764 | 0.82164461 |
| ENSMUSG00 Gm15851 | 659 | 0 | 0 | 0 | 0 | 0 | 0 | 0 | 0.11591017 | 0 | 0 | 0 | 0 | 0 | 0 | 0.06252133 | 0 | 0 | 0 |
| ENSMUSG00 Itga10 | 5157 | 0.02472669 | 0.09689614 | 0 | 0.0088348 | 0.01866864 | 0.01921766 | 0.03669783 | 0.05184154 | 0.1836901 | 0.0365722 | 0.09264099 | 0.05411641 | 0.03826404 | 0 | 0.06391554 | 0.00742498 | 0.05202328 | 0 |
| ENSMUSG00 Gm16050 | 2361 | 0 | 0.04232896 | 0.02048858 | 0.0771894 | 0 | 0 | 0 | 0 | 0 | 0.0399413 | 0.0404701 | 0 | 0.0208945 | 0 | 0 | 0 | 0 | 0 |
| ENSMUSG00 Gm15647 | 689 | 0 | 0 | 0 | 0 | 0 | 0 | 0 | 0 | 0 | 0 | 0 | 0 | 0 | 0 | 0 | 0 | 0 | 0 |
| ENSMUSG00 Tmem189 | 4212 | 5.75212496 | 6.70686949 | 5.9375867 | 6.71733346 | 5.49654684 | 7.28232243 | 5.84107147 | 6.77343889 | 7.71513792 | 7.44425871 | 4.94536607 | 6.50652667 | 8.01116605 | 11.2361171 | 9.11677353 | 7.79084771 | 7.20665412 | 6.93490875 |
| ENSMUSG00 Gm15657 | 1004 | 0 | 0 | 0 | 0 | 0 | 0 | 0 | 0 | 0 | 0 | 0 | 0 | 0 | 0 | 0.04103741 | 0 | 0 | 0 |
| ENSMUSG00 Trim34b | 2037 | 0.1460659 | 0.163539 | 0.28496925 | 0.13420043 | 0.23628829 | 0.09730532 | 0.37162634 | 0.35623743 | 0.04043841 | 0.27776556 | 0.51597893 | 0.05480183 | 0.41170476 | 0.04560048 | 0.26294562 | 0.63911667 | 0.84667813 | 0.45908435 |
| ENSMUSG00 Gm16549 | 501 | 0.33936268 | 0.0664928 | 0 | 0 | 0 | 0 | 0.84992836 | 0.68609106 | 0.73987753 | 0.09411318 | 0.76287359 | 0.33422552 | 0.1969338 | 0.55621662 | 0.82238635 | 0.07642836 | 0 | 0.10979862 |
| ENSMUSG00 Gm16116 | 428 | 0.09931116 | 0 | 0 | 0.10645104 | 0.11245777 | 0 | 0 | 0.08923459 | 0 | 0 | 0 | 0 | 0 | 0.10851422 | 0 | 0 | 0 | 0 |
| ENSMUSG00 4930516K23I | 1070 | 0 | 0 | 0 | 0 | 0 | 0 | 0 | 0 | 0 | 0 | 0 | 0 | 0 | 0 | 0 | 0 | 0 | 0 |
| ENSMUSG00 4930568K20I | 1612 | 0 | 0 | 0 | 0 | 0 | 0 | 0 | 0 | 0.07664984 | 0 | 0 | 0 | 0 | 0 | 0.02555928 | 0 | 0 | 0.13649902 |
| ENSMUSG00 Gm4245 | 1753 | 0 | 0 | 0 | 0 | 0 | 0 | 0 | 0 | 0 | 0 | 0 | 0 | 0 | 0 | 0 | 0 | 0 | 0 |
| ENSMUSG00 Gm16340 | 2157 | 0.25617399 | 0.09266452 | 0.0224263 | 0.06336724 | 0 | 0.13783792 | 0.3728862 | 0.46036275 | 0.17184916 | 0.19673451 | 0.33223198 | 0.31051829 | 0.297318 | 0.21531798 | 0.38202648 | 0.3017804 | 0.21322001 | 0.1785182 |
| ENSMUSG00 Pcp4 | 638 | 1.06596053 | 0.88764765 | 1.44059103 | 1.14259672 | 1.81060528 | 1.0096952 | 2.52136533 | 3.59176198 | 3.3568947 | 2.95615694 | 2.9952952 | 3.23695849 | 2.93826467 | 1.60152025 | 4.26222992 | 4.68129683 | 4.62558756 | 2.75907745 |
| ENSMUSG00 Gm15974 | 530 | 0 | 0.06285452 | 0 | 0.08596423 | 0.45407475 | 0 | 0.08926921 | 0 | 0.23313122 | 0 | 0 | 0 | 0.18615818 | 0 | 0.15547757 | 0.14449286 | 0.14462754 | 0 |
| ENSMUSG00 Gm11559 | 1095 | 0 | 0 | 0 | 0 | 0 | 0 | 0 | 0 | 0 | 0 | 0 | 0 | 0 | 0 | 0 | 0 | 0 | 0 |
| ENSMUSG00 Gm16110 | 890 | 0 | 0 | 0 | 0 | 0 | 0 | 0 | 0 | 0 | 0 | 0 | 0 | 0 | 0 | 0 | 0 | 0 | 0 |
| ENSMUSG00 Gm16554 | 458 | 0 | 0 | 0 | 0 | 0 | 0 | 0 | 0 | 0 | 0 | 0 | 0 | 0 | 0 | 0 | 0 | 0 | 0 |
| ENSMUSG00 Gm16289 | 182 | 0 | 0 | 0 | 0 | 0 | 0 | 0 | 0 | 0 | 0 | 0 | 0 | 0 | 0 | 0 | 0 | 0 | 0 |
| ENSMUSG00 Gm16315 | 781 | 0.05442404 | 0.63981231 | 0.06193794 | 0.23334722 | 0.43140008 | 0.25379121 | 0.06057961 | 0.24450962 | 0.63282739 | 0.3018611 | 0.18351457 | 0.21440075 | 0.12633013 | 0.71360954 | 0.5803036 | 0 | 0.14722009 | 1.19738132 |
| ENSMUSG00 Cfb | 3011 | 0.04234989 | 0 | 0 | 0 | 0 | 0 | 0 | 0 | 0 | 0.01565948 | 0 | 0 | 0 | 0 | 0.06841839 | 0.03815072 | 0 | 0 |
| ENSMUSG00 Gm16132 | 596 | 0 | 0.27947059 | 0 | 0.53511294 | 0.32303304 | 0.08314217 | 0 | 0.19224364 | 0.20731468 | 0 | 0.56111528 | 0 | 0 | 0.31170528 | 0.62217115 | 0.25698394 | 0.25722347 | 0 |
| ENSMUSG00 Gm16042 | 1246 | 0 | 0 | 0 | 0 | 0 | 0 | 0 | 0 | 0 | 0 | 0 | 0 | 0 | 0 | 0 | 0 | 0 | 0 |
| ENSMUSG00 Gm16244 | 2474 | 0.32643425 | 0.39049068 | 0.74300491 | 0.18415944 | 0.62256328 | 0.36052919 | 0.57371882 | 0.84906311 | 0.41619357 | 0.59081318 | 0.88829753 | 0.45121797 | 0.79760578 | 0.56318618 | 0.53292231 | 0.24763529 | 1.02244769 | 0.48916748 |
| ENSMUSG00 Car15 | 1536 | 0 | 0 | 0 | 0 | 0 | 0 | 0 | 0 | 0.05362828 | 0 | 0 | 0 | 0 | 0.06047407 | 0 | 0 | 0.02495202 | 0 |
| ENSMUSG00 Gm16308 | 936 | 0 | 0 | 0 | 0 | 0 | 0 | 0 | 0 | 0 | 0 | 0 | 0 | 0 | 0 | 0 | 0.16363507 | 0 | 0 |
| ENSMUSG00 Gm16207 | 728 | 0 | 0 | 0 | 0 | 0 | 0 | 0 | 0 | 0.22629954 | 0 | 0 | 0 | 0 | 0 | 0 | 0 | 0 | 0 |
| ENSMUSG00 Gm16103 | 1278 | 0.06651827 | 0.05213285 | 0.3028077 | 0.03565027 | 0.3012953 | 0 | 0.33318788 | 0.50803665 | 0.09668196 | 0.03689413 | 0 | 0.17469691 | 0.07720175 | 0 | 0 | 0.26965216 | 0.29989278 | 0.04304312 |
| ENSMUSG00 Gm16572 | 277 | 0 | 0.12026316 | 0 | 0 | 0 | 0 | 0 | 0.13787871 | 0 | 0 | 0.86236387 | 0 | 0 | 0 | 0 | 0.6911662 | 0.27672417 | 0 |
| ENSMUSG00 Gm17017 | 2403 | 0 | 0 | 0.10065237 | 0 | 0.14020952 | 0.0206212 | 0 | 0 | 0 | 0 | 0.03976276 | 0 | 0.06158791 | 0.03865509 | 0 | 0.14341051 | 0.07974677 | 0.02289185 |
| ENSMUSG00 Bloc1s1 | 1281 | 2.8867684 | 2.02841979 | 2.75664927 | 2.45410777 | 2.17927523 | 1.97282546 | 1.84670878 | 1.04350826 | 1.73619973 | 2.06123293 | 1.71557228 | 2.26574118 | 1.27084564 | 2.28413545 | 2.0906332 | 1.58423278 | 1.01724754 | 1.67475032 |
| ENSMUSG00 Gm14027 | 617 | 0.20667022 | 0.8638676 | 1.25441894 | 0.44305716 | 2.26227842 | 1.4456227 | 1.30359083 | 1.29990349 | 0.8677872 | 0.84061221 | 1.47118997 | 0.45231493 | 1.19931728 | 1.65602905 | 0.26710895 | 0.55853398 | 0.86964045 | 0.71324612 |
| ENSMUSG00 Gm3287 | 1323 | 0 | 0 | 0 | 0 | 0 | 0 | 0 | 0 | 0 | 0 | 0 | 0 | 0 | 0 | 0 | 0 | 0 | 0 |
| ENSMUSG00 Gm16028 | 888 | 0 | 0 | 0 | 0 | 0 | 0 | 0 | 0 | 0 | 0 | 0 | 0 | 0 | 0 | 0 | 0 | 0 | 0 |
| ENSMUSG00 Krtap1-4 | 520 | 0 | 0 | 0 | 0 | 0 | 0 | 0 | 0 | 0 | 0 | 0 | 0 | 0 | 0 | 0 | 0 | 0 | 0 |
| ENSMUSG00 Gm1965 | 2434 | 0 | 0 | 0 | 0 | 0 | 0 | 0 | 0 | 0 | 0 | 0 | 0 | 0 | 0 | 0 | 0 | 0 | 0 |
| ENSMUSG00 4921534H16 | 3949 | 0 | 0 | 0 | 0.05768681 | 0 | 0 | 0 | 0 | 0 | 0 | 0 | 0 | 0 | 0 | 0.19823489 | 0 | 0 | 0 |
| ENSMUSG00 Alms1-ps1 | 2157 | 0 | 0 | 0 | 0 | 0 | 0 | 0 | 0 | 0 | 0 | 0 | 0 | 0 | 0 | 0 | 0 | 0 | 0.0255026 |
| ENSMUSG00 Gm4524 | 2150 | 0 | 0 | 0 | 0 | 0.02238694 | 0 | 0 | 0 | 0 | 0.04386112 | 0 | 0.15576464 | 0 | 0 | 0 | 0.01780958 | 0 | 0 |
| ENSMUSG00 Churc1 | 5772 | 6.92218736 | 6.30243945 | 6.9224768 | 6.90677647 | 6.93793494 | 6.58471009 | 8.37726228 | 7.44394538 | 6.52904746 | 8.38125806 | 8.20252665 | 6.25653737 | 8.69205819 | 9.10857708 | 8.48729209 | 9.00214037 | 7.97468513 | 8.21515073 |
| ENSMUSG00 Gm16545 | 462 | 0 | 0 | 0 | 0 | 0 | 0 | 0 | 0 | 0 | 0 | 0 | 0 | 0 | 0 | 0 | 0 | 0 | 0 |
| ENSMUSG00 2810442N19I | 1087 | 0 | 0 | 0 | 0 | 0 | 0.04558669 | 0.13057777 | 0 | 0 | 0 | 0 | 0.20539342 | 0 | 0 | 0.03790392 | 0 | 0 | 0 |
| ENSMUSG00 Vmn2r-ps10 | 743 | 0 | 0 | 0 | 0 | 0 | 0 | 0 | 0 | 0 | 0 | 0 | 0 | 0 | 0 | 0 | 0 | 0 | 0 |
| ENSMUSG00 Mpv17 | 4399 | 32.0600532 | 27.7922987 | 29.074702 | 35.2867647 | 30.1877648 | 30.7860016 | 30.8892962 | 29.3887888 | 30.1853437 | 27.9645794 | 30.7132491 | 29.5255846 | 29.1348764 | 26.8064419 | 26.9650556 | 32.8068423 | 29.7793323 | 27.5983402 |
| ENSMUSG00 D730045A05I | 1266 | 0 | 0.10525401 | 0 | 0.03598819 | 0 | 0 | 0 | 0 | 0.03253279 | 0 | 0 | 0 | 0 | 0 | 0 | 0.09073604 | 0.27246183 | 0 |
| ENSMUSG00 Eif4ebp3 | 741 | 0.68834293 | 0.08991334 | 0.13056283 | 0.18445767 | 0.19486669 | 0.26749114 | 0.19154931 | 0 | 0.83373514 | 0.38178707 | 0.19342088 | 0.60259825 | 0.26629915 | 0.06267758 | 0.22241056 | 0.51674234 | 0.1034448 | 0 |
| ENSMUSG00 Gm9550 | 861 | 0 | 0 | 0 | 0 | 0 | 0 | 0 | 0 | 0 | 0 | 0 | 0 | 0 | 0 | 0 | 0 | 0 | 0 |
| ENSMUSG00 1110005A03I | 2163 | 11.7316644 | 8.5784938 | 10.6900358 | 8.97318765 | 10.5698861 | 9.82807337 | 11.0680608 | 11.0710293 | 9.95864436 | 9.13367761 | 12.3689213 | 9.65097448 | 10.4456857 | 10.6286746 | 9.75274924 | 9.71869786 | 12.3147371 | 9.18090038 |
| ENSMUSG00 Gm5286 | 1591 | 0 | 0 | 0 | 0 | 0 | 0 | 0 | 0 | 0 | 0 | 0 | 0 | 0 | 0 | 0 | 0 | 0 | 0 |
| ENSMUSG00 Gm17226 | 1165 | 0 | 0 | 0 | 0 | 0 | 0 | 0 | 0 | 0 | 0 | 0 | 0 | 0 | 0 | 0 | 0 | 0 | 0 |
| ENSMUSG00 Mndal | 1897 | 0 | 0 | 0 | 0 | 0 | 0 | 0.19952632 | 0.04026611 | 0.54278487 | 0 | 0.05036896 | 0 | 0.02600523 | 0 | 0.54298308 | 0.42388126 | 0 | 0 |
| ENSMUSG00 Prr22 | 1958 | 0.0651254 | 0.10208241 | 0.22235024 | 0.23269175 | 0.36873282 | 0.37961747 | 0 | 0 | 0.18931493 | 0.38529686 | 0 | 0.14253234 | 0.20156044 | 0.52184368 | 0.18938407 | 0.39111959 | 0.2740389 | 0.61807985 |
| ENSMUSG00 Gm17671 | 52 | 0 | 0 | 0 | 0.87617393 | 3.70245567 | 0 | 0 | 3.67234639 | 0 | 0 | 0 | 0 | 0 | 0 | 0 | 0 | 0 | 0 |
| ENSMUSG00 Cbln1 | 1815 | 0 | 0 | 0 | 0 | 0 | 0 | 0 | 0 | 0 | 0 | 0 | 0 | 0 | 0 | 0 | 0 | 0 | 0 |
| ENSMUSG00 Gm8338 | 362 | 0 | 0 | 0 | 0 | 0 | 0 | 0 | 0 | 0 | 0 | 0 | 0 | 0 | 0 | 0 | 0 | 0 | 0 |
| ENSMUSG00 D130062J21I | 2742 | 0.32553198 | 0.63175438 | 0.61745936 | 0.78095153 | 0.47394673 | 0.41565021 | 0.34509613 | 0.57107531 | 1.06646339 | 0.67063363 | 0.76662953 | 0.48853972 | 0.44978043 | 0.8638397 | 0.49586118 | 0.55857925 | 0.76876234 | 1.44444044 |
| ENSMUSG00 Gm9603 | 2058 | 0 | 0 | 0 | 0 | 0 | 0 | 0 | 0 | 0 | 0 | 0 | 0 | 0 | 0 | 0 | 0 | 0 | 0 |
| ENSMUSG00 Vmn2r20 | 2556 | 0 | 0 | 0 | 0 | 0 | 0 | 0 | 0 | 0 | 0 | 0 | 0 | 0 | 0 | 0 | 0 | 0 | 0 |
| ENSMUSG00 Gm17423 | 2166 | 0 | 0 | 0 | 0 | 0 | 0 | 0 | 0 | 0 | 0 | 0 | 0 | 0 | 0 | 0 | 0 | 0 | 0 |
| ENSMUSG00 Gm17688 | 1595 | 0.05329803 | 0 | 0 | 0 | 0 | 0 | 0 | 0.02394508 | 0 | 0.02956157 | 0.08985886 | 0 | 0 | 0.1164742 | 0 | 0 | 0.07208708 | 0 |
| ENSMUSG00 Gm17613 | 409 | 0 | 0 | 0 | 0 | 0 | 0 | 0 | 0 | 0 | 0 | 0 | 0 | 0 | 0 | 0 | 0 | 0 | 0 |
| ENSMUSG00 Gm17615 | 2274 | 0 | 0 | 0 | 0 | 0 | 0 | 0 | 0 | 0 | 0 | 0 | 0 | 0 | 0 | 0 | 0 | 0 | 0 |
| ENSMUSG00 Gm17614 | 1212 | 0 | 0 | 0 | 0 | 0 | 0 | 0 | 0 | 0 | 0 | 0 | 0 | 0 | 0 | 0 | 0 | 0 | 0 |
| ENSMUSG00 Gm17296 | 5030 | 0.57462266 | 0.37087914 | 0.14425506 | 0.71557107 | 0.30620707 | 0.76841217 | 0.52674155 | 1.18449598 | 0.73693567 | 0.65617281 | 0.11397604 | 0.521538 | 0.26480353 | 0.73867336 | 0.47508752 | 0.13702404 | 0.47241161 | 0.13123445 |
| ENSMUSG00 Lrrc10b | 1358 | 0.06259967 | 0.02453085 | 0.14248463 | 0.06710021 | 0 | 0 | 0 | 0 | 0 | 0 | 0 | 0.16440549 | 0 | 0 | 0 | 0 | 0 | 0 |
| ENSMUSG00 Gm17152 | 534 | 0 | 0 | 0 | 0 | 0 | 0 | 0 | 0 | 0 | 0 | 0 | 0 | 0 | 0 | 0 | 0 | 0 | 0 |
| ENSMUSG00 Gm17034 | 2227 | 0.07634517 | 0.2542969 | 0 | 0.10229242 | 0 | 0.08900356 | 0.02124503 | 0 | 0.14795336 | 0.23289526 | 0 | 0.15037897 | 0.68670383 | 0.16684001 | 0.12950646 | 0.20632568 | 0.15488849 | 0.14820595 |
| ENSMUSG00 Gm17208 | 2769 | 0 | 0 | 0 | 0 | 0.0869121 | 0.01789553 | 0.10251935 | 0.12413565 | 0 | 0 | 0.17253506 | 0 | 0 | 0 | 0 | 0 | 0 | 0 |
| ENSMUSG00 Gm6541 | 476 | 0 | 0 | 0 | 0 | 0 | 0 | 0 | 0 | 0 | 0 | 0 | 0 | 0 | 0 | 0 | 0 | 0 | 0 |
| ENSMUSG00 Gm17297 | 172 | 0 | 0 | 0 | 0 | 0 | 0 | 0 | 0 | 0 | 0 | 0 | 0 | 0 | 0 | 0 | 0 | 0 | 0 |
| ENSMUSG00 Gm4794 | 876 | 0 | 0 | 0 | 0 | 0 | 0 | 0 | 0 | 0 | 0 | 0 | 0 | 0 | 0 | 0 | 0 | 0 | 0 |
| ENSMUSG00 Gm17295 | 1499 | 0 | 0 | 0 | 0 | 0 | 0 | 0 | 0 | 0 | 0 | 0 | 0 | 0 | 0 | 0 | 0 | 0 | 0 |
| ENSMUSG00 Gm17119 | 279 | 0 | 0 | 0 | 0 | 0 | 0 | 0 | 0 | 0 | 0 | 0 | 0 | 0 | 0 | 0 | 0 | 0 | 0 |
| ENSMUSG00 Gm7714 | 682 | 0 | 0 | 0 | 0 | 0 | 0 | 0 | 0 | 0 | 0 | 0 | 0 | 0 | 0 | 0 | 0 | 0 | 0 |
| ENSMUSG00 Gm9479 | 1572 | 0 | 0 | 0.21540376 | 0.17389712 | 0.03061827 | 0.0315221 | 0.39126261 | 0 | 0.05240015 | 0.35992903 | 0 | 0.03550615 | 0 | 0.08863375 | 0 | 0.14614736 | 0.09752238 | 0.13997228 |
| ENSMUSG00 Vmn2r99 | 2565 | 0 | 0 | 0 | 0 | 0 | 0 | 0 | 0 | 0 | 0 | 0 | 0 | 0 | 0 | 0 | 0 | 0 | 0 |
| ENSMUSG00 Gm6459 | 812 | 0.47111648 | 0.41025732 | 0.05957331 | 0.16832898 | 0.05927577 | 0.1830766 | 0.23306738 | 0.14110493 | 0.20288924 | 0.05806737 | 0.11767231 | 0.2062155 | 0.06075359 | 0.17159146 | 0.05074083 | 0.2357796 | 0.14159962 | 0.20323562 |
| ENSMUSG00 Adh6-ps1 | 2785 | 0 | 0 | 0 | 0 | 0 | 0 | 0.25482592 | 0.02742722 | 0 | 0 | 0 | 0 | 0.08856718 | 0 | 0 | 0 | 0 | 0 |
| ENSMUSG00 1700071M16 | 3298 | 0.02577633 | 0 | 0 | 0 | 0 | 0 | 0 | 0 | 0 | 0 | 0 | 0 | 0 | 0 | 0 | 0 | 0 | 0 |
| ENSMUSG00 Gm5174 | 1442 | 0 | 0 | 0 | 0 | 0 | 0 | 0 | 0.05297143 | 0 | 0 | 0 | 0 | 0 | 0 | 0 | 0 | 0 | 0 |
| ENSMUSG00 Gm17506 | 443 | 0 | 0 | 0.21839066 | 0.1028466 | 0 | 0 | 0 | 0 | 0 | 0 | 0 | 0 | 0 | 0 | 0 | 0 | 0 | 0 |
| ENSMUSG00 Gm5156 | 1333 | 0 | 0 | 0 | 0 | 0 | 0 | 0 | 0 | 0 | 0 | 0 | 0 | 0 | 0 | 0 | 0 | 0 | 0 |
| ENSMUSG00 Gm17327 | 5282 | 1.33583098 | 0.97756505 | 0.74181294 | 0.83669468 | 1.49444064 | 1.09762776 | 2.79469061 | 1.53290218 | 1.27879375 | 1.16939457 | 2.36070885 | 0.60232729 | 0.80320804 | 0.92325429 | 1.45866924 | 1.08738946 | 1.828517 | 1.03103021 |
| ENSMUSG00 Gm17326 | 172 | 0 | 0 | 0 | 0 | 0 | 0 | 0 | 0 | 0 | 0 | 0 | 0 | 0 | 0 | 0 | 0 | 0 | 0 |
| ENSMUSG00 Gm17325 | 2655 | 0 | 0 | 0 | 0 | 0 | 0 | 0 | 0 | 0 | 0 | 0 | 0 | 0 | 0 | 0 | 0 | 0 | 0 |
| ENSMUSG00 2310050C09I | 1065 | 0 | 0 | 0 | 0 | 0 | 0 | 0.13327515 | 0 | 0 | 0.2213648 | 0 | 0 | 0 | 0 | 0 | 0.07190724 | 0 | 0 |
| ENSMUSG00 Vmn2r104 | 2577 | 0 | 0 | 0 | 0 | 0 | 0 | 0 | 0 | 0 | 0 | 0 | 0 | 0 | 0 | 0 | 0 | 0 | 0 |
| ENSMUSG00 Gm3662 | 1865 | 0 | 0 | 0 | 0 | 0 | 0 | 0 | 0 | 0 | 0 | 0 | 0 | 0 | 0 | 0 | 0 | 0 | 0 |
| ENSMUSG00 Gm17324 | 921 | 0.13845334 | 0.25319246 | 0.36765984 | 0.09893821 | 0.47034453 | 0 | 0.35959691 | 0.29027885 | 0.13415803 | 0.15358535 | 0.31123751 | 0.06060333 | 0.05356343 | 0.30256734 | 0.26841404 | 0.33260028 | 0.45775165 | 0.05972759 |
| ENSMUSG00 Gm17223 | 480 | 0 | 0 | 0 | 0 | 0 | 0 | 0 | 0 | 0 | 0 | 0 | 0 | 0 | 0 | 0 | 0 | 0 | 0 |
| ENSMUSG00 Gm4462 | 491 | 0.25970576 | 0.06784704 | 0 | 0 | 0 | 0.10092206 | 0.09635983 | 0 | 0 | 0 | 0.09730134 | 0 | 0.20094467 | 0 | 0 | 0.15611526 | 0 | 0 |
| ENSMUSG00 Gm17385 | 1499 | 0 | 0 | 0 | 0 | 0 | 0 | 0 | 0 | 0 | 0 | 0 | 0 | 0 | 0 | 0 | 0 | 0 | 0 |
| ENSMUSG00 Gm17090 | 487 | 0 | 0.0684043 | 0.59597779 | 0 | 0 | 0.10175099 | 0 | 0 | 0.0845719 | 0 | 0 | 0.11461122 | 0 | 0.28610321 | 0 | 0.07862548 | 0 | 0 |
| ENSMUSG00 Gm5263 | 361 | 0 | 0 | 0 | 0 | 0 | 0 | 0 | 0 | 0 | 0 | 0 | 0 | 0 | 0 | 0 | 0 | 0 | 0 |
| ENSMUSG00 4930505N22I | 943 | 0.40566976 | 0.45924457 | 0.4103799 | 0.43483499 | 0.20416511 | 0.10509594 | 0.80276019 | 0.40500957 | 0.91719706 | 1.00001492 | 0.81060375 | 1.18378924 | 0.15694141 | 0.24625709 | 0.43692 | 0.24363059 | 0.44707239 | 0.23333661 |
| ENSMUSG00 Gm17384 | 2418 | 0 | 0 | 0 | 0.0376849 | 0 | 0 | 0 | 0 | 0 | 0 | 0 | 0 | 0 | 0 | 0 | 0 | 0 | 0 |
| ENSMUSG00 Gm17111 | 409 | 1.4549449 | 0.97739543 | 0.23654538 | 0 | 0.35304589 | 0.24231166 | 0 | 0.6536597 | 0.70490369 | 0.11528289 | 0.70085514 | 0.95528029 | 2.7741665 | 0 | 0.6044238 | 1.49792107 | 0.56224396 | 0.26899319 |
| ENSMUSG00 Gm17073 | 555 | 0 | 0 | 0 | 0 | 0 | 0 | 0 | 0 | 0 | 0 | 0 | 0 | 0 | 0 | 0 | 0 | 0 | 0 |
| ENSMUSG00 Gm17160 | 258 | 0 | 0 | 0 | 0 | 0.55967353 | 0 | 0 | 0 | 0 | 0 | 0 | 0 | 0 | 0 | 0 | 0.29682641 | 0 | 0.21321359 |
| ENSMUSG00 9130221H12I | 1463 | 0.55201527 | 0.88804024 | 1.38871379 | 0.96540832 | 0.59219045 | 0.67741263 | 0.90550582 | 0.78316615 | 0.53488982 | 0.51566046 | 0.45717664 | 1.29715142 | 1.71970457 | 1.17459414 | 0.95752078 | 0.91604324 | 0.8907 | 1.27840713 |
| ENSMUSG00 Gm17199 | 415 | 0 | 0 | 0 | 0 | 0.23196108 | 0.23880835 | 0 | 0 | 0 | 0 | 0 | 0 | 0 | 0.33574039 | 0 | 0 | 0.64646766 | 0 |
| ENSMUSG00 Gm3216 | 483 | 0 | 0 | 0 | 0 | 0 | 0 | 0 | 0 | 0 | 0 | 0 | 0 | 0 | 0 | 0 | 0 | 0 | 0 |
| ENSMUSG00 Gm3025 | 486 | 0 | 0 | 0 | 0 | 0 | 0 | 0 | 0 | 0 | 0 | 0 | 0 | 0 | 0 | 0 | 0 | 0 | 0 |
| ENSMUSG00 Gm17149 | 374 | 0 | 0 | 0.25868198 | 0.24364195 | 0.51477993 | 0 | 0.37951347 | 0 | 0 | 0 | 0 | 0.14923974 | 0 | 0 | 0 | 0 | 0 | 0 |
| ENSMUSG00 Gm17698 | 1656 | 0 | 0.02011648 | 0.02921107 | 0.02751271 | 0 | 0 | 0 | 0 | 0 | 0 | 0 | 0 | 0 | 0 | 0 | 0 | 0 | 0 |
| ENSMUSG00 4932443I19R | 777 | 0 | 0.04287374 | 0 | 0 | 0 | 0.06377443 | 0 | 0 | 0 | 0 | 0 | 0 | 0 | 0 | 0 | 0 | 0 | 0 |
| ENSMUSG00 Gm8122 | 1399 | 0 | 0 | 0 | 0 | 0 | 0 | 0 | 0 | 0 | 0 | 0 | 0 | 0 | 0 | 0 | 0 | 0 | 0 |
| ENSMUSG00 Gm17081 | 477 | 0 | 0 | 0 | 0 | 0 | 0 | 0 | 0 | 0 | 0 | 0 | 0 | 0 | 0 | 0 | 0 | 0 | 0 |
| ENSMUSG00 Gm6669 | 1004 | 0 | 0 | 0 | 0 | 0 | 0 | 0 | 0 | 0.04102243 | 0 | 0 | 0 | 0 | 0 | 0 | 0 | 0 | 0 |
| ENSMUSG00 C030048H21 | 374 | 0.1136502 | 0 | 0 | 0 | 0.25738997 | 0.26498788 | 0.12650449 | 0 | 0 | 0 | 0 | 0 | 0 | 0 | 0 | 0 | 0 | 0 |
| ENSMUSG00 Gm17217 | 468 | 0 | 0 | 0 | 0 | 0 | 0 | 0 | 0 | 0 | 0 | 0 | 0 | 0 | 0 | 0 | 0 | 0 | 0 |
| ENSMUSG00 Vmn2r46 | 2424 | 0 | 0 | 0 | 0 | 0 | 0 | 0 | 0 | 0 | 0 | 0 | 0 | 0 | 0 | 0 | 0 | 0 | 0 |
| ENSMUSG00 Olfr723 | 1005 | 0 | 0 | 0 | 0 | 0 | 0 | 0 | 0 | 0 | 0 | 0 | 0 | 0 | 0 | 0 | 0 | 0 | 0 |
| ENSMUSG00 Gm17237 | 61 | 0 | 1.09222604 | 0 | 0.74690237 | 1.57809586 | 0.81233989 | 0 | 0 | 0 | 0 | 0 | 0 | 4.04359978 | 0 | 0 | 0.62829995 | 0 | 0 |
| ENSMUSG00 Gm5798 | 1746 | 0 | 0 | 0 | 0 | 0 | 0 | 0 | 0 | 0 | 0 | 0 | 0 | 0 | 0 | 0 | 0 | 0 | 0 |
| ENSMUSG00 Vmn1r5 | 1055 | 0 | 0 | 0 | 0 | 0 | 0 | 0 | 0 | 0 | 0 | 0 | 0 | 0 | 0 | 0 | 0 | 0 | 0 |
| ENSMUSG00 Gm17236 | 3653 | 0 | 0 | 0 | 0 | 0 | 0 | 0 | 0 | 0 | 0 | 0 | 0 | 0 | 0 | 0 | 0 | 0 | 0 |
| ENSMUSG00 Gm17261 | 43 | 0 | 0 | 0 | 0 | 0 | 0 | 0 | 0 | 0 | 0 | 0 | 0 | 0 | 0 | 0 | 0 | 0 | 0 |
| ENSMUSG00 Gm17148 | 248 | 0 | 0 | 0 | 0 | 0 | 0.59942823 | 0 | 0 | 0.16607466 | 0 | 0 | 0 | 0 | 0 | 0 | 0 | 0 | 0.66543274 |
| ENSMUSG00 Gm17204 | 475 | 0 | 0 | 0 | 0 | 0 | 0 | 0 | 0 | 0 | 0 | 0 | 0 | 0 | 0 | 0 | 0 | 0 | 0 |
| ENSMUSG00 Gm3972 | 2869 | 0 | 0 | 0 | 0 | 0 | 0 | 0 | 0 | 0 | 0 | 0 | 0 | 0 | 0 | 0 | 0 | 0 | 0 |
| ENSMUSG00 Gm17555 | 1499 | 0 | 0 | 0 | 0 | 0 | 0 | 0 | 0 | 0 | 0 | 0 | 0 | 0 | 0 | 0 | 0 | 0 | 0 |
| ENSMUSG00 Gm17556 | 3023 | 0 | 0 | 0 | 0 | 0 | 0 | 0 | 0 | 0 | 0 | 0 | 0 | 0 | 0 | 0 | 0 | 0 | 0 |
| ENSMUSG00 Vmn2r-ps128 | 2567 | 0 | 0 | 0 | 0 | 0 | 0 | 0 | 0 | 0 | 0 | 0 | 0 | 0 | 0 | 0 | 0 | 0 | 0 |
| ENSMUSG00 2310042E22I | 1247 | 0 | 0 | 0 | 0 | 0 | 0 | 0 | 0 | 0 | 0 | 0 | 0 | 0 | 0 | 0 | 0 | 0 | 0 |
| ENSMUSG00 Gm17077 | 277 | 0 | 0.12026316 | 0 | 0 | 0 | 0 | 0 | 0 | 0 | 0 | 0 | 0 | 0.17809356 | 0 | 0 | 0.27646648 | 0 | 0.39717767 |
| ENSMUSG00 Gm2822 | 2908 | 0 | 0 | 0 | 0 | 0.11586089 | 0 | 0 | 0 | 0 | 0 | 0 | 0 | 0 | 0 | 0 | 0 | 0.01317961 | 0 |
| ENSMUSG00 Gm17621 | 1130 | 0 | 0 | 0 | 0 | 0 | 0 | 0 | 0 | 0 | 0 | 0 | 0 | 0 | 0 | 0 | 0 | 0 | 0 |
| ENSMUSG00 Vmn2r79 | 2550 | 0 | 0 | 0 | 0 | 0 | 0 | 0 | 0 | 0 | 0 | 0 | 0 | 0 | 0 | 0 | 0 | 0 | 0 |
| ENSMUSG00 Gm3512 | 1855 | 0 | 0.0538753 | 0 | 0 | 0 | 0.02671306 | 0 | 0.08235558 | 0 | 0 | 0 | 0 | 0 | 0 | 0 | 0.02064184 | 0 | 0 |
| ENSMUSG00 Gm3043 | 816 | 0 | 0 | 0 | 0 | 0 | 0 | 0 | 0 | 0 | 0 | 0 | 0 | 0 | 0 | 0 | 0 | 0 | 0 |
| ENSMUSG00 Gm17070 | 783 | 0 | 0 | 0 | 0 | 0 | 0 | 0 | 0 | 0 | 0 | 0 | 0 | 0 | 0 | 0 | 0 | 0 | 0 |
| ENSMUSG00 Gm3623 | 4849 | 0 | 0 | 0 | 0 | 0 | 0 | 0 | 0 | 0 | 0 | 0 | 0 | 0 | 0 | 0 | 0 | 0 | 0 |
| ENSMUSG00 4933411K16I | 1379 | 0.33905507 | 0.16910099 | 0.56125924 | 0.13215676 | 0.20942099 | 0.07186763 | 0.20585647 | 0.19387006 | 0.29866944 | 0.13676781 | 0.45038032 | 0 | 0.35773689 | 0 | 0 | 0 | 0.25013537 | 0 |
| ENSMUSG00 Gm20533 | 1606 | 0 | 0 | 0 | 0 | 0 | 0 | 0 | 0 | 0 | 0 | 0 | 0 | 0 | 0 | 0 | 0 | 0 | 0 |
| ENSMUSG00 Gm17435 | 2992 | 0.08523765 | 0.11133989 | 0.2748496 | 0.41114579 | 0.40217182 | 0.29811137 | 0.17394367 | 0.63824202 | 0.19271765 | 0.09455355 | 0.36725403 | 0.35444438 | 0.54410203 | 0 | 0.08262344 | 0.12797663 | 0.06404796 | 0.73541587 |
| ENSMUSG00 Gm17232 | 449 | 0 | 0 | 0 | 0 | 0 | 0 | 0 | 0 | 0 | 0 | 0 | 0 | 0 | 0 | 0 | 0 | 0 | 0 |
| ENSMUSG00 Gm6882 | 1355 | 0 | 0 | 0 | 0 | 0 | 0 | 0 | 0 | 0 | 0 | 0 | 0 | 0 | 0 | 0 | 0 | 0 | 0 |
| ENSMUSG00 Gm8281 | 1947 | 0 | 0 | 0 | 0 | 0 | 0 | 0 | 0 | 0 | 0 | 0 | 0 | 0 | 0 | 0 | 0 | 0 | 0 |
| ENSMUSG00 Gm10925 | 770 | 1.2144336 | 1.94685745 | 3.32960663 | 1.59759506 | 2.3128327 | 1.60885499 | 3.25658699 | 2.92643083 | 2.94189398 | 3.85778481 | 3.53658784 | 2.89951494 | 2.17829245 | 4.28250676 | 4.33418965 | 2.88422757 | 3.68330646 | 4.78650669 |
| ENSMUSG00 Gm8229 | 1398 | 0 | 0 | 0 | 0 | 0 | 0 | 0 | 0 | 0 | 0 | 0 | 0 | 0 | 0 | 0 | 0 | 0 | 0 |
| ENSMUSG00 2610020F03I | 586 | 0 | 0.22739177 | 0 | 0.07774922 | 0.24640917 | 0 | 0 | 0 | 0 | 0.08046195 | 0 | 0.57149143 | 0.25255248 | 0 | 0.07030982 | 0 | 0 | 0 |
| ENSMUSG00 Gm6158 | 563 | 0.37748824 | 0.23668131 | 0.25776304 | 0.40462739 | 0.25647561 | 0.26404654 | 0.08403673 | 0.20351191 | 0 | 0 | 0.25457349 | 0.29741916 | 0 | 0.16498768 | 0 | 0.13602347 | 0.13615026 | 0 |
| ENSMUSG00 Gm17078 | 476 | 0 | 0 | 0 | 0 | 0 | 0 | 0 | 0 | 0 | 0 | 0 | 0 | 0 | 0 | 0 | 0 | 0 | 0 |
| ENSMUSG00 Vmn2r58 | 2583 | 0 | 0 | 0 | 0 | 0 | 0 | 0 | 0 | 0 | 0 | 0 | 0 | 0 | 0 | 0 | 0 | 0 | 0 |
| ENSMUSG00 Gm17492 | 6842 | 0 | 0 | 0 | 0 | 0 | 0 | 0 | 0 | 0 | 0 | 0 | 0 | 0 | 0 | 0 | 0 | 0 | 0 |
| ENSMUSG00 Gm7316 | 442 | 0 | 0 | 0 | 0 | 0 | 0 | 0 | 0 | 0 | 0 | 0 | 0 | 0 | 0 | 0 | 0 | 0 | 0 |
| ENSMUSG00 2810055G20I | 2171 | 0.76356604 | 0.70584668 | 0.40107027 | 0.65057226 | 0.46557826 | 0.13694906 | 0.41406766 | 0.59813067 | 0.53119412 | 0.45608695 | 0.37410147 | 0.95125726 | 0.34084696 | 0.51343072 | 0.5313881 | 0.31747164 | 0.56492008 | 0.43074842 |
| ENSMUSG00 Gm17056 | 2297 | 0.59214873 | 0.13052505 | 0.35801046 | 0.07934009 | 0.29335957 | 0.28044647 | 0.5767327 | 0.03325416 | 0.51998649 | 0.49264993 | 0.06239655 | 0.6560831 | 1.67518047 | 3.67994075 | 0.26905674 | 1.33358667 | 2.58623249 | 4.28673492 |
| ENSMUSG00 Mir706 | 2231 | 0.2857811 | 0.43302283 | 0.3035542 | 0.12253082 | 0.25888977 | 0.19989897 | 0.08482775 | 0.32526026 | 0.75690146 | 0.46495539 | 0.40686876 | 0.35025517 | 0.04422404 | 0.60371068 | 0 | 0.20595575 | 0.46383237 | 0.49313408 |
| ENSMUSG00 Gm7236 | 329 | 0.12919506 | 0 | 0 | 0 | 0 | 0 | 0.28761507 | 0 | 0 | 0.14331521 | 0 | 0 | 0 | 0 | 0.25046539 | 0.11638482 | 0 | 0 |
| ENSMUSG00 Gm7360 | 506 | 0 | 0 | 0 | 0 | 0 | 0 | 0 | 0 | 0 | 0 | 0 | 0 | 0 | 0 | 0 | 0 | 0 | 0 |
| ENSMUSG00 Gm14478 | 499 | 0 | 0 | 0 | 0 | 0 | 0 | 0 | 0 | 0 | 0 | 0 | 0 | 0 | 0 | 0 | 0 | 0 | 0 |
| ENSMUSG00 Gm13278 | 1555 | 0 | 0 | 0 | 0 | 0 | 0 | 0 | 0 | 0 | 0 | 0 | 0 | 0 | 0 | 0 | 0 | 0 | 0 |
| ENSMUSG00 Vmn1r91 | 923 | 0 | 0 | 0 | 0 | 0 | 0 | 0 | 0 | 0 | 0 | 0 | 0 | 0 | 0 | 0 | 0 | 0 | 0 |
| ENSMUSG00 4930523C07I | 5773 | 3.4678569 | 6.52062541 | 5.74817977 | 4.38800288 | 5.41098536 | 4.74669352 | 3.25361745 | 3.03660362 | 3.38167477 | 2.27871925 | 4.09641511 | 4.48613675 | 4.15300741 | 4.18342724 | 3.66125033 | 5.21993903 | 4.4746101 | 2.95389279 |
| ENSMUSG00 Gm17309 | 277 | 0 | 0 | 0 | 0 | 0 | 0 | 0 | 0 | 0 | 0 | 0 | 0 | 0 | 0 | 0 | 0 | 0 | 0 |
| ENSMUSG00 Gm6536 | 611 | 0 | 0 | 0 | 0 | 0 | 0 | 0 | 0 | 0 | 0 | 0 | 0 | 0 | 0 | 0 | 0 | 0 | 0 |
| ENSMUSG00 Gm17096 | 731 | 1.97698493 | 0.68357512 | 0.86026796 | 1.43352123 | 0.52675156 | 0.47451318 | 0.77667872 | 1.20167614 | 3.15519136 | 2.77357078 | 2.61422481 | 0.45813129 | 1.01228284 | 1.39777007 | 1.40908194 | 2.40953207 | 0.99616914 | 2.5585631 |
| ENSMUSG00 Gm17609 | 1089 | 0 | 0 | 0 | 0 | 0 | 0 | 0 | 0 | 0.03782049 | 0 | 0 | 0 | 0 | 0 | 0 | 0 | 0.07038806 | 0 |
| ENSMUSG00 Nav1 | 1643 | 0.1552228 | 0.62854517 | 0.08832659 | 0.0277304 | 0.02929515 | 0.30159911 | 0.37435473 | 0 | 0 | 0 | 0 | 0 | 0 | 0.22614285 | 0.25077027 | 0.0233053 | 0 | 0.53569428 |
| ENSMUSG00 A330094K24I | 1025 | 0 | 0 | 0 | 0 | 0 | 0 | 0.36926969 | 0.26082616 | 0 | 0 | 0 | 0 | 0 | 0.81560349 | 0 | 0 | 0 | 0.37567195 |
| ENSMUSG00 0610007N19I | 1480 | 3.56124448 | 3.48885041 | 3.33385142 | 2.58589711 | 3.96763155 | 3.21423137 | 5.5944046 | 2.2708996 | 3.89602176 | 5.32038341 | 5.87502868 | 2.3382237 | 7.29979046 | 9.03776836 | 6.45862227 | 5.17440641 | 3.62546052 | 9.10623726 |
| ENSMUSG00 Gm17400 | 850 | 0 | 0.03919164 | 0.51219032 | 0.05360123 | 0.05662579 | 0.05829733 | 0.11132395 | 0.31452567 | 0.04845472 | 0.38829991 | 0.56205834 | 0 | 0.17411265 | 0.10928021 | 0.04847242 | 0.36038219 | 0 | 0 |
| ENSMUSG00 Lca5 | 58 | 0 | 0.57436025 | 0 | 0 | 0 | 0 | 0 | 0 | 0 | 0 | 0 | 0 | 0 | 0 | 0 | 0 | 0 | 0 |
| ENSMUSG00 Gm8362 | 2139 | 0 | 0 | 0 | 0 | 0 | 0 | 0 | 0 | 0 | 0 | 0 | 0 | 0 | 0 | 0 | 0 | 0 | 0 |
| ENSMUSG00 Gm16501 | 335 | 0 | 0 | 0 | 0 | 0 | 0 | 0 | 0 | 0 | 0 | 0 | 0 | 0 | 0 | 0 | 0 | 0 | 0 |
| ENSMUSG00 Gm17058 | 926 | 0 | 0.50365067 | 0.52239234 | 0.24600996 | 0 | 0 | 0.05109361 | 0.20622248 | 0.26686727 | 0 | 0.56752111 | 0.12055219 | 0.10654842 | 0 | 0.13348236 | 0 | 0.08277818 | 0.11881017 |
| ENSMUSG00 Gm16042 | 1250 | 0 | 0 | 0 | 0 | 0 | 0 | 0 | 0 | 0 | 0 | 0 | 0 | 0 | 0 | 0 | 0 | 0 | 0 |
| ENSMUSG00 Gm17402 | 226 | 0 | 0 | 0 | 0 | 0 | 0 | 0 | 0 | 0 | 0 | 0 | 0 | 0 | 0 | 0 | 0 | 0 | 0 |
| ENSMUSG00 Gm17401 | 2835 | 1.09448954 | 0.81079002 | 0.66545597 | 0.67497843 | 0.9337763 | 0.99629835 | 0.60079592 | 0.79483307 | 0.88620016 | 0.78168714 | 1.04481391 | 0.78752258 | 1.04406174 | 0.80273731 | 0.56679389 | 0.7293449 | 0.68946777 | 0.62091408 |
| ENSMUSG00 Gm17582 | 341 | 0 | 0 | 0 | 0 | 0 | 0 | 0 | 0 | 0 | 0 | 0 | 0 | 0 | 0 | 0 | 0 | 0 | 0 |
| ENSMUSG00 Vmn1r174 | 941 | 0 | 0 | 0 | 0 | 0 | 0 | 0 | 0 | 0 | 0 | 0 | 0 | 0 | 0 | 0 | 0 | 0 | 0 |
| ENSMUSG00 Gm16484 | 326 | 0 | 0 | 0 | 0 | 0 | 0 | 0 | 0 | 0 | 0 | 0 | 0 | 0 | 0 | 0 | 0 | 0 | 0 |
| ENSMUSG00 Gm17583 | 66 | 0 | 0 | 0 | 0 | 0 | 1.50159799 | 0 | 0 | 0 | 0 | 0 | 0 | 0 | 0 | 0 | 0 | 0 | 0.83347131 |
| ENSMUSG00 Gm6713 | 1445 | 0 | 0 | 0 | 0 | 0 | 0 | 0 | 0 | 0 | 0 | 0 | 0 | 0 | 0 | 0.08553956 | 0 | 0 | 0 |
| ENSMUSG00 Gm17584 | 172 | 0 | 0 | 0 | 0 | 0 | 0 | 0 | 0 | 0 | 0 | 0 | 0 | 0 | 0 | 0 | 0 | 0 | 0 |
| ENSMUSG00 Gm4242 | 107 | 0 | 0 | 0 | 0 | 0 | 0 | 0 | 0 | 0 | 0 | 0 | 0 | 0 | 0 | 0 | 0 | 0 | 0 |
| ENSMUSG00 Vmn2r94 | 2451 | 0 | 0 | 0 | 0 | 0 | 0 | 0 | 0 | 0 | 0 | 0 | 0 | 0 | 0 | 0 | 0 | 0 | 0 |
| ENSMUSG00 Gm17585 | 2978 | 0 | 0 | 0.09746178 | 0.01529921 | 0.06465 | 0.11647721 | 0 | 0.06412425 | 0.19362365 | 0.07916505 | 0.08021316 | 0.01874267 | 0 | 0 | 0.01383531 | 0 | 0.01286981 | 0 |
| ENSMUSG00 Gm7980 | 1398 | 0 | 0 | 0 | 0 | 0 | 0 | 0 | 0 | 0 | 0 | 0 | 0 | 0 | 0 | 0 | 0 | 0 | 0 |
| ENSMUSG00 1700040D17 | 960 | 0.48703848 | 0.10410279 | 0.20155638 | 0.33221595 | 0.35096194 | 0.41293945 | 0 | 0.11935126 | 0.2145131 | 0.24557658 | 1.14460838 | 0.46513052 | 0.87358604 | 1.54813625 | 0.17167315 | 0.87749309 | 1.0779271 | 0 |
| ENSMUSG00 Gm4303 | 869 | 0 | 0 | 0 | 0 | 0 | 0 | 0 | 0 | 0 | 0 | 0 | 0 | 0 | 0 | 0 | 0 | 0 | 0 |
| ENSMUSG00 Gm17272 | 2213 | 0 | 0 | 0 | 0 | 0 | 0 | 0 | 0.29338944 | 0 | 0 | 0.04317665 | 0 | 0 | 0 | 0 | 0 | 0 | 0 |
| ENSMUSG00 Gm2871 | 1203 | 0 | 0 | 0 | 0 | 0 | 0 | 0 | 0 | 0 | 0 | 0 | 0 | 0 | 0 | 0 | 0 | 0 | 0 |
| ENSMUSG00 Efna5 | 2415 | 0 | 0 | 0 | 0 | 0 | 0 | 0 | 0 | 0 | 0 | 0 | 0 | 0 | 0 | 0 | 0 | 0 | 0 |
| ENSMUSG00 Gm17392 | 1506 | 0 | 0 | 0 | 0 | 0 | 0 | 0 | 0 | 0 | 0 | 0 | 0 | 0 | 0 | 0 | 0 | 0 | 0 |
| ENSMUSG00 Gm17225 | 495 | 0 | 0 | 0 | 0 | 0 | 0 | 0 | 0 | 0 | 0 | 0 | 0 | 0.49830219 | 0.09382644 | 0 | 0 | 0 | 0 |
| ENSMUSG00 Gm17271 | 376 | 0 | 0 | 0 | 0 | 0 | 0 | 0 | 0 | 0 | 0 | 0 | 0 | 0 | 0 | 0 | 0 | 0 | 0 |
| ENSMUSG00 Snhg4 | 1147 | 16.8241935 | 12.4887049 | 13.9174063 | 10.168812 | 12.7988114 | 10.5844985 | 23.9244584 | 15.7830852 | 24.1661073 | 15.6209828 | 17.7854466 | 17.8590655 | 14.5372171 | 17.9378646 | 20.3313693 | 20.998075 | 14.2679414 | 19.6152787 |
| ENSMUSG00 Gm17456 | 3375 | 0 | 0 | 0 | 0 | 0 | 0 | 0 | 0 | 0 | 0 | 0 | 0 | 0 | 0 | 0 | 0 | 0 | 0 |
| ENSMUSG00 9430037G07I | 4892 | 0.06950969 | 0.02723867 | 0.00988829 | 0.04656689 | 0.02951672 | 0 | 0.00967144 | 0.00780711 | 0.01683831 | 0 | 0 | 0.04563832 | 0.08067362 | 0.02848166 | 0 | 0.10958064 | 0.16452417 | 0.13493648 |
| ENSMUSG00 Gm6410 | 1103 | 0 | 0.03020208 | 0.04385633 | 0.12391943 | 0.17454913 | 0 | 0.04289454 | 0.03462593 | 0.2987236 | 0.12824307 | 0.0866273 | 0 | 0.62615308 | 0.4210706 | 0.93385213 | 0.27771973 | 0.41696787 | 0.29923358 |
| ENSMUSG00 Vmn2r75 | 2556 | 0 | 0 | 0 | 0 | 0 | 0 | 0 | 0 | 0 | 0 | 0 | 0 | 0 | 0 | 0 | 0 | 0 | 0 |
| ENSMUSG00 Gm17454 | 617 | 0 | 0 | 0 | 0 | 0 | 0 | 0 | 0 | 0 | 0 | 0 | 0 | 0 | 0 | 0 | 0 | 0 | 0 |
| ENSMUSG00 Gm17455 | 853 | 0 | 0 | 0 | 0 | 0 | 0 | 0 | 0 | 0 | 0 | 0 | 0 | 0 | 0 | 0 | 0 | 0 | 0 |
| ENSMUSG00 Gm9732 | 1462 | 0 | 0 | 0 | 0 | 0 | 0 | 0 | 0 | 0 | 0 | 0 | 0 | 0 | 0 | 0 | 0 | 0 | 0 |
| ENSMUSG00 Gm17651 | 34 | 0 | 0 | 0 | 0 | 0 | 0 | 0 | 0 | 0 | 0 | 0 | 0 | 0 | 0 | 0 | 0 | 0 | 0 |
| ENSMUSG00 Psg-ps1 | 1917 | 0 | 0 | 0 | 0 | 0 | 0 | 0 | 0 | 0 | 0 | 0 | 0 | 0 | 0 | 0 | 0 | 0 | 0 |
| ENSMUSG00 Gm16478 | 395 | 0 | 0 | 0 | 0 | 0 | 0 | 0 | 0 | 0 | 0 | 0 | 0 | 0 | 0 | 0 | 0 | 0 | 0 |
| ENSMUSG00 D930048G16 | 2092 | 1.36130344 | 3.05739756 | 1.9192175 | 3.98550244 | 1.97865461 | 2.08443621 | 1.19864818 | 1.82564065 | 1.10250711 | 2.47924348 | 1.41589265 | 0.56029107 | 1.55636068 | 0.95463468 | 0.98474082 | 2.04997516 | 1.70379809 | 0.81514451 |
| ENSMUSG00 Gm17653 | 2139 | 0 | 0.01557405 | 0.04523004 | 0 | 0.04500414 | 0 | 0 | 0.07142104 | 0.50063086 | 0.2204334 | 0 | 0 | 0 | 0.06513897 | 0.01926206 | 0.16111055 | 0 | 0 |
| ENSMUSG00 A930017M01 | 836 | 0 | 0 | 0 | 0 | 0 | 0 | 0 | 0 | 0 | 0 | 0 | 0 | 0 | 0 | 0 | 0 | 0 | 0 |
| ENSMUSG00 Gm17652 | 2359 | 0.6666772 | 0.26831072 | 0.59467248 | 0.46352906 | 0.44887763 | 0.52514554 | 0.16044995 | 0.33999171 | 0.97772144 | 0.47970194 | 0.95185377 | 1.39598308 | 0.27185881 | 0.49220101 | 0.80342161 | 1.07129296 | 0.27619629 | 0.88611533 |
| ENSMUSG00 Gm17192 | 510 | 0 | 0 | 0 | 0 | 0 | 0 | 0 | 0 | 0 | 0 | 0 | 0 | 0 | 0 | 0 | 0 | 0 | 0 |
| ENSMUSG00 Gm17076 | 1400 | 0 | 0 | 0 | 0 | 0 | 0 | 0 | 0 | 0 | 0 | 0 | 0 | 0 | 0 | 0 | 0 | 0 | 0 |
| ENSMUSG00 Trav5d-3 | 269 | 0 | 0 | 0 | 0 | 0 | 0 | 0 | 0 | 0 | 0 | 0 | 0 | 0 | 0 | 0 | 0 | 0 | 0 |
| ENSMUSG00 Gm6133 | 947 | 0 | 0.03517729 | 0 | 0 | 0 | 0 | 0 | 0.04032989 | 0 | 0.0995791 | 0 | 0 | 0.57302121 | 0.44139048 | 0.52208941 | 0.40433588 | 0.04047127 | 0 |
| ENSMUSG00 1700022N22I | 707 | 0.66132523 | 0.65966127 | 1.29999587 | 1.03108445 | 1.2935029 | 0.70088732 | 0.06692034 | 0.64824445 | 0.64080859 | 0.73360359 | 1.82450336 | 1.10526065 | 0.27910561 | 1.57660268 | 0.93242559 | 0.97486695 | 0.59630731 | 1.08928924 |
| ENSMUSG00 Gm17343 | 52 | 0 | 0 | 0 | 0 | 0 | 0 | 0 | 0 | 0 | 0 | 0 | 0 | 0 | 0 | 0 | 0 | 0 | 1.05786744 |
| ENSMUSG00 Gm17341 | 1955 | 0 | 0 | 0 | 0 | 0 | 0 | 0 | 0 | 0 | 0 | 0 | 0 | 0 | 0 | 0 | 0 | 0 | 0 |
| ENSMUSG00 Gm2431 | 514 | 0 | 0 | 0 | 0 | 0 | 0 | 0 | 0 | 0 | 0 | 0 | 0 | 0 | 0 | 0 | 0 | 0 | 0 |
| ENSMUSG00 Vmn2r-ps100 | 660 | 0 | 0 | 0 | 0 | 0 | 0 | 0 | 0 | 0 | 0 | 0 | 0 | 0 | 0 | 0 | 0 | 0 | 0 |
| ENSMUSG00 4930571K23I | 2152 | 0 | 0 | 0 | 0 | 0 | 0 | 0 | 0 | 0 | 0 | 0 | 0 | 0 | 0 | 0 | 0 | 0 | 0 |
| ENSMUSG00 Gm17122 | 370 | 0 | 0 | 0 | 0 | 0 | 0 | 0 | 0 | 0 | 0 | 0 | 0 | 0 | 0 | 0 | 0 | 0 | 0 |
| ENSMUSG00 Gm17210 | 389 | 2.62242731 | 2.99730411 | 1.61659613 | 2.57671716 | 3.71197355 | 2.16554363 | 0.4865057 | 2.15998163 | 0.52938966 | 1.33331037 | 1.10533323 | 0.14348499 | 0.50726907 | 2.02969019 | 0.84733277 | 1.08276782 | 0.09852518 | 1.41411585 |
| ENSMUSG00 Gm17511 | 496 | 0.4284796 | 0.26865237 | 0.39010912 | 0.27557083 | 0.09704017 | 0.19980941 | 0.38155386 | 0.61600649 | 0.08303733 | 0.47530951 | 0.57792288 | 0.45012631 | 0.79567609 | 0.28091182 | 0.58147357 | 1.15798208 | 0.7727076 | 0.33271637 |
| ENSMUSG00 Il1f6 | 1106 | 0 | 0 | 0 | 0 | 0 | 0 | 0 | 0 | 0 | 0 | 0 | 0 | 0 | 0 | 0 | 0 | 0 | 0 |
| ENSMUSG00 Gm3578 | 433 | 0 | 0 | 0 | 0 | 0 | 0 | 0 | 0 | 0 | 0 | 0 | 0 | 0 | 0 | 0 | 0 | 0 | 0 |
| ENSMUSG00 Vmn1r-ps73 | 4211 | 0 | 0 | 0 | 0 | 0 | 0 | 0 | 0 | 0 | 0 | 0 | 0 | 0 | 0 | 0 | 0 | 0 | 0 |
| ENSMUSG00 Gm17140 | 625 | 0 | 0 | 0 | 0 | 0 | 0 | 0 | 0 | 0 | 0 | 0 | 0 | 0 | 0 | 0 | 0 | 0 | 0 |
| ENSMUSG00 Gm3091 | 483 | 0 | 0 | 0 | 0 | 0 | 0 | 0 | 0 | 0 | 0 | 0 | 0 | 0 | 0 | 0 | 0 | 0 | 0 |
| ENSMUSG00 Gm17043 | 1279 | 0 | 0 | 0 | 0 | 0 | 0 | 0 | 0 | 0 | 0 | 0 | 0 | 0 | 0 | 0 | 0 | 0 | 0 |
| ENSMUSG00 Gm4613 | 348 | 0.12214131 | 0.19145342 | 0 | 0.26184508 | 0.13831013 | 0.14239291 | 0.40786792 | 0.10974828 | 0.47340823 | 0 | 0 | 0.16038984 | 0.28351677 | 0 | 0.23679055 | 0.22006096 | 0.22026807 | 0.94843287 |
| ENSMUSG00 Gm5331 | 398 | 0 | 0 | 0 | 0 | 0 | 0.12450436 | 0.11887608 | 0 | 0.10348371 | 0 | 0 | 0.14024036 | 0 | 0 | 0.1035215 | 0 | 0 | 0 |
| ENSMUSG00 AC166053.1 | 846 | 0 | 0 | 0 | 0 | 0 | 0 | 0 | 0 | 0 | 0 | 0 | 0 | 0 | 0 | 0 | 0 | 0 | 0 |
| ENSMUSG00 2410012M07 | 1813 | 0 | 0 | 0 | 0 | 0 | 0 | 0 | 0 | 0 | 0 | 0 | 0 | 0 | 0 | 0 | 0 | 0 | 0 |

| | | | | | | | | | | | | | | | | | | | |
|---|---|---|---|---|---|---|---|---|---|---|---|---|---|---|---|---|---|---|---|
| ENSMUSG00 | Gm4553 | 713 | 0 | 0 | 0 | 0 | 0 | 0 | 0 | 0 | 0 | 0 | 0 | 0 | 0 | 0 | 0 | 0 | 0 | 0 |
| ENSMUSG00 | Gm3047 | 817 | 0 | 0 | 0 | 0 | 0 | 0 | 0 | 0 | 0 | 0 | 0 | 0 | 0 | 0 | 0 | 0 | 0 | 0 |
| ENSMUSG00 | Gm17262 | 1648 | 1.31539076 | 1.87991454 | 1.14476194 | 0.80174168 | 0.75936287 | 1.11253103 | 0.54547385 | 1.39050009 | 0.39986909 | 0.34333036 | 0.86968978 | 0.88058691 | 1.1375078 | 0.84546276 | 1.10004154 | 0.53439561 | 0.27907498 | 0.20027587 |
| ENSMUSG00 | Gm9372 | 446 | 0 | 0.14938518 | 0 | 0.10215481 | 0 | 0 | 0.10608224 | 0.17126638 | 0 | 0 | 0.10711874 | 0 | 0 | 0.10413473 | 0 | 0.17170676 | 0.0859334 | 0 |
| ENSMUSG00 | Gm6245 | 767 | 0.05541744 | 0.21716359 | 0 | 0.05940162 | 0 | 0 | 0.06168537 | 0 | 0.10739639 | 0 | 0 | 0.0727714 | 0.06431801 | 0.30276459 | 0.16115341 | 0 | 0.09993819 | 0 |
| ENSMUSG00 | Gm17166 | 428 | 0 | 0 | 0 | 0 | 0 | 0 | 0 | 0 | 0 | 0 | 0 | 0 | 0 | 0 | 0 | 0 | 0 | 0 |
| ENSMUSG00 | Gm8068 | 1397 | 0 | 0 | 0 | 0 | 0 | 0 | 0 | 0 | 0 | 0 | 0 | 0 | 0 | 0 | 0 | 0 | 0 | 0 |
| ENSMUSG00 | Gm5656 | 1092 | 0.70063477 | 0.70164521 | 0.70876968 | 0.08344514 | 0.48484538 | 0.86218126 | 5.24252212 | 4.40681566 | 3.62079259 | 2.15891498 | 0.56874951 | 2.50454896 | 5.73732005 | 7.27283786 | 4.71629532 | 0.73635783 | 1.08801759 | 5.28933718 |
| ENSMUSG00 | Gm2683 | 865 | 0 | 0 | 0 | 0 | 0 | 0 | 0 | 0 | 0 | 0 | 0 | 0 | 0 | 0 | 0 | 0 | 0 | 0 |
| ENSMUSG00 | Gm2966 | 478 | 0 | 0 | 0 | 0 | 0 | 0 | 0 | 0 | 0 | 0 | 0 | 0 | 0 | 0 | 0 | 0 | 0 | 0 |
| ENSMUSG00 | Gm3675 | 1648 | 0 | 0 | 0 | 0 | 0 | 0 | 0 | 0 | 0 | 0 | 0 | 0 | 0 | 0 | 0 | 0 | 0 | 0 |
| ENSMUSG00 | Tmem29 | 1253 | 0 | 0 | 0 | 0 | 0 | 0 | 0 | 0 | 0 | 0 | 0 | 0 | 0 | 0 | 0 | 0 | 0 | 0 |
| ENSMUSG00 | Gm17417 | 3282 | 0.05180399 | 0.28420507 | 0.10317328 | 0.16658517 | 0.19065052 | 0.45295003 | 0.14415807 | 0.13964315 | 0.28863189 | 0.11493163 | 0.01455686 | 0.44217161 | 0.16534159 | 0.15566269 | 0.22596831 | 0.39667296 | 0.19852134 | 0.10056509 |
| ENSMUSG00 | Gm17416 | 200 | 183.62236 | 252.511738 | 198.573342 | 138.049964 | 119.84849 | 143.455164 | 15.1400573 | 9.93002463 | 6.38390994 | 2.12178165 | 6.2107446 | 5.86064457 | 2.95991504 | 1.8577635 | 5.56221005 | 0.95726519 | 1.34142039 | 0.27504553 |
| ENSMUSG00 | BC035947 | 2086 | 0 | 0.07984874 | 0 | 0 | 0.02307379 | 0 | 0 | 0 | 0 | 0 | 0 | 0 | 0 | 0 | 0 | 0 | 0 | 0 |
| ENSMUSG00 | Gm2888 | 1753 | 0 | 0 | 0 | 0 | 0 | 0 | 0 | 0 | 0 | 0 | 0 | 0 | 0 | 0 | 0 | 0 | 0 | 0 |
| ENSMUSG00 | A430105I06R | 1273 | 1.63609868 | 1.51779094 | 0.91199115 | 0.46527382 | 1.13429514 | 1.20670443 | 1.18932107 | 1.98012456 | 1.94123405 | 1.81491317 | 1.05082391 | 1.57844765 | 1.16257464 | 1.86068221 | 0.97097147 | 0.63165967 | 0.51182015 | 1.64206289 |
| ENSMUSG00 | Gm17415 | 800 | 0 | 0 | 0.06046691 | 0.05695131 | 0 | 0 | 0 | 0 | 0 | 0 | 0.0597187 | 0 | 0.0616649 | 0 | 0 | 0.09572652 | 0 | 0.13752277 |
| ENSMUSG00 | Gm17553 | 28 | 0 | 0 | 3.45525217 | 0 | 5.15699182 | 0 | 1.68973853 | 4.09204311 | 0 | 0 | 3.41249703 | 7.97366608 | 3.52370838 | 0 | 4.41445242 | 0 | 1.36879632 | 0 |
| ENSMUSG00 | Fryl | 1637 | 0 | 0 | 0 | 0 | 0 | 0 | 0 | 0 | 0 | 0 | 0 | 0 | 0 | 0 | 0 | 0 | 0 | 0 |
| ENSMUSG00 | Gm17176 | 164 | 0 | 0 | 0 | 0 | 0 | 0 | 0 | 0 | 0 | 0 | 0 | 0 | 0 | 0 | 0 | 0 | 0 | 0 |
| ENSMUSG00 | Gm3089 | 1845 | 0 | 0 | 0 | 0 | 0 | 0 | 0 | 0 | 0 | 0 | 0 | 0 | 0 | 0 | 0 | 0 | 0 | 0 |
| ENSMUSG00 | Traj36 | 62 | 0 | 0 | 0 | 0 | 0 | 0 | 0 | 0 | 0 | 0 | 0 | 0 | 0 | 0 | 0 | 0 | 0 | 0 |
| ENSMUSG00 | Vmn1r151 | 917 | 0 | 0 | 0 | 0 | 0 | 0 | 0 | 0 | 0 | 0 | 0 | 0 | 0 | 0 | 0 | 0 | 0 | 0 |
| ENSMUSG00 | Gm13693 | 2725 | 0 | 0 | 0 | 0 | 0 | 0 | 0 | 0 | 0 | 0 | 0 | 0 | 0 | 0 | 0 | 0 | 0 | 0 |
| ENSMUSG00 | Gm17611 | 230 | 0 | 0 | 0 | 0 | 0 | 0 | 0 | 0 | 0 | 0 | 0 | 0 | 0 | 0 | 0 | 0 | 0 | 0 |
| ENSMUSG00 | Kcnb2 | 4064 | 0 | 0 | 0 | 0 | 0 | 0 | 0 | 0 | 0 | 0 | 0 | 0 | 0 | 0 | 0 | 0 | 0 | 0 |
| ENSMUSG00 | Gm17057 | 589 | 0.64948486 | 1.75331022 | 1.39617999 | 1.23765146 | 0.98061644 | 0.84130278 | 1.20490693 | 1.29685577 | 0.13985235 | 0.96062553 | 1.21667976 | 0.85287091 | 0.33502151 | 0 | 0.20985512 | 0.78011424 | 0.52056091 | 1.58769918 |
| ENSMUSG00 | Gm3233 | 692 | 0 | 0 | 0 | 0 | 0 | 0 | 0 | 0 | 0 | 0 | 0 | 0 | 0 | 0 | 0 | 0 | 0 | 0 |
| ENSMUSG00 | Gm17735 | 1097 | 0 | 0 | 0 | 0 | 0 | 0 | 0 | 0 | 0 | 0 | 0 | 0 | 0 | 0 | 0 | 0 | 0 | 0 |
| ENSMUSG00 | Gm7352 | 1191 | 0 | 0 | 0 | 0 | 0 | 0 | 0 | 0 | 0 | 0 | 0 | 0 | 0 | 0 | 0 | 0 | 0 | 0 |
| ENSMUSG00 | Gm3486 | 878 | 0 | 0 | 0 | 0 | 0 | 0 | 0 | 0 | 0 | 0 | 0 | 0 | 0 | 0 | 0 | 0 | 0 | 0 |
| ENSMUSG00 | Gm17734 | 948 | 0 | 0 | 0 | 0 | 0.15231621 | 0 | 0 | 0 | 0 | 0 | 0 | 0 | 0 | 0 | 0 | 0 | 0 | 0 |
| ENSMUSG00 | Gm6476 | 795 | 0.16039689 | 0.16761205 | 0.30423604 | 0.22923796 | 0.2421732 | 0.37398289 | 0.35707682 | 0.48040758 | 0.10361388 | 0.05930906 | 0.12018857 | 0.28083352 | 0.18615818 | 0.23368094 | 0.10365171 | 0.86695715 | 0.33746425 | 0.27677538 |
| ENSMUSG00 | Gm7957 | 395 | 0 | 0 | 0 | 0 | 0 | 0 | 0 | 0 | 0 | 0 | 0 | 0 | 0 | 0 | 0 | 0 | 0 | 0 |
| ENSMUSG00 | Sfta2 | 1281 | 0.03318125 | 0 | 0 | 0.07113356 | 0.18786855 | 0.0773657 | 0 | 0.05962904 | 0 | 0.07361546 | 0 | 0 | 0 | 0.25379283 | 0.16081794 | 0.02989118 | 0 | 0 |
| ENSMUSG00 | Gm17554 | 3524 | 0 | 0 | 0 | 0 | 0 | 0 | 0.1476843 | 0 | 0 | 0 | 0 | 0 | 0.11199073 | 0 | 0.0233834 | 0 | 0 | 0 |
| ENSMUSG00 | Gm3510 | 1453 | 0 | 0 | 0 | 0 | 0 | 0 | 0 | 0 | 0 | 0 | 0 | 0 | 0 | 0 | 0 | 0 | 0 | 0 |
| ENSMUSG00 | Gm3424 | 1753 | 0 | 0 | 0 | 0 | 0 | 0 | 0 | 0 | 0 | 0 | 0 | 0 | 0 | 0 | 0 | 0 | 0 | 0 |
| ENSMUSG00 | Gm9140 | 305 | 0 | 0 | 0 | 0 | 0 | 0 | 0 | 0 | 0 | 0 | 0 | 0 | 0 | 0 | 0 | 0 | 0 | 0 |
| ENSMUSG00 | Gm7782 | 517 | 0 | 0.064435 | 0 | 0 | 0 | 0 | 0 | 0 | 0 | 0 | 0 | 0 | 0 | 0 | 0 | 0 | 0 | 0 |
| ENSMUSG00 | Krtap27-1 | 608 | 0 | 0 | 0 | 0 | 0 | 0 | 0 | 0 | 0 | 0 | 0 | 0 | 0 | 0 | 0 | 0 | 0 | 0 |
| ENSMUSG00 | Gm6202 | 474 | 5.7390955 | 2.8112147 | 3.87804674 | 2.11464763 | 3.35095671 | 3.24079059 | 4.5915258 | 3.14241286 | 3.64943811 | 3.9789623 | 3.22531365 | 3.29712775 | 6.55677382 | 5.09513195 | 4.52000191 | 9.12835155 | 4.36628699 | 4.17790684 |
| ENSMUSG00 | 6330403N20R | 1187 | 0 | 0 | 0.04075276 | 0.03838336 | 0 | 0.0417462 | 0.11957712 | 0 | 0.17348996 | 0.07944516 | 0.32198793 | 0.09404492 | 0.41560166 | 0 | 0.10413199 | 0.12903322 | 0 | 0.23171486 |
| ENSMUSG00 | Vmn2r35 | 2580 | 0 | 0 | 0 | 0 | 0 | 0 | 0 | 0 | 0 | 0 | 0 | 0 | 0 | 0 | 0 | 0 | 0 | 0 |
| ENSMUSG00 | Gm17061 | 496 | 0 | 0 | 0 | 0 | 0 | 0 | 0 | 0 | 0 | 0 | 0 | 0 | 0 | 0 | 0 | 0 | 0 | 0 |
| ENSMUSG00 | Gm17639 | 1226 | 0 | 0 | 0 | 0 | 0 | 0 | 0 | 0 | 0 | 0 | 0 | 0 | 0 | 0 | 0 | 0 | 0 | 0 |
| ENSMUSG00 | Gm5919 | 3836 | 0 | 0 | 0 | 0 | 0 | 0 | 0 | 0 | 0 | 0 | 0 | 0 | 0 | 0 | 0 | 0 | 0 | 0 |
| ENSMUSG00 | Gypc | 4203 | 0 | 0 | 0.04603715 | 0 | 0 | 0.01178985 | 0.03377065 | 0.02726082 | 0.01959863 | 0 | 0.5456098 | 0.22575928 | 0 | 0.35360714 | 0.02940868 | 0 | 0 | 0.18323281 |
| ENSMUSG00 | Gm13691 | 2725 | 0 | 0 | 0 | 0 | 0 | 0 | 0 | 0 | 0 | 0 | 0 | 0 | 0 | 0 | 0 | 0 | 0 | 0 |
| ENSMUSG00 | Gm7742 | 1526 | 0 | 0 | 0 | 0 | 0 | 0 | 0 | 0 | 0 | 0 | 0 | 0 | 0 | 0 | 0 | 0 | 0 | 0 |
| ENSMUSG00 | Gm17380 | 48 | 0 | 0 | 0 | 0 | 0 | 0 | 0 | 1.5913501 | 0 | 0 | 0 | 0 | 1.02774828 | 0 | 0 | 0 | 0 | 0 |
| ENSMUSG00 | Gm5538 | 1298 | 0 | 0 | 0 | 0 | 0 | 0 | 0 | 0 | 0 | 0 | 0 | 0 | 0 | 0 | 0 | 0 | 0 | 0 |
| ENSMUSG00 | Gm17381 | 1352 | 0 | 0 | 0 | 0 | 0 | 0 | 0 | 0 | 0 | 0 | 0 | 0 | 0 | 0 | 0 | 0 | 0 | 0 |
| ENSMUSG00 | Gm6803 | 493 | 0 | 0 | 0 | 0 | 0 | 0 | 0 | 0 | 0.08354263 | 0 | 0 | 0 | 0 | 0 | 0 | 0.07766857 | 0 | 0 |
| ENSMUSG00 | Gm17697 | 281 | 0 | 0 | 0 | 0 | 0 | 0 | 0 | 0 | 0 | 0 | 0 | 0 | 0 | 0 | 0 | 0.13626551 | 0 | 0 |
| ENSMUSG00 | Gm17696 | 2261 | 0 | 0 | 0 | 0 | 0 | 0 | 0 | 0 | 0 | 0 | 0 | 0 | 0 | 0 | 0 | 0 | 0 | 0 |
| ENSMUSG00 | Gm8214 | 416 | 0 | 0 | 0 | 0 | 0 | 0 | 0 | 0 | 0 | 0 | 0 | 0 | 0.35575902 | 0 | 0 | 0 | 0 | 0 |
| ENSMUSG00 | Gm4675 | 3126 | 0.29914071 | 0.21313432 | 0.17022036 | 0.33522202 | 0.06158915 | 0.12681442 | 0 | 0 | 0 | 0.07541699 | 0.04584929 | 0 | 0 | 0.13371618 | 0 | 0 | 0 | 0 |
| ENSMUSG00 | Gm8540 | 727 | 0 | 0 | 0 | 0 | 0 | 0 | 0 | 0 | 0 | 0 | 0 | 0 | 0 | 0 | 0 | 0 | 0 | 0 |
| ENSMUSG00 | Scn8a | 1069 | 0.23856974 | 0.15581335 | 0.18100479 | 0.21310124 | 0.09005037 | 0.3708343 | 0 | 0.28581779 | 0 | 0.1323219 | 0 | 0.30885187 | 0.23073862 | 0.69514069 | 0 | 0.42982908 | 0.17926238 | 0.46312625 |
| ENSMUSG00 | Gm17222 | 846 | 0 | 0 | 0 | 0 | 0 | 0 | 0 | 0 | 0 | 0 | 0 | 0 | 0 | 0 | 0 | 0 | 0 | 0 |
| ENSMUSG00 | Gm4775 | 404 | 0 | 0 | 0 | 0 | 0 | 0 | 0 | 0 | 0 | 0 | 0 | 0 | 0 | 0 | 0 | 0 | 0 | 0 |
| ENSMUSG00 | Gm9602 | 2057 | 0 | 0 | 0 | 0 | 0 | 0 | 0 | 0 | 0 | 0 | 0 | 0 | 0 | 0 | 0 | 0 | 0 | 0 |
| ENSMUSG00 | Gm17487 | 3072 | 0.51194385 | 0.40122952 | 0.37791821 | 0.51908742 | 0.43870243 | 0.38713073 | 1.12429218 | 0.74594536 | 1.05915845 | 0.92091217 | 0.34213837 | 0.56324399 | 0.69051837 | 0.69545183 | 0.87177771 | 0.6481483 | 0.88579658 | 0.26859915 |
| ENSMUSG00 | Gm10814 | 1928 | 0.39683256 | 0.24189861 | 0.25090005 | 0.18904998 | 0.24964691 | 0.33412113 | 0.39263634 | 0.27733072 | 0.32043451 | 0.39129214 | 0.42125223 | 0.57900065 | 0.69085154 | 0.28907108 | 0.70521335 | 0.19860274 | 0.53672719 | 0.22825355 |
| ENSMUSG00 | Gm17694 | 782 | 0 | 0 | 0 | 0 | 0 | 0 | 0 | 0 | 0 | 0 | 0 | 0 | 0 | 0 | 0 | 0 | 0 | 0 |
| ENSMUSG00 | Gm6576 | 954 | 0 | 0.03491918 | 0.05070601 | 0 | 0.15135825 | 0.05194207 | 0.09918801 | 0.04003396 | 0 | 0 | 0.10015715 | 0 | 0 | 0 | 0.04318821 | 0.04013691 | 0.12052295 | 0.28830769 |
| ENSMUSG00 | Gm17488 | 57 | 0 | 0 | 0 | 0 | 0 | 0 | 0 | 0 | 0 | 0 | 0 | 0 | 0 | 4.07404275 | 0 | 0 | 0 | 0 |
| ENSMUSG00 | Cdr1 | 2312 | 0 | 0.21613037 | 0 | 0 | 0 | 0 | 0 | 0 | 0 | 0 | 0 | 0 | 0 | 0 | 0 | 0 | 0 | 0 |
| ENSMUSG00 | Gm8265 | 471 | 0 | 0 | 0 | 0 | 0 | 0 | 0 | 0 | 0 | 0 | 0 | 0 | 0 | 0 | 0 | 0 | 0 | 0 |
| ENSMUSG00 | Gm17489 | 2859 | 0.13380433 | 0.20973491 | 0.11843816 | 0.35059216 | 0.20202276 | 0.48530134 | 0.52955779 | 0.06679329 | 0.46098933 | 0.19790432 | 0.15039336 | 0.09761396 | 0.53490362 | 0.29240769 | 0.01441118 | 0.41518322 | 0.50940863 | 0.01924068 |
| ENSMUSG00 | Rpl28-ps4 | 392 | 0 | 0 | 0 | 0 | 0 | 0 | 0 | 0 | 0 | 0 | 0 | 0 | 0 | 0 | 0 | 0 | 0 | 0 |
| ENSMUSG00 | Prlh | 259 | 0.49233795 | 0.77172728 | 0 | 0.17591137 | 0.55751263 | 0 | 0 | 0.14746101 | 0.79510648 | 0.36409809 | 0.7378372 | 0 | 1.52376578 | 1.61388721 | 0.15907937 | 0.29568037 | 0.44393394 | 0.84956149 |
| ENSMUSG00 | A730015C16R | 1220 | 0.31356277 | 0.54611302 | 0.23790261 | 0.29876095 | 0.31561917 | 0.20308497 | 0.5041515 | 0.65741021 | 0.43887271 | 0.30918494 | 0.31327842 | 0.41175489 | 0.36392398 | 1.06592987 | 0.57412004 | 0.18831446 | 0.25131998 | 0.09017886 |
| ENSMUSG00 | Gm17158 | 492 | 0 | 0 | 0 | 0 | 0 | 0 | 0 | 0 | 0 | 0 | 0 | 0 | 0 | 0 | 0 | 0 | 0 | 0 |
| ENSMUSG00 | Snrpe | 1329 | 43.4646608 | 33.8643492 | 42.0402013 | 36.784801 | 45.125942 | 37.2484431 | 47.3126789 | 37.272796 | 45.49421 | 40.9774735 | 41.4481017 | 35.1945261 | 41.5739258 | 48.5059393 | 42.4106309 | 44.1105492 | 41.0947879 | 43.2539628 |
| ENSMUSG00 | Gm17565 | 1316 | 0.29068889 | 0.02531375 | 0.18379001 | 0.10386256 | 0.07314882 | 0.07530811 | 0.17975942 | 0.05804316 | 0.21907721 | 0.10748641 | 0 | 0 | 0.03748626 | 0.10587558 | 0.09392452 | 0.02909621 | 0.02912333 | 0.08360047 |
| ENSMUSG00 | Gm8893 | 1238 | 0 | 0 | 0 | 0 | 0 | 0 | 0 | 0 | 0 | 0 | 0 | 0 | 0 | 0 | 0 | 0 | 0.03095824 | 0 |
| ENSMUSG00 | Olfr753-ps1 | 1801 | 0 | 0 | 0 | 0 | 0 | 0.05502802 | 0 | 0 | 0 | 0 | 0 | 0 | 0 | 0 | 0 | 0 | 0 | 0 |
| ENSMUSG00 | Gm17202 | 948 | 0 | 0 | 0 | 0 | 0 | 0 | 0.04990789 | 0 | 0 | 0 | 0 | 0 | 0 | 0.34294157 | 0 | 0 | 0 | 0 |
| ENSMUSG00 | Gm17137 | 5326 | 0.01596139 | 0.25019072 | 0.29972334 | 0.25663375 | 0.21689188 | 0.24190219 | 0.15101681 | 0.21512806 | 0.17786141 | 0.10623516 | 0.08073125 | 0.47159309 | 0.31492399 | 0.07848231 | 0.06962336 | 0.43855183 | 0.07915683 | 0.3408375 |
| ENSMUSG00 | Gm17190 | 848 | 0 | 0 | 0.22817703 | 0 | 0 | 0.11686965 | 0 | 0.09007642 | 0 | 0.05560224 | 0 | 0.06582036 | 0 | 0 | 0.09717348 | 0.4966942 | 0.40676494 | 0.12973846 |
| ENSMUSG00 | Gm17207 | 681 | 0 | 0.04891761 | 0 | 0 | 0 | 0 | 0 | 0 | 0.06047947 | 0 | 0 | 0 | 0 | 0.06819983 | 0 | 0 | 0.16883831 | 0 |
| ENSMUSG00 | Gm7358 | 1526 | 0 | 0 | 0 | 0 | 0 | 0 | 0 | 0 | 0 | 0 | 0 | 0 | 0 | 0 | 0 | 0 | 0 | 0 |
| ENSMUSG00 | Ighd6-1 | 28 | 0 | 0 | 0 | 0 | 0 | 0 | 0 | 0 | 0 | 0 | 0 | 0 | 0 | 0 | 0 | 0 | 0 | 0 |
| ENSMUSG00 | A430057M04R | 5891 | 0.01443055 | 0.03958416 | 0.14780573 | 0.05413806 | 0.0490225 | 0 | 0.12850159 | 0.07779814 | 0.03495715 | 0.01600771 | 0 | 0.05684841 | 0.18423055 | 0.1419103 | 0.06294585 | 0.10399758 | 0.06505907 | 0.09337822 |
| ENSMUSG00 | 4930405O22R | 1008 | 0.54818183 | 0.26438805 | 0.33592729 | 0.45199449 | 0.1909997 | 0.04915946 | 0.28162309 | 0.15155715 | 0.28601747 | 0.04677649 | 0.56874951 | 0.16611804 | 0.24470197 | 0.09215097 | 0.24524736 | 0.15194685 | 0.03802212 | 0.98230548 |
| ENSMUSG00 | Gm17465 | 2399 | 0 | 0 | 0 | 0 | 0 | 0 | 0 | 0.11144094 | 0 | 0 | 0 | 0 | 0 | 0 | 0 | 0 | 0 | 0 |
| ENSMUSG00 | Gm17466 | 1280 | 0 | 0 | 0 | 0 | 0 | 0 | 0 | 0 | 0 | 0 | 0 | 0 | 0 | 0 | 0 | 0 | 0 | 0 |
| ENSMUSG00 | Gm17041 | 477 | 0 | 0 | 0 | 0 | 0 | 0 | 0 | 0 | 0 | 0 | 0 | 0 | 0 | 0 | 0 | 0 | 0 | 0 |
| ENSMUSG00 | Gm17434 | 1685 | 0 | 0 | 0 | 0 | 0 | 0 | 0 | 0 | 0 | 0 | 0 | 0 | 0 | 0 | 0 | 0 | 0 | 0 |
| ENSMUSG00 | Vmn2r109 | 2571 | 0 | 0 | 0 | 0 | 0 | 0 | 0 | 0 | 0 | 0 | 0 | 0 | 0 | 0 | 0 | 0 | 0 | 0 |
| ENSMUSG00 | Gm2636 | 645 | 0 | 0 | 0 | 0 | 0 | 0 | 0 | 0 | 0 | 0 | 0 | 0 | 0 | 0 | 0 | 0 | 0 | 0 |
| ENSMUSG00 | Traj61 | 71 | 0 | 0 | 0 | 0 | 0 | 0 | 0 | 0 | 0 | 0 | 0 | 0 | 0 | 0 | 0 | 0 | 0 | 0 |
| ENSMUSG00 | Gm13289 | 1555 | 0 | 0 | 0 | 0 | 0 | 0 | 0 | 0 | 0 | 0 | 0 | 0 | 0 | 0 | 0 | 0 | 0 | 0 |
| ENSMUSG00 | Gm17055 | 2230 | 0.895849 | 1.46397472 | 0.91107098 | 0.65379077 | 1.33819698 | 0.93328018 | 0.67892633 | 0.44529258 | 0.72030229 | 0.63431439 | 0.94264492 | 1.15135447 | 1.48216971 | 0.7081613 | 0.9792298 | 0.37775487 | 0.87652069 | 0.51802298 |
| ENSMUSG00 | Gm3027 | 448 | 0 | 0 | 0 | 0 | 0 | 0 | 0 | 0 | 0 | 0 | 0 | 0 | 0 | 0 | 0 | 0 | 0 | 0 |
| ENSMUSG00 | Gm8323 | 1200 | 0 | 0 | 0 | 0 | 0 | 0 | 0 | 0 | 0 | 0 | 0 | 0 | 0 | 0 | 0 | 0 | 0 | 0 |
| ENSMUSG00 | Gm7137 | 690 | 0 | 0 | 0 | 0 | 0 | 0 | 0 | 0 | 0 | 0 | 0 | 0 | 0 | 0 | 0 | 0 | 0 | 0 |
| ENSMUSG00 | Gm4144 | 920 | 0 | 0 | 0 | 0 | 0 | 0 | 0 | 0 | 0 | 0 | 0 | 0 | 0 | 0 | 0 | 0 | 0 | 0 |
| ENSMUSG00 | Vmn2r6 | 2736 | 0 | 0 | 0 | 0 | 0 | 0 | 0 | 0 | 0 | 0 | 0 | 0 | 0 | 0 | 0 | 0 | 0 | 0 |
| ENSMUSG00 | Gm17024 | 632 | 0.47078518 | 0.31626165 | 0.91848475 | 1.00926364 | 0.15231621 | 0.78406224 | 1.27265117 | 1.87336151 | 0.71685391 | 0.89526652 | 1.28508591 | 0.97147514 | 0.62445465 | 2.49857432 | 1.95577006 | 1.4540737 | 2.60764362 | 1.30559589 |
| ENSMUSG00 | Gm17169 | 350 | 0 | 0 | 0 | 0 | 0 | 0 | 0 | 0 | 0 | 0 | 0 | 0 | 0 | 0 | 0 | 0 | 0 | 0 |
| ENSMUSG00 | 4933406F09R | 1291 | 0 | 0 | 0 | 0 | 0 | 0 | 0 | 0 | 0 | 0 | 0 | 0 | 0 | 0 | 0 | 0 | 0 | 0 |
| ENSMUSG00 | Gm17260 | 631 | 0.40416966 | 0.05279381 | 0.15332339 | 0.79424958 | 0.1525576 | 0 | 0.59984379 | 0.24210715 | 0.26108726 | 0.29889511 | 0 | 0.88455884 | 0 | 0 | 1.04473042 | 0 | 0.06073898 | 1.04613198 |
| ENSMUSG00 | Olfr1381 | 937 | 0 | 0 | 0 | 0 | 0 | 0 | 0 | 0 | 0 | 0 | 0 | 0 | 0 | 0 | 0 | 0 | 0 | 0 |
| ENSMUSG00 | Gm9573 | 5518 | 0 | 0 | 0 | 0 | 0 | 0 | 0.02572273 | 0.17995695 | 0.02239209 | 0.00854489 | 0.00865802 | 0.03034559 | 0 | 0.00841683 | 0 | 0 | 0 | 0 |
| ENSMUSG00 | Gm17180 | 431 | 0 | 0 | 0 | 0 | 0 | 0 | 0.32932259 | 0 | 0.19112072 | 0 | 0 | 0.1295027 | 0 | 0 | 0 | 0 | 0.17784825 | 0 |
| ENSMUSG00 | Gm17570 | 1274 | 0 | 0 | 0 | 0 | 0 | 0 | 0 | 0 | 0 | 0.07401994 | 0 | 0 | 0 | 0 | 0 | 0 | 0 | 0 |
| ENSMUSG00 | Gm17569 | 1514 | 0 | 0 | 0 | 0 | 0 | 0 | 0 | 0 | 0 | 0 | 0 | 0 | 0 | 0 | 0 | 0 | 0 | 0 |
| ENSMUSG00 | Gm17571 | 73 | 0 | 0 | 0 | 0 | 0 | 0 | 0 | 0 | 0 | 0 | 0 | 0 | 0 | 0 | 0 | 0 | 0 | 0 |
| ENSMUSG00 | 4930539J05R | 1404 | 0.21192039 | 0.40336125 | 0.58571938 | 0.12980355 | 0.41138396 | 0.21176382 | 0.23588942 | 0.02720257 | 1.08539963 | 0.47016371 | 0.64652722 | 0.75534016 | 0.77300725 | 2.01786989 | 0.44018756 | 0.24545261 | 0.46406485 | 0.07836055 |
| ENSMUSG00 | Gm3327 | 1384 | 0 | 0 | 0 | 0 | 0 | 0 | 0 | 0 | 0 | 0 | 0 | 0 | 0 | 0 | 0 | 0 | 0 | 0 |
| ENSMUSG00 | Gm17145 | 477 | 0 | 0 | 0 | 0 | 0 | 0 | 0 | 0 | 0 | 0 | 0 | 0 | 0 | 0 | 0 | 0 | 0 | 0 |
| ENSMUSG00 | Gm17121 | 846 | 0 | 0 | 0 | 0 | 0 | 0 | 0 | 0 | 0 | 0 | 0 | 0 | 0 | 0 | 0 | 0 | 0 | 0 |
| ENSMUSG00 | Gm8164 | 1191 | 0 | 0 | 0 | 0 | 0 | 0 | 0 | 0 | 0 | 0 | 0 | 0 | 0 | 0 | 0 | 0 | 0 | 0 |
| ENSMUSG00 | AL626783.1 | 2287 | 0.0929278 | 0.30589015 | 0.12690913 | 0.25898276 | 0.08418351 | 0.30333986 | 0.08275064 | 0.23379695 | 0.10805382 | 0.14431785 | 0.3342367 | 0.26846187 | 0.06471174 | 0.08123146 | 0.07206219 | 0.05022817 | 0.16758328 | 0.12026477 |
| ENSMUSG00 | Gm3395 | 2695 | 0 | 0 | 0 | 0 | 0 | 0 | 0 | 0 | 0 | 0 | 0 | 0 | 0 | 0 | 0 | 0.01420802 | 0 | 0 |
| ENSMUSG00 | Gm2877 | 1203 | 0 | 0 | 0 | 0 | 0 | 0 | 0 | 0 | 0 | 0 | 0 | 0 | 0 | 0 | 0 | 0 | 0 | 0 |
| ENSMUSG00 | Gm5611 | 854 | 0.04977187 | 0.07801615 | 0 | 0.05335017 | 0 | 0 | 0 | 0.22360891 | 0 | 0 | 0.05594257 | 0 | 0.05776571 | 0 | 0 | 0 | 0 | 0 |
| ENSMUSG00 | Gm17186 | 453 | 0 | 0 | 0 | 0 | 0 | 0 | 0 | 0 | 0 | 0 | 0 | 0 | 0 | 0 | 0 | 0 | 0 | 0 |
| ENSMUSG00 | Gm17378 | 925 | 0 | 0 | 0 | 0 | 0 | 0 | 0 | 0 | 0 | 0 | 0 | 0 | 0 | 0 | 0 | 0 | 0 | 0 |
| ENSMUSG00 | E230013L22R | 1854 | 2.22384146 | 1.45541771 | 2.32213819 | 2.28542455 | 2.02496766 | 2.03128789 | 2.60296292 | 2.26600014 | 2.17706502 | 1.83109527 | 2.86031305 | 2.61918158 | 3.56552153 | 4.13337347 | 3.26679003 | 2.54031538 | 4.81662739 | 2.16594649 |
| ENSMUSG00 | Gm17052 | 288 | 0 | 0 | 0 | 0 | 0 | 0 | 0 | 0 | 0 | 0 | 0 | 0 | 0 | 0 | 0 | 0 | 0 | 0 |
| ENSMUSG00 | Gm17200 | 2019 | 0.1894733 | 0.0494991 | 0 | 0.13539686 | 0 | 0 | 0.04686744 | 0.01891649 | 0 | 0 | 0.02366268 | 0 | 0.39094139 | 0.13802106 | 0.10203456 | 0 | 0.09491406 | 0.08173716 |
| ENSMUSG00 | Gm3571 | 1140 | 0.11185573 | 0.35066204 | 0.04243292 | 0.35969246 | 0.08444197 | 0.04346731 | 0.4150235 | 0.36852318 | 0.25289966 | 0.08272053 | 0.12572357 | 0.04896111 | 0.30291528 | 0.77406812 | 0.43370059 | 0.10076476 | 0.30257603 | 0.82031124 |
| ENSMUSG00 | C330013E15R | 876 | 5.5800174 | 4.18312599 | 2.208837 | 4.73293954 | 5.10989601 | 4.46879675 | 6.85925825 | 4.97024415 | 5.87707131 | 5.27485036 | 4.90838615 | 8.4742958 | 8.61619103 | 8.64199343 | 7.10209468 | 6.42547864 | 7.43775168 | 6.21678261 |
| ENSMUSG00 | D930030I03R | 918 | 1.15754837 | 0.72577112 | 1.05388955 | 0.69483074 | 0.41945032 | 0.75570618 | 1.18539392 | 0.54085101 | 0.71784777 | 0.87316117 | 0.57246682 | 0.60801375 | 0.53738472 | 0.65770494 | 0.71810991 | 0.91763983 | 0.66799646 | 0.95876439 |
| ENSMUSG00 | Gm17630 | 4427 | 0.52807424 | 0.48159594 | 0.65561595 | 0.51458148 | 0.73932026 | 0.81711081 | 0.30993171 | 1.0093768 | 0.55820893 | 0.61774131 | 0.71225373 | 1.21036901 | 0.45688019 | 0.68192132 | 0.16752383 | 0.71789483 | 0.75319355 | 0.23609059 |
| ENSMUSG00 | Gm17629 | 2536 | 0.20112859 | 0.10508799 | 0.17167262 | 0.01796571 | 0.09489733 | 0.09769861 | 0.03731284 | 0.0150601 | 0.09744444 | 0 | 0.13187094 | 0 | 0.01945265 | 0.01831391 | 0.04874001 | 0.2264823 | 0.19646761 | 0.08676515 |
| ENSMUSG00 | Gm17628 | 1594 | 0 | 0 | 0 | 0 | 0 | 0 | 0 | 0 | 0 | 0 | 0 | 0 | 0 | 0.43705227 | 0 | 0 | 0 | 0 |
| ENSMUSG00 | Gm17395 | 2970 | 0 | 0 | 0.01628738 | 0 | 0 | 0 | 0 | 0.03857818 | 0 | 0.01587566 | 0 | 0.01879315 | 0 | 0 | 0.05549031 | 0.02578492 | 0.01290448 | 0.01852158 |
| ENSMUSG00 | Gm8379 | 1191 | 0 | 0 | 0 | 0 | 0 | 0 | 0 | 0 | 0 | 0 | 0 | 0 | 0 | 0 | 0 | 0 | 0 | 0 |
| ENSMUSG00 | Vmn2r60 | 2556 | 0 | 0 | 0 | 0 | 0 | 0 | 0 | 0 | 0 | 0 | 0 | 0 | 0 | 0 | 0 | 0 | 0 | 0 |
| ENSMUSG00 | Gm6724 | 1109 | 3.67943814 | 2.10270748 | 2.96609564 | 2.79364384 | 2.51726923 | 3.08308261 | 4.90618402 | 3.13391219 | 3.75098115 | 5.27203535 | 3.01555193 | 6.14022618 | 4.00349193 | 4.77423441 | 4.86691057 | 5.11001794 | 2.83386506 | 5.90269044 |
| ENSMUSG00 | Gm4419 | 1727 | 1.55056519 | 1.35026207 | 0.81229437 | 0.73868514 | 1.47712331 | 0.71732388 | 1.12323094 | 1.87976503 | 1.24012671 | 1.28319806 | 0.88523374 | 1.97148548 | 1.25686413 | 1.18328885 | 1.14515037 | 1.13075911 | 1.48689166 | 0.63704814 |
| ENSMUSG00 | A930033H14R | 2967 | 0.71629889 | 1.19014722 | 1.22278894 | 1.47416928 | 1.44379549 | 1.35280466 | 1.05245595 | 1.40309129 | 1.34650894 | 1.00117772 | 1.14324976 | 1.39209944 | 1.03086581 | 0.78267758 | 0.55546418 | 0.85176275 | 1.40801023 | 1.7984103 |
| ENSMUSG00 | Cfhr3 | 965 | 0 | 0 | 0 | 0 | 0 | 0 | 0 | 0 | 0 | 0 | 0 | 0 | 0 | 0 | 0 | 0 | 0 | 0 |
| ENSMUSG00 | Gm17060 | 823 | 0 | 0 | 0 | 0 | 0.11696701 | 0 | 0 | 0.04640632 | 0 | 0 | 0 | 0 | 0 | 0 | 0 | 0 | 0 | 0.13367948 |
| ENSMUSG00 | AL593857.1 | 1471 | 0 | 0.02264643 | 0 | 0 | 0 | 0 | 0.09649085 | 0.10385426 | 0.05599798 | 0.06410701 | 0.12991151 | 0.07588805 | 0.03353631 | 0.06314628 | 0.02800922 | 0.07809097 | 0.15632752 | 0.07479144 |
| ENSMUSG00 | Tex9 | 1467 | 0 | 0 | 0 | 0 | 0 | 0 | 0 | 0 | 0 | 0 | 0 | 0 | 0 | 0 | 0 | 0 | 0 | 0 |
| ENSMUSG00 | Gm8356 | 2368 | 0 | 0 | 0 | 0 | 0 | 0 | 0 | 0 | 0 | 0 | 0 | 0 | 0 | 0 | 0 | 0 | 0 | 0 |
| ENSMUSG00 | Gm17083 | 1365 | 0 | 0.07321515 | 0.2126309 | 0.10013416 | 0 | 0.10890711 | 0.79720997 | 0.08393935 | 0 | 0.10362792 | 0.3499997 | 0 | 0 | 0.40829967 | 0.03018429 | 0 | 0.16846724 | 0.28209798 |
| ENSMUSG00 | Olfr180 | 1140 | 0 | 0 | 0 | 0 | 0 | 0 | 0 | 0 | 0 | 0 | 0 | 0 | 0 | 0 | 0 | 0 | 0 | 0 |
| ENSMUSG00 | Gm17403 | 3268 | 0.57228511 | 0.43832756 | 0.54768073 | 0.93408506 | 0.35347802 | 0.90978091 | 1.47671152 | 0.66614655 | 0.64275162 | 0.44726799 | 1.16952163 | 0.32450967 | 0.31700436 | 0.42635331 | 0.63037876 | 0.23433664 | 0.45738237 | 0.90896321 |
| ENSMUSG00 | Olfr456 | 1051 | 0.0404426 | 0 | 0.09205239 | 0.39015166 | 0 | 0.04714818 | 0 | 0 | 0 | 0.04486271 | 0 | 0 | 0 | 0.04419038 | 0 | 0 | 0 | 0 |
| ENSMUSG00 | Gm17404 | 170 | 0 | 0 | 0 | 0 | 0 | 0 | 0 | 0 | 0 | 0 | 0 | 0 | 0 | 0 | 0 | 0 | 0 | 0 |
| ENSMUSG00 | Vmn1r-ps79 | 3054 | 0 | 0 | 0 | 0 | 0 | 0 | 0 | 0 | 0 | 0 | 0 | 0 | 0 | 0 | 0 | 0 | 0 | 0 |
| ENSMUSG00 | Gm8126 | 1759 | 0 | 0 | 0 | 0 | 0 | 0 | 0 | 0 | 0 | 0 | 0 | 0 | 0.2524089 | 0 | 0.11711642 | 0 | 0 | 0.06254589 |
| ENSMUSG00 | Gm2951 | 487 | 0 | 0 | 0 | 0 | 0 | 0 | 0 | 0 | 0 | 0 | 0 | 0 | 0 | 0 | 0 | 0 | 0 | 0 |
| ENSMUSG00 | Gm17069 | 489 | 0 | 0 | 0 | 0 | 0 | 0 | 0 | 0 | 0 | 0 | 0 | 0 | 0 | 0 | 0 | 0 | 0 | 0 |
| ENSMUSG00 | Gm6189 | 2084 | 0 | 0 | 0 | 0 | 0.09238373 | 0 | 0 | 0.03665298 | 0 | 0 | 0 | 0.08034884 | 0.02367175 | 0 | 0 | 0.4593403 | 0 | 0 |
| ENSMUSG00 | Gm17028 | 702 | 0 | 0 | 0 | 0 | 0 | 0 | 0 | 0 | 0 | 0 | 0 | 0 | 0 | 0 | 0 | 0 | 0 | 0 |
| ENSMUSG00 | Gm20425 | 3005 | 0 | 0 | 0 | 0 | 0 | 0 | 0 | 0 | 0 | 0 | 0 | 0 | 0 | 0 | 0 | 0 | 0 | 0 |
| ENSMUSG00 | Gm17447 | 1946 | 0 | 0 | 0 | 0 | 0 | 0 | 0 | 0 | 0 | 0 | 0 | 0 | 0 | 0 | 0 | 0 | 0 | 0 |
| ENSMUSG00 | Zfp712 | 3966 | 0.92169857 | 1.78071951 | 1.36607045 | 1.04539966 | 1.67478706 | 2.23649504 | 0.27438013 | 0.2792687 | 1.30849748 | 1.188873 | 1.44553581 | 0.90070661 | 1.06972892 | 1.3115829 | 1.29508837 | 1.0234 | 0.5508318 | 2.67694342 |
| ENSMUSG00 | Gm17182 | 857 | 0.44637874 | 1.01065957 | 0.39511635 | 0.15949024 | 0.05616327 | 0.34692696 | 2.81557366 | 0.93586984 | 0.81700206 | 0.22007329 | 0.16724023 | 1.43284081 | 0.63319847 | 0.54193801 | 0.19230598 | 0.89359644 | 0.1341644 | 1.66888772 |
| ENSMUSG00 | Gm3453 | 1751 | 0 | 0 | 0 | 0 | 0 | 0 | 0 | 0 | 0 | 0 | 0 | 0 | 0 | 0 | 0 | 0 | 0 | 0 |
| ENSMUSG00 | B230364G03 | 1372 | 0 | 0 | 0 | 0 | 0 | 0 | 0 | 0 | 0 | 0 | 0 | 0 | 0 | 0 | 0 | 0 | 0 | 0 |
| ENSMUSG00 | Gm17448 | 43 | 0 | 0 | 0 | 0 | 0 | 0 | 0 | 0 | 0 | 0 | 0 | 0 | 0 | 0 | 0 | 0 | 0 | 0 |
| ENSMUSG00 | 2310010G23 | 1018 | 0 | 0 | 0 | 0 | 0 | 0 | 0 | 0 | 0 | 0 | 0 | 0 | 0 | 0 | 0 | 0 | 0 | 0 |
| ENSMUSG00 | Gm17446 | 3803 | 0 | 0 | 0 | 0 | 0 | 0 | 0 | 0 | 0 | 0 | 0 | 0 | 0 | 0 | 0 | 0 | 0 | 0 |
| ENSMUSG00 | Psmd14 | 336 | 0 | 0 | 0 | 0 | 0 | 0 | 0 | 0 | 0 | 0 | 0 | 0 | 0 | 0 | 0 | 0 | 0 | 0 |
| ENSMUSG00 | Gm11168 | 668 | 0 | 0 | 0 | 0 | 0 | 0 | 0.07082736 | 0 | 0 | 0 | 0 | 0.08355638 | 0 | 0 | 0 | 0 | 0 | 0 |
| ENSMUSG00 | Vmn1r117 | 923 | 0 | 0 | 0 | 0 | 0 | 0 | 0 | 0 | 0 | 0 | 0 | 0 | 0 | 0 | 0 | 0 | 0 | 0 |
| ENSMUSG00 | Gm4064 | 1721 | 0 | 0 | 0 | 0 | 0 | 0 | 0 | 0 | 0 | 0 | 0 | 0 | 0 | 0 | 0 | 0 | 0 | 0 |
| ENSMUSG00 | Gm17645 | 1162 | 0.18289663 | 0.43002875 | 0.24977727 | 0.74497405 | 0.16568648 | 0.1705774 | 0 | 0.49301724 | 0 | 0.56808076 | 0.69894518 | 0.04803413 | 0.04245432 | 0 | 0.31911704 | 0.29657097 | 0.26386435 | 0 |
| ENSMUSG00 | Vmn2r120 | 2564 | 0 | 0 | 0 | 0 | 0 | 0 | 0 | 0 | 0 | 0 | 0 | 0 | 0 | 0 | 0 | 0 | 0 | 0 |
| ENSMUSG00 | Gm17559 | 1785 | 0.45243605 | 0.78383281 | 0.9485006 | 0.61258547 | 0.59322259 | 0.49969143 | 0.4505694 | 0.66328542 | 0.96909449 | 0.89810863 | 0.53529365 | 0.78173197 | 0.91201864 | 0.57242012 | 0.41547787 | 0.90095547 | 0.5797255 | 0.73961824 |
| ENSMUSG00 | Gm8114 | 481 | 0 | 0 | 0 | 0 | 0 | 0 | 0 | 0 | 0 | 0 | 0 | 0 | 0 | 0 | 0 | 0 | 0 | 0 |
| ENSMUSG00 | Prss47 | 868 | 0 | 0 | 0 | 0 | 0 | 0 | 0 | 0 | 0 | 0 | 0 | 0 | 0 | 0 | 0 | 0 | 0 | 0 |
| ENSMUSG00 | Zfp493 | 3270 | 0.16898082 | 0.67237034 | 0.48817324 | 0.27866082 | 0.70652365 | 0.37884353 | 0.92599739 | 0.65405949 | 0.49121533 | 0.70653959 | 1.05192569 | 0.59741535 | 0.13577592 | 0.09942159 | 0.07559919 | 0.42154797 | 0.09376464 | 0.45420363 |
| ENSMUSG00 | Gm17030 | 390 | 0 | 0 | 0 | 0 | 0 | 0 | 0 | 0 | 0 | 0 | 0 | 0 | 0 | 0 | 0 | 0 | 0 | 0 |
| ENSMUSG00 | Gm17724 | 1831 | 1.32320865 | 1.0734357 | 0.63406047 | 1.46810574 | 1.52465952 | 1.00133869 | 0.9302329 | 1.08465588 | 0.87726604 | 2.24036657 | 0.70449147 | 1.31079929 | 0.99687654 | 1.49655989 | 1.23762183 | 2.17488977 | 0.9628671 | 1.62233302 |
| ENSMUSG00 | Vmn2r45 | 3613 | 0 | 0 | 0 | 0 | 0 | 0 | 0 | 0 | 0 | 0 | 0 | 0 | 0 | 0 | 0 | 0 | 0 | 0 |
| ENSMUSG00 | Srcap | 3051 | 6.29706312 | 4.8697315 | 5.13701207 | 4.73380893 | 5.53730095 | 4.92116626 | 3.93884643 | 5.11985991 | 4.79226912 | 5.68713825 | 4.68197725 | 4.86626229 | 5.69152241 | 4.74944453 | 5.30718173 | 5.28362691 | 3.7685641 | 4.95821185 |
| ENSMUSG00 | Gm17568 | 2380 | 0.9108252 | 0.75583878 | 0.46747529 | 0.80401843 | 0.99095137 | 1.39497191 | 0.61625758 | 0.67398357 | 0.69221035 | 0.85188245 | 0.52191131 | 0.5159431 | 1.09856791 | 0.81960154 | 0.81364417 | 0.48265472 | 0.85348476 | 0.76273131 |
| ENSMUSG00 | Gad1-ps | 3277 | 0 | 0 | 0 | 0 | 0 | 0 | 0 | 0 | 0 | 0 | 0 | 0 | 0 | 0.01417275 | 0 | 0.01169554 | 0 | 0 |
| ENSMUSG00 | Gm17567 | 65 | 0.65392579 | 0.51250606 | 0 | 0 | 0 | 1.52469949 | 0 | 0 | 0 | 0 | 0 | 0 | 0 | 0 | 0 | 0 | 0 | 0 |
| ENSMUSG00 | Gm765 | 822 | 0 | 0 | 0 | 0 | 0 | 0 | 0 | 0 | 0 | 0 | 0 | 0 | 0 | 0 | 0 | 0 | 0 | 0 |
| ENSMUSG00 | 4930448E22R | 1581 | 0 | 0 | 0 | 0 | 0 | 0.03134265 | 0 | 0 | 0 | 0 | 0 | 0 | 0 | 0 | 0 | 0 | 0 | 0 |
| ENSMUSG00 | Gm2990 | 1107 | 0 | 0 | 0 | 0 | 0 | 0 | 0 | 0 | 0 | 0 | 0 | 0 | 0 | 0 | 0 | 0 | 0 | 0 |
| ENSMUSG00 | Gm5067 | 1008 | 0 | 0 | 0 | 0 | 0.04774992 | 0 | 0 | 0.41678217 | 0.32687711 | 0 | 0 | 0.16611804 | 0.14682118 | 0 | 0 | 0.03798671 | 0 | 0.05457253 |
| ENSMUSG00 | Gm17256 | 2594 | 0.45880683 | 0.60358752 | 0.41026124 | 0.42153626 | 0.38965705 | 0.03820565 | 0.5106997 | 0.16195699 | 0.42869542 | 0.25447565 | 0.49727212 | 0.30124105 | 0.51347794 | 0.44761071 | 0.36531834 | 0.75282227 | 0.87172379 | 0.50895087 |
| ENSMUSG00 | Gm340 | 3727 | 1.22029886 | 1.32286245 | 1.24600454 | 0.75792454 | 1.05897981 | 1.18330917 | 1.07903883 | 0.81979935 | 0.78461031 | 0.5313468 | 0.94857712 | 0.9584659 | 1.00596343 | 1.38322879 | 0.88439079 | 0.81163348 | 0.74040606 | 1.04793308 |
| ENSMUSG00 | Gm17082 | 1090 | 0 | 0 | 0.17751754 | 0.04179912 | 0 | 0 | 0 | 0 | 0.03778579 | 0 | 0.21915119 | 0 | 0 | 0 | 0 | 0 | 0.03516174 | 0 |
| ENSMUSG00 | Olfr111 | 1040 | 0 | 0 | 0 | 0 | 0 | 0 | 0 | 0 | 0 | 0 | 0 | 0 | 0 | 0 | 0 | 0 | 0 | 0 |
| ENSMUSG00 | Gm6721 | 1430 | 0 | 0 | 0 | 0 | 0 | 0 | 0.03308579 | 0.02670797 | 0.31681935 | 0 | 0 | 0.03903193 | 0 | 0 | 0 | 0 | 0 | 0 |
| ENSMUSG00 | D930019O06R | 3661 | 0.01161026 | 0.08189458 | 0.44924885 | 0.1368947 | 0.11832486 | 0 | 0.07754058 | 0 | 0 | 0 | 0 | 0 | 0 | 0 | 0 | 0 | 0 | 0 |
| ENSMUSG00 | 6030442K20R | 1538 | 0.16581993 | 0 | 0.18871338 | 0.0888707 | 0.12518056 | 0.12887577 | 0 | 0.12416256 | 0.56236465 | 0.3372287 | 0.21744129 | 0.39920175 | 0.32075369 | 0 | 0.08036714 | 0.04979273 | 0.37379353 | 0 |
| ENSMUSG00 | Gm17255 | 622 | 0 | 0 | 0 | 0 | 0 | 0 | 0 | 0 | 0 | 0 | 0 | 0 | 0 | 0 | 0 | 0 | 0 | 0 |
| ENSMUSG00 | Vmn2r101 | 2568 | 0 | 0 | 0 | 0 | 0 | 0 | 0 | 0 | 0 | 0 | 0 | 0 | 0 | 0 | 0 | 0 | 0 | 0 |
| ENSMUSG00 | Gm3221 | 437 | 0 | 0 | 0 | 0 | 0 | 0 | 0 | 0 | 0 | 0 | 0 | 0 | 0 | 0 | 0 | 0 | 0 | 0 |
| ENSMUSG00 | Gm3852 | 263 | 0 | 0 | 0 | 0 | 0 | 0 | 0 | 0 | 0 | 0 | 0 | 0 | 0 | 0 | 0 | 0 | 0 | 0 |
| ENSMUSG00 | Gm4169 | 1151 | 0 | 0 | 0 | 0 | 0 | 0 | 0 | 0 | 0 | 0 | 0 | 0 | 0 | 0 | 0 | 0 | 0 | 0 |
| ENSMUSG00 | Gm17221 | 480 | 0 | 0 | 0 | 0 | 0 | 0 | 0 | 0 | 0 | 0 | 0 | 0 | 0 | 0 | 0 | 0 | 0 | 0 |
| ENSMUSG00 | Gm9047 | 680 | 0.06250761 | 0 | 0.07113754 | 0.40200921 | 0.07078224 | 0.14574333 | 0 | 0 | 0 | 0 | 0.14051458 | 0 | 0.07254694 | 0.34150064 | 0 | 0.05630972 | 0 | 0 |
| ENSMUSG00 | Gm3015 | 1398 | 0 | 0 | 0 | 0 | 0 | 0 | 0 | 0 | 0 | 0 | 0 | 0 | 0 | 0 | 0 | 0 | 0 | 0 |
| ENSMUSG00 | 9330151L19R | 2544 | 4.0600463 | 6.46877742 | 6.19880931 | 6.01749642 | 5.07050139 | 5.57078687 | 10.2101654 | 7.80662313 | 7.30154033 | 5.13394057 | 6.91084305 | 6.77949675 | 4.61517151 | 3.88859694 | 2.93140001 | 5.62920093 | 7.4573573 | 6.07608451 |
| ENSMUSG00 | Vmn2r12 | 2490 | 0 | 0 | 0 | 0 | 0 | 0 | 0 | 0 | 0 | 0 | 0 | 0 | 0 | 0 | 0 | 0 | 0 | 0 |
| ENSMUSG00 | Gm17695 | 1685 | 0 | 0 | 0 | 0 | 0 | 0 | 0 | 0 | 0 | 0 | 0 | 0 | 0 | 0 | 0 | 0 | 0 | 0 |
| ENSMUSG00 | Gm8020 | 1401 | 0 | 0 | 0 | 0 | 0 | 0 | 0 | 0 | 0 | 0 | 0 | 0 | 0 | 0 | 0 | 0 | 0 | 0 |
| ENSMUSG00 | Gm3667 | 2182 | 0.01947992 | 0 | 0 | 0 | 0 | 0 | 0.04336634 | 0 | 0 | 0 | 0 | 0 | 0 | 0 | 0 | 0 | 0 | 0 |
| ENSMUSG00 | Gm17172 | 188 | 0 | 0 | 0 | 0 | 0 | 0 | 0 | 0 | 0 | 0 | 0 | 0 | 0 | 0 | 0 | 0 | 0 | 0 |
| ENSMUSG00 | Trim43c | 1832 | 0 | 0 | 0 | 0 | 0.05254577 | 0 | 0 | 0.08338952 | 0 | 0 | 0 | 0 | 0 | 0 | 0 | 0 | 0 | 0 |
| ENSMUSG00 | Gm17379 | 40 | 0 | 0 | 2.41867652 | 1.13902611 | 0 | 0 | 0 | 0 | 0 | 0 | 0 | 0 | 0 | 0 | 2.06007779 | 0 | 0 | 0 |
| ENSMUSG00 | Nsun4 | 3914 | 0.0542989 | 0.03404486 | 0.0123591 | 0.01164053 | 0.01229737 | 0 | 0.01208806 | 0.04878948 | 0.03156861 | 0 | 0 | 0.02852103 | 0 | 0.07119686 | 0.03158014 | 0 | 0 | 0 |
| ENSMUSG00 | Apold1 | 3282 | 0 | 0 | 0.10317328 | 0 | 0 | 0 | 0.07207904 | 0.08145851 | 0.22588583 | 0.01436645 | 0.20379324 | 0.0680264 | 0.13527948 | 0.09905808 | 0 | 0 | 0.10509953 | 0 |
| ENSMUSG00 | Gm9071 | 471 | 0 | 0 | 0 | 0 | 0 | 0 | 0 | 0 | 0 | 0 | 0 | 0 | 0 | 0 | 0 | 0 | 0 | 0 |
| ENSMUSG00 | Cyp4f40 | 1846 | 0 | 0 | 0 | 0 | 0 | 0 | 0.15377902 | 0 | 0.22311222 | 0 | 0 | 0.06047201 | 0 | 0.02515931 | 0 | 0 | 0 | 0 |
| ENSMUSG00 | Gm6486 | 431 | 0 | 0 | 0 | 0 | 0 | 0 | 0 | 0 | 0 | 0 | 0 | 0 | 0 | 0 | 0 | 0 | 0 | 0 |
| ENSMUSG00 | Gm17552 | 3875 | 0.02193816 | 0.13755001 | 0 | 0 | 0 | 0 | 0 | 0.03942442 | 0 | 0.03650377 | 0.01232902 | 0.10082829 | 0.06365409 | 0 | 0 | 0 | 0.09890657 | 0.05678359 |
| ENSMUSG00 | Gm17048 | 846 | 0 | 0 | 0 | 0 | 0 | 0 | 0 | 0 | 0 | 0 | 0 | 0 | 0 | 0 | 0 | 0 | 0 | 0 |
| ENSMUSG00 | Trp53-ps | 140 | 0 | 0 | 0 | 0.65087206 | 0 | 0 | 0 | 0.27280287 | 0 | 0 | 0 | 0 | 0 | 0 | 0.29429683 | 0 | 0 | 0 |
| ENSMUSG00 | Gm17233 | 696 | 0 | 0 | 0.0695022 | 0 | 0 | 0 | 0 | 0 | 0 | 0 | 0 | 0 | 0 | 0 | 0 | 0 | 0 | 0 |
| ENSMUSG00 | Gm8237 | 2180 | 0 | 0 | 0 | 0.18809605 | 0 | 0 | 0 | 0 | 0 | 0 | 0 | 0 | 0 | 0 | 0 | 0 | 0 | 0 |
| ENSMUSG00 | Gm17196 | 342 | 0 | 0 | 0 | 0 | 0.28147324 | 0 | 0.13834117 | 0.44669476 | 0 | 0 | 0 | 0 | 0 | 0 | 0 | 0.22392168 | 0 | 0.32169068 |
| ENSMUSG00 | Gm17173 | 2862 | 0 | 0 | 0 | 0 | 0 | 0 | 0 | 0 | 0 | 0 | 0 | 0 | 0 | 0 | 0 | 0 | 0 | 0 |
| ENSMUSG00 | Gm9513 | 297 | 0 | 0 | 0 | 0 | 0 | 0 | 0 | 0 | 0 | 0 | 0 | 0 | 0 | 0 | 0 | 0 | 0.25808954 | 0 |
| ENSMUSG00 | Gm17027 | 1396 | 0 | 0 | 0 | 0 | 0 | 0 | 0 | 0 | 0 | 0 | 0 | 0 | 0 | 0 | 0 | 0 | 0 | 0 |
| ENSMUSG00 | Gm8159 | 1945 | 0 | 0 | 0 | 0 | 0 | 0 | 0 | 0 | 0 | 0 | 0 | 0 | 0 | 0 | 0 | 0 | 0 | 0 |
| ENSMUSG00 | Gm8127 | 1398 | 0 | 0 | 0 | 0 | 0 | 0 | 0 | 0 | 0 | 0 | 0 | 0 | 0 | 0 | 0 | 0 | 0 | 0 |
| ENSMUSG00 | Zscan4d | 1988 | 0 | 0 | 0 | 0 | 0 | 0 | 0 | 0 | 0 | 0 | 0 | 0 | 0 | 0 | 0 | 0 | 0 | 0 |
| ENSMUSG00 | Vmn1r167 | 947 | 0 | 0 | 0 | 0 | 0 | 0 | 0 | 0 | 0 | 0 | 0 | 0 | 0 | 0 | 0 | 0 | 0 | 0 |
| ENSMUSG00 | Gm3371 | 1434 | 0 | 0 | 0 | 0 | 0 | 0 | 0 | 0 | 0 | 0 | 0 | 0 | 0 | 0 | 0 | 0 | 0 | 0 |
| ENSMUSG00 | Gm17376 | 82 | 0 | 0 | 0 | 0 | 0 | 0 | 0 | 0 | 0 | 0 | 0 | 0 | 0 | 0 | 0 | 0 | 0 | 0 |
| ENSMUSG00 | Gm14476 | 499 | 0 | 0 | 0 | 0 | 0 | 0 | 0 | 0 | 0 | 0 | 0 | 0 | 0 | 0 | 0 | 0 | 0 | 0 |
| ENSMUSG00 | Gm17632 | 3939 | 0.51796102 | 0.1945155 | 0.12280663 | 0.25446636 | 0.18328988 | 0.06290014 | 0.36034028 | 0.09695964 | 0.06273651 | 0.02394044 | 0.09702962 | 0.35425021 | 0.13776367 | 0.0353725 | 0.65897386 | 0.17497612 | 0.36973833 | 0.26533969 |
| ENSMUSG00 | Gm8378 | 2103 | 0.08084674 | 0.39601634 | 0.23002154 | 0.58494921 | 0.36619628 | 0.51838333 | 0.02249771 | 0.30873554 | 0.13709254 | 0.20178618 | 0.06815258 | 0.31849166 | 0.4222418 | 0.61837111 | 0.2546934 | 0.14566089 | 0.43738998 | 0.81088079 |
| ENSMUSG00 | Gm9625 | 1096 | 3.25769597 | 2.21883328 | 3.00127743 | 2.61892864 | 2.63499932 | 2.57710384 | 4.10100775 | 3.380167 | 2.93115714 | 2.53822216 | 3.05132034 | 2.75004177 | 4.32104385 | 3.30532739 | 3.45852476 | 2.8997449 | 3.07729391 | 3.11182903 |
| ENSMUSG00 | 4930542C12R | 1202 | 0 | 0 | 0 | 0 | 0 | 0 | 0 | 0 | 0 | 0 | 0 | 0 | 0 | 0 | 0 | 0 | 0 | 0 |
| ENSMUSG00 | Gm17633 | 1249 | 0 | 0 | 0 | 0 | 0 | 0 | 0 | 0 | 0 | 0 | 0 | 0 | 0 | 0 | 0 | 0 | 0 | 0 |
| ENSMUSG00 | Gm17421 | 267 | 0 | 0 | 0 | 0 | 0 | 0 | 0 | 0 | 0 | 0 | 0 | 0 | 0 | 0 | 0 | 0 | 0 | 0 |
| ENSMUSG00 | Gm3956 | 2554 | 0 | 0 | 0 | 0 | 0 | 0 | 0 | 0 | 0 | 0 | 0 | 0 | 0 | 0 | 0 | 0 | 0 | 0 |
| ENSMUSG00 | Gm17420 | 150 | 0 | 0 | 0 | 0 | 0 | 0 | 0 | 0 | 0 | 0 | 0 | 0 | 0 | 0 | 0 | 0 | 0 | 0 |
| ENSMUSG00 | Gm2959 | 617 | 0 | 0 | 0 | 0 | 0 | 0 | 0 | 0 | 0 | 0 | 0 | 0 | 0 | 0 | 0 | 0 | 0 | 0 |
| ENSMUSG00 | 4930579O11R | 1293 | 0 | 0 | 0 | 0 | 0 | 0 | 0 | 0 | 0 | 0 | 0 | 0 | 0 | 0 | 0 | 0 | 0 | 0 |
| ENSMUSG00 | Rps27 | 341 | 307.009527 | 218.145727 | 291.376045 | 247.445906 | 304.882566 | 256.482624 | 326.471359 | 237.778505 | 275.381982 | 269.906665 | 257.788632 | 262.710083 | 482.469044 | 440.605932 | 354.865014 | 303.854497 | 306.835163 | 412.810276 |
| ENSMUSG00 | Gm9593 | 466 | 0 | 0 | 0 | 0 | 0 | 0 | 0 | 0 | 0 | 0 | 0 | 0 | 0 | 0 | 0 | 0 | 0 | 0 |
| ENSMUSG00 | Gm6788 | 269 | 0 | 0.37151927 | 0.17982725 | 0.84685956 | 0 | 0.18421091 | 0.35176713 | 0.14197919 | 0.15310972 | 0.17528142 | 0 | 0 | 0.18339003 | 0.1726546 | 0.15316564 | 0 | 0 | 0.61348446 |
| ENSMUSG00 | Gm17727 | 327 | 14.9483035 | 17.7261272 | 18.9352045 | 26.8907693 | 20.6069398 | 17.8814146 | 2.02561928 | 2.68631577 | 1.63738442 | 0.57676701 | 0.43830237 | 0.3413802 | 89.6121066 | 197.706941 | 112.768784 | 27.9861014 | 28.9498328 | 68.4669922 |
| ENSMUSG00 | D930016D06R | 4504 | 2.15168298 | 1.26476574 | 2.52392976 | 2.34683887 | 3.13113979 | 2.13437618 | 2.71018454 | 2.69653285 | 2.84391794 | 2.38684732 | 3.22459755 | 3.06093887 | 2.38773489 | 1.86642536 | 1.948475 | 2.09136088 | 1.94014114 | 1.58774065 |
| ENSMUSG00 | Gm17079 | 1347 | 0 | 0 | 0 | 0 | 0 | 0 | 0 | 0 | 0 | 0 | 0 | 0 | 0 | 0 | 0 | 0 | 0 | 0 |
| ENSMUSG00 | 1700091H14R | 1397 | 0 | 0 | 0 | 0 | 0 | 0 | 0 | 0 | 0 | 0 | 0 | 0 | 0 | 0 | 0 | 0 | 0 | 0 |
| ENSMUSG00 | Gm3008 | 567 | 0 | 0 | 0 | 0 | 0 | 0 | 0 | 0 | 0 | 0 | 0 | 0 | 0 | 0 | 0 | 0 | 0 | 0 |
| ENSMUSG00 | Gm17213 | 1359 | 0 | 0 | 0 | 0 | 0 | 0.07292529 | 0 | 0 | 0 | 0 | 0 | 0 | 0 | 0 | 0 | 0 | 0 | 0 |
| ENSMUSG00 | Gm6871 | 2273 | 0.03740007 | 0 | 0.1915362 | 0.16035563 | 0.10587753 | 0 | 0.22896589 | 0.0840132 | 0.7247957 | 0 | 0.18916614 | 0.39289512 | 0 | 0 | 0 | 0 | 0 | 0 |
| ENSMUSG00 | Vmn1r126 | 965 | 0 | 0 | 0 | 0 | 0 | 0 | 0 | 0 | 0 | 0 | 0 | 0 | 0 | 0 | 0 | 0 | 0 | 0 |
| ENSMUSG00 | Gm17654 | 1398 | 0 | 0 | 0 | 0 | 0 | 0 | 0 | 0 | 0 | 0 | 0 | 0 | 0 | 0 | 0 | 0 | 0 | 0 |
| ENSMUSG00 | Gm17495 | 616 | 0 | 0 | 0 | 0 | 0 | 0 | 0 | 0 | 0 | 0 | 0 | 0 | 0 | 0 | 0 | 0 | 0 | 0 |
| ENSMUSG00 | Olfr247 | 962 | 0 | 0 | 0 | 0 | 0 | 0 | 0 | 0 | 0 | 0 | 0 | 0 | 0 | 0 | 0 | 0 | 0 | 0 |
| ENSMUSG00 | Gm3495 | 385 | 0 | 0 | 0 | 0 | 0 | 0 | 0 | 0 | 0 | 0 | 0 | 0 | 0 | 0 | 0 | 0 | 0 | 0 |
| ENSMUSG00 | Vmn2r63 | 2574 | 0 | 0 | 0 | 0 | 0 | 0 | 0 | 0 | 0 | 0 | 0 | 0 | 0 | 0 | 0 | 0 | 0 | 0 |
| ENSMUSG00 | Gm6401 | 1399 | 0 | 0 | 0 | 0 | 0 | 0 | 0 | 0 | 0 | 0 | 0 | 0 | 0 | 0 | 0 | 0 | 0 | 0 |
| ENSMUSG00 | Rpl31-ps4 | 389 | 0 | 0 | 0 | 0.11712351 | 0 | 0 | 0 | 0 | 0 | 0.24242007 | 0 | 0 | 0 | 0 | 0.1059166 | 0 | 0 | 0 |
| ENSMUSG00 | C230091D08R | 52 | 0 | 0.64063258 | 1.8605204 | 0.87617393 | 0 | 0.95293718 | 8.18873289 | 4.40681566 | 0.79204838 | 1.81348859 | 0 | 1.07337813 | 0 | 0 | 0.79233761 | 0.73635783 | 2.21113251 | 1.05786744 |
| ENSMUSG00 | Gm17635 | 167 | 10.9444465 | 16.9556647 | 13.0347836 | 15.0051344 | 14.1225405 | 12.7590871 | 16.4319484 | 16.2374884 | 19.7300674 | 12.4229398 | 16.8785781 | 18.7166294 | 22.4504534 | 13.6273071 | 17.7635451 | 18.572091 | 19.0483991 | 17.45798 |
| ENSMUSG00 | Gm13277 | 1555 | 0.02733452 | 0 | 0.09332514 | 0.08789912 | 0 | 0.03186671 | 0.06085232 | 0 | 0.23837855 | 0.030322 | 0 | 0.28715453 | 0.03172471 | 0.02986758 | 0.05299235 | 0.14774511 | 0 | 0.03537563 |
| ENSMUSG00 | Gm17178 | 354 | 0 | 0 | 0.68324195 | 0 | 0 | 0 | 0 | 0 | 0 | 0 | 0 | 0 | 0 | 0 | 0 | 0 | 0 | 0 |
| ENSMUSG00 | Gm17634 | 1277 | 3.92765135 | 4.06954698 | 3.67441852 | 3.49646229 | 5.05064822 | 3.6863819 | 5.00173191 | 4.57591039 | 5.28941941 | 4.72614723 | 5.20024998 | 4.50196809 | 7.99663812 | 6.00099798 | 6.61418869 | 5.30731207 | 5.10216952 | 6.54767754 |
| ENSMUSG00 | Gm10634 | 3668 | 0 | 0 | 0 | 0 | 0 | 0 | 0 | 0 | 0 | 0 | 0 | 0 | 0 | 0 | 0 | 0 | 0 | 0 |
| ENSMUSG00 | Gm17201 | 388 | 1.31459307 | 0.60100582 | 1.62076261 | 0.58712686 | 0.62025675 | 0.25542646 | 0.48775958 | 0.88590624 | 1.37996058 | 0.97217945 | 0.49252535 | 1.72625761 | 0 | 0.83790879 | 0.31856873 | 1.48030699 | 0.88901204 | 0.14177605 |
| ENSMUSG00 | Vmn2r56 | 2334 | 0 | 0 | 0 | 0 | 0 | 0 | 0 | 0 | 0 | 0 | 0 | 0 | 0 | 0 | 0.01765277 | 0 | 0 | 0 |
| ENSMUSG00 | Gm3127 | 1944 | 0 | 0 | 0 | 0 | 0 | 0 | 0 | 0 | 0 | 0 | 0 | 0 | 0 | 0 | 0 | 0 | 0 | 0 |
| ENSMUSG00 | Gm17686 | 293 | 0 | 0 | 0 | 0 | 0 | 0 | 0 | 0 | 0 | 0 | 0 | 0 | 0 | 0 | 0 | 0 | 0 | 0 |
| ENSMUSG00 | Gm17684 | 250 | 0 | 0 | 0 | 0 | 0 | 0 | 0 | 0 | 0 | 0 | 0 | 0 | 0 | 0 | 0 | 0 | 0 | 0 |
| ENSMUSG00 | Gm17685 | 2730 | 0.49822917 | 0.36607576 | 0.67333119 | 0.30040249 | 0.44076853 | 0.72604738 | 0.29462108 | 0.02797978 | 0.96554469 | 0.6044962 | 0.75249035 | 1.12449137 | 0.90351497 | 0.90166177 | 0.61877795 | 1.31843117 | 1.02484237 | 1.43063977 |
| ENSMUSG00 | Gm2824 | 5459 | 0 | 0 | 0 | 0 | 0 | 0 | 0 | 0 | 0 | 0 | 0 | 0 | 0 | 0 | 0.06792709 | 0 | 0.00702075 | 0 |
| ENSMUSG00 | Gm7970 | 1398 | 0 | 0 | 0 | 0 | 0 | 0 | 0 | 0 | 0 | 0 | 0 | 0 | 0 | 0 | 0 | 0 | 0 | 0 |
| ENSMUSG00 | Gm17501 | 1318 | 0 | 0.05055067 | 0 | 0.03456832 | 0.47474583 | 0 | 0 | 0 | 0.31249253 | 0 | 0 | 0 | 0 | 0 | 0 | 0 | 0 | 0 |
| ENSMUSG00 | Gm2930 | 471 | 0 | 0 | 0 | 0 | 0 | 0 | 0 | 0 | 0 | 0 | 0 | 0 | 0 | 0 | 0 | 0 | 0 | 0 |
| ENSMUSG00 | Vmn2r-ps69 | 2658 | 0 | 0 | 0 | 0 | 0 | 0 | 0 | 0 | 0 | 0 | 0 | 0 | 0 | 0 | 0 | 0 | 0 | 0 |
| ENSMUSG00 | Vmn2r74 | 2556 | 0 | 0 | 0 | 0 | 0 | 0 | 0 | 0 | 0 | 0 | 0 | 0 | 0 | 0 | 0 | 0 | 0 | 0 |
| ENSMUSG00 | Vmn1r61 | 902 | 0 | 0 | 0 | 0 | 0 | 0 | 0 | 0 | 0 | 0 | 0 | 0 | 0 | 0 | 0 | 0 | 0 | 0 |
| ENSMUSG00 | Gm7873 | 742 | 0.11456921 | 0 | 0 | 0 | 0 | 0 | 0 | 0.05147224 | 0 | 0 | 0 | 0 | 0 | 0 | 0 | 0 | 0 | 0 |
| ENSMUSG00 | Gm3235 | 693 | 0.55201527 | 1.00948164 | 0.97724304 | 0.13148931 | 0.20836331 | 0.85805599 | 0.34136132 | 0.16533508 | 0.41602541 | 0.13607707 | 0.68939334 | 1.7719258 | 0 | 0.06701889 | 1.248532 | 1.10506803 | 1.54853725 | 0.47626932 |
| ENSMUSG00 | Gm17110 | 732 | 3.77436672 | 2.86709335 | 3.63462318 | 2.30294896 | 5.32607352 | 3.1139696 | 4.26589728 | 4.90448883 | 4.72632148 | 3.22067645 | 5.67817129 | 4.19379978 | 5.99800634 | 4.18758165 | 3.26460416 | 5.75405302 | 3.66508304 | 5.86162613 |
| ENSMUSG00 | Gm17532 | 58 | 3.66423931 | 2.29744098 | 0.83402639 | 3.92767624 | 0.82986075 | 2.56307243 | 0.81573584 | 1.97546909 | 4.97078639 | 3.25177264 | 0 | 2.88701703 | 12.7582545 | 0 | 1.42074331 | 5.28146309 | 1.32159644 | 0 |
| ENSMUSG00 | Igkv2-95-2 | 122 | 0 | 0 | 0 | 0 | 0 | 0 | 0 | 0 | 0 | 0 | 0 | 0 | 0 | 0 | 0 | 0 | 0 | 0 |
| ENSMUSG00 | Traj60 | 53 | 0 | 0 | 0 | 0 | 0 | 0 | 0 | 0 | 0 | 0 | 0 | 0 | 0 | 0 | 0 | 0 | 0 | 0 |
| ENSMUSG00 | Gm17485 | 40 | 0 | 0 | 0 | 0 | 0 | 0 | 0 | 0 | 0 | 0 | 1.19437396 | 0 | 0 | 0 | 0 | 0 | 0 | 0 |
| ENSMUSG00 | Gm17531 | 1416 | 0.36021336 | 0.04705211 | 0.0683242 | 0.03217588 | 0 | 0 | 0 | 0.51246868 | 0 | 0 | 0.23617564 | 0.23650704 | 0.06967785 | 0.09839849 | 0 | 0.10816556 | 0.24359934 | 0.15539296 |
| ENSMUSG00 | Gm17116 | 1009 | 0 | 0 | 0 | 0 | 0.0477026 | 0 | 0.04689066 | 0 | 0 | 0 | 0 | 0 | 0 | 0 | 0 | 0 | 0 | 0 |
| ENSMUSG00 | Gm9597 | 598 | 0 | 0 | 0 | 0 | 0 | 0.16572821 | 0 | 0.31933447 | 0 | 0 | 0 | 0 | 0 | 0 | 0 | 0 | 0 | 0 |
| ENSMUSG00 | Gm17377 | 206 | 0 | 0.80856539 | 0 | 0 | 0.23365011 | 0.72164175 | 0 | 0 | 0.39986909 | 0.45777382 | 0 | 0 | 0.23947533 | 0.22545673 | 0.40001511 | 0 | 0.18604998 | 0 |
| ENSMUSG00 | Vmn1r158 | 886 | 0 | 0 | 0 | 0 | 0 | 0 | 0 | 0 | 0 | 0 | 0 | 0 | 0 | 0 | 0 | 0 | 0 | 0 |
| ENSMUSG00 | Gm8221 | 481 | 0 | 0 | 0 | 0 | 0 | 0 | 0 | 0 | 0 | 0 | 0 | 0 | 0 | 0 | 0 | 0 | 0 | 0 |
| ENSMUSG00 | Gm6650 | 767 | 0 | 0 | 0 | 0 | 0 | 0 | 0 | 0 | 0 | 0 | 0 | 0 | 0 | 0 | 0 | 0 | 0 | 0 |
| ENSMUSG00 | Vmn1r60 | 1834 | 0 | 0 | 0 | 0 | 0 | 0 | 0 | 0 | 0 | 0 | 0 | 0 | 0 | 0 | 0 | 0 | 0 | 0 |
| ENSMUSG00 | Gm7346 | 473 | 0 | 0 | 0 | 0 | 0 | 0 | 0 | 0 | 0 | 0 | 0 | 0 | 0 | 0 | 0 | 0 | 0 | 0 |
| ENSMUSG00 | Gm3141 | 496 | 0 | 0 | 0 | 0 | 0 | 0 | 0 | 0 | 0 | 0 | 0 | 0 | 0 | 0 | 0 | 0 | 0 | 0 |
| ENSMUSG00 | Gm17335 | 932 | 0.09121282 | 0 | 0 | 0 | 0 | 0 | 0.05076468 | 0 | 0 | 0 | 0 | 0 | 0 | 0 | 0 | 0 | 0 | 0 |
| ENSMUSG00 | Gm8046 | 612 | 0 | 0 | 0 | 0 | 0 | 0 | 0 | 0 | 0 | 0.07704363 | 0 | 0 | 0.16121542 | 0 | 0.0673228 | 0 | 0 | 0 |
| ENSMUSG00 | Klhl33 | 1599 | 0.18607644 | 0 | 0 | 0 | 0.18060759 | 0.18593896 | 0 | 0.07165554 | 0 | 0 | 0 | 0 | 0 | 0 | 0 | 0 | 0 | 0 |
| ENSMUSG00 | 4833407H14R | 1483 | 0 | 0 | 0 | 0.12288886 | 0 | 0 | 0.2233235 | 0.1030139 | 0 | 0.09538241 | 0.38658092 | 0 | 1.09774327 | 0.65767083 | 0.27782573 | 0.05163939 | 0 | 0 |
| ENSMUSG00 | Gm4862 | 437 | 0 | 0.07623088 | 0 | 0 | 0 | 0 | 0 | 0 | 0 | 0 | 0 | 0 | 0 | 0 | 0 | 0 | 0 | 0 |
| ENSMUSG00 | Gm8074 | 256 | 0 | 0 | 0 | 0 | 0 | 0 | 0 | 0 | 0 | 0 | 0 | 0 | 0 | 0 | 0 | 0 | 0 | 0 |
| ENSMUSG00 | Gm17413 | 70 | 0 | 0 | 0 | 0 | 0 | 0 | 0 | 0 | 0 | 0 | 0 | 0 | 0 | 0 | 0 | 0 | 0 | 0 |
| ENSMUSG00 | Gm17729 | 57 | 0 | 0 | 0 | 0 | 0 | 0 | 0 | 0 | 0 | 0 | 0 | 0 | 0 | 0 | 0 | 0 | 0 | 0 |
| ENSMUSG00 | Vmn2r70 | 2559 | 0 | 0 | 0 | 0 | 0 | 0 | 0 | 0 | 0 | 0 | 0 | 0 | 0 | 0 | 0 | 0 | 0 | 0 |
| ENSMUSG00 | Gm1758 | 468 | 0 | 0 | 0 | 0 | 0 | 0 | 0 | 0 | 0 | 0 | 0 | 0 | 0 | 0 | 0 | 0 | 0 | 0 |
| ENSMUSG00 | Gm17412 | 172 | 0 | 0 | 0 | 0 | 0 | 0 | 0 | 0 | 0 | 0 | 0 | 0 | 0 | 0 | 0 | 0 | 0 | 0 |
| ENSMUSG00 | Gm3261 | 480 | 0 | 0 | 0 | 0 | 0 | 0 | 0 | 0 | 0 | 0 | 0 | 0 | 0 | 0 | 0 | 0 | 0 | 0 |
| ENSMUSG00 | Samd15 | 1193 | 0.10688644 | 0.05584727 | 0 | 0.03819031 | 0 | 0 | 0 | 0.03201375 | 0 | 0.11856841 | 0.08009213 | 0.37428776 | 0.08270229 | 0.31144401 | 0 | 0.12838427 | 0.06425196 | 0 |
| ENSMUSG00 | Gm4651 | 1281 | 0.92907489 | 1.3002691 | 1.39720579 | 1.0314366 | 1.08963762 | 1.46994838 | 2.91779987 | 2.3255327 | 2.0577182 | 0.62573142 | 1.60368713 | 1.83002172 | 3.04232745 | 1.84906203 | 1.86548809 | 0.71738843 | 1.82506176 | 1.46003874 |
| ENSMUSG00 | Vmn1r88 | 950 | 0 | 0 | 0 | 0 | 0 | 0 | 0 | 0 | 0 | 0 | 0 | 0 | 0 | 0 | 0 | 0 | 0 | 0 |
| ENSMUSG00 | Gm17601 | 80 | 0 | 0 | 0 | 0 | 0 | 0 | 0 | 0 | 0 | 0 | 0 | 0 | 0 | 0 | 0 | 0 | 0 | 0 |
| ENSMUSG00 | Gm6729 | 1451 | 0 | 0 | 0 | 0 | 0 | 0 | 0 | 0 | 0 | 0 | 0 | 0 | 0 | 0 | 0 | 0 | 0 | 0 |
| ENSMUSG00 | Gm4450 | 1684 | 0 | 0 | 0 | 0 | 0 | 0 | 0 | 0 | 0 | 0 | 0 | 0 | 0 | 0 | 0 | 0 | 0 | 0 |
| ENSMUSG00 | Trbj2-6 | 47 | 0 | 0 | 0 | 0 | 0 | 0 | 0 | 0 | 0 | 0 | 0 | 0 | 0 | 0 | 0 | 0 | 0 | 0 |
| ENSMUSG00 | Gm17603 | 1170 | 0.21797526 | 0.02847256 | 0.20672449 | 0 | 0.08227679 | 0.21176382 | 0.48525825 | 0 | 0 | 0.28209822 | 0.12249989 | 0 | 0 | 0.27787061 | 0.28172004 | 0.03272701 | 0.16378759 | 0 |
| ENSMUSG00 | Gm8161 | 478 | 0 | 0 | 0 | 0 | 0 | 0 | 0 | 0 | 0 | 0 | 0 | 0 | 0 | 0 | 0 | 0 | 0 | 0 |
| ENSMUSG00 | 2700012I20R | 682 | 0.93486458 | 1.46537658 | 2.41158362 | 1.73693131 | 1.12919469 | 1.52581731 | 2.49744346 | 1.40001475 | 1.57016042 | 0.62222336 | 0.84061511 | 0.40920574 | 0.79567609 | 1.49819637 | 1.51032096 | 0.61759044 | 0.674363 | 0.48395109 |
| ENSMUSG00 | Olfr1344 | 1060 | 0 | 0 | 0 | 0 | 0 | 0 | 0 | 0 | 0 | 0 | 0 | 0 | 0 | 0 | 0 | 0 | 0 | 0 |
| ENSMUSG00 | Gm2221 | 445 | 0 | 0 | 0 | 0 | 0 | 0.11135446 | 0 | 0 | 0 | 0 | 0 | 0 | 0 | 0 | 0 | 0 | 0 | 0 |
| ENSMUSG00 | Gm17285 | 37 | 0 | 0 | 0 | 0 | 0 | 0 | 0 | 0 | 0 | 0 | 0 | 0 | 0 | 0 | 0 | 0 | 0 | 0 |
| ENSMUSG00 | Gm8297 | 1945 | 0 | 0 | 0 | 0 | 0 | 0 | 0 | 0 | 0 | 0 | 0 | 0 | 0 | 0 | 0 | 0 | 0 | 0 |
| ENSMUSG00 | Sox2ot | 8689 | 0.05870205 | 0.23003495 | 0.16144923 | 0.24644598 | 0.11632759 | 0.22811708 | 0.31581717 | 0.1582376 | 0.26070415 | 0.18450039 | 0.54433432 | 0.24410118 | 0.08516271 | 0.00534516 | 0.04741806 | 0 | 0.12350516 | 0.22158116 |
| ENSMUSG00 | Vmn1r135 | 923 | 0 | 0 | 0 | 0 | 0 | 0 | 0 | 0 | 0 | 0 | 0 | 0 | 0 | 0 | 0 | 0 | 0 | 0 |
| ENSMUSG00 | Vmn1r163 | 923 | 0 | 0 | 0 | 0 | 0 | 0 | 0 | 0 | 0 | 0 | 0 | 0 | 0 | 0 | 0 | 0 | 0 | 0 |
| ENSMUSG00 | Gm17474 | 57 | 0 | 0 | 0 | 0 | 0 | 0 | 0 | 0 | 0 | 0 | 0 | 0 | 0 | 0 | 0 | 0 | 0 | 0 |
| ENSMUSG00 | Gm17476 | 3613 | 0 | 0.02766086 | 0.01338874 | 0.06305154 | 0 | 0.01371512 | 0.09166586 | 0.01057083 | 0.04559814 | 0.03915087 | 0.03966922 | 0 | 0.013654 | 0.02570943 | 0.03421109 | 0.04239204 | 0.01060789 | 0.01522533 |
| ENSMUSG00 | Vmn1r116 | 923 | 0 | 0 | 0 | 0 | 0 | 0 | 0 | 0 | 0 | 0 | 0 | 0 | 0 | 0 | 0 | 0 | 0 | 0 |
| ENSMUSG00 | Gm17475 | 3214 | 0 | 0 | 0 | 0 | 0 | 0 | 0 | 0 | 0 | 0 | 0 | 0 | 0 | 0 | 0 | 0 | 0 | 0 |
| ENSMUSG00 | Gm7233 | 1756 | 0 | 0 | 0 | 0 | 0 | 0 | 0 | 0 | 0 | 0 | 0 | 0 | 0 | 0 | 0 | 0 | 0 | 0 |
| ENSMUSG00 | Gm7084 | 317 | 0 | 0 | 0 | 0.14372569 | 0 | 0 | 0 | 0.24096153 | 0 | 0.14874039 | 0 | 0 | 0 | 0 | 0 | 0.12079056 | 0 | 0.34706061 |
| ENSMUSG00 | Gm17094 | 3323 | 0 | 0 | 0 | 0 | 0 | 0 | 0 | 0 | 0 | 0 | 0 | 0 | 0 | 0 | 0 | 0 | 0 | 0 |
| ENSMUSG00 | 1700092M07 | 378 | 1.12447556 | 1.0575522 | 0.25594461 | 0.36159559 | 0.63666566 | 0.26218378 | 0.37549745 | 2.32387634 | 0.98063133 | 1.87105965 | 0.12638878 | 2.80554918 | 0.13050772 | 0.61433978 | 0.65399295 | 0.40519161 | 0.20278464 | 0.14552674 |
| ENSMUSG00 | Myl6 | 659 | 437.306667 | 365.430623 | 380.161629 | 339.391755 | 407.697114 | 356.945108 | 375.845029 | 343.615711 | 353.429054 | 342.504426 | 332.177329 | 319.05554 | 388.741497 | 373.385091 | 359.3726 | 374.597187 | 346.623262 | 354.178513 |
| ENSMUSG00 | Gm17674 | 86 | 0 | 0 | 0 | 0 | 0 | 0 | 0 | 0 | 0 | 1.09652798 | 0 | 0 | 0 | 0 | 0 | 0 | 0 | 0 |
| ENSMUSG00 | Gm17673 | 1696 | 0 | 0 | 0 | 0 | 0 | 0 | 0 | 0 | 0 | 0 | 0 | 0 | 0 | 0 | 0 | 0 | 0 | 0 |
| ENSMUSG00 | Gm3302 | 1845 | 0 | 0 | 0 | 0 | 0 | 0 | 0 | 0 | 0 | 0 | 0 | 0 | 0 | 0 | 0 | 0 | 0 | 0 |
| ENSMUSG00 | Lsm3 | 627 | 71.994413 | 58.2311516 | 61.7206129 | 60.0214077 | 61.3355774 | 51.5285204 | 83.8347469 | 74.7401559 | 77.1179737 | 81.9685272 | 77.2628515 | 70.2369343 | 97.4049659 | 89.1844995 | 84.045917 | 86.169134 | 82.0317232 | 91.0677078 |
| ENSMUSG00 | Gm17672 | 1868 | 0 | 0 | 0 | 0 | 0 | 0 | 0 | 0 | 0 | 0 | 0 | 0 | 0 | 0 | 0 | 0 | 0 | 0 |
| ENSMUSG00 | Vmn1r165 | 923 | 0 | 0 | 0 | 0 | 0 | 0 | 0 | 0 | 0 | 0 | 0 | 0 | 0 | 0 | 0 | 0 | 0 | 0 |
| ENSMUSG00 | Vmn2r-ps36 | 3460 | 0 | 0 | 0 | 0 | 0 | 0 | 0 | 0 | 0 | 0 | 0 | 0 | 0 | 0 | 0 | 0 | 0 | 0 |
| ENSMUSG00 | Orai2-ps | 3033 | 0 | 0 | 0 | 0.03004355 | 0 | 0 | 0 | 0 | 0 | 0 | 0 | 0 | 0 | 0 | 0 | 0 | 0 | 0 |
| ENSMUSG00 | Vmn2r98 | 2568 | 0 | 0 | 0 | 0 | 0 | 0 | 0 | 0 | 0 | 0 | 0 | 0 | 0 | 0 | 0 | 0 | 0 | 0 |
| ENSMUSG00 | Tnfrsf22 | 1779 | 0 | 0 | 0 | 0 | 0 | 0 | 0 | 0 | 0 | 0 | 0 | 0 | 0 | 0 | 0 | 0 | 0 | 0 |
| ENSMUSG00 | Gm17364 | 70 | 0 | 0 | 0 | 0 | 0 | 0 | 0 | 0 | 0 | 0 | 0 | 0 | 0 | 0 | 0 | 0 | 0 | 0 |
| ENSMUSG00 | Gm4340 | 797 | 0 | 0 | 0 | 0 | 0 | 0 | 0 | 0 | 0 | 0 | 0 | 0 | 0 | 0 | 0 | 0 | 0 | 0 |
| ENSMUSG00 | Gm17219 | 592 | 0 | 0 | 0.08171204 | 0 | 0.08130393 | 0 | 0 | 0 | 0 | 0.15929292 | 0 | 0 | 0 | 0 | 0 | 0 | 0 | 0.09292079 |
| ENSMUSG00 | Gm17365 | 294 | 0 | 0 | 0 | 0 | 0.32742805 | 0.16854671 | 2.73576715 | 3.76727779 | 2.24144303 | 2.72640121 | 1.78749845 | 3.60713466 | 0 | 0 | 0.14014135 | 0 | 0 | 0 |
| ENSMUSG00 | Gm3238 | 692 | 0 | 0 | 0 | 0 | 0 | 0 | 0 | 0 | 0 | 0 | 0 | 0 | 0 | 0 | 0 | 0 | 0 | 0 |
| ENSMUSG00 | Gm17134 | 435 | 0 | 0 | 0 | 0 | 0 | 0 | 0 | 0 | 0 | 0 | 0 | 0 | 0 | 0 | 0 | 0 | 0 | 0 |
| ENSMUSG00 | 4930570N19R | 564 | 0.52754651 | 0.05906542 | 0.08576867 | 0 | 0.8534029 | 0.52715674 | 0.33555091 | 0.88032133 | 0.80328311 | 0.08360054 | 0.59295161 | 0.19792788 | 0.26240382 | 0.65878138 | 0.51136683 | 0.1357823 | 0.67954427 | 0.48766939 |
| ENSMUSG00 | Rps13 | 1443 | 58.2347422 | 47.4183539 | 52.1616169 | 48.4342633 | 46.7309945 | 45.1572035 | 49.1160035 | 48.6734775 | 51.2048574 | 42.4454356 | 46.2850949 | 42.8577645 | 62.4939326 | 59.6080734 | 51.509083 | 50.9745369 | 49.9596427 | 59.8123967 |
| ENSMUSG00 | A530084C06R | 3199 | 0.33217549 | 0.51026315 | 0.58973669 | 0.62666019 | 0.67706676 | 0.71254321 | 0.63596286 | 0.60888169 | 0.91411148 | 0.45691522 | 1.04540391 | 1.04687082 | 0.4626313 | 1.13242851 | 0.78565017 | 0.75408198 | 0.50318989 | 0.6190459 |
| ENSMUSG00 | Vmn2r40 | 3066 | 0 | 0 | 0 | 0 | 0 | 0 | 0 | 0 | 0 | 0 | 0 | 0 | 0 | 0 | 0 | 0 | 0 | 0 |
| ENSMUSG00 | Gm17074 | 555 | 0 | 0 | 0 | 0 | 0 | 0 | 0 | 0 | 0 | 0 | 0 | 0 | 0 | 0 | 0 | 0 | 0 | 0 |
| ENSMUSG00 | Snhg1 | 3501 | 3.27803414 | 2.43590429 | 2.94303398 | 3.09726608 | 3.21704374 | 2.40615959 | 4.37855126 | 3.54542439 | 3.84689822 | 2.97637973 | 5.14457565 | 3.74655262 | 4.32587792 | 5.12065631 | 4.66033023 | 3.62016311 | 4.63068369 | 4.63515752 |
| ENSMUSG00 | Trav9d-4 | 305 | 0 | 0 | 0 | 0 | 0 | 0 | 0 | 0 | 0 | 0 | 0 | 0 | 0 | 0 | 0 | 0 | 0 | 0 |
| ENSMUSG00 | Gm17540 | 1526 | 0 | 0 | 0 | 0 | 0 | 0 | 0 | 0 | 0 | 0 | 0 | 0 | 0 | 0 | 0 | 0 | 0 | 0 |
| ENSMUSG00 | Gm17722 | 50 | 0 | 1.33251577 | 0 | 0 | 0 | 0.99105467 | 0.94625358 | 0 | 1.64746063 | 0.94301407 | 0 | 2.2326265 | 3.94655338 | 2.78664524 | 0.82403112 | 3.06324859 | 0.76652594 | 4.40072854 |
| ENSMUSG00 | Gm3239 | 1746 | 0 | 0 | 0 | 0 | 0 | 0 | 0 | 0 | 0 | 0 | 0 | 0 | 0 | 0 | 0 | 0 | 0 | 0 |
| ENSMUSG00 | Gm17112 | 428 | 0.09931116 | 0.07783386 | 0 | 0.85160831 | 0.22491553 | 0 | 0.11054364 | 0 | 0.57738106 | 0 | 0.3348712 | 0.39123128 | 0.34578447 | 0.21702845 | 0 | 0.08946404 | 0 | 0 |
| ENSMUSG00 | Olfr733 | 993 | 0 | 0 | 0 | 0 | 0 | 0 | 0 | 0 | 0 | 0 | 0 | 0 | 0 | 0 | 0 | 0 | 0 | 0 |
| ENSMUSG00 | Gm14482 | 499 | 0 | 0 | 0 | 0 | 0 | 0 | 0 | 0 | 0 | 0 | 0 | 0 | 0 | 0 | 0 | 0 | 0 | 0 |
| ENSMUSG00 | Hspa1b | 5006 | 25.4980111 | 28.4018043 | 24.4186798 | 41.0559071 | 28.4887914 | 28.7853275 | 30.9715639 | 33.2714876 | 31.5439675 | 54.3937897 | 25.9011661 | 34.2520496 | 89.3611318 | 121.769606 | 114.674646 | 87.5727455 | 104.053676 | 170.071104 |
| ENSMUSG00 | Gm17640 | 1194 | 0.07119795 | 0.05580049 | 0.24308307 | 0 | 0.04031149 | 0.04150145 | 0.1981268 | 0.12794775 | 0.17247285 | 0.1579588 | 0 | 0.04674679 | 0.04131651 | 0 | 0.03450717 | 0.09620756 | 0 | 0.23035639 |
| ENSMUSG00 | Gm17721 | 622 | 0 | 0 | 0 | 0 | 0 | 0 | 0 | 0.30701288 | 0 | 0 | 0 | 0 | 0 | 0 | 0 | 0 | 0 | 0 |
| ENSMUSG00 | Vmn1r129 | 923 | 0 | 0 | 0 | 0 | 0 | 0 | 0 | 0 | 0 | 0 | 0 | 0 | 0 | 0 | 0 | 0 | 0 | 0 |
| ENSMUSG00 | Gm6904 | 892 | 0.09530309 | 0.07469259 | 0.05423042 | 0 | 0.16187867 | 0.16665718 | 0 | 0 | 0 | 0.05285953 | 0 | 0 | 0.05530484 | 0 | 0.09238017 | 0.04292669 | 0.0429667 | 0 |
| ENSMUSG00 | Gm17065 | 238 | 0 | 0 | 0 | 0 | 0 | 0 | 0 | 0 | 0 | 0 | 0 | 0 | 0 | 0 | 0 | 0 | 0 | 0 |
| ENSMUSG00 | Gm17301 | 76 | 0 | 0 | 0 | 0 | 0 | 0 | 0 | 0 | 0 | 0 | 0 | 0 | 0 | 0 | 0 | 0 | 0 | 0 |
| ENSMUSG00 | Vmn2r102 | 2568 | 0 | 0 | 0 | 0 | 0 | 0 | 0 | 0 | 0 | 0 | 0 | 0 | 0 | 0 | 0 | 0 | 0 | 0 |
| ENSMUSG00 | Gm3944 | 335 | 0 | 0 | 0 | 0.13600312 | 0 | 0 | 0 | 0 | 0 | 0 | 0 | 0 | 0.14725945 | 0 | 0 | 0 | 0 | 0 |
| ENSMUSG00 | Gm17427 | 3194 | 0 | 0 | 0 | 0 | 0 | 0 | 0 | 0 | 0 | 0 | 0 | 0 | 0 | 0 | 0 | 0 | 0 | 0 |
| ENSMUSG00 | Gm17426 | 34 | 0 | 0 | 0 | 0 | 0 | 0 | 0 | 0 | 0 | 0 | 0 | 0 | 0 | 0 | 0 | 0 | 0 | 0 |
| ENSMUSG00 | Gm17429 | 3218 | 0 | 0 | 0 | 0 | 0 | 0 | 0 | 0 | 0 | 0 | 0 | 0 | 0 | 0 | 0 | 0 | 0 | 0 |
| ENSMUSG00 | Gm17428 | 92 | 0 | 0 | 0 | 0 | 0 | 0 | 0 | 0 | 0 | 0 | 0 | 0 | 0 | 0 | 0 | 0 | 0 | 0 |
| ENSMUSG00 | D6Ertd527e | 1403 | 0 | 0 | 0.10343592 | 0 | 0 | 0 | 0.4046701 | 0.10888782 | 0 | 0.03360706 | 0 | 0.43761389 | 0 | 0 | 0 | 0.0545839 | 0.16390433 | 0.54891482 |
| ENSMUSG00 | Vmn2r41 | 3611 | 0 | 0 | 0 | 0 | 0 | 0 | 0 | 0 | 0 | 0 | 0 | 0 | 0 | 0 | 0 | 0 | 0 | 0 |
| ENSMUSG00 | Gm4840 | 1574 | 0 | 0.25397378 | 0.0921986 | 0.54997449 | 0.06115873 | 0.03148204 | 0.21041217 | 0.07279365 | 0 | 0.20969182 | 0.03035258 | 0.17730515 | 0 | 0 | 0.02617634 | 0 | 0.51134195 | 0 |
| ENSMUSG00 | Olfr110 | 1101 | 0 | 0 | 0 | 0 | 0 | 0 | 0 | 0 | 0 | 0 | 0 | 0 | 0 | 0 | 0 | 0 | 0 | 0 |
| ENSMUSG00 | Vmn2r33 | 2580 | 0 | 0 | 0 | 0 | 0 | 0 | 0 | 0 | 0 | 0 | 0 | 0 | 0 | 0 | 0 | 0 | 0 | 0 |
| ENSMUSG00 | Gm17616 | 70 | 0 | 0 | 0 | 0 | 0 | 0 | 0 | 0 | 0 | 0 | 0 | 0 | 0 | 0 | 0 | 0 | 0 | 0 |
| ENSMUSG00 | Gm5494 | 1224 | 0 | 0 | 0 | 0 | 0 | 0 | 0 | 0 | 0 | 0 | 0 | 0 | 0 | 0 | 0 | 0 | 0 | 0 |
| ENSMUSG00 | Gm7732 | 1526 | 0 | 0 | 0 | 0 | 0 | 0 | 0 | 0 | 0 | 0 | 0 | 0 | 0 | 0 | 0 | 0 | 0 | 0 |
| ENSMUSG00 | Gm3326 | 1042 | 0 | 0 | 0 | 0 | 0 | 0 | 0 | 0 | 0 | 0 | 0 | 0 | 0 | 0 | 0 | 0 | 0 | 0 |
| ENSMUSG00 | Abl1 | 483 | 0 | 0 | 0 | 0 | 0 | 0 | 0 | 0 | 0 | 0 | 0 | 0 | 0 | 0 | 0 | 0 | 0 | 0 |
| ENSMUSG00 | Gm17246 | 1455 | 0.49662405 | 0.36632736 | 0 | 0 | 0 | 0.13622745 | 0 | 0.18374352 | 0.31137572 | 0.25924785 | 0.03283502 | 0.11508384 | 0.20343059 | 0 | 0.19822054 | 0 | 0.15804659 | 0 |
| ENSMUSG00 | Gm17247 | 43 | 0 | 0 | 0 | 0 | 0 | 0 | 0 | 0 | 0 | 0 | 0 | 0 | 0 | 0 | 0 | 0 | 0 | 0 |
| ENSMUSG00 | Gm17248 | 2597 | 0.26187247 | 0.39765103 | 0.22352036 | 0.45613675 | 0.50040891 | 0.30529216 | 0.12752744 | 0.44119063 | 0.36476314 | 0.34496086 | 0.349528 | 0.49432431 | 0.15196586 | 0.07153498 | 0.20624576 | 0.07372085 | 0.26564241 | 0.12709074 |
| ENSMUSG00 | Gm17249 | 2507 | 0.91554827 | 0.70426143 | 1.00336002 | 0.94502366 | 0.82555753 | 0.71156698 | 0.73601694 | 1.47772758 | 1.38000292 | 0.41376764 | 1.06717099 | 0.24490319 | 0.23613203 | 0.57429865 | 0.41086514 | 0.27492259 | 0.15287713 | 0.59243952 |
| ENSMUSG00 | Mcart1 | 572 | 0 | 0.11647865 | 0 | 0 | 0 | 0 | 0 | 0.06676993 | 0.21601319 | 0.0824313 | 0.33409062 | 0 | 0 | 0.40597978 | 0 | 0.06694162 | 0.06700402 | 0 |
| ENSMUSG00 | C030017D09 | 921 | 0.13845334 | 0 | 0 | 0 | 0 | 0 | 0 | 0 | 0 | 0 | 0 | 0 | 0 | 0 | 0 | 0 | 0 | 0 |
| ENSMUSG00 | Gm17323 | 64 | 0 | 0 | 0 | 0 | 0 | 0 | 0 | 0 | 0 | 0 | 0 | 0 | 0 | 0 | 0 | 0 | 0 | 0 |
| ENSMUSG00 | Gm17706 | 34 | 7.50091342 | 11.7574921 | 2.84550179 | 8.04018429 | 14.1564481 | 8.74460004 | 15.3070432 | 17.9728952 | 14.5364173 | 12.4810685 | 5.62058335 | 1.64163713 | 5.80375497 | 9.56201799 | 13.3299151 | 5.63097168 | 9.01795221 | 3.23582981 |
| ENSMUSG00 | 4931403G20R | 1697 | 0.15028347 | 0.23556555 | 0.02850532 | 0 | 0.14181474 | 0.05840039 | 0.05576038 | 0.36009336 | 0 | 0 | 0.39413637 | 0.19734471 | 0.26163068 | 0.0547367 | 0 | 0.04512741 | 0.33877104 | 0 |
| ENSMUSG00 | 9530052C20R | 3093 | 0.508468 | 1.10935277 | 0.75070464 | 0.45664157 | 0.7780783 | 0.8811511 | 0.79542816 | 0.48157248 | 0.06658021 | 0.68599471 | 0.92676932 | 0.64964884 | 0.66988054 | 1.00606332 | 0.95910508 | 0.48281076 | 0.85499983 | 0.51576595 |
| ENSMUSG00 | Gm17563 | 1352 | 0 | 0 | 0 | 0 | 0 | 0 | 0 | 0 | 0 | 0 | 0 | 0 | 0 | 0 | 0 | 0 | 0 | 0 |
| ENSMUSG00 | Gm17562 | 28 | 0 | 0 | 0 | 0 | 0 | 0 | 0 | 0 | 0 | 0 | 1.70624852 | 0 | 0 | 0 | 0 | 0 | 0 | 0 |
| ENSMUSG00 | 1700007L15R | 735 | 9.25282743 | 6.39063685 | 5.98910376 | 4.58709834 | 7.00696032 | 5.79800692 | 8.30385793 | 8.83361688 | 10.0864937 | 8.66033325 | 8.57999254 | 8.80900253 | 7.91995407 | 9.98389906 | 9.1372158 | 6.25152774 | 8.60386257 | 12.4986678 |
| ENSMUSG00 | Pabpc4l | 5216 | 0.14668197 | 0.0510934 | 0 | 0 | 0 | 0 | 0.01814136 | 0.08786596 | 0.03158475 | 0.0632774 | 0.25646067 | 0.10700856 | 0 | 0.12465821 | 0 | 0.05138693 | 0 | 0 |
| ENSMUSG00 | 9130213A22R | 1579 | 0 | 0 | 0.0612711 | 0.02885437 | 0.03048254 | 0.43935293 | 0.20974589 | 0.09675086 | 0.28692316 | 0 | 0 | 0.84836979 | 0 | 0 | 0 | 0 | 0.04854502 | 0.1741897 |
| ENSMUSG00 | Gm17391 | 335 | 0 | 0 | 0 | 0 | 0 | 0 | 0 | 0 | 0 | 0 | 0 | 0 | 0 | 0 | 0 | 0 | 0 | 0 |
| ENSMUSG00 | Gm17390 | 2234 | 0.95132444 | 0.43244133 | 0.30314656 | 0.79538081 | 1.03416846 | 0.6654351 | 0.97420915 | 0.61545501 | 0.79275747 | 0.99197988 | 0.51324933 | 0.84947741 | 0.48581118 | 0.51974135 | 0.64550334 | 0.18853925 | 0.12009135 | 1.45279199 |
| ENSMUSG00 | 1810064F22R | 1205 | 0.17637003 | 0.08293667 | 0 | 0 | 0.19971753 | 0.0411226 | 1.09938175 | 0.41203422 | 0 | 0 | 0 | 0.04632005 | 0.3275148 | 0.53959936 | 0 | 0.03177644 | 0 | 0.04565071 |
| ENSMUSG00 | Gm17047 | 429 | 0 | 0 | 0 | 0 | 0 | 0 | 0 | 0 | 0 | 0 | 0 | 0 | 0 | 0 | 0 | 0 | 0 | 0 |
| ENSMUSG00 | C530050E15R | 3835 | 0.07758442 | 0 | 0.08829588 | 0 | 0.07530418 | 0.05168473 | 0 | 0.04979453 | 0.01073964 | 0.03688451 | 0 | 0 | 0 | 0.12110583 | 0.06446137 | 0.09984513 | 0.23985166 | 0.45900689 |
| ENSMUSG00 | Gm17704 | 956 | 0.04446148 | 0 | 0 | 0 | 0 | 0 | 0 | 0.03995021 | 0 | 0 | 0.24986903 | 0 | 0.25801212 | 0 | 0 | 0 | 0.08018054 | 0 |
| ENSMUSG00 | 4632411P08R | 3235 | 0.64381874 | 0.30892947 | 0.59812711 | 0.2816757 | 0.46123327 | 0.5820723 | 0.9652664 | 0.86183783 | 0.73842903 | 0.65588304 | 0.60549406 | 0.53486415 | 1.23520411 | 1.26339403 | 0.84058816 | 1.3848535 | 0.41465854 | 1.10528344 |
| ENSMUSG00 | Vmn1r101 | 923 | 0 | 0 | 0 | 0 | 0 | 0 | 0 | 0 | 0 | 0 | 0 | 0 | 0 | 0 | 0 | 0 | 0 | 0 |
| ENSMUSG00 | Gm17703 | 2237 | 0.11400584 | 0.13402595 | 0.1729943 | 0.04073406 | 0.06454885 | 0.06645427 | 0.04230012 | 0.22194959 | 0.0184115 | 0.1264659 | 0.12814026 | 0.09980449 | 0.19847441 | 0.04152355 | 0.01841822 | 0.13693557 | 0.0856645 | 0.04918114 |
| ENSMUSG00 | 4930598A11R | 625 | 0 | 0 | 0 | 0 | 0 | 0.07928437 | 0 | 0 | 0.39539055 | 0 | 0 | 0 | 0 | 0.22293162 | 0 | 0 | 0.2452883 | 0.70411657 |
| ENSMUSG00 | Synj2bp | 6020 | 11.021691 | 8.17881356 | 9.76309624 | 10.0052659 | 10.7697178 | 9.44959105 | 10.0441202 | 8.50129887 | 11.0081568 | 9.02285883 | 10.213683 | 9.85582215 | 7.08019544 | 8.62533052 | 7.67224986 | 8.46591336 | 8.05361555 | 9.37530623 |
| ENSMUSG00 | Gm17705 | 1553 | 1.9979896 | 1.54444841 | 1.96235184 | 0.70409856 | 1.08475037 | 1.2444022 | 0.67023756 | 1.77067159 | 2.30729354 | 0.81974822 | 0.89212736 | 0.57504868 | 0.44471786 | 1.43549014 | 1.0081514 | 0.34518255 | 0.93779735 | 0.92095092 |
| ENSMUSG00 | Gm5895 | 279 | 0 | 0 | 0 | 0 | 0 | 0 | 0 | 0 | 0 | 0 | 0 | 0 | 0 | 0 | 0 | 0 | 0 | 0 |
| ENSMUSG00 | Gm6905 | 512 | 0 | 0 | 0 | 0 | 0.18801533 | 0 | 0 | 0.29837814 | 0 | 0.09209122 | 0 | 0.3270449 | 0.0963514 | 0.09071111 | 0.24141537 | 0 | 0.07485605 | 0.10743966 |
| ENSMUSG00 | Gm17147 | 431 | 0 | 0.38646049 | 0 | 0 | 0 | 0.22994308 | 0 | 0 | 0 | 0 | 0 | 0 | 0 | 0 | 0.09559526 | 0 | 0 | 0 |
| ENSMUSG00 | AW121686 | 2198 | 0.05801434 | 0 | 0 | 0 | 0 | 0 | 0 | 0.0868799 | 0 | 0 | 0 | 0 | 0 | 0 | 0 | 0 | 0 | 0 |
| ENSMUSG00 | Gm17212 | 258 | 0.16474874 | 0.38735924 | 0.37498861 | 0 | 0 | 0.76825944 | 2.75073715 | 0.74016284 | 0.95782595 | 1.64479197 | 0.92587129 | 2.37973755 | 1.33846287 | 0.90007921 | 0.31939191 | 1.18730566 | 0.44565462 | 0 |
| ENSMUSG00 | F830016B08R | 3295 | 0 | 0 | 0 | 0 | 0 | 0.21054272 | 0 | 0 | 0 | 0 | 0 | 0 | 0 | 0 | 0 | 0 | 0 | 0 |
| ENSMUSG00 | Gm3594 | 2136 | 0 | 0 | 0 | 0 | 0 | 0 | 0 | 0 | 0 | 0 | 0 | 0 | 0 | 0 | 0 | 0 | 0 | 0 |
| ENSMUSG00 | Gm17513 | 3521 | 0 | 0 | 0.01373858 | 0 | 0.02733992 | 0 | 0 | 0.0325411 | 0.02339478 | 0.02678256 | 0.04070573 | 0.03170444 | 0 | 0 | 0 | 0.02174985 | 0.03265518 | 0.12498519 |
| ENSMUSG00 | Gm17514 | 1098 | 0 | 0 | 0 | 0 | 0 | 0 | 0 | 0 | 0 | 0 | 0 | 0 | 0 | 0 | 0 | 0 | 0 | 0 |
| ENSMUSG00 | 2010109K11R | 4239 | 3.14853156 | 3.60712867 | 2.85288573 | 3.36414411 | 2.75915486 | 3.53029619 | 1.99786967 | 2.87411568 | 3.90586915 | 3.0477218 | 3.61777817 | 3.67364234 | 3.42146348 | 3.4731719 | 4.3738382 | 3.29702093 | 3.4899624 | 3.16636519 |
| ENSMUSG00 | Gm3102 | 486 | 0 | 0 | 0 | 0 | 0 | 0 | 0 | 0 | 0 | 0 | 0 | 0 | 0 | 0 | 0 | 0 | 0 | 0 |
| ENSMUSG00 | Gm17512 | 1438 | 0.32514391 | 0.34749194 | 0.47095231 | 0.38020343 | 0 | 0.24121637 | 0 | 0.05311878 | 0.45826443 | 0.16394542 | 0.09966959 | 0 | 0.51458885 | 0.35527466 | 0 | 0 | 0.45309252 | 0 |
| ENSMUSG00 | Vmn2r77 | 2559 | 0 | 0 | 0 | 0 | 0 | 0 | 0 | 0 | 0 | 0 | 0 | 0 | 0 | 0 | 0 | 0 | 0 | 0 |
| ENSMUSG00 | Olfr181 | 1389 | 0 | 0 | 0 | 0 | 0 | 0 | 0 | 0 | 0 | 0 | 0 | 0 | 0 | 0 | 0 | 0 | 0 | 0 |
| ENSMUSG00 | Gm17251 | 923 | 12.2956468 | 7.651499 | 10.1149419 | 11.3038777 | 9.17791827 | 10.8983802 | 11.9947637 | 12.3721867 | 11.8695701 | 10.4211739 | 12.2672429 | 12.0339294 | 9.67397294 | 10.4159546 | 10.9811297 | 10.2467823 | 10.0071913 | 12.1580258 |
| ENSMUSG00 | Gm17250 | 1006 | 0 | 0 | 0 | 0 | 0 | 0.04915946 | 0 | 0 | 0 | 0.14032947 | 0 | 0 | 0 | 0.09215097 | 0 | 0.22792028 | 0.03802212 | 0 |
| ENSMUSG00 | Gm17345 | 92 | 0 | 0 | 0 | 0 | 0 | 0 | 0 | 0 | 0.44767952 | 0 | 0 | 0 | 0 | 2.01930815 | 0 | 0 | 0 | 0.59792507 |
| ENSMUSG00 | Gm17097 | 712 | 0.11939656 | 0.37430218 | 0.47558246 | 0.31995115 | 0.47320711 | 0.27838614 | 0 | 0.26820507 | 0.69415476 | 0.0662229 | 1.20779389 | 0.39196392 | 0.4157184 | 0.9784569 | 0.92587766 | 3.01162081 | 2.09933368 | 0 |
| ENSMUSG00 | Gm7535 | 908 | 0.23405934 | 0.14675284 | 0 | 0 | 0 | 0.0545735 | 0.05210647 | 0.04206212 | 0.0907292 | 0.05192809 | 0.05261559 | 0.12294199 | 0.16299092 | 0.15344983 | 0.09075233 | 0.04217027 | 0 | 0 |
| ENSMUSG00 | Lrrc32 | 3815 | 0 | 0 | 0 | 0 | 0 | 0 | 0 | 0 | 0 | 0 | 0 | 0 | 0 | 0 | 0 | 0 | 0 | 0 |
| ENSMUSG00 | Ighv1-62-1 | 89 | 0 | 0 | 0 | 0 | 0 | 0 | 0 | 0 | 0 | 0 | 0 | 0 | 0 | 0 | 0 | 0 | 0 | 0 |
| ENSMUSG00 | Vmn2r21 | 2556 | 0 | 0 | 0 | 0 | 0 | 0 | 0 | 0 | 0 | 0 | 0 | 0 | 0 | 0 | 0 | 0 | 0 | 0 |

| | | | | | | | | | | | | | | | | | | | |
|---|---|---|---|---|---|---|---|---|---|---|---|---|---|---|---|---|---|---|---|
| ENSMUSG00 Gm17656 | 1092 | 0 | 0 | 0 | 0 | 0 | 0 | 0 | 0 | 0 | 0 | 0 | 0 | 0 | 0 | 0 | 0 | 0 | 0 |
| ENSMUSG00 Gm3209 | 254 | 0 | 0 | 0 | 0 | 0 | 0 | 0 | 0 | 0 | 0 | 0 | 0 | 0 | 0 | 0 | 0 | 0 | 0 |
| ENSMUSG00 Vmn1r142 | 917 | 0 | 0 | 0 | 0 | 0 | 0 | 0 | 0 | 0 | 0 | 0 | 0 | 0 | 0 | 0 | 0 | 0 | 0 |
| ENSMUSG00 Slco6b1 | 2477 | 0 | 0 | 0 | 0 | 0 | 0 | 0 | 0 | 0 | 0 | 0 | 0 | 0 | 0 | 0 | 0 | 0 | 0 |
| ENSMUSG00 Gm17162 | 470 | 0 | 0 | 0 | 0 | 0 | 0 | 0 | 0 | 0 | 0 | 0 | 0 | 0 | 0 | 0 | 0 | 0 | 0 |
| ENSMUSG00 Gm14479 | 499 | 0 | 0 | 0 | 0 | 0 | 0 | 0 | 0 | 0 | 0 | 0 | 0 | 0 | 0 | 0 | 0 | 0 | 0 |
| ENSMUSG00 Gm17459 | 448 | 0 | 0 | 0 | 0 | 0 | 0 | 0 | 0 | 0 | 0 | 0 | 0 | 0 | 0 | 0 | 0 | 0 | 0 |
| ENSMUSG00 Gm8922 | 1119 | 0 | 0 | 0 | 0 | 0 | 0 | 0 | 0 | 0 | 0 | 0 | 0 | 0 | 0 | 0 | 0 | 0 | 0 |
| ENSMUSG00 Gm17274 | 3682 | 0.02308809 | 0.018095 | 0.03941352 | 0.01237399 | 0 | 0 | 0.01284972 | 0.02074547 | 0 | 0.05122293 | 0 | 0.01515906 | 0.13398131 | 0.01261382 | 0.02237999 | 0 | 0.13531827 | 0 |
| ENSMUSG00 Olfr718-ps1 | 989 | 0 | 0 | 0 | 0 | 0 | 0 | 0 | 0 | 0 | 0 | 0 | 0 | 0 | 0 | 0 | 0 | 0 | 0 |
| ENSMUSG00 9430076C15i | 610 | 0.20904185 | 0.27305651 | 0.23790261 | 0.14938047 | 0.15780959 | 0.16246798 | 0 | 0 | 0 | 0 | 0.0783196 | 0.18300217 | 0 | 0 | 0 | 0 | 0 | 0 |
| ENSMUSG00 AL451076.1 | 628 | 0 | 0 | 0 | 0 | 0 | 0 | 0 | 0 | 0 | 0 | 0 | 0 | 0 | 0 | 0 | 0 | 0 | 0 |
| ENSMUSG00 Gm17276 | 2780 | 0.09173779 | 0.01198306 | 0.0696022 | 0.11472205 | 0.05194092 | 0.10694835 | 0.22124634 | 0.16485929 | 0.14815293 | 0.10176411 | 0.08592618 | 0.04015515 | 0.5855947 | 0.5346082 | 0.04446211 | 0.3167928 | 0.06893219 | 0.35617407 |
| ENSMUSG00 Vmn1r170 | 914 | 0 | 0 | 0 | 0 | 0 | 0 | 0 | 0 | 0 | 0 | 0 | 0 | 0 | 0 | 0 | 0 | 0 | 0 |
| ENSMUSG00 Gm17275 | 2548 | 0.51713519 | 1.01978248 | 0.58853196 | 0.53643302 | 0.62337264 | 0.89459409 | 0.98413343 | 1.28906853 | 1.27697596 | 0.70318945 | 0.46874959 | 0.67907596 | 1.20038417 | 0.67442356 | 1.73020662 | 1.81835302 | 0.46629325 | 0.56131742 |
| ENSMUSG00 Gm4781 | 785 | 0 | 0 | 0 | 0 | 0 | 0 | 0 | 0 | 0 | 0 | 0 | 0 | 0 | 0 | 0 | 0 | 0 | 0 |
| ENSMUSG00 Gm8693 | 920 | 0 | 0 | 0 | 0 | 0 | 0 | 0 | 0 | 0 | 0 | 0 | 0 | 0 | 0 | 0 | 0 | 0 | 0 |
| ENSMUSG00 Gm17042 | 1212 | 0 | 0 | 0 | 0 | 0 | 0 | 0 | 0 | 0 | 0 | 0 | 0 | 0 | 0 | 0 | 0 | 0 | 0 |
| ENSMUSG00 Gm17197 | 449 | 0 | 0.22258058 | 0.2154723 | 0.20294452 | 0.32159414 | 0 | 0 | 0.42530515 | 0.36691774 | 0.31503811 | 0.53201513 | 0 | 0.10987064 | 0 | 0.27528879 | 0.42639875 | 0.08535924 | 0.24502943 |
| ENSMUSG00 Gm6647 | 637 | 0 | 0 | 0 | 0 | 0 | 0 | 0 | 0 | 0 | 0 | 0 | 0 | 0 | 0 | 0 | 0 | 0 | 0 |
| ENSMUSG00 Gm17588 | 4243 | 0 | 0 | 0 | 0 | 0 | 0 | 0 | 0 | 0 | 0 | 0 | 0 | 0 | 0 | 0 | 0 | 0 | 0 |
| ENSMUSG00 Gm17587 | 1814 | 0 | 0.03672866 | 0 | 0 | 0 | 0 | 0 | 0.06316274 | 0 | 0.05198534 | 0.13168401 | 0 | 0 | 0 | 0.0227131 | 0 | 0 | 0.06064951 |
| ENSMUSG00 Gm6153 | 1057 | 0 | 0 | 0 | 0 | 0 | 0 | 0 | 0 | 0 | 0 | 0 | 0 | 0 | 0 | 0 | 0.03622574 | 0 | 0 |
| ENSMUSG00 Gm17589 | 2142 | 0.15874949 | 0 | 0.02258335 | 0.02127033 | 0 | 0.02313386 | 0.06626426 | 0 | 0.3845613 | 0.22012467 | 0 | 0.20846186 | 0.76001553 | 0.47701677 | 0.442407 | 0.89380503 | 0.53678287 | 2.15721987 |
| ENSMUSG00 Gm20458 | 1363 | 0 | 0 | 0 | 0 | 0 | 0 | 0 | 0 | 0 | 0 | 0 | 0 | 0 | 0 | 0 | 0 | 0 | 0 |
| ENSMUSG00 Gm17586 | 906 | 12.3856142 | 11.3616825 | 9.77081242 | 10.962812 | 8.55324471 | 7.60246133 | 7.78100349 | 12.477871 | 9.04648635 | 11.5014409 | 13.2356673 | 9.30261043 | 11.8701523 | 13.4821137 | 11.5509881 | 12.2141121 | 9.05278978 | 9.04675155 |
| ENSMUSG00 Gm8266 | 1191 | 0 | 0 | 0 | 0 | 0 | 0 | 0 | 0 | 0 | 0 | 0 | 0 | 0 | 0 | 0 | 0 | 0 | 0 |
| ENSMUSG00 Tcerg1l | 3973 | 0 | 0 | 0 | 0 | 0 | 0.02494474 | 0.16671973 | 0.09612988 | 0.3006315 | 0.17801675 | 0.01202491 | 0.11238996 | 0 | 0 | 0.15555584 | 0.13492789 | 0.12540696 | 0.2353775 |
| ENSMUSG00 Gm7244 | 377 | 0.33823748 | 0.6185418 | 0.12831175 | 0 | 0.12767089 | 0.26287922 | 0 | 0.20261221 | 0 | 0.25013636 | 0 | 0 | 0 | 0 | 0 | 0.1015666 | 0 | 0 |
| ENSMUSG00 Gm6872 | 699 | 0 | 0 | 0 | 0 | 0 | 0 | 0 | 0 | 0 | 0.06745451 | 0 | 0 | 0 | 0 | 0 | 0 | 0 | 0 |
| ENSMUSG00 Traj55 | 62 | 0 | 0 | 0 | 0 | 0 | 0 | 0 | 0 | 0 | 0 | 0 | 0 | 0 | 0 | 0 | 0 | 0 | 0 |
| ENSMUSG00 Vmn2r69 | 2547 | 0 | 0 | 0 | 0 | 0 | 0 | 0 | 0 | 0 | 0 | 0 | 0 | 0 | 0 | 0 | 0 | 0 | 0 |
| ENSMUSG00 D630036H23 | 939 | 0.45266428 | 0.49667787 | 0.66970809 | 0.63077058 | 0.66636316 | 0.63326177 | 1.05811103 | 0.56942879 | 1.35972523 | 0.70299238 | 1.37372085 | 0.7132992 | 0.73551314 | 1.33545299 | 1.53573425 | 1.63112279 | 1.06121802 | 0.99590502 |
| ENSMUSG00 Gm17472 | 350 | 0 | 0 | 0 | 0 | 0 | 0 | 0 | 0 | 0 | 0 | 0 | 0 | 0 | 0 | 0 | 0 | 0 | 0 |
| ENSMUSG00 Gm17473 | 2195 | 0 | 0 | 0 | 0 | 0 | 0 | 0 | 0 | 0 | 0 | 0 | 0 | 0 | 0 | 0 | 0 | 0 | 0 |
| ENSMUSG00 Gm17107 | 552 | 0 | 0 | 0 | 0 | 0 | 0 | 0 | 0 | 0 | 0 | 0 | 0 | 0 | 0 | 0 | 0 | 0 | 0 |
| ENSMUSG00 Gm17187 | 408 | 0 | 0 | 0 | 0 | 0 | 0 | 0 | 0 | 0 | 0 | 0 | 0 | 0 | 0 | 0 | 0 | 0 | 0 |
| ENSMUSG00 Vmn1r34 | 929 | 0 | 0 | 0 | 0 | 0 | 0 | 0 | 0 | 0 | 0 | 0 | 0 | 0 | 0 | 0 | 0 | 0 | 0 |
| ENSMUSG00 Vmn1r1 | 1175 | 0 | 0 | 0 | 0 | 0 | 0 | 0 | 0 | 0 | 0 | 0 | 0 | 0 | 0 | 0 | 0 | 0 | 0 |
| ENSMUSG00 Gm5244 | 500 | 0 | 0 | 0 | 0 | 0 | 0 | 0 | 0 | 0 | 0 | 0 | 0 | 0 | 0 | 0 | 0 | 0 | 0 |
| ENSMUSG00 Gm17284 | 848 | 0.90223251 | 1.53207886 | 1.02679663 | 1.77301234 | 1.30546491 | 1.11026172 | 0.94848531 | 1.17099347 | 0.92281108 | 0.77843142 | 0.95775271 | 1.25058678 | 0.7562676 | 1.09537942 | 0.58304089 | 1.76100671 | 0.85872599 | 0.58382307 |
| ENSMUSG00 Gm2878 | 483 | 0 | 0 | 0 | 0 | 0 | 0 | 0 | 0 | 0 | 0 | 0 | 0 | 0 | 0 | 0 | 0 | 0 | 0 |
| ENSMUSG00 Fam71a | 2233 | 0.28552514 | 0.07459224 | 0 | 0 | 0 | 0.0221911 | 0.50851066 | 0 | 0.18444476 | 0.42230813 | 0.87719359 | 0 | 2.91617245 | 4.18059183 | 0.97791422 | 2.12630332 | 1.66486825 | 3.6705584 |
| ENSMUSG00 Rplp2-ps1 | 441 | 3.95172838 | 4.607906 | 4.60700289 | 5.57890339 | 5.23884883 | 6.62950404 | 7.72451901 | 6.58191062 | 5.23003375 | 6.52197936 | 6.39166111 | 4.17668223 | 7.04741675 | 8.21460049 | 7.28735002 | 6.85931516 | 4.34538514 | 7.85844382 |
| ENSMUSG00 Gm7502 | 732 | 0 | 0 | 0 | 0.06224186 | 0.26301598 | 0 | 0 | 0 | 0 | 0 | 0 | 0 | 0.60653997 | 0.76137848 | 0.22514511 | 0.88926274 | 0.62829995 | 0 |
| ENSMUSG00 Gm6828 | 3144 | 1.17619285 | 1.35625015 | 1.55398428 | 1.68100545 | 1.913642 | 1.38697219 | 1.6252447 | 1.53061155 | 1.45410409 | 1.43971613 | 2.21855723 | 1.02967825 | 1.33371914 | 1.25564486 | 1.37600616 | 1.69287355 | 2.06016036 | 1.57468817 |
| ENSMUSG00 Gm17300 | 1525 | 0.19510573 | 0.4150459 | 0.60268661 | 0.5676458 | 0.41030492 | 0.51989753 | 0.37229649 | 0.65114916 | 0.13503776 | 0.34010343 | 0.28195057 | 0.40260478 | 0.29113918 | 0.30455139 | 0.48631345 | 0.72814926 | 0.45237596 | 0.793574 |
| ENSMUSG00 Gm3115 | 1792 | 0 | 0 | 0 | 0 | 0 | 0 | 0 | 0 | 0 | 0 | 0 | 0 | 0 | 0 | 0 | 0 | 0 | 0 |
| ENSMUSG00 Gm17351 | 1764 | 0.26505495 | 0.0944243 | 0 | 0 | 0.08185701 | 0.08427336 | 0.13410623 | 0 | 0.16343855 | 0 | 0.05416662 | 0 | 0 | 0 | 0.04671378 | 0 | 0 | 0.46776451 |
| ENSMUSG00 Gm8305 | 1194 | 0 | 0 | 0 | 0 | 0 | 0 | 0 | 0 | 0 | 0 | 0 | 0 | 0 | 0 | 0 | 0 | 0 | 0 |
| ENSMUSG00 Gm6900 | 692 | 0.12284733 | 0.2407001 | 0.20971184 | 0 | 0.13910961 | 0.071608 | 0.20511277 | 0.22076533 | 0.53566278 | 0.13627371 | 0.20711687 | 0.08065847 | 0.0712889 | 0.06711573 | 0.05953982 | 0.38733273 | 0.55384822 | 0.07949293 |
| ENSMUSG00 Gm11107 | 504 | 0 | 0 | 0 | 0 | 0 | 0 | 0 | 0 | 0 | 0 | 0 | 0 | 0 | 0 | 0 | 0 | 0 | 0 |
| ENSMUSG00 Gm10722 | 647 | 0 | 0.05148824 | 0.14953178 | 0 | 0 | 0.22976538 | 0 | 0.23611995 | 0 | 0.07287589 | 0 | 0 | 0 | 0.07178375 | 0 | 0 | 0 | 0 |
| ENSMUSG00 Gm8138 | 1399 | 0 | 0 | 0 | 0 | 0 | 0 | 0 | 0.02729979 | 0 | 0 | 0 | 0 | 0 | 0 | 0 | 0 | 0 | 0 |
| ENSMUSG00 Gm17063 | 453 | 0 | 0 | 0 | 0 | 0 | 0 | 0 | 0 | 0 | 0 | 0 | 0 | 0 | 0 | 0 | 0 | 0 | 0 |
| ENSMUSG00 Gm17299 | 40 | 2.1252588 | 3.33128942 | 0 | 0 | 0 | 0 | 2.36563395 | 1.90962012 | 0 | 3.53630274 | 7.16624377 | 5.58156626 | 2.46659586 | 2.32220437 | 2.06007779 | 3.82906074 | 1.91631485 | 5.50091067 |
| ENSMUSG00 4930477E14i | 619 | 0 | 0 | 0 | 0 | 0 | 0 | 0 | 0 | 0 | 0 | 0 | 0 | 0 | 0 | 0 | 0 | 0 | 0 |
| ENSMUSG00 Gm17660 | 510 | 0 | 0 | 0 | 0 | 0 | 0 | 0 | 0 | 0 | 0 | 0 | 0 | 0.5803755 | 0 | 0 | 0 | 0 | 0.21572199 |
| ENSMUSG00 Gm17659 | 1535 | 1.60605877 | 1.06340835 | 0.81935621 | 1.0982141 | 0.81526385 | 0.54879249 | 0.67809703 | 1.14452802 | 0.96593783 | 1.65872181 | 1.36944506 | 1.16358385 | 1.4140745 | 0.48410775 | 1.82521551 | 1.14747097 | 1.62293765 | 1.54097172 |
| ENSMUSG00 Gm17174 | 476 | 0 | 0 | 0 | 0 | 0 | 0 | 0 | 0 | 0 | 0 | 0 | 0 | 0 | 0 | 0 | 0 | 0 | 0 |
| ENSMUSG00 Gm17661 | 268 | 0 | 0 | 0 | 0 | 0 | 0 | 0 | 0 | 0 | 0 | 0 | 0 | 0 | 0 | 0 | 0 | 0 | 0 |
| ENSMUSG00 Gm17662 | 22 | 0 | 0 | 0 | 0 | 0 | 0 | 0 | 0 | 0 | 0 | 0 | 0 | 0 | 0 | 0 | 0 | 0 | 0 |
| ENSMUSG00 Krtap20-2 | 527 | 0 | 0 | 0 | 0 | 0 | 0 | 0 | 0 | 0 | 0 | 0 | 0 | 0 | 0 | 0 | 0 | 0 | 0 |
| ENSMUSG00 Gm9359 | 1220 | 0.4180837 | 0.13652825 | 0.1189513 | 0.11203535 | 0.27616678 | 0.16246798 | 0.7368368 | 0.31305248 | 0.10127832 | 0.07729623 | 0.31327842 | 0 | 0.80871996 | 0.03806892 | 0.13508707 | 0.69048636 | 0.18848998 | 0.04508943 |
| ENSMUSG00 Gm17720 | 25 | 13.6016563 | 17.322705 | 7.73976486 | 12.7570924 | 15.4022156 | 7.92843737 | 9.46253579 | 12.2215688 | 14.8271457 | 13.2021969 | 34.3979701 | 13.395759 | 3.94655338 | 5.57329049 | 4.94418671 | 29.1008616 | 24.52883 | 17.6029142 |
| ENSMUSG00 Gm3727 | 990 | 0 | 0 | 0 | 0 | 0 | 0 | 0 | 0 | 0 | 0 | 0 | 0 | 0 | 0 | 0 | 0 | 0 | 0 |
| ENSMUSG00 Glyatl3 | 868 | 0 | 0 | 0 | 0 | 0 | 0 | 0 | 0 | 0 | 0 | 0 | 0 | 0 | 0 | 0 | 0 | 0 | 0 |
| ENSMUSG00 Gm3170 | 843 | 0 | 0 | 0 | 0 | 0 | 0 | 0 | 0 | 0 | 0 | 0 | 0 | 0 | 0 | 0 | 0 | 0 | 0 |
| ENSMUSG00 Vmn2r55 | 2341 | 0 | 0 | 0 | 0 | 0 | 0 | 0 | 0 | 0 | 0 | 0 | 0 | 0 | 0 | 0 | 0 | 0 | 0 |
| ENSMUSG00 4930532G15i | 697 | 0.06098304 | 0.43015215 | 0 | 0.26146941 | 0.06905584 | 0.21328293 | 0.33940229 | 0.65754495 | 0.88636691 | 0.60883261 | 0.1370874 | 0 | 1.13243999 | 0.99951408 | 2.0689447 | 0.76910833 | 0.65985016 | 1.42060821 |
| ENSMUSG00 Gm8207 | 499 | 0 | 0 | 0 | 0 | 0 | 0 | 0 | 0 | 0 | 0 | 0 | 0 | 0 | 0 | 0 | 0 | 0 | 0 |
| ENSMUSG00 Gm1979 | 4317 | 0 | 0 | 0 | 0 | 0 | 0 | 0 | 0 | 0 | 0 | 0 | 0 | 0 | 0 | 0 | 0 | 0 | 0 |
| ENSMUSG00 9330020H09i | 2145 | 0.79263732 | 1.04054262 | 0.69910463 | 1.63552467 | 1.00976064 | 1.06266934 | 1.43371754 | 0.71221263 | 0.90245512 | 1.29691911 | 1.78181663 | 1.89955402 | 1.37991377 | 1.84044169 | 1.97844301 | 1.23172583 | 1.87611244 | 1.6925879 |
| ENSMUSG00 Ighd5-8 | 28 | 0 | 0 | 0 | 0 | 0 | 0 | 0 | 0 | 0 | 0 | 0 | 0 | 0 | 0 | 0 | 0 | 0 | 0 |
| ENSMUSG00 Gm5551 | 617 | 0 | 0 | 0.07840118 | 0 | 0 | 0 | 0 | 0 | 0 | 0 | 0 | 0 | 0 | 0 | 0.06677724 | 0.06205933 | 0 | 0 |
| ENSMUSG00 Gm9742 | 587 | 0 | 0 | 0 | 0 | 0 | 0 | 0 | 0 | 0 | 0 | 0 | 0 | 0 | 0 | 0 | 0 | 0 | 0 |
| ENSMUSG00 Siglec15 | 1150 | 26.168404 | 27.5193474 | 28.8137985 | 23.7709797 | 26.744608 | 25.9828681 | 14.3583695 | 15.1773286 | 18.0146239 | 15.3342287 | 13.7093359 | 21.3070225 | 1.24402226 | 0.44424779 | 4.12015559 | 1.49832812 | 1.76634238 | 3.73105246 |
| ENSMUSG00 Gm17536 | 3633 | 0.32759288 | 0.18339055 | 0.19972556 | 0.12540888 | 0.30471628 | 0.32735084 | 0.44278312 | 0.34691695 | 0.12470456 | 0.23361207 | 0.15780333 | 0.12290815 | 0.19010373 | 0.23011109 | 0.05670459 | 0.16863466 | 0.36923215 | 0.19683963 |
| ENSMUSG00 Gm17535 | 668 | 0 | 0.0498696 | 0.21724639 | 0.20461547 | 0 | 0 | 0.14165473 | 0 | 0 | 0 | 0 | 0 | 0.14770035 | 0 | 0 | 0 | 0 | 0 |
| ENSMUSG00 Gm17538 | 326 | 0.26076795 | 0 | 0 | 0.27951561 | 0.44293181 | 0.15200225 | 0.14513092 | 0 | 0 | 0 | 0.14654895 | 0 | 0 | 0.14246653 | 0.12638514 | 0 | 0.11756533 | 0.16873959 |
| ENSMUSG00 Vmn2r14 | 2541 | 0 | 0 | 0 | 0 | 0 | 0 | 0 | 0 | 0 | 0 | 0 | 0 | 0 | 0 | 0 | 0 | 0 | 0 |
| ENSMUSG00 7420461P10i | 731 | 0 | 0 | 0 | 0 | 0 | 0 | 0 | 0 | 0 | 0 | 0 | 0 | 0 | 0 | 0 | 0 | 0 | 0 |
| ENSMUSG00 4930445B16i | 1118 | 0 | 0 | 0 | 0 | 0 | 0 | 0 | 0 | 0 | 0 | 0 | 0 | 0 | 0 | 0 | 0 | 0 | 0 |
| ENSMUSG00 Ighd5-6 | 11 | 0 | 0 | 0 | 0 | 0 | 0 | 0 | 0 | 0 | 0 | 0 | 0 | 0 | 0 | 0 | 0 | 0 | 0 |
| ENSMUSG00 Gm13698 | 2725 | 0 | 0 | 0 | 0 | 0 | 0 | 0 | 0 | 0 | 0 | 0 | 0 | 0 | 0 | 0 | 0 | 0 | 0 |
| ENSMUSG00 Gm996 | 2824 | 0 | 0 | 0 | 0 | 0 | 0 | 0 | 0 | 0 | 0 | 0 | 0 | 0 | 0 | 0 | 0 | 0 | 0 |
| ENSMUSG00 Gm17599 | 1272 | 0 | 0 | 0 | 0 | 0 | 0 | 0 | 0.12010189 | 0.12951735 | 0 | 0 | 0.17552095 | 0 | 0.10953794 | 0 | 0.27092411 | 0.09039221 | 0 |
| ENSMUSG00 Gm17598 | 994 | 0 | 0 | 0 | 0.04583606 | 0 | 0 | 0.42838442 | 0 | 0 | 0.04743532 | 0.04806334 | 0 | 0.09925939 | 0.04672443 | 0.04145026 | 0 | 0.30846114 | 0.22136461 |
| ENSMUSG00 Gm7889 | 424 | 0 | 0 | 0 | 0 | 0 | 0 | 0 | 0 | 0 | 0 | 0 | 0 | 0 | 0 | 0 | 0 | 0 | 0 |
| ENSMUSG00 Vmn1r175 | 914 | 0 | 0 | 0 | 0 | 0 | 0 | 0 | 0 | 0 | 0 | 0 | 0 | 0 | 0 | 0 | 0 | 0 | 0 |
| ENSMUSG00 Gm3764 | 1086 | 0 | 0 | 0 | 0 | 0 | 0.04562867 | 0 | 0 | 0 | 0 | 0 | 0 | 0 | 0 | 0 | 0 | 0 | 0 |
| ENSMUSG00 1700030C10i | 1470 | 0.14457543 | 0.2039565 | 0.52651462 | 0.18596345 | 0.22919964 | 0.13483737 | 0.25748397 | 0.33775594 | 0.08405411 | 0.12830123 | 0.16249986 | 0.03796984 | 0.10067738 | 0.37913541 | 0.02802827 | 0.83353703 | 0 | 0.74842322 |
| ENSMUSG00 Gm17411 | 34 | 0 | 0 | 0 | 0 | 0 | 0 | 0 | 0 | 0 | 0 | 0 | 0 | 0 | 0 | 0 | 0 | 0 | 0 |
| ENSMUSG00 Gm17410 | 3372 | 0 | 0.00987927 | 0 | 0 | 0 | 0 | 0 | 0 | 0 | 0.01398301 | 0 | 0 | 0 | 0 | 0 | 0 | 0 | 0 |
| ENSMUSG00 Vmn2r115 | 2568 | 0 | 0 | 0 | 0 | 0 | 0 | 0 | 0 | 0 | 0 | 0 | 0 | 0 | 0 | 0 | 0 | 0 | 0 |
| ENSMUSG00 Gm17218 | 406 | 0 | 0.08164925 | 0.35568772 | 0.11166923 | 0 | 0.12145278 | 0.69577469 | 0.09360883 | 0.7066314 | 0.2311309 | 0 | 0.41040928 | 0.12091156 | 0.68300129 | 0 | 0.09384953 | 0 | 0 |
| ENSMUSG00 Gm17105 | 650 | 0.13078516 | 0 | 0 | 0 | 0.22214734 | 0.45740985 | 0 | 0 | 0 | 0.07253954 | 0.07349994 | 0 | 0 | 0 | 0.19016103 | 0.05890863 | 0.23585413 | 0 |
| ENSMUSG00 Prr23a | 1058 | 0 | 0 | 0 | 0 | 0 | 0 | 0 | 0 | 0 | 0 | 0 | 0 | 0 | 0 | 0 | 0 | 0 | 0 |
| ENSMUSG00 Gm17679 | 172 | 0 | 0 | 0 | 0 | 0 | 0 | 0 | 0 | 0 | 0 | 0 | 0 | 0 | 0 | 0 | 0 | 0 | 0 |
| ENSMUSG00 Gm17130 | 2680 | 0 | 0.07458111 | 0 | 0 | 0 | 0 | 0.07061594 | 0 | 0.03073621 | 0.10556128 | 0 | 0.16661392 | 0 | 0 | 0.07686857 | 0.1428754 | 0 | 0 |
| ENSMUSG00 Gm4535 | 557 | 0 | 0 | 0.17369311 | 0.08179721 | 0.08641279 | 0 | 0 | 0 | 0 | 0 | 0 | 0 | 0 | 0.33353025 | 0 | 0 | 0 | 0 |
| ENSMUSG00 BC065403 | 1018 | 0 | 0.03272386 | 0.23759101 | 0 | 0 | 0 | 0 | 0 | 0 | 0.046317 | 0.3285115 | 0 | 0.04845964 | 0.13686863 | 0 | 0 | 0.07529724 | 0 |
| ENSMUSG00 Gm5428 | 887 | 0.23960077 | 0.07511363 | 0.05453611 | 0 | 0.05426372 | 0.11173108 | 0 | 0 | 0.0464335 | 0.1594725 | 0 | 0.06292634 | 0 | 0.10472173 | 0.09290092 | 0.129506 | 0.08641781 | 0 |
| ENSMUSG00 Ighv8-14 | 119 | 0 | 0 | 0 | 0 | 0 | 0 | 0 | 0 | 0 | 0 | 0 | 0 | 0 | 0 | 0 | 0 | 0 | 0 |
| ENSMUSG00 4921514A10i | 2114 | 0 | 0 | 0.11441232 | 0 | 0.02276818 | 0 | 0.08952257 | 0 | 0.07793097 | 0.04460804 | 0 | 0 | 0 | 0 | 0 | 0.03622574 | 0 | 0 |
| ENSMUSG00 A930018M24 | 1420 | 1.22726212 | 0.42227612 | 0.3065928 | 0.73796058 | 0.10168716 | 0.83751099 | 0.2665503 | 0.37654481 | 0.02900459 | 0.36525193 | 0 | 0.98267012 | 0 | 0 | 0.02901518 | 0.40447825 | 0.02699035 | 0.5036045 |
| ENSMUSG00 Gm5269 | 748 | 0 | 0 | 0 | 0 | 0 | 0 | 0 | 0 | 0 | 0 | 0 | 0 | 0 | 0 | 0 | 0.1023813 | 0 | 0 |
| ENSMUSG00 Kcnmb3 | 716 | 0 | 0 | 0.13512159 | 0.12726549 | 0 | 0 | 0 | 0.10668269 | 0 | 0.32926469 | 0 | 0.23386451 | 0 | 0.12973209 | 0 | 0.21391401 | 0.05352835 | 0.30731345 |
| ENSMUSG00 Gm17302 | 785 | 0 | 0 | 0 | 0 | 0 | 0 | 0 | 0 | 0 | 0 | 0 | 0 | 0 | 0 | 0 | 0 | 0 | 0 |
| ENSMUSG00 Gm9222 | 335 | 9.64296531 | 9.94414753 | 10.2523005 | 10.0642307 | 7.75857874 | 7.6917676 | 10.309927 | 8.20851634 | 8.4831928 | 8.30415371 | 9.98282714 | 13.4957274 | 8.83556727 | 7.34787054 | 8.60928034 | 7.7724218 | 8.23729366 | 12.3154717 |
| ENSMUSG00 Gm17305 | 501 | 0.08484067 | 0.19947841 | 0 | 0.63658146 | 0 | 0.29672296 | 0.18887297 | 0.07623234 | 0.08220861 | 0.09411318 | 0.0953592 | 0 | 0.0984669 | 0.27810831 | 0.16447727 | 0.22928507 | 0.38249797 | 0.10979862 |
| ENSMUSG00 Gm17304 | 67 | 0 | 0 | 0 | 2.72006235 | 0 | 0 | 0 | 1.71010757 | 0 | 0 | 0 | 0 | 0 | 0 | 0 | 0 | 0 | 0 |
| ENSMUSG00 Gm17303 | 74 | 0 | 0 | 0 | 0 | 1.9512942 | 0 | 3.83616316 | 0 | 4.4525963 | 1.27434333 | 2.58243019 | 0.75426571 | 0.66664753 | 3.13811401 | 0 | 0 | 6.21507517 | 2.23009892 |
| ENSMUSG00 Cldn21 | 653 | 0 | 0 | 0 | 0 | 0 | 0 | 0 | 0 | 0 | 0 | 0 | 0 | 0 | 0 | 0 | 0 | 0 | 0 |
| ENSMUSG00 Gm17164 | 503 | 0 | 0 | 0 | 0 | 0 | 0 | 0 | 0 | 0 | 0 | 0 | 0 | 0 | 0 | 0 | 0 | 0 | 0 |
| ENSMUSG00 Gm17626 | 3795 | 0.04480124 | 0.13167152 | 0.10197319 | 0.12005545 | 0.03804895 | 0.06528687 | 0.02493422 | 0.24153298 | 0.06511702 | 0.04969771 | 0.10071137 | 0.1911999 | 0.03899756 | 0.15909701 | 0.0108568 | 0.04035901 | 0.13128903 | 0 |
| ENSMUSG00 Gm4302 | 884 | 0 | 0 | 0 | 0 | 0 | 0 | 0 | 0 | 0 | 0 | 0 | 0 | 0 | 0 | 0 | 0 | 0 | 0 |
| ENSMUSG00 5830462I19R | 3382 | 0.28906536 | 0.09850057 | 0.07151616 | 0.13471628 | 0.12808614 | 0.16117093 | 0.12590601 | 0.30490682 | 0.40187907 | 0.12547496 | 0.09888371 | 0.31357114 | 0.21879916 | 0.04119819 | 0.14619121 | 0.2604033 | 0.16998653 | 0.34157044 |
| ENSMUSG00 Gm17623 | 58 | 0 | 0 | 0 | 0 | 0 | 2.56307243 | 0 | 0 | 0 | 0 | 0 | 2.88701703 | 0 | 1.60152025 | 0 | 3.30091443 | 2.64319289 | 0 |
| ENSMUSG00 Gm5633 | 548 | 0 | 0 | 0 | 0 | 0 | 0 | 0 | 0 | 0.07515788 | 0.08604143 | 0.34872232 | 0 | 0.09002175 | 0 | 0.07518532 | 0 | 0 | 0 |
| ENSMUSG00 Gm17625 | 3197 | 0 | 0 | 0 | 0 | 0 | 0 | 0 | 0.01194633 | 0 | 0 | 0 | 0 | 0.01543069 | 0 | 0 | 0 | 0 | 0 |
| ENSMUSG00 Gm17624 | 199 | 0.64078155 | 0 | 0 | 0.22894997 | 0.24186896 | 0 | 0 | 0 | 0 | 0 | 0 | 0.28048072 | 0 | 0 | 0.41408599 | 0.19241511 | 0 | 0 |
| ENSMUSG00 Gm13276 | 1555 | 0 | 0 | 0 | 0 | 0 | 0 | 0 | 0 | 0 | 0 | 0 | 0 | 0 | 0 | 0 | 0 | 0 | 0 |
| ENSMUSG00 Gm17622 | 43 | 0 | 0 | 0 | 1.05955917 | 1.11934706 | 2.30477831 | 1.10029486 | 0 | 0.95782595 | 0 | 0 | 0 | 0 | 0 | 0 | 0 | 0 | 0 |
| ENSMUSG00 Gm2832 | 1755 | 0 | 0 | 0 | 0 | 0 | 0 | 0 | 0 | 0 | 0 | 0 | 0 | 0 | 0 | 0 | 0 | 0 | 0 |
| ENSMUSG00 Gm8324 | 1203 | 0 | 0 | 0 | 0 | 0 | 0 | 0 | 0 | 0 | 0 | 0 | 0 | 0 | 0 | 0 | 0 | 0 | 0 |
| ENSMUSG00 Gm17310 | 40 | 0 | 0 | 0 | 0 | 0 | 0 | 0 | 0 | 0 | 0 | 0 | 0 | 0 | 0 | 0 | 0 | 0 | 0 |
| ENSMUSG00 Gm17311 | 1743 | 0 | 0 | 0 | 0.1306972 | 0 | 0 | 0 | 0 | 0 | 0 | 0 | 0 | 0 | 0 | 0 | 0 | 0 | 0 |
| ENSMUSG00 Gm3138 | 1755 | 0 | 0 | 0 | 0 | 0 | 0 | 0 | 0 | 0 | 0 | 0 | 0 | 0 | 0 | 0 | 0 | 0 | 0 |
| ENSMUSG00 BC027582 | 873 | 0 | 0 | 0 | 0 | 0 | 0 | 0 | 0 | 0 | 0 | 0 | 0 | 0.16952549 | 0.21280223 | 0 | 0 | 0.21950915 | 0 |
| ENSMUSG00 Gm7991 | 587 | 0 | 0 | 0 | 0 | 0 | 0 | 0 | 0 | 0 | 0 | 0 | 0 | 0 | 0 | 0 | 0 | 0 | 0 |
| ENSMUSG00 Gm9064 | 565 | 0 | 0.11792175 | 0 | 0 | 0 | 0 | 0 | 0 | 0 | 0 | 0 | 0 | 0.17462626 | 0 | 0 | 0 | 0 | 0 |
| ENSMUSG00 Gm20603 | 3483 | 0 | 0.00956443 | 0 | 0 | 0 | 0 | 0 | 0.06579225 | 0 | 0 | 0.01371661 | 0 | 0.02832726 | 0 | 0.03548799 | 0.076955 | 0.04401527 | 0 |
| ENSMUSG00 Ccdc152 | 789 | 0.37710549 | 0.71776832 | 0.42916947 | 0.40421712 | 0.61003706 | 0.75365374 | 0.53968835 | 0.53246695 | 0.62641089 | 0.2988004 | 0.48441023 | 0.49519599 | 0.56272149 | 0.64756946 | 0 | 0.72795832 | 0.63148525 | 0.55776027 |
| ENSMUSG00 Gm4545 | 1510 | 0 | 0 | 0 | 0 | 0 | 0 | 0 | 0 | 0 | 0 | 0 | 0 | 0 | 0 | 0 | 0 | 0 | 0 |
| ENSMUSG00 Vmn1r104 | 923 | 0 | 0 | 0 | 0 | 0 | 0 | 0 | 0 | 0 | 0 | 0 | 0 | 0 | 0 | 0 | 0 | 0 | 0 |
| ENSMUSG00 Gm8247 | 1400 | 0 | 0 | 0 | 0 | 0 | 0 | 0 | 0 | 0 | 0 | 0 | 0 | 0 | 0 | 0 | 0 | 0 | 0 |
| ENSMUSG00 Gm17181 | 122 | 0 | 0 | 0 | 0 | 0 | 0 | 0 | 0 | 0 | 0 | 0 | 0 | 0 | 0 | 0 | 0 | 0 | 0 |
| ENSMUSG00 Gm17239 | 49 | 0 | 0 | 1.97442981 | 0 | 0 | 0 | 0 | 0 | 0 | 0 | 0.97499915 | 4.55638062 | 0 | 0 | 0 | 0 | 0 | 0 |
| ENSMUSG00 Gm17238 | 696 | 9.89344614 | 11.2957515 | 13.1359156 | 8.05173628 | 7.88367715 | 6.69246689 | 11.8281697 | 14.9806406 | 10.9475653 | 13.4813074 | 8.85484144 | 9.6233901 | 12.4038585 | 11.6777519 | 11.3067488 | 7.75714891 | 9.85690682 | 8.85204016 |
| ENSMUSG00 Iqcf6 | 472 | 0 | 0 | 0 | 0 | 0 | 0 | 0 | 0 | 0 | 0 | 0 | 0 | 0 | 0 | 0 | 0 | 0 | 0 |
| ENSMUSG00 Gm14474 | 524 | 0 | 0 | 0 | 0 | 0 | 0 | 0 | 0 | 0 | 0 | 0 | 0 | 0 | 0 | 0 | 0 | 0 | 0 |
| ENSMUSG00 Gm7945 | 1423 | 0 | 0 | 0 | 0 | 0 | 0 | 0 | 0 | 0 | 0 | 0 | 0 | 0 | 0 | 0 | 0 | 0 | 0 |
| ENSMUSG00 Gm17109 | 357 | 0 | 0 | 0 | 0 | 0 | 0 | 0 | 0 | 0 | 0 | 0 | 0 | 0 | 0 | 0 | 0 | 0 | 0 |
| ENSMUSG00 Gm17108 | 2875 | 0.20698173 | 0.23174187 | 0.13460461 | 0.11093124 | 0.05022462 | 0.0517072 | 0.0987395 | 0.37196079 | 0.20056042 | 0.18040269 | 0.13293901 | 0.21355558 | 0.24022499 | 0.3230893 | 0.3009505 | 0.26636944 | 0.07998532 | 0.22960323 |
| ENSMUSG00 Gm17436 | 59 | 0 | 0 | 0 | 0 | 0 | 0 | 0 | 0 | 0 | 0 | 0 | 0 | 0 | 0 | 0 | 0 | 0 | 0 |
| ENSMUSG00 Gm17209 | 518 | 0 | 0 | 0 | 0 | 0 | 0 | 0 | 0 | 0 | 0 | 0 | 0 | 0 | 0 | 0 | 0 | 0 | 0 |
| ENSMUSG00 Gm17437 | 1966 | 0.15134091 | 0.0169445 | 0.07381515 | 0 | 0 | 0 | 0 | 0 | 0 | 0 | 0 | 0 | 0 | 0.04724729 | 0 | 0 | 0 | 0 |
| ENSMUSG00 Gm1153 | 506 | 0 | 0 | 0 | 0 | 0 | 0 | 0 | 0 | 0 | 0 | 0 | 0 | 0 | 0 | 0 | 0 | 0 | 0 |
| ENSMUSG00 E530011L22i | 2065 | 4.11272027 | 2.90760922 | 2.5574036 | 2.74956926 | 1.9444817 | 2.91632048 | 2.87887622 | 3.0096756 | 3.04020166 | 2.98660315 | 2.54958631 | 3.31275005 | 1.99296025 | 2.45539814 | 2.26043449 | 2.69275593 | 2.40853537 | 1.94795341 |
| ENSMUSG00 Gm6190 | 772 | 0 | 0 | 0.06266001 | 0.0590169 | 0 | 0 | 0.18385756 | 0.04947202 | 0.05335041 | 0.18322812 | 0.18565398 | 0 | 0.06390145 | 0 | 0.10673978 | 0.29759539 | 0.19858185 | 0.07125532 |
| ENSMUSG00 Gm17124 | 1374 | 0 | 0 | 0 | 0 | 0 | 0 | 0 | 0 | 0 | 0 | 0 | 0 | 0 | 0 | 0 | 0 | 0 | 0 |
| ENSMUSG00 Gm17367 | 58 | 0 | 0 | 0 | 0 | 0 | 0 | 0 | 0 | 0 | 0 | 0 | 0 | 0 | 0 | 0 | 0 | 0 | 0 |
| ENSMUSG00 Gm17175 | 1398 | 0 | 0 | 0 | 0 | 0 | 0 | 0 | 0 | 0 | 0 | 0 | 0 | 0 | 0 | 0 | 0 | 0 | 0 |
| ENSMUSG00 Gm6907 | 3448 | 0.43146205 | 0.34781444 | 0.14029446 | 0.11892384 | 0.19543124 | 0.02874288 | 0.13721775 | 0.23261034 | 0.07167027 | 0.16409758 | 0.180126 | 0.14569053 | 0.40060722 | 0.21551781 | 0.19119051 | 0.16657747 | 0.42238958 | 0.12763134 |
| ENSMUSG00 Gm4308 | 915 | 0 | 0 | 0 | 0 | 0 | 0 | 0 | 0 | 0 | 0 | 0 | 0 | 0 | 0 | 0 | 0 | 0 | 0 |
| ENSMUSG00 Gm3077 | 624 | 0 | 0 | 0 | 0 | 0 | 0 | 0 | 0 | 0 | 0 | 0 | 0 | 0 | 0 | 0 | 0 | 0 | 0 |
| ENSMUSG00 4921509A06i | 390 | 0 | 0 | 0 | 0 | 0.24683038 | 0 | 0 | 0 | 0 | 0 | 0.36749968 | 0 | 0 | 0 | 0 | 0.39272418 | 0 | 1.55153891 |
| ENSMUSG00 Gm3182 | 1939 | 0 | 0 | 0 | 0 | 0 | 0 | 0 | 0 | 0 | 0 | 0 | 0 | 0 | 0 | 0 | 0 | 0 | 0 |
| ENSMUSG00 Gm17020 | 273 | 0 | 0 | 0 | 0 | 0 | 0 | 0 | 0.27979782 | 0 | 0 | 0.3499997 | 0.40890595 | 0.90351497 | 0 | 0 | 0.98181045 | 0.14038937 | 0.60449568 |
| ENSMUSG00 Gm17544 | 73 | 0 | 0 | 0 | 0 | 0 | 0 | 0 | 0 | 0 | 0 | 0 | 0 | 0 | 0 | 0 | 0 | 0 | 0 |
| ENSMUSG00 Vmn1r224 | 896 | 0 | 0 | 0 | 0 | 0 | 0 | 0 | 0 | 0 | 0 | 0 | 0 | 0 | 0 | 0 | 0 | 0 | 0 |
| ENSMUSG00 Gm13696 | 2725 | 0 | 0 | 0 | 0 | 0 | 0 | 0 | 0 | 0 | 0 | 0 | 0 | 0 | 0 | 0 | 0 | 0 | 0 |
| ENSMUSG00 Olfr92 | 938 | 0 | 0 | 0 | 0 | 0 | 0 | 0 | 0 | 0 | 0 | 0 | 0 | 0 | 0 | 0 | 0 | 0 | 0 |
| ENSMUSG00 1700024N20i | 615 | 0.8984834 | 1.13751346 | 2.04505982 | 0.66674699 | 0.78263291 | 0.40286775 | 2.23100437 | 1.05572494 | 1.205459 | 1.99336306 | 1.16524289 | 0.54454305 | 1.44386099 | 2.71867829 | 3.14873679 | 2.17914026 | 1.55797955 | 2.14669685 |
| ENSMUSG00 Serpine3 | 1393 | 0.27462066 | 0.14348698 | 0.24308307 | 0.26165711 | 0.27642167 | 0.35572673 | 0.40757512 | 0.38384324 | 0.23653419 | 0.20308989 | 0.13718581 | 0.04006867 | 0.28331324 | 0.53345685 | 0.53239627 | 0.19241511 | 0.38518891 | 0.03948967 |
| ENSMUSG00 Gm3500 | 2040 | 0 | 0 | 0 | 0 | 0 | 0 | 0 | 0 | 0 | 0 | 0 | 0 | 0 | 0 | 0 | 0 | 0 | 0 |
| ENSMUSG00 Serpina3l-ps | 1260 | 0 | 0 | 0 | 0 | 0 | 0 | 0 | 0 | 0 | 0 | 0 | 0 | 0 | 0 | 0 | 0 | 0 | 0 |
| ENSMUSG00 Mirg | 3166 | 0 | 0 | 0 | 0 | 0 | 0 | 0 | 0.15682288 | 0 | 0 | 0 | 0 | 0 | 0 | 0 | 0 | 0 | 0 |
| ENSMUSG00 Gm17545 | 70 | 0 | 1.42769547 | 1.38210087 | 0 | 0 | 0 | 0 | 2.18242299 | 0.5883788 | 0 | 0 | 0 | 0 | 1.32697393 | 0 | 0.54700868 | 0 | 0 |
| ENSMUSG00 Gm17133 | 287 | 0 | 0 | 0 | 0 | 0 | 0 | 0.16485254 | 0 | 0 | 0 | 0 | 0 | 0 | 0 | 0 | 0 | 0 | 0 |
| ENSMUSG00 Gm6164 | 923 | 0 | 0 | 0 | 0 | 0 | 0 | 0 | 0 | 0 | 0 | 0 | 0 | 0 | 0 | 0 | 0 | 0 | 0 |
| ENSMUSG00 Gm7677 | 1782 | 0 | 0 | 0 | 0 | 0 | 0 | 0 | 0 | 0 | 0 | 0 | 0 | 0 | 0 | 0 | 0 | 0 | 0.24695446 |
| ENSMUSG00 Gm3962 | 2611 | 0 | 0 | 0 | 0 | 0 | 0 | 0 | 0 | 0 | 0 | 0 | 0 | 0 | 0 | 0 | 0 | 0 | 0 |
| ENSMUSG00 Gm17716 | 2541 | 0.35128245 | 0.20976242 | 0.53304166 | 0.16137324 | 0.47355297 | 0.50703309 | 0.446873 | 0.12024369 | 0.8266479 | 0.2041156 | 0.01880164 | 0.1976942 | 0.44653054 | 0.14622302 | 0.51887044 | 0.19589842 | 0.07541578 | 0.30308048 |
| ENSMUSG00 Gm17036 | 252 | 0 | 0 | 0 | 0.18079779 | 0.1909997 | 0 | 0 | 0.75778576 | 1.14406988 | 0 | 0.56874951 | 0.44298145 | 0.39152315 | 0.18430193 | 0 | 0.30389371 | 0.15208848 | 0.43658021 |
| ENSMUSG00 Tstd1 | 547 | 1.63182577 | 0.60901086 | 2.12242181 | 0.41646293 | 0.70394038 | 0.81531006 | 1.38391748 | 0.97750207 | 0.6023622 | 0.60339108 | 0.34935984 | 0.51019801 | 1.89391273 | 1.27360386 | 2.86226531 | 2.45003886 | 2.17205705 | 0.10056589 |
| ENSMUSG00 Gm14477 | 499 | 0 | 0 | 0 | 0 | 0 | 0 | 0 | 0 | 0 | 0 | 0 | 0 | 0 | 0 | 0 | 0 | 0 | 0 |
| ENSMUSG00 1700026J14R | 792 | 0 | 0.21030868 | 0.36646614 | 0 | 0 | 0.06256658 | 0.11947646 | 0 | 0 | 0.05953372 | 0.42225342 | 0 | 0.18686332 | 0.4691322 | 0.10404433 | 0.62850744 | 0.33874252 | 0 |
| ENSMUSG00 Gm17493 | 1132 | 0.22529245 | 0 | 0.47006081 | 0.08049654 | 0.08503873 | 0.83171549 | 2.08978264 | 0.84347178 | 0.21830309 | 0.58313591 | 0.42204027 | 0.64099259 | 1.52528013 | 1.47702045 | 0.90992835 | 0.43973312 | 0.44014299 | 0.58313541 |
| ENSMUSG00 Gm4307 | 869 | 0 | 0 | 0 | 0 | 0 | 0 | 0 | 0 | 0 | 0 | 0 | 0 | 0 | 0 | 0 | 0 | 0 | 0 |
| ENSMUSG00 Gm10060 | 4008 | 0.07423559 | 0.041558 | 0.06034622 | 0.05683763 | 0.02401793 | 0.01236346 | 0.14165473 | 0.03811617 | 0.14386508 | 0.05882074 | 0.0357597 | 0.11140851 | 0.06154181 | 0.03476354 | 0.06167898 | 0.04776772 | 0.06693715 | 0.19214758 |
| ENSMUSG00 Gm17120 | 741 | 0 | 0 | 0 | 0 | 0 | 0 | 0 | 0 | 0 | 0 | 0 | 0 | 0 | 0.06267758 | 0 | 0 | 0 | 0 |
| ENSMUSG00 Gm7354 | 506 | 0 | 0 | 0 | 0 | 0 | 0 | 0 | 0 | 0 | 0 | 0 | 0 | 0 | 0 | 0 | 0 | 0 | 0 |
| ENSMUSG00 Gm17677 | 348 | 0.24428262 | 2.39316769 | 0 | 1.70199304 | 0.69155063 | 2.84785825 | 0 | 0 | 0 | 0 | 0 | 0 | 0 | 0 | 0 | 0 | 0 | 0 |
| ENSMUSG00 9130204L05i | 488 | 0 | 0 | 0 | 0 | 0 | 0 | 0 | 0 | 0 | 0 | 0 | 0 | 0 | 0 | 0 | 0 | 0 | 0 |
| ENSMUSG00 Gm17678 | 242 | 0 | 0 | 0 | 0.56480633 | 0 | 0 | 0 | 0 | 0 | 0.38967523 | 0 | 0 | 0.61155269 | 0.76767087 | 0.17025436 | 0.15822565 | 0.63349251 | 0 |
| ENSMUSG00 Gm15494 | 449 | 0 | 0 | 0 | 0 | 0 | 0.22072487 | 0 | 0.08506103 | 0.73383547 | 0 | 0.10640303 | 0.37293316 | 0 | 0 | 0 | 0.1705595 | 0 | 0 |
| ENSMUSG00 Vmn2r44 | 2580 | 0 | 0 | 0 | 0 | 0 | 0 | 0 | 0 | 0 | 0 | 0 | 0 | 0 | 0 | 0 | 0 | 0 | 0 |
| ENSMUSG00 Gm3597 | 1754 | 0 | 0 | 0 | 0 | 0 | 0 | 0 | 0 | 0 | 0 | 0 | 0 | 0 | 0 | 0 | 0 | 0 | 0 |
| ENSMUSG00 Trav5n-3 | 269 | 0 | 0 | 0 | 0 | 0 | 0 | 0 | 0 | 0 | 0 | 0 | 0 | 0 | 0 | 0 | 0 | 0 | 0 |
| ENSMUSG00 Gm17647 | 34 | 0 | 0 | 0 | 0 | 0 | 0 | 1.39154938 | 0 | 1.21136811 | 0 | 0 | 0 | 0 | 0 | 0 | 0 | 0 | 0 |
| ENSMUSG00 Gm17649 | 49 | 0 | 0 | 0 | 0 | 0 | 0 | 0 | 0 | 0 | 0 | 0 | 0 | 0 | 0 | 0 | 0 | 0 | 0 |
| ENSMUSG00 Gm3604 | 1674 | 1.44730886 | 0.89550791 | 0.8091152 | 0.81650617 | 1.38012684 | 1.12485297 | 1.04574022 | 2.09898508 | 2.23893246 | 0.81682819 | 1.45550949 | 1.73381987 | 1.03143196 | 1.60917388 | 2.78122809 | 2.10438225 | 1.144752 | 1.5116003 |
| ENSMUSG00 Gm17059 | 517 | 0.57550528 | 0 | 0 | 0 | 0 | 0.09584668 | 0 | 0.44323871 | 0 | 0 | 0.6468563 | 0 | 0.09541957 | 0 | 0.39846766 | 0 | 0 | 0 |
| ENSMUSG00 Gm3278 | 1495 | 0 | 0 | 0 | 0 | 0 | 0 | 0 | 0 | 0 | 0 | 0 | 0 | 0 | 0 | 0 | 0 | 0 | 0 |
| ENSMUSG00 Gm2078 | 8379 | 0 | 0.00795152 | 0 | 0.00543753 | 0.00574435 | 0 | 0 | 0.00455811 | 0 | 0 | 0 | 0 | 0 | 0 | 0 | 0 | 0 | 0.01313023 |
| ENSMUSG00 Gm8082 | 1398 | 0 | 0 | 0 | 0 | 0 | 0 | 0 | 0 | 0 | 0 | 0 | 0 | 0 | 0 | 0 | 0 | 0 | 0 |
| ENSMUSG00 2700016F22i | 3603 | 0.23594325 | 0.22190104 | 0.17453675 | 0.30348739 | 0.34733001 | 0.17879143 | 0.22323495 | 0.23320368 | 0.49154043 | 0.18321117 | 0.29171498 | 0.18589729 | 0.20537851 | 0.06445197 | 0.35449576 | 0.17003878 | 0.20210925 | 0 |
| ENSMUSG00 Ear-ps3 | 444 | 0 | 0 | 0 | 0 | 0 | 0 | 0 | 0 | 0 | 0 | 0 | 0 | 0 | 0 | 0 | 0 | 0 | 0 |
| ENSMUSG00 2310031A07i | 988 | 0.17208573 | 0.10115251 | 0 | 0.18445767 | 0 | 0 | 0.38309861 | 0.23193767 | 0.16674703 | 0.09544677 | 0.09671044 | 0.05649359 | 0.24965545 | 0.37606548 | 0.16680792 | 0.3100454 | 0.0387918 | 0 |
| ENSMUSG00 Gm17334 | 906 | 0.23457603 | 0 | 0.05339242 | 0.10057626 | 0.31875446 | 0.38285776 | 0.104443 | 0.12646491 | 0.50005703 | 0.26021359 | 0.31639045 | 0.18482007 | 0.65340288 | 0.92273021 | 0.7276213 | 0.84526727 | 0.59223858 | 1.09289616 |
| ENSMUSG00 Gm17071 | 263 | 0 | 0 | 0 | 0.69294364 | 0 | 0 | 0 | 0 | 0.62641089 | 0 | 0 | 0 | 0 | 0.52978046 | 0 | 0 | 0 | 0 |
| ENSMUSG00 Gm17333 | 376 | 0 | 0 | 0 | 0 | 0 | 0 | 0 | 0 | 0 | 0 | 0 | 0 | 0 | 0 | 0 | 0 | 0 | 0 |
| ENSMUSG00 Gm19840 | 880 | 0.09660267 | 0.03785556 | 0 | 0 | 0 | 0 | 0 | 0 | 0 | 0 | 0 | 0 | 0 | 0 | 0 | 0 | 0 | 0 |
| ENSMUSG00 Gm17332 | 86 | 0 | 0 | 0 | 0 | 0 | 0 | 0 | 0 | 0 | 0 | 0 | 0 | 0 | 0 | 0 | 0 | 0 | 0 |
| ENSMUSG00 Gm17330 | 335 | 0 | 0 | 0 | 0.40800935 | 0 | 0.29583722 | 0 | 0.11400717 | 0 | 0 | 0 | 0 | 0 | 0 | 0 | 0 | 0 | 0 |
| ENSMUSG00 Vmn1r95 | 923 | 0 | 0 | 0 | 0 | 0 | 0 | 0 | 0 | 0 | 0 | 0 | 0 | 0 | 0 | 0 | 0 | 0 | 0 |
| ENSMUSG00 Gm5218 | 464 | 0 | 0 | 0 | 0 | 0 | 0 | 0 | 0 | 0 | 0 | 0 | 0 | 0 | 0 | 0 | 0 | 0 | 0 |
| ENSMUSG00 Gm2619 | 490 | 0 | 0 | 0 | 0 | 0 | 0 | 0 | 0 | 0 | 0 | 0 | 0 | 0 | 0.09478385 | 0 | 0 | 0 | 0 |
| ENSMUSG00 Vmn1r77 | 920 | 0 | 0 | 0 | 0 | 0 | 0 | 0 | 0 | 0 | 0 | 0 | 0 | 0 | 0 | 0 | 0 | 0 | 0 |
| ENSMUSG00 Gm3685 | 979 | 0 | 0 | 0 | 0 | 0 | 0 | 0 | 0 | 0 | 0 | 0 | 0 | 0 | 0 | 0 | 0 | 0 | 0 |
| ENSMUSG00 Gm8011 | 1746 | 0 | 0 | 0 | 0 | 0 | 0 | 0 | 0 | 0 | 0 | 0 | 0 | 0 | 0 | 0 | 0 | 0 | 0 |
| ENSMUSG00 Gm7271 | 516 | 0 | 0 | 0 | 0 | 0 | 0 | 0 | 0 | 0 | 0 | 0 | 0 | 0 | 0 | 0 | 0 | 0 | 0 |
| ENSMUSG00 Vmn2r71 | 2562 | 0 | 0 | 0 | 0 | 0 | 0 | 0 | 0 | 0 | 0 | 0 | 0 | 0 | 0 | 0 | 0 | 0 | 0 |
| ENSMUSG00 Vmn2r91 | 2565 | 0 | 0 | 0 | 0 | 0 | 0 | 0 | 0 | 0 | 0 | 0 | 0 | 0 | 0 | 0 | 0 | 0 | 0 |
| ENSMUSG00 Purb | 1125 | 33.0973637 | 35.5337538 | 38.3978334 | 31.4269959 | 33.2858992 | 36.6029525 | 29.1866659 | 35.5443958 | 30.2400551 | 32.8168895 | 34.7801698 | 36.1189354 | 35.6066816 | 36.4947318 | 31.825913 | 29.7815835 | 32.2962929 | 35.6459012 |
| ENSMUSG00 Gm3494 | 887 | 0 | 0 | 0 | 0 | 0 | 0 | 0 | 0 | 0 | 0 | 0 | 0 | 0 | 0 | 0 | 0 | 0 | 0 |
| ENSMUSG00 Gm3264 | 1004 | 0 | 0 | 0 | 0 | 0 | 0 | 0 | 0 | 0 | 0 | 0 | 0 | 0 | 0 | 0 | 0 | 0 | 0 |
| ENSMUSG00 1110054M08 | 1601 | 7.14171915 | 5.70126984 | 7.73493053 | 7.37058743 | 7.81655225 | 7.14970734 | 9.10200194 | 8.37322501 | 8.77239343 | 8.57017779 | 8.08682932 | 7.39095595 | 8.19630855 | 7.22334651 | 6.43372203 | 8.27517187 | 6.17625522 | 9.07083334 |
| ENSMUSG00 Gm4211 | 2218 | 0 | 0 | 0 | 0 | 0 | 0 | 0 | 0 | 0 | 0 | 0.51695176 | 0.02516486 | 0 | 0 | 0 | 0 | 0 | 0 |
| ENSMUSG00 Krtap11-1 | 1007 | 0 | 0 | 0 | 0 | 0 | 0 | 0 | 0 | 0 | 0 | 0 | 0 | 0 | 0 | 0 | 0 | 0 | 0 |
| ENSMUSG00 Gm17253 | 64 | 11.2904374 | 7.80770958 | 6.80252771 | 0.71189132 | 9.02473569 | 4.64556877 | 6.65334547 | 4.7740503 | 10.9401682 | 2.21018922 | 3.73241863 | 1.74423946 | 8.47892328 | 8.70826639 | 1.93132293 | 11.9658148 | 7.78502906 | 6.01662105 |
| ENSMUSG00 4732491K20i | 3140 | 6.01028604 | 4.62561206 | 4.80654187 | 5.74591515 | 5.11976513 | 5.94948425 | 2.41083714 | 2.82185903 | 2.41347736 | 2.53772893 | 4.04717801 | 3.99952996 | 2.30948785 | 2.27783104 | 2.34875112 | 3.26811554 | 2.28248966 | 3.87166006 |
| ENSMUSG00 Gm17252 | 383 | 4.43918287 | 1.82655556 | 1.01041317 | 0.23791668 | 1.13103737 | 0.38814152 | 153.426494 | 126.543495 | 110.332546 | 68.4485405 | 6.11220095 | 75.9267891 | 79.8584562 | 79.306614 | 33.0257902 | 27.6932069 | 7.40508086 | 36.1939814 |
| ENSMUSG00 Gm17208 | 527 | 0 | 0 | 0 | 0 | 0 | 0 | 0 | 0 | 0 | 0 | 0 | 0 | 0 | 0 | 0 | 0 | 0 | 0 |
| ENSMUSG00 5430400D12i | 1656 | 0.74435393 | 1.42827022 | 2.57057408 | 2.06345309 | 1.27886754 | 1.58592686 | 0.97139558 | 1.40684574 | 0.77100362 | 1.11043323 | 1.52902947 | 1.95489639 | 0.08936942 | 0.75724056 | 0.99520666 | 1.3642185 | 0.0925756 | 0.43183477 |
| ENSMUSG00 Gm17254 | 4474 | 0.38951994 | 0.20103892 | 0.43248574 | 0.06110109 | 0.04303256 | 0.08860569 | 0.61335167 | 0.12804784 | 0.2301437 | 0.17916003 | 0.10678355 | 0.13723118 | 0.48515967 | 0.05190443 | 0.11971842 | 0.35945586 | 0.1199303 | 0.06147643 |
| ENSMUSG00 Olfr724 | 1005 | 0 | 0 | 0 | 0 | 0 | 0 | 0 | 0 | 0 | 0 | 0 | 0 | 0 | 0 | 0 | 0 | 0 | 0 |
| ENSMUSG00 Gm3084 | 597 | 0 | 0 | 0 | 0 | 0 | 0 | 0 | 0 | 0 | 0 | 0 | 0 | 0 | 0 | 0 | 0 | 0 | 0 |
| ENSMUSG00 Gm17723 | 66 | 0 | 0 | 0 | 0 | 0 | 0 | 0 | 0 | 0 | 0 | 0 | 0 | 0 | 0 | 0 | 0 | 0 | 0 |
| ENSMUSG00 Gm8775 | 512 | 0 | 0.06506425 | 0 | 0 | 0 | 0 | 0 | 0.29837814 | 0 | 0 | 0 | 0.10901497 | 0 | 0 | 0.08047179 | 0 | 0 | 0.21487932 |
| ENSMUSG00 Vmn1r122 | 917 | 0 | 0 | 0 | 0 | 0 | 0 | 0 | 0 | 0 | 0 | 0 | 0 | 0 | 0 | 0 | 0 | 0 | 0 |
| ENSMUSG00 Vmn1r176 | 914 | 0 | 0 | 0 | 0 | 0 | 0 | 0 | 0 | 0 | 0 | 0 | 0 | 0 | 0 | 0 | 0 | 0 | 0 |
| ENSMUSG00 Gm3755 | 1457 | 0 | 0 | 0 | 0 | 0 | 0 | 0 | 0 | 0 | 0 | 0 | 0 | 0 | 0 | 0 | 0 | 0 | 0 |
| ENSMUSG00 Gm20390 | 983 | 0 | 0 | 0 | 0 | 0 | 0 | 0 | 0 | 0 | 0 | 0 | 0 | 0 | 0 | 0 | 0 | 0 | 0 |
| ENSMUSG00 Gm7177 | 539 | 0 | 0 | 0.08974681 | 0.08452884 | 0 | 0 | 0 | 0 | 0 | 0 | 0 | 0 | 0 | 0 | 0 | 0 | 0 | 0 |
| ENSMUSG00 Gm6970 | 665 | 0.51134046 | 1.45274276 | 2.10952238 | 1.37025697 | 2.31612264 | 2.83158478 | 1.77867214 | 0.63175402 | 0.49547688 | 1.20535632 | 0.71842043 | 2.01439985 | 2.29968336 | 0.48888513 | 0.37174336 | 1.03643749 | 3.34274469 | 2.31617291 |
| ENSMUSG00 Gm4033 | 2552 | 0 | 0 | 0 | 0 | 0 | 0 | 0 | 0 | 0 | 0 | 0 | 0 | 0 | 0 | 0 | 0 | 0 | 0 |
| ENSMUSG00 Gm17394 | 43 | 0 | 0 | 0 | 0 | 1.11934706 | 0 | 0 | 0 | 0 | 0 | 0 | 0 | 0 | 0 | 0 | 0 | 0 | 0 |
| ENSMUSG00 Gm6434 | 407 | 0 | 0 | 0 | 0 | 0.11826025 | 0 | 0.11624737 | 0.28151648 | 0 | 0 | 0.11738319 | 0.13713922 | 0.12120864 | 0 | 0 | 0 | 0.09416781 | 0.13515751 |
| ENSMUSG00 Gm17393 | 2797 | 1.3980966 | 1.42922678 | 1.07227704 | 1.35200811 | 1.34225601 | 1.66534035 | 1.13333911 | 1.11969145 | 1.29582173 | 2.15777262 | 1.8276441 | 0.7782663 | 1.35808281 | 0.99629791 | 0.76599246 | 0.62973469 | 0.80845603 | 1.94705097 |
| ENSMUSG00 Gm4207 | 1869 | 0 | 0 | 0 | 0 | 0 | 0 | 0 | 0 | 0 | 0 | 0 | 0 | 0 | 0 | 0 | 0 | 0 | 0 |
| ENSMUSG00 Vmn1r159 | 917 | 0 | 0 | 0 | 0 | 0 | 0 | 0 | 0 | 0 | 0 | 0 | 0 | 0 | 0 | 0 | 0 | 0 | 0 |
| ENSMUSG00 Gm17114 | 509 | 0 | 0 | 0 | 0 | 0 | 0 | 0 | 0 | 0 | 0 | 0 | 0 | 0 | 0 | 0 | 0 | 0 | 0 |
| ENSMUSG00 Gm17103 | 547 | 0 | 0 | 0 | 0 | 0 | 0.18118001 | 0 | 0 | 0 | 0 | 0 | 0 | 0 | 0 | 0 | 0 | 0 | 0 |
| ENSMUSG00 Vmn2r76 | 2556 | 0 | 0 | 0 | 0 | 0 | 0 | 0 | 0 | 0 | 0 | 0 | 0 | 0 | 0 | 0 | 0 | 0 | 0 |
| ENSMUSG00 Gm9182 | 1040 | 0.36783325 | 0.35234792 | 0.27907806 | 0.52570436 | 0.74049113 | 0.28588116 | 0.40943664 | 0.62429689 | 0.39602419 | 0.58938379 | 0.50531206 | 0.53668906 | 0.56921443 | 1.92028438 | 0.99042202 | 0.92044729 | 1.03186184 | 1.11076081 |
| ENSMUSG00 Ighv1-62-2 | 301 | 0 | 0 | 0 | 0 | 0 | 0 | 0 | 0 | 0 | 0 | 0 | 0 | 0 | 0 | 0 | 0 | 0 | 0 |
| ENSMUSG00 Vgll3 | 3318 | 0 | 0 | 0 | 0 | 0 | 0 | 0 | 0 | 0 | 0 | 0 | 0 | 0 | 0 | 0 | 0 | 0 | 0 |
| ENSMUSG00 Gm17690 | 3465 | 0.82188941 | 1.00948164 | 1.20061287 | 1.30174412 | 1.05570742 | 0.91525973 | 1.13331958 | 1.05814448 | 1.52146436 | 0.77563927 | 1.28227161 | 0.96650498 | 0.96812998 | 0.89805306 | 0.83235466 | 0.51938198 | 0.42031725 | 1.14304637 |
| ENSMUSG00 Gm3250 | 692 | 0 | 0 | 0 | 0 | 0 | 0 | 0 | 0 | 0 | 0 | 0 | 0 | 0 | 0 | 0 | 0 | 0 | 0 |
| ENSMUSG00 Gm3834 | 2375 | 0 | 0 | 0 | 0 | 0 | 0 | 0 | 0 | 0 | 0 | 0 | 0 | 0 | 0 | 0 | 0 | 0 | 0 |
| ENSMUSG00 Gm16529 | 1685 | 0.05045125 | 0 | 0 | 0 | 0 | 0 | 0 | 0 | 0 | 0 | 0.02835309 | 0 | 0 | 0 | 0 | 0 | 0 | 0 |
| ENSMUSG00 Gm17689 | 327 | 0 | 0 | 0 | 0 | 0 | 0 | 0 | 0 | 0 | 0 | 0 | 0 | 0 | 0 | 0.37799593 | 0 | 0 | 0 |
| ENSMUSG00 Kiss1 | 776 | 1.69801605 | 0.77272177 | 2.18179583 | 0.58712686 | 0.9924108 | 1.46870216 | 0.97551915 | 1.18120832 | 0.37152785 | 0.66837337 | 0.49252535 | 1.2946932 | 1.01715293 | 0.53865565 | 0.58404267 | 2.12177335 | 0.39511646 | 1.13420839 |
| ENSMUSG00 Gm6346 | 1443 | 0 | 0 | 0 | 0 | 0 | 0 | 0 | 0 | 0 | 0 | 0 | 0 | 0 | 0 | 0 | 0 | 0 | 0 |
| ENSMUSG00 Gm17505 | 82 | 0 | 0 | 0 | 0 | 0 | 0 | 0 | 0.465761 | 0 | 0 | 0 | 0 | 0.60160875 | 0 | 0 | 0 | 0 | 0 |
| ENSMUSG00 Speer4e | 1598 | 0 | 0.02084662 | 0 | 0 | 0 | 0 | 0 | 0 | 0 | 0 | 0 | 0 | 0 | 0 | 0 | 0 | 0 | 0 |
| ENSMUSG00 Gm3348 | 846 | 0 | 0 | 0 | 0 | 0 | 0 | 0 | 0 | 0 | 0 | 0 | 0 | 0 | 0 | 0 | 0 | 0 | 0 |
| ENSMUSG00 Vmn2r110 | 2571 | 0 | 0 | 0 | 0 | 0 | 0 | 0 | 0 | 0 | 0 | 0 | 0 | 0 | 0 | 0 | 0 | 0 | 0 |
| ENSMUSG00 Vmn2r19 | 2559 | 0 | 0 | 0 | 0 | 0 | 0 | 0 | 0 | 0 | 0 | 0 | 0 | 0 | 0 | 0 | 0 | 0 | 0 |
| ENSMUSG00 Gm8453 | 920 | 0 | 0 | 0 | 0 | 0 | 0 | 0 | 0 | 0 | 0 | 0 | 0 | 0 | 0 | 0 | 0 | 0 | 0 |
| ENSMUSG00 D430020J02i | 37 | 0 | 0 | 0 | 0 | 1.3008628 | 0 | 0 | 0 | 0 | 0 | 0 | 0 | 3.99988518 | 0 | 0 | 0 | 1.03584586 | 0 |
| ENSMUSG00 BC024659 | 4462 | 2.58155596 | 1.44838671 | 2.20076797 | 2.20555481 | 2.15741478 | 2.83190208 | 2.0888834 | 2.14842963 | 1.43995887 | 3.27582206 | 2.51616209 | 2.16407656 | 3.36102708 | 3.54940247 | 2.55778373 | 2.55728395 | 2.54248854 | 1.09722333 |
| ENSMUSG00 Gm17320 | 93 | 0 | 0 | 0 | 0 | 0 | 0 | 0 | 0 | 0 | 0 | 0 | 0 | 0 | 0 | 0 | 0 | 0 | 0 |
| ENSMUSG00 Gm17321 | 2775 | 0.41356387 | 0.3601394 | 0.20918283 | 0.2955311 | 0.39893126 | 0.64284627 | 0.34099228 | 0.35783873 | 0.1187359 | 0.49274609 | 0.58535084 | 0.36204754 | 0.1066636 | 0.2175759 | 0.10393185 | 0.35875884 | 0.22098045 | 0.19823102 |
| ENSMUSG00 Gm17322 | 2167 | 0.72574597 | 0.07686408 | 0.78129837 | 0.52562349 | 0.62191687 | 0.84607805 | 0.82966396 | 1.49808685 | 0.22807485 | 0.39165328 | 0.26455907 | 0.4121138 | 0.20488567 | 0.3643514 | 0.30421084 | 0.10601922 | 0.8135716 | 1.04078144 |
| ENSMUSG00 Igkv17-134 | 344 | 0 | 0 | 0 | 0 | 0 | 0 | 0 | 0 | 0 | 0 | 0 | 0 | 0 | 0 | 0 | 0 | 0 | 0 |
| ENSMUSG00 Gm6682 | 296 | 0.4307957 | 0.11254356 | 0 | 0.46176734 | 0.81303925 | 0.33481577 | 0.15984013 | 0.12902839 | 0.13914363 | 0.15929292 | 0.48420566 | 0.18856643 | 0.33332377 | 0.3138114 | 0.13919445 | 0.12936016 | 0.51792293 | 0.55752473 |
| ENSMUSG00 Gm17642 | 2446 | 2.72825537 | 2.07013897 | 1.40413764 | 2.27246419 | 1.90874759 | 1.05345141 | 1.35400144 | 2.10792082 | 1.56596319 | 2.06260231 | 2.94931265 | 2.0537243 | 1.61347236 | 1.53808994 | 1.97080214 | 1.01753454 | 0.78344842 | 1.05700246 |
| ENSMUSG00 Gm17104 | 321 | 0.13241488 | 0.62267092 | 0.15069636 | 0 | 0.14994369 | 0 | 0 | 0 | 0.25661381 | 0.14688693 | 0 | 0 | 0.46104596 | 0 | 0 | 0.23857076 | 0.11939656 | 0 |
| ENSMUSG00 Gm17641 | 2349 | 0 | 0 | 0 | 0 | 0 | 0 | 0 | 0 | 0 | 0 | 0 | 0 | 0 | 0 | 0 | 0 | 0 | 0 |
| ENSMUSG00 Gm6673 | 695 | 0 | 0 | 0 | 0 | 0 | 0 | 0 | 0 | 0 | 0 | 0 | 0 | 0 | 0.46778218 | 0 | 0 | 0 | 0 |
| ENSMUSG00 Igkv4-53 | 291 | 0 | 0 | 0 | 0 | 0 | 0 | 0 | 0 | 0.28306884 | 0 | 0 | 0 | 0 | 0.47880502 | 0 | 0 | 0 | 0 |
| ENSMUSG00 Gm3248 | 2048 | 0 | 0 | 0 | 0 | 0 | 0 | 0 | 0 | 0 | 0 | 0 | 0 | 0 | 0 | 0 | 0 | 0 | 0 |
| ENSMUSG00 Vmn1r149 | 923 | 0 | 0 | 0 | 0 | 0 | 0 | 0 | 0 | 0 | 0 | 0 | 0 | 0 | 0 | 0 | 0 | 0 | 0 |
| ENSMUSG00 Gm1818 | 2397 | 0.01773266 | 0.20846617 | 0.04036173 | 0.01900753 | 0.04016014 | 0.04134563 | 0.1776446 | 0.12746734 | 0.10309516 | 0.15736572 | 0.01993115 | 0.11642817 | 0.06174207 | 0.11625554 | 0.085944 | 0.03194878 | 0.23983915 | 0.0458983 |
| ENSMUSG00 Gm17643 | 1592 | 2.45632927 | 0.92070813 | 2.03582069 | 1.25922484 | 1.57214826 | 1.52517836 | 3.74459645 | 1.46340235 | 1.81096489 | 1.18469104 | 1.23038524 | 2.38408609 | 2.32405389 | 2.421394 | 2.30335331 | 2.74191535 | 1.32408689 | 1.17481761 |
| ENSMUSG00 Gm17032 | 336 | 0 | 0 | 0 | 0 | 0 | 0 | 0 | 0 | 0 | 0 | 0 | 0 | 0 | 0 | 0 | 0 | 0 | 0 |
| ENSMUSG00 Gm8065 | 601 | 0 | 0 | 0 | 0 | 0 | 0 | 0 | 0 | 0 | 0 | 0 | 0 | 0 | 0 | 0 | 0 | 0 | 0 |
| ENSMUSG00 Gm17431 | 2024 | 0 | 0 | 0 | 0 | 0 | 0 | 0 | 0 | 0 | 0 | 0 | 0 | 0 | 0 | 0 | 0 | 0 | 0 |
| ENSMUSG00 Gm17234 | 390 | 0 | 0 | 0 | 0 | 0 | 0 | 0 | 0 | 0 | 0 | 0 | 0 | 0 | 0 | 0 | 0 | 0 | 0 |
| ENSMUSG00 Gm17430 | 473 | 0 | 0 | 0 | 0 | 0 | 0 | 0 | 0 | 0 | 0 | 0 | 0 | 0 | 0 | 0 | 0 | 0 | 0 |
| ENSMUSG00 Gm3111 | 487 | 0 | 0 | 0 | 0 | 0 | 0 | 0 | 0 | 0 | 0 | 0 | 0 | 0 | 0 | 0 | 0 | 0 | 0 |
| ENSMUSG00 Gm17075 | 669 | 0 | 0 | 0 | 0 | 0 | 0 | 0 | 0 | 0 | 0 | 0 | 0 | 0 | 0 | 0 | 0 | 0 | 0 |
| ENSMUSG00 Gm17051 | 486 | 0 | 0 | 0 | 0 | 0 | 0 | 0 | 0 | 0 | 0 | 0 | 0 | 0 | 0 | 0 | 0 | 0 | 0 |
| ENSMUSG00 Gm3275 | 486 | 0 | 0 | 0 | 0 | 0 | 0 | 0 | 0 | 0 | 0 | 0 | 0 | 0 | 0 | 0 | 0 | 0 | 0 |
| ENSMUSG00 Gm17731 | 34 | 0 | 0 | 0 | 1.34003072 | 0 | 0 | 0 | 1.12330595 | 0 | 0 | 0 | 0 | 0 | 0 | 1.21181047 | 1.12619434 | 0 | 0 |
| ENSMUSG00 Gm3104 | 470 | 0 | 0 | 0 | 0 | 0 | 0 | 0 | 0 | 0 | 0 | 0 | 0 | 0 | 0 | 0 | 0 | 0 | 0 |
| ENSMUSG00 Gm6047 | 573 | 0 | 0 | 0.08442152 | 0 | 0 | 0 | 0 | 0 | 0 | 0 | 0 | 0 | 0 | 0 | 0 | 0 | 0 | 0 |
| ENSMUSG00 Glis3 | 1271 | 0 | 0 | 0 | 0 | 0 | 0 | 0 | 0 | 0 | 0 | 0 | 0 | 0 | 0 | 0 | 0 | 0 | 0 |
| ENSMUSG00 Flg | 935 | 0 | 0 | 0 | 0 | 0 | 0 | 0 | 0 | 0 | 0 | 0 | 0 | 0 | 0 | 0 | 0 | 0 | 0 |
| ENSMUSG00 Gm4181 | 893 | 0 | 0 | 0 | 0 | 0 | 0 | 0 | 0 | 0 | 0 | 0 | 0 | 0 | 0 | 0 | 0 | 0 | 0 |
| ENSMUSG00 Gm8439 | 417 | 3.26178809 | 4.95299626 | 2.90009175 | 2.29444108 | 2.77018266 | 2.97078738 | 6.92103936 | 3.20559732 | 6.12365462 | 2.939852 | 2.63506965 | 4.68476784 | 5.91509799 | 9.13289009 | 5.03903921 | 4.86667192 | 2.02201087 | 6.8596488 |
| ENSMUSG00 Gm17730 | 2757 | 0 | 0 | 0 | 0 | 0 | 0 | 0 | 0 | 0 | 0 | 0 | 0 | 0 | 0 | 0 | 0 | 0 | 0 |
| ENSMUSG00 Snhg7 | 1529 | 1.91815379 | 2.83235857 | 3.57502219 | 1.57929061 | 1.82580221 | 1.58802089 | 2.04227391 | 2.77263353 | 2.34351005 | 1.91193172 | 1.49980249 | 3.13940287 | 2.22622779 | 2.15665808 | 1.58985729 | 1.40240289 | 1.85490253 | 1.61897306 |
| ENSMUSG00 Ighv1-71 | 301 | 0 | 0 | 0 | 0 | 0 | 0 | 0 | 0 | 0 | 0 | 0 | 0 | 0 | 0 | 0 | 0 | 0 | 0 |
| ENSMUSG00 2310043M15 | 1551 | 0.05481003 | 0.79469831 | 0.21832025 | 0.2937527 | 0.77582082 | 0.63897787 | 0.94564349 | 0.51711183 | 1.11530217 | 0.76000489 | 0.15401341 | 0.50381643 | 1.14503483 | 0.65878138 | 0.47816119 | 0.66656763 | 0.39537121 | 0.31920178 |
| ENSMUSG00 4831440E17i | 1026 | 2.11283039 | 1.33121702 | 2.16879376 | 1.64303961 | 1.82957605 | 1.64209838 | 4.61137222 | 3.12686335 | 2.48885378 | 2.34374841 | 3.67857867 | 3.10087014 | 4.56777012 | 2.39915851 | 4.45747827 | 1.34353008 | 2.42807924 | 3.27052194 |
| ENSMUSG00 Speer6-ps1 | 541 | 0 | 0 | 0 | 0 | 0 | 0 | 0 | 0 | 0 | 0 | 0 | 0 | 0 | 0 | 0 | 0 | 0 | 0 |
| ENSMUSG00 Gm17100 | 271 | 0 | 0 | 0 | 0 | 0 | 0 | 0 | 0.14093137 | 0 | 0 | 0 | 0 | 0 | 0 | 0 | 0 | 0 | 0 |
| ENSMUSG00 Gm17606 | 156 | 0 | 1.28126516 | 0 | 0.58411595 | 0.30853797 | 0 | 0.3032864 | 0 | 0.52803225 | 0.3022481 | 0 | 0 | 0 | 0.59543702 | 0 | 0.98181045 | 0 | 3.52622479 |
| ENSMUSG00 Gm8895 | 1219 | 0 | 0 | 0 | 0 | 0 | 0 | 0 | 0 | 0 | 0 | 0 | 0 | 0 | 0 | 0 | 0 | 0 | 0 |
| ENSMUSG00 Olfr1380 | 937 | 0 | 0 | 0 | 0 | 0 | 0 | 0 | 0 | 0 | 0 | 0 | 0 | 0 | 0 | 0 | 0 | 0 | 0 |
| ENSMUSG00 Gm17605 | 709 | 0 | 0 | 0 | 0 | 0 | 0 | 0 | 0 | 0 | 0 | 0 | 0 | 0 | 0 | 0 | 0 | 0 | 0 |
| ENSMUSG00 Gm17288 | 1634 | 0 | 0 | 0 | 0 | 0 | 0 | 0 | 0 | 0 | 0 | 0 | 0 | 0 | 0 | 0 | 0 | 0 | 0 |
| ENSMUSG00 Gm4934 | 4347 | 0 | 0 | 0 | 0 | 0 | 0 | 0 | 0 | 0 | 0 | 0 | 0 | 0 | 0 | 0 | 0 | 0 | 0 |
| ENSMUSG00 Gm17490 | 61 | 0 | 0 | 0 | 0 | 0 | 0 | 0 | 0 | 0 | 0 | 0 | 0 | 0 | 0 | 0 | 0 | 0 | 0 |
| ENSMUSG00 Fth-ps2 | 515 | 1.40308348 | 0.51748185 | 0.75143348 | 0.44234024 | 0.65422032 | 0.4810945 | 0 | 0.88992006 | 1.03965962 | 0.09155476 | 0.64936837 | 1.62569891 | 1.82001248 | 1.80365388 | 0.48001813 | 1.63571527 | 1.56281987 | 1.28176559 |
| ENSMUSG00 Gm17368 | 158 | 0 | 0 | 0 | 0 | 0 | 0 | 0 | 0 | 0 | 0 | 0 | 0 | 0 | 0 | 0 | 0 | 0 | 0 |
| ENSMUSG00 Gm17214 | 540 | 0.07871329 | 0 | 0 | 0.0843723 | 0 | 0 | 0 | 0.07072667 | 0 | 0 | 0.08847215 | 0.10336234 | 0 | 0 | 0.07629918 | 0.49635973 | 0.14194925 | 0 |
| ENSMUSG00 Gm17289 | 1850 | 0.41356387 | 0.75629273 | 1.22894915 | 0.73882775 | 0.46831061 | 0.45534944 | 1.02297684 | 1.23867251 | 1.40256783 | 0.89204033 | 0.41318883 | 0.81460697 | 0.18666131 | 0.75314736 | 0.64586223 | 0.12418575 | 0.39362143 | 0.38655048 |
| ENSMUSG00 Gm5415 | 3747 | 0 | 0 | 0 | 0 | 0 | 0 | 0 | 0 | 0 | 0 | 0 | 0 | 0 | 0 | 0 | 0.010219 | 0 | 0 |
| ENSMUSG00 Gm17287 | 1053 | 0 | 0 | 0 | 0 | 0 | 0 | 0 | 0 | 0 | 0 | 0 | 0 | 0 | 0 | 0 | 0 | 0 | 0 |
| ENSMUSG00 Vmn1r123 | 923 | 0 | 0 | 0 | 0 | 0 | 0 | 0 | 0 | 0 | 0 | 0 | 0 | 0 | 0 | 0 | 0 | 0 | 0 |
| ENSMUSG00 Gm17480 | 1900 | 0 | 0.0350662 | 0 | 0 | 0 | 0 | 0 | 0.04020253 | 0 | 0 | 0 | 0.14688332 | 0 | 0 | 0 | 0 | 0 | 0 |
| ENSMUSG00 4930480E11i | 1468 | 0 | 0 | 0 | 0 | 0 | 0 | 0 | 0 | 0 | 0 | 0 | 0 | 0 | 0 | 0 | 0 | 0 | 0 |
| ENSMUSG00 Gm12166 | 811 | 0 | 0 | 0 | 0 | 0 | 0 | 0 | 0 | 0 | 0 | 0 | 0 | 0 | 0 | 0 | 0 | 0 | 0 |
| ENSMUSG00 Gm4673 | 3658 | 0 | 0 | 0 | 0 | 0 | 0 | 0 | 0 | 0 | 0 | 0 | 0 | 0 | 0 | 0 | 0 | 0 | 0 |
| ENSMUSG00 Gm3760 | 1750 | 0 | 0 | 0 | 0 | 0 | 0 | 0 | 0 | 0 | 0 | 0 | 0 | 0 | 0 | 0 | 0 | 0 | 0 |
| ENSMUSG00 Traj54 | 53 | 0 | 0 | 0 | 0 | 0 | 0 | 0 | 0 | 0 | 0 | 0 | 0 | 0 | 0 | 0 | 0 | 0 | 0 |
| ENSMUSG00 Raxos1 | 2545 | 0 | 0 | 0 | 0 | 0 | 0 | 0 | 0 | 0 | 0 | 0 | 0 | 0 | 0 | 0 | 0 | 0.01505945 | 0 |
| ENSMUSG00 Gm3542 | 2058 | 0 | 0 | 0 | 0 | 0 | 0 | 0 | 0 | 0 | 0 | 0 | 0 | 0 | 0 | 0 | 0 | 0 | 0 |
| ENSMUSG00 1700112D23 | 543 | 0 | 0.06134971 | 0.44542846 | 0.67124927 | 0.44320372 | 0.09125734 | 0.87132005 | 0.63302324 | 0.91019924 | 0.17366742 | 0.1759667 | 0.20558255 | 0.72680541 | 0.51319434 | 0.30351054 | 0.63465095 | 0.56465999 | 0.81044724 |
| ENSMUSG00 Gm20520 | 588 | 0 | 0 | 0 | 0 | 0 | 0 | 0 | 0 | 0 | 0 | 0 | 0 | 0 | 0 | 0 | 0 | 0 | 0 |
| ENSMUSG00 4930500F10i | 716 | 0.11872954 | 0.27915833 | 0 | 0.06363274 | 0 | 0.20762319 | 0 | 0.05334134 | 0.5177076 | 0 | 0 | 0.15590967 | 0 | 0.25946418 | 0 | 0.16043551 | 0.05352835 | 0 |
| ENSMUSG00 BC051077 | 1934 | 0.13186714 | 0.01722487 | 0 | 0.02355793 | 0 | 0.05124378 | 0 | 0.05924364 | 0.06388808 | 0 | 0 | 0.02886022 | 0 | 0 | 0.0213038 | 0.07919464 | 0.07926845 | 0 |
| ENSMUSG00 Ighd5-3 | 9 | 0 | 0 | 0 | 0 | 0 | 0 | 0 | 0 | 0 | 0 | 0 | 0 | 0 | 0 | 0 | 0 | 0 | 0 |
| ENSMUSG00 Gm8766 | 785 | 0 | 0 | 0 | 0 | 0 | 0 | 0 | 0 | 0 | 0 | 0 | 0 | 0 | 0 | 0 | 0 | 0 | 0 |
| ENSMUSG00 Gm17157 | 347 | 0 | 0 | 0 | 0 | 0 | 0 | 0 | 0 | 0 | 0 | 0.68839998 | 0 | 0 | 0 | 0 | 0 | 0 | 0 |
| ENSMUSG00 Eid1 | 1690 | 24.8743308 | 19.7906188 | 26.7914937 | 21.8639094 | 21.8729689 | 21.9028947 | 32.9788969 | 26.6216864 | 30.9995551 | 25.41674 | 25.1595935 | 24.6381564 | 22.7977677 | 27.5641536 | 21.1615092 | 21.4336773 | 19.4352878 | 18.1953199 |
| ENSMUSG00 Ighv1-78 | 293 | 0 | 0 | 0 | 0 | 0 | 0 | 0 | 0 | 0 | 0 | 0 | 0 | 0 | 0 | 0 | 0 | 0 | 0 |
| ENSMUSG00 Gm2866 | 480 | 0 | 0 | 0 | 0 | 0 | 0 | 0 | 0 | 0 | 0 | 0 | 0 | 0 | 0 | 0 | 0 | 0 | 0 |
| ENSMUSG00 Flg | 1487 | 0 | 0 | 0 | 0 | 0 | 0 | 0 | 0 | 0 | 0 | 0 | 0 | 0 | 0 | 0 | 0 | 0 | 0 |
| ENSMUSG00 Gm8104 | 1756 | 0 | 0 | 0 | 0 | 0 | 0 | 0 | 0 | 0 | 0 | 0 | 0 | 0 | 0 | 0 | 0 | 0 | 0 |
| ENSMUSG00 Gm17329 | 138 | 0 | 0 | 0 | 0 | 0 | 0 | 0 | 0 | 0 | 0 | 0 | 0 | 0 | 0 | 0 | 0 | 0 | 0 |
| ENSMUSG00 Gm17328 | 1395 | 0.3046966 | 0.50148443 | 0.17338183 | 0.06532049 | 0.17251586 | 0.10656502 | 0.16957949 | 0.32853679 | 0.11809754 | 0.06759957 | 0.10274185 | 0.40011228 | 0.35363382 | 0.56598529 | 0.0886055 | 0.41172696 | 0.32968858 | 0.35489746 |
| ENSMUSG00 1700037F24i | 971 | 0 | 0 | 0 | 0 | 0 | 0 | 0 | 0 | 0 | 0 | 0 | 0 | 0 | 0 | 0 | 0 | 0.03947095 | 0 |
| ENSMUSG00 Col6a5 | 9252 | 0.03215913 | 0.02520431 | 0 | 0 | 0 | 0 | 0 | 0 | 0 | 0 | 0 | 0 | 0.02666014 | 0.05521887 | 0.08906519 | 0.05794082 | 0.0331399 | 0 |
| ENSMUSG00 Gm17353 | 254 | 0 | 0 | 0 | 0 | 0 | 0 | 0 | 0 | 0 | 0 | 0 | 0 | 0 | 0 | 0 | 0 | 0 | 0 |
| ENSMUSG00 Gm7349 | 506 | 0 | 0 | 0 | 0 | 0 | 0 | 0 | 0 | 0 | 0 | 0 | 0 | 0 | 0 | 0 | 0 | 0 | 0 |
| ENSMUSG00 Vmn2r92 | 2577 | 0 | 0 | 0 | 0 | 0 | 0 | 0 | 0 | 0 | 0 | 0 | 0 | 0 | 0 | 0 | 0 | 0 | 0 |
| ENSMUSG00 Gm17135 | 561 | 0 | 0 | 0 | 0 | 0 | 0.08832929 | 0.59035428 | 0 | 0 | 0 | 0 | 0 | 0 | 0 | 0 | 0 | 0.61486038 | 0 |
| ENSMUSG00 Gm17519 | 1735 | 0.39197857 | 0.09600258 | 0.16728598 | 0 | 0.30515917 | 0.08568196 | 0 | 0 | 0 | 0 | 0.027536 | 0 | 0.17060029 | 0 | 0 | 0 | 0 | 0 |
| ENSMUSG00 Gm13285 | 1555 | 0.16400711 | 0 | 0.03110838 | 0.08789912 | 0 | 0.09560013 | 0.18255696 | 0 | 0.23837855 | 0.12128798 | 0 | 0.14357727 | 0.15862353 | 0 | 0.13248089 | 0.41861114 | 0 | 0.17687816 |
| ENSMUSG00 6030419C18 | 535 | 0 | 0 | 0 | 0 | 0.08998621 | 0 | 0 | 0.28555067 | 0 | 0 | 0 | 0 | 0 | 0 | 0 | 0 | 0 | 0 |
| ENSMUSG00 Gm17142 | 642 | 0 | 0 | 0 | 0 | 0 | 0 | 0 | 0 | 0 | 0.07344346 | 0 | 0 | 0.46104596 | 0.14468563 | 0 | 0 | 0 | 0 |
| ENSMUSG00 Speer5-ps1 | 772 | 0 | 0 | 0 | 0 | 0 | 0 | 0 | 0 | 0 | 0 | 0 | 0 | 0 | 0 | 0 | 0 | 0 | 0 |
| ENSMUSG00 Gm3129 | 1865 | 0 | 0 | 0.02593755 | 0 | 0 | 0 | 0 | 0 | 0 | 0 | 0 | 0 | 0 | 0 | 0.04418397 | 0 | 0 | 0 |
| ENSMUSG00 Gm5277 | 587 | 0 | 0 | 0 | 0 | 0 | 0 | 0.16120163 | 0 | 0 | 0 | 0 | 0 | 0 | 0 | 0 | 0 | 0 | 0 |
| ENSMUSG00 Gm7147 | 282 | 0 | 0 | 0 | 0 | 0 | 0 | 0 | 0 | 0 | 0 | 0 | 0 | 0 | 0 | 0 | 0 | 0 | 0 |
| ENSMUSG00 Gm7701 | 1478 | 1.29413594 | 1.48758526 | 1.30916185 | 0.98643668 | 1.53058215 | 1.13991403 | 0.76827084 | 0.54265254 | 1.25398729 | 1.05275589 | 0.74345335 | 1.13292955 | 1.20158932 | 0.43993046 | 0.89204992 | 0.41451266 | 0.62234853 | 1.07933971 |
| ENSMUSG00 Gm17045 | 1691 | 0 | 0 | 0 | 0 | 0 | 0 | 0 | 0 | 0 | 0 | 0 | 0 | 0 | 0 | 0 | 0.06799461 | 0 | 0 |
| ENSMUSG00 Gm17040 | 1434 | 0.32605086 | 0 | 0.26986628 | 0 | 0.1006944 | 0.0691112 | 0 | 0.69247034 | 0.20104994 | 0.06576109 | 0.23321109 | 0 | 0.34401616 | 0.7773069 | 0.05746382 | 0.08010587 | 0.56126376 | 0 |
| ENSMUSG00 Gm17711 | 235 | 0 | 0 | 0 | 0 | 0 | 0 | 0 | 0 | 0 | 0 | 0 | 0 | 0 | 0 | 0 | 0 | 0 | 0 |
| ENSMUSG00 Gm17085 | 501 | 0 | 0 | 0 | 0 | 0 | 0 | 0 | 0 | 0 | 0 | 0 | 0 | 0 | 0 | 0 | 0 | 0 | 0 |
| ENSMUSG00 5730435O14 | 694 | 0 | 0.09600258 | 0 | 0 | 0 | 0 | 0 | 0.11006456 | 0 | 0 | 0 | 0 | 0 | 0.06692232 | 0 | 0 | 0 | 0 |
| ENSMUSG00 Gm17397 | 1262 | 0 | 0.02639691 | 0.22998509 | 0 | 0 | 0 | 0 | 0 | 0 | 0 | 0.07571309 | 0 | 0.50817347 | 0 | 0 | 0 | 0 | 0 |
| ENSMUSG00 Gm14475 | 499 | 0 | 0 | 0 | 0 | 0 | 0 | 0 | 0 | 0 | 0 | 0 | 0 | 0 | 0 | 0 | 0 | 0 | 0 |
| ENSMUSG00 Gm8810 | 1031 | 0.04122713 | 0 | 0 | 0.04419112 | 0 | 0.14418836 | 0 | 0 | 0.03994812 | 0 | 0 | 0 | 0.09569722 | 0 | 0 | 0 | 0 | 0 |
| ENSMUSG00 Vmn2r15 | 2562 | 0 | 0 | 0 | 0 | 0 | 0 | 0 | 0 | 0 | 0 | 0 | 0 | 0 | 0 | 0 | 0 | 0 | 0 |
| ENSMUSG00 Aadacl2 | 1585 | 0 | 0 | 0 | 0 | 0 | 0 | 0 | 0 | 0 | 0 | 0 | 0 | 0 | 0 | 0 | 0 | 0 | 0 |
| ENSMUSG00 Gm17398 | 1475 | 0 | 0 | 0 | 0 | 0.06526363 | 0 | 0 | 0.18125208 | 0.02792306 | 0 | 0 | 0 | 0.1003361 | 0 | 0 | 0.18171814 | 0.18188751 | 0.44753172 |
| ENSMUSG00 Gm4219 | 2141 | 0 | 0.031119 | 0.22593896 | 0.04256053 | 0.0449621 | 0 | 0.37567284 | 0.07135432 | 0.01923705 | 0 | 0.1115716 | 0 | 0.27649837 | 0.04338542 | 0.21168478 | 0 | 0.42962687 | 0 |
| ENSMUSG00 Gm3159 | 1984 | 0 | 0 | 0 | 0 | 0 | 0 | 0 | 0 | 0 | 0 | 0 | 0 | 0 | 0 | 0 | 0 | 0 | 0 |
| ENSMUSG00 Gm17575 | 3238 | 0 | 0 | 0 | 0 | 0 | 0 | 0 | 0 | 0 | 0 | 0 | 0 | 0 | 0 | 0 | 0 | 0 | 0 |
| ENSMUSG00 Vmn2r83 | 2604 | 0 | 0 | 0 | 0 | 0 | 0 | 0 | 0 | 0 | 0 | 0 | 0 | 0 | 0 | 0 | 0 | 0 | 0 |
| ENSMUSG00 Vmn1r18 | 1086 | 0 | 0 | 0 | 0 | 0 | 0 | 0 | 0 | 0 | 0 | 0 | 0 | 0 | 0 | 0 | 0 | 0 | 0 |
| ENSMUSG00 Hist1h2al | 377 | 5.29905378 | 6.09705491 | 10.3932519 | 7.25109459 | 2.5534177 | 1.57727534 | 17.8206907 | 13.8789367 | 19.0091611 | 8.75477249 | 5.70258125 | 3.10909526 | 18.5812527 | 37.2045475 | 34.862855 | 13.4067909 | 10.369449 | 9.77615425 |
| ENSMUSG00 Gm17574 | 5749 | 0.05175443 | 0.04635644 | 0.00841425 | 0.03170015 | 0.07535003 | 0 | 0.08229723 | 0.03321656 | 0.07164118 | 0.0164031 | 0.09141147 | 0.06796132 | 0.18878104 | 0.05655046 | 0.18633509 | 0.07992473 | 0.01999981 | 0.01913693 |
| ENSMUSG00 Gm17573 | 245 | 0 | 0 | 0 | 0 | 0 | 0 | 0 | 0 | 0 | 0 | 0 | 0 | 0 | 0 | 0 | 0 | 0 | 0 |
| ENSMUSG00 Olfr443-ps1 | 914 | 0 | 0 | 0 | 0 | 0 | 0 | 0 | 0 | 0 | 0 | 0 | 0 | 0 | 0 | 0 | 0 | 0 | 0 |
| ENSMUSG00 Gcnt4 | 5090 | 0.43423755 | 0.94899208 | 0.73178032 | 0.53706536 | 0.34987842 | 1.11956078 | 0.36251365 | 0.54024616 | 1.03573163 | 0.39832618 | 0.88228803 | 0.39476697 | 0.07753543 | 0.65696941 | 0.3966358 | 0.91777094 | 1.00898306 | 0.57278638 |
| ENSMUSG00 Gm17576 | 190 | 0 | 0 | 0 | 0 | 0 | 0 | 0 | 0 | 0 | 0 | 0 | 0 | 0 | 0 | 0 | 0 | 0 | 0 |
| ENSMUSG00 Speer1-ps1 | 1093 | 0 | 0 | 0 | 0 | 0 | 0 | 0 | 0 | 0 | 0 | 0 | 0 | 0 | 0 | 0 | 0 | 0 | 0 |
| ENSMUSG00 Gm17168 | 341 | 0.62324305 | 0.09769177 | 0 | 0 | 0 | 0 | 0 | 0 | 0.24156314 | 0 | 0 | 0 | 0.14466838 | 0 | 0 | 0 | 0.22478767 | 0 |
| ENSMUSG00 Gm5373 | 471 | 0 | 0 | 0 | 0.19346516 | 0.10219092 | 0.1052075 | 0 | 0 | 0 | 0 | 0 | 0 | 0 | 0.29582221 | 0 | 0.32518563 | 0 | 0 |
| ENSMUSG00 Gm17263 | 2459 | 0 | 0 | 0 | 0 | 0 | 0 | 0 | 0 | 0 | 0 | 0 | 0 | 0 | 0 | 0 | 0 | 0 | 0 |
| ENSMUSG00 5330438I03R | 2844 | 0 | 0 | 0 | 0 | 0 | 0 | 0.01663596 | 0 | 0 | 0 | 0 | 0 | 0 | 0 | 0 | 0 | 0 | 0 |
| ENSMUSG00 Ighd5-2 | 9 | 0 | 0 | 0 | 0 | 0 | 0 | 0 | 0 | 0 | 0 | 0 | 0 | 0 | 0 | 0 | 0 | 0 | 0 |
| ENSMUSG00 Gm6297 | 2144 | 2.4979721 | 2.01990497 | 2.27879038 | 2.18880017 | 2.73885013 | 2.21877912 | 2.20674808 | 2.24451619 | 1.97864325 | 1.8253304 | 2.85223633 | 3.09797754 | 3.65847707 | 2.5994825 | 2.88257154 | 1.89309906 | 2.57415427 | 2.206522 |
| ENSMUSG00 Sparcn4 | 422 | 0 | 0 | 0 | 0 | 0 | 0 | 0 | 0 | 0 | 0 | 0 | 0 | 0 | 0 | 0 | 0 | 0 | 0 |
| ENSMUSG00 Gm17265 | 2718 | 4.14417647 | 3.76271469 | 4.16460857 | 4.22420279 | 5.40111726 | 5.17769549 | 5.15252133 | 3.96256714 | 5.62185333 | 4.75323499 | 4.90405203 | 5.50353112 | 2.75881215 | 2.39226352 | 4.45668044 | 4.00093175 | 4.37128478 | 3.50131548 |
| ENSMUSG00 Gm3164 | 2191 | 0 | 0 | 0 | 0 | 0 | 0 | 0 | 0 | 0 | 0 | 0 | 0 | 0 | 0 | 0 | 0 | 0 | 0 |
| ENSMUSG00 Gm6356 | 1752 | 0 | 0 | 0 | 0 | 0 | 0 | 0 | 0 | 0 | 0 | 0 | 0 | 0 | 0 | 0 | 0 | 0 | 0 |
| ENSMUSG00 Gm9252 | 1601 | 0 | 0 | 0 | 0 | 0 | 0 | 0 | 0 | 0 | 0 | 0 | 0 | 0 | 0 | 0 | 0 | 0 | 0 |
| ENSMUSG00 BC048943 | 758 | 0 | 0 | 0 | 0 | 0 | 0 | 0 | 0 | 0 | 0 | 0 | 0 | 0 | 0 | 0 | 0 | 0 | 0 |
| ENSMUSG00 9530085L02i | 3626 | 0 | 0 | 0 | 0 | 0 | 0 | 0 | 0 | 0 | 0 | 0 | 0 | 0 | 0 | 0 | 0.16896021 | 0 | 0 |
| ENSMUSG00 Gm3140 | 480 | 0 | 0 | 0 | 0 | 0 | 0 | 0 | 0 | 0 | 0 | 0 | 0 | 0 | 0 | 0 | 0 | 0 | 0 |
| ENSMUSG00 Hist2h4 | 1607 | 0.74060045 | 0.39386745 | 1.44488454 | 0.96395489 | 0.5990283 | 0.64754661 | 0.97157337 | 1.2833788 | 2.43479714 | 2.31792505 | 0.08918785 | 0.38206116 | 2.73213481 | 1.44505561 | 3.20485033 | 1.21519663 | 1.81256911 | 2.08808681 |
| ENSMUSG00 Gm17650 | 64 | 0 | 0 | 0 | 0 | 0 | 0 | 0 | 0 | 0 | 0 | 0 | 0 | 0 | 0 | 0 | 0 | 0 | 0 |
| ENSMUSG00 Vmn2r117 | 2568 | 0 | 0 | 0 | 0 | 0 | 0 | 0 | 0 | 0 | 0 | 0 | 0 | 0 | 0.0180857 | 0 | 0 | 0 | 0 |
| ENSMUSG00 Gm6728 | 1096 | 0.58173142 | 0.75987441 | 1.14754725 | 0.9976871 | 0.3952439 | 1.2207334 | 0.34534802 | 0.87117706 | 0.601263 | 0.98947644 | 0.39231262 | 0.56019369 | 0.72017397 | 0.76276786 | 0.41351927 | 0.76860708 | 0.0699385 | 0.45171712 |
| ENSMUSG00 Gm17150 | 380 | 0.22371145 | 0 | 0 | 0 | 0.12666296 | 0 | 0 | 0 | 0 | 0 | 0.5028943 | 0 | 0.12982083 | 0.12222128 | 0 | 0.10076476 | 0 | 0.14476081 |
| ENSMUSG00 Gm5916 | 1139 | 2.35103256 | 13.9218065 | 1.31657545 | 3.40007794 | 2.40870909 | 9.48419308 | 0 | 0 | 0 | 0 | 0 | 0 | 0 | 0 | 0 | 0 | 0 | 0 |
| ENSMUSG00 Gm2895 | 3192 | 0.19974237 | 0.05218185 | 0.54556613 | 0.25692318 | 0.21110493 | 0.15524039 | 0.07411134 | 0.05982519 | 0.07741826 | 0.13294371 | 0.17960511 | 0.08743055 | 0 | 0.04365046 | 0.14198531 | 0.04798322 | 0.02401397 | 0.17233429 |
| ENSMUSG00 Olfr772 | 1003 | 0 | 0 | 0 | 0 | 0 | 0 | 0 | 0 | 0 | 0 | 0 | 0 | 0 | 0 | 0 | 0 | 0 | 0 |
| ENSMUSG00 Akd1 | 5637 | 0.1432674 | 0.11228401 | 0.04290716 | 0.15356747 | 0.05977 | 0.01758124 | 0 | 0.03387653 | 0 | 0 | 0.02542574 | 0 | 0 | 0 | 0 | 0 | 0 | 0 |
| ENSMUSG00 Gm6327 | 1409 | 0 | 0 | 0 | 0 | 0 | 0 | 0 | 0 | 0 | 0 | 0 | 0 | 0 | 0 | 0 | 0 | 0 | 0 |
| ENSMUSG00 Gpr173 | 398 | 0 | 0 | 0 | 0 | 0 | 0 | 0 | 0 | 0 | 0 | 0 | 0 | 0 | 0 | 0 | 0 | 0 | 0 |
| ENSMUSG00 Gm17143 | 1948 | 0 | 0 | 0 | 0 | 0 | 0 | 0 | 0 | 0 | 0 | 0 | 0 | 0 | 0 | 0 | 0 | 0 | 0 |
| ENSMUSG00 Gm17450 | 463 | 0.45901918 | 0.50365067 | 0.10447847 | 0 | 0 | 0 | 0.20437442 | 0.08248899 | 0.17791152 | 0 | 0.20637131 | 0 | 0 | 0.1003112 | 0 | 0 | 0 | 0 |
| ENSMUSG00 Gm17510 | 338 | 6.53925785 | 9.46165043 | 6.01091206 | 5.12224759 | 6.40809635 | 9.52937184 | 4.61928522 | 5.87575422 | 4.0211687 | 5.99846225 | 7.06730155 | 4.6237827 | 6.71381123 | 8.3819211 | 5.72921043 | 6.45729178 | 9.07131288 | 4.71971626 |
| ENSMUSG00 4933402N22i | 1314 | 0 | 0 | 0 | 0 | 0 | 0 | 0 | 0 | 0 | 0 | 0 | 0 | 0 | 0 | 0 | 0 | 0 | 0 |
| ENSMUSG00 Gm17509 | 2374 | 0 | 0 | 0.46865678 | 0.13434175 | 0.10137305 | 0.12523859 | 0.65767414 | 0.09652671 | 0.83275179 | 0.27805806 | 0.04024849 | 0 | 0.10390042 | 0.46952742 | 0 | 0.19354983 | 0.1130093 | 0 |
| ENSMUSG00 Gm17118 | 333 | 0 | 0 | 0 | 0 | 0 | 0 | 0 | 0 | 0 | 0 | 0 | 0 | 0 | 0 | 0 | 0 | 0 | 0 |
| ENSMUSG00 Gm17228 | 490 | 0.34698103 | 0 | 0 | 0.09298172 | 0 | 0 | 0 | 0 | 0.33621646 | 0.28867778 | 0 | 0 | 0 | 0 | 0.16816962 | 0 | 0 | 0 |
| ENSMUSG00 Gm17508 | 679 | 0 | 0 | 0 | 0 | 0 | 0 | 0 | 0 | 0 | 0 | 0 | 0.16440549 | 0 | 0 | 0 | 0.05639265 | 0 | 0 |
| ENSMUSG00 Gm17507 | 34 | 0 | 0 | 0 | 0 | 0 | 0 | 1.39154938 | 0 | 0 | 0 | 0 | 0 | 0 | 4.09800771 | 0 | 2.25238867 | 2.25448805 | 0 |
| ENSMUSG00 Gm17093 | 1399 | 0 | 0 | 0 | 0 | 0 | 0 | 0 | 0 | 0 | 0 | 0 | 0 | 0 | 0 | 0 | 0 | 0 | 0 |
| ENSMUSG00 Gm17337 | 1464 | 0 | 0 | 0 | 0 | 0 | 0 | 0 | 0 | 0 | 0 | 0 | 0 | 0 | 0 | 0 | 0 | 0 | 0 |
| ENSMUSG00 Gm17336 | 2385 | 0 | 0.08380602 | 0.0202824 | 0.01910316 | 0 | 0 | 0.0198376 | 0.16013586 | 0.01726898 | 0.01976969 | 0.02003143 | 0.02340279 | 0 | 0 | 0 | 0 | 0.04820918 | 0 |
| ENSMUSG00 Gm17338 | 4274 | 0.41769242 | 0.35074409 | 0.10186284 | 0.30914139 | 0.31532379 | 0.27825587 | 0.18818801 | 0.69700688 | 0.33727844 | 0.19857573 | 0.30180718 | 0.50931466 | 0.20776194 | 0.11953322 | 0.14460069 | 0.17022029 | 0.17934627 | 0.60491999 |
| ENSMUSG00 Vmn1r119 | 923 | 0 | 0 | 0 | 0 | 0 | 0 | 0 | 0 | 0 | 0 | 0 | 0 | 0 | 0 | 0 | 0 | 0 | 0 |
| ENSMUSG00 1010001B22i | 587 | 4.56188431 | 3.74557243 | 4.20281098 | 4.65700624 | 3.36185498 | 3.3766769 | 9.18849301 | 5.40028859 | 3.99937206 | 6.58663998 | 6.10412587 | 5.61009215 | 5.29456693 | 5.69671941 | 8.70356547 | 4.24001615 | 5.28863722 | 3.65477881 |
| ENSMUSG00 Vmn1r115 | 887 | 0 | 0 | 0 | 0 | 0 | 0 | 0 | 0 | 0 | 0 | 0 | 0 | 0 | 0 | 0 | 0 | 0 | 0 |
| ENSMUSG00 Vmn1r155 | 923 | 0 | 0 | 0 | 0 | 0 | 0 | 0 | 0 | 0 | 0 | 0 | 0 | 0 | 0 | 0 | 0 | 0 | 0 |
| ENSMUSG00 C1galt1c1 | 533 | 1.3556998 | 1.18751405 | 1.2705993 | 0.34192153 | 0.36121519 | 1.39454222 | 2.48546906 | 0.78821093 | 0.77273013 | 0.70770286 | 0.89634068 | 1.67551708 | 0.37022077 | 0.87137125 | 0.54110861 | 0.21551937 | 0.07190675 | 1.75451185 |
| ENSMUSG00 Gm17088 | 579 | 0 | 0 | 0.25064005 | 0 | 0.1662588 | 0 | 0 | 0 | 0 | 0 | 0.08251288 | 0 | 0 | 0.08021431 | 0.07115985 | 0 | 0.06619395 | 0 |
| ENSMUSG00 Gm17387 | 70 | 0 | 0 | 0 | 0 | 0 | 0 | 0 | 0 | 0 | 0 | 0 | 0 | 0 | 0 | 0 | 0 | 0 | 0 |
| ENSMUSG00 Gm17031 | 1668 | 0.02548272 | 0 | 0 | 0.02731477 | 0 | 0 | 0 | 0 | 0 | 0 | 0 | 0 | 0 | 0.02784418 | 0 | 0 | 0.0229774 | 0 |
| ENSMUSG00 Gm17023 | 445 | 0.382069 | 0.22458131 | 0 | 0.20476874 | 0 | 0 | 0.31896188 | 0 | 0 | 0.21191327 | 0.10735946 | 0.37628537 | 0.77600769 | 0.20873747 | 0.09258777 | 0.34418524 | 0.08612651 | 0.49446388 |

| | | | | | | | | | | | | | | | | | | | | |
|---|---|---|---|---|---|---|---|---|---|---|---|---|---|---|---|---|---|---|---|---|
| ENSMUSG00 | 2010300F17I | 2804 | 0.10611135 | 0.19008784 | 0.05175485 | 0.04874577 | 0 | 0.14137727 | 0.30371905 | 0 | 0 | 0 | 0.03407629 | 0 | 0.03518682 | 0.04969054 | 0.0881631 | 0 | 0 | 0.05885425 |
| ENSMUSG00 | Gm17386 | 739 | 0 | 0 | 0 | 0 | 0 | 0 | 0 | 0 | 0 | 0 | 0 | 0 | 0 | 0 | 0 | 0 | 0 | 0 |
| ENSMUSG00 | Gm17388 | 1917 | 0.06651827 | 0.45181807 | 0.37850963 | 0.33273585 | 0.22597147 | 0.10339642 | 0.51829226 | 0.49807515 | 0.04296976 | 0.49192179 | 0.29906077 | 0.40762612 | 0.61761399 | 1.06600931 | 0.0859709 | 0.33956199 | 0.09996426 | 0.25825872 |
| ENSMUSG00 | Gm10269 | 444 | 0.28719714 | 0.07502904 | 0.21789879 | 0.10261496 | 0.3252157 | 0.33481577 | 0.21312018 | 0.17203785 | 0.18552485 | 0.63717167 | 0.53800629 | 0.2514219 | 0.44443169 | 0.6276228 | 0.37118519 | 0.34496043 | 0.43160244 | 0.24778877 |
| ENSMUSG00 | Vmn2r11 | 2580 | 0 | 0 | 0 | 0 | 0 | 0 | 0 | 0 | 0 | 0 | 0 | 0 | 0 | 0 | 0 | 0 | 0 | 0 |
| ENSMUSG00 | C430039J16I | 1031 | 0 | 0 | 0 | 0 | 0 | 0 | 0 | 0 | 0 | 0 | 0 | 0 | 0 | 0 | 0 | 0 | 0 | 0 |
| ENSMUSG00 | Gm6541 | 718 | 0 | 0 | 0 | 0 | 0 | 0 | 0 | 0 | 0 | 0 | 0 | 0 | 0 | 0 | 0 | 0 | 0 | 0 |
| ENSMUSG00 | Dynll1b | 802 | 2.17295788 | 3.32298197 | 2.95549001 | 1.5906599 | 2.70066903 | 1.23572902 | 3.12664836 | 2.38107247 | 2.56773789 | 3.17473563 | 2.20408163 | 1.53110296 | 2.70648923 | 2.54805467 | 3.13378418 | 2.86463397 | 2.67615041 | 2.53782662 |
| ENSMUSG00 | Otogl | 6919 | 0 | 0 | 0 | 0 | 0 | 0 | 0 | 0 | 0 | 0 | 0.07595383 | 0 | 0 | 0 | 0 | 0 | 0.01107857 | 0 |
| ENSMUSG00 | A930011O12I | 4049 | 0 | 0 | 0 | 0 | 0 | 0 | 0 | 0 | 0 | 0 | 0 | 0 | 0 | 0 | 0 | 0 | 0 | 0 |
| ENSMUSG00 | Gm17171 | 529 | 1.76770108 | 0.62973335 | 1.3716502 | 0.25838021 | 0.54591974 | 0.09367246 | 0.35775183 | 0 | 1.71286077 | 0.08913176 | 0.36124732 | 0.8440932 | 0 | 0.08779601 | 0.15577148 | 0 | 0 | 0 |
| ENSMUSG00 | Gm17444 | 841 | 0 | 0 | 0 | 0 | 0 | 0 | 0 | 0 | 0 | 0 | 0 | 0 | 0 | 0 | 0 | 0 | 0 | 0 |
| ENSMUSG00 | Gm17054 | 258 | 0 | 0 | 0 | 0 | 0 | 0 | 0 | 0 | 0 | 0 | 0 | 0 | 0 | 0 | 0 | 0 | 0 | 0 |
| ENSMUSG00 | Gm7808 | 158 | 0 | 0 | 0 | 0 | 0 | 0 | 0 | 0 | 0 | 0 | 0.60474631 | 1.05979106 | 0.31222732 | 0.29394992 | 0 | 0 | 0 | 0 |
| ENSMUSG00 | 2210008F06I | 1080 | 0.62970631 | 0.37014327 | 0.26874184 | 0.63279228 | 0.35653277 | 0.41293945 | 0.91996876 | 0.14145334 | 0.15254265 | 0.78584505 | 0 | 0.36176818 | 0.31974391 | 0.5590492 | 0.07629918 | 0.56726826 | 0.03548731 | 0 |
| ENSMUSG00 | Gm17084 | 4640 | 0.10076658 | 0 | 0.02085066 | 0 | 0.0518663 | 0.01067947 | 0.0203934 | 0 | 0 | 0 | 0.04118531 | 0.18043856 | 0 | 0.0100095 | 0.00887965 | 0.04951372 | 0 | 0 |
| ENSMUSG00 | Ms4a12 | 783 | 0 | 0 | 0 | 0 | 0 | 0 | 0 | 0 | 0 | 0 | 0 | 0 | 0 | 0 | 0 | 0 | 0 | 0 |
| ENSMUSG00 | Gm17183 | 490 | 0 | 0 | 0 | 0 | 0 | 0 | 0 | 0 | 0 | 0 | 0 | 0 | 0 | 0 | 0 | 0 | 0 | 0 |
| ENSMUSG00 | Gm17240 | 2027 | 0 | 0 | 0 | 0 | 0 | 0 | 0 | 0 | 0 | 0 | 0 | 0 | 0.29204884 | 0 | 0 | 0 | 0 | 0 |
| ENSMUSG00 | Vmn2r82 | 3039 | 0 | 0 | 0 | 0 | 0 | 0 | 0 | 0 | 0 | 0 | 0 | 0 | 0 | 0 | 0 | 0 | 0 | 0 |
| ENSMUSG00 | 4930525G20I | 2866 | 0.3707709 | 0.13948176 | 0.03375683 | 0.28614752 | 0.10076467 | 0.17289858 | 0.33016524 | 0.10660824 | 0.44549267 | 0 | 0.23337384 | 0.25317642 | 0.3098306 | 0 | 0.01437598 | 0.06680148 | 0.01337275 | 0 |
| ENSMUSG00 | Gm17159 | 479 | 0 | 0 | 0 | 0 | 0 | 0 | 0 | 0 | 0 | 0 | 0 | 0 | 0 | 0 | 0 | 0 | 0 | 0 |
| ENSMUSG00 | Gm20538 | 223 | 0 | 0 | 0 | 0 | 0 | 0 | 0 | 0 | 0 | 0 | 0 | 0 | 0 | 0 | 0 | 0 | 0 | 0 |
| ENSMUSG00 | Gm3739 | 3181 | 0.01336221 | 0 | 0 | 0 | 0 | 0 | 0 | 0 | 0 | 0 | 0 | 0 | 0 | 0.0438014 | 0 | 0 | 0 | 0 |
| ENSMUSG00 | 1700045I11R | 490 | 0 | 0 | 0 | 0 | 0 | 0 | 0 | 0 | 0 | 0 | 0 | 0 | 0 | 0 | 0 | 0 | 0 | 0 |
| ENSMUSG00 | 2610021A01I | 3998 | 3.89116919 | 2.39139586 | 2.54088078 | 1.84614537 | 2.48003409 | 2.34253793 | 3.17153526 | 2.52195954 | 2.90510191 | 2.77148956 | 3.15472462 | 3.47626313 | 2.17169021 | 2.23043141 | 1.85499751 | 2.36562782 | 2.5595601 | 1.14200997 |
| ENSMUSG00 | 2810468N07I | 1233 | 0 | 0 | 0 | 0 | 0 | 0.08037751 | 0 | 0 | 0 | 0 | 0 | 0 | 0 | 0 | 0 | 0 | 0 | 0 |
| ENSMUSG00 | Gm16432 | 1655 | 0 | 0.08051455 | 0 | 0 | 0 | 0 | 0 | 0 | 0 | 0.02848985 | 0.14433522 | 0 | 0 | 0 | 0 | 0 | 0.02315788 | 0 |
| ENSMUSG00 | Gm5799 | 1310 | 0 | 0 | 0 | 0 | 0 | 0 | 0 | 0 | 0 | 0.14397161 | 0 | 0.04260738 | 1.16739652 | 0.42544202 | 0.44032197 | 0 | 0.08777015 | 0.67186695 |
| ENSMUSG00 | Gm10039 | 943 | 10.7277114 | 9.99740092 | 11.6445296 | 8.11691988 | 9.0853472 | 10.2468537 | 15.5534788 | 10.9352584 | 9.60873114 | 12.5501872 | 12.9189973 | 10.5949137 | 11.0905265 | 10.9338148 | 7.90825198 | 9.78582861 | 10.323308 | 9.10012794 |
| ENSMUSG00 | Gm17035 | 417 | 0 | 0 | 0 | 0 | 0 | 0 | 0 | 0 | 0 | 0 | 0 | 0 | 0 | 0 | 0 | 0 | 0 | 0 |
| ENSMUSG00 | Gm17665 | 58 | 0 | 0 | 0 | 0 | 0 | 0 | 0 | 0 | 0 | 0 | 0 | 0 | 0 | 0 | 0 | 0 | 0 | 0 |
| ENSMUSG00 | Gm17154 | 428 | 0 | 0 | 0 | 0 | 0 | 0 | 0 | 0 | 0 | 0 | 0 | 0 | 0 | 0 | 0 | 0 | 0 | 0 |
| ENSMUSG00 | Gm8223 | 261 | 0 | 0 | 0 | 0 | 0 | 0 | 0 | 0 | 0 | 0 | 0 | 0 | 0.18901118 | 0 | 0 | 0 | 0 | 0 |
| ENSMUSG00 | Gm8715 | 527 | 0 | 0 | 0 | 0 | 0 | 0 | 0 | 0 | 0 | 0 | 0 | 0 | 0 | 0 | 0 | 0 | 0 | 0 |
| ENSMUSG00 | Gm7953 | 466 | 0 | 0 | 0 | 0 | 0 | 0 | 0 | 0 | 0 | 0 | 0 | 0 | 0 | 0 | 0 | 0 | 0 | 0 |
| ENSMUSG00 | Gm6586 | 639 | 0 | 0 | 0 | 0 | 0 | 0 | 0 | 0 | 0 | 0 | 0 | 0 | 0 | 0 | 0 | 0 | 0 | 0 |
| ENSMUSG00 | Gm6225 | 1698 | 0.60077987 | 0.13733231 | 0.19941974 | 0.37565054 | 0.68030988 | 0.204281 | 0.39009276 | 0.44985162 | 0.24255899 | 0.38875727 | 0.19695213 | 0.19722849 | 0.05810591 | 0.43763569 | 0.41250085 | 0.65396208 | 0.36114296 | 0.09718923 |
| ENSMUSG00 | Gm17664 | 54 | 0 | 0 | 0 | 0 | 0 | 0 | 0 | 0 | 0 | 0 | 0 | 0 | 0 | 0 | 0 | 0 | 0 | 0 |
| ENSMUSG00 | Triml2 | 1531 | 0 | 0 | 0 | 0.05951802 | 0 | 0 | 0 | 0 | 0 | 0 | 0 | 0 | 0 | 0 | 0 | 0 | 0 | 0 |
| ENSMUSG00 | Vmn2r97 | 2580 | 0 | 0 | 0 | 0 | 0 | 0 | 0 | 0 | 0 | 0 | 0 | 0 | 0 | 0 | 0 | 0 | 0 | 0 |
| ENSMUSG00 | Gm17025 | 578 | 1.02953714 | 0.63398242 | 2.25966318 | 0.23647601 | 0.3330929 | 0.42865686 | 1.71897276 | 1.85015098 | 1.28262506 | 0.97890733 | 0.57858946 | 0.48283445 | 1.19489073 | 0 | 1.06924453 | 0 | 0.99462708 | 0.38068586 |
| ENSMUSG00 | Gm4498 | 920 | 0 | 0 | 0 | 0 | 0 | 0 | 0 | 0 | 0 | 0 | 0 | 0 | 0 | 0 | 0 | 0 | 0 | 0 |
| ENSMUSG00 | Gm3269 | 1753 | 0 | 0 | 0 | 0 | 0 | 0 | 0 | 0 | 0 | 0 | 0 | 0 | 0 | 0 | 0 | 0 | 0 | 0 |
| ENSMUSG00 | Gm8398 | 363 | 0 | 0 | 0 | 0 | 0 | 0 | 0 | 0 | 0 | 0 | 0 | 0 | 0 | 0 | 0 | 0 | 0 | 0 |
| ENSMUSG00 | Gm44 | 992 | 0 | 0 | 0 | 0 | 0 | 0 | 0 | 0 | 0 | 0 | 0 | 0 | 0 | 0 | 0 | 0 | 0 | 0 |
| ENSMUSG00 | Gm3452 | 329 | 31.5235956 | 26.9338294 | 34.9936177 | 28.2506172 | 29.5521233 | 32.5331017 | 40.2661097 | 35.6385032 | 44.5665642 | 35.9721171 | 44.5802804 | 36.475281 | 50.3815325 | 41.7855619 | 39.5735309 | 39.919995 | 37.6273371 | 39.1250182 |
| ENSMUSG00 | Gm5327 | 1004 | 0 | 0.03318017 | 0 | 0 | 0 | 0 | 0 | 0 | 0 | 0 | 0 | 0 | 0 | 0 | 0 | 0 | 0 | 0 |
| ENSMUSG00 | Gm17292 | 2107 | 0.02017332 | 0.04743174 | 0 | 0 | 0.04568764 | 0.14110888 | 0 | 0.01812644 | 0.09773734 | 0 | 0.0906976 | 0 | 0 | 0.04408551 | 0.01955461 | 0.01817305 | 0.05456995 | 0 |
| ENSMUSG00 | Gm17293 | 506 | 0 | 0 | 0 | 0 | 0 | 0 | 0 | 0 | 0 | 0 | 0 | 0 | 0 | 0 | 0 | 0 | 0 | 0 |
| ENSMUSG00 | Gm7138 | 692 | 0 | 0 | 0 | 0 | 0 | 0 | 0 | 0 | 0 | 0 | 0 | 0 | 0 | 0 | 0 | 0 | 0 | 0 |
| ENSMUSG00 | Vmn2r118 | 2556 | 0 | 0 | 0 | 0 | 0 | 0 | 0 | 0 | 0 | 0 | 0 | 0 | 0 | 0 | 0 | 0 | 0 | 0 |
| ENSMUSG00 | Gm8720 | 923 | 0 | 0 | 0 | 0 | 0 | 0 | 0 | 0 | 0 | 0 | 0 | 0 | 0 | 0 | 0 | 0 | 0 | 0 |
| ENSMUSG00 | Gm17294 | 3549 | 0.03592999 | 0.04693279 | 0 | 0.03851314 | 0.09493476 | 0.09773715 | 0.1333127 | 0.118376 | 0.2669174 | 0.1727132 | 0.05384611 | 0 | 0.06950115 | 0.03925958 | 0.16253079 | 0.23736077 | 0 | 0 |
| ENSMUSG00 | Gm17194 | 347 | 0.61246651 | 0.76802062 | 0.55761995 | 0.26259968 | 0 | 0.14280327 | 0 | 0 | 0.35607939 | 0.13588099 | 0.13768 | 0.32170411 | 0 | 0.4015339 | 0.35620942 | 0.44139029 | 0.55225212 | 0.47558306 |
| ENSMUSG00 | Gm7951 | 1403 | 0 | 0 | 0 | 0 | 0 | 0 | 0 | 0 | 0 | 0 | 0 | 0 | 0 | 0 | 0 | 0 | 0 | 0 |
| ENSMUSG00 | Gm17066 | 1502 | 1.04706492 | 1.77432193 | 1.12721276 | 1.69868075 | 1.98680377 | 1.74853188 | 1.73248824 | 1.85622195 | 2.02916256 | 1.38124563 | 1.6858008 | 1.96952737 | 2.10202577 | 1.88621127 | 2.60595726 | 1.68254333 | 2.6027179 | 1.90444311 |
| ENSMUSG00 | Mtag2 | 711 | 0 | 0 | 0 | 0 | 0 | 0.06969442 | 0 | 0 | 0.05792759 | 0 | 0 | 0 | 0.48568695 | 0 | 0 | 0.05385458 | 0.05390478 | 0 |
| ENSMUSG00 | Vmn2r87 | 3030 | 0 | 0 | 0 | 0 | 0 | 0 | 0 | 0 | 0 | 0 | 0 | 0 | 0 | 0 | 0 | 0.02527433 | 0 | 0 |
| ENSMUSG00 | Lamtor3 | 1288 | 75.17608 | 65.9791872 | 68.8045867 | 65.8300493 | 68.5730434 | 57.9782372 | 80.8503155 | 73.2712937 | 70.3176616 | 68.6760243 | 80.1194956 | 63.3559773 | 70.6656734 | 68.15165 | 69.1597545 | 74.9164058 | 75.0159895 | 59.6643805 |
| ENSMUSG00 | 4731419I09R | 2818 | 0.03016691 | 0.1063932 | 0.03433182 | 0.0485036 | 0.13664137 | 0.19342799 | 0.11752617 | 0.25750733 | 0.14615513 | 0.18405172 | 0.0508605 | 0.21787519 | 0.14004803 | 0.03296245 | 0.292417 | 0.14946653 | 0.08160319 | 0.0976031 |
| ENSMUSG00 | Gm17484 | 2291 | 0.50093398 | 0.17448919 | 0.65455235 | 0.11932181 | 0.50421919 | 0.58399119 | 0.82606161 | 0.46677751 | 0.39550561 | 0.41161679 | 0.62559963 | 0.41417122 | 0.66752049 | 0.16217927 | 0.44960231 | 0.43455076 | 0.18401976 | 0.52824109 |
| ENSMUSG00 | Zfp422-rs1 | 432 | 0.29517483 | 0.30845272 | 0.11197576 | 0.21093076 | 0.77991543 | 0.91764321 | 0.65712054 | 1.23771674 | 0.28601747 | 0.43658059 | 0.33177054 | 0.51681169 | 0.68516552 | 0.86007569 | 0.85836575 | 0.53181399 | 0.4435914 | 0.25467179 |
| ENSMUSG00 | Ighd5-1 | 9 | 0 | 0 | 0 | 0 | 0 | 0 | 0 | 0 | 0 | 0 | 0 | 0 | 0 | 0 | 0 | 0 | 0 | 0 |
| ENSMUSG00 | Gm4066 | 239 | 0 | 0 | 0 | 0 | 0 | 0 | 0 | 0 | 0 | 0 | 0 | 0 | 0 | 0 | 0 | 0 | 0 | 0 |
| ENSMUSG00 | Skor2 | 3132 | 0 | 0 | 0 | 0 | 0 | 0 | 0 | 0 | 0 | 0 | 0 | 0 | 0 | 0.03946509 | 0 | 0 | 0 | 0 |
| ENSMUSG00 | Gm17419 | 2647 | 0 | 0.03775545 | 0 | 0 | 0 | 0 | 0 | 0 | 0 | 0 | 0 | 0 | 0 | 0 | 0 | 0 | 0 | 0 |
| ENSMUSG00 | AU040972 | 1038 | 0 | 0.16046674 | 0.13980789 | 0.04389311 | 0.09273974 | 0.09547733 | 0.04558062 | 0.18397111 | 0 | 0 | 0 | 0.26886157 | 0 | 0 | 0 | 0 | 0 | 0 |
| ENSMUSG00 | 2610020C07I | 2923 | 0.21812441 | 0.17095225 | 0.62887244 | 0.26498042 | 0.51046515 | 0.33905394 | 0.48559027 | 0.56184513 | 0.49316731 | 0.30648764 | 0.22882293 | 0.19095334 | 0.57382319 | 0.39722962 | 0.31010408 | 0.72048697 | 0.485143 | 1.03506701 |
| ENSMUSG00 | Gm6494 | 284 | 2.24499169 | 1.17298923 | 1.70329332 | 2.88767182 | 2.03374325 | 2.44274039 | 0 | 0 | 0 | 0 | 0 | 0.39306805 | 0 | 0 | 0 | 0 | 0 | 0 |
| ENSMUSG00 | Gm8330 | 3272 | 0.29878333 | 0.67195936 | 0.22176129 | 0.29241502 | 0.64725081 | 0.25745613 | 0.59285447 | 0.99216204 | 0.21398862 | 0.23056579 | 0.10220804 | 0.05117573 | 0.45230975 | 0.46841531 | 0.22665893 | 0.18724013 | 0.56224396 | 0.18493282 |
| ENSMUSG00 | Gm9268 | 2587 | 0 | 0 | 0 | 0 | 0 | 0 | 0 | 0 | 0 | 0 | 0 | 0 | 0 | 0 | 0 | 0 | 0 | 0 |
| ENSMUSG00 | Gm3485 | 446 | 0 | 0 | 0 | 0 | 0 | 0 | 0 | 0 | 0 | 0 | 0 | 0 | 0 | 0 | 0 | 0 | 0 | 0 |
| ENSMUSG00 | Cldn20 | 659 | 0 | 0.05055067 | 0.58723557 | 0 | 0 | 0.45116298 | 0 | 0 | 0.31249253 | 0.07154887 | 0.36248072 | 0 | 0 | 0.14095322 | 0.12504266 | 0.11620822 | 0 | 0 |
| ENSMUSG00 | Olfr102 | 1026 | 0 | 0 | 0 | 0 | 0 | 0 | 0 | 0 | 0 | 0 | 0 | 0 | 0 | 0 | 0 | 0 | 0 | 0 |
| ENSMUSG00 | Gm6465 | 1093 | 0 | 0 | 0 | 0 | 0 | 0 | 0 | 0 | 0 | 0 | 0 | 0 | 0 | 0 | 0 | 0 | 0 | 0 |
| ENSMUSG00 | Gm17195 | 3528 | 1.61442562 | 1.19918865 | 1.53566763 | 0.82650421 | 1.82813996 | 1.32028258 | 1.3812942 | 1.45061846 | 1.10904733 | 0.88207098 | 1.59791528 | 1.7244635 | 0.53135285 | 0.47391926 | 0.60727917 | 2.1815227 | 1.33620593 | 2.44796762 |
| ENSMUSG00 | Gm17612 | 3613 | 0.01176451 | 0 | 0 | 0 | 0 | 0.06857561 | 0.14404635 | 0 | 0 | 0 | 0 | 0.03089713 | 0.04096201 | 0 | 0.06842218 | 0.01059801 | 0 | 0 |
| ENSMUSG00 | Ccdc72 | 633 | 59.695263 | 58.4686026 | 55.4805419 | 44.6253515 | 60.5260841 | 48.6133453 | 79.4524134 | 70.4115855 | 68.9044553 | 59.888097 | 63.0961537 | 52.9940177 | 62.5806154 | 88.6326344 | 66.3911327 | 80.5736004 | 77.7424411 | 80.6452623 |
| ENSMUSG00 | Vmn1r238 | 1010 | 0 | 0 | 0 | 0 | 0 | 0 | 0 | 0 | 0 | 0 | 0 | 0 | 0 | 0 | 0 | 0 | 0 | 0 |
| ENSMUSG00 | Gm17548 | 2667 | 0.01593745 | 0.01249077 | 0 | 0.01708326 | 0 | 0 | 0 | 0.02864072 | 0.03088603 | 0.0176793 | 0 | 0.02092826 | 0 | 0.01741436 | 0 | 0.01435718 | 0.02874113 | 0 |
| ENSMUSG00 | Vmn1r9 | 1050 | 0 | 0 | 0 | 0 | 0 | 0 | 0 | 0 | 0 | 0 | 0 | 0 | 0 | 0 | 0 | 0 | 0 | 0 |
| ENSMUSG00 | Gm17167 | 5052 | 0.05889474 | 0.026376 | 0.02872537 | 0.04509209 | 0.09527301 | 0.01961708 | 0.04682569 | 0 | 0 | 0 | 0 | 0.05524115 | 0.00976483 | 0.01838642 | 0.01631099 | 0 | 0.05310453 | 0.1088858 |
| ENSMUSG00 | Gm17549 | 1207 | 1.86642447 | 2.40117796 | 1.28247968 | 1.43439908 | 1.79447934 | 0.98530705 | 4.03745313 | 3.79709055 | 2.32036709 | 1.83602573 | 2.05824179 | 2.17343508 | 1.4713745 | 2.23177885 | 2.52602745 | 1.87170326 | 2.7625417 | 4.28405636 |
| ENSMUSG00 | Vmn2r48 | 3115 | 0 | 0 | 0 | 0 | 0 | 0 | 0 | 0 | 0 | 0 | 0 | 0 | 0 | 0 | 0 | 0 | 0 | 0 |
| ENSMUSG00 | Vmn2r47 | 2553 | 0 | 0 | 0 | 0 | 0 | 0 | 0 | 0 | 0 | 0 | 0 | 0 | 0 | 0 | 0 | 0 | 0 | 0 |
| ENSMUSG00 | Gm2366 | 3980 | 0.01067969 | 0.07533067 | 0.06077077 | 0.12592248 | 0.07256069 | 0.08715305 | 0 | 0.08636473 | 0 | 0.03554073 | 0.01200376 | 0.08414421 | 0 | 0 | 0 | 0 | 0.11555667 | 0 |
| ENSMUSG00 | Gm17547 | 2009 | 0 | 0 | 0 | 0 | 0 | 0 | 0 | 0 | 0 | 0 | 0 | 0 | 0 | 0 | 0 | 0 | 0 | 0 |
| ENSMUSG00 | Gm17572 | 419 | 0 | 0 | 0 | 0 | 0 | 0 | 0 | 0 | 0 | 0 | 0 | 0 | 0 | 0 | 0 | 0 | 0 | 0 |
| ENSMUSG00 | Gm6548 | 1164 | 3.03086736 | 3.17674507 | 3.19996721 | 3.20962683 | 3.43208734 | 4.59767631 | 3.90207661 | 2.88739812 | 3.67989488 | 2.83552339 | 2.7088894 | 3.78817641 | 2.2885941 | 1.39651466 | 3.32727341 | 1.97374265 | 2.73288887 | 2.59922755 |
| ENSMUSG00 | Gm6811 | 1532 | 0 | 0.06523413 | 0 | 0 | 0.12567082 | 0.03234513 | 0 | 0 | 0 | 0 | 0 | 0 | 0.09094795 | 0.18825776 | 0 | 0 | 0 | 0.21544037 |
| ENSMUSG00 | Gm8711 | 877 | 0 | 0 | 0 | 0 | 0 | 0 | 0 | 0 | 0 | 0 | 0 | 0 | 0 | 0 | 0 | 0 | 0 | 0 |
| ENSMUSG00 | Serpina3e-ps | 1224 | 10.3832088 | 9.03585039 | 11.6191323 | 11.3902611 | 10.6173361 | 10.4449389 | 8.19467968 | 8.83043291 | 10.2293307 | 6.89540513 | 8.46990685 | 12.7226878 | 1.41063489 | 0.30355613 | 2.05334551 | 1.53287562 | 0.90805769 | 1.21343618 |
| ENSMUSG00 | Gm8653 | 923 | 0 | 0 | 0 | 0 | 0 | 0 | 0 | 0 | 0 | 0 | 0 | 0 | 0 | 0 | 0 | 0 | 0 | 0 |
| ENSMUSG00 | Gm17725 | 1670 | 0 | 0.49869602 | 0.14483093 | 0.24553856 | 0 | 0 | 0 | 0.43452434 | 0 | 0 | 0.02860776 | 0.0668451 | 0 | 0.22248665 | 0.34540226 | 0 | 0.13769927 | 0 |
| ENSMUSG00 | Gm14569 | 5711 | 0 | 0 | 0 | 0 | 0 | 0 | 0 | 0 | 0 | 0 | 0 | 0 | 0 | 0 | 0 | 0 | 0 | 0 |
| ENSMUSG00 | Gm8584 | 738 | 0 | 0 | 0 | 0 | 0 | 0 | 0 | 0 | 0 | 0 | 0 | 0 | 0 | 0 | 0 | 0 | 0 | 0 |
| ENSMUSG00 | Gm17227 | 1122 | 0 | 0.17814382 | 0.08622733 | 0 | 0.08579666 | 0.13249394 | 0.42168163 | 0.20423745 | 0 | 0.2521428 | 0.21290088 | 0.4974658 | 0.08793568 | 0.33115214 | 0 | 0.20476261 | 0.27327128 | 0 |
| ENSMUSG00 | Gm17726 | 3345 | 0.2287274 | 0.45811454 | 0.57845776 | 0.69465269 | 0.43167645 | 0.23702354 | 0.07072149 | 0.14843086 | 0.52945297 | 0.22553401 | 0.21423748 | 0.83431484 | 0.26546323 | 0.26380797 | 0.35720332 | 0.19460099 | 0.21769795 | 0.54269074 |
| ENSMUSG00 | Gm6665 | 252 | 3.20475534 | 2.90826854 | 3.45525217 | 2.53116913 | 2.29199637 | 2.94956747 | 3.00397961 | 5.15294318 | 4.57627953 | 4.11633124 | 4.36041288 | 4.87279594 | 5.67708572 | 8.6621909 | 7.03042422 | 3.64672451 | 3.65012351 | 3.92922191 |
| ENSMUSG00 | Gm9581 | 6258 | 0.58412354 | 0.72928516 | 0.71887797 | 0.88093422 | 0.66913988 | 0.88684982 | 0.2948537 | 0.5797824 | 0.66472325 | 0.69317109 | 0.89320392 | 0.55298355 | 0.74888657 | 0.34881306 | 0.43453223 | 0.67305318 | 0.81454099 | 1.11635611 |
| ENSMUSG00 | Gm8108 | 2082 | 0 | 0 | 0 | 0 | 0 | 0 | 0 | 0 | 0 | 0 | 0 | 0.05361735 | 0 | 0.02230744 | 0.05936824 | 0 | 0 | 0 |
| ENSMUSG00 | Gm17683 | 781 | 0.10884808 | 0.04265415 | 0 | 0 | 0 | 0 | 0 | 0 | 0 | 0 | 0 | 0 | 0 | 0 | 0 | 0 | 0 | 0 |
| ENSMUSG00 | Gm3317 | 685 | 0 | 0 | 0 | 0 | 0 | 0 | 0 | 0 | 0 | 0 | 0 | 0 | 0 | 0 | 0 | 0 | 0 | 0 |
| ENSMUSG00 | Gm13279 | 1555 | 0 | 0 | 0 | 0 | 0 | 0 | 0 | 0 | 0 | 0 | 0 | 0 | 0 | 0 | 0 | 0 | 0 | 0 |
| ENSMUSG00 | Gm8989 | 2132 | 0 | 0 | 0 | 0 | 0 | 0 | 0 | 0 | 0 | 0 | 0 | 0 | 0 | 0 | 0 | 0 | 0 | 0 |
| ENSMUSG00 | Gm8206 | 2332 | 0 | 0 | 0 | 0 | 0 | 0 | 0 | 0 | 0 | 0 | 0 | 0 | 0 | 0 | 0 | 0 | 0 | 0 |
| ENSMUSG00 | Gm8232 | 1397 | 0 | 0 | 0 | 0 | 0 | 0 | 0 | 0 | 0 | 0 | 0 | 0 | 0 | 0 | 0 | 0 | 0 | 0 |
| ENSMUSG00 | Gm7707 | 1155 | 0 | 0 | 0.12564553 | 0.07889358 | 0.08334532 | 0 | 0 | 0.06613403 | 0 | 0 | 0 | 0 | 0 | 0.04021133 | 0 | 0.03315204 | 0 | 0.04762693 |
| ENSMUSG00 | Gm6026 | 335 | 0 | 0 | 0 | 0 | 0 | 0 | 0 | 0 | 0 | 0 | 0 | 0 | 0 | 0 | 0 | 0 | 0 | 0 |
| ENSMUSG00 | Vmn2r3 | 2742 | 0 | 0 | 0 | 0 | 0 | 0 | 0 | 0 | 0 | 0 | 0 | 0 | 0 | 0 | 0 | 0 | 0 | 0 |
| ENSMUSG00 | Serpina3d-ps | 1268 | 0 | 0 | 0 | 0 | 0 | 0 | 0 | 0 | 0 | 0 | 0 | 0 | 0 | 0 | 0 | 0 | 0 | 0 |
| ENSMUSG00 | Gm14296 | 1264 | 0 | 0 | 0 | 0 | 0 | 0 | 0 | 0 | 0 | 0 | 0 | 0 | 0 | 0 | 0 | 0 | 0 | 0 |
| ENSMUSG00 | 2010016I18R | 1424 | 3.7609917 | 2.62011528 | 2.68364389 | 2.71958481 | 2.60263913 | 3.79301121 | 2.99026763 | 4.90815284 | 4.25169791 | 2.48335867 | 4.79762575 | 6.62419029 | 3.25646083 | 1.79383764 | 2.66189827 | 3.95275231 | 2.7452825 | 2.93587929 |
| ENSMUSG00 | Trim30b | 2769 | 0 | 0 | 0 | 0 | 0 | 0 | 0 | 0 | 0 | 0 | 0.03450701 | 0 | 0.07126315 | 0 | 0 | 0 | 0 | 0 |
| ENSMUSG00 | Gm6211 | 5205 | 0 | 0 | 0 | 0.02625997 | 0.02774174 | 0.12376283 | 0 | 0 | 0 | 0 | 0 | 0 | 0 | 0.00892298 | 0.01583153 | 0 | 0.01472672 | 0 |
| ENSMUSG00 | Gstm2-ps1 | 255 | 4.83392198 | 3.78852522 | 3.9837025 | 5.1814521 | 7.73885832 | 3.49784002 | 6.86497694 | 7.1891581 | 6.13759843 | 2.58866606 | 5.24587779 | 5.25323883 | 7.35142297 | 4.5533419 | 4.52409241 | 2.7028664 | 3.60718088 | 5.82449365 |
| ENSMUSG00 | CT030161.2 | 152 | 0 | 0 | 0 | 0 | 0 | 0 | 0 | 0 | 0 | 0 | 0 | 0 | 0 | 0 | 0 | 0 | 0 | 0 |
| ENSMUSG00 | Gm6467 | 607 | 1.68060004 | 1.86596113 | 1.83293443 | 2.55201896 | 1.66519011 | 3.10214807 | 1.71479232 | 1.76175827 | 2.03557739 | 0.85446085 | 2.2824939 | 2.39078621 | 0.81271692 | 1.37725465 | 2.10419808 | 2.08169694 | 1.45223201 | 2.44686307 |
| ENSMUSG00 | Gm17277 | 982 | 0 | 0 | 0 | 0 | 0 | 0 | 0 | 0 | 0 | 0 | 0 | 0 | 0 | 0 | 0 | 0 | 0 | 0 |
| ENSMUSG00 | Gm3187 | 596 | 0 | 0 | 0 | 0 | 0 | 0 | 0 | 0 | 0 | 0 | 0 | 0 | 0 | 0 | 0 | 0 | 0 | 0 |
| ENSMUSG00 | Gm17033 | 590 | 0 | 0 | 0 | 0 | 0 | 0 | 0 | 0 | 0 | 0 | 0 | 0 | 0 | 0 | 0 | 0 | 0 | 0 |
| ENSMUSG00 | Gm8180 | 1387 | 0 | 0 | 0 | 0 | 0 | 0 | 0 | 0 | 0 | 0 | 0 | 0 | 0 | 0 | 0 | 0 | 0 | 0 |
| ENSMUSG00 | Cyp4f17 | 1680 | 1.8975525 | 2.63727079 | 1.58365724 | 1.57294082 | 2.14874659 | 1.82873183 | 0.59140849 | 1.97782084 | 1.49546277 | 1.15070169 | 1.79156094 | 1.59473322 | 2.26104621 | 1.54813625 | 2.96749301 | 1.98290645 | 0.84409106 | 2.48850721 |
| ENSMUSG00 | Gm17191 | 415 | 3.07266333 | 1.20408051 | 0.46625089 | 1.64678474 | 0.11598054 | 1.31344595 | 0.45602582 | 0.55217931 | 0.19848923 | 1.13616152 | 0.57560191 | 1.21046015 | 1.90195344 | 0.11191346 | 0.09280858 | 1.38399786 | 0.92352523 | 0.92786445 |
| ENSMUSG00 | Gm17279 | 1015 | 0.08375404 | 0.03282059 | 0.04765865 | 0.08977546 | 0.04742061 | 0 | 0.09322695 | 0 | 0 | 0.04645389 | 0 | 0.2199632 | 0.04860287 | 0 | 0 | 0.0377599 | 0 | 0 |
| ENSMUSG00 | Gm17278 | 517 | 0 | 0 | 0 | 0 | 0 | 0 | 0 | 0 | 0 | 0 | 0 | 0 | 0 | 0 | 0 | 0 | 0 | 0 |
| ENSMUSG00 | Gm17461 | 1744 | 0.12186117 | 0 | 0.05547423 | 0 | 0.02759858 | 0.02841326 | 0.16277298 | 0 | 0.09446449 | 0.13517977 | 0.0273939 | 0.25603515 | 0.19800655 | 0.05326157 | 0.70874236 | 0.61475746 | 0.3516174 | 0.37850303 |
| ENSMUSG00 | Gm17460 | 2925 | 0 | 0 | 0 | 0 | 0 | 0 | 0 | 0 | 0 | 0 | 0 | 0 | 0 | 0 | 0 | 0 | 0 | 0 |
| ENSMUSG00 | 4631405K08I | 1739 | 0.2688654 | 0.21071986 | 0.41725299 | 0.41919305 | 0.22142346 | 0.82635381 | 1.08827324 | 0.10981139 | 0.42631241 | 0.46093269 | 0.3571446 | 0.67402468 | 1.50350294 | 1.14841619 | 1.06617022 | 2.55417508 | 0.88157095 | 0.28469348 |
| ENSMUSG00 | Gm17067 | 3669 | 0 | 0 | 0 | 0 | 0 | 0 | 0 | 0 | 0 | 0 | 0 | 0.10648941 | 0 | 0 | 0 | 0 | 0 | 0 |
| ENSMUSG00 | Gm3344 | 599 | 0 | 0 | 0 | 0 | 0 | 0 | 0 | 0 | 0 | 0 | 0 | 0 | 0 | 0 | 0 | 0 | 0 | 0 |
| ENSMUSG00 | 3110083C13I | 854 | 2.33927784 | 2.22346015 | 2.54895652 | 1.86725592 | 0.73268736 | 0.69629134 | 2.16064927 | 1.78887131 | 2.79721067 | 2.37409864 | 1.90204753 | 3.52902761 | 0.80871996 | 1.35960443 | 0.67543534 | 0.40353099 | 1.48099274 | 0.77296169 |
| ENSMUSG00 | Gm10310 | 962 | 0 | 0 | 0 | 0 | 0 | 0 | 0 | 0 | 0 | 0 | 0 | 0 | 0 | 0 | 0 | 0 | 0 | 0 |
| ENSMUSG00 | Gm10716 | 665 | 0 | 0 | 0.07274215 | 0 | 0 | 0 | 0 | 0 | 0 | 0 | 0 | 0.08393333 | 0.14836667 | 0 | 0 | 0 | 0 | 0 |
| ENSMUSG00 | Gm3991 | 2554 | 0 | 0 | 0 | 0 | 0 | 0 | 0 | 0 | 0 | 0 | 0 | 0 | 0 | 0 | 0 | 0 | 0 | 0 |
| ENSMUSG00 | Olfr93 | 1044 | 0 | 0 | 0 | 0 | 0 | 0 | 0 | 0 | 0 | 0 | 0 | 0 | 0 | 0 | 0 | 0 | 0 | 0 |
| ENSMUSG00 | Gm8879 | 1120 | 0 | 0 | 0 | 0 | 0 | 0 | 0 | 0 | 0 | 0 | 0 | 0 | 0 | 0 | 0 | 0 | 0 | 0 |
| ENSMUSG00 | Gm17346 | 546 | 0 | 0 | 0 | 0 | 0 | 0 | 0 | 0 | 0 | 0 | 0 | 0 | 0 | 0 | 0 | 0 | 0 | 0 |
| ENSMUSG00 | Gm17349 | 293 | 0.14506886 | 0.45478354 | 0.16509737 | 0.31099689 | 1.31418222 | 0.50736587 | 0.16147672 | 0.39104849 | 0.70284157 | 0.3218478 | 1.95665359 | 0.76198857 | 0.16836832 | 0.79256122 | 0.42185893 | 0.52273867 | 1.04645179 | 0.56323317 |
| ENSMUSG00 | Gm8256 | 450 | 0 | 0 | 0 | 0 | 0 | 0 | 0 | 0 | 0 | 0 | 0 | 0 | 0 | 0 | 0 | 0 | 0 | 0 |
| ENSMUSG00 | Ighv2-1 | 351 | 0 | 0 | 0 | 0 | 0 | 0 | 0 | 0 | 0 | 0 | 0 | 0 | 0 | 0 | 0 | 0 | 0 | 0 |
| ENSMUSG00 | Gm17348 | 1523 | 0 | 0 | 0 | 0 | 0 | 0 | 0 | 0 | 0 | 0 | 0 | 0 | 0 | 0 | 0 | 0 | 0 | 0 |
| ENSMUSG00 | Vmn1r111 | 920 | 0 | 0 | 0 | 0 | 0 | 0 | 0 | 0 | 0 | 0 | 0 | 0 | 0 | 0 | 0 | 0 | 0 | 0 |
| ENSMUSG00 | Acer2 | 2939 | 0 | 0 | 0 | 0 | 0 | 0 | 0 | 0 | 0 | 0 | 0 | 0 | 0 | 0 | 0 | 0 | 0 | 0 |
| ENSMUSG00 | Gm9364 | 787 | 0 | 0 | 0 | 0 | 0 | 0 | 0 | 0.0970582 | 0 | 0.05991195 | 0 | 0 | 0 | 0 | 0 | 0 | 0 | 0 |
| ENSMUSG00 | Gm17087 | 1098 | 0 | 0 | 0 | 0.12448373 | 0 | 0 | 0 | 0 | 0 | 0 | 0 | 0.05083394 | 0.04492889 | 0.16919522 | 0 | 0 | 0 | 0 |
| ENSMUSG00 | Gm17046 | 849 | 0 | 0 | 0 | 0 | 0 | 0 | 0.05572754 | 0 | 0 | 0 | 0 | 0 | 0 | 0 | 0 | 0 | 0 | 0 |
| ENSMUSG00 | Gm17657 | 509 | 0 | 0 | 0 | 0 | 0.28368521 | 0 | 0 | 0 | 0 | 0 | 0 | 0 | 0 | 0 | 0 | 0 | 0 | 0 |
| ENSMUSG00 | Gm17044 | 739 | 0 | 0 | 0 | 0 | 0 | 0 | 0 | 0 | 0 | 0 | 0 | 0 | 0 | 0 | 0 | 0 | 0 | 0 |
| ENSMUSG00 | Vmn1r169 | 914 | 0 | 0 | 0 | 0 | 0 | 0 | 0 | 0 | 0 | 0 | 0 | 0 | 0 | 0 | 0 | 0 | 0 | 0 |
| ENSMUSG00 | Gm3752 | 1946 | 0 | 0 | 0 | 0 | 0 | 0 | 0 | 0 | 0 | 0 | 0 | 0 | 0 | 0 | 0 | 0 | 0 | 0 |
| ENSMUSG00 | H60c | 1068 | 0 | 0 | 0 | 0 | 0 | 0 | 0 | 0 | 0 | 0 | 0 | 0 | 0 | 0 | 0 | 0 | 0 | 0 |
| ENSMUSG00 | Gm3625 | 101 | 8.41686654 | 8.90542717 | 9.09997106 | 9.02198898 | 7.1483055 | 13.7373915 | 14.053271 | 12.1005631 | 11.8258313 | 8.40309563 | 7.09529086 | 12.1578671 | 14.6530447 | 8.73700654 | 14.6857031 | 12.1316776 | 12.1429852 | 21.2411402 |
| ENSMUSG00 | Vmn2r23 | 2596 | 0 | 0 | 0 | 0 | 0 | 0 | 0 | 0 | 0 | 0 | 0 | 0 | 0 | 0 | 0 | 0 | 0 | 0 |
| ENSMUSG00 | Gm17463 | 1358 | 0 | 0.07359255 | 0 | 0 | 0 | 0 | 0 | 0 | 0 | 0 | 0 | 0 | 0 | 0 | 0 | 0 | 0 | 0 |
| ENSMUSG00 | Gm17464 | 1347 | 0 | 0 | 0 | 0 | 0 | 0 | 0 | 0 | 0 | 0 | 0 | 0 | 0 | 0 | 0 | 0 | 0.05690616 | 0 |
| ENSMUSG00 | Gm17092 | 696 | 0.48856524 | 0.19145342 | 0.2085066 | 0.72007398 | 0.41493038 | 0.21358937 | 0.13595597 | 0.10974828 | 0 | 0.60970737 | 0 | 0 | 0.99230868 | 0.53384008 | 0.11839528 | 0.22006096 | 0.16519956 | 0 |
| ENSMUSG00 | Vmn2r9 | 2541 | 0 | 0 | 0 | 0 | 0 | 0 | 0 | 0 | 0 | 0 | 0 | 0 | 0 | 0 | 0 | 0 | 0 | 0 |
| ENSMUSG00 | Lsm5 | 994 | 16.4632724 | 9.75256762 | 12.361043 | 10.542294 | 16.269946 | 9.97036893 | 13.9462927 | 16.3297495 | 16.1182642 | 13.5665001 | 15.1399516 | 16.901926 | 19.6037297 | 18.5496003 | 18.2795635 | 15.6398256 | 16.5026711 | 16.6023461 |
| ENSMUSG00 | Gm17462 | 1865 | 0 | 0 | 0 | 0 | 0 | 0 | 0 | 0 | 0.02208392 | 0 | 0 | 0 | 0 | 0.024903 | 0 | 0 | 0 | 0 |
| ENSMUSG00 | Gm5929 | 484 | 0 | 0 | 0 | 0 | 0 | 0 | 0 | 0 | 0 | 0 | 0 | 0 | 0 | 0 | 0 | 0 | 0 | 0 |
| ENSMUSG00 | Ighd5-4 | 9 | 0 | 0 | 0 | 0 | 0 | 0 | 0 | 0 | 0 | 0 | 0 | 0 | 0 | 0 | 0 | 0 | 0 | 0 |
| ENSMUSG00 | Vmn2r95 | 2532 | 0 | 0 | 0 | 0 | 0 | 0 | 0 | 0 | 0 | 0 | 0 | 0 | 0 | 0 | 0 | 0 | 0 | 0 |
| ENSMUSG00 | Gm17280 | 1489 | 0.31400734 | 0.46982591 | 0.61725794 | 0.30598418 | 0.19394999 | 0.33279203 | 0.25419841 | 0.25649699 | 0.19362365 | 0.34832622 | 0.32085264 | 0.41233868 | 0.33130905 | 0.37429755 | 0.2490356 | 0.18000957 | 0.33461508 | 0.33249292 |
| ENSMUSG00 | Gm13290 | 1555 | 0 | 0 | 0 | 0 | 0 | 0 | 0 | 0 | 0 | 0 | 0 | 0 | 0 | 0 | 0 | 0 | 0 | 0 |
| ENSMUSG00 | Vmn2r13 | 2490 | 0 | 0 | 0 | 0 | 0 | 0 | 0 | 0 | 0 | 0 | 0 | 0 | 0 | 0 | 0 | 0 | 0 | 0 |
| ENSMUSG00 | A330050F15I | 2294 | 0 | 0 | 0 | 0 | 0 | 0 | 0 | 0 | 0 | 0 | 0 | 0 | 0 | 0 | 0 | 0 | 0 | 0 |
| ENSMUSG00 | Gm17281 | 3587 | 0 | 0 | 0 | 0.1016137 | 0 | 0 | 0 | 0 | 0 | 0 | 0 | 0.1867265 | 0.16503569 | 0.05179157 | 0 | 0.16012242 | 0 | 0 |
| ENSMUSG00 | Vmn1r113 | 1561 | 0 | 0 | 0 | 0 | 0 | 0 | 0 | 0 | 0 | 0 | 0 | 0 | 0 | 0 | 0 | 0 | 0 | 0 |
| ENSMUSG00 | Gm3756 | 296 | 0 | 0.22508712 | 0.49027227 | 0 | 0.3252157 | 0.66963153 | 0.15984013 | 0 | 0 | 0 | 0.32280377 | 0.37713286 | 0.16666188 | 0 | 0.83516667 | 0 | 0.64740366 | 0.37168315 |
| ENSMUSG00 | Gm17592 | 199 | 0.4271877 | 0.16740148 | 1.4584984 | 0.22894997 | 0.48373793 | 1.24504356 | 0 | 1.72729458 | 0 | 0.47387641 | 0 | 2.24384573 | 0.74369725 | 0.46677475 | 0.20704299 | 0.57724534 | 0.57778337 | 0.82928302 |
| ENSMUSG00 | Gm17593 | 1408 | 0 | 0 | 0 | 0 | 0 | 0 | 0 | 0 | 0 | 0 | 0 | 0 | 0 | 0 | 0 | 0 | 0 | 0 |
| ENSMUSG00 | Mocs3 | 1056 | 0 | 0 | 0 | 0 | 0 | 0 | 0 | 0 | 0 | 0 | 0 | 0 | 0 | 0 | 0 | 0 | 0 | 0 |
| ENSMUSG00 | Zscan4b | 1741 | 0 | 0 | 0 | 0 | 0 | 0 | 0 | 0 | 0 | 0 | 0 | 0 | 0 | 0 | 0 | 0 | 0 | 0 |
| ENSMUSG00 | Gm17590 | 107 | 0 | 0 | 0 | 0 | 0 | 0 | 0 | 0 | 0 | 0 | 0 | 0 | 0 | 0 | 0 | 0 | 0 | 0 |
| ENSMUSG00 | Gm17591 | 1806 | 0.0706066 | 0.29513085 | 0.2142792 | 0.1009104 | 0.1332556 | 0.08231351 | 0.13098748 | 0.23262261 | 0.1140269 | 0.07832343 | 0.0793604 | 0.24724546 | 0.13657784 | 0.23144894 | 0.27376449 | 0.16961509 | 0.40321132 | 0.21321359 |
| ENSMUSG00 | Olfr819 | 962 | 0 | 0 | 0 | 0 | 0 | 0 | 0 | 0 | 0 | 0 | 0 | 0 | 0 | 0 | 0 | 0 | 0 | 0 |
| ENSMUSG00 | Gm4971 | 1049 | 0 | 0 | 0 | 0 | 0 | 0 | 0 | 0 | 0 | 0 | 0 | 0 | 0 | 0 | 0 | 0 | 0 | 0 |
| ENSMUSG00 | C2cd4d | 1314 | 0.06469585 | 0.3549319 | 0.40495345 | 0.10402065 | 0.18315039 | 0.11313409 | 0.54009907 | 0.43598633 | 0 | 0.21530001 | 0.43630099 | 0.38229906 | 0.30034653 | 0.38880134 | 0.37626992 | 0 | 0.23334123 | 0 |
| ENSMUSG00 | Gm4902 | 1154 | 0 | 0 | 0 | 0 | 0 | 0 | 0.08199771 | 0 | 0.24983155 | 0.36772646 | 0.28979611 | 0 | 1.23971023 | 0 | 0.21421953 | 0.03318077 | 0 | 0 |
| ENSMUSG00 | Apol11a | 1128 | 0 | 0 | 0 | 0 | 0 | 0 | 0 | 0 | 0 | 0 | 0 | 0 | 0 | 0 | 0 | 0 | 0 | 0 |
| ENSMUSG00 | Vmn2r36 | 2555 | 0 | 0 | 0 | 0 | 0 | 0 | 0 | 0 | 0 | 0 | 0 | 0 | 0 | 0 | 0 | 0 | 0 | 0 |
| ENSMUSG00 | Vmn1r57 | 932 | 0 | 0 | 0 | 0 | 0 | 0 | 0 | 0 | 0 | 0 | 0 | 0 | 0 | 0 | 0 | 0 | 0 | 0 |
| ENSMUSG00 | Gm17406 | 1840 | 0 | 0.01810483 | 0 | 0.07428431 | 0 | 0.0807925 | 0.07714024 | 0 | 0 | 0 | 0 | 0 | 0 | 0 | 0.0895686 | 0 | 0 | 0 |
| ENSMUSG00 | Gm8741 | 769 | 0 | 0.04331976 | 0.18871338 | 0 | 0.12518056 | 0 | 0 | 0.14899507 | 0.10711708 | 0 | 0 | 0.14516427 | 0.38490443 | 0 | 0.10715619 | 0.09958545 | 0.19935655 | 0.1430666 |
| ENSMUSG00 | Vmn2r106 | 2583 | 0 | 0 | 0 | 0 | 0 | 0 | 0 | 0 | 0 | 0 | 0 | 0 | 0 | 0 | 0 | 0 | 0 | 0 |
| ENSMUSG00 | Gm3072 | 1390 | 0 | 0 | 0 | 0 | 0 | 0 | 0 | 0 | 0 | 0 | 0 | 0 | 0 | 0 | 0 | 0 | 0 | 0 |
| ENSMUSG00 | Gm17405 | 103 | 0 | 0 | 0 | 0 | 0 | 0 | 0 | 0 | 0 | 0 | 0 | 0 | 0 | 0 | 0 | 0 | 0 | 0 |
| ENSMUSG00 | Gm17099 | 410 | 0 | 0 | 0 | 0 | 0 | 0 | 0 | 0 | 0 | 0 | 0 | 0 | 0 | 0 | 0 | 0 | 0 | 0 |
| ENSMUSG00 | Gm17021 | 459 | 0 | 0 | 0 | 0.19852307 | 0 | 0 | 0 | 0 | 0 | 0 | 0 | 0 | 0 | 0 | 0 | 0 | 0 | 0 |
| ENSMUSG00 | Vmn1r69 | 2389 | 0 | 0 | 0 | 0 | 0 | 0 | 0 | 0 | 0 | 0 | 0 | 0 | 0 | 0 | 0 | 0 | 0 | 0 |
| ENSMUSG00 | Gm2783 | 1865 | 0 | 0 | 0 | 0 | 0 | 0 | 0 | 0 | 0 | 0 | 0 | 0 | 0 | 0 | 0 | 0 | 0 | 0 |
| ENSMUSG00 | Gm10662 | 1355 | 0 | 0 | 0 | 0 | 0 | 0 | 0 | 0 | 0 | 0 | 0 | 0 | 0 | 0 | 0 | 0 | 0 | 0 |
| ENSMUSG00 | Gm17101 | 1587 | 0 | 0 | 0 | 0 | 0 | 0 | 0 | 0 | 0 | 0 | 0 | 0 | 0 | 0 | 0 | 0 | 0 | 0 |
| ENSMUSG00 | Gm17717 | 1055 | 0.28202486 | 0 | 0.04585169 | 0 | 0.27373606 | 0.23484708 | 0 | 0.18100665 | 0 | 0.04469261 | 0 | 0.05290584 | 1.07548256 | 0.22011416 | 0.35148247 | 0.03629441 | 0 | 0 |
| ENSMUSG00 | Gm17718 | 2437 | 0 | 0 | 0 | 0 | 0 | 0 | 0.01941431 | 0 | 0 | 0 | 0.019604 | 0 | 0 | 0 | 0 | 0 | 0 | 0 |
| ENSMUSG00 | Gm9307 | 728 | 0 | 0 | 0.06644716 | 0 | 0 | 0.06806694 | 0 | 0 | 0 | 0 | 0 | 0 | 0 | 0 | 0 | 0 | 0 | 0 |
| ENSMUSG00 | Gm17117 | 298 | 0 | 0 | 0 | 0 | 0 | 0 | 0 | 0 | 0 | 0 | 0 | 0 | 0 | 0 | 0 | 0 | 0 | 0 |
| ENSMUSG00 | Vmn2r105 | 2577 | 0 | 0 | 0 | 0 | 0 | 0 | 0 | 0 | 0 | 0 | 0 | 0 | 0 | 0 | 0 | 0 | 0 | 0 |
| ENSMUSG00 | Trbv10 | 288 | 0 | 0 | 0 | 0 | 0 | 0 | 0 | 0 | 0 | 0 | 0 | 0 | 0 | 0 | 0 | 0 | 0 | 0 |
| ENSMUSG00 | Gm17129 | 504 | 0 | 0 | 0 | 0 | 0 | 0 | 0 | 0 | 0 | 0 | 0 | 0 | 0 | 0 | 0 | 0 | 0 | 0 |
| ENSMUSG00 | Gm17525 | 1046 | 0.60953885 | 0.31847891 | 0.18498482 | 0.17422962 | 0.18406089 | 0.33161485 | 0 | 0.18256407 | 0.19687627 | 0.09015431 | 0.27404374 | 0.32016632 | 0.51878689 | 0.44401613 | 0.19694816 | 0.47588709 | 0.21984492 | 0.21035987 |
| ENSMUSG00 | C630004L07I | 1268 | 0.1005643 | 0.052544 | 0 | 0 | 0.03795893 | 0 | 0.11193851 | 0 | 0 | 0 | 0 | 0 | 0 | 0 | 0 | 0 | 0 | 0 |
| ENSMUSG00 | Gm8374 | 668 | 0 | 0 | 0 | 0 | 0 | 0 | 0 | 0 | 0 | 0 | 0 | 0 | 0 | 0 | 0 | 0 | 0 | 0 |
| ENSMUSG00 | Gm7356 | 1463 | 0 | 0 | 0 | 0 | 0 | 0 | 0 | 0 | 0 | 0 | 0 | 0 | 0 | 0 | 0 | 0 | 0 | 0 |
| ENSMUSG00 | Gm17527 | 193 | 0.44046815 | 0.51781701 | 0.50128011 | 0.47213517 | 0.49877641 | 0.51349983 | 0 | 0.39577619 | 0.42680327 | 0.48860832 | 0.24753864 | 0 | 0.76681737 | 0.48128588 | 0.21347956 | 0 | 0.19858185 | 0 |
| ENSMUSG00 | Vmn2r96 | 2338 | 0 | 0 | 0 | 0 | 0 | 0 | 0 | 0 | 0 | 0 | 0 | 0 | 0 | 0 | 0.12335795 | 0 | 0 | 0 |
| ENSMUSG00 | Klhdc7b | 1817 | 0 | 0 | 0 | 0 | 0 | 0 | 0 | 0 | 0 | 0 | 0 | 0 | 0 | 0 | 0 | 0 | 0 | 0 |
| ENSMUSG00 | Gm17037 | 395 | 0 | 0 | 0 | 0 | 0 | 0 | 0 | 0 | 0 | 0 | 0 | 0 | 0 | 0 | 0 | 0 | 0 | 0 |
| ENSMUSG00 | Gm17356 | 516 | 0 | 0 | 0 | 0 | 0 | 0 | 0 | 0 | 0 | 0 | 0 | 0 | 0 | 0 | 0 | 0 | 0 | 0 |
| ENSMUSG00 | Gm17357 | 2524 | 0.35364845 | 0.55433501 | 0.49830103 | 0.37907367 | 0.28604551 | 0.47118289 | 0.33741213 | 0.63553126 | 0.32635908 | 0.13076661 | 0.45427853 | 0.17691177 | 0.33226727 | 0.34961872 | 0.40609782 | 0.21238847 | 0.18221694 | 0.52306599 |
| ENSMUSG00 | Gm17358 | 2912 | 0.05838623 | 0.01143987 | 0.13289431 | 0.06258385 | 0.04958646 | 0.2892845 | 0 | 0.1705018 | 0.01414372 | 0.01619186 | 0.09843741 | 0.01916747 | 0 | 0 | 0.04244666 | 0.07889548 | 0.0131615 | 0 |
| ENSMUSG00 | Gm17359 | 773 | 0.71483478 | 1.20667663 | 1.50189486 | 1.59139482 | 1.86799186 | 0.57693998 | 0 | 0.24704012 | 0 | 0.18299109 | 0.0618046 | 0.1444131 | 0.06381878 | 0.36049744 | 0 | 0.04953507 | 0.09916248 | 0.85395767 |
| ENSMUSG00 | 4930471C04 | 861 | 0 | 0 | 0 | 0.10583286 | 0 | 0 | 0 | 0 | 0 | 0 | 0 | 0 | 0 | 0 | 0 | 0 | 0 | 0 |
| ENSMUSG00 | Gm4983 | 335 | 0.88816786 | 0.99441475 | 0.86639159 | 0.40800935 | 0.14367738 | 0.44375582 | 0 | 0 | 0.36883447 | 0.14074837 | 0.14261182 | 0 | 0.44177836 | 0.97047347 | 0.12298972 | 0.5715016 | 0.22881371 | 0 |
| ENSMUSG00 | Hivep3 | 1147 | 0 | 0 | 0 | 0 | 0 | 0 | 0 | 0 | 0 | 0 | 0 | 0 | 0 | 0 | 0 | 0 | 0 | 0 |
| ENSMUSG00 | Gm17355 | 53 | 0 | 0 | 0 | 0 | 0 | 0 | 0 | 0 | 0 | 0 | 0 | 0 | 0 | 0 | 0 | 0 | 0 | 0 |
| ENSMUSG00 | Gm17668 | 1603 | 0 | 0 | 0 | 0 | 0 | 0 | 0 | 0 | 0 | 0 | 0 | 0 | 0 | 0 | 0 | 0 | 0 | 0 |
| ENSMUSG00 | Gm17667 | 1814 | 0.02343174 | 0 | 0 | 0 | 0 | 0.02731683 | 0 | 0 | 0 | 0 | 0 | 0 | 0 | 0 | 0 | 0 | 0 | 0 |
| ENSMUSG00 | 4930433I11R | 2422 | 0 | 0 | 0 | 0 | 0 | 0 | 0 | 0 | 0 | 0 | 0 | 0 | 0 | 0 | 0 | 0 | 0 | 0 |
| ENSMUSG00 | Apol11b | 2032 | 0 | 0 | 0 | 0 | 0 | 0 | 0 | 0 | 0 | 0 | 0 | 0 | 0 | 0 | 0 | 0 | 0 | 0 |
| ENSMUSG00 | Gm17669 | 464 | 0.09160598 | 0.14359006 | 0.1042533 | 0.58915144 | 0.20746519 | 0.32038405 | 0.10196698 | 0.08231121 | 0.26629213 | 0 | 0 | 0.60146188 | 0.21263757 | 0.70066511 | 0.35518583 | 0.33009144 | 0.24779933 | 0.35566233 |
| ENSMUSG00 | Gm17670 | 3427 | 0.45891203 | 0.87486445 | 1.03042536 | 0.41213667 | 0.29494321 | 0.40486622 | 1.27013903 | 0.79126366 | 1.06962355 | 0.85303286 | 0.75280063 | 0.87949979 | 1.00765515 | 1.20616393 | 0.18033946 | 1.49721079 | 0.72693589 | 1.05941087 |
| ENSMUSG00 | Eif3s6-ps2 | 218 | 1.16986723 | 0 | 0.22189693 | 0.41799123 | 1.54552048 | 0 | 1.95327574 | 0 | 0 | 0.43257526 | 1.5340583 | 0 | 0 | 0.42609254 | 0 | 0.35128998 | 0.87904351 | 0 |
| ENSMUSG00 | Gm6526 | 1718 | 0 | 0 | 0 | 0 | 0 | 0 | 0 | 0.04446147 | 0 | 0 | 0 | 0 | 0 | 0 | 0 | 0 | 0 | 0 |
| ENSMUSG00 | Gm7876 | 1931 | 0 | 0 | 0 | 0 | 0 | 0 | 0 | 0 | 0 | 0 | 0 | 0 | 0 | 0 | 0 | 0 | 0 | 0 |
| ENSMUSG00 | Vmn1r94 | 917 | 0 | 0 | 0 | 0 | 0 | 0 | 0 | 0 | 0 | 0 | 0 | 0 | 0 | 0 | 0 | 0 | 0 | 0 |
| ENSMUSG00 | Gm14483 | 524 | 0 | 0 | 0 | 0 | 0 | 0 | 0 | 0 | 0 | 0 | 0 | 0 | 0 | 0 | 0 | 0 | 0 | 0 |
| ENSMUSG00 | Gm4222 | 428 | 0 | 0 | 0 | 0 | 0 | 0 | 0 | 0 | 0 | 0 | 0 | 0 | 0 | 0 | 0 | 0 | 0 | 0 |
| ENSMUSG00 | Ighd6-2 | 28 | 0 | 0 | 0 | 0 | 0 | 0 | 0 | 0 | 0 | 0 | 0 | 0 | 0 | 0 | 0 | 0 | 0 | 0 |
| ENSMUSG00 | H2-Q2 | 2173 | 17.8197033 | 17.9671938 | 18.4767741 | 18.0105555 | 17.0997906 | 21.4812126 | 20.1400037 | 19.2455962 | 25.8150182 | 24.1503602 | 18.0063005 | 23.1173936 | 32.2598502 | 30.1362923 | 37.5990268 | 40.9161484 | 36.3861714 | 33.0611566 |
| ENSMUSG00 | AA543186 | 1694 | 0.52692367 | 0.92426566 | 0.85667409 | 0.4034331 | 0.6535031 | 0.43877863 | 1.340619 | 1.35274153 | 1.21565867 | 0.83501836 | 0.95888346 | 0.95552197 | 0.2038509 | 1.09667267 | 0.6566954 | 0.58769527 | 1.17648609 | 1.07160598 |
| ENSMUSG00 | Gm8224 | 494 | 0 | 0 | 0 | 0 | 0 | 0 | 0 | 0 | 0 | 0 | 0 | 0 | 0 | 0 | 0 | 0 | 0 | 0 |
| ENSMUSG00 | Gm17189 | 1627 | 0 | 0 | 0.11892693 | 0 | 0.11833294 | 0.09136951 | 0.26171734 | 0.21126713 | 0 | 0.2898015 | 0 | 0.20583526 | 0.51545335 | 0 | 0.17726545 | 0.4706897 | 0.21200779 | 0.06762029 |
| ENSMUSG00 | 4930426I24R | 635 | 1.07099656 | 0.94430251 | 1.75211212 | 0.86099611 | 1.28857118 | 0.46821481 | 1.04311418 | 2.04494753 | 0.90804917 | 0.74253076 | 0.75236155 | 0.17579736 | 1.63144923 | 1.97478797 | 0.9732651 | 1.02510288 | 0.96570197 | 1.12617069 |
| ENSMUSG00 | Gm2885 | 1440 | 0.02951748 | 0.37014327 | 0.16796365 | 0.03163961 | 0.20054968 | 0.82587889 | 0.26284822 | 0.29174752 | 0.05720349 | 0.06548709 | 0.43130171 | 0.38760877 | 0.03425828 | 1.22560786 | 0.77252917 | 0.34567909 | 0.23953936 | 1.83363689 |
| ENSMUSG00 | Sec14l5 | 2321 | 0 | 0 | 0 | 0 | 0 | 0 | 0.22423071 | 0 | 0.14196128 | 0.04062965 | 0 | 0 | 0 | 0 | 0 | 0 | 0 | 0 |
| ENSMUSG00 | Gm17126 | 522 | 0 | 0 | 0 | 0 | 0 | 0 | 0 | 0 | 0 | 0 | 0 | 0 | 0 | 0 | 0 | 0 | 0 | 0 |
| ENSMUSG00 | Gm17675 | 3326 | 0 | 0 | 0 | 0 | 0 | 0 | 0 | 0 | 0 | 0 | 0 | 0 | 0 | 0 | 0 | 0 | 0 | 0 |
| ENSMUSG00 | Gm4305 | 869 | 0 | 0 | 0 | 0 | 0 | 0 | 0 | 0 | 0 | 0 | 0 | 0 | 0 | 0 | 0 | 0 | 0 | 0 |
| ENSMUSG00 | Gm17676 | 37 | 0 | 0 | 1.30739271 | 0 | 0 | 0 | 5.11488421 | 0 | 3.33944722 | 1.27434333 | 2.58243019 | 6.03412569 | 0 | 5.02098242 | 1.11355556 | 1.03488128 | 0 | 2.97346523 |
| ENSMUSG00 | Olfr773 | 1003 | 0 | 0 | 0 | 0 | 0 | 0 | 0 | 0 | 0 | 0 | 0 | 0 | 0 | 0 | 0 | 0 | 0 | 0 |
| ENSMUSG00 | Gm16506 | 1400 | 0 | 0 | 0 | 0 | 0 | 0 | 0 | 0 | 0 | 0 | 0 | 0 | 0 | 0 | 0 | 0 | 0 | 0 |
| ENSMUSG00 | Vmn2r22 | 2556 | 0 | 0 | 0 | 0 | 0 | 0 | 0 | 0 | 0 | 0 | 0 | 0 | 0 | 0 | 0 | 0 | 0 | 0 |
| ENSMUSG00 | Gm8890 | 1120 | 0 | 0 | 0 | 0 | 0 | 0 | 0 | 0 | 0 | 0 | 0 | 0 | 0 | 0 | 0 | 0 | 0 | 0 |
| ENSMUSG00 | Gm6549 | 652 | 0 | 0 | 0 | 0 | 0 | 0 | 0 | 0 | 0 | 0 | 0 | 0 | 0 | 0.28493305 | 0 | 0 | 0 | 0 |
| ENSMUSG00 | Siah3 | 903 | 2.9654774 | 3.02509117 | 2.46421085 | 2.11911834 | 2.23869412 | 3.12791342 | 0.31436996 | 0.80360537 | 0.95782595 | 0.93988113 | 0.95232475 | 1.79252958 | 0.4370491 | 0 | 0.18250966 | 0.04240377 | 0.12732989 | 0 |
| ENSMUSG00 | Gm17366 | 187 | 0 | 0 | 0 | 0 | 0 | 0 | 0 | 0 | 0 | 0 | 0 | 0 | 0 | 0 | 0 | 0 | 0 | 0 |
| ENSMUSG00 | A430010J10R | 2035 | 0.12532239 | 0.04910992 | 0.04754155 | 0 | 0.0473041 | 0.14610143 | 0.04649895 | 0.0563033 | 0.10119537 | 0.04633976 | 0.04695328 | 0.27427844 | 0.12120864 | 0.04564529 | 0 | 0 | 0.03766712 | 0 |
| ENSMUSG00 | Gm8220 | 1560 | 0 | 0 | 0 | 0 | 0 | 0 | 0 | 0 | 0 | 0 | 0 | 0 | 0 | 0 | 0 | 0 | 0 | 0 |
| ENSMUSG00 | 1810019N24 | 1472 | 0.0288758 | 0.09052417 | 0.06572491 | 0 | 0 | 0.10099063 | 0.61069355 | 0.12972963 | 0.22383976 | 0.32031728 | 0.32455814 | 0.03791825 | 0 | 0 | 0.02799019 | 0.05202528 | 0 | 0 |
| ENSMUSG00 | Gm8705 | 810 | 0 | 0 | 0 | 0 | 0 | 0 | 0 | 0 | 0 | 0 | 0 | 0 | 0 | 0 | 0 | 0 | 0 | 0 |
| ENSMUSG00 | Gm13697 | 2725 | 0 | 0 | 0 | 0 | 0 | 0 | 0 | 0 | 0 | 0 | 0 | 0 | 0 | 0 | 0 | 0 | 0 | 0 |
| ENSMUSG00 | Gm17230 | 3941 | 0 | 0.01690581 | 0 | 0 | 0.02442625 | 0.02514729 | 0 | 0 | 0 | 0 | 0 | 0 | 0 | 0 | 0 | 0.26233098 | 0 | 0 |
| ENSMUSG00 | Gm17542 | 97 | 0 | 0 | 0 | 0 | 0 | 0 | 0 | 0 | 0 | 0 | 0 | 0 | 0 | 0 | 0 | 0 | 0 | 0 |
| ENSMUSG00 | Gm17541 | 524 | 0.1622335 | 0.06357423 | 0.27694769 | 0 | 0 | 0 | 0 | 0.07288626 | 0 | 0 | 0 | 0 | 0.37657952 | 0.26590126 | 0.15725785 | 0.21922103 | 0.07314179 | 0 |
| ENSMUSG00 | Gm8094 | 1403 | 0 | 0 | 0 | 0 | 0 | 0 | 0 | 0 | 0 | 0 | 0 | 0 | 0 | 0 | 0 | 0 | 0 | 0 |
| ENSMUSG00 | Vmn1r29 | 1022 | 0 | 0 | 0 | 0 | 0 | 0 | 0 | 0 | 0 | 0 | 0 | 0 | 0 | 0 | 0 | 0 | 0 | 0 |
| ENSMUSG00 | Gpr62 | 1967 | 0 | 0 | 0 | 0.06948812 | 0 | 0 | 0 | 0 | 0 | 0 | 0 | 0 | 0.07523932 | 0 | 0.04189279 | 0.0194665 | 0.09742323 | 0 |
| ENSMUSG00 | Yy2 | 2866 | 0 | 0 | 0 | 0 | 0 | 0 | 0 | 0 | 0 | 0 | 0 | 0 | 0 | 0 | 0 | 0 | 0 | 0 |
| ENSMUSG00 | Gm17543 | 1475 | 0.05763414 | 0.02258501 | 0 | 0 | 0 | 0.03359507 | 0 | 0.07767946 | 0.19546143 | 0.06393316 | 0.0323898 | 0.03784113 | 0 | 0.09446255 | 0.27933258 | 0.25959734 | 0.18188751 | 0 |
| ENSMUSG00 | Gm8723 | 871 | 0 | 0 | 0 | 0 | 0 | 0 | 0 | 0 | 0 | 0 | 0 | 0 | 0 | 0 | 0 | 0 | 0 | 0 |
| ENSMUSG00 | Gm8701 | 914 | 0 | 0 | 0 | 0 | 0 | 0 | 0 | 0 | 0 | 0 | 0 | 0 | 0 | 0 | 0 | 0 | 0 | 0 |
| ENSMUSG00 | Gm7929 | 620 | 0 | 0 | 0 | 0 | 0 | 0 | 0 | 0 | 0 | 0 | 0 | 0 | 0 | 0 | 0 | 0 | 0 | 0 |
| ENSMUSG00 | Gm17161 | 521 | 0 | 0 | 0 | 0 | 0 | 0 | 0 | 0 | 0 | 0 | 0 | 0 | 0 | 0 | 0 | 0 | 0 | 0 |
| ENSMUSG00 | Gm5093 | 521 | 0.08158383 | 0 | 0 | 0 | 0 | 0.09511081 | 0 | 0.07330595 | 0 | 0.09050039 | 0 | 0 | 0 | 0 | 0 | 0.29397779 | 0 | 0 |
| ENSMUSG00 | Gm5705 | 626 | 0 | 0 | 0 | 0 | 0 | 0 | 0 | 0 | 0 | 0 | 0 | 0 | 0 | 0 | 0 | 0 | 0 | 0.17574794 |
| ENSMUSG00 | Gm17732 | 820 | 1.81424532 | 2.43752885 | 1.35681853 | 0.44449799 | 0.93915949 | 0.96688261 | 13.789915 | 13.9728301 | 13.8627785 | 7.99261921 | 2.27222363 | 10.6866573 | 48.5498259 | 79.577979 | 45.4222031 | 10.9268319 | 14.1620341 | 33.6092225 |
| ENSMUSG00 | Gm17098 | 626 | 0 | 0 | 0 | 0 | 0.30755223 | 0 | 0 | 0 | 0.39475894 | 0.07532061 | 0 | 0 | 0 | 0.2225755 | 0.723989 | 0 | 0 | 0 |
| ENSMUSG00 | Gm4511 | 1865 | 0.02279098 | 0 | 0.02593755 | 0 | 0 | 0.05313966 | 0 | 0 | 0.04416785 | 0 | 0 | 0 | 0 | 0.04980599 | 0 | 0.06159347 | 0.02055029 | 0 |
| ENSMUSG00 | D17H6S56E- | 5355 | 366.981197 | 370.647162 | 420.718731 | 419.544474 | 396.31763 | 441.338576 | 253.862783 | 300.404106 | 316.624705 | 272.690435 | 266.192612 | 309.555437 | 202.643172 | 211.630659 | 271.691754 | 303.936612 | 294.550653 | 231.757322 |
| ENSMUSG00 | Gm17733 | 981 | 1.60315139 | 2.54685736 | 2.02172757 | 1.95062575 | 2.25695055 | 1.31332423 | 1.49509995 | 3.62068647 | 2.72897403 | 2.21094021 | 3.45771871 | 2.3327647 | 4.32471446 | 8.37982005 | 3.1079665 | 1.95161098 | 5.03984944 | 4.48596181 |
| ENSMUSG00 | Tspy-ps | 698 | 0 | 0 | 0 | 0 | 0 | 0 | 0 | 0 | 0 | 0 | 0 | 0 | 0.56540879 | 0 | 0.11805603 | 0 | 0 | 0 |
| ENSMUSG00 | 2810029C07I | 1948 | 1.30919433 | 3.02689029 | 1.83759818 | 0.3508294 | 1.23541899 | 1.75520463 | 1.04437638 | 2.29389686 | 1.98743967 | 1.96057852 | 1.32435717 | 2.43548836 | 1.72205871 | 3.02792562 | 1.98816543 | 1.57250954 | 0.78698762 | 1.60960939 |
| ENSMUSG00 | C030010L15I | 713 | 0.29807276 | 0.88772088 | 0.88198583 | 0.57510435 | 0.87758066 | 0.2779957 | 3.31785967 | 1.12488142 | 0.40435569 | 0.26452007 | 1.40711659 | 0.23484851 | 1.45297372 | 0.32569486 | 0.40450335 | 0 | 1.61260716 | 1.08012271 |
| ENSMUSG00 | Gm3208 | 426 | 0 | 0 | 0 | 0 | 0 | 0 | 0 | 0 | 0 | 0 | 0 | 0 | 0 | 0 | 0 | 0 | 0 | 0 |
| ENSMUSG00 | Gm17432 | 1722 | 0.44430498 | 0.36756387 | 0.61801258 | 0.47624785 | 0.97829113 | 0.6330779 | 0.85173812 | 0.82062653 | 0.98063133 | 0.68453402 | 0.80457247 | 0.8427452 | 1.57564196 | 0.64730435 | 1.45952085 | 1.245223 | 0.48965072 | 0.35139383 |
| ENSMUSG00 | Gm3636 | 1945 | 0 | 0.0342549 | 0.04974142 | 0.09369881 | 0.09898596 | 0 | 0 | 0.03927239 | 0 | 0.02424201 | 0 | 0.086091 | 0 | 0 | 0 | 0.03937338 | 0 | 0.02828232 |
| ENSMUSG00 | Vmn1r152 | 923 | 0 | 0 | 0 | 0 | 0 | 0 | 0 | 0 | 0 | 0 | 0 | 0 | 0 | 0 | 0 | 0 | 0 | 0 |
| ENSMUSG00 | Gm3095 | 1564 | 0 | 0 | 0 | 0 | 0 | 0 | 0 | 0 | 0 | 0 | 0 | 0 | 0 | 0 | 0 | 0 | 0 | 0 |
| ENSMUSG00 | Gm17064 | 449 | 0 | 0 | 0 | 0 | 0 | 0 | 0 | 0 | 0 | 0 | 0 | 0 | 0 | 0 | 0 | 0 | 0 | 0 |
| ENSMUSG00 | Rgs21 | 456 | 0 | 0 | 0 | 0 | 0 | 0 | 0 | 0 | 0 | 0 | 0 | 0 | 0 | 0 | 0 | 0 | 0 | 0 |
| ENSMUSG00 | Gm17433 | 3681 | 0.01154718 | 0.41629805 | 0.2628282 | 0.09901884 | 0.06537887 | 0.24231166 | 0.02570643 | 0.14525771 | 0.45874685 | 0.38427631 | 0.11680919 | 0.45489538 | 0.14741948 | 0.20187596 | 0.31340493 | 0.52011148 | 0.33318161 | 0.14944066 |
| ENSMUSG00 | Gm17619 | 1447 | 1.76248138 | 0.64461717 | 1.30377863 | 1.41689495 | 1.53010953 | 0.71914817 | 0.26157666 | 0.39591295 | 0.39848737 | 0.2932663 | 0.36318213 | 0.2314402 | 0.61366587 | 0.22467769 | 0.14236889 | 0.42339303 | 0.84757533 | 0.68428744 |
| ENSMUSG00 | Gm4349 | 2557 | 0 | 0 | 0 | 0 | 0 | 0 | 0 | 0 | 0 | 0 | 0 | 0 | 0 | 0 | 0.06445296 | 0.0898489 | 0 | 0 |
| ENSMUSG00 | Gm3191 | 345 | 0 | 0 | 0 | 0 | 0 | 0 | 0 | 0 | 0 | 0 | 0 | 0 | 0 | 0 | 0 | 0 | 0 | 0 |
| ENSMUSG00 | Vmn1r107 | 923 | 0 | 0 | 0 | 0 | 0 | 0 | 0 | 0 | 0 | 0 | 0 | 0 | 0 | 0 | 0 | 0 | 0 | 0 |
| ENSMUSG00 | Zfp964 | 1413 | 4.60247129 | 3.96076874 | 4.07392082 | 3.38564024 | 2.89540942 | 3.61212425 | 5.25696433 | 3.24351613 | 5.45072785 | 3.80409071 | 2.19771571 | 2.44909489 | 3.03742166 | 4.27298752 | 6.18169133 | 1.86981735 | 3.22776315 | 5.21671642 |
| ENSMUSG00 | Gm5891 | 1355 | 0 | 0 | 0 | 0 | 0 | 0 | 0 | 0 | 0 | 0 | 0 | 0 | 0 | 0 | 0 | 0 | 0 | 0 |
| ENSMUSG00 | Gm4220 | 920 | 0 | 0 | 0 | 0 | 0 | 0 | 0 | 0 | 0 | 0 | 0 | 0 | 0 | 0 | 0 | 0 | 0 | 0 |
| ENSMUSG00 | Gm17618 | 633 | 0 | 0 | 0 | 0 | 0 | 0 | 0 | 0 | 0 | 0 | 0 | 0 | 0 | 0 | 0 | 0 | 0 | 0 |
| ENSMUSG00 | Gm17617 | 37 | 0 | 0 | 0 | 0 | 0 | 0 | 0 | 0 | 0 | 0 | 0 | 0 | 0 | 0 | 0 | 0 | 0 | 0 |
| ENSMUSG00 | 4921507G05I | 1651 | 0.2574511 | 0 | 0.02929953 | 0.08278809 | 0.02915319 | 0 | 0 | 0.04626578 | 0.4490353 | 0 | 0.40511776 | 0.06761437 | 0 | 0.02813088 | 0.09982206 | 0 | 0.02321399 | 0 |
| ENSMUSG00 | Gm17095 | 578 | 0 | 0 | 0 | 0 | 0 | 0 | 0 | 0 | 0 | 0 | 0 | 0 | 0 | 0 | 0 | 0 | 0 | 0 |
| ENSMUSG00 | Vmn2r103 | 2565 | 0 | 0 | 0 | 0 | 0 | 0 | 0 | 0 | 0 | 0 | 0 | 0 | 0 | 0 | 0 | 0 | 0 | 0 |
| ENSMUSG00 | Gm17307 | 101 | 0 | 0 | 0 | 0.45109945 | 0 | 0 | 0 | 0.7562852 | 0 | 1.40051594 | 0 | 0 | 0 | 0 | 0 | 1.13734477 | 0.75893657 | 0 |
| ENSMUSG00 | Gm17308 | 1995 | 0 | 0 | 0 | 0 | 0 | 0 | 0 | 0 | 0 | 0 | 0 | 0 | 0 | 0 | 0 | 0 | 0 | 0 |
| ENSMUSG00 | Gm7954 | 1398 | 0 | 0 | 0 | 0 | 0 | 0 | 0 | 0 | 0 | 0 | 0 | 0 | 0 | 0 | 0 | 0 | 0 | 0 |
| ENSMUSG00 | Gm16494 | 260 | 0 | 0 | 0.18605204 | 0 | 0 | 0 | 0 | 0 | 0.15840968 | 0 | 0 | 0 | 0 | 0.17863111 | 0 | 0 | 0 | 0 |
| ENSMUSG00 | Gm8340 | 1190 | 0 | 0 | 0 | 0 | 0 | 0 | 0 | 0 | 0 | 0 | 0 | 0 | 0 | 0 | 0 | 0 | 0 | 0 |
| ENSMUSG00 | Gm17564 | 82 | 0 | 0 | 0 | 0 | 0 | 0 | 0 | 0 | 0 | 0 | 0 | 0 | 0 | 0 | 0 | 0 | 0 | 0 |
| ENSMUSG00 | Gm17306 | 3971 | 0.06422338 | 0.05033426 | 0.1705438 | 0.13768132 | 0.07272514 | 0.13726519 | 0.08340185 | 0.23082792 | 0.06223095 | 0.18998017 | 0.26468121 | 0.09839074 | 0.06211523 | 0.19882888 | 0.16600979 | 0.12535329 | 0.23163715 | 0.40172856 |
| ENSMUSG00 | Gm6763 | 862 | 0 | 0 | 0 | 0 | 0 | 0 | 0 | 0 | 0 | 0 | 0 | 0 | 0 | 0 | 0 | 0 | 0 | 0 |
| ENSMUSG00 | Sco2 | 767 | 0 | 0 | 0 | 0 | 0 | 0 | 0 | 0 | 0 | 0 | 0 | 0 | 0 | 0 | 0 | 0 | 0 | 0 |
| ENSMUSG00 | Gm9590 | 323 | 0 | 0.1031359 | 0 | 0 | 0 | 0 | 0 | 0 | 0 | 0 | 0 | 0 | 0 | 0 | 0 | 0 | 0 | 0 |
| ENSMUSG00 | Gm8212 | 1399 | 0 | 0 | 0 | 0 | 0 | 0 | 0 | 0 | 0 | 0 | 0 | 0 | 0 | 0 | 0 | 0 | 0 | 0 |
| ENSMUSG00 | Gm17022 | 911 | 0.32660399 | 0.87761741 | 0.10619875 | 0.10002425 | 0.63400997 | 0.76151292 | 0.57128372 | 0.62885405 | 0.1808409 | 0.46581375 | 0.68175023 | 0.36761139 | 0 | 1.47846162 | 0.13568021 | 0.50437683 | 0.5889881 | 0.72459855 |
| ENSMUSG00 | Rpl29-ps5 | 476 | 0 | 0 | 0 | 0 | 0 | 0 | 0 | 0 | 0 | 0 | 0 | 0 | 0.20727696 | 0 | 0 | 0 | 0.08051743 | 0 |
| ENSMUSG00 | Gm8425 | 768 | 0 | 0 | 0.25194547 | 0 | 0 | 0 | 0 | 0 | 0 | 0 | 0 | 0 | 0 | 0 | 0 | 0 | 0 | 0.07162644 |
| ENSMUSG00 | Gm8032 | 570 | 0 | 0 | 0 | 0 | 0 | 0 | 0 | 0 | 0 | 0 | 0 | 0 | 0 | 0 | 0 | 0 | 0 | 0 |
| ENSMUSG00 | Vmn1r143 | 923 | 0 | 0 | 0 | 0 | 0 | 0 | 0 | 0 | 0 | 0 | 0 | 0 | 0 | 0 | 0 | 0 | 0 | 0 |
| ENSMUSG00 | Gm16500 | 359 | 0 | 0 | 0 | 0 | 0 | 0 | 0 | 0 | 0 | 0.13133901 | 0 | 0 | 0 | 0 | 0 | 0 | 0 | 0.15322871 |
| ENSMUSG00 | Gm3573 | 893 | 0 | 0 | 0 | 0 | 0 | 0 | 0 | 0 | 0 | 0 | 0 | 0 | 0 | 0 | 0 | 0 | 0 | 0 |
| ENSMUSG00 | Rian | 6489 | 0 | 0 | 0 | 0 | 0 | 0 | 0 | 0 | 0 | 0 | 0 | 0 | 0 | 0 | 0 | 0 | 0 | 0 |
| ENSMUSG00 | Vmn2r18 | 2380 | 0 | 0 | 0 | 0 | 0.22245847 | 0.3539481 | 0 | 0.04814168 | 0.08652629 | 0.05943366 | 0 | 0 | 0.04145539 | 0 | 0 | 0 | 0.16103486 | 0 |
| ENSMUSG00 | Gm17692 | 2465 | 0 | 0.13514359 | 0 | 0 | 0.01952614 | 0.08041012 | 0.26871298 | 0.10845713 | 0.01670853 | 0.15302459 | 0.03876264 | 0 | 0.18011653 | 0.11304849 | 0.05014388 | 0.03106743 | 0.12438555 | 0.04463214 |
| ENSMUSG00 | Gm17693 | 172 | 0 | 0 | 0 | 0 | 0 | 0 | 0 | 0 | 0 | 0 | 0 | 0 | 0 | 0 | 0 | 0 | 0 | 0 |
| ENSMUSG00 | Vmn2r53 | 2416 | 0 | 0 | 0 | 0 | 0 | 0 | 0 | 0 | 0 | 0 | 0 | 0 | 0 | 0 | 0 | 0 | 0 | 0 |
| ENSMUSG00 | Gm17691 | 1958 | 0.36904392 | 0.22117856 | 0.32117257 | 0.23269175 | 0.54080813 | 0.22777048 | 0.24163779 | 0.15604659 | 0.08413997 | 0.24081054 | 0.51239741 | 0.2565582 | 0.15117033 | 0.18976134 | 0.06312802 | 0.19555979 | 0.25446469 | 0.11237815 |
| ENSMUSG00 | Igkv1-115 | 341 | 0 | 0 | 0 | 0 | 0 | 0 | 0 | 0 | 0 | 0 | 0 | 0 | 0 | 0 | 0 | 0 | 0 | 0 |
| ENSMUSG00 | Gm17637 | 1968 | 0 | 0 | 0 | 0 | 0 | 0 | 0 | 0 | 0 | 0 | 0 | 0 | 0 | 0 | 0 | 0 | 0 | 0 |
| ENSMUSG00 | Gm17638 | 4341 | 0.42103722 | 0.54485498 | 0.53488354 | 0.49328935 | 0.27719376 | 0.29679131 | 0.22887958 | 0.43990328 | 0.55977987 | 0.2932663 | 0.88044153 | 0.54002714 | 0.35228966 | 0.57774262 | 0.36066785 | 0.38811028 | 0.37964312 | 0.22809581 |
| ENSMUSG00 | Cox16 | 3740 | 1.35243742 | 1.42515055 | 1.55209188 | 1.13293506 | 1.29981933 | 1.5501791 | 2.17587721 | 1.23563655 | 1.66287804 | 1.91628527 | 2.01830038 | 1.80580085 | 2.44021516 | 2.1483495 | 2.01601196 | 1.94524476 | 1.53715095 | 1.89737293 |
| ENSMUSG00 | Gm17636 | 103 | 0 | 0 | 0 | 0 | 0 | 0 | 0 | 0 | 0 | 0.45777382 | 0 | 0 | 0 | 0 | 0 | 0 | 0 | 0 |
| ENSMUSG00 | Vmn2r108 | 2562 | 0 | 0 | 0 | 0 | 0 | 0 | 0 | 0 | 0 | 0 | 0 | 0 | 0 | 0 | 0 | 0 | 0 | 0 |
| ENSMUSG00 | Gm5258 | 326 | 0 | 0 | 0 | 0 | 0 | 0 | 0 | 0 | 0 | 0 | 0 | 0 | 0 | 0 | 0 | 0 | 0 | 0 |
| ENSMUSG00 | Olfr1438-ps1 | 959 | 0 | 0 | 0 | 0 | 0 | 0 | 0 | 0 | 0 | 0 | 0 | 0 | 0 | 0 | 0 | 0 | 0 | 0 |
| ENSMUSG00 | Gm9671 | 544 | 0 | 0 | 0 | 0 | 0 | 0 | 0 | 0 | 0 | 0 | 0 | 0 | 0 | 0 | 0 | 0 | 0 | 0 |
| ENSMUSG00 | Olfr721-ps1 | 950 | 0 | 0 | 0 | 0 | 0 | 0 | 0 | 0 | 0 | 0 | 0 | 0 | 0 | 0 | 0 | 0 | 0 | 0 |
| ENSMUSG00 | Gm17442 | 1487 | 2.82986713 | 2.53151113 | 3.41575029 | 3.73802785 | 2.58947807 | 3.53234012 | 3.88173963 | 3.54441932 | 4.84710172 | 3.29769545 | 2.28111772 | 3.86618241 | 4.14693319 | 1.43673707 | 3.32494062 | 4.1457887 | 3.53107107 | 4.51318831 |
| ENSMUSG00 | Prr24 | 1678 | 7.2192939 | 6.13449602 | 7.06288137 | 6.84230225 | 6.167082 | 3.83900796 | 9.78396877 | 11.2665311 | 10.2843564 | 8.54220131 | 7.08937107 | 8.51538118 | 13.9352377 | 13.2301989 | 11.9332277 | 13.2123133 | 9.22754705 | 14.0637108 |
| ENSMUSG00 | Ear-ps5 | 462 | 0 | 0 | 0 | 0 | 0 | 0 | 0 | 0 | 0 | 0 | 0 | 0 | 0 | 0 | 0 | 0 | 0 | 0 |
| ENSMUSG00 | Ces2h | 1594 | 0 | 0 | 0 | 0 | 0 | 0 | 0 | 0 | 0 | 0 | 0 | 0 | 0 | 0 | 0 | 0 | 0 | 0 |
| ENSMUSG00 | Gm8050 | 1590 | 0 | 0 | 0 | 0 | 0 | 0 | 0 | 0 | 0 | 0 | 0 | 0 | 0 | 0 | 0 | 0 | 0 | 0 |
| ENSMUSG00 | Gm17153 | 487 | 0 | 0 | 0 | 0 | 0 | 0 | 0 | 0 | 0 | 0 | 0 | 0 | 0 | 0 | 0 | 0 | 0 | 0 |
| ENSMUSG00 | Gm17141 | 3143 | 0 | 0 | 0 | 0 | 0 | 0 | 0 | 0 | 0 | 0 | 0 | 0 | 0 | 0 | 0 | 0 | 0 | 0 |
| ENSMUSG00 | Gm17091 | 354 | 0 | 0 | 0 | 0 | 0 | 0 | 0 | 0 | 0 | 0 | 0 | 0 | 0 | 0 | 0 | 0 | 0 | 0 |
| ENSMUSG00 | Rpl19-ps7 | 681 | 4.68118679 | 4.40258514 | 4.83024973 | 5.3522519 | 4.52341133 | 5.02076155 | 5.48854866 | 3.86971478 | 4.71739828 | 5.33128363 | 4.77048044 | 3.68825964 | 5.57791135 | 6.13798512 | 5.8686504 | 5.0604327 | 3.54560456 | 6.38137949 |
| ENSMUSG00 | Gm17443 | 4689 | 0.00922221 | 0.00722779 | 0.0419818 | 0.00988523 | 0 | 0.0215026 | 0 | 0.12429725 | 0 | 0.01023014 | 0 | 0.01211015 | 0.03211017 | 0.06046095 | 0.05363622 | 0 | 0.0498932 | 0 |
| ENSMUSG00 | Gm1815 | 302 | 0 | 0 | 0.48053176 | 0 | 0.15937723 | 0.1640819 | 0 | 0 | 0.13637919 | 0 | 0 | 0 | 0 | 0 | 0 | 0 | 0 | 0 |
| ENSMUSG00 | Gm17503 | 1082 | 0 | 0 | 0 | 0 | 0 | 0 | 0 | 0 | 0 | 0 | 0 | 0 | 0 | 0 | 0 | 0 | 0 | 0 |
| ENSMUSG00 | Gm17502 | 1130 | 0.1504608 | 0.14740219 | 0.1284253 | 0 | 0 | 0.30696384 | 0 | 0.10139576 | 0.25513771 | 0.08345257 | 0.12683617 | 0.09678878 | 0.82947472 | 0.12330289 | 0 | 0.74548085 | 0 | 0.14604188 |
| ENSMUSG00 | Gm8897 | 1452 | 0 | 0 | 0 | 0 | 0 | 0 | 0 | 0 | 0 | 0 | 0 | 0 | 0 | 0 | 0 | 0 | 0 | 0 |
| ENSMUSG00 | Gm6553 | 917 | 0 | 0 | 0 | 0 | 0 | 0 | 0 | 0 | 0 | 0 | 0 | 0 | 0 | 0 | 0 | 0 | 0 | 0 |
| ENSMUSG00 | Gm5778 | 1071 | 0.07937475 | 0.09331343 | 0.13550009 | 0.04254066 | 0 | 0.09253545 | 0.04417617 | 0.35660506 | 0.34610517 | 0.22012467 | 0.26764683 | 0.05211546 | 0.09212309 | 0.13009548 | 0.19235087 | 0.0715044 | 0.21471315 | 0.05136238 |
| ENSMUSG00 | Gm17504 | 2035 | 0 | 0 | 0 | 0 | 0 | 0 | 0 | 0 | 0 | 0 | 0 | 0 | 0 | 0 | 0 | 0 | 0 | 0 |
| ENSMUSG00 | Gm3495 | 1000 | 0.80759834 | 1.06601262 | 0.53210883 | 0.4100494 | 0.19252769 | 0.2973164 | 0 | 0 | 0.20593258 | 0.0471507 | 0.14332488 | 0.61397229 | 0 | 0.04644409 | 0 | 0.07658121 | 0 | 0 |
| ENSMUSG00 | 5033417F24I | 1783 | 0.42910441 | 0.52314136 | 0.16278249 | 0.30663631 | 0.24295419 | 0.66700258 | 0.87566932 | 1.19953704 | 0.06929868 | 0.5024472 | 0.45550998 | 0.72000013 | 1.74307952 | 1.1721727 | 1.22472376 | 0.88049069 | 0.92430217 | 1.11067181 |
| ENSMUSG00 | 9330159M07 | 3902 | 0.06535906 | 0 | 0 | 0.05838166 | 0 | 0 | 0.18187857 | 0 | 0 | 0 | 0.01224371 | 0 | 0.11378453 | 0.16663691 | 0.01055909 | 0.14719608 | 0.01964444 | 0 |
| ENSMUSG00 | Gm4707 | 1598 | 1.06395935 | 0.77132483 | 0.27244166 | 0.76980488 | 1.2349242 | 0.4651383 | 0.94743788 | 1.19500633 | 1.77839148 | 0.32456679 | 0.38865736 | 0.69856899 | 1.20397045 | 0.87191653 | 0.74771284 | 1.05430959 | 0.47967831 | 0.20654233 |
| ENSMUSG00 | Gm17155 | 470 | 0 | 0 | 0 | 0 | 0 | 0 | 0 | 0 | 0 | 0 | 0 | 0 | 0 | 0 | 0 | 0 | 0 | 0 |
| ENSMUSG00 | Gm17317 | 3766 | 0.16929836 | 0.291908 | 0.30827529 | 0.08468596 | 0.17892908 | 0.1184213 | 0.03768934 | 0.50706854 | 0.24060099 | 0.05008041 | 0.07611518 | 0.38534446 | 0.09169501 | 0.09865977 | 0.13128483 | 0.15251171 | 0.12212309 | 0.07303386 |
| ENSMUSG00 | Gm17177 | 478 | 0 | 0 | 0 | 0 | 0 | 0 | 0 | 0 | 0 | 0 | 0 | 0 | 0 | 0 | 0 | 0 | 0 | 0 |
| ENSMUSG00 | Gm2075 | 995 | 0 | 0 | 0 | 0 | 0 | 0 | 0 | 0 | 0 | 0 | 0 | 0 | 0 | 0 | 0 | 0 | 0 | 0 |
| ENSMUSG00 | Gm17316 | 1499 | 0 | 0 | 0 | 0 | 0 | 0 | 0 | 0 | 0 | 0 | 0 | 0 | 0 | 0 | 0 | 0 | 0 | 0 |
| ENSMUSG00 | Gm17315 | 2502 | 0.03397696 | 0.06657253 | 0.09666972 | 0.12746895 | 0.05771214 | 0.079221 | 0.05672983 | 0.0305295 | 0.11523006 | 0.07538082 | 0.03818942 | 0.20077576 | 0 | 0.07425114 | 0 | 0 | 0.15318264 | 0.32979081 |
| ENSMUSG00 | Gm3005 | 1954 | 0 | 0 | 0 | 0 | 0 | 0 | 0 | 0 | 0 | 0 | 0 | 0 | 0 | 0 | 0 | 0 | 0 | 0 |
| ENSMUSG00 | 4930519A11I | 1394 | 0 | 0 | 0 | 0 | 0 | 0 | 0 | 0 | 0 | 0 | 0 | 0 | 0 | 0 | 0 | 0 | 0 | 0 |
| ENSMUSG00 | Gm16490 | 353 | 0.24082253 | 0.09437081 | 0.27407099 | 0.25813623 | 0.13635106 | 0.56150406 | 0.13403025 | 0.10819377 | 0.23335136 | 0.1335714 | 0 | 0.15811802 | 0.41925142 | 0 | 0.11671829 | 0.21694395 | 0.76001155 | 0.31166633 |
| ENSMUSG00 | Ighv1-76 | 293 | 0 | 0 | 0 | 0 | 0 | 0 | 0 | 0 | 0 | 0 | 0 | 0 | 0 | 0 | 0 | 0 | 0 | 0 |
| ENSMUSG00 | Gm8251 | 5984 | 0 | 0 | 0 | 0 | 0 | 0 | 0 | 0 | 0 | 0 | 0 | 0.04663742 | 0 | 0 | 0 | 0 | 0 | 0 |
| ENSMUSG00 | Gm4604 | 389 | 0.21853561 | 0.4281863 | 0.37306065 | 0.23424701 | 0.2474649 | 0.38215476 | 0.24325285 | 0 | 0.42351173 | 0 | 0.49125921 | 0 | 0.12681727 | 0.5969677 | 0.21183319 | 0.39373375 | 0.49262592 | 0.84846951 |
| ENSMUSG00 | Gm2855 | 453 | 0 | 0 | 0 | 0 | 0 | 0 | 0 | 0 | 0 | 0 | 0 | 0 | 0 | 0 | 0 | 0 | 0 | 0 |
| ENSMUSG00 | Gm7697 | 471 | 0 | 0 | 0 | 0 | 0 | 0 | 0 | 0 | 0 | 0 | 0 | 0 | 0 | 0 | 0 | 0 | 0 | 0 |
| ENSMUSG00 | Gm17188 | 2874 | 0 | 0 | 0 | 0 | 0 | 0 | 0 | 0 | 0 | 0 | 0 | 0 | 0 | 0 | 0 | 0 | 0 | 0 |
| ENSMUSG00 | Gm3460 | 1804 | 0 | 0 | 0 | 0 | 0 | 0 | 0 | 0 | 0 | 0 | 0 | 0 | 0 | 0 | 0 | 0 | 0 | 0 |
| ENSMUSG00 | Gm8024 | 1403 | 0 | 0 | 0 | 0 | 0 | 0 | 0 | 0 | 0 | 0 | 0 | 0 | 0 | 0 | 0 | 0 | 0 | 0 |
| ENSMUSG00 | 4930534D22I | 2713 | 0.01566722 | 0 | 0 | 0.0167936 | 0 | 0.03652984 | 0 | 0 | 0 | 0.03475909 | 0 | 0 | 0 | 0 | 0.03037343 | 0.16936502 | 0 | 0.04055223 |
| ENSMUSG00 | Gm17702 | 964 | 0.22046253 | 0.10367083 | 0.35126006 | 0.33083746 | 0.09985876 | 0 | 0.19631817 | 0.11885602 | 0.42724601 | 0.24455759 | 0.19823634 | 0.05790006 | 0.25587094 | 0.09635703 | 0.64110305 | 0.11916164 | 0.15903028 | 0.05706339 |
| ENSMUSG00 | Mier2 | 1397 | 0 | 0 | 0 | 0 | 0 | 0 | 0 | 0 | 0 | 0 | 0 | 0 | 0 | 0 | 0 | 0 | 0 | 0 |
| ENSMUSG00 | Gm17701 | 1348 | 0 | 0 | 0 | 0 | 0 | 0.03676019 | 0 | 0.11333057 | 0 | 0 | 0.10632409 | 0 | 0.10978913 | 0 | 0.03056495 | 0 | 0.14215985 | 0 |
| ENSMUSG00 | Ighd5-5 | 9 | 0 | 0 | 0 | 0 | 0 | 0 | 0 | 0 | 0 | 0 | 0 | 0 | 0 | 0 | 0 | 0 | 0 | 0 |
| ENSMUSG00 | Vmn2r100 | 2547 | 0 | 0 | 0 | 0 | 0 | 0 | 0 | 0 | 0 | 0 | 0 | 0 | 0 | 0 | 0 | 0 | 0 | 0 |
| ENSMUSG00 | Flg | 935 | 0 | 0 | 0 | 0 | 0 | 0 | 0 | 0 | 0 | 0 | 0 | 0 | 0 | 0 | 0 | 0 | 0 | 0 |
| ENSMUSG00 | C230036H16 | 1301 | 0 | 0 | 0 | 0 | 0 | 0 | 0 | 0 | 0 | 0 | 0 | 0 | 0 | 0 | 0 | 0 | 0 | 0 |
| ENSMUSG00 | Gm7157 | 635 | 0 | 0 | 0 | 0 | 0 | 0 | 0 | 0 | 0 | 0 | 0 | 0 | 0 | 0 | 0 | 0 | 0 | 0 |
| ENSMUSG00 | Gm17102 | 904 | 0.235095 | 0.03685055 | 0 | 0.15119816 | 0.15972984 | 0.27407485 | 0.05233703 | 0.29573763 | 0.27336183 | 0.365105 | 0.31709043 | 0.18522897 | 0 | 0.46238583 | 0.36461554 | 0.7624236 | 0.42396346 | 0 |
| ENSMUSG00 | Gm17244 | 1287 | 0.46237177 | 0.10353658 | 0.03758627 | 0.17700483 | 0.18699271 | 0.73154774 | 0.29409591 | 1.27604763 | 0.41602541 | 0.549542 | 0.63106006 | 0.56379457 | 0.42164032 | 0.07217418 | 0.38416369 | 0.83305129 | 0.26801606 | 0.25645271 |
| ENSMUSG00 | Gm6257 | 909 | 0.23380185 | 0.51306988 | 0.42572964 | 0.35085513 | 0.74130576 | 0.32708075 | 0.52049152 | 0.16806338 | 0.67964547 | 0.15561288 | 0.15767313 | 0.73684043 | 0.16281161 | 0.71531048 | 0.81587239 | 0.33699104 | 1.22273114 | 0.48412855 |
| ENSMUSG00 | Cyp2a22 | 2217 | 0 | 0 | 0 | 0 | 0 | 0 | 0 | 0 | 0 | 0 | 0 | 0 | 0 | 0 | 0 | 0 | 0 | 0 |
| ENSMUSG00 | 9330104G04 | 2233 | 1.16113557 | 1.13380204 | 1.06148813 | 0.71412295 | 0.68975439 | 1.10955516 | 0.69920215 | 0.46179797 | 1.30955782 | 0.6334622 | 1.19811808 | 0.99983274 | 1.06042635 | 0.45757722 | 0.49818272 | 0.25721411 | 0.72087079 | 0.2463462 |
| ENSMUSG00 | 1700101I11R | 882 | 0.24095905 | 0.60431554 | 0.21938109 | 0.10331303 | 0.6548561 | 0.95509804 | 0.10728499 | 0.56292657 | 0.74714768 | 0.10691769 | 0.43333296 | 0.69611371 | 0.89491006 | 1.73770395 | 1.54155481 | 0.17365355 | 1.21670784 | 2.18290106 |
| ENSMUSG00 | Gm17560 | 468 | 0.09082303 | 0.0711814 | 0.20672449 | 0.09735266 | 0 | 0 | 0 | 0.24482309 | 0.08800538 | 0.20149873 | 0 | 0.35779271 | 0 | 0 | 0 | 0.08181754 | 0 | 0 |
| ENSMUSG00 | Gm17561 | 2038 | 0 | 0 | 0 | 0 | 0 | 0 | 0 | 0 | 0 | 0 | 0 | 0 | 0 | 0 | 0 | 0 | 0 | 0 |
| ENSMUSG00 | Olfr732 | 1043 | 0 | 0 | 0 | 0 | 0 | 0 | 0 | 0 | 0 | 0 | 0 | 0 | 0 | 0 | 0 | 0 | 0 | 0 |
| ENSMUSG00 | Vmn1r56 | 1830 | 0 | 0 | 0 | 0 | 0 | 0 | 0 | 0 | 0 | 0 | 0 | 0 | 0 | 0 | 0 | 0 | 0 | 0 |
| ENSMUSG00 | Mir155 | 1259 | 0.6076991 | 0 | 0 | 0.07237656 | 0 | 0 | 0 | 0.18201304 | 0 | 0 | 0.0758935 | 0.48766663 | 0 | 0.55334496 | 0 | 0 | 0.12176742 | 0 |
| ENSMUSG00 | Gm3851 | 317 | 0 | 0 | 0 | 0 | 0 | 0 | 0 | 0 | 0 | 0.14874039 | 0 | 0.17607465 | 0.15562119 | 0 | 0 | 0 | 0 | 0 |
| ENSMUSG00 | Gm17699 | 3465 | 0 | 0 | 0 | 0 | 0 | 0 | 0 | 0 | 0 | 0 | 0 | 0 | 0 | 0 | 0 | 0 | 0 | 0 |
| ENSMUSG00 | C430002E04 | 984 | 0.21598159 | 0.16927284 | 0.49160092 | 0.13890562 | 0.39131645 | 0.60430163 | 0.76931185 | 0.07762683 | 0.20928108 | 0.19166953 | 0.04855179 | 0.45378587 | 0 | 0.42479348 | 0.502458 | 0.23347931 | 0.07789898 | 0.05590356 |
| ENSMUSG00 | Vmn2r17 | 2544 | 0 | 0 | 0 | 0 | 0 | 0 | 0 | 0 | 0 | 0 | 0 | 0 | 0 | 0 | 0 | 0 | 0 | 0 |
| ENSMUSG00 | Zscan4e | 1741 | 0 | 0 | 0 | 0 | 0 | 0 | 0 | 0 | 0 | 0 | 0 | 0 | 0 | 0 | 0 | 0 | 0 | 0 |
| ENSMUSG00 | Gm17146 | 370 | 0.45951542 | 0.45017425 | 0.13073927 | 0.49255183 | 0.13008628 | 0.13392631 | 0.25574421 | 0.30966813 | 0.89051926 | 0.50973733 | 0 | 0 | 1.06663605 | 0.75314736 | 0.22271111 | 0.41395251 | 0.62150752 | 1.04071283 |
| ENSMUSG00 | Gm6337 | 2135 | 0 | 0 | 0 | 0 | 0 | 0 | 0 | 0 | 0 | 0 | 0 | 0 | 0 | 0 | 0 | 0 | 0 | 0 |
| ENSMUSG00 | 1700039M10 | 520 | 0 | 0 | 0 | 0 | 0 | 0 | 0 | 0 | 0 | 0 | 0 | 0 | 0 | 0 | 0.39616881 | 0 | 0.07370442 | 0 |
| ENSMUSG00 | Vmn1r103 | 917 | 0 | 0 | 0 | 0 | 0 | 0 | 0 | 0 | 0 | 0 | 0 | 0 | 0 | 0 | 0 | 0 | 0 | 0 |
| ENSMUSG00 | Gm8926 | 1093 | 0 | 0 | 0 | 0 | 0 | 0 | 0 | 0 | 0 | 0 | 0 | 0 | 0 | 0 | 0 | 0 | 0 | 0 |
| ENSMUSG00 | AW047730 | 2577 | 0 | 0.03878102 | 0 | 0.19447865 | 0.14942002 | 0 | 0 | 0.02964098 | 0 | 0.01829674 | 0 | 0.02165916 | 0 | 0.05406762 | 0 | 0 | 0 | 0 |
| ENSMUSG00 | Gm17053 | 493 | 0 | 0 | 0 | 0 | 0 | 0 | 0 | 0 | 0 | 0 | 0 | 0 | 0 | 0 | 0 | 0 | 0 | 0 |
| ENSMUSG00 | Vmn2r80 | 2684 | 0 | 0 | 0 | 0 | 0 | 0 | 0 | 0 | 0 | 0 | 0 | 0 | 0 | 0 | 0 | 0 | 0 | 0 |
| ENSMUSG00 | Gm4216 | 917 | 0 | 0 | 0 | 0 | 0 | 0 | 0 | 0 | 0 | 0 | 0 | 0 | 0 | 0 | 0 | 0 | 0 | 0 |
| ENSMUSG00 | A830073O21I | 2024 | 0 | 0.06583576 | 0 | 0.13506238 | 0.11890297 | 0.04896515 | 0.60777157 | 0.37739528 | 0.10174535 | 0.37273283 | 0.68452263 | 0.71699962 | 0.29248172 | 0.04589337 | 0 | 0.47295711 | 0.15148734 | 0 |
| ENSMUSG00 | Gm17422 | 969 | 0 | 0.17189316 | 0.19968434 | 0.37614897 | 0 | 0.20455205 | 0 | 0.27589971 | 0.85008288 | 0.38927309 | 0.09860673 | 0.74881694 | 0.25455066 | 0.47929915 | 0.42519666 | 0.59273386 | 0.23731453 | 0.05676894 |
| ENSMUSG00 | Taar8c | 1108 | 0 | 0 | 0 | 0 | 0 | 0 | 0 | 0 | 0 | 0 | 0 | 0 | 0 | 0 | 0 | 0 | 0 | 0 |
| ENSMUSG00 | Gm3030 | 512 | 0 | 0 | 0 | 0 | 0 | 0 | 0 | 0 | 0 | 0 | 0 | 0 | 0 | 0 | 0 | 0 | 0 | 0 |
| ENSMUSG00 | Fgfr1 | 756 | 0 | 0 | 0 | 0 | 0 | 0 | 0 | 0 | 0 | 0 | 0 | 0 | 0 | 0 | 0 | 0 | 0 | 0 |
| ENSMUSG00 | Ube2d2 | 2877 | 25.1307975 | 25.6359916 | 25.8597462 | 26.0190462 | 24.5762238 | 24.7677549 | 26.8055845 | 28.0635171 | 22.2610469 | 23.0754919 | 24.6264385 | 23.4553827 | 23.6800062 | 26.7654838 | 23.4578201 | 25.2608873 | 25.2444674 | 26.0609706 |
| ENSMUSG00 | Gm17019 | 1157 | 0 | 0 | 0 | 0 | 0 | 0 | 0 | 0 | 0 | 0 | 0 | 0 | 0 | 0 | 0 | 0 | 0 | 0 |
| ENSMUSG00 | Tnnc1 | 1044 | 0.04071377 | 0 | 0 | 0 | 0 | 0 | 0 | 0.07316552 | 0 | 0 | 0 | 0 | 0 | 0 | 0 | 0 | 0 | 0 |
| ENSMUSG00 | Gm4353 | 1542 | 0.02756496 | 0.02160369 | 0 | 0.02954672 | 0 | 0.12854146 | 0.03068267 | 0.02476809 | 0.08012941 | 0 | 0 | 0.03619693 | 0 | 0 | 0.16031734 | 0.02483178 | 0.02485493 | 0.28539096 |
| ENSMUSG00 | Vmn2r67 | 2550 | 0 | 0 | 0 | 0 | 0 | 0 | 0 | 0 | 0 | 0 | 0 | 0 | 0 | 0 | 0 | 0 | 0 | 0 |
| ENSMUSG00 | Gm6918 | 1218 | 0 | 0 | 0 | 0 | 0 | 0 | 0 | 0 | 0 | 0 | 0 | 0 | 0 | 0 | 0 | 0 | 0 | 0 |
| ENSMUSG00 | Gm8871 | 1224 | 0 | 0 | 0 | 0 | 0 | 0 | 0 | 0 | 0 | 0 | 0 | 0 | 0 | 0 | 0 | 0 | 0 | 0 |
| ENSMUSG00 | Gm6395 | 721 | 0.29476544 | 0.32342616 | 0.2683691 | 0.31595731 | 0.13351435 | 0.41236671 | 0.06562091 | 0.1589143 | 0.28562078 | 0.3269813 | 0.13252416 | 0.30965694 | 0.20526457 | 0.51532968 | 0.62859517 | 0.42486111 | 0.53157139 | 0.38147786 |
| ENSMUSG00 | 1700099I09R | 874 | 0 | 0 | 0 | 0 | 0 | 0 | 0 | 0 | 0 | 0 | 0.16398727 | 0.12772463 | 0 | 0 | 0.04381076 | 0 | 0 | 0 |
| ENSMUSG00 | Gm9107 | 1022 | 0 | 0 | 0 | 0 | 0 | 0 | 0 | 0 | 0 | 0 | 0 | 0 | 0 | 0 | 0 | 0 | 0 | 0 |
| ENSMUSG00 | Gm17231 | 1100 | 0 | 0 | 0 | 0.04141913 | 0.13126888 | 0.04504794 | 0.08602305 | 0.31248329 | 0 | 0 | 0 | 0 | 0 | 0 | 0 | 0 | 0.06968418 | 0 |
| ENSMUSG00 | Gm17282 | 5126 | 1.15259841 | 1.06580465 | 1.34947617 | 1.19102223 | 1.2582282 | 0.95702704 | 0.7383953 | 1.11760834 | 1.31771139 | 1.22337954 | 1.87334504 | 1.08887364 | 0.58705559 | 1.8211591 | 0.93238012 | 1.39686765 | 1.85425705 | 2.60772785 |
| ENSMUSG00 | Gm17469 | 172 | 0 | 0 | 0 | 0 | 0 | 0 | 0 | 0 | 0 | 0 | 0 | 0 | 0 | 0 | 0 | 0 | 0 | 0 |
| ENSMUSG00 | Gm9587 | 451 | 0 | 0 | 0 | 0 | 0 | 0 | 0 | 0 | 0 | 0 | 0 | 0 | 0 | 0 | 0 | 0 | 0 | 0 |
| ENSMUSG00 | Gm17468 | 1995 | 0.36219949 | 0.48424759 | 0.26672122 | 0.45675232 | 0.2171365 | 0.29806156 | 0.28458754 | 0.21058467 | 0.8257948 | 0.23634438 | 0.11973674 | 0.53157774 | 0.24727778 | 0.39576415 | 0.30978613 | 0.07677315 | 0.23053412 | 0.05514697 |
| ENSMUSG00 | Gm17558 | 36 | 0 | 0 | 0 | 0 | 0 | 0 | 2.62848216 | 0 | 0 | 1.30974176 | 0 | 0 | 0 | 3.87034062 | 1.14448766 | 0 | 0 | 0 |
| ENSMUSG00 | 2010015M23 | 1197 | 0.99427312 | 1.11321284 | 0.6870092 | 1.10381812 | 1.52799758 | 0.66235901 | 1.18578143 | 0.79766922 | 1.68599772 | 0.94537751 | 1.71622658 | 0.74607402 | 1.48366669 | 0.38800407 | 1.54893067 | 0.51182098 | 0.32018627 | 1.51654179 |
| ENSMUSG00 | Gm4312 | 915 | 0 | 0 | 0 | 0 | 0 | 0 | 0 | 0 | 0 | 0 | 0 | 0 | 0 | 0 | 0 | 0 | 0 | 0 |
| ENSMUSG00 | Gm17467 | 172 | 0 | 0 | 0 | 0 | 0 | 0 | 0 | 0 | 0 | 0 | 0 | 0 | 0 | 0 | 0 | 0 | 0 | 0 |
| ENSMUSG00 | Gm4301 | 915 | 0 | 0 | 0 | 0 | 0 | 0 | 0 | 0 | 0 | 0 | 0 | 0 | 0 | 0 | 0 | 0 | 0 | 0 |
| ENSMUSG00 | Gm17470 | 1728 | 0.0245979 | 0 | 0 | 0.02636635 | 0 | 0.1147054 | 0.02738002 | 0 | 0.07150437 | 0 | 0 | 0.19380438 | 0 | 0 | 0.04768699 | 0.04431783 | 0 | 0 |
| ENSMUSG00 | Gm5861 | 1091 | 0 | 0 | 0 | 0 | 0 | 0 | 0 | 0 | 0 | 0 | 0 | 0 | 0 | 0 | 0 | 0 | 0 | 0 |
| ENSMUSG00 | B930041F14I | 2136 | 0 | 0 | 0 | 0 | 0 | 0 | 0 | 0 | 0 | 0 | 0 | 0 | 0 | 0 | 0 | 0 | 0 | 0 |
| ENSMUSG00 | Gm17409 | 961 | 0 | 0 | 0 | 0 | 0 | 0.05156372 | 0 | 0.03974235 | 0 | 0 | 0 | 0 | 0 | 0 | 0 | 0 | 0 | 0 |

| | | | | | | | | | | | | | | | | | | | |
|---|---|---|---|---|---|---|---|---|---|---|---|---|---|---|---|---|---|---|---|
| ENSMUSG00 Gm8267 | 2534 | 0 | 0 | 0 | 0 | 0 | 0 | 0 | 0 | 0 | 0 | 0 | 0 | 0 | 0 | 0 | 0 | 0 | 0 |
| ENSMUSG00 Vmn1r185 | 1143 | 0 | 0 | 0 | 0 | 0 | 0 | 0 | 0 | 0 | 0 | 0 | 0 | 0 | 0 | 0 | 0 | 0 | 0 |
| ENSMUSG00 Vmn2r62 | 2582 | 0 | 0 | 0 | 0 | 0 | 0 | 0 | 0 | 0 | 0 | 0 | 0 | 0 | 0 | 0 | 0 | 0 | 0 |
| ENSMUSG00 Gm17408 | 2437 | 0.06976639 | 0.27339265 | 0.01984962 | 0.09347773 | 0.27650674 | 0.10166749 | 0.19414312 | 0.09403136 | 0.23660698 | 0.03869569 | 0 | 0.29774461 | 0.04048578 | 0.15246315 | 0.10144002 | 0.23568285 | 0 | 0.02257247 |
| ENSMUSG00 Gm8979 | 2132 | 0 | 0 | 0 | 0 | 0 | 0 | 0 | 0 | 0 | 0 | 0 | 0 | 0 | 0 | 0 | 0 | 0 | 0 |
| ENSMUSG00 Gm8582 | 784 | 0 | 0 | 0 | 0 | 0 | 0 | 0 | 0 | 0 | 0 | 0 | 0 | 0 | 0 | 0 | 0 | 0 | 0 |
| ENSMUSG00 Vmn2r52 | 2553 | 0 | 0 | 0 | 0 | 0 | 0 | 0 | 0 | 0 | 0 | 0 | 0 | 0 | 0 | 0 | 0 | 0 | 0 |
| ENSMUSG00 Moap1 | 7408 | 1.43443494 | 1.00730134 | 0.90112678 | 1.06399307 | 0.83165311 | 0.84951366 | 1.05380562 | 1.02595681 | 1.15642485 | 0.91653635 | 0.98671283 | 1.17538382 | 1.5049964 | 1.02818984 | 1.0511736 | 1.2870358 | 0.68291998 | 0.8390968 |
| ENSMUSG00 Gm17596 | 2503 | 0.42454231 | 0.38598641 | 0.4638293 | 0.40045664 | 0.23075633 | 0.45533874 | 0.47255972 | 0.76293253 | 0.69110414 | 0.50861725 | 0.57261237 | 0.35679209 | 1.02487403 | 0.24121979 | 0.75719999 | 0.56602176 | 0.70435863 | 1.0988635 |
| ENSMUSG00 Gm8857 | 1172 | 0 | 0 | 0 | 0 | 0 | 0 | 0 | 0 | 0 | 0 | 0 | 0 | 0 | 0 | 0 | 0 | 0 | 0 |
| ENSMUSG00 Gm17597 | 1113 | 0.30551789 | 0 | 0.43462291 | 0.12280605 | 0.04324521 | 0.35617419 | 0.4676006 | 0.10294448 | 0.18502478 | 0.38127253 | 0.04292449 | 0.85253034 | 0.04432338 | 0.54247362 | 0.70335091 | 0.13761225 | 0.48209178 | 0.1976967 |
| ENSMUSG00 Vmn1r118 | 920 | 0 | 0 | 0 | 0 | 0 | 0 | 0 | 0 | 0 | 0 | 0 | 0 | 0 | 0 | 0 | 0 | 0 | 0 |
| ENSMUSG00 Gm3053 | 612 | 0 | 0 | 0 | 0 | 0 | 0 | 0 | 0 | 0 | 0 | 0 | 0 | 0 | 0 | 0 | 0 | 0 | 0 |
| ENSMUSG00 Gm3646 | 161 | 0 | 0 | 0 | 0 | 0 | 0 | 0 | 0 | 0 | 0 | 0 | 0 | 0 | 0 | 0 | 0 | 0 | 0 |
| ENSMUSG00 Gm2442 | 703 | 0 | 0 | 0 | 0 | 0 | 0 | 0 | 0 | 0 | 0 | 0 | 0 | 0 | 0 | 0 | 0 | 0 | 0 |
| ENSMUSG00 Gm17595 | 4310 | 0.00986199 | 0.0772921 | 0 | 0 | 0 | 0 | 0.03293226 | 0 | 0.22934487 | 0.07657887 | 0.25494757 | 0.19425405 | 0 | 0 | 0.04779763 | 0 | 0 | 0 |
| ENSMUSG00 Vmn1r-ps70 | 3054 | 0 | 0 | 0 | 0 | 0 | 0 | 0 | 0 | 0 | 0 | 0 | 0 | 0 | 0 | 0 | 0 | 0 | 0 |
| ENSMUSG00 Gm7399 | 857 | 0.04959764 | 0 | 0.05644519 | 0 | 0 | 0.11564232 | 0 | 0.04456523 | 0 | 0 | 0.27873371 | 0.26051651 | 0 | 0 | 0 | 0 | 0 | 0 |
| ENSMUSG00 Gm17138 | 458 | 0 | 0 | 0.10561906 | 0 | 0.10509154 | 0.21638748 | 0 | 0 | 0.17985378 | 0 | 0 | 0.12186826 | 0 | 0 | 0.08995973 | 0.16720789 | 0 | 0 |
| ENSMUSG00 A730099G02l | 2219 | 0 | 0 | 0 | 0 | 0 | 0.02233111 | 0 | 0 | 0 | 0 | 0 | 0 | 0 | 0 | 0 | 0 | 0 | 0 |
| ENSMUSG00 Gm17517 | 6618 | 0 | 0 | 0 | 0 | 0 | 0 | 0 | 0 | 0 | 0 | 0 | 0 | 0 | 0 | 0 | 0 | 0 | 0 |
| ENSMUSG00 Vmn2r114 | 2565 | 0 | 0 | 0 | 0 | 0 | 0.03863761 | 0.03689098 | 0 | 0 | 0 | 0 | 0 | 0 | 0 | 0 | 0 | 0 | 0.10723023 |
| ENSMUSG00 Gm17518 | 1344 | 1.80267488 | 0.37179569 | 0.39591431 | 0.40679504 | 0.39393688 | 0.14747837 | 0.77446349 | 0.99459381 | 0.30644729 | 1.43837711 | 1.38632692 | 0.16611804 | 0.44046355 | 0.79480209 | 1.62476374 | 1.79487222 | 1.62544563 | 0.20464697 |
| ENSMUSG00 6530401F13l | 1088 | 0 | 0.03061847 | 0 | 0.16750384 | 0 | 0 | 0 | 0 | 0 | 0.21668522 | 0 | 0.15390348 | 0 | 0.38418822 | 0 | 0.03519357 | 0.03522638 | 0 |
| ENSMUSG00 Olfr421 | 1049 | 0 | 0 | 0 | 0 | 0 | 0 | 0 | 0 | 0 | 0 | 0 | 0 | 0 | 0 | 0 | 0 | 0 | 0 |
| ENSMUSG00 Gm17709 | 1433 | 1.15680521 | 0.81364361 | 1.0802184 | 1.3353551 | 1.24276425 | 1.24486979 | 0.72636353 | 0.98612623 | 2.18435115 | 1.44775362 | 1.33356479 | 1.36325763 | 0.61966121 | 0.06482078 | 0.8913107 | 0.66801478 | 1.06981987 | 0.92129697 |
| ENSMUSG00 Gm17710 | 1553 | 0 | 0.02145067 | 0 | 0.02933744 | 0 | 0 | 0 | 0.04918532 | 0 | 0 | 0.06152603 | 0 | 0 | 0.50840276 | 0 | 0 | 0 | 0 |
| ENSMUSG00 Gm17179 | 392 | 0 | 0 | 0 | 0 | 0 | 0 | 0.12069561 | 0 | 0.10506764 | 0 | 0 | 0 | 0.50338691 | 0 | 0 | 0 | 0 | 0 |
| ENSMUSG00 Gm3788 | 474 | 59.3637691 | 44.8388745 | 40.3112753 | 43.5425171 | 50.2643507 | 42.9666108 | 30.0445493 | 30.6989564 | 28.0659169 | 23.9732479 | 24.2906434 | 32.6180138 | 25.3944891 | 23.4180103 | 15.4723142 | 17.1257569 | 15.6054331 | 18.6845278 |
| ENSMUSG00 C2cd4b | 1324 | 0.28893247 | 0.15096478 | 0.32882309 | 0.44735164 | 0.25447392 | 0.48654497 | 0 | 0.25961603 | 0.03110764 | 0.5341847 | 0 | 0.46372529 | 1.71394879 | 0.84188678 | 2.24056799 | 2.11118908 | 1.99736744 | 1.28797757 |
| ENSMUSG00 Gm8841 | 950 | 0.08948458 | 0 | 0 | 0 | 0 | 0 | 0.29881692 | 0 | 0 | 0.14889696 | 0 | 0.35251997 | 0.10385667 | 0.09777703 | 0.04337006 | 0 | 0.12103041 | 0.40533026 |
| ENSMUSG00 Gm17708 | 4867 | 0 | 0 | 0 | 0 | 0 | 0 | 0 | 0 | 0 | 0 | 0 | 0 | 0 | 0 | 0 | 0 | 0 | 0 |
| ENSMUSG00 Gm17038 | 491 | 0 | 0 | 0 | 0 | 0 | 0 | 0 | 0 | 0 | 0 | 0 | 0 | 0 | 0 | 0 | 0 | 0 | 0 |
| ENSMUSG00 Gm3123 | 477 | 0 | 0 | 0 | 0 | 0 | 0 | 0 | 0 | 0 | 0 | 0 | 0 | 0 | 0 | 0 | 0 | 0 | 0 |
| ENSMUSG00 Vmn2r78 | 2605 | 0 | 0 | 0 | 0 | 0 | 0 | 0 | 0 | 0 | 0 | 0 | 0 | 0 | 0 | 0 | 0 | 0 | 0 |
| ENSMUSG00 Gm17658 | 46 | 0 | 0 | 0 | 0 | 0 | 0 | 0 | 0 | 0 | 1.02501529 | 0 | 0 | 0 | 0 | 0 | 0 | 0 | 0 |
| ENSMUSG00 BC037034 | 3577 | 0 | 0 | 0 | 0 | 0 | 0 | 0 | 0 | 0 | 0 | 0 | 0 | 0 | 0 | 0 | 0 | 0 | 0 |
| ENSMUSG00 Gm2670 | 2802 | 0 | 0 | 0 | 0 | 0 | 0 | 0 | 0 | 0 | 0 | 0 | 0.03983987 | 0 | 0 | 0 | 0 | 0 | 0 |
| ENSMUSG00 Vmn2r-ps5 | 3197 | 0 | 0 | 0 | 0 | 0 | 0 | 0 | 0 | 0 | 0 | 0 | 0 | 0 | 0 | 0 | 0 | 0 | 0 |
| ENSMUSG00 Gm17115 | 883 | 0 | 0 | 0.10956632 | 0.05159801 | 0 | 0.11223722 | 0.05358174 | 0 | 0.27986307 | 0.53398305 | 0.75747386 | 0.12642279 | 0.11173707 | 0 | 0 | 0.21682111 | 0.2170232 | 0.18689391 |
| ENSMUSG00 Pcdha3 | 2437 | 0 | 0 | 0 | 0 | 0 | 0 | 0 | 0 | 0 | 0 | 0 | 0 | 0 | 0 | 0 | 0 | 0 | 0 |
| ENSMUSG00 Gm4868 | 788 | 0 | 0 | 0 | 0 | 0 | 0 | 0 | 0 | 0 | 0 | 0 | 0 | 0 | 0 | 0 | 0 | 0 | 0 |
| ENSMUSG00 Hspa1a | 2925 | 27.6392632 | 23.9511168 | 22.9050733 | 37.492456 | 28.1057524 | 28.5118806 | 34.4048096 | 31.4156992 | 28.0490733 | 56.725923 | 37.6319673 | 38.3362961 | 94.6835499 | 132.758637 | 109.265118 | 102.527192 | 93.9616667 | 187.237835 |
| ENSMUSG00 Tmem207 | 1533 | 0 | 0 | 0 | 0 | 0 | 0 | 0 | 0 | 0 | 0 | 0 | 0 | 0 | 0 | 0 | 0 | 0 | 0 |
| ENSMUSG00 Gm20474 | 1481 | 0 | 0 | 0 | 0 | 0 | 0 | 0 | 0 | 0 | 0 | 0 | 0 | 0 | 0 | 0 | 0.10341825 | 0 | 0 |
| ENSMUSG00 Gm7822 | 519 | 0 | 0 | 0 | 0 | 0 | 0 | 0 | 0 | 0 | 0 | 0 | 0 | 0 | 0 | 0 | 0 | 0 | 0 |
| ENSMUSG00 Gm2104 | 1649 | 0 | 0 | 0 | 0 | 0 | 0 | 0 | 0 | 0 | 0 | 0 | 0 | 0 | 0 | 0 | 0 | 0 | 0 |
| ENSMUSG00 Gm17350 | 22 | 0 | 0 | 0 | 0 | 0 | 0 | 0 | 0 | 0 | 0 | 0 | 0 | 0 | 0 | 0 | 0 | 0 | 0 |
| ENSMUSG00 Gm725 | 905 | 0 | 0 | 0 | 0 | 0 | 0 | 0 | 0 | 0 | 0 | 0 | 0 | 0 | 0 | 0 | 0 | 0 | 0 |
| ENSMUSG00 Gm3719 | 312 | 0 | 0 | 0 | 0 | 0 | 0 | 0 | 0 | 0 | 0 | 0 | 0 | 0 | 0 | 0 | 0 | 0 | 0 |
| ENSMUSG00 Gm8038 | 2063 | 0 | 0 | 0 | 0 | 0 | 0 | 0 | 0 | 0 | 0 | 0 | 0 | 0 | 0 | 0 | 0 | 0 | 0 |
| ENSMUSG00 Gm17259 | 4606 | 0.46141094 | 0.71601748 | 1.06073091 | 0.31653353 | 0.88823567 | 0.77459766 | 0.63686194 | 0.65505857 | 0.85842499 | 0.58349763 | 0.49787191 | 1.42992796 | 0.74972519 | 0.9680053 | 0.72456058 | 0.71493535 | 0.87369978 | 0.78823297 |
| ENSMUSG00 Olfr457 | 1038 | 0 | 0 | 0 | 0 | 0.13910961 | 0 | 0 | 0 | 0 | 0 | 0 | 0 | 0 | 0 | 0 | 0 | 0 | 0 |
| ENSMUSG00 Gm17258 | 1264 | 0 | 0 | 0 | 0 | 0 | 0 | 0 | 0 | 0 | 0 | 0 | 0 | 0 | 0 | 0 | 0 | 0 | 0 |
| ENSMUSG00 Gm17354 | 3037 | 1.55353129 | 2.11701962 | 0.9716119 | 1.39518509 | 1.60069947 | 1.25635841 | 1.51113923 | 1.35817565 | 1.12561107 | 1.30413536 | 1.5101732 | 1.3783914 | 0.95837442 | 1.34576216 | 1.4244858 | 1.2734117 | 0.97172369 | 0.77885795 |
| ENSMUSG00 Rps2-ps5 | 665 | 0 | 0 | 0.1454843 | 0.06851285 | 0 | 0.07451539 | 0.14229377 | 0.05743218 | 0 | 0.07090331 | 0.07184204 | 0.08393333 | 0.07418333 | 0.06984073 | 0 | 0.17273958 | 0.05763353 | 0.08272046 |
| ENSMUSG00 Gm3376 | 1496 | 0 | 0 | 0 | 0 | 0 | 0 | 0 | 0 | 0 | 0 | 0 | 0 | 0 | 0 | 0 | 0 | 0 | 0 |
| ENSMUSG00 Vmn1r120 | 917 | 0 | 0 | 0 | 0 | 0 | 0 | 0 | 0 | 0 | 0 | 0 | 0 | 0 | 0 | 0 | 0 | 0 | 0 |
| ENSMUSG00 Gm17257 | 467 | 0.09101751 | 0.07133382 | 0.41433431 | 0 | 0 | 0.10610864 | 0.40524779 | 0 | 0.08819382 | 0.60579062 | 0 | 0.11951962 | 0.1056358 | 0.09945201 | 0.52935618 | 0.40996368 | 0.16413832 | 0.23558504 |
| ENSMUSG00 Gm9672 | 422 | 0 | 0 | 0 | 0 | 0 | 0 | 0 | 0 | 0 | 0 | 0 | 0 | 0 | 0 | 0 | 0 | 0 | 0 |
| ENSMUSG00 Gm17458 | 1815 | 0 | 0.05506264 | 0.18656458 | 0 | 0 | 0 | 0.02606759 | 0.16834117 | 0 | 0.25978349 | 0.71070186 | 0.24601945 | 0.1359006 | 0.28147902 | 0.11350291 | 0 | 0 | 0 |
| ENSMUSG00 B930036N10 | 1315 | 0 | 0 | 0.11035786 | 0.03464718 | 0 | 0 | 0.17989612 | 0.11617461 | 0 | 0.17928024 | 0 | 0 | 0.1125443 | 0.31786828 | 0.21932387 | 0.23294666 | 0.20401831 | 0 |
| ENSMUSG00 E130317F20l | 3263 | 1.06816562 | 1.33741623 | 2.0013566 | 1.75932932 | 1.56358686 | 2.12607499 | 1.02948213 | 2.25900511 | 2.72641355 | 1.60396202 | 1.493425 | 1.89672465 | 1.76887353 | 2.04963181 | 1.41421215 | 1.49031785 | 2.36089049 | 1.92186275 |
| ENSMUSG00 Gm3339 | 471 | 0 | 0 | 0 | 0 | 0 | 0 | 0 | 0 | 0 | 0 | 0 | 0 | 0 | 0 | 0 | 0 | 0 | 0 |
| ENSMUSG00 BC049352 | 2678 | 0.23807978 | 0 | 0 | 0 | 0 | 0 | 0.03533434 | 0 | 0.04613874 | 0.03521337 | 0.05351937 | 0 | 0.07368472 | 0 | 0.04615559 | 0.08578926 | 0.05724615 | 0.04108223 |
| ENSMUSG00 Gm6611 | 1271 | 0.30098079 | 0.10483995 | 0 | 0.43015935 | 0.15147734 | 0.35088482 | 0.26057337 | 0.27044187 | 0.58328661 | 0.03709733 | 0.15035392 | 0.57089191 | 0 | 0.07308275 | 0.03241665 | 0.48202181 | 0.2110811 | 0 |
| ENSMUSG00 Gm17072 | 1351 | 0 | 0 | 0 | 0 | 0 | 0 | 0 | 0 | 0 | 0 | 0 | 0 | 0 | 0 | 0 | 0 | 0 | 0 |
| ENSMUSG00 Gm2858 | 245 | 0 | 0 | 0 | 0 | 0 | 0 | 0 | 0 | 0 | 0 | 0 | 0 | 0 | 0 | 0 | 0 | 0 | 0 |
| ENSMUSG00 Gm13272 | 1555 | 0.02733452 | 0 | 0.09332514 | 0.02929971 | 0.03095301 | 0.06373342 | 0.12170464 | 0.17192721 | 0.21189204 | 0.09096599 | 0.03072345 | 0.14357727 | 0.31724706 | 0.05973516 | 0.26496177 | 0.3201144 | 0.07394141 | 0.07075126 |
| ENSMUSG00 Tmem200a | 2883 | 0 | 0 | 0.01677889 | 0 | 0 | 0 | 0 | 0 | 0 | 0 | 0 | 0 | 0 | 0 | 0 | 0 | 0.01329389 | 0 |
| ENSMUSG00 Gm17375 | 3219 | 0.99033495 | 1.33499949 | 0.67623761 | 0.62276668 | 0.88219431 | 1.00059885 | 0.66140744 | 1.10341517 | 1.3562506 | 0.57125736 | 0.74207764 | 1.36981589 | 0.45975692 | 0.59155253 | 0.70397191 | 0.84455829 | 0.69056391 | 0.78608851 |
| ENSMUSG00 Gm3906 | 2136 | 0 | 0 | 0 | 0 | 0 | 0 | 0.04430026 | 0.17880338 | 0 | 0 | 0.06709966 | 0.13065464 | 0 | 0.08697395 | 0.17360206 | 0 | 0 | 0 |
| [illegible] | [illegible] | [illegible] | [illegible] | [illegible] | [illegible] | [illegible] | [illegible] | [illegible] | [illegible] | [illegible] | [illegible] | [illegible] | [illegible] | [illegible] | [illegible] | [illegible] | [illegible] | [illegible] | [illegible] |

| | | | | | | | | | | | | | | | | | | | |
|---|---|---|---|---|---|---|---|---|---|---|---|---|---|---|---|---|---|---|---|
| ENSMUSG00 Gm9566 | 528 | 0 | 0 | 0 | 0 | 0 | 0 | 0 | 0 | 0 | 0 | 0 | 0 | 0 | 0 | 0 | 0 | 0 | 0 |
| ENSMUSG00 Gm4189 | 428 | 0.79448927 | 0.54483705 | 1.24324494 | 1.49031453 | 0.89966212 | 0.57888707 | 0.7738055 | 0.53540751 | 0.96230177 | 0.66099117 | 1.22786108 | 0.65205213 | 2.0747068 | 3.14691246 | 1.34771445 | 0.89464036 | 0.89547423 | 2.0564152 |
| ENSMUSG00 Gm20401 | 209 | 0 | 0 | 0 | 0 | 0 | 0 | 0 | 0 | 0 | 0 | 0 | 0 | 0 | 0 | 0 | 0 | 0 | 0 |
| ENSMUSG00 Gm20536 | 404 | 0 | 0 | 0 | 0 | 0 | 0 | 0 | 0 | 0 | 0 | 0 | 0 | 0 | 0 | 0 | 0 | 0 | 0 |
| ENSMUSG00 Gm7648 | 1654 | 0 | 0 | 0 | 0 | 0 | 0 | 0 | 0 | 0 | 0 | 0 | 0 | 0 | 0 | 0 | 0 | 0 | 0 |
| ENSMUSG00 Olfr306-ps1 | 627 | 0 | 0 | 0 | 0 | 0 | 0 | 0 | 0 | 0 | 0 | 0 | 0 | 0 | 0 | 0 | 0 | 0 | 0 |
| ENSMUSG00 Gm20483 | 287 | 0 | 0 | 0 | 0 | 0 | 0 | 0 | 0 | 0 | 0 | 0 | 0 | 0 | 0 | 0 | 0 | 0 | 0 |
| ENSMUSG00 Gm20541 | 1403 | 0 | 0 | 0 | 0 | 0 | 0 | 0 | 0 | 0.14678017 | 0 | 0.034052 | 0 | 0 | 0 | 0.11746702 | 0.49125512 | 0 | 0 |
| ENSMUSG00 Gm4175 | 944 | 0 | 0 | 0 | 0 | 0 | 0 | 0 | 0 | 0 | 0 | 0 | 0 | 0 | 0 | 0 | 0 | 0 | 0 |
| ENSMUSG00 Gm20546 | 1885 | 0 | 0 | 0.0513247 | 0 | 0 | 0 | 0 | 0 | 0 | 0 | 0 | 0.08883129 | 0.07851234 | 0 | 0 | 0 | 0 | 0.26264295 |
| ENSMUSG00 Gm20399 | 137 | 0 | 0 | 0 | 0 | 0 | 0 | 0 | 0 | 0 | 0 | 0 | 0 | 0 | 0 | 0 | 0 | 0 | 0 |
| ENSMUSG00 Gm20445 | 564 | 0 | 0 | 0 | 0 | 0 | 0 | 0 | 0 | 0 | 0 | 0 | 0 | 0 | 0 | 0.36526202 | 0 | 0 | 0 |
| ENSMUSG00 Gm20463 | 364 | 0 | 0 | 0 | 0 | 0 | 0 | 0 | 0 | 0 | 0 | 0 | 0 | 0 | 0 | 0 | 0 | 0 | 0 |
| ENSMUSG00 Olfr758-ps1 | 838 | 0 | 0 | 0 | 0 | 0 | 0 | 0 | 0 | 0 | 0 | 0 | 0 | 0 | 0 | 0 | 0 | 0 | 0 |
| ENSMUSG00 C130080G10 | 2860 | 0 | 0 | 0 | 0 | 0 | 0 | 0 | 0 | 0 | 0 | 0 | 0 | 0 | 0 | 0 | 0 | 0 | 0 |
| ENSMUSG00 Vmn1r-ps59 | 198 | 0 | 0 | 0 | 0 | 0 | 0 | 0 | 0 | 0 | 0 | 0 | 0 | 0 | 0 | 0 | 0 | 0 | 0 |
| ENSMUSG00 4930556N13 | 1029 | 0 | 0.09712214 | 0 | 0 | 0 | 0 | 0 | 0 | 0 | 0 | 0 | 0 | 0 | 0 | 0 | 0 | 0 | 0 |
| ENSMUSG00 Gm20469 | 3838 | 0 | 0 | 0 | 0 | 0 | 0 | 0 | 0 | 0 | 0 | 0 | 0 | 0 | 0 | 0 | 0 | 0 | 0 |
| ENSMUSG00 Olfr300-ps1 | 905 | 0 | 0 | 0 | 0 | 0 | 0 | 0 | 0 | 0 | 0 | 0 | 0 | 0 | 0 | 0 | 0 | 0 | 0 |
| ENSMUSG00 Gm20485 | 392 | 0 | 0 | 0 | 0 | 0 | 0 | 0 | 0 | 0 | 0 | 0 | 0 | 0 | 0 | 0 | 0 | 0 | 0 |
| ENSMUSG00 Gm20497 | 668 | 0 | 0.14960881 | 0 | 0 | 0.07205378 | 0.14836148 | 0 | 0 | 0 | 0 | 0 | 0 | 0 | 0.06952708 | 0 | 0 | 0 | 0.08234896 |
| ENSMUSG00 Gm9921 | 4397 | 0 | 0 | 0 | 0 | 0 | 0 | 0 | 0 | 0 | 0 | 0 | 0 | 0 | 0 | 0 | 0 | 0 | 0 |
| ENSMUSG00 Gm20402 | 685 | 0 | 0.14589589 | 0 | 0 | 0 | 0 | 0 | 0 | 0 | 0 | 0 | 0 | 0 | 0.06780159 | 0 | 0 | 0 | 0 |
| ENSMUSG00 Gm20462 | 407 | 0 | 0.08184986 | 0 | 0 | 0 | 0 | 0 | 0 | 0.10119537 | 0 | 0 | 0.41141766 | 0 | 0.34233971 | 0 | 0 | 0 | 0 |
| ENSMUSG00 Gm4040 | 2355 | 0 | 0 | 0 | 0 | 0 | 0 | 0 | 0 | 0 | 0 | 0 | 0 | 0 | 0 | 0 | 0 | 0 | 0 |
| ENSMUSG00 Vhl-ps1 | 596 | 0 | 0 | 0 | 0 | 0 | 0 | 0 | 0 | 0 | 0 | 0 | 0 | 0 | 0 | 0 | 0 | 0 | 0 |

[illegible]

| | | | | | | | | | | | | | | | | | | | | |
|---|---|---|---|---|---|---|---|---|---|---|---|---|---|---|---|---|---|---|---|---|
| ENSMUSG00 | AC161765.1 | 108 | 0 | 0 | 0 | 0 | 0 | 0 | 0 | 0 | 0 | 0 | 0 | 0 | 0 | 0 | 0 | 0 | 0 | 0 |
| ENSMUSG00 | Mir3091 | 75 | 0 | 0.88834385 | 1.28996081 | 1.21496118 | 0 | 1.98210934 | 1.26167144 | 3.05539219 | 1.64746063 | 1.25735209 | 3.18499723 | 1.48841767 | 0.6577589 | 0 | 2.19741631 | 3.57379002 | 2.04406917 | 0 |
| ENSMUSG00 | AC154187.1 | 82 | 0 | 0 | 0 | 0 | 0 | 0 | 0 | 0 | 0 | 0 | 0 | 0 | 0 | 0 | 0 | 0 | 0 | 0 |
| ENSMUSG00 | AL954675.1 | 76 | 0 | 0 | 0 | 0 | 0 | 0 | 0 | 0 | 0 | 0 | 0 | 0 | 0 | 0 | 0 | 0 | 0 | 0 |
| ENSMUSG00 | Mir3963 | 70 | 0 | 0 | 0 | 0 | 0 | 0 | 0 | 0 | 0 | 0 | 0 | 0 | 0 | 0 | 0 | 0 | 0 | 0 |
| ENSMUSG00 | AC114668.1 | 288 | 0.59034967 | 0.11566977 | 0 | 0 | 0 | 0 | 0 | 0.26522502 | 0 | 0 | 0.16588527 | 0 | 0.17129138 | 0 | 0 | 0 | 0.26615484 | 0.19100384 |
| ENSMUSG00 | Mir3088 | 73 | 0 | 0 | 0 | 0 | 0 | 0 | 0 | 0 | 0 | 0 | 0 | 0 | 0 | 0 | 0 | 0 | 0 | 0 |
| ENSMUSG00 | AL808119.1 | 90 | 0 | 0 | 0 | 0 | 0 | 0 | 0 | 0 | 0 | 0 | 0 | 0 | 0 | 0 | 0 | 0 | 0 | 0 |
| ENSMUSG00 | AC114406.1 | 160 | 0 | 0 | 0 | 0 | 0 | 0 | 0 | 0 | 0 | 0 | 0 | 0 | 0 | 0 | 0 | 0 | 0 | 0 |
| ENSMUSG00 | AC154848.1 | 165 | 0 | 0 | 0 | 0 | 0 | 0 | 0 | 0 | 0 | 0 | 0 | 0 | 1.19592527 | 0 | 0 | 0 | 0 | 0 |
| ENSMUSG00 | AC103359.1 | 225 | 0 | 0 | 0 | 0 | 0 | 0 | 0 | 0 | 0 | 0 | 0 | 0 | 0 | 0 | 0 | 0 | 0 | 0 |
| ENSMUSG00 | AC027184.1 | 318 | 0 | 0 | 0 | 0 | 0 | 0 | 0.29756402 | 0 | 0 | 0.2965453 | 0 | 0.52656285 | 0.31026363 | 1.02235412 | 0 | 0.12041072 | 0.36156884 | 0 |
| ENSMUSG00 | AL671885.1 | 77 | 0 | 0 | 0 | 0 | 0 | 0 | 0 | 0 | 0 | 0 | 0 | 0 | 0 | 0 | 0 | 0 | 0 | 0 |
| ENSMUSG00 | AL606908.1 | 294 | 0 | 0 | 0 | 0 | 0.16371403 | 0 | 0 | 0 | 0.56036076 | 0.16037654 | 0.16249986 | 0.18984919 | 0 | 0 | 0 | 0 | 0 | 0 |
| ENSMUSG00 | CT033785.1 | 66 | 0 | 0 | 0 | 0 | 0 | 0 | 0 | 0 | 0 | 0 | 0 | 0 | 0 | 0 | 0 | 0 | 0 | 0 |
| ENSMUSG00 | AC124992.1 | 75 | 0 | 0 | 0.6449804 | 1.21496118 | 0.64175898 | 0.66070311 | 1.89250716 | 0 | 0 | 0.62867604 | 0 | 0.74420883 | 1.31551779 | 3.09627249 | 0.54935408 | 1.02108286 | 0.51101729 | 2.20036427 |
| ENSMUSG00 | AC102287.1 | 83 | 0 | 0 | 0 | 0 | 0 | 0 | 0 | 0 | 0 | 0 | 0 | 0 | 0 | 0 | 0 | 0 | 0 | 0 |
| ENSMUSG00 | AL935159.1 | 101 | 0 | 0 | 0 | 0 | 0 | 0 | 0 | 0 | 0 | 0 | 0 | 0 | 0 | 0 | 0 | 0 | 0 | 0 |
| ENSMUSG00 | AL592465.1 | 78 | 0 | 0 | 0 | 0 | 0 | 0 | 0 | 0 | 0 | 0 | 0 | 0 | 0 | 0 | 0 | 0 | 0 | 0 |
| ENSMUSG00 | AL732585.1 | 66 | 0 | 2.52370411 | 1.46586456 | 0 | 0 | 1.50159799 | 5.0180114 | 0 | 1.24807624 | 3.57202297 | 0 | 0 | 0 | 0.70369829 | 2.49706399 | 4.06112503 | 1.16140294 | 1.66694263 |
| ENSMUSG00 | AC107711.1 | 70 | 0 | 0 | 0 | 0 | 0 | 0 | 0 | 0 | 0 | 0 | 0 | 0 | 0 | 0 | 0 | 0 | 0 | 0 |
| ENSMUSG00 | Mir344d-1 | 68 | 0 | 0 | 0 | 0 | 0 | 0 | 0 | 0 | 0 | 0 | 0 | 0 | 0 | 0 | 0 | 0 | 0 | 0 |
| ENSMUSG00 | AL671011.1 | 75 | 0.56673568 | 0 | 0 | 0 | 0 | 0 | 0 | 0 | 0 | 0 | 0 | 0 | 0 | 0 | 0 | 0 | 0 | 0 |
| ENSMUSG00 | AL831766.1 | 77 | 0 | 0 | 0 | 0 | 0 | 0 | 0 | 0 | 0 | 0 | 0 | 0 | 0 | 0 | 0 | 0 | 0 | 0 |
| ENSMUSG00 | Mir344g | 70 | 0 | 0 | 0 | 0 | 0 | 0 | 0 | 0 | 0 | 0 | 0 | 0 | 0 | 0 | 0 | 0 | 0 | 0 |
| ENSMUSG00 | AL844840.1 | 124 | 0 | 0 | 0 | 0.36742778 | 0 | 0 | 0 | 0 | 0 | 0 | 0 | 0 | 0 | 0 | 0 | 0 | 0 | 0 |
| ENSMUSG00 | Mir5103 | 78 | 0 | 3.4167071 | 0 | 0 | 2.46830378 | 0 | 0 | 0 | 0 | 0 | 0 | 0 | 0 | 0 | 0 | 0.98181045 | 0 | 0 |
| ENSMUSG00 | AL772185.1 | 101 | 0 | 0 | 0 | 0 | 0 | 0 | 0 | 0 | 0 | 0 | 0 | 0 | 0 | 0 | 0 | 0 | 0 | 0 |
| ENSMUSG00 | AC124745.1 | 74 | 0 | 0 | 0 | 0 | 0 | 0 | 0 | 0 | 0 | 0 | 0 | 0 | 0 | 0 | 0 | 0 | 0 | 0 |
| ENSMUSG00 | AC152394.1 | 70 | 0 | 0 | 0 | 0 | 0 | 0 | 0 | 0 | 0 | 0 | 0 | 0 | 0 | 0 | 0 | 0 | 0 | 0 |
| ENSMUSG00 | AC099728.1 | 204 | 0 | 0 | 0 | 0 | 0 | 0 | 0 | 0 | 0 | 0 | 0 | 0 | 0 | 0 | 0 | 0 | 0 | 0 |
| ENSMUSG00 | AC158898.2 | 79 | 0 | 0 | 0 | 0 | 0 | 0 | 0 | 0 | 0 | 0 | 0 | 0 | 0 | 0 | 0 | 0 | 0 | 0 |
| ENSMUSG00 | AL844147.1 | 66 | 0 | 0 | 0 | 0 | 0 | 0 | 0 | 0 | 0 | 0 | 0 | 0 | 0 | 0 | 0 | 0 | 0 | 0 |
| ENSMUSG00 | AC139317.1 | 86 | 0 | 0 | 0 | 0 | 0 | 0 | 0 | 0 | 0 | 0 | 0 | 0 | 0 | 0 | 0 | 0 | 0 | 0 |
| ENSMUSG00 | CR956641.1 | 294 | 0 | 0 | 0 | 0 | 0 | 0 | 0.16092748 | 0 | 0 | 0 | 0 | 0 | 0 | 0 | 0 | 0 | 0 | 0 |
| ENSMUSG00 | Mir3094 | 63 | 0 | 0 | 1.53566763 | 3.6159559 | 0 | 3.93275663 | 0.7509949 | 1.81868583 | 0 | 0.74842386 | 0.75833267 | 1.7719258 | 0 | 0 | 0 | 0 | 1.21670784 | 0 |
| ENSMUSG00 | BX000694.1 | 271 | 0 | 0 | 0 | 0 | 0 | 0 | 0 | 0 | 0 | 0 | 0 | 0 | 0 | 0 | 0 | 0 | 0 | 0 |
| ENSMUSG00 | AC133502.1 | 70 | 0 | 0 | 0 | 0 | 0 | 0 | 0 | 0 | 0 | 0 | 0 | 0 | 0 | 0 | 0 | 0 | 0 | 0 |
| ENSMUSG00 | AL833775.1 | 82 | 0 | 0 | 1.1798422 | 0 | 0 | 0 | 0 | 1.39728301 | 0 | 0 | 1.16524289 | 0 | 0 | 0 | 0 | 0 | 0 | 0 |
| ENSMUSG00 | AC144794.1 | 184 | 0 | 0 | 0 | 0 | 0 | 0 | 0 | 0 | 0 | 0 | 0 | 0 | 0 | 0 | 0 | 0 | 0 | 0 |
| ENSMUSG00 | Mir3058 | 90 | 0 | 0 | 0 | 0 | 0 | 0 | 0 | 0 | 0 | 0 | 0 | 0 | 0 | 0 | 0 | 0 | 0 | 0 |
| ENSMUSG00 | Mir5113 | 81 | 0 | 0 | 0 | 0 | 0 | 0 | 0 | 0 | 0 | 0 | 0 | 0 | 0 | 0 | 0 | 0 | 0 | 0 |
| ENSMUSG00 | AL808027.2 | 84 | 0 | 3.17265659 | 0 | 2.16957354 | 0.57299909 | 1.17982699 | 3.37947707 | 0 | 0 | 0.5613179 | 0 | 0 | 0 | 0 | 0 | 0 | 1.36879632 | 2.61948127 |
| ENSMUSG00 | Mir3061 | 90 | 0 | 0 | 0 | 0 | 0 | 0 | 0 | 0 | 0 | 0 | 0 | 0 | 0 | 0 | 0 | 0 | 0 | 0 |
| ENSMUSG00 | AC138790.1 | 114 | 0 | 0 | 0 | 0 | 0 | 0 | 0 | 0 | 0 | 0 | 0 | 0 | 0 | 0 | 0 | 0 | 0 | 0 |
| ENSMUSG00 | AL672094.1 | 69 | 0 | 0 | 0 | 0 | 0 | 0 | 0 | 0 | 0 | 0 | 0 | 0 | 0 | 0 | 0 | 0 | 0 | 0 |
| ENSMUSG00 | AC110520.1 | 105 | 0 | 0 | 0 | 0 | 0 | 0 | 0 | 0 | 0 | 0 | 0 | 0 | 0 | 0 | 0 | 0 | 0 | 0 |
| ENSMUSG00 | AL645950.1 | 125 | 0 | 0 | 0 | 0 | 0 | 0 | 0 | 0 | 0 | 0 | 0 | 0 | 0 | 0 | 0 | 0 | 0 | 0 |
| ENSMUSG00 | AC149277.1 | 298 | 0 | 0 | 0 | 0 | 0 | 0 | 0 | 0 | 0 | 0 | 0 | 0 | 0 | 0 | 0 | 0 | 0 | 0 |
| ENSMUSG00 | AC044864.1 | 107 | 0 | 0 | 0 | 0 | 0 | 0 | 0 | 0 | 0 | 0 | 0 | 0 | 0 | 0 | 0 | 0 | 0 | 0 |
| ENSMUSG00 | AC166251.1 | 95 | 1.34226872 | 0 | 0 | 0 | 0 | 0 | 0 | 0 | 0.43354227 | 0 | 0 | 0 | 0 | 0 | 0 | 0 | 0 | 0 |
| ENSMUSG00 | AL590633.1 | 65 | 0 | 0 | 0 | 0 | 0 | 0 | 0 | 0 | 0 | 0 | 0 | 0 | 0 | 0 | 0 | 0 | 0 | 0 |
| ENSMUSG00 | BX005219.1 | 104 | 0 | 0 | 0 | 0 | 0 | 0 | 0 | 0 | 0 | 0 | 0 | 0 | 0 | 0 | 0 | 0 | 0 | 0 |
| ENSMUSG00 | Mir3471-1 | 130 | 0 | 0 | 0 | 0 | 0 | 0 | 0 | 0 | 0 | 0 | 0 | 0 | 0 | 0 | 0 | 0 | 0 | 0 |
| ENSMUSG00 | AC148768.1 | 100 | 0 | 0 | 0 | 0 | 0 | 0 | 0 | 0 | 0 | 0 | 0 | 0 | 0 | 0 | 0 | 0 | 0 | 0 |
| ENSMUSG00 | AL772216.41 | 126 | 0 | 0 | 0 | 0 | 0 | 0 | 0 | 0 | 0 | 0 | 0 | 0 | 0 | 0 | 0 | 0 | 0 | 0 |
| ENSMUSG00 | AC121802.1 | 70 | 0 | 0 | 0 | 0 | 0 | 0 | 0 | 0 | 0 | 0 | 0 | 0 | 0 | 0.66348696 | 0 | 0 | 0 | 0 |
| ENSMUSG00 | AC125484.1 | 70 | 0 | 0 | 0 | 0 | 0 | 0 | 0 | 0 | 0 | 0 | 0 | 0 | 0 | 0 | 0.58859366 | 0 | 0 | 0 |
| ENSMUSG00 | AL646020.1 | 52 | 10.626294 | 12.8126516 | 28.8380662 | 15.7711307 | 12.9585948 | 8.57643466 | 13.6478882 | 19.8306705 | 11.8807257 | 21.761863 | 17.4562348 | 11.8071594 | 18.9738143 | 8.03839974 | 7.92337613 | 8.09993618 | 7.37044171 | 9.52080693 |
| ENSMUSG00 | BX470220.1 | 69 | 0 | 0 | 0 | 0 | 0 | 0 | 0 | 0 | 0 | 0 | 0 | 0 | 0 | 0 | 0 | 0 | 0 | 0 |
| ENSMUSG00 | Snord53 | 77 | 1.10403055 | 0 | 0 | 0 | 0.62508992 | 0 | 0 | 0 | 0 | 0 | 0 | 0 | 0 | 0 | 1.07017028 | 0 | 0 | 0 |
| ENSMUSG00 | AC162464.1 | 199 | 0 | 0 | 0.24308307 | 0 | 0 | 0 | 0 | 0 | 0 | 0 | 0 | 0 | 0 | 0 | 0 | 0.19241511 | 0 | 0 |
| ENSMUSG00 | AL669944.1 | 53 | 8.01984453 | 10.0567228 | 12.7779137 | 6.01749642 | 4.54074752 | 3.73982895 | 12.4976888 | 8.64733639 | 4.66262443 | 6.22745137 | 12.6198004 | 8.42500567 | 0.93079089 | 11.391946 | 6.99649062 | 11.5594286 | 9.40078981 | 2.07581535 |
| ENSMUSG00 | AC120002.1 | 70 | 0 | 0 | 0 | 0 | 0 | 0 | 0 | 0 | 0 | 0 | 0 | 0 | 0 | 0 | 0 | 0 | 0 | 0 |
| ENSMUSG00 | AC163023.1 | 73 | 0 | 0 | 0 | 0 | 0 | 0 | 0 | 0 | 0 | 0 | 0 | 0 | 0 | 0 | 0 | 0 | 0 | 0 |
| ENSMUSG00 | Mir344d-3 | 79 | 0 | 0 | 0 | 0 | 0 | 0 | 0 | 0 | 0 | 0 | 0 | 0 | 0 | 0 | 0 | 0 | 0 | 0 |
| ENSMUSG00 | AC124489.1 | 259 | 0 | 0.38586364 | 0 | 0.70364547 | 0.55751263 | 0.57396989 | 0.36534887 | 0.14746101 | 0.31804259 | 1.27434333 | 0.1844593 | 0 | 0.76188289 | 0.5379624 | 0.31815873 | 0.29568037 | 0.14797798 | 0 |
| ENSMUSG00 | AL606479.2 | 76 | 0 | 0 | 0 | 0 | 0 | 0 | 0 | 0 | 0 | 0 | 0 | 0 | 0 | 0 | 0 | 0 | 0 | 0 |
| ENSMUSG00 | AC127086.1 | 100 | 0 | 0 | 0 | 0 | 0 | 0 | 0 | 0 | 0 | 0 | 0 | 0 | 0 | 0 | 0 | 0 | 0 | 0 |
| ENSMUSG00 | CT030685.1 | 80 | 0 | 0 | 0 | 0 | 0 | 0 | 0 | 0 | 0 | 0 | 0 | 0 | 0 | 0 | 0 | 0 | 0 | 0 |
| ENSMUSG00 | AC156551.1 | 83 | 0 | 0 | 0 | 0 | 0 | 0 | 0 | 0 | 0 | 0 | 0 | 0 | 0 | 0 | 0 | 0 | 0 | 0 |
| ENSMUSG00 | AL672284.1 | 108 | 0 | 0 | 0 | 0 | 0 | 0 | 0 | 0 | 0 | 0 | 0 | 0 | 0 | 0 | 0 | 0 | 0 | 0 |
| ENSMUSG00 | AC113276.1 | 90 | 0 | 0 | 0 | 0 | 0 | 0 | 0 | 0 | 0 | 0 | 0 | 0 | 0 | 0 | 0 | 0 | 0 | 0 |
| ENSMUSG00 | AC134897.1 | 101 | 0 | 0 | 0 | 0 | 0 | 0 | 0 | 0 | 0 | 0 | 0 | 0 | 0 | 0 | 0 | 0 | 0 | 0 |
| ENSMUSG00 | Mir5130 | 83 | 0 | 0 | 0 | 0 | 0 | 0 | 0 | 0 | 0 | 0 | 0 | 0 | 0 | 0 | 0 | 0 | 0 | 0 |
| ENSMUSG00 | AC132606.1 | 125 | 0 | 0 | 0 | 0 | 0 | 0 | 0 | 0 | 0.32949213 | 0 | 0 | 0 | 1.57862135 | 0 | 0 | 0 | 0 | 0 |
| ENSMUSG00 | AC142212.1 | 302 | 0 | 0 | 0 | 0 | 0 | 0 | 0.1566645 | 0 | 0 | 0 | 0 | 0 | 0 | 0 | 0 | 0 | 0 | 0 |
| ENSMUSG00 | AC124426.2 | 109 | 0 | 0 | 0 | 0 | 0 | 0 | 0 | 0 | 0 | 0 | 0 | 0 | 0 | 0 | 0 | 0 | 0 | 0 |
| ENSMUSG00 | AC122537.1 | 212 | 0 | 0 | 0 | 0 | 0 | 0 | 0 | 0 | 0 | 0 | 0 | 0 | 0 | 0 | 0 | 0 | 0 | 0 |
| ENSMUSG00 | AC098743.1 | 100 | 0 | 0 | 0 | 0 | 0 | 0 | 0 | 0 | 0 | 0 | 0 | 0 | 0 | 0 | 0 | 0 | 0 | 0 |
| ENSMUSG00 | Mir5132 | 70 | 0 | 0 | 0 | 0 | 0 | 0 | 0 | 0 | 0 | 0 | 0 | 0 | 0 | 0 | 0 | 0 | 0 | 0 |
| ENSMUSG00 | CT030729.1 | 124 | 0 | 0 | 0 | 0 | 0 | 0 | 0 | 0 | 0 | 0 | 0 | 0 | 0 | 0 | 0 | 0 | 0 | 0 |
| ENSMUSG00 | AL807811.1 | 59 | 0 | 0 | 0 | 0 | 0 | 0 | 0 | 0 | 0 | 0 | 0 | 0 | 0 | 0 | 0 | 0 | 0 | 0 |
| ENSMUSG00 | AC146595.1 | 101 | 0 | 0 | 0 | 0 | 0 | 0 | 0 | 0 | 0 | 0 | 0 | 0 | 0 | 0 | 0 | 0 | 0 | 0 |
| ENSMUSG00 | AC137107.1 | 72 | 1.18069933 | 0 | 0.67185459 | 0 | 0 | 0.68823241 | 0 | 0 | 0 | 0 | 1.99062327 | 0 | 0 | 3.22528385 | 0 | 0 | 0 | 0 |
| ENSMUSG00 | AC125186.1 | 97 | 0 | 0 | 0 | 0 | 0 | 0 | 0 | 0 | 0 | 0 | 0 | 0 | 0 | 0 | 0 | 0 | 0 | 0 |
| ENSMUSG00 | AL589678.1 | 115 | 0 | 0 | 0 | 0 | 0 | 0 | 0 | 0 | 0 | 0 | 0 | 0 | 0 | 0 | 0 | 0 | 0 | 0 |
| ENSMUSG00 | AC153950.1 | 136 | 0 | 0 | 0 | 0 | 0 | 0 | 0 | 0 | 0 | 0 | 0 | 0 | 0 | 0 | 0 | 0 | 0 | 0 |
| ENSMUSG00 | CT010508.1 | 119 | 1.07155906 | 1.95958201 | 0.40650026 | 1.14859776 | 0.40446995 | 1.6656381 | 1.19275661 | 0.96283367 | 0 | 1.98112199 | 0 | 0.46903918 | 2.07276963 | 1.17085935 | 4.50101031 | 1.93061886 | 1.28827889 | 0 |
| ENSMUSG00 | AC125115.1 | 82 | 0 | 0 | 0 | 0 | 0 | 0 | 0 | 0 | 0 | 0 | 0 | 0 | 0 | 0 | 0 | 0 | 0 | 0 |
| ENSMUSG00 | Mir3472 | 70 | 0 | 0 | 0 | 0 | 0 | 0 | 0 | 0 | 0 | 0 | 0 | 0 | 0 | 0 | 0 | 0 | 0 | 0 |
| ENSMUSG00 | AL928590.1 | 124 | 0 | 0 | 0 | 0 | 0 | 0.39961882 | 0 | 0 | 0 | 0 | 0 | 0 | 0 | 0 | 0 | 0 | 0 | 0 |
| ENSMUSG00 | AC124226.1 | 225 | 0 | 0 | 0 | 0 | 0 | 0 | 0 | 0 | 0 | 0 | 0 | 0 | 0 | 0 | 0 | 0 | 0 | 0 |
| ENSMUSG00 | AC110211.1 | 284 | 0 | 0 | 0 | 0 | 0 | 0 | 0 | 0 | 0 | 0 | 0 | 0 | 0 | 0 | 0 | 0 | 0 | 0 |
| ENSMUSG00 | Mir344b | 62 | 0 | 0 | 0 | 0 | 0 | 0 | 0 | 0 | 0 | 0 | 0 | 0 | 0 | 0 | 0 | 0 | 0 | 0 |
| ENSMUSG00 | Mir5101 | 82 | 3.11013483 | 0.81250962 | 0 | 0 | 1.17394936 | 0 | 0.57698389 | 0.465761 | 2.00909833 | 1.15001715 | 0 | 1.36135762 | 0 | 0 | 0 | 0 | 3.73915092 | 0 |
| ENSMUSG00 | BX005039.1 | 94 | 0 | 0.35439249 | 0 | 0 | 0 | 1.05431348 | 1.00665274 | 0.40630215 | 0.87630885 | 2.0064129 | 0.50824424 | 0.59378364 | 0 | 0.49408604 | 0 | 0 | 0 | 0 |
| ENSMUSG00 | AL806523.1 | 107 | 0.79448927 | 0 | 0 | 0 | 0 | 0 | 0 | 0 | 1.53968283 | 0.44066078 | 0 | 0 | 0 | 0 | 0 | 0 | 0 | 0 |
| ENSMUSG00 | Mir3960 | 72 | 0 | 0 | 0 | 0 | 0 | 0 | 0 | 0 | 0 | 0 | 0 | 0 | 0 | 0 | 0 | 0 | 0 | 0 |
| ENSMUSG00 | AC154229.1 | 125 | 0 | 0 | 0 | 0 | 0 | 0 | 0 | 0 | 0 | 0.75441125 | 0 | 0 | 0 | 0 | 0 | 0 | 0 | 0 |
| ENSMUSG00 | AC166290.1 | 92 | 0 | 0 | 0 | 0 | 0 | 0 | 0 | 0 | 0 | 0 | 0 | 0 | 0 | 0 | 0 | 0 | 0 | 0 |
| ENSMUSG00 | AC141881.3 | 118 | 0 | 0 | 0 | 0 | 0 | 0 | 0 | 0 | 0 | 0 | 0 | 0 | 0 | 0 | 0 | 0 | 0 | 0 |
| ENSMUSG00 | Mir3471-2 | 101 | 0 | 0 | 0 | 0 | 0 | 0 | 0 | 0 | 0 | 0.93367729 | 0 | 0 | 0 | 0 | 0 | 0 | 0 | 0 |
| ENSMUSG00 | AC132354.1 | 157 | 0 | 0 | 0 | 0 | 0 | 0 | 0 | 0 | 0 | 0 | 0 | 0 | 0 | 0 | 0 | 0 | 0 | 0 |
| ENSMUSG00 | AC161342.2 | 247 | 0 | 0 | 0 | 0 | 0 | 0 | 0 | 0 | 0 | 0 | 0 | 0 | 0 | 0 | 0 | 0 | 0 | 0 |
| ENSMUSG00 | AC114657.1 | 74 | 0 | 0 | 0 | 0 | 0 | 0 | 0 | 0 | 0 | 0 | 0 | 0 | 0 | 0 | 0 | 0 | 0 | 0 |
| ENSMUSG00 | AC134918.1 | 101 | 0 | 0 | 0 | 0 | 0 | 0 | 0 | 0 | 0 | 0 | 0 | 0 | 0 | 0 | 0 | 0 | 0 | 0 |
| ENSMUSG00 | AC125110.1 | 62 | 0 | 0 | 0 | 0 | 0 | 0 | 0 | 0 | 0 | 0 | 0 | 0 | 0 | 0 | 0 | 0 | 0 | 0 |
| ENSMUSG00 | AC153877.1 | 76 | 0 | 0 | 0 | 0 | 0 | 0 | 0 | 0 | 0 | 0 | 0 | 0 | 0 | 0 | 0 | 0 | 0 | 0 |
| ENSMUSG00 | AC158384.1 | 77 | 0.55201527 | 0 | 0 | 0 | 0 | 1.28708399 | 0 | 0 | 0 | 0 | 0 | 0 | 1.2813485 | 0 | 0 | 0.99456123 | 0 | 0 |
| ENSMUSG00 | CAAA010105[illegible] | 100 | 0 | 0 | 0 | 0 | 0 | 0 | 0 | 0 | 0 | 0 | 0 | 0 | 0 | 0 | 0 | 0 | 0 | 0 |
| ENSMUSG00 | AC015535.1 | 125 | 0 | 0 | 0 | 0 | 0 | 0 | 0 | 0 | 0 | 0 | 0 | 0 | 0 | 0 | 0 | 0 | 0 | 0 |
| ENSMUSG00 | AC117662.1 | 69 | 0 | 0 | 0 | 0 | 0 | 0 | 0 | 0 | 0 | 0 | 0 | 0 | 0 | 0 | 0 | 0 | 0 | 0 |
| ENSMUSG00 | AC093346.1 | 295 | 0 | 0 | 0.16397807 | 0 | 0 | 0 | 0 | 0 | 0 | 0 | 0 | 0 | 0 | 0 | 0 | 0 | 0 | 0 |
| ENSMUSG00 | AL591712.1 | 299 | 0 | 0 | 0 | 0 | 0 | 0 | 0 | 0 | 0 | 0 | 0 | 0 | 0 | 0 | 0 | 0 | 0 | 0 |
| ENSMUSG00 | AC159649.1 | 276 | 0 | 0 | 0 | 0 | 0 | 0 | 0 | 0 | 0 | 0 | 0 | 0 | 0 | 0 | 0 | 0 | 0 | 0 |
| ENSMUSG00 | AC119249.1 | 79 | 0 | 0 | 0 | 0 | 0 | 0 | 0 | 0 | 0 | 0 | 0 | 0 | 0 | 0 | 0 | 0 | 0 | 0 |
| ENSMUSG00 | AC140327.1 | 249 | 0 | 0 | 0 | 0 | 0 | 0 | 0 | 0 | 0 | 0 | 0 | 0 | 0 | 0.18652244 | 0 | 0 | 0 | 0 |
| ENSMUSG00 | Mir3070a | 88 | 0 | 0 | 0 | 0 | 0 | 0 | 0 | 0 | 0 | 0 | 0 | 0 | 0 | 0 | 0 | 0 | 0 | 0 |
| ENSMUSG00 | AC159140.1 | 259 | 0 | 0 | 0 | 0 | 0 | 0 | 0 | 0 | 0 | 0 | 0 | 0 | 0 | 0 | 0 | 0 | 0 | 0 |
| ENSMUSG00 | AC122189.1 | 71 | 0 | 0 | 0 | 0 | 0 | 0 | 0 | 0 | 0 | 0 | 0 | 0 | 0 | 0 | 0 | 0 | 0 | 0 |
| ENSMUSG00 | AC159624.1 | 90 | 0 | 0 | 0 | 0 | 0 | 0 | 0 | 0 | 0 | 0 | 0 | 0 | 0 | 0 | 0 | 0 | 0 | 0 |
| ENSMUSG00 | AL662901.1 | 80 | 0 | 0 | 0 | 0 | 0 | 0 | 0 | 0 | 0 | 0 | 0 | 0 | 0 | 0 | 0 | 0 | 0 | 0 |
| ENSMUSG00 | AL928889.1 | 287 | 0 | 0 | 0 | 0 | 0 | 0 | 0 | 0 | 0 | 0 | 0 | 0 | 0 | 0 | 0 | 0 | 0 | 0 |
| ENSMUSG00 | Mir3059 | 80 | 0 | 0 | 0 | 0 | 0 | 0 | 0 | 0 | 0 | 0 | 0 | 0 | 0 | 0 | 0 | 0 | 0 | 0 |
| ENSMUSG00 | AC162893.1 | 83 | 0 | 0 | 2.91406809 | 0 | 0 | 0 | 0 | 0.92029885 | 0 | 0 | 0 | 0 | 0 | 0 | 0 | 0 | 0.46176261 | 0 |
| ENSMUSG00 | AL808021.1 | 70 | 0 | 0 | 0.69105843 | 0 | 1.37519782 | 0 | 0 | 0 | 0 | 0.67358148 | 0 | 0 | 0.70474168 | 0 | 1.76578097 | 0.54700868 | 0 | 0 |
| ENSMUSG00 | AL732525.1 | 255 | 0 | 0 | 0 | 0 | 0 | 0 | 0 | 0 | 0 | 0 | 0 | 0 | 0 | 0 | 0 | 0 | 0.30059841 | 0 |
| ENSMUSG00 | AC044807.1 | 88 | 0 | 0 | 0 | 0 | 0 | 0 | 0 | 0 | 0 | 0 | 0 | 0 | 0 | 0 | 0 | 0 | 0 | 0 |
| ENSMUSG00 | AC084822.1 | 70 | 0 | 0 | 0 | 0 | 0 | 0 | 0 | 0 | 0 | 0 | 0 | 0 | 0 | 0 | 0 | 0 | 0 | 0 |
| ENSMUSG00 | Mir5128 | 83 | 0 | 0.80272034 | 0 | 0 | 0 | 2.38808355 | 0 | 0 | 0 | 0 | 1.72680573 | 0 | 0.59436045 | 0 | 0.49640429 | 0 | 0 | 0.66276032 |
| ENSMUSG00 | CT030685.2 | 66 | 0 | 0 | 0 | 0 | 0 | 0 | 0 | 0 | 0 | 0 | 0 | 0 | 0 | 0 | 0 | 0 | 0 | 0 |
| ENSMUSG00 | AC113329.1 | 70 | 0 | 0 | 0 | 0 | 0 | 0 | 0.67589541 | 0 | 0 | 0 | 0 | 0 | 0 | 0 | 0 | 0 | 0 | 0 |
| ENSMUSG00 | AL772216.42 | 102 | 0 | 0 | 0 | 0 | 0 | 0 | 0 | 0 | 0 | 0 | 0 | 0 | 0 | 0 | 0 | 0 | 0 | 0 |
| ENSMUSG00 | AC141881.4 | 118 | 0 | 0 | 0 | 0 | 0 | 0 | 0 | 0 | 0 | 0 | 0 | 0 | 0 | 0 | 0 | 0 | 0 | 0 |
| ENSMUSG00 | AL672070.1 | 87 | 0 | 0 | 0 | 0 | 0 | 0 | 0 | 0 | 0 | 0 | 0 | 0 | 0 | 0 | 0 | 0 | 0 | 0 |
| ENSMUSG00 | Mir3470a | 76 | 0 | 0 | 0 | 0 | 0 | 0 | 0 | 0 | 0 | 0 | 0 | 0 | 0 | 0 | 0 | 0 | 0 | 0 |
| ENSMUSG00 | AC091260.1 | 117 | 0 | 0 | 0 | 0 | 0 | 0 | 0 | 0 | 0 | 0 | 0 | 0 | 0 | 0 | 0 | 0 | 0 | 0 |
| ENSMUSG00 | AC126679.1 | 81 | 0 | 0 | 0 | 0 | 0 | 0 | 0 | 0 | 0 | 0 | 0 | 0 | 0 | 0 | 0 | 0 | 0 | 0 |
| ENSMUSG00 | AC167020.1 | 87 | 0 | 0 | 3.89212313 | 0 | 0.5532405 | 0 | 0 | 0 | 0.47340823 | 0 | 0 | 1.28311868 | 0 | 0 | 0 | 0 | 0.44053215 | 0 |
| ENSMUSG00 | AC118255.1 | 70 | 0 | 0 | 0 | 0 | 0 | 0 | 0 | 0 | 0 | 0 | 0 | 0 | 0 | 0 | 0.58859366 | 0 | 0 | 0 |
| ENSMUSG00 | AC160534.1 | 109 | 0 | 0 | 0 | 0 | 0 | 0 | 0 | 0 | 0 | 0 | 0 | 0 | 0 | 0 | 0 | 0 | 0 | 0 |
| ENSMUSG00 | AC091763.1 | 125 | 0 | 0.79950946 | 1.16096473 | 0 | 0 | 0 | 0.37850143 | 0 | 0 | 0 | 0 | 0.8930506 | 0.78931068 | 0.7431054 | 0.65922489 | 1.83794916 | 0.30661038 | 0.44007285 |
| ENSMUSG00 | AL833775.2 | 82 | 1.55506742 | 1.62501923 | 0 | 0 | 0 | 0 | 0 | 1.39728301 | 0 | 2.30003431 | 0 | 0 | 0 | 0 | 0 | 0 | 0 | 0 |
| ENSMUSG00 | Mir3078 | 86 | 0 | 0 | 0 | 0 | 0 | 0 | 0 | 0 | 0 | 0 | 0 | 0 | 0 | 0 | 0 | 0 | 0 | 0 |
| ENSMUSG00 | AC117949.1 | 69 | 0.61601704 | 0.48279557 | 0 | 1.32060998 | 0 | 0 | 0.685691 | 2.21405231 | 0.59690603 | 0 | 0 | 0.80892265 | 0 | 2.69241086 | 0.597124 | 0 | 0 | 0 |
| ENSMUSG00 | AL589670.1 | 232 | 0 | 0 | 0 | 0 | 0 | 0 | 0 | 0 | 0 | 0 | 0 | 0 | 0 | 0 | 0 | 0 | 0 | 0 |
| ENSMUSG00 | AC122252.1 | 125 | 0 | 0 | 0 | 0 | 0 | 0 | 0 | 0 | 0 | 0 | 0 | 0 | 0 | 0 | 0 | 0 | 0 | 0 |
| ENSMUSG00 | AC155948.1 | 100 | 0 | 0 | 0 | 0 | 0 | 0 | 0 | 0 | 0 | 0 | 0 | 0 | 0 | 0 | 0 | 0 | 0 | 0 |
| ENSMUSG00 | AC108434.1 | 251 | 0 | 0 | 0 | 0 | 0 | 0 | 0 | 0 | 0 | 0 | 0 | 0 | 0 | 0 | 0 | 0 | 0 | 0 |
| ENSMUSG00 | AC113178.1 | 101 | 0 | 0 | 0 | 0 | 0 | 0 | 0 | 0 | 0 | 0 | 0 | 0 | 0 | 0 | 0 | 0 | 0 | 0 |
| ENSMUSG00 | AC103598.1 | 85 | 0 | 0 | 0 | 0 | 0 | 0 | 0 | 0 | 0 | 0 | 0 | 0 | 0 | 0 | 0 | 0 | 0 | 0 |
| ENSMUSG00 | AC161246.1 | 295 | 0 | 0 | 0 | 0 | 0 | 0 | 0 | 0 | 0 | 0 | 0 | 0 | 0 | 0 | 0 | 0 | 0 | 0 |
| ENSMUSG00 | Mir5125 | 78 | 13.6234539 | 10.6772097 | 11.7832959 | 6.42527548 | 11.107367 | 10.7999548 | 3.63943684 | 5.38610803 | 13.2008063 | 13.2989163 | 7.96249308 | 10.7337813 | 3.79476287 | 11.9087404 | 6.86692598 | 10.3090097 | 9.33589284 | 24.6835735 |
| ENSMUSG00 | AC123853.1 | 55 | 0 | 0 | 0 | 0 | 0 | 0 | 0 | 0 | 0 | 0 | 0 | 0 | 0.89694395 | 0 | 1.4962384 | 0 | 0 | 0 |
| ENSMUSG00 | Mir5115 | 73 | 692.892596 | 681.773479 | 621.566733 | 473.085912 | 459.56097 | 539.649633 | 777.094548 | 671.244552 | 719.353529 | 762.162053 | 532.72351 | 660.612774 | 1533.34411 | 1510.38717 | 819.515879 | 792.56312 | 623.196088 | 1079.08275 |
| ENSMUSG00 | Mir378b | 91 | 1.86835939 | 0.36607576 | 0 | 0 | 0 | 1.6336066 | 3.63943684 | 3.77727057 | 0 | 0.5181396 | 2.09999817 | 3.06679465 | 3.79476287 | 1.02074917 | 0.9055287 | 1.26232772 | 1.26350429 | 0 |
| ENSMUSG00 | AC162799.1 | 246 | 0 | 0 | 0 | 0 | 0 | 0 | 0 | 0 | 0 | 0 | 0 | 0 | 0 | 0 | 0 | 0 | 0 | 0 |
| ENSMUSG00 | BX470216.1 | 206 | 0 | 0 | 0 | 0 | 0 | 0 | 0 | 0 | 0 | 0 | 0 | 0 | 0 | 0 | 0 | 0 | 0 | 0 |
| ENSMUSG00 | Mir3093 | 86 | 0 | 0 | 0 | 0 | 0 | 0 | 0 | 0 | 0 | 0 | 0 | 0 | 0 | 0 | 0 | 0 | 0 | 0 |
| ENSMUSG00 | AL596108.1 | 296 | 0 | 0 | 0 | 0 | 0 | 0 | 0 | 0 | 0 | 0 | 0 | 0 | 0 | 0 | 0 | 0 | 0.3884422 | 0 |
| ENSMUSG00 | Mir466q | 75 | 0 | 0 | 0 | 0 | 0.64175898 | 0 | 0 | 0 | 0 | 0 | 0 | 0 | 0 | 0 | 0 | 0 | 0 | 0 |
| ENSMUSG00 | AC158379.1 | 285 | 0 | 0 | 0 | 0 | 0 | 0.34773848 | 0 | 0 | 0 | 0 | 0 | 0 | 0 | 0 | 0 | 0 | 0 | 0 |
| ENSMUSG00 | AL731688.1 | 69 | 0 | 0 | 0 | 0 | 0 | 0 | 0 | 0 | 0 | 0 | 0 | 0 | 0 | 1.34620543 | 0 | 0 | 0 | 0 |
| ENSMUSG00 | AL772336.1 | 100 | 0 | 0 | 0 | 0 | 0 | 0 | 0 | 0 | 0 | 0 | 0 | 0 | 0 | 0 | 0 | 0 | 0 | 0 |
| ENSMUSG00 | AC113124.1 | 105 | 0 | 0.63453132 | 0 | 0 | 0 | 0 | 0 | 1.0912115 | 0 | 0 | 0 | 0 | 0 | 0 | 0 | 0 | 0 | 0 |
| ENSMUSG00 | Mir16-1 | 92 | 0 | 0 | 0 | 0 | 0 | 0 | 0 | 0 | 0 | 0 | 0 | 0 | 0 | 0 | 0 | 0 | 0 | 0 |
| ENSMUSG00 | AC119249.2 | 79 | 0 | 0 | 0 | 0 | 0 | 0 | 0 | 0 | 0 | 0 | 0 | 0 | 0 | 0 | 0 | 0 | 0 | 0 |
| ENSMUSG00 | AC091333.1 | 97 | 0 | 0 | 0 | 0 | 0 | 0 | 0 | 0 | 0 | 0 | 0 | 0 | 0 | 0 | 0 | 0 | 0 | 0 |
| ENSMUSG00 | Mir5099 | 73 | 0 | 0 | 0 | 0 | 0 | 0 | 0 | 0 | 0 | 0 | 0 | 0 | 0 | 0 | 0 | 0 | 0 | 0 |
| ENSMUSG00 | AC160471.1 | 106 | 0 | 0 | 0 | 0 | 0 | 0 | 0 | 0 | 0 | 0 | 0 | 0 | 0 | 0 | 0 | 0 | 0 | 0 |
| ENSMUSG00 | AC122183.1 | 109 | 0 | 0 | 0 | 0 | 0 | 0 | 0 | 0 | 0 | 0 | 0 | 0 | 0 | 0 | 0 | 0 | 0 | 0 |
| ENSMUSG00 | AL954388.2 | 312 | 0 | 0 | 0 | 0 | 0 | 0 | 0 | 0 | 0 | 0 | 0 | 0 | 0 | 0 | 0 | 0 | 0 | 0 |
| ENSMUSG00 | AC127694.1 | 129 | 0 | 0 | 0 | 0 | 0 | 0 | 0 | 0 | 0 | 0 | 0 | 0 | 0 | 0 | 0 | 0 | 0 | 0 |
| ENSMUSG00 | Mir3473c | 80 | 0 | 0 | 0 | 0 | 0 | 0 | 0 | 0 | 0 | 0 | 0 | 0 | 2.46659586 | 0 | 0 | 0 | 0 | 0 |
| ENSMUSG00 | AL596123.1 | 70 | 0 | 0 | 0 | 0 | 0 | 0 | 0 | 0 | 0 | 0 | 0 | 0 | 0 | 0 | 0 | 0 | 0 | 0 |
| ENSMUSG00 | AC138118.1 | 101 | 0 | 0 | 0 | 0 | 0 | 0 | 0 | 1.13442779 | 0 | 0 | 0 | 0 | 0 | 0 | 0 | 0 | 0 | 0 |
| ENSMUSG00 | AL935177.1 | 95 | 0.89484581 | 0 | 0 | 0 | 0 | 0 | 0 | 0 | 0 | 1.98529277 | 0 | 0 | 0 | 0 | 0 | 0 | 0 | 0 |
| ENSMUSG00 | Mir15a | 83 | 0 | 0.80272034 | 0 | 0 | 0 | 0 | 0 | 0 | 0 | 0 | 0 | 0 | 0 | 0 | 0 | 0 | 0 | 0 |
| ENSMUSG00 | Snord52 | 65 | 0 | 0 | 0 | 0 | 0 | 0 | 0 | 0 | 0 | 0 | 0 | 0 | 0 | 0 | 0 | 0 | 0 | 0 |
| ENSMUSG00 | AC139323.1 | 123 | 0 | 0 | 0 | 0 | 0 | 0 | 0 | 0 | 0 | 0 | 0 | 0 | 0 | 0 | 0 | 0 | 0 | 0 |
| ENSMUSG00 | Mir100 | 79 | 3.22824122 | 5.06018646 | 3.06161585 | 1.73016624 | 2.43705943 | 6.27249792 | 4.19226269 | 6.76827384 | 8.86292111 | 1.19368869 | 7.86170203 | 4.23916425 | 2.4978186 | 2.35159936 | 4.69384814 | 2.9081474 | 4.36628699 | 1.39263561 |
| ENSMUSG00 | AC122247.1 | 292 | 0 | 0 | 0 | 0 | 0 | 0 | 0 | 0 | 0 | 0 | 0 | 0 | 0 | 0 | 0 | 0 | 0 | 0 |
| ENSMUSG00 | AL591209.1 | 69 | 0 | 0 | 0 | 0 | 0 | 0 | 0 | 0 | 0 | 0 | 0 | 0 | 0 | 0 | 0 | 0 | 0 | 0 |
| ENSMUSG00 | AL732400.1 | 63 | 0 | 0 | 0 | 0 | 0 | 0 | 0 | 0 | 0 | 0 | 0 | 0 | 0 | 0 | 0 | 0 | 0 | 0 |
| ENSMUSG00 | AC103663.1 | 100 | 0 | 0 | 0 | 0 | 0 | 0 | 0 | 0 | 0 | 0 | 0 | 0 | 0 | 0 | 0 | 0 | 0 | 0 |
| ENSMUSG00 | AC122266.1 | 283 | 0.15019497 | 0 | 0 | 0 | 0 | 0 | 0 | 0 | 0.87321235 | 0 | 0.16881611 | 0.39445698 | 0.69727092 | 0 | 0.14558854 | 0.811815 | 0 | 0 |
| ENSMUSG00 | Mir5108 | 84 | 0 | 0 | 0 | 0 | 0 | 0 | 0 | 0 | 0 | 0 | 0 | 0 | 0 | 0 | 0 | 0 | 0 | 0 |
| ENSMUSG00 | AL672099.1 | 269 | 0 | 0 | 0 | 0 | 0 | 0 | 0 | 0 | 0 | 0 | 0 | 0 | 0 | 0 | 0 | 0 | 0 | 0 |
| ENSMUSG00 | AC119249.3 | 68 | 0 | 0 | 0 | 0 | 0 | 0 | 0 | 0 | 0 | 0 | 0 | 0 | 0 | 0 | 0 | 0 | 0 | 0 |
| ENSMUSG00 | AC149222.1 | 280 | 0 | 0 | 0 | 0 | 0 | 0 | 0 | 0 | 0 | 0 | 0 | 0 | 0 | 0 | 0 | 0 | 0 | 0 |
| ENSMUSG00 | AC138604.1 | 73 | 0 | 0 | 0 | 0 | 0 | 0 | 0.64811889 | 0 | 0 | 0 | 0 | 0 | 0 | 0 | 0 | 0 | 0 | 0 |
| ENSMUSG00 | AC127294.1 | 100 | 0 | 0 | 0 | 0 | 0 | 0 | 0 | 0 | 0 | 0 | 0 | 0 | 0 | 0 | 0 | 0 | 0 | 0 |
| ENSMUSG00 | AC142099.1 | 263 | 0 | 0.126665 | 0.73571909 | 0 | 0 | 0 | 0 | 0 | 0.78301361 | 0.53784072 | 0 | 0.63668056 | 0.56272149 | 0 | 0.31331982 | 0.29118333 | 0.58290946 | 0 |
| ENSMUSG00 | AL773580.1 | 101 | 0 | 0 | 0 | 0 | 0 | 0 | 0 | 0 | 0 | 0 | 0 | 0 | 0 | 0 | 0 | 0 | 0 | 0 |
| ENSMUSG00 | AC079245.1 | 83 | 0 | 0 | 0 | 0 | 0 | 0 | 0 | 0 | 0 | 0 | 0 | 0 | 0 | 0 | 0 | 0 | 0 | 0 |
| ENSMUSG00 | AC116595.1 | 112 | 0.3795105 | 0 | 1.72762608 | 1.22038512 | 0.42974932 | 0 | 0.84486927 | 0 | 0 | 0 | 0 | 0 | 0 | 0 | 0.36787103 | 0 | 0 | 0.49115274 |
| ENSMUSG00 | AC101777.1 | 120 | 0 | 0 | 0 | 0 | 0 | 0 | 0 | 0 | 0 | 0 | 0 | 0 | 0 | 0 | 0 | 0 | 0 | 0 |
| ENSMUSG00 | AL596246.1 | 229 | 0 | 0.43641346 | 0 | 0 | 0 | 0 | 0 | 0 | 0 | 0 | 0 | 0 | 0 | 0 | 0 | 0 | 0 | 0 |
| ENSMUSG00 | AC122043.1 | 116 | 0 | 0 | 0 | 0 | 0 | 0 | 0 | 0 | 0 | 0 | 0 | 0 | 0 | 0 | 0 | 0 | 0 | 0 |
| ENSMUSG00 | AC140381.1 | 124 | 0 | 0 | 0 | 0 | 0 | 0 | 0 | 0 | 0 | 0 | 0 | 0 | 0 | 0 | 0 | 0 | 0 | 0 |
| ENSMUSG00 | BX537327.1 | 253 | 0 | 0 | 0 | 0.36016636 | 0 | 0 | 0 | 0 | 0 | 0 | 0 | 0 | 0 | 0 | 0 | 0 | 0 | 0 |
| ENSMUSG00 | AC125372.1 | 122 | 0 | 0 | 0 | 0 | 0 | 0 | 0 | 0 | 0 | 0 | 0 | 0 | 0 | 0 | 0 | 0 | 0 | 0 |
| ENSMUSG00 | AL683878.1 | 103 | 0 | 0 | 0 | 0 | 0 | 0 | 0 | 0 | 0 | 0 | 0 | 0 | 0 | 0 | 0 | 0 | 0 | 0 |
| ENSMUSG00 | AC154327.1 | 118 | 0 | 0 | 0.40994517 | 0.38611055 | 0 | 0 | 0.40095491 | 0 | 0 | 0 | 0 | 0 | 0 | 0 | 0 | 0 | 0 | 0 |
| ENSMUSG00 | AC133597.1 | 70 | 0 | 0 | 0 | 0 | 0 | 0 | 0 | 0 | 0 | 0 | 0 | 0 | 0 | 0 | 0 | 0 | 0 | 0 |
| ENSMUSG00 | Mir5127 | 70 | 0 | 0 | 0 | 0 | 0 | 0 | 0 | 0 | 0 | 0 | 0 | 0 | 0 | 0 | 0 | 0 | 0 | 0 |
| ENSMUSG00 | Mir3066 | 82 | 0 | 0 | 0 | 0 | 0 | 0 | 0.57698389 | 0 | 0 | 0 | 0 | 0 | 0 | 0 | 0 | 1.40087588 | 0 | 0 |
| ENSMUSG00 | AC131596.1 | 100 | 0 | 0 | 0 | 0 | 0 | 0 | 0 | 0 | 0 | 0 | 0 | 0 | 0 | 0 | 0 | 0 | 0 | 0 |
| ENSMUSG00 | Mir1912 | 86 | 0 | 0 | 0 | 0 | 0 | 0 | 0 | 0 | 0 | 0 | 0 | 0 | 0 | 0 | 0 | 0 | 0 | 0 |
| ENSMUSG00 | AL663077.1 | 125 | 0 | 0 | 0 | 0 | 0 | 0 | 0 | 0 | 0 | 0 | 0 | 0 | 0 | 0 | 0 | 0 | 0 | 0 |
| ENSMUSG00 | AC132338.1 | 91 | 0 | 0 | 0 | 0 | 0 | 0 | 0 | 0 | 0 | 0 | 0 | 0 | 0 | 0 | 0 | 0 | 0 | 0 |
| ENSMUSG00 | Mir3109 | 87 | 0 | 0 | 0 | 0 | 0 | 0 | 0 | 0 | 0 | 0 | 0 | 0 | 0 | 0 | 0 | 0 | 0 | 0 |
| ENSMUSG00 | AC159379.1 | 311 | 0 | 0 | 0 | 0 | 0 | 0 | 0 | 0 | 0 | 0 | 0 | 0 | 0 | 0 | 0 | 0 | 0 | 0 |
| ENSMUSG00 | Snord8 | 109 | 4.67946892 | 3.97309748 | 2.66276314 | 5.01589479 | 3.09104097 | 5.00073458 | 3.90655147 | 6.65739124 | 3.02286354 | 4.32575259 | 11.834164 | 1.5362109 | 2.71551838 | 2.98264781 | 3.0239674 | 7.72837948 | 6.32911325 | 3.53269493 |
| ENSMUSG00 | AC162893.2 | 70 | 0 | 0 | 0 | 0 | 0 | 0 | 0 | 0 | 0 | 0 | 0 | 0 | 0 | 0 | 0 | 0 | 0 | 0 |
| ENSMUSG00 | Mir3081 | 83 | 0 | 0 | 0 | 0 | 0 | 0 | 0 | 0.46014943 | 0 | 0 | 0.57560191 | 0 | 0 | 0 | 0 | 0 | 0 | 1.32552064 |
| ENSMUSG00 | AL645683.1 | 100 | 0 | 0 | 0 | 0 | 0 | 0 | 0 | 0 | 0 | 0 | 0 | 0 | 0 | 0 | 0 | 0 | 0 | 0 |
| ENSMUSG00 | AC135118.1 | 103 | 0 | 0 | 0 | 0 | 0 | 0 | 0 | 0 | 0 | 0 | 0 | 0 | 0 | 0 | 0 | 0 | 0 | 0 |
| ENSMUSG00 | AC122782.1 | 74 | 0 | 0.45017425 | 0.65369636 | 0 | 0 | 0 | 0 | 0 | 0 | 0 | 0 | 0 | 0 | 0 | 0 | 0 | 0 | 0 |
| ENSMUSG00 | AC156026.1 | 69 | 0 | 0 | 0 | 0 | 0 | 0 | 0 | 0 | 0 | 0 | 0 | 0 | 0 | 0 | 0 | 0 | 0 | 0 |
| ENSMUSG00 | CT030727.1 | 83 | 0 | 0 | 0 | 0 | 0 | 0 | 0 | 0 | 1.48866924 | 0 | 0 | 0 | 0 | 0 | 0 | 0 | 0 | 0 |
| ENSMUSG00 | AC121954.1 | 148 | 0 | 0 | 0 | 0 | 0 | 0 | 0 | 0 | 0 | 0 | 0 | 0 | 0 | 0 | 0.55677778 | 1.03488128 | 0 | 0.74336631 |
| ENSMUSG00 | AC124134.1 | 69 | 0 | 0 | 0 | 0 | 0 | 0 | 0 | 0 | 0 | 0 | 0 | 0 | 0 | 0 | 0 | 0 | 0 | 0 |
| ENSMUSG00 | AC115933.1 | 99 | 0 | 0 | 0 | 0 | 0 | 0 | 0 | 0 | 0 | 0 | 0 | 0 | 0 | 0 | 0 | 0 | 0 | 0 |
| ENSMUSG00 | AL844869.1 | 277 | 0 | 0 | 0 | 0 | 0 | 0 | 0 | 0 | 0 | 0 | 0 | 0 | 0 | 0 | 0 | 0 | 0 | 0 |
| ENSMUSG00 | AC156988.1 | 99 | 0 | 0 | 0 | 0 | 0 | 0 | 0 | 0 | 0 | 0 | 0 | 0 | 0 | 0 | 0 | 0 | 0 | 0 |
| ENSMUSG00 | AC134453.1 | 69 | 0 | 0 | 0 | 0 | 0 | 0 | 0 | 0 | 0 | 0 | 0 | 0 | 0 | 0 | 0 | 0 | 0 | 0 |
| ENSMUSG00 | AL589767.1 | 267 | 0 | 0 | 0 | 0 | 0 | 0 | 0 | 0 | 0 | 0 | 0 | 0 | 0 | 0 | 0 | 0 | 0 | 0 |
| ENSMUSG00 | Mir5104 | 93 | 0 | 0 | 0 | 0 | 0 | 0 | 0.50873848 | 0 | 0 | 0 | 0 | 0 | 0 | 0 | 0 | 0 | 0 | 0 |
| ENSMUSG00 | Mir5098 | 81 | 0 | 0 | 0 | 0 | 0 | 0 | 0 | 0 | 0 | 0 | 0 | 0 | 0 | 0 | 0 | 0 | 0 | 0 |
| ENSMUSG00 | AL732357.1 | 115 | 0 | 0 | 0 | 0 | 0 | 0 | 0 | 0 | 0 | 0 | 0 | 0 | 0 | 0 | 0 | 0 | 0 | 0 |
| ENSMUSG00 | AC124613.1 | 285 | 0 | 0 | 0 | 0 | 0 | 0 | 0 | 0 | 0 | 0 | 0 | 0 | 0 | 0 | 0 | 0 | 0 | 0 |
| ENSMUSG00 | AL806517.1 | 64 | 0 | 0 | 0 | 0 | 0 | 0 | 0 | 0 | 0 | 0 | 0 | 0 | 0 | 0 | 0 | 0 | 0 | 0 |
| ENSMUSG00 | AL592465.2 | 78 | 0 | 0 | 0 | 0 | 0 | 0 | 0 | 0 | 0 | 0 | 0 | 0 | 0 | 0 | 0 | 0 | 0 | 0 |
| ENSMUSG00 | AC168054.1 | 85 | 0 | 0 | 0 | 0 | 0 | 0 | 0 | 0 | 0 | 0 | 0 | 0 | 0 | 0 | 0 | 0 | 0 | 0 |
| ENSMUSG00 | AC132118.1 | 78 | 0 | 0 | 0 | 0 | 0 | 0 | 0 | 0 | 0 | 0 | 0 | 0 | 0 | 0 | 0 | 0 | 0 | 0 |
| ENSMUSG00 | AC123530.1 | 125 | 0 | 0 | 0 | 0 | 0 | 0 | 0 | 0 | 0 | 0 | 0 | 0 | 0 | 0 | 0 | 0 | 0 | 0 |
| ENSMUSG00 | AC115293.1 | 73 | 2.91131343 | 0.45634102 | 0 | 0.6241239 | 0 | 0 | 0.64811889 | 0.52318359 | 2.82099423 | 3.22950022 | 0.65445149 | 3.82299059 | 1.35155938 | 2.5448815 | 0.56440488 | 1.57358661 | 2.10007106 | 4.52129644 |
| ENSMUSG00 | AL954710.1 | 63 | 0 | 0 | 0 | 0 | 0 | 0 | 0 | 0 | 0 | 0 | 0 | 0 | 0 | 0 | 0 | 0 | 0 | 0 |
| ENSMUSG00 | AC110817.1 | 99 | 0 | 0 | 0 | 0 | 0 | 0 | 0 | 0 | 0 | 0 | 0 | 0 | 0 | 0 | 0 | 0 | 0 | 0 |
| ENSMUSG00 | Mir124a-2 | 108 | 0 | 0 | 0 | 0 | 0 | 0 | 0 | 0 | 0 | 0 | 0 | 0 | 0 | 0 | 0 | 0 | 0 | 0 |
| ENSMUSG00 | CAAA010375 | 113 | 0 | 0 | 0 | 0 | 0 | 0 | 0 | 0 | 0 | 0 | 0 | 0 | 0 | 0 | 0 | 0 | 0 | 0 |
| ENSMUSG00 | AL807824.1 | 111 | 0 | 0.30011616 | 0 | 0 | 0.43362093 | 0 | 0.42624035 | 0.68815139 | 0 | 0 | 0 | 0 | 2.22215844 | 0 | 0.74237038 | 0 | 0.69056391 | 0 |
| ENSMUSG00 | Mir3082 | 63 | 0 | 0 | 0 | 0 | 0 | 0 | 0 | 0 | 0 | 0 | 0 | 0 | 0 | 0 | 0 | 0 | 0 | 0 |
| ENSMUSG00 | Mir5105 | 81 | 189.436649 | 265.269343 | 218.576693 | 231.180114 | 227.58675 | 312.610455 | 396.608753 | 284.321218 | 251.186899 | 239.246161 | 309.062694 | 257.716763 | 344.105349 | 629.575407 | 389.125806 | 209.416532 | 326.956434 | 484.895089 |
| ENSMUSG00 | AC150681.1 | 70 | 0 | 0 | 0 | 0 | 0 | 0 | 0 | 0 | 0 | 0 | 0 | 0 | 0 | 0 | 0 | 0 | 0 | 0 |
| ENSMUSG00 | AC140192.1 | 82 | 0 | 0 | 0 | 0 | 0 | 0 | 0 | 0 | 0 | 0 | 0 | 0 | 0 | 0 | 0 | 0 | 0 | 0 |
| ENSMUSG00 | Mir3060 | 82 | 1.03671161 | 2.43752885 | 0 | 0 | 0 | 1.81290489 | 0.57698389 | 0.93152201 | 0 | 2.30003431 | 3.49572867 | 1.36135762 | 1.20321749 | 0.56639131 | 3.51720599 | 0.49695863 | 2.80436319 | 0 |
| ENSMUSG00 | Mir3075 | 84 | 0 | 0 | 0 | 0 | 0 | 0 | 0.56324618 | 0 | 0 | 0 | 0 | 0 | 0 | 0 | 0 | 0 | 0 | 0 |
| ENSMUSG00 | AC105995.1 | 69 | 0 | 0 | 0 | 0 | 0 | 0 | 0 | 0 | 0 | 0 | 0 | 0 | 0 | 0 | 0 | 0 | 0 | 0 |
| ENSMUSG00 | AL627205.1 | 76 | 0 | 0 | 0 | 0 | 0 | 0 | 0 | 0 | 0 | 0 | 0 | 0 | 0 | 0 | 0 | 0 | 0 | 0 |
| ENSMUSG00 | Mir1186b | 63 | 0 | 0 | 0 | 0 | 0 | 0 | 0 | 0 | 0 | 0 | 0 | 0 | 0 | 0 | 0 | 0 | 0 | 0 |
| ENSMUSG00 | AC153815.1 | 73 | 0 | 0 | 0.6626511 | 0 | 0 | 0 | 1.94435667 | 0 | 0 | 0 | 0 | 0 | 0 | 0 | 0 | 0.52452887 | 0 | 0 |
| ENSMUSG00 | AC127419.1 | 80 | 0.5313147 | 1.24923353 | 0.60486913 | 1.70853916 | 0 | 0 | 0.59140849 | 0 | 0 | 2.35753516 | 0.59718698 | 0 | 0 | 2.90275546 | 1.54505835 | 0 | 1.43723613 | 0.68761383 |
| ENSMUSG00 | Mir3473d | 80 | 1.0626294 | 1.66564471 | 3.02334565 | 1.13902611 | 0 | 0 | 0 | 0.47740503 | 0 | 1.76815137 | 0 | 2.79078313 | 2.46659586 | 0 | 1.0300389 | 0 | 0 | 0 |
| ENSMUSG00 | AC160635.1 | 125 | 0 | 0 | 0 | 0 | 0 | 0 | 0 | 0 | 0 | 0.37720563 | 0 | 0 | 0 | 0 | 0 | 0 | 0 | 0 |
| ENSMUSG00 | AC121784.1 | 79 | 0 | 0 | 0 | 0 | 0 | 0 | 0 | 0 | 0 | 0 | 0 | 0 | 0 | 0 | 0 | 0 | 0 | 0 |
| ENSMUSG00 | Mir3073 | 84 | 0 | 0 | 0 | 0 | 0 | 0 | 0 | 0 | 0 | 0 | 0 | 0 | 0 | 0 | 0 | 0 | 0 | 0 |
| ENSMUSG00 | Mir3089 | 83 | 0 | 0 | 0 | 0 | 0 | 0 | 0 | 0 | 0 | 0 | 0 | 0 | 0 | 0 | 0 | 0 | 0 | 0 |
| ENSMUSG00 | AC169127.1 | 100 | 0 | 0 | 0 | 0 | 0 | 0 | 0 | 0 | 0 | 0 | 0 | 0 | 0 | 0 | 0 | 0 | 0 | 0 |
| ENSMUSG00 | AC132327.1 | 247 | 0 | 0 | 0 | 0 | 0 | 0 | 0 | 0 | 0 | 0 | 0 | 0 | 0 | 0 | 0 | 0 | 0 | 0 |
| ENSMUSG00 | AC140313.1 | 100 | 0 | 0 | 0 | 0 | 0 | 0 | 0 | 0 | 0 | 0 | 0 | 0 | 0 | 0 | 0 | 0 | 0 | 0 |
| ENSMUSG00 | AC170189.1 | 80 | 0 | 0 | 0 | 0 | 0 | 0 | 0 | 0 | 0 | 0 | 0 | 0 | 0 | 0 | 0 | 0 | 0 | 0 |
| ENSMUSG00 | Mir3968 | 99 | 0 | 0 | 0 | 0 | 0 | 0 | 0 | 0 | 0 | 0 | 0 | 0 | 0 | 0 | 0 | 0 | 0 | 0 |
| ENSMUSG00 | Mir3097 | 66 | 0.64401782 | 0 | 0 | 0 | 0 | 0 | 0 | 0 | 0 | 0 | 0 | 0 | 0 | 0 | 0 | 0 | 0 | 0 |
| ENSMUSG00 | Mir5102 | 86 | 259.479272 | 294.005659 | 213.181023 | 209.792716 | 232.264515 | 244.3065 | 311.383445 | 301.986438 | 343.859515 | 286.742067 | 204.43238 | 271.290081 | 586.246738 | 652.377414 | 311.886196 | 252.450865 | 266.947114 | 356.279912 |
| ENSMUSG00 | Rn5s | 118 | 0 | 0 | 0 | 0 | 0 | 0 | 0 | 0 | 0 | 0 | 0 | 0 | 0 | 0 | 0 | 0 | 0 | 0 |
| ENSMUSG00 | Mir3971 | 82 | 0 | 0.81250962 | 0 | 0 | 0 | 0 | 1.73095167 | 1.39728301 | 0 | 1.72502573 | 0.58262144 | 0.68067881 | 0 | 0 | 0 | 0 | 0 | 0 |
| ENSMUSG00 | AC119249.4 | 78 | 0 | 0 | 0 | 0 | 0 | 0 | 0 | 0 | 0 | 0 | 0 | 0 | 0 | 0 | 0 | 0 | 0 | 0 |
| ENSMUSG00 | Mir3475 | 64 | 0 | 0 | 0 | 0 | 0 | 0 | 0 | 0 | 0 | 0 | 0 | 0 | 0 | 0 | 0 | 0 | 0 | 0 |
| ENSMUSG00 | AC141881.5 | 118 | 0 | 0 | 0 | 0 | 0 | 0 | 0 | 0 | 0 | 0 | 0 | 0 | 0 | 0 | 0 | 0 | 0 | 0 |
| ENSMUSG00 | AL845455.1 | 152 | 1.95747521 | 0.43832756 | 0.31824691 | 0.29974371 | 0 | 0.32600483 | 1.55633812 | 0.50253161 | 1.08385568 | 2.79181796 | 0 | 0 | 0 | 1.52776603 | 0.54212574 | 0.50382378 | 0.25214669 | 0.72380404 |
| ENSMUSG00 | AC153918.1 | 124 | 0 | 0 | 0 | 0 | 1.16448202 | 0 | 0 | 0 | 0 | 0.76049521 | 0 | 0 | 0 | 0.37454969 | 0 | 0 | 0.30908304 | 0 |
| ENSMUSG00 | AC136976.1 | 124 | 0 | 0 | 0 | 0.36742778 | 0 | 0 | 0 | 0 | 0 | 0 | 0.77056385 | 0 | 0 | 0 | 0 | 0 | 0 | 0 |
| ENSMUSG00 | AC167122.1 | 126 | 2.36139867 | 1.5863283 | 4.9909198 | 3.6159559 | 1.14599818 | 2.75292964 | 3.75497452 | 5.15294318 | 3.26877109 | 0 | 2.27499802 | 5.75875884 | 1.95761576 | 0.73720774 | 2.94296828 | 1.82336226 | 1.21670784 | 3.4926417 |
| ENSMUSG00 | Mir344f | 67 | 0 | 0 | 0 | 0 | 0 | 0 | 0 | 0 | 0 | 0 | 0 | 0 | 0 | 0 | 0 | 0 | 0 | 0 |
| ENSMUSG00 | AC165159.1 | 284 | 0 | 0 | 0 | 0 | 0 | 0 | 0 | 0 | 0 | 0 | 0 | 0 | 0 | 0 | 0 | 0 | 0 | 0 |
| ENSMUSG00 | AC115767.1 | 85 | 0 | 1.17574921 | 0 | 0 | 0 | 0 | 0 | 0 | 0 | 0 | 0 | 0 | 0 | 0 | 0 | 0.90095547 | 0 | 0 |
| ENSMUSG00 | AC120136.1 | 190 | 1.11855726 | 0.35066204 | 0.50919506 | 0 | 1.01330366 | 1.56482317 | 0.7470423 | 0.80405058 | 0.86708454 | 0 | 0 | 1.76259987 | 0 | 0 | 1.73480235 | 0 | 0.80686941 | 2.89521614 |
| ENSMUSG00 | AC069308.1 | 275 | 0 | 0 | 0 | 0 | 0 | 0 | 0 | 0 | 0 | 0 | 0 | 0 | 0 | 0 | 0 | 0 | 0 | 0 |
| ENSMUSG00 | AL662785.1 | 68 | 0 | 0 | 0 | 0 | 0 | 0 | 0 | 0 | 0 | 0 | 0 | 0 | 0 | 0 | 0 | 0 | 0 | 0 |
| ENSMUSG00 | AC133174.1 | 67 | 0 | 0 | 0 | 0 | 0 | 0 | 0 | 0 | 0 | 0 | 0 | 0 | 0 | 0 | 0 | 0 | 0 | 0 |
| ENSMUSG00 | AL772159.1 | 100 | 0 | 0 | 0 | 0 | 0 | 0 | 0 | 0 | 0 | 0 | 0 | 0 | 0 | 0 | 0 | 0 | 0 | 0 |
| ENSMUSG00 | AC130536.1 | 107 | 0 | 0 | 0 | 0 | 0 | 0 | 0 | 0 | 0 | 0 | 0 | 0 | 0 | 0 | 0 | 0 | 0 | 0 |
| ENSMUSG00 | AC166574.1 | 104 | 0 | 0 | 0 | 0 | 0 | 0 | 0 | 0 | 0 | 0 | 0 | 0 | 0 | 0 | 0 | 0 | 0 | 0 |
| ENSMUSG00 | Mir3083 | 63 | 0 | 0 | 0 | 0 | 0 | 0 | 0 | 0 | 0 | 0 | 0 | 0 | 0 | 0 | 0 | 0 | 0 | 0 |
| ENSMUSG00 | AC122463.1 | 124 | 0 | 0 | 0.78021823 | 0 | 0 | 0 | 1.14466159 | 0 | 1.6607466 | 0 | 0 | 0 | 0 | 0 | 0 | 0 | 0 | 0 |
| ENSMUSG00 | Mir3474 | 57 | 0 | 0.58443674 | 0 | 0 | 0 | 0 | 0 | 0.67004215 | 1.4451409 | 0 | 0 | 0 | 0 | 0.81480855 | 0 | 1.34353008 | 0 | 0 |
| ENSMUSG00 | AC167242.1 | 237 | 0 | 0 | 0 | 0 | 0.20308829 | 0 | 0 | 0.16114908 | 0 | 0 | 0 | 0 | 0 | 0 | 0 | 0 | 0 | 0 |
| ENSMUSG00 | Mir1264 | 85 | 0 | 0 | 0 | 0 | 0 | 0 | 0 | 0 | 0 | 0 | 0 | 0 | 0 | 0 | 0 | 0 | 0 | 0 |
| ENSMUSG00 | AL671229.1 | 125 | 0 | 0 | 0 | 0 | 0 | 0 | 0 | 0 | 0 | 0 | 0 | 0 | 0 | 0 | 0 | 0 | 0 | 0 |
| ENSMUSG00 | AC114917.1 | 85 | 4.00048716 | 0.3919164 | 5.12190321 | 2.14404915 | 2.83128963 | 1.16594667 | 1.66985926 | 2.69593429 | 3.39183071 | 6.65656987 | 4.49646668 | 1.96996456 | 0.5803755 | 1.09280206 | 3.8777935 | 1.80191094 | 4.50897611 | 1.94149788 |
| ENSMUSG00 | AC130474.1 | 124 | 0 | 0 | 0 | 0 | 0.38816067 | 0 | 0 | 0 | 0 | 0 | 0 | 0 | 0 | 0 | 0 | 0 | 0 | 0 |
| ENSMUSG00 | AC161763.1 | 70 | 1.2144336 | 0.95179698 | 0 | 0.65087206 | 0.68759891 | 0 | 0.67589541 | 1.0912115 | 1.76513639 | 0 | 0 | 0 | 1.40948335 | 2.65394785 | 0.58859366 | 0.54700868 | 0.54751853 | 1.57168876 |
| ENSMUSG00 | AC167724.1 | 125 | 0 | 0 | 0 | 0 | 0 | 0 | 0 | 0 | 0 | 0 | 0 | 0 | 0 | 0 | 0 | 0 | 0 | 0 |
| ENSMUSG00 | AC134912.1 | 95 | 0 | 0 | 0 | 0 | 0 | 0 | 0 | 0 | 0 | 0 | 0 | 0 | 0 | 0 | 0 | 0 | 0 | 0 |
| ENSMUSG00 | AC107770.1 | 105 | 0 | 0 | 0 | 0 | 0 | 0 | 0 | 0 | 0 | 0 | 0 | 0 | 0 | 0 | 0 | 0 | 0 | 0 |
| ENSMUSG00 | AC140963.1 | 106 | 0 | 0 | 0.91270812 | 0 | 0 | 0 | 0 | 0 | 0 | 0 | 0 | 0 | 0 | 0 | 0 | 0 | 0 | 0 |
| ENSMUSG00 | AC144905.1 | 79 | 0 | 0 | 0 | 0 | 0.60926486 | 0 | 0 | 0 | 0 | 0 | 0 | 0 | 0 | 0 | 0 | 0 | 0 | 0 |
| ENSMUSG00 | AL603664.1 | 100 | 0 | 0 | 0 | 0 | 0 | 0 | 0 | 0 | 0 | 0 | 0 | 0 | 0 | 0 | 0 | 0 | 0 | 0 |
| ENSMUSG00 | CT025520.1 | 116 | 0 | 0 | 0 | 0 | 0 | 0 | 0 | 0 | 0 | 0 | 0 | 0 | 0 | 0 | 0 | 0 | 0 | 0 |
| ENSMUSG00 | AL671140.1 | 109 | 0 | 0 | 0 | 0.41799123 | 0 | 0.45461223 | 0 | 0 | 2.26714766 | 0 | 0.43830237 | 0 | 0.90517279 | 0 | 0 | 0 | 0 | 0 |
| ENSMUSG00 | AC122057.1 | 141 | 0 | 0 | 0 | 0 | 0 | 0 | 0 | 0 | 0 | 0 | 0 | 0 | 0 | 0 | 0 | 0 | 0 | 0 |
| ENSMUSG00 | AL845277.1 | 52 | 1.63481446 | 0.64063258 | 0 | 0.87617393 | 1.85122783 | 0.95293718 | 0 | 1.46893865 | 0.79204838 | 0.90674429 | 1.8374964 | 1.07337813 | 4.74345358 | 1.78631105 | 0 | 0 | 4.42226503 | 2.11573487 |
| ENSMUSG00 | AC161198.1 | 132 | 0 | 0 | 0 | 0 | 0.36463579 | 0.3753995 | 0 | 0 | 0 | 0 | 0 | 1.26853779 | 0 | 0 | 0 | 0 | 0 | 0 |
| ENSMUSG00 | Mir28b | 79 | 0 | 0 | 0 | 0 | 0 | 0 | 0 | 0 | 0 | 0 | 0 | 0 | 0 | 0 | 0 | 0 | 0 | 0 |
| ENSMUSG00 | AC163282.1 | 89 | 0 | 0 | 0 | 0 | 0 | 0 | 0 | 0 | 0 | 0 | 0 | 0 | 0 | 0 | 0 | 0 | 0 | 0 |
| ENSMUSG00 | Mir3106 | 81 | 0 | 0 | 0 | 0 | 0 | 0 | 0 | 0 | 0 | 0 | 0 | 0 | 0 | 0 | 0 | 0 | 0 | 0 |
| ENSMUSG00 | AC134910.1 | 274 | 0 | 0 | 0 | 0 | 0 | 0 | 0 | 0 | 0 | 0 | 0 | 0 | 0 | 0 | 0 | 0 | 0 | 0 |
| ENSMUSG00 | AC125186.2 | 99 | 0 | 0 | 0 | 0 | 0 | 0 | 0 | 0 | 0 | 0.47626973 | 0 | 0 | 0 | 0 | 0 | 0 | 0 | 0 |
| ENSMUSG00 | AC140457.1 | 73 | 0 | 0 | 0 | 0 | 0 | 0 | 0 | 0 | 0 | 0 | 0 | 0 | 0 | 0 | 0 | 0 | 0 | 0 |
| ENSMUSG00 | AC131793.1 | 129 | 0 | 0 | 0 | 0 | 0 | 0 | 0 | 0 | 0 | 0 | 0 | 0 | 0.76483593 | 0 | 0 | 0 | 0 | 0 |
| ENSMUSG00 | AL772271.1 | 104 | 0 | 0 | 0 | 0 | 0 | 0 | 0 | 0 | 0 | 0 | 0 | 0 | 0 | 0 | 0 | 0 | 0 | 0 |
| ENSMUSG00 | AC122359.1 | 82 | 0 | 0 | 0 | 0 | 0 | 0 | 0 | 0 | 0 | 0 | 0 | 0 | 0 | 0 | 0 | 0 | 0.46739386 | 0 |
| ENSMUSG00 | AC163694.1 | 83 | 0 | 0 | 0 | 0 | 0 | 1.79106266 | 0 | 0.46014943 | 1.48866924 | 0 | 0 | 0 | 1.1887209 | 0 | 0 | 0 | 0 | 0 |
| ENSMUSG00 | AC160932.1 | 70 | 0 | 0 | 0 | 0 | 0 | 0 | 0 | 0 | 0 | 0 | 0 | 0 | 0 | 0 | 0 | 0 | 0 | 0 |
| ENSMUSG00 | AC123662.1 | 94 | 0 | 0 | 0 | 0 | 0 | 0 | 0 | 0.81260431 | 0.43815442 | 0 | 0 | 0 | 0 | 0 | 0 | 0 | 0 | 0 |
| ENSMUSG00 | AC073947.1 | 83 | 0 | 0 | 0 | 0 | 0 | 0 | 0 | 0 | 0 | 0 | 0 | 0 | 0 | 0 | 0 | 0 | 0 | 0 |
| ENSMUSG00 | AC149058.1 | 69 | 0 | 0 | 0 | 0 | 0 | 0 | 0 | 0 | 0 | 0 | 0 | 0 | 0 | 0 | 0 | 0 | 0 | 0 |
| ENSMUSG00 | AC126942.1 | 57 | 2.98281937 | 4.67549393 | 3.39463371 | 6.39453254 | 4.22209857 | 3.47738481 | 3.32018799 | 4.69029503 | 2.16771136 | 5.79043724 | 10.057886 | 6.85455505 | 14.713028 | 8.96289406 | 9.39684608 | 8.73294555 | 8.06869408 | 16.4062248 |
| ENSMUSG00 | AC117610.1 | 76 | 0 | 0 | 0 | 0 | 0 | 0 | 1.86760575 | 0 | 0 | 0 | 0 | 0 | 0.64910417 | 0 | 0 | 1.00764756 | 0 | 0 |
| ENSMUSG00 | Mir3076 | 59 | 0 | 0 | 0 | 0 | 0 | 0 | 0 | 0 | 0 | 0 | 0 | 0 | 0 | 0 | 0 | 1.94698004 | 0 | 0 |
| ENSMUSG00 | Mir3098 | 86 | 2.47123116 | 1.93679618 | 0 | 3.17867751 | 4.47738825 | 0.57619458 | 0.55014743 | 4.88507473 | 0.47891297 | 0.54826399 | 0 | 5.19215466 | 0.57362695 | 3.78033269 | 1.91635144 | 1.78095848 | 3.11958231 | 0 |
| ENSMUSG00 | AC124466.1 | 49 | 0 | 1.35970997 | 0.98721491 | 0 | 0 | 0 | 0 | 0 | 0 | 0 | 0 | 0 | 0 | 0 | 0 | 0 | 0 | 0 |
| ENSMUSG00 | AC026231.1 | 109 | 0 | 0 | 0 | 0 | 0 | 0 | 0 | 0 | 0 | 0 | 0 | 0 | 0 | 0 | 0 | 0 | 0 | 0 |
| ENSMUSG00 | AL845460.1 | 77 | 0 | 0 | 0 | 0 | 0 | 0 | 0 | 0 | 0 | 0 | 0 | 0 | 0 | 0 | 0 | 0 | 0 | 0 |
| ENSMUSG00 | AC153489.1 | 72 | 0 | 0 | 0 | 0 | 0 | 0 | 0 | 0 | 0 | 0 | 0 | 0 | 0 | 0 | 0 | 0 | 0 | 0 |
| ENSMUSG00 | AC158662.1 | 61 | 0.69680616 | 0 | 0 | 0 | 0 | 0 | 0 | 0 | 0 | 0 | 0 | 0 | 0 | 3.04551393 | 0 | 0 | 0 | 0 |
| ENSMUSG00 | AL607123.1 | 81 | 0 | 0 | 0 | 0 | 0 | 0 | 0 | 0 | 0 | 1.74632234 | 0 | 0 | 0 | 0.57338379 | 0 | 0 | 0 | 0 |
| ENSMUSG00 | AL645971.1 | 251 | 0 | 0 | 0 | 0 | 0 | 0 | 0 | 0 | 0 | 0 | 0 | 0 | 0 | 0 | 0 | 0 | 0 | 0 |
| ENSMUSG00 | AC127366.1 | 125 | 0 | 0 | 0 | 0 | 0 | 0 | 0 | 0 | 0 | 0 | 0 | 0 | 0 | 0 | 0 | 0 | 0 | 0 |
| ENSMUSG00 | AC160757.1 | 320 | 0 | 0 | 0 | 0 | 0 | 0 | 0 | 0.23870252 | 0 | 0 | 0 | 0 | 0 | 0 | 0 | 0 | 0 | 0 |
| ENSMUSG00 | Mir5131 | 92 | 0 | 0 | 0 | 0 | 0 | 0 | 0 | 0 | 0 | 0 | 0 | 0 | 0 | 0 | 0 | 0 | 0 | 0 |
| ENSMUSG00 | AL663049.1 | 88 | 1.93205346 | 1.51422246 | 0 | 1.55321742 | 3.82867575 | 2.25239698 | 0.53764408 | 5.64205945 | 2.34014294 | 0 | 4.34317804 | 0 | 0.56058997 | 1.58332116 | 0.936399 | 5.656567 | 9.14604812 | 4.3757244 |
| ENSMUSG00 | Mir3079 | 89 | 0 | 0 | 0 | 0 | 0 | 0 | 0 | 0 | 0 | 0 | 0 | 0 | 0 | 0 | 0 | 0 | 0 | 0 |
| ENSMUSG00 | AC091333.2 | 69 | 0 | 0 | 0 | 0 | 0 | 0 | 0 | 0 | 0 | 0 | 0 | 0 | 0 | 0 | 0 | 0 | 0 | 0 |
| ENSMUSG00 | AC132452.1 | 72 | 0 | 0 | 0 | 0 | 0 | 0 | 0 | 0 | 0 | 0 | 0 | 0 | 0 | 0 | 0 | 0 | 0 | 0 |
| ENSMUSG00 | Mir3112 | 81 | 0 | 0 | 0 | 0 | 0 | 0 | 0 | 0 | 0 | 0 | 0 | 0 | 0 | 0 | 0 | 0 | 0 | 0 |
| ENSMUSG00 | AC139568.1 | 76 | 0 | 0 | 0 | 0 | 0 | 0 | 0 | 0 | 0 | 0 | 0 | 0 | 0 | 0 | 0 | 0 | 0 | 0 |
| ENSMUSG00 | BX323066.1 | 277 | 0.15344829 | 0 | 0 | 0 | 0 | 0 | 0 | 0 | 0 | 0 | 0 | 0 | 0 | 0 | 0 | 0 | 0 | 0 |
| ENSMUSG00 | AC133646.1 | 324 | 0 | 0 | 0 | 0 | 0 | 0 | 0 | 0 | 0 | 0.29105372 | 0 | 0 | 0 | 0 | 0 | 0 | 0 | 0 |
| ENSMUSG00 | Mir5119 | 56 | 4.554126 | 0 | 3.45525217 | 0 | 2.57849591 | 0 | 0 | 8.86609342 | 0 | 3.36790738 | 3.41249703 | 2.99012478 | 0 | 1.65871741 | 2.20722621 | 1.36752169 | 2.73759264 | 0 |
| ENSMUSG00 | AC110195.1 | 70 | 0 | 0 | 0 | 0 | 0 | 0 | 0 | 0 | 0 | 0 | 0 | 0 | 0 | 0 | 0 | 0 | 0 | 0 |
| ENSMUSG00 | Snord87 | 86 | 4.94246233 | 4.26095159 | 3.93738038 | 1.58933876 | 1.11934706 | 2.88097288 | 1.65044229 | 0 | 1.43673892 | 6.03090391 | 4.99970496 | 3.24509666 | 2.86813473 | 9.7208555 | 0.95817572 | 1.33571886 | 10.6957108 | 3.19820388 |
| ENSMUSG00 | AL672246.1 | 66 | 0 | 0 | 0 | 0 | 0 | 0 | 0 | 0 | 0 | 0 | 0 | 0 | 0 | 0 | 0 | 0 | 0 | 0 |
| ENSMUSG00 | AL670285.1 | 76 | 0 | 0 | 0 | 0 | 0 | 0 | 0 | 0 | 0 | 0 | 0 | 0 | 0 | 0 | 0 | 0 | 0 | 0 |
| ENSMUSG00 | AC157798.1 | 195 | 0 | 0 | 0 | 0 | 0 | 0 | 0 | 0 | 0 | 0 | 0 | 0 | 0 | 0 | 0 | 0 | 0 | 0 |
| ENSMUSG00 | AC136021.1 | 316 | 0 | 0 | 0 | 0 | 0 | 0 | 0.14972367 | 0 | 0.26067415 | 0.14921109 | 0 | 0 | 0 | 0 | 0 | 0 | 0 | 0 |
| ENSMUSG00 | AL772349.1 | 278 | 0 | 0 | 0 | 0 | 0 | 0 | 0 | 0 | 0 | 0 | 0 | 0 | 0 | 0 | 0 | 0 | 0 | 0 |
| ENSMUSG00 | AL808020.1 | 84 | 0 | 0 | 0 | 0 | 0 | 0 | 0 | 0 | 0 | 0 | 0 | 0 | 0 | 0 | 0 | 0 | 0 | 0 |
| ENSMUSG00 | AL772376.2 | 124 | 0 | 0 | 0 | 0 | 0 | 0 | 0 | 0 | 0 | 0 | 0 | 0 | 0 | 0 | 0 | 0 | 0 | 0 |
| ENSMUSG00 | AL713978.1 | 279 | 0 | 0 | 0 | 0 | 0 | 0 | 0 | 0 | 0 | 0 | 0 | 0 | 0 | 0 | 0 | 0 | 0 | 0 |
| ENSMUSG00 | Mir344-2 | 79 | 0 | 0 | 0 | 0 | 0 | 0 | 0 | 0 | 0 | 0 | 0 | 0 | 0 | 0 | 0 | 0 | 0 | 0 |
| ENSMUSG00 | AL732564.1 | 251 | 0 | 0 | 0 | 0 | 0 | 0 | 0 | 0 | 0.1640897 | 0 | 0 | 0.44474632 | 0 | 0 | 0 | 0.15255222 | 0 | 0 |
| ENSMUSG00 | AC161829.1 | 281 | 0 | 0.11855122 | 0 | 0.16213895 | 0 | 0 | 0 | 0 | 0.29314246 | 0 | 1.36014117 | 0.3972645 | 0.17555842 | 0.6611258 | 0.14662475 | 1.36265507 | 0.95474761 | 0.39152389 |
| ENSMUSG00 | AL732525.2 | 125 | 0 | 0 | 0 | 0 | 0 | 0 | 0 | 0 | 0 | 0 | 0 | 0 | 0 | 0 | 0 | 0 | 0.30661038 | 0 |
| ENSMUSG00 | AC113598.1 | 300 | 0 | 0 | 0 | 0 | 0 | 0 | 0 | 0 | 0 | 0 | 0 | 0 | 0 | 0 | 0 | 0 | 0 | 0 |
| ENSMUSG00 | AC125201.1 | 104 | 0 | 0 | 0 | 0 | 0 | 0 | 0 | 0 | 0 | 0 | 0 | 0 | 0 | 0 | 0 | 0 | 0 | 0 |
| ENSMUSG00 | AC105256.2 | 70 | 0 | 0 | 0 | 0 | 0 | 0 | 0 | 0 | 0 | 0 | 0 | 0 | 0 | 0 | 0 | 0 | 0 | 0 |
| ENSMUSG00 | AC126681.1 | 85 | 0 | 0 | 0 | 0 | 0 | 0 | 0 | 0 | 0 | 0 | 0 | 0 | 0 | 0 | 0 | 0 | 0 | 0 |
| ENSMUSG00 | AC115706.1 | 122 | 0 | 0 | 0 | 0 | 0 | 0 | 0 | 0 | 0 | 0 | 0 | 0 | 0 | 0 | 0 | 0 | 0 | 0 |
| ENSMUSG00 | AC156032.1 | 228 | 0.18642621 | 0.58443674 | 0 | 1.39880399 | 1.89994436 | 0.21733655 | 1.03755875 | 1.34008429 | 0.72257045 | 0 | 0.20953929 | 0 | 0.43273612 | 0.20370214 | 1.08425147 | 1.00764756 | 0.67239117 | 0.24126801 |
| ENSMUSG00 | AL929170.1 | 68 | 0 | 0 | 0 | 0 | 0 | 0 | 0 | 0 | 0 | 0 | 0 | 0 | 0 | 0 | 0 | 0 | 0 | 0 |
| ENSMUSG00 | Mir344d-2 | 79 | 0 | 0 | 0 | 0 | 0 | 0 | 0 | 0 | 0 | 0 | 0 | 0 | 0 | 0 | 0 | 0 | 0 | 0 |
| ENSMUSG00 | AC141881.6 | 118 | 0 | 0 | 0 | 0 | 0 | 0 | 0 | 0 | 0 | 0 | 0 | 0 | 0 | 0 | 0 | 0 | 0 | 0 |
| ENSMUSG00 | AC153589.1 | 327 | 0 | 0 | 0 | 0 | 0 | 0 | 0 | 0 | 0 | 0 | 0 | 0 | 0 | 0 | 0 | 0 | 0 | 0 |
| ENSMUSG00 | AC122454.1 | 109 | 0 | 0 | 0 | 0 | 0 | 0 | 0 | 0 | 0 | 0 | 0 | 0 | 0 | 0 | 0 | 0 | 0 | 0 |
| ENSMUSG00 | AL844481.1 | 125 | 0 | 0 | 0 | 0 | 0 | 0 | 0 | 0 | 0 | 0 | 0 | 0 | 0 | 0 | 0 | 0 | 0 | 0 |
| ENSMUSG00 | AC166574.2 | 106 | 0 | 0 | 0 | 0 | 0 | 0 | 0 | 0 | 0 | 0 | 0 | 0 | 0 | 0 | 0 | 0 | 0 | 0 |
| ENSMUSG00 | AC122417.1 | 268 | 0.95160842 | 0.12430184 | 0.90249124 | 0.34000779 | 0.71838692 | 0.18489826 | 0.17653985 | 0.99756275 | 0 | 0.52780638 | 1.7826477 | 0.4165348 | 0.18407432 | 0.69319533 | 0.6149486 | 0 | 1.43008571 | 0.61577358 |
| ENSMUSG00 | AL672290.1 | 90 | 0 | 0 | 0 | 0 | 0 | 0 | 0 | 0 | 0 | 0 | 0 | 0 | 0 | 0 | 0 | 0 | 0 | 0 |
| ENSMUSG00 | AC134475.1 | 100 | 0 | 0 | 0 | 0 | 0 | 0 | 0 | 0 | 0 | 0 | 0 | 0 | 0 | 0 | 0 | 0 | 0 | 0 |
| ENSMUSG00 | AC122860.1 | 61 | 0 | 0 | 0 | 0 | 0 | 0 | 0 | 0 | 0 | 0 | 0 | 0 | 0 | 0 | 0 | 0 | 0 | 0 |
| ENSMUSG00 | Mir3070b | 88 | 0 | 0 | 0 | 0 | 0 | 0 | 0 | 0 | 0 | 0 | 0 | 0 | 0 | 0 | 0 | 0 | 0 | 0 |
| ENSMUSG00 | AL662894.1 | 315 | 0 | 0 | 0 | 0 | 0 | 0 | 0 | 0.12124572 | 0 | 0 | 0 | 0 | 0 | 0 | 0 | 0 | 0 | 0 |
| ENSMUSG00 | AC157395.1 | 75 | 0 | 0 | 0 | 0 | 0 | 0 | 0 | 0 | 0 | 0 | 0 | 0 | 0 | 0 | 0 | 0 | 0 | 0 |
| ENSMUSG00 | AC123823.1 | 77 | 0 | 0 | 0 | 0 | 0 | 0 | 0 | 0 | 0 | 0 | 0 | 0 | 0 | 0 | 0 | 0 | 0 | 0 |
| ENSMUSG00 | AC101833.1 | 101 | 0 | 0 | 0 | 0 | 0 | 0 | 0 | 0 | 0 | 0 | 0 | 0 | 0 | 0 | 0 | 0 | 0 | 0 |
| ENSMUSG00 | AC154353.1 | 97 | 0 | 0 | 0 | 0 | 0 | 0 | 0 | 0 | 0 | 0 | 0 | 0 | 0 | 0 | 0 | 0 | 0 | 0 |
| ENSMUSG00 | Mir3103 | 66 | 0 | 0 | 0 | 0 | 0 | 0 | 0 | 0 | 0 | 0 | 0 | 0 | 0 | 0 | 0 | 0 | 0 | 0 |
| ENSMUSG00 | AC122376.1 | 89 | 0 | 0 | 0 | 0 | 0 | 0 | 0 | 0 | 0 | 0 | 0 | 0 | 0 | 0 | 0 | 0 | 0 | 0 |
| ENSMUSG00 | AL671117.1 | 68 | 0 | 0.97979101 | 1.42275089 | 0 | 0 | 0 | 0 | 0 | 0 | 0 | 0 | 0 | 0 | 0 | 0 | 0 | 0 | 0 |
| ENSMUSG00 | AC121784.2 | 119 | 0 | 0 | 0 | 0 | 0 | 0 | 0 | 0 | 0 | 0 | 0 | 0 | 0 | 0 | 0 | 0 | 0 | 0 |
| ENSMUSG00 | Mir3113 | 75 | 0 | 0 | 0 | 0 | 0 | 0 | 0 | 0 | 0 | 0 | 0 | 0 | 0 | 0 | 0 | 0 | 0 | 0 |
| ENSMUSG00 | AL627182.1 | 252 | 0 | 0 | 0 | 0 | 0 | 0 | 0 | 0 | 0 | 0 | 0 | 0 | 0 | 0 | 0 | 0 | 0 | 0 |
| ENSMUSG00 | Mir3105 | 84 | 0 | 0 | 0 | 0 | 0 | 0 | 0 | 0 | 0 | 0 | 0 | 0 | 0 | 0 | 0 | 0 | 0 | 0 |
| ENSMUSG00 | AC154683.1 | 106 | 0 | 0 | 0 | 0 | 0 | 0 | 1.33903808 | 0 | 0.77710407 | 0 | 0 | 0 | 0 | 0 | 0 | 0 | 0 | 0 |
| ENSMUSG00 | AC117806.1 | 125 | 0 | 0 | 0 | 0 | 0 | 0 | 0 | 0 | 0 | 0 | 0 | 0 | 0 | 0 | 0 | 0 | 0 | 0 |
| ENSMUSG00 | AC121784.3 | 111 | 0 | 0 | 0 | 0 | 0 | 0 | 0 | 0 | 0 | 0 | 0 | 0 | 0 | 0 | 0 | 0 | 0 | 0 |
| ENSMUSG00 | AL645531.1 | 73 | 0 | 0 | 0 | 0 | 0 | 0 | 0 | 0 | 0 | 0 | 0 | 0 | 0 | 0 | 0 | 0 | 0 | 0 |
| ENSMUSG00 | AC163349.1 | 110 | 0.38641069 | 0 | 0 | 0 | 0 | 0 | 0 | 0 | 0 | 0 | 0 | 0 | 0 | 0 | 0 | 0 | 0 | 0 |
| ENSMUSG00 | Mir1251 | 83 | 0 | 0 | 0 | 0 | 0 | 0 | 0 | 0 | 0 | 0 | 0 | 0 | 0 | 0 | 0 | 0 | 0 | 0 |
| ENSMUSG00 | AC167197.1 | 69 | 0 | 0 | 0 | 0 | 0 | 0 | 0 | 0 | 0 | 0 | 0 | 0 | 0 | 0 | 0 | 0 | 0 | 0 |
| ENSMUSG00 | AC139241.1 | 102 | 0 | 0 | 0 | 0 | 0 | 0 | 0 | 0 | 0 | 0 | 0 | 0 | 0 | 0 | 0 | 0 | 0 | 0 |
| ENSMUSG00 | AC161253.2 | 66 | 0 | 0 | 0 | 0 | 0 | 0 | 0 | 0 | 0 | 0 | 0 | 0 | 0 | 0 | 0 | 0 | 0 | 0 |
| ENSMUSG00 | AC147162.1 | 96 | 0 | 0 | 0 | 0 | 0 | 0.51617431 | 0.49284041 | 0 | 0 | 0 | 0 | 0 | 0 | 0 | 0 | 0 | 1.59692904 | 0 |
| ENSMUSG00 | AL591070.1 | 280 | 0 | 0 | 1.38210087 | 0.16271802 | 0 | 0 | 0 | 0 | 0 | 0.33679074 | 0 | 0 | 0 | 0.49761522 | 0 | 0 | 0.13687963 | 0 |
| ENSMUSG00 | Mir3085 | 88 | 0 | 0 | 0 | 0 | 0 | 0 | 0 | 0 | 0 | 0 | 0 | 0 | 0 | 0 | 0 | 0 | 0 | 0 |
| ENSMUSG00 | AC132271.1 | 77 | 0 | 0 | 0 | 0 | 0 | 0 | 0 | 0.99201045 | 0 | 0 | 0 | 0 | 0 | 0 | 0 | 0 | 0 | 0 |
| ENSMUSG00 | AC157986.1 | 76 | 0 | 0 | 0 | 0 | 0 | 0 | 0 | 0 | 0 | 0 | 0 | 0 | 0 | 0 | 0 | 0 | 0 | 0 |
| ENSMUSG00 | AC132575.1 | 275 | 0 | 0 | 0 | 0 | 0 | 0 | 0 | 0 | 0 | 0 | 0 | 0 | 0 | 0 | 0 | 0 | 0 | 0 |
| ENSMUSG00 | Mir9-3 | 89 | 0 | 0 | 0 | 0 | 0 | 0 | 0 | 0 | 0 | 0 | 0 | 0 | 0 | 0 | 0 | 0 | 0 | 0 |
| ENSMUSG00 | AC160118.1 | 126 | 0 | 0 | 0 | 0 | 0 | 0 | 0 | 0 | 0.32687711 | 0 | 0 | 0.44298145 | 0 | 0 | 0 | 0.60778742 | 0 | 0 |
| ENSMUSG00 | AL807771.1 | 111 | 0 | 0 | 0 | 0 | 0 | 0 | 0 | 0 | 0 | 0 | 0 | 0 | 0 | 0 | 0 | 0 | 0 | 0 |
| ENSMUSG00 | Mir3108 | 78 | 0 | 0 | 0 | 0 | 0 | 0 | 0 | 0 | 0 | 0 | 0 | 0 | 0 | 0 | 0 | 0 | 0 | 0 |
| ENSMUSG00 | AL772404.1 | 73 | 0 | 0 | 0 | 0 | 0 | 0 | 0 | 0 | 0 | 0 | 0 | 0 | 0 | 0 | 0 | 0 | 0 | 0 |
| ENSMUSG00 | AL583889.1 | 76 | 0 | 0 | 0 | 0 | 0 | 0 | 0 | 0 | 0 | 0 | 0 | 0 | 0 | 0 | 0 | 0 | 0 | 0 |
| ENSMUSG00 | AL645963.1 | 230 | 0 | 0 | 0 | 0 | 0 | 0 | 0 | 0 | 0 | 0 | 0 | 0 | 0 | 0 | 0 | 0 | 0 | 0 |
| ENSMUSG00 | AC131120.1 | 71 | 8.97996677 | 3.75356555 | 10.2197599 | 8.34216305 | 11.5245451 | 11.864739 | 11.9947637 | 25.2822945 | 19.1430285 | 24.5714932 | 16.1492817 | 18.0811301 | 18.0652091 | 11.1204153 | 13.3469829 | 8.62886927 | 21.5922799 | 7.74776151 |
| ENSMUSG00 | AC084822.2 | 87 | 0 | 0 | 0 | 0 | 0 | 0 | 0 | 0 | 0 | 0 | 0 | 0 | 0 | 0 | 0 | 0 | 0 | 0 |
| ENSMUSG00 | AC120136.2 | 190 | 0 | 0 | 0 | 0 | 0 | 0 | 2.2411269 | 0.40202529 | 0 | 0 | 0.25144715 | 1.46883323 | 0 | 0 | 0.43370059 | 0 | 0.4034347 | 0.28952161 |
| ENSMUSG00 | AC135292.1 | 100 | 0 | 0 | 0 | 0 | 0 | 0 | 0 | 0 | 0 | 0 | 0 | 0 | 1.97327669 | 0 | 0 | 0 | 0 | 0 |
| ENSMUSG00 | AC148990.1 | 242 | 0.17564122 | 0 | 0.39978124 | 0.18826878 | 0 | 0 | 0.19550694 | 0 | 0.34038443 | 0 | 0 | 0.46128647 | 0 | 0.19191772 | 0.17025436 | 0.3164513 | 0.31674626 | 0.22731036 |
| ENSMUSG00 | AC165248.1 | 69 | 0 | 0 | 0 | 0 | 0 | 0 | 0 | 0 | 0 | 0 | 0 | 0 | 0 | 0 | 0 | 0 | 0 | 0 |
| ENSMUSG00 | AC154311.1 | 71 | 0 | 0 | 0 | 0 | 0 | 0 | 0 | 0 | 0 | 0.66409441 | 0 | 0 | 3.47407868 | 0 | 0 | 0 | 0 | 0 |
| ENSMUSG00 | AC109245.1 | 82 | 0 | 0 | 0 | 0 | 0 | 0 | 0 | 0 | 0 | 0 | 0 | 0 | 0 | 0 | 0 | 0 | 0 | 0 |
| ENSMUSG00 | AL671211.1 | 98 | 0 | 0 | 0 | 0 | 0 | 0 | 0 | 0 | 0 | 0 | 0 | 0 | 0 | 0 | 0 | 0 | 0 | 0 |
| ENSMUSG00 | Mir5126 | 76 | 0 | 0 | 0 | 0 | 0 | 0 | 0 | 0 | 0 | 0 | 0 | 0 | 0 | 0 | 0 | 0 | 0 | 0 |
| ENSMUSG00 | AC132439.1 | 100 | 0 | 0 | 0 | 0 | 0 | 0 | 0 | 0 | 0 | 0 | 0 | 0 | 0 | 0 | 0 | 0 | 0 | 0 |
| ENSMUSG00 | AC134560.1 | 73 | 0.58226269 | 1.36902305 | 0 | 1.24824779 | 0.65934142 | 0 | 0.64811889 | 0 | 1.69259654 | 0 | 3.92670892 | 0.76459812 | 0 | 0 | 0 | 0.52452887 | 1.5750533 | 0 |
| ENSMUSG00 | CT010471.1 | 125 | 0 | 0 | 0 | 0 | 0 | 0 | 0 | 0 | 0 | 0 | 0 | 0 | 0 | 0 | 0 | 0 | 0 | 0 |
| ENSMUSG00 | AC132284.1 | 106 | 0 | 0 | 0 | 0 | 0 | 0 | 0 | 1.08091705 | 3.10841628 | 0 | 0 | 0.52656285 | 0 | 0 | 0 | 0 | 0 | 0 |
| ENSMUSG00 | AC133523.1 | 221 | 0 | 0 | 0 | 0 | 0 | 0 | 0 | 0 | 0 | 0 | 0 | 0 | 0 | 0 | 0 | 0 | 0 | 0 |
| ENSMUSG00 | AC122824.1 | 82 | 0 | 0 | 0 | 0 | 0 | 0 | 0 | 0 | 0 | 0 | 0 | 0 | 0 | 0 | 0 | 0 | 0 | 0 |
| ENSMUSG00 | AC142474.1 | 94 | 0 | 0 | 0 | 0 | 0 | 0 | 0 | 0 | 0 | 0 | 0 | 0 | 0 | 0 | 0 | 0 | 0 | 0 |
| ENSMUSG00 | AC132380.1 | 124 | 0 | 0 | 0 | 0 | 0.38816067 | 0 | 0 | 0.30800325 | 0 | 0 | 0 | 0.45012631 | 0 | 0 | 0 | 0 | 0.30908304 | 0 |
| ENSMUSG00 | AL645988.1 | 93 | 0 | 0 | 0 | 0 | 0 | 0 | 0 | 0 | 0 | 0 | 0 | 0 | 0 | 0 | 0 | 0 | 0 | 0 |
| ENSMUSG00 | AL669855.1 | 83 | 0 | 0 | 1.74844086 | 0 | 0 | 0 | 0.57003228 | 0.46014943 | 0 | 0 | 0 | 0 | 0.59436045 | 1.67870195 | 0.49640429 | 1.38399786 | 1.38528784 | 0.66276032 |
| ENSMUSG00 | AC127575.1 | 293 | 0 | 0 | 0 | 0 | 0 | 0 | 0 | 0 | 0 | 0 | 0 | 0 | 0 | 0 | 0 | 0 | 0 | 0 |
| ENSMUSG00 | AL671848.1 | 82 | 0 | 0 | 0 | 0 | 0 | 0 | 0 | 0 | 0 | 0 | 0 | 0 | 0 | 0 | 0 | 0 | 0 | 0 |

| | | | | | | | | | | | | | | | | | | | |
|---|---|---|---|---|---|---|---|---|---|---|---|---|---|---|---|---|---|---|---|
| ENSMUSG00 | AC122525.1 | 275 | 0 | 0.48455119 | 0.70361499 | 0.16567652 | 0 | 0 | 0 | 0 | 0 | 0 | 0.34745424 | 0 | 0 | 0 | 0.14982384 | 0 | 0 | 0 |
| ENSMUSG00 | AC154368.1 | 125 | 0 | 0 | 0 | 0 | 0 | 0 | 0 | 0 | 0 | 0 | 0 | 0 | 0 | 0 | 0 | 0 | 0 | 0 |
| ENSMUSG00 | Mir5124 | 71 | 0 | 0 | 0 | 0 | 0 | 2.7917033 | 0 | 0 | 0 | 0 | 0 | 0 | 0 | 0 | 0 | 0 | 0 | 0 |
| ENSMUSG00 | AC117577.2 | 77 | 0 | 0 | 0 | 0 | 0 | 0 | 0 | 0 | 0 | 0 | 0 | 0 | 0 | 0 | 0 | 0 | 0 | 0 |
| ENSMUSG00 | AC027654.1 | 69 | 0 | 0 | 0 | 0 | 0 | 0 | 0 | 0 | 0 | 0 | 0 | 0 | 0 | 0 | 0 | 0 | 0 | 0 |
| ENSMUSG00 | Mir3064 | 66 | 0 | 0 | 0 | 0 | 0 | 0 | 0 | 0 | 0 | 0 | 0 | 0 | 0 | 0 | 0 | 0 | 0 | 0 |
| ENSMUSG00 | AC122304.1 | 100 | 0 | 0 | 0 | 0 | 0 | 0 | 0 | 0 | 0 | 0 | 0 | 0 | 0 | 0 | 0 | 0 | 0 | 0 |
| ENSMUSG00 | AC158667.1 | 243 | 0 | 0 | 0 | 0 | 0 | 0 | 0 | 0 | 0 | 0 | 0 | 0 | 0.81204802 | 0 | 0 | 0 | 0 | 0 |
| ENSMUSG00 | AC171917.1 | 124 | 0 | 0.53730475 | 0 | 0 | 0.38816067 | 0 | 0 | 0 | 0 | 0 | 0 | 0 | 0 | 0 | 0 | 0 | 0 | 0 |
| ENSMUSG00 | AL772271.2 | 136 | 0 | 0 | 0 | 0 | 0 | 0 | 0 | 0 | 0 | 0 | 0 | 0 | 0 | 0 | 0 | 0 | 0 | 0 |
| ENSMUSG00 | Mir5110 | 86 | 0 | 0 | 0.56248291 | 0.52977959 | 2.23869412 | 0 | 0 | 0.8881954 | 0 | 0 | 0 | 0.64901933 | 0 | 0 | 0 | 1.33571886 | 0 | 0 |
| ENSMUSG00 | AC090443.1 | 116 | 0 | 0 | 0 | 0 | 0 | 0 | 0 | 0 | 0 | 0 | 0 | 0.48116951 | 0 | 0 | 0 | 0 | 0 | 0.94843287 |
| ENSMUSG00 | Mir2861 | 81 | 0 | 0 | 0 | 0 | 0 | 0 | 0 | 0 | 0 | 0 | 0 | 0 | 0 | 0 | 0 | 0 | 0 | 0 |
| ENSMUSG00 | AC134552.1 | 68 | 0 | 0 | 0 | 0 | 0 | 0 | 0 | 0 | 0 | 0 | 0 | 0 | 0 | 0 | 0 | 0 | 0 | 0 |
| ENSMUSG00 | AC153974.1 | 84 | 0 | 0 | 0 | 0.54239338 | 0 | 0 | 0 | 0 | 0 | 0 | 0 | 0 | 0 | 0 | 0 | 0 | 0 | 0 |
| ENSMUSG00 | AC102240.1 | 341 | 0 | 0 | 0 | 0 | 0 | 0 | 0 | 0 | 0 | 0 | 0 | 0 | 0 | 0 | 0 | 0 | 0 | 0 |
| ENSMUSG00 | AC131316.1 | 290 | 0 | 0 | 0 | 0 | 0 | 0 | 0 | 0 | 0 | 0 | 0 | 0 | 0 | 0 | 0 | 0 | 0 | 0 |
| ENSMUSG00 | AL807811.2 | 70 | 0 | 0 | 0 | 0 | 0 | 0 | 0 | 0 | 0 | 0 | 0 | 0 | 0 | 0 | 0 | 0 | 0 | 0.78584438 |
| ENSMUSG00 | AC116712.1 | 123 | 0 | 0 | 0 | 0 | 0 | 0 | 0 | 0.31050734 | 0 | 0 | 0 | 0 | 0 | 0 | 0 | 0 | 0 | 0 |
| ENSMUSG00 | AC153624.1 | 303 | 0 | 0 | 0 | 0 | 0 | 0 | 0 | 0 | 0 | 0 | 0 | 0 | 0 | 0 | 0 | 0 | 0 | 0 |
| ENSMUSG00 | AC153909.1 | 125 | 0 | 0 | 0 | 0 | 0 | 0 | 0 | 0.61107844 | 0 | 0 | 0 | 0 | 0 | 0 | 0 | 0 | 0 | 0 |
| ENSMUSG00 | Snord92 | 84 | 0 | 0 | 1.15175072 | 0 | 3.43799455 | 0 | 0.56324618 | 0 | 0 | 0.5613179 | 0 | 0 | 0 | 0 | 0 | 1.36752169 | 0 | 0.65487032 |
| ENSMUSG00 | Mir3572 | 85 | 0 | 0 | 0 | 0 | 0 | 0 | 0 | 0 | 0 | 0 | 0 | 0 | 0 | 0 | 0 | 0 | 0 | 0 |
| ENSMUSG00 | AC121784.4 | 116 | 0 | 0 | 0 | 0 | 0 | 0 | 0 | 0 | 0 | 0 | 0 | 0 | 0 | 0 | 0 | 0 | 0 | 0 |
| ENSMUSG00 | Mir5118 | 66 | 0 | 0 | 0 | 0 | 0 | 0 | 0 | 0 | 0 | 0 | 0 | 0 | 0 | 0 | 0 | 0 | 0 | 0 |
| ENSMUSG00 | Mir3095 | 86 | 0 | 0 | 0 | 0 | 0 | 0 | 0 | 0 | 0 | 0 | 0 | 0 | 0 | 0 | 0 | 0 | 0 | 0 |
| ENSMUSG00 | AL671011.2 | 75 | 0 | 0.44417192 | 0 | 0 | 0 | 0 | 0 | 0 | 0 | 0 | 0 | 0 | 1.97327669 | 0 | 0 | 0 | 1.53305188 | 0 |
| ENSMUSG00 | AC166252.2 | 100 | 0 | 0 | 0 | 0 | 0 | 0 | 0 | 0 | 0 | 0 | 0 | 0 | 0 | 0 | 0 | 0 | 0 | 0 |
| ENSMUSG00 | Mir3102 | 103 | 0 | 0 | 0 | 0 | 0 | 0 | 0 | 0 | 0 | 0 | 0 | 0 | 0 | 0 | 0 | 0 | 0 | 0 |
| ENSMUSG00 | AL645683.2 | 100 | 0 | 0 | 0 | 0 | 0 | 0 | 0 | 0 | 0 | 0 | 0 | 0 | 0 | 0 | 0 | 0 | 0 | 0 |
| ENSMUSG00 | Mir3473b | 54 | 0 | 0 | 0 | 0 | 0 | 0 | 0 | 0 | 0 | 0 | 0 | 0 | 0 | 0 | 0 | 0 | 0 | 0 |
| ENSMUSG00 | AL844901.1 | 74 | 0 | 0 | 0 | 0 | 0 | 0 | 0 | 0 | 0 | 0 | 0 | 0 | 0 | 0 | 0 | 0 | 0 | 0 |
| ENSMUSG00 | AC159245.1 | 119 | 0 | 0 | 0 | 0 | 0 | 0 | 0 | 0 | 0 | 0 | 0 | 0 | 0 | 0 | 0 | 0 | 0 | 0 |
| ENSMUSG00 | CAAA01039[illegible] | 101 | 0 | 0 | 0 | 0 | 0 | 0 | 0 | 0 | 0 | 0 | 0 | 0 | 0 | 0 | 0 | 0 | 0 | 0 |
| [illegible] | [illegible] | [illegible] | [illegible] | [illegible] | [illegible] | [illegible] | [illegible] | [illegible] | [illegible] | [illegible] | [illegible] | [illegible] | [illegible] | [illegible] | [illegible] | [illegible] | [illegible] | [illegible] | [illegible] | [illegible] |

| | | | | | | | | | | | | | | | | | | | |
|---|---|---|---|---|---|---|---|---|---|---|---|---|---|---|---|---|---|---|---|
| ENSMUSG00 Gm18397 | 586 | 0 | 0 | 0 | 0 | 0 | 0 | 0 | 0 | 0 | 0 | 0 | 0 | 0 | 0 | 0 | 0 | 0 | 0 |
| ENSMUSG00 AL691456.1 | 70 | 0 | 0 | 0 | 0 | 0 | 0 | 0 | 0 | 0 | 0 | 0 | 0 | 0 | 0 | 0 | 0 | 0 | 0 |
| ENSMUSG00 Vmn1r-ps142 | 851 | 0 | 0 | 0 | 0 | 0 | 0 | 0 | 0 | 0 | 0 | 0 | 0 | 0 | 0 | 0 | 0 | 0 | 0 |
| ENSMUSG00 AL591496.4 | 81 | 0 | 0 | 0 | 0 | 0 | 0 | 0 | 0 | 0 | 0 | 0 | 0 | 0 | 0 | 0 | 0 | 0 | 0 |
| ENSMUSG00 Gm3965 | 1133 | 0 | 0 | 0 | 0 | 0 | 0 | 0 | 0 | 0 | 0 | 0 | 0 | 0 | 0 | 0 | 0 | 0 | 0 |
| ENSMUSG00 RP24-91B1.4 | 258 | 0 | 0 | 0 | 0 | 0 | 0 | 0 | 0 | 0 | 0 | 0 | 0 | 0 | 0 | 0 | 0.74206603 | 0 | 0 |
| ENSMUSG00 RP23-278M1 | 2430 | 0 | 0 | 0 | 0 | 0 | 0 | 0 | 0 | 0 | 0 | 0 | 0 | 0 | 0 | 0 | 0 | 0 | 0 |
| ENSMUSG00 AC105298.1 | 79 | 0 | 0 | 0 | 0 | 0 | 0 | 0 | 0 | 0 | 0 | 0 | 0 | 0 | 0 | 0 | 0 | 0 | 0 |
| ENSMUSG00 Vmn2r-ps125 | 930 | 0 | 0 | 0 | 0 | 0 | 0 | 0 | 0 | 0 | 0 | 0 | 0 | 0 | 0 | 0 | 0 | 0 | 0 |
| ENSMUSG00 AC162914.1 | 70 | 0 | 0 | 0 | 0 | 0 | 0 | 0 | 0 | 0 | 0 | 0 | 0 | 0 | 0 | 0 | 0 | 0 | 0 |
| ENSMUSG00 Vmn1r235 | 986 | 0 | 0 | 0 | 0 | 0 | 0 | 0 | 0 | 0 | 0 | 0 | 0 | 0 | 0 | 0 | 0 | 0 | 0 |
| ENSMUSG00 Vmn2r-ps132 | 650 | 0 | 0 | 0 | 0 | 0 | 0 | 0 | 0 | 0 | 0 | 0 | 0 | 0 | 0 | 0 | 0 | 0 | 0 |
| ENSMUSG00 AC150683.1 | 68 | 0 | 0 | 0 | 0 | 0 | 0 | 0 | 0 | 0 | 0 | 0 | 0 | 0 | 0 | 0 | 0 | 0 | 0 |
| ENSMUSG00 Gm19815 | 334 | 0 | 0.79791383 | 0 | 0.40923094 | 0.72053778 | 0 | 0.28330945 | 0.45739404 | 0 | 0.42350931 | 0.1430388 | 0 | 1.18160281 | 0.13905415 | 0.49343181 | 0 | 0.22949878 | 0 |
| ENSMUSG00 RP23-345J21 | 1607 | 0 | 0 | 0 | 0 | 0 | 0 | 0 | 0 | 0 | 0 | 0 | 0 | 0 | 0 | 0 | 0 | 0 | 0 |
| ENSMUSG00 RP23-322O2 | 1529 | 0 | 0 | 0 | 0 | 0 | 0 | 0 | 0 | 0 | 0 | 0 | 0 | 0 | 0 | 0 | 0 | 0 | 0 |
| ENSMUSG00 Gm4016 | 947 | 0 | 0 | 0 | 0 | 0 | 0 | 0 | 0 | 0 | 0 | 0 | 0 | 0 | 0 | 0 | 0 | 0 | 0 |
| ENSMUSG00 RP23-372N1 | 1285 | 0.06615592 | 0.02592443 | 0.15057908 | 0.24819246 | 0.03745675 | 0.07712488 | 0 | 0.05944343 | 0 | 0.11007946 | 0.22307374 | 0 | 0.19195299 | 0.03614326 | 0 | 0.05959628 | 0 | 0.12842593 |
| ENSMUSG00 Gm3896 | 4092 | 0 | 0 | 0 | 0 | 0 | 0 | 0 | 0 | 0 | 0 | 0 | 0 | 0 | 0 | 0 | 0 | 0 | 0 |
| ENSMUSG00 RP24-166B2. | 235 | 0 | 0 | 0 | 0 | 0 | 0 | 0 | 0 | 0 | 0 | 0 | 0 | 0 | 0 | 0 | 0 | 0 | 0 |
| ENSMUSG00 Gm3987 | 1134 | 0 | 0 | 0 | 0 | 0 | 0 | 0 | 0 | 0 | 0 | 0 | 0 | 0 | 0 | 0 | 0 | 0 | 0 |
| ENSMUSG00 Vmn2r-ps32 | 756 | 0 | 0 | 0 | 0 | 0 | 0 | 0 | 0 | 0 | 0 | 0 | 0 | 0 | 0 | 0 | 0 | 0 | 0 |
| ENSMUSG00 Gm18595 | 1066 | 0 | 0 | 0 | 0 | 0 | 0 | 0 | 0 | 0 | 0 | 0 | 0 | 0 | 0 | 0 | 0 | 0 | 0 |
| ENSMUSG00 RP23-358B2 | 1929 | 0 | 0 | 0 | 0 | 0 | 0 | 0 | 0 | 0 | 0 | 0.22290027 | 0 | 0 | 0 | 0 | 0 | 0 | 0 |
| ENSMUSG00 Gm8053 | 1000 | 0 | 0 | 0 | 0 | 0 | 0 | 0 | 0 | 0 | 0 | 0 | 0 | 0 | 0 | 0 | 0 | 0 | 0 |
| ENSMUSG00 Vmn1r-ps8 | 837 | 0 | 0 | 0 | 0 | 0 | 0 | 0 | 0 | 0 | 0 | 0 | 0 | 0 | 0 | 0 | 0 | 0 | 0 |
| ENSMUSG00 Vmn1r-ps141 | 752 | 0 | 0 | 0 | 0 | 0 | 0 | 0 | 0 | 0 | 0 | 0 | 0 | 0 | 0 | 0 | 0 | 0 | 0 |
| ENSMUSG00 RP24-399B3. | 314 | 0 | 0 | 0 | 0 | 0 | 0 | 0 | 0 | 0 | 0 | 0 | 0 | 0 | 0 | 0 | 0 | 0 | 0 |
| ENSMUSG00 Vmn1r-ps18 | 874 | 0 | 0 | 0 | 0 | 0 | 0 | 0 | 0 | 0 | 0 | 0 | 0 | 0 | 0 | 0 | 0 | 0 | 0 |
| ENSMUSG00 RP23-60H16. | 1523 | 0 | 0 | 0 | 0 | 0 | 0 | 0 | 0 | 0 | 0 | 0 | 0 | 0 | 0 | 0 | 0 | 0 | 0 |
| ENSMUSG00 AL663090.2 | 109 | 0 | 0 | 0 | 0 | 0 | 0 | 0 | 0 | 0 | 0 | 0 | 0 | 0 | 0 | 0 | 0 | 0 | 0 |
| ENSMUSG00 AL645907.1 | 254 | 2.6774914 | 3.67228755 | 2.09491667 | 2.86998704 | 3.78991525 | 5.26741656 | 1.49016312 | 2.85691199 | 2.91872946 | 2.04195959 | 2.82135582 | 2.85670714 | 2.13642161 | 2.74276107 | 2.5953736 | 2.56275719 | 2.26336399 | 3.89828315 |
| ENSMUSG00 Vmn2r-ps33 | 903 | 0 | 0 | 0 | 0 | 0 | 0 | 0 | 0 | 0 | 0 | 0 | 0 | 0 | 0 | 0 | 0 | 0 | 0 |
| ENSMUSG00 RP23-349I13. | 723 | 0 | 0 | 0 | 0 | 0 | 0 | 0 | 0 | 0 | 0 | 0 | 0 | 0 | 0 | 0 | 0 | 0 | 0 |
| ENSMUSG00 RP24-356D2 | 663 | 0 | 0 | 0 | 0 | 0 | 0 | 0 | 0 | 0 | 0 | 0 | 0 | 0 | 0 | 0 | 0 | 0 | 0 |
| ENSMUSG00 AI606473 | 1728 | 0 | 0 | 0 | 0 | 0 | 0 | 0 | 0 | 0 | 0 | 0 | 0 | 0 | 0 | 0 | 0 | 0 | 0 |
| ENSMUSG00 RP23-329D1 | 679 | 0.31299835 | 1.0793574 | 0.78366544 | 0.33550106 | 0.70886486 | 0.43787393 | 1.04519909 | 0.84372023 | 0.7885489 | 0.41664834 | 0.70360764 | 0.16440549 | 0.50857647 | 0.34200359 | 1.09223565 | 0.05639265 | 1.01601376 | 0 |
| ENSMUSG00 Gm18417 | 647 | 0 | 0 | 0 | 0.07041892 | 0 | 0 | 0 | 0 | 0 | 0 | 0 | 0.43134206 | 0 | 0 | 0 | 0.23672709 | 0 | 1.27532705 |
| ENSMUSG00 RP24-97C18. | 260 | 0 | 0 | 0 | 0 | 0 | 0 | 0 | 0 | 0 | 0 | 0 | 0 | 0 | 0 | 0 | 0 | 0 | 0 |
| ENSMUSG00 Gm3977 | 1359 | 0 | 0 | 0 | 0 | 0 | 0 | 0 | 0 | 0 | 0 | 0 | 0 | 0 | 0 | 0 | 0 | 0 | 0 |
| ENSMUSG00 Vmn1r-ps17 | 893 | 0 | 0 | 0 | 0 | 0 | 0 | 0 | 0 | 0 | 0 | 0 | 0 | 0 | 0 | 0 | 0 | 0 | 0 |
| ENSMUSG00 RP23-177N1 | 336 | 0 | 0 | 0 | 0 | 0 | 0 | 0 | 0 | 0 | 0 | 0 | 0 | 0 | 0 | 0 | 0 | 0 | 0 |
| ENSMUSG00 AC136903.1 | 58 | 0 | 0 | 0 | 0 | 0 | 0 | 0 | 0 | 0 | 0 | 0 | 0 | 0 | 0 | 0 | 0 | 0 | 0 |
| ENSMUSG00 AL833801.1 | 59 | 0 | 0 | 0 | 0 | 0 | 0 | 0 | 0 | 0 | 0 | 0 | 0 | 0 | 0 | 0 | 0 | 0 | 0 |
| ENSMUSG00 RP24-97C18. | 429 | 0 | 0 | 0 | 0 | 0 | 0 | 0 | 0 | 0 | 0 | 0 | 0 | 0 | 0 | 0 | 0 | 0 | 0 |
| ENSMUSG00 Gm19031 | 1364 | 0 | 0 | 0 | 0 | 0 | 0 | 0 | 0 | 0 | 0 | 0 | 0 | 0 | 0 | 0 | 0 | 0 | 0 |
| ENSMUSG00 4921523L03R | 1184 | 0 | 0 | 0 | 0 | 0 | 0.08370394 | 0 | 0 | 0 | 0 | 0 | 0.04714161 | 0 | 0 | 0 | 0 | 0.16185092 | 0 |
| ENSMUSG00 AC139849.1 | 1617 | 0 | 0 | 0 | 0 | 0 | 0 | 0 | 0 | 0 | 0 | 0 | 0 | 0 | 0 | 0 | 0 | 0 | 0 |
| ENSMUSG00 Gm3307 | 631 | 0.06736161 | 0 | 0.0766617 | 0.07220451 | 0 | 0 | 0 | 0.24210715 | 0 | 0 | 0.15142618 | 0 | 0.07818053 | 0 | 0.06529565 | 0.12136484 | 0.18221694 | 0 |
| ENSMUSG00 RP23-391M1 | 5276 | 0 | 0 | 0 | 0 | 0 | 0 | 0 | 0 | 0 | 0 | 0 | 0 | 0 | 0 | 0 | 0 | 0 | 0 |
| ENSMUSG00 RP23-309A11 | 738 | 0 | 0 | 0 | 0 | 0 | 0 | 0 | 0 | 0 | 0 | 0 | 0 | 0 | 0 | 0 | 0 | 0 | 0 |
| ENSMUSG00 Vmn2r-ps130 | 2562 | 0 | 0 | 0 | 0 | 0 | 0 | 0 | 0 | 0 | 0 | 0 | 0 | 0 | 0 | 0 | 0 | 0 | 0 |
| ENSMUSG00 Vmn1r39 | 1231 | 0 | 0 | 0 | 0 | 0 | 0 | 0 | 0 | 0 | 0 | 0 | 0 | 0 | 0 | 0 | 0 | 0 | 0 |
| ENSMUSG00 Gm18594 | 1402 | 0 | 0 | 0 | 0 | 0 | 0 | 0 | 0 | 0 | 0 | 0 | 0 | 0 | 0 | 0 | 0 | 0 | 0 |
| ENSMUSG00 AC140469.1 | 69 | 0 | 0 | 0 | 0 | 0 | 0 | 0 | 0 | 0 | 0 | 0 | 0 | 0 | 0 | 0 | 0 | 0 | 0 |
| ENSMUSG00 RP24-262G2 | 346 | 0 | 0.19256008 | 0 | 0 | 0 | 0.14321599 | 0 | 0 | 0 | 0 | 0 | 0 | 0 | 0 | 0.11907964 | 0.1106665 | 0 | 0 |
| ENSMUSG00 2600006L11R | 302 | 0 | 0 | 0 | 0 | 0.31875446 | 0 | 0 | 0 | 0 | 0 | 0 | 0 | 0 | 0 | 0 | 0 | 0 | 0 |
| ENSMUSG00 RP23-152I2.1 | 1049 | 0 | 0.28581129 | 0.50725342 | 0.30402985 | 0.04588363 | 0.04723807 | 0.0902053 | 0.10922517 | 0 | 0.08989648 | 0.04554334 | 0.1064169 | 0 | 0.04427463 | 0 | 0.03650201 | 0 | 0 |
| ENSMUSG00 Gm5496 | 1041 | 0 | 0 | 0 | 0 | 0 | 0 | 0 | 0 | 0 | 0 | 0 | 0 | 0 | 0 | 0 | 0 | 0 | 0 |
| ENSMUSG00 AC147634.1 | 93 | 0 | 0 | 0 | 0 | 0 | 0 | 0 | 0 | 0 | 0 | 0 | 0 | 0 | 0 | 0 | 0 | 0 | 1.18299154 |
| ENSMUSG00 AC157086.1 | 59 | 0 | 0 | 0 | 0 | 0 | 0 | 0 | 0 | 0 | 0 | 0 | 0 | 0.83613419 | 0 | 0 | 0 | 0 | 0 |
| ENSMUSG00 Vmn1r36 | 1028 | 0 | 0 | 0 | 0 | 0 | 0 | 0 | 0 | 0 | 0 | 0 | 0 | 0 | 0 | 0 | 0 | 0 | 0 |
| ENSMUSG00 RP24-386J17 | 3151 | 0 | 0 | 0.0153518 | 0 | 0 | 0 | 0.06006053 | 0.01212072 | 0 | 0.02992745 | 0 | 0.14170908 | 0.01565596 | 0 | 0 | 0 | 0 | 0.43644166 |
| ENSMUSG00 RP24-347O5. | 1398 | 0 | 0 | 0 | 0 | 0 | 0 | 0 | 0 | 0 | 0 | 0 | 0 | 0 | 0 | 0 | 0 | 0 | 0 |
| ENSMUSG00 Vmn2r-ps13 | 605 | 0 | 0 | 0 | 0 | 0 | 0 | 0 | 0 | 0 | 0 | 0 | 0 | 0 | 0 | 0 | 0 | 0 | 0 |
| ENSMUSG00 RP23-142A14 | 196 | 0 | 0 | 0 | 0 | 0 | 0 | 0 | 0 | 0 | 0 | 0 | 0 | 0 | 0 | 0 | 0 | 0 | 0 |
| ENSMUSG00 Hist2h3c1 | 2364 | 0 | 0 | 0 | 0 | 0 | 0 | 0 | 0 | 0 | 0 | 0 | 0 | 0 | 0 | 0 | 0 | 0 | 0 |
| ENSMUSG00 Gm8072 | 1002 | 0 | 0 | 0 | 0 | 0 | 0 | 0 | 0 | 0 | 0 | 0 | 0 | 0 | 0 | 0 | 0 | 0 | 0 |
| ENSMUSG00 AU023070 | 385 | 0 | 0 | 0 | 0 | 0 | 0 | 0 | 0 | 0 | 0 | 0 | 0 | 0 | 0 | 0 | 0 | 0 | 0 |
| ENSMUSG00 4931403E22R | 664 | 0 | 0 | 0 | 0.06861603 | 0 | 0.07462761 | 0.07125403 | 0.40263075 | 0 | 0.14202019 | 0.07195024 | 0 | 0.07429506 | 0.20983774 | 0.24820214 | 0 | 0 | 0 |
| ENSMUSG00 Gm18355 | 408 | 0 | 0 | 0 | 0 | 0 | 0 | 0 | 0 | 0 | 0 | 0 | 0 | 0 | 0 | 0 | 0 | 0 | 0 |
| ENSMUSG00 Vmn1r-ps151 | 981 | 0 | 0 | 0 | 0 | 0 | 0.10102494 | 0 | 0 | 0 | 0 | 0 | 0 | 0 | 0 | 0 | 0 | 0 | 0 |
| ENSMUSG00 Vmn2r-ps114 | 1279 | 0 | 0 | 0 | 0 | 0 | 0 | 0 | 0 | 0 | 0 | 0 | 0 | 0 | 0 | 0 | 0 | 0 | 0 |
| ENSMUSG00 Vmn2r-ps120 | 2403 | 0 | 0 | 0 | 0 | 0 | 0 | 0 | 0 | 0 | 0 | 0 | 0 | 0 | 0 | 0 | 0 | 0 | 0 |
| ENSMUSG00 AC130205.1 | 68 | 0 | 0 | 0 | 0 | 0 | 0 | 0 | 0 | 0 | 0 | 0 | 0 | 0 | 0 | 0 | 0 | 0 | 0 |
| ENSMUSG00 RP23-71K9.1 | 1628 | 7.28436371 | 6.11827726 | 6.38839621 | 6.49272868 | 7.03648516 | 6.02668381 | 6.77140921 | 4.03506954 | 5.23686042 | 5.30010976 | 5.25289777 | 5.99984088 | 6.15133858 | 4.70717102 | 5.21346469 | 5.12736635 | 4.37880296 | 5.50766855 |
| ENSMUSG00 RP24-63G9.3 | 327 | 0 | 0.10187429 | 0 | 0 | 0 | 0 | 0 | 0.11679634 | 0 | 0 | 0 | 1.0241406 | 0 | 0 | 0 | 0 | 0.1172058 | 0 |
| ENSMUSG00 AC132271.2 | 95 | 0 | 0 | 0 | 0 | 0 | 0 | 0 | 0 | 0 | 0 | 0 | 0 | 0 | 0 | 0 | 0 | 0 | 0 |
| ENSMUSG00 Vmn1r-ps47 | 906 | 0 | 0 | 0 | 0 | 0 | 0 | 0 | 0 | 0 | 0 | 0 | 0 | 0 | 0 | 0 | 0 | 0 | 0 |
| ENSMUSG00 AC138214.1 | 52 | 0 | 0 | 0 | 0 | 0 | 0 | 0 | 0 | 0 | 0 | 0 | 0 | 0 | 0 | 0 | 0 | 0 | 0 |
| ENSMUSG00 AC132307.1 | 70 | 0 | 0 | 0 | 0 | 0 | 0 | 0 | 0 | 0 | 0 | 0 | 0 | 0 | 0 | 0 | 0 | 0 | 0 |
| ENSMUSG00 AC158612.2 | 75 | 0 | 0 | 0 | 1.82244177 | 0 | 0 | 0 | 0 | 0 | 0 | 0 | 0 | 0 | 0 | 0 | 0 | 0 | 0 |
| ENSMUSG00 Vmn1r10 | 971 | 0 | 0 | 0 | 0 | 0 | 0 | 0 | 0 | 0 | 0 | 0 | 0 | 0 | 0 | 0 | 0 | 0 | 0 |
| ENSMUSG00 AL591496.5 | 81 | 0 | 0 | 0 | 0 | 0 | 0 | 0 | 0 | 0 | 0 | 0 | 0 | 0 | 0 | 0 | 0 | 0 | 0 |
| ENSMUSG00 RP23-286K8. | 147 | 6.65046972 | 2.26618328 | 1.31628654 | 1.54969539 | 3.27428052 | 0.33709343 | 2.89669463 | 0.25981226 | 0.84054114 | 0.64150617 | 0 | 1.13909515 | 0 | 0 | 0 | 0 | 0.26072311 | 1.12263483 |
| ENSMUSG00 Vmn1r-ps147 | 512 | 0 | 0 | 0 | 0 | 0 | 0 | 0 | 0 | 0 | 0 | 0 | 0 | 0 | 0 | 0 | 0 | 0 | 0 |
| ENSMUSG00 RP24-73K9.3 | 1109 | 0 | 0 | 0 | 0 | 0 | 0 | 0 | 0 | 0 | 0 | 0 | 0 | 0 | 0 | 0 | 0 | 0 | 0 |

**immune-related genes**

| **Seg 1,2,3** | **Seg 4,5,6** | **Seg 8,9,10** | **Seg 7** |
|---|---|---|---|
| Abl1 | Abl1 | Abl1 | Abl1 |
| Actr1a | Actr1a | Actr1a | Actr1a |
| Adora2b | Adam8 | Adora2b | Adam8 |
| Ap1g1 | Adora2b | Ap1g1 | Adora2b |
| B2m | Ap1g1 | B2m | Ap1g1 |
| Bax | B2m | Bax | B2m |
| Bcar1 | Bax | Bcar1 | Bax |
| Bcl10 | Bcar1 | Bcl10 | Bcar1 |
| Bcl2 | Bcl10 | Bcl2 | Bcl10 |
| Blk | Bcl2 | Blk | Bcl2 |
| Braf | Braf | Braf | Blk |
| Btk | C1qb | C1qa | Braf |
| C1qa | C1qbp | C1qb | C1qb |
| C1qb | C2 | C1qbp | C1qbp |
| C1qbp | C9 | C2 | C2 |
| C2 | Cav1 | C3 | C3ar1 |
| C3ar1 | Cd14 | C3ar1 | C9 |
| Cav1 | Cd19 | Cav1 | Cav1 |
| Cd14 | Cd1d1 | Cd14 | Cd14 |
| Cd24a | Cd24a | Cd19 | Cd19 |
| Cd3e | Cd3e | Cd1d1 | Cd24a |
| Cd44 | Cd4 | Cd24a | Cd3e |
| Cfi | Cd44 | Cd3e | Cd4 |
| Colec12 | Cfi | Cd4 | Cd44 |
| Crhr1 | Cnr1 | Cd44 | Cfi |
| Cst11 | Colec12 | Cd79a | Cnr1 |
| Cyld | Crhr1 | Cfi | Coch |
| Ddx58 | Crtam | Cnr1 | Colec12 |
| Dhx15 | Cst11 | Colec12 | Crhr1 |
| Exosc3 | Cyld | Crhr1 | Crp |
| Exosc6 | Ddx58 | Crtam | Crtam |
| Fadd | Dhx15 | Cyba | Cst11 |
| Fcgr1 | Exosc3 | Cyld | Cyld |
| Fcna | F2rl1 | Ddx58 | Ddx58 |
| Fcnb | Fadd | Dhx15 | Dhx15 |
| Fgr | Fcer1a | Exosc3 | Exosc3 |
| Flot1 | Fcgr1 | F2rl1 | F2rl1 |
| Fpr-rs4 | Fcna | Fadd | Fadd |
| Gata2 | Fcnb | Fcer1a | Fcer1a |
| Gpr33 | Fgr | Fcgr1 | Fcgr1 |
| Gpx4 | Flot1 | Fcna | Fcna |
| H2-M3 | Gab2 | Fcnb | Fcnb |
| Hmgb1 | Gata2 | Fgr | Fgr |
| Hmgb2 | Gc | Flot1 | Flot1 |
| Hspa8 | Gpr33 | Fpr-rs4 | Foxp3 |
| Il4ra | Gpx4 | Gata2 | Fpr-rs4 |

| | | | |
|---|---|---|---|
| Irak1 | H2-M3 | Gata3 | Gata2 |
| Irak2 | Havcr2 | Gpr33 | Gc |
| Irak3 | Hmgb1 | Gpx4 | Gpr33 |
| Irak4 | Hmgb2 | H2-M3 | Gpx4 |
| Irf3 | Hspa8 | Havcr2 | H2-M3 |
| Irf7 | Il4ra | Hmgb1 | Havcr2 |
| Kars | Irak1 | Hmgb2 | Hmgb1 |
| Klrb1c | Irak2 | Hspa8 | Hmgb2 |
| Klre1 | Irak3 | Ifih1 | Hspa8 |
| Klrk1 | Irak4 | Il18 | Ifih1 |
| Lamp1 | Irf3 | Il4 | Il12b |
| Lbp | Irf7 | Irak1 | Il18 |
| Lck | Kars | Irak2 | Il4 |
| Lcp1 | Klk5 | Irak4 | Irak1 |
| Lgals9 | Klrb1c | Irf3 | Irak2 |
| Lrrfip2 | Klre1 | Irf7 | Irak4 |
| Ltf | Lamp1 | Kars | Irf3 |
| Ly96 | Lbp | Klk5 | Irf7 |
| Malt1 | Lck | Klrb1c | Kars |
| Mapk1 | Lcp1 | Klre1 | Klk5 |
| Mapk3 | Lgals9 | Klrk1 | Klrb1c |
| Mapkapk3 | Lrrfip2 | Lamp1 | Klre1 |
| Masp1 | Ly96 | Lbp | Klrk1 |
| Mef2c | Malt1 | Lck | Lag3 |
| Mif | Mapk1 | Lcp1 | Lamp1 |
| Mlh1 | Mapk3 | Lcp2 | Lbp |
| Mmp2 | Mapkapk3 | Lgals9 | Lck |
| Msh2 | Masp1 | Lrrfip2 | Lcp1 |
| Myd88 | Mef2c | Ltf | Lcp2 |
| Myo1g | Mif | Ly96 | Lgals9 |
| Nfatc2 | Mlh1 | Malt1 | Lrrfip2 |
| Nfkbia | Mmp12 | Mapk1 | Ltf |
| Nras | Mmp2 | Mapk3 | Ly96 |
| P2rx7 | Msh2 | Mapkapk3 | Malt1 |
| Park7 | Myb | Masp1 | Mapk1 |
| Pde4b | Myd88 | Mef2c | Mapk3 |
| Peli1 | Nfatc2 | Mif | Mapkapk3 |
| Phb | Nfkbia | Mlh1 | Masp1 |
| Pik3ap1 | Nr4a3 | Mmp12 | Mef2c |
| Plcg1 | Nras | Mmp2 | Mif |
| Plcg2 | P2rx7 | Msh2 | Mmp12 |
| Pms2 | Park7 | Myb | Mmp2 |
| Pnp | Pde4b | Myd88 | Msh2 |
| Polr3b | Peli1 | Nfatc2 | Myb |
| Polr3c | Phb | Nfkbia | Myd88 |
| Polr3d | Pik3ap1 | Nr4a3 | Myo1g |
| Polr3f | Plcg1 | Nras | Nfatc2 |
| Prkch | Plcg2 | P2rx7 | Nfkbia |

| | | | |
|---|---|---|---|
| Prkd2 | Pld2 | Park7 | Nr1h3 |
| Psen1 | Pms2 | Pde4b | Nr4a3 |
| Psen2 | Pnp | Peli1 | Nras |
| Pum2 | Polr3b | Phb | P2rx7 |
| Pycard | Polr3c | Pik3ap1 | Park7 |
| Rap1a | Polr3d | Plcg1 | Pde4b |
| Rapgef1 | Polr3f | Plcg2 | Peli1 |
| Rasgrp1 | Prkch | Pld2 | Phb |
| Rela | Prkd2 | Plscr1 | Pik3ap1 |
| Rnf31 | Psen1 | Pms2 | Plcg1 |
| Rps19 | Pum2 | Pnp | Plcg2 |
| Rps3 | Pycard | Polr3b | Pms2 |
| Rsad2 | Rap1a | Polr3c | Pnp |
| Serping1 | Rapgef1 | Polr3d | Polr3b |
| Shb | Rasgrp1 | Polr3f | Polr3c |
| Sin3a | Rela | Prkch | Polr3d |
| Slamf6 | Rnf31 | Prkd2 | Polr3f |
| Slc11a1 | Rps19 | Psen1 | Prkch |
| Slc39a10 | Rps3 | Psen2 | Prkd2 |
| Snx4 | Rsad2 | Pum2 | Psen1 |
| Spon2 | Serping1 | Pycard | Psen2 |
| Stat5b | Shb | Rap1a | Ptafr |
| Stat6 | Sin3a | Rapgef1 | Pum1 |
| Stk11 | Slamf6 | Rasgrp1 | Pum2 |
| Stoml2 | Slc11a1 | Rela | Pycard |
| Stx4a | Slc39a10 | Rnf31 | Rap1a |
| Stx7 | Snx4 | Rps19 | Rapgef1 |
| Tap2 | Spon2 | Rps3 | Rasgrp1 |
| Tec | Stat5a | Rsad2 | Rela |
| Tfrc | Stat5b | S100a14 | Rnf31 |
| Tgfb1 | Stat6 | Serping1 | Rps19 |
| Thy1 | Stk11 | Shb | Rps3 |
| Tlr12 | Stoml2 | Sin3a | Rsad2 |
| Tlr13 | Stx4a | Slamf6 | Serping1 |
| Tlr2 | Stx7 | Slc11a1 | Shb |
| Tlr5 | Syk | Slc39a10 | Sin3a |
| Tnfrsf21 | Tap2 | Snx4 | Sirt1 |
| Tnfsf13b | Tec | Socs5 | Slamf6 |
| Tnfsf4 | Tfrc | Spon2 | Slc11a1 |
| Tnip1 | Tgfb1 | Stat5a | Slc39a10 |
| Traf2 | Thy1 | Stat5b | Snx4 |
| Traf3 | Tlr12 | Stat6 | Spon2 |
| Traf6 | Tlr2 | Stk11 | Stat5a |
| Ube2n | Tlr5 | Stoml2 | Stat5b |
| Unc93b1 | Tlr9 | Stx4a | Stat6 |
| Ung | Tnfrsf21 | Stx7 | Stk11 |
| Usp12 | Tnfsf13b | Syk | Stoml2 |
| Vamp8 | Tnfsf4 | Tap2 | Stx4a |

| | | | |
|---|---|---|---|
| Vav1 | Tnip1 | Tec | Stx7 |
| Wdfy1 | Tnip2 | Tfrc | Syk |
| Wnt5a | Traf3 | Thy1 | Tap2 |
| Xbp1 | Traf6 | Tlr12 | Tec |
| Zap70 | Ube2n | Tlr13 | Tfrc |
| Zbtb1 | Ung | Tlr2 | Tgfb1 |
| Abl1 | Usp12 | Tlr3 | Thy1 |
| Actr1a | Vamp8 | Tlr5 | Tlr12 |
| Adora2b | Vav1 | Tlr8 | Tlr2 |
| Ap1g1 | Wdfy1 | Tlr9 | Tlr5 |
| B2m | Wnt5a | Tnfrsf21 | Tlr8 |
| Bax | Xbp1 | Tnfsf13b | Tlr9 |
| Bcar1 | Zap70 | Tnfsf4 | Tnfrsf21 |
| Bcl10 | Zbtb1 | Tnip1 | Tnfsf13b |
| Bcl2 | Abl1 | Tnip2 | Tnfsf4 |
| Braf | Actr1a | Traf2 | Tnip1 |
| C1qa | Adam8 | Traf3 | Tnip2 |
| C1qb | Adora2b | Traf6 | Traf3 |
| C1qbp | Ap1g1 | Ube2n | Traf6 |
| C2 | B2m | Unc93b1 | Txk |
| C3ar1 | Bax | Ung | Ube2n |
| Cav1 | Bcar1 | Usp12 | Ung |
| Cd14 | Bcl10 | Vamp8 | Usp12 |
| Cd19 | Bcl2 | Vav1 | Vamp8 |
| Cd24a | Braf | Wdfy1 | Vav1 |
| Cd3e | C1qb | Wnt5a | Wdfy1 |
| Cd4 | C1qbp | Xbp1 | Wnt5a |
| Cd44 | C2 | Zap70 | Xbp1 |
| Cfi | Cav1 | Zbtb1 | Zap70 |
| Colec12 | Cd14 | Abl1 | Zbtb1 |
| Crhr1 | Cd19 | Actr1a | Acp5 |
| Crtam | Cd1d1 | Adora2b | Adora1 |
| Cst11 | Cd24a | Ap1g1 | Adora2a |
| Cyld | Cd3e | B2m | Adrb2 |
| Ddx58 | Cd4 | Bax | Apod |
| Dhx15 | Cd44 | Bcar1 | Apoe |
| Exosc3 | Cfi | Bcl10 | Ash1l |
| Exosc6 | Colec12 | Bcl2 | Bcr |
| F2rl1 | Crhr1 | Braf | Calcrl |
| Fadd | Crtam | C1qa | Cd44 |
| Fcer1a | Cst11 | C1qb | Chrna7 |
| Fcgr1 | Cyld | C1qbp | Cnr2 |
| Fcna | Ddx58 | C2 | Cuedc2 |
| Fcnb | Dhx15 | C3 | Cx3cr1 |
| Fgr | Exosc3 | C3ar1 | Dusp10 |
| Flot1 | F2rl1 | Cav1 | Ets1 |
| Gab2 | Fadd | Cd14 | Fcgr2b |
| Gata2 | Fcer1a | Cd19 | Fem1a |

| | | | |
|---|---|---|---|
| Gpr33 | Fcgr1 | Cd1d1 | Fn1 |
| Gpx4 | Fcna | Cd24a | Foxp3 |
| H2-M3 | Fcnb | Cd3e | Gpx1 |
| Havcr2 | Fgr | Cd4 | Hamp |
| Hmgb1 | Flot1 | Cd44 | Ier3 |
| Hmgb2 | Gab2 | Cd79a | Il12b |
| Hspa8 | Gata2 | Cfi | Il22ra2 |
| Il4ra | Gc | Cnr1 | Il4 |
| Irak1 | Gpr33 | Colec12 | Ins2 |
| Irak2 | Gpx4 | Crhr1 | Lgals9 |
| Irak3 | H2-M3 | Crtam | Mefv |
| Irak4 | Havcr2 | Cst11 | Metrnl |
| Irf3 | Hmgb1 | Cyba | Ndfip1 |
| Irf7 | Hmgb2 | Cyld | Nr1h3 |
| Kars | Hspa8 | Ddx58 | Nt5e |
| Klk5 | Il4ra | Dhx15 | Pbk |
| Klrb1c | Irak1 | Exosc3 | Pparg |
| Klre1 | Irak2 | F2rl1 | Proc |
| Klrk1 | Irak3 | Fadd | Psma1 |
| Lamp1 | Irak4 | Fcer1a | Psmb4 |
| Lbp | Irf3 | Fcgr1 | Ptgds |
| Lck | Irf7 | Fcna | Ptgis |
| Lcp1 | Itk | Fcnb | Ptpn2 |
| Lcp2 | Kars | Fgr | Rabgef1 |
| Lgals9 | Klk5 | Flot1 | Rps19 |
| Lrrfip2 | Klrb1c | Gata2 | Serpinb1a |
| Ly96 | Klre1 | Gata3 | Serpinf1 |
| Malt1 | Lamp1 | Gpr33 | Slamf8 |
| Mapk1 | Lbp | Gpx4 | Slit2 |
| Mapk3 | Lck | H2-M3 | Smad3 |
| Mapkapk3 | Lcp1 | Havcr2 | Smpdl3b |
| Masp1 | Lgals9 | Hmgb1 | Spn |
| Mef2c | Lrrfip2 | Hmgb2 | Tff2 |
| Mif | Ly96 | Hspa8 | Tnfaip3 |
| Mlh1 | Malt1 | Ifih1 | Tnfrsf1a |
| Mmp12 | Mapk1 | Il18 | Tnfrsf1b |
| Mmp2 | Mapk3 | Irak1 | Uaca |
| Msh2 | Mapkapk3 | Irak2 | Zfp36 |
| Myb | Masp1 | Irak4 | Acp5 |
| Myd88 | Mef2c | Irf3 | Adora1 |
| Nfatc2 | Mif | Irf7 | Adora2a |
| Nfkbia | Mlh1 | Kars | Adrb2 |
| Nr4a3 | Mmp12 | Klk5 | Apod |
| Nras | Mmp2 | Klrb1c | Apoe |
| P2rx7 | Msh2 | Klre1 | Ash1l |
| Park7 | Myb | Lamp1 | Bax |
| Pde4b | Myd88 | Lbp | Bcr |
| Peli1 | Nfatc2 | Lck | Calcrl |

| | | | |
|---|---|---|---|
| Phb | Nfkbia | Lcp1 | Ccl2 |
| Pik3ap1 | Nr4a3 | Lgals9 | Cd22 |
| Plcg1 | Nras | Lrrfip2 | Cd4 |
| Plcg2 | P2rx7 | Ltf | Cd44 |
| Pms2 | Park7 | Ly96 | Cd5 |
| Pnp | Pde4b | Malt1 | Cd80 |
| Polr3b | Peli1 | Mapk1 | Chrna7 |
| Polr3c | Phb | Mapk3 | Cnr2 |
| Polr3d | Pik3ap1 | Mapkapk3 | Cuedc2 |
| Polr3f | Plcg1 | Masp1 | Cx3cr1 |
| Prkch | Plcg2 | Mef2c | Cxcl16 |
| Prkd2 | Pms2 | Mif | Cybb |
| Psen1 | Pnp | Mmp12 | Dusp10 |
| Pum2 | Polr3b | Mmp2 | Ets1 |
| Pycard | Polr3c | Msh2 | Fcgr2b |
| Rap1a | Polr3d | Myb | Fem1a |
| Rapgef1 | Polr3f | Myd88 | Foxp3 |
| Rasgrp1 | Prkch | Nfatc2 | Gpx1 |
| Rela | Prkd2 | Nfkbia | Hamp |
| Rnf31 | Psen1 | Nr4a3 | Havcr2 |
| Rps19 | Pum2 | Nras | Hmgb1 |
| Rps3 | Pycard | P2rx7 | Ier3 |
| Rsad2 | Rap1a | Park7 | Il12b |
| Serping1 | Rapgef1 | Pde4b | Il22ra2 |
| Shb | Rasgrp1 | Peli1 | Il4 |
| Sin3a | Rela | Phb | Ins2 |
| Slamf6 | Rnf31 | Pik3ap1 | Itgal |
| Slc11a1 | Rps19 | Plcg1 | Lgals9 |
| Slc39a10 | Rps3 | Plcg2 | Ltbr |
| Snx4 | Rsad2 | Pld2 | Mapk1 |
| Spon2 | Serping1 | Plscr1 | Mapk3 |
| Stat5a | Shb | Pms2 | Mefv |
| Stat5b | Sin3a | Pnp | Metrnl |
| Stat6 | Slamf6 | Polr3b | Msn |
| Stk11 | Slc11a1 | Polr3d | Myd88 |
| Stoml2 | Slc39a10 | Polr3f | Ndfip1 |
| Stx4a | Snx4 | Prkch | Nr1h3 |
| Stx7 | Spon2 | Prkd2 | Nt5e |
| Syk | Stat5a | Psen1 | Pbk |
| Tap2 | Stat5b | Psen2 | Pparg |
| Tec | Stat6 | Pum2 | Prf1 |
| Tfrc | Stk11 | Pycard | Prnp |
| Tgfb1 | Stoml2 | Rap1a | Proc |
| Thy1 | Stx4a | Rapgef1 | Psma1 |
| Tlr12 | Stx7 | Rasgrp1 | Psmb4 |
| Tlr13 | Syk | Rela | Pten |
| Tlr2 | Tap2 | Rnf31 | Ptgis |
| Tlr5 | Tec | Rps19 | Ptpn2 |

| | | | |
|---|---|---|---|
| Tlr9 | Tfrc | Rps3 | Rabgef1 |
| Tnfrsf21 | Thy1 | Rsad2 | Rap1a |
| Tnfsf13b | Tlr12 | S100a14 | Rps19 |
| Tnip1 | Tlr13 | Serping1 | Serpinf1 |
| Tnip2 | Tlr2 | Shb | Slamf8 |
| Traf3 | Tlr4 | Sin3a | Slit2 |
| Traf6 | Tlr5 | Slamf6 | Smad3 |
| Ube2n | Tlr9 | Slc11a1 | Smpdl3b |
| Unc93b1 | Tnfrsf21 | Slc39a10 | Spn |
| Ung | Tnfsf13b | Snx4 | Stat1 |
| Usp12 | Tnfsf4 | Socs5 | Stat3 |
| Vamp8 | Tnip1 | Spon2 | Stat5a |
| Vav1 | Tnip2 | Stat5a | Stat5b |
| Wdfy1 | Traf3 | Stat5b | Stk11 |
| Wnt5a | Traf6 | Stat6 | Tff2 |
| Xbp1 | Ube2n | Stk11 | Tgfb1 |
| Zap70 | Ung | Stoml2 | Tnfaip3 |
| Zbtb1 | Usp12 | Stx4a | Tnfrsf1a |
| Abl1 | Vamp8 | Stx7 | Tnfrsf1b |
| Actr1a | Vav1 | Syk | Trp53 |
| Adora2b | Wdfy1 | Tap2 | Uaca |
| Ap1g1 | Wnt5a | Tec | Usp18 |
| B2m | Xbp1 | Tfrc | Zap70 |
| Bax | Zap70 | Thy1 | Zfp36 |
| Bcar1 | Zbtb1 | Tlr12 | |
| Bcl10 | Abl1 | Tlr13 | |
| Bcl2 | Actr1a | Tlr2 | |
| Braf | Adora2b | Tlr3 | |
| C1qb | Ap1g1 | Tlr5 | |
| C1qbp | B2m | Tlr8 | |
| C2 | Bax | Tlr9 | |
| C3ar1 | Bcar1 | Tnfrsf21 | |
| Cav1 | Bcl10 | Tnfsf13b | |
| Cd14 | Bcl2 | Tnfsf4 | |
| Cd19 | Braf | Tnip1 | |
| Cd1d1 | C1qb | Tnip2 | |
| Cd24a | C1qbp | Traf3 | |
| Cd3e | C2 | Traf6 | |
| Cd4 | Cav1 | Ube2n | |
| Cd44 | Cd14 | Ung | |
| Cd79a | Cd19 | Usp12 | |
| Cfi | Cd1d1 | Vamp8 | |
| Cnr1 | Cd24a | Vav1 | |
| Colec12 | Cd3e | Wdfy1 | |
| Crhr1 | Cd4 | Wnt5a | |
| Crtam | Cd44 | Xbp1 | |
| Cst11 | Cfi | Zap70 | |
| Cyld | Colec12 | Zbtb1 | |

| | | |
|---|---|---|
| Ddx58 | Crhr1 | Abl1 |
| Dhx15 | Crtam | Actr1a |
| Exosc3 | Cst11 | Adora2b |
| F2rl1 | Cyld | Ap1g1 |
| Fadd | Ddx58 | B2m |
| Fcer1a | Dhx15 | Bax |
| Fcgr1 | Exosc3 | Bcar1 |
| Fcna | F2rl1 | Bcl10 |
| Fcnb | Fadd | Bcl2 |
| Fgr | Fcer1a | Braf |
| Flot1 | Fcgr1 | C1qa |
| Gab2 | Fcna | C1qb |
| Gata2 | Fcnb | C1qbp |
| Gc | Fgr | C2 |
| Gpr33 | Flot1 | C3 |
| Gpx4 | Gata2 | C3ar1 |
| H2-M3 | Gc | Cav1 |
| Havcr2 | Gpr33 | Cd14 |
| Hmgb1 | Gpx4 | Cd19 |
| Hmgb2 | H2-M3 | Cd1d1 |
| Hspa8 | Havcr2 | Cd24a |
| Il4ra | Hmgb1 | Cd3e |
| Irak1 | Hmgb2 | Cd4 |
| Irak2 | Hspa8 | Cd44 |
| Irak3 | Il4ra | Cfi |
| Irak4 | Irak1 | Cnr1 |
| Irf3 | Irak2 | Colec12 |
| Irf7 | Irak3 | Crhr1 |
| Kars | Irak4 | Crtam |
| Klk5 | Irf3 | Cyba |
| Klrb1c | Irf7 | Cyld |
| Klre1 | Kars | Ddx58 |
| Klrk1 | Klk5 | Dhx15 |
| Lamp1 | Klrb1c | Exosc3 |
| Lbp | Klre1 | F2rl1 |
| Lck | Lamp1 | Fadd |
| Lcp1 | Lbp | Fcer1a |
| Lcp2 | Lck | Fcgr1 |
| Lgals9 | Lcp1 | Fcna |
| Lrrfip2 | Lgals9 | Fcnb |
| Ltf | Lrrfip2 | Fgr |
| Ly96 | Ltf | Flot1 |
| Malt1 | Ly96 | Gata2 |
| Mapk1 | Malt1 | Gata3 |
| Mapk3 | Mapk1 | Gpr33 |
| Mapkapk3 | Mapk3 | Gpx4 |
| Masp1 | Mapkapk3 | H2-M3 |
| Mef2c | Masp1 | Havcr2 |

| | | |
|---|---|---|
| Mif | Mef2c | Hmgb1 |
| Mlh1 | Mif | Hmgb2 |
| Mmp12 | Mlh1 | Hspa8 |
| Mmp2 | Mmp12 | Ifih1 |
| Msh2 | Mmp2 | Irak1 |
| Myb | Msh2 | Irak2 |
| Myd88 | Myb | Irak4 |
| Nfatc2 | Myd88 | Irf3 |
| Nfkbia | Nfatc2 | Irf7 |
| Nr4a3 | Nfkbia | Kars |
| Nras | Nr4a3 | Klk5 |
| P2rx7 | Nras | Klrb1c |
| Park7 | P2rx7 | Klre1 |
| Pde4b | Park7 | Lamp1 |
| Peli1 | Pde4b | Lbp |
| Phb | Peli1 | Lcp1 |
| Pik3ap1 | Phb | Lgals9 |
| Plcg1 | Pik3ap1 | Lrrfip2 |
| Plcg2 | Plcg1 | Ltf |
| Pld2 | Plcg2 | Ly96 |
| Pms2 | Pms2 | Malt1 |
| Pnp | Pnp | Mapk1 |
| Polr3b | Polr3b | Mapk3 |
| Polr3c | Polr3c | Mapkapk3 |
| Polr3d | Polr3d | Masp1 |
| Polr3f | Polr3f | Mef2c |
| Prkch | Prkch | Mif |
| Prkd2 | Prkd2 | Mlh1 |
| Psen1 | Psen1 | Mmp12 |
| Pum2 | Psen2 | Mmp2 |
| Pycard | Pum2 | Msh2 |
| Rap1a | Pycard | Myb |
| Rapgef1 | Rap1a | Myd88 |
| Rasgrp1 | Rapgef1 | Nfatc2 |
| Rela | Rasgrp1 | Nfkbia |
| Rnf31 | Rela | Nr4a3 |
| Rps19 | Rnf31 | Nras |
| Rps3 | Rps19 | P2rx7 |
| Rsad2 | Rps3 | Park7 |
| Serping1 | Rsad2 | Pde4b |
| Shb | Serping1 | Peli1 |
| Sin3a | Shb | Phb |
| Slamf6 | Sin3a | Pik3ap1 |
| Slc11a1 | Slamf6 | Plcg1 |
| Slc39a10 | Slc11a1 | Plcg2 |
| Snx4 | Slc39a10 | Pld2 |
| Spon2 | Snx4 | Pms2 |
| Stat5a | Socs5 | Pnp |

| | | |
|---|---|---|
| Stat5b | Spon2 | Polr3b |
| Stat6 | Stat5a | Polr3d |
| Stk11 | Stat5b | Polr3f |
| Stoml2 | Stat6 | Prkch |
| Stx4a | Stk11 | Prkd2 |
| Stx7 | Stoml2 | Psen1 |
| Syk | Stx4a | Psen2 |
| Tap2 | Stx7 | Pum2 |
| Tec | Syk | Pycard |
| Tfrc | Tap2 | Rap1a |
| Tgfb1 | Tec | Rapgef1 |
| Thy1 | Tfrc | Rasgrp1 |
| Tlr12 | Tgfb1 | Rela |
| Tlr2 | Thy1 | Rnf31 |
| Tlr4 | Tlr12 | Rps19 |
| Tlr5 | Tlr13 | Rps3 |
| Tlr9 | Tlr2 | Rsad2 |
| Tnfrsf21 | Tlr5 | S100a14 |
| Tnfsf13b | Tlr9 | Serping1 |
| Tnfsf4 | Tnfrsf21 | Shb |
| Tnip1 | Tnfsf13b | Sin3a |
| Tnip2 | Tnip1 | Slamf6 |
| Traf3 | Tnip2 | Slc11a1 |
| Traf6 | Traf3 | Slc39a10 |
| Ube2n | Traf6 | Snx4 |
| Unc93b1 | Ube2n | Spon2 |
| Ung | Unc93b1 | Stat5a |
| Usp12 | Ung | Stat5b |
| Vamp8 | Usp12 | Stat6 |
| Vav1 | Vamp8 | Stk11 |
| Wdfy1 | Vav1 | Stoml2 |
| Wnt5a | Wdfy1 | Stx4a |
| Xbp1 | Wnt5a | Stx7 |
| Zap70 | Xbp1 | Syk |
| Zbtb1 | Zap70 | Tap2 |
| Adora1 | Zbtb1 | Tec |
| Adora2a | Acp5 | Tfrc |
| Apod | Adora2a | Thy1 |
| Apoe | Adrb2 | Tlr12 |
| Ash1l | Apod | Tlr13 |
| Bcr | Apoe | Tlr2 |
| Cd44 | Ash1l | Tlr3 |
| Cuedc2 | Bax | Tlr5 |
| Fcgr2b | Bcr | Tlr9 |
| Fem1a | Calcrl | Tnfrsf21 |
| Fn1 | Cd22 | Tnfsf13b |
| Gpx1 | Cd4 | Tnfsf4 |
| Ier3 | Cd44 | Tnip1 |

| | | |
|---|---|---|
| Il22ra2 | Cd80 | Tnip2 |
| Lgals9 | Chrna7 | Traf3 |
| Mefv | Cnr2 | Traf6 |
| Metrnl | Cuedc2 | Ube2n |
| Ndfip1 | Cxcl16 | Ung |
| Nt5e | Cybb | Usp12 |
| Pbk | Dusp10 | Vamp8 |
| Psma1 | Ets1 | Vav1 |
| Psmb4 | Fcgr2b | Wdfy1 |
| Ptgds | Fem1a | Wnt5a |
| Ptgis | Gpx1 | Xbp1 |
| Ptpn2 | Hamp | Zap70 |
| Rabgef1 | Havcr2 | Zbtb1 |
| Rps19 | Hmgb1 | Acp5 |
| Serpinf1 | Ier3 | Adora1 |
| Slit2 | Itgal | Adora2a |
| Smad3 | Lgals9 | Adrb2 |
| Spn | Ltbr | Apod |
| Tnfaip3 | Mapk1 | Apoe |
| Tnfrsf1a | Mapk3 | Ash1l |
| Tnfrsf1b | Mefv | Bcr |
| Uaca | Metrnl | Calcrl |
| Wfdc1 | Msn | Cd44 |
| Zfp36 | Myd88 | Chrna7 |
| Acp5 | Ndfip1 | Cnr2 |
| Adora2a | Nt5e | Cuedc2 |
| Adrb2 | Pbk | Dusp10 |
| Apod | Pparg | Ets1 |
| Apoe | Prf1 | Fcgr2b |
| Ash1l | Prnp | Fem1a |
| Bcr | Psma1 | Fn1 |
| Calcrl | Psmb4 | Gata3 |
| Cd44 | Pten | Gpx1 |
| Chrna7 | Ptgis | Hamp |
| Cnr2 | Ptpn2 | Ier3 |
| Cuedc2 | Rabgef1 | Il22ra2 |
| Dusp10 | Rap1a | Il4 |
| Ets1 | Rps19 | Lgals9 |
| Fcgr2b | Serpinf1 | Mefv |
| Fem1a | Slit2 | Metrnl |
| Fn1 | Smad3 | Ndfip1 |
| Gpx1 | Smpdl3b | Nt5e |
| Hamp | Spn | Pbk |
| Ier3 | Stat1 | Pparg |
| Il22ra2 | Stat3 | Psma1 |
| Lgals9 | Stat5a | Psmb4 |
| Mefv | Stat5b | Ptgds |
| Metrnl | Stk11 | Ptgis |

| | | |
|---|---|---|
| Ndfip1 | Tff2 | Ptpn2 |
| Nt5e | Tgfb1 | Rabgef1 |
| Pbk | Tnfaip3 | Rps19 |
| Pparg | Tnfrsf1a | Serpina3k |
| Psma1 | Trp53 | Serpinb1a |
| Psmb4 | Wfdc1 | Serpinf1 |
| Ptgds | Zap70 | Slit2 |
| Ptgis | Zfp36 | Smad3 |
| Ptpn2 | Acp5 | Socs5 |
| Rabgef1 | Adora2a | Spn |
| Rps19 | Adrb2 | Tff2 |
| Serpina3k | Apod | Tnfaip3 |
| Serpinb1a | Apoe | Tnfrsf1a |
| Serpinf1 | Ash1l | Tnfrsf1b |
| Slit2 | Bcr | Uaca |
| Smad3 | Calcrl | Wfdc1 |
| Smpdl3b | Cd44 | Zfp36 |
| Spn | Chrna7 | Acp5 |
| Tff2 | Cnr2 | Adora1 |
| Tnfaip3 | Cuedc2 | Adora2a |
| Tnfrsf1a | Dusp10 | Adrb2 |
| Tnfrsf1b | Ets1 | Apod |
| Uaca | Fcgr2b | Apoe |
| Wfdc1 | Fem1a | Ash1l |
| Zfp36 | Fn1 | Bcr |
| Acp5 | Gpx1 | Calcrl |
| Adora2a | Hamp | Cd44 |
| Adrb2 | Lgals9 | Chrna7 |
| Apod | Mefv | Cnr2 |
| Apoe | Metrnl | Cuedc2 |
| Ash1l | Ndfip1 | Dusp10 |
| Bcr | Nt5e | Ets1 |
| Calcrl | Pbk | Fcgr2b |
| Cd44 | Pparg | Fem1a |
| Chrna7 | Psma1 | Fn1 |
| Cnr2 | Psmb4 | Gata3 |
| Cuedc2 | Ptgds | Gpx1 |
| Dusp10 | Ptgis | Ier3 |
| Ets1 | Ptpn2 | Il22ra2 |
| Fcgr2b | Rabgef1 | Lgals9 |
| Fem1a | Rps19 | Mefv |
| Fn1 | Serpina3k | Metrnl |
| Gpx1 | Serpinb1a | Ndfip1 |
| Hamp | Serpinf1 | Nt5e |
| Ier3 | Slit2 | Pbk |
| Il22ra2 | Smad3 | Pparg |
| Lgals9 | Smpdl3b | Psma1 |
| Mefv | Spn | Psmb4 |

| | | |
|---|---|---|
| Metrnl | Tff2 | Ptgds |
| Ndfip1 | Tnfaip3 | Ptgis |
| Nt5e | Tnfrsf1a | Ptpn2 |
| Pbk | Uaca | Rabgef1 |
| Pparg | Wfdc1 | Rps19 |
| Psma1 | Acp5 | Serpinb1a |
| Psmb4 | Adora2a | Serpinf1 |
| Ptgds | Adrb2 | Slit2 |
| Ptgis | Apod | Smad3 |
| Ptpn2 | Apoe | Socs5 |
| Rabgef1 | Ash1l | Spn |
| Rps19 | Bcr | Tff2 |
| Serpina3k | Calcrl | Tnfaip3 |
| Serpinb1a | Cd44 | Tnfrsf1a |
| Serpinf1 | Chrna7 | Tnfrsf1b |
| Slit2 | Cnr2 | Wfdc1 |
| Smad3 | Cuedc2 | Zfp36 |
| Smpdl3b | Dusp10 | Acp5 |
| Spn | Ets1 | Adora1 |
| Tff2 | Fcgr2b | Adora2a |
| Tnfaip3 | Fem1a | Adrb2 |
| Tnfrsf1a | Fn1 | Apod |
| Tnfrsf1b | Gpx1 | Apoe |
| Uaca | Hamp | Ash1l |
| Wfdc1 | Ier3 | Bcr |
| Zfp36 | Il22ra2 | Calcrl |
| Adora1 | Lgals9 | Cd44 |
| Adora2a | Mefv | Chrna7 |
| Apod | Metrnl | Cnr2 |
| Apoe | Ndfip1 | Cuedc2 |
| Ash1l | Nt5e | D1Ertd622e |
| Bax | Pbk | Dusp10 |
| Bcr | Pparg | Ets1 |
| Ccl2 | Psma1 | Fcgr2b |
| Cd22 | Psmb4 | Fem1a |
| Cd44 | Ptgds | Fn1 |
| Cd80 | Ptgis | Gata3 |
| Cuedc2 | Ptpn2 | Gpx1 |
| Cxcl16 | Rabgef1 | Ier3 |
| Cybb | Rps19 | Il22ra2 |
| Fcgr2b | Serpina3k | Lgals9 |
| Fem1a | Serpinb1a | Mefv |
| Gpx1 | Serpinf1 | Metrnl |
| Hmgb1 | Slit2 | Ndfip1 |
| Ier3 | Smad3 | Nt5e |
| Il22ra2 | Smpdl3b | Pbk |
| Itgal | Socs5 | Pparg |
| Lgals9 | Tff2 | Psma1 |

| | | |
|---|---|---|
| Ltbr | Tnfaip3 | Psmb4 |
| Mapk1 | Tnfrsf1a | Ptgds |
| Mapk3 | Tnfrsf1b | Ptgis |
| Mefv | Uaca | Ptpn2 |
| Metrnl | Wfdc1 | Rabgef1 |
| Msn | Zfp36 | Rps19 |
| Myd88 | Acp5 | Serpinb1a |
| Ndfip1 | Adora2a | Serpinf1 |
| Nt5e | Adrb2 | Slit2 |
| Pbk | Apod | Smad3 |
| Prf1 | Apoe | Spn |
| Prnp | Ash1l | Tff2 |
| Psma1 | Bax | Tnfaip3 |
| Psmb4 | Bcr | Tnfrsf1a |
| Pten | Calcrl | Uaca |
| Ptgis | Cd22 | Wfdc1 |
| Ptpn2 | Cd4 | Zfp36 |
| Rabgef1 | Cd44 | Acp5 |
| Rap1a | Cd80 | Adora1 |
| Rps19 | Chrna7 | Adora2a |
| Serpinf1 | Cnr2 | Adrb2 |
| Slit2 | Cuedc2 | Apod |
| Smad3 | Cxcl16 | Apoe |
| Spn | Cybb | Ash1l |
| Stat1 | Dusp10 | Bax |
| Stat3 | Ets1 | Bcr |
| Stat5b | Fcgr2b | Calcrl |
| Stk11 | Fem1a | Cd22 |
| Tgfb1 | Gpx1 | Cd4 |
| Tnfaip3 | Hamp | Cd44 |
| Tnfrsf1a | Havcr2 | Cd80 |
| Tnfrsf1b | Hmgb1 | Chrna7 |
| Trp53 | Ier3 | Cnr2 |
| Uaca | Itgal | Cuedc2 |
| Usp18 | Lgals9 | Cxcl16 |
| Vip | Ltbr | Cybb |
| Wfdc1 | Mapk1 | Dusp10 |
| Zap70 | Mapk3 | Ets1 |
| Zfp36 | Mefv | Fcgr2b |
| Acp5 | Metrnl | Fem1a |
| Adora2a | Msn | Gata3 |
| Adrb2 | Myd88 | Gpx1 |
| Apod | Ndfip1 | Hamp |
| Apoe | Nt5e | Havcr2 |
| Ash1l | Pbk | Hmgb1 |
| Bax | Pparg | Ier3 |
| Bcr | Prf1 | Il22ra2 |
| Calcrl | Prnp | Il4 |

| | | |
|---|---|---|
| Cd22 | Psma1 | Itgal |
| Cd4 | Psmb4 | Lgals9 |
| Cd44 | Pten | Ltbr |
| Cd80 | Ptgis | Mapk1 |
| Chrna7 | Ptpn2 | Mapk3 |
| Cnr2 | Rabgef1 | Mefv |
| Cuedc2 | Rap1a | Metrnl |
| Cxcl16 | Rps19 | Msn |
| Cybb | Serpinf1 | Myd88 |
| Dusp10 | Slit2 | Ndfip1 |
| Ets1 | Smad3 | Nt5e |
| Fcgr2b | Smpdl3b | Pbk |
| Fem1a | Spn | Pparg |
| Gpx1 | Stat1 | Prf1 |
| Hamp | Stat3 | Prnp |
| Havcr2 | Stat5a | Psma1 |
| Hmgb1 | Stat5b | Psmb4 |
| Ier3 | Stk11 | Pten |
| Il22ra2 | Tff2 | Ptgis |
| Itgal | Tgfb1 | Ptpn2 |
| Lgals9 | Tnfaip3 | Rabgef1 |
| Ltbr | Tnfrsf1a | Rap1a |
| Mapk1 | Trp53 | Rps19 |
| Mapk3 | Wfdc1 | Serpinf1 |
| Mefv | Zap70 | Slit2 |
| Metrnl | Zfp36 | Smad3 |
| Msn | Acp5 | Socs5 |
| Myd88 | Adora2a | Spn |
| Ndfip1 | Adrb2 | Stat1 |
| Nt5e | Apod | Stat3 |
| Pbk | Apoe | Stat5a |
| Pparg | Ash1l | Stat5b |
| Prnp | Bax | Stk11 |
| Psma1 | Bcr | Tff2 |
| Psmb4 | Calcrl | Tnfaip3 |
| Pten | Cd22 | Tnfrsf1a |
| Ptgis | Cd4 | Tnfrsf1b |
| Ptpn2 | Cd44 | Trp53 |
| Rabgef1 | Cd80 | Uaca |
| Rap1a | Chrna7 | Usp18 |
| Rps19 | Cnr2 | Wfdc1 |
| Serpinf1 | Cuedc2 | Zap70 |
| Slit2 | Cxcl16 | Zfp36 |
| Smad3 | Cybb | Acp5 |
| Smpdl3b | Dusp10 | Adora1 |
| Spn | Ets1 | Adora2a |
| Stat1 | Fcgr2b | Adrb2 |
| Stat3 | Fem1a | Apod |

| | | |
|---|---|---|
| Stat5a | Gpx1 | Apoe |
| Stat5b | Hamp | Ash1l |
| Stk11 | Havcr2 | Bax |
| Tff2 | Hmgb1 | Bcr |
| Tgfb1 | Itgal | Calcrl |
| Tnfaip3 | Lgals9 | Cd22 |
| Tnfrsf1a | Ltbr | Cd4 |
| Tnfrsf1b | Mapk1 | Cd44 |
| Trp53 | Mapk3 | Cd80 |
| Uaca | Mefv | Chrna7 |
| Usp18 | Metrnl | Cnr2 |
| Wfdc1 | Msn | Cuedc2 |
| Zap70 | Myd88 | Cxcl16 |
| Zfp36 | Ndfip1 | Cybb |
| Acp5 | Nt5e | Dusp10 |
| Adora2a | Pbk | Ets1 |
| Adrb2 | Pparg | Fcgr2b |
| Apod | Prf1 | Fem1a |
| Apoe | Prnp | Gata3 |
| Ash1l | Psma1 | Gpx1 |
| Bax | Psmb4 | Havcr2 |
| Bcr | Pten | Hmgb1 |
| Calcrl | Ptgis | Ier3 |
| Cd22 | Ptpn2 | Il22ra2 |
| Cd4 | Rabgef1 | Itgal |
| Cd44 | Rap1a | Lgals9 |
| Cd80 | Rps19 | Ltbr |
| Chrna7 | Serpinf1 | Mapk1 |
| Cnr2 | Slit2 | Mapk3 |
| Cuedc2 | Smad3 | Mefv |
| Cxcl16 | Smpdl3b | Metrnl |
| Cybb | Spn | Msn |
| Dusp10 | Stat1 | Myd88 |
| Ets1 | Stat3 | Ndfip1 |
| Fcgr2b | Stat5a | Nt5e |
| Fem1a | Stat5b | Pbk |
| Gpx1 | Stk11 | Pparg |
| Hamp | Tff2 | Prnp |
| Havcr2 | Tlr4 | Psma1 |
| Hmgb1 | Tnfaip3 | Psmb4 |
| Ier3 | Tnfrsf1a | Pten |
| Il22ra2 | Trp53 | Ptgis |
| Itgal | Uaca | Ptpn2 |
| Lgals9 | Usp18 | Rabgef1 |
| Ltbr | Wfdc1 | Rap1a |
| Mapk1 | Zap70 | Rps19 |
| Mapk3 | Zfp36 | Serpinf1 |
| Mefv | Acp5 | Slit2 |

| | | |
|---|---|---|
| Metrnl | Adora2a | Smad3 |
| Msn | Adrb2 | Socs5 |
| Myd88 | Apod | Spn |
| Ndfip1 | Apoe | Stat3 |
| Nt5e | Ash1l | Stat5a |
| Pbk | Bax | Stat5b |
| Pparg | Bcr | Stk11 |
| Prf1 | Calcrl | Tff2 |
| Prnp | Cd22 | Tnfaip3 |
| Psma1 | Cd4 | Tnfrsf1a |
| Psmb4 | Cd44 | Tnfrsf1b |
| Pten | Cd80 | Usp18 |
| Ptgis | Chrna7 | Wfdc1 |
| Ptpn2 | Cnr2 | Zap70 |
| Rabgef1 | Cuedc2 | Zfp36 |
| Rap1a | Cxcl16 | Acp5 |
| Rps19 | Cybb | Adora1 |
| Serpinf1 | Dusp10 | Adora2a |
| Slit2 | Ets1 | Adrb2 |
| Smad3 | Fcgr2b | Apod |
| Smpdl3b | Fem1a | Apoe |
| Spn | Gpx1 | Ash1l |
| Stat1 | Hamp | Bax |
| Stat3 | Havcr2 | Bcr |
| Stat5a | Hmgb1 | Calcrl |
| Stat5b | Ier3 | Cd22 |
| Stk11 | Il22ra2 | Cd4 |
| Tff2 | Itgal | Cd44 |
| Tgfb1 | Lgals9 | Cd80 |
| Tlr4 | Ltbr | Chrna7 |
| Tnfaip3 | Mapk1 | Cnr2 |
| Tnfrsf1a | Mapk3 | Cuedc2 |
| Tnfrsf1b | Mefv | Cxcl16 |
| Trp53 | Metrnl | Cybb |
| Uaca | Msn | D1Ertd622e |
| Usp18 | Myd88 | Dusp10 |
| Wfdc1 | Ndfip1 | Ets1 |
| Zap70 | Nt5e | Fcgr2b |
| Zfp36 | Pbk | Fem1a |
| | Pparg | Gata3 |
| | Prf1 | Gpx1 |
| | Prnp | Havcr2 |
| | Psma1 | Hmgb1 |
| | Psmb4 | Ier3 |
| | Pten | Il22ra2 |
| | Ptgis | Itgal |
| | Ptpn2 | Lgals9 |
| | Rabgef1 | Ltbr |

| | |
|---|---|
| Rap1a | Mapk1 |
| Rps19 | Mapk3 |
| Serpinf1 | Mefv |
| Slit2 | Metrnl |
| Smad3 | Msn |
| Smpdl3b | Myd88 |
| Socs5 | Ndfip1 |
| Stat1 | Nt5e |
| Stat3 | Pbk |
| Stat5a | Pparg |
| Stat5b | Prnp |
| Stk11 | Psma1 |
| Tff2 | Psmb4 |
| Tgfb1 | Pten |
| Tnfaip3 | Ptgis |
| Tnfrsf1a | Ptpn2 |
| Tnfrsf1b | Rabgef1 |
| Trp53 | Rap1a |
| Uaca | Rps19 |
| Usp18 | Serpinf1 |
| Wfdc1 | Slit2 |
| Zap70 | Smad3 |
| Zfp36 | Spn |
| | Stat1 |
| | Stat3 |
| | Stat5a |
| | Stat5b |
| | Stk11 |
| | Tff2 |
| | Tnfaip3 |
| | Tnfrsf1a |
| | Trp53 |
| | Uaca |
| | Wfdc1 |
| | Zap70 |
| | Zfp36 |

**sperm maturation-related genes**

| Seg 1,2,3 | Seg 4,5,6 | Seg 8,9,10 | Seg 7 |
|---|---|---|---|
| 1600014C10Rik | 1600014C10Rik | 1600014C10Rik | 1600014C10Rik |
| 1700009P17Rik | 1700009P17Rik | 1700009P17Rik | 1700009P17Rik |
| 1700034H14Rik | 1700012A03Rik | 1700012A03Rik | 1700012A03Rik |
| 2300009A05Rik | 1700034H14Rik | 1700034H14Rik | 1700034H14Rik |
| 4930415O20Rik | 2300009A05Rik | 2300009A05Rik | 2300009A05Rik |
| 4930471M23Rik | 4921507P07Rik | 4732415M23Rik | 4921507P07Rik |
| 5730469M10Rik | 4921530D09Rik | 4921507P07Rik | 4921530D09Rik |
| AI314180 | 4930415O20Rik | 4921530D09Rik | 4930415O20Rik |
| Abca3 | 4930471M23Rik | 4930415O20Rik | 4930471M23Rik |
| Abcc3 | 5730469M10Rik | 4930471M23Rik | 5730469M10Rik |
| Acat2 | 9430016H08Rik | 5730469M10Rik | 8430406I07Rik |
| Acbd5 | AI314180 | 8430406I07Rik | 9430016H08Rik |
| Aco1 | Abca3 | 9430016H08Rik | AI314180 |
| Acp1 | Abcc3 | AI314180 | Abca3 |
| Actn1 | Acat2 | Abca3 | Abcc3 |
| Actr3 | Acbd5 | Abcc3 | Acat2 |
| Acyp1 | Aco1 | Acat2 | Acbd5 |
| Adck5 | Acp1 | Acbd5 | Aco1 |
| Adh5 | Actn1 | Aco1 | Acp1 |
| Akap4 | Actr3 | Acp1 | Actn1 |
| Anp32b | Acyp1 | Actn1 | Actr3 |
| Ap2a1 | Adck5 | Actr3 | Acyp1 |
| Ap2s1 | Adh5 | Acyp1 | Adck5 |
| Aprt | Akap4 | Adck5 | Adh5 |
| Arl3 | Anp32b | Adh5 | Akap4 |
| Armc1 | Ap2a1 | Akap4 | Anp32b |
| Arpc4 | Ap2s1 | Anp32b | Anxa13 |
| Arpc5l | Aprt | Anxa13 | Ap2a1 |
| Atp11a | Arl3 | Ap2a1 | Ap2s1 |
| Atp13a4 | Armc1 | Ap2s1 | Aprt |
| Atp6v0c | Arpc4 | Aprt | Arl3 |
| BC051142 | Arpc5l | Arl3 | Armc1 |
| Bloc1s1 | Atp11a | Armc1 | Arpc4 |
| Bsg | Atp13a4 | Arpc4 | Arpc5l |
| Calm3 | Atp6v0c | Arpc5l | Atp11a |
| Cap1 | BC051142 | Atp11a | Atp13a4 |
| Capza1 | Bloc1s1 | Atp13a4 | Atp6v0c |
| Car2 | Bsg | Atp6v0c | BC051142 |
| Cbr3 | C030006K11Rik | BC051142 | Bloc1s1 |
| Cct7 | Calm3 | Bloc1s1 | Bsg |
| Cct8 | Cap1 | Bsg | C030006K11Rik |
| Chchd1 | Capza1 | C030006K11Rik | Cabyr |
| Clca2 | Car2 | Cabyr | Calm3 |
| Clic4 | Cbr3 | Calm3 | Cap1 |
| Coasy | Cct7 | Cap1 | Capza1 |
| Cops4 | Cct8 | Capza1 | Car2 |

| | | | |
|---|---|---|---|
| Cops8 | Chchd1 | Car2 | Car4 |
| Crot | Ckmt1 | Car4 | Cbr3 |
| Csnk2a1 | Clca2 | Cbr3 | Cct7 |
| Ctsf | Clic3 | Cct7 | Cct8 |
| Cutc | Clic4 | Cct8 | Chchd1 |
| Cyfip1 | Coasy | Chchd1 | Ckmt1 |
| Ddb1 | Cops4 | Ckmt1 | Clca2 |
| Ddt | Cops8 | Clca2 | Clic3 |
| Dnaja4 | Cox6b2 | Clic3 | Clic4 |
| Dnajb6 | Crisp2 | Clic4 | Coasy |
| Dnm2 | Crot | Coasy | Cops8 |
| Dock7 | Csnk2a1 | Cops4 | Cox6b2 |
| Dpysl2 | Ctsf | Cops8 | Crisp2 |
| Dtymk | Cyfip1 | Cox6b2 | Crot |
| Dusp3 | Cyp17a1 | Crisp2 | Csnk2a1 |
| Ebp | Ddb1 | Crot | Ctsf |
| Eef1d | Ddt | Csnk2a1 | Cyfip1 |
| Eef1g | Defb19 | Ctsf | Cyp17a1 |
| Ehd2 | Defb2 | Cyfip1 | Ddb1 |
| Eif2s1 | Dnaja4 | Cyp17a1 | Ddt |
| Eif5a | Dnajb6 | Ddb1 | Defb11 |
| Endog | Dnm2 | Ddt | Defb2 |
| Eprs | Dock7 | Defb11 | Dnaja4 |
| Erbb2 | Dpysl2 | Defb2 | Dnajb6 |
| Fdx1 | Dtymk | Dnajb6 | Dnm2 |
| Fhl1 | Dusp3 | Dnm2 | Dock7 |
| Fmo2 | Ebp | Dock7 | Dpysl2 |
| Fundc1 | Eef1d | Dpysl2 | Dtymk |
| Fxc1 | Eef1g | Dtymk | Dusp3 |
| Gaa | Ehd2 | Dusp3 | Ebp |
| Galntl4 | Eif2s1 | Ebp | Eef1d |
| Gba | Eif5a | Eef1d | Eef1g |
| Gclm | Eprs | Eef1g | Ehd2 |
| Ghitm | Erbb2 | Ehd2 | Eif2s1 |
| Gm2a | Fdx1 | Eif2s1 | Eif5a |
| Gpi1 | Fhl1 | Eif5a | Eno1 |
| Gpnmb | Fhl4 | Eprs | Eprs |
| Gpx1 | Fmo2 | Erbb2 | Erbb2 |
| Gpx3 | Fundc1 | Fdx1 | Fdx1 |
| Gpx4 | Fxc1 | Fhl1 | Fhl1 |
| Gpx5 | Gaa | Fhl4 | Fhl4 |
| Gsr | Galntl4 | Fmo2 | Fmo2 |
| Gstm1 | Gba | Fundc1 | Fundc1 |
| Gstm2 | Gclm | Fxc1 | Fxc1 |
| Gtl3 | Ghitm | Fxn | Gaa |
| Hdhd2 | Gm2a | Gaa | Galntl4 |
| Hint1 | Gpi1 | Galntl4 | Gba |
| Hrsp12 | Gpnmb | Gba | Gclm |

| | | | |
|---|---|---|---|
| Hsd3b7 | Gpx1 | Gclm | Ghitm |
| Htatip2 | Gpx3 | Ghitm | Gm2a |
| Idi1 | Gpx4 | Gm2a | Gpi1 |
| Ipo7 | Gpx5 | Gpi1 | Gpnmb |
| Isg20 | Gsr | Gpnmb | Gpx1 |
| Itpr1 | Gstm1 | Gpx1 | Gpx3 |
| Itpr3 | Gstm2 | Gpx3 | Gpx4 |
| Lactb | Gtl3 | Gpx4 | Gpx5 |
| Lamb2 | Hdhd2 | Gpx5 | Gsr |
| Lamc1 | Hint1 | Gsr | Gstm1 |
| Lamp2 | Hrsp12 | Gstm1 | Gstm2 |
| Lancl1 | Hsd3b7 | Gstm2 | Gtl3 |
| Lancl2 | Htatip2 | Gtl3 | Hdhd2 |
| Lars | Idi1 | Hdhd2 | Hint1 |
| Lass2 | Ipo7 | Hint1 | Hrsp12 |
| Lcn12 | Isg20 | Hrsp12 | Hsd3b7 |
| Lcn8 | Itpr1 | Hsd3b7 | Htatip2 |
| Lgals1 | Itpr3 | Htatip2 | Idi1 |
| Lipe | Klhl10 | Idi1 | Immp2l |
| Lipg | Lactb | Ipo7 | Ipo7 |
| Ly6a | Lamb2 | Isg20 | Isg20 |
| Lypla1 | Lamc1 | Itpr1 | Itpr1 |
| Man2a2 | Lamp2 | Itpr3 | Itpr3 |
| Manba | Lancl1 | Klhl10 | Klhl10 |
| Mdk | Lancl2 | Lactb | Lactb |
| Mipep | Lars | Lamc1 | Lamc1 |
| Mrpl11 | Lass2 | Lamp2 | Lamp2 |
| Mrpl17 | Lcn12 | Lancl1 | Lancl1 |
| Mrpl18 | Lcn8 | Lancl2 | Lancl2 |
| Mrpl43 | Lgals1 | Lars | Lars |
| Mrpl48 | Lipe | Lass2 | Lass2 |
| Mrpl52 | Lipg | Lcn12 | Lcn12 |
| Mrpl55 | Ly6a | Lcn8 | Lcn8 |
| Mrps10 | Lypla1 | Lgals1 | Lgals1 |
| Mrps14 | Man2a2 | Lipe | Lipe |
| Mrps16 | Manba | Lipg | Lipg |
| Mrps18a | Mdk | Ly6a | Ly6a |
| Mrps18b | Mipep | Lypla1 | Lypla1 |
| Mrps21 | Mlf1 | Man2a2 | Man2a2 |
| Mrps24 | Mrpl11 | Manba | Manba |
| Mrps31 | Mrpl17 | Mdk | Mdk |
| Mrps36 | Mrpl18 | Mipep | Mipep |
| Mrps6 | Mrpl43 | Mlf1 | Mrpl11 |
| Mrrf | Mrpl48 | Mrpl11 | Mrpl17 |
| Mtpn | Mrpl52 | Mrpl17 | Mrpl18 |
| Ncam1 | Mrpl55 | Mrpl18 | Mrpl43 |
| Ndufc2 | Mrps10 | Mrpl43 | Mrpl48 |
| Ndufs6 | Mrps14 | Mrpl48 | Mrpl52 |

| | | | |
|---|---|---|---|
| Ndufv2 | Mrps16 | Mrpl52 | Mrpl55 |
| Nif3l1 | Mrps18a | Mrpl55 | Mrps10 |
| Nit1 | Mrps18b | Mrps10 | Mrps14 |
| Nqo1 | Mrps21 | Mrps14 | Mrps16 |
| Oplah | Mrps24 | Mrps16 | Mrps18a |
| Pcbp1 | Mrps31 | Mrps18a | Mrps18b |
| Pex11c | Mrps36 | Mrps18b | Mrps21 |
| Pfkm | Mrps6 | Mrps21 | Mrps24 |
| Pgls | Mrrf | Mrps24 | Mrps31 |
| Phb | Mtpn | Mrps31 | Mrps36 |
| Phpt1 | Ncam1 | Mrps36 | Mrps6 |
| Pmm2 | Ndufc2 | Mrps6 | Mrrf |
| Ppap2a | Ndufs6 | Mrrf | Mtpn |
| Ppil1 | Ndufv2 | Mtpn | Ncam1 |
| Ppp2r1b | Nif3l1 | Ncam1 | Ndufc2 |
| Ppp2r4 | Nit1 | Ndufc2 | Ndufs6 |
| Ppp6c | Nqo1 | Ndufs6 | Ndufv2 |
| Prkacb | Oplah | Ndufv2 | Nif3l1 |
| Psen1 | Pcbp1 | Nif3l1 | Nit1 |
| Psma1 | Pex11c | Nit1 | Nme5 |
| Psma2 | Pfkm | Nqo1 | Nqo1 |
| Psma6 | Pgk2 | Oplah | Oplah |
| Psma7 | Pgls | Pcbp1 | Pcbp1 |
| Psmb1 | Phb | Pex11c | Pex11c |
| Psmb5 | Phpt1 | Pfkm | Pfkm |
| Psmb6 | Pmm2 | Pgk2 | Pgk2 |
| Psmd7 | Ppap2a | Pgls | Pgls |
| Psme1 | Ppil1 | Phb | Phb |
| Psme4 | Ppp2r1b | Phpt1 | Phpt1 |
| Ptcd2 | Ppp2r4 | Pmm2 | Pmm2 |
| Ptges | Ppp6c | Ppap2a | Ppap2a |
| Ptp4a2 | Prkacb | Ppil1 | Ppil1 |
| Rab24 | Psen1 | Ppp2r1b | Ppp2r1b |
| Rab28 | Psma1 | Ppp2r4 | Ppp2r4 |
| Rab5b | Psma2 | Ppp6c | Ppp6c |
| Rac2 | Psma6 | Prkacb | Prkacb |
| Rdh13 | Psma7 | Psen1 | Psen1 |
| Recql | Psmb1 | Psma1 | Psma1 |
| Rpia | Psmb5 | Psma2 | Psma2 |
| Rpl37a | Psmb6 | Psma6 | Psma6 |
| Rpl5 | Psmd7 | Psma7 | Psma7 |
| Rplp2 | Psme1 | Psmb1 | Psmb1 |
| Rps10 | Psme4 | Psmb5 | Psmb5 |
| Rps13 | Ptcd2 | Psmb6 | Psmb6 |
| Rps19 | Ptges | Psmd7 | Psmd7 |
| Rps27 | Ptp4a2 | Psme1 | Psme1 |
| Rrbp1 | Rab24 | Psme4 | Psme4 |
| S100a16 | Rab28 | Ptcd2 | Ptcd2 |

| | | | |
|---|---|---|---|
| Sbds | Rab2b | Ptges | Ptges |
| Scarb2 | Rab5b | Ptp4a2 | Ptp4a2 |
| Sco1 | Rac2 | Rab24 | Rab24 |
| Sdf2l1 | Recql | Rab28 | Rab28 |
| Sdhc | Ribc1 | Rab2b | Rab2b |
| Sdhd | Rpia | Rab5b | Rab5b |
| Sec23ip | Rpl37a | Rac2 | Rac2 |
| Slc12a9 | Rpl5 | Rdh13 | Rdh13 |
| Slc2a5 | Rplp2 | Recql | Rdh9 |
| Slc35f2 | Rps10 | Ropn1l | Recql |
| Slco4c1 | Rps13 | Rpia | Rpia |
| Snx1 | Rps19 | Rpl37a | Rpl37a |
| Spata6 | Rps27 | Rpl5 | Rpl5 |
| Spnb3 | Rrbp1 | Rplp2 | Rplp2 |
| Stx17 | S100a16 | Rps10 | Rps10 |
| Syne2 | Sbds | Rps13 | Rps13 |
| Tcp1 | Scarb2 | Rps19 | Rps19 |
| Tcte1 | Sco1 | Rps27 | Rps27 |
| Tpm3 | Sdf2l1 | Rrbp1 | Rrbp1 |
| Tpmt | Sdhc | S100a16 | S100a16 |
| Tpp2 | Sdhd | Sbds | Sbds |
| Tram1 | Sec23ip | Scarb2 | Scarb2 |
| Trappc3 | Slc12a9 | Sco1 | Sco1 |
| Trappc4 | Slc16a1 | Sdf2l1 | Sdf2l1 |
| Tsta3 | Slc2a5 | Sdhc | Sdhc |
| Txn1 | Slc35f2 | Sdhd | Sdhd |
| Ube2d3 | Slco4c1 | Sec23ip | Sec23ip |
| Ube2j1 | Snx1 | Slc12a9 | Slc12a9 |
| Ube2n | Spata6 | Slc16a1 | Slc16a1 |
| Usp7 | Spg7 | Slc2a5 | Slc2a5 |
| Usp9x | Spnb3 | Slc35f2 | Slc35f2 |
| Vamp3 | Stx17 | Slco4c1 | Slco4c1 |
| Vars2 | Syne2 | Snx1 | Snx1 |
| Vcl | Tcp1 | Spata6 | Spata6 |
| Vim | Tcte1 | Spg7 | Spnb3 |
| Vkorc1 | Tpm3 | Spnb3 | Stx17 |
| Wrb | Tpmt | Stx17 | Syne2 |
| Xpo7 | Tpp2 | Syne2 | Tcp1 |
| Ywhab | Tram1 | Tcp1 | Tcte1 |
| Zfp185 | Trappc3 | Tcte1 | Tpm3 |
| 1600014C10Rik | Trappc4 | Tpm3 | Tpmt |
| 1700009P17Rik | Tsta3 | Tpmt | Tpp2 |
| 1700012A03Rik | Txn1 | Tpp2 | Tram1 |
| 1700034H14Rik | Ube2d3 | Tram1 | Trappc3 |
| 2300009A05Rik | Ube2j1 | Trappc3 | Trappc4 |
| 4921507P07Rik | Ube2n | Trappc4 | Tsta3 |
| 4921530D09Rik | Usp7 | Tsta3 | Txn1 |
| 4930415O20Rik | Usp9x | Txn1 | Ube2d3 |

| | | | |
|---|---|---|---|
| 4930471M23Rik | Vamp3 | Ube2d3 | Ube2j1 |
| 5730469M10Rik | Vars2 | Ube2j1 | Ube2n |
| 9430016H08Rik | Vcl | Ube2n | Usp7 |
| AI314180 | Vim | Usp7 | Usp9x |
| Abca3 | Vkorc1 | Usp9x | Vamp3 |
| Abcc3 | Wbscr16 | Vamp3 | Vars2 |
| Acat2 | Wrb | Vars2 | Vcl |
| Acbd5 | Xpo7 | Vcl | Vim |
| Aco1 | Ywhab | Vim | Vkorc1 |
| Acp1 | Zfp185 | Vkorc1 | Wbscr16 |
| Actn1 | 1600014C10Rik | Wrb | Wrb |
| Actr3 | 1700009P17Rik | Xpo7 | Xpo7 |
| Acyp1 | 1700012A03Rik | Ywhab | Ywhab |
| Adck5 | 1700034H14Rik | Zfp185 | Zfp185 |
| Adh5 | 2300009A05Rik | 1600014C10Rik | |
| Akap4 | 4921507P07Rik | 1700009P17Rik | |
| Anp32b | 4921530D09Rik | 1700012A03Rik | |
| Ap2a1 | 4930415O20Rik | 1700034H14Rik | |
| Ap2s1 | 4930471M23Rik | 2300009A05Rik | |
| Aprt | 5730469M10Rik | 4732415M23Rik | |
| Arl3 | 8430406I07Rik | 4921507P07Rik | |
| Armc1 | 9430016H08Rik | 4921530D09Rik | |
| Arpc4 | AI314180 | 4930415O20Rik | |
| Arpc5l | Abca3 | 4930471M23Rik | |
| Atp11a | Abcc3 | 5730469M10Rik | |
| Atp13a4 | Acat2 | 8430406I07Rik | |
| Atp6v0c | Acbd5 | 9430016H08Rik | |
| BC051142 | Aco1 | AI314180 | |
| Bloc1s1 | Acp1 | Abca3 | |
| Bsg | Actn1 | Abcc3 | |
| C030006K11Rik | Actr3 | Acat2 | |
| Calm3 | Acyp1 | Acbd5 | |
| Cap1 | Adck5 | Aco1 | |
| Capza1 | Adh5 | Acp1 | |
| Car2 | Akap4 | Actn1 | |
| Cbr3 | Anp32b | Actr3 | |
| Cct7 | Ap2a1 | Acyp1 | |
| Cct8 | Ap2s1 | Adck5 | |
| Chchd1 | Aprt | Adh5 | |
| Ckmt1 | Arl3 | Akap4 | |
| Clca2 | Armc1 | Anp32b | |
| Clic3 | Arpc4 | Anxa13 | |
| Clic4 | Arpc5l | Ap2a1 | |
| Coasy | Atp11a | Ap2s1 | |
| Cops4 | Atp13a4 | Aprt | |
| Cops8 | Atp6v0c | Arl3 | |
| Cox6b2 | BC051142 | Armc1 | |
| Crisp2 | Bloc1s1 | Arpc4 | |

| | | |
|---|---|---|
| Crot | Bsg | Arpc5l |
| Csnk2a1 | C030006K11Rik | Atp11a |
| Ctsf | Calm3 | Atp13a4 |
| Cutc | Cap1 | Atp6v0c |
| Cyfip1 | Capza1 | BC051142 |
| Ddb1 | Car2 | Bloc1s1 |
| Ddt | Cbr3 | Bsg |
| Defb2 | Cct7 | C030006K11Rik |
| Dnaja4 | Cct8 | Cabyr |
| Dnajb6 | Chchd1 | Calm3 |
| Dnm2 | Ckmt1 | Cap1 |
| Dock7 | Clca2 | Capza1 |
| Dpysl2 | Clic3 | Car2 |
| Dtymk | Clic4 | Cbr3 |
| Dusp3 | Coasy | Cct7 |
| Ebp | Cops4 | Cct8 |
| Eef1d | Cops8 | Chchd1 |
| Eef1g | Cox6b2 | Ckmt1 |
| Ehd2 | Crisp2 | Clca2 |
| Eif2s1 | Crot | Clic3 |
| Eif5a | Csnk2a1 | Clic4 |
| Endog | Ctsf | Coasy |
| Eprs | Cyfip1 | Cops4 |
| Erbb2 | Cyp17a1 | Cops8 |
| Fdx1 | Ddb1 | Cox6b2 |
| Fhl1 | Ddt | Crisp2 |
| Fhl4 | Defb11 | Crot |
| Fmo2 | Defb19 | Csnk2a1 |
| Fundc1 | Defb2 | Ctsf |
| Fxc1 | Dnaja4 | Cyfip1 |
| Gaa | Dnajb6 | Ddb1 |
| Galntl4 | Dnm2 | Ddt |
| Gba | Dock7 | Defb11 |
| Gclm | Dpysl2 | Defb2 |
| Ghitm | Dtymk | Dnajb6 |
| Gm2a | Dusp3 | Dnm2 |
| Gpi1 | Ebp | Dock7 |
| Gpnmb | Eef1d | Dpysl2 |
| Gpx1 | Eef1g | Dtymk |
| Gpx3 | Ehd2 | Dusp3 |
| Gpx4 | Eif2s1 | Ebp |
| Gpx5 | Eif5a | Eef1d |
| Gsr | Eprs | Eef1g |
| Gstm1 | Erbb2 | Ehd2 |
| Gstm2 | Fdx1 | Eif2s1 |
| Gtl3 | Fhl1 | Eif5a |
| Hdhd2 | Fhl4 | Eprs |
| Hint1 | Fmo2 | Erbb2 |

| | | |
|---|---|---|
| Hrsp12 | Fundc1 | Fdx1 |
| Hsd3b7 | Fxc1 | Fhl1 |
| Htatip2 | Gaa | Fhl4 |
| Idi1 | Galntl4 | Fmo2 |
| Immp2l | Gba | Fundc1 |
| Ipo7 | Gclm | Fxc1 |
| Isg20 | Ghitm | Fxn |
| Itpr1 | Gm2a | Gaa |
| Itpr3 | Gpi1 | Galntl4 |
| Klhl10 | Gpnmb | Gba |
| Lactb | Gpx1 | Gclm |
| Lamb2 | Gpx3 | Ghitm |
| Lamc1 | Gpx4 | Gm2a |
| Lamp2 | Gpx5 | Gpi1 |
| Lancl1 | Gsr | Gpnmb |
| Lancl2 | Gstm1 | Gpx1 |
| Lars | Gstm2 | Gpx3 |
| Lass2 | Gtl3 | Gpx4 |
| Lcn12 | Hdhd2 | Gpx5 |
| Lcn8 | Hint1 | Gsr |
| Lgals1 | Hrsp12 | Gstm1 |
| Lipe | Hsd3b7 | Gstm2 |
| Lipg | Htatip2 | Gtl3 |
| Ly6a | Idi1 | Hdhd2 |
| Lypla1 | Ipo7 | Hint1 |
| Man2a2 | Isg20 | Hrsp12 |
| Manba | Itpr1 | Hsd3b7 |
| Mdk | Itpr3 | Htatip2 |
| Mipep | Klhl10 | Idi1 |
| Mlf1 | Lamc1 | Ipo7 |
| Mrpl11 | Lamp2 | Isg20 |
| Mrpl17 | Lancl1 | Itpr1 |
| Mrpl18 | Lancl2 | Itpr3 |
| Mrpl43 | Lars | Klhl10 |
| Mrpl48 | Lass2 | Lactb |
| Mrpl52 | Lcn12 | Lamc1 |
| Mrpl55 | Lcn8 | Lamp2 |
| Mrps10 | Lgals1 | Lancl1 |
| Mrps14 | Lipe | Lancl2 |
| Mrps16 | Lipg | Lars |
| Mrps18a | Ly6a | Lass2 |
| Mrps18b | Lypla1 | Lcn12 |
| Mrps21 | Man2a2 | Lcn8 |
| Mrps24 | Manba | Lgals1 |
| Mrps31 | Mdk | Lipe |
| Mrps36 | Mipep | Lipg |
| Mrps6 | Mlf1 | Lpp |
| Mrrf | Mrpl11 | Ly6a |

| | | |
|---|---|---|
| Mtpn | Mrpl17 | Lypla1 |
| Ncam1 | Mrpl18 | Man2a2 |
| Ndufc2 | Mrpl43 | Manba |
| Ndufs6 | Mrpl48 | Mdk |
| Ndufv2 | Mrpl52 | Mipep |
| Nif3l1 | Mrpl55 | Mlf1 |
| Nit1 | Mrps10 | Mrpl11 |
| Nqo1 | Mrps14 | Mrpl17 |
| Oplah | Mrps16 | Mrpl18 |
| Pcbp1 | Mrps18a | Mrpl43 |
| Pex11c | Mrps18b | Mrpl48 |
| Pfkm | Mrps21 | Mrpl52 |
| Pgk2 | Mrps24 | Mrpl55 |
| Pgls | Mrps31 | Mrps10 |
| Phb | Mrps36 | Mrps14 |
| Phpt1 | Mrps6 | Mrps16 |
| Pmm2 | Mrrf | Mrps18a |
| Ppap2a | Mtpn | Mrps18b |
| Ppil1 | Ncam1 | Mrps21 |
| Ppp2r1b | Ndufc2 | Mrps24 |
| Ppp2r4 | Ndufs6 | Mrps31 |
| Ppp6c | Ndufv2 | Mrps36 |
| Prkacb | Nif3l1 | Mrps6 |
| Psen1 | Nit1 | Mrrf |
| Psma1 | Nqo1 | Mtpn |
| Psma2 | Oplah | Ncam1 |
| Psma6 | Pcbp1 | Ndufc2 |
| Psma7 | Pex11c | Ndufs6 |
| Psmb1 | Pfkm | Ndufv2 |
| Psmb5 | Pgk2 | Nif3l1 |
| Psmb6 | Pgls | Nit1 |
| Psmd7 | Phb | Nqo1 |
| Psme1 | Phpt1 | Oplah |
| Psme4 | Pmm2 | Pcbp1 |
| Ptcd2 | Ppap2a | Pex11c |
| Ptges | Ppil1 | Pfkm |
| Ptp4a2 | Ppp2r1b | Pgk2 |
| Rab24 | Ppp2r4 | Pgls |
| Rab28 | Ppp6c | Phb |
| Rab2b | Prkacb | Phpt1 |
| Rab5b | Psen1 | Pmm2 |
| Rac2 | Psma1 | Ppap2a |
| Rdh13 | Psma2 | Ppil1 |
| Recql | Psma6 | Ppp2r1b |
| Rpia | Psma7 | Ppp2r4 |
| Rpl37a | Psmb1 | Ppp6c |
| Rpl5 | Psmb5 | Prkacb |
| Rplp2 | Psmb6 | Psen1 |

| | | |
|---|---|---|
| Rps10 | Psmd7 | Psma1 |
| Rps13 | Psme1 | Psma2 |
| Rps19 | Psme4 | Psma6 |
| Rps27 | Ptcd2 | Psma7 |
| Rrbp1 | Ptges | Psmb1 |
| S100a16 | Ptp4a2 | Psmb5 |
| Sbds | Rab24 | Psmb6 |
| Scarb2 | Rab28 | Psmd7 |
| Sco1 | Rab2b | Psme1 |
| Sdf2l1 | Rab5b | Psme4 |
| Sdhc | Rac2 | Ptcd2 |
| Sdhd | Rdh13 | Ptges |
| Sec23ip | Recql | Ptp4a2 |
| Slc12a9 | Rpia | Rab24 |
| Slc16a1 | Rpl37a | Rab28 |
| Slc2a5 | Rpl5 | Rab2b |
| Slc35f2 | Rplp2 | Rab5b |
| Slco4c1 | Rps10 | Rac2 |
| Snx1 | Rps13 | Rdh13 |
| Spata6 | Rps19 | Recql |
| Spg7 | Rps27 | Ropn1l |
| Spnb3 | Rrbp1 | Rpia |
| Stx17 | S100a16 | Rpl37a |
| Syne2 | Sbds | Rpl5 |
| Tcp1 | Scarb2 | Rplp2 |
| Tcte1 | Sco1 | Rps10 |
| Tpm3 | Sdf2l1 | Rps13 |
| Tpmt | Sdhc | Rps19 |
| Tpp2 | Sdhd | Rps27 |
| Tram1 | Sec23ip | Rrbp1 |
| Trappc3 | Slc12a9 | S100a16 |
| Trappc4 | Slc16a1 | Sbds |
| Tsta3 | Slc2a5 | Scarb2 |
| Txn1 | Slc35f2 | Sco1 |
| Ube2d3 | Slco4c1 | Sdf2l1 |
| Ube2j1 | Snx1 | Sdhc |
| Ube2n | Spata6 | Sdhd |
| Usp7 | Spnb3 | Sec23ip |
| Usp9x | Stx17 | Slc12a9 |
| Vamp3 | Syne2 | Slc16a1 |
| Vars2 | Tcp1 | Slc2a5 |
| Vcl | Tcte1 | Slc35f2 |
| Vim | Tpm3 | Slco4c1 |
| Vkorc1 | Tpmt | Snx1 |
| Wbscr16 | Tpp2 | Spata6 |
| Wrb | Tram1 | Spg7 |
| Xpo7 | Trappc3 | Spnb3 |
| Ywhab | Trappc4 | Stx17 |

| | | |
|---|---|---|
| Zfp185 | Tsta3 | Syne2 |
| 1600014C10Rik | Txn1 | Tcp1 |
| 1700009P17Rik | Ube2d3 | Tcte1 |
| 1700012A03Rik | Ube2j1 | Tpm3 |
| 1700034H14Rik | Ube2n | Tpmt |
| 2300009A05Rik | Usp7 | Tpp2 |
| 4921507P07Rik | Usp9x | Tram1 |
| 4921530D09Rik | Vamp3 | Trappc3 |
| 4930415O20Rik | Vars2 | Trappc4 |
| 4930471M23Rik | Vcl | Tsta3 |
| 5730469M10Rik | Vim | Txn1 |
| 9430016H08Rik | Vkorc1 | Ube2d3 |
| AI314180 | Wbscr16 | Ube2j1 |
| Abca3 | Wrb | Ube2n |
| Abcc3 | Xpo7 | Usp7 |
| Acat2 | Ywhab | Usp9x |
| Acbd5 | Zfp185 | Vamp3 |
| Aco1 | 1600014C10Rik | Vars2 |
| Acp1 | 1700009P17Rik | Vcl |
| Actn1 | 1700012A03Rik | Vim |
| Actr3 | 1700034H14Rik | Vkorc1 |
| Acyp1 | 2300009A05Rik | Wrb |
| Adck5 | 4921507P07Rik | Xpo7 |
| Adh5 | 4921530D09Rik | Ywhab |
| Akap4 | 4930415O20Rik | Zfp185 |
| Anp32b | 4930471M23Rik | 1600014C10Rik |
| Ap2a1 | 5730469M10Rik | 1700009P17Rik |
| Ap2s1 | 8430406I07Rik | 1700012A03Rik |
| Aprt | 9430016H08Rik | 1700034H14Rik |
| Arl3 | AI314180 | 2300009A05Rik |
| Armc1 | Abca3 | 4921507P07Rik |
| Arpc4 | Abcc3 | 4921530D09Rik |
| Arpc5l | Acat2 | 4930415O20Rik |
| Atp11a | Acbd5 | 4930471M23Rik |
| Atp13a4 | Aco1 | 5730469M10Rik |
| Atp6v0c | Acp1 | 8430406I07Rik |
| BC051142 | Actn1 | 9430016H08Rik |
| Bloc1s1 | Actr3 | AI314180 |
| Bsg | Acyp1 | Abca3 |
| C030006K11Rik | Adck5 | Abcc3 |
| Calm3 | Adh5 | Acat2 |
| Cap1 | Akap4 | Acbd5 |
| Capza1 | Anp32b | Aco1 |
| Car2 | Ap2a1 | Acp1 |
| Cbr3 | Ap2s1 | Actn1 |
| Cct7 | Aprt | Actr3 |
| Cct8 | Arl3 | Acyp1 |
| Chchd1 | Armc1 | Adck5 |

| | | |
|---|---|---|
| Ckmt1 | Arpc4 | Adh5 |
| Clca2 | Arpc5l | Akap4 |
| Clic3 | Atp11a | Anp32b |
| Clic4 | Atp13a4 | Anxa13 |
| Coasy | Atp6v0c | Ap2a1 |
| Cops4 | BC051142 | Ap2s1 |
| Cops8 | Bloc1s1 | Aprt |
| Cox6b2 | Bsg | Arl3 |
| Crisp2 | C030006K11Rik | Armc1 |
| Crot | Calm3 | Arpc4 |
| Csnk2a1 | Cap1 | Arpc5l |
| Ctsf | Capza1 | Atp11a |
| Cutc | Car2 | Atp13a4 |
| Cyfip1 | Car4 | Atp6v0c |
| Ddb1 | Cbr3 | BC051142 |
| Ddt | Cct7 | Bloc1s1 |
| Defb2 | Cct8 | Bsg |
| Dnaja4 | Chchd1 | C030006K11Rik |
| Dnajb6 | Ckmt1 | Cabyr |
| Dnm2 | Clca2 | Calm3 |
| Dock7 | Clic3 | Cap1 |
| Dpysl2 | Clic4 | Capza1 |
| Dtymk | Coasy | Car2 |
| Dusp3 | Cops4 | Cbr3 |
| Ebp | Cops8 | Cct7 |
| Eef1d | Cox6b2 | Cct8 |
| Eef1g | Crisp2 | Chchd1 |
| Ehd2 | Crot | Ckmt1 |
| Eif2s1 | Csnk2a1 | Clca2 |
| Eif5a | Ctsf | Clic3 |
| Endog | Cyfip1 | Clic4 |
| Eprs | Cyp17a1 | Coasy |
| Erbb2 | Ddb1 | Cops4 |
| Fdx1 | Ddt | Cops8 |
| Fhl1 | Defb11 | Cox6b2 |
| Fhl4 | Defb19 | Crisp2 |
| Fmo2 | Defb2 | Crot |
| Fundc1 | Dnaja4 | Csnk2a1 |
| Fxc1 | Dnajb6 | Ctsf |
| Gaa | Dnm2 | Cutc |
| Galntl4 | Dock7 | Cyfip1 |
| Gba | Dpysl2 | Ddb1 |
| Gclm | Dtymk | Ddt |
| Ghitm | Dusp3 | Defb11 |
| Gm2a | Ebp | Defb2 |
| Gpi1 | Eef1d | Dnajb6 |
| Gpnmb | Eef1g | Dnm2 |
| Gpx1 | Ehd2 | Dock7 |

| | | |
|---|---|---|
| Gpx3 | Eif2s1 | Dpysl2 |
| Gpx4 | Eif5a | Dtymk |
| Gpx5 | Eno1 | Dusp3 |
| Gsr | Eprs | Ebp |
| Gstm1 | Erbb2 | Eef1d |
| Gstm2 | Fdx1 | Eef1g |
| Gtl3 | Fhl1 | Ehd2 |
| Hdhd2 | Fhl4 | Eif2s1 |
| Hint1 | Fmo2 | Eif5a |
| Hrsp12 | Fundc1 | Eprs |
| Hsd3b7 | Fxc1 | Erbb2 |
| Htatip2 | Gaa | Fdx1 |
| Idi1 | Galntl4 | Fhl1 |
| Immp2l | Gba | Fhl4 |
| Ipo7 | Gclm | Fmo2 |
| Isg20 | Ghitm | Fundc1 |
| Itpr1 | Gm2a | Fxc1 |
| Itpr3 | Gpi1 | Fxn |
| Klhl10 | Gpnmb | Gaa |
| Lactb | Gpx1 | Galntl4 |
| Lamb2 | Gpx3 | Gba |
| Lamc1 | Gpx4 | Gclm |
| Lamp2 | Gpx5 | Ghitm |
| Lancl1 | Gsr | Gm2a |
| Lancl2 | Gstm1 | Gpi1 |
| Lars | Gstm2 | Gpnmb |
| Lass2 | Gtl3 | Gpx1 |
| Lcn12 | Hdhd2 | Gpx3 |
| Lcn8 | Hint1 | Gpx4 |
| Lgals1 | Hrsp12 | Gpx5 |
| Lipe | Hsd3b7 | Gsr |
| Lipg | Htatip2 | Gstm1 |
| Ly6a | Idi1 | Gstm2 |
| Lypla1 | Ipo7 | Gtl3 |
| Man2a2 | Isg20 | Hdhd2 |
| Manba | Itpr1 | Hint1 |
| Mdk | Itpr3 | Hrsp12 |
| Mipep | Klhl10 | Hsd3b7 |
| Mlf1 | Lamc1 | Htatip2 |
| Mrpl11 | Lamp2 | Idi1 |
| Mrpl17 | Lancl1 | Ipo7 |
| Mrpl18 | Lancl2 | Isg20 |
| Mrpl43 | Lars | Itpr1 |
| Mrpl48 | Lass2 | Itpr3 |
| Mrpl52 | Lcn12 | Klhl10 |
| Mrpl55 | Lcn8 | Lactb |
| Mrps10 | Lgals1 | Lamb2 |
| Mrps14 | Lipe | Lamc1 |

| | | |
|---|---|---|
| Mrps16 | Lipg | Lamp2 |
| Mrps18a | Ly6a | Lancl1 |
| Mrps18b | Lypla1 | Lancl2 |
| Mrps21 | Man2a2 | Lars |
| Mrps24 | Manba | Lass2 |
| Mrps31 | Mdk | Lcn12 |
| Mrps36 | Mipep | Lcn8 |
| Mrps6 | Mlf1 | Lgals1 |
| Mrrf | Mrpl11 | Lipe |
| Mtpn | Mrpl17 | Lipg |
| Ncam1 | Mrpl18 | Lpp |
| Ndufc2 | Mrpl43 | Ly6a |
| Ndufs6 | Mrpl48 | Lypla1 |
| Ndufv2 | Mrpl52 | Man2a2 |
| Nif3l1 | Mrpl55 | Manba |
| Nit1 | Mrps10 | Mdk |
| Nme5 | Mrps14 | Mipep |
| Nqo1 | Mrps16 | Mrpl11 |
| Oplah | Mrps18a | Mrpl17 |
| Pcbp1 | Mrps18b | Mrpl18 |
| Pex11c | Mrps21 | Mrpl43 |
| Pfkm | Mrps24 | Mrpl48 |
| Pgk2 | Mrps31 | Mrpl52 |
| Pgls | Mrps36 | Mrpl55 |
| Phb | Mrps6 | Mrps10 |
| Phpt1 | Mrrf | Mrps14 |
| Pmm2 | Mtpn | Mrps16 |
| Ppap2a | Ncam1 | Mrps18a |
| Ppil1 | Ndufc2 | Mrps18b |
| Ppp2r1b | Ndufs6 | Mrps21 |
| Ppp2r4 | Ndufv2 | Mrps24 |
| Ppp6c | Nif3l1 | Mrps31 |
| Prkacb | Nit1 | Mrps36 |
| Psen1 | Nqo1 | Mrps6 |
| Psma1 | Pcbp1 | Mrrf |
| Psma2 | Pex11c | Mtpn |
| Psma6 | Pfkm | Ncam1 |
| Psma7 | Pgk2 | Ndufc2 |
| Psmb1 | Pgls | Ndufs6 |
| Psmb5 | Phb | Ndufv2 |
| Psmb6 | Phpt1 | Nif3l1 |
| Psmd7 | Pmm2 | Nit1 |
| Psme1 | Ppap2a | Nqo1 |
| Psme4 | Ppil1 | Oplah |
| Ptcd2 | Ppp2r1b | Pcbp1 |
| Ptges | Ppp2r4 | Pex11c |
| Ptp4a2 | Ppp6c | Pfkm |
| Rab24 | Prkacb | Pgk2 |

| | | |
|---|---|---|
| Rab28 | Psen1 | Pgls |
| Rab2b | Psma1 | Phb |
| Rab5b | Psma2 | Phpt1 |
| Rac2 | Psma6 | Pmm2 |
| Rdh13 | Psma7 | Ppap2a |
| Recql | Psmb1 | Ppil1 |
| Rpia | Psmb5 | Ppp2r1b |
| Rpl37a | Psmb6 | Ppp2r4 |
| Rpl5 | Psmd7 | Ppp6c |
| Rplp2 | Psme1 | Prkacb |
| Rps10 | Psme4 | Psen1 |
| Rps13 | Ptcd2 | Psma1 |
| Rps19 | Ptges | Psma2 |
| Rps27 | Ptp4a2 | Psma6 |
| Rrbp1 | Rab24 | Psma7 |
| S100a16 | Rab28 | Psmb1 |
| Sbds | Rab2b | Psmb5 |
| Scarb2 | Rab5b | Psmb6 |
| Sco1 | Rac2 | Psmd7 |
| Sdf2l1 | Rdh13 | Psme1 |
| Sdhc | Recql | Psme4 |
| Sdhd | Rpia | Ptcd2 |
| Sec23ip | Rpl37a | Ptges |
| Slc12a9 | Rpl5 | Ptp4a2 |
| Slc16a1 | Rplp2 | Rab24 |
| Slc2a5 | Rps10 | Rab28 |
| Slc35f2 | Rps13 | Rab2b |
| Slco4c1 | Rps19 | Rab5b |
| Snx1 | Rps27 | Rac2 |
| Spata6 | Rrbp1 | Rdh13 |
| Spg7 | S100a16 | Recql |
| Spnb3 | Sbds | Ribc1 |
| Stx17 | Scarb2 | Ropn1l |
| Syne2 | Sco1 | Rpia |
| Tcp1 | Sdf2l1 | Rpl37a |
| Tcte1 | Sdhc | Rpl5 |
| Tpm3 | Sdhd | Rplp2 |
| Tpmt | Sec23ip | Rps10 |
| Tpp2 | Slc12a9 | Rps13 |
| Tram1 | Slc16a1 | Rps19 |
| Trappc3 | Slc2a5 | Rps27 |
| Trappc4 | Slc35f2 | Rrbp1 |
| Tsta3 | Slco4c1 | S100a16 |
| Txn1 | Snx1 | Sbds |
| Ube2d3 | Spata6 | Scarb2 |
| Ube2j1 | Spg7 | Sco1 |
| Ube2n | Spnb3 | Sdf2l1 |
| Usp7 | Stx17 | Sdhc |

Usp9x
Vamp3
Vars2
Vcl
Vim
Vkorc1
Wbscr16
Wrb
Xpo7
Ywhab
Zfp185

Syne2
Tcp1
Tcte1
Tpm3
Tpmt
Tpp2
Tram1
Trappc3
Trappc4
Tsta3
Txn1
Ube2d3
Ube2j1
Ube2n
Usp7
Usp9x
Vamp3
Vars2
Vcl
Vim
Vkorc1
Wbscr16
Wrb
Xpo7
Ywhab
Zfp185

Sdhd
Sec23ip
Slc12a9
Slc16a1
Slc2a5
Slc35f2
Slco4c1
Snx1
Spata6
Spg7
Spnb3
Stx17
Syne2
Tcp1
Tcte1
Tpm3
Tpmt
Tpp2
Tram1
Trappc3
Trappc4
Tsta3
Txn1
Ube2d3
Ube2j1
Ube2n
Usp7
Usp9x
Vamp3
Vars2
Vcl
Vim
Vkorc1
Wbscr16
Wrb
Xpo7
Ywhab
Zfp185
